# Protein Interaction Patterns in *Arabidopsis thaliana* Leaf Mitochondria Change in Response to Illumination

**DOI:** 10.1101/2020.07.30.222745

**Authors:** Nils Rugen, Frank Schaarschmidt, Hans-Peter Braun, Holger Eubel

## Abstract

Mitochondrial biology is underpinned by the presence and activity of large protein complexes participating in the organelle-located parts of cellular respiration, the TCA cycle and oxidative phosphorylation. While the enzymatic roles of these complexes are undisputed, little is known about the interactions of the subunits beyond their presence in the monomeric protein complexes and their functions in regulating mitochondria metabolism. By applying one of the most important regulatory cues for plant metabolism, the presence or absence of light, we here assess the changes in the composition and molecular mass of known mitochondrial protein complexes by employing a differential complexome profiling strategy. Covering a mass range up to 25 MDa, we demonstrate dynamic associations of TCA-cycle enzymes and of OXPHOS components. The data presented here form the basis for future studies aiming to advance our understanding of the role of protein:protein interactions in the regulation of plant mitochondrial functions.

## Introduction

Plant mitochondria produce ATP to support metabolic and transport processes throughout the cell. For this, they rely on two essential metabolic pathways, the TCA-cycle (located in the mitochondrial matrix) and oxidative phosphorylation (OXPHOS, taking place within the inner mitochondrial membrane). In plants, flux through these two pathways responds to photosynthetic activity. Electrons fed into the respiratory chain are mostly supplied by reduction equivalents produced by the stepwise oxidation of the dicarboxylates pyruvate and malate in the TCA-cycle. During the day, however, this pathway only partially operates in the cyclic mode since its early intermediates, citrate and 2-oxoglutarate, are exported from mitochondria to serve as substrates for ammonium assimilation in chloroplasts. The reduced rate of NADH-production by the TCA cycle is compensated or even over-compensated by the oxidation of glycine in the mitochondrial matrix as part of the photorespiratory (C2) pathway. These electrons are injected into the respiratory chain of mitochondria using either the ubiquitous respiratory enzymes NADH dehydrogenase complex (complex I) reductase,) or the alternative enzymes NDA/B and alternative oxidase (AOX) and are transferred onto molecular oxygen by the components of the cytochrome c pathway (complex III; cytochrome c oxidase, complex IV) or the alternative oxidase (AOX).

Flux through the TCA cycle and the respiratory chain needs to be regulated tightly to prevent over-reduction of the NAD pool and the pools of the mobile respiratory electron transporter ubiquinone and cytochrome c, which would lead to an increased production of reactive oxygen species (ROS) damaging a wide spectrum of cellular components, including DNA, lipids, and proteins. As such, the activities of TCA-cycle and respiratory enzymes need to be synchronized with the prevailing light conditions to reconcile their sometimes opposing roles in ATP production and photorespiration, as well as in supporting photosynthesis (Braun 2020). Considering the costs associated with a periodic production and degradation of proteins, changes in protein abundance unsurprisingly do not seem to play an important role in regulating mitochondrial energy metabolism (Lee et al. 2010). Instead, several other mechanisms regulating TCA cycle, OXPHOS, and alternative respiratory enzyme functions have been described. Particularly the TCA-cycle is constrained by a network of regulatory events. Entry of carbon (in the form of pyruvate) to the TCA-cycle by the pyruvate dehydrogenase complex (PDC) is controlled by phosphorylation/dephosphorylation of the E1α-subunit (Budde und Randall 1987). Acetylation of lysine residues is another, experimentally verified post-translational modification (PTM) taking place in plant mitochondria. Acetyl-moieties attached to the ∊-amino group of lysine residues were discovered in enzymes of the OXPHOS system and in matrix proteins, also including TCA-cycle enzymes (König et al. 2014a). For many of these proteins a possible impact of the acetylation event on enzymatic properties awaits confirmation. However, the activity of the ATP-synthase complex reacts to a loss-of-function mutant for a SRT2, a mitochondria located sirtuin deacetylase (König et al. 2014b), thus strongly suggesting similar responses for other enzymes with confirmed acetylation sites. TCA cycle proteins are also affected by the NAD^+^/NADH ratio as well as ADP and ATP concentrations (Tovar-Méndez et al. 2003; Oestreicher et al. 1973; Igamberdiev und Gardeström 2003; Affourtit et al. 2001; Bunik und Strumilo 2009). They are also subject to allosteric downstream metabolite inhibition or upstream metabolite activation (Nunes-Nesi et al. 2013; Tronconi et al. 2010; Studart-Guimarães et al. 2005). In addition, second messengers such as calcium and reactive oxygen species (ROS) affect the activity of TCA enzymes (Tovar-Méndez et al. 2003; Sweetlove et al. 2002, 2002; Morgan et al. 2008). Redox regulation, either directly or indirectly, is expected to have a big impact on the regulation of mitochondrial functions (Møller et al. 2020; Daloso et al. 2015). Recently, the protein kinase target of rapamycin (TOR) has also been suggested to play a role in regulating respiratory activity but the mechanisms by which this is achieved still await discovery (O’Leary et al. 2019).

The regulation of OXPHOS enzymes in plant mitochondria is even less well understood. Calcium, inorganic phosphate (P_i_), as well as ADP/ATP ratio seem to influence respiration (Vinnakota et al. 2016; Gellerich et al. 2010) but most likely, these parameters rather regulate the provision of reducing equivalents than having an impact on OXOHOS enzymes themselves. In mammals, several OXPHOS targets for PTMs, such as phosphorylation, acetylation, succinylation, ubiquitination, and redox signals have already been identified (Stram und Payne 2016; Hofer und Wenz 2014). For plant mitochondria, however, data on PTMs of OXPHOS complexes focus on acetylation (König et al. 2014a; Møller et al. 2020) since phosphorylation of plant mitochondrial proteins seems to be a scarce event (Ito et al. 2009; Reiland et al. 2009). In the branched respiratory chain of plants, alternative NADH dehydrogenases and AOX possess important functions under stress. The alternative NADH dehydrogenases are regulated by calcium concentrations as well as pH (Rasmusson et al. 2008). AOX is regulated by redox events in which oxidation of a cysteine residue near the N-terminus of the protein induces inactivation by dimerization. This process is reversed by thioredoxin-mediated reduction of the cysteine residue (Gelhaye et al. 2004).

It is widely accepted that mitochondrial proteins interact to form, for example, the large dehydrogenase complexes in the mitochondrial matrix and the OXPHOS protein (super)complexes in the inner mitochondrial membrane, and that the enzymatic functions provided by these complexes are only accomplished by the protein:protein interactions (PPIs) taking place among the subunits. It is therefore even more surprising that a comprehensive overview of the plant mitochondrial PPIs is still missing and that the role of these PPIs in regulation plant mitochondrial metabolism remains unexplored. Recently, differential complexome profiling analysis of human mitochondria from patients suffering from Barth and Leigh syndrome has revealed altered abundance of inner membrane or inner membrane associated protein complexes (van Strien et al. 2019) and complex I assembly intermediates (Alston et al. 2020), respectively, indicating that dynamic PPIs have an impact on mitochondrial metabolism.

Complexome profiling has successfully been employed to characterize the general architecture of the plant mitochondrial proteome (Senkler et al. 2017) and, after considerably extending the upper molecular mass limit to ~30 MDa with large pore (lp) BN gels (Strecker et al. 2010), for elucidating the protein composition of Arabidopsis mitochondrial ribosomes (Rugen et al. 2019). Using a closely related complexome profiling strategy, we here assess changes in the PPI patterns of TCA-cycle and respiratory enzymes of mitochondria in response to a major cue for plant metabolism, the presence and absence of light. For this, mitochondria were isolated two hours before the end of the night (for the reminder of the manuscript referred to as ‘dark’ mitochondria) and two hours before the end of the day (for the reminder of the manuscript referred to as ‘light’ mitochondria). Results highlight the presence of TCA-cycle proteins in high molecular mass protein assemblies which respond to the presence or absence of light. Likewise, OXPHOS components, particularly the complexes I and III, as well as the ATP synthase complex also show altered abundance at different positions in the lpBN gel. Together, the data presented here are painting a picture of plant mitochondria as organelles with dynamic PPI patterns and provide new targets for future studies aiming to unravel the intricate regulatory network of this organelle.

## Results and discussion

### Assessment of the quality of isolated organelles and the complexome profiling workflow

Mitochondria from rosettes of five week old plants (from now on referred to as ‘isolates’) were produced in five replicates for both conditions (light, dark). From each freshly prepared isolate 125 μg of protein were solubilized in 2.5% digitonin, supplemented with Coomassie G250, and loaded onto lpBN gels (Suppl. Fig. S1). After separation, gel lanes were cut into 46 – 48 slices (Figs. S2, S3; from now on referred to as ‘fractions’), each of which would then be submitted to MS analysis for protein identification and quantitation using a label-free approach. The complexome profiling workflow was finalized by producing protein abundance profiles, which were then clustered hierarchically for each isolate using the NOVA software (Giese et al. 2015), and producing in a clustered protein abundance heatmap. An isolate processed this way will be referred to as ‘sample’ in the following.

Within the 469 MS/MS runs performed for this study, between 1102 (sample dark I) and 1304 (sample dark V) unique protein groups were identified for the dark samples, closely matched by 1019 (sample light V) and 1300 (sample light IV) unique protein groups for the light samples (Tab. 1). On average, fractions contain more than 350 unique protein groups, each of which being identified by an average of more than eight unique peptides. Most protein groupss are present in 13 to 15 fractions and less than eight percent of the identified protein groups are limited to a single fraction.

**Table 1:** Identification of protein groups in light and dark replicates.

| Light condition | dark |  |  |  |  | light |  |  |  |  |
| --- | --- | --- | --- | --- | --- | --- | --- | --- | --- | --- |
| sample | I | II | III | IV | V | I | II | III | IV | V |
| no. of prot. groups | 1102 | 1170 | 1178 | 1239 | 1304 | 1206 | 1055 | 1124 | 1300 | 1019 |
| no. of fractions | 47 | 46 | 46 | 47 | 47 | 48 | 48 | 46 | 47 | 47 |
| av. no. of prot. groups per fraction | 313 | 384 | 337 | 381 | 403 | 381 | 286 | 332 | 405 | 322 |
| prot. groups identif. in only one fraction | 82 | 65 | 91 | 76 | 87 | 81 | 66 | 73 | 88 | 67 |
| av. no. of fractions per prot. group | 13 | 15 | 13 | 14 | 15 | 14 | 13 | 14 | 15 | 15 |
| av. no. of unique pept. per prot. group | 8.5 | 9.5 | 8.9 | 10.0 | 9.4 | 9.2 | 9.0 | 8.9 | 8.6 | 9.1 |

Purity of the mitochondrial isolates was assessed by tallying the intensity Based Absolute Quantitation (iBAQ) values (Schwanhäusser et al. 2011) for each protein across all gel fractions. After querying the complete set of identified proteins against the SUBA4 database (Hooper et al. 2017) using the SUBAcon algorithm (Hooper et al. 2014) which assigns each protein to its most likely cellular compartment, the contribution of each of these compartments towards the total protein content of the isolates was calculated. Approximately 90% of the protein abundance in the isolates is of mitochondrial origin (88% – 92% in dark mitochondria, 86% to 92% in light mitochondria; Fig. S4). This value is in concordance with purities for mitochondria isolated from Arabidopsis leaves reported earlier (Senkler et al. 2017; Rugen et al. 2019).

### MS analysis reveals no alterations in protein abundance in ‘dark’ and ‘light’ mitochondria

Previously, diurnal variation of protein abundance in plant mitochondria was found to be limited (Lee et al. 2010). This follows the notion that a daily repeating synthesis and degradation of plant mitochondrial proteins is an energetically wasteful process. The MS data produced during this study lend themselves to a verification of the original data. To this end, MS raw data for all the gel fractions of each individual isolate were processed as a single sample, allowing a direct comparison of protein abundance in ‘dark’ and ‘light’ mitochondria by utilizing Label-Free Quantitation (LFQ) values (Cox et al. 2014). Within the 1562 proteins covered by this approach, only 49 were found to possess a p-value ≤0.05 (Tab. S1) but none of them passed multi-hypotheses testing by the permutation-based FDR correction (FDR ≤0.05). These results therefore largely match previous results by Lee at al. (2010) and clearly show, that the composition of the mitochondrial proteome in the light does not differ from that in the dark. Hence, protein abundance as a means to regulate mitochondrial functions can be largely excluded, at least for those proteins, which were amenable to MS detection by our approach. This opens the path for further research on the mechanisms regulating enzymatic activity in plant mitochondria, for example by elucidating their PPI patterns.

### Technical variation in the heatmaps requires alignment prior to further analysis

As reported previously (Rugen et al. 2019), cumulated intensities of mitochondrial proteins (based on iBAQ values) separated in lpBN gels drop sharply at molecular masses above 3 MDa to remain relatively constant at a low level. In contrast, the number of proteins identified in each fraction remains constant until approximately 7 MDa from where they slowly decline towards the upper end of the gel lane (Fig. S5). In terms of the number of identified proteins, variation within the two datasets is discernible but low across all fraction. In contrast, the cumulated iBAQ values display a higher degree of variation, particularly in the dark replicates between 0.4 MDa and 2 MDa (Fig. S5). This interval spans an area of closely spaced, highly abundant respiratory protein complexes, such as dimeric complex III (C-III_2_), monomeric ATP synthase (C-V_1_), monomeric complex I (C-I_1_) and a supercomplex made up by C-I_1_ and C-III_2_ (SC-I_1_III_2_). The resolution of the gel raster and technical variation between the individual samples in this hotspot of protein abundance are most likely to blame for the observed variation, also showcasing the need for heatmap alignment prior to further processing. The production of heatmaps based on lpBN gels is a manual process and liable to technical variation. Casting the gels and fractionating the soft upper parts of gel lanes are particular prone to introducing variance between replicates. Both effects also contribute to the unequal numbers of fractions obtained for the ten samples. In order to enable comparison of the dark and light heatmaps among and between each other, alignment is thus a critical and necessary step in the analytical procedure. For this, cumulated iBAQ values (Schwanhäusser et al. 2011) of each fraction were used as landmarks. To accommodate the different numbers of fractions for some of the gel lanes, the natural numbering system for the fractions had to be converted into a rational numbering system to enable alignment.

### A differential heatmap reveals differences in the abundance of mitochondrial protein complexes in response to changing light conditions

From the aligned dark and light heatmaps, average heatmaps for both conditions were produced (Fig. S6), in which log_10_ transformation of iBAQ values aids the visualization of low abundant protein complexes. Since the technical variation between the heatmaps could not be completely alleviated, the resolution of the average heatmaps was found to suffer in comparison with that of the primary heatmaps. Average heatmaps can, however, be used to build a differential heatmap of dark and light mitochondria (Fig. 1) in which only the (log_10_ transformed) differences between the dark and light mitochondria are shown. This heatmap looks different to the average heatmaps as some of the known protein complexes seem to be missing (indicating the absence of any change), while other potential complexes seem to appear (indicating abundance differences between dark and light mitochondria) at positions, in which no complex bands are discernible in the original heatmaps. A major problem associated with such heatmaps is the statistical verification. When viewed on a protein-by-protein basis, 152 proteins (in 900 positions) make the cut of p ≤ 0.05 (for student’s t-test at aligned positions; Tab. S2, Fig. S7), whereas another 230 proteins (in 374 positions) yield p-values of >0.05, ≤0.1. These numbers seem low when considering, that a total of 24322 positions entered the calculation. However, it can be anticipated that the subunits of a protein complex change uniformly at the position within the gel, representing the molecular mass of the holo-complex. Therefore, even in the absence of statistical significance for the individual subunits, a uniform change in protein abundance at such positions is a strong indicator for a *bona fide* change in complex abundance. We here mainly focus on investigating the changes of the (multi-) protein complexes and supercomplexes involved in mitochondrial ATP production, for which subunit compositions are known. These protein complexes include those of the TCA cycle, such as the 2-oxoglutarate dehydrogenase complex (OGDC) and the succinate dehydrogenase complex (SDH) as well as those of the mitochondrial electron transfer chain, and the ATP synthase.

**Figure 1:**
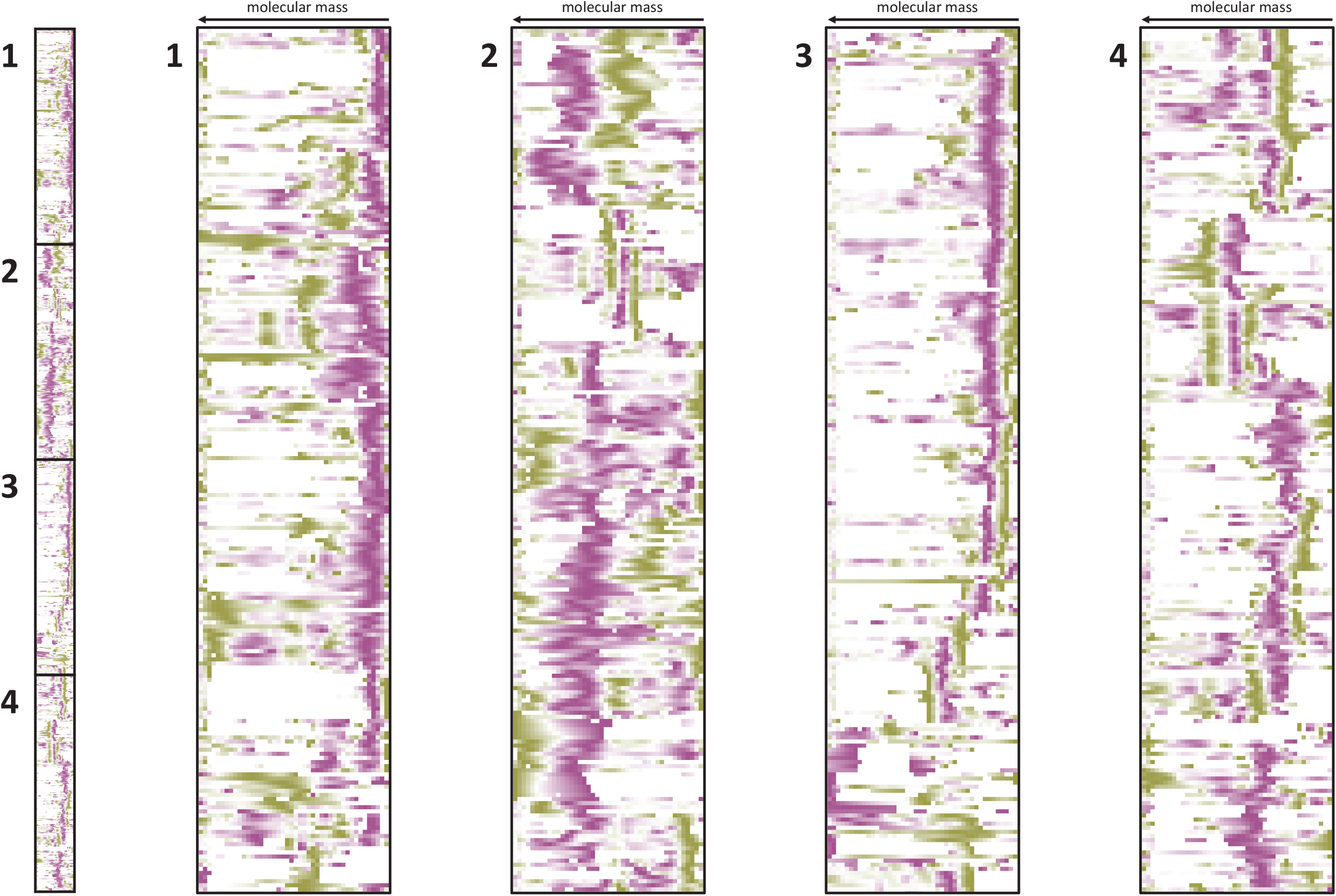
Differential complexome profile of mitochondria isolated from dark- and light-adapted plants. Results are displayed as a heatmap. Left panel shows relative differences in abundance profiles of 843 proteins present in mitochondria isolated from dark- and from light-adapted plants. For both conditions (‘dark’, ‘light’), log_10_-transformed abundance profiles of five biological replicates were merged into a single, average profile (see Figs. S2 and S3 for the ten individual complexome profiling experiments). Values of the ‘light’ profile were subsequently subtracted from values of the ‘dark’ profile. Purple color indicates higher abundance in mitochondria isolated from dark-adapted plants, whereas green indicates higher abundance in mitochondria isolated from light-adapted plants. The four panels to the right are cutouts of the left panel as indicated by the numbers.

### Light conditions trigger individual and common responses in the interaction patterns of TCA-cycle enzymes

The eight enzymes of the TCA-cycle as well as those closely associated with it (the pyruvate dehydrogenase complex, PDC; and the malic enzyme, ME) serve in producing reduction equivalents for the respiratory chain but also precursors for nitrogen assimilation in plastids. By large, these two processes are temporally separated, as the latter occurs mainly during the day when photosynthesis takes place, whereas the first is prevailing in the dark. As such, different TCA cycle flux modes have been proposed: mostly cyclic in the night and primarily linear during the day (Sweetlove et al. 2010). Transition between the two flux modes requires tight regulation of TCA-cycle enzymes. Given that most enzymes of the TCA cycle do not act as individual proteins but as heteromers or homomers of up to eight subunits, dynamics in the association and dissociation patterns of these enzymes may serve in regulating activity of the enzymes involved in this pathway in the absence of altered total protein abundance. Moreover, the interaction of TCA-cycle enzyme to form mitochondrial metabolons has been proposed decades ago (Robinson und Srere 1985) and coherent evidence for their presence across all lineages is beginning to emerge (Vélot et al. 1997; Wu und Minteer 2015; Zhang et al. 2017), particularly for the interaction between mitochondrial malate dehydrogenase (mMDH), citrate synthase (CSY), and aconitase (ACO).

The Pyruvate dehydrogenase complex (PDC) and mitochondrial malate dehydrogenase (mMDH) provide the substrates for aconitase as the first enzyme of the TCA cycle, acetyl-CoA and oxaloacetate. In plants malic enzyme (ME) is also indirectly involved in this, since it converts malate to CO_2_ and pyruvate, the latter of which then being the substrate of PDC. Two isoforms of NAD-dependent, mitochondrial ME are encoded by the Arabidopsis genome, NAD-ME1 and NAD-ME2. The two isoforms associate to form either homo-dimers or hetero-dimers (NAD-MEH). Depending on their composition, the dimers have molecular masses of 117 to 125 kDa. Distribution of the three different NAD-ME dimers is tissue-specific (Tronconi et al. 2008). Since the three complexes not only differ in respect to their composition but also regarding their activation (fumarate for NAD-ME1, CoA for NAD-ME2, and both, fumarate and CoA, for NAD-MEH) it has been proposed that the allocation of NAD-ME isoforms into homodimers and heterodimers is an act of adjusting NAD-ME activity to mitochondrial functions (Tronconi et al. 2010). In lpBN gels, abundance of the two NAD-ME isoforms concentrates at 50 kDa and therefore slightly below the expected molecular mass of approximately 60 kDa (Fig. 2A, upper panel). Most likely, this is due to a lack of raster resolution owing to the complexome profiling strategy. MEH is expected to produce a strong peak in the range of 120 kDa, which could not be observed in the individual heatmaps (Fig. 2A, upper panel). This lack could be due to organelle isolation procedure used here being much longer than the shorter protocol for crude extracts used elsewhere (Tronconi et al. 2008) and may therefore promote dissociation of the dimer. However, higher molecular mass complexes of low abundance seem to respond to light conditions (Fig. 2A, lower panel).

**Figure 2:**
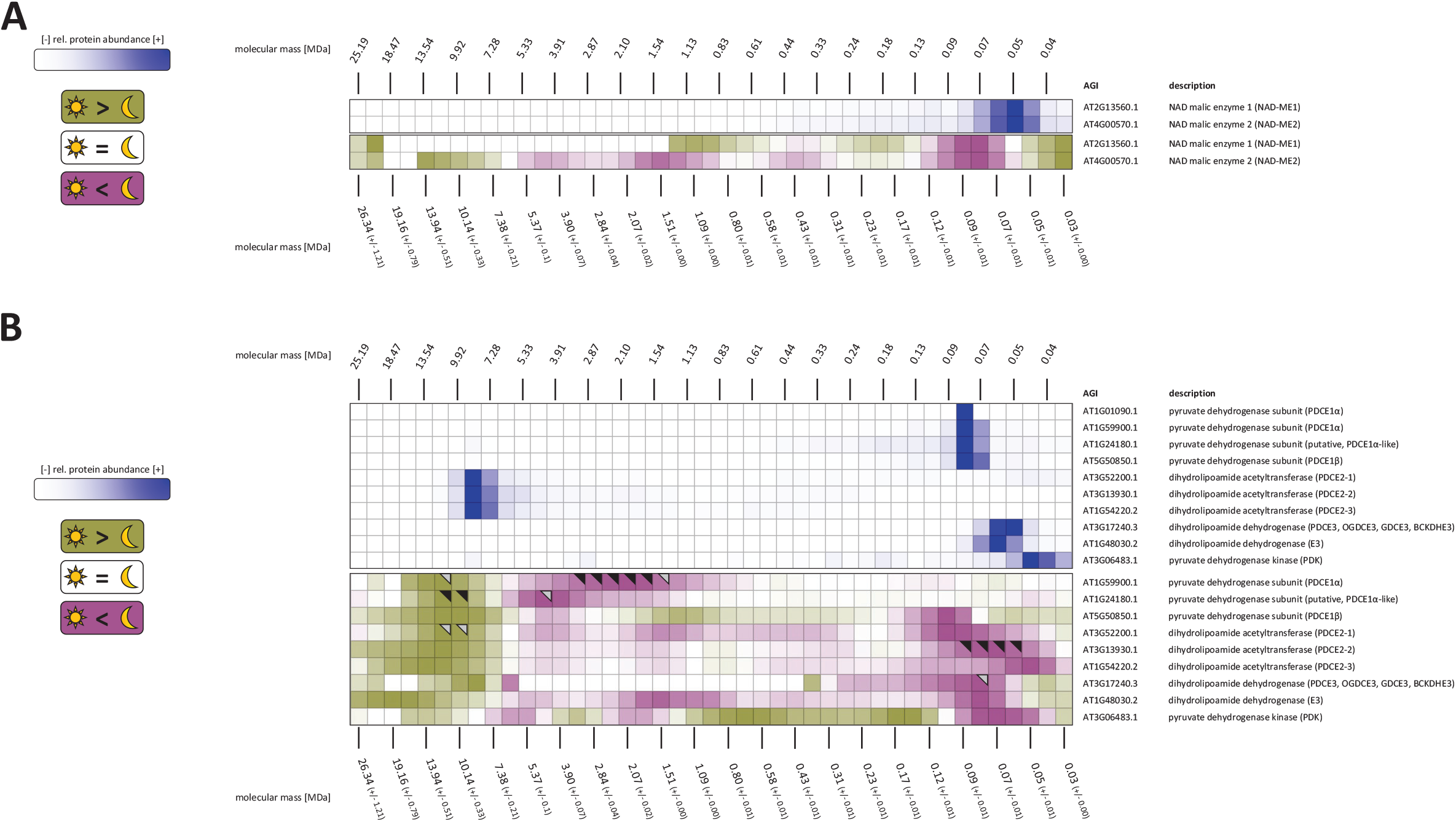
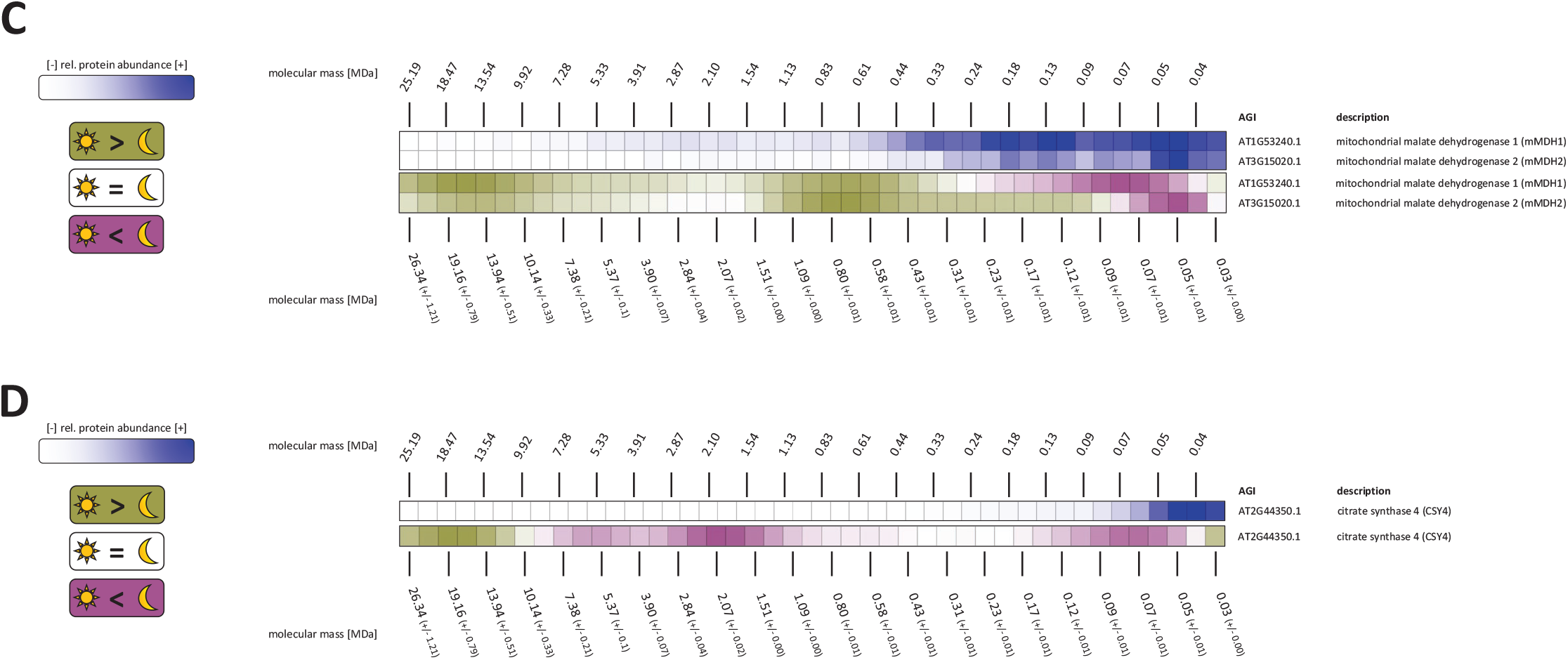
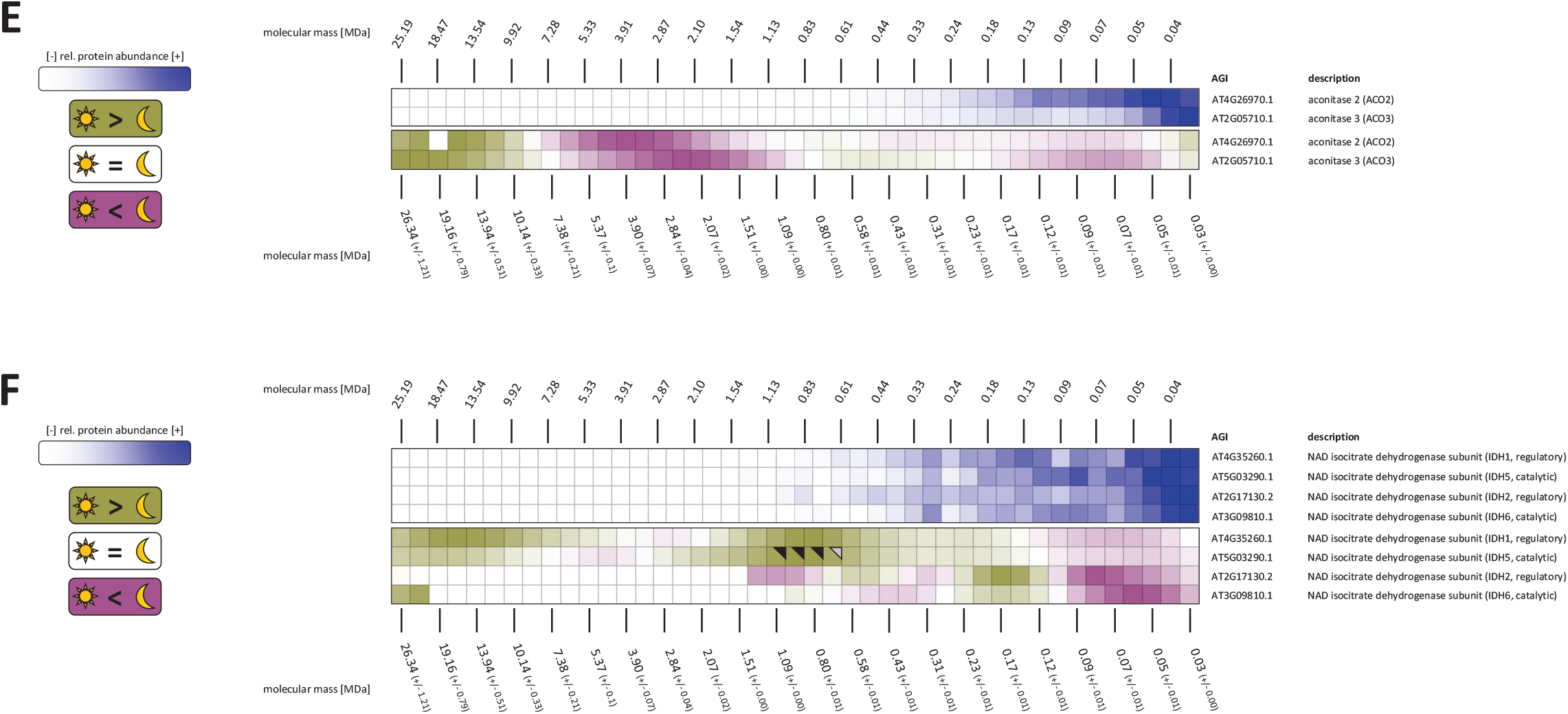
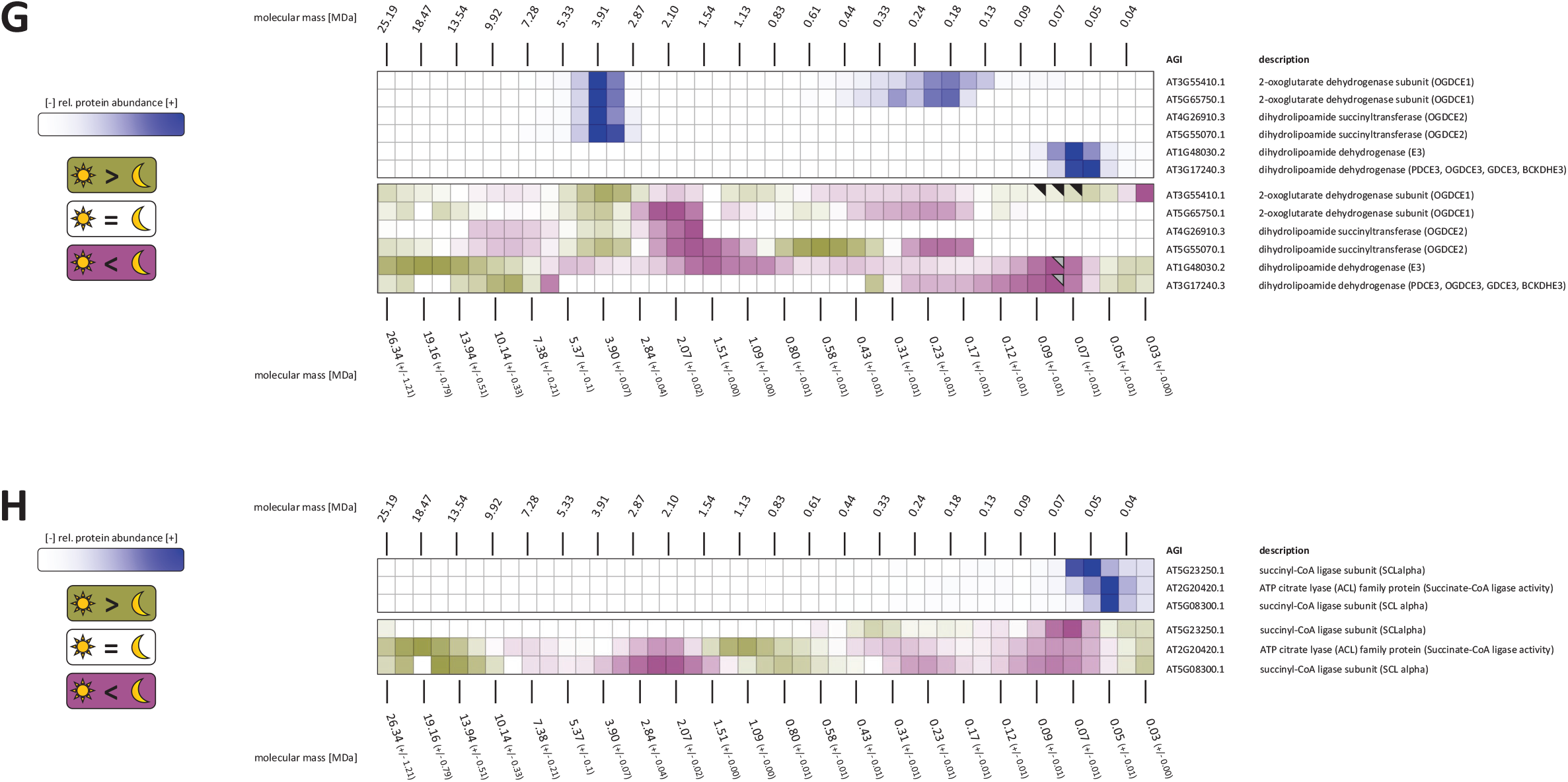
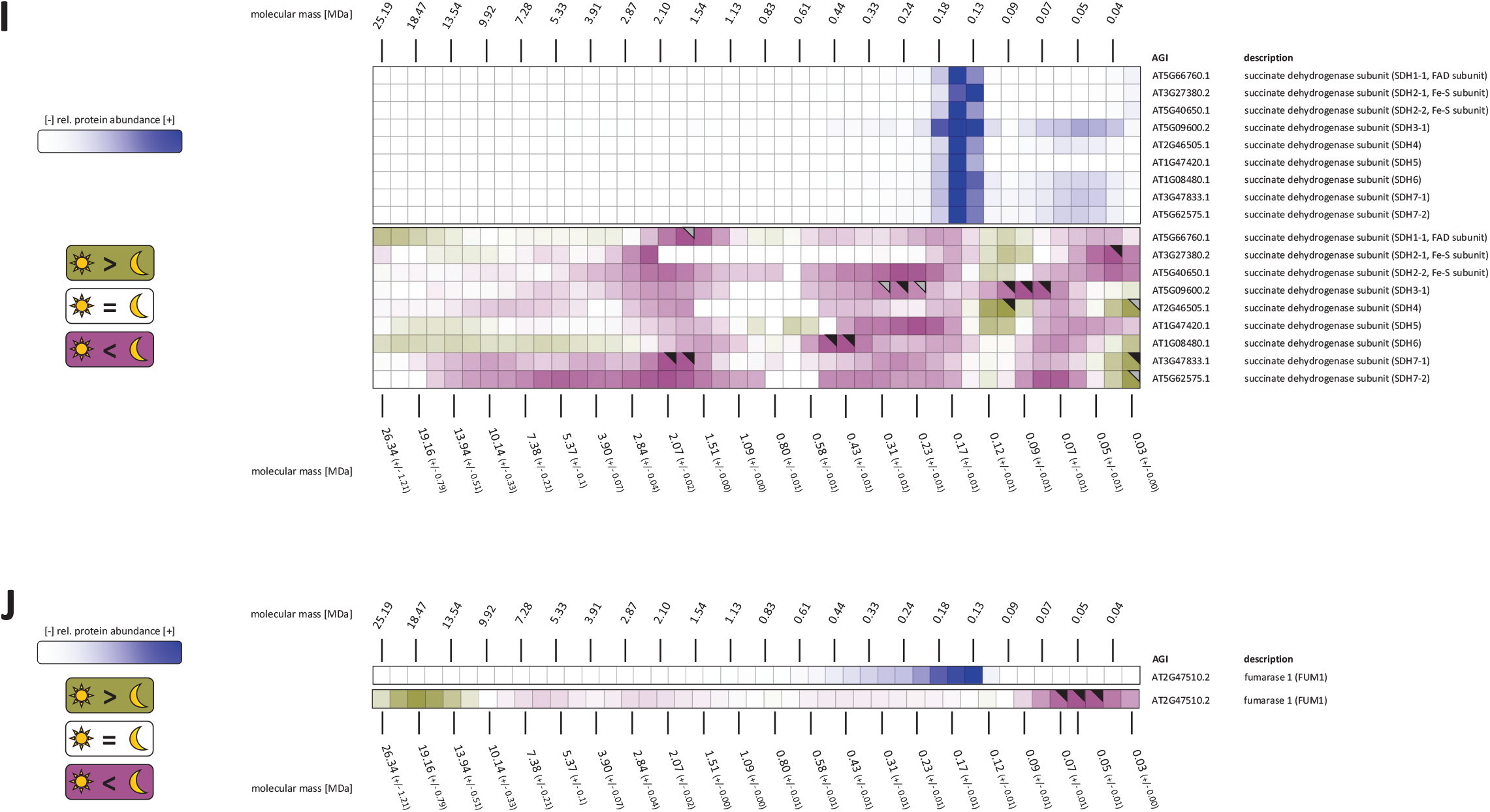
Differential heatmaps of TCA-cycle enzymes. A, mitochondrial malate dehydrogenase (mMDH); B, pyruvate dehydrogenase complex (PDC); C, NAD-malic enzyme (MAD-ME); D, citrate synthase (CSY); E, aconitose (ACO); F, Isocitrate dehydrogenase (ICDH); G, 2-oxoglutarate dehydrogenase complex (2-OGDC); H, succinyl-CoA-ligase (SCL); I, succinate dehydrogenase (SDH); J, fumarase (FUM). Upper panels in all figure parts show abundance profiles of TCA-cycle enzymes in a replicate of mitochondria isolated from dark-adapted plants (dark II, Figure S2). Color indicates relative protein abundance (dark blue, high relative abundance; light blue, low abundance; white, very little relative abundance or no detection). Lower panels in all parts of the figure show differential heatmaps of the TCA cycle enzymes. For this approach, log_10_-transformed average abundance profiles of five biological replicates for each condition (‘dark’, ‘light’) were merged into a single, average profile. Values of the ‘dark’ profiles were then subtracted from values of the ‘light’ profiles. Purple color indicates higher abundance in dark-adapted mitochondria, whereas green indicates higher abundance in light-adapted mitochondria. Black and grey triangles in the top right corners of tiles indicate p-values of ≤0.05 and >0.05 ≤0.1, respectively. Fractions 3 to 45 after alignment are shown. Please note that the number of fractions in the abundance heatmaps (upper panels of all parts of the figure) does not match the number of fractions in the differential heatmaps (lower panels of all parts of the figure) due to warping and alignment of the individual replicates, in which the original rasters were lost and replaced by a virtual raster containing a higher number of fractions. AGI, Arabidopsis genome identifier.

PDC is a protein complex composed of three different subunits (E1, heterotetramer; E2, trimer; E3, trimer) in a 22:20:20 stoichiometry with a molecular mass of ~9 MDa (Zhou et al. 2001). It is inhibited by phosphorylation and activated by dephosphorylation of the E1α subunit (Linn et al. 1969; Rubin und Randall 1977). The enzymes responsible for this (PDC kinase, PDK; PDC phosphorylase, PDP) are reported to be part of the holo-complex (Tovar-Méndez et al. 2003). In the lp-BN gels, PDC-E1α/β abundance concentrates at a molecular mass of approximately 80 kDa, indicating the presence of PDC-E1 dimers (Fig. 2B, upper panel, Fig. S8). PDC-E3 behaves similar, albeit at a slightly lower molecular mass of ~60 kDa, which matches the molecular mass of the protein. In contrast, PDC-E2 is mostly absent in the low molecular mass region of the gel but forms a sharp peak at ~8.5 MDa. PDC-E1 isoforms and PDC-3 also have minor peaks in this region. Together, this demonstrates that PDC-E2 abundance in plant mitochondria is mostly tied up in the PDC holo-complex whereas for PDC-E1 and PDC-E3 dimeric and monomeric forms dominate. Interestingly, the iBAQ values at the position of the holo-complex seem to suggest an unusual stoichiometry of the PDC complex, largely dominated by E2 subunits (Fig. S8). It is currently not clear if the dominance of the E2 SU is an artifact of the complexome profiling process, if it is caused by a differing amenability of the PDC SU for MS identification (which may distort iBAQ ratios), or if it indeed represents the true PDC stoichiometry. Changes in the diurnal pattern of the PDC subunits are mostly inconsistent, except for a diffuse accumulation of signal high up in the mass range (> 9 MDa) in the light (Fig. 2B, lower panel). Interestingly, the reactions of a PCD-E3 isoform (AT1G48030), all PDC-E2 isoforms and that of NAD-ME2 are remarkably similar in the mass range below 10 MDa. These proteins are of higher abundance at ~5 MDa, ~1,5 MDa and ~0.4 MDa in the dark and of higher abundance in the light at a molecular mass >10 MDa. Interactions of these proteins are supported by recent observations (Zhang et al. 2017). An assembly containing PDC and NAD-ME2 would be able to use both, malate and pyruvate, to provide acetyl-CoA for the TCA-cycle.

Mitochondrial MDHs are enzymatically active as homo-dimers with a molecular masses of around 70kDa (Dasika et al. 2015). For both isoforms of mitochondrial malate dehydrogenase, mMDH1 and mMDH2, a hotspot of protein abundance is found in the molecular mass range below 0.5 MDa (Fig 2C, upper panel). Interestingly, both isoforms behave similarly in the differential heat map (Fig. 2C, lower panel) since assemblies at 0.8 MDa and at 16 MDa are more abundant in the light whereas in the dark, the monomeric version of mMDH1 and a putative breakdown product of mMDH2 are of higher abundance.

Citrate synthase (CSY) is present in a single isoform in our dataset (CSY4). The enzyme is redox-regulated (Schmidtmann et al. 2014) and the active form seems to be a homodimer, which forms a single active site. In addition to dimers, monomers, tetramers and higher molecular mass complexes have also been reported (Schmidtmann et al. 2014). In our heatmaps, the predominant form is the monomer at ~45 kDa (Fig 2D, upper panel), which is lower than the calculated mass of the precursor protein (~53 kDa). Assemblies at ~2 MDa, ~6 MDa, and ~19 MDa seem to respond to light conditions (Fig. 2D, lower panel), comparably to another TCA cycle enzyme, fumarase (FUM1, Fig. 2J, lower panel). A physical interaction of CSY and FUM has been reported previously (Beeckmans und Kanarek 1981).

The iron-sulfur containing protein Aconitase (ACO), is represented by two isoforms in our heatmaps (ACO2, ACO3) and has a molecular mass of ~108 kDa (Fig. 2E). It catalyzes the second step of the TCA cycle and is a key TCA-cycle enzyme, due to its amenability to ROS (Sweetlove et al. 2002), its importance for balancing carbon (sucrose) flux (Carrari et al. 2003), and its role in regulating the expression of several genes (Moeder et al. 2007; Hirling et al. 1994). For both isoforms, a clear signal for the monomeric protein is missing. Instead, intensities are highest at lower molecular masses, indicative of the presence of degradation products (Fig. 2E, upper panel). In the differential heatmap, both isoforms are of increased abundance in the dark at molecular masses of ~2.1 MDa (ACO3) and ~3.9 MDa (ACO2), repectively. Simultaneously, both are also of increased abundance in the light at ~19 MDa (Fig. 2E, lower panel).

The NAD-dependent isocitrate dehydrogenase (IDH) of mitochondria is a branching point of the TCA-cycle in plants since it produces 2-oxoglutarate, which not only is a TCA cycle intermediate but also a substrate for chloroplast nitrogen fixation. Due to the presence of cytosolic isoforms of this enzyme, T-DNA-knockdown mutants of IDH genes do not have a discernible phenotype, rendering this enzyme not essential for both, TCA and nitrogen fixation (Lemaitre et al. 2007). In yeast, the functional enzyme is suggested to be a octamer with a mass of approximately 300 KDa, consisting of four regulatory units combined with the same amount of catalytic units (Keys und McAlister-Henn 1990). In porcine mitochondria the enzyme seems to be a hetero-tetramer containing three different proteins (Huang et al. 1997). Little information is available for the plant NAD-IDH. In contrast to earlier observations, the enzyme also seems to be a heteromer of unknown molecular mass (Lancien et al. 1998). Mitochondrial pea NAD-IDH is instable and the molecular mass of the complex was determined to be 1.4 MDa in fresh mitochondrial extracts. With increasing age of the extract, this declines continuously from 1.4 MDa down to 0.69 MDa, 0.3 MDa, 0.21 MDa, and finally to 0.094 MDa. The loss in molecular mass is accompanied by a loss of its activity (McIntosh und Oliver 1992). Complexes of all sizes appear to contain the same two subunits migrating at 45 kDa and 47 kDa in SDS gels. For plants, a matrix IDH as well as a membrane associated IDH have been described (Tezuka und Laties 1983). Arabidopsis contains a total of six NAD-IDH genes. One of them is missing its N-terminus and is considered inactive (IDH4), two are of the catalytic type (containing the binding sites for its regulators citrate and AMP), and the remaining three are of the regulatory type (containing the binding sites for its substrates isocitrate and NADH) (Lin und McAlister-Henn 2003). Of the latter, only two are present in vegetative tissue while the other one seems to be pollen specific (IDH3) (Lemaitre und Hodges 2006). All four isoforms present in vegetative tissue (IDH1 and IDH2, regulatory; IDH5 and IDH6, catalytic) are featured in the differential heatmap (Fig. 2F) but for IDH2 and IDH6 large gaps in the molecular mass range above 1MDa exist (Fig 2F, lower panel). This is explained by a recent estimation of the copy numbers of proteins present in a single mitochondrion. IDH1 is more than five times more abundant than IDH2, which is matched by a similar ratio of IDH5 over IDH6 (Fuchs et al. 2020). As such, protein abundance of IDH2 and IDH 6 in some areas of the upper part of the gel will be too low for a reliable identification and quantitation. All four isoforms have their main peak at approximately 40 kDa, which corresponds to the monomeric mass of the proteins (Fig 2F, upper panel) but assemblies with higher molecular masses, for example at ~280 kDa, can also be observed. The differential heatmap (Fig. 2F, lower panel) suggests an assembly of increased abundance at ~0.9 MDa in the light in which only the higher-abundant isoforms (NAD-IDH1 and NAD-IDH5) participate but not NAD-IDH2 and NAD-IDH6. A second, light-specific peak can be observed at ~16 MDa.

The 2-oxoglutarate decarboxylase (2-OGDC) catalyzes the conversion of 2-oxoglutarate, NAD^+^, and CoA to succinyl-CoA, NADH, H^+^, and CO_2_. The complex resembles the PDC in functional terms and is composed of three different proteins with similar functions and the E3-SU is shared with the PDC. However, with a molecular mass of ~ 3.9 MDa, it is considerably smaller than the PDC (~8.5 MDa). While considerable amounts of the two E1 isoforms are present in their monomeric forms (~118 kDa; Fig. 2G, upper panel), they are of highest abundance at the position of the holo-complex. This situation therefore differs considerably from the one described for the PDC since OGDC-E1 amounts compare with those of the OGDC-E2 SUs at this position (Fig. S9). However, PDC and OGDC both share the low amount of E3 subunits at the position of the expected holo-complexes. Judging from the patterns of OGDC-E1 and OGDC-E2 subunits the differential heatmap, the 2-OGDC holo-complex is more abundant in the light, while assemblies at smaller (~2 MDa) and larger (~8 MDa) molecular masses are more abundant in the dark (Fig. 2G, lower panel). Apart from OGDC and PDC the E3 subunits is also part the glycine decarboxylase complex (GDC), and the branch-chain alpha-ketoacid dehydrogenase (BCKDH), which may explain its divergence from the pattern of the other 2-OGDC subunits. Given that 2-OG (and also citrate) is exported from mitochondria to fuel nitrogen fixation in chloroplasts during the day, the data obtained for the holo-complex are surprising since its abundance is expected to be reduced in light and to be higher in the dark in order to promote a cyclic flux mode of the TCA metabolic pathway. This sparks the question about the functions of the other 2-OGDC assemblies with higher and lower molecular masses in the context of cyclic flux mode TCA-cycle activity in the dark.

The heterodimeric succinyl-CoA ligase enzyme consists of SCLα and SCLβ subunits (SUs) and catalyzes the reaction of succinyl-CoA, ADP and P_i_ to succinate and ATP. It is activated by metabolites upstream of its position in the TCA-cycle and is inactivated by downstream TCA cycle intermediates (Studart-Guimarães et al. 2005). *Arabidopsis thaliana* contains three SCL genes, two of which are encoding the α-SU and a single one the β-SU. The two α-SUs are not of similar abundance within mitochondria since one isoform dominates over the other one (AT5G08300, 10612 copies per mitochondrion; AT5G23250, 1054 copies per mitochondrion; (Fuchs et al. 2020)). The copy number of the β-SU (AT2G20240, 11290 copies per mitochondrion) is slightly higher than that of the dominant α-SU (Fuchs et al. 2020). The bulk of the more abundant α-SU and the β-SU can be found in the low molecular mass area of the gel, matching the monomeric molecular mass for AT2G20420 (45 kDa) but not that for the other two subunits (36-36 kDa; Fig. 2H, upper panel). In the differential heatmap (Fig. 2H, lower panel) the two more abundant proteins produce a matching pattern at higher molecular masses, in which an assembly at ~2.5 MDa is more abundant in the dark, while another one at ~19 MDa is present in higher amounts in light mitochondria.

Converting succinate and ubiquinone into fumarate and ubiquinol, the plant version of the FAD-containing succinate dehydrogenase (SDH) complex (C-II_1_) is composed of eight subunits and thus contains four subunits more than its bacterial, fungal, and mammalian counterparts (Eubel et al. 2003; Millar et al. 2004). This higher number of subunits in plants can partly be explained by the truncated membrane anchor subunits of the plant complex (SDH3 and SDH4), which are lacking their transmembrane helices. SDH6 and SDH7 show homology to the lost transmembrane domains of non-plant SDH3 and SDH4 subunits and, together with SDH5, compensate for the missing domains in plant SDH3 and SDH4 to form a functional membrane anchor for the succinate oxidizing SDH1 and SDH2 subunits (Schikowsky et al. 2017). The role of SDH8 remains unclear, as is its *bona fide* participation in the complex (Huang et al. 2019). It has been speculated that SDH8 may introduce side reactions to this respiratory chain component similar to those reported for the other respiratory chain enzymes (Eubel et al. 2003; Millar et al. 2004). Besides its role as a TCA-cycle and respiratory chain enzyme, reactive oxygen species (ROS) produced by SDH in response to salicylic acid are also involved in plant signaling events (Gleason et al. 2011; Belt et al. 2017). Given the diverse functions of SDH, its regulation is complex. Besides succinate, it is also activated by both, ADP and ATP, as well as oligomycin, but inhibited by uncouplers, potassium, and nitric oxide (Affourtit et al. 2001; Simonin und Galina 2013). SDH1, SDH2, and SDH7 are each present in two isoforms in Arabidopsis whereas the rest of the SUs is encoded by a single gene. SDH1 shows an extreme preference of one isoform over the other one with 7723 copies for SDH1-1 (AT5G66760) and only 2 for SDH1-2 (AT2G18450) in a single mitochondrion (Fuchs et al. 2020). Highest intensities of the SDH subunits are detected in the region of the native C-II_1_ at ~150 kDa but small portions of the signal are also found further up and down in the mass range (Fig. 2I, upper panel). The distribution of SDH subunits over the mass range is rather erratic except for a broad signal near 2 MDa, at which all subunits are of higher abundance in the dark.

Mitochondrial fumarase (FUM) is encoded by a single gene, the product of which is forming homo-tetramers with a molecular mass of approximately 200 kDa (Weaver et al. 1995). The enzyme catalyzes the reversible addition of water to fumarate to produce malate. It is inhibited by pyruvate, 2-OG, and uncomplexed adenylates (AMP, ADP, and ATP) (Behal und Oliver 1997). In a previous complexome profiling approach (Senkler et al. 2017) the bulk of fumarase migrates close to the expected molecular mass of 200 kDa while we here observe a peak at a lower molecular mass (Fig. 2J, upper panel). In the differential heatmap, several minor peaks in the mass range between 200 kDa and 10 MDa show elevated abundance, mostly in the dark. This is accompanied by a major differential peak for the monomeric protein at ~50 kDa, which is also of increased abundance in the dark, indicative of an increased degradation of homo-tetramers in this condition or an increased biosynthesis of fumarase subunits waiting for their assembly. Fumarase also displays a sharp increase at ~19 kDa in the light, minor peaks of high abundance in the dark can be observed at molecular masses of ~0.4 MDa, ~2.5 MDa, and ~6.3 MDa.

A common denominator in the differential heatmaps for mMDH, NAD-ME, CSY, ACO, IDH, SCL, and FUM is the higher abundance of these proteins in the light at molecular mass of ~19 MDa, pointing towards the formation of a high molecular mass protein assembly during the day. PDC does not seem to take part in this and the participation of NAD-ME is uncertain due to a lack of signal at this position. At the same time, CSY, ACO3, OGDC, SCL, SDH, and FUM are of increased abundance in the dark at a molecular mass slightly higher than 2 MDa, also potentially forming a stable assembly of TCA-cycle enzymes. Since the OGDC holo-enzyme has a molecular mass of ~3.9 MDa, only sub-stoichiometric assemblies of this protein complex can be expected to participate in this proposed assembly. The presence of FUM, CSY, and ACO in both assemblies would allow the channeling of citrate and fumarate as deduced from isotope dilution experiments carried out on isolated potato tuber mitochondria (Zhang et al. 2017). The considerable molecular mass of the 19 MDa-assembly indicates the formation of a TCA-cycle microdomain, which is expected to contain unknown but large numbers of each participating enzyme and which is most likely less well-defined than classical metabolons.

### Assembly of complexes I and III into a high-molecular mass supercomplex increases in the light

In spinach and potato mitochondria, monomeric complex I (C-I_1_) and dimeric complex III (C-III_2_) form a supercomplex also involving C-IV in varying stoichiometries (Krause et al. 2004; Eubel et al. 2004). In contrast, the predominant supercomplex in Arabidopsis lacks C-IV and contains only C-I and C-III in a SC-I_1_III_2_ stoichiometry (Eubel et al. 2003; Dudkina et al. 2005; Braun 2020). As such, the difference profiles of C-I and C-III subunits are here presented in the same figure (Fig. 3A, B). C-I_1_ migrates at 1.1 MDa (Fig. 3A, upper panel) which is higher than the estimated 1 MDa reported previously (Eubel et al. 2003; Senkler et al. 2017). Most likely, this is owed to a lack of raster resolution in the lp-BN gels which is further exacerbated by the matching process of the dark and light replicates, and ultimately merges C-I_1_ with SC-I_1_/III_2_ (1.5 MDa). For the sake of simplicity, this broad band in the differential heatmap (Fig. 3A and B, lower panels) will be referred to as ‘extended complex I area’ (ECIA) in the following. ECIA and C-III_2_, are of higher abundance in the dark whereas a supercomplex consisting of two C-I_1_ subunits associated with C-III_2_ (SC-I_2_/III_2_) and a molecular mass of 2.5 MDa is of higher abundance in the light. This suggests a shift in ECIA abundance towards the SC-I_2_/III_2_ in the light. Inspection of the pattern for C-III_2_ (Fig. 3B, upper and lower panel) reveals that both, C-III_2_ (~0.5 MDa) and SC-I_1_-III_2_ (~1.5 MDa), are of lower abundance in the light and therefore contribute towards the higher amounts of SC-I_2_/III_2_ in the light. C-III subunits also increase in abundance at ~0.9 MDa in the light, most likely as part of a supercomplex consisting of C-III_2_ and C-IV_1_ (SC-III_2_/IV_1_). The occurrence of this supercomplex was also observed earlier (Senkler et al. 2017) but is here not clearly supported by an increase of C-IV in this region (Fig. 4).

**Figure 3:**
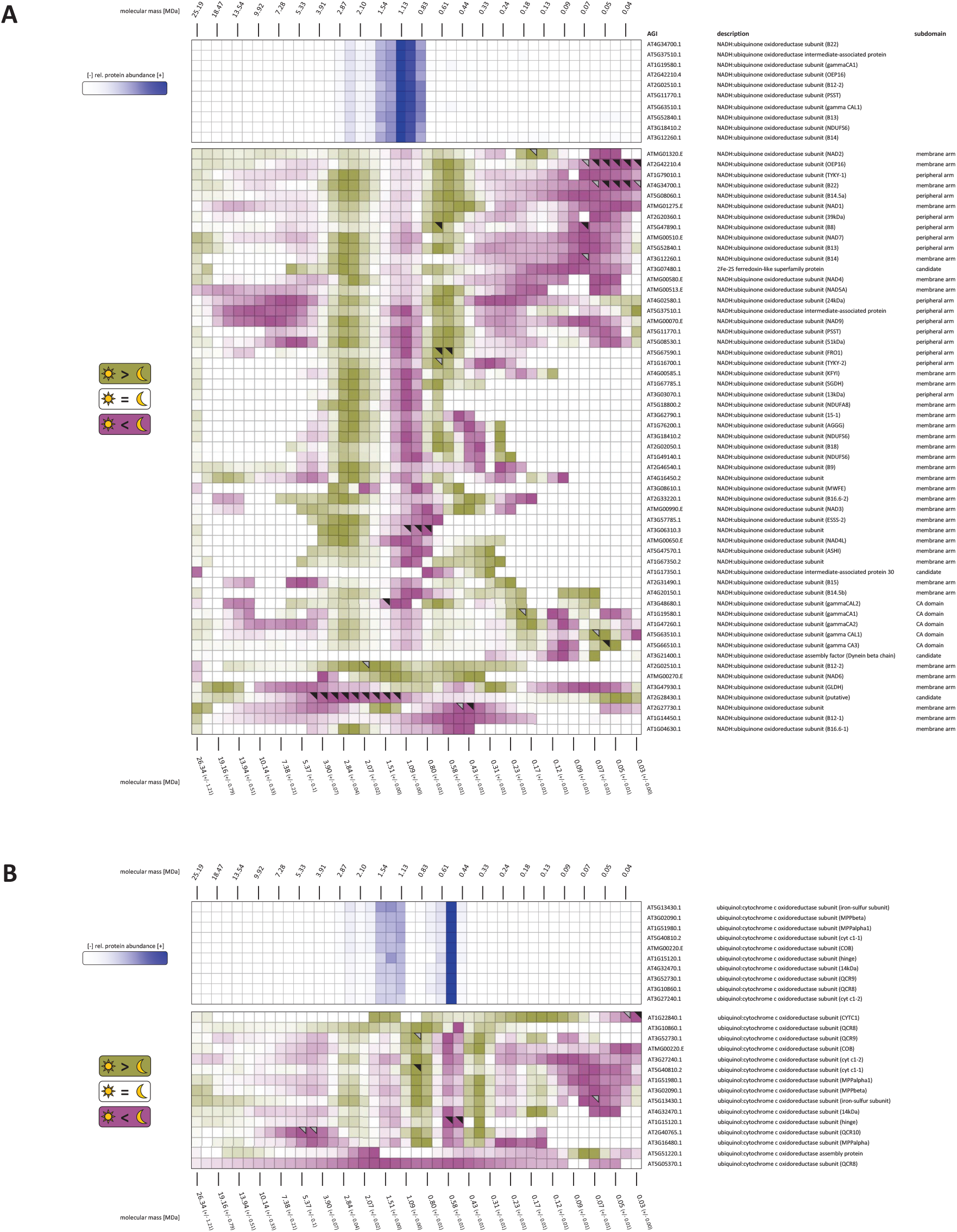
Differential heatmaps for complexes I and III. A, complex I (C-I); B complex III (C-III). Upper panels in the two figure parts show abundance profiles of C-I and C-III, respectively, in a replicate of mitochondria isolated from dark-adapted plants (dark II, Figure S2). Color indicates relative protein abundance (dark blue, high relative abundance; light blue, low abundance; white, very little relative abundance or no detection). Lower panels in the two figure parts show differential heatmaps of C-I and C-III. For this approach, log_10_-transformed abundance profiles of five biological replicates for each condition (‘dark’, ‘light’), were merged into a single, average profile. Values of the ‘dark’ profile were then subtracted from values of the ‘light’ profile. Purple color indicates higher abundance in dark-adapted mitochondria, whereas green indicates higher abundance in light-adapted mitochondria. Black and grey triangles in the top right corners of tiles indicate p-values of ≤0.05 and >0.05 ≤0.1, respectively. Fractions 3 to 45 after alignment are shown. Please note that the number of fractions in the abundance heatmaps (upper panels of the two parts of the figure) does not match the number of fractions in the differential heatmaps (lower panels of the two parts of the figure) due to warping and alignment of the individual replicates, in which the original rasters were lost and replaced by a virtual raster containing a higher number of fractions. AGI, Arabidopsis genome identifier.

**Figure 4:**
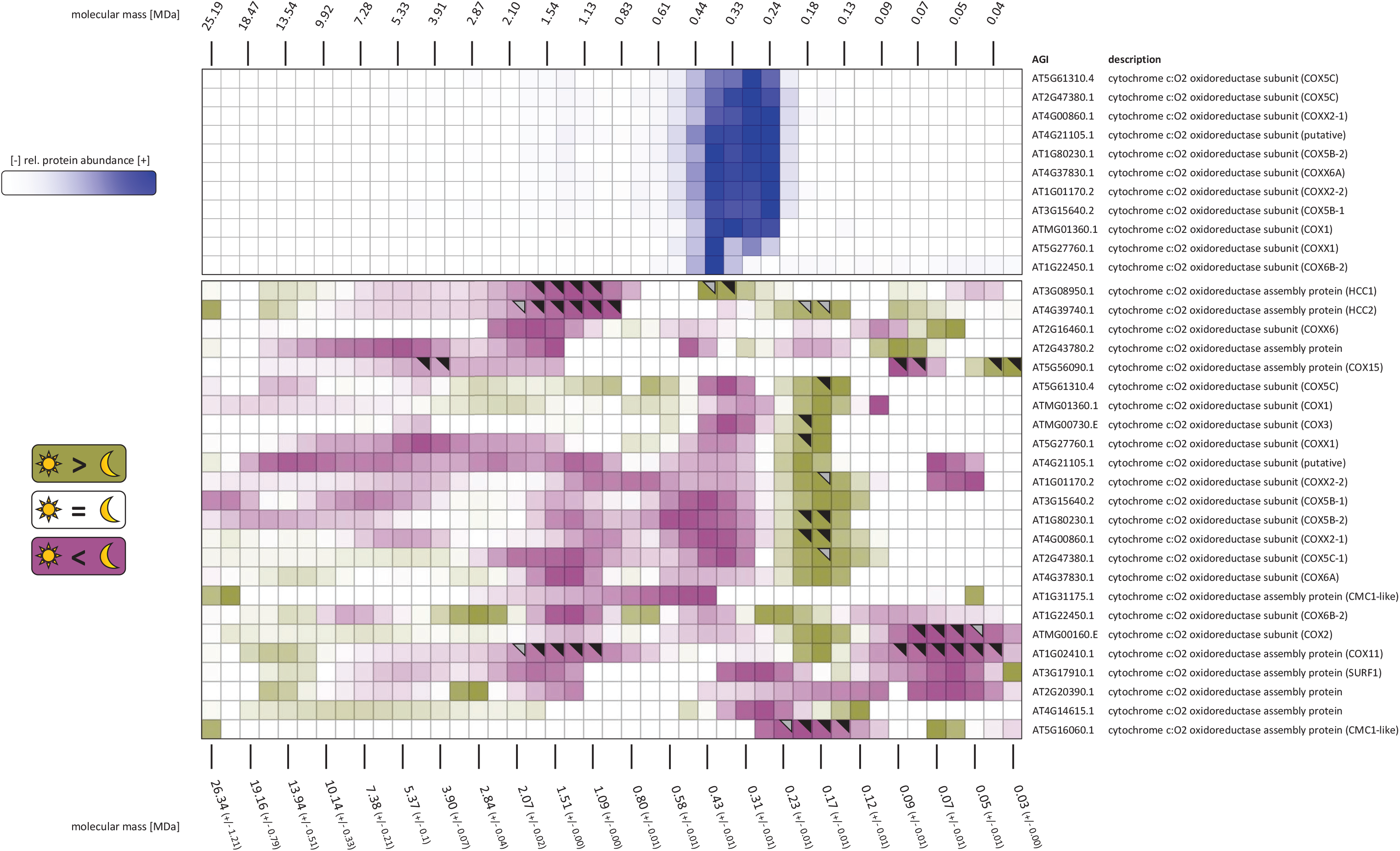
Differential heatmap for complex IV. Upper panel shows abundance profiles of complex IV (C-IV) subunits in a replicate of mitochondria isolated from dark-adapted plants (dark II, Figure S2). Color indicates relative protein abundance (dark blue, high relative abundance; light blue, low abundance; white, very little relative abundance or no detection). Lower panel shows a differential heatmap of the subunits of complex VI. For this approach, log_10_-transformed abundance profiles of five biological replicates for each condition (‘dark’, ‘light’) were merged into a single, average profile. Values of the ‘dark’ profile were then subtracted from values of the ‘light’ profile. Purple color indicates higher abundance in dark-adapted mitochondria, whereas green indicates higher abundance in light-adapted mitochondria. Black and grey triangles in the top right corners of tiles indicate p-values of ≤0.05 and >0.05 ≤0.1, respectively. Fractions 3 to 45 after alignment are shown. Please note that the number of fractions in the abundance heatmap (upper panel of the figure) does not match the number of fractions in the differential heatmap (lower panel of the figure) due to warping and alignment of the individual replicates, in which the original rasters were lost and replaced by a virtual raster containing a higher number of fractions. AGI, Arabidopsis genome identifier.

Together, these results indicate an increase in the amount of SC-I_2_/III_2_ in response to photosynthetic activity which comes at the expense of EICA and C-III_2_ abundance. Its benefit for daytime mitochondrial respiration is currently unknown but is unlikely to foster faster transfer of electrons from C-I to the cytochrome c pathway due to structural considerations (Milenkovic et al. 2017).

Plant C-I_1_ is composed of approximately 50 subunits (Braun et al. 2014). In the differential heatmap (Fig. 3A, lower panel) the vast majority of these proteins behave in a similar fashion within ECIA and in SC-I_2_/III_2_. There is, however, also non-uniform behavior of C-I subunits in other areas of the gel, particularly in the mass range below 1 MDa. This area harbors assembly intermediates as well as breakdown products of C-I_1_ (Meyer et al. 2011; Klodmann et al. 2011; Ligas et al. 2019). Particularly the five γ-carbonic anhydrases (γ-CA) and γ-CA-like (γ-CAL) proteins are interesting, since they form a domain which is assembled early in C-I_1_ biogenesis (Meyer et al. 2011; Fromm et al. 2016). In the differential heatmap, these subunits cluster tightly and show a distinct dark/light pattern, indicative of an altered assembly of C-I under these conditions. Another group containing off-pattern C-I subunits in the differential heatmap is composed of eight proteins (AT3G21400, dynein β-chain; AT1G14450, B12-1; AT2G02510, B12-2; AT1G04630, B16.6-1; ATMG00270, NAD6; AT3G47930, GLDH; AT2G28430, putative SU; AT2G27730, unknown function SU), most of them being located in the membrane arm. Interestingly, half of these eight proteins (AT2G02510, B12-2; ATMG00270, NAD6; AT2G27730, unknown function SU; AT1G04630, B16.6-1) are running in the normal C-I_1_ and SC-I_1_ positions in the individual heatmaps (Figs S2, S3). Their behavior in the differential heatmap is therefore indicative of specialized structural functions in C-I_1_. Among those behaving differently from the patterns in the differential heatmap as well as in the individual heatmaps (AT3G21400, dynein β-chain; AT3G47930, GLDH; AT2G28430, putative subunit; AT1G14450, B12-1;) are two proteins with predicted (dynein β-chain) and well documented (GLDH) functions in C-I_1_ assembly (Pineau et al. 2008; Schertl et al. 2012; Schimmeyer et al. 2016). GLDH is also the final enzyme of ascorbate synthesis by the Smirnoff-Wheeler pathway and has been found in several C-I_1_ subcomplexes but does not seem to be part of the holo-complex (Millar et al. 2003). An isoform of the B12 subunit (B12-1; AT1G14450) is only found in SC-I_2_/III_2_ in the individual heatmaps but not in the C-I_1_ holo-complex or in SC-I_1_/III_2_, suggesting a specific role for this protein in this supercomplex.

### Complex IV is of increased abundance in the dark

The highest intensity signals for C-IV subunits are found at the molecular mass of intact C-IV_1_ (240 kDa - 380 kDa, Fig. 4, upper panel). In BN/SDS-gels two versions of complex IV can be found, which also produce in-gel activity stains in BN gels (Eubel et al. 2004). Whereas the two versions of C-IV_1_ remain discernible to some degree in the individual heatmaps, this resolution is lost in the differential map. Here, the molecular mass range of the C-IV holo-complexes contains a cluster, which is of higher abundance in the dark and contains catalytic subunits COX1, COX2, and COX3, as well as the structurally important COX5B/C subunits (Mansilla et al. 2018), and the plant specific subunits (COXX1, COXX2-1, COXX2-2). Another cluster containing the same essential and plant specific subunits migrates at a much lower molecular mass of ~160 kDa, albeit this time being of stronger abundance in the light. The added masses of these proteins (~224 kDa) is above the apparent molecular mass in the gel. It is therefore likely that at least two sub-complexes with similar masses co-migrate at the same position and that all subunits of the dark-induced complex are found across these subcomplexes. Two assembly factors of C-IV_1_, the copper chaperone HCC2 (AT4G39740) and COX11 (AT1G02410) are behaving in a similar fashion to the complexes’ established subunits, whereas other assembly factors have little in common with HCC2 and COX11. Together, the data suggest a formation of C-IV_1_ subcomplexes late in the day in order to have increased amounts of functional C-IV_1_ ready for the night, when the dissipation of surplus reduction equivalents of photosynthetic origin by alternative oxidoreductases ceases to contribute to the electron flow along the plant respiratory chain and a tighter coupling becomes necessary to meet the ATP demands of the cell.

### Protein abundance profiles of alternative respiratory enzymes lack a common response to light conditions

Plant possess alternative oxidoreductases as part of a branched respiratory chain. A set of eight type-II NAD(P)H dehydrogenases (NDs; NDA1-3, NDB1-4, NDC1), facing either the matrix or the intermembrane space, locate in the inner mitochondria membrane (IMM) and bypass C-I. Oxidizing either NADH or NADPH, they transfer electrons onto ubiquinone. In addition, five isoforms of an alternative oxidase (AOX; AOX1a-d, AOX2) transfer electrons from ubiquinol to molecular oxygen, thus bypassing the cytochrome-c pathway (C-III and C-IV). Together, these enzymes form the alternative pathway (AP). In contrast to the classical respiratory complexes, none of the enzymes of the AP contributes towards ATP-synthesis by translocating protons from the mitochondrial matrix into the crista lumen. The expression of these enzymes responds to environmental stimuli, particularly to stresses such as high light. The expression of NDA1 and NDC1, for example, are light regulated (Michalecka et al. 2003; Escobar et al. 2004) and AOX expression is also adjusted over the diurnal cycle (Dutilleul et al. 2003). Since these enzymes also react to a wealth of additional stimuli, often in an individual fashion, their expression patterns are difficult to interpret. However, NDA2, NDB2, and AOX1a seem to be co-expressed (Clifton et al. 2005; Elhafez et al. 2006). For the four NDAs and NDBs found here, highest intensities are detected at a molecular mass of ~110 kDa (Fig. 5A, upper panel). Owing to molecular masses of 57 kDa (NDA1, 2) and 63-69 kDa (NDB1, NDB2), this indicates the presence of ND dimers. The dynamic patterns of these enzymes (Fig. 5A, lower panel) are almost devoid of a common response, which underlines the individual functions of these proteins in the context of photosynthesis and respiration. The two AOX isoforms identified in this study, AOX1a (~40 kDa) and AOX1c (~38 kDa) run at a molecular mass of ~70 kDa (Fig 5B, upper panel), which conforms to the dimeric nature of the enzyme. Higher molecular assemblies have already been described previously (Senkler et al. 2017) which are here partly confirmed by dynamic alterations of the AOX proteins (Fig 5B, lower panel). Similar to the situation found for the NDAs and NDBs, no common response among the AOX isoforms could be detected.

**Figure 5:**
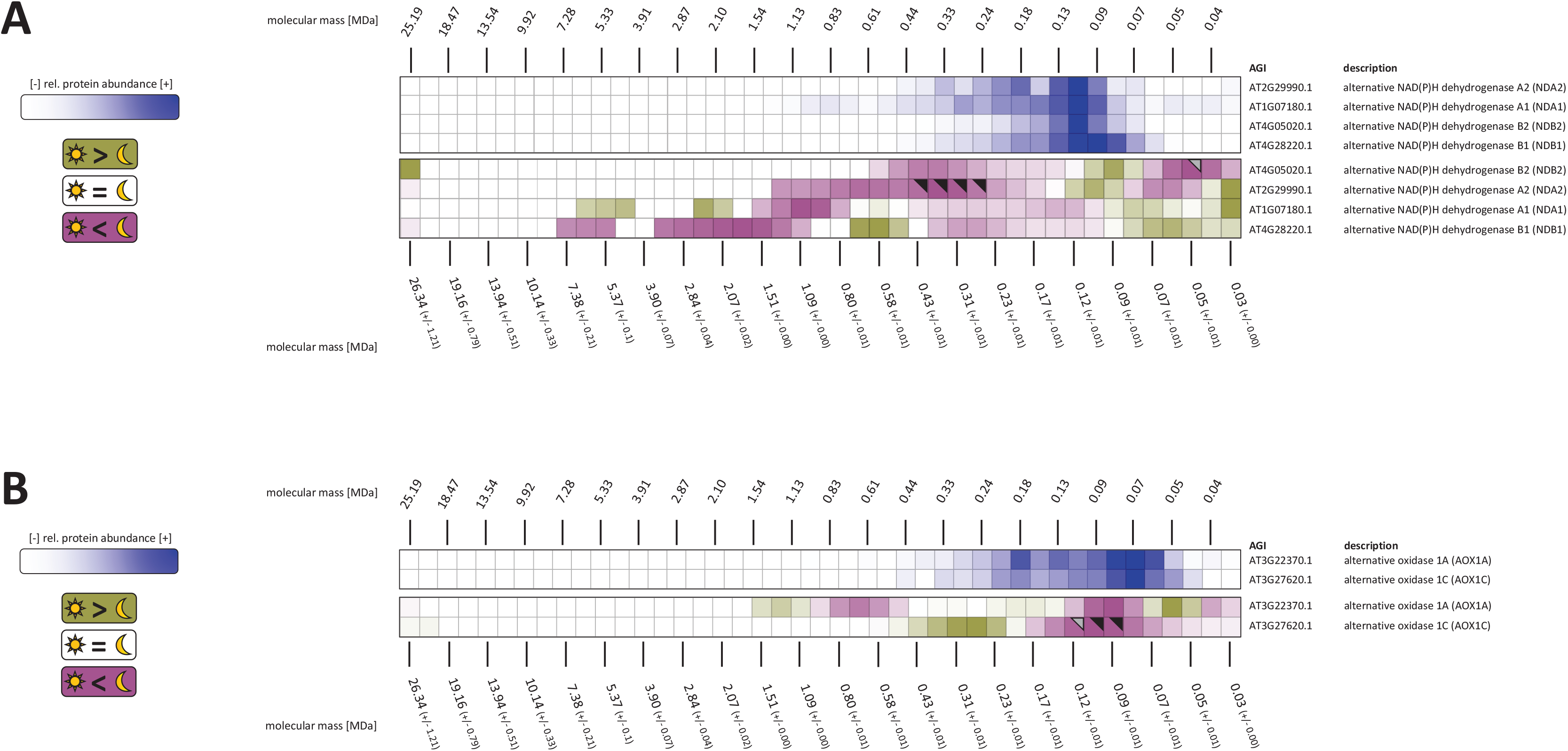
Differential heatmaps for alternative respiratory oxidoreductases. A, alternative NADH dehydrogenases (NDA, NDB); B, alternative oxidase (AOX). Upper panels in the two figure parts show abundance profiles of NDA and NDB isoforms and AOX isoforms, respectively, in a replicate of mitochondria isolated from dark-adapted plants (dark II, Figure S2). Color indicates relative protein abundance (dark blue, high relative abundance; light blue, low abundance; white, very little relative abundance or no detection). Lower panels in the two figure parts of the figure show differential heatmaps for NDA, NDB and AOX. For this approach, log_10_-transformed abundance profiles of five biological replicates for each condition (‘dark’, ‘light’) were merged into a single, average profile. Values of the ‘dark’ profile were then subtracted from values of the ‘light’ profile. Purple color indicates higher abundance in dark-adapted mitochondria, whereas green indicates higher abundance in light-adapted mitochondria. Black and grey triangles in the top right corners of tiles indicate p-values of ≤0.05 and >0.05 ≤0.1, respectively. Fractions 3 to 45 after alignment are shown. Please note that the number of fractions in the abundance heatmaps (upper panels of all parts of the figure) does not match the number of fractions in the differential heatmaps (lower panels of all parts of the figure) due to warping and alignment of the individual replicates, in which the original rasters were lost and replaced by a virtual raster containing a higher number of fractions. AGI, Arabidopsis genome identifier.

### The ATP synthase complex and its building blocks are induced in the dark

The ATP-synthase complex (C-V) uses the energy stored in the proton gradient formed by C-I, C-III, and C-IV to produce ATP from ADP and inorganic phosphate (P_i_). Structurally, it can be divided into a hydrophobic membrane part (F_O_) containing the rotor assembly from which the soluble part (F_1_) with the stalk and the three ADP binding sites protrudes into the mitochondrial matrix. Both parts are connected by the stator proteins. The complex exists in monomeric (C-V_1_), as well as dimeric (C-V_2_) form. In the latter, the monomers are arranged in a ‘V’ form, thus bending the inner mitochondrial membrane in this area (Dudkina et al. 2006). Rows of dimeric ATP-synthase dimers at the crista rims contribute towards crista formation and thus the inner structure of mitochondria. The bulk of C-V protein abundance is found at the position of the monomeric complex at ~700 kDa (Fig. 6, upper panel). C-V_1_ reacts to the light conditions by being of higher abundance in the dark (Fig. 6, lower panel). Other C-V assemblies at ~310 kDa, and at ~4 MDa react in a similar fashion. In contrast, signals at ~500 kDa and at ~1.3 MDa are of higher abundance in the light. However, in these cases, the signals are not uniform across all subunits but are interspersed with proteins either not responding at all or, in rare cases, responding diametrically. Apart from C-V_1_ at 700 kDa, it is therefore difficult to tell which subcomplex or supercomplex some the differential signals belong to. However, a common degradation product of C-V_1_ is the F_1_ subcomplex migrating at ~300 kDa in BN gels. We here observe a dark-induced pattern at a similar position. Most of the dark-induced subunits at this position are indeed components of the F_1_ part, but two F_0_ subunits (ATP8, ATMG00480 and ATP9, ATMG01080) behave in a similar fashion. Since F_O_ subunits are often not well detected in proteomic studies due to their hydrophobic nature, it is not entirely clear if the band at ~310 kDa represents the F_1_ subcomplex or another subcomplex containing both F_1_ and F_O_ subunits. The strongest response in the C-V differential heatmap is the increased dark abundance of either individual subunits or low molecular mass assemblies. However, these difference do not seem to translate into high absolute numbers since the dark II heatmap (Fig. 6, upper panel) does not indicate high protein intensities in this area of the gel.

**Figure 6:**
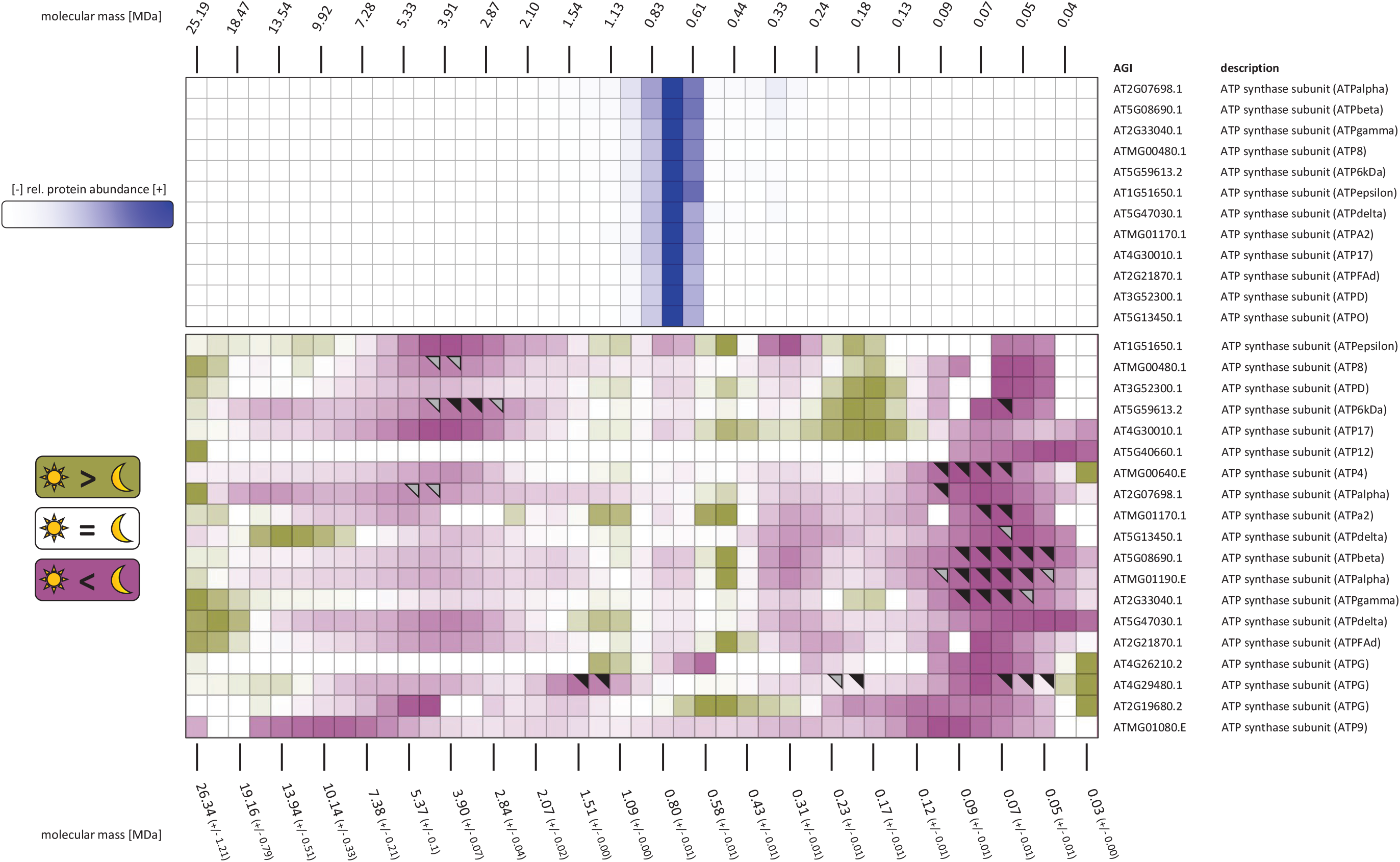
Differential heatmap for complex V. Upper panel shows abundance profiles of complex V (ATP synthase, C-V) subunits in a replicate of mitochondria isolated from dark-adapted plants (dark II, Figure S2). Color indicates relative protein abundance (dark blue, high relative abundance; light blue, low abundance; white, very little relative abundance or no detection). Lower panel shows a differential heatmap for complex V. For this approach, log_10_-transformed abundance profiles of five biological replicates for each condition (‘dark’, ‘light’), were merged into a single, average profile. Values of the ‘dark’ profile were then subtracted from values of the ‘light’ profile. Purple color indicates higher abundance in dark-adapted mitochondria, whereas green indicates higher abundance in light-adapted mitochondria. Black and grey triangles in the top right corners of tiles indicate p-values of ≤0.05 and >0.05 ≤0.1, respectively. Fractions 3 to 45 after alignment are shown. Please note that the number of fractions in the abundance heatmaps (upper panels of all parts of the figure) does not match the number of fractions in the differential heatmaps (lower panels in all parts of the figure) due to warping and alignment of the individual replicates, in which the original rasters were lost and replaced by a virtual raster containing a higher number of fractions. AGI, Arabidopsis genome identifier.

### Assessing the significance and biological meaning of altered abundance profiles of TCA-cycle and OXPHOS subunits and enzymes in the dark and in the light

All proteins covered in this study reacted in some way or another to the two conditions (dark and light) tested in this study. Due to the four-dimensional output of the differential complexome profiling approach undertaken here (light condition, protein ID, electrophoretic mobility, protein abundance), which is further challenged by technical limitations, such as variation in the migration patterns of proteins, it is difficult to assess the significance of the data presented here. Particularly in the high molecular mass region of lpBN gels, protein abundance is generally low and here, changes in protein abundance may only affect a very small portion of the total protein abundance (< 1%). The resulting low signal-to-noise (S/N) ratio further compromizes statistical significance testing. A statistical analysis of the abundance of all 843 proteins at all positions in the lpBN gels is found in Fig. S7 and Tab. S2. In total, 152 proteins show a change in at least one position in the lpBN gels with a p-value ≤0.05. This number is extended by another 230 proteins when the limit is extended to ≤0.1.

However, while the assessment of the statistical significance of individual proteins is a common feature of shotgun MS data it largely ignores the very nature of multiprotein assemblies. By definition, these assemblies consist of more than one protein species. In extreme cases (such as C-I), up to 50 different proteins associate to form a functional enzymatic entity. A uniform response of the subunits towards the light condition at a given position within the lpBN gel can thus be considered a strong signal in terms of significance. Hence (although provided), the work described here omits the results of individual statistical significance for individual proteins at individual gel positions and focusses solely on the group dynamics of protein complexes with known subunit compositions. Since the composition of TCA-cycle protein and OXPHOS protein complexes is well established for the model plant Arabidopsis, the data can be summarized as follows:

i. Many TCA-cycle enzymes increase their abundance in the high-molecular mass range at ~19 MDa in the light, whereas a shift towards 2 MDa assembly prevails in the dark. Considering previously published results documenting the dynamics of the TCA cycle and the interactions of its enzymes (Zhang et al. 2017; Sweetlove et al. 2010), it is safe to assume that the changes observed here represent dynamic adaptations of the TCA-cycle PPIs respinding to alterations in light conditions.
ii. The association of C-I and C-III is shifted from individual complexes or small supercomplexes towards the larger SC-I_2_/III_2_ in the light. The consequences this may have for electron transport remain unknown.
iii. The ATP-synthase complex reacts to illumination changes by altering the abundance of several assemblies with differing protein composition, which are difficult to interpret. The clearest signals originate from the monomeric complex and its subunits, both of which are of increased abundance in the dark.

In many cases, the observed changes take place in the high molecular mass range of the lpBN-gel where only minor amounts of the proteins are found, raising the question about the functional impact of the observed assemblies and their dynamic changes in abundance, particularly in case of the TCA-cycle enzymes. The minute abundances could be attributed to either biological or technical reasons, or the combination of both. The classical isolation procedure for mitochondria using centrifugation-based techniques takes hours from plant harvest and cell disruption to the final isolate. During this time, PPIs in organelles separated from their natural environment and depleted from their supply of metabolites can be expected to reduce in their intensity. It should also be considered, that leaf material was used for this study. These plant organs are composed of different cell types (epidermis cells, parenchyma cell, guard cells, cells of the vascular system), each of which being present in different amounts and reacting differently to changing light conditions. As such, the picture which emerges from the differential heatmaps presented here is a picture integrating all of these cell types and their dynamic adjustments. Finally, the association of, for example, TCA-cycle enzymes may improve flux through a pathway (or a part of said pathway) so efficiently that even small amounts can produce a considerable difference. Indeed, enzyme clusters catalyzing three subsequent reaction steps are reported to be more than 100 times more efficient than evenly distributed enzymes (Castellana et al. 2014).

## Outlook

The data presented merely grant a first glimpse at the dynamics of the Arabidopsis mitochondrial PPI landscape. Although some valuable conclusions can already be drawn from these data, they are clearly still limited by technical shortcomings. As outlined above, one of these shortcomings is the assumed loss of PPIs during the isolation procedure and, potentially, also during sample preparation and the separation process itself. New isolation techniques employing fast affinity purification based approaches are beginning to emerge (Niehaus et al. 2020; Kuhnert et al. 2020), considerably reducing the time required for obtaining high quality mitochondrial isolates. Another shortcoming is the composition of plant leaves from different cell types. By using an affinity purification procedure, in which the tag is expressed under a cell-type specific promotor, cell-type specific mitochondria can be isolated, which reduce interference from other cell types in a given plant organ (Boussardon et al. 2020). The organellar yields provided by such approaches are low but compatible with the complexome profiling workflow, given that highly sensitive mass spectrometers are used. This improved workflow, possibly combined with a cross-linking strategy, may also allow the investigation of highly unstable protein complexes, such as the glycine decarboxylase complex (GDC).

Another limitation is our current knowledge of the composition of mitochondrial protein complexes apart from the highly abundant complexes constituting the TCA-cycle, the OXPHOS system, and the protein import machinery. Non-differential complexome profiling is seeking to provide answers to these questions but co-migration of protein complexes is clouding our view on the interaction patterns of proteins in mitochondria (or other organelles). Pending the improvements to the (differential) complexome profiling procedure, this technique has the potential to detect co-migration of protein complexes when these complexes react differently to individual treatments, stresses, conditions, or gene knock-outs and are thus an important tool not only for investigating the dynamics of protein complexes, but also for elucidating their composition.

## Materials & Methods

### Plant material and plant cultivation conditions

*Arabidopsis thaliana* (ecotype Columbia) plants were grown under short-day conditions (10 h of light / 14 h of dark). Day conditions were set to 120 μmol sec^−1^ m^−2^ radiation at 22° C and 65% humidity. Night conditions were set to 20° C and 65% humidity. Rosette leaves were harvested 2 h before changes in the light conditions took place. For both conditions, five biological replicates were produced.

### Mitochondria Isolation

Organelle isolation from Arabidopsis leaf material was performed as described previously (Senkler et al. 2017) with minor adjustments. In brief, differential and isopycnic centrifugation was followed by four washing steps to remove Percoll from the isolated organelles. Isolation of dark mitochondria was achieved by performing plant harvesting and cell disruption under low-intensity green light. The concentration of mitochondrial isolates was adjusted to 3 mg protein/ml (according to Bradford). Only freshly isolated mitochondria were used for complexome profiling analysis.

### Large-Pore Blue-Native PAGE (lpBN-PAGE)

lpBN-PAGE was performed according to Strecker et al. (Strecker et al. 2010) with minor modifications as introduced by Rugen et al. (2019). To stabilize the gel matrix in the high molecular-mass area, gels were firmly attached to one of the two glass gel plates via binding silane (GE Healthcare, Uppsala, Sweden) whereas the opposite plate was coated with a releasing agent (Blueslick, SERVA electrophoresis GmbH, Heidelberg, Germany) according to the manufacturer’s directions. For sample preparation, 42 μl of mitochondria suspension (corresponding to 125 ug of protein) were treated with 42 ul of digitonin solubilization buffer (50 mM imidazole, 50 mM NaCl, 2 mM aminocaproic acid (ACA), 1 mM EDTA and 5% [w/v] digitonin, pH 7.0) and incubated for 20 min on ice. Separation of protein complexes in lpBN-PAGE was performed as described previously (Rugen et al. 2019). In brief, protein complexes were loaded onto an acrylamide gradient gel (2% T, 20% C to 13% T, 3% C, in 0.5 M 6-aminohexanoic acid, 25 mM imidazole/HCL, pH 7.4) overlaid with a 2.5% T, 25% C (in 0.5 M 6-aminohexanoic acid, 25 mM imidazole/HCL, pH 7.4) sample gel. After 1 h at 100 V, separation of protein complexes continued for another 20 h at 500 V and a maximum of 15 mA. After Coomassie staining, each gel lane was cut into 46 – 48 fractions of 4 mm each from bottom to top and subsequently subjected to tryptic in-gel digestion as described by Senkler et al. (2017). To avoid blocking the UPLC capillaries and columns with remnants of soft gel pieces from the top of the gel, peptide extraction was reduced to two steps, first using 50% [v/v] ACN, 2.5% [v/v] formic acid, and then using 100% [v/v] acetonitrile (ACN), 1% [v/v] formic acid.

### Liquid Chromatography Coupled Tandem Mass Spectrometry (LC-MS/MS)

LC-MS/MS was performed according to (Thal et al. 2018) with minor modifications. Extracted peptides were resuspended in 20 μl of 5% [v/v] ACN and 0.1% [v/v] TFA and 5 μl of peptide solution were loaded onto a 2 cm C18 reversed phase trap column (Acclaim PepMap100, diameter: 100 μm, granulometry: 5 μm, pore size: 100 Å; Thermo Fisher Scientific, Waltham, MA, USA) to be further separated on a 50 cm C18 reversed phase analytical column (Acclaim PepMap100, diameter: 75 μm, granulometry: 3 μm, pore size: 100 Å; Thermo Fisher Scientific). Peptides were eluted by a non-linear 5-36% [v/v] acetonitrile gradient in 0.1% [v/v] formic acid at a flow rate of 250 nl min-1 over a period of 60 min at 33° C. Transfer of eluting peptides into the mass spectrometer (Q-Exactive, Thermo Fisher Scientific, Dreieich, Germany) was achieved via electrospray ionization (ESI) using a NSI source (Thermo Fisher Scientific, Dreieich, Germany) equipped with a stainless steel nano-bore emitter (Thermo Fisher Scientific, Dreieich, Germany). Spray voltage was set to 2.2 kV, capillary temperature to 275° C, and S-lens RF level to 50%. The MS was run in positive ion mode, MS/MS spectra (top 10) were recorded from 20 to 100 min. Full MS scans were performed at a resolution of 70,000 whereas 17,500 was used for MS/MS scans. Automatic gain control (AGC) targets for MS and MS/MS were set to 1E6 and 1E5, respectively. Only peptides with 2, 3, or 4 positive charges were considered.

### Processing of MS Data

LC-MS/MS spectra were analyzed according to Fuchs et al. 2020 with minor adjustments. MaxQuant version 1.6.4.0; (Tyanova et al. 2016) was used to query acquired LC-MS/MS spectra against an in-house modified TAIR10 database (Berardini et al. 2015) incorporating RNA-edited mitochondrial and chloroplast protein models, as well as common contaminants. RNA editing site information was derived from the PREPACT database (Lenz et al. 2018) and a published RNA-seq analysis (Bentolila et al. 2013). Only sites that are edited at more than 50% efficiency were considered.

The search parameters were set to: carbamidomethyl (C) as fixed modification, oxidation at methionine (M) and acetylation at the protein N-terminus as variable modifications. The specific digestions mode was set to trypsin (P) and two missed cleavage sites were allowed. A FDR of 1% was applied on the PSM and protein level. Protein groups containing identified contaminants or false-positive identifications were manually removed before further data analysis, the iBAQ (Schwanhäusser et al. 2011) function of MaxQuant was enabled, “log fit” disabled. For building individual complexome maps, protein abundance profiles based on the iBAQ value were hierarchically clustered by the NOVA software (version 0.5.9.1) (Giese et al. 2015). Average linkage was applied based on the Pearson correlation distance. No normalization was performed before clustering.

### Alignment of Complexome Maps and Statistical Analysis

To identify significantly altered protein abundance between light and dark samples, cubic regression splines for the log-transformed relative protein abundances were fitted, depending on the normalized positions. To account for repeated measurements from the same biological replication, generalized additive mixed models were used for fitting the splines. Based on these fitted splines, significance tests between the light and dark treatment at different values of the normalized gel positions were performed using methods by Herberich et al. (Herberich et al. 2014). A detailed description of the statistical procedure is given in the supplemental methods section.

### Building a Differential Abundance Heatmap

From the aligned, normalized sample heatmaps, average abundance profiles were produced for each protein present in at least four of the five dark and light samples. At each (aligned) position, the values of average light abundance were subtracted from those of the average dark abundance. These values were loaded (in form of an Excel spreadsheet) into the NOVA software (Giese et al. 2015) to be displayed in a three color code with white indicating either an unchanged signal (value=0) or the absence of data at this position and purple and green indicating higher abundance in the dark and the light samples, respectively.

## Acknowledgements

We thank Marianna Langer for expert technical assistance and Michael Senkler for most helpful IT support. We also thank Mareike Schallenberg-Rüdinger and Etienne Meyer for critically reading and discussing the manuscript. This work is founded by a DFG grant (EU 54/4-1) to HE.

---

**Figure S1: Complexome profiling workflow.**1, *Arabidopsis thaliana* (Col-0) plants were grown for 5 weeks in a 10 h light / 14 h dark regime. 2, plants were harvested 2h before dawn or 2h before dusk and used for isolation of mitochondria by differential and isopycinic centrifugation. Five independent isolates were produced for each condition (dark, light) and processed individually as follows. 3, after solubilization, mitochondrial protein complexes were separated according to molecular mass in large-pore blue-native (lpBN) gels. 4, gel lanes of individual isolates were dissected into 46-48 gel fractions followed by MS analysis of each fraction. Heatmaps for each sample were produced on the basis of abundance profiles of proteins along the gel lane using the NOVA software (Giese et al. 2015)(Figs S2, 3). 5, after alignment (not shown) the heatmaps for all five samples produced from the ‘dark’ mitochondria or the ‘light’ mitochondria are summarized in average ‘dark’ and ‘light’ heatmaps (Fig. S6). 6, parallel to the production of the average heatmaps, statistics was performed to identify positions (molecular masses) at which protein abundance differs significantly (t-test) between ‘dark’ and ‘light’ mitochondria. Results are provided in graphical form (Fig. S7) and in tabular form (Tab. S2). 7, statistics results as well as the average heatmaps were used to produce a differential heatmap in which the average light-adapted intensities were subtracted from the corresponding dark-adapted intensities followed by cluster analysis as outlined for the individual heatmaps (Fig. 1).

**Figure S2: Dark replicate heatmaps.** Individual heatmaps produced from five replicates of mitochondria isolated form rosette leaves 2h hours before the end of the dark period (Fig. S1). Blue color indicates relative protein abundance (dark blue: high relative abundance; light blue: low abundance; white: no detection).

**Figure S3: Light replicate heatmaps.** Individual heatmaps produced from five replicates of mitochondria isolated from rosette leaves 2h before the light period (Fig. S1). Blue color indicates relative protein abundance (dark blue: high relative abundance; light blue: low abundance; white: no detection).

**Figure S4: Purity of mitochondrial fractions isolated from dark- and light-adapted plants.**All identified proteins of the ten samples (five mitochondrial fractions isolated from light-adapted plants, five mitochondrial fractions isolated from dark-adapted plants) were assigned to subcellular compartments in accordance with SUBAcon (Hooper et al. 2014). Next, accumulated iBAQ values were calculated for all subcellular compartments in all ten samples. Finally, the proportion of the accumulated iBAQ values for the subcellular compartments with respect to the total accumulated iBAQ value was displayed for all ten samples (in %).

**Figure S5: Average accumulated protein abundance and protein complexity profiles for mitochondria isolated in the dark and the light.** A, mitochondria isolated from light-adapted plants (5 samples); B, mitochondria isolated from dark-adapted plants (5 samples). Average protein intensities according to the cumulated iBAQ values in each fraction (top; yellow graphs) are shown above a representative lpBN gel lane (middle) and the number of protein species identified in each fraction (bottom; green graphs). Grey areas indicate variation, expressed as standard deviation.

**Figure S6: Average light and dark heatmaps produced after alignment of the individual heatmaps.** A, light; B, dark. These average heatmaps form the basis of the differential heatmap shown in Fig. 1. Numbers and bars on top of the heatmaps indicate molecular mass expressed in megadalton (MDa).

**Figure S7: Protein abundance profiles of 843 proteins passing the selection criteria for quantitation.** Horizontal axis, normalized gel fraction (1, top of the gel; 45 bottom of the gel); vertical axis, relative, log10-transformed protein intensity; bold graphs, average protein abundance; thin and dotted lines, data from individual samples. black frame indicates proteins for which the dark vs light t-test in at least one fraction has a p-value ≤0.05; grey frame indicates proteins for which the t-test in at least one fraction has a p-value >0.05 and ≤0.1. Fractions marked by “*”, “**”, or “***” have p-values of ≤0.05, ≤0.03, and ≤0.01, respectively. Fractions marked by “.” have p-values of >0.05 and ≤0.1.

**Figure S8:** Abundance of PDC subunits across the lpBN-gel. Top and bottom panel, absolute abundance of cumulated PDC-E1α (including PDC-E1 α-like), PDC-E1β, cumulated PDC-E2, and cumulated E3 subunits within the lpBN gel of the dark II replicate on a linear and logarithmic (log_10_) scale, respectively. Middle panel, relative abundance of individual PDC subunits as derived from the dark II heatmap.

**Figure S9:** Abundance of OGDC subunits across the lpBN-gel. Top and bottom panel, absolute abundance of cumulated OGDC-E1, cumulated OGDC-E2, and cumulated E3 subunits within the lpBN gel of the dark II replicate on a linear and logarithmic (log_10_) scale, respectively. Middle panel, relative abundance of individual OGDC subunits as derived from the dark II heatmap.

## References

Affourtit, C.; Krab, K.; Leach, G. R.; Whitehouse, D. G.; Moore, A. L. (2001): New insights into the regulation of plant succinate dehydrogenase. On the role of the protonmotive force. In: The Journal of biological chemistry 276 (35), S. 32567–32574. DOI: 10.1074/jbc.M103111200.

Alston, Charlotte L.; Veling, Mike T.; Heidler, Juliana; Taylor, Lucie S.; Alaimo, Joseph T.; Sung, Andrew Y. et al. (2020): Pathogenic Bi-allelic Mutations in NDUFAF8 Cause Leigh Syndrome with an Isolated Complex I Deficiency. In: American journal of human genetics 106 (1), S. 92–101. DOI: 10.1016/j.ajhg.2019.12.001.

Beeckmans, S.; Kanarek, L. (1981): Demonstration of physical interactions between consecutive enzymes of the citric acid cycle and of the aspartate-malate shuttle. A study involving fumarase, malate dehydrogenase, citrate synthesis and aspartate aminotransferase. In: European journal of biochemistry 117 (3), S. 527–535. DOI: 10.1111/j.1432-1033.1981.tb06369.x.

Behal, R. H.; Oliver, D. J. (1997): Biochemical and molecular characterization of fumarase from plants: purification and characterization of the enzyme--cloning, sequencing, and expression of the gene. In: Archives of biochemistry and biophysics 348 (1), S. 65–74. DOI: 10.1006/abbi.1997.0359.

Belt, Katharina; Huang, Shaobai; Thatcher, Louise F.; Casarotto, Hayley; Singh, Karam B.; van Aken, Olivier; Millar, A. Harvey (2017): Salicylic Acid-Dependent Plant Stress Signaling via Mitochondrial Succinate Dehydrogenase. In: Plant physiology 173 (4), S. 2029–2040. DOI: 10.1104/pp.16.00060.

Bentolila, Stéphane; Oh, Julyun; Hanson, Maureen R.; Bukowski, Robert (2013): Comprehensive high-resolution analysis of the role of an Arabidopsis gene family in RNA editing. In: PLoS genetics 9 (6), e1003584. DOI: 10.1371/journal.pgen.1003584.

Berardini, Tanya Z.; Reiser, Leonore; Li, Donghui; Mezheritsky, Yarik; Muller, Robert; Strait, Emily; Huala, Eva (2015): The Arabidopsis information resource: Making and mining the “gold standard” annotated reference plant genome. In: Genesis (New York, N.Y. : 2000) 53 (8), S. 474–485. DOI: 10.1002/dvg.22877.

Boussardon, Clément; Przybyla-Toscano, Jonathan; Carrie, Chris; Keech, Olivier (2020): Tissue-specific isolation of Arabidopsis/plant mitochondria - IMTACT (isolation of mitochondria tagged in specific cell types). In: The Plant journal : for cell and molecular biology. DOI: 10.1111/tpj.14723.

Braun, Hans-Peter (2020): The Oxidative Phosphorylation system of the mitochondria in plants. In: Mitochondrion. DOI: 10.1016/j.mito.2020.04.007.

Braun, Hans-Peter; Binder, Stefan; Brennicke, Axel; Eubel, Holger; Fernie, Alisdair R.; Finkemeier, Iris et al. (2014): The life of plant mitochondrial complex I. In: Mitochondrion 19 Pt B, S. 295–313. DOI: 10.1016/j.mito.2014.02.006.

Budde, Raymond J.A.; Randall, Douglas D. (1987): Regulation of pea mitochondrial pyruvate dehydrogenase complex activity: Inhibition of ATP-dependent inactivation. In: Archives of biochemistry and biophysics 258 (2), S. 600–606. DOI: 10.1016/0003-9861(87)90382-1.

Bunik, Victoria; Strumilo, Slawomir (2009): Regulation of Catalysis Within Cellular Network: Metabolic and Signaling Implications of the 2-Oxoglutarate Oxidative Decarboxylation. In: CCB 3 (3), S. 279–290. DOI: 10.2174/2212796810903030279.

Carrari, Fernando; Nunes-Nesi, Adriano; Gibon, Yves; Lytovchenko, Anna; Loureiro, Marcelo Ehlers; Fernie, Alisdair R. (2003): Reduced expression of aconitase results in an enhanced rate of photosynthesis and marked shifts in carbon partitioning in illuminated leaves of wild species tomato. In: Plant physiology 133 (3), S. 1322–1335. DOI: 10.1104/pp.103.026716.

Castellana, Michele; Wilson, Maxwell Z.; Xu, Yifan; Joshi, Preeti; Cristea, Ileana M.; Rabinowitz, Joshua D. et al. (2014): Enzyme clustering accelerates processing of intermediates through metabolic channeling. In: Nature biotechnology 32 (10), S. 1011–1018. DOI: 10.1038/nbt.3018.

Clifton, Rachel; Lister, Ryan; Parker, Karen L.; Sappl, Pia G.; Elhafez, Dina; Millar, A. Harvey et al. (2005): Stress-induced co-expression of alternative respiratory chain components in Arabidopsis thaliana. In: Plant molecular biology 58 (2), S. 193–212. DOI: 10.1007/s11103-005-5514-7.

Cox, Jürgen; Hein, Marco Y.; Luber, Christian A.; Paron, Igor; Nagaraj, Nagarjuna; Mann, Matthias (2014): Accurate proteome-wide label-free quantification by delayed normalization and maximal peptide ratio extraction, termed MaxLFQ. In: Molecular & cellular proteomics : MCP 13 (9), S. 2513–2526. DOI: 10.1074/mcp.M113.031591.

Daloso, Danilo M.; Müller, Karolin; Obata, Toshihiro; Florian, Alexandra; Tohge, Takayuki; Bottcher, Alexandra et al. (2015): Thioredoxin, a master regulator of the tricarboxylic acid cycle in plant mitochondria. In: Proceedings of the National Academy of Sciences of the United States of America 112 (11), E1392–400. DOI: 10.1073/pnas.1424840112.

Dasika, Santosh K.; Vinnakota, Kalyan C.; Beard, Daniel A. (2015): Determination of the catalytic mechanism for mitochondrial malate dehydrogenase. In: Biophysical journal 108 (2), S. 408–419. DOI: 10.1016/j.bpj.2014.11.3467.

Dudkina, Natalia V.; Eubel, Holger; Keegstra, Wilko; Boekema, Egbert J.; Braun, Hans-Peter (2005): Structure of a mitochondrial supercomplex formed by respiratory-chain complexes I and III. In: Proceedings of the National Academy of Sciences of the United States of America 102 (9), S. 3225–3229. DOI: 10.1073/pnas.0408870102.

Dudkina, Natalya V.; Sunderhaus, Stephanie; Braun, Hans-Peter; Boekema, Egbert J. (2006): Characterization of dimeric ATP synthase and cristae membrane ultrastructure from Saccharomyces and Polytomella mitochondria. In: FEBS Letters 580 (14), S. 3427–3432. DOI: 10.1016/j.febslet.2006.04.097.

Dutilleul, Christelle; Garmier, Marie; Noctor, Graham; Mathieu, Chantal; Chétrit, Philippe; Foyer, Christine H.; Paepe, Rosine de (2003): Leaf mitochondria modulate whole cell redox homeostasis, set antioxidant capacity, and determine stress resistance through altered signaling and diurnal regulation. In: The Plant cell 15 (5), S. 1212–1226. DOI: 10.1105/tpc.009464.

Elhafez, Dina; Murcha, Monika W.; Clifton, Rachel; Soole, Kathleen L.; Day, David A.; Whelan, James (2006): Characterization of mitochondrial alternative NAD(P)H dehydrogenases in Arabidopsis: intraorganelle location and expression. In: Plant & cell physiology 47 (1), S. 43–54. DOI: 10.1093/pcp/pci221.

Escobar, Matthew A.; Franklin, Keara A.; Svensson, A. Staffan; Salter, Michael G.; Whitelam, Garry C.; Rasmusson, Allan G. (2004): Light regulation of the Arabidopsis respiratory chain. Multiple discrete photoreceptor responses contribute to induction of type II NAD(P)H dehydrogenase genes. In: Plant physiology 136 (1), S. 2710–2721. DOI: 10.1104/pp.104.046698.

Eubel, Holger; Heinemeyer, Jesco; Braun, Hans-Peter (2004): Identification and characterization of respirasomes in potato mitochondria. In: Plant physiology 134 (4), S. 1450–1459. DOI: 10.1104/pp.103.038018.

Eubel, Holger; Jänsch, Lothar; Braun, Hans-Peter (2003): New insights into the respiratory chain of plant mitochondria. Supercomplexes and a unique composition of complex II. In: Plant physiology 133 (1), S. 274–286. DOI: 10.1104/pp.103.024620.

Fromm, Steffanie; Braun, Hans-Peter; Peterhansel, Christoph (2016): Mitochondrial gamma carbonic anhydrases are required for complex I assembly and plant reproductive development. In: The New phytologist 211 (1), S. 194–207. DOI: 10.1111/nph.13886.

Fuchs, Philippe; Rugen, Nils; Carrie, Chris; Elsässer, Marlene; Finkemeier, Iris; Giese, Jonas et al. (2020): Single organelle function and organization as estimated from Arabidopsis mitochondrial proteomics. In: The Plant journal : for cell and molecular biology 101 (2), S. 420–441. DOI: 10.1111/tpj.14534.

Gelhaye, Eric; Rouhier, Nicolas; Gérard, Joelle; Jolivet, Yves; Gualberto, José; Navrot, Nicolas et al. (2004): A specific form of thioredoxin h occurs in plant mitochondria and regulates the alternative oxidase. In: Proceedings of the National Academy of Sciences of the United States of America 101 (40), S. 14545–14550. DOI: 10.1073/pnas.0405282101.

Gellerich, Frank N.; Gizatullina, Zemfira; Trumbeckaite, Sonata; Nguyen, Huu P.; Pallas, Thilo; Arandarcikaite, Odeta et al. (2010): The regulation of OXPHOS by extramitochondrial calcium. In: Biochimica et biophysica acta 1797 (6-7), S. 1018–1027. DOI: 10.1016/j.bbabio.2010.02.005.

Giese, Heiko; Ackermann, Jörg; Heide, Heinrich; Bleier, Lea; Dröse, Stefan; Wittig, Ilka et al. (2015): NOVA: a software to analyze complexome profiling data. In: Bioinformatics (Oxford, England) 31 (3), S. 440–441. DOI: 10.1093/bioinformatics/btu623.

Gleason, Cynthia; Huang, Shaobai; Thatcher, Louise F.; Foley, Rhonda C.; Anderson, Carol R.; Carroll, Adam J. et al. (2011): Mitochondrial complex II has a key role in mitochondrial-derived reactive oxygen species influence on plant stress gene regulation and defense. In: Proceedings of the National Academy of Sciences of the United States of America 108 (26), S. 10768–10773. DOI: 10.1073/pnas.1016060108.

Herberich, Esther; Hassler, Christine; Hothorn, Torsten (2014): Multiple curve comparisons with an application to the formation of the dorsal funiculus of mutant mice. In: The international journal of biostatistics 10 (2), S. 289–302. DOI: 10.1515/ijb-2013-0003.

Hirling, H.; Henderson, B. R.; Kühn, L. C. (1994): Mutational analysis of the 4Fe-4S-cluster converting iron regulatory factor from its RNA-binding form to cytoplasmic aconitase. In: The EMBO journal 13 (2), S. 453–461.

Hofer, Annette; Wenz, Tina (2014): Post-translational modification of mitochondria as a novel mode of regulation. In: Experimental gerontology 56, S. 202–220. DOI: 10.1016/j.exger.2014.03.006.

Hooper, Cornelia M.; Castleden, Ian R.; Tanz, Sandra K.; Aryamanesh, Nader; Millar, A. Harvey (2017): SUBA4: the interactive data analysis centre for Arabidopsis subcellular protein locations. In: Nucleic acids research 45 (D1), D1064–D1074. DOI: 10.1093/nar/gkw1041.

Hooper, Cornelia M.; Tanz, Sandra K.; Castleden, Ian R.; Vacher, Michael A.; Small, Ian D.; Millar, A. Harvey (2014): SUBAcon: a consensus algorithm for unifying the subcellular localization data of the Arabidopsis proteome. In: Bioinformatics (Oxford, England) 30 (23), S. 3356–3364. DOI: 10.1093/bioinformatics/btu550.

Huang, Shaobai; Braun, Hans-Peter; Gawryluk, Ryan M. R.; Millar, A. Harvey (2019): Mitochondrial complex II of plants: subunit composition, assembly, and function in respiration and signaling. In: The Plant journal : for cell and molecular biology 98 (3), S. 405–417. DOI: 10.1111/tpj.14227.

Huang, Y. C.; Kumar, A.; Colman, R. F. (1997): Identification of the subunits and target peptides of pig heart NAD-specific isocitrate dehydrogenase modified by the affinity label 8-(4-bromo-2,3-dioxobutylthio)NAD. In: Archives of biochemistry and biophysics 348 (1), S. 207–218. DOI: 10.1006/abbi.1997.0392.

Igamberdiev, Abir U.; Gardeström, Per (2003): Regulation of NAD- and NADP-dependent isocitrate dehydrogenases by reduction levels of pyridine nucleotides in mitochondria and cytosol of pea leaves. In: Biochimica et Biophysica Acta (BBA) - Bioenergetics 1606 (1-3), S. 117–125. DOI: 10.1016/S0005-2728(03)00106-3.

Ito, Jun; Taylor, Nicolas L.; Castleden, Ian; Weckwerth, Wolfram; Millar, A. Harvey; Heazlewood, Joshua L. (2009): A survey of the Arabidopsis thaliana mitochondrial phosphoproteome. In: Proteomics 9 (17), S. 4229–4240. DOI: 10.1002/pmic.200900064.

Keys, D. A.; McAlister-Henn, L. (1990): Subunit structure, expression, and function of NAD(H)-specific isocitrate dehydrogenase in Saccharomyces cerevisiae. In: Journal of bacteriology 172 (8), S. 4280–4287. DOI: 10.1128/jb.172.8.4280-4287.1990.

Klodmann, Jennifer; Lewejohann, Dagmar; Braun, Hans-Peter (2011): Low-SDS Blue native PAGE. In: Proteomics 11 (9), S. 1834–1839. DOI: 10.1002/pmic.201000638.

König, Ann-Christine; Hartl, Markus; Boersema, Paul J.; Mann, Matthias; Finkemeier, Iris (2014a): The mitochondrial lysine acetylome of Arabidopsis. In: Mitochondrion 19 Pt B, S. 252–260. DOI: 10.1016/j.mito.2014.03.004.

König, Ann-Christine; Hartl, Markus; Pham, Phuong Anh; Laxa, Miriam; Boersema, Paul J.; Orwat, Anne et al. (2014b): The Arabidopsis class II sirtuin is a lysine deacetylase and interacts with mitochondrial energy metabolism. In: Plant physiology 164 (3), S. 1401–1414. DOI: 10.1104/pp.113.232496.

Krause, Frank; Reifschneider, Nicole H.; Vocke, Dirk; Seelert, Holger; Rexroth, Sascha; Dencher, Norbert A. (2004): “Respirasome”-like supercomplexes in green leaf mitochondria of spinach. In: The Journal of biological chemistry 279 (46), S. 48369–48375. DOI: 10.1074/jbc.M406085200.

Kuhnert, Franziska; Stefanski, Anja; Overbeck, Nina; Drews, Leonie; Reichert, Andreas S.; Stühler, Kai; Weber, Andreas P. M. (2020): Rapid Single-Step Affinity Purification of HA-Tagged Plant Mitochondria. In: Plant physiology 182 (2), S. 692–706. DOI: 10.1104/pp.19.00732.

Lancien, M.; Gadal, P.; Hodges, M. (1998): Molecular characterization of higher plant NAD-dependent isocitrate dehydrogenase: evidence for a heteromeric structure by the complementation of yeast mutants. In: The Plant journal : for cell and molecular biology 16 (3), S. 325–333. DOI: 10.1046/j.1365-313x.1998.00305.x.

Lee, Chun Pong; Eubel, Holger; Millar, A. Harvey (2010): Diurnal changes in mitochondrial function reveal daily optimization of light and dark respiratory metabolism in Arabidopsis. In: Molecular & cellular proteomics : MCP 9 (10), S. 2125–2139. DOI: 10.1074/mcp.M110.001214.

Lemaitre, Thomas; Hodges, Michael (2006): Expression analysis of Arabidopsis thaliana NAD-dependent isocitrate dehydrogenase genes shows the presence of a functional subunit that is mainly expressed in the pollen and absent from vegetative organs. In: Plant & cell physiology 47 (5), S. 634–643. DOI: 10.1093/pcp/pcj030.

Lemaitre, Thomas; Urbanczyk-Wochniak, Ewa; Flesch, Valerie; Bismuth, Evelyne; Fernie, Alisdair R.; Hodges, Michael (2007): NAD-dependent isocitrate dehydrogenase mutants of Arabidopsis suggest the enzyme is not limiting for nitrogen assimilation. In: Plant physiology 144 (3), S. 1546–1558. DOI: 10.1104/pp.107.100677.

Lenz, Henning; Hein, Anke; Knoop, Volker (2018): Plant organelle RNA editing and its specificity factors: enhancements of analyses and new database features in PREPACT 3.0. In: BMC bioinformatics 19 (1), S. 255. DOI: 10.1186/s12859-018-2244-9.

Ligas, Joanna; Pineau, Emmanuelle; Bock, Ralph; Huynen, Martijn A.; Meyer, Etienne H. (2019): The assembly pathway of complex I in Arabidopsis thaliana. In: The Plant journal : for cell and molecular biology 97 (3), S. 447–459. DOI: 10.1111/tpj.14133.

Lin, An-Ping; McAlister-Henn, Lee (2003): Homologous binding sites in yeast isocitrate dehydrogenase for cofactor (NAD+) and allosteric activator (AMP). In: The Journal of biological chemistry 278 (15), S. 12864–12872. DOI: 10.1074/jbc.M300154200.

Linn, T. C.; Pettit, F. H.; Reed, L. J. (1969): Alpha-keto acid dehydrogenase complexes. X. Regulation of the activity of the pyruvate dehydrogenase complex from beef kidney mitochondria by phosphorylation and dephosphorylation. In: Proceedings of the National Academy of Sciences of the United States of America 62 (1), S. 234–241. DOI: 10.1073/pnas.62.1.234.

Mansilla, Natanael; Racca, Sofia; Gras, Diana E.; Gonzalez, Daniel H.; Welchen, Elina (2018): The Complexity of Mitochondrial Complex IV: An Update of Cytochrome c Oxidase Biogenesis in Plants. In: International journal of molecular sciences 19 (3). DOI: 10.3390/ijms19030662.

McIntosh, C. A.; Oliver, D. J. (1992): NAD-Linked Isocitrate Dehydrogenase: Isolation, Purification, and Characterization of the Protein from Pea Mitochondria. In: Plant physiology 100 (1), S. 69–75. DOI: 10.1104/pp.100.1.69.

Meyer, Etienne H.; Solheim, Cory; Tanz, Sandra K.; Bonnard, Géraldine; Millar, A. Harvey (2011): Insights into the composition and assembly of the membrane arm of plant complex I through analysis of subcomplexes in Arabidopsis mutant lines. In: The Journal of biological chemistry 286 (29), S. 26081–26092. DOI: 10.1074/jbc.M110.209601.

Michalecka, Agnieszka M.; Svensson, Å. Staffan; Johansson, Fredrik I.; Agius, Stephanie C.; Johanson, Urban; Brennicke, Axel et al. (2003): Arabidopsis Genes Encoding Mitochondrial Type II NAD(P)H Dehydrogenases Have Different Evolutionary Origin and Show Distinct Responses to Light1. In: Plant physiology 133 (2), S. 642–652. DOI: 10.1104/pp.103.024208.

Milenkovic, Dusanka; Blaza, James N.; Larsson, Nils-Göran; Hirst, Judy (2017): The Enigma of the Respiratory Chain Supercomplex. In: Cell metabolism 25 (4), S. 765–776. DOI: 10.1016/j.cmet.2017.03.009.

Millar, A. Harvey; Eubel, Holger; Jänsch, Lothar; Kruft, Volker; Heazlewood, Joshua L.; Braun, Hans-Peter (2004): Mitochondrial cytochrome c oxidase and succinate dehydrogenase complexes contain plant specific subunits. In: Plant molecular biology 56 (1), S. 77–90. DOI: 10.1007/s11103-004-2316-2.

Millar, A. Harvey; Mittova, Valentina; Kiddle, Guy; Heazlewood, Joshua L.; Bartoli, Carlos G.; Theodoulou, Frederica L.; Foyer, Christine H. (2003): Control of ascorbate synthesis by respiration and its implications for stress responses. In: Plant physiology 133 (2), S. 443–447. DOI: 10.1104/pp.103.028399.

Moeder, Wolfgang; Del Pozo, Olga; Navarre, Duroy A.; Martin, Gregory B.; Klessig, Daniel F. (2007): Aconitase plays a role in regulating resistance to oxidative stress and cell death in Arabidopsis and Nicotiana benthamiana. In: Plant molecular biology 63 (2), S. 273–287. DOI: 10.1007/s11103-006-9087-x.

Møller, Ian Max; Igamberdiev, Abir U.; Bykova, Natalia V.; Finkemeier, Iris; Rasmusson, Allan G.; Schwarzländer, Markus (2020): Matrix Redox Physiology Governs the Regulation of Plant Mitochondrial Metabolism through Posttranslational Protein Modifications. In: The Plant cell 32 (3), S. 573–594. DOI: 10.1105/tpc.19.00535.

Morgan, Megan J.; Lehmann, Martin; Schwarzländer, Markus; Baxter, Charles J.; Sienkiewicz-Porzucek, Agata; Williams, Thomas C. R. et al. (2008): Decrease in manganese superoxide dismutase leads to reduced root growth and affects tricarboxylic acid cycle flux and mitochondrial redox homeostasis. In: Plant physiology 147 (1), S. 101–114. DOI: 10.1104/pp.107.113613.

Niehaus, Markus; Straube, Henryk; Künzler, Patrick; Rugen, Nils; Hegermann, Jan; Giavalisco, Patrick et al. (2020): Rapid Affinity Purification of Tagged Plant Mitochondria (Mito-AP) for Metabolome and Proteome Analyses. In: Plant physiology 182 (3), S. 1194–1210. DOI: 10.1104/pp.19.00736.

Nunes-Nesi, Adriano; Araújo, Wagner L.; Obata, Toshihiro; Fernie, Alisdair R. (2013): Regulation of the mitochondrial tricarboxylic acid cycle. In: Current opinion in plant biology 16 (3), S. 335–343. DOI: 10.1016/j.pbi.2013.01.004.

Oestreicher, Guillermo; Hogue, Patricia; Singer, Thomas P. (1973): Regulation of Succinate Dehydrogenase in Higher Plants: II. Activation by Substrates, Reduced Coenzyme Q, Nucleotides, and Anions 1. In: Plant physiology 52 (6), S. 622–626.

O’Leary, Brendan M.; Oh, Glenda Guek Khim; Lee, Chun Pong; Millar, A. Harvey (2019): Metabolite regulatory interactions control plant respiratory metabolism via Target of Rapamycin (TOR) kinase activation. In: The Plant cell. DOI: 10.1105/tpc.19.00157.

Pineau, Bernard; Layoune, Ouardia; Danon, Antoine; Paepe, Rosine de (2008): L-galactono-1,4-lactone dehydrogenase is required for the accumulation of plant respiratory complex I. In: The Journal of biological chemistry 283 (47), S. 32500–32505. DOI: 10.1074/jbc.M805320200.

Rasmusson, Allan G.; Geisler, Daniela A.; Møller, Ian M. (2008): The multiplicity of dehydrogenases in the electron transport chain of plant mitochondria. In: Mitochondrion 8 (1), S. 47–60. DOI: 10.1016/j.mito.2007.10.004.

Reiland, Sonja; Messerli, Gaëlle; Baerenfaller, Katja; Gerrits, Bertran; Endler, Anne; Grossmann, Jonas et al. (2009): Large-scale Arabidopsis phosphoproteome profiling reveals novel chloroplast kinase substrates and phosphorylation networks. In: Plant physiology 150 (2), S. 889–903. DOI: 10.1104/pp.109.138677.

Robinson, J. B.; Srere, P. A. (1985): Organization of Krebs tricarboxylic acid cycle enzymes in mitochondria. In: The Journal of biological chemistry 260 (19), S. 10800–10805.

Rubin, P. M.; Randall, D. D. (1977): Regulation of plant pyruvate dehydrogenase complex by phosphorylation. In: Plant physiology 60 (1), S. 34–39. DOI: 10.1104/pp.60.1.34.

Rugen, Nils; Straube, Henryk; Franken, Linda E.; Braun, Hans-Peter; Eubel, Holger (2019): Complexome Profiling Reveals Association of PPR Proteins with Ribosomes in the Mitochondria of Plants. In: Molecular & cellular proteomics : MCP 18 (7), S. 1345–1362. DOI: 10.1074/mcp.RA119.001396.

Schertl, Peter; Sunderhaus, Stephanie; Klodmann, Jennifer; Grozeff, Gustavo E. Gergoff; Bartoli, Carlos G.; Braun, Hans-Peter (2012): L-galactono-1,4-lactone dehydrogenase (GLDH) forms part of three subcomplexes of mitochondrial complex I in Arabidopsis thaliana. In: The Journal of biological chemistry 287 (18), S. 14412–14419. DOI: 10.1074/jbc.M111.305144.

Schikowsky, Christine; Senkler, Jennifer; Braun, Hans-Peter (2017): SDH6 and SDH7 Contribute to Anchoring Succinate Dehydrogenase to the Inner Mitochondrial Membrane in Arabidopsis thaliana. In: Plant physiology 173 (2), S. 1094–1108. DOI: 10.1104/pp.16.01675.

Schimmeyer, Joram; Bock, Ralph; Meyer, Etienne H. (2016): L-Galactono-1,4-lactone dehydrogenase is an assembly factor of the membrane arm of mitochondrial complex I in Arabidopsis. In: Plant molecular biology 90 (1-2), S. 117–126. DOI: 10.1007/s11103-015-0400-4.

Schmidtmann, Elisabeth; König, Ann-Christine; Orwat, Anne; Leister, Dario; Hartl, Markus; Finkemeier, Iris (2014): Redox regulation of Arabidopsis mitochondrial citrate synthase. In: Molecular plant 7 (1), S. 156–169. DOI: 10.1093/mp/sst144.

Schwanhäusser, Björn; Busse, Dorothea; Li, Na; Dittmar, Gunnar; Schuchhardt, Johannes; Wolf, Jana et al. (2011): Global quantification of mammalian gene expression control. In: Nature 473 (7347), S. 337–342. DOI: 10.1038/nature10098.

Senkler, Jennifer; Senkler, Michael; Eubel, Holger; Hildebrandt, Tatjana; Lengwenus, Christian; Schertl, Peter et al. (2017): The mitochondrial complexome of Arabidopsis thaliana. In: The Plant journal : for cell and molecular biology 89 (6), S. 1079–1092. DOI: 10.1111/tpj.13448.

Simonin, Vagner; Galina, Antonio (2013): Nitric oxide inhibits succinate dehydrogenase-driven oxygen consumption in potato tuber mitochondria in an oxygen tension-independent manner. In: The Biochemical journal 449 (1), S. 263–273. DOI: 10.1042/BJ20120396.

Stram, Amanda R.; Payne, R. Mark (2016): Post-translational modifications in mitochondria: protein signaling in the powerhouse. In: Cellular and molecular life sciences : CMLS 73 (21), S. 4063–4073. DOI: 10.1007/s00018-016-2280-4.

Strecker, Valentina; Wumaier, Zibiernisha; Wittig, Ilka; Schägger, Hermann (2010): Large pore gels to separate mega protein complexes larger than 10 MDa by blue native electrophoresis: isolation of putative respiratory strings or patches. In: Proteomics 10 (18), S. 3379–3387. DOI: 10.1002/pmic.201000343.

Studart-Guimarães, Claudia; Gibon, Yves; Frankel, Nicolás; Wood, Craig C.; Zanor, María Inés; Fernie, Alisdair R.; Carrari, Fernando (2005): Identification and characterisation of the alpha and beta subunits of succinyl CoA ligase of tomato. In: Plant molecular biology 59 (5), S. 781–791. DOI: 10.1007/s11103-005-1004-1.

Sweetlove, L. J.; Heazlewood, J. L.; Herald, V.; Holtzapffel, R.; Day, D. A.; Leaver, C. J.; Millar, A. H. (2002): The impact of oxidative stress on Arabidopsis mitochondria. In: The Plant journal : for cell and molecular biology 32 (6), S. 891–904. DOI: 10.1046/j.1365-313x.2002.01474.x.

Sweetlove, Lee J.; Beard, Katherine F. M.; Nunes-Nesi, Adriano; Fernie, Alisdair R.; Ratcliffe, R. George (2010): Not just a circle: flux modes in the plant TCA cycle. In: Trends in plant science 15 (8), S. 462–470. DOI: 10.1016/j.tplants.2010.05.006.

Tezuka, T.; Laties, G. G. (1983): Isolation and Characterization of Inner Membrane-Associated and Matrix NAD-Specific Isocitrate Dehydrogenase in Potato Mitochondria. In: Plant physiology 72 (4), S. 959–963. DOI: 10.1104/pp.72.4.959.

Thal, Beate; Braun, Hans-Peter; Eubel, Holger (2018): Proteomic analysis dissects the impact of nodulation and biological nitrogen fixation on Vicia faba root nodule physiology. In: Plant molecular biology 97 (3), S. 233–251. DOI: 10.1007/s11103-018-0736-7.

Tovar-Méndez, Alejandro; Miernyk, Jan A.; Randall, Douglas D. (2003): Regulation of pyruvate dehydrogenase complex activity in plant cells. In: European journal of biochemistry 270 (6), S. 1043–1049. DOI: 10.1046/j.1432-1033.2003.03469.x.

Tronconi, Marcos A.; Fahnenstich, Holger; Gerrard Weehler, Mariel C.; Andreo, Carlos S.; Flügge, Ulf-Ingo; Drincovich, María F.; Maurino, Verónica G. (2008): Arabidopsis NAD-malic enzyme functions as a homodimer and heterodimer and has a major impact on nocturnal metabolism. In: Plant physiology 146 (4), S. 1540–1552. DOI: 10.1104/pp.107.114975.

Tronconi, Marcos A.; Maurino, Verónica G.; Andreo, Carlos S.; Drincovich, María F. (2010): Three different and tissue-specific NAD-malic enzymes generated by alternative subunit association in Arabidopsis thaliana. In: The Journal of biological chemistry 285 (16), S. 11870–11879. DOI: 10.1074/jbc.M109.097477.

Tyanova, Stefka; Temu, Tikira; Cox, Juergen (2016): The MaxQuant computational platform for mass spectrometry-based shotgun proteomics. In: Nature protocols 11 (12), S. 2301–2319. DOI: 10.1038/nprot.2016.136.

van Strien, Joeri; Guerrero-Castillo, Sergio; Chatzispyrou, Iliana A.; Houtkooper, Riekelt H.; Brandt, Ulrich; Huynen, Martijn A. (2019): COmplexome Profiling ALignment (COPAL) reveals remodeling of mitochondrial protein complexes in Barth syndrome. In: Bioinformatics (Oxford, England) 35 (17), S. 3083–3091. DOI: 10.1093/bioinformatics/btz025.

Vélot, C.; Mixon, M. B.; Teige, M.; Srere, P. A. (1997): Model of a quinary structure between Krebs TCA cycle enzymes: a model for the metabolon. In: Biochemistry 36 (47), S. 14271–14276. DOI: 10.1021/bi972011j.

Vinnakota, Kalyan C.; Bazil, Jason N.; van den Bergh, Françoise; Wiseman, Robert W.; Beard, Daniel A. (2016): Feedback Regulation and Time Hierarchy of Oxidative Phosphorylation in Cardiac Mitochondria. In: Biophysical journal 110 (4), S. 972–980. DOI: 10.1016/j.bpj.2016.01.003.

Weaver, T. M.; Levitt, D. G.; Donnelly, M. I.; Stevens, P. P.; Banaszak, L. J. (1995): The multisubunit active site of fumarase C from Escherichia coli. In: Nature structural biology 2 (8), S. 654–662. DOI: 10.1038/nsb0895-654.

Wu, Fei; Minteer, Shelley (2015): Krebs cycle metabolon: structural evidence of substrate channeling revealed by cross-linking and mass spectrometry. In: Angewandte Chemie (International ed. in English) 54 (6), S. 1851–1854. DOI: 10.1002/anie.201409336.

Zhang, Youjun; Beard, Katherine F. M.; Swart, Corné; Bergmann, Susan; Krahnert, Ina; Nikoloski, Zoran et al. (2017): Protein-protein interactions and metabolite channelling in the plant tricarboxylic acid cycle. In: Nature communications 8, S. 15212. DOI: 10.1038/ncomms15212.

Zhou, Z. H.; McCarthy, D. B.; O’Connor, C. M.; Reed, L. J.; Stoops, J. K. (2001): The remarkable structural and functional organization of the eukaryotic pyruvate dehydrogenase complexes. In: Proceedings of the National Academy of Sciences of the United States of America 98 (26), S. 14802–14807. DOI: 10.1073/pnas.011597698.

