## Supplemental Figure 7 for "Protein Interaction Patterns in *Arabidopsis thaliana* Leaf Mitochondria Change in Response to Illumination"

### Chromosome 1 (AT1G01170 – AT1G17290)

Figure S7

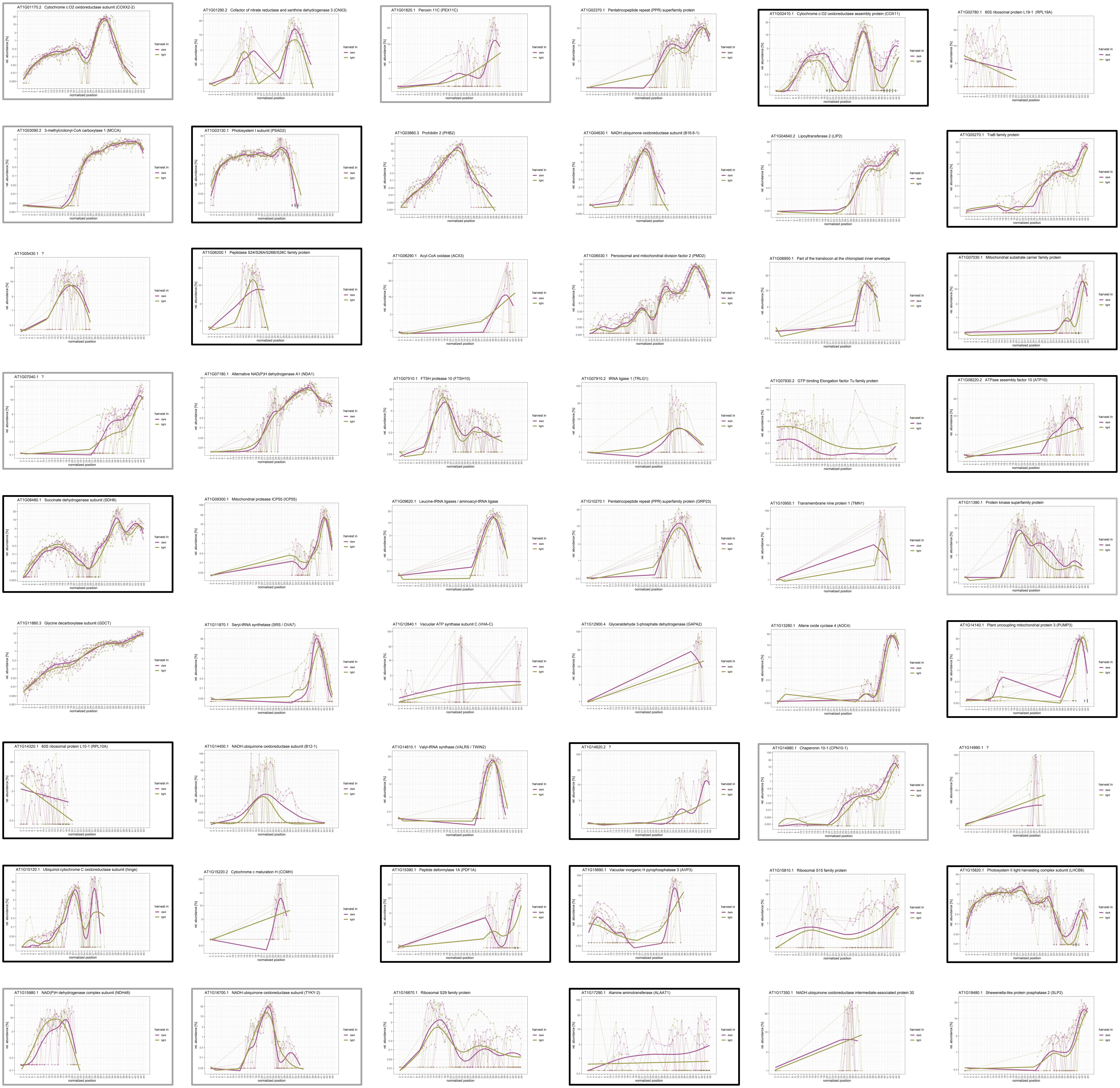

### Chromosome 1 (AT1G8600 – AT1G50200)

Figure S7

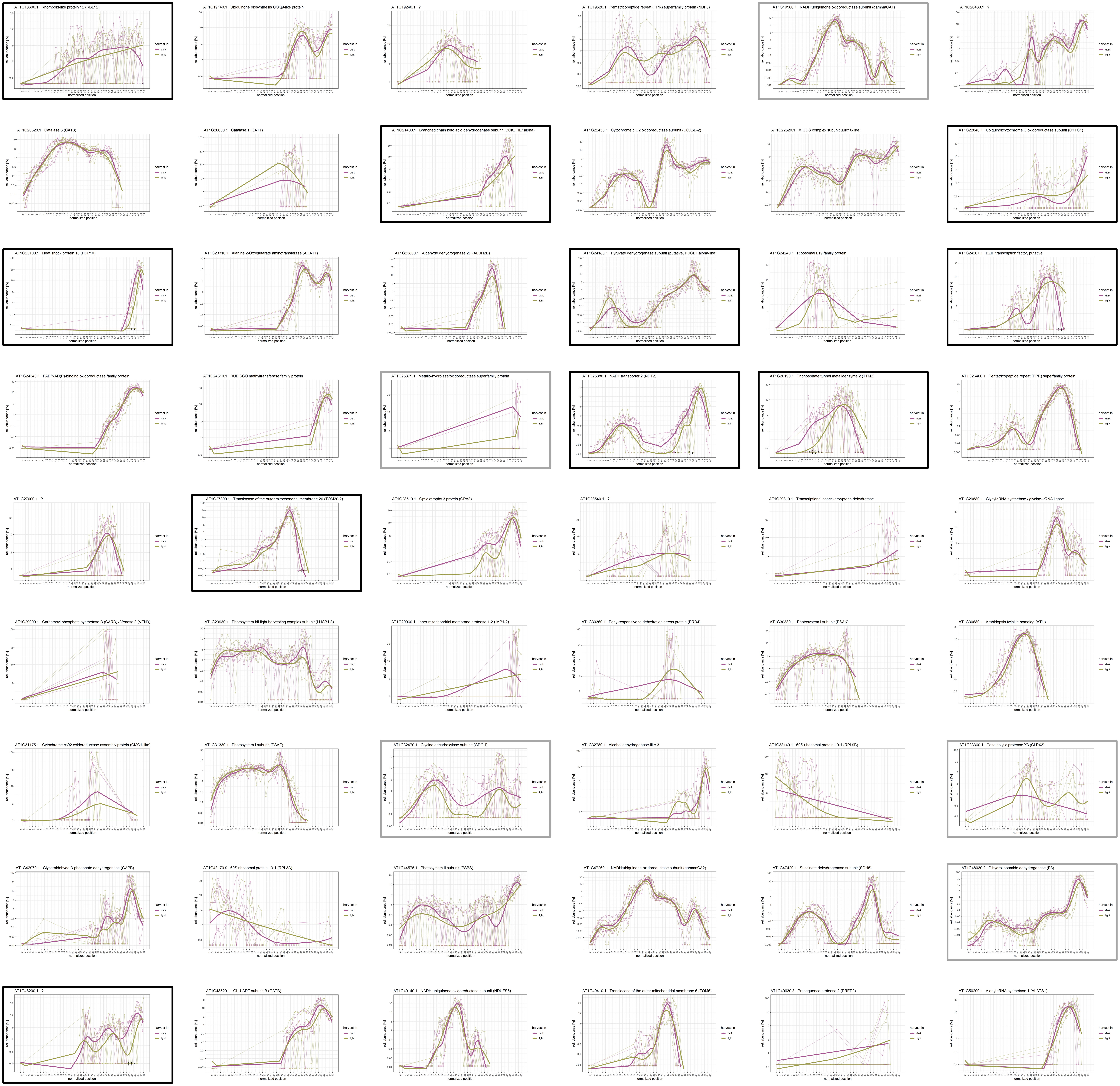

### **Chromosome 1** (AT1G50380 – AT1G72750)

Figure S7

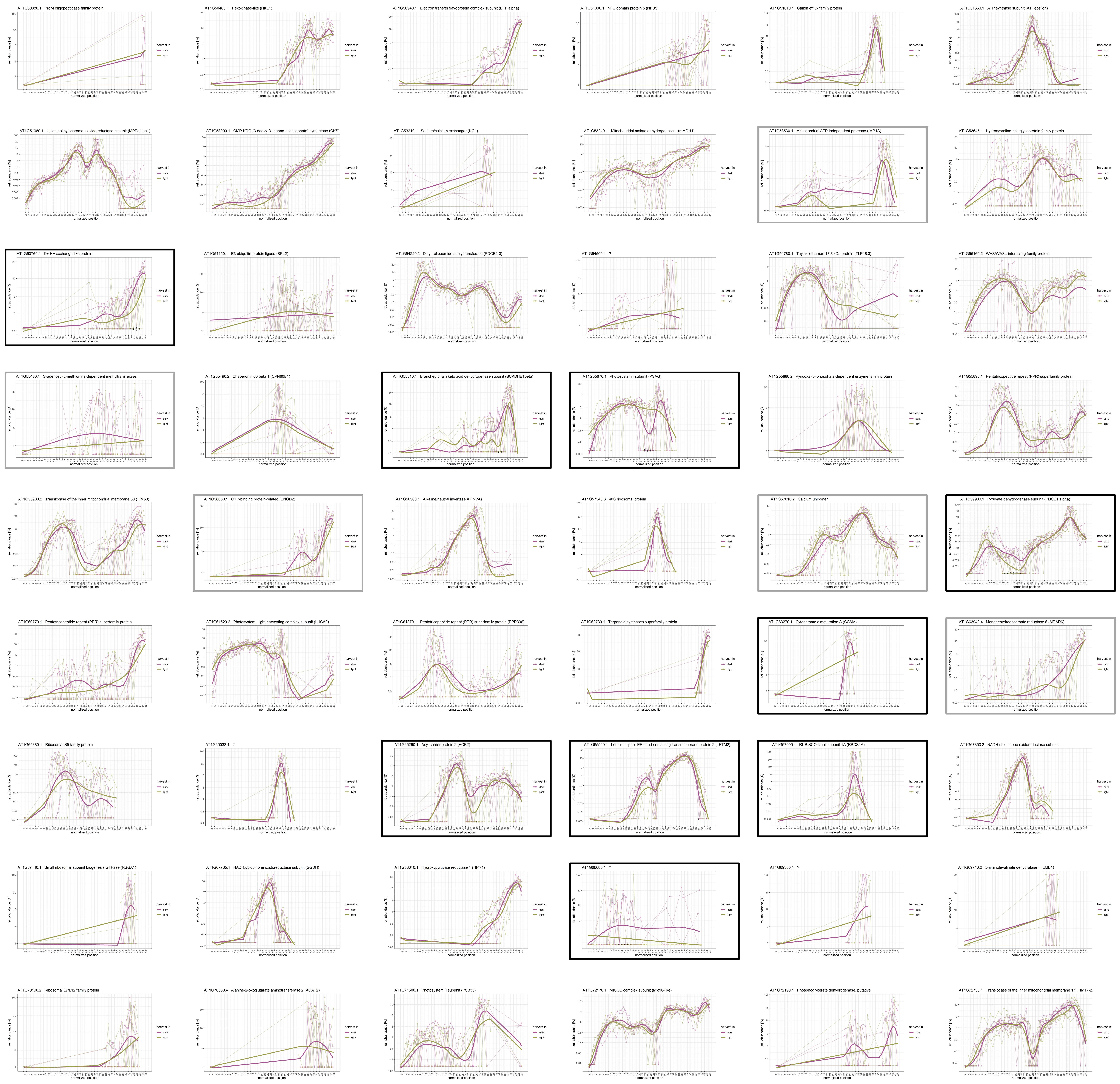

### Chromosome 1 (AT1G72820 – AT1G80620)

Figure S7

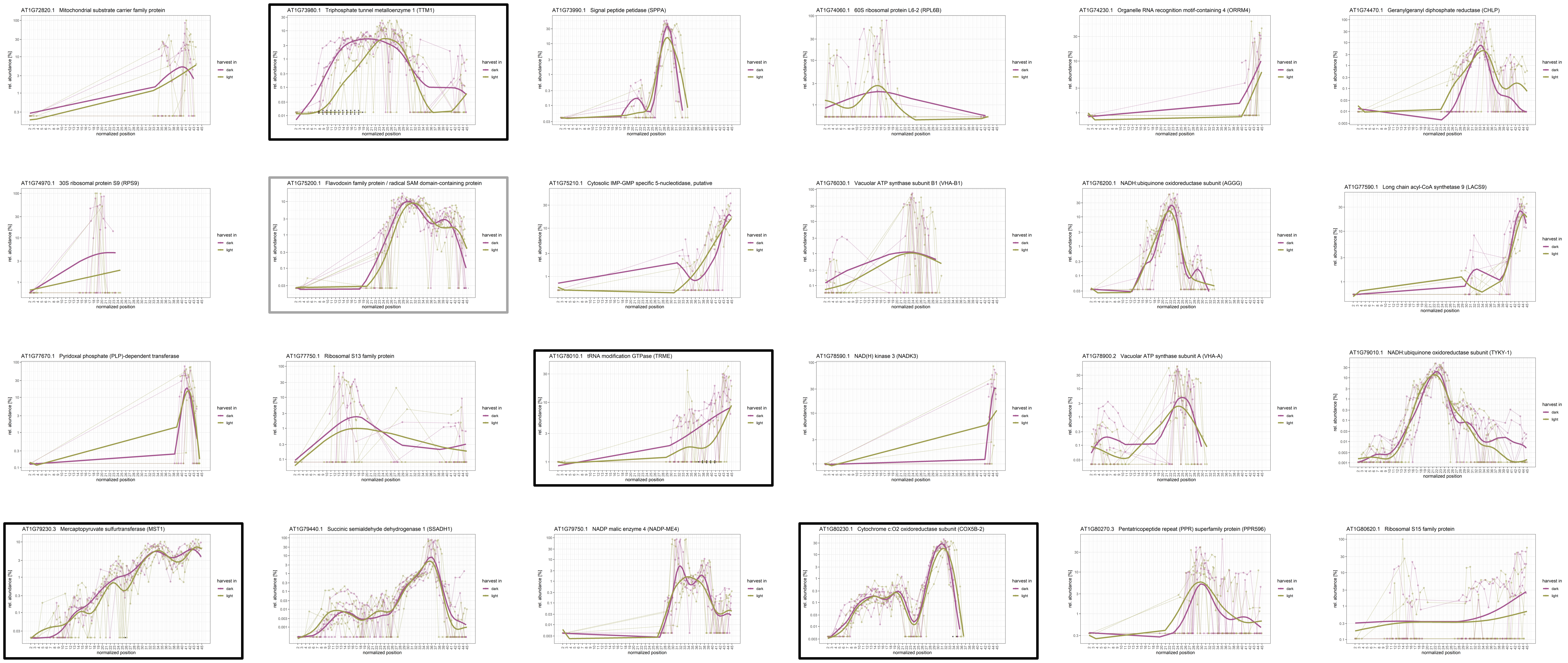

### **Chromosome 2** (AT101170 – AT1G17290)

Figure S7

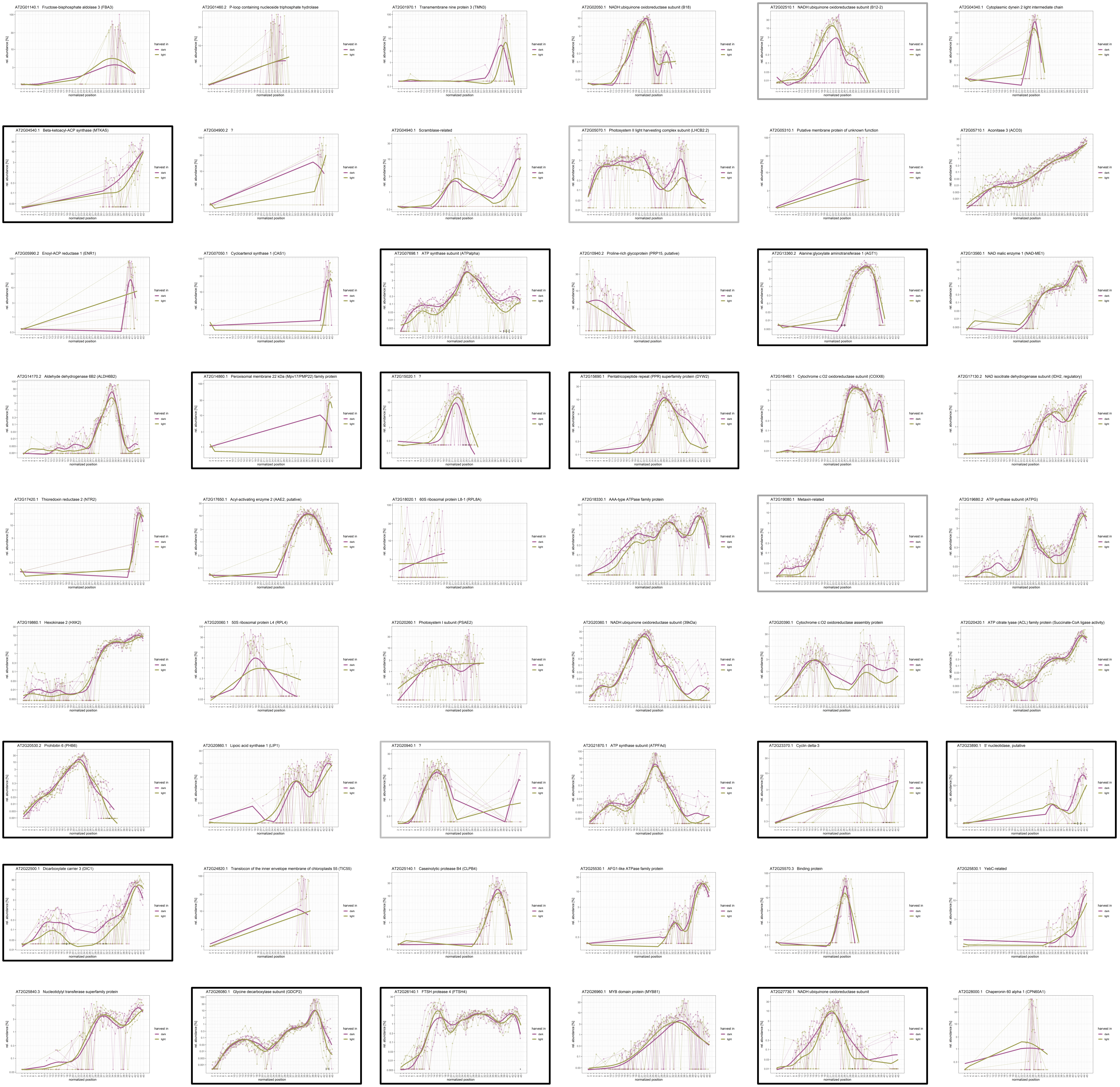

### Chromosome 2 (AT101170 – AT1G17290)

Figure S7

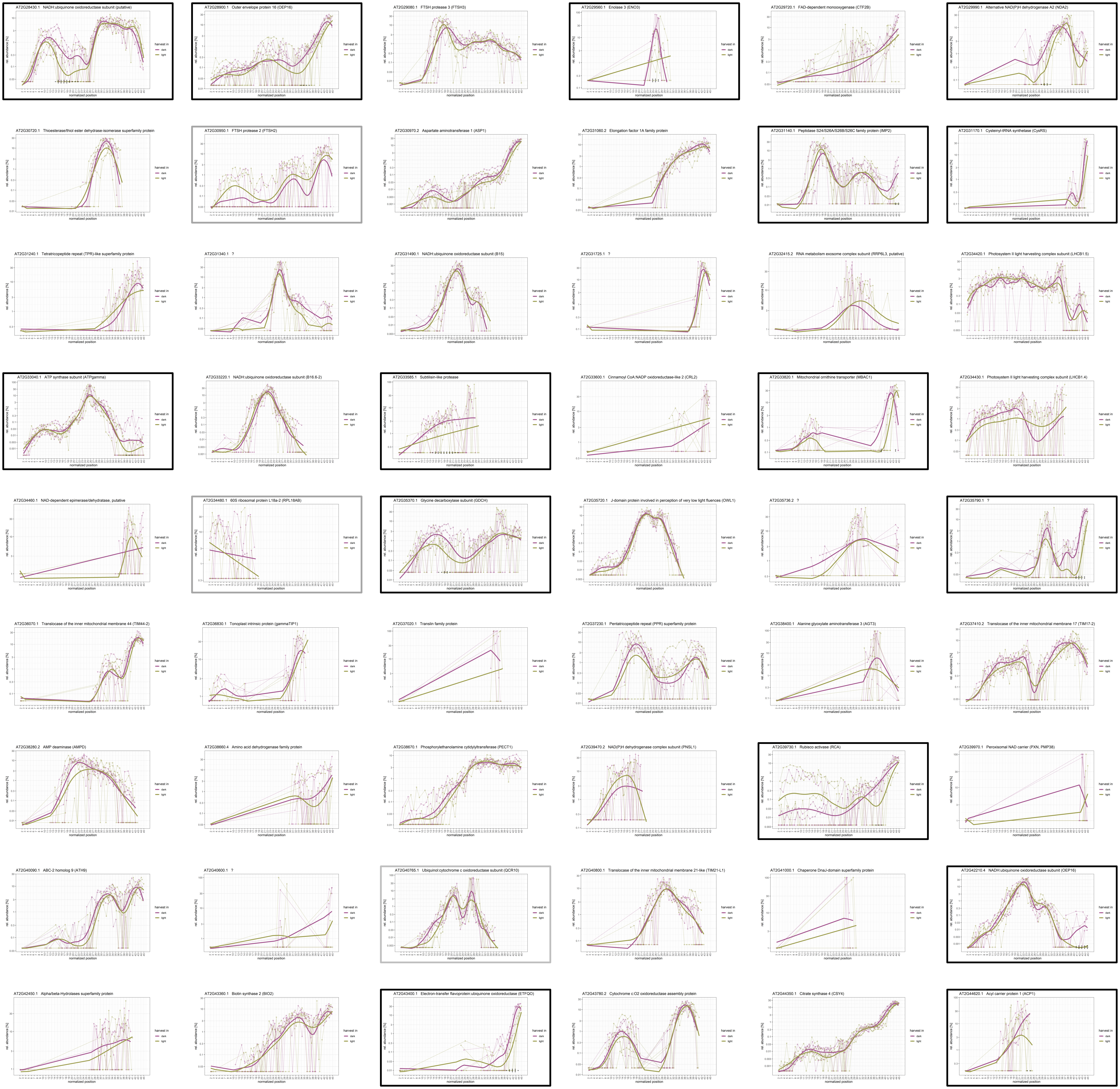

### **Chromosome 2** (AT2G45030 – AT2G47830)

Figure S7

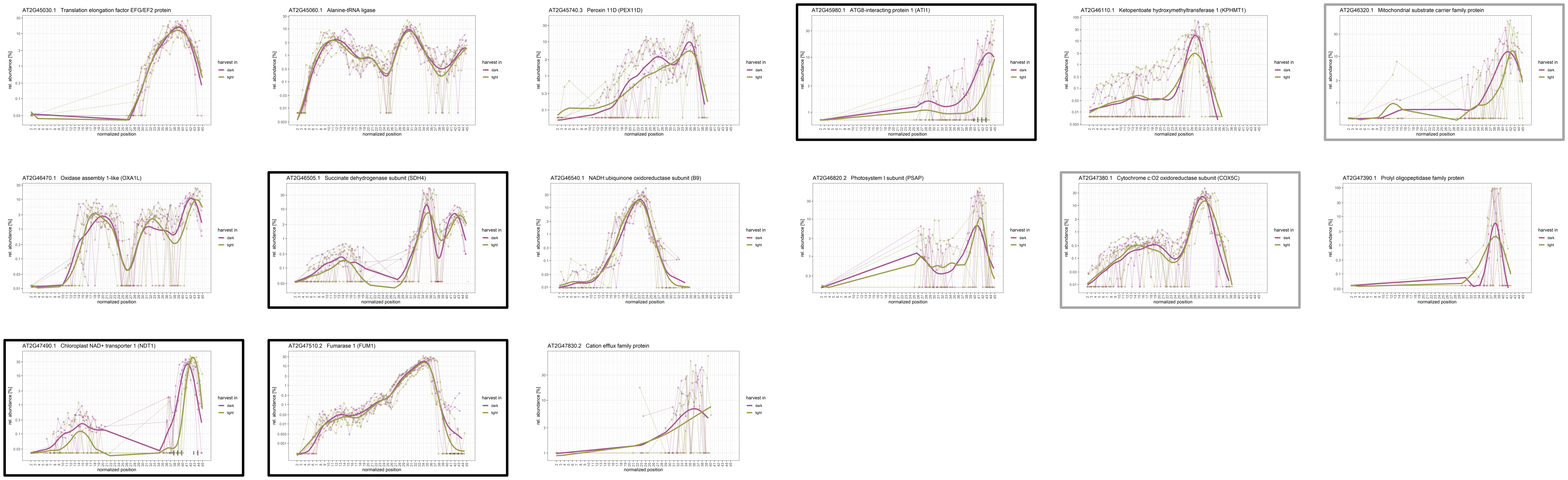

### **Chromosome 3** (AT3G01120 – AT3G13930)

Figure S7

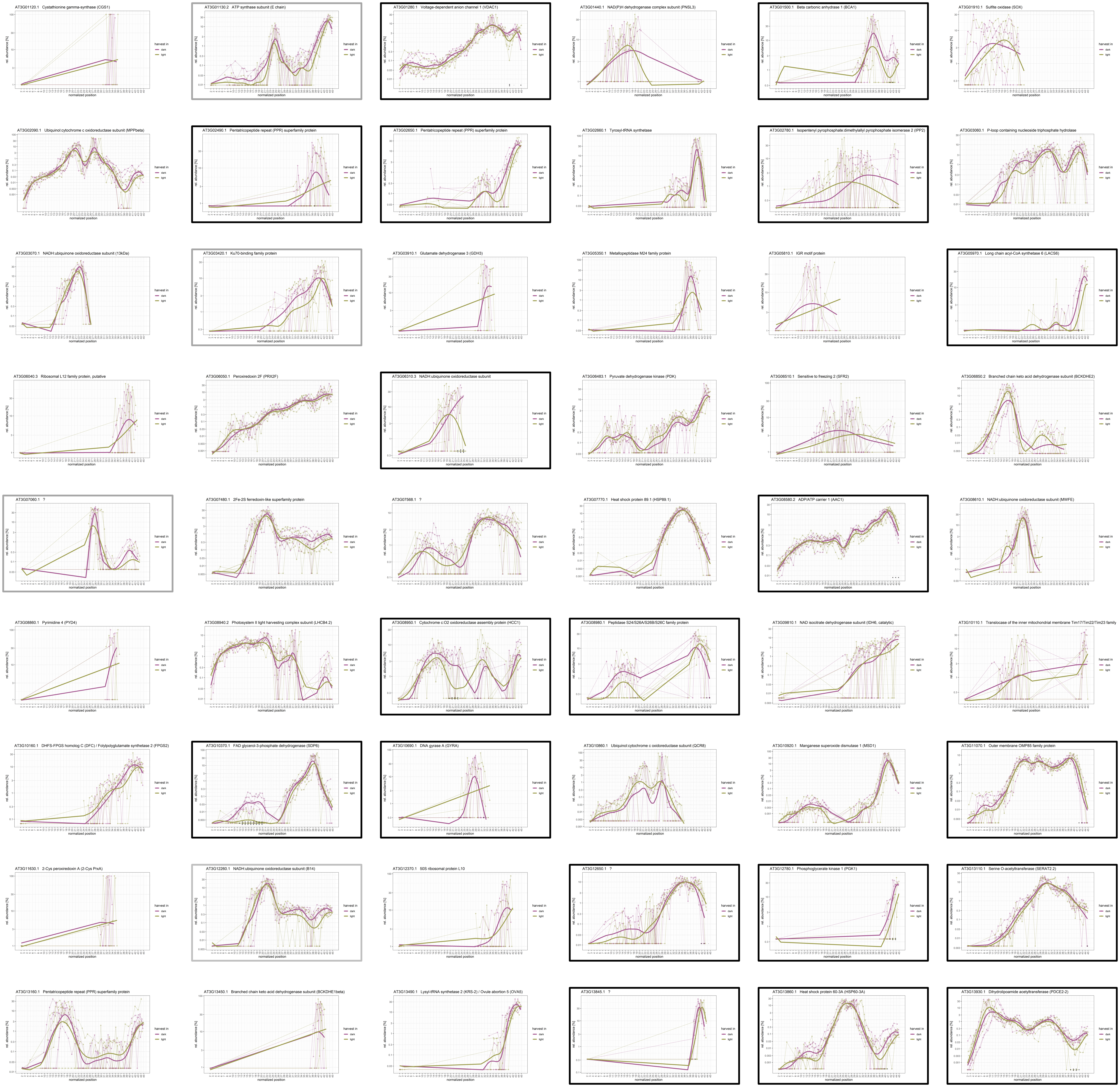

### **Chromosome 3** (AT3G14210 – AT1G44370)

Figure S7

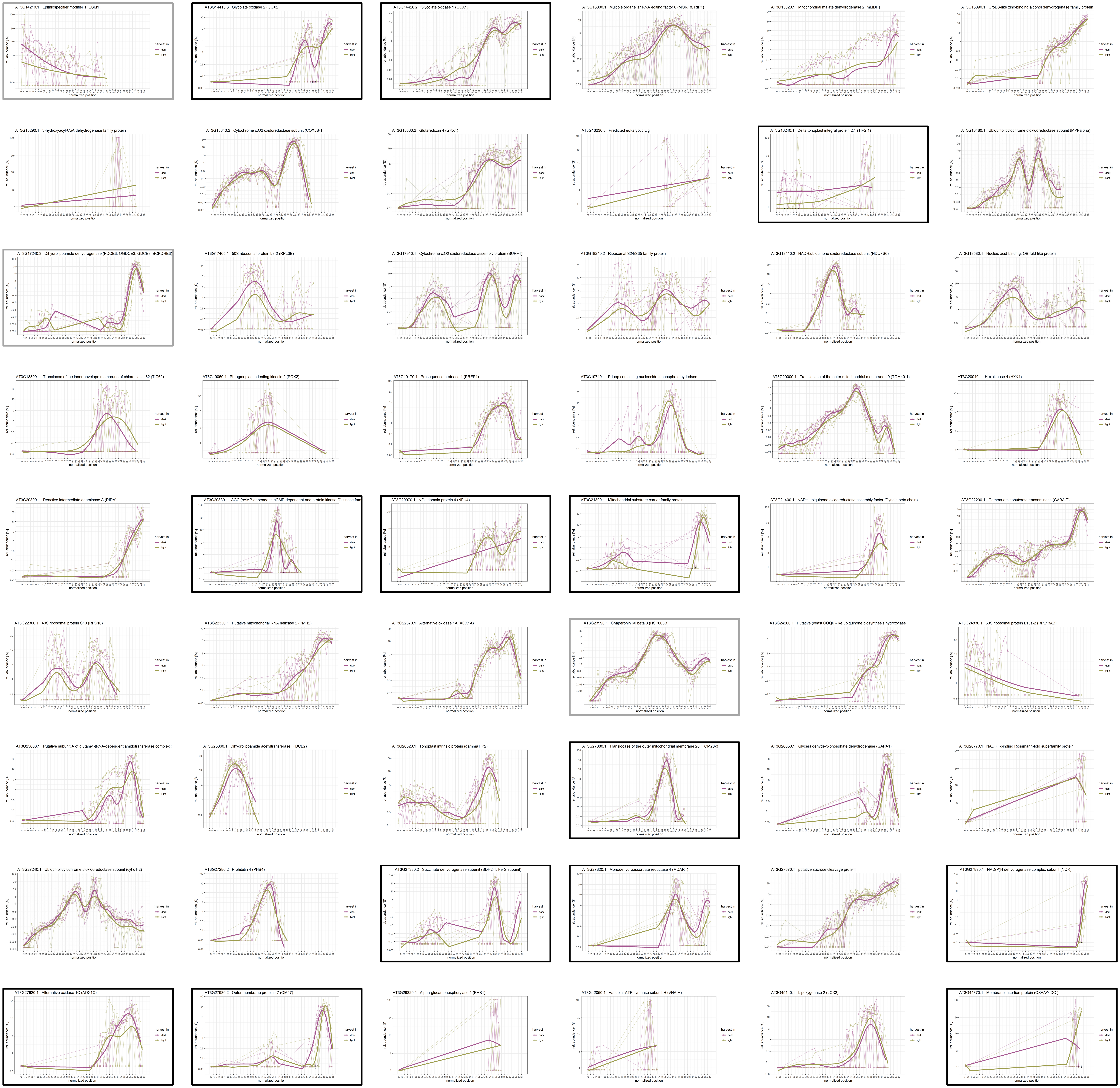

### Chromosome 3 (AT3G45300 – AT3G60980)

Figure S7

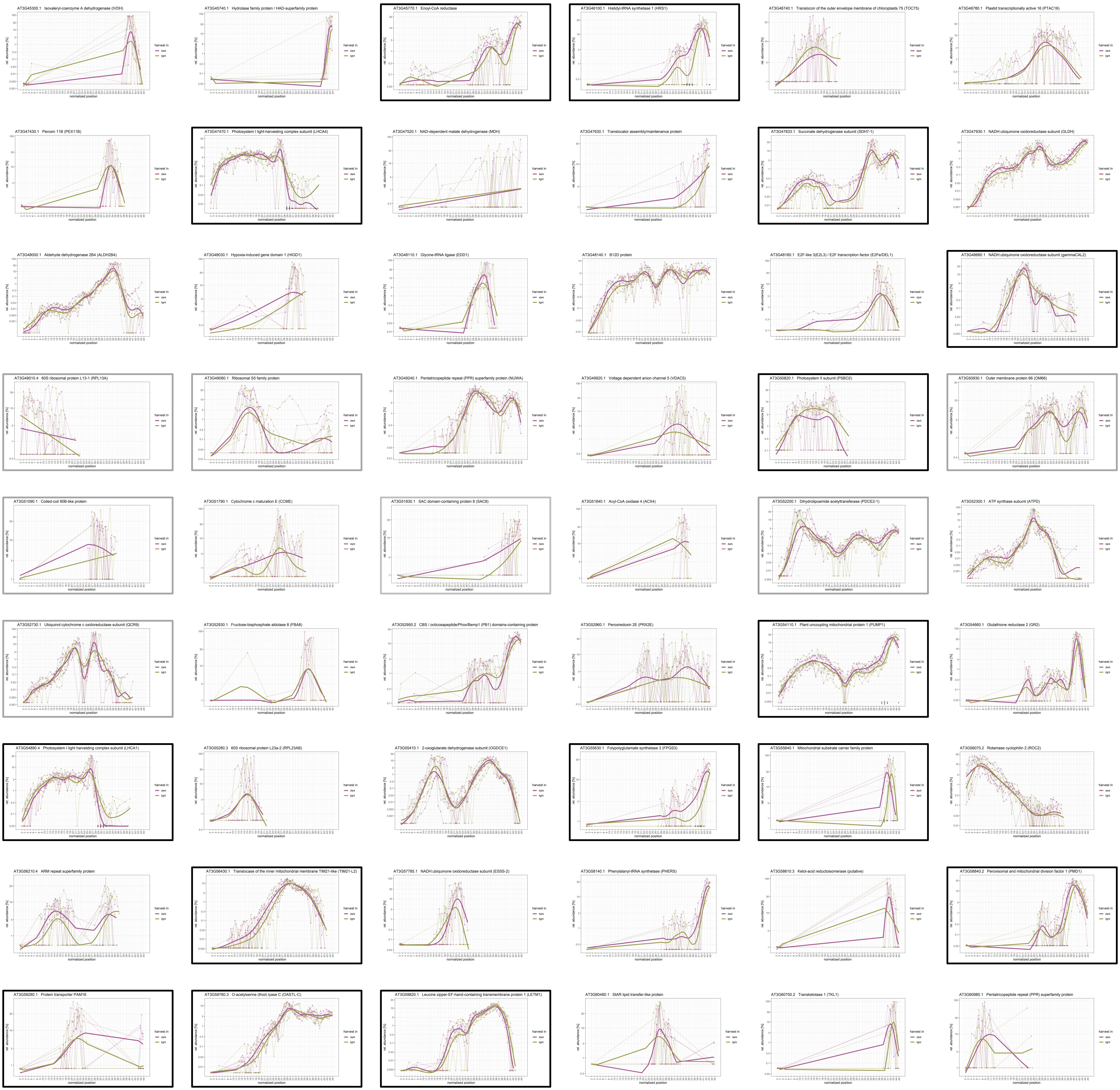

### Chromosome 3 (AT3G61070 – AT3G63520)

Figure S7

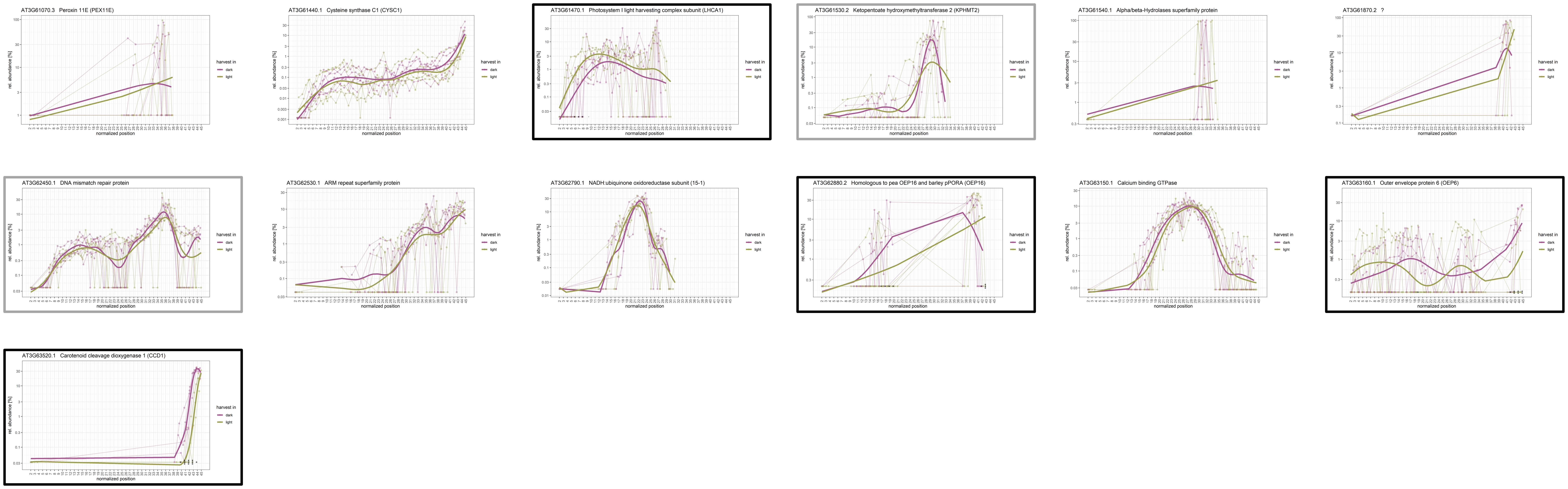

### Chromosome 4 (AT4G00026 – AT4G17340)

Figure S7

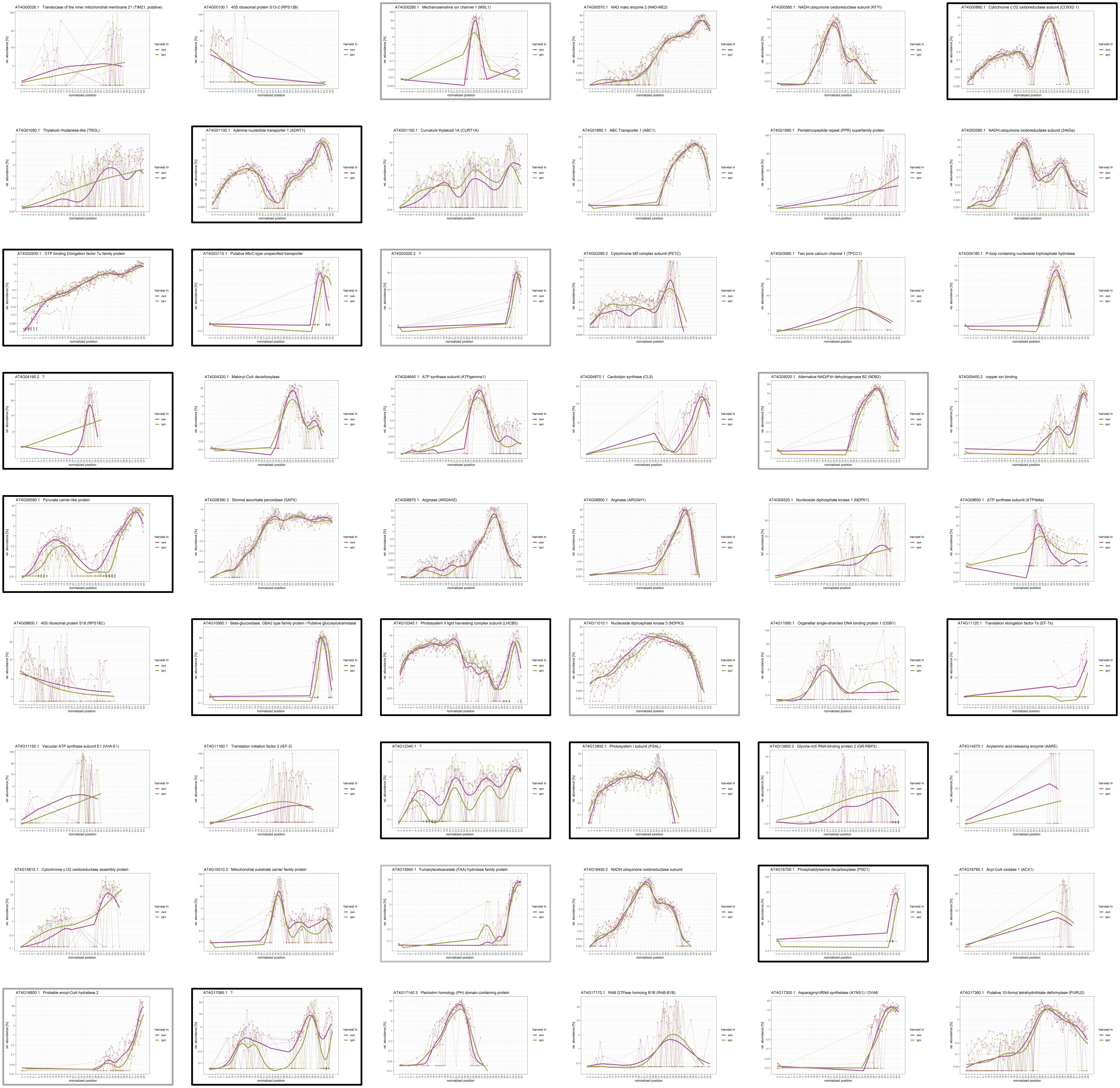

### **Chromosome 4** (AT4G18100 – AT4G35090)

Figure S7

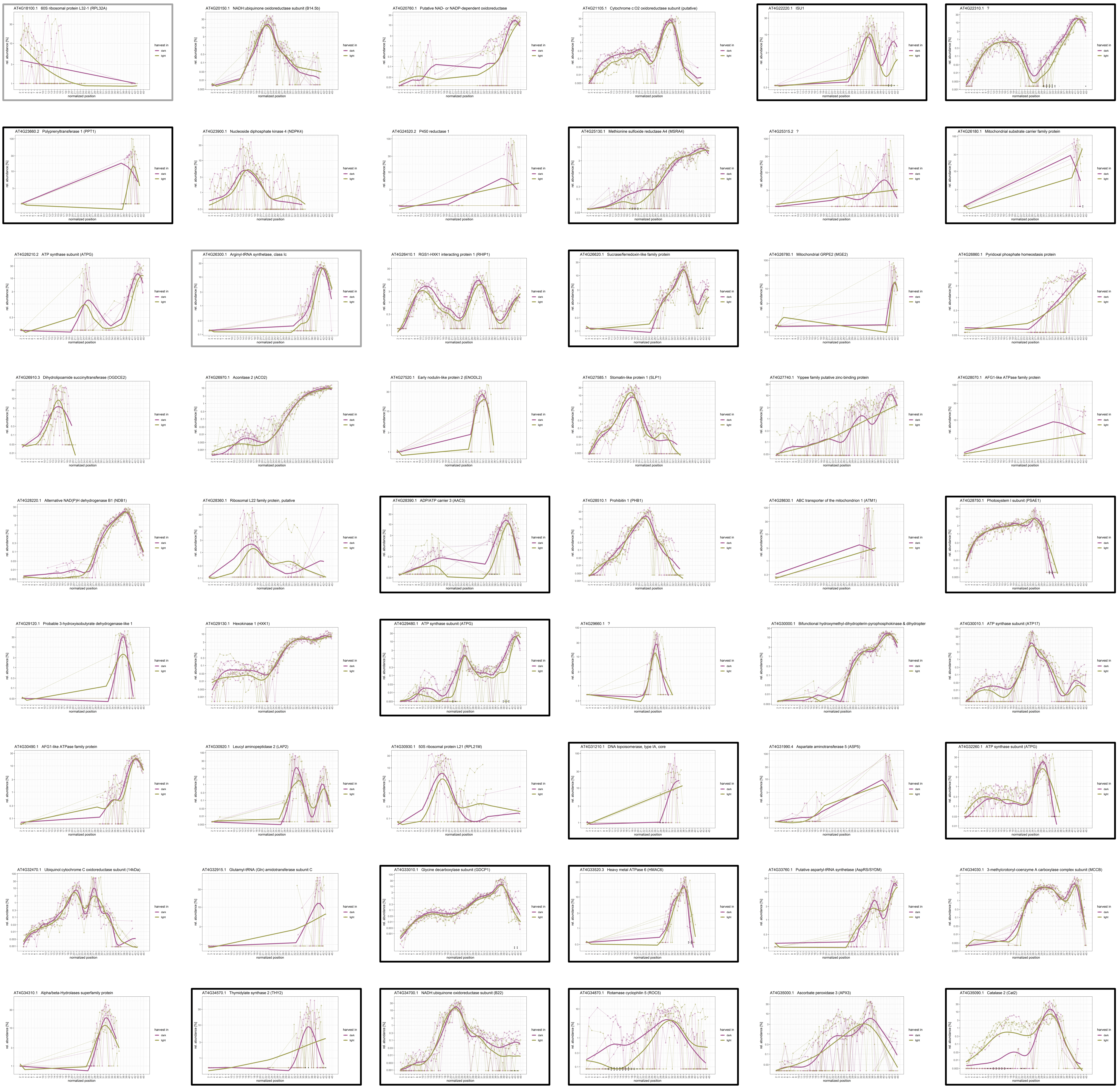

### **Chromosome 4** (AT4G352100 – AT4G39740)

Figure S7

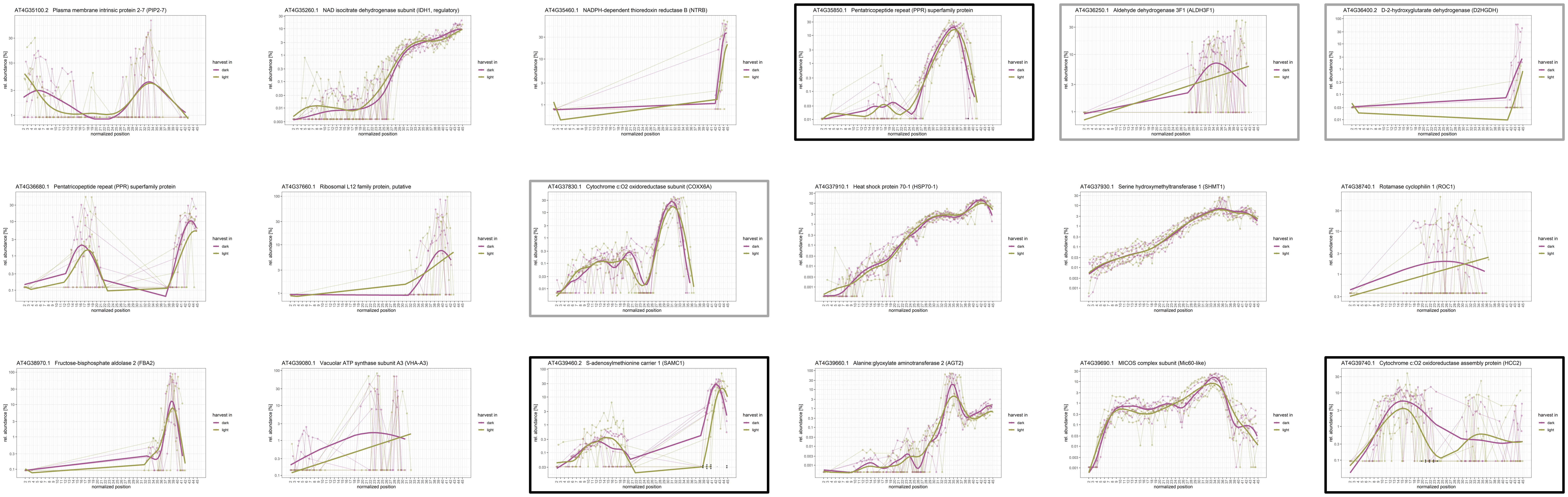

### **Chromosome 5** (AT5G01530 – AT5G15980)

Figure S7

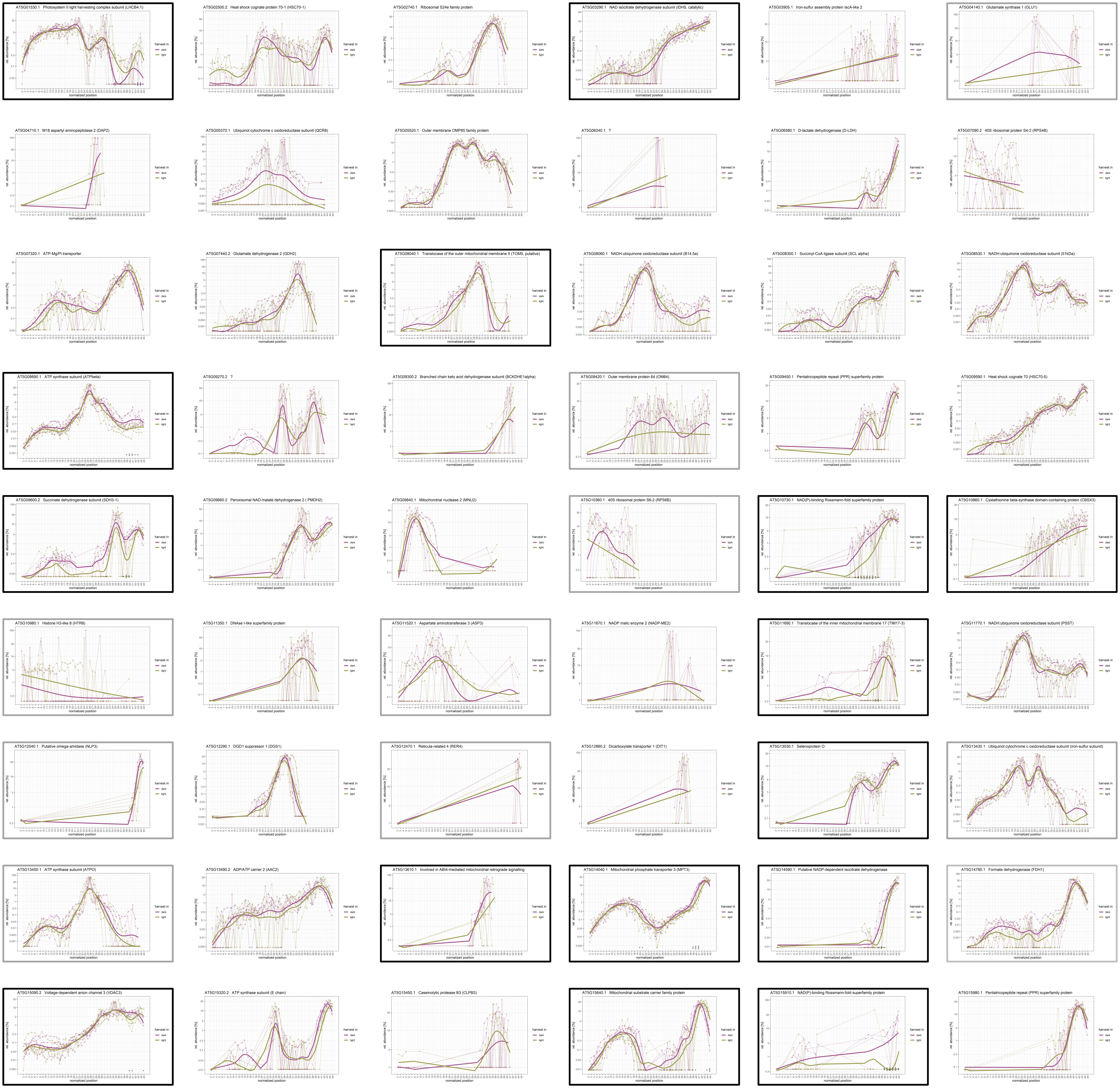

### **Chromosome 5** (AT5G16060 – AT5G42650)

Figure S7

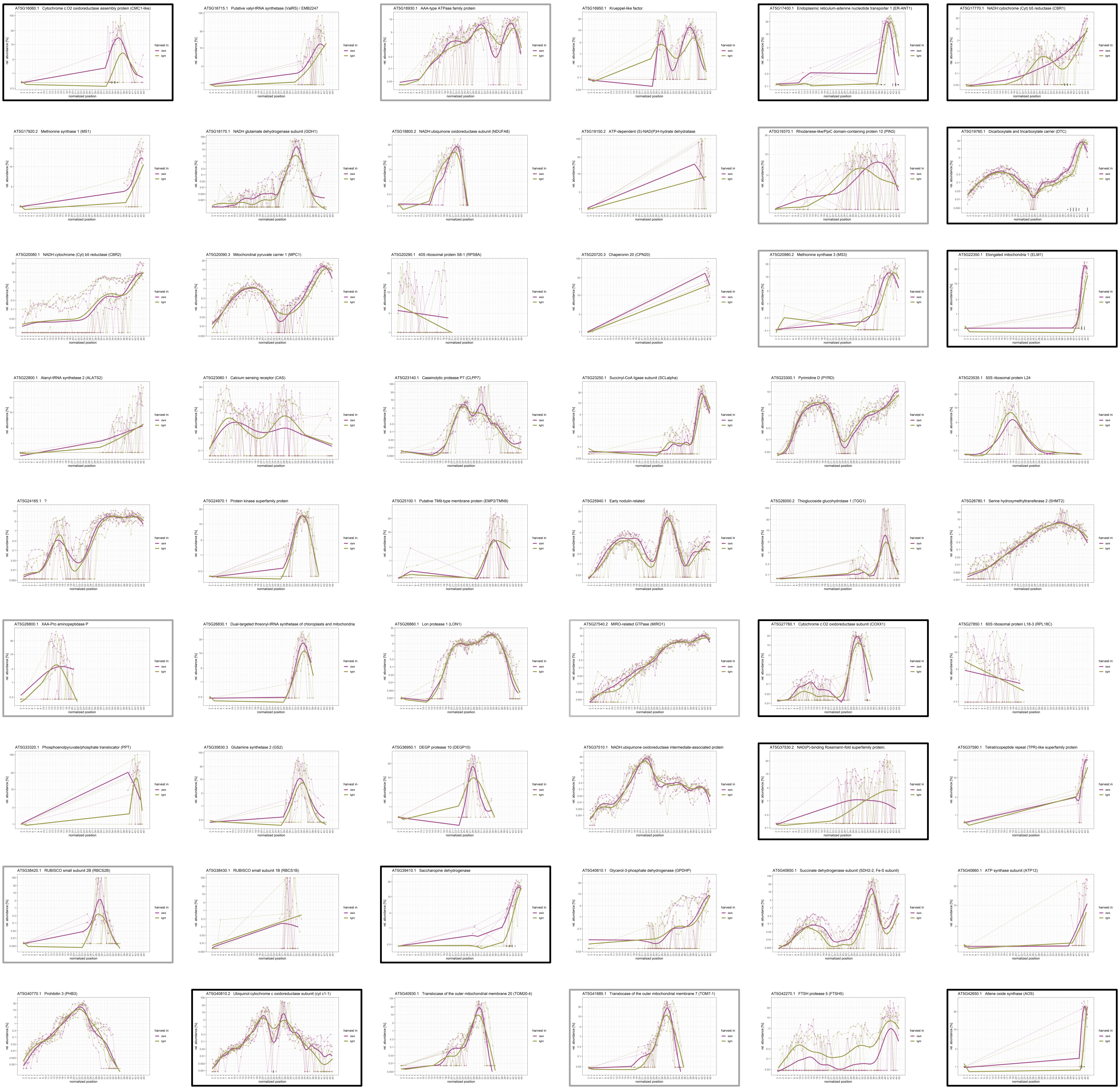

### Chromosome 5 (AT5G43430 – AT5G61790)

Figure S7

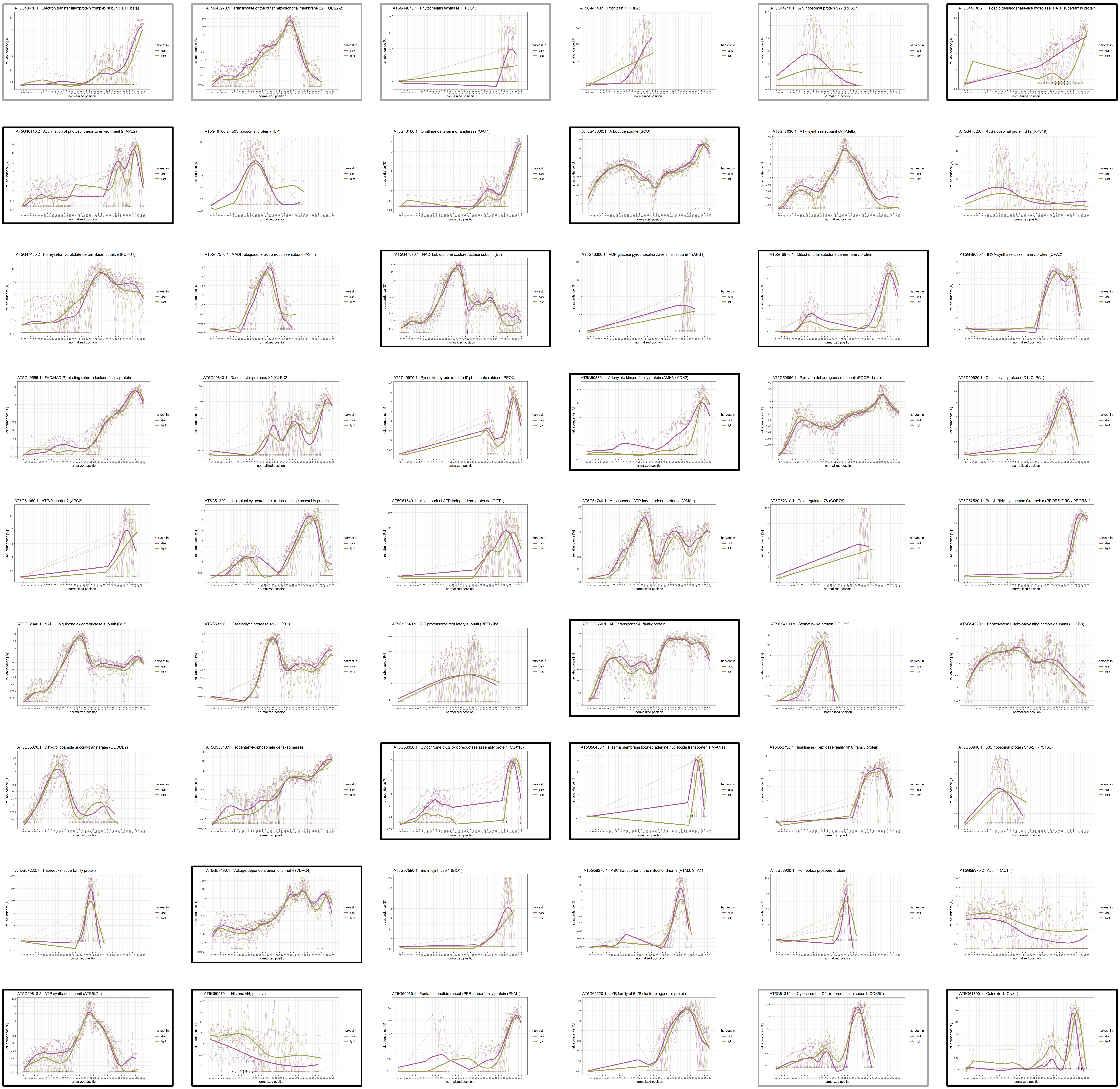

### Chromosome 5 (AT5G61810 – AT1G67590)

Figure S7

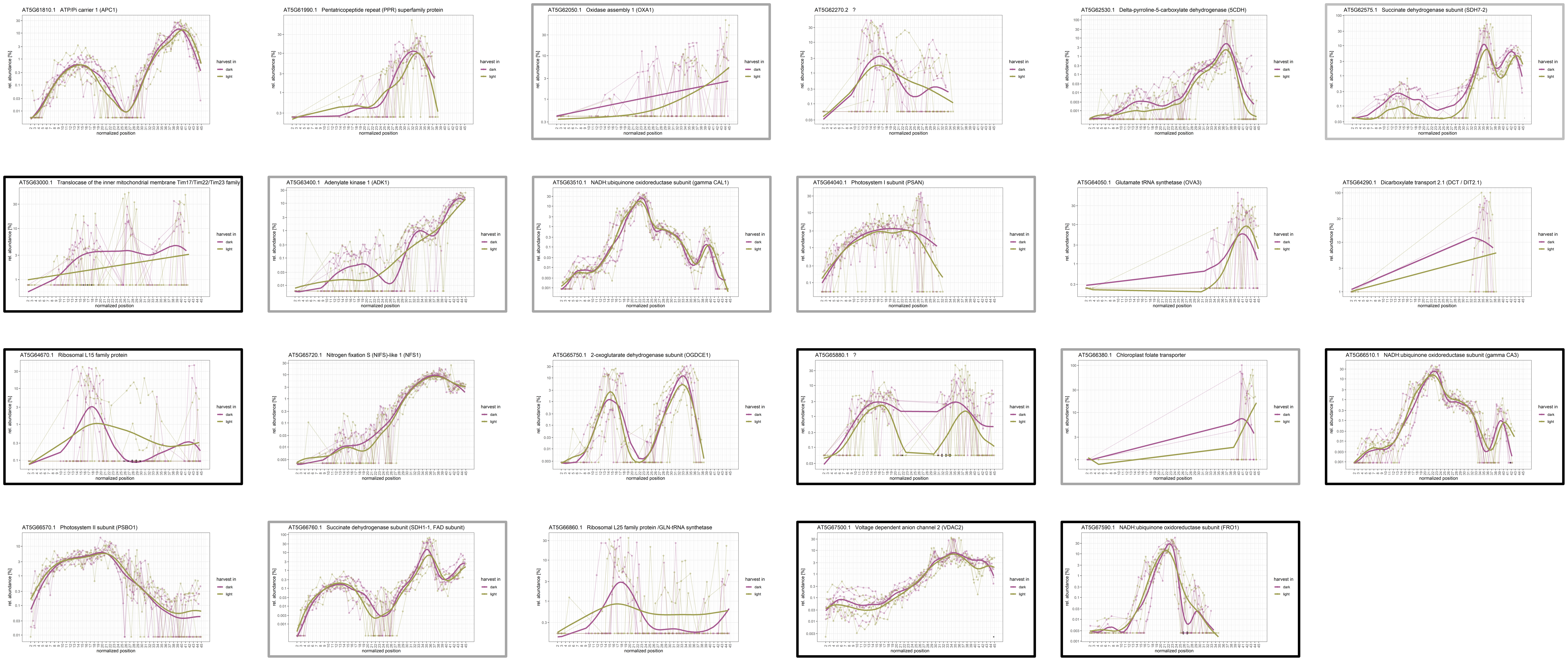

### **Mitochondrial and Plastid Genome** (ATMG00070 – ATMG01360; ATCG00020 – ATCG01110)

Figure S8

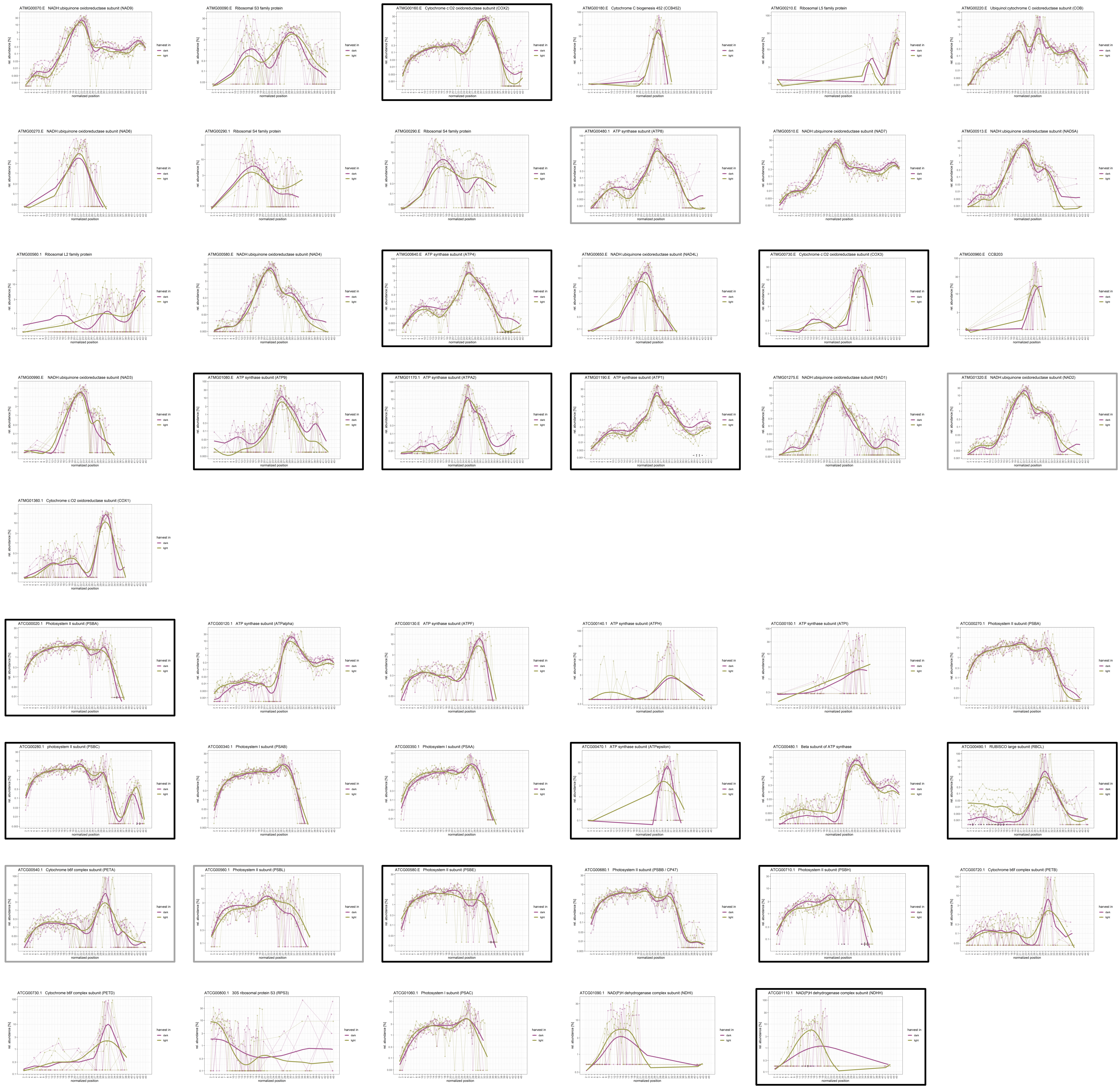
