## Supplemental Figure 9 for "Protein Interaction Patterns in *Arabidopsis thaliana* Leaf Mitochondria Change in Response to Illumination"

Figure S9

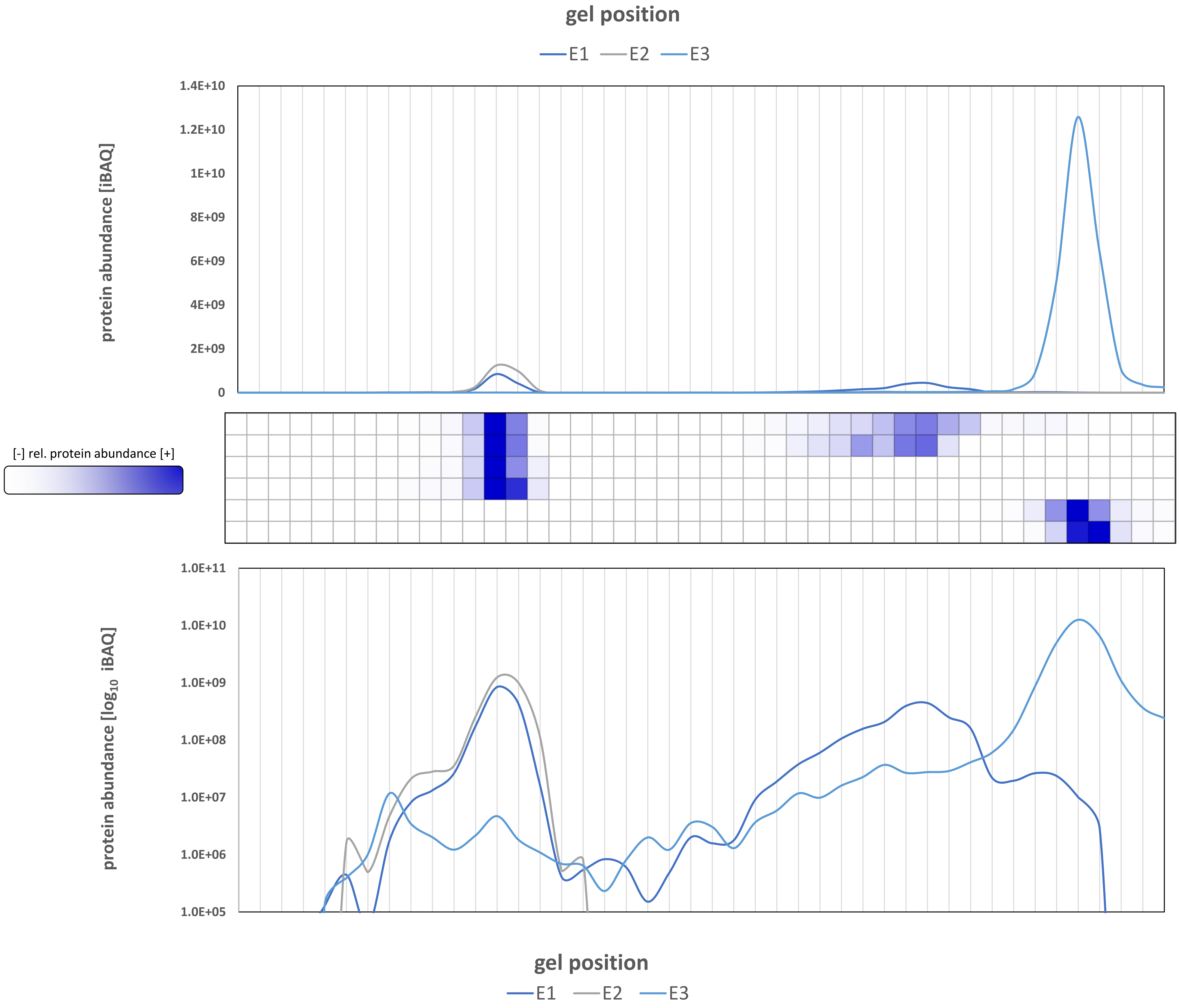


| AGI | description |
|---|---|
| AT3G55410.1 | 2-oxoglutarate dehydrogenase subunit (OGDCE1) |
| AT5G65750.1 | 2-oxoglutarate dehydrogenase subunit (OGDCE1) |
| AT4G26910.3 | dihydrolipoamide succinyltransferase (OGDCE2) |
| AT5G55070.1 | dihydrolipoamide succinyltransferase (OGDCE2) |
| AT1G48030.2 | dihydrolipoamide dehydrogenase (E3) |
| AT3G17240.3 | dihydrolipoamide dehydrogenase (PDCE3, OGDCE3, GDCE3, BCKDHE3) |
