## Supplemental Table 1 for "Protein Interaction Patterns in *Arabidopsis thaliana* Leaf Mitochondria Change in Response to Illumination"

| Table S1: LFQ-based assessment of altered protein abundance in dark and light mitochondria | | | | | |
|---|---|---|---|---|---|
| **majority protein IDs** | **primary protein ID** | **fasta headers** | **Welch's T-test p-value** | **Welch's T-test q-value** | **Welch's T-test Difference** |
| AT1G07030.1 | AT1G07030.1 | Mitochondrial substrate carrier family protein | 0.022 | 0.935 | -0.763 |
| AT5G59970.1; AT5G59690.1; AT3G53730.1; AT3G46320.1; AT3G45930.1; AT2G28740.1; AT1G07820.2; AT1G07820.1; AT1G07660.1; AT1G07660.2 | AT5G59970.1 | Histone superfamily protein | 0.031 | 0.880 | -1.798 |
| AT5G60390.3; AT5G60390.1; AT1G07940.2; AT1G07940.1; AT1G07930.1; AT1G07920.1; AT1G07930.2; AT5G60390.2 | AT5G60390.3 | GTP binding Elongation factor Tu family protein (EF1ALPHA) | 0.038 | 0.697 | -1.389 |
| AT1G14140.1 | AT1G14140.1 | Uncoupling protein 3 (UCP3) | 0.004 | 0.865 | -0.574 |
| AT1G22530.1 | AT1G22530.1 | PATL2 PATELLIN 2 | 0.035 | 0.739 | -1.844 |
| AT1G24267.1; AT1G24267.2 | AT1G24267.1 | bZIP transcription factor, putative;Protein of unknown function (DUF1664) | 0.032 | 0.823 | -0.797 |
| AT1G29880.1 | AT1G29880.1 | Glycyl-tRNA synthetase / glycine--tRNA ligase | 0.032 | 0.859 | -0.384 |
| AT1G56050.1 | AT1G56050.1 | GTP-binding protein-related | 0.009 | 0.973 | -0.824 |

| | | | | | |
|---|---|---|---|---|---|
| AT1G57540.3; AT1G57540.2; AT1G57540.1 | AT1G57540.3 | Ribosomal protein | 0.049 | 0.788 | 0.735 |
| AT1G71310.2; AT1G71310.1; AT1G71310.3 | AT1G71310.2 | Organellar DNA-binding protein 1 (ODB1) | 0.018 | 0.984 | -1.508 |
| AT1G72150.1 | AT1G72150.1 | PATL1 PATELLIN 1 | 0.034 | 0.776 | -1.437 |
| AT1G73980.1 | AT1G73980.1 | Triphosphate Tunnel Metalloenzyme (TTM1) | 0.033 | 0.795 | 0.660 |
| AT1G74470.1 | AT1G74470.1 | Pyridine nucleotide-disulphide oxidoreductase family protein | 0.027 | 1.000 | -1.366 |
| AT1G79440.1 | AT1G79440.1 | Succinic semialdehyde dehydrogenase 1 (SSADH1) | 0.025 | 0.977 | 0.455 |
| AT1G80270.3; AT1G80270.2; AT1G80270.1 | AT1G80270.3 | Pentatricopeptide repeat (PPR) superfamily protein (PPR596) | 0.031 | 0.913 | -1.294 |
| AT1G80700.1; AT1G80980.1 | AT1G80700.1 | unknown protein | 0.031 | 0.950 | 0.978 |
| AT2G02510.1 | AT2G02510.1 | NADH:ubiquinone oxidoreductase subunit (B12-2) | 0.039 | 0.685 | -0.510 |
| AT2G04900.2; AT2G04900.1 | AT2G04900.2 | unknown protein | 0.005 | 0.825 | 0.757 |
| ATMG00480.1; AT2G07707.1 | ATMG00480.1 | ATP synthase subunit (ATP8);Plant mitochondrial ATPase, F0 complex, subunit 8 protein | 0.032 | 0.848 | -0.448 |
| AT2G15020.1 | AT2G15020.1 | unknown protein;BEST Arabidopsis thaliana protein match is: unknown protein (TAIR:AT5G64190.1);Has 72 Blast hits to 72 proteins in 10 species: Archae - 0;Bacteria - 0;Metazoa - 0;Fungi - 0;Plants - 72;Viruses - 0;Other Eukaryotes - 0 (source: NCBI | 0.023 | 0.939 | -1.663 |

| | | | | | |
|---|---|---|---|---|---|
| AT2G40800.1 | AT2G40800.1 | Translocase of the inner mitochondrial membrane 21-like (TIM21-L1) | 0.030 | 0.968 | 0.506 |
| AT3G62250.1;<br>AT2G47110.2;<br>AT2G47110.1;<br>AT1G23410.1;<br>AT3G52590.1;<br>AT2G36170.1;<br>AT4G05050.4;<br>AT2G35635.1;<br>AT1G31340.1;<br>AT4G05050.3;<br>AT4G05050.2;<br>AT4G05050.1;<br>AT4G02890.2;<br>AT4G02890.1;<br>AT1G55060.1;<br>AT4G05320.5;<br>AT4G02890.4;<br>AT4G02890.3;<br>AT5G03240.3;<br>AT5G03240.2;<br>AT5G03240.1;<br>AT1G65350.1;<br>AT5G37640.1;<br>AT4G05320.6;<br>AT4G05320.3;<br>AT4G05320.1;<br>AT5G20620.1;<br>AT4G05320.4; | AT3G62250.1 | UBQ5 ubiquitin 5 | 0.044 | 0.765 | -0.886 |
| AT2G47490.1 | AT2G47490.1 | Chloroplast NAD+ transporter 1 (NDT1) | 0.037 | 0.716 | -0.250 |
| AT3G15590.1 | AT3G15590.1 | Pentatricopeptide repeat (PPR) superfamily protein | 0.035 | 0.764 | -1.030 |

| | | | | | |
|---|---|---|---|---|---|
| AT3G16000.1 | AT3G16000.1 | MAR binding filament-like protein 1 (MFP1) | 0.011 | 0.769 | -2.250 |
| AT3G21390.1 | AT3G21390.1 | Mitochondrial thiamin diphosphate carrier. | 0.004 | 1.000 | -0.970 |
| AT3G47930.1; AT3G47930.2 | AT3G47930.1 | NADH:ubiquinone oxidoreductase subunit (GLDH);ATGLDH, GLDH L-galactono-1,4-lactone dehydrogenase | 0.029 | 0.969 | -0.325 |
| AT3G52730.1 | AT3G52730.1 | Ubiquinol:cytochrome c oxidoreductase subunit (QCR9) | 0.003 | 1.000 | -0.825 |
| AT3G52950.2; AT3G52950.1 | AT3G52950.2 | CBS / octicosapeptide/Phox/Bemp1 (PB1) domains-containing protein;CBS / octicosapeptide/Phox/Bemp1 (PB1) domains-containing protein | 0.009 | 0.850 | -1.211 |
| AT3G58140.1 | AT3G58140.1 | Phenylalanyl-tRNA synthetase (PHERS) | 0.046 | 0.757 | -0.453 |
| AT3G60480.1 | AT3G60480.1 | StAR lipid transfer-like protein | 0.009 | 0.747 | -1.199 |
| AT4G17300.1 | AT4G17300.1 | NS1, OVA8, ATNS1 Class II aminoacyl-tRNA and biotin synthetases superfamily protein | 0.034 | 0.784 | -0.508 |
| AT4G28510.1 | AT4G28510.1 | Prohibitin 1 (PHB1) | 0.047 | 0.770 | 0.416 |
| AT4G32060.1; AT4G32060.2 | AT4G32060.1 | Calcium-binding EF hand family protein (MICU);Calcium-binding EF hand family protein (MICU) | 0.018 | 0.951 | 0.533 |
| AT5G10980.1; AT4G40040.2; AT4G40040.1; AT4G40030.3; AT4G40030.1; AT4G40030.2; AT1G75600.1 | AT5G10980.1 | Histone 3.3 (H3.3) | 0.006 | 0.832 | -1.734 |

| | | | | | |
|---|---|---|---|---|---|
| AT5G09300.2; AT5G09300.1 | AT5G09300.2 | Branched chain keto acid dehydrogenase subunit (BCKDHE1alpha);Branched chain keto acid dehydrogenase subunit (BCKDHE1alpha) | 0.015 | 1.000 | 0.690 |
| AT5G09450.1 | AT5G09450.1 | Tetratricopeptide repeat (TPR)-like superfamily protein | 0.027 | 0.984 | -0.713 |
| AT5G18170.1 | AT5G18170.1 | NADH glutamate dehydrogenase subunit (GDH1) | 0.038 | 0.714 | 0.530 |
| AT5G46180.1 | AT5G46180.1 | Ornithine delta-aminotransferase (OAT1) | 0.036 | 0.733 | 0.292 |
| AT5G46800.1 | AT5G46800.1 | A bout de souffle (BOU) | 0.011 | 0.813 | -0.436 |
| AT5G47570.1 | AT5G47570.1 | NADH:ubiquinone oxidoreductase subunit (ASHI) | 0.044 | 0.752 | 0.591 |
| AT5G49840.1 | AT5G49840.1 | ATP-dependent Clp protease | 0.035 | 0.749 | -0.777 |
| AT5G51220.1 | AT5G51220.1 | Ubiquinol:cytochrome c oxidoreductase assembly protein | 0.033 | 0.808 | -0.455 |
| AT5G53170.1 | AT5G53170.1 | FTSH protease 11 (FTSH11) | 0.029 | 0.993 | -0.680 |
| AT5G53350.1 | AT5G53350.1 | Caseinolytic protease X (CLPX) | 0.021 | 0.969 | -0.492 |
| AT5G63000.1 | AT5G63000.1 | Translocase of the inner mitochondrial membrane Tim17/Tim22/Tim23 family protein | 0.038 | 0.689 | -0.603 |
| ATCG00340.1 | ATCG00340.1 | PSAB Photosystem I, PsaA/PsaB protein | 0.037 | 0.735 | -0.714 |
| ATMG00220.E; ATMG00220.1; AT2G07727.1 | ATMG00220.E | RNA Edit COB apocytochrome b | 0.020 | 0.943 | -0.553 |
| ATMG00730.E; ATMG00730.1; AT2G07687.1 | ATMG00730.E | RNA Edit COX3 cytochrome c oxidase subunit 3 | 0.016 | 0.981 | 1.401 |
