## Supplemental Methods for "Protein Interaction Patterns in *Arabidopsis thaliana* Leaf Mitochondria Change in Response to Illumination"

### Statistical comparison of relative abundance patterns of proteins along a gel gradient

Frank Schaarschmidt

3 4 2020

#### Notation

Index $i = 1, 2$ denotes the 2 experimental treatments (dark, light). For each treatment, there were five independent biological replications (with index $j = 1....J_i$), i.e. $J_i = 5$ plants being subject to the light and 5 plants subject to the dark conditions. Although equal number of replications were used here, the method can in principle deal with different number of replications per treatment condition. For each biological replication, protein complexes were separated by size in a gel. Subsequently gels were manually cut into up to 48 slices. The index of the slices from the same gel (and biological replicate) is $k = 1, 2, ..., K_{ij}$. The order $k = 1, 2, 3, ..., K_{ij}$ represents decreasing size of protein complexes or proteins in downstream direction of the gel. I.e. largest complexes are expected slice $k = 1$ and smallest fragments are expected in slices with large $k$. The total number of slices in a given treatment $i$ and biological replication $j$ is denoted $K_{ij}$, and differed between individual gels from $K_{ij} = 46$ to $K_{ij} = 48$. Finally, from each slice $k$ of a given treatments biological replication $ij$, indidual proteins were detected and raw abundance was measured. The index of the proteins is $m = 1, ..., M$, where $M$ is the total number of proteins detected and quantified across all slices and replicates. The raw abundance measurement of protein $m$ from treatment $i$, biological replicate $j$, slice $k$ is denoted $y_{ijkm}$.

#### Normalization and preprocessing

##### Total protein proportions

Index $k = 1, ...., K_{ij}$ represents the downstream position of gel slices from the same biological replicate, and thus increasing $k$ represents decreasing size of protein complexes or their fragments. However, differences between biological replicates, gel preparation, complex fragmentation, the manual cutting process of the slices may lead to shifts along the size gradients represented by $k = 1, ..., K_{ij}$. That is, under the same treatment conditions, peak abundances of given proteins may appear in slightly different gel positions $k$. Note further, that complex size is a continuous variable mapped to the 48 gel positions that result from the cutting process. To remove these potential sources of variation from the data, and align the protein profiles across the gel positions of different replicates, we computed a normalized position $x_{ijk}$ based on the relative abundance of all proteins in a given replicate $ij$:

Denote the sum of abundances of all proteins ($m = 1, ..., M$) in a given slice $k$ of a given biological replicate ($ij$) by

$$y_{ijk.} = \sum_{m=1}^{M} y_{ijkm}$$

and the total protein abundance of all $k$ slices of given biological replicate $ij$ by

$$y_{ij..} = \sum_{k=1}^{K_{ij}} y_{ijk.}$$

The proportion of the total protein abundance found in slice $k$ of biological replicate $ij$ then is

$$p_{ijk.} = y_{ijk.}/y_{ij..} = \frac{\sum_{m=1}^{M} y_{ijkm}}{\sum_{m=1}^{M} \sum_{k=1}^{K_{ij}} y_{ijkm}}$$

$p_{ijk.}$ is a number between 0 and 1, sums up 1 for all $k = 1, ..., K_{ij}$ of a given replicate $ij$.

#### Identifying extreme values

Visual assessment of total protein proportions per slice(heat maps, line plots) showed a number of slices with extremely low total protein contents. To formally assess whether these values deviate significantly from the remaining values, the following procedure was performed: Within each treatment $i = 1, 2$ separately, the mean function of log-transformed proportions of total protein was fitted using a cubic regression spline (denoted $s(x_{ijk}, n)$, with $n = 40$ nodes). $x_{ijk}$ denotes the original gel positions of each slice ($x_{ij1} = 1, x_{ij2} = 2, ...x_{ijK_{ij}} = K_{ij}$) and additive shift parameters $d_{ij}$ for the replicates:

$$\log(p_{ijk.}) = d_{ij} + s(x_{ijk}, n) + \epsilon_{ijk}, \quad \epsilon_{ijk} \sim N(0, \sigma_i^2).$$

The residuals of the fitted model, $\hat{\epsilon}_{ijk} = \log_e(p_{ijk.}) - \left[\hat{d}_{ij} + \hat{s}(x_{ijk}, n)\right]$, and the residual standard deviation, $\hat{\sigma}_i$, obtained from the model fit were used to perform an approximate outlier test including a Bonferroni-correction for the total number of residuals: Let $z$ be the quantile of a standard normal distribution with $z = z_{1-0.05/2K_{i.}}$, where $K_{i.}$ is the number of observations in the above models, i.e., the total number of slices in treatment $i$, $K_{i.} = \sum_j K_{ij}$. Residuals that deviate more extremely from 0 than $z * \hat{\sigma}_i$ are treated as outliers and were omitted from all following steps, i.e. values with $|\hat{\epsilon}_{ijk}| > z\hat{\sigma}_i$ were omitted. Applying this rule led to omission of two slices (out of $K_{i.} = 236$ slices, both with extremely small $p_{ijk.}$) in the light treatment and no omission of slices in the dark treatment.

#### Normalization of gel position to align protein curves from different replicates

The cumulated sum of the proportions $p_{ijk.}$ in downstream direction in the gel (from $k = 1$ to $K_{ij}$) gives the cumulated distribution of total proteins across the slices:

$$c_{ijk} = \sum_{l \leq k} p_{ijl.}, \quad l = 1, ..., K_{ij}$$

The $c_{ijk}$ are again numbers in the range $[0, 1]$, that have values very close to zero at position $k = 1$ and approach 1 at the last gel position $K_{ij}$ in each replicate $ij$. The normalization procedure is now based on the assumption, that within a given treatment $i$, the pattern of cumulated proportions $c_{ijk}$ along the gradient of the gel slices should be equal between replicates. Shifts in the pattern of cumulated proportions between replicates of the same treatment are treated as technical artifacts. For each treatment separately, additive models are fitted to predict the observed position $x_{ijk}$ based on the $c_{ijk}$, including an additive shift $a_{ij}$ for each replicate and a cubic regression spline $s(c_{ijk})_{ij}$ specific for each replicate $ij$. In this step, the number nodes of the regression spline is chosen by the methods default (R-package `mgcv`, Wood, 2004).

For the purpose of obtaining a new, normalized position variable, denote the original position $x_{ijk}$ within each replicate $ij$, as the number of the slices, $k = 1, ..., K_{ij}$, i.e. $x_{ij1} = 1$, $x_{ij2} = 2$, ... , $x_{ijK_{ij}} = K_{ij}$. Note that

- the length of these vectors may differ between replicates of the same treatment because the total number of slices obtained from each gel, $K_{ij}$, may differ between replicates
- aim is to shift the profiles of proteins such that replicates of the same treatment match, but there should be no average shift of values in one treatment , i.e., for all replicates form the same treatment, there should be an average shift of 0, such that the normalization procedure does not induce treatment effects

In the first normalization step, an overall cubic regression spline $s(c_{ijk})$ for all replicates within a treatment is fitted, with the number of nodes chosen by the implementations default:

$$x_{ijk} = s(c_{ijk}) + r_{ijk}, \quad r_{ijk} \sim N(0, \sigma^2)$$

This step yields the residuals $\hat{r}_{ijk} = x_{ijk} - \hat{s}(c_{ijk})$, from which the overall trend is removed and which sum to 0. In a second step, a generalized additive model with an additive shift for each replicate ($a_{ij}$) and an individual cubic regression splines $s(c_{ijk})_{ij}$ for each replicate are fitted to the residuals of the first step:

$$\hat{r}_{ijk} = a_{ij} + s(c_{ijk})_{ij} + \epsilon_{ijk}, \quad \epsilon_{ijk} \sim N(0, \sigma^2)$$

,

Subtracting both the additive shifts, $\hat{a}_{ij}$, as well as the spline fits, $\hat{s}(c_{ijk})_{ij}$, from the original positions, $x_{ijk}$, yields a normalized position:

$$x^*_{ijk} = x_{ijk} - \hat{a}_{ij} - \hat{s}(c_{ijk})_{ij}$$

These normalized position values are not integer values (1,2,. . . ,48) anymore, but are numbers on a continuous scale. For first and last gel positions of some replicates they may have values below 1 or above 48. We believe that these normalized gel positions are better proxies for the actual size patterns of protein complexes than the original gel positions.

### Comparison of protein profiles between treatments

The statistical comparison of protein profiles was restricted to those proteins which were detected in all 5 replicates of each treatment.

#### Proportion of individual proteins per slice

Separately for each biological replicate $ij$, the proportion of the $m$th protein in each slice $k$ was computed: The total abundance of protein $m$ in replicate $ij$ is the sum across all its slices $k = 1, ..., K_{ij}$ and is denoted $y_{ij.m} = \sum_{k=1}^{K_{ij}} y_{ijkm}$. Dividing protein $m$s abundance in individual slices $k$ by this total, normalizes for differences in protein load between replicates and leads to the proportions:

$$p_{ijkm} = \frac{y_{ijkm}}{y_{ij.m}} = \frac{y_{ijkm}}{\sum_{k=1}^{K_{ij}} y_{ijkm}}.$$

These proportions are numbers between 0 and 1, are extremely right-skewed, and contain many 0-values for the majority of identified proteins. Before statistical analysis, they needed log-transformed after adding a small constant to deal with the zeros.

##### Dealing with zero observations in model fitting and treatment comparisons

In many proteins, a high proportion of positions $k$ with zero abundance in many or all replications occurred. Zero abundance ($y_{ijkm} = 0$, and thus $p_{ijkm} = 0$) is taken into consideration for the log-transformation, and the definition of the position variables in the model fitting and treatment comparisons. All these steps are performed separately for each protein, $m$.

For the log-transformation, the minimal observed non-zero proportion, $\tilde{p}_m = min_{i,j,k}(p_{ijkm}|y_{ijkm} > 0)$ was used to shift all observed proportions to values above 0: $\log_{10}(p_{ijkm} + \tilde{p}_m)$.

In model fitting, all those positions $x^*_{ijk}$ were omitted from the fitting process, where less than four observations $y_{ijkm} > 0$ were observed for a given protein $m$ and a given position index $k$.

For the purpose of treatment comparisons, we define a set of position values, $x^{**}_{qm}$, with index $q = 1, ..., Q_m$ at which the relative abundance of the $m$th protein is to be compared between the treatments. These values do not have to be values occurring in the normalized position data $x^*_{ijk}$. Practically, we chose that subset of $k = 1, ..., 48$, where at least four non-zero protein abundances of protein $m$ are observed. Thus, the total number of position values ($Q_m$), as well as the positions $x^{**}_{qm}$ where treatment comparisons are performed, may differ between proteins, $m = 1, ..., M$.

##### Fitting cubic regression splines

Separately for each protein $m$, treatment specific cubic regression splines were fitted to estimate the mean patterns of the log-transformed proportional abundance $p_{ijkm}$ in dependence of the normalized position, $x^*_{ijk}$. The splines were fitted in a joint model for both treatments in order to test differential abundance between treatments using methods of Herberich et al. (2014). In detail, a generalized additive mixed model was fitted with treatment specific shift effect, $d_{im}$ and treatment specific spline, $s(x, n)_{im}$, depending on the position $x$ and number of nodes, $n$. Mean differences between replicates $j$ were included as random effects. The model equation is

$$\log_{10}(p_{ijkm} + \tilde{p}_m) = d_{im} + s\left(x^*_{ijk}, n\right)_{im} + b_{ijm} + \epsilon_{ijkm},$$

with

$$b_{ijm} \sim N(0, \sigma^2_b)$$

representing the random effects for replicates and the residual error is

$$\epsilon_{ijkm} \sim N(0, \sigma^2_e).$$

Adding further terms to the model equation (especially serial correlation structures for residuals along the gel gradient) led to convergence problems for many proteins and were thus generally omitted from the model.

Fitting this model for each protein $m$ yields estimates $\hat{d}_{im} + \hat{s}\left(x^*_{ijk}, n\right)_{im}$ for each treatment $i$. Quantity $n$ denotes the number of nodes for the cubic regression spline, and was chosen to be $Q_m$ (number of positions with at least 4 non-zero abundance values) for the $m$th protein. The functions $\hat{d}_{im} + \hat{s}\left(x^*_{ijk}, n\right)_{im}$ estimate the mean pattern of log-proportions of protein $m$ along the potential positions on the gel, summarizing the 5 replicates of each treatment $i$. Note that this approach allows the normalized positions $x^*_{ijk}$ to differ between replicates $j$ of the same treatment $i$ and comparable slices $k$, as is generally allowed for the independent variable in regression models. However, these functions can be used to compare the mean functions of the two treatments, $i = 1, 2$ at prespecified position values $x_{qm}$ (Herberich et al. 2014).

#### Tests for differences between two curves of one protein

The model prediction for protein $m$ at position $x_{qm}^{**}$ in treatment $i$ can be computed by inserting the values $x_{qm}^{**}$ in to the fitted model equation:

$$\hat{\theta}_{iqm} = \hat{d}_{im} + \hat{s}\left(x_{qm}^{**}\right)_{im}$$

The corresponding differential abundance on the log-scale, at position $x_{qm}^{**}$ is then

$$\hat{\delta}_{qm} = \hat{\theta}_{1qm} - \hat{\theta}_{2qm}$$

Positive values of $\hat{\delta}_{qm}$ correspond to relative protein abundance that is higher in dark than in light treatment, and negative values $\hat{\delta}_{qm}$ indicate that relative abundance in light exceeds that in the dark treatment.

Standard errors of quantities like $\hat{\delta}_{qm}$ follow straightforwardly from theory of generalized additive models and Herberich et al. 2014, and are denoted $\widehat{s.e.}(\hat{\delta})_{qm}$.

Herberich et al. (2014) describe how to perform multiple tests for at least one significant difference between two such curves, $i = 1, 2$. We apply this method to test whether there is at least one gel position $(x_q^{**}, q = 1, ..., Q_m)$ where the two relative abundance profiles differ significantly: For protein $m$ the null hypothesis of equal relative abundance curves is

$$H_0 : \delta_{qm} = 0, \quad \text{for all} \quad q = 1, ..., Q_m,$$

that is, there is no difference of the log-scaled relative abundance between dark and light treatment at all gel positions considered for protein $m$.

The alternative hypothesis is that for at least one (or several) positions, the (log-scaled) relative abundance differs between dark and light treatment.

$$H_A : \delta_{qm} \neq 0, \quad \text{for at least one} \quad q = 1, ..., Q_m$$

The test statistics for protein $m$ at position $q$ resembles that of the t-Test,

$$t_{qm} = \frac{\hat{\delta}_{qm}}{\widehat{s.e.}(\hat{\delta})_{qm}}.$$

Herberich et al. (2014) describe how to compute adjusted p-values that asymptotically control the family wise error rate (FWER) among all $q = 1, ..., Q_m$ tests w.r.t. the $m$th protein. Deviating from their approach, we used probabilities of the multivariate t-distribution with residual degree of freedom, for computing adjusted p-values, but otherwise used their implementation (R package `multcomp`, Hothorn et al. 2008). For all positions within a given protein, the minimal p-value can be compared with $\alpha = 0.05$ to control the FWER at level 5%. For an error control across all proteins that have been considered in significance tests, we extracted the minimal p-value obtained for each protein, and then computed adjusted p-values to control the false discovery rate (FDR) for all proteins.

### Graphics for relative abundance profiles and heat maps

#### Line graphs of selected protein profiles

For selected proteins $m$, graphs show the observed relative abundance $p_{ijkm}$ in transparent dots and lines, where lines join observations from the same biological replicate $(ij)$. Bold solid lines interpolate the treatment specific estimates of the cubic regression splines, $\hat{\theta}_{iqm} = \hat{d}_{im} + \hat{s}(x^{**}_{qm}, n)_{im}$. The vertical difference between the two bold solid lines corresponds to the estimated difference $\hat{\delta}_{qm}$. Stars show significant p-values after FWER adjustment within protein ($* : 0.05 > p > 0.01$; $** : 0.01 > p > 0.001$; $*** : 0.001 > p$).

##### Heat maps

The treatment specific estimates of the cubic regression splines, $\hat{\theta}_{iqm} = \hat{d}_{im} + \hat{s}(x^{**}_{qm}, n)_{im}$, are used to show the mean relative abundance of proteins as a consensus map along the positions $x^{**}_{qm}$. Differential heat maps show the estimated difference between cubic regression splines, $\hat{\delta}_{qm}$ along the positions $x^{**}_{qm}$.

#### Software

Statistical analysis was performed in R (version 3.6.3, R Core Team, 2020). Summarizing and restructuring of data before model fitting was done using packages `plyr` (version 1.8.4, Wickham, 2011) and `reshape2` (version 1.4.3, Wickham, 2007). All fits of splines in generalized additive (mixed) models were performed using package `mgcv` (version 1.8-31, Wood, 2004). Multiple comparisons between treatments at given positions were performed using package `multcomp` (version 1.4-12, Hothorn et al. 2008). Graphical display of mean relative abundance curves were created using `ggplot2` (version 3.3.0, Wickham, 2016).

#### References


Herberich, E., Hassler, C., Hothorn, T. (2014): Multiple Curve Comparisons with an Application to the Formation of the Dorsal Funiculus of Mutant Mice. The International Journal of Biostatistics 10(2), 1-14.

Hothorn, T., Bretz, F., Westfall, P. (2008). Simultaneous Inference in General Parametric Models. Biometrical Journal 50(3), 346–363.

R Core Team (2020). R: A Language and Environment for Statistical Computing. R Foundation for Statistical Computing, Vienna, Austria. URL https://www.R-project.org/.

Wickham, H.(2016): ggplot2: Elegant Graphics for Data Analysis. Springer-Verlag New York.

Wood, S.N. (2004): Stable and Efficient Multiple Smoothing Parameter Estimation for Generalized Additive Models. Journal of the American Statistical Association 99:673-686.

Wickham, H. (2007). Reshaping Data with the reshape Package. Journal of Statistical Software, 21(12), 1-20. URL http://www.jstatsoft.org/v21/i12/.

Wickham, H. (2011). The Split-Apply-Combine Strategy for Data Analysis. Journal of Statistical Software, 40(1), 1-29. URL http://www.jstatsoft.org/v40/i01/.
