## Supplemental Table 2 for "Protein Interaction Patterns in *Arabidopsis thaliana* Leaf Mitochondria Change in Response to Illumination"

| Table S2: t-test statistics for 843 proteins commonly observed in dark and light samples across aligned lpBN fractions | | | | | | |
|---|---|---|---|---|---|---|
| **Arabidopsis genome** | **description** | **fraction** | **fold change** | **std. error** | **pvalue** | **significance** |
| AT1G01170.2 | Cytochrome c:O2 oxidoreductase subunit (COXX2-2) | 2 | 1.07 | 0.276 | 1.000 | |
| AT1G01170.2 | Cytochrome c:O2 oxidoreductase subunit (COXX2-2) | 3 | 1.09 | 0.163 | 1.000 | |
| AT1G01170.2 | Cytochrome c:O2 oxidoreductase subunit (COXX2-2) | 4 | 1.13 | 0.156 | 1.000 | |
| AT1G01170.2 | Cytochrome c:O2 oxidoreductase subunit (COXX2-2) | 5 | 1.16 | 0.155 | 1.000 | |
| AT1G01170.2 | Cytochrome c:O2 oxidoreductase subunit (COXX2-2) | 6 | 1.14 | 0.154 | 1.000 | |
| AT1G01170.2 | Cytochrome c:O2 oxidoreductase subunit (COXX2-2) | 7 | 1.11 | 0.155 | 1.000 | |
| AT1G01170.2 | Cytochrome c:O2 oxidoreductase subunit (COXX2-2) | 8 | 1.12 | 0.152 | 1.000 | |
| AT1G01170.2 | Cytochrome c:O2 oxidoreductase subunit (COXX2-2) | 9 | 1.13 | 0.150 | 1.000 | |
| AT1G01170.2 | Cytochrome c:O2 oxidoreductase subunit (COXX2-2) | 10 | 1.16 | 0.151 | 1.000 | |
| AT1G01170.2 | Cytochrome c:O2 oxidoreductase subunit (COXX2-2) | 11 | 1.26 | 0.153 | 1.000 | |
| AT1G01170.2 | Cytochrome c:O2 oxidoreductase subunit (COXX2-2) | 12 | 1.35 | 0.152 | 1.000 | |
| AT1G01170.2 | Cytochrome c:O2 oxidoreductase subunit (COXX2-2) | 13 | 1.41 | 0.150 | 1.000 | |
| AT1G01170.2 | Cytochrome c:O2 oxidoreductase subunit (COXX2-2) | 14 | 1.48 | 0.148 | 1.000 | |
| AT1G01170.2 | Cytochrome c:O2 oxidoreductase subunit (COXX2-2) | 15 | 1.37 | 0.147 | 1.000 | |
| AT1G01170.2 | Cytochrome c:O2 oxidoreductase subunit (COXX2-2) | 16 | 1.14 | 0.146 | 1.000 | |
| AT1G01170.2 | Cytochrome c:O2 oxidoreductase subunit (COXX2-2) | 17 | 1.05 | 0.145 | 1.000 | |
| AT1G01170.2 | Cytochrome c:O2 oxidoreductase subunit (COXX2-2) | 18 | 1.04 | 0.148 | 1.000 | |
| AT1G01170.2 | Cytochrome c:O2 oxidoreductase subunit (COXX2-2) | 19 | 1.02 | 0.152 | 1.000 | |
| AT1G01170.2 | Cytochrome c:O2 oxidoreductase subunit (COXX2-2) | 20 | 1.05 | 0.155 | 1.000 | |

| | | | | | | |
|---|---|---|---|---|---|---|
| AT1G01170.2 | Cytochrome c:O2 oxidoreductase subunit (COXX2-2) | 21 | 1.25 | 0.157 | 1.000 | |
| AT1G01170.2 | Cytochrome c:O2 oxidoreductase subunit (COXX2-2) | 22 | 1.70 | 0.157 | 0.984 | |
| AT1G01170.2 | Cytochrome c:O2 oxidoreductase subunit (COXX2-2) | 23 | 2.02 | 0.156 | 0.781 | |
| AT1G01170.2 | Cytochrome c:O2 oxidoreductase subunit (COXX2-2) | 24 | 1.93 | 0.152 | 0.833 | |
| AT1G01170.2 | Cytochrome c:O2 oxidoreductase subunit (COXX2-2) | 25 | 2.07 | 0.149 | 0.654 | |
| AT1G01170.2 | Cytochrome c:O2 oxidoreductase subunit (COXX2-2) | 26 | 2.08 | 0.150 | 0.644 | |
| AT1G01170.2 | Cytochrome c:O2 oxidoreductase subunit (COXX2-2) | 27 | 1.70 | 0.151 | 0.976 | |
| AT1G01170.2 | Cytochrome c:O2 oxidoreductase subunit (COXX2-2) | 28 | 1.50 | 0.151 | 0.999 | |
| AT1G01170.2 | Cytochrome c:O2 oxidoreductase subunit (COXX2-2) | 29 | 1.57 | 0.150 | 0.997 | |
| AT1G01170.2 | Cytochrome c:O2 oxidoreductase subunit (COXX2-2) | 30 | 1.61 | 0.150 | 0.994 | |
| AT1G01170.2 | Cytochrome c:O2 oxidoreductase subunit (COXX2-2) | 31 | 1.41 | 0.150 | 1.000 | |
| AT1G01170.2 | Cytochrome c:O2 oxidoreductase subunit (COXX2-2) | 32 | 1.09 | 0.151 | 1.000 | |
| AT1G01170.2 | Cytochrome c:O2 oxidoreductase subunit (COXX2-2) | 33 | 0.65 | 0.151 | 0.998 | |
| AT1G01170.2 | Cytochrome c:O2 oxidoreductase subunit (COXX2-2) | 34 | 0.37 | 0.152 | 0.148 | |
| AT1G01170.2 | Cytochrome c:O2 oxidoreductase subunit (COXX2-2) | 35 | 0.34 | 0.154 | 0.080 | . |
| AT1G01170.2 | Cytochrome c:O2 oxidoreductase subunit (COXX2-2) | 36 | 0.42 | 0.162 | 0.476 | |
| AT1G01170.2 | Cytochrome c:O2 oxidoreductase subunit (COXX2-2) | 37 | 0.68 | 0.187 | 1.000 | |
| AT1G01170.2 | Cytochrome c:O2 oxidoreductase subunit (COXX2-2) | 38 | 1.10 | 0.246 | 1.000 | |
| AT1G01170.2 | Cytochrome c:O2 oxidoreductase subunit (COXX2-2) | 41 | 2.14 | 0.245 | 0.995 | |
| AT1G01170.2 | Cytochrome c:O2 oxidoreductase subunit (COXX2-2) | 42 | 2.34 | 0.254 | 0.987 | |

| | | | | | |
|---|---|---|---|---|---|
| AT1G01170.2 | Cytochrome c:O2 oxidoreductase subunit (COXX2-2) | 43 | 2.51 | 0.333 | 0.999 |
| AT1G01170.2 | Cytochrome c:O2 oxidoreductase subunit (COXX2-2) | 46 | 2.93 | 1.202 | 1.000 |
| AT1G01170.2 | Cytochrome c:O2 oxidoreductase subunit (COXX2-2) | 47 | 3.06 | 1.588 | 1.000 |
| AT1G01290.2 | Cofactor of nitrate reductase and xanthine dehydrogenase 3 (CNX3) | 2 | 0.91 | 0.491 | 1.000 |
| AT1G01290.2 | Cofactor of nitrate reductase and xanthine dehydrogenase 3 (CNX3) | 3 | 0.96 | 0.440 | 1.000 |
| AT1G01290.2 | Cofactor of nitrate reductase and xanthine dehydrogenase 3 (CNX3) | 12 | 0.68 | 0.289 | 1.000 |
| AT1G01290.2 | Cofactor of nitrate reductase and xanthine dehydrogenase 3 (CNX3) | 13 | 0.58 | 0.247 | 0.995 |
| AT1G01290.2 | Cofactor of nitrate reductase and xanthine dehydrogenase 3 (CNX3) | 14 | 0.54 | 0.232 | 0.978 |
| AT1G01290.2 | Cofactor of nitrate reductase and xanthine dehydrogenase 3 (CNX3) | 15 | 0.68 | 0.220 | 1.000 |
| AT1G01290.2 | Cofactor of nitrate reductase and xanthine dehydrogenase 3 (CNX3) | 16 | 1.38 | 0.215 | 1.000 |
| AT1G01290.2 | Cofactor of nitrate reductase and xanthine dehydrogenase 3 (CNX3) | 17 | 3.57 | 0.244 | 0.317 |
| AT1G01290.2 | Cofactor of nitrate reductase and xanthine dehydrogenase 3 (CNX3) | 18 | 6.88 | 0.328 | 0.169 |
| AT1G01290.2 | Cofactor of nitrate reductase and xanthine dehydrogenase 3 (CNX3) | 27 | 0.34 | 0.321 | 0.882 |
| AT1G01290.2 | Cofactor of nitrate reductase and xanthine dehydrogenase 3 (CNX3) | 28 | 0.42 | 0.240 | 0.821 |
| AT1G01290.2 | Cofactor of nitrate reductase and xanthine dehydrogenase 3 (CNX3) | 29 | 0.72 | 0.211 | 1.000 |
| AT1G01290.2 | Cofactor of nitrate reductase and xanthine dehydrogenase 3 (CNX3) | 30 | 1.34 | 0.199 | 1.000 |
| AT1G01290.2 | Cofactor of nitrate reductase and xanthine dehydrogenase 3 (CNX3) | 31 | 2.07 | 0.196 | 0.791 |
| AT1G01290.2 | Cofactor of nitrate reductase and xanthine dehydrogenase 3 (CNX3) | 32 | 2.33 | 0.197 | 0.609 |
| AT1G01290.2 | Cofactor of nitrate reductase and xanthine dehydrogenase 3 (CNX3) | 33 | 2.11 | 0.200 | 0.784 |
| AT1G01290.2 | Cofactor of nitrate reductase and xanthine dehydrogenase 3 (CNX3) | 34 | 1.96 | 0.209 | 0.910 |

| | | | | | | |
|---|---|---|---|---|---|---|
| AT1G01290.2 | Cofactor of nitrate reductase and xanthine dehydrogenase 3 (CNX3) | 35 | 2.07 | 0.224 | 0.905 | |
| AT1G01290.2 | Cofactor of nitrate reductase and xanthine dehydrogenase 3 (CNX3) | 36 | 2.56 | 0.253 | 0.792 | |
| AT1G01290.2 | Cofactor of nitrate reductase and xanthine dehydrogenase 3 (CNX3) | 37 | 3.13 | 0.311 | 0.804 | |
| AT1G01290.2 | Cofactor of nitrate reductase and xanthine dehydrogenase 3 (CNX3) | 38 | 3.93 | 0.444 | 0.933 | |
| AT1G01290.2 | Cofactor of nitrate reductase and xanthine dehydrogenase 3 (CNX3) | 46 | 353.70 | 2.938 | 0.998 | |
| AT1G01290.2 | Cofactor of nitrate reductase and xanthine dehydrogenase 3 (CNX3) | 47 | 732.76 | 3.414 | 0.999 | |
| AT1G01820.1 | Peroxin 11C (PEX11C) | 2 | 1.14 | 0.339 | 1.000 | |
| AT1G01820.1 | Peroxin 11C (PEX11C) | 3 | 1.10 | 0.304 | 1.000 | |
| AT1G01820.1 | Peroxin 11C (PEX11C) | 4 | 1.06 | 0.397 | 1.000 | |
| AT1G01820.1 | Peroxin 11C (PEX11C) | 19 | 1.23 | 0.226 | 1.000 | |
| AT1G01820.1 | Peroxin 11C (PEX11C) | 20 | 1.30 | 0.207 | 1.000 | |
| AT1G01820.1 | Peroxin 11C (PEX11C) | 21 | 1.37 | 0.189 | 1.000 | |
| AT1G01820.1 | Peroxin 11C (PEX11C) | 22 | 1.42 | 0.176 | 0.997 | |
| AT1G01820.1 | Peroxin 11C (PEX11C) | 23 | 1.43 | 0.163 | 0.993 | |
| AT1G01820.1 | Peroxin 11C (PEX11C) | 24 | 1.32 | 0.153 | 0.999 | |
| AT1G01820.1 | Peroxin 11C (PEX11C) | 25 | 1.14 | 0.148 | 1.000 | |
| AT1G01820.1 | Peroxin 11C (PEX11C) | 26 | 1.03 | 0.145 | 1.000 | |
| AT1G01820.1 | Peroxin 11C (PEX11C) | 27 | 0.91 | 0.146 | 1.000 | |
| AT1G01820.1 | Peroxin 11C (PEX11C) | 28 | 0.77 | 0.150 | 0.999 | |
| AT1G01820.1 | Peroxin 11C (PEX11C) | 29 | 0.63 | 0.156 | 0.931 | |
| AT1G01820.1 | Peroxin 11C (PEX11C) | 30 | 0.53 | 0.156 | 0.644 | |
| AT1G01820.1 | Peroxin 11C (PEX11C) | 31 | 0.51 | 0.152 | 0.538 | |
| AT1G01820.1 | Peroxin 11C (PEX11C) | 32 | 0.64 | 0.149 | 0.925 | |
| AT1G01820.1 | Peroxin 11C (PEX11C) | 33 | 1.03 | 0.151 | 1.000 | |
| AT1G01820.1 | Peroxin 11C (PEX11C) | 34 | 1.91 | 0.155 | 0.611 | |
| AT1G01820.1 | Peroxin 11C (PEX11C) | 35 | 2.94 | 0.164 | 0.070 | . |
| AT1G01820.1 | Peroxin 11C (PEX11C) | 36 | 2.93 | 0.184 | 0.159 | |
| AT1G01820.1 | Peroxin 11C (PEX11C) | 37 | 2.01 | 0.231 | 0.921 | |
| AT1G01820.1 | Peroxin 11C (PEX11C) | 38 | 1.21 | 0.309 | 1.000 | |
| AT1G01820.1 | Peroxin 11C (PEX11C) | 46 | 0.04 | 0.455 | 0.035 | * |
| AT1G01820.1 | Peroxin 11C (PEX11C) | 47 | 0.03 | 0.501 | 0.032 | * |
| AT1G02370.1 | Pentatricopeptide repeat (PPR) superfamily protein | 2 | 1.04 | 0.333 | 1.000 | |
| AT1G02370.1 | Pentatricopeptide repeat (PPR) superfamily protein | 3 | 0.89 | 0.313 | 1.000 | |
| AT1G02370.1 | Pentatricopeptide repeat (PPR) superfamily protein | 23 | 0.53 | 0.278 | 0.996 | |
| AT1G02370.1 | Pentatricopeptide repeat (PPR) superfamily protein | 24 | 0.68 | 0.213 | 1.000 | |
| AT1G02370.1 | Pentatricopeptide repeat (PPR) superfamily protein | 25 | 0.88 | 0.187 | 1.000 | |
| AT1G02370.1 | Pentatricopeptide repeat (PPR) superfamily protein | 26 | 1.16 | 0.166 | 1.000 | |
| AT1G02370.1 | Pentatricopeptide repeat (PPR) superfamily protein | 27 | 1.43 | 0.153 | 0.995 | |
| AT1G02370.1 | Pentatricopeptide repeat (PPR) superfamily protein | 28 | 1.60 | 0.140 | 0.897 | |
| AT1G02370.1 | Pentatricopeptide repeat (PPR) superfamily protein | 29 | 1.82 | 0.132 | 0.556 | |

| | | | | | |
|---|---|---|---|---|---|
| AT1G02370.1 | Pentatricopeptide repeat (PPR) superfamily protein | 30 | 1.74 | 0.129 | 0.637 |
| AT1G02370.1 | Pentatricopeptide repeat (PPR) superfamily protein | 31 | 1.48 | 0.129 | 0.949 |
| AT1G02370.1 | Pentatricopeptide repeat (PPR) superfamily protein | 32 | 1.32 | 0.129 | 0.998 |
| AT1G02370.1 | Pentatricopeptide repeat (PPR) superfamily protein | 33 | 1.28 | 0.130 | 1.000 |
| AT1G02370.1 | Pentatricopeptide repeat (PPR) superfamily protein | 34 | 1.28 | 0.130 | 0.999 |
| AT1G02370.1 | Pentatricopeptide repeat (PPR) superfamily protein | 35 | 1.32 | 0.130 | 0.998 |
| AT1G02370.1 | Pentatricopeptide repeat (PPR) superfamily protein | 36 | 1.42 | 0.129 | 0.981 |
| AT1G02370.1 | Pentatricopeptide repeat (PPR) superfamily protein | 37 | 1.61 | 0.129 | 0.823 |
| AT1G02370.1 | Pentatricopeptide repeat (PPR) superfamily protein | 38 | 1.71 | 0.129 | 0.677 |
| AT1G02370.1 | Pentatricopeptide repeat (PPR) superfamily protein | 39 | 1.64 | 0.129 | 0.789 |
| AT1G02370.1 | Pentatricopeptide repeat (PPR) superfamily protein | 40 | 1.48 | 0.129 | 0.955 |
| AT1G02370.1 | Pentatricopeptide repeat (PPR) superfamily protein | 41 | 1.30 | 0.130 | 0.999 |
| AT1G02370.1 | Pentatricopeptide repeat (PPR) superfamily protein | 42 | 1.13 | 0.132 | 1.000 |
| AT1G02370.1 | Pentatricopeptide repeat (PPR) superfamily protein | 43 | 0.96 | 0.133 | 1.000 |
| AT1G02370.1 | Pentatricopeptide repeat (PPR) superfamily protein | 44 | 0.76 | 0.133 | 0.999 |
| AT1G02370.1 | Pentatricopeptide repeat (PPR) superfamily protein | 45 | 0.56 | 0.134 | 0.651 |
| AT1G02370.1 | Pentatricopeptide repeat (PPR) superfamily protein | 46 | 0.39 | 0.168 | 0.245 |
| AT1G02370.1 | Pentatricopeptide repeat (PPR) superfamily protein | 47 | 0.26 | 0.281 | 0.474 |
| AT1G02410.1 | Cytochrome c:O2 oxidoreductase assembly protein (COX11) | 2 | 1.04 | 0.385 | 1.000 |
| AT1G02410.1 | Cytochrome c:O2 oxidoreductase assembly protein (COX11) | 3 | 0.97 | 0.276 | 1.000 |
| AT1G02410.1 | Cytochrome c:O2 oxidoreductase assembly protein (COX11) | 5 | 0.82 | 0.248 | 1.000 |
| AT1G02410.1 | Cytochrome c:O2 oxidoreductase assembly protein (COX11) | 6 | 0.75 | 0.202 | 1.000 |
| AT1G02410.1 | Cytochrome c:O2 oxidoreductase assembly protein (COX11) | 7 | 0.75 | 0.179 | 1.000 |
| AT1G02410.1 | Cytochrome c:O2 oxidoreductase assembly protein (COX11) | 8 | 0.84 | 0.169 | 1.000 |
| AT1G02410.1 | Cytochrome c:O2 oxidoreductase assembly protein (COX11) | 9 | 0.96 | 0.164 | 1.000 |
| AT1G02410.1 | Cytochrome c:O2 oxidoreductase assembly protein (COX11) | 10 | 1.09 | 0.164 | 1.000 |

| | | | | | | |
|---|---|---|---|---|---|---|
| AT1G02410.1 | Cytochrome c:O2 oxidoreductase assembly protein (COX11) | 11 | 1.24 | 0.166 | 1.000 | |
| AT1G02410.1 | Cytochrome c:O2 oxidoreductase assembly protein (COX11) | 12 | 1.40 | 0.165 | 1.000 | |
| AT1G02410.1 | Cytochrome c:O2 oxidoreductase assembly protein (COX11) | 13 | 1.50 | 0.162 | 1.000 | |
| AT1G02410.1 | Cytochrome c:O2 oxidoreductase assembly protein (COX11) | 14 | 1.59 | 0.159 | 0.999 | |
| AT1G02410.1 | Cytochrome c:O2 oxidoreductase assembly protein (COX11) | 15 | 1.67 | 0.160 | 0.993 | |
| AT1G02410.1 | Cytochrome c:O2 oxidoreductase assembly protein (COX11) | 16 | 1.64 | 0.158 | 0.996 | |
| AT1G02410.1 | Cytochrome c:O2 oxidoreductase assembly protein (COX11) | 17 | 1.79 | 0.157 | 0.963 | |
| AT1G02410.1 | Cytochrome c:O2 oxidoreductase assembly protein (COX11) | 18 | 2.30 | 0.159 | 0.532 | |
| AT1G02410.1 | Cytochrome c:O2 oxidoreductase assembly protein (COX11) | 19 | 3.40 | 0.169 | 0.063 | . |
| AT1G02410.1 | Cytochrome c:O2 oxidoreductase assembly protein (COX11) | 20 | 5.06 | 0.175 | 0.002 | ** |
| AT1G02410.1 | Cytochrome c:O2 oxidoreductase assembly protein (COX11) | 21 | 5.83 | 0.176 | 0.001 | *** |
| AT1G02410.1 | Cytochrome c:O2 oxidoreductase assembly protein (COX11) | 22 | 5.80 | 0.188 | 0.002 | ** |
| AT1G02410.1 | Cytochrome c:O2 oxidoreductase assembly protein (COX11) | 23 | 5.16 | 0.203 | 0.020 | * |
| AT1G02410.1 | Cytochrome c:O2 oxidoreductase assembly protein (COX11) | 24 | 3.98 | 0.221 | 0.215 | |
| AT1G02410.1 | Cytochrome c:O2 oxidoreductase assembly protein (COX11) | 25 | 2.78 | 0.212 | 0.686 | |
| AT1G02410.1 | Cytochrome c:O2 oxidoreductase assembly protein (COX11) | 26 | 1.83 | 0.210 | 0.999 | |
| AT1G02410.1 | Cytochrome c:O2 oxidoreductase assembly protein (COX11) | 27 | 1.21 | 0.187 | 1.000 | |
| AT1G02410.1 | Cytochrome c:O2 oxidoreductase assembly protein (COX11) | 28 | 0.95 | 0.173 | 1.000 | |
| AT1G02410.1 | Cytochrome c:O2 oxidoreductase assembly protein (COX11) | 29 | 0.99 | 0.164 | 1.000 | |
| AT1G02410.1 | Cytochrome c:O2 oxidoreductase assembly protein (COX11) | 30 | 1.15 | 0.164 | 1.000 | |

| | | | | | | |
|---|---|---|---|---|---|---|
| AT1G02410.1 | Cytochrome c:O2 oxidoreductase assembly protein (COX11) | 31 | 1.28 | 0.164 | 1.000 | |
| AT1G02410.1 | Cytochrome c:O2 oxidoreductase assembly protein (COX11) | 32 | 1.25 | 0.164 | 1.000 | |
| AT1G02410.1 | Cytochrome c:O2 oxidoreductase assembly protein (COX11) | 33 | 0.97 | 0.164 | 1.000 | |
| AT1G02410.1 | Cytochrome c:O2 oxidoreductase assembly protein (COX11) | 34 | 0.68 | 0.164 | 1.000 | |
| AT1G02410.1 | Cytochrome c:O2 oxidoreductase assembly protein (COX11) | 35 | 0.60 | 0.166 | 0.996 | |
| AT1G02410.1 | Cytochrome c:O2 oxidoreductase assembly protein (COX11) | 36 | 0.90 | 0.170 | 1.000 | |
| AT1G02410.1 | Cytochrome c:O2 oxidoreductase assembly protein (COX11) | 37 | 1.99 | 0.178 | 0.945 | |
| AT1G02410.1 | Cytochrome c:O2 oxidoreductase assembly protein (COX11) | 38 | 3.49 | 0.184 | 0.113 | |
| AT1G02410.1 | Cytochrome c:O2 oxidoreductase assembly protein (COX11) | 39 | 4.42 | 0.181 | 0.015 | * |
| AT1G02410.1 | Cytochrome c:O2 oxidoreductase assembly protein (COX11) | 40 | 5.89 | 0.169 | 0.000 | *** |
| AT1G02410.1 | Cytochrome c:O2 oxidoreductase assembly protein (COX11) | 41 | 7.78 | 0.164 | 0.000 | *** |
| AT1G02410.1 | Cytochrome c:O2 oxidoreductase assembly protein (COX11) | 42 | 8.14 | 0.166 | 0.000 | *** |
| AT1G02410.1 | Cytochrome c:O2 oxidoreductase assembly protein (COX11) | 43 | 6.64 | 0.167 | 0.000 | *** |
| AT1G02410.1 | Cytochrome c:O2 oxidoreductase assembly protein (COX11) | 44 | 4.39 | 0.167 | 0.006 | ** |
| AT1G02410.1 | Cytochrome c:O2 oxidoreductase assembly protein (COX11) | 45 | 2.21 | 0.167 | 0.714 | |
| AT1G02410.1 | Cytochrome c:O2 oxidoreductase assembly protein (COX11) | 46 | 0.82 | 0.194 | 1.000 | |
| AT1G02410.1 | Cytochrome c:O2 oxidoreductase assembly protein (COX11) | 47 | 0.24 | 0.366 | 0.939 | |
| AT1G02780.1 | 60S ribosomal protein L19-1 (RPL19A) | 2 | 0.81 | 0.193 | 0.890 | |
| AT1G02780.1 | 60S ribosomal protein L19-1 (RPL19A) | 3 | 0.85 | 0.183 | 0.921 | |
| AT1G02780.1 | 60S ribosomal protein L19-1 (RPL19A) | 4 | 0.88 | 0.173 | 0.951 | |
| AT1G02780.1 | 60S ribosomal protein L19-1 (RPL19A) | 5 | 0.92 | 0.164 | 0.976 | |
| AT1G02780.1 | 60S ribosomal protein L19-1 (RPL19A) | 6 | 0.96 | 0.156 | 0.994 | |

| | | | | | | |
|---|---|---|---|---|---|---|
| AT1G02780.1 | 60S ribosomal protein L19-1 (RPL19A) | 7 | 1.01 | 0.150 | 1.000 | |
| AT1G02780.1 | 60S ribosomal protein L19-1 (RPL19A) | 8 | 1.05 | 0.145 | 0.989 | |
| AT1G02780.1 | 60S ribosomal protein L19-1 (RPL19A) | 9 | 1.10 | 0.141 | 0.959 | |
| AT1G02780.1 | 60S ribosomal protein L19-1 (RPL19A) | 10 | 1.14 | 0.139 | 0.911 | |
| AT1G02780.1 | 60S ribosomal protein L19-1 (RPL19A) | 11 | 1.20 | 0.138 | 0.849 | |
| AT1G02780.1 | 60S ribosomal protein L19-1 (RPL19A) | 12 | 1.25 | 0.139 | 0.780 | |
| AT1G02780.1 | 60S ribosomal protein L19-1 (RPL19A) | 13 | 1.30 | 0.142 | 0.712 | |
| AT1G02780.1 | 60S ribosomal protein L19-1 (RPL19A) | 14 | 1.36 | 0.146 | 0.649 | |
| AT1G02780.1 | 60S ribosomal protein L19-1 (RPL19A) | 15 | 1.42 | 0.152 | 0.595 | |
| AT1G02780.1 | 60S ribosomal protein L19-1 (RPL19A) | 16 | 1.48 | 0.159 | 0.549 | |
| AT1G02780.1 | 60S ribosomal protein L19-1 (RPL19A) | 17 | 1.55 | 0.167 | 0.514 | |
| AT1G02780.1 | 60S ribosomal protein L19-1 (RPL19A) | 18 | 1.62 | 0.176 | 0.485 | |
| AT1G02780.1 | 60S ribosomal protein L19-1 (RPL19A) | 19 | 1.69 | 0.186 | 0.462 | |
| AT1G02780.1 | 60S ribosomal protein L19-1 (RPL19A) | 46 | 5.43 | 0.556 | 0.407 | |
| AT1G02780.1 | 60S ribosomal protein L19-1 (RPL19A) | 47 | 5.67 | 0.571 | 0.408 | |
| AT1G03090.2 | 3-methylcrotonyl-CoA carboxylase 1 (MCCA) | 2 | 0.99 | 0.248 | 1.000 | |
| AT1G03090.2 | 3-methylcrotonyl-CoA carboxylase 1 (MCCA) | 3 | 0.94 | 0.226 | 1.000 | |
| AT1G03090.2 | 3-methylcrotonyl-CoA carboxylase 1 (MCCA) | 18 | 1.86 | 0.154 | 0.806 | |
| AT1G03090.2 | 3-methylcrotonyl-CoA carboxylase 1 (MCCA) | 19 | 2.20 | 0.139 | 0.278 | |
| AT1G03090.2 | 3-methylcrotonyl-CoA carboxylase 1 (MCCA) | 20 | 2.36 | 0.125 | 0.071 | . |
| AT1G03090.2 | 3-methylcrotonyl-CoA carboxylase 1 (MCCA) | 21 | 1.87 | 0.121 | 0.416 | |
| AT1G03090.2 | 3-methylcrotonyl-CoA carboxylase 1 (MCCA) | 22 | 1.19 | 0.119 | 1.000 | |
| AT1G03090.2 | 3-methylcrotonyl-CoA carboxylase 1 (MCCA) | 23 | 0.86 | 0.118 | 1.000 | |
| AT1G03090.2 | 3-methylcrotonyl-CoA carboxylase 1 (MCCA) | 24 | 0.81 | 0.115 | 1.000 | |
| AT1G03090.2 | 3-methylcrotonyl-CoA carboxylase 1 (MCCA) | 25 | 0.90 | 0.114 | 1.000 | |
| AT1G03090.2 | 3-methylcrotonyl-CoA carboxylase 1 (MCCA) | 26 | 1.06 | 0.114 | 1.000 | |
| AT1G03090.2 | 3-methylcrotonyl-CoA carboxylase 1 (MCCA) | 27 | 1.16 | 0.115 | 1.000 | |
| AT1G03090.2 | 3-methylcrotonyl-CoA carboxylase 1 (MCCA) | 28 | 1.13 | 0.115 | 1.000 | |
| AT1G03090.2 | 3-methylcrotonyl-CoA carboxylase 1 (MCCA) | 29 | 1.07 | 0.115 | 1.000 | |
| AT1G03090.2 | 3-methylcrotonyl-CoA carboxylase 1 (MCCA) | 30 | 1.04 | 0.115 | 1.000 | |
| AT1G03090.2 | 3-methylcrotonyl-CoA carboxylase 1 (MCCA) | 31 | 1.06 | 0.115 | 1.000 | |

| | | | | | | |
|---|---|---|---|---|---|---|
| AT1G03090.2 | 3-methylcrotonyl-CoA carboxylase 1 (MCCA) | 32 | 1.08 | 0.115 | 1.000 | |
| AT1G03090.2 | 3-methylcrotonyl-CoA carboxylase 1 (MCCA) | 33 | 0.95 | 0.115 | 1.000 | |
| AT1G03090.2 | 3-methylcrotonyl-CoA carboxylase 1 (MCCA) | 34 | 0.80 | 0.115 | 1.000 | |
| AT1G03090.2 | 3-methylcrotonyl-CoA carboxylase 1 (MCCA) | 35 | 0.77 | 0.115 | 1.000 | |
| AT1G03090.2 | 3-methylcrotonyl-CoA carboxylase 1 (MCCA) | 36 | 0.86 | 0.114 | 1.000 | |
| AT1G03090.2 | 3-methylcrotonyl-CoA carboxylase 1 (MCCA) | 37 | 1.02 | 0.114 | 1.000 | |
| AT1G03090.2 | 3-methylcrotonyl-CoA carboxylase 1 (MCCA) | 38 | 1.18 | 0.115 | 1.000 | |
| AT1G03090.2 | 3-methylcrotonyl-CoA carboxylase 1 (MCCA) | 39 | 1.29 | 0.115 | 1.000 | |
| AT1G03090.2 | 3-methylcrotonyl-CoA carboxylase 1 (MCCA) | 40 | 1.35 | 0.115 | 0.996 | |
| AT1G03090.2 | 3-methylcrotonyl-CoA carboxylase 1 (MCCA) | 41 | 1.29 | 0.116 | 1.000 | |
| AT1G03090.2 | 3-methylcrotonyl-CoA carboxylase 1 (MCCA) | 42 | 1.15 | 0.117 | 1.000 | |
| AT1G03090.2 | 3-methylcrotonyl-CoA carboxylase 1 (MCCA) | 43 | 0.97 | 0.118 | 1.000 | |
| AT1G03090.2 | 3-methylcrotonyl-CoA carboxylase 1 (MCCA) | 44 | 0.73 | 0.118 | 0.996 | |
| AT1G03090.2 | 3-methylcrotonyl-CoA carboxylase 1 (MCCA) | 45 | 0.52 | 0.118 | 0.294 | |
| AT1G03090.2 | 3-methylcrotonyl-CoA carboxylase 1 (MCCA) | 46 | 0.33 | 0.134 | 0.011 | * |
| AT1G03090.2 | 3-methylcrotonyl-CoA carboxylase 1 (MCCA) | 47 | 0.18 | 0.247 | 0.076 | . |
| AT1G03130.1 | Photosystem I subunit (PSAD2) | 2 | 0.30 | 0.249 | 0.580 | |
| AT1G03130.1 | Photosystem I subunit (PSAD2) | 3 | 0.35 | 0.150 | 0.078 | . |
| AT1G03130.1 | Photosystem I subunit (PSAD2) | 4 | 0.56 | 0.139 | 0.819 | |
| AT1G03130.1 | Photosystem I subunit (PSAD2) | 5 | 0.81 | 0.139 | 1.000 | |
| AT1G03130.1 | Photosystem I subunit (PSAD2) | 6 | 0.89 | 0.140 | 1.000 | |
| AT1G03130.1 | Photosystem I subunit (PSAD2) | 7 | 0.85 | 0.141 | 1.000 | |
| AT1G03130.1 | Photosystem I subunit (PSAD2) | 8 | 0.93 | 0.136 | 1.000 | |
| AT1G03130.1 | Photosystem I subunit (PSAD2) | 9 | 1.04 | 0.133 | 1.000 | |
| AT1G03130.1 | Photosystem I subunit (PSAD2) | 10 | 0.99 | 0.134 | 1.000 | |
| AT1G03130.1 | Photosystem I subunit (PSAD2) | 11 | 0.95 | 0.135 | 1.000 | |
| AT1G03130.1 | Photosystem I subunit (PSAD2) | 12 | 1.02 | 0.135 | 1.000 | |
| AT1G03130.1 | Photosystem I subunit (PSAD2) | 13 | 1.13 | 0.133 | 1.000 | |
| AT1G03130.1 | Photosystem I subunit (PSAD2) | 14 | 1.13 | 0.131 | 1.000 | |
| AT1G03130.1 | Photosystem I subunit (PSAD2) | 15 | 0.95 | 0.131 | 1.000 | |
| AT1G03130.1 | Photosystem I subunit (PSAD2) | 16 | 0.84 | 0.130 | 1.000 | |
| AT1G03130.1 | Photosystem I subunit (PSAD2) | 17 | 0.89 | 0.129 | 1.000 | |
| AT1G03130.1 | Photosystem I subunit (PSAD2) | 18 | 1.04 | 0.129 | 1.000 | |
| AT1G03130.1 | Photosystem I subunit (PSAD2) | 19 | 1.30 | 0.134 | 1.000 | |
| AT1G03130.1 | Photosystem I subunit (PSAD2) | 20 | 1.48 | 0.139 | 0.996 | |
| AT1G03130.1 | Photosystem I subunit (PSAD2) | 21 | 1.35 | 0.140 | 1.000 | |
| AT1G03130.1 | Photosystem I subunit (PSAD2) | 22 | 1.09 | 0.140 | 1.000 | |
| AT1G03130.1 | Photosystem I subunit (PSAD2) | 23 | 0.84 | 0.140 | 1.000 | |
| AT1G03130.1 | Photosystem I subunit (PSAD2) | 24 | 0.65 | 0.136 | 0.985 | |
| AT1G03130.1 | Photosystem I subunit (PSAD2) | 25 | 0.66 | 0.133 | 0.983 | |
| AT1G03130.1 | Photosystem I subunit (PSAD2) | 26 | 0.98 | 0.132 | 1.000 | |
| AT1G03130.1 | Photosystem I subunit (PSAD2) | 27 | 1.29 | 0.133 | 1.000 | |
| AT1G03130.1 | Photosystem I subunit (PSAD2) | 28 | 1.42 | 0.134 | 0.998 | |
| AT1G03130.1 | Photosystem I subunit (PSAD2) | 29 | 1.67 | 0.136 | 0.914 | |
| AT1G03130.1 | Photosystem I subunit (PSAD2) | 30 | 0.98 | 0.139 | 1.000 | |
| AT1G03130.1 | Photosystem I subunit (PSAD2) | 31 | 0.26 | 0.148 | 0.003 | ** |

| | | | | | | |
|---|---|---|---|---|---|---|
| AT1G03130.1 | Photosystem I subunit (PSAD2) | 32 | 0.08 | 0.186 | 0.000 | *** |
| AT1G03130.1 | Photosystem I subunit (PSAD2) | 33 | 0.04 | 0.310 | 0.000 | *** |
| AT1G03130.1 | Photosystem I subunit (PSAD2) | 46 | 1813525.38 | 3.061 | 0.642 | |
| AT1G03130.1 | Photosystem I subunit (PSAD2) | 47 | 20115384.23 | 3.390 | 0.548 | |
| AT1G03860.3 | Prohibitin 2 (PHB2) | 2 | 1.04 | 0.356 | 1.000 | |
| AT1G03860.3 | Prohibitin 2 (PHB2) | 3 | 0.99 | 0.237 | 1.000 | |
| AT1G03860.3 | Prohibitin 2 (PHB2) | 4 | 0.93 | 0.209 | 1.000 | |
| AT1G03860.3 | Prohibitin 2 (PHB2) | 5 | 0.90 | 0.207 | 1.000 | |
| AT1G03860.3 | Prohibitin 2 (PHB2) | 6 | 0.87 | 0.208 | 1.000 | |
| AT1G03860.3 | Prohibitin 2 (PHB2) | 7 | 0.87 | 0.207 | 1.000 | |
| AT1G03860.3 | Prohibitin 2 (PHB2) | 8 | 0.96 | 0.203 | 1.000 | |
| AT1G03860.3 | Prohibitin 2 (PHB2) | 9 | 1.10 | 0.201 | 1.000 | |
| AT1G03860.3 | Prohibitin 2 (PHB2) | 10 | 1.26 | 0.201 | 1.000 | |
| AT1G03860.3 | Prohibitin 2 (PHB2) | 11 | 1.44 | 0.201 | 1.000 | |
| AT1G03860.3 | Prohibitin 2 (PHB2) | 12 | 1.56 | 0.201 | 0.999 | |
| AT1G03860.3 | Prohibitin 2 (PHB2) | 13 | 1.55 | 0.199 | 0.999 | |
| AT1G03860.3 | Prohibitin 2 (PHB2) | 14 | 1.30 | 0.196 | 1.000 | |
| AT1G03860.3 | Prohibitin 2 (PHB2) | 15 | 0.92 | 0.196 | 1.000 | |
| AT1G03860.3 | Prohibitin 2 (PHB2) | 16 | 0.67 | 0.195 | 1.000 | |
| AT1G03860.3 | Prohibitin 2 (PHB2) | 17 | 0.61 | 0.195 | 0.996 | |
| AT1G03860.3 | Prohibitin 2 (PHB2) | 18 | 0.68 | 0.197 | 1.000 | |
| AT1G03860.3 | Prohibitin 2 (PHB2) | 19 | 0.80 | 0.201 | 1.000 | |
| AT1G03860.3 | Prohibitin 2 (PHB2) | 20 | 0.96 | 0.206 | 1.000 | |
| AT1G03860.3 | Prohibitin 2 (PHB2) | 21 | 1.19 | 0.208 | 1.000 | |
| AT1G03860.3 | Prohibitin 2 (PHB2) | 22 | 1.51 | 0.208 | 1.000 | |
| AT1G03860.3 | Prohibitin 2 (PHB2) | 23 | 1.70 | 0.207 | 0.995 | |
| AT1G03860.3 | Prohibitin 2 (PHB2) | 24 | 1.41 | 0.203 | 1.000 | |
| AT1G03860.3 | Prohibitin 2 (PHB2) | 25 | 0.81 | 0.200 | 1.000 | |
| AT1G03860.3 | Prohibitin 2 (PHB2) | 26 | 0.43 | 0.199 | 0.700 | |
| AT1G03860.3 | Prohibitin 2 (PHB2) | 27 | 0.29 | 0.199 | 0.133 | |
| AT1G03860.3 | Prohibitin 2 (PHB2) | 28 | 0.29 | 0.199 | 0.149 | |
| AT1G03860.3 | Prohibitin 2 (PHB2) | 29 | 0.45 | 0.200 | 0.780 | |
| AT1G03860.3 | Prohibitin 2 (PHB2) | 30 | 0.87 | 0.202 | 1.000 | |
| AT1G03860.3 | Prohibitin 2 (PHB2) | 31 | 1.68 | 0.204 | 0.996 | |
| AT1G03860.3 | Prohibitin 2 (PHB2) | 32 | 2.70 | 0.208 | 0.535 | |
| AT1G03860.3 | Prohibitin 2 (PHB2) | 33 | 3.55 | 0.225 | 0.268 | |
| AT1G03860.3 | Prohibitin 2 (PHB2) | 34 | 4.29 | 0.281 | 0.396 | |
| AT1G03860.3 | Prohibitin 2 (PHB2) | 35 | 5.26 | 0.401 | 0.748 | |
| AT1G03860.3 | Prohibitin 2 (PHB2) | 46 | 685.20 | 2.801 | 0.998 | |
| AT1G03860.3 | Prohibitin 2 (PHB2) | 47 | 1199.11 | 3.135 | 0.999 | |
| AT1G04630.1 | NADH:ubiquinone oxidoreductase subunit (B16.6-1) | 2 | 0.83 | 0.679 | 1.000 | |
| AT1G04630.1 | NADH:ubiquinone oxidoreductase subunit (B16.6-1) | 3 | 0.84 | 0.610 | 1.000 | |
| AT1G04630.1 | NADH:ubiquinone oxidoreductase subunit (B16.6-1) | 14 | 1.65 | 0.391 | 1.000 | |
| AT1G04630.1 | NADH:ubiquinone oxidoreductase subunit (B16.6-1) | 15 | 1.36 | 0.335 | 1.000 | |
| AT1G04630.1 | NADH:ubiquinone oxidoreductase subunit (B16.6-1) | 16 | 0.91 | 0.304 | 1.000 | |
| AT1G04630.1 | NADH:ubiquinone oxidoreductase subunit (B16.6-1) | 17 | 0.60 | 0.275 | 0.998 | |
| AT1G04630.1 | NADH:ubiquinone oxidoreductase subunit (B16.6-1) | 18 | 0.57 | 0.267 | 0.993 | |

| | | | | | |
|---|---|---|---|---|---|
| AT1G04630.1 | NADH:ubiquinone oxidoreductase subunit (B16.6-1) | 19 | 0.72 | 0.274 | 1.000 |
| AT1G04630.1 | NADH:ubiquinone oxidoreductase subunit (B16.6-1) | 20 | 0.91 | 0.280 | 1.000 |
| AT1G04630.1 | NADH:ubiquinone oxidoreductase subunit (B16.6-1) | 21 | 1.14 | 0.282 | 1.000 |
| AT1G04630.1 | NADH:ubiquinone oxidoreductase subunit (B16.6-1) | 22 | 1.46 | 0.283 | 1.000 |
| AT1G04630.1 | NADH:ubiquinone oxidoreductase subunit (B16.6-1) | 23 | 1.81 | 0.280 | 0.993 |
| AT1G04630.1 | NADH:ubiquinone oxidoreductase subunit (B16.6-1) | 24 | 1.90 | 0.276 | 0.985 |
| AT1G04630.1 | NADH:ubiquinone oxidoreductase subunit (B16.6-1) | 25 | 1.93 | 0.273 | 0.981 |
| AT1G04630.1 | NADH:ubiquinone oxidoreductase subunit (B16.6-1) | 26 | 2.64 | 0.278 | 0.800 |
| AT1G04630.1 | NADH:ubiquinone oxidoreductase subunit (B16.6-1) | 27 | 4.24 | 0.302 | 0.394 |
| AT1G04630.1 | NADH:ubiquinone oxidoreductase subunit (B16.6-1) | 28 | 4.99 | 0.381 | 0.575 |
| AT1G04630.1 | NADH:ubiquinone oxidoreductase subunit (B16.6-1) | 29 | 4.71 | 0.566 | 0.952 |
| AT1G04630.1 | NADH:ubiquinone oxidoreductase subunit (B16.6-1) | 46 | 5706.05 | 9.895 | 1.000 |
| AT1G04630.1 | NADH:ubiquinone oxidoreductase subunit (B16.6-1) | 47 | 11172.26 | 10.734 | 1.000 |
| AT1G04640.2 | Lipoyltransferase 2 (LIP2) | 2 | 1.00 | 0.442 | 1.000 |
| AT1G04640.2 | Lipoyltransferase 2 (LIP2) | 3 | 1.05 | 0.415 | 1.000 |
| AT1G04640.2 | Lipoyltransferase 2 (LIP2) | 22 | 1.41 | 0.376 | 1.000 |
| AT1G04640.2 | Lipoyltransferase 2 (LIP2) | 23 | 1.35 | 0.324 | 1.000 |
| AT1G04640.2 | Lipoyltransferase 2 (LIP2) | 24 | 1.27 | 0.303 | 1.000 |
| AT1G04640.2 | Lipoyltransferase 2 (LIP2) | 27 | 0.96 | 0.252 | 1.000 |
| AT1G04640.2 | Lipoyltransferase 2 (LIP2) | 28 | 0.88 | 0.226 | 1.000 |
| AT1G04640.2 | Lipoyltransferase 2 (LIP2) | 29 | 0.93 | 0.208 | 1.000 |
| AT1G04640.2 | Lipoyltransferase 2 (LIP2) | 30 | 1.12 | 0.199 | 1.000 |
| AT1G04640.2 | Lipoyltransferase 2 (LIP2) | 31 | 1.49 | 0.195 | 0.995 |
| AT1G04640.2 | Lipoyltransferase 2 (LIP2) | 32 | 1.77 | 0.195 | 0.919 |
| AT1G04640.2 | Lipoyltransferase 2 (LIP2) | 33 | 1.73 | 0.195 | 0.938 |
| AT1G04640.2 | Lipoyltransferase 2 (LIP2) | 34 | 1.52 | 0.195 | 0.993 |
| AT1G04640.2 | Lipoyltransferase 2 (LIP2) | 35 | 1.37 | 0.195 | 1.000 |
| AT1G04640.2 | Lipoyltransferase 2 (LIP2) | 36 | 1.38 | 0.195 | 0.999 |
| AT1G04640.2 | Lipoyltransferase 2 (LIP2) | 37 | 1.51 | 0.194 | 0.993 |
| AT1G04640.2 | Lipoyltransferase 2 (LIP2) | 38 | 1.65 | 0.194 | 0.966 |
| AT1G04640.2 | Lipoyltransferase 2 (LIP2) | 39 | 1.83 | 0.194 | 0.885 |
| AT1G04640.2 | Lipoyltransferase 2 (LIP2) | 40 | 2.07 | 0.195 | 0.723 |
| AT1G04640.2 | Lipoyltransferase 2 (LIP2) | 41 | 2.13 | 0.196 | 0.675 |
| AT1G04640.2 | Lipoyltransferase 2 (LIP2) | 42 | 1.93 | 0.197 | 0.836 |
| AT1G04640.2 | Lipoyltransferase 2 (LIP2) | 43 | 1.51 | 0.199 | 0.994 |
| AT1G04640.2 | Lipoyltransferase 2 (LIP2) | 44 | 1.01 | 0.198 | 1.000 |

| | | | | | | |
|---|---|---|---|---|---|---|
| AT1G04640.2 | Lipoyltransferase 2 (LIP2) | 45 | 0.56 | 0.202 | 0.928 | |
| AT1G04640.2 | Lipoyltransferase 2 (LIP2) | 46 | 0.26 | 0.238 | 0.180 | |
| AT1G04640.2 | Lipoyltransferase 2 (LIP2) | 47 | 0.11 | 0.353 | 0.092 | . |
| AT1G05270.1 | TraB family protein | 2 | 0.87 | 0.378 | 1.000 | |
| AT1G05270.1 | TraB family protein | 3 | 0.93 | 0.344 | 1.000 | |
| AT1G05270.1 | TraB family protein | 15 | 0.71 | 0.245 | 1.000 | |
| AT1G05270.1 | TraB family protein | 16 | 0.69 | 0.224 | 1.000 | |
| AT1G05270.1 | TraB family protein | 17 | 0.78 | 0.208 | 1.000 | |
| AT1G05270.1 | TraB family protein | 18 | 0.98 | 0.203 | 1.000 | |
| AT1G05270.1 | TraB family protein | 19 | 1.26 | 0.209 | 1.000 | |
| AT1G05270.1 | TraB family protein | 20 | 1.44 | 0.238 | 1.000 | |
| AT1G05270.1 | TraB family protein | 21 | 1.32 | 0.283 | 1.000 | |
| AT1G05270.1 | TraB family protein | 26 | 0.31 | 0.254 | 0.506 | |
| AT1G05270.1 | TraB family protein | 27 | 0.36 | 0.208 | 0.415 | |
| AT1G05270.1 | TraB family protein | 28 | 0.54 | 0.188 | 0.911 | |
| AT1G05270.1 | TraB family protein | 29 | 0.80 | 0.181 | 1.000 | |
| AT1G05270.1 | TraB family protein | 30 | 0.90 | 0.179 | 1.000 | |
| AT1G05270.1 | TraB family protein | 31 | 0.91 | 0.178 | 1.000 | |
| AT1G05270.1 | TraB family protein | 32 | 0.93 | 0.178 | 1.000 | |
| AT1G05270.1 | TraB family protein | 33 | 0.91 | 0.178 | 1.000 | |
| AT1G05270.1 | TraB family protein | 34 | 0.91 | 0.179 | 1.000 | |
| AT1G05270.1 | TraB family protein | 35 | 1.00 | 0.179 | 1.000 | |
| AT1G05270.1 | TraB family protein | 36 | 1.17 | 0.179 | 1.000 | |
| AT1G05270.1 | TraB family protein | 37 | 1.45 | 0.178 | 0.999 | |
| AT1G05270.1 | TraB family protein | 38 | 1.71 | 0.178 | 0.950 | |
| AT1G05270.1 | TraB family protein | 39 | 1.99 | 0.178 | 0.760 | |
| AT1G05270.1 | TraB family protein | 40 | 2.55 | 0.178 | 0.323 | |
| AT1G05270.1 | TraB family protein | 41 | 3.36 | 0.179 | 0.065 | . |
| AT1G05270.1 | TraB family protein | 42 | 3.78 | 0.180 | 0.032 | * |
| AT1G05270.1 | TraB family protein | 43 | 3.34 | 0.182 | 0.077 | . |
| AT1G05270.1 | TraB family protein | 44 | 2.09 | 0.181 | 0.703 | |
| AT1G05270.1 | TraB family protein | 45 | 0.87 | 0.183 | 1.000 | |
| AT1G05270.1 | TraB family protein | 46 | 0.24 | 0.209 | 0.066 | . |
| AT1G05270.1 | TraB family protein | 47 | 0.05 | 0.327 | 0.002 | ** |
| AT1G05430.1 | ? | 2 | 0.91 | 0.549 | 1.000 | |
| AT1G05430.1 | ? | 3 | 0.91 | 0.487 | 1.000 | |
| AT1G05430.1 | ? | 13 | 1.01 | 0.286 | 1.000 | |
| AT1G05430.1 | ? | 14 | 1.04 | 0.241 | 1.000 | |
| AT1G05430.1 | ? | 15 | 0.99 | 0.213 | 1.000 | |
| AT1G05430.1 | ? | 16 | 0.88 | 0.194 | 1.000 | |
| AT1G05430.1 | ? | 17 | 0.81 | 0.185 | 1.000 | |
| AT1G05430.1 | ? | 18 | 0.82 | 0.184 | 1.000 | |
| AT1G05430.1 | ? | 19 | 0.90 | 0.187 | 1.000 | |
| AT1G05430.1 | ? | 20 | 1.03 | 0.192 | 1.000 | |
| AT1G05430.1 | ? | 21 | 1.20 | 0.195 | 1.000 | |
| AT1G05430.1 | ? | 22 | 1.38 | 0.197 | 0.996 | |
| AT1G05430.1 | ? | 23 | 1.51 | 0.202 | 0.981 | |
| AT1G05430.1 | ? | 24 | 1.53 | 0.222 | 0.987 | |
| AT1G05430.1 | ? | 25 | 1.54 | 0.277 | 0.997 | |
| AT1G05430.1 | ? | 26 | 1.65 | 0.381 | 0.999 | |
| AT1G05430.1 | ? | 46 | 247138.32 | 5.715 | 0.971 | |
| AT1G05430.1 | ? | 47 | 579407.01 | 6.109 | 0.972 | |
| AT1G06200.1 | Peptidase S24/S26A/S26B/S26C family protein | 2 | 0.81 | 0.455 | 1.000 | |
| AT1G06200.1 | Peptidase S24/S26A/S26B/S26C family protein | 3 | 1.08 | 0.402 | 1.000 | |
| AT1G06200.1 | Peptidase S24/S26A/S26B/S26C family protein | 13 | 1.63 | 0.242 | 0.967 | |
| AT1G06200.1 | Peptidase S24/S26A/S26B/S26C family protein | 14 | 1.17 | 0.210 | 1.000 | |
| AT1G06200.1 | Peptidase S24/S26A/S26B/S26C family protein | 15 | 0.81 | 0.186 | 0.999 | |

| | | | | | | |
|---|---|---|---|---|---|---|
| AT1G06200.1 | Peptidase S24/S26A/S26B/S26C family protein | 16 | 0.58 | 0.168 | 0.712 | |
| AT1G06200.1 | Peptidase S24/S26A/S26B/S26C family protein | 17 | 0.48 | 0.160 | 0.322 | |
| AT1G06200.1 | Peptidase S24/S26A/S26B/S26C family protein | 18 | 0.55 | 0.163 | 0.603 | |
| AT1G06200.1 | Peptidase S24/S26A/S26B/S26C family protein | 19 | 0.87 | 0.176 | 1.000 | |
| AT1G06200.1 | Peptidase S24/S26A/S26B/S26C family protein | 20 | 1.76 | 0.194 | 0.809 | |
| AT1G06200.1 | Peptidase S24/S26A/S26B/S26C family protein | 21 | 3.80 | 0.226 | 0.094 | . |
| AT1G06200.1 | Peptidase S24/S26A/S26B/S26C family protein | 22 | 7.84 | 0.297 | 0.029 | * |
| AT1G06200.1 | Peptidase S24/S26A/S26B/S26C family protein | 46 | 28060852.84 | 11.330 | 0.994 | |
| AT1G06200.1 | Peptidase S24/S26A/S26B/S26C family protein | 47 | 50334094.61 | 12.010 | 0.995 | |
| AT1G06290.1 | Acyl-CoA oxidase (ACX3) | 2 | 1.11 | 0.397 | 1.000 | |
| AT1G06290.1 | Acyl-CoA oxidase (ACX3) | 3 | 0.99 | 0.361 | 1.000 | |
| AT1G06290.1 | Acyl-CoA oxidase (ACX3) | 4 | 0.88 | 0.485 | 1.000 | |
| AT1G06290.1 | Acyl-CoA oxidase (ACX3) | 32 | 0.48 | 0.401 | 0.971 | |
| AT1G06290.1 | Acyl-CoA oxidase (ACX3) | 33 | 0.61 | 0.364 | 0.994 | |
| AT1G06290.1 | Acyl-CoA oxidase (ACX3) | 34 | 0.78 | 0.349 | 1.000 | |
| AT1G06290.1 | Acyl-CoA oxidase (ACX3) | 39 | 1.61 | 0.258 | 0.968 | |
| AT1G06290.1 | Acyl-CoA oxidase (ACX3) | 40 | 1.31 | 0.235 | 0.998 | |
| AT1G06290.1 | Acyl-CoA oxidase (ACX3) | 41 | 0.85 | 0.239 | 1.000 | |
| AT1G06290.1 | Acyl-CoA oxidase (ACX3) | 42 | 0.49 | 0.271 | 0.850 | |
| AT1G06290.1 | Acyl-CoA oxidase (ACX3) | 43 | 0.28 | 0.312 | 0.436 | |
| AT1G06290.1 | Acyl-CoA oxidase (ACX3) | 46 | 0.05 | 0.435 | 0.036 | * |
| AT1G06290.1 | Acyl-CoA oxidase (ACX3) | 47 | 0.03 | 0.509 | 0.031 | * |
| AT1G06530.1 | Peroxisomal and mitochondrial division factor 2 (PMD2) | 2 | 0.62 | 0.395 | 1.000 | |
| AT1G06530.1 | Peroxisomal and mitochondrial division factor 2 (PMD2) | 3 | 0.64 | 0.248 | 1.000 | |
| AT1G06530.1 | Peroxisomal and mitochondrial division factor 2 (PMD2) | 4 | 0.74 | 0.231 | 1.000 | |
| AT1G06530.1 | Peroxisomal and mitochondrial division factor 2 (PMD2) | 5 | 0.99 | 0.227 | 1.000 | |
| AT1G06530.1 | Peroxisomal and mitochondrial division factor 2 (PMD2) | 6 | 1.40 | 0.219 | 1.000 | |
| AT1G06530.1 | Peroxisomal and mitochondrial division factor 2 (PMD2) | 7 | 1.69 | 0.223 | 1.000 | |
| AT1G06530.1 | Peroxisomal and mitochondrial division factor 2 (PMD2) | 8 | 1.67 | 0.221 | 1.000 | |
| AT1G06530.1 | Peroxisomal and mitochondrial division factor 2 (PMD2) | 9 | 1.54 | 0.213 | 1.000 | |
| AT1G06530.1 | Peroxisomal and mitochondrial division factor 2 (PMD2) | 10 | 1.39 | 0.214 | 1.000 | |
| AT1G06530.1 | Peroxisomal and mitochondrial division factor 2 (PMD2) | 11 | 1.25 | 0.216 | 1.000 | |

| | | | | | |
|---|---|---|---|---|---|
| AT1G06530.1 | Peroxisomal and mitochondrial division factor 2 (PMD2) | 12 | 1.19 | 0.208 | 1.000 |
| AT1G06530.1 | Peroxisomal and mitochondrial division factor 2 (PMD2) | 13 | 1.25 | 0.200 | 1.000 |
| AT1G06530.1 | Peroxisomal and mitochondrial division factor 2 (PMD2) | 14 | 1.56 | 0.196 | 1.000 |
| AT1G06530.1 | Peroxisomal and mitochondrial division factor 2 (PMD2) | 15 | 1.66 | 0.195 | 1.000 |
| AT1G06530.1 | Peroxisomal and mitochondrial division factor 2 (PMD2) | 16 | 1.49 | 0.193 | 1.000 |
| AT1G06530.1 | Peroxisomal and mitochondrial division factor 2 (PMD2) | 17 | 1.43 | 0.192 | 1.000 |
| AT1G06530.1 | Peroxisomal and mitochondrial division factor 2 (PMD2) | 18 | 1.56 | 0.196 | 1.000 |
| AT1G06530.1 | Peroxisomal and mitochondrial division factor 2 (PMD2) | 19 | 1.58 | 0.203 | 1.000 |
| AT1G06530.1 | Peroxisomal and mitochondrial division factor 2 (PMD2) | 20 | 1.41 | 0.208 | 1.000 |
| AT1G06530.1 | Peroxisomal and mitochondrial division factor 2 (PMD2) | 21 | 1.26 | 0.211 | 1.000 |
| AT1G06530.1 | Peroxisomal and mitochondrial division factor 2 (PMD2) | 22 | 1.08 | 0.211 | 1.000 |
| AT1G06530.1 | Peroxisomal and mitochondrial division factor 2 (PMD2) | 23 | 0.90 | 0.209 | 1.000 |
| AT1G06530.1 | Peroxisomal and mitochondrial division factor 2 (PMD2) | 24 | 0.76 | 0.204 | 1.000 |
| AT1G06530.1 | Peroxisomal and mitochondrial division factor 2 (PMD2) | 25 | 0.68 | 0.200 | 1.000 |
| AT1G06530.1 | Peroxisomal and mitochondrial division factor 2 (PMD2) | 26 | 0.92 | 0.199 | 1.000 |
| AT1G06530.1 | Peroxisomal and mitochondrial division factor 2 (PMD2) | 27 | 1.62 | 0.200 | 1.000 |
| AT1G06530.1 | Peroxisomal and mitochondrial division factor 2 (PMD2) | 28 | 2.26 | 0.200 | 0.917 |
| AT1G06530.1 | Peroxisomal and mitochondrial division factor 2 (PMD2) | 29 | 2.51 | 0.199 | 0.777 |
| AT1G06530.1 | Peroxisomal and mitochondrial division factor 2 (PMD2) | 30 | 1.71 | 0.199 | 1.000 |
| AT1G06530.1 | Peroxisomal and mitochondrial division factor 2 (PMD2) | 31 | 0.94 | 0.200 | 1.000 |

| | | | | | |
|---|---|---|---|---|---|
| AT1G06530.1 | Peroxisomal and mitochondrial division factor 2 (PMD2) | 32 | 0.75 | 0.200 | 1.000 |
| AT1G06530.1 | Peroxisomal and mitochondrial division factor 2 (PMD2) | 33 | 0.86 | 0.200 | 1.000 |
| AT1G06530.1 | Peroxisomal and mitochondrial division factor 2 (PMD2) | 34 | 1.18 | 0.201 | 1.000 |
| AT1G06530.1 | Peroxisomal and mitochondrial division factor 2 (PMD2) | 35 | 1.74 | 0.200 | 1.000 |
| AT1G06530.1 | Peroxisomal and mitochondrial division factor 2 (PMD2) | 36 | 2.51 | 0.199 | 0.776 |
| AT1G06530.1 | Peroxisomal and mitochondrial division factor 2 (PMD2) | 37 | 3.02 | 0.199 | 0.432 |
| AT1G06530.1 | Peroxisomal and mitochondrial division factor 2 (PMD2) | 38 | 2.67 | 0.200 | 0.675 |
| AT1G06530.1 | Peroxisomal and mitochondrial division factor 2 (PMD2) | 39 | 1.76 | 0.201 | 0.999 |
| AT1G06530.1 | Peroxisomal and mitochondrial division factor 2 (PMD2) | 40 | 1.00 | 0.201 | 1.000 |
| AT1G06530.1 | Peroxisomal and mitochondrial division factor 2 (PMD2) | 41 | 0.57 | 0.202 | 0.999 |
| AT1G06530.1 | Peroxisomal and mitochondrial division factor 2 (PMD2) | 42 | 0.40 | 0.205 | 0.825 |
| AT1G06530.1 | Peroxisomal and mitochondrial division factor 2 (PMD2) | 43 | 0.42 | 0.207 | 0.886 |
| AT1G06530.1 | Peroxisomal and mitochondrial division factor 2 (PMD2) | 44 | 0.50 | 0.207 | 0.989 |
| AT1G06530.1 | Peroxisomal and mitochondrial division factor 2 (PMD2) | 45 | 0.53 | 0.206 | 0.997 |
| AT1G06530.1 | Peroxisomal and mitochondrial division factor 2 (PMD2) | 46 | 0.54 | 0.237 | 1.000 |
| AT1G06530.1 | Peroxisomal and mitochondrial division factor 2 (PMD2) | 47 | 0.57 | 0.442 | 1.000 |
| AT1G06950.1 | Part of the translocon at the chloroplast inner envelope | 2 | 0.80 | 0.376 | 1.000 |
| AT1G06950.1 | Part of the translocon at the chloroplast inner envelope | 3 | 0.64 | 0.419 | 1.000 |
| AT1G06950.1 | Part of the translocon at the chloroplast inner envelope | 30 | 0.41 | 0.256 | 0.703 |
| AT1G06950.1 | Part of the translocon at the chloroplast inner envelope | 31 | 0.60 | 0.184 | 0.894 |
| AT1G06950.1 | Part of the translocon at the chloroplast inner envelope | 32 | 0.88 | 0.173 | 1.000 |
| AT1G06950.1 | Part of the translocon at the chloroplast inner envelope | 33 | 0.99 | 0.164 | 1.000 |
| AT1G06950.1 | Part of the translocon at the chloroplast inner envelope | 34 | 0.78 | 0.161 | 0.998 |

| | | | | | | |
|---|---|---|---|---|---|---|
| AT1G06950.1 | Part of the translocon at the chloroplast inner envelope | 35 | 0.72 | 0.163 | 0.980 | |
| AT1G06950.1 | Part of the translocon at the chloroplast inner envelope | 36 | 0.94 | 0.172 | 1.000 | |
| AT1G06950.1 | Part of the translocon at the chloroplast inner envelope | 37 | 1.65 | 0.214 | 0.957 | |
| AT1G06950.1 | Part of the translocon at the chloroplast inner envelope | 38 | 3.61 | 0.345 | 0.635 | |
| AT1G06950.1 | Part of the translocon at the chloroplast inner envelope | 46 | 3858.96 | 2.886 | 0.872 | |
| AT1G06950.1 | Part of the translocon at the chloroplast inner envelope | 47 | 9546.52 | 3.371 | 0.901 | |
| AT1G07030.1 | Mitochondrial substrate carrier family protein | 2 | 1.15 | 0.494 | 1.000 | |
| AT1G07030.1 | Mitochondrial substrate carrier family protein | 3 | 1.20 | 0.482 | 1.000 | |
| AT1G07030.1 | Mitochondrial substrate carrier family protein | 34 | 1.14 | 0.378 | 1.000 | |
| AT1G07030.1 | Mitochondrial substrate carrier family protein | 35 | 1.04 | 0.299 | 1.000 | |
| AT1G07030.1 | Mitochondrial substrate carrier family protein | 36 | 0.95 | 0.258 | 1.000 | |
| AT1G07030.1 | Mitochondrial substrate carrier family protein | 37 | 0.90 | 0.254 | 1.000 | |
| AT1G07030.1 | Mitochondrial substrate carrier family protein | 38 | 0.77 | 0.265 | 1.000 | |
| AT1G07030.1 | Mitochondrial substrate carrier family protein | 39 | 1.10 | 0.273 | 1.000 | |
| AT1G07030.1 | Mitochondrial substrate carrier family protein | 40 | 3.06 | 0.262 | 0.548 | |
| AT1G07030.1 | Mitochondrial substrate carrier family protein | 41 | 6.82 | 0.247 | 0.013 | * |
| AT1G07030.1 | Mitochondrial substrate carrier family protein | 42 | 7.39 | 0.245 | 0.008 | ** |
| AT1G07030.1 | Mitochondrial substrate carrier family protein | 43 | 3.06 | 0.246 | 0.456 | |
| AT1G07030.1 | Mitochondrial substrate carrier family protein | 44 | 0.78 | 0.244 | 1.000 | |
| AT1G07030.1 | Mitochondrial substrate carrier family protein | 45 | 0.32 | 0.243 | 0.423 | |
| AT1G07030.1 | Mitochondrial substrate carrier family protein | 46 | 0.24 | 0.269 | 0.243 | |
| AT1G07030.1 | Mitochondrial substrate carrier family protein | 47 | 0.26 | 0.556 | 0.974 | |
| AT1G07040.1 | ? | 2 | 0.96 | 0.408 | 1.000 | |
| AT1G07040.1 | ? | 3 | 0.86 | 0.390 | 1.000 | |
| AT1G07040.1 | ? | 27 | 0.62 | 0.276 | 0.999 | |
| AT1G07040.1 | ? | 28 | 0.73 | 0.198 | 1.000 | |
| AT1G07040.1 | ? | 29 | 0.89 | 0.168 | 1.000 | |
| AT1G07040.1 | ? | 30 | 1.11 | 0.158 | 1.000 | |
| AT1G07040.1 | ? | 31 | 1.47 | 0.155 | 0.981 | |
| AT1G07040.1 | ? | 32 | 2.02 | 0.154 | 0.488 | |
| AT1G07040.1 | ? | 33 | 2.53 | 0.154 | 0.130 | |
| AT1G07040.1 | ? | 34 | 2.78 | 0.153 | 0.065 | . |
| AT1G07040.1 | ? | 35 | 2.83 | 0.153 | 0.056 | . |
| AT1G07040.1 | ? | 36 | 2.74 | 0.153 | 0.070 | . |
| AT1G07040.1 | ? | 37 | 2.68 | 0.152 | 0.080 | . |
| AT1G07040.1 | ? | 38 | 2.73 | 0.152 | 0.068 | . |
| AT1G07040.1 | ? | 39 | 2.81 | 0.152 | 0.054 | . |
| AT1G07040.1 | ? | 40 | 2.85 | 0.152 | 0.051 | . |
| AT1G07040.1 | ? | 41 | 2.74 | 0.154 | 0.074 | . |
| AT1G07040.1 | ? | 42 | 2.53 | 0.156 | 0.143 | |

| | | | | | | |
|---|---|---|---|---|---|---|
| AT1G07040.1 | ? | 43 | 2.10 | 0.157 | 0.430 | |
| AT1G07040.1 | ? | 44 | 1.38 | 0.157 | 0.996 | |
| AT1G07040.1 | ? | 45 | 0.67 | 0.163 | 0.985 | |
| AT1G07040.1 | ? | 46 | 0.26 | 0.202 | 0.068 | . |
| AT1G07040.1 | ? | 47 | 0.09 | 0.316 | 0.021 | * |
| AT1G07180.1 | Alternative NAD(P)H dehydrogenase A1 (NDA1) | 2 | 1.02 | 0.419 | 1.000 | |
| AT1G07180.1 | Alternative NAD(P)H dehydrogenase A1 (NDA1) | 3 | 1.00 | 0.369 | 1.000 | |
| AT1G07180.1 | Alternative NAD(P)H dehydrogenase A1 (NDA1) | 12 | 0.81 | 0.326 | 1.000 | |
| AT1G07180.1 | Alternative NAD(P)H dehydrogenase A1 (NDA1) | 13 | 0.77 | 0.285 | 1.000 | |
| AT1G07180.1 | Alternative NAD(P)H dehydrogenase A1 (NDA1) | 14 | 0.74 | 0.263 | 1.000 | |
| AT1G07180.1 | Alternative NAD(P)H dehydrogenase A1 (NDA1) | 18 | 0.68 | 0.202 | 1.000 | |
| AT1G07180.1 | Alternative NAD(P)H dehydrogenase A1 (NDA1) | 19 | 0.79 | 0.183 | 1.000 | |
| AT1G07180.1 | Alternative NAD(P)H dehydrogenase A1 (NDA1) | 20 | 1.00 | 0.169 | 1.000 | |
| AT1G07180.1 | Alternative NAD(P)H dehydrogenase A1 (NDA1) | 21 | 1.27 | 0.161 | 1.000 | |
| AT1G07180.1 | Alternative NAD(P)H dehydrogenase A1 (NDA1) | 22 | 1.51 | 0.157 | 0.993 | |
| AT1G07180.1 | Alternative NAD(P)H dehydrogenase A1 (NDA1) | 23 | 1.74 | 0.154 | 0.898 | |
| AT1G07180.1 | Alternative NAD(P)H dehydrogenase A1 (NDA1) | 24 | 1.68 | 0.151 | 0.923 | |
| AT1G07180.1 | Alternative NAD(P)H dehydrogenase A1 (NDA1) | 25 | 1.30 | 0.148 | 1.000 | |
| AT1G07180.1 | Alternative NAD(P)H dehydrogenase A1 (NDA1) | 26 | 0.98 | 0.146 | 1.000 | |
| AT1G07180.1 | Alternative NAD(P)H dehydrogenase A1 (NDA1) | 27 | 0.93 | 0.146 | 1.000 | |
| AT1G07180.1 | Alternative NAD(P)H dehydrogenase A1 (NDA1) | 28 | 0.99 | 0.147 | 1.000 | |
| AT1G07180.1 | Alternative NAD(P)H dehydrogenase A1 (NDA1) | 29 | 1.09 | 0.147 | 1.000 | |
| AT1G07180.1 | Alternative NAD(P)H dehydrogenase A1 (NDA1) | 30 | 1.14 | 0.148 | 1.000 | |
| AT1G07180.1 | Alternative NAD(P)H dehydrogenase A1 (NDA1) | 31 | 1.17 | 0.148 | 1.000 | |
| AT1G07180.1 | Alternative NAD(P)H dehydrogenase A1 (NDA1) | 32 | 1.18 | 0.148 | 1.000 | |
| AT1G07180.1 | Alternative NAD(P)H dehydrogenase A1 (NDA1) | 33 | 1.16 | 0.148 | 1.000 | |
| AT1G07180.1 | Alternative NAD(P)H dehydrogenase A1 (NDA1) | 34 | 1.16 | 0.148 | 1.000 | |
| AT1G07180.1 | Alternative NAD(P)H dehydrogenase A1 (NDA1) | 35 | 1.21 | 0.148 | 1.000 | |
| AT1G07180.1 | Alternative NAD(P)H dehydrogenase A1 (NDA1) | 36 | 1.28 | 0.148 | 1.000 | |
| AT1G07180.1 | Alternative NAD(P)H dehydrogenase A1 (NDA1) | 37 | 1.26 | 0.148 | 1.000 | |
| AT1G07180.1 | Alternative NAD(P)H dehydrogenase A1 (NDA1) | 38 | 1.10 | 0.148 | 1.000 | |
| AT1G07180.1 | Alternative NAD(P)H dehydrogenase A1 (NDA1) | 39 | 0.91 | 0.148 | 1.000 | |
| AT1G07180.1 | Alternative NAD(P)H dehydrogenase A1 (NDA1) | 40 | 0.81 | 0.148 | 1.000 | |

| | | | | | |
|---|---|---|---|---|---|
| AT1G07180.1 | Alternative NAD(P)H dehydrogenase A1 (NDA1) | 41 | 0.83 | 0.150 | 1.000 |
| AT1G07180.1 | Alternative NAD(P)H dehydrogenase A1 (NDA1) | 42 | 0.91 | 0.151 | 1.000 |
| AT1G07180.1 | Alternative NAD(P)H dehydrogenase A1 (NDA1) | 43 | 0.97 | 0.152 | 1.000 |
| AT1G07180.1 | Alternative NAD(P)H dehydrogenase A1 (NDA1) | 44 | 0.89 | 0.153 | 1.000 |
| AT1G07180.1 | Alternative NAD(P)H dehydrogenase A1 (NDA1) | 45 | 0.64 | 0.161 | 0.987 |
| AT1G07180.1 | Alternative NAD(P)H dehydrogenase A1 (NDA1) | 46 | 0.37 | 0.201 | 0.450 |
| AT1G07180.1 | Alternative NAD(P)H dehydrogenase A1 (NDA1) | 47 | 0.18 | 0.307 | 0.303 |
| AT1G07510.1 | FTSH protease 10 (FTSH10) | 2 | 0.59 | 0.611 | 1.000 |
| AT1G07510.1 | FTSH protease 10 (FTSH10) | 3 | 0.70 | 0.536 | 1.000 |
| AT1G07510.1 | FTSH protease 10 (FTSH10) | 12 | 1.52 | 0.369 | 1.000 |
| AT1G07510.1 | FTSH protease 10 (FTSH10) | 13 | 1.38 | 0.324 | 1.000 |
| AT1G07510.1 | FTSH protease 10 (FTSH10) | 14 | 1.13 | 0.294 | 1.000 |
| AT1G07510.1 | FTSH protease 10 (FTSH10) | 15 | 0.76 | 0.272 | 1.000 |
| AT1G07510.1 | FTSH protease 10 (FTSH10) | 16 | 0.52 | 0.259 | 0.990 |
| AT1G07510.1 | FTSH protease 10 (FTSH10) | 17 | 0.51 | 0.256 | 0.987 |
| AT1G07510.1 | FTSH protease 10 (FTSH10) | 18 | 0.71 | 0.258 | 1.000 |
| AT1G07510.1 | FTSH protease 10 (FTSH10) | 19 | 1.15 | 0.264 | 1.000 |
| AT1G07510.1 | FTSH protease 10 (FTSH10) | 20 | 1.89 | 0.268 | 0.995 |
| AT1G07510.1 | FTSH protease 10 (FTSH10) | 21 | 2.64 | 0.271 | 0.852 |
| AT1G07510.1 | FTSH protease 10 (FTSH10) | 22 | 2.80 | 0.271 | 0.791 |
| AT1G07510.1 | FTSH protease 10 (FTSH10) | 23 | 2.61 | 0.268 | 0.852 |
| AT1G07510.1 | FTSH protease 10 (FTSH10) | 24 | 2.18 | 0.265 | 0.964 |
| AT1G07510.1 | FTSH protease 10 (FTSH10) | 25 | 1.67 | 0.262 | 0.999 |
| AT1G07510.1 | FTSH protease 10 (FTSH10) | 26 | 1.35 | 0.260 | 1.000 |
| AT1G07510.1 | FTSH protease 10 (FTSH10) | 27 | 1.29 | 0.260 | 1.000 |
| AT1G07510.1 | FTSH protease 10 (FTSH10) | 28 | 1.44 | 0.260 | 1.000 |
| AT1G07510.1 | FTSH protease 10 (FTSH10) | 29 | 1.68 | 0.261 | 0.999 |
| AT1G07510.1 | FTSH protease 10 (FTSH10) | 30 | 1.60 | 0.261 | 1.000 |
| AT1G07510.1 | FTSH protease 10 (FTSH10) | 31 | 1.20 | 0.262 | 1.000 |
| AT1G07510.1 | FTSH protease 10 (FTSH10) | 32 | 0.89 | 0.262 | 1.000 |
| AT1G07510.1 | FTSH protease 10 (FTSH10) | 33 | 0.81 | 0.263 | 1.000 |
| AT1G07510.1 | FTSH protease 10 (FTSH10) | 34 | 0.86 | 0.265 | 1.000 |
| AT1G07510.1 | FTSH protease 10 (FTSH10) | 35 | 1.00 | 0.270 | 1.000 |
| AT1G07510.1 | FTSH protease 10 (FTSH10) | 36 | 1.19 | 0.288 | 1.000 |
| AT1G07510.1 | FTSH protease 10 (FTSH10) | 37 | 1.46 | 0.344 | 1.000 |
| AT1G07510.1 | FTSH protease 10 (FTSH10) | 38 | 1.78 | 0.477 | 1.000 |
| AT1G07510.1 | FTSH protease 10 (FTSH10) | 46 | 2.84 | 3.547 | 1.000 |
| AT1G07510.1 | FTSH protease 10 (FTSH10) | 47 | 2.82 | 4.120 | 1.000 |
| AT1G07910.2 | tRNA ligase 1 (TRLG1) | 2 | 1.02 | 0.408 | 1.000 |
| AT1G07910.2 | tRNA ligase 1 (TRLG1) | 3 | 0.99 | 0.369 | 1.000 |
| AT1G07910.2 | tRNA ligase 1 (TRLG1) | 4 | 0.96 | 0.377 | 1.000 |
| AT1G07910.2 | tRNA ligase 1 (TRLG1) | 22 | 0.57 | 0.379 | 0.996 |
| AT1G07910.2 | tRNA ligase 1 (TRLG1) | 23 | 0.57 | 0.346 | 0.993 |
| AT1G07910.2 | tRNA ligase 1 (TRLG1) | 24 | 0.57 | 0.313 | 0.987 |
| AT1G07910.2 | tRNA ligase 1 (TRLG1) | 28 | 0.66 | 0.212 | 0.978 |
| AT1G07910.2 | tRNA ligase 1 (TRLG1) | 29 | 0.72 | 0.191 | 0.990 |
| AT1G07910.2 | tRNA ligase 1 (TRLG1) | 30 | 0.80 | 0.174 | 0.999 |
| AT1G07910.2 | tRNA ligase 1 (TRLG1) | 31 | 0.89 | 0.164 | 1.000 |
| AT1G07910.2 | tRNA ligase 1 (TRLG1) | 32 | 0.98 | 0.161 | 1.000 |
| AT1G07910.2 | tRNA ligase 1 (TRLG1) | 33 | 1.03 | 0.163 | 1.000 |
| AT1G07910.2 | tRNA ligase 1 (TRLG1) | 34 | 1.04 | 0.172 | 1.000 |
| AT1G07910.2 | tRNA ligase 1 (TRLG1) | 35 | 1.02 | 0.187 | 1.000 |
| AT1G07910.2 | tRNA ligase 1 (TRLG1) | 36 | 0.98 | 0.207 | 1.000 |
| AT1G07910.2 | tRNA ligase 1 (TRLG1) | 37 | 0.95 | 0.230 | 1.000 |
| AT1G07910.2 | tRNA ligase 1 (TRLG1) | 41 | 0.86 | 0.321 | 1.000 |
| AT1G07910.2 | tRNA ligase 1 (TRLG1) | 42 | 0.87 | 0.353 | 1.000 |

| | | | | | |
|---|---|---|---|---|---|
| AT1G07910.2 | tRNA ligase 1 (TRLG1) | 43 | 0.90 | 0.396 | 1.000 |
| AT1G07910.2 | tRNA ligase 1 (TRLG1) | 46 | 1.08 | 0.591 | 1.000 |
| AT1G07910.2 | tRNA ligase 1 (TRLG1) | 47 | 1.16 | 0.683 | 1.000 |
| AT1G07930.2 | GTP binding Elongation factor Tu family protein | 2 | 0.24 | 0.384 | 0.560 |
| AT1G07930.2 | GTP binding Elongation factor Tu family protein | 3 | 0.25 | 0.332 | 0.408 |
| AT1G07930.2 | GTP binding Elongation factor Tu family protein | 4 | 0.25 | 0.300 | 0.311 |
| AT1G07930.2 | GTP binding Elongation factor Tu family protein | 5 | 0.26 | 0.282 | 0.260 |
| AT1G07930.2 | GTP binding Elongation factor Tu family protein | 6 | 0.26 | 0.274 | 0.249 |
| AT1G07930.2 | GTP binding Elongation factor Tu family protein | 7 | 0.27 | 0.271 | 0.251 |
| AT1G07930.2 | GTP binding Elongation factor Tu family protein | 8 | 0.27 | 0.269 | 0.254 |
| AT1G07930.2 | GTP binding Elongation factor Tu family protein | 9 | 0.27 | 0.268 | 0.244 |
| AT1G07930.2 | GTP binding Elongation factor Tu family protein | 10 | 0.26 | 0.267 | 0.222 |
| AT1G07930.2 | GTP binding Elongation factor Tu family protein | 11 | 0.25 | 0.267 | 0.191 |
| AT1G07930.2 | GTP binding Elongation factor Tu family protein | 12 | 0.24 | 0.266 | 0.158 |
| AT1G07930.2 | GTP binding Elongation factor Tu family protein | 13 | 0.23 | 0.265 | 0.128 |
| AT1G07930.2 | GTP binding Elongation factor Tu family protein | 14 | 0.22 | 0.265 | 0.109 |
| AT1G07930.2 | GTP binding Elongation factor Tu family protein | 15 | 0.21 | 0.266 | 0.101 |
| AT1G07930.2 | GTP binding Elongation factor Tu family protein | 16 | 0.21 | 0.268 | 0.109 |
| AT1G07930.2 | GTP binding Elongation factor Tu family protein | 17 | 0.22 | 0.272 | 0.134 |
| AT1G07930.2 | GTP binding Elongation factor Tu family protein | 18 | 0.24 | 0.278 | 0.183 |
| AT1G07930.2 | GTP binding Elongation factor Tu family protein | 19 | 0.25 | 0.287 | 0.262 |
| AT1G07930.2 | GTP binding Elongation factor Tu family protein | 20 | 0.27 | 0.299 | 0.372 |
| AT1G07930.2 | GTP binding Elongation factor Tu family protein | 21 | 0.30 | 0.312 | 0.514 |
| AT1G07930.2 | GTP binding Elongation factor Tu family protein | 22 | 0.33 | 0.324 | 0.660 |
| AT1G07930.2 | GTP binding Elongation factor Tu family protein | 26 | 0.51 | 0.359 | 0.980 |
| AT1G07930.2 | GTP binding Elongation factor Tu family protein | 27 | 0.56 | 0.364 | 0.993 |
| AT1G07930.2 | GTP binding Elongation factor Tu family protein | 28 | 0.59 | 0.371 | 0.997 |
| AT1G07930.2 | GTP binding Elongation factor Tu family protein | 30 | 0.62 | 0.388 | 0.999 |
| AT1G07930.2 | GTP binding Elongation factor Tu family protein | 31 | 0.60 | 0.401 | 0.999 |
| AT1G07930.2 | GTP binding Elongation factor Tu family protein | 32 | 0.58 | 0.414 | 0.998 |
| AT1G07930.2 | GTP binding Elongation factor Tu family protein | 41 | 0.45 | 0.382 | 0.958 |
| AT1G07930.2 | GTP binding Elongation factor Tu family protein | 42 | 0.51 | 0.362 | 0.980 |

| | | | | | | |
|---|---|---|---|---|---|---|
| AT1G07930.2 | GTP binding Elongation factor Tu family protein | 43 | 0.61 | 0.347 | 0.997 | |
| AT1G07930.2 | GTP binding Elongation factor Tu family protein | 44 | 0.79 | 0.343 | 1.000 | |
| AT1G07930.2 | GTP binding Elongation factor Tu family protein | 45 | 1.08 | 0.353 | 1.000 | |
| AT1G07930.2 | GTP binding Elongation factor Tu family protein | 46 | 1.54 | 0.384 | 1.000 | |
| AT1G07930.2 | GTP binding Elongation factor Tu family protein | 47 | 2.24 | 0.438 | 0.982 | |
| AT1G08220.2 | ATPase assembly factor 10 (ATP10) | 2 | 1.29 | 0.302 | 1.000 | |
| AT1G08220.2 | ATPase assembly factor 10 (ATP10) | 3 | 1.11 | 0.281 | 1.000 | |
| AT1G08220.2 | ATPase assembly factor 10 (ATP10) | 4 | 0.95 | 0.352 | 1.000 | |
| AT1G08220.2 | ATPase assembly factor 10 (ATP10) | 29 | 0.52 | 0.198 | 0.759 | |
| AT1G08220.2 | ATPase assembly factor 10 (ATP10) | 30 | 0.68 | 0.154 | 0.937 | |
| AT1G08220.2 | ATPase assembly factor 10 (ATP10) | 31 | 0.90 | 0.133 | 1.000 | |
| AT1G08220.2 | ATPase assembly factor 10 (ATP10) | 32 | 1.13 | 0.126 | 1.000 | |
| AT1G08220.2 | ATPase assembly factor 10 (ATP10) | 33 | 1.31 | 0.123 | 0.975 | |
| AT1G08220.2 | ATPase assembly factor 10 (ATP10) | 34 | 1.40 | 0.121 | 0.895 | |
| AT1G08220.2 | ATPase assembly factor 10 (ATP10) | 35 | 1.53 | 0.120 | 0.694 | |
| AT1G08220.2 | ATPase assembly factor 10 (ATP10) | 36 | 1.81 | 0.119 | 0.278 | |
| AT1G08220.2 | ATPase assembly factor 10 (ATP10) | 37 | 2.17 | 0.119 | 0.060 | . |
| AT1G08220.2 | ATPase assembly factor 10 (ATP10) | 38 | 2.46 | 0.120 | 0.018 | * |
| AT1G08220.2 | ATPase assembly factor 10 (ATP10) | 39 | 2.52 | 0.122 | 0.016 | * |
| AT1G08220.2 | ATPase assembly factor 10 (ATP10) | 40 | 2.31 | 0.127 | 0.054 | . |
| AT1G08220.2 | ATPase assembly factor 10 (ATP10) | 41 | 1.87 | 0.135 | 0.368 | |
| AT1G08220.2 | ATPase assembly factor 10 (ATP10) | 42 | 1.32 | 0.149 | 0.992 | |
| AT1G08220.2 | ATPase assembly factor 10 (ATP10) | 43 | 0.86 | 0.169 | 1.000 | |
| AT1G08220.2 | ATPase assembly factor 10 (ATP10) | 44 | 0.54 | 0.186 | 0.769 | |
| AT1G08220.2 | ATPase assembly factor 10 (ATP10) | 46 | 0.20 | 0.225 | 0.028 | * |
| AT1G08220.2 | ATPase assembly factor 10 (ATP10) | 47 | 0.12 | 0.271 | 0.012 | * |
| AT1G08480.1 | Succinate dehydrogenase subunit (SDH6) | 3 | 0.62 | 0.192 | 1.000 | |
| AT1G08480.1 | Succinate dehydrogenase subunit (SDH6) | 4 | 0.55 | 0.180 | 0.990 | |
| AT1G08480.1 | Succinate dehydrogenase subunit (SDH6) | 5 | 0.58 | 0.180 | 0.998 | |
| AT1G08480.1 | Succinate dehydrogenase subunit (SDH6) | 6 | 0.61 | 0.181 | 1.000 | |
| AT1G08480.1 | Succinate dehydrogenase subunit (SDH6) | 7 | 0.57 | 0.181 | 0.997 | |

| | | | | | | |
|---|---|---|---|---|---|---|
| AT1G08480.1 | Succinate dehydrogenase subunit (SDH6) | 8 | 0.59 | 0.177 | 0.998 | |
| AT1G08480.1 | Succinate dehydrogenase subunit (SDH6) | 9 | 0.65 | 0.174 | 1.000 | |
| AT1G08480.1 | Succinate dehydrogenase subunit (SDH6) | 10 | 0.65 | 0.174 | 1.000 | |
| AT1G08480.1 | Succinate dehydrogenase subunit (SDH6) | 11 | 0.61 | 0.175 | 0.999 | |
| AT1G08480.1 | Succinate dehydrogenase subunit (SDH6) | 12 | 0.57 | 0.175 | 0.993 | |
| AT1G08480.1 | Succinate dehydrogenase subunit (SDH6) | 13 | 0.58 | 0.173 | 0.995 | |
| AT1G08480.1 | Succinate dehydrogenase subunit (SDH6) | 14 | 0.67 | 0.170 | 1.000 | |
| AT1G08480.1 | Succinate dehydrogenase subunit (SDH6) | 15 | 0.71 | 0.170 | 1.000 | |
| AT1G08480.1 | Succinate dehydrogenase subunit (SDH6) | 16 | 0.73 | 0.169 | 1.000 | |
| AT1G08480.1 | Succinate dehydrogenase subunit (SDH6) | 17 | 0.83 | 0.168 | 1.000 | |
| AT1G08480.1 | Succinate dehydrogenase subunit (SDH6) | 18 | 1.12 | 0.171 | 1.000 | |
| AT1G08480.1 | Succinate dehydrogenase subunit (SDH6) | 19 | 1.55 | 0.176 | 1.000 | |
| AT1G08480.1 | Succinate dehydrogenase subunit (SDH6) | 20 | 1.99 | 0.181 | 0.952 | |
| AT1G08480.1 | Succinate dehydrogenase subunit (SDH6) | 21 | 2.17 | 0.183 | 0.874 | |
| AT1G08480.1 | Succinate dehydrogenase subunit (SDH6) | 22 | 1.88 | 0.183 | 0.984 | |
| AT1G08480.1 | Succinate dehydrogenase subunit (SDH6) | 23 | 1.48 | 0.182 | 1.000 | |
| AT1G08480.1 | Succinate dehydrogenase subunit (SDH6) | 24 | 1.08 | 0.178 | 1.000 | |
| AT1G08480.1 | Succinate dehydrogenase subunit (SDH6) | 25 | 0.86 | 0.174 | 1.000 | |
| AT1G08480.1 | Succinate dehydrogenase subunit (SDH6) | 26 | 1.09 | 0.174 | 1.000 | |
| AT1G08480.1 | Succinate dehydrogenase subunit (SDH6) | 27 | 2.15 | 0.174 | 0.829 | |
| AT1G08480.1 | Succinate dehydrogenase subunit (SDH6) | 28 | 3.78 | 0.173 | 0.036 | * |
| AT1G08480.1 | Succinate dehydrogenase subunit (SDH6) | 29 | 4.09 | 0.173 | 0.019 | * |
| AT1G08480.1 | Succinate dehydrogenase subunit (SDH6) | 30 | 2.89 | 0.173 | 0.246 | |
| AT1G08480.1 | Succinate dehydrogenase subunit (SDH6) | 31 | 1.94 | 0.174 | 0.950 | |
| AT1G08480.1 | Succinate dehydrogenase subunit (SDH6) | 32 | 1.67 | 0.174 | 0.998 | |
| AT1G08480.1 | Succinate dehydrogenase subunit (SDH6) | 33 | 1.77 | 0.174 | 0.992 | |
| AT1G08480.1 | Succinate dehydrogenase subunit (SDH6) | 34 | 2.06 | 0.175 | 0.894 | |
| AT1G08480.1 | Succinate dehydrogenase subunit (SDH6) | 35 | 2.08 | 0.175 | 0.878 | |
| AT1G08480.1 | Succinate dehydrogenase subunit (SDH6) | 36 | 1.45 | 0.174 | 1.000 | |
| AT1G08480.1 | Succinate dehydrogenase subunit (SDH6) | 37 | 0.93 | 0.174 | 1.000 | |
| AT1G08480.1 | Succinate dehydrogenase subunit (SDH6) | 38 | 0.93 | 0.174 | 1.000 | |

| | | | | | | |
|---|---|---|---|---|---|---|
| AT1G08480.1 | Succinate dehydrogenase subunit (SDH6) | 39 | 1.44 | 0.175 | 1.000 | |
| AT1G08480.1 | Succinate dehydrogenase subunit (SDH6) | 40 | 2.28 | 0.175 | 0.730 | |
| AT1G08480.1 | Succinate dehydrogenase subunit (SDH6) | 41 | 2.57 | 0.176 | 0.482 | |
| AT1G08480.1 | Succinate dehydrogenase subunit (SDH6) | 42 | 2.02 | 0.178 | 0.931 | |
| AT1G08480.1 | Succinate dehydrogenase subunit (SDH6) | 43 | 1.32 | 0.180 | 1.000 | |
| AT1G08480.1 | Succinate dehydrogenase subunit (SDH6) | 44 | 0.78 | 0.180 | 1.000 | |
| AT1G08480.1 | Succinate dehydrogenase subunit (SDH6) | 45 | 0.43 | 0.180 | 0.731 | |
| AT1G08480.1 | Succinate dehydrogenase subunit (SDH6) | 46 | 0.21 | 0.206 | 0.039 | * |
| AT1G08480.1 | Succinate dehydrogenase subunit (SDH6) | 47 | 0.09 | 0.372 | 0.154 | |
| AT1G08480.1 | Succinate dehydrogenase subunit (SDH6) | 48 | 0.04 | 0.639 | 0.541 | |
| AT1G09300.1 | Mitochondrial protease ICP55 (ICP55) | 2 | 1.10 | 0.561 | 1.000 | |
| AT1G09300.1 | Mitochondrial protease ICP55 (ICP55) | 3 | 1.05 | 0.551 | 1.000 | |
| AT1G09300.1 | Mitochondrial protease ICP55 (ICP55) | 31 | 0.50 | 0.412 | 1.000 | |
| AT1G09300.1 | Mitochondrial protease ICP55 (ICP55) | 32 | 0.51 | 0.335 | 0.998 | |
| AT1G09300.1 | Mitochondrial protease ICP55 (ICP55) | 33 | 0.51 | 0.309 | 0.996 | |
| AT1G09300.1 | Mitochondrial protease ICP55 (ICP55) | 34 | 0.44 | 0.298 | 0.970 | |
| AT1G09300.1 | Mitochondrial protease ICP55 (ICP55) | 35 | 0.46 | 0.297 | 0.978 | |
| AT1G09300.1 | Mitochondrial protease ICP55 (ICP55) | 36 | 0.68 | 0.291 | 1.000 | |
| AT1G09300.1 | Mitochondrial protease ICP55 (ICP55) | 37 | 1.02 | 0.276 | 1.000 | |
| AT1G09300.1 | Mitochondrial protease ICP55 (ICP55) | 38 | 1.30 | 0.266 | 1.000 | |
| AT1G09300.1 | Mitochondrial protease ICP55 (ICP55) | 39 | 1.79 | 0.266 | 0.996 | |
| AT1G09300.1 | Mitochondrial protease ICP55 (ICP55) | 40 | 2.76 | 0.268 | 0.769 | |
| AT1G09300.1 | Mitochondrial protease ICP55 (ICP55) | 41 | 2.65 | 0.266 | 0.803 | |
| AT1G09300.1 | Mitochondrial protease ICP55 (ICP55) | 42 | 1.38 | 0.270 | 1.000 | |
| AT1G09300.1 | Mitochondrial protease ICP55 (ICP55) | 43 | 0.67 | 0.272 | 1.000 | |
| AT1G09300.1 | Mitochondrial protease ICP55 (ICP55) | 44 | 0.46 | 0.270 | 0.956 | |
| AT1G09300.1 | Mitochondrial protease ICP55 (ICP55) | 45 | 0.48 | 0.271 | 0.971 | |
| AT1G09300.1 | Mitochondrial protease ICP55 (ICP55) | 46 | 0.67 | 0.302 | 1.000 | |
| AT1G09300.1 | Mitochondrial protease ICP55 (ICP55) | 47 | 1.16 | 0.632 | 1.000 | |
| AT1G09620.1 | Leucine-tRNA ligases / aminoacyl-tRNA ligase | 2 | 0.79 | 0.453 | 1.000 | |
| AT1G09620.1 | Leucine-tRNA ligases / aminoacyl-tRNA ligase | 3 | 0.87 | 0.437 | 1.000 | |

| | | | | | |
|---|---|---|---|---|---|
| AT1G09620.1 | Leucine-tRNA ligases / aminoacyl-tRNA ligase | 28 | 1.71 | 0.318 | 0.998 |
| AT1G09620.1 | Leucine-tRNA ligases / aminoacyl-tRNA ligase | 29 | 1.51 | 0.232 | 0.997 |
| AT1G09620.1 | Leucine-tRNA ligases / aminoacyl-tRNA ligase | 30 | 1.40 | 0.204 | 0.998 |
| AT1G09620.1 | Leucine-tRNA ligases / aminoacyl-tRNA ligase | 31 | 1.56 | 0.192 | 0.976 |
| AT1G09620.1 | Leucine-tRNA ligases / aminoacyl-tRNA ligase | 32 | 1.67 | 0.188 | 0.933 |
| AT1G09620.1 | Leucine-tRNA ligases / aminoacyl-tRNA ligase | 33 | 1.52 | 0.187 | 0.981 |
| AT1G09620.1 | Leucine-tRNA ligases / aminoacyl-tRNA ligase | 34 | 1.33 | 0.188 | 0.999 |
| AT1G09620.1 | Leucine-tRNA ligases / aminoacyl-tRNA ligase | 35 | 1.14 | 0.188 | 1.000 |
| AT1G09620.1 | Leucine-tRNA ligases / aminoacyl-tRNA ligase | 36 | 0.89 | 0.188 | 1.000 |
| AT1G09620.1 | Leucine-tRNA ligases / aminoacyl-tRNA ligase | 37 | 0.66 | 0.191 | 0.983 |
| AT1G09620.1 | Leucine-tRNA ligases / aminoacyl-tRNA ligase | 38 | 0.56 | 0.204 | 0.911 |
| AT1G09620.1 | Leucine-tRNA ligases / aminoacyl-tRNA ligase | 39 | 0.69 | 0.260 | 1.000 |
| AT1G09620.1 | Leucine-tRNA ligases / aminoacyl-tRNA ligase | 40 | 1.25 | 0.391 | 1.000 |
| AT1G09620.1 | Leucine-tRNA ligases / aminoacyl-tRNA ligase | 46 | 864.52 | 2.177 | 0.861 |
| AT1G09620.1 | Leucine-tRNA ligases / aminoacyl-tRNA ligase | 47 | 3261.81 | 2.626 | 0.867 |
| AT1G10270.1 | Pentatricopeptide repeat (PPR) superfamily protein (GRP23) | 2 | 0.95 | 0.424 | 1.000 |
| AT1G10270.1 | Pentatricopeptide repeat (PPR) superfamily protein (GRP23) | 3 | 0.81 | 0.405 | 1.000 |
| AT1G10270.1 | Pentatricopeptide repeat (PPR) superfamily protein (GRP23) | 26 | 0.57 | 0.280 | 0.991 |
| AT1G10270.1 | Pentatricopeptide repeat (PPR) superfamily protein (GRP23) | 27 | 0.75 | 0.205 | 1.000 |
| AT1G10270.1 | Pentatricopeptide repeat (PPR) superfamily protein (GRP23) | 28 | 0.99 | 0.169 | 1.000 |
| AT1G10270.1 | Pentatricopeptide repeat (PPR) superfamily protein (GRP23) | 29 | 1.22 | 0.157 | 1.000 |
| AT1G10270.1 | Pentatricopeptide repeat (PPR) superfamily protein (GRP23) | 30 | 1.33 | 0.153 | 0.995 |
| AT1G10270.1 | Pentatricopeptide repeat (PPR) superfamily protein (GRP23) | 31 | 1.37 | 0.152 | 0.988 |
| AT1G10270.1 | Pentatricopeptide repeat (PPR) superfamily protein (GRP23) | 32 | 1.36 | 0.152 | 0.991 |
| AT1G10270.1 | Pentatricopeptide repeat (PPR) superfamily protein (GRP23) | 33 | 1.34 | 0.152 | 0.994 |
| AT1G10270.1 | Pentatricopeptide repeat (PPR) superfamily protein (GRP23) | 34 | 1.37 | 0.152 | 0.990 |
| AT1G10270.1 | Pentatricopeptide repeat (PPR) superfamily protein (GRP23) | 35 | 1.42 | 0.153 | 0.977 |
| AT1G10270.1 | Pentatricopeptide repeat (PPR) superfamily protein (GRP23) | 36 | 1.42 | 0.157 | 0.982 |
| AT1G10270.1 | Pentatricopeptide repeat (PPR) superfamily protein (GRP23) | 37 | 1.34 | 0.167 | 0.997 |
| AT1G10270.1 | Pentatricopeptide repeat (PPR) superfamily protein (GRP23) | 38 | 1.19 | 0.184 | 1.000 |
| AT1G10270.1 | Pentatricopeptide repeat (PPR) superfamily protein (GRP23) | 39 | 1.01 | 0.204 | 1.000 |

| | | | | | |
|---|---|---|---|---|---|
| AT1G10270.1 | Pentatricopeptide repeat (PPR) superfamily protein (GRP23) | 40 | 0.89 | 0.231 | 1.000 |
| AT1G10270.1 | Pentatricopeptide repeat (PPR) superfamily protein (GRP23) | 41 | 0.86 | 0.275 | 1.000 |
| AT1G10270.1 | Pentatricopeptide repeat (PPR) superfamily protein (GRP23) | 42 | 0.95 | 0.341 | 1.000 |
| AT1G10270.1 | Pentatricopeptide repeat (PPR) superfamily protein (GRP23) | 46 | 3.67 | 0.819 | 0.999 |
| AT1G10270.1 | Pentatricopeptide repeat (PPR) superfamily protein (GRP23) | 47 | 5.78 | 1.005 | 0.997 |
| AT1G10950.1 | Transmembrane nine protein 1 (TMN1) | 2 | 0.89 | 0.541 | 1.000 |
| AT1G10950.1 | Transmembrane nine protein 1 (TMN1) | 3 | 1.09 | 0.373 | 1.000 |
| AT1G10950.1 | Transmembrane nine protein 1 (TMN1) | 4 | 1.36 | 0.432 | 1.000 |
| AT1G10950.1 | Transmembrane nine protein 1 (TMN1) | 37 | 2.14 | 0.383 | 0.960 |
| AT1G10950.1 | Transmembrane nine protein 1 (TMN1) | 38 | 0.73 | 0.288 | 0.999 |
| AT1G10950.1 | Transmembrane nine protein 1 (TMN1) | 39 | 0.38 | 0.251 | 0.512 |
| AT1G10950.1 | Transmembrane nine protein 1 (TMN1) | 40 | 0.42 | 0.241 | 0.598 |
| AT1G10950.1 | Transmembrane nine protein 1 (TMN1) | 41 | 0.83 | 0.271 | 1.000 |
| AT1G10950.1 | Transmembrane nine protein 1 (TMN1) | 42 | 1.96 | 0.417 | 0.987 |
| AT1G10950.1 | Transmembrane nine protein 1 (TMN1) | 46 | 68.28 | 2.150 | 0.963 |
| AT1G10950.1 | Transmembrane nine protein 1 (TMN1) | 47 | 166.28 | 2.765 | 0.973 |
| AT1G11390.1 | Protein kinase superfamily protein | 2 | 0.95 | 0.502 | 1.000 |
| AT1G11390.1 | Protein kinase superfamily protein | 3 | 0.89 | 0.457 | 1.000 |
| AT1G11390.1 | Protein kinase superfamily protein | 17 | 1.89 | 0.272 | 0.996 |
| AT1G11390.1 | Protein kinase superfamily protein | 18 | 2.09 | 0.233 | 0.944 |
| AT1G11390.1 | Protein kinase superfamily protein | 19 | 2.03 | 0.210 | 0.912 |
| AT1G11390.1 | Protein kinase superfamily protein | 20 | 1.81 | 0.201 | 0.967 |
| AT1G11390.1 | Protein kinase superfamily protein | 21 | 1.49 | 0.198 | 0.999 |
| AT1G11390.1 | Protein kinase superfamily protein | 22 | 1.13 | 0.196 | 1.000 |
| AT1G11390.1 | Protein kinase superfamily protein | 23 | 0.82 | 0.194 | 1.000 |
| AT1G11390.1 | Protein kinase superfamily protein | 24 | 0.65 | 0.190 | 0.997 |
| AT1G11390.1 | Protein kinase superfamily protein | 25 | 0.67 | 0.187 | 0.999 |
| AT1G11390.1 | Protein kinase superfamily protein | 26 | 0.97 | 0.186 | 1.000 |
| AT1G11390.1 | Protein kinase superfamily protein | 27 | 1.60 | 0.186 | 0.992 |
| AT1G11390.1 | Protein kinase superfamily protein | 28 | 2.39 | 0.186 | 0.530 |
| AT1G11390.1 | Protein kinase superfamily protein | 29 | 3.15 | 0.187 | 0.144 |

| | | | | | | |
|---|---|---|---|---|---|---|
| AT1G11390.1 | Protein kinase superfamily protein | 30 | 3.60 | 0.187 | 0.061 | . |
| AT1G11390.1 | Protein kinase superfamily protein | 31 | 3.71 | 0.187 | 0.051 | . |
| AT1G11390.1 | Protein kinase superfamily protein | 32 | 3.56 | 0.188 | 0.071 | . |
| AT1G11390.1 | Protein kinase superfamily protein | 33 | 3.12 | 0.188 | 0.159 | |
| AT1G11390.1 | Protein kinase superfamily protein | 34 | 2.46 | 0.188 | 0.486 | |
| AT1G11390.1 | Protein kinase superfamily protein | 35 | 1.87 | 0.188 | 0.916 | |
| AT1G11390.1 | Protein kinase superfamily protein | 36 | 1.46 | 0.188 | 0.999 | |
| AT1G11390.1 | Protein kinase superfamily protein | 37 | 1.24 | 0.188 | 1.000 | |
| AT1G11390.1 | Protein kinase superfamily protein | 38 | 1.21 | 0.188 | 1.000 | |
| AT1G11390.1 | Protein kinase superfamily protein | 39 | 1.35 | 0.188 | 1.000 | |
| AT1G11390.1 | Protein kinase superfamily protein | 40 | 1.48 | 0.189 | 0.999 | |
| AT1G11390.1 | Protein kinase superfamily protein | 41 | 1.47 | 0.194 | 0.999 | |
| AT1G11390.1 | Protein kinase superfamily protein | 42 | 1.37 | 0.214 | 1.000 | |
| AT1G11390.1 | Protein kinase superfamily protein | 43 | 1.24 | 0.270 | 1.000 | |
| AT1G11390.1 | Protein kinase superfamily protein | 44 | 1.11 | 0.353 | 1.000 | |
| AT1G11390.1 | Protein kinase superfamily protein | 46 | 0.85 | 0.630 | 1.000 | |
| AT1G11390.1 | Protein kinase superfamily protein | 47 | 0.74 | 0.835 | 1.000 | |
| AT1G11860.3 | Glycine decarboxylase subunit (GDCT) | 3 | 0.96 | 0.106 | 1.000 | |
| AT1G11860.3 | Glycine decarboxylase subunit (GDCT) | 4 | 0.90 | 0.092 | 1.000 | |
| AT1G11860.3 | Glycine decarboxylase subunit (GDCT) | 5 | 0.89 | 0.090 | 1.000 | |
| AT1G11860.3 | Glycine decarboxylase subunit (GDCT) | 6 | 0.91 | 0.090 | 1.000 | |
| AT1G11860.3 | Glycine decarboxylase subunit (GDCT) | 7 | 0.96 | 0.090 | 1.000 | |
| AT1G11860.3 | Glycine decarboxylase subunit (GDCT) | 8 | 1.04 | 0.089 | 1.000 | |
| AT1G11860.3 | Glycine decarboxylase subunit (GDCT) | 9 | 1.14 | 0.088 | 1.000 | |
| AT1G11860.3 | Glycine decarboxylase subunit (GDCT) | 10 | 1.26 | 0.088 | 0.985 | |
| AT1G11860.3 | Glycine decarboxylase subunit (GDCT) | 11 | 1.39 | 0.088 | 0.792 | |
| AT1G11860.3 | Glycine decarboxylase subunit (GDCT) | 12 | 1.49 | 0.087 | 0.504 | |
| AT1G11860.3 | Glycine decarboxylase subunit (GDCT) | 13 | 1.56 | 0.087 | 0.345 | |
| AT1G11860.3 | Glycine decarboxylase subunit (GDCT) | 14 | 1.58 | 0.086 | 0.304 | |
| AT1G11860.3 | Glycine decarboxylase subunit (GDCT) | 15 | 1.56 | 0.086 | 0.331 | |
| AT1G11860.3 | Glycine decarboxylase subunit (GDCT) | 16 | 1.55 | 0.086 | 0.349 | |

| | | | | | |
|---|---|---|---|---|---|
| AT1G11860.3 | Glycine decarboxylase subunit (GDCT) | 17 | 1.55 | 0.086 | 0.340 |
| AT1G11860.3 | Glycine decarboxylase subunit (GDCT) | 18 | 1.54 | 0.087 | 0.388 |
| AT1G11860.3 | Glycine decarboxylase subunit (GDCT) | 19 | 1.47 | 0.088 | 0.571 |
| AT1G11860.3 | Glycine decarboxylase subunit (GDCT) | 20 | 1.36 | 0.089 | 0.860 |
| AT1G11860.3 | Glycine decarboxylase subunit (GDCT) | 21 | 1.23 | 0.090 | 0.996 |
| AT1G11860.3 | Glycine decarboxylase subunit (GDCT) | 22 | 1.10 | 0.090 | 1.000 |
| AT1G11860.3 | Glycine decarboxylase subunit (GDCT) | 23 | 1.00 | 0.089 | 1.000 |
| AT1G11860.3 | Glycine decarboxylase subunit (GDCT) | 24 | 0.92 | 0.088 | 1.000 |
| AT1G11860.3 | Glycine decarboxylase subunit (GDCT) | 25 | 0.88 | 0.088 | 1.000 |
| AT1G11860.3 | Glycine decarboxylase subunit (GDCT) | 26 | 0.86 | 0.087 | 1.000 |
| AT1G11860.3 | Glycine decarboxylase subunit (GDCT) | 27 | 0.88 | 0.087 | 1.000 |
| AT1G11860.3 | Glycine decarboxylase subunit (GDCT) | 28 | 0.92 | 0.087 | 1.000 |
| AT1G11860.3 | Glycine decarboxylase subunit (GDCT) | 29 | 0.97 | 0.087 | 1.000 |
| AT1G11860.3 | Glycine decarboxylase subunit (GDCT) | 30 | 1.01 | 0.087 | 1.000 |
| AT1G11860.3 | Glycine decarboxylase subunit (GDCT) | 31 | 1.04 | 0.087 | 1.000 |
| AT1G11860.3 | Glycine decarboxylase subunit (GDCT) | 32 | 1.06 | 0.087 | 1.000 |
| AT1G11860.3 | Glycine decarboxylase subunit (GDCT) | 33 | 1.05 | 0.088 | 1.000 |
| AT1G11860.3 | Glycine decarboxylase subunit (GDCT) | 34 | 1.05 | 0.088 | 1.000 |
| AT1G11860.3 | Glycine decarboxylase subunit (GDCT) | 35 | 1.07 | 0.088 | 1.000 |
| AT1G11860.3 | Glycine decarboxylase subunit (GDCT) | 36 | 1.12 | 0.087 | 1.000 |
| AT1G11860.3 | Glycine decarboxylase subunit (GDCT) | 37 | 1.18 | 0.087 | 1.000 |
| AT1G11860.3 | Glycine decarboxylase subunit (GDCT) | 38 | 1.25 | 0.087 | 0.989 |
| AT1G11860.3 | Glycine decarboxylase subunit (GDCT) | 39 | 1.30 | 0.088 | 0.955 |
| AT1G11860.3 | Glycine decarboxylase subunit (GDCT) | 40 | 1.31 | 0.088 | 0.934 |
| AT1G11860.3 | Glycine decarboxylase subunit (GDCT) | 41 | 1.29 | 0.088 | 0.968 |
| AT1G11860.3 | Glycine decarboxylase subunit (GDCT) | 42 | 1.22 | 0.089 | 0.998 |
| AT1G11860.3 | Glycine decarboxylase subunit (GDCT) | 43 | 1.12 | 0.089 | 1.000 |
| AT1G11860.3 | Glycine decarboxylase subunit (GDCT) | 44 | 1.00 | 0.089 | 1.000 |
| AT1G11860.3 | Glycine decarboxylase subunit (GDCT) | 45 | 0.86 | 0.093 | 1.000 |
| AT1G11860.3 | Glycine decarboxylase subunit (GDCT) | 46 | 0.74 | 0.111 | 0.975 |
| AT1G11860.3 | Glycine decarboxylase subunit (GDCT) | 47 | 0.62 | 0.154 | 0.936 |

| | | | | | |
|---|---|---|---|---|---|
| AT1G11860.3 | Glycine decarboxylase subunit (GDCT) | 48 | 0.52 | 0.215 | 0.948 |
| AT1G11870.1 | Seryl-tRNA synthetase (SRS / OVA7) | 2 | 0.82 | 0.585 | 1.000 |
| AT1G11870.1 | Seryl-tRNA synthetase (SRS / OVA7) | 3 | 0.85 | 0.579 | 1.000 |
| AT1G11870.1 | Seryl-tRNA synthetase (SRS / OVA7) | 31 | 0.35 | 0.452 | 0.984 |
| AT1G11870.1 | Seryl-tRNA synthetase (SRS / OVA7) | 32 | 0.31 | 0.364 | 0.872 |
| AT1G11870.1 | Seryl-tRNA synthetase (SRS / OVA7) | 33 | 0.33 | 0.334 | 0.844 |
| AT1G11870.1 | Seryl-tRNA synthetase (SRS / OVA7) | 34 | 0.40 | 0.321 | 0.929 |
| AT1G11870.1 | Seryl-tRNA synthetase (SRS / OVA7) | 35 | 0.46 | 0.317 | 0.979 |
| AT1G11870.1 | Seryl-tRNA synthetase (SRS / OVA7) | 36 | 0.53 | 0.314 | 0.995 |
| AT1G11870.1 | Seryl-tRNA synthetase (SRS / OVA7) | 37 | 1.22 | 0.312 | 1.000 |
| AT1G11870.1 | Seryl-tRNA synthetase (SRS / OVA7) | 38 | 4.33 | 0.313 | 0.415 |
| AT1G11870.1 | Seryl-tRNA synthetase (SRS / OVA7) | 39 | 5.63 | 0.315 | 0.212 |
| AT1G11870.1 | Seryl-tRNA synthetase (SRS / OVA7) | 40 | 2.38 | 0.314 | 0.948 |
| AT1G11870.1 | Seryl-tRNA synthetase (SRS / OVA7) | 41 | 0.83 | 0.314 | 1.000 |
| AT1G11870.1 | Seryl-tRNA synthetase (SRS / OVA7) | 42 | 0.39 | 0.318 | 0.911 |
| AT1G11870.1 | Seryl-tRNA synthetase (SRS / OVA7) | 43 | 0.26 | 0.320 | 0.584 |
| AT1G11870.1 | Seryl-tRNA synthetase (SRS / OVA7) | 44 | 0.37 | 0.321 | 0.892 |
| AT1G11870.1 | Seryl-tRNA synthetase (SRS / OVA7) | 45 | 0.71 | 0.377 | 1.000 |
| AT1G11870.1 | Seryl-tRNA synthetase (SRS / OVA7) | 46 | 1.67 | 0.539 | 1.000 |
| AT1G11870.1 | Seryl-tRNA synthetase (SRS / OVA7) | 47 | 4.59 | 0.932 | 0.999 |
| AT1G12840.1 | Vacuolar ATP synthase subunit C (VHA-C) | 2 | 1.33 | 0.237 | 0.973 |
| AT1G12840.1 | Vacuolar ATP synthase subunit C (VHA-C) | 3 | 1.35 | 0.223 | 0.959 |
| AT1G12840.1 | Vacuolar ATP synthase subunit C (VHA-C) | 4 | 1.37 | 0.211 | 0.942 |
| AT1G12840.1 | Vacuolar ATP synthase subunit C (VHA-C) | 5 | 1.39 | 0.201 | 0.921 |
| AT1G12840.1 | Vacuolar ATP synthase subunit C (VHA-C) | 6 | 1.40 | 0.193 | 0.898 |
| AT1G12840.1 | Vacuolar ATP synthase subunit C (VHA-C) | 7 | 1.42 | 0.188 | 0.875 |
| AT1G12840.1 | Vacuolar ATP synthase subunit C (VHA-C) | 8 | 1.44 | 0.184 | 0.853 |
| AT1G12840.1 | Vacuolar ATP synthase subunit C (VHA-C) | 21 | 1.59 | 0.191 | 0.731 |
| AT1G12840.1 | Vacuolar ATP synthase subunit C (VHA-C) | 22 | 1.60 | 0.192 | 0.727 |
| AT1G12840.1 | Vacuolar ATP synthase subunit C (VHA-C) | 23 | 1.61 | 0.193 | 0.726 |
| AT1G12840.1 | Vacuolar ATP synthase subunit C (VHA-C) | 24 | 1.61 | 0.194 | 0.726 |

| | | | | | | |
|---|---|---|---|---|---|---|
| AT1G12840.1 | Vacuolar ATP synthase subunit C (VHA-C) | 25 | 1.61 | 0.195 | 0.730 | |
| AT1G12840.1 | Vacuolar ATP synthase subunit C (VHA-C) | 26 | 1.61 | 0.196 | 0.735 | |
| AT1G12840.1 | Vacuolar ATP synthase subunit C (VHA-C) | 42 | 1.36 | 0.205 | 0.940 | |
| AT1G12840.1 | Vacuolar ATP synthase subunit C (VHA-C) | 43 | 1.33 | 0.210 | 0.958 | |
| AT1G12840.1 | Vacuolar ATP synthase subunit C (VHA-C) | 44 | 1.30 | 0.217 | 0.973 | |
| AT1G12840.1 | Vacuolar ATP synthase subunit C (VHA-C) | 45 | 1.27 | 0.226 | 0.984 | |
| AT1G12840.1 | Vacuolar ATP synthase subunit C (VHA-C) | 46 | 1.24 | 0.238 | 0.992 | |
| AT1G12840.1 | Vacuolar ATP synthase subunit C (VHA-C) | 47 | 1.21 | 0.251 | 0.996 | |
| AT1G12900.4 | Glyceraldehyde 3-phosphate dehydrogenase (GAPA2) | 2 | 1.07 | 0.557 | 1.000 | |
| AT1G12900.4 | Glyceraldehyde 3-phosphate dehydrogenase (GAPA2) | 3 | 1.25 | 0.564 | 1.000 | |
| AT1G12900.4 | Glyceraldehyde 3-phosphate dehydrogenase (GAPA2) | 39 | 2.07 | 0.281 | 0.699 | |
| AT1G12900.4 | Glyceraldehyde 3-phosphate dehydrogenase (GAPA2) | 40 | 1.54 | 0.250 | 0.907 | |
| AT1G12900.4 | Glyceraldehyde 3-phosphate dehydrogenase (GAPA2) | 41 | 1.08 | 0.242 | 1.000 | |
| AT1G12900.4 | Glyceraldehyde 3-phosphate dehydrogenase (GAPA2) | 42 | 0.71 | 0.250 | 0.960 | |
| AT1G12900.4 | Glyceraldehyde 3-phosphate dehydrogenase (GAPA2) | 43 | 0.44 | 0.266 | 0.565 | |
| AT1G12900.4 | Glyceraldehyde 3-phosphate dehydrogenase (GAPA2) | 46 | 0.09 | 0.326 | 0.015 | * |
| AT1G12900.4 | Glyceraldehyde 3-phosphate dehydrogenase (GAPA2) | 47 | 0.05 | 0.370 | 0.007 | ** |
| AT1G13280.1 | Allene oxide cyclase 4 (AOC4) | 2 | 1.32 | 0.343 | 1.000 | |
| AT1G13280.1 | Allene oxide cyclase 4 (AOC4) | 3 | 0.86 | 0.254 | 1.000 | |
| AT1G13280.1 | Allene oxide cyclase 4 (AOC4) | 4 | 0.59 | 0.490 | 1.000 | |
| AT1G13280.1 | Allene oxide cyclase 4 (AOC4) | 31 | 1.04 | 0.256 | 1.000 | |
| AT1G13280.1 | Allene oxide cyclase 4 (AOC4) | 32 | 1.08 | 0.199 | 1.000 | |
| AT1G13280.1 | Allene oxide cyclase 4 (AOC4) | 33 | 1.03 | 0.180 | 1.000 | |
| AT1G13280.1 | Allene oxide cyclase 4 (AOC4) | 34 | 0.93 | 0.171 | 1.000 | |
| AT1G13280.1 | Allene oxide cyclase 4 (AOC4) | 35 | 0.85 | 0.169 | 1.000 | |
| AT1G13280.1 | Allene oxide cyclase 4 (AOC4) | 36 | 0.86 | 0.168 | 1.000 | |
| AT1G13280.1 | Allene oxide cyclase 4 (AOC4) | 37 | 1.19 | 0.169 | 1.000 | |
| AT1G13280.1 | Allene oxide cyclase 4 (AOC4) | 38 | 1.63 | 0.169 | 0.964 | |
| AT1G13280.1 | Allene oxide cyclase 4 (AOC4) | 39 | 2.01 | 0.167 | 0.672 | |
| AT1G13280.1 | Allene oxide cyclase 4 (AOC4) | 40 | 2.66 | 0.166 | 0.174 | |
| AT1G13280.1 | Allene oxide cyclase 4 (AOC4) | 41 | 2.32 | 0.168 | 0.379 | |
| AT1G13280.1 | Allene oxide cyclase 4 (AOC4) | 42 | 1.45 | 0.170 | 0.997 | |
| AT1G13280.1 | Allene oxide cyclase 4 (AOC4) | 43 | 0.93 | 0.171 | 1.000 | |
| AT1G13280.1 | Allene oxide cyclase 4 (AOC4) | 44 | 0.68 | 0.170 | 0.996 | |
| AT1G13280.1 | Allene oxide cyclase 4 (AOC4) | 45 | 0.55 | 0.171 | 0.855 | |
| AT1G13280.1 | Allene oxide cyclase 4 (AOC4) | 46 | 0.39 | 0.190 | 0.419 | |
| AT1G13280.1 | Allene oxide cyclase 4 (AOC4) | 47 | 0.23 | 0.377 | 0.762 | |
| AT1G14140.1 | Plant uncoupling mitochondrial protein 3 (PUMP3) | 2 | 1.04 | 0.465 | 1.000 | |
| AT1G14140.1 | Plant uncoupling mitochondrial protein 3 (PUMP3) | 3 | 1.12 | 0.446 | 1.000 | |

| | | | | | | |
|---|---|---|---|---|---|---|
| AT1G14140.1 | Plant uncoupling mitochondrial protein 3 (PUMP3) | 10 | 0.89 | 0.321 | 1.000 | |
| AT1G14140.1 | Plant uncoupling mitochondrial protein 3 (PUMP3) | 11 | 0.98 | 0.287 | 1.000 | |
| AT1G14140.1 | Plant uncoupling mitochondrial protein 3 (PUMP3) | 12 | 1.46 | 0.307 | 1.000 | |
| AT1G14140.1 | Plant uncoupling mitochondrial protein 3 (PUMP3) | 13 | 3.23 | 0.290 | 0.678 | |
| AT1G14140.1 | Plant uncoupling mitochondrial protein 3 (PUMP3) | 14 | 5.99 | 0.284 | 0.104 | |
| AT1G14140.1 | Plant uncoupling mitochondrial protein 3 (PUMP3) | 36 | 2.09 | 0.405 | 0.999 | |
| AT1G14140.1 | Plant uncoupling mitochondrial protein 3 (PUMP3) | 37 | 2.37 | 0.303 | 0.956 | |
| AT1G14140.1 | Plant uncoupling mitochondrial protein 3 (PUMP3) | 38 | 3.13 | 0.314 | 0.805 | |
| AT1G14140.1 | Plant uncoupling mitochondrial protein 3 (PUMP3) | 39 | 5.22 | 0.282 | 0.170 | |
| AT1G14140.1 | Plant uncoupling mitochondrial protein 3 (PUMP3) | 40 | 7.58 | 0.258 | 0.015 | * |
| AT1G14140.1 | Plant uncoupling mitochondrial protein 3 (PUMP3) | 41 | 5.32 | 0.227 | 0.028 | * |
| AT1G14140.1 | Plant uncoupling mitochondrial protein 3 (PUMP3) | 42 | 1.92 | 0.226 | 0.953 | |
| AT1G14140.1 | Plant uncoupling mitochondrial protein 3 (PUMP3) | 43 | 0.56 | 0.226 | 0.978 | |
| AT1G14140.1 | Plant uncoupling mitochondrial protein 3 (PUMP3) | 44 | 0.15 | 0.224 | 0.007 | ** |
| AT1G14140.1 | Plant uncoupling mitochondrial protein 3 (PUMP3) | 45 | 0.07 | 0.226 | 0.000 | *** |
| AT1G14140.1 | Plant uncoupling mitochondrial protein 3 (PUMP3) | 46 | 0.12 | 0.243 | 0.005 | ** |
| AT1G14140.1 | Plant uncoupling mitochondrial protein 3 (PUMP3) | 47 | 0.51 | 0.506 | 1.000 | |
| AT1G14320.1 | 60S ribosomal protein L10-1 (RPL10A) | 2 | 0.60 | 0.214 | 0.583 | |
| AT1G14320.1 | 60S ribosomal protein L10-1 (RPL10A) | 3 | 0.67 | 0.199 | 0.675 | |
| AT1G14320.1 | 60S ribosomal protein L10-1 (RPL10A) | 4 | 0.74 | 0.185 | 0.781 | |
| AT1G14320.1 | 60S ribosomal protein L10-1 (RPL10A) | 5 | 0.82 | 0.173 | 0.887 | |
| AT1G14320.1 | 60S ribosomal protein L10-1 (RPL10A) | 6 | 0.92 | 0.161 | 0.972 | |

| | | | | | | |
|---|---|---|---|---|---|---|
| AT1G14320.1 | 60S ribosomal protein L10-1 (RPL10A) | 7 | 1.02 | 0.151 | 0.999 | |
| AT1G14320.1 | 60S ribosomal protein L10-1 (RPL10A) | 8 | 1.13 | 0.143 | 0.932 | |
| AT1G14320.1 | 60S ribosomal protein L10-1 (RPL10A) | 9 | 1.26 | 0.137 | 0.766 | |
| AT1G14320.1 | 60S ribosomal protein L10-1 (RPL10A) | 10 | 1.39 | 0.133 | 0.551 | |
| AT1G14320.1 | 60S ribosomal protein L10-1 (RPL10A) | 11 | 1.55 | 0.133 | 0.353 | |
| AT1G14320.1 | 60S ribosomal protein L10-1 (RPL10A) | 12 | 1.72 | 0.134 | 0.212 | |
| AT1G14320.1 | 60S ribosomal protein L10-1 (RPL10A) | 13 | 1.91 | 0.139 | 0.126 | |
| AT1G14320.1 | 60S ribosomal protein L10-1 (RPL10A) | 14 | 2.12 | 0.145 | 0.079 | . |
| AT1G14320.1 | 60S ribosomal protein L10-1 (RPL10A) | 15 | 2.36 | 0.154 | 0.054 | . |
| AT1G14320.1 | 60S ribosomal protein L10-1 (RPL10A) | 16 | 2.62 | 0.165 | 0.040 | * |
| AT1G14320.1 | 60S ribosomal protein L10-1 (RPL10A) | 17 | 2.91 | 0.177 | 0.032 | * |
| AT1G14320.1 | 60S ribosomal protein L10-1 (RPL10A) | 18 | 3.23 | 0.190 | 0.028 | * |
| AT1G14320.1 | 60S ribosomal protein L10-1 (RPL10A) | 19 | 3.59 | 0.204 | 0.026 | * |
| AT1G14320.1 | 60S ribosomal protein L10-1 (RPL10A) | 46 | 61.16 | 0.680 | 0.033 | * |
| AT1G14320.1 | 60S ribosomal protein L10-1 (RPL10A) | 47 | 67.93 | 0.699 | 0.033 | * |
| AT1G14450.1 | NADH:ubiquinone oxidoreductase subunit (B12-1) | 2 | 0.78 | 0.592 | 1.000 | |
| AT1G14450.1 | NADH:ubiquinone oxidoreductase subunit (B12-1) | 3 | 0.88 | 0.484 | 1.000 | |
| AT1G14450.1 | NADH:ubiquinone oxidoreductase subunit (B12-1) | 4 | 0.99 | 0.427 | 1.000 | |
| AT1G14450.1 | NADH:ubiquinone oxidoreductase subunit (B12-1) | 5 | 1.10 | 0.404 | 1.000 | |
| AT1G14450.1 | NADH:ubiquinone oxidoreductase subunit (B12-1) | 6 | 1.23 | 0.396 | 1.000 | |
| AT1G14450.1 | NADH:ubiquinone oxidoreductase subunit (B12-1) | 7 | 1.37 | 0.393 | 1.000 | |
| AT1G14450.1 | NADH:ubiquinone oxidoreductase subunit (B12-1) | 8 | 1.53 | 0.392 | 1.000 | |
| AT1G14450.1 | NADH:ubiquinone oxidoreductase subunit (B12-1) | 9 | 1.70 | 0.389 | 1.000 | |
| AT1G14450.1 | NADH:ubiquinone oxidoreductase subunit (B12-1) | 10 | 1.86 | 0.384 | 0.999 | |
| AT1G14450.1 | NADH:ubiquinone oxidoreductase subunit (B12-1) | 11 | 2.01 | 0.379 | 0.996 | |
| AT1G14450.1 | NADH:ubiquinone oxidoreductase subunit (B12-1) | 12 | 2.09 | 0.375 | 0.992 | |
| AT1G14450.1 | NADH:ubiquinone oxidoreductase subunit (B12-1) | 13 | 2.07 | 0.372 | 0.993 | |
| AT1G14450.1 | NADH:ubiquinone oxidoreductase subunit (B12-1) | 14 | 1.91 | 0.370 | 0.997 | |
| AT1G14450.1 | NADH:ubiquinone oxidoreductase subunit (B12-1) | 15 | 1.65 | 0.368 | 1.000 | |
| AT1G14450.1 | NADH:ubiquinone oxidoreductase subunit (B12-1) | 16 | 1.40 | 0.368 | 1.000 | |
| AT1G14450.1 | NADH:ubiquinone oxidoreductase subunit (B12-1) | 17 | 1.23 | 0.368 | 1.000 | |

| | | | | | |
|---|---|---|---|---|---|
| AT1G14450.1 | NADH:ubiquinone oxidoreductase subunit (B12-1) | 18 | 1.17 | 0.368 | 1.000 |
| AT1G14450.1 | NADH:ubiquinone oxidoreductase subunit (B12-1) | 19 | 1.18 | 0.368 | 1.000 |
| AT1G14450.1 | NADH:ubiquinone oxidoreductase subunit (B12-1) | 20 | 1.25 | 0.369 | 1.000 |
| AT1G14450.1 | NADH:ubiquinone oxidoreductase subunit (B12-1) | 21 | 1.44 | 0.369 | 1.000 |
| AT1G14450.1 | NADH:ubiquinone oxidoreductase subunit (B12-1) | 22 | 1.81 | 0.370 | 0.999 |
| AT1G14450.1 | NADH:ubiquinone oxidoreductase subunit (B12-1) | 23 | 2.46 | 0.370 | 0.960 |
| AT1G14450.1 | NADH:ubiquinone oxidoreductase subunit (B12-1) | 24 | 3.38 | 0.369 | 0.781 |
| AT1G14450.1 | NADH:ubiquinone oxidoreductase subunit (B12-1) | 25 | 4.59 | 0.368 | 0.518 |
| AT1G14450.1 | NADH:ubiquinone oxidoreductase subunit (B12-1) | 26 | 5.85 | 0.367 | 0.322 |
| AT1G14450.1 | NADH:ubiquinone oxidoreductase subunit (B12-1) | 27 | 6.71 | 0.367 | 0.235 |
| AT1G14450.1 | NADH:ubiquinone oxidoreductase subunit (B12-1) | 28 | 6.92 | 0.366 | 0.218 |
| AT1G14450.1 | NADH:ubiquinone oxidoreductase subunit (B12-1) | 29 | 6.49 | 0.367 | 0.256 |
| AT1G14450.1 | NADH:ubiquinone oxidoreductase subunit (B12-1) | 30 | 5.63 | 0.370 | 0.357 |
| AT1G14450.1 | NADH:ubiquinone oxidoreductase subunit (B12-1) | 31 | 4.62 | 0.377 | 0.540 |
| AT1G14450.1 | NADH:ubiquinone oxidoreductase subunit (B12-1) | 32 | 3.68 | 0.390 | 0.771 |
| AT1G14450.1 | NADH:ubiquinone oxidoreductase subunit (B12-1) | 33 | 2.92 | 0.414 | 0.940 |
| AT1G14450.1 | NADH:ubiquinone oxidoreductase subunit (B12-1) | 34 | 2.38 | 0.450 | 0.994 |
| AT1G14450.1 | NADH:ubiquinone oxidoreductase subunit (B12-1) | 35 | 1.99 | 0.494 | 1.000 |
| AT1G14450.1 | NADH:ubiquinone oxidoreductase subunit (B12-1) | 41 | 1.12 | 0.622 | 1.000 |
| AT1G14450.1 | NADH:ubiquinone oxidoreductase subunit (B12-1) | 42 | 1.08 | 0.668 | 1.000 |
| AT1G14450.1 | NADH:ubiquinone oxidoreductase subunit (B12-1) | 43 | 1.05 | 0.749 | 1.000 |
| AT1G14450.1 | NADH:ubiquinone oxidoreductase subunit (B12-1) | 46 | 1.00 | 1.293 | 1.000 |
| AT1G14450.1 | NADH:ubiquinone oxidoreductase subunit (B12-1) | 47 | 0.99 | 1.546 | 1.000 |
| AT1G14610.1 | Valyl-tRNA synthase (VALRS / TWIN2) | 2 | 0.91 | 0.360 | 1.000 |
| AT1G14610.1 | Valyl-tRNA synthase (VALRS / TWIN2) | 3 | 0.74 | 0.361 | 1.000 |
| AT1G14610.1 | Valyl-tRNA synthase (VALRS / TWIN2) | 29 | 0.61 | 0.264 | 0.996 |
| AT1G14610.1 | Valyl-tRNA synthase (VALRS / TWIN2) | 30 | 0.98 | 0.197 | 1.000 |
| AT1G14610.1 | Valyl-tRNA synthase (VALRS / TWIN2) | 31 | 1.75 | 0.180 | 0.879 |
| AT1G14610.1 | Valyl-tRNA synthase (VALRS / TWIN2) | 32 | 2.39 | 0.170 | 0.285 |
| AT1G14610.1 | Valyl-tRNA synthase (VALRS / TWIN2) | 33 | 1.99 | 0.167 | 0.588 |
| AT1G14610.1 | Valyl-tRNA synthase (VALRS / TWIN2) | 34 | 1.40 | 0.168 | 0.994 |

| | | | | | | |
|---|---|---|---|---|---|---|
| AT1G14610.1 | Valyl-tRNA synthase (VALRS / TWIN2) | 35 | 1.14 | 0.167 | 1.000 | |
| AT1G14610.1 | Valyl-tRNA synthase (VALRS / TWIN2) | 36 | 0.95 | 0.167 | 1.000 | |
| AT1G14610.1 | Valyl-tRNA synthase (VALRS / TWIN2) | 37 | 0.62 | 0.169 | 0.925 | |
| AT1G14610.1 | Valyl-tRNA synthase (VALRS / TWIN2) | 38 | 0.36 | 0.176 | 0.150 | |
| AT1G14610.1 | Valyl-tRNA synthase (VALRS / TWIN2) | 39 | 0.27 | 0.223 | 0.143 | |
| AT1G14610.1 | Valyl-tRNA synthase (VALRS / TWIN2) | 40 | 0.32 | 0.375 | 0.887 | |
| AT1G14610.1 | Valyl-tRNA synthase (VALRS / TWIN2) | 46 | 144.34 | 2.220 | 0.985 | |
| AT1G14610.1 | Valyl-tRNA synthase (VALRS / TWIN2) | 47 | 606.43 | 2.695 | 0.977 | |
| AT1G14620.2 | ? | 2 | 1.08 | 0.370 | 1.000 | |
| AT1G14620.2 | ? | 3 | 1.05 | 0.300 | 1.000 | |
| AT1G14620.2 | ? | 4 | 1.02 | 0.292 | 1.000 | |
| AT1G14620.2 | ? | 6 | 0.96 | 0.317 | 1.000 | |
| AT1G14620.2 | ? | 7 | 0.94 | 0.350 | 1.000 | |
| AT1G14620.2 | ? | 8 | 0.94 | 0.395 | 1.000 | |
| AT1G14620.2 | ? | 22 | 1.02 | 0.340 | 1.000 | |
| AT1G14620.2 | ? | 23 | 0.98 | 0.330 | 1.000 | |
| AT1G14620.2 | ? | 24 | 0.92 | 0.316 | 1.000 | |
| AT1G14620.2 | ? | 28 | 0.71 | 0.243 | 1.000 | |
| AT1G14620.2 | ? | 29 | 0.70 | 0.206 | 1.000 | |
| AT1G14620.2 | ? | 30 | 0.71 | 0.189 | 0.999 | |
| AT1G14620.2 | ? | 31 | 0.76 | 0.184 | 1.000 | |
| AT1G14620.2 | ? | 32 | 0.89 | 0.183 | 1.000 | |
| AT1G14620.2 | ? | 33 | 1.11 | 0.182 | 1.000 | |
| AT1G14620.2 | ? | 34 | 1.35 | 0.184 | 1.000 | |
| AT1G14620.2 | ? | 35 | 1.39 | 0.192 | 1.000 | |
| AT1G14620.2 | ? | 36 | 1.20 | 0.204 | 1.000 | |
| AT1G14620.2 | ? | 37 | 0.94 | 0.213 | 1.000 | |
| AT1G14620.2 | ? | 38 | 0.77 | 0.205 | 1.000 | |
| AT1G14620.2 | ? | 39 | 0.73 | 0.193 | 1.000 | |
| AT1G14620.2 | ? | 40 | 0.84 | 0.183 | 1.000 | |
| AT1G14620.2 | ? | 41 | 1.32 | 0.182 | 1.000 | |
| AT1G14620.2 | ? | 42 | 2.63 | 0.184 | 0.295 | |
| AT1G14620.2 | ? | 43 | 4.56 | 0.188 | 0.011 | * |
| AT1G14620.2 | ? | 44 | 4.60 | 0.193 | 0.014 | * |
| AT1G14620.2 | ? | 45 | 2.53 | 0.199 | 0.462 | |
| AT1G14620.2 | ? | 46 | 0.97 | 0.223 | 1.000 | |
| AT1G14620.2 | ? | 47 | 0.33 | 0.317 | 0.838 | |
| AT1G14980.1 | Chaperonin 10-1 (CPN10-1) | 2 | 0.82 | 0.502 | 1.000 | |
| AT1G14980.1 | Chaperonin 10-1 (CPN10-1) | 3 | 0.50 | 0.392 | 1.000 | |
| AT1G14980.1 | Chaperonin 10-1 (CPN10-1) | 4 | 0.36 | 0.459 | 0.997 | |
| AT1G14980.1 | Chaperonin 10-1 (CPN10-1) | 9 | 0.51 | 0.471 | 1.000 | |
| AT1G14980.1 | Chaperonin 10-1 (CPN10-1) | 10 | 0.60 | 0.462 | 1.000 | |
| AT1G14980.1 | Chaperonin 10-1 (CPN10-1) | 11 | 0.69 | 0.463 | 1.000 | |
| AT1G14980.1 | Chaperonin 10-1 (CPN10-1) | 22 | 1.55 | 0.458 | 1.000 | |
| AT1G14980.1 | Chaperonin 10-1 (CPN10-1) | 23 | 1.93 | 0.396 | 1.000 | |
| AT1G14980.1 | Chaperonin 10-1 (CPN10-1) | 24 | 2.71 | 0.334 | 0.955 | |
| AT1G14980.1 | Chaperonin 10-1 (CPN10-1) | 25 | 3.65 | 0.307 | 0.646 | |
| AT1G14980.1 | Chaperonin 10-1 (CPN10-1) | 26 | 3.66 | 0.290 | 0.556 | |
| AT1G14980.1 | Chaperonin 10-1 (CPN10-1) | 27 | 3.19 | 0.282 | 0.685 | |
| AT1G14980.1 | Chaperonin 10-1 (CPN10-1) | 28 | 2.55 | 0.280 | 0.898 | |
| AT1G14980.1 | Chaperonin 10-1 (CPN10-1) | 29 | 1.92 | 0.279 | 0.995 | |
| AT1G14980.1 | Chaperonin 10-1 (CPN10-1) | 30 | 1.47 | 0.280 | 1.000 | |
| AT1G14980.1 | Chaperonin 10-1 (CPN10-1) | 31 | 1.25 | 0.281 | 1.000 | |
| AT1G14980.1 | Chaperonin 10-1 (CPN10-1) | 32 | 1.20 | 0.281 | 1.000 | |
| AT1G14980.1 | Chaperonin 10-1 (CPN10-1) | 33 | 1.20 | 0.281 | 1.000 | |

| | | | | | | |
|---|---|---|---|---|---|---|
| AT1G14980.1 | Chaperonin 10-1 (CPN10-1) | 34 | 1.29 | 0.281 | 1.000 | |
| AT1G14980.1 | Chaperonin 10-1 (CPN10-1) | 35 | 1.63 | 0.281 | 1.000 | |
| AT1G14980.1 | Chaperonin 10-1 (CPN10-1) | 36 | 2.37 | 0.281 | 0.944 | |
| AT1G14980.1 | Chaperonin 10-1 (CPN10-1) | 37 | 3.65 | 0.281 | 0.512 | |
| AT1G14980.1 | Chaperonin 10-1 (CPN10-1) | 38 | 5.16 | 0.280 | 0.181 | |
| AT1G14980.1 | Chaperonin 10-1 (CPN10-1) | 39 | 6.21 | 0.280 | 0.090 | . |
| AT1G14980.1 | Chaperonin 10-1 (CPN10-1) | 40 | 6.05 | 0.281 | 0.101 | |
| AT1G14980.1 | Chaperonin 10-1 (CPN10-1) | 41 | 5.22 | 0.282 | 0.180 | |
| AT1G14980.1 | Chaperonin 10-1 (CPN10-1) | 42 | 4.25 | 0.284 | 0.360 | |
| AT1G14980.1 | Chaperonin 10-1 (CPN10-1) | 43 | 2.85 | 0.286 | 0.822 | |
| AT1G14980.1 | Chaperonin 10-1 (CPN10-1) | 44 | 1.31 | 0.286 | 1.000 | |
| AT1G14980.1 | Chaperonin 10-1 (CPN10-1) | 45 | 0.44 | 0.287 | 0.969 | |
| AT1G14980.1 | Chaperonin 10-1 (CPN10-1) | 46 | 0.12 | 0.331 | 0.111 | |
| AT1G14980.1 | Chaperonin 10-1 (CPN10-1) | 47 | 0.03 | 0.516 | 0.061 | . |
| AT1G14990.1 | ? | 2 | 1.06 | 0.587 | 1.000 | |
| AT1G14990.1 | ? | 3 | 1.05 | 0.560 | 1.000 | |
| AT1G14990.1 | ? | 23 | 0.74 | 0.232 | 0.943 | |
| AT1G14990.1 | ? | 24 | 0.71 | 0.226 | 0.916 | |
| AT1G14990.1 | ? | 25 | 0.69 | 0.224 | 0.884 | |
| AT1G14990.1 | ? | 26 | 0.66 | 0.226 | 0.848 | |
| AT1G14990.1 | ? | 27 | 0.63 | 0.231 | 0.807 | |
| AT1G14990.1 | ? | 28 | 0.59 | 0.240 | 0.759 | |
| AT1G14990.1 | ? | 29 | 0.55 | 0.251 | 0.708 | |
| AT1G14990.1 | ? | 46 | 0.06 | 0.497 | 0.067 | . |
| AT1G14990.1 | ? | 47 | 0.05 | 0.519 | 0.063 | . |
| AT1G15120.1 | Ubiquinol:cytochrome C oxidoreductase subunit (hinge) | 2 | 1.01 | 0.629 | 1.000 | |
| AT1G15120.1 | Ubiquinol:cytochrome C oxidoreductase subunit (hinge) | 3 | 0.65 | 0.401 | 1.000 | |
| AT1G15120.1 | Ubiquinol:cytochrome C oxidoreductase subunit (hinge) | 4 | 0.46 | 0.391 | 1.000 | |
| AT1G15120.1 | Ubiquinol:cytochrome C oxidoreductase subunit (hinge) | 5 | 0.47 | 0.363 | 1.000 | |
| AT1G15120.1 | Ubiquinol:cytochrome C oxidoreductase subunit (hinge) | 6 | 0.59 | 0.347 | 1.000 | |
| AT1G15120.1 | Ubiquinol:cytochrome C oxidoreductase subunit (hinge) | 7 | 0.66 | 0.349 | 1.000 | |
| AT1G15120.1 | Ubiquinol:cytochrome C oxidoreductase subunit (hinge) | 8 | 0.79 | 0.334 | 1.000 | |
| AT1G15120.1 | Ubiquinol:cytochrome C oxidoreductase subunit (hinge) | 9 | 1.07 | 0.327 | 1.000 | |
| AT1G15120.1 | Ubiquinol:cytochrome C oxidoreductase subunit (hinge) | 10 | 1.36 | 0.329 | 1.000 | |
| AT1G15120.1 | Ubiquinol:cytochrome C oxidoreductase subunit (hinge) | 11 | 1.38 | 0.335 | 1.000 | |
| AT1G15120.1 | Ubiquinol:cytochrome C oxidoreductase subunit (hinge) | 12 | 1.52 | 0.335 | 1.000 | |
| AT1G15120.1 | Ubiquinol:cytochrome C oxidoreductase subunit (hinge) | 13 | 1.85 | 0.331 | 1.000 | |
| AT1G15120.1 | Ubiquinol:cytochrome C oxidoreductase subunit (hinge) | 14 | 1.97 | 0.324 | 1.000 | |
| AT1G15120.1 | Ubiquinol:cytochrome C oxidoreductase subunit (hinge) | 15 | 1.34 | 0.322 | 1.000 | |
| AT1G15120.1 | Ubiquinol:cytochrome C oxidoreductase subunit (hinge) | 16 | 0.72 | 0.319 | 1.000 | |
| AT1G15120.1 | Ubiquinol:cytochrome C oxidoreductase subunit (hinge) | 17 | 0.29 | 0.319 | 0.887 | |
| AT1G15120.1 | Ubiquinol:cytochrome C oxidoreductase subunit (hinge) | 18 | 0.27 | 0.325 | 0.863 | |
| AT1G15120.1 | Ubiquinol:cytochrome C oxidoreductase subunit (hinge) | 19 | 0.50 | 0.335 | 1.000 | |

| | | | | | | |
|---|---|---|---|---|---|---|
| AT1G15120.1 | Ubiquinol:cytochrome C oxidoreductase subunit (hinge) | 20 | 1.00 | 0.342 | 1.000 | |
| AT1G15120.1 | Ubiquinol:cytochrome C oxidoreductase subunit (hinge) | 21 | 2.23 | 0.345 | 1.000 | |
| AT1G15120.1 | Ubiquinol:cytochrome C oxidoreductase subunit (hinge) | 22 | 3.22 | 0.345 | 0.964 | |
| AT1G15120.1 | Ubiquinol:cytochrome C oxidoreductase subunit (hinge) | 23 | 1.43 | 0.348 | 1.000 | |
| AT1G15120.1 | Ubiquinol:cytochrome C oxidoreductase subunit (hinge) | 24 | 0.39 | 0.333 | 0.996 | |
| AT1G15120.1 | Ubiquinol:cytochrome C oxidoreductase subunit (hinge) | 25 | 0.27 | 0.325 | 0.839 | |
| AT1G15120.1 | Ubiquinol:cytochrome C oxidoreductase subunit (hinge) | 26 | 1.05 | 0.328 | 1.000 | |
| AT1G15120.1 | Ubiquinol:cytochrome C oxidoreductase subunit (hinge) | 27 | 13.69 | 0.336 | 0.023 | * |
| AT1G15120.1 | Ubiquinol:cytochrome C oxidoreductase subunit (hinge) | 28 | 13.98 | 0.359 | 0.042 | * |
| AT1G15120.1 | Ubiquinol:cytochrome C oxidoreductase subunit (hinge) | 29 | 1.01 | 0.424 | 1.000 | |
| AT1G15120.1 | Ubiquinol:cytochrome C oxidoreductase subunit (hinge) | 30 | 0.04 | 0.734 | 0.714 | |
| AT1G15120.1 | Ubiquinol:cytochrome C oxidoreductase subunit (hinge) | 46 | 333882.00 | 13.219 | 1.000 | |
| AT1G15120.1 | Ubiquinol:cytochrome C oxidoreductase subunit (hinge) | 47 | 7839387.90 | 14.428 | 1.000 | |
| AT1G15220.2 | Cytochrome c maturation H (CCMH) | 2 | 1.08 | 0.556 | 1.000 | |
| AT1G15220.2 | Cytochrome c maturation H (CCMH) | 3 | 0.84 | 0.806 | 1.000 | |
| AT1G15220.2 | Cytochrome c maturation H (CCMH) | 23 | 0.16 | 0.321 | 0.112 | |
| AT1G15220.2 | Cytochrome c maturation H (CCMH) | 24 | 0.37 | 0.237 | 0.421 | |
| AT1G15220.2 | Cytochrome c maturation H (CCMH) | 25 | 1.07 | 0.222 | 1.000 | |
| AT1G15220.2 | Cytochrome c maturation H (CCMH) | 26 | 2.94 | 0.222 | 0.262 | |
| AT1G15220.2 | Cytochrome c maturation H (CCMH) | 27 | 3.75 | 0.238 | 0.143 | |
| AT1G15220.2 | Cytochrome c maturation H (CCMH) | 28 | 1.75 | 0.310 | 0.982 | |
| AT1G15220.2 | Cytochrome c maturation H (CCMH) | 29 | 0.59 | 0.488 | 0.999 | |
| AT1G15220.2 | Cytochrome c maturation H (CCMH) | 46 | 0.01 | 0.457 | 0.002 | ** |
| AT1G15220.2 | Cytochrome c maturation H (CCMH) | 47 | 0.02 | 0.470 | 0.005 | ** |
| AT1G15390.1 | Peptide deformylase 1A (PDF1A) | 2 | 0.81 | 0.483 | 1.000 | |
| AT1G15390.1 | Peptide deformylase 1A (PDF1A) | 3 | 1.23 | 0.470 | 1.000 | |
| AT1G15390.1 | Peptide deformylase 1A (PDF1A) | 33 | 3.18 | 0.345 | 0.772 | |
| AT1G15390.1 | Peptide deformylase 1A (PDF1A) | 34 | 1.59 | 0.258 | 0.995 | |
| AT1G15390.1 | Peptide deformylase 1A (PDF1A) | 35 | 0.84 | 0.229 | 1.000 | |
| AT1G15390.1 | Peptide deformylase 1A (PDF1A) | 36 | 0.52 | 0.225 | 0.888 | |
| AT1G15390.1 | Peptide deformylase 1A (PDF1A) | 37 | 0.42 | 0.228 | 0.619 | |

| | | | | | | |
|---|---|---|---|---|---|---|
| AT1G15390.1 | Peptide deformylase 1A (PDF1A) | 38 | 0.41 | 0.230 | 0.601 | |
| AT1G15390.1 | Peptide deformylase 1A (PDF1A) | 39 | 0.49 | 0.223 | 0.821 | |
| AT1G15390.1 | Peptide deformylase 1A (PDF1A) | 40 | 0.80 | 0.215 | 1.000 | |
| AT1G15390.1 | Peptide deformylase 1A (PDF1A) | 41 | 1.73 | 0.212 | 0.938 | |
| AT1G15390.1 | Peptide deformylase 1A (PDF1A) | 42 | 4.10 | 0.213 | 0.053 | . |
| AT1G15390.1 | Peptide deformylase 1A (PDF1A) | 43 | 7.05 | 0.214 | 0.002 | ** |
| AT1G15390.1 | Peptide deformylase 1A (PDF1A) | 44 | 6.42 | 0.214 | 0.003 | ** |
| AT1G15390.1 | Peptide deformylase 1A (PDF1A) | 45 | 3.24 | 0.217 | 0.192 | |
| AT1G15390.1 | Peptide deformylase 1A (PDF1A) | 46 | 1.10 | 0.255 | 1.000 | |
| AT1G15390.1 | Peptide deformylase 1A (PDF1A) | 47 | 0.32 | 0.403 | 0.891 | |
| AT1G15690.1 | Vacuolar inorganic H pyrophosphatase 3 (AVP3) | 2 | 0.33 | 0.416 | 0.966 | |
| AT1G15690.1 | Vacuolar inorganic H pyrophosphatase 3 (AVP3) | 3 | 0.74 | 0.304 | 1.000 | |
| AT1G15690.1 | Vacuolar inorganic H pyrophosphatase 3 (AVP3) | 4 | 1.55 | 0.289 | 1.000 | |
| AT1G15690.1 | Vacuolar inorganic H pyrophosphatase 3 (AVP3) | 5 | 2.45 | 0.284 | 0.887 | |
| AT1G15690.1 | Vacuolar inorganic H pyrophosphatase 3 (AVP3) | 6 | 2.64 | 0.285 | 0.826 | |
| AT1G15690.1 | Vacuolar inorganic H pyrophosphatase 3 (AVP3) | 7 | 2.27 | 0.288 | 0.943 | |
| AT1G15690.1 | Vacuolar inorganic H pyrophosphatase 3 (AVP3) | 8 | 2.00 | 0.281 | 0.982 | |
| AT1G15690.1 | Vacuolar inorganic H pyrophosphatase 3 (AVP3) | 9 | 1.78 | 0.278 | 0.997 | |
| AT1G15690.1 | Vacuolar inorganic H pyrophosphatase 3 (AVP3) | 10 | 1.47 | 0.280 | 1.000 | |
| AT1G15690.1 | Vacuolar inorganic H pyrophosphatase 3 (AVP3) | 11 | 1.19 | 0.284 | 1.000 | |
| AT1G15690.1 | Vacuolar inorganic H pyrophosphatase 3 (AVP3) | 12 | 1.19 | 0.284 | 1.000 | |
| AT1G15690.1 | Vacuolar inorganic H pyrophosphatase 3 (AVP3) | 13 | 1.48 | 0.285 | 1.000 | |
| AT1G15690.1 | Vacuolar inorganic H pyrophosphatase 3 (AVP3) | 14 | 1.57 | 0.287 | 1.000 | |
| AT1G15690.1 | Vacuolar inorganic H pyrophosphatase 3 (AVP3) | 15 | 1.13 | 0.300 | 1.000 | |
| AT1G15690.1 | Vacuolar inorganic H pyrophosphatase 3 (AVP3) | 16 | 0.74 | 0.307 | 1.000 | |
| AT1G15690.1 | Vacuolar inorganic H pyrophosphatase 3 (AVP3) | 17 | 0.55 | 0.343 | 0.999 | |
| AT1G15690.1 | Vacuolar inorganic H pyrophosphatase 3 (AVP3) | 18 | 0.43 | 0.417 | 0.997 | |
| AT1G15690.1 | Vacuolar inorganic H pyrophosphatase 3 (AVP3) | 28 | 0.18 | 0.375 | 0.465 | |
| AT1G15690.1 | Vacuolar inorganic H pyrophosphatase 3 (AVP3) | 29 | 0.26 | 0.308 | 0.534 | |
| AT1G15690.1 | Vacuolar inorganic H pyrophosphatase 3 (AVP3) | 30 | 0.73 | 0.293 | 1.000 | |
| AT1G15690.1 | Vacuolar inorganic H pyrophosphatase 3 (AVP3) | 31 | 3.66 | 0.286 | 0.463 | |

| | | | | | | |
|---|---|---|---|---|---|---|
| AT1G15690.1 | Vacuolar inorganic H pyrophosphatase 3 (AVP3) | 32 | 9.56 | 0.286 | 0.014 | * |
| AT1G15690.1 | Vacuolar inorganic H pyrophosphatase 3 (AVP3) | 33 | 5.12 | 0.297 | 0.216 | |
| AT1G15690.1 | Vacuolar inorganic H pyrophosphatase 3 (AVP3) | 34 | 0.70 | 0.355 | 1.000 | |
| AT1G15690.1 | Vacuolar inorganic H pyrophosphatase 3 (AVP3) | 35 | 0.06 | 0.541 | 0.305 | |
| AT1G15690.1 | Vacuolar inorganic H pyrophosphatase 3 (AVP3) | 46 | 0.00 | 1.441 | 0.013 | * |
| AT1G15690.1 | Vacuolar inorganic H pyrophosphatase 3 (AVP3) | 47 | 0.00 | 1.589 | 0.033 | * |
| AT1G15810.1 | Ribosomal S15 family protein | 2 | 2.51 | 0.522 | 0.983 | |
| AT1G15810.1 | Ribosomal S15 family protein | 3 | 2.34 | 0.473 | 0.981 | |
| AT1G15810.1 | Ribosomal S15 family protein | 11 | 1.42 | 0.270 | 0.998 | |
| AT1G15810.1 | Ribosomal S15 family protein | 12 | 1.35 | 0.255 | 0.999 | |
| AT1G15810.1 | Ribosomal S15 family protein | 13 | 1.30 | 0.243 | 1.000 | |
| AT1G15810.1 | Ribosomal S15 family protein | 14 | 1.27 | 0.235 | 1.000 | |
| AT1G15810.1 | Ribosomal S15 family protein | 15 | 1.27 | 0.232 | 1.000 | |
| AT1G15810.1 | Ribosomal S15 family protein | 16 | 1.29 | 0.233 | 1.000 | |
| AT1G15810.1 | Ribosomal S15 family protein | 17 | 1.32 | 0.239 | 0.999 | |
| AT1G15810.1 | Ribosomal S15 family protein | 18 | 1.34 | 0.248 | 0.999 | |
| AT1G15810.1 | Ribosomal S15 family protein | 19 | 1.36 | 0.258 | 0.999 | |
| AT1G15810.1 | Ribosomal S15 family protein | 26 | 1.42 | 0.265 | 0.998 | |
| AT1G15810.1 | Ribosomal S15 family protein | 27 | 1.43 | 0.255 | 0.996 | |
| AT1G15810.1 | Ribosomal S15 family protein | 28 | 1.44 | 0.245 | 0.994 | |
| AT1G15810.1 | Ribosomal S15 family protein | 29 | 1.46 | 0.235 | 0.991 | |
| AT1G15810.1 | Ribosomal S15 family protein | 30 | 1.46 | 0.228 | 0.988 | |
| AT1G15810.1 | Ribosomal S15 family protein | 31 | 1.47 | 0.223 | 0.984 | |
| AT1G15810.1 | Ribosomal S15 family protein | 32 | 1.48 | 0.219 | 0.982 | |
| AT1G15810.1 | Ribosomal S15 family protein | 33 | 1.48 | 0.218 | 0.980 | |
| AT1G15810.1 | Ribosomal S15 family protein | 34 | 1.50 | 0.217 | 0.976 | |
| AT1G15810.1 | Ribosomal S15 family protein | 35 | 1.54 | 0.218 | 0.965 | |
| AT1G15810.1 | Ribosomal S15 family protein | 36 | 1.60 | 0.219 | 0.945 | |
| AT1G15810.1 | Ribosomal S15 family protein | 37 | 1.68 | 0.220 | 0.913 | |
| AT1G15810.1 | Ribosomal S15 family protein | 38 | 1.75 | 0.221 | 0.875 | |
| AT1G15810.1 | Ribosomal S15 family protein | 39 | 1.80 | 0.220 | 0.846 | |
| AT1G15810.1 | Ribosomal S15 family protein | 40 | 1.80 | 0.220 | 0.843 | |
| AT1G15810.1 | Ribosomal S15 family protein | 41 | 1.75 | 0.219 | 0.875 | |
| AT1G15810.1 | Ribosomal S15 family protein | 42 | 1.63 | 0.219 | 0.935 | |
| AT1G15810.1 | Ribosomal S15 family protein | 43 | 1.46 | 0.223 | 0.987 | |
| AT1G15810.1 | Ribosomal S15 family protein | 44 | 1.26 | 0.232 | 1.000 | |
| AT1G15810.1 | Ribosomal S15 family protein | 45 | 1.06 | 0.250 | 1.000 | |
| AT1G15810.1 | Ribosomal S15 family protein | 46 | 0.88 | 0.280 | 1.000 | |
| AT1G15810.1 | Ribosomal S15 family protein | 47 | 0.72 | 0.325 | 1.000 | |
| AT1G15820.1 | Photosystem II light harvesting complex subunit (LHCB6) | 2 | 0.70 | 0.308 | 1.000 | |
| AT1G15820.1 | Photosystem II light harvesting complex subunit (LHCB6) | 3 | 0.55 | 0.180 | 0.995 | |
| AT1G15820.1 | Photosystem II light harvesting complex subunit (LHCB6) | 4 | 0.65 | 0.174 | 1.000 | |
| AT1G15820.1 | Photosystem II light harvesting complex subunit (LHCB6) | 5 | 0.90 | 0.173 | 1.000 | |
| AT1G15820.1 | Photosystem II light harvesting complex subunit (LHCB6) | 6 | 0.97 | 0.174 | 1.000 | |
| AT1G15820.1 | Photosystem II light harvesting complex subunit (LHCB6) | 7 | 0.97 | 0.175 | 1.000 | |
| AT1G15820.1 | Photosystem II light harvesting complex subunit (LHCB6) | 8 | 1.02 | 0.171 | 1.000 | |
| AT1G15820.1 | Photosystem II light harvesting complex subunit (LHCB6) | 9 | 1.04 | 0.168 | 1.000 | |

| | | | | | | |
|---|---|---|---|---|---|---|
| AT1G15820.1 | Photosystem II light harvesting complex subunit (LHCB6) | 10 | 0.94 | 0.167 | 1.000 | |
| AT1G15820.1 | Photosystem II light harvesting complex subunit (LHCB6) | 11 | 0.88 | 0.168 | 1.000 | |
| AT1G15820.1 | Photosystem II light harvesting complex subunit (LHCB6) | 12 | 0.89 | 0.168 | 1.000 | |
| AT1G15820.1 | Photosystem II light harvesting complex subunit (LHCB6) | 13 | 0.95 | 0.166 | 1.000 | |
| AT1G15820.1 | Photosystem II light harvesting complex subunit (LHCB6) | 14 | 1.07 | 0.163 | 1.000 | |
| AT1G15820.1 | Photosystem II light harvesting complex subunit (LHCB6) | 15 | 1.08 | 0.164 | 1.000 | |
| AT1G15820.1 | Photosystem II light harvesting complex subunit (LHCB6) | 16 | 0.96 | 0.162 | 1.000 | |
| AT1G15820.1 | Photosystem II light harvesting complex subunit (LHCB6) | 17 | 0.97 | 0.162 | 1.000 | |
| AT1G15820.1 | Photosystem II light harvesting complex subunit (LHCB6) | 18 | 1.15 | 0.164 | 1.000 | |
| AT1G15820.1 | Photosystem II light harvesting complex subunit (LHCB6) | 19 | 1.37 | 0.169 | 1.000 | |
| AT1G15820.1 | Photosystem II light harvesting complex subunit (LHCB6) | 20 | 1.52 | 0.173 | 1.000 | |
| AT1G15820.1 | Photosystem II light harvesting complex subunit (LHCB6) | 21 | 1.58 | 0.176 | 1.000 | |
| AT1G15820.1 | Photosystem II light harvesting complex subunit (LHCB6) | 22 | 1.53 | 0.176 | 1.000 | |
| AT1G15820.1 | Photosystem II light harvesting complex subunit (LHCB6) | 23 | 1.23 | 0.176 | 1.000 | |
| AT1G15820.1 | Photosystem II light harvesting complex subunit (LHCB6) | 24 | 0.93 | 0.172 | 1.000 | |
| AT1G15820.1 | Photosystem II light harvesting complex subunit (LHCB6) | 25 | 0.81 | 0.167 | 1.000 | |
| AT1G15820.1 | Photosystem II light harvesting complex subunit (LHCB6) | 26 | 1.00 | 0.166 | 1.000 | |
| AT1G15820.1 | Photosystem II light harvesting complex subunit (LHCB6) | 27 | 1.31 | 0.166 | 1.000 | |
| AT1G15820.1 | Photosystem II light harvesting complex subunit (LHCB6) | 28 | 1.07 | 0.166 | 1.000 | |
| AT1G15820.1 | Photosystem II light harvesting complex subunit (LHCB6) | 29 | 0.74 | 0.166 | 1.000 | |
| AT1G15820.1 | Photosystem II light harvesting complex subunit (LHCB6) | 30 | 0.83 | 0.166 | 1.000 | |
| AT1G15820.1 | Photosystem II light harvesting complex subunit (LHCB6) | 31 | 1.51 | 0.167 | 1.000 | |
| AT1G15820.1 | Photosystem II light harvesting complex subunit (LHCB6) | 32 | 1.66 | 0.167 | 0.999 | |
| AT1G15820.1 | Photosystem II light harvesting complex subunit (LHCB6) | 33 | 0.72 | 0.168 | 1.000 | |
| AT1G15820.1 | Photosystem II light harvesting complex subunit (LHCB6) | 34 | 0.49 | 0.169 | 0.893 | |
| AT1G15820.1 | Photosystem II light harvesting complex subunit (LHCB6) | 35 | 0.69 | 0.170 | 1.000 | |
| AT1G15820.1 | Photosystem II light harvesting complex subunit (LHCB6) | 36 | 1.18 | 0.177 | 1.000 | |
| AT1G15820.1 | Photosystem II light harvesting complex subunit (LHCB6) | 37 | 1.89 | 0.194 | 0.995 | |
| AT1G15820.1 | Photosystem II light harvesting complex subunit (LHCB6) | 38 | 2.75 | 0.224 | 0.828 | |
| AT1G15820.1 | Photosystem II light harvesting complex subunit (LHCB6) | 39 | 3.85 | 0.212 | 0.205 | |
| AT1G15820.1 | Photosystem II light harvesting complex subunit (LHCB6) | 40 | 5.63 | 0.197 | 0.007 | ** |

| | | | | | | |
|---|---|---|---|---|---|---|
| AT1G15820.1 | Photosystem II light harvesting complex subunit (LHCB6) | 41 | 7.57 | 0.175 | 0.000 | *** |
| AT1G15820.1 | Photosystem II light harvesting complex subunit (LHCB6) | 42 | 5.08 | 0.172 | 0.002 | ** |
| AT1G15820.1 | Photosystem II light harvesting complex subunit (LHCB6) | 43 | 1.76 | 0.173 | 0.996 | |
| AT1G15820.1 | Photosystem II light harvesting complex subunit (LHCB6) | 44 | 0.48 | 0.173 | 0.906 | |
| AT1G15820.1 | Photosystem II light harvesting complex subunit (LHCB6) | 45 | 0.24 | 0.173 | 0.014 | * |
| AT1G15820.1 | Photosystem II light harvesting complex subunit (LHCB6) | 46 | 0.24 | 0.192 | 0.059 | . |
| AT1G15820.1 | Photosystem II light harvesting complex subunit (LHCB6) | 47 | 0.38 | 0.382 | 1.000 | |
| AT1G15980.1 | NAD(P)H dehydrogenase complex subunit (NDH48) | 2 | 1.59 | 0.376 | 1.000 | |
| AT1G15980.1 | NAD(P)H dehydrogenase complex subunit (NDH48) | 3 | 1.08 | 0.256 | 1.000 | |
| AT1G15980.1 | NAD(P)H dehydrogenase complex subunit (NDH48) | 4 | 0.74 | 0.215 | 1.000 | |
| AT1G15980.1 | NAD(P)H dehydrogenase complex subunit (NDH48) | 5 | 0.52 | 0.202 | 0.871 | |
| AT1G15980.1 | NAD(P)H dehydrogenase complex subunit (NDH48) | 6 | 0.38 | 0.194 | 0.346 | |
| AT1G15980.1 | NAD(P)H dehydrogenase complex subunit (NDH48) | 7 | 0.31 | 0.190 | 0.113 | |
| AT1G15980.1 | NAD(P)H dehydrogenase complex subunit (NDH48) | 8 | 0.29 | 0.185 | 0.062 | . |
| AT1G15980.1 | NAD(P)H dehydrogenase complex subunit (NDH48) | 9 | 0.30 | 0.181 | 0.060 | . |
| AT1G15980.1 | NAD(P)H dehydrogenase complex subunit (NDH48) | 10 | 0.31 | 0.181 | 0.074 | . |
| AT1G15980.1 | NAD(P)H dehydrogenase complex subunit (NDH48) | 11 | 0.34 | 0.181 | 0.139 | |
| AT1G15980.1 | NAD(P)H dehydrogenase complex subunit (NDH48) | 12 | 0.38 | 0.181 | 0.247 | |
| AT1G15980.1 | NAD(P)H dehydrogenase complex subunit (NDH48) | 13 | 0.40 | 0.178 | 0.297 | |
| AT1G15980.1 | NAD(P)H dehydrogenase complex subunit (NDH48) | 14 | 0.42 | 0.176 | 0.374 | |
| AT1G15980.1 | NAD(P)H dehydrogenase complex subunit (NDH48) | 15 | 0.49 | 0.176 | 0.625 | |
| AT1G15980.1 | NAD(P)H dehydrogenase complex subunit (NDH48) | 16 | 0.72 | 0.175 | 0.999 | |
| AT1G15980.1 | NAD(P)H dehydrogenase complex subunit (NDH48) | 17 | 1.28 | 0.177 | 1.000 | |
| AT1G15980.1 | NAD(P)H dehydrogenase complex subunit (NDH48) | 18 | 2.18 | 0.184 | 0.590 | |
| AT1G15980.1 | NAD(P)H dehydrogenase complex subunit (NDH48) | 19 | 2.77 | 0.195 | 0.284 | |
| AT1G15980.1 | NAD(P)H dehydrogenase complex subunit (NDH48) | 20 | 2.47 | 0.213 | 0.582 | |
| AT1G15980.1 | NAD(P)H dehydrogenase complex subunit (NDH48) | 21 | 1.82 | 0.275 | 0.993 | |
| AT1G15980.1 | NAD(P)H dehydrogenase complex subunit (NDH48) | 22 | 1.35 | 0.415 | 1.000 | |
| AT1G15980.1 | NAD(P)H dehydrogenase complex subunit (NDH48) | 46 | ########### | 7.214 | 0.971 | |
| AT1G15980.1 | NAD(P)H dehydrogenase complex subunit (NDH48) | 47 | ########### | 7.636 | 0.969 | |

| | | | | | | |
|---|---|---|---|---|---|---|
| AT1G16700.1 | NADH:ubiquinone oxidoreductase subunit (TYKY-2) | 2 | 0.85 | 0.505 | 1.000 | |
| AT1G16700.1 | NADH:ubiquinone oxidoreductase subunit (TYKY-2) | 3 | 0.75 | 0.469 | 1.000 | |
| AT1G16700.1 | NADH:ubiquinone oxidoreductase subunit (TYKY-2) | 14 | 0.93 | 0.313 | 1.000 | |
| AT1G16700.1 | NADH:ubiquinone oxidoreductase subunit (TYKY-2) | 15 | 0.92 | 0.272 | 1.000 | |
| AT1G16700.1 | NADH:ubiquinone oxidoreductase subunit (TYKY-2) | 16 | 0.63 | 0.252 | 1.000 | |
| AT1G16700.1 | NADH:ubiquinone oxidoreductase subunit (TYKY-2) | 17 | 0.34 | 0.226 | 0.479 | |
| AT1G16700.1 | NADH:ubiquinone oxidoreductase subunit (TYKY-2) | 18 | 0.33 | 0.223 | 0.431 | |
| AT1G16700.1 | NADH:ubiquinone oxidoreductase subunit (TYKY-2) | 19 | 0.52 | 0.227 | 0.979 | |
| AT1G16700.1 | NADH:ubiquinone oxidoreductase subunit (TYKY-2) | 20 | 0.74 | 0.236 | 1.000 | |
| AT1G16700.1 | NADH:ubiquinone oxidoreductase subunit (TYKY-2) | 21 | 1.11 | 0.240 | 1.000 | |
| AT1G16700.1 | NADH:ubiquinone oxidoreductase subunit (TYKY-2) | 22 | 1.85 | 0.238 | 0.991 | |
| AT1G16700.1 | NADH:ubiquinone oxidoreductase subunit (TYKY-2) | 23 | 2.33 | 0.238 | 0.881 | |
| AT1G16700.1 | NADH:ubiquinone oxidoreductase subunit (TYKY-2) | 24 | 1.35 | 0.229 | 1.000 | |
| AT1G16700.1 | NADH:ubiquinone oxidoreductase subunit (TYKY-2) | 25 | 0.39 | 0.226 | 0.712 | |
| AT1G16700.1 | NADH:ubiquinone oxidoreductase subunit (TYKY-2) | 26 | 0.21 | 0.225 | 0.051 | . |
| AT1G16700.1 | NADH:ubiquinone oxidoreductase subunit (TYKY-2) | 27 | 0.34 | 0.225 | 0.512 | |
| AT1G16700.1 | NADH:ubiquinone oxidoreductase subunit (TYKY-2) | 28 | 0.77 | 0.230 | 1.000 | |
| AT1G16700.1 | NADH:ubiquinone oxidoreductase subunit (TYKY-2) | 29 | 1.80 | 0.244 | 0.996 | |
| AT1G16700.1 | NADH:ubiquinone oxidoreductase subunit (TYKY-2) | 30 | 3.33 | 0.283 | 0.679 | |
| AT1G16700.1 | NADH:ubiquinone oxidoreductase subunit (TYKY-2) | 31 | 3.86 | 0.310 | 0.643 | |

| | | | | | |
|---|---|---|---|---|---|
| AT1G16700.1 | NADH:ubiquinone oxidoreductase subunit (TYKY-2) | 32 | 3.00 | 0.324 | 0.913 |
| AT1G16700.1 | NADH:ubiquinone oxidoreductase subunit (TYKY-2) | 33 | 1.85 | 0.358 | 1.000 |
| AT1G16700.1 | NADH:ubiquinone oxidoreductase subunit (TYKY-2) | 34 | 1.14 | 0.541 | 1.000 |
| AT1G16700.1 | NADH:ubiquinone oxidoreductase subunit (TYKY-2) | 46 | 1.74 | 5.020 | 1.000 |
| AT1G16700.1 | NADH:ubiquinone oxidoreductase subunit (TYKY-2) | 47 | 2.34 | 5.597 | 1.000 |
| AT1G16870.1 | Ribosomal S29 family protein | 2 | 1.01 | 0.789 | 1.000 |
| AT1G16870.1 | Ribosomal S29 family protein | 3 | 0.96 | 0.683 | 1.000 |
| AT1G16870.1 | Ribosomal S29 family protein | 10 | 1.37 | 0.439 | 1.000 |
| AT1G16870.1 | Ribosomal S29 family protein | 11 | 1.67 | 0.385 | 1.000 |
| AT1G16870.1 | Ribosomal S29 family protein | 12 | 1.94 | 0.347 | 0.999 |
| AT1G16870.1 | Ribosomal S29 family protein | 13 | 2.00 | 0.323 | 0.998 |
| AT1G16870.1 | Ribosomal S29 family protein | 14 | 1.99 | 0.308 | 0.996 |
| AT1G16870.1 | Ribosomal S29 family protein | 15 | 2.16 | 0.301 | 0.986 |
| AT1G16870.1 | Ribosomal S29 family protein | 16 | 2.57 | 0.299 | 0.925 |
| AT1G16870.1 | Ribosomal S29 family protein | 17 | 3.03 | 0.299 | 0.802 |
| AT1G16870.1 | Ribosomal S29 family protein | 18 | 3.12 | 0.302 | 0.782 |
| AT1G16870.1 | Ribosomal S29 family protein | 19 | 2.70 | 0.307 | 0.913 |
| AT1G16870.1 | Ribosomal S29 family protein | 20 | 1.95 | 0.314 | 0.998 |
| AT1G16870.1 | Ribosomal S29 family protein | 21 | 1.22 | 0.322 | 1.000 |
| AT1G16870.1 | Ribosomal S29 family protein | 22 | 0.75 | 0.331 | 1.000 |
| AT1G16870.1 | Ribosomal S29 family protein | 23 | 0.50 | 0.339 | 0.999 |
| AT1G16870.1 | Ribosomal S29 family protein | 24 | 0.38 | 0.345 | 0.967 |
| AT1G16870.1 | Ribosomal S29 family protein | 25 | 0.32 | 0.343 | 0.889 |
| AT1G16870.1 | Ribosomal S29 family protein | 26 | 0.29 | 0.336 | 0.819 |
| AT1G16870.1 | Ribosomal S29 family protein | 27 | 0.30 | 0.326 | 0.799 |
| AT1G16870.1 | Ribosomal S29 family protein | 28 | 0.34 | 0.317 | 0.875 |
| AT1G16870.1 | Ribosomal S29 family protein | 29 | 0.42 | 0.310 | 0.971 |
| AT1G16870.1 | Ribosomal S29 family protein | 30 | 0.53 | 0.306 | 0.998 |
| AT1G16870.1 | Ribosomal S29 family protein | 31 | 0.64 | 0.305 | 1.000 |
| AT1G16870.1 | Ribosomal S29 family protein | 32 | 0.74 | 0.305 | 1.000 |
| AT1G16870.1 | Ribosomal S29 family protein | 33 | 0.78 | 0.306 | 1.000 |
| AT1G16870.1 | Ribosomal S29 family protein | 34 | 0.75 | 0.309 | 1.000 |
| AT1G16870.1 | Ribosomal S29 family protein | 35 | 0.69 | 0.315 | 1.000 |
| AT1G16870.1 | Ribosomal S29 family protein | 36 | 0.67 | 0.326 | 1.000 |
| AT1G16870.1 | Ribosomal S29 family protein | 37 | 0.69 | 0.340 | 1.000 |
| AT1G16870.1 | Ribosomal S29 family protein | 38 | 0.78 | 0.350 | 1.000 |
| AT1G16870.1 | Ribosomal S29 family protein | 39 | 0.96 | 0.347 | 1.000 |
| AT1G16870.1 | Ribosomal S29 family protein | 40 | 1.21 | 0.338 | 1.000 |
| AT1G16870.1 | Ribosomal S29 family protein | 41 | 1.46 | 0.326 | 1.000 |
| AT1G16870.1 | Ribosomal S29 family protein | 42 | 1.60 | 0.318 | 1.000 |
| AT1G16870.1 | Ribosomal S29 family protein | 43 | 1.58 | 0.314 | 1.000 |
| AT1G16870.1 | Ribosomal S29 family protein | 44 | 1.41 | 0.314 | 1.000 |
| AT1G16870.1 | Ribosomal S29 family protein | 45 | 1.13 | 0.331 | 1.000 |
| AT1G16870.1 | Ribosomal S29 family protein | 46 | 0.85 | 0.401 | 1.000 |
| AT1G16870.1 | Ribosomal S29 family protein | 47 | 0.62 | 0.555 | 1.000 |
| AT1G17290.1 | Alanine aminotransferase (ALAAT1) | 2 | 0.60 | 0.255 | 0.893 |
| AT1G17290.1 | Alanine aminotransferase (ALAAT1) | 3 | 0.64 | 0.238 | 0.929 |
| AT1G17290.1 | Alanine aminotransferase (ALAAT1) | 4 | 0.70 | 0.224 | 0.962 |

| | | | | | | |
|---|---|---|---|---|---|---|
| AT1G17290.1 | Alanine aminotransferase (ALAAT1) | 6 | 0.82 | 0.203 | 0.998 | |
| AT1G17290.1 | Alanine aminotransferase (ALAAT1) | 7 | 0.89 | 0.195 | 1.000 | |
| AT1G17290.1 | Alanine aminotransferase (ALAAT1) | 8 | 0.96 | 0.188 | 1.000 | |
| AT1G17290.1 | Alanine aminotransferase (ALAAT1) | 10 | 1.13 | 0.179 | 1.000 | |
| AT1G17290.1 | Alanine aminotransferase (ALAAT1) | 11 | 1.22 | 0.175 | 0.995 | |
| AT1G17290.1 | Alanine aminotransferase (ALAAT1) | 12 | 1.31 | 0.173 | 0.968 | |
| AT1G17290.1 | Alanine aminotransferase (ALAAT1) | 13 | 1.40 | 0.171 | 0.908 | |
| AT1G17290.1 | Alanine aminotransferase (ALAAT1) | 14 | 1.48 | 0.169 | 0.825 | |
| AT1G17290.1 | Alanine aminotransferase (ALAAT1) | 15 | 1.56 | 0.169 | 0.738 | |
| AT1G17290.1 | Alanine aminotransferase (ALAAT1) | 16 | 1.63 | 0.168 | 0.662 | |
| AT1G17290.1 | Alanine aminotransferase (ALAAT1) | 20 | 1.82 | 0.168 | 0.458 | |
| AT1G17290.1 | Alanine aminotransferase (ALAAT1) | 21 | 1.86 | 0.167 | 0.422 | |
| AT1G17290.1 | Alanine aminotransferase (ALAAT1) | 22 | 1.89 | 0.167 | 0.394 | |
| AT1G17290.1 | Alanine aminotransferase (ALAAT1) | 23 | 1.91 | 0.166 | 0.370 | |
| AT1G17290.1 | Alanine aminotransferase (ALAAT1) | 24 | 1.93 | 0.165 | 0.354 | |
| AT1G17290.1 | Alanine aminotransferase (ALAAT1) | 25 | 1.93 | 0.165 | 0.344 | |
| AT1G17290.1 | Alanine aminotransferase (ALAAT1) | 26 | 1.93 | 0.164 | 0.340 | |
| AT1G17290.1 | Alanine aminotransferase (ALAAT1) | 29 | 1.89 | 0.160 | 0.350 | |
| AT1G17290.1 | Alanine aminotransferase (ALAAT1) | 30 | 1.87 | 0.158 | 0.355 | |
| AT1G17290.1 | Alanine aminotransferase (ALAAT1) | 31 | 1.86 | 0.157 | 0.357 | |
| AT1G17290.1 | Alanine aminotransferase (ALAAT1) | 32 | 1.85 | 0.156 | 0.358 | |
| AT1G17290.1 | Alanine aminotransferase (ALAAT1) | 33 | 1.85 | 0.155 | 0.354 | |
| AT1G17290.1 | Alanine aminotransferase (ALAAT1) | 34 | 1.86 | 0.155 | 0.342 | |
| AT1G17290.1 | Alanine aminotransferase (ALAAT1) | 35 | 1.89 | 0.155 | 0.323 | |
| AT1G17290.1 | Alanine aminotransferase (ALAAT1) | 36 | 1.93 | 0.155 | 0.292 | |
| AT1G17290.1 | Alanine aminotransferase (ALAAT1) | 37 | 1.99 | 0.156 | 0.251 | |
| AT1G17290.1 | Alanine aminotransferase (ALAAT1) | 38 | 2.08 | 0.157 | 0.203 | |
| AT1G17290.1 | Alanine aminotransferase (ALAAT1) | 39 | 2.19 | 0.157 | 0.154 | |
| AT1G17290.1 | Alanine aminotransferase (ALAAT1) | 40 | 2.34 | 0.159 | 0.110 | |
| AT1G17290.1 | Alanine aminotransferase (ALAAT1) | 41 | 2.52 | 0.160 | 0.072 | . |
| AT1G17290.1 | Alanine aminotransferase (ALAAT1) | 42 | 2.74 | 0.163 | 0.046 | * |

| | | | | | | |
|---|---|---|---|---|---|---|
| AT1G17290.1 | Alanine aminotransferase (ALAAT1) | 43 | 2.99 | 0.166 | 0.029 | * |
| AT1G17290.1 | Alanine aminotransferase (ALAAT1) | 44 | 3.27 | 0.172 | 0.020 | * |
| AT1G17290.1 | Alanine aminotransferase (ALAAT1) | 45 | 3.59 | 0.180 | 0.015 | * |
| AT1G17290.1 | Alanine aminotransferase (ALAAT1) | 46 | 3.94 | 0.191 | 0.014 | * |
| AT1G17290.1 | Alanine aminotransferase (ALAAT1) | 47 | 4.33 | 0.205 | 0.014 | * |
| AT1G17350.1 | NADH:ubiquinone oxidoreductase intermediate-associated protein 30 | 2 | 1.15 | 0.580 | 1.000 | |
| AT1G17350.1 | NADH:ubiquinone oxidoreductase intermediate-associated protein 30 | 3 | 1.18 | 0.555 | 1.000 | |
| AT1G17350.1 | NADH:ubiquinone oxidoreductase intermediate-associated protein 30 | 25 | 1.21 | 0.210 | 0.989 | |
| AT1G17350.1 | NADH:ubiquinone oxidoreductase intermediate-associated protein 30 | 26 | 1.15 | 0.201 | 0.996 | |
| AT1G17350.1 | NADH:ubiquinone oxidoreductase intermediate-associated protein 30 | 27 | 1.09 | 0.197 | 1.000 | |
| AT1G17350.1 | NADH:ubiquinone oxidoreductase intermediate-associated protein 30 | 28 | 1.01 | 0.198 | 1.000 | |
| AT1G17350.1 | NADH:ubiquinone oxidoreductase intermediate-associated protein 30 | 29 | 0.93 | 0.205 | 1.000 | |
| AT1G17350.1 | NADH:ubiquinone oxidoreductase intermediate-associated protein 30 | 30 | 0.83 | 0.216 | 0.992 | |
| AT1G17350.1 | NADH:ubiquinone oxidoreductase intermediate-associated protein 30 | 31 | 0.74 | 0.231 | 0.959 | |
| AT1G17350.1 | NADH:ubiquinone oxidoreductase intermediate-associated protein 30 | 46 | 0.06 | 0.435 | 0.026 | * |
| AT1G17350.1 | NADH:ubiquinone oxidoreductase intermediate-associated protein 30 | 47 | 0.05 | 0.457 | 0.023 | * |
| AT1G18480.1 | Shewenella-like protein posphatase 2 (SLP2) | 2 | 1.00 | 0.366 | 1.000 | |
| AT1G18480.1 | Shewenella-like protein posphatase 2 (SLP2) | 3 | 0.91 | 0.351 | 1.000 | |
| AT1G18480.1 | Shewenella-like protein posphatase 2 (SLP2) | 27 | 0.78 | 0.315 | 1.000 | |
| AT1G18480.1 | Shewenella-like protein posphatase 2 (SLP2) | 28 | 0.92 | 0.257 | 1.000 | |
| AT1G18480.1 | Shewenella-like protein posphatase 2 (SLP2) | 29 | 1.06 | 0.249 | 1.000 | |
| AT1G18480.1 | Shewenella-like protein posphatase 2 (SLP2) | 31 | 1.23 | 0.209 | 1.000 | |
| AT1G18480.1 | Shewenella-like protein posphatase 2 (SLP2) | 32 | 1.24 | 0.189 | 1.000 | |
| AT1G18480.1 | Shewenella-like protein posphatase 2 (SLP2) | 33 | 1.19 | 0.176 | 1.000 | |
| AT1G18480.1 | Shewenella-like protein posphatase 2 (SLP2) | 34 | 1.14 | 0.170 | 1.000 | |

| | | | | | | |
|---|---|---|---|---|---|---|
| AT1G18480.1 | Shewenella-like protein posphatase 2 (SLP2) | 35 | 1.17 | 0.167 | 1.000 | |
| AT1G18480.1 | Shewenella-like protein posphatase 2 (SLP2) | 36 | 1.27 | 0.166 | 1.000 | |
| AT1G18480.1 | Shewenella-like protein posphatase 2 (SLP2) | 37 | 1.31 | 0.166 | 0.999 | |
| AT1G18480.1 | Shewenella-like protein posphatase 2 (SLP2) | 38 | 1.29 | 0.166 | 1.000 | |
| AT1G18480.1 | Shewenella-like protein posphatase 2 (SLP2) | 39 | 1.29 | 0.166 | 1.000 | |
| AT1G18480.1 | Shewenella-like protein posphatase 2 (SLP2) | 40 | 1.46 | 0.166 | 0.985 | |
| AT1G18480.1 | Shewenella-like protein posphatase 2 (SLP2) | 41 | 1.86 | 0.167 | 0.716 | |
| AT1G18480.1 | Shewenella-like protein posphatase 2 (SLP2) | 42 | 2.15 | 0.169 | 0.444 | |
| AT1G18480.1 | Shewenella-like protein posphatase 2 (SLP2) | 43 | 2.03 | 0.170 | 0.561 | |
| AT1G18480.1 | Shewenella-like protein posphatase 2 (SLP2) | 44 | 1.47 | 0.170 | 0.986 | |
| AT1G18480.1 | Shewenella-like protein posphatase 2 (SLP2) | 45 | 0.80 | 0.172 | 1.000 | |
| AT1G18480.1 | Shewenella-like protein posphatase 2 (SLP2) | 46 | 0.34 | 0.199 | 0.223 | |
| AT1G18480.1 | Shewenella-like protein posphatase 2 (SLP2) | 47 | 0.13 | 0.305 | 0.050 | * |
| AT1G18600.1 | Rhomboid-like protein 12 (RBL12) | 2 | 0.91 | 0.341 | 1.000 | |
| AT1G18600.1 | Rhomboid-like protein 12 (RBL12) | 3 | 0.83 | 0.305 | 1.000 | |
| AT1G18600.1 | Rhomboid-like protein 12 (RBL12) | 4 | 0.76 | 0.288 | 1.000 | |
| AT1G18600.1 | Rhomboid-like protein 12 (RBL12) | 12 | 0.57 | 0.189 | 0.925 | |
| AT1G18600.1 | Rhomboid-like protein 12 (RBL12) | 13 | 0.60 | 0.168 | 0.923 | |
| AT1G18600.1 | Rhomboid-like protein 12 (RBL12) | 14 | 0.65 | 0.152 | 0.955 | |
| AT1G18600.1 | Rhomboid-like protein 12 (RBL12) | 15 | 0.72 | 0.140 | 0.990 | |
| AT1G18600.1 | Rhomboid-like protein 12 (RBL12) | 16 | 0.82 | 0.132 | 1.000 | |
| AT1G18600.1 | Rhomboid-like protein 12 (RBL12) | 17 | 0.93 | 0.128 | 1.000 | |
| AT1G18600.1 | Rhomboid-like protein 12 (RBL12) | 18 | 1.06 | 0.127 | 1.000 | |
| AT1G18600.1 | Rhomboid-like protein 12 (RBL12) | 19 | 1.19 | 0.127 | 1.000 | |
| AT1G18600.1 | Rhomboid-like protein 12 (RBL12) | 20 | 1.30 | 0.129 | 0.996 | |
| AT1G18600.1 | Rhomboid-like protein 12 (RBL12) | 21 | 1.37 | 0.130 | 0.984 | |
| AT1G18600.1 | Rhomboid-like protein 12 (RBL12) | 22 | 1.38 | 0.130 | 0.983 | |
| AT1G18600.1 | Rhomboid-like protein 12 (RBL12) | 23 | 1.31 | 0.129 | 0.996 | |
| AT1G18600.1 | Rhomboid-like protein 12 (RBL12) | 24 | 1.20 | 0.127 | 1.000 | |
| AT1G18600.1 | Rhomboid-like protein 12 (RBL12) | 25 | 1.11 | 0.126 | 1.000 | |
| AT1G18600.1 | Rhomboid-like protein 12 (RBL12) | 26 | 1.10 | 0.124 | 1.000 | |

| | | | | | | |
|---|---|---|---|---|---|---|
| AT1G18600.1 | Rhomboid-like protein 12 (RBL12) | 27 | 1.16 | 0.123 | 1.000 | |
| AT1G18600.1 | Rhomboid-like protein 12 (RBL12) | 28 | 1.28 | 0.123 | 0.998 | |
| AT1G18600.1 | Rhomboid-like protein 12 (RBL12) | 29 | 1.40 | 0.123 | 0.959 | |
| AT1G18600.1 | Rhomboid-like protein 12 (RBL12) | 30 | 1.50 | 0.123 | 0.876 | |
| AT1G18600.1 | Rhomboid-like protein 12 (RBL12) | 31 | 1.52 | 0.124 | 0.850 | |
| AT1G18600.1 | Rhomboid-like protein 12 (RBL12) | 32 | 1.49 | 0.124 | 0.889 | |
| AT1G18600.1 | Rhomboid-like protein 12 (RBL12) | 33 | 1.44 | 0.124 | 0.940 | |
| AT1G18600.1 | Rhomboid-like protein 12 (RBL12) | 34 | 1.38 | 0.125 | 0.974 | |
| AT1G18600.1 | Rhomboid-like protein 12 (RBL12) | 35 | 1.35 | 0.125 | 0.987 | |
| AT1G18600.1 | Rhomboid-like protein 12 (RBL12) | 36 | 1.34 | 0.125 | 0.990 | |
| AT1G18600.1 | Rhomboid-like protein 12 (RBL12) | 37 | 1.32 | 0.126 | 0.993 | |
| AT1G18600.1 | Rhomboid-like protein 12 (RBL12) | 38 | 1.27 | 0.127 | 0.998 | |
| AT1G18600.1 | Rhomboid-like protein 12 (RBL12) | 39 | 1.18 | 0.129 | 1.000 | |
| AT1G18600.1 | Rhomboid-like protein 12 (RBL12) | 40 | 1.05 | 0.131 | 1.000 | |
| AT1G18600.1 | Rhomboid-like protein 12 (RBL12) | 41 | 0.89 | 0.134 | 1.000 | |
| AT1G18600.1 | Rhomboid-like protein 12 (RBL12) | 42 | 0.69 | 0.136 | 0.963 | |
| AT1G18600.1 | Rhomboid-like protein 12 (RBL12) | 43 | 0.49 | 0.140 | 0.342 | |
| AT1G18600.1 | Rhomboid-like protein 12 (RBL12) | 44 | 0.32 | 0.146 | 0.017 | * |
| AT1G18600.1 | Rhomboid-like protein 12 (RBL12) | 45 | 0.20 | 0.162 | 0.001 | *** |
| AT1G18600.1 | Rhomboid-like protein 12 (RBL12) | 46 | 0.12 | 0.193 | 0.000 | *** |
| AT1G18600.1 | Rhomboid-like protein 12 (RBL12) | 47 | 0.07 | 0.244 | 0.000 | *** |
| AT1G19140.1 | Ubiquinone biosynthesis COQ9-like protein | 2 | 0.82 | 0.371 | 1.000 | |
| AT1G19140.1 | Ubiquinone biosynthesis COQ9-like protein | 3 | 0.91 | 0.353 | 1.000 | |
| AT1G19140.1 | Ubiquinone biosynthesis COQ9-like protein | 26 | 1.53 | 0.278 | 1.000 | |
| AT1G19140.1 | Ubiquinone biosynthesis COQ9-like protein | 27 | 1.33 | 0.198 | 1.000 | |
| AT1G19140.1 | Ubiquinone biosynthesis COQ9-like protein | 28 | 1.07 | 0.173 | 1.000 | |
| AT1G19140.1 | Ubiquinone biosynthesis COQ9-like protein | 29 | 0.81 | 0.163 | 1.000 | |
| AT1G19140.1 | Ubiquinone biosynthesis COQ9-like protein | 30 | 0.68 | 0.161 | 0.994 | |
| AT1G19140.1 | Ubiquinone biosynthesis COQ9-like protein | 31 | 0.70 | 0.161 | 0.997 | |
| AT1G19140.1 | Ubiquinone biosynthesis COQ9-like protein | 32 | 0.71 | 0.161 | 0.997 | |
| AT1G19140.1 | Ubiquinone biosynthesis COQ9-like protein | 33 | 0.69 | 0.161 | 0.994 | |

| | | | | | |
|---|---|---|---|---|---|
| AT1G19140.1 | Ubiquinone biosynthesis COQ9-like protein | 34 | 0.72 | 0.162 | 0.999 |
| AT1G19140.1 | Ubiquinone biosynthesis COQ9-like protein | 35 | 0.77 | 0.162 | 1.000 |
| AT1G19140.1 | Ubiquinone biosynthesis COQ9-like protein | 36 | 0.77 | 0.161 | 1.000 |
| AT1G19140.1 | Ubiquinone biosynthesis COQ9-like protein | 37 | 0.81 | 0.160 | 1.000 |
| AT1G19140.1 | Ubiquinone biosynthesis COQ9-like protein | 38 | 1.01 | 0.160 | 1.000 |
| AT1G19140.1 | Ubiquinone biosynthesis COQ9-like protein | 39 | 1.34 | 0.160 | 1.000 |
| AT1G19140.1 | Ubiquinone biosynthesis COQ9-like protein | 40 | 1.54 | 0.160 | 0.979 |
| AT1G19140.1 | Ubiquinone biosynthesis COQ9-like protein | 41 | 1.45 | 0.161 | 0.995 |
| AT1G19140.1 | Ubiquinone biosynthesis COQ9-like protein | 42 | 1.36 | 0.164 | 0.999 |
| AT1G19140.1 | Ubiquinone biosynthesis COQ9-like protein | 43 | 1.32 | 0.165 | 1.000 |
| AT1G19140.1 | Ubiquinone biosynthesis COQ9-like protein | 44 | 1.32 | 0.165 | 1.000 |
| AT1G19140.1 | Ubiquinone biosynthesis COQ9-like protein | 45 | 1.36 | 0.165 | 0.999 |
| AT1G19140.1 | Ubiquinone biosynthesis COQ9-like protein | 46 | 1.25 | 0.196 | 1.000 |
| AT1G19140.1 | Ubiquinone biosynthesis COQ9-like protein | 47 | 0.97 | 0.342 | 1.000 |
| AT1G19240.1 | ? | 2 | 1.06 | 0.335 | 1.000 |
| AT1G19240.1 | ? | 3 | 0.97 | 0.301 | 1.000 |
| AT1G19240.1 | ? | 14 | 0.70 | 0.171 | 0.977 |
| AT1G19240.1 | ? | 15 | 0.75 | 0.148 | 0.986 |
| AT1G19240.1 | ? | 16 | 0.82 | 0.132 | 0.997 |
| AT1G19240.1 | ? | 17 | 0.89 | 0.122 | 1.000 |
| AT1G19240.1 | ? | 18 | 0.99 | 0.118 | 1.000 |
| AT1G19240.1 | ? | 19 | 1.12 | 0.118 | 1.000 |
| AT1G19240.1 | ? | 20 | 1.23 | 0.119 | 0.992 |
| AT1G19240.1 | ? | 21 | 1.31 | 0.120 | 0.959 |
| AT1G19240.1 | ? | 22 | 1.35 | 0.120 | 0.925 |
| AT1G19240.1 | ? | 23 | 1.40 | 0.119 | 0.877 |
| AT1G19240.1 | ? | 24 | 1.45 | 0.118 | 0.793 |
| AT1G19240.1 | ? | 25 | 1.52 | 0.118 | 0.677 |
| AT1G19240.1 | ? | 26 | 1.59 | 0.119 | 0.553 |
| AT1G19240.1 | ? | 27 | 1.66 | 0.124 | 0.508 |
| AT1G19240.1 | ? | 28 | 1.65 | 0.139 | 0.650 |
| AT1G19240.1 | ? | 29 | 1.57 | 0.168 | 0.896 |
| AT1G19240.1 | ? | 30 | 1.45 | 0.212 | 0.992 |
| AT1G19240.1 | ? | 46 | 0.42 | 1.755 | 1.000 |
| AT1G19240.1 | ? | 47 | 0.38 | 1.900 | 1.000 |
| AT1G19520.1 | Pentatricopeptide repeat (PPR) superfamily protein (NDF5) | 2 | 1.02 | 0.624 | 1.000 |
| AT1G19520.1 | Pentatricopeptide repeat (PPR) superfamily protein (NDF5) | 3 | 1.03 | 0.535 | 1.000 |
| AT1G19520.1 | Pentatricopeptide repeat (PPR) superfamily protein (NDF5) | 10 | 1.31 | 0.340 | 1.000 |
| AT1G19520.1 | Pentatricopeptide repeat (PPR) superfamily protein (NDF5) | 11 | 1.42 | 0.291 | 1.000 |
| AT1G19520.1 | Pentatricopeptide repeat (PPR) superfamily protein (NDF5) | 12 | 1.52 | 0.257 | 1.000 |
| AT1G19520.1 | Pentatricopeptide repeat (PPR) superfamily protein (NDF5) | 13 | 1.51 | 0.236 | 1.000 |

| | | | | | |
|---|---|---|---|---|---|
| AT1G19520.1 | Pentatricopeptide repeat (PPR) superfamily protein (NDF5) | 14 | 1.49 | 0.223 | 1.000 |
| AT1G19520.1 | Pentatricopeptide repeat (PPR) superfamily protein (NDF5) | 15 | 1.63 | 0.217 | 0.999 |
| AT1G19520.1 | Pentatricopeptide repeat (PPR) superfamily protein (NDF5) | 16 | 1.93 | 0.215 | 0.969 |
| AT1G19520.1 | Pentatricopeptide repeat (PPR) superfamily protein (NDF5) | 17 | 2.06 | 0.217 | 0.935 |
| AT1G19520.1 | Pentatricopeptide repeat (PPR) superfamily protein (NDF5) | 18 | 1.82 | 0.222 | 0.990 |
| AT1G19520.1 | Pentatricopeptide repeat (PPR) superfamily protein (NDF5) | 19 | 1.33 | 0.230 | 1.000 |
| AT1G19520.1 | Pentatricopeptide repeat (PPR) superfamily protein (NDF5) | 20 | 0.88 | 0.243 | 1.000 |
| AT1G19520.1 | Pentatricopeptide repeat (PPR) superfamily protein (NDF5) | 21 | 0.55 | 0.263 | 0.999 |
| AT1G19520.1 | Pentatricopeptide repeat (PPR) superfamily protein (NDF5) | 22 | 0.35 | 0.280 | 0.856 |
| AT1G19520.1 | Pentatricopeptide repeat (PPR) superfamily protein (NDF5) | 23 | 0.24 | 0.290 | 0.480 |
| AT1G19520.1 | Pentatricopeptide repeat (PPR) superfamily protein (NDF5) | 24 | 0.19 | 0.287 | 0.227 |
| AT1G19520.1 | Pentatricopeptide repeat (PPR) superfamily protein (NDF5) | 25 | 0.18 | 0.274 | 0.131 |
| AT1G19520.1 | Pentatricopeptide repeat (PPR) superfamily protein (NDF5) | 26 | 0.20 | 0.255 | 0.137 |
| AT1G19520.1 | Pentatricopeptide repeat (PPR) superfamily protein (NDF5) | 27 | 0.28 | 0.237 | 0.339 |
| AT1G19520.1 | Pentatricopeptide repeat (PPR) superfamily protein (NDF5) | 28 | 0.45 | 0.226 | 0.900 |
| AT1G19520.1 | Pentatricopeptide repeat (PPR) superfamily protein (NDF5) | 29 | 0.75 | 0.222 | 1.000 |
| AT1G19520.1 | Pentatricopeptide repeat (PPR) superfamily protein (NDF5) | 30 | 1.19 | 0.220 | 1.000 |
| AT1G19520.1 | Pentatricopeptide repeat (PPR) superfamily protein (NDF5) | 31 | 1.66 | 0.220 | 0.998 |
| AT1G19520.1 | Pentatricopeptide repeat (PPR) superfamily protein (NDF5) | 32 | 1.95 | 0.221 | 0.972 |
| AT1G19520.1 | Pentatricopeptide repeat (PPR) superfamily protein (NDF5) | 33 | 1.95 | 0.221 | 0.972 |
| AT1G19520.1 | Pentatricopeptide repeat (PPR) superfamily protein (NDF5) | 34 | 1.74 | 0.221 | 0.996 |
| AT1G19520.1 | Pentatricopeptide repeat (PPR) superfamily protein (NDF5) | 35 | 1.49 | 0.221 | 1.000 |
| AT1G19520.1 | Pentatricopeptide repeat (PPR) superfamily protein (NDF5) | 36 | 1.35 | 0.221 | 1.000 |
| AT1G19520.1 | Pentatricopeptide repeat (PPR) superfamily protein (NDF5) | 37 | 1.37 | 0.220 | 1.000 |
| AT1G19520.1 | Pentatricopeptide repeat (PPR) superfamily protein (NDF5) | 38 | 1.51 | 0.221 | 1.000 |
| AT1G19520.1 | Pentatricopeptide repeat (PPR) superfamily protein (NDF5) | 39 | 1.57 | 0.221 | 1.000 |
| AT1G19520.1 | Pentatricopeptide repeat (PPR) superfamily protein (NDF5) | 40 | 1.43 | 0.223 | 1.000 |
| AT1G19520.1 | Pentatricopeptide repeat (PPR) superfamily protein (NDF5) | 41 | 1.10 | 0.224 | 1.000 |
| AT1G19520.1 | Pentatricopeptide repeat (PPR) superfamily protein (NDF5) | 42 | 0.76 | 0.226 | 1.000 |
| AT1G19520.1 | Pentatricopeptide repeat (PPR) superfamily protein (NDF5) | 43 | 0.51 | 0.231 | 0.979 |
| AT1G19520.1 | Pentatricopeptide repeat (PPR) superfamily protein (NDF5) | 44 | 0.36 | 0.247 | 0.757 |

| | | | | | |
|---|---|---|---|---|---|
| AT1G19520.1 | Pentatricopeptide repeat (PPR) superfamily protein (NDF5) | 45 | 0.30 | 0.298 | 0.760 |
| AT1G19520.1 | Pentatricopeptide repeat (PPR) superfamily protein (NDF5) | 46 | 0.26 | 0.403 | 0.941 |
| AT1G19520.1 | Pentatricopeptide repeat (PPR) superfamily protein (NDF5) | 47 | 0.25 | 0.574 | 0.997 |
| AT1G19580.1 | NADH:ubiquinone oxidoreductase subunit (gammaCA1) | 2 | 1.09 | 0.533 | 1.000 |
| AT1G19580.1 | NADH:ubiquinone oxidoreductase subunit (gammaCA1) | 3 | 0.90 | 0.367 | 1.000 |
| AT1G19580.1 | NADH:ubiquinone oxidoreductase subunit (gammaCA1) | 4 | 0.85 | 0.416 | 1.000 |
| AT1G19580.1 | NADH:ubiquinone oxidoreductase subunit (gammaCA1) | 6 | 1.11 | 0.483 | 1.000 |
| AT1G19580.1 | NADH:ubiquinone oxidoreductase subunit (gammaCA1) | 7 | 1.45 | 0.450 | 1.000 |
| AT1G19580.1 | NADH:ubiquinone oxidoreductase subunit (gammaCA1) | 8 | 1.79 | 0.424 | 1.000 |
| AT1G19580.1 | NADH:ubiquinone oxidoreductase subunit (gammaCA1) | 11 | 0.79 | 0.355 | 1.000 |
| AT1G19580.1 | NADH:ubiquinone oxidoreductase subunit (gammaCA1) | 12 | 0.66 | 0.335 | 1.000 |
| AT1G19580.1 | NADH:ubiquinone oxidoreductase subunit (gammaCA1) | 13 | 0.98 | 0.313 | 1.000 |
| AT1G19580.1 | NADH:ubiquinone oxidoreductase subunit (gammaCA1) | 14 | 1.43 | 0.282 | 1.000 |
| AT1G19580.1 | NADH:ubiquinone oxidoreductase subunit (gammaCA1) | 15 | 1.38 | 0.268 | 1.000 |
| AT1G19580.1 | NADH:ubiquinone oxidoreductase subunit (gammaCA1) | 16 | 0.77 | 0.262 | 1.000 |
| AT1G19580.1 | NADH:ubiquinone oxidoreductase subunit (gammaCA1) | 17 | 0.46 | 0.261 | 0.997 |
| AT1G19580.1 | NADH:ubiquinone oxidoreductase subunit (gammaCA1) | 18 | 0.51 | 0.265 | 1.000 |
| AT1G19580.1 | NADH:ubiquinone oxidoreductase subunit (gammaCA1) | 19 | 0.75 | 0.272 | 1.000 |
| AT1G19580.1 | NADH:ubiquinone oxidoreductase subunit (gammaCA1) | 20 | 0.90 | 0.280 | 1.000 |
| AT1G19580.1 | NADH:ubiquinone oxidoreductase subunit (gammaCA1) | 21 | 1.05 | 0.283 | 1.000 |
| AT1G19580.1 | NADH:ubiquinone oxidoreductase subunit (gammaCA1) | 22 | 1.30 | 0.284 | 1.000 |

| | | | | | | |
|---|---|---|---|---|---|---|
| AT1G19580.1 | NADH:ubiquinone oxidoreductase subunit (gammaCA1) | 23 | 1.50 | 0.283 | 1.000 | |
| AT1G19580.1 | NADH:ubiquinone oxidoreductase subunit (gammaCA1) | 24 | 1.34 | 0.277 | 1.000 | |
| AT1G19580.1 | NADH:ubiquinone oxidoreductase subunit (gammaCA1) | 25 | 0.83 | 0.269 | 1.000 | |
| AT1G19580.1 | NADH:ubiquinone oxidoreductase subunit (gammaCA1) | 26 | 0.64 | 0.268 | 1.000 | |
| AT1G19580.1 | NADH:ubiquinone oxidoreductase subunit (gammaCA1) | 27 | 0.76 | 0.270 | 1.000 | |
| AT1G19580.1 | NADH:ubiquinone oxidoreductase subunit (gammaCA1) | 28 | 0.87 | 0.271 | 1.000 | |
| AT1G19580.1 | NADH:ubiquinone oxidoreductase subunit (gammaCA1) | 29 | 0.91 | 0.269 | 1.000 | |
| AT1G19580.1 | NADH:ubiquinone oxidoreductase subunit (gammaCA1) | 30 | 1.01 | 0.269 | 1.000 | |
| AT1G19580.1 | NADH:ubiquinone oxidoreductase subunit (gammaCA1) | 31 | 1.02 | 0.272 | 1.000 | |
| AT1G19580.1 | NADH:ubiquinone oxidoreductase subunit (gammaCA1) | 32 | 0.69 | 0.274 | 1.000 | |
| AT1G19580.1 | NADH:ubiquinone oxidoreductase subunit (gammaCA1) | 33 | 0.27 | 0.273 | 0.684 | |
| AT1G19580.1 | NADH:ubiquinone oxidoreductase subunit (gammaCA1) | 34 | 0.14 | 0.277 | 0.084 | . |
| AT1G19580.1 | NADH:ubiquinone oxidoreductase subunit (gammaCA1) | 35 | 0.23 | 0.285 | 0.521 | |
| AT1G19580.1 | NADH:ubiquinone oxidoreductase subunit (gammaCA1) | 36 | 0.81 | 0.294 | 1.000 | |
| AT1G19580.1 | NADH:ubiquinone oxidoreductase subunit (gammaCA1) | 37 | 2.30 | 0.303 | 0.999 | |
| AT1G19580.1 | NADH:ubiquinone oxidoreductase subunit (gammaCA1) | 38 | 2.05 | 0.301 | 1.000 | |
| AT1G19580.1 | NADH:ubiquinone oxidoreductase subunit (gammaCA1) | 39 | 0.76 | 0.281 | 1.000 | |
| AT1G19580.1 | NADH:ubiquinone oxidoreductase subunit (gammaCA1) | 40 | 0.51 | 0.272 | 1.000 | |
| AT1G19580.1 | NADH:ubiquinone oxidoreductase subunit (gammaCA1) | 41 | 0.93 | 0.274 | 1.000 | |
| AT1G19580.1 | NADH:ubiquinone oxidoreductase subunit (gammaCA1) | 42 | 2.00 | 0.291 | 1.000 | |

| | | | | | | |
|---|---|---|---|---|---|---|
| AT1G19580.1 | NADH:ubiquinone oxidoreductase subunit (gammaCA1) | 43 | 2.05 | 0.374 | 1.000 | |
| AT1G19580.1 | NADH:ubiquinone oxidoreductase subunit (gammaCA1) | 44 | 1.54 | 0.576 | 1.000 | |
| AT1G19580.1 | NADH:ubiquinone oxidoreductase subunit (gammaCA1) | 46 | 0.64 | 1.502 | 1.000 | |
| AT1G19580.1 | NADH:ubiquinone oxidoreductase subunit (gammaCA1) | 47 | 0.39 | 2.173 | 1.000 | |
| AT1G20430.1 | ? | 2 | 1.07 | 0.448 | 1.000 | |
| AT1G20430.1 | ? | 3 | 1.11 | 0.369 | 1.000 | |
| AT1G20430.1 | ? | 8 | 1.47 | 0.380 | 1.000 | |
| AT1G20430.1 | ? | 9 | 1.57 | 0.297 | 1.000 | |
| AT1G20430.1 | ? | 10 | 1.59 | 0.275 | 1.000 | |
| AT1G20430.1 | ? | 11 | 1.47 | 0.238 | 1.000 | |
| AT1G20430.1 | ? | 12 | 1.42 | 0.246 | 1.000 | |
| AT1G20430.1 | ? | 13 | 1.66 | 0.258 | 1.000 | |
| AT1G20430.1 | ? | 14 | 2.26 | 0.237 | 0.977 | |
| AT1G20430.1 | ? | 15 | 3.12 | 0.233 | 0.627 | |
| AT1G20430.1 | ? | 16 | 3.54 | 0.277 | 0.740 | |
| AT1G20430.1 | ? | 22 | 0.28 | 0.328 | 0.927 | |
| AT1G20430.1 | ? | 23 | 0.36 | 0.262 | 0.921 | |
| AT1G20430.1 | ? | 24 | 1.06 | 0.213 | 1.000 | |
| AT1G20430.1 | ? | 25 | 3.23 | 0.205 | 0.335 | |
| AT1G20430.1 | ? | 26 | 2.38 | 0.204 | 0.840 | |
| AT1G20430.1 | ? | 27 | 0.90 | 0.203 | 1.000 | |
| AT1G20430.1 | ? | 28 | 1.12 | 0.202 | 1.000 | |
| AT1G20430.1 | ? | 29 | 1.93 | 0.203 | 0.989 | |
| AT1G20430.1 | ? | 30 | 1.63 | 0.205 | 1.000 | |
| AT1G20430.1 | ? | 31 | 1.18 | 0.205 | 1.000 | |
| AT1G20430.1 | ? | 32 | 1.12 | 0.205 | 1.000 | |
| AT1G20430.1 | ? | 33 | 1.31 | 0.204 | 1.000 | |
| AT1G20430.1 | ? | 34 | 1.50 | 0.205 | 1.000 | |
| AT1G20430.1 | ? | 35 | 1.72 | 0.206 | 0.999 | |
| AT1G20430.1 | ? | 36 | 1.70 | 0.206 | 1.000 | |
| AT1G20430.1 | ? | 37 | 1.43 | 0.205 | 1.000 | |
| AT1G20430.1 | ? | 38 | 1.39 | 0.203 | 1.000 | |
| AT1G20430.1 | ? | 39 | 1.88 | 0.203 | 0.993 | |
| AT1G20430.1 | ? | 40 | 3.10 | 0.206 | 0.407 | |
| AT1G20430.1 | ? | 41 | 3.65 | 0.205 | 0.178 | |
| AT1G20430.1 | ? | 42 | 2.66 | 0.207 | 0.683 | |
| AT1G20430.1 | ? | 43 | 1.65 | 0.209 | 1.000 | |
| AT1G20430.1 | ? | 44 | 0.86 | 0.207 | 1.000 | |
| AT1G20430.1 | ? | 45 | 0.39 | 0.209 | 0.745 | |
| AT1G20430.1 | ? | 46 | 0.14 | 0.235 | 0.012 | * |
| AT1G20430.1 | ? | 47 | 0.04 | 0.448 | 0.059 | . |
| AT1G20620.1 | Catalase 3 (CAT3) | 2 | 0.70 | 0.244 | 1.000 | |
| AT1G20620.1 | Catalase 3 (CAT3) | 3 | 0.74 | 0.164 | 1.000 | |
| AT1G20620.1 | Catalase 3 (CAT3) | 4 | 0.83 | 0.141 | 1.000 | |
| AT1G20620.1 | Catalase 3 (CAT3) | 5 | 0.97 | 0.139 | 1.000 | |
| AT1G20620.1 | Catalase 3 (CAT3) | 6 | 1.09 | 0.140 | 1.000 | |
| AT1G20620.1 | Catalase 3 (CAT3) | 7 | 1.16 | 0.139 | 1.000 | |
| AT1G20620.1 | Catalase 3 (CAT3) | 8 | 1.20 | 0.137 | 1.000 | |
| AT1G20620.1 | Catalase 3 (CAT3) | 9 | 1.22 | 0.135 | 1.000 | |
| AT1G20620.1 | Catalase 3 (CAT3) | 10 | 1.21 | 0.135 | 1.000 | |
| AT1G20620.1 | Catalase 3 (CAT3) | 11 | 1.16 | 0.135 | 1.000 | |
| AT1G20620.1 | Catalase 3 (CAT3) | 12 | 1.06 | 0.135 | 1.000 | |
| AT1G20620.1 | Catalase 3 (CAT3) | 13 | 0.95 | 0.133 | 1.000 | |
| AT1G20620.1 | Catalase 3 (CAT3) | 14 | 0.86 | 0.132 | 1.000 | |
| AT1G20620.1 | Catalase 3 (CAT3) | 15 | 0.81 | 0.131 | 1.000 | |

| | | | | | |
|---|---|---|---|---|---|
| AT1G20620.1 | Catalase 3 (CAT3) | 16 | 0.83 | 0.131 | 1.000 |
| AT1G20620.1 | Catalase 3 (CAT3) | 17 | 0.91 | 0.131 | 1.000 |
| AT1G20620.1 | Catalase 3 (CAT3) | 18 | 1.02 | 0.132 | 1.000 |
| AT1G20620.1 | Catalase 3 (CAT3) | 19 | 1.14 | 0.135 | 1.000 |
| AT1G20620.1 | Catalase 3 (CAT3) | 20 | 1.19 | 0.138 | 1.000 |
| AT1G20620.1 | Catalase 3 (CAT3) | 21 | 1.17 | 0.140 | 1.000 |
| AT1G20620.1 | Catalase 3 (CAT3) | 22 | 1.08 | 0.140 | 1.000 |
| AT1G20620.1 | Catalase 3 (CAT3) | 23 | 0.95 | 0.139 | 1.000 |
| AT1G20620.1 | Catalase 3 (CAT3) | 24 | 0.84 | 0.136 | 1.000 |
| AT1G20620.1 | Catalase 3 (CAT3) | 25 | 0.76 | 0.134 | 1.000 |
| AT1G20620.1 | Catalase 3 (CAT3) | 26 | 0.76 | 0.133 | 1.000 |
| AT1G20620.1 | Catalase 3 (CAT3) | 27 | 0.83 | 0.133 | 1.000 |
| AT1G20620.1 | Catalase 3 (CAT3) | 28 | 0.94 | 0.133 | 1.000 |
| AT1G20620.1 | Catalase 3 (CAT3) | 29 | 1.06 | 0.134 | 1.000 |
| AT1G20620.1 | Catalase 3 (CAT3) | 30 | 1.27 | 0.135 | 1.000 |
| AT1G20620.1 | Catalase 3 (CAT3) | 31 | 1.63 | 0.135 | 0.898 |
| AT1G20620.1 | Catalase 3 (CAT3) | 32 | 2.06 | 0.136 | 0.369 |
| AT1G20620.1 | Catalase 3 (CAT3) | 33 | 2.24 | 0.137 | 0.218 |
| AT1G20620.1 | Catalase 3 (CAT3) | 34 | 2.02 | 0.140 | 0.459 |
| AT1G20620.1 | Catalase 3 (CAT3) | 35 | 1.63 | 0.153 | 0.958 |
| AT1G20620.1 | Catalase 3 (CAT3) | 36 | 1.48 | 0.191 | 1.000 |
| AT1G20620.1 | Catalase 3 (CAT3) | 37 | 1.60 | 0.266 | 1.000 |
| AT1G20620.1 | Catalase 3 (CAT3) | 46 | 671.52 | 1.470 | 0.662 |
| AT1G20620.1 | Catalase 3 (CAT3) | 47 | 1752.76 | 1.672 | 0.650 |
| AT1G20630.1 | Catalase 1 (CAT1) | 2 | 1.33 | 0.675 | 1.000 |
| AT1G20630.1 | Catalase 1 (CAT1) | 3 | 1.13 | 0.642 | 1.000 |
| AT1G20630.1 | Catalase 1 (CAT1) | 27 | 0.24 | 0.313 | 0.248 |
| AT1G20630.1 | Catalase 1 (CAT1) | 28 | 0.28 | 0.268 | 0.199 |
| AT1G20630.1 | Catalase 1 (CAT1) | 29 | 0.32 | 0.233 | 0.188 |
| AT1G20630.1 | Catalase 1 (CAT1) | 30 | 0.38 | 0.209 | 0.239 |
| AT1G20630.1 | Catalase 1 (CAT1) | 31 | 0.47 | 0.196 | 0.411 |
| AT1G20630.1 | Catalase 1 (CAT1) | 32 | 0.59 | 0.194 | 0.732 |
| AT1G20630.1 | Catalase 1 (CAT1) | 33 | 0.74 | 0.205 | 0.973 |
| AT1G20630.1 | Catalase 1 (CAT1) | 34 | 0.92 | 0.229 | 1.000 |
| AT1G20630.1 | Catalase 1 (CAT1) | 35 | 1.15 | 0.267 | 1.000 |
| AT1G20630.1 | Catalase 1 (CAT1) | 36 | 1.43 | 0.320 | 0.993 |
| AT1G20630.1 | Catalase 1 (CAT1) | 46 | 9.57 | 1.402 | 0.962 |
| AT1G20630.1 | Catalase 1 (CAT1) | 47 | 11.44 | 1.552 | 0.966 |
| AT1G21400.1 | Branched chain keto acid dehydrogenase subunit (BCKDHE1alpha) | 2 | 1.06 | 0.575 | 1.000 |
| AT1G21400.1 | Branched chain keto acid dehydrogenase subunit (BCKDHE1alpha) | 3 | 0.80 | 0.561 | 1.000 |
| AT1G21400.1 | Branched chain keto acid dehydrogenase subunit (BCKDHE1alpha) | 31 | 0.86 | 0.346 | 1.000 |
| AT1G21400.1 | Branched chain keto acid dehydrogenase subunit (BCKDHE1alpha) | 32 | 1.40 | 0.256 | 0.999 |
| AT1G21400.1 | Branched chain keto acid dehydrogenase subunit (BCKDHE1alpha) | 33 | 1.95 | 0.222 | 0.842 |
| AT1G21400.1 | Branched chain keto acid dehydrogenase subunit (BCKDHE1alpha) | 34 | 2.13 | 0.210 | 0.672 |
| AT1G21400.1 | Branched chain keto acid dehydrogenase subunit (BCKDHE1alpha) | 35 | 2.11 | 0.202 | 0.645 |
| AT1G21400.1 | Branched chain keto acid dehydrogenase subunit (BCKDHE1alpha) | 36 | 2.20 | 0.198 | 0.544 |

| | | | | | | |
|---|---|---|---|---|---|---|
| AT1G21400.1 | Branched chain keto acid dehydrogenase subunit (BCKDHE1alpha) | 37 | 2.56 | 0.195 | 0.306 | |
| AT1G21400.1 | Branched chain keto acid dehydrogenase subunit (BCKDHE1alpha) | 38 | 3.16 | 0.195 | 0.112 | |
| AT1G21400.1 | Branched chain keto acid dehydrogenase subunit (BCKDHE1alpha) | 39 | 3.31 | 0.198 | 0.099 | . |
| AT1G21400.1 | Branched chain keto acid dehydrogenase subunit (BCKDHE1alpha) | 40 | 2.20 | 0.208 | 0.614 | |
| AT1G21400.1 | Branched chain keto acid dehydrogenase subunit (BCKDHE1alpha) | 41 | 0.84 | 0.240 | 1.000 | |
| AT1G21400.1 | Branched chain keto acid dehydrogenase subunit (BCKDHE1alpha) | 42 | 0.24 | 0.307 | 0.364 | |
| AT1G21400.1 | Branched chain keto acid dehydrogenase subunit (BCKDHE1alpha) | 43 | 0.07 | 0.375 | 0.033 | * |
| AT1G21400.1 | Branched chain keto acid dehydrogenase subunit (BCKDHE1alpha) | 46 | 0.00 | 0.483 | 0.000 | *** |
| AT1G21400.1 | Branched chain keto acid dehydrogenase subunit (BCKDHE1alpha) | 47 | 0.00 | 0.578 | 0.000 | *** |
| AT1G22450.1 | Cytochrome c:O2 oxidoreductase subunit (COX6B-2) | 2 | 1.04 | 0.340 | 1.000 | |
| AT1G22450.1 | Cytochrome c:O2 oxidoreductase subunit (COX6B-2) | 3 | 1.05 | 0.209 | 1.000 | |
| AT1G22450.1 | Cytochrome c:O2 oxidoreductase subunit (COX6B-2) | 4 | 0.99 | 0.203 | 1.000 | |
| AT1G22450.1 | Cytochrome c:O2 oxidoreductase subunit (COX6B-2) | 5 | 0.92 | 0.202 | 1.000 | |
| AT1G22450.1 | Cytochrome c:O2 oxidoreductase subunit (COX6B-2) | 6 | 0.92 | 0.203 | 1.000 | |
| AT1G22450.1 | Cytochrome c:O2 oxidoreductase subunit (COX6B-2) | 7 | 0.78 | 0.204 | 1.000 | |
| AT1G22450.1 | Cytochrome c:O2 oxidoreductase subunit (COX6B-2) | 8 | 0.82 | 0.200 | 1.000 | |
| AT1G22450.1 | Cytochrome c:O2 oxidoreductase subunit (COX6B-2) | 9 | 1.39 | 0.196 | 1.000 | |
| AT1G22450.1 | Cytochrome c:O2 oxidoreductase subunit (COX6B-2) | 10 | 2.03 | 0.196 | 0.971 | |
| AT1G22450.1 | Cytochrome c:O2 oxidoreductase subunit (COX6B-2) | 11 | 1.87 | 0.197 | 0.994 | |
| AT1G22450.1 | Cytochrome c:O2 oxidoreductase subunit (COX6B-2) | 12 | 1.39 | 0.197 | 1.000 | |

| | | | | | |
|---|---|---|---|---|---|
| AT1G22450.1 | Cytochrome c:O2 oxidoreductase subunit (COX6B-2) | 13 | 1.12 | 0.195 | 1.000 |
| AT1G22450.1 | Cytochrome c:O2 oxidoreductase subunit (COX6B-2) | 14 | 0.95 | 0.192 | 1.000 |
| AT1G22450.1 | Cytochrome c:O2 oxidoreductase subunit (COX6B-2) | 15 | 0.74 | 0.193 | 1.000 |
| AT1G22450.1 | Cytochrome c:O2 oxidoreductase subunit (COX6B-2) | 16 | 0.57 | 0.191 | 0.999 |
| AT1G22450.1 | Cytochrome c:O2 oxidoreductase subunit (COX6B-2) | 17 | 0.50 | 0.191 | 0.972 |
| AT1G22450.1 | Cytochrome c:O2 oxidoreductase subunit (COX6B-2) | 18 | 0.56 | 0.193 | 0.998 |
| AT1G22450.1 | Cytochrome c:O2 oxidoreductase subunit (COX6B-2) | 19 | 0.90 | 0.198 | 1.000 |
| AT1G22450.1 | Cytochrome c:O2 oxidoreductase subunit (COX6B-2) | 20 | 1.91 | 0.203 | 0.994 |
| AT1G22450.1 | Cytochrome c:O2 oxidoreductase subunit (COX6B-2) | 21 | 3.94 | 0.210 | 0.158 |
| AT1G22450.1 | Cytochrome c:O2 oxidoreductase subunit (COX6B-2) | 22 | 4.28 | 0.219 | 0.137 |
| AT1G22450.1 | Cytochrome c:O2 oxidoreductase subunit (COX6B-2) | 23 | 2.83 | 0.231 | 0.782 |
| AT1G22450.1 | Cytochrome c:O2 oxidoreductase subunit (COX6B-2) | 24 | 1.31 | 0.247 | 1.000 |
| AT1G22450.1 | Cytochrome c:O2 oxidoreductase subunit (COX6B-2) | 25 | 0.63 | 0.239 | 1.000 |
| AT1G22450.1 | Cytochrome c:O2 oxidoreductase subunit (COX6B-2) | 26 | 0.60 | 0.221 | 1.000 |
| AT1G22450.1 | Cytochrome c:O2 oxidoreductase subunit (COX6B-2) | 27 | 1.06 | 0.206 | 1.000 |
| AT1G22450.1 | Cytochrome c:O2 oxidoreductase subunit (COX6B-2) | 28 | 1.67 | 0.197 | 1.000 |
| AT1G22450.1 | Cytochrome c:O2 oxidoreductase subunit (COX6B-2) | 29 | 2.22 | 0.196 | 0.902 |
| AT1G22450.1 | Cytochrome c:O2 oxidoreductase subunit (COX6B-2) | 30 | 1.85 | 0.195 | 0.995 |
| AT1G22450.1 | Cytochrome c:O2 oxidoreductase subunit (COX6B-2) | 31 | 0.91 | 0.196 | 1.000 |
| AT1G22450.1 | Cytochrome c:O2 oxidoreductase subunit (COX6B-2) | 32 | 0.53 | 0.196 | 0.991 |

| | | | | | |
|---|---|---|---|---|---|
| AT1G22450.1 | Cytochrome c:O2 oxidoreductase subunit (COX6B-2) | 33 | 0.53 | 0.196 | 0.992 |
| AT1G22450.1 | Cytochrome c:O2 oxidoreductase subunit (COX6B-2) | 34 | 0.65 | 0.197 | 1.000 |
| AT1G22450.1 | Cytochrome c:O2 oxidoreductase subunit (COX6B-2) | 35 | 0.80 | 0.196 | 1.000 |
| AT1G22450.1 | Cytochrome c:O2 oxidoreductase subunit (COX6B-2) | 36 | 1.12 | 0.195 | 1.000 |
| AT1G22450.1 | Cytochrome c:O2 oxidoreductase subunit (COX6B-2) | 37 | 1.67 | 0.195 | 1.000 |
| AT1G22450.1 | Cytochrome c:O2 oxidoreductase subunit (COX6B-2) | 38 | 2.12 | 0.196 | 0.944 |
| AT1G22450.1 | Cytochrome c:O2 oxidoreductase subunit (COX6B-2) | 39 | 2.17 | 0.197 | 0.930 |
| AT1G22450.1 | Cytochrome c:O2 oxidoreductase subunit (COX6B-2) | 40 | 1.96 | 0.198 | 0.986 |
| AT1G22450.1 | Cytochrome c:O2 oxidoreductase subunit (COX6B-2) | 41 | 1.58 | 0.198 | 1.000 |
| AT1G22450.1 | Cytochrome c:O2 oxidoreductase subunit (COX6B-2) | 42 | 1.34 | 0.201 | 1.000 |
| AT1G22450.1 | Cytochrome c:O2 oxidoreductase subunit (COX6B-2) | 43 | 1.31 | 0.202 | 1.000 |
| AT1G22450.1 | Cytochrome c:O2 oxidoreductase subunit (COX6B-2) | 44 | 1.21 | 0.202 | 1.000 |
| AT1G22450.1 | Cytochrome c:O2 oxidoreductase subunit (COX6B-2) | 45 | 1.00 | 0.202 | 1.000 |
| AT1G22450.1 | Cytochrome c:O2 oxidoreductase subunit (COX6B-2) | 46 | 0.63 | 0.220 | 1.000 |
| AT1G22450.1 | Cytochrome c:O2 oxidoreductase subunit (COX6B-2) | 47 | 0.27 | 0.415 | 0.996 |
| AT1G22520.1 | MICOS complex subunit (Mic10-like) | 2 | 1.04 | 0.249 | 1.000 |
| AT1G22520.1 | MICOS complex subunit (Mic10-like) | 3 | 0.82 | 0.165 | 1.000 |
| AT1G22520.1 | MICOS complex subunit (Mic10-like) | 4 | 0.68 | 0.145 | 0.999 |
| AT1G22520.1 | MICOS complex subunit (Mic10-like) | 5 | 0.59 | 0.142 | 0.943 |
| AT1G22520.1 | MICOS complex subunit (Mic10-like) | 6 | 0.51 | 0.136 | 0.632 |
| AT1G22520.1 | MICOS complex subunit (Mic10-like) | 7 | 0.47 | 0.133 | 0.346 |
| AT1G22520.1 | MICOS complex subunit (Mic10-like) | 8 | 0.49 | 0.129 | 0.420 |
| AT1G22520.1 | MICOS complex subunit (Mic10-like) | 9 | 0.57 | 0.126 | 0.761 |

| | | | | | |
|---|---|---|---|---|---|
| AT1G22520.1 | MICOS complex subunit (Mic10-like) | 10 | 0.66 | 0.126 | 0.987 |
| AT1G22520.1 | MICOS complex subunit (Mic10-like) | 11 | 0.82 | 0.127 | 1.000 |
| AT1G22520.1 | MICOS complex subunit (Mic10-like) | 12 | 1.03 | 0.127 | 1.000 |
| AT1G22520.1 | MICOS complex subunit (Mic10-like) | 13 | 1.21 | 0.125 | 1.000 |
| AT1G22520.1 | MICOS complex subunit (Mic10-like) | 14 | 1.36 | 0.123 | 1.000 |
| AT1G22520.1 | MICOS complex subunit (Mic10-like) | 15 | 1.39 | 0.122 | 0.999 |
| AT1G22520.1 | MICOS complex subunit (Mic10-like) | 16 | 1.32 | 0.122 | 1.000 |
| AT1G22520.1 | MICOS complex subunit (Mic10-like) | 17 | 1.27 | 0.121 | 1.000 |
| AT1G22520.1 | MICOS complex subunit (Mic10-like) | 18 | 1.26 | 0.123 | 1.000 |
| AT1G22520.1 | MICOS complex subunit (Mic10-like) | 19 | 1.38 | 0.127 | 1.000 |
| AT1G22520.1 | MICOS complex subunit (Mic10-like) | 20 | 1.53 | 0.130 | 0.988 |
| AT1G22520.1 | MICOS complex subunit (Mic10-like) | 21 | 1.56 | 0.131 | 0.980 |
| AT1G22520.1 | MICOS complex subunit (Mic10-like) | 22 | 1.49 | 0.131 | 0.994 |
| AT1G22520.1 | MICOS complex subunit (Mic10-like) | 23 | 1.42 | 0.130 | 0.999 |
| AT1G22520.1 | MICOS complex subunit (Mic10-like) | 24 | 1.34 | 0.127 | 1.000 |
| AT1G22520.1 | MICOS complex subunit (Mic10-like) | 25 | 1.26 | 0.125 | 1.000 |
| AT1G22520.1 | MICOS complex subunit (Mic10-like) | 26 | 1.29 | 0.125 | 1.000 |
| AT1G22520.1 | MICOS complex subunit (Mic10-like) | 27 | 1.39 | 0.124 | 0.999 |
| AT1G22520.1 | MICOS complex subunit (Mic10-like) | 28 | 1.42 | 0.125 | 0.998 |
| AT1G22520.1 | MICOS complex subunit (Mic10-like) | 29 | 1.40 | 0.125 | 0.999 |
| AT1G22520.1 | MICOS complex subunit (Mic10-like) | 30 | 1.33 | 0.126 | 1.000 |
| AT1G22520.1 | MICOS complex subunit (Mic10-like) | 31 | 1.23 | 0.126 | 1.000 |
| AT1G22520.1 | MICOS complex subunit (Mic10-like) | 32 | 1.13 | 0.126 | 1.000 |
| AT1G22520.1 | MICOS complex subunit (Mic10-like) | 33 | 1.02 | 0.126 | 1.000 |
| AT1G22520.1 | MICOS complex subunit (Mic10-like) | 34 | 0.90 | 0.127 | 1.000 |
| AT1G22520.1 | MICOS complex subunit (Mic10-like) | 35 | 0.74 | 0.126 | 1.000 |
| AT1G22520.1 | MICOS complex subunit (Mic10-like) | 36 | 0.62 | 0.126 | 0.933 |
| AT1G22520.1 | MICOS complex subunit (Mic10-like) | 37 | 0.60 | 0.126 | 0.892 |
| AT1G22520.1 | MICOS complex subunit (Mic10-like) | 38 | 0.71 | 0.126 | 0.999 |
| AT1G22520.1 | MICOS complex subunit (Mic10-like) | 39 | 0.91 | 0.126 | 1.000 |
| AT1G22520.1 | MICOS complex subunit (Mic10-like) | 40 | 1.22 | 0.126 | 1.000 |

| | | | | | | |
|---|---|---|---|---|---|---|
| AT1G22520.1 | MICOS complex subunit (Mic10-like) | 41 | 1.63 | 0.127 | 0.926 | |
| AT1G22520.1 | MICOS complex subunit (Mic10-like) | 42 | 1.93 | 0.128 | 0.538 | |
| AT1G22520.1 | MICOS complex subunit (Mic10-like) | 43 | 1.78 | 0.130 | 0.785 | |
| AT1G22520.1 | MICOS complex subunit (Mic10-like) | 44 | 1.13 | 0.130 | 1.000 | |
| AT1G22520.1 | MICOS complex subunit (Mic10-like) | 45 | 0.50 | 0.131 | 0.474 | |
| AT1G22520.1 | MICOS complex subunit (Mic10-like) | 46 | 0.16 | 0.158 | 0.000 | *** |
| AT1G22520.1 | MICOS complex subunit (Mic10-like) | 47 | 0.05 | 0.262 | 0.000 | *** |
| AT1G22840.1 | Ubiquinol:cytochrome C oxidoreductase subunit (CYTC1) | 2 | 0.95 | 0.413 | 1.000 | |
| AT1G22840.1 | Ubiquinol:cytochrome C oxidoreductase subunit (CYTC1) | 3 | 0.90 | 0.382 | 1.000 | |
| AT1G22840.1 | Ubiquinol:cytochrome C oxidoreductase subunit (CYTC1) | 4 | 0.85 | 0.397 | 1.000 | |
| AT1G22840.1 | Ubiquinol:cytochrome C oxidoreductase subunit (CYTC1) | 20 | 0.50 | 0.369 | 0.987 | |
| AT1G22840.1 | Ubiquinol:cytochrome C oxidoreductase subunit (CYTC1) | 21 | 0.52 | 0.343 | 0.984 | |
| AT1G22840.1 | Ubiquinol:cytochrome C oxidoreductase subunit (CYTC1) | 22 | 0.54 | 0.319 | 0.984 | |
| AT1G22840.1 | Ubiquinol:cytochrome C oxidoreductase subunit (CYTC1) | 25 | 0.66 | 0.259 | 0.996 | |
| AT1G22840.1 | Ubiquinol:cytochrome C oxidoreductase subunit (CYTC1) | 26 | 0.73 | 0.242 | 0.999 | |
| AT1G22840.1 | Ubiquinol:cytochrome C oxidoreductase subunit (CYTC1) | 27 | 0.79 | 0.229 | 1.000 | |
| AT1G22840.1 | Ubiquinol:cytochrome C oxidoreductase subunit (CYTC1) | 28 | 0.83 | 0.222 | 1.000 | |
| AT1G22840.1 | Ubiquinol:cytochrome C oxidoreductase subunit (CYTC1) | 29 | 0.82 | 0.219 | 1.000 | |
| AT1G22840.1 | Ubiquinol:cytochrome C oxidoreductase subunit (CYTC1) | 30 | 0.77 | 0.221 | 1.000 | |
| AT1G22840.1 | Ubiquinol:cytochrome C oxidoreductase subunit (CYTC1) | 31 | 0.70 | 0.224 | 0.996 | |
| AT1G22840.1 | Ubiquinol:cytochrome C oxidoreductase subunit (CYTC1) | 32 | 0.62 | 0.228 | 0.973 | |
| AT1G22840.1 | Ubiquinol:cytochrome C oxidoreductase subunit (CYTC1) | 33 | 0.55 | 0.232 | 0.915 | |
| AT1G22840.1 | Ubiquinol:cytochrome C oxidoreductase subunit (CYTC1) | 34 | 0.50 | 0.236 | 0.841 | |
| AT1G22840.1 | Ubiquinol:cytochrome C oxidoreductase subunit (CYTC1) | 35 | 0.47 | 0.239 | 0.780 | |
| AT1G22840.1 | Ubiquinol:cytochrome C oxidoreductase subunit (CYTC1) | 36 | 0.46 | 0.241 | 0.759 | |
| AT1G22840.1 | Ubiquinol:cytochrome C oxidoreductase subunit (CYTC1) | 37 | 0.48 | 0.240 | 0.793 | |
| AT1G22840.1 | Ubiquinol:cytochrome C oxidoreductase subunit (CYTC1) | 38 | 0.53 | 0.235 | 0.881 | |
| AT1G22840.1 | Ubiquinol:cytochrome C oxidoreductase subunit (CYTC1) | 39 | 0.63 | 0.230 | 0.977 | |
| AT1G22840.1 | Ubiquinol:cytochrome C oxidoreductase subunit (CYTC1) | 40 | 0.81 | 0.224 | 1.000 | |
| AT1G22840.1 | Ubiquinol:cytochrome C oxidoreductase subunit (CYTC1) | 41 | 1.12 | 0.220 | 1.000 | |
| AT1G22840.1 | Ubiquinol:cytochrome C oxidoreductase subunit (CYTC1) | 42 | 1.65 | 0.219 | 0.954 | |

| | | | | | | |
|---|---|---|---|---|---|---|
| AT1G22840.1 | Ubiquinol:cytochrome C oxidoreductase subunit (CYTC1) | 43 | 2.53 | 0.220 | 0.444 | |
| AT1G22840.1 | Ubiquinol:cytochrome C oxidoreductase subunit (CYTC1) | 44 | 3.96 | 0.226 | 0.086 | . |
| AT1G22840.1 | Ubiquinol:cytochrome C oxidoreductase subunit (CYTC1) | 45 | 6.15 | 0.243 | 0.016 | * |
| AT1G22840.1 | Ubiquinol:cytochrome C oxidoreductase subunit (CYTC1) | 46 | 9.40 | 0.279 | 0.007 | ** |
| AT1G22840.1 | Ubiquinol:cytochrome C oxidoreductase subunit (CYTC1) | 47 | 14.25 | 0.341 | 0.011 | * |
| AT1G23100.1 | Heat shock protein 10 (HSP10) | 2 | 0.94 | 0.480 | 1.000 | |
| AT1G23100.1 | Heat shock protein 10 (HSP10) | 3 | 0.73 | 0.466 | 1.000 | |
| AT1G23100.1 | Heat shock protein 10 (HSP10) | 38 | 1.70 | 0.414 | 1.000 | |
| AT1G23100.1 | Heat shock protein 10 (HSP10) | 39 | 3.13 | 0.298 | 0.626 | |
| AT1G23100.1 | Heat shock protein 10 (HSP10) | 40 | 6.22 | 0.255 | 0.028 | * |
| AT1G23100.1 | Heat shock protein 10 (HSP10) | 41 | 9.67 | 0.249 | 0.002 | ** |
| AT1G23100.1 | Heat shock protein 10 (HSP10) | 42 | 11.02 | 0.254 | 0.001 | ** |
| AT1G23100.1 | Heat shock protein 10 (HSP10) | 43 | 3.66 | 0.254 | 0.257 | |
| AT1G23100.1 | Heat shock protein 10 (HSP10) | 44 | 0.51 | 0.248 | 0.914 | |
| AT1G23100.1 | Heat shock protein 10 (HSP10) | 45 | 0.14 | 0.252 | 0.010 | * |
| AT1G23100.1 | Heat shock protein 10 (HSP10) | 46 | 0.15 | 0.266 | 0.028 | * |
| AT1G23100.1 | Heat shock protein 10 (HSP10) | 47 | 0.34 | 0.631 | 0.996 | |
| AT1G23310.1 | Alanine:2-Oxoglutarate aminotransferase (AOAT1) | 2 | 0.88 | 0.351 | 1.000 | |
| AT1G23310.1 | Alanine:2-Oxoglutarate aminotransferase (AOAT1) | 3 | 0.82 | 0.339 | 1.000 | |
| AT1G23310.1 | Alanine:2-Oxoglutarate aminotransferase (AOAT1) | 26 | 0.99 | 0.283 | 1.000 | |
| AT1G23310.1 | Alanine:2-Oxoglutarate aminotransferase (AOAT1) | 27 | 1.17 | 0.200 | 1.000 | |
| AT1G23310.1 | Alanine:2-Oxoglutarate aminotransferase (AOAT1) | 28 | 1.30 | 0.180 | 1.000 | |
| AT1G23310.1 | Alanine:2-Oxoglutarate aminotransferase (AOAT1) | 29 | 1.14 | 0.171 | 1.000 | |
| AT1G23310.1 | Alanine:2-Oxoglutarate aminotransferase (AOAT1) | 30 | 0.80 | 0.171 | 1.000 | |
| AT1G23310.1 | Alanine:2-Oxoglutarate aminotransferase (AOAT1) | 31 | 0.71 | 0.172 | 1.000 | |
| AT1G23310.1 | Alanine:2-Oxoglutarate aminotransferase (AOAT1) | 32 | 1.06 | 0.173 | 1.000 | |
| AT1G23310.1 | Alanine:2-Oxoglutarate aminotransferase (AOAT1) | 33 | 1.71 | 0.172 | 0.957 | |
| AT1G23310.1 | Alanine:2-Oxoglutarate aminotransferase (AOAT1) | 34 | 1.92 | 0.173 | 0.826 | |
| AT1G23310.1 | Alanine:2-Oxoglutarate aminotransferase (AOAT1) | 35 | 1.42 | 0.173 | 1.000 | |
| AT1G23310.1 | Alanine:2-Oxoglutarate aminotransferase (AOAT1) | 36 | 1.02 | 0.172 | 1.000 | |
| AT1G23310.1 | Alanine:2-Oxoglutarate aminotransferase (AOAT1) | 37 | 0.93 | 0.171 | 1.000 | |
| AT1G23310.1 | Alanine:2-Oxoglutarate aminotransferase (AOAT1) | 38 | 0.93 | 0.170 | 1.000 | |
| AT1G23310.1 | Alanine:2-Oxoglutarate aminotransferase (AOAT1) | 39 | 0.95 | 0.170 | 1.000 | |
| AT1G23310.1 | Alanine:2-Oxoglutarate aminotransferase (AOAT1) | 40 | 1.09 | 0.170 | 1.000 | |
| AT1G23310.1 | Alanine:2-Oxoglutarate aminotransferase (AOAT1) | 41 | 1.25 | 0.172 | 1.000 | |
| AT1G23310.1 | Alanine:2-Oxoglutarate aminotransferase (AOAT1) | 42 | 1.31 | 0.174 | 1.000 | |
| AT1G23310.1 | Alanine:2-Oxoglutarate aminotransferase (AOAT1) | 43 | 1.10 | 0.175 | 1.000 | |

| | | | | | | |
|---|---|---|---|---|---|---|
| AT1G23310.1 | Alanine:2-Oxoglutarate aminotransferase (AOAT1) | 44 | 0.64 | 0.175 | 0.992 | |
| AT1G23310.1 | Alanine:2-Oxoglutarate aminotransferase (AOAT1) | 45 | 0.34 | 0.175 | 0.156 | |
| AT1G23310.1 | Alanine:2-Oxoglutarate aminotransferase (AOAT1) | 46 | 0.23 | 0.195 | 0.024 | * |
| AT1G23310.1 | Alanine:2-Oxoglutarate aminotransferase (AOAT1) | 47 | 0.18 | 0.369 | 0.522 | |
| AT1G23800.1 | Aldehyde dehydrogenase 2B (ALDH2B) | 2 | 0.81 | 0.603 | 1.000 | |
| AT1G23800.1 | Aldehyde dehydrogenase 2B (ALDH2B) | 3 | 0.75 | 0.599 | 1.000 | |
| AT1G23800.1 | Aldehyde dehydrogenase 2B (ALDH2B) | 26 | 0.50 | 0.475 | 1.000 | |
| AT1G23800.1 | Aldehyde dehydrogenase 2B (ALDH2B) | 27 | 0.63 | 0.336 | 1.000 | |
| AT1G23800.1 | Aldehyde dehydrogenase 2B (ALDH2B) | 28 | 1.10 | 0.297 | 1.000 | |
| AT1G23800.1 | Aldehyde dehydrogenase 2B (ALDH2B) | 29 | 1.73 | 0.279 | 0.997 | |
| AT1G23800.1 | Aldehyde dehydrogenase 2B (ALDH2B) | 30 | 1.64 | 0.275 | 0.999 | |
| AT1G23800.1 | Aldehyde dehydrogenase 2B (ALDH2B) | 31 | 1.34 | 0.278 | 1.000 | |
| AT1G23800.1 | Aldehyde dehydrogenase 2B (ALDH2B) | 32 | 1.36 | 0.277 | 1.000 | |
| AT1G23800.1 | Aldehyde dehydrogenase 2B (ALDH2B) | 33 | 1.89 | 0.277 | 0.990 | |
| AT1G23800.1 | Aldehyde dehydrogenase 2B (ALDH2B) | 34 | 2.93 | 0.278 | 0.709 | |
| AT1G23800.1 | Aldehyde dehydrogenase 2B (ALDH2B) | 35 | 3.07 | 0.280 | 0.664 | |
| AT1G23800.1 | Aldehyde dehydrogenase 2B (ALDH2B) | 36 | 1.60 | 0.283 | 0.999 | |
| AT1G23800.1 | Aldehyde dehydrogenase 2B (ALDH2B) | 37 | 0.55 | 0.295 | 0.996 | |
| AT1G23800.1 | Aldehyde dehydrogenase 2B (ALDH2B) | 38 | 0.26 | 0.368 | 0.778 | |
| AT1G23800.1 | Aldehyde dehydrogenase 2B (ALDH2B) | 39 | 0.29 | 0.630 | 0.997 | |
| AT1G23800.1 | Aldehyde dehydrogenase 2B (ALDH2B) | 46 | 1748378.50 | 4.015 | 0.796 | |
| AT1G23800.1 | Aldehyde dehydrogenase 2B (ALDH2B) | 47 | 45633012.97 | 4.764 | 0.761 | |
| AT1G24180.1 | Pyruvate dehydrogenase subunit (putative, PDCE1 alpha-like) | 2 | 1.30 | 0.381 | 1.000 | |
| AT1G24180.1 | Pyruvate dehydrogenase subunit (putative, PDCE1 alpha-like) | 3 | 1.14 | 0.276 | 1.000 | |
| AT1G24180.1 | Pyruvate dehydrogenase subunit (putative, PDCE1 alpha-like) | 4 | 0.93 | 0.265 | 1.000 | |
| AT1G24180.1 | Pyruvate dehydrogenase subunit (putative, PDCE1 alpha-like) | 6 | 0.45 | 0.268 | 0.991 | |
| AT1G24180.1 | Pyruvate dehydrogenase subunit (putative, PDCE1 alpha-like) | 7 | 0.27 | 0.239 | 0.391 | |
| AT1G24180.1 | Pyruvate dehydrogenase subunit (putative, PDCE1 alpha-like) | 8 | 0.19 | 0.215 | 0.032 | * |

| | | | | | | |
|---|---|---|---|---|---|---|
| AT1G24180.1 | Pyruvate dehydrogenase subunit (putative, PDCE1 alpha-like) | 9 | 0.21 | 0.200 | 0.022 | * |
| AT1G24180.1 | Pyruvate dehydrogenase subunit (putative, PDCE1 alpha-like) | 10 | 0.33 | 0.194 | 0.305 | |
| AT1G24180.1 | Pyruvate dehydrogenase subunit (putative, PDCE1 alpha-like) | 11 | 0.72 | 0.193 | 1.000 | |
| AT1G24180.1 | Pyruvate dehydrogenase subunit (putative, PDCE1 alpha-like) | 12 | 1.66 | 0.193 | 0.998 | |
| AT1G24180.1 | Pyruvate dehydrogenase subunit (putative, PDCE1 alpha-like) | 13 | 2.91 | 0.191 | 0.336 | |
| AT1G24180.1 | Pyruvate dehydrogenase subunit (putative, PDCE1 alpha-like) | 14 | 3.61 | 0.190 | 0.099 | . |
| AT1G24180.1 | Pyruvate dehydrogenase subunit (putative, PDCE1 alpha-like) | 15 | 3.40 | 0.190 | 0.141 | |
| AT1G24180.1 | Pyruvate dehydrogenase subunit (putative, PDCE1 alpha-like) | 16 | 2.64 | 0.189 | 0.480 | |
| AT1G24180.1 | Pyruvate dehydrogenase subunit (putative, PDCE1 alpha-like) | 17 | 2.00 | 0.190 | 0.933 | |
| AT1G24180.1 | Pyruvate dehydrogenase subunit (putative, PDCE1 alpha-like) | 18 | 1.73 | 0.195 | 0.996 | |
| AT1G24180.1 | Pyruvate dehydrogenase subunit (putative, PDCE1 alpha-like) | 19 | 1.71 | 0.202 | 0.998 | |
| AT1G24180.1 | Pyruvate dehydrogenase subunit (putative, PDCE1 alpha-like) | 20 | 1.68 | 0.211 | 0.999 | |
| AT1G24180.1 | Pyruvate dehydrogenase subunit (putative, PDCE1 alpha-like) | 21 | 1.49 | 0.217 | 1.000 | |
| AT1G24180.1 | Pyruvate dehydrogenase subunit (putative, PDCE1 alpha-like) | 22 | 1.21 | 0.215 | 1.000 | |
| AT1G24180.1 | Pyruvate dehydrogenase subunit (putative, PDCE1 alpha-like) | 23 | 0.98 | 0.207 | 1.000 | |
| AT1G24180.1 | Pyruvate dehydrogenase subunit (putative, PDCE1 alpha-like) | 24 | 0.89 | 0.198 | 1.000 | |
| AT1G24180.1 | Pyruvate dehydrogenase subunit (putative, PDCE1 alpha-like) | 25 | 0.90 | 0.193 | 1.000 | |
| AT1G24180.1 | Pyruvate dehydrogenase subunit (putative, PDCE1 alpha-like) | 26 | 0.97 | 0.192 | 1.000 | |
| AT1G24180.1 | Pyruvate dehydrogenase subunit (putative, PDCE1 alpha-like) | 27 | 0.99 | 0.191 | 1.000 | |
| AT1G24180.1 | Pyruvate dehydrogenase subunit (putative, PDCE1 alpha-like) | 28 | 0.93 | 0.192 | 1.000 | |

| | | | | | |
|---|---|---|---|---|---|
| AT1G24180.1 | Pyruvate dehydrogenase subunit (putative, PDCE1 alpha-like) | 29 | 0.87 | 0.192 | 1.000 |
| AT1G24180.1 | Pyruvate dehydrogenase subunit (putative, PDCE1 alpha-like) | 30 | 0.88 | 0.193 | 1.000 |
| AT1G24180.1 | Pyruvate dehydrogenase subunit (putative, PDCE1 alpha-like) | 31 | 0.94 | 0.193 | 1.000 |
| AT1G24180.1 | Pyruvate dehydrogenase subunit (putative, PDCE1 alpha-like) | 32 | 0.97 | 0.193 | 1.000 |
| AT1G24180.1 | Pyruvate dehydrogenase subunit (putative, PDCE1 alpha-like) | 33 | 0.93 | 0.193 | 1.000 |
| AT1G24180.1 | Pyruvate dehydrogenase subunit (putative, PDCE1 alpha-like) | 34 | 0.88 | 0.194 | 1.000 |
| AT1G24180.1 | Pyruvate dehydrogenase subunit (putative, PDCE1 alpha-like) | 35 | 0.88 | 0.194 | 1.000 |
| AT1G24180.1 | Pyruvate dehydrogenase subunit (putative, PDCE1 alpha-like) | 36 | 0.92 | 0.193 | 1.000 |
| AT1G24180.1 | Pyruvate dehydrogenase subunit (putative, PDCE1 alpha-like) | 37 | 0.96 | 0.192 | 1.000 |
| AT1G24180.1 | Pyruvate dehydrogenase subunit (putative, PDCE1 alpha-like) | 38 | 0.95 | 0.192 | 1.000 |
| AT1G24180.1 | Pyruvate dehydrogenase subunit (putative, PDCE1 alpha-like) | 39 | 0.89 | 0.193 | 1.000 |
| AT1G24180.1 | Pyruvate dehydrogenase subunit (putative, PDCE1 alpha-like) | 40 | 0.85 | 0.193 | 1.000 |
| AT1G24180.1 | Pyruvate dehydrogenase subunit (putative, PDCE1 alpha-like) | 41 | 0.92 | 0.194 | 1.000 |
| AT1G24180.1 | Pyruvate dehydrogenase subunit (putative, PDCE1 alpha-like) | 42 | 1.12 | 0.197 | 1.000 |
| AT1G24180.1 | Pyruvate dehydrogenase subunit (putative, PDCE1 alpha-like) | 43 | 1.26 | 0.199 | 1.000 |
| AT1G24180.1 | Pyruvate dehydrogenase subunit (putative, PDCE1 alpha-like) | 44 | 1.17 | 0.198 | 1.000 |
| AT1G24180.1 | Pyruvate dehydrogenase subunit (putative, PDCE1 alpha-like) | 45 | 0.89 | 0.199 | 1.000 |
| AT1G24180.1 | Pyruvate dehydrogenase subunit (putative, PDCE1 alpha-like) | 46 | 0.57 | 0.243 | 1.000 |
| AT1G24180.1 | Pyruvate dehydrogenase subunit (putative, PDCE1 alpha-like) | 47 | 0.33 | 0.400 | 0.997 |
| AT1G24240.1 | Ribosomal L19 family protein | 2 | 1.12 | 0.537 | 1.000 |
| AT1G24240.1 | Ribosomal L19 family protein | 3 | 1.42 | 0.475 | 1.000 |
| AT1G24240.1 | Ribosomal L19 family protein | 12 | 2.65 | 0.283 | 0.734 |
| AT1G24240.1 | Ribosomal L19 family protein | 13 | 2.06 | 0.248 | 0.874 |
| AT1G24240.1 | Ribosomal L19 family protein | 14 | 1.53 | 0.223 | 0.990 |

| | | | | | | |
|---|---|---|---|---|---|---|
| AT1G24240.1 | Ribosomal L19 family protein | 15 | 1.12 | 0.204 | 1.000 | |
| AT1G24240.1 | Ribosomal L19 family protein | 16 | 0.86 | 0.194 | 1.000 | |
| AT1G24240.1 | Ribosomal L19 family protein | 17 | 0.76 | 0.190 | 0.999 | |
| AT1G24240.1 | Ribosomal L19 family protein | 18 | 0.78 | 0.193 | 1.000 | |
| AT1G24240.1 | Ribosomal L19 family protein | 19 | 0.88 | 0.199 | 1.000 | |
| AT1G24240.1 | Ribosomal L19 family protein | 20 | 1.10 | 0.209 | 1.000 | |
| AT1G24240.1 | Ribosomal L19 family protein | 21 | 1.47 | 0.226 | 0.996 | |
| AT1G24240.1 | Ribosomal L19 family protein | 22 | 1.94 | 0.251 | 0.923 | |
| AT1G24240.1 | Ribosomal L19 family protein | 23 | 2.38 | 0.285 | 0.843 | |
| AT1G24240.1 | Ribosomal L19 family protein | 31 | 1.09 | 0.374 | 1.000 | |
| AT1G24240.1 | Ribosomal L19 family protein | 32 | 0.92 | 0.385 | 1.000 | |
| AT1G24240.1 | Ribosomal L19 family protein | 33 | 0.80 | 0.397 | 1.000 | |
| AT1G24240.1 | Ribosomal L19 family protein | 43 | 0.52 | 0.384 | 0.996 | |
| AT1G24240.1 | Ribosomal L19 family protein | 44 | 0.48 | 0.355 | 0.984 | |
| AT1G24240.1 | Ribosomal L19 family protein | 45 | 0.44 | 0.405 | 0.986 | |
| AT1G24240.1 | Ribosomal L19 family protein | 46 | 0.41 | 0.536 | 0.996 | |
| AT1G24240.1 | Ribosomal L19 family protein | 47 | 0.37 | 0.721 | 0.999 | |
| AT1G24267.1 | BZIP transcription factor, putative | 2 | 1.03 | 0.477 | 1.000 | |
| AT1G24267.1 | BZIP transcription factor, putative | 3 | 1.03 | 0.437 | 1.000 | |
| AT1G24267.1 | BZIP transcription factor, putative | 15 | 1.01 | 0.257 | 1.000 | |
| AT1G24267.1 | BZIP transcription factor, putative | 16 | 0.99 | 0.222 | 1.000 | |
| AT1G24267.1 | BZIP transcription factor, putative | 17 | 1.03 | 0.202 | 1.000 | |
| AT1G24267.1 | BZIP transcription factor, putative | 18 | 1.26 | 0.187 | 1.000 | |
| AT1G24267.1 | BZIP transcription factor, putative | 19 | 1.66 | 0.187 | 0.979 | |
| AT1G24267.1 | BZIP transcription factor, putative | 20 | 1.74 | 0.192 | 0.963 | |
| AT1G24267.1 | BZIP transcription factor, putative | 21 | 1.24 | 0.198 | 1.000 | |
| AT1G24267.1 | BZIP transcription factor, putative | 22 | 0.65 | 0.202 | 0.998 | |
| AT1G24267.1 | BZIP transcription factor, putative | 23 | 0.35 | 0.207 | 0.353 | |
| AT1G24267.1 | BZIP transcription factor, putative | 24 | 0.24 | 0.206 | 0.057 | . |
| AT1G24267.1 | BZIP transcription factor, putative | 25 | 0.23 | 0.196 | 0.022 | * |
| AT1G24267.1 | BZIP transcription factor, putative | 26 | 0.28 | 0.186 | 0.056 | . |
| AT1G24267.1 | BZIP transcription factor, putative | 27 | 0.43 | 0.183 | 0.531 | |
| AT1G24267.1 | BZIP transcription factor, putative | 28 | 0.83 | 0.182 | 1.000 | |
| AT1G24267.1 | BZIP transcription factor, putative | 29 | 1.47 | 0.182 | 0.998 | |
| AT1G24267.1 | BZIP transcription factor, putative | 30 | 1.68 | 0.182 | 0.968 | |
| AT1G24267.1 | BZIP transcription factor, putative | 31 | 1.55 | 0.184 | 0.994 | |
| AT1G24267.1 | BZIP transcription factor, putative | 32 | 1.32 | 0.185 | 1.000 | |
| AT1G24267.1 | BZIP transcription factor, putative | 33 | 0.92 | 0.186 | 1.000 | |
| AT1G24267.1 | BZIP transcription factor, putative | 34 | 0.48 | 0.190 | 0.761 | |

| | | | | | | |
|---|---|---|---|---|---|---|
| AT1G24267.1 | BZIP transcription factor, putative | 35 | 0.18 | 0.207 | 0.007 | ** |
| AT1G24267.1 | BZIP transcription factor, putative | 36 | 0.07 | 0.269 | 0.000 | *** |
| AT1G24267.1 | BZIP transcription factor, putative | 37 | 0.03 | 0.397 | 0.004 | ** |
| AT1G24267.1 | BZIP transcription factor, putative | 46 | 0.50 | 1.264 | 1.000 | |
| AT1G24267.1 | BZIP transcription factor, putative | 47 | 1.11 | 1.423 | 1.000 | |
| AT1G24340.1 | FAD/NAD(P)-binding oxidoreductase family protein | 2 | 0.84 | 0.244 | 1.000 | |
| AT1G24340.1 | FAD/NAD(P)-binding oxidoreductase family protein | 3 | 0.86 | 0.236 | 1.000 | |
| AT1G24340.1 | FAD/NAD(P)-binding oxidoreductase family protein | 28 | 1.53 | 0.202 | 0.994 | |
| AT1G24340.1 | FAD/NAD(P)-binding oxidoreductase family protein | 29 | 1.56 | 0.156 | 0.931 | |
| AT1G24340.1 | FAD/NAD(P)-binding oxidoreductase family protein | 30 | 1.54 | 0.142 | 0.900 | |
| AT1G24340.1 | FAD/NAD(P)-binding oxidoreductase family protein | 31 | 1.44 | 0.132 | 0.949 | |
| AT1G24340.1 | FAD/NAD(P)-binding oxidoreductase family protein | 32 | 1.31 | 0.127 | 0.993 | |
| AT1G24340.1 | FAD/NAD(P)-binding oxidoreductase family protein | 33 | 1.15 | 0.124 | 1.000 | |
| AT1G24340.1 | FAD/NAD(P)-binding oxidoreductase family protein | 34 | 0.90 | 0.123 | 1.000 | |
| AT1G24340.1 | FAD/NAD(P)-binding oxidoreductase family protein | 35 | 0.68 | 0.122 | 0.877 | |
| AT1G24340.1 | FAD/NAD(P)-binding oxidoreductase family protein | 36 | 0.66 | 0.122 | 0.804 | |
| AT1G24340.1 | FAD/NAD(P)-binding oxidoreductase family protein | 37 | 0.82 | 0.122 | 1.000 | |
| AT1G24340.1 | FAD/NAD(P)-binding oxidoreductase family protein | 38 | 1.08 | 0.122 | 1.000 | |
| AT1G24340.1 | FAD/NAD(P)-binding oxidoreductase family protein | 39 | 1.31 | 0.122 | 0.991 | |
| AT1G24340.1 | FAD/NAD(P)-binding oxidoreductase family protein | 40 | 1.38 | 0.122 | 0.961 | |
| AT1G24340.1 | FAD/NAD(P)-binding oxidoreductase family protein | 41 | 1.31 | 0.122 | 0.990 | |
| AT1G24340.1 | FAD/NAD(P)-binding oxidoreductase family protein | 42 | 1.21 | 0.123 | 1.000 | |
| AT1G24340.1 | FAD/NAD(P)-binding oxidoreductase family protein | 43 | 1.12 | 0.124 | 1.000 | |
| AT1G24340.1 | FAD/NAD(P)-binding oxidoreductase family protein | 44 | 0.91 | 0.124 | 1.000 | |
| AT1G24340.1 | FAD/NAD(P)-binding oxidoreductase family protein | 45 | 0.60 | 0.125 | 0.576 | |
| AT1G24340.1 | FAD/NAD(P)-binding oxidoreductase family protein | 46 | 0.31 | 0.140 | 0.006 | ** |
| AT1G24340.1 | FAD/NAD(P)-binding oxidoreductase family protein | 47 | 0.13 | 0.224 | 0.002 | ** |
| AT1G24610.1 | RUBISCO methyltransferase family protein | 2 | 0.82 | 0.305 | 1.000 | |
| AT1G24610.1 | RUBISCO methyltransferase family protein | 3 | 0.88 | 0.295 | 1.000 | |
| AT1G24610.1 | RUBISCO methyltransferase family protein | 38 | 2.27 | 0.248 | 0.734 | |
| AT1G24610.1 | RUBISCO methyltransferase family protein | 39 | 2.07 | 0.168 | 0.419 | |

| | | | | | | |
|---|---|---|---|---|---|---|
| AT1G24610.1 | RUBISCO methyltransferase family protein | 40 | 1.70 | 0.140 | 0.586 | |
| AT1G24610.1 | RUBISCO methyltransferase family protein | 41 | 1.23 | 0.134 | 0.996 | |
| AT1G24610.1 | RUBISCO methyltransferase family protein | 42 | 0.95 | 0.135 | 1.000 | |
| AT1G24610.1 | RUBISCO methyltransferase family protein | 43 | 0.84 | 0.136 | 0.999 | |
| AT1G24610.1 | RUBISCO methyltransferase family protein | 44 | 0.71 | 0.135 | 0.906 | |
| AT1G24610.1 | RUBISCO methyltransferase family protein | 45 | 0.57 | 0.134 | 0.477 | |
| AT1G24610.1 | RUBISCO methyltransferase family protein | 46 | 0.50 | 0.159 | 0.394 | |
| AT1G24610.1 | RUBISCO methyltransferase family protein | 47 | 0.43 | 0.300 | 0.859 | |
| AT1G25375.1 | Metallo-hydrolase/oxidoreductase superfamily protein | 2 | 0.84 | 0.551 | 1.000 | |
| AT1G25375.1 | Metallo-hydrolase/oxidoreductase superfamily protein | 3 | 1.42 | 0.529 | 0.998 | |
| AT1G25375.1 | Metallo-hydrolase/oxidoreductase superfamily protein | 42 | 13.50 | 0.459 | 0.079 | . |
| AT1G25375.1 | Metallo-hydrolase/oxidoreductase superfamily protein | 43 | 5.29 | 0.333 | 0.146 | |
| AT1G25375.1 | Metallo-hydrolase/oxidoreductase superfamily protein | 44 | 1.90 | 0.237 | 0.664 | |
| AT1G25375.1 | Metallo-hydrolase/oxidoreductase superfamily protein | 45 | 0.64 | 0.213 | 0.830 | |
| AT1G25375.1 | Metallo-hydrolase/oxidoreductase superfamily protein | 46 | 0.21 | 0.284 | 0.092 | . |
| AT1G25375.1 | Metallo-hydrolase/oxidoreductase superfamily protein | 47 | 0.07 | 0.422 | 0.038 | * |
| AT1G25380.1 | NAD+ transporter 2 (NDT2) | 2 | 1.06 | 0.381 | 1.000 | |
| AT1G25380.1 | NAD+ transporter 2 (NDT2) | 3 | 1.14 | 0.246 | 1.000 | |
| AT1G25380.1 | NAD+ transporter 2 (NDT2) | 4 | 1.24 | 0.235 | 1.000 | |
| AT1G25380.1 | NAD+ transporter 2 (NDT2) | 5 | 1.28 | 0.228 | 1.000 | |
| AT1G25380.1 | NAD+ transporter 2 (NDT2) | 6 | 1.28 | 0.218 | 1.000 | |
| AT1G25380.1 | NAD+ transporter 2 (NDT2) | 7 | 1.23 | 0.213 | 1.000 | |
| AT1G25380.1 | NAD+ transporter 2 (NDT2) | 8 | 1.17 | 0.209 | 1.000 | |
| AT1G25380.1 | NAD+ transporter 2 (NDT2) | 9 | 1.32 | 0.207 | 1.000 | |
| AT1G25380.1 | NAD+ transporter 2 (NDT2) | 10 | 1.55 | 0.207 | 1.000 | |
| AT1G25380.1 | NAD+ transporter 2 (NDT2) | 11 | 1.45 | 0.209 | 1.000 | |
| AT1G25380.1 | NAD+ transporter 2 (NDT2) | 12 | 1.24 | 0.208 | 1.000 | |
| AT1G25380.1 | NAD+ transporter 2 (NDT2) | 13 | 1.17 | 0.206 | 1.000 | |
| AT1G25380.1 | NAD+ transporter 2 (NDT2) | 14 | 1.27 | 0.204 | 1.000 | |
| AT1G25380.1 | NAD+ transporter 2 (NDT2) | 15 | 1.46 | 0.204 | 1.000 | |
| AT1G25380.1 | NAD+ transporter 2 (NDT2) | 16 | 1.61 | 0.203 | 1.000 | |
| AT1G25380.1 | NAD+ transporter 2 (NDT2) | 17 | 2.32 | 0.202 | 0.823 | |
| AT1G25380.1 | NAD+ transporter 2 (NDT2) | 18 | 3.67 | 0.206 | 0.166 | |
| AT1G25380.1 | NAD+ transporter 2 (NDT2) | 19 | 4.13 | 0.215 | 0.121 | |
| AT1G25380.1 | NAD+ transporter 2 (NDT2) | 20 | 3.66 | 0.228 | 0.304 | |
| AT1G25380.1 | NAD+ transporter 2 (NDT2) | 21 | 2.89 | 0.264 | 0.859 | |
| AT1G25380.1 | NAD+ transporter 2 (NDT2) | 22 | 2.11 | 0.346 | 1.000 | |
| AT1G25380.1 | NAD+ transporter 2 (NDT2) | 28 | 2.57 | 0.338 | 0.997 | |

| | | | | | | |
|---|---|---|---|---|---|---|
| AT1G25380.1 | NAD+ transporter 2 (NDT2) | 29 | 3.24 | 0.304 | 0.892 | |
| AT1G25380.1 | NAD+ transporter 2 (NDT2) | 30 | 4.09 | 0.256 | 0.364 | |
| AT1G25380.1 | NAD+ transporter 2 (NDT2) | 31 | 4.97 | 0.244 | 0.122 | |
| AT1G25380.1 | NAD+ transporter 2 (NDT2) | 32 | 5.55 | 0.231 | 0.042 | * |
| AT1G25380.1 | NAD+ transporter 2 (NDT2) | 33 | 5.78 | 0.222 | 0.021 | * |
| AT1G25380.1 | NAD+ transporter 2 (NDT2) | 34 | 5.17 | 0.212 | 0.027 | * |
| AT1G25380.1 | NAD+ transporter 2 (NDT2) | 35 | 3.16 | 0.208 | 0.350 | |
| AT1G25380.1 | NAD+ transporter 2 (NDT2) | 36 | 2.70 | 0.206 | 0.601 | |
| AT1G25380.1 | NAD+ transporter 2 (NDT2) | 37 | 5.00 | 0.207 | 0.025 | * |
| AT1G25380.1 | NAD+ transporter 2 (NDT2) | 38 | 8.62 | 0.208 | 0.000 | *** |
| AT1G25380.1 | NAD+ transporter 2 (NDT2) | 39 | 6.68 | 0.208 | 0.003 | ** |
| AT1G25380.1 | NAD+ transporter 2 (NDT2) | 40 | 2.68 | 0.207 | 0.622 | |
| AT1G25380.1 | NAD+ transporter 2 (NDT2) | 41 | 0.82 | 0.207 | 1.000 | |
| AT1G25380.1 | NAD+ transporter 2 (NDT2) | 42 | 0.33 | 0.210 | 0.437 | |
| AT1G25380.1 | NAD+ transporter 2 (NDT2) | 43 | 0.23 | 0.211 | 0.082 | . |
| AT1G25380.1 | NAD+ transporter 2 (NDT2) | 44 | 0.29 | 0.211 | 0.246 | |
| AT1G25380.1 | NAD+ transporter 2 (NDT2) | 45 | 0.51 | 0.210 | 0.981 | |
| AT1G25380.1 | NAD+ transporter 2 (NDT2) | 46 | 1.05 | 0.237 | 1.000 | |
| AT1G25380.1 | NAD+ transporter 2 (NDT2) | 47 | 2.43 | 0.445 | 1.000 | |
| AT1G26190.1 | Triphosphate tunnel metalloenzyme 2 (TTM2) | 2 | 1.08 | 0.455 | 1.000 | |
| AT1G26190.1 | Triphosphate tunnel metalloenzyme 2 (TTM2) | 3 | 1.10 | 0.397 | 1.000 | |
| AT1G26190.1 | Triphosphate tunnel metalloenzyme 2 (TTM2) | 11 | 2.34 | 0.246 | 0.809 | |
| AT1G26190.1 | Triphosphate tunnel metalloenzyme 2 (TTM2) | 12 | 2.98 | 0.213 | 0.290 | |
| AT1G26190.1 | Triphosphate tunnel metalloenzyme 2 (TTM2) | 13 | 3.88 | 0.189 | 0.032 | * |
| AT1G26190.1 | Triphosphate tunnel metalloenzyme 2 (TTM2) | 14 | 4.79 | 0.174 | 0.002 | ** |
| AT1G26190.1 | Triphosphate tunnel metalloenzyme 2 (TTM2) | 15 | 5.29 | 0.165 | 0.000 | *** |
| AT1G26190.1 | Triphosphate tunnel metalloenzyme 2 (TTM2) | 16 | 5.15 | 0.161 | 0.000 | *** |
| AT1G26190.1 | Triphosphate tunnel metalloenzyme 2 (TTM2) | 17 | 4.48 | 0.160 | 0.001 | ** |
| AT1G26190.1 | Triphosphate tunnel metalloenzyme 2 (TTM2) | 18 | 3.64 | 0.162 | 0.010 | * |
| AT1G26190.1 | Triphosphate tunnel metalloenzyme 2 (TTM2) | 19 | 2.91 | 0.165 | 0.074 | . |
| AT1G26190.1 | Triphosphate tunnel metalloenzyme 2 (TTM2) | 20 | 2.30 | 0.168 | 0.332 | |
| AT1G26190.1 | Triphosphate tunnel metalloenzyme 2 (TTM2) | 21 | 1.84 | 0.169 | 0.769 | |
| AT1G26190.1 | Triphosphate tunnel metalloenzyme 2 (TTM2) | 22 | 1.50 | 0.169 | 0.980 | |
| AT1G26190.1 | Triphosphate tunnel metalloenzyme 2 (TTM2) | 23 | 1.28 | 0.168 | 1.000 | |
| AT1G26190.1 | Triphosphate tunnel metalloenzyme 2 (TTM2) | 24 | 1.11 | 0.166 | 1.000 | |
| AT1G26190.1 | Triphosphate tunnel metalloenzyme 2 (TTM2) | 25 | 0.94 | 0.164 | 1.000 | |
| AT1G26190.1 | Triphosphate tunnel metalloenzyme 2 (TTM2) | 26 | 0.79 | 0.164 | 1.000 | |
| AT1G26190.1 | Triphosphate tunnel metalloenzyme 2 (TTM2) | 27 | 0.66 | 0.164 | 0.974 | |
| AT1G26190.1 | Triphosphate tunnel metalloenzyme 2 (TTM2) | 28 | 0.53 | 0.165 | 0.671 | |
| AT1G26190.1 | Triphosphate tunnel metalloenzyme 2 (TTM2) | 29 | 0.40 | 0.167 | 0.201 | |

| | | | | | | |
|---|---|---|---|---|---|---|
| AT1G26190.1 | Triphosphate tunnel metalloenzyme 2 (TTM2) | 30 | 0.30 | 0.174 | 0.038 | * |
| AT1G26190.1 | Triphosphate tunnel metalloenzyme 2 (TTM2) | 31 | 0.24 | 0.194 | 0.023 | * |
| AT1G26190.1 | Triphosphate tunnel metalloenzyme 2 (TTM2) | 32 | 0.21 | 0.239 | 0.067 | . |
| AT1G26190.1 | Triphosphate tunnel metalloenzyme 2 (TTM2) | 33 | 0.20 | 0.315 | 0.293 | |
| AT1G26190.1 | Triphosphate tunnel metalloenzyme 2 (TTM2) | 46 | 41.06 | 2.165 | 0.999 | |
| AT1G26190.1 | Triphosphate tunnel metalloenzyme 2 (TTM2) | 47 | 77.16 | 2.381 | 0.998 | |
| AT1G26460.1 | Pentatricopeptide repeat (PPR) superfamily protein | 2 | 1.04 | 0.431 | 1.000 | |
| AT1G26460.1 | Pentatricopeptide repeat (PPR) superfamily protein | 3 | 0.91 | 0.325 | 1.000 | |
| AT1G26460.1 | Pentatricopeptide repeat (PPR) superfamily protein | 4 | 0.84 | 0.434 | 1.000 | |
| AT1G26460.1 | Pentatricopeptide repeat (PPR) superfamily protein | 11 | 1.37 | 0.310 | 1.000 | |
| AT1G26460.1 | Pentatricopeptide repeat (PPR) superfamily protein | 12 | 1.38 | 0.272 | 1.000 | |
| AT1G26460.1 | Pentatricopeptide repeat (PPR) superfamily protein | 13 | 1.33 | 0.249 | 1.000 | |
| AT1G26460.1 | Pentatricopeptide repeat (PPR) superfamily protein | 14 | 1.28 | 0.228 | 1.000 | |
| AT1G26460.1 | Pentatricopeptide repeat (PPR) superfamily protein | 15 | 1.20 | 0.217 | 1.000 | |
| AT1G26460.1 | Pentatricopeptide repeat (PPR) superfamily protein | 16 | 0.97 | 0.213 | 1.000 | |
| AT1G26460.1 | Pentatricopeptide repeat (PPR) superfamily protein | 17 | 0.79 | 0.212 | 1.000 | |
| AT1G26460.1 | Pentatricopeptide repeat (PPR) superfamily protein | 18 | 0.82 | 0.215 | 1.000 | |
| AT1G26460.1 | Pentatricopeptide repeat (PPR) superfamily protein | 19 | 1.15 | 0.221 | 1.000 | |
| AT1G26460.1 | Pentatricopeptide repeat (PPR) superfamily protein | 20 | 1.79 | 0.227 | 0.998 | |
| AT1G26460.1 | Pentatricopeptide repeat (PPR) superfamily protein | 21 | 2.21 | 0.234 | 0.959 | |
| AT1G26460.1 | Pentatricopeptide repeat (PPR) superfamily protein | 22 | 1.84 | 0.244 | 0.999 | |
| AT1G26460.1 | Pentatricopeptide repeat (PPR) superfamily protein | 23 | 1.20 | 0.261 | 1.000 | |
| AT1G26460.1 | Pentatricopeptide repeat (PPR) superfamily protein | 24 | 0.70 | 0.277 | 1.000 | |
| AT1G26460.1 | Pentatricopeptide repeat (PPR) superfamily protein | 25 | 0.43 | 0.267 | 0.980 | |
| AT1G26460.1 | Pentatricopeptide repeat (PPR) superfamily protein | 26 | 0.36 | 0.251 | 0.842 | |
| AT1G26460.1 | Pentatricopeptide repeat (PPR) superfamily protein | 27 | 0.47 | 0.231 | 0.974 | |
| AT1G26460.1 | Pentatricopeptide repeat (PPR) superfamily protein | 28 | 0.72 | 0.221 | 1.000 | |
| AT1G26460.1 | Pentatricopeptide repeat (PPR) superfamily protein | 29 | 1.06 | 0.218 | 1.000 | |
| AT1G26460.1 | Pentatricopeptide repeat (PPR) superfamily protein | 30 | 1.21 | 0.218 | 1.000 | |
| AT1G26460.1 | Pentatricopeptide repeat (PPR) superfamily protein | 31 | 1.16 | 0.220 | 1.000 | |
| AT1G26460.1 | Pentatricopeptide repeat (PPR) superfamily protein | 32 | 1.14 | 0.220 | 1.000 | |

| | | | | | |
|---|---|---|---|---|---|
| AT1G26460.1 | Pentatricopeptide repeat (PPR) superfamily protein | 33 | 1.17 | 0.220 | 1.000 |
| AT1G26460.1 | Pentatricopeptide repeat (PPR) superfamily protein | 34 | 1.22 | 0.220 | 1.000 |
| AT1G26460.1 | Pentatricopeptide repeat (PPR) superfamily protein | 35 | 1.24 | 0.220 | 1.000 |
| AT1G26460.1 | Pentatricopeptide repeat (PPR) superfamily protein | 36 | 1.14 | 0.219 | 1.000 |
| AT1G26460.1 | Pentatricopeptide repeat (PPR) superfamily protein | 37 | 0.91 | 0.219 | 1.000 |
| AT1G26460.1 | Pentatricopeptide repeat (PPR) superfamily protein | 38 | 0.65 | 0.219 | 1.000 |
| AT1G26460.1 | Pentatricopeptide repeat (PPR) superfamily protein | 39 | 0.44 | 0.220 | 0.905 |
| AT1G26460.1 | Pentatricopeptide repeat (PPR) superfamily protein | 40 | 0.35 | 0.222 | 0.634 |
| AT1G26460.1 | Pentatricopeptide repeat (PPR) superfamily protein | 41 | 0.37 | 0.237 | 0.799 |
| AT1G26460.1 | Pentatricopeptide repeat (PPR) superfamily protein | 42 | 0.56 | 0.313 | 1.000 |
| AT1G26460.1 | Pentatricopeptide repeat (PPR) superfamily protein | 43 | 1.36 | 0.454 | 1.000 |
| AT1G26460.1 | Pentatricopeptide repeat (PPR) superfamily protein | 46 | 166.07 | 1.263 | 0.836 |
| AT1G26460.1 | Pentatricopeptide repeat (PPR) superfamily protein | 47 | 1143.37 | 1.656 | 0.777 |
| AT1G27000.1 | ? | 2 | 0.96 | 0.450 | 1.000 |
| AT1G27000.1 | ? | 3 | 0.88 | 0.431 | 1.000 |
| AT1G27000.1 | ? | 26 | 0.75 | 0.310 | 1.000 |
| AT1G27000.1 | ? | 27 | 0.86 | 0.223 | 1.000 |
| AT1G27000.1 | ? | 28 | 1.01 | 0.186 | 1.000 |
| AT1G27000.1 | ? | 29 | 1.13 | 0.173 | 1.000 |
| AT1G27000.1 | ? | 30 | 1.19 | 0.170 | 1.000 |
| AT1G27000.1 | ? | 31 | 1.27 | 0.169 | 0.999 |
| AT1G27000.1 | ? | 32 | 1.27 | 0.169 | 0.999 |
| AT1G27000.1 | ? | 33 | 1.10 | 0.170 | 1.000 |
| AT1G27000.1 | ? | 34 | 0.89 | 0.171 | 1.000 |
| AT1G27000.1 | ? | 35 | 0.69 | 0.175 | 0.979 |
| AT1G27000.1 | ? | 36 | 0.55 | 0.191 | 0.838 |
| AT1G27000.1 | ? | 37 | 0.51 | 0.245 | 0.913 |
| AT1G27000.1 | ? | 38 | 0.55 | 0.352 | 0.996 |
| AT1G27000.1 | ? | 46 | 47.82 | 1.831 | 0.981 |
| AT1G27000.1 | ? | 47 | 105.56 | 2.112 | 0.975 |
| AT1G27390.1 | Translocase of the outer mitochondrial membrane 20 (TOM20-2) | 2 | 0.98 | 0.523 | 1.000 |
| AT1G27390.1 | Translocase of the outer mitochondrial membrane 20 (TOM20-2) | 3 | 0.72 | 0.415 | 1.000 |
| AT1G27390.1 | Translocase of the outer mitochondrial membrane 20 (TOM20-2) | 4 | 0.56 | 0.627 | 1.000 |
| AT1G27390.1 | Translocase of the outer mitochondrial membrane 20 (TOM20-2) | 15 | 0.96 | 0.376 | 1.000 |
| AT1G27390.1 | Translocase of the outer mitochondrial membrane 20 (TOM20-2) | 16 | 1.15 | 0.348 | 1.000 |
| AT1G27390.1 | Translocase of the outer mitochondrial membrane 20 (TOM20-2) | 17 | 1.30 | 0.319 | 1.000 |

| | | | | | | |
|---|---|---|---|---|---|---|
| AT1G27390.1 | Translocase of the outer mitochondrial membrane 20 (TOM20-2) | 18 | 1.62 | 0.304 | 1.000 | |
| AT1G27390.1 | Translocase of the outer mitochondrial membrane 20 (TOM20-2) | 19 | 2.23 | 0.304 | 0.977 | |
| AT1G27390.1 | Translocase of the outer mitochondrial membrane 20 (TOM20-2) | 20 | 2.24 | 0.309 | 0.979 | |
| AT1G27390.1 | Translocase of the outer mitochondrial membrane 20 (TOM20-2) | 21 | 1.77 | 0.311 | 0.999 | |
| AT1G27390.1 | Translocase of the outer mitochondrial membrane 20 (TOM20-2) | 22 | 1.57 | 0.311 | 1.000 | |
| AT1G27390.1 | Translocase of the outer mitochondrial membrane 20 (TOM20-2) | 23 | 1.42 | 0.310 | 1.000 | |
| AT1G27390.1 | Translocase of the outer mitochondrial membrane 20 (TOM20-2) | 24 | 1.06 | 0.306 | 1.000 | |
| AT1G27390.1 | Translocase of the outer mitochondrial membrane 20 (TOM20-2) | 25 | 0.79 | 0.302 | 1.000 | |
| AT1G27390.1 | Translocase of the outer mitochondrial membrane 20 (TOM20-2) | 26 | 0.77 | 0.301 | 1.000 | |
| AT1G27390.1 | Translocase of the outer mitochondrial membrane 20 (TOM20-2) | 27 | 0.84 | 0.300 | 1.000 | |
| AT1G27390.1 | Translocase of the outer mitochondrial membrane 20 (TOM20-2) | 28 | 1.04 | 0.301 | 1.000 | |
| AT1G27390.1 | Translocase of the outer mitochondrial membrane 20 (TOM20-2) | 29 | 1.79 | 0.302 | 0.999 | |
| AT1G27390.1 | Translocase of the outer mitochondrial membrane 20 (TOM20-2) | 30 | 2.50 | 0.302 | 0.931 | |
| AT1G27390.1 | Translocase of the outer mitochondrial membrane 20 (TOM20-2) | 31 | 1.40 | 0.302 | 1.000 | |
| AT1G27390.1 | Translocase of the outer mitochondrial membrane 20 (TOM20-2) | 32 | 0.32 | 0.304 | 0.744 | |
| AT1G27390.1 | Translocase of the outer mitochondrial membrane 20 (TOM20-2) | 33 | 0.07 | 0.308 | 0.003 | ** |
| AT1G27390.1 | Translocase of the outer mitochondrial membrane 20 (TOM20-2) | 34 | 0.04 | 0.320 | 0.000 | *** |
| AT1G27390.1 | Translocase of the outer mitochondrial membrane 20 (TOM20-2) | 35 | 0.06 | 0.372 | 0.023 | * |
| AT1G27390.1 | Translocase of the outer mitochondrial membrane 20 (TOM20-2) | 36 | 0.16 | 0.562 | 0.900 | |
| AT1G27390.1 | Translocase of the outer mitochondrial membrane 20 (TOM20-2) | 46 | 14829832.04 | 4.185 | 0.695 | |

| | | | | | |
|---|---|---|---|---|---|
| AT1G27390.1 | Translocase of the outer mitochondrial membrane 20 (TOM20-2) | 47 | ########### | 4.750 | 0.695 |
| AT1G28510.1 | Optic atrophy 3 protein (OPA3) | 2 | 0.81 | 0.486 | 1.000 |
| AT1G28510.1 | Optic atrophy 3 protein (OPA3) | 3 | 0.93 | 0.464 | 1.000 |
| AT1G28510.1 | Optic atrophy 3 protein (OPA3) | 28 | 4.54 | 0.350 | 0.570 |
| AT1G28510.1 | Optic atrophy 3 protein (OPA3) | 29 | 4.13 | 0.262 | 0.249 |
| AT1G28510.1 | Optic atrophy 3 protein (OPA3) | 30 | 3.51 | 0.235 | 0.263 |
| AT1G28510.1 | Optic atrophy 3 protein (OPA3) | 31 | 2.70 | 0.224 | 0.531 |
| AT1G28510.1 | Optic atrophy 3 protein (OPA3) | 32 | 2.17 | 0.220 | 0.823 |
| AT1G28510.1 | Optic atrophy 3 protein (OPA3) | 33 | 2.28 | 0.219 | 0.756 |
| AT1G28510.1 | Optic atrophy 3 protein (OPA3) | 34 | 2.90 | 0.220 | 0.401 |
| AT1G28510.1 | Optic atrophy 3 protein (OPA3) | 35 | 3.37 | 0.220 | 0.223 |
| AT1G28510.1 | Optic atrophy 3 protein (OPA3) | 36 | 3.71 | 0.219 | 0.142 |
| AT1G28510.1 | Optic atrophy 3 protein (OPA3) | 37 | 3.60 | 0.218 | 0.160 |
| AT1G28510.1 | Optic atrophy 3 protein (OPA3) | 38 | 2.80 | 0.218 | 0.439 |
| AT1G28510.1 | Optic atrophy 3 protein (OPA3) | 39 | 2.07 | 0.218 | 0.864 |
| AT1G28510.1 | Optic atrophy 3 protein (OPA3) | 40 | 1.77 | 0.217 | 0.975 |
| AT1G28510.1 | Optic atrophy 3 protein (OPA3) | 41 | 1.36 | 0.219 | 1.000 |
| AT1G28510.1 | Optic atrophy 3 protein (OPA3) | 42 | 0.90 | 0.222 | 1.000 |
| AT1G28510.1 | Optic atrophy 3 protein (OPA3) | 43 | 0.59 | 0.224 | 0.989 |
| AT1G28510.1 | Optic atrophy 3 protein (OPA3) | 44 | 0.44 | 0.224 | 0.777 |
| AT1G28510.1 | Optic atrophy 3 protein (OPA3) | 45 | 0.42 | 0.223 | 0.709 |
| AT1G28510.1 | Optic atrophy 3 protein (OPA3) | 46 | 0.47 | 0.262 | 0.952 |
| AT1G28510.1 | Optic atrophy 3 protein (OPA3) | 47 | 0.56 | 0.459 | 1.000 |
| AT1G28540.1 | ? | 2 | 0.97 | 0.365 | 1.000 |
| AT1G28540.1 | ? | 3 | 1.00 | 0.329 | 1.000 |
| AT1G28540.1 | ? | 11 | 1.27 | 0.165 | 0.996 |
| AT1G28540.1 | ? | 12 | 1.32 | 0.152 | 0.983 |
| AT1G28540.1 | ? | 13 | 1.38 | 0.141 | 0.941 |
| AT1G28540.1 | ? | 14 | 1.45 | 0.133 | 0.851 |
| AT1G28540.1 | ? | 15 | 1.52 | 0.129 | 0.721 |
| AT1G28540.1 | ? | 16 | 1.60 | 0.127 | 0.591 |
| AT1G28540.1 | ? | 17 | 1.67 | 0.129 | 0.496 |
| AT1G28540.1 | ? | 18 | 1.73 | 0.133 | 0.452 |
| AT1G28540.1 | ? | 19 | 1.77 | 0.139 | 0.453 |
| AT1G28540.1 | ? | 20 | 1.77 | 0.145 | 0.501 |
| AT1G28540.1 | ? | 21 | 1.74 | 0.150 | 0.580 |
| AT1G28540.1 | ? | 27 | 1.21 | 0.148 | 0.998 |
| AT1G28540.1 | ? | 28 | 1.13 | 0.143 | 1.000 |
| AT1G28540.1 | ? | 29 | 1.07 | 0.138 | 1.000 |
| AT1G28540.1 | ? | 30 | 1.03 | 0.134 | 1.000 |
| AT1G28540.1 | ? | 31 | 1.01 | 0.132 | 1.000 |
| AT1G28540.1 | ? | 32 | 1.02 | 0.131 | 1.000 |
| AT1G28540.1 | ? | 33 | 1.04 | 0.133 | 1.000 |
| AT1G28540.1 | ? | 34 | 1.09 | 0.136 | 1.000 |
| AT1G28540.1 | ? | 35 | 1.14 | 0.141 | 1.000 |
| AT1G28540.1 | ? | 36 | 1.21 | 0.147 | 0.999 |
| AT1G28540.1 | ? | 37 | 1.27 | 0.155 | 0.994 |
| AT1G28540.1 | ? | 38 | 1.34 | 0.164 | 0.984 |
| AT1G28540.1 | ? | 45 | 1.78 | 0.245 | 0.928 |
| AT1G28540.1 | ? | 46 | 1.84 | 0.268 | 0.939 |
| AT1G28540.1 | ? | 47 | 1.90 | 0.298 | 0.954 |
| AT1G29810.1 | Transcriptional coactivator/pterin dehydratase | 2 | 1.11 | 0.312 | 1.000 |
| AT1G29810.1 | Transcriptional coactivator/pterin dehydratase | 3 | 1.08 | 0.303 | 1.000 |
| AT1G29810.1 | Transcriptional coactivator/pterin dehydratase | 4 | 1.04 | 0.298 | 1.000 |
| AT1G29810.1 | Transcriptional coactivator/pterin dehydratase | 35 | 0.93 | 0.215 | 1.000 |
| AT1G29810.1 | Transcriptional coactivator/pterin dehydratase | 36 | 0.97 | 0.204 | 1.000 |

| | | | | | |
|---|---|---|---|---|---|
| AT1G29810.1 | Transcriptional coactivator/pterin dehydratase | 37 | 1.02 | 0.194 | 1.000 |
| AT1G29810.1 | Transcriptional coactivator/pterin dehydratase | 38 | 1.08 | 0.186 | 1.000 |
| AT1G29810.1 | Transcriptional coactivator/pterin dehydratase | 39 | 1.14 | 0.179 | 0.994 |
| AT1G29810.1 | Transcriptional coactivator/pterin dehydratase | 40 | 1.22 | 0.175 | 0.968 |
| AT1G29810.1 | Transcriptional coactivator/pterin dehydratase | 41 | 1.31 | 0.172 | 0.905 |
| AT1G29810.1 | Transcriptional coactivator/pterin dehydratase | 42 | 1.41 | 0.171 | 0.806 |
| AT1G29810.1 | Transcriptional coactivator/pterin dehydratase | 43 | 1.53 | 0.174 | 0.685 |
| AT1G29810.1 | Transcriptional coactivator/pterin dehydratase | 44 | 1.67 | 0.179 | 0.569 |
| AT1G29810.1 | Transcriptional coactivator/pterin dehydratase | 45 | 1.82 | 0.188 | 0.478 |
| AT1G29810.1 | Transcriptional coactivator/pterin dehydratase | 46 | 1.98 | 0.201 | 0.417 |
| AT1G29810.1 | Transcriptional coactivator/pterin dehydratase | 47 | 2.16 | 0.219 | 0.383 |
| AT1G29880.1 | Glycyl-tRNA synthetase / glycine--tRNA ligase | 2 | 0.83 | 0.373 | 1.000 |
| AT1G29880.1 | Glycyl-tRNA synthetase / glycine--tRNA ligase | 3 | 1.04 | 0.365 | 1.000 |
| AT1G29880.1 | Glycyl-tRNA synthetase / glycine--tRNA ligase | 28 | 1.03 | 0.281 | 1.000 |
| AT1G29880.1 | Glycyl-tRNA synthetase / glycine--tRNA ligase | 29 | 0.72 | 0.210 | 1.000 |
| AT1G29880.1 | Glycyl-tRNA synthetase / glycine--tRNA ligase | 30 | 0.69 | 0.190 | 0.998 |
| AT1G29880.1 | Glycyl-tRNA synthetase / glycine--tRNA ligase | 31 | 0.89 | 0.182 | 1.000 |
| AT1G29880.1 | Glycyl-tRNA synthetase / glycine--tRNA ligase | 32 | 1.16 | 0.179 | 1.000 |
| AT1G29880.1 | Glycyl-tRNA synthetase / glycine--tRNA ligase | 33 | 1.48 | 0.178 | 0.994 |
| AT1G29880.1 | Glycyl-tRNA synthetase / glycine--tRNA ligase | 34 | 1.68 | 0.178 | 0.945 |
| AT1G29880.1 | Glycyl-tRNA synthetase / glycine--tRNA ligase | 35 | 1.37 | 0.178 | 0.999 |
| AT1G29880.1 | Glycyl-tRNA synthetase / glycine--tRNA ligase | 36 | 0.80 | 0.178 | 1.000 |
| AT1G29880.1 | Glycyl-tRNA synthetase / glycine--tRNA ligase | 37 | 0.51 | 0.177 | 0.725 |
| AT1G29880.1 | Glycyl-tRNA synthetase / glycine--tRNA ligase | 38 | 0.54 | 0.177 | 0.826 |
| AT1G29880.1 | Glycyl-tRNA synthetase / glycine--tRNA ligase | 39 | 0.81 | 0.178 | 1.000 |
| AT1G29880.1 | Glycyl-tRNA synthetase / glycine--tRNA ligase | 40 | 1.08 | 0.178 | 1.000 |
| AT1G29880.1 | Glycyl-tRNA synthetase / glycine--tRNA ligase | 41 | 1.18 | 0.179 | 1.000 |
| AT1G29880.1 | Glycyl-tRNA synthetase / glycine--tRNA ligase | 42 | 1.15 | 0.181 | 1.000 |
| AT1G29880.1 | Glycyl-tRNA synthetase / glycine--tRNA ligase | 43 | 1.01 | 0.188 | 1.000 |
| AT1G29880.1 | Glycyl-tRNA synthetase / glycine--tRNA ligase | 44 | 0.90 | 0.235 | 1.000 |
| AT1G29880.1 | Glycyl-tRNA synthetase / glycine--tRNA ligase | 45 | 0.85 | 0.322 | 1.000 |

| | | | | | | |
|---|---|---|---|---|---|---|
| AT1G29880.1 | Glycyl-tRNA synthetase / glycine--tRNA ligase | 46 | 0.85 | 0.461 | 1.000 | |
| AT1G29880.1 | Glycyl-tRNA synthetase / glycine--tRNA ligase | 47 | 0.86 | 0.687 | 1.000 | |
| AT1G29900.1 | Carbamoyl phosphate synthetase B (CARB) / Venosa 3 (VEN3) | 2 | 1.00 | 0.404 | 1.000 | |
| AT1G29900.1 | Carbamoyl phosphate synthetase B (CARB) / Venosa 3 (VEN3) | 3 | 1.07 | 0.388 | 1.000 | |
| AT1G29900.1 | Carbamoyl phosphate synthetase B (CARB) / Venosa 3 (VEN3) | 4 | 1.15 | 0.382 | 1.000 | |
| AT1G29900.1 | Carbamoyl phosphate synthetase B (CARB) / Venosa 3 (VEN3) | 31 | 1.23 | 0.237 | 0.991 | |
| AT1G29900.1 | Carbamoyl phosphate synthetase B (CARB) / Venosa 3 (VEN3) | 32 | 1.08 | 0.229 | 1.000 | |
| AT1G29900.1 | Carbamoyl phosphate synthetase B (CARB) / Venosa 3 (VEN3) | 33 | 0.93 | 0.227 | 1.000 | |
| AT1G29900.1 | Carbamoyl phosphate synthetase B (CARB) / Venosa 3 (VEN3) | 34 | 0.80 | 0.231 | 0.985 | |
| AT1G29900.1 | Carbamoyl phosphate synthetase B (CARB) / Venosa 3 (VEN3) | 35 | 0.68 | 0.238 | 0.907 | |
| AT1G29900.1 | Carbamoyl phosphate synthetase B (CARB) / Venosa 3 (VEN3) | 36 | 0.58 | 0.249 | 0.770 | |
| AT1G29900.1 | Carbamoyl phosphate synthetase B (CARB) / Venosa 3 (VEN3) | 46 | 0.10 | 0.387 | 0.055 | . |
| AT1G29900.1 | Carbamoyl phosphate synthetase B (CARB) / Venosa 3 (VEN3) | 47 | 0.09 | 0.411 | 0.050 | . |
| AT1G29930.1 | Photosystem I/II light harvesting complex subunit (LHCB1.3) | 3 | 0.64 | 0.270 | 1.000 | |
| AT1G29930.1 | Photosystem I/II light harvesting complex subunit (LHCB1.3) | 4 | 0.66 | 0.248 | 1.000 | |
| AT1G29930.1 | Photosystem I/II light harvesting complex subunit (LHCB1.3) | 5 | 0.71 | 0.249 | 1.000 | |
| AT1G29930.1 | Photosystem I/II light harvesting complex subunit (LHCB1.3) | 6 | 0.65 | 0.249 | 1.000 | |
| AT1G29930.1 | Photosystem I/II light harvesting complex subunit (LHCB1.3) | 7 | 0.55 | 0.249 | 0.999 | |
| AT1G29930.1 | Photosystem I/II light harvesting complex subunit (LHCB1.3) | 8 | 0.49 | 0.245 | 0.991 | |
| AT1G29930.1 | Photosystem I/II light harvesting complex subunit (LHCB1.3) | 9 | 0.48 | 0.242 | 0.987 | |
| AT1G29930.1 | Photosystem I/II light harvesting complex subunit (LHCB1.3) | 10 | 0.55 | 0.242 | 0.999 | |

| | | | | | |
|---|---|---|---|---|---|
| AT1G29930.1 | Photosystem I/II light harvesting complex subunit (LHCB1.3) | 11 | 0.68 | 0.243 | 1.000 |
| AT1G29930.1 | Photosystem I/II light harvesting complex subunit (LHCB1.3) | 12 | 0.80 | 0.243 | 1.000 |
| AT1G29930.1 | Photosystem I/II light harvesting complex subunit (LHCB1.3) | 13 | 0.97 | 0.240 | 1.000 |
| AT1G29930.1 | Photosystem I/II light harvesting complex subunit (LHCB1.3) | 14 | 1.13 | 0.238 | 1.000 |
| AT1G29930.1 | Photosystem I/II light harvesting complex subunit (LHCB1.3) | 15 | 1.14 | 0.237 | 1.000 |
| AT1G29930.1 | Photosystem I/II light harvesting complex subunit (LHCB1.3) | 16 | 1.02 | 0.236 | 1.000 |
| AT1G29930.1 | Photosystem I/II light harvesting complex subunit (LHCB1.3) | 17 | 0.98 | 0.236 | 1.000 |
| AT1G29930.1 | Photosystem I/II light harvesting complex subunit (LHCB1.3) | 18 | 1.14 | 0.239 | 1.000 |
| AT1G29930.1 | Photosystem I/II light harvesting complex subunit (LHCB1.3) | 19 | 1.44 | 0.243 | 1.000 |
| AT1G29930.1 | Photosystem I/II light harvesting complex subunit (LHCB1.3) | 20 | 1.75 | 0.248 | 1.000 |
| AT1G29930.1 | Photosystem I/II light harvesting complex subunit (LHCB1.3) | 21 | 1.89 | 0.251 | 0.998 |
| AT1G29930.1 | Photosystem I/II light harvesting complex subunit (LHCB1.3) | 22 | 1.74 | 0.251 | 1.000 |
| AT1G29930.1 | Photosystem I/II light harvesting complex subunit (LHCB1.3) | 23 | 1.34 | 0.249 | 1.000 |
| AT1G29930.1 | Photosystem I/II light harvesting complex subunit (LHCB1.3) | 24 | 1.03 | 0.245 | 1.000 |
| AT1G29930.1 | Photosystem I/II light harvesting complex subunit (LHCB1.3) | 25 | 0.91 | 0.242 | 1.000 |
| AT1G29930.1 | Photosystem I/II light harvesting complex subunit (LHCB1.3) | 26 | 0.86 | 0.241 | 1.000 |
| AT1G29930.1 | Photosystem I/II light harvesting complex subunit (LHCB1.3) | 27 | 0.86 | 0.241 | 1.000 |
| AT1G29930.1 | Photosystem I/II light harvesting complex subunit (LHCB1.3) | 28 | 0.91 | 0.241 | 1.000 |
| AT1G29930.1 | Photosystem I/II light harvesting complex subunit (LHCB1.3) | 29 | 0.99 | 0.241 | 1.000 |
| AT1G29930.1 | Photosystem I/II light harvesting complex subunit (LHCB1.3) | 30 | 1.19 | 0.241 | 1.000 |

| | | | | | |
|---|---|---|---|---|---|
| AT1G29930.1 | Photosystem I/II light harvesting complex subunit (LHCB1.3) | 31 | 1.50 | 0.241 | 1.000 |
| AT1G29930.1 | Photosystem I/II light harvesting complex subunit (LHCB1.3) | 32 | 1.74 | 0.242 | 1.000 |
| AT1G29930.1 | Photosystem I/II light harvesting complex subunit (LHCB1.3) | 33 | 2.13 | 0.242 | 0.980 |
| AT1G29930.1 | Photosystem I/II light harvesting complex subunit (LHCB1.3) | 34 | 2.80 | 0.242 | 0.752 |
| AT1G29930.1 | Photosystem I/II light harvesting complex subunit (LHCB1.3) | 35 | 3.16 | 0.242 | 0.574 |
| AT1G29930.1 | Photosystem I/II light harvesting complex subunit (LHCB1.3) | 36 | 2.41 | 0.241 | 0.912 |
| AT1G29930.1 | Photosystem I/II light harvesting complex subunit (LHCB1.3) | 37 | 1.41 | 0.241 | 1.000 |
| AT1G29930.1 | Photosystem I/II light harvesting complex subunit (LHCB1.3) | 38 | 1.03 | 0.242 | 1.000 |
| AT1G29930.1 | Photosystem I/II light harvesting complex subunit (LHCB1.3) | 39 | 1.18 | 0.242 | 1.000 |
| AT1G29930.1 | Photosystem I/II light harvesting complex subunit (LHCB1.3) | 40 | 1.73 | 0.242 | 1.000 |
| AT1G29930.1 | Photosystem I/II light harvesting complex subunit (LHCB1.3) | 41 | 2.30 | 0.243 | 0.948 |
| AT1G29930.1 | Photosystem I/II light harvesting complex subunit (LHCB1.3) | 42 | 2.22 | 0.246 | 0.970 |
| AT1G29930.1 | Photosystem I/II light harvesting complex subunit (LHCB1.3) | 43 | 1.57 | 0.248 | 1.000 |
| AT1G29930.1 | Photosystem I/II light harvesting complex subunit (LHCB1.3) | 44 | 1.04 | 0.248 | 1.000 |
| AT1G29930.1 | Photosystem I/II light harvesting complex subunit (LHCB1.3) | 45 | 0.77 | 0.249 | 1.000 |
| AT1G29930.1 | Photosystem I/II light harvesting complex subunit (LHCB1.3) | 46 | 0.66 | 0.287 | 1.000 |
| AT1G29930.1 | Photosystem I/II light harvesting complex subunit (LHCB1.3) | 47 | 0.60 | 0.456 | 1.000 |
| AT1G29930.1 | Photosystem I/II light harvesting complex subunit (LHCB1.3) | 48 | 0.55 | 0.722 | 1.000 |
| AT1G29960.1 | Inner mitochondrial membrane protease 1-2 (IMP1-2) | 2 | 1.24 | 0.382 | 1.000 |
| AT1G29960.1 | Inner mitochondrial membrane protease 1-2 (IMP1-2) | 3 | 1.17 | 0.357 | 1.000 |

| | | | | | |
|---|---|---|---|---|---|
| AT1G29960.1 | Inner mitochondrial membrane protease 1-2 (IMP1-2) | 4 | 1.11 | 0.348 | 1.000 |
| AT1G29960.1 | Inner mitochondrial membrane protease 1-2 (IMP1-2) | 16 | 0.70 | 0.298 | 0.996 |
| AT1G29960.1 | Inner mitochondrial membrane protease 1-2 (IMP1-2) | 17 | 0.69 | 0.286 | 0.994 |
| AT1G29960.1 | Inner mitochondrial membrane protease 1-2 (IMP1-2) | 18 | 0.69 | 0.277 | 0.992 |
| AT1G29960.1 | Inner mitochondrial membrane protease 1-2 (IMP1-2) | 22 | 0.74 | 0.297 | 0.999 |
| AT1G29960.1 | Inner mitochondrial membrane protease 1-2 (IMP1-2) | 23 | 0.78 | 0.313 | 1.000 |
| AT1G29960.1 | Inner mitochondrial membrane protease 1-2 (IMP1-2) | 24 | 0.83 | 0.330 | 1.000 |
| AT1G29960.1 | Inner mitochondrial membrane protease 1-2 (IMP1-2) | 40 | 1.65 | 0.224 | 0.908 |
| AT1G29960.1 | Inner mitochondrial membrane protease 1-2 (IMP1-2) | 41 | 1.48 | 0.200 | 0.948 |
| AT1G29960.1 | Inner mitochondrial membrane protease 1-2 (IMP1-2) | 42 | 1.27 | 0.184 | 0.994 |
| AT1G29960.1 | Inner mitochondrial membrane protease 1-2 (IMP1-2) | 43 | 1.03 | 0.176 | 1.000 |
| AT1G29960.1 | Inner mitochondrial membrane protease 1-2 (IMP1-2) | 44 | 0.78 | 0.177 | 0.990 |
| AT1G29960.1 | Inner mitochondrial membrane protease 1-2 (IMP1-2) | 45 | 0.56 | 0.191 | 0.725 |
| AT1G29960.1 | Inner mitochondrial membrane protease 1-2 (IMP1-2) | 46 | 0.40 | 0.220 | 0.387 |
| AT1G29960.1 | Inner mitochondrial membrane protease 1-2 (IMP1-2) | 47 | 0.29 | 0.265 | 0.247 |
| AT1G30360.1 | Early-responsive to dehydration stress protein (ERD4) | 2 | 1.00 | 0.325 | 1.000 |
| AT1G30360.1 | Early-responsive to dehydration stress protein (ERD4) | 3 | 1.08 | 0.257 | 1.000 |
| AT1G30360.1 | Early-responsive to dehydration stress protein (ERD4) | 4 | 1.17 | 0.222 | 1.000 |
| AT1G30360.1 | Early-responsive to dehydration stress protein (ERD4) | 5 | 1.26 | 0.220 | 1.000 |
| AT1G30360.1 | Early-responsive to dehydration stress protein (ERD4) | 6 | 1.37 | 0.242 | 0.998 |

| | | | | | |
|---|---|---|---|---|---|
| AT1G30360.1 | Early-responsive to dehydration stress protein (ERD4) | 7 | 1.49 | 0.277 | 0.997 |
| AT1G30360.1 | Early-responsive to dehydration stress protein (ERD4) | 22 | 2.73 | 0.360 | 0.857 |
| AT1G30360.1 | Early-responsive to dehydration stress protein (ERD4) | 23 | 2.46 | 0.332 | 0.874 |
| AT1G30360.1 | Early-responsive to dehydration stress protein (ERD4) | 24 | 2.15 | 0.304 | 0.910 |
| AT1G30360.1 | Early-responsive to dehydration stress protein (ERD4) | 28 | 0.93 | 0.223 | 1.000 |
| AT1G30360.1 | Early-responsive to dehydration stress protein (ERD4) | 29 | 0.72 | 0.208 | 0.993 |
| AT1G30360.1 | Early-responsive to dehydration stress protein (ERD4) | 30 | 0.57 | 0.199 | 0.839 |
| AT1G30360.1 | Early-responsive to dehydration stress protein (ERD4) | 31 | 0.48 | 0.199 | 0.605 |
| AT1G30360.1 | Early-responsive to dehydration stress protein (ERD4) | 32 | 0.44 | 0.208 | 0.534 |
| AT1G30360.1 | Early-responsive to dehydration stress protein (ERD4) | 33 | 0.43 | 0.225 | 0.601 |
| AT1G30360.1 | Early-responsive to dehydration stress protein (ERD4) | 34 | 0.45 | 0.248 | 0.740 |
| AT1G30360.1 | Early-responsive to dehydration stress protein (ERD4) | 41 | 1.11 | 0.366 | 1.000 |
| AT1G30360.1 | Early-responsive to dehydration stress protein (ERD4) | 42 | 1.32 | 0.398 | 1.000 |
| AT1G30360.1 | Early-responsive to dehydration stress protein (ERD4) | 43 | 1.56 | 0.452 | 1.000 |
| AT1G30360.1 | Early-responsive to dehydration stress protein (ERD4) | 46 | 2.59 | 0.742 | 0.999 |
| AT1G30360.1 | Early-responsive to dehydration stress protein (ERD4) | 47 | 3.08 | 0.872 | 0.999 |
| AT1G30380.1 | Photosystem I subunit (PSAK) | 2 | 0.77 | 0.276 | 1.000 |
| AT1G30380.1 | Photosystem I subunit (PSAK) | 3 | 0.75 | 0.198 | 1.000 |
| AT1G30380.1 | Photosystem I subunit (PSAK) | 4 | 0.74 | 0.160 | 0.999 |
| AT1G30380.1 | Photosystem I subunit (PSAK) | 5 | 0.76 | 0.149 | 0.999 |
| AT1G30380.1 | Photosystem I subunit (PSAK) | 6 | 0.77 | 0.148 | 0.999 |
| AT1G30380.1 | Photosystem I subunit (PSAK) | 7 | 0.80 | 0.147 | 1.000 |
| AT1G30380.1 | Photosystem I subunit (PSAK) | 8 | 0.85 | 0.146 | 1.000 |
| AT1G30380.1 | Photosystem I subunit (PSAK) | 9 | 0.92 | 0.144 | 1.000 |
| AT1G30380.1 | Photosystem I subunit (PSAK) | 10 | 0.99 | 0.143 | 1.000 |
| AT1G30380.1 | Photosystem I subunit (PSAK) | 11 | 1.07 | 0.142 | 1.000 |
| AT1G30380.1 | Photosystem I subunit (PSAK) | 12 | 1.13 | 0.141 | 1.000 |
| AT1G30380.1 | Photosystem I subunit (PSAK) | 13 | 1.18 | 0.140 | 1.000 |
| AT1G30380.1 | Photosystem I subunit (PSAK) | 14 | 1.20 | 0.140 | 1.000 |
| AT1G30380.1 | Photosystem I subunit (PSAK) | 15 | 1.18 | 0.139 | 1.000 |

| | | | | | |
|---|---|---|---|---|---|
| AT1G30380.1 | Photosystem I subunit (PSAK) | 16 | 1.12 | 0.138 | 1.000 |
| AT1G30380.1 | Photosystem I subunit (PSAK) | 17 | 1.07 | 0.138 | 1.000 |
| AT1G30380.1 | Photosystem I subunit (PSAK) | 18 | 1.06 | 0.140 | 1.000 |
| AT1G30380.1 | Photosystem I subunit (PSAK) | 19 | 1.08 | 0.143 | 1.000 |
| AT1G30380.1 | Photosystem I subunit (PSAK) | 20 | 1.11 | 0.145 | 1.000 |
| AT1G30380.1 | Photosystem I subunit (PSAK) | 21 | 1.11 | 0.147 | 1.000 |
| AT1G30380.1 | Photosystem I subunit (PSAK) | 22 | 1.08 | 0.147 | 1.000 |
| AT1G30380.1 | Photosystem I subunit (PSAK) | 23 | 1.02 | 0.146 | 1.000 |
| AT1G30380.1 | Photosystem I subunit (PSAK) | 24 | 0.94 | 0.145 | 1.000 |
| AT1G30380.1 | Photosystem I subunit (PSAK) | 25 | 0.88 | 0.145 | 1.000 |
| AT1G30380.1 | Photosystem I subunit (PSAK) | 26 | 0.89 | 0.148 | 1.000 |
| AT1G30380.1 | Photosystem I subunit (PSAK) | 27 | 1.01 | 0.156 | 1.000 |
| AT1G30380.1 | Photosystem I subunit (PSAK) | 28 | 1.28 | 0.176 | 1.000 |
| AT1G30380.1 | Photosystem I subunit (PSAK) | 29 | 1.83 | 0.211 | 0.950 |
| AT1G30380.1 | Photosystem I subunit (PSAK) | 30 | 2.83 | 0.271 | 0.734 |
| AT1G30380.1 | Photosystem I subunit (PSAK) | 46 | 211816.26 | 3.159 | 0.723 |
| AT1G30380.1 | Photosystem I subunit (PSAK) | 47 | 486414.74 | 3.428 | 0.743 |
| AT1G30680.1 | Arabidopsis twinkle homolog (ATH) | 2 | 1.36 | 0.344 | 1.000 |
| AT1G30680.1 | Arabidopsis twinkle homolog (ATH) | 3 | 1.31 | 0.292 | 1.000 |
| AT1G30680.1 | Arabidopsis twinkle homolog (ATH) | 9 | 1.03 | 0.247 | 1.000 |
| AT1G30680.1 | Arabidopsis twinkle homolog (ATH) | 10 | 0.99 | 0.207 | 1.000 |
| AT1G30680.1 | Arabidopsis twinkle homolog (ATH) | 11 | 0.99 | 0.183 | 1.000 |
| AT1G30680.1 | Arabidopsis twinkle homolog (ATH) | 12 | 1.05 | 0.169 | 1.000 |
| AT1G30680.1 | Arabidopsis twinkle homolog (ATH) | 13 | 1.21 | 0.162 | 1.000 |
| AT1G30680.1 | Arabidopsis twinkle homolog (ATH) | 14 | 1.49 | 0.156 | 0.978 |
| AT1G30680.1 | Arabidopsis twinkle homolog (ATH) | 15 | 1.70 | 0.151 | 0.824 |
| AT1G30680.1 | Arabidopsis twinkle homolog (ATH) | 16 | 1.68 | 0.149 | 0.832 |
| AT1G30680.1 | Arabidopsis twinkle homolog (ATH) | 17 | 1.54 | 0.149 | 0.945 |
| AT1G30680.1 | Arabidopsis twinkle homolog (ATH) | 18 | 1.40 | 0.150 | 0.993 |
| AT1G30680.1 | Arabidopsis twinkle homolog (ATH) | 19 | 1.33 | 0.154 | 0.999 |
| AT1G30680.1 | Arabidopsis twinkle homolog (ATH) | 20 | 1.29 | 0.157 | 1.000 |
| AT1G30680.1 | Arabidopsis twinkle homolog (ATH) | 21 | 1.24 | 0.158 | 1.000 |
| AT1G30680.1 | Arabidopsis twinkle homolog (ATH) | 22 | 1.16 | 0.158 | 1.000 |
| AT1G30680.1 | Arabidopsis twinkle homolog (ATH) | 23 | 1.08 | 0.157 | 1.000 |
| AT1G30680.1 | Arabidopsis twinkle homolog (ATH) | 24 | 1.03 | 0.154 | 1.000 |
| AT1G30680.1 | Arabidopsis twinkle homolog (ATH) | 25 | 0.99 | 0.153 | 1.000 |
| AT1G30680.1 | Arabidopsis twinkle homolog (ATH) | 26 | 0.92 | 0.153 | 1.000 |
| AT1G30680.1 | Arabidopsis twinkle homolog (ATH) | 27 | 0.77 | 0.156 | 1.000 |
| AT1G30680.1 | Arabidopsis twinkle homolog (ATH) | 28 | 0.63 | 0.168 | 0.965 |

| | | | | | |
|---|---|---|---|---|---|
| AT1G30680.1 | Arabidopsis twinkle homolog (ATH) | 29 | 0.61 | 0.203 | 0.987 |
| AT1G30680.1 | Arabidopsis twinkle homolog (ATH) | 30 | 0.67 | 0.286 | 1.000 |
| AT1G30680.1 | Arabidopsis twinkle homolog (ATH) | 46 | 812100.33 | 4.218 | 0.890 |
| AT1G30680.1 | Arabidopsis twinkle homolog (ATH) | 47 | 2856080.98 | 4.596 | 0.888 |
| AT1G31175.1 | Cytochrome c:O2 oxidoreductase assembly protein (CMC1-like) | 2 | 0.99 | 0.364 | 1.000 |
| AT1G31175.1 | Cytochrome c:O2 oxidoreductase assembly protein (CMC1-like) | 3 | 0.97 | 0.331 | 1.000 |
| AT1G31175.1 | Cytochrome c:O2 oxidoreductase assembly protein (CMC1-like) | 4 | 0.96 | 0.320 | 1.000 |
| AT1G31175.1 | Cytochrome c:O2 oxidoreductase assembly protein (CMC1-like) | 17 | 1.12 | 0.231 | 1.000 |
| AT1G31175.1 | Cytochrome c:O2 oxidoreductase assembly protein (CMC1-like) | 18 | 1.18 | 0.210 | 1.000 |
| AT1G31175.1 | Cytochrome c:O2 oxidoreductase assembly protein (CMC1-like) | 19 | 1.26 | 0.191 | 0.999 |
| AT1G31175.1 | Cytochrome c:O2 oxidoreductase assembly protein (CMC1-like) | 20 | 1.34 | 0.177 | 0.988 |
| AT1G31175.1 | Cytochrome c:O2 oxidoreductase assembly protein (CMC1-like) | 21 | 1.42 | 0.167 | 0.954 |
| AT1G31175.1 | Cytochrome c:O2 oxidoreductase assembly protein (CMC1-like) | 22 | 1.50 | 0.160 | 0.887 |
| AT1G31175.1 | Cytochrome c:O2 oxidoreductase assembly protein (CMC1-like) | 23 | 1.59 | 0.156 | 0.786 |
| AT1G31175.1 | Cytochrome c:O2 oxidoreductase assembly protein (CMC1-like) | 24 | 1.70 | 0.156 | 0.655 |
| AT1G31175.1 | Cytochrome c:O2 oxidoreductase assembly protein (CMC1-like) | 25 | 1.85 | 0.159 | 0.511 |
| AT1G31175.1 | Cytochrome c:O2 oxidoreductase assembly protein (CMC1-like) | 26 | 2.03 | 0.167 | 0.405 |
| AT1G31175.1 | Cytochrome c:O2 oxidoreductase assembly protein (CMC1-like) | 27 | 2.21 | 0.183 | 0.377 |
| AT1G31175.1 | Cytochrome c:O2 oxidoreductase assembly protein (CMC1-like) | 28 | 2.31 | 0.205 | 0.445 |
| AT1G31175.1 | Cytochrome c:O2 oxidoreductase assembly protein (CMC1-like) | 29 | 2.34 | 0.231 | 0.571 |
| AT1G31175.1 | Cytochrome c:O2 oxidoreductase assembly protein (CMC1-like) | 41 | 1.05 | 0.362 | 1.000 |
| AT1G31175.1 | Cytochrome c:O2 oxidoreductase assembly protein (CMC1-like) | 42 | 1.00 | 0.372 | 1.000 |

| | | | | | |
|---|---|---|---|---|---|
| AT1G31175.1 | Cytochrome c:O2 oxidoreductase assembly protein (CMC1-like) | 43 | 0.97 | 0.390 | 1.000 |
| AT1G31175.1 | Cytochrome c:O2 oxidoreductase assembly protein (CMC1-like) | 46 | 0.93 | 0.509 | 1.000 |
| AT1G31175.1 | Cytochrome c:O2 oxidoreductase assembly protein (CMC1-like) | 47 | 0.92 | 0.571 | 1.000 |
| AT1G31330.1 | Photosystem I subunit (PSAF) | 2 | 0.37 | 0.260 | 0.905 |
| AT1G31330.1 | Photosystem I subunit (PSAF) | 3 | 0.41 | 0.160 | 0.344 |
| AT1G31330.1 | Photosystem I subunit (PSAF) | 4 | 0.51 | 0.147 | 0.696 |
| AT1G31330.1 | Photosystem I subunit (PSAF) | 5 | 0.67 | 0.147 | 0.997 |
| AT1G31330.1 | Photosystem I subunit (PSAF) | 6 | 0.76 | 0.147 | 1.000 |
| AT1G31330.1 | Photosystem I subunit (PSAF) | 7 | 0.76 | 0.147 | 1.000 |
| AT1G31330.1 | Photosystem I subunit (PSAF) | 8 | 0.78 | 0.144 | 1.000 |
| AT1G31330.1 | Photosystem I subunit (PSAF) | 9 | 0.80 | 0.142 | 1.000 |
| AT1G31330.1 | Photosystem I subunit (PSAF) | 10 | 0.80 | 0.142 | 1.000 |
| AT1G31330.1 | Photosystem I subunit (PSAF) | 11 | 0.82 | 0.142 | 1.000 |
| AT1G31330.1 | Photosystem I subunit (PSAF) | 12 | 0.85 | 0.142 | 1.000 |
| AT1G31330.1 | Photosystem I subunit (PSAF) | 13 | 0.89 | 0.140 | 1.000 |
| AT1G31330.1 | Photosystem I subunit (PSAF) | 14 | 0.93 | 0.138 | 1.000 |
| AT1G31330.1 | Photosystem I subunit (PSAF) | 15 | 0.94 | 0.138 | 1.000 |
| AT1G31330.1 | Photosystem I subunit (PSAF) | 16 | 0.94 | 0.137 | 1.000 |
| AT1G31330.1 | Photosystem I subunit (PSAF) | 17 | 1.00 | 0.137 | 1.000 |
| AT1G31330.1 | Photosystem I subunit (PSAF) | 18 | 1.15 | 0.138 | 1.000 |
| AT1G31330.1 | Photosystem I subunit (PSAF) | 19 | 1.34 | 0.142 | 1.000 |
| AT1G31330.1 | Photosystem I subunit (PSAF) | 20 | 1.46 | 0.146 | 0.998 |
| AT1G31330.1 | Photosystem I subunit (PSAF) | 21 | 1.42 | 0.148 | 1.000 |
| AT1G31330.1 | Photosystem I subunit (PSAF) | 22 | 1.22 | 0.148 | 1.000 |
| AT1G31330.1 | Photosystem I subunit (PSAF) | 23 | 0.97 | 0.147 | 1.000 |
| AT1G31330.1 | Photosystem I subunit (PSAF) | 24 | 0.80 | 0.143 | 1.000 |
| AT1G31330.1 | Photosystem I subunit (PSAF) | 25 | 0.79 | 0.141 | 1.000 |
| AT1G31330.1 | Photosystem I subunit (PSAF) | 26 | 1.01 | 0.140 | 1.000 |
| AT1G31330.1 | Photosystem I subunit (PSAF) | 27 | 1.27 | 0.140 | 1.000 |
| AT1G31330.1 | Photosystem I subunit (PSAF) | 28 | 1.29 | 0.140 | 1.000 |
| AT1G31330.1 | Photosystem I subunit (PSAF) | 29 | 1.08 | 0.141 | 1.000 |
| AT1G31330.1 | Photosystem I subunit (PSAF) | 30 | 0.72 | 0.142 | 1.000 |
| AT1G31330.1 | Photosystem I subunit (PSAF) | 31 | 0.53 | 0.143 | 0.723 |
| AT1G31330.1 | Photosystem I subunit (PSAF) | 32 | 0.62 | 0.144 | 0.976 |
| AT1G31330.1 | Photosystem I subunit (PSAF) | 33 | 0.86 | 0.147 | 1.000 |
| AT1G31330.1 | Photosystem I subunit (PSAF) | 34 | 1.03 | 0.157 | 1.000 |
| AT1G31330.1 | Photosystem I subunit (PSAF) | 35 | 0.95 | 0.193 | 1.000 |
| AT1G31330.1 | Photosystem I subunit (PSAF) | 36 | 0.81 | 0.294 | 1.000 |
| AT1G31330.1 | Photosystem I subunit (PSAF) | 46 | 0.85 | 2.675 | 1.000 |
| AT1G31330.1 | Photosystem I subunit (PSAF) | 47 | 0.93 | 3.037 | 1.000 |
| AT1G32470.1 | Glycine decarboxylase subunit (GDCH) | 2 | 0.66 | 0.528 | 1.000 |
| AT1G32470.1 | Glycine decarboxylase subunit (GDCH) | 3 | 0.66 | 0.398 | 1.000 |
| AT1G32470.1 | Glycine decarboxylase subunit (GDCH) | 4 | 0.67 | 0.329 | 1.000 |
| AT1G32470.1 | Glycine decarboxylase subunit (GDCH) | 5 | 0.67 | 0.292 | 1.000 |
| AT1G32470.1 | Glycine decarboxylase subunit (GDCH) | 6 | 0.67 | 0.270 | 1.000 |
| AT1G32470.1 | Glycine decarboxylase subunit (GDCH) | 7 | 0.66 | 0.250 | 1.000 |
| AT1G32470.1 | Glycine decarboxylase subunit (GDCH) | 8 | 0.69 | 0.234 | 1.000 |
| AT1G32470.1 | Glycine decarboxylase subunit (GDCH) | 9 | 0.78 | 0.222 | 1.000 |

| | | | | | | |
|---|---|---|---|---|---|---|
| AT1G32470.1 | Glycine decarboxylase subunit (GDCH) | 10 | 0.96 | 0.215 | 1.000 | |
| AT1G32470.1 | Glycine decarboxylase subunit (GDCH) | 11 | 1.21 | 0.212 | 1.000 | |
| AT1G32470.1 | Glycine decarboxylase subunit (GDCH) | 12 | 1.45 | 0.211 | 1.000 | |
| AT1G32470.1 | Glycine decarboxylase subunit (GDCH) | 13 | 1.58 | 0.210 | 1.000 | |
| AT1G32470.1 | Glycine decarboxylase subunit (GDCH) | 14 | 1.66 | 0.208 | 0.998 | |
| AT1G32470.1 | Glycine decarboxylase subunit (GDCH) | 15 | 1.85 | 0.207 | 0.983 | |
| AT1G32470.1 | Glycine decarboxylase subunit (GDCH) | 16 | 2.27 | 0.206 | 0.826 | |
| AT1G32470.1 | Glycine decarboxylase subunit (GDCH) | 17 | 2.97 | 0.207 | 0.403 | |
| AT1G32470.1 | Glycine decarboxylase subunit (GDCH) | 18 | 3.90 | 0.213 | 0.132 | |
| AT1G32470.1 | Glycine decarboxylase subunit (GDCH) | 19 | 4.70 | 0.221 | 0.065 | . |
| AT1G32470.1 | Glycine decarboxylase subunit (GDCH) | 20 | 4.83 | 0.233 | 0.084 | . |
| AT1G32470.1 | Glycine decarboxylase subunit (GDCH) | 21 | 4.31 | 0.247 | 0.221 | |
| AT1G32470.1 | Glycine decarboxylase subunit (GDCH) | 22 | 3.47 | 0.264 | 0.585 | |
| AT1G32470.1 | Glycine decarboxylase subunit (GDCH) | 23 | 2.68 | 0.277 | 0.919 | |
| AT1G32470.1 | Glycine decarboxylase subunit (GDCH) | 24 | 2.26 | 0.291 | 0.990 | |
| AT1G32470.1 | Glycine decarboxylase subunit (GDCH) | 25 | 2.22 | 0.300 | 0.995 | |
| AT1G32470.1 | Glycine decarboxylase subunit (GDCH) | 26 | 2.48 | 0.316 | 0.987 | |
| AT1G32470.1 | Glycine decarboxylase subunit (GDCH) | 28 | 3.94 | 0.320 | 0.731 | |
| AT1G32470.1 | Glycine decarboxylase subunit (GDCH) | 29 | 4.93 | 0.312 | 0.450 | |
| AT1G32470.1 | Glycine decarboxylase subunit (GDCH) | 30 | 5.65 | 0.290 | 0.207 | |
| AT1G32470.1 | Glycine decarboxylase subunit (GDCH) | 31 | 5.65 | 0.270 | 0.131 | |
| AT1G32470.1 | Glycine decarboxylase subunit (GDCH) | 32 | 4.78 | 0.250 | 0.154 | |
| AT1G32470.1 | Glycine decarboxylase subunit (GDCH) | 33 | 3.52 | 0.234 | 0.358 | |
| AT1G32470.1 | Glycine decarboxylase subunit (GDCH) | 34 | 2.45 | 0.222 | 0.805 | |
| AT1G32470.1 | Glycine decarboxylase subunit (GDCH) | 35 | 1.77 | 0.215 | 0.995 | |
| AT1G32470.1 | Glycine decarboxylase subunit (GDCH) | 36 | 1.36 | 0.212 | 1.000 | |
| AT1G32470.1 | Glycine decarboxylase subunit (GDCH) | 37 | 1.18 | 0.211 | 1.000 | |
| AT1G32470.1 | Glycine decarboxylase subunit (GDCH) | 38 | 1.19 | 0.212 | 1.000 | |
| AT1G32470.1 | Glycine decarboxylase subunit (GDCH) | 39 | 1.44 | 0.212 | 1.000 | |
| AT1G32470.1 | Glycine decarboxylase subunit (GDCH) | 40 | 2.04 | 0.213 | 0.950 | |
| AT1G32470.1 | Glycine decarboxylase subunit (GDCH) | 41 | 2.89 | 0.216 | 0.513 | |

| | | | | | |
|---|---|---|---|---|---|
| AT1G32470.1 | Glycine decarboxylase subunit (GDCH) | 42 | 3.44 | 0.218 | 0.284 |
| AT1G32470.1 | Glycine decarboxylase subunit (GDCH) | 43 | 3.32 | 0.219 | 0.330 |
| AT1G32470.1 | Glycine decarboxylase subunit (GDCH) | 44 | 2.55 | 0.219 | 0.733 |
| AT1G32470.1 | Glycine decarboxylase subunit (GDCH) | 45 | 1.66 | 0.234 | 1.000 |
| AT1G32470.1 | Glycine decarboxylase subunit (GDCH) | 46 | 1.05 | 0.302 | 1.000 |
| AT1G32470.1 | Glycine decarboxylase subunit (GDCH) | 47 | 0.66 | 0.452 | 1.000 |
| AT1G32780.1 | Alcohol dehydrogenase-like 3 | 2 | 1.02 | 0.389 | 1.000 |
| AT1G32780.1 | Alcohol dehydrogenase-like 3 | 3 | 0.91 | 0.274 | 1.000 |
| AT1G32780.1 | Alcohol dehydrogenase-like 3 | 4 | 0.84 | 0.507 | 1.000 |
| AT1G32780.1 | Alcohol dehydrogenase-like 3 | 31 | 0.87 | 0.260 | 1.000 |
| AT1G32780.1 | Alcohol dehydrogenase-like 3 | 32 | 0.62 | 0.212 | 0.997 |
| AT1G32780.1 | Alcohol dehydrogenase-like 3 | 33 | 0.44 | 0.197 | 0.666 |
| AT1G32780.1 | Alcohol dehydrogenase-like 3 | 34 | 0.39 | 0.191 | 0.410 |
| AT1G32780.1 | Alcohol dehydrogenase-like 3 | 35 | 0.44 | 0.189 | 0.617 |
| AT1G32780.1 | Alcohol dehydrogenase-like 3 | 36 | 0.49 | 0.188 | 0.808 |
| AT1G32780.1 | Alcohol dehydrogenase-like 3 | 37 | 0.64 | 0.189 | 0.994 |
| AT1G32780.1 | Alcohol dehydrogenase-like 3 | 38 | 0.97 | 0.189 | 1.000 |
| AT1G32780.1 | Alcohol dehydrogenase-like 3 | 39 | 1.52 | 0.188 | 0.997 |
| AT1G32780.1 | Alcohol dehydrogenase-like 3 | 40 | 1.71 | 0.187 | 0.968 |
| AT1G32780.1 | Alcohol dehydrogenase-like 3 | 41 | 1.36 | 0.187 | 1.000 |
| AT1G32780.1 | Alcohol dehydrogenase-like 3 | 42 | 1.23 | 0.191 | 1.000 |
| AT1G32780.1 | Alcohol dehydrogenase-like 3 | 43 | 0.94 | 0.190 | 1.000 |
| AT1G32780.1 | Alcohol dehydrogenase-like 3 | 44 | 0.44 | 0.188 | 0.625 |
| AT1G32780.1 | Alcohol dehydrogenase-like 3 | 45 | 0.34 | 0.190 | 0.229 |
| AT1G32780.1 | Alcohol dehydrogenase-like 3 | 46 | 0.64 | 0.207 | 0.998 |
| AT1G32780.1 | Alcohol dehydrogenase-like 3 | 47 | 1.84 | 0.440 | 1.000 |
| AT1G33140.1 | 60S ribosomal protein L9-1 (RPL9B) | 2 | 0.45 | 0.162 | 0.175 |
| AT1G33140.1 | 60S ribosomal protein L9-1 (RPL9B) | 3 | 0.49 | 0.145 | 0.184 |
| AT1G33140.1 | 60S ribosomal protein L9-1 (RPL9B) | 4 | 0.54 | 0.130 | 0.217 |
| AT1G33140.1 | 60S ribosomal protein L9-1 (RPL9B) | 5 | 0.59 | 0.119 | 0.294 |
| AT1G33140.1 | 60S ribosomal protein L9-1 (RPL9B) | 6 | 0.66 | 0.110 | 0.437 |
| AT1G33140.1 | 60S ribosomal protein L9-1 (RPL9B) | 7 | 0.73 | 0.104 | 0.657 |
| AT1G33140.1 | 60S ribosomal protein L9-1 (RPL9B) | 8 | 0.80 | 0.100 | 0.886 |
| AT1G33140.1 | 60S ribosomal protein L9-1 (RPL9B) | 9 | 0.89 | 0.096 | 1.000 |
| AT1G33140.1 | 60S ribosomal protein L9-1 (RPL9B) | 10 | 0.98 | 0.094 | 1.000 |
| AT1G33140.1 | 60S ribosomal protein L9-1 (RPL9B) | 11 | 1.08 | 0.093 | 1.000 |
| AT1G33140.1 | 60S ribosomal protein L9-1 (RPL9B) | 12 | 1.18 | 0.092 | 0.945 |
| AT1G33140.1 | 60S ribosomal protein L9-1 (RPL9B) | 13 | 1.29 | 0.091 | 0.738 |
| AT1G33140.1 | 60S ribosomal protein L9-1 (RPL9B) | 14 | 1.40 | 0.091 | 0.474 |
| AT1G33140.1 | 60S ribosomal protein L9-1 (RPL9B) | 15 | 1.50 | 0.091 | 0.273 |
| AT1G33140.1 | 60S ribosomal protein L9-1 (RPL9B) | 16 | 1.61 | 0.093 | 1.000 |

| | | | | | | |
|---|---|---|---|---|---|---|
| AT1G33140.1 | 60S ribosomal protein L9-1 (RPL9B) | 17 | 1.71 | 0.095 | 1.000 | |
| AT1G33140.1 | 60S ribosomal protein L9-1 (RPL9B) | 18 | 1.79 | 0.098 | 1.000 | |
| AT1G33140.1 | 60S ribosomal protein L9-1 (RPL9B) | 19 | 1.87 | 0.103 | 1.000 | |
| AT1G33140.1 | 60S ribosomal protein L9-1 (RPL9B) | 20 | 1.93 | 0.108 | 1.000 | |
| AT1G33140.1 | 60S ribosomal protein L9-1 (RPL9B) | 21 | 1.98 | 0.115 | 1.000 | |
| AT1G33140.1 | 60S ribosomal protein L9-1 (RPL9B) | 22 | 2.01 | 0.123 | 1.000 | |
| AT1G33140.1 | 60S ribosomal protein L9-1 (RPL9B) | 23 | 2.03 | 0.132 | 1.000 | |
| AT1G33140.1 | 60S ribosomal protein L9-1 (RPL9B) | 24 | 2.04 | 0.142 | 1.000 | |
| AT1G33140.1 | 60S ribosomal protein L9-1 (RPL9B) | 41 | 1.18 | 0.301 | 1.000 | |
| AT1G33140.1 | 60S ribosomal protein L9-1 (RPL9B) | 42 | 1.12 | 0.317 | 1.000 | |
| AT1G33140.1 | 60S ribosomal protein L9-1 (RPL9B) | 43 | 1.07 | 0.334 | 1.000 | |
| AT1G33140.1 | 60S ribosomal protein L9-1 (RPL9B) | 46 | 0.92 | 0.403 | 1.000 | |
| AT1G33140.1 | 60S ribosomal protein L9-1 (RPL9B) | 47 | 0.87 | 0.430 | 1.000 | |
| AT1G33360.1 | Caseinolytic protease X3 (CLPX3) | 2 | 2.44 | 0.774 | 1.000 | |
| AT1G33360.1 | Caseinolytic protease X3 (CLPX3) | 3 | 3.28 | 0.705 | 0.999 | |
| AT1G33360.1 | Caseinolytic protease X3 (CLPX3) | 20 | 0.84 | 0.428 | 1.000 | |
| AT1G33360.1 | Caseinolytic protease X3 (CLPX3) | 21 | 0.45 | 0.355 | 0.985 | |
| AT1G33360.1 | Caseinolytic protease X3 (CLPX3) | 22 | 0.25 | 0.313 | 0.476 | |
| AT1G33360.1 | Caseinolytic protease X3 (CLPX3) | 23 | 0.17 | 0.290 | 0.115 | |
| AT1G33360.1 | Caseinolytic protease X3 (CLPX3) | 24 | 0.17 | 0.282 | 0.081 | . |
| AT1G33360.1 | Caseinolytic protease X3 (CLPX3) | 25 | 0.23 | 0.280 | 0.262 | |
| AT1G33360.1 | Caseinolytic protease X3 (CLPX3) | 26 | 0.40 | 0.281 | 0.835 | |
| AT1G33360.1 | Caseinolytic protease X3 (CLPX3) | 27 | 0.67 | 0.283 | 1.000 | |
| AT1G33360.1 | Caseinolytic protease X3 (CLPX3) | 28 | 1.01 | 0.287 | 1.000 | |
| AT1G33360.1 | Caseinolytic protease X3 (CLPX3) | 29 | 1.36 | 0.292 | 1.000 | |
| AT1G33360.1 | Caseinolytic protease X3 (CLPX3) | 30 | 1.55 | 0.297 | 1.000 | |
| AT1G33360.1 | Caseinolytic protease X3 (CLPX3) | 31 | 1.52 | 0.305 | 1.000 | |
| AT1G33360.1 | Caseinolytic protease X3 (CLPX3) | 32 | 1.35 | 0.321 | 1.000 | |
| AT1G33360.1 | Caseinolytic protease X3 (CLPX3) | 33 | 1.09 | 0.344 | 1.000 | |
| AT1G33360.1 | Caseinolytic protease X3 (CLPX3) | 34 | 0.80 | 0.367 | 1.000 | |
| AT1G33360.1 | Caseinolytic protease X3 (CLPX3) | 35 | 0.54 | 0.389 | 0.999 | |

| | | | | | |
|---|---|---|---|---|---|
| AT1G33360.1 | Caseinolytic protease X3 (CLPX3) | 38 | 0.20 | 0.391 | 0.584 |
| AT1G33360.1 | Caseinolytic protease X3 (CLPX3) | 39 | 0.20 | 0.370 | 0.523 |
| AT1G33360.1 | Caseinolytic protease X3 (CLPX3) | 40 | 0.25 | 0.352 | 0.654 |
| AT1G33360.1 | Caseinolytic protease X3 (CLPX3) | 41 | 0.34 | 0.343 | 0.860 |
| AT1G33360.1 | Caseinolytic protease X3 (CLPX3) | 42 | 0.41 | 0.341 | 0.957 |
| AT1G33360.1 | Caseinolytic protease X3 (CLPX3) | 43 | 0.42 | 0.343 | 0.965 |
| AT1G33360.1 | Caseinolytic protease X3 (CLPX3) | 44 | 0.35 | 0.360 | 0.910 |
| AT1G33360.1 | Caseinolytic protease X3 (CLPX3) | 45 | 0.25 | 0.421 | 0.832 |
| AT1G33360.1 | Caseinolytic protease X3 (CLPX3) | 46 | 0.17 | 0.562 | 0.869 |
| AT1G33360.1 | Caseinolytic protease X3 (CLPX3) | 47 | 0.12 | 0.785 | 0.943 |
| AT1G42970.1 | Glyceraldehyde-3-phosphate dehydrogenase (GAPB) | 2 | 1.28 | 0.627 | 1.000 |
| AT1G42970.1 | Glyceraldehyde-3-phosphate dehydrogenase (GAPB) | 3 | 0.95 | 0.490 | 1.000 |
| AT1G42970.1 | Glyceraldehyde-3-phosphate dehydrogenase (GAPB) | 6 | 0.43 | 0.432 | 1.000 |
| AT1G42970.1 | Glyceraldehyde-3-phosphate dehydrogenase (GAPB) | 7 | 0.36 | 0.483 | 1.000 |
| AT1G42970.1 | Glyceraldehyde-3-phosphate dehydrogenase (GAPB) | 8 | 0.31 | 0.539 | 0.999 |
| AT1G42970.1 | Glyceraldehyde-3-phosphate dehydrogenase (GAPB) | 26 | 0.94 | 0.473 | 1.000 |
| AT1G42970.1 | Glyceraldehyde-3-phosphate dehydrogenase (GAPB) | 27 | 0.97 | 0.345 | 1.000 |
| AT1G42970.1 | Glyceraldehyde-3-phosphate dehydrogenase (GAPB) | 28 | 0.99 | 0.300 | 1.000 |
| AT1G42970.1 | Glyceraldehyde-3-phosphate dehydrogenase (GAPB) | 29 | 1.01 | 0.282 | 1.000 |
| AT1G42970.1 | Glyceraldehyde-3-phosphate dehydrogenase (GAPB) | 30 | 1.17 | 0.278 | 1.000 |
| AT1G42970.1 | Glyceraldehyde-3-phosphate dehydrogenase (GAPB) | 31 | 2.78 | 0.278 | 0.876 |
| AT1G42970.1 | Glyceraldehyde-3-phosphate dehydrogenase (GAPB) | 32 | 5.30 | 0.278 | 0.183 |
| AT1G42970.1 | Glyceraldehyde-3-phosphate dehydrogenase (GAPB) | 33 | 3.62 | 0.279 | 0.587 |
| AT1G42970.1 | Glyceraldehyde-3-phosphate dehydrogenase (GAPB) | 34 | 1.53 | 0.281 | 1.000 |
| AT1G42970.1 | Glyceraldehyde-3-phosphate dehydrogenase (GAPB) | 35 | 0.93 | 0.281 | 1.000 |
| AT1G42970.1 | Glyceraldehyde-3-phosphate dehydrogenase (GAPB) | 36 | 0.98 | 0.279 | 1.000 |
| AT1G42970.1 | Glyceraldehyde-3-phosphate dehydrogenase (GAPB) | 37 | 1.83 | 0.276 | 0.999 |
| AT1G42970.1 | Glyceraldehyde-3-phosphate dehydrogenase (GAPB) | 38 | 3.27 | 0.277 | 0.705 |
| AT1G42970.1 | Glyceraldehyde-3-phosphate dehydrogenase (GAPB) | 39 | 3.71 | 0.279 | 0.554 |
| AT1G42970.1 | Glyceraldehyde-3-phosphate dehydrogenase (GAPB) | 40 | 2.39 | 0.279 | 0.965 |
| AT1G42970.1 | Glyceraldehyde-3-phosphate dehydrogenase (GAPB) | 41 | 0.97 | 0.278 | 1.000 |

| | | | | | |
|---|---|---|---|---|---|
| AT1G42970.1 | Glyceraldehyde-3-phosphate dehydrogenase (GAPB) | 42 | 0.42 | 0.283 | 0.973 |
| AT1G42970.1 | Glyceraldehyde-3-phosphate dehydrogenase (GAPB) | 43 | 0.32 | 0.285 | 0.805 |
| AT1G42970.1 | Glyceraldehyde-3-phosphate dehydrogenase (GAPB) | 44 | 0.36 | 0.284 | 0.896 |
| AT1G42970.1 | Glyceraldehyde-3-phosphate dehydrogenase (GAPB) | 45 | 0.38 | 0.284 | 0.921 |
| AT1G42970.1 | Glyceraldehyde-3-phosphate dehydrogenase (GAPB) | 46 | 0.31 | 0.327 | 0.903 |
| AT1G42970.1 | Glyceraldehyde-3-phosphate dehydrogenase (GAPB) | 47 | 0.24 | 0.639 | 0.999 |
| AT1G43170.9 | 60S ribosomal protein L3-1 (RPL3A) | 2 | 0.33 | 0.224 | 0.268 |
| AT1G43170.9 | 60S ribosomal protein L3-1 (RPL3A) | 3 | 0.44 | 0.187 | 0.418 |
| AT1G43170.9 | 60S ribosomal protein L3-1 (RPL3A) | 4 | 0.58 | 0.163 | 0.764 |
| AT1G43170.9 | 60S ribosomal protein L3-1 (RPL3A) | 5 | 0.76 | 0.150 | 0.993 |
| AT1G43170.9 | 60S ribosomal protein L3-1 (RPL3A) | 6 | 0.95 | 0.143 | 1.000 |
| AT1G43170.9 | 60S ribosomal protein L3-1 (RPL3A) | 7 | 1.15 | 0.139 | 1.000 |
| AT1G43170.9 | 60S ribosomal protein L3-1 (RPL3A) | 8 | 1.32 | 0.137 | 0.986 |
| AT1G43170.9 | 60S ribosomal protein L3-1 (RPL3A) | 9 | 1.44 | 0.135 | 0.911 |
| AT1G43170.9 | 60S ribosomal protein L3-1 (RPL3A) | 10 | 1.50 | 0.132 | 0.843 |
| AT1G43170.9 | 60S ribosomal protein L3-1 (RPL3A) | 11 | 1.49 | 0.131 | 0.843 |
| AT1G43170.9 | 60S ribosomal protein L3-1 (RPL3A) | 12 | 1.42 | 0.129 | 0.904 |
| AT1G43170.9 | 60S ribosomal protein L3-1 (RPL3A) | 13 | 1.32 | 0.127 | 0.975 |
| AT1G43170.9 | 60S ribosomal protein L3-1 (RPL3A) | 14 | 1.19 | 0.126 | 1.000 |
| AT1G43170.9 | 60S ribosomal protein L3-1 (RPL3A) | 15 | 1.03 | 0.127 | 1.000 |
| AT1G43170.9 | 60S ribosomal protein L3-1 (RPL3A) | 16 | 0.88 | 0.128 | 1.000 |
| AT1G43170.9 | 60S ribosomal protein L3-1 (RPL3A) | 17 | 0.74 | 0.132 | 0.964 |
| AT1G43170.9 | 60S ribosomal protein L3-1 (RPL3A) | 18 | 0.62 | 0.138 | 0.729 |
| AT1G43170.9 | 60S ribosomal protein L3-1 (RPL3A) | 19 | 0.53 | 0.146 | 0.457 |
| AT1G43170.9 | 60S ribosomal protein L3-1 (RPL3A) | 20 | 0.47 | 0.154 | 0.302 |
| AT1G43170.9 | 60S ribosomal protein L3-1 (RPL3A) | 21 | 0.43 | 0.162 | 0.225 |
| AT1G43170.9 | 60S ribosomal protein L3-1 (RPL3A) | 25 | 0.38 | 0.165 | 0.127 |
| AT1G43170.9 | 60S ribosomal protein L3-1 (RPL3A) | 26 | 0.39 | 0.163 | 0.137 |
| AT1G43170.9 | 60S ribosomal protein L3-1 (RPL3A) | 27 | 0.41 | 0.164 | 0.180 |
| AT1G43170.9 | 60S ribosomal protein L3-1 (RPL3A) | 28 | 0.43 | 0.170 | 0.275 |
| AT1G43170.9 | 60S ribosomal protein L3-1 (RPL3A) | 29 | 0.45 | 0.179 | 0.426 |

| | | | | | |
|---|---|---|---|---|---|
| AT1G43170.9 | 60S ribosomal protein L3-1 (RPL3A) | 30 | 0.48 | 0.192 | 0.612 |
| AT1G43170.9 | 60S ribosomal protein L3-1 (RPL3A) | 31 | 0.51 | 0.207 | 0.787 |
| AT1G43170.9 | 60S ribosomal protein L3-1 (RPL3A) | 32 | 0.54 | 0.223 | 0.904 |
| AT1G43170.9 | 60S ribosomal protein L3-1 (RPL3A) | 41 | 1.11 | 0.227 | 1.000 |
| AT1G43170.9 | 60S ribosomal protein L3-1 (RPL3A) | 42 | 1.25 | 0.216 | 1.000 |
| AT1G43170.9 | 60S ribosomal protein L3-1 (RPL3A) | 43 | 1.41 | 0.209 | 0.997 |
| AT1G43170.9 | 60S ribosomal protein L3-1 (RPL3A) | 44 | 1.61 | 0.210 | 0.969 |
| AT1G43170.9 | 60S ribosomal protein L3-1 (RPL3A) | 45 | 1.87 | 0.221 | 0.888 |
| AT1G43170.9 | 60S ribosomal protein L3-1 (RPL3A) | 46 | 2.20 | 0.246 | 0.798 |
| AT1G43170.9 | 60S ribosomal protein L3-1 (RPL3A) | 47 | 2.60 | 0.284 | 0.755 |
| AT1G44575.1 | Photosystem II subunit (PSBS) | 2 | 0.29 | 0.450 | 0.982 |
| AT1G44575.1 | Photosystem II subunit (PSBS) | 3 | 0.46 | 0.335 | 0.998 |
| AT1G44575.1 | Photosystem II subunit (PSBS) | 4 | 0.73 | 0.297 | 1.000 |
| AT1G44575.1 | Photosystem II subunit (PSBS) | 5 | 1.16 | 0.288 | 1.000 |
| AT1G44575.1 | Photosystem II subunit (PSBS) | 6 | 1.61 | 0.286 | 1.000 |
| AT1G44575.1 | Photosystem II subunit (PSBS) | 7 | 2.00 | 0.284 | 0.995 |
| AT1G44575.1 | Photosystem II subunit (PSBS) | 8 | 2.40 | 0.281 | 0.946 |
| AT1G44575.1 | Photosystem II subunit (PSBS) | 9 | 2.69 | 0.278 | 0.861 |
| AT1G44575.1 | Photosystem II subunit (PSBS) | 10 | 2.76 | 0.279 | 0.836 |
| AT1G44575.1 | Photosystem II subunit (PSBS) | 11 | 2.75 | 0.279 | 0.841 |
| AT1G44575.1 | Photosystem II subunit (PSBS) | 12 | 2.70 | 0.278 | 0.853 |
| AT1G44575.1 | Photosystem II subunit (PSBS) | 13 | 2.58 | 0.275 | 0.884 |
| AT1G44575.1 | Photosystem II subunit (PSBS) | 14 | 2.35 | 0.272 | 0.940 |
| AT1G44575.1 | Photosystem II subunit (PSBS) | 15 | 1.98 | 0.270 | 0.992 |
| AT1G44575.1 | Photosystem II subunit (PSBS) | 16 | 1.62 | 0.270 | 1.000 |
| AT1G44575.1 | Photosystem II subunit (PSBS) | 17 | 1.48 | 0.270 | 1.000 |
| AT1G44575.1 | Photosystem II subunit (PSBS) | 18 | 1.60 | 0.273 | 1.000 |
| AT1G44575.1 | Photosystem II subunit (PSBS) | 19 | 1.75 | 0.278 | 1.000 |
| AT1G44575.1 | Photosystem II subunit (PSBS) | 20 | 1.57 | 0.283 | 1.000 |
| AT1G44575.1 | Photosystem II subunit (PSBS) | 21 | 1.18 | 0.286 | 1.000 |
| AT1G44575.1 | Photosystem II subunit (PSBS) | 22 | 0.89 | 0.290 | 1.000 |
| AT1G44575.1 | Photosystem II subunit (PSBS) | 23 | 0.70 | 0.293 | 1.000 |
| AT1G44575.1 | Photosystem II subunit (PSBS) | 24 | 0.58 | 0.295 | 1.000 |
| AT1G44575.1 | Photosystem II subunit (PSBS) | 25 | 0.50 | 0.294 | 0.997 |
| AT1G44575.1 | Photosystem II subunit (PSBS) | 26 | 0.49 | 0.287 | 0.995 |
| AT1G44575.1 | Photosystem II subunit (PSBS) | 27 | 0.55 | 0.281 | 0.999 |
| AT1G44575.1 | Photosystem II subunit (PSBS) | 28 | 0.70 | 0.278 | 1.000 |
| AT1G44575.1 | Photosystem II subunit (PSBS) | 29 | 0.94 | 0.277 | 1.000 |
| AT1G44575.1 | Photosystem II subunit (PSBS) | 30 | 1.14 | 0.277 | 1.000 |
| AT1G44575.1 | Photosystem II subunit (PSBS) | 31 | 1.29 | 0.277 | 1.000 |
| AT1G44575.1 | Photosystem II subunit (PSBS) | 32 | 1.32 | 0.277 | 1.000 |
| AT1G44575.1 | Photosystem II subunit (PSBS) | 33 | 1.01 | 0.277 | 1.000 |
| AT1G44575.1 | Photosystem II subunit (PSBS) | 34 | 0.56 | 0.277 | 0.999 |
| AT1G44575.1 | Photosystem II subunit (PSBS) | 35 | 0.28 | 0.277 | 0.504 |
| AT1G44575.1 | Photosystem II subunit (PSBS) | 36 | 0.18 | 0.277 | 0.120 |
| AT1G44575.1 | Photosystem II subunit (PSBS) | 37 | 0.19 | 0.277 | 0.151 |
| AT1G44575.1 | Photosystem II subunit (PSBS) | 38 | 0.29 | 0.277 | 0.556 |
| AT1G44575.1 | Photosystem II subunit (PSBS) | 39 | 0.54 | 0.277 | 0.999 |
| AT1G44575.1 | Photosystem II subunit (PSBS) | 40 | 1.09 | 0.278 | 1.000 |
| AT1G44575.1 | Photosystem II subunit (PSBS) | 41 | 1.90 | 0.278 | 0.997 |
| AT1G44575.1 | Photosystem II subunit (PSBS) | 42 | 2.56 | 0.280 | 0.904 |
| AT1G44575.1 | Photosystem II subunit (PSBS) | 43 | 2.51 | 0.283 | 0.922 |
| AT1G44575.1 | Photosystem II subunit (PSBS) | 44 | 1.64 | 0.288 | 1.000 |

| | | | | | | |
|---|---|---|---|---|---|---|
| AT1G44575.1 | Photosystem II subunit (PSBS) | 45 | 0.67 | 0.301 | 1.000 | |
| AT1G44575.1 | Photosystem II subunit (PSBS) | 46 | 0.18 | 0.346 | 0.433 | |
| AT1G44575.1 | Photosystem II subunit (PSBS) | 47 | 0.04 | 0.469 | 0.068 | . |
| AT1G47260.1 | NADH:ubiquinone oxidoreductase subunit (gammaCA2) | 2 | 0.90 | 0.409 | 1.000 | |
| AT1G47260.1 | NADH:ubiquinone oxidoreductase subunit (gammaCA2) | 3 | 0.72 | 0.252 | 1.000 | |
| AT1G47260.1 | NADH:ubiquinone oxidoreductase subunit (gammaCA2) | 4 | 0.72 | 0.241 | 1.000 | |
| AT1G47260.1 | NADH:ubiquinone oxidoreductase subunit (gammaCA2) | 5 | 0.89 | 0.241 | 1.000 | |
| AT1G47260.1 | NADH:ubiquinone oxidoreductase subunit (gammaCA2) | 6 | 1.13 | 0.241 | 1.000 | |
| AT1G47260.1 | NADH:ubiquinone oxidoreductase subunit (gammaCA2) | 7 | 1.32 | 0.243 | 1.000 | |
| AT1G47260.1 | NADH:ubiquinone oxidoreductase subunit (gammaCA2) | 8 | 1.73 | 0.238 | 1.000 | |
| AT1G47260.1 | NADH:ubiquinone oxidoreductase subunit (gammaCA2) | 9 | 1.88 | 0.235 | 0.999 | |
| AT1G47260.1 | NADH:ubiquinone oxidoreductase subunit (gammaCA2) | 10 | 1.73 | 0.235 | 1.000 | |
| AT1G47260.1 | NADH:ubiquinone oxidoreductase subunit (gammaCA2) | 11 | 1.81 | 0.235 | 1.000 | |
| AT1G47260.1 | NADH:ubiquinone oxidoreductase subunit (gammaCA2) | 12 | 1.95 | 0.236 | 0.999 | |
| AT1G47260.1 | NADH:ubiquinone oxidoreductase subunit (gammaCA2) | 13 | 1.89 | 0.233 | 0.999 | |
| AT1G47260.1 | NADH:ubiquinone oxidoreductase subunit (gammaCA2) | 14 | 1.64 | 0.230 | 1.000 | |
| AT1G47260.1 | NADH:ubiquinone oxidoreductase subunit (gammaCA2) | 15 | 1.17 | 0.229 | 1.000 | |
| AT1G47260.1 | NADH:ubiquinone oxidoreductase subunit (gammaCA2) | 16 | 0.68 | 0.227 | 1.000 | |
| AT1G47260.1 | NADH:ubiquinone oxidoreductase subunit (gammaCA2) | 17 | 0.46 | 0.225 | 0.978 | |
| AT1G47260.1 | NADH:ubiquinone oxidoreductase subunit (gammaCA2) | 18 | 0.51 | 0.229 | 0.997 | |
| AT1G47260.1 | NADH:ubiquinone oxidoreductase subunit (gammaCA2) | 19 | 0.73 | 0.237 | 1.000 | |
| AT1G47260.1 | NADH:ubiquinone oxidoreductase subunit (gammaCA2) | 20 | 0.91 | 0.242 | 1.000 | |

| | | | | | |
|---|---|---|---|---|---|
| AT1G47260.1 | NADH:ubiquinone oxidoreductase subunit (gammaCA2) | 21 | 1.08 | 0.241 | 1.000 |
| AT1G47260.1 | NADH:ubiquinone oxidoreductase subunit (gammaCA2) | 22 | 1.33 | 0.242 | 1.000 |
| AT1G47260.1 | NADH:ubiquinone oxidoreductase subunit (gammaCA2) | 23 | 1.57 | 0.240 | 1.000 |
| AT1G47260.1 | NADH:ubiquinone oxidoreductase subunit (gammaCA2) | 24 | 1.38 | 0.234 | 1.000 |
| AT1G47260.1 | NADH:ubiquinone oxidoreductase subunit (gammaCA2) | 25 | 0.95 | 0.231 | 1.000 |
| AT1G47260.1 | NADH:ubiquinone oxidoreductase subunit (gammaCA2) | 26 | 0.77 | 0.231 | 1.000 |
| AT1G47260.1 | NADH:ubiquinone oxidoreductase subunit (gammaCA2) | 27 | 0.78 | 0.231 | 1.000 |
| AT1G47260.1 | NADH:ubiquinone oxidoreductase subunit (gammaCA2) | 28 | 0.80 | 0.232 | 1.000 |
| AT1G47260.1 | NADH:ubiquinone oxidoreductase subunit (gammaCA2) | 29 | 0.83 | 0.234 | 1.000 |
| AT1G47260.1 | NADH:ubiquinone oxidoreductase subunit (gammaCA2) | 30 | 0.96 | 0.234 | 1.000 |
| AT1G47260.1 | NADH:ubiquinone oxidoreductase subunit (gammaCA2) | 31 | 1.15 | 0.235 | 1.000 |
| AT1G47260.1 | NADH:ubiquinone oxidoreductase subunit (gammaCA2) | 32 | 1.14 | 0.236 | 1.000 |
| AT1G47260.1 | NADH:ubiquinone oxidoreductase subunit (gammaCA2) | 33 | 0.72 | 0.235 | 1.000 |
| AT1G47260.1 | NADH:ubiquinone oxidoreductase subunit (gammaCA2) | 34 | 0.37 | 0.236 | 0.858 |
| AT1G47260.1 | NADH:ubiquinone oxidoreductase subunit (gammaCA2) | 35 | 0.30 | 0.236 | 0.534 |
| AT1G47260.1 | NADH:ubiquinone oxidoreductase subunit (gammaCA2) | 36 | 0.55 | 0.235 | 1.000 |
| AT1G47260.1 | NADH:ubiquinone oxidoreductase subunit (gammaCA2) | 37 | 1.49 | 0.235 | 1.000 |
| AT1G47260.1 | NADH:ubiquinone oxidoreductase subunit (gammaCA2) | 38 | 2.50 | 0.236 | 0.930 |
| AT1G47260.1 | NADH:ubiquinone oxidoreductase subunit (gammaCA2) | 39 | 1.80 | 0.235 | 1.000 |
| AT1G47260.1 | NADH:ubiquinone oxidoreductase subunit (gammaCA2) | 40 | 0.92 | 0.234 | 1.000 |

| | | | | | |
|---|---|---|---|---|---|
| AT1G47260.1 | NADH:ubiquinone oxidoreductase subunit (gammaCA2) | 41 | 0.61 | 0.235 | 1.000 |
| AT1G47260.1 | NADH:ubiquinone oxidoreductase subunit (gammaCA2) | 42 | 0.69 | 0.239 | 1.000 |
| AT1G47260.1 | NADH:ubiquinone oxidoreductase subunit (gammaCA2) | 43 | 1.11 | 0.242 | 1.000 |
| AT1G47260.1 | NADH:ubiquinone oxidoreductase subunit (gammaCA2) | 44 | 1.41 | 0.244 | 1.000 |
| AT1G47260.1 | NADH:ubiquinone oxidoreductase subunit (gammaCA2) | 45 | 1.20 | 0.284 | 1.000 |
| AT1G47260.1 | NADH:ubiquinone oxidoreductase subunit (gammaCA2) | 46 | 0.97 | 0.438 | 1.000 |
| AT1G47260.1 | NADH:ubiquinone oxidoreductase subunit (gammaCA2) | 47 | 0.80 | 0.778 | 1.000 |
| AT1G47420.1 | Succinate dehydrogenase subunit (SDH5) | 2 | 1.23 | 0.438 | 1.000 |
| AT1G47420.1 | Succinate dehydrogenase subunit (SDH5) | 3 | 0.95 | 0.263 | 1.000 |
| AT1G47420.1 | Succinate dehydrogenase subunit (SDH5) | 4 | 0.75 | 0.251 | 1.000 |
| AT1G47420.1 | Succinate dehydrogenase subunit (SDH5) | 5 | 0.68 | 0.250 | 1.000 |
| AT1G47420.1 | Succinate dehydrogenase subunit (SDH5) | 6 | 0.68 | 0.249 | 1.000 |
| AT1G47420.1 | Succinate dehydrogenase subunit (SDH5) | 7 | 0.68 | 0.250 | 1.000 |
| AT1G47420.1 | Succinate dehydrogenase subunit (SDH5) | 8 | 0.73 | 0.245 | 1.000 |
| AT1G47420.1 | Succinate dehydrogenase subunit (SDH5) | 9 | 0.85 | 0.242 | 1.000 |
| AT1G47420.1 | Succinate dehydrogenase subunit (SDH5) | 10 | 1.03 | 0.242 | 1.000 |
| AT1G47420.1 | Succinate dehydrogenase subunit (SDH5) | 11 | 1.11 | 0.244 | 1.000 |
| AT1G47420.1 | Succinate dehydrogenase subunit (SDH5) | 12 | 1.09 | 0.244 | 1.000 |
| AT1G47420.1 | Succinate dehydrogenase subunit (SDH5) | 13 | 1.05 | 0.242 | 1.000 |
| AT1G47420.1 | Succinate dehydrogenase subunit (SDH5) | 14 | 1.05 | 0.239 | 1.000 |
| AT1G47420.1 | Succinate dehydrogenase subunit (SDH5) | 15 | 1.07 | 0.238 | 1.000 |
| AT1G47420.1 | Succinate dehydrogenase subunit (SDH5) | 16 | 1.08 | 0.235 | 1.000 |
| AT1G47420.1 | Succinate dehydrogenase subunit (SDH5) | 17 | 1.24 | 0.235 | 1.000 |
| AT1G47420.1 | Succinate dehydrogenase subunit (SDH5) | 18 | 1.44 | 0.243 | 1.000 |
| AT1G47420.1 | Succinate dehydrogenase subunit (SDH5) | 19 | 1.63 | 0.252 | 1.000 |
| AT1G47420.1 | Succinate dehydrogenase subunit (SDH5) | 20 | 1.85 | 0.263 | 1.000 |
| AT1G47420.1 | Succinate dehydrogenase subunit (SDH5) | 21 | 1.76 | 0.290 | 1.000 |

| | | | | | |
|---|---|---|---|---|---|
| AT1G47420.1 | Succinate dehydrogenase subunit (SDH5) | 22 | 1.35 | 0.311 | 1.000 |
| AT1G47420.1 | Succinate dehydrogenase subunit (SDH5) | 23 | 0.95 | 0.356 | 1.000 |
| AT1G47420.1 | Succinate dehydrogenase subunit (SDH5) | 24 | 0.69 | 0.361 | 1.000 |
| AT1G47420.1 | Succinate dehydrogenase subunit (SDH5) | 26 | 0.53 | 0.334 | 1.000 |
| AT1G47420.1 | Succinate dehydrogenase subunit (SDH5) | 27 | 0.61 | 0.282 | 1.000 |
| AT1G47420.1 | Succinate dehydrogenase subunit (SDH5) | 28 | 1.02 | 0.258 | 1.000 |
| AT1G47420.1 | Succinate dehydrogenase subunit (SDH5) | 29 | 2.15 | 0.244 | 0.995 |
| AT1G47420.1 | Succinate dehydrogenase subunit (SDH5) | 30 | 3.01 | 0.241 | 0.770 |
| AT1G47420.1 | Succinate dehydrogenase subunit (SDH5) | 31 | 3.06 | 0.243 | 0.758 |
| AT1G47420.1 | Succinate dehydrogenase subunit (SDH5) | 32 | 3.75 | 0.245 | 0.464 |
| AT1G47420.1 | Succinate dehydrogenase subunit (SDH5) | 33 | 4.31 | 0.245 | 0.282 |
| AT1G47420.1 | Succinate dehydrogenase subunit (SDH5) | 34 | 3.72 | 0.245 | 0.482 |
| AT1G47420.1 | Succinate dehydrogenase subunit (SDH5) | 35 | 2.31 | 0.245 | 0.985 |
| AT1G47420.1 | Succinate dehydrogenase subunit (SDH5) | 36 | 0.90 | 0.244 | 1.000 |
| AT1G47420.1 | Succinate dehydrogenase subunit (SDH5) | 37 | 0.39 | 0.243 | 0.936 |
| AT1G47420.1 | Succinate dehydrogenase subunit (SDH5) | 38 | 0.43 | 0.242 | 0.983 |
| AT1G47420.1 | Succinate dehydrogenase subunit (SDH5) | 39 | 0.95 | 0.241 | 1.000 |
| AT1G47420.1 | Succinate dehydrogenase subunit (SDH5) | 40 | 1.85 | 0.242 | 1.000 |
| AT1G47420.1 | Succinate dehydrogenase subunit (SDH5) | 41 | 2.32 | 0.245 | 0.983 |
| AT1G47420.1 | Succinate dehydrogenase subunit (SDH5) | 42 | 2.50 | 0.256 | 0.974 |
| AT1G47420.1 | Succinate dehydrogenase subunit (SDH5) | 43 | 2.33 | 0.284 | 0.998 |
| AT1G47420.1 | Succinate dehydrogenase subunit (SDH5) | 44 | 1.97 | 0.292 | 1.000 |
| AT1G47420.1 | Succinate dehydrogenase subunit (SDH5) | 45 | 1.61 | 0.275 | 1.000 |
| AT1G47420.1 | Succinate dehydrogenase subunit (SDH5) | 46 | 1.32 | 0.283 | 1.000 |
| AT1G47420.1 | Succinate dehydrogenase subunit (SDH5) | 47 | 1.07 | 0.539 | 1.000 |
| AT1G48030.2 | Dihydrolipoamide dehydrogenase (E3) | 3 | 0.34 | 0.219 | 0.650 |
| AT1G48030.2 | Dihydrolipoamide dehydrogenase (E3) | 4 | 0.28 | 0.207 | 0.234 |
| AT1G48030.2 | Dihydrolipoamide dehydrogenase (E3) | 5 | 0.26 | 0.207 | 0.150 |
| AT1G48030.2 | Dihydrolipoamide dehydrogenase (E3) | 6 | 0.26 | 0.207 | 0.171 |
| AT1G48030.2 | Dihydrolipoamide dehydrogenase (E3) | 7 | 0.31 | 0.207 | 0.381 |
| AT1G48030.2 | Dihydrolipoamide dehydrogenase (E3) | 8 | 0.41 | 0.203 | 0.818 |

| | | | | | |
|---|---|---|---|---|---|
| AT1G48030.2 | Dihydrolipoamide dehydrogenase (E3) | 9 | 0.52 | 0.200 | 0.991 |
| AT1G48030.2 | Dihydrolipoamide dehydrogenase (E3) | 10 | 0.60 | 0.200 | 1.000 |
| AT1G48030.2 | Dihydrolipoamide dehydrogenase (E3) | 11 | 0.80 | 0.200 | 1.000 |
| AT1G48030.2 | Dihydrolipoamide dehydrogenase (E3) | 12 | 1.27 | 0.201 | 1.000 |
| AT1G48030.2 | Dihydrolipoamide dehydrogenase (E3) | 13 | 1.69 | 0.199 | 1.000 |
| AT1G48030.2 | Dihydrolipoamide dehydrogenase (E3) | 14 | 1.61 | 0.196 | 1.000 |
| AT1G48030.2 | Dihydrolipoamide dehydrogenase (E3) | 15 | 1.31 | 0.195 | 1.000 |
| AT1G48030.2 | Dihydrolipoamide dehydrogenase (E3) | 16 | 1.20 | 0.193 | 1.000 |
| AT1G48030.2 | Dihydrolipoamide dehydrogenase (E3) | 17 | 1.32 | 0.193 | 1.000 |
| AT1G48030.2 | Dihydrolipoamide dehydrogenase (E3) | 18 | 1.51 | 0.197 | 1.000 |
| AT1G48030.2 | Dihydrolipoamide dehydrogenase (E3) | 19 | 2.07 | 0.202 | 0.973 |
| AT1G48030.2 | Dihydrolipoamide dehydrogenase (E3) | 20 | 3.11 | 0.207 | 0.436 |
| AT1G48030.2 | Dihydrolipoamide dehydrogenase (E3) | 21 | 3.51 | 0.210 | 0.282 |
| AT1G48030.2 | Dihydrolipoamide dehydrogenase (E3) | 22 | 2.86 | 0.210 | 0.612 |
| AT1G48030.2 | Dihydrolipoamide dehydrogenase (E3) | 23 | 2.72 | 0.208 | 0.690 |
| AT1G48030.2 | Dihydrolipoamide dehydrogenase (E3) | 24 | 2.60 | 0.204 | 0.732 |
| AT1G48030.2 | Dihydrolipoamide dehydrogenase (E3) | 25 | 1.80 | 0.200 | 0.998 |
| AT1G48030.2 | Dihydrolipoamide dehydrogenase (E3) | 26 | 1.29 | 0.199 | 1.000 |
| AT1G48030.2 | Dihydrolipoamide dehydrogenase (E3) | 27 | 1.34 | 0.199 | 1.000 |
| AT1G48030.2 | Dihydrolipoamide dehydrogenase (E3) | 28 | 1.52 | 0.199 | 1.000 |
| AT1G48030.2 | Dihydrolipoamide dehydrogenase (E3) | 29 | 1.62 | 0.199 | 1.000 |
| AT1G48030.2 | Dihydrolipoamide dehydrogenase (E3) | 30 | 1.44 | 0.199 | 1.000 |
| AT1G48030.2 | Dihydrolipoamide dehydrogenase (E3) | 31 | 1.34 | 0.199 | 1.000 |
| AT1G48030.2 | Dihydrolipoamide dehydrogenase (E3) | 32 | 1.39 | 0.200 | 1.000 |
| AT1G48030.2 | Dihydrolipoamide dehydrogenase (E3) | 33 | 1.35 | 0.200 | 1.000 |
| AT1G48030.2 | Dihydrolipoamide dehydrogenase (E3) | 34 | 1.24 | 0.201 | 1.000 |
| AT1G48030.2 | Dihydrolipoamide dehydrogenase (E3) | 35 | 1.21 | 0.200 | 1.000 |
| AT1G48030.2 | Dihydrolipoamide dehydrogenase (E3) | 36 | 1.27 | 0.199 | 1.000 |
| AT1G48030.2 | Dihydrolipoamide dehydrogenase (E3) | 37 | 1.47 | 0.199 | 1.000 |
| AT1G48030.2 | Dihydrolipoamide dehydrogenase (E3) | 38 | 2.02 | 0.200 | 0.979 |
| AT1G48030.2 | Dihydrolipoamide dehydrogenase (E3) | 39 | 3.14 | 0.200 | 0.360 |

| | | | | | | |
|---|---|---|---|---|---|---|
| AT1G48030.2 | Dihydrolipoamide dehydrogenase (E3) | 40 | 4.05 | 0.201 | 0.093 | . |
| AT1G48030.2 | Dihydrolipoamide dehydrogenase (E3) | 41 | 3.12 | 0.202 | 0.382 | |
| AT1G48030.2 | Dihydrolipoamide dehydrogenase (E3) | 42 | 1.51 | 0.204 | 1.000 | |
| AT1G48030.2 | Dihydrolipoamide dehydrogenase (E3) | 43 | 0.75 | 0.206 | 1.000 | |
| AT1G48030.2 | Dihydrolipoamide dehydrogenase (E3) | 44 | 0.58 | 0.206 | 1.000 | |
| AT1G48030.2 | Dihydrolipoamide dehydrogenase (E3) | 45 | 0.58 | 0.206 | 1.000 | |
| AT1G48030.2 | Dihydrolipoamide dehydrogenase (E3) | 46 | 0.46 | 0.234 | 0.989 | |
| AT1G48030.2 | Dihydrolipoamide dehydrogenase (E3) | 47 | 0.26 | 0.430 | 0.996 | |
| AT1G48030.2 | Dihydrolipoamide dehydrogenase (E3) | 48 | 0.14 | 0.742 | 1.000 | |
| AT1G48200.1 | ? | 2 | 1.09 | 0.573 | 1.000 | |
| AT1G48200.1 | ? | 3 | 1.00 | 0.534 | 1.000 | |
| AT1G48200.1 | ? | 22 | 0.41 | 0.406 | 0.998 | |
| AT1G48200.1 | ? | 23 | 0.47 | 0.320 | 0.996 | |
| AT1G48200.1 | ? | 24 | 0.77 | 0.276 | 1.000 | |
| AT1G48200.1 | ? | 25 | 1.51 | 0.264 | 1.000 | |
| AT1G48200.1 | ? | 26 | 2.15 | 0.268 | 0.977 | |
| AT1G48200.1 | ? | 27 | 2.35 | 0.271 | 0.947 | |
| AT1G48200.1 | ? | 28 | 2.54 | 0.267 | 0.892 | |
| AT1G48200.1 | ? | 29 | 2.74 | 0.261 | 0.798 | |
| AT1G48200.1 | ? | 30 | 2.85 | 0.253 | 0.711 | |
| AT1G48200.1 | ? | 31 | 3.15 | 0.250 | 0.560 | |
| AT1G48200.1 | ? | 32 | 3.05 | 0.250 | 0.605 | |
| AT1G48200.1 | ? | 33 | 2.15 | 0.250 | 0.957 | |
| AT1G48200.1 | ? | 34 | 1.30 | 0.251 | 1.000 | |
| AT1G48200.1 | ? | 35 | 0.89 | 0.251 | 1.000 | |
| AT1G48200.1 | ? | 36 | 0.94 | 0.250 | 1.000 | |
| AT1G48200.1 | ? | 37 | 1.59 | 0.250 | 1.000 | |
| AT1G48200.1 | ? | 38 | 3.44 | 0.250 | 0.442 | |
| AT1G48200.1 | ? | 39 | 7.67 | 0.250 | 0.010 | * |
| AT1G48200.1 | ? | 40 | 13.22 | 0.249 | 0.000 | *** |
| AT1G48200.1 | ? | 41 | 11.97 | 0.251 | 0.001 | *** |
| AT1G48200.1 | ? | 42 | 6.79 | 0.254 | 0.026 | * |
| AT1G48200.1 | ? | 43 | 3.12 | 0.256 | 0.610 | |
| AT1G48200.1 | ? | 44 | 1.31 | 0.256 | 1.000 | |
| AT1G48200.1 | ? | 45 | 0.58 | 0.256 | 0.999 | |
| AT1G48200.1 | ? | 46 | 0.25 | 0.303 | 0.578 | |
| AT1G48200.1 | ? | 47 | 0.10 | 0.525 | 0.641 | |
| AT1G48520.1 | GLU-ADT subunit B (GATB) | 2 | 0.54 | 0.683 | 1.000 | |
| AT1G48520.1 | GLU-ADT subunit B (GATB) | 3 | 0.45 | 0.654 | 1.000 | |
| AT1G48520.1 | GLU-ADT subunit B (GATB) | 27 | 0.55 | 0.512 | 1.000 | |
| AT1G48520.1 | GLU-ADT subunit B (GATB) | 28 | 0.82 | 0.412 | 1.000 | |
| AT1G48520.1 | GLU-ADT subunit B (GATB) | 29 | 1.16 | 0.377 | 1.000 | |
| AT1G48520.1 | GLU-ADT subunit B (GATB) | 30 | 1.47 | 0.365 | 1.000 | |
| AT1G48520.1 | GLU-ADT subunit B (GATB) | 31 | 1.74 | 0.362 | 0.998 | |
| AT1G48520.1 | GLU-ADT subunit B (GATB) | 32 | 1.87 | 0.361 | 0.995 | |
| AT1G48520.1 | GLU-ADT subunit B (GATB) | 33 | 1.73 | 0.361 | 0.998 | |
| AT1G48520.1 | GLU-ADT subunit B (GATB) | 34 | 1.54 | 0.361 | 1.000 | |
| AT1G48520.1 | GLU-ADT subunit B (GATB) | 35 | 1.50 | 0.361 | 1.000 | |
| AT1G48520.1 | GLU-ADT subunit B (GATB) | 36 | 1.77 | 0.360 | 0.998 | |
| AT1G48520.1 | GLU-ADT subunit B (GATB) | 37 | 2.38 | 0.360 | 0.942 | |
| AT1G48520.1 | GLU-ADT subunit B (GATB) | 38 | 3.08 | 0.360 | 0.783 | |
| AT1G48520.1 | GLU-ADT subunit B (GATB) | 39 | 3.49 | 0.359 | 0.674 | |
| AT1G48520.1 | GLU-ADT subunit B (GATB) | 40 | 3.03 | 0.360 | 0.795 | |
| AT1G48520.1 | GLU-ADT subunit B (GATB) | 41 | 1.92 | 0.361 | 0.993 | |

| | | | | | |
|---|---|---|---|---|---|
| AT1G48520.1 | GLU-ADT subunit B (GATB) | 42 | 1.13 | 0.364 | 1.000 |
| AT1G48520.1 | GLU-ADT subunit B (GATB) | 43 | 0.77 | 0.365 | 1.000 |
| AT1G48520.1 | GLU-ADT subunit B (GATB) | 44 | 0.61 | 0.365 | 0.999 |
| AT1G48520.1 | GLU-ADT subunit B (GATB) | 45 | 0.47 | 0.368 | 0.980 |
| AT1G48520.1 | GLU-ADT subunit B (GATB) | 46 | 0.31 | 0.412 | 0.864 |
| AT1G48520.1 | GLU-ADT subunit B (GATB) | 47 | 0.18 | 0.576 | 0.836 |
| AT1G49140.1 | NADH:ubiquinone oxidoreductase subunit (NDUFS6) | 2 | 0.96 | 0.596 | 1.000 |
| AT1G49140.1 | NADH:ubiquinone oxidoreductase subunit (NDUFS6) | 3 | 0.86 | 0.540 | 1.000 |
| AT1G49140.1 | NADH:ubiquinone oxidoreductase subunit (NDUFS6) | 13 | 1.25 | 0.365 | 1.000 |
| AT1G49140.1 | NADH:ubiquinone oxidoreductase subunit (NDUFS6) | 14 | 1.36 | 0.331 | 1.000 |
| AT1G49140.1 | NADH:ubiquinone oxidoreductase subunit (NDUFS6) | 15 | 1.11 | 0.310 | 1.000 |
| AT1G49140.1 | NADH:ubiquinone oxidoreductase subunit (NDUFS6) | 16 | 0.62 | 0.276 | 1.000 |
| AT1G49140.1 | NADH:ubiquinone oxidoreductase subunit (NDUFS6) | 17 | 0.36 | 0.263 | 0.818 |
| AT1G49140.1 | NADH:ubiquinone oxidoreductase subunit (NDUFS6) | 18 | 0.40 | 0.263 | 0.907 |
| AT1G49140.1 | NADH:ubiquinone oxidoreductase subunit (NDUFS6) | 19 | 0.64 | 0.276 | 1.000 |
| AT1G49140.1 | NADH:ubiquinone oxidoreductase subunit (NDUFS6) | 20 | 0.79 | 0.284 | 1.000 |
| AT1G49140.1 | NADH:ubiquinone oxidoreductase subunit (NDUFS6) | 21 | 0.99 | 0.287 | 1.000 |
| AT1G49140.1 | NADH:ubiquinone oxidoreductase subunit (NDUFS6) | 22 | 1.57 | 0.285 | 1.000 |
| AT1G49140.1 | NADH:ubiquinone oxidoreductase subunit (NDUFS6) | 23 | 2.57 | 0.285 | 0.933 |
| AT1G49140.1 | NADH:ubiquinone oxidoreductase subunit (NDUFS6) | 24 | 2.83 | 0.274 | 0.831 |
| AT1G49140.1 | NADH:ubiquinone oxidoreductase subunit (NDUFS6) | 25 | 1.59 | 0.268 | 1.000 |
| AT1G49140.1 | NADH:ubiquinone oxidoreductase subunit (NDUFS6) | 26 | 0.70 | 0.269 | 1.000 |
| AT1G49140.1 | NADH:ubiquinone oxidoreductase subunit (NDUFS6) | 27 | 0.43 | 0.271 | 0.963 |
| AT1G49140.1 | NADH:ubiquinone oxidoreductase subunit (NDUFS6) | 28 | 0.75 | 0.271 | 1.000 |

| | | | | | |
|---|---|---|---|---|---|
| AT1G49140.1 | NADH:ubiquinone oxidoreductase subunit (NDUFS6) | 29 | 2.17 | 0.271 | 0.981 |
| AT1G49140.1 | NADH:ubiquinone oxidoreductase subunit (NDUFS6) | 30 | 2.32 | 0.277 | 0.967 |
| AT1G49140.1 | NADH:ubiquinone oxidoreductase subunit (NDUFS6) | 31 | 0.88 | 0.298 | 1.000 |
| AT1G49140.1 | NADH:ubiquinone oxidoreductase subunit (NDUFS6) | 32 | 0.35 | 0.368 | 0.979 |
| AT1G49140.1 | NADH:ubiquinone oxidoreductase subunit (NDUFS6) | 33 | 0.20 | 0.620 | 0.993 |
| AT1G49140.1 | NADH:ubiquinone oxidoreductase subunit (NDUFS6) | 46 | 19360.06 | 7.647 | 1.000 |
| AT1G49140.1 | NADH:ubiquinone oxidoreductase subunit (NDUFS6) | 47 | 92849.39 | 8.486 | 1.000 |
| AT1G49410.1 | Translocase of the outer mitochondrial membrane 6 (TOM6) | 2 | 1.00 | 0.521 | 1.000 |
| AT1G49410.1 | Translocase of the outer mitochondrial membrane 6 (TOM6) | 3 | 0.95 | 0.476 | 1.000 |
| AT1G49410.1 | Translocase of the outer mitochondrial membrane 6 (TOM6) | 16 | 0.41 | 0.338 | 0.979 |
| AT1G49410.1 | Translocase of the outer mitochondrial membrane 6 (TOM6) | 17 | 0.49 | 0.292 | 0.989 |
| AT1G49410.1 | Translocase of the outer mitochondrial membrane 6 (TOM6) | 18 | 0.75 | 0.267 | 1.000 |
| AT1G49410.1 | Translocase of the outer mitochondrial membrane 6 (TOM6) | 19 | 1.29 | 0.258 | 1.000 |
| AT1G49410.1 | Translocase of the outer mitochondrial membrane 6 (TOM6) | 20 | 1.81 | 0.260 | 0.995 |
| AT1G49410.1 | Translocase of the outer mitochondrial membrane 6 (TOM6) | 21 | 1.89 | 0.261 | 0.990 |
| AT1G49410.1 | Translocase of the outer mitochondrial membrane 6 (TOM6) | 22 | 1.75 | 0.261 | 0.997 |
| AT1G49410.1 | Translocase of the outer mitochondrial membrane 6 (TOM6) | 23 | 1.88 | 0.260 | 0.990 |
| AT1G49410.1 | Translocase of the outer mitochondrial membrane 6 (TOM6) | 24 | 1.74 | 0.255 | 0.997 |
| AT1G49410.1 | Translocase of the outer mitochondrial membrane 6 (TOM6) | 25 | 1.08 | 0.252 | 1.000 |
| AT1G49410.1 | Translocase of the outer mitochondrial membrane 6 (TOM6) | 26 | 0.79 | 0.250 | 1.000 |

| | | | | | |
|---|---|---|---|---|---|
| AT1G49410.1 | Translocase of the outer mitochondrial membrane 6 (TOM6) | 27 | 0.88 | 0.250 | 1.000 |
| AT1G49410.1 | Translocase of the outer mitochondrial membrane 6 (TOM6) | 28 | 1.03 | 0.251 | 1.000 |
| AT1G49410.1 | Translocase of the outer mitochondrial membrane 6 (TOM6) | 29 | 1.52 | 0.252 | 1.000 |
| AT1G49410.1 | Translocase of the outer mitochondrial membrane 6 (TOM6) | 30 | 2.20 | 0.251 | 0.922 |
| AT1G49410.1 | Translocase of the outer mitochondrial membrane 6 (TOM6) | 31 | 1.87 | 0.253 | 0.988 |
| AT1G49410.1 | Translocase of the outer mitochondrial membrane 6 (TOM6) | 32 | 0.97 | 0.258 | 1.000 |
| AT1G49410.1 | Translocase of the outer mitochondrial membrane 6 (TOM6) | 33 | 0.42 | 0.272 | 0.910 |
| AT1G49410.1 | Translocase of the outer mitochondrial membrane 6 (TOM6) | 34 | 0.24 | 0.326 | 0.569 |
| AT1G49410.1 | Translocase of the outer mitochondrial membrane 6 (TOM6) | 35 | 0.22 | 0.509 | 0.945 |
| AT1G49410.1 | Translocase of the outer mitochondrial membrane 6 (TOM6) | 46 | 23849745.65 | 5.136 | 0.889 |
| AT1G49410.1 | Translocase of the outer mitochondrial membrane 6 (TOM6) | 47 | ########### | 5.786 | 0.874 |
| AT1G49630.3 | Presequence protease 2 (PREP2) | 2 | 1.99 | 0.577 | 0.954 |
| AT1G49630.3 | Presequence protease 2 (PREP2) | 3 | 2.00 | 0.559 | 0.948 |
| AT1G49630.3 | Presequence protease 2 (PREP2) | 22 | 1.80 | 0.329 | 0.843 |
| AT1G49630.3 | Presequence protease 2 (PREP2) | 23 | 1.76 | 0.319 | 0.847 |
| AT1G49630.3 | Presequence protease 2 (PREP2) | 24 | 1.72 | 0.310 | 0.853 |
| AT1G49630.3 | Presequence protease 2 (PREP2) | 33 | 1.22 | 0.240 | 0.988 |
| AT1G49630.3 | Presequence protease 2 (PREP2) | 34 | 1.16 | 0.236 | 0.996 |
| AT1G49630.3 | Presequence protease 2 (PREP2) | 35 | 1.10 | 0.232 | 0.999 |
| AT1G49630.3 | Presequence protease 2 (PREP2) | 39 | 0.89 | 0.232 | 0.999 |
| AT1G49630.3 | Presequence protease 2 (PREP2) | 40 | 0.84 | 0.237 | 0.993 |
| AT1G49630.3 | Presequence protease 2 (PREP2) | 41 | 0.80 | 0.244 | 0.982 |
| AT1G49630.3 | Presequence protease 2 (PREP2) | 42 | 0.76 | 0.253 | 0.965 |
| AT1G49630.3 | Presequence protease 2 (PREP2) | 45 | 0.64 | 0.292 | 0.901 |
| AT1G49630.3 | Presequence protease 2 (PREP2) | 46 | 0.61 | 0.309 | 0.882 |

| | | | | | |
|---|---|---|---|---|---|
| AT1G49630.3 | Presequence protease 2 (PREP2) | 47 | 0.57 | 0.328 | 0.866 |
| AT1G50200.1 | Alanyl-tRNA synthetase 1 (ALATS1) | 2 | 0.69 | 0.383 | 1.000 |
| AT1G50200.1 | Alanyl-tRNA synthetase 1 (ALATS1) | 3 | 0.63 | 0.369 | 1.000 |
| AT1G50200.1 | Alanyl-tRNA synthetase 1 (ALATS1) | 30 | 0.71 | 0.269 | 1.000 |
| AT1G50200.1 | Alanyl-tRNA synthetase 1 (ALATS1) | 31 | 0.86 | 0.207 | 1.000 |
| AT1G50200.1 | Alanyl-tRNA synthetase 1 (ALATS1) | 32 | 1.12 | 0.185 | 1.000 |
| AT1G50200.1 | Alanyl-tRNA synthetase 1 (ALATS1) | 33 | 1.56 | 0.175 | 0.954 |
| AT1G50200.1 | Alanyl-tRNA synthetase 1 (ALATS1) | 34 | 1.95 | 0.172 | 0.626 |
| AT1G50200.1 | Alanyl-tRNA synthetase 1 (ALATS1) | 35 | 1.94 | 0.171 | 0.623 |
| AT1G50200.1 | Alanyl-tRNA synthetase 1 (ALATS1) | 36 | 1.67 | 0.170 | 0.876 |
| AT1G50200.1 | Alanyl-tRNA synthetase 1 (ALATS1) | 37 | 1.39 | 0.169 | 0.993 |
| AT1G50200.1 | Alanyl-tRNA synthetase 1 (ALATS1) | 38 | 1.14 | 0.170 | 1.000 |
| AT1G50200.1 | Alanyl-tRNA synthetase 1 (ALATS1) | 39 | 0.89 | 0.171 | 1.000 |
| AT1G50200.1 | Alanyl-tRNA synthetase 1 (ALATS1) | 40 | 0.67 | 0.172 | 0.975 |
| AT1G50200.1 | Alanyl-tRNA synthetase 1 (ALATS1) | 41 | 0.56 | 0.184 | 0.839 |
| AT1G50200.1 | Alanyl-tRNA synthetase 1 (ALATS1) | 42 | 0.61 | 0.237 | 0.989 |
| AT1G50200.1 | Alanyl-tRNA synthetase 1 (ALATS1) | 43 | 0.82 | 0.343 | 1.000 |
| AT1G50200.1 | Alanyl-tRNA synthetase 1 (ALATS1) | 46 | 3.36 | 0.989 | 1.000 |
| AT1G50200.1 | Alanyl-tRNA synthetase 1 (ALATS1) | 47 | 5.88 | 1.298 | 1.000 |
| AT1G50380.1 | Prolyl oligopeptidase family protein | 2 | 0.96 | 0.689 | 1.000 |
| AT1G50380.1 | Prolyl oligopeptidase family protein | 3 | 0.89 | 0.672 | 1.000 |
| AT1G50380.1 | Prolyl oligopeptidase family protein | 43 | 0.80 | 0.395 | 0.996 |
| AT1G50380.1 | Prolyl oligopeptidase family protein | 44 | 0.92 | 0.316 | 1.000 |
| AT1G50380.1 | Prolyl oligopeptidase family protein | 45 | 1.07 | 0.280 | 1.000 |
| AT1G50380.1 | Prolyl oligopeptidase family protein | 46 | 1.23 | 0.308 | 0.992 |
| AT1G50380.1 | Prolyl oligopeptidase family protein | 47 | 1.41 | 0.394 | 0.981 |
| AT1G50460.1 | Hexokinase-like (HKL1) | 2 | 0.95 | 0.342 | 1.000 |
| AT1G50460.1 | Hexokinase-like (HKL1) | 3 | 0.98 | 0.329 | 1.000 |
| AT1G50460.1 | Hexokinase-like (HKL1) | 26 | 1.02 | 0.246 | 1.000 |
| AT1G50460.1 | Hexokinase-like (HKL1) | 27 | 0.96 | 0.171 | 1.000 |
| AT1G50460.1 | Hexokinase-like (HKL1) | 28 | 0.97 | 0.148 | 1.000 |
| AT1G50460.1 | Hexokinase-like (HKL1) | 29 | 0.95 | 0.140 | 1.000 |
| AT1G50460.1 | Hexokinase-like (HKL1) | 30 | 0.77 | 0.137 | 0.999 |
| AT1G50460.1 | Hexokinase-like (HKL1) | 31 | 0.57 | 0.137 | 0.698 |
| AT1G50460.1 | Hexokinase-like (HKL1) | 32 | 0.47 | 0.137 | 0.267 |
| AT1G50460.1 | Hexokinase-like (HKL1) | 33 | 0.49 | 0.137 | 0.324 |

| | | | | | | |
|---|---|---|---|---|---|---|
| AT1G50460.1 | Hexokinase-like (HKL1) | 34 | 0.71 | 0.137 | 0.990 | |
| AT1G50460.1 | Hexokinase-like (HKL1) | 35 | 1.26 | 0.137 | 1.000 | |
| AT1G50460.1 | Hexokinase-like (HKL1) | 36 | 1.71 | 0.137 | 0.738 | |
| AT1G50460.1 | Hexokinase-like (HKL1) | 37 | 1.55 | 0.136 | 0.916 | |
| AT1G50460.1 | Hexokinase-like (HKL1) | 38 | 1.15 | 0.136 | 1.000 | |
| AT1G50460.1 | Hexokinase-like (HKL1) | 39 | 0.96 | 0.136 | 1.000 | |
| AT1G50460.1 | Hexokinase-like (HKL1) | 40 | 0.95 | 0.136 | 1.000 | |
| AT1G50460.1 | Hexokinase-like (HKL1) | 41 | 1.05 | 0.138 | 1.000 | |
| AT1G50460.1 | Hexokinase-like (HKL1) | 42 | 1.27 | 0.139 | 1.000 | |
| AT1G50460.1 | Hexokinase-like (HKL1) | 43 | 1.48 | 0.139 | 0.970 | |
| AT1G50460.1 | Hexokinase-like (HKL1) | 44 | 1.30 | 0.140 | 0.999 | |
| AT1G50460.1 | Hexokinase-like (HKL1) | 45 | 0.77 | 0.146 | 1.000 | |
| AT1G50460.1 | Hexokinase-like (HKL1) | 46 | 0.34 | 0.172 | 0.106 | |
| AT1G50460.1 | Hexokinase-like (HKL1) | 47 | 0.13 | 0.275 | 0.030 | * |
| AT1G50940.1 | Electron transfer flavoprotein complex subunit (ETF alpha) | 2 | 0.80 | 0.418 | 1.000 | |
| AT1G50940.1 | Electron transfer flavoprotein complex subunit (ETF alpha) | 3 | 0.81 | 0.404 | 1.000 | |
| AT1G50940.1 | Electron transfer flavoprotein complex subunit (ETF alpha) | 28 | 0.98 | 0.306 | 1.000 | |
| AT1G50940.1 | Electron transfer flavoprotein complex subunit (ETF alpha) | 29 | 0.98 | 0.244 | 1.000 | |
| AT1G50940.1 | Electron transfer flavoprotein complex subunit (ETF alpha) | 30 | 0.98 | 0.223 | 1.000 | |
| AT1G50940.1 | Electron transfer flavoprotein complex subunit (ETF alpha) | 31 | 1.05 | 0.218 | 1.000 | |
| AT1G50940.1 | Electron transfer flavoprotein complex subunit (ETF alpha) | 32 | 1.19 | 0.219 | 1.000 | |
| AT1G50940.1 | Electron transfer flavoprotein complex subunit (ETF alpha) | 33 | 1.37 | 0.223 | 0.999 | |
| AT1G50940.1 | Electron transfer flavoprotein complex subunit (ETF alpha) | 34 | 1.65 | 0.225 | 0.970 | |
| AT1G50940.1 | Electron transfer flavoprotein complex subunit (ETF alpha) | 35 | 2.08 | 0.223 | 0.756 | |
| AT1G50940.1 | Electron transfer flavoprotein complex subunit (ETF alpha) | 36 | 2.37 | 0.222 | 0.565 | |
| AT1G50940.1 | Electron transfer flavoprotein complex subunit (ETF alpha) | 37 | 2.18 | 0.223 | 0.691 | |
| AT1G50940.1 | Electron transfer flavoprotein complex subunit (ETF alpha) | 38 | 1.85 | 0.224 | 0.896 | |
| AT1G50940.1 | Electron transfer flavoprotein complex subunit (ETF alpha) | 39 | 1.77 | 0.220 | 0.923 | |
| AT1G50940.1 | Electron transfer flavoprotein complex subunit (ETF alpha) | 40 | 2.09 | 0.215 | 0.714 | |
| AT1G50940.1 | Electron transfer flavoprotein complex subunit (ETF alpha) | 41 | 2.68 | 0.213 | 0.347 | |
| AT1G50940.1 | Electron transfer flavoprotein complex subunit (ETF alpha) | 42 | 3.01 | 0.214 | 0.223 | |
| AT1G50940.1 | Electron transfer flavoprotein complex subunit (ETF alpha) | 43 | 2.69 | 0.214 | 0.348 | |
| AT1G50940.1 | Electron transfer flavoprotein complex subunit (ETF alpha) | 44 | 1.86 | 0.214 | 0.858 | |
| AT1G50940.1 | Electron transfer flavoprotein complex subunit (ETF alpha) | 45 | 1.04 | 0.217 | 1.000 | |
| AT1G50940.1 | Electron transfer flavoprotein complex subunit (ETF alpha) | 46 | 0.50 | 0.244 | 0.879 | |
| AT1G50940.1 | Electron transfer flavoprotein complex subunit (ETF alpha) | 47 | 0.22 | 0.343 | 0.415 | |
| AT1G51390.1 | NFU domain protein 5 (NFU5) | 2 | 1.00 | 0.316 | 1.000 | |
| AT1G51390.1 | NFU domain protein 5 (NFU5) | 3 | 0.98 | 0.304 | 1.000 | |
| AT1G51390.1 | NFU domain protein 5 (NFU5) | 30 | 0.98 | 0.158 | 1.000 | |
| AT1G51390.1 | NFU domain protein 5 (NFU5) | 31 | 1.03 | 0.135 | 1.000 | |

| | | | | | |
|---|---|---|---|---|---|
| AT1G51390.1 | NFU domain protein 5 (NFU5) | 32 | 1.08 | 0.118 | 1.000 |
| AT1G51390.1 | NFU domain protein 5 (NFU5) | 33 | 1.16 | 0.108 | 0.993 |
| AT1G51390.1 | NFU domain protein 5 (NFU5) | 34 | 1.26 | 0.103 | 0.908 |
| AT1G51390.1 | NFU domain protein 5 (NFU5) | 35 | 1.38 | 0.103 | 0.692 |
| AT1G51390.1 | NFU domain protein 5 (NFU5) | 36 | 1.49 | 0.107 | 0.504 |
| AT1G51390.1 | NFU domain protein 5 (NFU5) | 37 | 1.56 | 0.114 | 0.462 |
| AT1G51390.1 | NFU domain protein 5 (NFU5) | 38 | 1.54 | 0.123 | 0.564 |
| AT1G51390.1 | NFU domain protein 5 (NFU5) | 42 | 0.97 | 0.130 | 1.000 |
| AT1G51390.1 | NFU domain protein 5 (NFU5) | 43 | 0.82 | 0.125 | 0.984 |
| AT1G51390.1 | NFU domain protein 5 (NFU5) | 44 | 0.71 | 0.122 | 0.771 |
| AT1G51390.1 | NFU domain protein 5 (NFU5) | 45 | 0.62 | 0.128 | 0.511 |
| AT1G51390.1 | NFU domain protein 5 (NFU5) | 46 | 0.55 | 0.150 | 0.441 |
| AT1G51390.1 | NFU domain protein 5 (NFU5) | 47 | 0.49 | 0.189 | 0.502 |
| AT1G51610.1 | Cation efflux family protein | 2 | 0.99 | 0.515 | 1.000 |
| AT1G51610.1 | Cation efflux family protein | 3 | 0.87 | 0.480 | 1.000 |
| AT1G51610.1 | Cation efflux family protein | 12 | 0.57 | 0.394 | 1.000 |
| AT1G51610.1 | Cation efflux family protein | 13 | 0.62 | 0.367 | 1.000 |
| AT1G51610.1 | Cation efflux family protein | 14 | 0.71 | 0.410 | 1.000 |
| AT1G51610.1 | Cation efflux family protein | 32 | 1.74 | 0.363 | 1.000 |
| AT1G51610.1 | Cation efflux family protein | 33 | 1.15 | 0.292 | 1.000 |
| AT1G51610.1 | Cation efflux family protein | 34 | 1.16 | 0.260 | 1.000 |
| AT1G51610.1 | Cation efflux family protein | 35 | 1.77 | 0.246 | 0.983 |
| AT1G51610.1 | Cation efflux family protein | 36 | 2.92 | 0.241 | 0.489 |
| AT1G51610.1 | Cation efflux family protein | 37 | 1.76 | 0.243 | 0.983 |
| AT1G51610.1 | Cation efflux family protein | 38 | 0.49 | 0.258 | 0.947 |
| AT1G51610.1 | Cation efflux family protein | 39 | 0.21 | 0.314 | 0.354 |
| AT1G51610.1 | Cation efflux family protein | 40 | 0.34 | 0.569 | 0.997 |
| AT1G51610.1 | Cation efflux family protein | 46 | 534725.00 | 4.173 | 0.874 |
| AT1G51610.1 | Cation efflux family protein | 47 | 13875307.17 | 5.111 | 0.861 |
| AT1G51650.1 | ATP synthase subunit (ATPepsilon) | 2 | 1.06 | 0.402 | 1.000 |
| AT1G51650.1 | ATP synthase subunit (ATPepsilon) | 3 | 0.92 | 0.264 | 1.000 |
| AT1G51650.1 | ATP synthase subunit (ATPepsilon) | 4 | 0.85 | 0.251 | 1.000 |
| AT1G51650.1 | ATP synthase subunit (ATPepsilon) | 5 | 0.88 | 0.235 | 1.000 |
| AT1G51650.1 | ATP synthase subunit (ATPepsilon) | 6 | 0.92 | 0.225 | 1.000 |
| AT1G51650.1 | ATP synthase subunit (ATPepsilon) | 7 | 0.87 | 0.223 | 1.000 |
| AT1G51650.1 | ATP synthase subunit (ATPepsilon) | 8 | 0.78 | 0.217 | 1.000 |
| AT1G51650.1 | ATP synthase subunit (ATPepsilon) | 9 | 0.80 | 0.213 | 1.000 |
| AT1G51650.1 | ATP synthase subunit (ATPepsilon) | 10 | 0.95 | 0.215 | 1.000 |
| AT1G51650.1 | ATP synthase subunit (ATPepsilon) | 11 | 1.22 | 0.217 | 1.000 |
| AT1G51650.1 | ATP synthase subunit (ATPepsilon) | 12 | 1.68 | 0.216 | 1.000 |
| AT1G51650.1 | ATP synthase subunit (ATPepsilon) | 13 | 2.57 | 0.213 | 0.757 |
| AT1G51650.1 | ATP synthase subunit (ATPepsilon) | 14 | 3.54 | 0.208 | 0.222 |
| AT1G51650.1 | ATP synthase subunit (ATPepsilon) | 15 | 3.65 | 0.209 | 0.195 |
| AT1G51650.1 | ATP synthase subunit (ATPepsilon) | 16 | 3.16 | 0.208 | 0.370 |
| AT1G51650.1 | ATP synthase subunit (ATPepsilon) | 17 | 2.65 | 0.207 | 0.661 |

| | | | | | |
|---|---|---|---|---|---|
| AT1G51650.1 | ATP synthase subunit (ATPepsilon) | 18 | 2.04 | 0.208 | 0.971 |
| AT1G51650.1 | ATP synthase subunit (ATPepsilon) | 19 | 1.91 | 0.217 | 0.993 |
| AT1G51650.1 | ATP synthase subunit (ATPepsilon) | 20 | 1.71 | 0.223 | 1.000 |
| AT1G51650.1 | ATP synthase subunit (ATPepsilon) | 21 | 1.20 | 0.225 | 1.000 |
| AT1G51650.1 | ATP synthase subunit (ATPepsilon) | 22 | 0.81 | 0.224 | 1.000 |
| AT1G51650.1 | ATP synthase subunit (ATPepsilon) | 23 | 0.77 | 0.224 | 1.000 |
| AT1G51650.1 | ATP synthase subunit (ATPepsilon) | 24 | 1.19 | 0.217 | 1.000 |
| AT1G51650.1 | ATP synthase subunit (ATPepsilon) | 25 | 2.03 | 0.212 | 0.977 |
| AT1G51650.1 | ATP synthase subunit (ATPepsilon) | 26 | 1.82 | 0.211 | 0.997 |
| AT1G51650.1 | ATP synthase subunit (ATPepsilon) | 27 | 0.80 | 0.212 | 1.000 |
| AT1G51650.1 | ATP synthase subunit (ATPepsilon) | 28 | 0.54 | 0.214 | 0.997 |
| AT1G51650.1 | ATP synthase subunit (ATPepsilon) | 29 | 1.05 | 0.214 | 1.000 |
| AT1G51650.1 | ATP synthase subunit (ATPepsilon) | 30 | 2.67 | 0.213 | 0.694 |
| AT1G51650.1 | ATP synthase subunit (ATPepsilon) | 31 | 3.52 | 0.213 | 0.264 |
| AT1G51650.1 | ATP synthase subunit (ATPepsilon) | 32 | 1.86 | 0.215 | 0.996 |
| AT1G51650.1 | ATP synthase subunit (ATPepsilon) | 33 | 0.81 | 0.218 | 1.000 |
| AT1G51650.1 | ATP synthase subunit (ATPepsilon) | 34 | 0.62 | 0.231 | 1.000 |
| AT1G51650.1 | ATP synthase subunit (ATPepsilon) | 35 | 0.73 | 0.276 | 1.000 |
| AT1G51650.1 | ATP synthase subunit (ATPepsilon) | 36 | 0.99 | 0.395 | 1.000 |
| AT1G51650.1 | ATP synthase subunit (ATPepsilon) | 41 | 2.76 | 0.356 | 0.997 |
| AT1G51650.1 | ATP synthase subunit (ATPepsilon) | 42 | 2.72 | 0.361 | 0.998 |
| AT1G51650.1 | ATP synthase subunit (ATPepsilon) | 43 | 2.42 | 0.432 | 1.000 |
| AT1G51650.1 | ATP synthase subunit (ATPepsilon) | 46 | 1.14 | 1.197 | 1.000 |
| AT1G51650.1 | ATP synthase subunit (ATPepsilon) | 47 | 0.84 | 1.548 | 1.000 |
| AT1G51980.1 | Ubiquinol:cytochrome c oxidoreductase subunit (MPPalpha1) | 3 | 0.79 | 0.286 | 1.000 |
| AT1G51980.1 | Ubiquinol:cytochrome c oxidoreductase subunit (MPPalpha1) | 4 | 0.76 | 0.268 | 1.000 |
| AT1G51980.1 | Ubiquinol:cytochrome c oxidoreductase subunit (MPPalpha1) | 5 | 0.93 | 0.268 | 1.000 |
| AT1G51980.1 | Ubiquinol:cytochrome c oxidoreductase subunit (MPPalpha1) | 6 | 1.11 | 0.269 | 1.000 |

| | | | | | |
|---|---|---|---|---|---|
| AT1G51980.1 | Ubiquinol:cytochrome c oxidoreductase subunit (MPPalpha1) | 7 | 1.12 | 0.270 | 1.000 |
| AT1G51980.1 | Ubiquinol:cytochrome c oxidoreductase subunit (MPPalpha1) | 8 | 1.19 | 0.264 | 1.000 |
| AT1G51980.1 | Ubiquinol:cytochrome c oxidoreductase subunit (MPPalpha1) | 9 | 1.33 | 0.259 | 1.000 |
| AT1G51980.1 | Ubiquinol:cytochrome c oxidoreductase subunit (MPPalpha1) | 10 | 1.41 | 0.260 | 1.000 |
| AT1G51980.1 | Ubiquinol:cytochrome c oxidoreductase subunit (MPPalpha1) | 11 | 1.51 | 0.261 | 1.000 |
| AT1G51980.1 | Ubiquinol:cytochrome c oxidoreductase subunit (MPPalpha1) | 12 | 1.62 | 0.261 | 1.000 |
| AT1G51980.1 | Ubiquinol:cytochrome c oxidoreductase subunit (MPPalpha1) | 13 | 1.73 | 0.258 | 1.000 |
| AT1G51980.1 | Ubiquinol:cytochrome c oxidoreductase subunit (MPPalpha1) | 14 | 1.82 | 0.253 | 1.000 |
| AT1G51980.1 | Ubiquinol:cytochrome c oxidoreductase subunit (MPPalpha1) | 15 | 1.60 | 0.253 | 1.000 |
| AT1G51980.1 | Ubiquinol:cytochrome c oxidoreductase subunit (MPPalpha1) | 16 | 1.10 | 0.251 | 1.000 |
| AT1G51980.1 | Ubiquinol:cytochrome c oxidoreductase subunit (MPPalpha1) | 17 | 0.75 | 0.251 | 1.000 |
| AT1G51980.1 | Ubiquinol:cytochrome c oxidoreductase subunit (MPPalpha1) | 18 | 0.72 | 0.255 | 1.000 |
| AT1G51980.1 | Ubiquinol:cytochrome c oxidoreductase subunit (MPPalpha1) | 19 | 0.94 | 0.262 | 1.000 |
| AT1G51980.1 | Ubiquinol:cytochrome c oxidoreductase subunit (MPPalpha1) | 20 | 1.31 | 0.269 | 1.000 |
| AT1G51980.1 | Ubiquinol:cytochrome c oxidoreductase subunit (MPPalpha1) | 21 | 1.65 | 0.273 | 1.000 |
| AT1G51980.1 | Ubiquinol:cytochrome c oxidoreductase subunit (MPPalpha1) | 22 | 1.39 | 0.272 | 1.000 |
| AT1G51980.1 | Ubiquinol:cytochrome c oxidoreductase subunit (MPPalpha1) | 23 | 0.72 | 0.271 | 1.000 |
| AT1G51980.1 | Ubiquinol:cytochrome c oxidoreductase subunit (MPPalpha1) | 24 | 0.44 | 0.265 | 0.995 |
| AT1G51980.1 | Ubiquinol:cytochrome c oxidoreductase subunit (MPPalpha1) | 25 | 0.56 | 0.259 | 1.000 |
| AT1G51980.1 | Ubiquinol:cytochrome c oxidoreductase subunit (MPPalpha1) | 26 | 1.20 | 0.258 | 1.000 |

| | | | | | |
|---|---|---|---|---|---|
| AT1G51980.1 | Ubiquinol:cytochrome c oxidoreductase subunit (MPPalpha1) | 27 | 2.25 | 0.259 | 0.994 |
| AT1G51980.1 | Ubiquinol:cytochrome c oxidoreductase subunit (MPPalpha1) | 28 | 1.66 | 0.258 | 1.000 |
| AT1G51980.1 | Ubiquinol:cytochrome c oxidoreductase subunit (MPPalpha1) | 29 | 0.63 | 0.258 | 1.000 |
| AT1G51980.1 | Ubiquinol:cytochrome c oxidoreductase subunit (MPPalpha1) | 30 | 0.46 | 0.258 | 0.996 |
| AT1G51980.1 | Ubiquinol:cytochrome c oxidoreductase subunit (MPPalpha1) | 31 | 0.77 | 0.259 | 1.000 |
| AT1G51980.1 | Ubiquinol:cytochrome c oxidoreductase subunit (MPPalpha1) | 32 | 1.43 | 0.260 | 1.000 |
| AT1G51980.1 | Ubiquinol:cytochrome c oxidoreductase subunit (MPPalpha1) | 33 | 1.64 | 0.260 | 1.000 |
| AT1G51980.1 | Ubiquinol:cytochrome c oxidoreductase subunit (MPPalpha1) | 34 | 1.21 | 0.260 | 1.000 |
| AT1G51980.1 | Ubiquinol:cytochrome c oxidoreductase subunit (MPPalpha1) | 35 | 0.79 | 0.260 | 1.000 |
| AT1G51980.1 | Ubiquinol:cytochrome c oxidoreductase subunit (MPPalpha1) | 36 | 0.75 | 0.259 | 1.000 |
| AT1G51980.1 | Ubiquinol:cytochrome c oxidoreductase subunit (MPPalpha1) | 37 | 1.28 | 0.258 | 1.000 |
| AT1G51980.1 | Ubiquinol:cytochrome c oxidoreductase subunit (MPPalpha1) | 38 | 2.90 | 0.259 | 0.887 |
| AT1G51980.1 | Ubiquinol:cytochrome c oxidoreductase subunit (MPPalpha1) | 39 | 5.14 | 0.260 | 0.194 |
| AT1G51980.1 | Ubiquinol:cytochrome c oxidoreductase subunit (MPPalpha1) | 40 | 5.99 | 0.261 | 0.101 |
| AT1G51980.1 | Ubiquinol:cytochrome c oxidoreductase subunit (MPPalpha1) | 41 | 5.04 | 0.262 | 0.222 |
| AT1G51980.1 | Ubiquinol:cytochrome c oxidoreductase subunit (MPPalpha1) | 42 | 3.49 | 0.265 | 0.703 |
| AT1G51980.1 | Ubiquinol:cytochrome c oxidoreductase subunit (MPPalpha1) | 43 | 3.49 | 0.267 | 0.716 |
| AT1G51980.1 | Ubiquinol:cytochrome c oxidoreductase subunit (MPPalpha1) | 44 | 3.37 | 0.267 | 0.762 |
| AT1G51980.1 | Ubiquinol:cytochrome c oxidoreductase subunit (MPPalpha1) | 45 | 2.07 | 0.268 | 0.999 |
| AT1G51980.1 | Ubiquinol:cytochrome c oxidoreductase subunit (MPPalpha1) | 46 | 0.85 | 0.306 | 1.000 |

| | | | | | |
|---|---|---|---|---|---|
| AT1G51980.1 | Ubiquinol:cytochrome c oxidoreductase subunit (MPPalpha1) | 47 | 0.27 | 0.532 | 1.000 |
| AT1G51980.1 | Ubiquinol:cytochrome c oxidoreductase subunit (MPPalpha1) | 48 | 0.08 | 0.906 | 0.999 |
| AT1G53000.1 | CMP-KDO (3-deoxy-D-manno-octulosonate) synthetase (CKS) | 2 | 0.67 | 0.244 | 1.000 |
| AT1G53000.1 | CMP-KDO (3-deoxy-D-manno-octulosonate) synthetase (CKS) | 3 | 0.74 | 0.186 | 1.000 |
| AT1G53000.1 | CMP-KDO (3-deoxy-D-manno-octulosonate) synthetase (CKS) | 4 | 0.82 | 0.162 | 1.000 |
| AT1G53000.1 | CMP-KDO (3-deoxy-D-manno-octulosonate) synthetase (CKS) | 5 | 0.91 | 0.154 | 1.000 |
| AT1G53000.1 | CMP-KDO (3-deoxy-D-manno-octulosonate) synthetase (CKS) | 6 | 1.02 | 0.146 | 1.000 |
| AT1G53000.1 | CMP-KDO (3-deoxy-D-manno-octulosonate) synthetase (CKS) | 7 | 1.12 | 0.142 | 1.000 |
| AT1G53000.1 | CMP-KDO (3-deoxy-D-manno-octulosonate) synthetase (CKS) | 8 | 1.18 | 0.141 | 1.000 |
| AT1G53000.1 | CMP-KDO (3-deoxy-D-manno-octulosonate) synthetase (CKS) | 9 | 1.21 | 0.141 | 1.000 |
| AT1G53000.1 | CMP-KDO (3-deoxy-D-manno-octulosonate) synthetase (CKS) | 10 | 1.23 | 0.141 | 1.000 |
| AT1G53000.1 | CMP-KDO (3-deoxy-D-manno-octulosonate) synthetase (CKS) | 11 | 1.28 | 0.141 | 1.000 |
| AT1G53000.1 | CMP-KDO (3-deoxy-D-manno-octulosonate) synthetase (CKS) | 12 | 1.38 | 0.139 | 0.999 |
| AT1G53000.1 | CMP-KDO (3-deoxy-D-manno-octulosonate) synthetase (CKS) | 13 | 1.51 | 0.136 | 0.976 |
| AT1G53000.1 | CMP-KDO (3-deoxy-D-manno-octulosonate) synthetase (CKS) | 14 | 1.66 | 0.133 | 0.851 |
| AT1G53000.1 | CMP-KDO (3-deoxy-D-manno-octulosonate) synthetase (CKS) | 15 | 1.79 | 0.130 | 0.650 |
| AT1G53000.1 | CMP-KDO (3-deoxy-D-manno-octulosonate) synthetase (CKS) | 16 | 1.89 | 0.128 | 0.473 |
| AT1G53000.1 | CMP-KDO (3-deoxy-D-manno-octulosonate) synthetase (CKS) | 17 | 2.03 | 0.128 | 0.303 |
| AT1G53000.1 | CMP-KDO (3-deoxy-D-manno-octulosonate) synthetase (CKS) | 18 | 2.16 | 0.128 | 0.187 |
| AT1G53000.1 | CMP-KDO (3-deoxy-D-manno-octulosonate) synthetase (CKS) | 19 | 2.18 | 0.128 | 0.178 |
| AT1G53000.1 | CMP-KDO (3-deoxy-D-manno-octulosonate) synthetase (CKS) | 20 | 1.95 | 0.128 | 0.392 |
| AT1G53000.1 | CMP-KDO (3-deoxy-D-manno-octulosonate) synthetase (CKS) | 21 | 1.58 | 0.127 | 0.901 |
| AT1G53000.1 | CMP-KDO (3-deoxy-D-manno-octulosonate) synthetase (CKS) | 22 | 1.23 | 0.126 | 1.000 |
| AT1G53000.1 | CMP-KDO (3-deoxy-D-manno-octulosonate) synthetase (CKS) | 23 | 0.98 | 0.125 | 1.000 |
| AT1G53000.1 | CMP-KDO (3-deoxy-D-manno-octulosonate) synthetase (CKS) | 24 | 0.83 | 0.122 | 1.000 |
| AT1G53000.1 | CMP-KDO (3-deoxy-D-manno-octulosonate) synthetase (CKS) | 25 | 0.77 | 0.121 | 1.000 |
| AT1G53000.1 | CMP-KDO (3-deoxy-D-manno-octulosonate) synthetase (CKS) | 26 | 0.82 | 0.120 | 1.000 |
| AT1G53000.1 | CMP-KDO (3-deoxy-D-manno-octulosonate) synthetase (CKS) | 27 | 0.96 | 0.120 | 1.000 |
| AT1G53000.1 | CMP-KDO (3-deoxy-D-manno-octulosonate) synthetase (CKS) | 28 | 1.10 | 0.120 | 1.000 |
| AT1G53000.1 | CMP-KDO (3-deoxy-D-manno-octulosonate) synthetase (CKS) | 29 | 1.24 | 0.120 | 1.000 |

| | | | | | | |
|---|---|---|---|---|---|---|
| AT1G53000.1 | CMP-KDO (3-deoxy-D-manno-octulosonate) synthetase (CKS) | 30 | 1.31 | 0.120 | 0.999 | |
| AT1G53000.1 | CMP-KDO (3-deoxy-D-manno-octulosonate) synthetase (CKS) | 31 | 1.38 | 0.120 | 0.994 | |
| AT1G53000.1 | CMP-KDO (3-deoxy-D-manno-octulosonate) synthetase (CKS) | 32 | 1.43 | 0.121 | 0.980 | |
| AT1G53000.1 | CMP-KDO (3-deoxy-D-manno-octulosonate) synthetase (CKS) | 33 | 1.44 | 0.121 | 0.979 | |
| AT1G53000.1 | CMP-KDO (3-deoxy-D-manno-octulosonate) synthetase (CKS) | 34 | 1.37 | 0.121 | 0.995 | |
| AT1G53000.1 | CMP-KDO (3-deoxy-D-manno-octulosonate) synthetase (CKS) | 35 | 1.29 | 0.121 | 1.000 | |
| AT1G53000.1 | CMP-KDO (3-deoxy-D-manno-octulosonate) synthetase (CKS) | 36 | 1.23 | 0.121 | 1.000 | |
| AT1G53000.1 | CMP-KDO (3-deoxy-D-manno-octulosonate) synthetase (CKS) | 37 | 1.25 | 0.121 | 1.000 | |
| AT1G53000.1 | CMP-KDO (3-deoxy-D-manno-octulosonate) synthetase (CKS) | 38 | 1.36 | 0.121 | 0.997 | |
| AT1G53000.1 | CMP-KDO (3-deoxy-D-manno-octulosonate) synthetase (CKS) | 39 | 1.56 | 0.121 | 0.889 | |
| AT1G53000.1 | CMP-KDO (3-deoxy-D-manno-octulosonate) synthetase (CKS) | 40 | 1.82 | 0.121 | 0.498 | |
| AT1G53000.1 | CMP-KDO (3-deoxy-D-manno-octulosonate) synthetase (CKS) | 41 | 2.04 | 0.122 | 0.231 | |
| AT1G53000.1 | CMP-KDO (3-deoxy-D-manno-octulosonate) synthetase (CKS) | 42 | 2.10 | 0.123 | 0.192 | |
| AT1G53000.1 | CMP-KDO (3-deoxy-D-manno-octulosonate) synthetase (CKS) | 43 | 1.86 | 0.124 | 0.467 | |
| AT1G53000.1 | CMP-KDO (3-deoxy-D-manno-octulosonate) synthetase (CKS) | 44 | 1.34 | 0.126 | 0.999 | |
| AT1G53000.1 | CMP-KDO (3-deoxy-D-manno-octulosonate) synthetase (CKS) | 45 | 0.78 | 0.134 | 1.000 | |
| AT1G53000.1 | CMP-KDO (3-deoxy-D-manno-octulosonate) synthetase (CKS) | 46 | 0.39 | 0.163 | 0.242 | |
| AT1G53000.1 | CMP-KDO (3-deoxy-D-manno-octulosonate) synthetase (CKS) | 47 | 0.18 | 0.230 | 0.037 | * |
| AT1G53210.1 | Sodium/calcium exchanger (NCL) | 2 | 1.32 | 0.266 | 0.991 | |
| AT1G53210.1 | Sodium/calcium exchanger (NCL) | 3 | 1.45 | 0.239 | 0.952 | |
| AT1G53210.1 | Sodium/calcium exchanger (NCL) | 4 | 1.58 | 0.223 | 0.862 | |
| AT1G53210.1 | Sodium/calcium exchanger (NCL) | 5 | 1.73 | 0.218 | 0.750 | |
| AT1G53210.1 | Sodium/calcium exchanger (NCL) | 6 | 1.88 | 0.223 | 0.659 | |
| AT1G53210.1 | Sodium/calcium exchanger (NCL) | 7 | 2.03 | 0.235 | 0.606 | |
| AT1G53210.1 | Sodium/calcium exchanger (NCL) | 8 | 2.18 | 0.252 | 0.582 | |
| AT1G53210.1 | Sodium/calcium exchanger (NCL) | 31 | 1.44 | 0.227 | 0.942 | |
| AT1G53210.1 | Sodium/calcium exchanger (NCL) | 32 | 1.27 | 0.223 | 0.990 | |
| AT1G53210.1 | Sodium/calcium exchanger (NCL) | 33 | 1.09 | 0.225 | 1.000 | |
| AT1G53210.1 | Sodium/calcium exchanger (NCL) | 34 | 0.92 | 0.231 | 1.000 | |
| AT1G53210.1 | Sodium/calcium exchanger (NCL) | 35 | 0.77 | 0.241 | 0.990 | |
| AT1G53210.1 | Sodium/calcium exchanger (NCL) | 46 | 0.06 | 0.354 | 0.006 | ** |

| | | | | | | |
|---|---|---|---|---|---|---|
| AT1G53210.1 | Sodium/calcium exchanger (NCL) | 47 | 0.04 | 0.374 | 0.004 | ** |
| AT1G53240.1 | Mitochondrial malate dehydrogenase 1 (mMDH1) | 3 | 0.65 | 0.260 | 0.996 | |
| AT1G53240.1 | Mitochondrial malate dehydrogenase 1 (mMDH1) | 4 | 0.57 | 0.238 | 0.938 | |
| AT1G53240.1 | Mitochondrial malate dehydrogenase 1 (mMDH1) | 5 | 0.53 | 0.233 | 0.844 | |
| AT1G53240.1 | Mitochondrial malate dehydrogenase 1 (mMDH1) | 6 | 0.52 | 0.232 | 0.812 | |
| AT1G53240.1 | Mitochondrial malate dehydrogenase 1 (mMDH1) | 7 | 0.53 | 0.232 | 0.850 | |
| AT1G53240.1 | Mitochondrial malate dehydrogenase 1 (mMDH1) | 8 | 0.57 | 0.231 | 0.919 | |
| AT1G53240.1 | Mitochondrial malate dehydrogenase 1 (mMDH1) | 9 | 0.62 | 0.230 | 0.974 | |
| AT1G53240.1 | Mitochondrial malate dehydrogenase 1 (mMDH1) | 10 | 0.68 | 0.230 | 0.995 | |
| AT1G53240.1 | Mitochondrial malate dehydrogenase 1 (mMDH1) | 11 | 0.72 | 0.229 | 0.999 | |
| AT1G53240.1 | Mitochondrial malate dehydrogenase 1 (mMDH1) | 12 | 0.76 | 0.229 | 1.000 | |
| AT1G53240.1 | Mitochondrial malate dehydrogenase 1 (mMDH1) | 13 | 0.77 | 0.228 | 1.000 | |
| AT1G53240.1 | Mitochondrial malate dehydrogenase 1 (mMDH1) | 14 | 0.78 | 0.227 | 1.000 | |
| AT1G53240.1 | Mitochondrial malate dehydrogenase 1 (mMDH1) | 15 | 0.78 | 0.227 | 1.000 | |
| AT1G53240.1 | Mitochondrial malate dehydrogenase 1 (mMDH1) | 16 | 0.80 | 0.227 | 1.000 | |
| AT1G53240.1 | Mitochondrial malate dehydrogenase 1 (mMDH1) | 17 | 0.84 | 0.227 | 1.000 | |
| AT1G53240.1 | Mitochondrial malate dehydrogenase 1 (mMDH1) | 18 | 0.88 | 0.228 | 1.000 | |
| AT1G53240.1 | Mitochondrial malate dehydrogenase 1 (mMDH1) | 19 | 0.90 | 0.229 | 1.000 | |
| AT1G53240.1 | Mitochondrial malate dehydrogenase 1 (mMDH1) | 20 | 0.87 | 0.231 | 1.000 | |
| AT1G53240.1 | Mitochondrial malate dehydrogenase 1 (mMDH1) | 21 | 0.80 | 0.231 | 1.000 | |
| AT1G53240.1 | Mitochondrial malate dehydrogenase 1 (mMDH1) | 22 | 0.72 | 0.231 | 0.999 | |
| AT1G53240.1 | Mitochondrial malate dehydrogenase 1 (mMDH1) | 23 | 0.65 | 0.231 | 0.989 | |
| AT1G53240.1 | Mitochondrial malate dehydrogenase 1 (mMDH1) | 24 | 0.60 | 0.230 | 0.955 | |
| AT1G53240.1 | Mitochondrial malate dehydrogenase 1 (mMDH1) | 25 | 0.57 | 0.229 | 0.912 | |
| AT1G53240.1 | Mitochondrial malate dehydrogenase 1 (mMDH1) | 26 | 0.55 | 0.229 | 0.882 | |
| AT1G53240.1 | Mitochondrial malate dehydrogenase 1 (mMDH1) | 27 | 0.57 | 0.228 | 0.908 | |
| AT1G53240.1 | Mitochondrial malate dehydrogenase 1 (mMDH1) | 28 | 0.62 | 0.228 | 0.968 | |
| AT1G53240.1 | Mitochondrial malate dehydrogenase 1 (mMDH1) | 29 | 0.70 | 0.228 | 0.998 | |
| AT1G53240.1 | Mitochondrial malate dehydrogenase 1 (mMDH1) | 30 | 0.80 | 0.229 | 1.000 | |
| AT1G53240.1 | Mitochondrial malate dehydrogenase 1 (mMDH1) | 31 | 0.89 | 0.229 | 1.000 | |
| AT1G53240.1 | Mitochondrial malate dehydrogenase 1 (mMDH1) | 32 | 0.98 | 0.229 | 1.000 | |

| | | | | | |
|---|---|---|---|---|---|
| AT1G53240.1 | Mitochondrial malate dehydrogenase 1 (mMDH1) | 33 | 1.03 | 0.229 | 1.000 |
| AT1G53240.1 | Mitochondrial malate dehydrogenase 1 (mMDH1) | 34 | 1.07 | 0.229 | 1.000 |
| AT1G53240.1 | Mitochondrial malate dehydrogenase 1 (mMDH1) | 35 | 1.09 | 0.229 | 1.000 |
| AT1G53240.1 | Mitochondrial malate dehydrogenase 1 (mMDH1) | 36 | 1.12 | 0.229 | 1.000 |
| AT1G53240.1 | Mitochondrial malate dehydrogenase 1 (mMDH1) | 37 | 1.16 | 0.229 | 1.000 |
| AT1G53240.1 | Mitochondrial malate dehydrogenase 1 (mMDH1) | 38 | 1.22 | 0.229 | 1.000 |
| AT1G53240.1 | Mitochondrial malate dehydrogenase 1 (mMDH1) | 39 | 1.28 | 0.229 | 1.000 |
| AT1G53240.1 | Mitochondrial malate dehydrogenase 1 (mMDH1) | 40 | 1.32 | 0.230 | 1.000 |
| AT1G53240.1 | Mitochondrial malate dehydrogenase 1 (mMDH1) | 41 | 1.30 | 0.231 | 1.000 |
| AT1G53240.1 | Mitochondrial malate dehydrogenase 1 (mMDH1) | 42 | 1.24 | 0.231 | 1.000 |
| AT1G53240.1 | Mitochondrial malate dehydrogenase 1 (mMDH1) | 43 | 1.15 | 0.231 | 1.000 |
| AT1G53240.1 | Mitochondrial malate dehydrogenase 1 (mMDH1) | 44 | 1.03 | 0.233 | 1.000 |
| AT1G53240.1 | Mitochondrial malate dehydrogenase 1 (mMDH1) | 45 | 0.89 | 0.241 | 1.000 |
| AT1G53240.1 | Mitochondrial malate dehydrogenase 1 (mMDH1) | 46 | 0.75 | 0.267 | 1.000 |
| AT1G53240.1 | Mitochondrial malate dehydrogenase 1 (mMDH1) | 47 | 0.63 | 0.324 | 0.999 |
| AT1G53240.1 | Mitochondrial malate dehydrogenase 1 (mMDH1) | 48 | 0.52 | 0.407 | 0.997 |
| AT1G53530.1 | Mitochondrial ATP-independent protease (IMP1A) | 2 | 0.97 | 0.471 | 1.000 |
| AT1G53530.1 | Mitochondrial ATP-independent protease (IMP1A) | 3 | 0.92 | 0.423 | 1.000 |
| AT1G53530.1 | Mitochondrial ATP-independent protease (IMP1A) | 11 | 1.24 | 0.284 | 1.000 |
| AT1G53530.1 | Mitochondrial ATP-independent protease (IMP1A) | 12 | 1.32 | 0.238 | 1.000 |
| AT1G53530.1 | Mitochondrial ATP-independent protease (IMP1A) | 13 | 1.34 | 0.221 | 1.000 |
| AT1G53530.1 | Mitochondrial ATP-independent protease (IMP1A) | 14 | 1.34 | 0.206 | 1.000 |
| AT1G53530.1 | Mitochondrial ATP-independent protease (IMP1A) | 15 | 1.28 | 0.197 | 1.000 |
| AT1G53530.1 | Mitochondrial ATP-independent protease (IMP1A) | 16 | 1.34 | 0.199 | 1.000 |
| AT1G53530.1 | Mitochondrial ATP-independent protease (IMP1A) | 17 | 1.70 | 0.213 | 0.988 |
| AT1G53530.1 | Mitochondrial ATP-independent protease (IMP1A) | 18 | 2.27 | 0.238 | 0.866 |
| AT1G53530.1 | Mitochondrial ATP-independent protease (IMP1A) | 19 | 2.90 | 0.302 | 0.848 |
| AT1G53530.1 | Mitochondrial ATP-independent protease (IMP1A) | 20 | 3.47 | 0.448 | 0.970 |
| AT1G53530.1 | Mitochondrial ATP-independent protease (IMP1A) | 37 | 2.34 | 0.345 | 0.989 |
| AT1G53530.1 | Mitochondrial ATP-independent protease (IMP1A) | 38 | 2.40 | 0.239 | 0.813 |
| AT1G53530.1 | Mitochondrial ATP-independent protease (IMP1A) | 39 | 1.94 | 0.204 | 0.905 |

| | | | | | | |
|---|---|---|---|---|---|---|
| AT1G53530.1 | Mitochondrial ATP-independent protease (IMP1A) | 40 | 0.97 | 0.195 | 1.000 | |
| AT1G53530.1 | Mitochondrial ATP-independent protease (IMP1A) | 41 | 0.42 | 0.194 | 0.552 | |
| AT1G53530.1 | Mitochondrial ATP-independent protease (IMP1A) | 42 | 0.26 | 0.197 | 0.066 | . |
| AT1G53530.1 | Mitochondrial ATP-independent protease (IMP1A) | 43 | 0.27 | 0.208 | 0.106 | |
| AT1G53530.1 | Mitochondrial ATP-independent protease (IMP1A) | 44 | 0.41 | 0.260 | 0.869 | |
| AT1G53530.1 | Mitochondrial ATP-independent protease (IMP1A) | 45 | 0.83 | 0.381 | 1.000 | |
| AT1G53530.1 | Mitochondrial ATP-independent protease (IMP1A) | 46 | 2.02 | 0.583 | 1.000 | |
| AT1G53530.1 | Mitochondrial ATP-independent protease (IMP1A) | 47 | 5.32 | 0.882 | 0.999 | |
| AT1G53645.1 | Hydroxyproline-rich glycoprotein family protein | 2 | 1.43 | 0.778 | 1.000 | |
| AT1G53645.1 | Hydroxyproline-rich glycoprotein family protein | 3 | 1.54 | 0.679 | 1.000 | |
| AT1G53645.1 | Hydroxyproline-rich glycoprotein family protein | 11 | 2.50 | 0.405 | 0.996 | |
| AT1G53645.1 | Hydroxyproline-rich glycoprotein family protein | 12 | 2.54 | 0.353 | 0.983 | |
| AT1G53645.1 | Hydroxyproline-rich glycoprotein family protein | 13 | 2.49 | 0.313 | 0.961 | |
| AT1G53645.1 | Hydroxyproline-rich glycoprotein family protein | 14 | 2.40 | 0.287 | 0.945 | |
| AT1G53645.1 | Hydroxyproline-rich glycoprotein family protein | 15 | 2.43 | 0.272 | 0.911 | |
| AT1G53645.1 | Hydroxyproline-rich glycoprotein family protein | 16 | 2.57 | 0.265 | 0.845 | |
| AT1G53645.1 | Hydroxyproline-rich glycoprotein family protein | 17 | 2.58 | 0.263 | 0.837 | |
| AT1G53645.1 | Hydroxyproline-rich glycoprotein family protein | 18 | 2.32 | 0.265 | 0.925 | |
| AT1G53645.1 | Hydroxyproline-rich glycoprotein family protein | 19 | 1.92 | 0.270 | 0.992 | |
| AT1G53645.1 | Hydroxyproline-rich glycoprotein family protein | 20 | 1.53 | 0.274 | 1.000 | |
| AT1G53645.1 | Hydroxyproline-rich glycoprotein family protein | 21 | 1.24 | 0.276 | 1.000 | |
| AT1G53645.1 | Hydroxyproline-rich glycoprotein family protein | 22 | 1.07 | 0.277 | 1.000 | |
| AT1G53645.1 | Hydroxyproline-rich glycoprotein family protein | 23 | 0.98 | 0.275 | 1.000 | |
| AT1G53645.1 | Hydroxyproline-rich glycoprotein family protein | 24 | 0.92 | 0.272 | 1.000 | |
| AT1G53645.1 | Hydroxyproline-rich glycoprotein family protein | 25 | 0.88 | 0.269 | 1.000 | |
| AT1G53645.1 | Hydroxyproline-rich glycoprotein family protein | 26 | 0.88 | 0.267 | 1.000 | |
| AT1G53645.1 | Hydroxyproline-rich glycoprotein family protein | 27 | 0.91 | 0.267 | 1.000 | |
| AT1G53645.1 | Hydroxyproline-rich glycoprotein family protein | 28 | 0.92 | 0.267 | 1.000 | |
| AT1G53645.1 | Hydroxyproline-rich glycoprotein family protein | 29 | 0.91 | 0.267 | 1.000 | |
| AT1G53645.1 | Hydroxyproline-rich glycoprotein family protein | 30 | 0.90 | 0.267 | 1.000 | |
| AT1G53645.1 | Hydroxyproline-rich glycoprotein family protein | 31 | 0.89 | 0.268 | 1.000 | |

| | | | | | | |
|---|---|---|---|---|---|---|
| AT1G53645.1 | Hydroxyproline-rich glycoprotein family protein | 32 | 0.89 | 0.268 | 1.000 | |
| AT1G53645.1 | Hydroxyproline-rich glycoprotein family protein | 33 | 0.89 | 0.269 | 1.000 | |
| AT1G53645.1 | Hydroxyproline-rich glycoprotein family protein | 34 | 0.93 | 0.269 | 1.000 | |
| AT1G53645.1 | Hydroxyproline-rich glycoprotein family protein | 35 | 1.03 | 0.269 | 1.000 | |
| AT1G53645.1 | Hydroxyproline-rich glycoprotein family protein | 36 | 1.24 | 0.269 | 1.000 | |
| AT1G53645.1 | Hydroxyproline-rich glycoprotein family protein | 37 | 1.51 | 0.270 | 1.000 | |
| AT1G53645.1 | Hydroxyproline-rich glycoprotein family protein | 38 | 1.77 | 0.270 | 0.998 | |
| AT1G53645.1 | Hydroxyproline-rich glycoprotein family protein | 39 | 1.91 | 0.271 | 0.993 | |
| AT1G53645.1 | Hydroxyproline-rich glycoprotein family protein | 40 | 1.83 | 0.274 | 0.997 | |
| AT1G53645.1 | Hydroxyproline-rich glycoprotein family protein | 41 | 1.51 | 0.286 | 1.000 | |
| AT1G53645.1 | Hydroxyproline-rich glycoprotein family protein | 42 | 1.11 | 0.317 | 1.000 | |
| AT1G53645.1 | Hydroxyproline-rich glycoprotein family protein | 43 | 0.74 | 0.380 | 1.000 | |
| AT1G53645.1 | Hydroxyproline-rich glycoprotein family protein | 44 | 0.45 | 0.477 | 1.000 | |
| AT1G53645.1 | Hydroxyproline-rich glycoprotein family protein | 46 | 0.13 | 0.790 | 0.988 | |
| AT1G53645.1 | Hydroxyproline-rich glycoprotein family protein | 47 | 0.07 | 1.007 | 0.983 | |
| AT1G53760.1 | K+-H+ exchange-like protein | 2 | 1.30 | 0.347 | 1.000 | |
| AT1G53760.1 | K+-H+ exchange-like protein | 3 | 1.20 | 0.329 | 1.000 | |
| AT1G53760.1 | K+-H+ exchange-like protein | 23 | 0.69 | 0.271 | 1.000 | |
| AT1G53760.1 | K+-H+ exchange-like protein | 24 | 0.75 | 0.222 | 1.000 | |
| AT1G53760.1 | K+-H+ exchange-like protein | 25 | 0.83 | 0.200 | 1.000 | |
| AT1G53760.1 | K+-H+ exchange-like protein | 26 | 0.93 | 0.182 | 1.000 | |
| AT1G53760.1 | K+-H+ exchange-like protein | 27 | 1.07 | 0.171 | 1.000 | |
| AT1G53760.1 | K+-H+ exchange-like protein | 28 | 1.27 | 0.160 | 1.000 | |
| AT1G53760.1 | K+-H+ exchange-like protein | 29 | 1.52 | 0.150 | 0.947 | |
| AT1G53760.1 | K+-H+ exchange-like protein | 30 | 1.72 | 0.144 | 0.718 | |
| AT1G53760.1 | K+-H+ exchange-like protein | 31 | 1.78 | 0.141 | 0.617 | |
| AT1G53760.1 | K+-H+ exchange-like protein | 32 | 1.67 | 0.139 | 0.743 | |
| AT1G53760.1 | K+-H+ exchange-like protein | 33 | 1.48 | 0.139 | 0.940 | |
| AT1G53760.1 | K+-H+ exchange-like protein | 34 | 1.26 | 0.138 | 0.999 | |
| AT1G53760.1 | K+-H+ exchange-like protein | 35 | 1.12 | 0.138 | 1.000 | |
| AT1G53760.1 | K+-H+ exchange-like protein | 36 | 1.15 | 0.138 | 1.000 | |
| AT1G53760.1 | K+-H+ exchange-like protein | 37 | 1.31 | 0.138 | 0.997 | |
| AT1G53760.1 | K+-H+ exchange-like protein | 38 | 1.63 | 0.138 | 0.789 | |
| AT1G53760.1 | K+-H+ exchange-like protein | 39 | 2.18 | 0.138 | 0.185 | |
| AT1G53760.1 | K+-H+ exchange-like protein | 40 | 2.92 | 0.138 | 0.016 | * |
| AT1G53760.1 | K+-H+ exchange-like protein | 41 | 3.57 | 0.139 | 0.002 | ** |
| AT1G53760.1 | K+-H+ exchange-like protein | 42 | 3.89 | 0.140 | 0.001 | *** |
| AT1G53760.1 | K+-H+ exchange-like protein | 43 | 3.63 | 0.141 | 0.002 | ** |
| AT1G53760.1 | K+-H+ exchange-like protein | 44 | 2.48 | 0.142 | 0.083 | . |
| AT1G53760.1 | K+-H+ exchange-like protein | 45 | 1.19 | 0.147 | 1.000 | |
| AT1G53760.1 | K+-H+ exchange-like protein | 46 | 0.43 | 0.177 | 0.382 | |
| AT1G53760.1 | K+-H+ exchange-like protein | 47 | 0.13 | 0.257 | 0.013 | * |
| AT1G54150.1 | E3 ubiquitin-protein ligase (SPL2) | 2 | 1.96 | 0.270 | 0.801 | |
| AT1G54150.1 | E3 ubiquitin-protein ligase (SPL2) | 3 | 1.89 | 0.258 | 0.807 | |

| | | | | | |
|---|---|---|---|---|---|
| AT1G54150.1 | E3 ubiquitin-protein ligase (SPL2) | 20 | 1.00 | 0.128 | 1.000 |
| AT1G54150.1 | E3 ubiquitin-protein ligase (SPL2) | 21 | 0.97 | 0.120 | 1.000 |
| AT1G54150.1 | E3 ubiquitin-protein ligase (SPL2) | 22 | 0.93 | 0.113 | 1.000 |
| AT1G54150.1 | E3 ubiquitin-protein ligase (SPL2) | 23 | 0.90 | 0.106 | 0.997 |
| AT1G54150.1 | E3 ubiquitin-protein ligase (SPL2) | 24 | 0.87 | 0.099 | 0.979 |
| AT1G54150.1 | E3 ubiquitin-protein ligase (SPL2) | 25 | 0.84 | 0.093 | 0.928 |
| AT1G54150.1 | E3 ubiquitin-protein ligase (SPL2) | 26 | 0.81 | 0.089 | 0.838 |
| AT1G54150.1 | E3 ubiquitin-protein ligase (SPL2) | 27 | 0.79 | 0.085 | 0.730 |
| AT1G54150.1 | E3 ubiquitin-protein ligase (SPL2) | 28 | 0.78 | 0.082 | 0.636 |
| AT1G54150.1 | E3 ubiquitin-protein ligase (SPL2) | 29 | 0.77 | 0.079 | 0.577 |
| AT1G54150.1 | E3 ubiquitin-protein ligase (SPL2) | 30 | 0.77 | 0.078 | 0.557 |
| AT1G54150.1 | E3 ubiquitin-protein ligase (SPL2) | 31 | 0.78 | 0.077 | 0.577 |
| AT1G54150.1 | E3 ubiquitin-protein ligase (SPL2) | 32 | 0.79 | 0.077 | 0.632 |
| AT1G54150.1 | E3 ubiquitin-protein ligase (SPL2) | 33 | 0.80 | 0.078 | 0.710 |
| AT1G54150.1 | E3 ubiquitin-protein ligase (SPL2) | 34 | 0.82 | 0.079 | 0.800 |
| AT1G54150.1 | E3 ubiquitin-protein ligase (SPL2) | 35 | 0.84 | 0.080 | 0.885 |
| AT1G54150.1 | E3 ubiquitin-protein ligase (SPL2) | 36 | 0.87 | 0.082 | 0.950 |
| AT1G54150.1 | E3 ubiquitin-protein ligase (SPL2) | 37 | 0.89 | 0.084 | 0.986 |
| AT1G54150.1 | E3 ubiquitin-protein ligase (SPL2) | 38 | 0.93 | 0.087 | 0.999 |
| AT1G54150.1 | E3 ubiquitin-protein ligase (SPL2) | 39 | 0.96 | 0.089 | 1.000 |
| AT1G54150.1 | E3 ubiquitin-protein ligase (SPL2) | 40 | 1.00 | 0.093 | 1.000 |
| AT1G54150.1 | E3 ubiquitin-protein ligase (SPL2) | 41 | 1.04 | 0.097 | 1.000 |
| AT1G54150.1 | E3 ubiquitin-protein ligase (SPL2) | 42 | 1.08 | 0.101 | 0.999 |
| AT1G54150.1 | E3 ubiquitin-protein ligase (SPL2) | 43 | 1.13 | 0.108 | 0.994 |
| AT1G54150.1 | E3 ubiquitin-protein ligase (SPL2) | 44 | 1.18 | 0.115 | 0.982 |
| AT1G54150.1 | E3 ubiquitin-protein ligase (SPL2) | 45 | 1.22 | 0.125 | 0.963 |
| AT1G54150.1 | E3 ubiquitin-protein ligase (SPL2) | 46 | 1.28 | 0.137 | 0.944 |
| AT1G54150.1 | E3 ubiquitin-protein ligase (SPL2) | 47 | 1.33 | 0.151 | 0.928 |
| AT1G54220.2 | Dihydrolipoamide acetyltransferase (PDCE2-3) | 3 | 0.85 | 0.246 | 1.000 |
| AT1G54220.2 | Dihydrolipoamide acetyltransferase (PDCE2-3) | 4 | 0.52 | 0.223 | 0.993 |
| AT1G54220.2 | Dihydrolipoamide acetyltransferase (PDCE2-3) | 5 | 0.38 | 0.224 | 0.787 |

| | | | | | |
|---|---|---|---|---|---|
| AT1G54220.2 | Dihydrolipoamide acetyltransferase (PDCE2-3) | 6 | 0.32 | 0.224 | 0.506 |
| AT1G54220.2 | Dihydrolipoamide acetyltransferase (PDCE2-3) | 7 | 0.29 | 0.223 | 0.361 |
| AT1G54220.2 | Dihydrolipoamide acetyltransferase (PDCE2-3) | 8 | 0.30 | 0.220 | 0.379 |
| AT1G54220.2 | Dihydrolipoamide acetyltransferase (PDCE2-3) | 9 | 0.35 | 0.217 | 0.616 |
| AT1G54220.2 | Dihydrolipoamide acetyltransferase (PDCE2-3) | 10 | 0.44 | 0.217 | 0.911 |
| AT1G54220.2 | Dihydrolipoamide acetyltransferase (PDCE2-3) | 11 | 0.61 | 0.217 | 1.000 |
| AT1G54220.2 | Dihydrolipoamide acetyltransferase (PDCE2-3) | 12 | 0.91 | 0.217 | 1.000 |
| AT1G54220.2 | Dihydrolipoamide acetyltransferase (PDCE2-3) | 13 | 1.24 | 0.215 | 1.000 |
| AT1G54220.2 | Dihydrolipoamide acetyltransferase (PDCE2-3) | 14 | 1.41 | 0.213 | 1.000 |
| AT1G54220.2 | Dihydrolipoamide acetyltransferase (PDCE2-3) | 15 | 1.36 | 0.212 | 1.000 |
| AT1G54220.2 | Dihydrolipoamide acetyltransferase (PDCE2-3) | 16 | 1.21 | 0.210 | 1.000 |
| AT1G54220.2 | Dihydrolipoamide acetyltransferase (PDCE2-3) | 17 | 1.08 | 0.210 | 1.000 |
| AT1G54220.2 | Dihydrolipoamide acetyltransferase (PDCE2-3) | 18 | 1.01 | 0.213 | 1.000 |
| AT1G54220.2 | Dihydrolipoamide acetyltransferase (PDCE2-3) | 19 | 1.09 | 0.218 | 1.000 |
| AT1G54220.2 | Dihydrolipoamide acetyltransferase (PDCE2-3) | 20 | 1.29 | 0.223 | 1.000 |
| AT1G54220.2 | Dihydrolipoamide acetyltransferase (PDCE2-3) | 21 | 1.41 | 0.225 | 1.000 |
| AT1G54220.2 | Dihydrolipoamide acetyltransferase (PDCE2-3) | 22 | 1.24 | 0.225 | 1.000 |
| AT1G54220.2 | Dihydrolipoamide acetyltransferase (PDCE2-3) | 23 | 0.99 | 0.224 | 1.000 |
| AT1G54220.2 | Dihydrolipoamide acetyltransferase (PDCE2-3) | 24 | 0.85 | 0.220 | 1.000 |
| AT1G54220.2 | Dihydrolipoamide acetyltransferase (PDCE2-3) | 25 | 0.82 | 0.217 | 1.000 |
| AT1G54220.2 | Dihydrolipoamide acetyltransferase (PDCE2-3) | 26 | 0.94 | 0.215 | 1.000 |
| AT1G54220.2 | Dihydrolipoamide acetyltransferase (PDCE2-3) | 27 | 1.18 | 0.216 | 1.000 |
| AT1G54220.2 | Dihydrolipoamide acetyltransferase (PDCE2-3) | 28 | 1.33 | 0.215 | 1.000 |
| AT1G54220.2 | Dihydrolipoamide acetyltransferase (PDCE2-3) | 29 | 1.36 | 0.215 | 1.000 |
| AT1G54220.2 | Dihydrolipoamide acetyltransferase (PDCE2-3) | 30 | 1.33 | 0.215 | 1.000 |
| AT1G54220.2 | Dihydrolipoamide acetyltransferase (PDCE2-3) | 31 | 1.31 | 0.216 | 1.000 |
| AT1G54220.2 | Dihydrolipoamide acetyltransferase (PDCE2-3) | 32 | 1.25 | 0.217 | 1.000 |
| AT1G54220.2 | Dihydrolipoamide acetyltransferase (PDCE2-3) | 33 | 1.16 | 0.217 | 1.000 |
| AT1G54220.2 | Dihydrolipoamide acetyltransferase (PDCE2-3) | 34 | 1.19 | 0.217 | 1.000 |
| AT1G54220.2 | Dihydrolipoamide acetyltransferase (PDCE2-3) | 35 | 1.38 | 0.217 | 1.000 |
| AT1G54220.2 | Dihydrolipoamide acetyltransferase (PDCE2-3) | 36 | 1.72 | 0.216 | 0.999 |

| | | | | | | |
|---|---|---|---|---|---|---|
| AT1G54220.2 | Dihydrolipoamide acetyltransferase (PDCE2-3) | 37 | 2.04 | 0.216 | 0.976 | |
| AT1G54220.2 | Dihydrolipoamide acetyltransferase (PDCE2-3) | 38 | 2.10 | 0.216 | 0.966 | |
| AT1G54220.2 | Dihydrolipoamide acetyltransferase (PDCE2-3) | 39 | 2.04 | 0.217 | 0.978 | |
| AT1G54220.2 | Dihydrolipoamide acetyltransferase (PDCE2-3) | 40 | 2.15 | 0.217 | 0.955 | |
| AT1G54220.2 | Dihydrolipoamide acetyltransferase (PDCE2-3) | 41 | 2.73 | 0.218 | 0.681 | |
| AT1G54220.2 | Dihydrolipoamide acetyltransferase (PDCE2-3) | 42 | 3.80 | 0.221 | 0.221 | |
| AT1G54220.2 | Dihydrolipoamide acetyltransferase (PDCE2-3) | 43 | 4.39 | 0.223 | 0.117 | |
| AT1G54220.2 | Dihydrolipoamide acetyltransferase (PDCE2-3) | 44 | 3.22 | 0.223 | 0.450 | |
| AT1G54220.2 | Dihydrolipoamide acetyltransferase (PDCE2-3) | 45 | 1.50 | 0.223 | 1.000 | |
| AT1G54220.2 | Dihydrolipoamide acetyltransferase (PDCE2-3) | 46 | 0.52 | 0.263 | 1.000 | |
| AT1G54220.2 | Dihydrolipoamide acetyltransferase (PDCE2-3) | 47 | 0.15 | 0.435 | 0.775 | |
| AT1G54220.2 | Dihydrolipoamide acetyltransferase (PDCE2-3) | 48 | 0.04 | 0.696 | 0.716 | |
| AT1G54500.1 | ? | 2 | 0.92 | 0.219 | 1.000 | |
| AT1G54500.1 | ? | 3 | 0.94 | 0.199 | 1.000 | |
| AT1G54500.1 | ? | 4 | 0.97 | 0.182 | 1.000 | |
| AT1G54500.1 | ? | 6 | 1.03 | 0.155 | 1.000 | |
| AT1G54500.1 | ? | 7 | 1.06 | 0.145 | 1.000 | |
| AT1G54500.1 | ? | 8 | 1.09 | 0.137 | 1.000 | |
| AT1G54500.1 | ? | 9 | 1.13 | 0.131 | 0.998 | |
| AT1G54500.1 | ? | 10 | 1.17 | 0.126 | 0.989 | |
| AT1G54500.1 | ? | 11 | 1.21 | 0.124 | 0.968 | |
| AT1G54500.1 | ? | 12 | 1.25 | 0.123 | 0.932 | |
| AT1G54500.1 | ? | 13 | 1.29 | 0.124 | 0.889 | |
| AT1G54500.1 | ? | 14 | 1.33 | 0.125 | 0.848 | |
| AT1G54500.1 | ? | 15 | 1.36 | 0.128 | 0.818 | |
| AT1G54500.1 | ? | 16 | 1.38 | 0.131 | 0.802 | |
| AT1G54500.1 | ? | 28 | 0.97 | 0.137 | 1.000 | |
| AT1G54500.1 | ? | 29 | 0.90 | 0.137 | 0.999 | |
| AT1G54500.1 | ? | 30 | 0.83 | 0.139 | 0.982 | |
| AT1G54500.1 | ? | 31 | 0.75 | 0.142 | 0.906 | |
| AT1G54500.1 | ? | 32 | 0.68 | 0.146 | 0.764 | |
| AT1G54500.1 | ? | 33 | 0.62 | 0.152 | 0.595 | |
| AT1G54500.1 | ? | 34 | 0.55 | 0.159 | 0.443 | |
| AT1G54500.1 | ? | 35 | 0.50 | 0.167 | 0.326 | |
| AT1G54500.1 | ? | 46 | 0.15 | 0.282 | 0.029 | * |
| AT1G54500.1 | ? | 47 | 0.14 | 0.299 | 0.028 | * |
| AT1G54780.1 | Thylakoid lumen 18.3 kDa protein (TLP18.3) | 2 | 0.41 | 0.354 | 0.986 | |
| AT1G54780.1 | Thylakoid lumen 18.3 kDa protein (TLP18.3) | 3 | 0.49 | 0.258 | 0.969 | |
| AT1G54780.1 | Thylakoid lumen 18.3 kDa protein (TLP18.3) | 4 | 0.59 | 0.215 | 0.990 | |
| AT1G54780.1 | Thylakoid lumen 18.3 kDa protein (TLP18.3) | 5 | 0.71 | 0.204 | 1.000 | |
| AT1G54780.1 | Thylakoid lumen 18.3 kDa protein (TLP18.3) | 6 | 0.79 | 0.204 | 1.000 | |
| AT1G54780.1 | Thylakoid lumen 18.3 kDa protein (TLP18.3) | 7 | 0.82 | 0.203 | 1.000 | |
| AT1G54780.1 | Thylakoid lumen 18.3 kDa protein (TLP18.3) | 8 | 0.84 | 0.201 | 1.000 | |

| | | | | | |
|---|---|---|---|---|---|
| AT1G54780.1 | Thylakoid lumen 18.3 kDa protein (TLP18.3) | 9 | 0.88 | 0.199 | 1.000 |
| AT1G54780.1 | Thylakoid lumen 18.3 kDa protein (TLP18.3) | 10 | 0.92 | 0.198 | 1.000 |
| AT1G54780.1 | Thylakoid lumen 18.3 kDa protein (TLP18.3) | 11 | 0.97 | 0.197 | 1.000 |
| AT1G54780.1 | Thylakoid lumen 18.3 kDa protein (TLP18.3) | 12 | 1.03 | 0.197 | 1.000 |
| AT1G54780.1 | Thylakoid lumen 18.3 kDa protein (TLP18.3) | 13 | 1.08 | 0.195 | 1.000 |
| AT1G54780.1 | Thylakoid lumen 18.3 kDa protein (TLP18.3) | 14 | 1.13 | 0.193 | 1.000 |
| AT1G54780.1 | Thylakoid lumen 18.3 kDa protein (TLP18.3) | 15 | 1.18 | 0.192 | 1.000 |
| AT1G54780.1 | Thylakoid lumen 18.3 kDa protein (TLP18.3) | 16 | 1.22 | 0.192 | 1.000 |
| AT1G54780.1 | Thylakoid lumen 18.3 kDa protein (TLP18.3) | 17 | 1.24 | 0.192 | 1.000 |
| AT1G54780.1 | Thylakoid lumen 18.3 kDa protein (TLP18.3) | 18 | 1.20 | 0.194 | 1.000 |
| AT1G54780.1 | Thylakoid lumen 18.3 kDa protein (TLP18.3) | 19 | 1.06 | 0.198 | 1.000 |
| AT1G54780.1 | Thylakoid lumen 18.3 kDa protein (TLP18.3) | 20 | 0.83 | 0.201 | 1.000 |
| AT1G54780.1 | Thylakoid lumen 18.3 kDa protein (TLP18.3) | 21 | 0.61 | 0.203 | 0.990 |
| AT1G54780.1 | Thylakoid lumen 18.3 kDa protein (TLP18.3) | 22 | 0.45 | 0.204 | 0.723 |
| AT1G54780.1 | Thylakoid lumen 18.3 kDa protein (TLP18.3) | 23 | 0.35 | 0.204 | 0.325 |
| AT1G54780.1 | Thylakoid lumen 18.3 kDa protein (TLP18.3) | 24 | 0.29 | 0.205 | 0.150 |
| AT1G54780.1 | Thylakoid lumen 18.3 kDa protein (TLP18.3) | 25 | 0.27 | 0.207 | 0.117 |
| AT1G54780.1 | Thylakoid lumen 18.3 kDa protein (TLP18.3) | 26 | 0.29 | 0.211 | 0.171 |
| AT1G54780.1 | Thylakoid lumen 18.3 kDa protein (TLP18.3) | 27 | 0.34 | 0.215 | 0.372 |
| AT1G54780.1 | Thylakoid lumen 18.3 kDa protein (TLP18.3) | 28 | 0.43 | 0.223 | 0.786 |
| AT1G54780.1 | Thylakoid lumen 18.3 kDa protein (TLP18.3) | 29 | 0.58 | 0.244 | 0.996 |
| AT1G54780.1 | Thylakoid lumen 18.3 kDa protein (TLP18.3) | 30 | 0.77 | 0.287 | 1.000 |
| AT1G54780.1 | Thylakoid lumen 18.3 kDa protein (TLP18.3) | 31 | 1.02 | 0.351 | 1.000 |
| AT1G54780.1 | Thylakoid lumen 18.3 kDa protein (TLP18.3) | 43 | 8.25 | 0.357 | 0.169 |
| AT1G54780.1 | Thylakoid lumen 18.3 kDa protein (TLP18.3) | 44 | 6.12 | 0.284 | 0.102 |
| AT1G54780.1 | Thylakoid lumen 18.3 kDa protein (TLP18.3) | 45 | 3.99 | 0.242 | 0.200 |
| AT1G54780.1 | Thylakoid lumen 18.3 kDa protein (TLP18.3) | 46 | 2.31 | 0.278 | 0.943 |
| AT1G54780.1 | Thylakoid lumen 18.3 kDa protein (TLP18.3) | 47 | 1.27 | 0.399 | 1.000 |
| AT1G55160.2 | WAS/WASL-interacting family protein | 3 | 0.57 | 0.354 | 0.998 |
| AT1G55160.2 | WAS/WASL-interacting family protein | 4 | 0.46 | 0.332 | 0.943 |
| AT1G55160.2 | WAS/WASL-interacting family protein | 5 | 0.40 | 0.330 | 0.838 |

| | | | | | |
|---|---|---|---|---|---|
| AT1G55160.2 | WAS/WASL-interacting family protein | 6 | 0.36 | 0.330 | 0.744 |
| AT1G55160.2 | WAS/WASL-interacting family protein | 7 | 0.34 | 0.330 | 0.686 |
| AT1G55160.2 | WAS/WASL-interacting family protein | 8 | 0.34 | 0.328 | 0.680 |
| AT1G55160.2 | WAS/WASL-interacting family protein | 9 | 0.35 | 0.327 | 0.715 |
| AT1G55160.2 | WAS/WASL-interacting family protein | 10 | 0.37 | 0.327 | 0.771 |
| AT1G55160.2 | WAS/WASL-interacting family protein | 11 | 0.41 | 0.326 | 0.851 |
| AT1G55160.2 | WAS/WASL-interacting family protein | 12 | 0.46 | 0.326 | 0.933 |
| AT1G55160.2 | WAS/WASL-interacting family protein | 13 | 0.53 | 0.325 | 0.987 |
| AT1G55160.2 | WAS/WASL-interacting family protein | 14 | 0.63 | 0.324 | 1.000 |
| AT1G55160.2 | WAS/WASL-interacting family protein | 15 | 0.72 | 0.324 | 1.000 |
| AT1G55160.2 | WAS/WASL-interacting family protein | 16 | 0.78 | 0.323 | 1.000 |
| AT1G55160.2 | WAS/WASL-interacting family protein | 17 | 0.89 | 0.324 | 1.000 |
| AT1G55160.2 | WAS/WASL-interacting family protein | 18 | 1.09 | 0.325 | 1.000 |
| AT1G55160.2 | WAS/WASL-interacting family protein | 19 | 1.29 | 0.327 | 1.000 |
| AT1G55160.2 | WAS/WASL-interacting family protein | 20 | 1.32 | 0.329 | 1.000 |
| AT1G55160.2 | WAS/WASL-interacting family protein | 21 | 1.16 | 0.330 | 1.000 |
| AT1G55160.2 | WAS/WASL-interacting family protein | 22 | 0.94 | 0.330 | 1.000 |
| AT1G55160.2 | WAS/WASL-interacting family protein | 23 | 0.78 | 0.329 | 1.000 |
| AT1G55160.2 | WAS/WASL-interacting family protein | 24 | 0.72 | 0.328 | 1.000 |
| AT1G55160.2 | WAS/WASL-interacting family protein | 25 | 0.77 | 0.327 | 1.000 |
| AT1G55160.2 | WAS/WASL-interacting family protein | 26 | 0.89 | 0.326 | 1.000 |
| AT1G55160.2 | WAS/WASL-interacting family protein | 27 | 1.02 | 0.326 | 1.000 |
| AT1G55160.2 | WAS/WASL-interacting family protein | 28 | 1.06 | 0.326 | 1.000 |
| AT1G55160.2 | WAS/WASL-interacting family protein | 29 | 0.96 | 0.326 | 1.000 |
| AT1G55160.2 | WAS/WASL-interacting family protein | 30 | 0.82 | 0.326 | 1.000 |
| AT1G55160.2 | WAS/WASL-interacting family protein | 31 | 0.67 | 0.326 | 1.000 |
| AT1G55160.2 | WAS/WASL-interacting family protein | 32 | 0.54 | 0.326 | 0.991 |
| AT1G55160.2 | WAS/WASL-interacting family protein | 33 | 0.45 | 0.326 | 0.917 |
| AT1G55160.2 | WAS/WASL-interacting family protein | 34 | 0.38 | 0.327 | 0.787 |
| AT1G55160.2 | WAS/WASL-interacting family protein | 35 | 0.34 | 0.326 | 0.684 |
| AT1G55160.2 | WAS/WASL-interacting family protein | 36 | 0.34 | 0.326 | 0.683 |

| | | | | | | |
|---|---|---|---|---|---|---|
| AT1G55160.2 | WAS/WASL-interacting family protein | 37 | 0.38 | 0.326 | 0.794 | |
| AT1G55160.2 | WAS/WASL-interacting family protein | 38 | 0.43 | 0.326 | 0.889 | |
| AT1G55160.2 | WAS/WASL-interacting family protein | 39 | 0.44 | 0.326 | 0.901 | |
| AT1G55160.2 | WAS/WASL-interacting family protein | 40 | 0.42 | 0.327 | 0.867 | |
| AT1G55160.2 | WAS/WASL-interacting family protein | 41 | 0.37 | 0.328 | 0.775 | |
| AT1G55160.2 | WAS/WASL-interacting family protein | 42 | 0.33 | 0.329 | 0.662 | |
| AT1G55160.2 | WAS/WASL-interacting family protein | 43 | 0.30 | 0.330 | 0.561 | |
| AT1G55160.2 | WAS/WASL-interacting family protein | 44 | 0.27 | 0.329 | 0.472 | |
| AT1G55160.2 | WAS/WASL-interacting family protein | 45 | 0.25 | 0.334 | 0.400 | |
| AT1G55160.2 | WAS/WASL-interacting family protein | 46 | 0.22 | 0.365 | 0.420 | |
| AT1G55160.2 | WAS/WASL-interacting family protein | 47 | 0.20 | 0.458 | 0.603 | |
| AT1G55160.2 | WAS/WASL-interacting family protein | 48 | 0.18 | 0.607 | 0.823 | |
| AT1G55450.1 | S-adenosyl-L-methionine-dependent methyltransferase | 2 | 0.88 | 0.312 | 1.000 | |
| AT1G55450.1 | S-adenosyl-L-methionine-dependent methyltransferase | 3 | 0.93 | 0.298 | 1.000 | |
| AT1G55450.1 | S-adenosyl-L-methionine-dependent methyltransferase | 4 | 0.98 | 0.285 | 1.000 | |
| AT1G55450.1 | S-adenosyl-L-methionine-dependent methyltransferase | 19 | 1.81 | 0.174 | 0.513 | |
| AT1G55450.1 | S-adenosyl-L-methionine-dependent methyltransferase | 20 | 1.85 | 0.167 | 0.427 | |
| AT1G55450.1 | S-adenosyl-L-methionine-dependent methyltransferase | 21 | 1.90 | 0.160 | 0.346 | |
| AT1G55450.1 | S-adenosyl-L-methionine-dependent methyltransferase | 22 | 1.94 | 0.153 | 0.271 | |
| AT1G55450.1 | S-adenosyl-L-methionine-dependent methyltransferase | 23 | 1.98 | 0.147 | 0.210 | |
| AT1G55450.1 | S-adenosyl-L-methionine-dependent methyltransferase | 24 | 2.01 | 0.141 | 0.161 | |
| AT1G55450.1 | S-adenosyl-L-methionine-dependent methyltransferase | 26 | 2.05 | 0.131 | 0.097 | . |
| AT1G55450.1 | S-adenosyl-L-methionine-dependent methyltransferase | 27 | 2.04 | 0.126 | 0.082 | . |
| AT1G55450.1 | S-adenosyl-L-methionine-dependent methyltransferase | 28 | 2.02 | 0.123 | 0.074 | . |
| AT1G55450.1 | S-adenosyl-L-methionine-dependent methyltransferase | 29 | 1.99 | 0.120 | 0.076 | . |
| AT1G55450.1 | S-adenosyl-L-methionine-dependent methyltransferase | 30 | 1.94 | 0.118 | 0.089 | . |
| AT1G55450.1 | S-adenosyl-L-methionine-dependent methyltransferase | 31 | 1.87 | 0.118 | 0.112 | |
| AT1G55450.1 | S-adenosyl-L-methionine-dependent methyltransferase | 32 | 1.80 | 0.117 | 0.153 | |
| AT1G55450.1 | S-adenosyl-L-methionine-dependent methyltransferase | 33 | 1.72 | 0.118 | 0.214 | |
| AT1G55450.1 | S-adenosyl-L-methionine-dependent methyltransferase | 34 | 1.65 | 0.119 | 0.302 | |
| AT1G55450.1 | S-adenosyl-L-methionine-dependent methyltransferase | 35 | 1.57 | 0.120 | 0.415 | |

| | | | | | |
|---|---|---|---|---|---|
| AT1G55450.1 | S-adenosyl-L-methionine-dependent methyltransferase | 36 | 1.49 | 0.122 | 0.548 |
| AT1G55450.1 | S-adenosyl-L-methionine-dependent methyltransferase | 37 | 1.42 | 0.124 | 0.687 |
| AT1G55450.1 | S-adenosyl-L-methionine-dependent methyltransferase | 38 | 1.35 | 0.127 | 0.813 |
| AT1G55450.1 | S-adenosyl-L-methionine-dependent methyltransferase | 39 | 1.28 | 0.130 | 0.909 |
| AT1G55450.1 | S-adenosyl-L-methionine-dependent methyltransferase | 40 | 1.22 | 0.133 | 0.968 |
| AT1G55450.1 | S-adenosyl-L-methionine-dependent methyltransferase | 41 | 1.16 | 0.138 | 0.993 |
| AT1G55450.1 | S-adenosyl-L-methionine-dependent methyltransferase | 42 | 1.11 | 0.144 | 0.999 |
| AT1G55450.1 | S-adenosyl-L-methionine-dependent methyltransferase | 43 | 1.05 | 0.151 | 1.000 |
| AT1G55450.1 | S-adenosyl-L-methionine-dependent methyltransferase | 44 | 1.00 | 0.161 | 1.000 |
| AT1G55450.1 | S-adenosyl-L-methionine-dependent methyltransferase | 45 | 0.95 | 0.172 | 1.000 |
| AT1G55450.1 | S-adenosyl-L-methionine-dependent methyltransferase | 46 | 0.91 | 0.185 | 1.000 |
| AT1G55450.1 | S-adenosyl-L-methionine-dependent methyltransferase | 47 | 0.86 | 0.201 | 0.999 |
| AT1G55490.2 | Chaperonin 60 beta 1 (CPN60B1) | 2 | 1.25 | 0.688 | 1.000 |
| AT1G55490.2 | Chaperonin 60 beta 1 (CPN60B1) | 3 | 1.23 | 0.649 | 1.000 |
| AT1G55490.2 | Chaperonin 60 beta 1 (CPN60B1) | 23 | 1.17 | 0.306 | 1.000 |
| AT1G55490.2 | Chaperonin 60 beta 1 (CPN60B1) | 24 | 1.20 | 0.266 | 1.000 |
| AT1G55490.2 | Chaperonin 60 beta 1 (CPN60B1) | 25 | 1.25 | 0.232 | 0.999 |
| AT1G55490.2 | Chaperonin 60 beta 1 (CPN60B1) | 26 | 1.29 | 0.208 | 0.995 |
| AT1G55490.2 | Chaperonin 60 beta 1 (CPN60B1) | 27 | 1.33 | 0.196 | 0.987 |
| AT1G55490.2 | Chaperonin 60 beta 1 (CPN60B1) | 28 | 1.36 | 0.197 | 0.982 |
| AT1G55490.2 | Chaperonin 60 beta 1 (CPN60B1) | 29 | 1.38 | 0.208 | 0.982 |
| AT1G55490.2 | Chaperonin 60 beta 1 (CPN60B1) | 30 | 1.40 | 0.229 | 0.986 |
| AT1G55490.2 | Chaperonin 60 beta 1 (CPN60B1) | 31 | 1.41 | 0.255 | 0.991 |
| AT1G55490.2 | Chaperonin 60 beta 1 (CPN60B1) | 32 | 1.43 | 0.284 | 0.994 |
| AT1G55490.2 | Chaperonin 60 beta 1 (CPN60B1) | 43 | 1.03 | 0.324 | 1.000 |
| AT1G55490.2 | Chaperonin 60 beta 1 (CPN60B1) | 44 | 0.97 | 0.321 | 1.000 |
| AT1G55490.2 | Chaperonin 60 beta 1 (CPN60B1) | 45 | 0.90 | 0.339 | 1.000 |
| AT1G55490.2 | Chaperonin 60 beta 1 (CPN60B1) | 46 | 0.85 | 0.381 | 1.000 |
| AT1G55490.2 | Chaperonin 60 beta 1 (CPN60B1) | 47 | 0.79 | 0.447 | 1.000 |
| AT1G55510.1 | Branched chain keto acid dehydrogenase subunit (BCKDHE1beta) | 2 | 0.94 | 0.540 | 1.000 |

| | | | | | | |
|---|---|---|---|---|---|---|
| AT1G55510.1 | Branched chain keto acid dehydrogenase subunit (BCKDHE1beta) | 3 | 1.11 | 0.480 | 1.000 | |
| AT1G55510.1 | Branched chain keto acid dehydrogenase subunit (BCKDHE1beta) | 14 | 0.67 | 0.438 | 1.000 | |
| AT1G55510.1 | Branched chain keto acid dehydrogenase subunit (BCKDHE1beta) | 15 | 0.49 | 0.412 | 1.000 | |
| AT1G55510.1 | Branched chain keto acid dehydrogenase subunit (BCKDHE1beta) | 16 | 0.39 | 0.443 | 1.000 | |
| AT1G55510.1 | Branched chain keto acid dehydrogenase subunit (BCKDHE1beta) | 22 | 0.53 | 0.390 | 1.000 | |
| AT1G55510.1 | Branched chain keto acid dehydrogenase subunit (BCKDHE1beta) | 23 | 0.48 | 0.306 | 0.998 | |
| AT1G55510.1 | Branched chain keto acid dehydrogenase subunit (BCKDHE1beta) | 24 | 0.43 | 0.277 | 0.977 | |
| AT1G55510.1 | Branched chain keto acid dehydrogenase subunit (BCKDHE1beta) | 25 | 0.50 | 0.275 | 0.997 | |
| AT1G55510.1 | Branched chain keto acid dehydrogenase subunit (BCKDHE1beta) | 26 | 0.66 | 0.288 | 1.000 | |
| AT1G55510.1 | Branched chain keto acid dehydrogenase subunit (BCKDHE1beta) | 27 | 0.82 | 0.288 | 1.000 | |
| AT1G55510.1 | Branched chain keto acid dehydrogenase subunit (BCKDHE1beta) | 28 | 0.95 | 0.274 | 1.000 | |
| AT1G55510.1 | Branched chain keto acid dehydrogenase subunit (BCKDHE1beta) | 29 | 0.92 | 0.252 | 1.000 | |
| AT1G55510.1 | Branched chain keto acid dehydrogenase subunit (BCKDHE1beta) | 30 | 0.85 | 0.237 | 1.000 | |
| AT1G55510.1 | Branched chain keto acid dehydrogenase subunit (BCKDHE1beta) | 31 | 1.32 | 0.234 | 1.000 | |
| AT1G55510.1 | Branched chain keto acid dehydrogenase subunit (BCKDHE1beta) | 32 | 2.43 | 0.236 | 0.871 | |
| AT1G55510.1 | Branched chain keto acid dehydrogenase subunit (BCKDHE1beta) | 33 | 2.92 | 0.235 | 0.629 | |
| AT1G55510.1 | Branched chain keto acid dehydrogenase subunit (BCKDHE1beta) | 34 | 2.29 | 0.235 | 0.922 | |
| AT1G55510.1 | Branched chain keto acid dehydrogenase subunit (BCKDHE1beta) | 35 | 2.26 | 0.236 | 0.931 | |
| AT1G55510.1 | Branched chain keto acid dehydrogenase subunit (BCKDHE1beta) | 36 | 4.13 | 0.235 | 0.185 | |
| AT1G55510.1 | Branched chain keto acid dehydrogenase subunit (BCKDHE1beta) | 37 | 7.72 | 0.233 | 0.005 | ** |

| | | | | | | |
|---|---|---|---|---|---|---|
| AT1G55510.1 | Branched chain keto acid dehydrogenase subunit (BCKDHE1beta) | 38 | 7.60 | 0.236 | 0.006 | ** |
| AT1G55510.1 | Branched chain keto acid dehydrogenase subunit (BCKDHE1beta) | 39 | 3.64 | 0.234 | 0.310 | |
| AT1G55510.1 | Branched chain keto acid dehydrogenase subunit (BCKDHE1beta) | 40 | 1.14 | 0.236 | 1.000 | |
| AT1G55510.1 | Branched chain keto acid dehydrogenase subunit (BCKDHE1beta) | 41 | 0.46 | 0.238 | 0.959 | |
| AT1G55510.1 | Branched chain keto acid dehydrogenase subunit (BCKDHE1beta) | 42 | 0.53 | 0.256 | 0.998 | |
| AT1G55510.1 | Branched chain keto acid dehydrogenase subunit (BCKDHE1beta) | 43 | 1.58 | 0.314 | 1.000 | |
| AT1G55510.1 | Branched chain keto acid dehydrogenase subunit (BCKDHE1beta) | 44 | 7.17 | 0.505 | 0.838 | |
| AT1G55510.1 | Branched chain keto acid dehydrogenase subunit (BCKDHE1beta) | 46 | 280.44 | 1.435 | 0.831 | |
| AT1G55510.1 | Branched chain keto acid dehydrogenase subunit (BCKDHE1beta) | 47 | 2027.59 | 2.090 | 0.898 | |
| AT1G55670.1 | Photosystem I subunit (PSAG) | 2 | 0.14 | 0.350 | 0.312 | |
| AT1G55670.1 | Photosystem I subunit (PSAG) | 3 | 0.22 | 0.228 | 0.095 | . |
| AT1G55670.1 | Photosystem I subunit (PSAG) | 4 | 0.39 | 0.203 | 0.602 | |
| AT1G55670.1 | Photosystem I subunit (PSAG) | 5 | 0.67 | 0.197 | 1.000 | |
| AT1G55670.1 | Photosystem I subunit (PSAG) | 6 | 0.79 | 0.197 | 1.000 | |
| AT1G55670.1 | Photosystem I subunit (PSAG) | 7 | 0.74 | 0.198 | 1.000 | |
| AT1G55670.1 | Photosystem I subunit (PSAG) | 8 | 0.80 | 0.193 | 1.000 | |
| AT1G55670.1 | Photosystem I subunit (PSAG) | 9 | 0.97 | 0.189 | 1.000 | |
| AT1G55670.1 | Photosystem I subunit (PSAG) | 10 | 1.09 | 0.190 | 1.000 | |
| AT1G55670.1 | Photosystem I subunit (PSAG) | 11 | 1.18 | 0.192 | 1.000 | |
| AT1G55670.1 | Photosystem I subunit (PSAG) | 12 | 1.25 | 0.191 | 1.000 | |
| AT1G55670.1 | Photosystem I subunit (PSAG) | 13 | 1.28 | 0.187 | 1.000 | |
| AT1G55670.1 | Photosystem I subunit (PSAG) | 14 | 1.22 | 0.184 | 1.000 | |
| AT1G55670.1 | Photosystem I subunit (PSAG) | 15 | 1.08 | 0.183 | 1.000 | |
| AT1G55670.1 | Photosystem I subunit (PSAG) | 16 | 1.02 | 0.183 | 1.000 | |
| AT1G55670.1 | Photosystem I subunit (PSAG) | 17 | 1.09 | 0.182 | 1.000 | |
| AT1G55670.1 | Photosystem I subunit (PSAG) | 18 | 1.11 | 0.183 | 1.000 | |
| AT1G55670.1 | Photosystem I subunit (PSAG) | 19 | 0.89 | 0.188 | 1.000 | |
| AT1G55670.1 | Photosystem I subunit (PSAG) | 20 | 0.57 | 0.196 | 0.990 | |
| AT1G55670.1 | Photosystem I subunit (PSAG) | 21 | 0.31 | 0.198 | 0.210 | |
| AT1G55670.1 | Photosystem I subunit (PSAG) | 22 | 0.15 | 0.199 | 0.001 | ** |
| AT1G55670.1 | Photosystem I subunit (PSAG) | 23 | 0.10 | 0.197 | 0.000 | *** |
| AT1G55670.1 | Photosystem I subunit (PSAG) | 24 | 0.10 | 0.192 | 0.000 | *** |
| AT1G55670.1 | Photosystem I subunit (PSAG) | 25 | 0.21 | 0.188 | 0.011 | * |
| AT1G55670.1 | Photosystem I subunit (PSAG) | 26 | 0.92 | 0.188 | 1.000 | |
| AT1G55670.1 | Photosystem I subunit (PSAG) | 27 | 2.70 | 0.189 | 0.397 | |
| AT1G55670.1 | Photosystem I subunit (PSAG) | 28 | 3.46 | 0.192 | 0.118 | |
| AT1G55670.1 | Photosystem I subunit (PSAG) | 29 | 2.43 | 0.197 | 0.651 | |
| AT1G55670.1 | Photosystem I subunit (PSAG) | 30 | 1.12 | 0.215 | 1.000 | |
| AT1G55670.1 | Photosystem I subunit (PSAG) | 31 | 0.51 | 0.286 | 0.999 | |
| AT1G55670.1 | Photosystem I subunit (PSAG) | 32 | 0.27 | 0.450 | 0.986 | |
| AT1G55670.1 | Photosystem I subunit (PSAG) | 46 | 276.07 | 1.970 | 0.990 | |
| AT1G55670.1 | Photosystem I subunit (PSAG) | 47 | 789.93 | 2.149 | 0.976 | |

| | | | | | |
|---|---|---|---|---|---|
| AT1G55880.2 | Pyridoxal-5'-phosphate-dependent enzyme family protein | 2 | 0.96 | 0.325 | 1.000 |
| AT1G55880.2 | Pyridoxal-5'-phosphate-dependent enzyme family protein | 3 | 1.00 | 0.287 | 1.000 |
| AT1G55880.2 | Pyridoxal-5'-phosphate-dependent enzyme family protein | 4 | 1.03 | 0.296 | 1.000 |
| AT1G55880.2 | Pyridoxal-5'-phosphate-dependent enzyme family protein | 18 | 1.46 | 0.278 | 1.000 |
| AT1G55880.2 | Pyridoxal-5'-phosphate-dependent enzyme family protein | 19 | 1.45 | 0.244 | 0.999 |
| AT1G55880.2 | Pyridoxal-5'-phosphate-dependent enzyme family protein | 20 | 1.42 | 0.216 | 0.999 |
| AT1G55880.2 | Pyridoxal-5'-phosphate-dependent enzyme family protein | 21 | 1.37 | 0.194 | 0.999 |
| AT1G55880.2 | Pyridoxal-5'-phosphate-dependent enzyme family protein | 22 | 1.30 | 0.174 | 0.999 |
| AT1G55880.2 | Pyridoxal-5'-phosphate-dependent enzyme family protein | 23 | 1.23 | 0.158 | 1.000 |
| AT1G55880.2 | Pyridoxal-5'-phosphate-dependent enzyme family protein | 24 | 1.15 | 0.146 | 1.000 |
| AT1G55880.2 | Pyridoxal-5'-phosphate-dependent enzyme family protein | 25 | 1.08 | 0.139 | 1.000 |
| AT1G55880.2 | Pyridoxal-5'-phosphate-dependent enzyme family protein | 26 | 1.05 | 0.135 | 1.000 |
| AT1G55880.2 | Pyridoxal-5'-phosphate-dependent enzyme family protein | 27 | 1.04 | 0.133 | 1.000 |
| AT1G55880.2 | Pyridoxal-5'-phosphate-dependent enzyme family protein | 28 | 1.05 | 0.132 | 1.000 |
| AT1G55880.2 | Pyridoxal-5'-phosphate-dependent enzyme family protein | 29 | 1.06 | 0.132 | 1.000 |
| AT1G55880.2 | Pyridoxal-5'-phosphate-dependent enzyme family protein | 30 | 1.04 | 0.132 | 1.000 |
| AT1G55880.2 | Pyridoxal-5'-phosphate-dependent enzyme family protein | 31 | 1.00 | 0.132 | 1.000 |
| AT1G55880.2 | Pyridoxal-5'-phosphate-dependent enzyme family protein | 32 | 0.94 | 0.133 | 1.000 |
| AT1G55880.2 | Pyridoxal-5'-phosphate-dependent enzyme family protein | 33 | 0.86 | 0.135 | 1.000 |
| AT1G55880.2 | Pyridoxal-5'-phosphate-dependent enzyme family protein | 34 | 0.78 | 0.138 | 0.998 |

| | | | | | |
|---|---|---|---|---|---|
| AT1G55880.2 | Pyridoxal-5'-phosphate-dependent enzyme family protein | 35 | 0.72 | 0.143 | 0.981 |
| AT1G55880.2 | Pyridoxal-5'-phosphate-dependent enzyme family protein | 36 | 0.69 | 0.150 | 0.965 |
| AT1G55880.2 | Pyridoxal-5'-phosphate-dependent enzyme family protein | 37 | 0.68 | 0.157 | 0.969 |
| AT1G55880.2 | Pyridoxal-5'-phosphate-dependent enzyme family protein | 38 | 0.69 | 0.164 | 0.985 |
| AT1G55880.2 | Pyridoxal-5'-phosphate-dependent enzyme family protein | 39 | 0.73 | 0.174 | 0.997 |
| AT1G55880.2 | Pyridoxal-5'-phosphate-dependent enzyme family protein | 40 | 0.78 | 0.187 | 1.000 |
| AT1G55880.2 | Pyridoxal-5'-phosphate-dependent enzyme family protein | 41 | 0.86 | 0.208 | 1.000 |
| AT1G55880.2 | Pyridoxal-5'-phosphate-dependent enzyme family protein | 42 | 0.96 | 0.240 | 1.000 |
| AT1G55880.2 | Pyridoxal-5'-phosphate-dependent enzyme family protein | 43 | 1.10 | 0.284 | 1.000 |
| AT1G55880.2 | Pyridoxal-5'-phosphate-dependent enzyme family protein | 46 | 1.78 | 0.493 | 1.000 |
| AT1G55880.2 | Pyridoxal-5'-phosphate-dependent enzyme family protein | 47 | 2.12 | 0.591 | 1.000 |
| AT1G55890.1 | Pentatricopeptide repeat (PPR) superfamily protein | 2 | 1.08 | 0.745 | 1.000 |
| AT1G55890.1 | Pentatricopeptide repeat (PPR) superfamily protein | 3 | 0.94 | 0.642 | 1.000 |
| AT1G55890.1 | Pentatricopeptide repeat (PPR) superfamily protein | 10 | 1.00 | 0.439 | 1.000 |
| AT1G55890.1 | Pentatricopeptide repeat (PPR) superfamily protein | 11 | 1.27 | 0.379 | 1.000 |
| AT1G55890.1 | Pentatricopeptide repeat (PPR) superfamily protein | 12 | 1.55 | 0.342 | 1.000 |
| AT1G55890.1 | Pentatricopeptide repeat (PPR) superfamily protein | 13 | 1.47 | 0.318 | 1.000 |
| AT1G55890.1 | Pentatricopeptide repeat (PPR) superfamily protein | 14 | 1.28 | 0.303 | 1.000 |
| AT1G55890.1 | Pentatricopeptide repeat (PPR) superfamily protein | 15 | 1.39 | 0.297 | 1.000 |
| AT1G55890.1 | Pentatricopeptide repeat (PPR) superfamily protein | 16 | 1.94 | 0.295 | 0.999 |
| AT1G55890.1 | Pentatricopeptide repeat (PPR) superfamily protein | 17 | 2.74 | 0.294 | 0.916 |
| AT1G55890.1 | Pentatricopeptide repeat (PPR) superfamily protein | 18 | 3.23 | 0.298 | 0.796 |
| AT1G55890.1 | Pentatricopeptide repeat (PPR) superfamily protein | 19 | 3.04 | 0.303 | 0.866 |
| AT1G55890.1 | Pentatricopeptide repeat (PPR) superfamily protein | 20 | 2.29 | 0.310 | 0.991 |
| AT1G55890.1 | Pentatricopeptide repeat (PPR) superfamily protein | 21 | 1.41 | 0.315 | 1.000 |

| | | | | | |
|---|---|---|---|---|---|
| AT1G55890.1 | Pentatricopeptide repeat (PPR) superfamily protein | 22 | 0.84 | 0.321 | 1.000 |
| AT1G55890.1 | Pentatricopeptide repeat (PPR) superfamily protein | 23 | 0.60 | 0.331 | 1.000 |
| AT1G55890.1 | Pentatricopeptide repeat (PPR) superfamily protein | 24 | 0.50 | 0.346 | 1.000 |
| AT1G55890.1 | Pentatricopeptide repeat (PPR) superfamily protein | 25 | 0.46 | 0.358 | 0.999 |
| AT1G55890.1 | Pentatricopeptide repeat (PPR) superfamily protein | 26 | 0.45 | 0.349 | 0.998 |
| AT1G55890.1 | Pentatricopeptide repeat (PPR) superfamily protein | 27 | 0.45 | 0.334 | 0.997 |
| AT1G55890.1 | Pentatricopeptide repeat (PPR) superfamily protein | 28 | 0.49 | 0.318 | 0.999 |
| AT1G55890.1 | Pentatricopeptide repeat (PPR) superfamily protein | 29 | 0.60 | 0.307 | 1.000 |
| AT1G55890.1 | Pentatricopeptide repeat (PPR) superfamily protein | 30 | 0.83 | 0.303 | 1.000 |
| AT1G55890.1 | Pentatricopeptide repeat (PPR) superfamily protein | 31 | 1.40 | 0.303 | 1.000 |
| AT1G55890.1 | Pentatricopeptide repeat (PPR) superfamily protein | 32 | 2.17 | 0.305 | 0.995 |
| AT1G55890.1 | Pentatricopeptide repeat (PPR) superfamily protein | 33 | 2.35 | 0.312 | 0.988 |
| AT1G55890.1 | Pentatricopeptide repeat (PPR) superfamily protein | 34 | 1.88 | 0.330 | 1.000 |
| AT1G55890.1 | Pentatricopeptide repeat (PPR) superfamily protein | 35 | 1.32 | 0.361 | 1.000 |
| AT1G55890.1 | Pentatricopeptide repeat (PPR) superfamily protein | 36 | 0.95 | 0.392 | 1.000 |
| AT1G55890.1 | Pentatricopeptide repeat (PPR) superfamily protein | 38 | 0.75 | 0.376 | 1.000 |
| AT1G55890.1 | Pentatricopeptide repeat (PPR) superfamily protein | 39 | 0.88 | 0.348 | 1.000 |
| AT1G55890.1 | Pentatricopeptide repeat (PPR) superfamily protein | 40 | 1.21 | 0.321 | 1.000 |
| AT1G55890.1 | Pentatricopeptide repeat (PPR) superfamily protein | 41 | 1.77 | 0.310 | 1.000 |
| AT1G55890.1 | Pentatricopeptide repeat (PPR) superfamily protein | 42 | 2.32 | 0.309 | 0.989 |
| AT1G55890.1 | Pentatricopeptide repeat (PPR) superfamily protein | 43 | 2.31 | 0.309 | 0.990 |
| AT1G55890.1 | Pentatricopeptide repeat (PPR) superfamily protein | 44 | 1.57 | 0.309 | 1.000 |
| AT1G55890.1 | Pentatricopeptide repeat (PPR) superfamily protein | 45 | 0.75 | 0.319 | 1.000 |
| AT1G55890.1 | Pentatricopeptide repeat (PPR) superfamily protein | 46 | 0.28 | 0.382 | 0.933 |
| AT1G55890.1 | Pentatricopeptide repeat (PPR) superfamily protein | 47 | 0.10 | 0.568 | 0.732 |
| AT1G55900.2 | Translocase of the inner mitochondrial membrane 50 (TIM50) | 2 | 0.98 | 0.391 | 1.000 |
| AT1G55900.2 | Translocase of the inner mitochondrial membrane 50 (TIM50) | 3 | 0.95 | 0.281 | 1.000 |
| AT1G55900.2 | Translocase of the inner mitochondrial membrane 50 (TIM50) | 4 | 0.92 | 0.248 | 1.000 |
| AT1G55900.2 | Translocase of the inner mitochondrial membrane 50 (TIM50) | 5 | 0.89 | 0.245 | 1.000 |

| | | | | | |
|---|---|---|---|---|---|
| AT1G55900.2 | Translocase of the inner mitochondrial membrane 50 (TIM50) | 6 | 0.88 | 0.228 | 1.000 |
| AT1G55900.2 | Translocase of the inner mitochondrial membrane 50 (TIM50) | 7 | 0.88 | 0.213 | 1.000 |
| AT1G55900.2 | Translocase of the inner mitochondrial membrane 50 (TIM50) | 8 | 0.90 | 0.200 | 1.000 |
| AT1G55900.2 | Translocase of the inner mitochondrial membrane 50 (TIM50) | 9 | 0.91 | 0.193 | 1.000 |
| AT1G55900.2 | Translocase of the inner mitochondrial membrane 50 (TIM50) | 10 | 0.87 | 0.191 | 1.000 |
| AT1G55900.2 | Translocase of the inner mitochondrial membrane 50 (TIM50) | 11 | 0.81 | 0.192 | 1.000 |
| AT1G55900.2 | Translocase of the inner mitochondrial membrane 50 (TIM50) | 12 | 0.74 | 0.191 | 1.000 |
| AT1G55900.2 | Translocase of the inner mitochondrial membrane 50 (TIM50) | 13 | 0.70 | 0.189 | 1.000 |
| AT1G55900.2 | Translocase of the inner mitochondrial membrane 50 (TIM50) | 14 | 0.69 | 0.188 | 1.000 |
| AT1G55900.2 | Translocase of the inner mitochondrial membrane 50 (TIM50) | 15 | 0.71 | 0.186 | 1.000 |
| AT1G55900.2 | Translocase of the inner mitochondrial membrane 50 (TIM50) | 16 | 0.76 | 0.185 | 1.000 |
| AT1G55900.2 | Translocase of the inner mitochondrial membrane 50 (TIM50) | 17 | 0.93 | 0.185 | 1.000 |
| AT1G55900.2 | Translocase of the inner mitochondrial membrane 50 (TIM50) | 18 | 1.29 | 0.188 | 1.000 |
| AT1G55900.2 | Translocase of the inner mitochondrial membrane 50 (TIM50) | 19 | 1.89 | 0.192 | 0.965 |
| AT1G55900.2 | Translocase of the inner mitochondrial membrane 50 (TIM50) | 20 | 2.52 | 0.199 | 0.641 |
| AT1G55900.2 | Translocase of the inner mitochondrial membrane 50 (TIM50) | 21 | 2.68 | 0.207 | 0.604 |
| AT1G55900.2 | Translocase of the inner mitochondrial membrane 50 (TIM50) | 22 | 2.38 | 0.224 | 0.873 |
| AT1G55900.2 | Translocase of the inner mitochondrial membrane 50 (TIM50) | 23 | 2.04 | 0.255 | 0.994 |
| AT1G55900.2 | Translocase of the inner mitochondrial membrane 50 (TIM50) | 24 | 1.79 | 0.296 | 1.000 |
| AT1G55900.2 | Translocase of the inner mitochondrial membrane 50 (TIM50) | 28 | 1.16 | 0.281 | 1.000 |

| | | | | | |
|---|---|---|---|---|---|
| AT1G55900.2 | Translocase of the inner mitochondrial membrane 50 (TIM50) | 29 | 1.04 | 0.253 | 1.000 |
| AT1G55900.2 | Translocase of the inner mitochondrial membrane 50 (TIM50) | 30 | 0.94 | 0.229 | 1.000 |
| AT1G55900.2 | Translocase of the inner mitochondrial membrane 50 (TIM50) | 31 | 0.84 | 0.216 | 1.000 |
| AT1G55900.2 | Translocase of the inner mitochondrial membrane 50 (TIM50) | 32 | 0.79 | 0.207 | 1.000 |
| AT1G55900.2 | Translocase of the inner mitochondrial membrane 50 (TIM50) | 33 | 0.84 | 0.200 | 1.000 |
| AT1G55900.2 | Translocase of the inner mitochondrial membrane 50 (TIM50) | 34 | 0.99 | 0.195 | 1.000 |
| AT1G55900.2 | Translocase of the inner mitochondrial membrane 50 (TIM50) | 35 | 1.24 | 0.193 | 1.000 |
| AT1G55900.2 | Translocase of the inner mitochondrial membrane 50 (TIM50) | 36 | 1.62 | 0.191 | 0.998 |
| AT1G55900.2 | Translocase of the inner mitochondrial membrane 50 (TIM50) | 37 | 1.96 | 0.191 | 0.939 |
| AT1G55900.2 | Translocase of the inner mitochondrial membrane 50 (TIM50) | 38 | 2.13 | 0.191 | 0.855 |
| AT1G55900.2 | Translocase of the inner mitochondrial membrane 50 (TIM50) | 39 | 2.27 | 0.191 | 0.763 |
| AT1G55900.2 | Translocase of the inner mitochondrial membrane 50 (TIM50) | 40 | 2.36 | 0.191 | 0.697 |
| AT1G55900.2 | Translocase of the inner mitochondrial membrane 50 (TIM50) | 41 | 2.19 | 0.193 | 0.826 |
| AT1G55900.2 | Translocase of the inner mitochondrial membrane 50 (TIM50) | 42 | 1.80 | 0.195 | 0.987 |
| AT1G55900.2 | Translocase of the inner mitochondrial membrane 50 (TIM50) | 43 | 1.31 | 0.197 | 1.000 |
| AT1G55900.2 | Translocase of the inner mitochondrial membrane 50 (TIM50) | 44 | 0.85 | 0.196 | 1.000 |
| AT1G55900.2 | Translocase of the inner mitochondrial membrane 50 (TIM50) | 45 | 0.47 | 0.201 | 0.900 |
| AT1G55900.2 | Translocase of the inner mitochondrial membrane 50 (TIM50) | 46 | 0.22 | 0.252 | 0.210 |
| AT1G55900.2 | Translocase of the inner mitochondrial membrane 50 (TIM50) | 47 | 0.09 | 0.395 | 0.224 |
| AT1G56050.1 | GTP-binding protein-related (ENGD2) | 2 | 1.00 | 0.289 | 1.000 |
| AT1G56050.1 | GTP-binding protein-related (ENGD2) | 3 | 0.92 | 0.281 | 1.000 |

| | | | | | | |
|---|---|---|---|---|---|---|
| AT1G56050.1 | GTP-binding protein-related (ENGD2) | 28 | 0.93 | 0.210 | 1.000 | |
| AT1G56050.1 | GTP-binding protein-related (ENGD2) | 29 | 1.09 | 0.163 | 1.000 | |
| AT1G56050.1 | GTP-binding protein-related (ENGD2) | 30 | 1.28 | 0.145 | 0.999 | |
| AT1G56050.1 | GTP-binding protein-related (ENGD2) | 31 | 1.52 | 0.137 | 0.903 | |
| AT1G56050.1 | GTP-binding protein-related (ENGD2) | 32 | 1.80 | 0.129 | 0.452 | |
| AT1G56050.1 | GTP-binding protein-related (ENGD2) | 33 | 2.03 | 0.121 | 0.152 | |
| AT1G56050.1 | GTP-binding protein-related (ENGD2) | 34 | 2.12 | 0.123 | 0.112 | |
| AT1G56050.1 | GTP-binding protein-related (ENGD2) | 35 | 1.89 | 0.136 | 0.427 | |
| AT1G56050.1 | GTP-binding protein-related (ENGD2) | 36 | 1.47 | 0.151 | 0.967 | |
| AT1G56050.1 | GTP-binding protein-related (ENGD2) | 38 | 0.93 | 0.139 | 1.000 | |
| AT1G56050.1 | GTP-binding protein-related (ENGD2) | 39 | 0.89 | 0.123 | 1.000 | |
| AT1G56050.1 | GTP-binding protein-related (ENGD2) | 40 | 1.04 | 0.112 | 1.000 | |
| AT1G56050.1 | GTP-binding protein-related (ENGD2) | 41 | 1.40 | 0.109 | 0.895 | |
| AT1G56050.1 | GTP-binding protein-related (ENGD2) | 42 | 1.84 | 0.107 | 0.179 | |
| AT1G56050.1 | GTP-binding protein-related (ENGD2) | 43 | 2.01 | 0.107 | 0.073 | . |
| AT1G56050.1 | GTP-binding protein-related (ENGD2) | 44 | 1.66 | 0.110 | 0.442 | |
| AT1G56050.1 | GTP-binding protein-related (ENGD2) | 45 | 1.06 | 0.116 | 1.000 | |
| AT1G56050.1 | GTP-binding protein-related (ENGD2) | 46 | 0.54 | 0.137 | 0.477 | |
| AT1G56050.1 | GTP-binding protein-related (ENGD2) | 47 | 0.24 | 0.211 | 0.054 | . |
| AT1G56560.1 | Alkaline/neutral invertase A (INVA) | 2 | 1.28 | 0.514 | 1.000 | |
| AT1G56560.1 | Alkaline/neutral invertase A (INVA) | 3 | 1.16 | 0.455 | 1.000 | |
| AT1G56560.1 | Alkaline/neutral invertase A (INVA) | 12 | 0.71 | 0.315 | 1.000 | |
| AT1G56560.1 | Alkaline/neutral invertase A (INVA) | 13 | 0.81 | 0.269 | 1.000 | |
| AT1G56560.1 | Alkaline/neutral invertase A (INVA) | 14 | 1.02 | 0.248 | 1.000 | |
| AT1G56560.1 | Alkaline/neutral invertase A (INVA) | 15 | 1.33 | 0.228 | 1.000 | |
| AT1G56560.1 | Alkaline/neutral invertase A (INVA) | 16 | 1.58 | 0.215 | 1.000 | |
| AT1G56560.1 | Alkaline/neutral invertase A (INVA) | 17 | 1.55 | 0.213 | 1.000 | |
| AT1G56560.1 | Alkaline/neutral invertase A (INVA) | 18 | 1.37 | 0.215 | 1.000 | |
| AT1G56560.1 | Alkaline/neutral invertase A (INVA) | 19 | 1.30 | 0.221 | 1.000 | |
| AT1G56560.1 | Alkaline/neutral invertase A (INVA) | 20 | 1.30 | 0.226 | 1.000 | |
| AT1G56560.1 | Alkaline/neutral invertase A (INVA) | 21 | 1.23 | 0.227 | 1.000 | |

| | | | | | |
|---|---|---|---|---|---|
| AT1G56560.1 | Alkaline/neutral invertase A (INVA) | 22 | 1.11 | 0.228 | 1.000 |
| AT1G56560.1 | Alkaline/neutral invertase A (INVA) | 23 | 0.98 | 0.226 | 1.000 |
| AT1G56560.1 | Alkaline/neutral invertase A (INVA) | 24 | 0.88 | 0.222 | 1.000 |
| AT1G56560.1 | Alkaline/neutral invertase A (INVA) | 25 | 0.86 | 0.218 | 1.000 |
| AT1G56560.1 | Alkaline/neutral invertase A (INVA) | 26 | 1.03 | 0.217 | 1.000 |
| AT1G56560.1 | Alkaline/neutral invertase A (INVA) | 27 | 1.44 | 0.218 | 1.000 |
| AT1G56560.1 | Alkaline/neutral invertase A (INVA) | 28 | 2.00 | 0.218 | 0.960 |
| AT1G56560.1 | Alkaline/neutral invertase A (INVA) | 29 | 2.04 | 0.218 | 0.952 |
| AT1G56560.1 | Alkaline/neutral invertase A (INVA) | 30 | 1.34 | 0.219 | 1.000 |
| AT1G56560.1 | Alkaline/neutral invertase A (INVA) | 31 | 0.88 | 0.220 | 1.000 |
| AT1G56560.1 | Alkaline/neutral invertase A (INVA) | 32 | 0.91 | 0.225 | 1.000 |
| AT1G56560.1 | Alkaline/neutral invertase A (INVA) | 33 | 1.35 | 0.238 | 1.000 |
| AT1G56560.1 | Alkaline/neutral invertase A (INVA) | 34 | 2.10 | 0.260 | 0.986 |
| AT1G56560.1 | Alkaline/neutral invertase A (INVA) | 35 | 2.94 | 0.282 | 0.845 |
| AT1G56560.1 | Alkaline/neutral invertase A (INVA) | 36 | 3.70 | 0.311 | 0.738 |
| AT1G56560.1 | Alkaline/neutral invertase A (INVA) | 37 | 4.28 | 0.321 | 0.627 |
| AT1G56560.1 | Alkaline/neutral invertase A (INVA) | 38 | 4.63 | 0.348 | 0.669 |
| AT1G56560.1 | Alkaline/neutral invertase A (INVA) | 40 | 4.64 | 0.330 | 0.580 |
| AT1G56560.1 | Alkaline/neutral invertase A (INVA) | 41 | 4.21 | 0.367 | 0.823 |
| AT1G56560.1 | Alkaline/neutral invertase A (INVA) | 42 | 3.51 | 0.455 | 0.990 |
| AT1G56560.1 | Alkaline/neutral invertase A (INVA) | 46 | 0.91 | 1.498 | 1.000 |
| AT1G56560.1 | Alkaline/neutral invertase A (INVA) | 47 | 0.60 | 1.891 | 1.000 |
| AT1G57540.3 | 40S ribosomal protein | 2 | 0.78 | 0.452 | 1.000 |
| AT1G57540.3 | 40S ribosomal protein | 3 | 1.03 | 0.440 | 1.000 |
| AT1G57540.3 | 40S ribosomal protein | 23 | 0.25 | 0.345 | 0.621 |
| AT1G57540.3 | 40S ribosomal protein | 24 | 0.23 | 0.266 | 0.195 |
| AT1G57540.3 | 40S ribosomal protein | 25 | 0.52 | 0.224 | 0.912 |
| AT1G57540.3 | 40S ribosomal protein | 26 | 1.86 | 0.213 | 0.910 |
| AT1G57540.3 | 40S ribosomal protein | 27 | 2.26 | 0.212 | 0.678 |
| AT1G57540.3 | 40S ribosomal protein | 28 | 1.00 | 0.212 | 1.000 |
| AT1G57540.3 | 40S ribosomal protein | 29 | 0.51 | 0.218 | 0.876 |
| AT1G57540.3 | 40S ribosomal protein | 30 | 0.53 | 0.231 | 0.933 |
| AT1G57540.3 | 40S ribosomal protein | 31 | 0.84 | 0.286 | 1.000 |
| AT1G57540.3 | 40S ribosomal protein | 32 | 1.39 | 0.453 | 1.000 |
| AT1G57540.3 | 40S ribosomal protein | 45 | 0.72 | 0.566 | 1.000 |
| AT1G57540.3 | 40S ribosomal protein | 46 | 0.44 | 0.297 | 0.940 |
| AT1G57540.3 | 40S ribosomal protein | 47 | 0.28 | 0.522 | 0.971 |
| AT1G57610.2 | Calcium uniporter | 2 | 0.83 | 0.444 | 1.000 |
| AT1G57610.2 | Calcium uniporter | 3 | 0.90 | 0.357 | 1.000 |
| AT1G57610.2 | Calcium uniporter | 7 | 1.10 | 0.310 | 1.000 |

| | | | | | | |
|---|---|---|---|---|---|---|
| AT1G57610.2 | Calcium uniporter | 8 | 1.08 | 0.262 | 1.000 | |
| AT1G57610.2 | Calcium uniporter | 9 | 1.00 | 0.236 | 1.000 | |
| AT1G57610.2 | Calcium uniporter | 10 | 0.88 | 0.214 | 1.000 | |
| AT1G57610.2 | Calcium uniporter | 11 | 0.74 | 0.204 | 1.000 | |
| AT1G57610.2 | Calcium uniporter | 12 | 0.63 | 0.196 | 0.999 | |
| AT1G57610.2 | Calcium uniporter | 13 | 0.57 | 0.190 | 0.989 | |
| AT1G57610.2 | Calcium uniporter | 14 | 0.53 | 0.182 | 0.951 | |
| AT1G57610.2 | Calcium uniporter | 15 | 0.46 | 0.176 | 0.720 | |
| AT1G57610.2 | Calcium uniporter | 16 | 0.35 | 0.173 | 0.193 | |
| AT1G57610.2 | Calcium uniporter | 17 | 0.30 | 0.172 | 0.071 | . |
| AT1G57610.2 | Calcium uniporter | 18 | 0.34 | 0.175 | 0.190 | |
| AT1G57610.2 | Calcium uniporter | 19 | 0.49 | 0.179 | 0.864 | |
| AT1G57610.2 | Calcium uniporter | 20 | 0.80 | 0.182 | 1.000 | |
| AT1G57610.2 | Calcium uniporter | 21 | 1.24 | 0.184 | 1.000 | |
| AT1G57610.2 | Calcium uniporter | 22 | 1.57 | 0.184 | 0.999 | |
| AT1G57610.2 | Calcium uniporter | 23 | 1.75 | 0.183 | 0.986 | |
| AT1G57610.2 | Calcium uniporter | 24 | 2.07 | 0.179 | 0.835 | |
| AT1G57610.2 | Calcium uniporter | 25 | 2.57 | 0.177 | 0.405 | |
| AT1G57610.2 | Calcium uniporter | 26 | 2.88 | 0.176 | 0.219 | |
| AT1G57610.2 | Calcium uniporter | 27 | 2.60 | 0.176 | 0.373 | |
| AT1G57610.2 | Calcium uniporter | 28 | 2.04 | 0.177 | 0.839 | |
| AT1G57610.2 | Calcium uniporter | 29 | 1.59 | 0.177 | 0.998 | |
| AT1G57610.2 | Calcium uniporter | 30 | 1.30 | 0.177 | 1.000 | |
| AT1G57610.2 | Calcium uniporter | 31 | 1.10 | 0.177 | 1.000 | |
| AT1G57610.2 | Calcium uniporter | 32 | 0.97 | 0.177 | 1.000 | |
| AT1G57610.2 | Calcium uniporter | 33 | 0.87 | 0.177 | 1.000 | |
| AT1G57610.2 | Calcium uniporter | 34 | 0.81 | 0.178 | 1.000 | |
| AT1G57610.2 | Calcium uniporter | 35 | 0.82 | 0.178 | 1.000 | |
| AT1G57610.2 | Calcium uniporter | 36 | 0.98 | 0.178 | 1.000 | |
| AT1G57610.2 | Calcium uniporter | 37 | 1.30 | 0.177 | 1.000 | |
| AT1G57610.2 | Calcium uniporter | 38 | 1.57 | 0.177 | 0.999 | |
| AT1G57610.2 | Calcium uniporter | 39 | 1.63 | 0.177 | 0.996 | |
| AT1G57610.2 | Calcium uniporter | 40 | 1.69 | 0.177 | 0.991 | |
| AT1G57610.2 | Calcium uniporter | 41 | 1.62 | 0.179 | 0.997 | |
| AT1G57610.2 | Calcium uniporter | 42 | 1.37 | 0.182 | 1.000 | |
| AT1G57610.2 | Calcium uniporter | 43 | 1.15 | 0.185 | 1.000 | |
| AT1G57610.2 | Calcium uniporter | 44 | 0.95 | 0.189 | 1.000 | |
| AT1G57610.2 | Calcium uniporter | 45 | 0.85 | 0.224 | 1.000 | |
| AT1G57610.2 | Calcium uniporter | 46 | 0.75 | 0.342 | 1.000 | |
| AT1G57610.2 | Calcium uniporter | 47 | 0.66 | 0.552 | 1.000 | |
| AT1G59900.1 | Pyruvate dehydrogenase subunit (PDCE1 alpha) | 2 | 1.45 | 0.477 | 1.000 | |
| AT1G59900.1 | Pyruvate dehydrogenase subunit (PDCE1 alpha) | 3 | 0.98 | 0.345 | 1.000 | |
| AT1G59900.1 | Pyruvate dehydrogenase subunit (PDCE1 alpha) | 4 | 0.61 | 0.345 | 1.000 | |
| AT1G59900.1 | Pyruvate dehydrogenase subunit (PDCE1 alpha) | 6 | 0.23 | 0.345 | 0.805 | |
| AT1G59900.1 | Pyruvate dehydrogenase subunit (PDCE1 alpha) | 7 | 0.16 | 0.305 | 0.248 | |
| AT1G59900.1 | Pyruvate dehydrogenase subunit (PDCE1 alpha) | 8 | 0.15 | 0.272 | 0.074 | . |
| AT1G59900.1 | Pyruvate dehydrogenase subunit (PDCE1 alpha) | 9 | 0.20 | 0.255 | 0.165 | |
| AT1G59900.1 | Pyruvate dehydrogenase subunit (PDCE1 alpha) | 10 | 0.33 | 0.249 | 0.759 | |
| AT1G59900.1 | Pyruvate dehydrogenase subunit (PDCE1 alpha) | 11 | 0.64 | 0.247 | 1.000 | |
| AT1G59900.1 | Pyruvate dehydrogenase subunit (PDCE1 alpha) | 12 | 1.27 | 0.247 | 1.000 | |
| AT1G59900.1 | Pyruvate dehydrogenase subunit (PDCE1 alpha) | 13 | 2.23 | 0.245 | 0.980 | |

| | | | | | | |
|---|---|---|---|---|---|---|
| AT1G59900.1 | Pyruvate dehydrogenase subunit (PDCE1 alpha) | 14 | 3.32 | 0.243 | 0.564 | |
| AT1G59900.1 | Pyruvate dehydrogenase subunit (PDCE1 alpha) | 15 | 4.57 | 0.241 | 0.172 | |
| AT1G59900.1 | Pyruvate dehydrogenase subunit (PDCE1 alpha) | 16 | 5.81 | 0.239 | 0.047 | * |
| AT1G59900.1 | Pyruvate dehydrogenase subunit (PDCE1 alpha) | 17 | 7.32 | 0.239 | 0.012 | * |
| AT1G59900.1 | Pyruvate dehydrogenase subunit (PDCE1 alpha) | 18 | 8.66 | 0.243 | 0.005 | ** |
| AT1G59900.1 | Pyruvate dehydrogenase subunit (PDCE1 alpha) | 19 | 9.40 | 0.249 | 0.004 | ** |
| AT1G59900.1 | Pyruvate dehydrogenase subunit (PDCE1 alpha) | 20 | 8.71 | 0.254 | 0.009 | ** |
| AT1G59900.1 | Pyruvate dehydrogenase subunit (PDCE1 alpha) | 21 | 6.40 | 0.257 | 0.057 | . |
| AT1G59900.1 | Pyruvate dehydrogenase subunit (PDCE1 alpha) | 22 | 4.22 | 0.256 | 0.339 | |
| AT1G59900.1 | Pyruvate dehydrogenase subunit (PDCE1 alpha) | 23 | 2.93 | 0.254 | 0.811 | |
| AT1G59900.1 | Pyruvate dehydrogenase subunit (PDCE1 alpha) | 24 | 2.14 | 0.250 | 0.991 | |
| AT1G59900.1 | Pyruvate dehydrogenase subunit (PDCE1 alpha) | 25 | 1.68 | 0.247 | 1.000 | |
| AT1G59900.1 | Pyruvate dehydrogenase subunit (PDCE1 alpha) | 26 | 1.27 | 0.246 | 1.000 | |
| AT1G59900.1 | Pyruvate dehydrogenase subunit (PDCE1 alpha) | 27 | 0.94 | 0.246 | 1.000 | |
| AT1G59900.1 | Pyruvate dehydrogenase subunit (PDCE1 alpha) | 28 | 0.80 | 0.246 | 1.000 | |
| AT1G59900.1 | Pyruvate dehydrogenase subunit (PDCE1 alpha) | 29 | 0.89 | 0.245 | 1.000 | |
| AT1G59900.1 | Pyruvate dehydrogenase subunit (PDCE1 alpha) | 30 | 1.01 | 0.245 | 1.000 | |
| AT1G59900.1 | Pyruvate dehydrogenase subunit (PDCE1 alpha) | 31 | 1.03 | 0.246 | 1.000 | |
| AT1G59900.1 | Pyruvate dehydrogenase subunit (PDCE1 alpha) | 32 | 1.02 | 0.247 | 1.000 | |
| AT1G59900.1 | Pyruvate dehydrogenase subunit (PDCE1 alpha) | 33 | 0.99 | 0.247 | 1.000 | |
| AT1G59900.1 | Pyruvate dehydrogenase subunit (PDCE1 alpha) | 34 | 0.99 | 0.247 | 1.000 | |
| AT1G59900.1 | Pyruvate dehydrogenase subunit (PDCE1 alpha) | 35 | 1.09 | 0.247 | 1.000 | |
| AT1G59900.1 | Pyruvate dehydrogenase subunit (PDCE1 alpha) | 36 | 1.23 | 0.246 | 1.000 | |
| AT1G59900.1 | Pyruvate dehydrogenase subunit (PDCE1 alpha) | 37 | 1.29 | 0.246 | 1.000 | |
| AT1G59900.1 | Pyruvate dehydrogenase subunit (PDCE1 alpha) | 38 | 1.25 | 0.246 | 1.000 | |
| AT1G59900.1 | Pyruvate dehydrogenase subunit (PDCE1 alpha) | 39 | 1.04 | 0.247 | 1.000 | |
| AT1G59900.1 | Pyruvate dehydrogenase subunit (PDCE1 alpha) | 40 | 0.80 | 0.247 | 1.000 | |
| AT1G59900.1 | Pyruvate dehydrogenase subunit (PDCE1 alpha) | 41 | 0.71 | 0.248 | 1.000 | |
| AT1G59900.1 | Pyruvate dehydrogenase subunit (PDCE1 alpha) | 42 | 0.80 | 0.252 | 1.000 | |
| AT1G59900.1 | Pyruvate dehydrogenase subunit (PDCE1 alpha) | 43 | 1.01 | 0.254 | 1.000 | |
| AT1G59900.1 | Pyruvate dehydrogenase subunit (PDCE1 alpha) | 44 | 1.11 | 0.254 | 1.000 | |

| | | | | | |
|---|---|---|---|---|---|
| AT1G59900.1 | Pyruvate dehydrogenase subunit (PDCE1 alpha) | 45 | 0.97 | 0.254 | 1.000 |
| AT1G59900.1 | Pyruvate dehydrogenase subunit (PDCE1 alpha) | 46 | 0.69 | 0.302 | 1.000 |
| AT1G59900.1 | Pyruvate dehydrogenase subunit (PDCE1 alpha) | 47 | 0.42 | 0.506 | 1.000 |
| AT1G60770.1 | Pentatricopeptide repeat (PPR) superfamily protein | 2 | 0.97 | 0.442 | 1.000 |
| AT1G60770.1 | Pentatricopeptide repeat (PPR) superfamily protein | 3 | 0.93 | 0.381 | 1.000 |
| AT1G60770.1 | Pentatricopeptide repeat (PPR) superfamily protein | 4 | 0.89 | 0.426 | 1.000 |
| AT1G60770.1 | Pentatricopeptide repeat (PPR) superfamily protein | 12 | 1.24 | 0.305 | 1.000 |
| AT1G60770.1 | Pentatricopeptide repeat (PPR) superfamily protein | 13 | 1.41 | 0.264 | 1.000 |
| AT1G60770.1 | Pentatricopeptide repeat (PPR) superfamily protein | 14 | 1.54 | 0.242 | 1.000 |
| AT1G60770.1 | Pentatricopeptide repeat (PPR) superfamily protein | 15 | 1.62 | 0.226 | 0.999 |
| AT1G60770.1 | Pentatricopeptide repeat (PPR) superfamily protein | 16 | 1.80 | 0.218 | 0.986 |
| AT1G60770.1 | Pentatricopeptide repeat (PPR) superfamily protein | 17 | 2.17 | 0.218 | 0.877 |
| AT1G60770.1 | Pentatricopeptide repeat (PPR) superfamily protein | 18 | 2.71 | 0.224 | 0.613 |
| AT1G60770.1 | Pentatricopeptide repeat (PPR) superfamily protein | 19 | 3.26 | 0.239 | 0.443 |
| AT1G60770.1 | Pentatricopeptide repeat (PPR) superfamily protein | 20 | 3.57 | 0.263 | 0.478 |
| AT1G60770.1 | Pentatricopeptide repeat (PPR) superfamily protein | 21 | 3.64 | 0.282 | 0.565 |
| AT1G60770.1 | Pentatricopeptide repeat (PPR) superfamily protein | 23 | 3.03 | 0.288 | 0.801 |
| AT1G60770.1 | Pentatricopeptide repeat (PPR) superfamily protein | 24 | 2.40 | 0.272 | 0.939 |
| AT1G60770.1 | Pentatricopeptide repeat (PPR) superfamily protein | 25 | 1.75 | 0.268 | 0.999 |
| AT1G60770.1 | Pentatricopeptide repeat (PPR) superfamily protein | 26 | 1.28 | 0.272 | 1.000 |
| AT1G60770.1 | Pentatricopeptide repeat (PPR) superfamily protein | 27 | 1.05 | 0.268 | 1.000 |
| AT1G60770.1 | Pentatricopeptide repeat (PPR) superfamily protein | 28 | 1.05 | 0.260 | 1.000 |
| AT1G60770.1 | Pentatricopeptide repeat (PPR) superfamily protein | 29 | 1.19 | 0.247 | 1.000 |
| AT1G60770.1 | Pentatricopeptide repeat (PPR) superfamily protein | 30 | 1.30 | 0.233 | 1.000 |
| AT1G60770.1 | Pentatricopeptide repeat (PPR) superfamily protein | 31 | 1.20 | 0.224 | 1.000 |
| AT1G60770.1 | Pentatricopeptide repeat (PPR) superfamily protein | 32 | 1.00 | 0.221 | 1.000 |
| AT1G60770.1 | Pentatricopeptide repeat (PPR) superfamily protein | 33 | 0.83 | 0.225 | 1.000 |
| AT1G60770.1 | Pentatricopeptide repeat (PPR) superfamily protein | 34 | 0.68 | 0.238 | 1.000 |
| AT1G60770.1 | Pentatricopeptide repeat (PPR) superfamily protein | 35 | 0.53 | 0.259 | 0.994 |
| AT1G60770.1 | Pentatricopeptide repeat (PPR) superfamily protein | 36 | 0.43 | 0.275 | 0.953 |
| AT1G60770.1 | Pentatricopeptide repeat (PPR) superfamily protein | 38 | 0.50 | 0.268 | 0.990 |

| | | | | | |
|---|---|---|---|---|---|
| AT1G60770.1 | Pentatricopeptide repeat (PPR) superfamily protein | 39 | 0.81 | 0.244 | 1.000 |
| AT1G60770.1 | Pentatricopeptide repeat (PPR) superfamily protein | 40 | 1.50 | 0.228 | 1.000 |
| AT1G60770.1 | Pentatricopeptide repeat (PPR) superfamily protein | 41 | 2.51 | 0.221 | 0.703 |
| AT1G60770.1 | Pentatricopeptide repeat (PPR) superfamily protein | 42 | 3.40 | 0.219 | 0.259 |
| AT1G60770.1 | Pentatricopeptide repeat (PPR) superfamily protein | 43 | 3.47 | 0.221 | 0.247 |
| AT1G60770.1 | Pentatricopeptide repeat (PPR) superfamily protein | 44 | 2.43 | 0.225 | 0.779 |
| AT1G60770.1 | Pentatricopeptide repeat (PPR) superfamily protein | 45 | 1.12 | 0.239 | 1.000 |
| AT1G60770.1 | Pentatricopeptide repeat (PPR) superfamily protein | 46 | 0.37 | 0.284 | 0.882 |
| AT1G60770.1 | Pentatricopeptide repeat (PPR) superfamily protein | 47 | 0.10 | 0.399 | 0.221 |
| AT1G61520.2 | Photosystem I light harvesting complex subunit (LHCA3) | 2 | 0.17 | 0.400 | 0.711 |
| AT1G61520.2 | Photosystem I light harvesting complex subunit (LHCA3) | 3 | 0.28 | 0.258 | 0.541 |
| AT1G61520.2 | Photosystem I light harvesting complex subunit (LHCA3) | 4 | 0.46 | 0.229 | 0.960 |
| AT1G61520.2 | Photosystem I light harvesting complex subunit (LHCA3) | 5 | 0.66 | 0.227 | 1.000 |
| AT1G61520.2 | Photosystem I light harvesting complex subunit (LHCA3) | 6 | 0.75 | 0.227 | 1.000 |
| AT1G61520.2 | Photosystem I light harvesting complex subunit (LHCA3) | 7 | 0.76 | 0.227 | 1.000 |
| AT1G61520.2 | Photosystem I light harvesting complex subunit (LHCA3) | 8 | 0.79 | 0.223 | 1.000 |
| AT1G61520.2 | Photosystem I light harvesting complex subunit (LHCA3) | 9 | 0.79 | 0.220 | 1.000 |
| AT1G61520.2 | Photosystem I light harvesting complex subunit (LHCA3) | 10 | 0.76 | 0.221 | 1.000 |
| AT1G61520.2 | Photosystem I light harvesting complex subunit (LHCA3) | 11 | 0.80 | 0.223 | 1.000 |
| AT1G61520.2 | Photosystem I light harvesting complex subunit (LHCA3) | 12 | 0.88 | 0.222 | 1.000 |
| AT1G61520.2 | Photosystem I light harvesting complex subunit (LHCA3) | 13 | 0.96 | 0.219 | 1.000 |
| AT1G61520.2 | Photosystem I light harvesting complex subunit (LHCA3) | 14 | 0.99 | 0.215 | 1.000 |
| AT1G61520.2 | Photosystem I light harvesting complex subunit (LHCA3) | 15 | 0.96 | 0.214 | 1.000 |
| AT1G61520.2 | Photosystem I light harvesting complex subunit (LHCA3) | 16 | 0.97 | 0.213 | 1.000 |
| AT1G61520.2 | Photosystem I light harvesting complex subunit (LHCA3) | 17 | 1.14 | 0.213 | 1.000 |
| AT1G61520.2 | Photosystem I light harvesting complex subunit (LHCA3) | 18 | 1.54 | 0.215 | 1.000 |
| AT1G61520.2 | Photosystem I light harvesting complex subunit (LHCA3) | 19 | 2.03 | 0.220 | 0.972 |
| AT1G61520.2 | Photosystem I light harvesting complex subunit (LHCA3) | 20 | 2.09 | 0.227 | 0.970 |
| AT1G61520.2 | Photosystem I light harvesting complex subunit (LHCA3) | 21 | 1.52 | 0.229 | 1.000 |
| AT1G61520.2 | Photosystem I light harvesting complex subunit (LHCA3) | 22 | 0.95 | 0.230 | 1.000 |
| AT1G61520.2 | Photosystem I light harvesting complex subunit (LHCA3) | 23 | 0.66 | 0.228 | 1.000 |

| | | | | | |
|---|---|---|---|---|---|
| AT1G61520.2 | Photosystem I light harvesting complex subunit (LHCA3) | 24 | 0.53 | 0.223 | 0.993 |
| AT1G61520.2 | Photosystem I light harvesting complex subunit (LHCA3) | 25 | 0.56 | 0.220 | 0.998 |
| AT1G61520.2 | Photosystem I light harvesting complex subunit (LHCA3) | 26 | 0.88 | 0.220 | 1.000 |
| AT1G61520.2 | Photosystem I light harvesting complex subunit (LHCA3) | 27 | 0.95 | 0.224 | 1.000 |
| AT1G61520.2 | Photosystem I light harvesting complex subunit (LHCA3) | 28 | 0.51 | 0.239 | 0.993 |
| AT1G61520.2 | Photosystem I light harvesting complex subunit (LHCA3) | 29 | 0.34 | 0.268 | 0.836 |
| AT1G61520.2 | Photosystem I light harvesting complex subunit (LHCA3) | 30 | 0.35 | 0.274 | 0.873 |
| AT1G61520.2 | Photosystem I light harvesting complex subunit (LHCA3) | 31 | 0.42 | 0.306 | 0.993 |
| AT1G61520.2 | Photosystem I light harvesting complex subunit (LHCA3) | 32 | 0.49 | 0.363 | 1.000 |
| AT1G61520.2 | Photosystem I light harvesting complex subunit (LHCA3) | 33 | 0.54 | 0.424 | 1.000 |
| AT1G61520.2 | Photosystem I light harvesting complex subunit (LHCA3) | 42 | 2.34 | 0.392 | 1.000 |
| AT1G61520.2 | Photosystem I light harvesting complex subunit (LHCA3) | 43 | 2.85 | 0.290 | 0.923 |
| AT1G61520.2 | Photosystem I light harvesting complex subunit (LHCA3) | 44 | 2.63 | 0.242 | 0.838 |
| AT1G61520.2 | Photosystem I light harvesting complex subunit (LHCA3) | 45 | 1.51 | 0.234 | 1.000 |
| AT1G61520.2 | Photosystem I light harvesting complex subunit (LHCA3) | 46 | 0.44 | 0.278 | 0.989 |
| AT1G61520.2 | Photosystem I light harvesting complex subunit (LHCA3) | 47 | 0.08 | 0.449 | 0.332 |
| AT1G61870.1 | Pentatricopeptide repeat (PPR) superfamily protein (PPR336) | 2 | 0.94 | 0.499 | 1.000 |
| AT1G61870.1 | Pentatricopeptide repeat (PPR) superfamily protein (PPR336) | 3 | 1.05 | 0.430 | 1.000 |
| AT1G61870.1 | Pentatricopeptide repeat (PPR) superfamily protein (PPR336) | 10 | 1.38 | 0.259 | 1.000 |
| AT1G61870.1 | Pentatricopeptide repeat (PPR) superfamily protein (PPR336) | 11 | 1.27 | 0.225 | 1.000 |
| AT1G61870.1 | Pentatricopeptide repeat (PPR) superfamily protein (PPR336) | 12 | 1.14 | 0.199 | 1.000 |
| AT1G61870.1 | Pentatricopeptide repeat (PPR) superfamily protein (PPR336) | 13 | 1.01 | 0.181 | 1.000 |
| AT1G61870.1 | Pentatricopeptide repeat (PPR) superfamily protein (PPR336) | 14 | 0.93 | 0.170 | 1.000 |
| AT1G61870.1 | Pentatricopeptide repeat (PPR) superfamily protein (PPR336) | 15 | 0.96 | 0.165 | 1.000 |
| AT1G61870.1 | Pentatricopeptide repeat (PPR) superfamily protein (PPR336) | 16 | 1.14 | 0.164 | 1.000 |
| AT1G61870.1 | Pentatricopeptide repeat (PPR) superfamily protein (PPR336) | 17 | 1.48 | 0.165 | 0.992 |
| AT1G61870.1 | Pentatricopeptide repeat (PPR) superfamily protein (PPR336) | 18 | 1.90 | 0.171 | 0.795 |
| AT1G61870.1 | Pentatricopeptide repeat (PPR) superfamily protein (PPR336) | 19 | 2.24 | 0.181 | 0.561 |
| AT1G61870.1 | Pentatricopeptide repeat (PPR) superfamily protein (PPR336) | 20 | 2.43 | 0.194 | 0.531 |
| AT1G61870.1 | Pentatricopeptide repeat (PPR) superfamily protein (PPR336) | 21 | 2.41 | 0.212 | 0.670 |
| AT1G61870.1 | Pentatricopeptide repeat (PPR) superfamily protein (PPR336) | 22 | 2.21 | 0.226 | 0.856 |

| | | | | | |
|---|---|---|---|---|---|
| AT1G61870.1 | Pentatricopeptide repeat (PPR) superfamily protein (PPR336) | 25 | 1.47 | 0.237 | 1.000 |
| AT1G61870.1 | Pentatricopeptide repeat (PPR) superfamily protein (PPR336) | 26 | 1.33 | 0.228 | 1.000 |
| AT1G61870.1 | Pentatricopeptide repeat (PPR) superfamily protein (PPR336) | 27 | 1.25 | 0.221 | 1.000 |
| AT1G61870.1 | Pentatricopeptide repeat (PPR) superfamily protein (PPR336) | 28 | 1.22 | 0.219 | 1.000 |
| AT1G61870.1 | Pentatricopeptide repeat (PPR) superfamily protein (PPR336) | 29 | 1.19 | 0.222 | 1.000 |
| AT1G61870.1 | Pentatricopeptide repeat (PPR) superfamily protein (PPR336) | 30 | 1.17 | 0.223 | 1.000 |
| AT1G61870.1 | Pentatricopeptide repeat (PPR) superfamily protein (PPR336) | 32 | 1.12 | 0.218 | 1.000 |
| AT1G61870.1 | Pentatricopeptide repeat (PPR) superfamily protein (PPR336) | 33 | 1.11 | 0.215 | 1.000 |
| AT1G61870.1 | Pentatricopeptide repeat (PPR) superfamily protein (PPR336) | 34 | 1.12 | 0.214 | 1.000 |
| AT1G61870.1 | Pentatricopeptide repeat (PPR) superfamily protein (PPR336) | 35 | 1.14 | 0.221 | 1.000 |
| AT1G61870.1 | Pentatricopeptide repeat (PPR) superfamily protein (PPR336) | 36 | 1.16 | 0.230 | 1.000 |
| AT1G61870.1 | Pentatricopeptide repeat (PPR) superfamily protein (PPR336) | 39 | 1.17 | 0.225 | 1.000 |
| AT1G61870.1 | Pentatricopeptide repeat (PPR) superfamily protein (PPR336) | 40 | 1.16 | 0.209 | 1.000 |
| AT1G61870.1 | Pentatricopeptide repeat (PPR) superfamily protein (PPR336) | 41 | 1.13 | 0.192 | 1.000 |
| AT1G61870.1 | Pentatricopeptide repeat (PPR) superfamily protein (PPR336) | 42 | 1.06 | 0.180 | 1.000 |
| AT1G61870.1 | Pentatricopeptide repeat (PPR) superfamily protein (PPR336) | 43 | 0.91 | 0.174 | 1.000 |
| AT1G61870.1 | Pentatricopeptide repeat (PPR) superfamily protein (PPR336) | 44 | 0.71 | 0.174 | 0.999 |
| AT1G61870.1 | Pentatricopeptide repeat (PPR) superfamily protein (PPR336) | 45 | 0.50 | 0.190 | 0.831 |
| AT1G61870.1 | Pentatricopeptide repeat (PPR) superfamily protein (PPR336) | 46 | 0.34 | 0.242 | 0.572 |
| AT1G61870.1 | Pentatricopeptide repeat (PPR) superfamily protein (PPR336) | 47 | 0.23 | 0.343 | 0.609 |
| AT1G62730.1 | Terpenoid synthases superfamily protein | 2 | 0.75 | 0.333 | 1.000 |
| AT1G62730.1 | Terpenoid synthases superfamily protein | 3 | 0.95 | 0.330 | 1.000 |
| AT1G62730.1 | Terpenoid synthases superfamily protein | 40 | 2.25 | 0.313 | 0.892 |
| AT1G62730.1 | Terpenoid synthases superfamily protein | 41 | 1.55 | 0.199 | 0.950 |
| AT1G62730.1 | Terpenoid synthases superfamily protein | 42 | 1.24 | 0.170 | 0.999 |
| AT1G62730.1 | Terpenoid synthases superfamily protein | 43 | 1.53 | 0.166 | 0.896 |
| AT1G62730.1 | Terpenoid synthases superfamily protein | 44 | 1.67 | 0.164 | 0.763 |
| AT1G62730.1 | Terpenoid synthases superfamily protein | 45 | 1.11 | 0.163 | 1.000 |
| AT1G62730.1 | Terpenoid synthases superfamily protein | 46 | 0.46 | 0.182 | 0.414 |
| AT1G62730.1 | Terpenoid synthases superfamily protein | 47 | 0.15 | 0.364 | 0.184 |
| AT1G63270.1 | Cytochrome c maturation A (CCMA) | 2 | 1.20 | 0.483 | 1.000 |

| | | | | | | |
|---|---|---|---|---|---|---|
| AT1G63270.1 | Cytochrome c maturation A (CCMA) | 3 | 0.61 | 0.895 | 1.000 | |
| AT1G63270.1 | Cytochrome c maturation A (CCMA) | 25 | 0.12 | 0.305 | 0.033 | * |
| AT1G63270.1 | Cytochrome c maturation A (CCMA) | 26 | 0.42 | 0.214 | 0.477 | |
| AT1G63270.1 | Cytochrome c maturation A (CCMA) | 27 | 1.45 | 0.209 | 0.982 | |
| AT1G63270.1 | Cytochrome c maturation A (CCMA) | 28 | 2.80 | 0.219 | 0.297 | |
| AT1G63270.1 | Cytochrome c maturation A (CCMA) | 29 | 1.80 | 0.282 | 0.959 | |
| AT1G63270.1 | Cytochrome c maturation A (CCMA) | 30 | 0.65 | 0.465 | 1.000 | |
| AT1G63270.1 | Cytochrome c maturation A (CCMA) | 46 | 0.01 | 0.397 | 0.000 | *** |
| AT1G63270.1 | Cytochrome c maturation A (CCMA) | 47 | 0.01 | 0.403 | 0.000 | *** |
| AT1G63940.4 | Monodehydroascorbate reductase 6 (MDAR6) | 2 | 0.70 | 0.388 | 1.000 | |
| AT1G63940.4 | Monodehydroascorbate reductase 6 (MDAR6) | 3 | 0.71 | 0.322 | 1.000 | |
| AT1G63940.4 | Monodehydroascorbate reductase 6 (MDAR6) | 4 | 0.73 | 0.287 | 1.000 | |
| AT1G63940.4 | Monodehydroascorbate reductase 6 (MDAR6) | 6 | 0.85 | 0.268 | 1.000 | |
| AT1G63940.4 | Monodehydroascorbate reductase 6 (MDAR6) | 7 | 0.98 | 0.259 | 1.000 | |
| AT1G63940.4 | Monodehydroascorbate reductase 6 (MDAR6) | 8 | 1.14 | 0.250 | 1.000 | |
| AT1G63940.4 | Monodehydroascorbate reductase 6 (MDAR6) | 9 | 1.28 | 0.237 | 1.000 | |
| AT1G63940.4 | Monodehydroascorbate reductase 6 (MDAR6) | 10 | 1.39 | 0.226 | 1.000 | |
| AT1G63940.4 | Monodehydroascorbate reductase 6 (MDAR6) | 11 | 1.43 | 0.216 | 1.000 | |
| AT1G63940.4 | Monodehydroascorbate reductase 6 (MDAR6) | 12 | 1.40 | 0.208 | 1.000 | |
| AT1G63940.4 | Monodehydroascorbate reductase 6 (MDAR6) | 13 | 1.33 | 0.203 | 1.000 | |
| AT1G63940.4 | Monodehydroascorbate reductase 6 (MDAR6) | 14 | 1.25 | 0.200 | 1.000 | |
| AT1G63940.4 | Monodehydroascorbate reductase 6 (MDAR6) | 15 | 1.16 | 0.202 | 1.000 | |
| AT1G63940.4 | Monodehydroascorbate reductase 6 (MDAR6) | 16 | 1.08 | 0.208 | 1.000 | |
| AT1G63940.4 | Monodehydroascorbate reductase 6 (MDAR6) | 17 | 1.04 | 0.216 | 1.000 | |
| AT1G63940.4 | Monodehydroascorbate reductase 6 (MDAR6) | 18 | 1.01 | 0.224 | 1.000 | |
| AT1G63940.4 | Monodehydroascorbate reductase 6 (MDAR6) | 19 | 0.99 | 0.230 | 1.000 | |
| AT1G63940.4 | Monodehydroascorbate reductase 6 (MDAR6) | 20 | 0.95 | 0.234 | 1.000 | |
| AT1G63940.4 | Monodehydroascorbate reductase 6 (MDAR6) | 21 | 0.90 | 0.230 | 1.000 | |
| AT1G63940.4 | Monodehydroascorbate reductase 6 (MDAR6) | 22 | 0.84 | 0.223 | 1.000 | |
| AT1G63940.4 | Monodehydroascorbate reductase 6 (MDAR6) | 23 | 0.78 | 0.212 | 1.000 | |
| AT1G63940.4 | Monodehydroascorbate reductase 6 (MDAR6) | 24 | 0.73 | 0.202 | 1.000 | |

| | | | | | | |
|---|---|---|---|---|---|---|
| AT1G63940.4 | Monodehydroascorbate reductase 6 (MDAR6) | 25 | 0.69 | 0.196 | 0.999 | |
| AT1G63940.4 | Monodehydroascorbate reductase 6 (MDAR6) | 26 | 0.65 | 0.192 | 0.993 | |
| AT1G63940.4 | Monodehydroascorbate reductase 6 (MDAR6) | 27 | 0.63 | 0.190 | 0.986 | |
| AT1G63940.4 | Monodehydroascorbate reductase 6 (MDAR6) | 28 | 0.63 | 0.189 | 0.988 | |
| AT1G63940.4 | Monodehydroascorbate reductase 6 (MDAR6) | 29 | 0.67 | 0.189 | 0.996 | |
| AT1G63940.4 | Monodehydroascorbate reductase 6 (MDAR6) | 30 | 0.74 | 0.188 | 1.000 | |
| AT1G63940.4 | Monodehydroascorbate reductase 6 (MDAR6) | 31 | 0.89 | 0.188 | 1.000 | |
| AT1G63940.4 | Monodehydroascorbate reductase 6 (MDAR6) | 32 | 1.16 | 0.188 | 1.000 | |
| AT1G63940.4 | Monodehydroascorbate reductase 6 (MDAR6) | 33 | 1.63 | 0.188 | 0.977 | |
| AT1G63940.4 | Monodehydroascorbate reductase 6 (MDAR6) | 34 | 2.38 | 0.188 | 0.483 | |
| AT1G63940.4 | Monodehydroascorbate reductase 6 (MDAR6) | 35 | 3.37 | 0.188 | 0.090 | . |
| AT1G63940.4 | Monodehydroascorbate reductase 6 (MDAR6) | 36 | 4.43 | 0.188 | 1.000 | |
| AT1G63940.4 | Monodehydroascorbate reductase 6 (MDAR6) | 37 | 5.23 | 0.189 | 1.000 | |
| AT1G63940.4 | Monodehydroascorbate reductase 6 (MDAR6) | 38 | 5.28 | 0.189 | 1.000 | |
| AT1G63940.4 | Monodehydroascorbate reductase 6 (MDAR6) | 39 | 4.56 | 0.190 | 1.000 | |
| AT1G63940.4 | Monodehydroascorbate reductase 6 (MDAR6) | 40 | 3.49 | 0.191 | 0.078 | . |
| AT1G63940.4 | Monodehydroascorbate reductase 6 (MDAR6) | 41 | 2.53 | 0.192 | 0.411 | |
| AT1G63940.4 | Monodehydroascorbate reductase 6 (MDAR6) | 42 | 1.82 | 0.193 | 0.916 | |
| AT1G63940.4 | Monodehydroascorbate reductase 6 (MDAR6) | 43 | 1.35 | 0.194 | 1.000 | |
| AT1G63940.4 | Monodehydroascorbate reductase 6 (MDAR6) | 44 | 1.05 | 0.200 | 1.000 | |
| AT1G63940.4 | Monodehydroascorbate reductase 6 (MDAR6) | 45 | 0.86 | 0.221 | 1.000 | |
| AT1G63940.4 | Monodehydroascorbate reductase 6 (MDAR6) | 46 | 0.73 | 0.269 | 1.000 | |
| AT1G63940.4 | Monodehydroascorbate reductase 6 (MDAR6) | 47 | 0.62 | 0.353 | 1.000 | |
| AT1G64880.1 | Ribosomal S5 family protein | 2 | 0.93 | 0.841 | 1.000 | |
| AT1G64880.1 | Ribosomal S5 family protein | 3 | 0.75 | 0.733 | 1.000 | |
| AT1G64880.1 | Ribosomal S5 family protein | 10 | 0.76 | 0.459 | 1.000 | |
| AT1G64880.1 | Ribosomal S5 family protein | 11 | 1.10 | 0.409 | 1.000 | |
| AT1G64880.1 | Ribosomal S5 family protein | 12 | 1.59 | 0.374 | 1.000 | |
| AT1G64880.1 | Ribosomal S5 family protein | 13 | 2.01 | 0.351 | 0.994 | |
| AT1G64880.1 | Ribosomal S5 family protein | 14 | 2.29 | 0.336 | 0.966 | |
| AT1G64880.1 | Ribosomal S5 family protein | 15 | 2.58 | 0.329 | 0.908 | |
| AT1G64880.1 | Ribosomal S5 family protein | 16 | 2.85 | 0.326 | 0.835 | |
| AT1G64880.1 | Ribosomal S5 family protein | 17 | 2.76 | 0.325 | 0.858 | |
| AT1G64880.1 | Ribosomal S5 family protein | 18 | 2.18 | 0.328 | 0.974 | |
| AT1G64880.1 | Ribosomal S5 family protein | 19 | 1.44 | 0.333 | 1.000 | |
| AT1G64880.1 | Ribosomal S5 family protein | 20 | 0.82 | 0.337 | 1.000 | |
| AT1G64880.1 | Ribosomal S5 family protein | 21 | 0.44 | 0.340 | 0.972 | |
| AT1G64880.1 | Ribosomal S5 family protein | 22 | 0.25 | 0.340 | 0.571 | |
| AT1G64880.1 | Ribosomal S5 family protein | 23 | 0.16 | 0.339 | 0.224 | |

| | | | | | | |
|---|---|---|---|---|---|---|
| AT1G64880.1 | Ribosomal S5 family protein | 24 | 0.14 | 0.335 | 0.124 | |
| AT1G64880.1 | Ribosomal S5 family protein | 25 | 0.15 | 0.332 | 0.144 | |
| AT1G64880.1 | Ribosomal S5 family protein | 26 | 0.19 | 0.331 | 0.312 | |
| AT1G64880.1 | Ribosomal S5 family protein | 27 | 0.29 | 0.331 | 0.684 | |
| AT1G64880.1 | Ribosomal S5 family protein | 28 | 0.42 | 0.332 | 0.953 | |
| AT1G64880.1 | Ribosomal S5 family protein | 29 | 0.55 | 0.333 | 0.997 | |
| AT1G64880.1 | Ribosomal S5 family protein | 30 | 0.59 | 0.337 | 0.999 | |
| AT1G64880.1 | Ribosomal S5 family protein | 31 | 0.54 | 0.349 | 0.998 | |
| AT1G64880.1 | Ribosomal S5 family protein | 32 | 0.43 | 0.386 | 0.986 | |
| AT1G64880.1 | Ribosomal S5 family protein | 33 | 0.33 | 0.465 | 0.971 | |
| AT1G64880.1 | Ribosomal S5 family protein | 34 | 0.25 | 0.590 | 0.977 | |
| AT1G64880.1 | Ribosomal S5 family protein | 46 | 0.17 | 2.415 | 1.000 | |
| AT1G64880.1 | Ribosomal S5 family protein | 47 | 0.18 | 2.653 | 1.000 | |
| AT1G65032.1 | ? | 2 | 0.85 | 0.517 | 1.000 | |
| AT1G65032.1 | ? | 3 | 2.01 | 0.566 | 1.000 | |
| AT1G65032.1 | ? | 22 | 0.47 | 0.397 | 0.992 | |
| AT1G65032.1 | ? | 23 | 0.47 | 0.291 | 0.940 | |
| AT1G65032.1 | ? | 24 | 0.76 | 0.254 | 1.000 | |
| AT1G65032.1 | ? | 25 | 1.67 | 0.237 | 0.978 | |
| AT1G65032.1 | ? | 26 | 2.32 | 0.235 | 0.704 | |
| AT1G65032.1 | ? | 27 | 2.13 | 0.240 | 0.824 | |
| AT1G65032.1 | ? | 28 | 0.85 | 0.258 | 1.000 | |
| AT1G65032.1 | ? | 29 | 0.26 | 0.321 | 0.506 | |
| AT1G65032.1 | ? | 30 | 0.14 | 0.543 | 0.689 | |
| AT1G65032.1 | ? | 46 | 62027884.18 | 10.660 | 0.997 | |
| AT1G65032.1 | ? | 47 | ########### | 11.652 | 0.996 | |
| AT1G65290.1 | Acyl carrier protein 2 (ACP2) | 2 | 0.78 | 0.610 | 1.000 | |
| AT1G65290.1 | Acyl carrier protein 2 (ACP2) | 3 | 0.83 | 0.544 | 1.000 | |
| AT1G65290.1 | Acyl carrier protein 2 (ACP2) | 14 | 0.94 | 0.355 | 1.000 | |
| AT1G65290.1 | Acyl carrier protein 2 (ACP2) | 15 | 0.88 | 0.321 | 1.000 | |
| AT1G65290.1 | Acyl carrier protein 2 (ACP2) | 16 | 0.82 | 0.285 | 1.000 | |
| AT1G65290.1 | Acyl carrier protein 2 (ACP2) | 17 | 0.78 | 0.261 | 1.000 | |
| AT1G65290.1 | Acyl carrier protein 2 (ACP2) | 18 | 0.92 | 0.255 | 1.000 | |
| AT1G65290.1 | Acyl carrier protein 2 (ACP2) | 19 | 1.20 | 0.261 | 1.000 | |
| AT1G65290.1 | Acyl carrier protein 2 (ACP2) | 20 | 1.46 | 0.266 | 1.000 | |
| AT1G65290.1 | Acyl carrier protein 2 (ACP2) | 21 | 1.58 | 0.269 | 1.000 | |
| AT1G65290.1 | Acyl carrier protein 2 (ACP2) | 22 | 1.49 | 0.268 | 1.000 | |
| AT1G65290.1 | Acyl carrier protein 2 (ACP2) | 23 | 1.23 | 0.265 | 1.000 | |
| AT1G65290.1 | Acyl carrier protein 2 (ACP2) | 24 | 1.12 | 0.260 | 1.000 | |
| AT1G65290.1 | Acyl carrier protein 2 (ACP2) | 25 | 1.37 | 0.257 | 1.000 | |
| AT1G65290.1 | Acyl carrier protein 2 (ACP2) | 26 | 2.43 | 0.256 | 0.938 | |
| AT1G65290.1 | Acyl carrier protein 2 (ACP2) | 27 | 4.79 | 0.256 | 0.182 | |
| AT1G65290.1 | Acyl carrier protein 2 (ACP2) | 28 | 8.23 | 0.257 | 0.012 | * |
| AT1G65290.1 | Acyl carrier protein 2 (ACP2) | 29 | 10.02 | 0.256 | 0.003 | ** |
| AT1G65290.1 | Acyl carrier protein 2 (ACP2) | 30 | 7.68 | 0.256 | 0.018 | * |
| AT1G65290.1 | Acyl carrier protein 2 (ACP2) | 31 | 4.31 | 0.257 | 0.277 | |
| AT1G65290.1 | Acyl carrier protein 2 (ACP2) | 32 | 2.14 | 0.257 | 0.986 | |
| AT1G65290.1 | Acyl carrier protein 2 (ACP2) | 33 | 1.15 | 0.257 | 1.000 | |
| AT1G65290.1 | Acyl carrier protein 2 (ACP2) | 34 | 0.74 | 0.257 | 1.000 | |
| AT1G65290.1 | Acyl carrier protein 2 (ACP2) | 35 | 0.64 | 0.257 | 1.000 | |
| AT1G65290.1 | Acyl carrier protein 2 (ACP2) | 36 | 0.76 | 0.256 | 1.000 | |
| AT1G65290.1 | Acyl carrier protein 2 (ACP2) | 37 | 1.12 | 0.256 | 1.000 | |
| AT1G65290.1 | Acyl carrier protein 2 (ACP2) | 38 | 1.51 | 0.257 | 1.000 | |
| AT1G65290.1 | Acyl carrier protein 2 (ACP2) | 39 | 1.71 | 0.257 | 1.000 | |
| AT1G65290.1 | Acyl carrier protein 2 (ACP2) | 40 | 1.70 | 0.258 | 1.000 | |
| AT1G65290.1 | Acyl carrier protein 2 (ACP2) | 41 | 1.53 | 0.259 | 1.000 | |
| AT1G65290.1 | Acyl carrier protein 2 (ACP2) | 42 | 1.37 | 0.263 | 1.000 | |
| AT1G65290.1 | Acyl carrier protein 2 (ACP2) | 43 | 1.22 | 0.265 | 1.000 | |
| AT1G65290.1 | Acyl carrier protein 2 (ACP2) | 44 | 1.20 | 0.265 | 1.000 | |
| AT1G65290.1 | Acyl carrier protein 2 (ACP2) | 45 | 1.27 | 0.265 | 1.000 | |
| AT1G65290.1 | Acyl carrier protein 2 (ACP2) | 46 | 1.27 | 0.317 | 1.000 | |
| AT1G65290.1 | Acyl carrier protein 2 (ACP2) | 47 | 1.25 | 0.538 | 1.000 | |

| | | | | | | |
|---|---|---|---|---|---|---|
| AT1G65540.1 | Leucine zipper-EF-hand-containing transmembrane protein 2 (LETM2) | 2 | 0.92 | 0.457 | 1.000 | |
| AT1G65540.1 | Leucine zipper-EF-hand-containing transmembrane protein 2 (LETM2) | 3 | 0.86 | 0.414 | 1.000 | |
| AT1G65540.1 | Leucine zipper-EF-hand-containing transmembrane protein 2 (LETM2) | 15 | 1.44 | 0.275 | 1.000 | |
| AT1G65540.1 | Leucine zipper-EF-hand-containing transmembrane protein 2 (LETM2) | 16 | 1.72 | 0.249 | 1.000 | |
| AT1G65540.1 | Leucine zipper-EF-hand-containing transmembrane protein 2 (LETM2) | 17 | 1.95 | 0.221 | 0.987 | |
| AT1G65540.1 | Leucine zipper-EF-hand-containing transmembrane protein 2 (LETM2) | 18 | 2.14 | 0.206 | 0.911 | |
| AT1G65540.1 | Leucine zipper-EF-hand-containing transmembrane protein 2 (LETM2) | 19 | 2.65 | 0.206 | 0.605 | |
| AT1G65540.1 | Leucine zipper-EF-hand-containing transmembrane protein 2 (LETM2) | 20 | 3.59 | 0.211 | 0.202 | |
| AT1G65540.1 | Leucine zipper-EF-hand-containing transmembrane protein 2 (LETM2) | 21 | 4.17 | 0.213 | 0.097 | . |
| AT1G65540.1 | Leucine zipper-EF-hand-containing transmembrane protein 2 (LETM2) | 22 | 3.98 | 0.213 | 0.126 | |
| AT1G65540.1 | Leucine zipper-EF-hand-containing transmembrane protein 2 (LETM2) | 23 | 3.06 | 0.211 | 0.408 | |
| AT1G65540.1 | Leucine zipper-EF-hand-containing transmembrane protein 2 (LETM2) | 24 | 1.65 | 0.206 | 0.999 | |
| AT1G65540.1 | Leucine zipper-EF-hand-containing transmembrane protein 2 (LETM2) | 25 | 0.87 | 0.204 | 1.000 | |
| AT1G65540.1 | Leucine zipper-EF-hand-containing transmembrane protein 2 (LETM2) | 26 | 0.89 | 0.203 | 1.000 | |
| AT1G65540.1 | Leucine zipper-EF-hand-containing transmembrane protein 2 (LETM2) | 27 | 1.50 | 0.202 | 1.000 | |
| AT1G65540.1 | Leucine zipper-EF-hand-containing transmembrane protein 2 (LETM2) | 28 | 1.92 | 0.202 | 0.974 | |
| AT1G65540.1 | Leucine zipper-EF-hand-containing transmembrane protein 2 (LETM2) | 29 | 1.76 | 0.203 | 0.995 | |
| AT1G65540.1 | Leucine zipper-EF-hand-containing transmembrane protein 2 (LETM2) | 30 | 1.45 | 0.204 | 1.000 | |
| AT1G65540.1 | Leucine zipper-EF-hand-containing transmembrane protein 2 (LETM2) | 31 | 1.29 | 0.204 | 1.000 | |
| AT1G65540.1 | Leucine zipper-EF-hand-containing transmembrane protein 2 (LETM2) | 32 | 1.21 | 0.205 | 1.000 | |

| | | | | | | |
|---|---|---|---|---|---|---|
| AT1G65540.1 | Leucine zipper-EF-hand-containing transmembrane protein 2 (LETM2) | 33 | 1.11 | 0.205 | 1.000 | |
| AT1G65540.1 | Leucine zipper-EF-hand-containing transmembrane protein 2 (LETM2) | 34 | 1.07 | 0.205 | 1.000 | |
| AT1G65540.1 | Leucine zipper-EF-hand-containing transmembrane protein 2 (LETM2) | 35 | 1.09 | 0.205 | 1.000 | |
| AT1G65540.1 | Leucine zipper-EF-hand-containing transmembrane protein 2 (LETM2) | 36 | 1.07 | 0.205 | 1.000 | |
| AT1G65540.1 | Leucine zipper-EF-hand-containing transmembrane protein 2 (LETM2) | 37 | 0.94 | 0.205 | 1.000 | |
| AT1G65540.1 | Leucine zipper-EF-hand-containing transmembrane protein 2 (LETM2) | 38 | 0.75 | 0.204 | 1.000 | |
| AT1G65540.1 | Leucine zipper-EF-hand-containing transmembrane protein 2 (LETM2) | 39 | 0.52 | 0.203 | 0.973 | |
| AT1G65540.1 | Leucine zipper-EF-hand-containing transmembrane protein 2 (LETM2) | 40 | 0.29 | 0.204 | 0.202 | |
| AT1G65540.1 | Leucine zipper-EF-hand-containing transmembrane protein 2 (LETM2) | 41 | 0.16 | 0.215 | 0.007 | ** |
| AT1G65540.1 | Leucine zipper-EF-hand-containing transmembrane protein 2 (LETM2) | 42 | 0.19 | 0.240 | 0.085 | . |
| AT1G65540.1 | Leucine zipper-EF-hand-containing transmembrane protein 2 (LETM2) | 43 | 0.44 | 0.253 | 0.971 | |
| AT1G65540.1 | Leucine zipper-EF-hand-containing transmembrane protein 2 (LETM2) | 44 | 1.23 | 0.304 | 1.000 | |
| AT1G65540.1 | Leucine zipper-EF-hand-containing transmembrane protein 2 (LETM2) | 45 | 3.40 | 0.389 | 0.980 | |
| AT1G65540.1 | Leucine zipper-EF-hand-containing transmembrane protein 2 (LETM2) | 46 | 9.10 | 0.561 | 0.861 | |
| AT1G65540.1 | Leucine zipper-EF-hand-containing transmembrane protein 2 (LETM2) | 47 | 23.99 | 0.832 | 0.888 | |
| AT1G67090.1 | RUBISCO small subunit 1A (RBCS1A) | 2 | 0.80 | 0.984 | 1.000 | |
| AT1G67090.1 | RUBISCO small subunit 1A (RBCS1A) | 3 | 0.88 | 0.866 | 1.000 | |
| AT1G67090.1 | RUBISCO small subunit 1A (RBCS1A) | 11 | 0.67 | 0.778 | 1.000 | |
| AT1G67090.1 | RUBISCO small subunit 1A (RBCS1A) | 12 | 0.58 | 0.731 | 1.000 | |
| AT1G67090.1 | RUBISCO small subunit 1A (RBCS1A) | 13 | 0.52 | 0.780 | 1.000 | |
| AT1G67090.1 | RUBISCO small subunit 1A (RBCS1A) | 24 | 0.69 | 0.662 | 1.000 | |
| AT1G67090.1 | RUBISCO small subunit 1A (RBCS1A) | 25 | 0.58 | 0.500 | 1.000 | |
| AT1G67090.1 | RUBISCO small subunit 1A (RBCS1A) | 26 | 0.60 | 0.434 | 1.000 | |

| | | | | | | |
|---|---|---|---|---|---|---|
| AT1G67090.1 | RUBISCO small subunit 1A (RBCS1A) | 27 | 1.26 | 0.410 | 1.000 | |
| AT1G67090.1 | RUBISCO small subunit 1A (RBCS1A) | 28 | 4.92 | 0.404 | 0.628 | |
| AT1G67090.1 | RUBISCO small subunit 1A (RBCS1A) | 29 | 16.33 | 0.402 | 0.040 | * |
| AT1G67090.1 | RUBISCO small subunit 1A (RBCS1A) | 30 | 16.55 | 0.404 | 0.040 | * |
| AT1G67090.1 | RUBISCO small subunit 1A (RBCS1A) | 31 | 5.98 | 0.407 | 0.481 | |
| AT1G67090.1 | RUBISCO small subunit 1A (RBCS1A) | 32 | 2.13 | 0.417 | 0.997 | |
| AT1G67090.1 | RUBISCO small subunit 1A (RBCS1A) | 33 | 1.22 | 0.448 | 1.000 | |
| AT1G67090.1 | RUBISCO small subunit 1A (RBCS1A) | 34 | 0.94 | 0.574 | 1.000 | |
| AT1G67090.1 | RUBISCO small subunit 1A (RBCS1A) | 35 | 0.92 | 0.866 | 1.000 | |
| AT1G67090.1 | RUBISCO small subunit 1A (RBCS1A) | 46 | 18856.20 | 4.488 | 0.986 | |
| AT1G67090.1 | RUBISCO small subunit 1A (RBCS1A) | 47 | 72601.56 | 5.010 | 0.984 | |
| AT1G67350.2 | NADH:ubiquinone oxidoreductase subunit | 2 | 1.16 | 0.580 | 1.000 | |
| AT1G67350.2 | NADH:ubiquinone oxidoreductase subunit | 3 | 0.91 | 0.430 | 1.000 | |
| AT1G67350.2 | NADH:ubiquinone oxidoreductase subunit | 4 | 0.71 | 0.671 | 1.000 | |
| AT1G67350.2 | NADH:ubiquinone oxidoreductase subunit | 13 | 0.44 | 0.391 | 0.999 | |
| AT1G67350.2 | NADH:ubiquinone oxidoreductase subunit | 14 | 0.60 | 0.353 | 1.000 | |
| AT1G67350.2 | NADH:ubiquinone oxidoreductase subunit | 15 | 0.77 | 0.329 | 1.000 | |
| AT1G67350.2 | NADH:ubiquinone oxidoreductase subunit | 16 | 0.73 | 0.294 | 1.000 | |
| AT1G67350.2 | NADH:ubiquinone oxidoreductase subunit | 17 | 0.59 | 0.280 | 1.000 | |
| AT1G67350.2 | NADH:ubiquinone oxidoreductase subunit | 18 | 0.60 | 0.281 | 1.000 | |
| AT1G67350.2 | NADH:ubiquinone oxidoreductase subunit | 19 | 0.72 | 0.293 | 1.000 | |
| AT1G67350.2 | NADH:ubiquinone oxidoreductase subunit | 20 | 0.79 | 0.300 | 1.000 | |
| AT1G67350.2 | NADH:ubiquinone oxidoreductase subunit | 21 | 1.02 | 0.303 | 1.000 | |
| AT1G67350.2 | NADH:ubiquinone oxidoreductase subunit | 22 | 1.76 | 0.301 | 1.000 | |
| AT1G67350.2 | NADH:ubiquinone oxidoreductase subunit | 23 | 2.92 | 0.300 | 0.880 | |
| AT1G67350.2 | NADH:ubiquinone oxidoreductase subunit | 24 | 2.48 | 0.291 | 0.955 | |
| AT1G67350.2 | NADH:ubiquinone oxidoreductase subunit | 25 | 0.91 | 0.286 | 1.000 | |
| AT1G67350.2 | NADH:ubiquinone oxidoreductase subunit | 26 | 0.46 | 0.287 | 0.989 | |
| AT1G67350.2 | NADH:ubiquinone oxidoreductase subunit | 27 | 0.45 | 0.288 | 0.986 | |
| AT1G67350.2 | NADH:ubiquinone oxidoreductase subunit | 28 | 0.50 | 0.288 | 0.996 | |
| AT1G67350.2 | NADH:ubiquinone oxidoreductase subunit | 29 | 0.52 | 0.294 | 0.998 | |

| | | | | | | |
|---|---|---|---|---|---|---|
| AT1G67350.2 | NADH:ubiquinone oxidoreductase subunit | 30 | 0.42 | 0.316 | 0.984 | |
| AT1G67350.2 | NADH:ubiquinone oxidoreductase subunit | 31 | 0.30 | 0.392 | 0.960 | |
| AT1G67350.2 | NADH:ubiquinone oxidoreductase subunit | 32 | 0.25 | 0.630 | 0.998 | |
| AT1G67350.2 | NADH:ubiquinone oxidoreductase subunit | 46 | 6120.06 | 9.338 | 1.000 | |
| AT1G67350.2 | NADH:ubiquinone oxidoreductase subunit | 47 | 19755.51 | 10.305 | 1.000 | |
| AT1G67440.1 | Small ribosomal subunit biogenesis GTPase (RSGA1) | 2 | 1.10 | 0.537 | 1.000 | |
| AT1G67440.1 | Small ribosomal subunit biogenesis GTPase (RSGA1) | 3 | 0.75 | 0.647 | 1.000 | |
| AT1G67440.1 | Small ribosomal subunit biogenesis GTPase (RSGA1) | 36 | 0.33 | 0.287 | 0.470 | |
| AT1G67440.1 | Small ribosomal subunit biogenesis GTPase (RSGA1) | 37 | 0.66 | 0.218 | 0.957 | |
| AT1G67440.1 | Small ribosomal subunit biogenesis GTPase (RSGA1) | 38 | 1.29 | 0.198 | 0.995 | |
| AT1G67440.1 | Small ribosomal subunit biogenesis GTPase (RSGA1) | 39 | 2.00 | 0.199 | 0.586 | |
| AT1G67440.1 | Small ribosomal subunit biogenesis GTPase (RSGA1) | 40 | 2.11 | 0.218 | 0.599 | |
| AT1G67440.1 | Small ribosomal subunit biogenesis GTPase (RSGA1) | 41 | 1.62 | 0.262 | 0.964 | |
| AT1G67440.1 | Small ribosomal subunit biogenesis GTPase (RSGA1) | 42 | 1.10 | 0.319 | 1.000 | |
| AT1G67440.1 | Small ribosomal subunit biogenesis GTPase (RSGA1) | 46 | 0.17 | 0.303 | 0.093 | . |
| AT1G67440.1 | Small ribosomal subunit biogenesis GTPase (RSGA1) | 47 | 0.10 | 0.350 | 0.047 | * |
| AT1G67785.1 | NADH:ubiquinone oxidoreductase subunit (SGDH) | 2 | 0.90 | 0.542 | 1.000 | |
| AT1G67785.1 | NADH:ubiquinone oxidoreductase subunit (SGDH) | 3 | 0.76 | 0.501 | 1.000 | |
| AT1G67785.1 | NADH:ubiquinone oxidoreductase subunit (SGDH) | 15 | 0.95 | 0.339 | 1.000 | |
| AT1G67785.1 | NADH:ubiquinone oxidoreductase subunit (SGDH) | 16 | 0.68 | 0.289 | 1.000 | |
| AT1G67785.1 | NADH:ubiquinone oxidoreductase subunit (SGDH) | 17 | 0.41 | 0.262 | 0.875 | |
| AT1G67785.1 | NADH:ubiquinone oxidoreductase subunit (SGDH) | 18 | 0.40 | 0.240 | 0.744 | |
| AT1G67785.1 | NADH:ubiquinone oxidoreductase subunit (SGDH) | 19 | 0.60 | 0.246 | 0.997 | |
| AT1G67785.1 | NADH:ubiquinone oxidoreductase subunit (SGDH) | 20 | 0.77 | 0.254 | 1.000 | |
| AT1G67785.1 | NADH:ubiquinone oxidoreductase subunit (SGDH) | 21 | 1.02 | 0.257 | 1.000 | |
| AT1G67785.1 | NADH:ubiquinone oxidoreductase subunit (SGDH) | 22 | 1.74 | 0.254 | 0.996 | |
| AT1G67785.1 | NADH:ubiquinone oxidoreductase subunit (SGDH) | 23 | 2.55 | 0.254 | 0.794 | |
| AT1G67785.1 | NADH:ubiquinone oxidoreductase subunit (SGDH) | 24 | 1.52 | 0.244 | 1.000 | |
| AT1G67785.1 | NADH:ubiquinone oxidoreductase subunit (SGDH) | 25 | 0.50 | 0.241 | 0.958 | |
| AT1G67785.1 | NADH:ubiquinone oxidoreductase subunit (SGDH) | 26 | 0.40 | 0.242 | 0.761 | |
| AT1G67785.1 | NADH:ubiquinone oxidoreductase subunit (SGDH) | 27 | 0.71 | 0.245 | 1.000 | |

| | | | | | |
|---|---|---|---|---|---|
| AT1G67785.1 | NADH:ubiquinone oxidoreductase subunit (SGDH) | 28 | 0.82 | 0.262 | 1.000 |
| AT1G67785.1 | NADH:ubiquinone oxidoreductase subunit (SGDH) | 29 | 0.72 | 0.330 | 1.000 |
| AT1G67785.1 | NADH:ubiquinone oxidoreductase subunit (SGDH) | 30 | 0.69 | 0.558 | 1.000 |
| AT1G67785.1 | NADH:ubiquinone oxidoreductase subunit (SGDH) | 46 | 623.01 | 8.236 | 1.000 |
| AT1G67785.1 | NADH:ubiquinone oxidoreductase subunit (SGDH) | 47 | 1203.94 | 8.983 | 1.000 |
| AT1G68010.1 | Hydroxypyruvate reductase 1 (HPR1) | 2 | 0.88 | 0.423 | 1.000 |
| AT1G68010.1 | Hydroxypyruvate reductase 1 (HPR1) | 3 | 0.91 | 0.407 | 1.000 |
| AT1G68010.1 | Hydroxypyruvate reductase 1 (HPR1) | 28 | 0.65 | 0.322 | 1.000 |
| AT1G68010.1 | Hydroxypyruvate reductase 1 (HPR1) | 29 | 0.59 | 0.260 | 0.984 |
| AT1G68010.1 | Hydroxypyruvate reductase 1 (HPR1) | 30 | 0.55 | 0.241 | 0.939 |
| AT1G68010.1 | Hydroxypyruvate reductase 1 (HPR1) | 31 | 0.56 | 0.235 | 0.935 |
| AT1G68010.1 | Hydroxypyruvate reductase 1 (HPR1) | 32 | 0.60 | 0.233 | 0.972 |
| AT1G68010.1 | Hydroxypyruvate reductase 1 (HPR1) | 33 | 0.67 | 0.233 | 0.996 |
| AT1G68010.1 | Hydroxypyruvate reductase 1 (HPR1) | 34 | 0.78 | 0.233 | 1.000 |
| AT1G68010.1 | Hydroxypyruvate reductase 1 (HPR1) | 35 | 0.95 | 0.232 | 1.000 |
| AT1G68010.1 | Hydroxypyruvate reductase 1 (HPR1) | 36 | 1.23 | 0.232 | 1.000 |
| AT1G68010.1 | Hydroxypyruvate reductase 1 (HPR1) | 37 | 1.69 | 0.232 | 0.966 |
| AT1G68010.1 | Hydroxypyruvate reductase 1 (HPR1) | 38 | 2.07 | 0.231 | 0.782 |
| AT1G68010.1 | Hydroxypyruvate reductase 1 (HPR1) | 39 | 2.13 | 0.231 | 0.748 |
| AT1G68010.1 | Hydroxypyruvate reductase 1 (HPR1) | 40 | 1.93 | 0.231 | 0.865 |
| AT1G68010.1 | Hydroxypyruvate reductase 1 (HPR1) | 41 | 1.64 | 0.232 | 0.978 |
| AT1G68010.1 | Hydroxypyruvate reductase 1 (HPR1) | 42 | 1.41 | 0.234 | 0.999 |
| AT1G68010.1 | Hydroxypyruvate reductase 1 (HPR1) | 43 | 1.19 | 0.235 | 1.000 |
| AT1G68010.1 | Hydroxypyruvate reductase 1 (HPR1) | 44 | 0.87 | 0.235 | 1.000 |
| AT1G68010.1 | Hydroxypyruvate reductase 1 (HPR1) | 45 | 0.51 | 0.236 | 0.867 |
| AT1G68010.1 | Hydroxypyruvate reductase 1 (HPR1) | 46 | 0.27 | 0.260 | 0.253 |
| AT1G68010.1 | Hydroxypyruvate reductase 1 (HPR1) | 47 | 0.14 | 0.375 | 0.210 |
| AT1G68680.1 | ? | 2 | 0.51 | 0.257 | 0.848 |
| AT1G68680.1 | ? | 3 | 0.62 | 0.230 | 0.942 |
| AT1G68680.1 | ? | 4 | 0.74 | 0.208 | 0.994 |
| AT1G68680.1 | ? | 6 | 1.07 | 0.179 | 1.000 |
| AT1G68680.1 | ? | 7 | 1.27 | 0.168 | 0.994 |
| AT1G68680.1 | ? | 8 | 1.50 | 0.160 | 0.859 |
| AT1G68680.1 | ? | 9 | 1.75 | 0.153 | 0.539 |
| AT1G68680.1 | ? | 10 | 2.00 | 0.148 | 0.260 |

| | | | | | | |
|---|---|---|---|---|---|---|
| AT1G68680.1 | ? | 11 | 2.24 | 0.143 | 0.112 | |
| AT1G68680.1 | ? | 12 | 2.43 | 0.139 | 0.050 | * |
| AT1G68680.1 | ? | 13 | 2.57 | 0.136 | 0.025 | * |
| AT1G68680.1 | ? | 14 | 2.66 | 0.134 | 0.016 | * |
| AT1G68680.1 | ? | 15 | 2.69 | 0.133 | 0.013 | * |
| AT1G68680.1 | ? | 16 | 2.67 | 0.133 | 0.014 | * |
| AT1G68680.1 | ? | 17 | 2.62 | 0.135 | 0.021 | * |
| AT1G68680.1 | ? | 18 | 2.56 | 0.139 | 0.032 | * |
| AT1G68680.1 | ? | 19 | 2.50 | 0.145 | 0.053 | . |
| AT1G68680.1 | ? | 20 | 2.45 | 0.152 | 0.085 | . |
| AT1G68680.1 | ? | 21 | 2.43 | 0.161 | 0.124 | |
| AT1G68680.1 | ? | 22 | 2.42 | 0.170 | 0.169 | |
| AT1G68680.1 | ? | 29 | 2.89 | 0.211 | 0.193 | |
| AT1G68680.1 | ? | 30 | 3.01 | 0.213 | 0.175 | |
| AT1G68680.1 | ? | 31 | 3.12 | 0.216 | 0.156 | |
| AT1G68680.1 | ? | 34 | 3.40 | 0.225 | 0.136 | |
| AT1G68680.1 | ? | 35 | 3.43 | 0.229 | 0.143 | |
| AT1G68680.1 | ? | 36 | 3.42 | 0.233 | 0.158 | |
| AT1G68680.1 | ? | 40 | 2.99 | 0.251 | 0.337 | |
| AT1G68680.1 | ? | 41 | 2.81 | 0.256 | 0.428 | |
| AT1G68680.1 | ? | 42 | 2.61 | 0.262 | 0.541 | |
| AT1G68680.1 | ? | 46 | 1.76 | 0.312 | 0.973 | |
| AT1G68680.1 | ? | 47 | 1.58 | 0.335 | 0.996 | |
| AT1G69380.1 | ? | 2 | 1.10 | 0.592 | 1.000 | |
| AT1G69380.1 | ? | 3 | 0.92 | 0.587 | 1.000 | |
| AT1G69380.1 | ? | 27 | 0.47 | 0.357 | 0.882 | |
| AT1G69380.1 | ? | 28 | 0.62 | 0.324 | 0.971 | |
| AT1G69380.1 | ? | 29 | 0.82 | 0.304 | 1.000 | |
| AT1G69380.1 | ? | 32 | 1.75 | 0.286 | 0.911 | |
| AT1G69380.1 | ? | 33 | 2.01 | 0.295 | 0.826 | |
| AT1G69380.1 | ? | 34 | 2.11 | 0.318 | 0.831 | |
| AT1G69380.1 | ? | 35 | 2.06 | 0.355 | 0.898 | |
| AT1G69380.1 | ? | 46 | 0.12 | 0.447 | 0.204 | |
| AT1G69380.1 | ? | 47 | 0.08 | 0.471 | 0.130 | |
| AT1G69740.2 | 5-aminolevulinate dehydratase (HEMB1) | 2 | 1.28 | 0.656 | 0.999 | |
| AT1G69740.2 | 5-aminolevulinate dehydratase (HEMB1) | 3 | 1.32 | 0.633 | 0.999 | |
| AT1G69740.2 | 5-aminolevulinate dehydratase (HEMB1) | 31 | 0.95 | 0.288 | 1.000 | |
| AT1G69740.2 | 5-aminolevulinate dehydratase (HEMB1) | 32 | 0.86 | 0.284 | 0.997 | |
| AT1G69740.2 | 5-aminolevulinate dehydratase (HEMB1) | 33 | 0.76 | 0.283 | 0.978 | |
| AT1G69740.2 | 5-aminolevulinate dehydratase (HEMB1) | 34 | 0.67 | 0.287 | 0.927 | |
| AT1G69740.2 | 5-aminolevulinate dehydratase (HEMB1) | 35 | 0.59 | 0.295 | 0.841 | |
| AT1G69740.2 | 5-aminolevulinate dehydratase (HEMB1) | 46 | 0.08 | 0.459 | 0.080 | . |
| AT1G69740.2 | 5-aminolevulinate dehydratase (HEMB1) | 47 | 0.07 | 0.485 | 0.072 | . |
| AT1G70190.2 | Ribosomal L7/L12 family protein | 2 | 0.98 | 0.340 | 1.000 | |
| AT1G70190.2 | Ribosomal L7/L12 family protein | 3 | 0.97 | 0.313 | 1.000 | |
| AT1G70190.2 | Ribosomal L7/L12 family protein | 4 | 0.96 | 0.376 | 1.000 | |
| AT1G70190.2 | Ribosomal L7/L12 family protein | 29 | 0.91 | 0.239 | 1.000 | |
| AT1G70190.2 | Ribosomal L7/L12 family protein | 30 | 0.93 | 0.195 | 1.000 | |

| | | | | | | |
|---|---|---|---|---|---|---|
| AT1G70190.2 | Ribosomal L7/L12 family protein | 31 | 0.96 | 0.171 | 1.000 | |
| AT1G70190.2 | Ribosomal L7/L12 family protein | 32 | 0.99 | 0.161 | 1.000 | |
| AT1G70190.2 | Ribosomal L7/L12 family protein | 33 | 1.02 | 0.158 | 1.000 | |
| AT1G70190.2 | Ribosomal L7/L12 family protein | 34 | 1.07 | 0.157 | 1.000 | |
| AT1G70190.2 | Ribosomal L7/L12 family protein | 35 | 1.15 | 0.156 | 1.000 | |
| AT1G70190.2 | Ribosomal L7/L12 family protein | 36 | 1.29 | 0.153 | 0.993 | |
| AT1G70190.2 | Ribosomal L7/L12 family protein | 37 | 1.49 | 0.150 | 0.891 | |
| AT1G70190.2 | Ribosomal L7/L12 family protein | 38 | 1.70 | 0.149 | 0.655 | |
| AT1G70190.2 | Ribosomal L7/L12 family protein | 39 | 1.79 | 0.153 | 0.570 | |
| AT1G70190.2 | Ribosomal L7/L12 family protein | 40 | 1.63 | 0.163 | 0.815 | |
| AT1G70190.2 | Ribosomal L7/L12 family protein | 41 | 1.28 | 0.182 | 0.999 | |
| AT1G70190.2 | Ribosomal L7/L12 family protein | 42 | 0.89 | 0.209 | 1.000 | |
| AT1G70190.2 | Ribosomal L7/L12 family protein | 43 | 0.58 | 0.237 | 0.946 | |
| AT1G70190.2 | Ribosomal L7/L12 family protein | 46 | 0.12 | 0.334 | 0.066 | . |
| AT1G70190.2 | Ribosomal L7/L12 family protein | 47 | 0.07 | 0.394 | 0.039 | * |
| AT1G70580.4 | Alanine-2-oxoglutarate aminotransferase 2 (AOAT2) | 2 | 1.07 | 0.352 | 1.000 | |
| AT1G70580.4 | Alanine-2-oxoglutarate aminotransferase 2 (AOAT2) | 3 | 0.93 | 0.325 | 1.000 | |
| AT1G70580.4 | Alanine-2-oxoglutarate aminotransferase 2 (AOAT2) | 4 | 0.81 | 0.373 | 1.000 | |
| AT1G70580.4 | Alanine-2-oxoglutarate aminotransferase 2 (AOAT2) | 34 | 0.58 | 0.243 | 0.928 | |
| AT1G70580.4 | Alanine-2-oxoglutarate aminotransferase 2 (AOAT2) | 35 | 0.75 | 0.200 | 0.993 | |
| AT1G70580.4 | Alanine-2-oxoglutarate aminotransferase 2 (AOAT2) | 36 | 0.96 | 0.172 | 1.000 | |
| AT1G70580.4 | Alanine-2-oxoglutarate aminotransferase 2 (AOAT2) | 37 | 1.19 | 0.157 | 0.999 | |
| AT1G70580.4 | Alanine-2-oxoglutarate aminotransferase 2 (AOAT2) | 38 | 1.36 | 0.149 | 0.952 | |
| AT1G70580.4 | Alanine-2-oxoglutarate aminotransferase 2 (AOAT2) | 39 | 1.44 | 0.146 | 0.879 | |
| AT1G70580.4 | Alanine-2-oxoglutarate aminotransferase 2 (AOAT2) | 40 | 1.44 | 0.144 | 0.881 | |
| AT1G70580.4 | Alanine-2-oxoglutarate aminotransferase 2 (AOAT2) | 41 | 1.36 | 0.144 | 0.944 | |
| AT1G70580.4 | Alanine-2-oxoglutarate aminotransferase 2 (AOAT2) | 42 | 1.22 | 0.146 | 0.995 | |
| AT1G70580.4 | Alanine-2-oxoglutarate aminotransferase 2 (AOAT2) | 43 | 1.04 | 0.150 | 1.000 | |
| AT1G70580.4 | Alanine-2-oxoglutarate aminotransferase 2 (AOAT2) | 44 | 0.85 | 0.158 | 1.000 | |
| AT1G70580.4 | Alanine-2-oxoglutarate aminotransferase 2 (AOAT2) | 45 | 0.68 | 0.177 | 0.935 | |
| AT1G70580.4 | Alanine-2-oxoglutarate aminotransferase 2 (AOAT2) | 46 | 0.53 | 0.214 | 0.780 | |

| | | | | | |
|---|---|---|---|---|---|
| AT1G70580.4 | Alanine-2-oxoglutarate aminotransferase 2 (AOAT2) | 47 | 0.42 | 0.275 | 0.719 |
| AT1G71500.1 | Photosystem II subunit (PSB33) | 2 | 0.74 | 0.481 | 1.000 |
| AT1G71500.1 | Photosystem II subunit (PSB33) | 3 | 0.66 | 0.366 | 1.000 |
| AT1G71500.1 | Photosystem II subunit (PSB33) | 4 | 0.60 | 0.329 | 1.000 |
| AT1G71500.1 | Photosystem II subunit (PSB33) | 5 | 0.60 | 0.316 | 1.000 |
| AT1G71500.1 | Photosystem II subunit (PSB33) | 6 | 0.62 | 0.310 | 1.000 |
| AT1G71500.1 | Photosystem II subunit (PSB33) | 7 | 0.65 | 0.306 | 1.000 |
| AT1G71500.1 | Photosystem II subunit (PSB33) | 8 | 0.67 | 0.301 | 1.000 |
| AT1G71500.1 | Photosystem II subunit (PSB33) | 9 | 0.70 | 0.297 | 1.000 |
| AT1G71500.1 | Photosystem II subunit (PSB33) | 10 | 0.71 | 0.297 | 1.000 |
| AT1G71500.1 | Photosystem II subunit (PSB33) | 11 | 0.70 | 0.298 | 1.000 |
| AT1G71500.1 | Photosystem II subunit (PSB33) | 12 | 0.68 | 0.297 | 1.000 |
| AT1G71500.1 | Photosystem II subunit (PSB33) | 13 | 0.68 | 0.295 | 1.000 |
| AT1G71500.1 | Photosystem II subunit (PSB33) | 14 | 0.71 | 0.292 | 1.000 |
| AT1G71500.1 | Photosystem II subunit (PSB33) | 15 | 0.75 | 0.292 | 1.000 |
| AT1G71500.1 | Photosystem II subunit (PSB33) | 16 | 0.80 | 0.291 | 1.000 |
| AT1G71500.1 | Photosystem II subunit (PSB33) | 17 | 0.93 | 0.293 | 1.000 |
| AT1G71500.1 | Photosystem II subunit (PSB33) | 18 | 1.22 | 0.297 | 1.000 |
| AT1G71500.1 | Photosystem II subunit (PSB33) | 19 | 1.55 | 0.302 | 1.000 |
| AT1G71500.1 | Photosystem II subunit (PSB33) | 20 | 1.77 | 0.312 | 0.999 |
| AT1G71500.1 | Photosystem II subunit (PSB33) | 21 | 1.66 | 0.328 | 1.000 |
| AT1G71500.1 | Photosystem II subunit (PSB33) | 22 | 1.30 | 0.342 | 1.000 |
| AT1G71500.1 | Photosystem II subunit (PSB33) | 23 | 0.91 | 0.349 | 1.000 |
| AT1G71500.1 | Photosystem II subunit (PSB33) | 24 | 0.60 | 0.350 | 1.000 |
| AT1G71500.1 | Photosystem II subunit (PSB33) | 25 | 0.39 | 0.354 | 0.971 |
| AT1G71500.1 | Photosystem II subunit (PSB33) | 26 | 0.28 | 0.345 | 0.748 |
| AT1G71500.1 | Photosystem II subunit (PSB33) | 27 | 0.27 | 0.337 | 0.693 |
| AT1G71500.1 | Photosystem II subunit (PSB33) | 28 | 0.44 | 0.320 | 0.976 |
| AT1G71500.1 | Photosystem II subunit (PSB33) | 29 | 1.03 | 0.310 | 1.000 |
| AT1G71500.1 | Photosystem II subunit (PSB33) | 30 | 2.24 | 0.308 | 0.972 |
| AT1G71500.1 | Photosystem II subunit (PSB33) | 31 | 3.09 | 0.324 | 0.812 |
| AT1G71500.1 | Photosystem II subunit (PSB33) | 32 | 2.77 | 0.380 | 0.966 |
| AT1G71500.1 | Photosystem II subunit (PSB33) | 33 | 2.43 | 0.469 | 0.999 |
| AT1G71500.1 | Photosystem II subunit (PSB33) | 44 | 1.75 | 0.486 | 1.000 |
| AT1G71500.1 | Photosystem II subunit (PSB33) | 45 | 1.31 | 0.374 | 1.000 |
| AT1G71500.1 | Photosystem II subunit (PSB33) | 46 | 0.89 | 0.364 | 1.000 |
| AT1G71500.1 | Photosystem II subunit (PSB33) | 47 | 0.57 | 0.508 | 1.000 |
| AT1G72170.1 | MICOS complex subunit (Mic10-like) | 3 | 0.59 | 0.136 | 0.942 |
| AT1G72170.1 | MICOS complex subunit (Mic10-like) | 4 | 0.48 | 0.126 | 0.318 |
| AT1G72170.1 | MICOS complex subunit (Mic10-like) | 5 | 0.55 | 0.126 | 0.742 |
| AT1G72170.1 | MICOS complex subunit (Mic10-like) | 6 | 0.65 | 0.126 | 0.991 |
| AT1G72170.1 | MICOS complex subunit (Mic10-like) | 7 | 0.68 | 0.126 | 0.998 |
| AT1G72170.1 | MICOS complex subunit (Mic10-like) | 8 | 0.76 | 0.123 | 1.000 |
| AT1G72170.1 | MICOS complex subunit (Mic10-like) | 9 | 0.91 | 0.121 | 1.000 |
| AT1G72170.1 | MICOS complex subunit (Mic10-like) | 10 | 1.01 | 0.121 | 1.000 |
| AT1G72170.1 | MICOS complex subunit (Mic10-like) | 11 | 1.10 | 0.122 | 1.000 |
| AT1G72170.1 | MICOS complex subunit (Mic10-like) | 12 | 1.28 | 0.122 | 1.000 |
| AT1G72170.1 | MICOS complex subunit (Mic10-like) | 13 | 1.54 | 0.120 | 0.978 |
| AT1G72170.1 | MICOS complex subunit (Mic10-like) | 14 | 1.73 | 0.118 | 0.766 |

| | | | | | |
|---|---|---|---|---|---|
| AT1G72170.1 | MICOS complex subunit (Mic10-like) | 15 | 1.71 | 0.118 | 0.800 |
| AT1G72170.1 | MICOS complex subunit (Mic10-like) | 16 | 1.49 | 0.117 | 0.988 |
| AT1G72170.1 | MICOS complex subunit (Mic10-like) | 17 | 1.26 | 0.117 | 1.000 |
| AT1G72170.1 | MICOS complex subunit (Mic10-like) | 18 | 1.08 | 0.119 | 1.000 |
| AT1G72170.1 | MICOS complex subunit (Mic10-like) | 19 | 1.03 | 0.122 | 1.000 |
| AT1G72170.1 | MICOS complex subunit (Mic10-like) | 20 | 1.11 | 0.126 | 1.000 |
| AT1G72170.1 | MICOS complex subunit (Mic10-like) | 21 | 1.29 | 0.128 | 1.000 |
| AT1G72170.1 | MICOS complex subunit (Mic10-like) | 22 | 1.53 | 0.128 | 0.991 |
| AT1G72170.1 | MICOS complex subunit (Mic10-like) | 23 | 1.67 | 0.127 | 0.925 |
| AT1G72170.1 | MICOS complex subunit (Mic10-like) | 24 | 1.52 | 0.124 | 0.991 |
| AT1G72170.1 | MICOS complex subunit (Mic10-like) | 25 | 1.16 | 0.121 | 1.000 |
| AT1G72170.1 | MICOS complex subunit (Mic10-like) | 26 | 0.78 | 0.121 | 1.000 |
| AT1G72170.1 | MICOS complex subunit (Mic10-like) | 27 | 0.69 | 0.121 | 0.998 |
| AT1G72170.1 | MICOS complex subunit (Mic10-like) | 28 | 0.85 | 0.120 | 1.000 |
| AT1G72170.1 | MICOS complex subunit (Mic10-like) | 29 | 1.05 | 0.120 | 1.000 |
| AT1G72170.1 | MICOS complex subunit (Mic10-like) | 30 | 1.21 | 0.120 | 1.000 |
| AT1G72170.1 | MICOS complex subunit (Mic10-like) | 31 | 1.20 | 0.121 | 1.000 |
| AT1G72170.1 | MICOS complex subunit (Mic10-like) | 32 | 1.08 | 0.121 | 1.000 |
| AT1G72170.1 | MICOS complex subunit (Mic10-like) | 33 | 1.00 | 0.121 | 1.000 |
| AT1G72170.1 | MICOS complex subunit (Mic10-like) | 34 | 1.04 | 0.122 | 1.000 |
| AT1G72170.1 | MICOS complex subunit (Mic10-like) | 35 | 1.15 | 0.121 | 1.000 |
| AT1G72170.1 | MICOS complex subunit (Mic10-like) | 36 | 1.13 | 0.121 | 1.000 |
| AT1G72170.1 | MICOS complex subunit (Mic10-like) | 37 | 0.97 | 0.121 | 1.000 |
| AT1G72170.1 | MICOS complex subunit (Mic10-like) | 38 | 0.83 | 0.121 | 1.000 |
| AT1G72170.1 | MICOS complex subunit (Mic10-like) | 39 | 0.81 | 0.121 | 1.000 |
| AT1G72170.1 | MICOS complex subunit (Mic10-like) | 40 | 1.00 | 0.122 | 1.000 |
| AT1G72170.1 | MICOS complex subunit (Mic10-like) | 41 | 1.46 | 0.122 | 0.997 |
| AT1G72170.1 | MICOS complex subunit (Mic10-like) | 42 | 1.94 | 0.124 | 0.510 |
| AT1G72170.1 | MICOS complex subunit (Mic10-like) | 43 | 1.86 | 0.125 | 0.651 |
| AT1G72170.1 | MICOS complex subunit (Mic10-like) | 44 | 1.16 | 0.125 | 1.000 |
| AT1G72170.1 | MICOS complex subunit (Mic10-like) | 45 | 0.50 | 0.125 | 0.428 |

| | | | | | | |
|---|---|---|---|---|---|---|
| AT1G72170.1 | MICOS complex subunit (Mic10-like) | 46 | 0.16 | 0.146 | 0.000 | *** |
| AT1G72170.1 | MICOS complex subunit (Mic10-like) | 47 | 0.04 | 0.266 | 0.000 | *** |
| AT1G72170.1 | MICOS complex subunit (Mic10-like) | 48 | 0.01 | 0.457 | 0.001 | *** |
| AT1G72190.1 | Phosphoglycerate dehydrogenase, putative | 2 | 1.26 | 0.537 | 1.000 | |
| AT1G72190.1 | Phosphoglycerate dehydrogenase, putative | 3 | 1.09 | 0.486 | 1.000 | |
| AT1G72190.1 | Phosphoglycerate dehydrogenase, putative | 4 | 0.94 | 0.741 | 1.000 | |
| AT1G72190.1 | Phosphoglycerate dehydrogenase, putative | 26 | 0.87 | 0.387 | 1.000 | |
| AT1G72190.1 | Phosphoglycerate dehydrogenase, putative | 27 | 1.15 | 0.286 | 1.000 | |
| AT1G72190.1 | Phosphoglycerate dehydrogenase, putative | 28 | 1.61 | 0.259 | 0.998 | |
| AT1G72190.1 | Phosphoglycerate dehydrogenase, putative | 29 | 2.09 | 0.262 | 0.934 | |
| AT1G72190.1 | Phosphoglycerate dehydrogenase, putative | 30 | 1.96 | 0.269 | 0.971 | |
| AT1G72190.1 | Phosphoglycerate dehydrogenase, putative | 31 | 1.49 | 0.269 | 1.000 | |
| AT1G72190.1 | Phosphoglycerate dehydrogenase, putative | 32 | 1.12 | 0.264 | 1.000 | |
| AT1G72190.1 | Phosphoglycerate dehydrogenase, putative | 33 | 0.83 | 0.273 | 1.000 | |
| AT1G72190.1 | Phosphoglycerate dehydrogenase, putative | 34 | 0.58 | 0.316 | 0.999 | |
| AT1G72190.1 | Phosphoglycerate dehydrogenase, putative | 35 | 0.43 | 0.384 | 0.989 | |
| AT1G72190.1 | Phosphoglycerate dehydrogenase, putative | 39 | 0.48 | 0.326 | 0.986 | |
| AT1G72190.1 | Phosphoglycerate dehydrogenase, putative | 40 | 0.75 | 0.280 | 1.000 | |
| AT1G72190.1 | Phosphoglycerate dehydrogenase, putative | 41 | 1.40 | 0.260 | 1.000 | |
| AT1G72190.1 | Phosphoglycerate dehydrogenase, putative | 42 | 2.85 | 0.259 | 0.616 | |
| AT1G72190.1 | Phosphoglycerate dehydrogenase, putative | 43 | 4.74 | 0.264 | 0.139 | |
| AT1G72190.1 | Phosphoglycerate dehydrogenase, putative | 44 | 3.91 | 0.267 | 0.294 | |
| AT1G72190.1 | Phosphoglycerate dehydrogenase, putative | 45 | 1.51 | 0.270 | 1.000 | |
| AT1G72190.1 | Phosphoglycerate dehydrogenase, putative | 46 | 0.42 | 0.296 | 0.911 | |
| AT1G72190.1 | Phosphoglycerate dehydrogenase, putative | 47 | 0.11 | 0.422 | 0.285 | |
| AT1G72750.1 | Translocase of the inner mitochondrial membrane 17 (TIM17-2) | 2 | 1.16 | 0.311 | 1.000 | |
| AT1G72750.1 | Translocase of the inner mitochondrial membrane 17 (TIM17-2) | 3 | 0.98 | 0.202 | 1.000 | |
| AT1G72750.1 | Translocase of the inner mitochondrial membrane 17 (TIM17-2) | 4 | 0.84 | 0.192 | 1.000 | |
| AT1G72750.1 | Translocase of the inner mitochondrial membrane 17 (TIM17-2) | 5 | 0.72 | 0.188 | 1.000 | |

| | | | | | |
|---|---|---|---|---|---|
| AT1G72750.1 | Translocase of the inner mitochondrial membrane 17 (TIM17-2) | 6 | 0.57 | 0.180 | 0.995 |
| AT1G72750.1 | Translocase of the inner mitochondrial membrane 17 (TIM17-2) | 7 | 0.48 | 0.179 | 0.885 |
| AT1G72750.1 | Translocase of the inner mitochondrial membrane 17 (TIM17-2) | 8 | 0.60 | 0.173 | 0.998 |
| AT1G72750.1 | Translocase of the inner mitochondrial membrane 17 (TIM17-2) | 9 | 0.88 | 0.170 | 1.000 |
| AT1G72750.1 | Translocase of the inner mitochondrial membrane 17 (TIM17-2) | 10 | 1.04 | 0.170 | 1.000 |
| AT1G72750.1 | Translocase of the inner mitochondrial membrane 17 (TIM17-2) | 11 | 1.12 | 0.170 | 1.000 |
| AT1G72750.1 | Translocase of the inner mitochondrial membrane 17 (TIM17-2) | 12 | 1.21 | 0.171 | 1.000 |
| AT1G72750.1 | Translocase of the inner mitochondrial membrane 17 (TIM17-2) | 13 | 1.24 | 0.169 | 1.000 |
| AT1G72750.1 | Translocase of the inner mitochondrial membrane 17 (TIM17-2) | 14 | 1.22 | 0.166 | 1.000 |
| AT1G72750.1 | Translocase of the inner mitochondrial membrane 17 (TIM17-2) | 15 | 1.09 | 0.166 | 1.000 |
| AT1G72750.1 | Translocase of the inner mitochondrial membrane 17 (TIM17-2) | 16 | 0.93 | 0.164 | 1.000 |
| AT1G72750.1 | Translocase of the inner mitochondrial membrane 17 (TIM17-2) | 17 | 0.97 | 0.164 | 1.000 |
| AT1G72750.1 | Translocase of the inner mitochondrial membrane 17 (TIM17-2) | 18 | 1.10 | 0.167 | 1.000 |
| AT1G72750.1 | Translocase of the inner mitochondrial membrane 17 (TIM17-2) | 19 | 1.05 | 0.171 | 1.000 |
| AT1G72750.1 | Translocase of the inner mitochondrial membrane 17 (TIM17-2) | 20 | 0.91 | 0.175 | 1.000 |
| AT1G72750.1 | Translocase of the inner mitochondrial membrane 17 (TIM17-2) | 21 | 0.98 | 0.178 | 1.000 |
| AT1G72750.1 | Translocase of the inner mitochondrial membrane 17 (TIM17-2) | 22 | 1.41 | 0.177 | 1.000 |
| AT1G72750.1 | Translocase of the inner mitochondrial membrane 17 (TIM17-2) | 23 | 1.99 | 0.177 | 0.930 |
| AT1G72750.1 | Translocase of the inner mitochondrial membrane 17 (TIM17-2) | 24 | 1.67 | 0.173 | 0.998 |
| AT1G72750.1 | Translocase of the inner mitochondrial membrane 17 (TIM17-2) | 25 | 0.83 | 0.170 | 1.000 |

| | | | | | |
|---|---|---|---|---|---|
| AT1G72750.1 | Translocase of the inner mitochondrial membrane 17 (TIM17-2) | 26 | 0.61 | 0.169 | 0.998 |
| AT1G72750.1 | Translocase of the inner mitochondrial membrane 17 (TIM17-2) | 27 | 0.68 | 0.169 | 1.000 |
| AT1G72750.1 | Translocase of the inner mitochondrial membrane 17 (TIM17-2) | 28 | 1.02 | 0.169 | 1.000 |
| AT1G72750.1 | Translocase of the inner mitochondrial membrane 17 (TIM17-2) | 29 | 1.60 | 0.169 | 0.999 |
| AT1G72750.1 | Translocase of the inner mitochondrial membrane 17 (TIM17-2) | 30 | 1.44 | 0.168 | 1.000 |
| AT1G72750.1 | Translocase of the inner mitochondrial membrane 17 (TIM17-2) | 31 | 1.12 | 0.169 | 1.000 |
| AT1G72750.1 | Translocase of the inner mitochondrial membrane 17 (TIM17-2) | 32 | 1.10 | 0.170 | 1.000 |
| AT1G72750.1 | Translocase of the inner mitochondrial membrane 17 (TIM17-2) | 33 | 1.31 | 0.170 | 1.000 |
| AT1G72750.1 | Translocase of the inner mitochondrial membrane 17 (TIM17-2) | 34 | 1.45 | 0.170 | 1.000 |
| AT1G72750.1 | Translocase of the inner mitochondrial membrane 17 (TIM17-2) | 35 | 1.51 | 0.170 | 1.000 |
| AT1G72750.1 | Translocase of the inner mitochondrial membrane 17 (TIM17-2) | 36 | 1.65 | 0.169 | 0.998 |
| AT1G72750.1 | Translocase of the inner mitochondrial membrane 17 (TIM17-2) | 37 | 1.73 | 0.169 | 0.991 |
| AT1G72750.1 | Translocase of the inner mitochondrial membrane 17 (TIM17-2) | 38 | 1.71 | 0.170 | 0.994 |
| AT1G72750.1 | Translocase of the inner mitochondrial membrane 17 (TIM17-2) | 39 | 1.45 | 0.170 | 1.000 |
| AT1G72750.1 | Translocase of the inner mitochondrial membrane 17 (TIM17-2) | 40 | 1.02 | 0.171 | 1.000 |
| AT1G72750.1 | Translocase of the inner mitochondrial membrane 17 (TIM17-2) | 41 | 0.61 | 0.171 | 0.999 |
| AT1G72750.1 | Translocase of the inner mitochondrial membrane 17 (TIM17-2) | 42 | 0.40 | 0.173 | 0.506 |
| AT1G72750.1 | Translocase of the inner mitochondrial membrane 17 (TIM17-2) | 43 | 0.35 | 0.175 | 0.281 |
| AT1G72750.1 | Translocase of the inner mitochondrial membrane 17 (TIM17-2) | 44 | 0.45 | 0.176 | 0.755 |
| AT1G72750.1 | Translocase of the inner mitochondrial membrane 17 (TIM17-2) | 45 | 0.63 | 0.208 | 1.000 |

| | | | | | | |
|---|---|---|---|---|---|---|
| AT1G72750.1 | Translocase of the inner mitochondrial membrane 17 (TIM17-2) | 46 | 0.89 | 0.306 | 1.000 | |
| AT1G72750.1 | Translocase of the inner mitochondrial membrane 17 (TIM17-2) | 47 | 1.27 | 0.534 | 1.000 | |
| AT1G72820.1 | Mitochondrial substrate carrier family protein | 2 | 1.53 | 0.697 | 1.000 | |
| AT1G72820.1 | Mitochondrial substrate carrier family protein | 3 | 1.44 | 0.717 | 1.000 | |
| AT1G72820.1 | Mitochondrial substrate carrier family protein | 34 | 1.28 | 0.412 | 1.000 | |
| AT1G72820.1 | Mitochondrial substrate carrier family protein | 35 | 1.44 | 0.359 | 0.998 | |
| AT1G72820.1 | Mitochondrial substrate carrier family protein | 36 | 1.58 | 0.333 | 0.986 | |
| AT1G72820.1 | Mitochondrial substrate carrier family protein | 37 | 1.67 | 0.325 | 0.971 | |
| AT1G72820.1 | Mitochondrial substrate carrier family protein | 38 | 1.73 | 0.322 | 0.960 | |
| AT1G72820.1 | Mitochondrial substrate carrier family protein | 39 | 1.72 | 0.317 | 0.959 | |
| AT1G72820.1 | Mitochondrial substrate carrier family protein | 40 | 1.59 | 0.314 | 0.980 | |
| AT1G72820.1 | Mitochondrial substrate carrier family protein | 41 | 1.26 | 0.318 | 1.000 | |
| AT1G72820.1 | Mitochondrial substrate carrier family protein | 42 | 0.82 | 0.332 | 1.000 | |
| AT1G72820.1 | Mitochondrial substrate carrier family protein | 43 | 0.45 | 0.357 | 0.870 | |
| AT1G72820.1 | Mitochondrial substrate carrier family protein | 44 | 0.23 | 0.387 | 0.420 | |
| AT1G72820.1 | Mitochondrial substrate carrier family protein | 46 | 0.05 | 0.463 | 0.038 | * |
| AT1G72820.1 | Mitochondrial substrate carrier family protein | 47 | 0.02 | 0.534 | 0.019 | * |
| AT1G73980.1 | Triphosphate tunnel metalloenzyme 1 (TTM1) | 2 | 0.59 | 0.567 | 1.000 | |
| AT1G73980.1 | Triphosphate tunnel metalloenzyme 1 (TTM1) | 3 | 0.88 | 0.454 | 1.000 | |
| AT1G73980.1 | Triphosphate tunnel metalloenzyme 1 (TTM1) | 6 | 3.08 | 0.302 | 0.853 | |
| AT1G73980.1 | Triphosphate tunnel metalloenzyme 1 (TTM1) | 7 | 4.76 | 0.265 | 0.200 | |
| AT1G73980.1 | Triphosphate tunnel metalloenzyme 1 (TTM1) | 8 | 7.36 | 0.235 | 0.007 | ** |
| AT1G73980.1 | Triphosphate tunnel metalloenzyme 1 (TTM1) | 9 | 11.13 | 0.215 | 0.000 | *** |
| AT1G73980.1 | Triphosphate tunnel metalloenzyme 1 (TTM1) | 10 | 15.64 | 0.204 | 0.000 | *** |
| AT1G73980.1 | Triphosphate tunnel metalloenzyme 1 (TTM1) | 11 | 19.32 | 0.200 | 0.000 | *** |
| AT1G73980.1 | Triphosphate tunnel metalloenzyme 1 (TTM1) | 12 | 20.82 | 0.198 | 0.000 | *** |
| AT1G73980.1 | Triphosphate tunnel metalloenzyme 1 (TTM1) | 13 | 20.28 | 0.196 | 0.000 | *** |
| AT1G73980.1 | Triphosphate tunnel metalloenzyme 1 (TTM1) | 14 | 18.58 | 0.195 | 0.000 | *** |
| AT1G73980.1 | Triphosphate tunnel metalloenzyme 1 (TTM1) | 15 | 16.24 | 0.193 | 0.000 | *** |
| AT1G73980.1 | Triphosphate tunnel metalloenzyme 1 (TTM1) | 16 | 13.27 | 0.192 | 0.000 | *** |

| | | | | | | |
|---|---|---|---|---|---|---|
| AT1G73980.1 | Triphosphate tunnel metalloenzyme 1 (TTM1) | 17 | 10.07 | 0.193 | 0.000 | *** |
| AT1G73980.1 | Triphosphate tunnel metalloenzyme 1 (TTM1) | 18 | 7.24 | 0.195 | 0.000 | *** |
| AT1G73980.1 | Triphosphate tunnel metalloenzyme 1 (TTM1) | 19 | 4.99 | 0.199 | 0.012 | * |
| AT1G73980.1 | Triphosphate tunnel metalloenzyme 1 (TTM1) | 20 | 3.33 | 0.202 | 0.191 | |
| AT1G73980.1 | Triphosphate tunnel metalloenzyme 1 (TTM1) | 21 | 2.18 | 0.204 | 0.828 | |
| AT1G73980.1 | Triphosphate tunnel metalloenzyme 1 (TTM1) | 22 | 1.44 | 0.204 | 1.000 | |
| AT1G73980.1 | Triphosphate tunnel metalloenzyme 1 (TTM1) | 23 | 1.00 | 0.203 | 1.000 | |
| AT1G73980.1 | Triphosphate tunnel metalloenzyme 1 (TTM1) | 24 | 0.79 | 0.201 | 1.000 | |
| AT1G73980.1 | Triphosphate tunnel metalloenzyme 1 (TTM1) | 25 | 0.69 | 0.199 | 1.000 | |
| AT1G73980.1 | Triphosphate tunnel metalloenzyme 1 (TTM1) | 26 | 0.65 | 0.198 | 0.999 | |
| AT1G73980.1 | Triphosphate tunnel metalloenzyme 1 (TTM1) | 27 | 0.61 | 0.197 | 0.995 | |
| AT1G73980.1 | Triphosphate tunnel metalloenzyme 1 (TTM1) | 28 | 0.59 | 0.197 | 0.989 | |
| AT1G73980.1 | Triphosphate tunnel metalloenzyme 1 (TTM1) | 29 | 0.61 | 0.197 | 0.995 | |
| AT1G73980.1 | Triphosphate tunnel metalloenzyme 1 (TTM1) | 30 | 0.73 | 0.198 | 1.000 | |
| AT1G73980.1 | Triphosphate tunnel metalloenzyme 1 (TTM1) | 31 | 1.01 | 0.199 | 1.000 | |
| AT1G73980.1 | Triphosphate tunnel metalloenzyme 1 (TTM1) | 32 | 1.54 | 0.203 | 0.999 | |
| AT1G73980.1 | Triphosphate tunnel metalloenzyme 1 (TTM1) | 33 | 2.34 | 0.214 | 0.785 | |
| AT1G73980.1 | Triphosphate tunnel metalloenzyme 1 (TTM1) | 34 | 3.42 | 0.234 | 0.362 | |
| AT1G73980.1 | Triphosphate tunnel metalloenzyme 1 (TTM1) | 35 | 4.77 | 0.267 | 0.206 | |
| AT1G73980.1 | Triphosphate tunnel metalloenzyme 1 (TTM1) | 36 | 6.35 | 0.301 | 0.156 | |
| AT1G73980.1 | Triphosphate tunnel metalloenzyme 1 (TTM1) | 41 | 7.03 | 0.322 | 0.171 | |
| AT1G73980.1 | Triphosphate tunnel metalloenzyme 1 (TTM1) | 42 | 4.99 | 0.281 | 0.238 | |
| AT1G73980.1 | Triphosphate tunnel metalloenzyme 1 (TTM1) | 43 | 3.11 | 0.243 | 0.555 | |
| AT1G73980.1 | Triphosphate tunnel metalloenzyme 1 (TTM1) | 44 | 1.72 | 0.235 | 0.998 | |
| AT1G73980.1 | Triphosphate tunnel metalloenzyme 1 (TTM1) | 45 | 0.86 | 0.285 | 1.000 | |
| AT1G73980.1 | Triphosphate tunnel metalloenzyme 1 (TTM1) | 46 | 0.40 | 0.404 | 0.998 | |
| AT1G73980.1 | Triphosphate tunnel metalloenzyme 1 (TTM1) | 47 | 0.18 | 0.580 | 0.975 | |
| AT1G73990.1 | Signal peptide petidase (SPPA) | 2 | 1.06 | 0.523 | 1.000 | |
| AT1G73990.1 | Signal peptide petidase (SPPA) | 3 | 0.54 | 0.608 | 1.000 | |
| AT1G73990.1 | Signal peptide petidase (SPPA) | 20 | 2.82 | 0.360 | 0.946 | |
| AT1G73990.1 | Signal peptide petidase (SPPA) | 21 | 3.01 | 0.289 | 0.738 | |
| AT1G73990.1 | Signal peptide petidase (SPPA) | 22 | 2.38 | 0.274 | 0.900 | |
| AT1G73990.1 | Signal peptide petidase (SPPA) | 23 | 1.36 | 0.290 | 1.000 | |
| AT1G73990.1 | Signal peptide petidase (SPPA) | 24 | 0.64 | 0.318 | 1.000 | |
| AT1G73990.1 | Signal peptide petidase (SPPA) | 25 | 0.36 | 0.308 | 0.879 | |

| | | | | | |
|---|---|---|---|---|---|
| AT1G73990.1 | Signal peptide petidase (SPPA) | 26 | 0.43 | 0.271 | 0.907 |
| AT1G73990.1 | Signal peptide petidase (SPPA) | 27 | 0.92 | 0.247 | 1.000 |
| AT1G73990.1 | Signal peptide petidase (SPPA) | 28 | 1.87 | 0.235 | 0.969 |
| AT1G73990.1 | Signal peptide petidase (SPPA) | 29 | 2.30 | 0.235 | 0.815 |
| AT1G73990.1 | Signal peptide petidase (SPPA) | 30 | 1.36 | 0.241 | 1.000 |
| AT1G73990.1 | Signal peptide petidase (SPPA) | 31 | 0.42 | 0.257 | 0.867 |
| AT1G73990.1 | Signal peptide petidase (SPPA) | 32 | 0.14 | 0.320 | 0.134 |
| AT1G73990.1 | Signal peptide petidase (SPPA) | 33 | 0.10 | 0.560 | 0.627 |
| AT1G73990.1 | Signal peptide petidase (SPPA) | 46 | ########### | 8.198 | 0.985 |
| AT1G73990.1 | Signal peptide petidase (SPPA) | 47 | ########### | 9.123 | 0.983 |
| AT1G74060.1 | 60S ribosomal protein L6-2 (RPL6B) | 2 | 0.66 | 0.289 | 0.999 |
| AT1G74060.1 | 60S ribosomal protein L6-2 (RPL6B) | 3 | 0.76 | 0.226 | 1.000 |
| AT1G74060.1 | 60S ribosomal protein L6-2 (RPL6B) | 4 | 0.88 | 0.191 | 1.000 |
| AT1G74060.1 | 60S ribosomal protein L6-2 (RPL6B) | 5 | 1.05 | 0.174 | 1.000 |
| AT1G74060.1 | 60S ribosomal protein L6-2 (RPL6B) | 6 | 1.26 | 0.168 | 0.999 |
| AT1G74060.1 | 60S ribosomal protein L6-2 (RPL6B) | 7 | 1.47 | 0.166 | 0.968 |
| AT1G74060.1 | 60S ribosomal protein L6-2 (RPL6B) | 8 | 1.61 | 0.164 | 0.885 |
| AT1G74060.1 | 60S ribosomal protein L6-2 (RPL6B) | 9 | 1.64 | 0.164 | 0.863 |
| AT1G74060.1 | 60S ribosomal protein L6-2 (RPL6B) | 10 | 1.52 | 0.163 | 0.941 |
| AT1G74060.1 | 60S ribosomal protein L6-2 (RPL6B) | 11 | 1.30 | 0.161 | 0.997 |
| AT1G74060.1 | 60S ribosomal protein L6-2 (RPL6B) | 12 | 1.07 | 0.159 | 1.000 |
| AT1G74060.1 | 60S ribosomal protein L6-2 (RPL6B) | 13 | 0.88 | 0.157 | 1.000 |
| AT1G74060.1 | 60S ribosomal protein L6-2 (RPL6B) | 14 | 0.77 | 0.156 | 0.997 |
| AT1G74060.1 | 60S ribosomal protein L6-2 (RPL6B) | 15 | 0.73 | 0.157 | 0.987 |
| AT1G74060.1 | 60S ribosomal protein L6-2 (RPL6B) | 16 | 0.77 | 0.160 | 0.997 |
| AT1G74060.1 | 60S ribosomal protein L6-2 (RPL6B) | 17 | 0.90 | 0.166 | 1.000 |
| AT1G74060.1 | 60S ribosomal protein L6-2 (RPL6B) | 18 | 1.13 | 0.176 | 1.000 |
| AT1G74060.1 | 60S ribosomal protein L6-2 (RPL6B) | 19 | 1.41 | 0.189 | 0.994 |
| AT1G74060.1 | 60S ribosomal protein L6-2 (RPL6B) | 20 | 1.72 | 0.206 | 0.932 |
| AT1G74060.1 | 60S ribosomal protein L6-2 (RPL6B) | 21 | 2.03 | 0.225 | 0.833 |
| AT1G74060.1 | 60S ribosomal protein L6-2 (RPL6B) | 22 | 2.31 | 0.250 | 0.781 |
| AT1G74060.1 | 60S ribosomal protein L6-2 (RPL6B) | 23 | 2.54 | 0.280 | 0.780 |
| AT1G74060.1 | 60S ribosomal protein L6-2 (RPL6B) | 24 | 2.73 | 0.317 | 0.824 |
| AT1G74060.1 | 60S ribosomal protein L6-2 (RPL6B) | 41 | 1.23 | 0.406 | 1.000 |
| AT1G74060.1 | 60S ribosomal protein L6-2 (RPL6B) | 42 | 1.12 | 0.404 | 1.000 |
| AT1G74060.1 | 60S ribosomal protein L6-2 (RPL6B) | 43 | 1.01 | 0.443 | 1.000 |

| | | | | | |
|---|---|---|---|---|---|
| AT1G74060.1 | 60S ribosomal protein L6-2 (RPL6B) | 46 | 0.76 | 0.811 | 1.000 |
| AT1G74060.1 | 60S ribosomal protein L6-2 (RPL6B) | 47 | 0.70 | 0.981 | 1.000 |
| AT1G74230.1 | Organelle RNA recognition motif-containing 4 (ORRM4) | 2 | 0.86 | 0.473 | 1.000 |
| AT1G74230.1 | Organelle RNA recognition motif-containing 4 (ORRM4) | 3 | 0.91 | 0.456 | 1.000 |
| AT1G74230.1 | Organelle RNA recognition motif-containing 4 (ORRM4) | 40 | 2.11 | 0.296 | 0.745 |
| AT1G74230.1 | Organelle RNA recognition motif-containing 4 (ORRM4) | 41 | 2.03 | 0.232 | 0.597 |
| AT1G74230.1 | Organelle RNA recognition motif-containing 4 (ORRM4) | 42 | 1.93 | 0.185 | 0.448 |
| AT1G74230.1 | Organelle RNA recognition motif-containing 4 (ORRM4) | 43 | 1.86 | 0.158 | 0.353 |
| AT1G74230.1 | Organelle RNA recognition motif-containing 4 (ORRM4) | 44 | 1.80 | 0.152 | 0.369 |
| AT1G74230.1 | Organelle RNA recognition motif-containing 4 (ORRM4) | 45 | 1.76 | 0.172 | 0.529 |
| AT1G74230.1 | Organelle RNA recognition motif-containing 4 (ORRM4) | 46 | 1.74 | 0.222 | 0.756 |
| AT1G74230.1 | Organelle RNA recognition motif-containing 4 (ORRM4) | 47 | 1.72 | 0.301 | 0.912 |
| AT1G74470.1 | Geranylgeranyl diphosphate reductase (CHLP) | 2 | 0.76 | 0.749 | 1.000 |
| AT1G74470.1 | Geranylgeranyl diphosphate reductase (CHLP) | 3 | 0.88 | 0.716 | 1.000 |
| AT1G74470.1 | Geranylgeranyl diphosphate reductase (CHLP) | 24 | 0.22 | 0.591 | 0.955 |
| AT1G74470.1 | Geranylgeranyl diphosphate reductase (CHLP) | 25 | 0.14 | 0.462 | 0.523 |
| AT1G74470.1 | Geranylgeranyl diphosphate reductase (CHLP) | 26 | 0.11 | 0.414 | 0.237 |
| AT1G74470.1 | Geranylgeranyl diphosphate reductase (CHLP) | 27 | 0.13 | 0.401 | 0.269 |
| AT1G74470.1 | Geranylgeranyl diphosphate reductase (CHLP) | 28 | 0.21 | 0.397 | 0.618 |
| AT1G74470.1 | Geranylgeranyl diphosphate reductase (CHLP) | 29 | 0.38 | 0.396 | 0.970 |
| AT1G74470.1 | Geranylgeranyl diphosphate reductase (CHLP) | 30 | 0.63 | 0.396 | 1.000 |
| AT1G74470.1 | Geranylgeranyl diphosphate reductase (CHLP) | 31 | 0.98 | 0.397 | 1.000 |
| AT1G74470.1 | Geranylgeranyl diphosphate reductase (CHLP) | 32 | 1.52 | 0.398 | 1.000 |
| AT1G74470.1 | Geranylgeranyl diphosphate reductase (CHLP) | 33 | 1.87 | 0.398 | 0.999 |
| AT1G74470.1 | Geranylgeranyl diphosphate reductase (CHLP) | 34 | 1.17 | 0.399 | 1.000 |
| AT1G74470.1 | Geranylgeranyl diphosphate reductase (CHLP) | 35 | 0.45 | 0.401 | 0.995 |
| AT1G74470.1 | Geranylgeranyl diphosphate reductase (CHLP) | 36 | 0.24 | 0.407 | 0.754 |
| AT1G74470.1 | Geranylgeranyl diphosphate reductase (CHLP) | 37 | 0.24 | 0.415 | 0.754 |
| AT1G74470.1 | Geranylgeranyl diphosphate reductase (CHLP) | 38 | 0.34 | 0.420 | 0.955 |
| AT1G74470.1 | Geranylgeranyl diphosphate reductase (CHLP) | 39 | 0.48 | 0.416 | 0.998 |
| AT1G74470.1 | Geranylgeranyl diphosphate reductase (CHLP) | 40 | 0.44 | 0.405 | 0.993 |

| | | | | | |
|---|---|---|---|---|---|
| AT1G74470.1 | Geranylgeranyl diphosphate reductase (CHLP) | 41 | 0.26 | 0.400 | 0.794 |
| AT1G74470.1 | Geranylgeranyl diphosphate reductase (CHLP) | 42 | 0.14 | 0.402 | 0.298 |
| AT1G74470.1 | Geranylgeranyl diphosphate reductase (CHLP) | 43 | 0.09 | 0.404 | 0.116 |
| AT1G74470.1 | Geranylgeranyl diphosphate reductase (CHLP) | 44 | 0.09 | 0.411 | 0.131 |
| AT1G74470.1 | Geranylgeranyl diphosphate reductase (CHLP) | 45 | 0.13 | 0.458 | 0.420 |
| AT1G74470.1 | Geranylgeranyl diphosphate reductase (CHLP) | 46 | 0.21 | 0.599 | 0.945 |
| AT1G74470.1 | Geranylgeranyl diphosphate reductase (CHLP) | 47 | 0.35 | 0.899 | 1.000 |
| AT1G74970.1 | 30S ribosomal protein S9 (RPS9) | 2 | 0.89 | 0.420 | 1.000 |
| AT1G74970.1 | 30S ribosomal protein S9 (RPS9) | 3 | 0.98 | 0.392 | 1.000 |
| AT1G74970.1 | 30S ribosomal protein S9 (RPS9) | 4 | 1.07 | 0.370 | 1.000 |
| AT1G74970.1 | 30S ribosomal protein S9 (RPS9) | 17 | 2.84 | 0.220 | 0.161 |
| AT1G74970.1 | 30S ribosomal protein S9 (RPS9) | 18 | 2.94 | 0.218 | 0.136 |
| AT1G74970.1 | 30S ribosomal protein S9 (RPS9) | 19 | 2.98 | 0.221 | 0.135 |
| AT1G74970.1 | 30S ribosomal protein S9 (RPS9) | 20 | 2.98 | 0.230 | 0.161 |
| AT1G74970.1 | 30S ribosomal protein S9 (RPS9) | 21 | 2.92 | 0.243 | 0.215 |
| AT1G74970.1 | 30S ribosomal protein S9 (RPS9) | 22 | 2.81 | 0.261 | 0.302 |
| AT1G74970.1 | 30S ribosomal protein S9 (RPS9) | 23 | 2.67 | 0.282 | 0.417 |
| AT1G74970.1 | 30S ribosomal protein S9 (RPS9) | 24 | 2.51 | 0.306 | 0.546 |
| AT1G74970.1 | 30S ribosomal protein S9 (RPS9) | 46 | 0.14 | 0.654 | 0.543 |
| AT1G74970.1 | 30S ribosomal protein S9 (RPS9) | 47 | 0.12 | 0.678 | 0.502 |
| AT1G75200.1 | Flavodoxin family protein / radical SAM domain-containing protein | 2 | 1.04 | 0.387 | 1.000 |
| AT1G75200.1 | Flavodoxin family protein / radical SAM domain-containing protein | 3 | 0.87 | 0.303 | 1.000 |
| AT1G75200.1 | Flavodoxin family protein / radical SAM domain-containing protein | 4 | 0.75 | 0.418 | 1.000 |
| AT1G75200.1 | Flavodoxin family protein / radical SAM domain-containing protein | 21 | 2.00 | 0.293 | 0.996 |
| AT1G75200.1 | Flavodoxin family protein / radical SAM domain-containing protein | 22 | 2.38 | 0.235 | 0.846 |
| AT1G75200.1 | Flavodoxin family protein / radical SAM domain-containing protein | 23 | 2.77 | 0.199 | 0.392 |
| AT1G75200.1 | Flavodoxin family protein / radical SAM domain-containing protein | 24 | 2.99 | 0.181 | 0.164 |

| | | | | | | |
|---|---|---|---|---|---|---|
| AT1G75200.1 | Flavodoxin family protein / radical SAM domain-containing protein | 25 | 2.93 | 0.174 | 0.141 | |
| AT1G75200.1 | Flavodoxin family protein / radical SAM domain-containing protein | 26 | 2.68 | 0.172 | 0.224 | |
| AT1G75200.1 | Flavodoxin family protein / radical SAM domain-containing protein | 27 | 2.32 | 0.172 | 0.457 | |
| AT1G75200.1 | Flavodoxin family protein / radical SAM domain-containing protein | 28 | 1.93 | 0.172 | 0.811 | |
| AT1G75200.1 | Flavodoxin family protein / radical SAM domain-containing protein | 29 | 1.60 | 0.172 | 0.984 | |
| AT1G75200.1 | Flavodoxin family protein / radical SAM domain-containing protein | 30 | 1.31 | 0.172 | 1.000 | |
| AT1G75200.1 | Flavodoxin family protein / radical SAM domain-containing protein | 31 | 1.07 | 0.173 | 1.000 | |
| AT1G75200.1 | Flavodoxin family protein / radical SAM domain-containing protein | 32 | 0.89 | 0.173 | 1.000 | |
| AT1G75200.1 | Flavodoxin family protein / radical SAM domain-containing protein | 33 | 0.75 | 0.173 | 1.000 | |
| AT1G75200.1 | Flavodoxin family protein / radical SAM domain-containing protein | 34 | 0.67 | 0.174 | 0.997 | |
| AT1G75200.1 | Flavodoxin family protein / radical SAM domain-containing protein | 35 | 0.67 | 0.174 | 0.997 | |
| AT1G75200.1 | Flavodoxin family protein / radical SAM domain-containing protein | 36 | 0.76 | 0.173 | 1.000 | |
| AT1G75200.1 | Flavodoxin family protein / radical SAM domain-containing protein | 37 | 1.03 | 0.173 | 1.000 | |
| AT1G75200.1 | Flavodoxin family protein / radical SAM domain-containing protein | 38 | 1.45 | 0.173 | 0.999 | |
| AT1G75200.1 | Flavodoxin family protein / radical SAM domain-containing protein | 39 | 1.76 | 0.173 | 0.931 | |
| AT1G75200.1 | Flavodoxin family protein / radical SAM domain-containing protein | 40 | 1.69 | 0.173 | 0.963 | |
| AT1G75200.1 | Flavodoxin family protein / radical SAM domain-containing protein | 41 | 1.21 | 0.175 | 1.000 | |
| AT1G75200.1 | Flavodoxin family protein / radical SAM domain-containing protein | 42 | 0.70 | 0.178 | 0.999 | |
| AT1G75200.1 | Flavodoxin family protein / radical SAM domain-containing protein | 43 | 0.42 | 0.180 | 0.470 | |
| AT1G75200.1 | Flavodoxin family protein / radical SAM domain-containing protein | 44 | 0.28 | 0.189 | 0.070 | . |

| | | | | | | |
|---|---|---|---|---|---|---|
| AT1G75200.1 | Flavodoxin family protein / radical SAM domain-containing protein | 45 | 0.22 | 0.231 | 0.096 | . |
| AT1G75200.1 | Flavodoxin family protein / radical SAM domain-containing protein | 46 | 0.20 | 0.334 | 0.474 | |
| AT1G75200.1 | Flavodoxin family protein / radical SAM domain-containing protein | 47 | 0.19 | 0.511 | 0.927 | |
| AT1G75210.1 | Cytosolic IMP-GMP specific 5-nucleotidase, putative | 2 | 1.19 | 0.349 | 1.000 | |
| AT1G75210.1 | Cytosolic IMP-GMP specific 5-nucleotidase, putative | 3 | 1.47 | 0.350 | 1.000 | |
| AT1G75210.1 | Cytosolic IMP-GMP specific 5-nucleotidase, putative | 32 | 3.15 | 0.265 | 0.465 | |
| AT1G75210.1 | Cytosolic IMP-GMP specific 5-nucleotidase, putative | 33 | 2.17 | 0.212 | 0.681 | |
| AT1G75210.1 | Cytosolic IMP-GMP specific 5-nucleotidase, putative | 34 | 1.39 | 0.194 | 0.997 | |
| AT1G75210.1 | Cytosolic IMP-GMP specific 5-nucleotidase, putative | 35 | 0.91 | 0.184 | 1.000 | |
| AT1G75210.1 | Cytosolic IMP-GMP specific 5-nucleotidase, putative | 36 | 0.67 | 0.187 | 0.984 | |
| AT1G75210.1 | Cytosolic IMP-GMP specific 5-nucleotidase, putative | 37 | 0.56 | 0.199 | 0.884 | |
| AT1G75210.1 | Cytosolic IMP-GMP specific 5-nucleotidase, putative | 38 | 0.50 | 0.188 | 0.671 | |
| AT1G75210.1 | Cytosolic IMP-GMP specific 5-nucleotidase, putative | 39 | 0.48 | 0.173 | 0.484 | |
| AT1G75210.1 | Cytosolic IMP-GMP specific 5-nucleotidase, putative | 40 | 0.48 | 0.163 | 0.419 | |
| AT1G75210.1 | Cytosolic IMP-GMP specific 5-nucleotidase, putative | 41 | 0.50 | 0.162 | 0.488 | |
| AT1G75210.1 | Cytosolic IMP-GMP specific 5-nucleotidase, putative | 42 | 0.67 | 0.161 | 0.949 | |
| AT1G75210.1 | Cytosolic IMP-GMP specific 5-nucleotidase, putative | 43 | 1.10 | 0.161 | 1.000 | |
| AT1G75210.1 | Cytosolic IMP-GMP specific 5-nucleotidase, putative | 44 | 1.45 | 0.164 | 0.974 | |
| AT1G75210.1 | Cytosolic IMP-GMP specific 5-nucleotidase, putative | 45 | 1.00 | 0.169 | 1.000 | |
| AT1G75210.1 | Cytosolic IMP-GMP specific 5-nucleotidase, putative | 46 | 0.34 | 0.185 | 0.131 | |
| AT1G75210.1 | Cytosolic IMP-GMP specific 5-nucleotidase, putative | 47 | 0.08 | 0.272 | 0.001 | ** |
| AT1G76030.1 | Vacuolar ATP synthase subunit B1 (VHA-B1) | 2 | 1.59 | 0.403 | 0.996 | |
| AT1G76030.1 | Vacuolar ATP synthase subunit B1 (VHA-B1) | 3 | 1.75 | 0.343 | 0.974 | |
| AT1G76030.1 | Vacuolar ATP synthase subunit B1 (VHA-B1) | 4 | 1.94 | 0.304 | 0.907 | |
| AT1G76030.1 | Vacuolar ATP synthase subunit B1 (VHA-B1) | 5 | 2.13 | 0.287 | 0.809 | |
| AT1G76030.1 | Vacuolar ATP synthase subunit B1 (VHA-B1) | 6 | 2.31 | 0.289 | 0.740 | |
| AT1G76030.1 | Vacuolar ATP synthase subunit B1 (VHA-B1) | 7 | 2.46 | 0.306 | 0.726 | |
| AT1G76030.1 | Vacuolar ATP synthase subunit B1 (VHA-B1) | 8 | 2.55 | 0.329 | 0.753 | |
| AT1G76030.1 | Vacuolar ATP synthase subunit B1 (VHA-B1) | 21 | 1.19 | 0.322 | 1.000 | |

| | | | | | |
|---|---|---|---|---|---|
| AT1G76030.1 | Vacuolar ATP synthase subunit B1 (VHA-B1) | 22 | 1.13 | 0.296 | 1.000 |
| AT1G76030.1 | Vacuolar ATP synthase subunit B1 (VHA-B1) | 23 | 1.08 | 0.273 | 1.000 |
| AT1G76030.1 | Vacuolar ATP synthase subunit B1 (VHA-B1) | 24 | 1.06 | 0.255 | 1.000 |
| AT1G76030.1 | Vacuolar ATP synthase subunit B1 (VHA-B1) | 25 | 1.05 | 0.245 | 1.000 |
| AT1G76030.1 | Vacuolar ATP synthase subunit B1 (VHA-B1) | 26 | 1.05 | 0.245 | 1.000 |
| AT1G76030.1 | Vacuolar ATP synthase subunit B1 (VHA-B1) | 27 | 1.06 | 0.257 | 1.000 |
| AT1G76030.1 | Vacuolar ATP synthase subunit B1 (VHA-B1) | 28 | 1.08 | 0.281 | 1.000 |
| AT1G76030.1 | Vacuolar ATP synthase subunit B1 (VHA-B1) | 29 | 1.09 | 0.318 | 1.000 |
| AT1G76030.1 | Vacuolar ATP synthase subunit B1 (VHA-B1) | 30 | 1.12 | 0.366 | 1.000 |
| AT1G76030.1 | Vacuolar ATP synthase subunit B1 (VHA-B1) | 46 | 1.18 | 1.945 | 1.000 |
| AT1G76030.1 | Vacuolar ATP synthase subunit B1 (VHA-B1) | 47 | 1.17 | 2.084 | 1.000 |
| AT1G76200.1 | NADH:ubiquinone oxidoreductase subunit (AGGG) | 2 | 0.94 | 0.521 | 1.000 |
| AT1G76200.1 | NADH:ubiquinone oxidoreductase subunit (AGGG) | 3 | 0.84 | 0.479 | 1.000 |
| AT1G76200.1 | NADH:ubiquinone oxidoreductase subunit (AGGG) | 14 | 1.25 | 0.324 | 1.000 |
| AT1G76200.1 | NADH:ubiquinone oxidoreductase subunit (AGGG) | 15 | 1.18 | 0.282 | 1.000 |
| AT1G76200.1 | NADH:ubiquinone oxidoreductase subunit (AGGG) | 16 | 0.81 | 0.262 | 1.000 |
| AT1G76200.1 | NADH:ubiquinone oxidoreductase subunit (AGGG) | 17 | 0.46 | 0.234 | 0.907 |
| AT1G76200.1 | NADH:ubiquinone oxidoreductase subunit (AGGG) | 18 | 0.42 | 0.229 | 0.812 |
| AT1G76200.1 | NADH:ubiquinone oxidoreductase subunit (AGGG) | 19 | 0.61 | 0.237 | 0.999 |
| AT1G76200.1 | NADH:ubiquinone oxidoreductase subunit (AGGG) | 20 | 0.75 | 0.242 | 1.000 |
| AT1G76200.1 | NADH:ubiquinone oxidoreductase subunit (AGGG) | 21 | 0.95 | 0.245 | 1.000 |
| AT1G76200.1 | NADH:ubiquinone oxidoreductase subunit (AGGG) | 22 | 1.54 | 0.246 | 1.000 |
| AT1G76200.1 | NADH:ubiquinone oxidoreductase subunit (AGGG) | 23 | 2.29 | 0.246 | 0.905 |
| AT1G76200.1 | NADH:ubiquinone oxidoreductase subunit (AGGG) | 24 | 1.76 | 0.242 | 0.996 |
| AT1G76200.1 | NADH:ubiquinone oxidoreductase subunit (AGGG) | 25 | 0.76 | 0.242 | 1.000 |
| AT1G76200.1 | NADH:ubiquinone oxidoreductase subunit (AGGG) | 26 | 0.54 | 0.257 | 0.994 |
| AT1G76200.1 | NADH:ubiquinone oxidoreductase subunit (AGGG) | 27 | 0.84 | 0.260 | 1.000 |
| AT1G76200.1 | NADH:ubiquinone oxidoreductase subunit (AGGG) | 28 | 1.86 | 0.251 | 0.993 |
| AT1G76200.1 | NADH:ubiquinone oxidoreductase subunit (AGGG) | 29 | 2.58 | 0.250 | 0.803 |
| AT1G76200.1 | NADH:ubiquinone oxidoreductase subunit (AGGG) | 30 | 1.75 | 0.260 | 0.998 |
| AT1G76200.1 | NADH:ubiquinone oxidoreductase subunit (AGGG) | 31 | 0.87 | 0.325 | 1.000 |

| | | | | | | |
|---|---|---|---|---|---|---|
| AT1G76200.1 | NADH:ubiquinone oxidoreductase subunit (AGGG) | 32 | 0.47 | 0.545 | 1.000 | |
| AT1G76200.1 | NADH:ubiquinone oxidoreductase subunit (AGGG) | 46 | 3.60 | 7.214 | 1.000 | |
| AT1G76200.1 | NADH:ubiquinone oxidoreductase subunit (AGGG) | 47 | 6.04 | 7.960 | 1.000 | |
| AT1G77590.1 | Long chain acyl-CoA synthetase 9 (LACS9) | 2 | 1.13 | 0.274 | 1.000 | |
| AT1G77590.1 | Long chain acyl-CoA synthetase 9 (LACS9) | 3 | 0.77 | 0.279 | 1.000 | |
| AT1G77590.1 | Long chain acyl-CoA synthetase 9 (LACS9) | 30 | 0.99 | 0.211 | 1.000 | |
| AT1G77590.1 | Long chain acyl-CoA synthetase 9 (LACS9) | 31 | 1.70 | 0.176 | 0.891 | |
| AT1G77590.1 | Long chain acyl-CoA synthetase 9 (LACS9) | 32 | 2.34 | 0.185 | 0.437 | |
| AT1G77590.1 | Long chain acyl-CoA synthetase 9 (LACS9) | 33 | 2.65 | 0.252 | 0.666 | |
| AT1G77590.1 | Long chain acyl-CoA synthetase 9 (LACS9) | 39 | 1.09 | 0.224 | 1.000 | |
| AT1G77590.1 | Long chain acyl-CoA synthetase 9 (LACS9) | 40 | 1.06 | 0.158 | 1.000 | |
| AT1G77590.1 | Long chain acyl-CoA synthetase 9 (LACS9) | 41 | 1.32 | 0.134 | 0.991 | |
| AT1G77590.1 | Long chain acyl-CoA synthetase 9 (LACS9) | 42 | 1.74 | 0.131 | 0.555 | |
| AT1G77590.1 | Long chain acyl-CoA synthetase 9 (LACS9) | 43 | 1.66 | 0.132 | 0.678 | |
| AT1G77590.1 | Long chain acyl-CoA synthetase 9 (LACS9) | 44 | 1.07 | 0.132 | 1.000 | |
| AT1G77590.1 | Long chain acyl-CoA synthetase 9 (LACS9) | 45 | 0.62 | 0.132 | 0.746 | |
| AT1G77590.1 | Long chain acyl-CoA synthetase 9 (LACS9) | 46 | 0.39 | 0.147 | 0.082 | . |
| AT1G77590.1 | Long chain acyl-CoA synthetase 9 (LACS9) | 47 | 0.25 | 0.293 | 0.377 | |
| AT1G77670.1 | Pyridoxal phosphate (PLP)-dependent transferase | 2 | 1.03 | 0.789 | 1.000 | |
| AT1G77670.1 | Pyridoxal phosphate (PLP)-dependent transferase | 3 | 0.37 | 1.646 | 1.000 | |
| AT1G77670.1 | Pyridoxal phosphate (PLP)-dependent transferase | 39 | 0.46 | 0.615 | 0.999 | |
| AT1G77670.1 | Pyridoxal phosphate (PLP)-dependent transferase | 40 | 0.93 | 0.382 | 1.000 | |
| AT1G77670.1 | Pyridoxal phosphate (PLP)-dependent transferase | 41 | 1.38 | 0.343 | 1.000 | |
| AT1G77670.1 | Pyridoxal phosphate (PLP)-dependent transferase | 42 | 0.83 | 0.338 | 1.000 | |
| AT1G77670.1 | Pyridoxal phosphate (PLP)-dependent transferase | 43 | 0.54 | 0.348 | 0.988 | |
| AT1G77670.1 | Pyridoxal phosphate (PLP)-dependent transferase | 44 | 0.99 | 0.442 | 1.000 | |
| AT1G77670.1 | Pyridoxal phosphate (PLP)-dependent transferase | 45 | 4.40 | 0.716 | 0.968 | |
| AT1G77670.1 | Pyridoxal phosphate (PLP)-dependent transferase | 46 | 30.54 | 1.170 | 0.827 | |
| AT1G77670.1 | Pyridoxal phosphate (PLP)-dependent transferase | 47 | 257.58 | 1.876 | 0.818 | |
| AT1G77750.1 | Ribosomal S13 family protein | 2 | 1.47 | 0.701 | 1.000 | |
| AT1G77750.1 | Ribosomal S13 family protein | 3 | 1.45 | 0.635 | 1.000 | |
| AT1G77750.1 | Ribosomal S13 family protein | 11 | 1.61 | 0.363 | 0.996 | |
| AT1G77750.1 | Ribosomal S13 family protein | 12 | 1.73 | 0.341 | 0.985 | |

| | | | | | | |
|---|---|---|---|---|---|---|
| AT1G77750.1 | Ribosomal S13 family protein | 13 | 1.87 | 0.325 | 0.960 | |
| AT1G77750.1 | Ribosomal S13 family protein | 14 | 2.01 | 0.315 | 0.920 | |
| AT1G77750.1 | Ribosomal S13 family protein | 15 | 2.16 | 0.313 | 0.872 | |
| AT1G77750.1 | Ribosomal S13 family protein | 16 | 2.30 | 0.317 | 0.834 | |
| AT1G77750.1 | Ribosomal S13 family protein | 17 | 2.38 | 0.328 | 0.831 | |
| AT1G77750.1 | Ribosomal S13 family protein | 18 | 2.35 | 0.345 | 0.868 | |
| AT1G77750.1 | Ribosomal S13 family protein | 19 | 2.21 | 0.366 | 0.927 | |
| AT1G77750.1 | Ribosomal S13 family protein | 27 | 0.63 | 0.468 | 0.999 | |
| AT1G77750.1 | Ribosomal S13 family protein | 28 | 0.56 | 0.477 | 0.998 | |
| AT1G77750.1 | Ribosomal S13 family protein | 29 | 0.52 | 0.483 | 0.995 | |
| AT1G77750.1 | Ribosomal S13 family protein | 39 | 0.88 | 0.399 | 1.000 | |
| AT1G77750.1 | Ribosomal S13 family protein | 40 | 0.98 | 0.373 | 1.000 | |
| AT1G77750.1 | Ribosomal S13 family protein | 41 | 1.09 | 0.350 | 1.000 | |
| AT1G77750.1 | Ribosomal S13 family protein | 42 | 1.21 | 0.335 | 1.000 | |
| AT1G77750.1 | Ribosomal S13 family protein | 43 | 1.35 | 0.330 | 1.000 | |
| AT1G77750.1 | Ribosomal S13 family protein | 44 | 1.52 | 0.341 | 0.997 | |
| AT1G77750.1 | Ribosomal S13 family protein | 45 | 1.71 | 0.371 | 0.992 | |
| AT1G77750.1 | Ribosomal S13 family protein | 46 | 1.94 | 0.422 | 0.987 | |
| AT1G77750.1 | Ribosomal S13 family protein | 47 | 2.20 | 0.494 | 0.986 | |
| AT1G78010.1 | tRNA modification GTPase (TRME) | 2 | 0.87 | 0.354 | 1.000 | |
| AT1G78010.1 | tRNA modification GTPase (TRME) | 3 | 0.89 | 0.337 | 1.000 | |
| AT1G78010.1 | tRNA modification GTPase (TRME) | 30 | 1.47 | 0.173 | 0.943 | |
| AT1G78010.1 | tRNA modification GTPase (TRME) | 31 | 1.47 | 0.146 | 0.872 | |
| AT1G78010.1 | tRNA modification GTPase (TRME) | 32 | 1.48 | 0.127 | 0.762 | |
| AT1G78010.1 | tRNA modification GTPase (TRME) | 33 | 1.51 | 0.115 | 0.612 | |
| AT1G78010.1 | tRNA modification GTPase (TRME) | 34 | 1.58 | 0.109 | 0.415 | |
| AT1G78010.1 | tRNA modification GTPase (TRME) | 35 | 1.74 | 0.106 | 0.190 | |
| AT1G78010.1 | tRNA modification GTPase (TRME) | 36 | 1.98 | 0.106 | 0.052 | . |
| AT1G78010.1 | tRNA modification GTPase (TRME) | 37 | 2.26 | 0.106 | 0.011 | * |
| AT1G78010.1 | tRNA modification GTPase (TRME) | 38 | 2.50 | 0.107 | 0.002 | ** |
| AT1G78010.1 | tRNA modification GTPase (TRME) | 39 | 2.63 | 0.107 | 0.001 | ** |
| AT1G78010.1 | tRNA modification GTPase (TRME) | 40 | 2.56 | 0.106 | 0.002 | ** |
| AT1G78010.1 | tRNA modification GTPase (TRME) | 41 | 2.32 | 0.105 | 0.006 | ** |
| AT1G78010.1 | tRNA modification GTPase (TRME) | 42 | 1.94 | 0.105 | 0.059 | . |
| AT1G78010.1 | tRNA modification GTPase (TRME) | 43 | 1.54 | 0.107 | 0.467 | |
| AT1G78010.1 | tRNA modification GTPase (TRME) | 44 | 1.19 | 0.112 | 0.993 | |
| AT1G78010.1 | tRNA modification GTPase (TRME) | 45 | 0.93 | 0.126 | 1.000 | |
| AT1G78010.1 | tRNA modification GTPase (TRME) | 46 | 0.73 | 0.156 | 0.966 | |
| AT1G78010.1 | tRNA modification GTPase (TRME) | 47 | 0.57 | 0.206 | 0.857 | |
| AT1G78590.1 | NAD(H) kinase 3 (NADK3) | 2 | 1.05 | 0.448 | 1.000 | |
| AT1G78590.1 | NAD(H) kinase 3 (NADK3) | 3 | 0.51 | 0.464 | 0.988 | |
| AT1G78590.1 | NAD(H) kinase 3 (NADK3) | 42 | 0.23 | 0.395 | 0.520 | |

| | | | | | | |
|---|---|---|---|---|---|---|
| AT1G78590.1 | NAD(H) kinase 3 (NADK3) | 43 | 0.91 | 0.253 | 1.000 | |
| AT1G78590.1 | NAD(H) kinase 3 (NADK3) | 44 | 2.84 | 0.220 | 0.250 | |
| AT1G78590.1 | NAD(H) kinase 3 (NADK3) | 45 | 2.31 | 0.214 | 0.448 | |
| AT1G78590.1 | NAD(H) kinase 3 (NADK3) | 46 | 0.52 | 0.238 | 0.800 | |
| AT1G78590.1 | NAD(H) kinase 3 (NADK3) | 47 | 0.07 | 0.402 | 0.040 | * |
| AT1G78900.2 | Vacuolar ATP synthase subunit A (VHA-A) | 2 | 0.56 | 0.531 | 1.000 | |
| AT1G78900.2 | Vacuolar ATP synthase subunit A (VHA-A) | 3 | 1.08 | 0.362 | 1.000 | |
| AT1G78900.2 | Vacuolar ATP synthase subunit A (VHA-A) | 4 | 1.89 | 0.303 | 0.990 | |
| AT1G78900.2 | Vacuolar ATP synthase subunit A (VHA-A) | 5 | 2.85 | 0.295 | 0.758 | |
| AT1G78900.2 | Vacuolar ATP synthase subunit A (VHA-A) | 6 | 3.62 | 0.299 | 0.517 | |
| AT1G78900.2 | Vacuolar ATP synthase subunit A (VHA-A) | 7 | 3.99 | 0.309 | 0.461 | |
| AT1G78900.2 | Vacuolar ATP synthase subunit A (VHA-A) | 8 | 4.05 | 0.332 | 0.544 | |
| AT1G78900.2 | Vacuolar ATP synthase subunit A (VHA-A) | 9 | 3.94 | 0.382 | 0.747 | |
| AT1G78900.2 | Vacuolar ATP synthase subunit A (VHA-A) | 10 | 3.50 | 0.461 | 0.940 | |
| AT1G78900.2 | Vacuolar ATP synthase subunit A (VHA-A) | 21 | 0.50 | 0.458 | 0.999 | |
| AT1G78900.2 | Vacuolar ATP synthase subunit A (VHA-A) | 22 | 0.73 | 0.377 | 1.000 | |
| AT1G78900.2 | Vacuolar ATP synthase subunit A (VHA-A) | 23 | 1.21 | 0.329 | 1.000 | |
| AT1G78900.2 | Vacuolar ATP synthase subunit A (VHA-A) | 24 | 1.81 | 0.301 | 0.994 | |
| AT1G78900.2 | Vacuolar ATP synthase subunit A (VHA-A) | 25 | 2.26 | 0.289 | 0.925 | |
| AT1G78900.2 | Vacuolar ATP synthase subunit A (VHA-A) | 26 | 2.65 | 0.287 | 0.799 | |
| AT1G78900.2 | Vacuolar ATP synthase subunit A (VHA-A) | 27 | 2.40 | 0.295 | 0.898 | |
| AT1G78900.2 | Vacuolar ATP synthase subunit A (VHA-A) | 28 | 1.39 | 0.331 | 1.000 | |
| AT1G78900.2 | Vacuolar ATP synthase subunit A (VHA-A) | 29 | 0.69 | 0.428 | 1.000 | |
| AT1G78900.2 | Vacuolar ATP synthase subunit A (VHA-A) | 30 | 0.39 | 0.605 | 0.999 | |
| AT1G78900.2 | Vacuolar ATP synthase subunit A (VHA-A) | 46 | 3958.98 | 4.697 | 0.998 | |
| AT1G78900.2 | Vacuolar ATP synthase subunit A (VHA-A) | 47 | 12486.68 | 5.091 | 0.996 | |
| AT1G79010.1 | NADH:ubiquinone oxidoreductase subunit (TYKY-1) | 2 | 0.68 | 0.499 | 1.000 | |
| AT1G79010.1 | NADH:ubiquinone oxidoreductase subunit (TYKY-1) | 3 | 0.74 | 0.344 | 1.000 | |
| AT1G79010.1 | NADH:ubiquinone oxidoreductase subunit (TYKY-1) | 4 | 0.82 | 0.308 | 1.000 | |
| AT1G79010.1 | NADH:ubiquinone oxidoreductase subunit (TYKY-1) | 5 | 0.98 | 0.300 | 1.000 | |
| AT1G79010.1 | NADH:ubiquinone oxidoreductase subunit (TYKY-1) | 6 | 1.22 | 0.298 | 1.000 | |

| | | | | | |
|---|---|---|---|---|---|
| AT1G79010.1 | NADH:ubiquinone oxidoreductase subunit (TYKY-1) | 7 | 1.45 | 0.299 | 1.000 |
| AT1G79010.1 | NADH:ubiquinone oxidoreductase subunit (TYKY-1) | 8 | 1.57 | 0.295 | 1.000 |
| AT1G79010.1 | NADH:ubiquinone oxidoreductase subunit (TYKY-1) | 9 | 1.61 | 0.289 | 1.000 |
| AT1G79010.1 | NADH:ubiquinone oxidoreductase subunit (TYKY-1) | 10 | 1.66 | 0.282 | 1.000 |
| AT1G79010.1 | NADH:ubiquinone oxidoreductase subunit (TYKY-1) | 11 | 1.77 | 0.277 | 1.000 |
| AT1G79010.1 | NADH:ubiquinone oxidoreductase subunit (TYKY-1) | 12 | 1.86 | 0.272 | 1.000 |
| AT1G79010.1 | NADH:ubiquinone oxidoreductase subunit (TYKY-1) | 13 | 1.80 | 0.267 | 1.000 |
| AT1G79010.1 | NADH:ubiquinone oxidoreductase subunit (TYKY-1) | 14 | 1.42 | 0.264 | 1.000 |
| AT1G79010.1 | NADH:ubiquinone oxidoreductase subunit (TYKY-1) | 15 | 0.89 | 0.263 | 1.000 |
| AT1G79010.1 | NADH:ubiquinone oxidoreductase subunit (TYKY-1) | 16 | 0.51 | 0.262 | 0.998 |
| AT1G79010.1 | NADH:ubiquinone oxidoreductase subunit (TYKY-1) | 17 | 0.37 | 0.261 | 0.892 |
| AT1G79010.1 | NADH:ubiquinone oxidoreductase subunit (TYKY-1) | 18 | 0.39 | 0.264 | 0.936 |
| AT1G79010.1 | NADH:ubiquinone oxidoreductase subunit (TYKY-1) | 19 | 0.53 | 0.271 | 0.999 |
| AT1G79010.1 | NADH:ubiquinone oxidoreductase subunit (TYKY-1) | 20 | 0.74 | 0.276 | 1.000 |
| AT1G79010.1 | NADH:ubiquinone oxidoreductase subunit (TYKY-1) | 21 | 1.08 | 0.279 | 1.000 |
| AT1G79010.1 | NADH:ubiquinone oxidoreductase subunit (TYKY-1) | 22 | 1.50 | 0.279 | 1.000 |
| AT1G79010.1 | NADH:ubiquinone oxidoreductase subunit (TYKY-1) | 23 | 1.73 | 0.277 | 1.000 |
| AT1G79010.1 | NADH:ubiquinone oxidoreductase subunit (TYKY-1) | 24 | 1.44 | 0.272 | 1.000 |
| AT1G79010.1 | NADH:ubiquinone oxidoreductase subunit (TYKY-1) | 25 | 0.87 | 0.268 | 1.000 |
| AT1G79010.1 | NADH:ubiquinone oxidoreductase subunit (TYKY-1) | 26 | 0.51 | 0.266 | 0.999 |

| | | | | | |
|---|---|---|---|---|---|
| AT1G79010.1 | NADH:ubiquinone oxidoreductase subunit (TYKY-1) | 27 | 0.39 | 0.267 | 0.943 |
| AT1G79010.1 | NADH:ubiquinone oxidoreductase subunit (TYKY-1) | 28 | 0.48 | 0.268 | 0.996 |
| AT1G79010.1 | NADH:ubiquinone oxidoreductase subunit (TYKY-1) | 29 | 0.86 | 0.268 | 1.000 |
| AT1G79010.1 | NADH:ubiquinone oxidoreductase subunit (TYKY-1) | 30 | 1.48 | 0.268 | 1.000 |
| AT1G79010.1 | NADH:ubiquinone oxidoreductase subunit (TYKY-1) | 31 | 1.98 | 0.268 | 0.998 |
| AT1G79010.1 | NADH:ubiquinone oxidoreductase subunit (TYKY-1) | 32 | 2.07 | 0.269 | 0.997 |
| AT1G79010.1 | NADH:ubiquinone oxidoreductase subunit (TYKY-1) | 33 | 1.76 | 0.270 | 1.000 |
| AT1G79010.1 | NADH:ubiquinone oxidoreductase subunit (TYKY-1) | 34 | 1.55 | 0.271 | 1.000 |
| AT1G79010.1 | NADH:ubiquinone oxidoreductase subunit (TYKY-1) | 35 | 1.59 | 0.275 | 1.000 |
| AT1G79010.1 | NADH:ubiquinone oxidoreductase subunit (TYKY-1) | 36 | 2.01 | 0.290 | 0.999 |
| AT1G79010.1 | NADH:ubiquinone oxidoreductase subunit (TYKY-1) | 37 | 3.14 | 0.324 | 0.939 |
| AT1G79010.1 | NADH:ubiquinone oxidoreductase subunit (TYKY-1) | 38 | 5.30 | 0.351 | 0.598 |
| AT1G79010.1 | NADH:ubiquinone oxidoreductase subunit (TYKY-1) | 40 | 9.61 | 0.359 | 0.158 |
| AT1G79010.1 | NADH:ubiquinone oxidoreductase subunit (TYKY-1) | 41 | 8.89 | 0.331 | 0.115 |
| AT1G79010.1 | NADH:ubiquinone oxidoreductase subunit (TYKY-1) | 42 | 7.91 | 0.320 | 0.133 |
| AT1G79010.1 | NADH:ubiquinone oxidoreductase subunit (TYKY-1) | 43 | 6.99 | 0.315 | 0.181 |
| AT1G79010.1 | NADH:ubiquinone oxidoreductase subunit (TYKY-1) | 44 | 5.38 | 0.309 | 0.360 |
| AT1G79010.1 | NADH:ubiquinone oxidoreductase subunit (TYKY-1) | 45 | 3.11 | 0.342 | 0.966 |
| AT1G79010.1 | NADH:ubiquinone oxidoreductase subunit (TYKY-1) | 46 | 1.49 | 0.465 | 1.000 |
| AT1G79010.1 | NADH:ubiquinone oxidoreductase subunit (TYKY-1) | 47 | 0.66 | 0.714 | 1.000 |
| AT1G79230.3 | Mercaptopyruvate sulfurtransferase (MST1) | 2 | 1.06 | 0.288 | 1.000 |

| | | | | | | |
|---|---|---|---|---|---|---|
| AT1G79230.3 | Mercaptopyruvate sulfurtransferase (MST1) | 3 | 0.83 | 0.214 | 1.000 | |
| AT1G79230.3 | Mercaptopyruvate sulfurtransferase (MST1) | 4 | 0.67 | 0.209 | 1.000 | |
| AT1G79230.3 | Mercaptopyruvate sulfurtransferase (MST1) | 7 | 0.53 | 0.194 | 0.981 | |
| AT1G79230.3 | Mercaptopyruvate sulfurtransferase (MST1) | 8 | 0.57 | 0.169 | 0.972 | |
| AT1G79230.3 | Mercaptopyruvate sulfurtransferase (MST1) | 9 | 0.65 | 0.151 | 0.995 | |
| AT1G79230.3 | Mercaptopyruvate sulfurtransferase (MST1) | 10 | 0.78 | 0.141 | 1.000 | |
| AT1G79230.3 | Mercaptopyruvate sulfurtransferase (MST1) | 11 | 0.94 | 0.136 | 1.000 | |
| AT1G79230.3 | Mercaptopyruvate sulfurtransferase (MST1) | 12 | 1.12 | 0.134 | 1.000 | |
| AT1G79230.3 | Mercaptopyruvate sulfurtransferase (MST1) | 13 | 1.19 | 0.133 | 1.000 | |
| AT1G79230.3 | Mercaptopyruvate sulfurtransferase (MST1) | 14 | 1.16 | 0.131 | 1.000 | |
| AT1G79230.3 | Mercaptopyruvate sulfurtransferase (MST1) | 15 | 1.26 | 0.130 | 1.000 | |
| AT1G79230.3 | Mercaptopyruvate sulfurtransferase (MST1) | 16 | 1.62 | 0.129 | 0.921 | |
| AT1G79230.3 | Mercaptopyruvate sulfurtransferase (MST1) | 17 | 2.10 | 0.128 | 0.295 | |
| AT1G79230.3 | Mercaptopyruvate sulfurtransferase (MST1) | 18 | 2.24 | 0.131 | 0.195 | |
| AT1G79230.3 | Mercaptopyruvate sulfurtransferase (MST1) | 19 | 2.01 | 0.134 | 0.470 | |
| AT1G79230.3 | Mercaptopyruvate sulfurtransferase (MST1) | 20 | 1.65 | 0.137 | 0.935 | |
| AT1G79230.3 | Mercaptopyruvate sulfurtransferase (MST1) | 21 | 1.35 | 0.139 | 1.000 | |
| AT1G79230.3 | Mercaptopyruvate sulfurtransferase (MST1) | 22 | 1.16 | 0.139 | 1.000 | |
| AT1G79230.3 | Mercaptopyruvate sulfurtransferase (MST1) | 23 | 1.29 | 0.138 | 1.000 | |
| AT1G79230.3 | Mercaptopyruvate sulfurtransferase (MST1) | 24 | 1.80 | 0.136 | 0.780 | |
| AT1G79230.3 | Mercaptopyruvate sulfurtransferase (MST1) | 25 | 2.50 | 0.134 | 0.090 | . |
| AT1G79230.3 | Mercaptopyruvate sulfurtransferase (MST1) | 26 | 2.67 | 0.133 | 0.044 | * |
| AT1G79230.3 | Mercaptopyruvate sulfurtransferase (MST1) | 27 | 2.21 | 0.133 | 0.239 | |
| AT1G79230.3 | Mercaptopyruvate sulfurtransferase (MST1) | 28 | 1.64 | 0.133 | 0.926 | |
| AT1G79230.3 | Mercaptopyruvate sulfurtransferase (MST1) | 29 | 1.26 | 0.132 | 1.000 | |
| AT1G79230.3 | Mercaptopyruvate sulfurtransferase (MST1) | 30 | 1.08 | 0.132 | 1.000 | |
| AT1G79230.3 | Mercaptopyruvate sulfurtransferase (MST1) | 31 | 1.05 | 0.133 | 1.000 | |
| AT1G79230.3 | Mercaptopyruvate sulfurtransferase (MST1) | 32 | 1.06 | 0.133 | 1.000 | |
| AT1G79230.3 | Mercaptopyruvate sulfurtransferase (MST1) | 33 | 1.00 | 0.134 | 1.000 | |
| AT1G79230.3 | Mercaptopyruvate sulfurtransferase (MST1) | 34 | 0.92 | 0.134 | 1.000 | |
| AT1G79230.3 | Mercaptopyruvate sulfurtransferase (MST1) | 35 | 0.90 | 0.134 | 1.000 | |

| | | | | | |
|---|---|---|---|---|---|
| AT1G79230.3 | Mercaptopyruvate sulfurtransferase (MST1) | 36 | 0.94 | 0.133 | 1.000 |
| AT1G79230.3 | Mercaptopyruvate sulfurtransferase (MST1) | 37 | 1.06 | 0.133 | 1.000 |
| AT1G79230.3 | Mercaptopyruvate sulfurtransferase (MST1) | 38 | 1.25 | 0.133 | 1.000 |
| AT1G79230.3 | Mercaptopyruvate sulfurtransferase (MST1) | 39 | 1.45 | 0.134 | 0.997 |
| AT1G79230.3 | Mercaptopyruvate sulfurtransferase (MST1) | 40 | 1.54 | 0.134 | 0.981 |
| AT1G79230.3 | Mercaptopyruvate sulfurtransferase (MST1) | 41 | 1.41 | 0.135 | 0.999 |
| AT1G79230.3 | Mercaptopyruvate sulfurtransferase (MST1) | 42 | 1.15 | 0.137 | 1.000 |
| AT1G79230.3 | Mercaptopyruvate sulfurtransferase (MST1) | 43 | 0.91 | 0.139 | 1.000 |
| AT1G79230.3 | Mercaptopyruvate sulfurtransferase (MST1) | 44 | 0.71 | 0.138 | 0.999 |
| AT1G79230.3 | Mercaptopyruvate sulfurtransferase (MST1) | 45 | 0.56 | 0.139 | 0.812 |
| AT1G79230.3 | Mercaptopyruvate sulfurtransferase (MST1) | 46 | 0.41 | 0.173 | 0.519 |
| AT1G79230.3 | Mercaptopyruvate sulfurtransferase (MST1) | 47 | 0.29 | 0.286 | 0.778 |
| AT1G79440.1 | Succinic semialdehyde dehydrogenase 1 (SSADH1) | 2 | 1.31 | 0.596 | 1.000 |
| AT1G79440.1 | Succinic semialdehyde dehydrogenase 1 (SSADH1) | 3 | 1.00 | 0.437 | 1.000 |
| AT1G79440.1 | Succinic semialdehyde dehydrogenase 1 (SSADH1) | 4 | 0.80 | 0.440 | 1.000 |
| AT1G79440.1 | Succinic semialdehyde dehydrogenase 1 (SSADH1) | 6 | 0.54 | 0.437 | 1.000 |
| AT1G79440.1 | Succinic semialdehyde dehydrogenase 1 (SSADH1) | 7 | 0.42 | 0.388 | 1.000 |
| AT1G79440.1 | Succinic semialdehyde dehydrogenase 1 (SSADH1) | 8 | 0.31 | 0.348 | 0.965 |
| AT1G79440.1 | Succinic semialdehyde dehydrogenase 1 (SSADH1) | 9 | 0.26 | 0.328 | 0.812 |
| AT1G79440.1 | Succinic semialdehyde dehydrogenase 1 (SSADH1) | 10 | 0.28 | 0.322 | 0.859 |
| AT1G79440.1 | Succinic semialdehyde dehydrogenase 1 (SSADH1) | 11 | 0.39 | 0.320 | 0.992 |
| AT1G79440.1 | Succinic semialdehyde dehydrogenase 1 (SSADH1) | 12 | 0.59 | 0.320 | 1.000 |
| AT1G79440.1 | Succinic semialdehyde dehydrogenase 1 (SSADH1) | 13 | 0.82 | 0.317 | 1.000 |
| AT1G79440.1 | Succinic semialdehyde dehydrogenase 1 (SSADH1) | 14 | 1.05 | 0.314 | 1.000 |
| AT1G79440.1 | Succinic semialdehyde dehydrogenase 1 (SSADH1) | 15 | 1.18 | 0.312 | 1.000 |
| AT1G79440.1 | Succinic semialdehyde dehydrogenase 1 (SSADH1) | 16 | 1.18 | 0.310 | 1.000 |
| AT1G79440.1 | Succinic semialdehyde dehydrogenase 1 (SSADH1) | 17 | 1.18 | 0.310 | 1.000 |
| AT1G79440.1 | Succinic semialdehyde dehydrogenase 1 (SSADH1) | 18 | 1.18 | 0.314 | 1.000 |
| AT1G79440.1 | Succinic semialdehyde dehydrogenase 1 (SSADH1) | 19 | 1.08 | 0.321 | 1.000 |
| AT1G79440.1 | Succinic semialdehyde dehydrogenase 1 (SSADH1) | 20 | 0.85 | 0.328 | 1.000 |
| AT1G79440.1 | Succinic semialdehyde dehydrogenase 1 (SSADH1) | 21 | 0.61 | 0.331 | 1.000 |

| | | | | | |
|---|---|---|---|---|---|
| AT1G79440.1 | Succinic semialdehyde dehydrogenase 1 (SSADH1) | 22 | 0.49 | 0.330 | 1.000 |
| AT1G79440.1 | Succinic semialdehyde dehydrogenase 1 (SSADH1) | 23 | 0.52 | 0.328 | 1.000 |
| AT1G79440.1 | Succinic semialdehyde dehydrogenase 1 (SSADH1) | 24 | 0.62 | 0.323 | 1.000 |
| AT1G79440.1 | Succinic semialdehyde dehydrogenase 1 (SSADH1) | 25 | 0.71 | 0.319 | 1.000 |
| AT1G79440.1 | Succinic semialdehyde dehydrogenase 1 (SSADH1) | 26 | 0.86 | 0.317 | 1.000 |
| AT1G79440.1 | Succinic semialdehyde dehydrogenase 1 (SSADH1) | 27 | 1.17 | 0.318 | 1.000 |
| AT1G79440.1 | Succinic semialdehyde dehydrogenase 1 (SSADH1) | 28 | 1.39 | 0.317 | 1.000 |
| AT1G79440.1 | Succinic semialdehyde dehydrogenase 1 (SSADH1) | 29 | 1.35 | 0.317 | 1.000 |
| AT1G79440.1 | Succinic semialdehyde dehydrogenase 1 (SSADH1) | 30 | 1.15 | 0.317 | 1.000 |
| AT1G79440.1 | Succinic semialdehyde dehydrogenase 1 (SSADH1) | 31 | 0.97 | 0.318 | 1.000 |
| AT1G79440.1 | Succinic semialdehyde dehydrogenase 1 (SSADH1) | 32 | 0.87 | 0.319 | 1.000 |
| AT1G79440.1 | Succinic semialdehyde dehydrogenase 1 (SSADH1) | 33 | 0.86 | 0.319 | 1.000 |
| AT1G79440.1 | Succinic semialdehyde dehydrogenase 1 (SSADH1) | 34 | 0.98 | 0.319 | 1.000 |
| AT1G79440.1 | Succinic semialdehyde dehydrogenase 1 (SSADH1) | 35 | 1.33 | 0.319 | 1.000 |
| AT1G79440.1 | Succinic semialdehyde dehydrogenase 1 (SSADH1) | 36 | 1.78 | 0.318 | 1.000 |
| AT1G79440.1 | Succinic semialdehyde dehydrogenase 1 (SSADH1) | 37 | 1.93 | 0.318 | 1.000 |
| AT1G79440.1 | Succinic semialdehyde dehydrogenase 1 (SSADH1) | 38 | 1.87 | 0.318 | 1.000 |
| AT1G79440.1 | Succinic semialdehyde dehydrogenase 1 (SSADH1) | 39 | 2.11 | 0.319 | 1.000 |
| AT1G79440.1 | Succinic semialdehyde dehydrogenase 1 (SSADH1) | 40 | 2.62 | 0.319 | 0.989 |
| AT1G79440.1 | Succinic semialdehyde dehydrogenase 1 (SSADH1) | 41 | 2.98 | 0.321 | 0.961 |
| AT1G79440.1 | Succinic semialdehyde dehydrogenase 1 (SSADH1) | 42 | 3.18 | 0.324 | 0.939 |
| AT1G79440.1 | Succinic semialdehyde dehydrogenase 1 (SSADH1) | 43 | 2.77 | 0.327 | 0.985 |
| AT1G79440.1 | Succinic semialdehyde dehydrogenase 1 (SSADH1) | 44 | 1.88 | 0.327 | 1.000 |
| AT1G79440.1 | Succinic semialdehyde dehydrogenase 1 (SSADH1) | 45 | 1.26 | 0.327 | 1.000 |
| AT1G79440.1 | Succinic semialdehyde dehydrogenase 1 (SSADH1) | 46 | 0.85 | 0.384 | 1.000 |
| AT1G79440.1 | Succinic semialdehyde dehydrogenase 1 (SSADH1) | 47 | 0.55 | 0.629 | 1.000 |
| AT1G79750.1 | NADP malic enzyme 4 (NADP-ME4) | 2 | 0.72 | 0.755 | 1.000 |
| AT1G79750.1 | NADP malic enzyme 4 (NADP-ME4) | 3 | 0.72 | 0.730 | 1.000 |
| AT1G79750.1 | NADP malic enzyme 4 (NADP-ME4) | 27 | 0.52 | 0.561 | 1.000 |
| AT1G79750.1 | NADP malic enzyme 4 (NADP-ME4) | 28 | 0.50 | 0.388 | 1.000 |
| AT1G79750.1 | NADP malic enzyme 4 (NADP-ME4) | 29 | 0.53 | 0.344 | 1.000 |

| | | | | | |
|---|---|---|---|---|---|
| AT1G79750.1 | NADP malic enzyme 4 (NADP-ME4) | 30 | 1.11 | 0.329 | 1.000 |
| AT1G79750.1 | NADP malic enzyme 4 (NADP-ME4) | 31 | 3.98 | 0.326 | 0.658 |
| AT1G79750.1 | NADP malic enzyme 4 (NADP-ME4) | 32 | 4.63 | 0.326 | 0.500 |
| AT1G79750.1 | NADP malic enzyme 4 (NADP-ME4) | 33 | 1.42 | 0.326 | 1.000 |
| AT1G79750.1 | NADP malic enzyme 4 (NADP-ME4) | 34 | 0.51 | 0.326 | 0.999 |
| AT1G79750.1 | NADP malic enzyme 4 (NADP-ME4) | 35 | 0.51 | 0.326 | 0.999 |
| AT1G79750.1 | NADP malic enzyme 4 (NADP-ME4) | 36 | 0.97 | 0.325 | 1.000 |
| AT1G79750.1 | NADP malic enzyme 4 (NADP-ME4) | 37 | 2.11 | 0.323 | 0.996 |
| AT1G79750.1 | NADP malic enzyme 4 (NADP-ME4) | 38 | 3.00 | 0.321 | 0.893 |
| AT1G79750.1 | NADP malic enzyme 4 (NADP-ME4) | 39 | 2.10 | 0.320 | 0.996 |
| AT1G79750.1 | NADP malic enzyme 4 (NADP-ME4) | 40 | 1.12 | 0.321 | 1.000 |
| AT1G79750.1 | NADP malic enzyme 4 (NADP-ME4) | 41 | 0.85 | 0.325 | 1.000 |
| AT1G79750.1 | NADP malic enzyme 4 (NADP-ME4) | 42 | 0.81 | 0.329 | 1.000 |
| AT1G79750.1 | NADP malic enzyme 4 (NADP-ME4) | 43 | 0.78 | 0.331 | 1.000 |
| AT1G79750.1 | NADP malic enzyme 4 (NADP-ME4) | 44 | 0.72 | 0.331 | 1.000 |
| AT1G79750.1 | NADP malic enzyme 4 (NADP-ME4) | 45 | 0.56 | 0.336 | 1.000 |
| AT1G79750.1 | NADP malic enzyme 4 (NADP-ME4) | 46 | 0.33 | 0.394 | 0.972 |
| AT1G79750.1 | NADP malic enzyme 4 (NADP-ME4) | 47 | 0.18 | 0.685 | 0.990 |
| AT1G80230.1 | Cytochrome c:O2 oxidoreductase subunit (COX5B-2) | 2 | 0.98 | 0.396 | 1.000 |
| AT1G80230.1 | Cytochrome c:O2 oxidoreductase subunit (COX5B-2) | 3 | 1.13 | 0.252 | 1.000 |
| AT1G80230.1 | Cytochrome c:O2 oxidoreductase subunit (COX5B-2) | 4 | 1.33 | 0.244 | 1.000 |
| AT1G80230.1 | Cytochrome c:O2 oxidoreductase subunit (COX5B-2) | 5 | 1.49 | 0.236 | 1.000 |
| AT1G80230.1 | Cytochrome c:O2 oxidoreductase subunit (COX5B-2) | 6 | 1.51 | 0.226 | 1.000 |
| AT1G80230.1 | Cytochrome c:O2 oxidoreductase subunit (COX5B-2) | 7 | 1.45 | 0.223 | 1.000 |
| AT1G80230.1 | Cytochrome c:O2 oxidoreductase subunit (COX5B-2) | 8 | 1.38 | 0.216 | 1.000 |
| AT1G80230.1 | Cytochrome c:O2 oxidoreductase subunit (COX5B-2) | 9 | 1.31 | 0.213 | 1.000 |

| | | | | | |
|---|---|---|---|---|---|
| AT1G80230.1 | Cytochrome c:O2 oxidoreductase subunit (COX5B-2) | 10 | 1.30 | 0.212 | 1.000 |
| AT1G80230.1 | Cytochrome c:O2 oxidoreductase subunit (COX5B-2) | 11 | 1.31 | 0.214 | 1.000 |
| AT1G80230.1 | Cytochrome c:O2 oxidoreductase subunit (COX5B-2) | 12 | 1.20 | 0.215 | 1.000 |
| AT1G80230.1 | Cytochrome c:O2 oxidoreductase subunit (COX5B-2) | 13 | 1.07 | 0.213 | 1.000 |
| AT1G80230.1 | Cytochrome c:O2 oxidoreductase subunit (COX5B-2) | 14 | 1.02 | 0.211 | 1.000 |
| AT1G80230.1 | Cytochrome c:O2 oxidoreductase subunit (COX5B-2) | 15 | 0.99 | 0.211 | 1.000 |
| AT1G80230.1 | Cytochrome c:O2 oxidoreductase subunit (COX5B-2) | 16 | 0.92 | 0.208 | 1.000 |
| AT1G80230.1 | Cytochrome c:O2 oxidoreductase subunit (COX5B-2) | 17 | 0.85 | 0.206 | 1.000 |
| AT1G80230.1 | Cytochrome c:O2 oxidoreductase subunit (COX5B-2) | 18 | 0.90 | 0.209 | 1.000 |
| AT1G80230.1 | Cytochrome c:O2 oxidoreductase subunit (COX5B-2) | 19 | 1.01 | 0.218 | 1.000 |
| AT1G80230.1 | Cytochrome c:O2 oxidoreductase subunit (COX5B-2) | 20 | 1.25 | 0.223 | 1.000 |
| AT1G80230.1 | Cytochrome c:O2 oxidoreductase subunit (COX5B-2) | 21 | 1.69 | 0.225 | 1.000 |
| AT1G80230.1 | Cytochrome c:O2 oxidoreductase subunit (COX5B-2) | 22 | 1.87 | 0.224 | 0.997 |
| AT1G80230.1 | Cytochrome c:O2 oxidoreductase subunit (COX5B-2) | 23 | 1.53 | 0.222 | 1.000 |
| AT1G80230.1 | Cytochrome c:O2 oxidoreductase subunit (COX5B-2) | 24 | 1.32 | 0.218 | 1.000 |
| AT1G80230.1 | Cytochrome c:O2 oxidoreductase subunit (COX5B-2) | 25 | 1.36 | 0.214 | 1.000 |
| AT1G80230.1 | Cytochrome c:O2 oxidoreductase subunit (COX5B-2) | 26 | 1.57 | 0.213 | 1.000 |
| AT1G80230.1 | Cytochrome c:O2 oxidoreductase subunit (COX5B-2) | 27 | 2.20 | 0.212 | 0.932 |
| AT1G80230.1 | Cytochrome c:O2 oxidoreductase subunit (COX5B-2) | 28 | 2.21 | 0.212 | 0.924 |
| AT1G80230.1 | Cytochrome c:O2 oxidoreductase subunit (COX5B-2) | 29 | 2.07 | 0.212 | 0.966 |

| | | | | | | |
|---|---|---|---|---|---|---|
| AT1G80230.1 | Cytochrome c:O2 oxidoreductase subunit (COX5B-2) | 30 | 1.95 | 0.212 | 0.988 | |
| AT1G80230.1 | Cytochrome c:O2 oxidoreductase subunit (COX5B-2) | 31 | 1.64 | 0.214 | 1.000 | |
| AT1G80230.1 | Cytochrome c:O2 oxidoreductase subunit (COX5B-2) | 32 | 1.15 | 0.216 | 1.000 | |
| AT1G80230.1 | Cytochrome c:O2 oxidoreductase subunit (COX5B-2) | 33 | 0.50 | 0.220 | 0.988 | |
| AT1G80230.1 | Cytochrome c:O2 oxidoreductase subunit (COX5B-2) | 34 | 0.17 | 0.233 | 0.033 | * |
| AT1G80230.1 | Cytochrome c:O2 oxidoreductase subunit (COX5B-2) | 35 | 0.12 | 0.269 | 0.026 | * |
| AT1G80230.1 | Cytochrome c:O2 oxidoreductase subunit (COX5B-2) | 36 | 0.20 | 0.432 | 0.919 | |
| AT1G80230.1 | Cytochrome c:O2 oxidoreductase subunit (COX5B-2) | 46 | ########### | 5.601 | 0.655 | |
| AT1G80230.1 | Cytochrome c:O2 oxidoreductase subunit (COX5B-2) | 47 | ########### | 6.388 | 0.642 | |
| AT1G80270.3 | Pentatricopeptide repeat (PPR) superfamily protein (PPR596) | 2 | 1.15 | 0.519 | 1.000 | |
| AT1G80270.3 | Pentatricopeptide repeat (PPR) superfamily protein (PPR596) | 3 | 1.24 | 0.487 | 1.000 | |
| AT1G80270.3 | Pentatricopeptide repeat (PPR) superfamily protein (PPR596) | 22 | 0.58 | 0.394 | 1.000 | |
| AT1G80270.3 | Pentatricopeptide repeat (PPR) superfamily protein (PPR596) | 23 | 0.47 | 0.335 | 0.987 | |
| AT1G80270.3 | Pentatricopeptide repeat (PPR) superfamily protein (PPR596) | 24 | 0.41 | 0.292 | 0.886 | |
| AT1G80270.3 | Pentatricopeptide repeat (PPR) superfamily protein (PPR596) | 25 | 0.38 | 0.255 | 0.691 | |
| AT1G80270.3 | Pentatricopeptide repeat (PPR) superfamily protein (PPR596) | 26 | 0.39 | 0.232 | 0.606 | |
| AT1G80270.3 | Pentatricopeptide repeat (PPR) superfamily protein (PPR596) | 27 | 0.45 | 0.216 | 0.721 | |
| AT1G80270.3 | Pentatricopeptide repeat (PPR) superfamily protein (PPR596) | 28 | 0.58 | 0.207 | 0.960 | |
| AT1G80270.3 | Pentatricopeptide repeat (PPR) superfamily protein (PPR596) | 29 | 0.78 | 0.204 | 1.000 | |
| AT1G80270.3 | Pentatricopeptide repeat (PPR) superfamily protein (PPR596) | 30 | 0.90 | 0.203 | 1.000 | |
| AT1G80270.3 | Pentatricopeptide repeat (PPR) superfamily protein (PPR596) | 31 | 0.89 | 0.203 | 1.000 | |
| AT1G80270.3 | Pentatricopeptide repeat (PPR) superfamily protein (PPR596) | 32 | 0.78 | 0.203 | 1.000 | |
| AT1G80270.3 | Pentatricopeptide repeat (PPR) superfamily protein (PPR596) | 33 | 0.67 | 0.203 | 0.995 | |
| AT1G80270.3 | Pentatricopeptide repeat (PPR) superfamily protein (PPR596) | 34 | 0.62 | 0.203 | 0.981 | |
| AT1G80270.3 | Pentatricopeptide repeat (PPR) superfamily protein (PPR596) | 35 | 0.63 | 0.204 | 0.987 | |
| AT1G80270.3 | Pentatricopeptide repeat (PPR) superfamily protein (PPR596) | 36 | 0.66 | 0.206 | 0.995 | |

| | | | | | |
|---|---|---|---|---|---|
| AT1G80270.3 | Pentatricopeptide repeat (PPR) superfamily protein (PPR596) | 37 | 0.73 | 0.211 | 1.000 |
| AT1G80270.3 | Pentatricopeptide repeat (PPR) superfamily protein (PPR596) | 38 | 0.88 | 0.216 | 1.000 |
| AT1G80270.3 | Pentatricopeptide repeat (PPR) superfamily protein (PPR596) | 39 | 1.08 | 0.219 | 1.000 |
| AT1G80270.3 | Pentatricopeptide repeat (PPR) superfamily protein (PPR596) | 40 | 1.27 | 0.220 | 1.000 |
| AT1G80270.3 | Pentatricopeptide repeat (PPR) superfamily protein (PPR596) | 41 | 1.34 | 0.222 | 1.000 |
| AT1G80270.3 | Pentatricopeptide repeat (PPR) superfamily protein (PPR596) | 42 | 1.23 | 0.227 | 1.000 |
| AT1G80270.3 | Pentatricopeptide repeat (PPR) superfamily protein (PPR596) | 43 | 1.02 | 0.230 | 1.000 |
| AT1G80270.3 | Pentatricopeptide repeat (PPR) superfamily protein (PPR596) | 44 | 0.82 | 0.238 | 1.000 |
| AT1G80270.3 | Pentatricopeptide repeat (PPR) superfamily protein (PPR596) | 45 | 0.68 | 0.268 | 1.000 |
| AT1G80270.3 | Pentatricopeptide repeat (PPR) superfamily protein (PPR596) | 46 | 0.56 | 0.341 | 0.999 |
| AT1G80270.3 | Pentatricopeptide repeat (PPR) superfamily protein (PPR596) | 47 | 0.46 | 0.470 | 0.999 |
| AT1G80620.1 | Ribosomal S15 family protein | 2 | 1.72 | 0.521 | 0.991 |
| AT1G80620.1 | Ribosomal S15 family protein | 3 | 1.65 | 0.487 | 0.991 |
| AT1G80620.1 | Ribosomal S15 family protein | 11 | 1.21 | 0.327 | 1.000 |
| AT1G80620.1 | Ribosomal S15 family protein | 12 | 1.17 | 0.319 | 1.000 |
| AT1G80620.1 | Ribosomal S15 family protein | 13 | 1.14 | 0.313 | 1.000 |
| AT1G80620.1 | Ribosomal S15 family protein | 14 | 1.11 | 0.310 | 1.000 |
| AT1G80620.1 | Ribosomal S15 family protein | 15 | 1.09 | 0.308 | 1.000 |
| AT1G80620.1 | Ribosomal S15 family protein | 16 | 1.07 | 0.308 | 1.000 |
| AT1G80620.1 | Ribosomal S15 family protein | 17 | 1.05 | 0.310 | 1.000 |
| AT1G80620.1 | Ribosomal S15 family protein | 18 | 1.04 | 0.312 | 1.000 |
| AT1G80620.1 | Ribosomal S15 family protein | 19 | 1.03 | 0.315 | 1.000 |
| AT1G80620.1 | Ribosomal S15 family protein | 28 | 1.05 | 0.311 | 1.000 |
| AT1G80620.1 | Ribosomal S15 family protein | 29 | 1.08 | 0.307 | 1.000 |
| AT1G80620.1 | Ribosomal S15 family protein | 30 | 1.12 | 0.304 | 1.000 |
| AT1G80620.1 | Ribosomal S15 family protein | 31 | 1.17 | 0.301 | 1.000 |
| AT1G80620.1 | Ribosomal S15 family protein | 32 | 1.23 | 0.299 | 0.999 |
| AT1G80620.1 | Ribosomal S15 family protein | 33 | 1.30 | 0.297 | 0.996 |
| AT1G80620.1 | Ribosomal S15 family protein | 34 | 1.38 | 0.296 | 0.988 |
| AT1G80620.1 | Ribosomal S15 family protein | 35 | 1.48 | 0.295 | 0.968 |
| AT1G80620.1 | Ribosomal S15 family protein | 36 | 1.60 | 0.294 | 0.932 |
| AT1G80620.1 | Ribosomal S15 family protein | 37 | 1.74 | 0.294 | 0.874 |
| AT1G80620.1 | Ribosomal S15 family protein | 38 | 1.90 | 0.294 | 0.793 |
| AT1G80620.1 | Ribosomal S15 family protein | 39 | 2.10 | 0.294 | 0.694 |
| AT1G80620.1 | Ribosomal S15 family protein | 40 | 2.33 | 0.294 | 0.584 |
| AT1G80620.1 | Ribosomal S15 family protein | 41 | 2.61 | 0.296 | 0.474 |
| AT1G80620.1 | Ribosomal S15 family protein | 42 | 2.93 | 0.298 | 0.378 |
| AT1G80620.1 | Ribosomal S15 family protein | 43 | 3.27 | 0.303 | 0.304 |
| AT1G80620.1 | Ribosomal S15 family protein | 44 | 3.64 | 0.310 | 0.253 |
| AT1G80620.1 | Ribosomal S15 family protein | 45 | 4.01 | 0.321 | 0.221 |
| AT1G80620.1 | Ribosomal S15 family protein | 46 | 4.39 | 0.336 | 0.208 |
| AT1G80620.1 | Ribosomal S15 family protein | 47 | 4.78 | 0.356 | 0.209 |
| AT2G01140.1 | Fructose-bisphosphate aldolase 3 (FBA3) | 2 | 0.96 | 0.348 | 1.000 |
| AT2G01140.1 | Fructose-bisphosphate aldolase 3 (FBA3) | 3 | 0.98 | 0.304 | 1.000 |
| AT2G01140.1 | Fructose-bisphosphate aldolase 3 (FBA3) | 4 | 1.00 | 0.281 | 1.000 |
| AT2G01140.1 | Fructose-bisphosphate aldolase 3 (FBA3) | 6 | 1.04 | 0.287 | 1.000 |

| | | | | | | |
|---|---|---|---|---|---|---|
| AT2G01140.1 | Fructose-bisphosphate aldolase 3 (FBA3) | 7 | 1.06 | 0.306 | 1.000 | |
| AT2G01140.1 | Fructose-bisphosphate aldolase 3 (FBA3) | 8 | 1.08 | 0.331 | 1.000 | |
| AT2G01140.1 | Fructose-bisphosphate aldolase 3 (FBA3) | 22 | 1.08 | 0.372 | 1.000 | |
| AT2G01140.1 | Fructose-bisphosphate aldolase 3 (FBA3) | 23 | 1.04 | 0.357 | 1.000 | |
| AT2G01140.1 | Fructose-bisphosphate aldolase 3 (FBA3) | 24 | 0.99 | 0.344 | 1.000 | |
| AT2G01140.1 | Fructose-bisphosphate aldolase 3 (FBA3) | 32 | 0.64 | 0.217 | 0.957 | |
| AT2G01140.1 | Fructose-bisphosphate aldolase 3 (FBA3) | 33 | 0.64 | 0.198 | 0.927 | |
| AT2G01140.1 | Fructose-bisphosphate aldolase 3 (FBA3) | 34 | 0.65 | 0.183 | 0.911 | |
| AT2G01140.1 | Fructose-bisphosphate aldolase 3 (FBA3) | 35 | 0.67 | 0.173 | 0.917 | |
| AT2G01140.1 | Fructose-bisphosphate aldolase 3 (FBA3) | 36 | 0.69 | 0.171 | 0.943 | |
| AT2G01140.1 | Fructose-bisphosphate aldolase 3 (FBA3) | 37 | 0.72 | 0.176 | 0.974 | |
| AT2G01140.1 | Fructose-bisphosphate aldolase 3 (FBA3) | 38 | 0.76 | 0.189 | 0.993 | |
| AT2G01140.1 | Fructose-bisphosphate aldolase 3 (FBA3) | 39 | 0.80 | 0.210 | 0.999 | |
| AT2G01140.1 | Fructose-bisphosphate aldolase 3 (FBA3) | 41 | 0.90 | 0.274 | 1.000 | |
| AT2G01140.1 | Fructose-bisphosphate aldolase 3 (FBA3) | 42 | 0.95 | 0.319 | 1.000 | |
| AT2G01140.1 | Fructose-bisphosphate aldolase 3 (FBA3) | 43 | 1.01 | 0.373 | 1.000 | |
| AT2G01140.1 | Fructose-bisphosphate aldolase 3 (FBA3) | 46 | 1.19 | 0.586 | 1.000 | |
| AT2G01140.1 | Fructose-bisphosphate aldolase 3 (FBA3) | 47 | 1.25 | 0.673 | 1.000 | |
| AT2G01460.2 | P-loop containing nucleoside triphosphate hydrolase | 2 | 0.96 | 0.532 | 1.000 | |
| AT2G01460.2 | P-loop containing nucleoside triphosphate hydrolase | 3 | 0.96 | 0.504 | 1.000 | |
| AT2G01460.2 | P-loop containing nucleoside triphosphate hydrolase | 21 | 0.99 | 0.181 | 1.000 | |
| AT2G01460.2 | P-loop containing nucleoside triphosphate hydrolase | 22 | 1.00 | 0.170 | 1.000 | |
| AT2G01460.2 | P-loop containing nucleoside triphosphate hydrolase | 23 | 1.00 | 0.163 | 1.000 | |
| AT2G01460.2 | P-loop containing nucleoside triphosphate hydrolase | 24 | 1.00 | 0.160 | 1.000 | |
| AT2G01460.2 | P-loop containing nucleoside triphosphate hydrolase | 25 | 0.99 | 0.161 | 1.000 | |
| AT2G01460.2 | P-loop containing nucleoside triphosphate hydrolase | 26 | 0.97 | 0.167 | 1.000 | |
| AT2G01460.2 | P-loop containing nucleoside triphosphate hydrolase | 27 | 0.94 | 0.177 | 1.000 | |
| AT2G01460.2 | P-loop containing nucleoside triphosphate hydrolase | 28 | 0.90 | 0.191 | 0.999 | |
| AT2G01460.2 | P-loop containing nucleoside triphosphate hydrolase | 29 | 0.85 | 0.206 | 0.995 | |
| AT2G01460.2 | P-loop containing nucleoside triphosphate hydrolase | 46 | 0.10 | 0.439 | 0.107 | |
| AT2G01460.2 | P-loop containing nucleoside triphosphate hydrolase | 47 | 0.09 | 0.460 | 0.097 | . |

| | | | | | |
|---|---|---|---|---|---|
| AT2G01970.1 | Transmembrane nine protein 3 (TMN3) | 2 | 1.03 | 0.548 | 1.000 |
| AT2G01970.1 | Transmembrane nine protein 3 (TMN3) | 3 | 1.00 | 0.339 | 1.000 |
| AT2G01970.1 | Transmembrane nine protein 3 (TMN3) | 4 | 0.96 | 0.345 | 1.000 |
| AT2G01970.1 | Transmembrane nine protein 3 (TMN3) | 5 | 0.91 | 0.397 | 1.000 |
| AT2G01970.1 | Transmembrane nine protein 3 (TMN3) | 6 | 0.87 | 0.450 | 1.000 |
| AT2G01970.1 | Transmembrane nine protein 3 (TMN3) | 7 | 0.88 | 0.477 | 1.000 |
| AT2G01970.1 | Transmembrane nine protein 3 (TMN3) | 8 | 0.91 | 0.561 | 1.000 |
| AT2G01970.1 | Transmembrane nine protein 3 (TMN3) | 22 | 0.98 | 0.578 | 1.000 |
| AT2G01970.1 | Transmembrane nine protein 3 (TMN3) | 23 | 0.95 | 0.548 | 1.000 |
| AT2G01970.1 | Transmembrane nine protein 3 (TMN3) | 24 | 0.92 | 0.643 | 1.000 |
| AT2G01970.1 | Transmembrane nine protein 3 (TMN3) | 33 | 2.09 | 0.511 | 1.000 |
| AT2G01970.1 | Transmembrane nine protein 3 (TMN3) | 34 | 3.23 | 0.452 | 0.975 |
| AT2G01970.1 | Transmembrane nine protein 3 (TMN3) | 35 | 5.07 | 0.458 | 0.816 |
| AT2G01970.1 | Transmembrane nine protein 3 (TMN3) | 37 | 6.63 | 0.377 | 0.365 |
| AT2G01970.1 | Transmembrane nine protein 3 (TMN3) | 38 | 2.92 | 0.326 | 0.880 |
| AT2G01970.1 | Transmembrane nine protein 3 (TMN3) | 39 | 0.69 | 0.290 | 1.000 |
| AT2G01970.1 | Transmembrane nine protein 3 (TMN3) | 40 | 0.22 | 0.281 | 0.277 |
| AT2G01970.1 | Transmembrane nine protein 3 (TMN3) | 41 | 0.22 | 0.306 | 0.391 |
| AT2G01970.1 | Transmembrane nine protein 3 (TMN3) | 42 | 0.49 | 0.410 | 0.999 |
| AT2G01970.1 | Transmembrane nine protein 3 (TMN3) | 43 | 1.57 | 0.634 | 1.000 |
| AT2G01970.1 | Transmembrane nine protein 3 (TMN3) | 46 | 194.27 | 1.912 | 0.961 |
| AT2G01970.1 | Transmembrane nine protein 3 (TMN3) | 47 | 1192.73 | 2.540 | 0.957 |
| AT2G02050.1 | NADH:ubiquinone oxidoreductase subunit (B18) | 2 | 0.92 | 0.655 | 1.000 |
| AT2G02050.1 | NADH:ubiquinone oxidoreductase subunit (B18) | 3 | 0.89 | 0.589 | 1.000 |
| AT2G02050.1 | NADH:ubiquinone oxidoreductase subunit (B18) | 12 | 0.60 | 0.402 | 1.000 |
| AT2G02050.1 | NADH:ubiquinone oxidoreductase subunit (B18) | 13 | 0.76 | 0.366 | 1.000 |
| AT2G02050.1 | NADH:ubiquinone oxidoreductase subunit (B18) | 14 | 1.13 | 0.342 | 1.000 |
| AT2G02050.1 | NADH:ubiquinone oxidoreductase subunit (B18) | 15 | 0.92 | 0.309 | 1.000 |
| AT2G02050.1 | NADH:ubiquinone oxidoreductase subunit (B18) | 16 | 0.55 | 0.292 | 0.999 |
| AT2G02050.1 | NADH:ubiquinone oxidoreductase subunit (B18) | 17 | 0.38 | 0.286 | 0.924 |
| AT2G02050.1 | NADH:ubiquinone oxidoreductase subunit (B18) | 18 | 0.46 | 0.291 | 0.992 |

| | | | | | |
|---|---|---|---|---|---|
| AT2G02050.1 | NADH:ubiquinone oxidoreductase subunit (B18) | 19 | 0.68 | 0.303 | 1.000 |
| AT2G02050.1 | NADH:ubiquinone oxidoreductase subunit (B18) | 20 | 0.78 | 0.311 | 1.000 |
| AT2G02050.1 | NADH:ubiquinone oxidoreductase subunit (B18) | 21 | 0.95 | 0.314 | 1.000 |
| AT2G02050.1 | NADH:ubiquinone oxidoreductase subunit (B18) | 22 | 1.58 | 0.312 | 1.000 |
| AT2G02050.1 | NADH:ubiquinone oxidoreductase subunit (B18) | 23 | 2.85 | 0.312 | 0.930 |
| AT2G02050.1 | NADH:ubiquinone oxidoreductase subunit (B18) | 24 | 2.42 | 0.301 | 0.977 |
| AT2G02050.1 | NADH:ubiquinone oxidoreductase subunit (B18) | 25 | 0.62 | 0.297 | 1.000 |
| AT2G02050.1 | NADH:ubiquinone oxidoreductase subunit (B18) | 26 | 0.21 | 0.298 | 0.357 |
| AT2G02050.1 | NADH:ubiquinone oxidoreductase subunit (B18) | 27 | 0.26 | 0.298 | 0.592 |
| AT2G02050.1 | NADH:ubiquinone oxidoreductase subunit (B18) | 28 | 0.74 | 0.297 | 1.000 |
| AT2G02050.1 | NADH:ubiquinone oxidoreductase subunit (B18) | 29 | 2.54 | 0.303 | 0.965 |
| AT2G02050.1 | NADH:ubiquinone oxidoreductase subunit (B18) | 30 | 3.38 | 0.326 | 0.853 |
| AT2G02050.1 | NADH:ubiquinone oxidoreductase subunit (B18) | 31 | 1.16 | 0.405 | 1.000 |
| AT2G02050.1 | NADH:ubiquinone oxidoreductase subunit (B18) | 32 | 0.24 | 0.696 | 1.000 |
| AT2G02050.1 | NADH:ubiquinone oxidoreductase subunit (B18) | 46 | 0.03 | 8.907 | 1.000 |
| AT2G02050.1 | NADH:ubiquinone oxidoreductase subunit (B18) | 47 | 0.05 | 9.825 | 1.000 |
| AT2G02510.1 | NADH:ubiquinone oxidoreductase subunit (B12-2) | 2 | 2.03 | 0.546 | 1.000 |
| AT2G02510.1 | NADH:ubiquinone oxidoreductase subunit (B12-2) | 3 | 1.31 | 0.439 | 1.000 |
| AT2G02510.1 | NADH:ubiquinone oxidoreductase subunit (B12-2) | 4 | 0.85 | 0.432 | 1.000 |
| AT2G02510.1 | NADH:ubiquinone oxidoreductase subunit (B12-2) | 6 | 0.51 | 0.440 | 1.000 |
| AT2G02510.1 | NADH:ubiquinone oxidoreductase subunit (B12-2) | 7 | 0.54 | 0.412 | 1.000 |
| AT2G02510.1 | NADH:ubiquinone oxidoreductase subunit (B12-2) | 8 | 0.64 | 0.395 | 1.000 |
| AT2G02510.1 | NADH:ubiquinone oxidoreductase subunit (B12-2) | 9 | 0.73 | 0.385 | 1.000 |
| AT2G02510.1 | NADH:ubiquinone oxidoreductase subunit (B12-2) | 10 | 0.78 | 0.380 | 1.000 |
| AT2G02510.1 | NADH:ubiquinone oxidoreductase subunit (B12-2) | 11 | 0.81 | 0.374 | 1.000 |
| AT2G02510.1 | NADH:ubiquinone oxidoreductase subunit (B12-2) | 12 | 0.80 | 0.367 | 1.000 |
| AT2G02510.1 | NADH:ubiquinone oxidoreductase subunit (B12-2) | 13 | 0.69 | 0.361 | 1.000 |
| AT2G02510.1 | NADH:ubiquinone oxidoreductase subunit (B12-2) | 14 | 0.53 | 0.355 | 0.999 |
| AT2G02510.1 | NADH:ubiquinone oxidoreductase subunit (B12-2) | 15 | 0.35 | 0.349 | 0.897 |
| AT2G02510.1 | NADH:ubiquinone oxidoreductase subunit (B12-2) | 16 | 0.22 | 0.347 | 0.485 |
| AT2G02510.1 | NADH:ubiquinone oxidoreductase subunit (B12-2) | 17 | 0.15 | 0.346 | 0.210 |

| | | | | | | |
|---|---|---|---|---|---|---|
| AT2G02510.1 | NADH:ubiquinone oxidoreductase subunit (B12-2) | 18 | 0.13 | 0.348 | 0.124 | |
| AT2G02510.1 | NADH:ubiquinone oxidoreductase subunit (B12-2) | 19 | 0.11 | 0.352 | 0.096 | . |
| AT2G02510.1 | NADH:ubiquinone oxidoreductase subunit (B12-2) | 20 | 0.11 | 0.356 | 0.103 | |
| AT2G02510.1 | NADH:ubiquinone oxidoreductase subunit (B12-2) | 21 | 0.12 | 0.358 | 0.138 | |
| AT2G02510.1 | NADH:ubiquinone oxidoreductase subunit (B12-2) | 22 | 0.17 | 0.358 | 0.319 | |
| AT2G02510.1 | NADH:ubiquinone oxidoreductase subunit (B12-2) | 23 | 0.31 | 0.356 | 0.828 | |
| AT2G02510.1 | NADH:ubiquinone oxidoreductase subunit (B12-2) | 24 | 0.47 | 0.353 | 0.991 | |
| AT2G02510.1 | NADH:ubiquinone oxidoreductase subunit (B12-2) | 25 | 0.40 | 0.351 | 0.961 | |
| AT2G02510.1 | NADH:ubiquinone oxidoreductase subunit (B12-2) | 26 | 0.28 | 0.350 | 0.718 | |
| AT2G02510.1 | NADH:ubiquinone oxidoreductase subunit (B12-2) | 27 | 0.23 | 0.350 | 0.513 | |
| AT2G02510.1 | NADH:ubiquinone oxidoreductase subunit (B12-2) | 28 | 0.23 | 0.351 | 0.516 | |
| AT2G02510.1 | NADH:ubiquinone oxidoreductase subunit (B12-2) | 29 | 0.26 | 0.351 | 0.633 | |
| AT2G02510.1 | NADH:ubiquinone oxidoreductase subunit (B12-2) | 30 | 0.28 | 0.353 | 0.720 | |
| AT2G02510.1 | NADH:ubiquinone oxidoreductase subunit (B12-2) | 31 | 0.28 | 0.358 | 0.732 | |
| AT2G02510.1 | NADH:ubiquinone oxidoreductase subunit (B12-2) | 32 | 0.29 | 0.374 | 0.812 | |
| AT2G02510.1 | NADH:ubiquinone oxidoreductase subunit (B12-2) | 33 | 0.35 | 0.427 | 0.971 | |
| AT2G02510.1 | NADH:ubiquinone oxidoreductase subunit (B12-2) | 34 | 0.47 | 0.554 | 1.000 | |
| AT2G02510.1 | NADH:ubiquinone oxidoreductase subunit (B12-2) | 46 | 14313.91 | 4.505 | 0.993 | |
| AT2G02510.1 | NADH:ubiquinone oxidoreductase subunit (B12-2) | 47 | 44064.73 | 5.014 | 0.993 | |
| AT2G04340.1 | Cytoplasmic dynein 2 light intermediate chain | 2 | 0.86 | 0.687 | 1.000 | |
| AT2G04340.1 | Cytoplasmic dynein 2 light intermediate chain | 3 | 0.99 | 1.273 | 1.000 | |
| AT2G04340.1 | Cytoplasmic dynein 2 light intermediate chain | 23 | 0.37 | 0.485 | 0.976 | |
| AT2G04340.1 | Cytoplasmic dynein 2 light intermediate chain | 24 | 0.35 | 0.343 | 0.801 | |
| AT2G04340.1 | Cytoplasmic dynein 2 light intermediate chain | 25 | 0.56 | 0.313 | 0.988 | |
| AT2G04340.1 | Cytoplasmic dynein 2 light intermediate chain | 26 | 1.60 | 0.303 | 0.996 | |
| AT2G04340.1 | Cytoplasmic dynein 2 light intermediate chain | 27 | 2.09 | 0.324 | 0.955 | |
| AT2G04340.1 | Cytoplasmic dynein 2 light intermediate chain | 28 | 0.87 | 0.403 | 1.000 | |
| AT2G04340.1 | Cytoplasmic dynein 2 light intermediate chain | 29 | 0.32 | 0.651 | 0.991 | |
| AT2G04340.1 | Cytoplasmic dynein 2 light intermediate chain | 46 | ########### | 15.610 | 0.983 | |
| AT2G04340.1 | Cytoplasmic dynein 2 light intermediate chain | 47 | ########### | 16.985 | 0.981 | |
| AT2G04540.1 | Beta-ketoacyl-ACP synthase (MTKAS) | 2 | 0.89 | 0.634 | 1.000 | |

| | | | | | | |
|---|---|---|---|---|---|---|
| AT2G04540.1 | Beta-ketoacyl-ACP synthase (MTKAS) | 3 | 0.85 | 0.605 | 1.000 | |
| AT2G04540.1 | Beta-ketoacyl-ACP synthase (MTKAS) | 31 | 1.42 | 0.323 | 0.999 | |
| AT2G04540.1 | Beta-ketoacyl-ACP synthase (MTKAS) | 32 | 1.65 | 0.274 | 0.973 | |
| AT2G04540.1 | Beta-ketoacyl-ACP synthase (MTKAS) | 33 | 1.98 | 0.240 | 0.799 | |
| AT2G04540.1 | Beta-ketoacyl-ACP synthase (MTKAS) | 34 | 2.43 | 0.220 | 0.447 | |
| AT2G04540.1 | Beta-ketoacyl-ACP synthase (MTKAS) | 35 | 3.01 | 0.212 | 0.177 | |
| AT2G04540.1 | Beta-ketoacyl-ACP synthase (MTKAS) | 36 | 3.62 | 0.210 | 0.071 | . |
| AT2G04540.1 | Beta-ketoacyl-ACP synthase (MTKAS) | 37 | 4.09 | 0.211 | 0.037 | * |
| AT2G04540.1 | Beta-ketoacyl-ACP synthase (MTKAS) | 38 | 4.25 | 0.211 | 0.030 | * |
| AT2G04540.1 | Beta-ketoacyl-ACP synthase (MTKAS) | 39 | 4.05 | 0.209 | 0.037 | * |
| AT2G04540.1 | Beta-ketoacyl-ACP synthase (MTKAS) | 40 | 3.60 | 0.203 | 0.057 | . |
| AT2G04540.1 | Beta-ketoacyl-ACP synthase (MTKAS) | 41 | 3.06 | 0.196 | 0.110 | |
| AT2G04540.1 | Beta-ketoacyl-ACP synthase (MTKAS) | 42 | 2.51 | 0.190 | 0.242 | |
| AT2G04540.1 | Beta-ketoacyl-ACP synthase (MTKAS) | 43 | 1.99 | 0.188 | 0.558 | |
| AT2G04540.1 | Beta-ketoacyl-ACP synthase (MTKAS) | 44 | 1.52 | 0.195 | 0.938 | |
| AT2G04540.1 | Beta-ketoacyl-ACP synthase (MTKAS) | 45 | 1.14 | 0.222 | 1.000 | |
| AT2G04540.1 | Beta-ketoacyl-ACP synthase (MTKAS) | 46 | 0.85 | 0.278 | 1.000 | |
| AT2G04540.1 | Beta-ketoacyl-ACP synthase (MTKAS) | 47 | 0.63 | 0.365 | 0.997 | |
| AT2G04900.2 | ? | 2 | 0.89 | 0.552 | 1.000 | |
| AT2G04900.2 | ? | 3 | 1.30 | 0.545 | 1.000 | |
| AT2G04900.2 | ? | 39 | 5.92 | 0.381 | 0.215 | |
| AT2G04900.2 | ? | 40 | 2.81 | 0.279 | 0.422 | |
| AT2G04900.2 | ? | 41 | 1.22 | 0.234 | 0.997 | |
| AT2G04900.2 | ? | 42 | 0.48 | 0.273 | 0.712 | |
| AT2G04900.2 | ? | 43 | 0.18 | 0.376 | 0.231 | |
| AT2G04900.2 | ? | 46 | 0.01 | 0.842 | 0.077 | . |
| AT2G04900.2 | ? | 47 | 0.00 | 1.035 | 0.078 | . |
| AT2G04940.1 | Scramblase-related | 2 | 1.00 | 0.493 | 1.000 | |
| AT2G04940.1 | Scramblase-related | 3 | 0.98 | 0.448 | 1.000 | |
| AT2G04940.1 | Scramblase-related | 16 | 0.77 | 0.255 | 1.000 | |
| AT2G04940.1 | Scramblase-related | 17 | 0.78 | 0.216 | 1.000 | |
| AT2G04940.1 | Scramblase-related | 18 | 0.83 | 0.192 | 1.000 | |
| AT2G04940.1 | Scramblase-related | 19 | 0.90 | 0.183 | 1.000 | |
| AT2G04940.1 | Scramblase-related | 20 | 0.94 | 0.181 | 1.000 | |
| AT2G04940.1 | Scramblase-related | 21 | 0.90 | 0.182 | 1.000 | |
| AT2G04940.1 | Scramblase-related | 22 | 0.85 | 0.188 | 1.000 | |
| AT2G04940.1 | Scramblase-related | 23 | 0.78 | 0.206 | 1.000 | |
| AT2G04940.1 | Scramblase-related | 24 | 0.71 | 0.252 | 1.000 | |
| AT2G04940.1 | Scramblase-related | 25 | 0.65 | 0.326 | 1.000 | |
| AT2G04940.1 | Scramblase-related | 36 | 1.59 | 0.294 | 1.000 | |
| AT2G04940.1 | Scramblase-related | 37 | 1.80 | 0.237 | 0.975 | |
| AT2G04940.1 | Scramblase-related | 38 | 1.98 | 0.216 | 0.883 | |
| AT2G04940.1 | Scramblase-related | 39 | 2.21 | 0.227 | 0.800 | |
| AT2G04940.1 | Scramblase-related | 40 | 2.65 | 0.240 | 0.629 | |

| | | | | | | |
|---|---|---|---|---|---|---|
| AT2G04940.1 | Scramblase-related | 42 | 3.88 | 0.236 | 0.169 | |
| AT2G04940.1 | Scramblase-related | 43 | 4.12 | 0.210 | 0.056 | . |
| AT2G04940.1 | Scramblase-related | 44 | 3.10 | 0.191 | 0.141 | |
| AT2G04940.1 | Scramblase-related | 45 | 1.46 | 0.191 | 0.996 | |
| AT2G04940.1 | Scramblase-related | 46 | 0.47 | 0.235 | 0.880 | |
| AT2G04940.1 | Scramblase-related | 47 | 0.13 | 0.353 | 0.170 | |
| AT2G05070.1 | Photosystem II light harvesting complex subunit (LHCB2.2) | 2 | 1.07 | 0.502 | 1.000 | |
| AT2G05070.1 | Photosystem II light harvesting complex subunit (LHCB2.2) | 3 | 1.45 | 0.367 | 1.000 | |
| AT2G05070.1 | Photosystem II light harvesting complex subunit (LHCB2.2) | 4 | 1.83 | 0.339 | 1.000 | |
| AT2G05070.1 | Photosystem II light harvesting complex subunit (LHCB2.2) | 5 | 1.85 | 0.337 | 1.000 | |
| AT2G05070.1 | Photosystem II light harvesting complex subunit (LHCB2.2) | 6 | 1.42 | 0.338 | 1.000 | |
| AT2G05070.1 | Photosystem II light harvesting complex subunit (LHCB2.2) | 7 | 1.00 | 0.337 | 1.000 | |
| AT2G05070.1 | Photosystem II light harvesting complex subunit (LHCB2.2) | 8 | 0.83 | 0.334 | 1.000 | |
| AT2G05070.1 | Photosystem II light harvesting complex subunit (LHCB2.2) | 9 | 0.83 | 0.331 | 1.000 | |
| AT2G05070.1 | Photosystem II light harvesting complex subunit (LHCB2.2) | 10 | 0.91 | 0.331 | 1.000 | |
| AT2G05070.1 | Photosystem II light harvesting complex subunit (LHCB2.2) | 11 | 1.08 | 0.332 | 1.000 | |
| AT2G05070.1 | Photosystem II light harvesting complex subunit (LHCB2.2) | 12 | 1.29 | 0.332 | 1.000 | |
| AT2G05070.1 | Photosystem II light harvesting complex subunit (LHCB2.2) | 13 | 1.51 | 0.329 | 1.000 | |
| AT2G05070.1 | Photosystem II light harvesting complex subunit (LHCB2.2) | 14 | 1.68 | 0.327 | 1.000 | |
| AT2G05070.1 | Photosystem II light harvesting complex subunit (LHCB2.2) | 15 | 1.76 | 0.326 | 1.000 | |
| AT2G05070.1 | Photosystem II light harvesting complex subunit (LHCB2.2) | 16 | 1.83 | 0.325 | 1.000 | |
| AT2G05070.1 | Photosystem II light harvesting complex subunit (LHCB2.2) | 17 | 2.12 | 0.325 | 0.996 | |
| AT2G05070.1 | Photosystem II light harvesting complex subunit (LHCB2.2) | 18 | 2.69 | 0.327 | 0.943 | |
| AT2G05070.1 | Photosystem II light harvesting complex subunit (LHCB2.2) | 19 | 3.41 | 0.332 | 0.787 | |
| AT2G05070.1 | Photosystem II light harvesting complex subunit (LHCB2.2) | 20 | 4.09 | 0.336 | 0.628 | |
| AT2G05070.1 | Photosystem II light harvesting complex subunit (LHCB2.2) | 21 | 4.50 | 0.339 | 0.540 | |
| AT2G05070.1 | Photosystem II light harvesting complex subunit (LHCB2.2) | 22 | 4.04 | 0.339 | 0.650 | |
| AT2G05070.1 | Photosystem II light harvesting complex subunit (LHCB2.2) | 23 | 2.66 | 0.338 | 0.960 | |
| AT2G05070.1 | Photosystem II light harvesting complex subunit (LHCB2.2) | 24 | 1.44 | 0.333 | 1.000 | |
| AT2G05070.1 | Photosystem II light harvesting complex subunit (LHCB2.2) | 25 | 0.79 | 0.330 | 1.000 | |
| AT2G05070.1 | Photosystem II light harvesting complex subunit (LHCB2.2) | 26 | 0.58 | 0.329 | 1.000 | |
| AT2G05070.1 | Photosystem II light harvesting complex subunit (LHCB2.2) | 27 | 0.60 | 0.329 | 1.000 | |
| AT2G05070.1 | Photosystem II light harvesting complex subunit (LHCB2.2) | 28 | 0.78 | 0.330 | 1.000 | |
| AT2G05070.1 | Photosystem II light harvesting complex subunit (LHCB2.2) | 29 | 1.21 | 0.330 | 1.000 | |

| | | | | | | |
|---|---|---|---|---|---|---|
| AT2G05070.1 | Photosystem II light harvesting complex subunit (LHCB2.2) | 30 | 1.90 | 0.330 | 1.000 | |
| AT2G05070.1 | Photosystem II light harvesting complex subunit (LHCB2.2) | 31 | 2.97 | 0.331 | 0.893 | |
| AT2G05070.1 | Photosystem II light harvesting complex subunit (LHCB2.2) | 32 | 4.49 | 0.331 | 0.509 | |
| AT2G05070.1 | Photosystem II light harvesting complex subunit (LHCB2.2) | 33 | 6.93 | 0.331 | 0.175 | |
| AT2G05070.1 | Photosystem II light harvesting complex subunit (LHCB2.2) | 34 | 9.65 | 0.331 | 0.059 | . |
| AT2G05070.1 | Photosystem II light harvesting complex subunit (LHCB2.2) | 35 | 8.65 | 0.332 | 0.088 | . |
| AT2G05070.1 | Photosystem II light harvesting complex subunit (LHCB2.2) | 36 | 3.95 | 0.336 | 0.659 | |
| AT2G05070.1 | Photosystem II light harvesting complex subunit (LHCB2.2) | 37 | 1.29 | 0.351 | 1.000 | |
| AT2G05070.1 | Photosystem II light harvesting complex subunit (LHCB2.2) | 38 | 0.62 | 0.390 | 1.000 | |
| AT2G05070.1 | Photosystem II light harvesting complex subunit (LHCB2.2) | 39 | 0.51 | 0.448 | 1.000 | |
| AT2G05070.1 | Photosystem II light harvesting complex subunit (LHCB2.2) | 42 | 1.17 | 0.455 | 1.000 | |
| AT2G05070.1 | Photosystem II light harvesting complex subunit (LHCB2.2) | 43 | 1.52 | 0.483 | 1.000 | |
| AT2G05070.1 | Photosystem II light harvesting complex subunit (LHCB2.2) | 44 | 1.59 | 0.533 | 1.000 | |
| AT2G05070.1 | Photosystem II light harvesting complex subunit (LHCB2.2) | 46 | 1.09 | 0.840 | 1.000 | |
| AT2G05070.1 | Photosystem II light harvesting complex subunit (LHCB2.2) | 47 | 0.81 | 1.090 | 1.000 | |
| AT2G05310.1 | [MISSING] | 2 | 1.08 | 0.439 | 1.000 | |
| AT2G05310.1 | [MISSING] | 3 | 1.13 | 0.421 | 1.000 | |
| AT2G05310.1 | [MISSING] | 4 | 1.18 | 0.412 | 1.000 | |
| AT2G05310.1 | [MISSING] | 30 | 1.35 | 0.284 | 0.974 | |
| AT2G05310.1 | [MISSING] | 31 | 1.25 | 0.281 | 0.991 | |
| AT2G05310.1 | [MISSING] | 32 | 1.15 | 0.282 | 0.999 | |
| AT2G05310.1 | [MISSING] | 33 | 1.04 | 0.289 | 1.000 | |
| AT2G05310.1 | [MISSING] | 34 | 0.93 | 0.299 | 1.000 | |
| AT2G05310.1 | [MISSING] | 46 | 0.11 | 0.464 | 0.158 | |
| AT2G05310.1 | [MISSING] | 47 | 0.09 | 0.488 | 0.134 | |
| AT2G05710.1 | Aconitase 3 (ACO3) | 2 | 0.47 | 0.278 | 0.940 | |
| AT2G05710.1 | Aconitase 3 (ACO3) | 3 | 0.45 | 0.223 | 0.744 | |
| AT2G05710.1 | Aconitase 3 (ACO3) | 4 | 0.45 | 0.192 | 0.534 | |
| AT2G05710.1 | Aconitase 3 (ACO3) | 5 | 0.46 | 0.180 | 0.476 | |
| AT2G05710.1 | Aconitase 3 (ACO3) | 6 | 0.49 | 0.175 | 0.569 | |
| AT2G05710.1 | Aconitase 3 (ACO3) | 7 | 0.55 | 0.172 | 0.759 | |
| AT2G05710.1 | Aconitase 3 (ACO3) | 8 | 0.63 | 0.170 | 0.939 | |
| AT2G05710.1 | Aconitase 3 (ACO3) | 9 | 0.73 | 0.168 | 0.998 | |
| AT2G05710.1 | Aconitase 3 (ACO3) | 10 | 0.86 | 0.167 | 1.000 | |
| AT2G05710.1 | Aconitase 3 (ACO3) | 11 | 1.01 | 0.166 | 1.000 | |
| AT2G05710.1 | Aconitase 3 (ACO3) | 12 | 1.18 | 0.166 | 1.000 | |
| AT2G05710.1 | Aconitase 3 (ACO3) | 13 | 1.35 | 0.165 | 0.999 | |
| AT2G05710.1 | Aconitase 3 (ACO3) | 14 | 1.51 | 0.164 | 0.971 | |
| AT2G05710.1 | Aconitase 3 (ACO3) | 15 | 1.67 | 0.164 | 0.868 | |
| AT2G05710.1 | Aconitase 3 (ACO3) | 16 | 1.83 | 0.164 | 0.699 | |
| AT2G05710.1 | Aconitase 3 (ACO3) | 17 | 1.98 | 0.164 | 0.539 | |
| AT2G05710.1 | Aconitase 3 (ACO3) | 18 | 2.07 | 0.165 | 0.469 | |
| AT2G05710.1 | Aconitase 3 (ACO3) | 19 | 2.04 | 0.166 | 0.507 | |
| AT2G05710.1 | Aconitase 3 (ACO3) | 20 | 1.90 | 0.168 | 0.663 | |
| AT2G05710.1 | Aconitase 3 (ACO3) | 21 | 1.68 | 0.169 | 0.875 | |
| AT2G05710.1 | Aconitase 3 (ACO3) | 22 | 1.45 | 0.169 | 0.989 | |
| AT2G05710.1 | Aconitase 3 (ACO3) | 23 | 1.25 | 0.168 | 1.000 | |

| | | | | | | |
|---|---|---|---|---|---|---|
| AT2G05710.1 | Aconitase 3 (ACO3) | 24 | 1.10 | 0.167 | 1.000 | |
| AT2G05710.1 | Aconitase 3 (ACO3) | 25 | 0.99 | 0.166 | 1.000 | |
| AT2G05710.1 | Aconitase 3 (ACO3) | 26 | 0.91 | 0.166 | 1.000 | |
| AT2G05710.1 | Aconitase 3 (ACO3) | 27 | 0.86 | 0.165 | 1.000 | |
| AT2G05710.1 | Aconitase 3 (ACO3) | 28 | 0.85 | 0.165 | 1.000 | |
| AT2G05710.1 | Aconitase 3 (ACO3) | 29 | 0.86 | 0.166 | 1.000 | |
| AT2G05710.1 | Aconitase 3 (ACO3) | 30 | 0.88 | 0.166 | 1.000 | |
| AT2G05710.1 | Aconitase 3 (ACO3) | 31 | 0.90 | 0.166 | 1.000 | |
| AT2G05710.1 | Aconitase 3 (ACO3) | 32 | 0.93 | 0.166 | 1.000 | |
| AT2G05710.1 | Aconitase 3 (ACO3) | 33 | 0.97 | 0.166 | 1.000 | |
| AT2G05710.1 | Aconitase 3 (ACO3) | 34 | 1.02 | 0.166 | 1.000 | |
| AT2G05710.1 | Aconitase 3 (ACO3) | 35 | 1.08 | 0.166 | 1.000 | |
| AT2G05710.1 | Aconitase 3 (ACO3) | 36 | 1.16 | 0.166 | 1.000 | |
| AT2G05710.1 | Aconitase 3 (ACO3) | 37 | 1.25 | 0.166 | 1.000 | |
| AT2G05710.1 | Aconitase 3 (ACO3) | 38 | 1.34 | 0.166 | 0.999 | |
| AT2G05710.1 | Aconitase 3 (ACO3) | 39 | 1.43 | 0.167 | 0.992 | |
| AT2G05710.1 | Aconitase 3 (ACO3) | 40 | 1.49 | 0.168 | 0.980 | |
| AT2G05710.1 | Aconitase 3 (ACO3) | 41 | 1.49 | 0.168 | 0.980 | |
| AT2G05710.1 | Aconitase 3 (ACO3) | 42 | 1.42 | 0.169 | 0.994 | |
| AT2G05710.1 | Aconitase 3 (ACO3) | 43 | 1.27 | 0.169 | 1.000 | |
| AT2G05710.1 | Aconitase 3 (ACO3) | 44 | 1.07 | 0.171 | 1.000 | |
| AT2G05710.1 | Aconitase 3 (ACO3) | 45 | 0.84 | 0.182 | 1.000 | |
| AT2G05710.1 | Aconitase 3 (ACO3) | 46 | 0.64 | 0.213 | 0.993 | |
| AT2G05710.1 | Aconitase 3 (ACO3) | 47 | 0.47 | 0.273 | 0.942 | |
| AT2G05990.2 | Enoyl-ACP reductase 1 (ENR1) | 2 | 1.13 | 0.609 | 1.000 | |
| AT2G05990.2 | Enoyl-ACP reductase 1 (ENR1) | 3 | 0.55 | 1.324 | 1.000 | |
| AT2G05990.2 | Enoyl-ACP reductase 1 (ENR1) | 38 | 0.21 | 0.362 | 0.379 | |
| AT2G05990.2 | Enoyl-ACP reductase 1 (ENR1) | 39 | 0.90 | 0.271 | 1.000 | |
| AT2G05990.2 | Enoyl-ACP reductase 1 (ENR1) | 40 | 2.69 | 0.263 | 0.550 | |
| AT2G05990.2 | Enoyl-ACP reductase 1 (ENR1) | 41 | 2.21 | 0.276 | 0.810 | |
| AT2G05990.2 | Enoyl-ACP reductase 1 (ENR1) | 42 | 0.71 | 0.364 | 1.000 | |
| AT2G05990.2 | Enoyl-ACP reductase 1 (ENR1) | 43 | 0.26 | 0.520 | 0.868 | |
| AT2G05990.2 | Enoyl-ACP reductase 1 (ENR1) | 46 | 0.05 | 0.365 | 0.006 | ** |
| AT2G05990.2 | Enoyl-ACP reductase 1 (ENR1) | 47 | 0.03 | 0.417 | 0.007 | ** |
| AT2G07050.1 | Cycloartenol synthase 1 (CAS1) | 2 | 0.85 | 0.453 | 1.000 | |
| AT2G07050.1 | Cycloartenol synthase 1 (CAS1) | 3 | 0.62 | 0.445 | 0.999 | |
| AT2G07050.1 | Cycloartenol synthase 1 (CAS1) | 41 | 3.25 | 0.486 | 0.913 | |
| AT2G07050.1 | Cycloartenol synthase 1 (CAS1) | 42 | 5.47 | 0.289 | 0.102 | |
| AT2G07050.1 | Cycloartenol synthase 1 (CAS1) | 43 | 3.55 | 0.244 | 0.196 | |
| AT2G07050.1 | Cycloartenol synthase 1 (CAS1) | 44 | 0.82 | 0.233 | 1.000 | |
| AT2G07050.1 | Cycloartenol synthase 1 (CAS1) | 45 | 0.26 | 0.234 | 0.110 | |
| AT2G07050.1 | Cycloartenol synthase 1 (CAS1) | 46 | 0.25 | 0.247 | 0.127 | |
| AT2G07050.1 | Cycloartenol synthase 1 (CAS1) | 47 | 0.38 | 0.593 | 0.989 | |
| AT2G07698.1 | ATP synthase subunit (ATPalpha) | 3 | 0.81 | 0.209 | 1.000 | |
| AT2G07698.1 | ATP synthase subunit (ATPalpha) | 4 | 1.52 | 0.187 | 1.000 | |
| AT2G07698.1 | ATP synthase subunit (ATPalpha) | 5 | 2.53 | 0.188 | 0.539 | |
| AT2G07698.1 | ATP synthase subunit (ATPalpha) | 6 | 3.20 | 0.188 | 0.181 | |
| AT2G07698.1 | ATP synthase subunit (ATPalpha) | 7 | 3.16 | 0.187 | 0.187 | |
| AT2G07698.1 | ATP synthase subunit (ATPalpha) | 8 | 2.67 | 0.184 | 0.407 | |
| AT2G07698.1 | ATP synthase subunit (ATPalpha) | 9 | 2.34 | 0.182 | 0.642 | |
| AT2G07698.1 | ATP synthase subunit (ATPalpha) | 10 | 2.33 | 0.182 | 0.641 | |
| AT2G07698.1 | ATP synthase subunit (ATPalpha) | 11 | 2.66 | 0.182 | 0.389 | |
| AT2G07698.1 | ATP synthase subunit (ATPalpha) | 12 | 3.15 | 0.182 | 0.162 | |

| | | | | | | |
|---|---|---|---|---|---|---|
| AT2G07698.1 | ATP synthase subunit (ATPalpha) | 13 | 3.51 | 0.180 | 0.076 | . |
| AT2G07698.1 | ATP synthase subunit (ATPalpha) | 14 | 3.48 | 0.179 | 0.073 | . |
| AT2G07698.1 | ATP synthase subunit (ATPalpha) | 15 | 2.99 | 0.178 | 0.190 | |
| AT2G07698.1 | ATP synthase subunit (ATPalpha) | 16 | 2.40 | 0.177 | 0.532 | |
| AT2G07698.1 | ATP synthase subunit (ATPalpha) | 17 | 2.11 | 0.177 | 0.784 | |
| AT2G07698.1 | ATP synthase subunit (ATPalpha) | 18 | 1.93 | 0.179 | 0.921 | |
| AT2G07698.1 | ATP synthase subunit (ATPalpha) | 19 | 1.83 | 0.183 | 0.970 | |
| AT2G07698.1 | ATP synthase subunit (ATPalpha) | 20 | 1.78 | 0.186 | 0.986 | |
| AT2G07698.1 | ATP synthase subunit (ATPalpha) | 21 | 1.71 | 0.189 | 0.994 | |
| AT2G07698.1 | ATP synthase subunit (ATPalpha) | 22 | 1.66 | 0.189 | 0.998 | |
| AT2G07698.1 | ATP synthase subunit (ATPalpha) | 23 | 1.49 | 0.187 | 1.000 | |
| AT2G07698.1 | ATP synthase subunit (ATPalpha) | 24 | 1.29 | 0.184 | 1.000 | |
| AT2G07698.1 | ATP synthase subunit (ATPalpha) | 25 | 1.20 | 0.182 | 1.000 | |
| AT2G07698.1 | ATP synthase subunit (ATPalpha) | 26 | 1.12 | 0.181 | 1.000 | |
| AT2G07698.1 | ATP synthase subunit (ATPalpha) | 27 | 0.96 | 0.181 | 1.000 | |
| AT2G07698.1 | ATP synthase subunit (ATPalpha) | 28 | 0.92 | 0.181 | 1.000 | |
| AT2G07698.1 | ATP synthase subunit (ATPalpha) | 29 | 1.14 | 0.181 | 1.000 | |
| AT2G07698.1 | ATP synthase subunit (ATPalpha) | 30 | 1.57 | 0.181 | 0.999 | |
| AT2G07698.1 | ATP synthase subunit (ATPalpha) | 31 | 1.90 | 0.181 | 0.945 | |
| AT2G07698.1 | ATP synthase subunit (ATPalpha) | 32 | 1.85 | 0.182 | 0.961 | |
| AT2G07698.1 | ATP synthase subunit (ATPalpha) | 33 | 1.73 | 0.182 | 0.989 | |
| AT2G07698.1 | ATP synthase subunit (ATPalpha) | 34 | 1.68 | 0.182 | 0.995 | |
| AT2G07698.1 | ATP synthase subunit (ATPalpha) | 35 | 1.75 | 0.182 | 0.987 | |
| AT2G07698.1 | ATP synthase subunit (ATPalpha) | 36 | 2.08 | 0.181 | 0.838 | |
| AT2G07698.1 | ATP synthase subunit (ATPalpha) | 37 | 2.84 | 0.181 | 0.283 | |
| AT2G07698.1 | ATP synthase subunit (ATPalpha) | 38 | 3.95 | 0.182 | 0.033 | * |
| AT2G07698.1 | ATP synthase subunit (ATPalpha) | 39 | 5.53 | 0.182 | 0.002 | ** |
| AT2G07698.1 | ATP synthase subunit (ATPalpha) | 40 | 6.81 | 0.182 | 0.000 | *** |
| AT2G07698.1 | ATP synthase subunit (ATPalpha) | 41 | 6.16 | 0.183 | 0.001 | *** |
| AT2G07698.1 | ATP synthase subunit (ATPalpha) | 42 | 4.60 | 0.185 | 0.013 | * |
| AT2G07698.1 | ATP synthase subunit (ATPalpha) | 43 | 3.19 | 0.187 | 0.180 | |

| | | | | | | |
|---|---|---|---|---|---|---|
| AT2G07698.1 | ATP synthase subunit (ATPalpha) | 44 | 2.11 | 0.187 | 0.850 | |
| AT2G07698.1 | ATP synthase subunit (ATPalpha) | 45 | 1.30 | 0.188 | 1.000 | |
| AT2G07698.1 | ATP synthase subunit (ATPalpha) | 46 | 0.70 | 0.224 | 1.000 | |
| AT2G07698.1 | ATP synthase subunit (ATPalpha) | 47 | 0.34 | 0.362 | 0.991 | |
| AT2G07698.1 | ATP synthase subunit (ATPalpha) | 48 | 0.16 | 0.566 | 0.980 | |
| AT2G10940.2 | Proline-rich glycoprotein (PRP15, putative) | 2 | 0.88 | 0.264 | 1.000 | |
| AT2G10940.2 | Proline-rich glycoprotein (PRP15, putative) | 3 | 1.06 | 0.235 | 1.000 | |
| AT2G10940.2 | Proline-rich glycoprotein (PRP15, putative) | 4 | 1.27 | 0.217 | 0.990 | |
| AT2G10940.2 | Proline-rich glycoprotein (PRP15, putative) | 5 | 1.49 | 0.208 | 0.888 | |
| AT2G10940.2 | Proline-rich glycoprotein (PRP15, putative) | 6 | 1.67 | 0.203 | 0.729 | |
| AT2G10940.2 | Proline-rich glycoprotein (PRP15, putative) | 7 | 1.80 | 0.200 | 0.611 | |
| AT2G10940.2 | Proline-rich glycoprotein (PRP15, putative) | 8 | 1.85 | 0.199 | 0.564 | |
| AT2G10940.2 | Proline-rich glycoprotein (PRP15, putative) | 9 | 1.81 | 0.198 | 0.587 | |
| AT2G10940.2 | Proline-rich glycoprotein (PRP15, putative) | 10 | 1.71 | 0.197 | 0.670 | |
| AT2G10940.2 | Proline-rich glycoprotein (PRP15, putative) | 11 | 1.60 | 0.198 | 0.779 | |
| AT2G10940.2 | Proline-rich glycoprotein (PRP15, putative) | 12 | 1.48 | 0.200 | 0.879 | |
| AT2G10940.2 | Proline-rich glycoprotein (PRP15, putative) | 13 | 1.37 | 0.203 | 0.949 | |
| AT2G10940.2 | Proline-rich glycoprotein (PRP15, putative) | 14 | 1.28 | 0.208 | 0.986 | |
| AT2G10940.2 | Proline-rich glycoprotein (PRP15, putative) | 15 | 1.20 | 0.217 | 0.998 | |
| AT2G10940.2 | Proline-rich glycoprotein (PRP15, putative) | 16 | 1.12 | 0.229 | 1.000 | |
| AT2G10940.2 | Proline-rich glycoprotein (PRP15, putative) | 17 | 1.06 | 0.248 | 1.000 | |
| AT2G10940.2 | Proline-rich glycoprotein (PRP15, putative) | 18 | 1.01 | 0.273 | 1.000 | |
| AT2G10940.2 | Proline-rich glycoprotein (PRP15, putative) | 46 | 20.98 | 0.541 | 0.076 | . |
| AT2G10940.2 | Proline-rich glycoprotein (PRP15, putative) | 47 | 25.25 | 0.557 | 0.062 | . |
| AT2G13360.2 | Alanine:glyoxylate aminotransferase 1 (AGT1) | 2 | 0.83 | 0.470 | 1.000 | |
| AT2G13360.2 | Alanine:glyoxylate aminotransferase 1 (AGT1) | 3 | 0.88 | 0.456 | 1.000 | |
| AT2G13360.2 | Alanine:glyoxylate aminotransferase 1 (AGT1) | 24 | 0.12 | 0.365 | 0.194 | |
| AT2G13360.2 | Alanine:glyoxylate aminotransferase 1 (AGT1) | 25 | 0.11 | 0.258 | 0.005 | ** |
| AT2G13360.2 | Alanine:glyoxylate aminotransferase 1 (AGT1) | 26 | 0.16 | 0.218 | 0.007 | ** |
| AT2G13360.2 | Alanine:glyoxylate aminotransferase 1 (AGT1) | 27 | 0.38 | 0.209 | 0.479 | |
| AT2G13360.2 | Alanine:glyoxylate aminotransferase 1 (AGT1) | 28 | 0.77 | 0.205 | 1.000 | |

| | | | | | | |
|---|---|---|---|---|---|---|
| AT2G13360.2 | Alanine:glyoxylate aminotransferase 1 (AGT1) | 29 | 1.05 | 0.205 | 1.000 | |
| AT2G13360.2 | Alanine:glyoxylate aminotransferase 1 (AGT1) | 30 | 1.11 | 0.207 | 1.000 | |
| AT2G13360.2 | Alanine:glyoxylate aminotransferase 1 (AGT1) | 31 | 1.12 | 0.206 | 1.000 | |
| AT2G13360.2 | Alanine:glyoxylate aminotransferase 1 (AGT1) | 32 | 1.17 | 0.207 | 1.000 | |
| AT2G13360.2 | Alanine:glyoxylate aminotransferase 1 (AGT1) | 33 | 1.18 | 0.206 | 1.000 | |
| AT2G13360.2 | Alanine:glyoxylate aminotransferase 1 (AGT1) | 34 | 1.12 | 0.207 | 1.000 | |
| AT2G13360.2 | Alanine:glyoxylate aminotransferase 1 (AGT1) | 35 | 1.07 | 0.208 | 1.000 | |
| AT2G13360.2 | Alanine:glyoxylate aminotransferase 1 (AGT1) | 36 | 0.93 | 0.208 | 1.000 | |
| AT2G13360.2 | Alanine:glyoxylate aminotransferase 1 (AGT1) | 37 | 0.57 | 0.210 | 0.978 | |
| AT2G13360.2 | Alanine:glyoxylate aminotransferase 1 (AGT1) | 38 | 0.24 | 0.221 | 0.096 | . |
| AT2G13360.2 | Alanine:glyoxylate aminotransferase 1 (AGT1) | 39 | 0.16 | 0.285 | 0.082 | . |
| AT2G13360.2 | Alanine:glyoxylate aminotransferase 1 (AGT1) | 40 | 0.20 | 0.468 | 0.866 | |
| AT2G13360.2 | Alanine:glyoxylate aminotransferase 1 (AGT1) | 46 | 7467.40 | 2.109 | 0.631 | |
| AT2G13360.2 | Alanine:glyoxylate aminotransferase 1 (AGT1) | 47 | 88350.33 | 2.539 | 0.543 | |
| AT2G13560.1 | NAD malic enzyme 1 (NAD-ME1) | 2 | 1.32 | 0.348 | 1.000 | |
| AT2G13560.1 | NAD malic enzyme 1 (NAD-ME1) | 3 | 0.73 | 0.262 | 1.000 | |
| AT2G13560.1 | NAD malic enzyme 1 (NAD-ME1) | 4 | 0.44 | 0.421 | 1.000 | |
| AT2G13560.1 | NAD malic enzyme 1 (NAD-ME1) | 22 | 0.55 | 0.278 | 0.999 | |
| AT2G13560.1 | NAD malic enzyme 1 (NAD-ME1) | 23 | 0.51 | 0.213 | 0.954 | |
| AT2G13560.1 | NAD malic enzyme 1 (NAD-ME1) | 24 | 0.54 | 0.187 | 0.936 | |
| AT2G13560.1 | NAD malic enzyme 1 (NAD-ME1) | 25 | 0.67 | 0.173 | 0.998 | |
| AT2G13560.1 | NAD malic enzyme 1 (NAD-ME1) | 26 | 0.76 | 0.170 | 1.000 | |
| AT2G13560.1 | NAD malic enzyme 1 (NAD-ME1) | 27 | 0.84 | 0.170 | 1.000 | |
| AT2G13560.1 | NAD malic enzyme 1 (NAD-ME1) | 28 | 0.95 | 0.170 | 1.000 | |
| AT2G13560.1 | NAD malic enzyme 1 (NAD-ME1) | 29 | 1.09 | 0.171 | 1.000 | |
| AT2G13560.1 | NAD malic enzyme 1 (NAD-ME1) | 30 | 1.07 | 0.172 | 1.000 | |
| AT2G13560.1 | NAD malic enzyme 1 (NAD-ME1) | 31 | 0.91 | 0.172 | 1.000 | |
| AT2G13560.1 | NAD malic enzyme 1 (NAD-ME1) | 32 | 0.80 | 0.172 | 1.000 | |
| AT2G13560.1 | NAD malic enzyme 1 (NAD-ME1) | 33 | 0.71 | 0.172 | 1.000 | |
| AT2G13560.1 | NAD malic enzyme 1 (NAD-ME1) | 34 | 0.66 | 0.173 | 0.996 | |
| AT2G13560.1 | NAD malic enzyme 1 (NAD-ME1) | 35 | 0.70 | 0.173 | 0.999 | |
| AT2G13560.1 | NAD malic enzyme 1 (NAD-ME1) | 36 | 0.86 | 0.172 | 1.000 | |
| AT2G13560.1 | NAD malic enzyme 1 (NAD-ME1) | 37 | 1.14 | 0.171 | 1.000 | |
| AT2G13560.1 | NAD malic enzyme 1 (NAD-ME1) | 38 | 1.49 | 0.171 | 0.998 | |
| AT2G13560.1 | NAD malic enzyme 1 (NAD-ME1) | 39 | 1.80 | 0.171 | 0.912 | |
| AT2G13560.1 | NAD malic enzyme 1 (NAD-ME1) | 40 | 1.84 | 0.171 | 0.887 | |
| AT2G13560.1 | NAD malic enzyme 1 (NAD-ME1) | 41 | 1.46 | 0.173 | 0.999 | |
| AT2G13560.1 | NAD malic enzyme 1 (NAD-ME1) | 42 | 1.00 | 0.175 | 1.000 | |
| AT2G13560.1 | NAD malic enzyme 1 (NAD-ME1) | 43 | 0.71 | 0.176 | 1.000 | |
| AT2G13560.1 | NAD malic enzyme 1 (NAD-ME1) | 44 | 0.56 | 0.176 | 0.935 | |
| AT2G13560.1 | NAD malic enzyme 1 (NAD-ME1) | 45 | 0.46 | 0.176 | 0.620 | |
| AT2G13560.1 | NAD malic enzyme 1 (NAD-ME1) | 46 | 0.32 | 0.205 | 0.283 | |
| AT2G13560.1 | NAD malic enzyme 1 (NAD-ME1) | 47 | 0.19 | 0.354 | 0.556 | |
| AT2G14170.2 | Aldehyde dehydrogenase 6B2 (ALDH6B2) | 2 | 1.23 | 0.532 | 1.000 | |
| AT2G14170.2 | Aldehyde dehydrogenase 6B2 (ALDH6B2) | 3 | 0.77 | 0.391 | 1.000 | |

| | | | | | |
|---|---|---|---|---|---|
| AT2G14170.2 | Aldehyde dehydrogenase 6B2 (ALDH6B2) | 4 | 0.54 | 0.569 | 1.000 |
| AT2G14170.2 | Aldehyde dehydrogenase 6B2 (ALDH6B2) | 11 | 0.62 | 0.399 | 1.000 |
| AT2G14170.2 | Aldehyde dehydrogenase 6B2 (ALDH6B2) | 12 | 0.80 | 0.375 | 1.000 |
| AT2G14170.2 | Aldehyde dehydrogenase 6B2 (ALDH6B2) | 13 | 1.04 | 0.349 | 1.000 |
| AT2G14170.2 | Aldehyde dehydrogenase 6B2 (ALDH6B2) | 14 | 1.33 | 0.338 | 1.000 |
| AT2G14170.2 | Aldehyde dehydrogenase 6B2 (ALDH6B2) | 15 | 1.58 | 0.329 | 1.000 |
| AT2G14170.2 | Aldehyde dehydrogenase 6B2 (ALDH6B2) | 16 | 1.55 | 0.336 | 1.000 |
| AT2G14170.2 | Aldehyde dehydrogenase 6B2 (ALDH6B2) | 18 | 1.53 | 0.337 | 1.000 |
| AT2G14170.2 | Aldehyde dehydrogenase 6B2 (ALDH6B2) | 19 | 2.07 | 0.307 | 1.000 |
| AT2G14170.2 | Aldehyde dehydrogenase 6B2 (ALDH6B2) | 20 | 2.80 | 0.287 | 0.954 |
| AT2G14170.2 | Aldehyde dehydrogenase 6B2 (ALDH6B2) | 21 | 2.81 | 0.278 | 0.937 |
| AT2G14170.2 | Aldehyde dehydrogenase 6B2 (ALDH6B2) | 22 | 2.33 | 0.279 | 0.993 |
| AT2G14170.2 | Aldehyde dehydrogenase 6B2 (ALDH6B2) | 23 | 1.80 | 0.286 | 1.000 |
| AT2G14170.2 | Aldehyde dehydrogenase 6B2 (ALDH6B2) | 24 | 1.24 | 0.290 | 1.000 |
| AT2G14170.2 | Aldehyde dehydrogenase 6B2 (ALDH6B2) | 25 | 0.79 | 0.293 | 1.000 |
| AT2G14170.2 | Aldehyde dehydrogenase 6B2 (ALDH6B2) | 26 | 0.71 | 0.286 | 1.000 |
| AT2G14170.2 | Aldehyde dehydrogenase 6B2 (ALDH6B2) | 27 | 1.12 | 0.272 | 1.000 |
| AT2G14170.2 | Aldehyde dehydrogenase 6B2 (ALDH6B2) | 28 | 1.68 | 0.262 | 1.000 |
| AT2G14170.2 | Aldehyde dehydrogenase 6B2 (ALDH6B2) | 29 | 1.69 | 0.261 | 1.000 |
| AT2G14170.2 | Aldehyde dehydrogenase 6B2 (ALDH6B2) | 30 | 1.19 | 0.260 | 1.000 |
| AT2G14170.2 | Aldehyde dehydrogenase 6B2 (ALDH6B2) | 31 | 0.97 | 0.261 | 1.000 |
| AT2G14170.2 | Aldehyde dehydrogenase 6B2 (ALDH6B2) | 32 | 1.25 | 0.262 | 1.000 |
| AT2G14170.2 | Aldehyde dehydrogenase 6B2 (ALDH6B2) | 33 | 2.16 | 0.262 | 0.995 |
| AT2G14170.2 | Aldehyde dehydrogenase 6B2 (ALDH6B2) | 34 | 2.54 | 0.263 | 0.961 |
| AT2G14170.2 | Aldehyde dehydrogenase 6B2 (ALDH6B2) | 35 | 1.43 | 0.262 | 1.000 |
| AT2G14170.2 | Aldehyde dehydrogenase 6B2 (ALDH6B2) | 36 | 0.66 | 0.262 | 1.000 |
| AT2G14170.2 | Aldehyde dehydrogenase 6B2 (ALDH6B2) | 37 | 0.49 | 0.261 | 0.998 |
| AT2G14170.2 | Aldehyde dehydrogenase 6B2 (ALDH6B2) | 38 | 0.66 | 0.263 | 1.000 |
| AT2G14170.2 | Aldehyde dehydrogenase 6B2 (ALDH6B2) | 39 | 1.15 | 0.275 | 1.000 |
| AT2G14170.2 | Aldehyde dehydrogenase 6B2 (ALDH6B2) | 40 | 1.84 | 0.296 | 1.000 |
| AT2G14170.2 | Aldehyde dehydrogenase 6B2 (ALDH6B2) | 41 | 2.51 | 0.310 | 0.995 |

| | | | | | | |
|---|---|---|---|---|---|---|
| AT2G14170.2 | Aldehyde dehydrogenase 6B2 (ALDH6B2) | 42 | 2.59 | 0.306 | 0.990 | |
| AT2G14170.2 | Aldehyde dehydrogenase 6B2 (ALDH6B2) | 43 | 1.87 | 0.378 | 1.000 | |
| AT2G14170.2 | Aldehyde dehydrogenase 6B2 (ALDH6B2) | 44 | 1.26 | 0.557 | 1.000 | |
| AT2G14170.2 | Aldehyde dehydrogenase 6B2 (ALDH6B2) | 46 | 0.62 | 1.220 | 1.000 | |
| AT2G14170.2 | Aldehyde dehydrogenase 6B2 (ALDH6B2) | 47 | 0.45 | 1.727 | 1.000 | |
| AT2G14860.1 | Peroxisomal membrane 22 kDa (Mpv17/PMP22) family protein | 2 | 0.81 | 0.562 | 1.000 | |
| AT2G14860.1 | Peroxisomal membrane 22 kDa (Mpv17/PMP22) family protein | 3 | 1.33 | 0.524 | 1.000 | |
| AT2G14860.1 | Peroxisomal membrane 22 kDa (Mpv17/PMP22) family protein | 41 | 18.05 | 0.550 | 0.174 | |
| AT2G14860.1 | Peroxisomal membrane 22 kDa (Mpv17/PMP22) family protein | 42 | 6.66 | 0.336 | 0.121 | |
| AT2G14860.1 | Peroxisomal membrane 22 kDa (Mpv17/PMP22) family protein | 43 | 1.03 | 0.264 | 1.000 | |
| AT2G14860.1 | Peroxisomal membrane 22 kDa (Mpv17/PMP22) family protein | 44 | 0.17 | 0.244 | 0.022 | * |
| AT2G14860.1 | Peroxisomal membrane 22 kDa (Mpv17/PMP22) family protein | 45 | 0.17 | 0.248 | 0.026 | * |
| AT2G14860.1 | Peroxisomal membrane 22 kDa (Mpv17/PMP22) family protein | 46 | 0.42 | 0.284 | 0.738 | |
| AT2G14860.1 | Peroxisomal membrane 22 kDa (Mpv17/PMP22) family protein | 47 | 1.19 | 0.671 | 1.000 | |
| AT2G15020.1 | ? | 2 | 1.35 | 0.512 | 1.000 | |
| AT2G15020.1 | ? | 3 | 1.36 | 0.466 | 1.000 | |
| AT2G15020.1 | ? | 15 | 0.60 | 0.310 | 0.998 | |
| AT2G15020.1 | ? | 16 | 0.46 | 0.266 | 0.890 | |
| AT2G15020.1 | ? | 17 | 0.37 | 0.240 | 0.532 | |
| AT2G15020.1 | ? | 18 | 0.36 | 0.226 | 0.407 | |
| AT2G15020.1 | ? | 19 | 0.42 | 0.225 | 0.632 | |
| AT2G15020.1 | ? | 20 | 0.52 | 0.229 | 0.905 | |
| AT2G15020.1 | ? | 21 | 0.63 | 0.231 | 0.991 | |
| AT2G15020.1 | ? | 22 | 0.65 | 0.231 | 0.996 | |
| AT2G15020.1 | ? | 23 | 0.51 | 0.230 | 0.897 | |
| AT2G15020.1 | ? | 24 | 0.33 | 0.228 | 0.317 | |
| AT2G15020.1 | ? | 25 | 0.20 | 0.234 | 0.042 | * |
| AT2G15020.1 | ? | 26 | 0.15 | 0.248 | 0.015 | * |
| AT2G15020.1 | ? | 27 | 0.16 | 0.274 | 0.047 | * |
| AT2G15020.1 | ? | 28 | 0.19 | 0.334 | 0.298 | |
| AT2G15020.1 | ? | 29 | 0.25 | 0.473 | 0.892 | |
| AT2G15020.1 | ? | 46 | 11450.06 | 5.984 | 0.999 | |
| AT2G15020.1 | ? | 47 | 25622.77 | 6.483 | 0.999 | |
| AT2G15690.1 | Pentatricopeptide repeat (PPR) superfamily protein (DYW2) | 2 | 0.96 | 0.447 | 1.000 | |
| AT2G15690.1 | Pentatricopeptide repeat (PPR) superfamily protein (DYW2) | 3 | 1.05 | 0.410 | 1.000 | |
| AT2G15690.1 | Pentatricopeptide repeat (PPR) superfamily protein (DYW2) | 19 | 1.55 | 0.274 | 1.000 | |
| AT2G15690.1 | Pentatricopeptide repeat (PPR) superfamily protein (DYW2) | 20 | 1.39 | 0.233 | 1.000 | |
| AT2G15690.1 | Pentatricopeptide repeat (PPR) superfamily protein (DYW2) | 21 | 1.23 | 0.211 | 1.000 | |
| AT2G15690.1 | Pentatricopeptide repeat (PPR) superfamily protein (DYW2) | 22 | 1.11 | 0.200 | 1.000 | |
| AT2G15690.1 | Pentatricopeptide repeat (PPR) superfamily protein (DYW2) | 23 | 1.06 | 0.191 | 1.000 | |

| | | | | | | |
|---|---|---|---|---|---|---|
| AT2G15690.1 | Pentatricopeptide repeat (PPR) superfamily protein (DYW2) | 24 | 1.09 | 0.180 | 1.000 | |
| AT2G15690.1 | Pentatricopeptide repeat (PPR) superfamily protein (DYW2) | 25 | 1.18 | 0.172 | 1.000 | |
| AT2G15690.1 | Pentatricopeptide repeat (PPR) superfamily protein (DYW2) | 26 | 1.30 | 0.168 | 1.000 | |
| AT2G15690.1 | Pentatricopeptide repeat (PPR) superfamily protein (DYW2) | 27 | 1.36 | 0.168 | 0.999 | |
| AT2G15690.1 | Pentatricopeptide repeat (PPR) superfamily protein (DYW2) | 28 | 1.34 | 0.168 | 1.000 | |
| AT2G15690.1 | Pentatricopeptide repeat (PPR) superfamily protein (DYW2) | 29 | 1.25 | 0.168 | 1.000 | |
| AT2G15690.1 | Pentatricopeptide repeat (PPR) superfamily protein (DYW2) | 30 | 1.14 | 0.168 | 1.000 | |
| AT2G15690.1 | Pentatricopeptide repeat (PPR) superfamily protein (DYW2) | 31 | 1.05 | 0.169 | 1.000 | |
| AT2G15690.1 | Pentatricopeptide repeat (PPR) superfamily protein (DYW2) | 32 | 1.07 | 0.169 | 1.000 | |
| AT2G15690.1 | Pentatricopeptide repeat (PPR) superfamily protein (DYW2) | 33 | 1.24 | 0.170 | 1.000 | |
| AT2G15690.1 | Pentatricopeptide repeat (PPR) superfamily protein (DYW2) | 34 | 1.67 | 0.171 | 0.942 | |
| AT2G15690.1 | Pentatricopeptide repeat (PPR) superfamily protein (DYW2) | 35 | 2.50 | 0.175 | 0.309 | |
| AT2G15690.1 | Pentatricopeptide repeat (PPR) superfamily protein (DYW2) | 36 | 3.77 | 0.190 | 0.047 | * |
| AT2G15690.1 | Pentatricopeptide repeat (PPR) superfamily protein (DYW2) | 37 | 5.04 | 0.221 | 0.031 | * |
| AT2G15690.1 | Pentatricopeptide repeat (PPR) superfamily protein (DYW2) | 38 | 5.91 | 0.261 | 0.060 | . |
| AT2G15690.1 | Pentatricopeptide repeat (PPR) superfamily protein (DYW2) | 42 | 3.34 | 0.273 | 0.563 | |
| AT2G15690.1 | Pentatricopeptide repeat (PPR) superfamily protein (DYW2) | 43 | 2.12 | 0.233 | 0.903 | |
| AT2G15690.1 | Pentatricopeptide repeat (PPR) superfamily protein (DYW2) | 44 | 1.28 | 0.238 | 1.000 | |
| AT2G15690.1 | Pentatricopeptide repeat (PPR) superfamily protein (DYW2) | 45 | 0.79 | 0.316 | 1.000 | |
| AT2G15690.1 | Pentatricopeptide repeat (PPR) superfamily protein (DYW2) | 46 | 0.51 | 0.461 | 1.000 | |
| AT2G15690.1 | Pentatricopeptide repeat (PPR) superfamily protein (DYW2) | 47 | 0.34 | 0.659 | 1.000 | |
| AT2G16460.1 | Cytochrome c:O2 oxidoreductase subunit (COXX6) | 2 | 0.98 | 0.480 | 1.000 | |
| AT2G16460.1 | Cytochrome c:O2 oxidoreductase subunit (COXX6) | 3 | 1.04 | 0.366 | 1.000 | |
| AT2G16460.1 | Cytochrome c:O2 oxidoreductase subunit (COXX6) | 6 | 1.09 | 0.364 | 1.000 | |
| AT2G16460.1 | Cytochrome c:O2 oxidoreductase subunit (COXX6) | 7 | 1.02 | 0.374 | 1.000 | |
| AT2G16460.1 | Cytochrome c:O2 oxidoreductase subunit (COXX6) | 8 | 0.94 | 0.385 | 1.000 | |
| AT2G16460.1 | Cytochrome c:O2 oxidoreductase subunit (COXX6) | 18 | 2.94 | 0.309 | 0.946 | |
| AT2G16460.1 | Cytochrome c:O2 oxidoreductase subunit (COXX6) | 19 | 3.69 | 0.253 | 0.451 | |
| AT2G16460.1 | Cytochrome c:O2 oxidoreductase subunit (COXX6) | 20 | 4.07 | 0.235 | 0.219 | |
| AT2G16460.1 | Cytochrome c:O2 oxidoreductase subunit (COXX6) | 21 | 3.66 | 0.228 | 0.289 | |
| AT2G16460.1 | Cytochrome c:O2 oxidoreductase subunit (COXX6) | 22 | 2.32 | 0.225 | 0.907 | |

| | | | | | |
|---|---|---|---|---|---|
| AT2G16460.1 | Cytochrome c:O2 oxidoreductase subunit (COXX6) | 23 | 1.36 | 0.224 | 1.000 |
| AT2G16460.1 | Cytochrome c:O2 oxidoreductase subunit (COXX6) | 24 | 0.87 | 0.217 | 1.000 |
| AT2G16460.1 | Cytochrome c:O2 oxidoreductase subunit (COXX6) | 25 | 0.66 | 0.214 | 1.000 |
| AT2G16460.1 | Cytochrome c:O2 oxidoreductase subunit (COXX6) | 26 | 0.84 | 0.212 | 1.000 |
| AT2G16460.1 | Cytochrome c:O2 oxidoreductase subunit (COXX6) | 27 | 1.34 | 0.212 | 1.000 |
| AT2G16460.1 | Cytochrome c:O2 oxidoreductase subunit (COXX6) | 28 | 1.52 | 0.215 | 1.000 |
| AT2G16460.1 | Cytochrome c:O2 oxidoreductase subunit (COXX6) | 29 | 1.34 | 0.214 | 1.000 |
| AT2G16460.1 | Cytochrome c:O2 oxidoreductase subunit (COXX6) | 30 | 1.12 | 0.213 | 1.000 |
| AT2G16460.1 | Cytochrome c:O2 oxidoreductase subunit (COXX6) | 31 | 0.99 | 0.214 | 1.000 |
| AT2G16460.1 | Cytochrome c:O2 oxidoreductase subunit (COXX6) | 32 | 0.92 | 0.217 | 1.000 |
| AT2G16460.1 | Cytochrome c:O2 oxidoreductase subunit (COXX6) | 33 | 1.02 | 0.216 | 1.000 |
| AT2G16460.1 | Cytochrome c:O2 oxidoreductase subunit (COXX6) | 34 | 1.32 | 0.217 | 1.000 |
| AT2G16460.1 | Cytochrome c:O2 oxidoreductase subunit (COXX6) | 35 | 1.22 | 0.215 | 1.000 |
| AT2G16460.1 | Cytochrome c:O2 oxidoreductase subunit (COXX6) | 36 | 0.97 | 0.215 | 1.000 |
| AT2G16460.1 | Cytochrome c:O2 oxidoreductase subunit (COXX6) | 37 | 1.45 | 0.216 | 1.000 |
| AT2G16460.1 | Cytochrome c:O2 oxidoreductase subunit (COXX6) | 38 | 2.58 | 0.216 | 0.733 |
| AT2G16460.1 | Cytochrome c:O2 oxidoreductase subunit (COXX6) | 39 | 1.91 | 0.216 | 0.988 |
| AT2G16460.1 | Cytochrome c:O2 oxidoreductase subunit (COXX6) | 40 | 0.54 | 0.226 | 0.996 |
| AT2G16460.1 | Cytochrome c:O2 oxidoreductase subunit (COXX6) | 41 | 0.17 | 0.305 | 0.258 |
| AT2G16460.1 | Cytochrome c:O2 oxidoreductase subunit (COXX6) | 42 | 0.13 | 0.497 | 0.807 |
| AT2G16460.1 | Cytochrome c:O2 oxidoreductase subunit (COXX6) | 46 | 45.09 | 1.645 | 1.000 |
| AT2G16460.1 | Cytochrome c:O2 oxidoreductase subunit (COXX6) | 47 | 449.12 | 2.103 | 0.992 |
| AT2G17130.2 | NAD isocitrate dehydrogenase subunit (IDH2, regulatory) | 2 | 0.97 | 0.309 | 1.000 |
| AT2G17130.2 | NAD isocitrate dehydrogenase subunit (IDH2, regulatory) | 3 | 0.97 | 0.288 | 1.000 |
| AT2G17130.2 | NAD isocitrate dehydrogenase subunit (IDH2, regulatory) | 22 | 1.34 | 0.243 | 1.000 |
| AT2G17130.2 | NAD isocitrate dehydrogenase subunit (IDH2, regulatory) | 23 | 1.38 | 0.185 | 1.000 |
| AT2G17130.2 | NAD isocitrate dehydrogenase subunit (IDH2, regulatory) | 24 | 1.35 | 0.158 | 1.000 |
| AT2G17130.2 | NAD isocitrate dehydrogenase subunit (IDH2, regulatory) | 25 | 1.15 | 0.140 | 1.000 |
| AT2G17130.2 | NAD isocitrate dehydrogenase subunit (IDH2, regulatory) | 26 | 0.92 | 0.130 | 1.000 |
| AT2G17130.2 | NAD isocitrate dehydrogenase subunit (IDH2, regulatory) | 27 | 0.80 | 0.125 | 1.000 |
| AT2G17130.2 | NAD isocitrate dehydrogenase subunit (IDH2, regulatory) | 28 | 0.80 | 0.124 | 1.000 |

| | | | | | |
|---|---|---|---|---|---|
| AT2G17130.2 | NAD isocitrate dehydrogenase subunit (IDH2, regulatory) | 29 | 0.91 | 0.124 | 1.000 |
| AT2G17130.2 | NAD isocitrate dehydrogenase subunit (IDH2, regulatory) | 30 | 1.06 | 0.124 | 1.000 |
| AT2G17130.2 | NAD isocitrate dehydrogenase subunit (IDH2, regulatory) | 31 | 1.17 | 0.124 | 1.000 |
| AT2G17130.2 | NAD isocitrate dehydrogenase subunit (IDH2, regulatory) | 32 | 1.08 | 0.124 | 1.000 |
| AT2G17130.2 | NAD isocitrate dehydrogenase subunit (IDH2, regulatory) | 33 | 0.84 | 0.124 | 1.000 |
| AT2G17130.2 | NAD isocitrate dehydrogenase subunit (IDH2, regulatory) | 34 | 0.66 | 0.125 | 0.901 |
| AT2G17130.2 | NAD isocitrate dehydrogenase subunit (IDH2, regulatory) | 35 | 0.59 | 0.125 | 0.633 |
| AT2G17130.2 | NAD isocitrate dehydrogenase subunit (IDH2, regulatory) | 36 | 0.62 | 0.124 | 0.785 |
| AT2G17130.2 | NAD isocitrate dehydrogenase subunit (IDH2, regulatory) | 37 | 0.80 | 0.124 | 1.000 |
| AT2G17130.2 | NAD isocitrate dehydrogenase subunit (IDH2, regulatory) | 38 | 1.12 | 0.124 | 1.000 |
| AT2G17130.2 | NAD isocitrate dehydrogenase subunit (IDH2, regulatory) | 39 | 1.50 | 0.124 | 0.918 |
| AT2G17130.2 | NAD isocitrate dehydrogenase subunit (IDH2, regulatory) | 40 | 1.73 | 0.124 | 0.584 |
| AT2G17130.2 | NAD isocitrate dehydrogenase subunit (IDH2, regulatory) | 41 | 1.65 | 0.125 | 0.732 |
| AT2G17130.2 | NAD isocitrate dehydrogenase subunit (IDH2, regulatory) | 42 | 1.50 | 0.127 | 0.926 |
| AT2G17130.2 | NAD isocitrate dehydrogenase subunit (IDH2, regulatory) | 43 | 1.40 | 0.128 | 0.986 |
| AT2G17130.2 | NAD isocitrate dehydrogenase subunit (IDH2, regulatory) | 44 | 1.26 | 0.127 | 1.000 |
| AT2G17130.2 | NAD isocitrate dehydrogenase subunit (IDH2, regulatory) | 45 | 1.12 | 0.129 | 1.000 |
| AT2G17130.2 | NAD isocitrate dehydrogenase subunit (IDH2, regulatory) | 46 | 0.99 | 0.159 | 1.000 |
| AT2G17130.2 | NAD isocitrate dehydrogenase subunit (IDH2, regulatory) | 47 | 0.88 | 0.259 | 1.000 |
| AT2G17420.1 | Thioredoxin reductase 2 (NTR2) | 2 | 0.79 | 0.412 | 1.000 |
| AT2G17420.1 | Thioredoxin reductase 2 (NTR2) | 3 | 0.64 | 0.407 | 1.000 |
| AT2G17420.1 | Thioredoxin reductase 2 (NTR2) | 40 | 0.88 | 0.406 | 1.000 |
| AT2G17420.1 | Thioredoxin reductase 2 (NTR2) | 41 | 1.50 | 0.258 | 0.995 |
| AT2G17420.1 | Thioredoxin reductase 2 (NTR2) | 42 | 3.37 | 0.224 | 0.172 |
| AT2G17420.1 | Thioredoxin reductase 2 (NTR2) | 43 | 3.37 | 0.222 | 0.164 |
| AT2G17420.1 | Thioredoxin reductase 2 (NTR2) | 44 | 1.09 | 0.216 | 1.000 |
| AT2G17420.1 | Thioredoxin reductase 2 (NTR2) | 45 | 0.50 | 0.216 | 0.770 |
| AT2G17420.1 | Thioredoxin reductase 2 (NTR2) | 46 | 0.35 | 0.231 | 0.366 |
| AT2G17420.1 | Thioredoxin reductase 2 (NTR2) | 47 | 0.21 | 0.530 | 0.825 |
| AT2G17650.1 | Acyl-activating enzyme 2 (AAE2, putative) | 2 | 0.88 | 0.296 | 1.000 |
| AT2G17650.1 | Acyl-activating enzyme 2 (AAE2, putative) | 3 | 0.80 | 0.281 | 1.000 |
| AT2G17650.1 | Acyl-activating enzyme 2 (AAE2, putative) | 26 | 0.50 | 0.209 | 0.887 |
| AT2G17650.1 | Acyl-activating enzyme 2 (AAE2, putative) | 27 | 0.57 | 0.148 | 0.769 |
| AT2G17650.1 | Acyl-activating enzyme 2 (AAE2, putative) | 28 | 0.71 | 0.125 | 0.964 |
| AT2G17650.1 | Acyl-activating enzyme 2 (AAE2, putative) | 29 | 0.89 | 0.117 | 1.000 |
| AT2G17650.1 | Acyl-activating enzyme 2 (AAE2, putative) | 30 | 1.03 | 0.115 | 1.000 |

| | | | | | |
|---|---|---|---|---|---|
| AT2G17650.1 | Acyl-activating enzyme 2 (AAE2, putative) | 31 | 1.12 | 0.115 | 1.000 |
| AT2G17650.1 | Acyl-activating enzyme 2 (AAE2, putative) | 32 | 1.16 | 0.115 | 1.000 |
| AT2G17650.1 | Acyl-activating enzyme 2 (AAE2, putative) | 33 | 1.15 | 0.115 | 1.000 |
| AT2G17650.1 | Acyl-activating enzyme 2 (AAE2, putative) | 34 | 1.11 | 0.115 | 1.000 |
| AT2G17650.1 | Acyl-activating enzyme 2 (AAE2, putative) | 35 | 1.09 | 0.115 | 1.000 |
| AT2G17650.1 | Acyl-activating enzyme 2 (AAE2, putative) | 36 | 1.07 | 0.114 | 1.000 |
| AT2G17650.1 | Acyl-activating enzyme 2 (AAE2, putative) | 37 | 1.07 | 0.114 | 1.000 |
| AT2G17650.1 | Acyl-activating enzyme 2 (AAE2, putative) | 38 | 1.09 | 0.114 | 1.000 |
| AT2G17650.1 | Acyl-activating enzyme 2 (AAE2, putative) | 39 | 1.11 | 0.114 | 1.000 |
| AT2G17650.1 | Acyl-activating enzyme 2 (AAE2, putative) | 40 | 1.14 | 0.114 | 1.000 |
| AT2G17650.1 | Acyl-activating enzyme 2 (AAE2, putative) | 41 | 1.18 | 0.115 | 1.000 |
| AT2G17650.1 | Acyl-activating enzyme 2 (AAE2, putative) | 42 | 1.30 | 0.117 | 0.994 |
| AT2G17650.1 | Acyl-activating enzyme 2 (AAE2, putative) | 43 | 1.46 | 0.118 | 0.905 |
| AT2G17650.1 | Acyl-activating enzyme 2 (AAE2, putative) | 44 | 1.43 | 0.117 | 0.932 |
| AT2G17650.1 | Acyl-activating enzyme 2 (AAE2, putative) | 45 | 1.19 | 0.119 | 1.000 |
| AT2G17650.1 | Acyl-activating enzyme 2 (AAE2, putative) | 46 | 0.88 | 0.149 | 1.000 |
| AT2G17650.1 | Acyl-activating enzyme 2 (AAE2, putative) | 47 | 0.61 | 0.248 | 0.998 |
| AT2G18020.1 | 60S ribosomal protein L8-1 (RPL8A) | 2 | 0.63 | 0.245 | 0.859 |
| AT2G18020.1 | 60S ribosomal protein L8-1 (RPL8A) | 3 | 0.68 | 0.219 | 0.893 |
| AT2G18020.1 | 60S ribosomal protein L8-1 (RPL8A) | 4 | 0.74 | 0.197 | 0.932 |
| AT2G18020.1 | 60S ribosomal protein L8-1 (RPL8A) | 5 | 0.80 | 0.178 | 0.970 |
| AT2G18020.1 | 60S ribosomal protein L8-1 (RPL8A) | 6 | 0.87 | 0.162 | 0.993 |
| AT2G18020.1 | 60S ribosomal protein L8-1 (RPL8A) | 7 | 0.94 | 0.149 | 1.000 |
| AT2G18020.1 | 60S ribosomal protein L8-1 (RPL8A) | 8 | 1.01 | 0.140 | 1.000 |
| AT2G18020.1 | 60S ribosomal protein L8-1 (RPL8A) | 10 | 1.16 | 0.130 | 0.979 |
| AT2G18020.1 | 60S ribosomal protein L8-1 (RPL8A) | 11 | 1.24 | 0.129 | 0.911 |
| AT2G18020.1 | 60S ribosomal protein L8-1 (RPL8A) | 12 | 1.32 | 0.131 | 0.806 |
| AT2G18020.1 | 60S ribosomal protein L8-1 (RPL8A) | 13 | 1.41 | 0.137 | 0.702 |
| AT2G18020.1 | 60S ribosomal protein L8-1 (RPL8A) | 14 | 1.50 | 0.146 | 0.625 |
| AT2G18020.1 | 60S ribosomal protein L8-1 (RPL8A) | 15 | 1.58 | 0.158 | 0.582 |
| AT2G18020.1 | 60S ribosomal protein L8-1 (RPL8A) | 16 | 1.67 | 0.173 | 0.568 |

| | | | | | |
|---|---|---|---|---|---|
| AT2G18020.1 | 60S ribosomal protein L8-1 (RPL8A) | 17 | 1.75 | 0.190 | 0.577 |
| AT2G18020.1 | 60S ribosomal protein L8-1 (RPL8A) | 18 | 1.81 | 0.210 | 0.605 |
| AT2G18020.1 | 60S ribosomal protein L8-1 (RPL8A) | 46 | 0.42 | 0.660 | 0.960 |
| AT2G18020.1 | 60S ribosomal protein L8-1 (RPL8A) | 47 | 0.38 | 0.680 | 0.947 |
| AT2G18330.1 | AAA-type ATPase family protein | 2 | 1.16 | 0.481 | 1.000 |
| AT2G18330.1 | AAA-type ATPase family protein | 3 | 1.08 | 0.357 | 1.000 |
| AT2G18330.1 | AAA-type ATPase family protein | 4 | 1.11 | 0.397 | 1.000 |
| AT2G18330.1 | AAA-type ATPase family protein | 8 | 2.06 | 0.367 | 1.000 |
| AT2G18330.1 | AAA-type ATPase family protein | 9 | 2.26 | 0.305 | 0.998 |
| AT2G18330.1 | AAA-type ATPase family protein | 10 | 2.36 | 0.275 | 0.985 |
| AT2G18330.1 | AAA-type ATPase family protein | 11 | 2.56 | 0.258 | 0.933 |
| AT2G18330.1 | AAA-type ATPase family protein | 12 | 2.88 | 0.251 | 0.799 |
| AT2G18330.1 | AAA-type ATPase family protein | 13 | 2.83 | 0.247 | 0.801 |
| AT2G18330.1 | AAA-type ATPase family protein | 14 | 2.51 | 0.245 | 0.912 |
| AT2G18330.1 | AAA-type ATPase family protein | 15 | 2.26 | 0.244 | 0.970 |
| AT2G18330.1 | AAA-type ATPase family protein | 16 | 2.12 | 0.242 | 0.986 |
| AT2G18330.1 | AAA-type ATPase family protein | 17 | 2.02 | 0.242 | 0.994 |
| AT2G18330.1 | AAA-type ATPase family protein | 18 | 2.07 | 0.244 | 0.992 |
| AT2G18330.1 | AAA-type ATPase family protein | 19 | 2.32 | 0.249 | 0.967 |
| AT2G18330.1 | AAA-type ATPase family protein | 20 | 2.68 | 0.255 | 0.889 |
| AT2G18330.1 | AAA-type ATPase family protein | 21 | 2.82 | 0.257 | 0.849 |
| AT2G18330.1 | AAA-type ATPase family protein | 22 | 2.35 | 0.257 | 0.972 |
| AT2G18330.1 | AAA-type ATPase family protein | 23 | 1.71 | 0.255 | 1.000 |
| AT2G18330.1 | AAA-type ATPase family protein | 24 | 1.49 | 0.251 | 1.000 |
| AT2G18330.1 | AAA-type ATPase family protein | 25 | 1.42 | 0.248 | 1.000 |
| AT2G18330.1 | AAA-type ATPase family protein | 26 | 1.32 | 0.247 | 1.000 |
| AT2G18330.1 | AAA-type ATPase family protein | 27 | 1.17 | 0.247 | 1.000 |
| AT2G18330.1 | AAA-type ATPase family protein | 28 | 1.02 | 0.248 | 1.000 |
| AT2G18330.1 | AAA-type ATPase family protein | 29 | 0.96 | 0.248 | 1.000 |
| AT2G18330.1 | AAA-type ATPase family protein | 30 | 0.99 | 0.248 | 1.000 |
| AT2G18330.1 | AAA-type ATPase family protein | 31 | 1.08 | 0.249 | 1.000 |
| AT2G18330.1 | AAA-type ATPase family protein | 32 | 1.12 | 0.249 | 1.000 |
| AT2G18330.1 | AAA-type ATPase family protein | 33 | 0.98 | 0.249 | 1.000 |
| AT2G18330.1 | AAA-type ATPase family protein | 34 | 0.77 | 0.250 | 1.000 |
| AT2G18330.1 | AAA-type ATPase family protein | 35 | 0.65 | 0.249 | 1.000 |
| AT2G18330.1 | AAA-type ATPase family protein | 36 | 0.72 | 0.248 | 1.000 |
| AT2G18330.1 | AAA-type ATPase family protein | 37 | 1.03 | 0.248 | 1.000 |
| AT2G18330.1 | AAA-type ATPase family protein | 38 | 1.90 | 0.248 | 0.999 |
| AT2G18330.1 | AAA-type ATPase family protein | 39 | 3.24 | 0.248 | 0.625 |
| AT2G18330.1 | AAA-type ATPase family protein | 40 | 3.17 | 0.249 | 0.661 |
| AT2G18330.1 | AAA-type ATPase family protein | 41 | 1.85 | 0.251 | 0.999 |
| AT2G18330.1 | AAA-type ATPase family protein | 42 | 0.92 | 0.255 | 1.000 |
| AT2G18330.1 | AAA-type ATPase family protein | 43 | 0.51 | 0.258 | 0.999 |
| AT2G18330.1 | AAA-type ATPase family protein | 44 | 0.41 | 0.263 | 0.968 |
| AT2G18330.1 | AAA-type ATPase family protein | 45 | 0.51 | 0.305 | 1.000 |
| AT2G18330.1 | AAA-type ATPase family protein | 46 | 0.73 | 0.463 | 1.000 |
| AT2G18330.1 | AAA-type ATPase family protein | 47 | 1.10 | 0.766 | 1.000 |
| AT2G19080.1 | Metaxin-related | 2 | 0.78 | 0.293 | 1.000 |
| AT2G19080.1 | Metaxin-related | 3 | 1.19 | 0.194 | 1.000 |
| AT2G19080.1 | Metaxin-related | 4 | 1.73 | 0.176 | 0.988 |
| AT2G19080.1 | Metaxin-related | 5 | 2.15 | 0.168 | 0.708 |
| AT2G19080.1 | Metaxin-related | 6 | 2.15 | 0.161 | 0.632 |
| AT2G19080.1 | Metaxin-related | 7 | 2.05 | 0.159 | 0.722 |
| AT2G19080.1 | Metaxin-related | 8 | 2.18 | 0.153 | 0.516 |
| AT2G19080.1 | Metaxin-related | 9 | 2.43 | 0.150 | 0.253 |
| AT2G19080.1 | Metaxin-related | 10 | 2.36 | 0.150 | 0.306 |
| AT2G19080.1 | Metaxin-related | 11 | 1.94 | 0.152 | 0.770 |
| AT2G19080.1 | Metaxin-related | 12 | 1.54 | 0.152 | 0.996 |
| AT2G19080.1 | Metaxin-related | 13 | 1.25 | 0.149 | 1.000 |

| | | | | | | |
|---|---|---|---|---|---|---|
| AT2G19080.1 | Metaxin-related | 14 | 1.07 | 0.147 | 1.000 | |
| AT2G19080.1 | Metaxin-related | 15 | 0.92 | 0.147 | 1.000 | |
| AT2G19080.1 | Metaxin-related | 16 | 0.83 | 0.146 | 1.000 | |
| AT2G19080.1 | Metaxin-related | 17 | 0.85 | 0.145 | 1.000 | |
| AT2G19080.1 | Metaxin-related | 18 | 0.93 | 0.146 | 1.000 | |
| AT2G19080.1 | Metaxin-related | 19 | 1.05 | 0.152 | 1.000 | |
| AT2G19080.1 | Metaxin-related | 20 | 1.18 | 0.157 | 1.000 | |
| AT2G19080.1 | Metaxin-related | 21 | 1.25 | 0.158 | 1.000 | |
| AT2G19080.1 | Metaxin-related | 22 | 1.24 | 0.158 | 1.000 | |
| AT2G19080.1 | Metaxin-related | 23 | 1.09 | 0.157 | 1.000 | |
| AT2G19080.1 | Metaxin-related | 24 | 0.80 | 0.153 | 1.000 | |
| AT2G19080.1 | Metaxin-related | 25 | 0.61 | 0.150 | 0.979 | |
| AT2G19080.1 | Metaxin-related | 26 | 0.73 | 0.150 | 1.000 | |
| AT2G19080.1 | Metaxin-related | 27 | 1.17 | 0.150 | 1.000 | |
| AT2G19080.1 | Metaxin-related | 28 | 1.50 | 0.150 | 0.998 | |
| AT2G19080.1 | Metaxin-related | 29 | 1.33 | 0.150 | 1.000 | |
| AT2G19080.1 | Metaxin-related | 30 | 0.91 | 0.150 | 1.000 | |
| AT2G19080.1 | Metaxin-related | 31 | 0.66 | 0.150 | 0.998 | |
| AT2G19080.1 | Metaxin-related | 32 | 0.63 | 0.151 | 0.989 | |
| AT2G19080.1 | Metaxin-related | 33 | 0.76 | 0.152 | 1.000 | |
| AT2G19080.1 | Metaxin-related | 34 | 1.33 | 0.153 | 1.000 | |
| AT2G19080.1 | Metaxin-related | 35 | 2.54 | 0.156 | 0.236 | |
| AT2G19080.1 | Metaxin-related | 36 | 3.12 | 0.165 | 0.083 | . |
| AT2G19080.1 | Metaxin-related | 37 | 2.00 | 0.209 | 0.978 | |
| AT2G19080.1 | Metaxin-related | 38 | 1.18 | 0.325 | 1.000 | |
| AT2G19080.1 | Metaxin-related | 46 | 5.61 | 1.633 | 1.000 | |
| AT2G19080.1 | Metaxin-related | 47 | 9.71 | 1.888 | 1.000 | |
| AT2G19680.2 | ATP synthase subunit (ATPG) | 2 | 1.11 | 0.514 | 1.000 | |
| AT2G19680.2 | ATP synthase subunit (ATPG) | 3 | 0.99 | 0.391 | 1.000 | |
| AT2G19680.2 | ATP synthase subunit (ATPG) | 6 | 0.88 | 0.353 | 1.000 | |
| AT2G19680.2 | ATP synthase subunit (ATPG) | 7 | 0.99 | 0.276 | 1.000 | |
| AT2G19680.2 | ATP synthase subunit (ATPG) | 8 | 1.16 | 0.239 | 1.000 | |
| AT2G19680.2 | ATP synthase subunit (ATPG) | 9 | 1.35 | 0.224 | 1.000 | |
| AT2G19680.2 | ATP synthase subunit (ATPG) | 10 | 1.50 | 0.222 | 1.000 | |
| AT2G19680.2 | ATP synthase subunit (ATPG) | 11 | 1.84 | 0.233 | 0.999 | |
| AT2G19680.2 | ATP synthase subunit (ATPG) | 12 | 2.85 | 0.249 | 0.857 | |
| AT2G19680.2 | ATP synthase subunit (ATPG) | 13 | 4.42 | 0.265 | 0.379 | |
| AT2G19680.2 | ATP synthase subunit (ATPG) | 14 | 4.85 | 0.263 | 0.257 | |
| AT2G19680.2 | ATP synthase subunit (ATPG) | 17 | 1.59 | 0.290 | 1.000 | |
| AT2G19680.2 | ATP synthase subunit (ATPG) | 18 | 1.30 | 0.258 | 1.000 | |
| AT2G19680.2 | ATP synthase subunit (ATPG) | 19 | 1.38 | 0.236 | 1.000 | |
| AT2G19680.2 | ATP synthase subunit (ATPG) | 20 | 1.63 | 0.230 | 1.000 | |
| AT2G19680.2 | ATP synthase subunit (ATPG) | 21 | 1.59 | 0.228 | 1.000 | |
| AT2G19680.2 | ATP synthase subunit (ATPG) | 22 | 1.26 | 0.228 | 1.000 | |
| AT2G19680.2 | ATP synthase subunit (ATPG) | 23 | 1.03 | 0.226 | 1.000 | |
| AT2G19680.2 | ATP synthase subunit (ATPG) | 24 | 0.99 | 0.220 | 1.000 | |
| AT2G19680.2 | ATP synthase subunit (ATPG) | 25 | 0.94 | 0.218 | 1.000 | |
| AT2G19680.2 | ATP synthase subunit (ATPG) | 26 | 0.70 | 0.217 | 1.000 | |
| AT2G19680.2 | ATP synthase subunit (ATPG) | 27 | 0.49 | 0.218 | 0.990 | |
| AT2G19680.2 | ATP synthase subunit (ATPG) | 28 | 0.49 | 0.218 | 0.989 | |
| AT2G19680.2 | ATP synthase subunit (ATPG) | 29 | 0.63 | 0.217 | 1.000 | |
| AT2G19680.2 | ATP synthase subunit (ATPG) | 30 | 0.71 | 0.217 | 1.000 | |
| AT2G19680.2 | ATP synthase subunit (ATPG) | 31 | 0.77 | 0.218 | 1.000 | |
| AT2G19680.2 | ATP synthase subunit (ATPG) | 32 | 1.10 | 0.218 | 1.000 | |
| AT2G19680.2 | ATP synthase subunit (ATPG) | 33 | 2.06 | 0.219 | 0.987 | |
| AT2G19680.2 | ATP synthase subunit (ATPG) | 34 | 2.81 | 0.220 | 0.708 | |
| AT2G19680.2 | ATP synthase subunit (ATPG) | 35 | 2.60 | 0.220 | 0.820 | |
| AT2G19680.2 | ATP synthase subunit (ATPG) | 36 | 3.07 | 0.220 | 0.554 | |
| AT2G19680.2 | ATP synthase subunit (ATPG) | 37 | 3.63 | 0.219 | 0.288 | |
| AT2G19680.2 | ATP synthase subunit (ATPG) | 38 | 3.61 | 0.218 | 0.293 | |
| AT2G19680.2 | ATP synthase subunit (ATPG) | 39 | 3.32 | 0.218 | 0.408 | |
| AT2G19680.2 | ATP synthase subunit (ATPG) | 40 | 3.43 | 0.217 | 0.354 | |
| AT2G19680.2 | ATP synthase subunit (ATPG) | 41 | 3.32 | 0.219 | 0.417 | |

| | | | | | |
|---|---|---|---|---|---|
| AT2G19680.2 | ATP synthase subunit (ATPG) | 42 | 2.71 | 0.223 | 0.780 |
| AT2G19680.2 | ATP synthase subunit (ATPG) | 43 | 1.87 | 0.224 | 0.999 |
| AT2G19680.2 | ATP synthase subunit (ATPG) | 44 | 1.02 | 0.224 | 1.000 |
| AT2G19680.2 | ATP synthase subunit (ATPG) | 45 | 0.49 | 0.224 | 0.991 |
| AT2G19680.2 | ATP synthase subunit (ATPG) | 46 | 0.22 | 0.261 | 0.333 |
| AT2G19680.2 | ATP synthase subunit (ATPG) | 47 | 0.09 | 0.478 | 0.580 |
| AT2G19860.1 | Hexokinase 2 (HXK2) | 2 | 0.70 | 0.280 | 1.000 |
| AT2G19860.1 | Hexokinase 2 (HXK2) | 3 | 0.86 | 0.184 | 1.000 |
| AT2G19860.1 | Hexokinase 2 (HXK2) | 4 | 1.08 | 0.156 | 1.000 |
| AT2G19860.1 | Hexokinase 2 (HXK2) | 5 | 1.39 | 0.153 | 1.000 |
| AT2G19860.1 | Hexokinase 2 (HXK2) | 6 | 1.68 | 0.154 | 0.970 |
| AT2G19860.1 | Hexokinase 2 (HXK2) | 7 | 1.89 | 0.153 | 0.828 |
| AT2G19860.1 | Hexokinase 2 (HXK2) | 8 | 1.96 | 0.150 | 0.723 |
| AT2G19860.1 | Hexokinase 2 (HXK2) | 9 | 1.88 | 0.148 | 0.796 |
| AT2G19860.1 | Hexokinase 2 (HXK2) | 10 | 1.73 | 0.147 | 0.923 |
| AT2G19860.1 | Hexokinase 2 (HXK2) | 11 | 1.63 | 0.148 | 0.974 |
| AT2G19860.1 | Hexokinase 2 (HXK2) | 12 | 1.64 | 0.148 | 0.970 |
| AT2G19860.1 | Hexokinase 2 (HXK2) | 13 | 1.82 | 0.146 | 0.843 |
| AT2G19860.1 | Hexokinase 2 (HXK2) | 14 | 2.00 | 0.146 | 0.638 |
| AT2G19860.1 | Hexokinase 2 (HXK2) | 15 | 1.80 | 0.148 | 0.881 |
| AT2G19860.1 | Hexokinase 2 (HXK2) | 16 | 1.39 | 0.153 | 1.000 |
| AT2G19860.1 | Hexokinase 2 (HXK2) | 17 | 1.13 | 0.162 | 1.000 |
| AT2G19860.1 | Hexokinase 2 (HXK2) | 18 | 1.07 | 0.183 | 1.000 |
| AT2G19860.1 | Hexokinase 2 (HXK2) | 19 | 1.09 | 0.211 | 1.000 |
| AT2G19860.1 | Hexokinase 2 (HXK2) | 22 | 0.96 | 0.247 | 1.000 |
| AT2G19860.1 | Hexokinase 2 (HXK2) | 23 | 0.77 | 0.217 | 1.000 |
| AT2G19860.1 | Hexokinase 2 (HXK2) | 24 | 0.59 | 0.190 | 0.997 |
| AT2G19860.1 | Hexokinase 2 (HXK2) | 25 | 0.50 | 0.168 | 0.841 |
| AT2G19860.1 | Hexokinase 2 (HXK2) | 26 | 0.53 | 0.155 | 0.833 |
| AT2G19860.1 | Hexokinase 2 (HXK2) | 27 | 0.71 | 0.148 | 1.000 |
| AT2G19860.1 | Hexokinase 2 (HXK2) | 28 | 1.03 | 0.147 | 1.000 |
| AT2G19860.1 | Hexokinase 2 (HXK2) | 29 | 1.31 | 0.147 | 1.000 |
| AT2G19860.1 | Hexokinase 2 (HXK2) | 30 | 1.34 | 0.147 | 1.000 |
| AT2G19860.1 | Hexokinase 2 (HXK2) | 31 | 1.22 | 0.147 | 1.000 |
| AT2G19860.1 | Hexokinase 2 (HXK2) | 32 | 1.09 | 0.148 | 1.000 |
| AT2G19860.1 | Hexokinase 2 (HXK2) | 33 | 0.97 | 0.148 | 1.000 |
| AT2G19860.1 | Hexokinase 2 (HXK2) | 34 | 0.89 | 0.148 | 1.000 |
| AT2G19860.1 | Hexokinase 2 (HXK2) | 35 | 0.87 | 0.148 | 1.000 |
| AT2G19860.1 | Hexokinase 2 (HXK2) | 36 | 0.88 | 0.147 | 1.000 |
| AT2G19860.1 | Hexokinase 2 (HXK2) | 37 | 0.95 | 0.147 | 1.000 |
| AT2G19860.1 | Hexokinase 2 (HXK2) | 38 | 1.08 | 0.147 | 1.000 |
| AT2G19860.1 | Hexokinase 2 (HXK2) | 39 | 1.29 | 0.147 | 1.000 |
| AT2G19860.1 | Hexokinase 2 (HXK2) | 40 | 1.50 | 0.147 | 0.997 |
| AT2G19860.1 | Hexokinase 2 (HXK2) | 41 | 1.62 | 0.149 | 0.979 |
| AT2G19860.1 | Hexokinase 2 (HXK2) | 42 | 1.59 | 0.151 | 0.988 |
| AT2G19860.1 | Hexokinase 2 (HXK2) | 43 | 1.41 | 0.152 | 1.000 |
| AT2G19860.1 | Hexokinase 2 (HXK2) | 44 | 1.06 | 0.152 | 1.000 |
| AT2G19860.1 | Hexokinase 2 (HXK2) | 45 | 0.67 | 0.158 | 0.999 |
| AT2G19860.1 | Hexokinase 2 (HXK2) | 46 | 0.37 | 0.197 | 0.520 |
| AT2G19860.1 | Hexokinase 2 (HXK2) | 47 | 0.19 | 0.307 | 0.382 |
| AT2G20060.1 | 50S ribosomal protein L4 (RPL4) | 2 | 1.33 | 0.813 | 1.000 |
| AT2G20060.1 | 50S ribosomal protein L4 (RPL4) | 3 | 1.09 | 0.729 | 1.000 |
| AT2G20060.1 | 50S ribosomal protein L4 (RPL4) | 11 | 0.61 | 0.419 | 1.000 |
| AT2G20060.1 | 50S ribosomal protein L4 (RPL4) | 12 | 0.74 | 0.372 | 1.000 |
| AT2G20060.1 | 50S ribosomal protein L4 (RPL4) | 13 | 0.96 | 0.335 | 1.000 |
| AT2G20060.1 | 50S ribosomal protein L4 (RPL4) | 14 | 1.34 | 0.307 | 1.000 |
| AT2G20060.1 | 50S ribosomal protein L4 (RPL4) | 15 | 1.95 | 0.289 | 0.957 |
| AT2G20060.1 | 50S ribosomal protein L4 (RPL4) | 16 | 2.69 | 0.279 | 0.676 |
| AT2G20060.1 | 50S ribosomal protein L4 (RPL4) | 17 | 3.27 | 0.276 | 0.443 |
| AT2G20060.1 | 50S ribosomal protein L4 (RPL4) | 18 | 3.30 | 0.282 | 0.461 |
| AT2G20060.1 | 50S ribosomal protein L4 (RPL4) | 19 | 2.75 | 0.296 | 0.716 |
| AT2G20060.1 | 50S ribosomal protein L4 (RPL4) | 20 | 2.02 | 0.319 | 0.967 |
| AT2G20060.1 | 50S ribosomal protein L4 (RPL4) | 21 | 1.39 | 0.350 | 1.000 |

| | | | | | |
|---|---|---|---|---|---|
| AT2G20060.1 | 50S ribosomal protein L4 (RPL4) | 22 | 0.95 | 0.382 | 1.000 |
| AT2G20060.1 | 50S ribosomal protein L4 (RPL4) | 27 | 0.20 | 0.365 | 0.421 |
| AT2G20060.1 | 50S ribosomal protein L4 (RPL4) | 28 | 0.17 | 0.341 | 0.227 |
| AT2G20060.1 | 50S ribosomal protein L4 (RPL4) | 29 | 0.15 | 0.333 | 0.145 |
| AT2G20060.1 | 50S ribosomal protein L4 (RPL4) | 30 | 0.14 | 0.348 | 0.150 |
| AT2G20060.1 | 50S ribosomal protein L4 (RPL4) | 31 | 0.14 | 0.390 | 0.241 |
| AT2G20060.1 | 50S ribosomal protein L4 (RPL4) | 32 | 0.14 | 0.459 | 0.424 |
| AT2G20060.1 | 50S ribosomal protein L4 (RPL4) | 33 | 0.14 | 0.551 | 0.656 |
| AT2G20060.1 | 50S ribosomal protein L4 (RPL4) | 46 | 0.66 | 1.636 | 1.000 |
| AT2G20060.1 | 50S ribosomal protein L4 (RPL4) | 47 | 0.79 | 1.770 | 1.000 |
| AT2G20260.1 | Photosystem I subunit (PSAE2) | 2 | 0.31 | 0.310 | 0.538 |
| AT2G20260.1 | Photosystem I subunit (PSAE2) | 3 | 0.35 | 0.259 | 0.448 |
| AT2G20260.1 | Photosystem I subunit (PSAE2) | 4 | 0.39 | 0.233 | 0.461 |
| AT2G20260.1 | Photosystem I subunit (PSAE2) | 5 | 0.43 | 0.223 | 0.555 |
| AT2G20260.1 | Photosystem I subunit (PSAE2) | 6 | 0.48 | 0.219 | 0.691 |
| AT2G20260.1 | Photosystem I subunit (PSAE2) | 7 | 0.55 | 0.217 | 0.848 |
| AT2G20260.1 | Photosystem I subunit (PSAE2) | 8 | 0.64 | 0.215 | 0.972 |
| AT2G20260.1 | Photosystem I subunit (PSAE2) | 9 | 0.79 | 0.214 | 1.000 |
| AT2G20260.1 | Photosystem I subunit (PSAE2) | 10 | 0.98 | 0.213 | 1.000 |
| AT2G20260.1 | Photosystem I subunit (PSAE2) | 11 | 1.20 | 0.213 | 1.000 |
| AT2G20260.1 | Photosystem I subunit (PSAE2) | 12 | 1.44 | 0.212 | 0.992 |
| AT2G20260.1 | Photosystem I subunit (PSAE2) | 13 | 1.64 | 0.211 | 0.937 |
| AT2G20260.1 | Photosystem I subunit (PSAE2) | 14 | 1.77 | 0.210 | 0.861 |
| AT2G20260.1 | Photosystem I subunit (PSAE2) | 15 | 1.82 | 0.210 | 0.825 |
| AT2G20260.1 | Photosystem I subunit (PSAE2) | 16 | 1.79 | 0.210 | 0.847 |
| AT2G20260.1 | Photosystem I subunit (PSAE2) | 17 | 1.68 | 0.210 | 0.915 |
| AT2G20260.1 | Photosystem I subunit (PSAE2) | 18 | 1.51 | 0.211 | 0.981 |
| AT2G20260.1 | Photosystem I subunit (PSAE2) | 19 | 1.29 | 0.213 | 1.000 |
| AT2G20260.1 | Photosystem I subunit (PSAE2) | 20 | 1.05 | 0.215 | 1.000 |
| AT2G20260.1 | Photosystem I subunit (PSAE2) | 21 | 0.82 | 0.216 | 1.000 |
| AT2G20260.1 | Photosystem I subunit (PSAE2) | 22 | 0.64 | 0.216 | 0.972 |
| AT2G20260.1 | Photosystem I subunit (PSAE2) | 23 | 0.53 | 0.216 | 0.808 |
| AT2G20260.1 | Photosystem I subunit (PSAE2) | 24 | 0.48 | 0.215 | 0.663 |
| AT2G20260.1 | Photosystem I subunit (PSAE2) | 25 | 0.50 | 0.214 | 0.718 |
| AT2G20260.1 | Photosystem I subunit (PSAE2) | 26 | 0.60 | 0.215 | 0.928 |
| AT2G20260.1 | Photosystem I subunit (PSAE2) | 27 | 0.74 | 0.218 | 0.999 |
| AT2G20260.1 | Photosystem I subunit (PSAE2) | 28 | 0.87 | 0.227 | 1.000 |
| AT2G20260.1 | Photosystem I subunit (PSAE2) | 29 | 0.95 | 0.246 | 1.000 |
| AT2G20260.1 | Photosystem I subunit (PSAE2) | 30 | 0.96 | 0.281 | 1.000 |
| AT2G20260.1 | Photosystem I subunit (PSAE2) | 31 | 0.92 | 0.330 | 1.000 |
| AT2G20260.1 | Photosystem I subunit (PSAE2) | 46 | 0.03 | 0.906 | 0.499 |
| AT2G20260.1 | Photosystem I subunit (PSAE2) | 47 | 0.02 | 0.971 | 0.477 |
| AT2G20360.1 | NADH:ubiquinone oxidoreductase subunit (39kDa) | 2 | 0.68 | 0.496 | 1.000 |
| AT2G20360.1 | NADH:ubiquinone oxidoreductase subunit (39kDa) | 3 | 0.82 | 0.339 | 1.000 |
| AT2G20360.1 | NADH:ubiquinone oxidoreductase subunit (39kDa) | 4 | 1.03 | 0.306 | 1.000 |
| AT2G20360.1 | NADH:ubiquinone oxidoreductase subunit (39kDa) | 5 | 1.27 | 0.305 | 1.000 |
| AT2G20360.1 | NADH:ubiquinone oxidoreductase subunit (39kDa) | 6 | 1.32 | 0.306 | 1.000 |
| AT2G20360.1 | NADH:ubiquinone oxidoreductase subunit (39kDa) | 7 | 1.18 | 0.305 | 1.000 |
| AT2G20360.1 | NADH:ubiquinone oxidoreductase subunit (39kDa) | 8 | 1.06 | 0.301 | 1.000 |
| AT2G20360.1 | NADH:ubiquinone oxidoreductase subunit (39kDa) | 9 | 1.03 | 0.297 | 1.000 |
| AT2G20360.1 | NADH:ubiquinone oxidoreductase subunit (39kDa) | 10 | 1.05 | 0.297 | 1.000 |
| AT2G20360.1 | NADH:ubiquinone oxidoreductase subunit (39kDa) | 11 | 1.18 | 0.298 | 1.000 |

| | | | | | |
|---|---|---|---|---|---|
| AT2G20360.1 | NADH:ubiquinone oxidoreductase subunit (39kDa) | 12 | 1.38 | 0.297 | 1.000 |
| AT2G20360.1 | NADH:ubiquinone oxidoreductase subunit (39kDa) | 13 | 1.44 | 0.295 | 1.000 |
| AT2G20360.1 | NADH:ubiquinone oxidoreductase subunit (39kDa) | 14 | 1.25 | 0.292 | 1.000 |
| AT2G20360.1 | NADH:ubiquinone oxidoreductase subunit (39kDa) | 15 | 0.93 | 0.291 | 1.000 |
| AT2G20360.1 | NADH:ubiquinone oxidoreductase subunit (39kDa) | 16 | 0.66 | 0.289 | 1.000 |
| AT2G20360.1 | NADH:ubiquinone oxidoreductase subunit (39kDa) | 17 | 0.53 | 0.290 | 1.000 |
| AT2G20360.1 | NADH:ubiquinone oxidoreductase subunit (39kDa) | 18 | 0.56 | 0.292 | 1.000 |
| AT2G20360.1 | NADH:ubiquinone oxidoreductase subunit (39kDa) | 19 | 0.72 | 0.298 | 1.000 |
| AT2G20360.1 | NADH:ubiquinone oxidoreductase subunit (39kDa) | 20 | 0.97 | 0.304 | 1.000 |
| AT2G20360.1 | NADH:ubiquinone oxidoreductase subunit (39kDa) | 21 | 1.32 | 0.307 | 1.000 |
| AT2G20360.1 | NADH:ubiquinone oxidoreductase subunit (39kDa) | 22 | 1.67 | 0.307 | 1.000 |
| AT2G20360.1 | NADH:ubiquinone oxidoreductase subunit (39kDa) | 23 | 1.67 | 0.306 | 1.000 |
| AT2G20360.1 | NADH:ubiquinone oxidoreductase subunit (39kDa) | 24 | 1.10 | 0.301 | 1.000 |
| AT2G20360.1 | NADH:ubiquinone oxidoreductase subunit (39kDa) | 25 | 0.51 | 0.296 | 1.000 |
| AT2G20360.1 | NADH:ubiquinone oxidoreductase subunit (39kDa) | 26 | 0.27 | 0.295 | 0.651 |
| AT2G20360.1 | NADH:ubiquinone oxidoreductase subunit (39kDa) | 27 | 0.24 | 0.295 | 0.520 |
| AT2G20360.1 | NADH:ubiquinone oxidoreductase subunit (39kDa) | 28 | 0.38 | 0.295 | 0.959 |
| AT2G20360.1 | NADH:ubiquinone oxidoreductase subunit (39kDa) | 29 | 0.81 | 0.295 | 1.000 |
| AT2G20360.1 | NADH:ubiquinone oxidoreductase subunit (39kDa) | 30 | 1.53 | 0.296 | 1.000 |
| AT2G20360.1 | NADH:ubiquinone oxidoreductase subunit (39kDa) | 31 | 2.33 | 0.296 | 0.990 |
| AT2G20360.1 | NADH:ubiquinone oxidoreductase subunit (39kDa) | 32 | 2.96 | 0.297 | 0.897 |
| AT2G20360.1 | NADH:ubiquinone oxidoreductase subunit (39kDa) | 33 | 2.68 | 0.297 | 0.954 |
| AT2G20360.1 | NADH:ubiquinone oxidoreductase subunit (39kDa) | 34 | 1.76 | 0.298 | 1.000 |
| AT2G20360.1 | NADH:ubiquinone oxidoreductase subunit (39kDa) | 35 | 1.20 | 0.299 | 1.000 |
| AT2G20360.1 | NADH:ubiquinone oxidoreductase subunit (39kDa) | 36 | 1.14 | 0.302 | 1.000 |
| AT2G20360.1 | NADH:ubiquinone oxidoreductase subunit (39kDa) | 37 | 1.46 | 0.318 | 1.000 |
| AT2G20360.1 | NADH:ubiquinone oxidoreductase subunit (39kDa) | 38 | 2.47 | 0.353 | 0.997 |
| AT2G20360.1 | NADH:ubiquinone oxidoreductase subunit (39kDa) | 39 | 4.67 | 0.395 | 0.835 |
| AT2G20360.1 | NADH:ubiquinone oxidoreductase subunit (39kDa) | 41 | 10.38 | 0.374 | 0.154 |
| AT2G20360.1 | NADH:ubiquinone oxidoreductase subunit (39kDa) | 42 | 9.23 | 0.346 | 0.128 |
| AT2G20360.1 | NADH:ubiquinone oxidoreductase subunit (39kDa) | 43 | 5.34 | 0.319 | 0.389 |

| | | | | | |
|---|---|---|---|---|---|
| AT2G20360.1 | NADH:ubiquinone oxidoreductase subunit (39kDa) | 44 | 2.57 | 0.307 | 0.978 |
| AT2G20360.1 | NADH:ubiquinone oxidoreductase subunit (39kDa) | 45 | 1.00 | 0.308 | 1.000 |
| AT2G20360.1 | NADH:ubiquinone oxidoreductase subunit (39kDa) | 46 | 0.34 | 0.360 | 0.982 |
| AT2G20360.1 | NADH:ubiquinone oxidoreductase subunit (39kDa) | 47 | 0.11 | 0.549 | 0.811 |
| AT2G20390.1 | Cytochrome c:O2 oxidoreductase assembly protein | 2 | 1.04 | 0.579 | 1.000 |
| AT2G20390.1 | Cytochrome c:O2 oxidoreductase assembly protein | 3 | 1.00 | 0.468 | 1.000 |
| AT2G20390.1 | Cytochrome c:O2 oxidoreductase assembly protein | 6 | 0.94 | 0.347 | 1.000 |
| AT2G20390.1 | Cytochrome c:O2 oxidoreductase assembly protein | 7 | 0.95 | 0.310 | 1.000 |
| AT2G20390.1 | Cytochrome c:O2 oxidoreductase assembly protein | 8 | 0.98 | 0.280 | 1.000 |
| AT2G20390.1 | Cytochrome c:O2 oxidoreductase assembly protein | 9 | 1.05 | 0.256 | 1.000 |
| AT2G20390.1 | Cytochrome c:O2 oxidoreductase assembly protein | 10 | 1.15 | 0.239 | 1.000 |
| AT2G20390.1 | Cytochrome c:O2 oxidoreductase assembly protein | 11 | 1.27 | 0.229 | 1.000 |
| AT2G20390.1 | Cytochrome c:O2 oxidoreductase assembly protein | 12 | 1.33 | 0.223 | 1.000 |
| AT2G20390.1 | Cytochrome c:O2 oxidoreductase assembly protein | 13 | 1.30 | 0.219 | 1.000 |
| AT2G20390.1 | Cytochrome c:O2 oxidoreductase assembly protein | 14 | 1.18 | 0.217 | 1.000 |
| AT2G20390.1 | Cytochrome c:O2 oxidoreductase assembly protein | 15 | 1.03 | 0.216 | 1.000 |
| AT2G20390.1 | Cytochrome c:O2 oxidoreductase assembly protein | 16 | 0.92 | 0.216 | 1.000 |
| AT2G20390.1 | Cytochrome c:O2 oxidoreductase assembly protein | 17 | 0.90 | 0.218 | 1.000 |
| AT2G20390.1 | Cytochrome c:O2 oxidoreductase assembly protein | 18 | 1.02 | 0.224 | 1.000 |
| AT2G20390.1 | Cytochrome c:O2 oxidoreductase assembly protein | 19 | 1.29 | 0.236 | 1.000 |
| AT2G20390.1 | Cytochrome c:O2 oxidoreductase assembly protein | 20 | 1.68 | 0.261 | 0.999 |
| AT2G20390.1 | Cytochrome c:O2 oxidoreductase assembly protein | 21 | 2.09 | 0.302 | 0.994 |

| | | | | | |
|---|---|---|---|---|---|
| AT2G20390.1 | Cytochrome c:O2 oxidoreductase assembly protein | 22 | 2.37 | 0.357 | 0.994 |
| AT2G20390.1 | Cytochrome c:O2 oxidoreductase assembly protein | 29 | 1.32 | 0.325 | 1.000 |
| AT2G20390.1 | Cytochrome c:O2 oxidoreductase assembly protein | 30 | 1.29 | 0.278 | 1.000 |
| AT2G20390.1 | Cytochrome c:O2 oxidoreductase assembly protein | 31 | 1.38 | 0.247 | 1.000 |
| AT2G20390.1 | Cytochrome c:O2 oxidoreductase assembly protein | 32 | 1.57 | 0.230 | 0.999 |
| AT2G20390.1 | Cytochrome c:O2 oxidoreductase assembly protein | 33 | 1.78 | 0.224 | 0.990 |
| AT2G20390.1 | Cytochrome c:O2 oxidoreductase assembly protein | 34 | 2.01 | 0.225 | 0.947 |
| AT2G20390.1 | Cytochrome c:O2 oxidoreductase assembly protein | 35 | 2.22 | 0.231 | 0.887 |
| AT2G20390.1 | Cytochrome c:O2 oxidoreductase assembly protein | 36 | 2.32 | 0.244 | 0.890 |
| AT2G20390.1 | Cytochrome c:O2 oxidoreductase assembly protein | 37 | 2.41 | 0.263 | 0.911 |
| AT2G20390.1 | Cytochrome c:O2 oxidoreductase assembly protein | 38 | 2.58 | 0.282 | 0.909 |
| AT2G20390.1 | Cytochrome c:O2 oxidoreductase assembly protein | 40 | 3.07 | 0.277 | 0.733 |
| AT2G20390.1 | Cytochrome c:O2 oxidoreductase assembly protein | 41 | 3.31 | 0.263 | 0.564 |
| AT2G20390.1 | Cytochrome c:O2 oxidoreductase assembly protein | 42 | 3.41 | 0.245 | 0.412 |
| AT2G20390.1 | Cytochrome c:O2 oxidoreductase assembly protein | 43 | 3.07 | 0.233 | 0.476 |
| AT2G20390.1 | Cytochrome c:O2 oxidoreductase assembly protein | 44 | 2.19 | 0.228 | 0.892 |
| AT2G20390.1 | Cytochrome c:O2 oxidoreductase assembly protein | 45 | 1.27 | 0.240 | 1.000 |
| AT2G20390.1 | Cytochrome c:O2 oxidoreductase assembly protein | 46 | 0.66 | 0.300 | 1.000 |
| AT2G20390.1 | Cytochrome c:O2 oxidoreductase assembly protein | 47 | 0.32 | 0.432 | 0.988 |
| AT2G20420.1 | ATP citrate lyase (ACL) family protein (Succinate-CoA ligase activity) | 3 | 0.48 | 0.220 | 0.967 |

| | | | | | |
|---|---|---|---|---|---|
| AT2G20420.1 | ATP citrate lyase (ACL) family protein (Succinate-CoA ligase activity) | 4 | 0.32 | 0.201 | 0.314 |
| AT2G20420.1 | ATP citrate lyase (ACL) family protein (Succinate-CoA ligase activity) | 5 | 0.28 | 0.201 | 0.169 |
| AT2G20420.1 | ATP citrate lyase (ACL) family protein (Succinate-CoA ligase activity) | 6 | 0.34 | 0.201 | 0.405 |
| AT2G20420.1 | ATP citrate lyase (ACL) family protein (Succinate-CoA ligase activity) | 7 | 0.48 | 0.200 | 0.920 |
| AT2G20420.1 | ATP citrate lyase (ACL) family protein (Succinate-CoA ligase activity) | 8 | 0.72 | 0.198 | 1.000 |
| AT2G20420.1 | ATP citrate lyase (ACL) family protein (Succinate-CoA ligase activity) | 9 | 1.05 | 0.196 | 1.000 |
| AT2G20420.1 | ATP citrate lyase (ACL) family protein (Succinate-CoA ligase activity) | 10 | 1.27 | 0.195 | 1.000 |
| AT2G20420.1 | ATP citrate lyase (ACL) family protein (Succinate-CoA ligase activity) | 11 | 1.30 | 0.196 | 1.000 |
| AT2G20420.1 | ATP citrate lyase (ACL) family protein (Succinate-CoA ligase activity) | 12 | 1.19 | 0.196 | 1.000 |
| AT2G20420.1 | ATP citrate lyase (ACL) family protein (Succinate-CoA ligase activity) | 13 | 1.03 | 0.194 | 1.000 |
| AT2G20420.1 | ATP citrate lyase (ACL) family protein (Succinate-CoA ligase activity) | 14 | 0.97 | 0.192 | 1.000 |
| AT2G20420.1 | ATP citrate lyase (ACL) family protein (Succinate-CoA ligase activity) | 15 | 1.01 | 0.191 | 1.000 |
| AT2G20420.1 | ATP citrate lyase (ACL) family protein (Succinate-CoA ligase activity) | 16 | 1.22 | 0.190 | 1.000 |
| AT2G20420.1 | ATP citrate lyase (ACL) family protein (Succinate-CoA ligase activity) | 17 | 1.66 | 0.190 | 0.998 |
| AT2G20420.1 | ATP citrate lyase (ACL) family protein (Succinate-CoA ligase activity) | 18 | 2.16 | 0.193 | 0.843 |
| AT2G20420.1 | ATP citrate lyase (ACL) family protein (Succinate-CoA ligase activity) | 19 | 2.07 | 0.197 | 0.912 |
| AT2G20420.1 | ATP citrate lyase (ACL) family protein (Succinate-CoA ligase activity) | 20 | 1.28 | 0.200 | 1.000 |
| AT2G20420.1 | ATP citrate lyase (ACL) family protein (Succinate-CoA ligase activity) | 21 | 0.65 | 0.202 | 1.000 |
| AT2G20420.1 | ATP citrate lyase (ACL) family protein (Succinate-CoA ligase activity) | 22 | 0.38 | 0.202 | 0.604 |
| AT2G20420.1 | ATP citrate lyase (ACL) family protein (Succinate-CoA ligase activity) | 23 | 0.34 | 0.201 | 0.393 |

| | | | | | |
|---|---|---|---|---|---|
| AT2G20420.1 | ATP citrate lyase (ACL) family protein (Succinate-CoA ligase activity) | 24 | 0.44 | 0.198 | 0.800 |
| AT2G20420.1 | ATP citrate lyase (ACL) family protein (Succinate-CoA ligase activity) | 25 | 0.59 | 0.195 | 0.997 |
| AT2G20420.1 | ATP citrate lyase (ACL) family protein (Succinate-CoA ligase activity) | 26 | 0.67 | 0.194 | 1.000 |
| AT2G20420.1 | ATP citrate lyase (ACL) family protein (Succinate-CoA ligase activity) | 27 | 0.81 | 0.195 | 1.000 |
| AT2G20420.1 | ATP citrate lyase (ACL) family protein (Succinate-CoA ligase activity) | 28 | 1.05 | 0.194 | 1.000 |
| AT2G20420.1 | ATP citrate lyase (ACL) family protein (Succinate-CoA ligase activity) | 29 | 1.27 | 0.194 | 1.000 |
| AT2G20420.1 | ATP citrate lyase (ACL) family protein (Succinate-CoA ligase activity) | 30 | 1.45 | 0.194 | 1.000 |
| AT2G20420.1 | ATP citrate lyase (ACL) family protein (Succinate-CoA ligase activity) | 31 | 1.57 | 0.195 | 1.000 |
| AT2G20420.1 | ATP citrate lyase (ACL) family protein (Succinate-CoA ligase activity) | 32 | 1.55 | 0.195 | 1.000 |
| AT2G20420.1 | ATP citrate lyase (ACL) family protein (Succinate-CoA ligase activity) | 33 | 1.37 | 0.195 | 1.000 |
| AT2G20420.1 | ATP citrate lyase (ACL) family protein (Succinate-CoA ligase activity) | 34 | 1.19 | 0.196 | 1.000 |
| AT2G20420.1 | ATP citrate lyase (ACL) family protein (Succinate-CoA ligase activity) | 35 | 1.11 | 0.196 | 1.000 |
| AT2G20420.1 | ATP citrate lyase (ACL) family protein (Succinate-CoA ligase activity) | 36 | 1.14 | 0.195 | 1.000 |
| AT2G20420.1 | ATP citrate lyase (ACL) family protein (Succinate-CoA ligase activity) | 37 | 1.29 | 0.195 | 1.000 |
| AT2G20420.1 | ATP citrate lyase (ACL) family protein (Succinate-CoA ligase activity) | 38 | 1.56 | 0.195 | 1.000 |
| AT2G20420.1 | ATP citrate lyase (ACL) family protein (Succinate-CoA ligase activity) | 39 | 1.91 | 0.195 | 0.968 |
| AT2G20420.1 | ATP citrate lyase (ACL) family protein (Succinate-CoA ligase activity) | 40 | 2.18 | 0.196 | 0.849 |
| AT2G20420.1 | ATP citrate lyase (ACL) family protein (Succinate-CoA ligase activity) | 41 | 2.13 | 0.196 | 0.881 |
| AT2G20420.1 | ATP citrate lyase (ACL) family protein (Succinate-CoA ligase activity) | 42 | 1.72 | 0.199 | 0.997 |
| AT2G20420.1 | ATP citrate lyase (ACL) family protein (Succinate-CoA ligase activity) | 43 | 1.19 | 0.201 | 1.000 |

| | | | | | | |
|---|---|---|---|---|---|---|
| AT2G20420.1 | ATP citrate lyase (ACL) family protein (Succinate-CoA ligase activity) | 44 | 0.79 | 0.200 | 1.000 | |
| AT2G20420.1 | ATP citrate lyase (ACL) family protein (Succinate-CoA ligase activity) | 45 | 0.59 | 0.200 | 0.998 | |
| AT2G20420.1 | ATP citrate lyase (ACL) family protein (Succinate-CoA ligase activity) | 46 | 0.48 | 0.234 | 0.984 | |
| AT2G20420.1 | ATP citrate lyase (ACL) family protein (Succinate-CoA ligase activity) | 47 | 0.40 | 0.382 | 1.000 | |
| AT2G20420.1 | ATP citrate lyase (ACL) family protein (Succinate-CoA ligase activity) | 48 | 0.33 | 0.609 | 1.000 | |
| AT2G20530.2 | Prohibitin 6 (PHB6) | 2 | 0.62 | 0.376 | 1.000 | |
| AT2G20530.2 | Prohibitin 6 (PHB6) | 3 | 0.59 | 0.243 | 1.000 | |
| AT2G20530.2 | Prohibitin 6 (PHB6) | 4 | 0.58 | 0.208 | 0.996 | |
| AT2G20530.2 | Prohibitin 6 (PHB6) | 5 | 0.63 | 0.207 | 1.000 | |
| AT2G20530.2 | Prohibitin 6 (PHB6) | 6 | 0.71 | 0.208 | 1.000 | |
| AT2G20530.2 | Prohibitin 6 (PHB6) | 7 | 0.76 | 0.206 | 1.000 | |
| AT2G20530.2 | Prohibitin 6 (PHB6) | 8 | 0.83 | 0.202 | 1.000 | |
| AT2G20530.2 | Prohibitin 6 (PHB6) | 9 | 0.95 | 0.200 | 1.000 | |
| AT2G20530.2 | Prohibitin 6 (PHB6) | 10 | 1.10 | 0.199 | 1.000 | |
| AT2G20530.2 | Prohibitin 6 (PHB6) | 11 | 1.28 | 0.200 | 1.000 | |
| AT2G20530.2 | Prohibitin 6 (PHB6) | 12 | 1.46 | 0.199 | 1.000 | |
| AT2G20530.2 | Prohibitin 6 (PHB6) | 13 | 1.53 | 0.197 | 1.000 | |
| AT2G20530.2 | Prohibitin 6 (PHB6) | 14 | 1.34 | 0.195 | 1.000 | |
| AT2G20530.2 | Prohibitin 6 (PHB6) | 15 | 0.99 | 0.194 | 1.000 | |
| AT2G20530.2 | Prohibitin 6 (PHB6) | 16 | 0.74 | 0.193 | 1.000 | |
| AT2G20530.2 | Prohibitin 6 (PHB6) | 17 | 0.68 | 0.193 | 1.000 | |
| AT2G20530.2 | Prohibitin 6 (PHB6) | 18 | 0.76 | 0.195 | 1.000 | |
| AT2G20530.2 | Prohibitin 6 (PHB6) | 19 | 0.90 | 0.200 | 1.000 | |
| AT2G20530.2 | Prohibitin 6 (PHB6) | 20 | 1.07 | 0.205 | 1.000 | |
| AT2G20530.2 | Prohibitin 6 (PHB6) | 21 | 1.28 | 0.207 | 1.000 | |
| AT2G20530.2 | Prohibitin 6 (PHB6) | 22 | 1.55 | 0.208 | 1.000 | |
| AT2G20530.2 | Prohibitin 6 (PHB6) | 23 | 1.77 | 0.206 | 0.991 | |
| AT2G20530.2 | Prohibitin 6 (PHB6) | 24 | 1.65 | 0.202 | 0.998 | |
| AT2G20530.2 | Prohibitin 6 (PHB6) | 25 | 1.10 | 0.199 | 1.000 | |
| AT2G20530.2 | Prohibitin 6 (PHB6) | 26 | 0.63 | 0.197 | 0.999 | |
| AT2G20530.2 | Prohibitin 6 (PHB6) | 27 | 0.43 | 0.197 | 0.731 | |
| AT2G20530.2 | Prohibitin 6 (PHB6) | 28 | 0.38 | 0.198 | 0.538 | |
| AT2G20530.2 | Prohibitin 6 (PHB6) | 29 | 0.51 | 0.199 | 0.941 | |
| AT2G20530.2 | Prohibitin 6 (PHB6) | 30 | 1.00 | 0.200 | 1.000 | |
| AT2G20530.2 | Prohibitin 6 (PHB6) | 31 | 2.32 | 0.203 | 0.775 | |
| AT2G20530.2 | Prohibitin 6 (PHB6) | 32 | 4.41 | 0.209 | 0.054 | . |
| AT2G20530.2 | Prohibitin 6 (PHB6) | 33 | 5.86 | 0.229 | 0.024 | * |
| AT2G20530.2 | Prohibitin 6 (PHB6) | 34 | 5.90 | 0.287 | 0.163 | |
| AT2G20530.2 | Prohibitin 6 (PHB6) | 35 | 5.71 | 0.408 | 0.737 | |
| AT2G20530.2 | Prohibitin 6 (PHB6) | 46 | 170.48 | 3.520 | 1.000 | |
| AT2G20530.2 | Prohibitin 6 (PHB6) | 47 | 274.80 | 3.947 | 1.000 | |
| AT2G20860.1 | Lipoic acid synthase 1 (LIP1) | 2 | 1.16 | 0.602 | 1.000 | |
| AT2G20860.1 | Lipoic acid synthase 1 (LIP1) | 3 | 1.51 | 0.559 | 1.000 | |
| AT2G20860.1 | Lipoic acid synthase 1 (LIP1) | 20 | 1.87 | 0.459 | 1.000 | |
| AT2G20860.1 | Lipoic acid synthase 1 (LIP1) | 21 | 1.22 | 0.413 | 1.000 | |
| AT2G20860.1 | Lipoic acid synthase 1 (LIP1) | 22 | 0.83 | 0.402 | 1.000 | |
| AT2G20860.1 | Lipoic acid synthase 1 (LIP1) | 25 | 0.43 | 0.314 | 0.975 | |
| AT2G20860.1 | Lipoic acid synthase 1 (LIP1) | 26 | 0.42 | 0.278 | 0.925 | |
| AT2G20860.1 | Lipoic acid synthase 1 (LIP1) | 27 | 0.44 | 0.248 | 0.887 | |
| AT2G20860.1 | Lipoic acid synthase 1 (LIP1) | 28 | 0.48 | 0.234 | 0.917 | |
| AT2G20860.1 | Lipoic acid synthase 1 (LIP1) | 29 | 0.54 | 0.228 | 0.970 | |
| AT2G20860.1 | Lipoic acid synthase 1 (LIP1) | 30 | 0.61 | 0.228 | 0.996 | |
| AT2G20860.1 | Lipoic acid synthase 1 (LIP1) | 31 | 0.75 | 0.228 | 1.000 | |

| | | | | | | |
|---|---|---|---|---|---|---|
| AT2G20860.1 | Lipoic acid synthase 1 (LIP1) | 32 | 0.91 | 0.228 | 1.000 | |
| AT2G20860.1 | Lipoic acid synthase 1 (LIP1) | 33 | 0.94 | 0.228 | 1.000 | |
| AT2G20860.1 | Lipoic acid synthase 1 (LIP1) | 34 | 0.81 | 0.230 | 1.000 | |
| AT2G20860.1 | Lipoic acid synthase 1 (LIP1) | 35 | 0.68 | 0.233 | 1.000 | |
| AT2G20860.1 | Lipoic acid synthase 1 (LIP1) | 36 | 0.63 | 0.239 | 0.999 | |
| AT2G20860.1 | Lipoic acid synthase 1 (LIP1) | 37 | 0.66 | 0.246 | 1.000 | |
| AT2G20860.1 | Lipoic acid synthase 1 (LIP1) | 38 | 0.80 | 0.249 | 1.000 | |
| AT2G20860.1 | Lipoic acid synthase 1 (LIP1) | 39 | 1.10 | 0.243 | 1.000 | |
| AT2G20860.1 | Lipoic acid synthase 1 (LIP1) | 40 | 1.54 | 0.237 | 0.999 | |
| AT2G20860.1 | Lipoic acid synthase 1 (LIP1) | 41 | 1.99 | 0.233 | 0.946 | |
| AT2G20860.1 | Lipoic acid synthase 1 (LIP1) | 42 | 2.26 | 0.233 | 0.842 | |
| AT2G20860.1 | Lipoic acid synthase 1 (LIP1) | 43 | 2.16 | 0.234 | 0.888 | |
| AT2G20860.1 | Lipoic acid synthase 1 (LIP1) | 44 | 1.56 | 0.234 | 0.999 | |
| AT2G20860.1 | Lipoic acid synthase 1 (LIP1) | 45 | 0.86 | 0.244 | 1.000 | |
| AT2G20860.1 | Lipoic acid synthase 1 (LIP1) | 46 | 0.39 | 0.299 | 0.921 | |
| AT2G20860.1 | Lipoic acid synthase 1 (LIP1) | 47 | 0.17 | 0.449 | 0.696 | |
| AT2G20940.1 | ? | 2 | 1.00 | 0.468 | 1.000 | |
| AT2G20940.1 | ? | 3 | 0.94 | 0.364 | 1.000 | |
| AT2G20940.1 | ? | 6 | 0.85 | 0.287 | 1.000 | |
| AT2G20940.1 | ? | 7 | 0.89 | 0.237 | 1.000 | |
| AT2G20940.1 | ? | 8 | 0.99 | 0.203 | 1.000 | |
| AT2G20940.1 | ? | 9 | 1.15 | 0.187 | 1.000 | |
| AT2G20940.1 | ? | 10 | 1.27 | 0.183 | 1.000 | |
| AT2G20940.1 | ? | 11 | 1.22 | 0.182 | 1.000 | |
| AT2G20940.1 | ? | 12 | 1.09 | 0.182 | 1.000 | |
| AT2G20940.1 | ? | 13 | 0.94 | 0.179 | 1.000 | |
| AT2G20940.1 | ? | 14 | 0.88 | 0.176 | 1.000 | |
| AT2G20940.1 | ? | 15 | 0.83 | 0.176 | 1.000 | |
| AT2G20940.1 | ? | 16 | 0.80 | 0.175 | 1.000 | |
| AT2G20940.1 | ? | 17 | 0.87 | 0.176 | 1.000 | |
| AT2G20940.1 | ? | 18 | 1.19 | 0.183 | 1.000 | |
| AT2G20940.1 | ? | 19 | 1.77 | 0.191 | 0.957 | |
| AT2G20940.1 | ? | 20 | 2.37 | 0.208 | 0.689 | |
| AT2G20940.1 | ? | 21 | 2.50 | 0.270 | 0.894 | |
| AT2G20940.1 | ? | 22 | 2.23 | 0.408 | 0.999 | |
| AT2G20940.1 | ? | 40 | 0.45 | 0.382 | 0.998 | |
| AT2G20940.1 | ? | 41 | 0.93 | 0.320 | 1.000 | |
| AT2G20940.1 | ? | 42 | 2.10 | 0.293 | 0.989 | |
| AT2G20940.1 | ? | 44 | 6.04 | 0.284 | 0.114 | |
| AT2G20940.1 | ? | 45 | 4.80 | 0.231 | 0.065 | . |
| AT2G20940.1 | ? | 46 | 1.89 | 0.239 | 0.983 | |
| AT2G20940.1 | ? | 47 | 0.50 | 0.390 | 1.000 | |
| AT2G21870.1 | ATP synthase subunit (ATPFAd) | 2 | 0.81 | 0.395 | 1.000 | |
| AT2G21870.1 | ATP synthase subunit (ATPFAd) | 3 | 0.77 | 0.258 | 1.000 | |
| AT2G21870.1 | ATP synthase subunit (ATPFAd) | 4 | 0.80 | 0.236 | 1.000 | |
| AT2G21870.1 | ATP synthase subunit (ATPFAd) | 5 | 0.92 | 0.236 | 1.000 | |
| AT2G21870.1 | ATP synthase subunit (ATPFAd) | 6 | 1.05 | 0.236 | 1.000 | |
| AT2G21870.1 | ATP synthase subunit (ATPFAd) | 7 | 1.10 | 0.235 | 1.000 | |
| AT2G21870.1 | ATP synthase subunit (ATPFAd) | 8 | 1.18 | 0.232 | 1.000 | |
| AT2G21870.1 | ATP synthase subunit (ATPFAd) | 9 | 1.33 | 0.229 | 1.000 | |
| AT2G21870.1 | ATP synthase subunit (ATPFAd) | 10 | 1.48 | 0.229 | 1.000 | |
| AT2G21870.1 | ATP synthase subunit (ATPFAd) | 11 | 1.71 | 0.230 | 1.000 | |
| AT2G21870.1 | ATP synthase subunit (ATPFAd) | 12 | 2.05 | 0.230 | 0.984 | |
| AT2G21870.1 | ATP synthase subunit (ATPFAd) | 13 | 2.43 | 0.228 | 0.875 | |
| AT2G21870.1 | ATP synthase subunit (ATPFAd) | 14 | 2.75 | 0.226 | 0.701 | |
| AT2G21870.1 | ATP synthase subunit (ATPFAd) | 15 | 2.76 | 0.225 | 0.694 | |
| AT2G21870.1 | ATP synthase subunit (ATPFAd) | 16 | 2.46 | 0.224 | 0.846 | |
| AT2G21870.1 | ATP synthase subunit (ATPFAd) | 17 | 2.08 | 0.223 | 0.973 | |
| AT2G21870.1 | ATP synthase subunit (ATPFAd) | 18 | 1.69 | 0.225 | 1.000 | |
| AT2G21870.1 | ATP synthase subunit (ATPFAd) | 19 | 1.45 | 0.231 | 1.000 | |
| AT2G21870.1 | ATP synthase subunit (ATPFAd) | 20 | 1.31 | 0.235 | 1.000 | |
| AT2G21870.1 | ATP synthase subunit (ATPFAd) | 21 | 1.15 | 0.237 | 1.000 | |
| AT2G21870.1 | ATP synthase subunit (ATPFAd) | 22 | 0.99 | 0.237 | 1.000 | |

| | | | | | | |
|---|---|---|---|---|---|---|
| AT2G21870.1 | ATP synthase subunit (ATPFAd) | 23 | 0.94 | 0.235 | 1.000 | |
| AT2G21870.1 | ATP synthase subunit (ATPFAd) | 24 | 1.09 | 0.231 | 1.000 | |
| AT2G21870.1 | ATP synthase subunit (ATPFAd) | 25 | 1.36 | 0.229 | 1.000 | |
| AT2G21870.1 | ATP synthase subunit (ATPFAd) | 26 | 1.34 | 0.228 | 1.000 | |
| AT2G21870.1 | ATP synthase subunit (ATPFAd) | 27 | 0.95 | 0.229 | 1.000 | |
| AT2G21870.1 | ATP synthase subunit (ATPFAd) | 28 | 0.75 | 0.229 | 1.000 | |
| AT2G21870.1 | ATP synthase subunit (ATPFAd) | 29 | 0.89 | 0.229 | 1.000 | |
| AT2G21870.1 | ATP synthase subunit (ATPFAd) | 30 | 1.33 | 0.229 | 1.000 | |
| AT2G21870.1 | ATP synthase subunit (ATPFAd) | 31 | 1.93 | 0.229 | 0.994 | |
| AT2G21870.1 | ATP synthase subunit (ATPFAd) | 32 | 2.41 | 0.230 | 0.893 | |
| AT2G21870.1 | ATP synthase subunit (ATPFAd) | 33 | 2.38 | 0.230 | 0.906 | |
| AT2G21870.1 | ATP synthase subunit (ATPFAd) | 34 | 1.87 | 0.231 | 0.997 | |
| AT2G21870.1 | ATP synthase subunit (ATPFAd) | 35 | 1.35 | 0.234 | 1.000 | |
| AT2G21870.1 | ATP synthase subunit (ATPFAd) | 36 | 1.27 | 0.245 | 1.000 | |
| AT2G21870.1 | ATP synthase subunit (ATPFAd) | 37 | 1.83 | 0.274 | 1.000 | |
| AT2G21870.1 | ATP synthase subunit (ATPFAd) | 38 | 3.17 | 0.314 | 0.924 | |
| AT2G21870.1 | ATP synthase subunit (ATPFAd) | 40 | 5.32 | 0.297 | 0.318 | |
| AT2G21870.1 | ATP synthase subunit (ATPFAd) | 41 | 4.54 | 0.280 | 0.382 | |
| AT2G21870.1 | ATP synthase subunit (ATPFAd) | 42 | 2.99 | 0.272 | 0.844 | |
| AT2G21870.1 | ATP synthase subunit (ATPFAd) | 43 | 2.05 | 0.265 | 0.997 | |
| AT2G21870.1 | ATP synthase subunit (ATPFAd) | 44 | 1.50 | 0.255 | 1.000 | |
| AT2G21870.1 | ATP synthase subunit (ATPFAd) | 45 | 1.10 | 0.241 | 1.000 | |
| AT2G21870.1 | ATP synthase subunit (ATPFAd) | 46 | 0.84 | 0.276 | 1.000 | |
| AT2G21870.1 | ATP synthase subunit (ATPFAd) | 47 | 0.63 | 0.451 | 1.000 | |
| AT2G22500.1 | Dicarboxylate carrier 3 (DIC1) | 2 | 0.84 | 0.483 | 1.000 | |
| AT2G22500.1 | Dicarboxylate carrier 3 (DIC1) | 3 | 0.86 | 0.377 | 1.000 | |
| AT2G22500.1 | Dicarboxylate carrier 3 (DIC1) | 6 | 1.20 | 0.297 | 1.000 | |
| AT2G22500.1 | Dicarboxylate carrier 3 (DIC1) | 7 | 1.58 | 0.253 | 1.000 | |
| AT2G22500.1 | Dicarboxylate carrier 3 (DIC1) | 8 | 2.21 | 0.223 | 0.907 | |
| AT2G22500.1 | Dicarboxylate carrier 3 (DIC1) | 9 | 2.98 | 0.207 | 0.376 | |
| AT2G22500.1 | Dicarboxylate carrier 3 (DIC1) | 10 | 3.36 | 0.202 | 0.193 | |
| AT2G22500.1 | Dicarboxylate carrier 3 (DIC1) | 11 | 3.18 | 0.202 | 0.247 | |
| AT2G22500.1 | Dicarboxylate carrier 3 (DIC1) | 12 | 2.79 | 0.201 | 0.425 | |
| AT2G22500.1 | Dicarboxylate carrier 3 (DIC1) | 13 | 2.54 | 0.199 | 0.574 | |
| AT2G22500.1 | Dicarboxylate carrier 3 (DIC1) | 14 | 2.69 | 0.198 | 0.465 | |
| AT2G22500.1 | Dicarboxylate carrier 3 (DIC1) | 15 | 3.24 | 0.200 | 0.213 | |
| AT2G22500.1 | Dicarboxylate carrier 3 (DIC1) | 16 | 3.93 | 0.202 | 0.078 | . |
| AT2G22500.1 | Dicarboxylate carrier 3 (DIC1) | 17 | 5.10 | 0.209 | 0.020 | * |
| AT2G22500.1 | Dicarboxylate carrier 3 (DIC1) | 18 | 6.64 | 0.225 | 0.008 | ** |
| AT2G22500.1 | Dicarboxylate carrier 3 (DIC1) | 19 | 7.28 | 0.250 | 0.016 | * |
| AT2G22500.1 | Dicarboxylate carrier 3 (DIC1) | 20 | 6.99 | 0.303 | 0.120 | |
| AT2G22500.1 | Dicarboxylate carrier 3 (DIC1) | 21 | 6.44 | 0.354 | 0.375 | |
| AT2G22500.1 | Dicarboxylate carrier 3 (DIC1) | 28 | 3.68 | 0.302 | 0.699 | |
| AT2G22500.1 | Dicarboxylate carrier 3 (DIC1) | 29 | 3.88 | 0.245 | 0.300 | |
| AT2G22500.1 | Dicarboxylate carrier 3 (DIC1) | 30 | 4.16 | 0.216 | 0.099 | . |
| AT2G22500.1 | Dicarboxylate carrier 3 (DIC1) | 31 | 4.47 | 0.206 | 0.042 | * |
| AT2G22500.1 | Dicarboxylate carrier 3 (DIC1) | 32 | 4.43 | 0.202 | 0.038 | * |
| AT2G22500.1 | Dicarboxylate carrier 3 (DIC1) | 33 | 3.89 | 0.201 | 0.082 | . |
| AT2G22500.1 | Dicarboxylate carrier 3 (DIC1) | 34 | 3.32 | 0.201 | 0.197 | |
| AT2G22500.1 | Dicarboxylate carrier 3 (DIC1) | 35 | 2.89 | 0.200 | 0.367 | |
| AT2G22500.1 | Dicarboxylate carrier 3 (DIC1) | 36 | 2.53 | 0.200 | 0.586 | |
| AT2G22500.1 | Dicarboxylate carrier 3 (DIC1) | 37 | 2.34 | 0.200 | 0.721 | |
| AT2G22500.1 | Dicarboxylate carrier 3 (DIC1) | 38 | 2.35 | 0.200 | 0.716 | |
| AT2G22500.1 | Dicarboxylate carrier 3 (DIC1) | 39 | 2.41 | 0.200 | 0.673 | |
| AT2G22500.1 | Dicarboxylate carrier 3 (DIC1) | 40 | 2.38 | 0.200 | 0.694 | |
| AT2G22500.1 | Dicarboxylate carrier 3 (DIC1) | 41 | 2.02 | 0.201 | 0.920 | |
| AT2G22500.1 | Dicarboxylate carrier 3 (DIC1) | 42 | 1.36 | 0.204 | 1.000 | |
| AT2G22500.1 | Dicarboxylate carrier 3 (DIC1) | 43 | 0.79 | 0.205 | 1.000 | |
| AT2G22500.1 | Dicarboxylate carrier 3 (DIC1) | 44 | 0.45 | 0.205 | 0.816 | |
| AT2G22500.1 | Dicarboxylate carrier 3 (DIC1) | 45 | 0.27 | 0.208 | 0.130 | |
| AT2G22500.1 | Dicarboxylate carrier 3 (DIC1) | 46 | 0.17 | 0.252 | 0.051 | . |
| AT2G22500.1 | Dicarboxylate carrier 3 (DIC1) | 47 | 0.10 | 0.393 | 0.221 | |
| AT2G23370.1 | Cyclin delta-3 | 2 | 1.09 | 0.430 | 1.000 | |

| | | | | | | |
|---|---|---|---|---|---|---|
| AT2G23370.1 | Cyclin delta-3 | 3 | 1.07 | 0.396 | 1.000 | |
| AT2G23370.1 | Cyclin delta-3 | 12 | 1.01 | 0.363 | 1.000 | |
| AT2G23370.1 | Cyclin delta-3 | 13 | 1.02 | 0.358 | 1.000 | |
| AT2G23370.1 | Cyclin delta-3 | 14 | 1.04 | 0.362 | 1.000 | |
| AT2G23370.1 | Cyclin delta-3 | 31 | 1.94 | 0.257 | 0.872 | |
| AT2G23370.1 | Cyclin delta-3 | 32 | 2.06 | 0.231 | 0.732 | |
| AT2G23370.1 | Cyclin delta-3 | 33 | 2.24 | 0.218 | 0.556 | |
| AT2G23370.1 | Cyclin delta-3 | 34 | 2.54 | 0.218 | 0.383 | |
| AT2G23370.1 | Cyclin delta-3 | 35 | 3.03 | 0.228 | 0.239 | |
| AT2G23370.1 | Cyclin delta-3 | 40 | 5.63 | 0.235 | 0.016 | * |
| AT2G23370.1 | Cyclin delta-3 | 41 | 5.01 | 0.220 | 0.017 | * |
| AT2G23370.1 | Cyclin delta-3 | 42 | 3.92 | 0.205 | 0.039 | * |
| AT2G23370.1 | Cyclin delta-3 | 43 | 2.70 | 0.196 | 0.204 | |
| AT2G23370.1 | Cyclin delta-3 | 44 | 1.67 | 0.194 | 0.859 | |
| AT2G23370.1 | Cyclin delta-3 | 45 | 0.97 | 0.206 | 1.000 | |
| AT2G23370.1 | Cyclin delta-3 | 46 | 0.56 | 0.242 | 0.906 | |
| AT2G23370.1 | Cyclin delta-3 | 47 | 0.32 | 0.311 | 0.561 | |
| AT2G23890.1 | 5' nucleotidase, putative | 2 | 1.00 | 0.346 | 1.000 | |
| AT2G23890.1 | 5' nucleotidase, putative | 3 | 0.88 | 0.340 | 1.000 | |
| AT2G23890.1 | 5' nucleotidase, putative | 31 | 0.66 | 0.242 | 0.997 | |
| AT2G23890.1 | 5' nucleotidase, putative | 32 | 0.82 | 0.180 | 1.000 | |
| AT2G23890.1 | 5' nucleotidase, putative | 33 | 0.95 | 0.172 | 1.000 | |
| AT2G23890.1 | 5' nucleotidase, putative | 34 | 0.92 | 0.197 | 1.000 | |
| AT2G23890.1 | 5' nucleotidase, putative | 35 | 0.88 | 0.233 | 1.000 | |
| AT2G23890.1 | 5' nucleotidase, putative | 39 | 1.46 | 0.208 | 0.995 | |
| AT2G23890.1 | 5' nucleotidase, putative | 40 | 2.05 | 0.172 | 0.523 | |
| AT2G23890.1 | 5' nucleotidase, putative | 41 | 2.94 | 0.152 | 0.033 | * |
| AT2G23890.1 | 5' nucleotidase, putative | 42 | 3.69 | 0.147 | 0.002 | ** |
| AT2G23890.1 | 5' nucleotidase, putative | 43 | 3.56 | 0.146 | 0.004 | ** |
| AT2G23890.1 | 5' nucleotidase, putative | 44 | 2.48 | 0.146 | 0.086 | . |
| AT2G23890.1 | 5' nucleotidase, putative | 45 | 1.21 | 0.149 | 1.000 | |
| AT2G23890.1 | 5' nucleotidase, putative | 46 | 0.46 | 0.177 | 0.450 | |
| AT2G23890.1 | 5' nucleotidase, putative | 47 | 0.16 | 0.283 | 0.062 | . |
| AT2G24820.1 | Translocon of the inner envelope membrane of chloroplasts 55 (TIC55) | 2 | 1.17 | 0.620 | 1.000 | |
| AT2G24820.1 | Translocon of the inner envelope membrane of chloroplasts 55 (TIC55) | 3 | 1.27 | 0.601 | 1.000 | |
| AT2G24820.1 | Translocon of the inner envelope membrane of chloroplasts 55 (TIC55) | 33 | 1.47 | 0.281 | 0.942 | |
| AT2G24820.1 | Translocon of the inner envelope membrane of chloroplasts 55 (TIC55) | 34 | 1.27 | 0.269 | 0.988 | |
| AT2G24820.1 | Translocon of the inner envelope membrane of chloroplasts 55 (TIC55) | 35 | 1.07 | 0.267 | 1.000 | |
| AT2G24820.1 | Translocon of the inner envelope membrane of chloroplasts 55 (TIC55) | 36 | 0.89 | 0.271 | 0.999 | |
| AT2G24820.1 | Translocon of the inner envelope membrane of chloroplasts 55 (TIC55) | 37 | 0.73 | 0.282 | 0.970 | |
| AT2G24820.1 | Translocon of the inner envelope membrane of chloroplasts 55 (TIC55) | 46 | 0.08 | 0.411 | 0.036 | * |
| AT2G24820.1 | Translocon of the inner envelope membrane of chloroplasts 55 (TIC55) | 47 | 0.06 | 0.440 | 0.029 | * |
| AT2G25140.1 | Caseinolytic protease B4 (CLPB4) | 2 | 1.16 | 0.375 | 1.000 | |

| | | | | | |
|---|---|---|---|---|---|
| AT2G25140.1 | Caseinolytic protease B4 (CLPB4) | 3 | 0.89 | 0.280 | 1.000 |
| AT2G25140.1 | Caseinolytic protease B4 (CLPB4) | 4 | 0.70 | 0.500 | 1.000 |
| AT2G25140.1 | Caseinolytic protease B4 (CLPB4) | 29 | 0.98 | 0.279 | 1.000 |
| AT2G25140.1 | Caseinolytic protease B4 (CLPB4) | 30 | 0.99 | 0.206 | 1.000 |
| AT2G25140.1 | Caseinolytic protease B4 (CLPB4) | 31 | 0.99 | 0.183 | 1.000 |
| AT2G25140.1 | Caseinolytic protease B4 (CLPB4) | 32 | 1.01 | 0.173 | 1.000 |
| AT2G25140.1 | Caseinolytic protease B4 (CLPB4) | 33 | 0.92 | 0.169 | 1.000 |
| AT2G25140.1 | Caseinolytic protease B4 (CLPB4) | 34 | 0.91 | 0.168 | 1.000 |
| AT2G25140.1 | Caseinolytic protease B4 (CLPB4) | 35 | 1.13 | 0.168 | 1.000 |
| AT2G25140.1 | Caseinolytic protease B4 (CLPB4) | 36 | 1.39 | 0.167 | 0.997 |
| AT2G25140.1 | Caseinolytic protease B4 (CLPB4) | 37 | 1.37 | 0.167 | 0.998 |
| AT2G25140.1 | Caseinolytic protease B4 (CLPB4) | 38 | 1.12 | 0.169 | 1.000 |
| AT2G25140.1 | Caseinolytic protease B4 (CLPB4) | 39 | 0.97 | 0.170 | 1.000 |
| AT2G25140.1 | Caseinolytic protease B4 (CLPB4) | 40 | 1.03 | 0.182 | 1.000 |
| AT2G25140.1 | Caseinolytic protease B4 (CLPB4) | 41 | 1.29 | 0.237 | 1.000 |
| AT2G25140.1 | Caseinolytic protease B4 (CLPB4) | 42 | 2.09 | 0.356 | 0.995 |
| AT2G25140.1 | Caseinolytic protease B4 (CLPB4) | 46 | 107.34 | 1.284 | 0.770 |
| AT2G25140.1 | Caseinolytic protease B4 (CLPB4) | 47 | 365.35 | 1.635 | 0.780 |
| AT2G25530.1 | AFG1-like ATPase family protein | 2 | 0.92 | 0.323 | 1.000 |
| AT2G25530.1 | AFG1-like ATPase family protein | 3 | 1.11 | 0.312 | 1.000 |
| AT2G25530.1 | AFG1-like ATPase family protein | 28 | 1.64 | 0.268 | 1.000 |
| AT2G25530.1 | AFG1-like ATPase family protein | 29 | 1.23 | 0.199 | 1.000 |
| AT2G25530.1 | AFG1-like ATPase family protein | 30 | 1.01 | 0.179 | 1.000 |
| AT2G25530.1 | AFG1-like ATPase family protein | 31 | 1.01 | 0.163 | 1.000 |
| AT2G25530.1 | AFG1-like ATPase family protein | 32 | 1.15 | 0.155 | 1.000 |
| AT2G25530.1 | AFG1-like ATPase family protein | 33 | 1.06 | 0.150 | 1.000 |
| AT2G25530.1 | AFG1-like ATPase family protein | 34 | 0.80 | 0.148 | 1.000 |
| AT2G25530.1 | AFG1-like ATPase family protein | 35 | 0.80 | 0.147 | 1.000 |
| AT2G25530.1 | AFG1-like ATPase family protein | 36 | 1.10 | 0.146 | 1.000 |
| AT2G25530.1 | AFG1-like ATPase family protein | 37 | 1.50 | 0.145 | 0.975 |
| AT2G25530.1 | AFG1-like ATPase family protein | 38 | 1.72 | 0.146 | 0.825 |

| | | | | | |
|---|---|---|---|---|---|
| AT2G25530.1 | AFG1-like ATPase family protein | 39 | 1.76 | 0.147 | 0.780 |
| AT2G25530.1 | AFG1-like ATPase family protein | 40 | 1.72 | 0.146 | 0.822 |
| AT2G25530.1 | AFG1-like ATPase family protein | 41 | 1.44 | 0.146 | 0.991 |
| AT2G25530.1 | AFG1-like ATPase family protein | 42 | 1.13 | 0.148 | 1.000 |
| AT2G25530.1 | AFG1-like ATPase family protein | 43 | 0.89 | 0.149 | 1.000 |
| AT2G25530.1 | AFG1-like ATPase family protein | 44 | 0.70 | 0.149 | 0.993 |
| AT2G25530.1 | AFG1-like ATPase family protein | 45 | 0.52 | 0.149 | 0.593 |
| AT2G25530.1 | AFG1-like ATPase family protein | 46 | 0.37 | 0.172 | 0.196 |
| AT2G25530.1 | AFG1-like ATPase family protein | 47 | 0.25 | 0.333 | 0.697 |
| AT2G25570.3 | Binding protein | 2 | 0.85 | 0.476 | 1.000 |
| AT2G25570.3 | Binding protein | 3 | 1.52 | 0.517 | 1.000 |
| AT2G25570.3 | Binding protein | 22 | 0.78 | 0.364 | 1.000 |
| AT2G25570.3 | Binding protein | 23 | 0.68 | 0.267 | 0.999 |
| AT2G25570.3 | Binding protein | 24 | 0.72 | 0.233 | 0.999 |
| AT2G25570.3 | Binding protein | 25 | 0.97 | 0.217 | 1.000 |
| AT2G25570.3 | Binding protein | 26 | 1.80 | 0.216 | 0.915 |
| AT2G25570.3 | Binding protein | 27 | 1.76 | 0.220 | 0.938 |
| AT2G25570.3 | Binding protein | 28 | 0.68 | 0.237 | 0.998 |
| AT2G25570.3 | Binding protein | 29 | 0.27 | 0.295 | 0.422 |
| AT2G25570.3 | Binding protein | 30 | 0.17 | 0.494 | 0.700 |
| AT2G25570.3 | Binding protein | 46 | 55199764.44 | 10.243 | 0.996 |
| AT2G25570.3 | Binding protein | 47 | ########### | 11.198 | 0.995 |
| AT2G25830.1 | YebC-related | 2 | 1.30 | 0.354 | 1.000 |
| AT2G25830.1 | YebC-related | 3 | 1.36 | 0.345 | 1.000 |
| AT2G25830.1 | YebC-related | 31 | 1.08 | 0.252 | 1.000 |
| AT2G25830.1 | YebC-related | 32 | 0.97 | 0.202 | 1.000 |
| AT2G25830.1 | YebC-related | 33 | 0.89 | 0.186 | 1.000 |
| AT2G25830.1 | YebC-related | 34 | 0.86 | 0.181 | 1.000 |
| AT2G25830.1 | YebC-related | 35 | 0.89 | 0.178 | 1.000 |
| AT2G25830.1 | YebC-related | 36 | 0.97 | 0.177 | 1.000 |
| AT2G25830.1 | YebC-related | 37 | 0.99 | 0.177 | 1.000 |
| AT2G25830.1 | YebC-related | 38 | 0.85 | 0.176 | 1.000 |
| AT2G25830.1 | YebC-related | 39 | 0.65 | 0.176 | 0.919 |
| AT2G25830.1 | YebC-related | 40 | 0.56 | 0.176 | 0.706 |
| AT2G25830.1 | YebC-related | 41 | 0.67 | 0.177 | 0.953 |
| AT2G25830.1 | YebC-related | 42 | 1.06 | 0.177 | 1.000 |
| AT2G25830.1 | YebC-related | 43 | 1.74 | 0.178 | 0.764 |
| AT2G25830.1 | YebC-related | 44 | 2.13 | 0.180 | 0.428 |
| AT2G25830.1 | YebC-related | 45 | 1.80 | 0.185 | 0.749 |
| AT2G25830.1 | YebC-related | 46 | 1.12 | 0.202 | 1.000 |
| AT2G25830.1 | YebC-related | 47 | 0.56 | 0.274 | 0.970 |
| AT2G25840.3 | Nucleotidylyl transferase superfamily protein | 2 | 0.99 | 0.440 | 1.000 |
| AT2G25840.3 | Nucleotidylyl transferase superfamily protein | 3 | 1.01 | 0.382 | 1.000 |
| AT2G25840.3 | Nucleotidylyl transferase superfamily protein | 11 | 1.29 | 0.277 | 1.000 |
| AT2G25840.3 | Nucleotidylyl transferase superfamily protein | 12 | 1.35 | 0.252 | 1.000 |
| AT2G25840.3 | Nucleotidylyl transferase superfamily protein | 13 | 1.39 | 0.252 | 1.000 |
| AT2G25840.3 | Nucleotidylyl transferase superfamily protein | 14 | 1.40 | 0.256 | 1.000 |

| | | | | | |
|---|---|---|---|---|---|
| AT2G25840.3 | Nucleotidylyl transferase superfamily protein | 21 | 0.83 | 0.240 | 1.000 |
| AT2G25840.3 | Nucleotidylyl transferase superfamily protein | 22 | 0.74 | 0.206 | 1.000 |
| AT2G25840.3 | Nucleotidylyl transferase superfamily protein | 23 | 0.68 | 0.178 | 0.999 |
| AT2G25840.3 | Nucleotidylyl transferase superfamily protein | 24 | 0.63 | 0.162 | 0.979 |
| AT2G25840.3 | Nucleotidylyl transferase superfamily protein | 25 | 0.58 | 0.155 | 0.900 |
| AT2G25840.3 | Nucleotidylyl transferase superfamily protein | 26 | 0.56 | 0.152 | 0.828 |
| AT2G25840.3 | Nucleotidylyl transferase superfamily protein | 27 | 0.60 | 0.152 | 0.919 |
| AT2G25840.3 | Nucleotidylyl transferase superfamily protein | 28 | 0.70 | 0.152 | 0.997 |
| AT2G25840.3 | Nucleotidylyl transferase superfamily protein | 29 | 0.83 | 0.152 | 1.000 |
| AT2G25840.3 | Nucleotidylyl transferase superfamily protein | 30 | 0.90 | 0.153 | 1.000 |
| AT2G25840.3 | Nucleotidylyl transferase superfamily protein | 31 | 0.90 | 0.153 | 1.000 |
| AT2G25840.3 | Nucleotidylyl transferase superfamily protein | 32 | 0.86 | 0.153 | 1.000 |
| AT2G25840.3 | Nucleotidylyl transferase superfamily protein | 33 | 0.79 | 0.153 | 1.000 |
| AT2G25840.3 | Nucleotidylyl transferase superfamily protein | 34 | 0.73 | 0.153 | 0.999 |
| AT2G25840.3 | Nucleotidylyl transferase superfamily protein | 35 | 0.72 | 0.153 | 0.999 |
| AT2G25840.3 | Nucleotidylyl transferase superfamily protein | 36 | 0.74 | 0.153 | 0.999 |
| AT2G25840.3 | Nucleotidylyl transferase superfamily protein | 37 | 0.78 | 0.153 | 1.000 |
| AT2G25840.3 | Nucleotidylyl transferase superfamily protein | 38 | 0.91 | 0.153 | 1.000 |
| AT2G25840.3 | Nucleotidylyl transferase superfamily protein | 39 | 1.21 | 0.153 | 1.000 |
| AT2G25840.3 | Nucleotidylyl transferase superfamily protein | 40 | 1.58 | 0.154 | 0.969 |
| AT2G25840.3 | Nucleotidylyl transferase superfamily protein | 41 | 1.85 | 0.155 | 0.772 |
| AT2G25840.3 | Nucleotidylyl transferase superfamily protein | 42 | 1.91 | 0.157 | 0.731 |
| AT2G25840.3 | Nucleotidylyl transferase superfamily protein | 43 | 1.69 | 0.158 | 0.926 |
| AT2G25840.3 | Nucleotidylyl transferase superfamily protein | 44 | 1.23 | 0.158 | 1.000 |
| AT2G25840.3 | Nucleotidylyl transferase superfamily protein | 45 | 0.73 | 0.167 | 1.000 |
| AT2G25840.3 | Nucleotidylyl transferase superfamily protein | 46 | 0.35 | 0.210 | 0.451 |
| AT2G25840.3 | Nucleotidylyl transferase superfamily protein | 47 | 0.16 | 0.319 | 0.204 |
| AT2G26080.1 | Glycine decarboxylase subunit (GDCP2) | 3 | 1.10 | 0.217 | 1.000 |
| AT2G26080.1 | Glycine decarboxylase subunit (GDCP2) | 4 | 1.16 | 0.204 | 1.000 |
| AT2G26080.1 | Glycine decarboxylase subunit (GDCP2) | 5 | 1.08 | 0.204 | 1.000 |
| AT2G26080.1 | Glycine decarboxylase subunit (GDCP2) | 6 | 0.91 | 0.204 | 1.000 |

| | | | | | |
|---|---|---|---|---|---|
| AT2G26080.1 | Glycine decarboxylase subunit (GDCP2) | 7 | 0.81 | 0.203 | 1.000 |
| AT2G26080.1 | Glycine decarboxylase subunit (GDCP2) | 8 | 0.82 | 0.201 | 1.000 |
| AT2G26080.1 | Glycine decarboxylase subunit (GDCP2) | 9 | 0.84 | 0.198 | 1.000 |
| AT2G26080.1 | Glycine decarboxylase subunit (GDCP2) | 10 | 0.82 | 0.197 | 1.000 |
| AT2G26080.1 | Glycine decarboxylase subunit (GDCP2) | 11 | 0.84 | 0.197 | 1.000 |
| AT2G26080.1 | Glycine decarboxylase subunit (GDCP2) | 12 | 0.89 | 0.198 | 1.000 |
| AT2G26080.1 | Glycine decarboxylase subunit (GDCP2) | 13 | 0.93 | 0.196 | 1.000 |
| AT2G26080.1 | Glycine decarboxylase subunit (GDCP2) | 14 | 0.98 | 0.194 | 1.000 |
| AT2G26080.1 | Glycine decarboxylase subunit (GDCP2) | 15 | 1.03 | 0.194 | 1.000 |
| AT2G26080.1 | Glycine decarboxylase subunit (GDCP2) | 16 | 1.09 | 0.192 | 1.000 |
| AT2G26080.1 | Glycine decarboxylase subunit (GDCP2) | 17 | 1.39 | 0.192 | 1.000 |
| AT2G26080.1 | Glycine decarboxylase subunit (GDCP2) | 18 | 1.85 | 0.195 | 0.992 |
| AT2G26080.1 | Glycine decarboxylase subunit (GDCP2) | 19 | 2.04 | 0.200 | 0.964 |
| AT2G26080.1 | Glycine decarboxylase subunit (GDCP2) | 20 | 1.95 | 0.203 | 0.986 |
| AT2G26080.1 | Glycine decarboxylase subunit (GDCP2) | 21 | 1.57 | 0.206 | 1.000 |
| AT2G26080.1 | Glycine decarboxylase subunit (GDCP2) | 22 | 1.03 | 0.206 | 1.000 |
| AT2G26080.1 | Glycine decarboxylase subunit (GDCP2) | 23 | 0.68 | 0.204 | 1.000 |
| AT2G26080.1 | Glycine decarboxylase subunit (GDCP2) | 24 | 0.69 | 0.201 | 1.000 |
| AT2G26080.1 | Glycine decarboxylase subunit (GDCP2) | 25 | 0.87 | 0.198 | 1.000 |
| AT2G26080.1 | Glycine decarboxylase subunit (GDCP2) | 26 | 1.05 | 0.197 | 1.000 |
| AT2G26080.1 | Glycine decarboxylase subunit (GDCP2) | 27 | 1.14 | 0.197 | 1.000 |
| AT2G26080.1 | Glycine decarboxylase subunit (GDCP2) | 28 | 1.12 | 0.197 | 1.000 |
| AT2G26080.1 | Glycine decarboxylase subunit (GDCP2) | 29 | 1.09 | 0.196 | 1.000 |
| AT2G26080.1 | Glycine decarboxylase subunit (GDCP2) | 30 | 1.13 | 0.196 | 1.000 |
| AT2G26080.1 | Glycine decarboxylase subunit (GDCP2) | 31 | 1.20 | 0.197 | 1.000 |
| AT2G26080.1 | Glycine decarboxylase subunit (GDCP2) | 32 | 1.12 | 0.197 | 1.000 |
| AT2G26080.1 | Glycine decarboxylase subunit (GDCP2) | 33 | 0.95 | 0.198 | 1.000 |
| AT2G26080.1 | Glycine decarboxylase subunit (GDCP2) | 34 | 0.92 | 0.198 | 1.000 |
| AT2G26080.1 | Glycine decarboxylase subunit (GDCP2) | 35 | 1.06 | 0.198 | 1.000 |
| AT2G26080.1 | Glycine decarboxylase subunit (GDCP2) | 36 | 1.29 | 0.197 | 1.000 |
| AT2G26080.1 | Glycine decarboxylase subunit (GDCP2) | 37 | 1.48 | 0.197 | 1.000 |

| | | | | | | |
|---|---|---|---|---|---|---|
| AT2G26080.1 | Glycine decarboxylase subunit (GDCP2) | 38 | 1.41 | 0.198 | 1.000 | |
| AT2G26080.1 | Glycine decarboxylase subunit (GDCP2) | 39 | 1.04 | 0.198 | 1.000 | |
| AT2G26080.1 | Glycine decarboxylase subunit (GDCP2) | 40 | 0.89 | 0.199 | 1.000 | |
| AT2G26080.1 | Glycine decarboxylase subunit (GDCP2) | 41 | 1.30 | 0.199 | 1.000 | |
| AT2G26080.1 | Glycine decarboxylase subunit (GDCP2) | 42 | 2.91 | 0.202 | 0.477 | |
| AT2G26080.1 | Glycine decarboxylase subunit (GDCP2) | 43 | 6.40 | 0.204 | 0.004 | ** |
| AT2G26080.1 | Glycine decarboxylase subunit (GDCP2) | 44 | 7.04 | 0.203 | 0.001 | ** |
| AT2G26080.1 | Glycine decarboxylase subunit (GDCP2) | 45 | 4.26 | 0.202 | 0.067 | . |
| AT2G26080.1 | Glycine decarboxylase subunit (GDCP2) | 46 | 2.28 | 0.231 | 0.966 | |
| AT2G26080.1 | Glycine decarboxylase subunit (GDCP2) | 47 | 1.19 | 0.424 | 1.000 | |
| AT2G26080.1 | Glycine decarboxylase subunit (GDCP2) | 48 | 0.62 | 0.722 | 1.000 | |
| AT2G26140.1 | FTSH protease 4 (FTSH4) | 2 | 0.77 | 0.372 | 1.000 | |
| AT2G26140.1 | FTSH protease 4 (FTSH4) | 3 | 0.58 | 0.251 | 1.000 | |
| AT2G26140.1 | FTSH protease 4 (FTSH4) | 4 | 0.47 | 0.236 | 0.993 | |
| AT2G26140.1 | FTSH protease 4 (FTSH4) | 5 | 0.43 | 0.238 | 0.971 | |
| AT2G26140.1 | FTSH protease 4 (FTSH4) | 6 | 0.44 | 0.221 | 0.951 | |
| AT2G26140.1 | FTSH protease 4 (FTSH4) | 7 | 0.51 | 0.202 | 0.989 | |
| AT2G26140.1 | FTSH protease 4 (FTSH4) | 8 | 0.71 | 0.190 | 1.000 | |
| AT2G26140.1 | FTSH protease 4 (FTSH4) | 9 | 1.05 | 0.184 | 1.000 | |
| AT2G26140.1 | FTSH protease 4 (FTSH4) | 10 | 1.24 | 0.183 | 1.000 | |
| AT2G26140.1 | FTSH protease 4 (FTSH4) | 11 | 1.18 | 0.182 | 1.000 | |
| AT2G26140.1 | FTSH protease 4 (FTSH4) | 12 | 0.93 | 0.182 | 1.000 | |
| AT2G26140.1 | FTSH protease 4 (FTSH4) | 13 | 0.62 | 0.181 | 1.000 | |
| AT2G26140.1 | FTSH protease 4 (FTSH4) | 14 | 0.43 | 0.178 | 0.697 | |
| AT2G26140.1 | FTSH protease 4 (FTSH4) | 15 | 0.38 | 0.177 | 0.434 | |
| AT2G26140.1 | FTSH protease 4 (FTSH4) | 16 | 0.48 | 0.176 | 0.862 | |
| AT2G26140.1 | FTSH protease 4 (FTSH4) | 17 | 0.87 | 0.175 | 1.000 | |
| AT2G26140.1 | FTSH protease 4 (FTSH4) | 18 | 1.78 | 0.179 | 0.992 | |
| AT2G26140.1 | FTSH protease 4 (FTSH4) | 19 | 2.72 | 0.184 | 0.451 | |
| AT2G26140.1 | FTSH protease 4 (FTSH4) | 20 | 2.68 | 0.188 | 0.525 | |
| AT2G26140.1 | FTSH protease 4 (FTSH4) | 21 | 1.94 | 0.191 | 0.979 | |
| AT2G26140.1 | FTSH protease 4 (FTSH4) | 22 | 1.35 | 0.190 | 1.000 | |
| AT2G26140.1 | FTSH protease 4 (FTSH4) | 23 | 1.07 | 0.189 | 1.000 | |
| AT2G26140.1 | FTSH protease 4 (FTSH4) | 24 | 1.01 | 0.185 | 1.000 | |
| AT2G26140.1 | FTSH protease 4 (FTSH4) | 25 | 1.05 | 0.182 | 1.000 | |
| AT2G26140.1 | FTSH protease 4 (FTSH4) | 26 | 1.12 | 0.181 | 1.000 | |
| AT2G26140.1 | FTSH protease 4 (FTSH4) | 27 | 1.15 | 0.181 | 1.000 | |
| AT2G26140.1 | FTSH protease 4 (FTSH4) | 28 | 1.12 | 0.181 | 1.000 | |
| AT2G26140.1 | FTSH protease 4 (FTSH4) | 29 | 1.10 | 0.181 | 1.000 | |
| AT2G26140.1 | FTSH protease 4 (FTSH4) | 30 | 1.07 | 0.181 | 1.000 | |
| AT2G26140.1 | FTSH protease 4 (FTSH4) | 31 | 1.03 | 0.181 | 1.000 | |
| AT2G26140.1 | FTSH protease 4 (FTSH4) | 32 | 0.95 | 0.182 | 1.000 | |
| AT2G26140.1 | FTSH protease 4 (FTSH4) | 33 | 0.85 | 0.182 | 1.000 | |
| AT2G26140.1 | FTSH protease 4 (FTSH4) | 34 | 0.78 | 0.183 | 1.000 | |
| AT2G26140.1 | FTSH protease 4 (FTSH4) | 35 | 0.79 | 0.182 | 1.000 | |
| AT2G26140.1 | FTSH protease 4 (FTSH4) | 36 | 0.87 | 0.181 | 1.000 | |
| AT2G26140.1 | FTSH protease 4 (FTSH4) | 37 | 1.05 | 0.181 | 1.000 | |
| AT2G26140.1 | FTSH protease 4 (FTSH4) | 38 | 1.32 | 0.182 | 1.000 | |
| AT2G26140.1 | FTSH protease 4 (FTSH4) | 39 | 1.61 | 0.182 | 1.000 | |
| AT2G26140.1 | FTSH protease 4 (FTSH4) | 40 | 1.72 | 0.183 | 0.997 | |
| AT2G26140.1 | FTSH protease 4 (FTSH4) | 41 | 1.53 | 0.183 | 1.000 | |
| AT2G26140.1 | FTSH protease 4 (FTSH4) | 42 | 1.15 | 0.186 | 1.000 | |

| | | | | | | |
|---|---|---|---|---|---|---|
| AT2G26140.1 | FTSH protease 4 (FTSH4) | 43 | 0.78 | 0.189 | 1.000 | |
| AT2G26140.1 | FTSH protease 4 (FTSH4) | 44 | 0.46 | 0.188 | 0.901 | |
| AT2G26140.1 | FTSH protease 4 (FTSH4) | 45 | 0.22 | 0.187 | 0.019 | * |
| AT2G26140.1 | FTSH protease 4 (FTSH4) | 46 | 0.08 | 0.226 | 0.000 | *** |
| AT2G26140.1 | FTSH protease 4 (FTSH4) | 47 | 0.03 | 0.406 | 0.006 | ** |
| AT2G26960.1 | MYB domain protein (MYB81) | 2 | 0.86 | 0.429 | 1.000 | |
| AT2G26960.1 | MYB domain protein (MYB81) | 3 | 0.85 | 0.363 | 1.000 | |
| AT2G26960.1 | MYB domain protein (MYB81) | 4 | 0.84 | 0.323 | 1.000 | |
| AT2G26960.1 | MYB domain protein (MYB81) | 6 | 0.83 | 0.291 | 1.000 | |
| AT2G26960.1 | MYB domain protein (MYB81) | 7 | 0.82 | 0.288 | 1.000 | |
| AT2G26960.1 | MYB domain protein (MYB81) | 8 | 0.82 | 0.282 | 1.000 | |
| AT2G26960.1 | MYB domain protein (MYB81) | 11 | 0.80 | 0.246 | 1.000 | |
| AT2G26960.1 | MYB domain protein (MYB81) | 12 | 0.80 | 0.231 | 1.000 | |
| AT2G26960.1 | MYB domain protein (MYB81) | 13 | 0.80 | 0.218 | 1.000 | |
| AT2G26960.1 | MYB domain protein (MYB81) | 14 | 0.80 | 0.206 | 1.000 | |
| AT2G26960.1 | MYB domain protein (MYB81) | 15 | 0.81 | 0.198 | 1.000 | |
| AT2G26960.1 | MYB domain protein (MYB81) | 16 | 0.81 | 0.193 | 1.000 | |
| AT2G26960.1 | MYB domain protein (MYB81) | 17 | 0.83 | 0.191 | 1.000 | |
| AT2G26960.1 | MYB domain protein (MYB81) | 18 | 0.84 | 0.191 | 1.000 | |
| AT2G26960.1 | MYB domain protein (MYB81) | 19 | 0.84 | 0.192 | 1.000 | |
| AT2G26960.1 | MYB domain protein (MYB81) | 20 | 0.84 | 0.194 | 1.000 | |
| AT2G26960.1 | MYB domain protein (MYB81) | 21 | 0.81 | 0.195 | 1.000 | |
| AT2G26960.1 | MYB domain protein (MYB81) | 22 | 0.76 | 0.195 | 1.000 | |
| AT2G26960.1 | MYB domain protein (MYB81) | 23 | 0.70 | 0.194 | 0.998 | |
| AT2G26960.1 | MYB domain protein (MYB81) | 24 | 0.64 | 0.193 | 0.989 | |
| AT2G26960.1 | MYB domain protein (MYB81) | 25 | 0.61 | 0.191 | 0.968 | |
| AT2G26960.1 | MYB domain protein (MYB81) | 26 | 0.58 | 0.190 | 0.943 | |
| AT2G26960.1 | MYB domain protein (MYB81) | 27 | 0.57 | 0.190 | 0.925 | |
| AT2G26960.1 | MYB domain protein (MYB81) | 28 | 0.57 | 0.189 | 0.929 | |
| AT2G26960.1 | MYB domain protein (MYB81) | 29 | 0.61 | 0.190 | 0.966 | |
| AT2G26960.1 | MYB domain protein (MYB81) | 30 | 0.67 | 0.190 | 0.995 | |
| AT2G26960.1 | MYB domain protein (MYB81) | 31 | 0.76 | 0.191 | 1.000 | |
| AT2G26960.1 | MYB domain protein (MYB81) | 32 | 0.88 | 0.192 | 1.000 | |
| AT2G26960.1 | MYB domain protein (MYB81) | 33 | 1.02 | 0.192 | 1.000 | |
| AT2G26960.1 | MYB domain protein (MYB81) | 34 | 1.17 | 0.192 | 1.000 | |
| AT2G26960.1 | MYB domain protein (MYB81) | 35 | 1.29 | 0.193 | 1.000 | |
| AT2G26960.1 | MYB domain protein (MYB81) | 36 | 1.37 | 0.193 | 1.000 | |
| AT2G26960.1 | MYB domain protein (MYB81) | 37 | 1.39 | 0.194 | 0.999 | |
| AT2G26960.1 | MYB domain protein (MYB81) | 38 | 1.35 | 0.194 | 1.000 | |
| AT2G26960.1 | MYB domain protein (MYB81) | 39 | 1.28 | 0.195 | 1.000 | |
| AT2G26960.1 | MYB domain protein (MYB81) | 40 | 1.20 | 0.195 | 1.000 | |
| AT2G26960.1 | MYB domain protein (MYB81) | 41 | 1.12 | 0.196 | 1.000 | |
| AT2G26960.1 | MYB domain protein (MYB81) | 42 | 1.05 | 0.197 | 1.000 | |
| AT2G26960.1 | MYB domain protein (MYB81) | 43 | 0.99 | 0.199 | 1.000 | |
| AT2G26960.1 | MYB domain protein (MYB81) | 44 | 0.95 | 0.209 | 1.000 | |
| AT2G26960.1 | MYB domain protein (MYB81) | 45 | 0.90 | 0.234 | 1.000 | |
| AT2G26960.1 | MYB domain protein (MYB81) | 46 | 0.83 | 0.282 | 1.000 | |
| AT2G26960.1 | MYB domain protein (MYB81) | 47 | 0.76 | 0.356 | 1.000 | |
| AT2G27730.1 | NADH:ubiquinone oxidoreductase subunit | 2 | 0.80 | 0.482 | 1.000 | |
| AT2G27730.1 | NADH:ubiquinone oxidoreductase subunit | 3 | 0.80 | 0.365 | 1.000 | |
| AT2G27730.1 | NADH:ubiquinone oxidoreductase subunit | 4 | 0.84 | 0.330 | 1.000 | |
| AT2G27730.1 | NADH:ubiquinone oxidoreductase subunit | 6 | 1.12 | 0.323 | 1.000 | |
| AT2G27730.1 | NADH:ubiquinone oxidoreductase subunit | 7 | 1.41 | 0.296 | 1.000 | |
| AT2G27730.1 | NADH:ubiquinone oxidoreductase subunit | 8 | 1.70 | 0.272 | 1.000 | |
| AT2G27730.1 | NADH:ubiquinone oxidoreductase subunit | 9 | 1.82 | 0.254 | 0.998 | |

| | | | | | | |
|---|---|---|---|---|---|---|
| AT2G27730.1 | NADH:ubiquinone oxidoreductase subunit | 10 | 1.86 | 0.245 | 0.994 | |
| AT2G27730.1 | NADH:ubiquinone oxidoreductase subunit | 11 | 2.02 | 0.242 | 0.978 | |
| AT2G27730.1 | NADH:ubiquinone oxidoreductase subunit | 12 | 2.40 | 0.241 | 0.872 | |
| AT2G27730.1 | NADH:ubiquinone oxidoreductase subunit | 13 | 3.12 | 0.240 | 0.526 | |
| AT2G27730.1 | NADH:ubiquinone oxidoreductase subunit | 14 | 3.80 | 0.238 | 0.267 | |
| AT2G27730.1 | NADH:ubiquinone oxidoreductase subunit | 15 | 3.85 | 0.237 | 0.244 | |
| AT2G27730.1 | NADH:ubiquinone oxidoreductase subunit | 16 | 3.07 | 0.235 | 0.516 | |
| AT2G27730.1 | NADH:ubiquinone oxidoreductase subunit | 17 | 2.03 | 0.236 | 0.970 | |
| AT2G27730.1 | NADH:ubiquinone oxidoreductase subunit | 18 | 1.31 | 0.238 | 1.000 | |
| AT2G27730.1 | NADH:ubiquinone oxidoreductase subunit | 19 | 0.97 | 0.242 | 1.000 | |
| AT2G27730.1 | NADH:ubiquinone oxidoreductase subunit | 20 | 0.81 | 0.246 | 1.000 | |
| AT2G27730.1 | NADH:ubiquinone oxidoreductase subunit | 21 | 0.78 | 0.248 | 1.000 | |
| AT2G27730.1 | NADH:ubiquinone oxidoreductase subunit | 22 | 0.85 | 0.249 | 1.000 | |
| AT2G27730.1 | NADH:ubiquinone oxidoreductase subunit | 23 | 1.02 | 0.247 | 1.000 | |
| AT2G27730.1 | NADH:ubiquinone oxidoreductase subunit | 24 | 1.27 | 0.244 | 1.000 | |
| AT2G27730.1 | NADH:ubiquinone oxidoreductase subunit | 25 | 1.69 | 0.242 | 0.999 | |
| AT2G27730.1 | NADH:ubiquinone oxidoreductase subunit | 26 | 2.50 | 0.241 | 0.828 | |
| AT2G27730.1 | NADH:ubiquinone oxidoreductase subunit | 27 | 3.89 | 0.240 | 0.253 | |
| AT2G27730.1 | NADH:ubiquinone oxidoreductase subunit | 28 | 5.45 | 0.240 | 0.053 | . |
| AT2G27730.1 | NADH:ubiquinone oxidoreductase subunit | 29 | 5.62 | 0.241 | 0.046 | * |
| AT2G27730.1 | NADH:ubiquinone oxidoreductase subunit | 30 | 4.21 | 0.242 | 0.191 | |
| AT2G27730.1 | NADH:ubiquinone oxidoreductase subunit | 31 | 2.75 | 0.245 | 0.730 | |
| AT2G27730.1 | NADH:ubiquinone oxidoreductase subunit | 32 | 1.86 | 0.253 | 0.996 | |
| AT2G27730.1 | NADH:ubiquinone oxidoreductase subunit | 33 | 1.31 | 0.274 | 1.000 | |
| AT2G27730.1 | NADH:ubiquinone oxidoreductase subunit | 34 | 1.05 | 0.316 | 1.000 | |
| AT2G27730.1 | NADH:ubiquinone oxidoreductase subunit | 35 | 0.99 | 0.374 | 1.000 | |
| AT2G27730.1 | NADH:ubiquinone oxidoreductase subunit | 42 | 2.66 | 0.414 | 0.997 | |
| AT2G27730.1 | NADH:ubiquinone oxidoreductase subunit | 43 | 2.50 | 0.334 | 0.987 | |
| AT2G27730.1 | NADH:ubiquinone oxidoreductase subunit | 44 | 2.15 | 0.280 | 0.988 | |
| AT2G27730.1 | NADH:ubiquinone oxidoreductase subunit | 45 | 1.74 | 0.315 | 1.000 | |
| AT2G27730.1 | NADH:ubiquinone oxidoreductase subunit | 46 | 1.36 | 0.466 | 1.000 | |

| | | | | | |
|---|---|---|---|---|---|
| AT2G27730.1 | NADH:ubiquinone oxidoreductase subunit | 47 | 1.04 | 0.712 | 1.000 |
| AT2G28000.1 | Chaperonin 60 alpha 1 (CPN60A1) | 2 | 1.69 | 0.824 | 0.999 |
| AT2G28000.1 | Chaperonin 60 alpha 1 (CPN60A1) | 3 | 1.54 | 0.780 | 0.999 |
| AT2G28000.1 | Chaperonin 60 alpha 1 (CPN60A1) | 23 | 0.56 | 0.371 | 0.942 |
| AT2G28000.1 | Chaperonin 60 alpha 1 (CPN60A1) | 24 | 0.59 | 0.335 | 0.940 |
| AT2G28000.1 | Chaperonin 60 alpha 1 (CPN60A1) | 25 | 0.63 | 0.307 | 0.952 |
| AT2G28000.1 | Chaperonin 60 alpha 1 (CPN60A1) | 26 | 0.70 | 0.293 | 0.977 |
| AT2G28000.1 | Chaperonin 60 alpha 1 (CPN60A1) | 27 | 0.79 | 0.296 | 0.996 |
| AT2G28000.1 | Chaperonin 60 alpha 1 (CPN60A1) | 28 | 0.90 | 0.317 | 1.000 |
| AT2G28000.1 | Chaperonin 60 alpha 1 (CPN60A1) | 29 | 1.05 | 0.355 | 1.000 |
| AT2G28000.1 | Chaperonin 60 alpha 1 (CPN60A1) | 30 | 1.22 | 0.409 | 1.000 |
| AT2G28000.1 | Chaperonin 60 alpha 1 (CPN60A1) | 46 | 7.26 | 2.278 | 0.995 |
| AT2G28000.1 | Chaperonin 60 alpha 1 (CPN60A1) | 47 | 7.96 | 2.436 | 0.995 |
| AT2G28430.1 | NADH:ubiquinone oxidoreductase subunit (putative) | 2 | 1.03 | 0.381 | 1.000 |
| AT2G28430.1 | NADH:ubiquinone oxidoreductase subunit (putative) | 3 | 0.86 | 0.256 | 1.000 |
| AT2G28430.1 | NADH:ubiquinone oxidoreductase subunit (putative) | 4 | 0.73 | 0.227 | 1.000 |
| AT2G28430.1 | NADH:ubiquinone oxidoreductase subunit (putative) | 5 | 0.64 | 0.198 | 1.000 |
| AT2G28430.1 | NADH:ubiquinone oxidoreductase subunit (putative) | 6 | 0.65 | 0.182 | 1.000 |
| AT2G28430.1 | NADH:ubiquinone oxidoreductase subunit (putative) | 7 | 0.76 | 0.175 | 1.000 |
| AT2G28430.1 | NADH:ubiquinone oxidoreductase subunit (putative) | 8 | 0.94 | 0.170 | 1.000 |
| AT2G28430.1 | NADH:ubiquinone oxidoreductase subunit (putative) | 9 | 1.15 | 0.167 | 1.000 |
| AT2G28430.1 | NADH:ubiquinone oxidoreductase subunit (putative) | 10 | 1.34 | 0.166 | 1.000 |
| AT2G28430.1 | NADH:ubiquinone oxidoreductase subunit (putative) | 11 | 1.57 | 0.168 | 0.999 |
| AT2G28430.1 | NADH:ubiquinone oxidoreductase subunit (putative) | 12 | 1.98 | 0.168 | 0.900 |
| AT2G28430.1 | NADH:ubiquinone oxidoreductase subunit (putative) | 13 | 2.63 | 0.166 | 0.315 |

| | | | | | | |
|---|---|---|---|---|---|---|
| AT2G28430.1 | NADH:ubiquinone oxidoreductase subunit (putative) | 14 | 3.80 | 0.164 | 0.017 | * |
| AT2G28430.1 | NADH:ubiquinone oxidoreductase subunit (putative) | 15 | 5.11 | 0.164 | 0.001 | *** |
| AT2G28430.1 | NADH:ubiquinone oxidoreductase subunit (putative) | 16 | 5.83 | 0.162 | 0.000 | *** |
| AT2G28430.1 | NADH:ubiquinone oxidoreductase subunit (putative) | 17 | 5.92 | 0.161 | 0.000 | *** |
| AT2G28430.1 | NADH:ubiquinone oxidoreductase subunit (putative) | 18 | 5.43 | 0.162 | 0.000 | *** |
| AT2G28430.1 | NADH:ubiquinone oxidoreductase subunit (putative) | 19 | 4.53 | 0.168 | 0.004 | ** |
| AT2G28430.1 | NADH:ubiquinone oxidoreductase subunit (putative) | 20 | 4.24 | 0.172 | 0.011 | * |
| AT2G28430.1 | NADH:ubiquinone oxidoreductase subunit (putative) | 21 | 4.34 | 0.173 | 0.011 | * |
| AT2G28430.1 | NADH:ubiquinone oxidoreductase subunit (putative) | 22 | 3.92 | 0.174 | 0.026 | * |
| AT2G28430.1 | NADH:ubiquinone oxidoreductase subunit (putative) | 23 | 2.88 | 0.172 | 0.229 | |
| AT2G28430.1 | NADH:ubiquinone oxidoreductase subunit (putative) | 24 | 1.88 | 0.168 | 0.951 | |
| AT2G28430.1 | NADH:ubiquinone oxidoreductase subunit (putative) | 25 | 1.39 | 0.166 | 1.000 | |
| AT2G28430.1 | NADH:ubiquinone oxidoreductase subunit (putative) | 26 | 1.30 | 0.166 | 1.000 | |
| AT2G28430.1 | NADH:ubiquinone oxidoreductase subunit (putative) | 27 | 1.40 | 0.166 | 1.000 | |
| AT2G28430.1 | NADH:ubiquinone oxidoreductase subunit (putative) | 28 | 1.38 | 0.167 | 1.000 | |
| AT2G28430.1 | NADH:ubiquinone oxidoreductase subunit (putative) | 29 | 1.27 | 0.167 | 1.000 | |
| AT2G28430.1 | NADH:ubiquinone oxidoreductase subunit (putative) | 30 | 1.24 | 0.167 | 1.000 | |
| AT2G28430.1 | NADH:ubiquinone oxidoreductase subunit (putative) | 31 | 1.22 | 0.167 | 1.000 | |
| AT2G28430.1 | NADH:ubiquinone oxidoreductase subunit (putative) | 32 | 1.14 | 0.167 | 1.000 | |
| AT2G28430.1 | NADH:ubiquinone oxidoreductase subunit (putative) | 33 | 1.00 | 0.168 | 1.000 | |

| | | | | | |
|---|---|---|---|---|---|
| AT2G28430.1 | NADH:ubiquinone oxidoreductase subunit (putative) | 34 | 0.93 | 0.167 | 1.000 |
| AT2G28430.1 | NADH:ubiquinone oxidoreductase subunit (putative) | 35 | 0.91 | 0.167 | 1.000 |
| AT2G28430.1 | NADH:ubiquinone oxidoreductase subunit (putative) | 36 | 0.87 | 0.166 | 1.000 |
| AT2G28430.1 | NADH:ubiquinone oxidoreductase subunit (putative) | 37 | 0.83 | 0.166 | 1.000 |
| AT2G28430.1 | NADH:ubiquinone oxidoreductase subunit (putative) | 38 | 0.84 | 0.167 | 1.000 |
| AT2G28430.1 | NADH:ubiquinone oxidoreductase subunit (putative) | 39 | 0.88 | 0.167 | 1.000 |
| AT2G28430.1 | NADH:ubiquinone oxidoreductase subunit (putative) | 40 | 0.86 | 0.168 | 1.000 |
| AT2G28430.1 | NADH:ubiquinone oxidoreductase subunit (putative) | 41 | 0.66 | 0.168 | 1.000 |
| AT2G28430.1 | NADH:ubiquinone oxidoreductase subunit (putative) | 42 | 0.46 | 0.171 | 0.773 |
| AT2G28430.1 | NADH:ubiquinone oxidoreductase subunit (putative) | 43 | 0.38 | 0.174 | 0.412 |
| AT2G28430.1 | NADH:ubiquinone oxidoreductase subunit (putative) | 44 | 0.42 | 0.177 | 0.634 |
| AT2G28430.1 | NADH:ubiquinone oxidoreductase subunit (putative) | 45 | 0.55 | 0.209 | 0.998 |
| AT2G28430.1 | NADH:ubiquinone oxidoreductase subunit (putative) | 46 | 0.74 | 0.323 | 1.000 |
| AT2G28430.1 | NADH:ubiquinone oxidoreductase subunit (putative) | 47 | 1.02 | 0.555 | 1.000 |
| AT2G28900.1 | Outer envelope protein 16 (OEP16) | 2 | 0.62 | 0.322 | 1.000 |
| AT2G28900.1 | Outer envelope protein 16 (OEP16) | 3 | 0.66 | 0.229 | 1.000 |
| AT2G28900.1 | Outer envelope protein 16 (OEP16) | 4 | 0.71 | 0.194 | 1.000 |
| AT2G28900.1 | Outer envelope protein 16 (OEP16) | 5 | 0.80 | 0.188 | 1.000 |
| AT2G28900.1 | Outer envelope protein 16 (OEP16) | 6 | 0.91 | 0.188 | 1.000 |
| AT2G28900.1 | Outer envelope protein 16 (OEP16) | 7 | 1.04 | 0.187 | 1.000 |
| AT2G28900.1 | Outer envelope protein 16 (OEP16) | 8 | 1.17 | 0.185 | 1.000 |
| AT2G28900.1 | Outer envelope protein 16 (OEP16) | 9 | 1.30 | 0.183 | 1.000 |
| AT2G28900.1 | Outer envelope protein 16 (OEP16) | 10 | 1.43 | 0.182 | 1.000 |
| AT2G28900.1 | Outer envelope protein 16 (OEP16) | 11 | 1.54 | 0.182 | 0.998 |

| | | | | | | |
|---|---|---|---|---|---|---|
| AT2G28900.1 | Outer envelope protein 16 (OEP16) | 12 | 1.62 | 0.181 | 0.994 | |
| AT2G28900.1 | Outer envelope protein 16 (OEP16) | 13 | 1.65 | 0.180 | 0.989 | |
| AT2G28900.1 | Outer envelope protein 16 (OEP16) | 14 | 1.61 | 0.178 | 0.993 | |
| AT2G28900.1 | Outer envelope protein 16 (OEP16) | 15 | 1.48 | 0.177 | 0.999 | |
| AT2G28900.1 | Outer envelope protein 16 (OEP16) | 16 | 1.29 | 0.176 | 1.000 | |
| AT2G28900.1 | Outer envelope protein 16 (OEP16) | 17 | 1.16 | 0.177 | 1.000 | |
| AT2G28900.1 | Outer envelope protein 16 (OEP16) | 18 | 1.10 | 0.179 | 1.000 | |
| AT2G28900.1 | Outer envelope protein 16 (OEP16) | 19 | 1.05 | 0.182 | 1.000 | |
| AT2G28900.1 | Outer envelope protein 16 (OEP16) | 20 | 0.97 | 0.186 | 1.000 | |
| AT2G28900.1 | Outer envelope protein 16 (OEP16) | 21 | 0.88 | 0.187 | 1.000 | |
| AT2G28900.1 | Outer envelope protein 16 (OEP16) | 22 | 0.83 | 0.187 | 1.000 | |
| AT2G28900.1 | Outer envelope protein 16 (OEP16) | 23 | 0.87 | 0.186 | 1.000 | |
| AT2G28900.1 | Outer envelope protein 16 (OEP16) | 24 | 0.99 | 0.183 | 1.000 | |
| AT2G28900.1 | Outer envelope protein 16 (OEP16) | 25 | 1.22 | 0.181 | 1.000 | |
| AT2G28900.1 | Outer envelope protein 16 (OEP16) | 26 | 1.59 | 0.181 | 0.996 | |
| AT2G28900.1 | Outer envelope protein 16 (OEP16) | 27 | 1.92 | 0.181 | 0.900 | |
| AT2G28900.1 | Outer envelope protein 16 (OEP16) | 28 | 2.06 | 0.184 | 0.821 | |
| AT2G28900.1 | Outer envelope protein 16 (OEP16) | 29 | 2.01 | 0.189 | 0.876 | |
| AT2G28900.1 | Outer envelope protein 16 (OEP16) | 30 | 1.93 | 0.193 | 0.935 | |
| AT2G28900.1 | Outer envelope protein 16 (OEP16) | 31 | 2.03 | 0.194 | 0.888 | |
| AT2G28900.1 | Outer envelope protein 16 (OEP16) | 32 | 2.46 | 0.192 | 0.567 | |
| AT2G28900.1 | Outer envelope protein 16 (OEP16) | 33 | 3.13 | 0.188 | 0.180 | |
| AT2G28900.1 | Outer envelope protein 16 (OEP16) | 34 | 3.76 | 0.185 | 0.050 | * |
| AT2G28900.1 | Outer envelope protein 16 (OEP16) | 35 | 4.06 | 0.183 | 0.025 | * |
| AT2G28900.1 | Outer envelope protein 16 (OEP16) | 36 | 3.99 | 0.182 | 0.027 | * |
| AT2G28900.1 | Outer envelope protein 16 (OEP16) | 37 | 3.75 | 0.182 | 0.044 | * |
| AT2G28900.1 | Outer envelope protein 16 (OEP16) | 38 | 3.44 | 0.182 | 0.077 | . |
| AT2G28900.1 | Outer envelope protein 16 (OEP16) | 39 | 3.16 | 0.182 | 0.137 | |
| AT2G28900.1 | Outer envelope protein 16 (OEP16) | 40 | 3.03 | 0.182 | 0.177 | |
| AT2G28900.1 | Outer envelope protein 16 (OEP16) | 41 | 2.93 | 0.184 | 0.223 | |
| AT2G28900.1 | Outer envelope protein 16 (OEP16) | 42 | 2.55 | 0.185 | 0.443 | |

| | | | | | | |
|---|---|---|---|---|---|---|
| AT2G28900.1 | Outer envelope protein 16 (OEP16) | 43 | 1.86 | 0.186 | 0.944 | |
| AT2G28900.1 | Outer envelope protein 16 (OEP16) | 44 | 1.06 | 0.187 | 1.000 | |
| AT2G28900.1 | Outer envelope protein 16 (OEP16) | 45 | 0.48 | 0.197 | 0.870 | |
| AT2G28900.1 | Outer envelope protein 16 (OEP16) | 46 | 0.18 | 0.242 | 0.051 | . |
| AT2G28900.1 | Outer envelope protein 16 (OEP16) | 47 | 0.06 | 0.346 | 0.013 | * |
| AT2G29080.1 | FTSH protease 3 (FTSH3) | 2 | 0.73 | 0.514 | 1.000 | |
| AT2G29080.1 | FTSH protease 3 (FTSH3) | 3 | 0.69 | 0.443 | 1.000 | |
| AT2G29080.1 | FTSH protease 3 (FTSH3) | 11 | 1.32 | 0.308 | 1.000 | |
| AT2G29080.1 | FTSH protease 3 (FTSH3) | 12 | 1.71 | 0.269 | 1.000 | |
| AT2G29080.1 | FTSH protease 3 (FTSH3) | 13 | 1.81 | 0.240 | 0.998 | |
| AT2G29080.1 | FTSH protease 3 (FTSH3) | 14 | 1.35 | 0.223 | 1.000 | |
| AT2G29080.1 | FTSH protease 3 (FTSH3) | 15 | 0.74 | 0.212 | 1.000 | |
| AT2G29080.1 | FTSH protease 3 (FTSH3) | 16 | 0.46 | 0.208 | 0.900 | |
| AT2G29080.1 | FTSH protease 3 (FTSH3) | 17 | 0.46 | 0.207 | 0.876 | |
| AT2G29080.1 | FTSH protease 3 (FTSH3) | 18 | 0.67 | 0.211 | 1.000 | |
| AT2G29080.1 | FTSH protease 3 (FTSH3) | 19 | 1.26 | 0.215 | 1.000 | |
| AT2G29080.1 | FTSH protease 3 (FTSH3) | 20 | 2.41 | 0.219 | 0.820 | |
| AT2G29080.1 | FTSH protease 3 (FTSH3) | 21 | 3.72 | 0.221 | 0.218 | |
| AT2G29080.1 | FTSH protease 3 (FTSH3) | 22 | 3.82 | 0.221 | 0.192 | |
| AT2G29080.1 | FTSH protease 3 (FTSH3) | 23 | 2.90 | 0.219 | 0.535 | |
| AT2G29080.1 | FTSH protease 3 (FTSH3) | 24 | 1.96 | 0.216 | 0.976 | |
| AT2G29080.1 | FTSH protease 3 (FTSH3) | 25 | 1.39 | 0.214 | 1.000 | |
| AT2G29080.1 | FTSH protease 3 (FTSH3) | 26 | 1.14 | 0.213 | 1.000 | |
| AT2G29080.1 | FTSH protease 3 (FTSH3) | 27 | 1.00 | 0.213 | 1.000 | |
| AT2G29080.1 | FTSH protease 3 (FTSH3) | 28 | 0.93 | 0.213 | 1.000 | |
| AT2G29080.1 | FTSH protease 3 (FTSH3) | 29 | 0.92 | 0.212 | 1.000 | |
| AT2G29080.1 | FTSH protease 3 (FTSH3) | 30 | 0.93 | 0.213 | 1.000 | |
| AT2G29080.1 | FTSH protease 3 (FTSH3) | 31 | 0.92 | 0.213 | 1.000 | |
| AT2G29080.1 | FTSH protease 3 (FTSH3) | 32 | 0.91 | 0.213 | 1.000 | |
| AT2G29080.1 | FTSH protease 3 (FTSH3) | 33 | 0.88 | 0.213 | 1.000 | |
| AT2G29080.1 | FTSH protease 3 (FTSH3) | 34 | 0.83 | 0.214 | 1.000 | |
| AT2G29080.1 | FTSH protease 3 (FTSH3) | 35 | 0.84 | 0.214 | 1.000 | |
| AT2G29080.1 | FTSH protease 3 (FTSH3) | 36 | 0.95 | 0.213 | 1.000 | |
| AT2G29080.1 | FTSH protease 3 (FTSH3) | 37 | 1.16 | 0.213 | 1.000 | |
| AT2G29080.1 | FTSH protease 3 (FTSH3) | 38 | 1.30 | 0.213 | 1.000 | |
| AT2G29080.1 | FTSH protease 3 (FTSH3) | 39 | 1.26 | 0.214 | 1.000 | |
| AT2G29080.1 | FTSH protease 3 (FTSH3) | 40 | 1.14 | 0.214 | 1.000 | |
| AT2G29080.1 | FTSH protease 3 (FTSH3) | 41 | 1.01 | 0.215 | 1.000 | |
| AT2G29080.1 | FTSH protease 3 (FTSH3) | 42 | 0.93 | 0.218 | 1.000 | |
| AT2G29080.1 | FTSH protease 3 (FTSH3) | 43 | 0.94 | 0.220 | 1.000 | |
| AT2G29080.1 | FTSH protease 3 (FTSH3) | 44 | 0.88 | 0.219 | 1.000 | |
| AT2G29080.1 | FTSH protease 3 (FTSH3) | 45 | 0.67 | 0.220 | 1.000 | |
| AT2G29080.1 | FTSH protease 3 (FTSH3) | 46 | 0.38 | 0.267 | 0.911 | |
| AT2G29080.1 | FTSH protease 3 (FTSH3) | 47 | 0.19 | 0.440 | 0.874 | |
| AT2G29560.1 | Enolase 3 (ENO3) | 2 | 1.02 | 0.530 | 1.000 | |
| AT2G29560.1 | Enolase 3 (ENO3) | 3 | 0.73 | 0.778 | 1.000 | |
| AT2G29560.1 | Enolase 3 (ENO3) | 23 | 0.53 | 0.315 | 0.981 | |
| AT2G29560.1 | Enolase 3 (ENO3) | 24 | 2.15 | 0.230 | 0.745 | |
| AT2G29560.1 | Enolase 3 (ENO3) | 25 | 9.26 | 0.215 | 0.000 | *** |
| AT2G29560.1 | Enolase 3 (ENO3) | 26 | 17.59 | 0.217 | 0.000 | *** |
| AT2G29560.1 | Enolase 3 (ENO3) | 27 | 6.99 | 0.219 | 0.003 | ** |
| AT2G29560.1 | Enolase 3 (ENO3) | 28 | 1.11 | 0.235 | 1.000 | |
| AT2G29560.1 | Enolase 3 (ENO3) | 29 | 0.26 | 0.307 | 0.415 | |
| AT2G29560.1 | Enolase 3 (ENO3) | 30 | 0.08 | 0.532 | 0.314 | |
| AT2G29560.1 | Enolase 3 (ENO3) | 46 | 0.03 | 0.428 | 0.009 | ** |
| AT2G29560.1 | Enolase 3 (ENO3) | 47 | 0.05 | 0.435 | 0.041 | * |
| AT2G29720.1 | FAD-dependent monooxygenase (CTF2B) | 2 | 1.24 | 0.479 | 1.000 | |

| | | | | | |
|---|---|---|---|---|---|
| AT2G29720.1 | FAD-dependent monooxygenase (CTF2B) | 3 | 1.20 | 0.430 | 1.000 |
| AT2G29720.1 | FAD-dependent monooxygenase (CTF2B) | 9 | 0.97 | 0.256 | 1.000 |
| AT2G29720.1 | FAD-dependent monooxygenase (CTF2B) | 10 | 0.95 | 0.244 | 1.000 |
| AT2G29720.1 | FAD-dependent monooxygenase (CTF2B) | 11 | 0.92 | 0.235 | 1.000 |
| AT2G29720.1 | FAD-dependent monooxygenase (CTF2B) | 12 | 0.89 | 0.231 | 1.000 |
| AT2G29720.1 | FAD-dependent monooxygenase (CTF2B) | 13 | 0.86 | 0.231 | 1.000 |
| AT2G29720.1 | FAD-dependent monooxygenase (CTF2B) | 14 | 0.83 | 0.233 | 1.000 |
| AT2G29720.1 | FAD-dependent monooxygenase (CTF2B) | 15 | 0.78 | 0.237 | 1.000 |
| AT2G29720.1 | FAD-dependent monooxygenase (CTF2B) | 22 | 0.49 | 0.241 | 0.760 |
| AT2G29720.1 | FAD-dependent monooxygenase (CTF2B) | 23 | 0.47 | 0.235 | 0.685 |
| AT2G29720.1 | FAD-dependent monooxygenase (CTF2B) | 24 | 0.46 | 0.229 | 0.621 |
| AT2G29720.1 | FAD-dependent monooxygenase (CTF2B) | 25 | 0.45 | 0.222 | 0.579 |
| AT2G29720.1 | FAD-dependent monooxygenase (CTF2B) | 26 | 0.46 | 0.216 | 0.564 |
| AT2G29720.1 | FAD-dependent monooxygenase (CTF2B) | 27 | 0.47 | 0.210 | 0.578 |
| AT2G29720.1 | FAD-dependent monooxygenase (CTF2B) | 28 | 0.49 | 0.205 | 0.618 |
| AT2G29720.1 | FAD-dependent monooxygenase (CTF2B) | 29 | 0.52 | 0.202 | 0.680 |
| AT2G29720.1 | FAD-dependent monooxygenase (CTF2B) | 30 | 0.55 | 0.201 | 0.759 |
| AT2G29720.1 | FAD-dependent monooxygenase (CTF2B) | 31 | 0.59 | 0.200 | 0.846 |
| AT2G29720.1 | FAD-dependent monooxygenase (CTF2B) | 32 | 0.64 | 0.202 | 0.928 |
| AT2G29720.1 | FAD-dependent monooxygenase (CTF2B) | 33 | 0.70 | 0.204 | 0.981 |
| AT2G29720.1 | FAD-dependent monooxygenase (CTF2B) | 34 | 0.77 | 0.208 | 0.998 |
| AT2G29720.1 | FAD-dependent monooxygenase (CTF2B) | 35 | 0.86 | 0.211 | 1.000 |
| AT2G29720.1 | FAD-dependent monooxygenase (CTF2B) | 36 | 0.97 | 0.214 | 1.000 |
| AT2G29720.1 | FAD-dependent monooxygenase (CTF2B) | 39 | 1.33 | 0.214 | 0.997 |
| AT2G29720.1 | FAD-dependent monooxygenase (CTF2B) | 40 | 1.46 | 0.211 | 0.978 |
| AT2G29720.1 | FAD-dependent monooxygenase (CTF2B) | 41 | 1.59 | 0.208 | 0.929 |
| AT2G29720.1 | FAD-dependent monooxygenase (CTF2B) | 42 | 1.72 | 0.207 | 0.855 |
| AT2G29720.1 | FAD-dependent monooxygenase (CTF2B) | 43 | 1.83 | 0.210 | 0.788 |
| AT2G29720.1 | FAD-dependent monooxygenase (CTF2B) | 44 | 1.91 | 0.219 | 0.762 |
| AT2G29720.1 | FAD-dependent monooxygenase (CTF2B) | 45 | 1.96 | 0.236 | 0.790 |
| AT2G29720.1 | FAD-dependent monooxygenase (CTF2B) | 46 | 1.99 | 0.263 | 0.854 |

| | | | | | | |
|---|---|---|---|---|---|---|
| AT2G29720.1 | FAD-dependent monooxygenase (CTF2B) | 47 | 2.00 | 0.300 | 0.916 | |
| AT2G29990.1 | Alternative NAD(P)H dehydrogenase A2 (NDA2) | 2 | 1.10 | 0.467 | 1.000 | |
| AT2G29990.1 | Alternative NAD(P)H dehydrogenase A2 (NDA2) | 3 | 1.21 | 0.433 | 1.000 | |
| AT2G29990.1 | Alternative NAD(P)H dehydrogenase A2 (NDA2) | 22 | 2.96 | 0.370 | 0.975 | |
| AT2G29990.1 | Alternative NAD(P)H dehydrogenase A2 (NDA2) | 23 | 2.93 | 0.295 | 0.872 | |
| AT2G29990.1 | Alternative NAD(P)H dehydrogenase A2 (NDA2) | 24 | 3.16 | 0.274 | 0.715 | |
| AT2G29990.1 | Alternative NAD(P)H dehydrogenase A2 (NDA2) | 25 | 3.85 | 0.248 | 0.310 | |
| AT2G29990.1 | Alternative NAD(P)H dehydrogenase A2 (NDA2) | 26 | 4.55 | 0.247 | 0.153 | |
| AT2G29990.1 | Alternative NAD(P)H dehydrogenase A2 (NDA2) | 27 | 4.26 | 0.233 | 0.139 | |
| AT2G29990.1 | Alternative NAD(P)H dehydrogenase A2 (NDA2) | 28 | 3.87 | 0.224 | 0.171 | |
| AT2G29990.1 | Alternative NAD(P)H dehydrogenase A2 (NDA2) | 29 | 4.52 | 0.205 | 0.035 | * |
| AT2G29990.1 | Alternative NAD(P)H dehydrogenase A2 (NDA2) | 30 | 5.87 | 0.191 | 0.002 | ** |
| AT2G29990.1 | Alternative NAD(P)H dehydrogenase A2 (NDA2) | 31 | 5.43 | 0.188 | 0.003 | ** |
| AT2G29990.1 | Alternative NAD(P)H dehydrogenase A2 (NDA2) | 32 | 4.01 | 0.189 | 0.034 | * |
| AT2G29990.1 | Alternative NAD(P)H dehydrogenase A2 (NDA2) | 33 | 2.62 | 0.188 | 0.402 | |
| AT2G29990.1 | Alternative NAD(P)H dehydrogenase A2 (NDA2) | 34 | 1.93 | 0.189 | 0.905 | |
| AT2G29990.1 | Alternative NAD(P)H dehydrogenase A2 (NDA2) | 35 | 1.51 | 0.189 | 0.999 | |
| AT2G29990.1 | Alternative NAD(P)H dehydrogenase A2 (NDA2) | 36 | 1.03 | 0.188 | 1.000 | |
| AT2G29990.1 | Alternative NAD(P)H dehydrogenase A2 (NDA2) | 37 | 0.64 | 0.188 | 0.997 | |
| AT2G29990.1 | Alternative NAD(P)H dehydrogenase A2 (NDA2) | 38 | 0.47 | 0.188 | 0.781 | |
| AT2G29990.1 | Alternative NAD(P)H dehydrogenase A2 (NDA2) | 39 | 0.65 | 0.188 | 0.998 | |
| AT2G29990.1 | Alternative NAD(P)H dehydrogenase A2 (NDA2) | 40 | 1.62 | 0.188 | 0.993 | |
| AT2G29990.1 | Alternative NAD(P)H dehydrogenase A2 (NDA2) | 41 | 3.22 | 0.189 | 0.147 | |
| AT2G29990.1 | Alternative NAD(P)H dehydrogenase A2 (NDA2) | 42 | 3.40 | 0.193 | 0.124 | |
| AT2G29990.1 | Alternative NAD(P)H dehydrogenase A2 (NDA2) | 43 | 2.08 | 0.195 | 0.843 | |
| AT2G29990.1 | Alternative NAD(P)H dehydrogenase A2 (NDA2) | 44 | 0.81 | 0.193 | 1.000 | |
| AT2G29990.1 | Alternative NAD(P)H dehydrogenase A2 (NDA2) | 45 | 0.38 | 0.194 | 0.445 | |
| AT2G29990.1 | Alternative NAD(P)H dehydrogenase A2 (NDA2) | 46 | 0.27 | 0.238 | 0.280 | |
| AT2G29990.1 | Alternative NAD(P)H dehydrogenase A2 (NDA2) | 47 | 0.21 | 0.450 | 0.910 | |
| AT2G30720.1 | Thioesterase/thiol ester dehydrase-isomerase superfamily protein | 2 | 0.90 | 0.589 | 1.000 | |

| | | | | | |
|---|---|---|---|---|---|
| AT2G30720.1 | Thioesterase/thiol ester dehydrase-isomerase superfamily protein | 3 | 1.04 | 0.554 | 1.000 |
| AT2G30720.1 | Thioesterase/thiol ester dehydrase-isomerase superfamily protein | 22 | 0.76 | 0.457 | 1.000 |
| AT2G30720.1 | Thioesterase/thiol ester dehydrase-isomerase superfamily protein | 23 | 0.57 | 0.374 | 1.000 |
| AT2G30720.1 | Thioesterase/thiol ester dehydrase-isomerase superfamily protein | 24 | 0.42 | 0.300 | 0.931 |
| AT2G30720.1 | Thioesterase/thiol ester dehydrase-isomerase superfamily protein | 25 | 0.39 | 0.265 | 0.773 |
| AT2G30720.1 | Thioesterase/thiol ester dehydrase-isomerase superfamily protein | 26 | 0.55 | 0.243 | 0.974 |
| AT2G30720.1 | Thioesterase/thiol ester dehydrase-isomerase superfamily protein | 27 | 0.97 | 0.234 | 1.000 |
| AT2G30720.1 | Thioesterase/thiol ester dehydrase-isomerase superfamily protein | 28 | 1.48 | 0.232 | 0.999 |
| AT2G30720.1 | Thioesterase/thiol ester dehydrase-isomerase superfamily protein | 29 | 1.85 | 0.231 | 0.958 |
| AT2G30720.1 | Thioesterase/thiol ester dehydrase-isomerase superfamily protein | 30 | 2.02 | 0.233 | 0.904 |
| AT2G30720.1 | Thioesterase/thiol ester dehydrase-isomerase superfamily protein | 31 | 2.13 | 0.233 | 0.859 |
| AT2G30720.1 | Thioesterase/thiol ester dehydrase-isomerase superfamily protein | 32 | 2.12 | 0.234 | 0.866 |
| AT2G30720.1 | Thioesterase/thiol ester dehydrase-isomerase superfamily protein | 33 | 1.87 | 0.237 | 0.959 |
| AT2G30720.1 | Thioesterase/thiol ester dehydrase-isomerase superfamily protein | 34 | 1.41 | 0.242 | 1.000 |
| AT2G30720.1 | Thioesterase/thiol ester dehydrase-isomerase superfamily protein | 35 | 0.84 | 0.259 | 1.000 |
| AT2G30720.1 | Thioesterase/thiol ester dehydrase-isomerase superfamily protein | 36 | 0.44 | 0.334 | 0.977 |
| AT2G30720.1 | Thioesterase/thiol ester dehydrase-isomerase superfamily protein | 37 | 0.28 | 0.511 | 0.974 |
| AT2G30720.1 | Thioesterase/thiol ester dehydrase-isomerase superfamily protein | 46 | 659.94 | 3.106 | 0.993 |
| AT2G30720.1 | Thioesterase/thiol ester dehydrase-isomerase superfamily protein | 47 | 2982.86 | 3.550 | 0.987 |
| AT2G30950.1 | FTSH protease 2 (FTSH2) | 2 | 1.22 | 0.485 | 1.000 |
| AT2G30950.1 | FTSH protease 2 (FTSH2) | 3 | 1.01 | 0.354 | 1.000 |
| AT2G30950.1 | FTSH protease 2 (FTSH2) | 4 | 0.85 | 0.316 | 1.000 |
| AT2G30950.1 | FTSH protease 2 (FTSH2) | 5 | 0.70 | 0.312 | 1.000 |
| AT2G30950.1 | FTSH protease 2 (FTSH2) | 6 | 0.52 | 0.312 | 0.999 |

| | | | | | | |
|---|---|---|---|---|---|---|
| AT2G30950.1 | FTSH protease 2 (FTSH2) | 7 | 0.38 | 0.312 | 0.930 | |
| AT2G30950.1 | FTSH protease 2 (FTSH2) | 8 | 0.29 | 0.308 | 0.710 | |
| AT2G30950.1 | FTSH protease 2 (FTSH2) | 9 | 0.25 | 0.306 | 0.520 | |
| AT2G30950.1 | FTSH protease 2 (FTSH2) | 10 | 0.23 | 0.305 | 0.437 | |
| AT2G30950.1 | FTSH protease 2 (FTSH2) | 11 | 0.24 | 0.306 | 0.493 | |
| AT2G30950.1 | FTSH protease 2 (FTSH2) | 12 | 0.29 | 0.305 | 0.666 | |
| AT2G30950.1 | FTSH protease 2 (FTSH2) | 13 | 0.34 | 0.303 | 0.848 | |
| AT2G30950.1 | FTSH protease 2 (FTSH2) | 14 | 0.41 | 0.301 | 0.953 | |
| AT2G30950.1 | FTSH protease 2 (FTSH2) | 15 | 0.44 | 0.300 | 0.977 | |
| AT2G30950.1 | FTSH protease 2 (FTSH2) | 16 | 0.45 | 0.301 | 0.986 | |
| AT2G30950.1 | FTSH protease 2 (FTSH2) | 17 | 0.50 | 0.303 | 0.997 | |
| AT2G30950.1 | FTSH protease 2 (FTSH2) | 18 | 0.61 | 0.306 | 1.000 | |
| AT2G30950.1 | FTSH protease 2 (FTSH2) | 19 | 0.76 | 0.314 | 1.000 | |
| AT2G30950.1 | FTSH protease 2 (FTSH2) | 20 | 0.87 | 0.325 | 1.000 | |
| AT2G30950.1 | FTSH protease 2 (FTSH2) | 21 | 0.88 | 0.334 | 1.000 | |
| AT2G30950.1 | FTSH protease 2 (FTSH2) | 22 | 0.78 | 0.328 | 1.000 | |
| AT2G30950.1 | FTSH protease 2 (FTSH2) | 23 | 0.66 | 0.317 | 1.000 | |
| AT2G30950.1 | FTSH protease 2 (FTSH2) | 24 | 0.57 | 0.309 | 1.000 | |
| AT2G30950.1 | FTSH protease 2 (FTSH2) | 25 | 0.52 | 0.304 | 0.999 | |
| AT2G30950.1 | FTSH protease 2 (FTSH2) | 26 | 0.52 | 0.302 | 0.999 | |
| AT2G30950.1 | FTSH protease 2 (FTSH2) | 27 | 0.53 | 0.301 | 0.999 | |
| AT2G30950.1 | FTSH protease 2 (FTSH2) | 28 | 0.53 | 0.302 | 0.999 | |
| AT2G30950.1 | FTSH protease 2 (FTSH2) | 29 | 0.54 | 0.303 | 0.999 | |
| AT2G30950.1 | FTSH protease 2 (FTSH2) | 30 | 0.55 | 0.304 | 1.000 | |
| AT2G30950.1 | FTSH protease 2 (FTSH2) | 31 | 0.60 | 0.305 | 1.000 | |
| AT2G30950.1 | FTSH protease 2 (FTSH2) | 32 | 0.58 | 0.306 | 1.000 | |
| AT2G30950.1 | FTSH protease 2 (FTSH2) | 33 | 0.46 | 0.306 | 0.991 | |
| AT2G30950.1 | FTSH protease 2 (FTSH2) | 34 | 0.32 | 0.307 | 0.803 | |
| AT2G30950.1 | FTSH protease 2 (FTSH2) | 35 | 0.22 | 0.306 | 0.395 | |
| AT2G30950.1 | FTSH protease 2 (FTSH2) | 36 | 0.18 | 0.306 | 0.230 | |
| AT2G30950.1 | FTSH protease 2 (FTSH2) | 37 | 0.21 | 0.306 | 0.344 | |
| AT2G30950.1 | FTSH protease 2 (FTSH2) | 38 | 0.31 | 0.306 | 0.763 | |
| AT2G30950.1 | FTSH protease 2 (FTSH2) | 39 | 0.49 | 0.306 | 0.997 | |
| AT2G30950.1 | FTSH protease 2 (FTSH2) | 40 | 0.72 | 0.307 | 1.000 | |
| AT2G30950.1 | FTSH protease 2 (FTSH2) | 41 | 0.82 | 0.308 | 1.000 | |
| AT2G30950.1 | FTSH protease 2 (FTSH2) | 42 | 0.67 | 0.310 | 1.000 | |
| AT2G30950.1 | FTSH protease 2 (FTSH2) | 43 | 0.46 | 0.311 | 0.993 | |
| AT2G30950.1 | FTSH protease 2 (FTSH2) | 44 | 0.27 | 0.311 | 0.641 | |
| AT2G30950.1 | FTSH protease 2 (FTSH2) | 45 | 0.12 | 0.318 | 0.072 | . |
| AT2G30950.1 | FTSH protease 2 (FTSH2) | 46 | 0.04 | 0.367 | 0.005 | ** |
| AT2G30950.1 | FTSH protease 2 (FTSH2) | 47 | 0.01 | 0.515 | 0.011 | * |
| AT2G30970.2 | Aspartate aminotransferase 1 (ASP1) | 2 | 0.97 | 0.360 | 1.000 | |
| AT2G30970.2 | Aspartate aminotransferase 1 (ASP1) | 3 | 0.80 | 0.277 | 1.000 | |
| AT2G30970.2 | Aspartate aminotransferase 1 (ASP1) | 4 | 0.69 | 0.256 | 1.000 | |
| AT2G30970.2 | Aspartate aminotransferase 1 (ASP1) | 6 | 0.62 | 0.244 | 1.000 | |
| AT2G30970.2 | Aspartate aminotransferase 1 (ASP1) | 7 | 0.69 | 0.222 | 1.000 | |
| AT2G30970.2 | Aspartate aminotransferase 1 (ASP1) | 8 | 0.87 | 0.201 | 1.000 | |
| AT2G30970.2 | Aspartate aminotransferase 1 (ASP1) | 9 | 1.19 | 0.188 | 1.000 | |
| AT2G30970.2 | Aspartate aminotransferase 1 (ASP1) | 10 | 1.62 | 0.183 | 0.996 | |
| AT2G30970.2 | Aspartate aminotransferase 1 (ASP1) | 11 | 2.07 | 0.181 | 0.821 | |
| AT2G30970.2 | Aspartate aminotransferase 1 (ASP1) | 12 | 2.44 | 0.180 | 0.507 | |
| AT2G30970.2 | Aspartate aminotransferase 1 (ASP1) | 13 | 2.67 | 0.178 | 0.324 | |

| | | | | | |
|---|---|---|---|---|---|
| AT2G30970.2 | Aspartate aminotransferase 1 (ASP1) | 14 | 2.71 | 0.176 | 0.284 |
| AT2G30970.2 | Aspartate aminotransferase 1 (ASP1) | 15 | 2.54 | 0.175 | 0.385 |
| AT2G30970.2 | Aspartate aminotransferase 1 (ASP1) | 16 | 2.24 | 0.174 | 0.618 |
| AT2G30970.2 | Aspartate aminotransferase 1 (ASP1) | 17 | 1.98 | 0.174 | 0.847 |
| AT2G30970.2 | Aspartate aminotransferase 1 (ASP1) | 18 | 1.76 | 0.176 | 0.971 |
| AT2G30970.2 | Aspartate aminotransferase 1 (ASP1) | 19 | 1.48 | 0.180 | 1.000 |
| AT2G30970.2 | Aspartate aminotransferase 1 (ASP1) | 20 | 1.19 | 0.184 | 1.000 |
| AT2G30970.2 | Aspartate aminotransferase 1 (ASP1) | 21 | 0.94 | 0.186 | 1.000 |
| AT2G30970.2 | Aspartate aminotransferase 1 (ASP1) | 22 | 0.79 | 0.186 | 1.000 |
| AT2G30970.2 | Aspartate aminotransferase 1 (ASP1) | 23 | 0.76 | 0.185 | 1.000 |
| AT2G30970.2 | Aspartate aminotransferase 1 (ASP1) | 24 | 0.81 | 0.182 | 1.000 |
| AT2G30970.2 | Aspartate aminotransferase 1 (ASP1) | 25 | 0.86 | 0.179 | 1.000 |
| AT2G30970.2 | Aspartate aminotransferase 1 (ASP1) | 26 | 0.90 | 0.178 | 1.000 |
| AT2G30970.2 | Aspartate aminotransferase 1 (ASP1) | 27 | 0.99 | 0.178 | 1.000 |
| AT2G30970.2 | Aspartate aminotransferase 1 (ASP1) | 28 | 1.18 | 0.178 | 1.000 |
| AT2G30970.2 | Aspartate aminotransferase 1 (ASP1) | 29 | 1.43 | 0.178 | 1.000 |
| AT2G30970.2 | Aspartate aminotransferase 1 (ASP1) | 30 | 1.65 | 0.179 | 0.992 |
| AT2G30970.2 | Aspartate aminotransferase 1 (ASP1) | 31 | 1.88 | 0.179 | 0.928 |
| AT2G30970.2 | Aspartate aminotransferase 1 (ASP1) | 32 | 2.10 | 0.179 | 0.789 |
| AT2G30970.2 | Aspartate aminotransferase 1 (ASP1) | 33 | 2.16 | 0.179 | 0.741 |
| AT2G30970.2 | Aspartate aminotransferase 1 (ASP1) | 34 | 2.04 | 0.180 | 0.833 |
| AT2G30970.2 | Aspartate aminotransferase 1 (ASP1) | 35 | 1.84 | 0.179 | 0.950 |
| AT2G30970.2 | Aspartate aminotransferase 1 (ASP1) | 36 | 1.65 | 0.179 | 0.993 |
| AT2G30970.2 | Aspartate aminotransferase 1 (ASP1) | 37 | 1.54 | 0.179 | 0.999 |
| AT2G30970.2 | Aspartate aminotransferase 1 (ASP1) | 38 | 1.61 | 0.179 | 0.996 |
| AT2G30970.2 | Aspartate aminotransferase 1 (ASP1) | 39 | 1.88 | 0.179 | 0.932 |
| AT2G30970.2 | Aspartate aminotransferase 1 (ASP1) | 40 | 2.26 | 0.180 | 0.657 |
| AT2G30970.2 | Aspartate aminotransferase 1 (ASP1) | 41 | 2.51 | 0.181 | 0.468 |
| AT2G30970.2 | Aspartate aminotransferase 1 (ASP1) | 42 | 2.43 | 0.183 | 0.548 |
| AT2G30970.2 | Aspartate aminotransferase 1 (ASP1) | 43 | 2.05 | 0.184 | 0.857 |
| AT2G30970.2 | Aspartate aminotransferase 1 (ASP1) | 44 | 1.48 | 0.185 | 1.000 |

| | | | | | | |
|---|---|---|---|---|---|---|
| AT2G30970.2 | Aspartate aminotransferase 1 (ASP1) | 45 | 0.91 | 0.193 | 1.000 | |
| AT2G30970.2 | Aspartate aminotransferase 1 (ASP1) | 46 | 0.49 | 0.236 | 0.985 | |
| AT2G30970.2 | Aspartate aminotransferase 1 (ASP1) | 47 | 0.25 | 0.349 | 0.841 | |
| AT2G31060.2 | Elongation factor 1A family protein | 2 | 0.90 | 0.328 | 1.000 | |
| AT2G31060.2 | Elongation factor 1A family protein | 3 | 0.75 | 0.312 | 1.000 | |
| AT2G31060.2 | Elongation factor 1A family protein | 25 | 0.46 | 0.227 | 0.841 | |
| AT2G31060.2 | Elongation factor 1A family protein | 26 | 0.61 | 0.165 | 0.926 | |
| AT2G31060.2 | Elongation factor 1A family protein | 27 | 0.75 | 0.138 | 0.995 | |
| AT2G31060.2 | Elongation factor 1A family protein | 28 | 0.83 | 0.129 | 1.000 | |
| AT2G31060.2 | Elongation factor 1A family protein | 29 | 0.88 | 0.126 | 1.000 | |
| AT2G31060.2 | Elongation factor 1A family protein | 30 | 0.93 | 0.126 | 1.000 | |
| AT2G31060.2 | Elongation factor 1A family protein | 31 | 0.98 | 0.126 | 1.000 | |
| AT2G31060.2 | Elongation factor 1A family protein | 32 | 0.99 | 0.126 | 1.000 | |
| AT2G31060.2 | Elongation factor 1A family protein | 33 | 0.96 | 0.126 | 1.000 | |
| AT2G31060.2 | Elongation factor 1A family protein | 34 | 0.95 | 0.126 | 1.000 | |
| AT2G31060.2 | Elongation factor 1A family protein | 35 | 0.97 | 0.125 | 1.000 | |
| AT2G31060.2 | Elongation factor 1A family protein | 36 | 1.01 | 0.125 | 1.000 | |
| AT2G31060.2 | Elongation factor 1A family protein | 37 | 1.09 | 0.125 | 1.000 | |
| AT2G31060.2 | Elongation factor 1A family protein | 38 | 1.23 | 0.125 | 1.000 | |
| AT2G31060.2 | Elongation factor 1A family protein | 39 | 1.35 | 0.125 | 0.985 | |
| AT2G31060.2 | Elongation factor 1A family protein | 40 | 1.43 | 0.125 | 0.950 | |
| AT2G31060.2 | Elongation factor 1A family protein | 41 | 1.44 | 0.126 | 0.944 | |
| AT2G31060.2 | Elongation factor 1A family protein | 42 | 1.39 | 0.128 | 0.974 | |
| AT2G31060.2 | Elongation factor 1A family protein | 43 | 1.24 | 0.128 | 1.000 | |
| AT2G31060.2 | Elongation factor 1A family protein | 44 | 0.94 | 0.129 | 1.000 | |
| AT2G31060.2 | Elongation factor 1A family protein | 45 | 0.59 | 0.134 | 0.680 | |
| AT2G31060.2 | Elongation factor 1A family protein | 46 | 0.31 | 0.164 | 0.033 | * |
| AT2G31060.2 | Elongation factor 1A family protein | 47 | 0.15 | 0.248 | 0.016 | * |
| AT2G31140.1 | Peptidase S24/S26A/S26B/S26C family protein (IMP2) | 2 | 0.90 | 0.689 | 1.000 | |
| AT2G31140.1 | Peptidase S24/S26A/S26B/S26C family protein (IMP2) | 3 | 0.99 | 0.587 | 1.000 | |
| AT2G31140.1 | Peptidase S24/S26A/S26B/S26C family protein (IMP2) | 10 | 1.55 | 0.414 | 1.000 | |

| | | | | | |
|---|---|---|---|---|---|
| AT2G31140.1 | Peptidase S24/S26A/S26B/S26C family protein (IMP2) | 11 | 1.52 | 0.354 | 1.000 |
| AT2G31140.1 | Peptidase S24/S26A/S26B/S26C family protein (IMP2) | 12 | 1.38 | 0.315 | 1.000 |
| AT2G31140.1 | Peptidase S24/S26A/S26B/S26C family protein (IMP2) | 13 | 1.16 | 0.292 | 1.000 |
| AT2G31140.1 | Peptidase S24/S26A/S26B/S26C family protein (IMP2) | 14 | 0.87 | 0.278 | 1.000 |
| AT2G31140.1 | Peptidase S24/S26A/S26B/S26C family protein (IMP2) | 15 | 0.58 | 0.273 | 1.000 |
| AT2G31140.1 | Peptidase S24/S26A/S26B/S26C family protein (IMP2) | 16 | 0.43 | 0.271 | 0.974 |
| AT2G31140.1 | Peptidase S24/S26A/S26B/S26C family protein (IMP2) | 17 | 0.44 | 0.270 | 0.984 |
| AT2G31140.1 | Peptidase S24/S26A/S26B/S26C family protein (IMP2) | 18 | 0.60 | 0.273 | 1.000 |
| AT2G31140.1 | Peptidase S24/S26A/S26B/S26C family protein (IMP2) | 19 | 0.87 | 0.280 | 1.000 |
| AT2G31140.1 | Peptidase S24/S26A/S26B/S26C family protein (IMP2) | 20 | 1.22 | 0.286 | 1.000 |
| AT2G31140.1 | Peptidase S24/S26A/S26B/S26C family protein (IMP2) | 21 | 1.53 | 0.288 | 1.000 |
| AT2G31140.1 | Peptidase S24/S26A/S26B/S26C family protein (IMP2) | 22 | 1.48 | 0.289 | 1.000 |
| AT2G31140.1 | Peptidase S24/S26A/S26B/S26C family protein (IMP2) | 23 | 1.17 | 0.286 | 1.000 |
| AT2G31140.1 | Peptidase S24/S26A/S26B/S26C family protein (IMP2) | 24 | 0.97 | 0.282 | 1.000 |
| AT2G31140.1 | Peptidase S24/S26A/S26B/S26C family protein (IMP2) | 25 | 0.93 | 0.278 | 1.000 |
| AT2G31140.1 | Peptidase S24/S26A/S26B/S26C family protein (IMP2) | 26 | 0.99 | 0.277 | 1.000 |
| AT2G31140.1 | Peptidase S24/S26A/S26B/S26C family protein (IMP2) | 27 | 1.03 | 0.276 | 1.000 |
| AT2G31140.1 | Peptidase S24/S26A/S26B/S26C family protein (IMP2) | 28 | 1.05 | 0.276 | 1.000 |
| AT2G31140.1 | Peptidase S24/S26A/S26B/S26C family protein (IMP2) | 29 | 1.19 | 0.276 | 1.000 |
| AT2G31140.1 | Peptidase S24/S26A/S26B/S26C family protein (IMP2) | 30 | 1.21 | 0.277 | 1.000 |
| AT2G31140.1 | Peptidase S24/S26A/S26B/S26C family protein (IMP2) | 31 | 1.08 | 0.278 | 1.000 |
| AT2G31140.1 | Peptidase S24/S26A/S26B/S26C family protein (IMP2) | 32 | 0.94 | 0.279 | 1.000 |
| AT2G31140.1 | Peptidase S24/S26A/S26B/S26C family protein (IMP2) | 33 | 0.80 | 0.279 | 1.000 |
| AT2G31140.1 | Peptidase S24/S26A/S26B/S26C family protein (IMP2) | 34 | 0.73 | 0.280 | 1.000 |
| AT2G31140.1 | Peptidase S24/S26A/S26B/S26C family protein (IMP2) | 35 | 0.72 | 0.280 | 1.000 |
| AT2G31140.1 | Peptidase S24/S26A/S26B/S26C family protein (IMP2) | 36 | 0.84 | 0.285 | 1.000 |
| AT2G31140.1 | Peptidase S24/S26A/S26B/S26C family protein (IMP2) | 37 | 1.08 | 0.301 | 1.000 |
| AT2G31140.1 | Peptidase S24/S26A/S26B/S26C family protein (IMP2) | 38 | 1.34 | 0.339 | 1.000 |
| AT2G31140.1 | Peptidase S24/S26A/S26B/S26C family protein (IMP2) | 39 | 1.67 | 0.377 | 1.000 |
| AT2G31140.1 | Peptidase S24/S26A/S26B/S26C family protein (IMP2) | 41 | 2.73 | 0.376 | 0.996 |
| AT2G31140.1 | Peptidase S24/S26A/S26B/S26C family protein (IMP2) | 42 | 4.16 | 0.341 | 0.774 |

| | | | | | | |
|---|---|---|---|---|---|---|
| AT2G31140.1 | Peptidase S24/S26A/S26B/S26C family protein (IMP2) | 43 | 7.94 | 0.305 | 0.085 | . |
| AT2G31140.1 | Peptidase S24/S26A/S26B/S26C family protein (IMP2) | 44 | 13.40 | 0.289 | 0.004 | ** |
| AT2G31140.1 | Peptidase S24/S26A/S26B/S26C family protein (IMP2) | 45 | 13.97 | 0.288 | 0.003 | ** |
| AT2G31140.1 | Peptidase S24/S26A/S26B/S26C family protein (IMP2) | 46 | 8.61 | 0.352 | 0.183 | |
| AT2G31140.1 | Peptidase S24/S26A/S26B/S26C family protein (IMP2) | 47 | 4.08 | 0.572 | 0.998 | |
| AT2G31170.1 | Cysteinyl-tRNA synthetase (CysRS) | 2 | 1.10 | 0.524 | 1.000 | |
| AT2G31170.1 | Cysteinyl-tRNA synthetase (CysRS) | 3 | 0.74 | 0.530 | 1.000 | |
| AT2G31170.1 | Cysteinyl-tRNA synthetase (CysRS) | 38 | 1.07 | 0.441 | 1.000 | |
| AT2G31170.1 | Cysteinyl-tRNA synthetase (CysRS) | 39 | 2.15 | 0.331 | 0.968 | |
| AT2G31170.1 | Cysteinyl-tRNA synthetase (CysRS) | 40 | 2.96 | 0.313 | 0.737 | |
| AT2G31170.1 | Cysteinyl-tRNA synthetase (CysRS) | 41 | 1.54 | 0.319 | 0.999 | |
| AT2G31170.1 | Cysteinyl-tRNA synthetase (CysRS) | 42 | 1.08 | 0.288 | 1.000 | |
| AT2G31170.1 | Cysteinyl-tRNA synthetase (CysRS) | 43 | 2.38 | 0.274 | 0.822 | |
| AT2G31170.1 | Cysteinyl-tRNA synthetase (CysRS) | 44 | 6.72 | 0.269 | 0.032 | * |
| AT2G31170.1 | Cysteinyl-tRNA synthetase (CysRS) | 45 | 3.84 | 0.267 | 0.270 | |
| AT2G31170.1 | Cysteinyl-tRNA synthetase (CysRS) | 46 | 0.57 | 0.296 | 0.992 | |
| AT2G31170.1 | Cysteinyl-tRNA synthetase (CysRS) | 47 | 0.03 | 0.586 | 0.103 | |
| AT2G31240.1 | Tetratricopeptide repeat (TPR)-like superfamily protein | 2 | 1.13 | 0.524 | 1.000 | |
| AT2G31240.1 | Tetratricopeptide repeat (TPR)-like superfamily protein | 3 | 1.15 | 0.501 | 1.000 | |
| AT2G31240.1 | Tetratricopeptide repeat (TPR)-like superfamily protein | 29 | 0.76 | 0.356 | 1.000 | |
| AT2G31240.1 | Tetratricopeptide repeat (TPR)-like superfamily protein | 30 | 0.69 | 0.294 | 0.999 | |
| AT2G31240.1 | Tetratricopeptide repeat (TPR)-like superfamily protein | 31 | 0.63 | 0.257 | 0.991 | |
| AT2G31240.1 | Tetratricopeptide repeat (TPR)-like superfamily protein | 32 | 0.57 | 0.229 | 0.935 | |
| AT2G31240.1 | Tetratricopeptide repeat (TPR)-like superfamily protein | 33 | 0.52 | 0.208 | 0.792 | |
| AT2G31240.1 | Tetratricopeptide repeat (TPR)-like superfamily protein | 34 | 0.50 | 0.193 | 0.641 | |
| AT2G31240.1 | Tetratricopeptide repeat (TPR)-like superfamily protein | 35 | 0.50 | 0.185 | 0.607 | |
| AT2G31240.1 | Tetratricopeptide repeat (TPR)-like superfamily protein | 36 | 0.54 | 0.181 | 0.709 | |
| AT2G31240.1 | Tetratricopeptide repeat (TPR)-like superfamily protein | 37 | 0.62 | 0.179 | 0.901 | |
| AT2G31240.1 | Tetratricopeptide repeat (TPR)-like superfamily protein | 38 | 0.76 | 0.178 | 0.997 | |
| AT2G31240.1 | Tetratricopeptide repeat (TPR)-like superfamily protein | 39 | 0.96 | 0.178 | 1.000 | |
| AT2G31240.1 | Tetratricopeptide repeat (TPR)-like superfamily protein | 40 | 1.23 | 0.178 | 1.000 | |

| | | | | | |
|---|---|---|---|---|---|
| AT2G31240.1 | Tetratricopeptide repeat (TPR)-like superfamily protein | 41 | 1.56 | 0.179 | 0.930 |
| AT2G31240.1 | Tetratricopeptide repeat (TPR)-like superfamily protein | 42 | 1.84 | 0.181 | 0.716 |
| AT2G31240.1 | Tetratricopeptide repeat (TPR)-like superfamily protein | 43 | 1.88 | 0.183 | 0.701 |
| AT2G31240.1 | Tetratricopeptide repeat (TPR)-like superfamily protein | 44 | 1.55 | 0.188 | 0.948 |
| AT2G31240.1 | Tetratricopeptide repeat (TPR)-like superfamily protein | 45 | 1.08 | 0.203 | 1.000 |
| AT2G31240.1 | Tetratricopeptide repeat (TPR)-like superfamily protein | 46 | 0.68 | 0.243 | 0.996 |
| AT2G31240.1 | Tetratricopeptide repeat (TPR)-like superfamily protein | 47 | 0.42 | 0.328 | 0.896 |
| AT2G31340.1 | ? | 2 | 1.02 | 0.479 | 1.000 |
| AT2G31340.1 | ? | 3 | 0.80 | 0.405 | 1.000 |
| AT2G31340.1 | ? | 10 | 0.97 | 0.417 | 1.000 |
| AT2G31340.1 | ? | 11 | 1.51 | 0.361 | 1.000 |
| AT2G31340.1 | ? | 12 | 2.37 | 0.380 | 0.999 |
| AT2G31340.1 | ? | 22 | 1.52 | 0.352 | 1.000 |
| AT2G31340.1 | ? | 23 | 0.94 | 0.259 | 1.000 |
| AT2G31340.1 | ? | 24 | 0.73 | 0.227 | 1.000 |
| AT2G31340.1 | ? | 25 | 0.94 | 0.211 | 1.000 |
| AT2G31340.1 | ? | 26 | 1.43 | 0.209 | 1.000 |
| AT2G31340.1 | ? | 27 | 1.86 | 0.208 | 0.976 |
| AT2G31340.1 | ? | 28 | 1.60 | 0.210 | 0.999 |
| AT2G31340.1 | ? | 29 | 1.18 | 0.208 | 1.000 |
| AT2G31340.1 | ? | 30 | 1.23 | 0.209 | 1.000 |
| AT2G31340.1 | ? | 31 | 1.84 | 0.211 | 0.983 |
| AT2G31340.1 | ? | 32 | 2.83 | 0.211 | 0.479 |
| AT2G31340.1 | ? | 33 | 3.11 | 0.215 | 0.367 |
| AT2G31340.1 | ? | 34 | 3.10 | 0.230 | 0.479 |
| AT2G31340.1 | ? | 35 | 3.53 | 0.276 | 0.603 |
| AT2G31340.1 | ? | 36 | 4.17 | 0.376 | 0.848 |
| AT2G31340.1 | ? | 41 | 2.82 | 0.363 | 0.985 |
| AT2G31340.1 | ? | 42 | 2.38 | 0.270 | 0.955 |
| AT2G31340.1 | ? | 43 | 2.24 | 0.228 | 0.906 |
| AT2G31340.1 | ? | 44 | 2.06 | 0.221 | 0.948 |
| AT2G31340.1 | ? | 45 | 1.64 | 0.265 | 1.000 |
| AT2G31340.1 | ? | 46 | 1.15 | 0.422 | 1.000 |
| AT2G31340.1 | ? | 47 | 0.74 | 0.757 | 1.000 |
| AT2G31490.1 | NADH:ubiquinone oxidoreductase subunit (B15) | 2 | 0.80 | 0.609 | 1.000 |
| AT2G31490.1 | NADH:ubiquinone oxidoreductase subunit (B15) | 3 | 0.91 | 0.472 | 1.000 |
| AT2G31490.1 | NADH:ubiquinone oxidoreductase subunit (B15) | 6 | 1.29 | 0.487 | 1.000 |
| AT2G31490.1 | NADH:ubiquinone oxidoreductase subunit (B15) | 7 | 1.42 | 0.450 | 1.000 |
| AT2G31490.1 | NADH:ubiquinone oxidoreductase subunit (B15) | 8 | 1.55 | 0.444 | 1.000 |
| AT2G31490.1 | NADH:ubiquinone oxidoreductase subunit (B15) | 12 | 2.04 | 0.350 | 1.000 |
| AT2G31490.1 | NADH:ubiquinone oxidoreductase subunit (B15) | 13 | 2.16 | 0.310 | 0.995 |
| AT2G31490.1 | NADH:ubiquinone oxidoreductase subunit (B15) | 14 | 1.96 | 0.290 | 0.998 |
| AT2G31490.1 | NADH:ubiquinone oxidoreductase subunit (B15) | 15 | 1.34 | 0.266 | 1.000 |
| AT2G31490.1 | NADH:ubiquinone oxidoreductase subunit (B15) | 16 | 0.83 | 0.251 | 1.000 |

| | | | | | |
|---|---|---|---|---|---|
| AT2G31490.1 | NADH:ubiquinone oxidoreductase subunit (B15) | 17 | 0.60 | 0.248 | 1.000 |
| AT2G31490.1 | NADH:ubiquinone oxidoreductase subunit (B15) | 18 | 0.65 | 0.250 | 1.000 |
| AT2G31490.1 | NADH:ubiquinone oxidoreductase subunit (B15) | 19 | 0.83 | 0.259 | 1.000 |
| AT2G31490.1 | NADH:ubiquinone oxidoreductase subunit (B15) | 20 | 0.93 | 0.266 | 1.000 |
| AT2G31490.1 | NADH:ubiquinone oxidoreductase subunit (B15) | 21 | 1.05 | 0.269 | 1.000 |
| AT2G31490.1 | NADH:ubiquinone oxidoreductase subunit (B15) | 22 | 1.35 | 0.268 | 1.000 |
| AT2G31490.1 | NADH:ubiquinone oxidoreductase subunit (B15) | 23 | 1.67 | 0.266 | 1.000 |
| AT2G31490.1 | NADH:ubiquinone oxidoreductase subunit (B15) | 24 | 1.45 | 0.259 | 1.000 |
| AT2G31490.1 | NADH:ubiquinone oxidoreductase subunit (B15) | 25 | 0.95 | 0.256 | 1.000 |
| AT2G31490.1 | NADH:ubiquinone oxidoreductase subunit (B15) | 26 | 0.63 | 0.254 | 1.000 |
| AT2G31490.1 | NADH:ubiquinone oxidoreductase subunit (B15) | 27 | 0.48 | 0.254 | 0.980 |
| AT2G31490.1 | NADH:ubiquinone oxidoreductase subunit (B15) | 28 | 0.60 | 0.256 | 1.000 |
| AT2G31490.1 | NADH:ubiquinone oxidoreductase subunit (B15) | 29 | 1.00 | 0.258 | 1.000 |
| AT2G31490.1 | NADH:ubiquinone oxidoreductase subunit (B15) | 30 | 0.95 | 0.264 | 1.000 |
| AT2G31490.1 | NADH:ubiquinone oxidoreductase subunit (B15) | 31 | 0.42 | 0.283 | 0.965 |
| AT2G31490.1 | NADH:ubiquinone oxidoreductase subunit (B15) | 32 | 0.17 | 0.359 | 0.488 |
| AT2G31490.1 | NADH:ubiquinone oxidoreductase subunit (B15) | 33 | 0.10 | 0.560 | 0.758 |
| AT2G31490.1 | NADH:ubiquinone oxidoreductase subunit (B15) | 46 | 11151.63 | 5.808 | 1.000 |
| AT2G31490.1 | NADH:ubiquinone oxidoreductase subunit (B15) | 47 | 54460.11 | 6.435 | 1.000 |
| AT2G31725.1 | ? | 2 | 0.84 | 0.446 | 1.000 |
| AT2G31725.1 | ? | 3 | 0.87 | 0.437 | 1.000 |
| AT2G31725.1 | ? | 38 | 1.11 | 0.379 | 1.000 |
| AT2G31725.1 | ? | 39 | 1.16 | 0.263 | 1.000 |
| AT2G31725.1 | ? | 40 | 1.68 | 0.226 | 0.964 |
| AT2G31725.1 | ? | 41 | 3.09 | 0.219 | 0.242 |
| AT2G31725.1 | ? | 42 | 2.96 | 0.222 | 0.300 |
| AT2G31725.1 | ? | 43 | 1.36 | 0.222 | 0.999 |
| AT2G31725.1 | ? | 44 | 0.71 | 0.220 | 0.998 |
| AT2G31725.1 | ? | 45 | 0.49 | 0.221 | 0.797 |
| AT2G31725.1 | ? | 46 | 0.34 | 0.244 | 0.421 |
| AT2G31725.1 | ? | 47 | 0.21 | 0.508 | 0.841 |
| AT2G32415.2 | RNA metabolism exosome complex subunit (RRP6L3, putative) | 2 | 0.97 | 0.287 | 1.000 |
| AT2G32415.2 | RNA metabolism exosome complex subunit (RRP6L3, putative) | 3 | 0.98 | 0.241 | 1.000 |
| AT2G32415.2 | RNA metabolism exosome complex subunit (RRP6L3, putative) | 4 | 0.98 | 0.222 | 1.000 |
| AT2G32415.2 | RNA metabolism exosome complex subunit (RRP6L3, putative) | 6 | 0.99 | 0.239 | 1.000 |

| | | | | | |
|---|---|---|---|---|---|
| AT2G32415.2 | RNA metabolism exosome complex subunit (RRP6L3, putative) | 7 | 1.00 | 0.262 | 1.000 |
| AT2G32415.2 | RNA metabolism exosome complex subunit (RRP6L3, putative) | 8 | 1.01 | 0.287 | 1.000 |
| AT2G32415.2 | RNA metabolism exosome complex subunit (RRP6L3, putative) | 21 | 1.27 | 0.204 | 1.000 |
| AT2G32415.2 | RNA metabolism exosome complex subunit (RRP6L3, putative) | 22 | 1.26 | 0.179 | 1.000 |
| AT2G32415.2 | RNA metabolism exosome complex subunit (RRP6L3, putative) | 23 | 1.25 | 0.157 | 1.000 |
| AT2G32415.2 | RNA metabolism exosome complex subunit (RRP6L3, putative) | 24 | 1.22 | 0.141 | 1.000 |
| AT2G32415.2 | RNA metabolism exosome complex subunit (RRP6L3, putative) | 25 | 1.17 | 0.131 | 1.000 |
| AT2G32415.2 | RNA metabolism exosome complex subunit (RRP6L3, putative) | 26 | 1.11 | 0.126 | 1.000 |
| AT2G32415.2 | RNA metabolism exosome complex subunit (RRP6L3, putative) | 27 | 1.05 | 0.124 | 1.000 |
| AT2G32415.2 | RNA metabolism exosome complex subunit (RRP6L3, putative) | 28 | 0.98 | 0.123 | 1.000 |
| AT2G32415.2 | RNA metabolism exosome complex subunit (RRP6L3, putative) | 29 | 0.90 | 0.122 | 1.000 |
| AT2G32415.2 | RNA metabolism exosome complex subunit (RRP6L3, putative) | 30 | 0.80 | 0.122 | 0.996 |
| AT2G32415.2 | RNA metabolism exosome complex subunit (RRP6L3, putative) | 31 | 0.71 | 0.123 | 0.930 |
| AT2G32415.2 | RNA metabolism exosome complex subunit (RRP6L3, putative) | 32 | 0.64 | 0.124 | 0.720 |
| AT2G32415.2 | RNA metabolism exosome complex subunit (RRP6L3, putative) | 33 | 0.57 | 0.127 | 0.481 |
| AT2G32415.2 | RNA metabolism exosome complex subunit (RRP6L3, putative) | 34 | 0.53 | 0.132 | 0.360 |
| AT2G32415.2 | RNA metabolism exosome complex subunit (RRP6L3, putative) | 35 | 0.51 | 0.140 | 0.354 |
| AT2G32415.2 | RNA metabolism exosome complex subunit (RRP6L3, putative) | 36 | 0.51 | 0.149 | 0.431 |
| AT2G32415.2 | RNA metabolism exosome complex subunit (RRP6L3, putative) | 37 | 0.52 | 0.159 | 0.558 |
| AT2G32415.2 | RNA metabolism exosome complex subunit (RRP6L3, putative) | 39 | 0.55 | 0.172 | 0.781 |

| | | | | | |
|---|---|---|---|---|---|
| AT2G32415.2 | RNA metabolism exosome complex subunit (RRP6L3, putative) | 40 | 0.58 | 0.176 | 0.857 |
| AT2G32415.2 | RNA metabolism exosome complex subunit (RRP6L3, putative) | 41 | 0.60 | 0.179 | 0.906 |
| AT2G32415.2 | RNA metabolism exosome complex subunit (RRP6L3, putative) | 42 | 0.62 | 0.183 | 0.944 |
| AT2G32415.2 | RNA metabolism exosome complex subunit (RRP6L3, putative) | 43 | 0.63 | 0.191 | 0.970 |
| AT2G32415.2 | RNA metabolism exosome complex subunit (RRP6L3, putative) | 44 | 0.65 | 0.208 | 0.989 |
| AT2G32415.2 | RNA metabolism exosome complex subunit (RRP6L3, putative) | 45 | 0.67 | 0.237 | 0.998 |
| AT2G32415.2 | RNA metabolism exosome complex subunit (RRP6L3, putative) | 46 | 0.68 | 0.284 | 1.000 |
| AT2G32415.2 | RNA metabolism exosome complex subunit (RRP6L3, putative) | 47 | 0.70 | 0.347 | 1.000 |
| AT2G33040.1 | ATP synthase subunit (ATPgamma) | 3 | 0.51 | 0.272 | 0.998 |
| AT2G33040.1 | ATP synthase subunit (ATPgamma) | 4 | 0.62 | 0.250 | 1.000 |
| AT2G33040.1 | ATP synthase subunit (ATPgamma) | 5 | 0.77 | 0.250 | 1.000 |
| AT2G33040.1 | ATP synthase subunit (ATPgamma) | 6 | 0.89 | 0.250 | 1.000 |
| AT2G33040.1 | ATP synthase subunit (ATPgamma) | 7 | 0.93 | 0.250 | 1.000 |
| AT2G33040.1 | ATP synthase subunit (ATPgamma) | 8 | 0.93 | 0.247 | 1.000 |
| AT2G33040.1 | ATP synthase subunit (ATPgamma) | 9 | 0.94 | 0.244 | 1.000 |
| AT2G33040.1 | ATP synthase subunit (ATPgamma) | 10 | 0.95 | 0.244 | 1.000 |
| AT2G33040.1 | ATP synthase subunit (ATPgamma) | 11 | 1.00 | 0.244 | 1.000 |
| AT2G33040.1 | ATP synthase subunit (ATPgamma) | 12 | 1.12 | 0.244 | 1.000 |
| AT2G33040.1 | ATP synthase subunit (ATPgamma) | 13 | 1.28 | 0.243 | 1.000 |
| AT2G33040.1 | ATP synthase subunit (ATPgamma) | 14 | 1.47 | 0.240 | 1.000 |
| AT2G33040.1 | ATP synthase subunit (ATPgamma) | 15 | 1.56 | 0.240 | 1.000 |
| AT2G33040.1 | ATP synthase subunit (ATPgamma) | 16 | 1.55 | 0.238 | 1.000 |
| AT2G33040.1 | ATP synthase subunit (ATPgamma) | 17 | 1.49 | 0.239 | 1.000 |
| AT2G33040.1 | ATP synthase subunit (ATPgamma) | 18 | 1.35 | 0.241 | 1.000 |
| AT2G33040.1 | ATP synthase subunit (ATPgamma) | 19 | 1.19 | 0.245 | 1.000 |
| AT2G33040.1 | ATP synthase subunit (ATPgamma) | 20 | 1.09 | 0.249 | 1.000 |
| AT2G33040.1 | ATP synthase subunit (ATPgamma) | 21 | 0.98 | 0.252 | 1.000 |

| | | | | | | |
|---|---|---|---|---|---|---|
| AT2G33040.1 | ATP synthase subunit (ATPgamma) | 22 | 0.91 | 0.251 | 1.000 | |
| AT2G33040.1 | ATP synthase subunit (ATPgamma) | 23 | 0.93 | 0.250 | 1.000 | |
| AT2G33040.1 | ATP synthase subunit (ATPgamma) | 24 | 1.07 | 0.247 | 1.000 | |
| AT2G33040.1 | ATP synthase subunit (ATPgamma) | 25 | 1.27 | 0.244 | 1.000 | |
| AT2G33040.1 | ATP synthase subunit (ATPgamma) | 26 | 1.26 | 0.243 | 1.000 | |
| AT2G33040.1 | ATP synthase subunit (ATPgamma) | 27 | 1.00 | 0.243 | 1.000 | |
| AT2G33040.1 | ATP synthase subunit (ATPgamma) | 28 | 0.90 | 0.243 | 1.000 | |
| AT2G33040.1 | ATP synthase subunit (ATPgamma) | 29 | 1.15 | 0.243 | 1.000 | |
| AT2G33040.1 | ATP synthase subunit (ATPgamma) | 30 | 1.66 | 0.243 | 1.000 | |
| AT2G33040.1 | ATP synthase subunit (ATPgamma) | 31 | 1.91 | 0.243 | 0.994 | |
| AT2G33040.1 | ATP synthase subunit (ATPgamma) | 32 | 1.52 | 0.244 | 1.000 | |
| AT2G33040.1 | ATP synthase subunit (ATPgamma) | 33 | 1.02 | 0.244 | 1.000 | |
| AT2G33040.1 | ATP synthase subunit (ATPgamma) | 34 | 0.75 | 0.244 | 1.000 | |
| AT2G33040.1 | ATP synthase subunit (ATPgamma) | 35 | 0.73 | 0.244 | 1.000 | |
| AT2G33040.1 | ATP synthase subunit (ATPgamma) | 36 | 1.01 | 0.244 | 1.000 | |
| AT2G33040.1 | ATP synthase subunit (ATPgamma) | 37 | 1.88 | 0.243 | 0.996 | |
| AT2G33040.1 | ATP synthase subunit (ATPgamma) | 38 | 3.81 | 0.244 | 0.295 | |
| AT2G33040.1 | ATP synthase subunit (ATPgamma) | 39 | 6.32 | 0.244 | 0.030 | * |
| AT2G33040.1 | ATP synthase subunit (ATPgamma) | 40 | 7.34 | 0.244 | 0.013 | * |
| AT2G33040.1 | ATP synthase subunit (ATPgamma) | 41 | 6.66 | 0.245 | 0.023 | * |
| AT2G33040.1 | ATP synthase subunit (ATPgamma) | 42 | 5.28 | 0.247 | 0.087 | . |
| AT2G33040.1 | ATP synthase subunit (ATPgamma) | 43 | 4.07 | 0.249 | 0.264 | |
| AT2G33040.1 | ATP synthase subunit (ATPgamma) | 44 | 2.65 | 0.249 | 0.807 | |
| AT2G33040.1 | ATP synthase subunit (ATPgamma) | 45 | 1.46 | 0.250 | 1.000 | |
| AT2G33040.1 | ATP synthase subunit (ATPgamma) | 46 | 0.79 | 0.288 | 1.000 | |
| AT2G33040.1 | ATP synthase subunit (ATPgamma) | 47 | 0.45 | 0.443 | 1.000 | |
| AT2G33040.1 | ATP synthase subunit (ATPgamma) | 48 | 0.26 | 0.686 | 1.000 | |
| AT2G33220.1 | NADH:ubiquinone oxidoreductase subunit (B16.6-2) | 2 | 0.76 | 0.630 | 1.000 | |
| AT2G33220.1 | NADH:ubiquinone oxidoreductase subunit (B16.6-2) | 3 | 1.07 | 0.480 | 1.000 | |

| | | | | | |
|---|---|---|---|---|---|
| AT2G33220.1 | NADH:ubiquinone oxidoreductase subunit (B16.6-2) | 5 | 1.70 | 0.436 | 1.000 |
| AT2G33220.1 | NADH:ubiquinone oxidoreductase subunit (B16.6-2) | 6 | 1.76 | 0.432 | 1.000 |
| AT2G33220.1 | NADH:ubiquinone oxidoreductase subunit (B16.6-2) | 7 | 1.62 | 0.439 | 1.000 |
| AT2G33220.1 | NADH:ubiquinone oxidoreductase subunit (B16.6-2) | 11 | 0.66 | 0.367 | 1.000 |
| AT2G33220.1 | NADH:ubiquinone oxidoreductase subunit (B16.6-2) | 12 | 0.62 | 0.331 | 1.000 |
| AT2G33220.1 | NADH:ubiquinone oxidoreductase subunit (B16.6-2) | 13 | 0.65 | 0.307 | 1.000 |
| AT2G33220.1 | NADH:ubiquinone oxidoreductase subunit (B16.6-2) | 14 | 0.67 | 0.285 | 1.000 |
| AT2G33220.1 | NADH:ubiquinone oxidoreductase subunit (B16.6-2) | 15 | 0.60 | 0.272 | 1.000 |
| AT2G33220.1 | NADH:ubiquinone oxidoreductase subunit (B16.6-2) | 16 | 0.48 | 0.267 | 0.975 |
| AT2G33220.1 | NADH:ubiquinone oxidoreductase subunit (B16.6-2) | 17 | 0.41 | 0.266 | 0.907 |
| AT2G33220.1 | NADH:ubiquinone oxidoreductase subunit (B16.6-2) | 18 | 0.48 | 0.269 | 0.978 |
| AT2G33220.1 | NADH:ubiquinone oxidoreductase subunit (B16.6-2) | 19 | 0.65 | 0.275 | 1.000 |
| AT2G33220.1 | NADH:ubiquinone oxidoreductase subunit (B16.6-2) | 20 | 0.85 | 0.280 | 1.000 |
| AT2G33220.1 | NADH:ubiquinone oxidoreductase subunit (B16.6-2) | 21 | 1.06 | 0.282 | 1.000 |
| AT2G33220.1 | NADH:ubiquinone oxidoreductase subunit (B16.6-2) | 22 | 1.29 | 0.282 | 1.000 |
| AT2G33220.1 | NADH:ubiquinone oxidoreductase subunit (B16.6-2) | 23 | 1.48 | 0.280 | 1.000 |
| AT2G33220.1 | NADH:ubiquinone oxidoreductase subunit (B16.6-2) | 24 | 1.54 | 0.276 | 1.000 |
| AT2G33220.1 | NADH:ubiquinone oxidoreductase subunit (B16.6-2) | 25 | 1.43 | 0.273 | 1.000 |
| AT2G33220.1 | NADH:ubiquinone oxidoreductase subunit (B16.6-2) | 26 | 1.21 | 0.272 | 1.000 |
| AT2G33220.1 | NADH:ubiquinone oxidoreductase subunit (B16.6-2) | 27 | 0.90 | 0.271 | 1.000 |

| | | | | | | |
|---|---|---|---|---|---|---|
| AT2G33220.1 | NADH:ubiquinone oxidoreductase subunit (B16.6-2) | 28 | 0.58 | 0.271 | 0.999 | |
| AT2G33220.1 | NADH:ubiquinone oxidoreductase subunit (B16.6-2) | 29 | 0.42 | 0.273 | 0.932 | |
| AT2G33220.1 | NADH:ubiquinone oxidoreductase subunit (B16.6-2) | 30 | 0.47 | 0.274 | 0.974 | |
| AT2G33220.1 | NADH:ubiquinone oxidoreductase subunit (B16.6-2) | 31 | 0.74 | 0.276 | 1.000 | |
| AT2G33220.1 | NADH:ubiquinone oxidoreductase subunit (B16.6-2) | 32 | 1.30 | 0.283 | 1.000 | |
| AT2G33220.1 | NADH:ubiquinone oxidoreductase subunit (B16.6-2) | 33 | 1.98 | 0.305 | 0.997 | |
| AT2G33220.1 | NADH:ubiquinone oxidoreductase subunit (B16.6-2) | 34 | 2.45 | 0.370 | 0.993 | |
| AT2G33220.1 | NADH:ubiquinone oxidoreductase subunit (B16.6-2) | 35 | 2.86 | 0.508 | 0.999 | |
| AT2G33220.1 | NADH:ubiquinone oxidoreductase subunit (B16.6-2) | 46 | 150.44 | 4.213 | 1.000 | |
| AT2G33220.1 | NADH:ubiquinone oxidoreductase subunit (B16.6-2) | 47 | 239.49 | 4.726 | 1.000 | |
| AT2G33585.1 | Subtilisin-like protease | 2 | 0.58 | 0.302 | 0.961 | |
| AT2G33585.1 | Subtilisin-like protease | 3 | 0.65 | 0.272 | 0.981 | |
| AT2G33585.1 | Subtilisin-like protease | 4 | 0.73 | 0.246 | 0.995 | |
| AT2G33585.1 | Subtilisin-like protease | 8 | 1.22 | 0.181 | 0.998 | |
| AT2G33585.1 | Subtilisin-like protease | 9 | 1.40 | 0.169 | 0.943 | |
| AT2G33585.1 | Subtilisin-like protease | 10 | 1.61 | 0.160 | 0.720 | |
| AT2G33585.1 | Subtilisin-like protease | 11 | 1.84 | 0.152 | 0.404 | |
| AT2G33585.1 | Subtilisin-like protease | 12 | 2.10 | 0.146 | 0.175 | |
| AT2G33585.1 | Subtilisin-like protease | 13 | 2.35 | 0.141 | 0.065 | . |
| AT2G33585.1 | Subtilisin-like protease | 14 | 2.60 | 0.138 | 0.023 | * |
| AT2G33585.1 | Subtilisin-like protease | 15 | 2.83 | 0.136 | 0.009 | ** |
| AT2G33585.1 | Subtilisin-like protease | 16 | 3.02 | 0.135 | 0.004 | ** |
| AT2G33585.1 | Subtilisin-like protease | 17 | 3.15 | 0.136 | 0.003 | ** |
| AT2G33585.1 | Subtilisin-like protease | 18 | 3.22 | 0.137 | 0.002 | ** |
| AT2G33585.1 | Subtilisin-like protease | 19 | 3.22 | 0.139 | 0.003 | ** |
| AT2G33585.1 | Subtilisin-like protease | 20 | 3.16 | 0.143 | 0.005 | ** |
| AT2G33585.1 | Subtilisin-like protease | 21 | 3.06 | 0.147 | 0.009 | ** |
| AT2G33585.1 | Subtilisin-like protease | 22 | 2.94 | 0.152 | 0.019 | * |
| AT2G33585.1 | Subtilisin-like protease | 23 | 2.81 | 0.158 | 0.037 | * |
| AT2G33585.1 | Subtilisin-like protease | 24 | 2.71 | 0.166 | 0.067 | . |
| AT2G33585.1 | Subtilisin-like protease | 25 | 2.62 | 0.175 | 0.114 | |
| AT2G33585.1 | Subtilisin-like protease | 26 | 2.54 | 0.186 | 0.185 | |
| AT2G33585.1 | Subtilisin-like protease | 27 | 2.44 | 0.200 | 0.290 | |
| AT2G33585.1 | Subtilisin-like protease | 28 | 2.32 | 0.216 | 0.443 | |
| AT2G33585.1 | Subtilisin-like protease | 29 | 2.16 | 0.235 | 0.628 | |
| AT2G33585.1 | Subtilisin-like protease | 46 | 0.16 | 0.490 | 0.499 | |
| AT2G33585.1 | Subtilisin-like protease | 47 | 0.13 | 0.516 | 0.443 | |
| AT2G33600.1 | Cinnamoyl CoA:NADP oxidoreductase-like 2 (CRL2) | 2 | 0.82 | 0.584 | 1.000 | |
| AT2G33600.1 | Cinnamoyl CoA:NADP oxidoreductase-like 2 (CRL2) | 3 | 0.76 | 0.565 | 1.000 | |

| | | | | | | |
|---|---|---|---|---|---|---|
| AT2G33600.1 | Cinnamoyl CoA:NADP oxidoreductase-like 2 (CRL2) | 32 | 0.28 | 0.252 | 0.125 | |
| AT2G33600.1 | Cinnamoyl CoA:NADP oxidoreductase-like 2 (CRL2) | 33 | 0.29 | 0.239 | 0.117 | |
| AT2G33600.1 | Cinnamoyl CoA:NADP oxidoreductase-like 2 (CRL2) | 34 | 0.31 | 0.227 | 0.116 | |
| AT2G33600.1 | Cinnamoyl CoA:NADP oxidoreductase-like 2 (CRL2) | 35 | 0.33 | 0.217 | 0.124 | |
| AT2G33600.1 | Cinnamoyl CoA:NADP oxidoreductase-like 2 (CRL2) | 36 | 0.36 | 0.208 | 0.140 | |
| AT2G33600.1 | Cinnamoyl CoA:NADP oxidoreductase-like 2 (CRL2) | 41 | 0.54 | 0.177 | 0.443 | |
| AT2G33600.1 | Cinnamoyl CoA:NADP oxidoreductase-like 2 (CRL2) | 42 | 0.59 | 0.176 | 0.567 | |
| AT2G33600.1 | Cinnamoyl CoA:NADP oxidoreductase-like 2 (CRL2) | 43 | 0.64 | 0.177 | 0.701 | |
| AT2G33600.1 | Cinnamoyl CoA:NADP oxidoreductase-like 2 (CRL2) | 44 | 0.68 | 0.184 | 0.821 | |
| AT2G33600.1 | Cinnamoyl CoA:NADP oxidoreductase-like 2 (CRL2) | 45 | 0.72 | 0.195 | 0.910 | |
| AT2G33600.1 | Cinnamoyl CoA:NADP oxidoreductase-like 2 (CRL2) | 46 | 0.76 | 0.212 | 0.963 | |
| AT2G33600.1 | Cinnamoyl CoA:NADP oxidoreductase-like 2 (CRL2) | 47 | 0.80 | 0.235 | 0.988 | |
| AT2G33820.1 | Mitochondrial ornithine transporter (MBAC1) | 2 | 1.05 | 0.413 | 1.000 | |
| AT2G33820.1 | Mitochondrial ornithine transporter (MBAC1) | 3 | 1.07 | 0.345 | 1.000 | |
| AT2G33820.1 | Mitochondrial ornithine transporter (MBAC1) | 8 | 1.06 | 0.291 | 1.000 | |
| AT2G33820.1 | Mitochondrial ornithine transporter (MBAC1) | 9 | 1.06 | 0.239 | 1.000 | |
| AT2G33820.1 | Mitochondrial ornithine transporter (MBAC1) | 10 | 1.12 | 0.216 | 1.000 | |
| AT2G33820.1 | Mitochondrial ornithine transporter (MBAC1) | 11 | 1.24 | 0.201 | 1.000 | |
| AT2G33820.1 | Mitochondrial ornithine transporter (MBAC1) | 12 | 1.34 | 0.195 | 1.000 | |
| AT2G33820.1 | Mitochondrial ornithine transporter (MBAC1) | 13 | 1.21 | 0.193 | 1.000 | |
| AT2G33820.1 | Mitochondrial ornithine transporter (MBAC1) | 14 | 1.37 | 0.197 | 1.000 | |
| AT2G33820.1 | Mitochondrial ornithine transporter (MBAC1) | 15 | 1.76 | 0.203 | 0.986 | |
| AT2G33820.1 | Mitochondrial ornithine transporter (MBAC1) | 16 | 1.83 | 0.205 | 0.977 | |
| AT2G33820.1 | Mitochondrial ornithine transporter (MBAC1) | 17 | 2.50 | 0.215 | 0.701 | |
| AT2G33820.1 | Mitochondrial ornithine transporter (MBAC1) | 18 | 3.96 | 0.311 | 0.640 | |
| AT2G33820.1 | Mitochondrial ornithine transporter (MBAC1) | 36 | 2.12 | 0.356 | 0.999 | |
| AT2G33820.1 | Mitochondrial ornithine transporter (MBAC1) | 37 | 3.32 | 0.285 | 0.716 | |
| AT2G33820.1 | Mitochondrial ornithine transporter (MBAC1) | 38 | 5.87 | 0.273 | 0.106 | |
| AT2G33820.1 | Mitochondrial ornithine transporter (MBAC1) | 39 | 11.12 | 0.247 | 0.001 | *** |
| AT2G33820.1 | Mitochondrial ornithine transporter (MBAC1) | 40 | 18.95 | 0.228 | 1.000 | |
| AT2G33820.1 | Mitochondrial ornithine transporter (MBAC1) | 41 | 21.32 | 0.199 | 1.000 | |

| | | | | | | |
|---|---|---|---|---|---|---|
| AT2G33820.1 | Mitochondrial ornithine transporter (MBAC1) | 42 | 10.22 | 0.198 | 1.000 | |
| AT2G33820.1 | Mitochondrial ornithine transporter (MBAC1) | 43 | 1.53 | 0.200 | 0.999 | |
| AT2G33820.1 | Mitochondrial ornithine transporter (MBAC1) | 44 | 0.17 | 0.197 | 0.003 | ** |
| AT2G33820.1 | Mitochondrial ornithine transporter (MBAC1) | 45 | 0.06 | 0.197 | 1.000 | |
| AT2G33820.1 | Mitochondrial ornithine transporter (MBAC1) | 46 | 0.09 | 0.215 | 1.000 | |
| AT2G33820.1 | Mitochondrial ornithine transporter (MBAC1) | 47 | 0.24 | 0.452 | 0.953 | |
| AT2G34420.1 | Photosystem II light harvesting complex subunit (LHCB1.5) | 2 | 0.34 | 0.472 | 1.000 | |
| AT2G34420.1 | Photosystem II light harvesting complex subunit (LHCB1.5) | 3 | 0.40 | 0.298 | 0.995 | |
| AT2G34420.1 | Photosystem II light harvesting complex subunit (LHCB1.5) | 4 | 0.53 | 0.277 | 1.000 | |
| AT2G34420.1 | Photosystem II light harvesting complex subunit (LHCB1.5) | 5 | 0.75 | 0.277 | 1.000 | |
| AT2G34420.1 | Photosystem II light harvesting complex subunit (LHCB1.5) | 6 | 1.12 | 0.278 | 1.000 | |
| AT2G34420.1 | Photosystem II light harvesting complex subunit (LHCB1.5) | 7 | 1.15 | 0.280 | 1.000 | |
| AT2G34420.1 | Photosystem II light harvesting complex subunit (LHCB1.5) | 8 | 0.95 | 0.273 | 1.000 | |
| AT2G34420.1 | Photosystem II light harvesting complex subunit (LHCB1.5) | 9 | 0.70 | 0.269 | 1.000 | |
| AT2G34420.1 | Photosystem II light harvesting complex subunit (LHCB1.5) | 10 | 0.43 | 0.271 | 0.992 | |
| AT2G34420.1 | Photosystem II light harvesting complex subunit (LHCB1.5) | 11 | 0.38 | 0.271 | 0.970 | |
| AT2G34420.1 | Photosystem II light harvesting complex subunit (LHCB1.5) | 12 | 0.49 | 0.271 | 1.000 | |
| AT2G34420.1 | Photosystem II light harvesting complex subunit (LHCB1.5) | 13 | 0.77 | 0.268 | 1.000 | |
| AT2G34420.1 | Photosystem II light harvesting complex subunit (LHCB1.5) | 14 | 0.89 | 0.264 | 1.000 | |
| AT2G34420.1 | Photosystem II light harvesting complex subunit (LHCB1.5) | 15 | 0.67 | 0.262 | 1.000 | |
| AT2G34420.1 | Photosystem II light harvesting complex subunit (LHCB1.5) | 16 | 0.47 | 0.261 | 0.997 | |
| AT2G34420.1 | Photosystem II light harvesting complex subunit (LHCB1.5) | 17 | 0.53 | 0.259 | 1.000 | |
| AT2G34420.1 | Photosystem II light harvesting complex subunit (LHCB1.5) | 18 | 0.66 | 0.262 | 1.000 | |
| AT2G34420.1 | Photosystem II light harvesting complex subunit (LHCB1.5) | 19 | 0.64 | 0.271 | 1.000 | |
| AT2G34420.1 | Photosystem II light harvesting complex subunit (LHCB1.5) | 20 | 0.72 | 0.278 | 1.000 | |
| AT2G34420.1 | Photosystem II light harvesting complex subunit (LHCB1.5) | 21 | 1.05 | 0.278 | 1.000 | |
| AT2G34420.1 | Photosystem II light harvesting complex subunit (LHCB1.5) | 22 | 1.28 | 0.279 | 1.000 | |
| AT2G34420.1 | Photosystem II light harvesting complex subunit (LHCB1.5) | 23 | 1.01 | 0.277 | 1.000 | |
| AT2G34420.1 | Photosystem II light harvesting complex subunit (LHCB1.5) | 24 | 0.65 | 0.270 | 1.000 | |
| AT2G34420.1 | Photosystem II light harvesting complex subunit (LHCB1.5) | 25 | 0.56 | 0.266 | 1.000 | |
| AT2G34420.1 | Photosystem II light harvesting complex subunit (LHCB1.5) | 26 | 0.77 | 0.266 | 1.000 | |

| | | | | | |
|---|---|---|---|---|---|
| AT2G34420.1 | Photosystem II light harvesting complex subunit (LHCB1.5) | 27 | 1.03 | 0.266 | 1.000 |
| AT2G34420.1 | Photosystem II light harvesting complex subunit (LHCB1.5) | 28 | 1.01 | 0.267 | 1.000 |
| AT2G34420.1 | Photosystem II light harvesting complex subunit (LHCB1.5) | 29 | 0.81 | 0.268 | 1.000 |
| AT2G34420.1 | Photosystem II light harvesting complex subunit (LHCB1.5) | 30 | 0.84 | 0.269 | 1.000 |
| AT2G34420.1 | Photosystem II light harvesting complex subunit (LHCB1.5) | 31 | 1.15 | 0.270 | 1.000 |
| AT2G34420.1 | Photosystem II light harvesting complex subunit (LHCB1.5) | 32 | 1.27 | 0.270 | 1.000 |
| AT2G34420.1 | Photosystem II light harvesting complex subunit (LHCB1.5) | 33 | 1.35 | 0.270 | 1.000 |
| AT2G34420.1 | Photosystem II light harvesting complex subunit (LHCB1.5) | 34 | 2.00 | 0.271 | 1.000 |
| AT2G34420.1 | Photosystem II light harvesting complex subunit (LHCB1.5) | 35 | 2.51 | 0.270 | 0.980 |
| AT2G34420.1 | Photosystem II light harvesting complex subunit (LHCB1.5) | 36 | 1.30 | 0.270 | 1.000 |
| AT2G34420.1 | Photosystem II light harvesting complex subunit (LHCB1.5) | 37 | 0.50 | 0.270 | 1.000 |
| AT2G34420.1 | Photosystem II light harvesting complex subunit (LHCB1.5) | 38 | 0.38 | 0.270 | 0.961 |
| AT2G34420.1 | Photosystem II light harvesting complex subunit (LHCB1.5) | 39 | 0.67 | 0.269 | 1.000 |
| AT2G34420.1 | Photosystem II light harvesting complex subunit (LHCB1.5) | 40 | 1.72 | 0.269 | 1.000 |
| AT2G34420.1 | Photosystem II light harvesting complex subunit (LHCB1.5) | 41 | 3.50 | 0.270 | 0.717 |
| AT2G34420.1 | Photosystem II light harvesting complex subunit (LHCB1.5) | 42 | 3.40 | 0.275 | 0.775 |
| AT2G34420.1 | Photosystem II light harvesting complex subunit (LHCB1.5) | 43 | 1.20 | 0.276 | 1.000 |
| AT2G34420.1 | Photosystem II light harvesting complex subunit (LHCB1.5) | 44 | 0.44 | 0.283 | 0.998 |
| AT2G34420.1 | Photosystem II light harvesting complex subunit (LHCB1.5) | 45 | 0.25 | 0.338 | 0.884 |
| AT2G34420.1 | Photosystem II light harvesting complex subunit (LHCB1.5) | 46 | 0.18 | 0.460 | 0.946 |
| AT2G34420.1 | Photosystem II light harvesting complex subunit (LHCB1.5) | 47 | 0.14 | 0.736 | 0.999 |
| AT2G34430.1 | Photosystem II light harvesting complex subunit (LHCB1.4) | 2 | 1.00 | 0.501 | 1.000 |
| AT2G34430.1 | Photosystem II light harvesting complex subunit (LHCB1.4) | 3 | 1.33 | 0.389 | 1.000 |
| AT2G34430.1 | Photosystem II light harvesting complex subunit (LHCB1.4) | 4 | 1.72 | 0.329 | 1.000 |
| AT2G34430.1 | Photosystem II light harvesting complex subunit (LHCB1.4) | 5 | 2.04 | 0.302 | 0.989 |
| AT2G34430.1 | Photosystem II light harvesting complex subunit (LHCB1.4) | 6 | 2.16 | 0.290 | 0.970 |
| AT2G34430.1 | Photosystem II light harvesting complex subunit (LHCB1.4) | 7 | 2.10 | 0.283 | 0.973 |
| AT2G34430.1 | Photosystem II light harvesting complex subunit (LHCB1.4) | 8 | 2.00 | 0.278 | 0.982 |
| AT2G34430.1 | Photosystem II light harvesting complex subunit (LHCB1.4) | 9 | 1.92 | 0.275 | 0.988 |
| AT2G34430.1 | Photosystem II light harvesting complex subunit (LHCB1.4) | 10 | 1.90 | 0.273 | 0.989 |
| AT2G34430.1 | Photosystem II light harvesting complex subunit (LHCB1.4) | 11 | 1.95 | 0.272 | 0.984 |

| | | | | | |
|---|---|---|---|---|---|
| AT2G34430.1 | Photosystem II light harvesting complex subunit (LHCB1.4) | 12 | 2.09 | 0.271 | 0.964 |
| AT2G34430.1 | Photosystem II light harvesting complex subunit (LHCB1.4) | 13 | 2.31 | 0.268 | 0.909 |
| AT2G34430.1 | Photosystem II light harvesting complex subunit (LHCB1.4) | 14 | 2.48 | 0.266 | 0.840 |
| AT2G34430.1 | Photosystem II light harvesting complex subunit (LHCB1.4) | 15 | 2.53 | 0.264 | 0.816 |
| AT2G34430.1 | Photosystem II light harvesting complex subunit (LHCB1.4) | 16 | 2.47 | 0.264 | 0.841 |
| AT2G34430.1 | Photosystem II light harvesting complex subunit (LHCB1.4) | 17 | 2.48 | 0.265 | 0.837 |
| AT2G34430.1 | Photosystem II light harvesting complex subunit (LHCB1.4) | 18 | 2.66 | 0.267 | 0.775 |
| AT2G34430.1 | Photosystem II light harvesting complex subunit (LHCB1.4) | 19 | 2.80 | 0.271 | 0.731 |
| AT2G34430.1 | Photosystem II light harvesting complex subunit (LHCB1.4) | 20 | 2.69 | 0.275 | 0.792 |
| AT2G34430.1 | Photosystem II light harvesting complex subunit (LHCB1.4) | 21 | 2.21 | 0.277 | 0.947 |
| AT2G34430.1 | Photosystem II light harvesting complex subunit (LHCB1.4) | 22 | 1.49 | 0.278 | 1.000 |
| AT2G34430.1 | Photosystem II light harvesting complex subunit (LHCB1.4) | 23 | 0.86 | 0.276 | 1.000 |
| AT2G34430.1 | Photosystem II light harvesting complex subunit (LHCB1.4) | 24 | 0.48 | 0.274 | 0.967 |
| AT2G34430.1 | Photosystem II light harvesting complex subunit (LHCB1.4) | 25 | 0.29 | 0.272 | 0.470 |
| AT2G34430.1 | Photosystem II light harvesting complex subunit (LHCB1.4) | 26 | 0.21 | 0.272 | 0.168 |
| AT2G34430.1 | Photosystem II light harvesting complex subunit (LHCB1.4) | 27 | 0.19 | 0.275 | 0.130 |
| AT2G34430.1 | Photosystem II light harvesting complex subunit (LHCB1.4) | 28 | 0.21 | 0.281 | 0.214 |
| AT2G34430.1 | Photosystem II light harvesting complex subunit (LHCB1.4) | 29 | 0.27 | 0.286 | 0.463 |
| AT2G34430.1 | Photosystem II light harvesting complex subunit (LHCB1.4) | 30 | 0.36 | 0.287 | 0.800 |
| AT2G34430.1 | Photosystem II light harvesting complex subunit (LHCB1.4) | 31 | 0.45 | 0.284 | 0.957 |
| AT2G34430.1 | Photosystem II light harvesting complex subunit (LHCB1.4) | 32 | 0.52 | 0.281 | 0.989 |
| AT2G34430.1 | Photosystem II light harvesting complex subunit (LHCB1.4) | 33 | 0.56 | 0.282 | 0.996 |
| AT2G34430.1 | Photosystem II light harvesting complex subunit (LHCB1.4) | 34 | 0.61 | 0.293 | 1.000 |
| AT2G34430.1 | Photosystem II light harvesting complex subunit (LHCB1.4) | 35 | 0.64 | 0.326 | 1.000 |
| AT2G34430.1 | Photosystem II light harvesting complex subunit (LHCB1.4) | 36 | 0.57 | 0.390 | 1.000 |
| AT2G34430.1 | Photosystem II light harvesting complex subunit (LHCB1.4) | 37 | 0.43 | 0.483 | 0.999 |
| AT2G34430.1 | Photosystem II light harvesting complex subunit (LHCB1.4) | 46 | 0.00 | 1.422 | 0.215 |
| AT2G34430.1 | Photosystem II light harvesting complex subunit (LHCB1.4) | 47 | 0.00 | 1.587 | 0.205 |
| AT2G34460.1 | NAD-dependent epimerase/dehydratase, putative | 2 | 0.65 | 0.453 | 1.000 |
| AT2G34460.1 | NAD-dependent epimerase/dehydratase, putative | 3 | 0.91 | 0.433 | 1.000 |

| | | | | | |
|---|---|---|---|---|---|
| AT2G34460.1 | NAD-dependent epimerase/dehydratase, putative | 36 | 4.90 | 0.402 | 0.519 |
| AT2G34460.1 | NAD-dependent epimerase/dehydratase, putative | 37 | 2.94 | 0.269 | 0.501 |
| AT2G34460.1 | NAD-dependent epimerase/dehydratase, putative | 38 | 1.62 | 0.196 | 0.921 |
| AT2G34460.1 | NAD-dependent epimerase/dehydratase, putative | 39 | 0.81 | 0.171 | 0.999 |
| AT2G34460.1 | NAD-dependent epimerase/dehydratase, putative | 40 | 0.48 | 0.167 | 0.388 |
| AT2G34460.1 | NAD-dependent epimerase/dehydratase, putative | 41 | 0.45 | 0.167 | 0.278 |
| AT2G34460.1 | NAD-dependent epimerase/dehydratase, putative | 42 | 0.62 | 0.170 | 0.844 |
| AT2G34460.1 | NAD-dependent epimerase/dehydratase, putative | 43 | 1.00 | 0.173 | 1.000 |
| AT2G34460.1 | NAD-dependent epimerase/dehydratase, putative | 44 | 1.52 | 0.173 | 0.926 |
| AT2G34460.1 | NAD-dependent epimerase/dehydratase, putative | 45 | 2.15 | 0.176 | 0.404 |
| AT2G34460.1 | NAD-dependent epimerase/dehydratase, putative | 46 | 2.93 | 0.221 | 0.272 |
| AT2G34460.1 | NAD-dependent epimerase/dehydratase, putative | 47 | 3.92 | 0.369 | 0.597 |
| AT2G34480.1 | 60S ribosomal protein L18a-2 (RPL18AB) | 2 | 0.58 | 0.219 | 0.544 |
| AT2G34480.1 | 60S ribosomal protein L18a-2 (RPL18AB) | 3 | 0.64 | 0.202 | 0.624 |
| AT2G34480.1 | 60S ribosomal protein L18a-2 (RPL18AB) | 4 | 0.71 | 0.186 | 0.721 |
| AT2G34480.1 | 60S ribosomal protein L18a-2 (RPL18AB) | 5 | 0.79 | 0.172 | 0.829 |
| AT2G34480.1 | 60S ribosomal protein L18a-2 (RPL18AB) | 6 | 0.87 | 0.159 | 0.932 |
| AT2G34480.1 | 60S ribosomal protein L18a-2 (RPL18AB) | 7 | 0.97 | 0.148 | 0.996 |
| AT2G34480.1 | 60S ribosomal protein L18a-2 (RPL18AB) | 8 | 1.07 | 0.140 | 0.975 |
| AT2G34480.1 | 60S ribosomal protein L18a-2 (RPL18AB) | 9 | 1.19 | 0.135 | 0.851 |
| AT2G34480.1 | 60S ribosomal protein L18a-2 (RPL18AB) | 10 | 1.32 | 0.134 | 0.659 |
| AT2G34480.1 | 60S ribosomal protein L18a-2 (RPL18AB) | 11 | 1.47 | 0.136 | 0.467 |
| AT2G34480.1 | 60S ribosomal protein L18a-2 (RPL18AB) | 12 | 1.63 | 0.141 | 0.321 |
| AT2G34480.1 | 60S ribosomal protein L18a-2 (RPL18AB) | 13 | 1.80 | 0.149 | 0.225 |
| AT2G34480.1 | 60S ribosomal protein L18a-2 (RPL18AB) | 14 | 2.00 | 0.160 | 0.168 |

| | | | | | | |
|---|---|---|---|---|---|---|
| AT2G34480.1 | 60S ribosomal protein L18a-2 (RPL18AB) | 15 | 2.22 | 0.173 | 0.133 | |
| AT2G34480.1 | 60S ribosomal protein L18a-2 (RPL18AB) | 16 | 2.46 | 0.187 | 0.112 | |
| AT2G34480.1 | 60S ribosomal protein L18a-2 (RPL18AB) | 17 | 2.73 | 0.203 | 0.099 | . |
| AT2G34480.1 | 60S ribosomal protein L18a-2 (RPL18AB) | 18 | 3.03 | 0.220 | 0.092 | . |
| AT2G34480.1 | 60S ribosomal protein L18a-2 (RPL18AB) | 46 | 55.49 | 0.794 | 0.088 | . |
| AT2G34480.1 | 60S ribosomal protein L18a-2 (RPL18AB) | 47 | 61.56 | 0.815 | 0.090 | . |
| AT2G35370.1 | Glycine decarboxylase subunit (GDCH) | 2 | 0.48 | 0.486 | 1.000 | |
| AT2G35370.1 | Glycine decarboxylase subunit (GDCH) | 3 | 0.48 | 0.370 | 1.000 | |
| AT2G35370.1 | Glycine decarboxylase subunit (GDCH) | 4 | 0.49 | 0.295 | 0.996 | |
| AT2G35370.1 | Glycine decarboxylase subunit (GDCH) | 5 | 0.52 | 0.251 | 0.991 | |
| AT2G35370.1 | Glycine decarboxylase subunit (GDCH) | 6 | 0.58 | 0.229 | 0.997 | |
| AT2G35370.1 | Glycine decarboxylase subunit (GDCH) | 7 | 0.70 | 0.218 | 1.000 | |
| AT2G35370.1 | Glycine decarboxylase subunit (GDCH) | 8 | 0.87 | 0.212 | 1.000 | |
| AT2G35370.1 | Glycine decarboxylase subunit (GDCH) | 9 | 1.11 | 0.208 | 1.000 | |
| AT2G35370.1 | Glycine decarboxylase subunit (GDCH) | 10 | 1.43 | 0.206 | 1.000 | |
| AT2G35370.1 | Glycine decarboxylase subunit (GDCH) | 11 | 1.80 | 0.204 | 0.978 | |
| AT2G35370.1 | Glycine decarboxylase subunit (GDCH) | 12 | 2.22 | 0.203 | 0.793 | |
| AT2G35370.1 | Glycine decarboxylase subunit (GDCH) | 13 | 2.66 | 0.201 | 0.486 | |
| AT2G35370.1 | Glycine decarboxylase subunit (GDCH) | 14 | 3.08 | 0.200 | 0.258 | |
| AT2G35370.1 | Glycine decarboxylase subunit (GDCH) | 15 | 3.52 | 0.199 | 0.124 | |
| AT2G35370.1 | Glycine decarboxylase subunit (GDCH) | 16 | 4.08 | 0.198 | 0.049 | * |
| AT2G35370.1 | Glycine decarboxylase subunit (GDCH) | 17 | 4.80 | 0.199 | 0.017 | * |
| AT2G35370.1 | Glycine decarboxylase subunit (GDCH) | 18 | 5.43 | 0.201 | 0.007 | ** |
| AT2G35370.1 | Glycine decarboxylase subunit (GDCH) | 19 | 5.49 | 0.204 | 0.009 | ** |
| AT2G35370.1 | Glycine decarboxylase subunit (GDCH) | 20 | 4.81 | 0.207 | 0.026 | * |
| AT2G35370.1 | Glycine decarboxylase subunit (GDCH) | 21 | 3.77 | 0.209 | 0.126 | |
| AT2G35370.1 | Glycine decarboxylase subunit (GDCH) | 22 | 2.84 | 0.210 | 0.445 | |
| AT2G35370.1 | Glycine decarboxylase subunit (GDCH) | 23 | 2.22 | 0.209 | 0.825 | |
| AT2G35370.1 | Glycine decarboxylase subunit (GDCH) | 24 | 1.85 | 0.206 | 0.972 | |
| AT2G35370.1 | Glycine decarboxylase subunit (GDCH) | 25 | 1.65 | 0.204 | 0.996 | |
| AT2G35370.1 | Glycine decarboxylase subunit (GDCH) | 26 | 1.55 | 0.203 | 0.999 | |

| | | | | | |
|---|---|---|---|---|---|
| AT2G35370.1 | Glycine decarboxylase subunit (GDCH) | 27 | 1.52 | 0.202 | 0.999 |
| AT2G35370.1 | Glycine decarboxylase subunit (GDCH) | 28 | 1.54 | 0.202 | 0.999 |
| AT2G35370.1 | Glycine decarboxylase subunit (GDCH) | 29 | 1.55 | 0.202 | 0.999 |
| AT2G35370.1 | Glycine decarboxylase subunit (GDCH) | 30 | 1.50 | 0.203 | 1.000 |
| AT2G35370.1 | Glycine decarboxylase subunit (GDCH) | 31 | 1.37 | 0.203 | 1.000 |
| AT2G35370.1 | Glycine decarboxylase subunit (GDCH) | 32 | 1.17 | 0.203 | 1.000 |
| AT2G35370.1 | Glycine decarboxylase subunit (GDCH) | 33 | 0.95 | 0.204 | 1.000 |
| AT2G35370.1 | Glycine decarboxylase subunit (GDCH) | 34 | 0.79 | 0.204 | 1.000 |
| AT2G35370.1 | Glycine decarboxylase subunit (GDCH) | 35 | 0.73 | 0.204 | 1.000 |
| AT2G35370.1 | Glycine decarboxylase subunit (GDCH) | 36 | 0.75 | 0.203 | 1.000 |
| AT2G35370.1 | Glycine decarboxylase subunit (GDCH) | 37 | 0.80 | 0.203 | 1.000 |
| AT2G35370.1 | Glycine decarboxylase subunit (GDCH) | 38 | 0.84 | 0.204 | 1.000 |
| AT2G35370.1 | Glycine decarboxylase subunit (GDCH) | 39 | 0.88 | 0.205 | 1.000 |
| AT2G35370.1 | Glycine decarboxylase subunit (GDCH) | 40 | 0.95 | 0.206 | 1.000 |
| AT2G35370.1 | Glycine decarboxylase subunit (GDCH) | 41 | 1.03 | 0.208 | 1.000 |
| AT2G35370.1 | Glycine decarboxylase subunit (GDCH) | 42 | 1.10 | 0.209 | 1.000 |
| AT2G35370.1 | Glycine decarboxylase subunit (GDCH) | 43 | 1.09 | 0.209 | 1.000 |
| AT2G35370.1 | Glycine decarboxylase subunit (GDCH) | 44 | 0.95 | 0.213 | 1.000 |
| AT2G35370.1 | Glycine decarboxylase subunit (GDCH) | 45 | 0.72 | 0.233 | 1.000 |
| AT2G35370.1 | Glycine decarboxylase subunit (GDCH) | 46 | 0.49 | 0.290 | 0.995 |
| AT2G35370.1 | Glycine decarboxylase subunit (GDCH) | 47 | 0.32 | 0.399 | 0.980 |
| AT2G35720.1 | J-domain protein involved in perception of very low light fluences (OWL1) | 2 | 1.04 | 0.282 | 1.000 |
| AT2G35720.1 | J-domain protein involved in perception of very low light fluences (OWL1) | 3 | 1.09 | 0.176 | 1.000 |
| AT2G35720.1 | J-domain protein involved in perception of very low light fluences (OWL1) | 4 | 1.17 | 0.159 | 1.000 |
| AT2G35720.1 | J-domain protein involved in perception of very low light fluences (OWL1) | 5 | 1.36 | 0.160 | 1.000 |
| AT2G35720.1 | J-domain protein involved in perception of very low light fluences (OWL1) | 6 | 1.57 | 0.160 | 0.993 |
| AT2G35720.1 | J-domain protein involved in perception of very low light fluences (OWL1) | 7 | 1.68 | 0.159 | 0.969 |

| | | | | | |
|---|---|---|---|---|---|
| AT2G35720.1 | J-domain protein involved in perception of very low light fluences (OWL1) | 8 | 1.62 | 0.156 | 0.983 |
| AT2G35720.1 | J-domain protein involved in perception of very low light fluences (OWL1) | 9 | 1.45 | 0.154 | 0.999 |
| AT2G35720.1 | J-domain protein involved in perception of very low light fluences (OWL1) | 10 | 1.29 | 0.154 | 1.000 |
| AT2G35720.1 | J-domain protein involved in perception of very low light fluences (OWL1) | 11 | 1.18 | 0.155 | 1.000 |
| AT2G35720.1 | J-domain protein involved in perception of very low light fluences (OWL1) | 12 | 1.14 | 0.155 | 1.000 |
| AT2G35720.1 | J-domain protein involved in perception of very low light fluences (OWL1) | 13 | 1.17 | 0.153 | 1.000 |
| AT2G35720.1 | J-domain protein involved in perception of very low light fluences (OWL1) | 14 | 1.37 | 0.151 | 1.000 |
| AT2G35720.1 | J-domain protein involved in perception of very low light fluences (OWL1) | 15 | 1.62 | 0.151 | 0.972 |
| AT2G35720.1 | J-domain protein involved in perception of very low light fluences (OWL1) | 16 | 1.62 | 0.150 | 0.973 |
| AT2G35720.1 | J-domain protein involved in perception of very low light fluences (OWL1) | 17 | 1.49 | 0.149 | 0.996 |
| AT2G35720.1 | J-domain protein involved in perception of very low light fluences (OWL1) | 18 | 1.38 | 0.151 | 1.000 |
| AT2G35720.1 | J-domain protein involved in perception of very low light fluences (OWL1) | 19 | 1.28 | 0.154 | 1.000 |
| AT2G35720.1 | J-domain protein involved in perception of very low light fluences (OWL1) | 20 | 1.21 | 0.159 | 1.000 |
| AT2G35720.1 | J-domain protein involved in perception of very low light fluences (OWL1) | 21 | 1.15 | 0.160 | 1.000 |
| AT2G35720.1 | J-domain protein involved in perception of very low light fluences (OWL1) | 22 | 1.11 | 0.160 | 1.000 |
| AT2G35720.1 | J-domain protein involved in perception of very low light fluences (OWL1) | 23 | 1.05 | 0.159 | 1.000 |
| AT2G35720.1 | J-domain protein involved in perception of very low light fluences (OWL1) | 24 | 0.94 | 0.156 | 1.000 |
| AT2G35720.1 | J-domain protein involved in perception of very low light fluences (OWL1) | 25 | 0.84 | 0.153 | 1.000 |
| AT2G35720.1 | J-domain protein involved in perception of very low light fluences (OWL1) | 26 | 0.90 | 0.153 | 1.000 |
| AT2G35720.1 | J-domain protein involved in perception of very low light fluences (OWL1) | 27 | 1.11 | 0.153 | 1.000 |

| | | | | | |
|---|---|---|---|---|---|
| AT2G35720.1 | J-domain protein involved in perception of very low light fluences (OWL1) | 28 | 1.26 | 0.153 | 1.000 |
| AT2G35720.1 | J-domain protein involved in perception of very low light fluences (OWL1) | 29 | 1.29 | 0.154 | 1.000 |
| AT2G35720.1 | J-domain protein involved in perception of very low light fluences (OWL1) | 30 | 1.43 | 0.155 | 0.999 |
| AT2G35720.1 | J-domain protein involved in perception of very low light fluences (OWL1) | 31 | 1.78 | 0.156 | 0.913 |
| AT2G35720.1 | J-domain protein involved in perception of very low light fluences (OWL1) | 32 | 1.93 | 0.160 | 0.811 |
| AT2G35720.1 | J-domain protein involved in perception of very low light fluences (OWL1) | 33 | 1.50 | 0.172 | 0.999 |
| AT2G35720.1 | J-domain protein involved in perception of very low light fluences (OWL1) | 34 | 1.08 | 0.212 | 1.000 |
| AT2G35720.1 | J-domain protein involved in perception of very low light fluences (OWL1) | 35 | 0.91 | 0.316 | 1.000 |
| AT2G35720.1 | J-domain protein involved in perception of very low light fluences (OWL1) | 46 | 8456.41 | 3.156 | 0.992 |
| AT2G35720.1 | J-domain protein involved in perception of very low light fluences (OWL1) | 47 | 31628.07 | 3.549 | 0.990 |
| AT2G35736.2 | ? | 2 | 0.87 | 0.525 | 1.000 |
| AT2G35736.2 | ? | 3 | 0.97 | 0.483 | 1.000 |
| AT2G35736.2 | ? | 17 | 2.72 | 0.285 | 0.648 |
| AT2G35736.2 | ? | 18 | 2.72 | 0.268 | 0.579 |
| AT2G35736.2 | ? | 19 | 2.66 | 0.258 | 0.558 |
| AT2G35736.2 | ? | 20 | 2.55 | 0.253 | 0.588 |
| AT2G35736.2 | ? | 26 | 1.37 | 0.207 | 0.995 |
| AT2G35736.2 | ? | 27 | 1.22 | 0.190 | 1.000 |
| AT2G35736.2 | ? | 28 | 1.11 | 0.175 | 1.000 |
| AT2G35736.2 | ? | 29 | 1.06 | 0.165 | 1.000 |
| AT2G35736.2 | ? | 30 | 1.04 | 0.163 | 1.000 |
| AT2G35736.2 | ? | 31 | 1.06 | 0.169 | 1.000 |
| AT2G35736.2 | ? | 32 | 1.09 | 0.182 | 1.000 |
| AT2G35736.2 | ? | 33 | 1.14 | 0.202 | 1.000 |
| AT2G35736.2 | ? | 34 | 1.22 | 0.226 | 1.000 |
| AT2G35736.2 | ? | 43 | 2.39 | 0.239 | 0.606 |
| AT2G35736.2 | ? | 44 | 2.44 | 0.220 | 0.474 |
| AT2G35736.2 | ? | 45 | 2.44 | 0.216 | 0.454 |
| AT2G35736.2 | ? | 46 | 2.40 | 0.244 | 0.619 |
| AT2G35736.2 | ? | 47 | 2.35 | 0.303 | 0.840 |
| AT2G35790.1 | ? | 2 | 1.08 | 0.593 | 1.000 |
| AT2G35790.1 | ? | 3 | 1.18 | 0.510 | 1.000 |
| AT2G35790.1 | ? | 11 | 1.47 | 0.512 | 1.000 |
| AT2G35790.1 | ? | 12 | 1.37 | 0.446 | 1.000 |
| AT2G35790.1 | ? | 13 | 1.27 | 0.471 | 1.000 |
| AT2G35790.1 | ? | 22 | 1.70 | 0.513 | 1.000 |
| AT2G35790.1 | ? | 23 | 1.85 | 0.436 | 1.000 |
| AT2G35790.1 | ? | 24 | 1.99 | 0.354 | 1.000 |
| AT2G35790.1 | ? | 25 | 1.89 | 0.345 | 1.000 |
| AT2G35790.1 | ? | 26 | 1.32 | 0.307 | 1.000 |
| AT2G35790.1 | ? | 27 | 0.83 | 0.282 | 1.000 |
| AT2G35790.1 | ? | 28 | 0.87 | 0.272 | 1.000 |

| | | | | | | |
|---|---|---|---|---|---|---|
| AT2G35790.1 | ? | 29 | 1.50 | 0.269 | 1.000 | |
| AT2G35790.1 | ? | 30 | 2.68 | 0.268 | 0.903 | |
| AT2G35790.1 | ? | 31 | 3.67 | 0.271 | 0.565 | |
| AT2G35790.1 | ? | 32 | 1.94 | 0.275 | 0.999 | |
| AT2G35790.1 | ? | 33 | 0.69 | 0.285 | 1.000 | |
| AT2G35790.1 | ? | 34 | 0.69 | 0.298 | 1.000 | |
| AT2G35790.1 | ? | 35 | 1.36 | 0.306 | 1.000 | |
| AT2G35790.1 | ? | 36 | 3.14 | 0.296 | 0.865 | |
| AT2G35790.1 | ? | 37 | 5.42 | 0.279 | 0.190 | |
| AT2G35790.1 | ? | 38 | 5.97 | 0.271 | 0.104 | |
| AT2G35790.1 | ? | 39 | 5.32 | 0.271 | 0.169 | |
| AT2G35790.1 | ? | 40 | 6.51 | 0.270 | 0.068 | . |
| AT2G35790.1 | ? | 41 | 10.87 | 0.270 | 0.004 | ** |
| AT2G35790.1 | ? | 42 | 22.91 | 0.275 | 0.000 | *** |
| AT2G35790.1 | ? | 43 | 25.14 | 0.277 | 0.000 | *** |
| AT2G35790.1 | ? | 44 | 10.21 | 0.276 | 0.008 | ** |
| AT2G35790.1 | ? | 45 | 2.17 | 0.275 | 0.992 | |
| AT2G35790.1 | ? | 46 | 0.29 | 0.312 | 0.841 | |
| AT2G35790.1 | ? | 47 | 0.02 | 0.586 | 0.141 | |
| AT2G36070.1 | Translocase of the inner mitochondrial membrane 44 (TIM44-2) | 2 | 0.87 | 0.405 | 1.000 | |
| AT2G36070.1 | Translocase of the inner mitochondrial membrane 44 (TIM44-2) | 3 | 0.83 | 0.390 | 1.000 | |
| AT2G36070.1 | Translocase of the inner mitochondrial membrane 44 (TIM44-2) | 27 | 0.92 | 0.319 | 1.000 | |
| AT2G36070.1 | Translocase of the inner mitochondrial membrane 44 (TIM44-2) | 28 | 1.00 | 0.245 | 1.000 | |
| AT2G36070.1 | Translocase of the inner mitochondrial membrane 44 (TIM44-2) | 29 | 1.01 | 0.226 | 1.000 | |
| AT2G36070.1 | Translocase of the inner mitochondrial membrane 44 (TIM44-2) | 30 | 1.05 | 0.219 | 1.000 | |
| AT2G36070.1 | Translocase of the inner mitochondrial membrane 44 (TIM44-2) | 31 | 1.23 | 0.217 | 1.000 | |
| AT2G36070.1 | Translocase of the inner mitochondrial membrane 44 (TIM44-2) | 32 | 1.28 | 0.217 | 1.000 | |
| AT2G36070.1 | Translocase of the inner mitochondrial membrane 44 (TIM44-2) | 33 | 0.97 | 0.217 | 1.000 | |
| AT2G36070.1 | Translocase of the inner mitochondrial membrane 44 (TIM44-2) | 34 | 0.71 | 0.218 | 1.000 | |
| AT2G36070.1 | Translocase of the inner mitochondrial membrane 44 (TIM44-2) | 35 | 0.70 | 0.218 | 0.999 | |
| AT2G36070.1 | Translocase of the inner mitochondrial membrane 44 (TIM44-2) | 36 | 0.87 | 0.217 | 1.000 | |
| AT2G36070.1 | Translocase of the inner mitochondrial membrane 44 (TIM44-2) | 37 | 1.10 | 0.217 | 1.000 | |
| AT2G36070.1 | Translocase of the inner mitochondrial membrane 44 (TIM44-2) | 38 | 1.38 | 0.216 | 1.000 | |

| | | | | | |
|---|---|---|---|---|---|
| AT2G36070.1 | Translocase of the inner mitochondrial membrane 44 (TIM44-2) | 39 | 1.90 | 0.216 | 0.913 |
| AT2G36070.1 | Translocase of the inner mitochondrial membrane 44 (TIM44-2) | 40 | 2.12 | 0.216 | 0.792 |
| AT2G36070.1 | Translocase of the inner mitochondrial membrane 44 (TIM44-2) | 41 | 1.66 | 0.217 | 0.986 |
| AT2G36070.1 | Translocase of the inner mitochondrial membrane 44 (TIM44-2) | 42 | 1.26 | 0.219 | 1.000 |
| AT2G36070.1 | Translocase of the inner mitochondrial membrane 44 (TIM44-2) | 43 | 1.09 | 0.220 | 1.000 |
| AT2G36070.1 | Translocase of the inner mitochondrial membrane 44 (TIM44-2) | 44 | 0.91 | 0.220 | 1.000 |
| AT2G36070.1 | Translocase of the inner mitochondrial membrane 44 (TIM44-2) | 45 | 0.62 | 0.220 | 0.992 |
| AT2G36070.1 | Translocase of the inner mitochondrial membrane 44 (TIM44-2) | 46 | 0.34 | 0.242 | 0.489 |
| AT2G36070.1 | Translocase of the inner mitochondrial membrane 44 (TIM44-2) | 47 | 0.16 | 0.399 | 0.409 |
| AT2G36830.1 | Tonoplast intrinsic protein (gammaTIP1) | 2 | 0.67 | 0.231 | 0.998 |
| AT2G36830.1 | Tonoplast intrinsic protein (gammaTIP1) | 3 | 0.77 | 0.159 | 0.999 |
| AT2G36830.1 | Tonoplast intrinsic protein (gammaTIP1) | 4 | 0.90 | 0.134 | 1.000 |
| AT2G36830.1 | Tonoplast intrinsic protein (gammaTIP1) | 5 | 1.08 | 0.133 | 1.000 |
| AT2G36830.1 | Tonoplast intrinsic protein (gammaTIP1) | 6 | 1.29 | 0.143 | 0.998 |
| AT2G36830.1 | Tonoplast intrinsic protein (gammaTIP1) | 7 | 1.47 | 0.160 | 0.976 |
| AT2G36830.1 | Tonoplast intrinsic protein (gammaTIP1) | 8 | 1.54 | 0.178 | 0.976 |
| AT2G36830.1 | Tonoplast intrinsic protein (gammaTIP1) | 11 | 1.22 | 0.185 | 1.000 |
| AT2G36830.1 | Tonoplast intrinsic protein (gammaTIP1) | 12 | 1.17 | 0.186 | 1.000 |
| AT2G36830.1 | Tonoplast intrinsic protein (gammaTIP1) | 13 | 1.18 | 0.198 | 1.000 |
| AT2G36830.1 | Tonoplast intrinsic protein (gammaTIP1) | 14 | 1.24 | 0.211 | 1.000 |
| AT2G36830.1 | Tonoplast intrinsic protein (gammaTIP1) | 28 | 1.31 | 0.197 | 1.000 |
| AT2G36830.1 | Tonoplast intrinsic protein (gammaTIP1) | 29 | 1.24 | 0.154 | 1.000 |
| AT2G36830.1 | Tonoplast intrinsic protein (gammaTIP1) | 30 | 1.21 | 0.135 | 1.000 |
| AT2G36830.1 | Tonoplast intrinsic protein (gammaTIP1) | 31 | 1.32 | 0.128 | 0.988 |
| AT2G36830.1 | Tonoplast intrinsic protein (gammaTIP1) | 32 | 1.58 | 0.127 | 0.760 |
| AT2G36830.1 | Tonoplast intrinsic protein (gammaTIP1) | 33 | 1.67 | 0.130 | 0.654 |

| | | | | | | |
|---|---|---|---|---|---|---|
| AT2G36830.1 | Tonoplast intrinsic protein (gammaTIP1) | 34 | 1.33 | 0.146 | 0.996 | |
| AT2G36830.1 | Tonoplast intrinsic protein (gammaTIP1) | 35 | 0.83 | 0.188 | 1.000 | |
| AT2G36830.1 | Tonoplast intrinsic protein (gammaTIP1) | 36 | 0.46 | 0.263 | 0.913 | |
| AT2G36830.1 | Tonoplast intrinsic protein (gammaTIP1) | 46 | 0.00 | 1.071 | 0.062 | . |
| AT2G36830.1 | Tonoplast intrinsic protein (gammaTIP1) | 47 | 0.00 | 1.195 | 0.073 | . |
| AT2G37020.1 | Translin family protein | 2 | 1.17 | 0.745 | 1.000 | |
| AT2G37020.1 | Translin family protein | 3 | 1.47 | 0.732 | 0.999 | |
| AT2G37020.1 | Translin family protein | 35 | 4.99 | 0.409 | 0.306 | |
| AT2G37020.1 | Translin family protein | 36 | 3.55 | 0.392 | 0.477 | |
| AT2G37020.1 | Translin family protein | 37 | 2.45 | 0.393 | 0.746 | |
| AT2G37020.1 | Translin family protein | 38 | 1.66 | 0.406 | 0.959 | |
| AT2G37020.1 | Translin family protein | 46 | 0.05 | 0.524 | 0.078 | . |
| AT2G37020.1 | Translin family protein | 47 | 0.03 | 0.560 | 0.053 | . |
| AT2G37230.1 | Pentatricopeptide repeat (PPR) superfamily protein | 2 | 0.71 | 0.709 | 1.000 | |
| AT2G37230.1 | Pentatricopeptide repeat (PPR) superfamily protein | 3 | 0.73 | 0.626 | 1.000 | |
| AT2G37230.1 | Pentatricopeptide repeat (PPR) superfamily protein | 12 | 1.86 | 0.406 | 1.000 | |
| AT2G37230.1 | Pentatricopeptide repeat (PPR) superfamily protein | 13 | 2.30 | 0.354 | 0.995 | |
| AT2G37230.1 | Pentatricopeptide repeat (PPR) superfamily protein | 14 | 2.69 | 0.319 | 0.939 | |
| AT2G37230.1 | Pentatricopeptide repeat (PPR) superfamily protein | 15 | 2.82 | 0.298 | 0.869 | |
| AT2G37230.1 | Pentatricopeptide repeat (PPR) superfamily protein | 16 | 2.89 | 0.286 | 0.808 | |
| AT2G37230.1 | Pentatricopeptide repeat (PPR) superfamily protein | 17 | 3.16 | 0.282 | 0.695 | |
| AT2G37230.1 | Pentatricopeptide repeat (PPR) superfamily protein | 18 | 3.44 | 0.283 | 0.600 | |
| AT2G37230.1 | Pentatricopeptide repeat (PPR) superfamily protein | 19 | 3.58 | 0.287 | 0.572 | |
| AT2G37230.1 | Pentatricopeptide repeat (PPR) superfamily protein | 20 | 3.55 | 0.292 | 0.609 | |
| AT2G37230.1 | Pentatricopeptide repeat (PPR) superfamily protein | 21 | 3.24 | 0.295 | 0.725 | |
| AT2G37230.1 | Pentatricopeptide repeat (PPR) superfamily protein | 22 | 2.59 | 0.295 | 0.920 | |
| AT2G37230.1 | Pentatricopeptide repeat (PPR) superfamily protein | 23 | 1.87 | 0.293 | 0.998 | |
| AT2G37230.1 | Pentatricopeptide repeat (PPR) superfamily protein | 24 | 1.30 | 0.289 | 1.000 | |
| AT2G37230.1 | Pentatricopeptide repeat (PPR) superfamily protein | 25 | 0.94 | 0.286 | 1.000 | |
| AT2G37230.1 | Pentatricopeptide repeat (PPR) superfamily protein | 26 | 0.75 | 0.285 | 1.000 | |
| AT2G37230.1 | Pentatricopeptide repeat (PPR) superfamily protein | 27 | 0.63 | 0.285 | 1.000 | |
| AT2G37230.1 | Pentatricopeptide repeat (PPR) superfamily protein | 28 | 0.57 | 0.285 | 0.999 | |
| AT2G37230.1 | Pentatricopeptide repeat (PPR) superfamily protein | 29 | 0.53 | 0.285 | 0.997 | |
| AT2G37230.1 | Pentatricopeptide repeat (PPR) superfamily protein | 30 | 0.50 | 0.285 | 0.993 | |
| AT2G37230.1 | Pentatricopeptide repeat (PPR) superfamily protein | 31 | 0.48 | 0.286 | 0.986 | |

| | | | | | |
|---|---|---|---|---|---|
| AT2G37230.1 | Pentatricopeptide repeat (PPR) superfamily protein | 32 | 0.46 | 0.286 | 0.979 |
| AT2G37230.1 | Pentatricopeptide repeat (PPR) superfamily protein | 33 | 0.45 | 0.287 | 0.975 |
| AT2G37230.1 | Pentatricopeptide repeat (PPR) superfamily protein | 34 | 0.45 | 0.287 | 0.978 |
| AT2G37230.1 | Pentatricopeptide repeat (PPR) superfamily protein | 35 | 0.50 | 0.287 | 0.993 |
| AT2G37230.1 | Pentatricopeptide repeat (PPR) superfamily protein | 36 | 0.62 | 0.287 | 1.000 |
| AT2G37230.1 | Pentatricopeptide repeat (PPR) superfamily protein | 37 | 0.82 | 0.286 | 1.000 |
| AT2G37230.1 | Pentatricopeptide repeat (PPR) superfamily protein | 38 | 1.04 | 0.286 | 1.000 |
| AT2G37230.1 | Pentatricopeptide repeat (PPR) superfamily protein | 39 | 1.19 | 0.287 | 1.000 |
| AT2G37230.1 | Pentatricopeptide repeat (PPR) superfamily protein | 40 | 1.21 | 0.288 | 1.000 |
| AT2G37230.1 | Pentatricopeptide repeat (PPR) superfamily protein | 41 | 1.08 | 0.291 | 1.000 |
| AT2G37230.1 | Pentatricopeptide repeat (PPR) superfamily protein | 42 | 0.89 | 0.296 | 1.000 |
| AT2G37230.1 | Pentatricopeptide repeat (PPR) superfamily protein | 43 | 0.73 | 0.316 | 1.000 |
| AT2G37230.1 | Pentatricopeptide repeat (PPR) superfamily protein | 44 | 0.66 | 0.371 | 1.000 |
| AT2G37230.1 | Pentatricopeptide repeat (PPR) superfamily protein | 45 | 0.64 | 0.472 | 1.000 |
| AT2G37230.1 | Pentatricopeptide repeat (PPR) superfamily protein | 46 | 0.63 | 0.639 | 1.000 |
| AT2G37230.1 | Pentatricopeptide repeat (PPR) superfamily protein | 47 | 0.64 | 0.872 | 1.000 |
| AT2G37410.2 | Translocase of the inner mitochondrial membrane 17 (TIM17-2) | 2 | 1.24 | 0.322 | 1.000 |
| AT2G37410.2 | Translocase of the inner mitochondrial membrane 17 (TIM17-2) | 3 | 1.08 | 0.225 | 1.000 |
| AT2G37410.2 | Translocase of the inner mitochondrial membrane 17 (TIM17-2) | 4 | 0.97 | 0.220 | 1.000 |
| AT2G37410.2 | Translocase of the inner mitochondrial membrane 17 (TIM17-2) | 5 | 0.92 | 0.215 | 1.000 |
| AT2G37410.2 | Translocase of the inner mitochondrial membrane 17 (TIM17-2) | 6 | 0.92 | 0.195 | 1.000 |
| AT2G37410.2 | Translocase of the inner mitochondrial membrane 17 (TIM17-2) | 7 | 0.97 | 0.181 | 1.000 |
| AT2G37410.2 | Translocase of the inner mitochondrial membrane 17 (TIM17-2) | 8 | 1.04 | 0.171 | 1.000 |
| AT2G37410.2 | Translocase of the inner mitochondrial membrane 17 (TIM17-2) | 9 | 1.13 | 0.166 | 1.000 |
| AT2G37410.2 | Translocase of the inner mitochondrial membrane 17 (TIM17-2) | 10 | 1.23 | 0.166 | 1.000 |
| AT2G37410.2 | Translocase of the inner mitochondrial membrane 17 (TIM17-2) | 11 | 1.41 | 0.166 | 1.000 |

| | | | | | |
|---|---|---|---|---|---|
| AT2G37410.2 | Translocase of the inner mitochondrial membrane 17 (TIM17-2) | 12 | 1.63 | 0.166 | 0.998 |
| AT2G37410.2 | Translocase of the inner mitochondrial membrane 17 (TIM17-2) | 13 | 1.77 | 0.164 | 0.978 |
| AT2G37410.2 | Translocase of the inner mitochondrial membrane 17 (TIM17-2) | 14 | 1.75 | 0.161 | 0.979 |
| AT2G37410.2 | Translocase of the inner mitochondrial membrane 17 (TIM17-2) | 15 | 1.54 | 0.160 | 0.999 |
| AT2G37410.2 | Translocase of the inner mitochondrial membrane 17 (TIM17-2) | 16 | 1.35 | 0.159 | 1.000 |
| AT2G37410.2 | Translocase of the inner mitochondrial membrane 17 (TIM17-2) | 17 | 1.27 | 0.159 | 1.000 |
| AT2G37410.2 | Translocase of the inner mitochondrial membrane 17 (TIM17-2) | 18 | 1.21 | 0.161 | 1.000 |
| AT2G37410.2 | Translocase of the inner mitochondrial membrane 17 (TIM17-2) | 19 | 1.22 | 0.166 | 1.000 |
| AT2G37410.2 | Translocase of the inner mitochondrial membrane 17 (TIM17-2) | 20 | 1.26 | 0.171 | 1.000 |
| AT2G37410.2 | Translocase of the inner mitochondrial membrane 17 (TIM17-2) | 21 | 1.46 | 0.174 | 1.000 |
| AT2G37410.2 | Translocase of the inner mitochondrial membrane 17 (TIM17-2) | 22 | 1.95 | 0.173 | 0.938 |
| AT2G37410.2 | Translocase of the inner mitochondrial membrane 17 (TIM17-2) | 23 | 2.28 | 0.172 | 0.679 |
| AT2G37410.2 | Translocase of the inner mitochondrial membrane 17 (TIM17-2) | 24 | 1.57 | 0.168 | 1.000 |
| AT2G37410.2 | Translocase of the inner mitochondrial membrane 17 (TIM17-2) | 25 | 0.69 | 0.165 | 1.000 |
| AT2G37410.2 | Translocase of the inner mitochondrial membrane 17 (TIM17-2) | 26 | 0.50 | 0.164 | 0.850 |
| AT2G37410.2 | Translocase of the inner mitochondrial membrane 17 (TIM17-2) | 27 | 0.74 | 0.164 | 1.000 |
| AT2G37410.2 | Translocase of the inner mitochondrial membrane 17 (TIM17-2) | 28 | 1.30 | 0.164 | 1.000 |
| AT2G37410.2 | Translocase of the inner mitochondrial membrane 17 (TIM17-2) | 29 | 1.77 | 0.164 | 0.979 |
| AT2G37410.2 | Translocase of the inner mitochondrial membrane 17 (TIM17-2) | 30 | 1.76 | 0.164 | 0.982 |
| AT2G37410.2 | Translocase of the inner mitochondrial membrane 17 (TIM17-2) | 31 | 1.75 | 0.164 | 0.983 |

| | | | | | |
|---|---|---|---|---|---|
| AT2G37410.2 | Translocase of the inner mitochondrial membrane 17 (TIM17-2) | 32 | 1.90 | 0.165 | 0.929 |
| AT2G37410.2 | Translocase of the inner mitochondrial membrane 17 (TIM17-2) | 33 | 2.24 | 0.165 | 0.653 |
| AT2G37410.2 | Translocase of the inner mitochondrial membrane 17 (TIM17-2) | 34 | 2.37 | 0.165 | 0.520 |
| AT2G37410.2 | Translocase of the inner mitochondrial membrane 17 (TIM17-2) | 35 | 2.18 | 0.165 | 0.705 |
| AT2G37410.2 | Translocase of the inner mitochondrial membrane 17 (TIM17-2) | 36 | 1.90 | 0.164 | 0.930 |
| AT2G37410.2 | Translocase of the inner mitochondrial membrane 17 (TIM17-2) | 37 | 1.57 | 0.164 | 0.999 |
| AT2G37410.2 | Translocase of the inner mitochondrial membrane 17 (TIM17-2) | 38 | 1.28 | 0.164 | 1.000 |
| AT2G37410.2 | Translocase of the inner mitochondrial membrane 17 (TIM17-2) | 39 | 1.05 | 0.165 | 1.000 |
| AT2G37410.2 | Translocase of the inner mitochondrial membrane 17 (TIM17-2) | 40 | 0.84 | 0.165 | 1.000 |
| AT2G37410.2 | Translocase of the inner mitochondrial membrane 17 (TIM17-2) | 41 | 0.72 | 0.166 | 1.000 |
| AT2G37410.2 | Translocase of the inner mitochondrial membrane 17 (TIM17-2) | 42 | 0.75 | 0.168 | 1.000 |
| AT2G37410.2 | Translocase of the inner mitochondrial membrane 17 (TIM17-2) | 43 | 0.82 | 0.169 | 1.000 |
| AT2G37410.2 | Translocase of the inner mitochondrial membrane 17 (TIM17-2) | 44 | 0.78 | 0.170 | 1.000 |
| AT2G37410.2 | Translocase of the inner mitochondrial membrane 17 (TIM17-2) | 45 | 0.61 | 0.171 | 0.998 |
| AT2G37410.2 | Translocase of the inner mitochondrial membrane 17 (TIM17-2) | 46 | 0.41 | 0.199 | 0.790 |
| AT2G37410.2 | Translocase of the inner mitochondrial membrane 17 (TIM17-2) | 47 | 0.27 | 0.339 | 0.939 |
| AT2G38280.2 | AMP deaminase (AMPD) | 2 | 0.88 | 0.662 | 1.000 |
| AT2G38280.2 | AMP deaminase (AMPD) | 3 | 0.87 | 0.593 | 1.000 |
| AT2G38280.2 | AMP deaminase (AMPD) | 14 | 1.73 | 0.326 | 1.000 |
| AT2G38280.2 | AMP deaminase (AMPD) | 15 | 2.05 | 0.281 | 0.979 |
| AT2G38280.2 | AMP deaminase (AMPD) | 16 | 2.37 | 0.249 | 0.843 |
| AT2G38280.2 | AMP deaminase (AMPD) | 17 | 2.66 | 0.230 | 0.598 |
| AT2G38280.2 | AMP deaminase (AMPD) | 18 | 2.96 | 0.221 | 0.387 |
| AT2G38280.2 | AMP deaminase (AMPD) | 19 | 3.27 | 0.220 | 0.259 |
| AT2G38280.2 | AMP deaminase (AMPD) | 20 | 3.50 | 0.221 | 0.203 |
| AT2G38280.2 | AMP deaminase (AMPD) | 21 | 3.49 | 0.223 | 0.209 |
| AT2G38280.2 | AMP deaminase (AMPD) | 22 | 3.16 | 0.223 | 0.317 |
| AT2G38280.2 | AMP deaminase (AMPD) | 23 | 2.60 | 0.221 | 0.577 |
| AT2G38280.2 | AMP deaminase (AMPD) | 24 | 2.04 | 0.219 | 0.889 |
| AT2G38280.2 | AMP deaminase (AMPD) | 25 | 1.56 | 0.216 | 0.997 |

| | | | | | |
|---|---|---|---|---|---|
| AT2G38280.2 | AMP deaminase (AMPD) | 26 | 1.25 | 0.215 | 1.000 |
| AT2G38280.2 | AMP deaminase (AMPD) | 27 | 1.07 | 0.214 | 1.000 |
| AT2G38280.2 | AMP deaminase (AMPD) | 28 | 0.98 | 0.214 | 1.000 |
| AT2G38280.2 | AMP deaminase (AMPD) | 29 | 0.93 | 0.214 | 1.000 |
| AT2G38280.2 | AMP deaminase (AMPD) | 30 | 0.89 | 0.215 | 1.000 |
| AT2G38280.2 | AMP deaminase (AMPD) | 31 | 0.86 | 0.215 | 1.000 |
| AT2G38280.2 | AMP deaminase (AMPD) | 32 | 0.83 | 0.216 | 1.000 |
| AT2G38280.2 | AMP deaminase (AMPD) | 33 | 0.78 | 0.216 | 1.000 |
| AT2G38280.2 | AMP deaminase (AMPD) | 34 | 0.75 | 0.217 | 1.000 |
| AT2G38280.2 | AMP deaminase (AMPD) | 35 | 0.76 | 0.217 | 1.000 |
| AT2G38280.2 | AMP deaminase (AMPD) | 36 | 0.81 | 0.217 | 1.000 |
| AT2G38280.2 | AMP deaminase (AMPD) | 37 | 0.94 | 0.217 | 1.000 |
| AT2G38280.2 | AMP deaminase (AMPD) | 38 | 1.16 | 0.218 | 1.000 |
| AT2G38280.2 | AMP deaminase (AMPD) | 39 | 1.44 | 0.221 | 1.000 |
| AT2G38280.2 | AMP deaminase (AMPD) | 40 | 1.67 | 0.233 | 0.994 |
| AT2G38280.2 | AMP deaminase (AMPD) | 41 | 1.82 | 0.263 | 0.993 |
| AT2G38280.2 | AMP deaminase (AMPD) | 42 | 1.91 | 0.321 | 0.998 |
| AT2G38280.2 | AMP deaminase (AMPD) | 43 | 2.02 | 0.399 | 0.999 |
| AT2G38280.2 | AMP deaminase (AMPD) | 46 | 2.49 | 0.780 | 1.000 |
| AT2G38280.2 | AMP deaminase (AMPD) | 47 | 2.69 | 0.957 | 1.000 |
| AT2G38400.1 | Alanine:glyoxylate aminotransferase 3 (AGT3) | 2 | 1.06 | 0.738 | 1.000 |
| AT2G38400.1 | Alanine:glyoxylate aminotransferase 3 (AGT3) | 3 | 0.73 | 0.730 | 1.000 |
| AT2G38400.1 | Alanine:glyoxylate aminotransferase 3 (AGT3) | 33 | 0.42 | 0.423 | 0.943 |
| AT2G38400.1 | Alanine:glyoxylate aminotransferase 3 (AGT3) | 34 | 0.77 | 0.326 | 1.000 |
| AT2G38400.1 | Alanine:glyoxylate aminotransferase 3 (AGT3) | 35 | 1.45 | 0.277 | 0.995 |
| AT2G38400.1 | Alanine:glyoxylate aminotransferase 3 (AGT3) | 36 | 2.43 | 0.267 | 0.650 |
| AT2G38400.1 | Alanine:glyoxylate aminotransferase 3 (AGT3) | 37 | 3.27 | 0.286 | 0.405 |
| AT2G38400.1 | Alanine:glyoxylate aminotransferase 3 (AGT3) | 38 | 3.49 | 0.330 | 0.508 |
| AT2G38400.1 | Alanine:glyoxylate aminotransferase 3 (AGT3) | 39 | 3.20 | 0.387 | 0.746 |
| AT2G38400.1 | Alanine:glyoxylate aminotransferase 3 (AGT3) | 44 | 0.83 | 0.377 | 1.000 |
| AT2G38400.1 | Alanine:glyoxylate aminotransferase 3 (AGT3) | 45 | 0.63 | 0.343 | 0.994 |
| AT2G38400.1 | Alanine:glyoxylate aminotransferase 3 (AGT3) | 46 | 0.50 | 0.367 | 0.962 |
| AT2G38400.1 | Alanine:glyoxylate aminotransferase 3 (AGT3) | 47 | 0.40 | 0.487 | 0.965 |
| AT2G38660.4 | Amino acid dehydrogenase family protein | 2 | 0.88 | 0.742 | 1.000 |
| AT2G38660.4 | Amino acid dehydrogenase family protein | 3 | 0.86 | 0.709 | 1.000 |
| AT2G38660.4 | Amino acid dehydrogenase family protein | 31 | 0.71 | 0.443 | 1.000 |
| AT2G38660.4 | Amino acid dehydrogenase family protein | 32 | 0.73 | 0.376 | 1.000 |
| AT2G38660.4 | Amino acid dehydrogenase family protein | 33 | 0.82 | 0.335 | 1.000 |
| AT2G38660.4 | Amino acid dehydrogenase family protein | 34 | 0.97 | 0.314 | 1.000 |
| AT2G38660.4 | Amino acid dehydrogenase family protein | 35 | 1.21 | 0.305 | 1.000 |
| AT2G38660.4 | Amino acid dehydrogenase family protein | 36 | 1.56 | 0.302 | 0.992 |

| | | | | | |
|---|---|---|---|---|---|
| AT2G38660.4 | Amino acid dehydrogenase family protein | 37 | 2.03 | 0.301 | 0.900 |
| AT2G38660.4 | Amino acid dehydrogenase family protein | 38 | 2.59 | 0.301 | 0.691 |
| AT2G38660.4 | Amino acid dehydrogenase family protein | 39 | 3.08 | 0.301 | 0.514 |
| AT2G38660.4 | Amino acid dehydrogenase family protein | 40 | 3.29 | 0.302 | 0.450 |
| AT2G38660.4 | Amino acid dehydrogenase family protein | 41 | 3.27 | 0.303 | 0.463 |
| AT2G38660.4 | Amino acid dehydrogenase family protein | 42 | 3.10 | 0.304 | 0.518 |
| AT2G38660.4 | Amino acid dehydrogenase family protein | 43 | 2.75 | 0.305 | 0.643 |
| AT2G38660.4 | Amino acid dehydrogenase family protein | 44 | 2.12 | 0.309 | 0.882 |
| AT2G38660.4 | Amino acid dehydrogenase family protein | 45 | 1.41 | 0.328 | 0.999 |
| AT2G38660.4 | Amino acid dehydrogenase family protein | 46 | 0.88 | 0.386 | 1.000 |
| AT2G38660.4 | Amino acid dehydrogenase family protein | 47 | 0.54 | 0.501 | 0.998 |
| AT2G38670.1 | Phosphorylethanolamine cytidylyltransferase (PECT1) | 2 | 1.19 | 0.218 | 1.000 |
| AT2G38670.1 | Phosphorylethanolamine cytidylyltransferase (PECT1) | 3 | 1.17 | 0.168 | 1.000 |
| AT2G38670.1 | Phosphorylethanolamine cytidylyltransferase (PECT1) | 4 | 1.14 | 0.146 | 1.000 |
| AT2G38670.1 | Phosphorylethanolamine cytidylyltransferase (PECT1) | 5 | 1.09 | 0.141 | 1.000 |
| AT2G38670.1 | Phosphorylethanolamine cytidylyltransferase (PECT1) | 6 | 1.02 | 0.139 | 1.000 |
| AT2G38670.1 | Phosphorylethanolamine cytidylyltransferase (PECT1) | 7 | 0.97 | 0.139 | 1.000 |
| AT2G38670.1 | Phosphorylethanolamine cytidylyltransferase (PECT1) | 8 | 0.95 | 0.138 | 1.000 |
| AT2G38670.1 | Phosphorylethanolamine cytidylyltransferase (PECT1) | 9 | 0.97 | 0.137 | 1.000 |
| AT2G38670.1 | Phosphorylethanolamine cytidylyltransferase (PECT1) | 10 | 1.06 | 0.136 | 1.000 |
| AT2G38670.1 | Phosphorylethanolamine cytidylyltransferase (PECT1) | 11 | 1.20 | 0.135 | 1.000 |
| AT2G38670.1 | Phosphorylethanolamine cytidylyltransferase (PECT1) | 12 | 1.39 | 0.133 | 0.986 |
| AT2G38670.1 | Phosphorylethanolamine cytidylyltransferase (PECT1) | 13 | 1.57 | 0.131 | 0.827 |
| AT2G38670.1 | Phosphorylethanolamine cytidylyltransferase (PECT1) | 14 | 1.72 | 0.129 | 0.590 |
| AT2G38670.1 | Phosphorylethanolamine cytidylyltransferase (PECT1) | 15 | 1.85 | 0.128 | 0.396 |
| AT2G38670.1 | Phosphorylethanolamine cytidylyltransferase (PECT1) | 16 | 1.97 | 0.128 | 0.268 |
| AT2G38670.1 | Phosphorylethanolamine cytidylyltransferase (PECT1) | 17 | 2.08 | 0.128 | 0.183 |
| AT2G38670.1 | Phosphorylethanolamine cytidylyltransferase (PECT1) | 18 | 2.13 | 0.129 | 0.160 |
| AT2G38670.1 | Phosphorylethanolamine cytidylyltransferase (PECT1) | 19 | 2.08 | 0.131 | 0.205 |
| AT2G38670.1 | Phosphorylethanolamine cytidylyltransferase (PECT1) | 20 | 1.90 | 0.132 | 0.378 |
| AT2G38670.1 | Phosphorylethanolamine cytidylyltransferase (PECT1) | 21 | 1.66 | 0.133 | 0.724 |

| | | | | | |
|---|---|---|---|---|---|
| AT2G38670.1 | Phosphorylethanolamine cytidylyltransferase (PECT1) | 22 | 1.41 | 0.133 | 0.979 |
| AT2G38670.1 | Phosphorylethanolamine cytidylyltransferase (PECT1) | 23 | 1.21 | 0.133 | 1.000 |
| AT2G38670.1 | Phosphorylethanolamine cytidylyltransferase (PECT1) | 24 | 1.08 | 0.132 | 1.000 |
| AT2G38670.1 | Phosphorylethanolamine cytidylyltransferase (PECT1) | 25 | 1.01 | 0.130 | 1.000 |
| AT2G38670.1 | Phosphorylethanolamine cytidylyltransferase (PECT1) | 26 | 1.00 | 0.130 | 1.000 |
| AT2G38670.1 | Phosphorylethanolamine cytidylyltransferase (PECT1) | 27 | 1.03 | 0.130 | 1.000 |
| AT2G38670.1 | Phosphorylethanolamine cytidylyltransferase (PECT1) | 28 | 1.03 | 0.130 | 1.000 |
| AT2G38670.1 | Phosphorylethanolamine cytidylyltransferase (PECT1) | 29 | 0.99 | 0.130 | 1.000 |
| AT2G38670.1 | Phosphorylethanolamine cytidylyltransferase (PECT1) | 30 | 0.92 | 0.130 | 1.000 |
| AT2G38670.1 | Phosphorylethanolamine cytidylyltransferase (PECT1) | 31 | 0.87 | 0.130 | 1.000 |
| AT2G38670.1 | Phosphorylethanolamine cytidylyltransferase (PECT1) | 32 | 0.86 | 0.130 | 1.000 |
| AT2G38670.1 | Phosphorylethanolamine cytidylyltransferase (PECT1) | 33 | 0.87 | 0.130 | 1.000 |
| AT2G38670.1 | Phosphorylethanolamine cytidylyltransferase (PECT1) | 34 | 0.89 | 0.130 | 1.000 |
| AT2G38670.1 | Phosphorylethanolamine cytidylyltransferase (PECT1) | 35 | 0.91 | 0.130 | 1.000 |
| AT2G38670.1 | Phosphorylethanolamine cytidylyltransferase (PECT1) | 36 | 0.94 | 0.130 | 1.000 |
| AT2G38670.1 | Phosphorylethanolamine cytidylyltransferase (PECT1) | 37 | 0.99 | 0.130 | 1.000 |
| AT2G38670.1 | Phosphorylethanolamine cytidylyltransferase (PECT1) | 38 | 1.03 | 0.130 | 1.000 |
| AT2G38670.1 | Phosphorylethanolamine cytidylyltransferase (PECT1) | 39 | 1.09 | 0.130 | 1.000 |
| AT2G38670.1 | Phosphorylethanolamine cytidylyltransferase (PECT1) | 40 | 1.16 | 0.131 | 1.000 |
| AT2G38670.1 | Phosphorylethanolamine cytidylyltransferase (PECT1) | 41 | 1.21 | 0.132 | 1.000 |
| AT2G38670.1 | Phosphorylethanolamine cytidylyltransferase (PECT1) | 42 | 1.23 | 0.132 | 1.000 |
| AT2G38670.1 | Phosphorylethanolamine cytidylyltransferase (PECT1) | 43 | 1.18 | 0.133 | 1.000 |
| AT2G38670.1 | Phosphorylethanolamine cytidylyltransferase (PECT1) | 44 | 1.03 | 0.133 | 1.000 |
| AT2G38670.1 | Phosphorylethanolamine cytidylyltransferase (PECT1) | 45 | 0.81 | 0.138 | 1.000 |
| AT2G38670.1 | Phosphorylethanolamine cytidylyltransferase (PECT1) | 46 | 0.57 | 0.161 | 0.808 |
| AT2G38670.1 | Phosphorylethanolamine cytidylyltransferase (PECT1) | 47 | 0.37 | 0.217 | 0.468 |
| AT2G39470.2 | NAD(P)H dehydrogenase complex subunit (PNSL1) | 2 | 1.02 | 0.430 | 1.000 |
| AT2G39470.2 | NAD(P)H dehydrogenase complex subunit (PNSL1) | 3 | 0.97 | 0.328 | 1.000 |
| AT2G39470.2 | NAD(P)H dehydrogenase complex subunit (PNSL1) | 4 | 0.92 | 0.268 | 1.000 |
| AT2G39470.2 | NAD(P)H dehydrogenase complex subunit (PNSL1) | 5 | 0.86 | 0.240 | 1.000 |
| AT2G39470.2 | NAD(P)H dehydrogenase complex subunit (PNSL1) | 6 | 0.78 | 0.223 | 1.000 |

| | | | | | |
|---|---|---|---|---|---|
| AT2G39470.2 | NAD(P)H dehydrogenase complex subunit (PNSL1) | 7 | 0.70 | 0.212 | 0.995 |
| AT2G39470.2 | NAD(P)H dehydrogenase complex subunit (PNSL1) | 8 | 0.62 | 0.203 | 0.960 |
| AT2G39470.2 | NAD(P)H dehydrogenase complex subunit (PNSL1) | 9 | 0.56 | 0.196 | 0.864 |
| AT2G39470.2 | NAD(P)H dehydrogenase complex subunit (PNSL1) | 10 | 0.52 | 0.191 | 0.743 |
| AT2G39470.2 | NAD(P)H dehydrogenase complex subunit (PNSL1) | 11 | 0.50 | 0.189 | 0.645 |
| AT2G39470.2 | NAD(P)H dehydrogenase complex subunit (PNSL1) | 12 | 0.48 | 0.187 | 0.568 |
| AT2G39470.2 | NAD(P)H dehydrogenase complex subunit (PNSL1) | 13 | 0.46 | 0.187 | 0.502 |
| AT2G39470.2 | NAD(P)H dehydrogenase complex subunit (PNSL1) | 14 | 0.44 | 0.186 | 0.444 |
| AT2G39470.2 | NAD(P)H dehydrogenase complex subunit (PNSL1) | 15 | 0.43 | 0.186 | 1.000 |
| AT2G39470.2 | NAD(P)H dehydrogenase complex subunit (PNSL1) | 16 | 0.42 | 0.186 | 1.000 |
| AT2G39470.2 | NAD(P)H dehydrogenase complex subunit (PNSL1) | 17 | 0.45 | 0.189 | 0.491 |
| AT2G39470.2 | NAD(P)H dehydrogenase complex subunit (PNSL1) | 18 | 0.58 | 0.197 | 0.906 |
| AT2G39470.2 | NAD(P)H dehydrogenase complex subunit (PNSL1) | 19 | 0.92 | 0.214 | 1.000 |
| AT2G39470.2 | NAD(P)H dehydrogenase complex subunit (PNSL1) | 20 | 1.68 | 0.242 | 0.977 |
| AT2G39470.2 | NAD(P)H dehydrogenase complex subunit (PNSL1) | 21 | 3.27 | 0.290 | 0.533 |
| AT2G39470.2 | NAD(P)H dehydrogenase complex subunit (PNSL1) | 22 | 6.46 | 0.371 | 1.000 |
| AT2G39470.2 | NAD(P)H dehydrogenase complex subunit (PNSL1) | 46 | ########### | 7.463 | 0.941 |
| AT2G39470.2 | NAD(P)H dehydrogenase complex subunit (PNSL1) | 47 | ########### | 7.895 | 0.947 |
| AT2G39730.1 | Rubisco activase (RCA) | 3 | 0.13 | 0.450 | 0.297 |
| AT2G39730.1 | Rubisco activase (RCA) | 4 | 0.11 | 0.417 | 0.175 |
| AT2G39730.1 | Rubisco activase (RCA) | 5 | 0.11 | 0.407 | 0.138 |
| AT2G39730.1 | Rubisco activase (RCA) | 6 | 0.11 | 0.404 | 0.142 |
| AT2G39730.1 | Rubisco activase (RCA) | 7 | 0.12 | 0.403 | 0.161 |
| AT2G39730.1 | Rubisco activase (RCA) | 8 | 0.13 | 0.402 | 0.193 |
| AT2G39730.1 | Rubisco activase (RCA) | 9 | 0.14 | 0.401 | 0.238 |
| AT2G39730.1 | Rubisco activase (RCA) | 10 | 0.15 | 0.400 | 0.275 |
| AT2G39730.1 | Rubisco activase (RCA) | 11 | 0.16 | 0.399 | 0.289 |
| AT2G39730.1 | Rubisco activase (RCA) | 12 | 0.15 | 0.398 | 0.271 |
| AT2G39730.1 | Rubisco activase (RCA) | 13 | 0.14 | 0.397 | 0.228 |
| AT2G39730.1 | Rubisco activase (RCA) | 14 | 0.13 | 0.397 | 0.180 |
| AT2G39730.1 | Rubisco activase (RCA) | 15 | 0.11 | 0.396 | 0.142 |
| AT2G39730.1 | Rubisco activase (RCA) | 16 | 0.11 | 0.395 | 0.119 |
| AT2G39730.1 | Rubisco activase (RCA) | 17 | 0.11 | 0.395 | 0.119 |
| AT2G39730.1 | Rubisco activase (RCA) | 18 | 0.12 | 0.396 | 0.148 |
| AT2G39730.1 | Rubisco activase (RCA) | 19 | 0.14 | 0.398 | 0.214 |
| AT2G39730.1 | Rubisco activase (RCA) | 20 | 0.17 | 0.400 | 0.319 |
| AT2G39730.1 | Rubisco activase (RCA) | 21 | 0.20 | 0.401 | 0.449 |
| AT2G39730.1 | Rubisco activase (RCA) | 22 | 0.24 | 0.401 | 0.584 |
| AT2G39730.1 | Rubisco activase (RCA) | 23 | 0.29 | 0.401 | 0.753 |
| AT2G39730.1 | Rubisco activase (RCA) | 24 | 0.38 | 0.400 | 0.927 |
| AT2G39730.1 | Rubisco activase (RCA) | 25 | 0.50 | 0.399 | 0.994 |
| AT2G39730.1 | Rubisco activase (RCA) | 26 | 0.58 | 0.399 | 1.000 |
| AT2G39730.1 | Rubisco activase (RCA) | 27 | 0.59 | 0.399 | 1.000 |
| AT2G39730.1 | Rubisco activase (RCA) | 28 | 0.56 | 0.399 | 0.999 |

| | | | | | | |
|---|---|---|---|---|---|---|
| AT2G39730.1 | Rubisco activase (RCA) | 29 | 0.54 | 0.398 | 0.998 | |
| AT2G39730.1 | Rubisco activase (RCA) | 30 | 0.54 | 0.398 | 0.998 | |
| AT2G39730.1 | Rubisco activase (RCA) | 31 | 0.61 | 0.398 | 1.000 | |
| AT2G39730.1 | Rubisco activase (RCA) | 32 | 0.82 | 0.398 | 1.000 | |
| AT2G39730.1 | Rubisco activase (RCA) | 33 | 1.27 | 0.398 | 1.000 | |
| AT2G39730.1 | Rubisco activase (RCA) | 34 | 2.31 | 0.398 | 0.970 | |
| AT2G39730.1 | Rubisco activase (RCA) | 35 | 4.63 | 0.398 | 0.497 | |
| AT2G39730.1 | Rubisco activase (RCA) | 36 | 8.87 | 0.398 | 0.139 | |
| AT2G39730.1 | Rubisco activase (RCA) | 37 | 14.15 | 0.398 | 0.042 | * |
| AT2G39730.1 | Rubisco activase (RCA) | 38 | 17.19 | 0.398 | 0.023 | * |
| AT2G39730.1 | Rubisco activase (RCA) | 39 | 15.21 | 0.399 | 0.035 | * |
| AT2G39730.1 | Rubisco activase (RCA) | 40 | 10.08 | 0.399 | 0.104 | |
| AT2G39730.1 | Rubisco activase (RCA) | 41 | 5.69 | 0.401 | 0.358 | |
| AT2G39730.1 | Rubisco activase (RCA) | 42 | 3.25 | 0.402 | 0.793 | |
| AT2G39730.1 | Rubisco activase (RCA) | 43 | 2.03 | 0.403 | 0.993 | |
| AT2G39730.1 | Rubisco activase (RCA) | 44 | 1.41 | 0.406 | 1.000 | |
| AT2G39730.1 | Rubisco activase (RCA) | 45 | 1.08 | 0.422 | 1.000 | |
| AT2G39730.1 | Rubisco activase (RCA) | 46 | 0.87 | 0.471 | 1.000 | |
| AT2G39730.1 | Rubisco activase (RCA) | 47 | 0.71 | 0.579 | 1.000 | |
| AT2G39730.1 | Rubisco activase (RCA) | 48 | 0.57 | 0.735 | 1.000 | |
| AT2G39970.1 | Peroxisomal NAD carrier (PXN, PMP38) | 2 | 0.74 | 0.453 | 0.999 | |
| AT2G39970.1 | Peroxisomal NAD carrier (PXN, PMP38) | 3 | 1.14 | 0.396 | 1.000 | |
| AT2G39970.1 | Peroxisomal NAD carrier (PXN, PMP38) | 4 | 1.78 | 0.505 | 0.990 | |
| AT2G39970.1 | Peroxisomal NAD carrier (PXN, PMP38) | 42 | 8.16 | 0.412 | 0.157 | |
| AT2G39970.1 | Peroxisomal NAD carrier (PXN, PMP38) | 43 | 3.11 | 0.301 | 0.429 | |
| AT2G39970.1 | Peroxisomal NAD carrier (PXN, PMP38) | 44 | 1.05 | 0.227 | 1.000 | |
| AT2G39970.1 | Peroxisomal NAD carrier (PXN, PMP38) | 45 | 0.34 | 0.215 | 0.158 | |
| AT2G39970.1 | Peroxisomal NAD carrier (PXN, PMP38) | 46 | 0.11 | 0.281 | 0.008 | ** |
| AT2G39970.1 | Peroxisomal NAD carrier (PXN, PMP38) | 47 | 0.04 | 0.407 | 0.005 | ** |
| AT2G40090.1 | ABC-2 homolog 9 (ATH9) | 2 | 0.99 | 0.451 | 1.000 | |
| AT2G40090.1 | ABC-2 homolog 9 (ATH9) | 3 | 0.97 | 0.395 | 1.000 | |
| AT2G40090.1 | ABC-2 homolog 9 (ATH9) | 12 | 1.07 | 0.353 | 1.000 | |
| AT2G40090.1 | ABC-2 homolog 9 (ATH9) | 13 | 1.11 | 0.311 | 1.000 | |
| AT2G40090.1 | ABC-2 homolog 9 (ATH9) | 14 | 1.09 | 0.300 | 1.000 | |
| AT2G40090.1 | ABC-2 homolog 9 (ATH9) | 16 | 0.87 | 0.260 | 1.000 | |
| AT2G40090.1 | ABC-2 homolog 9 (ATH9) | 17 | 0.81 | 0.260 | 1.000 | |
| AT2G40090.1 | ABC-2 homolog 9 (ATH9) | 18 | 0.85 | 0.252 | 1.000 | |
| AT2G40090.1 | ABC-2 homolog 9 (ATH9) | 19 | 1.00 | 0.251 | 1.000 | |
| AT2G40090.1 | ABC-2 homolog 9 (ATH9) | 20 | 1.24 | 0.234 | 1.000 | |
| AT2G40090.1 | ABC-2 homolog 9 (ATH9) | 21 | 1.41 | 0.231 | 1.000 | |
| AT2G40090.1 | ABC-2 homolog 9 (ATH9) | 22 | 1.49 | 0.240 | 1.000 | |
| AT2G40090.1 | ABC-2 homolog 9 (ATH9) | 23 | 1.66 | 0.235 | 1.000 | |
| AT2G40090.1 | ABC-2 homolog 9 (ATH9) | 24 | 1.89 | 0.225 | 0.993 | |
| AT2G40090.1 | ABC-2 homolog 9 (ATH9) | 25 | 2.15 | 0.210 | 0.918 | |
| AT2G40090.1 | ABC-2 homolog 9 (ATH9) | 26 | 2.57 | 0.200 | 0.610 | |
| AT2G40090.1 | ABC-2 homolog 9 (ATH9) | 27 | 2.96 | 0.193 | 0.311 | |
| AT2G40090.1 | ABC-2 homolog 9 (ATH9) | 28 | 2.77 | 0.186 | 0.350 | |
| AT2G40090.1 | ABC-2 homolog 9 (ATH9) | 29 | 2.19 | 0.179 | 0.732 | |
| AT2G40090.1 | ABC-2 homolog 9 (ATH9) | 30 | 1.67 | 0.178 | 0.991 | |
| AT2G40090.1 | ABC-2 homolog 9 (ATH9) | 31 | 1.40 | 0.178 | 1.000 | |
| AT2G40090.1 | ABC-2 homolog 9 (ATH9) | 32 | 1.23 | 0.178 | 1.000 | |
| AT2G40090.1 | ABC-2 homolog 9 (ATH9) | 33 | 1.06 | 0.178 | 1.000 | |
| AT2G40090.1 | ABC-2 homolog 9 (ATH9) | 34 | 0.89 | 0.179 | 1.000 | |
| AT2G40090.1 | ABC-2 homolog 9 (ATH9) | 35 | 0.82 | 0.178 | 1.000 | |

| | | | | | | |
|---|---|---|---|---|---|---|
| AT2G40090.1 | ABC-2 homolog 9 (ATH9) | 36 | 0.86 | 0.178 | 1.000 | |
| AT2G40090.1 | ABC-2 homolog 9 (ATH9) | 37 | 1.04 | 0.178 | 1.000 | |
| AT2G40090.1 | ABC-2 homolog 9 (ATH9) | 38 | 1.42 | 0.177 | 1.000 | |
| AT2G40090.1 | ABC-2 homolog 9 (ATH9) | 39 | 1.88 | 0.177 | 0.934 | |
| AT2G40090.1 | ABC-2 homolog 9 (ATH9) | 40 | 2.10 | 0.178 | 0.791 | |
| AT2G40090.1 | ABC-2 homolog 9 (ATH9) | 41 | 1.80 | 0.178 | 0.966 | |
| AT2G40090.1 | ABC-2 homolog 9 (ATH9) | 42 | 1.38 | 0.181 | 1.000 | |
| AT2G40090.1 | ABC-2 homolog 9 (ATH9) | 43 | 1.00 | 0.184 | 1.000 | |
| AT2G40090.1 | ABC-2 homolog 9 (ATH9) | 44 | 0.62 | 0.183 | 0.997 | |
| AT2G40090.1 | ABC-2 homolog 9 (ATH9) | 45 | 0.31 | 0.184 | 0.132 | |
| AT2G40090.1 | ABC-2 homolog 9 (ATH9) | 46 | 0.12 | 0.225 | 0.002 | ** |
| AT2G40090.1 | ABC-2 homolog 9 (ATH9) | 47 | 0.05 | 0.388 | 0.020 | * |
| AT2G40600.1 | ? | 2 | 0.95 | 0.400 | 1.000 | |
| AT2G40600.1 | ? | 3 | 0.93 | 0.354 | 1.000 | |
| AT2G40600.1 | ? | 4 | 0.90 | 0.355 | 1.000 | |
| AT2G40600.1 | ? | 25 | 0.67 | 0.301 | 0.993 | |
| AT2G40600.1 | ? | 26 | 0.70 | 0.266 | 0.993 | |
| AT2G40600.1 | ? | 27 | 0.75 | 0.248 | 0.997 | |
| AT2G40600.1 | ? | 28 | 0.83 | 0.248 | 1.000 | |
| AT2G40600.1 | ? | 29 | 0.94 | 0.265 | 1.000 | |
| AT2G40600.1 | ? | 42 | 4.19 | 0.282 | 0.186 | |
| AT2G40600.1 | ? | 43 | 3.56 | 0.240 | 0.152 | |
| AT2G40600.1 | ? | 44 | 2.81 | 0.210 | 0.215 | |
| AT2G40600.1 | ? | 45 | 2.08 | 0.210 | 0.580 | |
| AT2G40600.1 | ? | 46 | 1.49 | 0.250 | 0.982 | |
| AT2G40600.1 | ? | 47 | 1.05 | 0.326 | 1.000 | |
| AT2G40765.1 | Ubiquinol:cytochrome c oxidoreductase subunit (QCR10) | 2 | 0.92 | 0.540 | 1.000 | |
| AT2G40765.1 | Ubiquinol:cytochrome c oxidoreductase subunit (QCR10) | 3 | 0.91 | 0.417 | 1.000 | |
| AT2G40765.1 | Ubiquinol:cytochrome c oxidoreductase subunit (QCR10) | 6 | 0.90 | 0.397 | 1.000 | |
| AT2G40765.1 | Ubiquinol:cytochrome c oxidoreductase subunit (QCR10) | 7 | 0.93 | 0.298 | 1.000 | |
| AT2G40765.1 | Ubiquinol:cytochrome c oxidoreductase subunit (QCR10) | 8 | 1.08 | 0.260 | 1.000 | |
| AT2G40765.1 | Ubiquinol:cytochrome c oxidoreductase subunit (QCR10) | 9 | 1.39 | 0.246 | 1.000 | |
| AT2G40765.1 | Ubiquinol:cytochrome c oxidoreductase subunit (QCR10) | 10 | 1.95 | 0.243 | 0.998 | |
| AT2G40765.1 | Ubiquinol:cytochrome c oxidoreductase subunit (QCR10) | 11 | 2.96 | 0.245 | 0.765 | |
| AT2G40765.1 | Ubiquinol:cytochrome c oxidoreductase subunit (QCR10) | 12 | 4.23 | 0.246 | 0.272 | |
| AT2G40765.1 | Ubiquinol:cytochrome c oxidoreductase subunit (QCR10) | 13 | 5.23 | 0.243 | 0.094 | . |
| AT2G40765.1 | Ubiquinol:cytochrome c oxidoreductase subunit (QCR10) | 14 | 5.12 | 0.238 | 0.088 | . |
| AT2G40765.1 | Ubiquinol:cytochrome c oxidoreductase subunit (QCR10) | 15 | 3.77 | 0.237 | 0.350 | |
| AT2G40765.1 | Ubiquinol:cytochrome c oxidoreductase subunit (QCR10) | 16 | 2.14 | 0.234 | 0.984 | |
| AT2G40765.1 | Ubiquinol:cytochrome c oxidoreductase subunit (QCR10) | 17 | 1.05 | 0.234 | 1.000 | |
| AT2G40765.1 | Ubiquinol:cytochrome c oxidoreductase subunit (QCR10) | 18 | 0.69 | 0.238 | 1.000 | |
| AT2G40765.1 | Ubiquinol:cytochrome c oxidoreductase subunit (QCR10) | 19 | 0.72 | 0.245 | 1.000 | |
| AT2G40765.1 | Ubiquinol:cytochrome c oxidoreductase subunit (QCR10) | 20 | 0.93 | 0.252 | 1.000 | |
| AT2G40765.1 | Ubiquinol:cytochrome c oxidoreductase subunit (QCR10) | 21 | 1.31 | 0.254 | 1.000 | |

| | | | | | |
|---|---|---|---|---|---|
| AT2G40765.1 | Ubiquinol:cytochrome c oxidoreductase subunit (QCR10) | 22 | 1.25 | 0.254 | 1.000 |
| AT2G40765.1 | Ubiquinol:cytochrome c oxidoreductase subunit (QCR10) | 23 | 0.53 | 0.254 | 1.000 |
| AT2G40765.1 | Ubiquinol:cytochrome c oxidoreductase subunit (QCR10) | 24 | 0.27 | 0.245 | 0.428 |
| AT2G40765.1 | Ubiquinol:cytochrome c oxidoreductase subunit (QCR10) | 25 | 0.39 | 0.240 | 0.910 |
| AT2G40765.1 | Ubiquinol:cytochrome c oxidoreductase subunit (QCR10) | 26 | 1.28 | 0.242 | 1.000 |
| AT2G40765.1 | Ubiquinol:cytochrome c oxidoreductase subunit (QCR10) | 27 | 3.63 | 0.244 | 0.453 |
| AT2G40765.1 | Ubiquinol:cytochrome c oxidoreductase subunit (QCR10) | 28 | 2.10 | 0.244 | 0.993 |
| AT2G40765.1 | Ubiquinol:cytochrome c oxidoreductase subunit (QCR10) | 29 | 0.40 | 0.242 | 0.931 |
| AT2G40765.1 | Ubiquinol:cytochrome c oxidoreductase subunit (QCR10) | 30 | 0.23 | 0.241 | 0.227 |
| AT2G40765.1 | Ubiquinol:cytochrome c oxidoreductase subunit (QCR10) | 31 | 0.46 | 0.242 | 0.986 |
| AT2G40765.1 | Ubiquinol:cytochrome c oxidoreductase subunit (QCR10) | 32 | 1.20 | 0.244 | 1.000 |
| AT2G40765.1 | Ubiquinol:cytochrome c oxidoreductase subunit (QCR10) | 33 | 1.38 | 0.251 | 1.000 |
| AT2G40765.1 | Ubiquinol:cytochrome c oxidoreductase subunit (QCR10) | 34 | 1.02 | 0.266 | 1.000 |
| AT2G40765.1 | Ubiquinol:cytochrome c oxidoreductase subunit (QCR10) | 35 | 1.39 | 0.316 | 1.000 |
| AT2G40765.1 | Ubiquinol:cytochrome c oxidoreductase subunit (QCR10) | 36 | 2.68 | 0.555 | 1.000 |
| AT2G40765.1 | Ubiquinol:cytochrome c oxidoreductase subunit (QCR10) | 46 | 156.14 | 5.218 | 1.000 |
| AT2G40765.1 | Ubiquinol:cytochrome c oxidoreductase subunit (QCR10) | 47 | 205.17 | 5.939 | 1.000 |
| AT2G40800.1 | Translocase of the inner mitochondrial membrane 21-like (TIM21-L1) | 2 | 0.96 | 0.455 | 1.000 |
| AT2G40800.1 | Translocase of the inner mitochondrial membrane 21-like (TIM21-L1) | 3 | 0.84 | 0.418 | 1.000 |
| AT2G40800.1 | Translocase of the inner mitochondrial membrane 21-like (TIM21-L1) | 19 | 0.84 | 0.247 | 1.000 |
| AT2G40800.1 | Translocase of the inner mitochondrial membrane 21-like (TIM21-L1) | 20 | 1.00 | 0.207 | 1.000 |
| AT2G40800.1 | Translocase of the inner mitochondrial membrane 21-like (TIM21-L1) | 21 | 1.14 | 0.181 | 1.000 |
| AT2G40800.1 | Translocase of the inner mitochondrial membrane 21-like (TIM21-L1) | 22 | 1.20 | 0.167 | 1.000 |
| AT2G40800.1 | Translocase of the inner mitochondrial membrane 21-like (TIM21-L1) | 23 | 1.19 | 0.161 | 1.000 |
| AT2G40800.1 | Translocase of the inner mitochondrial membrane 21-like (TIM21-L1) | 24 | 1.13 | 0.157 | 1.000 |
| AT2G40800.1 | Translocase of the inner mitochondrial membrane 21-like (TIM21-L1) | 25 | 1.07 | 0.154 | 1.000 |

| | | | | | |
|---|---|---|---|---|---|
| AT2G40800.1 | Translocase of the inner mitochondrial membrane 21-like (TIM21-L1) | 26 | 1.04 | 0.153 | 1.000 |
| AT2G40800.1 | Translocase of the inner mitochondrial membrane 21-like (TIM21-L1) | 27 | 1.05 | 0.152 | 1.000 |
| AT2G40800.1 | Translocase of the inner mitochondrial membrane 21-like (TIM21-L1) | 28 | 1.05 | 0.152 | 1.000 |
| AT2G40800.1 | Translocase of the inner mitochondrial membrane 21-like (TIM21-L1) | 29 | 1.02 | 0.153 | 1.000 |
| AT2G40800.1 | Translocase of the inner mitochondrial membrane 21-like (TIM21-L1) | 30 | 0.96 | 0.153 | 1.000 |
| AT2G40800.1 | Translocase of the inner mitochondrial membrane 21-like (TIM21-L1) | 31 | 0.91 | 0.153 | 1.000 |
| AT2G40800.1 | Translocase of the inner mitochondrial membrane 21-like (TIM21-L1) | 32 | 0.91 | 0.154 | 1.000 |
| AT2G40800.1 | Translocase of the inner mitochondrial membrane 21-like (TIM21-L1) | 33 | 0.98 | 0.154 | 1.000 |
| AT2G40800.1 | Translocase of the inner mitochondrial membrane 21-like (TIM21-L1) | 34 | 1.13 | 0.154 | 1.000 |
| AT2G40800.1 | Translocase of the inner mitochondrial membrane 21-like (TIM21-L1) | 35 | 1.34 | 0.154 | 0.999 |
| AT2G40800.1 | Translocase of the inner mitochondrial membrane 21-like (TIM21-L1) | 36 | 1.50 | 0.154 | 0.976 |
| AT2G40800.1 | Translocase of the inner mitochondrial membrane 21-like (TIM21-L1) | 37 | 1.51 | 0.155 | 0.972 |
| AT2G40800.1 | Translocase of the inner mitochondrial membrane 21-like (TIM21-L1) | 38 | 1.37 | 0.155 | 0.998 |
| AT2G40800.1 | Translocase of the inner mitochondrial membrane 21-like (TIM21-L1) | 39 | 1.16 | 0.156 | 1.000 |
| AT2G40800.1 | Translocase of the inner mitochondrial membrane 21-like (TIM21-L1) | 40 | 0.97 | 0.162 | 1.000 |
| AT2G40800.1 | Translocase of the inner mitochondrial membrane 21-like (TIM21-L1) | 41 | 0.86 | 0.183 | 1.000 |
| AT2G40800.1 | Translocase of the inner mitochondrial membrane 21-like (TIM21-L1) | 42 | 0.84 | 0.232 | 1.000 |
| AT2G40800.1 | Translocase of the inner mitochondrial membrane 21-like (TIM21-L1) | 43 | 0.89 | 0.308 | 1.000 |
| AT2G40800.1 | Translocase of the inner mitochondrial membrane 21-like (TIM21-L1) | 46 | 1.57 | 0.711 | 1.000 |
| AT2G40800.1 | Translocase of the inner mitochondrial membrane 21-like (TIM21-L1) | 47 | 2.01 | 0.897 | 1.000 |
| AT2G41000.1 | [MISSING] | 2 | 1.47 | 0.623 | 0.994 |
| AT2G41000.1 | [MISSING] | 3 | 1.57 | 0.597 | 0.988 |

| | | | | | |
|---|---|---|---|---|---|
| AT2G41000.1 | [MISSING] | 26 | 1.97 | 0.325 | 0.757 |
| AT2G41000.1 | [MISSING] | 27 | 1.80 | 0.327 | 0.831 |
| AT2G41000.1 | [MISSING] | 28 | 1.62 | 0.333 | 0.903 |
| AT2G41000.1 | [MISSING] | 29 | 1.46 | 0.342 | 0.959 |
| AT2G41000.1 | [MISSING] | 46 | 0.12 | 0.573 | 0.310 |
| AT2G41000.1 | [MISSING] | 47 | 0.10 | 0.596 | 0.282 |
| AT2G42210.4 | NADH:ubiquinone oxidoreductase subunit (OEP16) | 2 | 0.94 | 0.392 | 1.000 |
| AT2G42210.4 | NADH:ubiquinone oxidoreductase subunit (OEP16) | 3 | 0.79 | 0.248 | 1.000 |
| AT2G42210.4 | NADH:ubiquinone oxidoreductase subunit (OEP16) | 4 | 0.72 | 0.227 | 1.000 |
| AT2G42210.4 | NADH:ubiquinone oxidoreductase subunit (OEP16) | 5 | 0.73 | 0.227 | 1.000 |
| AT2G42210.4 | NADH:ubiquinone oxidoreductase subunit (OEP16) | 6 | 0.78 | 0.226 | 1.000 |
| AT2G42210.4 | NADH:ubiquinone oxidoreductase subunit (OEP16) | 7 | 0.79 | 0.226 | 1.000 |
| AT2G42210.4 | NADH:ubiquinone oxidoreductase subunit (OEP16) | 8 | 0.77 | 0.223 | 1.000 |
| AT2G42210.4 | NADH:ubiquinone oxidoreductase subunit (OEP16) | 9 | 0.91 | 0.220 | 1.000 |
| AT2G42210.4 | NADH:ubiquinone oxidoreductase subunit (OEP16) | 10 | 1.15 | 0.221 | 1.000 |
| AT2G42210.4 | NADH:ubiquinone oxidoreductase subunit (OEP16) | 11 | 1.59 | 0.223 | 1.000 |
| AT2G42210.4 | NADH:ubiquinone oxidoreductase subunit (OEP16) | 12 | 2.13 | 0.222 | 0.977 |
| AT2G42210.4 | NADH:ubiquinone oxidoreductase subunit (OEP16) | 13 | 2.29 | 0.219 | 0.933 |
| AT2G42210.4 | NADH:ubiquinone oxidoreductase subunit (OEP16) | 14 | 2.11 | 0.216 | 0.971 |
| AT2G42210.4 | NADH:ubiquinone oxidoreductase subunit (OEP16) | 15 | 1.63 | 0.215 | 1.000 |
| AT2G42210.4 | NADH:ubiquinone oxidoreductase subunit (OEP16) | 16 | 1.08 | 0.214 | 1.000 |
| AT2G42210.4 | NADH:ubiquinone oxidoreductase subunit (OEP16) | 17 | 0.73 | 0.213 | 1.000 |
| AT2G42210.4 | NADH:ubiquinone oxidoreductase subunit (OEP16) | 18 | 0.65 | 0.215 | 1.000 |
| AT2G42210.4 | NADH:ubiquinone oxidoreductase subunit (OEP16) | 19 | 0.72 | 0.221 | 1.000 |
| AT2G42210.4 | NADH:ubiquinone oxidoreductase subunit (OEP16) | 20 | 0.85 | 0.227 | 1.000 |
| AT2G42210.4 | NADH:ubiquinone oxidoreductase subunit (OEP16) | 21 | 1.11 | 0.229 | 1.000 |
| AT2G42210.4 | NADH:ubiquinone oxidoreductase subunit (OEP16) | 22 | 1.52 | 0.230 | 1.000 |
| AT2G42210.4 | NADH:ubiquinone oxidoreductase subunit (OEP16) | 23 | 1.74 | 0.228 | 1.000 |
| AT2G42210.4 | NADH:ubiquinone oxidoreductase subunit (OEP16) | 24 | 1.28 | 0.223 | 1.000 |
| AT2G42210.4 | NADH:ubiquinone oxidoreductase subunit (OEP16) | 25 | 0.66 | 0.220 | 1.000 |
| AT2G42210.4 | NADH:ubiquinone oxidoreductase subunit (OEP16) | 26 | 0.39 | 0.219 | 0.816 |
| AT2G42210.4 | NADH:ubiquinone oxidoreductase subunit (OEP16) | 27 | 0.34 | 0.219 | 0.582 |
| AT2G42210.4 | NADH:ubiquinone oxidoreductase subunit (OEP16) | 28 | 0.43 | 0.219 | 0.926 |
| AT2G42210.4 | NADH:ubiquinone oxidoreductase subunit (OEP16) | 29 | 0.68 | 0.219 | 1.000 |

| | | | | | | |
|---|---|---|---|---|---|---|
| AT2G42210.4 | NADH:ubiquinone oxidoreductase subunit (OEP16) | 30 | 0.95 | 0.220 | 1.000 | |
| AT2G42210.4 | NADH:ubiquinone oxidoreductase subunit (OEP16) | 31 | 1.10 | 0.221 | 1.000 | |
| AT2G42210.4 | NADH:ubiquinone oxidoreductase subunit (OEP16) | 32 | 1.08 | 0.222 | 1.000 | |
| AT2G42210.4 | NADH:ubiquinone oxidoreductase subunit (OEP16) | 33 | 0.91 | 0.226 | 1.000 | |
| AT2G42210.4 | NADH:ubiquinone oxidoreductase subunit (OEP16) | 34 | 0.79 | 0.238 | 1.000 | |
| AT2G42210.4 | NADH:ubiquinone oxidoreductase subunit (OEP16) | 35 | 0.72 | 0.259 | 1.000 | |
| AT2G42210.4 | NADH:ubiquinone oxidoreductase subunit (OEP16) | 36 | 0.96 | 0.273 | 1.000 | |
| AT2G42210.4 | NADH:ubiquinone oxidoreductase subunit (OEP16) | 37 | 1.46 | 0.281 | 1.000 | |
| AT2G42210.4 | NADH:ubiquinone oxidoreductase subunit (OEP16) | 38 | 2.16 | 0.266 | 0.997 | |
| AT2G42210.4 | NADH:ubiquinone oxidoreductase subunit (OEP16) | 39 | 3.98 | 0.272 | 0.542 | |
| AT2G42210.4 | NADH:ubiquinone oxidoreductase subunit (OEP16) | 40 | 8.61 | 0.296 | 0.057 | . |
| AT2G42210.4 | NADH:ubiquinone oxidoreductase subunit (OEP16) | 41 | 15.13 | 0.297 | 0.004 | ** |
| AT2G42210.4 | NADH:ubiquinone oxidoreductase subunit (OEP16) | 42 | 21.35 | 0.292 | 0.000 | *** |
| AT2G42210.4 | NADH:ubiquinone oxidoreductase subunit (OEP16) | 43 | 27.89 | 0.276 | 0.000 | *** |
| AT2G42210.4 | NADH:ubiquinone oxidoreductase subunit (OEP16) | 44 | 28.89 | 0.249 | 0.000 | *** |
| AT2G42210.4 | NADH:ubiquinone oxidoreductase subunit (OEP16) | 45 | 13.64 | 0.232 | 0.000 | *** |
| AT2G42210.4 | NADH:ubiquinone oxidoreductase subunit (OEP16) | 46 | 3.26 | 0.266 | 0.766 | |
| AT2G42210.4 | NADH:ubiquinone oxidoreductase subunit (OEP16) | 47 | 0.63 | 0.448 | 1.000 | |
| AT2G42450.1 | Alpha/beta-Hydrolases superfamily protein | 2 | 1.11 | 0.311 | 1.000 | |
| AT2G42450.1 | Alpha/beta-Hydrolases superfamily protein | 3 | 1.12 | 0.293 | 1.000 | |
| AT2G42450.1 | Alpha/beta-Hydrolases superfamily protein | 4 | 1.13 | 0.302 | 1.000 | |
| AT2G42450.1 | Alpha/beta-Hydrolases superfamily protein | 27 | 1.57 | 0.194 | 0.913 | |
| AT2G42450.1 | Alpha/beta-Hydrolases superfamily protein | 28 | 1.60 | 0.165 | 0.801 | |
| AT2G42450.1 | Alpha/beta-Hydrolases superfamily protein | 29 | 1.61 | 0.144 | 0.673 | |
| AT2G42450.1 | Alpha/beta-Hydrolases superfamily protein | 30 | 1.60 | 0.131 | 0.589 | |
| AT2G42450.1 | Alpha/beta-Hydrolases superfamily protein | 31 | 1.56 | 0.123 | 0.581 | |
| AT2G42450.1 | Alpha/beta-Hydrolases superfamily protein | 32 | 1.48 | 0.119 | 0.672 | |
| AT2G42450.1 | Alpha/beta-Hydrolases superfamily protein | 33 | 1.39 | 0.116 | 0.812 | |
| AT2G42450.1 | Alpha/beta-Hydrolases superfamily protein | 34 | 1.30 | 0.116 | 0.921 | |
| AT2G42450.1 | Alpha/beta-Hydrolases superfamily protein | 35 | 1.24 | 0.116 | 0.972 | |
| AT2G42450.1 | Alpha/beta-Hydrolases superfamily protein | 36 | 1.21 | 0.120 | 0.990 | |

| | | | | | | |
|---|---|---|---|---|---|---|
| AT2G42450.1 | Alpha/beta-Hydrolases superfamily protein | 37 | 1.17 | 0.127 | 0.998 | |
| AT2G42450.1 | Alpha/beta-Hydrolases superfamily protein | 38 | 1.11 | 0.139 | 1.000 | |
| AT2G42450.1 | Alpha/beta-Hydrolases superfamily protein | 39 | 1.00 | 0.156 | 1.000 | |
| AT2G42450.1 | Alpha/beta-Hydrolases superfamily protein | 40 | 0.85 | 0.177 | 1.000 | |
| AT2G42450.1 | Alpha/beta-Hydrolases superfamily protein | 41 | 0.69 | 0.201 | 0.974 | |
| AT2G42450.1 | Alpha/beta-Hydrolases superfamily protein | 46 | 0.14 | 0.340 | 0.097 | . |
| AT2G42450.1 | Alpha/beta-Hydrolases superfamily protein | 47 | 0.10 | 0.383 | 0.069 | . |
| AT2G43360.1 | Biotin synthase 2 (BIO2) | 2 | 1.25 | 0.532 | 1.000 | |
| AT2G43360.1 | Biotin synthase 2 (BIO2) | 3 | 1.24 | 0.472 | 1.000 | |
| AT2G43360.1 | Biotin synthase 2 (BIO2) | 11 | 1.15 | 0.350 | 1.000 | |
| AT2G43360.1 | Biotin synthase 2 (BIO2) | 12 | 1.12 | 0.307 | 1.000 | |
| AT2G43360.1 | Biotin synthase 2 (BIO2) | 13 | 1.05 | 0.284 | 1.000 | |
| AT2G43360.1 | Biotin synthase 2 (BIO2) | 14 | 0.95 | 0.266 | 1.000 | |
| AT2G43360.1 | Biotin synthase 2 (BIO2) | 15 | 0.85 | 0.256 | 1.000 | |
| AT2G43360.1 | Biotin synthase 2 (BIO2) | 16 | 0.78 | 0.250 | 1.000 | |
| AT2G43360.1 | Biotin synthase 2 (BIO2) | 17 | 0.76 | 0.249 | 1.000 | |
| AT2G43360.1 | Biotin synthase 2 (BIO2) | 18 | 0.78 | 0.250 | 1.000 | |
| AT2G43360.1 | Biotin synthase 2 (BIO2) | 19 | 0.89 | 0.251 | 1.000 | |
| AT2G43360.1 | Biotin synthase 2 (BIO2) | 20 | 1.04 | 0.251 | 1.000 | |
| AT2G43360.1 | Biotin synthase 2 (BIO2) | 21 | 1.20 | 0.250 | 1.000 | |
| AT2G43360.1 | Biotin synthase 2 (BIO2) | 22 | 1.34 | 0.249 | 1.000 | |
| AT2G43360.1 | Biotin synthase 2 (BIO2) | 23 | 1.47 | 0.247 | 1.000 | |
| AT2G43360.1 | Biotin synthase 2 (BIO2) | 24 | 1.53 | 0.244 | 1.000 | |
| AT2G43360.1 | Biotin synthase 2 (BIO2) | 25 | 1.44 | 0.242 | 1.000 | |
| AT2G43360.1 | Biotin synthase 2 (BIO2) | 26 | 1.30 | 0.241 | 1.000 | |
| AT2G43360.1 | Biotin synthase 2 (BIO2) | 27 | 1.29 | 0.241 | 1.000 | |
| AT2G43360.1 | Biotin synthase 2 (BIO2) | 28 | 1.42 | 0.241 | 1.000 | |
| AT2G43360.1 | Biotin synthase 2 (BIO2) | 29 | 1.52 | 0.242 | 1.000 | |
| AT2G43360.1 | Biotin synthase 2 (BIO2) | 30 | 1.47 | 0.242 | 1.000 | |
| AT2G43360.1 | Biotin synthase 2 (BIO2) | 31 | 1.44 | 0.242 | 1.000 | |
| AT2G43360.1 | Biotin synthase 2 (BIO2) | 32 | 1.55 | 0.242 | 0.999 | |
| AT2G43360.1 | Biotin synthase 2 (BIO2) | 33 | 1.62 | 0.242 | 0.998 | |
| AT2G43360.1 | Biotin synthase 2 (BIO2) | 34 | 1.44 | 0.242 | 1.000 | |
| AT2G43360.1 | Biotin synthase 2 (BIO2) | 35 | 1.10 | 0.242 | 1.000 | |
| AT2G43360.1 | Biotin synthase 2 (BIO2) | 36 | 0.77 | 0.242 | 1.000 | |
| AT2G43360.1 | Biotin synthase 2 (BIO2) | 37 | 0.55 | 0.242 | 0.983 | |
| AT2G43360.1 | Biotin synthase 2 (BIO2) | 38 | 0.49 | 0.242 | 0.934 | |
| AT2G43360.1 | Biotin synthase 2 (BIO2) | 39 | 0.65 | 0.242 | 0.999 | |
| AT2G43360.1 | Biotin synthase 2 (BIO2) | 40 | 1.08 | 0.243 | 1.000 | |
| AT2G43360.1 | Biotin synthase 2 (BIO2) | 41 | 1.59 | 0.244 | 0.999 | |
| AT2G43360.1 | Biotin synthase 2 (BIO2) | 42 | 1.82 | 0.245 | 0.985 | |
| AT2G43360.1 | Biotin synthase 2 (BIO2) | 43 | 1.68 | 0.246 | 0.996 | |
| AT2G43360.1 | Biotin synthase 2 (BIO2) | 44 | 1.22 | 0.248 | 1.000 | |
| AT2G43360.1 | Biotin synthase 2 (BIO2) | 45 | 0.66 | 0.256 | 1.000 | |
| AT2G43360.1 | Biotin synthase 2 (BIO2) | 46 | 0.27 | 0.291 | 0.489 | |
| AT2G43360.1 | Biotin synthase 2 (BIO2) | 47 | 0.10 | 0.393 | 0.142 | |
| AT2G43400.1 | Electron-transfer flavoprotein:ubiquinone oxidoreductase (ETFQO) | 2 | 1.52 | 0.514 | 1.000 | |
| AT2G43400.1 | Electron-transfer flavoprotein:ubiquinone oxidoreductase (ETFQO) | 3 | 1.31 | 0.494 | 1.000 | |
| AT2G43400.1 | Electron-transfer flavoprotein:ubiquinone oxidoreductase (ETFQO) | 23 | 0.42 | 0.438 | 0.997 | |

| | | | | | | |
|---|---|---|---|---|---|---|
| AT2G43400.1 | Electron-transfer flavoprotein:ubiquinone oxidoreductase (ETFQO) | 24 | 0.46 | 0.392 | 0.997 | |
| AT2G43400.1 | Electron-transfer flavoprotein:ubiquinone oxidoreductase (ETFQO) | 25 | 0.49 | 0.449 | 1.000 | |
| AT2G43400.1 | Electron-transfer flavoprotein:ubiquinone oxidoreductase (ETFQO) | 32 | 0.59 | 0.365 | 1.000 | |
| AT2G43400.1 | Electron-transfer flavoprotein:ubiquinone oxidoreductase (ETFQO) | 33 | 0.80 | 0.298 | 1.000 | |
| AT2G43400.1 | Electron-transfer flavoprotein:ubiquinone oxidoreductase (ETFQO) | 34 | 1.13 | 0.292 | 1.000 | |
| AT2G43400.1 | Electron-transfer flavoprotein:ubiquinone oxidoreductase (ETFQO) | 35 | 1.61 | 0.304 | 1.000 | |
| AT2G43400.1 | Electron-transfer flavoprotein:ubiquinone oxidoreductase (ETFQO) | 36 | 2.41 | 0.290 | 0.915 | |
| AT2G43400.1 | Electron-transfer flavoprotein:ubiquinone oxidoreductase (ETFQO) | 37 | 3.27 | 0.262 | 0.487 | |
| AT2G43400.1 | Electron-transfer flavoprotein:ubiquinone oxidoreductase (ETFQO) | 38 | 3.54 | 0.244 | 0.293 | |
| AT2G43400.1 | Electron-transfer flavoprotein:ubiquinone oxidoreductase (ETFQO) | 39 | 3.53 | 0.238 | 0.264 | |
| AT2G43400.1 | Electron-transfer flavoprotein:ubiquinone oxidoreductase (ETFQO) | 40 | 5.08 | 0.235 | 0.047 | * |
| AT2G43400.1 | Electron-transfer flavoprotein:ubiquinone oxidoreductase (ETFQO) | 41 | 9.02 | 0.236 | 0.002 | ** |
| AT2G43400.1 | Electron-transfer flavoprotein:ubiquinone oxidoreductase (ETFQO) | 42 | 14.47 | 0.238 | 0.000 | *** |
| AT2G43400.1 | Electron-transfer flavoprotein:ubiquinone oxidoreductase (ETFQO) | 43 | 14.70 | 0.240 | 0.000 | *** |
| AT2G43400.1 | Electron-transfer flavoprotein:ubiquinone oxidoreductase (ETFQO) | 44 | 6.55 | 0.240 | 0.014 | * |
| AT2G43400.1 | Electron-transfer flavoprotein:ubiquinone oxidoreductase (ETFQO) | 45 | 1.28 | 0.243 | 1.000 | |
| AT2G43400.1 | Electron-transfer flavoprotein:ubiquinone oxidoreductase (ETFQO) | 46 | 0.13 | 0.277 | 0.025 | * |
| AT2G43400.1 | Electron-transfer flavoprotein:ubiquinone oxidoreductase (ETFQO) | 47 | 0.01 | 0.448 | 0.000 | *** |
| AT2G43780.2 | Cytochrome c:O2 oxidoreductase assembly protein | 2 | 0.92 | 0.534 | 1.000 | |
| AT2G43780.2 | Cytochrome c:O2 oxidoreductase assembly protein | 3 | 0.95 | 0.412 | 1.000 | |

| | | | | | |
|---|---|---|---|---|---|
| AT2G43780.2 | Cytochrome c:O2 oxidoreductase assembly protein | 6 | 1.15 | 0.330 | 1.000 |
| AT2G43780.2 | Cytochrome c:O2 oxidoreductase assembly protein | 7 | 1.27 | 0.275 | 1.000 |
| AT2G43780.2 | Cytochrome c:O2 oxidoreductase assembly protein | 8 | 1.41 | 0.238 | 1.000 |
| AT2G43780.2 | Cytochrome c:O2 oxidoreductase assembly protein | 9 | 1.54 | 0.220 | 1.000 |
| AT2G43780.2 | Cytochrome c:O2 oxidoreductase assembly protein | 10 | 1.63 | 0.215 | 0.999 |
| AT2G43780.2 | Cytochrome c:O2 oxidoreductase assembly protein | 11 | 1.67 | 0.213 | 0.998 |
| AT2G43780.2 | Cytochrome c:O2 oxidoreductase assembly protein | 12 | 1.77 | 0.212 | 0.993 |
| AT2G43780.2 | Cytochrome c:O2 oxidoreductase assembly protein | 13 | 1.84 | 0.210 | 0.985 |
| AT2G43780.2 | Cytochrome c:O2 oxidoreductase assembly protein | 14 | 1.73 | 0.208 | 0.995 |
| AT2G43780.2 | Cytochrome c:O2 oxidoreductase assembly protein | 15 | 1.44 | 0.208 | 1.000 |
| AT2G43780.2 | Cytochrome c:O2 oxidoreductase assembly protein | 16 | 1.21 | 0.208 | 1.000 |
| AT2G43780.2 | Cytochrome c:O2 oxidoreductase assembly protein | 17 | 1.17 | 0.211 | 1.000 |
| AT2G43780.2 | Cytochrome c:O2 oxidoreductase assembly protein | 18 | 1.31 | 0.220 | 1.000 |
| AT2G43780.2 | Cytochrome c:O2 oxidoreductase assembly protein | 19 | 1.50 | 0.241 | 1.000 |
| AT2G43780.2 | Cytochrome c:O2 oxidoreductase assembly protein | 20 | 1.68 | 0.295 | 1.000 |
| AT2G43780.2 | Cytochrome c:O2 oxidoreductase assembly protein | 21 | 1.88 | 0.371 | 1.000 |
| AT2G43780.2 | Cytochrome c:O2 oxidoreductase assembly protein | 28 | 1.77 | 0.332 | 1.000 |
| AT2G43780.2 | Cytochrome c:O2 oxidoreductase assembly protein | 29 | 1.35 | 0.265 | 1.000 |
| AT2G43780.2 | Cytochrome c:O2 oxidoreductase assembly protein | 30 | 1.01 | 0.232 | 1.000 |
| AT2G43780.2 | Cytochrome c:O2 oxidoreductase assembly protein | 31 | 0.87 | 0.219 | 1.000 |

| | | | | | |
|---|---|---|---|---|---|
| AT2G43780.2 | Cytochrome c:O2 oxidoreductase assembly protein | 32 | 0.95 | 0.214 | 1.000 |
| AT2G43780.2 | Cytochrome c:O2 oxidoreductase assembly protein | 33 | 1.18 | 0.213 | 1.000 |
| AT2G43780.2 | Cytochrome c:O2 oxidoreductase assembly protein | 34 | 1.37 | 0.213 | 1.000 |
| AT2G43780.2 | Cytochrome c:O2 oxidoreductase assembly protein | 35 | 1.37 | 0.213 | 1.000 |
| AT2G43780.2 | Cytochrome c:O2 oxidoreductase assembly protein | 36 | 1.20 | 0.213 | 1.000 |
| AT2G43780.2 | Cytochrome c:O2 oxidoreductase assembly protein | 37 | 0.99 | 0.214 | 1.000 |
| AT2G43780.2 | Cytochrome c:O2 oxidoreductase assembly protein | 38 | 0.81 | 0.217 | 1.000 |
| AT2G43780.2 | Cytochrome c:O2 oxidoreductase assembly protein | 39 | 0.70 | 0.235 | 1.000 |
| AT2G43780.2 | Cytochrome c:O2 oxidoreductase assembly protein | 40 | 0.74 | 0.296 | 1.000 |
| AT2G43780.2 | Cytochrome c:O2 oxidoreductase assembly protein | 41 | 0.94 | 0.422 | 1.000 |
| AT2G43780.2 | Cytochrome c:O2 oxidoreductase assembly protein | 46 | 13.53 | 1.728 | 1.000 |
| AT2G43780.2 | Cytochrome c:O2 oxidoreductase assembly protein | 47 | 26.53 | 2.120 | 1.000 |
| AT2G44350.1 | Citrate synthase 4 (CSY4) | 3 | 0.64 | 0.168 | 0.999 |
| AT2G44350.1 | Citrate synthase 4 (CSY4) | 4 | 0.56 | 0.146 | 0.888 |
| AT2G44350.1 | Citrate synthase 4 (CSY4) | 5 | 0.51 | 0.145 | 0.705 |
| AT2G44350.1 | Citrate synthase 4 (CSY4) | 6 | 0.52 | 0.146 | 0.749 |
| AT2G44350.1 | Citrate synthase 4 (CSY4) | 7 | 0.59 | 0.145 | 0.940 |
| AT2G44350.1 | Citrate synthase 4 (CSY4) | 8 | 0.71 | 0.142 | 1.000 |
| AT2G44350.1 | Citrate synthase 4 (CSY4) | 9 | 0.90 | 0.140 | 1.000 |
| AT2G44350.1 | Citrate synthase 4 (CSY4) | 10 | 1.10 | 0.140 | 1.000 |
| AT2G44350.1 | Citrate synthase 4 (CSY4) | 11 | 1.33 | 0.140 | 1.000 |
| AT2G44350.1 | Citrate synthase 4 (CSY4) | 12 | 1.54 | 0.140 | 0.990 |
| AT2G44350.1 | Citrate synthase 4 (CSY4) | 13 | 1.56 | 0.139 | 0.983 |
| AT2G44350.1 | Citrate synthase 4 (CSY4) | 14 | 1.44 | 0.137 | 0.999 |
| AT2G44350.1 | Citrate synthase 4 (CSY4) | 15 | 1.33 | 0.136 | 1.000 |
| AT2G44350.1 | Citrate synthase 4 (CSY4) | 16 | 1.39 | 0.135 | 1.000 |
| AT2G44350.1 | Citrate synthase 4 (CSY4) | 17 | 1.65 | 0.135 | 0.932 |
| AT2G44350.1 | Citrate synthase 4 (CSY4) | 18 | 2.01 | 0.137 | 0.521 |
| AT2G44350.1 | Citrate synthase 4 (CSY4) | 19 | 2.22 | 0.141 | 0.325 |
| AT2G44350.1 | Citrate synthase 4 (CSY4) | 20 | 2.15 | 0.144 | 0.437 |
| AT2G44350.1 | Citrate synthase 4 (CSY4) | 21 | 1.87 | 0.146 | 0.800 |
| AT2G44350.1 | Citrate synthase 4 (CSY4) | 22 | 1.55 | 0.146 | 0.992 |
| AT2G44350.1 | Citrate synthase 4 (CSY4) | 23 | 1.29 | 0.145 | 1.000 |
| AT2G44350.1 | Citrate synthase 4 (CSY4) | 24 | 1.16 | 0.142 | 1.000 |
| AT2G44350.1 | Citrate synthase 4 (CSY4) | 25 | 1.11 | 0.140 | 1.000 |
| AT2G44350.1 | Citrate synthase 4 (CSY4) | 26 | 1.10 | 0.139 | 1.000 |
| AT2G44350.1 | Citrate synthase 4 (CSY4) | 27 | 1.11 | 0.139 | 1.000 |
| AT2G44350.1 | Citrate synthase 4 (CSY4) | 28 | 1.08 | 0.139 | 1.000 |

| | | | | | | |
|---|---|---|---|---|---|---|
| AT2G44350.1 | Citrate synthase 4 (CSY4) | 29 | 1.03 | 0.139 | 1.000 | |
| AT2G44350.1 | Citrate synthase 4 (CSY4) | 30 | 1.01 | 0.139 | 1.000 | |
| AT2G44350.1 | Citrate synthase 4 (CSY4) | 31 | 1.02 | 0.140 | 1.000 | |
| AT2G44350.1 | Citrate synthase 4 (CSY4) | 32 | 1.03 | 0.140 | 1.000 | |
| AT2G44350.1 | Citrate synthase 4 (CSY4) | 33 | 1.01 | 0.140 | 1.000 | |
| AT2G44350.1 | Citrate synthase 4 (CSY4) | 34 | 1.01 | 0.140 | 1.000 | |
| AT2G44350.1 | Citrate synthase 4 (CSY4) | 35 | 1.06 | 0.140 | 1.000 | |
| AT2G44350.1 | Citrate synthase 4 (CSY4) | 36 | 1.17 | 0.140 | 1.000 | |
| AT2G44350.1 | Citrate synthase 4 (CSY4) | 37 | 1.30 | 0.139 | 1.000 | |
| AT2G44350.1 | Citrate synthase 4 (CSY4) | 38 | 1.49 | 0.140 | 0.996 | |
| AT2G44350.1 | Citrate synthase 4 (CSY4) | 39 | 1.72 | 0.140 | 0.902 | |
| AT2G44350.1 | Citrate synthase 4 (CSY4) | 40 | 1.92 | 0.140 | 0.681 | |
| AT2G44350.1 | Citrate synthase 4 (CSY4) | 41 | 1.95 | 0.141 | 0.648 | |
| AT2G44350.1 | Citrate synthase 4 (CSY4) | 42 | 1.79 | 0.143 | 0.856 | |
| AT2G44350.1 | Citrate synthase 4 (CSY4) | 43 | 1.49 | 0.144 | 0.998 | |
| AT2G44350.1 | Citrate synthase 4 (CSY4) | 44 | 1.08 | 0.144 | 1.000 | |
| AT2G44350.1 | Citrate synthase 4 (CSY4) | 45 | 0.67 | 0.147 | 0.998 | |
| AT2G44350.1 | Citrate synthase 4 (CSY4) | 46 | 0.36 | 0.181 | 0.352 | |
| AT2G44350.1 | Citrate synthase 4 (CSY4) | 47 | 0.18 | 0.289 | 0.272 | |
| AT2G44350.1 | Citrate synthase 4 (CSY4) | 48 | 0.09 | 0.448 | 0.425 | |
| AT2G44620.1 | Acyl carrier protein 1 (ACP1) | 2 | 0.99 | 0.637 | 1.000 | |
| AT2G44620.1 | Acyl carrier protein 1 (ACP1) | 3 | 1.06 | 0.575 | 1.000 | |
| AT2G44620.1 | Acyl carrier protein 1 (ACP1) | 15 | 0.96 | 0.316 | 1.000 | |
| AT2G44620.1 | Acyl carrier protein 1 (ACP1) | 16 | 0.86 | 0.272 | 1.000 | |
| AT2G44620.1 | Acyl carrier protein 1 (ACP1) | 17 | 0.82 | 0.242 | 1.000 | |
| AT2G44620.1 | Acyl carrier protein 1 (ACP1) | 18 | 0.87 | 0.225 | 1.000 | |
| AT2G44620.1 | Acyl carrier protein 1 (ACP1) | 19 | 1.07 | 0.219 | 1.000 | |
| AT2G44620.1 | Acyl carrier protein 1 (ACP1) | 20 | 1.43 | 0.219 | 0.983 | |
| AT2G44620.1 | Acyl carrier protein 1 (ACP1) | 21 | 2.06 | 0.224 | 0.681 | |
| AT2G44620.1 | Acyl carrier protein 1 (ACP1) | 22 | 3.14 | 0.236 | 0.232 | |
| AT2G44620.1 | Acyl carrier protein 1 (ACP1) | 23 | 5.09 | 0.264 | 0.063 | . |
| AT2G44620.1 | Acyl carrier protein 1 (ACP1) | 24 | 8.58 | 0.320 | 0.034 | * |
| AT2G44620.1 | Acyl carrier protein 1 (ACP1) | 25 | 14.33 | 0.409 | 0.043 | * |
| AT2G44620.1 | Acyl carrier protein 1 (ACP1) | 46 | 136.32 | 5.191 | 0.999 | |
| AT2G44620.1 | Acyl carrier protein 1 (ACP1) | 47 | 124.01 | 5.525 | 1.000 | |
| AT2G45030.1 | Translation elongation factor EFG/EF2 protein | 2 | 0.89 | 0.304 | 1.000 | |
| AT2G45030.1 | Translation elongation factor EFG/EF2 protein | 3 | 0.85 | 0.291 | 1.000 | |
| AT2G45030.1 | Translation elongation factor EFG/EF2 protein | 27 | 0.78 | 0.221 | 1.000 | |
| AT2G45030.1 | Translation elongation factor EFG/EF2 protein | 28 | 0.85 | 0.160 | 1.000 | |
| AT2G45030.1 | Translation elongation factor EFG/EF2 protein | 29 | 0.90 | 0.141 | 1.000 | |
| AT2G45030.1 | Translation elongation factor EFG/EF2 protein | 30 | 0.97 | 0.134 | 1.000 | |
| AT2G45030.1 | Translation elongation factor EFG/EF2 protein | 31 | 1.04 | 0.132 | 1.000 | |
| AT2G45030.1 | Translation elongation factor EFG/EF2 protein | 32 | 1.02 | 0.132 | 1.000 | |
| AT2G45030.1 | Translation elongation factor EFG/EF2 protein | 33 | 0.91 | 0.132 | 1.000 | |
| AT2G45030.1 | Translation elongation factor EFG/EF2 protein | 34 | 0.82 | 0.132 | 1.000 | |
| AT2G45030.1 | Translation elongation factor EFG/EF2 protein | 35 | 0.81 | 0.132 | 1.000 | |
| AT2G45030.1 | Translation elongation factor EFG/EF2 protein | 36 | 0.91 | 0.131 | 1.000 | |
| AT2G45030.1 | Translation elongation factor EFG/EF2 protein | 37 | 1.06 | 0.131 | 1.000 | |

| | | | | | |
|---|---|---|---|---|---|
| AT2G45030.1 | Translation elongation factor EFG/EF2 protein | 38 | 1.19 | 0.131 | 1.000 |
| AT2G45030.1 | Translation elongation factor EFG/EF2 protein | 39 | 1.26 | 0.130 | 0.999 |
| AT2G45030.1 | Translation elongation factor EFG/EF2 protein | 40 | 1.24 | 0.130 | 1.000 |
| AT2G45030.1 | Translation elongation factor EFG/EF2 protein | 41 | 1.07 | 0.131 | 1.000 |
| AT2G45030.1 | Translation elongation factor EFG/EF2 protein | 42 | 0.84 | 0.133 | 1.000 |
| AT2G45030.1 | Translation elongation factor EFG/EF2 protein | 43 | 0.64 | 0.134 | 0.882 |
| AT2G45030.1 | Translation elongation factor EFG/EF2 protein | 44 | 0.52 | 0.134 | 0.402 |
| AT2G45030.1 | Translation elongation factor EFG/EF2 protein | 45 | 0.47 | 0.135 | 0.208 |
| AT2G45030.1 | Translation elongation factor EFG/EF2 protein | 46 | 0.46 | 0.161 | 0.426 |
| AT2G45030.1 | Translation elongation factor EFG/EF2 protein | 47 | 0.50 | 0.270 | 0.979 |
| AT2G45060.1 | Alanine-tRNA ligase | 3 | 0.80 | 0.188 | 1.000 |
| AT2G45060.1 | Alanine-tRNA ligase | 4 | 0.68 | 0.173 | 1.000 |
| AT2G45060.1 | Alanine-tRNA ligase | 5 | 0.65 | 0.173 | 1.000 |
| AT2G45060.1 | Alanine-tRNA ligase | 6 | 0.65 | 0.173 | 1.000 |
| AT2G45060.1 | Alanine-tRNA ligase | 7 | 0.65 | 0.173 | 1.000 |
| AT2G45060.1 | Alanine-tRNA ligase | 8 | 0.72 | 0.169 | 1.000 |
| AT2G45060.1 | Alanine-tRNA ligase | 9 | 0.82 | 0.167 | 1.000 |
| AT2G45060.1 | Alanine-tRNA ligase | 10 | 0.92 | 0.166 | 1.000 |
| AT2G45060.1 | Alanine-tRNA ligase | 11 | 1.03 | 0.167 | 1.000 |
| AT2G45060.1 | Alanine-tRNA ligase | 12 | 1.16 | 0.167 | 1.000 |
| AT2G45060.1 | Alanine-tRNA ligase | 13 | 1.27 | 0.165 | 1.000 |
| AT2G45060.1 | Alanine-tRNA ligase | 14 | 1.40 | 0.163 | 1.000 |
| AT2G45060.1 | Alanine-tRNA ligase | 15 | 1.40 | 0.162 | 1.000 |
| AT2G45060.1 | Alanine-tRNA ligase | 16 | 1.26 | 0.160 | 1.000 |
| AT2G45060.1 | Alanine-tRNA ligase | 17 | 1.14 | 0.160 | 1.000 |
| AT2G45060.1 | Alanine-tRNA ligase | 18 | 1.03 | 0.163 | 1.000 |
| AT2G45060.1 | Alanine-tRNA ligase | 19 | 0.94 | 0.168 | 1.000 |
| AT2G45060.1 | Alanine-tRNA ligase | 20 | 0.94 | 0.173 | 1.000 |
| AT2G45060.1 | Alanine-tRNA ligase | 21 | 1.01 | 0.175 | 1.000 |
| AT2G45060.1 | Alanine-tRNA ligase | 22 | 1.03 | 0.175 | 1.000 |
| AT2G45060.1 | Alanine-tRNA ligase | 23 | 0.93 | 0.174 | 1.000 |
| AT2G45060.1 | Alanine-tRNA ligase | 24 | 0.81 | 0.170 | 1.000 |
| AT2G45060.1 | Alanine-tRNA ligase | 25 | 0.86 | 0.167 | 1.000 |
| AT2G45060.1 | Alanine-tRNA ligase | 26 | 1.05 | 0.166 | 1.000 |
| AT2G45060.1 | Alanine-tRNA ligase | 27 | 1.16 | 0.166 | 1.000 |
| AT2G45060.1 | Alanine-tRNA ligase | 28 | 1.16 | 0.166 | 1.000 |
| AT2G45060.1 | Alanine-tRNA ligase | 29 | 1.18 | 0.166 | 1.000 |
| AT2G45060.1 | Alanine-tRNA ligase | 30 | 1.17 | 0.165 | 1.000 |
| AT2G45060.1 | Alanine-tRNA ligase | 31 | 1.07 | 0.166 | 1.000 |
| AT2G45060.1 | Alanine-tRNA ligase | 32 | 0.89 | 0.167 | 1.000 |
| AT2G45060.1 | Alanine-tRNA ligase | 33 | 0.77 | 0.167 | 1.000 |
| AT2G45060.1 | Alanine-tRNA ligase | 34 | 0.86 | 0.167 | 1.000 |
| AT2G45060.1 | Alanine-tRNA ligase | 35 | 1.14 | 0.167 | 1.000 |
| AT2G45060.1 | Alanine-tRNA ligase | 36 | 1.37 | 0.166 | 1.000 |
| AT2G45060.1 | Alanine-tRNA ligase | 37 | 1.61 | 0.166 | 0.999 |
| AT2G45060.1 | Alanine-tRNA ligase | 38 | 1.87 | 0.166 | 0.958 |
| AT2G45060.1 | Alanine-tRNA ligase | 39 | 1.80 | 0.167 | 0.980 |
| AT2G45060.1 | Alanine-tRNA ligase | 40 | 1.51 | 0.167 | 1.000 |
| AT2G45060.1 | Alanine-tRNA ligase | 41 | 1.30 | 0.168 | 1.000 |
| AT2G45060.1 | Alanine-tRNA ligase | 42 | 1.18 | 0.170 | 1.000 |
| AT2G45060.1 | Alanine-tRNA ligase | 43 | 1.27 | 0.172 | 1.000 |
| AT2G45060.1 | Alanine-tRNA ligase | 44 | 1.30 | 0.172 | 1.000 |
| AT2G45060.1 | Alanine-tRNA ligase | 45 | 0.98 | 0.172 | 1.000 |

| | | | | | |
|---|---|---|---|---|---|
| AT2G45060.1 | Alanine-tRNA ligase | 46 | 0.57 | 0.203 | 0.999 |
| AT2G45060.1 | Alanine-tRNA ligase | 47 | 0.30 | 0.372 | 0.993 |
| AT2G45060.1 | Alanine-tRNA ligase | 48 | 0.16 | 0.629 | 0.998 |
| AT2G45740.3 | Peroxin 11D (PEX11D) | 2 | 0.73 | 0.421 | 1.000 |
| AT2G45740.3 | Peroxin 11D (PEX11D) | 3 | 0.58 | 0.343 | 1.000 |
| AT2G45740.3 | Peroxin 11D (PEX11D) | 4 | 0.50 | 0.441 | 1.000 |
| AT2G45740.3 | Peroxin 11D (PEX11D) | 12 | 0.71 | 0.315 | 1.000 |
| AT2G45740.3 | Peroxin 11D (PEX11D) | 13 | 0.74 | 0.284 | 1.000 |
| AT2G45740.3 | Peroxin 11D (PEX11D) | 14 | 0.75 | 0.263 | 1.000 |
| AT2G45740.3 | Peroxin 11D (PEX11D) | 15 | 0.74 | 0.248 | 1.000 |
| AT2G45740.3 | Peroxin 11D (PEX11D) | 16 | 0.78 | 0.239 | 1.000 |
| AT2G45740.3 | Peroxin 11D (PEX11D) | 17 | 0.97 | 0.237 | 1.000 |
| AT2G45740.3 | Peroxin 11D (PEX11D) | 18 | 1.30 | 0.238 | 1.000 |
| AT2G45740.3 | Peroxin 11D (PEX11D) | 19 | 1.66 | 0.242 | 0.999 |
| AT2G45740.3 | Peroxin 11D (PEX11D) | 20 | 1.85 | 0.247 | 0.990 |
| AT2G45740.3 | Peroxin 11D (PEX11D) | 21 | 1.75 | 0.248 | 0.997 |
| AT2G45740.3 | Peroxin 11D (PEX11D) | 22 | 1.65 | 0.249 | 0.999 |
| AT2G45740.3 | Peroxin 11D (PEX11D) | 23 | 1.81 | 0.247 | 0.993 |
| AT2G45740.3 | Peroxin 11D (PEX11D) | 24 | 2.01 | 0.244 | 0.966 |
| AT2G45740.3 | Peroxin 11D (PEX11D) | 25 | 2.26 | 0.241 | 0.881 |
| AT2G45740.3 | Peroxin 11D (PEX11D) | 26 | 2.63 | 0.240 | 0.694 |
| AT2G45740.3 | Peroxin 11D (PEX11D) | 27 | 2.51 | 0.240 | 0.759 |
| AT2G45740.3 | Peroxin 11D (PEX11D) | 28 | 1.94 | 0.240 | 0.974 |
| AT2G45740.3 | Peroxin 11D (PEX11D) | 29 | 1.40 | 0.241 | 1.000 |
| AT2G45740.3 | Peroxin 11D (PEX11D) | 30 | 1.02 | 0.241 | 1.000 |
| AT2G45740.3 | Peroxin 11D (PEX11D) | 31 | 0.84 | 0.242 | 1.000 |
| AT2G45740.3 | Peroxin 11D (PEX11D) | 32 | 0.89 | 0.242 | 1.000 |
| AT2G45740.3 | Peroxin 11D (PEX11D) | 33 | 1.19 | 0.242 | 1.000 |
| AT2G45740.3 | Peroxin 11D (PEX11D) | 34 | 1.67 | 0.243 | 0.998 |
| AT2G45740.3 | Peroxin 11D (PEX11D) | 35 | 1.83 | 0.244 | 0.990 |
| AT2G45740.3 | Peroxin 11D (PEX11D) | 36 | 1.28 | 0.246 | 1.000 |
| AT2G45740.3 | Peroxin 11D (PEX11D) | 37 | 0.65 | 0.258 | 1.000 |
| AT2G45740.3 | Peroxin 11D (PEX11D) | 38 | 0.39 | 0.310 | 0.941 |
| AT2G45740.3 | Peroxin 11D (PEX11D) | 39 | 0.35 | 0.429 | 0.992 |
| AT2G45740.3 | Peroxin 11D (PEX11D) | 46 | 797.60 | 1.774 | 0.778 |
| AT2G45740.3 | Peroxin 11D (PEX11D) | 47 | 4308.73 | 2.075 | 0.694 |
| AT2G45980.1 | ATG8-interacting protein 1 (ATI1) | 2 | 1.00 | 0.347 | 1.000 |
| AT2G45980.1 | ATG8-interacting protein 1 (ATI1) | 3 | 0.97 | 0.333 | 1.000 |
| AT2G45980.1 | ATG8-interacting protein 1 (ATI1) | 26 | 1.24 | 0.251 | 1.000 |
| AT2G45980.1 | ATG8-interacting protein 1 (ATI1) | 27 | 1.34 | 0.192 | 1.000 |
| AT2G45980.1 | ATG8-interacting protein 1 (ATI1) | 28 | 1.44 | 0.157 | 0.983 |
| AT2G45980.1 | ATG8-interacting protein 1 (ATI1) | 29 | 1.47 | 0.142 | 0.947 |
| AT2G45980.1 | ATG8-interacting protein 1 (ATI1) | 30 | 1.43 | 0.137 | 0.961 |
| AT2G45980.1 | ATG8-interacting protein 1 (ATI1) | 31 | 1.38 | 0.137 | 0.981 |
| AT2G45980.1 | ATG8-interacting protein 1 (ATI1) | 32 | 1.35 | 0.136 | 0.989 |
| AT2G45980.1 | ATG8-interacting protein 1 (ATI1) | 33 | 1.35 | 0.135 | 0.988 |
| AT2G45980.1 | ATG8-interacting protein 1 (ATI1) | 34 | 1.37 | 0.135 | 0.981 |
| AT2G45980.1 | ATG8-interacting protein 1 (ATI1) | 35 | 1.43 | 0.137 | 0.961 |
| AT2G45980.1 | ATG8-interacting protein 1 (ATI1) | 36 | 1.57 | 0.141 | 0.868 |

| | | | | | | |
|---|---|---|---|---|---|---|
| AT2G45980.1 | ATG8-interacting protein 1 (ATI1) | 37 | 1.84 | 0.145 | 0.575 | |
| AT2G45980.1 | ATG8-interacting protein 1 (ATI1) | 38 | 2.21 | 0.150 | 0.260 | |
| AT2G45980.1 | ATG8-interacting protein 1 (ATI1) | 39 | 2.79 | 0.153 | 0.059 | . |
| AT2G45980.1 | ATG8-interacting protein 1 (ATI1) | 40 | 3.62 | 0.150 | 0.005 | ** |
| AT2G45980.1 | ATG8-interacting protein 1 (ATI1) | 41 | 4.47 | 0.141 | 0.000 | *** |
| AT2G45980.1 | ATG8-interacting protein 1 (ATI1) | 42 | 4.55 | 0.133 | 0.000 | *** |
| AT2G45980.1 | ATG8-interacting protein 1 (ATI1) | 43 | 3.61 | 0.128 | 0.000 | *** |
| AT2G45980.1 | ATG8-interacting protein 1 (ATI1) | 44 | 2.14 | 0.126 | 0.122 | |
| AT2G45980.1 | ATG8-interacting protein 1 (ATI1) | 45 | 0.99 | 0.132 | 1.000 | |
| AT2G45980.1 | ATG8-interacting protein 1 (ATI1) | 46 | 0.39 | 0.167 | 0.190 | |
| AT2G45980.1 | ATG8-interacting protein 1 (ATI1) | 47 | 0.15 | 0.251 | 0.018 | * |
| AT2G46110.1 | Ketopentoate hydroxymethyltransferase 1 (KPHMT1) | 2 | 1.19 | 0.567 | 1.000 | |
| AT2G46110.1 | Ketopentoate hydroxymethyltransferase 1 (KPHMT1) | 3 | 1.13 | 0.432 | 1.000 | |
| AT2G46110.1 | Ketopentoate hydroxymethyltransferase 1 (KPHMT1) | 4 | 1.05 | 0.407 | 1.000 | |
| AT2G46110.1 | Ketopentoate hydroxymethyltransferase 1 (KPHMT1) | 5 | 0.95 | 0.406 | 1.000 | |
| AT2G46110.1 | Ketopentoate hydroxymethyltransferase 1 (KPHMT1) | 6 | 0.88 | 0.407 | 1.000 | |
| AT2G46110.1 | Ketopentoate hydroxymethyltransferase 1 (KPHMT1) | 7 | 0.85 | 0.406 | 1.000 | |
| AT2G46110.1 | Ketopentoate hydroxymethyltransferase 1 (KPHMT1) | 8 | 0.82 | 0.402 | 1.000 | |
| AT2G46110.1 | Ketopentoate hydroxymethyltransferase 1 (KPHMT1) | 9 | 0.80 | 0.400 | 1.000 | |
| AT2G46110.1 | Ketopentoate hydroxymethyltransferase 1 (KPHMT1) | 10 | 0.81 | 0.400 | 1.000 | |
| AT2G46110.1 | Ketopentoate hydroxymethyltransferase 1 (KPHMT1) | 11 | 0.84 | 0.401 | 1.000 | |
| AT2G46110.1 | Ketopentoate hydroxymethyltransferase 1 (KPHMT1) | 12 | 0.89 | 0.401 | 1.000 | |
| AT2G46110.1 | Ketopentoate hydroxymethyltransferase 1 (KPHMT1) | 13 | 0.91 | 0.398 | 1.000 | |
| AT2G46110.1 | Ketopentoate hydroxymethyltransferase 1 (KPHMT1) | 14 | 0.84 | 0.396 | 1.000 | |

| | | | | | |
|---|---|---|---|---|---|
| AT2G46110.1 | Ketopentoate hydroxymethyltransferase 1 (KPHMT1) | 15 | 0.74 | 0.395 | 1.000 |
| AT2G46110.1 | Ketopentoate hydroxymethyltransferase 1 (KPHMT1) | 16 | 0.77 | 0.395 | 1.000 |
| AT2G46110.1 | Ketopentoate hydroxymethyltransferase 1 (KPHMT1) | 17 | 0.92 | 0.394 | 1.000 |
| AT2G46110.1 | Ketopentoate hydroxymethyltransferase 1 (KPHMT1) | 18 | 1.06 | 0.396 | 1.000 |
| AT2G46110.1 | Ketopentoate hydroxymethyltransferase 1 (KPHMT1) | 19 | 1.11 | 0.400 | 1.000 |
| AT2G46110.1 | Ketopentoate hydroxymethyltransferase 1 (KPHMT1) | 20 | 1.04 | 0.405 | 1.000 |
| AT2G46110.1 | Ketopentoate hydroxymethyltransferase 1 (KPHMT1) | 21 | 0.84 | 0.407 | 1.000 |
| AT2G46110.1 | Ketopentoate hydroxymethyltransferase 1 (KPHMT1) | 22 | 0.61 | 0.407 | 1.000 |
| AT2G46110.1 | Ketopentoate hydroxymethyltransferase 1 (KPHMT1) | 23 | 0.48 | 0.406 | 0.999 |
| AT2G46110.1 | Ketopentoate hydroxymethyltransferase 1 (KPHMT1) | 24 | 0.48 | 0.402 | 0.998 |
| AT2G46110.1 | Ketopentoate hydroxymethyltransferase 1 (KPHMT1) | 25 | 0.65 | 0.399 | 1.000 |
| AT2G46110.1 | Ketopentoate hydroxymethyltransferase 1 (KPHMT1) | 26 | 1.23 | 0.398 | 1.000 |
| AT2G46110.1 | Ketopentoate hydroxymethyltransferase 1 (KPHMT1) | 27 | 2.63 | 0.398 | 0.973 |
| AT2G46110.1 | Ketopentoate hydroxymethyltransferase 1 (KPHMT1) | 28 | 4.67 | 0.399 | 0.618 |
| AT2G46110.1 | Ketopentoate hydroxymethyltransferase 1 (KPHMT1) | 29 | 5.98 | 0.400 | 0.426 |
| AT2G46110.1 | Ketopentoate hydroxymethyltransferase 1 (KPHMT1) | 30 | 4.77 | 0.401 | 0.609 |
| AT2G46110.1 | Ketopentoate hydroxymethyltransferase 1 (KPHMT1) | 31 | 2.26 | 0.403 | 0.995 |
| AT2G46110.1 | Ketopentoate hydroxymethyltransferase 1 (KPHMT1) | 32 | 0.88 | 0.407 | 1.000 |
| AT2G46110.1 | Ketopentoate hydroxymethyltransferase 1 (KPHMT1) | 33 | 0.40 | 0.422 | 0.989 |
| AT2G46110.1 | Ketopentoate hydroxymethyltransferase 1 (KPHMT1) | 34 | 0.24 | 0.481 | 0.887 |

| | | | | | | |
|---|---|---|---|---|---|---|
| AT2G46110.1 | Ketopentoate hydroxymethyltransferase 1 (KPHMT1) | 35 | 0.20 | 0.630 | 0.959 | |
| AT2G46110.1 | Ketopentoate hydroxymethyltransferase 1 (KPHMT1) | 46 | 63074.92 | 3.870 | 0.911 | |
| AT2G46110.1 | Ketopentoate hydroxymethyltransferase 1 (KPHMT1) | 47 | 382030.10 | 4.330 | 0.887 | |
| AT2G46320.1 | Mitochondrial substrate carrier family protein | 2 | 0.97 | 0.357 | 1.000 | |
| AT2G46320.1 | Mitochondrial substrate carrier family protein | 3 | 0.97 | 0.269 | 1.000 | |
| AT2G46320.1 | Mitochondrial substrate carrier family protein | 4 | 0.99 | 0.306 | 1.000 | |
| AT2G46320.1 | Mitochondrial substrate carrier family protein | 11 | 0.72 | 0.285 | 1.000 | |
| AT2G46320.1 | Mitochondrial substrate carrier family protein | 12 | 0.63 | 0.268 | 1.000 | |
| AT2G46320.1 | Mitochondrial substrate carrier family protein | 13 | 0.62 | 0.262 | 1.000 | |
| AT2G46320.1 | Mitochondrial substrate carrier family protein | 14 | 0.73 | 0.256 | 1.000 | |
| AT2G46320.1 | Mitochondrial substrate carrier family protein | 30 | 1.42 | 0.257 | 1.000 | |
| AT2G46320.1 | Mitochondrial substrate carrier family protein | 31 | 1.17 | 0.203 | 1.000 | |
| AT2G46320.1 | Mitochondrial substrate carrier family protein | 32 | 1.01 | 0.179 | 1.000 | |
| AT2G46320.1 | Mitochondrial substrate carrier family protein | 33 | 0.95 | 0.168 | 1.000 | |
| AT2G46320.1 | Mitochondrial substrate carrier family protein | 34 | 1.02 | 0.165 | 1.000 | |
| AT2G46320.1 | Mitochondrial substrate carrier family protein | 35 | 1.34 | 0.166 | 1.000 | |
| AT2G46320.1 | Mitochondrial substrate carrier family protein | 36 | 1.84 | 0.170 | 0.847 | |
| AT2G46320.1 | Mitochondrial substrate carrier family protein | 37 | 2.36 | 0.175 | 0.425 | |
| AT2G46320.1 | Mitochondrial substrate carrier family protein | 38 | 2.88 | 0.179 | 0.171 | |
| AT2G46320.1 | Mitochondrial substrate carrier family protein | 39 | 3.38 | 0.176 | 0.055 | . |
| AT2G46320.1 | Mitochondrial substrate carrier family protein | 40 | 3.17 | 0.169 | 0.061 | . |
| AT2G46320.1 | Mitochondrial substrate carrier family protein | 41 | 1.86 | 0.165 | 0.793 | |
| AT2G46320.1 | Mitochondrial substrate carrier family protein | 42 | 0.97 | 0.166 | 1.000 | |
| AT2G46320.1 | Mitochondrial substrate carrier family protein | 43 | 0.77 | 0.168 | 1.000 | |
| AT2G46320.1 | Mitochondrial substrate carrier family protein | 44 | 0.90 | 0.167 | 1.000 | |
| AT2G46320.1 | Mitochondrial substrate carrier family protein | 45 | 1.09 | 0.167 | 1.000 | |
| AT2G46320.1 | Mitochondrial substrate carrier family protein | 46 | 0.96 | 0.208 | 1.000 | |
| AT2G46320.1 | Mitochondrial substrate carrier family protein | 47 | 0.74 | 0.362 | 1.000 | |
| AT2G46470.1 | Oxidase assembly 1-like (OXA1L) | 2 | 0.90 | 0.639 | 1.000 | |

| | | | | | |
|---|---|---|---|---|---|
| AT2G46470.1 | Oxidase assembly 1-like (OXA1L) | 3 | 0.81 | 0.552 | 1.000 |
| AT2G46470.1 | Oxidase assembly 1-like (OXA1L) | 11 | 1.34 | 0.384 | 1.000 |
| AT2G46470.1 | Oxidase assembly 1-like (OXA1L) | 12 | 1.91 | 0.338 | 1.000 |
| AT2G46470.1 | Oxidase assembly 1-like (OXA1L) | 13 | 2.64 | 0.302 | 0.985 |
| AT2G46470.1 | Oxidase assembly 1-like (OXA1L) | 14 | 2.44 | 0.280 | 0.986 |
| AT2G46470.1 | Oxidase assembly 1-like (OXA1L) | 15 | 1.17 | 0.264 | 1.000 |
| AT2G46470.1 | Oxidase assembly 1-like (OXA1L) | 16 | 0.52 | 0.259 | 0.999 |
| AT2G46470.1 | Oxidase assembly 1-like (OXA1L) | 17 | 0.42 | 0.258 | 0.974 |
| AT2G46470.1 | Oxidase assembly 1-like (OXA1L) | 18 | 0.53 | 0.262 | 1.000 |
| AT2G46470.1 | Oxidase assembly 1-like (OXA1L) | 19 | 0.83 | 0.269 | 1.000 |
| AT2G46470.1 | Oxidase assembly 1-like (OXA1L) | 20 | 1.19 | 0.276 | 1.000 |
| AT2G46470.1 | Oxidase assembly 1-like (OXA1L) | 21 | 1.39 | 0.279 | 1.000 |
| AT2G46470.1 | Oxidase assembly 1-like (OXA1L) | 22 | 1.42 | 0.280 | 1.000 |
| AT2G46470.1 | Oxidase assembly 1-like (OXA1L) | 23 | 1.37 | 0.282 | 1.000 |
| AT2G46470.1 | Oxidase assembly 1-like (OXA1L) | 24 | 1.21 | 0.284 | 1.000 |
| AT2G46470.1 | Oxidase assembly 1-like (OXA1L) | 25 | 1.04 | 0.292 | 1.000 |
| AT2G46470.1 | Oxidase assembly 1-like (OXA1L) | 26 | 1.04 | 0.293 | 1.000 |
| AT2G46470.1 | Oxidase assembly 1-like (OXA1L) | 27 | 0.90 | 0.289 | 1.000 |
| AT2G46470.1 | Oxidase assembly 1-like (OXA1L) | 28 | 0.85 | 0.279 | 1.000 |
| AT2G46470.1 | Oxidase assembly 1-like (OXA1L) | 29 | 0.92 | 0.270 | 1.000 |
| AT2G46470.1 | Oxidase assembly 1-like (OXA1L) | 30 | 0.93 | 0.268 | 1.000 |
| AT2G46470.1 | Oxidase assembly 1-like (OXA1L) | 31 | 0.78 | 0.269 | 1.000 |
| AT2G46470.1 | Oxidase assembly 1-like (OXA1L) | 32 | 0.55 | 0.269 | 1.000 |
| AT2G46470.1 | Oxidase assembly 1-like (OXA1L) | 33 | 0.50 | 0.269 | 0.999 |
| AT2G46470.1 | Oxidase assembly 1-like (OXA1L) | 34 | 0.62 | 0.270 | 1.000 |
| AT2G46470.1 | Oxidase assembly 1-like (OXA1L) | 35 | 0.89 | 0.270 | 1.000 |
| AT2G46470.1 | Oxidase assembly 1-like (OXA1L) | 36 | 1.30 | 0.268 | 1.000 |
| AT2G46470.1 | Oxidase assembly 1-like (OXA1L) | 37 | 1.69 | 0.266 | 1.000 |
| AT2G46470.1 | Oxidase assembly 1-like (OXA1L) | 38 | 2.03 | 0.266 | 0.999 |
| AT2G46470.1 | Oxidase assembly 1-like (OXA1L) | 39 | 2.89 | 0.266 | 0.886 |
| AT2G46470.1 | Oxidase assembly 1-like (OXA1L) | 40 | 4.03 | 0.267 | 0.481 |

| | | | | | |
|---|---|---|---|---|---|
| AT2G46470.1 | Oxidase assembly 1-like (OXA1L) | 41 | 3.58 | 0.269 | 0.656 |
| AT2G46470.1 | Oxidase assembly 1-like (OXA1L) | 42 | 2.10 | 0.273 | 0.998 |
| AT2G46470.1 | Oxidase assembly 1-like (OXA1L) | 43 | 1.04 | 0.276 | 1.000 |
| AT2G46470.1 | Oxidase assembly 1-like (OXA1L) | 44 | 0.47 | 0.276 | 0.998 |
| AT2G46470.1 | Oxidase assembly 1-like (OXA1L) | 45 | 0.22 | 0.275 | 0.383 |
| AT2G46470.1 | Oxidase assembly 1-like (OXA1L) | 46 | 0.12 | 0.326 | 0.134 |
| AT2G46470.1 | Oxidase assembly 1-like (OXA1L) | 47 | 0.07 | 0.590 | 0.744 |
| AT2G46505.1 | Succinate dehydrogenase subunit (SDH4) | 2 | 1.03 | 0.451 | 1.000 |
| AT2G46505.1 | Succinate dehydrogenase subunit (SDH4) | 3 | 1.09 | 0.307 | 1.000 |
| AT2G46505.1 | Succinate dehydrogenase subunit (SDH4) | 4 | 1.12 | 0.293 | 1.000 |
| AT2G46505.1 | Succinate dehydrogenase subunit (SDH4) | 5 | 1.10 | 0.244 | 1.000 |
| AT2G46505.1 | Succinate dehydrogenase subunit (SDH4) | 6 | 1.11 | 0.223 | 1.000 |
| AT2G46505.1 | Succinate dehydrogenase subunit (SDH4) | 7 | 1.25 | 0.220 | 1.000 |
| AT2G46505.1 | Succinate dehydrogenase subunit (SDH4) | 8 | 1.49 | 0.213 | 1.000 |
| AT2G46505.1 | Succinate dehydrogenase subunit (SDH4) | 9 | 1.67 | 0.210 | 1.000 |
| AT2G46505.1 | Succinate dehydrogenase subunit (SDH4) | 10 | 1.76 | 0.213 | 0.999 |
| AT2G46505.1 | Succinate dehydrogenase subunit (SDH4) | 11 | 1.66 | 0.214 | 1.000 |
| AT2G46505.1 | Succinate dehydrogenase subunit (SDH4) | 12 | 1.58 | 0.214 | 1.000 |
| AT2G46505.1 | Succinate dehydrogenase subunit (SDH4) | 13 | 1.44 | 0.210 | 1.000 |
| AT2G46505.1 | Succinate dehydrogenase subunit (SDH4) | 14 | 1.28 | 0.205 | 1.000 |
| AT2G46505.1 | Succinate dehydrogenase subunit (SDH4) | 15 | 1.01 | 0.208 | 1.000 |
| AT2G46505.1 | Succinate dehydrogenase subunit (SDH4) | 16 | 1.07 | 0.213 | 1.000 |
| AT2G46505.1 | Succinate dehydrogenase subunit (SDH4) | 17 | 1.66 | 0.217 | 1.000 |
| AT2G46505.1 | Succinate dehydrogenase subunit (SDH4) | 18 | 2.22 | 0.222 | 0.965 |
| AT2G46505.1 | Succinate dehydrogenase subunit (SDH4) | 19 | 2.61 | 0.272 | 0.971 |
| AT2G46505.1 | Succinate dehydrogenase subunit (SDH4) | 20 | 2.77 | 0.385 | 0.999 |
| AT2G46505.1 | Succinate dehydrogenase subunit (SDH4) | 28 | 1.76 | 0.348 | 1.000 |
| AT2G46505.1 | Succinate dehydrogenase subunit (SDH4) | 29 | 1.81 | 0.282 | 1.000 |
| AT2G46505.1 | Succinate dehydrogenase subunit (SDH4) | 30 | 1.82 | 0.245 | 1.000 |
| AT2G46505.1 | Succinate dehydrogenase subunit (SDH4) | 31 | 1.61 | 0.231 | 1.000 |
| AT2G46505.1 | Succinate dehydrogenase subunit (SDH4) | 32 | 1.21 | 0.221 | 1.000 |

| | | | | | | |
|---|---|---|---|---|---|---|
| AT2G46505.1 | Succinate dehydrogenase subunit (SDH4) | 33 | 1.17 | 0.215 | 1.000 | |
| AT2G46505.1 | Succinate dehydrogenase subunit (SDH4) | 34 | 1.78 | 0.214 | 0.999 | |
| AT2G46505.1 | Succinate dehydrogenase subunit (SDH4) | 35 | 1.96 | 0.211 | 0.991 | |
| AT2G46505.1 | Succinate dehydrogenase subunit (SDH4) | 36 | 0.82 | 0.210 | 1.000 | |
| AT2G46505.1 | Succinate dehydrogenase subunit (SDH4) | 37 | 0.25 | 0.210 | 0.121 | |
| AT2G46505.1 | Succinate dehydrogenase subunit (SDH4) | 38 | 0.20 | 0.212 | 0.034 | * |
| AT2G46505.1 | Succinate dehydrogenase subunit (SDH4) | 39 | 0.57 | 0.213 | 0.999 | |
| AT2G46505.1 | Succinate dehydrogenase subunit (SDH4) | 40 | 1.83 | 0.211 | 0.998 | |
| AT2G46505.1 | Succinate dehydrogenase subunit (SDH4) | 41 | 2.44 | 0.211 | 0.845 | |
| AT2G46505.1 | Succinate dehydrogenase subunit (SDH4) | 42 | 1.72 | 0.214 | 1.000 | |
| AT2G46505.1 | Succinate dehydrogenase subunit (SDH4) | 43 | 0.88 | 0.215 | 1.000 | |
| AT2G46505.1 | Succinate dehydrogenase subunit (SDH4) | 44 | 0.39 | 0.214 | 0.784 | |
| AT2G46505.1 | Succinate dehydrogenase subunit (SDH4) | 45 | 0.22 | 0.216 | 0.072 | . |
| AT2G46505.1 | Succinate dehydrogenase subunit (SDH4) | 46 | 0.16 | 0.240 | 0.031 | * |
| AT2G46505.1 | Succinate dehydrogenase subunit (SDH4) | 47 | 0.11 | 0.453 | 0.597 | |
| AT2G46540.1 | NADH:ubiquinone oxidoreductase subunit (B9) | 2 | 0.86 | 0.341 | 1.000 | |
| AT2G46540.1 | NADH:ubiquinone oxidoreductase subunit (B9) | 3 | 0.84 | 0.221 | 1.000 | |
| AT2G46540.1 | NADH:ubiquinone oxidoreductase subunit (B9) | 4 | 0.81 | 0.204 | 1.000 | |
| AT2G46540.1 | NADH:ubiquinone oxidoreductase subunit (B9) | 5 | 0.80 | 0.199 | 1.000 | |
| AT2G46540.1 | NADH:ubiquinone oxidoreductase subunit (B9) | 6 | 0.86 | 0.199 | 1.000 | |
| AT2G46540.1 | NADH:ubiquinone oxidoreductase subunit (B9) | 7 | 0.91 | 0.206 | 1.000 | |
| AT2G46540.1 | NADH:ubiquinone oxidoreductase subunit (B9) | 8 | 0.90 | 0.214 | 1.000 | |
| AT2G46540.1 | NADH:ubiquinone oxidoreductase subunit (B9) | 9 | 0.87 | 0.221 | 1.000 | |
| AT2G46540.1 | NADH:ubiquinone oxidoreductase subunit (B9) | 10 | 0.84 | 0.217 | 1.000 | |
| AT2G46540.1 | NADH:ubiquinone oxidoreductase subunit (B9) | 11 | 0.85 | 0.206 | 1.000 | |
| AT2G46540.1 | NADH:ubiquinone oxidoreductase subunit (B9) | 12 | 0.95 | 0.201 | 1.000 | |
| AT2G46540.1 | NADH:ubiquinone oxidoreductase subunit (B9) | 13 | 1.15 | 0.193 | 1.000 | |
| AT2G46540.1 | NADH:ubiquinone oxidoreductase subunit (B9) | 14 | 1.28 | 0.180 | 1.000 | |
| AT2G46540.1 | NADH:ubiquinone oxidoreductase subunit (B9) | 15 | 1.08 | 0.172 | 1.000 | |
| AT2G46540.1 | NADH:ubiquinone oxidoreductase subunit (B9) | 16 | 0.71 | 0.168 | 1.000 | |
| AT2G46540.1 | NADH:ubiquinone oxidoreductase subunit (B9) | 17 | 0.50 | 0.167 | 0.767 | |

| | | | | | |
|---|---|---|---|---|---|
| AT2G46540.1 | NADH:ubiquinone oxidoreductase subunit (B9) | 18 | 0.49 | 0.170 | 0.757 |
| AT2G46540.1 | NADH:ubiquinone oxidoreductase subunit (B9) | 19 | 0.61 | 0.175 | 0.991 |
| AT2G46540.1 | NADH:ubiquinone oxidoreductase subunit (B9) | 20 | 0.75 | 0.180 | 1.000 |
| AT2G46540.1 | NADH:ubiquinone oxidoreductase subunit (B9) | 21 | 0.89 | 0.182 | 1.000 |
| AT2G46540.1 | NADH:ubiquinone oxidoreductase subunit (B9) | 22 | 1.09 | 0.181 | 1.000 |
| AT2G46540.1 | NADH:ubiquinone oxidoreductase subunit (B9) | 23 | 1.29 | 0.179 | 1.000 |
| AT2G46540.1 | NADH:ubiquinone oxidoreductase subunit (B9) | 24 | 1.28 | 0.176 | 1.000 |
| AT2G46540.1 | NADH:ubiquinone oxidoreductase subunit (B9) | 25 | 1.13 | 0.173 | 1.000 |
| AT2G46540.1 | NADH:ubiquinone oxidoreductase subunit (B9) | 26 | 1.11 | 0.173 | 1.000 |
| AT2G46540.1 | NADH:ubiquinone oxidoreductase subunit (B9) | 27 | 1.13 | 0.177 | 1.000 |
| AT2G46540.1 | NADH:ubiquinone oxidoreductase subunit (B9) | 28 | 1.12 | 0.191 | 1.000 |
| AT2G46540.1 | NADH:ubiquinone oxidoreductase subunit (B9) | 29 | 1.25 | 0.220 | 1.000 |
| AT2G46540.1 | NADH:ubiquinone oxidoreductase subunit (B9) | 30 | 1.55 | 0.243 | 1.000 |
| AT2G46540.1 | NADH:ubiquinone oxidoreductase subunit (B9) | 32 | 1.72 | 0.265 | 1.000 |
| AT2G46540.1 | NADH:ubiquinone oxidoreductase subunit (B9) | 33 | 1.52 | 0.298 | 1.000 |
| AT2G46540.1 | NADH:ubiquinone oxidoreductase subunit (B9) | 34 | 1.33 | 0.392 | 1.000 |
| AT2G46540.1 | NADH:ubiquinone oxidoreductase subunit (B9) | 46 | 3.45 | 4.353 | 1.000 |
| AT2G46540.1 | NADH:ubiquinone oxidoreductase subunit (B9) | 47 | 4.15 | 4.854 | 1.000 |
| AT2G46820.2 | Photosystem I subunit (PSAP) | 2 | 0.84 | 0.585 | 1.000 |
| AT2G46820.2 | Photosystem I subunit (PSAP) | 3 | 1.15 | 0.554 | 1.000 |
| AT2G46820.2 | Photosystem I subunit (PSAP) | 26 | 1.37 | 0.442 | 1.000 |
| AT2G46820.2 | Photosystem I subunit (PSAP) | 27 | 0.91 | 0.341 | 1.000 |
| AT2G46820.2 | Photosystem I subunit (PSAP) | 28 | 0.97 | 0.295 | 1.000 |
| AT2G46820.2 | Photosystem I subunit (PSAP) | 29 | 1.05 | 0.276 | 1.000 |
| AT2G46820.2 | Photosystem I subunit (PSAP) | 30 | 0.75 | 0.273 | 1.000 |
| AT2G46820.2 | Photosystem I subunit (PSAP) | 31 | 0.62 | 0.274 | 1.000 |
| AT2G46820.2 | Photosystem I subunit (PSAP) | 32 | 0.68 | 0.281 | 1.000 |
| AT2G46820.2 | Photosystem I subunit (PSAP) | 33 | 0.65 | 0.299 | 1.000 |
| AT2G46820.2 | Photosystem I subunit (PSAP) | 34 | 0.61 | 0.328 | 1.000 |
| AT2G46820.2 | Photosystem I subunit (PSAP) | 35 | 0.77 | 0.376 | 1.000 |
| AT2G46820.2 | Photosystem I subunit (PSAP) | 38 | 2.61 | 0.400 | 0.989 |
| AT2G46820.2 | Photosystem I subunit (PSAP) | 39 | 2.32 | 0.329 | 0.981 |
| AT2G46820.2 | Photosystem I subunit (PSAP) | 40 | 1.47 | 0.286 | 1.000 |
| AT2G46820.2 | Photosystem I subunit (PSAP) | 41 | 0.65 | 0.275 | 1.000 |
| AT2G46820.2 | Photosystem I subunit (PSAP) | 42 | 0.40 | 0.279 | 0.878 |
| AT2G46820.2 | Photosystem I subunit (PSAP) | 43 | 0.69 | 0.281 | 1.000 |
| AT2G46820.2 | Photosystem I subunit (PSAP) | 44 | 1.48 | 0.279 | 1.000 |
| AT2G46820.2 | Photosystem I subunit (PSAP) | 45 | 1.72 | 0.277 | 0.999 |
| AT2G46820.2 | Photosystem I subunit (PSAP) | 46 | 1.30 | 0.318 | 1.000 |
| AT2G46820.2 | Photosystem I subunit (PSAP) | 47 | 0.85 | 0.591 | 1.000 |
| AT2G47380.1 | Cytochrome c:O2 oxidoreductase subunit (COX5C) | 2 | 0.91 | 0.395 | 1.000 |
| AT2G47380.1 | Cytochrome c:O2 oxidoreductase subunit (COX5C) | 3 | 0.82 | 0.257 | 1.000 |

| | | | | | |
|---|---|---|---|---|---|
| AT2G47380.1 | Cytochrome c:O2 oxidoreductase subunit (COX5C) | 4 | 0.79 | 0.236 | 1.000 |
| AT2G47380.1 | Cytochrome c:O2 oxidoreductase subunit (COX5C) | 5 | 0.82 | 0.237 | 1.000 |
| AT2G47380.1 | Cytochrome c:O2 oxidoreductase subunit (COX5C) | 6 | 0.85 | 0.237 | 1.000 |
| AT2G47380.1 | Cytochrome c:O2 oxidoreductase subunit (COX5C) | 7 | 0.84 | 0.236 | 1.000 |
| AT2G47380.1 | Cytochrome c:O2 oxidoreductase subunit (COX5C) | 8 | 0.84 | 0.232 | 1.000 |
| AT2G47380.1 | Cytochrome c:O2 oxidoreductase subunit (COX5C) | 9 | 0.81 | 0.230 | 1.000 |
| AT2G47380.1 | Cytochrome c:O2 oxidoreductase subunit (COX5C) | 10 | 0.74 | 0.230 | 1.000 |
| AT2G47380.1 | Cytochrome c:O2 oxidoreductase subunit (COX5C) | 11 | 0.72 | 0.230 | 1.000 |
| AT2G47380.1 | Cytochrome c:O2 oxidoreductase subunit (COX5C) | 12 | 0.70 | 0.230 | 1.000 |
| AT2G47380.1 | Cytochrome c:O2 oxidoreductase subunit (COX5C) | 13 | 0.65 | 0.227 | 1.000 |
| AT2G47380.1 | Cytochrome c:O2 oxidoreductase subunit (COX5C) | 14 | 0.65 | 0.225 | 1.000 |
| AT2G47380.1 | Cytochrome c:O2 oxidoreductase subunit (COX5C) | 15 | 0.69 | 0.225 | 1.000 |
| AT2G47380.1 | Cytochrome c:O2 oxidoreductase subunit (COX5C) | 16 | 0.87 | 0.224 | 1.000 |
| AT2G47380.1 | Cytochrome c:O2 oxidoreductase subunit (COX5C) | 17 | 1.13 | 0.223 | 1.000 |
| AT2G47380.1 | Cytochrome c:O2 oxidoreductase subunit (COX5C) | 18 | 1.40 | 0.226 | 1.000 |
| AT2G47380.1 | Cytochrome c:O2 oxidoreductase subunit (COX5C) | 19 | 1.62 | 0.230 | 1.000 |
| AT2G47380.1 | Cytochrome c:O2 oxidoreductase subunit (COX5C) | 20 | 1.65 | 0.235 | 1.000 |
| AT2G47380.1 | Cytochrome c:O2 oxidoreductase subunit (COX5C) | 21 | 1.70 | 0.237 | 1.000 |
| AT2G47380.1 | Cytochrome c:O2 oxidoreductase subunit (COX5C) | 22 | 1.78 | 0.238 | 0.999 |
| AT2G47380.1 | Cytochrome c:O2 oxidoreductase subunit (COX5C) | 23 | 1.57 | 0.237 | 1.000 |
| AT2G47380.1 | Cytochrome c:O2 oxidoreductase subunit (COX5C) | 24 | 1.35 | 0.232 | 1.000 |
| AT2G47380.1 | Cytochrome c:O2 oxidoreductase subunit (COX5C) | 25 | 1.23 | 0.228 | 1.000 |
| AT2G47380.1 | Cytochrome c:O2 oxidoreductase subunit (COX5C) | 26 | 1.18 | 0.228 | 1.000 |
| AT2G47380.1 | Cytochrome c:O2 oxidoreductase subunit (COX5C) | 27 | 1.19 | 0.228 | 1.000 |
| AT2G47380.1 | Cytochrome c:O2 oxidoreductase subunit (COX5C) | 28 | 1.37 | 0.228 | 1.000 |
| AT2G47380.1 | Cytochrome c:O2 oxidoreductase subunit (COX5C) | 29 | 1.70 | 0.228 | 0.999 |
| AT2G47380.1 | Cytochrome c:O2 oxidoreductase subunit (COX5C) | 30 | 1.83 | 0.229 | 0.996 |
| AT2G47380.1 | Cytochrome c:O2 oxidoreductase subunit (COX5C) | 31 | 1.55 | 0.230 | 1.000 |
| AT2G47380.1 | Cytochrome c:O2 oxidoreductase subunit (COX5C) | 32 | 1.00 | 0.231 | 1.000 |
| AT2G47380.1 | Cytochrome c:O2 oxidoreductase subunit (COX5C) | 33 | 0.49 | 0.231 | 0.980 |
| AT2G47380.1 | Cytochrome c:O2 oxidoreductase subunit (COX5C) | 34 | 0.24 | 0.233 | 0.168 |

| | | | | | | |
|---|---|---|---|---|---|---|
| AT2G47380.1 | Cytochrome c:O2 oxidoreductase subunit (COX5C) | 35 | 0.19 | 0.236 | 0.062 | . |
| AT2G47380.1 | Cytochrome c:O2 oxidoreductase subunit (COX5C) | 36 | 0.23 | 0.249 | 0.226 | |
| AT2G47380.1 | Cytochrome c:O2 oxidoreductase subunit (COX5C) | 37 | 0.33 | 0.302 | 0.905 | |
| AT2G47380.1 | Cytochrome c:O2 oxidoreductase subunit (COX5C) | 38 | 0.60 | 0.437 | 1.000 | |
| AT2G47380.1 | Cytochrome c:O2 oxidoreductase subunit (COX5C) | 46 | 80736.38 | 2.902 | 0.849 | |
| AT2G47380.1 | Cytochrome c:O2 oxidoreductase subunit (COX5C) | 47 | 539722.52 | 3.368 | 0.842 | |
| AT2G47390.1 | Prolyl oligopeptidase family protein | 2 | 0.95 | 0.869 | 1.000 | |
| AT2G47390.1 | Prolyl oligopeptidase family protein | 3 | 1.29 | 1.598 | 1.000 | |
| AT2G47390.1 | Prolyl oligopeptidase family protein | 31 | 0.99 | 0.685 | 1.000 | |
| AT2G47390.1 | Prolyl oligopeptidase family protein | 32 | 0.51 | 0.547 | 1.000 | |
| AT2G47390.1 | Prolyl oligopeptidase family protein | 33 | 0.26 | 0.563 | 0.966 | |
| AT2G47390.1 | Prolyl oligopeptidase family protein | 35 | 0.19 | 0.487 | 0.775 | |
| AT2G47390.1 | Prolyl oligopeptidase family protein | 36 | 0.41 | 0.412 | 0.983 | |
| AT2G47390.1 | Prolyl oligopeptidase family protein | 37 | 1.45 | 0.367 | 1.000 | |
| AT2G47390.1 | Prolyl oligopeptidase family protein | 38 | 2.94 | 0.355 | 0.865 | |
| AT2G47390.1 | Prolyl oligopeptidase family protein | 39 | 1.52 | 0.352 | 1.000 | |
| AT2G47390.1 | Prolyl oligopeptidase family protein | 40 | 0.39 | 0.373 | 0.952 | |
| AT2G47390.1 | Prolyl oligopeptidase family protein | 41 | 0.17 | 0.512 | 0.767 | |
| AT2G47390.1 | Prolyl oligopeptidase family protein | 42 | 0.18 | 0.821 | 0.985 | |
| AT2G47390.1 | Prolyl oligopeptidase family protein | 46 | 22.72 | 1.927 | 0.998 | |
| AT2G47390.1 | Prolyl oligopeptidase family protein | 47 | 133.42 | 2.410 | 0.989 | |
| AT2G47490.1 | Chloroplast NAD+ transporter 1 (NDT1) | 2 | 1.05 | 0.494 | 1.000 | |
| AT2G47490.1 | Chloroplast NAD+ transporter 1 (NDT1) | 3 | 1.07 | 0.373 | 1.000 | |
| AT2G47490.1 | Chloroplast NAD+ transporter 1 (NDT1) | 6 | 1.18 | 0.357 | 1.000 | |
| AT2G47490.1 | Chloroplast NAD+ transporter 1 (NDT1) | 7 | 1.30 | 0.279 | 1.000 | |
| AT2G47490.1 | Chloroplast NAD+ transporter 1 (NDT1) | 8 | 1.74 | 0.241 | 1.000 | |
| AT2G47490.1 | Chloroplast NAD+ transporter 1 (NDT1) | 9 | 2.46 | 0.229 | 0.872 | |
| AT2G47490.1 | Chloroplast NAD+ transporter 1 (NDT1) | 10 | 2.83 | 0.226 | 0.663 | |
| AT2G47490.1 | Chloroplast NAD+ transporter 1 (NDT1) | 11 | 2.57 | 0.227 | 0.808 | |
| AT2G47490.1 | Chloroplast NAD+ transporter 1 (NDT1) | 12 | 2.02 | 0.226 | 0.984 | |
| AT2G47490.1 | Chloroplast NAD+ transporter 1 (NDT1) | 13 | 1.63 | 0.223 | 1.000 | |

| | | | | | | |
|---|---|---|---|---|---|---|
| AT2G47490.1 | Chloroplast NAD+ transporter 1 (NDT1) | 14 | 1.68 | 0.221 | 1.000 | |
| AT2G47490.1 | Chloroplast NAD+ transporter 1 (NDT1) | 15 | 1.93 | 0.221 | 0.991 | |
| AT2G47490.1 | Chloroplast NAD+ transporter 1 (NDT1) | 16 | 2.10 | 0.219 | 0.963 | |
| AT2G47490.1 | Chloroplast NAD+ transporter 1 (NDT1) | 17 | 2.80 | 0.220 | 0.634 | |
| AT2G47490.1 | Chloroplast NAD+ transporter 1 (NDT1) | 18 | 4.25 | 0.231 | 0.163 | |
| AT2G47490.1 | Chloroplast NAD+ transporter 1 (NDT1) | 19 | 4.97 | 0.249 | 0.132 | |
| AT2G47490.1 | Chloroplast NAD+ transporter 1 (NDT1) | 20 | 5.85 | 0.293 | 0.212 | |
| AT2G47490.1 | Chloroplast NAD+ transporter 1 (NDT1) | 21 | 6.52 | 0.453 | 0.815 | |
| AT2G47490.1 | Chloroplast NAD+ transporter 1 (NDT1) | 35 | 1.62 | 0.363 | 1.000 | |
| AT2G47490.1 | Chloroplast NAD+ transporter 1 (NDT1) | 36 | 2.95 | 0.269 | 0.848 | |
| AT2G47490.1 | Chloroplast NAD+ transporter 1 (NDT1) | 37 | 5.78 | 0.235 | 0.036 | * |
| AT2G47490.1 | Chloroplast NAD+ transporter 1 (NDT1) | 38 | 15.00 | 0.225 | 0.000 | *** |
| AT2G47490.1 | Chloroplast NAD+ transporter 1 (NDT1) | 39 | 36.69 | 0.226 | 0.000 | *** |
| AT2G47490.1 | Chloroplast NAD+ transporter 1 (NDT1) | 40 | 22.85 | 0.225 | 0.000 | *** |
| AT2G47490.1 | Chloroplast NAD+ transporter 1 (NDT1) | 41 | 3.80 | 0.224 | 0.225 | |
| AT2G47490.1 | Chloroplast NAD+ transporter 1 (NDT1) | 42 | 0.56 | 0.229 | 0.999 | |
| AT2G47490.1 | Chloroplast NAD+ transporter 1 (NDT1) | 43 | 0.14 | 0.231 | 0.007 | ** |
| AT2G47490.1 | Chloroplast NAD+ transporter 1 (NDT1) | 44 | 0.09 | 0.234 | 0.000 | *** |
| AT2G47490.1 | Chloroplast NAD+ transporter 1 (NDT1) | 45 | 0.27 | 0.267 | 0.545 | |
| AT2G47490.1 | Chloroplast NAD+ transporter 1 (NDT1) | 46 | 2.10 | 0.491 | 1.000 | |
| AT2G47490.1 | Chloroplast NAD+ transporter 1 (NDT1) | 47 | 22.64 | 0.993 | 0.982 | |
| AT2G47510.2 | Fumarase 1 (FUM1) | 2 | 1.19 | 0.413 | 1.000 | |
| AT2G47510.2 | Fumarase 1 (FUM1) | 3 | 0.74 | 0.283 | 1.000 | |
| AT2G47510.2 | Fumarase 1 (FUM1) | 4 | 0.49 | 0.271 | 0.999 | |
| AT2G47510.2 | Fumarase 1 (FUM1) | 5 | 0.40 | 0.276 | 0.982 | |
| AT2G47510.2 | Fumarase 1 (FUM1) | 6 | 0.44 | 0.249 | 0.986 | |
| AT2G47510.2 | Fumarase 1 (FUM1) | 7 | 0.59 | 0.230 | 1.000 | |
| AT2G47510.2 | Fumarase 1 (FUM1) | 8 | 0.79 | 0.217 | 1.000 | |
| AT2G47510.2 | Fumarase 1 (FUM1) | 9 | 1.02 | 0.210 | 1.000 | |
| AT2G47510.2 | Fumarase 1 (FUM1) | 10 | 1.19 | 0.209 | 1.000 | |
| AT2G47510.2 | Fumarase 1 (FUM1) | 11 | 1.41 | 0.210 | 1.000 | |
| AT2G47510.2 | Fumarase 1 (FUM1) | 12 | 1.60 | 0.210 | 1.000 | |
| AT2G47510.2 | Fumarase 1 (FUM1) | 13 | 1.50 | 0.208 | 1.000 | |
| AT2G47510.2 | Fumarase 1 (FUM1) | 14 | 1.26 | 0.206 | 1.000 | |
| AT2G47510.2 | Fumarase 1 (FUM1) | 15 | 1.16 | 0.205 | 1.000 | |
| AT2G47510.2 | Fumarase 1 (FUM1) | 16 | 1.24 | 0.203 | 1.000 | |
| AT2G47510.2 | Fumarase 1 (FUM1) | 17 | 1.45 | 0.202 | 1.000 | |
| AT2G47510.2 | Fumarase 1 (FUM1) | 18 | 1.51 | 0.205 | 1.000 | |
| AT2G47510.2 | Fumarase 1 (FUM1) | 19 | 1.39 | 0.212 | 1.000 | |
| AT2G47510.2 | Fumarase 1 (FUM1) | 20 | 1.29 | 0.217 | 1.000 | |
| AT2G47510.2 | Fumarase 1 (FUM1) | 21 | 1.22 | 0.219 | 1.000 | |

| | | | | | | |
|---|---|---|---|---|---|---|
| AT2G47510.2 | Fumarase 1 (FUM1) | 22 | 1.15 | 0.219 | 1.000 | |
| AT2G47510.2 | Fumarase 1 (FUM1) | 23 | 1.07 | 0.217 | 1.000 | |
| AT2G47510.2 | Fumarase 1 (FUM1) | 24 | 0.99 | 0.213 | 1.000 | |
| AT2G47510.2 | Fumarase 1 (FUM1) | 25 | 0.96 | 0.209 | 1.000 | |
| AT2G47510.2 | Fumarase 1 (FUM1) | 26 | 1.05 | 0.208 | 1.000 | |
| AT2G47510.2 | Fumarase 1 (FUM1) | 27 | 1.19 | 0.208 | 1.000 | |
| AT2G47510.2 | Fumarase 1 (FUM1) | 28 | 1.27 | 0.207 | 1.000 | |
| AT2G47510.2 | Fumarase 1 (FUM1) | 29 | 1.28 | 0.208 | 1.000 | |
| AT2G47510.2 | Fumarase 1 (FUM1) | 30 | 1.26 | 0.209 | 1.000 | |
| AT2G47510.2 | Fumarase 1 (FUM1) | 31 | 1.22 | 0.210 | 1.000 | |
| AT2G47510.2 | Fumarase 1 (FUM1) | 32 | 1.19 | 0.211 | 1.000 | |
| AT2G47510.2 | Fumarase 1 (FUM1) | 33 | 1.17 | 0.211 | 1.000 | |
| AT2G47510.2 | Fumarase 1 (FUM1) | 34 | 1.15 | 0.211 | 1.000 | |
| AT2G47510.2 | Fumarase 1 (FUM1) | 35 | 1.11 | 0.211 | 1.000 | |
| AT2G47510.2 | Fumarase 1 (FUM1) | 36 | 1.01 | 0.210 | 1.000 | |
| AT2G47510.2 | Fumarase 1 (FUM1) | 37 | 0.92 | 0.209 | 1.000 | |
| AT2G47510.2 | Fumarase 1 (FUM1) | 38 | 1.00 | 0.209 | 1.000 | |
| AT2G47510.2 | Fumarase 1 (FUM1) | 39 | 1.55 | 0.209 | 1.000 | |
| AT2G47510.2 | Fumarase 1 (FUM1) | 40 | 2.93 | 0.210 | 0.544 | |
| AT2G47510.2 | Fumarase 1 (FUM1) | 41 | 4.84 | 0.212 | 0.046 | * |
| AT2G47510.2 | Fumarase 1 (FUM1) | 42 | 6.03 | 0.217 | 0.013 | * |
| AT2G47510.2 | Fumarase 1 (FUM1) | 43 | 5.61 | 0.227 | 0.037 | * |
| AT2G47510.2 | Fumarase 1 (FUM1) | 44 | 4.03 | 0.275 | 0.562 | |
| AT2G47510.2 | Fumarase 1 (FUM1) | 45 | 2.82 | 0.397 | 0.999 | |
| AT2G47510.2 | Fumarase 1 (FUM1) | 46 | 2.00 | 0.633 | 1.000 | |
| AT2G47510.2 | Fumarase 1 (FUM1) | 47 | 1.42 | 0.961 | 1.000 | |
| AT2G47830.2 | Cation efflux family protein | 2 | 1.10 | 0.451 | 1.000 | |
| AT2G47830.2 | Cation efflux family protein | 3 | 1.09 | 0.435 | 1.000 | |
| AT2G47830.2 | Cation efflux family protein | 22 | 0.94 | 0.335 | 1.000 | |
| AT2G47830.2 | Cation efflux family protein | 23 | 0.95 | 0.309 | 1.000 | |
| AT2G47830.2 | Cation efflux family protein | 24 | 0.95 | 0.283 | 1.000 | |
| AT2G47830.2 | Cation efflux family protein | 29 | 1.09 | 0.187 | 1.000 | |
| AT2G47830.2 | Cation efflux family protein | 30 | 1.17 | 0.167 | 1.000 | |
| AT2G47830.2 | Cation efflux family protein | 31 | 1.27 | 0.151 | 0.989 | |
| AT2G47830.2 | Cation efflux family protein | 32 | 1.39 | 0.140 | 0.912 | |
| AT2G47830.2 | Cation efflux family protein | 33 | 1.50 | 0.133 | 0.757 | |
| AT2G47830.2 | Cation efflux family protein | 34 | 1.56 | 0.130 | 0.649 | |
| AT2G47830.2 | Cation efflux family protein | 35 | 1.54 | 0.131 | 0.680 | |
| AT2G47830.2 | Cation efflux family protein | 36 | 1.43 | 0.136 | 0.854 | |
| AT2G47830.2 | Cation efflux family protein | 37 | 1.24 | 0.146 | 0.994 | |
| AT2G47830.2 | Cation efflux family protein | 38 | 0.99 | 0.163 | 1.000 | |
| AT2G47830.2 | Cation efflux family protein | 39 | 0.75 | 0.187 | 0.990 | |
| AT2G47830.2 | Cation efflux family protein | 40 | 0.56 | 0.214 | 0.832 | |
| AT2G47830.2 | Cation efflux family protein | 46 | 0.08 | 0.383 | 0.049 | * |
| AT2G47830.2 | Cation efflux family protein | 47 | 0.06 | 0.431 | 0.046 | * |
| AT3G01120.1 | Cystathionine gamma-synthase (CGS1) | 2 | 1.05 | 0.452 | 1.000 | |
| AT3G01120.1 | Cystathionine gamma-synthase (CGS1) | 3 | 1.09 | 0.435 | 1.000 | |
| AT3G01120.1 | Cystathionine gamma-synthase (CGS1) | 4 | 1.13 | 0.425 | 1.000 | |
| AT3G01120.1 | Cystathionine gamma-synthase (CGS1) | 32 | 1.20 | 0.295 | 0.996 | |
| AT3G01120.1 | Cystathionine gamma-synthase (CGS1) | 33 | 1.11 | 0.292 | 1.000 | |
| AT3G01120.1 | Cystathionine gamma-synthase (CGS1) | 34 | 1.03 | 0.292 | 1.000 | |
| AT3G01120.1 | Cystathionine gamma-synthase (CGS1) | 35 | 0.93 | 0.297 | 1.000 | |
| AT3G01120.1 | Cystathionine gamma-synthase (CGS1) | 36 | 0.84 | 0.304 | 0.997 | |
| AT3G01120.1 | Cystathionine gamma-synthase (CGS1) | 46 | 0.18 | 0.453 | 0.311 | |

| | | | | | | |
|---|---|---|---|---|---|---|
| AT3G01120.1 | Cystathionine gamma-synthase (CGS1) | 47 | 0.15 | 0.478 | 0.277 | |
| AT3G01130.2 | ATP synthase subunit (E chain) | 2 | 1.03 | 0.485 | 1.000 | |
| AT3G01130.2 | ATP synthase subunit (E chain) | 3 | 1.20 | 0.396 | 1.000 | |
| AT3G01130.2 | ATP synthase subunit (E chain) | 8 | 1.84 | 0.380 | 1.000 | |
| AT3G01130.2 | ATP synthase subunit (E chain) | 9 | 1.75 | 0.293 | 1.000 | |
| AT3G01130.2 | ATP synthase subunit (E chain) | 10 | 1.63 | 0.271 | 1.000 | |
| AT3G01130.2 | ATP synthase subunit (E chain) | 11 | 1.55 | 0.263 | 1.000 | |
| AT3G01130.2 | ATP synthase subunit (E chain) | 12 | 1.58 | 0.263 | 1.000 | |
| AT3G01130.2 | ATP synthase subunit (E chain) | 13 | 1.66 | 0.258 | 1.000 | |
| AT3G01130.2 | ATP synthase subunit (E chain) | 14 | 1.67 | 0.245 | 1.000 | |
| AT3G01130.2 | ATP synthase subunit (E chain) | 15 | 1.61 | 0.245 | 1.000 | |
| AT3G01130.2 | ATP synthase subunit (E chain) | 16 | 1.61 | 0.248 | 1.000 | |
| AT3G01130.2 | ATP synthase subunit (E chain) | 17 | 1.70 | 0.233 | 1.000 | |
| AT3G01130.2 | ATP synthase subunit (E chain) | 18 | 1.92 | 0.220 | 0.997 | |
| AT3G01130.2 | ATP synthase subunit (E chain) | 19 | 2.90 | 0.225 | 0.702 | |
| AT3G01130.2 | ATP synthase subunit (E chain) | 20 | 4.03 | 0.229 | 0.244 | |
| AT3G01130.2 | ATP synthase subunit (E chain) | 21 | 4.01 | 0.233 | 0.272 | |
| AT3G01130.2 | ATP synthase subunit (E chain) | 22 | 2.65 | 0.231 | 0.863 | |
| AT3G01130.2 | ATP synthase subunit (E chain) | 23 | 1.42 | 0.229 | 1.000 | |
| AT3G01130.2 | ATP synthase subunit (E chain) | 24 | 1.26 | 0.225 | 1.000 | |
| AT3G01130.2 | ATP synthase subunit (E chain) | 25 | 2.12 | 0.221 | 0.983 | |
| AT3G01130.2 | ATP synthase subunit (E chain) | 26 | 3.19 | 0.219 | 0.489 | |
| AT3G01130.2 | ATP synthase subunit (E chain) | 27 | 2.01 | 0.219 | 0.992 | |
| AT3G01130.2 | ATP synthase subunit (E chain) | 28 | 1.02 | 0.219 | 1.000 | |
| AT3G01130.2 | ATP synthase subunit (E chain) | 29 | 0.85 | 0.221 | 1.000 | |
| AT3G01130.2 | ATP synthase subunit (E chain) | 30 | 0.89 | 0.221 | 1.000 | |
| AT3G01130.2 | ATP synthase subunit (E chain) | 31 | 1.18 | 0.222 | 1.000 | |
| AT3G01130.2 | ATP synthase subunit (E chain) | 32 | 1.68 | 0.222 | 1.000 | |
| AT3G01130.2 | ATP synthase subunit (E chain) | 33 | 1.48 | 0.222 | 1.000 | |
| AT3G01130.2 | ATP synthase subunit (E chain) | 34 | 1.02 | 0.222 | 1.000 | |
| AT3G01130.2 | ATP synthase subunit (E chain) | 35 | 1.27 | 0.222 | 1.000 | |
| AT3G01130.2 | ATP synthase subunit (E chain) | 36 | 2.32 | 0.221 | 0.943 | |
| AT3G01130.2 | ATP synthase subunit (E chain) | 37 | 4.30 | 0.222 | 0.141 | |
| AT3G01130.2 | ATP synthase subunit (E chain) | 38 | 5.18 | 0.223 | 0.051 | . |
| AT3G01130.2 | ATP synthase subunit (E chain) | 39 | 4.34 | 0.222 | 0.134 | |
| AT3G01130.2 | ATP synthase subunit (E chain) | 40 | 3.83 | 0.221 | 0.246 | |
| AT3G01130.2 | ATP synthase subunit (E chain) | 41 | 3.10 | 0.221 | 0.555 | |
| AT3G01130.2 | ATP synthase subunit (E chain) | 42 | 2.21 | 0.225 | 0.974 | |
| AT3G01130.2 | ATP synthase subunit (E chain) | 43 | 1.30 | 0.226 | 1.000 | |
| AT3G01130.2 | ATP synthase subunit (E chain) | 44 | 0.64 | 0.225 | 1.000 | |
| AT3G01130.2 | ATP synthase subunit (E chain) | 45 | 0.36 | 0.225 | 0.756 | |
| AT3G01130.2 | ATP synthase subunit (E chain) | 46 | 0.23 | 0.253 | 0.316 | |
| AT3G01130.2 | ATP synthase subunit (E chain) | 47 | 0.14 | 0.506 | 0.924 | |
| AT3G01280.1 | Voltage-dependent anion channel 1 (VDAC1) | 3 | 0.72 | 0.140 | 1.000 | |
| AT3G01280.1 | Voltage-dependent anion channel 1 (VDAC1) | 4 | 0.89 | 0.133 | 1.000 | |
| AT3G01280.1 | Voltage-dependent anion channel 1 (VDAC1) | 5 | 1.23 | 0.133 | 1.000 | |
| AT3G01280.1 | Voltage-dependent anion channel 1 (VDAC1) | 6 | 1.50 | 0.133 | 0.991 | |
| AT3G01280.1 | Voltage-dependent anion channel 1 (VDAC1) | 7 | 1.56 | 0.133 | 0.973 | |
| AT3G01280.1 | Voltage-dependent anion channel 1 (VDAC1) | 8 | 1.48 | 0.132 | 0.993 | |
| AT3G01280.1 | Voltage-dependent anion channel 1 (VDAC1) | 9 | 1.32 | 0.130 | 1.000 | |
| AT3G01280.1 | Voltage-dependent anion channel 1 (VDAC1) | 10 | 1.21 | 0.130 | 1.000 | |
| AT3G01280.1 | Voltage-dependent anion channel 1 (VDAC1) | 11 | 1.23 | 0.130 | 1.000 | |

| | | | | | | |
|---|---|---|---|---|---|---|
| AT3G01280.1 | Voltage-dependent anion channel 1 (VDAC1) | 12 | 1.32 | 0.130 | 1.000 | |
| AT3G01280.1 | Voltage-dependent anion channel 1 (VDAC1) | 13 | 1.47 | 0.129 | 0.993 | |
| AT3G01280.1 | Voltage-dependent anion channel 1 (VDAC1) | 14 | 1.56 | 0.128 | 0.960 | |
| AT3G01280.1 | Voltage-dependent anion channel 1 (VDAC1) | 15 | 1.46 | 0.128 | 0.994 | |
| AT3G01280.1 | Voltage-dependent anion channel 1 (VDAC1) | 16 | 1.25 | 0.127 | 1.000 | |
| AT3G01280.1 | Voltage-dependent anion channel 1 (VDAC1) | 17 | 1.22 | 0.127 | 1.000 | |
| AT3G01280.1 | Voltage-dependent anion channel 1 (VDAC1) | 18 | 1.33 | 0.128 | 1.000 | |
| AT3G01280.1 | Voltage-dependent anion channel 1 (VDAC1) | 19 | 1.46 | 0.131 | 0.995 | |
| AT3G01280.1 | Voltage-dependent anion channel 1 (VDAC1) | 20 | 1.58 | 0.133 | 0.964 | |
| AT3G01280.1 | Voltage-dependent anion channel 1 (VDAC1) | 21 | 1.60 | 0.135 | 0.955 | |
| AT3G01280.1 | Voltage-dependent anion channel 1 (VDAC1) | 22 | 1.53 | 0.135 | 0.986 | |
| AT3G01280.1 | Voltage-dependent anion channel 1 (VDAC1) | 23 | 1.34 | 0.134 | 1.000 | |
| AT3G01280.1 | Voltage-dependent anion channel 1 (VDAC1) | 24 | 1.15 | 0.132 | 1.000 | |
| AT3G01280.1 | Voltage-dependent anion channel 1 (VDAC1) | 25 | 0.99 | 0.130 | 1.000 | |
| AT3G01280.1 | Voltage-dependent anion channel 1 (VDAC1) | 26 | 0.95 | 0.130 | 1.000 | |
| AT3G01280.1 | Voltage-dependent anion channel 1 (VDAC1) | 27 | 0.95 | 0.130 | 1.000 | |
| AT3G01280.1 | Voltage-dependent anion channel 1 (VDAC1) | 28 | 0.97 | 0.130 | 1.000 | |
| AT3G01280.1 | Voltage-dependent anion channel 1 (VDAC1) | 29 | 1.11 | 0.129 | 1.000 | |
| AT3G01280.1 | Voltage-dependent anion channel 1 (VDAC1) | 30 | 1.28 | 0.129 | 1.000 | |
| AT3G01280.1 | Voltage-dependent anion channel 1 (VDAC1) | 31 | 1.22 | 0.130 | 1.000 | |
| AT3G01280.1 | Voltage-dependent anion channel 1 (VDAC1) | 32 | 1.06 | 0.130 | 1.000 | |
| AT3G01280.1 | Voltage-dependent anion channel 1 (VDAC1) | 33 | 1.02 | 0.130 | 1.000 | |
| AT3G01280.1 | Voltage-dependent anion channel 1 (VDAC1) | 34 | 1.06 | 0.130 | 1.000 | |
| AT3G01280.1 | Voltage-dependent anion channel 1 (VDAC1) | 35 | 1.08 | 0.130 | 1.000 | |
| AT3G01280.1 | Voltage-dependent anion channel 1 (VDAC1) | 36 | 1.04 | 0.130 | 1.000 | |
| AT3G01280.1 | Voltage-dependent anion channel 1 (VDAC1) | 37 | 1.04 | 0.129 | 1.000 | |
| AT3G01280.1 | Voltage-dependent anion channel 1 (VDAC1) | 38 | 1.13 | 0.130 | 1.000 | |
| AT3G01280.1 | Voltage-dependent anion channel 1 (VDAC1) | 39 | 1.45 | 0.130 | 0.996 | |
| AT3G01280.1 | Voltage-dependent anion channel 1 (VDAC1) | 40 | 2.27 | 0.131 | 0.171 | |
| AT3G01280.1 | Voltage-dependent anion channel 1 (VDAC1) | 41 | 3.02 | 0.131 | 0.010 | ** |
| AT3G01280.1 | Voltage-dependent anion channel 1 (VDAC1) | 42 | 2.51 | 0.132 | 0.077 | . |

| | | | | | | |
|---|---|---|---|---|---|---|
| AT3G01280.1 | Voltage-dependent anion channel 1 (VDAC1) | 43 | 1.45 | 0.133 | 0.997 | |
| AT3G01280.1 | Voltage-dependent anion channel 1 (VDAC1) | 44 | 0.69 | 0.133 | 0.997 | |
| AT3G01280.1 | Voltage-dependent anion channel 1 (VDAC1) | 45 | 0.34 | 0.133 | 0.017 | * |
| AT3G01280.1 | Voltage-dependent anion channel 1 (VDAC1) | 46 | 0.19 | 0.148 | 0.000 | *** |
| AT3G01280.1 | Voltage-dependent anion channel 1 (VDAC1) | 47 | 0.11 | 0.259 | 0.011 | * |
| AT3G01280.1 | Voltage-dependent anion channel 1 (VDAC1) | 48 | 0.07 | 0.439 | 0.194 | |
| AT3G01440.1 | NAD(P)H dehydrogenase complex subunit (PNSL3) | 2 | 1.02 | 0.318 | 1.000 | |
| AT3G01440.1 | NAD(P)H dehydrogenase complex subunit (PNSL3) | 3 | 1.04 | 0.258 | 1.000 | |
| AT3G01440.1 | NAD(P)H dehydrogenase complex subunit (PNSL3) | 4 | 1.05 | 0.220 | 1.000 | |
| AT3G01440.1 | NAD(P)H dehydrogenase complex subunit (PNSL3) | 7 | 1.00 | 0.172 | 1.000 | |
| AT3G01440.1 | NAD(P)H dehydrogenase complex subunit (PNSL3) | 8 | 0.95 | 0.159 | 1.000 | |
| AT3G01440.1 | NAD(P)H dehydrogenase complex subunit (PNSL3) | 9 | 0.90 | 0.147 | 1.000 | |
| AT3G01440.1 | NAD(P)H dehydrogenase complex subunit (PNSL3) | 10 | 0.87 | 0.137 | 1.000 | |
| AT3G01440.1 | NAD(P)H dehydrogenase complex subunit (PNSL3) | 11 | 0.86 | 0.130 | 1.000 | |
| AT3G01440.1 | NAD(P)H dehydrogenase complex subunit (PNSL3) | 12 | 0.84 | 0.126 | 0.999 | |
| AT3G01440.1 | NAD(P)H dehydrogenase complex subunit (PNSL3) | 13 | 0.80 | 0.123 | 0.996 | |
| AT3G01440.1 | NAD(P)H dehydrogenase complex subunit (PNSL3) | 14 | 0.76 | 0.122 | 0.979 | |
| AT3G01440.1 | NAD(P)H dehydrogenase complex subunit (PNSL3) | 15 | 0.73 | 0.122 | 0.943 | |
| AT3G01440.1 | NAD(P)H dehydrogenase complex subunit (PNSL3) | 16 | 0.72 | 0.122 | 0.922 | |
| AT3G01440.1 | NAD(P)H dehydrogenase complex subunit (PNSL3) | 17 | 0.76 | 0.124 | 0.980 | |
| AT3G01440.1 | NAD(P)H dehydrogenase complex subunit (PNSL3) | 18 | 0.90 | 0.128 | 1.000 | |
| AT3G01440.1 | NAD(P)H dehydrogenase complex subunit (PNSL3) | 19 | 1.16 | 0.136 | 1.000 | |
| AT3G01440.1 | NAD(P)H dehydrogenase complex subunit (PNSL3) | 20 | 1.57 | 0.146 | 0.853 | |
| AT3G01440.1 | NAD(P)H dehydrogenase complex subunit (PNSL3) | 21 | 2.13 | 0.162 | 0.378 | |
| AT3G01440.1 | NAD(P)H dehydrogenase complex subunit (PNSL3) | 22 | 2.77 | 0.184 | 0.182 | |
| AT3G01440.1 | NAD(P)H dehydrogenase complex subunit (PNSL3) | 23 | 3.41 | 0.213 | 0.145 | |
| AT3G01440.1 | NAD(P)H dehydrogenase complex subunit (PNSL3) | 24 | 3.99 | 0.251 | 0.188 | |
| AT3G01440.1 | NAD(P)H dehydrogenase complex subunit (PNSL3) | 41 | 1.31 | 0.323 | 1.000 | |
| AT3G01440.1 | NAD(P)H dehydrogenase complex subunit (PNSL3) | 42 | 1.13 | 0.316 | 1.000 | |
| AT3G01440.1 | NAD(P)H dehydrogenase complex subunit (PNSL3) | 43 | 0.98 | 0.349 | 1.000 | |
| AT3G01440.1 | NAD(P)H dehydrogenase complex subunit (PNSL3) | 46 | 0.65 | 0.686 | 1.000 | |

| | | | | | | |
|---|---|---|---|---|---|---|
| AT3G01440.1 | NAD(P)H dehydrogenase complex subunit (PNSL3) | 47 | 0.56 | 0.839 | 1.000 | |
| AT3G01500.1 | Beta carbonic anhydrase 1 (BCA1) | 2 | 1.33 | 0.385 | 1.000 | |
| AT3G01500.1 | Beta carbonic anhydrase 1 (BCA1) | 3 | 0.68 | 0.288 | 1.000 | |
| AT3G01500.1 | Beta carbonic anhydrase 1 (BCA1) | 4 | 0.37 | 0.524 | 0.998 | |
| AT3G01500.1 | Beta carbonic anhydrase 1 (BCA1) | 30 | 0.59 | 0.285 | 0.999 | |
| AT3G01500.1 | Beta carbonic anhydrase 1 (BCA1) | 31 | 0.72 | 0.212 | 1.000 | |
| AT3G01500.1 | Beta carbonic anhydrase 1 (BCA1) | 32 | 0.80 | 0.190 | 1.000 | |
| AT3G01500.1 | Beta carbonic anhydrase 1 (BCA1) | 33 | 0.90 | 0.179 | 1.000 | |
| AT3G01500.1 | Beta carbonic anhydrase 1 (BCA1) | 34 | 1.26 | 0.176 | 1.000 | |
| AT3G01500.1 | Beta carbonic anhydrase 1 (BCA1) | 35 | 2.07 | 0.177 | 0.654 | |
| AT3G01500.1 | Beta carbonic anhydrase 1 (BCA1) | 36 | 2.86 | 0.188 | 0.219 | |
| AT3G01500.1 | Beta carbonic anhydrase 1 (BCA1) | 37 | 2.60 | 0.217 | 0.552 | |
| AT3G01500.1 | Beta carbonic anhydrase 1 (BCA1) | 38 | 2.19 | 0.243 | 0.893 | |
| AT3G01500.1 | Beta carbonic anhydrase 1 (BCA1) | 40 | 2.36 | 0.248 | 0.843 | |
| AT3G01500.1 | Beta carbonic anhydrase 1 (BCA1) | 41 | 3.30 | 0.209 | 0.191 | |
| AT3G01500.1 | Beta carbonic anhydrase 1 (BCA1) | 42 | 4.19 | 0.186 | 0.018 | * |
| AT3G01500.1 | Beta carbonic anhydrase 1 (BCA1) | 43 | 3.60 | 0.180 | 0.039 | * |
| AT3G01500.1 | Beta carbonic anhydrase 1 (BCA1) | 44 | 1.94 | 0.178 | 0.776 | |
| AT3G01500.1 | Beta carbonic anhydrase 1 (BCA1) | 45 | 0.91 | 0.178 | 1.000 | |
| AT3G01500.1 | Beta carbonic anhydrase 1 (BCA1) | 46 | 0.46 | 0.211 | 0.781 | |
| AT3G01500.1 | Beta carbonic anhydrase 1 (BCA1) | 47 | 0.25 | 0.376 | 0.768 | |
| AT3G01910.1 | Sulfite oxidase (SOX) | 2 | 1.25 | 0.298 | 1.000 | |
| AT3G01910.1 | Sulfite oxidase (SOX) | 3 | 1.37 | 0.225 | 0.999 | |
| AT3G01910.1 | Sulfite oxidase (SOX) | 4 | 1.50 | 0.182 | 0.966 | |
| AT3G01910.1 | Sulfite oxidase (SOX) | 5 | 1.65 | 0.164 | 0.818 | |
| AT3G01910.1 | Sulfite oxidase (SOX) | 6 | 1.82 | 0.158 | 0.614 | |
| AT3G01910.1 | Sulfite oxidase (SOX) | 7 | 1.94 | 0.158 | 0.473 | |
| AT3G01910.1 | Sulfite oxidase (SOX) | 8 | 1.99 | 0.157 | 0.427 | |
| AT3G01910.1 | Sulfite oxidase (SOX) | 9 | 1.92 | 0.156 | 0.489 | |
| AT3G01910.1 | Sulfite oxidase (SOX) | 10 | 1.74 | 0.156 | 0.679 | |
| AT3G01910.1 | Sulfite oxidase (SOX) | 11 | 1.50 | 0.155 | 0.916 | |
| AT3G01910.1 | Sulfite oxidase (SOX) | 12 | 1.24 | 0.155 | 0.999 | |
| AT3G01910.1 | Sulfite oxidase (SOX) | 13 | 1.00 | 0.154 | 1.000 | |
| AT3G01910.1 | Sulfite oxidase (SOX) | 14 | 0.82 | 0.155 | 0.999 | |
| AT3G01910.1 | Sulfite oxidase (SOX) | 15 | 0.70 | 0.155 | 0.955 | |
| AT3G01910.1 | Sulfite oxidase (SOX) | 16 | 0.64 | 0.156 | 0.856 | |
| AT3G01910.1 | Sulfite oxidase (SOX) | 17 | 0.63 | 0.159 | 0.857 | |
| AT3G01910.1 | Sulfite oxidase (SOX) | 18 | 0.69 | 0.165 | 0.960 | |
| AT3G01910.1 | Sulfite oxidase (SOX) | 19 | 0.82 | 0.178 | 1.000 | |
| AT3G01910.1 | Sulfite oxidase (SOX) | 20 | 1.07 | 0.205 | 1.000 | |
| AT3G01910.1 | Sulfite oxidase (SOX) | 21 | 1.49 | 0.250 | 0.996 | |

| | | | | | |
|---|---|---|---|---|---|
| AT3G01910.1 | Sulfite oxidase (SOX) | 22 | 2.14 | 0.317 | 0.946 |
| AT3G01910.1 | Sulfite oxidase (SOX) | 46 | 36656.75 | 4.779 | 0.967 |
| AT3G01910.1 | Sulfite oxidase (SOX) | 47 | 56234.14 | 5.047 | 0.970 |
| AT3G02090.1 | Ubiquinol:cytochrome c oxidoreductase subunit (MPPbeta) | 3 | 0.65 | 0.278 | 1.000 |
| AT3G02090.1 | Ubiquinol:cytochrome c oxidoreductase subunit (MPPbeta) | 4 | 0.68 | 0.265 | 1.000 |
| AT3G02090.1 | Ubiquinol:cytochrome c oxidoreductase subunit (MPPbeta) | 5 | 0.94 | 0.265 | 1.000 |
| AT3G02090.1 | Ubiquinol:cytochrome c oxidoreductase subunit (MPPbeta) | 6 | 1.08 | 0.266 | 1.000 |
| AT3G02090.1 | Ubiquinol:cytochrome c oxidoreductase subunit (MPPbeta) | 7 | 0.97 | 0.268 | 1.000 |
| AT3G02090.1 | Ubiquinol:cytochrome c oxidoreductase subunit (MPPbeta) | 8 | 0.94 | 0.261 | 1.000 |
| AT3G02090.1 | Ubiquinol:cytochrome c oxidoreductase subunit (MPPbeta) | 9 | 1.01 | 0.257 | 1.000 |
| AT3G02090.1 | Ubiquinol:cytochrome c oxidoreductase subunit (MPPbeta) | 10 | 1.07 | 0.257 | 1.000 |
| AT3G02090.1 | Ubiquinol:cytochrome c oxidoreductase subunit (MPPbeta) | 11 | 1.18 | 0.258 | 1.000 |
| AT3G02090.1 | Ubiquinol:cytochrome c oxidoreductase subunit (MPPbeta) | 12 | 1.32 | 0.259 | 1.000 |
| AT3G02090.1 | Ubiquinol:cytochrome c oxidoreductase subunit (MPPbeta) | 13 | 1.53 | 0.255 | 1.000 |
| AT3G02090.1 | Ubiquinol:cytochrome c oxidoreductase subunit (MPPbeta) | 14 | 1.71 | 0.251 | 1.000 |
| AT3G02090.1 | Ubiquinol:cytochrome c oxidoreductase subunit (MPPbeta) | 15 | 1.53 | 0.251 | 1.000 |
| AT3G02090.1 | Ubiquinol:cytochrome c oxidoreductase subunit (MPPbeta) | 16 | 1.04 | 0.249 | 1.000 |
| AT3G02090.1 | Ubiquinol:cytochrome c oxidoreductase subunit (MPPbeta) | 17 | 0.70 | 0.249 | 1.000 |
| AT3G02090.1 | Ubiquinol:cytochrome c oxidoreductase subunit (MPPbeta) | 18 | 0.67 | 0.253 | 1.000 |
| AT3G02090.1 | Ubiquinol:cytochrome c oxidoreductase subunit (MPPbeta) | 19 | 0.90 | 0.259 | 1.000 |
| AT3G02090.1 | Ubiquinol:cytochrome c oxidoreductase subunit (MPPbeta) | 20 | 1.28 | 0.266 | 1.000 |
| AT3G02090.1 | Ubiquinol:cytochrome c oxidoreductase subunit (MPPbeta) | 21 | 1.73 | 0.270 | 1.000 |

| | | | | | |
|---|---|---|---|---|---|
| AT3G02090.1 | Ubiquinol:cytochrome c oxidoreductase subunit (MPPbeta) | 22 | 1.61 | 0.269 | 1.000 |
| AT3G02090.1 | Ubiquinol:cytochrome c oxidoreductase subunit (MPPbeta) | 23 | 0.81 | 0.269 | 1.000 |
| AT3G02090.1 | Ubiquinol:cytochrome c oxidoreductase subunit (MPPbeta) | 24 | 0.47 | 0.263 | 0.999 |
| AT3G02090.1 | Ubiquinol:cytochrome c oxidoreductase subunit (MPPbeta) | 25 | 0.58 | 0.257 | 1.000 |
| AT3G02090.1 | Ubiquinol:cytochrome c oxidoreductase subunit (MPPbeta) | 26 | 1.19 | 0.256 | 1.000 |
| AT3G02090.1 | Ubiquinol:cytochrome c oxidoreductase subunit (MPPbeta) | 27 | 2.25 | 0.256 | 0.995 |
| AT3G02090.1 | Ubiquinol:cytochrome c oxidoreductase subunit (MPPbeta) | 28 | 1.70 | 0.256 | 1.000 |
| AT3G02090.1 | Ubiquinol:cytochrome c oxidoreductase subunit (MPPbeta) | 29 | 0.65 | 0.256 | 1.000 |
| AT3G02090.1 | Ubiquinol:cytochrome c oxidoreductase subunit (MPPbeta) | 30 | 0.49 | 0.255 | 0.999 |
| AT3G02090.1 | Ubiquinol:cytochrome c oxidoreductase subunit (MPPbeta) | 31 | 0.84 | 0.256 | 1.000 |
| AT3G02090.1 | Ubiquinol:cytochrome c oxidoreductase subunit (MPPbeta) | 32 | 1.56 | 0.257 | 1.000 |
| AT3G02090.1 | Ubiquinol:cytochrome c oxidoreductase subunit (MPPbeta) | 33 | 1.71 | 0.257 | 1.000 |
| AT3G02090.1 | Ubiquinol:cytochrome c oxidoreductase subunit (MPPbeta) | 34 | 1.18 | 0.258 | 1.000 |
| AT3G02090.1 | Ubiquinol:cytochrome c oxidoreductase subunit (MPPbeta) | 35 | 0.78 | 0.257 | 1.000 |
| AT3G02090.1 | Ubiquinol:cytochrome c oxidoreductase subunit (MPPbeta) | 36 | 0.68 | 0.256 | 1.000 |
| AT3G02090.1 | Ubiquinol:cytochrome c oxidoreductase subunit (MPPbeta) | 37 | 0.89 | 0.256 | 1.000 |
| AT3G02090.1 | Ubiquinol:cytochrome c oxidoreductase subunit (MPPbeta) | 38 | 1.37 | 0.257 | 1.000 |
| AT3G02090.1 | Ubiquinol:cytochrome c oxidoreductase subunit (MPPbeta) | 39 | 1.95 | 0.258 | 1.000 |
| AT3G02090.1 | Ubiquinol:cytochrome c oxidoreductase subunit (MPPbeta) | 40 | 2.84 | 0.259 | 0.912 |
| AT3G02090.1 | Ubiquinol:cytochrome c oxidoreductase subunit (MPPbeta) | 41 | 3.54 | 0.259 | 0.660 |

| | | | | | | |
|---|---|---|---|---|---|---|
| AT3G02090.1 | Ubiquinol:cytochrome c oxidoreductase subunit (MPPbeta) | 42 | 3.00 | 0.262 | 0.878 | |
| AT3G02090.1 | Ubiquinol:cytochrome c oxidoreductase subunit (MPPbeta) | 43 | 2.45 | 0.264 | 0.986 | |
| AT3G02090.1 | Ubiquinol:cytochrome c oxidoreductase subunit (MPPbeta) | 44 | 1.68 | 0.264 | 1.000 | |
| AT3G02090.1 | Ubiquinol:cytochrome c oxidoreductase subunit (MPPbeta) | 45 | 0.80 | 0.265 | 1.000 | |
| AT3G02090.1 | Ubiquinol:cytochrome c oxidoreductase subunit (MPPbeta) | 46 | 0.28 | 0.297 | 0.839 | |
| AT3G02090.1 | Ubiquinol:cytochrome c oxidoreductase subunit (MPPbeta) | 47 | 0.07 | 0.538 | 0.670 | |
| AT3G02090.1 | Ubiquinol:cytochrome c oxidoreductase subunit (MPPbeta) | 48 | 0.02 | 0.941 | 0.873 | |
| AT3G02490.1 | Pentatricopeptide repeat (PPR) superfamily protein | 2 | 1.11 | 0.332 | 1.000 | |
| AT3G02490.1 | Pentatricopeptide repeat (PPR) superfamily protein | 3 | 1.09 | 0.302 | 1.000 | |
| AT3G02490.1 | Pentatricopeptide repeat (PPR) superfamily protein | 4 | 1.06 | 0.427 | 1.000 | |
| AT3G02490.1 | Pentatricopeptide repeat (PPR) superfamily protein | 30 | 0.70 | 0.248 | 0.999 | |
| AT3G02490.1 | Pentatricopeptide repeat (PPR) superfamily protein | 31 | 0.74 | 0.195 | 0.998 | |
| AT3G02490.1 | Pentatricopeptide repeat (PPR) superfamily protein | 32 | 0.78 | 0.171 | 0.999 | |
| AT3G02490.1 | Pentatricopeptide repeat (PPR) superfamily protein | 33 | 0.82 | 0.162 | 1.000 | |
| AT3G02490.1 | Pentatricopeptide repeat (PPR) superfamily protein | 34 | 0.85 | 0.157 | 1.000 | |
| AT3G02490.1 | Pentatricopeptide repeat (PPR) superfamily protein | 35 | 0.94 | 0.152 | 1.000 | |
| AT3G02490.1 | Pentatricopeptide repeat (PPR) superfamily protein | 36 | 1.19 | 0.149 | 1.000 | |
| AT3G02490.1 | Pentatricopeptide repeat (PPR) superfamily protein | 37 | 1.68 | 0.147 | 0.718 | |
| AT3G02490.1 | Pentatricopeptide repeat (PPR) superfamily protein | 38 | 2.37 | 0.146 | 0.121 | |
| AT3G02490.1 | Pentatricopeptide repeat (PPR) superfamily protein | 39 | 2.72 | 0.147 | 0.043 | * |
| AT3G02490.1 | Pentatricopeptide repeat (PPR) superfamily protein | 40 | 2.34 | 0.149 | 0.147 | |
| AT3G02490.1 | Pentatricopeptide repeat (PPR) superfamily protein | 41 | 1.60 | 0.154 | 0.847 | |
| AT3G02490.1 | Pentatricopeptide repeat (PPR) superfamily protein | 42 | 0.96 | 0.164 | 1.000 | |
| AT3G02490.1 | Pentatricopeptide repeat (PPR) superfamily protein | 43 | 0.58 | 0.180 | 0.859 | |
| AT3G02490.1 | Pentatricopeptide repeat (PPR) superfamily protein | 44 | 0.37 | 0.205 | 0.321 | |
| AT3G02490.1 | Pentatricopeptide repeat (PPR) superfamily protein | 45 | 0.25 | 0.229 | 0.103 | |
| AT3G02490.1 | Pentatricopeptide repeat (PPR) superfamily protein | 46 | 0.18 | 0.259 | 0.048 | * |

| | | | | | | |
|---|---|---|---|---|---|---|
| AT3G02490.1 | Pentatricopeptide repeat (PPR) superfamily protein | 47 | 0.13 | 0.322 | 0.067 | . |
| AT3G02650.1 | Pentatricopeptide repeat (PPR) superfamily protein | 2 | 1.09 | 0.467 | 1.000 | |
| AT3G02650.1 | Pentatricopeptide repeat (PPR) superfamily protein | 3 | 1.13 | 0.416 | 1.000 | |
| AT3G02650.1 | Pentatricopeptide repeat (PPR) superfamily protein | 13 | 1.83 | 0.326 | 0.999 | |
| AT3G02650.1 | Pentatricopeptide repeat (PPR) superfamily protein | 14 | 1.96 | 0.309 | 0.997 | |
| AT3G02650.1 | Pentatricopeptide repeat (PPR) superfamily protein | 15 | 2.11 | 0.291 | 0.984 | |
| AT3G02650.1 | Pentatricopeptide repeat (PPR) superfamily protein | 16 | 2.28 | 0.319 | 0.982 | |
| AT3G02650.1 | Pentatricopeptide repeat (PPR) superfamily protein | 27 | 1.71 | 0.323 | 1.000 | |
| AT3G02650.1 | Pentatricopeptide repeat (PPR) superfamily protein | 28 | 1.30 | 0.251 | 1.000 | |
| AT3G02650.1 | Pentatricopeptide repeat (PPR) superfamily protein | 29 | 0.98 | 0.224 | 1.000 | |
| AT3G02650.1 | Pentatricopeptide repeat (PPR) superfamily protein | 30 | 0.84 | 0.214 | 1.000 | |
| AT3G02650.1 | Pentatricopeptide repeat (PPR) superfamily protein | 31 | 0.89 | 0.211 | 1.000 | |
| AT3G02650.1 | Pentatricopeptide repeat (PPR) superfamily protein | 32 | 0.98 | 0.211 | 1.000 | |
| AT3G02650.1 | Pentatricopeptide repeat (PPR) superfamily protein | 33 | 1.12 | 0.214 | 1.000 | |
| AT3G02650.1 | Pentatricopeptide repeat (PPR) superfamily protein | 34 | 1.42 | 0.221 | 1.000 | |
| AT3G02650.1 | Pentatricopeptide repeat (PPR) superfamily protein | 35 | 1.97 | 0.231 | 0.950 | |
| AT3G02650.1 | Pentatricopeptide repeat (PPR) superfamily protein | 36 | 3.03 | 0.241 | 0.496 | |
| AT3G02650.1 | Pentatricopeptide repeat (PPR) superfamily protein | 37 | 4.59 | 0.242 | 0.110 | |
| AT3G02650.1 | Pentatricopeptide repeat (PPR) superfamily protein | 38 | 5.57 | 0.238 | 0.035 | * |
| AT3G02650.1 | Pentatricopeptide repeat (PPR) superfamily protein | 39 | 5.34 | 0.228 | 0.030 | * |
| AT3G02650.1 | Pentatricopeptide repeat (PPR) superfamily protein | 40 | 4.82 | 0.217 | 0.034 | * |
| AT3G02650.1 | Pentatricopeptide repeat (PPR) superfamily protein | 41 | 3.85 | 0.212 | 0.102 | |
| AT3G02650.1 | Pentatricopeptide repeat (PPR) superfamily protein | 42 | 2.92 | 0.214 | 0.368 | |
| AT3G02650.1 | Pentatricopeptide repeat (PPR) superfamily protein | 43 | 2.15 | 0.216 | 0.833 | |
| AT3G02650.1 | Pentatricopeptide repeat (PPR) superfamily protein | 44 | 1.37 | 0.215 | 1.000 | |
| AT3G02650.1 | Pentatricopeptide repeat (PPR) superfamily protein | 45 | 0.71 | 0.215 | 1.000 | |
| AT3G02650.1 | Pentatricopeptide repeat (PPR) superfamily protein | 46 | 0.28 | 0.251 | 0.370 | |
| AT3G02650.1 | Pentatricopeptide repeat (PPR) superfamily protein | 47 | 0.09 | 0.404 | 0.170 | |
| AT3G02660.1 | Tyrosyl-tRNA synthetase | 2 | 0.87 | 0.657 | 1.000 | |
| AT3G02660.1 | Tyrosyl-tRNA synthetase | 3 | 1.17 | 0.663 | 1.000 | |
| AT3G02660.1 | Tyrosyl-tRNA synthetase | 29 | 0.74 | 0.500 | 1.000 | |
| AT3G02660.1 | Tyrosyl-tRNA synthetase | 30 | 0.49 | 0.391 | 1.000 | |
| AT3G02660.1 | Tyrosyl-tRNA synthetase | 31 | 0.58 | 0.358 | 1.000 | |
| AT3G02660.1 | Tyrosyl-tRNA synthetase | 32 | 0.91 | 0.338 | 1.000 | |

| | | | | | |
|---|---|---|---|---|---|
| AT3G02660.1 | Tyrosyl-tRNA synthetase | 33 | 1.11 | 0.331 | 1.000 |
| AT3G02660.1 | Tyrosyl-tRNA synthetase | 34 | 1.60 | 0.333 | 1.000 |
| AT3G02660.1 | Tyrosyl-tRNA synthetase | 35 | 2.54 | 0.334 | 0.971 |
| AT3G02660.1 | Tyrosyl-tRNA synthetase | 36 | 2.68 | 0.333 | 0.955 |
| AT3G02660.1 | Tyrosyl-tRNA synthetase | 37 | 1.84 | 0.328 | 0.999 |
| AT3G02660.1 | Tyrosyl-tRNA synthetase | 38 | 1.54 | 0.330 | 1.000 |
| AT3G02660.1 | Tyrosyl-tRNA synthetase | 39 | 2.86 | 0.334 | 0.929 |
| AT3G02660.1 | Tyrosyl-tRNA synthetase | 40 | 3.48 | 0.332 | 0.792 |
| AT3G02660.1 | Tyrosyl-tRNA synthetase | 41 | 1.13 | 0.334 | 1.000 |
| AT3G02660.1 | Tyrosyl-tRNA synthetase | 42 | 0.29 | 0.347 | 0.842 |
| AT3G02660.1 | Tyrosyl-tRNA synthetase | 43 | 0.23 | 0.470 | 0.933 |
| AT3G02660.1 | Tyrosyl-tRNA synthetase | 44 | 0.81 | 0.742 | 1.000 |
| AT3G02660.1 | Tyrosyl-tRNA synthetase | 46 | 238.99 | 1.807 | 0.946 |
| AT3G02660.1 | Tyrosyl-tRNA synthetase | 47 | 8424.60 | 2.624 | 0.873 |
| AT3G02780.1 | Isopentenyl pyrophosphate:dimethylallyl pyrophosphate isomerase 2 (IPP2) | 2 | 1.20 | 0.400 | 1.000 |
| AT3G02780.1 | Isopentenyl pyrophosphate:dimethylallyl pyrophosphate isomerase 2 (IPP2) | 3 | 1.16 | 0.364 | 1.000 |
| AT3G02780.1 | Isopentenyl pyrophosphate:dimethylallyl pyrophosphate isomerase 2 (IPP2) | 4 | 1.12 | 0.339 | 1.000 |
| AT3G02780.1 | Isopentenyl pyrophosphate:dimethylallyl pyrophosphate isomerase 2 (IPP2) | 12 | 0.76 | 0.260 | 1.000 |
| AT3G02780.1 | Isopentenyl pyrophosphate:dimethylallyl pyrophosphate isomerase 2 (IPP2) | 13 | 0.70 | 0.247 | 0.998 |
| AT3G02780.1 | Isopentenyl pyrophosphate:dimethylallyl pyrophosphate isomerase 2 (IPP2) | 14 | 0.64 | 0.233 | 0.983 |
| AT3G02780.1 | Isopentenyl pyrophosphate:dimethylallyl pyrophosphate isomerase 2 (IPP2) | 15 | 0.58 | 0.220 | 0.924 |
| AT3G02780.1 | Isopentenyl pyrophosphate:dimethylallyl pyrophosphate isomerase 2 (IPP2) | 16 | 0.52 | 0.209 | 0.785 |
| AT3G02780.1 | Isopentenyl pyrophosphate:dimethylallyl pyrophosphate isomerase 2 (IPP2) | 17 | 0.47 | 0.200 | 0.583 |
| AT3G02780.1 | Isopentenyl pyrophosphate:dimethylallyl pyrophosphate isomerase 2 (IPP2) | 18 | 0.43 | 0.194 | 0.392 |
| AT3G02780.1 | Isopentenyl pyrophosphate:dimethylallyl pyrophosphate isomerase 2 (IPP2) | 19 | 0.40 | 0.189 | 0.263 |
| AT3G02780.1 | Isopentenyl pyrophosphate:dimethylallyl pyrophosphate isomerase 2 (IPP2) | 20 | 0.38 | 0.187 | 0.195 |

| | | | | | |
|---|---|---|---|---|---|
| AT3G02780.1 | Isopentenyl pyrophosphate:dimethylallyl pyrophosphate isomerase 2 (IPP2) | 21 | 0.37 | 0.185 | 0.177 |
| AT3G02780.1 | Isopentenyl pyrophosphate:dimethylallyl pyrophosphate isomerase 2 (IPP2) | 22 | 0.38 | 0.184 | 0.199 |
| AT3G02780.1 | Isopentenyl pyrophosphate:dimethylallyl pyrophosphate isomerase 2 (IPP2) | 23 | 0.41 | 0.183 | 0.270 |
| AT3G02780.1 | Isopentenyl pyrophosphate:dimethylallyl pyrophosphate isomerase 2 (IPP2) | 24 | 0.46 | 0.182 | 0.417 |
| AT3G02780.1 | Isopentenyl pyrophosphate:dimethylallyl pyrophosphate isomerase 2 (IPP2) | 25 | 0.53 | 0.181 | 0.648 |
| AT3G02780.1 | Isopentenyl pyrophosphate:dimethylallyl pyrophosphate isomerase 2 (IPP2) | 26 | 0.62 | 0.181 | 0.891 |
| AT3G02780.1 | Isopentenyl pyrophosphate:dimethylallyl pyrophosphate isomerase 2 (IPP2) | 27 | 0.75 | 0.180 | 0.995 |
| AT3G02780.1 | Isopentenyl pyrophosphate:dimethylallyl pyrophosphate isomerase 2 (IPP2) | 28 | 0.91 | 0.181 | 1.000 |
| AT3G02780.1 | Isopentenyl pyrophosphate:dimethylallyl pyrophosphate isomerase 2 (IPP2) | 29 | 1.12 | 0.181 | 1.000 |
| AT3G02780.1 | Isopentenyl pyrophosphate:dimethylallyl pyrophosphate isomerase 2 (IPP2) | 30 | 1.36 | 0.182 | 0.993 |
| AT3G02780.1 | Isopentenyl pyrophosphate:dimethylallyl pyrophosphate isomerase 2 (IPP2) | 31 | 1.64 | 0.185 | 0.882 |
| AT3G02780.1 | Isopentenyl pyrophosphate:dimethylallyl pyrophosphate isomerase 2 (IPP2) | 32 | 1.96 | 0.188 | 0.640 |
| AT3G02780.1 | Isopentenyl pyrophosphate:dimethylallyl pyrophosphate isomerase 2 (IPP2) | 33 | 2.28 | 0.193 | 0.421 |
| AT3G02780.1 | Isopentenyl pyrophosphate:dimethylallyl pyrophosphate isomerase 2 (IPP2) | 34 | 2.61 | 0.200 | 0.282 |
| AT3G02780.1 | Isopentenyl pyrophosphate:dimethylallyl pyrophosphate isomerase 2 (IPP2) | 35 | 2.92 | 0.207 | 0.204 |

| | | | | | | |
|---|---|---|---|---|---|---|
| AT3G02780.1 | Isopentenyl pyrophosphate:dimethylallyl pyrophosphate isomerase 2 (IPP2) | 36 | 3.23 | 0.215 | 0.158 | |
| AT3G02780.1 | Isopentenyl pyrophosphate:dimethylallyl pyrophosphate isomerase 2 (IPP2) | 37 | 3.55 | 0.223 | 0.126 | |
| AT3G02780.1 | Isopentenyl pyrophosphate:dimethylallyl pyrophosphate isomerase 2 (IPP2) | 38 | 3.87 | 0.229 | 0.099 | . |
| AT3G02780.1 | Isopentenyl pyrophosphate:dimethylallyl pyrophosphate isomerase 2 (IPP2) | 42 | 5.01 | 0.235 | 0.033 | * |
| AT3G02780.1 | Isopentenyl pyrophosphate:dimethylallyl pyrophosphate isomerase 2 (IPP2) | 43 | 5.14 | 0.239 | 0.034 | * |
| AT3G02780.1 | Isopentenyl pyrophosphate:dimethylallyl pyrophosphate isomerase 2 (IPP2) | 44 | 5.18 | 0.251 | 0.047 | * |
| AT3G02780.1 | Isopentenyl pyrophosphate:dimethylallyl pyrophosphate isomerase 2 (IPP2) | 45 | 5.12 | 0.274 | 0.093 | . |
| AT3G02780.1 | Isopentenyl pyrophosphate:dimethylallyl pyrophosphate isomerase 2 (IPP2) | 46 | 4.99 | 0.309 | 0.201 | |
| AT3G02780.1 | Isopentenyl pyrophosphate:dimethylallyl pyrophosphate isomerase 2 (IPP2) | 47 | 4.82 | 0.358 | 0.388 | |
| AT3G03060.1 | P-loop containing nucleoside triphosphate hydrolase | 2 | 0.87 | 0.621 | 1.000 | |
| AT3G03060.1 | P-loop containing nucleoside triphosphate hydrolase | 3 | 0.73 | 0.537 | 1.000 | |
| AT3G03060.1 | P-loop containing nucleoside triphosphate hydrolase | 11 | 1.07 | 0.373 | 1.000 | |
| AT3G03060.1 | P-loop containing nucleoside triphosphate hydrolase | 12 | 1.70 | 0.324 | 1.000 | |
| AT3G03060.1 | P-loop containing nucleoside triphosphate hydrolase | 13 | 2.90 | 0.290 | 0.918 | |
| AT3G03060.1 | P-loop containing nucleoside triphosphate hydrolase | 14 | 4.01 | 0.270 | 0.463 | |
| AT3G03060.1 | P-loop containing nucleoside triphosphate hydrolase | 15 | 3.24 | 0.256 | 0.659 | |
| AT3G03060.1 | P-loop containing nucleoside triphosphate hydrolase | 16 | 1.82 | 0.251 | 0.999 | |
| AT3G03060.1 | P-loop containing nucleoside triphosphate hydrolase | 17 | 1.14 | 0.250 | 1.000 | |
| AT3G03060.1 | P-loop containing nucleoside triphosphate hydrolase | 18 | 1.01 | 0.254 | 1.000 | |
| AT3G03060.1 | P-loop containing nucleoside triphosphate hydrolase | 19 | 1.34 | 0.261 | 1.000 | |
| AT3G03060.1 | P-loop containing nucleoside triphosphate hydrolase | 20 | 2.15 | 0.267 | 0.993 | |
| AT3G03060.1 | P-loop containing nucleoside triphosphate hydrolase | 21 | 3.41 | 0.270 | 0.677 | |

| | | | | | |
|---|---|---|---|---|---|
| AT3G03060.1 | P-loop containing nucleoside triphosphate hydrolase | 22 | 4.43 | 0.269 | 0.341 |
| AT3G03060.1 | P-loop containing nucleoside triphosphate hydrolase | 23 | 4.38 | 0.267 | 0.337 |
| AT3G03060.1 | P-loop containing nucleoside triphosphate hydrolase | 24 | 3.08 | 0.262 | 0.766 |
| AT3G03060.1 | P-loop containing nucleoside triphosphate hydrolase | 25 | 2.00 | 0.258 | 0.997 |
| AT3G03060.1 | P-loop containing nucleoside triphosphate hydrolase | 26 | 1.53 | 0.257 | 1.000 |
| AT3G03060.1 | P-loop containing nucleoside triphosphate hydrolase | 27 | 1.40 | 0.257 | 1.000 |
| AT3G03060.1 | P-loop containing nucleoside triphosphate hydrolase | 28 | 1.38 | 0.257 | 1.000 |
| AT3G03060.1 | P-loop containing nucleoside triphosphate hydrolase | 29 | 1.42 | 0.257 | 1.000 |
| AT3G03060.1 | P-loop containing nucleoside triphosphate hydrolase | 30 | 1.41 | 0.257 | 1.000 |
| AT3G03060.1 | P-loop containing nucleoside triphosphate hydrolase | 31 | 1.28 | 0.258 | 1.000 |
| AT3G03060.1 | P-loop containing nucleoside triphosphate hydrolase | 32 | 1.05 | 0.258 | 1.000 |
| AT3G03060.1 | P-loop containing nucleoside triphosphate hydrolase | 33 | 0.74 | 0.258 | 1.000 |
| AT3G03060.1 | P-loop containing nucleoside triphosphate hydrolase | 34 | 0.55 | 0.259 | 1.000 |
| AT3G03060.1 | P-loop containing nucleoside triphosphate hydrolase | 35 | 0.58 | 0.258 | 1.000 |
| AT3G03060.1 | P-loop containing nucleoside triphosphate hydrolase | 36 | 0.84 | 0.257 | 1.000 |
| AT3G03060.1 | P-loop containing nucleoside triphosphate hydrolase | 37 | 1.28 | 0.257 | 1.000 |
| AT3G03060.1 | P-loop containing nucleoside triphosphate hydrolase | 38 | 1.83 | 0.258 | 1.000 |
| AT3G03060.1 | P-loop containing nucleoside triphosphate hydrolase | 39 | 2.37 | 0.258 | 0.965 |
| AT3G03060.1 | P-loop containing nucleoside triphosphate hydrolase | 40 | 2.61 | 0.259 | 0.913 |
| AT3G03060.1 | P-loop containing nucleoside triphosphate hydrolase | 41 | 2.34 | 0.260 | 0.972 |
| AT3G03060.1 | P-loop containing nucleoside triphosphate hydrolase | 42 | 1.80 | 0.263 | 1.000 |
| AT3G03060.1 | P-loop containing nucleoside triphosphate hydrolase | 43 | 1.35 | 0.266 | 1.000 |
| AT3G03060.1 | P-loop containing nucleoside triphosphate hydrolase | 44 | 0.90 | 0.266 | 1.000 |
| AT3G03060.1 | P-loop containing nucleoside triphosphate hydrolase | 45 | 0.49 | 0.267 | 0.998 |
| AT3G03060.1 | P-loop containing nucleoside triphosphate hydrolase | 46 | 0.26 | 0.319 | 0.783 |
| AT3G03060.1 | P-loop containing nucleoside triphosphate hydrolase | 47 | 0.15 | 0.533 | 0.929 |
| AT3G03070.1 | NADH:ubiquinone oxidoreductase subunit (13kDa) | 2 | 1.11 | 0.533 | 1.000 |
| AT3G03070.1 | NADH:ubiquinone oxidoreductase subunit (13kDa) | 3 | 0.85 | 0.491 | 1.000 |
| AT3G03070.1 | NADH:ubiquinone oxidoreductase subunit (13kDa) | 14 | 0.91 | 0.355 | 1.000 |
| AT3G03070.1 | NADH:ubiquinone oxidoreductase subunit (13kDa) | 15 | 1.07 | 0.310 | 1.000 |
| AT3G03070.1 | NADH:ubiquinone oxidoreductase subunit (13kDa) | 16 | 0.86 | 0.287 | 1.000 |

| | | | | | | |
|---|---|---|---|---|---|---|
| AT3G03070.1 | NADH:ubiquinone oxidoreductase subunit (13kDa) | 17 | 0.49 | 0.262 | 0.940 | |
| AT3G03070.1 | NADH:ubiquinone oxidoreductase subunit (13kDa) | 18 | 0.42 | 0.259 | 0.799 | |
| AT3G03070.1 | NADH:ubiquinone oxidoreductase subunit (13kDa) | 19 | 0.54 | 0.264 | 0.976 | |
| AT3G03070.1 | NADH:ubiquinone oxidoreductase subunit (13kDa) | 20 | 0.73 | 0.271 | 1.000 | |
| AT3G03070.1 | NADH:ubiquinone oxidoreductase subunit (13kDa) | 21 | 1.08 | 0.275 | 1.000 | |
| AT3G03070.1 | NADH:ubiquinone oxidoreductase subunit (13kDa) | 22 | 1.74 | 0.274 | 0.992 | |
| AT3G03070.1 | NADH:ubiquinone oxidoreductase subunit (13kDa) | 23 | 2.39 | 0.277 | 0.852 | |
| AT3G03070.1 | NADH:ubiquinone oxidoreductase subunit (13kDa) | 24 | 1.92 | 0.281 | 0.977 | |
| AT3G03070.1 | NADH:ubiquinone oxidoreductase subunit (13kDa) | 25 | 0.97 | 0.324 | 1.000 | |
| AT3G03070.1 | NADH:ubiquinone oxidoreductase subunit (13kDa) | 26 | 0.60 | 0.508 | 1.000 | |
| AT3G03070.1 | NADH:ubiquinone oxidoreductase subunit (13kDa) | 46 | ########### | 13.020 | 0.901 | |
| AT3G03070.1 | NADH:ubiquinone oxidoreductase subunit (13kDa) | 47 | ########### | 13.961 | 0.896 | |
| AT3G03420.1 | Ku70-binding family protein | 2 | 1.00 | 0.398 | 1.000 | |
| AT3G03420.1 | Ku70-binding family protein | 3 | 0.94 | 0.371 | 1.000 | |
| AT3G03420.1 | Ku70-binding family protein | 22 | 1.00 | 0.305 | 1.000 | |
| AT3G03420.1 | Ku70-binding family protein | 23 | 1.15 | 0.247 | 1.000 | |
| AT3G03420.1 | Ku70-binding family protein | 24 | 1.35 | 0.225 | 1.000 | |
| AT3G03420.1 | Ku70-binding family protein | 25 | 1.62 | 0.212 | 0.995 | |
| AT3G03420.1 | Ku70-binding family protein | 26 | 1.89 | 0.214 | 0.952 | |
| AT3G03420.1 | Ku70-binding family protein | 28 | 1.98 | 0.194 | 0.857 | |
| AT3G03420.1 | Ku70-binding family protein | 29 | 1.86 | 0.178 | 0.860 | |
| AT3G03420.1 | Ku70-binding family protein | 30 | 1.81 | 0.162 | 0.817 | |
| AT3G03420.1 | Ku70-binding family protein | 31 | 1.94 | 0.152 | 0.594 | |
| AT3G03420.1 | Ku70-binding family protein | 32 | 2.17 | 0.148 | 0.318 | |
| AT3G03420.1 | Ku70-binding family protein | 33 | 2.33 | 0.147 | 0.200 | |
| AT3G03420.1 | Ku70-binding family protein | 34 | 2.40 | 0.147 | 0.165 | |
| AT3G03420.1 | Ku70-binding family protein | 35 | 2.38 | 0.147 | 0.170 | |
| AT3G03420.1 | Ku70-binding family protein | 36 | 2.45 | 0.146 | 0.134 | |
| AT3G03420.1 | Ku70-binding family protein | 37 | 2.65 | 0.147 | 0.074 | . |
| AT3G03420.1 | Ku70-binding family protein | 38 | 2.76 | 0.147 | 0.053 | . |
| AT3G03420.1 | Ku70-binding family protein | 39 | 2.41 | 0.147 | 0.152 | |
| AT3G03420.1 | Ku70-binding family protein | 40 | 1.67 | 0.147 | 0.856 | |
| AT3G03420.1 | Ku70-binding family protein | 41 | 1.02 | 0.148 | 1.000 | |
| AT3G03420.1 | Ku70-binding family protein | 42 | 0.72 | 0.151 | 0.997 | |
| AT3G03420.1 | Ku70-binding family protein | 43 | 0.68 | 0.152 | 0.985 | |
| AT3G03420.1 | Ku70-binding family protein | 44 | 0.77 | 0.151 | 1.000 | |
| AT3G03420.1 | Ku70-binding family protein | 45 | 0.99 | 0.154 | 1.000 | |
| AT3G03420.1 | Ku70-binding family protein | 46 | 1.46 | 0.198 | 0.999 | |
| AT3G03420.1 | Ku70-binding family protein | 47 | 2.26 | 0.325 | 0.988 | |
| AT3G03910.1 | Glutamate dehydrogenase 3 (GDH3) | 2 | 1.04 | 0.597 | 1.000 | |
| AT3G03910.1 | Glutamate dehydrogenase 3 (GDH3) | 3 | 0.62 | 0.739 | 1.000 | |
| AT3G03910.1 | Glutamate dehydrogenase 3 (GDH3) | 30 | 0.23 | 0.336 | 0.337 | |
| AT3G03910.1 | Glutamate dehydrogenase 3 (GDH3) | 31 | 0.57 | 0.255 | 0.916 | |
| AT3G03910.1 | Glutamate dehydrogenase 3 (GDH3) | 32 | 1.29 | 0.238 | 0.998 | |

| | | | | | | |
|---|---|---|---|---|---|---|
| AT3G03910.1 | Glutamate dehydrogenase 3 (GDH3) | 33 | 1.95 | 0.256 | 0.828 | |
| AT3G03910.1 | Glutamate dehydrogenase 3 (GDH3) | 34 | 1.93 | 0.320 | 0.934 | |
| AT3G03910.1 | Glutamate dehydrogenase 3 (GDH3) | 35 | 1.63 | 0.441 | 0.998 | |
| AT3G03910.1 | Glutamate dehydrogenase 3 (GDH3) | 46 | 0.05 | 0.409 | 0.016 | * |
| AT3G03910.1 | Glutamate dehydrogenase 3 (GDH3) | 47 | 0.03 | 0.435 | 0.009 | ** |
| AT3G05350.1 | Metallopeptidase M24 family protein | 2 | 0.93 | 0.787 | 1.000 | |
| AT3G05350.1 | Metallopeptidase M24 family protein | 3 | 0.80 | 0.788 | 1.000 | |
| AT3G05350.1 | Metallopeptidase M24 family protein | 33 | 0.35 | 0.563 | 0.989 | |
| AT3G05350.1 | Metallopeptidase M24 family protein | 34 | 0.46 | 0.409 | 0.989 | |
| AT3G05350.1 | Metallopeptidase M24 family protein | 35 | 0.73 | 0.358 | 1.000 | |
| AT3G05350.1 | Metallopeptidase M24 family protein | 36 | 1.76 | 0.339 | 0.995 | |
| AT3G05350.1 | Metallopeptidase M24 family protein | 37 | 4.84 | 0.334 | 0.333 | |
| AT3G05350.1 | Metallopeptidase M24 family protein | 38 | 6.96 | 0.332 | 0.121 | |
| AT3G05350.1 | Metallopeptidase M24 family protein | 39 | 3.94 | 0.335 | 0.515 | |
| AT3G05350.1 | Metallopeptidase M24 family protein | 40 | 1.38 | 0.357 | 1.000 | |
| AT3G05350.1 | Metallopeptidase M24 family protein | 41 | 0.52 | 0.487 | 0.999 | |
| AT3G05350.1 | Metallopeptidase M24 family protein | 42 | 0.33 | 0.736 | 0.998 | |
| AT3G05350.1 | Metallopeptidase M24 family protein | 46 | 3.40 | 1.873 | 1.000 | |
| AT3G05350.1 | Metallopeptidase M24 family protein | 47 | 9.93 | 2.345 | 1.000 | |
| AT3G05810.1 | IGR motif protein | 2 | 0.77 | 0.362 | 0.999 | |
| AT3G05810.1 | IGR motif protein | 3 | 0.85 | 0.330 | 1.000 | |
| AT3G05810.1 | IGR motif protein | 4 | 0.93 | 0.304 | 1.000 | |
| AT3G05810.1 | IGR motif protein | 11 | 1.57 | 0.188 | 0.795 | |
| AT3G05810.1 | IGR motif protein | 12 | 1.61 | 0.176 | 0.708 | |
| AT3G05810.1 | IGR motif protein | 13 | 1.61 | 0.168 | 0.667 | |
| AT3G05810.1 | IGR motif protein | 14 | 1.57 | 0.164 | 0.703 | |
| AT3G05810.1 | IGR motif protein | 15 | 1.48 | 0.166 | 0.808 | |
| AT3G05810.1 | IGR motif protein | 16 | 1.36 | 0.174 | 0.927 | |
| AT3G05810.1 | IGR motif protein | 17 | 1.23 | 0.187 | 0.992 | |
| AT3G05810.1 | IGR motif protein | 18 | 1.08 | 0.204 | 1.000 | |
| AT3G05810.1 | IGR motif protein | 22 | 0.55 | 0.299 | 0.887 | |
| AT3G05810.1 | IGR motif protein | 23 | 0.46 | 0.327 | 0.802 | |
| AT3G05810.1 | IGR motif protein | 24 | 0.39 | 0.356 | 0.725 | |
| AT3G05810.1 | IGR motif protein | 46 | 0.03 | 0.684 | 0.158 | |
| AT3G05810.1 | IGR motif protein | 47 | 0.03 | 0.707 | 0.165 | |
| AT3G05970.1 | Long chain acyl-CoA synthetase 6 (LACS6) | 2 | 1.08 | 0.309 | 1.000 | |
| AT3G05970.1 | Long chain acyl-CoA synthetase 6 (LACS6) | 3 | 1.03 | 0.218 | 1.000 | |
| AT3G05970.1 | Long chain acyl-CoA synthetase 6 (LACS6) | 4 | 1.01 | 0.256 | 1.000 | |
| AT3G05970.1 | Long chain acyl-CoA synthetase 6 (LACS6) | 6 | 1.00 | 0.274 | 1.000 | |

| | | | | | | |
|---|---|---|---|---|---|---|
| AT3G05970.1 | Long chain acyl-CoA synthetase 6 (LACS6) | 7 | 1.00 | 0.268 | 1.000 | |
| AT3G05970.1 | Long chain acyl-CoA synthetase 6 (LACS6) | 8 | 1.01 | 0.319 | 1.000 | |
| AT3G05970.1 | Long chain acyl-CoA synthetase 6 (LACS6) | 22 | 0.98 | 0.346 | 1.000 | |
| AT3G05970.1 | Long chain acyl-CoA synthetase 6 (LACS6) | 23 | 0.88 | 0.231 | 1.000 | |
| AT3G05970.1 | Long chain acyl-CoA synthetase 6 (LACS6) | 24 | 0.77 | 0.199 | 1.000 | |
| AT3G05970.1 | Long chain acyl-CoA synthetase 6 (LACS6) | 25 | 0.68 | 0.211 | 1.000 | |
| AT3G05970.1 | Long chain acyl-CoA synthetase 6 (LACS6) | 26 | 0.67 | 0.205 | 1.000 | |
| AT3G05970.1 | Long chain acyl-CoA synthetase 6 (LACS6) | 27 | 0.74 | 0.195 | 1.000 | |
| AT3G05970.1 | Long chain acyl-CoA synthetase 6 (LACS6) | 28 | 0.88 | 0.190 | 1.000 | |
| AT3G05970.1 | Long chain acyl-CoA synthetase 6 (LACS6) | 29 | 1.00 | 0.195 | 1.000 | |
| AT3G05970.1 | Long chain acyl-CoA synthetase 6 (LACS6) | 30 | 1.05 | 0.204 | 1.000 | |
| AT3G05970.1 | Long chain acyl-CoA synthetase 6 (LACS6) | 31 | 1.01 | 0.199 | 1.000 | |
| AT3G05970.1 | Long chain acyl-CoA synthetase 6 (LACS6) | 32 | 0.94 | 0.188 | 1.000 | |
| AT3G05970.1 | Long chain acyl-CoA synthetase 6 (LACS6) | 33 | 0.89 | 0.177 | 1.000 | |
| AT3G05970.1 | Long chain acyl-CoA synthetase 6 (LACS6) | 34 | 0.87 | 0.168 | 1.000 | |
| AT3G05970.1 | Long chain acyl-CoA synthetase 6 (LACS6) | 35 | 0.97 | 0.165 | 1.000 | |
| AT3G05970.1 | Long chain acyl-CoA synthetase 6 (LACS6) | 36 | 1.10 | 0.169 | 1.000 | |
| AT3G05970.1 | Long chain acyl-CoA synthetase 6 (LACS6) | 37 | 1.21 | 0.178 | 1.000 | |
| AT3G05970.1 | Long chain acyl-CoA synthetase 6 (LACS6) | 38 | 1.40 | 0.185 | 1.000 | |
| AT3G05970.1 | Long chain acyl-CoA synthetase 6 (LACS6) | 39 | 1.69 | 0.178 | 0.985 | |
| AT3G05970.1 | Long chain acyl-CoA synthetase 6 (LACS6) | 40 | 2.31 | 0.167 | 0.476 | |
| AT3G05970.1 | Long chain acyl-CoA synthetase 6 (LACS6) | 41 | 3.20 | 0.162 | 0.054 | . |
| AT3G05970.1 | Long chain acyl-CoA synthetase 6 (LACS6) | 42 | 4.15 | 0.163 | 0.006 | ** |
| AT3G05970.1 | Long chain acyl-CoA synthetase 6 (LACS6) | 43 | 3.93 | 0.164 | 0.010 | * |
| AT3G05970.1 | Long chain acyl-CoA synthetase 6 (LACS6) | 44 | 2.11 | 0.167 | 0.660 | |
| AT3G05970.1 | Long chain acyl-CoA synthetase 6 (LACS6) | 45 | 1.05 | 0.166 | 1.000 | |
| AT3G05970.1 | Long chain acyl-CoA synthetase 6 (LACS6) | 46 | 0.70 | 0.190 | 1.000 | |
| AT3G05970.1 | Long chain acyl-CoA synthetase 6 (LACS6) | 47 | 0.48 | 0.349 | 1.000 | |
| AT3G06040.3 | Ribosomal L12 family protein, putative | 2 | 1.03 | 0.490 | 1.000 | |
| AT3G06040.3 | Ribosomal L12 family protein, putative | 3 | 0.92 | 0.477 | 1.000 | |
| AT3G06040.3 | Ribosomal L12 family protein, putative | 34 | 0.77 | 0.277 | 0.999 | |

| | | | | | | |
|---|---|---|---|---|---|---|
| AT3G06040.3 | Ribosomal L12 family protein, putative | 35 | 0.94 | 0.216 | 1.000 | |
| AT3G06040.3 | Ribosomal L12 family protein, putative | 36 | 1.17 | 0.181 | 1.000 | |
| AT3G06040.3 | Ribosomal L12 family protein, putative | 37 | 1.47 | 0.164 | 0.891 | |
| AT3G06040.3 | Ribosomal L12 family protein, putative | 38 | 1.77 | 0.156 | 0.531 | |
| AT3G06040.3 | Ribosomal L12 family protein, putative | 39 | 1.92 | 0.156 | 0.387 | |
| AT3G06040.3 | Ribosomal L12 family protein, putative | 40 | 1.74 | 0.167 | 0.638 | |
| AT3G06040.3 | Ribosomal L12 family protein, putative | 41 | 1.30 | 0.193 | 0.992 | |
| AT3G06040.3 | Ribosomal L12 family protein, putative | 42 | 0.86 | 0.233 | 1.000 | |
| AT3G06040.3 | Ribosomal L12 family protein, putative | 43 | 0.53 | 0.275 | 0.902 | |
| AT3G06040.3 | Ribosomal L12 family protein, putative | 46 | 0.10 | 0.400 | 0.094 | . |
| AT3G06040.3 | Ribosomal L12 family protein, putative | 47 | 0.05 | 0.475 | 0.064 | . |
| AT3G06050.1 | Peroxiredoxin 2F (PRX2F) | 2 | 1.25 | 0.259 | 1.000 | |
| AT3G06050.1 | Peroxiredoxin 2F (PRX2F) | 3 | 1.03 | 0.190 | 1.000 | |
| AT3G06050.1 | Peroxiredoxin 2F (PRX2F) | 4 | 0.84 | 0.166 | 1.000 | |
| AT3G06050.1 | Peroxiredoxin 2F (PRX2F) | 5 | 0.68 | 0.161 | 0.999 | |
| AT3G06050.1 | Peroxiredoxin 2F (PRX2F) | 6 | 0.56 | 0.154 | 0.890 | |
| AT3G06050.1 | Peroxiredoxin 2F (PRX2F) | 7 | 0.47 | 0.145 | 0.425 | |
| AT3G06050.1 | Peroxiredoxin 2F (PRX2F) | 8 | 0.42 | 0.137 | 0.146 | |
| AT3G06050.1 | Peroxiredoxin 2F (PRX2F) | 9 | 0.43 | 0.133 | 0.140 | |
| AT3G06050.1 | Peroxiredoxin 2F (PRX2F) | 10 | 0.49 | 0.131 | 0.361 | |
| AT3G06050.1 | Peroxiredoxin 2F (PRX2F) | 11 | 0.63 | 0.131 | 0.923 | |
| AT3G06050.1 | Peroxiredoxin 2F (PRX2F) | 12 | 0.83 | 0.131 | 1.000 | |
| AT3G06050.1 | Peroxiredoxin 2F (PRX2F) | 13 | 0.96 | 0.130 | 1.000 | |
| AT3G06050.1 | Peroxiredoxin 2F (PRX2F) | 14 | 0.95 | 0.129 | 1.000 | |
| AT3G06050.1 | Peroxiredoxin 2F (PRX2F) | 15 | 0.88 | 0.128 | 1.000 | |
| AT3G06050.1 | Peroxiredoxin 2F (PRX2F) | 16 | 0.87 | 0.127 | 1.000 | |
| AT3G06050.1 | Peroxiredoxin 2F (PRX2F) | 17 | 0.95 | 0.127 | 1.000 | |
| AT3G06050.1 | Peroxiredoxin 2F (PRX2F) | 18 | 1.07 | 0.129 | 1.000 | |
| AT3G06050.1 | Peroxiredoxin 2F (PRX2F) | 19 | 1.15 | 0.132 | 1.000 | |
| AT3G06050.1 | Peroxiredoxin 2F (PRX2F) | 20 | 1.18 | 0.134 | 1.000 | |
| AT3G06050.1 | Peroxiredoxin 2F (PRX2F) | 21 | 1.19 | 0.135 | 1.000 | |
| AT3G06050.1 | Peroxiredoxin 2F (PRX2F) | 22 | 1.21 | 0.135 | 1.000 | |
| AT3G06050.1 | Peroxiredoxin 2F (PRX2F) | 23 | 1.23 | 0.134 | 1.000 | |
| AT3G06050.1 | Peroxiredoxin 2F (PRX2F) | 24 | 1.27 | 0.133 | 1.000 | |
| AT3G06050.1 | Peroxiredoxin 2F (PRX2F) | 25 | 1.32 | 0.131 | 1.000 | |
| AT3G06050.1 | Peroxiredoxin 2F (PRX2F) | 26 | 1.35 | 0.130 | 0.999 | |
| AT3G06050.1 | Peroxiredoxin 2F (PRX2F) | 27 | 1.39 | 0.130 | 0.998 | |
| AT3G06050.1 | Peroxiredoxin 2F (PRX2F) | 28 | 1.41 | 0.130 | 0.996 | |
| AT3G06050.1 | Peroxiredoxin 2F (PRX2F) | 29 | 1.39 | 0.130 | 0.998 | |
| AT3G06050.1 | Peroxiredoxin 2F (PRX2F) | 30 | 1.32 | 0.130 | 1.000 | |
| AT3G06050.1 | Peroxiredoxin 2F (PRX2F) | 31 | 1.26 | 0.131 | 1.000 | |
| AT3G06050.1 | Peroxiredoxin 2F (PRX2F) | 32 | 1.21 | 0.131 | 1.000 | |
| AT3G06050.1 | Peroxiredoxin 2F (PRX2F) | 33 | 1.14 | 0.131 | 1.000 | |
| AT3G06050.1 | Peroxiredoxin 2F (PRX2F) | 34 | 1.10 | 0.131 | 1.000 | |
| AT3G06050.1 | Peroxiredoxin 2F (PRX2F) | 35 | 1.11 | 0.131 | 1.000 | |
| AT3G06050.1 | Peroxiredoxin 2F (PRX2F) | 36 | 1.14 | 0.131 | 1.000 | |
| AT3G06050.1 | Peroxiredoxin 2F (PRX2F) | 37 | 1.19 | 0.131 | 1.000 | |
| AT3G06050.1 | Peroxiredoxin 2F (PRX2F) | 38 | 1.25 | 0.131 | 1.000 | |
| AT3G06050.1 | Peroxiredoxin 2F (PRX2F) | 39 | 1.33 | 0.131 | 1.000 | |
| AT3G06050.1 | Peroxiredoxin 2F (PRX2F) | 40 | 1.41 | 0.131 | 0.997 | |
| AT3G06050.1 | Peroxiredoxin 2F (PRX2F) | 41 | 1.40 | 0.132 | 0.997 | |
| AT3G06050.1 | Peroxiredoxin 2F (PRX2F) | 42 | 1.33 | 0.134 | 1.000 | |

| | | | | | | |
|---|---|---|---|---|---|---|
| AT3G06050.1 | Peroxiredoxin 2F (PRX2F) | 43 | 1.22 | 0.135 | 1.000 | |
| AT3G06050.1 | Peroxiredoxin 2F (PRX2F) | 44 | 1.07 | 0.134 | 1.000 | |
| AT3G06050.1 | Peroxiredoxin 2F (PRX2F) | 45 | 0.85 | 0.139 | 1.000 | |
| AT3G06050.1 | Peroxiredoxin 2F (PRX2F) | 46 | 0.61 | 0.171 | 0.991 | |
| AT3G06050.1 | Peroxiredoxin 2F (PRX2F) | 47 | 0.41 | 0.258 | 0.944 | |
| AT3G06310.3 | NADH:ubiquinone oxidoreductase subunit | 2 | 0.81 | 0.527 | 1.000 | |
| AT3G06310.3 | NADH:ubiquinone oxidoreductase subunit | 3 | 0.85 | 0.474 | 1.000 | |
| AT3G06310.3 | NADH:ubiquinone oxidoreductase subunit | 15 | 0.61 | 0.262 | 0.973 | |
| AT3G06310.3 | NADH:ubiquinone oxidoreductase subunit | 16 | 0.56 | 0.223 | 0.872 | |
| AT3G06310.3 | NADH:ubiquinone oxidoreductase subunit | 17 | 0.56 | 0.196 | 0.777 | |
| AT3G06310.3 | NADH:ubiquinone oxidoreductase subunit | 18 | 0.61 | 0.181 | 0.846 | |
| AT3G06310.3 | NADH:ubiquinone oxidoreductase subunit | 19 | 0.76 | 0.175 | 0.990 | |
| AT3G06310.3 | NADH:ubiquinone oxidoreductase subunit | 20 | 1.06 | 0.176 | 1.000 | |
| AT3G06310.3 | NADH:ubiquinone oxidoreductase subunit | 21 | 1.66 | 0.181 | 0.822 | |
| AT3G06310.3 | NADH:ubiquinone oxidoreductase subunit | 22 | 2.93 | 0.191 | 0.118 | |
| AT3G06310.3 | NADH:ubiquinone oxidoreductase subunit | 23 | 5.84 | 0.213 | 0.005 | ** |
| AT3G06310.3 | NADH:ubiquinone oxidoreductase subunit | 24 | 12.61 | 0.257 | 0.000 | *** |
| AT3G06310.3 | NADH:ubiquinone oxidoreductase subunit | 25 | 27.62 | 0.336 | 0.001 | *** |
| AT3G06310.3 | NADH:ubiquinone oxidoreductase subunit | 46 | 284844.60 | 6.014 | 0.952 | |
| AT3G06310.3 | NADH:ubiquinone oxidoreductase subunit | 47 | 372377.73 | 6.410 | 0.962 | |
| AT3G06483.1 | Pyruvate dehydrogenase kinase (PDK) | 2 | 1.04 | 0.399 | 1.000 | |
| AT3G06483.1 | Pyruvate dehydrogenase kinase (PDK) | 3 | 0.92 | 0.308 | 1.000 | |
| AT3G06483.1 | Pyruvate dehydrogenase kinase (PDK) | 6 | 0.69 | 0.286 | 1.000 | |
| AT3G06483.1 | Pyruvate dehydrogenase kinase (PDK) | 7 | 0.67 | 0.234 | 1.000 | |
| AT3G06483.1 | Pyruvate dehydrogenase kinase (PDK) | 8 | 0.71 | 0.203 | 1.000 | |
| AT3G06483.1 | Pyruvate dehydrogenase kinase (PDK) | 9 | 0.85 | 0.181 | 1.000 | |
| AT3G06483.1 | Pyruvate dehydrogenase kinase (PDK) | 10 | 1.01 | 0.171 | 1.000 | |
| AT3G06483.1 | Pyruvate dehydrogenase kinase (PDK) | 11 | 1.20 | 0.165 | 1.000 | |
| AT3G06483.1 | Pyruvate dehydrogenase kinase (PDK) | 12 | 1.43 | 0.162 | 1.000 | |
| AT3G06483.1 | Pyruvate dehydrogenase kinase (PDK) | 13 | 1.42 | 0.159 | 1.000 | |
| AT3G06483.1 | Pyruvate dehydrogenase kinase (PDK) | 14 | 1.07 | 0.157 | 1.000 | |
| AT3G06483.1 | Pyruvate dehydrogenase kinase (PDK) | 15 | 0.75 | 0.157 | 1.000 | |
| AT3G06483.1 | Pyruvate dehydrogenase kinase (PDK) | 16 | 0.65 | 0.156 | 0.998 | |

| | | | | | |
|---|---|---|---|---|---|
| AT3G06483.1 | Pyruvate dehydrogenase kinase (PDK) | 17 | 0.80 | 0.158 | 1.000 |
| AT3G06483.1 | Pyruvate dehydrogenase kinase (PDK) | 18 | 1.15 | 0.164 | 1.000 |
| AT3G06483.1 | Pyruvate dehydrogenase kinase (PDK) | 19 | 1.37 | 0.174 | 1.000 |
| AT3G06483.1 | Pyruvate dehydrogenase kinase (PDK) | 20 | 1.38 | 0.191 | 1.000 |
| AT3G06483.1 | Pyruvate dehydrogenase kinase (PDK) | 21 | 1.17 | 0.206 | 1.000 |
| AT3G06483.1 | Pyruvate dehydrogenase kinase (PDK) | 22 | 0.89 | 0.210 | 1.000 |
| AT3G06483.1 | Pyruvate dehydrogenase kinase (PDK) | 23 | 0.68 | 0.197 | 1.000 |
| AT3G06483.1 | Pyruvate dehydrogenase kinase (PDK) | 24 | 0.57 | 0.176 | 0.988 |
| AT3G06483.1 | Pyruvate dehydrogenase kinase (PDK) | 25 | 0.50 | 0.162 | 0.840 |
| AT3G06483.1 | Pyruvate dehydrogenase kinase (PDK) | 26 | 0.48 | 0.159 | 0.716 |
| AT3G06483.1 | Pyruvate dehydrogenase kinase (PDK) | 27 | 0.49 | 0.159 | 0.771 |
| AT3G06483.1 | Pyruvate dehydrogenase kinase (PDK) | 28 | 0.53 | 0.159 | 0.904 |
| AT3G06483.1 | Pyruvate dehydrogenase kinase (PDK) | 29 | 0.59 | 0.160 | 0.985 |
| AT3G06483.1 | Pyruvate dehydrogenase kinase (PDK) | 30 | 0.59 | 0.160 | 0.987 |
| AT3G06483.1 | Pyruvate dehydrogenase kinase (PDK) | 31 | 0.60 | 0.160 | 0.990 |
| AT3G06483.1 | Pyruvate dehydrogenase kinase (PDK) | 32 | 0.63 | 0.160 | 0.997 |
| AT3G06483.1 | Pyruvate dehydrogenase kinase (PDK) | 33 | 0.61 | 0.160 | 0.994 |
| AT3G06483.1 | Pyruvate dehydrogenase kinase (PDK) | 34 | 0.55 | 0.161 | 0.944 |
| AT3G06483.1 | Pyruvate dehydrogenase kinase (PDK) | 35 | 0.48 | 0.161 | 0.741 |
| AT3G06483.1 | Pyruvate dehydrogenase kinase (PDK) | 36 | 0.48 | 0.160 | 0.743 |
| AT3G06483.1 | Pyruvate dehydrogenase kinase (PDK) | 37 | 0.63 | 0.161 | 0.997 |
| AT3G06483.1 | Pyruvate dehydrogenase kinase (PDK) | 38 | 0.97 | 0.161 | 1.000 |
| AT3G06483.1 | Pyruvate dehydrogenase kinase (PDK) | 39 | 1.47 | 0.160 | 1.000 |
| AT3G06483.1 | Pyruvate dehydrogenase kinase (PDK) | 40 | 1.85 | 0.160 | 0.926 |
| AT3G06483.1 | Pyruvate dehydrogenase kinase (PDK) | 41 | 1.87 | 0.161 | 0.916 |
| AT3G06483.1 | Pyruvate dehydrogenase kinase (PDK) | 42 | 1.80 | 0.163 | 0.958 |
| AT3G06483.1 | Pyruvate dehydrogenase kinase (PDK) | 43 | 1.63 | 0.165 | 0.996 |
| AT3G06483.1 | Pyruvate dehydrogenase kinase (PDK) | 44 | 1.20 | 0.165 | 1.000 |
| AT3G06483.1 | Pyruvate dehydrogenase kinase (PDK) | 45 | 0.72 | 0.165 | 1.000 |
| AT3G06483.1 | Pyruvate dehydrogenase kinase (PDK) | 46 | 0.37 | 0.200 | 0.573 |
| AT3G06483.1 | Pyruvate dehydrogenase kinase (PDK) | 47 | 0.16 | 0.351 | 0.496 |

| | | | | | |
|---|---|---|---|---|---|
| AT3G06510.1 | Sensitive to freezing 2 (SFR2) | 2 | 1.03 | 0.358 | 1.000 |
| AT3G06510.1 | Sensitive to freezing 2 (SFR2) | 3 | 1.06 | 0.329 | 1.000 |
| AT3G06510.1 | Sensitive to freezing 2 (SFR2) | 15 | 1.61 | 0.135 | 0.653 |
| AT3G06510.1 | Sensitive to freezing 2 (SFR2) | 16 | 1.66 | 0.124 | 0.474 |
| AT3G06510.1 | Sensitive to freezing 2 (SFR2) | 17 | 1.70 | 0.114 | 0.314 |
| AT3G06510.1 | Sensitive to freezing 2 (SFR2) | 18 | 1.73 | 0.106 | 0.204 |
| AT3G06510.1 | Sensitive to freezing 2 (SFR2) | 19 | 1.74 | 0.100 | 0.143 |
| AT3G06510.1 | Sensitive to freezing 2 (SFR2) | 20 | 1.73 | 0.097 | 0.121 |
| AT3G06510.1 | Sensitive to freezing 2 (SFR2) | 21 | 1.71 | 0.094 | 0.125 |
| AT3G06510.1 | Sensitive to freezing 2 (SFR2) | 22 | 1.66 | 0.093 | 0.155 |
| AT3G06510.1 | Sensitive to freezing 2 (SFR2) | 23 | 1.60 | 0.092 | 0.217 |
| AT3G06510.1 | Sensitive to freezing 2 (SFR2) | 24 | 1.53 | 0.092 | 0.323 |
| AT3G06510.1 | Sensitive to freezing 2 (SFR2) | 25 | 1.45 | 0.092 | 0.489 |
| AT3G06510.1 | Sensitive to freezing 2 (SFR2) | 26 | 1.36 | 0.093 | 0.708 |
| AT3G06510.1 | Sensitive to freezing 2 (SFR2) | 27 | 1.27 | 0.095 | 0.911 |
| AT3G06510.1 | Sensitive to freezing 2 (SFR2) | 28 | 1.16 | 0.098 | 0.994 |
| AT3G06510.1 | Sensitive to freezing 2 (SFR2) | 29 | 1.06 | 0.101 | 1.000 |
| AT3G06510.1 | Sensitive to freezing 2 (SFR2) | 30 | 0.97 | 0.105 | 1.000 |
| AT3G06510.1 | Sensitive to freezing 2 (SFR2) | 31 | 0.89 | 0.110 | 1.000 |
| AT3G06510.1 | Sensitive to freezing 2 (SFR2) | 32 | 0.83 | 0.116 | 0.994 |
| AT3G06510.1 | Sensitive to freezing 2 (SFR2) | 33 | 0.80 | 0.121 | 0.981 |
| AT3G06510.1 | Sensitive to freezing 2 (SFR2) | 34 | 0.77 | 0.126 | 0.970 |
| AT3G06510.1 | Sensitive to freezing 2 (SFR2) | 35 | 0.76 | 0.131 | 0.966 |
| AT3G06510.1 | Sensitive to freezing 2 (SFR2) | 40 | 0.80 | 0.140 | 0.994 |
| AT3G06510.1 | Sensitive to freezing 2 (SFR2) | 41 | 0.81 | 0.144 | 0.997 |
| AT3G06510.1 | Sensitive to freezing 2 (SFR2) | 42 | 0.82 | 0.151 | 0.998 |
| AT3G06510.1 | Sensitive to freezing 2 (SFR2) | 43 | 0.82 | 0.164 | 0.999 |
| AT3G06510.1 | Sensitive to freezing 2 (SFR2) | 44 | 0.82 | 0.183 | 1.000 |
| AT3G06510.1 | Sensitive to freezing 2 (SFR2) | 46 | 0.81 | 0.236 | 1.000 |
| AT3G06510.1 | Sensitive to freezing 2 (SFR2) | 47 | 0.80 | 0.270 | 1.000 |
| AT3G06850.2 | Branched chain keto acid dehydrogenase subunit (BCKDHE2) | 2 | 0.92 | 0.550 | 1.000 |
| AT3G06850.2 | Branched chain keto acid dehydrogenase subunit (BCKDHE2) | 3 | 0.82 | 0.386 | 1.000 |
| AT3G06850.2 | Branched chain keto acid dehydrogenase subunit (BCKDHE2) | 4 | 0.72 | 0.392 | 1.000 |
| AT3G06850.2 | Branched chain keto acid dehydrogenase subunit (BCKDHE2) | 6 | 0.53 | 0.396 | 1.000 |
| AT3G06850.2 | Branched chain keto acid dehydrogenase subunit (BCKDHE2) | 7 | 0.46 | 0.344 | 1.000 |
| AT3G06850.2 | Branched chain keto acid dehydrogenase subunit (BCKDHE2) | 8 | 0.50 | 0.302 | 0.999 |
| AT3G06850.2 | Branched chain keto acid dehydrogenase subunit (BCKDHE2) | 9 | 0.68 | 0.282 | 1.000 |
| AT3G06850.2 | Branched chain keto acid dehydrogenase subunit (BCKDHE2) | 10 | 0.90 | 0.275 | 1.000 |
| AT3G06850.2 | Branched chain keto acid dehydrogenase subunit (BCKDHE2) | 11 | 1.03 | 0.273 | 1.000 |
| AT3G06850.2 | Branched chain keto acid dehydrogenase subunit (BCKDHE2) | 12 | 0.95 | 0.273 | 1.000 |

| | | | | | |
|---|---|---|---|---|---|
| AT3G06850.2 | Branched chain keto acid dehydrogenase subunit (BCKDHE2) | 13 | 0.73 | 0.271 | 1.000 |
| AT3G06850.2 | Branched chain keto acid dehydrogenase subunit (BCKDHE2) | 14 | 0.53 | 0.268 | 0.999 |
| AT3G06850.2 | Branched chain keto acid dehydrogenase subunit (BCKDHE2) | 15 | 0.37 | 0.267 | 0.895 |
| AT3G06850.2 | Branched chain keto acid dehydrogenase subunit (BCKDHE2) | 16 | 0.27 | 0.265 | 0.513 |
| AT3G06850.2 | Branched chain keto acid dehydrogenase subunit (BCKDHE2) | 17 | 0.31 | 0.265 | 0.689 |
| AT3G06850.2 | Branched chain keto acid dehydrogenase subunit (BCKDHE2) | 18 | 0.65 | 0.273 | 1.000 |
| AT3G06850.2 | Branched chain keto acid dehydrogenase subunit (BCKDHE2) | 19 | 1.54 | 0.288 | 1.000 |
| AT3G06850.2 | Branched chain keto acid dehydrogenase subunit (BCKDHE2) | 20 | 2.71 | 0.310 | 0.968 |
| AT3G06850.2 | Branched chain keto acid dehydrogenase subunit (BCKDHE2) | 21 | 3.17 | 0.359 | 0.969 |
| AT3G06850.2 | Branched chain keto acid dehydrogenase subunit (BCKDHE2) | 22 | 2.66 | 0.419 | 0.999 |
| AT3G06850.2 | Branched chain keto acid dehydrogenase subunit (BCKDHE2) | 26 | 0.33 | 0.411 | 0.994 |
| AT3G06850.2 | Branched chain keto acid dehydrogenase subunit (BCKDHE2) | 27 | 0.22 | 0.342 | 0.683 |
| AT3G06850.2 | Branched chain keto acid dehydrogenase subunit (BCKDHE2) | 28 | 0.19 | 0.296 | 0.295 |
| AT3G06850.2 | Branched chain keto acid dehydrogenase subunit (BCKDHE2) | 29 | 0.22 | 0.275 | 0.332 |
| AT3G06850.2 | Branched chain keto acid dehydrogenase subunit (BCKDHE2) | 30 | 0.30 | 0.272 | 0.688 |
| AT3G06850.2 | Branched chain keto acid dehydrogenase subunit (BCKDHE2) | 31 | 0.48 | 0.275 | 0.996 |
| AT3G06850.2 | Branched chain keto acid dehydrogenase subunit (BCKDHE2) | 32 | 0.76 | 0.280 | 1.000 |
| AT3G06850.2 | Branched chain keto acid dehydrogenase subunit (BCKDHE2) | 33 | 0.94 | 0.291 | 1.000 |
| AT3G06850.2 | Branched chain keto acid dehydrogenase subunit (BCKDHE2) | 34 | 0.85 | 0.310 | 1.000 |
| AT3G06850.2 | Branched chain keto acid dehydrogenase subunit (BCKDHE2) | 35 | 0.67 | 0.348 | 1.000 |

| | | | | | | |
|---|---|---|---|---|---|---|
| AT3G06850.2 | Branched chain keto acid dehydrogenase subunit (BCKDHE2) | 36 | 0.54 | 0.415 | 1.000 | |
| AT3G06850.2 | Branched chain keto acid dehydrogenase subunit (BCKDHE2) | 37 | 0.46 | 0.581 | 1.000 | |
| AT3G06850.2 | Branched chain keto acid dehydrogenase subunit (BCKDHE2) | 46 | 0.40 | 5.764 | 1.000 | |
| AT3G06850.2 | Branched chain keto acid dehydrogenase subunit (BCKDHE2) | 47 | 0.43 | 6.631 | 1.000 | |
| AT3G07060.1 | ? | 2 | 1.23 | 0.664 | 1.000 | |
| AT3G07060.1 | ? | 3 | 0.61 | 1.145 | 1.000 | |
| AT3G07060.1 | ? | 25 | 0.03 | 0.506 | 0.074 | . |
| AT3G07060.1 | ? | 26 | 0.28 | 0.347 | 0.845 | |
| AT3G07060.1 | ? | 27 | 2.26 | 0.317 | 0.989 | |
| AT3G07060.1 | ? | 28 | 4.74 | 0.304 | 0.371 | |
| AT3G07060.1 | ? | 29 | 1.44 | 0.302 | 1.000 | |
| AT3G07060.1 | ? | 30 | 0.34 | 0.301 | 0.852 | |
| AT3G07060.1 | ? | 31 | 0.34 | 0.308 | 0.870 | |
| AT3G07060.1 | ? | 32 | 0.67 | 0.324 | 1.000 | |
| AT3G07060.1 | ? | 33 | 0.88 | 0.374 | 1.000 | |
| AT3G07060.1 | ? | 34 | 0.98 | 0.461 | 1.000 | |
| AT3G07060.1 | ? | 36 | 1.35 | 0.473 | 1.000 | |
| AT3G07060.1 | ? | 37 | 1.70 | 0.422 | 1.000 | |
| AT3G07060.1 | ? | 38 | 2.28 | 0.431 | 1.000 | |
| AT3G07060.1 | ? | 39 | 3.12 | 0.386 | 0.964 | |
| AT3G07060.1 | ? | 40 | 3.27 | 0.342 | 0.884 | |
| AT3G07060.1 | ? | 41 | 2.28 | 0.315 | 0.987 | |
| AT3G07060.1 | ? | 42 | 1.49 | 0.317 | 1.000 | |
| AT3G07060.1 | ? | 43 | 1.18 | 0.436 | 1.000 | |
| AT3G07060.1 | ? | 44 | 1.01 | 0.678 | 1.000 | |
| AT3G07060.1 | ? | 46 | 0.77 | 1.136 | 1.000 | |
| AT3G07060.1 | ? | 47 | 0.69 | 1.571 | 1.000 | |
| AT3G07480.1 | 2Fe-2S ferredoxin-like superfamily protein | 2 | 1.07 | 0.451 | 1.000 | |
| AT3G07480.1 | 2Fe-2S ferredoxin-like superfamily protein | 3 | 0.81 | 0.359 | 1.000 | |
| AT3G07480.1 | 2Fe-2S ferredoxin-like superfamily protein | 4 | 0.63 | 0.462 | 1.000 | |
| AT3G07480.1 | 2Fe-2S ferredoxin-like superfamily protein | 12 | 0.53 | 0.337 | 1.000 | |
| AT3G07480.1 | 2Fe-2S ferredoxin-like superfamily protein | 13 | 0.66 | 0.303 | 1.000 | |
| AT3G07480.1 | 2Fe-2S ferredoxin-like superfamily protein | 14 | 0.79 | 0.281 | 1.000 | |
| AT3G07480.1 | 2Fe-2S ferredoxin-like superfamily protein | 15 | 0.78 | 0.268 | 1.000 | |
| AT3G07480.1 | 2Fe-2S ferredoxin-like superfamily protein | 16 | 0.59 | 0.258 | 1.000 | |
| AT3G07480.1 | 2Fe-2S ferredoxin-like superfamily protein | 17 | 0.45 | 0.255 | 0.953 | |
| AT3G07480.1 | 2Fe-2S ferredoxin-like superfamily protein | 18 | 0.51 | 0.258 | 0.992 | |
| AT3G07480.1 | 2Fe-2S ferredoxin-like superfamily protein | 19 | 0.73 | 0.263 | 1.000 | |
| AT3G07480.1 | 2Fe-2S ferredoxin-like superfamily protein | 20 | 0.98 | 0.267 | 1.000 | |
| AT3G07480.1 | 2Fe-2S ferredoxin-like superfamily protein | 21 | 1.25 | 0.269 | 1.000 | |

| | | | | | |
|---|---|---|---|---|---|
| AT3G07480.1 | 2Fe-2S ferredoxin-like superfamily protein | 22 | 1.44 | 0.268 | 1.000 |
| AT3G07480.1 | 2Fe-2S ferredoxin-like superfamily protein | 23 | 1.28 | 0.266 | 1.000 |
| AT3G07480.1 | 2Fe-2S ferredoxin-like superfamily protein | 24 | 0.89 | 0.263 | 1.000 |
| AT3G07480.1 | 2Fe-2S ferredoxin-like superfamily protein | 25 | 0.61 | 0.261 | 1.000 |
| AT3G07480.1 | 2Fe-2S ferredoxin-like superfamily protein | 26 | 0.58 | 0.260 | 0.999 |
| AT3G07480.1 | 2Fe-2S ferredoxin-like superfamily protein | 27 | 0.69 | 0.260 | 1.000 |
| AT3G07480.1 | 2Fe-2S ferredoxin-like superfamily protein | 28 | 0.81 | 0.260 | 1.000 |
| AT3G07480.1 | 2Fe-2S ferredoxin-like superfamily protein | 29 | 0.92 | 0.260 | 1.000 |
| AT3G07480.1 | 2Fe-2S ferredoxin-like superfamily protein | 30 | 1.07 | 0.260 | 1.000 |
| AT3G07480.1 | 2Fe-2S ferredoxin-like superfamily protein | 31 | 1.25 | 0.261 | 1.000 |
| AT3G07480.1 | 2Fe-2S ferredoxin-like superfamily protein | 32 | 1.40 | 0.261 | 1.000 |
| AT3G07480.1 | 2Fe-2S ferredoxin-like superfamily protein | 33 | 1.62 | 0.261 | 1.000 |
| AT3G07480.1 | 2Fe-2S ferredoxin-like superfamily protein | 34 | 2.11 | 0.261 | 0.976 |
| AT3G07480.1 | 2Fe-2S ferredoxin-like superfamily protein | 35 | 2.65 | 0.261 | 0.818 |
| AT3G07480.1 | 2Fe-2S ferredoxin-like superfamily protein | 36 | 2.89 | 0.260 | 0.709 |
| AT3G07480.1 | 2Fe-2S ferredoxin-like superfamily protein | 37 | 3.02 | 0.260 | 0.653 |
| AT3G07480.1 | 2Fe-2S ferredoxin-like superfamily protein | 38 | 3.20 | 0.260 | 0.575 |
| AT3G07480.1 | 2Fe-2S ferredoxin-like superfamily protein | 39 | 3.30 | 0.261 | 0.537 |
| AT3G07480.1 | 2Fe-2S ferredoxin-like superfamily protein | 40 | 3.03 | 0.261 | 0.652 |
| AT3G07480.1 | 2Fe-2S ferredoxin-like superfamily protein | 41 | 2.63 | 0.262 | 0.831 |
| AT3G07480.1 | 2Fe-2S ferredoxin-like superfamily protein | 42 | 2.43 | 0.264 | 0.909 |
| AT3G07480.1 | 2Fe-2S ferredoxin-like superfamily protein | 43 | 2.29 | 0.266 | 0.948 |
| AT3G07480.1 | 2Fe-2S ferredoxin-like superfamily protein | 44 | 1.75 | 0.266 | 0.999 |
| AT3G07480.1 | 2Fe-2S ferredoxin-like superfamily protein | 45 | 1.03 | 0.267 | 1.000 |
| AT3G07480.1 | 2Fe-2S ferredoxin-like superfamily protein | 46 | 0.48 | 0.301 | 0.996 |
| AT3G07480.1 | 2Fe-2S ferredoxin-like superfamily protein | 47 | 0.19 | 0.459 | 0.850 |
| AT3G07568.1 | ? | 2 | 0.86 | 0.348 | 1.000 |
| AT3G07568.1 | ? | 3 | 0.73 | 0.275 | 1.000 |
| AT3G07568.1 | ? | 6 | 0.51 | 0.201 | 0.957 |
| AT3G07568.1 | ? | 7 | 0.50 | 0.172 | 0.830 |
| AT3G07568.1 | ? | 8 | 0.55 | 0.152 | 0.850 |
| AT3G07568.1 | ? | 9 | 0.68 | 0.141 | 0.993 |
| AT3G07568.1 | ? | 10 | 0.88 | 0.137 | 1.000 |
| AT3G07568.1 | ? | 11 | 1.13 | 0.136 | 1.000 |
| AT3G07568.1 | ? | 12 | 1.39 | 0.135 | 0.999 |
| AT3G07568.1 | ? | 13 | 1.60 | 0.133 | 0.928 |

| | | | | | |
|---|---|---|---|---|---|
| AT3G07568.1 | ? | 14 | 1.83 | 0.132 | 0.644 |
| AT3G07568.1 | ? | 15 | 2.00 | 0.131 | 0.393 |
| AT3G07568.1 | ? | 16 | 2.04 | 0.130 | 0.350 |
| AT3G07568.1 | ? | 17 | 1.95 | 0.130 | 0.460 |
| AT3G07568.1 | ? | 18 | 1.84 | 0.132 | 0.638 |
| AT3G07568.1 | ? | 19 | 1.72 | 0.135 | 0.827 |
| AT3G07568.1 | ? | 20 | 1.54 | 0.138 | 0.978 |
| AT3G07568.1 | ? | 21 | 1.25 | 0.140 | 1.000 |
| AT3G07568.1 | ? | 22 | 0.92 | 0.143 | 1.000 |
| AT3G07568.1 | ? | 23 | 0.66 | 0.145 | 0.992 |
| AT3G07568.1 | ? | 24 | 0.51 | 0.146 | 0.640 |
| AT3G07568.1 | ? | 25 | 0.46 | 0.146 | 0.386 |
| AT3G07568.1 | ? | 26 | 0.52 | 0.143 | 0.640 |
| AT3G07568.1 | ? | 27 | 0.70 | 0.139 | 0.997 |
| AT3G07568.1 | ? | 28 | 0.96 | 0.136 | 1.000 |
| AT3G07568.1 | ? | 29 | 1.17 | 0.135 | 1.000 |
| AT3G07568.1 | ? | 30 | 1.22 | 0.134 | 1.000 |
| AT3G07568.1 | ? | 31 | 1.12 | 0.134 | 1.000 |
| AT3G07568.1 | ? | 32 | 0.98 | 0.134 | 1.000 |
| AT3G07568.1 | ? | 33 | 0.88 | 0.135 | 1.000 |
| AT3G07568.1 | ? | 34 | 0.85 | 0.135 | 1.000 |
| AT3G07568.1 | ? | 35 | 0.90 | 0.134 | 1.000 |
| AT3G07568.1 | ? | 36 | 0.99 | 0.134 | 1.000 |
| AT3G07568.1 | ? | 37 | 1.10 | 0.134 | 1.000 |
| AT3G07568.1 | ? | 38 | 1.21 | 0.134 | 1.000 |
| AT3G07568.1 | ? | 39 | 1.36 | 0.134 | 0.999 |
| AT3G07568.1 | ? | 40 | 1.46 | 0.135 | 0.993 |
| AT3G07568.1 | ? | 41 | 1.37 | 0.136 | 0.999 |
| AT3G07568.1 | ? | 42 | 1.14 | 0.137 | 1.000 |
| AT3G07568.1 | ? | 43 | 0.88 | 0.138 | 1.000 |
| AT3G07568.1 | ? | 44 | 0.65 | 0.138 | 0.979 |
| AT3G07568.1 | ? | 45 | 0.49 | 0.144 | 0.519 |
| AT3G07568.1 | ? | 46 | 0.38 | 0.177 | 0.326 |
| AT3G07568.1 | ? | 47 | 0.29 | 0.265 | 0.616 |
| AT3G07770.1 | Heat shock protein 89.1 (HSP89.1) | 2 | 1.09 | 0.368 | 1.000 |
| AT3G07770.1 | Heat shock protein 89.1 (HSP89.1) | 3 | 0.55 | 0.287 | 0.999 |
| AT3G07770.1 | Heat shock protein 89.1 (HSP89.1) | 4 | 0.33 | 0.395 | 0.974 |
| AT3G07770.1 | Heat shock protein 89.1 (HSP89.1) | 15 | 0.70 | 0.347 | 1.000 |
| AT3G07770.1 | Heat shock protein 89.1 (HSP89.1) | 16 | 0.65 | 0.330 | 1.000 |
| AT3G07770.1 | Heat shock protein 89.1 (HSP89.1) | 17 | 0.55 | 0.329 | 1.000 |
| AT3G07770.1 | Heat shock protein 89.1 (HSP89.1) | 25 | 0.31 | 0.290 | 0.722 |
| AT3G07770.1 | Heat shock protein 89.1 (HSP89.1) | 26 | 0.50 | 0.234 | 0.964 |
| AT3G07770.1 | Heat shock protein 89.1 (HSP89.1) | 27 | 0.93 | 0.202 | 1.000 |
| AT3G07770.1 | Heat shock protein 89.1 (HSP89.1) | 28 | 1.38 | 0.192 | 1.000 |
| AT3G07770.1 | Heat shock protein 89.1 (HSP89.1) | 29 | 1.53 | 0.189 | 0.997 |
| AT3G07770.1 | Heat shock protein 89.1 (HSP89.1) | 30 | 1.41 | 0.189 | 1.000 |
| AT3G07770.1 | Heat shock protein 89.1 (HSP89.1) | 31 | 1.25 | 0.189 | 1.000 |
| AT3G07770.1 | Heat shock protein 89.1 (HSP89.1) | 32 | 1.16 | 0.189 | 1.000 |

| | | | | | |
|---|---|---|---|---|---|
| AT3G07770.1 | Heat shock protein 89.1 (HSP89.1) | 33 | 1.12 | 0.190 | 1.000 |
| AT3G07770.1 | Heat shock protein 89.1 (HSP89.1) | 34 | 1.10 | 0.190 | 1.000 |
| AT3G07770.1 | Heat shock protein 89.1 (HSP89.1) | 35 | 1.07 | 0.190 | 1.000 |
| AT3G07770.1 | Heat shock protein 89.1 (HSP89.1) | 36 | 1.00 | 0.189 | 1.000 |
| AT3G07770.1 | Heat shock protein 89.1 (HSP89.1) | 37 | 0.92 | 0.189 | 1.000 |
| AT3G07770.1 | Heat shock protein 89.1 (HSP89.1) | 38 | 0.85 | 0.189 | 1.000 |
| AT3G07770.1 | Heat shock protein 89.1 (HSP89.1) | 39 | 0.83 | 0.189 | 1.000 |
| AT3G07770.1 | Heat shock protein 89.1 (HSP89.1) | 40 | 0.88 | 0.189 | 1.000 |
| AT3G07770.1 | Heat shock protein 89.1 (HSP89.1) | 41 | 1.05 | 0.190 | 1.000 |
| AT3G07770.1 | Heat shock protein 89.1 (HSP89.1) | 42 | 1.36 | 0.192 | 1.000 |
| AT3G07770.1 | Heat shock protein 89.1 (HSP89.1) | 43 | 1.67 | 0.193 | 0.986 |
| AT3G07770.1 | Heat shock protein 89.1 (HSP89.1) | 44 | 1.72 | 0.193 | 0.977 |
| AT3G07770.1 | Heat shock protein 89.1 (HSP89.1) | 45 | 1.46 | 0.195 | 1.000 |
| AT3G07770.1 | Heat shock protein 89.1 (HSP89.1) | 46 | 1.22 | 0.230 | 1.000 |
| AT3G07770.1 | Heat shock protein 89.1 (HSP89.1) | 47 | 1.14 | 0.370 | 1.000 |
| AT3G08580.2 | ADP/ATP carrier 1 (AAC1) | 3 | 0.79 | 0.104 | 1.000 |
| AT3G08580.2 | ADP/ATP carrier 1 (AAC1) | 4 | 0.79 | 0.102 | 1.000 |
| AT3G08580.2 | ADP/ATP carrier 1 (AAC1) | 5 | 0.95 | 0.101 | 1.000 |
| AT3G08580.2 | ADP/ATP carrier 1 (AAC1) | 6 | 0.99 | 0.102 | 1.000 |
| AT3G08580.2 | ADP/ATP carrier 1 (AAC1) | 7 | 0.87 | 0.103 | 1.000 |
| AT3G08580.2 | ADP/ATP carrier 1 (AAC1) | 8 | 0.86 | 0.101 | 1.000 |
| AT3G08580.2 | ADP/ATP carrier 1 (AAC1) | 9 | 0.93 | 0.099 | 1.000 |
| AT3G08580.2 | ADP/ATP carrier 1 (AAC1) | 10 | 0.92 | 0.099 | 1.000 |
| AT3G08580.2 | ADP/ATP carrier 1 (AAC1) | 11 | 0.93 | 0.099 | 1.000 |
| AT3G08580.2 | ADP/ATP carrier 1 (AAC1) | 12 | 0.94 | 0.099 | 1.000 |
| AT3G08580.2 | ADP/ATP carrier 1 (AAC1) | 13 | 0.99 | 0.099 | 1.000 |
| AT3G08580.2 | ADP/ATP carrier 1 (AAC1) | 14 | 1.07 | 0.097 | 1.000 |
| AT3G08580.2 | ADP/ATP carrier 1 (AAC1) | 15 | 1.06 | 0.097 | 1.000 |
| AT3G08580.2 | ADP/ATP carrier 1 (AAC1) | 16 | 0.93 | 0.096 | 1.000 |
| AT3G08580.2 | ADP/ATP carrier 1 (AAC1) | 17 | 0.94 | 0.096 | 1.000 |
| AT3G08580.2 | ADP/ATP carrier 1 (AAC1) | 18 | 1.12 | 0.098 | 1.000 |
| AT3G08580.2 | ADP/ATP carrier 1 (AAC1) | 19 | 1.26 | 0.100 | 1.000 |
| AT3G08580.2 | ADP/ATP carrier 1 (AAC1) | 20 | 1.27 | 0.102 | 1.000 |
| AT3G08580.2 | ADP/ATP carrier 1 (AAC1) | 21 | 1.23 | 0.104 | 1.000 |
| AT3G08580.2 | ADP/ATP carrier 1 (AAC1) | 22 | 1.30 | 0.103 | 1.000 |
| AT3G08580.2 | ADP/ATP carrier 1 (AAC1) | 23 | 1.34 | 0.103 | 0.998 |
| AT3G08580.2 | ADP/ATP carrier 1 (AAC1) | 24 | 1.19 | 0.101 | 1.000 |
| AT3G08580.2 | ADP/ATP carrier 1 (AAC1) | 25 | 0.97 | 0.099 | 1.000 |
| AT3G08580.2 | ADP/ATP carrier 1 (AAC1) | 26 | 1.08 | 0.099 | 1.000 |
| AT3G08580.2 | ADP/ATP carrier 1 (AAC1) | 27 | 1.44 | 0.099 | 0.952 |
| AT3G08580.2 | ADP/ATP carrier 1 (AAC1) | 28 | 1.67 | 0.099 | 0.517 |
| AT3G08580.2 | ADP/ATP carrier 1 (AAC1) | 29 | 1.70 | 0.099 | 0.447 |
| AT3G08580.2 | ADP/ATP carrier 1 (AAC1) | 30 | 1.56 | 0.098 | 0.746 |
| AT3G08580.2 | ADP/ATP carrier 1 (AAC1) | 31 | 1.45 | 0.099 | 0.940 |
| AT3G08580.2 | ADP/ATP carrier 1 (AAC1) | 32 | 1.39 | 0.099 | 0.987 |
| AT3G08580.2 | ADP/ATP carrier 1 (AAC1) | 33 | 1.47 | 0.099 | 0.920 |
| AT3G08580.2 | ADP/ATP carrier 1 (AAC1) | 34 | 1.70 | 0.099 | 0.463 |

| | | | | | | |
|---|---|---|---|---|---|---|
| AT3G08580.2 | ADP/ATP carrier 1 (AAC1) | 35 | 1.72 | 0.099 | 0.413 | |
| AT3G08580.2 | ADP/ATP carrier 1 (AAC1) | 36 | 1.45 | 0.098 | 0.942 | |
| AT3G08580.2 | ADP/ATP carrier 1 (AAC1) | 37 | 1.33 | 0.098 | 0.998 | |
| AT3G08580.2 | ADP/ATP carrier 1 (AAC1) | 38 | 1.51 | 0.099 | 0.857 | |
| AT3G08580.2 | ADP/ATP carrier 1 (AAC1) | 39 | 1.68 | 0.099 | 0.502 | |
| AT3G08580.2 | ADP/ATP carrier 1 (AAC1) | 40 | 1.39 | 0.100 | 0.987 | |
| AT3G08580.2 | ADP/ATP carrier 1 (AAC1) | 41 | 0.87 | 0.100 | 1.000 | |
| AT3G08580.2 | ADP/ATP carrier 1 (AAC1) | 42 | 0.57 | 0.101 | 0.365 | |
| AT3G08580.2 | ADP/ATP carrier 1 (AAC1) | 43 | 0.47 | 0.102 | 0.046 | * |
| AT3G08580.2 | ADP/ATP carrier 1 (AAC1) | 44 | 0.46 | 0.101 | 0.031 | * |
| AT3G08580.2 | ADP/ATP carrier 1 (AAC1) | 45 | 0.47 | 0.102 | 0.048 | * |
| AT3G08580.2 | ADP/ATP carrier 1 (AAC1) | 46 | 0.46 | 0.110 | 0.079 | . |
| AT3G08580.2 | ADP/ATP carrier 1 (AAC1) | 47 | 0.42 | 0.206 | 0.852 | |
| AT3G08580.2 | ADP/ATP carrier 1 (AAC1) | 48 | 0.38 | 0.371 | 1.000 | |
| AT3G08610.1 | NADH:ubiquinone oxidoreductase subunit (MWFE) | 2 | 0.71 | 0.567 | 1.000 | |
| AT3G08610.1 | NADH:ubiquinone oxidoreductase subunit (MWFE) | 3 | 1.20 | 0.514 | 1.000 | |
| AT3G08610.1 | NADH:ubiquinone oxidoreductase subunit (MWFE) | 16 | 0.27 | 0.386 | 0.822 | |
| AT3G08610.1 | NADH:ubiquinone oxidoreductase subunit (MWFE) | 17 | 0.34 | 0.316 | 0.818 | |
| AT3G08610.1 | NADH:ubiquinone oxidoreductase subunit (MWFE) | 18 | 0.84 | 0.281 | 1.000 | |
| AT3G08610.1 | NADH:ubiquinone oxidoreductase subunit (MWFE) | 19 | 1.72 | 0.275 | 0.996 | |
| AT3G08610.1 | NADH:ubiquinone oxidoreductase subunit (MWFE) | 20 | 1.39 | 0.275 | 1.000 | |
| AT3G08610.1 | NADH:ubiquinone oxidoreductase subunit (MWFE) | 21 | 1.00 | 0.278 | 1.000 | |
| AT3G08610.1 | NADH:ubiquinone oxidoreductase subunit (MWFE) | 22 | 1.05 | 0.275 | 1.000 | |
| AT3G08610.1 | NADH:ubiquinone oxidoreductase subunit (MWFE) | 23 | 1.36 | 0.273 | 1.000 | |
| AT3G08610.1 | NADH:ubiquinone oxidoreductase subunit (MWFE) | 24 | 1.79 | 0.269 | 0.991 | |
| AT3G08610.1 | NADH:ubiquinone oxidoreductase subunit (MWFE) | 25 | 1.52 | 0.268 | 1.000 | |
| AT3G08610.1 | NADH:ubiquinone oxidoreductase subunit (MWFE) | 26 | 0.79 | 0.283 | 1.000 | |
| AT3G08610.1 | NADH:ubiquinone oxidoreductase subunit (MWFE) | 27 | 0.39 | 0.345 | 0.948 | |
| AT3G08610.1 | NADH:ubiquinone oxidoreductase subunit (MWFE) | 28 | 0.23 | 0.516 | 0.939 | |
| AT3G08610.1 | NADH:ubiquinone oxidoreductase subunit (MWFE) | 46 | 0.00 | 20.856 | 1.000 | |
| AT3G08610.1 | NADH:ubiquinone oxidoreductase subunit (MWFE) | 47 | 0.00 | 22.584 | 1.000 | |
| AT3G08860.1 | Pyrimidine 4 (PYD4) | 2 | 1.03 | 0.563 | 1.000 | |
| AT3G08860.1 | Pyrimidine 4 (PYD4) | 3 | 0.69 | 0.662 | 1.000 | |
| AT3G08860.1 | Pyrimidine 4 (PYD4) | 32 | 0.43 | 0.335 | 0.806 | |
| AT3G08860.1 | Pyrimidine 4 (PYD4) | 33 | 0.88 | 0.268 | 1.000 | |
| AT3G08860.1 | Pyrimidine 4 (PYD4) | 34 | 1.65 | 0.266 | 0.937 | |
| AT3G08860.1 | Pyrimidine 4 (PYD4) | 35 | 2.56 | 0.317 | 0.687 | |
| AT3G08860.1 | Pyrimidine 4 (PYD4) | 36 | 3.28 | 0.408 | 0.701 | |
| AT3G08860.1 | Pyrimidine 4 (PYD4) | 46 | 0.07 | 0.409 | 0.046 | * |
| AT3G08860.1 | Pyrimidine 4 (PYD4) | 47 | 0.04 | 0.431 | 0.012 | * |
| AT3G08940.2 | Photosystem II light harvesting complex subunit (LHCB4.2) | 2 | 0.32 | 0.387 | 0.996 | |
| AT3G08940.2 | Photosystem II light harvesting complex subunit (LHCB4.2) | 3 | 0.50 | 0.242 | 0.998 | |

| | | | | | | |
|---|---|---|---|---|---|---|
| AT3G08940.2 | Photosystem II light harvesting complex subunit (LHCB4.2) | 4 | 0.83 | 0.224 | 1.000 | |
| AT3G08940.2 | Photosystem II light harvesting complex subunit (LHCB4.2) | 5 | 1.11 | 0.223 | 1.000 | |
| AT3G08940.2 | Photosystem II light harvesting complex subunit (LHCB4.2) | 6 | 1.07 | 0.224 | 1.000 | |
| AT3G08940.2 | Photosystem II light harvesting complex subunit (LHCB4.2) | 7 | 0.90 | 0.225 | 1.000 | |
| AT3G08940.2 | Photosystem II light harvesting complex subunit (LHCB4.2) | 8 | 0.86 | 0.219 | 1.000 | |
| AT3G08940.2 | Photosystem II light harvesting complex subunit (LHCB4.2) | 9 | 0.87 | 0.215 | 1.000 | |
| AT3G08940.2 | Photosystem II light harvesting complex subunit (LHCB4.2) | 10 | 0.82 | 0.216 | 1.000 | |
| AT3G08940.2 | Photosystem II light harvesting complex subunit (LHCB4.2) | 11 | 0.81 | 0.218 | 1.000 | |
| AT3G08940.2 | Photosystem II light harvesting complex subunit (LHCB4.2) | 12 | 0.85 | 0.218 | 1.000 | |
| AT3G08940.2 | Photosystem II light harvesting complex subunit (LHCB4.2) | 13 | 0.95 | 0.215 | 1.000 | |
| AT3G08940.2 | Photosystem II light harvesting complex subunit (LHCB4.2) | 14 | 1.05 | 0.212 | 1.000 | |
| AT3G08940.2 | Photosystem II light harvesting complex subunit (LHCB4.2) | 15 | 1.01 | 0.212 | 1.000 | |
| AT3G08940.2 | Photosystem II light harvesting complex subunit (LHCB4.2) | 16 | 0.90 | 0.210 | 1.000 | |
| AT3G08940.2 | Photosystem II light harvesting complex subunit (LHCB4.2) | 17 | 0.88 | 0.209 | 1.000 | |
| AT3G08940.2 | Photosystem II light harvesting complex subunit (LHCB4.2) | 18 | 0.99 | 0.211 | 1.000 | |
| AT3G08940.2 | Photosystem II light harvesting complex subunit (LHCB4.2) | 19 | 1.20 | 0.219 | 1.000 | |
| AT3G08940.2 | Photosystem II light harvesting complex subunit (LHCB4.2) | 20 | 1.46 | 0.225 | 1.000 | |
| AT3G08940.2 | Photosystem II light harvesting complex subunit (LHCB4.2) | 21 | 1.78 | 0.227 | 1.000 | |
| AT3G08940.2 | Photosystem II light harvesting complex subunit (LHCB4.2) | 22 | 1.93 | 0.227 | 0.997 | |
| AT3G08940.2 | Photosystem II light harvesting complex subunit (LHCB4.2) | 23 | 1.53 | 0.225 | 1.000 | |
| AT3G08940.2 | Photosystem II light harvesting complex subunit (LHCB4.2) | 24 | 0.91 | 0.220 | 1.000 | |
| AT3G08940.2 | Photosystem II light harvesting complex subunit (LHCB4.2) | 25 | 0.49 | 0.216 | 0.985 | |
| AT3G08940.2 | Photosystem II light harvesting complex subunit (LHCB4.2) | 26 | 0.45 | 0.216 | 0.947 | |
| AT3G08940.2 | Photosystem II light harvesting complex subunit (LHCB4.2) | 27 | 0.67 | 0.216 | 1.000 | |
| AT3G08940.2 | Photosystem II light harvesting complex subunit (LHCB4.2) | 28 | 0.65 | 0.216 | 1.000 | |
| AT3G08940.2 | Photosystem II light harvesting complex subunit (LHCB4.2) | 29 | 0.58 | 0.216 | 1.000 | |
| AT3G08940.2 | Photosystem II light harvesting complex subunit (LHCB4.2) | 30 | 0.85 | 0.215 | 1.000 | |
| AT3G08940.2 | Photosystem II light harvesting complex subunit (LHCB4.2) | 31 | 1.62 | 0.217 | 1.000 | |
| AT3G08940.2 | Photosystem II light harvesting complex subunit (LHCB4.2) | 32 | 1.21 | 0.221 | 1.000 | |
| AT3G08940.2 | Photosystem II light harvesting complex subunit (LHCB4.2) | 33 | 0.35 | 0.233 | 0.733 | |
| AT3G08940.2 | Photosystem II light harvesting complex subunit (LHCB4.2) | 34 | 0.13 | 0.285 | 0.075 | . |

| | | | | | |
|---|---|---|---|---|---|
| AT3G08940.2 | Photosystem II light harvesting complex subunit (LHCB4.2) | 35 | 0.11 | 0.405 | 0.375 |
| AT3G08940.2 | Photosystem II light harvesting complex subunit (LHCB4.2) | 40 | 1.08 | 0.327 | 1.000 |
| AT3G08940.2 | Photosystem II light harvesting complex subunit (LHCB4.2) | 41 | 1.66 | 0.261 | 1.000 |
| AT3G08940.2 | Photosystem II light harvesting complex subunit (LHCB4.2) | 42 | 1.96 | 0.234 | 0.997 |
| AT3G08940.2 | Photosystem II light harvesting complex subunit (LHCB4.2) | 43 | 1.55 | 0.225 | 1.000 |
| AT3G08940.2 | Photosystem II light harvesting complex subunit (LHCB4.2) | 44 | 0.99 | 0.230 | 1.000 |
| AT3G08940.2 | Photosystem II light harvesting complex subunit (LHCB4.2) | 45 | 0.71 | 0.279 | 1.000 |
| AT3G08940.2 | Photosystem II light harvesting complex subunit (LHCB4.2) | 46 | 0.56 | 0.371 | 1.000 |
| AT3G08940.2 | Photosystem II light harvesting complex subunit (LHCB4.2) | 47 | 0.47 | 0.585 | 1.000 |
| AT3G08950.1 | Cytochrome c:O2 oxidoreductase assembly protein (HCC1) | 2 | 1.06 | 0.415 | 1.000 |
| AT3G08950.1 | Cytochrome c:O2 oxidoreductase assembly protein (HCC1) | 3 | 0.88 | 0.321 | 1.000 |
| AT3G08950.1 | Cytochrome c:O2 oxidoreductase assembly protein (HCC1) | 6 | 0.59 | 0.266 | 1.000 |
| AT3G08950.1 | Cytochrome c:O2 oxidoreductase assembly protein (HCC1) | 7 | 0.58 | 0.213 | 0.999 |
| AT3G08950.1 | Cytochrome c:O2 oxidoreductase assembly protein (HCC1) | 8 | 0.65 | 0.182 | 1.000 |
| AT3G08950.1 | Cytochrome c:O2 oxidoreductase assembly protein (HCC1) | 9 | 0.83 | 0.170 | 1.000 |
| AT3G08950.1 | Cytochrome c:O2 oxidoreductase assembly protein (HCC1) | 10 | 1.07 | 0.167 | 1.000 |
| AT3G08950.1 | Cytochrome c:O2 oxidoreductase assembly protein (HCC1) | 11 | 1.35 | 0.167 | 1.000 |
| AT3G08950.1 | Cytochrome c:O2 oxidoreductase assembly protein (HCC1) | 12 | 1.58 | 0.166 | 0.997 |
| AT3G08950.1 | Cytochrome c:O2 oxidoreductase assembly protein (HCC1) | 13 | 1.62 | 0.163 | 0.992 |
| AT3G08950.1 | Cytochrome c:O2 oxidoreductase assembly protein (HCC1) | 14 | 1.56 | 0.161 | 0.997 |
| AT3G08950.1 | Cytochrome c:O2 oxidoreductase assembly protein (HCC1) | 15 | 1.42 | 0.160 | 1.000 |
| AT3G08950.1 | Cytochrome c:O2 oxidoreductase assembly protein (HCC1) | 16 | 1.38 | 0.159 | 1.000 |
| AT3G08950.1 | Cytochrome c:O2 oxidoreductase assembly protein (HCC1) | 17 | 1.61 | 0.158 | 0.990 |

| | | | | | | |
|---|---|---|---|---|---|---|
| AT3G08950.1 | Cytochrome c:O2 oxidoreductase assembly protein (HCC1) | 18 | 2.13 | 0.161 | 0.643 | |
| AT3G08950.1 | Cytochrome c:O2 oxidoreductase assembly protein (HCC1) | 19 | 2.88 | 0.166 | 0.152 | |
| AT3G08950.1 | Cytochrome c:O2 oxidoreductase assembly protein (HCC1) | 20 | 3.79 | 0.171 | 0.024 | * |
| AT3G08950.1 | Cytochrome c:O2 oxidoreductase assembly protein (HCC1) | 21 | 4.77 | 0.173 | 0.004 | ** |
| AT3G08950.1 | Cytochrome c:O2 oxidoreductase assembly protein (HCC1) | 22 | 5.46 | 0.177 | 0.001 | ** |
| AT3G08950.1 | Cytochrome c:O2 oxidoreductase assembly protein (HCC1) | 23 | 5.09 | 0.187 | 0.006 | ** |
| AT3G08950.1 | Cytochrome c:O2 oxidoreductase assembly protein (HCC1) | 24 | 3.70 | 0.218 | 0.224 | |
| AT3G08950.1 | Cytochrome c:O2 oxidoreductase assembly protein (HCC1) | 25 | 2.15 | 0.280 | 0.997 | |
| AT3G08950.1 | Cytochrome c:O2 oxidoreductase assembly protein (HCC1) | 29 | 0.17 | 0.243 | 0.058 | . |
| AT3G08950.1 | Cytochrome c:O2 oxidoreductase assembly protein (HCC1) | 30 | 0.20 | 0.200 | 0.017 | * |
| AT3G08950.1 | Cytochrome c:O2 oxidoreductase assembly protein (HCC1) | 31 | 0.35 | 0.185 | 0.319 | |
| AT3G08950.1 | Cytochrome c:O2 oxidoreductase assembly protein (HCC1) | 32 | 0.62 | 0.189 | 0.999 | |
| AT3G08950.1 | Cytochrome c:O2 oxidoreductase assembly protein (HCC1) | 33 | 0.85 | 0.228 | 1.000 | |
| AT3G08950.1 | Cytochrome c:O2 oxidoreductase assembly protein (HCC1) | 34 | 0.97 | 0.301 | 1.000 | |
| AT3G08950.1 | Cytochrome c:O2 oxidoreductase assembly protein (HCC1) | 39 | 0.71 | 0.261 | 1.000 | |
| AT3G08950.1 | Cytochrome c:O2 oxidoreductase assembly protein (HCC1) | 40 | 0.78 | 0.206 | 1.000 | |
| AT3G08950.1 | Cytochrome c:O2 oxidoreductase assembly protein (HCC1) | 41 | 1.01 | 0.177 | 1.000 | |
| AT3G08950.1 | Cytochrome c:O2 oxidoreductase assembly protein (HCC1) | 42 | 1.42 | 0.170 | 1.000 | |
| AT3G08950.1 | Cytochrome c:O2 oxidoreductase assembly protein (HCC1) | 43 | 1.86 | 0.170 | 0.932 | |
| AT3G08950.1 | Cytochrome c:O2 oxidoreductase assembly protein (HCC1) | 44 | 1.66 | 0.170 | 0.991 | |

| | | | | | | |
|---|---|---|---|---|---|---|
| AT3G08950.1 | Cytochrome c:O2 oxidoreductase assembly protein (HCC1) | 45 | 0.98 | 0.171 | 1.000 | |
| AT3G08950.1 | Cytochrome c:O2 oxidoreductase assembly protein (HCC1) | 46 | 0.41 | 0.206 | 0.754 | |
| AT3G08950.1 | Cytochrome c:O2 oxidoreductase assembly protein (HCC1) | 47 | 0.14 | 0.353 | 0.319 | |
| AT3G08980.1 | Peptidase S24/S26A/S26B/S26C family protein | 2 | 0.89 | 0.413 | 1.000 | |
| AT3G08980.1 | Peptidase S24/S26A/S26B/S26C family protein | 3 | 0.98 | 0.339 | 1.000 | |
| AT3G08980.1 | Peptidase S24/S26A/S26B/S26C family protein | 8 | 1.85 | 0.240 | 0.983 | |
| AT3G08980.1 | Peptidase S24/S26A/S26B/S26C family protein | 9 | 2.18 | 0.200 | 0.739 | |
| AT3G08980.1 | Peptidase S24/S26A/S26B/S26C family protein | 10 | 2.46 | 0.174 | 0.326 | |
| AT3G08980.1 | Peptidase S24/S26A/S26B/S26C family protein | 11 | 2.57 | 0.160 | 0.166 | |
| AT3G08980.1 | Peptidase S24/S26A/S26B/S26C family protein | 12 | 2.50 | 0.154 | 0.155 | |
| AT3G08980.1 | Peptidase S24/S26A/S26B/S26C family protein | 13 | 2.35 | 0.151 | 0.209 | |
| AT3G08980.1 | Peptidase S24/S26A/S26B/S26C family protein | 14 | 2.26 | 0.149 | 0.255 | |
| AT3G08980.1 | Peptidase S24/S26A/S26B/S26C family protein | 15 | 2.24 | 0.149 | 0.264 | |
| AT3G08980.1 | Peptidase S24/S26A/S26B/S26C family protein | 16 | 2.29 | 0.149 | 0.235 | |
| AT3G08980.1 | Peptidase S24/S26A/S26B/S26C family protein | 17 | 2.51 | 0.152 | 0.139 | |
| AT3G08980.1 | Peptidase S24/S26A/S26B/S26C family protein | 18 | 2.97 | 0.159 | 0.056 | . |
| AT3G08980.1 | Peptidase S24/S26A/S26B/S26C family protein | 19 | 3.55 | 0.178 | 0.040 | * |
| AT3G08980.1 | Peptidase S24/S26A/S26B/S26C family protein | 20 | 4.12 | 0.224 | 0.106 | |
| AT3G08980.1 | Peptidase S24/S26A/S26B/S26C family protein | 21 | 4.72 | 0.300 | 0.325 | |
| AT3G08980.1 | Peptidase S24/S26A/S26B/S26C family protein | 38 | 2.02 | 0.284 | 0.987 | |
| AT3G08980.1 | Peptidase S24/S26A/S26B/S26C family protein | 39 | 1.46 | 0.208 | 0.999 | |
| AT3G08980.1 | Peptidase S24/S26A/S26B/S26C family protein | 40 | 0.99 | 0.169 | 1.000 | |
| AT3G08980.1 | Peptidase S24/S26A/S26B/S26C family protein | 41 | 0.66 | 0.158 | 0.976 | |
| AT3G08980.1 | Peptidase S24/S26A/S26B/S26C family protein | 42 | 0.46 | 0.158 | 0.421 | |
| AT3G08980.1 | Peptidase S24/S26A/S26B/S26C family protein | 43 | 0.36 | 0.159 | 0.092 | . |
| AT3G08980.1 | Peptidase S24/S26A/S26B/S26C family protein | 44 | 0.28 | 0.165 | 0.021 | * |
| AT3G08980.1 | Peptidase S24/S26A/S26B/S26C family protein | 45 | 0.24 | 0.201 | 0.037 | * |
| AT3G08980.1 | Peptidase S24/S26A/S26B/S26C family protein | 46 | 0.20 | 0.297 | 0.282 | |
| AT3G08980.1 | Peptidase S24/S26A/S26B/S26C family protein | 47 | 0.18 | 0.460 | 0.775 | |

| | | | | | |
|---|---|---|---|---|---|
| AT3G09810.1 | NAD isocitrate dehydrogenase subunit (IDH6, catalytic) | 2 | 0.57 | 0.602 | 1.000 |
| AT3G09810.1 | NAD isocitrate dehydrogenase subunit (IDH6, catalytic) | 3 | 0.47 | 0.553 | 0.998 |
| AT3G09810.1 | NAD isocitrate dehydrogenase subunit (IDH6, catalytic) | 4 | 0.40 | 0.594 | 0.994 |
| AT3G09810.1 | NAD isocitrate dehydrogenase subunit (IDH6, catalytic) | 24 | 0.83 | 0.492 | 1.000 |
| AT3G09810.1 | NAD isocitrate dehydrogenase subunit (IDH6, catalytic) | 25 | 0.93 | 0.431 | 1.000 |
| AT3G09810.1 | NAD isocitrate dehydrogenase subunit (IDH6, catalytic) | 26 | 1.03 | 0.391 | 1.000 |
| AT3G09810.1 | NAD isocitrate dehydrogenase subunit (IDH6, catalytic) | 27 | 1.15 | 0.369 | 1.000 |
| AT3G09810.1 | NAD isocitrate dehydrogenase subunit (IDH6, catalytic) | 28 | 1.26 | 0.359 | 1.000 |
| AT3G09810.1 | NAD isocitrate dehydrogenase subunit (IDH6, catalytic) | 29 | 1.34 | 0.356 | 1.000 |
| AT3G09810.1 | NAD isocitrate dehydrogenase subunit (IDH6, catalytic) | 30 | 1.32 | 0.355 | 1.000 |
| AT3G09810.1 | NAD isocitrate dehydrogenase subunit (IDH6, catalytic) | 31 | 1.20 | 0.355 | 1.000 |
| AT3G09810.1 | NAD isocitrate dehydrogenase subunit (IDH6, catalytic) | 32 | 1.00 | 0.355 | 1.000 |
| AT3G09810.1 | NAD isocitrate dehydrogenase subunit (IDH6, catalytic) | 33 | 0.81 | 0.355 | 1.000 |
| AT3G09810.1 | NAD isocitrate dehydrogenase subunit (IDH6, catalytic) | 34 | 0.67 | 0.355 | 1.000 |
| AT3G09810.1 | NAD isocitrate dehydrogenase subunit (IDH6, catalytic) | 35 | 0.61 | 0.355 | 0.997 |
| AT3G09810.1 | NAD isocitrate dehydrogenase subunit (IDH6, catalytic) | 36 | 0.64 | 0.355 | 0.999 |
| AT3G09810.1 | NAD isocitrate dehydrogenase subunit (IDH6, catalytic) | 37 | 0.78 | 0.355 | 1.000 |
| AT3G09810.1 | NAD isocitrate dehydrogenase subunit (IDH6, catalytic) | 38 | 1.05 | 0.355 | 1.000 |
| AT3G09810.1 | NAD isocitrate dehydrogenase subunit (IDH6, catalytic) | 39 | 1.47 | 0.356 | 1.000 |
| AT3G09810.1 | NAD isocitrate dehydrogenase subunit (IDH6, catalytic) | 40 | 2.00 | 0.357 | 0.971 |
| AT3G09810.1 | NAD isocitrate dehydrogenase subunit (IDH6, catalytic) | 41 | 2.50 | 0.357 | 0.870 |
| AT3G09810.1 | NAD isocitrate dehydrogenase subunit (IDH6, catalytic) | 42 | 2.79 | 0.358 | 0.794 |
| AT3G09810.1 | NAD isocitrate dehydrogenase subunit (IDH6, catalytic) | 43 | 2.69 | 0.360 | 0.826 |
| AT3G09810.1 | NAD isocitrate dehydrogenase subunit (IDH6, catalytic) | 44 | 2.18 | 0.366 | 0.949 |
| AT3G09810.1 | NAD isocitrate dehydrogenase subunit (IDH6, catalytic) | 45 | 1.53 | 0.387 | 1.000 |
| AT3G09810.1 | NAD isocitrate dehydrogenase subunit (IDH6, catalytic) | 46 | 0.97 | 0.436 | 1.000 |
| AT3G09810.1 | NAD isocitrate dehydrogenase subunit (IDH6, catalytic) | 47 | 0.58 | 0.526 | 1.000 |
| AT3G10110.1 | Translocase of the inner mitochondrial membrane Tim17/Tim22/Tim23 family protein (MEE67) | 2 | 1.07 | 0.461 | 1.000 |
| AT3G10110.1 | Translocase of the inner mitochondrial membrane Tim17/Tim22/Tim23 family protein (MEE67) | 3 | 1.20 | 0.413 | 1.000 |

| | | | | | |
|---|---|---|---|---|---|
| AT3G10110.1 | Translocase of the inner mitochondrial membrane Tim17/Tim22/Tim23 family protein (MEE67) | 11 | 2.09 | 0.250 | 0.841 |
| AT3G10110.1 | Translocase of the inner mitochondrial membrane Tim17/Tim22/Tim23 family protein (MEE67) | 12 | 2.04 | 0.226 | 0.787 |
| AT3G10110.1 | Translocase of the inner mitochondrial membrane Tim17/Tim22/Tim23 family protein (MEE67) | 13 | 1.94 | 0.204 | 0.765 |
| AT3G10110.1 | Translocase of the inner mitochondrial membrane Tim17/Tim22/Tim23 family protein (MEE67) | 14 | 1.78 | 0.186 | 0.804 |
| AT3G10110.1 | Translocase of the inner mitochondrial membrane Tim17/Tim22/Tim23 family protein (MEE67) | 15 | 1.60 | 0.174 | 0.895 |
| AT3G10110.1 | Translocase of the inner mitochondrial membrane Tim17/Tim22/Tim23 family protein (MEE67) | 16 | 1.40 | 0.165 | 0.978 |
| AT3G10110.1 | Translocase of the inner mitochondrial membrane Tim17/Tim22/Tim23 family protein (MEE67) | 17 | 1.21 | 0.161 | 1.000 |
| AT3G10110.1 | Translocase of the inner mitochondrial membrane Tim17/Tim22/Tim23 family protein (MEE67) | 18 | 1.06 | 0.159 | 1.000 |
| AT3G10110.1 | Translocase of the inner mitochondrial membrane Tim17/Tim22/Tim23 family protein (MEE67) | 19 | 0.96 | 0.160 | 1.000 |
| AT3G10110.1 | Translocase of the inner mitochondrial membrane Tim17/Tim22/Tim23 family protein (MEE67) | 20 | 0.92 | 0.162 | 1.000 |
| AT3G10110.1 | Translocase of the inner mitochondrial membrane Tim17/Tim22/Tim23 family protein (MEE67) | 21 | 0.93 | 0.167 | 1.000 |
| AT3G10110.1 | Translocase of the inner mitochondrial membrane Tim17/Tim22/Tim23 family protein (MEE67) | 22 | 1.03 | 0.176 | 1.000 |
| AT3G10110.1 | Translocase of the inner mitochondrial membrane Tim17/Tim22/Tim23 family protein (MEE67) | 23 | 1.21 | 0.190 | 1.000 |
| AT3G10110.1 | Translocase of the inner mitochondrial membrane Tim17/Tim22/Tim23 family protein (MEE67) | 24 | 1.50 | 0.213 | 0.986 |
| AT3G10110.1 | Translocase of the inner mitochondrial membrane Tim17/Tim22/Tim23 family protein (MEE67) | 25 | 1.89 | 0.246 | 0.913 |

| | | | | | |
|---|---|---|---|---|---|
| AT3G10110.1 | Translocase of the inner mitochondrial membrane Tim17/Tim22/Tim23 family protein (MEE67) | 41 | 2.29 | 0.291 | 0.863 |
| AT3G10110.1 | Translocase of the inner mitochondrial membrane Tim17/Tim22/Tim23 family protein (MEE67) | 42 | 1.59 | 0.249 | 0.988 |
| AT3G10110.1 | Translocase of the inner mitochondrial membrane Tim17/Tim22/Tim23 family protein (MEE67) | 43 | 1.04 | 0.214 | 1.000 |
| AT3G10110.1 | Translocase of the inner mitochondrial membrane Tim17/Tim22/Tim23 family protein (MEE67) | 44 | 0.67 | 0.197 | 0.977 |
| AT3G10110.1 | Translocase of the inner mitochondrial membrane Tim17/Tim22/Tim23 family protein (MEE67) | 45 | 0.43 | 0.208 | 0.497 |
| AT3G10110.1 | Translocase of the inner mitochondrial membrane Tim17/Tim22/Tim23 family protein (MEE67) | 46 | 0.27 | 0.252 | 0.217 |
| AT3G10110.1 | Translocase of the inner mitochondrial membrane Tim17/Tim22/Tim23 family protein (MEE67) | 47 | 0.17 | 0.327 | 0.179 |
| AT3G10160.1 | DHFS-FPGS homolog C (DFC) / Folylpolyglutamate synthetase 2 (FPGS2) | 2 | 0.95 | 0.401 | 1.000 |
| AT3G10160.1 | DHFS-FPGS homolog C (DFC) / Folylpolyglutamate synthetase 2 (FPGS2) | 3 | 0.86 | 0.380 | 1.000 |
| AT3G10160.1 | DHFS-FPGS homolog C (DFC) / Folylpolyglutamate synthetase 2 (FPGS2) | 25 | 0.50 | 0.280 | 0.958 |
| AT3G10160.1 | DHFS-FPGS homolog C (DFC) / Folylpolyglutamate synthetase 2 (FPGS2) | 26 | 0.56 | 0.219 | 0.937 |
| AT3G10160.1 | DHFS-FPGS homolog C (DFC) / Folylpolyglutamate synthetase 2 (FPGS2) | 27 | 0.62 | 0.189 | 0.948 |
| AT3G10160.1 | DHFS-FPGS homolog C (DFC) / Folylpolyglutamate synthetase 2 (FPGS2) | 28 | 0.65 | 0.178 | 0.961 |
| AT3G10160.1 | DHFS-FPGS homolog C (DFC) / Folylpolyglutamate synthetase 2 (FPGS2) | 29 | 0.64 | 0.175 | 0.952 |
| AT3G10160.1 | DHFS-FPGS homolog C (DFC) / Folylpolyglutamate synthetase 2 (FPGS2) | 30 | 0.61 | 0.175 | 0.900 |
| AT3G10160.1 | DHFS-FPGS homolog C (DFC) / Folylpolyglutamate synthetase 2 (FPGS2) | 31 | 0.58 | 0.175 | 0.844 |
| AT3G10160.1 | DHFS-FPGS homolog C (DFC) / Folylpolyglutamate synthetase 2 (FPGS2) | 32 | 0.59 | 0.175 | 0.868 |
| AT3G10160.1 | DHFS-FPGS homolog C (DFC) / Folylpolyglutamate synthetase 2 (FPGS2) | 33 | 0.66 | 0.175 | 0.966 |

| | | | | | | |
|---|---|---|---|---|---|---|
| AT3G10160.1 | DHFS-FPGS homolog C (DFC) / Folylpolyglutamate synthetase 2 (FPGS2) | 34 | 0.80 | 0.175 | 1.000 | |
| AT3G10160.1 | DHFS-FPGS homolog C (DFC) / Folylpolyglutamate synthetase 2 (FPGS2) | 35 | 1.03 | 0.175 | 1.000 | |
| AT3G10160.1 | DHFS-FPGS homolog C (DFC) / Folylpolyglutamate synthetase 2 (FPGS2) | 36 | 1.37 | 0.175 | 0.996 | |
| AT3G10160.1 | DHFS-FPGS homolog C (DFC) / Folylpolyglutamate synthetase 2 (FPGS2) | 37 | 1.80 | 0.174 | 0.772 | |
| AT3G10160.1 | DHFS-FPGS homolog C (DFC) / Folylpolyglutamate synthetase 2 (FPGS2) | 38 | 2.18 | 0.174 | 0.417 | |
| AT3G10160.1 | DHFS-FPGS homolog C (DFC) / Folylpolyglutamate synthetase 2 (FPGS2) | 39 | 2.36 | 0.174 | 0.295 | |
| AT3G10160.1 | DHFS-FPGS homolog C (DFC) / Folylpolyglutamate synthetase 2 (FPGS2) | 40 | 2.20 | 0.175 | 0.413 | |
| AT3G10160.1 | DHFS-FPGS homolog C (DFC) / Folylpolyglutamate synthetase 2 (FPGS2) | 41 | 1.80 | 0.176 | 0.775 | |
| AT3G10160.1 | DHFS-FPGS homolog C (DFC) / Folylpolyglutamate synthetase 2 (FPGS2) | 42 | 1.42 | 0.178 | 0.991 | |
| AT3G10160.1 | DHFS-FPGS homolog C (DFC) / Folylpolyglutamate synthetase 2 (FPGS2) | 43 | 1.11 | 0.178 | 1.000 | |
| AT3G10160.1 | DHFS-FPGS homolog C (DFC) / Folylpolyglutamate synthetase 2 (FPGS2) | 44 | 0.82 | 0.178 | 1.000 | |
| AT3G10160.1 | DHFS-FPGS homolog C (DFC) / Folylpolyglutamate synthetase 2 (FPGS2) | 45 | 0.57 | 0.183 | 0.848 | |
| AT3G10160.1 | DHFS-FPGS homolog C (DFC) / Folylpolyglutamate synthetase 2 (FPGS2) | 46 | 0.37 | 0.217 | 0.388 | |
| AT3G10160.1 | DHFS-FPGS homolog C (DFC) / Folylpolyglutamate synthetase 2 (FPGS2) | 47 | 0.23 | 0.306 | 0.334 | |
| AT3G10370.1 | FAD glycerol-3-phosphate dehydrogenase (SDP6) | 2 | 1.02 | 0.390 | 1.000 | |
| AT3G10370.1 | FAD glycerol-3-phosphate dehydrogenase (SDP6) | 3 | 0.77 | 0.290 | 1.000 | |
| AT3G10370.1 | FAD glycerol-3-phosphate dehydrogenase (SDP6) | 4 | 0.67 | 0.355 | 1.000 | |
| AT3G10370.1 | FAD glycerol-3-phosphate dehydrogenase (SDP6) | 8 | 1.34 | 0.339 | 1.000 | |
| AT3G10370.1 | FAD glycerol-3-phosphate dehydrogenase (SDP6) | 9 | 1.96 | 0.275 | 0.999 | |
| AT3G10370.1 | FAD glycerol-3-phosphate dehydrogenase (SDP6) | 10 | 2.90 | 0.249 | 0.770 | |
| AT3G10370.1 | FAD glycerol-3-phosphate dehydrogenase (SDP6) | 11 | 4.25 | 0.233 | 0.175 | |
| AT3G10370.1 | FAD glycerol-3-phosphate dehydrogenase (SDP6) | 12 | 6.37 | 0.228 | 0.015 | * |
| AT3G10370.1 | FAD glycerol-3-phosphate dehydrogenase (SDP6) | 13 | 9.15 | 0.217 | 0.000 | *** |
| AT3G10370.1 | FAD glycerol-3-phosphate dehydrogenase (SDP6) | 14 | 11.95 | 0.209 | 0.000 | *** |

| | | | | | | |
|---|---|---|---|---|---|---|
| AT3G10370.1 | FAD glycerol-3-phosphate dehydrogenase (SDP6) | 15 | 12.97 | 0.208 | 0.000 | *** |
| AT3G10370.1 | FAD glycerol-3-phosphate dehydrogenase (SDP6) | 16 | 12.66 | 0.206 | 0.000 | *** |
| AT3G10370.1 | FAD glycerol-3-phosphate dehydrogenase (SDP6) | 17 | 14.71 | 0.206 | 0.000 | *** |
| AT3G10370.1 | FAD glycerol-3-phosphate dehydrogenase (SDP6) | 18 | 19.42 | 0.212 | 0.000 | *** |
| AT3G10370.1 | FAD glycerol-3-phosphate dehydrogenase (SDP6) | 19 | 19.86 | 0.221 | 0.000 | *** |
| AT3G10370.1 | FAD glycerol-3-phosphate dehydrogenase (SDP6) | 20 | 13.09 | 0.233 | 0.000 | *** |
| AT3G10370.1 | FAD glycerol-3-phosphate dehydrogenase (SDP6) | 21 | 6.53 | 0.267 | 0.068 | . |
| AT3G10370.1 | FAD glycerol-3-phosphate dehydrogenase (SDP6) | 22 | 3.38 | 0.337 | 0.932 | |
| AT3G10370.1 | FAD glycerol-3-phosphate dehydrogenase (SDP6) | 26 | 1.17 | 0.305 | 1.000 | |
| AT3G10370.1 | FAD glycerol-3-phosphate dehydrogenase (SDP6) | 27 | 1.15 | 0.246 | 1.000 | |
| AT3G10370.1 | FAD glycerol-3-phosphate dehydrogenase (SDP6) | 28 | 1.51 | 0.223 | 1.000 | |
| AT3G10370.1 | FAD glycerol-3-phosphate dehydrogenase (SDP6) | 29 | 2.20 | 0.212 | 0.917 | |
| AT3G10370.1 | FAD glycerol-3-phosphate dehydrogenase (SDP6) | 30 | 2.11 | 0.211 | 0.945 | |
| AT3G10370.1 | FAD glycerol-3-phosphate dehydrogenase (SDP6) | 31 | 1.79 | 0.212 | 0.997 | |
| AT3G10370.1 | FAD glycerol-3-phosphate dehydrogenase (SDP6) | 32 | 1.57 | 0.212 | 1.000 | |
| AT3G10370.1 | FAD glycerol-3-phosphate dehydrogenase (SDP6) | 33 | 1.37 | 0.212 | 1.000 | |
| AT3G10370.1 | FAD glycerol-3-phosphate dehydrogenase (SDP6) | 34 | 1.37 | 0.212 | 1.000 | |
| AT3G10370.1 | FAD glycerol-3-phosphate dehydrogenase (SDP6) | 35 | 1.55 | 0.211 | 1.000 | |
| AT3G10370.1 | FAD glycerol-3-phosphate dehydrogenase (SDP6) | 36 | 1.80 | 0.210 | 0.996 | |
| AT3G10370.1 | FAD glycerol-3-phosphate dehydrogenase (SDP6) | 37 | 1.91 | 0.211 | 0.987 | |
| AT3G10370.1 | FAD glycerol-3-phosphate dehydrogenase (SDP6) | 38 | 1.53 | 0.212 | 1.000 | |
| AT3G10370.1 | FAD glycerol-3-phosphate dehydrogenase (SDP6) | 39 | 0.94 | 0.212 | 1.000 | |
| AT3G10370.1 | FAD glycerol-3-phosphate dehydrogenase (SDP6) | 40 | 0.70 | 0.211 | 1.000 | |
| AT3G10370.1 | FAD glycerol-3-phosphate dehydrogenase (SDP6) | 41 | 0.72 | 0.211 | 1.000 | |
| AT3G10370.1 | FAD glycerol-3-phosphate dehydrogenase (SDP6) | 42 | 0.77 | 0.214 | 1.000 | |
| AT3G10370.1 | FAD glycerol-3-phosphate dehydrogenase (SDP6) | 43 | 0.70 | 0.215 | 1.000 | |
| AT3G10370.1 | FAD glycerol-3-phosphate dehydrogenase (SDP6) | 44 | 0.56 | 0.215 | 0.997 | |
| AT3G10370.1 | FAD glycerol-3-phosphate dehydrogenase (SDP6) | 45 | 0.52 | 0.215 | 0.986 | |
| AT3G10370.1 | FAD glycerol-3-phosphate dehydrogenase (SDP6) | 46 | 0.57 | 0.244 | 1.000 | |
| AT3G10370.1 | FAD glycerol-3-phosphate dehydrogenase (SDP6) | 47 | 0.68 | 0.422 | 1.000 | |
| AT3G10690.1 | DNA gyrase A (GYRA) | 2 | 1.14 | 0.460 | 1.000 | |
| AT3G10690.1 | DNA gyrase A (GYRA) | 3 | 1.13 | 0.408 | 1.000 | |

| | | | | | | |
|---|---|---|---|---|---|---|
| AT3G10690.1 | DNA gyrase A (GYRA) | 4 | 1.12 | 0.661 | 1.000 | |
| AT3G10690.1 | DNA gyrase A (GYRA) | 22 | 0.14 | 0.353 | 0.189 | |
| AT3G10690.1 | DNA gyrase A (GYRA) | 23 | 0.16 | 0.280 | 0.057 | . |
| AT3G10690.1 | DNA gyrase A (GYRA) | 24 | 0.18 | 0.246 | 0.042 | * |
| AT3G10690.1 | DNA gyrase A (GYRA) | 25 | 0.30 | 0.224 | 0.203 | |
| AT3G10690.1 | DNA gyrase A (GYRA) | 26 | 0.71 | 0.211 | 0.998 | |
| AT3G10690.1 | DNA gyrase A (GYRA) | 27 | 1.81 | 0.210 | 0.907 | |
| AT3G10690.1 | DNA gyrase A (GYRA) | 28 | 3.40 | 0.213 | 0.149 | |
| AT3G10690.1 | DNA gyrase A (GYRA) | 29 | 3.06 | 0.216 | 0.255 | |
| AT3G10690.1 | DNA gyrase A (GYRA) | 30 | 1.23 | 0.221 | 1.000 | |
| AT3G10690.1 | DNA gyrase A (GYRA) | 31 | 0.37 | 0.242 | 0.551 | |
| AT3G10690.1 | DNA gyrase A (GYRA) | 32 | 0.11 | 0.320 | 0.049 | * |
| AT3G10690.1 | DNA gyrase A (GYRA) | 33 | 0.04 | 0.494 | 0.079 | . |
| AT3G10690.1 | DNA gyrase A (GYRA) | 46 | 0.02 | 0.405 | 0.000 | *** |
| AT3G10690.1 | DNA gyrase A (GYRA) | 47 | 0.02 | 0.424 | 0.003 | ** |
| AT3G10860.1 | Ubiquinol:cytochrome c oxidoreductase subunit (QCR8) | 2 | 0.97 | 0.675 | 1.000 | |
| AT3G10860.1 | Ubiquinol:cytochrome c oxidoreductase subunit (QCR8) | 3 | 0.86 | 0.521 | 1.000 | |
| AT3G10860.1 | Ubiquinol:cytochrome c oxidoreductase subunit (QCR8) | 4 | 0.71 | 0.502 | 1.000 | |
| AT3G10860.1 | Ubiquinol:cytochrome c oxidoreductase subunit (QCR8) | 5 | 0.64 | 0.501 | 1.000 | |
| AT3G10860.1 | Ubiquinol:cytochrome c oxidoreductase subunit (QCR8) | 6 | 0.63 | 0.501 | 1.000 | |
| AT3G10860.1 | Ubiquinol:cytochrome c oxidoreductase subunit (QCR8) | 7 | 0.65 | 0.502 | 1.000 | |
| AT3G10860.1 | Ubiquinol:cytochrome c oxidoreductase subunit (QCR8) | 8 | 0.72 | 0.496 | 1.000 | |
| AT3G10860.1 | Ubiquinol:cytochrome c oxidoreductase subunit (QCR8) | 9 | 0.77 | 0.491 | 1.000 | |
| AT3G10860.1 | Ubiquinol:cytochrome c oxidoreductase subunit (QCR8) | 10 | 0.78 | 0.492 | 1.000 | |
| AT3G10860.1 | Ubiquinol:cytochrome c oxidoreductase subunit (QCR8) | 11 | 0.79 | 0.494 | 1.000 | |
| AT3G10860.1 | Ubiquinol:cytochrome c oxidoreductase subunit (QCR8) | 12 | 0.78 | 0.494 | 1.000 | |
| AT3G10860.1 | Ubiquinol:cytochrome c oxidoreductase subunit (QCR8) | 13 | 0.66 | 0.492 | 1.000 | |
| AT3G10860.1 | Ubiquinol:cytochrome c oxidoreductase subunit (QCR8) | 14 | 0.50 | 0.490 | 1.000 | |
| AT3G10860.1 | Ubiquinol:cytochrome c oxidoreductase subunit (QCR8) | 15 | 0.43 | 0.490 | 0.999 | |
| AT3G10860.1 | Ubiquinol:cytochrome c oxidoreductase subunit (QCR8) | 16 | 0.39 | 0.490 | 0.998 | |
| AT3G10860.1 | Ubiquinol:cytochrome c oxidoreductase subunit (QCR8) | 17 | 0.31 | 0.490 | 0.978 | |
| AT3G10860.1 | Ubiquinol:cytochrome c oxidoreductase subunit (QCR8) | 18 | 0.23 | 0.492 | 0.881 | |
| AT3G10860.1 | Ubiquinol:cytochrome c oxidoreductase subunit (QCR8) | 19 | 0.20 | 0.498 | 0.829 | |
| AT3G10860.1 | Ubiquinol:cytochrome c oxidoreductase subunit (QCR8) | 20 | 0.25 | 0.506 | 0.943 | |
| AT3G10860.1 | Ubiquinol:cytochrome c oxidoreductase subunit (QCR8) | 21 | 0.42 | 0.507 | 1.000 | |
| AT3G10860.1 | Ubiquinol:cytochrome c oxidoreductase subunit (QCR8) | 22 | 0.43 | 0.504 | 1.000 | |
| AT3G10860.1 | Ubiquinol:cytochrome c oxidoreductase subunit (QCR8) | 23 | 0.20 | 0.501 | 0.835 | |
| AT3G10860.1 | Ubiquinol:cytochrome c oxidoreductase subunit (QCR8) | 24 | 0.10 | 0.498 | 0.377 | |

| | | | | | |
|---|---|---|---|---|---|
| AT3G10860.1 | Ubiquinol:cytochrome c oxidoreductase subunit (QCR8) | 25 | 0.09 | 0.495 | 0.308 |
| AT3G10860.1 | Ubiquinol:cytochrome c oxidoreductase subunit (QCR8) | 26 | 0.17 | 0.494 | 0.701 |
| AT3G10860.1 | Ubiquinol:cytochrome c oxidoreductase subunit (QCR8) | 27 | 0.64 | 0.493 | 1.000 |
| AT3G10860.1 | Ubiquinol:cytochrome c oxidoreductase subunit (QCR8) | 28 | 1.30 | 0.493 | 1.000 |
| AT3G10860.1 | Ubiquinol:cytochrome c oxidoreductase subunit (QCR8) | 29 | 0.64 | 0.494 | 1.000 |
| AT3G10860.1 | Ubiquinol:cytochrome c oxidoreductase subunit (QCR8) | 30 | 0.26 | 0.496 | 0.945 |
| AT3G10860.1 | Ubiquinol:cytochrome c oxidoreductase subunit (QCR8) | 31 | 0.26 | 0.497 | 0.944 |
| AT3G10860.1 | Ubiquinol:cytochrome c oxidoreductase subunit (QCR8) | 32 | 0.44 | 0.502 | 1.000 |
| AT3G10860.1 | Ubiquinol:cytochrome c oxidoreductase subunit (QCR8) | 33 | 0.65 | 0.516 | 1.000 |
| AT3G10860.1 | Ubiquinol:cytochrome c oxidoreductase subunit (QCR8) | 34 | 0.76 | 0.576 | 1.000 |
| AT3G10860.1 | Ubiquinol:cytochrome c oxidoreductase subunit (QCR8) | 35 | 0.89 | 0.769 | 1.000 |
| AT3G10860.1 | Ubiquinol:cytochrome c oxidoreductase subunit (QCR8) | 46 | 3496.92 | 4.712 | 0.999 |
| AT3G10860.1 | Ubiquinol:cytochrome c oxidoreductase subunit (QCR8) | 47 | 9908.58 | 5.278 | 0.999 |
| AT3G10920.1 | Manganese superoxide dismutase 1 (MSD1) | 2 | 0.78 | 0.404 | 1.000 |
| AT3G10920.1 | Manganese superoxide dismutase 1 (MSD1) | 3 | 0.57 | 0.245 | 1.000 |
| AT3G10920.1 | Manganese superoxide dismutase 1 (MSD1) | 4 | 0.43 | 0.231 | 0.951 |
| AT3G10920.1 | Manganese superoxide dismutase 1 (MSD1) | 5 | 0.37 | 0.231 | 0.816 |
| AT3G10920.1 | Manganese superoxide dismutase 1 (MSD1) | 6 | 0.45 | 0.231 | 0.973 |
| AT3G10920.1 | Manganese superoxide dismutase 1 (MSD1) | 7 | 0.60 | 0.231 | 1.000 |
| AT3G10920.1 | Manganese superoxide dismutase 1 (MSD1) | 8 | 0.79 | 0.226 | 1.000 |
| AT3G10920.1 | Manganese superoxide dismutase 1 (MSD1) | 9 | 0.95 | 0.223 | 1.000 |
| AT3G10920.1 | Manganese superoxide dismutase 1 (MSD1) | 10 | 1.16 | 0.223 | 1.000 |
| AT3G10920.1 | Manganese superoxide dismutase 1 (MSD1) | 11 | 1.35 | 0.224 | 1.000 |
| AT3G10920.1 | Manganese superoxide dismutase 1 (MSD1) | 12 | 1.50 | 0.223 | 1.000 |
| AT3G10920.1 | Manganese superoxide dismutase 1 (MSD1) | 13 | 1.65 | 0.221 | 1.000 |
| AT3G10920.1 | Manganese superoxide dismutase 1 (MSD1) | 14 | 1.79 | 0.218 | 0.999 |
| AT3G10920.1 | Manganese superoxide dismutase 1 (MSD1) | 15 | 1.70 | 0.219 | 1.000 |
| AT3G10920.1 | Manganese superoxide dismutase 1 (MSD1) | 16 | 1.45 | 0.218 | 1.000 |
| AT3G10920.1 | Manganese superoxide dismutase 1 (MSD1) | 17 | 1.65 | 0.223 | 1.000 |
| AT3G10920.1 | Manganese superoxide dismutase 1 (MSD1) | 18 | 2.50 | 0.231 | 0.906 |
| AT3G10920.1 | Manganese superoxide dismutase 1 (MSD1) | 19 | 3.66 | 0.247 | 0.490 |

| | | | | | |
|---|---|---|---|---|---|
| AT3G10920.1 | Manganese superoxide dismutase 1 (MSD1) | 20 | 4.94 | 0.298 | 0.454 |
| AT3G10920.1 | Manganese superoxide dismutase 1 (MSD1) | 21 | 5.18 | 0.401 | 0.875 |
| AT3G10920.1 | Manganese superoxide dismutase 1 (MSD1) | 27 | 0.58 | 0.346 | 1.000 |
| AT3G10920.1 | Manganese superoxide dismutase 1 (MSD1) | 28 | 0.58 | 0.262 | 1.000 |
| AT3G10920.1 | Manganese superoxide dismutase 1 (MSD1) | 29 | 0.65 | 0.239 | 1.000 |
| AT3G10920.1 | Manganese superoxide dismutase 1 (MSD1) | 30 | 0.65 | 0.226 | 1.000 |
| AT3G10920.1 | Manganese superoxide dismutase 1 (MSD1) | 31 | 0.67 | 0.223 | 1.000 |
| AT3G10920.1 | Manganese superoxide dismutase 1 (MSD1) | 32 | 0.93 | 0.223 | 1.000 |
| AT3G10920.1 | Manganese superoxide dismutase 1 (MSD1) | 33 | 1.16 | 0.223 | 1.000 |
| AT3G10920.1 | Manganese superoxide dismutase 1 (MSD1) | 34 | 1.05 | 0.223 | 1.000 |
| AT3G10920.1 | Manganese superoxide dismutase 1 (MSD1) | 35 | 1.00 | 0.223 | 1.000 |
| AT3G10920.1 | Manganese superoxide dismutase 1 (MSD1) | 36 | 1.08 | 0.222 | 1.000 |
| AT3G10920.1 | Manganese superoxide dismutase 1 (MSD1) | 37 | 1.50 | 0.222 | 1.000 |
| AT3G10920.1 | Manganese superoxide dismutase 1 (MSD1) | 38 | 2.60 | 0.223 | 0.825 |
| AT3G10920.1 | Manganese superoxide dismutase 1 (MSD1) | 39 | 3.38 | 0.223 | 0.416 |
| AT3G10920.1 | Manganese superoxide dismutase 1 (MSD1) | 40 | 2.68 | 0.224 | 0.792 |
| AT3G10920.1 | Manganese superoxide dismutase 1 (MSD1) | 41 | 1.43 | 0.225 | 1.000 |
| AT3G10920.1 | Manganese superoxide dismutase 1 (MSD1) | 42 | 0.80 | 0.228 | 1.000 |
| AT3G10920.1 | Manganese superoxide dismutase 1 (MSD1) | 43 | 0.64 | 0.230 | 1.000 |
| AT3G10920.1 | Manganese superoxide dismutase 1 (MSD1) | 44 | 0.68 | 0.230 | 1.000 |
| AT3G10920.1 | Manganese superoxide dismutase 1 (MSD1) | 45 | 0.73 | 0.229 | 1.000 |
| AT3G10920.1 | Manganese superoxide dismutase 1 (MSD1) | 46 | 0.62 | 0.263 | 1.000 |
| AT3G10920.1 | Manganese superoxide dismutase 1 (MSD1) | 47 | 0.43 | 0.485 | 1.000 |
| AT3G11070.1 | Outer membrane OMP85 family protein | 3 | 0.55 | 0.178 | 0.983 |
| AT3G11070.1 | Outer membrane OMP85 family protein | 4 | 0.67 | 0.162 | 1.000 |
| AT3G11070.1 | Outer membrane OMP85 family protein | 5 | 1.09 | 0.163 | 1.000 |
| AT3G11070.1 | Outer membrane OMP85 family protein | 6 | 1.75 | 0.163 | 0.979 |
| AT3G11070.1 | Outer membrane OMP85 family protein | 7 | 2.12 | 0.162 | 0.731 |
| AT3G11070.1 | Outer membrane OMP85 family protein | 8 | 2.05 | 0.160 | 0.777 |
| AT3G11070.1 | Outer membrane OMP85 family protein | 9 | 1.81 | 0.157 | 0.946 |
| AT3G11070.1 | Outer membrane OMP85 family protein | 10 | 1.60 | 0.157 | 0.997 |

| | | | | | |
|---|---|---|---|---|---|
| AT3G11070.1 | Outer membrane OMP85 family protein | 11 | 1.73 | 0.157 | 0.978 |
| AT3G11070.1 | Outer membrane OMP85 family protein | 12 | 2.21 | 0.158 | 0.591 |
| AT3G11070.1 | Outer membrane OMP85 family protein | 13 | 2.47 | 0.156 | 0.319 |
| AT3G11070.1 | Outer membrane OMP85 family protein | 14 | 2.18 | 0.154 | 0.575 |
| AT3G11070.1 | Outer membrane OMP85 family protein | 15 | 1.64 | 0.153 | 0.991 |
| AT3G11070.1 | Outer membrane OMP85 family protein | 16 | 1.18 | 0.152 | 1.000 |
| AT3G11070.1 | Outer membrane OMP85 family protein | 17 | 0.99 | 0.152 | 1.000 |
| AT3G11070.1 | Outer membrane OMP85 family protein | 18 | 0.96 | 0.154 | 1.000 |
| AT3G11070.1 | Outer membrane OMP85 family protein | 19 | 0.99 | 0.158 | 1.000 |
| AT3G11070.1 | Outer membrane OMP85 family protein | 20 | 1.04 | 0.162 | 1.000 |
| AT3G11070.1 | Outer membrane OMP85 family protein | 21 | 1.12 | 0.165 | 1.000 |
| AT3G11070.1 | Outer membrane OMP85 family protein | 22 | 1.19 | 0.164 | 1.000 |
| AT3G11070.1 | Outer membrane OMP85 family protein | 23 | 1.19 | 0.163 | 1.000 |
| AT3G11070.1 | Outer membrane OMP85 family protein | 24 | 1.06 | 0.160 | 1.000 |
| AT3G11070.1 | Outer membrane OMP85 family protein | 25 | 0.90 | 0.157 | 1.000 |
| AT3G11070.1 | Outer membrane OMP85 family protein | 26 | 0.92 | 0.156 | 1.000 |
| AT3G11070.1 | Outer membrane OMP85 family protein | 27 | 1.08 | 0.156 | 1.000 |
| AT3G11070.1 | Outer membrane OMP85 family protein | 28 | 1.16 | 0.156 | 1.000 |
| AT3G11070.1 | Outer membrane OMP85 family protein | 29 | 1.05 | 0.156 | 1.000 |
| AT3G11070.1 | Outer membrane OMP85 family protein | 30 | 0.93 | 0.156 | 1.000 |
| AT3G11070.1 | Outer membrane OMP85 family protein | 31 | 0.96 | 0.157 | 1.000 |
| AT3G11070.1 | Outer membrane OMP85 family protein | 32 | 1.11 | 0.157 | 1.000 |
| AT3G11070.1 | Outer membrane OMP85 family protein | 33 | 1.23 | 0.157 | 1.000 |
| AT3G11070.1 | Outer membrane OMP85 family protein | 34 | 1.30 | 0.158 | 1.000 |
| AT3G11070.1 | Outer membrane OMP85 family protein | 35 | 1.36 | 0.157 | 1.000 |
| AT3G11070.1 | Outer membrane OMP85 family protein | 36 | 1.38 | 0.157 | 1.000 |
| AT3G11070.1 | Outer membrane OMP85 family protein | 37 | 1.33 | 0.156 | 1.000 |
| AT3G11070.1 | Outer membrane OMP85 family protein | 38 | 1.16 | 0.157 | 1.000 |
| AT3G11070.1 | Outer membrane OMP85 family protein | 39 | 0.89 | 0.157 | 1.000 |
| AT3G11070.1 | Outer membrane OMP85 family protein | 40 | 0.61 | 0.158 | 0.993 |
| AT3G11070.1 | Outer membrane OMP85 family protein | 41 | 0.38 | 0.158 | 0.241 |

| | | | | | | |
|---|---|---|---|---|---|---|
| AT3G11070.1 | Outer membrane OMP85 family protein | 42 | 0.27 | 0.160 | 0.015 | * |
| AT3G11070.1 | Outer membrane OMP85 family protein | 43 | 0.27 | 0.162 | 0.018 | * |
| AT3G11070.1 | Outer membrane OMP85 family protein | 44 | 0.41 | 0.162 | 0.434 | |
| AT3G11070.1 | Outer membrane OMP85 family protein | 45 | 0.72 | 0.162 | 1.000 | |
| AT3G11070.1 | Outer membrane OMP85 family protein | 46 | 1.19 | 0.191 | 1.000 | |
| AT3G11070.1 | Outer membrane OMP85 family protein | 47 | 1.99 | 0.334 | 1.000 | |
| AT3G11070.1 | Outer membrane OMP85 family protein | 48 | 3.32 | 0.552 | 1.000 | |
| AT3G11630.1 | 2-Cys peroxiredoxin A (2-Cys PrxA) | 2 | 1.26 | 0.601 | 1.000 | |
| AT3G11630.1 | 2-Cys peroxiredoxin A (2-Cys PrxA) | 3 | 1.29 | 0.579 | 0.999 | |
| AT3G11630.1 | 2-Cys peroxiredoxin A (2-Cys PrxA) | 30 | 1.16 | 0.235 | 0.996 | |
| AT3G11630.1 | 2-Cys peroxiredoxin A (2-Cys PrxA) | 31 | 1.10 | 0.228 | 0.999 | |
| AT3G11630.1 | 2-Cys peroxiredoxin A (2-Cys PrxA) | 32 | 1.03 | 0.226 | 1.000 | |
| AT3G11630.1 | 2-Cys peroxiredoxin A (2-Cys PrxA) | 33 | 0.96 | 0.228 | 1.000 | |
| AT3G11630.1 | 2-Cys peroxiredoxin A (2-Cys PrxA) | 34 | 0.88 | 0.234 | 0.998 | |
| AT3G11630.1 | 2-Cys peroxiredoxin A (2-Cys PrxA) | 35 | 0.80 | 0.243 | 0.983 | |
| AT3G11630.1 | 2-Cys peroxiredoxin A (2-Cys PrxA) | 46 | 0.16 | 0.404 | 0.187 | |
| AT3G11630.1 | 2-Cys peroxiredoxin A (2-Cys PrxA) | 47 | 0.14 | 0.428 | 0.168 | |
| AT3G12260.1 | NADH:ubiquinone oxidoreductase subunit (B14) | 2 | 0.68 | 0.532 | 1.000 | |
| AT3G12260.1 | NADH:ubiquinone oxidoreductase subunit (B14) | 3 | 0.68 | 0.472 | 1.000 | |
| AT3G12260.1 | NADH:ubiquinone oxidoreductase subunit (B14) | 13 | 1.28 | 0.320 | 1.000 | |
| AT3G12260.1 | NADH:ubiquinone oxidoreductase subunit (B14) | 14 | 1.21 | 0.290 | 1.000 | |
| AT3G12260.1 | NADH:ubiquinone oxidoreductase subunit (B14) | 15 | 0.82 | 0.262 | 1.000 | |
| AT3G12260.1 | NADH:ubiquinone oxidoreductase subunit (B14) | 16 | 0.46 | 0.243 | 0.967 | |
| AT3G12260.1 | NADH:ubiquinone oxidoreductase subunit (B14) | 17 | 0.32 | 0.233 | 0.496 | |
| AT3G12260.1 | NADH:ubiquinone oxidoreductase subunit (B14) | 18 | 0.38 | 0.235 | 0.783 | |
| AT3G12260.1 | NADH:ubiquinone oxidoreductase subunit (B14) | 19 | 0.61 | 0.241 | 1.000 | |
| AT3G12260.1 | NADH:ubiquinone oxidoreductase subunit (B14) | 20 | 0.87 | 0.246 | 1.000 | |
| AT3G12260.1 | NADH:ubiquinone oxidoreductase subunit (B14) | 21 | 1.09 | 0.248 | 1.000 | |
| AT3G12260.1 | NADH:ubiquinone oxidoreductase subunit (B14) | 22 | 1.27 | 0.248 | 1.000 | |
| AT3G12260.1 | NADH:ubiquinone oxidoreductase subunit (B14) | 23 | 1.33 | 0.245 | 1.000 | |
| AT3G12260.1 | NADH:ubiquinone oxidoreductase subunit (B14) | 24 | 1.17 | 0.241 | 1.000 | |

| | | | | | | |
|---|---|---|---|---|---|---|
| AT3G12260.1 | NADH:ubiquinone oxidoreductase subunit (B14) | 25 | 1.00 | 0.238 | 1.000 | |
| AT3G12260.1 | NADH:ubiquinone oxidoreductase subunit (B14) | 26 | 0.98 | 0.237 | 1.000 | |
| AT3G12260.1 | NADH:ubiquinone oxidoreductase subunit (B14) | 27 | 1.11 | 0.238 | 1.000 | |
| AT3G12260.1 | NADH:ubiquinone oxidoreductase subunit (B14) | 28 | 1.27 | 0.238 | 1.000 | |
| AT3G12260.1 | NADH:ubiquinone oxidoreductase subunit (B14) | 29 | 1.47 | 0.237 | 1.000 | |
| AT3G12260.1 | NADH:ubiquinone oxidoreductase subunit (B14) | 30 | 1.62 | 0.237 | 1.000 | |
| AT3G12260.1 | NADH:ubiquinone oxidoreductase subunit (B14) | 31 | 1.81 | 0.238 | 0.998 | |
| AT3G12260.1 | NADH:ubiquinone oxidoreductase subunit (B14) | 32 | 2.05 | 0.238 | 0.981 | |
| AT3G12260.1 | NADH:ubiquinone oxidoreductase subunit (B14) | 33 | 2.22 | 0.238 | 0.951 | |
| AT3G12260.1 | NADH:ubiquinone oxidoreductase subunit (B14) | 34 | 2.23 | 0.239 | 0.949 | |
| AT3G12260.1 | NADH:ubiquinone oxidoreductase subunit (B14) | 35 | 2.16 | 0.238 | 0.963 | |
| AT3G12260.1 | NADH:ubiquinone oxidoreductase subunit (B14) | 36 | 2.10 | 0.238 | 0.974 | |
| AT3G12260.1 | NADH:ubiquinone oxidoreductase subunit (B14) | 37 | 2.27 | 0.238 | 0.935 | |
| AT3G12260.1 | NADH:ubiquinone oxidoreductase subunit (B14) | 38 | 3.18 | 0.238 | 0.523 | |
| AT3G12260.1 | NADH:ubiquinone oxidoreductase subunit (B14) | 39 | 4.65 | 0.238 | 0.122 | |
| AT3G12260.1 | NADH:ubiquinone oxidoreductase subunit (B14) | 40 | 5.06 | 0.239 | 0.082 | . |
| AT3G12260.1 | NADH:ubiquinone oxidoreductase subunit (B14) | 41 | 3.71 | 0.240 | 0.330 | |
| AT3G12260.1 | NADH:ubiquinone oxidoreductase subunit (B14) | 42 | 2.53 | 0.243 | 0.861 | |
| AT3G12260.1 | NADH:ubiquinone oxidoreductase subunit (B14) | 43 | 1.91 | 0.245 | 0.996 | |
| AT3G12260.1 | NADH:ubiquinone oxidoreductase subunit (B14) | 44 | 1.47 | 0.245 | 1.000 | |
| AT3G12260.1 | NADH:ubiquinone oxidoreductase subunit (B14) | 45 | 1.09 | 0.245 | 1.000 | |
| AT3G12260.1 | NADH:ubiquinone oxidoreductase subunit (B14) | 46 | 0.69 | 0.285 | 1.000 | |
| AT3G12260.1 | NADH:ubiquinone oxidoreductase subunit (B14) | 47 | 0.37 | 0.472 | 1.000 | |
| AT3G12370.1 | 50S ribosomal protein L10 | 2 | 1.08 | 0.421 | 1.000 | |
| AT3G12370.1 | 50S ribosomal protein L10 | 3 | 1.01 | 0.428 | 1.000 | |
| AT3G12370.1 | 50S ribosomal protein L10 | 29 | 0.60 | 0.286 | 0.989 | |
| AT3G12370.1 | 50S ribosomal protein L10 | 30 | 0.64 | 0.231 | 0.982 | |
| AT3G12370.1 | 50S ribosomal protein L10 | 31 | 0.69 | 0.200 | 0.985 | |
| AT3G12370.1 | 50S ribosomal protein L10 | 32 | 0.74 | 0.182 | 0.993 | |
| AT3G12370.1 | 50S ribosomal protein L10 | 33 | 0.79 | 0.170 | 0.998 | |
| AT3G12370.1 | 50S ribosomal protein L10 | 34 | 0.87 | 0.161 | 1.000 | |
| AT3G12370.1 | 50S ribosomal protein L10 | 35 | 1.02 | 0.156 | 1.000 | |
| AT3G12370.1 | 50S ribosomal protein L10 | 36 | 1.28 | 0.153 | 0.995 | |
| AT3G12370.1 | 50S ribosomal protein L10 | 37 | 1.65 | 0.152 | 0.734 | |
| AT3G12370.1 | 50S ribosomal protein L10 | 38 | 2.03 | 0.153 | 0.330 | |
| AT3G12370.1 | 50S ribosomal protein L10 | 39 | 2.16 | 0.159 | 0.279 | |
| AT3G12370.1 | 50S ribosomal protein L10 | 40 | 1.82 | 0.178 | 0.710 | |
| AT3G12370.1 | 50S ribosomal protein L10 | 41 | 1.24 | 0.217 | 1.000 | |
| AT3G12370.1 | 50S ribosomal protein L10 | 42 | 0.73 | 0.270 | 1.000 | |

| | | | | | | |
|---|---|---|---|---|---|---|
| AT3G12370.1 | 50S ribosomal protein L10 | 46 | 0.04 | 0.494 | 0.056 | . |
| AT3G12370.1 | 50S ribosomal protein L10 | 47 | 0.02 | 0.589 | 0.037 | * |
| AT3G12650.1 | ? | 2 | 0.97 | 0.471 | 1.000 | |
| AT3G12650.1 | ? | 3 | 1.13 | 0.379 | 1.000 | |
| AT3G12650.1 | ? | 6 | 1.65 | 0.311 | 1.000 | |
| AT3G12650.1 | ? | 7 | 1.79 | 0.284 | 1.000 | |
| AT3G12650.1 | ? | 8 | 1.85 | 0.265 | 0.999 | |
| AT3G12650.1 | ? | 9 | 1.80 | 0.244 | 0.998 | |
| AT3G12650.1 | ? | 10 | 1.69 | 0.227 | 0.999 | |
| AT3G12650.1 | ? | 11 | 1.64 | 0.211 | 0.999 | |
| AT3G12650.1 | ? | 12 | 1.69 | 0.199 | 0.995 | |
| AT3G12650.1 | ? | 13 | 1.82 | 0.190 | 0.967 | |
| AT3G12650.1 | ? | 14 | 2.08 | 0.184 | 0.816 | |
| AT3G12650.1 | ? | 15 | 2.41 | 0.180 | 0.507 | |
| AT3G12650.1 | ? | 16 | 2.77 | 0.178 | 0.259 | |
| AT3G12650.1 | ? | 17 | 3.01 | 0.181 | 0.177 | |
| AT3G12650.1 | ? | 18 | 3.04 | 0.185 | 0.193 | |
| AT3G12650.1 | ? | 19 | 2.79 | 0.187 | 0.320 | |
| AT3G12650.1 | ? | 20 | 2.14 | 0.187 | 0.790 | |
| AT3G12650.1 | ? | 21 | 1.38 | 0.186 | 1.000 | |
| AT3G12650.1 | ? | 22 | 0.89 | 0.184 | 1.000 | |
| AT3G12650.1 | ? | 23 | 0.65 | 0.182 | 0.998 | |
| AT3G12650.1 | ? | 24 | 0.57 | 0.178 | 0.963 | |
| AT3G12650.1 | ? | 25 | 0.56 | 0.176 | 0.955 | |
| AT3G12650.1 | ? | 26 | 0.64 | 0.175 | 0.996 | |
| AT3G12650.1 | ? | 27 | 0.85 | 0.175 | 1.000 | |
| AT3G12650.1 | ? | 28 | 1.16 | 0.175 | 1.000 | |
| AT3G12650.1 | ? | 29 | 1.39 | 0.175 | 1.000 | |
| AT3G12650.1 | ? | 30 | 1.41 | 0.175 | 1.000 | |
| AT3G12650.1 | ? | 31 | 1.33 | 0.176 | 1.000 | |
| AT3G12650.1 | ? | 32 | 1.26 | 0.176 | 1.000 | |
| AT3G12650.1 | ? | 33 | 1.18 | 0.176 | 1.000 | |
| AT3G12650.1 | ? | 34 | 1.11 | 0.177 | 1.000 | |
| AT3G12650.1 | ? | 35 | 1.05 | 0.177 | 1.000 | |
| AT3G12650.1 | ? | 36 | 0.99 | 0.176 | 1.000 | |
| AT3G12650.1 | ? | 37 | 0.96 | 0.176 | 1.000 | |
| AT3G12650.1 | ? | 38 | 0.95 | 0.177 | 1.000 | |
| AT3G12650.1 | ? | 39 | 0.87 | 0.177 | 1.000 | |
| AT3G12650.1 | ? | 40 | 0.65 | 0.178 | 0.998 | |
| AT3G12650.1 | ? | 41 | 0.40 | 0.184 | 0.469 | |
| AT3G12650.1 | ? | 42 | 0.23 | 0.205 | 0.045 | * |
| AT3G12650.1 | ? | 43 | 0.14 | 0.252 | 0.025 | * |
| AT3G12650.1 | ? | 44 | 0.11 | 0.320 | 0.062 | . |
| AT3G12650.1 | ? | 46 | 0.08 | 0.532 | 0.590 | |
| AT3G12650.1 | ? | 47 | 0.08 | 0.690 | 0.890 | |
| AT3G12780.1 | Phosphoglycerate kinase 1 (PGK1) | 2 | 0.84 | 0.376 | 1.000 | |
| AT3G12780.1 | Phosphoglycerate kinase 1 (PGK1) | 3 | 0.90 | 0.371 | 1.000 | |
| AT3G12780.1 | Phosphoglycerate kinase 1 (PGK1) | 39 | 2.99 | 0.296 | 0.571 | |
| AT3G12780.1 | Phosphoglycerate kinase 1 (PGK1) | 40 | 2.89 | 0.211 | 0.221 | |
| AT3G12780.1 | Phosphoglycerate kinase 1 (PGK1) | 41 | 2.90 | 0.182 | 0.101 | |
| AT3G12780.1 | Phosphoglycerate kinase 1 (PGK1) | 42 | 3.35 | 0.175 | 0.031 | * |
| AT3G12780.1 | Phosphoglycerate kinase 1 (PGK1) | 43 | 4.69 | 0.174 | 0.002 | ** |
| AT3G12780.1 | Phosphoglycerate kinase 1 (PGK1) | 44 | 4.08 | 0.173 | 0.007 | ** |
| AT3G12780.1 | Phosphoglycerate kinase 1 (PGK1) | 45 | 1.76 | 0.177 | 0.728 | |

| | | | | | | |
|---|---|---|---|---|---|---|
| AT3G12780.1 | Phosphoglycerate kinase 1 (PGK1) | 46 | 0.43 | 0.201 | 0.417 | |
| AT3G12780.1 | Phosphoglycerate kinase 1 (PGK1) | 47 | 0.07 | 0.328 | 0.008 | ** |
| AT3G13110.1 | Serine O-acetyltransferase (SERAT2.2) | 2 | 0.94 | 0.361 | 1.000 | |
| AT3G13110.1 | Serine O-acetyltransferase (SERAT2.2) | 3 | 0.97 | 0.309 | 1.000 | |
| AT3G13110.1 | Serine O-acetyltransferase (SERAT2.2) | 10 | 1.22 | 0.200 | 1.000 | |
| AT3G13110.1 | Serine O-acetyltransferase (SERAT2.2) | 11 | 1.27 | 0.171 | 1.000 | |
| AT3G13110.1 | Serine O-acetyltransferase (SERAT2.2) | 12 | 1.33 | 0.151 | 1.000 | |
| AT3G13110.1 | Serine O-acetyltransferase (SERAT2.2) | 13 | 1.41 | 0.139 | 0.996 | |
| AT3G13110.1 | Serine O-acetyltransferase (SERAT2.2) | 14 | 1.52 | 0.132 | 0.957 | |
| AT3G13110.1 | Serine O-acetyltransferase (SERAT2.2) | 15 | 1.65 | 0.129 | 0.809 | |
| AT3G13110.1 | Serine O-acetyltransferase (SERAT2.2) | 16 | 1.77 | 0.128 | 0.633 | |
| AT3G13110.1 | Serine O-acetyltransferase (SERAT2.2) | 17 | 1.80 | 0.128 | 0.578 | |
| AT3G13110.1 | Serine O-acetyltransferase (SERAT2.2) | 18 | 1.75 | 0.129 | 0.676 | |
| AT3G13110.1 | Serine O-acetyltransferase (SERAT2.2) | 19 | 1.67 | 0.132 | 0.813 | |
| AT3G13110.1 | Serine O-acetyltransferase (SERAT2.2) | 20 | 1.66 | 0.134 | 0.847 | |
| AT3G13110.1 | Serine O-acetyltransferase (SERAT2.2) | 21 | 1.68 | 0.135 | 0.829 | |
| AT3G13110.1 | Serine O-acetyltransferase (SERAT2.2) | 22 | 1.68 | 0.136 | 0.826 | |
| AT3G13110.1 | Serine O-acetyltransferase (SERAT2.2) | 23 | 1.64 | 0.134 | 0.869 | |
| AT3G13110.1 | Serine O-acetyltransferase (SERAT2.2) | 24 | 1.54 | 0.133 | 0.948 | |
| AT3G13110.1 | Serine O-acetyltransferase (SERAT2.2) | 25 | 1.42 | 0.131 | 0.991 | |
| AT3G13110.1 | Serine O-acetyltransferase (SERAT2.2) | 26 | 1.35 | 0.130 | 0.998 | |
| AT3G13110.1 | Serine O-acetyltransferase (SERAT2.2) | 27 | 1.28 | 0.130 | 1.000 | |
| AT3G13110.1 | Serine O-acetyltransferase (SERAT2.2) | 28 | 1.21 | 0.130 | 1.000 | |
| AT3G13110.1 | Serine O-acetyltransferase (SERAT2.2) | 29 | 1.15 | 0.130 | 1.000 | |
| AT3G13110.1 | Serine O-acetyltransferase (SERAT2.2) | 30 | 1.08 | 0.130 | 1.000 | |
| AT3G13110.1 | Serine O-acetyltransferase (SERAT2.2) | 31 | 1.01 | 0.131 | 1.000 | |
| AT3G13110.1 | Serine O-acetyltransferase (SERAT2.2) | 32 | 0.94 | 0.131 | 1.000 | |
| AT3G13110.1 | Serine O-acetyltransferase (SERAT2.2) | 33 | 0.91 | 0.131 | 1.000 | |
| AT3G13110.1 | Serine O-acetyltransferase (SERAT2.2) | 34 | 0.94 | 0.131 | 1.000 | |
| AT3G13110.1 | Serine O-acetyltransferase (SERAT2.2) | 35 | 1.03 | 0.131 | 1.000 | |
| AT3G13110.1 | Serine O-acetyltransferase (SERAT2.2) | 36 | 1.15 | 0.131 | 1.000 | |

| | | | | | | |
|---|---|---|---|---|---|---|
| AT3G13110.1 | Serine O-acetyltransferase (SERAT2.2) | 37 | 1.25 | 0.131 | 1.000 | |
| AT3G13110.1 | Serine O-acetyltransferase (SERAT2.2) | 38 | 1.29 | 0.131 | 1.000 | |
| AT3G13110.1 | Serine O-acetyltransferase (SERAT2.2) | 39 | 1.24 | 0.131 | 1.000 | |
| AT3G13110.1 | Serine O-acetyltransferase (SERAT2.2) | 40 | 1.15 | 0.132 | 1.000 | |
| AT3G13110.1 | Serine O-acetyltransferase (SERAT2.2) | 41 | 1.02 | 0.133 | 1.000 | |
| AT3G13110.1 | Serine O-acetyltransferase (SERAT2.2) | 42 | 0.84 | 0.134 | 1.000 | |
| AT3G13110.1 | Serine O-acetyltransferase (SERAT2.2) | 43 | 0.65 | 0.135 | 0.959 | |
| AT3G13110.1 | Serine O-acetyltransferase (SERAT2.2) | 44 | 0.48 | 0.135 | 0.309 | |
| AT3G13110.1 | Serine O-acetyltransferase (SERAT2.2) | 45 | 0.35 | 0.142 | 0.033 | * |
| AT3G13110.1 | Serine O-acetyltransferase (SERAT2.2) | 46 | 0.26 | 0.178 | 0.030 | * |
| AT3G13110.1 | Serine O-acetyltransferase (SERAT2.2) | 47 | 0.20 | 0.262 | 0.159 | |
| AT3G13160.1 | Pentatricopeptide repeat (PPR) superfamily protein | 2 | 1.16 | 0.521 | 1.000 | |
| AT3G13160.1 | Pentatricopeptide repeat (PPR) superfamily protein | 3 | 1.06 | 0.415 | 1.000 | |
| AT3G13160.1 | Pentatricopeptide repeat (PPR) superfamily protein | 4 | 0.98 | 0.470 | 1.000 | |
| AT3G13160.1 | Pentatricopeptide repeat (PPR) superfamily protein | 10 | 1.24 | 0.386 | 1.000 | |
| AT3G13160.1 | Pentatricopeptide repeat (PPR) superfamily protein | 11 | 1.48 | 0.333 | 1.000 | |
| AT3G13160.1 | Pentatricopeptide repeat (PPR) superfamily protein | 12 | 1.62 | 0.300 | 1.000 | |
| AT3G13160.1 | Pentatricopeptide repeat (PPR) superfamily protein | 13 | 1.46 | 0.278 | 1.000 | |
| AT3G13160.1 | Pentatricopeptide repeat (PPR) superfamily protein | 14 | 1.22 | 0.265 | 1.000 | |
| AT3G13160.1 | Pentatricopeptide repeat (PPR) superfamily protein | 15 | 1.26 | 0.260 | 1.000 | |
| AT3G13160.1 | Pentatricopeptide repeat (PPR) superfamily protein | 16 | 1.68 | 0.258 | 1.000 | |
| AT3G13160.1 | Pentatricopeptide repeat (PPR) superfamily protein | 17 | 2.31 | 0.257 | 0.962 | |
| AT3G13160.1 | Pentatricopeptide repeat (PPR) superfamily protein | 18 | 2.69 | 0.260 | 0.871 | |
| AT3G13160.1 | Pentatricopeptide repeat (PPR) superfamily protein | 19 | 2.64 | 0.267 | 0.905 | |
| AT3G13160.1 | Pentatricopeptide repeat (PPR) superfamily protein | 20 | 2.23 | 0.273 | 0.986 | |
| AT3G13160.1 | Pentatricopeptide repeat (PPR) superfamily protein | 21 | 1.57 | 0.277 | 1.000 | |
| AT3G13160.1 | Pentatricopeptide repeat (PPR) superfamily protein | 22 | 0.92 | 0.283 | 1.000 | |
| AT3G13160.1 | Pentatricopeptide repeat (PPR) superfamily protein | 23 | 0.55 | 0.293 | 1.000 | |
| AT3G13160.1 | Pentatricopeptide repeat (PPR) superfamily protein | 24 | 0.43 | 0.307 | 0.993 | |
| AT3G13160.1 | Pentatricopeptide repeat (PPR) superfamily protein | 25 | 0.44 | 0.313 | 0.997 | |
| AT3G13160.1 | Pentatricopeptide repeat (PPR) superfamily protein | 26 | 0.54 | 0.313 | 1.000 | |

| | | | | | |
|---|---|---|---|---|---|
| AT3G13160.1 | Pentatricopeptide repeat (PPR) superfamily protein | 27 | 0.64 | 0.295 | 1.000 |
| AT3G13160.1 | Pentatricopeptide repeat (PPR) superfamily protein | 28 | 0.71 | 0.279 | 1.000 |
| AT3G13160.1 | Pentatricopeptide repeat (PPR) superfamily protein | 29 | 0.69 | 0.270 | 1.000 |
| AT3G13160.1 | Pentatricopeptide repeat (PPR) superfamily protein | 30 | 0.65 | 0.266 | 1.000 |
| AT3G13160.1 | Pentatricopeptide repeat (PPR) superfamily protein | 31 | 0.64 | 0.265 | 1.000 |
| AT3G13160.1 | Pentatricopeptide repeat (PPR) superfamily protein | 32 | 0.63 | 0.265 | 1.000 |
| AT3G13160.1 | Pentatricopeptide repeat (PPR) superfamily protein | 33 | 0.56 | 0.265 | 1.000 |
| AT3G13160.1 | Pentatricopeptide repeat (PPR) superfamily protein | 34 | 0.49 | 0.266 | 0.995 |
| AT3G13160.1 | Pentatricopeptide repeat (PPR) superfamily protein | 35 | 0.47 | 0.270 | 0.992 |
| AT3G13160.1 | Pentatricopeptide repeat (PPR) superfamily protein | 36 | 0.51 | 0.276 | 0.999 |
| AT3G13160.1 | Pentatricopeptide repeat (PPR) superfamily protein | 37 | 0.54 | 0.284 | 1.000 |
| AT3G13160.1 | Pentatricopeptide repeat (PPR) superfamily protein | 38 | 0.51 | 0.288 | 0.999 |
| AT3G13160.1 | Pentatricopeptide repeat (PPR) superfamily protein | 39 | 0.53 | 0.283 | 1.000 |
| AT3G13160.1 | Pentatricopeptide repeat (PPR) superfamily protein | 40 | 0.70 | 0.274 | 1.000 |
| AT3G13160.1 | Pentatricopeptide repeat (PPR) superfamily protein | 41 | 1.00 | 0.269 | 1.000 |
| AT3G13160.1 | Pentatricopeptide repeat (PPR) superfamily protein | 42 | 1.24 | 0.270 | 1.000 |
| AT3G13160.1 | Pentatricopeptide repeat (PPR) superfamily protein | 43 | 1.23 | 0.271 | 1.000 |
| AT3G13160.1 | Pentatricopeptide repeat (PPR) superfamily protein | 44 | 1.00 | 0.271 | 1.000 |
| AT3G13160.1 | Pentatricopeptide repeat (PPR) superfamily protein | 45 | 0.64 | 0.278 | 1.000 |
| AT3G13160.1 | Pentatricopeptide repeat (PPR) superfamily protein | 46 | 0.33 | 0.337 | 0.958 |
| AT3G13160.1 | Pentatricopeptide repeat (PPR) superfamily protein | 47 | 0.15 | 0.516 | 0.885 |
| AT3G13450.1 | Branched chain keto acid dehydrogenase subunit (BCKDHE1beta) | 2 | 1.10 | 0.559 | 1.000 |
| AT3G13450.1 | Branched chain keto acid dehydrogenase subunit (BCKDHE1beta) | 3 | 1.05 | 0.585 | 1.000 |
| AT3G13450.1 | Branched chain keto acid dehydrogenase subunit (BCKDHE1beta) | 38 | 0.99 | 0.277 | 1.000 |
| AT3G13450.1 | Branched chain keto acid dehydrogenase subunit (BCKDHE1beta) | 39 | 1.02 | 0.230 | 1.000 |
| AT3G13450.1 | Branched chain keto acid dehydrogenase subunit (BCKDHE1beta) | 40 | 0.99 | 0.215 | 1.000 |
| AT3G13450.1 | Branched chain keto acid dehydrogenase subunit (BCKDHE1beta) | 41 | 0.85 | 0.223 | 0.999 |

| | | | | | | |
|---|---|---|---|---|---|---|
| AT3G13450.1 | Branched chain keto acid dehydrogenase subunit (BCKDHE1beta) | 42 | 0.63 | 0.244 | 0.917 | |
| AT3G13450.1 | Branched chain keto acid dehydrogenase subunit (BCKDHE1beta) | 43 | 0.44 | 0.267 | 0.599 | |
| AT3G13450.1 | Branched chain keto acid dehydrogenase subunit (BCKDHE1beta) | 46 | 0.11 | 0.312 | 0.017 | * |
| AT3G13450.1 | Branched chain keto acid dehydrogenase subunit (BCKDHE1beta) | 47 | 0.06 | 0.360 | 0.010 | ** |
| AT3G13490.1 | Lysyl-tRNA synthetase 2 (KRS-2) / Ovule abortion 5 (OVA5) | 2 | 1.01 | 0.505 | 1.000 | |
| AT3G13490.1 | Lysyl-tRNA synthetase 2 (KRS-2) / Ovule abortion 5 (OVA5) | 3 | 0.84 | 0.495 | 1.000 | |
| AT3G13490.1 | Lysyl-tRNA synthetase 2 (KRS-2) / Ovule abortion 5 (OVA5) | 32 | 0.55 | 0.432 | 1.000 | |
| AT3G13490.1 | Lysyl-tRNA synthetase 2 (KRS-2) / Ovule abortion 5 (OVA5) | 33 | 0.68 | 0.352 | 1.000 | |
| AT3G13490.1 | Lysyl-tRNA synthetase 2 (KRS-2) / Ovule abortion 5 (OVA5) | 34 | 0.71 | 0.343 | 1.000 | |
| AT3G13490.1 | Lysyl-tRNA synthetase 2 (KRS-2) / Ovule abortion 5 (OVA5) | 35 | 0.59 | 0.307 | 0.998 | |
| AT3G13490.1 | Lysyl-tRNA synthetase 2 (KRS-2) / Ovule abortion 5 (OVA5) | 36 | 0.48 | 0.284 | 0.951 | |
| AT3G13490.1 | Lysyl-tRNA synthetase 2 (KRS-2) / Ovule abortion 5 (OVA5) | 37 | 0.48 | 0.261 | 0.912 | |
| AT3G13490.1 | Lysyl-tRNA synthetase 2 (KRS-2) / Ovule abortion 5 (OVA5) | 38 | 0.58 | 0.253 | 0.985 | |
| AT3G13490.1 | Lysyl-tRNA synthetase 2 (KRS-2) / Ovule abortion 5 (OVA5) | 39 | 1.07 | 0.252 | 1.000 | |
| AT3G13490.1 | Lysyl-tRNA synthetase 2 (KRS-2) / Ovule abortion 5 (OVA5) | 40 | 2.28 | 0.252 | 0.818 | |
| AT3G13490.1 | Lysyl-tRNA synthetase 2 (KRS-2) / Ovule abortion 5 (OVA5) | 41 | 2.71 | 0.252 | 0.610 | |
| AT3G13490.1 | Lysyl-tRNA synthetase 2 (KRS-2) / Ovule abortion 5 (OVA5) | 42 | 2.07 | 0.255 | 0.911 | |
| AT3G13490.1 | Lysyl-tRNA synthetase 2 (KRS-2) / Ovule abortion 5 (OVA5) | 43 | 1.53 | 0.257 | 0.999 | |
| AT3G13490.1 | Lysyl-tRNA synthetase 2 (KRS-2) / Ovule abortion 5 (OVA5) | 44 | 1.06 | 0.256 | 1.000 | |
| AT3G13490.1 | Lysyl-tRNA synthetase 2 (KRS-2) / Ovule abortion 5 (OVA5) | 45 | 0.60 | 0.256 | 0.992 | |
| AT3G13490.1 | Lysyl-tRNA synthetase 2 (KRS-2) / Ovule abortion 5 (OVA5) | 46 | 0.25 | 0.287 | 0.335 | |
| AT3G13490.1 | Lysyl-tRNA synthetase 2 (KRS-2) / Ovule abortion 5 (OVA5) | 47 | 0.08 | 0.479 | 0.220 | |
| AT3G13845.1 | ? | 2 | 1.00 | 0.421 | 1.000 | |
| AT3G13845.1 | ? | 3 | 0.94 | 1.181 | 1.000 | |
| AT3G13845.1 | ? | 38 | 1.53 | 0.320 | 0.999 | |
| AT3G13845.1 | ? | 39 | 2.56 | 0.211 | 0.401 | |
| AT3G13845.1 | ? | 40 | 4.02 | 0.196 | 0.031 | * |
| AT3G13845.1 | ? | 41 | 2.47 | 0.190 | 0.317 | |
| AT3G13845.1 | ? | 42 | 0.61 | 0.199 | 0.939 | |
| AT3G13845.1 | ? | 43 | 0.29 | 0.263 | 0.328 | |
| AT3G13845.1 | ? | 44 | 0.36 | 0.492 | 0.976 | |
| AT3G13845.1 | ? | 46 | 1.26 | 1.389 | 1.000 | |
| AT3G13845.1 | ? | 47 | 2.82 | 2.092 | 1.000 | |
| AT3G13860.1 | Heat shock protein 60-3A (HSP60-3A) | 2 | 1.06 | 0.432 | 1.000 | |

| | | | | | |
|---|---|---|---|---|---|
| AT3G13860.1 | Heat shock protein 60-3A (HSP60-3A) | 3 | 0.74 | 0.295 | 1.000 |
| AT3G13860.1 | Heat shock protein 60-3A (HSP60-3A) | 4 | 0.55 | 0.275 | 1.000 |
| AT3G13860.1 | Heat shock protein 60-3A (HSP60-3A) | 5 | 0.46 | 0.274 | 0.998 |
| AT3G13860.1 | Heat shock protein 60-3A (HSP60-3A) | 6 | 0.48 | 0.254 | 0.998 |
| AT3G13860.1 | Heat shock protein 60-3A (HSP60-3A) | 7 | 0.61 | 0.233 | 1.000 |
| AT3G13860.1 | Heat shock protein 60-3A (HSP60-3A) | 8 | 0.80 | 0.218 | 1.000 |
| AT3G13860.1 | Heat shock protein 60-3A (HSP60-3A) | 9 | 1.09 | 0.211 | 1.000 |
| AT3G13860.1 | Heat shock protein 60-3A (HSP60-3A) | 10 | 1.51 | 0.210 | 1.000 |
| AT3G13860.1 | Heat shock protein 60-3A (HSP60-3A) | 11 | 2.13 | 0.209 | 0.967 |
| AT3G13860.1 | Heat shock protein 60-3A (HSP60-3A) | 12 | 2.84 | 0.209 | 0.608 |
| AT3G13860.1 | Heat shock protein 60-3A (HSP60-3A) | 13 | 3.31 | 0.207 | 0.330 |
| AT3G13860.1 | Heat shock protein 60-3A (HSP60-3A) | 14 | 3.53 | 0.204 | 0.223 |
| AT3G13860.1 | Heat shock protein 60-3A (HSP60-3A) | 15 | 3.54 | 0.203 | 0.209 |
| AT3G13860.1 | Heat shock protein 60-3A (HSP60-3A) | 16 | 3.34 | 0.201 | 0.265 |
| AT3G13860.1 | Heat shock protein 60-3A (HSP60-3A) | 17 | 2.77 | 0.201 | 0.571 |
| AT3G13860.1 | Heat shock protein 60-3A (HSP60-3A) | 18 | 1.88 | 0.205 | 0.995 |
| AT3G13860.1 | Heat shock protein 60-3A (HSP60-3A) | 19 | 1.41 | 0.211 | 1.000 |
| AT3G13860.1 | Heat shock protein 60-3A (HSP60-3A) | 20 | 1.27 | 0.216 | 1.000 |
| AT3G13860.1 | Heat shock protein 60-3A (HSP60-3A) | 21 | 1.16 | 0.219 | 1.000 |
| AT3G13860.1 | Heat shock protein 60-3A (HSP60-3A) | 22 | 1.03 | 0.219 | 1.000 |
| AT3G13860.1 | Heat shock protein 60-3A (HSP60-3A) | 23 | 0.94 | 0.217 | 1.000 |
| AT3G13860.1 | Heat shock protein 60-3A (HSP60-3A) | 24 | 0.93 | 0.213 | 1.000 |
| AT3G13860.1 | Heat shock protein 60-3A (HSP60-3A) | 25 | 0.97 | 0.209 | 1.000 |
| AT3G13860.1 | Heat shock protein 60-3A (HSP60-3A) | 26 | 1.01 | 0.208 | 1.000 |
| AT3G13860.1 | Heat shock protein 60-3A (HSP60-3A) | 27 | 1.08 | 0.208 | 1.000 |
| AT3G13860.1 | Heat shock protein 60-3A (HSP60-3A) | 28 | 1.24 | 0.208 | 1.000 |
| AT3G13860.1 | Heat shock protein 60-3A (HSP60-3A) | 29 | 1.33 | 0.207 | 1.000 |
| AT3G13860.1 | Heat shock protein 60-3A (HSP60-3A) | 30 | 1.02 | 0.208 | 1.000 |
| AT3G13860.1 | Heat shock protein 60-3A (HSP60-3A) | 31 | 0.63 | 0.208 | 1.000 |
| AT3G13860.1 | Heat shock protein 60-3A (HSP60-3A) | 32 | 0.49 | 0.209 | 0.980 |
| AT3G13860.1 | Heat shock protein 60-3A (HSP60-3A) | 33 | 0.54 | 0.209 | 0.998 |

| | | | | | | |
|---|---|---|---|---|---|---|
| AT3G13860.1 | Heat shock protein 60-3A (HSP60-3A) | 34 | 0.62 | 0.209 | 1.000 | |
| AT3G13860.1 | Heat shock protein 60-3A (HSP60-3A) | 35 | 0.70 | 0.209 | 1.000 | |
| AT3G13860.1 | Heat shock protein 60-3A (HSP60-3A) | 36 | 1.01 | 0.208 | 1.000 | |
| AT3G13860.1 | Heat shock protein 60-3A (HSP60-3A) | 37 | 1.68 | 0.208 | 1.000 | |
| AT3G13860.1 | Heat shock protein 60-3A (HSP60-3A) | 38 | 2.93 | 0.209 | 0.543 | |
| AT3G13860.1 | Heat shock protein 60-3A (HSP60-3A) | 39 | 4.39 | 0.209 | 0.075 | . |
| AT3G13860.1 | Heat shock protein 60-3A (HSP60-3A) | 40 | 5.20 | 0.209 | 0.025 | * |
| AT3G13860.1 | Heat shock protein 60-3A (HSP60-3A) | 41 | 5.18 | 0.210 | 0.028 | * |
| AT3G13860.1 | Heat shock protein 60-3A (HSP60-3A) | 42 | 4.51 | 0.214 | 0.080 | . |
| AT3G13860.1 | Heat shock protein 60-3A (HSP60-3A) | 43 | 3.37 | 0.216 | 0.380 | |
| AT3G13860.1 | Heat shock protein 60-3A (HSP60-3A) | 44 | 2.02 | 0.216 | 0.989 | |
| AT3G13860.1 | Heat shock protein 60-3A (HSP60-3A) | 45 | 1.05 | 0.216 | 1.000 | |
| AT3G13860.1 | Heat shock protein 60-3A (HSP60-3A) | 46 | 0.46 | 0.264 | 0.998 | |
| AT3G13860.1 | Heat shock protein 60-3A (HSP60-3A) | 47 | 0.18 | 0.458 | 0.946 | |
| AT3G13930.1 | Dihydrolipoamide acetyltransferase (PDCE2-2) | 3 | 0.53 | 0.201 | 0.991 | |
| AT3G13930.1 | Dihydrolipoamide acetyltransferase (PDCE2-2) | 4 | 0.46 | 0.182 | 0.834 | |
| AT3G13930.1 | Dihydrolipoamide acetyltransferase (PDCE2-2) | 5 | 0.42 | 0.183 | 0.703 | |
| AT3G13930.1 | Dihydrolipoamide acetyltransferase (PDCE2-2) | 6 | 0.39 | 0.183 | 0.543 | |
| AT3G13930.1 | Dihydrolipoamide acetyltransferase (PDCE2-2) | 7 | 0.35 | 0.182 | 0.314 | |
| AT3G13930.1 | Dihydrolipoamide acetyltransferase (PDCE2-2) | 8 | 0.33 | 0.179 | 0.209 | |
| AT3G13930.1 | Dihydrolipoamide acetyltransferase (PDCE2-2) | 9 | 0.36 | 0.177 | 0.318 | |
| AT3G13930.1 | Dihydrolipoamide acetyltransferase (PDCE2-2) | 10 | 0.43 | 0.176 | 0.677 | |
| AT3G13930.1 | Dihydrolipoamide acetyltransferase (PDCE2-2) | 11 | 0.60 | 0.176 | 0.997 | |
| AT3G13930.1 | Dihydrolipoamide acetyltransferase (PDCE2-2) | 12 | 0.89 | 0.176 | 1.000 | |
| AT3G13930.1 | Dihydrolipoamide acetyltransferase (PDCE2-2) | 13 | 1.22 | 0.175 | 1.000 | |
| AT3G13930.1 | Dihydrolipoamide acetyltransferase (PDCE2-2) | 14 | 1.42 | 0.173 | 1.000 | |
| AT3G13930.1 | Dihydrolipoamide acetyltransferase (PDCE2-2) | 15 | 1.42 | 0.172 | 1.000 | |
| AT3G13930.1 | Dihydrolipoamide acetyltransferase (PDCE2-2) | 16 | 1.28 | 0.171 | 1.000 | |
| AT3G13930.1 | Dihydrolipoamide acetyltransferase (PDCE2-2) | 17 | 1.18 | 0.170 | 1.000 | |
| AT3G13930.1 | Dihydrolipoamide acetyltransferase (PDCE2-2) | 18 | 1.17 | 0.173 | 1.000 | |
| AT3G13930.1 | Dihydrolipoamide acetyltransferase (PDCE2-2) | 19 | 1.30 | 0.178 | 1.000 | |

| | | | | | | |
|---|---|---|---|---|---|---|
| AT3G13930.1 | Dihydrolipoamide acetyltransferase (PDCE2-2) | 20 | 1.51 | 0.182 | 1.000 | |
| AT3G13930.1 | Dihydrolipoamide acetyltransferase (PDCE2-2) | 21 | 1.54 | 0.184 | 1.000 | |
| AT3G13930.1 | Dihydrolipoamide acetyltransferase (PDCE2-2) | 22 | 1.28 | 0.184 | 1.000 | |
| AT3G13930.1 | Dihydrolipoamide acetyltransferase (PDCE2-2) | 23 | 1.00 | 0.183 | 1.000 | |
| AT3G13930.1 | Dihydrolipoamide acetyltransferase (PDCE2-2) | 24 | 0.85 | 0.179 | 1.000 | |
| AT3G13930.1 | Dihydrolipoamide acetyltransferase (PDCE2-2) | 25 | 0.85 | 0.176 | 1.000 | |
| AT3G13930.1 | Dihydrolipoamide acetyltransferase (PDCE2-2) | 26 | 0.97 | 0.175 | 1.000 | |
| AT3G13930.1 | Dihydrolipoamide acetyltransferase (PDCE2-2) | 27 | 1.11 | 0.175 | 1.000 | |
| AT3G13930.1 | Dihydrolipoamide acetyltransferase (PDCE2-2) | 28 | 1.18 | 0.175 | 1.000 | |
| AT3G13930.1 | Dihydrolipoamide acetyltransferase (PDCE2-2) | 29 | 1.22 | 0.175 | 1.000 | |
| AT3G13930.1 | Dihydrolipoamide acetyltransferase (PDCE2-2) | 30 | 1.31 | 0.175 | 1.000 | |
| AT3G13930.1 | Dihydrolipoamide acetyltransferase (PDCE2-2) | 31 | 1.42 | 0.176 | 1.000 | |
| AT3G13930.1 | Dihydrolipoamide acetyltransferase (PDCE2-2) | 32 | 1.37 | 0.176 | 1.000 | |
| AT3G13930.1 | Dihydrolipoamide acetyltransferase (PDCE2-2) | 33 | 1.10 | 0.176 | 1.000 | |
| AT3G13930.1 | Dihydrolipoamide acetyltransferase (PDCE2-2) | 34 | 0.88 | 0.177 | 1.000 | |
| AT3G13930.1 | Dihydrolipoamide acetyltransferase (PDCE2-2) | 35 | 0.91 | 0.177 | 1.000 | |
| AT3G13930.1 | Dihydrolipoamide acetyltransferase (PDCE2-2) | 36 | 1.26 | 0.176 | 1.000 | |
| AT3G13930.1 | Dihydrolipoamide acetyltransferase (PDCE2-2) | 37 | 1.98 | 0.176 | 0.923 | |
| AT3G13930.1 | Dihydrolipoamide acetyltransferase (PDCE2-2) | 38 | 2.83 | 0.176 | 0.283 | |
| AT3G13930.1 | Dihydrolipoamide acetyltransferase (PDCE2-2) | 39 | 3.76 | 0.176 | 0.043 | * |
| AT3G13930.1 | Dihydrolipoamide acetyltransferase (PDCE2-2) | 40 | 4.86 | 0.177 | 0.005 | ** |
| AT3G13930.1 | Dihydrolipoamide acetyltransferase (PDCE2-2) | 41 | 5.14 | 0.177 | 0.003 | ** |
| AT3G13930.1 | Dihydrolipoamide acetyltransferase (PDCE2-2) | 42 | 4.26 | 0.180 | 0.019 | * |
| AT3G13930.1 | Dihydrolipoamide acetyltransferase (PDCE2-2) | 43 | 3.13 | 0.182 | 0.196 | |
| AT3G13930.1 | Dihydrolipoamide acetyltransferase (PDCE2-2) | 44 | 1.92 | 0.182 | 0.965 | |
| AT3G13930.1 | Dihydrolipoamide acetyltransferase (PDCE2-2) | 45 | 0.98 | 0.181 | 1.000 | |
| AT3G13930.1 | Dihydrolipoamide acetyltransferase (PDCE2-2) | 46 | 0.41 | 0.217 | 0.881 | |
| AT3G13930.1 | Dihydrolipoamide acetyltransferase (PDCE2-2) | 47 | 0.14 | 0.378 | 0.539 | |
| AT3G13930.1 | Dihydrolipoamide acetyltransferase (PDCE2-2) | 48 | 0.05 | 0.619 | 0.638 | |
| AT3G14210.1 | Epithiospecifier modifier 1 (ESM1) | 2 | 4.62 | 0.298 | 0.088 | . |
| AT3G14210.1 | Epithiospecifier modifier 1 (ESM1) | 3 | 4.15 | 0.288 | 0.106 | |

| | | | | | |
|---|---|---|---|---|---|
| AT3G14210.1 | Epithiospecifier modifier 1 (ESM1) | 4 | 3.73 | 0.281 | 0.134 |
| AT3G14210.1 | Epithiospecifier modifier 1 (ESM1) | 5 | 3.35 | 0.276 | 0.176 |
| AT3G14210.1 | Epithiospecifier modifier 1 (ESM1) | 6 | 3.00 | 0.273 | 0.233 |
| AT3G14210.1 | Epithiospecifier modifier 1 (ESM1) | 7 | 2.69 | 0.271 | 0.308 |
| AT3G14210.1 | Epithiospecifier modifier 1 (ESM1) | 8 | 2.42 | 0.269 | 0.399 |
| AT3G14210.1 | Epithiospecifier modifier 1 (ESM1) | 9 | 2.18 | 0.269 | 0.501 |
| AT3G14210.1 | Epithiospecifier modifier 1 (ESM1) | 10 | 1.98 | 0.268 | 0.609 |
| AT3G14210.1 | Epithiospecifier modifier 1 (ESM1) | 11 | 1.80 | 0.268 | 0.715 |
| AT3G14210.1 | Epithiospecifier modifier 1 (ESM1) | 12 | 1.64 | 0.268 | 0.813 |
| AT3G14210.1 | Epithiospecifier modifier 1 (ESM1) | 13 | 1.51 | 0.268 | 0.894 |
| AT3G14210.1 | Epithiospecifier modifier 1 (ESM1) | 14 | 1.39 | 0.268 | 0.951 |
| AT3G14210.1 | Epithiospecifier modifier 1 (ESM1) | 15 | 1.30 | 0.268 | 0.983 |
| AT3G14210.1 | Epithiospecifier modifier 1 (ESM1) | 16 | 1.21 | 0.268 | 0.996 |
| AT3G14210.1 | Epithiospecifier modifier 1 (ESM1) | 17 | 1.15 | 0.268 | 1.000 |
| AT3G14210.1 | Epithiospecifier modifier 1 (ESM1) | 18 | 1.09 | 0.268 | 1.000 |
| AT3G14210.1 | Epithiospecifier modifier 1 (ESM1) | 19 | 1.04 | 0.269 | 1.000 |
| AT3G14210.1 | Epithiospecifier modifier 1 (ESM1) | 20 | 1.01 | 0.270 | 1.000 |
| AT3G14210.1 | Epithiospecifier modifier 1 (ESM1) | 21 | 0.98 | 0.270 | 1.000 |
| AT3G14210.1 | Epithiospecifier modifier 1 (ESM1) | 22 | 0.95 | 0.271 | 1.000 |
| AT3G14210.1 | Epithiospecifier modifier 1 (ESM1) | 23 | 0.94 | 0.273 | 1.000 |
| AT3G14210.1 | Epithiospecifier modifier 1 (ESM1) | 24 | 0.93 | 0.274 | 1.000 |
| AT3G14210.1 | Epithiospecifier modifier 1 (ESM1) | 25 | 0.94 | 0.276 | 1.000 |
| AT3G14210.1 | Epithiospecifier modifier 1 (ESM1) | 27 | 0.96 | 0.281 | 1.000 |
| AT3G14210.1 | Epithiospecifier modifier 1 (ESM1) | 28 | 0.97 | 0.285 | 1.000 |
| AT3G14210.1 | Epithiospecifier modifier 1 (ESM1) | 29 | 0.99 | 0.290 | 1.000 |
| AT3G14210.1 | Epithiospecifier modifier 1 (ESM1) | 30 | 1.00 | 0.295 | 1.000 |
| AT3G14210.1 | Epithiospecifier modifier 1 (ESM1) | 31 | 1.01 | 0.302 | 1.000 |
| AT3G14210.1 | Epithiospecifier modifier 1 (ESM1) | 46 | 0.93 | 0.503 | 1.000 |
| AT3G14210.1 | Epithiospecifier modifier 1 (ESM1) | 47 | 0.92 | 0.526 | 1.000 |
| AT3G14415.3 | Glycolate oxidase 2 (GOX2) | 2 | 1.00 | 0.483 | 1.000 |
| AT3G14415.3 | Glycolate oxidase 2 (GOX2) | 3 | 0.90 | 0.380 | 1.000 |
| AT3G14415.3 | Glycolate oxidase 2 (GOX2) | 4 | 0.80 | 0.719 | 1.000 |
| AT3G14415.3 | Glycolate oxidase 2 (GOX2) | 30 | 0.31 | 0.363 | 0.916 |

| | | | | | | |
|---|---|---|---|---|---|---|
| AT3G14415.3 | Glycolate oxidase 2 (GOX2) | 31 | 0.42 | 0.259 | 0.882 | |
| AT3G14415.3 | Glycolate oxidase 2 (GOX2) | 32 | 0.66 | 0.232 | 1.000 | |
| AT3G14415.3 | Glycolate oxidase 2 (GOX2) | 33 | 1.21 | 0.223 | 1.000 | |
| AT3G14415.3 | Glycolate oxidase 2 (GOX2) | 34 | 1.87 | 0.219 | 0.962 | |
| AT3G14415.3 | Glycolate oxidase 2 (GOX2) | 35 | 2.05 | 0.217 | 0.895 | |
| AT3G14415.3 | Glycolate oxidase 2 (GOX2) | 36 | 1.34 | 0.217 | 1.000 | |
| AT3G14415.3 | Glycolate oxidase 2 (GOX2) | 37 | 0.48 | 0.216 | 0.870 | |
| AT3G14415.3 | Glycolate oxidase 2 (GOX2) | 38 | 0.17 | 0.215 | 0.009 | ** |
| AT3G14415.3 | Glycolate oxidase 2 (GOX2) | 39 | 0.15 | 0.214 | 0.004 | ** |
| AT3G14415.3 | Glycolate oxidase 2 (GOX2) | 40 | 0.36 | 0.214 | 0.461 | |
| AT3G14415.3 | Glycolate oxidase 2 (GOX2) | 41 | 1.13 | 0.217 | 1.000 | |
| AT3G14415.3 | Glycolate oxidase 2 (GOX2) | 42 | 2.34 | 0.219 | 0.747 | |
| AT3G14415.3 | Glycolate oxidase 2 (GOX2) | 43 | 3.07 | 0.219 | 0.346 | |
| AT3G14415.3 | Glycolate oxidase 2 (GOX2) | 44 | 2.47 | 0.220 | 0.680 | |
| AT3G14415.3 | Glycolate oxidase 2 (GOX2) | 45 | 1.24 | 0.229 | 1.000 | |
| AT3G14415.3 | Glycolate oxidase 2 (GOX2) | 46 | 0.39 | 0.265 | 0.853 | |
| AT3G14415.3 | Glycolate oxidase 2 (GOX2) | 47 | 0.09 | 0.438 | 0.222 | |
| AT3G14420.2 | Glycolate oxidase 1 (GOX1) | 2 | 0.95 | 0.500 | 1.000 | |
| AT3G14420.2 | Glycolate oxidase 1 (GOX1) | 3 | 0.99 | 0.402 | 1.000 | |
| AT3G14420.2 | Glycolate oxidase 1 (GOX1) | 6 | 1.14 | 0.380 | 1.000 | |
| AT3G14420.2 | Glycolate oxidase 1 (GOX1) | 7 | 1.21 | 0.373 | 1.000 | |
| AT3G14420.2 | Glycolate oxidase 1 (GOX1) | 8 | 1.27 | 0.371 | 1.000 | |
| AT3G14420.2 | Glycolate oxidase 1 (GOX1) | 13 | 1.60 | 0.321 | 1.000 | |
| AT3G14420.2 | Glycolate oxidase 1 (GOX1) | 14 | 1.59 | 0.299 | 1.000 | |
| AT3G14420.2 | Glycolate oxidase 1 (GOX1) | 15 | 1.53 | 0.275 | 1.000 | |
| AT3G14420.2 | Glycolate oxidase 1 (GOX1) | 16 | 1.40 | 0.264 | 1.000 | |
| AT3G14420.2 | Glycolate oxidase 1 (GOX1) | 17 | 1.18 | 0.260 | 1.000 | |
| AT3G14420.2 | Glycolate oxidase 1 (GOX1) | 18 | 0.87 | 0.263 | 1.000 | |
| AT3G14420.2 | Glycolate oxidase 1 (GOX1) | 19 | 0.61 | 0.263 | 1.000 | |
| AT3G14420.2 | Glycolate oxidase 1 (GOX1) | 20 | 0.46 | 0.260 | 0.960 | |
| AT3G14420.2 | Glycolate oxidase 1 (GOX1) | 21 | 0.38 | 0.257 | 0.816 | |
| AT3G14420.2 | Glycolate oxidase 1 (GOX1) | 22 | 0.35 | 0.255 | 0.698 | |
| AT3G14420.2 | Glycolate oxidase 1 (GOX1) | 23 | 0.34 | 0.248 | 0.604 | |
| AT3G14420.2 | Glycolate oxidase 1 (GOX1) | 24 | 0.33 | 0.236 | 0.482 | |
| AT3G14420.2 | Glycolate oxidase 1 (GOX1) | 25 | 0.27 | 0.228 | 0.227 | |
| AT3G14420.2 | Glycolate oxidase 1 (GOX1) | 26 | 0.21 | 0.226 | 0.062 | . |
| AT3G14420.2 | Glycolate oxidase 1 (GOX1) | 27 | 0.20 | 0.225 | 0.049 | * |
| AT3G14420.2 | Glycolate oxidase 1 (GOX1) | 28 | 0.26 | 0.225 | 0.170 | |
| AT3G14420.2 | Glycolate oxidase 1 (GOX1) | 29 | 0.41 | 0.225 | 0.731 | |
| AT3G14420.2 | Glycolate oxidase 1 (GOX1) | 30 | 0.65 | 0.226 | 1.000 | |
| AT3G14420.2 | Glycolate oxidase 1 (GOX1) | 31 | 0.94 | 0.226 | 1.000 | |
| AT3G14420.2 | Glycolate oxidase 1 (GOX1) | 32 | 1.17 | 0.226 | 1.000 | |
| AT3G14420.2 | Glycolate oxidase 1 (GOX1) | 33 | 1.31 | 0.226 | 1.000 | |
| AT3G14420.2 | Glycolate oxidase 1 (GOX1) | 34 | 1.43 | 0.227 | 1.000 | |
| AT3G14420.2 | Glycolate oxidase 1 (GOX1) | 35 | 1.49 | 0.227 | 1.000 | |
| AT3G14420.2 | Glycolate oxidase 1 (GOX1) | 36 | 1.37 | 0.226 | 1.000 | |
| AT3G14420.2 | Glycolate oxidase 1 (GOX1) | 37 | 1.18 | 0.226 | 1.000 | |
| AT3G14420.2 | Glycolate oxidase 1 (GOX1) | 38 | 1.14 | 0.226 | 1.000 | |
| AT3G14420.2 | Glycolate oxidase 1 (GOX1) | 39 | 1.38 | 0.226 | 1.000 | |
| AT3G14420.2 | Glycolate oxidase 1 (GOX1) | 40 | 1.79 | 0.226 | 0.990 | |
| AT3G14420.2 | Glycolate oxidase 1 (GOX1) | 41 | 2.04 | 0.228 | 0.942 | |
| AT3G14420.2 | Glycolate oxidase 1 (GOX1) | 42 | 1.92 | 0.229 | 0.974 | |
| AT3G14420.2 | Glycolate oxidase 1 (GOX1) | 43 | 1.51 | 0.230 | 1.000 | |
| AT3G14420.2 | Glycolate oxidase 1 (GOX1) | 44 | 1.02 | 0.230 | 1.000 | |
| AT3G14420.2 | Glycolate oxidase 1 (GOX1) | 45 | 0.56 | 0.235 | 0.994 | |
| AT3G14420.2 | Glycolate oxidase 1 (GOX1) | 46 | 0.24 | 0.273 | 0.343 | |
| AT3G14420.2 | Glycolate oxidase 1 (GOX1) | 47 | 0.09 | 0.396 | 0.141 | |
| AT3G15000.1 | Multiple organellar RNA editing factor 8 (MORF8, RIP1) | 2 | 0.66 | 0.331 | 1.000 | |
| AT3G15000.1 | Multiple organellar RNA editing factor 8 (MORF8, RIP1) | 3 | 0.65 | 0.257 | 1.000 | |
| AT3G15000.1 | Multiple organellar RNA editing factor 8 (MORF8, RIP1) | 4 | 0.65 | 0.221 | 0.999 | |

| | | | | | |
|---|---|---|---|---|---|
| AT3G15000.1 | Multiple organellar RNA editing factor 8 (MORF8, RIP1) | 5 | 0.67 | 0.208 | 0.999 |
| AT3G15000.1 | Multiple organellar RNA editing factor 8 (MORF8, RIP1) | 6 | 0.70 | 0.205 | 1.000 |
| AT3G15000.1 | Multiple organellar RNA editing factor 8 (MORF8, RIP1) | 7 | 0.73 | 0.201 | 1.000 |
| AT3G15000.1 | Multiple organellar RNA editing factor 8 (MORF8, RIP1) | 8 | 0.77 | 0.197 | 1.000 |
| AT3G15000.1 | Multiple organellar RNA editing factor 8 (MORF8, RIP1) | 9 | 0.80 | 0.194 | 1.000 |
| AT3G15000.1 | Multiple organellar RNA editing factor 8 (MORF8, RIP1) | 10 | 0.82 | 0.191 | 1.000 |
| AT3G15000.1 | Multiple organellar RNA editing factor 8 (MORF8, RIP1) | 11 | 0.86 | 0.190 | 1.000 |
| AT3G15000.1 | Multiple organellar RNA editing factor 8 (MORF8, RIP1) | 12 | 0.91 | 0.189 | 1.000 |
| AT3G15000.1 | Multiple organellar RNA editing factor 8 (MORF8, RIP1) | 13 | 0.97 | 0.187 | 1.000 |
| AT3G15000.1 | Multiple organellar RNA editing factor 8 (MORF8, RIP1) | 14 | 1.06 | 0.186 | 1.000 |
| AT3G15000.1 | Multiple organellar RNA editing factor 8 (MORF8, RIP1) | 15 | 1.17 | 0.186 | 1.000 |
| AT3G15000.1 | Multiple organellar RNA editing factor 8 (MORF8, RIP1) | 16 | 1.27 | 0.185 | 1.000 |
| AT3G15000.1 | Multiple organellar RNA editing factor 8 (MORF8, RIP1) | 17 | 1.35 | 0.186 | 1.000 |
| AT3G15000.1 | Multiple organellar RNA editing factor 8 (MORF8, RIP1) | 18 | 1.40 | 0.187 | 1.000 |
| AT3G15000.1 | Multiple organellar RNA editing factor 8 (MORF8, RIP1) | 19 | 1.44 | 0.190 | 0.999 |
| AT3G15000.1 | Multiple organellar RNA editing factor 8 (MORF8, RIP1) | 20 | 1.51 | 0.192 | 0.997 |
| AT3G15000.1 | Multiple organellar RNA editing factor 8 (MORF8, RIP1) | 21 | 1.59 | 0.193 | 0.992 |
| AT3G15000.1 | Multiple organellar RNA editing factor 8 (MORF8, RIP1) | 22 | 1.63 | 0.193 | 0.987 |
| AT3G15000.1 | Multiple organellar RNA editing factor 8 (MORF8, RIP1) | 23 | 1.64 | 0.192 | 0.985 |
| AT3G15000.1 | Multiple organellar RNA editing factor 8 (MORF8, RIP1) | 24 | 1.63 | 0.191 | 0.985 |
| AT3G15000.1 | Multiple organellar RNA editing factor 8 (MORF8, RIP1) | 25 | 1.62 | 0.189 | 0.986 |
| AT3G15000.1 | Multiple organellar RNA editing factor 8 (MORF8, RIP1) | 26 | 1.58 | 0.188 | 0.991 |
| AT3G15000.1 | Multiple organellar RNA editing factor 8 (MORF8, RIP1) | 27 | 1.50 | 0.188 | 0.997 |
| AT3G15000.1 | Multiple organellar RNA editing factor 8 (MORF8, RIP1) | 28 | 1.38 | 0.188 | 1.000 |
| AT3G15000.1 | Multiple organellar RNA editing factor 8 (MORF8, RIP1) | 29 | 1.26 | 0.188 | 1.000 |
| AT3G15000.1 | Multiple organellar RNA editing factor 8 (MORF8, RIP1) | 30 | 1.16 | 0.188 | 1.000 |
| AT3G15000.1 | Multiple organellar RNA editing factor 8 (MORF8, RIP1) | 31 | 1.09 | 0.189 | 1.000 |
| AT3G15000.1 | Multiple organellar RNA editing factor 8 (MORF8, RIP1) | 32 | 1.06 | 0.189 | 1.000 |
| AT3G15000.1 | Multiple organellar RNA editing factor 8 (MORF8, RIP1) | 33 | 1.05 | 0.189 | 1.000 |
| AT3G15000.1 | Multiple organellar RNA editing factor 8 (MORF8, RIP1) | 34 | 1.05 | 0.189 | 1.000 |
| AT3G15000.1 | Multiple organellar RNA editing factor 8 (MORF8, RIP1) | 35 | 1.07 | 0.189 | 1.000 |

| | | | | | |
|---|---|---|---|---|---|
| AT3G15000.1 | Multiple organellar RNA editing factor 8 (MORF8, RIP1) | 36 | 1.11 | 0.189 | 1.000 |
| AT3G15000.1 | Multiple organellar RNA editing factor 8 (MORF8, RIP1) | 37 | 1.19 | 0.189 | 1.000 |
| AT3G15000.1 | Multiple organellar RNA editing factor 8 (MORF8, RIP1) | 38 | 1.31 | 0.189 | 1.000 |
| AT3G15000.1 | Multiple organellar RNA editing factor 8 (MORF8, RIP1) | 39 | 1.46 | 0.190 | 0.999 |
| AT3G15000.1 | Multiple organellar RNA editing factor 8 (MORF8, RIP1) | 40 | 1.58 | 0.191 | 0.993 |
| AT3G15000.1 | Multiple organellar RNA editing factor 8 (MORF8, RIP1) | 41 | 1.60 | 0.192 | 0.991 |
| AT3G15000.1 | Multiple organellar RNA editing factor 8 (MORF8, RIP1) | 42 | 1.44 | 0.193 | 0.999 |
| AT3G15000.1 | Multiple organellar RNA editing factor 8 (MORF8, RIP1) | 43 | 1.15 | 0.193 | 1.000 |
| AT3G15000.1 | Multiple organellar RNA editing factor 8 (MORF8, RIP1) | 44 | 0.85 | 0.196 | 1.000 |
| AT3G15000.1 | Multiple organellar RNA editing factor 8 (MORF8, RIP1) | 45 | 0.59 | 0.212 | 0.987 |
| AT3G15000.1 | Multiple organellar RNA editing factor 8 (MORF8, RIP1) | 46 | 0.40 | 0.258 | 0.842 |
| AT3G15000.1 | Multiple organellar RNA editing factor 8 (MORF8, RIP1) | 47 | 0.28 | 0.344 | 0.788 |
| AT3G15020.1 | Mitochondrial malate dehydrogenase 2 (mMDH) | 2 | 0.88 | 0.583 | 1.000 |
| AT3G15020.1 | Mitochondrial malate dehydrogenase 2 (mMDH) | 3 | 0.74 | 0.526 | 1.000 |
| AT3G15020.1 | Mitochondrial malate dehydrogenase 2 (mMDH) | 4 | 0.62 | 0.509 | 1.000 |
| AT3G15020.1 | Mitochondrial malate dehydrogenase 2 (mMDH) | 5 | 0.55 | 0.503 | 0.999 |
| AT3G15020.1 | Mitochondrial malate dehydrogenase 2 (mMDH) | 6 | 0.51 | 0.495 | 0.996 |
| AT3G15020.1 | Mitochondrial malate dehydrogenase 2 (mMDH) | 7 | 0.51 | 0.492 | 0.996 |
| AT3G15020.1 | Mitochondrial malate dehydrogenase 2 (mMDH) | 8 | 0.54 | 0.493 | 0.999 |
| AT3G15020.1 | Mitochondrial malate dehydrogenase 2 (mMDH) | 9 | 0.60 | 0.491 | 1.000 |
| AT3G15020.1 | Mitochondrial malate dehydrogenase 2 (mMDH) | 10 | 0.67 | 0.490 | 1.000 |
| AT3G15020.1 | Mitochondrial malate dehydrogenase 2 (mMDH) | 11 | 0.72 | 0.488 | 1.000 |
| AT3G15020.1 | Mitochondrial malate dehydrogenase 2 (mMDH) | 12 | 0.75 | 0.483 | 1.000 |
| AT3G15020.1 | Mitochondrial malate dehydrogenase 2 (mMDH) | 13 | 0.77 | 0.479 | 1.000 |
| AT3G15020.1 | Mitochondrial malate dehydrogenase 2 (mMDH) | 14 | 0.81 | 0.475 | 1.000 |
| AT3G15020.1 | Mitochondrial malate dehydrogenase 2 (mMDH) | 15 | 0.86 | 0.473 | 1.000 |
| AT3G15020.1 | Mitochondrial malate dehydrogenase 2 (mMDH) | 16 | 0.91 | 0.472 | 1.000 |
| AT3G15020.1 | Mitochondrial malate dehydrogenase 2 (mMDH) | 17 | 0.97 | 0.472 | 1.000 |
| AT3G15020.1 | Mitochondrial malate dehydrogenase 2 (mMDH) | 18 | 1.04 | 0.472 | 1.000 |
| AT3G15020.1 | Mitochondrial malate dehydrogenase 2 (mMDH) | 19 | 1.05 | 0.474 | 1.000 |
| AT3G15020.1 | Mitochondrial malate dehydrogenase 2 (mMDH) | 20 | 0.98 | 0.477 | 1.000 |

| | | | | | | |
|---|---|---|---|---|---|---|
| AT3G15020.1 | Mitochondrial malate dehydrogenase 2 (mMDH) | 21 | 0.82 | 0.478 | 1.000 | |
| AT3G15020.1 | Mitochondrial malate dehydrogenase 2 (mMDH) | 22 | 0.64 | 0.480 | 1.000 | |
| AT3G15020.1 | Mitochondrial malate dehydrogenase 2 (mMDH) | 23 | 0.51 | 0.481 | 0.995 | |
| AT3G15020.1 | Mitochondrial malate dehydrogenase 2 (mMDH) | 24 | 0.43 | 0.482 | 0.966 | |
| AT3G15020.1 | Mitochondrial malate dehydrogenase 2 (mMDH) | 25 | 0.39 | 0.481 | 0.930 | |
| AT3G15020.1 | Mitochondrial malate dehydrogenase 2 (mMDH) | 26 | 0.38 | 0.479 | 0.924 | |
| AT3G15020.1 | Mitochondrial malate dehydrogenase 2 (mMDH) | 27 | 0.42 | 0.477 | 0.956 | |
| AT3G15020.1 | Mitochondrial malate dehydrogenase 2 (mMDH) | 28 | 0.48 | 0.475 | 0.989 | |
| AT3G15020.1 | Mitochondrial malate dehydrogenase 2 (mMDH) | 29 | 0.54 | 0.475 | 0.998 | |
| AT3G15020.1 | Mitochondrial malate dehydrogenase 2 (mMDH) | 30 | 0.57 | 0.475 | 0.999 | |
| AT3G15020.1 | Mitochondrial malate dehydrogenase 2 (mMDH) | 31 | 0.58 | 0.474 | 0.999 | |
| AT3G15020.1 | Mitochondrial malate dehydrogenase 2 (mMDH) | 32 | 0.58 | 0.474 | 0.999 | |
| AT3G15020.1 | Mitochondrial malate dehydrogenase 2 (mMDH) | 33 | 0.59 | 0.474 | 1.000 | |
| AT3G15020.1 | Mitochondrial malate dehydrogenase 2 (mMDH) | 34 | 0.60 | 0.474 | 1.000 | |
| AT3G15020.1 | Mitochondrial malate dehydrogenase 2 (mMDH) | 35 | 0.61 | 0.474 | 1.000 | |
| AT3G15020.1 | Mitochondrial malate dehydrogenase 2 (mMDH) | 36 | 0.60 | 0.474 | 1.000 | |
| AT3G15020.1 | Mitochondrial malate dehydrogenase 2 (mMDH) | 37 | 0.59 | 0.474 | 1.000 | |
| AT3G15020.1 | Mitochondrial malate dehydrogenase 2 (mMDH) | 38 | 0.64 | 0.474 | 1.000 | |
| AT3G15020.1 | Mitochondrial malate dehydrogenase 2 (mMDH) | 39 | 0.82 | 0.474 | 1.000 | |
| AT3G15020.1 | Mitochondrial malate dehydrogenase 2 (mMDH) | 40 | 1.29 | 0.474 | 1.000 | |
| AT3G15020.1 | Mitochondrial malate dehydrogenase 2 (mMDH) | 41 | 2.37 | 0.475 | 0.958 | |
| AT3G15020.1 | Mitochondrial malate dehydrogenase 2 (mMDH) | 42 | 4.03 | 0.475 | 0.645 | |
| AT3G15020.1 | Mitochondrial malate dehydrogenase 2 (mMDH) | 43 | 4.86 | 0.476 | 0.522 | |
| AT3G15020.1 | Mitochondrial malate dehydrogenase 2 (mMDH) | 44 | 3.24 | 0.478 | 0.798 | |
| AT3G15020.1 | Mitochondrial malate dehydrogenase 2 (mMDH) | 45 | 1.09 | 0.485 | 1.000 | |
| AT3G15020.1 | Mitochondrial malate dehydrogenase 2 (mMDH) | 46 | 0.22 | 0.506 | 0.619 | |
| AT3G15020.1 | Mitochondrial malate dehydrogenase 2 (mMDH) | 47 | 0.03 | 0.573 | 0.069 | . |
| AT3G15090.1 | GroES-like zinc-binding alcohol dehydrogenase family protein | 2 | 1.49 | 0.312 | 1.000 | |
| AT3G15090.1 | GroES-like zinc-binding alcohol dehydrogenase family protein | 3 | 0.70 | 0.236 | 1.000 | |
| AT3G15090.1 | GroES-like zinc-binding alcohol dehydrogenase family protein | 4 | 0.36 | 0.368 | 0.964 | |
| AT3G15090.1 | GroES-like zinc-binding alcohol dehydrogenase family protein | 28 | 1.00 | 0.239 | 1.000 | |

| | | | | | |
|---|---|---|---|---|---|
| AT3G15090.1 | GroES-like zinc-binding alcohol dehydrogenase family protein | 29 | 1.01 | 0.175 | 1.000 |
| AT3G15090.1 | GroES-like zinc-binding alcohol dehydrogenase family protein | 30 | 1.00 | 0.151 | 1.000 |
| AT3G15090.1 | GroES-like zinc-binding alcohol dehydrogenase family protein | 31 | 1.03 | 0.143 | 1.000 |
| AT3G15090.1 | GroES-like zinc-binding alcohol dehydrogenase family protein | 32 | 1.08 | 0.139 | 1.000 |
| AT3G15090.1 | GroES-like zinc-binding alcohol dehydrogenase family protein | 33 | 1.07 | 0.139 | 1.000 |
| AT3G15090.1 | GroES-like zinc-binding alcohol dehydrogenase family protein | 34 | 1.16 | 0.139 | 1.000 |
| AT3G15090.1 | GroES-like zinc-binding alcohol dehydrogenase family protein | 35 | 1.44 | 0.138 | 0.976 |
| AT3G15090.1 | GroES-like zinc-binding alcohol dehydrogenase family protein | 36 | 1.77 | 0.138 | 0.652 |
| AT3G15090.1 | GroES-like zinc-binding alcohol dehydrogenase family protein | 37 | 2.00 | 0.138 | 0.362 |
| AT3G15090.1 | GroES-like zinc-binding alcohol dehydrogenase family protein | 38 | 1.95 | 0.137 | 0.413 |
| AT3G15090.1 | GroES-like zinc-binding alcohol dehydrogenase family protein | 39 | 1.62 | 0.137 | 0.846 |
| AT3G15090.1 | GroES-like zinc-binding alcohol dehydrogenase family protein | 40 | 1.31 | 0.137 | 0.999 |
| AT3G15090.1 | GroES-like zinc-binding alcohol dehydrogenase family protein | 41 | 1.16 | 0.138 | 1.000 |
| AT3G15090.1 | GroES-like zinc-binding alcohol dehydrogenase family protein | 42 | 1.16 | 0.141 | 1.000 |
| AT3G15090.1 | GroES-like zinc-binding alcohol dehydrogenase family protein | 43 | 1.24 | 0.142 | 1.000 |
| AT3G15090.1 | GroES-like zinc-binding alcohol dehydrogenase family protein | 44 | 1.23 | 0.142 | 1.000 |
| AT3G15090.1 | GroES-like zinc-binding alcohol dehydrogenase family protein | 45 | 1.03 | 0.143 | 1.000 |
| AT3G15090.1 | GroES-like zinc-binding alcohol dehydrogenase family protein | 46 | 0.74 | 0.180 | 1.000 |
| AT3G15090.1 | GroES-like zinc-binding alcohol dehydrogenase family protein | 47 | 0.49 | 0.301 | 0.990 |
| AT3G15290.1 | 3-hydroxyacyl-CoA dehydrogenase family protein | 2 | 1.24 | 0.304 | 0.951 |
| AT3G15290.1 | 3-hydroxyacyl-CoA dehydrogenase family protein | 3 | 1.21 | 0.295 | 0.958 |
| AT3G15290.1 | 3-hydroxyacyl-CoA dehydrogenase family protein | 4 | 1.19 | 0.286 | 0.965 |
| AT3G15290.1 | 3-hydroxyacyl-CoA dehydrogenase family protein | 6 | 1.13 | 0.268 | 0.978 |
| AT3G15290.1 | 3-hydroxyacyl-CoA dehydrogenase family protein | 7 | 1.11 | 0.260 | 0.984 |
| AT3G15290.1 | 3-hydroxyacyl-CoA dehydrogenase family protein | 8 | 1.08 | 0.251 | 0.990 |
| AT3G15290.1 | 3-hydroxyacyl-CoA dehydrogenase family protein | 22 | 0.79 | 0.164 | 0.820 |
| AT3G15290.1 | 3-hydroxyacyl-CoA dehydrogenase family protein | 23 | 0.78 | 0.161 | 0.781 |
| AT3G15290.1 | 3-hydroxyacyl-CoA dehydrogenase family protein | 24 | 0.76 | 0.159 | 0.742 |
| AT3G15290.1 | 3-hydroxyacyl-CoA dehydrogenase family protein | 33 | 0.62 | 0.171 | 0.464 |
| AT3G15290.1 | 3-hydroxyacyl-CoA dehydrogenase family protein | 34 | 0.61 | 0.175 | 0.450 |
| AT3G15290.1 | 3-hydroxyacyl-CoA dehydrogenase family protein | 35 | 0.59 | 0.180 | 0.439 |

| | | | | | |
|---|---|---|---|---|---|
| AT3G15290.1 | 3-hydroxyacyl-CoA dehydrogenase family protein | 36 | 0.58 | 0.186 | 0.430 |
| AT3G15290.1 | 3-hydroxyacyl-CoA dehydrogenase family protein | 37 | 0.57 | 0.192 | 0.423 |
| AT3G15290.1 | 3-hydroxyacyl-CoA dehydrogenase family protein | 38 | 0.56 | 0.198 | 0.419 |
| AT3G15290.1 | 3-hydroxyacyl-CoA dehydrogenase family protein | 41 | 0.52 | 0.219 | 0.413 |
| AT3G15290.1 | 3-hydroxyacyl-CoA dehydrogenase family protein | 42 | 0.51 | 0.226 | 0.413 |
| AT3G15290.1 | 3-hydroxyacyl-CoA dehydrogenase family protein | 43 | 0.50 | 0.234 | 0.415 |
| AT3G15290.1 | 3-hydroxyacyl-CoA dehydrogenase family protein | 46 | 0.46 | 0.258 | 0.420 |
| AT3G15290.1 | 3-hydroxyacyl-CoA dehydrogenase family protein | 47 | 0.45 | 0.267 | 0.422 |
| AT3G15640.2 | Cytochrome c:O2 oxidoreductase subunit (COX5B-1 | 2 | 1.49 | 0.374 | 1.000 |
| AT3G15640.2 | Cytochrome c:O2 oxidoreductase subunit (COX5B-1 | 3 | 1.63 | 0.231 | 1.000 |
| AT3G15640.2 | Cytochrome c:O2 oxidoreductase subunit (COX5B-1 | 4 | 1.66 | 0.220 | 1.000 |
| AT3G15640.2 | Cytochrome c:O2 oxidoreductase subunit (COX5B-1 | 5 | 1.35 | 0.219 | 1.000 |
| AT3G15640.2 | Cytochrome c:O2 oxidoreductase subunit (COX5B-1 | 6 | 1.02 | 0.220 | 1.000 |
| AT3G15640.2 | Cytochrome c:O2 oxidoreductase subunit (COX5B-1 | 7 | 0.93 | 0.220 | 1.000 |
| AT3G15640.2 | Cytochrome c:O2 oxidoreductase subunit (COX5B-1 | 8 | 1.14 | 0.216 | 1.000 |
| AT3G15640.2 | Cytochrome c:O2 oxidoreductase subunit (COX5B-1 | 9 | 1.38 | 0.213 | 1.000 |
| AT3G15640.2 | Cytochrome c:O2 oxidoreductase subunit (COX5B-1 | 10 | 1.53 | 0.213 | 1.000 |
| AT3G15640.2 | Cytochrome c:O2 oxidoreductase subunit (COX5B-1 | 11 | 1.68 | 0.214 | 1.000 |
| AT3G15640.2 | Cytochrome c:O2 oxidoreductase subunit (COX5B-1 | 12 | 1.61 | 0.214 | 1.000 |
| AT3G15640.2 | Cytochrome c:O2 oxidoreductase subunit (COX5B-1 | 13 | 1.48 | 0.211 | 1.000 |
| AT3G15640.2 | Cytochrome c:O2 oxidoreductase subunit (COX5B-1 | 14 | 1.30 | 0.208 | 1.000 |
| AT3G15640.2 | Cytochrome c:O2 oxidoreductase subunit (COX5B-1 | 15 | 1.08 | 0.209 | 1.000 |
| AT3G15640.2 | Cytochrome c:O2 oxidoreductase subunit (COX5B-1 | 16 | 0.92 | 0.207 | 1.000 |

| | | | | | |
|---|---|---|---|---|---|
| AT3G15640.2 | Cytochrome c:O2 oxidoreductase subunit (COX5B-1 | 17 | 0.89 | 0.207 | 1.000 |
| AT3G15640.2 | Cytochrome c:O2 oxidoreductase subunit (COX5B-1 | 18 | 0.93 | 0.208 | 1.000 |
| AT3G15640.2 | Cytochrome c:O2 oxidoreductase subunit (COX5B-1 | 19 | 1.04 | 0.213 | 1.000 |
| AT3G15640.2 | Cytochrome c:O2 oxidoreductase subunit (COX5B-1 | 20 | 1.25 | 0.219 | 1.000 |
| AT3G15640.2 | Cytochrome c:O2 oxidoreductase subunit (COX5B-1 | 21 | 1.41 | 0.221 | 1.000 |
| AT3G15640.2 | Cytochrome c:O2 oxidoreductase subunit (COX5B-1 | 22 | 1.35 | 0.221 | 1.000 |
| AT3G15640.2 | Cytochrome c:O2 oxidoreductase subunit (COX5B-1 | 23 | 1.19 | 0.221 | 1.000 |
| AT3G15640.2 | Cytochrome c:O2 oxidoreductase subunit (COX5B-1 | 24 | 1.07 | 0.215 | 1.000 |
| AT3G15640.2 | Cytochrome c:O2 oxidoreductase subunit (COX5B-1 | 25 | 1.04 | 0.211 | 1.000 |
| AT3G15640.2 | Cytochrome c:O2 oxidoreductase subunit (COX5B-1 | 26 | 1.20 | 0.211 | 1.000 |
| AT3G15640.2 | Cytochrome c:O2 oxidoreductase subunit (COX5B-1 | 27 | 1.59 | 0.211 | 1.000 |
| AT3G15640.2 | Cytochrome c:O2 oxidoreductase subunit (COX5B-1 | 28 | 1.92 | 0.210 | 0.987 |
| AT3G15640.2 | Cytochrome c:O2 oxidoreductase subunit (COX5B-1 | 29 | 2.03 | 0.211 | 0.968 |
| AT3G15640.2 | Cytochrome c:O2 oxidoreductase subunit (COX5B-1 | 30 | 1.80 | 0.213 | 0.997 |
| AT3G15640.2 | Cytochrome c:O2 oxidoreductase subunit (COX5B-1 | 31 | 1.42 | 0.214 | 1.000 |
| AT3G15640.2 | Cytochrome c:O2 oxidoreductase subunit (COX5B-1 | 32 | 1.06 | 0.215 | 1.000 |
| AT3G15640.2 | Cytochrome c:O2 oxidoreductase subunit (COX5B-1 | 33 | 0.67 | 0.215 | 1.000 |
| AT3G15640.2 | Cytochrome c:O2 oxidoreductase subunit (COX5B-1 | 34 | 0.37 | 0.220 | 0.713 |
| AT3G15640.2 | Cytochrome c:O2 oxidoreductase subunit (COX5B-1 | 35 | 0.26 | 0.231 | 0.286 |
| AT3G15640.2 | Cytochrome c:O2 oxidoreductase subunit (COX5B-1 | 36 | 0.28 | 0.275 | 0.646 |

| | | | | | | |
|---|---|---|---|---|---|---|
| AT3G15640.2 | Cytochrome c:O2 oxidoreductase subunit (COX5B-1 | 37 | 0.44 | 0.424 | 1.000 | |
| AT3G15640.2 | Cytochrome c:O2 oxidoreductase subunit (COX5B-1 | 46 | 15307531.83 | 4.120 | 0.858 | |
| AT3G15640.2 | Cytochrome c:O2 oxidoreductase subunit (COX5B-1 | 47 | ########### | 4.741 | 0.850 | |
| AT3G15660.2 | Glutaredoxin 4 (GRX4) | 2 | 0.99 | 0.372 | 1.000 | |
| AT3G15660.2 | Glutaredoxin 4 (GRX4) | 3 | 0.91 | 0.317 | 1.000 | |
| AT3G15660.2 | Glutaredoxin 4 (GRX4) | 4 | 0.85 | 0.324 | 1.000 | |
| AT3G15660.2 | Glutaredoxin 4 (GRX4) | 13 | 0.68 | 0.382 | 1.000 | |
| AT3G15660.2 | Glutaredoxin 4 (GRX4) | 14 | 0.67 | 0.368 | 1.000 | |
| AT3G15660.2 | Glutaredoxin 4 (GRX4) | 15 | 0.67 | 0.368 | 1.000 | |
| AT3G15660.2 | Glutaredoxin 4 (GRX4) | 23 | 0.69 | 0.283 | 1.000 | |
| AT3G15660.2 | Glutaredoxin 4 (GRX4) | 24 | 0.77 | 0.253 | 1.000 | |
| AT3G15660.2 | Glutaredoxin 4 (GRX4) | 25 | 0.91 | 0.220 | 1.000 | |
| AT3G15660.2 | Glutaredoxin 4 (GRX4) | 26 | 1.08 | 0.198 | 1.000 | |
| AT3G15660.2 | Glutaredoxin 4 (GRX4) | 27 | 1.25 | 0.183 | 1.000 | |
| AT3G15660.2 | Glutaredoxin 4 (GRX4) | 28 | 1.37 | 0.175 | 0.999 | |
| AT3G15660.2 | Glutaredoxin 4 (GRX4) | 29 | 1.43 | 0.172 | 0.996 | |
| AT3G15660.2 | Glutaredoxin 4 (GRX4) | 30 | 1.44 | 0.171 | 0.994 | |
| AT3G15660.2 | Glutaredoxin 4 (GRX4) | 31 | 1.41 | 0.171 | 0.996 | |
| AT3G15660.2 | Glutaredoxin 4 (GRX4) | 32 | 1.31 | 0.171 | 1.000 | |
| AT3G15660.2 | Glutaredoxin 4 (GRX4) | 33 | 1.16 | 0.172 | 1.000 | |
| AT3G15660.2 | Glutaredoxin 4 (GRX4) | 34 | 1.02 | 0.173 | 1.000 | |
| AT3G15660.2 | Glutaredoxin 4 (GRX4) | 35 | 0.92 | 0.175 | 1.000 | |
| AT3G15660.2 | Glutaredoxin 4 (GRX4) | 36 | 0.86 | 0.178 | 1.000 | |
| AT3G15660.2 | Glutaredoxin 4 (GRX4) | 37 | 0.83 | 0.182 | 1.000 | |
| AT3G15660.2 | Glutaredoxin 4 (GRX4) | 38 | 0.82 | 0.183 | 1.000 | |
| AT3G15660.2 | Glutaredoxin 4 (GRX4) | 39 | 0.81 | 0.182 | 1.000 | |
| AT3G15660.2 | Glutaredoxin 4 (GRX4) | 40 | 0.80 | 0.179 | 1.000 | |
| AT3G15660.2 | Glutaredoxin 4 (GRX4) | 41 | 0.78 | 0.178 | 1.000 | |
| AT3G15660.2 | Glutaredoxin 4 (GRX4) | 42 | 0.79 | 0.176 | 1.000 | |
| AT3G15660.2 | Glutaredoxin 4 (GRX4) | 43 | 0.81 | 0.176 | 1.000 | |
| AT3G15660.2 | Glutaredoxin 4 (GRX4) | 44 | 0.81 | 0.178 | 1.000 | |
| AT3G15660.2 | Glutaredoxin 4 (GRX4) | 45 | 0.74 | 0.195 | 1.000 | |
| AT3G15660.2 | Glutaredoxin 4 (GRX4) | 46 | 0.63 | 0.243 | 0.998 | |
| AT3G15660.2 | Glutaredoxin 4 (GRX4) | 47 | 0.52 | 0.334 | 0.997 | |
| AT3G16230.3 | Predicted eukaryotic LigT | 2 | 2.32 | 0.645 | 0.841 | |
| AT3G16230.3 | Predicted eukaryotic LigT | 3 | 2.27 | 0.629 | 0.840 | |
| AT3G16230.3 | Predicted eukaryotic LigT | 29 | 1.35 | 0.264 | 0.878 | |
| AT3G16230.3 | Predicted eukaryotic LigT | 30 | 1.32 | 0.257 | 0.886 | |
| AT3G16230.3 | Predicted eukaryotic LigT | 31 | 1.30 | 0.250 | 0.896 | |
| AT3G16230.3 | Predicted eukaryotic LigT | 32 | 1.27 | 0.244 | 0.907 | |
| AT3G16230.3 | Predicted eukaryotic LigT | 43 | 1.02 | 0.264 | 1.000 | |
| AT3G16230.3 | Predicted eukaryotic LigT | 44 | 1.00 | 0.273 | 1.000 | |
| AT3G16230.3 | Predicted eukaryotic LigT | 45 | 0.98 | 0.282 | 0.999 | |
| AT3G16230.3 | Predicted eukaryotic LigT | 46 | 0.96 | 0.293 | 0.998 | |
| AT3G16230.3 | Predicted eukaryotic LigT | 47 | 0.94 | 0.304 | 0.996 | |
| AT3G16240.1 | Delta tonoplast integral protein 2;1 (TIP2;1) | 2 | 2.18 | 0.193 | 0.404 | |
| AT3G16240.1 | Delta tonoplast integral protein 2;1 (TIP2;1) | 3 | 2.21 | 0.166 | 0.227 | |
| AT3G16240.1 | Delta tonoplast integral protein 2;1 (TIP2;1) | 4 | 2.24 | 0.144 | 0.106 | |
| AT3G16240.1 | Delta tonoplast integral protein 2;1 (TIP2;1) | 5 | 2.26 | 0.127 | 0.046 | * |
| AT3G16240.1 | Delta tonoplast integral protein 2;1 (TIP2;1) | 6 | 2.26 | 0.117 | 0.022 | * |
| AT3G16240.1 | Delta tonoplast integral protein 2;1 (TIP2;1) | 7 | 2.26 | 0.111 | 0.014 | * |

| | | | | | | |
|---|---|---|---|---|---|---|
| AT3G16240.1 | Delta tonoplast integral protein 2;1 (TIP2;1) | 8 | 2.24 | 0.110 | 0.014 | * |
| AT3G16240.1 | Delta tonoplast integral protein 2;1 (TIP2;1) | 9 | 2.22 | 0.113 | 0.019 | * |
| AT3G16240.1 | Delta tonoplast integral protein 2;1 (TIP2;1) | 10 | 2.19 | 0.117 | 0.031 | * |
| AT3G16240.1 | Delta tonoplast integral protein 2;1 (TIP2;1) | 11 | 2.17 | 0.123 | 0.051 | . |
| AT3G16240.1 | Delta tonoplast integral protein 2;1 (TIP2;1) | 12 | 2.15 | 0.130 | 0.081 | . |
| AT3G16240.1 | Delta tonoplast integral protein 2;1 (TIP2;1) | 13 | 2.13 | 0.139 | 0.125 | |
| AT3G16240.1 | Delta tonoplast integral protein 2;1 (TIP2;1) | 14 | 2.11 | 0.147 | 0.179 | |
| AT3G16240.1 | Delta tonoplast integral protein 2;1 (TIP2;1) | 15 | 2.08 | 0.156 | 0.247 | |
| AT3G16240.1 | Delta tonoplast integral protein 2;1 (TIP2;1) | 16 | 2.06 | 0.165 | 0.322 | |
| AT3G16240.1 | Delta tonoplast integral protein 2;1 (TIP2;1) | 17 | 2.03 | 0.174 | 0.400 | |
| AT3G16240.1 | Delta tonoplast integral protein 2;1 (TIP2;1) | 31 | 1.06 | 0.166 | 1.000 | |
| AT3G16240.1 | Delta tonoplast integral protein 2;1 (TIP2;1) | 32 | 0.95 | 0.163 | 1.000 | |
| AT3G16240.1 | Delta tonoplast integral protein 2;1 (TIP2;1) | 33 | 0.83 | 0.162 | 0.997 | |
| AT3G16240.1 | Delta tonoplast integral protein 2;1 (TIP2;1) | 34 | 0.72 | 0.167 | 0.942 | |
| AT3G16240.1 | Delta tonoplast integral protein 2;1 (TIP2;1) | 35 | 0.61 | 0.175 | 0.776 | |
| AT3G16240.1 | Delta tonoplast integral protein 2;1 (TIP2;1) | 36 | 0.52 | 0.188 | 0.568 | |
| AT3G16240.1 | Delta tonoplast integral protein 2;1 (TIP2;1) | 46 | 0.07 | 0.463 | 0.081 | . |
| AT3G16240.1 | Delta tonoplast integral protein 2;1 (TIP2;1) | 47 | 0.05 | 0.503 | 0.082 | . |
| AT3G16480.1 | Ubiquinol:cytochrome c oxidoreductase subunit (MPPalpha) | 2 | 0.97 | 0.775 | 1.000 | |
| AT3G16480.1 | Ubiquinol:cytochrome c oxidoreductase subunit (MPPalpha) | 3 | 0.94 | 0.556 | 1.000 | |
| AT3G16480.1 | Ubiquinol:cytochrome c oxidoreductase subunit (MPPalpha) | 5 | 0.85 | 0.541 | 1.000 | |
| AT3G16480.1 | Ubiquinol:cytochrome c oxidoreductase subunit (MPPalpha) | 6 | 0.77 | 0.417 | 1.000 | |
| AT3G16480.1 | Ubiquinol:cytochrome c oxidoreductase subunit (MPPalpha) | 7 | 0.75 | 0.378 | 1.000 | |
| AT3G16480.1 | Ubiquinol:cytochrome c oxidoreductase subunit (MPPalpha) | 8 | 0.94 | 0.359 | 1.000 | |
| AT3G16480.1 | Ubiquinol:cytochrome c oxidoreductase subunit (MPPalpha) | 9 | 1.20 | 0.348 | 1.000 | |
| AT3G16480.1 | Ubiquinol:cytochrome c oxidoreductase subunit (MPPalpha) | 10 | 1.17 | 0.348 | 1.000 | |

| | | | | | |
|---|---|---|---|---|---|
| AT3G16480.1 | Ubiquinol:cytochrome c oxidoreductase subunit (MPPalpha) | 11 | 1.27 | 0.355 | 1.000 |
| AT3G16480.1 | Ubiquinol:cytochrome c oxidoreductase subunit (MPPalpha) | 12 | 1.72 | 0.355 | 1.000 |
| AT3G16480.1 | Ubiquinol:cytochrome c oxidoreductase subunit (MPPalpha) | 13 | 2.73 | 0.350 | 0.997 |
| AT3G16480.1 | Ubiquinol:cytochrome c oxidoreductase subunit (MPPalpha) | 14 | 3.93 | 0.343 | 0.898 |
| AT3G16480.1 | Ubiquinol:cytochrome c oxidoreductase subunit (MPPalpha) | 15 | 3.19 | 0.341 | 0.978 |
| AT3G16480.1 | Ubiquinol:cytochrome c oxidoreductase subunit (MPPalpha) | 16 | 2.27 | 0.337 | 1.000 |
| AT3G16480.1 | Ubiquinol:cytochrome c oxidoreductase subunit (MPPalpha) | 17 | 1.84 | 0.337 | 1.000 |
| AT3G16480.1 | Ubiquinol:cytochrome c oxidoreductase subunit (MPPalpha) | 18 | 1.24 | 0.342 | 1.000 |
| AT3G16480.1 | Ubiquinol:cytochrome c oxidoreductase subunit (MPPalpha) | 19 | 0.85 | 0.349 | 1.000 |
| AT3G16480.1 | Ubiquinol:cytochrome c oxidoreductase subunit (MPPalpha) | 20 | 0.90 | 0.361 | 1.000 |
| AT3G16480.1 | Ubiquinol:cytochrome c oxidoreductase subunit (MPPalpha) | 21 | 1.46 | 0.364 | 1.000 |
| AT3G16480.1 | Ubiquinol:cytochrome c oxidoreductase subunit (MPPalpha) | 22 | 2.20 | 0.364 | 1.000 |
| AT3G16480.1 | Ubiquinol:cytochrome c oxidoreductase subunit (MPPalpha) | 23 | 1.15 | 0.363 | 1.000 |
| AT3G16480.1 | Ubiquinol:cytochrome c oxidoreductase subunit (MPPalpha) | 24 | 0.31 | 0.350 | 0.984 |
| AT3G16480.1 | Ubiquinol:cytochrome c oxidoreductase subunit (MPPalpha) | 25 | 0.29 | 0.346 | 0.963 |
| AT3G16480.1 | Ubiquinol:cytochrome c oxidoreductase subunit (MPPalpha) | 26 | 0.94 | 0.348 | 1.000 |
| AT3G16480.1 | Ubiquinol:cytochrome c oxidoreductase subunit (MPPalpha) | 27 | 3.08 | 0.351 | 0.989 |
| AT3G16480.1 | Ubiquinol:cytochrome c oxidoreductase subunit (MPPalpha) | 28 | 2.42 | 0.350 | 1.000 |
| AT3G16480.1 | Ubiquinol:cytochrome c oxidoreductase subunit (MPPalpha) | 29 | 0.56 | 0.348 | 1.000 |
| AT3G16480.1 | Ubiquinol:cytochrome c oxidoreductase subunit (MPPalpha) | 30 | 0.46 | 0.346 | 1.000 |

| | | | | | |
|---|---|---|---|---|---|
| AT3G16480.1 | Ubiquinol:cytochrome c oxidoreductase subunit (MPPalpha) | 31 | 1.60 | 0.348 | 1.000 |
| AT3G16480.1 | Ubiquinol:cytochrome c oxidoreductase subunit (MPPalpha) | 32 | 4.40 | 0.351 | 0.842 |
| AT3G16480.1 | Ubiquinol:cytochrome c oxidoreductase subunit (MPPalpha) | 33 | 4.26 | 0.360 | 0.889 |
| AT3G16480.1 | Ubiquinol:cytochrome c oxidoreductase subunit (MPPalpha) | 34 | 3.33 | 0.383 | 0.991 |
| AT3G16480.1 | Ubiquinol:cytochrome c oxidoreductase subunit (MPPalpha) | 35 | 3.63 | 0.449 | 0.997 |
| AT3G16480.1 | Ubiquinol:cytochrome c oxidoreductase subunit (MPPalpha) | 36 | 4.10 | 0.776 | 1.000 |
| AT3G16480.1 | Ubiquinol:cytochrome c oxidoreductase subunit (MPPalpha) | 46 | 0.00 | 9.657 | 1.000 |
| AT3G16480.1 | Ubiquinol:cytochrome c oxidoreductase subunit (MPPalpha) | 47 | 0.00 | 11.015 | 1.000 |
| AT3G17240.3 | Dihydrolipoamide dehydrogenase (PDCE3, OGDCE3, GDCE3, BCKDHE3) | 2 | 1.00 | 0.504 | 1.000 |
| AT3G17240.3 | Dihydrolipoamide dehydrogenase (PDCE3, OGDCE3, GDCE3, BCKDHE3) | 3 | 0.63 | 0.352 | 1.000 |
| AT3G17240.3 | Dihydrolipoamide dehydrogenase (PDCE3, OGDCE3, GDCE3, BCKDHE3) | 4 | 0.50 | 0.438 | 1.000 |
| AT3G17240.3 | Dihydrolipoamide dehydrogenase (PDCE3, OGDCE3, GDCE3, BCKDHE3) | 7 | 0.52 | 0.394 | 1.000 |
| AT3G17240.3 | Dihydrolipoamide dehydrogenase (PDCE3, OGDCE3, GDCE3, BCKDHE3) | 8 | 0.45 | 0.319 | 0.998 |
| AT3G17240.3 | Dihydrolipoamide dehydrogenase (PDCE3, OGDCE3, GDCE3, BCKDHE3) | 9 | 0.28 | 0.291 | 0.693 |
| AT3G17240.3 | Dihydrolipoamide dehydrogenase (PDCE3, OGDCE3, GDCE3, BCKDHE3) | 10 | 0.23 | 0.298 | 0.513 |
| AT3G17240.3 | Dihydrolipoamide dehydrogenase (PDCE3, OGDCE3, GDCE3, BCKDHE3) | 11 | 0.63 | 0.320 | 1.000 |
| AT3G17240.3 | Dihydrolipoamide dehydrogenase (PDCE3, OGDCE3, GDCE3, BCKDHE3) | 12 | 3.60 | 0.328 | 0.843 |
| AT3G17240.3 | Dihydrolipoamide dehydrogenase (PDCE3, OGDCE3, GDCE3, BCKDHE3) | 30 | 0.36 | 0.391 | 0.996 |
| AT3G17240.3 | Dihydrolipoamide dehydrogenase (PDCE3, OGDCE3, GDCE3, BCKDHE3) | 31 | 1.14 | 0.312 | 1.000 |
| AT3G17240.3 | Dihydrolipoamide dehydrogenase (PDCE3, OGDCE3, GDCE3, BCKDHE3) | 32 | 2.06 | 0.283 | 0.997 |

| | | | | | | |
|---|---|---|---|---|---|---|
| AT3G17240.3 | Dihydrolipoamide dehydrogenase (PDCE3, OGDCE3, GDCE3, BCKDHE3) | 33 | 2.16 | 0.268 | 0.989 | |
| AT3G17240.3 | Dihydrolipoamide dehydrogenase (PDCE3, OGDCE3, GDCE3, BCKDHE3) | 34 | 2.06 | 0.276 | 0.996 | |
| AT3G17240.3 | Dihydrolipoamide dehydrogenase (PDCE3, OGDCE3, GDCE3, BCKDHE3) | 35 | 2.13 | 0.283 | 0.995 | |
| AT3G17240.3 | Dihydrolipoamide dehydrogenase (PDCE3, OGDCE3, GDCE3, BCKDHE3) | 36 | 2.63 | 0.291 | 0.956 | |
| AT3G17240.3 | Dihydrolipoamide dehydrogenase (PDCE3, OGDCE3, GDCE3, BCKDHE3) | 37 | 3.46 | 0.289 | 0.730 | |
| AT3G17240.3 | Dihydrolipoamide dehydrogenase (PDCE3, OGDCE3, GDCE3, BCKDHE3) | 38 | 4.25 | 0.269 | 0.355 | |
| AT3G17240.3 | Dihydrolipoamide dehydrogenase (PDCE3, OGDCE3, GDCE3, BCKDHE3) | 39 | 4.91 | 0.259 | 0.167 | |
| AT3G17240.3 | Dihydrolipoamide dehydrogenase (PDCE3, OGDCE3, GDCE3, BCKDHE3) | 40 | 5.71 | 0.257 | 0.080 | . |
| AT3G17240.3 | Dihydrolipoamide dehydrogenase (PDCE3, OGDCE3, GDCE3, BCKDHE3) | 41 | 3.77 | 0.260 | 0.442 | |
| AT3G17240.3 | Dihydrolipoamide dehydrogenase (PDCE3, OGDCE3, GDCE3, BCKDHE3) | 42 | 1.49 | 0.265 | 1.000 | |
| AT3G17240.3 | Dihydrolipoamide dehydrogenase (PDCE3, OGDCE3, GDCE3, BCKDHE3) | 43 | 0.60 | 0.265 | 1.000 | |
| AT3G17240.3 | Dihydrolipoamide dehydrogenase (PDCE3, OGDCE3, GDCE3, BCKDHE3) | 44 | 0.46 | 0.263 | 0.984 | |
| AT3G17240.3 | Dihydrolipoamide dehydrogenase (PDCE3, OGDCE3, GDCE3, BCKDHE3) | 45 | 0.62 | 0.263 | 1.000 | |
| AT3G17240.3 | Dihydrolipoamide dehydrogenase (PDCE3, OGDCE3, GDCE3, BCKDHE3) | 46 | 0.74 | 0.289 | 1.000 | |
| AT3G17240.3 | Dihydrolipoamide dehydrogenase (PDCE3, OGDCE3, GDCE3, BCKDHE3) | 47 | 0.74 | 0.601 | 1.000 | |
| AT3G17465.1 | 50S ribosomal protein L3-2 (RPL3B) | 2 | 1.25 | 0.779 | 1.000 | |
| AT3G17465.1 | 50S ribosomal protein L3-2 (RPL3B) | 3 | 1.56 | 0.699 | 1.000 | |
| AT3G17465.1 | 50S ribosomal protein L3-2 (RPL3B) | 13 | 6.15 | 0.410 | 0.496 | |
| AT3G17465.1 | 50S ribosomal protein L3-2 (RPL3B) | 14 | 5.90 | 0.354 | 0.320 | |
| AT3G17465.1 | 50S ribosomal protein L3-2 (RPL3B) | 15 | 5.26 | 0.314 | 0.255 | |
| AT3G17465.1 | 50S ribosomal protein L3-2 (RPL3B) | 16 | 4.49 | 0.289 | 0.275 | |
| AT3G17465.1 | 50S ribosomal protein L3-2 (RPL3B) | 17 | 4.08 | 0.278 | 0.310 | |
| AT3G17465.1 | 50S ribosomal protein L3-2 (RPL3B) | 18 | 4.07 | 0.279 | 0.315 | |

| | | | | | |
|---|---|---|---|---|---|
| AT3G17465.1 | 50S ribosomal protein L3-2 (RPL3B) | 19 | 4.23 | 0.288 | 0.322 |
| AT3G17465.1 | 50S ribosomal protein L3-2 (RPL3B) | 20 | 4.31 | 0.304 | 0.381 |
| AT3G17465.1 | 50S ribosomal protein L3-2 (RPL3B) | 21 | 4.17 | 0.328 | 0.517 |
| AT3G17465.1 | 50S ribosomal protein L3-2 (RPL3B) | 22 | 3.72 | 0.354 | 0.726 |
| AT3G17465.1 | 50S ribosomal protein L3-2 (RPL3B) | 23 | 3.12 | 0.380 | 0.906 |
| AT3G17465.1 | 50S ribosomal protein L3-2 (RPL3B) | 24 | 2.52 | 0.397 | 0.983 |
| AT3G17465.1 | 50S ribosomal protein L3-2 (RPL3B) | 27 | 1.21 | 0.363 | 1.000 |
| AT3G17465.1 | 50S ribosomal protein L3-2 (RPL3B) | 28 | 0.96 | 0.332 | 1.000 |
| AT3G17465.1 | 50S ribosomal protein L3-2 (RPL3B) | 29 | 0.77 | 0.308 | 1.000 |
| AT3G17465.1 | 50S ribosomal protein L3-2 (RPL3B) | 30 | 0.63 | 0.296 | 1.000 |
| AT3G17465.1 | 50S ribosomal protein L3-2 (RPL3B) | 31 | 0.54 | 0.299 | 0.993 |
| AT3G17465.1 | 50S ribosomal protein L3-2 (RPL3B) | 32 | 0.49 | 0.313 | 0.987 |
| AT3G17465.1 | 50S ribosomal protein L3-2 (RPL3B) | 33 | 0.50 | 0.332 | 0.992 |
| AT3G17465.1 | 50S ribosomal protein L3-2 (RPL3B) | 34 | 0.55 | 0.356 | 0.999 |
| AT3G17465.1 | 50S ribosomal protein L3-2 (RPL3B) | 35 | 0.65 | 0.383 | 1.000 |
| AT3G17465.1 | 50S ribosomal protein L3-2 (RPL3B) | 36 | 0.77 | 0.425 | 1.000 |
| AT3G17465.1 | 50S ribosomal protein L3-2 (RPL3B) | 37 | 0.87 | 0.487 | 1.000 |
| AT3G17465.1 | 50S ribosomal protein L3-2 (RPL3B) | 38 | 0.92 | 0.590 | 1.000 |
| AT3G17465.1 | 50S ribosomal protein L3-2 (RPL3B) | 46 | 0.27 | 2.445 | 1.000 |
| AT3G17465.1 | 50S ribosomal protein L3-2 (RPL3B) | 47 | 0.21 | 2.792 | 1.000 |
| AT3G17910.1 | Cytochrome c:O2 oxidoreductase assembly protein (SURF1) | 2 | 0.98 | 0.490 | 1.000 |
| AT3G17910.1 | Cytochrome c:O2 oxidoreductase assembly protein (SURF1) | 3 | 0.99 | 0.373 | 1.000 |
| AT3G17910.1 | Cytochrome c:O2 oxidoreductase assembly protein (SURF1) | 6 | 0.93 | 0.344 | 1.000 |
| AT3G17910.1 | Cytochrome c:O2 oxidoreductase assembly protein (SURF1) | 7 | 0.88 | 0.268 | 1.000 |
| AT3G17910.1 | Cytochrome c:O2 oxidoreductase assembly protein (SURF1) | 8 | 0.89 | 0.231 | 1.000 |
| AT3G17910.1 | Cytochrome c:O2 oxidoreductase assembly protein (SURF1) | 9 | 1.01 | 0.218 | 1.000 |
| AT3G17910.1 | Cytochrome c:O2 oxidoreductase assembly protein (SURF1) | 10 | 1.18 | 0.214 | 1.000 |

| | | | | | |
|---|---|---|---|---|---|
| AT3G17910.1 | Cytochrome c:O2 oxidoreductase assembly protein (SURF1) | 11 | 1.32 | 0.216 | 1.000 |
| AT3G17910.1 | Cytochrome c:O2 oxidoreductase assembly protein (SURF1) | 12 | 1.59 | 0.215 | 1.000 |
| AT3G17910.1 | Cytochrome c:O2 oxidoreductase assembly protein (SURF1) | 13 | 1.73 | 0.212 | 0.999 |
| AT3G17910.1 | Cytochrome c:O2 oxidoreductase assembly protein (SURF1) | 14 | 1.65 | 0.209 | 1.000 |
| AT3G17910.1 | Cytochrome c:O2 oxidoreductase assembly protein (SURF1) | 15 | 1.42 | 0.208 | 1.000 |
| AT3G17910.1 | Cytochrome c:O2 oxidoreductase assembly protein (SURF1) | 16 | 1.21 | 0.206 | 1.000 |
| AT3G17910.1 | Cytochrome c:O2 oxidoreductase assembly protein (SURF1) | 17 | 1.34 | 0.207 | 1.000 |
| AT3G17910.1 | Cytochrome c:O2 oxidoreductase assembly protein (SURF1) | 18 | 1.88 | 0.212 | 0.994 |
| AT3G17910.1 | Cytochrome c:O2 oxidoreductase assembly protein (SURF1) | 19 | 2.60 | 0.223 | 0.809 |
| AT3G17910.1 | Cytochrome c:O2 oxidoreductase assembly protein (SURF1) | 20 | 3.06 | 0.237 | 0.661 |
| AT3G17910.1 | Cytochrome c:O2 oxidoreductase assembly protein (SURF1) | 21 | 2.82 | 0.290 | 0.955 |
| AT3G17910.1 | Cytochrome c:O2 oxidoreductase assembly protein (SURF1) | 22 | 2.23 | 0.430 | 1.000 |
| AT3G17910.1 | Cytochrome c:O2 oxidoreductase assembly protein (SURF1) | 30 | 3.10 | 0.324 | 0.966 |
| AT3G17910.1 | Cytochrome c:O2 oxidoreductase assembly protein (SURF1) | 31 | 3.70 | 0.244 | 0.429 |
| AT3G17910.1 | Cytochrome c:O2 oxidoreductase assembly protein (SURF1) | 32 | 3.89 | 0.232 | 0.277 |
| AT3G17910.1 | Cytochrome c:O2 oxidoreductase assembly protein (SURF1) | 33 | 2.86 | 0.219 | 0.637 |
| AT3G17910.1 | Cytochrome c:O2 oxidoreductase assembly protein (SURF1) | 34 | 1.57 | 0.215 | 1.000 |
| AT3G17910.1 | Cytochrome c:O2 oxidoreductase assembly protein (SURF1) | 35 | 0.99 | 0.214 | 1.000 |
| AT3G17910.1 | Cytochrome c:O2 oxidoreductase assembly protein (SURF1) | 36 | 0.89 | 0.215 | 1.000 |
| AT3G17910.1 | Cytochrome c:O2 oxidoreductase assembly protein (SURF1) | 37 | 1.22 | 0.222 | 1.000 |

| | | | | | |
|---|---|---|---|---|---|
| AT3G17910.1 | Cytochrome c:O2 oxidoreductase assembly protein (SURF1) | 38 | 1.76 | 0.234 | 1.000 |
| AT3G17910.1 | Cytochrome c:O2 oxidoreductase assembly protein (SURF1) | 39 | 2.21 | 0.238 | 0.979 |
| AT3G17910.1 | Cytochrome c:O2 oxidoreductase assembly protein (SURF1) | 40 | 2.69 | 0.234 | 0.822 |
| AT3G17910.1 | Cytochrome c:O2 oxidoreductase assembly protein (SURF1) | 41 | 3.43 | 0.220 | 0.347 |
| AT3G17910.1 | Cytochrome c:O2 oxidoreductase assembly protein (SURF1) | 42 | 3.93 | 0.217 | 0.174 |
| AT3G17910.1 | Cytochrome c:O2 oxidoreductase assembly protein (SURF1) | 43 | 2.90 | 0.218 | 0.602 |
| AT3G17910.1 | Cytochrome c:O2 oxidoreductase assembly protein (SURF1) | 44 | 1.49 | 0.218 | 1.000 |
| AT3G17910.1 | Cytochrome c:O2 oxidoreductase assembly protein (SURF1) | 45 | 0.68 | 0.218 | 1.000 |
| AT3G17910.1 | Cytochrome c:O2 oxidoreductase assembly protein (SURF1) | 46 | 0.34 | 0.249 | 0.777 |
| AT3G17910.1 | Cytochrome c:O2 oxidoreductase assembly protein (SURF1) | 47 | 0.15 | 0.479 | 0.887 |
| AT3G18240.2 | Ribosomal S24/S35 family protein | 2 | 1.12 | 0.610 | 1.000 |
| AT3G18240.2 | Ribosomal S24/S35 family protein | 3 | 1.24 | 0.531 | 1.000 |
| AT3G18240.2 | Ribosomal S24/S35 family protein | 11 | 2.06 | 0.315 | 0.997 |
| AT3G18240.2 | Ribosomal S24/S35 family protein | 12 | 1.99 | 0.270 | 0.991 |
| AT3G18240.2 | Ribosomal S24/S35 family protein | 13 | 1.79 | 0.238 | 0.994 |
| AT3G18240.2 | Ribosomal S24/S35 family protein | 14 | 1.55 | 0.217 | 0.999 |
| AT3G18240.2 | Ribosomal S24/S35 family protein | 15 | 1.45 | 0.206 | 1.000 |
| AT3G18240.2 | Ribosomal S24/S35 family protein | 16 | 1.47 | 0.203 | 1.000 |
| AT3G18240.2 | Ribosomal S24/S35 family protein | 17 | 1.41 | 0.204 | 1.000 |
| AT3G18240.2 | Ribosomal S24/S35 family protein | 18 | 1.22 | 0.209 | 1.000 |
| AT3G18240.2 | Ribosomal S24/S35 family protein | 19 | 0.98 | 0.218 | 1.000 |
| AT3G18240.2 | Ribosomal S24/S35 family protein | 20 | 0.78 | 0.223 | 1.000 |
| AT3G18240.2 | Ribosomal S24/S35 family protein | 21 | 0.65 | 0.224 | 1.000 |
| AT3G18240.2 | Ribosomal S24/S35 family protein | 22 | 0.59 | 0.219 | 0.996 |
| AT3G18240.2 | Ribosomal S24/S35 family protein | 23 | 0.60 | 0.213 | 0.995 |
| AT3G18240.2 | Ribosomal S24/S35 family protein | 24 | 0.65 | 0.207 | 0.999 |

| | | | | | |
|---|---|---|---|---|---|
| AT3G18240.2 | Ribosomal S24/S35 family protein | 25 | 0.76 | 0.203 | 1.000 |
| AT3G18240.2 | Ribosomal S24/S35 family protein | 26 | 0.95 | 0.201 | 1.000 |
| AT3G18240.2 | Ribosomal S24/S35 family protein | 27 | 1.22 | 0.201 | 1.000 |
| AT3G18240.2 | Ribosomal S24/S35 family protein | 28 | 1.43 | 0.201 | 1.000 |
| AT3G18240.2 | Ribosomal S24/S35 family protein | 29 | 1.49 | 0.201 | 0.999 |
| AT3G18240.2 | Ribosomal S24/S35 family protein | 30 | 1.43 | 0.202 | 1.000 |
| AT3G18240.2 | Ribosomal S24/S35 family protein | 31 | 1.34 | 0.206 | 1.000 |
| AT3G18240.2 | Ribosomal S24/S35 family protein | 32 | 1.21 | 0.216 | 1.000 |
| AT3G18240.2 | Ribosomal S24/S35 family protein | 33 | 1.02 | 0.236 | 1.000 |
| AT3G18240.2 | Ribosomal S24/S35 family protein | 34 | 0.88 | 0.267 | 1.000 |
| AT3G18240.2 | Ribosomal S24/S35 family protein | 35 | 0.85 | 0.304 | 1.000 |
| AT3G18240.2 | Ribosomal S24/S35 family protein | 39 | 1.95 | 0.290 | 0.997 |
| AT3G18240.2 | Ribosomal S24/S35 family protein | 40 | 2.45 | 0.258 | 0.893 |
| AT3G18240.2 | Ribosomal S24/S35 family protein | 41 | 2.80 | 0.232 | 0.611 |
| AT3G18240.2 | Ribosomal S24/S35 family protein | 42 | 2.77 | 0.218 | 0.525 |
| AT3G18240.2 | Ribosomal S24/S35 family protein | 43 | 2.31 | 0.211 | 0.766 |
| AT3G18240.2 | Ribosomal S24/S35 family protein | 44 | 1.59 | 0.211 | 0.998 |
| AT3G18240.2 | Ribosomal S24/S35 family protein | 45 | 0.94 | 0.228 | 1.000 |
| AT3G18240.2 | Ribosomal S24/S35 family protein | 46 | 0.52 | 0.290 | 0.997 |
| AT3G18240.2 | Ribosomal S24/S35 family protein | 47 | 0.28 | 0.417 | 0.953 |
| AT3G18410.2 | NADH:ubiquinone oxidoreductase subunit (NDUFS6) | 2 | 0.93 | 0.580 | 1.000 |
| AT3G18410.2 | NADH:ubiquinone oxidoreductase subunit (NDUFS6) | 3 | 0.84 | 0.522 | 1.000 |
| AT3G18410.2 | NADH:ubiquinone oxidoreductase subunit (NDUFS6) | 13 | 1.14 | 0.342 | 1.000 |
| AT3G18410.2 | NADH:ubiquinone oxidoreductase subunit (NDUFS6) | 14 | 1.30 | 0.306 | 1.000 |
| AT3G18410.2 | NADH:ubiquinone oxidoreductase subunit (NDUFS6) | 15 | 1.22 | 0.285 | 1.000 |
| AT3G18410.2 | NADH:ubiquinone oxidoreductase subunit (NDUFS6) | 16 | 0.81 | 0.255 | 1.000 |
| AT3G18410.2 | NADH:ubiquinone oxidoreductase subunit (NDUFS6) | 17 | 0.54 | 0.241 | 0.991 |

| | | | | | |
|---|---|---|---|---|---|
| AT3G18410.2 | NADH:ubiquinone oxidoreductase subunit (NDUFS6) | 18 | 0.57 | 0.242 | 0.996 |
| AT3G18410.2 | NADH:ubiquinone oxidoreductase subunit (NDUFS6) | 19 | 0.71 | 0.252 | 1.000 |
| AT3G18410.2 | NADH:ubiquinone oxidoreductase subunit (NDUFS6) | 20 | 0.79 | 0.260 | 1.000 |
| AT3G18410.2 | NADH:ubiquinone oxidoreductase subunit (NDUFS6) | 21 | 1.02 | 0.262 | 1.000 |
| AT3G18410.2 | NADH:ubiquinone oxidoreductase subunit (NDUFS6) | 22 | 1.61 | 0.261 | 1.000 |
| AT3G18410.2 | NADH:ubiquinone oxidoreductase subunit (NDUFS6) | 23 | 2.20 | 0.260 | 0.957 |
| AT3G18410.2 | NADH:ubiquinone oxidoreductase subunit (NDUFS6) | 24 | 1.77 | 0.252 | 0.997 |
| AT3G18410.2 | NADH:ubiquinone oxidoreductase subunit (NDUFS6) | 25 | 0.84 | 0.247 | 1.000 |
| AT3G18410.2 | NADH:ubiquinone oxidoreductase subunit (NDUFS6) | 26 | 0.54 | 0.247 | 0.993 |
| AT3G18410.2 | NADH:ubiquinone oxidoreductase subunit (NDUFS6) | 27 | 0.65 | 0.248 | 1.000 |
| AT3G18410.2 | NADH:ubiquinone oxidoreductase subunit (NDUFS6) | 28 | 1.13 | 0.249 | 1.000 |
| AT3G18410.2 | NADH:ubiquinone oxidoreductase subunit (NDUFS6) | 29 | 1.85 | 0.255 | 0.995 |
| AT3G18410.2 | NADH:ubiquinone oxidoreductase subunit (NDUFS6) | 30 | 1.81 | 0.275 | 0.998 |
| AT3G18410.2 | NADH:ubiquinone oxidoreductase subunit (NDUFS6) | 31 | 1.04 | 0.348 | 1.000 |
| AT3G18410.2 | NADH:ubiquinone oxidoreductase subunit (NDUFS6) | 32 | 0.52 | 0.566 | 1.000 |
| AT3G18410.2 | NADH:ubiquinone oxidoreductase subunit (NDUFS6) | 46 | 0.17 | 7.301 | 1.000 |
| AT3G18410.2 | NADH:ubiquinone oxidoreductase subunit (NDUFS6) | 47 | 0.22 | 8.050 | 1.000 |
| AT3G18580.1 | Nucleic acid-binding, OB-fold-like protein | 2 | 1.28 | 0.474 | 1.000 |
| AT3G18580.1 | Nucleic acid-binding, OB-fold-like protein | 3 | 1.18 | 0.400 | 1.000 |
| AT3G18580.1 | Nucleic acid-binding, OB-fold-like protein | 8 | 0.92 | 0.270 | 1.000 |
| AT3G18580.1 | Nucleic acid-binding, OB-fold-like protein | 9 | 0.92 | 0.239 | 1.000 |
| AT3G18580.1 | Nucleic acid-binding, OB-fold-like protein | 10 | 0.95 | 0.217 | 1.000 |

| | | | | | |
|---|---|---|---|---|---|
| AT3G18580.1 | Nucleic acid-binding, OB-fold-like protein | 11 | 1.00 | 0.205 | 1.000 |
| AT3G18580.1 | Nucleic acid-binding, OB-fold-like protein | 12 | 1.08 | 0.198 | 1.000 |
| AT3G18580.1 | Nucleic acid-binding, OB-fold-like protein | 13 | 1.19 | 0.194 | 1.000 |
| AT3G18580.1 | Nucleic acid-binding, OB-fold-like protein | 14 | 1.33 | 0.192 | 1.000 |
| AT3G18580.1 | Nucleic acid-binding, OB-fold-like protein | 15 | 1.49 | 0.191 | 0.998 |
| AT3G18580.1 | Nucleic acid-binding, OB-fold-like protein | 16 | 1.68 | 0.191 | 0.972 |
| AT3G18580.1 | Nucleic acid-binding, OB-fold-like protein | 17 | 1.93 | 0.191 | 0.850 |
| AT3G18580.1 | Nucleic acid-binding, OB-fold-like protein | 18 | 2.17 | 0.193 | 0.681 |
| AT3G18580.1 | Nucleic acid-binding, OB-fold-like protein | 19 | 2.25 | 0.196 | 0.643 |
| AT3G18580.1 | Nucleic acid-binding, OB-fold-like protein | 20 | 2.07 | 0.200 | 0.799 |
| AT3G18580.1 | Nucleic acid-binding, OB-fold-like protein | 21 | 1.67 | 0.205 | 0.985 |
| AT3G18580.1 | Nucleic acid-binding, OB-fold-like protein | 22 | 1.24 | 0.211 | 1.000 |
| AT3G18580.1 | Nucleic acid-binding, OB-fold-like protein | 23 | 0.93 | 0.220 | 1.000 |
| AT3G18580.1 | Nucleic acid-binding, OB-fold-like protein | 24 | 0.80 | 0.230 | 1.000 |
| AT3G18580.1 | Nucleic acid-binding, OB-fold-like protein | 25 | 0.76 | 0.241 | 1.000 |
| AT3G18580.1 | Nucleic acid-binding, OB-fold-like protein | 26 | 0.78 | 0.243 | 1.000 |
| AT3G18580.1 | Nucleic acid-binding, OB-fold-like protein | 28 | 0.91 | 0.233 | 1.000 |
| AT3G18580.1 | Nucleic acid-binding, OB-fold-like protein | 29 | 0.97 | 0.219 | 1.000 |
| AT3G18580.1 | Nucleic acid-binding, OB-fold-like protein | 30 | 0.99 | 0.208 | 1.000 |
| AT3G18580.1 | Nucleic acid-binding, OB-fold-like protein | 31 | 0.92 | 0.202 | 1.000 |
| AT3G18580.1 | Nucleic acid-binding, OB-fold-like protein | 32 | 0.81 | 0.204 | 1.000 |
| AT3G18580.1 | Nucleic acid-binding, OB-fold-like protein | 33 | 0.72 | 0.215 | 1.000 |
| AT3G18580.1 | Nucleic acid-binding, OB-fold-like protein | 34 | 0.70 | 0.232 | 1.000 |
| AT3G18580.1 | Nucleic acid-binding, OB-fold-like protein | 35 | 0.72 | 0.250 | 1.000 |
| AT3G18580.1 | Nucleic acid-binding, OB-fold-like protein | 38 | 0.86 | 0.245 | 1.000 |
| AT3G18580.1 | Nucleic acid-binding, OB-fold-like protein | 39 | 0.86 | 0.226 | 1.000 |
| AT3G18580.1 | Nucleic acid-binding, OB-fold-like protein | 40 | 0.80 | 0.211 | 1.000 |
| AT3G18580.1 | Nucleic acid-binding, OB-fold-like protein | 41 | 0.69 | 0.203 | 1.000 |
| AT3G18580.1 | Nucleic acid-binding, OB-fold-like protein | 42 | 0.55 | 0.203 | 0.953 |
| AT3G18580.1 | Nucleic acid-binding, OB-fold-like protein | 43 | 0.43 | 0.217 | 0.713 |
| AT3G18580.1 | Nucleic acid-binding, OB-fold-like protein | 44 | 0.33 | 0.250 | 0.533 |

| | | | | | |
|---|---|---|---|---|---|
| AT3G18580.1 | Nucleic acid-binding, OB-fold-like protein | 45 | 0.24 | 0.308 | 0.484 |
| AT3G18580.1 | Nucleic acid-binding, OB-fold-like protein | 46 | 0.17 | 0.396 | 0.545 |
| AT3G18580.1 | Nucleic acid-binding, OB-fold-like protein | 47 | 0.12 | 0.519 | 0.672 |
| AT3G18890.1 | Translocon of the inner envelope membrane of chloroplasts 62 (TIC62) | 2 | 1.22 | 0.705 | 1.000 |
| AT3G18890.1 | Translocon of the inner envelope membrane of chloroplasts 62 (TIC62) | 3 | 1.17 | 0.587 | 1.000 |
| AT3G18890.1 | Translocon of the inner envelope membrane of chloroplasts 62 (TIC62) | 4 | 1.12 | 0.531 | 1.000 |
| AT3G18890.1 | Translocon of the inner envelope membrane of chloroplasts 62 (TIC62) | 7 | 1.00 | 0.544 | 1.000 |
| AT3G18890.1 | Translocon of the inner envelope membrane of chloroplasts 62 (TIC62) | 8 | 0.98 | 0.560 | 1.000 |
| AT3G18890.1 | Translocon of the inner envelope membrane of chloroplasts 62 (TIC62) | 9 | 0.96 | 0.584 | 1.000 |
| AT3G18890.1 | Translocon of the inner envelope membrane of chloroplasts 62 (TIC62) | 16 | 0.80 | 0.589 | 1.000 |
| AT3G18890.1 | Translocon of the inner envelope membrane of chloroplasts 62 (TIC62) | 17 | 0.75 | 0.570 | 1.000 |
| AT3G18890.1 | Translocon of the inner envelope membrane of chloroplasts 62 (TIC62) | 18 | 0.70 | 0.557 | 1.000 |
| AT3G18890.1 | Translocon of the inner envelope membrane of chloroplasts 62 (TIC62) | 22 | 0.53 | 0.515 | 1.000 |
| AT3G18890.1 | Translocon of the inner envelope membrane of chloroplasts 62 (TIC62) | 23 | 0.52 | 0.497 | 1.000 |
| AT3G18890.1 | Translocon of the inner envelope membrane of chloroplasts 62 (TIC62) | 24 | 0.52 | 0.464 | 1.000 |
| AT3G18890.1 | Translocon of the inner envelope membrane of chloroplasts 62 (TIC62) | 26 | 0.64 | 0.389 | 1.000 |
| AT3G18890.1 | Translocon of the inner envelope membrane of chloroplasts 62 (TIC62) | 27 | 0.80 | 0.354 | 1.000 |
| AT3G18890.1 | Translocon of the inner envelope membrane of chloroplasts 62 (TIC62) | 28 | 1.10 | 0.328 | 1.000 |
| AT3G18890.1 | Translocon of the inner envelope membrane of chloroplasts 62 (TIC62) | 29 | 1.51 | 0.312 | 1.000 |
| AT3G18890.1 | Translocon of the inner envelope membrane of chloroplasts 62 (TIC62) | 30 | 1.86 | 0.305 | 0.994 |
| AT3G18890.1 | Translocon of the inner envelope membrane of chloroplasts 62 (TIC62) | 31 | 2.01 | 0.302 | 0.984 |

| | | | | | |
|---|---|---|---|---|---|
| AT3G18890.1 | Translocon of the inner envelope membrane of chloroplasts 62 (TIC62) | 32 | 1.80 | 0.302 | 0.996 |
| AT3G18890.1 | Translocon of the inner envelope membrane of chloroplasts 62 (TIC62) | 33 | 1.32 | 0.303 | 1.000 |
| AT3G18890.1 | Translocon of the inner envelope membrane of chloroplasts 62 (TIC62) | 34 | 0.86 | 0.304 | 1.000 |
| AT3G18890.1 | Translocon of the inner envelope membrane of chloroplasts 62 (TIC62) | 35 | 0.57 | 0.305 | 0.997 |
| AT3G18890.1 | Translocon of the inner envelope membrane of chloroplasts 62 (TIC62) | 36 | 0.40 | 0.306 | 0.900 |
| AT3G18890.1 | Translocon of the inner envelope membrane of chloroplasts 62 (TIC62) | 37 | 0.30 | 0.309 | 0.657 |
| AT3G18890.1 | Translocon of the inner envelope membrane of chloroplasts 62 (TIC62) | 38 | 0.24 | 0.317 | 0.478 |
| AT3G18890.1 | Translocon of the inner envelope membrane of chloroplasts 62 (TIC62) | 39 | 0.22 | 0.336 | 0.460 |
| AT3G18890.1 | Translocon of the inner envelope membrane of chloroplasts 62 (TIC62) | 40 | 0.22 | 0.371 | 0.630 |
| AT3G18890.1 | Translocon of the inner envelope membrane of chloroplasts 62 (TIC62) | 41 | 0.27 | 0.425 | 0.888 |
| AT3G18890.1 | Translocon of the inner envelope membrane of chloroplasts 62 (TIC62) | 42 | 0.36 | 0.501 | 0.994 |
| AT3G18890.1 | Translocon of the inner envelope membrane of chloroplasts 62 (TIC62) | 43 | 0.52 | 0.594 | 1.000 |
| AT3G18890.1 | Translocon of the inner envelope membrane of chloroplasts 62 (TIC62) | 46 | 2.14 | 0.992 | 1.000 |
| AT3G18890.1 | Translocon of the inner envelope membrane of chloroplasts 62 (TIC62) | 47 | 3.60 | 1.182 | 1.000 |
| AT3G19050.1 | Phragmoplast orienting kinesin 2 (POK2) | 2 | 1.08 | 0.348 | 1.000 |
| AT3G19050.1 | Phragmoplast orienting kinesin 2 (POK2) | 3 | 1.09 | 0.289 | 1.000 |
| AT3G19050.1 | Phragmoplast orienting kinesin 2 (POK2) | 4 | 1.10 | 0.253 | 1.000 |
| AT3G19050.1 | Phragmoplast orienting kinesin 2 (POK2) | 5 | 1.12 | 0.241 | 1.000 |
| AT3G19050.1 | Phragmoplast orienting kinesin 2 (POK2) | 6 | 1.13 | 0.245 | 1.000 |
| AT3G19050.1 | Phragmoplast orienting kinesin 2 (POK2) | 7 | 1.15 | 0.256 | 1.000 |
| AT3G19050.1 | Phragmoplast orienting kinesin 2 (POK2) | 8 | 1.16 | 0.270 | 1.000 |
| AT3G19050.1 | Phragmoplast orienting kinesin 2 (POK2) | 15 | 1.16 | 0.232 | 1.000 |
| AT3G19050.1 | Phragmoplast orienting kinesin 2 (POK2) | 16 | 1.15 | 0.211 | 1.000 |
| AT3G19050.1 | Phragmoplast orienting kinesin 2 (POK2) | 17 | 1.14 | 0.192 | 1.000 |

| | | | | | |
|---|---|---|---|---|---|
| AT3G19050.1 | Phragmoplast orienting kinesin 2 (POK2) | 18 | 1.15 | 0.178 | 1.000 |
| AT3G19050.1 | Phragmoplast orienting kinesin 2 (POK2) | 19 | 1.17 | 0.168 | 1.000 |
| AT3G19050.1 | Phragmoplast orienting kinesin 2 (POK2) | 20 | 1.20 | 0.163 | 1.000 |
| AT3G19050.1 | Phragmoplast orienting kinesin 2 (POK2) | 21 | 1.22 | 0.164 | 0.999 |
| AT3G19050.1 | Phragmoplast orienting kinesin 2 (POK2) | 22 | 1.22 | 0.170 | 0.999 |
| AT3G19050.1 | Phragmoplast orienting kinesin 2 (POK2) | 23 | 1.22 | 0.182 | 1.000 |
| AT3G19050.1 | Phragmoplast orienting kinesin 2 (POK2) | 24 | 1.21 | 0.203 | 1.000 |
| AT3G19050.1 | Phragmoplast orienting kinesin 2 (POK2) | 25 | 1.19 | 0.233 | 1.000 |
| AT3G19050.1 | Phragmoplast orienting kinesin 2 (POK2) | 26 | 1.18 | 0.268 | 1.000 |
| AT3G19050.1 | Phragmoplast orienting kinesin 2 (POK2) | 41 | 1.14 | 0.426 | 1.000 |
| AT3G19050.1 | Phragmoplast orienting kinesin 2 (POK2) | 42 | 1.17 | 0.431 | 1.000 |
| AT3G19050.1 | Phragmoplast orienting kinesin 2 (POK2) | 43 | 1.22 | 0.453 | 1.000 |
| AT3G19050.1 | Phragmoplast orienting kinesin 2 (POK2) | 46 | 1.42 | 0.635 | 1.000 |
| AT3G19050.1 | Phragmoplast orienting kinesin 2 (POK2) | 47 | 1.51 | 0.729 | 1.000 |
| AT3G19170.1 | Presequence protease 1 (PREP1) | 2 | 0.87 | 0.541 | 1.000 |
| AT3G19170.1 | Presequence protease 1 (PREP1) | 3 | 1.08 | 0.524 | 1.000 |
| AT3G19170.1 | Presequence protease 1 (PREP1) | 28 | 0.94 | 0.391 | 1.000 |
| AT3G19170.1 | Presequence protease 1 (PREP1) | 29 | 0.65 | 0.284 | 1.000 |
| AT3G19170.1 | Presequence protease 1 (PREP1) | 30 | 0.59 | 0.252 | 0.998 |
| AT3G19170.1 | Presequence protease 1 (PREP1) | 31 | 0.77 | 0.241 | 1.000 |
| AT3G19170.1 | Presequence protease 1 (PREP1) | 32 | 1.07 | 0.235 | 1.000 |
| AT3G19170.1 | Presequence protease 1 (PREP1) | 33 | 1.15 | 0.234 | 1.000 |
| AT3G19170.1 | Presequence protease 1 (PREP1) | 34 | 1.01 | 0.234 | 1.000 |
| AT3G19170.1 | Presequence protease 1 (PREP1) | 35 | 0.89 | 0.233 | 1.000 |
| AT3G19170.1 | Presequence protease 1 (PREP1) | 36 | 0.84 | 0.232 | 1.000 |
| AT3G19170.1 | Presequence protease 1 (PREP1) | 37 | 0.95 | 0.232 | 1.000 |
| AT3G19170.1 | Presequence protease 1 (PREP1) | 38 | 1.23 | 0.233 | 1.000 |
| AT3G19170.1 | Presequence protease 1 (PREP1) | 39 | 1.43 | 0.233 | 1.000 |
| AT3G19170.1 | Presequence protease 1 (PREP1) | 40 | 1.18 | 0.232 | 1.000 |
| AT3G19170.1 | Presequence protease 1 (PREP1) | 41 | 0.70 | 0.234 | 1.000 |
| AT3G19170.1 | Presequence protease 1 (PREP1) | 42 | 0.43 | 0.246 | 0.862 |

| | | | | | |
|---|---|---|---|---|---|
| AT3G19170.1 | Presequence protease 1 (PREP1) | 43 | 0.45 | 0.272 | 0.949 |
| AT3G19170.1 | Presequence protease 1 (PREP1) | 44 | 0.78 | 0.283 | 1.000 |
| AT3G19170.1 | Presequence protease 1 (PREP1) | 45 | 1.68 | 0.263 | 0.999 |
| AT3G19170.1 | Presequence protease 1 (PREP1) | 46 | 3.09 | 0.286 | 0.728 |
| AT3G19170.1 | Presequence protease 1 (PREP1) | 47 | 4.62 | 0.503 | 0.941 |
| AT3G19740.1 | P-loop containing nucleoside triphosphate hydrolase | 2 | 1.27 | 0.512 | 1.000 |
| AT3G19740.1 | P-loop containing nucleoside triphosphate hydrolase | 3 | 1.10 | 0.377 | 1.000 |
| AT3G19740.1 | P-loop containing nucleoside triphosphate hydrolase | 4 | 0.94 | 0.393 | 1.000 |
| AT3G19740.1 | P-loop containing nucleoside triphosphate hydrolase | 6 | 0.85 | 0.459 | 1.000 |
| AT3G19740.1 | P-loop containing nucleoside triphosphate hydrolase | 7 | 0.94 | 0.453 | 1.000 |
| AT3G19740.1 | P-loop containing nucleoside triphosphate hydrolase | 8 | 1.15 | 0.478 | 1.000 |
| AT3G19740.1 | P-loop containing nucleoside triphosphate hydrolase | 13 | 3.54 | 0.391 | 0.952 |
| AT3G19740.1 | P-loop containing nucleoside triphosphate hydrolase | 14 | 3.26 | 0.335 | 0.905 |
| AT3G19740.1 | P-loop containing nucleoside triphosphate hydrolase | 15 | 2.63 | 0.303 | 0.956 |
| AT3G19740.1 | P-loop containing nucleoside triphosphate hydrolase | 16 | 2.04 | 0.276 | 0.994 |
| AT3G19740.1 | P-loop containing nucleoside triphosphate hydrolase | 17 | 1.72 | 0.262 | 1.000 |
| AT3G19740.1 | P-loop containing nucleoside triphosphate hydrolase | 18 | 1.91 | 0.261 | 0.996 |
| AT3G19740.1 | P-loop containing nucleoside triphosphate hydrolase | 19 | 2.51 | 0.267 | 0.920 |
| AT3G19740.1 | P-loop containing nucleoside triphosphate hydrolase | 20 | 2.72 | 0.272 | 0.873 |
| AT3G19740.1 | P-loop containing nucleoside triphosphate hydrolase | 21 | 2.00 | 0.277 | 0.996 |
| AT3G19740.1 | P-loop containing nucleoside triphosphate hydrolase | 22 | 1.25 | 0.284 | 1.000 |
| AT3G19740.1 | P-loop containing nucleoside triphosphate hydrolase | 23 | 0.98 | 0.293 | 1.000 |
| AT3G19740.1 | P-loop containing nucleoside triphosphate hydrolase | 24 | 0.91 | 0.293 | 1.000 |
| AT3G19740.1 | P-loop containing nucleoside triphosphate hydrolase | 25 | 0.85 | 0.284 | 1.000 |
| AT3G19740.1 | P-loop containing nucleoside triphosphate hydrolase | 26 | 0.68 | 0.272 | 1.000 |
| AT3G19740.1 | P-loop containing nucleoside triphosphate hydrolase | 27 | 0.42 | 0.266 | 0.951 |
| AT3G19740.1 | P-loop containing nucleoside triphosphate hydrolase | 28 | 0.27 | 0.265 | 0.465 |
| AT3G19740.1 | P-loop containing nucleoside triphosphate hydrolase | 29 | 0.25 | 0.265 | 0.357 |
| AT3G19740.1 | P-loop containing nucleoside triphosphate hydrolase | 30 | 0.35 | 0.266 | 0.795 |
| AT3G19740.1 | P-loop containing nucleoside triphosphate hydrolase | 31 | 0.66 | 0.268 | 1.000 |
| AT3G19740.1 | P-loop containing nucleoside triphosphate hydrolase | 32 | 1.04 | 0.269 | 1.000 |

| | | | | | |
|---|---|---|---|---|---|
| AT3G19740.1 | P-loop containing nucleoside triphosphate hydrolase | 33 | 1.07 | 0.271 | 1.000 |
| AT3G19740.1 | P-loop containing nucleoside triphosphate hydrolase | 34 | 0.89 | 0.275 | 1.000 |
| AT3G19740.1 | P-loop containing nucleoside triphosphate hydrolase | 35 | 0.82 | 0.289 | 1.000 |
| AT3G19740.1 | P-loop containing nucleoside triphosphate hydrolase | 36 | 0.81 | 0.334 | 1.000 |
| AT3G19740.1 | P-loop containing nucleoside triphosphate hydrolase | 37 | 0.84 | 0.417 | 1.000 |
| AT3G19740.1 | P-loop containing nucleoside triphosphate hydrolase | 41 | 1.06 | 0.444 | 1.000 |
| AT3G19740.1 | P-loop containing nucleoside triphosphate hydrolase | 42 | 1.08 | 0.473 | 1.000 |
| AT3G19740.1 | P-loop containing nucleoside triphosphate hydrolase | 43 | 1.07 | 0.558 | 1.000 |
| AT3G19740.1 | P-loop containing nucleoside triphosphate hydrolase | 46 | 0.88 | 1.353 | 1.000 |
| AT3G19740.1 | P-loop containing nucleoside triphosphate hydrolase | 47 | 0.80 | 1.731 | 1.000 |
| AT3G20000.1 | Translocase of the outer mitochondrial membrane 40 (TOM40-1) | 2 | 0.68 | 0.409 | 1.000 |
| AT3G20000.1 | Translocase of the outer mitochondrial membrane 40 (TOM40-1) | 3 | 0.67 | 0.247 | 1.000 |
| AT3G20000.1 | Translocase of the outer mitochondrial membrane 40 (TOM40-1) | 4 | 0.85 | 0.238 | 1.000 |
| AT3G20000.1 | Translocase of the outer mitochondrial membrane 40 (TOM40-1) | 5 | 1.15 | 0.237 | 1.000 |
| AT3G20000.1 | Translocase of the outer mitochondrial membrane 40 (TOM40-1) | 6 | 1.45 | 0.237 | 1.000 |
| AT3G20000.1 | Translocase of the outer mitochondrial membrane 40 (TOM40-1) | 7 | 1.55 | 0.238 | 1.000 |
| AT3G20000.1 | Translocase of the outer mitochondrial membrane 40 (TOM40-1) | 8 | 1.65 | 0.233 | 1.000 |
| AT3G20000.1 | Translocase of the outer mitochondrial membrane 40 (TOM40-1) | 9 | 1.82 | 0.230 | 1.000 |
| AT3G20000.1 | Translocase of the outer mitochondrial membrane 40 (TOM40-1) | 10 | 2.01 | 0.229 | 0.997 |
| AT3G20000.1 | Translocase of the outer mitochondrial membrane 40 (TOM40-1) | 11 | 2.12 | 0.230 | 0.991 |
| AT3G20000.1 | Translocase of the outer mitochondrial membrane 40 (TOM40-1) | 12 | 2.27 | 0.230 | 0.974 |
| AT3G20000.1 | Translocase of the outer mitochondrial membrane 40 (TOM40-1) | 13 | 2.28 | 0.228 | 0.969 |
| AT3G20000.1 | Translocase of the outer mitochondrial membrane 40 (TOM40-1) | 14 | 2.18 | 0.225 | 0.981 |
| AT3G20000.1 | Translocase of the outer mitochondrial membrane 40 (TOM40-1) | 15 | 1.81 | 0.225 | 1.000 |

| | | | | | |
|---|---|---|---|---|---|
| AT3G20000.1 | Translocase of the outer mitochondrial membrane 40 (TOM40-1) | 16 | 1.36 | 0.223 | 1.000 |
| AT3G20000.1 | Translocase of the outer mitochondrial membrane 40 (TOM40-1) | 17 | 1.19 | 0.223 | 1.000 |
| AT3G20000.1 | Translocase of the outer mitochondrial membrane 40 (TOM40-1) | 18 | 1.42 | 0.226 | 1.000 |
| AT3G20000.1 | Translocase of the outer mitochondrial membrane 40 (TOM40-1) | 19 | 1.84 | 0.232 | 1.000 |
| AT3G20000.1 | Translocase of the outer mitochondrial membrane 40 (TOM40-1) | 20 | 2.22 | 0.237 | 0.987 |
| AT3G20000.1 | Translocase of the outer mitochondrial membrane 40 (TOM40-1) | 21 | 2.29 | 0.240 | 0.982 |
| AT3G20000.1 | Translocase of the outer mitochondrial membrane 40 (TOM40-1) | 22 | 2.07 | 0.240 | 0.997 |
| AT3G20000.1 | Translocase of the outer mitochondrial membrane 40 (TOM40-1) | 23 | 1.65 | 0.240 | 1.000 |
| AT3G20000.1 | Translocase of the outer mitochondrial membrane 40 (TOM40-1) | 24 | 1.32 | 0.234 | 1.000 |
| AT3G20000.1 | Translocase of the outer mitochondrial membrane 40 (TOM40-1) | 25 | 1.11 | 0.230 | 1.000 |
| AT3G20000.1 | Translocase of the outer mitochondrial membrane 40 (TOM40-1) | 26 | 1.07 | 0.228 | 1.000 |
| AT3G20000.1 | Translocase of the outer mitochondrial membrane 40 (TOM40-1) | 27 | 1.03 | 0.228 | 1.000 |
| AT3G20000.1 | Translocase of the outer mitochondrial membrane 40 (TOM40-1) | 28 | 1.05 | 0.228 | 1.000 |
| AT3G20000.1 | Translocase of the outer mitochondrial membrane 40 (TOM40-1) | 29 | 1.34 | 0.228 | 1.000 |
| AT3G20000.1 | Translocase of the outer mitochondrial membrane 40 (TOM40-1) | 30 | 1.73 | 0.228 | 1.000 |
| AT3G20000.1 | Translocase of the outer mitochondrial membrane 40 (TOM40-1) | 31 | 1.50 | 0.229 | 1.000 |
| AT3G20000.1 | Translocase of the outer mitochondrial membrane 40 (TOM40-1) | 32 | 0.91 | 0.229 | 1.000 |
| AT3G20000.1 | Translocase of the outer mitochondrial membrane 40 (TOM40-1) | 33 | 0.57 | 0.230 | 1.000 |
| AT3G20000.1 | Translocase of the outer mitochondrial membrane 40 (TOM40-1) | 34 | 0.51 | 0.231 | 0.998 |
| AT3G20000.1 | Translocase of the outer mitochondrial membrane 40 (TOM40-1) | 35 | 0.78 | 0.232 | 1.000 |

| | | | | | |
|---|---|---|---|---|---|
| AT3G20000.1 | Translocase of the outer mitochondrial membrane 40 (TOM40-1) | 36 | 1.74 | 0.230 | 1.000 |
| AT3G20000.1 | Translocase of the outer mitochondrial membrane 40 (TOM40-1) | 37 | 3.55 | 0.231 | 0.429 |
| AT3G20000.1 | Translocase of the outer mitochondrial membrane 40 (TOM40-1) | 38 | 4.71 | 0.232 | 0.128 |
| AT3G20000.1 | Translocase of the outer mitochondrial membrane 40 (TOM40-1) | 39 | 3.98 | 0.232 | 0.281 |
| AT3G20000.1 | Translocase of the outer mitochondrial membrane 40 (TOM40-1) | 40 | 1.90 | 0.231 | 0.999 |
| AT3G20000.1 | Translocase of the outer mitochondrial membrane 40 (TOM40-1) | 41 | 0.59 | 0.232 | 1.000 |
| AT3G20000.1 | Translocase of the outer mitochondrial membrane 40 (TOM40-1) | 42 | 0.31 | 0.246 | 0.700 |
| AT3G20000.1 | Translocase of the outer mitochondrial membrane 40 (TOM40-1) | 43 | 0.57 | 0.310 | 1.000 |
| AT3G20000.1 | Translocase of the outer mitochondrial membrane 40 (TOM40-1) | 44 | 2.38 | 0.466 | 1.000 |
| AT3G20000.1 | Translocase of the outer mitochondrial membrane 40 (TOM40-1) | 46 | 185.84 | 1.220 | 0.850 |
| AT3G20000.1 | Translocase of the outer mitochondrial membrane 40 (TOM40-1) | 47 | 2302.45 | 1.757 | 0.816 |
| AT3G20040.1 | Hexokinase 4 (HXK4) | 2 | 0.94 | 0.420 | 1.000 |
| AT3G20040.1 | Hexokinase 4 (HXK4) | 3 | 0.90 | 0.417 | 1.000 |
| AT3G20040.1 | Hexokinase 4 (HXK4) | 28 | 0.96 | 0.316 | 1.000 |
| AT3G20040.1 | Hexokinase 4 (HXK4) | 29 | 1.01 | 0.232 | 1.000 |
| AT3G20040.1 | Hexokinase 4 (HXK4) | 30 | 1.07 | 0.180 | 1.000 |
| AT3G20040.1 | Hexokinase 4 (HXK4) | 31 | 1.19 | 0.158 | 1.000 |
| AT3G20040.1 | Hexokinase 4 (HXK4) | 32 | 1.32 | 0.150 | 0.992 |
| AT3G20040.1 | Hexokinase 4 (HXK4) | 33 | 1.42 | 0.148 | 0.964 |
| AT3G20040.1 | Hexokinase 4 (HXK4) | 34 | 1.37 | 0.148 | 0.980 |
| AT3G20040.1 | Hexokinase 4 (HXK4) | 35 | 1.21 | 0.152 | 1.000 |
| AT3G20040.1 | Hexokinase 4 (HXK4) | 36 | 1.06 | 0.161 | 1.000 |
| AT3G20040.1 | Hexokinase 4 (HXK4) | 37 | 1.00 | 0.177 | 1.000 |
| AT3G20040.1 | Hexokinase 4 (HXK4) | 38 | 1.04 | 0.197 | 1.000 |
| AT3G20040.1 | Hexokinase 4 (HXK4) | 40 | 1.41 | 0.209 | 0.997 |
| AT3G20040.1 | Hexokinase 4 (HXK4) | 41 | 1.78 | 0.210 | 0.912 |
| AT3G20040.1 | Hexokinase 4 (HXK4) | 42 | 2.28 | 0.233 | 0.718 |
| AT3G20040.1 | Hexokinase 4 (HXK4) | 43 | 3.01 | 0.301 | 0.679 |
| AT3G20040.1 | Hexokinase 4 (HXK4) | 46 | 8.03 | 0.725 | 0.888 |
| AT3G20040.1 | Hexokinase 4 (HXK4) | 47 | 11.38 | 0.925 | 0.930 |
| AT3G20390.1 | Reactive intermediate deaminase A (RIDA) | 2 | 0.93 | 0.429 | 1.000 |
| AT3G20390.1 | Reactive intermediate deaminase A (RIDA) | 3 | 0.86 | 0.357 | 1.000 |
| AT3G20390.1 | Reactive intermediate deaminase A (RIDA) | 4 | 0.82 | 0.585 | 1.000 |
| AT3G20390.1 | Reactive intermediate deaminase A (RIDA) | 31 | 0.99 | 0.405 | 1.000 |
| AT3G20390.1 | Reactive intermediate deaminase A (RIDA) | 32 | 0.88 | 0.336 | 1.000 |

| | | | | | |
|---|---|---|---|---|---|
| AT3G20390.1 | Reactive intermediate deaminase A (RIDA) | 33 | 0.78 | 0.315 | 1.000 |
| AT3G20390.1 | Reactive intermediate deaminase A (RIDA) | 35 | 0.62 | 0.257 | 0.996 |
| AT3G20390.1 | Reactive intermediate deaminase A (RIDA) | 36 | 0.64 | 0.226 | 0.993 |
| AT3G20390.1 | Reactive intermediate deaminase A (RIDA) | 37 | 0.76 | 0.203 | 1.000 |
| AT3G20390.1 | Reactive intermediate deaminase A (RIDA) | 38 | 1.00 | 0.193 | 1.000 |
| AT3G20390.1 | Reactive intermediate deaminase A (RIDA) | 39 | 1.43 | 0.189 | 0.995 |
| AT3G20390.1 | Reactive intermediate deaminase A (RIDA) | 40 | 1.86 | 0.188 | 0.825 |
| AT3G20390.1 | Reactive intermediate deaminase A (RIDA) | 41 | 1.88 | 0.189 | 0.807 |
| AT3G20390.1 | Reactive intermediate deaminase A (RIDA) | 42 | 1.72 | 0.190 | 0.919 |
| AT3G20390.1 | Reactive intermediate deaminase A (RIDA) | 43 | 1.65 | 0.192 | 0.952 |
| AT3G20390.1 | Reactive intermediate deaminase A (RIDA) | 44 | 1.44 | 0.194 | 0.996 |
| AT3G20390.1 | Reactive intermediate deaminase A (RIDA) | 45 | 0.87 | 0.205 | 1.000 |
| AT3G20390.1 | Reactive intermediate deaminase A (RIDA) | 46 | 0.34 | 0.249 | 0.524 |
| AT3G20390.1 | Reactive intermediate deaminase A (RIDA) | 47 | 0.11 | 0.376 | 0.131 |
| AT3G20830.1 | AGC (cAMP-dependent, cGMP-dependent and protein kinase C) kinase family protein (AGC2-4) | 2 | 0.92 | 0.540 | 1.000 |
| AT3G20830.1 | AGC (cAMP-dependent, cGMP-dependent and protein kinase C) kinase family protein (AGC2-4) | 3 | 0.97 | 0.398 | 1.000 |
| AT3G20830.1 | AGC (cAMP-dependent, cGMP-dependent and protein kinase C) kinase family protein (AGC2-4) | 4 | 1.07 | 0.704 | 1.000 |
| AT3G20830.1 | AGC (cAMP-dependent, cGMP-dependent and protein kinase C) kinase family protein (AGC2-4) | 20 | 1.17 | 0.367 | 1.000 |
| AT3G20830.1 | AGC (cAMP-dependent, cGMP-dependent and protein kinase C) kinase family protein (AGC2-4) | 21 | 0.58 | 0.302 | 0.999 |
| AT3G20830.1 | AGC (cAMP-dependent, cGMP-dependent and protein kinase C) kinase family protein (AGC2-4) | 22 | 0.27 | 0.270 | 0.426 |
| AT3G20830.1 | AGC (cAMP-dependent, cGMP-dependent and protein kinase C) kinase family protein (AGC2-4) | 23 | 0.29 | 0.263 | 0.469 |
| AT3G20830.1 | AGC (cAMP-dependent, cGMP-dependent and protein kinase C) kinase family protein (AGC2-4) | 24 | 0.81 | 0.247 | 1.000 |

| | | | | | | |
|---|---|---|---|---|---|---|
| AT3G20830.1 | AGC (cAMP-dependent, cGMP-dependent and protein kinase C) kinase family protein (AGC2-4) | 25 | 3.02 | 0.238 | 0.485 | |
| AT3G20830.1 | AGC (cAMP-dependent, cGMP-dependent and protein kinase C) kinase family protein (AGC2-4) | 26 | 2.64 | 0.240 | 0.685 | |
| AT3G20830.1 | AGC (cAMP-dependent, cGMP-dependent and protein kinase C) kinase family protein (AGC2-4) | 27 | 0.42 | 0.238 | 0.790 | |
| AT3G20830.1 | AGC (cAMP-dependent, cGMP-dependent and protein kinase C) kinase family protein (AGC2-4) | 28 | 0.16 | 0.242 | 0.020 | * |
| AT3G20830.1 | AGC (cAMP-dependent, cGMP-dependent and protein kinase C) kinase family protein (AGC2-4) | 29 | 0.27 | 0.243 | 0.273 | |
| AT3G20830.1 | AGC (cAMP-dependent, cGMP-dependent and protein kinase C) kinase family protein (AGC2-4) | 30 | 0.75 | 0.248 | 1.000 | |
| AT3G20830.1 | AGC (cAMP-dependent, cGMP-dependent and protein kinase C) kinase family protein (AGC2-4) | 31 | 1.23 | 0.268 | 1.000 | |
| AT3G20830.1 | AGC (cAMP-dependent, cGMP-dependent and protein kinase C) kinase family protein (AGC2-4) | 32 | 0.88 | 0.335 | 1.000 | |
| AT3G20830.1 | AGC (cAMP-dependent, cGMP-dependent and protein kinase C) kinase family protein (AGC2-4) | 33 | 0.53 | 0.610 | 1.000 | |
| AT3G20830.1 | AGC (cAMP-dependent, cGMP-dependent and protein kinase C) kinase family protein (AGC2-4) | 46 | 34.96 | 5.145 | 1.000 | |
| AT3G20830.1 | AGC (cAMP-dependent, cGMP-dependent and protein kinase C) kinase family protein (AGC2-4) | 47 | 73.38 | 5.694 | 1.000 | |
| AT3G20970.1 | NFU domain protein 4 (NFU4) | 2 | 0.51 | 0.286 | 0.979 | |
| AT3G20970.1 | NFU domain protein 4 (NFU4) | 3 | 0.65 | 0.272 | 0.999 | |
| AT3G20970.1 | NFU domain protein 4 (NFU4) | 27 | 2.30 | 0.196 | 0.541 | |
| AT3G20970.1 | NFU domain protein 4 (NFU4) | 28 | 1.64 | 0.156 | 0.866 | |
| AT3G20970.1 | NFU domain protein 4 (NFU4) | 29 | 1.13 | 0.135 | 1.000 | |
| AT3G20970.1 | NFU domain protein 4 (NFU4) | 30 | 0.79 | 0.125 | 0.996 | |
| AT3G20970.1 | NFU domain protein 4 (NFU4) | 31 | 0.58 | 0.121 | 0.466 | |
| AT3G20970.1 | NFU domain protein 4 (NFU4) | 32 | 0.48 | 0.118 | 0.096 | . |
| AT3G20970.1 | NFU domain protein 4 (NFU4) | 33 | 0.45 | 0.117 | 0.044 | * |
| AT3G20970.1 | NFU domain protein 4 (NFU4) | 34 | 0.49 | 0.117 | 0.105 | |
| AT3G20970.1 | NFU domain protein 4 (NFU4) | 35 | 0.62 | 0.117 | 0.602 | |
| AT3G20970.1 | NFU domain protein 4 (NFU4) | 36 | 0.91 | 0.119 | 1.000 | |
| AT3G20970.1 | NFU domain protein 4 (NFU4) | 37 | 1.40 | 0.122 | 0.940 | |
| AT3G20970.1 | NFU domain protein 4 (NFU4) | 38 | 1.88 | 0.125 | 0.299 | |
| AT3G20970.1 | NFU domain protein 4 (NFU4) | 39 | 1.94 | 0.126 | 0.254 | |
| AT3G20970.1 | NFU domain protein 4 (NFU4) | 40 | 1.58 | 0.123 | 0.717 | |
| AT3G20970.1 | NFU domain protein 4 (NFU4) | 41 | 1.18 | 0.121 | 1.000 | |
| AT3G20970.1 | NFU domain protein 4 (NFU4) | 42 | 0.90 | 0.123 | 1.000 | |

| | | | | | | |
|---|---|---|---|---|---|---|
| AT3G20970.1 | NFU domain protein 4 (NFU4) | 43 | 0.71 | 0.125 | 0.943 | |
| AT3G20970.1 | NFU domain protein 4 (NFU4) | 44 | 0.62 | 0.126 | 0.684 | |
| AT3G20970.1 | NFU domain protein 4 (NFU4) | 45 | 0.64 | 0.130 | 0.805 | |
| AT3G20970.1 | NFU domain protein 4 (NFU4) | 46 | 0.76 | 0.160 | 0.999 | |
| AT3G20970.1 | NFU domain protein 4 (NFU4) | 47 | 0.91 | 0.252 | 1.000 | |
| AT3G21390.1 | Mitochondrial substrate carrier family protein | 2 | 0.85 | 0.484 | 1.000 | |
| AT3G21390.1 | Mitochondrial substrate carrier family protein | 3 | 0.98 | 0.369 | 1.000 | |
| AT3G21390.1 | Mitochondrial substrate carrier family protein | 6 | 1.39 | 0.332 | 1.000 | |
| AT3G21390.1 | Mitochondrial substrate carrier family protein | 7 | 1.48 | 0.271 | 1.000 | |
| AT3G21390.1 | Mitochondrial substrate carrier family protein | 8 | 1.52 | 0.235 | 1.000 | |
| AT3G21390.1 | Mitochondrial substrate carrier family protein | 9 | 1.42 | 0.221 | 1.000 | |
| AT3G21390.1 | Mitochondrial substrate carrier family protein | 10 | 1.22 | 0.217 | 1.000 | |
| AT3G21390.1 | Mitochondrial substrate carrier family protein | 11 | 1.21 | 0.216 | 1.000 | |
| AT3G21390.1 | Mitochondrial substrate carrier family protein | 12 | 1.56 | 0.219 | 0.999 | |
| AT3G21390.1 | Mitochondrial substrate carrier family protein | 13 | 2.11 | 0.222 | 0.911 | |
| AT3G21390.1 | Mitochondrial substrate carrier family protein | 14 | 2.79 | 0.228 | 0.576 | |
| AT3G21390.1 | Mitochondrial substrate carrier family protein | 15 | 3.61 | 0.240 | 0.313 | |
| AT3G21390.1 | Mitochondrial substrate carrier family protein | 16 | 3.76 | 0.251 | 0.328 | |
| AT3G21390.1 | Mitochondrial substrate carrier family protein | 17 | 2.97 | 0.286 | 0.804 | |
| AT3G21390.1 | Mitochondrial substrate carrier family protein | 18 | 2.09 | 0.398 | 1.000 | |
| AT3G21390.1 | Mitochondrial substrate carrier family protein | 38 | 6.86 | 0.382 | 0.401 | |
| AT3G21390.1 | Mitochondrial substrate carrier family protein | 39 | 7.98 | 0.259 | 0.013 | * |
| AT3G21390.1 | Mitochondrial substrate carrier family protein | 40 | 6.39 | 0.225 | 0.009 | ** |
| AT3G21390.1 | Mitochondrial substrate carrier family protein | 41 | 2.49 | 0.218 | 0.686 | |
| AT3G21390.1 | Mitochondrial substrate carrier family protein | 42 | 0.78 | 0.220 | 1.000 | |
| AT3G21390.1 | Mitochondrial substrate carrier family protein | 43 | 0.37 | 0.222 | 0.592 | |
| AT3G21390.1 | Mitochondrial substrate carrier family protein | 44 | 0.34 | 0.220 | 0.434 | |
| AT3G21390.1 | Mitochondrial substrate carrier family protein | 45 | 0.41 | 0.218 | 0.726 | |
| AT3G21390.1 | Mitochondrial substrate carrier family protein | 46 | 0.38 | 0.250 | 0.804 | |
| AT3G21390.1 | Mitochondrial substrate carrier family protein | 47 | 0.30 | 0.471 | 0.989 | |
| AT3G21400.1 | NADH:ubiquinone oxidoreductase assembly factor (Dynein beta chain) | 2 | 0.95 | 0.407 | 1.000 | |
| AT3G21400.1 | NADH:ubiquinone oxidoreductase assembly factor (Dynein beta chain) | 3 | 0.98 | 0.324 | 1.000 | |

| | | | | | |
|---|---|---|---|---|---|
| AT3G21400.1 | NADH:ubiquinone oxidoreductase assembly factor (Dynein beta chain) | 4 | 1.03 | 0.556 | 1.000 |
| AT3G21400.1 | NADH:ubiquinone oxidoreductase assembly factor (Dynein beta chain) | 31 | 1.40 | 0.321 | 1.000 |
| AT3G21400.1 | NADH:ubiquinone oxidoreductase assembly factor (Dynein beta chain) | 32 | 1.25 | 0.234 | 1.000 |
| AT3G21400.1 | NADH:ubiquinone oxidoreductase assembly factor (Dynein beta chain) | 33 | 1.12 | 0.204 | 1.000 |
| AT3G21400.1 | NADH:ubiquinone oxidoreductase assembly factor (Dynein beta chain) | 34 | 1.05 | 0.189 | 1.000 |
| AT3G21400.1 | NADH:ubiquinone oxidoreductase assembly factor (Dynein beta chain) | 35 | 1.06 | 0.183 | 1.000 |
| AT3G21400.1 | NADH:ubiquinone oxidoreductase assembly factor (Dynein beta chain) | 36 | 1.31 | 0.180 | 0.999 |
| AT3G21400.1 | NADH:ubiquinone oxidoreductase assembly factor (Dynein beta chain) | 37 | 1.96 | 0.179 | 0.682 |
| AT3G21400.1 | NADH:ubiquinone oxidoreductase assembly factor (Dynein beta chain) | 38 | 2.22 | 0.182 | 0.479 |
| AT3G21400.1 | NADH:ubiquinone oxidoreductase assembly factor (Dynein beta chain) | 39 | 1.57 | 0.200 | 0.980 |
| AT3G21400.1 | NADH:ubiquinone oxidoreductase assembly factor (Dynein beta chain) | 40 | 0.97 | 0.249 | 1.000 |
| AT3G21400.1 | NADH:ubiquinone oxidoreductase assembly factor (Dynein beta chain) | 41 | 0.74 | 0.313 | 1.000 |
| AT3G21400.1 | NADH:ubiquinone oxidoreductase assembly factor (Dynein beta chain) | 45 | 0.96 | 0.322 | 1.000 |
| AT3G21400.1 | NADH:ubiquinone oxidoreductase assembly factor (Dynein beta chain) | 46 | 1.09 | 0.301 | 1.000 |
| AT3G21400.1 | NADH:ubiquinone oxidoreductase assembly factor (Dynein beta chain) | 47 | 1.20 | 0.410 | 1.000 |
| AT3G22200.1 | Gamma-aminobutyrate transaminase (GABA-T) | 3 | 0.62 | 0.216 | 1.000 |
| AT3G22200.1 | Gamma-aminobutyrate transaminase (GABA-T) | 4 | 0.49 | 0.202 | 0.972 |
| AT3G22200.1 | Gamma-aminobutyrate transaminase (GABA-T) | 5 | 0.47 | 0.202 | 0.953 |
| AT3G22200.1 | Gamma-aminobutyrate transaminase (GABA-T) | 6 | 0.56 | 0.202 | 0.999 |
| AT3G22200.1 | Gamma-aminobutyrate transaminase (GABA-T) | 7 | 0.67 | 0.202 | 1.000 |
| AT3G22200.1 | Gamma-aminobutyrate transaminase (GABA-T) | 8 | 0.77 | 0.199 | 1.000 |
| AT3G22200.1 | Gamma-aminobutyrate transaminase (GABA-T) | 9 | 0.85 | 0.196 | 1.000 |
| AT3G22200.1 | Gamma-aminobutyrate transaminase (GABA-T) | 10 | 0.91 | 0.195 | 1.000 |

| | | | | | |
|---|---|---|---|---|---|
| AT3G22200.1 | Gamma-aminobutyrate transaminase (GABA-T) | 11 | 0.99 | 0.196 | 1.000 |
| AT3G22200.1 | Gamma-aminobutyrate transaminase (GABA-T) | 12 | 1.07 | 0.196 | 1.000 |
| AT3G22200.1 | Gamma-aminobutyrate transaminase (GABA-T) | 13 | 1.13 | 0.194 | 1.000 |
| AT3G22200.1 | Gamma-aminobutyrate transaminase (GABA-T) | 14 | 1.22 | 0.191 | 1.000 |
| AT3G22200.1 | Gamma-aminobutyrate transaminase (GABA-T) | 15 | 1.31 | 0.191 | 1.000 |
| AT3G22200.1 | Gamma-aminobutyrate transaminase (GABA-T) | 16 | 1.47 | 0.189 | 1.000 |
| AT3G22200.1 | Gamma-aminobutyrate transaminase (GABA-T) | 17 | 1.87 | 0.189 | 0.989 |
| AT3G22200.1 | Gamma-aminobutyrate transaminase (GABA-T) | 18 | 2.60 | 0.192 | 0.612 |
| AT3G22200.1 | Gamma-aminobutyrate transaminase (GABA-T) | 19 | 3.27 | 0.197 | 0.265 |
| AT3G22200.1 | Gamma-aminobutyrate transaminase (GABA-T) | 20 | 3.03 | 0.202 | 0.424 |
| AT3G22200.1 | Gamma-aminobutyrate transaminase (GABA-T) | 21 | 2.27 | 0.205 | 0.909 |
| AT3G22200.1 | Gamma-aminobutyrate transaminase (GABA-T) | 22 | 1.70 | 0.204 | 1.000 |
| AT3G22200.1 | Gamma-aminobutyrate transaminase (GABA-T) | 23 | 1.30 | 0.203 | 1.000 |
| AT3G22200.1 | Gamma-aminobutyrate transaminase (GABA-T) | 24 | 0.98 | 0.199 | 1.000 |
| AT3G22200.1 | Gamma-aminobutyrate transaminase (GABA-T) | 25 | 0.68 | 0.195 | 1.000 |
| AT3G22200.1 | Gamma-aminobutyrate transaminase (GABA-T) | 26 | 0.50 | 0.195 | 0.968 |
| AT3G22200.1 | Gamma-aminobutyrate transaminase (GABA-T) | 27 | 0.58 | 0.195 | 0.999 |
| AT3G22200.1 | Gamma-aminobutyrate transaminase (GABA-T) | 28 | 0.79 | 0.194 | 1.000 |
| AT3G22200.1 | Gamma-aminobutyrate transaminase (GABA-T) | 29 | 1.00 | 0.194 | 1.000 |
| AT3G22200.1 | Gamma-aminobutyrate transaminase (GABA-T) | 30 | 1.06 | 0.194 | 1.000 |
| AT3G22200.1 | Gamma-aminobutyrate transaminase (GABA-T) | 31 | 1.11 | 0.195 | 1.000 |
| AT3G22200.1 | Gamma-aminobutyrate transaminase (GABA-T) | 32 | 1.16 | 0.195 | 1.000 |
| AT3G22200.1 | Gamma-aminobutyrate transaminase (GABA-T) | 33 | 1.09 | 0.195 | 1.000 |
| AT3G22200.1 | Gamma-aminobutyrate transaminase (GABA-T) | 34 | 1.02 | 0.196 | 1.000 |
| AT3G22200.1 | Gamma-aminobutyrate transaminase (GABA-T) | 35 | 1.07 | 0.196 | 1.000 |
| AT3G22200.1 | Gamma-aminobutyrate transaminase (GABA-T) | 36 | 1.18 | 0.195 | 1.000 |
| AT3G22200.1 | Gamma-aminobutyrate transaminase (GABA-T) | 37 | 1.29 | 0.195 | 1.000 |
| AT3G22200.1 | Gamma-aminobutyrate transaminase (GABA-T) | 38 | 1.44 | 0.195 | 1.000 |
| AT3G22200.1 | Gamma-aminobutyrate transaminase (GABA-T) | 39 | 1.75 | 0.196 | 0.998 |
| AT3G22200.1 | Gamma-aminobutyrate transaminase (GABA-T) | 40 | 2.27 | 0.196 | 0.875 |
| AT3G22200.1 | Gamma-aminobutyrate transaminase (GABA-T) | 41 | 2.30 | 0.197 | 0.858 |

| | | | | | |
|---|---|---|---|---|---|
| AT3G22200.1 | Gamma-aminobutyrate transaminase (GABA-T) | 42 | 1.66 | 0.199 | 1.000 |
| AT3G22200.1 | Gamma-aminobutyrate transaminase (GABA-T) | 43 | 1.07 | 0.201 | 1.000 |
| AT3G22200.1 | Gamma-aminobutyrate transaminase (GABA-T) | 44 | 0.67 | 0.201 | 1.000 |
| AT3G22200.1 | Gamma-aminobutyrate transaminase (GABA-T) | 45 | 0.43 | 0.201 | 0.880 |
| AT3G22200.1 | Gamma-aminobutyrate transaminase (GABA-T) | 46 | 0.26 | 0.231 | 0.325 |
| AT3G22200.1 | Gamma-aminobutyrate transaminase (GABA-T) | 47 | 0.13 | 0.415 | 0.637 |
| AT3G22200.1 | Gamma-aminobutyrate transaminase (GABA-T) | 48 | 0.06 | 0.705 | 0.926 |
| AT3G22300.1 | 40S ribosomal protein S10 (RPS10) | 2 | 0.96 | 0.561 | 1.000 |
| AT3G22300.1 | 40S ribosomal protein S10 (RPS10) | 3 | 1.09 | 0.486 | 1.000 |
| AT3G22300.1 | 40S ribosomal protein S10 (RPS10) | 11 | 1.88 | 0.305 | 0.998 |
| AT3G22300.1 | 40S ribosomal protein S10 (RPS10) | 12 | 1.80 | 0.258 | 0.996 |
| AT3G22300.1 | 40S ribosomal protein S10 (RPS10) | 13 | 1.63 | 0.227 | 0.998 |
| AT3G22300.1 | 40S ribosomal protein S10 (RPS10) | 14 | 1.44 | 0.208 | 1.000 |
| AT3G22300.1 | 40S ribosomal protein S10 (RPS10) | 15 | 1.44 | 0.196 | 1.000 |
| AT3G22300.1 | 40S ribosomal protein S10 (RPS10) | 16 | 1.73 | 0.191 | 0.966 |
| AT3G22300.1 | 40S ribosomal protein S10 (RPS10) | 17 | 2.19 | 0.190 | 0.690 |
| AT3G22300.1 | 40S ribosomal protein S10 (RPS10) | 18 | 2.42 | 0.192 | 0.530 |
| AT3G22300.1 | 40S ribosomal protein S10 (RPS10) | 19 | 2.22 | 0.197 | 0.717 |
| AT3G22300.1 | 40S ribosomal protein S10 (RPS10) | 20 | 1.78 | 0.201 | 0.969 |
| AT3G22300.1 | 40S ribosomal protein S10 (RPS10) | 21 | 1.38 | 0.204 | 1.000 |
| AT3G22300.1 | 40S ribosomal protein S10 (RPS10) | 22 | 1.09 | 0.204 | 1.000 |
| AT3G22300.1 | 40S ribosomal protein S10 (RPS10) | 23 | 0.89 | 0.202 | 1.000 |
| AT3G22300.1 | 40S ribosomal protein S10 (RPS10) | 24 | 0.78 | 0.199 | 1.000 |
| AT3G22300.1 | 40S ribosomal protein S10 (RPS10) | 25 | 0.75 | 0.195 | 1.000 |
| AT3G22300.1 | 40S ribosomal protein S10 (RPS10) | 26 | 0.83 | 0.194 | 1.000 |
| AT3G22300.1 | 40S ribosomal protein S10 (RPS10) | 27 | 0.97 | 0.194 | 1.000 |
| AT3G22300.1 | 40S ribosomal protein S10 (RPS10) | 28 | 1.11 | 0.195 | 1.000 |
| AT3G22300.1 | 40S ribosomal protein S10 (RPS10) | 29 | 1.22 | 0.195 | 1.000 |
| AT3G22300.1 | 40S ribosomal protein S10 (RPS10) | 30 | 1.32 | 0.196 | 1.000 |
| AT3G22300.1 | 40S ribosomal protein S10 (RPS10) | 31 | 1.36 | 0.197 | 1.000 |
| AT3G22300.1 | 40S ribosomal protein S10 (RPS10) | 32 | 1.25 | 0.200 | 1.000 |

| | | | | | |
|---|---|---|---|---|---|
| AT3G22300.1 | 40S ribosomal protein S10 (RPS10) | 33 | 0.97 | 0.207 | 1.000 |
| AT3G22300.1 | 40S ribosomal protein S10 (RPS10) | 34 | 0.74 | 0.231 | 1.000 |
| AT3G22300.1 | 40S ribosomal protein S10 (RPS10) | 35 | 0.62 | 0.288 | 1.000 |
| AT3G22300.1 | 40S ribosomal protein S10 (RPS10) | 36 | 0.59 | 0.395 | 1.000 |
| AT3G22300.1 | 40S ribosomal protein S10 (RPS10) | 46 | 7.80 | 2.777 | 1.000 |
| AT3G22300.1 | 40S ribosomal protein S10 (RPS10) | 47 | 11.75 | 3.134 | 1.000 |
| AT3G22330.1 | Putative mitochondrial RNA helicase 2 (PMH2) | 2 | 0.92 | 0.341 | 1.000 |
| AT3G22330.1 | Putative mitochondrial RNA helicase 2 (PMH2) | 3 | 0.99 | 0.298 | 1.000 |
| AT3G22330.1 | Putative mitochondrial RNA helicase 2 (PMH2) | 4 | 1.05 | 0.331 | 1.000 |
| AT3G22330.1 | Putative mitochondrial RNA helicase 2 (PMH2) | 14 | 0.99 | 0.346 | 1.000 |
| AT3G22330.1 | Putative mitochondrial RNA helicase 2 (PMH2) | 15 | 0.93 | 0.339 | 1.000 |
| AT3G22330.1 | Putative mitochondrial RNA helicase 2 (PMH2) | 16 | 0.87 | 0.336 | 1.000 |
| AT3G22330.1 | Putative mitochondrial RNA helicase 2 (PMH2) | 26 | 0.95 | 0.256 | 1.000 |
| AT3G22330.1 | Putative mitochondrial RNA helicase 2 (PMH2) | 27 | 1.11 | 0.216 | 1.000 |
| AT3G22330.1 | Putative mitochondrial RNA helicase 2 (PMH2) | 28 | 1.30 | 0.191 | 1.000 |
| AT3G22330.1 | Putative mitochondrial RNA helicase 2 (PMH2) | 29 | 1.50 | 0.180 | 0.987 |
| AT3G22330.1 | Putative mitochondrial RNA helicase 2 (PMH2) | 30 | 1.66 | 0.175 | 0.920 |
| AT3G22330.1 | Putative mitochondrial RNA helicase 2 (PMH2) | 31 | 1.72 | 0.173 | 0.877 |
| AT3G22330.1 | Putative mitochondrial RNA helicase 2 (PMH2) | 32 | 1.68 | 0.172 | 0.904 |
| AT3G22330.1 | Putative mitochondrial RNA helicase 2 (PMH2) | 33 | 1.53 | 0.172 | 0.974 |
| AT3G22330.1 | Putative mitochondrial RNA helicase 2 (PMH2) | 34 | 1.32 | 0.172 | 0.999 |
| AT3G22330.1 | Putative mitochondrial RNA helicase 2 (PMH2) | 35 | 1.12 | 0.172 | 1.000 |
| AT3G22330.1 | Putative mitochondrial RNA helicase 2 (PMH2) | 36 | 0.99 | 0.172 | 1.000 |
| AT3G22330.1 | Putative mitochondrial RNA helicase 2 (PMH2) | 37 | 0.98 | 0.172 | 1.000 |
| AT3G22330.1 | Putative mitochondrial RNA helicase 2 (PMH2) | 38 | 1.11 | 0.172 | 1.000 |
| AT3G22330.1 | Putative mitochondrial RNA helicase 2 (PMH2) | 39 | 1.35 | 0.172 | 0.999 |
| AT3G22330.1 | Putative mitochondrial RNA helicase 2 (PMH2) | 40 | 1.63 | 0.173 | 0.935 |
| AT3G22330.1 | Putative mitochondrial RNA helicase 2 (PMH2) | 41 | 1.76 | 0.174 | 0.850 |
| AT3G22330.1 | Putative mitochondrial RNA helicase 2 (PMH2) | 42 | 1.64 | 0.175 | 0.935 |
| AT3G22330.1 | Putative mitochondrial RNA helicase 2 (PMH2) | 43 | 1.33 | 0.176 | 0.999 |
| AT3G22330.1 | Putative mitochondrial RNA helicase 2 (PMH2) | 44 | 0.94 | 0.179 | 1.000 |

| | | | | | |
|---|---|---|---|---|---|
| AT3G22330.1 | Putative mitochondrial RNA helicase 2 (PMH2) | 45 | 0.57 | 0.191 | 0.922 |
| AT3G22330.1 | Putative mitochondrial RNA helicase 2 (PMH2) | 46 | 0.31 | 0.227 | 0.276 |
| AT3G22330.1 | Putative mitochondrial RNA helicase 2 (PMH2) | 47 | 0.15 | 0.308 | 0.115 |
| AT3G22370.1 | Alternative oxidase 1A (AOX1A) | 2 | 0.86 | 0.255 | 1.000 |
| AT3G22370.1 | Alternative oxidase 1A (AOX1A) | 3 | 1.03 | 0.234 | 1.000 |
| AT3G22370.1 | Alternative oxidase 1A (AOX1A) | 21 | 0.78 | 0.185 | 1.000 |
| AT3G22370.1 | Alternative oxidase 1A (AOX1A) | 22 | 0.68 | 0.169 | 1.000 |
| AT3G22370.1 | Alternative oxidase 1A (AOX1A) | 23 | 0.81 | 0.141 | 1.000 |
| AT3G22370.1 | Alternative oxidase 1A (AOX1A) | 24 | 1.13 | 0.152 | 1.000 |
| AT3G22370.1 | Alternative oxidase 1A (AOX1A) | 25 | 1.40 | 0.153 | 1.000 |
| AT3G22370.1 | Alternative oxidase 1A (AOX1A) | 26 | 1.51 | 0.160 | 0.998 |
| AT3G22370.1 | Alternative oxidase 1A (AOX1A) | 27 | 1.40 | 0.149 | 1.000 |
| AT3G22370.1 | Alternative oxidase 1A (AOX1A) | 28 | 1.16 | 0.141 | 1.000 |
| AT3G22370.1 | Alternative oxidase 1A (AOX1A) | 29 | 1.00 | 0.129 | 1.000 |
| AT3G22370.1 | Alternative oxidase 1A (AOX1A) | 30 | 0.96 | 0.120 | 1.000 |
| AT3G22370.1 | Alternative oxidase 1A (AOX1A) | 31 | 0.96 | 0.122 | 1.000 |
| AT3G22370.1 | Alternative oxidase 1A (AOX1A) | 32 | 1.00 | 0.122 | 1.000 |
| AT3G22370.1 | Alternative oxidase 1A (AOX1A) | 33 | 0.92 | 0.121 | 1.000 |
| AT3G22370.1 | Alternative oxidase 1A (AOX1A) | 34 | 0.86 | 0.121 | 1.000 |
| AT3G22370.1 | Alternative oxidase 1A (AOX1A) | 35 | 0.86 | 0.121 | 1.000 |
| AT3G22370.1 | Alternative oxidase 1A (AOX1A) | 36 | 0.92 | 0.121 | 1.000 |
| AT3G22370.1 | Alternative oxidase 1A (AOX1A) | 37 | 1.24 | 0.121 | 1.000 |
| AT3G22370.1 | Alternative oxidase 1A (AOX1A) | 38 | 1.64 | 0.120 | 0.809 |
| AT3G22370.1 | Alternative oxidase 1A (AOX1A) | 39 | 1.73 | 0.120 | 0.644 |
| AT3G22370.1 | Alternative oxidase 1A (AOX1A) | 40 | 1.42 | 0.121 | 0.992 |
| AT3G22370.1 | Alternative oxidase 1A (AOX1A) | 41 | 0.78 | 0.121 | 1.000 |
| AT3G22370.1 | Alternative oxidase 1A (AOX1A) | 42 | 0.46 | 0.124 | 0.152 |
| AT3G22370.1 | Alternative oxidase 1A (AOX1A) | 43 | 0.66 | 0.124 | 0.954 |
| AT3G22370.1 | Alternative oxidase 1A (AOX1A) | 44 | 1.32 | 0.122 | 0.999 |
| AT3G22370.1 | Alternative oxidase 1A (AOX1A) | 45 | 1.16 | 0.123 | 1.000 |
| AT3G22370.1 | Alternative oxidase 1A (AOX1A) | 46 | 0.46 | 0.135 | 0.262 |
| AT3G22370.1 | Alternative oxidase 1A (AOX1A) | 47 | 0.16 | 0.310 | 0.244 |
| AT3G23990.1 | Chaperonin 60 beta 3 (HSP603B) | 3 | 0.58 | 0.165 | 0.992 |
| AT3G23990.1 | Chaperonin 60 beta 3 (HSP603B) | 4 | 0.44 | 0.152 | 0.439 |
| AT3G23990.1 | Chaperonin 60 beta 3 (HSP603B) | 5 | 0.44 | 0.152 | 0.494 |
| AT3G23990.1 | Chaperonin 60 beta 3 (HSP603B) | 6 | 0.57 | 0.152 | 0.968 |
| AT3G23990.1 | Chaperonin 60 beta 3 (HSP603B) | 7 | 0.75 | 0.152 | 1.000 |
| AT3G23990.1 | Chaperonin 60 beta 3 (HSP603B) | 8 | 0.95 | 0.149 | 1.000 |
| AT3G23990.1 | Chaperonin 60 beta 3 (HSP603B) | 9 | 1.15 | 0.147 | 1.000 |
| AT3G23990.1 | Chaperonin 60 beta 3 (HSP603B) | 10 | 1.26 | 0.146 | 1.000 |
| AT3G23990.1 | Chaperonin 60 beta 3 (HSP603B) | 11 | 1.43 | 0.147 | 1.000 |
| AT3G23990.1 | Chaperonin 60 beta 3 (HSP603B) | 12 | 1.67 | 0.147 | 0.981 |
| AT3G23990.1 | Chaperonin 60 beta 3 (HSP603B) | 13 | 1.91 | 0.145 | 0.813 |
| AT3G23990.1 | Chaperonin 60 beta 3 (HSP603B) | 14 | 2.16 | 0.143 | 0.473 |
| AT3G23990.1 | Chaperonin 60 beta 3 (HSP603B) | 15 | 2.29 | 0.142 | 0.334 |

| | | | | | | |
|---|---|---|---|---|---|---|
| AT3G23990.1 | Chaperonin 60 beta 3 (HSP603B) | 16 | 2.18 | 0.141 | 0.425 | |
| AT3G23990.1 | Chaperonin 60 beta 3 (HSP603B) | 17 | 1.88 | 0.141 | 0.809 | |
| AT3G23990.1 | Chaperonin 60 beta 3 (HSP603B) | 18 | 1.51 | 0.144 | 0.999 | |
| AT3G23990.1 | Chaperonin 60 beta 3 (HSP603B) | 19 | 1.32 | 0.148 | 1.000 | |
| AT3G23990.1 | Chaperonin 60 beta 3 (HSP603B) | 20 | 1.27 | 0.152 | 1.000 | |
| AT3G23990.1 | Chaperonin 60 beta 3 (HSP603B) | 21 | 1.17 | 0.154 | 1.000 | |
| AT3G23990.1 | Chaperonin 60 beta 3 (HSP603B) | 22 | 1.09 | 0.154 | 1.000 | |
| AT3G23990.1 | Chaperonin 60 beta 3 (HSP603B) | 23 | 1.04 | 0.153 | 1.000 | |
| AT3G23990.1 | Chaperonin 60 beta 3 (HSP603B) | 24 | 1.07 | 0.149 | 1.000 | |
| AT3G23990.1 | Chaperonin 60 beta 3 (HSP603B) | 25 | 1.10 | 0.146 | 1.000 | |
| AT3G23990.1 | Chaperonin 60 beta 3 (HSP603B) | 26 | 1.06 | 0.146 | 1.000 | |
| AT3G23990.1 | Chaperonin 60 beta 3 (HSP603B) | 27 | 1.03 | 0.146 | 1.000 | |
| AT3G23990.1 | Chaperonin 60 beta 3 (HSP603B) | 28 | 1.23 | 0.146 | 1.000 | |
| AT3G23990.1 | Chaperonin 60 beta 3 (HSP603B) | 29 | 1.46 | 0.146 | 1.000 | |
| AT3G23990.1 | Chaperonin 60 beta 3 (HSP603B) | 30 | 1.17 | 0.145 | 1.000 | |
| AT3G23990.1 | Chaperonin 60 beta 3 (HSP603B) | 31 | 0.75 | 0.146 | 1.000 | |
| AT3G23990.1 | Chaperonin 60 beta 3 (HSP603B) | 32 | 0.62 | 0.146 | 0.992 | |
| AT3G23990.1 | Chaperonin 60 beta 3 (HSP603B) | 33 | 0.73 | 0.147 | 1.000 | |
| AT3G23990.1 | Chaperonin 60 beta 3 (HSP603B) | 34 | 0.93 | 0.147 | 1.000 | |
| AT3G23990.1 | Chaperonin 60 beta 3 (HSP603B) | 35 | 1.10 | 0.147 | 1.000 | |
| AT3G23990.1 | Chaperonin 60 beta 3 (HSP603B) | 36 | 1.29 | 0.146 | 1.000 | |
| AT3G23990.1 | Chaperonin 60 beta 3 (HSP603B) | 37 | 1.56 | 0.146 | 0.997 | |
| AT3G23990.1 | Chaperonin 60 beta 3 (HSP603B) | 38 | 1.85 | 0.146 | 0.880 | |
| AT3G23990.1 | Chaperonin 60 beta 3 (HSP603B) | 39 | 2.26 | 0.147 | 0.412 | |
| AT3G23990.1 | Chaperonin 60 beta 3 (HSP603B) | 40 | 2.76 | 0.147 | 0.099 | . |
| AT3G23990.1 | Chaperonin 60 beta 3 (HSP603B) | 41 | 2.87 | 0.148 | 0.072 | . |
| AT3G23990.1 | Chaperonin 60 beta 3 (HSP603B) | 42 | 2.62 | 0.150 | 0.172 | |
| AT3G23990.1 | Chaperonin 60 beta 3 (HSP603B) | 43 | 2.30 | 0.151 | 0.432 | |
| AT3G23990.1 | Chaperonin 60 beta 3 (HSP603B) | 44 | 1.72 | 0.151 | 0.975 | |
| AT3G23990.1 | Chaperonin 60 beta 3 (HSP603B) | 45 | 0.99 | 0.151 | 1.000 | |
| AT3G23990.1 | Chaperonin 60 beta 3 (HSP603B) | 46 | 0.41 | 0.178 | 0.633 | |

| | | | | | |
|---|---|---|---|---|---|
| AT3G23990.1 | Chaperonin 60 beta 3 (HSP603B) | 47 | 0.13 | 0.317 | 0.175 |
| AT3G23990.1 | Chaperonin 60 beta 3 (HSP603B) | 48 | 0.04 | 0.533 | 0.260 |
| AT3G24200.1 | Putative (yeast COQ6)-like ubiquinone biosynthesis hydroxylase | 2 | 0.79 | 0.354 | 1.000 |
| AT3G24200.1 | Putative (yeast COQ6)-like ubiquinone biosynthesis hydroxylase | 3 | 0.74 | 0.340 | 1.000 |
| AT3G24200.1 | Putative (yeast COQ6)-like ubiquinone biosynthesis hydroxylase | 29 | 0.50 | 0.250 | 0.938 |
| AT3G24200.1 | Putative (yeast COQ6)-like ubiquinone biosynthesis hydroxylase | 30 | 0.56 | 0.192 | 0.896 |
| AT3G24200.1 | Putative (yeast COQ6)-like ubiquinone biosynthesis hydroxylase | 31 | 0.69 | 0.172 | 0.988 |
| AT3G24200.1 | Putative (yeast COQ6)-like ubiquinone biosynthesis hydroxylase | 32 | 0.85 | 0.165 | 1.000 |
| AT3G24200.1 | Putative (yeast COQ6)-like ubiquinone biosynthesis hydroxylase | 33 | 0.94 | 0.162 | 1.000 |
| AT3G24200.1 | Putative (yeast COQ6)-like ubiquinone biosynthesis hydroxylase | 34 | 0.90 | 0.162 | 1.000 |
| AT3G24200.1 | Putative (yeast COQ6)-like ubiquinone biosynthesis hydroxylase | 35 | 0.91 | 0.161 | 1.000 |
| AT3G24200.1 | Putative (yeast COQ6)-like ubiquinone biosynthesis hydroxylase | 36 | 1.14 | 0.161 | 1.000 |
| AT3G24200.1 | Putative (yeast COQ6)-like ubiquinone biosynthesis hydroxylase | 37 | 1.69 | 0.161 | 0.838 |
| AT3G24200.1 | Putative (yeast COQ6)-like ubiquinone biosynthesis hydroxylase | 38 | 2.33 | 0.160 | 0.244 |
| AT3G24200.1 | Putative (yeast COQ6)-like ubiquinone biosynthesis hydroxylase | 39 | 2.52 | 0.160 | 0.150 |
| AT3G24200.1 | Putative (yeast COQ6)-like ubiquinone biosynthesis hydroxylase | 40 | 2.13 | 0.160 | 0.385 |
| AT3G24200.1 | Putative (yeast COQ6)-like ubiquinone biosynthesis hydroxylase | 41 | 1.57 | 0.161 | 0.932 |
| AT3G24200.1 | Putative (yeast COQ6)-like ubiquinone biosynthesis hydroxylase | 42 | 1.22 | 0.163 | 1.000 |
| AT3G24200.1 | Putative (yeast COQ6)-like ubiquinone biosynthesis hydroxylase | 43 | 1.06 | 0.164 | 1.000 |
| AT3G24200.1 | Putative (yeast COQ6)-like ubiquinone biosynthesis hydroxylase | 44 | 0.90 | 0.164 | 1.000 |
| AT3G24200.1 | Putative (yeast COQ6)-like ubiquinone biosynthesis hydroxylase | 45 | 0.66 | 0.165 | 0.968 |

| | | | | | |
|---|---|---|---|---|---|
| AT3G24200.1 | Putative (yeast COQ6)-like ubiquinone biosynthesis hydroxylase | 46 | 0.40 | 0.193 | 0.386 |
| AT3G24200.1 | Putative (yeast COQ6)-like ubiquinone biosynthesis hydroxylase | 47 | 0.21 | 0.309 | 0.307 |
| AT3G24830.1 | 60S ribosomal protein L13a-2 (RPL13AB) | 2 | 1.40 | 0.223 | 0.963 |
| AT3G24830.1 | 60S ribosomal protein L13a-2 (RPL13AB) | 3 | 1.40 | 0.200 | 0.942 |
| AT3G24830.1 | 60S ribosomal protein L13a-2 (RPL13AB) | 4 | 1.40 | 0.179 | 0.912 |
| AT3G24830.1 | 60S ribosomal protein L13a-2 (RPL13AB) | 5 | 1.40 | 0.162 | 0.874 |
| AT3G24830.1 | 60S ribosomal protein L13a-2 (RPL13AB) | 6 | 1.40 | 0.148 | 0.833 |
| AT3G24830.1 | 60S ribosomal protein L13a-2 (RPL13AB) | 7 | 1.39 | 0.137 | 0.796 |
| AT3G24830.1 | 60S ribosomal protein L13a-2 (RPL13AB) | 8 | 1.39 | 0.130 | 0.771 |
| AT3G24830.1 | 60S ribosomal protein L13a-2 (RPL13AB) | 9 | 1.38 | 0.127 | 0.762 |
| AT3G24830.1 | 60S ribosomal protein L13a-2 (RPL13AB) | 10 | 1.37 | 0.126 | 0.768 |
| AT3G24830.1 | 60S ribosomal protein L13a-2 (RPL13AB) | 11 | 1.37 | 0.127 | 0.786 |
| AT3G24830.1 | 60S ribosomal protein L13a-2 (RPL13AB) | 12 | 1.36 | 0.131 | 0.809 |
| AT3G24830.1 | 60S ribosomal protein L13a-2 (RPL13AB) | 13 | 1.36 | 0.136 | 0.833 |
| AT3G24830.1 | 60S ribosomal protein L13a-2 (RPL13AB) | 14 | 1.36 | 0.143 | 0.854 |
| AT3G24830.1 | 60S ribosomal protein L13a-2 (RPL13AB) | 15 | 1.37 | 0.151 | 0.873 |
| AT3G24830.1 | 60S ribosomal protein L13a-2 (RPL13AB) | 16 | 1.38 | 0.160 | 0.889 |
| AT3G24830.1 | 60S ribosomal protein L13a-2 (RPL13AB) | 17 | 1.38 | 0.170 | 0.905 |
| AT3G24830.1 | 60S ribosomal protein L13a-2 (RPL13AB) | 22 | 1.40 | 0.221 | 0.960 |
| AT3G24830.1 | 60S ribosomal protein L13a-2 (RPL13AB) | 23 | 1.41 | 0.231 | 0.966 |
| AT3G24830.1 | 60S ribosomal protein L13a-2 (RPL13AB) | 24 | 1.41 | 0.241 | 0.971 |
| AT3G24830.1 | 60S ribosomal protein L13a-2 (RPL13AB) | 41 | 1.45 | 0.382 | 0.995 |
| AT3G24830.1 | 60S ribosomal protein L13a-2 (RPL13AB) | 42 | 1.46 | 0.398 | 0.996 |
| AT3G24830.1 | 60S ribosomal protein L13a-2 (RPL13AB) | 43 | 1.47 | 0.415 | 0.996 |
| AT3G24830.1 | 60S ribosomal protein L13a-2 (RPL13AB) | 46 | 1.49 | 0.479 | 0.998 |
| AT3G24830.1 | 60S ribosomal protein L13a-2 (RPL13AB) | 47 | 1.50 | 0.505 | 0.998 |
| AT3G25660.1 | Putative subunit A of glutamyl-tRNA-dependent amidotransferase complex (GATa) | 2 | 0.85 | 0.597 | 1.000 |
| AT3G25660.1 | Putative subunit A of glutamyl-tRNA-dependent amidotransferase complex (GATa) | 3 | 1.23 | 0.573 | 1.000 |

| | | | | | |
|---|---|---|---|---|---|
| AT3G25660.1 | Putative subunit A of glutamyl-tRNA-dependent amidotransferase complex (GATa) | 24 | 2.09 | 0.489 | 1.000 |
| AT3G25660.1 | Putative subunit A of glutamyl-tRNA-dependent amidotransferase complex (GATa) | 25 | 1.14 | 0.359 | 1.000 |
| AT3G25660.1 | Putative subunit A of glutamyl-tRNA-dependent amidotransferase complex (GATa) | 26 | 0.73 | 0.321 | 1.000 |
| AT3G25660.1 | Putative subunit A of glutamyl-tRNA-dependent amidotransferase complex (GATa) | 27 | 0.51 | 0.311 | 0.997 |
| AT3G25660.1 | Putative subunit A of glutamyl-tRNA-dependent amidotransferase complex (GATa) | 28 | 0.39 | 0.311 | 0.944 |
| AT3G25660.1 | Putative subunit A of glutamyl-tRNA-dependent amidotransferase complex (GATa) | 29 | 0.37 | 0.310 | 0.908 |
| AT3G25660.1 | Putative subunit A of glutamyl-tRNA-dependent amidotransferase complex (GATa) | 30 | 0.41 | 0.309 | 0.956 |
| AT3G25660.1 | Putative subunit A of glutamyl-tRNA-dependent amidotransferase complex (GATa) | 31 | 0.53 | 0.309 | 0.999 |
| AT3G25660.1 | Putative subunit A of glutamyl-tRNA-dependent amidotransferase complex (GATa) | 32 | 0.59 | 0.310 | 1.000 |
| AT3G25660.1 | Putative subunit A of glutamyl-tRNA-dependent amidotransferase complex (GATa) | 33 | 0.50 | 0.310 | 0.995 |
| AT3G25660.1 | Putative subunit A of glutamyl-tRNA-dependent amidotransferase complex (GATa) | 34 | 0.31 | 0.311 | 0.768 |
| AT3G25660.1 | Putative subunit A of glutamyl-tRNA-dependent amidotransferase complex (GATa) | 35 | 0.17 | 0.311 | 0.216 |
| AT3G25660.1 | Putative subunit A of glutamyl-tRNA-dependent amidotransferase complex (GATa) | 36 | 0.18 | 0.311 | 0.238 |
| AT3G25660.1 | Putative subunit A of glutamyl-tRNA-dependent amidotransferase complex (GATa) | 37 | 0.36 | 0.311 | 0.885 |
| AT3G25660.1 | Putative subunit A of glutamyl-tRNA-dependent amidotransferase complex (GATa) | 38 | 0.96 | 0.310 | 1.000 |

| | | | | | |
|---|---|---|---|---|---|
| AT3G25660.1 | Putative subunit A of glutamyl-tRNA-dependent amidotransferase complex (GATa) | 39 | 2.64 | 0.309 | 0.923 |
| AT3G25660.1 | Putative subunit A of glutamyl-tRNA-dependent amidotransferase complex (GATa) | 40 | 3.73 | 0.308 | 0.605 |
| AT3G25660.1 | Putative subunit A of glutamyl-tRNA-dependent amidotransferase complex (GATa) | 41 | 1.65 | 0.311 | 1.000 |
| AT3G25660.1 | Putative subunit A of glutamyl-tRNA-dependent amidotransferase complex (GATa) | 42 | 0.37 | 0.315 | 0.917 |
| AT3G25660.1 | Putative subunit A of glutamyl-tRNA-dependent amidotransferase complex (GATa) | 43 | 0.14 | 0.316 | 0.129 |
| AT3G25660.1 | Putative subunit A of glutamyl-tRNA-dependent amidotransferase complex (GATa) | 44 | 0.23 | 0.320 | 0.510 |
| AT3G25660.1 | Putative subunit A of glutamyl-tRNA-dependent amidotransferase complex (GATa) | 45 | 0.90 | 0.374 | 1.000 |
| AT3G25660.1 | Putative subunit A of glutamyl-tRNA-dependent amidotransferase complex (GATa) | 46 | 4.12 | 0.503 | 0.966 |
| AT3G25660.1 | Putative subunit A of glutamyl-tRNA-dependent amidotransferase complex (GATa) | 47 | 19.18 | 0.811 | 0.807 |
| AT3G25860.1 | Dihydrolipoamide acetyltransferase (PDCE2) | 2 | 1.08 | 0.374 | 1.000 |
| AT3G25860.1 | Dihydrolipoamide acetyltransferase (PDCE2) | 3 | 0.93 | 0.288 | 1.000 |
| AT3G25860.1 | Dihydrolipoamide acetyltransferase (PDCE2) | 6 | 0.63 | 0.226 | 0.986 |
| AT3G25860.1 | Dihydrolipoamide acetyltransferase (PDCE2) | 7 | 0.58 | 0.190 | 0.893 |
| AT3G25860.1 | Dihydrolipoamide acetyltransferase (PDCE2) | 8 | 0.57 | 0.163 | 0.743 |
| AT3G25860.1 | Dihydrolipoamide acetyltransferase (PDCE2) | 9 | 0.60 | 0.149 | 0.765 |
| AT3G25860.1 | Dihydrolipoamide acetyltransferase (PDCE2) | 10 | 0.64 | 0.145 | 0.858 |
| AT3G25860.1 | Dihydrolipoamide acetyltransferase (PDCE2) | 11 | 0.68 | 0.144 | 0.936 |
| AT3G25860.1 | Dihydrolipoamide acetyltransferase (PDCE2) | 12 | 0.73 | 0.144 | 0.979 |
| AT3G25860.1 | Dihydrolipoamide acetyltransferase (PDCE2) | 13 | 0.78 | 0.144 | 0.996 |
| AT3G25860.1 | Dihydrolipoamide acetyltransferase (PDCE2) | 14 | 0.91 | 0.147 | 1.000 |
| AT3G25860.1 | Dihydrolipoamide acetyltransferase (PDCE2) | 15 | 1.30 | 0.151 | 0.997 |
| AT3G25860.1 | Dihydrolipoamide acetyltransferase (PDCE2) | 16 | 2.01 | 0.158 | 0.446 |

| | | | | | |
|---|---|---|---|---|---|
| AT3G25860.1 | Dihydrolipoamide acetyltransferase (PDCE2) | 17 | 2.74 | 0.185 | 0.193 |
| AT3G25860.1 | Dihydrolipoamide acetyltransferase (PDCE2) | 18 | 3.11 | 0.260 | 0.463 |
| AT3G25860.1 | Dihydrolipoamide acetyltransferase (PDCE2) | 46 | 617213.41 | 11.761 | 1.000 |
| AT3G25860.1 | Dihydrolipoamide acetyltransferase (PDCE2) | 47 | 1135097.92 | 12.378 | 1.000 |
| AT3G26520.1 | Tonoplast intrinsic protein (gammaTIP2) | 2 | 0.80 | 0.325 | 1.000 |
| AT3G26520.1 | Tonoplast intrinsic protein (gammaTIP2) | 3 | 0.65 | 0.222 | 1.000 |
| AT3G26520.1 | Tonoplast intrinsic protein (gammaTIP2) | 4 | 0.59 | 0.203 | 0.990 |
| AT3G26520.1 | Tonoplast intrinsic protein (gammaTIP2) | 5 | 0.61 | 0.203 | 0.996 |
| AT3G26520.1 | Tonoplast intrinsic protein (gammaTIP2) | 6 | 0.71 | 0.203 | 1.000 |
| AT3G26520.1 | Tonoplast intrinsic protein (gammaTIP2) | 7 | 0.83 | 0.202 | 1.000 |
| AT3G26520.1 | Tonoplast intrinsic protein (gammaTIP2) | 8 | 0.95 | 0.200 | 1.000 |
| AT3G26520.1 | Tonoplast intrinsic protein (gammaTIP2) | 9 | 1.05 | 0.199 | 1.000 |
| AT3G26520.1 | Tonoplast intrinsic protein (gammaTIP2) | 10 | 1.16 | 0.199 | 1.000 |
| AT3G26520.1 | Tonoplast intrinsic protein (gammaTIP2) | 11 | 1.25 | 0.199 | 1.000 |
| AT3G26520.1 | Tonoplast intrinsic protein (gammaTIP2) | 12 | 1.25 | 0.199 | 1.000 |
| AT3G26520.1 | Tonoplast intrinsic protein (gammaTIP2) | 13 | 1.19 | 0.197 | 1.000 |
| AT3G26520.1 | Tonoplast intrinsic protein (gammaTIP2) | 14 | 1.11 | 0.196 | 1.000 |
| AT3G26520.1 | Tonoplast intrinsic protein (gammaTIP2) | 15 | 1.02 | 0.195 | 1.000 |
| AT3G26520.1 | Tonoplast intrinsic protein (gammaTIP2) | 16 | 0.95 | 0.194 | 1.000 |
| AT3G26520.1 | Tonoplast intrinsic protein (gammaTIP2) | 17 | 1.00 | 0.195 | 1.000 |
| AT3G26520.1 | Tonoplast intrinsic protein (gammaTIP2) | 18 | 1.19 | 0.199 | 1.000 |
| AT3G26520.1 | Tonoplast intrinsic protein (gammaTIP2) | 19 | 1.39 | 0.206 | 1.000 |
| AT3G26520.1 | Tonoplast intrinsic protein (gammaTIP2) | 20 | 1.52 | 0.218 | 1.000 |
| AT3G26520.1 | Tonoplast intrinsic protein (gammaTIP2) | 21 | 1.50 | 0.241 | 1.000 |
| AT3G26520.1 | Tonoplast intrinsic protein (gammaTIP2) | 22 | 1.33 | 0.265 | 1.000 |
| AT3G26520.1 | Tonoplast intrinsic protein (gammaTIP2) | 25 | 1.04 | 0.260 | 1.000 |
| AT3G26520.1 | Tonoplast intrinsic protein (gammaTIP2) | 26 | 1.22 | 0.234 | 1.000 |
| AT3G26520.1 | Tonoplast intrinsic protein (gammaTIP2) | 27 | 1.67 | 0.210 | 0.995 |
| AT3G26520.1 | Tonoplast intrinsic protein (gammaTIP2) | 28 | 2.26 | 0.200 | 0.722 |
| AT3G26520.1 | Tonoplast intrinsic protein (gammaTIP2) | 29 | 2.73 | 0.198 | 0.387 |
| AT3G26520.1 | Tonoplast intrinsic protein (gammaTIP2) | 30 | 2.96 | 0.198 | 0.274 |

| | | | | | |
|---|---|---|---|---|---|
| AT3G26520.1 | Tonoplast intrinsic protein (gammaTIP2) | 31 | 3.02 | 0.198 | 0.253 |
| AT3G26520.1 | Tonoplast intrinsic protein (gammaTIP2) | 32 | 2.70 | 0.199 | 0.407 |
| AT3G26520.1 | Tonoplast intrinsic protein (gammaTIP2) | 33 | 2.09 | 0.200 | 0.838 |
| AT3G26520.1 | Tonoplast intrinsic protein (gammaTIP2) | 34 | 1.56 | 0.204 | 0.999 |
| AT3G26520.1 | Tonoplast intrinsic protein (gammaTIP2) | 35 | 1.42 | 0.216 | 1.000 |
| AT3G26520.1 | Tonoplast intrinsic protein (gammaTIP2) | 36 | 1.74 | 0.252 | 0.999 |
| AT3G26520.1 | Tonoplast intrinsic protein (gammaTIP2) | 37 | 2.52 | 0.346 | 0.987 |
| AT3G26520.1 | Tonoplast intrinsic protein (gammaTIP2) | 46 | 809.82 | 2.911 | 0.998 |
| AT3G26520.1 | Tonoplast intrinsic protein (gammaTIP2) | 47 | 1749.35 | 3.340 | 0.998 |
| AT3G26650.1 | Glyceraldehyde-3-phosphate dehydrogenase (GAPA1) | 2 | 0.99 | 0.792 | 1.000 |
| AT3G26650.1 | Glyceraldehyde-3-phosphate dehydrogenase (GAPA1) | 3 | 0.89 | 1.122 | 1.000 |
| AT3G26650.1 | Glyceraldehyde-3-phosphate dehydrogenase (GAPA1) | 31 | 6.27 | 0.543 | 0.856 |
| AT3G26650.1 | Glyceraldehyde-3-phosphate dehydrogenase (GAPA1) | 32 | 3.65 | 0.420 | 0.916 |
| AT3G26650.1 | Glyceraldehyde-3-phosphate dehydrogenase (GAPA1) | 33 | 1.84 | 0.479 | 1.000 |
| AT3G26650.1 | Glyceraldehyde-3-phosphate dehydrogenase (GAPA1) | 34 | 0.94 | 0.505 | 1.000 |
| AT3G26650.1 | Glyceraldehyde-3-phosphate dehydrogenase (GAPA1) | 36 | 1.36 | 0.478 | 1.000 |
| AT3G26650.1 | Glyceraldehyde-3-phosphate dehydrogenase (GAPA1) | 37 | 2.29 | 0.405 | 0.997 |
| AT3G26650.1 | Glyceraldehyde-3-phosphate dehydrogenase (GAPA1) | 38 | 3.43 | 0.353 | 0.830 |
| AT3G26650.1 | Glyceraldehyde-3-phosphate dehydrogenase (GAPA1) | 39 | 5.63 | 0.340 | 0.331 |
| AT3G26650.1 | Glyceraldehyde-3-phosphate dehydrogenase (GAPA1) | 40 | 4.80 | 0.338 | 0.465 |
| AT3G26650.1 | Glyceraldehyde-3-phosphate dehydrogenase (GAPA1) | 41 | 1.25 | 0.337 | 1.000 |
| AT3G26650.1 | Glyceraldehyde-3-phosphate dehydrogenase (GAPA1) | 42 | 0.29 | 0.354 | 0.837 |
| AT3G26650.1 | Glyceraldehyde-3-phosphate dehydrogenase (GAPA1) | 43 | 0.23 | 0.406 | 0.806 |
| AT3G26650.1 | Glyceraldehyde-3-phosphate dehydrogenase (GAPA1) | 44 | 0.46 | 0.437 | 0.999 |
| AT3G26650.1 | Glyceraldehyde-3-phosphate dehydrogenase (GAPA1) | 45 | 1.02 | 0.460 | 1.000 |
| AT3G26650.1 | Glyceraldehyde-3-phosphate dehydrogenase (GAPA1) | 46 | 2.63 | 0.696 | 1.000 |
| AT3G26650.1 | Glyceraldehyde-3-phosphate dehydrogenase (GAPA1) | 47 | 8.35 | 1.294 | 1.000 |
| AT3G26770.1 | NAD(P)-binding Rossmann-fold superfamily protein | 2 | 1.05 | 0.389 | 1.000 |
| AT3G26770.1 | NAD(P)-binding Rossmann-fold superfamily protein | 3 | 0.93 | 0.335 | 1.000 |
| AT3G26770.1 | NAD(P)-binding Rossmann-fold superfamily protein | 4 | 0.83 | 0.416 | 1.000 |
| AT3G26770.1 | NAD(P)-binding Rossmann-fold superfamily protein | 41 | 0.94 | 0.357 | 1.000 |

| | | | | | |
|---|---|---|---|---|---|
| AT3G26770.1 | NAD(P)-binding Rossmann-fold superfamily protein | 42 | 0.95 | 0.275 | 1.000 |
| AT3G26770.1 | NAD(P)-binding Rossmann-fold superfamily protein | 43 | 0.94 | 0.219 | 1.000 |
| AT3G26770.1 | NAD(P)-binding Rossmann-fold superfamily protein | 44 | 0.92 | 0.200 | 1.000 |
| AT3G26770.1 | NAD(P)-binding Rossmann-fold superfamily protein | 45 | 0.89 | 0.222 | 1.000 |
| AT3G26770.1 | NAD(P)-binding Rossmann-fold superfamily protein | 46 | 0.86 | 0.284 | 1.000 |
| AT3G26770.1 | NAD(P)-binding Rossmann-fold superfamily protein | 47 | 0.82 | 0.382 | 1.000 |
| AT3G27080.1 | Translocase of the outer mitochondrial membrane 20 (TOM20-3) | 2 | 0.97 | 0.544 | 1.000 |
| AT3G27080.1 | Translocase of the outer mitochondrial membrane 20 (TOM20-3) | 3 | 0.98 | 0.500 | 1.000 |
| AT3G27080.1 | Translocase of the outer mitochondrial membrane 20 (TOM20-3) | 18 | 0.75 | 0.420 | 1.000 |
| AT3G27080.1 | Translocase of the outer mitochondrial membrane 20 (TOM20-3) | 19 | 0.80 | 0.404 | 1.000 |
| AT3G27080.1 | Translocase of the outer mitochondrial membrane 20 (TOM20-3) | 20 | 0.94 | 0.385 | 1.000 |
| AT3G27080.1 | Translocase of the outer mitochondrial membrane 20 (TOM20-3) | 22 | 1.56 | 0.402 | 1.000 |
| AT3G27080.1 | Translocase of the outer mitochondrial membrane 20 (TOM20-3) | 23 | 1.86 | 0.324 | 0.999 |
| AT3G27080.1 | Translocase of the outer mitochondrial membrane 20 (TOM20-3) | 24 | 1.69 | 0.274 | 0.999 |
| AT3G27080.1 | Translocase of the outer mitochondrial membrane 20 (TOM20-3) | 25 | 1.09 | 0.260 | 1.000 |
| AT3G27080.1 | Translocase of the outer mitochondrial membrane 20 (TOM20-3) | 26 | 0.71 | 0.257 | 1.000 |
| AT3G27080.1 | Translocase of the outer mitochondrial membrane 20 (TOM20-3) | 27 | 0.66 | 0.256 | 1.000 |
| AT3G27080.1 | Translocase of the outer mitochondrial membrane 20 (TOM20-3) | 28 | 0.88 | 0.258 | 1.000 |
| AT3G27080.1 | Translocase of the outer mitochondrial membrane 20 (TOM20-3) | 29 | 2.06 | 0.256 | 0.971 |
| AT3G27080.1 | Translocase of the outer mitochondrial membrane 20 (TOM20-3) | 30 | 3.37 | 0.259 | 0.490 |
| AT3G27080.1 | Translocase of the outer mitochondrial membrane 20 (TOM20-3) | 31 | 1.56 | 0.257 | 1.000 |
| AT3G27080.1 | Translocase of the outer mitochondrial membrane 20 (TOM20-3) | 32 | 0.33 | 0.260 | 0.633 |

| | | | | | | |
|---|---|---|---|---|---|---|
| AT3G27080.1 | Translocase of the outer mitochondrial membrane 20 (TOM20-3) | 33 | 0.16 | 0.265 | 0.046 | * |
| AT3G27080.1 | Translocase of the outer mitochondrial membrane 20 (TOM20-3) | 34 | 0.34 | 0.281 | 0.777 | |
| AT3G27080.1 | Translocase of the outer mitochondrial membrane 20 (TOM20-3) | 35 | 1.12 | 0.339 | 1.000 | |
| AT3G27080.1 | Translocase of the outer mitochondrial membrane 20 (TOM20-3) | 36 | 2.97 | 0.595 | 1.000 | |
| AT3G27080.1 | Translocase of the outer mitochondrial membrane 20 (TOM20-3) | 46 | 86.18 | 6.001 | 1.000 | |
| AT3G27080.1 | Translocase of the outer mitochondrial membrane 20 (TOM20-3) | 47 | 89.08 | 6.840 | 1.000 | |
| AT3G27240.1 | Ubiquinol:cytochrome c oxidoreductase subunit (cyt c1-2) | 3 | 1.28 | 0.224 | 1.000 | |
| AT3G27240.1 | Ubiquinol:cytochrome c oxidoreductase subunit (cyt c1-2) | 4 | 1.43 | 0.211 | 1.000 | |
| AT3G27240.1 | Ubiquinol:cytochrome c oxidoreductase subunit (cyt c1-2) | 5 | 1.43 | 0.210 | 1.000 | |
| AT3G27240.1 | Ubiquinol:cytochrome c oxidoreductase subunit (cyt c1-2) | 6 | 1.16 | 0.211 | 1.000 | |
| AT3G27240.1 | Ubiquinol:cytochrome c oxidoreductase subunit (cyt c1-2) | 7 | 0.93 | 0.213 | 1.000 | |
| AT3G27240.1 | Ubiquinol:cytochrome c oxidoreductase subunit (cyt c1-2) | 8 | 0.98 | 0.207 | 1.000 | |
| AT3G27240.1 | Ubiquinol:cytochrome c oxidoreductase subunit (cyt c1-2) | 9 | 1.09 | 0.203 | 1.000 | |
| AT3G27240.1 | Ubiquinol:cytochrome c oxidoreductase subunit (cyt c1-2) | 10 | 1.16 | 0.203 | 1.000 | |
| AT3G27240.1 | Ubiquinol:cytochrome c oxidoreductase subunit (cyt c1-2) | 11 | 1.31 | 0.204 | 1.000 | |
| AT3G27240.1 | Ubiquinol:cytochrome c oxidoreductase subunit (cyt c1-2) | 12 | 1.49 | 0.205 | 1.000 | |
| AT3G27240.1 | Ubiquinol:cytochrome c oxidoreductase subunit (cyt c1-2) | 13 | 1.60 | 0.202 | 1.000 | |
| AT3G27240.1 | Ubiquinol:cytochrome c oxidoreductase subunit (cyt c1-2) | 14 | 1.65 | 0.197 | 1.000 | |
| AT3G27240.1 | Ubiquinol:cytochrome c oxidoreductase subunit (cyt c1-2) | 15 | 1.43 | 0.197 | 1.000 | |
| AT3G27240.1 | Ubiquinol:cytochrome c oxidoreductase subunit (cyt c1-2) | 16 | 0.97 | 0.195 | 1.000 | |

| | | | | | |
|---|---|---|---|---|---|
| AT3G27240.1 | Ubiquinol:cytochrome c oxidoreductase subunit (cyt c1-2) | 17 | 0.67 | 0.196 | 1.000 |
| AT3G27240.1 | Ubiquinol:cytochrome c oxidoreductase subunit (cyt c1-2) | 18 | 0.66 | 0.199 | 1.000 |
| AT3G27240.1 | Ubiquinol:cytochrome c oxidoreductase subunit (cyt c1-2) | 19 | 0.91 | 0.205 | 1.000 |
| AT3G27240.1 | Ubiquinol:cytochrome c oxidoreductase subunit (cyt c1-2) | 20 | 1.35 | 0.211 | 1.000 |
| AT3G27240.1 | Ubiquinol:cytochrome c oxidoreductase subunit (cyt c1-2) | 21 | 1.86 | 0.215 | 0.999 |
| AT3G27240.1 | Ubiquinol:cytochrome c oxidoreductase subunit (cyt c1-2) | 22 | 1.66 | 0.214 | 1.000 |
| AT3G27240.1 | Ubiquinol:cytochrome c oxidoreductase subunit (cyt c1-2) | 23 | 0.79 | 0.214 | 1.000 |
| AT3G27240.1 | Ubiquinol:cytochrome c oxidoreductase subunit (cyt c1-2) | 24 | 0.44 | 0.208 | 0.941 |
| AT3G27240.1 | Ubiquinol:cytochrome c oxidoreductase subunit (cyt c1-2) | 25 | 0.50 | 0.203 | 0.991 |
| AT3G27240.1 | Ubiquinol:cytochrome c oxidoreductase subunit (cyt c1-2) | 26 | 1.13 | 0.202 | 1.000 |
| AT3G27240.1 | Ubiquinol:cytochrome c oxidoreductase subunit (cyt c1-2) | 27 | 2.58 | 0.202 | 0.768 |
| AT3G27240.1 | Ubiquinol:cytochrome c oxidoreductase subunit (cyt c1-2) | 28 | 2.13 | 0.202 | 0.970 |
| AT3G27240.1 | Ubiquinol:cytochrome c oxidoreductase subunit (cyt c1-2) | 29 | 0.80 | 0.202 | 1.000 |
| AT3G27240.1 | Ubiquinol:cytochrome c oxidoreductase subunit (cyt c1-2) | 30 | 0.61 | 0.201 | 1.000 |
| AT3G27240.1 | Ubiquinol:cytochrome c oxidoreductase subunit (cyt c1-2) | 31 | 1.04 | 0.202 | 1.000 |
| AT3G27240.1 | Ubiquinol:cytochrome c oxidoreductase subunit (cyt c1-2) | 32 | 1.83 | 0.203 | 0.999 |
| AT3G27240.1 | Ubiquinol:cytochrome c oxidoreductase subunit (cyt c1-2) | 33 | 2.02 | 0.203 | 0.989 |
| AT3G27240.1 | Ubiquinol:cytochrome c oxidoreductase subunit (cyt c1-2) | 34 | 1.68 | 0.204 | 1.000 |
| AT3G27240.1 | Ubiquinol:cytochrome c oxidoreductase subunit (cyt c1-2) | 35 | 1.70 | 0.203 | 1.000 |
| AT3G27240.1 | Ubiquinol:cytochrome c oxidoreductase subunit (cyt c1-2) | 36 | 2.18 | 0.202 | 0.956 |

| | | | | | |
|---|---|---|---|---|---|
| AT3G27240.1 | Ubiquinol:cytochrome c oxidoreductase subunit (cyt c1-2) | 37 | 2.83 | 0.202 | 0.595 |
| AT3G27240.1 | Ubiquinol:cytochrome c oxidoreductase subunit (cyt c1-2) | 38 | 3.20 | 0.203 | 0.375 |
| AT3G27240.1 | Ubiquinol:cytochrome c oxidoreductase subunit (cyt c1-2) | 39 | 3.14 | 0.203 | 0.416 |
| AT3G27240.1 | Ubiquinol:cytochrome c oxidoreductase subunit (cyt c1-2) | 40 | 2.82 | 0.204 | 0.621 |
| AT3G27240.1 | Ubiquinol:cytochrome c oxidoreductase subunit (cyt c1-2) | 41 | 2.31 | 0.205 | 0.923 |
| AT3G27240.1 | Ubiquinol:cytochrome c oxidoreductase subunit (cyt c1-2) | 42 | 1.77 | 0.208 | 1.000 |
| AT3G27240.1 | Ubiquinol:cytochrome c oxidoreductase subunit (cyt c1-2) | 43 | 1.81 | 0.210 | 1.000 |
| AT3G27240.1 | Ubiquinol:cytochrome c oxidoreductase subunit (cyt c1-2) | 44 | 2.18 | 0.210 | 0.971 |
| AT3G27240.1 | Ubiquinol:cytochrome c oxidoreductase subunit (cyt c1-2) | 45 | 1.79 | 0.211 | 1.000 |
| AT3G27240.1 | Ubiquinol:cytochrome c oxidoreductase subunit (cyt c1-2) | 46 | 0.78 | 0.240 | 1.000 |
| AT3G27240.1 | Ubiquinol:cytochrome c oxidoreductase subunit (cyt c1-2) | 47 | 0.26 | 0.439 | 0.998 |
| AT3G27240.1 | Ubiquinol:cytochrome c oxidoreductase subunit (cyt c1-2) | 48 | 0.09 | 0.772 | 0.997 |
| AT3G27280.2 | Prohibitin 4 (PHB4) | 2 | 0.96 | 0.555 | 1.000 |
| AT3G27280.2 | Prohibitin 4 (PHB4) | 3 | 0.94 | 0.508 | 1.000 |
| AT3G27280.2 | Prohibitin 4 (PHB4) | 14 | 2.65 | 0.339 | 0.944 |
| AT3G27280.2 | Prohibitin 4 (PHB4) | 15 | 1.88 | 0.295 | 0.995 |
| AT3G27280.2 | Prohibitin 4 (PHB4) | 16 | 1.12 | 0.273 | 1.000 |
| AT3G27280.2 | Prohibitin 4 (PHB4) | 17 | 0.88 | 0.244 | 1.000 |
| AT3G27280.2 | Prohibitin 4 (PHB4) | 18 | 0.95 | 0.239 | 1.000 |
| AT3G27280.2 | Prohibitin 4 (PHB4) | 19 | 0.96 | 0.246 | 1.000 |
| AT3G27280.2 | Prohibitin 4 (PHB4) | 20 | 0.91 | 0.252 | 1.000 |
| AT3G27280.2 | Prohibitin 4 (PHB4) | 21 | 1.01 | 0.255 | 1.000 |
| AT3G27280.2 | Prohibitin 4 (PHB4) | 22 | 1.66 | 0.256 | 0.997 |
| AT3G27280.2 | Prohibitin 4 (PHB4) | 23 | 3.34 | 0.255 | 0.425 |
| AT3G27280.2 | Prohibitin 4 (PHB4) | 24 | 4.08 | 0.250 | 0.200 |
| AT3G27280.2 | Prohibitin 4 (PHB4) | 25 | 1.70 | 0.247 | 0.995 |
| AT3G27280.2 | Prohibitin 4 (PHB4) | 26 | 0.50 | 0.262 | 0.970 |
| AT3G27280.2 | Prohibitin 4 (PHB4) | 27 | 0.28 | 0.325 | 0.686 |
| AT3G27280.2 | Prohibitin 4 (PHB4) | 28 | 0.23 | 0.525 | 0.949 |
| AT3G27280.2 | Prohibitin 4 (PHB4) | 46 | 54557.92 | 10.843 | 1.000 |
| AT3G27280.2 | Prohibitin 4 (PHB4) | 47 | 166135.02 | 11.719 | 1.000 |
| AT3G27380.2 | Succinate dehydrogenase subunit (SDH2-1, Fe-S subunit) | 2 | 2.47 | 0.730 | 1.000 |
| AT3G27380.2 | Succinate dehydrogenase subunit (SDH2-1, Fe-S subunit) | 3 | 1.64 | 0.579 | 1.000 |
| AT3G27380.2 | Succinate dehydrogenase subunit (SDH2-1, Fe-S subunit) | 6 | 0.64 | 0.554 | 1.000 |

| | | | | | | |
|---|---|---|---|---|---|---|
| AT3G27380.2 | Succinate dehydrogenase subunit (SDH2-1, Fe-S subunit) | 7 | 0.57 | 0.445 | 1.000 | |
| AT3G27380.2 | Succinate dehydrogenase subunit (SDH2-1, Fe-S subunit) | 8 | 0.59 | 0.396 | 1.000 | |
| AT3G27380.2 | Succinate dehydrogenase subunit (SDH2-1, Fe-S subunit) | 9 | 0.66 | 0.382 | 1.000 | |
| AT3G27380.2 | Succinate dehydrogenase subunit (SDH2-1, Fe-S subunit) | 10 | 0.86 | 0.380 | 1.000 | |
| AT3G27380.2 | Succinate dehydrogenase subunit (SDH2-1, Fe-S subunit) | 11 | 1.28 | 0.378 | 1.000 | |
| AT3G27380.2 | Succinate dehydrogenase subunit (SDH2-1, Fe-S subunit) | 12 | 1.82 | 0.378 | 1.000 | |
| AT3G27380.2 | Succinate dehydrogenase subunit (SDH2-1, Fe-S subunit) | 13 | 1.63 | 0.376 | 1.000 | |
| AT3G27380.2 | Succinate dehydrogenase subunit (SDH2-1, Fe-S subunit) | 14 | 1.15 | 0.381 | 1.000 | |
| AT3G27380.2 | Succinate dehydrogenase subunit (SDH2-1, Fe-S subunit) | 15 | 1.28 | 0.391 | 1.000 | |
| AT3G27380.2 | Succinate dehydrogenase subunit (SDH2-1, Fe-S subunit) | 16 | 2.49 | 0.397 | 0.999 | |
| AT3G27380.2 | Succinate dehydrogenase subunit (SDH2-1, Fe-S subunit) | 17 | 6.85 | 0.424 | 0.607 | |
| AT3G27380.2 | Succinate dehydrogenase subunit (SDH2-1, Fe-S subunit) | 18 | 16.44 | 0.575 | 0.492 | |
| AT3G27380.2 | Succinate dehydrogenase subunit (SDH2-1, Fe-S subunit) | 31 | 1.15 | 0.533 | 1.000 | |
| AT3G27380.2 | Succinate dehydrogenase subunit (SDH2-1, Fe-S subunit) | 32 | 0.98 | 0.425 | 1.000 | |
| AT3G27380.2 | Succinate dehydrogenase subunit (SDH2-1, Fe-S subunit) | 33 | 1.10 | 0.395 | 1.000 | |
| AT3G27380.2 | Succinate dehydrogenase subunit (SDH2-1, Fe-S subunit) | 34 | 2.15 | 0.383 | 1.000 | |
| AT3G27380.2 | Succinate dehydrogenase subunit (SDH2-1, Fe-S subunit) | 35 | 4.25 | 0.380 | 0.840 | |
| AT3G27380.2 | Succinate dehydrogenase subunit (SDH2-1, Fe-S subunit) | 36 | 2.14 | 0.380 | 1.000 | |
| AT3G27380.2 | Succinate dehydrogenase subunit (SDH2-1, Fe-S subunit) | 37 | 0.41 | 0.391 | 0.999 | |
| AT3G27380.2 | Succinate dehydrogenase subunit (SDH2-1, Fe-S subunit) | 38 | 0.18 | 0.440 | 0.821 | |
| AT3G27380.2 | Succinate dehydrogenase subunit (SDH2-1, Fe-S subunit) | 39 | 0.27 | 0.548 | 0.998 | |
| AT3G27380.2 | Succinate dehydrogenase subunit (SDH2-1, Fe-S subunit) | 41 | 1.43 | 0.491 | 1.000 | |
| AT3G27380.2 | Succinate dehydrogenase subunit (SDH2-1, Fe-S subunit) | 42 | 4.22 | 0.428 | 0.935 | |
| AT3G27380.2 | Succinate dehydrogenase subunit (SDH2-1, Fe-S subunit) | 43 | 11.85 | 0.392 | 0.134 | |
| AT3G27380.2 | Succinate dehydrogenase subunit (SDH2-1, Fe-S subunit) | 44 | 17.95 | 0.382 | 0.029 | * |
| AT3G27380.2 | Succinate dehydrogenase subunit (SDH2-1, Fe-S subunit) | 45 | 8.36 | 0.381 | 0.280 | |
| AT3G27380.2 | Succinate dehydrogenase subunit (SDH2-1, Fe-S subunit) | 46 | 1.11 | 0.414 | 1.000 | |
| AT3G27380.2 | Succinate dehydrogenase subunit (SDH2-1, Fe-S subunit) | 47 | 0.07 | 0.723 | 0.853 | |
| AT3G27570.1 | putative sucrose cleavage protein | 2 | 0.86 | 0.318 | 1.000 | |
| AT3G27570.1 | putative sucrose cleavage protein | 3 | 0.60 | 0.259 | 1.000 | |
| AT3G27570.1 | putative sucrose cleavage protein | 4 | 0.45 | 0.356 | 0.999 | |

| | | | | | | |
|---|---|---|---|---|---|---|
| AT3G27570.1 | putative sucrose cleavage protein | 17 | 1.19 | 0.211 | 1.000 | |
| AT3G27570.1 | putative sucrose cleavage protein | 18 | 1.55 | 0.185 | 0.998 | |
| AT3G27570.1 | putative sucrose cleavage protein | 19 | 1.90 | 0.167 | 0.838 | |
| AT3G27570.1 | putative sucrose cleavage protein | 20 | 2.05 | 0.160 | 0.648 | |
| AT3G27570.1 | putative sucrose cleavage protein | 21 | 1.89 | 0.158 | 0.794 | |
| AT3G27570.1 | putative sucrose cleavage protein | 22 | 1.55 | 0.157 | 0.989 | |
| AT3G27570.1 | putative sucrose cleavage protein | 23 | 1.19 | 0.155 | 1.000 | |
| AT3G27570.1 | putative sucrose cleavage protein | 24 | 0.92 | 0.152 | 1.000 | |
| AT3G27570.1 | putative sucrose cleavage protein | 25 | 0.79 | 0.149 | 1.000 | |
| AT3G27570.1 | putative sucrose cleavage protein | 26 | 0.86 | 0.149 | 1.000 | |
| AT3G27570.1 | putative sucrose cleavage protein | 27 | 1.07 | 0.149 | 1.000 | |
| AT3G27570.1 | putative sucrose cleavage protein | 28 | 1.13 | 0.149 | 1.000 | |
| AT3G27570.1 | putative sucrose cleavage protein | 29 | 1.00 | 0.149 | 1.000 | |
| AT3G27570.1 | putative sucrose cleavage protein | 30 | 0.83 | 0.150 | 1.000 | |
| AT3G27570.1 | putative sucrose cleavage protein | 31 | 0.71 | 0.150 | 0.999 | |
| AT3G27570.1 | putative sucrose cleavage protein | 32 | 0.63 | 0.150 | 0.968 | |
| AT3G27570.1 | putative sucrose cleavage protein | 33 | 0.58 | 0.150 | 0.885 | |
| AT3G27570.1 | putative sucrose cleavage protein | 34 | 0.59 | 0.150 | 0.916 | |
| AT3G27570.1 | putative sucrose cleavage protein | 35 | 0.66 | 0.150 | 0.991 | |
| AT3G27570.1 | putative sucrose cleavage protein | 36 | 0.80 | 0.150 | 1.000 | |
| AT3G27570.1 | putative sucrose cleavage protein | 37 | 1.03 | 0.150 | 1.000 | |
| AT3G27570.1 | putative sucrose cleavage protein | 38 | 1.33 | 0.149 | 1.000 | |
| AT3G27570.1 | putative sucrose cleavage protein | 39 | 1.59 | 0.150 | 0.968 | |
| AT3G27570.1 | putative sucrose cleavage protein | 40 | 1.74 | 0.150 | 0.883 | |
| AT3G27570.1 | putative sucrose cleavage protein | 41 | 1.79 | 0.152 | 0.845 | |
| AT3G27570.1 | putative sucrose cleavage protein | 42 | 1.85 | 0.153 | 0.795 | |
| AT3G27570.1 | putative sucrose cleavage protein | 43 | 1.86 | 0.154 | 0.795 | |
| AT3G27570.1 | putative sucrose cleavage protein | 44 | 1.53 | 0.155 | 0.991 | |
| AT3G27570.1 | putative sucrose cleavage protein | 45 | 0.89 | 0.161 | 1.000 | |
| AT3G27570.1 | putative sucrose cleavage protein | 46 | 0.36 | 0.196 | 0.380 | |
| AT3G27570.1 | putative sucrose cleavage protein | 47 | 0.11 | 0.304 | 0.048 | * |

| | | | | | | |
|---|---|---|---|---|---|---|
| AT3G27620.1 | Alternative oxidase 1C (AOX1C) | 2 | 0.97 | 0.358 | 1.000 | |
| AT3G27620.1 | Alternative oxidase 1C (AOX1C) | 3 | 0.95 | 0.268 | 1.000 | |
| AT3G27620.1 | Alternative oxidase 1C (AOX1C) | 4 | 0.94 | 0.372 | 1.000 | |
| AT3G27620.1 | Alternative oxidase 1C (AOX1C) | 27 | 1.04 | 0.284 | 1.000 | |
| AT3G27620.1 | Alternative oxidase 1C (AOX1C) | 28 | 0.93 | 0.213 | 1.000 | |
| AT3G27620.1 | Alternative oxidase 1C (AOX1C) | 29 | 0.80 | 0.178 | 1.000 | |
| AT3G27620.1 | Alternative oxidase 1C (AOX1C) | 30 | 0.66 | 0.161 | 0.977 | |
| AT3G27620.1 | Alternative oxidase 1C (AOX1C) | 31 | 0.57 | 0.153 | 0.774 | |
| AT3G27620.1 | Alternative oxidase 1C (AOX1C) | 32 | 0.55 | 0.149 | 0.687 | |
| AT3G27620.1 | Alternative oxidase 1C (AOX1C) | 33 | 0.66 | 0.147 | 0.951 | |
| AT3G27620.1 | Alternative oxidase 1C (AOX1C) | 34 | 0.94 | 0.146 | 1.000 | |
| AT3G27620.1 | Alternative oxidase 1C (AOX1C) | 35 | 1.47 | 0.146 | 0.973 | |
| AT3G27620.1 | Alternative oxidase 1C (AOX1C) | 36 | 2.12 | 0.146 | 0.315 | |
| AT3G27620.1 | Alternative oxidase 1C (AOX1C) | 37 | 2.71 | 0.145 | 0.052 | . |
| AT3G27620.1 | Alternative oxidase 1C (AOX1C) | 38 | 3.01 | 0.145 | 0.020 | * |
| AT3G27620.1 | Alternative oxidase 1C (AOX1C) | 39 | 2.88 | 0.145 | 0.030 | * |
| AT3G27620.1 | Alternative oxidase 1C (AOX1C) | 40 | 2.40 | 0.146 | 0.141 | |
| AT3G27620.1 | Alternative oxidase 1C (AOX1C) | 41 | 1.81 | 0.147 | 0.669 | |
| AT3G27620.1 | Alternative oxidase 1C (AOX1C) | 42 | 1.42 | 0.151 | 0.991 | |
| AT3G27620.1 | Alternative oxidase 1C (AOX1C) | 43 | 1.20 | 0.157 | 1.000 | |
| AT3G27620.1 | Alternative oxidase 1C (AOX1C) | 44 | 1.13 | 0.166 | 1.000 | |
| AT3G27620.1 | Alternative oxidase 1C (AOX1C) | 45 | 1.12 | 0.180 | 1.000 | |
| AT3G27620.1 | Alternative oxidase 1C (AOX1C) | 46 | 1.14 | 0.222 | 1.000 | |
| AT3G27620.1 | Alternative oxidase 1C (AOX1C) | 47 | 1.18 | 0.331 | 1.000 | |
| AT3G27820.1 | Monodehydroascorbate reductase 4 (MDAR4) | 2 | 0.99 | 0.579 | 1.000 | |
| AT3G27820.1 | Monodehydroascorbate reductase 4 (MDAR4) | 3 | 0.68 | 0.567 | 1.000 | |
| AT3G27820.1 | Monodehydroascorbate reductase 4 (MDAR4) | 27 | 0.11 | 0.432 | 0.329 | |
| AT3G27820.1 | Monodehydroascorbate reductase 4 (MDAR4) | 28 | 0.17 | 0.306 | 0.189 | |
| AT3G27820.1 | Monodehydroascorbate reductase 4 (MDAR4) | 29 | 0.25 | 0.270 | 0.365 | |
| AT3G27820.1 | Monodehydroascorbate reductase 4 (MDAR4) | 30 | 0.36 | 0.256 | 0.730 | |
| AT3G27820.1 | Monodehydroascorbate reductase 4 (MDAR4) | 31 | 0.60 | 0.251 | 0.999 | |
| AT3G27820.1 | Monodehydroascorbate reductase 4 (MDAR4) | 32 | 1.24 | 0.251 | 1.000 | |
| AT3G27820.1 | Monodehydroascorbate reductase 4 (MDAR4) | 33 | 1.80 | 0.252 | 0.994 | |
| AT3G27820.1 | Monodehydroascorbate reductase 4 (MDAR4) | 34 | 1.20 | 0.253 | 1.000 | |
| AT3G27820.1 | Monodehydroascorbate reductase 4 (MDAR4) | 35 | 0.75 | 0.256 | 1.000 | |
| AT3G27820.1 | Monodehydroascorbate reductase 4 (MDAR4) | 36 | 0.93 | 0.270 | 1.000 | |
| AT3G27820.1 | Monodehydroascorbate reductase 4 (MDAR4) | 37 | 1.30 | 0.314 | 1.000 | |
| AT3G27820.1 | Monodehydroascorbate reductase 4 (MDAR4) | 38 | 1.37 | 0.389 | 1.000 | |
| AT3G27820.1 | Monodehydroascorbate reductase 4 (MDAR4) | 40 | 2.02 | 0.345 | 0.999 | |
| AT3G27820.1 | Monodehydroascorbate reductase 4 (MDAR4) | 41 | 3.38 | 0.297 | 0.692 | |
| AT3G27820.1 | Monodehydroascorbate reductase 4 (MDAR4) | 42 | 6.83 | 0.268 | 0.037 | * |
| AT3G27820.1 | Monodehydroascorbate reductase 4 (MDAR4) | 43 | 10.59 | 0.259 | 0.003 | ** |
| AT3G27820.1 | Monodehydroascorbate reductase 4 (MDAR4) | 44 | 6.76 | 0.256 | 0.027 | * |

| | | | | | | |
|---|---|---|---|---|---|---|
| AT3G27820.1 | Monodehydroascorbate reductase 4 (MDAR4) | 45 | 2.08 | 0.258 | 0.969 | |
| AT3G27820.1 | Monodehydroascorbate reductase 4 (MDAR4) | 46 | 0.43 | 0.303 | 0.970 | |
| AT3G27820.1 | Monodehydroascorbate reductase 4 (MDAR4) | 47 | 0.07 | 0.539 | 0.376 | |
| AT3G27890.1 | NAD(P)H dehydrogenase complex subunit (NQR) | 2 | 0.75 | 0.589 | 1.000 | |
| AT3G27890.1 | NAD(P)H dehydrogenase complex subunit (NQR) | 3 | 0.63 | 0.590 | 1.000 | |
| AT3G27890.1 | NAD(P)H dehydrogenase complex subunit (NQR) | 41 | 1.74 | 0.642 | 1.000 | |
| AT3G27890.1 | NAD(P)H dehydrogenase complex subunit (NQR) | 42 | 3.21 | 0.388 | 0.796 | |
| AT3G27890.1 | NAD(P)H dehydrogenase complex subunit (NQR) | 43 | 9.44 | 0.333 | 0.042 | * |
| AT3G27890.1 | NAD(P)H dehydrogenase complex subunit (NQR) | 44 | 7.85 | 0.319 | 0.056 | . |
| AT3G27890.1 | NAD(P)H dehydrogenase complex subunit (NQR) | 45 | 1.25 | 0.319 | 1.000 | |
| AT3G27890.1 | NAD(P)H dehydrogenase complex subunit (NQR) | 46 | 0.45 | 0.336 | 0.924 | |
| AT3G27890.1 | NAD(P)H dehydrogenase complex subunit (NQR) | 47 | 0.26 | 0.808 | 0.989 | |
| AT3G27930.2 | Outer membrane protein 47 (OM47) | 2 | 0.97 | 0.512 | 1.000 | |
| AT3G27930.2 | Outer membrane protein 47 (OM47) | 3 | 1.00 | 0.346 | 1.000 | |
| AT3G27930.2 | Outer membrane protein 47 (OM47) | 4 | 1.04 | 0.377 | 1.000 | |
| AT3G27930.2 | Outer membrane protein 47 (OM47) | 6 | 1.19 | 0.423 | 1.000 | |
| AT3G27930.2 | Outer membrane protein 47 (OM47) | 7 | 1.30 | 0.443 | 1.000 | |
| AT3G27930.2 | Outer membrane protein 47 (OM47) | 8 | 1.41 | 0.557 | 1.000 | |
| AT3G27930.2 | Outer membrane protein 47 (OM47) | 20 | 1.15 | 0.350 | 1.000 | |
| AT3G27930.2 | Outer membrane protein 47 (OM47) | 21 | 1.09 | 0.312 | 1.000 | |
| AT3G27930.2 | Outer membrane protein 47 (OM47) | 22 | 1.05 | 0.273 | 1.000 | |
| AT3G27930.2 | Outer membrane protein 47 (OM47) | 23 | 0.97 | 0.264 | 1.000 | |
| AT3G27930.2 | Outer membrane protein 47 (OM47) | 24 | 0.83 | 0.270 | 1.000 | |
| AT3G27930.2 | Outer membrane protein 47 (OM47) | 25 | 0.68 | 0.307 | 1.000 | |
| AT3G27930.2 | Outer membrane protein 47 (OM47) | 26 | 0.56 | 0.437 | 1.000 | |
| AT3G27930.2 | Outer membrane protein 47 (OM47) | 35 | 0.97 | 0.391 | 1.000 | |
| AT3G27930.2 | Outer membrane protein 47 (OM47) | 36 | 1.56 | 0.326 | 1.000 | |
| AT3G27930.2 | Outer membrane protein 47 (OM47) | 37 | 3.29 | 0.289 | 0.756 | |
| AT3G27930.2 | Outer membrane protein 47 (OM47) | 38 | 9.83 | 0.278 | 0.011 | * |
| AT3G27930.2 | Outer membrane protein 47 (OM47) | 39 | 19.62 | 0.275 | 0.000 | *** |
| AT3G27930.2 | Outer membrane protein 47 (OM47) | 40 | 13.66 | 0.252 | 0.000 | *** |

| | | | | | | |
|---|---|---|---|---|---|---|
| AT3G27930.2 | Outer membrane protein 47 (OM47) | 41 | 2.91 | 0.245 | 0.677 | |
| AT3G27930.2 | Outer membrane protein 47 (OM47) | 42 | 0.58 | 0.249 | 0.999 | |
| AT3G27930.2 | Outer membrane protein 47 (OM47) | 43 | 0.21 | 0.253 | 0.144 | |
| AT3G27930.2 | Outer membrane protein 47 (OM47) | 44 | 0.19 | 0.257 | 0.119 | |
| AT3G27930.2 | Outer membrane protein 47 (OM47) | 45 | 0.71 | 0.291 | 1.000 | |
| AT3G27930.2 | Outer membrane protein 47 (OM47) | 46 | 6.06 | 0.533 | 0.932 | |
| AT3G27930.2 | Outer membrane protein 47 (OM47) | 47 | 67.91 | 1.066 | 0.804 | |
| AT3G29320.1 | Alpha-glucan phosphorylase 1 (PHS1) | 2 | 1.03 | 0.448 | 1.000 | |
| AT3G29320.1 | Alpha-glucan phosphorylase 1 (PHS1) | 3 | 1.10 | 0.432 | 1.000 | |
| AT3G29320.1 | Alpha-glucan phosphorylase 1 (PHS1) | 4 | 1.18 | 0.427 | 1.000 | |
| AT3G29320.1 | Alpha-glucan phosphorylase 1 (PHS1) | 34 | 1.62 | 0.298 | 0.896 | |
| AT3G29320.1 | Alpha-glucan phosphorylase 1 (PHS1) | 35 | 1.45 | 0.291 | 0.950 | |
| AT3G29320.1 | Alpha-glucan phosphorylase 1 (PHS1) | 36 | 1.29 | 0.289 | 0.988 | |
| AT3G29320.1 | Alpha-glucan phosphorylase 1 (PHS1) | 37 | 1.12 | 0.293 | 1.000 | |
| AT3G29320.1 | Alpha-glucan phosphorylase 1 (PHS1) | 38 | 0.96 | 0.300 | 1.000 | |
| AT3G29320.1 | Alpha-glucan phosphorylase 1 (PHS1) | 46 | 0.20 | 0.426 | 0.324 | |
| AT3G29320.1 | Alpha-glucan phosphorylase 1 (PHS1) | 47 | 0.16 | 0.455 | 0.273 | |
| AT3G42050.1 | Vacuolar ATP synthase subunit H (VHA-H) | 2 | 1.26 | 0.239 | 0.990 | |
| AT3G42050.1 | Vacuolar ATP synthase subunit H (VHA-H) | 3 | 1.28 | 0.220 | 0.982 | |
| AT3G42050.1 | Vacuolar ATP synthase subunit H (VHA-H) | 4 | 1.29 | 0.206 | 0.972 | |
| AT3G42050.1 | Vacuolar ATP synthase subunit H (VHA-H) | 5 | 1.31 | 0.197 | 0.961 | |
| AT3G42050.1 | Vacuolar ATP synthase subunit H (VHA-H) | 6 | 1.32 | 0.193 | 0.952 | |
| AT3G42050.1 | Vacuolar ATP synthase subunit H (VHA-H) | 7 | 1.33 | 0.192 | 0.946 | |
| AT3G42050.1 | Vacuolar ATP synthase subunit H (VHA-H) | 8 | 1.33 | 0.193 | 0.944 | |
| AT3G42050.1 | Vacuolar ATP synthase subunit H (VHA-H) | 21 | 1.18 | 0.196 | 0.995 | |
| AT3G42050.1 | Vacuolar ATP synthase subunit H (VHA-H) | 22 | 1.14 | 0.199 | 0.998 | |
| AT3G42050.1 | Vacuolar ATP synthase subunit H (VHA-H) | 23 | 1.10 | 0.204 | 1.000 | |
| AT3G42050.1 | Vacuolar ATP synthase subunit H (VHA-H) | 24 | 1.04 | 0.212 | 1.000 | |
| AT3G42050.1 | Vacuolar ATP synthase subunit H (VHA-H) | 25 | 0.98 | 0.223 | 1.000 | |
| AT3G42050.1 | Vacuolar ATP synthase subunit H (VHA-H) | 26 | 0.91 | 0.235 | 1.000 | |
| AT3G42050.1 | Vacuolar ATP synthase subunit H (VHA-H) | 46 | 0.05 | 0.459 | 0.034 | * |

| | | | | | | |
|---|---|---|---|---|---|---|
| AT3G42050.1 | Vacuolar ATP synthase subunit H (VHA-H) | 47 | 0.05 | 0.477 | 0.030 | * |
| AT3G44370.1 | Membrane insertion protein (OXAA/YIDC ) | 2 | 0.93 | 0.561 | 1.000 | |
| AT3G44370.1 | Membrane insertion protein (OXAA/YIDC ) | 3 | 1.34 | 0.533 | 1.000 | |
| AT3G44370.1 | Membrane insertion protein (OXAA/YIDC ) | 38 | 4.24 | 0.350 | 0.323 | |
| AT3G44370.1 | Membrane insertion protein (OXAA/YIDC ) | 39 | 1.95 | 0.268 | 0.768 | |
| AT3G44370.1 | Membrane insertion protein (OXAA/YIDC ) | 40 | 0.85 | 0.215 | 0.998 | |
| AT3G44370.1 | Membrane insertion protein (OXAA/YIDC ) | 41 | 0.35 | 0.209 | 0.151 | |
| AT3G44370.1 | Membrane insertion protein (OXAA/YIDC ) | 42 | 0.13 | 0.258 | 0.009 | ** |
| AT3G44370.1 | Membrane insertion protein (OXAA/YIDC ) | 43 | 0.05 | 0.353 | 0.004 | ** |
| AT3G44370.1 | Membrane insertion protein (OXAA/YIDC ) | 46 | 0.00 | 0.797 | 0.013 | * |
| AT3G44370.1 | Membrane insertion protein (OXAA/YIDC ) | 47 | 0.00 | 0.982 | 0.021 | * |
| AT3G45140.1 | Lipoxygenase 2 (LOX2) | 2 | 1.37 | 0.569 | 1.000 | |
| AT3G45140.1 | Lipoxygenase 2 (LOX2) | 3 | 1.22 | 0.456 | 1.000 | |
| AT3G45140.1 | Lipoxygenase 2 (LOX2) | 4 | 1.09 | 0.480 | 1.000 | |
| AT3G45140.1 | Lipoxygenase 2 (LOX2) | 11 | 0.85 | 0.438 | 1.000 | |
| AT3G45140.1 | Lipoxygenase 2 (LOX2) | 12 | 0.91 | 0.406 | 1.000 | |
| AT3G45140.1 | Lipoxygenase 2 (LOX2) | 13 | 0.99 | 0.382 | 1.000 | |
| AT3G45140.1 | Lipoxygenase 2 (LOX2) | 14 | 1.04 | 0.378 | 1.000 | |
| AT3G45140.1 | Lipoxygenase 2 (LOX2) | 15 | 1.05 | 0.385 | 1.000 | |
| AT3G45140.1 | Lipoxygenase 2 (LOX2) | 16 | 1.00 | 0.408 | 1.000 | |
| AT3G45140.1 | Lipoxygenase 2 (LOX2) | 17 | 0.92 | 0.440 | 1.000 | |
| AT3G45140.1 | Lipoxygenase 2 (LOX2) | 22 | 0.71 | 0.418 | 1.000 | |
| AT3G45140.1 | Lipoxygenase 2 (LOX2) | 23 | 0.71 | 0.376 | 1.000 | |
| AT3G45140.1 | Lipoxygenase 2 (LOX2) | 24 | 0.72 | 0.340 | 1.000 | |
| AT3G45140.1 | Lipoxygenase 2 (LOX2) | 25 | 0.73 | 0.316 | 1.000 | |
| AT3G45140.1 | Lipoxygenase 2 (LOX2) | 26 | 0.73 | 0.303 | 1.000 | |
| AT3G45140.1 | Lipoxygenase 2 (LOX2) | 27 | 0.72 | 0.294 | 1.000 | |
| AT3G45140.1 | Lipoxygenase 2 (LOX2) | 28 | 0.69 | 0.289 | 1.000 | |
| AT3G45140.1 | Lipoxygenase 2 (LOX2) | 29 | 0.66 | 0.287 | 1.000 | |
| AT3G45140.1 | Lipoxygenase 2 (LOX2) | 30 | 0.62 | 0.287 | 1.000 | |
| AT3G45140.1 | Lipoxygenase 2 (LOX2) | 31 | 0.61 | 0.288 | 1.000 | |
| AT3G45140.1 | Lipoxygenase 2 (LOX2) | 32 | 0.59 | 0.289 | 0.999 | |
| AT3G45140.1 | Lipoxygenase 2 (LOX2) | 33 | 0.57 | 0.290 | 0.999 | |
| AT3G45140.1 | Lipoxygenase 2 (LOX2) | 34 | 0.54 | 0.290 | 0.997 | |
| AT3G45140.1 | Lipoxygenase 2 (LOX2) | 35 | 0.53 | 0.291 | 0.995 | |
| AT3G45140.1 | Lipoxygenase 2 (LOX2) | 36 | 0.55 | 0.291 | 0.998 | |
| AT3G45140.1 | Lipoxygenase 2 (LOX2) | 37 | 0.59 | 0.292 | 0.999 | |
| AT3G45140.1 | Lipoxygenase 2 (LOX2) | 38 | 0.60 | 0.298 | 1.000 | |
| AT3G45140.1 | Lipoxygenase 2 (LOX2) | 39 | 0.58 | 0.321 | 1.000 | |
| AT3G45140.1 | Lipoxygenase 2 (LOX2) | 40 | 0.65 | 0.385 | 1.000 | |
| AT3G45140.1 | Lipoxygenase 2 (LOX2) | 41 | 0.90 | 0.505 | 1.000 | |
| AT3G45140.1 | Lipoxygenase 2 (LOX2) | 46 | 50.60 | 1.664 | 0.991 | |
| AT3G45140.1 | Lipoxygenase 2 (LOX2) | 47 | 142.03 | 2.012 | 0.987 | |
| AT3G45300.1 | Isovaleryl-coenzyme A dehydrogenase (IVDH) | 2 | 1.78 | 1.045 | 1.000 | |
| AT3G45300.1 | Isovaleryl-coenzyme A dehydrogenase (IVDH) | 3 | 0.70 | 0.782 | 1.000 | |
| AT3G45300.1 | Isovaleryl-coenzyme A dehydrogenase (IVDH) | 4 | 0.24 | 1.608 | 1.000 | |
| AT3G45300.1 | Isovaleryl-coenzyme A dehydrogenase (IVDH) | 38 | 0.16 | 0.853 | 0.968 | |

| | | | | | | |
|---|---|---|---|---|---|---|
| AT3G45300.1 | Isovaleryl-coenzyme A dehydrogenase (IVDH) | 39 | 1.06 | 0.571 | 1.000 | |
| AT3G45300.1 | Isovaleryl-coenzyme A dehydrogenase (IVDH) | 40 | 4.08 | 0.493 | 0.866 | |
| AT3G45300.1 | Isovaleryl-coenzyme A dehydrogenase (IVDH) | 41 | 3.47 | 0.473 | 0.910 | |
| AT3G45300.1 | Isovaleryl-coenzyme A dehydrogenase (IVDH) | 42 | 1.55 | 0.476 | 1.000 | |
| AT3G45300.1 | Isovaleryl-coenzyme A dehydrogenase (IVDH) | 43 | 1.31 | 0.497 | 1.000 | |
| AT3G45300.1 | Isovaleryl-coenzyme A dehydrogenase (IVDH) | 44 | 2.72 | 0.646 | 0.997 | |
| AT3G45300.1 | Isovaleryl-coenzyme A dehydrogenase (IVDH) | 45 | 10.98 | 0.929 | 0.918 | |
| AT3G45300.1 | Isovaleryl-coenzyme A dehydrogenase (IVDH) | 46 | 67.28 | 1.320 | 0.784 | |
| AT3G45300.1 | Isovaleryl-coenzyme A dehydrogenase (IVDH) | 47 | 495.83 | 1.971 | 0.794 | |
| AT3G45740.1 | Hydrolase family protein / HAD-superfamily protein | 2 | 0.77 | 0.639 | 1.000 | |
| AT3G45740.1 | Hydrolase family protein / HAD-superfamily protein | 3 | 0.61 | 0.635 | 1.000 | |
| AT3G45740.1 | Hydrolase family protein / HAD-superfamily protein | 41 | 0.94 | 0.687 | 1.000 | |
| AT3G45740.1 | Hydrolase family protein / HAD-superfamily protein | 42 | 1.75 | 0.411 | 0.997 | |
| AT3G45740.1 | Hydrolase family protein / HAD-superfamily protein | 43 | 4.23 | 0.349 | 0.450 | |
| AT3G45740.1 | Hydrolase family protein / HAD-superfamily protein | 44 | 3.15 | 0.335 | 0.665 | |
| AT3G45740.1 | Hydrolase family protein / HAD-superfamily protein | 45 | 0.93 | 0.335 | 1.000 | |
| AT3G45740.1 | Hydrolase family protein / HAD-superfamily protein | 46 | 0.40 | 0.356 | 0.887 | |
| AT3G45740.1 | Hydrolase family protein / HAD-superfamily protein | 47 | 0.19 | 0.825 | 0.965 | |
| AT3G45770.1 | Enoyl-CoA reductase | 2 | 1.22 | 0.484 | 1.000 | |
| AT3G45770.1 | Enoyl-CoA reductase | 3 | 0.93 | 0.370 | 1.000 | |
| AT3G45770.1 | Enoyl-CoA reductase | 5 | 0.67 | 0.329 | 1.000 | |
| AT3G45770.1 | Enoyl-CoA reductase | 6 | 0.68 | 0.299 | 1.000 | |
| AT3G45770.1 | Enoyl-CoA reductase | 7 | 0.76 | 0.281 | 1.000 | |
| AT3G45770.1 | Enoyl-CoA reductase | 8 | 0.86 | 0.273 | 1.000 | |
| AT3G45770.1 | Enoyl-CoA reductase | 9 | 0.94 | 0.269 | 1.000 | |
| AT3G45770.1 | Enoyl-CoA reductase | 10 | 1.03 | 0.272 | 1.000 | |
| AT3G45770.1 | Enoyl-CoA reductase | 11 | 1.17 | 0.279 | 1.000 | |
| AT3G45770.1 | Enoyl-CoA reductase | 12 | 1.31 | 0.284 | 1.000 | |
| AT3G45770.1 | Enoyl-CoA reductase | 13 | 1.47 | 0.286 | 1.000 | |
| AT3G45770.1 | Enoyl-CoA reductase | 14 | 1.60 | 0.315 | 1.000 | |
| AT3G45770.1 | Enoyl-CoA reductase | 27 | 0.57 | 0.356 | 1.000 | |
| AT3G45770.1 | Enoyl-CoA reductase | 28 | 0.85 | 0.290 | 1.000 | |
| AT3G45770.1 | Enoyl-CoA reductase | 29 | 1.36 | 0.269 | 1.000 | |
| AT3G45770.1 | Enoyl-CoA reductase | 30 | 1.85 | 0.260 | 0.991 | |
| AT3G45770.1 | Enoyl-CoA reductase | 31 | 2.31 | 0.257 | 0.880 | |
| AT3G45770.1 | Enoyl-CoA reductase | 32 | 2.45 | 0.257 | 0.823 | |
| AT3G45770.1 | Enoyl-CoA reductase | 33 | 1.88 | 0.257 | 0.988 | |
| AT3G45770.1 | Enoyl-CoA reductase | 34 | 1.26 | 0.258 | 1.000 | |
| AT3G45770.1 | Enoyl-CoA reductase | 35 | 0.95 | 0.257 | 1.000 | |
| AT3G45770.1 | Enoyl-CoA reductase | 36 | 0.91 | 0.257 | 1.000 | |
| AT3G45770.1 | Enoyl-CoA reductase | 37 | 1.12 | 0.256 | 1.000 | |
| AT3G45770.1 | Enoyl-CoA reductase | 38 | 1.91 | 0.256 | 0.984 | |
| AT3G45770.1 | Enoyl-CoA reductase | 39 | 3.88 | 0.257 | 0.275 | |
| AT3G45770.1 | Enoyl-CoA reductase | 40 | 6.66 | 0.257 | 0.028 | * |

| | | | | | | |
|---|---|---|---|---|---|---|
| AT3G45770.1 | Enoyl-CoA reductase | 41 | 6.44 | 0.257 | 0.033 | * |
| AT3G45770.1 | Enoyl-CoA reductase | 42 | 3.64 | 0.259 | 0.349 | |
| AT3G45770.1 | Enoyl-CoA reductase | 43 | 1.82 | 0.261 | 0.994 | |
| AT3G45770.1 | Enoyl-CoA reductase | 44 | 0.98 | 0.260 | 1.000 | |
| AT3G45770.1 | Enoyl-CoA reductase | 45 | 0.57 | 0.260 | 0.996 | |
| AT3G45770.1 | Enoyl-CoA reductase | 46 | 0.32 | 0.289 | 0.678 | |
| AT3G45770.1 | Enoyl-CoA reductase | 47 | 0.16 | 0.447 | 0.642 | |
| AT3G46100.1 | Histidyl-tRNA synthetase 1 (HRS1) | 2 | 0.90 | 0.424 | 1.000 | |
| AT3G46100.1 | Histidyl-tRNA synthetase 1 (HRS1) | 3 | 0.93 | 0.406 | 1.000 | |
| AT3G46100.1 | Histidyl-tRNA synthetase 1 (HRS1) | 28 | 1.58 | 0.360 | 1.000 | |
| AT3G46100.1 | Histidyl-tRNA synthetase 1 (HRS1) | 29 | 1.60 | 0.288 | 1.000 | |
| AT3G46100.1 | Histidyl-tRNA synthetase 1 (HRS1) | 30 | 1.63 | 0.265 | 0.999 | |
| AT3G46100.1 | Histidyl-tRNA synthetase 1 (HRS1) | 31 | 1.67 | 0.232 | 0.995 | |
| AT3G46100.1 | Histidyl-tRNA synthetase 1 (HRS1) | 32 | 1.68 | 0.217 | 0.989 | |
| AT3G46100.1 | Histidyl-tRNA synthetase 1 (HRS1) | 33 | 1.77 | 0.203 | 0.961 | |
| AT3G46100.1 | Histidyl-tRNA synthetase 1 (HRS1) | 34 | 2.23 | 0.193 | 0.645 | |
| AT3G46100.1 | Histidyl-tRNA synthetase 1 (HRS1) | 35 | 3.06 | 0.192 | 0.175 | |
| AT3G46100.1 | Histidyl-tRNA synthetase 1 (HRS1) | 36 | 4.22 | 0.198 | 0.033 | * |
| AT3G46100.1 | Histidyl-tRNA synthetase 1 (HRS1) | 37 | 5.65 | 0.205 | 0.006 | ** |
| AT3G46100.1 | Histidyl-tRNA synthetase 1 (HRS1) | 38 | 6.66 | 0.210 | 0.003 | ** |
| AT3G46100.1 | Histidyl-tRNA synthetase 1 (HRS1) | 39 | 5.53 | 0.208 | 0.008 | ** |
| AT3G46100.1 | Histidyl-tRNA synthetase 1 (HRS1) | 40 | 3.20 | 0.195 | 0.151 | |
| AT3G46100.1 | Histidyl-tRNA synthetase 1 (HRS1) | 41 | 1.76 | 0.190 | 0.940 | |
| AT3G46100.1 | Histidyl-tRNA synthetase 1 (HRS1) | 42 | 1.07 | 0.192 | 1.000 | |
| AT3G46100.1 | Histidyl-tRNA synthetase 1 (HRS1) | 43 | 0.62 | 0.194 | 0.985 | |
| AT3G46100.1 | Histidyl-tRNA synthetase 1 (HRS1) | 44 | 0.35 | 0.193 | 0.262 | |
| AT3G46100.1 | Histidyl-tRNA synthetase 1 (HRS1) | 45 | 0.27 | 0.192 | 0.050 | * |
| AT3G46100.1 | Histidyl-tRNA synthetase 1 (HRS1) | 46 | 0.27 | 0.225 | 0.161 | |
| AT3G46100.1 | Histidyl-tRNA synthetase 1 (HRS1) | 47 | 0.30 | 0.414 | 0.946 | |
| AT3G46740.1 | Translocon of the outer envelope membrane of chloroplasts 75 (TOC75) | 2 | 1.09 | 0.251 | 1.000 | |
| AT3G46740.1 | Translocon of the outer envelope membrane of chloroplasts 75 (TOC75) | 3 | 1.07 | 0.200 | 1.000 | |
| AT3G46740.1 | Translocon of the outer envelope membrane of chloroplasts 75 (TOC75) | 4 | 1.05 | 0.168 | 1.000 | |

| | | | | | |
|---|---|---|---|---|---|
| AT3G46740.1 | Translocon of the outer envelope membrane of chloroplasts 75 (TOC75) | 5 | 1.02 | 0.150 | 1.000 |
| AT3G46740.1 | Translocon of the outer envelope membrane of chloroplasts 75 (TOC75) | 6 | 0.98 | 0.142 | 1.000 |
| AT3G46740.1 | Translocon of the outer envelope membrane of chloroplasts 75 (TOC75) | 7 | 0.92 | 0.136 | 1.000 |
| AT3G46740.1 | Translocon of the outer envelope membrane of chloroplasts 75 (TOC75) | 8 | 0.86 | 0.132 | 1.000 |
| AT3G46740.1 | Translocon of the outer envelope membrane of chloroplasts 75 (TOC75) | 9 | 0.79 | 0.128 | 0.989 |
| AT3G46740.1 | Translocon of the outer envelope membrane of chloroplasts 75 (TOC75) | 10 | 0.72 | 0.125 | 0.919 |
| AT3G46740.1 | Translocon of the outer envelope membrane of chloroplasts 75 (TOC75) | 11 | 0.67 | 0.121 | 0.766 |
| AT3G46740.1 | Translocon of the outer envelope membrane of chloroplasts 75 (TOC75) | 12 | 0.63 | 0.118 | 0.586 |
| AT3G46740.1 | Translocon of the outer envelope membrane of chloroplasts 75 (TOC75) | 13 | 0.61 | 0.115 | 0.457 |
| AT3G46740.1 | Translocon of the outer envelope membrane of chloroplasts 75 (TOC75) | 14 | 0.61 | 0.113 | 0.419 |
| AT3G46740.1 | Translocon of the outer envelope membrane of chloroplasts 75 (TOC75) | 15 | 0.62 | 0.112 | 0.449 |
| AT3G46740.1 | Translocon of the outer envelope membrane of chloroplasts 75 (TOC75) | 16 | 0.64 | 0.112 | 0.560 |
| AT3G46740.1 | Translocon of the outer envelope membrane of chloroplasts 75 (TOC75) | 17 | 0.69 | 0.113 | 0.762 |
| AT3G46740.1 | Translocon of the outer envelope membrane of chloroplasts 75 (TOC75) | 18 | 0.75 | 0.116 | 0.937 |
| AT3G46740.1 | Translocon of the outer envelope membrane of chloroplasts 75 (TOC75) | 19 | 0.81 | 0.122 | 0.992 |
| AT3G46740.1 | Translocon of the outer envelope membrane of chloroplasts 75 (TOC75) | 20 | 0.83 | 0.132 | 0.999 |
| AT3G46740.1 | Translocon of the outer envelope membrane of chloroplasts 75 (TOC75) | 21 | 0.82 | 0.149 | 0.999 |
| AT3G46740.1 | Translocon of the outer envelope membrane of chloroplasts 75 (TOC75) | 22 | 0.76 | 0.174 | 0.997 |
| AT3G46740.1 | Translocon of the outer envelope membrane of chloroplasts 75 (TOC75) | 23 | 0.68 | 0.209 | 0.990 |
| AT3G46740.1 | Translocon of the outer envelope membrane of chloroplasts 75 (TOC75) | 24 | 0.60 | 0.257 | 0.985 |

| | | | | | |
|---|---|---|---|---|---|
| AT3G46740.1 | Translocon of the outer envelope membrane of chloroplasts 75 (TOC75) | 46 | 0.29 | 2.812 | 1.000 |
| AT3G46740.1 | Translocon of the outer envelope membrane of chloroplasts 75 (TOC75) | 47 | 0.30 | 2.981 | 1.000 |
| AT3G46780.1 | Plastid transcriptionally active 16 (PTAC16) | 2 | 0.95 | 0.554 | 1.000 |
| AT3G46780.1 | Plastid transcriptionally active 16 (PTAC16) | 3 | 0.98 | 0.469 | 1.000 |
| AT3G46780.1 | Plastid transcriptionally active 16 (PTAC16) | 6 | 1.02 | 0.373 | 1.000 |
| AT3G46780.1 | Plastid transcriptionally active 16 (PTAC16) | 7 | 1.02 | 0.366 | 1.000 |
| AT3G46780.1 | Plastid transcriptionally active 16 (PTAC16) | 8 | 1.02 | 0.361 | 1.000 |
| AT3G46780.1 | Plastid transcriptionally active 16 (PTAC16) | 11 | 1.01 | 0.334 | 1.000 |
| AT3G46780.1 | Plastid transcriptionally active 16 (PTAC16) | 12 | 1.01 | 0.319 | 1.000 |
| AT3G46780.1 | Plastid transcriptionally active 16 (PTAC16) | 13 | 1.00 | 0.305 | 1.000 |
| AT3G46780.1 | Plastid transcriptionally active 16 (PTAC16) | 14 | 0.99 | 0.291 | 1.000 |
| AT3G46780.1 | Plastid transcriptionally active 16 (PTAC16) | 15 | 0.98 | 0.280 | 1.000 |
| AT3G46780.1 | Plastid transcriptionally active 16 (PTAC16) | 16 | 0.98 | 0.273 | 1.000 |
| AT3G46780.1 | Plastid transcriptionally active 16 (PTAC16) | 17 | 1.00 | 0.270 | 1.000 |
| AT3G46780.1 | Plastid transcriptionally active 16 (PTAC16) | 18 | 1.06 | 0.272 | 1.000 |
| AT3G46780.1 | Plastid transcriptionally active 16 (PTAC16) | 19 | 1.12 | 0.278 | 1.000 |
| AT3G46780.1 | Plastid transcriptionally active 16 (PTAC16) | 20 | 1.17 | 0.286 | 1.000 |
| AT3G46780.1 | Plastid transcriptionally active 16 (PTAC16) | 21 | 1.17 | 0.294 | 1.000 |
| AT3G46780.1 | Plastid transcriptionally active 16 (PTAC16) | 22 | 1.14 | 0.298 | 1.000 |
| AT3G46780.1 | Plastid transcriptionally active 16 (PTAC16) | 23 | 1.11 | 0.294 | 1.000 |
| AT3G46780.1 | Plastid transcriptionally active 16 (PTAC16) | 24 | 1.10 | 0.286 | 1.000 |
| AT3G46780.1 | Plastid transcriptionally active 16 (PTAC16) | 25 | 1.11 | 0.273 | 1.000 |
| AT3G46780.1 | Plastid transcriptionally active 16 (PTAC16) | 26 | 1.19 | 0.259 | 1.000 |
| AT3G46780.1 | Plastid transcriptionally active 16 (PTAC16) | 27 | 1.31 | 0.248 | 1.000 |
| AT3G46780.1 | Plastid transcriptionally active 16 (PTAC16) | 28 | 1.44 | 0.240 | 1.000 |
| AT3G46780.1 | Plastid transcriptionally active 16 (PTAC16) | 29 | 1.50 | 0.235 | 0.999 |
| AT3G46780.1 | Plastid transcriptionally active 16 (PTAC16) | 30 | 1.45 | 0.234 | 1.000 |
| AT3G46780.1 | Plastid transcriptionally active 16 (PTAC16) | 31 | 1.30 | 0.233 | 1.000 |
| AT3G46780.1 | Plastid transcriptionally active 16 (PTAC16) | 32 | 1.10 | 0.234 | 1.000 |
| AT3G46780.1 | Plastid transcriptionally active 16 (PTAC16) | 33 | 0.91 | 0.235 | 1.000 |

| | | | | | |
|---|---|---|---|---|---|
| AT3G46780.1 | Plastid transcriptionally active 16 (PTAC16) | 34 | 0.81 | 0.236 | 1.000 |
| AT3G46780.1 | Plastid transcriptionally active 16 (PTAC16) | 35 | 0.77 | 0.239 | 1.000 |
| AT3G46780.1 | Plastid transcriptionally active 16 (PTAC16) | 36 | 0.79 | 0.242 | 1.000 |
| AT3G46780.1 | Plastid transcriptionally active 16 (PTAC16) | 37 | 0.80 | 0.245 | 1.000 |
| AT3G46780.1 | Plastid transcriptionally active 16 (PTAC16) | 38 | 0.77 | 0.246 | 1.000 |
| AT3G46780.1 | Plastid transcriptionally active 16 (PTAC16) | 39 | 0.73 | 0.248 | 1.000 |
| AT3G46780.1 | Plastid transcriptionally active 16 (PTAC16) | 40 | 0.70 | 0.253 | 1.000 |
| AT3G46780.1 | Plastid transcriptionally active 16 (PTAC16) | 41 | 0.69 | 0.273 | 1.000 |
| AT3G46780.1 | Plastid transcriptionally active 16 (PTAC16) | 42 | 0.73 | 0.312 | 1.000 |
| AT3G46780.1 | Plastid transcriptionally active 16 (PTAC16) | 43 | 0.81 | 0.371 | 1.000 |
| AT3G46780.1 | Plastid transcriptionally active 16 (PTAC16) | 46 | 1.44 | 0.657 | 1.000 |
| AT3G46780.1 | Plastid transcriptionally active 16 (PTAC16) | 47 | 1.81 | 0.795 | 1.000 |
| AT3G47430.1 | Peroxin 11B (PEX11B) | 2 | 0.89 | 0.497 | 1.000 |
| AT3G47430.1 | Peroxin 11B (PEX11B) | 3 | 0.68 | 0.538 | 1.000 |
| AT3G47430.1 | Peroxin 11B (PEX11B) | 30 | 0.18 | 0.335 | 0.231 |
| AT3G47430.1 | Peroxin 11B (PEX11B) | 31 | 0.31 | 0.238 | 0.278 |
| AT3G47430.1 | Peroxin 11B (PEX11B) | 32 | 0.59 | 0.216 | 0.933 |
| AT3G47430.1 | Peroxin 11B (PEX11B) | 33 | 0.97 | 0.207 | 1.000 |
| AT3G47430.1 | Peroxin 11B (PEX11B) | 34 | 1.00 | 0.203 | 1.000 |
| AT3G47430.1 | Peroxin 11B (PEX11B) | 35 | 0.82 | 0.205 | 1.000 |
| AT3G47430.1 | Peroxin 11B (PEX11B) | 36 | 0.78 | 0.219 | 1.000 |
| AT3G47430.1 | Peroxin 11B (PEX11B) | 37 | 0.85 | 0.280 | 1.000 |
| AT3G47430.1 | Peroxin 11B (PEX11B) | 38 | 1.07 | 0.434 | 1.000 |
| AT3G47430.1 | Peroxin 11B (PEX11B) | 46 | 402.16 | 2.533 | 0.942 |
| AT3G47430.1 | Peroxin 11B (PEX11B) | 47 | 1084.73 | 2.937 | 0.940 |
| AT3G47470.1 | Photosystem I light-harvesting complex subunit (LHCA4) | 2 | 0.26 | 0.334 | 0.890 |
| AT3G47470.1 | Photosystem I light-harvesting complex subunit (LHCA4) | 3 | 0.35 | 0.208 | 0.558 |
| AT3G47470.1 | Photosystem I light-harvesting complex subunit (LHCA4) | 4 | 0.51 | 0.194 | 0.973 |
| AT3G47470.1 | Photosystem I light-harvesting complex subunit (LHCA4) | 5 | 0.73 | 0.193 | 1.000 |
| AT3G47470.1 | Photosystem I light-harvesting complex subunit (LHCA4) | 6 | 0.71 | 0.193 | 1.000 |
| AT3G47470.1 | Photosystem I light-harvesting complex subunit (LHCA4) | 7 | 0.60 | 0.195 | 0.999 |
| AT3G47470.1 | Photosystem I light-harvesting complex subunit (LHCA4) | 8 | 0.63 | 0.190 | 1.000 |
| AT3G47470.1 | Photosystem I light-harvesting complex subunit (LHCA4) | 9 | 0.70 | 0.187 | 1.000 |
| AT3G47470.1 | Photosystem I light-harvesting complex subunit (LHCA4) | 10 | 0.71 | 0.189 | 1.000 |
| AT3G47470.1 | Photosystem I light-harvesting complex subunit (LHCA4) | 11 | 0.76 | 0.192 | 1.000 |
| AT3G47470.1 | Photosystem I light-harvesting complex subunit (LHCA4) | 12 | 0.84 | 0.191 | 1.000 |
| AT3G47470.1 | Photosystem I light-harvesting complex subunit (LHCA4) | 13 | 0.87 | 0.188 | 1.000 |

| | | | | | | |
|---|---|---|---|---|---|---|
| AT3G47470.1 | Photosystem I light-harvesting complex subunit (LHCA4) | 14 | 0.87 | 0.183 | 1.000 | |
| AT3G47470.1 | Photosystem I light-harvesting complex subunit (LHCA4) | 15 | 0.84 | 0.182 | 1.000 | |
| AT3G47470.1 | Photosystem I light-harvesting complex subunit (LHCA4) | 16 | 0.87 | 0.182 | 1.000 | |
| AT3G47470.1 | Photosystem I light-harvesting complex subunit (LHCA4) | 17 | 1.05 | 0.182 | 1.000 | |
| AT3G47470.1 | Photosystem I light-harvesting complex subunit (LHCA4) | 18 | 1.34 | 0.183 | 1.000 | |
| AT3G47470.1 | Photosystem I light-harvesting complex subunit (LHCA4) | 19 | 1.66 | 0.187 | 0.999 | |
| AT3G47470.1 | Photosystem I light-harvesting complex subunit (LHCA4) | 20 | 1.77 | 0.194 | 0.996 | |
| AT3G47470.1 | Photosystem I light-harvesting complex subunit (LHCA4) | 21 | 1.57 | 0.196 | 1.000 | |
| AT3G47470.1 | Photosystem I light-harvesting complex subunit (LHCA4) | 22 | 1.12 | 0.196 | 1.000 | |
| AT3G47470.1 | Photosystem I light-harvesting complex subunit (LHCA4) | 23 | 0.71 | 0.195 | 1.000 | |
| AT3G47470.1 | Photosystem I light-harvesting complex subunit (LHCA4) | 24 | 0.49 | 0.189 | 0.924 | |
| AT3G47470.1 | Photosystem I light-harvesting complex subunit (LHCA4) | 25 | 0.72 | 0.186 | 1.000 | |
| AT3G47470.1 | Photosystem I light-harvesting complex subunit (LHCA4) | 26 | 2.30 | 0.187 | 0.759 | |
| AT3G47470.1 | Photosystem I light-harvesting complex subunit (LHCA4) | 27 | 2.90 | 0.188 | 0.334 | |
| AT3G47470.1 | Photosystem I light-harvesting complex subunit (LHCA4) | 28 | 0.78 | 0.188 | 1.000 | |
| AT3G47470.1 | Photosystem I light-harvesting complex subunit (LHCA4) | 29 | 0.16 | 0.187 | 0.001 | *** |
| AT3G47470.1 | Photosystem I light-harvesting complex subunit (LHCA4) | 30 | 0.11 | 0.187 | 0.000 | *** |
| AT3G47470.1 | Photosystem I light-harvesting complex subunit (LHCA4) | 31 | 0.21 | 0.187 | 0.012 | * |
| AT3G47470.1 | Photosystem I light-harvesting complex subunit (LHCA4) | 32 | 0.34 | 0.189 | 0.331 | |
| AT3G47470.1 | Photosystem I light-harvesting complex subunit (LHCA4) | 33 | 0.39 | 0.192 | 0.619 | |
| AT3G47470.1 | Photosystem I light-harvesting complex subunit (LHCA4) | 34 | 0.43 | 0.202 | 0.842 | |
| AT3G47470.1 | Photosystem I light-harvesting complex subunit (LHCA4) | 35 | 0.51 | 0.228 | 0.995 | |
| AT3G47470.1 | Photosystem I light-harvesting complex subunit (LHCA4) | 36 | 0.51 | 0.289 | 1.000 | |
| AT3G47470.1 | Photosystem I light-harvesting complex subunit (LHCA4) | 38 | 0.18 | 0.263 | 0.146 | |
| AT3G47470.1 | Photosystem I light-harvesting complex subunit (LHCA4) | 39 | 0.09 | 0.304 | 0.019 | * |
| AT3G47470.1 | Photosystem I light-harvesting complex subunit (LHCA4) | 40 | 0.05 | 0.385 | 0.018 | * |
| AT3G47470.1 | Photosystem I light-harvesting complex subunit (LHCA4) | 46 | 0.00 | 1.469 | 0.879 | |
| AT3G47470.1 | Photosystem I light-harvesting complex subunit (LHCA4) | 47 | 0.00 | 1.748 | 0.955 | |
| AT3G47520.1 | NAD-dependent malate dehydrogenase (MDH) | 2 | 0.72 | 0.333 | 0.907 | |
| AT3G47520.1 | NAD-dependent malate dehydrogenase (MDH) | 3 | 0.72 | 0.324 | 0.906 | |
| AT3G47520.1 | NAD-dependent malate dehydrogenase (MDH) | 4 | 0.73 | 0.315 | 0.904 | |

| | | | | | |
|---|---|---|---|---|---|
| AT3G47520.1 | NAD-dependent malate dehydrogenase (MDH) | 15 | 0.78 | 0.221 | 0.886 |
| AT3G47520.1 | NAD-dependent malate dehydrogenase (MDH) | 16 | 0.79 | 0.213 | 0.884 |
| AT3G47520.1 | NAD-dependent malate dehydrogenase (MDH) | 17 | 0.79 | 0.205 | 0.882 |
| AT3G47520.1 | NAD-dependent malate dehydrogenase (MDH) | 18 | 0.80 | 0.198 | 0.881 |
| AT3G47520.1 | NAD-dependent malate dehydrogenase (MDH) | 19 | 0.80 | 0.191 | 0.880 |
| AT3G47520.1 | NAD-dependent malate dehydrogenase (MDH) | 27 | 0.85 | 0.149 | 0.886 |
| AT3G47520.1 | NAD-dependent malate dehydrogenase (MDH) | 28 | 0.85 | 0.147 | 0.891 |
| AT3G47520.1 | NAD-dependent malate dehydrogenase (MDH) | 29 | 0.86 | 0.144 | 0.896 |
| AT3G47520.1 | NAD-dependent malate dehydrogenase (MDH) | 30 | 0.86 | 0.143 | 0.903 |
| AT3G47520.1 | NAD-dependent malate dehydrogenase (MDH) | 31 | 0.87 | 0.142 | 0.910 |
| AT3G47520.1 | NAD-dependent malate dehydrogenase (MDH) | 32 | 0.88 | 0.142 | 0.918 |
| AT3G47520.1 | NAD-dependent malate dehydrogenase (MDH) | 33 | 0.88 | 0.143 | 0.926 |
| AT3G47520.1 | NAD-dependent malate dehydrogenase (MDH) | 34 | 0.89 | 0.144 | 0.935 |
| AT3G47520.1 | NAD-dependent malate dehydrogenase (MDH) | 35 | 0.89 | 0.146 | 0.943 |
| AT3G47520.1 | NAD-dependent malate dehydrogenase (MDH) | 36 | 0.90 | 0.149 | 0.951 |
| AT3G47520.1 | NAD-dependent malate dehydrogenase (MDH) | 37 | 0.91 | 0.152 | 0.959 |
| AT3G47520.1 | NAD-dependent malate dehydrogenase (MDH) | 38 | 0.91 | 0.156 | 0.966 |
| AT3G47520.1 | NAD-dependent malate dehydrogenase (MDH) | 39 | 0.92 | 0.161 | 0.972 |
| AT3G47520.1 | NAD-dependent malate dehydrogenase (MDH) | 40 | 0.92 | 0.166 | 0.978 |
| AT3G47520.1 | NAD-dependent malate dehydrogenase (MDH) | 41 | 0.93 | 0.171 | 0.982 |
| AT3G47520.1 | NAD-dependent malate dehydrogenase (MDH) | 42 | 0.94 | 0.177 | 0.986 |
| AT3G47520.1 | NAD-dependent malate dehydrogenase (MDH) | 43 | 0.94 | 0.183 | 0.990 |
| AT3G47520.1 | NAD-dependent malate dehydrogenase (MDH) | 44 | 0.95 | 0.190 | 0.992 |
| AT3G47520.1 | NAD-dependent malate dehydrogenase (MDH) | 45 | 0.96 | 0.197 | 0.995 |
| AT3G47520.1 | NAD-dependent malate dehydrogenase (MDH) | 46 | 0.96 | 0.204 | 0.996 |
| AT3G47520.1 | NAD-dependent malate dehydrogenase (MDH) | 47 | 0.97 | 0.211 | 0.998 |
| AT3G47630.1 | Translocator assembly/maintenance protein | 2 | 0.86 | 0.394 | 1.000 |
| AT3G47630.1 | Translocator assembly/maintenance protein | 3 | 0.87 | 0.376 | 1.000 |
| AT3G47630.1 | Translocator assembly/maintenance protein | 33 | 1.80 | 0.256 | 0.884 |
| AT3G47630.1 | Translocator assembly/maintenance protein | 34 | 1.87 | 0.221 | 0.751 |
| AT3G47630.1 | Translocator assembly/maintenance protein | 35 | 1.95 | 0.194 | 0.572 |

| | | | | | | |
|---|---|---|---|---|---|---|
| AT3G47630.1 | Translocator assembly/maintenance protein | 36 | 2.04 | 0.178 | 0.411 | |
| AT3G47630.1 | Translocator assembly/maintenance protein | 39 | 2.12 | 0.156 | 0.217 | |
| AT3G47630.1 | Translocator assembly/maintenance protein | 40 | 2.00 | 0.146 | 0.233 | |
| AT3G47630.1 | Translocator assembly/maintenance protein | 41 | 1.78 | 0.135 | 0.345 | |
| AT3G47630.1 | Translocator assembly/maintenance protein | 42 | 1.49 | 0.127 | 0.663 | |
| AT3G47630.1 | Translocator assembly/maintenance protein | 43 | 1.18 | 0.123 | 0.990 | |
| AT3G47630.1 | Translocator assembly/maintenance protein | 44 | 0.90 | 0.127 | 1.000 | |
| AT3G47630.1 | Translocator assembly/maintenance protein | 45 | 0.68 | 0.144 | 0.798 | |
| AT3G47630.1 | Translocator assembly/maintenance protein | 46 | 0.51 | 0.179 | 0.494 | |
| AT3G47630.1 | Translocator assembly/maintenance protein | 47 | 0.39 | 0.233 | 0.400 | |
| AT3G47833.1 | Succinate dehydrogenase subunit (SDH7-1) | 2 | 0.90 | 0.361 | 1.000 | |
| AT3G47833.1 | Succinate dehydrogenase subunit (SDH7-1) | 3 | 0.94 | 0.228 | 1.000 | |
| AT3G47833.1 | Succinate dehydrogenase subunit (SDH7-1) | 4 | 1.04 | 0.211 | 1.000 | |
| AT3G47833.1 | Succinate dehydrogenase subunit (SDH7-1) | 5 | 1.25 | 0.207 | 1.000 | |
| AT3G47833.1 | Succinate dehydrogenase subunit (SDH7-1) | 6 | 1.58 | 0.197 | 1.000 | |
| AT3G47833.1 | Succinate dehydrogenase subunit (SDH7-1) | 7 | 1.85 | 0.192 | 0.993 | |
| AT3G47833.1 | Succinate dehydrogenase subunit (SDH7-1) | 8 | 2.15 | 0.187 | 0.896 | |
| AT3G47833.1 | Succinate dehydrogenase subunit (SDH7-1) | 9 | 2.42 | 0.183 | 0.683 | |
| AT3G47833.1 | Succinate dehydrogenase subunit (SDH7-1) | 10 | 2.19 | 0.183 | 0.848 | |
| AT3G47833.1 | Succinate dehydrogenase subunit (SDH7-1) | 11 | 1.76 | 0.184 | 0.996 | |
| AT3G47833.1 | Succinate dehydrogenase subunit (SDH7-1) | 12 | 1.58 | 0.184 | 1.000 | |
| AT3G47833.1 | Succinate dehydrogenase subunit (SDH7-1) | 13 | 1.69 | 0.183 | 0.998 | |
| AT3G47833.1 | Succinate dehydrogenase subunit (SDH7-1) | 14 | 2.10 | 0.181 | 0.896 | |
| AT3G47833.1 | Succinate dehydrogenase subunit (SDH7-1) | 15 | 2.18 | 0.180 | 0.841 | |
| AT3G47833.1 | Succinate dehydrogenase subunit (SDH7-1) | 16 | 1.80 | 0.178 | 0.990 | |
| AT3G47833.1 | Succinate dehydrogenase subunit (SDH7-1) | 17 | 1.90 | 0.178 | 0.969 | |
| AT3G47833.1 | Succinate dehydrogenase subunit (SDH7-1) | 18 | 3.10 | 0.183 | 0.230 | |
| AT3G47833.1 | Succinate dehydrogenase subunit (SDH7-1) | 19 | 4.95 | 0.191 | 0.012 | * |
| AT3G47833.1 | Succinate dehydrogenase subunit (SDH7-1) | 20 | 4.77 | 0.198 | 0.026 | * |
| AT3G47833.1 | Succinate dehydrogenase subunit (SDH7-1) | 21 | 2.62 | 0.210 | 0.762 | |
| AT3G47833.1 | Succinate dehydrogenase subunit (SDH7-1) | 22 | 1.35 | 0.218 | 1.000 | |

| | | | | | | |
|---|---|---|---|---|---|---|
| AT3G47833.1 | Succinate dehydrogenase subunit (SDH7-1) | 23 | 0.97 | 0.231 | 1.000 | |
| AT3G47833.1 | Succinate dehydrogenase subunit (SDH7-1) | 24 | 1.09 | 0.234 | 1.000 | |
| AT3G47833.1 | Succinate dehydrogenase subunit (SDH7-1) | 25 | 1.26 | 0.255 | 1.000 | |
| AT3G47833.1 | Succinate dehydrogenase subunit (SDH7-1) | 27 | 1.25 | 0.249 | 1.000 | |
| AT3G47833.1 | Succinate dehydrogenase subunit (SDH7-1) | 28 | 1.41 | 0.216 | 1.000 | |
| AT3G47833.1 | Succinate dehydrogenase subunit (SDH7-1) | 29 | 1.66 | 0.197 | 1.000 | |
| AT3G47833.1 | Succinate dehydrogenase subunit (SDH7-1) | 30 | 1.95 | 0.186 | 0.971 | |
| AT3G47833.1 | Succinate dehydrogenase subunit (SDH7-1) | 31 | 2.60 | 0.184 | 0.536 | |
| AT3G47833.1 | Succinate dehydrogenase subunit (SDH7-1) | 32 | 2.98 | 0.184 | 0.288 | |
| AT3G47833.1 | Succinate dehydrogenase subunit (SDH7-1) | 33 | 2.57 | 0.185 | 0.575 | |
| AT3G47833.1 | Succinate dehydrogenase subunit (SDH7-1) | 34 | 2.03 | 0.186 | 0.945 | |
| AT3G47833.1 | Succinate dehydrogenase subunit (SDH7-1) | 35 | 1.56 | 0.185 | 1.000 | |
| AT3G47833.1 | Succinate dehydrogenase subunit (SDH7-1) | 36 | 1.08 | 0.185 | 1.000 | |
| AT3G47833.1 | Succinate dehydrogenase subunit (SDH7-1) | 37 | 0.88 | 0.184 | 1.000 | |
| AT3G47833.1 | Succinate dehydrogenase subunit (SDH7-1) | 38 | 1.07 | 0.184 | 1.000 | |
| AT3G47833.1 | Succinate dehydrogenase subunit (SDH7-1) | 39 | 1.58 | 0.183 | 1.000 | |
| AT3G47833.1 | Succinate dehydrogenase subunit (SDH7-1) | 40 | 2.04 | 0.183 | 0.936 | |
| AT3G47833.1 | Succinate dehydrogenase subunit (SDH7-1) | 41 | 1.94 | 0.184 | 0.972 | |
| AT3G47833.1 | Succinate dehydrogenase subunit (SDH7-1) | 42 | 1.39 | 0.188 | 1.000 | |
| AT3G47833.1 | Succinate dehydrogenase subunit (SDH7-1) | 43 | 0.80 | 0.189 | 1.000 | |
| AT3G47833.1 | Succinate dehydrogenase subunit (SDH7-1) | 44 | 0.41 | 0.189 | 0.714 | |
| AT3G47833.1 | Succinate dehydrogenase subunit (SDH7-1) | 45 | 0.24 | 0.188 | 0.036 | * |
| AT3G47833.1 | Succinate dehydrogenase subunit (SDH7-1) | 46 | 0.17 | 0.219 | 0.017 | * |
| AT3G47833.1 | Succinate dehydrogenase subunit (SDH7-1) | 47 | 0.12 | 0.409 | 0.567 | |
| AT3G47930.1 | NADH:ubiquinone oxidoreductase subunit (GLDH) | 2 | 1.39 | 0.248 | 1.000 | |
| AT3G47930.1 | NADH:ubiquinone oxidoreductase subunit (GLDH) | 3 | 0.94 | 0.163 | 1.000 | |
| AT3G47930.1 | NADH:ubiquinone oxidoreductase subunit (GLDH) | 4 | 0.66 | 0.149 | 0.999 | |
| AT3G47930.1 | NADH:ubiquinone oxidoreductase subunit (GLDH) | 5 | 0.54 | 0.146 | 0.860 | |
| AT3G47930.1 | NADH:ubiquinone oxidoreductase subunit (GLDH) | 6 | 0.55 | 0.140 | 0.855 | |
| AT3G47930.1 | NADH:ubiquinone oxidoreductase subunit (GLDH) | 7 | 0.66 | 0.138 | 0.997 | |
| AT3G47930.1 | NADH:ubiquinone oxidoreductase subunit (GLDH) | 8 | 1.02 | 0.134 | 1.000 | |

| | | | | | |
|---|---|---|---|---|---|
| AT3G47930.1 | NADH:ubiquinone oxidoreductase subunit (GLDH) | 9 | 1.48 | 0.131 | 0.997 |
| AT3G47930.1 | NADH:ubiquinone oxidoreductase subunit (GLDH) | 10 | 1.69 | 0.132 | 0.911 |
| AT3G47930.1 | NADH:ubiquinone oxidoreductase subunit (GLDH) | 11 | 1.81 | 0.133 | 0.787 |
| AT3G47930.1 | NADH:ubiquinone oxidoreductase subunit (GLDH) | 12 | 1.94 | 0.132 | 0.600 |
| AT3G47930.1 | NADH:ubiquinone oxidoreductase subunit (GLDH) | 13 | 2.03 | 0.130 | 0.435 |
| AT3G47930.1 | NADH:ubiquinone oxidoreductase subunit (GLDH) | 14 | 2.03 | 0.128 | 0.409 |
| AT3G47930.1 | NADH:ubiquinone oxidoreductase subunit (GLDH) | 15 | 1.79 | 0.128 | 0.756 |
| AT3G47930.1 | NADH:ubiquinone oxidoreductase subunit (GLDH) | 16 | 1.50 | 0.127 | 0.993 |
| AT3G47930.1 | NADH:ubiquinone oxidoreductase subunit (GLDH) | 17 | 1.39 | 0.126 | 1.000 |
| AT3G47930.1 | NADH:ubiquinone oxidoreductase subunit (GLDH) | 18 | 1.38 | 0.128 | 1.000 |
| AT3G47930.1 | NADH:ubiquinone oxidoreductase subunit (GLDH) | 19 | 1.37 | 0.132 | 1.000 |
| AT3G47930.1 | NADH:ubiquinone oxidoreductase subunit (GLDH) | 20 | 1.15 | 0.135 | 1.000 |
| AT3G47930.1 | NADH:ubiquinone oxidoreductase subunit (GLDH) | 21 | 0.90 | 0.136 | 1.000 |
| AT3G47930.1 | NADH:ubiquinone oxidoreductase subunit (GLDH) | 22 | 0.77 | 0.136 | 1.000 |
| AT3G47930.1 | NADH:ubiquinone oxidoreductase subunit (GLDH) | 23 | 0.75 | 0.135 | 1.000 |
| AT3G47930.1 | NADH:ubiquinone oxidoreductase subunit (GLDH) | 24 | 0.74 | 0.132 | 1.000 |
| AT3G47930.1 | NADH:ubiquinone oxidoreductase subunit (GLDH) | 25 | 0.72 | 0.130 | 1.000 |
| AT3G47930.1 | NADH:ubiquinone oxidoreductase subunit (GLDH) | 26 | 0.89 | 0.129 | 1.000 |
| AT3G47930.1 | NADH:ubiquinone oxidoreductase subunit (GLDH) | 27 | 1.34 | 0.129 | 1.000 |
| AT3G47930.1 | NADH:ubiquinone oxidoreductase subunit (GLDH) | 28 | 1.58 | 0.130 | 0.973 |
| AT3G47930.1 | NADH:ubiquinone oxidoreductase subunit (GLDH) | 29 | 1.24 | 0.130 | 1.000 |
| AT3G47930.1 | NADH:ubiquinone oxidoreductase subunit (GLDH) | 30 | 0.75 | 0.131 | 1.000 |
| AT3G47930.1 | NADH:ubiquinone oxidoreductase subunit (GLDH) | 31 | 0.52 | 0.131 | 0.624 |
| AT3G47930.1 | NADH:ubiquinone oxidoreductase subunit (GLDH) | 32 | 0.56 | 0.132 | 0.815 |
| AT3G47930.1 | NADH:ubiquinone oxidoreductase subunit (GLDH) | 33 | 0.76 | 0.132 | 1.000 |
| AT3G47930.1 | NADH:ubiquinone oxidoreductase subunit (GLDH) | 34 | 1.03 | 0.132 | 1.000 |
| AT3G47930.1 | NADH:ubiquinone oxidoreductase subunit (GLDH) | 35 | 1.31 | 0.132 | 1.000 |
| AT3G47930.1 | NADH:ubiquinone oxidoreductase subunit (GLDH) | 36 | 1.49 | 0.132 | 0.996 |
| AT3G47930.1 | NADH:ubiquinone oxidoreductase subunit (GLDH) | 37 | 1.67 | 0.131 | 0.930 |
| AT3G47930.1 | NADH:ubiquinone oxidoreductase subunit (GLDH) | 38 | 1.92 | 0.132 | 0.616 |
| AT3G47930.1 | NADH:ubiquinone oxidoreductase subunit (GLDH) | 39 | 2.16 | 0.131 | 0.301 |

| | | | | | | |
|---|---|---|---|---|---|---|
| AT3G47930.1 | NADH:ubiquinone oxidoreductase subunit (GLDH) | 40 | 2.31 | 0.131 | 0.176 | |
| AT3G47930.1 | NADH:ubiquinone oxidoreductase subunit (GLDH) | 41 | 2.25 | 0.132 | 0.222 | |
| AT3G47930.1 | NADH:ubiquinone oxidoreductase subunit (GLDH) | 42 | 2.08 | 0.133 | 0.425 | |
| AT3G47930.1 | NADH:ubiquinone oxidoreductase subunit (GLDH) | 43 | 1.86 | 0.135 | 0.743 | |
| AT3G47930.1 | NADH:ubiquinone oxidoreductase subunit (GLDH) | 44 | 1.42 | 0.134 | 1.000 | |
| AT3G47930.1 | NADH:ubiquinone oxidoreductase subunit (GLDH) | 45 | 0.83 | 0.135 | 1.000 | |
| AT3G47930.1 | NADH:ubiquinone oxidoreductase subunit (GLDH) | 46 | 0.36 | 0.156 | 0.138 | |
| AT3G47930.1 | NADH:ubiquinone oxidoreductase subunit (GLDH) | 47 | 0.12 | 0.266 | 0.024 | * |
| AT3G48000.1 | Aldehyde dehydrogenase 2B4 (ALDH2B4) | 3 | 0.51 | 0.275 | 1.000 | |
| AT3G48000.1 | Aldehyde dehydrogenase 2B4 (ALDH2B4) | 4 | 0.40 | 0.255 | 0.963 | |
| AT3G48000.1 | Aldehyde dehydrogenase 2B4 (ALDH2B4) | 5 | 0.38 | 0.255 | 0.930 | |
| AT3G48000.1 | Aldehyde dehydrogenase 2B4 (ALDH2B4) | 6 | 0.40 | 0.255 | 0.958 | |
| AT3G48000.1 | Aldehyde dehydrogenase 2B4 (ALDH2B4) | 7 | 0.40 | 0.254 | 0.965 | |
| AT3G48000.1 | Aldehyde dehydrogenase 2B4 (ALDH2B4) | 8 | 0.49 | 0.251 | 0.998 | |
| AT3G48000.1 | Aldehyde dehydrogenase 2B4 (ALDH2B4) | 9 | 0.71 | 0.248 | 1.000 | |
| AT3G48000.1 | Aldehyde dehydrogenase 2B4 (ALDH2B4) | 10 | 0.96 | 0.247 | 1.000 | |
| AT3G48000.1 | Aldehyde dehydrogenase 2B4 (ALDH2B4) | 11 | 1.24 | 0.247 | 1.000 | |
| AT3G48000.1 | Aldehyde dehydrogenase 2B4 (ALDH2B4) | 12 | 1.58 | 0.248 | 1.000 | |
| AT3G48000.1 | Aldehyde dehydrogenase 2B4 (ALDH2B4) | 13 | 1.89 | 0.245 | 0.999 | |
| AT3G48000.1 | Aldehyde dehydrogenase 2B4 (ALDH2B4) | 14 | 2.04 | 0.242 | 0.997 | |
| AT3G48000.1 | Aldehyde dehydrogenase 2B4 (ALDH2B4) | 15 | 1.97 | 0.241 | 0.998 | |
| AT3G48000.1 | Aldehyde dehydrogenase 2B4 (ALDH2B4) | 16 | 1.83 | 0.240 | 1.000 | |
| AT3G48000.1 | Aldehyde dehydrogenase 2B4 (ALDH2B4) | 17 | 1.75 | 0.239 | 1.000 | |
| AT3G48000.1 | Aldehyde dehydrogenase 2B4 (ALDH2B4) | 18 | 1.65 | 0.243 | 1.000 | |
| AT3G48000.1 | Aldehyde dehydrogenase 2B4 (ALDH2B4) | 19 | 1.34 | 0.249 | 1.000 | |
| AT3G48000.1 | Aldehyde dehydrogenase 2B4 (ALDH2B4) | 20 | 1.08 | 0.254 | 1.000 | |
| AT3G48000.1 | Aldehyde dehydrogenase 2B4 (ALDH2B4) | 21 | 0.94 | 0.258 | 1.000 | |
| AT3G48000.1 | Aldehyde dehydrogenase 2B4 (ALDH2B4) | 22 | 0.89 | 0.258 | 1.000 | |
| AT3G48000.1 | Aldehyde dehydrogenase 2B4 (ALDH2B4) | 23 | 0.86 | 0.256 | 1.000 | |
| AT3G48000.1 | Aldehyde dehydrogenase 2B4 (ALDH2B4) | 24 | 0.84 | 0.251 | 1.000 | |
| AT3G48000.1 | Aldehyde dehydrogenase 2B4 (ALDH2B4) | 25 | 0.81 | 0.247 | 1.000 | |

| | | | | | |
|---|---|---|---|---|---|
| AT3G48000.1 | Aldehyde dehydrogenase 2B4 (ALDH2B4) | 26 | 0.82 | 0.246 | 1.000 |
| AT3G48000.1 | Aldehyde dehydrogenase 2B4 (ALDH2B4) | 27 | 0.91 | 0.246 | 1.000 |
| AT3G48000.1 | Aldehyde dehydrogenase 2B4 (ALDH2B4) | 28 | 1.05 | 0.246 | 1.000 |
| AT3G48000.1 | Aldehyde dehydrogenase 2B4 (ALDH2B4) | 29 | 1.18 | 0.246 | 1.000 |
| AT3G48000.1 | Aldehyde dehydrogenase 2B4 (ALDH2B4) | 30 | 1.19 | 0.246 | 1.000 |
| AT3G48000.1 | Aldehyde dehydrogenase 2B4 (ALDH2B4) | 31 | 1.16 | 0.246 | 1.000 |
| AT3G48000.1 | Aldehyde dehydrogenase 2B4 (ALDH2B4) | 32 | 1.26 | 0.247 | 1.000 |
| AT3G48000.1 | Aldehyde dehydrogenase 2B4 (ALDH2B4) | 33 | 1.48 | 0.247 | 1.000 |
| AT3G48000.1 | Aldehyde dehydrogenase 2B4 (ALDH2B4) | 34 | 1.60 | 0.248 | 1.000 |
| AT3G48000.1 | Aldehyde dehydrogenase 2B4 (ALDH2B4) | 35 | 1.40 | 0.247 | 1.000 |
| AT3G48000.1 | Aldehyde dehydrogenase 2B4 (ALDH2B4) | 36 | 1.13 | 0.246 | 1.000 |
| AT3G48000.1 | Aldehyde dehydrogenase 2B4 (ALDH2B4) | 37 | 1.16 | 0.246 | 1.000 |
| AT3G48000.1 | Aldehyde dehydrogenase 2B4 (ALDH2B4) | 38 | 1.68 | 0.247 | 1.000 |
| AT3G48000.1 | Aldehyde dehydrogenase 2B4 (ALDH2B4) | 39 | 2.93 | 0.247 | 0.805 |
| AT3G48000.1 | Aldehyde dehydrogenase 2B4 (ALDH2B4) | 40 | 3.96 | 0.248 | 0.383 |
| AT3G48000.1 | Aldehyde dehydrogenase 2B4 (ALDH2B4) | 41 | 4.00 | 0.249 | 0.373 |
| AT3G48000.1 | Aldehyde dehydrogenase 2B4 (ALDH2B4) | 42 | 3.62 | 0.252 | 0.536 |
| AT3G48000.1 | Aldehyde dehydrogenase 2B4 (ALDH2B4) | 43 | 2.54 | 0.254 | 0.951 |
| AT3G48000.1 | Aldehyde dehydrogenase 2B4 (ALDH2B4) | 44 | 1.22 | 0.254 | 1.000 |
| AT3G48000.1 | Aldehyde dehydrogenase 2B4 (ALDH2B4) | 45 | 0.48 | 0.253 | 0.997 |
| AT3G48000.1 | Aldehyde dehydrogenase 2B4 (ALDH2B4) | 46 | 0.18 | 0.295 | 0.284 |
| AT3G48000.1 | Aldehyde dehydrogenase 2B4 (ALDH2B4) | 47 | 0.07 | 0.517 | 0.495 |
| AT3G48000.1 | Aldehyde dehydrogenase 2B4 (ALDH2B4) | 48 | 0.03 | 0.859 | 0.830 |
| AT3G48030.1 | Hypoxia-induced gene domain 1 (HIGD1) | 2 | 1.27 | 0.593 | 1.000 |
| AT3G48030.1 | Hypoxia-induced gene domain 1 (HIGD1) | 3 | 1.34 | 0.542 | 1.000 |
| AT3G48030.1 | Hypoxia-induced gene domain 1 (HIGD1) | 12 | 2.24 | 0.332 | 0.871 |
| AT3G48030.1 | Hypoxia-induced gene domain 1 (HIGD1) | 13 | 2.36 | 0.321 | 0.815 |
| AT3G48030.1 | Hypoxia-induced gene domain 1 (HIGD1) | 14 | 2.48 | 0.315 | 0.760 |
| AT3G48030.1 | Hypoxia-induced gene domain 1 (HIGD1) | 16 | 2.67 | 0.314 | 0.689 |
| AT3G48030.1 | Hypoxia-induced gene domain 1 (HIGD1) | 17 | 2.73 | 0.318 | 0.679 |
| AT3G48030.1 | Hypoxia-induced gene domain 1 (HIGD1) | 18 | 2.78 | 0.323 | 0.677 |

| | | | | | | |
|---|---|---|---|---|---|---|
| AT3G48030.1 | Hypoxia-induced gene domain 1 (HIGD1) | 28 | 2.39 | 0.236 | 0.519 | |
| AT3G48030.1 | Hypoxia-induced gene domain 1 (HIGD1) | 29 | 2.27 | 0.216 | 0.484 | |
| AT3G48030.1 | Hypoxia-induced gene domain 1 (HIGD1) | 30 | 2.11 | 0.201 | 0.510 | |
| AT3G48030.1 | Hypoxia-induced gene domain 1 (HIGD1) | 31 | 1.89 | 0.197 | 0.661 | |
| AT3G48030.1 | Hypoxia-induced gene domain 1 (HIGD1) | 32 | 1.59 | 0.204 | 0.902 | |
| AT3G48030.1 | Hypoxia-induced gene domain 1 (HIGD1) | 33 | 1.26 | 0.223 | 0.999 | |
| AT3G48030.1 | Hypoxia-induced gene domain 1 (HIGD1) | 34 | 0.96 | 0.251 | 1.000 | |
| AT3G48030.1 | Hypoxia-induced gene domain 1 (HIGD1) | 35 | 0.72 | 0.287 | 0.997 | |
| AT3G48030.1 | Hypoxia-induced gene domain 1 (HIGD1) | 46 | 0.01 | 0.760 | 0.075 | . |
| AT3G48030.1 | Hypoxia-induced gene domain 1 (HIGD1) | 47 | 0.01 | 0.825 | 0.071 | . |
| AT3G48110.1 | Glycine-tRNA ligase (EDD1) | 2 | 0.85 | 0.657 | 1.000 | |
| AT3G48110.1 | Glycine-tRNA ligase (EDD1) | 3 | 0.61 | 0.648 | 1.000 | |
| AT3G48110.1 | Glycine-tRNA ligase (EDD1) | 25 | 0.21 | 0.504 | 0.862 | |
| AT3G48110.1 | Glycine-tRNA ligase (EDD1) | 26 | 0.37 | 0.350 | 0.917 | |
| AT3G48110.1 | Glycine-tRNA ligase (EDD1) | 27 | 0.76 | 0.301 | 1.000 | |
| AT3G48110.1 | Glycine-tRNA ligase (EDD1) | 28 | 1.19 | 0.283 | 1.000 | |
| AT3G48110.1 | Glycine-tRNA ligase (EDD1) | 29 | 1.40 | 0.280 | 1.000 | |
| AT3G48110.1 | Glycine-tRNA ligase (EDD1) | 30 | 1.57 | 0.279 | 0.999 | |
| AT3G48110.1 | Glycine-tRNA ligase (EDD1) | 31 | 2.27 | 0.279 | 0.897 | |
| AT3G48110.1 | Glycine-tRNA ligase (EDD1) | 32 | 3.33 | 0.284 | 0.527 | |
| AT3G48110.1 | Glycine-tRNA ligase (EDD1) | 33 | 2.22 | 0.289 | 0.928 | |
| AT3G48110.1 | Glycine-tRNA ligase (EDD1) | 34 | 0.57 | 0.305 | 0.995 | |
| AT3G48110.1 | Glycine-tRNA ligase (EDD1) | 35 | 0.14 | 0.390 | 0.295 | |
| AT3G48110.1 | Glycine-tRNA ligase (EDD1) | 36 | 0.07 | 0.649 | 0.549 | |
| AT3G48110.1 | Glycine-tRNA ligase (EDD1) | 46 | 39034.51 | 4.373 | 0.968 | |
| AT3G48110.1 | Glycine-tRNA ligase (EDD1) | 47 | 400414.07 | 4.955 | 0.949 | |
| AT3G48140.1 | B12D protein | 3 | 1.16 | 0.150 | 1.000 | |
| AT3G48140.1 | B12D protein | 4 | 1.25 | 0.135 | 1.000 | |
| AT3G48140.1 | B12D protein | 5 | 1.33 | 0.135 | 1.000 | |
| AT3G48140.1 | B12D protein | 6 | 1.45 | 0.135 | 0.999 | |
| AT3G48140.1 | B12D protein | 7 | 1.64 | 0.135 | 0.960 | |
| AT3G48140.1 | B12D protein | 8 | 1.98 | 0.132 | 0.546 | |
| AT3G48140.1 | B12D protein | 9 | 2.15 | 0.130 | 0.298 | |
| AT3G48140.1 | B12D protein | 10 | 2.06 | 0.130 | 0.403 | |
| AT3G48140.1 | B12D protein | 11 | 1.89 | 0.131 | 0.658 | |
| AT3G48140.1 | B12D protein | 12 | 1.75 | 0.131 | 0.843 | |
| AT3G48140.1 | B12D protein | 13 | 1.70 | 0.129 | 0.890 | |
| AT3G48140.1 | B12D protein | 14 | 1.65 | 0.127 | 0.918 | |
| AT3G48140.1 | B12D protein | 15 | 1.52 | 0.126 | 0.987 | |
| AT3G48140.1 | B12D protein | 16 | 1.37 | 0.125 | 1.000 | |
| AT3G48140.1 | B12D protein | 17 | 1.23 | 0.125 | 1.000 | |
| AT3G48140.1 | B12D protein | 18 | 1.08 | 0.127 | 1.000 | |
| AT3G48140.1 | B12D protein | 19 | 0.99 | 0.131 | 1.000 | |
| AT3G48140.1 | B12D protein | 20 | 1.00 | 0.135 | 1.000 | |
| AT3G48140.1 | B12D protein | 21 | 1.11 | 0.137 | 1.000 | |
| AT3G48140.1 | B12D protein | 22 | 1.26 | 0.137 | 1.000 | |
| AT3G48140.1 | B12D protein | 23 | 1.24 | 0.136 | 1.000 | |
| AT3G48140.1 | B12D protein | 24 | 1.05 | 0.133 | 1.000 | |
| AT3G48140.1 | B12D protein | 25 | 0.82 | 0.130 | 1.000 | |
| AT3G48140.1 | B12D protein | 26 | 0.76 | 0.129 | 1.000 | |
| AT3G48140.1 | B12D protein | 27 | 0.87 | 0.129 | 1.000 | |
| AT3G48140.1 | B12D protein | 28 | 1.03 | 0.129 | 1.000 | |
| AT3G48140.1 | B12D protein | 29 | 1.13 | 0.129 | 1.000 | |

| | | | | | |
|---|---|---|---|---|---|
| AT3G48140.1 | B12D protein | 30 | 1.12 | 0.129 | 1.000 |
| AT3G48140.1 | B12D protein | 31 | 1.04 | 0.130 | 1.000 |
| AT3G48140.1 | B12D protein | 32 | 0.90 | 0.130 | 1.000 |
| AT3G48140.1 | B12D protein | 33 | 0.72 | 0.130 | 1.000 |
| AT3G48140.1 | B12D protein | 34 | 0.58 | 0.130 | 0.880 |
| AT3G48140.1 | B12D protein | 35 | 0.52 | 0.130 | 0.603 |
| AT3G48140.1 | B12D protein | 36 | 0.64 | 0.130 | 0.980 |
| AT3G48140.1 | B12D protein | 37 | 1.06 | 0.130 | 1.000 |
| AT3G48140.1 | B12D protein | 38 | 1.48 | 0.130 | 0.996 |
| AT3G48140.1 | B12D protein | 39 | 1.47 | 0.130 | 0.997 |
| AT3G48140.1 | B12D protein | 40 | 1.27 | 0.130 | 1.000 |
| AT3G48140.1 | B12D protein | 41 | 1.13 | 0.131 | 1.000 |
| AT3G48140.1 | B12D protein | 42 | 1.09 | 0.133 | 1.000 |
| AT3G48140.1 | B12D protein | 43 | 1.11 | 0.134 | 1.000 |
| AT3G48140.1 | B12D protein | 44 | 1.08 | 0.134 | 1.000 |
| AT3G48140.1 | B12D protein | 45 | 0.96 | 0.135 | 1.000 |
| AT3G48140.1 | B12D protein | 46 | 0.78 | 0.161 | 1.000 |
| AT3G48140.1 | B12D protein | 47 | 0.61 | 0.276 | 1.000 |
| AT3G48140.1 | B12D protein | 48 | 0.47 | 0.452 | 1.000 |
| AT3G48160.1 | E2F-like 3(E2L3) / E2F transcription factor (E2Fe/DEL1) | 2 | 0.97 | 0.492 | 1.000 |
| AT3G48160.1 | E2F-like 3(E2L3) / E2F transcription factor (E2Fe/DEL1) | 3 | 0.96 | 0.402 | 1.000 |
| AT3G48160.1 | E2F-like 3(E2L3) / E2F transcription factor (E2Fe/DEL1) | 4 | 0.95 | 0.403 | 1.000 |
| AT3G48160.1 | E2F-like 3(E2L3) / E2F transcription factor (E2Fe/DEL1) | 10 | 1.09 | 0.391 | 1.000 |
| AT3G48160.1 | E2F-like 3(E2L3) / E2F transcription factor (E2Fe/DEL1) | 11 | 1.20 | 0.350 | 1.000 |
| AT3G48160.1 | E2F-like 3(E2L3) / E2F transcription factor (E2Fe/DEL1) | 12 | 1.34 | 0.324 | 1.000 |
| AT3G48160.1 | E2F-like 3(E2L3) / E2F transcription factor (E2Fe/DEL1) | 13 | 1.54 | 0.305 | 1.000 |
| AT3G48160.1 | E2F-like 3(E2L3) / E2F transcription factor (E2Fe/DEL1) | 14 | 1.77 | 0.300 | 0.999 |
| AT3G48160.1 | E2F-like 3(E2L3) / E2F transcription factor (E2Fe/DEL1) | 15 | 2.01 | 0.305 | 0.993 |
| AT3G48160.1 | E2F-like 3(E2L3) / E2F transcription factor (E2Fe/DEL1) | 16 | 2.22 | 0.318 | 0.984 |
| AT3G48160.1 | E2F-like 3(E2L3) / E2F transcription factor (E2Fe/DEL1) | 19 | 2.61 | 0.380 | 0.983 |
| AT3G48160.1 | E2F-like 3(E2L3) / E2F transcription factor (E2Fe/DEL1) | 20 | 2.71 | 0.423 | 0.990 |
| AT3G48160.1 | E2F-like 3(E2L3) / E2F transcription factor (E2Fe/DEL1) | 21 | 2.82 | 0.460 | 0.994 |
| AT3G48160.1 | E2F-like 3(E2L3) / E2F transcription factor (E2Fe/DEL1) | 30 | 2.93 | 0.430 | 0.984 |
| AT3G48160.1 | E2F-like 3(E2L3) / E2F transcription factor (E2Fe/DEL1) | 31 | 2.65 | 0.385 | 0.983 |
| AT3G48160.1 | E2F-like 3(E2L3) / E2F transcription factor (E2Fe/DEL1) | 32 | 2.30 | 0.348 | 0.989 |
| AT3G48160.1 | E2F-like 3(E2L3) / E2F transcription factor (E2Fe/DEL1) | 34 | 1.71 | 0.285 | 0.999 |
| AT3G48160.1 | E2F-like 3(E2L3) / E2F transcription factor (E2Fe/DEL1) | 35 | 1.63 | 0.256 | 0.999 |
| AT3G48160.1 | E2F-like 3(E2L3) / E2F transcription factor (E2Fe/DEL1) | 36 | 1.64 | 0.232 | 0.996 |
| AT3G48160.1 | E2F-like 3(E2L3) / E2F transcription factor (E2Fe/DEL1) | 37 | 1.62 | 0.218 | 0.995 |
| AT3G48160.1 | E2F-like 3(E2L3) / E2F transcription factor (E2Fe/DEL1) | 38 | 1.42 | 0.211 | 1.000 |

| | | | | | |
|---|---|---|---|---|---|
| AT3G48160.1 | E2F-like 3(E2L3) / E2F transcription factor (E2Fe/DEL1) | 39 | 1.10 | 0.210 | 1.000 |
| AT3G48160.1 | E2F-like 3(E2L3) / E2F transcription factor (E2Fe/DEL1) | 40 | 0.82 | 0.211 | 1.000 |
| AT3G48160.1 | E2F-like 3(E2L3) / E2F transcription factor (E2Fe/DEL1) | 41 | 0.66 | 0.214 | 0.998 |
| AT3G48160.1 | E2F-like 3(E2L3) / E2F transcription factor (E2Fe/DEL1) | 42 | 0.64 | 0.216 | 0.997 |
| AT3G48160.1 | E2F-like 3(E2L3) / E2F transcription factor (E2Fe/DEL1) | 43 | 0.76 | 0.220 | 1.000 |
| AT3G48160.1 | E2F-like 3(E2L3) / E2F transcription factor (E2Fe/DEL1) | 44 | 1.05 | 0.236 | 1.000 |
| AT3G48160.1 | E2F-like 3(E2L3) / E2F transcription factor (E2Fe/DEL1) | 45 | 1.54 | 0.290 | 1.000 |
| AT3G48160.1 | E2F-like 3(E2L3) / E2F transcription factor (E2Fe/DEL1) | 46 | 2.29 | 0.404 | 0.997 |
| AT3G48160.1 | E2F-like 3(E2L3) / E2F transcription factor (E2Fe/DEL1) | 47 | 3.44 | 0.581 | 0.996 |
| AT3G48680.1 | NADH:ubiquinone oxidoreductase subunit (gammaCAL2) | 2 | 1.01 | 0.605 | 1.000 |
| AT3G48680.1 | NADH:ubiquinone oxidoreductase subunit (gammaCAL2) | 3 | 1.17 | 0.475 | 1.000 |
| AT3G48680.1 | NADH:ubiquinone oxidoreductase subunit (gammaCAL2) | 6 | 1.71 | 0.475 | 1.000 |
| AT3G48680.1 | NADH:ubiquinone oxidoreductase subunit (gammaCAL2) | 7 | 1.83 | 0.438 | 1.000 |
| AT3G48680.1 | NADH:ubiquinone oxidoreductase subunit (gammaCAL2) | 8 | 1.77 | 0.447 | 1.000 |
| AT3G48680.1 | NADH:ubiquinone oxidoreductase subunit (gammaCAL2) | 12 | 0.40 | 0.341 | 0.995 |
| AT3G48680.1 | NADH:ubiquinone oxidoreductase subunit (gammaCAL2) | 13 | 0.45 | 0.302 | 0.996 |
| AT3G48680.1 | NADH:ubiquinone oxidoreductase subunit (gammaCAL2) | 14 | 0.62 | 0.283 | 1.000 |
| AT3G48680.1 | NADH:ubiquinone oxidoreductase subunit (gammaCAL2) | 15 | 0.66 | 0.260 | 1.000 |
| AT3G48680.1 | NADH:ubiquinone oxidoreductase subunit (gammaCAL2) | 16 | 0.50 | 0.247 | 0.992 |
| AT3G48680.1 | NADH:ubiquinone oxidoreductase subunit (gammaCAL2) | 17 | 0.40 | 0.244 | 0.886 |
| AT3G48680.1 | NADH:ubiquinone oxidoreductase subunit (gammaCAL2) | 18 | 0.49 | 0.246 | 0.990 |
| AT3G48680.1 | NADH:ubiquinone oxidoreductase subunit (gammaCAL2) | 19 | 0.76 | 0.254 | 1.000 |
| AT3G48680.1 | NADH:ubiquinone oxidoreductase subunit (gammaCAL2) | 20 | 1.08 | 0.260 | 1.000 |

| | | | | | | |
|---|---|---|---|---|---|---|
| AT3G48680.1 | NADH:ubiquinone oxidoreductase subunit (gammaCAL2) | 21 | 1.57 | 0.264 | 1.000 | |
| AT3G48680.1 | NADH:ubiquinone oxidoreductase subunit (gammaCAL2) | 22 | 2.25 | 0.265 | 0.981 | |
| AT3G48680.1 | NADH:ubiquinone oxidoreductase subunit (gammaCAL2) | 23 | 2.21 | 0.264 | 0.985 | |
| AT3G48680.1 | NADH:ubiquinone oxidoreductase subunit (gammaCAL2) | 24 | 0.97 | 0.256 | 1.000 | |
| AT3G48680.1 | NADH:ubiquinone oxidoreductase subunit (gammaCAL2) | 25 | 0.24 | 0.250 | 0.281 | |
| AT3G48680.1 | NADH:ubiquinone oxidoreductase subunit (gammaCAL2) | 26 | 0.13 | 0.250 | 0.012 | * |
| AT3G48680.1 | NADH:ubiquinone oxidoreductase subunit (gammaCAL2) | 27 | 0.23 | 0.250 | 0.215 | |
| AT3G48680.1 | NADH:ubiquinone oxidoreductase subunit (gammaCAL2) | 28 | 0.57 | 0.250 | 1.000 | |
| AT3G48680.1 | NADH:ubiquinone oxidoreductase subunit (gammaCAL2) | 29 | 1.00 | 0.252 | 1.000 | |
| AT3G48680.1 | NADH:ubiquinone oxidoreductase subunit (gammaCAL2) | 30 | 1.19 | 0.253 | 1.000 | |
| AT3G48680.1 | NADH:ubiquinone oxidoreductase subunit (gammaCAL2) | 31 | 1.11 | 0.258 | 1.000 | |
| AT3G48680.1 | NADH:ubiquinone oxidoreductase subunit (gammaCAL2) | 32 | 0.74 | 0.278 | 1.000 | |
| AT3G48680.1 | NADH:ubiquinone oxidoreductase subunit (gammaCAL2) | 33 | 0.35 | 0.337 | 0.979 | |
| AT3G48680.1 | NADH:ubiquinone oxidoreductase subunit (gammaCAL2) | 34 | 0.23 | 0.430 | 0.945 | |
| AT3G48680.1 | NADH:ubiquinone oxidoreductase subunit (gammaCAL2) | 38 | 0.65 | 0.394 | 1.000 | |
| AT3G48680.1 | NADH:ubiquinone oxidoreductase subunit (gammaCAL2) | 39 | 0.58 | 0.327 | 1.000 | |
| AT3G48680.1 | NADH:ubiquinone oxidoreductase subunit (gammaCAL2) | 40 | 0.38 | 0.377 | 0.998 | |
| AT3G48680.1 | NADH:ubiquinone oxidoreductase subunit (gammaCAL2) | 41 | 0.25 | 0.562 | 0.998 | |
| AT3G48680.1 | NADH:ubiquinone oxidoreductase subunit (gammaCAL2) | 46 | 0.34 | 1.522 | 1.000 | |
| AT3G48680.1 | NADH:ubiquinone oxidoreductase subunit (gammaCAL2) | 47 | 0.45 | 1.842 | 1.000 | |
| AT3G49010.4 | 60S ribosomal protein L13-1 (RPL13A) | 2 | 0.40 | 0.187 | 0.109 | |

| | | | | | | |
|---|---|---|---|---|---|---|
| AT3G49010.4 | 60S ribosomal protein L13-1 (RPL13A) | 3 | 0.44 | 0.175 | 0.130 | |
| AT3G49010.4 | 60S ribosomal protein L13-1 (RPL13A) | 4 | 0.49 | 0.163 | 0.158 | |
| AT3G49010.4 | 60S ribosomal protein L13-1 (RPL13A) | 5 | 0.54 | 0.152 | 0.203 | |
| AT3G49010.4 | 60S ribosomal protein L13-1 (RPL13A) | 6 | 0.59 | 0.142 | 0.267 | |
| AT3G49010.4 | 60S ribosomal protein L13-1 (RPL13A) | 7 | 0.65 | 0.132 | 0.361 | |
| AT3G49010.4 | 60S ribosomal protein L13-1 (RPL13A) | 8 | 0.71 | 0.125 | 0.493 | |
| AT3G49010.4 | 60S ribosomal protein L13-1 (RPL13A) | 9 | 0.78 | 0.118 | 0.666 | |
| AT3G49010.4 | 60S ribosomal protein L13-1 (RPL13A) | 10 | 0.86 | 0.114 | 0.848 | |
| AT3G49010.4 | 60S ribosomal protein L13-1 (RPL13A) | 11 | 0.95 | 0.112 | 0.977 | |
| AT3G49010.4 | 60S ribosomal protein L13-1 (RPL13A) | 12 | 1.04 | 0.111 | 0.988 | |
| AT3G49010.4 | 60S ribosomal protein L13-1 (RPL13A) | 13 | 1.14 | 0.113 | 0.875 | |
| AT3G49010.4 | 60S ribosomal protein L13-1 (RPL13A) | 14 | 1.26 | 0.118 | 0.696 | |
| AT3G49010.4 | 60S ribosomal protein L13-1 (RPL13A) | 15 | 1.38 | 0.124 | 0.519 | |
| AT3G49010.4 | 60S ribosomal protein L13-1 (RPL13A) | 16 | 1.52 | 0.131 | 0.380 | |
| AT3G49010.4 | 60S ribosomal protein L13-1 (RPL13A) | 17 | 1.67 | 0.140 | 0.279 | |
| AT3G49010.4 | 60S ribosomal protein L13-1 (RPL13A) | 18 | 1.83 | 0.150 | 0.212 | |
| AT3G49010.4 | 60S ribosomal protein L13-1 (RPL13A) | 19 | 2.02 | 0.161 | 0.165 | |
| AT3G49010.4 | 60S ribosomal protein L13-1 (RPL13A) | 20 | 2.22 | 0.173 | 0.133 | |
| AT3G49010.4 | 60S ribosomal protein L13-1 (RPL13A) | 21 | 2.44 | 0.185 | 0.112 | |
| AT3G49010.4 | 60S ribosomal protein L13-1 (RPL13A) | 22 | 2.68 | 0.198 | 0.096 | . |
| AT3G49010.4 | 60S ribosomal protein L13-1 (RPL13A) | 46 | 25.91 | 0.552 | 0.037 | * |
| AT3G49010.4 | 60S ribosomal protein L13-1 (RPL13A) | 47 | 28.48 | 0.567 | 0.038 | * |
| AT3G49080.1 | Ribosomal S5 family protein | 2 | 1.02 | 0.730 | 1.000 | |
| AT3G49080.1 | Ribosomal S5 family protein | 3 | 1.05 | 0.630 | 1.000 | |
| AT3G49080.1 | Ribosomal S5 family protein | 10 | 1.52 | 0.384 | 1.000 | |
| AT3G49080.1 | Ribosomal S5 family protein | 11 | 1.69 | 0.333 | 1.000 | |
| AT3G49080.1 | Ribosomal S5 family protein | 12 | 1.81 | 0.296 | 0.999 | |
| AT3G49080.1 | Ribosomal S5 family protein | 13 | 1.80 | 0.271 | 0.997 | |
| AT3G49080.1 | Ribosomal S5 family protein | 14 | 1.74 | 0.256 | 0.997 | |
| AT3G49080.1 | Ribosomal S5 family protein | 15 | 1.76 | 0.248 | 0.996 | |
| AT3G49080.1 | Ribosomal S5 family protein | 16 | 1.85 | 0.246 | 0.989 | |
| AT3G49080.1 | Ribosomal S5 family protein | 17 | 1.87 | 0.246 | 0.987 | |
| AT3G49080.1 | Ribosomal S5 family protein | 18 | 1.66 | 0.249 | 0.999 | |
| AT3G49080.1 | Ribosomal S5 family protein | 19 | 1.29 | 0.255 | 1.000 | |
| AT3G49080.1 | Ribosomal S5 family protein | 20 | 0.88 | 0.263 | 1.000 | |
| AT3G49080.1 | Ribosomal S5 family protein | 21 | 0.57 | 0.273 | 0.998 | |
| AT3G49080.1 | Ribosomal S5 family protein | 22 | 0.36 | 0.284 | 0.842 | |
| AT3G49080.1 | Ribosomal S5 family protein | 23 | 0.25 | 0.294 | 0.464 | |
| AT3G49080.1 | Ribosomal S5 family protein | 24 | 0.18 | 0.299 | 0.223 | |
| AT3G49080.1 | Ribosomal S5 family protein | 25 | 0.16 | 0.297 | 0.117 | |

| | | | | | | |
|---|---|---|---|---|---|---|
| AT3G49080.1 | Ribosomal S5 family protein | 26 | 0.15 | 0.286 | 0.071 | . |
| AT3G49080.1 | Ribosomal S5 family protein | 27 | 0.16 | 0.277 | 0.072 | . |
| AT3G49080.1 | Ribosomal S5 family protein | 28 | 0.18 | 0.274 | 0.122 | |
| AT3G49080.1 | Ribosomal S5 family protein | 29 | 0.22 | 0.277 | 0.258 | |
| AT3G49080.1 | Ribosomal S5 family protein | 30 | 0.27 | 0.282 | 0.493 | |
| AT3G49080.1 | Ribosomal S5 family protein | 31 | 0.33 | 0.293 | 0.787 | |
| AT3G49080.1 | Ribosomal S5 family protein | 32 | 0.41 | 0.310 | 0.967 | |
| AT3G49080.1 | Ribosomal S5 family protein | 33 | 0.52 | 0.338 | 0.999 | |
| AT3G49080.1 | Ribosomal S5 family protein | 34 | 0.65 | 0.377 | 1.000 | |
| AT3G49080.1 | Ribosomal S5 family protein | 35 | 0.80 | 0.410 | 1.000 | |
| AT3G49080.1 | Ribosomal S5 family protein | 40 | 1.30 | 0.379 | 1.000 | |
| AT3G49080.1 | Ribosomal S5 family protein | 41 | 1.15 | 0.330 | 1.000 | |
| AT3G49080.1 | Ribosomal S5 family protein | 42 | 0.97 | 0.293 | 1.000 | |
| AT3G49080.1 | Ribosomal S5 family protein | 43 | 0.79 | 0.272 | 1.000 | |
| AT3G49080.1 | Ribosomal S5 family protein | 44 | 0.63 | 0.264 | 1.000 | |
| AT3G49080.1 | Ribosomal S5 family protein | 45 | 0.49 | 0.283 | 0.988 | |
| AT3G49080.1 | Ribosomal S5 family protein | 46 | 0.38 | 0.355 | 0.977 | |
| AT3G49080.1 | Ribosomal S5 family protein | 47 | 0.29 | 0.501 | 0.991 | |
| AT3G49240.1 | Pentatricopeptide repeat (PPR) superfamily protein (NUWA) | 2 | 1.01 | 0.454 | 1.000 | |
| AT3G49240.1 | Pentatricopeptide repeat (PPR) superfamily protein (NUWA) | 3 | 1.17 | 0.410 | 1.000 | |
| AT3G49240.1 | Pentatricopeptide repeat (PPR) superfamily protein (NUWA) | 16 | 1.59 | 0.254 | 1.000 | |
| AT3G49240.1 | Pentatricopeptide repeat (PPR) superfamily protein (NUWA) | 17 | 1.32 | 0.227 | 1.000 | |
| AT3G49240.1 | Pentatricopeptide repeat (PPR) superfamily protein (NUWA) | 18 | 1.11 | 0.200 | 1.000 | |
| AT3G49240.1 | Pentatricopeptide repeat (PPR) superfamily protein (NUWA) | 19 | 1.02 | 0.192 | 1.000 | |
| AT3G49240.1 | Pentatricopeptide repeat (PPR) superfamily protein (NUWA) | 20 | 1.09 | 0.191 | 1.000 | |
| AT3G49240.1 | Pentatricopeptide repeat (PPR) superfamily protein (NUWA) | 21 | 1.07 | 0.191 | 1.000 | |
| AT3G49240.1 | Pentatricopeptide repeat (PPR) superfamily protein (NUWA) | 22 | 0.96 | 0.192 | 1.000 | |
| AT3G49240.1 | Pentatricopeptide repeat (PPR) superfamily protein (NUWA) | 23 | 0.93 | 0.189 | 1.000 | |
| AT3G49240.1 | Pentatricopeptide repeat (PPR) superfamily protein (NUWA) | 24 | 1.08 | 0.185 | 1.000 | |
| AT3G49240.1 | Pentatricopeptide repeat (PPR) superfamily protein (NUWA) | 25 | 1.36 | 0.183 | 1.000 | |
| AT3G49240.1 | Pentatricopeptide repeat (PPR) superfamily protein (NUWA) | 26 | 1.60 | 0.182 | 0.997 | |
| AT3G49240.1 | Pentatricopeptide repeat (PPR) superfamily protein (NUWA) | 27 | 1.62 | 0.183 | 0.996 | |
| AT3G49240.1 | Pentatricopeptide repeat (PPR) superfamily protein (NUWA) | 28 | 1.48 | 0.183 | 1.000 | |
| AT3G49240.1 | Pentatricopeptide repeat (PPR) superfamily protein (NUWA) | 29 | 1.30 | 0.183 | 1.000 | |
| AT3G49240.1 | Pentatricopeptide repeat (PPR) superfamily protein (NUWA) | 30 | 1.05 | 0.183 | 1.000 | |
| AT3G49240.1 | Pentatricopeptide repeat (PPR) superfamily protein (NUWA) | 31 | 0.82 | 0.183 | 1.000 | |
| AT3G49240.1 | Pentatricopeptide repeat (PPR) superfamily protein (NUWA) | 32 | 0.64 | 0.184 | 0.999 | |
| AT3G49240.1 | Pentatricopeptide repeat (PPR) superfamily protein (NUWA) | 33 | 0.52 | 0.184 | 0.931 | |
| AT3G49240.1 | Pentatricopeptide repeat (PPR) superfamily protein (NUWA) | 34 | 0.50 | 0.184 | 0.876 | |
| AT3G49240.1 | Pentatricopeptide repeat (PPR) superfamily protein (NUWA) | 35 | 0.59 | 0.183 | 0.990 | |

| | | | | | | |
|---|---|---|---|---|---|---|
| AT3G49240.1 | Pentatricopeptide repeat (PPR) superfamily protein (NUWA) | 36 | 0.82 | 0.183 | 1.000 | |
| AT3G49240.1 | Pentatricopeptide repeat (PPR) superfamily protein (NUWA) | 37 | 1.26 | 0.183 | 1.000 | |
| AT3G49240.1 | Pentatricopeptide repeat (PPR) superfamily protein (NUWA) | 38 | 1.78 | 0.183 | 0.976 | |
| AT3G49240.1 | Pentatricopeptide repeat (PPR) superfamily protein (NUWA) | 39 | 1.93 | 0.183 | 0.922 | |
| AT3G49240.1 | Pentatricopeptide repeat (PPR) superfamily protein (NUWA) | 40 | 1.77 | 0.184 | 0.978 | |
| AT3G49240.1 | Pentatricopeptide repeat (PPR) superfamily protein (NUWA) | 41 | 1.47 | 0.185 | 1.000 | |
| AT3G49240.1 | Pentatricopeptide repeat (PPR) superfamily protein (NUWA) | 42 | 1.17 | 0.188 | 1.000 | |
| AT3G49240.1 | Pentatricopeptide repeat (PPR) superfamily protein (NUWA) | 43 | 0.88 | 0.190 | 1.000 | |
| AT3G49240.1 | Pentatricopeptide repeat (PPR) superfamily protein (NUWA) | 44 | 0.56 | 0.189 | 0.981 | |
| AT3G49240.1 | Pentatricopeptide repeat (PPR) superfamily protein (NUWA) | 45 | 0.31 | 0.190 | 0.171 | |
| AT3G49240.1 | Pentatricopeptide repeat (PPR) superfamily protein (NUWA) | 46 | 0.17 | 0.231 | 0.028 | * |
| AT3G49240.1 | Pentatricopeptide repeat (PPR) superfamily protein (NUWA) | 47 | 0.10 | 0.401 | 0.281 | |
| AT3G49920.1 | Voltage dependent anion channel 5 (VDAC5) | 2 | 1.07 | 0.416 | 1.000 | |
| AT3G49920.1 | Voltage dependent anion channel 5 (VDAC5) | 3 | 0.96 | 0.382 | 1.000 | |
| AT3G49920.1 | Voltage dependent anion channel 5 (VDAC5) | 4 | 0.86 | 0.406 | 1.000 | |
| AT3G49920.1 | Voltage dependent anion channel 5 (VDAC5) | 26 | 0.67 | 0.343 | 1.000 | |
| AT3G49920.1 | Voltage dependent anion channel 5 (VDAC5) | 27 | 0.80 | 0.295 | 1.000 | |
| AT3G49920.1 | Voltage dependent anion channel 5 (VDAC5) | 28 | 0.98 | 0.256 | 1.000 | |
| AT3G49920.1 | Voltage dependent anion channel 5 (VDAC5) | 29 | 1.19 | 0.227 | 1.000 | |
| AT3G49920.1 | Voltage dependent anion channel 5 (VDAC5) | 30 | 1.41 | 0.204 | 0.994 | |
| AT3G49920.1 | Voltage dependent anion channel 5 (VDAC5) | 31 | 1.60 | 0.189 | 0.925 | |
| AT3G49920.1 | Voltage dependent anion channel 5 (VDAC5) | 32 | 1.77 | 0.180 | 0.772 | |
| AT3G49920.1 | Voltage dependent anion channel 5 (VDAC5) | 33 | 1.92 | 0.175 | 0.605 | |
| AT3G49920.1 | Voltage dependent anion channel 5 (VDAC5) | 34 | 1.99 | 0.173 | 0.530 | |
| AT3G49920.1 | Voltage dependent anion channel 5 (VDAC5) | 35 | 1.93 | 0.173 | 0.584 | |
| AT3G49920.1 | Voltage dependent anion channel 5 (VDAC5) | 36 | 1.77 | 0.176 | 0.755 | |
| AT3G49920.1 | Voltage dependent anion channel 5 (VDAC5) | 37 | 1.55 | 0.181 | 0.932 | |
| AT3G49920.1 | Voltage dependent anion channel 5 (VDAC5) | 38 | 1.33 | 0.187 | 0.997 | |
| AT3G49920.1 | Voltage dependent anion channel 5 (VDAC5) | 39 | 1.11 | 0.193 | 1.000 | |
| AT3G49920.1 | Voltage dependent anion channel 5 (VDAC5) | 40 | 0.94 | 0.196 | 1.000 | |
| AT3G49920.1 | Voltage dependent anion channel 5 (VDAC5) | 41 | 0.80 | 0.197 | 1.000 | |

| | | | | | | |
|---|---|---|---|---|---|---|
| AT3G49920.1 | Voltage dependent anion channel 5 (VDAC5) | 42 | 0.71 | 0.198 | 0.992 | |
| AT3G49920.1 | Voltage dependent anion channel 5 (VDAC5) | 43 | 0.64 | 0.204 | 0.966 | |
| AT3G49920.1 | Voltage dependent anion channel 5 (VDAC5) | 44 | 0.60 | 0.221 | 0.950 | |
| AT3G49920.1 | Voltage dependent anion channel 5 (VDAC5) | 45 | 0.57 | 0.256 | 0.960 | |
| AT3G49920.1 | Voltage dependent anion channel 5 (VDAC5) | 46 | 0.54 | 0.311 | 0.979 | |
| AT3G49920.1 | Voltage dependent anion channel 5 (VDAC5) | 47 | 0.51 | 0.387 | 0.992 | |
| AT3G50820.1 | Photosystem II subunit (PSBO2) | 2 | 0.52 | 0.332 | 0.997 | |
| AT3G50820.1 | Photosystem II subunit (PSBO2) | 3 | 0.54 | 0.239 | 0.968 | |
| AT3G50820.1 | Photosystem II subunit (PSBO2) | 4 | 0.58 | 0.203 | 0.960 | |
| AT3G50820.1 | Photosystem II subunit (PSBO2) | 5 | 0.69 | 0.194 | 0.997 | |
| AT3G50820.1 | Photosystem II subunit (PSBO2) | 6 | 0.86 | 0.193 | 1.000 | |
| AT3G50820.1 | Photosystem II subunit (PSBO2) | 7 | 1.05 | 0.192 | 1.000 | |
| AT3G50820.1 | Photosystem II subunit (PSBO2) | 8 | 1.17 | 0.190 | 1.000 | |
| AT3G50820.1 | Photosystem II subunit (PSBO2) | 9 | 1.15 | 0.188 | 1.000 | |
| AT3G50820.1 | Photosystem II subunit (PSBO2) | 10 | 0.95 | 0.188 | 1.000 | |
| AT3G50820.1 | Photosystem II subunit (PSBO2) | 11 | 0.72 | 0.188 | 0.999 | |
| AT3G50820.1 | Photosystem II subunit (PSBO2) | 12 | 0.56 | 0.187 | 0.886 | |
| AT3G50820.1 | Photosystem II subunit (PSBO2) | 13 | 0.46 | 0.185 | 0.559 | |
| AT3G50820.1 | Photosystem II subunit (PSBO2) | 14 | 0.40 | 0.182 | 0.313 | |
| AT3G50820.1 | Photosystem II subunit (PSBO2) | 15 | 0.37 | 0.181 | 0.211 | |
| AT3G50820.1 | Photosystem II subunit (PSBO2) | 16 | 0.35 | 0.181 | 0.171 | |
| AT3G50820.1 | Photosystem II subunit (PSBO2) | 17 | 0.36 | 0.182 | 0.202 | |
| AT3G50820.1 | Photosystem II subunit (PSBO2) | 18 | 0.42 | 0.184 | 0.415 | |
| AT3G50820.1 | Photosystem II subunit (PSBO2) | 19 | 0.52 | 0.188 | 0.810 | |
| AT3G50820.1 | Photosystem II subunit (PSBO2) | 20 | 0.63 | 0.193 | 0.982 | |
| AT3G50820.1 | Photosystem II subunit (PSBO2) | 21 | 0.65 | 0.196 | 0.991 | |
| AT3G50820.1 | Photosystem II subunit (PSBO2) | 22 | 0.53 | 0.201 | 0.873 | |
| AT3G50820.1 | Photosystem II subunit (PSBO2) | 23 | 0.34 | 0.207 | 0.287 | |
| AT3G50820.1 | Photosystem II subunit (PSBO2) | 24 | 0.22 | 0.220 | 0.047 | * |
| AT3G50820.1 | Photosystem II subunit (PSBO2) | 25 | 0.16 | 0.250 | 0.027 | * |
| AT3G50820.1 | Photosystem II subunit (PSBO2) | 26 | 0.14 | 0.313 | 0.091 | . |
| AT3G50820.1 | Photosystem II subunit (PSBO2) | 27 | 0.13 | 0.406 | 0.319 | |
| AT3G50820.1 | Photosystem II subunit (PSBO2) | 46 | 266.52 | 2.805 | 0.996 | |
| AT3G50820.1 | Photosystem II subunit (PSBO2) | 47 | 503.00 | 2.994 | 0.994 | |
| AT3G50930.1 | Outer membrane protein 66 (OM66) | 2 | 0.89 | 0.392 | 1.000 | |
| AT3G50930.1 | Outer membrane protein 66 (OM66) | 3 | 0.99 | 0.365 | 1.000 | |
| AT3G50930.1 | Outer membrane protein 66 (OM66) | 22 | 1.28 | 0.266 | 1.000 | |
| AT3G50930.1 | Outer membrane protein 66 (OM66) | 23 | 1.10 | 0.203 | 1.000 | |
| AT3G50930.1 | Outer membrane protein 66 (OM66) | 24 | 1.00 | 0.171 | 1.000 | |
| AT3G50930.1 | Outer membrane protein 66 (OM66) | 25 | 1.04 | 0.156 | 1.000 | |
| AT3G50930.1 | Outer membrane protein 66 (OM66) | 26 | 1.21 | 0.152 | 1.000 | |
| AT3G50930.1 | Outer membrane protein 66 (OM66) | 27 | 1.30 | 0.151 | 1.000 | |
| AT3G50930.1 | Outer membrane protein 66 (OM66) | 28 | 1.32 | 0.151 | 1.000 | |
| AT3G50930.1 | Outer membrane protein 66 (OM66) | 29 | 1.32 | 0.151 | 1.000 | |
| AT3G50930.1 | Outer membrane protein 66 (OM66) | 30 | 1.16 | 0.152 | 1.000 | |

| | | | | | | |
|---|---|---|---|---|---|---|
| AT3G50930.1 | Outer membrane protein 66 (OM66) | 31 | 0.99 | 0.152 | 1.000 | |
| AT3G50930.1 | Outer membrane protein 66 (OM66) | 32 | 0.91 | 0.152 | 1.000 | |
| AT3G50930.1 | Outer membrane protein 66 (OM66) | 33 | 0.96 | 0.152 | 1.000 | |
| AT3G50930.1 | Outer membrane protein 66 (OM66) | 34 | 1.17 | 0.153 | 1.000 | |
| AT3G50930.1 | Outer membrane protein 66 (OM66) | 35 | 1.47 | 0.153 | 0.993 | |
| AT3G50930.1 | Outer membrane protein 66 (OM66) | 36 | 1.89 | 0.152 | 0.701 | |
| AT3G50930.1 | Outer membrane protein 66 (OM66) | 37 | 2.39 | 0.151 | 0.220 | |
| AT3G50930.1 | Outer membrane protein 66 (OM66) | 38 | 2.73 | 0.151 | 0.083 | . |
| AT3G50930.1 | Outer membrane protein 66 (OM66) | 39 | 2.56 | 0.152 | 0.136 | |
| AT3G50930.1 | Outer membrane protein 66 (OM66) | 40 | 1.94 | 0.152 | 0.640 | |
| AT3G50930.1 | Outer membrane protein 66 (OM66) | 41 | 1.29 | 0.153 | 1.000 | |
| AT3G50930.1 | Outer membrane protein 66 (OM66) | 42 | 0.87 | 0.156 | 1.000 | |
| AT3G50930.1 | Outer membrane protein 66 (OM66) | 43 | 0.66 | 0.158 | 0.989 | |
| AT3G50930.1 | Outer membrane protein 66 (OM66) | 44 | 0.59 | 0.157 | 0.921 | |
| AT3G50930.1 | Outer membrane protein 66 (OM66) | 45 | 0.62 | 0.158 | 0.965 | |
| AT3G50930.1 | Outer membrane protein 66 (OM66) | 46 | 0.62 | 0.198 | 0.996 | |
| AT3G50930.1 | Outer membrane protein 66 (OM66) | 47 | 0.59 | 0.343 | 1.000 | |
| AT3G51090.1 | Coiled-coil 90B-like protein | 2 | 1.16 | 0.506 | 1.000 | |
| AT3G51090.1 | Coiled-coil 90B-like protein | 3 | 1.26 | 0.483 | 1.000 | |
| AT3G51090.1 | Coiled-coil 90B-like protein | 26 | 2.56 | 0.175 | 0.100 | . |
| AT3G51090.1 | Coiled-coil 90B-like protein | 27 | 2.41 | 0.159 | 0.085 | . |
| AT3G51090.1 | Coiled-coil 90B-like protein | 28 | 2.23 | 0.147 | 0.090 | . |
| AT3G51090.1 | Coiled-coil 90B-like protein | 29 | 2.04 | 0.140 | 0.134 | |
| AT3G51090.1 | Coiled-coil 90B-like protein | 30 | 1.82 | 0.138 | 0.258 | |
| AT3G51090.1 | Coiled-coil 90B-like protein | 31 | 1.59 | 0.142 | 0.525 | |
| AT3G51090.1 | Coiled-coil 90B-like protein | 32 | 1.37 | 0.149 | 0.849 | |
| AT3G51090.1 | Coiled-coil 90B-like protein | 33 | 1.17 | 0.160 | 0.992 | |
| AT3G51090.1 | Coiled-coil 90B-like protein | 34 | 1.00 | 0.174 | 1.000 | |
| AT3G51090.1 | Coiled-coil 90B-like protein | 35 | 0.86 | 0.188 | 0.997 | |
| AT3G51090.1 | Coiled-coil 90B-like protein | 46 | 0.15 | 0.325 | 0.064 | . |
| AT3G51090.1 | Coiled-coil 90B-like protein | 47 | 0.13 | 0.347 | 0.059 | . |
| AT3G51790.1 | Cytochrome c maturation E (CCME) | 2 | 1.12 | 0.414 | 1.000 | |
| AT3G51790.1 | Cytochrome c maturation E (CCME) | 3 | 1.08 | 0.364 | 1.000 | |
| AT3G51790.1 | Cytochrome c maturation E (CCME) | 11 | 0.90 | 0.228 | 1.000 | |
| AT3G51790.1 | Cytochrome c maturation E (CCME) | 12 | 0.94 | 0.202 | 1.000 | |
| AT3G51790.1 | Cytochrome c maturation E (CCME) | 13 | 1.04 | 0.183 | 1.000 | |
| AT3G51790.1 | Cytochrome c maturation E (CCME) | 14 | 1.23 | 0.169 | 1.000 | |
| AT3G51790.1 | Cytochrome c maturation E (CCME) | 15 | 1.52 | 0.162 | 0.958 | |

| | | | | | | |
|---|---|---|---|---|---|---|
| AT3G51790.1 | Cytochrome c maturation E (CCME) | 16 | 1.88 | 0.159 | 0.622 | |
| AT3G51790.1 | Cytochrome c maturation E (CCME) | 17 | 2.28 | 0.160 | 0.271 | |
| AT3G51790.1 | Cytochrome c maturation E (CCME) | 18 | 2.63 | 0.163 | 0.131 | |
| AT3G51790.1 | Cytochrome c maturation E (CCME) | 19 | 2.82 | 0.168 | 0.101 | |
| AT3G51790.1 | Cytochrome c maturation E (CCME) | 20 | 2.78 | 0.174 | 0.137 | |
| AT3G51790.1 | Cytochrome c maturation E (CCME) | 21 | 2.48 | 0.180 | 0.298 | |
| AT3G51790.1 | Cytochrome c maturation E (CCME) | 23 | 1.48 | 0.182 | 0.989 | |
| AT3G51790.1 | Cytochrome c maturation E (CCME) | 24 | 1.06 | 0.176 | 1.000 | |
| AT3G51790.1 | Cytochrome c maturation E (CCME) | 25 | 0.81 | 0.167 | 1.000 | |
| AT3G51790.1 | Cytochrome c maturation E (CCME) | 26 | 0.70 | 0.161 | 0.986 | |
| AT3G51790.1 | Cytochrome c maturation E (CCME) | 27 | 0.73 | 0.160 | 0.995 | |
| AT3G51790.1 | Cytochrome c maturation E (CCME) | 28 | 0.88 | 0.162 | 1.000 | |
| AT3G51790.1 | Cytochrome c maturation E (CCME) | 29 | 1.15 | 0.166 | 1.000 | |
| AT3G51790.1 | Cytochrome c maturation E (CCME) | 30 | 1.56 | 0.170 | 0.956 | |
| AT3G51790.1 | Cytochrome c maturation E (CCME) | 31 | 2.01 | 0.175 | 0.619 | |
| AT3G51790.1 | Cytochrome c maturation E (CCME) | 32 | 2.34 | 0.179 | 0.381 | |
| AT3G51790.1 | Cytochrome c maturation E (CCME) | 33 | 2.49 | 0.188 | 0.343 | |
| AT3G51790.1 | Cytochrome c maturation E (CCME) | 34 | 2.55 | 0.207 | 0.447 | |
| AT3G51790.1 | Cytochrome c maturation E (CCME) | 35 | 2.60 | 0.252 | 0.685 | |
| AT3G51790.1 | Cytochrome c maturation E (CCME) | 46 | 3.11 | 2.275 | 1.000 | |
| AT3G51790.1 | Cytochrome c maturation E (CCME) | 47 | 3.16 | 2.543 | 1.000 | |
| AT3G51830.1 | SAC domain-containing protein 8 (SAC8) | 2 | 0.80 | 0.413 | 1.000 | |
| AT3G51830.1 | SAC domain-containing protein 8 (SAC8) | 3 | 0.85 | 0.397 | 1.000 | |
| AT3G51830.1 | SAC domain-containing protein 8 (SAC8) | 32 | 2.91 | 0.254 | 0.305 | |
| AT3G51830.1 | SAC domain-containing protein 8 (SAC8) | 33 | 2.89 | 0.233 | 0.233 | |
| AT3G51830.1 | SAC domain-containing protein 8 (SAC8) | 34 | 2.84 | 0.214 | 0.177 | |
| AT3G51830.1 | SAC domain-containing protein 8 (SAC8) | 38 | 2.42 | 0.151 | 0.067 | . |
| AT3G51830.1 | SAC domain-containing protein 8 (SAC8) | 39 | 2.25 | 0.137 | 0.064 | . |
| AT3G51830.1 | SAC domain-containing protein 8 (SAC8) | 40 | 2.07 | 0.126 | 0.073 | . |
| AT3G51830.1 | SAC domain-containing protein 8 (SAC8) | 41 | 1.87 | 0.117 | 0.115 | |
| AT3G51830.1 | SAC domain-containing protein 8 (SAC8) | 42 | 1.66 | 0.112 | 0.238 | |

| | | | | | | |
|---|---|---|---|---|---|---|
| AT3G51830.1 | SAC domain-containing protein 8 (SAC8) | 43 | 1.47 | 0.114 | 0.521 | |
| AT3G51830.1 | SAC domain-containing protein 8 (SAC8) | 44 | 1.29 | 0.123 | 0.878 | |
| AT3G51830.1 | SAC domain-containing protein 8 (SAC8) | 45 | 1.13 | 0.141 | 0.998 | |
| AT3G51830.1 | SAC domain-containing protein 8 (SAC8) | 46 | 0.99 | 0.168 | 1.000 | |
| AT3G51830.1 | SAC domain-containing protein 8 (SAC8) | 47 | 0.86 | 0.204 | 0.999 | |
| AT3G51840.1 | Acyl-CoA oxidase 4 (ACX4) | 2 | 1.06 | 0.527 | 1.000 | |
| AT3G51840.1 | Acyl-CoA oxidase 4 (ACX4) | 3 | 0.91 | 0.523 | 1.000 | |
| AT3G51840.1 | Acyl-CoA oxidase 4 (ACX4) | 33 | 0.64 | 0.299 | 0.968 | |
| AT3G51840.1 | Acyl-CoA oxidase 4 (ACX4) | 34 | 0.86 | 0.237 | 1.000 | |
| AT3G51840.1 | Acyl-CoA oxidase 4 (ACX4) | 35 | 1.15 | 0.203 | 0.999 | |
| AT3G51840.1 | Acyl-CoA oxidase 4 (ACX4) | 36 | 1.49 | 0.208 | 0.910 | |
| AT3G51840.1 | Acyl-CoA oxidase 4 (ACX4) | 37 | 1.81 | 0.247 | 0.810 | |
| AT3G51840.1 | Acyl-CoA oxidase 4 (ACX4) | 38 | 2.07 | 0.309 | 0.819 | |
| AT3G51840.1 | Acyl-CoA oxidase 4 (ACX4) | 46 | 1.64 | 0.996 | 1.000 | |
| AT3G51840.1 | Acyl-CoA oxidase 4 (ACX4) | 47 | 1.47 | 1.117 | 1.000 | |
| AT3G52200.1 | Dihydrolipoamide acetyltransferase (PDCE2-1) | 3 | 1.11 | 0.199 | 1.000 | |
| AT3G52200.1 | Dihydrolipoamide acetyltransferase (PDCE2-1) | 4 | 0.83 | 0.180 | 1.000 | |
| AT3G52200.1 | Dihydrolipoamide acetyltransferase (PDCE2-1) | 5 | 0.64 | 0.181 | 1.000 | |
| AT3G52200.1 | Dihydrolipoamide acetyltransferase (PDCE2-1) | 6 | 0.48 | 0.181 | 0.899 | |
| AT3G52200.1 | Dihydrolipoamide acetyltransferase (PDCE2-1) | 7 | 0.34 | 0.180 | 0.267 | |
| AT3G52200.1 | Dihydrolipoamide acetyltransferase (PDCE2-1) | 8 | 0.27 | 0.177 | 0.053 | . |
| AT3G52200.1 | Dihydrolipoamide acetyltransferase (PDCE2-1) | 9 | 0.29 | 0.175 | 0.069 | . |
| AT3G52200.1 | Dihydrolipoamide acetyltransferase (PDCE2-1) | 10 | 0.37 | 0.174 | 0.366 | |
| AT3G52200.1 | Dihydrolipoamide acetyltransferase (PDCE2-1) | 11 | 0.59 | 0.174 | 0.997 | |
| AT3G52200.1 | Dihydrolipoamide acetyltransferase (PDCE2-1) | 12 | 1.00 | 0.174 | 1.000 | |
| AT3G52200.1 | Dihydrolipoamide acetyltransferase (PDCE2-1) | 13 | 1.45 | 0.173 | 1.000 | |
| AT3G52200.1 | Dihydrolipoamide acetyltransferase (PDCE2-1) | 14 | 1.72 | 0.171 | 0.993 | |
| AT3G52200.1 | Dihydrolipoamide acetyltransferase (PDCE2-1) | 15 | 1.67 | 0.170 | 0.997 | |
| AT3G52200.1 | Dihydrolipoamide acetyltransferase (PDCE2-1) | 16 | 1.37 | 0.168 | 1.000 | |
| AT3G52200.1 | Dihydrolipoamide acetyltransferase (PDCE2-1) | 17 | 1.17 | 0.168 | 1.000 | |
| AT3G52200.1 | Dihydrolipoamide acetyltransferase (PDCE2-1) | 18 | 1.15 | 0.171 | 1.000 | |
| AT3G52200.1 | Dihydrolipoamide acetyltransferase (PDCE2-1) | 19 | 1.36 | 0.176 | 1.000 | |
| AT3G52200.1 | Dihydrolipoamide acetyltransferase (PDCE2-1) | 20 | 1.78 | 0.180 | 0.993 | |
| AT3G52200.1 | Dihydrolipoamide acetyltransferase (PDCE2-1) | 21 | 2.09 | 0.182 | 0.906 | |
| AT3G52200.1 | Dihydrolipoamide acetyltransferase (PDCE2-1) | 22 | 1.99 | 0.182 | 0.951 | |
| AT3G52200.1 | Dihydrolipoamide acetyltransferase (PDCE2-1) | 23 | 1.64 | 0.181 | 0.999 | |

| | | | | | |
|---|---|---|---|---|---|
| AT3G52200.1 | Dihydrolipoamide acetyltransferase (PDCE2-1) | 24 | 1.37 | 0.177 | 1.000 |
| AT3G52200.1 | Dihydrolipoamide acetyltransferase (PDCE2-1) | 25 | 1.36 | 0.174 | 1.000 |
| AT3G52200.1 | Dihydrolipoamide acetyltransferase (PDCE2-1) | 26 | 1.44 | 0.173 | 1.000 |
| AT3G52200.1 | Dihydrolipoamide acetyltransferase (PDCE2-1) | 27 | 1.51 | 0.173 | 1.000 |
| AT3G52200.1 | Dihydrolipoamide acetyltransferase (PDCE2-1) | 28 | 1.52 | 0.173 | 1.000 |
| AT3G52200.1 | Dihydrolipoamide acetyltransferase (PDCE2-1) | 29 | 1.56 | 0.173 | 1.000 |
| AT3G52200.1 | Dihydrolipoamide acetyltransferase (PDCE2-1) | 30 | 1.58 | 0.173 | 1.000 |
| AT3G52200.1 | Dihydrolipoamide acetyltransferase (PDCE2-1) | 31 | 1.44 | 0.174 | 1.000 |
| AT3G52200.1 | Dihydrolipoamide acetyltransferase (PDCE2-1) | 32 | 1.18 | 0.174 | 1.000 |
| AT3G52200.1 | Dihydrolipoamide acetyltransferase (PDCE2-1) | 33 | 0.97 | 0.174 | 1.000 |
| AT3G52200.1 | Dihydrolipoamide acetyltransferase (PDCE2-1) | 34 | 0.98 | 0.175 | 1.000 |
| AT3G52200.1 | Dihydrolipoamide acetyltransferase (PDCE2-1) | 35 | 1.19 | 0.174 | 1.000 |
| AT3G52200.1 | Dihydrolipoamide acetyltransferase (PDCE2-1) | 36 | 1.61 | 0.174 | 0.999 |
| AT3G52200.1 | Dihydrolipoamide acetyltransferase (PDCE2-1) | 37 | 2.38 | 0.173 | 0.610 |
| AT3G52200.1 | Dihydrolipoamide acetyltransferase (PDCE2-1) | 38 | 3.05 | 0.174 | 0.175 |
| AT3G52200.1 | Dihydrolipoamide acetyltransferase (PDCE2-1) | 39 | 2.99 | 0.174 | 0.200 |
| AT3G52200.1 | Dihydrolipoamide acetyltransferase (PDCE2-1) | 40 | 2.61 | 0.175 | 0.427 |
| AT3G52200.1 | Dihydrolipoamide acetyltransferase (PDCE2-1) | 41 | 2.14 | 0.175 | 0.835 |
| AT3G52200.1 | Dihydrolipoamide acetyltransferase (PDCE2-1) | 42 | 1.71 | 0.178 | 0.997 |
| AT3G52200.1 | Dihydrolipoamide acetyltransferase (PDCE2-1) | 43 | 1.41 | 0.180 | 1.000 |
| AT3G52200.1 | Dihydrolipoamide acetyltransferase (PDCE2-1) | 44 | 1.10 | 0.180 | 1.000 |
| AT3G52200.1 | Dihydrolipoamide acetyltransferase (PDCE2-1) | 45 | 0.76 | 0.179 | 1.000 |
| AT3G52200.1 | Dihydrolipoamide acetyltransferase (PDCE2-1) | 46 | 0.44 | 0.216 | 0.950 |
| AT3G52200.1 | Dihydrolipoamide acetyltransferase (PDCE2-1) | 47 | 0.22 | 0.387 | 0.937 |
| AT3G52200.1 | Dihydrolipoamide acetyltransferase (PDCE2-1) | 48 | 0.11 | 0.641 | 0.983 |
| AT3G52300.1 | ATP synthase subunit (ATPD) | 2 | 0.54 | 0.386 | 1.000 |
| AT3G52300.1 | ATP synthase subunit (ATPD) | 3 | 0.61 | 0.242 | 1.000 |
| AT3G52300.1 | ATP synthase subunit (ATPD) | 4 | 0.80 | 0.226 | 1.000 |
| AT3G52300.1 | ATP synthase subunit (ATPD) | 5 | 1.00 | 0.226 | 1.000 |
| AT3G52300.1 | ATP synthase subunit (ATPD) | 6 | 1.03 | 0.225 | 1.000 |
| AT3G52300.1 | ATP synthase subunit (ATPD) | 7 | 0.97 | 0.226 | 1.000 |
| AT3G52300.1 | ATP synthase subunit (ATPD) | 8 | 1.01 | 0.222 | 1.000 |
| AT3G52300.1 | ATP synthase subunit (ATPD) | 9 | 1.20 | 0.219 | 1.000 |
| AT3G52300.1 | ATP synthase subunit (ATPD) | 10 | 1.48 | 0.220 | 1.000 |
| AT3G52300.1 | ATP synthase subunit (ATPD) | 11 | 1.78 | 0.222 | 0.999 |
| AT3G52300.1 | ATP synthase subunit (ATPD) | 12 | 2.10 | 0.222 | 0.975 |
| AT3G52300.1 | ATP synthase subunit (ATPD) | 13 | 2.42 | 0.219 | 0.865 |

| | | | | | |
|---|---|---|---|---|---|
| AT3G52300.1 | ATP synthase subunit (ATPD) | 14 | 2.59 | 0.215 | 0.755 |
| AT3G52300.1 | ATP synthase subunit (ATPD) | 15 | 2.38 | 0.214 | 0.863 |
| AT3G52300.1 | ATP synthase subunit (ATPD) | 16 | 1.98 | 0.213 | 0.985 |
| AT3G52300.1 | ATP synthase subunit (ATPD) | 17 | 1.63 | 0.213 | 1.000 |
| AT3G52300.1 | ATP synthase subunit (ATPD) | 18 | 1.42 | 0.215 | 1.000 |
| AT3G52300.1 | ATP synthase subunit (ATPD) | 19 | 1.45 | 0.220 | 1.000 |
| AT3G52300.1 | ATP synthase subunit (ATPD) | 20 | 1.43 | 0.226 | 1.000 |
| AT3G52300.1 | ATP synthase subunit (ATPD) | 21 | 1.12 | 0.228 | 1.000 |
| AT3G52300.1 | ATP synthase subunit (ATPD) | 22 | 0.83 | 0.228 | 1.000 |
| AT3G52300.1 | ATP synthase subunit (ATPD) | 23 | 0.78 | 0.227 | 1.000 |
| AT3G52300.1 | ATP synthase subunit (ATPD) | 24 | 1.07 | 0.222 | 1.000 |
| AT3G52300.1 | ATP synthase subunit (ATPD) | 25 | 1.67 | 0.218 | 1.000 |
| AT3G52300.1 | ATP synthase subunit (ATPD) | 26 | 1.62 | 0.218 | 1.000 |
| AT3G52300.1 | ATP synthase subunit (ATPD) | 27 | 0.88 | 0.219 | 1.000 |
| AT3G52300.1 | ATP synthase subunit (ATPD) | 28 | 0.63 | 0.219 | 1.000 |
| AT3G52300.1 | ATP synthase subunit (ATPD) | 29 | 0.86 | 0.219 | 1.000 |
| AT3G52300.1 | ATP synthase subunit (ATPD) | 30 | 1.38 | 0.219 | 1.000 |
| AT3G52300.1 | ATP synthase subunit (ATPD) | 31 | 1.58 | 0.219 | 1.000 |
| AT3G52300.1 | ATP synthase subunit (ATPD) | 32 | 1.14 | 0.220 | 1.000 |
| AT3G52300.1 | ATP synthase subunit (ATPD) | 33 | 0.67 | 0.220 | 1.000 |
| AT3G52300.1 | ATP synthase subunit (ATPD) | 34 | 0.46 | 0.221 | 0.958 |
| AT3G52300.1 | ATP synthase subunit (ATPD) | 35 | 0.41 | 0.224 | 0.888 |
| AT3G52300.1 | ATP synthase subunit (ATPD) | 36 | 0.58 | 0.236 | 1.000 |
| AT3G52300.1 | ATP synthase subunit (ATPD) | 37 | 1.23 | 0.275 | 1.000 |
| AT3G52300.1 | ATP synthase subunit (ATPD) | 38 | 2.73 | 0.347 | 0.996 |
| AT3G52300.1 | ATP synthase subunit (ATPD) | 41 | 8.50 | 0.339 | 0.172 |
| AT3G52300.1 | ATP synthase subunit (ATPD) | 42 | 8.11 | 0.372 | 0.336 |
| AT3G52300.1 | ATP synthase subunit (ATPD) | 43 | 6.32 | 0.452 | 0.853 |
| AT3G52300.1 | ATP synthase subunit (ATPD) | 46 | 1.35 | 1.200 | 1.000 |
| AT3G52300.1 | ATP synthase subunit (ATPD) | 47 | 0.71 | 1.554 | 1.000 |
| AT3G52730.1 | Ubiquinol:cytochrome c oxidoreductase subunit (QCR9) | 2 | 1.15 | 0.470 | 1.000 |
| AT3G52730.1 | Ubiquinol:cytochrome c oxidoreductase subunit (QCR9) | 3 | 0.88 | 0.277 | 1.000 |
| AT3G52730.1 | Ubiquinol:cytochrome c oxidoreductase subunit (QCR9) | 4 | 0.95 | 0.265 | 1.000 |
| AT3G52730.1 | Ubiquinol:cytochrome c oxidoreductase subunit (QCR9) | 5 | 1.35 | 0.264 | 1.000 |
| AT3G52730.1 | Ubiquinol:cytochrome c oxidoreductase subunit (QCR9) | 6 | 1.37 | 0.263 | 1.000 |
| AT3G52730.1 | Ubiquinol:cytochrome c oxidoreductase subunit (QCR9) | 7 | 1.15 | 0.267 | 1.000 |
| AT3G52730.1 | Ubiquinol:cytochrome c oxidoreductase subunit (QCR9) | 8 | 1.16 | 0.258 | 1.000 |
| AT3G52730.1 | Ubiquinol:cytochrome c oxidoreductase subunit (QCR9) | 9 | 1.11 | 0.254 | 1.000 |
| AT3G52730.1 | Ubiquinol:cytochrome c oxidoreductase subunit (QCR9) | 10 | 1.06 | 0.256 | 1.000 |
| AT3G52730.1 | Ubiquinol:cytochrome c oxidoreductase subunit (QCR9) | 11 | 1.12 | 0.262 | 1.000 |
| AT3G52730.1 | Ubiquinol:cytochrome c oxidoreductase subunit (QCR9) | 12 | 1.25 | 0.261 | 1.000 |
| AT3G52730.1 | Ubiquinol:cytochrome c oxidoreductase subunit (QCR9) | 13 | 1.35 | 0.256 | 1.000 |
| AT3G52730.1 | Ubiquinol:cytochrome c oxidoreductase subunit (QCR9) | 14 | 1.67 | 0.251 | 1.000 |
| AT3G52730.1 | Ubiquinol:cytochrome c oxidoreductase subunit (QCR9) | 15 | 1.68 | 0.249 | 1.000 |
| AT3G52730.1 | Ubiquinol:cytochrome c oxidoreductase subunit (QCR9) | 16 | 1.08 | 0.247 | 1.000 |
| AT3G52730.1 | Ubiquinol:cytochrome c oxidoreductase subunit (QCR9) | 17 | 0.61 | 0.247 | 1.000 |

| | | | | | | |
|---|---|---|---|---|---|---|
| AT3G52730.1 | Ubiquinol:cytochrome c oxidoreductase subunit (QCR9) | 18 | 0.49 | 0.251 | 0.999 | |
| AT3G52730.1 | Ubiquinol:cytochrome c oxidoreductase subunit (QCR9) | 19 | 0.61 | 0.259 | 1.000 | |
| AT3G52730.1 | Ubiquinol:cytochrome c oxidoreductase subunit (QCR9) | 20 | 0.90 | 0.266 | 1.000 | |
| AT3G52730.1 | Ubiquinol:cytochrome c oxidoreductase subunit (QCR9) | 21 | 1.23 | 0.268 | 1.000 | |
| AT3G52730.1 | Ubiquinol:cytochrome c oxidoreductase subunit (QCR9) | 22 | 0.87 | 0.268 | 1.000 | |
| AT3G52730.1 | Ubiquinol:cytochrome c oxidoreductase subunit (QCR9) | 23 | 0.28 | 0.270 | 0.733 | |
| AT3G52730.1 | Ubiquinol:cytochrome c oxidoreductase subunit (QCR9) | 24 | 0.16 | 0.259 | 0.074 | . |
| AT3G52730.1 | Ubiquinol:cytochrome c oxidoreductase subunit (QCR9) | 25 | 0.41 | 0.252 | 0.983 | |
| AT3G52730.1 | Ubiquinol:cytochrome c oxidoreductase subunit (QCR9) | 26 | 1.46 | 0.253 | 1.000 | |
| AT3G52730.1 | Ubiquinol:cytochrome c oxidoreductase subunit (QCR9) | 27 | 3.61 | 0.255 | 0.616 | |
| AT3G52730.1 | Ubiquinol:cytochrome c oxidoreductase subunit (QCR9) | 28 | 2.13 | 0.258 | 0.999 | |
| AT3G52730.1 | Ubiquinol:cytochrome c oxidoreductase subunit (QCR9) | 29 | 0.40 | 0.257 | 0.976 | |
| AT3G52730.1 | Ubiquinol:cytochrome c oxidoreductase subunit (QCR9) | 30 | 0.18 | 0.256 | 0.135 | |
| AT3G52730.1 | Ubiquinol:cytochrome c oxidoreductase subunit (QCR9) | 31 | 0.39 | 0.256 | 0.967 | |
| AT3G52730.1 | Ubiquinol:cytochrome c oxidoreductase subunit (QCR9) | 32 | 1.12 | 0.256 | 1.000 | |
| AT3G52730.1 | Ubiquinol:cytochrome c oxidoreductase subunit (QCR9) | 33 | 1.37 | 0.256 | 1.000 | |
| AT3G52730.1 | Ubiquinol:cytochrome c oxidoreductase subunit (QCR9) | 34 | 0.71 | 0.260 | 1.000 | |
| AT3G52730.1 | Ubiquinol:cytochrome c oxidoreductase subunit (QCR9) | 35 | 0.31 | 0.268 | 0.828 | |
| AT3G52730.1 | Ubiquinol:cytochrome c oxidoreductase subunit (QCR9) | 36 | 0.36 | 0.284 | 0.974 | |
| AT3G52730.1 | Ubiquinol:cytochrome c oxidoreductase subunit (QCR9) | 37 | 0.98 | 0.315 | 1.000 | |
| AT3G52730.1 | Ubiquinol:cytochrome c oxidoreductase subunit (QCR9) | 38 | 2.24 | 0.324 | 1.000 | |
| AT3G52730.1 | Ubiquinol:cytochrome c oxidoreductase subunit (QCR9) | 39 | 2.20 | 0.315 | 1.000 | |
| AT3G52730.1 | Ubiquinol:cytochrome c oxidoreductase subunit (QCR9) | 40 | 1.17 | 0.393 | 1.000 | |
| AT3G52730.1 | Ubiquinol:cytochrome c oxidoreductase subunit (QCR9) | 41 | 0.60 | 0.557 | 1.000 | |
| AT3G52730.1 | Ubiquinol:cytochrome c oxidoreductase subunit (QCR9) | 46 | 0.03 | 2.483 | 1.000 | |
| AT3G52730.1 | Ubiquinol:cytochrome c oxidoreductase subunit (QCR9) | 47 | 0.02 | 3.067 | 1.000 | |
| AT3G52930.1 | Fructose-bisphosphate aldolase 8 (FBA8) | 2 | 1.09 | 0.422 | 1.000 | |
| AT3G52930.1 | Fructose-bisphosphate aldolase 8 (FBA8) | 3 | 1.01 | 0.345 | 1.000 | |
| AT3G52930.1 | Fructose-bisphosphate aldolase 8 (FBA8) | 4 | 0.93 | 0.403 | 1.000 | |
| AT3G52930.1 | Fructose-bisphosphate aldolase 8 (FBA8) | 15 | 0.42 | 0.394 | 0.988 | |
| AT3G52930.1 | Fructose-bisphosphate aldolase 8 (FBA8) | 16 | 0.46 | 0.362 | 0.991 | |

| | | | | | |
|---|---|---|---|---|---|
| AT3G52930.1 | Fructose-bisphosphate aldolase 8 (FBA8) | 17 | 0.52 | 0.360 | 0.998 |
| AT3G52930.1 | Fructose-bisphosphate aldolase 8 (FBA8) | 22 | 0.90 | 0.388 | 1.000 |
| AT3G52930.1 | Fructose-bisphosphate aldolase 8 (FBA8) | 23 | 0.93 | 0.393 | 1.000 |
| AT3G52930.1 | Fructose-bisphosphate aldolase 8 (FBA8) | 24 | 0.93 | 0.417 | 1.000 |
| AT3G52930.1 | Fructose-bisphosphate aldolase 8 (FBA8) | 32 | 0.65 | 0.272 | 0.999 |
| AT3G52930.1 | Fructose-bisphosphate aldolase 8 (FBA8) | 33 | 0.65 | 0.223 | 0.996 |
| AT3G52930.1 | Fructose-bisphosphate aldolase 8 (FBA8) | 34 | 0.70 | 0.196 | 0.998 |
| AT3G52930.1 | Fructose-bisphosphate aldolase 8 (FBA8) | 35 | 0.84 | 0.186 | 1.000 |
| AT3G52930.1 | Fructose-bisphosphate aldolase 8 (FBA8) | 36 | 0.99 | 0.189 | 1.000 |
| AT3G52930.1 | Fructose-bisphosphate aldolase 8 (FBA8) | 37 | 1.04 | 0.203 | 1.000 |
| AT3G52930.1 | Fructose-bisphosphate aldolase 8 (FBA8) | 38 | 0.97 | 0.228 | 1.000 |
| AT3G52930.1 | Fructose-bisphosphate aldolase 8 (FBA8) | 39 | 0.90 | 0.257 | 1.000 |
| AT3G52930.1 | Fructose-bisphosphate aldolase 8 (FBA8) | 41 | 0.92 | 0.311 | 1.000 |
| AT3G52930.1 | Fructose-bisphosphate aldolase 8 (FBA8) | 42 | 1.05 | 0.360 | 1.000 |
| AT3G52930.1 | Fructose-bisphosphate aldolase 8 (FBA8) | 43 | 1.33 | 0.441 | 1.000 |
| AT3G52930.1 | Fructose-bisphosphate aldolase 8 (FBA8) | 46 | 3.97 | 0.830 | 0.999 |
| AT3G52930.1 | Fructose-bisphosphate aldolase 8 (FBA8) | 47 | 6.08 | 1.027 | 0.998 |
| AT3G52950.2 | CBS / octicosapeptide/Phox/Bemp1 (PB1) domains-containing protein | 2 | 0.83 | 0.425 | 1.000 |
| AT3G52950.2 | CBS / octicosapeptide/Phox/Bemp1 (PB1) domains-containing protein | 3 | 0.83 | 0.336 | 1.000 |
| AT3G52950.2 | CBS / octicosapeptide/Phox/Bemp1 (PB1) domains-containing protein | 6 | 0.76 | 0.293 | 1.000 |
| AT3G52950.2 | CBS / octicosapeptide/Phox/Bemp1 (PB1) domains-containing protein | 7 | 0.70 | 0.327 | 1.000 |
| AT3G52950.2 | CBS / octicosapeptide/Phox/Bemp1 (PB1) domains-containing protein | 8 | 0.63 | 0.363 | 1.000 |
| AT3G52950.2 | CBS / octicosapeptide/Phox/Bemp1 (PB1) domains-containing protein | 25 | 0.48 | 0.300 | 0.994 |
| AT3G52950.2 | CBS / octicosapeptide/Phox/Bemp1 (PB1) domains-containing protein | 26 | 0.67 | 0.220 | 1.000 |

| | | | | | |
|---|---|---|---|---|---|
| AT3G52950.2 | CBS / octicosapeptide/Phox/Bemp1 (PB1) domains-containing protein | 27 | 0.89 | 0.186 | 1.000 |
| AT3G52950.2 | CBS / octicosapeptide/Phox/Bemp1 (PB1) domains-containing protein | 28 | 0.97 | 0.175 | 1.000 |
| AT3G52950.2 | CBS / octicosapeptide/Phox/Bemp1 (PB1) domains-containing protein | 29 | 0.87 | 0.171 | 1.000 |
| AT3G52950.2 | CBS / octicosapeptide/Phox/Bemp1 (PB1) domains-containing protein | 30 | 0.75 | 0.171 | 1.000 |
| AT3G52950.2 | CBS / octicosapeptide/Phox/Bemp1 (PB1) domains-containing protein | 31 | 0.82 | 0.172 | 1.000 |
| AT3G52950.2 | CBS / octicosapeptide/Phox/Bemp1 (PB1) domains-containing protein | 32 | 0.91 | 0.172 | 1.000 |
| AT3G52950.2 | CBS / octicosapeptide/Phox/Bemp1 (PB1) domains-containing protein | 33 | 0.80 | 0.172 | 1.000 |
| AT3G52950.2 | CBS / octicosapeptide/Phox/Bemp1 (PB1) domains-containing protein | 34 | 0.55 | 0.172 | 0.887 |
| AT3G52950.2 | CBS / octicosapeptide/Phox/Bemp1 (PB1) domains-containing protein | 35 | 0.41 | 0.175 | 0.359 |
| AT3G52950.2 | CBS / octicosapeptide/Phox/Bemp1 (PB1) domains-containing protein | 36 | 0.40 | 0.180 | 0.369 |
| AT3G52950.2 | CBS / octicosapeptide/Phox/Bemp1 (PB1) domains-containing protein | 37 | 0.46 | 0.187 | 0.695 |
| AT3G52950.2 | CBS / octicosapeptide/Phox/Bemp1 (PB1) domains-containing protein | 38 | 0.59 | 0.190 | 0.979 |
| AT3G52950.2 | CBS / octicosapeptide/Phox/Bemp1 (PB1) domains-containing protein | 39 | 0.85 | 0.184 | 1.000 |
| AT3G52950.2 | CBS / octicosapeptide/Phox/Bemp1 (PB1) domains-containing protein | 40 | 1.26 | 0.177 | 1.000 |
| AT3G52950.2 | CBS / octicosapeptide/Phox/Bemp1 (PB1) domains-containing protein | 41 | 1.62 | 0.174 | 0.979 |

| | | | | | |
|---|---|---|---|---|---|
| AT3G52950.2 | CBS / octicosapeptide/Phox/Bemp1 (PB1) domains-containing protein | 42 | 1.67 | 0.175 | 0.967 |
| AT3G52950.2 | CBS / octicosapeptide/Phox/Bemp1 (PB1) domains-containing protein | 43 | 1.48 | 0.175 | 0.997 |
| AT3G52950.2 | CBS / octicosapeptide/Phox/Bemp1 (PB1) domains-containing protein | 44 | 1.13 | 0.176 | 1.000 |
| AT3G52950.2 | CBS / octicosapeptide/Phox/Bemp1 (PB1) domains-containing protein | 45 | 0.71 | 0.179 | 1.000 |
| AT3G52950.2 | CBS / octicosapeptide/Phox/Bemp1 (PB1) domains-containing protein | 46 | 0.37 | 0.215 | 0.553 |
| AT3G52950.2 | CBS / octicosapeptide/Phox/Bemp1 (PB1) domains-containing protein | 47 | 0.17 | 0.348 | 0.402 |
| AT3G52960.1 | Peroxiredoxin 2E (PRX2E) | 2 | 0.99 | 0.509 | 1.000 |
| AT3G52960.1 | Peroxiredoxin 2E (PRX2E) | 3 | 0.98 | 0.473 | 1.000 |
| AT3G52960.1 | Peroxiredoxin 2E (PRX2E) | 19 | 1.54 | 0.210 | 0.987 |
| AT3G52960.1 | Peroxiredoxin 2E (PRX2E) | 20 | 1.63 | 0.188 | 0.938 |
| AT3G52960.1 | Peroxiredoxin 2E (PRX2E) | 21 | 1.68 | 0.173 | 0.863 |
| AT3G52960.1 | Peroxiredoxin 2E (PRX2E) | 22 | 1.66 | 0.164 | 0.842 |
| AT3G52960.1 | Peroxiredoxin 2E (PRX2E) | 23 | 1.57 | 0.160 | 0.904 |
| AT3G52960.1 | Peroxiredoxin 2E (PRX2E) | 24 | 1.41 | 0.158 | 0.979 |
| AT3G52960.1 | Peroxiredoxin 2E (PRX2E) | 25 | 1.25 | 0.155 | 0.999 |
| AT3G52960.1 | Peroxiredoxin 2E (PRX2E) | 26 | 1.11 | 0.151 | 1.000 |
| AT3G52960.1 | Peroxiredoxin 2E (PRX2E) | 27 | 1.02 | 0.146 | 1.000 |
| AT3G52960.1 | Peroxiredoxin 2E (PRX2E) | 28 | 0.97 | 0.142 | 1.000 |
| AT3G52960.1 | Peroxiredoxin 2E (PRX2E) | 29 | 0.94 | 0.138 | 1.000 |
| AT3G52960.1 | Peroxiredoxin 2E (PRX2E) | 30 | 0.93 | 0.136 | 1.000 |
| AT3G52960.1 | Peroxiredoxin 2E (PRX2E) | 31 | 0.97 | 0.135 | 1.000 |
| AT3G52960.1 | Peroxiredoxin 2E (PRX2E) | 32 | 1.08 | 0.135 | 1.000 |
| AT3G52960.1 | Peroxiredoxin 2E (PRX2E) | 33 | 1.26 | 0.136 | 0.997 |
| AT3G52960.1 | Peroxiredoxin 2E (PRX2E) | 34 | 1.51 | 0.139 | 0.872 |
| AT3G52960.1 | Peroxiredoxin 2E (PRX2E) | 35 | 1.82 | 0.144 | 0.524 |
| AT3G52960.1 | Peroxiredoxin 2E (PRX2E) | 36 | 2.11 | 0.150 | 0.290 |
| AT3G52960.1 | Peroxiredoxin 2E (PRX2E) | 37 | 2.29 | 0.157 | 0.218 |
| AT3G52960.1 | Peroxiredoxin 2E (PRX2E) | 38 | 2.32 | 0.161 | 0.231 |
| AT3G52960.1 | Peroxiredoxin 2E (PRX2E) | 40 | 2.02 | 0.161 | 0.454 |
| AT3G52960.1 | Peroxiredoxin 2E (PRX2E) | 41 | 1.78 | 0.157 | 0.677 |
| AT3G52960.1 | Peroxiredoxin 2E (PRX2E) | 42 | 1.52 | 0.153 | 0.916 |
| AT3G52960.1 | Peroxiredoxin 2E (PRX2E) | 43 | 1.26 | 0.153 | 0.999 |
| AT3G52960.1 | Peroxiredoxin 2E (PRX2E) | 44 | 1.01 | 0.159 | 1.000 |
| AT3G52960.1 | Peroxiredoxin 2E (PRX2E) | 45 | 0.79 | 0.176 | 1.000 |
| AT3G52960.1 | Peroxiredoxin 2E (PRX2E) | 46 | 0.62 | 0.210 | 0.969 |
| AT3G52960.1 | Peroxiredoxin 2E (PRX2E) | 47 | 0.48 | 0.264 | 0.903 |
| AT3G54110.1 | Plant uncoupling mitochondrial protein 1 (PUMP1) | 3 | 0.75 | 0.164 | 1.000 |
| AT3G54110.1 | Plant uncoupling mitochondrial protein 1 (PUMP1) | 4 | 0.70 | 0.160 | 1.000 |

| | | | | | |
|---|---|---|---|---|---|
| AT3G54110.1 | Plant uncoupling mitochondrial protein 1 (PUMP1) | 5 | 0.90 | 0.159 | 1.000 |
| AT3G54110.1 | Plant uncoupling mitochondrial protein 1 (PUMP1) | 6 | 1.10 | 0.159 | 1.000 |
| AT3G54110.1 | Plant uncoupling mitochondrial protein 1 (PUMP1) | 7 | 1.06 | 0.160 | 1.000 |
| AT3G54110.1 | Plant uncoupling mitochondrial protein 1 (PUMP1) | 8 | 1.02 | 0.158 | 1.000 |
| AT3G54110.1 | Plant uncoupling mitochondrial protein 1 (PUMP1) | 9 | 1.02 | 0.156 | 1.000 |
| AT3G54110.1 | Plant uncoupling mitochondrial protein 1 (PUMP1) | 10 | 1.00 | 0.155 | 1.000 |
| AT3G54110.1 | Plant uncoupling mitochondrial protein 1 (PUMP1) | 11 | 1.03 | 0.156 | 1.000 |
| AT3G54110.1 | Plant uncoupling mitochondrial protein 1 (PUMP1) | 12 | 1.08 | 0.156 | 1.000 |
| AT3G54110.1 | Plant uncoupling mitochondrial protein 1 (PUMP1) | 13 | 1.11 | 0.155 | 1.000 |
| AT3G54110.1 | Plant uncoupling mitochondrial protein 1 (PUMP1) | 14 | 1.17 | 0.153 | 1.000 |
| AT3G54110.1 | Plant uncoupling mitochondrial protein 1 (PUMP1) | 15 | 1.14 | 0.153 | 1.000 |
| AT3G54110.1 | Plant uncoupling mitochondrial protein 1 (PUMP1) | 16 | 1.00 | 0.152 | 1.000 |
| AT3G54110.1 | Plant uncoupling mitochondrial protein 1 (PUMP1) | 17 | 1.03 | 0.152 | 1.000 |
| AT3G54110.1 | Plant uncoupling mitochondrial protein 1 (PUMP1) | 18 | 1.25 | 0.154 | 1.000 |
| AT3G54110.1 | Plant uncoupling mitochondrial protein 1 (PUMP1) | 19 | 1.48 | 0.157 | 0.999 |
| AT3G54110.1 | Plant uncoupling mitochondrial protein 1 (PUMP1) | 20 | 1.63 | 0.159 | 0.986 |
| AT3G54110.1 | Plant uncoupling mitochondrial protein 1 (PUMP1) | 21 | 1.62 | 0.161 | 0.990 |
| AT3G54110.1 | Plant uncoupling mitochondrial protein 1 (PUMP1) | 22 | 1.43 | 0.161 | 1.000 |
| AT3G54110.1 | Plant uncoupling mitochondrial protein 1 (PUMP1) | 23 | 1.22 | 0.160 | 1.000 |
| AT3G54110.1 | Plant uncoupling mitochondrial protein 1 (PUMP1) | 24 | 1.21 | 0.158 | 1.000 |

| | | | | | | |
|---|---|---|---|---|---|---|
| AT3G54110.1 | Plant uncoupling mitochondrial protein 1 (PUMP1) | 25 | 1.43 | 0.156 | 1.000 | |
| AT3G54110.1 | Plant uncoupling mitochondrial protein 1 (PUMP1) | 26 | 1.72 | 0.155 | 0.946 | |
| AT3G54110.1 | Plant uncoupling mitochondrial protein 1 (PUMP1) | 27 | 1.67 | 0.156 | 0.969 | |
| AT3G54110.1 | Plant uncoupling mitochondrial protein 1 (PUMP1) | 28 | 1.51 | 0.155 | 0.998 | |
| AT3G54110.1 | Plant uncoupling mitochondrial protein 1 (PUMP1) | 29 | 1.50 | 0.155 | 0.999 | |
| AT3G54110.1 | Plant uncoupling mitochondrial protein 1 (PUMP1) | 30 | 1.46 | 0.155 | 0.999 | |
| AT3G54110.1 | Plant uncoupling mitochondrial protein 1 (PUMP1) | 31 | 1.57 | 0.155 | 0.993 | |
| AT3G54110.1 | Plant uncoupling mitochondrial protein 1 (PUMP1) | 32 | 1.79 | 0.156 | 0.897 | |
| AT3G54110.1 | Plant uncoupling mitochondrial protein 1 (PUMP1) | 33 | 1.90 | 0.156 | 0.793 | |
| AT3G54110.1 | Plant uncoupling mitochondrial protein 1 (PUMP1) | 34 | 2.07 | 0.156 | 0.614 | |
| AT3G54110.1 | Plant uncoupling mitochondrial protein 1 (PUMP1) | 35 | 2.17 | 0.156 | 0.504 | |
| AT3G54110.1 | Plant uncoupling mitochondrial protein 1 (PUMP1) | 36 | 2.01 | 0.155 | 0.671 | |
| AT3G54110.1 | Plant uncoupling mitochondrial protein 1 (PUMP1) | 37 | 1.97 | 0.155 | 0.711 | |
| AT3G54110.1 | Plant uncoupling mitochondrial protein 1 (PUMP1) | 38 | 2.49 | 0.156 | 0.241 | |
| AT3G54110.1 | Plant uncoupling mitochondrial protein 1 (PUMP1) | 39 | 3.82 | 0.156 | 0.007 | ** |
| AT3G54110.1 | Plant uncoupling mitochondrial protein 1 (PUMP1) | 40 | 5.06 | 0.157 | 0.000 | *** |
| AT3G54110.1 | Plant uncoupling mitochondrial protein 1 (PUMP1) | 41 | 3.99 | 0.157 | 0.005 | ** |
| AT3G54110.1 | Plant uncoupling mitochondrial protein 1 (PUMP1) | 42 | 2.04 | 0.158 | 0.667 | |
| AT3G54110.1 | Plant uncoupling mitochondrial protein 1 (PUMP1) | 43 | 0.95 | 0.159 | 1.000 | |
| AT3G54110.1 | Plant uncoupling mitochondrial protein 1 (PUMP1) | 44 | 0.46 | 0.159 | 0.544 | |

| | | | | | | |
|---|---|---|---|---|---|---|
| AT3G54110.1 | Plant uncoupling mitochondrial protein 1 (PUMP1) | 45 | 0.28 | 0.159 | 0.021 | * |
| AT3G54110.1 | Plant uncoupling mitochondrial protein 1 (PUMP1) | 46 | 0.21 | 0.171 | 0.003 | ** |
| AT3G54110.1 | Plant uncoupling mitochondrial protein 1 (PUMP1) | 47 | 0.16 | 0.308 | 0.243 | |
| AT3G54110.1 | Plant uncoupling mitochondrial protein 1 (PUMP1) | 48 | 0.13 | 0.541 | 0.883 | |
| AT3G54660.1 | Glutathione reductase 2 (GR2) | 2 | 0.96 | 0.519 | 1.000 | |
| AT3G54660.1 | Glutathione reductase 2 (GR2) | 3 | 0.87 | 0.484 | 1.000 | |
| AT3G54660.1 | Glutathione reductase 2 (GR2) | 22 | 1.26 | 0.399 | 1.000 | |
| AT3G54660.1 | Glutathione reductase 2 (GR2) | 23 | 1.79 | 0.344 | 1.000 | |
| AT3G54660.1 | Glutathione reductase 2 (GR2) | 24 | 2.74 | 0.289 | 0.940 | |
| AT3G54660.1 | Glutathione reductase 2 (GR2) | 25 | 1.93 | 0.298 | 1.000 | |
| AT3G54660.1 | Glutathione reductase 2 (GR2) | 26 | 1.17 | 0.292 | 1.000 | |
| AT3G54660.1 | Glutathione reductase 2 (GR2) | 27 | 0.96 | 0.290 | 1.000 | |
| AT3G54660.1 | Glutathione reductase 2 (GR2) | 28 | 0.80 | 0.282 | 1.000 | |
| AT3G54660.1 | Glutathione reductase 2 (GR2) | 29 | 0.71 | 0.271 | 1.000 | |
| AT3G54660.1 | Glutathione reductase 2 (GR2) | 30 | 0.68 | 0.266 | 1.000 | |
| AT3G54660.1 | Glutathione reductase 2 (GR2) | 31 | 1.20 | 0.268 | 1.000 | |
| AT3G54660.1 | Glutathione reductase 2 (GR2) | 32 | 1.87 | 0.273 | 1.000 | |
| AT3G54660.1 | Glutathione reductase 2 (GR2) | 33 | 1.27 | 0.270 | 1.000 | |
| AT3G54660.1 | Glutathione reductase 2 (GR2) | 34 | 1.12 | 0.270 | 1.000 | |
| AT3G54660.1 | Glutathione reductase 2 (GR2) | 35 | 1.99 | 0.273 | 0.998 | |
| AT3G54660.1 | Glutathione reductase 2 (GR2) | 36 | 2.04 | 0.286 | 0.998 | |
| AT3G54660.1 | Glutathione reductase 2 (GR2) | 37 | 1.29 | 0.296 | 1.000 | |
| AT3G54660.1 | Glutathione reductase 2 (GR2) | 38 | 0.98 | 0.312 | 1.000 | |
| AT3G54660.1 | Glutathione reductase 2 (GR2) | 39 | 1.48 | 0.295 | 1.000 | |
| AT3G54660.1 | Glutathione reductase 2 (GR2) | 40 | 3.87 | 0.278 | 0.540 | |
| AT3G54660.1 | Glutathione reductase 2 (GR2) | 41 | 2.73 | 0.269 | 0.895 | |
| AT3G54660.1 | Glutathione reductase 2 (GR2) | 42 | 0.52 | 0.275 | 0.999 | |
| AT3G54660.1 | Glutathione reductase 2 (GR2) | 43 | 0.23 | 0.279 | 0.410 | |
| AT3G54660.1 | Glutathione reductase 2 (GR2) | 44 | 0.56 | 0.363 | 1.000 | |
| AT3G54660.1 | Glutathione reductase 2 (GR2) | 45 | 4.61 | 0.602 | 0.998 | |
| AT3G54660.1 | Glutathione reductase 2 (GR2) | 46 | 77.64 | 1.046 | 0.787 | |
| AT3G54660.1 | Glutathione reductase 2 (GR2) | 47 | 1798.17 | 1.734 | 0.735 | |
| AT3G54890.4 | Photosystem I light harvesting complex subunit (LHCA1) | 2 | 0.84 | 0.256 | 1.000 | |
| AT3G54890.4 | Photosystem I light harvesting complex subunit (LHCA1) | 3 | 0.58 | 0.152 | 0.960 | |
| AT3G54890.4 | Photosystem I light harvesting complex subunit (LHCA1) | 4 | 0.48 | 0.142 | 0.522 | |
| AT3G54890.4 | Photosystem I light harvesting complex subunit (LHCA1) | 5 | 0.54 | 0.142 | 0.824 | |
| AT3G54890.4 | Photosystem I light harvesting complex subunit (LHCA1) | 6 | 0.64 | 0.141 | 0.991 | |
| AT3G54890.4 | Photosystem I light harvesting complex subunit (LHCA1) | 7 | 0.64 | 0.142 | 0.994 | |
| AT3G54890.4 | Photosystem I light harvesting complex subunit (LHCA1) | 8 | 0.71 | 0.139 | 1.000 | |
| AT3G54890.4 | Photosystem I light harvesting complex subunit (LHCA1) | 9 | 0.77 | 0.137 | 1.000 | |
| AT3G54890.4 | Photosystem I light harvesting complex subunit (LHCA1) | 10 | 0.74 | 0.138 | 1.000 | |
| AT3G54890.4 | Photosystem I light harvesting complex subunit (LHCA1) | 11 | 0.81 | 0.140 | 1.000 | |
| AT3G54890.4 | Photosystem I light harvesting complex subunit (LHCA1) | 12 | 0.98 | 0.139 | 1.000 | |

| | | | | | | |
|---|---|---|---|---|---|---|
| AT3G54890.4 | Photosystem I light harvesting complex subunit (LHCA1) | 13 | 1.07 | 0.137 | 1.000 | |
| AT3G54890.4 | Photosystem I light harvesting complex subunit (LHCA1) | 14 | 1.07 | 0.134 | 1.000 | |
| AT3G54890.4 | Photosystem I light harvesting complex subunit (LHCA1) | 15 | 0.93 | 0.133 | 1.000 | |
| AT3G54890.4 | Photosystem I light harvesting complex subunit (LHCA1) | 16 | 0.80 | 0.132 | 1.000 | |
| AT3G54890.4 | Photosystem I light harvesting complex subunit (LHCA1) | 17 | 0.82 | 0.133 | 1.000 | |
| AT3G54890.4 | Photosystem I light harvesting complex subunit (LHCA1) | 18 | 1.01 | 0.134 | 1.000 | |
| AT3G54890.4 | Photosystem I light harvesting complex subunit (LHCA1) | 19 | 1.34 | 0.137 | 1.000 | |
| AT3G54890.4 | Photosystem I light harvesting complex subunit (LHCA1) | 20 | 1.66 | 0.142 | 0.968 | |
| AT3G54890.4 | Photosystem I light harvesting complex subunit (LHCA1) | 21 | 1.76 | 0.144 | 0.916 | |
| AT3G54890.4 | Photosystem I light harvesting complex subunit (LHCA1) | 22 | 1.52 | 0.144 | 0.997 | |
| AT3G54890.4 | Photosystem I light harvesting complex subunit (LHCA1) | 23 | 1.04 | 0.143 | 1.000 | |
| AT3G54890.4 | Photosystem I light harvesting complex subunit (LHCA1) | 24 | 0.73 | 0.139 | 1.000 | |
| AT3G54890.4 | Photosystem I light harvesting complex subunit (LHCA1) | 25 | 0.81 | 0.136 | 1.000 | |
| AT3G54890.4 | Photosystem I light harvesting complex subunit (LHCA1) | 26 | 1.41 | 0.136 | 1.000 | |
| AT3G54890.4 | Photosystem I light harvesting complex subunit (LHCA1) | 27 | 1.48 | 0.137 | 0.998 | |
| AT3G54890.4 | Photosystem I light harvesting complex subunit (LHCA1) | 28 | 0.67 | 0.137 | 0.997 | |
| AT3G54890.4 | Photosystem I light harvesting complex subunit (LHCA1) | 29 | 0.31 | 0.137 | 0.007 | ** |
| AT3G54890.4 | Photosystem I light harvesting complex subunit (LHCA1) | 30 | 0.23 | 0.137 | 0.000 | *** |
| AT3G54890.4 | Photosystem I light harvesting complex subunit (LHCA1) | 31 | 0.26 | 0.137 | 0.001 | *** |
| AT3G54890.4 | Photosystem I light harvesting complex subunit (LHCA1) | 32 | 0.36 | 0.138 | 0.048 | * |
| AT3G54890.4 | Photosystem I light harvesting complex subunit (LHCA1) | 33 | 0.45 | 0.140 | 0.342 | |
| AT3G54890.4 | Photosystem I light harvesting complex subunit (LHCA1) | 34 | 0.45 | 0.149 | 0.445 | |
| AT3G54890.4 | Photosystem I light harvesting complex subunit (LHCA1) | 35 | 0.42 | 0.170 | 0.517 | |
| AT3G54890.4 | Photosystem I light harvesting complex subunit (LHCA1) | 36 | 0.38 | 0.213 | 0.735 | |
| AT3G54890.4 | Photosystem I light harvesting complex subunit (LHCA1) | 38 | 0.28 | 0.200 | 0.159 | |
| AT3G54890.4 | Photosystem I light harvesting complex subunit (LHCA1) | 39 | 0.22 | 0.230 | 0.147 | |
| AT3G54890.4 | Photosystem I light harvesting complex subunit (LHCA1) | 40 | 0.18 | 0.292 | 0.268 | |
| AT3G54890.4 | Photosystem I light harvesting complex subunit (LHCA1) | 46 | 0.03 | 1.499 | 1.000 | |
| AT3G54890.4 | Photosystem I light harvesting complex subunit (LHCA1) | 47 | 0.02 | 1.798 | 1.000 | |
| AT3G55280.3 | 60S ribosomal protein L23a-2 (RPL23AB) | 2 | 0.92 | 0.382 | 1.000 | |
| AT3G55280.3 | 60S ribosomal protein L23a-2 (RPL23AB) | 3 | 0.93 | 0.304 | 1.000 | |

| | | | | | |
|---|---|---|---|---|---|
| AT3G55280.3 | 60S ribosomal protein L23a-2 (RPL23AB) | 4 | 0.94 | 0.295 | 1.000 |
| AT3G55280.3 | 60S ribosomal protein L23a-2 (RPL23AB) | 9 | 1.14 | 0.272 | 1.000 |
| AT3G55280.3 | 60S ribosomal protein L23a-2 (RPL23AB) | 10 | 1.14 | 0.240 | 1.000 |
| AT3G55280.3 | 60S ribosomal protein L23a-2 (RPL23AB) | 11 | 1.10 | 0.213 | 1.000 |
| AT3G55280.3 | 60S ribosomal protein L23a-2 (RPL23AB) | 12 | 1.03 | 0.193 | 1.000 |
| AT3G55280.3 | 60S ribosomal protein L23a-2 (RPL23AB) | 13 | 0.97 | 0.180 | 1.000 |
| AT3G55280.3 | 60S ribosomal protein L23a-2 (RPL23AB) | 14 | 0.93 | 0.175 | 1.000 |
| AT3G55280.3 | 60S ribosomal protein L23a-2 (RPL23AB) | 15 | 0.93 | 0.173 | 1.000 |
| AT3G55280.3 | 60S ribosomal protein L23a-2 (RPL23AB) | 16 | 0.98 | 0.173 | 1.000 |
| AT3G55280.3 | 60S ribosomal protein L23a-2 (RPL23AB) | 17 | 1.08 | 0.177 | 1.000 |
| AT3G55280.3 | 60S ribosomal protein L23a-2 (RPL23AB) | 18 | 1.20 | 0.187 | 1.000 |
| AT3G55280.3 | 60S ribosomal protein L23a-2 (RPL23AB) | 19 | 1.29 | 0.212 | 1.000 |
| AT3G55280.3 | 60S ribosomal protein L23a-2 (RPL23AB) | 20 | 1.38 | 0.269 | 1.000 |
| AT3G55280.3 | 60S ribosomal protein L23a-2 (RPL23AB) | 21 | 1.50 | 0.367 | 1.000 |
| AT3G55280.3 | 60S ribosomal protein L23a-2 (RPL23AB) | 46 | 21210.07 | 7.771 | 0.999 |
| AT3G55280.3 | 60S ribosomal protein L23a-2 (RPL23AB) | 47 | 36073.02 | 8.208 | 0.999 |
| AT3G55410.1 | 2-oxoglutarate dehydrogenase subunit (OGDCE1) | 2 | 0.47 | 0.533 | 1.000 |
| AT3G55410.1 | 2-oxoglutarate dehydrogenase subunit (OGDCE1) | 3 | 0.60 | 0.350 | 1.000 |
| AT3G55410.1 | 2-oxoglutarate dehydrogenase subunit (OGDCE1) | 4 | 0.72 | 0.312 | 1.000 |
| AT3G55410.1 | 2-oxoglutarate dehydrogenase subunit (OGDCE1) | 5 | 0.80 | 0.307 | 1.000 |
| AT3G55410.1 | 2-oxoglutarate dehydrogenase subunit (OGDCE1) | 6 | 0.89 | 0.295 | 1.000 |
| AT3G55410.1 | 2-oxoglutarate dehydrogenase subunit (OGDCE1) | 7 | 0.99 | 0.286 | 1.000 |
| AT3G55410.1 | 2-oxoglutarate dehydrogenase subunit (OGDCE1) | 8 | 1.12 | 0.278 | 1.000 |
| AT3G55410.1 | 2-oxoglutarate dehydrogenase subunit (OGDCE1) | 9 | 1.22 | 0.274 | 1.000 |
| AT3G55410.1 | 2-oxoglutarate dehydrogenase subunit (OGDCE1) | 10 | 1.24 | 0.273 | 1.000 |
| AT3G55410.1 | 2-oxoglutarate dehydrogenase subunit (OGDCE1) | 11 | 1.16 | 0.274 | 1.000 |
| AT3G55410.1 | 2-oxoglutarate dehydrogenase subunit (OGDCE1) | 12 | 0.95 | 0.274 | 1.000 |
| AT3G55410.1 | 2-oxoglutarate dehydrogenase subunit (OGDCE1) | 13 | 0.67 | 0.271 | 1.000 |
| AT3G55410.1 | 2-oxoglutarate dehydrogenase subunit (OGDCE1) | 14 | 0.44 | 0.269 | 0.992 |
| AT3G55410.1 | 2-oxoglutarate dehydrogenase subunit (OGDCE1) | 15 | 0.31 | 0.268 | 0.803 |
| AT3G55410.1 | 2-oxoglutarate dehydrogenase subunit (OGDCE1) | 16 | 0.36 | 0.266 | 0.928 |

| | | | | | |
|---|---|---|---|---|---|
| AT3G55410.1 | 2-oxoglutarate dehydrogenase subunit (OGDCE1) | 17 | 0.70 | 0.265 | 1.000 |
| AT3G55410.1 | 2-oxoglutarate dehydrogenase subunit (OGDCE1) | 18 | 1.28 | 0.267 | 1.000 |
| AT3G55410.1 | 2-oxoglutarate dehydrogenase subunit (OGDCE1) | 19 | 1.55 | 0.274 | 1.000 |
| AT3G55410.1 | 2-oxoglutarate dehydrogenase subunit (OGDCE1) | 20 | 1.34 | 0.281 | 1.000 |
| AT3G55410.1 | 2-oxoglutarate dehydrogenase subunit (OGDCE1) | 21 | 0.98 | 0.284 | 1.000 |
| AT3G55410.1 | 2-oxoglutarate dehydrogenase subunit (OGDCE1) | 22 | 0.72 | 0.284 | 1.000 |
| AT3G55410.1 | 2-oxoglutarate dehydrogenase subunit (OGDCE1) | 23 | 0.63 | 0.281 | 1.000 |
| AT3G55410.1 | 2-oxoglutarate dehydrogenase subunit (OGDCE1) | 24 | 0.68 | 0.276 | 1.000 |
| AT3G55410.1 | 2-oxoglutarate dehydrogenase subunit (OGDCE1) | 25 | 0.84 | 0.272 | 1.000 |
| AT3G55410.1 | 2-oxoglutarate dehydrogenase subunit (OGDCE1) | 26 | 1.08 | 0.272 | 1.000 |
| AT3G55410.1 | 2-oxoglutarate dehydrogenase subunit (OGDCE1) | 27 | 1.31 | 0.272 | 1.000 |
| AT3G55410.1 | 2-oxoglutarate dehydrogenase subunit (OGDCE1) | 28 | 1.41 | 0.273 | 1.000 |
| AT3G55410.1 | 2-oxoglutarate dehydrogenase subunit (OGDCE1) | 29 | 1.40 | 0.273 | 1.000 |
| AT3G55410.1 | 2-oxoglutarate dehydrogenase subunit (OGDCE1) | 30 | 1.38 | 0.273 | 1.000 |
| AT3G55410.1 | 2-oxoglutarate dehydrogenase subunit (OGDCE1) | 31 | 1.40 | 0.274 | 1.000 |
| AT3G55410.1 | 2-oxoglutarate dehydrogenase subunit (OGDCE1) | 32 | 1.49 | 0.274 | 1.000 |
| AT3G55410.1 | 2-oxoglutarate dehydrogenase subunit (OGDCE1) | 33 | 1.54 | 0.274 | 1.000 |
| AT3G55410.1 | 2-oxoglutarate dehydrogenase subunit (OGDCE1) | 34 | 1.42 | 0.274 | 1.000 |
| AT3G55410.1 | 2-oxoglutarate dehydrogenase subunit (OGDCE1) | 35 | 1.18 | 0.274 | 1.000 |
| AT3G55410.1 | 2-oxoglutarate dehydrogenase subunit (OGDCE1) | 36 | 0.94 | 0.273 | 1.000 |
| AT3G55410.1 | 2-oxoglutarate dehydrogenase subunit (OGDCE1) | 37 | 0.82 | 0.273 | 1.000 |
| AT3G55410.1 | 2-oxoglutarate dehydrogenase subunit (OGDCE1) | 38 | 0.82 | 0.273 | 1.000 |
| AT3G55410.1 | 2-oxoglutarate dehydrogenase subunit (OGDCE1) | 39 | 0.84 | 0.274 | 1.000 |
| AT3G55410.1 | 2-oxoglutarate dehydrogenase subunit (OGDCE1) | 40 | 0.77 | 0.275 | 1.000 |
| AT3G55410.1 | 2-oxoglutarate dehydrogenase subunit (OGDCE1) | 41 | 0.62 | 0.277 | 1.000 |
| AT3G55410.1 | 2-oxoglutarate dehydrogenase subunit (OGDCE1) | 42 | 0.54 | 0.283 | 1.000 |
| AT3G55410.1 | 2-oxoglutarate dehydrogenase subunit (OGDCE1) | 43 | 0.68 | 0.297 | 1.000 |
| AT3G55410.1 | 2-oxoglutarate dehydrogenase subunit (OGDCE1) | 44 | 1.32 | 0.358 | 1.000 |
| AT3G55410.1 | 2-oxoglutarate dehydrogenase subunit (OGDCE1) | 45 | 3.95 | 0.514 | 0.999 |
| AT3G55410.1 | 2-oxoglutarate dehydrogenase subunit (OGDCE1) | 46 | 15.97 | 0.813 | 0.975 |
| AT3G55410.1 | 2-oxoglutarate dehydrogenase subunit (OGDCE1) | 47 | 73.71 | 1.222 | 0.964 |

| | | | | | | |
|---|---|---|---|---|---|---|
| AT3G55630.1 | Folypolyglutamate synthetase 3 (FPGS3) | 2 | 0.96 | 0.312 | 1.000 | |
| AT3G55630.1 | Folypolyglutamate synthetase 3 (FPGS3) | 3 | 1.01 | 0.301 | 1.000 | |
| AT3G55630.1 | Folypolyglutamate synthetase 3 (FPGS3) | 28 | 1.20 | 0.238 | 1.000 | |
| AT3G55630.1 | Folypolyglutamate synthetase 3 (FPGS3) | 29 | 1.15 | 0.182 | 1.000 | |
| AT3G55630.1 | Folypolyglutamate synthetase 3 (FPGS3) | 30 | 1.19 | 0.154 | 1.000 | |
| AT3G55630.1 | Folypolyglutamate synthetase 3 (FPGS3) | 31 | 1.32 | 0.141 | 0.996 | |
| AT3G55630.1 | Folypolyglutamate synthetase 3 (FPGS3) | 32 | 1.45 | 0.134 | 0.939 | |
| AT3G55630.1 | Folypolyglutamate synthetase 3 (FPGS3) | 33 | 1.49 | 0.132 | 0.903 | |
| AT3G55630.1 | Folypolyglutamate synthetase 3 (FPGS3) | 34 | 1.53 | 0.136 | 0.880 | |
| AT3G55630.1 | Folypolyglutamate synthetase 3 (FPGS3) | 35 | 1.64 | 0.143 | 0.799 | |
| AT3G55630.1 | Folypolyglutamate synthetase 3 (FPGS3) | 36 | 1.87 | 0.154 | 0.606 | |
| AT3G55630.1 | Folypolyglutamate synthetase 3 (FPGS3) | 37 | 2.14 | 0.163 | 0.410 | |
| AT3G55630.1 | Folypolyglutamate synthetase 3 (FPGS3) | 38 | 2.35 | 0.162 | 0.258 | |
| AT3G55630.1 | Folypolyglutamate synthetase 3 (FPGS3) | 39 | 2.51 | 0.158 | 0.155 | |
| AT3G55630.1 | Folypolyglutamate synthetase 3 (FPGS3) | 40 | 2.68 | 0.144 | 0.048 | * |
| AT3G55630.1 | Folypolyglutamate synthetase 3 (FPGS3) | 41 | 2.87 | 0.134 | 0.013 | * |
| AT3G55630.1 | Folypolyglutamate synthetase 3 (FPGS3) | 42 | 3.08 | 0.131 | 0.004 | ** |
| AT3G55630.1 | Folypolyglutamate synthetase 3 (FPGS3) | 43 | 2.83 | 0.130 | 0.010 | * |
| AT3G55630.1 | Folypolyglutamate synthetase 3 (FPGS3) | 44 | 1.86 | 0.130 | 0.392 | |
| AT3G55630.1 | Folypolyglutamate synthetase 3 (FPGS3) | 45 | 0.87 | 0.133 | 1.000 | |
| AT3G55630.1 | Folypolyglutamate synthetase 3 (FPGS3) | 46 | 0.32 | 0.160 | 0.034 | * |
| AT3G55630.1 | Folypolyglutamate synthetase 3 (FPGS3) | 47 | 0.11 | 0.247 | 0.002 | ** |
| AT3G55640.1 | Mitochondrial substrate carrier family protein | 2 | 1.09 | 0.499 | 1.000 | |
| AT3G55640.1 | Mitochondrial substrate carrier family protein | 3 | 0.93 | 0.378 | 1.000 | |
| AT3G55640.1 | Mitochondrial substrate carrier family protein | 4 | 0.75 | 0.832 | 1.000 | |
| AT3G55640.1 | Mitochondrial substrate carrier family protein | 39 | 8.39 | 0.407 | 0.229 | |
| AT3G55640.1 | Mitochondrial substrate carrier family protein | 40 | 15.24 | 0.264 | 0.001 | *** |
| AT3G55640.1 | Mitochondrial substrate carrier family protein | 41 | 5.68 | 0.236 | 0.024 | * |
| AT3G55640.1 | Mitochondrial substrate carrier family protein | 42 | 0.49 | 0.236 | 0.845 | |
| AT3G55640.1 | Mitochondrial substrate carrier family protein | 43 | 0.18 | 0.246 | 0.038 | * |
| AT3G55640.1 | Mitochondrial substrate carrier family protein | 44 | 0.45 | 0.299 | 0.910 | |

| | | | | | |
|---|---|---|---|---|---|
| AT3G55640.1 | Mitochondrial substrate carrier family protein | 45 | 3.03 | 0.551 | 0.984 |
| AT3G55640.1 | Mitochondrial substrate carrier family protein | 46 | 31.13 | 1.132 | 0.837 |
| AT3G55640.1 | Mitochondrial substrate carrier family protein | 47 | 383.25 | 1.978 | 0.844 |
| AT3G56070.2 | Rotamase cyclophilin 2 (ROC2) | 2 | 3.70 | 0.254 | 0.355 |
| AT3G56070.2 | Rotamase cyclophilin 2 (ROC2) | 3 | 2.35 | 0.181 | 0.490 |
| AT3G56070.2 | Rotamase cyclophilin 2 (ROC2) | 4 | 1.57 | 0.154 | 0.963 |
| AT3G56070.2 | Rotamase cyclophilin 2 (ROC2) | 5 | 1.14 | 0.149 | 1.000 |
| AT3G56070.2 | Rotamase cyclophilin 2 (ROC2) | 6 | 0.91 | 0.149 | 1.000 |
| AT3G56070.2 | Rotamase cyclophilin 2 (ROC2) | 7 | 0.80 | 0.148 | 1.000 |
| AT3G56070.2 | Rotamase cyclophilin 2 (ROC2) | 8 | 0.77 | 0.147 | 1.000 |
| AT3G56070.2 | Rotamase cyclophilin 2 (ROC2) | 9 | 0.79 | 0.146 | 1.000 |
| AT3G56070.2 | Rotamase cyclophilin 2 (ROC2) | 10 | 0.82 | 0.145 | 1.000 |
| AT3G56070.2 | Rotamase cyclophilin 2 (ROC2) | 11 | 0.86 | 0.144 | 1.000 |
| AT3G56070.2 | Rotamase cyclophilin 2 (ROC2) | 12 | 0.90 | 0.144 | 1.000 |
| AT3G56070.2 | Rotamase cyclophilin 2 (ROC2) | 13 | 0.93 | 0.143 | 1.000 |
| AT3G56070.2 | Rotamase cyclophilin 2 (ROC2) | 14 | 0.95 | 0.142 | 1.000 |
| AT3G56070.2 | Rotamase cyclophilin 2 (ROC2) | 15 | 0.96 | 0.142 | 1.000 |
| AT3G56070.2 | Rotamase cyclophilin 2 (ROC2) | 16 | 0.94 | 0.141 | 1.000 |
| AT3G56070.2 | Rotamase cyclophilin 2 (ROC2) | 17 | 0.94 | 0.141 | 1.000 |
| AT3G56070.2 | Rotamase cyclophilin 2 (ROC2) | 18 | 0.95 | 0.143 | 1.000 |
| AT3G56070.2 | Rotamase cyclophilin 2 (ROC2) | 19 | 0.96 | 0.145 | 1.000 |
| AT3G56070.2 | Rotamase cyclophilin 2 (ROC2) | 20 | 0.99 | 0.147 | 1.000 |
| AT3G56070.2 | Rotamase cyclophilin 2 (ROC2) | 21 | 1.03 | 0.148 | 1.000 |
| AT3G56070.2 | Rotamase cyclophilin 2 (ROC2) | 22 | 1.04 | 0.148 | 1.000 |
| AT3G56070.2 | Rotamase cyclophilin 2 (ROC2) | 23 | 1.00 | 0.147 | 1.000 |
| AT3G56070.2 | Rotamase cyclophilin 2 (ROC2) | 24 | 0.95 | 0.146 | 1.000 |
| AT3G56070.2 | Rotamase cyclophilin 2 (ROC2) | 25 | 0.89 | 0.144 | 1.000 |
| AT3G56070.2 | Rotamase cyclophilin 2 (ROC2) | 26 | 0.84 | 0.144 | 1.000 |
| AT3G56070.2 | Rotamase cyclophilin 2 (ROC2) | 27 | 0.84 | 0.143 | 1.000 |
| AT3G56070.2 | Rotamase cyclophilin 2 (ROC2) | 28 | 0.90 | 0.144 | 1.000 |
| AT3G56070.2 | Rotamase cyclophilin 2 (ROC2) | 29 | 1.01 | 0.144 | 1.000 |
| AT3G56070.2 | Rotamase cyclophilin 2 (ROC2) | 30 | 1.12 | 0.145 | 1.000 |
| AT3G56070.2 | Rotamase cyclophilin 2 (ROC2) | 31 | 1.21 | 0.146 | 1.000 |
| AT3G56070.2 | Rotamase cyclophilin 2 (ROC2) | 32 | 1.26 | 0.147 | 1.000 |
| AT3G56070.2 | Rotamase cyclophilin 2 (ROC2) | 33 | 1.28 | 0.153 | 1.000 |
| AT3G56070.2 | Rotamase cyclophilin 2 (ROC2) | 34 | 1.28 | 0.167 | 1.000 |
| AT3G56070.2 | Rotamase cyclophilin 2 (ROC2) | 35 | 1.33 | 0.199 | 1.000 |
| AT3G56070.2 | Rotamase cyclophilin 2 (ROC2) | 36 | 1.41 | 0.258 | 1.000 |
| AT3G56070.2 | Rotamase cyclophilin 2 (ROC2) | 46 | 3.02 | 1.880 | 1.000 |
| AT3G56070.2 | Rotamase cyclophilin 2 (ROC2) | 47 | 3.29 | 2.121 | 1.000 |
| AT3G56210.4 | ARM repeat superfamily protein | 2 | 0.82 | 0.410 | 1.000 |
| AT3G56210.4 | ARM repeat superfamily protein | 3 | 0.90 | 0.341 | 1.000 |
| AT3G56210.4 | ARM repeat superfamily protein | 8 | 1.52 | 0.224 | 0.999 |
| AT3G56210.4 | ARM repeat superfamily protein | 9 | 1.76 | 0.192 | 0.949 |
| AT3G56210.4 | ARM repeat superfamily protein | 10 | 2.05 | 0.169 | 1.000 |
| AT3G56210.4 | ARM repeat superfamily protein | 11 | 2.27 | 0.155 | 1.000 |
| AT3G56210.4 | ARM repeat superfamily protein | 12 | 2.28 | 0.148 | 0.225 |
| AT3G56210.4 | ARM repeat superfamily protein | 13 | 2.09 | 0.144 | 1.000 |
| AT3G56210.4 | ARM repeat superfamily protein | 14 | 1.86 | 0.143 | 1.000 |

| | | | | | |
|---|---|---|---|---|---|
| AT3G56210.4 | ARM repeat superfamily protein | 15 | 1.68 | 0.142 | 1.000 |
| AT3G56210.4 | ARM repeat superfamily protein | 16 | 1.60 | 0.141 | 0.889 |
| AT3G56210.4 | ARM repeat superfamily protein | 17 | 1.66 | 0.143 | 1.000 |
| AT3G56210.4 | ARM repeat superfamily protein | 18 | 1.85 | 0.147 | 1.000 |
| AT3G56210.4 | ARM repeat superfamily protein | 19 | 2.11 | 0.156 | 1.000 |
| AT3G56210.4 | ARM repeat superfamily protein | 20 | 2.33 | 0.173 | 1.000 |
| AT3G56210.4 | ARM repeat superfamily protein | 21 | 2.48 | 0.201 | 1.000 |
| AT3G56210.4 | ARM repeat superfamily protein | 22 | 2.55 | 0.237 | 1.000 |
| AT3G56210.4 | ARM repeat superfamily protein | 28 | 1.98 | 0.241 | 0.963 |
| AT3G56210.4 | ARM repeat superfamily protein | 29 | 1.85 | 0.211 | 0.954 |
| AT3G56210.4 | ARM repeat superfamily protein | 30 | 1.73 | 0.184 | 0.945 |
| AT3G56210.4 | ARM repeat superfamily protein | 31 | 1.55 | 0.167 | 1.000 |
| AT3G56210.4 | ARM repeat superfamily protein | 32 | 1.30 | 0.157 | 1.000 |
| AT3G56210.4 | ARM repeat superfamily protein | 33 | 1.04 | 0.158 | 1.000 |
| AT3G56210.4 | ARM repeat superfamily protein | 34 | 0.83 | 0.173 | 1.000 |
| AT3G56210.4 | ARM repeat superfamily protein | 35 | 0.69 | 0.209 | 1.000 |
| AT3G56210.4 | ARM repeat superfamily protein | 36 | 0.61 | 0.264 | 0.999 |
| AT3G56210.4 | ARM repeat superfamily protein | 45 | 0.77 | 0.289 | 1.000 |
| AT3G56210.4 | ARM repeat superfamily protein | 46 | 0.79 | 0.262 | 1.000 |
| AT3G56210.4 | ARM repeat superfamily protein | 47 | 0.80 | 0.329 | 1.000 |
| AT3G56430.1 | Translocase of the inner mitochondrial membrane TIM21-like (TIM21-L2) | 2 | 0.88 | 0.418 | 1.000 |
| AT3G56430.1 | Translocase of the inner mitochondrial membrane TIM21-like (TIM21-L2) | 3 | 0.95 | 0.333 | 1.000 |
| AT3G56430.1 | Translocase of the inner mitochondrial membrane TIM21-like (TIM21-L2) | 6 | 1.18 | 0.223 | 1.000 |
| AT3G56430.1 | Translocase of the inner mitochondrial membrane TIM21-like (TIM21-L2) | 7 | 1.28 | 0.194 | 1.000 |
| AT3G56430.1 | Translocase of the inner mitochondrial membrane TIM21-like (TIM21-L2) | 8 | 1.39 | 0.173 | 1.000 |
| AT3G56430.1 | Translocase of the inner mitochondrial membrane TIM21-like (TIM21-L2) | 9 | 1.50 | 0.158 | 0.995 |
| AT3G56430.1 | Translocase of the inner mitochondrial membrane TIM21-like (TIM21-L2) | 10 | 1.60 | 0.150 | 0.967 |

| | | | | | | |
|---|---|---|---|---|---|---|
| AT3G56430.1 | Translocase of the inner mitochondrial membrane TIM21-like (TIM21-L2) | 11 | 1.67 | 0.147 | 0.917 | |
| AT3G56430.1 | Translocase of the inner mitochondrial membrane TIM21-like (TIM21-L2) | 12 | 1.74 | 0.146 | 0.855 | |
| AT3G56430.1 | Translocase of the inner mitochondrial membrane TIM21-like (TIM21-L2) | 13 | 1.81 | 0.145 | 0.772 | |
| AT3G56430.1 | Translocase of the inner mitochondrial membrane TIM21-like (TIM21-L2) | 14 | 1.89 | 0.144 | 0.656 | |
| AT3G56430.1 | Translocase of the inner mitochondrial membrane TIM21-like (TIM21-L2) | 15 | 2.05 | 0.143 | 0.451 | |
| AT3G56430.1 | Translocase of the inner mitochondrial membrane TIM21-like (TIM21-L2) | 16 | 2.31 | 0.142 | 0.209 | |
| AT3G56430.1 | Translocase of the inner mitochondrial membrane TIM21-like (TIM21-L2) | 17 | 2.58 | 0.142 | 0.089 | . |
| AT3G56430.1 | Translocase of the inner mitochondrial membrane TIM21-like (TIM21-L2) | 18 | 2.75 | 0.144 | 0.057 | . |
| AT3G56430.1 | Translocase of the inner mitochondrial membrane TIM21-like (TIM21-L2) | 19 | 2.85 | 0.147 | 0.051 | . |
| AT3G56430.1 | Translocase of the inner mitochondrial membrane TIM21-like (TIM21-L2) | 20 | 2.92 | 0.150 | 0.050 | * |
| AT3G56430.1 | Translocase of the inner mitochondrial membrane TIM21-like (TIM21-L2) | 21 | 2.91 | 0.151 | 0.054 | . |
| AT3G56430.1 | Translocase of the inner mitochondrial membrane TIM21-like (TIM21-L2) | 22 | 2.81 | 0.152 | 0.075 | . |
| AT3G56430.1 | Translocase of the inner mitochondrial membrane TIM21-like (TIM21-L2) | 23 | 2.60 | 0.150 | 0.131 | |
| AT3G56430.1 | Translocase of the inner mitochondrial membrane TIM21-like (TIM21-L2) | 24 | 2.34 | 0.149 | 0.254 | |
| AT3G56430.1 | Translocase of the inner mitochondrial membrane TIM21-like (TIM21-L2) | 25 | 2.03 | 0.147 | 0.523 | |
| AT3G56430.1 | Translocase of the inner mitochondrial membrane TIM21-like (TIM21-L2) | 26 | 1.70 | 0.146 | 0.884 | |
| AT3G56430.1 | Translocase of the inner mitochondrial membrane TIM21-like (TIM21-L2) | 27 | 1.43 | 0.145 | 0.997 | |
| AT3G56430.1 | Translocase of the inner mitochondrial membrane TIM21-like (TIM21-L2) | 28 | 1.23 | 0.145 | 1.000 | |
| AT3G56430.1 | Translocase of the inner mitochondrial membrane TIM21-like (TIM21-L2) | 29 | 1.10 | 0.145 | 1.000 | |
| AT3G56430.1 | Translocase of the inner mitochondrial membrane TIM21-like (TIM21-L2) | 30 | 1.02 | 0.146 | 1.000 | |

| | | | | | |
|---|---|---|---|---|---|
| AT3G56430.1 | Translocase of the inner mitochondrial membrane TIM21-like (TIM21-L2) | 31 | 0.97 | 0.146 | 1.000 |
| AT3G56430.1 | Translocase of the inner mitochondrial membrane TIM21-like (TIM21-L2) | 32 | 0.94 | 0.146 | 1.000 |
| AT3G56430.1 | Translocase of the inner mitochondrial membrane TIM21-like (TIM21-L2) | 33 | 0.90 | 0.147 | 1.000 |
| AT3G56430.1 | Translocase of the inner mitochondrial membrane TIM21-like (TIM21-L2) | 34 | 0.87 | 0.147 | 1.000 |
| AT3G56430.1 | Translocase of the inner mitochondrial membrane TIM21-like (TIM21-L2) | 35 | 0.83 | 0.147 | 1.000 |
| AT3G56430.1 | Translocase of the inner mitochondrial membrane TIM21-like (TIM21-L2) | 36 | 0.80 | 0.147 | 1.000 |
| AT3G56430.1 | Translocase of the inner mitochondrial membrane TIM21-like (TIM21-L2) | 37 | 0.77 | 0.147 | 1.000 |
| AT3G56430.1 | Translocase of the inner mitochondrial membrane TIM21-like (TIM21-L2) | 38 | 0.75 | 0.148 | 1.000 |
| AT3G56430.1 | Translocase of the inner mitochondrial membrane TIM21-like (TIM21-L2) | 39 | 0.74 | 0.150 | 1.000 |
| AT3G56430.1 | Translocase of the inner mitochondrial membrane TIM21-like (TIM21-L2) | 40 | 0.74 | 0.156 | 1.000 |
| AT3G56430.1 | Translocase of the inner mitochondrial membrane TIM21-like (TIM21-L2) | 41 | 0.79 | 0.176 | 1.000 |
| AT3G56430.1 | Translocase of the inner mitochondrial membrane TIM21-like (TIM21-L2) | 42 | 0.93 | 0.221 | 1.000 |
| AT3G56430.1 | Translocase of the inner mitochondrial membrane TIM21-like (TIM21-L2) | 43 | 1.19 | 0.291 | 1.000 |
| AT3G56430.1 | Translocase of the inner mitochondrial membrane TIM21-like (TIM21-L2) | 46 | 3.55 | 0.657 | 1.000 |
| AT3G56430.1 | Translocase of the inner mitochondrial membrane TIM21-like (TIM21-L2) | 47 | 5.38 | 0.824 | 1.000 |
| AT3G57785.1 | NADH:ubiquinone oxidoreductase subunit (ESSS-2) | 2 | 0.82 | 0.714 | 1.000 |
| AT3G57785.1 | NADH:ubiquinone oxidoreductase subunit (ESSS-2) | 3 | 0.89 | 0.641 | 1.000 |
| AT3G57785.1 | NADH:ubiquinone oxidoreductase subunit (ESSS-2) | 14 | 0.62 | 0.421 | 1.000 |
| AT3G57785.1 | NADH:ubiquinone oxidoreductase subunit (ESSS-2) | 15 | 0.48 | 0.358 | 0.985 |
| AT3G57785.1 | NADH:ubiquinone oxidoreductase subunit (ESSS-2) | 16 | 0.38 | 0.325 | 0.862 |

| | | | | | |
|---|---|---|---|---|---|
| AT3G57785.1 | NADH:ubiquinone oxidoreductase subunit (ESSS-2) | 17 | 0.33 | 0.297 | 0.660 |
| AT3G57785.1 | NADH:ubiquinone oxidoreductase subunit (ESSS-2) | 18 | 0.34 | 0.290 | 0.667 |
| AT3G57785.1 | NADH:ubiquinone oxidoreductase subunit (ESSS-2) | 19 | 0.51 | 0.294 | 0.967 |
| AT3G57785.1 | NADH:ubiquinone oxidoreductase subunit (ESSS-2) | 20 | 0.89 | 0.301 | 1.000 |
| AT3G57785.1 | NADH:ubiquinone oxidoreductase subunit (ESSS-2) | 21 | 1.54 | 0.306 | 0.999 |
| AT3G57785.1 | NADH:ubiquinone oxidoreductase subunit (ESSS-2) | 22 | 2.54 | 0.308 | 0.854 |
| AT3G57785.1 | NADH:ubiquinone oxidoreductase subunit (ESSS-2) | 23 | 4.20 | 0.311 | 0.381 |
| AT3G57785.1 | NADH:ubiquinone oxidoreductase subunit (ESSS-2) | 24 | 6.98 | 0.325 | 0.109 |
| AT3G57785.1 | NADH:ubiquinone oxidoreductase subunit (ESSS-2) | 25 | 10.05 | 0.395 | 0.126 |
| AT3G57785.1 | NADH:ubiquinone oxidoreductase subunit (ESSS-2) | 26 | 13.54 | 0.581 | 0.415 |
| AT3G57785.1 | NADH:ubiquinone oxidoreductase subunit (ESSS-2) | 46 | ########### | 12.160 | 0.986 |
| AT3G57785.1 | NADH:ubiquinone oxidoreductase subunit (ESSS-2) | 47 | ########### | 13.023 | 0.987 |
| AT3G58140.1 | Phenylalanyl-tRNA synthetase (PHERS) | 2 | 1.12 | 0.356 | 1.000 |
| AT3G58140.1 | Phenylalanyl-tRNA synthetase (PHERS) | 3 | 1.18 | 0.348 | 1.000 |
| AT3G58140.1 | Phenylalanyl-tRNA synthetase (PHERS) | 29 | 1.34 | 0.260 | 1.000 |
| AT3G58140.1 | Phenylalanyl-tRNA synthetase (PHERS) | 30 | 1.27 | 0.198 | 1.000 |
| AT3G58140.1 | Phenylalanyl-tRNA synthetase (PHERS) | 31 | 1.17 | 0.179 | 1.000 |
| AT3G58140.1 | Phenylalanyl-tRNA synthetase (PHERS) | 32 | 0.94 | 0.169 | 1.000 |
| AT3G58140.1 | Phenylalanyl-tRNA synthetase (PHERS) | 33 | 0.79 | 0.168 | 1.000 |
| AT3G58140.1 | Phenylalanyl-tRNA synthetase (PHERS) | 34 | 0.78 | 0.175 | 1.000 |
| AT3G58140.1 | Phenylalanyl-tRNA synthetase (PHERS) | 35 | 0.72 | 0.185 | 0.999 |
| AT3G58140.1 | Phenylalanyl-tRNA synthetase (PHERS) | 36 | 0.72 | 0.203 | 1.000 |
| AT3G58140.1 | Phenylalanyl-tRNA synthetase (PHERS) | 37 | 0.92 | 0.213 | 1.000 |
| AT3G58140.1 | Phenylalanyl-tRNA synthetase (PHERS) | 38 | 1.40 | 0.226 | 1.000 |
| AT3G58140.1 | Phenylalanyl-tRNA synthetase (PHERS) | 39 | 2.23 | 0.209 | 0.742 |

| | | | | | |
|---|---|---|---|---|---|
| AT3G58140.1 | Phenylalanyl-tRNA synthetase (PHERS) | 40 | 2.95 | 0.192 | 0.206 |
| AT3G58140.1 | Phenylalanyl-tRNA synthetase (PHERS) | 41 | 2.75 | 0.171 | 0.159 |
| AT3G58140.1 | Phenylalanyl-tRNA synthetase (PHERS) | 42 | 2.16 | 0.168 | 0.495 |
| AT3G58140.1 | Phenylalanyl-tRNA synthetase (PHERS) | 43 | 1.84 | 0.169 | 0.811 |
| AT3G58140.1 | Phenylalanyl-tRNA synthetase (PHERS) | 44 | 1.45 | 0.169 | 0.995 |
| AT3G58140.1 | Phenylalanyl-tRNA synthetase (PHERS) | 45 | 0.94 | 0.168 | 1.000 |
| AT3G58140.1 | Phenylalanyl-tRNA synthetase (PHERS) | 46 | 0.47 | 0.191 | 0.710 |
| AT3G58140.1 | Phenylalanyl-tRNA synthetase (PHERS) | 47 | 0.18 | 0.359 | 0.453 |
| AT3G58610.3 | Ketol-acid reductoisomerase (putative) | 2 | 1.22 | 0.429 | 1.000 |
| AT3G58610.3 | Ketol-acid reductoisomerase (putative) | 3 | 0.94 | 0.356 | 1.000 |
| AT3G58610.3 | Ketol-acid reductoisomerase (putative) | 4 | 0.66 | 0.693 | 1.000 |
| AT3G58610.3 | Ketol-acid reductoisomerase (putative) | 40 | 0.43 | 0.353 | 0.927 |
| AT3G58610.3 | Ketol-acid reductoisomerase (putative) | 41 | 2.32 | 0.256 | 0.727 |
| AT3G58610.3 | Ketol-acid reductoisomerase (putative) | 42 | 3.16 | 0.220 | 0.204 |
| AT3G58610.3 | Ketol-acid reductoisomerase (putative) | 43 | 1.42 | 0.209 | 0.991 |
| AT3G58610.3 | Ketol-acid reductoisomerase (putative) | 44 | 0.59 | 0.219 | 0.928 |
| AT3G58610.3 | Ketol-acid reductoisomerase (putative) | 45 | 0.42 | 0.284 | 0.795 |
| AT3G58610.3 | Ketol-acid reductoisomerase (putative) | 46 | 0.52 | 0.331 | 0.975 |
| AT3G58610.3 | Ketol-acid reductoisomerase (putative) | 47 | 0.83 | 0.455 | 1.000 |
| AT3G58840.2 | Peroxisomal and mitochondrial division factor 1 (PMD1) | 2 | 0.90 | 0.509 | 1.000 |
| AT3G58840.2 | Peroxisomal and mitochondrial division factor 1 (PMD1) | 3 | 0.85 | 0.485 | 1.000 |
| AT3G58840.2 | Peroxisomal and mitochondrial division factor 1 (PMD1) | 25 | 0.44 | 0.396 | 0.999 |
| AT3G58840.2 | Peroxisomal and mitochondrial division factor 1 (PMD1) | 26 | 0.46 | 0.271 | 0.979 |
| AT3G58840.2 | Peroxisomal and mitochondrial division factor 1 (PMD1) | 27 | 0.59 | 0.235 | 0.998 |
| AT3G58840.2 | Peroxisomal and mitochondrial division factor 1 (PMD1) | 28 | 1.02 | 0.222 | 1.000 |
| AT3G58840.2 | Peroxisomal and mitochondrial division factor 1 (PMD1) | 29 | 1.73 | 0.219 | 0.994 |
| AT3G58840.2 | Peroxisomal and mitochondrial division factor 1 (PMD1) | 30 | 1.80 | 0.221 | 0.990 |

| | | | | | | |
|---|---|---|---|---|---|---|
| AT3G58840.2 | Peroxisomal and mitochondrial division factor 1 (PMD1) | 31 | 1.34 | 0.222 | 1.000 | |
| AT3G58840.2 | Peroxisomal and mitochondrial division factor 1 (PMD1) | 32 | 1.12 | 0.222 | 1.000 | |
| AT3G58840.2 | Peroxisomal and mitochondrial division factor 1 (PMD1) | 33 | 1.03 | 0.222 | 1.000 | |
| AT3G58840.2 | Peroxisomal and mitochondrial division factor 1 (PMD1) | 34 | 1.04 | 0.223 | 1.000 | |
| AT3G58840.2 | Peroxisomal and mitochondrial division factor 1 (PMD1) | 35 | 1.42 | 0.222 | 1.000 | |
| AT3G58840.2 | Peroxisomal and mitochondrial division factor 1 (PMD1) | 36 | 2.92 | 0.221 | 0.475 | |
| AT3G58840.2 | Peroxisomal and mitochondrial division factor 1 (PMD1) | 37 | 5.18 | 0.219 | 0.027 | * |
| AT3G58840.2 | Peroxisomal and mitochondrial division factor 1 (PMD1) | 38 | 5.02 | 0.219 | 0.033 | * |
| AT3G58840.2 | Peroxisomal and mitochondrial division factor 1 (PMD1) | 39 | 2.90 | 0.219 | 0.477 | |
| AT3G58840.2 | Peroxisomal and mitochondrial division factor 1 (PMD1) | 40 | 1.44 | 0.220 | 1.000 | |
| AT3G58840.2 | Peroxisomal and mitochondrial division factor 1 (PMD1) | 41 | 0.80 | 0.222 | 1.000 | |
| AT3G58840.2 | Peroxisomal and mitochondrial division factor 1 (PMD1) | 42 | 0.62 | 0.225 | 0.999 | |
| AT3G58840.2 | Peroxisomal and mitochondrial division factor 1 (PMD1) | 43 | 0.64 | 0.227 | 1.000 | |
| AT3G58840.2 | Peroxisomal and mitochondrial division factor 1 (PMD1) | 44 | 0.61 | 0.227 | 0.999 | |
| AT3G58840.2 | Peroxisomal and mitochondrial division factor 1 (PMD1) | 45 | 0.47 | 0.227 | 0.928 | |
| AT3G58840.2 | Peroxisomal and mitochondrial division factor 1 (PMD1) | 46 | 0.34 | 0.266 | 0.759 | |
| AT3G58840.2 | Peroxisomal and mitochondrial division factor 1 (PMD1) | 47 | 0.26 | 0.496 | 0.985 | |
| AT3G59280.1 | Protein transporter PAM16 | 2 | 0.79 | 0.470 | 1.000 | |
| AT3G59280.1 | Protein transporter PAM16 | 3 | 0.82 | 0.427 | 1.000 | |
| AT3G59280.1 | Protein transporter PAM16 | 16 | 0.91 | 0.219 | 1.000 | |
| AT3G59280.1 | Protein transporter PAM16 | 17 | 0.88 | 0.189 | 1.000 | |
| AT3G59280.1 | Protein transporter PAM16 | 18 | 0.86 | 0.167 | 1.000 | |
| AT3G59280.1 | Protein transporter PAM16 | 19 | 0.86 | 0.155 | 1.000 | |
| AT3G59280.1 | Protein transporter PAM16 | 20 | 0.89 | 0.150 | 1.000 | |
| AT3G59280.1 | Protein transporter PAM16 | 21 | 0.96 | 0.152 | 1.000 | |
| AT3G59280.1 | Protein transporter PAM16 | 22 | 1.09 | 0.162 | 1.000 | |
| AT3G59280.1 | Protein transporter PAM16 | 23 | 1.29 | 0.182 | 0.996 | |
| AT3G59280.1 | Protein transporter PAM16 | 24 | 1.59 | 0.214 | 0.953 | |

| | | | | | | |
|---|---|---|---|---|---|---|
| AT3G59280.1 | Protein transporter PAM16 | 25 | 1.98 | 0.259 | 0.876 | |
| AT3G59280.1 | Protein transporter PAM16 | 43 | 5.98 | 0.299 | 0.088 | . |
| AT3G59280.1 | Protein transporter PAM16 | 44 | 4.99 | 0.230 | 0.028 | * |
| AT3G59280.1 | Protein transporter PAM16 | 45 | 4.01 | 0.194 | 0.021 | * |
| AT3G59280.1 | Protein transporter PAM16 | 46 | 3.12 | 0.223 | 0.212 | |
| AT3G59280.1 | Protein transporter PAM16 | 47 | 2.39 | 0.314 | 0.846 | |
| AT3G59760.3 | O-acetylserine (thiol) lyase C (OASTL-C) | 2 | 0.98 | 0.224 | 1.000 | |
| AT3G59760.3 | O-acetylserine (thiol) lyase C (OASTL-C) | 3 | 0.90 | 0.179 | 1.000 | |
| AT3G59760.3 | O-acetylserine (thiol) lyase C (OASTL-C) | 4 | 0.85 | 0.212 | 1.000 | |
| AT3G59760.3 | O-acetylserine (thiol) lyase C (OASTL-C) | 12 | 0.99 | 0.161 | 1.000 | |
| AT3G59760.3 | O-acetylserine (thiol) lyase C (OASTL-C) | 13 | 1.07 | 0.139 | 1.000 | |
| AT3G59760.3 | O-acetylserine (thiol) lyase C (OASTL-C) | 14 | 1.17 | 0.124 | 1.000 | |
| AT3G59760.3 | O-acetylserine (thiol) lyase C (OASTL-C) | 15 | 1.33 | 0.116 | 0.998 | |
| AT3G59760.3 | O-acetylserine (thiol) lyase C (OASTL-C) | 16 | 1.60 | 0.110 | 0.730 | |
| AT3G59760.3 | O-acetylserine (thiol) lyase C (OASTL-C) | 17 | 1.93 | 0.108 | 0.180 | |
| AT3G59760.3 | O-acetylserine (thiol) lyase C (OASTL-C) | 18 | 2.22 | 0.109 | 0.041 | * |
| AT3G59760.3 | O-acetylserine (thiol) lyase C (OASTL-C) | 19 | 2.37 | 0.112 | 0.023 | * |
| AT3G59760.3 | O-acetylserine (thiol) lyase C (OASTL-C) | 20 | 2.24 | 0.114 | 0.056 | . |
| AT3G59760.3 | O-acetylserine (thiol) lyase C (OASTL-C) | 21 | 1.96 | 0.115 | 0.234 | |
| AT3G59760.3 | O-acetylserine (thiol) lyase C (OASTL-C) | 22 | 1.71 | 0.115 | 0.600 | |
| AT3G59760.3 | O-acetylserine (thiol) lyase C (OASTL-C) | 23 | 1.49 | 0.114 | 0.922 | |
| AT3G59760.3 | O-acetylserine (thiol) lyase C (OASTL-C) | 24 | 1.31 | 0.112 | 0.999 | |
| AT3G59760.3 | O-acetylserine (thiol) lyase C (OASTL-C) | 25 | 1.17 | 0.111 | 1.000 | |
| AT3G59760.3 | O-acetylserine (thiol) lyase C (OASTL-C) | 26 | 1.12 | 0.110 | 1.000 | |
| AT3G59760.3 | O-acetylserine (thiol) lyase C (OASTL-C) | 27 | 1.16 | 0.110 | 1.000 | |
| AT3G59760.3 | O-acetylserine (thiol) lyase C (OASTL-C) | 28 | 1.24 | 0.110 | 1.000 | |
| AT3G59760.3 | O-acetylserine (thiol) lyase C (OASTL-C) | 29 | 1.28 | 0.110 | 0.999 | |
| AT3G59760.3 | O-acetylserine (thiol) lyase C (OASTL-C) | 30 | 1.20 | 0.110 | 1.000 | |
| AT3G59760.3 | O-acetylserine (thiol) lyase C (OASTL-C) | 31 | 1.02 | 0.111 | 1.000 | |
| AT3G59760.3 | O-acetylserine (thiol) lyase C (OASTL-C) | 32 | 0.85 | 0.111 | 1.000 | |
| AT3G59760.3 | O-acetylserine (thiol) lyase C (OASTL-C) | 33 | 0.76 | 0.111 | 0.998 | |
| AT3G59760.3 | O-acetylserine (thiol) lyase C (OASTL-C) | 34 | 0.76 | 0.111 | 0.998 | |
| AT3G59760.3 | O-acetylserine (thiol) lyase C (OASTL-C) | 35 | 0.84 | 0.111 | 1.000 | |
| AT3G59760.3 | O-acetylserine (thiol) lyase C (OASTL-C) | 36 | 0.96 | 0.111 | 1.000 | |

| | | | | | |
|---|---|---|---|---|---|
| AT3G59760.3 | O-acetylserine (thiol) lyase C (OASTL-C) | 37 | 1.09 | 0.111 | 1.000 |
| AT3G59760.3 | O-acetylserine (thiol) lyase C (OASTL-C) | 38 | 1.17 | 0.111 | 1.000 |
| AT3G59760.3 | O-acetylserine (thiol) lyase C (OASTL-C) | 39 | 1.23 | 0.111 | 1.000 |
| AT3G59760.3 | O-acetylserine (thiol) lyase C (OASTL-C) | 40 | 1.30 | 0.111 | 0.999 |
| AT3G59760.3 | O-acetylserine (thiol) lyase C (OASTL-C) | 41 | 1.35 | 0.112 | 0.994 |
| AT3G59760.3 | O-acetylserine (thiol) lyase C (OASTL-C) | 42 | 1.35 | 0.113 | 0.996 |
| AT3G59760.3 | O-acetylserine (thiol) lyase C (OASTL-C) | 43 | 1.27 | 0.114 | 1.000 |
| AT3G59760.3 | O-acetylserine (thiol) lyase C (OASTL-C) | 44 | 1.10 | 0.114 | 1.000 |
| AT3G59760.3 | O-acetylserine (thiol) lyase C (OASTL-C) | 45 | 0.86 | 0.117 | 1.000 |
| AT3G59760.3 | O-acetylserine (thiol) lyase C (OASTL-C) | 46 | 0.62 | 0.143 | 0.945 |
| AT3G59760.3 | O-acetylserine (thiol) lyase C (OASTL-C) | 47 | 0.43 | 0.219 | 0.840 |
| AT3G59820.1 | Leucine zipper-EF-hand-containing transmembrane protein 1 (LETM1) | 2 | 1.00 | 0.373 | 1.000 |
| AT3G59820.1 | Leucine zipper-EF-hand-containing transmembrane protein 1 (LETM1) | 3 | 0.97 | 0.271 | 1.000 |
| AT3G59820.1 | Leucine zipper-EF-hand-containing transmembrane protein 1 (LETM1) | 4 | 1.02 | 0.300 | 1.000 |
| AT3G59820.1 | Leucine zipper-EF-hand-containing transmembrane protein 1 (LETM1) | 6 | 1.26 | 0.324 | 1.000 |
| AT3G59820.1 | Leucine zipper-EF-hand-containing transmembrane protein 1 (LETM1) | 7 | 1.37 | 0.332 | 1.000 |
| AT3G59820.1 | Leucine zipper-EF-hand-containing transmembrane protein 1 (LETM1) | 8 | 1.43 | 0.335 | 1.000 |
| AT3G59820.1 | Leucine zipper-EF-hand-containing transmembrane protein 1 (LETM1) | 14 | 1.03 | 0.267 | 1.000 |
| AT3G59820.1 | Leucine zipper-EF-hand-containing transmembrane protein 1 (LETM1) | 15 | 1.23 | 0.238 | 1.000 |
| AT3G59820.1 | Leucine zipper-EF-hand-containing transmembrane protein 1 (LETM1) | 16 | 1.66 | 0.221 | 1.000 |
| AT3G59820.1 | Leucine zipper-EF-hand-containing transmembrane protein 1 (LETM1) | 17 | 2.16 | 0.201 | 0.893 |
| AT3G59820.1 | Leucine zipper-EF-hand-containing transmembrane protein 1 (LETM1) | 18 | 2.88 | 0.196 | 0.384 |
| AT3G59820.1 | Leucine zipper-EF-hand-containing transmembrane protein 1 (LETM1) | 19 | 3.73 | 0.203 | 0.130 |
| AT3G59820.1 | Leucine zipper-EF-hand-containing transmembrane protein 1 (LETM1) | 20 | 3.90 | 0.208 | 0.122 |

| | | | | | |
|---|---|---|---|---|---|
| AT3G59820.1 | Leucine zipper-EF-hand-containing transmembrane protein 1 (LETM1) | 21 | 3.10 | 0.210 | 0.391 |
| AT3G59820.1 | Leucine zipper-EF-hand-containing transmembrane protein 1 (LETM1) | 22 | 2.35 | 0.209 | 0.828 |
| AT3G59820.1 | Leucine zipper-EF-hand-containing transmembrane protein 1 (LETM1) | 23 | 2.05 | 0.209 | 0.958 |
| AT3G59820.1 | Leucine zipper-EF-hand-containing transmembrane protein 1 (LETM1) | 24 | 1.78 | 0.202 | 0.994 |
| AT3G59820.1 | Leucine zipper-EF-hand-containing transmembrane protein 1 (LETM1) | 25 | 1.23 | 0.199 | 1.000 |
| AT3G59820.1 | Leucine zipper-EF-hand-containing transmembrane protein 1 (LETM1) | 26 | 1.12 | 0.199 | 1.000 |
| AT3G59820.1 | Leucine zipper-EF-hand-containing transmembrane protein 1 (LETM1) | 27 | 1.49 | 0.200 | 1.000 |
| AT3G59820.1 | Leucine zipper-EF-hand-containing transmembrane protein 1 (LETM1) | 28 | 1.67 | 0.200 | 0.999 |
| AT3G59820.1 | Leucine zipper-EF-hand-containing transmembrane protein 1 (LETM1) | 29 | 1.51 | 0.199 | 1.000 |
| AT3G59820.1 | Leucine zipper-EF-hand-containing transmembrane protein 1 (LETM1) | 30 | 1.27 | 0.199 | 1.000 |
| AT3G59820.1 | Leucine zipper-EF-hand-containing transmembrane protein 1 (LETM1) | 31 | 1.23 | 0.201 | 1.000 |
| AT3G59820.1 | Leucine zipper-EF-hand-containing transmembrane protein 1 (LETM1) | 32 | 1.21 | 0.201 | 1.000 |
| AT3G59820.1 | Leucine zipper-EF-hand-containing transmembrane protein 1 (LETM1) | 33 | 1.13 | 0.201 | 1.000 |
| AT3G59820.1 | Leucine zipper-EF-hand-containing transmembrane protein 1 (LETM1) | 34 | 1.07 | 0.201 | 1.000 |
| AT3G59820.1 | Leucine zipper-EF-hand-containing transmembrane protein 1 (LETM1) | 35 | 1.05 | 0.201 | 1.000 |
| AT3G59820.1 | Leucine zipper-EF-hand-containing transmembrane protein 1 (LETM1) | 36 | 0.96 | 0.201 | 1.000 |
| AT3G59820.1 | Leucine zipper-EF-hand-containing transmembrane protein 1 (LETM1) | 37 | 0.82 | 0.201 | 1.000 |
| AT3G59820.1 | Leucine zipper-EF-hand-containing transmembrane protein 1 (LETM1) | 38 | 0.67 | 0.201 | 1.000 |
| AT3G59820.1 | Leucine zipper-EF-hand-containing transmembrane protein 1 (LETM1) | 39 | 0.52 | 0.201 | 0.978 |
| AT3G59820.1 | Leucine zipper-EF-hand-containing transmembrane protein 1 (LETM1) | 40 | 0.34 | 0.202 | 0.396 |

| | | | | | | |
|---|---|---|---|---|---|---|
| AT3G59820.1 | Leucine zipper-EF-hand-containing transmembrane protein 1 (LETM1) | 41 | 0.18 | 0.212 | 0.014 | * |
| AT3G59820.1 | Leucine zipper-EF-hand-containing transmembrane protein 1 (LETM1) | 42 | 0.17 | 0.285 | 0.182 | |
| AT3G59820.1 | Leucine zipper-EF-hand-containing transmembrane protein 1 (LETM1) | 43 | 0.45 | 0.420 | 1.000 | |
| AT3G59820.1 | Leucine zipper-EF-hand-containing transmembrane protein 1 (LETM1) | 46 | 512.13 | 1.123 | 0.337 | |
| AT3G59820.1 | Leucine zipper-EF-hand-containing transmembrane protein 1 (LETM1) | 47 | 10267.07 | 1.477 | 0.169 | |
| AT3G60480.1 | StAR lipid transfer-like protein | 2 | 0.99 | 0.507 | 1.000 | |
| AT3G60480.1 | StAR lipid transfer-like protein | 3 | 0.96 | 0.487 | 1.000 | |
| AT3G60480.1 | StAR lipid transfer-like protein | 22 | 0.24 | 0.331 | 0.469 | |
| AT3G60480.1 | StAR lipid transfer-like protein | 23 | 0.35 | 0.261 | 0.566 | |
| AT3G60480.1 | StAR lipid transfer-like protein | 24 | 0.66 | 0.214 | 0.992 | |
| AT3G60480.1 | StAR lipid transfer-like protein | 25 | 1.37 | 0.194 | 0.998 | |
| AT3G60480.1 | StAR lipid transfer-like protein | 26 | 1.81 | 0.189 | 0.836 | |
| AT3G60480.1 | StAR lipid transfer-like protein | 27 | 1.59 | 0.188 | 0.958 | |
| AT3G60480.1 | StAR lipid transfer-like protein | 28 | 1.41 | 0.190 | 0.995 | |
| AT3G60480.1 | StAR lipid transfer-like protein | 29 | 1.30 | 0.195 | 1.000 | |
| AT3G60480.1 | StAR lipid transfer-like protein | 30 | 1.03 | 0.216 | 1.000 | |
| AT3G60480.1 | StAR lipid transfer-like protein | 31 | 0.78 | 0.282 | 1.000 | |
| AT3G60480.1 | StAR lipid transfer-like protein | 32 | 0.64 | 0.397 | 1.000 | |
| AT3G60480.1 | StAR lipid transfer-like protein | 44 | 1.55 | 0.420 | 1.000 | |
| AT3G60480.1 | StAR lipid transfer-like protein | 45 | 1.39 | 0.340 | 1.000 | |
| AT3G60480.1 | StAR lipid transfer-like protein | 46 | 1.11 | 0.371 | 1.000 | |
| AT3G60480.1 | StAR lipid transfer-like protein | 47 | 0.84 | 0.526 | 1.000 | |
| AT3G60750.2 | Transketolase 1 (TKL1) | 2 | 1.15 | 0.794 | 1.000 | |
| AT3G60750.2 | Transketolase 1 (TKL1) | 3 | 0.88 | 0.611 | 1.000 | |
| AT3G60750.2 | Transketolase 1 (TKL1) | 4 | 0.72 | 1.229 | 1.000 | |
| AT3G60750.2 | Transketolase 1 (TKL1) | 40 | 33.37 | 0.740 | 0.324 | |
| AT3G60750.2 | Transketolase 1 (TKL1) | 41 | 13.40 | 0.447 | 0.124 | |
| AT3G60750.2 | Transketolase 1 (TKL1) | 42 | 2.42 | 0.384 | 0.956 | |
| AT3G60750.2 | Transketolase 1 (TKL1) | 43 | 0.37 | 0.377 | 0.911 | |
| AT3G60750.2 | Transketolase 1 (TKL1) | 44 | 0.22 | 0.369 | 0.512 | |
| AT3G60750.2 | Transketolase 1 (TKL1) | 45 | 0.46 | 0.368 | 0.971 | |
| AT3G60750.2 | Transketolase 1 (TKL1) | 46 | 0.86 | 0.406 | 1.000 | |
| AT3G60750.2 | Transketolase 1 (TKL1) | 47 | 1.65 | 0.893 | 1.000 | |
| AT3G60980.1 | Pentatricopeptide repeat (PPR) superfamily protein | 2 | 0.85 | 0.350 | 1.000 | |
| AT3G60980.1 | Pentatricopeptide repeat (PPR) superfamily protein | 3 | 0.77 | 0.255 | 1.000 | |
| AT3G60980.1 | Pentatricopeptide repeat (PPR) superfamily protein | 4 | 0.71 | 0.208 | 0.987 | |
| AT3G60980.1 | Pentatricopeptide repeat (PPR) superfamily protein | 5 | 0.67 | 0.191 | 0.953 | |
| AT3G60980.1 | Pentatricopeptide repeat (PPR) superfamily protein | 6 | 0.69 | 0.185 | 0.963 | |
| AT3G60980.1 | Pentatricopeptide repeat (PPR) superfamily protein | 7 | 0.79 | 0.183 | 0.998 | |
| AT3G60980.1 | Pentatricopeptide repeat (PPR) superfamily protein | 8 | 1.00 | 0.185 | 1.000 | |
| AT3G60980.1 | Pentatricopeptide repeat (PPR) superfamily protein | 9 | 1.35 | 0.196 | 0.993 | |
| AT3G60980.1 | Pentatricopeptide repeat (PPR) superfamily protein | 10 | 1.86 | 0.219 | 0.821 | |

| | | | | | |
|---|---|---|---|---|---|
| AT3G60980.1 | Pentatricopeptide repeat (PPR) superfamily protein | 11 | 2.49 | 0.256 | 0.612 |
| AT3G60980.1 | Pentatricopeptide repeat (PPR) superfamily protein | 23 | 0.40 | 0.387 | 0.913 |
| AT3G60980.1 | Pentatricopeptide repeat (PPR) superfamily protein | 24 | 0.31 | 0.382 | 0.766 |
| AT3G60980.1 | Pentatricopeptide repeat (PPR) superfamily protein | 25 | 0.25 | 0.434 | 0.729 |
| AT3G60980.1 | Pentatricopeptide repeat (PPR) superfamily protein | 46 | 0.14 | 6.044 | 1.000 |
| AT3G60980.1 | Pentatricopeptide repeat (PPR) superfamily protein | 47 | 0.15 | 6.430 | 1.000 |
| AT3G61070.3 | Peroxin 11E (PEX11E) | 2 | 1.19 | 0.390 | 1.000 |
| AT3G61070.3 | Peroxin 11E (PEX11E) | 3 | 1.21 | 0.373 | 1.000 |
| AT3G61070.3 | Peroxin 11E (PEX11E) | 4 | 1.22 | 0.363 | 1.000 |
| AT3G61070.3 | Peroxin 11E (PEX11E) | 26 | 1.34 | 0.249 | 0.986 |
| AT3G61070.3 | Peroxin 11E (PEX11E) | 27 | 1.32 | 0.234 | 0.985 |
| AT3G61070.3 | Peroxin 11E (PEX11E) | 28 | 1.30 | 0.221 | 0.985 |
| AT3G61070.3 | Peroxin 11E (PEX11E) | 29 | 1.27 | 0.210 | 0.988 |
| AT3G61070.3 | Peroxin 11E (PEX11E) | 30 | 1.23 | 0.201 | 0.992 |
| AT3G61070.3 | Peroxin 11E (PEX11E) | 33 | 1.06 | 0.189 | 1.000 |
| AT3G61070.3 | Peroxin 11E (PEX11E) | 34 | 0.98 | 0.191 | 1.000 |
| AT3G61070.3 | Peroxin 11E (PEX11E) | 35 | 0.89 | 0.197 | 1.000 |
| AT3G61070.3 | Peroxin 11E (PEX11E) | 36 | 0.79 | 0.206 | 0.987 |
| AT3G61070.3 | Peroxin 11E (PEX11E) | 37 | 0.69 | 0.220 | 0.936 |
| AT3G61070.3 | Peroxin 11E (PEX11E) | 38 | 0.60 | 0.236 | 0.840 |
| AT3G61070.3 | Peroxin 11E (PEX11E) | 46 | 0.14 | 0.434 | 0.215 |
| AT3G61070.3 | Peroxin 11E (PEX11E) | 47 | 0.11 | 0.470 | 0.200 |
| AT3G61440.1 | Cysteine synthase C1 (CYSC1) | 3 | 0.49 | 0.242 | 0.732 |
| AT3G61440.1 | Cysteine synthase C1 (CYSC1) | 4 | 0.54 | 0.230 | 0.815 |
| AT3G61440.1 | Cysteine synthase C1 (CYSC1) | 5 | 0.63 | 0.229 | 0.969 |
| AT3G61440.1 | Cysteine synthase C1 (CYSC1) | 6 | 0.77 | 0.229 | 1.000 |
| AT3G61440.1 | Cysteine synthase C1 (CYSC1) | 7 | 0.91 | 0.229 | 1.000 |
| AT3G61440.1 | Cysteine synthase C1 (CYSC1) | 8 | 1.06 | 0.228 | 1.000 |
| AT3G61440.1 | Cysteine synthase C1 (CYSC1) | 9 | 1.22 | 0.227 | 1.000 |
| AT3G61440.1 | Cysteine synthase C1 (CYSC1) | 10 | 1.33 | 0.227 | 1.000 |
| AT3G61440.1 | Cysteine synthase C1 (CYSC1) | 11 | 1.37 | 0.227 | 0.999 |
| AT3G61440.1 | Cysteine synthase C1 (CYSC1) | 12 | 1.39 | 0.227 | 0.998 |
| AT3G61440.1 | Cysteine synthase C1 (CYSC1) | 13 | 1.42 | 0.226 | 0.997 |
| AT3G61440.1 | Cysteine synthase C1 (CYSC1) | 14 | 1.50 | 0.226 | 0.987 |
| AT3G61440.1 | Cysteine synthase C1 (CYSC1) | 15 | 1.60 | 0.225 | 0.955 |
| AT3G61440.1 | Cysteine synthase C1 (CYSC1) | 16 | 1.72 | 0.225 | 0.890 |
| AT3G61440.1 | Cysteine synthase C1 (CYSC1) | 17 | 1.89 | 0.225 | 0.771 |
| AT3G61440.1 | Cysteine synthase C1 (CYSC1) | 18 | 2.04 | 0.226 | 0.665 |
| AT3G61440.1 | Cysteine synthase C1 (CYSC1) | 19 | 2.01 | 0.227 | 0.696 |
| AT3G61440.1 | Cysteine synthase C1 (CYSC1) | 20 | 1.79 | 0.228 | 0.851 |
| AT3G61440.1 | Cysteine synthase C1 (CYSC1) | 21 | 1.52 | 0.229 | 0.984 |
| AT3G61440.1 | Cysteine synthase C1 (CYSC1) | 22 | 1.29 | 0.229 | 1.000 |
| AT3G61440.1 | Cysteine synthase C1 (CYSC1) | 23 | 1.14 | 0.228 | 1.000 |
| AT3G61440.1 | Cysteine synthase C1 (CYSC1) | 24 | 1.08 | 0.228 | 1.000 |
| AT3G61440.1 | Cysteine synthase C1 (CYSC1) | 25 | 1.10 | 0.227 | 1.000 |
| AT3G61440.1 | Cysteine synthase C1 (CYSC1) | 26 | 1.14 | 0.227 | 1.000 |
| AT3G61440.1 | Cysteine synthase C1 (CYSC1) | 27 | 1.22 | 0.226 | 1.000 |
| AT3G61440.1 | Cysteine synthase C1 (CYSC1) | 28 | 1.27 | 0.226 | 1.000 |
| AT3G61440.1 | Cysteine synthase C1 (CYSC1) | 29 | 1.31 | 0.226 | 1.000 |
| AT3G61440.1 | Cysteine synthase C1 (CYSC1) | 30 | 1.33 | 0.227 | 1.000 |
| AT3G61440.1 | Cysteine synthase C1 (CYSC1) | 31 | 1.33 | 0.227 | 1.000 |
| AT3G61440.1 | Cysteine synthase C1 (CYSC1) | 32 | 1.33 | 0.227 | 1.000 |
| AT3G61440.1 | Cysteine synthase C1 (CYSC1) | 33 | 1.28 | 0.227 | 1.000 |
| AT3G61440.1 | Cysteine synthase C1 (CYSC1) | 34 | 1.22 | 0.227 | 1.000 |
| AT3G61440.1 | Cysteine synthase C1 (CYSC1) | 35 | 1.20 | 0.227 | 1.000 |
| AT3G61440.1 | Cysteine synthase C1 (CYSC1) | 36 | 1.27 | 0.227 | 1.000 |
| AT3G61440.1 | Cysteine synthase C1 (CYSC1) | 37 | 1.41 | 0.227 | 0.998 |

| | | | | | | |
|---|---|---|---|---|---|---|
| AT3G61440.1 | Cysteine synthase C1 (CYSC1) | 38 | 1.61 | 0.227 | 0.954 | |
| AT3G61440.1 | Cysteine synthase C1 (CYSC1) | 39 | 1.83 | 0.227 | 0.823 | |
| AT3G61440.1 | Cysteine synthase C1 (CYSC1) | 40 | 2.04 | 0.227 | 0.668 | |
| AT3G61440.1 | Cysteine synthase C1 (CYSC1) | 41 | 2.21 | 0.228 | 0.554 | |
| AT3G61440.1 | Cysteine synthase C1 (CYSC1) | 42 | 2.33 | 0.228 | 0.488 | |
| AT3G61440.1 | Cysteine synthase C1 (CYSC1) | 43 | 2.33 | 0.229 | 0.494 | |
| AT3G61440.1 | Cysteine synthase C1 (CYSC1) | 44 | 2.04 | 0.228 | 0.676 | |
| AT3G61440.1 | Cysteine synthase C1 (CYSC1) | 45 | 1.53 | 0.231 | 0.984 | |
| AT3G61440.1 | Cysteine synthase C1 (CYSC1) | 46 | 1.02 | 0.249 | 1.000 | |
| AT3G61440.1 | Cysteine synthase C1 (CYSC1) | 47 | 0.64 | 0.304 | 0.999 | |
| AT3G61440.1 | Cysteine synthase C1 (CYSC1) | 48 | 0.41 | 0.395 | 0.919 | |
| AT3G61470.1 | Photosystem I light harvesting complex subunit (LHCA1) | 2 | 0.37 | 0.403 | 0.969 | |
| AT3G61470.1 | Photosystem I light harvesting complex subunit (LHCA1) | 3 | 0.30 | 0.306 | 0.644 | |
| AT3G61470.1 | Photosystem I light harvesting complex subunit (LHCA1) | 4 | 0.25 | 0.249 | 0.190 | |
| AT3G61470.1 | Photosystem I light harvesting complex subunit (LHCA1) | 5 | 0.22 | 0.225 | 0.051 | . |
| AT3G61470.1 | Photosystem I light harvesting complex subunit (LHCA1) | 6 | 0.20 | 0.219 | 0.026 | * |
| AT3G61470.1 | Photosystem I light harvesting complex subunit (LHCA1) | 7 | 0.20 | 0.217 | 0.024 | * |
| AT3G61470.1 | Photosystem I light harvesting complex subunit (LHCA1) | 8 | 0.22 | 0.216 | 0.037 | * |
| AT3G61470.1 | Photosystem I light harvesting complex subunit (LHCA1) | 9 | 0.25 | 0.215 | 0.078 | . |
| AT3G61470.1 | Photosystem I light harvesting complex subunit (LHCA1) | 10 | 0.30 | 0.213 | 0.182 | |
| AT3G61470.1 | Photosystem I light harvesting complex subunit (LHCA1) | 11 | 0.37 | 0.211 | 0.379 | |
| AT3G61470.1 | Photosystem I light harvesting complex subunit (LHCA1) | 12 | 0.44 | 0.210 | 0.635 | |
| AT3G61470.1 | Photosystem I light harvesting complex subunit (LHCA1) | 13 | 0.51 | 0.208 | 0.846 | |
| AT3G61470.1 | Photosystem I light harvesting complex subunit (LHCA1) | 14 | 0.58 | 0.207 | 0.953 | |
| AT3G61470.1 | Photosystem I light harvesting complex subunit (LHCA1) | 15 | 0.64 | 0.206 | 0.989 | |
| AT3G61470.1 | Photosystem I light harvesting complex subunit (LHCA1) | 16 | 0.70 | 0.205 | 0.998 | |
| AT3G61470.1 | Photosystem I light harvesting complex subunit (LHCA1) | 17 | 0.73 | 0.206 | 0.999 | |
| AT3G61470.1 | Photosystem I light harvesting complex subunit (LHCA1) | 18 | 0.75 | 0.208 | 1.000 | |
| AT3G61470.1 | Photosystem I light harvesting complex subunit (LHCA1) | 19 | 0.75 | 0.212 | 1.000 | |
| AT3G61470.1 | Photosystem I light harvesting complex subunit (LHCA1) | 20 | 0.73 | 0.214 | 1.000 | |
| AT3G61470.1 | Photosystem I light harvesting complex subunit (LHCA1) | 21 | 0.68 | 0.216 | 0.998 | |
| AT3G61470.1 | Photosystem I light harvesting complex subunit (LHCA1) | 22 | 0.62 | 0.217 | 0.986 | |
| AT3G61470.1 | Photosystem I light harvesting complex subunit (LHCA1) | 23 | 0.55 | 0.217 | 0.939 | |
| AT3G61470.1 | Photosystem I light harvesting complex subunit (LHCA1) | 24 | 0.49 | 0.218 | 0.834 | |
| AT3G61470.1 | Photosystem I light harvesting complex subunit (LHCA1) | 25 | 0.44 | 0.223 | 0.722 | |
| AT3G61470.1 | Photosystem I light harvesting complex subunit (LHCA1) | 26 | 0.43 | 0.237 | 0.744 | |

| | | | | | |
|---|---|---|---|---|---|
| AT3G61470.1 | Photosystem I light harvesting complex subunit (LHCA1) | 27 | 0.45 | 0.267 | 0.896 |
| AT3G61470.1 | Photosystem I light harvesting complex subunit (LHCA1) | 28 | 0.51 | 0.318 | 0.990 |
| AT3G61470.1 | Photosystem I light harvesting complex subunit (LHCA1) | 29 | 0.61 | 0.394 | 1.000 |
| AT3G61470.1 | Photosystem I light harvesting complex subunit (LHCA1) | 46 | 24.68 | 3.786 | 1.000 |
| AT3G61470.1 | Photosystem I light harvesting complex subunit (LHCA1) | 47 | 31.38 | 4.080 | 1.000 |
| AT3G61530.2 | Ketopentoate hydroxymethyltransferase 2 (KPHMT2) | 2 | 1.02 | 0.587 | 1.000 |
| AT3G61530.2 | Ketopentoate hydroxymethyltransferase 2 (KPHMT2) | 3 | 0.89 | 0.478 | 1.000 |
| AT3G61530.2 | Ketopentoate hydroxymethyltransferase 2 (KPHMT2) | 6 | 0.72 | 0.447 | 1.000 |
| AT3G61530.2 | Ketopentoate hydroxymethyltransferase 2 (KPHMT2) | 7 | 0.73 | 0.428 | 1.000 |
| AT3G61530.2 | Ketopentoate hydroxymethyltransferase 2 (KPHMT2) | 8 | 0.74 | 0.443 | 1.000 |
| AT3G61530.2 | Ketopentoate hydroxymethyltransferase 2 (KPHMT2) | 12 | 0.82 | 0.341 | 1.000 |
| AT3G61530.2 | Ketopentoate hydroxymethyltransferase 2 (KPHMT2) | 13 | 0.88 | 0.315 | 1.000 |
| AT3G61530.2 | Ketopentoate hydroxymethyltransferase 2 (KPHMT2) | 14 | 0.94 | 0.293 | 1.000 |
| AT3G61530.2 | Ketopentoate hydroxymethyltransferase 2 (KPHMT2) | 15 | 1.05 | 0.276 | 1.000 |
| AT3G61530.2 | Ketopentoate hydroxymethyltransferase 2 (KPHMT2) | 16 | 1.22 | 0.265 | 1.000 |
| AT3G61530.2 | Ketopentoate hydroxymethyltransferase 2 (KPHMT2) | 17 | 1.38 | 0.268 | 1.000 |
| AT3G61530.2 | Ketopentoate hydroxymethyltransferase 2 (KPHMT2) | 18 | 1.43 | 0.268 | 1.000 |
| AT3G61530.2 | Ketopentoate hydroxymethyltransferase 2 (KPHMT2) | 19 | 1.40 | 0.271 | 1.000 |
| AT3G61530.2 | Ketopentoate hydroxymethyltransferase 2 (KPHMT2) | 20 | 1.21 | 0.298 | 1.000 |
| AT3G61530.2 | Ketopentoate hydroxymethyltransferase 2 (KPHMT2) | 21 | 0.91 | 0.327 | 1.000 |
| AT3G61530.2 | Ketopentoate hydroxymethyltransferase 2 (KPHMT2) | 22 | 0.65 | 0.333 | 1.000 |
| AT3G61530.2 | Ketopentoate hydroxymethyltransferase 2 (KPHMT2) | 23 | 0.49 | 0.297 | 0.995 |

| | | | | | | |
|---|---|---|---|---|---|---|
| AT3G61530.2 | Ketopentoate hydroxymethyltransferase 2 (KPHMT2) | 24 | 0.42 | 0.268 | 0.928 | |
| AT3G61530.2 | Ketopentoate hydroxymethyltransferase 2 (KPHMT2) | 25 | 0.45 | 0.255 | 0.939 | |
| AT3G61530.2 | Ketopentoate hydroxymethyltransferase 2 (KPHMT2) | 26 | 0.73 | 0.252 | 1.000 | |
| AT3G61530.2 | Ketopentoate hydroxymethyltransferase 2 (KPHMT2) | 27 | 1.70 | 0.251 | 0.999 | |
| AT3G61530.2 | Ketopentoate hydroxymethyltransferase 2 (KPHMT2) | 28 | 3.68 | 0.253 | 0.356 | |
| AT3G61530.2 | Ketopentoate hydroxymethyltransferase 2 (KPHMT2) | 29 | 5.57 | 0.254 | 0.070 | . |
| AT3G61530.2 | Ketopentoate hydroxymethyltransferase 2 (KPHMT2) | 30 | 4.12 | 0.260 | 0.276 | |
| AT3G61530.2 | Ketopentoate hydroxymethyltransferase 2 (KPHMT2) | 31 | 1.25 | 0.278 | 1.000 | |
| AT3G61530.2 | Ketopentoate hydroxymethyltransferase 2 (KPHMT2) | 32 | 0.27 | 0.355 | 0.823 | |
| AT3G61530.2 | Ketopentoate hydroxymethyltransferase 2 (KPHMT2) | 33 | 0.07 | 0.543 | 0.473 | |
| AT3G61530.2 | Ketopentoate hydroxymethyltransferase 2 (KPHMT2) | 46 | 36.32 | 3.375 | 1.000 | |
| AT3G61530.2 | Ketopentoate hydroxymethyltransferase 2 (KPHMT2) | 47 | 139.39 | 3.722 | 1.000 | |
| AT3G61540.1 | Alpha/beta-Hydrolases superfamily protein | 2 | 1.27 | 0.756 | 1.000 | |
| AT3G61540.1 | Alpha/beta-Hydrolases superfamily protein | 3 | 1.30 | 0.727 | 1.000 | |
| AT3G61540.1 | Alpha/beta-Hydrolases superfamily protein | 29 | 1.05 | 0.293 | 1.000 | |
| AT3G61540.1 | Alpha/beta-Hydrolases superfamily protein | 30 | 0.98 | 0.286 | 1.000 | |
| AT3G61540.1 | Alpha/beta-Hydrolases superfamily protein | 31 | 0.90 | 0.285 | 1.000 | |
| AT3G61540.1 | Alpha/beta-Hydrolases superfamily protein | 32 | 0.82 | 0.288 | 0.994 | |
| AT3G61540.1 | Alpha/beta-Hydrolases superfamily protein | 33 | 0.73 | 0.296 | 0.973 | |
| AT3G61540.1 | Alpha/beta-Hydrolases superfamily protein | 34 | 0.65 | 0.307 | 0.927 | |
| AT3G61540.1 | Alpha/beta-Hydrolases superfamily protein | 46 | 0.08 | 0.524 | 0.149 | |
| AT3G61540.1 | Alpha/beta-Hydrolases superfamily protein | 47 | 0.07 | 0.553 | 0.136 | |
| AT3G61870.2 | ? | 2 | 0.89 | 0.691 | 1.000 | |
| AT3G61870.2 | ? | 3 | 0.84 | 0.950 | 1.000 | |
| AT3G61870.2 | ? | 39 | 2.69 | 0.519 | 0.938 | |
| AT3G61870.2 | ? | 40 | 2.22 | 0.357 | 0.881 | |
| AT3G61870.2 | ? | 41 | 1.20 | 0.313 | 1.000 | |
| AT3G61870.2 | ? | 42 | 0.37 | 0.404 | 0.828 | |

| | | | | | | |
|---|---|---|---|---|---|---|
| AT3G61870.2 | ? | 43 | 0.08 | 0.565 | 0.309 | |
| AT3G61870.2 | ? | 46 | 0.00 | 1.054 | 0.022 | * |
| AT3G61870.2 | ? | 47 | 0.00 | 1.297 | 0.021 | * |
| AT3G62450.1 | DNA mismatch repair protein | 2 | 1.24 | 0.461 | 1.000 | |
| AT3G62450.1 | DNA mismatch repair protein | 3 | 1.04 | 0.339 | 1.000 | |
| AT3G62450.1 | DNA mismatch repair protein | 5 | 0.85 | 0.280 | 1.000 | |
| AT3G62450.1 | DNA mismatch repair protein | 6 | 0.87 | 0.236 | 1.000 | |
| AT3G62450.1 | DNA mismatch repair protein | 7 | 0.98 | 0.214 | 1.000 | |
| AT3G62450.1 | DNA mismatch repair protein | 8 | 1.17 | 0.202 | 1.000 | |
| AT3G62450.1 | DNA mismatch repair protein | 9 | 1.30 | 0.195 | 1.000 | |
| AT3G62450.1 | DNA mismatch repair protein | 10 | 1.30 | 0.194 | 1.000 | |
| AT3G62450.1 | DNA mismatch repair protein | 11 | 1.29 | 0.195 | 1.000 | |
| AT3G62450.1 | DNA mismatch repair protein | 12 | 1.32 | 0.195 | 1.000 | |
| AT3G62450.1 | DNA mismatch repair protein | 13 | 1.34 | 0.192 | 1.000 | |
| AT3G62450.1 | DNA mismatch repair protein | 14 | 1.35 | 0.189 | 1.000 | |
| AT3G62450.1 | DNA mismatch repair protein | 15 | 1.25 | 0.189 | 1.000 | |
| AT3G62450.1 | DNA mismatch repair protein | 16 | 1.09 | 0.188 | 1.000 | |
| AT3G62450.1 | DNA mismatch repair protein | 17 | 1.05 | 0.188 | 1.000 | |
| AT3G62450.1 | DNA mismatch repair protein | 18 | 1.20 | 0.190 | 1.000 | |
| AT3G62450.1 | DNA mismatch repair protein | 19 | 1.28 | 0.195 | 1.000 | |
| AT3G62450.1 | DNA mismatch repair protein | 20 | 1.15 | 0.200 | 1.000 | |
| AT3G62450.1 | DNA mismatch repair protein | 21 | 1.01 | 0.203 | 1.000 | |
| AT3G62450.1 | DNA mismatch repair protein | 22 | 0.91 | 0.203 | 1.000 | |
| AT3G62450.1 | DNA mismatch repair protein | 23 | 0.71 | 0.202 | 1.000 | |
| AT3G62450.1 | DNA mismatch repair protein | 24 | 0.57 | 0.197 | 0.997 | |
| AT3G62450.1 | DNA mismatch repair protein | 25 | 0.59 | 0.193 | 0.998 | |
| AT3G62450.1 | DNA mismatch repair protein | 26 | 0.81 | 0.192 | 1.000 | |
| AT3G62450.1 | DNA mismatch repair protein | 27 | 1.21 | 0.192 | 1.000 | |
| AT3G62450.1 | DNA mismatch repair protein | 28 | 1.45 | 0.192 | 1.000 | |
| AT3G62450.1 | DNA mismatch repair protein | 29 | 1.43 | 0.193 | 1.000 | |
| AT3G62450.1 | DNA mismatch repair protein | 30 | 1.39 | 0.193 | 1.000 | |
| AT3G62450.1 | DNA mismatch repair protein | 31 | 1.52 | 0.193 | 1.000 | |
| AT3G62450.1 | DNA mismatch repair protein | 32 | 1.66 | 0.194 | 0.999 | |
| AT3G62450.1 | DNA mismatch repair protein | 33 | 1.71 | 0.194 | 0.998 | |
| AT3G62450.1 | DNA mismatch repair protein | 34 | 1.67 | 0.194 | 0.999 | |
| AT3G62450.1 | DNA mismatch repair protein | 35 | 1.62 | 0.194 | 1.000 | |
| AT3G62450.1 | DNA mismatch repair protein | 36 | 1.44 | 0.193 | 1.000 | |
| AT3G62450.1 | DNA mismatch repair protein | 37 | 1.05 | 0.193 | 1.000 | |
| AT3G62450.1 | DNA mismatch repair protein | 38 | 0.67 | 0.193 | 1.000 | |
| AT3G62450.1 | DNA mismatch repair protein | 39 | 0.49 | 0.194 | 0.950 | |
| AT3G62450.1 | DNA mismatch repair protein | 40 | 0.55 | 0.194 | 0.994 | |
| AT3G62450.1 | DNA mismatch repair protein | 41 | 0.87 | 0.195 | 1.000 | |
| AT3G62450.1 | DNA mismatch repair protein | 42 | 1.68 | 0.198 | 0.999 | |
| AT3G62450.1 | DNA mismatch repair protein | 43 | 3.20 | 0.199 | 0.290 | |
| AT3G62450.1 | DNA mismatch repair protein | 44 | 3.96 | 0.199 | 0.089 | . |
| AT3G62450.1 | DNA mismatch repair protein | 45 | 2.32 | 0.202 | 0.850 | |
| AT3G62450.1 | DNA mismatch repair protein | 46 | 0.65 | 0.239 | 1.000 | |
| AT3G62450.1 | DNA mismatch repair protein | 47 | 0.12 | 0.390 | 0.449 | |
| AT3G62530.1 | ARM repeat superfamily protein | 2 | 1.00 | 0.517 | 1.000 | |
| AT3G62530.1 | ARM repeat superfamily protein | 3 | 1.08 | 0.475 | 1.000 | |
| AT3G62530.1 | ARM repeat superfamily protein | 16 | 1.92 | 0.370 | 0.998 | |
| AT3G62530.1 | ARM repeat superfamily protein | 17 | 1.93 | 0.336 | 0.996 | |
| AT3G62530.1 | ARM repeat superfamily protein | 18 | 1.97 | 0.319 | 0.990 | |
| AT3G62530.1 | ARM repeat superfamily protein | 20 | 2.20 | 0.284 | 0.932 | |
| AT3G62530.1 | ARM repeat superfamily protein | 21 | 2.22 | 0.275 | 0.908 | |

| | | | | | |
|---|---|---|---|---|---|
| AT3G62530.1 | ARM repeat superfamily protein | 22 | 2.04 | 0.260 | 0.936 |
| AT3G62530.1 | ARM repeat superfamily protein | 23 | 1.70 | 0.247 | 0.990 |
| AT3G62530.1 | ARM repeat superfamily protein | 24 | 1.35 | 0.235 | 1.000 |
| AT3G62530.1 | ARM repeat superfamily protein | 25 | 1.10 | 0.227 | 1.000 |
| AT3G62530.1 | ARM repeat superfamily protein | 26 | 0.97 | 0.223 | 1.000 |
| AT3G62530.1 | ARM repeat superfamily protein | 27 | 0.91 | 0.222 | 1.000 |
| AT3G62530.1 | ARM repeat superfamily protein | 28 | 0.92 | 0.222 | 1.000 |
| AT3G62530.1 | ARM repeat superfamily protein | 29 | 1.01 | 0.222 | 1.000 |
| AT3G62530.1 | ARM repeat superfamily protein | 30 | 1.14 | 0.222 | 1.000 |
| AT3G62530.1 | ARM repeat superfamily protein | 31 | 1.30 | 0.223 | 1.000 |
| AT3G62530.1 | ARM repeat superfamily protein | 32 | 1.41 | 0.223 | 1.000 |
| AT3G62530.1 | ARM repeat superfamily protein | 33 | 1.49 | 0.223 | 0.998 |
| AT3G62530.1 | ARM repeat superfamily protein | 34 | 1.58 | 0.223 | 0.993 |
| AT3G62530.1 | ARM repeat superfamily protein | 35 | 1.60 | 0.223 | 0.991 |
| AT3G62530.1 | ARM repeat superfamily protein | 36 | 1.46 | 0.223 | 0.999 |
| AT3G62530.1 | ARM repeat superfamily protein | 37 | 1.23 | 0.223 | 1.000 |
| AT3G62530.1 | ARM repeat superfamily protein | 38 | 1.13 | 0.223 | 1.000 |
| AT3G62530.1 | ARM repeat superfamily protein | 39 | 1.24 | 0.223 | 1.000 |
| AT3G62530.1 | ARM repeat superfamily protein | 40 | 1.44 | 0.224 | 0.999 |
| AT3G62530.1 | ARM repeat superfamily protein | 41 | 1.48 | 0.225 | 0.999 |
| AT3G62530.1 | ARM repeat superfamily protein | 42 | 1.31 | 0.226 | 1.000 |
| AT3G62530.1 | ARM repeat superfamily protein | 43 | 1.04 | 0.227 | 1.000 |
| AT3G62530.1 | ARM repeat superfamily protein | 44 | 0.77 | 0.228 | 1.000 |
| AT3G62530.1 | ARM repeat superfamily protein | 45 | 0.51 | 0.238 | 0.914 |
| AT3G62530.1 | ARM repeat superfamily protein | 46 | 0.30 | 0.276 | 0.472 |
| AT3G62530.1 | ARM repeat superfamily protein | 47 | 0.16 | 0.367 | 0.320 |
| AT3G62790.1 | NADH:ubiquinone oxidoreductase subunit (15-1) | 2 | 0.96 | 0.505 | 1.000 |
| AT3G62790.1 | NADH:ubiquinone oxidoreductase subunit (15-1) | 3 | 0.73 | 0.462 | 1.000 |
| AT3G62790.1 | NADH:ubiquinone oxidoreductase subunit (15-1) | 14 | 1.05 | 0.308 | 1.000 |
| AT3G62790.1 | NADH:ubiquinone oxidoreductase subunit (15-1) | 15 | 1.17 | 0.269 | 1.000 |
| AT3G62790.1 | NADH:ubiquinone oxidoreductase subunit (15-1) | 16 | 0.84 | 0.249 | 1.000 |

| | | | | | | |
|---|---|---|---|---|---|---|
| AT3G62790.1 | NADH:ubiquinone oxidoreductase subunit (15-1) | 17 | 0.44 | 0.224 | 0.791 | |
| AT3G62790.1 | NADH:ubiquinone oxidoreductase subunit (15-1) | 18 | 0.38 | 0.218 | 0.562 | |
| AT3G62790.1 | NADH:ubiquinone oxidoreductase subunit (15-1) | 19 | 0.52 | 0.226 | 0.958 | |
| AT3G62790.1 | NADH:ubiquinone oxidoreductase subunit (15-1) | 20 | 0.70 | 0.232 | 1.000 | |
| AT3G62790.1 | NADH:ubiquinone oxidoreductase subunit (15-1) | 21 | 0.95 | 0.234 | 1.000 | |
| AT3G62790.1 | NADH:ubiquinone oxidoreductase subunit (15-1) | 22 | 1.40 | 0.233 | 1.000 | |
| AT3G62790.1 | NADH:ubiquinone oxidoreductase subunit (15-1) | 23 | 1.79 | 0.232 | 0.985 | |
| AT3G62790.1 | NADH:ubiquinone oxidoreductase subunit (15-1) | 24 | 1.36 | 0.225 | 1.000 | |
| AT3G62790.1 | NADH:ubiquinone oxidoreductase subunit (15-1) | 25 | 0.72 | 0.223 | 1.000 | |
| AT3G62790.1 | NADH:ubiquinone oxidoreductase subunit (15-1) | 26 | 0.67 | 0.223 | 0.999 | |
| AT3G62790.1 | NADH:ubiquinone oxidoreductase subunit (15-1) | 27 | 1.34 | 0.227 | 1.000 | |
| AT3G62790.1 | NADH:ubiquinone oxidoreductase subunit (15-1) | 28 | 2.19 | 0.242 | 0.899 | |
| AT3G62790.1 | NADH:ubiquinone oxidoreductase subunit (15-1) | 29 | 1.73 | 0.305 | 0.999 | |
| AT3G62790.1 | NADH:ubiquinone oxidoreductase subunit (15-1) | 30 | 0.97 | 0.500 | 1.000 | |
| AT3G62790.1 | NADH:ubiquinone oxidoreductase subunit (15-1) | 46 | 299.03 | 6.605 | 1.000 | |
| AT3G62790.1 | NADH:ubiquinone oxidoreductase subunit (15-1) | 47 | 693.23 | 7.198 | 1.000 | |
| AT3G62880.2 | Homologous to pea OEP16 and barley pPORA (OEP16) | 2 | 1.04 | 0.519 | 1.000 | |
| AT3G62880.2 | Homologous to pea OEP16 and barley pPORA (OEP16) | 3 | 1.00 | 0.463 | 1.000 | |
| AT3G62880.2 | Homologous to pea OEP16 and barley pPORA (OEP16) | 11 | 1.07 | 0.223 | 1.000 | |
| AT3G62880.2 | Homologous to pea OEP16 and barley pPORA (OEP16) | 12 | 1.21 | 0.198 | 1.000 | |
| AT3G62880.2 | Homologous to pea OEP16 and barley pPORA (OEP16) | 13 | 1.41 | 0.180 | 0.976 | |
| AT3G62880.2 | Homologous to pea OEP16 and barley pPORA (OEP16) | 14 | 1.70 | 0.169 | 1.000 | |
| AT3G62880.2 | Homologous to pea OEP16 and barley pPORA (OEP16) | 15 | 2.09 | 0.163 | 0.339 | |
| AT3G62880.2 | Homologous to pea OEP16 and barley pPORA (OEP16) | 16 | 2.58 | 0.165 | 0.111 | |
| AT3G62880.2 | Homologous to pea OEP16 and barley pPORA (OEP16) | 17 | 3.20 | 0.174 | 0.040 | * |
| AT3G62880.2 | Homologous to pea OEP16 and barley pPORA (OEP16) | 18 | 4.00 | 0.195 | 0.023 | * |
| AT3G62880.2 | Homologous to pea OEP16 and barley pPORA (OEP16) | 19 | 4.99 | 0.227 | 0.024 | * |
| AT3G62880.2 | Homologous to pea OEP16 and barley pPORA (OEP16) | 20 | 6.17 | 0.268 | 0.034 | * |
| AT3G62880.2 | Homologous to pea OEP16 and barley pPORA (OEP16) | 38 | 1.88 | 0.253 | 1.000 | |
| AT3G62880.2 | Homologous to pea OEP16 and barley pPORA (OEP16) | 39 | 1.15 | 0.227 | 1.000 | |
| AT3G62880.2 | Homologous to pea OEP16 and barley pPORA (OEP16) | 40 | 0.64 | 0.217 | 0.965 | |

| | | | | | | |
|---|---|---|---|---|---|---|
| AT3G62880.2 | Homologous to pea OEP16 and barley pPORA (OEP16) | 41 | 0.33 | 0.222 | 0.237 | |
| AT3G62880.2 | Homologous to pea OEP16 and barley pPORA (OEP16) | 42 | 0.17 | 0.238 | 0.015 | * |
| AT3G62880.2 | Homologous to pea OEP16 and barley pPORA (OEP16) | 43 | 0.09 | 0.258 | 0.001 | *** |
| AT3G62880.2 | Homologous to pea OEP16 and barley pPORA (OEP16) | 46 | 0.01 | 0.344 | 0.000 | *** |
| AT3G62880.2 | Homologous to pea OEP16 and barley pPORA (OEP16) | 47 | 0.01 | 0.396 | 0.000 | *** |
| AT3G63150.1 | Calcium binding GTPase | 2 | 1.00 | 0.421 | 1.000 | |
| AT3G63150.1 | Calcium binding GTPase | 3 | 0.89 | 0.377 | 1.000 | |
| AT3G63150.1 | Calcium binding GTPase | 14 | 1.01 | 0.233 | 1.000 | |
| AT3G63150.1 | Calcium binding GTPase | 15 | 1.28 | 0.202 | 1.000 | |
| AT3G63150.1 | Calcium binding GTPase | 16 | 1.65 | 0.180 | 0.982 | |
| AT3G63150.1 | Calcium binding GTPase | 17 | 2.02 | 0.167 | 0.695 | |
| AT3G63150.1 | Calcium binding GTPase | 18 | 2.25 | 0.162 | 0.427 | |
| AT3G63150.1 | Calcium binding GTPase | 19 | 2.24 | 0.164 | 0.452 | |
| AT3G63150.1 | Calcium binding GTPase | 20 | 2.01 | 0.166 | 0.700 | |
| AT3G63150.1 | Calcium binding GTPase | 21 | 1.68 | 0.167 | 0.956 | |
| AT3G63150.1 | Calcium binding GTPase | 22 | 1.41 | 0.168 | 0.999 | |
| AT3G63150.1 | Calcium binding GTPase | 23 | 1.26 | 0.166 | 1.000 | |
| AT3G63150.1 | Calcium binding GTPase | 24 | 1.20 | 0.164 | 1.000 | |
| AT3G63150.1 | Calcium binding GTPase | 25 | 1.17 | 0.162 | 1.000 | |
| AT3G63150.1 | Calcium binding GTPase | 26 | 1.16 | 0.161 | 1.000 | |
| AT3G63150.1 | Calcium binding GTPase | 27 | 1.13 | 0.161 | 1.000 | |
| AT3G63150.1 | Calcium binding GTPase | 28 | 1.06 | 0.161 | 1.000 | |
| AT3G63150.1 | Calcium binding GTPase | 29 | 0.96 | 0.161 | 1.000 | |
| AT3G63150.1 | Calcium binding GTPase | 30 | 0.85 | 0.161 | 1.000 | |
| AT3G63150.1 | Calcium binding GTPase | 31 | 0.74 | 0.161 | 1.000 | |
| AT3G63150.1 | Calcium binding GTPase | 32 | 0.64 | 0.162 | 0.984 | |
| AT3G63150.1 | Calcium binding GTPase | 33 | 0.55 | 0.162 | 0.845 | |
| AT3G63150.1 | Calcium binding GTPase | 34 | 0.47 | 0.162 | 0.560 | |
| AT3G63150.1 | Calcium binding GTPase | 35 | 0.43 | 0.162 | 0.375 | |
| AT3G63150.1 | Calcium binding GTPase | 36 | 0.45 | 0.162 | 0.458 | |
| AT3G63150.1 | Calcium binding GTPase | 37 | 0.56 | 0.162 | 0.883 | |
| AT3G63150.1 | Calcium binding GTPase | 38 | 0.77 | 0.162 | 1.000 | |
| AT3G63150.1 | Calcium binding GTPase | 39 | 1.04 | 0.162 | 1.000 | |
| AT3G63150.1 | Calcium binding GTPase | 40 | 1.26 | 0.163 | 1.000 | |
| AT3G63150.1 | Calcium binding GTPase | 41 | 1.33 | 0.165 | 1.000 | |
| AT3G63150.1 | Calcium binding GTPase | 42 | 1.29 | 0.168 | 1.000 | |
| AT3G63150.1 | Calcium binding GTPase | 43 | 1.22 | 0.180 | 1.000 | |
| AT3G63150.1 | Calcium binding GTPase | 44 | 1.11 | 0.216 | 1.000 | |
| AT3G63150.1 | Calcium binding GTPase | 45 | 0.98 | 0.285 | 1.000 | |
| AT3G63150.1 | Calcium binding GTPase | 46 | 0.84 | 0.402 | 1.000 | |
| AT3G63150.1 | Calcium binding GTPase | 47 | 0.72 | 0.562 | 1.000 | |
| AT3G63160.1 | Outer envelope protein 6 (OEP6) | 2 | 0.60 | 0.313 | 1.000 | |
| AT3G63160.1 | Outer envelope protein 6 (OEP6) | 3 | 0.54 | 0.244 | 0.977 | |
| AT3G63160.1 | Outer envelope protein 6 (OEP6) | 4 | 0.49 | 0.207 | 0.825 | |
| AT3G63160.1 | Outer envelope protein 6 (OEP6) | 5 | 0.46 | 0.193 | 0.653 | |
| AT3G63160.1 | Outer envelope protein 6 (OEP6) | 6 | 0.45 | 0.190 | 0.601 | |
| AT3G63160.1 | Outer envelope protein 6 (OEP6) | 7 | 0.46 | 0.189 | 0.633 | |
| AT3G63160.1 | Outer envelope protein 6 (OEP6) | 8 | 0.48 | 0.188 | 0.709 | |
| AT3G63160.1 | Outer envelope protein 6 (OEP6) | 9 | 0.52 | 0.187 | 0.803 | |

| | | | | | | |
|---|---|---|---|---|---|---|
| AT3G63160.1 | Outer envelope protein 6 (OEP6) | 10 | 0.56 | 0.186 | 0.900 | |
| AT3G63160.1 | Outer envelope protein 6 (OEP6) | 11 | 0.63 | 0.185 | 0.981 | |
| AT3G63160.1 | Outer envelope protein 6 (OEP6) | 12 | 0.75 | 0.184 | 1.000 | |
| AT3G63160.1 | Outer envelope protein 6 (OEP6) | 13 | 0.91 | 0.183 | 1.000 | |
| AT3G63160.1 | Outer envelope protein 6 (OEP6) | 14 | 1.17 | 0.183 | 1.000 | |
| AT3G63160.1 | Outer envelope protein 6 (OEP6) | 15 | 1.55 | 0.182 | 0.985 | |
| AT3G63160.1 | Outer envelope protein 6 (OEP6) | 16 | 2.04 | 0.182 | 0.685 | |
| AT3G63160.1 | Outer envelope protein 6 (OEP6) | 17 | 2.65 | 0.183 | 0.265 | |
| AT3G63160.1 | Outer envelope protein 6 (OEP6) | 18 | 3.28 | 0.186 | 0.090 | . |
| AT3G63160.1 | Outer envelope protein 6 (OEP6) | 19 | 3.72 | 0.191 | 0.049 | * |
| AT3G63160.1 | Outer envelope protein 6 (OEP6) | 20 | 3.77 | 0.196 | 0.057 | . |
| AT3G63160.1 | Outer envelope protein 6 (OEP6) | 21 | 3.42 | 0.203 | 0.126 | |
| AT3G63160.1 | Outer envelope protein 6 (OEP6) | 22 | 2.83 | 0.209 | 0.349 | |
| AT3G63160.1 | Outer envelope protein 6 (OEP6) | 23 | 2.21 | 0.215 | 0.763 | |
| AT3G63160.1 | Outer envelope protein 6 (OEP6) | 24 | 1.66 | 0.220 | 0.989 | |
| AT3G63160.1 | Outer envelope protein 6 (OEP6) | 25 | 1.23 | 0.222 | 1.000 | |
| AT3G63160.1 | Outer envelope protein 6 (OEP6) | 27 | 0.72 | 0.218 | 1.000 | |
| AT3G63160.1 | Outer envelope protein 6 (OEP6) | 28 | 0.61 | 0.210 | 0.987 | |
| AT3G63160.1 | Outer envelope protein 6 (OEP6) | 29 | 0.57 | 0.201 | 0.954 | |
| AT3G63160.1 | Outer envelope protein 6 (OEP6) | 30 | 0.61 | 0.195 | 0.974 | |
| AT3G63160.1 | Outer envelope protein 6 (OEP6) | 31 | 0.70 | 0.194 | 0.999 | |
| AT3G63160.1 | Outer envelope protein 6 (OEP6) | 32 | 0.88 | 0.199 | 1.000 | |
| AT3G63160.1 | Outer envelope protein 6 (OEP6) | 33 | 1.13 | 0.212 | 1.000 | |
| AT3G63160.1 | Outer envelope protein 6 (OEP6) | 34 | 1.49 | 0.232 | 0.999 | |
| AT3G63160.1 | Outer envelope protein 6 (OEP6) | 35 | 1.93 | 0.253 | 0.971 | |
| AT3G63160.1 | Outer envelope protein 6 (OEP6) | 41 | 5.82 | 0.270 | 0.076 | . |
| AT3G63160.1 | Outer envelope protein 6 (OEP6) | 42 | 6.22 | 0.242 | 0.021 | * |
| AT3G63160.1 | Outer envelope protein 6 (OEP6) | 43 | 6.39 | 0.217 | 0.005 | ** |
| AT3G63160.1 | Outer envelope protein 6 (OEP6) | 44 | 6.26 | 0.204 | 0.003 | ** |
| AT3G63160.1 | Outer envelope protein 6 (OEP6) | 45 | 5.79 | 0.213 | 0.008 | ** |
| AT3G63160.1 | Outer envelope protein 6 (OEP6) | 46 | 5.10 | 0.258 | 0.097 | . |

| | | | | | | |
|---|---|---|---|---|---|---|
| AT3G63160.1 | Outer envelope protein 6 (OEP6) | 47 | 4.43 | 0.342 | 0.543 | |
| AT3G63520.1 | Carotenoid cleavage dioxygenase 1 (CCD1) | 2 | 1.33 | 0.368 | 1.000 | |
| AT3G63520.1 | Carotenoid cleavage dioxygenase 1 (CCD1) | 3 | 0.97 | 0.359 | 1.000 | |
| AT3G63520.1 | Carotenoid cleavage dioxygenase 1 (CCD1) | 39 | 2.84 | 0.343 | 0.823 | |
| AT3G63520.1 | Carotenoid cleavage dioxygenase 1 (CCD1) | 40 | 5.43 | 0.244 | 0.035 | * |
| AT3G63520.1 | Carotenoid cleavage dioxygenase 1 (CCD1) | 41 | 8.95 | 0.215 | 0.000 | *** |
| AT3G63520.1 | Carotenoid cleavage dioxygenase 1 (CCD1) | 42 | 23.42 | 0.217 | 0.000 | *** |
| AT3G63520.1 | Carotenoid cleavage dioxygenase 1 (CCD1) | 43 | 24.21 | 0.217 | 0.000 | *** |
| AT3G63520.1 | Carotenoid cleavage dioxygenase 1 (CCD1) | 44 | 5.11 | 0.211 | 0.013 | * |
| AT3G63520.1 | Carotenoid cleavage dioxygenase 1 (CCD1) | 45 | 0.87 | 0.213 | 1.000 | |
| AT3G63520.1 | Carotenoid cleavage dioxygenase 1 (CCD1) | 46 | 0.19 | 0.222 | 0.017 | * |
| AT3G63520.1 | Carotenoid cleavage dioxygenase 1 (CCD1) | 47 | 0.02 | 0.492 | 0.017 | * |
| AT4G00026.1 | Translocase of the inner mitochondrial membrane 21 (TIM21, putative) | 2 | 1.08 | 0.349 | 1.000 | |
| AT4G00026.1 | Translocase of the inner mitochondrial membrane 21 (TIM21, putative) | 3 | 1.12 | 0.323 | 1.000 | |
| AT4G00026.1 | Translocase of the inner mitochondrial membrane 21 (TIM21, putative) | 10 | 1.38 | 0.197 | 0.951 | |
| AT4G00026.1 | Translocase of the inner mitochondrial membrane 21 (TIM21, putative) | 11 | 1.41 | 0.187 | 0.918 | |
| AT4G00026.1 | Translocase of the inner mitochondrial membrane 21 (TIM21, putative) | 12 | 1.45 | 0.180 | 0.879 | |
| AT4G00026.1 | Translocase of the inner mitochondrial membrane 21 (TIM21, putative) | 13 | 1.48 | 0.175 | 0.835 | |
| AT4G00026.1 | Translocase of the inner mitochondrial membrane 21 (TIM21, putative) | 14 | 1.52 | 0.171 | 0.794 | |
| AT4G00026.1 | Translocase of the inner mitochondrial membrane 21 (TIM21, putative) | 16 | 1.56 | 0.170 | 0.739 | |
| AT4G00026.1 | Translocase of the inner mitochondrial membrane 21 (TIM21, putative) | 17 | 1.58 | 0.170 | 0.726 | |
| AT4G00026.1 | Translocase of the inner mitochondrial membrane 21 (TIM21, putative) | 18 | 1.58 | 0.171 | 0.721 | |
| AT4G00026.1 | Translocase of the inner mitochondrial membrane 21 (TIM21, putative) | 31 | 1.20 | 0.155 | 0.989 | |
| AT4G00026.1 | Translocase of the inner mitochondrial membrane 21 (TIM21, putative) | 32 | 1.14 | 0.153 | 0.998 | |

| | | | | | |
|---|---|---|---|---|---|
| AT4G00026.1 | Translocase of the inner mitochondrial membrane 21 (TIM21, putative) | 33 | 1.08 | 0.153 | 1.000 |
| AT4G00026.1 | Translocase of the inner mitochondrial membrane 21 (TIM21, putative) | 34 | 1.02 | 0.154 | 1.000 |
| AT4G00026.1 | Translocase of the inner mitochondrial membrane 21 (TIM21, putative) | 35 | 0.95 | 0.158 | 1.000 |
| AT4G00026.1 | Translocase of the inner mitochondrial membrane 21 (TIM21, putative) | 36 | 0.88 | 0.163 | 0.999 |
| AT4G00026.1 | Translocase of the inner mitochondrial membrane 21 (TIM21, putative) | 37 | 0.81 | 0.170 | 0.985 |
| AT4G00026.1 | Translocase of the inner mitochondrial membrane 21 (TIM21, putative) | 38 | 0.74 | 0.178 | 0.943 |
| AT4G00026.1 | Translocase of the inner mitochondrial membrane 21 (TIM21, putative) | 46 | 0.30 | 0.287 | 0.293 |
| AT4G00026.1 | Translocase of the inner mitochondrial membrane 21 (TIM21, putative) | 47 | 0.26 | 0.307 | 0.266 |
| AT4G00100.1 | 40S ribosomal protein S13-2 (RPS13B) | 2 | 0.66 | 0.234 | 0.953 |
| AT4G00100.1 | 40S ribosomal protein S13-2 (RPS13B) | 3 | 0.71 | 0.202 | 0.960 |
| AT4G00100.1 | 40S ribosomal protein S13-2 (RPS13B) | 4 | 0.76 | 0.178 | 0.976 |
| AT4G00100.1 | 40S ribosomal protein S13-2 (RPS13B) | 5 | 0.82 | 0.161 | 0.992 |
| AT4G00100.1 | 40S ribosomal protein S13-2 (RPS13B) | 6 | 0.89 | 0.150 | 0.999 |
| AT4G00100.1 | 40S ribosomal protein S13-2 (RPS13B) | 7 | 0.96 | 0.144 | 1.000 |
| AT4G00100.1 | 40S ribosomal protein S13-2 (RPS13B) | 8 | 1.04 | 0.141 | 1.000 |
| AT4G00100.1 | 40S ribosomal protein S13-2 (RPS13B) | 9 | 1.12 | 0.140 | 0.999 |
| AT4G00100.1 | 40S ribosomal protein S13-2 (RPS13B) | 10 | 1.20 | 0.141 | 0.988 |
| AT4G00100.1 | 40S ribosomal protein S13-2 (RPS13B) | 11 | 1.29 | 0.144 | 0.951 |
| AT4G00100.1 | 40S ribosomal protein S13-2 (RPS13B) | 12 | 1.38 | 0.149 | 0.888 |
| AT4G00100.1 | 40S ribosomal protein S13-2 (RPS13B) | 13 | 1.47 | 0.156 | 0.823 |
| AT4G00100.1 | 40S ribosomal protein S13-2 (RPS13B) | 14 | 1.55 | 0.165 | 0.779 |
| AT4G00100.1 | 40S ribosomal protein S13-2 (RPS13B) | 15 | 1.61 | 0.178 | 0.763 |
| AT4G00100.1 | 40S ribosomal protein S13-2 (RPS13B) | 16 | 1.67 | 0.193 | 0.775 |
| AT4G00100.1 | 40S ribosomal protein S13-2 (RPS13B) | 17 | 1.71 | 0.211 | 0.803 |
| AT4G00100.1 | 40S ribosomal protein S13-2 (RPS13B) | 41 | 1.16 | 0.364 | 1.000 |
| AT4G00100.1 | 40S ribosomal protein S13-2 (RPS13B) | 42 | 1.12 | 0.372 | 1.000 |
| AT4G00100.1 | 40S ribosomal protein S13-2 (RPS13B) | 43 | 1.09 | 0.388 | 1.000 |

| | | | | | | |
|---|---|---|---|---|---|---|
| AT4G00100.1 | 40S ribosomal protein S13-2 (RPS13B) | 46 | 0.99 | 0.489 | 1.000 | |
| AT4G00100.1 | 40S ribosomal protein S13-2 (RPS13B) | 47 | 0.96 | 0.538 | 1.000 | |
| AT4G00290.1 | Mechanosensitive ion channel 1 (MSL1) | 2 | 0.98 | 0.808 | 1.000 | |
| AT4G00290.1 | Mechanosensitive ion channel 1 (MSL1) | 3 | 0.37 | 1.303 | 1.000 | |
| AT4G00290.1 | Mechanosensitive ion channel 1 (MSL1) | 25 | 0.02 | 0.597 | 0.062 | . |
| AT4G00290.1 | Mechanosensitive ion channel 1 (MSL1) | 26 | 0.13 | 0.400 | 0.298 | |
| AT4G00290.1 | Mechanosensitive ion channel 1 (MSL1) | 27 | 1.01 | 0.359 | 1.000 | |
| AT4G00290.1 | Mechanosensitive ion channel 1 (MSL1) | 28 | 4.59 | 0.343 | 0.498 | |
| AT4G00290.1 | Mechanosensitive ion channel 1 (MSL1) | 29 | 5.85 | 0.341 | 0.287 | |
| AT4G00290.1 | Mechanosensitive ion channel 1 (MSL1) | 30 | 1.64 | 0.345 | 1.000 | |
| AT4G00290.1 | Mechanosensitive ion channel 1 (MSL1) | 31 | 0.27 | 0.372 | 0.779 | |
| AT4G00290.1 | Mechanosensitive ion channel 1 (MSL1) | 32 | 0.10 | 0.474 | 0.371 | |
| AT4G00290.1 | Mechanosensitive ion channel 1 (MSL1) | 33 | 0.08 | 0.762 | 0.828 | |
| AT4G00290.1 | Mechanosensitive ion channel 1 (MSL1) | 42 | 3.66 | 0.664 | 0.996 | |
| AT4G00290.1 | Mechanosensitive ion channel 1 (MSL1) | 43 | 2.42 | 0.443 | 0.996 | |
| AT4G00290.1 | Mechanosensitive ion channel 1 (MSL1) | 44 | 1.16 | 0.497 | 1.000 | |
| AT4G00290.1 | Mechanosensitive ion channel 1 (MSL1) | 45 | 0.57 | 0.727 | 1.000 | |
| AT4G00290.1 | Mechanosensitive ion channel 1 (MSL1) | 46 | 0.30 | 0.995 | 1.000 | |
| AT4G00290.1 | Mechanosensitive ion channel 1 (MSL1) | 47 | 0.16 | 1.461 | 1.000 | |
| AT4G00570.1 | NAD malic enzyme 2 (NAD-ME2) | 2 | 0.98 | 0.315 | 1.000 | |
| AT4G00570.1 | NAD malic enzyme 2 (NAD-ME2) | 3 | 0.78 | 0.242 | 1.000 | |
| AT4G00570.1 | NAD malic enzyme 2 (NAD-ME2) | 4 | 0.65 | 0.239 | 1.000 | |
| AT4G00570.1 | NAD malic enzyme 2 (NAD-ME2) | 7 | 0.56 | 0.232 | 0.998 | |
| AT4G00570.1 | NAD malic enzyme 2 (NAD-ME2) | 8 | 0.60 | 0.204 | 0.998 | |
| AT4G00570.1 | NAD malic enzyme 2 (NAD-ME2) | 9 | 0.65 | 0.187 | 1.000 | |
| AT4G00570.1 | NAD malic enzyme 2 (NAD-ME2) | 10 | 0.72 | 0.177 | 1.000 | |
| AT4G00570.1 | NAD malic enzyme 2 (NAD-ME2) | 11 | 0.83 | 0.171 | 1.000 | |
| AT4G00570.1 | NAD malic enzyme 2 (NAD-ME2) | 12 | 0.99 | 0.169 | 1.000 | |
| AT4G00570.1 | NAD malic enzyme 2 (NAD-ME2) | 13 | 1.13 | 0.167 | 1.000 | |
| AT4G00570.1 | NAD malic enzyme 2 (NAD-ME2) | 14 | 1.21 | 0.164 | 1.000 | |
| AT4G00570.1 | NAD malic enzyme 2 (NAD-ME2) | 15 | 1.18 | 0.159 | 1.000 | |
| AT4G00570.1 | NAD malic enzyme 2 (NAD-ME2) | 16 | 1.11 | 0.157 | 1.000 | |
| AT4G00570.1 | NAD malic enzyme 2 (NAD-ME2) | 17 | 1.08 | 0.156 | 1.000 | |
| AT4G00570.1 | NAD malic enzyme 2 (NAD-ME2) | 18 | 1.08 | 0.158 | 1.000 | |
| AT4G00570.1 | NAD malic enzyme 2 (NAD-ME2) | 19 | 1.17 | 0.161 | 1.000 | |
| AT4G00570.1 | NAD malic enzyme 2 (NAD-ME2) | 20 | 1.32 | 0.164 | 1.000 | |
| AT4G00570.1 | NAD malic enzyme 2 (NAD-ME2) | 21 | 1.39 | 0.166 | 1.000 | |
| AT4G00570.1 | NAD malic enzyme 2 (NAD-ME2) | 22 | 1.32 | 0.166 | 1.000 | |
| AT4G00570.1 | NAD malic enzyme 2 (NAD-ME2) | 23 | 1.22 | 0.165 | 1.000 | |
| AT4G00570.1 | NAD malic enzyme 2 (NAD-ME2) | 24 | 1.14 | 0.162 | 1.000 | |
| AT4G00570.1 | NAD malic enzyme 2 (NAD-ME2) | 25 | 1.02 | 0.160 | 1.000 | |
| AT4G00570.1 | NAD malic enzyme 2 (NAD-ME2) | 26 | 0.97 | 0.159 | 1.000 | |
| AT4G00570.1 | NAD malic enzyme 2 (NAD-ME2) | 27 | 1.04 | 0.159 | 1.000 | |

| | | | | | |
|---|---|---|---|---|---|
| AT4G00570.1 | NAD malic enzyme 2 (NAD-ME2) | 28 | 1.13 | 0.160 | 1.000 |
| AT4G00570.1 | NAD malic enzyme 2 (NAD-ME2) | 29 | 1.18 | 0.160 | 1.000 |
| AT4G00570.1 | NAD malic enzyme 2 (NAD-ME2) | 30 | 1.15 | 0.160 | 1.000 |
| AT4G00570.1 | NAD malic enzyme 2 (NAD-ME2) | 31 | 1.06 | 0.161 | 1.000 |
| AT4G00570.1 | NAD malic enzyme 2 (NAD-ME2) | 32 | 0.96 | 0.161 | 1.000 |
| AT4G00570.1 | NAD malic enzyme 2 (NAD-ME2) | 33 | 0.90 | 0.161 | 1.000 |
| AT4G00570.1 | NAD malic enzyme 2 (NAD-ME2) | 34 | 0.88 | 0.161 | 1.000 |
| AT4G00570.1 | NAD malic enzyme 2 (NAD-ME2) | 35 | 0.92 | 0.161 | 1.000 |
| AT4G00570.1 | NAD malic enzyme 2 (NAD-ME2) | 36 | 1.02 | 0.161 | 1.000 |
| AT4G00570.1 | NAD malic enzyme 2 (NAD-ME2) | 37 | 1.16 | 0.160 | 1.000 |
| AT4G00570.1 | NAD malic enzyme 2 (NAD-ME2) | 38 | 1.31 | 0.160 | 1.000 |
| AT4G00570.1 | NAD malic enzyme 2 (NAD-ME2) | 39 | 1.44 | 0.160 | 1.000 |
| AT4G00570.1 | NAD malic enzyme 2 (NAD-ME2) | 40 | 1.48 | 0.161 | 0.999 |
| AT4G00570.1 | NAD malic enzyme 2 (NAD-ME2) | 41 | 1.34 | 0.162 | 1.000 |
| AT4G00570.1 | NAD malic enzyme 2 (NAD-ME2) | 42 | 1.07 | 0.164 | 1.000 |
| AT4G00570.1 | NAD malic enzyme 2 (NAD-ME2) | 43 | 0.82 | 0.165 | 1.000 |
| AT4G00570.1 | NAD malic enzyme 2 (NAD-ME2) | 44 | 0.65 | 0.165 | 0.996 |
| AT4G00570.1 | NAD malic enzyme 2 (NAD-ME2) | 45 | 0.53 | 0.168 | 0.877 |
| AT4G00570.1 | NAD malic enzyme 2 (NAD-ME2) | 46 | 0.43 | 0.205 | 0.809 |
| AT4G00570.1 | NAD malic enzyme 2 (NAD-ME2) | 47 | 0.36 | 0.315 | 0.963 |
| AT4G00585.1 | NADH:ubiquinone oxidoreductase subunit (KFYI) | 2 | 0.89 | 0.611 | 1.000 |
| AT4G00585.1 | NADH:ubiquinone oxidoreductase subunit (KFYI) | 3 | 0.84 | 0.545 | 1.000 |
| AT4G00585.1 | NADH:ubiquinone oxidoreductase subunit (KFYI) | 13 | 1.04 | 0.353 | 1.000 |
| AT4G00585.1 | NADH:ubiquinone oxidoreductase subunit (KFYI) | 14 | 1.14 | 0.309 | 1.000 |
| AT4G00585.1 | NADH:ubiquinone oxidoreductase subunit (KFYI) | 15 | 1.09 | 0.284 | 1.000 |
| AT4G00585.1 | NADH:ubiquinone oxidoreductase subunit (KFYI) | 16 | 0.80 | 0.256 | 1.000 |
| AT4G00585.1 | NADH:ubiquinone oxidoreductase subunit (KFYI) | 17 | 0.52 | 0.242 | 0.991 |
| AT4G00585.1 | NADH:ubiquinone oxidoreductase subunit (KFYI) | 18 | 0.47 | 0.243 | 0.959 |
| AT4G00585.1 | NADH:ubiquinone oxidoreductase subunit (KFYI) | 19 | 0.59 | 0.251 | 0.999 |
| AT4G00585.1 | NADH:ubiquinone oxidoreductase subunit (KFYI) | 20 | 0.76 | 0.258 | 1.000 |
| AT4G00585.1 | NADH:ubiquinone oxidoreductase subunit (KFYI) | 21 | 1.04 | 0.261 | 1.000 |
| AT4G00585.1 | NADH:ubiquinone oxidoreductase subunit (KFYI) | 22 | 1.46 | 0.260 | 1.000 |
| AT4G00585.1 | NADH:ubiquinone oxidoreductase subunit (KFYI) | 23 | 1.73 | 0.258 | 0.999 |
| AT4G00585.1 | NADH:ubiquinone oxidoreductase subunit (KFYI) | 24 | 1.46 | 0.251 | 1.000 |
| AT4G00585.1 | NADH:ubiquinone oxidoreductase subunit (KFYI) | 25 | 0.94 | 0.248 | 1.000 |
| AT4G00585.1 | NADH:ubiquinone oxidoreductase subunit (KFYI) | 26 | 0.66 | 0.247 | 1.000 |
| AT4G00585.1 | NADH:ubiquinone oxidoreductase subunit (KFYI) | 27 | 0.60 | 0.247 | 1.000 |
| AT4G00585.1 | NADH:ubiquinone oxidoreductase subunit (KFYI) | 28 | 0.74 | 0.246 | 1.000 |
| AT4G00585.1 | NADH:ubiquinone oxidoreductase subunit (KFYI) | 29 | 1.04 | 0.248 | 1.000 |
| AT4G00585.1 | NADH:ubiquinone oxidoreductase subunit (KFYI) | 30 | 1.17 | 0.248 | 1.000 |
| AT4G00585.1 | NADH:ubiquinone oxidoreductase subunit (KFYI) | 31 | 0.97 | 0.250 | 1.000 |

| | | | | | |
|---|---|---|---|---|---|
| AT4G00585.1 | NADH:ubiquinone oxidoreductase subunit (KFYI) | 32 | 0.93 | 0.255 | 1.000 |
| AT4G00585.1 | NADH:ubiquinone oxidoreductase subunit (KFYI) | 33 | 1.22 | 0.266 | 1.000 |
| AT4G00585.1 | NADH:ubiquinone oxidoreductase subunit (KFYI) | 34 | 1.46 | 0.285 | 1.000 |
| AT4G00585.1 | NADH:ubiquinone oxidoreductase subunit (KFYI) | 35 | 1.22 | 0.320 | 1.000 |
| AT4G00585.1 | NADH:ubiquinone oxidoreductase subunit (KFYI) | 36 | 0.83 | 0.393 | 1.000 |
| AT4G00585.1 | NADH:ubiquinone oxidoreductase subunit (KFYI) | 37 | 0.56 | 0.566 | 1.000 |
| AT4G00585.1 | NADH:ubiquinone oxidoreductase subunit (KFYI) | 46 | 0.28 | 4.105 | 1.000 |
| AT4G00585.1 | NADH:ubiquinone oxidoreductase subunit (KFYI) | 47 | 0.30 | 4.702 | 1.000 |
| AT4G00860.1 | Cytochrome c:O2 oxidoreductase subunit (COXX2-1) | 2 | 1.22 | 0.286 | 1.000 |
| AT4G00860.1 | Cytochrome c:O2 oxidoreductase subunit (COXX2-1) | 3 | 0.89 | 0.167 | 1.000 |
| AT4G00860.1 | Cytochrome c:O2 oxidoreductase subunit (COXX2-1) | 4 | 0.78 | 0.161 | 1.000 |
| AT4G00860.1 | Cytochrome c:O2 oxidoreductase subunit (COXX2-1) | 5 | 0.88 | 0.160 | 1.000 |
| AT4G00860.1 | Cytochrome c:O2 oxidoreductase subunit (COXX2-1) | 6 | 0.95 | 0.161 | 1.000 |
| AT4G00860.1 | Cytochrome c:O2 oxidoreductase subunit (COXX2-1) | 7 | 0.90 | 0.162 | 1.000 |
| AT4G00860.1 | Cytochrome c:O2 oxidoreductase subunit (COXX2-1) | 8 | 0.89 | 0.157 | 1.000 |
| AT4G00860.1 | Cytochrome c:O2 oxidoreductase subunit (COXX2-1) | 9 | 0.96 | 0.155 | 1.000 |
| AT4G00860.1 | Cytochrome c:O2 oxidoreductase subunit (COXX2-1) | 10 | 1.07 | 0.155 | 1.000 |
| AT4G00860.1 | Cytochrome c:O2 oxidoreductase subunit (COXX2-1) | 11 | 1.24 | 0.155 | 1.000 |
| AT4G00860.1 | Cytochrome c:O2 oxidoreductase subunit (COXX2-1) | 12 | 1.42 | 0.155 | 1.000 |
| AT4G00860.1 | Cytochrome c:O2 oxidoreductase subunit (COXX2-1) | 13 | 1.47 | 0.153 | 1.000 |
| AT4G00860.1 | Cytochrome c:O2 oxidoreductase subunit (COXX2-1) | 14 | 1.43 | 0.150 | 1.000 |
| AT4G00860.1 | Cytochrome c:O2 oxidoreductase subunit (COXX2-1) | 15 | 1.25 | 0.152 | 1.000 |
| AT4G00860.1 | Cytochrome c:O2 oxidoreductase subunit (COXX2-1) | 16 | 1.04 | 0.150 | 1.000 |

| | | | | | | |
|---|---|---|---|---|---|---|
| AT4G00860.1 | Cytochrome c:O2 oxidoreductase subunit (COXX2-1) | 17 | 0.99 | 0.150 | 1.000 | |
| AT4G00860.1 | Cytochrome c:O2 oxidoreductase subunit (COXX2-1) | 18 | 1.03 | 0.151 | 1.000 | |
| AT4G00860.1 | Cytochrome c:O2 oxidoreductase subunit (COXX2-1) | 19 | 1.06 | 0.155 | 1.000 | |
| AT4G00860.1 | Cytochrome c:O2 oxidoreductase subunit (COXX2-1) | 20 | 1.10 | 0.160 | 1.000 | |
| AT4G00860.1 | Cytochrome c:O2 oxidoreductase subunit (COXX2-1) | 21 | 1.20 | 0.161 | 1.000 | |
| AT4G00860.1 | Cytochrome c:O2 oxidoreductase subunit (COXX2-1) | 22 | 1.44 | 0.162 | 1.000 | |
| AT4G00860.1 | Cytochrome c:O2 oxidoreductase subunit (COXX2-1) | 23 | 1.64 | 0.163 | 0.996 | |
| AT4G00860.1 | Cytochrome c:O2 oxidoreductase subunit (COXX2-1) | 24 | 1.59 | 0.157 | 0.997 | |
| AT4G00860.1 | Cytochrome c:O2 oxidoreductase subunit (COXX2-1) | 25 | 1.22 | 0.153 | 1.000 | |
| AT4G00860.1 | Cytochrome c:O2 oxidoreductase subunit (COXX2-1) | 26 | 1.10 | 0.153 | 1.000 | |
| AT4G00860.1 | Cytochrome c:O2 oxidoreductase subunit (COXX2-1) | 27 | 1.48 | 0.153 | 1.000 | |
| AT4G00860.1 | Cytochrome c:O2 oxidoreductase subunit (COXX2-1) | 28 | 1.86 | 0.152 | 0.893 | |
| AT4G00860.1 | Cytochrome c:O2 oxidoreductase subunit (COXX2-1) | 29 | 2.06 | 0.153 | 0.697 | |
| AT4G00860.1 | Cytochrome c:O2 oxidoreductase subunit (COXX2-1) | 30 | 1.99 | 0.154 | 0.788 | |
| AT4G00860.1 | Cytochrome c:O2 oxidoreductase subunit (COXX2-1) | 31 | 1.65 | 0.156 | 0.992 | |
| AT4G00860.1 | Cytochrome c:O2 oxidoreductase subunit (COXX2-1) | 32 | 1.16 | 0.157 | 1.000 | |
| AT4G00860.1 | Cytochrome c:O2 oxidoreductase subunit (COXX2-1) | 33 | 0.58 | 0.157 | 0.980 | |
| AT4G00860.1 | Cytochrome c:O2 oxidoreductase subunit (COXX2-1) | 34 | 0.28 | 0.158 | 0.018 | * |
| AT4G00860.1 | Cytochrome c:O2 oxidoreductase subunit (COXX2-1) | 35 | 0.27 | 0.160 | 0.014 | * |
| AT4G00860.1 | Cytochrome c:O2 oxidoreductase subunit (COXX2-1) | 36 | 0.43 | 0.168 | 0.575 | |

| | | | | | |
|---|---|---|---|---|---|
| AT4G00860.1 | Cytochrome c:O2 oxidoreductase subunit (COXX2-1) | 37 | 0.56 | 0.200 | 0.998 |
| AT4G00860.1 | Cytochrome c:O2 oxidoreductase subunit (COXX2-1) | 38 | 0.90 | 0.329 | 1.000 |
| AT4G00860.1 | Cytochrome c:O2 oxidoreductase subunit (COXX2-1) | 46 | 2630201.23 | 3.265 | 0.771 |
| AT4G00860.1 | Cytochrome c:O2 oxidoreductase subunit (COXX2-1) | 47 | 33048145.66 | 3.818 | 0.768 |
| AT4G01050.1 | Thylakoid rhodanese-like (TROL) | 2 | 0.96 | 0.597 | 1.000 |
| AT4G01050.1 | Thylakoid rhodanese-like (TROL) | 3 | 0.90 | 0.525 | 1.000 |
| AT4G01050.1 | Thylakoid rhodanese-like (TROL) | 9 | 0.65 | 0.367 | 1.000 |
| AT4G01050.1 | Thylakoid rhodanese-like (TROL) | 10 | 0.64 | 0.330 | 1.000 |
| AT4G01050.1 | Thylakoid rhodanese-like (TROL) | 11 | 0.63 | 0.307 | 1.000 |
| AT4G01050.1 | Thylakoid rhodanese-like (TROL) | 12 | 0.63 | 0.288 | 1.000 |
| AT4G01050.1 | Thylakoid rhodanese-like (TROL) | 13 | 0.64 | 0.276 | 0.999 |
| AT4G01050.1 | Thylakoid rhodanese-like (TROL) | 14 | 0.65 | 0.268 | 0.999 |
| AT4G01050.1 | Thylakoid rhodanese-like (TROL) | 15 | 0.64 | 0.262 | 0.999 |
| AT4G01050.1 | Thylakoid rhodanese-like (TROL) | 16 | 0.61 | 0.258 | 0.997 |
| AT4G01050.1 | Thylakoid rhodanese-like (TROL) | 17 | 0.55 | 0.256 | 0.978 |
| AT4G01050.1 | Thylakoid rhodanese-like (TROL) | 18 | 0.47 | 0.258 | 0.894 |
| AT4G01050.1 | Thylakoid rhodanese-like (TROL) | 19 | 0.39 | 0.262 | 0.721 |
| AT4G01050.1 | Thylakoid rhodanese-like (TROL) | 20 | 0.32 | 0.266 | 0.506 |
| AT4G01050.1 | Thylakoid rhodanese-like (TROL) | 21 | 0.27 | 0.266 | 0.312 |
| AT4G01050.1 | Thylakoid rhodanese-like (TROL) | 22 | 0.25 | 0.262 | 0.212 |
| AT4G01050.1 | Thylakoid rhodanese-like (TROL) | 23 | 0.25 | 0.258 | 0.201 |
| AT4G01050.1 | Thylakoid rhodanese-like (TROL) | 24 | 0.27 | 0.254 | 0.258 |
| AT4G01050.1 | Thylakoid rhodanese-like (TROL) | 25 | 0.32 | 0.251 | 0.424 |
| AT4G01050.1 | Thylakoid rhodanese-like (TROL) | 26 | 0.43 | 0.250 | 0.778 |
| AT4G01050.1 | Thylakoid rhodanese-like (TROL) | 27 | 0.62 | 0.249 | 0.997 |
| AT4G01050.1 | Thylakoid rhodanese-like (TROL) | 28 | 0.95 | 0.250 | 1.000 |
| AT4G01050.1 | Thylakoid rhodanese-like (TROL) | 29 | 1.42 | 0.250 | 1.000 |
| AT4G01050.1 | Thylakoid rhodanese-like (TROL) | 30 | 1.90 | 0.250 | 0.956 |
| AT4G01050.1 | Thylakoid rhodanese-like (TROL) | 31 | 2.36 | 0.251 | 0.766 |

| | | | | | |
|---|---|---|---|---|---|
| AT4G01050.1 | Thylakoid rhodanese-like (TROL) | 32 | 2.64 | 0.251 | 0.624 |
| AT4G01050.1 | Thylakoid rhodanese-like (TROL) | 33 | 2.61 | 0.251 | 0.641 |
| AT4G01050.1 | Thylakoid rhodanese-like (TROL) | 34 | 2.39 | 0.251 | 0.756 |
| AT4G01050.1 | Thylakoid rhodanese-like (TROL) | 35 | 2.03 | 0.251 | 0.919 |
| AT4G01050.1 | Thylakoid rhodanese-like (TROL) | 36 | 1.51 | 0.251 | 0.999 |
| AT4G01050.1 | Thylakoid rhodanese-like (TROL) | 37 | 0.99 | 0.251 | 1.000 |
| AT4G01050.1 | Thylakoid rhodanese-like (TROL) | 38 | 0.65 | 0.252 | 0.999 |
| AT4G01050.1 | Thylakoid rhodanese-like (TROL) | 39 | 0.54 | 0.253 | 0.970 |
| AT4G01050.1 | Thylakoid rhodanese-like (TROL) | 40 | 0.57 | 0.254 | 0.988 |
| AT4G01050.1 | Thylakoid rhodanese-like (TROL) | 41 | 0.71 | 0.257 | 1.000 |
| AT4G01050.1 | Thylakoid rhodanese-like (TROL) | 42 | 0.91 | 0.260 | 1.000 |
| AT4G01050.1 | Thylakoid rhodanese-like (TROL) | 43 | 1.04 | 0.263 | 1.000 |
| AT4G01050.1 | Thylakoid rhodanese-like (TROL) | 44 | 0.92 | 0.267 | 1.000 |
| AT4G01050.1 | Thylakoid rhodanese-like (TROL) | 45 | 0.59 | 0.275 | 0.997 |
| AT4G01050.1 | Thylakoid rhodanese-like (TROL) | 46 | 0.30 | 0.305 | 0.584 |
| AT4G01050.1 | Thylakoid rhodanese-like (TROL) | 47 | 0.13 | 0.381 | 0.228 |
| AT4G01100.1 | Adenine nucleotide transporter 1 (ADNT1) | 2 | 1.29 | 0.294 | 1.000 |
| AT4G01100.1 | Adenine nucleotide transporter 1 (ADNT1) | 3 | 1.00 | 0.180 | 1.000 |
| AT4G01100.1 | Adenine nucleotide transporter 1 (ADNT1) | 4 | 0.88 | 0.170 | 1.000 |
| AT4G01100.1 | Adenine nucleotide transporter 1 (ADNT1) | 5 | 0.92 | 0.170 | 1.000 |
| AT4G01100.1 | Adenine nucleotide transporter 1 (ADNT1) | 6 | 0.93 | 0.170 | 1.000 |
| AT4G01100.1 | Adenine nucleotide transporter 1 (ADNT1) | 7 | 0.87 | 0.171 | 1.000 |
| AT4G01100.1 | Adenine nucleotide transporter 1 (ADNT1) | 8 | 0.90 | 0.168 | 1.000 |
| AT4G01100.1 | Adenine nucleotide transporter 1 (ADNT1) | 9 | 1.02 | 0.166 | 1.000 |
| AT4G01100.1 | Adenine nucleotide transporter 1 (ADNT1) | 10 | 1.06 | 0.166 | 1.000 |
| AT4G01100.1 | Adenine nucleotide transporter 1 (ADNT1) | 11 | 1.01 | 0.166 | 1.000 |
| AT4G01100.1 | Adenine nucleotide transporter 1 (ADNT1) | 12 | 1.00 | 0.166 | 1.000 |
| AT4G01100.1 | Adenine nucleotide transporter 1 (ADNT1) | 13 | 1.00 | 0.165 | 1.000 |
| AT4G01100.1 | Adenine nucleotide transporter 1 (ADNT1) | 14 | 0.98 | 0.163 | 1.000 |
| AT4G01100.1 | Adenine nucleotide transporter 1 (ADNT1) | 15 | 0.88 | 0.162 | 1.000 |
| AT4G01100.1 | Adenine nucleotide transporter 1 (ADNT1) | 16 | 0.75 | 0.160 | 1.000 |

| | | | | | | |
|---|---|---|---|---|---|---|
| AT4G01100.1 | Adenine nucleotide transporter 1 (ADNT1) | 17 | 0.78 | 0.159 | 1.000 | |
| AT4G01100.1 | Adenine nucleotide transporter 1 (ADNT1) | 18 | 1.07 | 0.161 | 1.000 | |
| AT4G01100.1 | Adenine nucleotide transporter 1 (ADNT1) | 19 | 1.85 | 0.167 | 0.963 | |
| AT4G01100.1 | Adenine nucleotide transporter 1 (ADNT1) | 20 | 3.14 | 0.171 | 0.124 | |
| AT4G01100.1 | Adenine nucleotide transporter 1 (ADNT1) | 21 | 3.55 | 0.171 | 0.050 | * |
| AT4G01100.1 | Adenine nucleotide transporter 1 (ADNT1) | 22 | 2.69 | 0.174 | 0.360 | |
| AT4G01100.1 | Adenine nucleotide transporter 1 (ADNT1) | 23 | 1.85 | 0.176 | 0.980 | |
| AT4G01100.1 | Adenine nucleotide transporter 1 (ADNT1) | 24 | 1.33 | 0.179 | 1.000 | |
| AT4G01100.1 | Adenine nucleotide transporter 1 (ADNT1) | 25 | 1.05 | 0.183 | 1.000 | |
| AT4G01100.1 | Adenine nucleotide transporter 1 (ADNT1) | 26 | 0.99 | 0.179 | 1.000 | |
| AT4G01100.1 | Adenine nucleotide transporter 1 (ADNT1) | 27 | 1.08 | 0.170 | 1.000 | |
| AT4G01100.1 | Adenine nucleotide transporter 1 (ADNT1) | 28 | 1.48 | 0.164 | 1.000 | |
| AT4G01100.1 | Adenine nucleotide transporter 1 (ADNT1) | 29 | 2.05 | 0.164 | 0.823 | |
| AT4G01100.1 | Adenine nucleotide transporter 1 (ADNT1) | 30 | 1.91 | 0.165 | 0.932 | |
| AT4G01100.1 | Adenine nucleotide transporter 1 (ADNT1) | 31 | 1.57 | 0.166 | 0.999 | |
| AT4G01100.1 | Adenine nucleotide transporter 1 (ADNT1) | 32 | 1.39 | 0.166 | 1.000 | |
| AT4G01100.1 | Adenine nucleotide transporter 1 (ADNT1) | 33 | 1.43 | 0.166 | 1.000 | |
| AT4G01100.1 | Adenine nucleotide transporter 1 (ADNT1) | 34 | 1.71 | 0.167 | 0.992 | |
| AT4G01100.1 | Adenine nucleotide transporter 1 (ADNT1) | 35 | 1.89 | 0.166 | 0.946 | |
| AT4G01100.1 | Adenine nucleotide transporter 1 (ADNT1) | 36 | 1.85 | 0.166 | 0.959 | |
| AT4G01100.1 | Adenine nucleotide transporter 1 (ADNT1) | 37 | 2.12 | 0.166 | 0.781 | |
| AT4G01100.1 | Adenine nucleotide transporter 1 (ADNT1) | 38 | 3.00 | 0.166 | 0.140 | |
| AT4G01100.1 | Adenine nucleotide transporter 1 (ADNT1) | 39 | 3.71 | 0.166 | 0.026 | * |
| AT4G01100.1 | Adenine nucleotide transporter 1 (ADNT1) | 40 | 2.80 | 0.166 | 0.219 | |
| AT4G01100.1 | Adenine nucleotide transporter 1 (ADNT1) | 41 | 1.32 | 0.166 | 1.000 | |
| AT4G01100.1 | Adenine nucleotide transporter 1 (ADNT1) | 42 | 0.55 | 0.169 | 0.977 | |
| AT4G01100.1 | Adenine nucleotide transporter 1 (ADNT1) | 43 | 0.29 | 0.170 | 0.068 | . |
| AT4G01100.1 | Adenine nucleotide transporter 1 (ADNT1) | 44 | 0.22 | 0.169 | 0.006 | ** |
| AT4G01100.1 | Adenine nucleotide transporter 1 (ADNT1) | 45 | 0.24 | 0.169 | 0.012 | * |
| AT4G01100.1 | Adenine nucleotide transporter 1 (ADNT1) | 46 | 0.29 | 0.192 | 0.177 | |
| AT4G01100.1 | Adenine nucleotide transporter 1 (ADNT1) | 47 | 0.35 | 0.355 | 0.998 | |

| | | | | | |
|---|---|---|---|---|---|
| AT4G01150.1 | Curvature thylakoid 1A (CURT1A) | 2 | 0.86 | 0.479 | 1.000 |
| AT4G01150.1 | Curvature thylakoid 1A (CURT1A) | 3 | 0.78 | 0.354 | 1.000 |
| AT4G01150.1 | Curvature thylakoid 1A (CURT1A) | 4 | 0.70 | 0.310 | 1.000 |
| AT4G01150.1 | Curvature thylakoid 1A (CURT1A) | 5 | 0.61 | 0.302 | 1.000 |
| AT4G01150.1 | Curvature thylakoid 1A (CURT1A) | 6 | 0.52 | 0.295 | 0.999 |
| AT4G01150.1 | Curvature thylakoid 1A (CURT1A) | 7 | 0.45 | 0.290 | 0.980 |
| AT4G01150.1 | Curvature thylakoid 1A (CURT1A) | 8 | 0.41 | 0.284 | 0.938 |
| AT4G01150.1 | Curvature thylakoid 1A (CURT1A) | 9 | 0.41 | 0.281 | 0.937 |
| AT4G01150.1 | Curvature thylakoid 1A (CURT1A) | 10 | 0.45 | 0.280 | 0.976 |
| AT4G01150.1 | Curvature thylakoid 1A (CURT1A) | 11 | 0.53 | 0.280 | 0.998 |
| AT4G01150.1 | Curvature thylakoid 1A (CURT1A) | 12 | 0.65 | 0.279 | 1.000 |
| AT4G01150.1 | Curvature thylakoid 1A (CURT1A) | 13 | 0.80 | 0.278 | 1.000 |
| AT4G01150.1 | Curvature thylakoid 1A (CURT1A) | 14 | 0.95 | 0.276 | 1.000 |
| AT4G01150.1 | Curvature thylakoid 1A (CURT1A) | 15 | 1.07 | 0.275 | 1.000 |
| AT4G01150.1 | Curvature thylakoid 1A (CURT1A) | 16 | 1.15 | 0.274 | 1.000 |
| AT4G01150.1 | Curvature thylakoid 1A (CURT1A) | 17 | 1.23 | 0.274 | 1.000 |
| AT4G01150.1 | Curvature thylakoid 1A (CURT1A) | 18 | 1.32 | 0.277 | 1.000 |
| AT4G01150.1 | Curvature thylakoid 1A (CURT1A) | 19 | 1.22 | 0.281 | 1.000 |
| AT4G01150.1 | Curvature thylakoid 1A (CURT1A) | 20 | 0.95 | 0.284 | 1.000 |
| AT4G01150.1 | Curvature thylakoid 1A (CURT1A) | 21 | 0.66 | 0.287 | 1.000 |
| AT4G01150.1 | Curvature thylakoid 1A (CURT1A) | 22 | 0.45 | 0.287 | 0.981 |
| AT4G01150.1 | Curvature thylakoid 1A (CURT1A) | 23 | 0.36 | 0.285 | 0.834 |
| AT4G01150.1 | Curvature thylakoid 1A (CURT1A) | 24 | 0.33 | 0.282 | 0.758 |
| AT4G01150.1 | Curvature thylakoid 1A (CURT1A) | 25 | 0.35 | 0.280 | 0.791 |
| AT4G01150.1 | Curvature thylakoid 1A (CURT1A) | 26 | 0.37 | 0.279 | 0.844 |
| AT4G01150.1 | Curvature thylakoid 1A (CURT1A) | 27 | 0.41 | 0.278 | 0.922 |
| AT4G01150.1 | Curvature thylakoid 1A (CURT1A) | 28 | 0.42 | 0.278 | 0.937 |
| AT4G01150.1 | Curvature thylakoid 1A (CURT1A) | 29 | 0.41 | 0.279 | 0.935 |
| AT4G01150.1 | Curvature thylakoid 1A (CURT1A) | 30 | 0.42 | 0.279 | 0.944 |
| AT4G01150.1 | Curvature thylakoid 1A (CURT1A) | 31 | 0.45 | 0.279 | 0.971 |
| AT4G01150.1 | Curvature thylakoid 1A (CURT1A) | 32 | 0.48 | 0.280 | 0.988 |

| | | | | | |
|---|---|---|---|---|---|
| AT4G01150.1 | Curvature thylakoid 1A (CURT1A) | 33 | 0.46 | 0.280 | 0.982 |
| AT4G01150.1 | Curvature thylakoid 1A (CURT1A) | 34 | 0.42 | 0.281 | 0.946 |
| AT4G01150.1 | Curvature thylakoid 1A (CURT1A) | 35 | 0.42 | 0.284 | 0.945 |
| AT4G01150.1 | Curvature thylakoid 1A (CURT1A) | 36 | 0.47 | 0.288 | 0.989 |
| AT4G01150.1 | Curvature thylakoid 1A (CURT1A) | 37 | 0.53 | 0.294 | 0.999 |
| AT4G01150.1 | Curvature thylakoid 1A (CURT1A) | 38 | 0.57 | 0.296 | 1.000 |
| AT4G01150.1 | Curvature thylakoid 1A (CURT1A) | 39 | 0.63 | 0.294 | 1.000 |
| AT4G01150.1 | Curvature thylakoid 1A (CURT1A) | 40 | 0.80 | 0.288 | 1.000 |
| AT4G01150.1 | Curvature thylakoid 1A (CURT1A) | 41 | 1.08 | 0.284 | 1.000 |
| AT4G01150.1 | Curvature thylakoid 1A (CURT1A) | 42 | 1.54 | 0.285 | 1.000 |
| AT4G01150.1 | Curvature thylakoid 1A (CURT1A) | 43 | 2.65 | 0.286 | 0.888 |
| AT4G01150.1 | Curvature thylakoid 1A (CURT1A) | 44 | 4.63 | 0.285 | 0.287 |
| AT4G01150.1 | Curvature thylakoid 1A (CURT1A) | 45 | 6.49 | 0.291 | 0.105 |
| AT4G01150.1 | Curvature thylakoid 1A (CURT1A) | 46 | 7.07 | 0.342 | 0.214 |
| AT4G01150.1 | Curvature thylakoid 1A (CURT1A) | 47 | 6.87 | 0.489 | 0.740 |
| AT4G01660.1 | ABC Transporter 1 (ABC1) | 2 | 0.84 | 0.286 | 1.000 |
| AT4G01660.1 | ABC Transporter 1 (ABC1) | 3 | 0.67 | 0.274 | 1.000 |
| AT4G01660.1 | ABC Transporter 1 (ABC1) | 27 | 0.57 | 0.212 | 0.984 |
| AT4G01660.1 | ABC Transporter 1 (ABC1) | 28 | 0.84 | 0.149 | 1.000 |
| AT4G01660.1 | ABC Transporter 1 (ABC1) | 29 | 1.10 | 0.132 | 1.000 |
| AT4G01660.1 | ABC Transporter 1 (ABC1) | 30 | 1.19 | 0.126 | 1.000 |
| AT4G01660.1 | ABC Transporter 1 (ABC1) | 31 | 1.17 | 0.124 | 1.000 |
| AT4G01660.1 | ABC Transporter 1 (ABC1) | 32 | 1.08 | 0.125 | 1.000 |
| AT4G01660.1 | ABC Transporter 1 (ABC1) | 33 | 0.93 | 0.125 | 1.000 |
| AT4G01660.1 | ABC Transporter 1 (ABC1) | 34 | 0.85 | 0.125 | 1.000 |
| AT4G01660.1 | ABC Transporter 1 (ABC1) | 35 | 0.85 | 0.125 | 1.000 |
| AT4G01660.1 | ABC Transporter 1 (ABC1) | 36 | 0.91 | 0.124 | 1.000 |
| AT4G01660.1 | ABC Transporter 1 (ABC1) | 37 | 1.04 | 0.124 | 1.000 |
| AT4G01660.1 | ABC Transporter 1 (ABC1) | 38 | 1.16 | 0.123 | 1.000 |
| AT4G01660.1 | ABC Transporter 1 (ABC1) | 39 | 1.20 | 0.123 | 1.000 |
| AT4G01660.1 | ABC Transporter 1 (ABC1) | 40 | 1.17 | 0.123 | 1.000 |
| AT4G01660.1 | ABC Transporter 1 (ABC1) | 41 | 1.05 | 0.124 | 1.000 |
| AT4G01660.1 | ABC Transporter 1 (ABC1) | 42 | 0.92 | 0.126 | 1.000 |
| AT4G01660.1 | ABC Transporter 1 (ABC1) | 43 | 0.85 | 0.127 | 1.000 |
| AT4G01660.1 | ABC Transporter 1 (ABC1) | 44 | 0.84 | 0.127 | 1.000 |
| AT4G01660.1 | ABC Transporter 1 (ABC1) | 45 | 0.86 | 0.127 | 1.000 |
| AT4G01660.1 | ABC Transporter 1 (ABC1) | 46 | 0.81 | 0.150 | 1.000 |
| AT4G01660.1 | ABC Transporter 1 (ABC1) | 47 | 0.73 | 0.278 | 1.000 |
| AT4G01990.1 | Pentatricopeptide repeat (PPR) superfamily protein | 2 | 1.00 | 0.279 | 1.000 |
| AT4G01990.1 | Pentatricopeptide repeat (PPR) superfamily protein | 3 | 1.04 | 0.267 | 1.000 |
| AT4G01990.1 | Pentatricopeptide repeat (PPR) superfamily protein | 4 | 1.08 | 0.258 | 1.000 |
| AT4G01990.1 | Pentatricopeptide repeat (PPR) superfamily protein | 26 | 1.66 | 0.151 | 0.494 |

| | | | | | |
|---|---|---|---|---|---|
| AT4G01990.1 | Pentatricopeptide repeat (PPR) superfamily protein | 27 | 1.63 | 0.144 | 0.482 |
| AT4G01990.1 | Pentatricopeptide repeat (PPR) superfamily protein | 28 | 1.59 | 0.137 | 0.483 |
| AT4G01990.1 | Pentatricopeptide repeat (PPR) superfamily protein | 29 | 1.55 | 0.132 | 0.505 |
| AT4G01990.1 | Pentatricopeptide repeat (PPR) superfamily protein | 30 | 1.50 | 0.127 | 0.550 |
| AT4G01990.1 | Pentatricopeptide repeat (PPR) superfamily protein | 31 | 1.44 | 0.124 | 0.620 |
| AT4G01990.1 | Pentatricopeptide repeat (PPR) superfamily protein | 32 | 1.37 | 0.121 | 0.713 |
| AT4G01990.1 | Pentatricopeptide repeat (PPR) superfamily protein | 33 | 1.31 | 0.120 | 0.817 |
| AT4G01990.1 | Pentatricopeptide repeat (PPR) superfamily protein | 34 | 1.24 | 0.119 | 0.912 |
| AT4G01990.1 | Pentatricopeptide repeat (PPR) superfamily protein | 35 | 1.17 | 0.118 | 0.976 |
| AT4G01990.1 | Pentatricopeptide repeat (PPR) superfamily protein | 38 | 0.95 | 0.118 | 1.000 |
| AT4G01990.1 | Pentatricopeptide repeat (PPR) superfamily protein | 39 | 0.89 | 0.118 | 0.992 |
| AT4G01990.1 | Pentatricopeptide repeat (PPR) superfamily protein | 40 | 0.82 | 0.119 | 0.934 |
| AT4G01990.1 | Pentatricopeptide repeat (PPR) superfamily protein | 41 | 0.76 | 0.121 | 0.805 |
| AT4G01990.1 | Pentatricopeptide repeat (PPR) superfamily protein | 42 | 0.70 | 0.124 | 0.645 |
| AT4G01990.1 | Pentatricopeptide repeat (PPR) superfamily protein | 43 | 0.65 | 0.129 | 0.503 |
| AT4G01990.1 | Pentatricopeptide repeat (PPR) superfamily protein | 44 | 0.61 | 0.136 | 0.404 |
| AT4G01990.1 | Pentatricopeptide repeat (PPR) superfamily protein | 45 | 0.57 | 0.145 | 0.342 |
| AT4G01990.1 | Pentatricopeptide repeat (PPR) superfamily protein | 46 | 0.53 | 0.157 | 0.312 |
| AT4G01990.1 | Pentatricopeptide repeat (PPR) superfamily protein | 47 | 0.50 | 0.172 | 0.303 |
| AT4G02580.1 | NADH:ubiquinone oxidoreductase subunit (24kDa) | 2 | 0.80 | 0.519 | 1.000 |
| AT4G02580.1 | NADH:ubiquinone oxidoreductase subunit (24kDa) | 3 | 0.97 | 0.342 | 1.000 |
| AT4G02580.1 | NADH:ubiquinone oxidoreductase subunit (24kDa) | 4 | 1.16 | 0.295 | 1.000 |
| AT4G02580.1 | NADH:ubiquinone oxidoreductase subunit (24kDa) | 5 | 1.37 | 0.283 | 1.000 |
| AT4G02580.1 | NADH:ubiquinone oxidoreductase subunit (24kDa) | 6 | 1.57 | 0.278 | 1.000 |
| AT4G02580.1 | NADH:ubiquinone oxidoreductase subunit (24kDa) | 7 | 1.71 | 0.275 | 1.000 |
| AT4G02580.1 | NADH:ubiquinone oxidoreductase subunit (24kDa) | 8 | 1.83 | 0.270 | 1.000 |
| AT4G02580.1 | NADH:ubiquinone oxidoreductase subunit (24kDa) | 9 | 2.12 | 0.267 | 0.997 |
| AT4G02580.1 | NADH:ubiquinone oxidoreductase subunit (24kDa) | 10 | 2.57 | 0.267 | 0.958 |
| AT4G02580.1 | NADH:ubiquinone oxidoreductase subunit (24kDa) | 11 | 2.86 | 0.267 | 0.895 |
| AT4G02580.1 | NADH:ubiquinone oxidoreductase subunit (24kDa) | 12 | 2.83 | 0.268 | 0.901 |
| AT4G02580.1 | NADH:ubiquinone oxidoreductase subunit (24kDa) | 13 | 2.55 | 0.265 | 0.960 |

| | | | | | |
|---|---|---|---|---|---|
| AT4G02580.1 | NADH:ubiquinone oxidoreductase subunit (24kDa) | 14 | 1.95 | 0.262 | 0.999 |
| AT4G02580.1 | NADH:ubiquinone oxidoreductase subunit (24kDa) | 15 | 1.26 | 0.260 | 1.000 |
| AT4G02580.1 | NADH:ubiquinone oxidoreductase subunit (24kDa) | 16 | 0.73 | 0.259 | 1.000 |
| AT4G02580.1 | NADH:ubiquinone oxidoreductase subunit (24kDa) | 17 | 0.49 | 0.258 | 0.998 |
| AT4G02580.1 | NADH:ubiquinone oxidoreductase subunit (24kDa) | 18 | 0.49 | 0.262 | 0.998 |
| AT4G02580.1 | NADH:ubiquinone oxidoreductase subunit (24kDa) | 19 | 0.63 | 0.270 | 1.000 |
| AT4G02580.1 | NADH:ubiquinone oxidoreductase subunit (24kDa) | 20 | 0.82 | 0.275 | 1.000 |
| AT4G02580.1 | NADH:ubiquinone oxidoreductase subunit (24kDa) | 21 | 1.10 | 0.276 | 1.000 |
| AT4G02580.1 | NADH:ubiquinone oxidoreductase subunit (24kDa) | 22 | 1.44 | 0.277 | 1.000 |
| AT4G02580.1 | NADH:ubiquinone oxidoreductase subunit (24kDa) | 23 | 1.61 | 0.274 | 1.000 |
| AT4G02580.1 | NADH:ubiquinone oxidoreductase subunit (24kDa) | 24 | 1.21 | 0.268 | 1.000 |
| AT4G02580.1 | NADH:ubiquinone oxidoreductase subunit (24kDa) | 25 | 0.64 | 0.265 | 1.000 |
| AT4G02580.1 | NADH:ubiquinone oxidoreductase subunit (24kDa) | 26 | 0.40 | 0.264 | 0.962 |
| AT4G02580.1 | NADH:ubiquinone oxidoreductase subunit (24kDa) | 27 | 0.42 | 0.264 | 0.978 |
| AT4G02580.1 | NADH:ubiquinone oxidoreductase subunit (24kDa) | 28 | 0.74 | 0.266 | 1.000 |
| AT4G02580.1 | NADH:ubiquinone oxidoreductase subunit (24kDa) | 29 | 1.65 | 0.266 | 1.000 |
| AT4G02580.1 | NADH:ubiquinone oxidoreductase subunit (24kDa) | 30 | 2.61 | 0.267 | 0.952 |
| AT4G02580.1 | NADH:ubiquinone oxidoreductase subunit (24kDa) | 31 | 2.84 | 0.268 | 0.899 |
| AT4G02580.1 | NADH:ubiquinone oxidoreductase subunit (24kDa) | 32 | 2.80 | 0.268 | 0.910 |
| AT4G02580.1 | NADH:ubiquinone oxidoreductase subunit (24kDa) | 33 | 2.73 | 0.268 | 0.929 |
| AT4G02580.1 | NADH:ubiquinone oxidoreductase subunit (24kDa) | 34 | 2.46 | 0.268 | 0.976 |
| AT4G02580.1 | NADH:ubiquinone oxidoreductase subunit (24kDa) | 35 | 2.02 | 0.268 | 0.999 |
| AT4G02580.1 | NADH:ubiquinone oxidoreductase subunit (24kDa) | 36 | 1.71 | 0.268 | 1.000 |
| AT4G02580.1 | NADH:ubiquinone oxidoreductase subunit (24kDa) | 37 | 1.60 | 0.267 | 1.000 |
| AT4G02580.1 | NADH:ubiquinone oxidoreductase subunit (24kDa) | 38 | 1.49 | 0.268 | 1.000 |
| AT4G02580.1 | NADH:ubiquinone oxidoreductase subunit (24kDa) | 39 | 1.34 | 0.267 | 1.000 |
| AT4G02580.1 | NADH:ubiquinone oxidoreductase subunit (24kDa) | 40 | 1.18 | 0.267 | 1.000 |
| AT4G02580.1 | NADH:ubiquinone oxidoreductase subunit (24kDa) | 41 | 0.91 | 0.268 | 1.000 |
| AT4G02580.1 | NADH:ubiquinone oxidoreductase subunit (24kDa) | 42 | 0.71 | 0.271 | 1.000 |
| AT4G02580.1 | NADH:ubiquinone oxidoreductase subunit (24kDa) | 43 | 0.66 | 0.274 | 1.000 |
| AT4G02580.1 | NADH:ubiquinone oxidoreductase subunit (24kDa) | 44 | 0.63 | 0.274 | 1.000 |

| | | | | | | |
|---|---|---|---|---|---|---|
| AT4G02580.1 | NADH:ubiquinone oxidoreductase subunit (24kDa) | 45 | 0.62 | 0.274 | 1.000 | |
| AT4G02580.1 | NADH:ubiquinone oxidoreductase subunit (24kDa) | 46 | 0.55 | 0.320 | 1.000 | |
| AT4G02580.1 | NADH:ubiquinone oxidoreductase subunit (24kDa) | 47 | 0.43 | 0.535 | 1.000 | |
| AT4G02930.1 | GTP binding Elongation factor Tu family protein | 3 | 0.07 | 0.128 | 0.000 | *** |
| AT4G02930.1 | GTP binding Elongation factor Tu family protein | 4 | 0.09 | 0.114 | 0.000 | *** |
| AT4G02930.1 | GTP binding Elongation factor Tu family protein | 5 | 0.13 | 0.112 | 0.000 | *** |
| AT4G02930.1 | GTP binding Elongation factor Tu family protein | 6 | 0.20 | 0.112 | 0.000 | *** |
| AT4G02930.1 | GTP binding Elongation factor Tu family protein | 7 | 0.30 | 0.112 | 0.000 | *** |
| AT4G02930.1 | GTP binding Elongation factor Tu family protein | 8 | 0.47 | 0.111 | 0.050 | . |
| AT4G02930.1 | GTP binding Elongation factor Tu family protein | 9 | 0.68 | 0.110 | 0.797 | |
| AT4G02930.1 | GTP binding Elongation factor Tu family protein | 10 | 0.90 | 0.110 | 1.000 | |
| AT4G02930.1 | GTP binding Elongation factor Tu family protein | 11 | 1.09 | 0.110 | 1.000 | |
| AT4G02930.1 | GTP binding Elongation factor Tu family protein | 12 | 1.24 | 0.109 | 0.998 | |
| AT4G02930.1 | GTP binding Elongation factor Tu family protein | 13 | 1.36 | 0.109 | 0.945 | |
| AT4G02930.1 | GTP binding Elongation factor Tu family protein | 14 | 1.43 | 0.108 | 0.837 | |
| AT4G02930.1 | GTP binding Elongation factor Tu family protein | 15 | 1.44 | 0.108 | 0.825 | |
| AT4G02930.1 | GTP binding Elongation factor Tu family protein | 16 | 1.40 | 0.107 | 0.882 | |
| AT4G02930.1 | GTP binding Elongation factor Tu family protein | 17 | 1.37 | 0.108 | 0.933 | |
| AT4G02930.1 | GTP binding Elongation factor Tu family protein | 18 | 1.35 | 0.108 | 0.956 | |
| AT4G02930.1 | GTP binding Elongation factor Tu family protein | 19 | 1.33 | 0.110 | 0.971 | |
| AT4G02930.1 | GTP binding Elongation factor Tu family protein | 20 | 1.29 | 0.111 | 0.990 | |
| AT4G02930.1 | GTP binding Elongation factor Tu family protein | 21 | 1.23 | 0.112 | 0.999 | |
| AT4G02930.1 | GTP binding Elongation factor Tu family protein | 22 | 1.14 | 0.112 | 1.000 | |
| AT4G02930.1 | GTP binding Elongation factor Tu family protein | 23 | 1.05 | 0.111 | 1.000 | |
| AT4G02930.1 | GTP binding Elongation factor Tu family protein | 24 | 0.97 | 0.110 | 1.000 | |
| AT4G02930.1 | GTP binding Elongation factor Tu family protein | 25 | 0.91 | 0.109 | 1.000 | |
| AT4G02930.1 | GTP binding Elongation factor Tu family protein | 26 | 0.90 | 0.109 | 1.000 | |
| AT4G02930.1 | GTP binding Elongation factor Tu family protein | 27 | 0.93 | 0.109 | 1.000 | |
| AT4G02930.1 | GTP binding Elongation factor Tu family protein | 28 | 0.97 | 0.109 | 1.000 | |
| AT4G02930.1 | GTP binding Elongation factor Tu family protein | 29 | 1.02 | 0.109 | 1.000 | |
| AT4G02930.1 | GTP binding Elongation factor Tu family protein | 30 | 1.05 | 0.109 | 1.000 | |

| | | | | | | |
|---|---|---|---|---|---|---|
| AT4G02930.1 | GTP binding Elongation factor Tu family protein | 31 | 1.06 | 0.109 | 1.000 | |
| AT4G02930.1 | GTP binding Elongation factor Tu family protein | 32 | 1.04 | 0.109 | 1.000 | |
| AT4G02930.1 | GTP binding Elongation factor Tu family protein | 33 | 1.00 | 0.109 | 1.000 | |
| AT4G02930.1 | GTP binding Elongation factor Tu family protein | 34 | 0.94 | 0.109 | 1.000 | |
| AT4G02930.1 | GTP binding Elongation factor Tu family protein | 35 | 0.89 | 0.109 | 1.000 | |
| AT4G02930.1 | GTP binding Elongation factor Tu family protein | 36 | 0.88 | 0.109 | 1.000 | |
| AT4G02930.1 | GTP binding Elongation factor Tu family protein | 37 | 0.91 | 0.109 | 1.000 | |
| AT4G02930.1 | GTP binding Elongation factor Tu family protein | 38 | 0.99 | 0.109 | 1.000 | |
| AT4G02930.1 | GTP binding Elongation factor Tu family protein | 39 | 1.12 | 0.109 | 1.000 | |
| AT4G02930.1 | GTP binding Elongation factor Tu family protein | 40 | 1.26 | 0.110 | 0.996 | |
| AT4G02930.1 | GTP binding Elongation factor Tu family protein | 41 | 1.39 | 0.110 | 0.921 | |
| AT4G02930.1 | GTP binding Elongation factor Tu family protein | 42 | 1.45 | 0.111 | 0.833 | |
| AT4G02930.1 | GTP binding Elongation factor Tu family protein | 43 | 1.40 | 0.111 | 0.906 | |
| AT4G02930.1 | GTP binding Elongation factor Tu family protein | 44 | 1.21 | 0.112 | 1.000 | |
| AT4G02930.1 | GTP binding Elongation factor Tu family protein | 45 | 0.94 | 0.116 | 1.000 | |
| AT4G02930.1 | GTP binding Elongation factor Tu family protein | 46 | 0.67 | 0.133 | 0.917 | |
| AT4G02930.1 | GTP binding Elongation factor Tu family protein | 47 | 0.47 | 0.175 | 0.510 | |
| AT4G02930.1 | GTP binding Elongation factor Tu family protein | 48 | 0.32 | 0.239 | 0.396 | |
| AT4G03115.1 | Putative MtcC-type unspecified transporter | 2 | 0.99 | 0.473 | 1.000 | |
| AT4G03115.1 | Putative MtcC-type unspecified transporter | 3 | 0.52 | 0.824 | 1.000 | |
| AT4G03115.1 | Putative MtcC-type unspecified transporter | 38 | 3.67 | 0.360 | 0.651 | |
| AT4G03115.1 | Putative MtcC-type unspecified transporter | 39 | 6.25 | 0.237 | 0.013 | * |
| AT4G03115.1 | Putative MtcC-type unspecified transporter | 40 | 6.87 | 0.213 | 0.002 | ** |
| AT4G03115.1 | Putative MtcC-type unspecified transporter | 41 | 2.63 | 0.207 | 0.335 | |
| AT4G03115.1 | Putative MtcC-type unspecified transporter | 42 | 0.64 | 0.210 | 0.968 | |
| AT4G03115.1 | Putative MtcC-type unspecified transporter | 43 | 0.17 | 0.216 | 0.006 | ** |
| AT4G03115.1 | Putative MtcC-type unspecified transporter | 44 | 0.09 | 0.277 | 0.005 | ** |
| AT4G03115.1 | Putative MtcC-type unspecified transporter | 45 | 0.09 | 0.423 | 0.136 | |
| AT4G03115.1 | Putative MtcC-type unspecified transporter | 46 | 0.11 | 0.638 | 0.715 | |
| AT4G03115.1 | Putative MtcC-type unspecified transporter | 47 | 0.16 | 0.994 | 0.988 | |
| AT4G03200.2 | ? | 2 | 0.86 | 0.313 | 1.000 | |
| AT4G03200.2 | ? | 3 | 0.79 | 0.308 | 1.000 | |

| | | | | | | |
|---|---|---|---|---|---|---|
| AT4G03200.2 | ? | 40 | 1.51 | 0.307 | 0.998 | |
| AT4G03200.2 | ? | 41 | 1.98 | 0.193 | 0.665 | |
| AT4G03200.2 | ? | 42 | 2.82 | 0.167 | 0.077 | . |
| AT4G03200.2 | ? | 43 | 2.28 | 0.166 | 0.255 | |
| AT4G03200.2 | ? | 44 | 0.85 | 0.161 | 1.000 | |
| AT4G03200.2 | ? | 45 | 0.43 | 0.162 | 0.203 | |
| AT4G03200.2 | ? | 46 | 0.37 | 0.173 | 0.123 | |
| AT4G03200.2 | ? | 47 | 0.37 | 0.396 | 0.918 | |
| AT4G03280.2 | Cytochrome b6f complex subunit (PETC) | 2 | 0.80 | 0.472 | 1.000 | |
| AT4G03280.2 | Cytochrome b6f complex subunit (PETC) | 3 | 0.81 | 0.319 | 1.000 | |
| AT4G03280.2 | Cytochrome b6f complex subunit (PETC) | 4 | 0.86 | 0.290 | 1.000 | |
| AT4G03280.2 | Cytochrome b6f complex subunit (PETC) | 5 | 0.87 | 0.288 | 1.000 | |
| AT4G03280.2 | Cytochrome b6f complex subunit (PETC) | 6 | 0.73 | 0.289 | 1.000 | |
| AT4G03280.2 | Cytochrome b6f complex subunit (PETC) | 7 | 0.51 | 0.289 | 0.999 | |
| AT4G03280.2 | Cytochrome b6f complex subunit (PETC) | 8 | 0.40 | 0.284 | 0.944 | |
| AT4G03280.2 | Cytochrome b6f complex subunit (PETC) | 9 | 0.37 | 0.280 | 0.887 | |
| AT4G03280.2 | Cytochrome b6f complex subunit (PETC) | 10 | 0.38 | 0.281 | 0.910 | |
| AT4G03280.2 | Cytochrome b6f complex subunit (PETC) | 11 | 0.42 | 0.283 | 0.970 | |
| AT4G03280.2 | Cytochrome b6f complex subunit (PETC) | 12 | 0.45 | 0.282 | 0.986 | |
| AT4G03280.2 | Cytochrome b6f complex subunit (PETC) | 13 | 0.43 | 0.279 | 0.971 | |
| AT4G03280.2 | Cytochrome b6f complex subunit (PETC) | 14 | 0.36 | 0.276 | 0.850 | |
| AT4G03280.2 | Cytochrome b6f complex subunit (PETC) | 15 | 0.27 | 0.275 | 0.503 | |
| AT4G03280.2 | Cytochrome b6f complex subunit (PETC) | 16 | 0.22 | 0.274 | 1.000 | |
| AT4G03280.2 | Cytochrome b6f complex subunit (PETC) | 17 | 0.26 | 0.273 | 0.463 | |
| AT4G03280.2 | Cytochrome b6f complex subunit (PETC) | 18 | 0.40 | 0.276 | 0.935 | |
| AT4G03280.2 | Cytochrome b6f complex subunit (PETC) | 19 | 0.59 | 0.283 | 1.000 | |
| AT4G03280.2 | Cytochrome b6f complex subunit (PETC) | 20 | 0.77 | 0.289 | 1.000 | |
| AT4G03280.2 | Cytochrome b6f complex subunit (PETC) | 21 | 0.92 | 0.292 | 1.000 | |
| AT4G03280.2 | Cytochrome b6f complex subunit (PETC) | 22 | 0.97 | 0.292 | 1.000 | |
| AT4G03280.2 | Cytochrome b6f complex subunit (PETC) | 23 | 0.92 | 0.290 | 1.000 | |
| AT4G03280.2 | Cytochrome b6f complex subunit (PETC) | 24 | 0.72 | 0.285 | 1.000 | |
| AT4G03280.2 | Cytochrome b6f complex subunit (PETC) | 25 | 0.52 | 0.280 | 0.999 | |
| AT4G03280.2 | Cytochrome b6f complex subunit (PETC) | 26 | 0.47 | 0.279 | 0.991 | |
| AT4G03280.2 | Cytochrome b6f complex subunit (PETC) | 27 | 0.59 | 0.279 | 1.000 | |
| AT4G03280.2 | Cytochrome b6f complex subunit (PETC) | 28 | 0.81 | 0.279 | 1.000 | |

| | | | | | |
|---|---|---|---|---|---|
| AT4G03280.2 | Cytochrome b6f complex subunit (PETC) | 29 | 1.32 | 0.279 | 1.000 |
| AT4G03280.2 | Cytochrome b6f complex subunit (PETC) | 30 | 2.52 | 0.282 | 0.942 |
| AT4G03280.2 | Cytochrome b6f complex subunit (PETC) | 31 | 3.36 | 0.287 | 0.706 |
| AT4G03280.2 | Cytochrome b6f complex subunit (PETC) | 32 | 1.70 | 0.298 | 1.000 |
| AT4G03280.2 | Cytochrome b6f complex subunit (PETC) | 33 | 0.47 | 0.315 | 0.998 |
| AT4G03280.2 | Cytochrome b6f complex subunit (PETC) | 34 | 0.17 | 0.340 | 0.353 |
| AT4G03280.2 | Cytochrome b6f complex subunit (PETC) | 35 | 0.10 | 0.419 | 1.000 |
| AT4G03280.2 | Cytochrome b6f complex subunit (PETC) | 36 | 0.09 | 0.562 | 0.664 |
| AT4G03280.2 | Cytochrome b6f complex subunit (PETC) | 46 | 21285.85 | 3.244 | 0.967 |
| AT4G03280.2 | Cytochrome b6f complex subunit (PETC) | 47 | 143049.08 | 3.654 | 0.946 |
| AT4G03560.1 | Two pore calcium channel 1 (TPCC1) | 2 | 0.96 | 0.386 | 1.000 |
| AT4G03560.1 | Two pore calcium channel 1 (TPCC1) | 3 | 0.99 | 0.344 | 1.000 |
| AT4G03560.1 | Two pore calcium channel 1 (TPCC1) | 4 | 1.01 | 0.317 | 1.000 |
| AT4G03560.1 | Two pore calcium channel 1 (TPCC1) | 6 | 1.07 | 0.304 | 1.000 |
| AT4G03560.1 | Two pore calcium channel 1 (TPCC1) | 7 | 1.10 | 0.312 | 1.000 |
| AT4G03560.1 | Two pore calcium channel 1 (TPCC1) | 8 | 1.13 | 0.325 | 1.000 |
| AT4G03560.1 | Two pore calcium channel 1 (TPCC1) | 22 | 1.44 | 0.345 | 0.998 |
| AT4G03560.1 | Two pore calcium channel 1 (TPCC1) | 23 | 1.44 | 0.330 | 0.997 |
| AT4G03560.1 | Two pore calcium channel 1 (TPCC1) | 24 | 1.43 | 0.316 | 0.997 |
| AT4G03560.1 | Two pore calcium channel 1 (TPCC1) | 30 | 1.20 | 0.238 | 1.000 |
| AT4G03560.1 | Two pore calcium channel 1 (TPCC1) | 31 | 1.14 | 0.232 | 1.000 |
| AT4G03560.1 | Two pore calcium channel 1 (TPCC1) | 32 | 1.09 | 0.232 | 1.000 |
| AT4G03560.1 | Two pore calcium channel 1 (TPCC1) | 33 | 1.04 | 0.236 | 1.000 |
| AT4G03560.1 | Two pore calcium channel 1 (TPCC1) | 34 | 1.00 | 0.243 | 1.000 |
| AT4G03560.1 | Two pore calcium channel 1 (TPCC1) | 41 | 0.85 | 0.351 | 1.000 |
| AT4G03560.1 | Two pore calcium channel 1 (TPCC1) | 42 | 0.85 | 0.380 | 1.000 |
| AT4G03560.1 | Two pore calcium channel 1 (TPCC1) | 43 | 0.84 | 0.415 | 1.000 |
| AT4G03560.1 | Two pore calcium channel 1 (TPCC1) | 46 | 0.83 | 0.565 | 1.000 |
| AT4G03560.1 | Two pore calcium channel 1 (TPCC1) | 47 | 0.83 | 0.629 | 1.000 |
| AT4G04180.1 | P-loop containing nucleoside triphosphate hydrolase | 2 | 0.87 | 0.373 | 1.000 |
| AT4G04180.1 | P-loop containing nucleoside triphosphate hydrolase | 3 | 0.80 | 0.365 | 1.000 |

| | | | | | | |
|---|---|---|---|---|---|---|
| AT4G04180.1 | P-loop containing nucleoside triphosphate hydrolase | 28 | 1.22 | 0.268 | 1.000 | |
| AT4G04180.1 | P-loop containing nucleoside triphosphate hydrolase | 29 | 1.39 | 0.194 | 0.998 | |
| AT4G04180.1 | P-loop containing nucleoside triphosphate hydrolase | 30 | 1.45 | 0.173 | 0.986 | |
| AT4G04180.1 | P-loop containing nucleoside triphosphate hydrolase | 31 | 1.52 | 0.164 | 0.954 | |
| AT4G04180.1 | P-loop containing nucleoside triphosphate hydrolase | 32 | 1.65 | 0.162 | 0.857 | |
| AT4G04180.1 | P-loop containing nucleoside triphosphate hydrolase | 33 | 1.75 | 0.160 | 0.758 | |
| AT4G04180.1 | P-loop containing nucleoside triphosphate hydrolase | 34 | 1.59 | 0.161 | 0.905 | |
| AT4G04180.1 | P-loop containing nucleoside triphosphate hydrolase | 35 | 1.21 | 0.162 | 1.000 | |
| AT4G04180.1 | P-loop containing nucleoside triphosphate hydrolase | 36 | 0.91 | 0.164 | 1.000 | |
| AT4G04180.1 | P-loop containing nucleoside triphosphate hydrolase | 37 | 0.79 | 0.173 | 1.000 | |
| AT4G04180.1 | P-loop containing nucleoside triphosphate hydrolase | 38 | 0.77 | 0.219 | 1.000 | |
| AT4G04180.1 | P-loop containing nucleoside triphosphate hydrolase | 39 | 0.88 | 0.346 | 1.000 | |
| AT4G04180.1 | P-loop containing nucleoside triphosphate hydrolase | 46 | 268.20 | 2.224 | 0.957 | |
| AT4G04180.1 | P-loop containing nucleoside triphosphate hydrolase | 47 | 842.15 | 2.634 | 0.952 | |
| AT4G04190.2 | ? | 2 | 1.06 | 0.480 | 1.000 | |
| AT4G04190.2 | ? | 3 | 1.51 | 0.605 | 1.000 | |
| AT4G04190.2 | ? | 22 | 0.22 | 0.276 | 0.163 | |
| AT4G04190.2 | ? | 23 | 0.35 | 0.216 | 0.266 | |
| AT4G04190.2 | ? | 24 | 0.77 | 0.195 | 0.998 | |
| AT4G04190.2 | ? | 25 | 2.36 | 0.189 | 0.357 | |
| AT4G04190.2 | ? | 26 | 4.03 | 0.192 | 0.022 | * |
| AT4G04190.2 | ? | 27 | 2.51 | 0.206 | 0.375 | |
| AT4G04190.2 | ? | 28 | 0.85 | 0.269 | 1.000 | |
| AT4G04190.2 | ? | 29 | 0.27 | 0.438 | 0.819 | |
| AT4G04190.2 | ? | 46 | 0.03 | 0.394 | 0.003 | ** |
| AT4G04190.2 | ? | 47 | 0.05 | 0.405 | 0.018 | * |
| AT4G04320.1 | Malonyl-CoA decarboxylase | 2 | 0.79 | 0.584 | 1.000 | |
| AT4G04320.1 | Malonyl-CoA decarboxylase | 3 | 0.81 | 0.561 | 1.000 | |
| AT4G04320.1 | Malonyl-CoA decarboxylase | 24 | 0.29 | 0.451 | 0.977 | |
| AT4G04320.1 | Malonyl-CoA decarboxylase | 25 | 0.29 | 0.318 | 0.746 | |
| AT4G04320.1 | Malonyl-CoA decarboxylase | 26 | 0.45 | 0.268 | 0.950 | |
| AT4G04320.1 | Malonyl-CoA decarboxylase | 27 | 0.94 | 0.256 | 1.000 | |
| AT4G04320.1 | Malonyl-CoA decarboxylase | 28 | 1.60 | 0.252 | 1.000 | |
| AT4G04320.1 | Malonyl-CoA decarboxylase | 29 | 2.20 | 0.251 | 0.933 | |
| AT4G04320.1 | Malonyl-CoA decarboxylase | 30 | 2.79 | 0.251 | 0.698 | |
| AT4G04320.1 | Malonyl-CoA decarboxylase | 31 | 3.34 | 0.254 | 0.472 | |
| AT4G04320.1 | Malonyl-CoA decarboxylase | 32 | 2.81 | 0.254 | 0.702 | |
| AT4G04320.1 | Malonyl-CoA decarboxylase | 33 | 1.53 | 0.254 | 1.000 | |
| AT4G04320.1 | Malonyl-CoA decarboxylase | 34 | 0.88 | 0.253 | 1.000 | |
| AT4G04320.1 | Malonyl-CoA decarboxylase | 35 | 0.80 | 0.254 | 1.000 | |
| AT4G04320.1 | Malonyl-CoA decarboxylase | 36 | 1.14 | 0.253 | 1.000 | |
| AT4G04320.1 | Malonyl-CoA decarboxylase | 37 | 1.85 | 0.254 | 0.992 | |
| AT4G04320.1 | Malonyl-CoA decarboxylase | 38 | 2.41 | 0.254 | 0.872 | |
| AT4G04320.1 | Malonyl-CoA decarboxylase | 39 | 1.93 | 0.255 | 0.986 | |
| AT4G04320.1 | Malonyl-CoA decarboxylase | 40 | 1.09 | 0.272 | 1.000 | |
| AT4G04320.1 | Malonyl-CoA decarboxylase | 41 | 0.56 | 0.361 | 1.000 | |
| AT4G04320.1 | Malonyl-CoA decarboxylase | 42 | 0.37 | 0.549 | 1.000 | |
| AT4G04320.1 | Malonyl-CoA decarboxylase | 46 | 0.79 | 1.802 | 1.000 | |

| | | | | | |
|---|---|---|---|---|---|
| AT4G04320.1 | Malonyl-CoA decarboxylase | 47 | 1.27 | 2.289 | 1.000 |
| AT4G04640.1 | ATP synthase subunit (ATPgamma1) | 2 | 0.89 | 0.676 | 1.000 |
| AT4G04640.1 | ATP synthase subunit (ATPgamma1) | 3 | 1.08 | 0.532 | 1.000 |
| AT4G04640.1 | ATP synthase subunit (ATPgamma1) | 6 | 1.32 | 0.474 | 1.000 |
| AT4G04640.1 | ATP synthase subunit (ATPgamma1) | 7 | 1.16 | 0.409 | 1.000 |
| AT4G04640.1 | ATP synthase subunit (ATPgamma1) | 8 | 0.98 | 0.386 | 1.000 |
| AT4G04640.1 | ATP synthase subunit (ATPgamma1) | 9 | 0.85 | 0.368 | 1.000 |
| AT4G04640.1 | ATP synthase subunit (ATPgamma1) | 10 | 0.80 | 0.363 | 1.000 |
| AT4G04640.1 | ATP synthase subunit (ATPgamma1) | 11 | 0.79 | 0.377 | 1.000 |
| AT4G04640.1 | ATP synthase subunit (ATPgamma1) | 12 | 0.76 | 0.392 | 1.000 |
| AT4G04640.1 | ATP synthase subunit (ATPgamma1) | 13 | 0.70 | 0.387 | 1.000 |
| AT4G04640.1 | ATP synthase subunit (ATPgamma1) | 14 | 0.57 | 0.413 | 1.000 |
| AT4G04640.1 | ATP synthase subunit (ATPgamma1) | 25 | 0.05 | 0.489 | 0.191 |
| AT4G04640.1 | ATP synthase subunit (ATPgamma1) | 26 | 0.21 | 0.376 | 0.724 |
| AT4G04640.1 | ATP synthase subunit (ATPgamma1) | 27 | 0.81 | 0.331 | 1.000 |
| AT4G04640.1 | ATP synthase subunit (ATPgamma1) | 28 | 1.62 | 0.317 | 1.000 |
| AT4G04640.1 | ATP synthase subunit (ATPgamma1) | 29 | 2.41 | 0.313 | 0.985 |
| AT4G04640.1 | ATP synthase subunit (ATPgamma1) | 30 | 3.17 | 0.312 | 0.854 |
| AT4G04640.1 | ATP synthase subunit (ATPgamma1) | 31 | 2.92 | 0.314 | 0.916 |
| AT4G04640.1 | ATP synthase subunit (ATPgamma1) | 32 | 1.57 | 0.315 | 1.000 |
| AT4G04640.1 | ATP synthase subunit (ATPgamma1) | 33 | 0.59 | 0.314 | 1.000 |
| AT4G04640.1 | ATP synthase subunit (ATPgamma1) | 34 | 0.25 | 0.314 | 0.651 |
| AT4G04640.1 | ATP synthase subunit (ATPgamma1) | 35 | 0.21 | 0.314 | 0.460 |
| AT4G04640.1 | ATP synthase subunit (ATPgamma1) | 36 | 0.36 | 0.314 | 0.934 |
| AT4G04640.1 | ATP synthase subunit (ATPgamma1) | 37 | 0.83 | 0.314 | 1.000 |
| AT4G04640.1 | ATP synthase subunit (ATPgamma1) | 38 | 2.03 | 0.313 | 0.999 |
| AT4G04640.1 | ATP synthase subunit (ATPgamma1) | 39 | 3.99 | 0.312 | 0.624 |
| AT4G04640.1 | ATP synthase subunit (ATPgamma1) | 40 | 4.84 | 0.313 | 0.421 |
| AT4G04640.1 | ATP synthase subunit (ATPgamma1) | 41 | 3.77 | 0.315 | 0.702 |
| AT4G04640.1 | ATP synthase subunit (ATPgamma1) | 42 | 2.49 | 0.318 | 0.981 |
| AT4G04640.1 | ATP synthase subunit (ATPgamma1) | 43 | 1.47 | 0.319 | 1.000 |

| | | | | | |
|---|---|---|---|---|---|
| AT4G04640.1 | ATP synthase subunit (ATPgamma1) | 44 | 0.98 | 0.319 | 1.000 |
| AT4G04640.1 | ATP synthase subunit (ATPgamma1) | 45 | 0.92 | 0.322 | 1.000 |
| AT4G04640.1 | ATP synthase subunit (ATPgamma1) | 46 | 0.97 | 0.367 | 1.000 |
| AT4G04640.1 | ATP synthase subunit (ATPgamma1) | 47 | 0.88 | 0.581 | 1.000 |
| AT4G04870.1 | Cardiolipin synthase (CLS) | 2 | 1.03 | 0.399 | 1.000 |
| AT4G04870.1 | Cardiolipin synthase (CLS) | 3 | 1.22 | 0.382 | 1.000 |
| AT4G04870.1 | Cardiolipin synthase (CLS) | 26 | 1.54 | 0.301 | 1.000 |
| AT4G04870.1 | Cardiolipin synthase (CLS) | 27 | 1.16 | 0.234 | 1.000 |
| AT4G04870.1 | Cardiolipin synthase (CLS) | 28 | 0.91 | 0.235 | 1.000 |
| AT4G04870.1 | Cardiolipin synthase (CLS) | 29 | 0.78 | 0.272 | 1.000 |
| AT4G04870.1 | Cardiolipin synthase (CLS) | 33 | 1.18 | 0.240 | 1.000 |
| AT4G04870.1 | Cardiolipin synthase (CLS) | 34 | 1.35 | 0.198 | 1.000 |
| AT4G04870.1 | Cardiolipin synthase (CLS) | 35 | 1.45 | 0.174 | 0.994 |
| AT4G04870.1 | Cardiolipin synthase (CLS) | 36 | 1.60 | 0.162 | 0.943 |
| AT4G04870.1 | Cardiolipin synthase (CLS) | 37 | 1.85 | 0.159 | 0.715 |
| AT4G04870.1 | Cardiolipin synthase (CLS) | 38 | 2.07 | 0.159 | 0.478 |
| AT4G04870.1 | Cardiolipin synthase (CLS) | 39 | 2.34 | 0.158 | 0.252 |
| AT4G04870.1 | Cardiolipin synthase (CLS) | 40 | 2.57 | 0.158 | 0.141 |
| AT4G04870.1 | Cardiolipin synthase (CLS) | 41 | 2.19 | 0.159 | 0.373 |
| AT4G04870.1 | Cardiolipin synthase (CLS) | 42 | 1.51 | 0.162 | 0.977 |
| AT4G04870.1 | Cardiolipin synthase (CLS) | 43 | 0.99 | 0.164 | 1.000 |
| AT4G04870.1 | Cardiolipin synthase (CLS) | 44 | 0.64 | 0.163 | 0.956 |
| AT4G04870.1 | Cardiolipin synthase (CLS) | 45 | 0.42 | 0.164 | 0.278 |
| AT4G04870.1 | Cardiolipin synthase (CLS) | 46 | 0.31 | 0.202 | 0.169 |
| AT4G04870.1 | Cardiolipin synthase (CLS) | 47 | 0.24 | 0.347 | 0.625 |
| AT4G05020.1 | Alternative NAD(P)H dehydrogenase B2 (NDB2) | 2 | 0.79 | 0.418 | 1.000 |
| AT4G05020.1 | Alternative NAD(P)H dehydrogenase B2 (NDB2) | 3 | 0.64 | 0.399 | 1.000 |
| AT4G05020.1 | Alternative NAD(P)H dehydrogenase B2 (NDB2) | 27 | 1.36 | 0.308 | 1.000 |
| AT4G05020.1 | Alternative NAD(P)H dehydrogenase B2 (NDB2) | 28 | 2.15 | 0.227 | 0.904 |
| AT4G05020.1 | Alternative NAD(P)H dehydrogenase B2 (NDB2) | 29 | 2.67 | 0.197 | 0.410 |
| AT4G05020.1 | Alternative NAD(P)H dehydrogenase B2 (NDB2) | 30 | 2.71 | 0.188 | 0.315 |
| AT4G05020.1 | Alternative NAD(P)H dehydrogenase B2 (NDB2) | 31 | 2.30 | 0.185 | 0.573 |
| AT4G05020.1 | Alternative NAD(P)H dehydrogenase B2 (NDB2) | 32 | 1.84 | 0.185 | 0.914 |
| AT4G05020.1 | Alternative NAD(P)H dehydrogenase B2 (NDB2) | 33 | 1.48 | 0.184 | 0.998 |
| AT4G05020.1 | Alternative NAD(P)H dehydrogenase B2 (NDB2) | 34 | 1.37 | 0.184 | 1.000 |
| AT4G05020.1 | Alternative NAD(P)H dehydrogenase B2 (NDB2) | 35 | 1.37 | 0.184 | 1.000 |
| AT4G05020.1 | Alternative NAD(P)H dehydrogenase B2 (NDB2) | 36 | 1.27 | 0.183 | 1.000 |
| AT4G05020.1 | Alternative NAD(P)H dehydrogenase B2 (NDB2) | 37 | 1.07 | 0.184 | 1.000 |
| AT4G05020.1 | Alternative NAD(P)H dehydrogenase B2 (NDB2) | 38 | 0.81 | 0.185 | 1.000 |
| AT4G05020.1 | Alternative NAD(P)H dehydrogenase B2 (NDB2) | 39 | 0.67 | 0.184 | 0.998 |
| AT4G05020.1 | Alternative NAD(P)H dehydrogenase B2 (NDB2) | 40 | 0.85 | 0.183 | 1.000 |

| | | | | | | |
|---|---|---|---|---|---|---|
| AT4G05020.1 | Alternative NAD(P)H dehydrogenase B2 (NDB2) | 41 | 1.65 | 0.184 | 0.981 | |
| AT4G05020.1 | Alternative NAD(P)H dehydrogenase B2 (NDB2) | 42 | 3.12 | 0.188 | 0.153 | |
| AT4G05020.1 | Alternative NAD(P)H dehydrogenase B2 (NDB2) | 43 | 3.72 | 0.191 | 0.057 | . |
| AT4G05020.1 | Alternative NAD(P)H dehydrogenase B2 (NDB2) | 44 | 3.04 | 0.194 | 0.212 | |
| AT4G05020.1 | Alternative NAD(P)H dehydrogenase B2 (NDB2) | 45 | 1.85 | 0.226 | 0.981 | |
| AT4G05020.1 | Alternative NAD(P)H dehydrogenase B2 (NDB2) | 46 | 1.00 | 0.392 | 1.000 | |
| AT4G05020.1 | Alternative NAD(P)H dehydrogenase B2 (NDB2) | 47 | 0.53 | 0.731 | 1.000 | |
| AT4G05400.2 | copper ion binding | 2 | 1.28 | 0.443 | 1.000 | |
| AT4G05400.2 | copper ion binding | 3 | 1.19 | 0.427 | 1.000 | |
| AT4G05400.2 | copper ion binding | 27 | 1.38 | 0.354 | 1.000 | |
| AT4G05400.2 | copper ion binding | 28 | 1.58 | 0.287 | 1.000 | |
| AT4G05400.2 | copper ion binding | 29 | 1.63 | 0.264 | 0.999 | |
| AT4G05400.2 | copper ion binding | 30 | 1.60 | 0.252 | 0.999 | |
| AT4G05400.2 | copper ion binding | 31 | 1.51 | 0.245 | 1.000 | |
| AT4G05400.2 | copper ion binding | 32 | 1.20 | 0.241 | 1.000 | |
| AT4G05400.2 | copper ion binding | 33 | 0.96 | 0.235 | 1.000 | |
| AT4G05400.2 | copper ion binding | 34 | 1.05 | 0.234 | 1.000 | |
| AT4G05400.2 | copper ion binding | 35 | 1.27 | 0.236 | 1.000 | |
| AT4G05400.2 | copper ion binding | 36 | 1.43 | 0.242 | 1.000 | |
| AT4G05400.2 | copper ion binding | 37 | 1.80 | 0.249 | 0.989 | |
| AT4G05400.2 | copper ion binding | 38 | 2.63 | 0.254 | 0.734 | |
| AT4G05400.2 | copper ion binding | 39 | 3.11 | 0.249 | 0.479 | |
| AT4G05400.2 | copper ion binding | 40 | 2.96 | 0.239 | 0.486 | |
| AT4G05400.2 | copper ion binding | 41 | 2.47 | 0.234 | 0.711 | |
| AT4G05400.2 | copper ion binding | 42 | 1.80 | 0.236 | 0.983 | |
| AT4G05400.2 | copper ion binding | 43 | 1.21 | 0.237 | 1.000 | |
| AT4G05400.2 | copper ion binding | 44 | 0.78 | 0.236 | 1.000 | |
| AT4G05400.2 | copper ion binding | 45 | 0.51 | 0.236 | 0.954 | |
| AT4G05400.2 | copper ion binding | 46 | 0.36 | 0.259 | 0.695 | |
| AT4G05400.2 | copper ion binding | 47 | 0.24 | 0.460 | 0.913 | |
| AT4G05590.1 | Pyruvate carrier-like protein | 2 | 0.89 | 0.322 | 1.000 | |
| AT4G05590.1 | Pyruvate carrier-like protein | 3 | 1.06 | 0.227 | 1.000 | |
| AT4G05590.1 | Pyruvate carrier-like protein | 4 | 1.27 | 0.211 | 1.000 | |
| AT4G05590.1 | Pyruvate carrier-like protein | 5 | 1.55 | 0.214 | 1.000 | |
| AT4G05590.1 | Pyruvate carrier-like protein | 6 | 2.00 | 0.198 | 0.952 | |
| AT4G05590.1 | Pyruvate carrier-like protein | 7 | 2.88 | 0.184 | 0.293 | |
| AT4G05590.1 | Pyruvate carrier-like protein | 8 | 4.44 | 0.174 | 0.009 | ** |
| AT4G05590.1 | Pyruvate carrier-like protein | 9 | 5.83 | 0.169 | 0.000 | *** |
| AT4G05590.1 | Pyruvate carrier-like protein | 10 | 5.63 | 0.168 | 0.000 | *** |
| AT4G05590.1 | Pyruvate carrier-like protein | 11 | 4.34 | 0.168 | 0.006 | ** |
| AT4G05590.1 | Pyruvate carrier-like protein | 12 | 3.35 | 0.168 | 0.057 | . |
| AT4G05590.1 | Pyruvate carrier-like protein | 13 | 2.82 | 0.167 | 0.187 | |
| AT4G05590.1 | Pyruvate carrier-like protein | 14 | 2.44 | 0.166 | 0.403 | |
| AT4G05590.1 | Pyruvate carrier-like protein | 15 | 1.98 | 0.165 | 0.824 | |
| AT4G05590.1 | Pyruvate carrier-like protein | 16 | 1.49 | 0.164 | 1.000 | |
| AT4G05590.1 | Pyruvate carrier-like protein | 17 | 1.37 | 0.163 | 1.000 | |
| AT4G05590.1 | Pyruvate carrier-like protein | 18 | 1.60 | 0.166 | 0.995 | |
| AT4G05590.1 | Pyruvate carrier-like protein | 19 | 1.98 | 0.169 | 0.850 | |
| AT4G05590.1 | Pyruvate carrier-like protein | 20 | 2.40 | 0.172 | 0.504 | |
| AT4G05590.1 | Pyruvate carrier-like protein | 21 | 2.56 | 0.174 | 0.398 | |
| AT4G05590.1 | Pyruvate carrier-like protein | 22 | 2.46 | 0.178 | 0.513 | |
| AT4G05590.1 | Pyruvate carrier-like protein | 23 | 2.30 | 0.184 | 0.700 | |
| AT4G05590.1 | Pyruvate carrier-like protein | 24 | 2.11 | 0.194 | 0.899 | |
| AT4G05590.1 | Pyruvate carrier-like protein | 25 | 1.84 | 0.207 | 0.993 | |
| AT4G05590.1 | Pyruvate carrier-like protein | 26 | 1.52 | 0.212 | 1.000 | |
| AT4G05590.1 | Pyruvate carrier-like protein | 27 | 1.28 | 0.207 | 1.000 | |

| | | | | | | |
|---|---|---|---|---|---|---|
| AT4G05590.1 | Pyruvate carrier-like protein | 28 | 1.22 | 0.196 | 1.000 | |
| AT4G05590.1 | Pyruvate carrier-like protein | 29 | 1.45 | 0.183 | 1.000 | |
| AT4G05590.1 | Pyruvate carrier-like protein | 30 | 2.10 | 0.176 | 0.802 | |
| AT4G05590.1 | Pyruvate carrier-like protein | 31 | 3.70 | 0.171 | 0.032 | * |
| AT4G05590.1 | Pyruvate carrier-like protein | 32 | 7.10 | 0.169 | 0.000 | *** |
| AT4G05590.1 | Pyruvate carrier-like protein | 33 | 11.33 | 0.168 | 0.000 | *** |
| AT4G05590.1 | Pyruvate carrier-like protein | 34 | 10.86 | 0.169 | 0.000 | *** |
| AT4G05590.1 | Pyruvate carrier-like protein | 35 | 6.27 | 0.169 | 0.000 | *** |
| AT4G05590.1 | Pyruvate carrier-like protein | 36 | 3.15 | 0.168 | 0.093 | . |
| AT4G05590.1 | Pyruvate carrier-like protein | 37 | 2.02 | 0.168 | 0.816 | |
| AT4G05590.1 | Pyruvate carrier-like protein | 38 | 1.79 | 0.169 | 0.959 | |
| AT4G05590.1 | Pyruvate carrier-like protein | 39 | 1.83 | 0.168 | 0.942 | |
| AT4G05590.1 | Pyruvate carrier-like protein | 40 | 1.70 | 0.168 | 0.984 | |
| AT4G05590.1 | Pyruvate carrier-like protein | 41 | 1.32 | 0.169 | 1.000 | |
| AT4G05590.1 | Pyruvate carrier-like protein | 42 | 0.95 | 0.171 | 1.000 | |
| AT4G05590.1 | Pyruvate carrier-like protein | 43 | 0.69 | 0.173 | 1.000 | |
| AT4G05590.1 | Pyruvate carrier-like protein | 44 | 0.48 | 0.172 | 0.778 | |
| AT4G05590.1 | Pyruvate carrier-like protein | 45 | 0.31 | 0.172 | 0.094 | . |
| AT4G05590.1 | Pyruvate carrier-like protein | 46 | 0.19 | 0.206 | 0.017 | * |
| AT4G05590.1 | Pyruvate carrier-like protein | 47 | 0.11 | 0.343 | 0.139 | |
| AT4G08390.3 | Stromal ascorbate peroxidase (SAPX) | 2 | 0.98 | 0.241 | 1.000 | |
| AT4G08390.3 | Stromal ascorbate peroxidase (SAPX) | 3 | 0.97 | 0.184 | 1.000 | |
| AT4G08390.3 | Stromal ascorbate peroxidase (SAPX) | 4 | 0.99 | 0.202 | 1.000 | |
| AT4G08390.3 | Stromal ascorbate peroxidase (SAPX) | 8 | 1.13 | 0.177 | 1.000 | |
| AT4G08390.3 | Stromal ascorbate peroxidase (SAPX) | 9 | 1.08 | 0.145 | 1.000 | |
| AT4G08390.3 | Stromal ascorbate peroxidase (SAPX) | 10 | 1.06 | 0.129 | 1.000 | |
| AT4G08390.3 | Stromal ascorbate peroxidase (SAPX) | 11 | 1.16 | 0.121 | 1.000 | |
| AT4G08390.3 | Stromal ascorbate peroxidase (SAPX) | 12 | 1.32 | 0.117 | 1.000 | |
| AT4G08390.3 | Stromal ascorbate peroxidase (SAPX) | 13 | 1.33 | 0.114 | 1.000 | |
| AT4G08390.3 | Stromal ascorbate peroxidase (SAPX) | 14 | 1.17 | 0.112 | 1.000 | |
| AT4G08390.3 | Stromal ascorbate peroxidase (SAPX) | 15 | 0.98 | 0.111 | 1.000 | |
| AT4G08390.3 | Stromal ascorbate peroxidase (SAPX) | 16 | 0.90 | 0.110 | 1.000 | |
| AT4G08390.3 | Stromal ascorbate peroxidase (SAPX) | 17 | 0.93 | 0.110 | 1.000 | |
| AT4G08390.3 | Stromal ascorbate peroxidase (SAPX) | 18 | 1.01 | 0.112 | 1.000 | |
| AT4G08390.3 | Stromal ascorbate peroxidase (SAPX) | 19 | 1.12 | 0.116 | 1.000 | |
| AT4G08390.3 | Stromal ascorbate peroxidase (SAPX) | 20 | 1.21 | 0.119 | 1.000 | |
| AT4G08390.3 | Stromal ascorbate peroxidase (SAPX) | 21 | 1.30 | 0.120 | 1.000 | |
| AT4G08390.3 | Stromal ascorbate peroxidase (SAPX) | 22 | 1.31 | 0.120 | 1.000 | |
| AT4G08390.3 | Stromal ascorbate peroxidase (SAPX) | 23 | 1.29 | 0.119 | 1.000 | |
| AT4G08390.3 | Stromal ascorbate peroxidase (SAPX) | 24 | 1.31 | 0.117 | 1.000 | |
| AT4G08390.3 | Stromal ascorbate peroxidase (SAPX) | 25 | 1.30 | 0.115 | 1.000 | |

| | | | | | | |
|---|---|---|---|---|---|---|
| AT4G08390.3 | Stromal ascorbate peroxidase (SAPX) | 26 | 1.15 | 0.114 | 1.000 | |
| AT4G08390.3 | Stromal ascorbate peroxidase (SAPX) | 27 | 0.99 | 0.114 | 1.000 | |
| AT4G08390.3 | Stromal ascorbate peroxidase (SAPX) | 28 | 0.95 | 0.114 | 1.000 | |
| AT4G08390.3 | Stromal ascorbate peroxidase (SAPX) | 29 | 1.01 | 0.114 | 1.000 | |
| AT4G08390.3 | Stromal ascorbate peroxidase (SAPX) | 30 | 1.05 | 0.114 | 1.000 | |
| AT4G08390.3 | Stromal ascorbate peroxidase (SAPX) | 31 | 1.03 | 0.114 | 1.000 | |
| AT4G08390.3 | Stromal ascorbate peroxidase (SAPX) | 32 | 0.96 | 0.115 | 1.000 | |
| AT4G08390.3 | Stromal ascorbate peroxidase (SAPX) | 33 | 0.88 | 0.115 | 1.000 | |
| AT4G08390.3 | Stromal ascorbate peroxidase (SAPX) | 34 | 0.85 | 0.115 | 1.000 | |
| AT4G08390.3 | Stromal ascorbate peroxidase (SAPX) | 35 | 0.88 | 0.115 | 1.000 | |
| AT4G08390.3 | Stromal ascorbate peroxidase (SAPX) | 36 | 0.97 | 0.114 | 1.000 | |
| AT4G08390.3 | Stromal ascorbate peroxidase (SAPX) | 37 | 1.09 | 0.114 | 1.000 | |
| AT4G08390.3 | Stromal ascorbate peroxidase (SAPX) | 38 | 1.23 | 0.114 | 1.000 | |
| AT4G08390.3 | Stromal ascorbate peroxidase (SAPX) | 39 | 1.36 | 0.115 | 0.998 | |
| AT4G08390.3 | Stromal ascorbate peroxidase (SAPX) | 40 | 1.47 | 0.115 | 0.978 | |
| AT4G08390.3 | Stromal ascorbate peroxidase (SAPX) | 41 | 1.50 | 0.116 | 0.960 | |
| AT4G08390.3 | Stromal ascorbate peroxidase (SAPX) | 42 | 1.48 | 0.117 | 0.977 | |
| AT4G08390.3 | Stromal ascorbate peroxidase (SAPX) | 43 | 1.38 | 0.119 | 0.998 | |
| AT4G08390.3 | Stromal ascorbate peroxidase (SAPX) | 44 | 1.11 | 0.119 | 1.000 | |
| AT4G08390.3 | Stromal ascorbate peroxidase (SAPX) | 45 | 0.72 | 0.121 | 0.998 | |
| AT4G08390.3 | Stromal ascorbate peroxidase (SAPX) | 46 | 0.36 | 0.150 | 0.095 | . |
| AT4G08390.3 | Stromal ascorbate peroxidase (SAPX) | 47 | 0.15 | 0.244 | 0.027 | * |
| AT4G08870.1 | Arginase (ARGAH2) | 2 | 1.16 | 0.588 | 1.000 | |
| AT4G08870.1 | Arginase (ARGAH2) | 3 | 1.04 | 0.454 | 1.000 | |
| AT4G08870.1 | Arginase (ARGAH2) | 6 | 0.85 | 0.389 | 1.000 | |
| AT4G08870.1 | Arginase (ARGAH2) | 7 | 0.87 | 0.317 | 1.000 | |
| AT4G08870.1 | Arginase (ARGAH2) | 8 | 0.94 | 0.277 | 1.000 | |
| AT4G08870.1 | Arginase (ARGAH2) | 9 | 1.05 | 0.260 | 1.000 | |
| AT4G08870.1 | Arginase (ARGAH2) | 10 | 1.19 | 0.254 | 1.000 | |
| AT4G08870.1 | Arginase (ARGAH2) | 11 | 1.49 | 0.254 | 1.000 | |
| AT4G08870.1 | Arginase (ARGAH2) | 12 | 1.94 | 0.253 | 0.999 | |
| AT4G08870.1 | Arginase (ARGAH2) | 13 | 2.22 | 0.251 | 0.986 | |
| AT4G08870.1 | Arginase (ARGAH2) | 14 | 2.27 | 0.248 | 0.979 | |
| AT4G08870.1 | Arginase (ARGAH2) | 15 | 2.15 | 0.247 | 0.989 | |
| AT4G08870.1 | Arginase (ARGAH2) | 16 | 2.09 | 0.245 | 0.993 | |
| AT4G08870.1 | Arginase (ARGAH2) | 17 | 2.45 | 0.245 | 0.940 | |
| AT4G08870.1 | Arginase (ARGAH2) | 18 | 2.89 | 0.248 | 0.806 | |
| AT4G08870.1 | Arginase (ARGAH2) | 19 | 3.22 | 0.254 | 0.699 | |
| AT4G08870.1 | Arginase (ARGAH2) | 20 | 2.95 | 0.260 | 0.838 | |
| AT4G08870.1 | Arginase (ARGAH2) | 21 | 1.93 | 0.264 | 0.999 | |

| | | | | | |
|---|---|---|---|---|---|
| AT4G08870.1 | Arginase (ARGAH2) | 22 | 1.16 | 0.263 | 1.000 |
| AT4G08870.1 | Arginase (ARGAH2) | 23 | 0.91 | 0.261 | 1.000 |
| AT4G08870.1 | Arginase (ARGAH2) | 24 | 0.87 | 0.257 | 1.000 |
| AT4G08870.1 | Arginase (ARGAH2) | 25 | 0.83 | 0.252 | 1.000 |
| AT4G08870.1 | Arginase (ARGAH2) | 26 | 0.93 | 0.251 | 1.000 |
| AT4G08870.1 | Arginase (ARGAH2) | 27 | 1.31 | 0.251 | 1.000 |
| AT4G08870.1 | Arginase (ARGAH2) | 28 | 1.65 | 0.251 | 1.000 |
| AT4G08870.1 | Arginase (ARGAH2) | 29 | 1.72 | 0.251 | 1.000 |
| AT4G08870.1 | Arginase (ARGAH2) | 30 | 1.40 | 0.251 | 1.000 |
| AT4G08870.1 | Arginase (ARGAH2) | 31 | 1.07 | 0.252 | 1.000 |
| AT4G08870.1 | Arginase (ARGAH2) | 32 | 0.96 | 0.253 | 1.000 |
| AT4G08870.1 | Arginase (ARGAH2) | 33 | 1.13 | 0.253 | 1.000 |
| AT4G08870.1 | Arginase (ARGAH2) | 34 | 1.58 | 0.253 | 1.000 |
| AT4G08870.1 | Arginase (ARGAH2) | 35 | 1.80 | 0.253 | 1.000 |
| AT4G08870.1 | Arginase (ARGAH2) | 36 | 1.33 | 0.252 | 1.000 |
| AT4G08870.1 | Arginase (ARGAH2) | 37 | 0.75 | 0.252 | 1.000 |
| AT4G08870.1 | Arginase (ARGAH2) | 38 | 0.46 | 0.252 | 0.992 |
| AT4G08870.1 | Arginase (ARGAH2) | 39 | 0.45 | 0.253 | 0.988 |
| AT4G08870.1 | Arginase (ARGAH2) | 40 | 0.72 | 0.253 | 1.000 |
| AT4G08870.1 | Arginase (ARGAH2) | 41 | 1.30 | 0.254 | 1.000 |
| AT4G08870.1 | Arginase (ARGAH2) | 42 | 1.81 | 0.257 | 1.000 |
| AT4G08870.1 | Arginase (ARGAH2) | 43 | 1.59 | 0.260 | 1.000 |
| AT4G08870.1 | Arginase (ARGAH2) | 44 | 1.00 | 0.260 | 1.000 |
| AT4G08870.1 | Arginase (ARGAH2) | 45 | 0.54 | 0.260 | 1.000 |
| AT4G08870.1 | Arginase (ARGAH2) | 46 | 0.30 | 0.306 | 0.896 |
| AT4G08870.1 | Arginase (ARGAH2) | 47 | 0.18 | 0.514 | 0.973 |
| AT4G08900.1 | Arginase (ARGAH1) | 2 | 0.88 | 0.652 | 1.000 |
| AT4G08900.1 | Arginase (ARGAH1) | 3 | 0.85 | 0.610 | 1.000 |
| AT4G08900.1 | Arginase (ARGAH1) | 22 | 2.41 | 0.562 | 1.000 |
| AT4G08900.1 | Arginase (ARGAH1) | 23 | 2.64 | 0.488 | 0.999 |
| AT4G08900.1 | Arginase (ARGAH1) | 24 | 2.56 | 0.436 | 0.997 |
| AT4G08900.1 | Arginase (ARGAH1) | 25 | 2.14 | 0.380 | 0.999 |
| AT4G08900.1 | Arginase (ARGAH1) | 26 | 1.81 | 0.354 | 1.000 |
| AT4G08900.1 | Arginase (ARGAH1) | 27 | 1.44 | 0.324 | 1.000 |
| AT4G08900.1 | Arginase (ARGAH1) | 28 | 1.08 | 0.310 | 1.000 |
| AT4G08900.1 | Arginase (ARGAH1) | 29 | 1.17 | 0.308 | 1.000 |
| AT4G08900.1 | Arginase (ARGAH1) | 30 | 1.16 | 0.307 | 1.000 |
| AT4G08900.1 | Arginase (ARGAH1) | 31 | 1.19 | 0.310 | 1.000 |
| AT4G08900.1 | Arginase (ARGAH1) | 32 | 1.20 | 0.310 | 1.000 |
| AT4G08900.1 | Arginase (ARGAH1) | 33 | 1.27 | 0.309 | 1.000 |
| AT4G08900.1 | Arginase (ARGAH1) | 34 | 1.58 | 0.311 | 1.000 |
| AT4G08900.1 | Arginase (ARGAH1) | 35 | 2.00 | 0.311 | 0.996 |
| AT4G08900.1 | Arginase (ARGAH1) | 36 | 1.85 | 0.309 | 0.999 |
| AT4G08900.1 | Arginase (ARGAH1) | 37 | 1.11 | 0.310 | 1.000 |
| AT4G08900.1 | Arginase (ARGAH1) | 38 | 0.48 | 0.314 | 0.995 |
| AT4G08900.1 | Arginase (ARGAH1) | 39 | 0.22 | 0.326 | 0.499 |
| AT4G08900.1 | Arginase (ARGAH1) | 40 | 0.19 | 0.412 | 0.728 |
| AT4G08900.1 | Arginase (ARGAH1) | 41 | 0.53 | 0.652 | 1.000 |
| AT4G08900.1 | Arginase (ARGAH1) | 46 | 2747124.23 | 2.907 | 0.357 |
| AT4G08900.1 | Arginase (ARGAH1) | 47 | ########### | 3.610 | 0.319 |
| AT4G09320.1 | Nucleoside diphosphate kinase 1 (NDPK1) | 2 | 1.15 | 0.497 | 1.000 |
| AT4G09320.1 | Nucleoside diphosphate kinase 1 (NDPK1) | 3 | 1.13 | 0.479 | 1.000 |
| AT4G09320.1 | Nucleoside diphosphate kinase 1 (NDPK1) | 22 | 0.87 | 0.239 | 1.000 |
| AT4G09320.1 | Nucleoside diphosphate kinase 1 (NDPK1) | 23 | 0.84 | 0.208 | 1.000 |
| AT4G09320.1 | Nucleoside diphosphate kinase 1 (NDPK1) | 24 | 0.81 | 0.186 | 0.999 |
| AT4G09320.1 | Nucleoside diphosphate kinase 1 (NDPK1) | 25 | 0.76 | 0.171 | 0.994 |

| | | | | | | |
|---|---|---|---|---|---|---|
| AT4G09320.1 | Nucleoside diphosphate kinase 1 (NDPK1) | 26 | 0.72 | 0.160 | 0.963 | |
| AT4G09320.1 | Nucleoside diphosphate kinase 1 (NDPK1) | 27 | 0.68 | 0.152 | 0.894 | |
| AT4G09320.1 | Nucleoside diphosphate kinase 1 (NDPK1) | 28 | 0.65 | 0.149 | 0.825 | |
| AT4G09320.1 | Nucleoside diphosphate kinase 1 (NDPK1) | 29 | 0.64 | 0.152 | 0.801 | |
| AT4G09320.1 | Nucleoside diphosphate kinase 1 (NDPK1) | 30 | 0.63 | 0.162 | 0.844 | |
| AT4G09320.1 | Nucleoside diphosphate kinase 1 (NDPK1) | 31 | 0.66 | 0.179 | 0.933 | |
| AT4G09320.1 | Nucleoside diphosphate kinase 1 (NDPK1) | 32 | 0.72 | 0.197 | 0.990 | |
| AT4G09320.1 | Nucleoside diphosphate kinase 1 (NDPK1) | 38 | 1.46 | 0.193 | 0.975 | |
| AT4G09320.1 | Nucleoside diphosphate kinase 1 (NDPK1) | 39 | 1.47 | 0.183 | 0.964 | |
| AT4G09320.1 | Nucleoside diphosphate kinase 1 (NDPK1) | 40 | 1.34 | 0.180 | 0.993 | |
| AT4G09320.1 | Nucleoside diphosphate kinase 1 (NDPK1) | 41 | 1.10 | 0.185 | 1.000 | |
| AT4G09320.1 | Nucleoside diphosphate kinase 1 (NDPK1) | 42 | 0.84 | 0.198 | 1.000 | |
| AT4G09320.1 | Nucleoside diphosphate kinase 1 (NDPK1) | 43 | 0.61 | 0.214 | 0.938 | |
| AT4G09320.1 | Nucleoside diphosphate kinase 1 (NDPK1) | 46 | 0.20 | 0.277 | 0.108 | |
| AT4G09320.1 | Nucleoside diphosphate kinase 1 (NDPK1) | 47 | 0.13 | 0.321 | 0.068 | . |
| AT4G09650.1 | ATP synthase subunit (ATPdelta) | 2 | 0.67 | 0.573 | 1.000 | |
| AT4G09650.1 | ATP synthase subunit (ATPdelta) | 3 | 0.39 | 0.561 | 1.000 | |
| AT4G09650.1 | ATP synthase subunit (ATPdelta) | 24 | 0.07 | 0.441 | 0.113 | |
| AT4G09650.1 | ATP synthase subunit (ATPdelta) | 25 | 0.23 | 0.342 | 0.543 | |
| AT4G09650.1 | ATP synthase subunit (ATPdelta) | 26 | 1.52 | 0.305 | 1.000 | |
| AT4G09650.1 | ATP synthase subunit (ATPdelta) | 27 | 5.16 | 0.294 | 0.195 | |
| AT4G09650.1 | ATP synthase subunit (ATPdelta) | 28 | 4.00 | 0.291 | 0.393 | |
| AT4G09650.1 | ATP synthase subunit (ATPdelta) | 29 | 1.92 | 0.290 | 0.991 | |
| AT4G09650.1 | ATP synthase subunit (ATPdelta) | 30 | 0.93 | 0.290 | 1.000 | |
| AT4G09650.1 | ATP synthase subunit (ATPdelta) | 31 | 0.50 | 0.291 | 0.984 | |
| AT4G09650.1 | ATP synthase subunit (ATPdelta) | 32 | 0.34 | 0.291 | 0.742 | |
| AT4G09650.1 | ATP synthase subunit (ATPdelta) | 33 | 0.29 | 0.291 | 0.536 | |
| AT4G09650.1 | ATP synthase subunit (ATPdelta) | 34 | 0.28 | 0.292 | 0.507 | |
| AT4G09650.1 | ATP synthase subunit (ATPdelta) | 35 | 0.24 | 0.296 | 0.386 | |
| AT4G09650.1 | ATP synthase subunit (ATPdelta) | 36 | 0.22 | 0.307 | 0.338 | |
| AT4G09650.1 | ATP synthase subunit (ATPdelta) | 37 | 0.26 | 0.338 | 0.645 | |

| | | | | | |
|---|---|---|---|---|---|
| AT4G09650.1 | ATP synthase subunit (ATPdelta) | 38 | 0.38 | 0.393 | 0.981 |
| AT4G09650.1 | ATP synthase subunit (ATPdelta) | 40 | 0.50 | 0.356 | 0.998 |
| AT4G09650.1 | ATP synthase subunit (ATPdelta) | 41 | 0.41 | 0.332 | 0.956 |
| AT4G09650.1 | ATP synthase subunit (ATPdelta) | 42 | 0.31 | 0.357 | 0.848 |
| AT4G09650.1 | ATP synthase subunit (ATPdelta) | 43 | 0.31 | 0.366 | 0.874 |
| AT4G09650.1 | ATP synthase subunit (ATPdelta) | 44 | 0.37 | 0.358 | 0.952 |
| AT4G09650.1 | ATP synthase subunit (ATPdelta) | 45 | 0.44 | 0.324 | 0.974 |
| AT4G09650.1 | ATP synthase subunit (ATPdelta) | 46 | 0.40 | 0.338 | 0.954 |
| AT4G09650.1 | ATP synthase subunit (ATPdelta) | 47 | 0.28 | 0.489 | 0.965 |
| AT4G09800.1 | 40S ribosomal protein S18 (RPS18C) | 2 | 0.86 | 0.167 | 0.999 |
| AT4G09800.1 | 40S ribosomal protein S18 (RPS18C) | 3 | 0.90 | 0.152 | 1.000 |
| AT4G09800.1 | 40S ribosomal protein S18 (RPS18C) | 4 | 0.94 | 0.140 | 1.000 |
| AT4G09800.1 | 40S ribosomal protein S18 (RPS18C) | 5 | 0.98 | 0.131 | 1.000 |
| AT4G09800.1 | 40S ribosomal protein S18 (RPS18C) | 6 | 1.02 | 0.124 | 1.000 |
| AT4G09800.1 | 40S ribosomal protein S18 (RPS18C) | 7 | 1.06 | 0.118 | 1.000 |
| AT4G09800.1 | 40S ribosomal protein S18 (RPS18C) | 8 | 1.10 | 0.114 | 0.999 |
| AT4G09800.1 | 40S ribosomal protein S18 (RPS18C) | 9 | 1.14 | 0.112 | 0.992 |
| AT4G09800.1 | 40S ribosomal protein S18 (RPS18C) | 10 | 1.18 | 0.111 | 0.975 |
| AT4G09800.1 | 40S ribosomal protein S18 (RPS18C) | 11 | 1.21 | 0.110 | 0.947 |
| AT4G09800.1 | 40S ribosomal protein S18 (RPS18C) | 12 | 1.24 | 0.110 | 0.913 |
| AT4G09800.1 | 40S ribosomal protein S18 (RPS18C) | 13 | 1.27 | 0.110 | 0.879 |
| AT4G09800.1 | 40S ribosomal protein S18 (RPS18C) | 14 | 1.29 | 0.111 | 0.848 |
| AT4G09800.1 | 40S ribosomal protein S18 (RPS18C) | 15 | 1.30 | 0.112 | 0.826 |
| AT4G09800.1 | 40S ribosomal protein S18 (RPS18C) | 16 | 1.32 | 0.113 | 0.813 |
| AT4G09800.1 | 40S ribosomal protein S18 (RPS18C) | 17 | 1.32 | 0.114 | 0.807 |
| AT4G09800.1 | 40S ribosomal protein S18 (RPS18C) | 18 | 1.33 | 0.116 | 0.805 |
| AT4G09800.1 | 40S ribosomal protein S18 (RPS18C) | 19 | 1.33 | 0.117 | 0.808 |
| AT4G09800.1 | 40S ribosomal protein S18 (RPS18C) | 20 | 1.33 | 0.119 | 0.814 |
| AT4G09800.1 | 40S ribosomal protein S18 (RPS18C) | 21 | 1.33 | 0.121 | 0.824 |
| AT4G09800.1 | 40S ribosomal protein S18 (RPS18C) | 22 | 1.33 | 0.122 | 0.837 |
| AT4G09800.1 | 40S ribosomal protein S18 (RPS18C) | 23 | 1.33 | 0.124 | 0.851 |

| | | | | | | |
|---|---|---|---|---|---|---|
| AT4G09800.1 | 40S ribosomal protein S18 (RPS18C) | 24 | 1.32 | 0.125 | 0.865 | |
| AT4G09800.1 | 40S ribosomal protein S18 (RPS18C) | 26 | 1.30 | 0.128 | 0.894 | |
| AT4G09800.1 | 40S ribosomal protein S18 (RPS18C) | 27 | 1.30 | 0.130 | 0.909 | |
| AT4G09800.1 | 40S ribosomal protein S18 (RPS18C) | 28 | 1.29 | 0.132 | 0.922 | |
| AT4G09800.1 | 40S ribosomal protein S18 (RPS18C) | 29 | 1.28 | 0.136 | 0.935 | |
| AT4G09800.1 | 40S ribosomal protein S18 (RPS18C) | 30 | 1.28 | 0.140 | 0.947 | |
| AT4G09800.1 | 40S ribosomal protein S18 (RPS18C) | 31 | 1.27 | 0.146 | 0.958 | |
| AT4G09800.1 | 40S ribosomal protein S18 (RPS18C) | 32 | 1.27 | 0.154 | 0.967 | |
| AT4G09800.1 | 40S ribosomal protein S18 (RPS18C) | 33 | 1.28 | 0.163 | 0.974 | |
| AT4G09800.1 | 40S ribosomal protein S18 (RPS18C) | 34 | 1.28 | 0.175 | 0.980 | |
| AT4G09800.1 | 40S ribosomal protein S18 (RPS18C) | 46 | 1.44 | 0.442 | 0.999 | |
| AT4G09800.1 | 40S ribosomal protein S18 (RPS18C) | 47 | 1.46 | 0.473 | 0.999 | |
| AT4G10060.1 | Beta-glucosidase, GBA2 type family protein / Putative glucosylceramidase | 2 | 0.83 | 0.446 | 1.000 | |
| AT4G10060.1 | Beta-glucosidase, GBA2 type family protein / Putative glucosylceramidase | 3 | 0.56 | 0.437 | 0.999 | |
| AT4G10060.1 | Beta-glucosidase, GBA2 type family protein / Putative glucosylceramidase | 38 | 4.56 | 0.383 | 0.567 | |
| AT4G10060.1 | Beta-glucosidase, GBA2 type family protein / Putative glucosylceramidase | 39 | 8.97 | 0.284 | 0.014 | * |
| AT4G10060.1 | Beta-glucosidase, GBA2 type family protein / Putative glucosylceramidase | 40 | 8.57 | 0.243 | 0.003 | ** |
| AT4G10060.1 | Beta-glucosidase, GBA2 type family protein / Putative glucosylceramidase | 41 | 2.23 | 0.235 | 0.742 | |
| AT4G10060.1 | Beta-glucosidase, GBA2 type family protein / Putative glucosylceramidase | 42 | 0.58 | 0.242 | 0.967 | |
| AT4G10060.1 | Beta-glucosidase, GBA2 type family protein / Putative glucosylceramidase | 43 | 0.26 | 0.244 | 0.171 | |
| AT4G10060.1 | Beta-glucosidase, GBA2 type family protein / Putative glucosylceramidase | 44 | 0.17 | 0.241 | 0.023 | * |
| AT4G10060.1 | Beta-glucosidase, GBA2 type family protein / Putative glucosylceramidase | 45 | 0.37 | 0.280 | 0.711 | |
| AT4G10060.1 | Beta-glucosidase, GBA2 type family protein / Putative glucosylceramidase | 46 | 1.95 | 0.513 | 0.999 | |
| AT4G10060.1 | Beta-glucosidase, GBA2 type family protein / Putative glucosylceramidase | 47 | 15.60 | 1.088 | 0.937 | |
| AT4G10340.1 | Photosystem II light harvesting complex subunit (LHCB5) | 3 | 0.61 | 0.193 | 1.000 | |

| | | | | | |
|---|---|---|---|---|---|
| AT4G10340.1 | Photosystem II light harvesting complex subunit (LHCB5) | 4 | 0.68 | 0.188 | 1.000 |
| AT4G10340.1 | Photosystem II light harvesting complex subunit (LHCB5) | 5 | 0.86 | 0.187 | 1.000 |
| AT4G10340.1 | Photosystem II light harvesting complex subunit (LHCB5) | 6 | 0.88 | 0.188 | 1.000 |
| AT4G10340.1 | Photosystem II light harvesting complex subunit (LHCB5) | 7 | 0.81 | 0.189 | 1.000 |
| AT4G10340.1 | Photosystem II light harvesting complex subunit (LHCB5) | 8 | 0.83 | 0.185 | 1.000 |
| AT4G10340.1 | Photosystem II light harvesting complex subunit (LHCB5) | 9 | 0.84 | 0.182 | 1.000 |
| AT4G10340.1 | Photosystem II light harvesting complex subunit (LHCB5) | 10 | 0.80 | 0.182 | 1.000 |
| AT4G10340.1 | Photosystem II light harvesting complex subunit (LHCB5) | 11 | 0.77 | 0.182 | 1.000 |
| AT4G10340.1 | Photosystem II light harvesting complex subunit (LHCB5) | 12 | 0.77 | 0.183 | 1.000 |
| AT4G10340.1 | Photosystem II light harvesting complex subunit (LHCB5) | 13 | 0.82 | 0.181 | 1.000 |
| AT4G10340.1 | Photosystem II light harvesting complex subunit (LHCB5) | 14 | 0.91 | 0.178 | 1.000 |
| AT4G10340.1 | Photosystem II light harvesting complex subunit (LHCB5) | 15 | 0.91 | 0.178 | 1.000 |
| AT4G10340.1 | Photosystem II light harvesting complex subunit (LHCB5) | 16 | 0.84 | 0.176 | 1.000 |
| AT4G10340.1 | Photosystem II light harvesting complex subunit (LHCB5) | 17 | 0.89 | 0.176 | 1.000 |
| AT4G10340.1 | Photosystem II light harvesting complex subunit (LHCB5) | 18 | 1.16 | 0.179 | 1.000 |
| AT4G10340.1 | Photosystem II light harvesting complex subunit (LHCB5) | 19 | 1.47 | 0.184 | 1.000 |
| AT4G10340.1 | Photosystem II light harvesting complex subunit (LHCB5) | 20 | 1.68 | 0.188 | 0.999 |
| AT4G10340.1 | Photosystem II light harvesting complex subunit (LHCB5) | 21 | 1.76 | 0.191 | 0.998 |
| AT4G10340.1 | Photosystem II light harvesting complex subunit (LHCB5) | 22 | 1.76 | 0.190 | 0.998 |
| AT4G10340.1 | Photosystem II light harvesting complex subunit (LHCB5) | 23 | 1.40 | 0.190 | 1.000 |
| AT4G10340.1 | Photosystem II light harvesting complex subunit (LHCB5) | 24 | 0.87 | 0.186 | 1.000 |
| AT4G10340.1 | Photosystem II light harvesting complex subunit (LHCB5) | 25 | 0.54 | 0.182 | 0.988 |
| AT4G10340.1 | Photosystem II light harvesting complex subunit (LHCB5) | 26 | 0.57 | 0.182 | 0.997 |
| AT4G10340.1 | Photosystem II light harvesting complex subunit (LHCB5) | 27 | 0.74 | 0.182 | 1.000 |
| AT4G10340.1 | Photosystem II light harvesting complex subunit (LHCB5) | 28 | 0.54 | 0.181 | 0.988 |
| AT4G10340.1 | Photosystem II light harvesting complex subunit (LHCB5) | 29 | 0.42 | 0.181 | 0.705 |
| AT4G10340.1 | Photosystem II light harvesting complex subunit (LHCB5) | 30 | 0.50 | 0.180 | 0.950 |
| AT4G10340.1 | Photosystem II light harvesting complex subunit (LHCB5) | 31 | 0.56 | 0.181 | 0.994 |
| AT4G10340.1 | Photosystem II light harvesting complex subunit (LHCB5) | 32 | 0.70 | 0.182 | 1.000 |
| AT4G10340.1 | Photosystem II light harvesting complex subunit (LHCB5) | 33 | 0.93 | 0.182 | 1.000 |
| AT4G10340.1 | Photosystem II light harvesting complex subunit (LHCB5) | 34 | 0.79 | 0.182 | 1.000 |

| | | | | | | |
|---|---|---|---|---|---|---|
| AT4G10340.1 | Photosystem II light harvesting complex subunit (LHCB5) | 35 | 0.72 | 0.182 | 1.000 | |
| AT4G10340.1 | Photosystem II light harvesting complex subunit (LHCB5) | 36 | 0.87 | 0.181 | 1.000 | |
| AT4G10340.1 | Photosystem II light harvesting complex subunit (LHCB5) | 37 | 1.45 | 0.181 | 1.000 | |
| AT4G10340.1 | Photosystem II light harvesting complex subunit (LHCB5) | 38 | 2.86 | 0.182 | 0.337 | |
| AT4G10340.1 | Photosystem II light harvesting complex subunit (LHCB5) | 39 | 4.76 | 0.183 | 0.010 | ** |
| AT4G10340.1 | Photosystem II light harvesting complex subunit (LHCB5) | 40 | 5.92 | 0.183 | 0.001 | ** |
| AT4G10340.1 | Photosystem II light harvesting complex subunit (LHCB5) | 41 | 4.89 | 0.183 | 0.008 | ** |
| AT4G10340.1 | Photosystem II light harvesting complex subunit (LHCB5) | 42 | 2.32 | 0.186 | 0.786 | |
| AT4G10340.1 | Photosystem II light harvesting complex subunit (LHCB5) | 43 | 0.73 | 0.187 | 1.000 | |
| AT4G10340.1 | Photosystem II light harvesting complex subunit (LHCB5) | 44 | 0.22 | 0.187 | 0.018 | * |
| AT4G10340.1 | Photosystem II light harvesting complex subunit (LHCB5) | 45 | 0.12 | 0.187 | 0.000 | *** |
| AT4G10340.1 | Photosystem II light harvesting complex subunit (LHCB5) | 46 | 0.14 | 0.204 | 0.001 | ** |
| AT4G10340.1 | Photosystem II light harvesting complex subunit (LHCB5) | 47 | 0.23 | 0.393 | 0.954 | |
| AT4G10340.1 | Photosystem II light harvesting complex subunit (LHCB5) | 48 | 0.38 | 0.708 | 1.000 | |
| AT4G11010.1 | Nucleoside diphosphate kinase 3 (NDPK3) | 2 | 1.24 | 0.307 | 1.000 | |
| AT4G11010.1 | Nucleoside diphosphate kinase 3 (NDPK3) | 3 | 1.05 | 0.214 | 1.000 | |
| AT4G11010.1 | Nucleoside diphosphate kinase 3 (NDPK3) | 4 | 0.93 | 0.190 | 1.000 | |
| AT4G11010.1 | Nucleoside diphosphate kinase 3 (NDPK3) | 5 | 0.91 | 0.189 | 1.000 | |
| AT4G11010.1 | Nucleoside diphosphate kinase 3 (NDPK3) | 6 | 0.96 | 0.190 | 1.000 | |
| AT4G11010.1 | Nucleoside diphosphate kinase 3 (NDPK3) | 7 | 1.05 | 0.189 | 1.000 | |
| AT4G11010.1 | Nucleoside diphosphate kinase 3 (NDPK3) | 8 | 1.21 | 0.187 | 1.000 | |
| AT4G11010.1 | Nucleoside diphosphate kinase 3 (NDPK3) | 9 | 1.40 | 0.185 | 1.000 | |
| AT4G11010.1 | Nucleoside diphosphate kinase 3 (NDPK3) | 10 | 1.60 | 0.185 | 0.996 | |
| AT4G11010.1 | Nucleoside diphosphate kinase 3 (NDPK3) | 11 | 1.87 | 0.185 | 0.926 | |
| AT4G11010.1 | Nucleoside diphosphate kinase 3 (NDPK3) | 12 | 2.17 | 0.185 | 0.709 | |
| AT4G11010.1 | Nucleoside diphosphate kinase 3 (NDPK3) | 13 | 2.27 | 0.183 | 0.615 | |
| AT4G11010.1 | Nucleoside diphosphate kinase 3 (NDPK3) | 14 | 2.14 | 0.181 | 0.719 | |
| AT4G11010.1 | Nucleoside diphosphate kinase 3 (NDPK3) | 15 | 2.08 | 0.181 | 0.759 | |
| AT4G11010.1 | Nucleoside diphosphate kinase 3 (NDPK3) | 16 | 2.37 | 0.180 | 0.504 | |
| AT4G11010.1 | Nucleoside diphosphate kinase 3 (NDPK3) | 17 | 2.90 | 0.180 | 0.199 | |
| AT4G11010.1 | Nucleoside diphosphate kinase 3 (NDPK3) | 18 | 3.29 | 0.182 | 0.101 | |

| | | | | | | |
|---|---|---|---|---|---|---|
| AT4G11010.1 | Nucleoside diphosphate kinase 3 (NDPK3) | 19 | 3.43 | 0.185 | 0.090 | . |
| AT4G11010.1 | Nucleoside diphosphate kinase 3 (NDPK3) | 20 | 3.29 | 0.188 | 0.131 | |
| AT4G11010.1 | Nucleoside diphosphate kinase 3 (NDPK3) | 21 | 2.90 | 0.190 | 0.268 | |
| AT4G11010.1 | Nucleoside diphosphate kinase 3 (NDPK3) | 22 | 2.41 | 0.190 | 0.564 | |
| AT4G11010.1 | Nucleoside diphosphate kinase 3 (NDPK3) | 23 | 1.90 | 0.189 | 0.925 | |
| AT4G11010.1 | Nucleoside diphosphate kinase 3 (NDPK3) | 24 | 1.43 | 0.187 | 1.000 | |
| AT4G11010.1 | Nucleoside diphosphate kinase 3 (NDPK3) | 25 | 1.08 | 0.184 | 1.000 | |
| AT4G11010.1 | Nucleoside diphosphate kinase 3 (NDPK3) | 26 | 0.85 | 0.183 | 1.000 | |
| AT4G11010.1 | Nucleoside diphosphate kinase 3 (NDPK3) | 27 | 0.73 | 0.183 | 1.000 | |
| AT4G11010.1 | Nucleoside diphosphate kinase 3 (NDPK3) | 28 | 0.66 | 0.183 | 0.999 | |
| AT4G11010.1 | Nucleoside diphosphate kinase 3 (NDPK3) | 29 | 0.64 | 0.183 | 0.997 | |
| AT4G11010.1 | Nucleoside diphosphate kinase 3 (NDPK3) | 30 | 0.64 | 0.184 | 0.997 | |
| AT4G11010.1 | Nucleoside diphosphate kinase 3 (NDPK3) | 31 | 0.66 | 0.184 | 0.999 | |
| AT4G11010.1 | Nucleoside diphosphate kinase 3 (NDPK3) | 32 | 0.70 | 0.184 | 1.000 | |
| AT4G11010.1 | Nucleoside diphosphate kinase 3 (NDPK3) | 33 | 0.75 | 0.185 | 1.000 | |
| AT4G11010.1 | Nucleoside diphosphate kinase 3 (NDPK3) | 34 | 0.80 | 0.185 | 1.000 | |
| AT4G11010.1 | Nucleoside diphosphate kinase 3 (NDPK3) | 35 | 0.85 | 0.185 | 1.000 | |
| AT4G11010.1 | Nucleoside diphosphate kinase 3 (NDPK3) | 36 | 0.87 | 0.185 | 1.000 | |
| AT4G11010.1 | Nucleoside diphosphate kinase 3 (NDPK3) | 37 | 0.82 | 0.185 | 1.000 | |
| AT4G11010.1 | Nucleoside diphosphate kinase 3 (NDPK3) | 38 | 0.71 | 0.186 | 1.000 | |
| AT4G11010.1 | Nucleoside diphosphate kinase 3 (NDPK3) | 39 | 0.55 | 0.186 | 0.955 | |
| AT4G11010.1 | Nucleoside diphosphate kinase 3 (NDPK3) | 40 | 0.43 | 0.188 | 0.599 | |
| AT4G11010.1 | Nucleoside diphosphate kinase 3 (NDPK3) | 41 | 0.39 | 0.203 | 0.578 | |
| AT4G11010.1 | Nucleoside diphosphate kinase 3 (NDPK3) | 42 | 0.55 | 0.253 | 0.999 | |
| AT4G11010.1 | Nucleoside diphosphate kinase 3 (NDPK3) | 43 | 1.23 | 0.329 | 1.000 | |
| AT4G11010.1 | Nucleoside diphosphate kinase 3 (NDPK3) | 46 | 84.81 | 0.743 | 0.186 | |
| AT4G11010.1 | Nucleoside diphosphate kinase 3 (NDPK3) | 47 | 462.44 | 0.946 | 0.108 | |
| AT4G11060.1 | Organellar single-stranded DNA binding protein 1 (OSB1) | 2 | 1.18 | 0.435 | 1.000 | |
| AT4G11060.1 | Organellar single-stranded DNA binding protein 1 (OSB1) | 3 | 1.10 | 0.363 | 1.000 | |
| AT4G11060.1 | Organellar single-stranded DNA binding protein 1 (OSB1) | 4 | 1.02 | 0.344 | 1.000 | |
| AT4G11060.1 | Organellar single-stranded DNA binding protein 1 (OSB1) | 8 | 0.80 | 0.327 | 1.000 | |

| | | | | | |
|---|---|---|---|---|---|
| AT4G11060.1 | Organellar single-stranded DNA binding protein 1 (OSB1) | 9 | 0.78 | 0.301 | 1.000 |
| AT4G11060.1 | Organellar single-stranded DNA binding protein 1 (OSB1) | 10 | 0.78 | 0.279 | 1.000 |
| AT4G11060.1 | Organellar single-stranded DNA binding protein 1 (OSB1) | 11 | 0.81 | 0.262 | 1.000 |
| AT4G11060.1 | Organellar single-stranded DNA binding protein 1 (OSB1) | 12 | 0.87 | 0.252 | 1.000 |
| AT4G11060.1 | Organellar single-stranded DNA binding protein 1 (OSB1) | 13 | 0.98 | 0.244 | 1.000 |
| AT4G11060.1 | Organellar single-stranded DNA binding protein 1 (OSB1) | 14 | 1.15 | 0.238 | 1.000 |
| AT4G11060.1 | Organellar single-stranded DNA binding protein 1 (OSB1) | 15 | 1.37 | 0.234 | 1.000 |
| AT4G11060.1 | Organellar single-stranded DNA binding protein 1 (OSB1) | 16 | 1.64 | 0.233 | 0.995 |
| AT4G11060.1 | Organellar single-stranded DNA binding protein 1 (OSB1) | 17 | 1.82 | 0.233 | 0.973 |
| AT4G11060.1 | Organellar single-stranded DNA binding protein 1 (OSB1) | 18 | 1.79 | 0.234 | 0.980 |
| AT4G11060.1 | Organellar single-stranded DNA binding protein 1 (OSB1) | 19 | 1.62 | 0.237 | 0.997 |
| AT4G11060.1 | Organellar single-stranded DNA binding protein 1 (OSB1) | 20 | 1.44 | 0.241 | 1.000 |
| AT4G11060.1 | Organellar single-stranded DNA binding protein 1 (OSB1) | 21 | 1.31 | 0.245 | 1.000 |
| AT4G11060.1 | Organellar single-stranded DNA binding protein 1 (OSB1) | 22 | 1.23 | 0.251 | 1.000 |
| AT4G11060.1 | Organellar single-stranded DNA binding protein 1 (OSB1) | 23 | 1.20 | 0.259 | 1.000 |
| AT4G11060.1 | Organellar single-stranded DNA binding protein 1 (OSB1) | 24 | 1.20 | 0.269 | 1.000 |
| AT4G11060.1 | Organellar single-stranded DNA binding protein 1 (OSB1) | 25 | 1.20 | 0.278 | 1.000 |
| AT4G11060.1 | Organellar single-stranded DNA binding protein 1 (OSB1) | 27 | 1.07 | 0.280 | 1.000 |
| AT4G11060.1 | Organellar single-stranded DNA binding protein 1 (OSB1) | 28 | 0.98 | 0.273 | 1.000 |
| AT4G11060.1 | Organellar single-stranded DNA binding protein 1 (OSB1) | 29 | 0.89 | 0.264 | 1.000 |
| AT4G11060.1 | Organellar single-stranded DNA binding protein 1 (OSB1) | 30 | 0.80 | 0.258 | 1.000 |
| AT4G11060.1 | Organellar single-stranded DNA binding protein 1 (OSB1) | 31 | 0.71 | 0.259 | 1.000 |
| AT4G11060.1 | Organellar single-stranded DNA binding protein 1 (OSB1) | 32 | 0.61 | 0.270 | 0.999 |
| AT4G11060.1 | Organellar single-stranded DNA binding protein 1 (OSB1) | 33 | 0.53 | 0.292 | 0.994 |
| AT4G11060.1 | Organellar single-stranded DNA binding protein 1 (OSB1) | 34 | 0.47 | 0.320 | 0.986 |
| AT4G11060.1 | Organellar single-stranded DNA binding protein 1 (OSB1) | 39 | 0.39 | 0.308 | 0.902 |
| AT4G11060.1 | Organellar single-stranded DNA binding protein 1 (OSB1) | 40 | 0.42 | 0.278 | 0.892 |
| AT4G11060.1 | Organellar single-stranded DNA binding protein 1 (OSB1) | 41 | 0.48 | 0.257 | 0.942 |
| AT4G11060.1 | Organellar single-stranded DNA binding protein 1 (OSB1) | 42 | 0.59 | 0.247 | 0.994 |
| AT4G11060.1 | Organellar single-stranded DNA binding protein 1 (OSB1) | 43 | 0.75 | 0.245 | 1.000 |
| AT4G11060.1 | Organellar single-stranded DNA binding protein 1 (OSB1) | 44 | 0.94 | 0.255 | 1.000 |

| | | | | | | |
|---|---|---|---|---|---|---|
| AT4G11060.1 | Organellar single-stranded DNA binding protein 1 (OSB1) | 45 | 1.10 | 0.289 | 1.000 | |
| AT4G11060.1 | Organellar single-stranded DNA binding protein 1 (OSB1) | 46 | 1.21 | 0.366 | 1.000 | |
| AT4G11060.1 | Organellar single-stranded DNA binding protein 1 (OSB1) | 47 | 1.31 | 0.493 | 1.000 | |
| AT4G11120.1 | Translation elongation factor Ts (EF-Ts) | 2 | 1.06 | 0.437 | 1.000 | |
| AT4G11120.1 | Translation elongation factor Ts (EF-Ts) | 3 | 1.11 | 0.425 | 1.000 | |
| AT4G11120.1 | Translation elongation factor Ts (EF-Ts) | 33 | 2.03 | 0.336 | 0.927 | |
| AT4G11120.1 | Translation elongation factor Ts (EF-Ts) | 34 | 2.07 | 0.303 | 0.878 | |
| AT4G11120.1 | Translation elongation factor Ts (EF-Ts) | 35 | 2.11 | 0.283 | 0.826 | |
| AT4G11120.1 | Translation elongation factor Ts (EF-Ts) | 41 | 2.68 | 0.250 | 0.447 | |
| AT4G11120.1 | Translation elongation factor Ts (EF-Ts) | 42 | 2.89 | 0.209 | 0.187 | |
| AT4G11120.1 | Translation elongation factor Ts (EF-Ts) | 43 | 3.02 | 0.177 | 0.060 | . |
| AT4G11120.1 | Translation elongation factor Ts (EF-Ts) | 44 | 2.89 | 0.160 | 0.039 | * |
| AT4G11120.1 | Translation elongation factor Ts (EF-Ts) | 45 | 2.45 | 0.165 | 0.138 | |
| AT4G11120.1 | Translation elongation factor Ts (EF-Ts) | 46 | 1.86 | 0.210 | 0.742 | |
| AT4G11120.1 | Translation elongation factor Ts (EF-Ts) | 47 | 1.34 | 0.309 | 0.999 | |
| AT4G11150.1 | Vacuolar ATP synthase subunit E1 (VHA-E1) | 2 | 1.51 | 0.374 | 0.993 | |
| AT4G11150.1 | Vacuolar ATP synthase subunit E1 (VHA-E1) | 3 | 1.64 | 0.338 | 0.971 | |
| AT4G11150.1 | Vacuolar ATP synthase subunit E1 (VHA-E1) | 4 | 1.78 | 0.311 | 0.923 | |
| AT4G11150.1 | Vacuolar ATP synthase subunit E1 (VHA-E1) | 5 | 1.93 | 0.295 | 0.848 | |
| AT4G11150.1 | Vacuolar ATP synthase subunit E1 (VHA-E1) | 6 | 2.09 | 0.287 | 0.767 | |
| AT4G11150.1 | Vacuolar ATP synthase subunit E1 (VHA-E1) | 7 | 2.24 | 0.287 | 0.695 | |
| AT4G11150.1 | Vacuolar ATP synthase subunit E1 (VHA-E1) | 8 | 2.39 | 0.292 | 0.642 | |
| AT4G11150.1 | Vacuolar ATP synthase subunit E1 (VHA-E1) | 21 | 2.39 | 0.248 | 0.487 | |
| AT4G11150.1 | Vacuolar ATP synthase subunit E1 (VHA-E1) | 22 | 2.19 | 0.239 | 0.555 | |
| AT4G11150.1 | Vacuolar ATP synthase subunit E1 (VHA-E1) | 23 | 1.96 | 0.233 | 0.670 | |
| AT4G11150.1 | Vacuolar ATP synthase subunit E1 (VHA-E1) | 24 | 1.71 | 0.232 | 0.830 | |
| AT4G11150.1 | Vacuolar ATP synthase subunit E1 (VHA-E1) | 25 | 1.45 | 0.237 | 0.962 | |
| AT4G11150.1 | Vacuolar ATP synthase subunit E1 (VHA-E1) | 26 | 1.20 | 0.247 | 0.999 | |
| AT4G11150.1 | Vacuolar ATP synthase subunit E1 (VHA-E1) | 27 | 0.98 | 0.263 | 1.000 | |
| AT4G11150.1 | Vacuolar ATP synthase subunit E1 (VHA-E1) | 28 | 0.79 | 0.283 | 0.999 | |
| AT4G11150.1 | Vacuolar ATP synthase subunit E1 (VHA-E1) | 29 | 0.64 | 0.305 | 0.970 | |

| | | | | | | |
|---|---|---|---|---|---|---|
| AT4G11150.1 | Vacuolar ATP synthase subunit E1 (VHA-E1) | 46 | 0.01 | 0.568 | 0.004 | ** |
| AT4G11150.1 | Vacuolar ATP synthase subunit E1 (VHA-E1) | 47 | 0.01 | 0.595 | 0.003 | ** |
| AT4G11160.1 | Translation initiation factor 2 (IEF-2) | 2 | 1.09 | 0.425 | 1.000 | |
| AT4G11160.1 | Translation initiation factor 2 (IEF-2) | 3 | 1.03 | 0.398 | 1.000 | |
| AT4G11160.1 | Translation initiation factor 2 (IEF-2) | 4 | 0.97 | 0.378 | 1.000 | |
| AT4G11160.1 | Translation initiation factor 2 (IEF-2) | 17 | 0.54 | 0.307 | 0.921 | |
| AT4G11160.1 | Translation initiation factor 2 (IEF-2) | 18 | 0.52 | 0.296 | 0.898 | |
| AT4G11160.1 | Translation initiation factor 2 (IEF-2) | 19 | 0.52 | 0.284 | 0.870 | |
| AT4G11160.1 | Translation initiation factor 2 (IEF-2) | 23 | 0.52 | 0.234 | 0.757 | |
| AT4G11160.1 | Translation initiation factor 2 (IEF-2) | 24 | 0.53 | 0.222 | 0.739 | |
| AT4G11160.1 | Translation initiation factor 2 (IEF-2) | 25 | 0.55 | 0.211 | 0.733 | |
| AT4G11160.1 | Translation initiation factor 2 (IEF-2) | 26 | 0.57 | 0.202 | 0.746 | |
| AT4G11160.1 | Translation initiation factor 2 (IEF-2) | 27 | 0.59 | 0.194 | 0.778 | |
| AT4G11160.1 | Translation initiation factor 2 (IEF-2) | 28 | 0.63 | 0.188 | 0.829 | |
| AT4G11160.1 | Translation initiation factor 2 (IEF-2) | 29 | 0.66 | 0.184 | 0.889 | |
| AT4G11160.1 | Translation initiation factor 2 (IEF-2) | 30 | 0.71 | 0.183 | 0.946 | |
| AT4G11160.1 | Translation initiation factor 2 (IEF-2) | 31 | 0.76 | 0.184 | 0.983 | |
| AT4G11160.1 | Translation initiation factor 2 (IEF-2) | 32 | 0.82 | 0.188 | 0.997 | |
| AT4G11160.1 | Translation initiation factor 2 (IEF-2) | 33 | 0.88 | 0.195 | 1.000 | |
| AT4G11160.1 | Translation initiation factor 2 (IEF-2) | 34 | 0.95 | 0.206 | 1.000 | |
| AT4G11160.1 | Translation initiation factor 2 (IEF-2) | 35 | 1.02 | 0.222 | 1.000 | |
| AT4G11160.1 | Translation initiation factor 2 (IEF-2) | 36 | 1.08 | 0.242 | 1.000 | |
| AT4G11160.1 | Translation initiation factor 2 (IEF-2) | 37 | 1.15 | 0.267 | 1.000 | |
| AT4G11160.1 | Translation initiation factor 2 (IEF-2) | 38 | 1.20 | 0.297 | 1.000 | |
| AT4G11160.1 | Translation initiation factor 2 (IEF-2) | 46 | 1.40 | 0.679 | 1.000 | |
| AT4G11160.1 | Translation initiation factor 2 (IEF-2) | 47 | 1.41 | 0.743 | 1.000 | |
| AT4G12340.1 | ? | 2 | 1.06 | 0.494 | 1.000 | |
| AT4G12340.1 | ? | 3 | 1.00 | 0.361 | 1.000 | |
| AT4G12340.1 | ? | 5 | 0.95 | 0.287 | 1.000 | |
| AT4G12340.1 | ? | 6 | 1.00 | 0.246 | 1.000 | |
| AT4G12340.1 | ? | 7 | 1.18 | 0.219 | 1.000 | |
| AT4G12340.1 | ? | 8 | 1.66 | 0.203 | 1.000 | |
| AT4G12340.1 | ? | 9 | 2.52 | 0.195 | 0.668 | |
| AT4G12340.1 | ? | 10 | 3.62 | 0.193 | 0.120 | |
| AT4G12340.1 | ? | 11 | 4.99 | 0.194 | 0.012 | * |
| AT4G12340.1 | ? | 12 | 6.45 | 0.193 | 0.001 | ** |

| | | | | | | |
|---|---|---|---|---|---|---|
| AT4G12340.1 | ? | 13 | 7.20 | 0.192 | 0.000 | *** |
| AT4G12340.1 | ? | 14 | 6.69 | 0.190 | 0.001 | *** |
| AT4G12340.1 | ? | 15 | 5.22 | 0.189 | 0.006 | ** |
| AT4G12340.1 | ? | 16 | 3.72 | 0.187 | 0.076 | . |
| AT4G12340.1 | ? | 17 | 2.69 | 0.186 | 0.455 | |
| AT4G12340.1 | ? | 18 | 2.08 | 0.190 | 0.918 | |
| AT4G12340.1 | ? | 19 | 1.85 | 0.195 | 0.990 | |
| AT4G12340.1 | ? | 20 | 1.90 | 0.200 | 0.987 | |
| AT4G12340.1 | ? | 21 | 1.98 | 0.202 | 0.977 | |
| AT4G12340.1 | ? | 22 | 2.23 | 0.202 | 0.898 | |
| AT4G12340.1 | ? | 23 | 2.68 | 0.200 | 0.604 | |
| AT4G12340.1 | ? | 24 | 3.00 | 0.198 | 0.377 | |
| AT4G12340.1 | ? | 25 | 3.05 | 0.197 | 0.347 | |
| AT4G12340.1 | ? | 26 | 3.18 | 0.202 | 0.325 | |
| AT4G12340.1 | ? | 27 | 3.29 | 0.209 | 0.333 | |
| AT4G12340.1 | ? | 28 | 3.14 | 0.211 | 0.416 | |
| AT4G12340.1 | ? | 29 | 2.84 | 0.208 | 0.563 | |
| AT4G12340.1 | ? | 30 | 2.60 | 0.202 | 0.669 | |
| AT4G12340.1 | ? | 31 | 2.53 | 0.198 | 0.684 | |
| AT4G12340.1 | ? | 32 | 2.39 | 0.195 | 0.767 | |
| AT4G12340.1 | ? | 33 | 1.96 | 0.195 | 0.971 | |
| AT4G12340.1 | ? | 34 | 1.63 | 0.196 | 1.000 | |
| AT4G12340.1 | ? | 35 | 1.62 | 0.198 | 1.000 | |
| AT4G12340.1 | ? | 36 | 1.91 | 0.204 | 0.989 | |
| AT4G12340.1 | ? | 37 | 2.39 | 0.212 | 0.863 | |
| AT4G12340.1 | ? | 38 | 2.76 | 0.216 | 0.681 | |
| AT4G12340.1 | ? | 39 | 3.04 | 0.211 | 0.471 | |
| AT4G12340.1 | ? | 40 | 2.99 | 0.202 | 0.419 | |
| AT4G12340.1 | ? | 41 | 2.38 | 0.197 | 0.785 | |
| AT4G12340.1 | ? | 42 | 1.78 | 0.198 | 0.996 | |
| AT4G12340.1 | ? | 43 | 1.31 | 0.200 | 1.000 | |
| AT4G12340.1 | ? | 44 | 0.95 | 0.199 | 1.000 | |
| AT4G12340.1 | ? | 45 | 0.74 | 0.201 | 1.000 | |
| AT4G12340.1 | ? | 46 | 0.60 | 0.252 | 1.000 | |
| AT4G12340.1 | ? | 47 | 0.45 | 0.425 | 1.000 | |
| AT4G12800.1 | Photosystem I subunit (PSAL) | 2 | 0.28 | 0.207 | 0.190 | |
| AT4G12800.1 | Photosystem I subunit (PSAL) | 3 | 0.43 | 0.126 | 0.092 | . |
| AT4G12800.1 | Photosystem I subunit (PSAL) | 4 | 0.69 | 0.115 | 0.975 | |
| AT4G12800.1 | Photosystem I subunit (PSAL) | 5 | 1.12 | 0.114 | 1.000 | |
| AT4G12800.1 | Photosystem I subunit (PSAL) | 6 | 1.29 | 0.115 | 1.000 | |
| AT4G12800.1 | Photosystem I subunit (PSAL) | 7 | 1.05 | 0.116 | 1.000 | |
| AT4G12800.1 | Photosystem I subunit (PSAL) | 8 | 0.88 | 0.112 | 1.000 | |
| AT4G12800.1 | Photosystem I subunit (PSAL) | 9 | 0.79 | 0.110 | 1.000 | |
| AT4G12800.1 | Photosystem I subunit (PSAL) | 10 | 0.74 | 0.110 | 0.996 | |
| AT4G12800.1 | Photosystem I subunit (PSAL) | 11 | 0.74 | 0.111 | 0.997 | |
| AT4G12800.1 | Photosystem I subunit (PSAL) | 12 | 0.80 | 0.111 | 1.000 | |
| AT4G12800.1 | Photosystem I subunit (PSAL) | 13 | 0.91 | 0.109 | 1.000 | |
| AT4G12800.1 | Photosystem I subunit (PSAL) | 14 | 1.00 | 0.108 | 1.000 | |
| AT4G12800.1 | Photosystem I subunit (PSAL) | 15 | 0.99 | 0.107 | 1.000 | |
| AT4G12800.1 | Photosystem I subunit (PSAL) | 16 | 0.94 | 0.107 | 1.000 | |
| AT4G12800.1 | Photosystem I subunit (PSAL) | 17 | 0.99 | 0.106 | 1.000 | |
| AT4G12800.1 | Photosystem I subunit (PSAL) | 18 | 1.13 | 0.107 | 1.000 | |
| AT4G12800.1 | Photosystem I subunit (PSAL) | 19 | 1.27 | 0.110 | 1.000 | |
| AT4G12800.1 | Photosystem I subunit (PSAL) | 20 | 1.28 | 0.114 | 1.000 | |
| AT4G12800.1 | Photosystem I subunit (PSAL) | 21 | 1.14 | 0.116 | 1.000 | |
| AT4G12800.1 | Photosystem I subunit (PSAL) | 22 | 0.94 | 0.116 | 1.000 | |
| AT4G12800.1 | Photosystem I subunit (PSAL) | 23 | 0.75 | 0.115 | 0.999 | |
| AT4G12800.1 | Photosystem I subunit (PSAL) | 24 | 0.65 | 0.112 | 0.900 | |
| AT4G12800.1 | Photosystem I subunit (PSAL) | 25 | 0.79 | 0.109 | 1.000 | |
| AT4G12800.1 | Photosystem I subunit (PSAL) | 26 | 1.26 | 0.109 | 1.000 | |
| AT4G12800.1 | Photosystem I subunit (PSAL) | 27 | 1.58 | 0.109 | 0.798 | |
| AT4G12800.1 | Photosystem I subunit (PSAL) | 28 | 1.56 | 0.111 | 0.846 | |
| AT4G12800.1 | Photosystem I subunit (PSAL) | 29 | 1.67 | 0.112 | 0.681 | |

| | | | | | | |
|---|---|---|---|---|---|---|
| AT4G12800.1 | Photosystem I subunit (PSAL) | 30 | 1.36 | 0.115 | 0.997 | |
| AT4G12800.1 | Photosystem I subunit (PSAL) | 31 | 0.68 | 0.123 | 0.984 | |
| AT4G12800.1 | Photosystem I subunit (PSAL) | 32 | 0.32 | 0.157 | 0.045 | * |
| AT4G12800.1 | Photosystem I subunit (PSAL) | 33 | 0.18 | 0.255 | 0.084 | . |
| AT4G12800.1 | Photosystem I subunit (PSAL) | 46 | 5905.96 | 2.264 | 0.892 | |
| AT4G12800.1 | Photosystem I subunit (PSAL) | 47 | 26096.99 | 2.504 | 0.838 | |
| AT4G13850.3 | Glycine-rich RNA-binding protein 2 (GR-RBP2) | 2 | 0.91 | 0.366 | 1.000 | |
| AT4G13850.3 | Glycine-rich RNA-binding protein 2 (GR-RBP2) | 3 | 0.85 | 0.331 | 1.000 | |
| AT4G13850.3 | Glycine-rich RNA-binding protein 2 (GR-RBP2) | 4 | 0.80 | 0.310 | 1.000 | |
| AT4G13850.3 | Glycine-rich RNA-binding protein 2 (GR-RBP2) | 11 | 0.53 | 0.245 | 0.951 | |
| AT4G13850.3 | Glycine-rich RNA-binding protein 2 (GR-RBP2) | 12 | 0.51 | 0.232 | 0.896 | |
| AT4G13850.3 | Glycine-rich RNA-binding protein 2 (GR-RBP2) | 13 | 0.48 | 0.217 | 0.796 | |
| AT4G13850.3 | Glycine-rich RNA-binding protein 2 (GR-RBP2) | 16 | 0.44 | 0.177 | 0.383 | |
| AT4G13850.3 | Glycine-rich RNA-binding protein 2 (GR-RBP2) | 17 | 0.43 | 0.168 | 0.296 | |
| AT4G13850.3 | Glycine-rich RNA-binding protein 2 (GR-RBP2) | 18 | 0.43 | 0.160 | 0.244 | |
| AT4G13850.3 | Glycine-rich RNA-binding protein 2 (GR-RBP2) | 19 | 0.44 | 0.155 | 0.233 | |
| AT4G13850.3 | Glycine-rich RNA-binding protein 2 (GR-RBP2) | 20 | 0.46 | 0.151 | 0.261 | |
| AT4G13850.3 | Glycine-rich RNA-binding protein 2 (GR-RBP2) | 21 | 0.49 | 0.148 | 0.336 | |
| AT4G13850.3 | Glycine-rich RNA-binding protein 2 (GR-RBP2) | 22 | 0.52 | 0.145 | 0.456 | |
| AT4G13850.3 | Glycine-rich RNA-binding protein 2 (GR-RBP2) | 23 | 0.57 | 0.142 | 0.614 | |
| AT4G13850.3 | Glycine-rich RNA-binding protein 2 (GR-RBP2) | 24 | 0.61 | 0.139 | 0.755 | |
| AT4G13850.3 | Glycine-rich RNA-binding protein 2 (GR-RBP2) | 25 | 0.64 | 0.137 | 0.832 | |
| AT4G13850.3 | Glycine-rich RNA-binding protein 2 (GR-RBP2) | 26 | 0.64 | 0.135 | 0.821 | |
| AT4G13850.3 | Glycine-rich RNA-binding protein 2 (GR-RBP2) | 27 | 0.61 | 0.134 | 0.710 | |
| AT4G13850.3 | Glycine-rich RNA-binding protein 2 (GR-RBP2) | 28 | 0.57 | 0.133 | 0.546 | |
| AT4G13850.3 | Glycine-rich RNA-binding protein 2 (GR-RBP2) | 29 | 0.54 | 0.133 | 0.403 | |
| AT4G13850.3 | Glycine-rich RNA-binding protein 2 (GR-RBP2) | 30 | 0.52 | 0.133 | 0.314 | |
| AT4G13850.3 | Glycine-rich RNA-binding protein 2 (GR-RBP2) | 31 | 0.51 | 0.133 | 0.286 | |
| AT4G13850.3 | Glycine-rich RNA-binding protein 2 (GR-RBP2) | 32 | 0.52 | 0.133 | 0.307 | |
| AT4G13850.3 | Glycine-rich RNA-binding protein 2 (GR-RBP2) | 33 | 0.53 | 0.133 | 0.374 | |
| AT4G13850.3 | Glycine-rich RNA-binding protein 2 (GR-RBP2) | 34 | 0.56 | 0.133 | 0.482 | |
| AT4G13850.3 | Glycine-rich RNA-binding protein 2 (GR-RBP2) | 35 | 0.60 | 0.134 | 0.637 | |
| AT4G13850.3 | Glycine-rich RNA-binding protein 2 (GR-RBP2) | 36 | 0.64 | 0.134 | 0.808 | |
| AT4G13850.3 | Glycine-rich RNA-binding protein 2 (GR-RBP2) | 37 | 0.68 | 0.134 | 0.908 | |

| | | | | | | |
|---|---|---|---|---|---|---|
| AT4G13850.3 | Glycine-rich RNA-binding protein 2 (GR-RBP2) | 38 | 0.69 | 0.135 | 0.930 | |
| AT4G13850.3 | Glycine-rich RNA-binding protein 2 (GR-RBP2) | 39 | 0.66 | 0.137 | 0.874 | |
| AT4G13850.3 | Glycine-rich RNA-binding protein 2 (GR-RBP2) | 40 | 0.59 | 0.139 | 0.667 | |
| AT4G13850.3 | Glycine-rich RNA-binding protein 2 (GR-RBP2) | 41 | 0.50 | 0.141 | 0.310 | |
| AT4G13850.3 | Glycine-rich RNA-binding protein 2 (GR-RBP2) | 42 | 0.39 | 0.145 | 0.070 | . |
| AT4G13850.3 | Glycine-rich RNA-binding protein 2 (GR-RBP2) | 43 | 0.30 | 0.150 | 0.009 | ** |
| AT4G13850.3 | Glycine-rich RNA-binding protein 2 (GR-RBP2) | 44 | 0.23 | 0.160 | 0.001 | ** |
| AT4G13850.3 | Glycine-rich RNA-binding protein 2 (GR-RBP2) | 45 | 0.17 | 0.177 | 0.000 | *** |
| AT4G13850.3 | Glycine-rich RNA-binding protein 2 (GR-RBP2) | 46 | 0.13 | 0.207 | 0.000 | *** |
| AT4G13850.3 | Glycine-rich RNA-binding protein 2 (GR-RBP2) | 47 | 0.09 | 0.252 | 0.001 | ** |
| AT4G14570.1 | Acylamino acid-releasing enzyme (AARE) | 2 | 1.27 | 0.640 | 1.000 | |
| AT4G14570.1 | Acylamino acid-releasing enzyme (AARE) | 3 | 1.43 | 0.620 | 0.997 | |
| AT4G14570.1 | Acylamino acid-releasing enzyme (AARE) | 32 | 3.46 | 0.341 | 0.353 | |
| AT4G14570.1 | Acylamino acid-releasing enzyme (AARE) | 33 | 2.97 | 0.336 | 0.450 | |
| AT4G14570.1 | Acylamino acid-releasing enzyme (AARE) | 34 | 2.50 | 0.338 | 0.597 | |
| AT4G14570.1 | Acylamino acid-releasing enzyme (AARE) | 35 | 2.08 | 0.347 | 0.766 | |
| AT4G14570.1 | Acylamino acid-releasing enzyme (AARE) | 46 | 0.17 | 0.485 | 0.360 | |
| AT4G14570.1 | Acylamino acid-releasing enzyme (AARE) | 47 | 0.14 | 0.512 | 0.295 | |
| AT4G14615.1 | Cytochrome c:O2 oxidoreductase assembly protein | 2 | 0.98 | 0.397 | 1.000 | |
| AT4G14615.1 | Cytochrome c:O2 oxidoreductase assembly protein | 3 | 0.88 | 0.310 | 1.000 | |
| AT4G14615.1 | Cytochrome c:O2 oxidoreductase assembly protein | 5 | 0.72 | 0.219 | 1.000 | |
| AT4G14615.1 | Cytochrome c:O2 oxidoreductase assembly protein | 6 | 0.65 | 0.191 | 0.994 | |
| AT4G14615.1 | Cytochrome c:O2 oxidoreductase assembly protein | 7 | 0.59 | 0.175 | 0.937 | |
| AT4G14615.1 | Cytochrome c:O2 oxidoreductase assembly protein | 8 | 0.54 | 0.166 | 0.792 | |
| AT4G14615.1 | Cytochrome c:O2 oxidoreductase assembly protein | 9 | 0.51 | 0.161 | 0.625 | |
| AT4G14615.1 | Cytochrome c:O2 oxidoreductase assembly protein | 10 | 0.49 | 0.158 | 0.522 | |

| | | | | | |
|---|---|---|---|---|---|
| AT4G14615.1 | Cytochrome c:O2 oxidoreductase assembly protein | 11 | 0.49 | 0.155 | 0.511 |
| AT4G14615.1 | Cytochrome c:O2 oxidoreductase assembly protein | 12 | 0.51 | 0.151 | 0.555 |
| AT4G14615.1 | Cytochrome c:O2 oxidoreductase assembly protein | 13 | 0.54 | 0.146 | 0.630 |
| AT4G14615.1 | Cytochrome c:O2 oxidoreductase assembly protein | 14 | 0.57 | 0.144 | 0.735 |
| AT4G14615.1 | Cytochrome c:O2 oxidoreductase assembly protein | 15 | 0.62 | 0.145 | 0.890 |
| AT4G14615.1 | Cytochrome c:O2 oxidoreductase assembly protein | 16 | 0.69 | 0.149 | 0.987 |
| AT4G14615.1 | Cytochrome c:O2 oxidoreductase assembly protein | 17 | 0.76 | 0.155 | 1.000 |
| AT4G14615.1 | Cytochrome c:O2 oxidoreductase assembly protein | 18 | 0.77 | 0.172 | 1.000 |
| AT4G14615.1 | Cytochrome c:O2 oxidoreductase assembly protein | 19 | 0.73 | 0.208 | 1.000 |
| AT4G14615.1 | Cytochrome c:O2 oxidoreductase assembly protein | 20 | 0.63 | 0.253 | 0.999 |
| AT4G14615.1 | Cytochrome c:O2 oxidoreductase assembly protein | 28 | 0.55 | 0.224 | 0.977 |
| AT4G14615.1 | Cytochrome c:O2 oxidoreductase assembly protein | 29 | 0.91 | 0.185 | 1.000 |
| AT4G14615.1 | Cytochrome c:O2 oxidoreductase assembly protein | 30 | 1.53 | 0.164 | 0.982 |
| AT4G14615.1 | Cytochrome c:O2 oxidoreductase assembly protein | 31 | 2.19 | 0.156 | 0.366 |
| AT4G14615.1 | Cytochrome c:O2 oxidoreductase assembly protein | 32 | 2.40 | 0.153 | 0.195 |
| AT4G14615.1 | Cytochrome c:O2 oxidoreductase assembly protein | 33 | 1.92 | 0.153 | 0.612 |
| AT4G14615.1 | Cytochrome c:O2 oxidoreductase assembly protein | 34 | 1.23 | 0.159 | 1.000 |
| AT4G14615.1 | Cytochrome c:O2 oxidoreductase assembly protein | 35 | 0.70 | 0.181 | 0.999 |
| AT4G14615.1 | Cytochrome c:O2 oxidoreductase assembly protein | 36 | 0.36 | 0.229 | 0.557 |
| AT4G14615.1 | Cytochrome c:O2 oxidoreductase assembly protein | 37 | 0.18 | 0.303 | 0.218 |

| | | | | | | |
|---|---|---|---|---|---|---|
| AT4G14615.1 | Cytochrome c:O2 oxidoreductase assembly protein | 46 | 0.00 | 0.693 | 0.001 | *** |
| AT4G14615.1 | Cytochrome c:O2 oxidoreductase assembly protein | 47 | 0.00 | 0.769 | 0.002 | ** |
| AT4G15010.3 | Mitochondrial substrate carrier family protein | 2 | 0.74 | 0.488 | 1.000 | |
| AT4G15010.3 | Mitochondrial substrate carrier family protein | 3 | 1.03 | 0.455 | 1.000 | |
| AT4G15010.3 | Mitochondrial substrate carrier family protein | 22 | 1.30 | 0.351 | 1.000 | |
| AT4G15010.3 | Mitochondrial substrate carrier family protein | 23 | 0.81 | 0.291 | 1.000 | |
| AT4G15010.3 | Mitochondrial substrate carrier family protein | 24 | 0.68 | 0.238 | 1.000 | |
| AT4G15010.3 | Mitochondrial substrate carrier family protein | 25 | 0.89 | 0.226 | 1.000 | |
| AT4G15010.3 | Mitochondrial substrate carrier family protein | 26 | 1.51 | 0.224 | 1.000 | |
| AT4G15010.3 | Mitochondrial substrate carrier family protein | 27 | 1.80 | 0.225 | 0.996 | |
| AT4G15010.3 | Mitochondrial substrate carrier family protein | 28 | 1.17 | 0.224 | 1.000 | |
| AT4G15010.3 | Mitochondrial substrate carrier family protein | 29 | 0.98 | 0.223 | 1.000 | |
| AT4G15010.3 | Mitochondrial substrate carrier family protein | 30 | 1.85 | 0.224 | 0.993 | |
| AT4G15010.3 | Mitochondrial substrate carrier family protein | 31 | 3.52 | 0.227 | 0.306 | |
| AT4G15010.3 | Mitochondrial substrate carrier family protein | 32 | 3.44 | 0.227 | 0.337 | |
| AT4G15010.3 | Mitochondrial substrate carrier family protein | 33 | 2.15 | 0.226 | 0.947 | |
| AT4G15010.3 | Mitochondrial substrate carrier family protein | 34 | 1.51 | 0.227 | 1.000 | |
| AT4G15010.3 | Mitochondrial substrate carrier family protein | 35 | 1.43 | 0.229 | 1.000 | |
| AT4G15010.3 | Mitochondrial substrate carrier family protein | 36 | 1.47 | 0.238 | 1.000 | |
| AT4G15010.3 | Mitochondrial substrate carrier family protein | 37 | 1.67 | 0.263 | 1.000 | |
| AT4G15010.3 | Mitochondrial substrate carrier family protein | 38 | 2.17 | 0.301 | 0.997 | |
| AT4G15010.3 | Mitochondrial substrate carrier family protein | 39 | 2.65 | 0.289 | 0.946 | |
| AT4G15010.3 | Mitochondrial substrate carrier family protein | 40 | 2.95 | 0.264 | 0.786 | |
| AT4G15010.3 | Mitochondrial substrate carrier family protein | 41 | 2.81 | 0.234 | 0.683 | |
| AT4G15010.3 | Mitochondrial substrate carrier family protein | 42 | 2.24 | 0.230 | 0.927 | |
| AT4G15010.3 | Mitochondrial substrate carrier family protein | 43 | 1.40 | 0.230 | 1.000 | |
| AT4G15010.3 | Mitochondrial substrate carrier family protein | 44 | 0.73 | 0.230 | 1.000 | |
| AT4G15010.3 | Mitochondrial substrate carrier family protein | 45 | 0.55 | 0.230 | 0.997 | |
| AT4G15010.3 | Mitochondrial substrate carrier family protein | 46 | 0.53 | 0.260 | 0.998 | |
| AT4G15010.3 | Mitochondrial substrate carrier family protein | 47 | 0.57 | 0.517 | 1.000 | |

| | | | | | | |
|---|---|---|---|---|---|---|
| AT4G15940.1 | Fumarylacetoacetate (FAA) hydrolase family protein | 2 | 0.84 | 0.410 | 1.000 | |
| AT4G15940.1 | Fumarylacetoacetate (FAA) hydrolase family protein | 3 | 0.99 | 0.409 | 1.000 | |
| AT4G15940.1 | Fumarylacetoacetate (FAA) hydrolase family protein | 31 | 0.28 | 0.311 | 0.673 | |
| AT4G15940.1 | Fumarylacetoacetate (FAA) hydrolase family protein | 32 | 0.21 | 0.253 | 0.116 | |
| AT4G15940.1 | Fumarylacetoacetate (FAA) hydrolase family protein | 33 | 0.21 | 0.233 | 0.062 | . |
| AT4G15940.1 | Fumarylacetoacetate (FAA) hydrolase family protein | 34 | 0.27 | 0.223 | 0.176 | |
| AT4G15940.1 | Fumarylacetoacetate (FAA) hydrolase family protein | 35 | 0.45 | 0.213 | 0.752 | |
| AT4G15940.1 | Fumarylacetoacetate (FAA) hydrolase family protein | 36 | 0.91 | 0.208 | 1.000 | |
| AT4G15940.1 | Fumarylacetoacetate (FAA) hydrolase family protein | 37 | 1.54 | 0.211 | 0.997 | |
| AT4G15940.1 | Fumarylacetoacetate (FAA) hydrolase family protein | 38 | 1.23 | 0.219 | 1.000 | |
| AT4G15940.1 | Fumarylacetoacetate (FAA) hydrolase family protein | 39 | 0.79 | 0.212 | 1.000 | |
| AT4G15940.1 | Fumarylacetoacetate (FAA) hydrolase family protein | 40 | 0.97 | 0.198 | 1.000 | |
| AT4G15940.1 | Fumarylacetoacetate (FAA) hydrolase family protein | 41 | 1.46 | 0.194 | 0.998 | |
| AT4G15940.1 | Fumarylacetoacetate (FAA) hydrolase family protein | 42 | 1.56 | 0.196 | 0.994 | |
| AT4G15940.1 | Fumarylacetoacetate (FAA) hydrolase family protein | 43 | 1.54 | 0.198 | 0.995 | |
| AT4G15940.1 | Fumarylacetoacetate (FAA) hydrolase family protein | 44 | 1.43 | 0.196 | 0.999 | |
| AT4G15940.1 | Fumarylacetoacetate (FAA) hydrolase family protein | 45 | 1.15 | 0.196 | 1.000 | |
| AT4G15940.1 | Fumarylacetoacetate (FAA) hydrolase family protein | 46 | 0.67 | 0.221 | 0.999 | |
| AT4G15940.1 | Fumarylacetoacetate (FAA) hydrolase family protein | 47 | 0.25 | 0.433 | 0.899 | |
| AT4G16450.2 | NADH:ubiquinone oxidoreductase subunit | 2 | 1.22 | 0.341 | 1.000 | |
| AT4G16450.2 | NADH:ubiquinone oxidoreductase subunit | 3 | 1.03 | 0.205 | 1.000 | |
| AT4G16450.2 | NADH:ubiquinone oxidoreductase subunit | 4 | 0.85 | 0.189 | 1.000 | |
| AT4G16450.2 | NADH:ubiquinone oxidoreductase subunit | 5 | 0.71 | 0.189 | 1.000 | |
| AT4G16450.2 | NADH:ubiquinone oxidoreductase subunit | 6 | 0.68 | 0.189 | 1.000 | |
| AT4G16450.2 | NADH:ubiquinone oxidoreductase subunit | 7 | 0.73 | 0.189 | 1.000 | |
| AT4G16450.2 | NADH:ubiquinone oxidoreductase subunit | 8 | 0.86 | 0.185 | 1.000 | |
| AT4G16450.2 | NADH:ubiquinone oxidoreductase subunit | 9 | 1.08 | 0.182 | 1.000 | |
| AT4G16450.2 | NADH:ubiquinone oxidoreductase subunit | 10 | 1.16 | 0.182 | 1.000 | |
| AT4G16450.2 | NADH:ubiquinone oxidoreductase subunit | 11 | 1.16 | 0.183 | 1.000 | |
| AT4G16450.2 | NADH:ubiquinone oxidoreductase subunit | 12 | 1.31 | 0.184 | 1.000 | |
| AT4G16450.2 | NADH:ubiquinone oxidoreductase subunit | 13 | 1.56 | 0.181 | 1.000 | |

| | | | | | | |
|---|---|---|---|---|---|---|
| AT4G16450.2 | NADH:ubiquinone oxidoreductase subunit | 14 | 1.68 | 0.179 | 0.996 | |
| AT4G16450.2 | NADH:ubiquinone oxidoreductase subunit | 15 | 1.42 | 0.178 | 1.000 | |
| AT4G16450.2 | NADH:ubiquinone oxidoreductase subunit | 16 | 0.93 | 0.176 | 1.000 | |
| AT4G16450.2 | NADH:ubiquinone oxidoreductase subunit | 17 | 0.62 | 0.175 | 0.998 | |
| AT4G16450.2 | NADH:ubiquinone oxidoreductase subunit | 18 | 0.59 | 0.178 | 0.994 | |
| AT4G16450.2 | NADH:ubiquinone oxidoreductase subunit | 19 | 0.71 | 0.185 | 1.000 | |
| AT4G16450.2 | NADH:ubiquinone oxidoreductase subunit | 20 | 0.86 | 0.190 | 1.000 | |
| AT4G16450.2 | NADH:ubiquinone oxidoreductase subunit | 21 | 1.07 | 0.193 | 1.000 | |
| AT4G16450.2 | NADH:ubiquinone oxidoreductase subunit | 22 | 1.41 | 0.192 | 1.000 | |
| AT4G16450.2 | NADH:ubiquinone oxidoreductase subunit | 23 | 1.63 | 0.190 | 0.999 | |
| AT4G16450.2 | NADH:ubiquinone oxidoreductase subunit | 24 | 1.32 | 0.186 | 1.000 | |
| AT4G16450.2 | NADH:ubiquinone oxidoreductase subunit | 25 | 0.85 | 0.182 | 1.000 | |
| AT4G16450.2 | NADH:ubiquinone oxidoreductase subunit | 26 | 0.72 | 0.182 | 1.000 | |
| AT4G16450.2 | NADH:ubiquinone oxidoreductase subunit | 27 | 0.85 | 0.181 | 1.000 | |
| AT4G16450.2 | NADH:ubiquinone oxidoreductase subunit | 28 | 1.00 | 0.181 | 1.000 | |
| AT4G16450.2 | NADH:ubiquinone oxidoreductase subunit | 29 | 1.02 | 0.181 | 1.000 | |
| AT4G16450.2 | NADH:ubiquinone oxidoreductase subunit | 30 | 1.10 | 0.181 | 1.000 | |
| AT4G16450.2 | NADH:ubiquinone oxidoreductase subunit | 31 | 1.38 | 0.182 | 1.000 | |
| AT4G16450.2 | NADH:ubiquinone oxidoreductase subunit | 32 | 1.65 | 0.184 | 0.998 | |
| AT4G16450.2 | NADH:ubiquinone oxidoreductase subunit | 33 | 1.47 | 0.187 | 1.000 | |
| AT4G16450.2 | NADH:ubiquinone oxidoreductase subunit | 34 | 1.02 | 0.196 | 1.000 | |
| AT4G16450.2 | NADH:ubiquinone oxidoreductase subunit | 35 | 0.78 | 0.209 | 1.000 | |
| AT4G16450.2 | NADH:ubiquinone oxidoreductase subunit | 36 | 0.90 | 0.225 | 1.000 | |
| AT4G16450.2 | NADH:ubiquinone oxidoreductase subunit | 37 | 1.38 | 0.284 | 1.000 | |
| AT4G16450.2 | NADH:ubiquinone oxidoreductase subunit | 38 | 1.98 | 0.400 | 1.000 | |
| AT4G16450.2 | NADH:ubiquinone oxidoreductase subunit | 46 | 1.60 | 2.779 | 1.000 | |
| AT4G16450.2 | NADH:ubiquinone oxidoreductase subunit | 47 | 1.28 | 3.225 | 1.000 | |
| AT4G16700.1 | Phosphatidylserine decarboxylase (PSD1) | 2 | 1.04 | 0.457 | 1.000 | |
| AT4G16700.1 | Phosphatidylserine decarboxylase (PSD1) | 3 | 0.84 | 0.451 | 1.000 | |
| AT4G16700.1 | Phosphatidylserine decarboxylase (PSD1) | 41 | 4.68 | 0.467 | 0.671 | |
| AT4G16700.1 | Phosphatidylserine decarboxylase (PSD1) | 42 | 7.44 | 0.289 | 0.030 | * |

| | | | | | | |
|---|---|---|---|---|---|---|
| AT4G16700.1 | Phosphatidylserine decarboxylase (PSD1) | 43 | 7.84 | 0.244 | 0.005 | ** |
| AT4G16700.1 | Phosphatidylserine decarboxylase (PSD1) | 44 | 3.51 | 0.236 | 0.164 | |
| AT4G16700.1 | Phosphatidylserine decarboxylase (PSD1) | 45 | 0.72 | 0.235 | 0.994 | |
| AT4G16700.1 | Phosphatidylserine decarboxylase (PSD1) | 46 | 0.16 | 0.255 | 0.020 | * |
| AT4G16700.1 | Phosphatidylserine decarboxylase (PSD1) | 47 | 0.04 | 0.505 | 0.053 | . |
| AT4G16760.1 | Acyl-CoA oxidase 1 (ACX1) | 2 | 1.18 | 0.598 | 1.000 | |
| AT4G16760.1 | Acyl-CoA oxidase 1 (ACX1) | 3 | 1.16 | 0.577 | 1.000 | |
| AT4G16760.1 | Acyl-CoA oxidase 1 (ACX1) | 34 | 0.71 | 0.309 | 0.984 | |
| AT4G16760.1 | Acyl-CoA oxidase 1 (ACX1) | 35 | 0.72 | 0.264 | 0.975 | |
| AT4G16760.1 | Acyl-CoA oxidase 1 (ACX1) | 36 | 0.74 | 0.230 | 0.966 | |
| AT4G16760.1 | Acyl-CoA oxidase 1 (ACX1) | 37 | 0.75 | 0.213 | 0.964 | |
| AT4G16760.1 | Acyl-CoA oxidase 1 (ACX1) | 38 | 0.76 | 0.218 | 0.973 | |
| AT4G16760.1 | Acyl-CoA oxidase 1 (ACX1) | 39 | 0.77 | 0.245 | 0.985 | |
| AT4G16760.1 | Acyl-CoA oxidase 1 (ACX1) | 40 | 0.77 | 0.290 | 0.993 | |
| AT4G16760.1 | Acyl-CoA oxidase 1 (ACX1) | 46 | 0.71 | 0.752 | 1.000 | |
| AT4G16760.1 | Acyl-CoA oxidase 1 (ACX1) | 47 | 0.69 | 0.851 | 1.000 | |
| AT4G16800.1 | Probable enoyl-CoA hydratase 2 | 2 | 0.95 | 0.408 | 1.000 | |
| AT4G16800.1 | Probable enoyl-CoA hydratase 2 | 3 | 1.00 | 0.401 | 1.000 | |
| AT4G16800.1 | Probable enoyl-CoA hydratase 2 | 28 | 0.86 | 0.350 | 1.000 | |
| AT4G16800.1 | Probable enoyl-CoA hydratase 2 | 29 | 0.80 | 0.287 | 1.000 | |
| AT4G16800.1 | Probable enoyl-CoA hydratase 2 | 30 | 0.86 | 0.280 | 1.000 | |
| AT4G16800.1 | Probable enoyl-CoA hydratase 2 | 31 | 1.10 | 0.244 | 1.000 | |
| AT4G16800.1 | Probable enoyl-CoA hydratase 2 | 32 | 1.38 | 0.222 | 1.000 | |
| AT4G16800.1 | Probable enoyl-CoA hydratase 2 | 33 | 1.49 | 0.210 | 0.998 | |
| AT4G16800.1 | Probable enoyl-CoA hydratase 2 | 34 | 1.44 | 0.206 | 0.999 | |
| AT4G16800.1 | Probable enoyl-CoA hydratase 2 | 35 | 1.59 | 0.206 | 0.991 | |
| AT4G16800.1 | Probable enoyl-CoA hydratase 2 | 36 | 2.37 | 0.211 | 0.632 | |
| AT4G16800.1 | Probable enoyl-CoA hydratase 2 | 37 | 3.08 | 0.217 | 0.299 | |
| AT4G16800.1 | Probable enoyl-CoA hydratase 2 | 38 | 3.14 | 0.224 | 0.316 | |
| AT4G16800.1 | Probable enoyl-CoA hydratase 2 | 39 | 2.54 | 0.215 | 0.551 | |
| AT4G16800.1 | Probable enoyl-CoA hydratase 2 | 40 | 2.13 | 0.206 | 0.763 | |
| AT4G16800.1 | Probable enoyl-CoA hydratase 2 | 41 | 1.82 | 0.204 | 0.934 | |
| AT4G16800.1 | Probable enoyl-CoA hydratase 2 | 42 | 2.00 | 0.205 | 0.847 | |
| AT4G16800.1 | Probable enoyl-CoA hydratase 2 | 43 | 3.08 | 0.206 | 0.232 | |
| AT4G16800.1 | Probable enoyl-CoA hydratase 2 | 44 | 3.92 | 0.205 | 0.065 | . |
| AT4G16800.1 | Probable enoyl-CoA hydratase 2 | 45 | 2.10 | 0.208 | 0.796 | |

| | | | | | | |
|---|---|---|---|---|---|---|
| AT4G16800.1 | Probable enoyl-CoA hydratase 2 | 46 | 0.40 | 0.231 | 0.677 | |
| AT4G16800.1 | Probable enoyl-CoA hydratase 2 | 47 | 0.04 | 0.372 | 0.003 | ** |
| AT4G17085.1 | ? | 2 | 0.89 | 0.533 | 1.000 | |
| AT4G17085.1 | ? | 3 | 0.91 | 0.408 | 1.000 | |
| AT4G17085.1 | ? | 6 | 0.98 | 0.390 | 1.000 | |
| AT4G17085.1 | ? | 7 | 1.04 | 0.320 | 1.000 | |
| AT4G17085.1 | ? | 8 | 1.18 | 0.275 | 1.000 | |
| AT4G17085.1 | ? | 9 | 1.41 | 0.252 | 1.000 | |
| AT4G17085.1 | ? | 10 | 1.52 | 0.237 | 1.000 | |
| AT4G17085.1 | ? | 11 | 1.43 | 0.227 | 1.000 | |
| AT4G17085.1 | ? | 12 | 1.28 | 0.221 | 1.000 | |
| AT4G17085.1 | ? | 13 | 1.29 | 0.217 | 1.000 | |
| AT4G17085.1 | ? | 14 | 1.66 | 0.216 | 1.000 | |
| AT4G17085.1 | ? | 15 | 2.27 | 0.216 | 0.922 | |
| AT4G17085.1 | ? | 16 | 2.19 | 0.216 | 0.947 | |
| AT4G17085.1 | ? | 17 | 1.57 | 0.220 | 1.000 | |
| AT4G17085.1 | ? | 18 | 1.44 | 0.228 | 1.000 | |
| AT4G17085.1 | ? | 19 | 1.94 | 0.247 | 0.998 | |
| AT4G17085.1 | ? | 20 | 3.03 | 0.272 | 0.861 | |
| AT4G17085.1 | ? | 21 | 4.63 | 0.292 | 0.466 | |
| AT4G17085.1 | ? | 22 | 6.09 | 0.306 | 0.262 | |
| AT4G17085.1 | ? | 23 | 6.94 | 0.347 | 0.357 | |
| AT4G17085.1 | ? | 24 | 7.13 | 0.418 | 0.665 | |
| AT4G17085.1 | ? | 30 | 3.47 | 0.334 | 0.934 | |
| AT4G17085.1 | ? | 31 | 3.21 | 0.270 | 0.791 | |
| AT4G17085.1 | ? | 32 | 3.35 | 0.241 | 0.546 | |
| AT4G17085.1 | ? | 33 | 3.86 | 0.226 | 0.243 | |
| AT4G17085.1 | ? | 34 | 4.58 | 0.222 | 0.091 | . |
| AT4G17085.1 | ? | 35 | 4.49 | 0.221 | 0.097 | . |
| AT4G17085.1 | ? | 36 | 2.95 | 0.219 | 0.580 | |
| AT4G17085.1 | ? | 37 | 1.54 | 0.220 | 1.000 | |
| AT4G17085.1 | ? | 38 | 0.84 | 0.222 | 1.000 | |
| AT4G17085.1 | ? | 39 | 0.72 | 0.229 | 1.000 | |
| AT4G17085.1 | ? | 40 | 1.24 | 0.241 | 1.000 | |
| AT4G17085.1 | ? | 41 | 2.91 | 0.253 | 0.820 | |
| AT4G17085.1 | ? | 42 | 6.54 | 0.250 | 0.038 | * |
| AT4G17085.1 | ? | 43 | 10.19 | 0.235 | 0.001 | *** |
| AT4G17085.1 | ? | 44 | 5.95 | 0.226 | 0.022 | * |
| AT4G17085.1 | ? | 45 | 1.92 | 0.224 | 0.995 | |
| AT4G17085.1 | ? | 46 | 0.58 | 0.267 | 1.000 | |
| AT4G17085.1 | ? | 47 | 0.17 | 0.508 | 0.958 | |
| AT4G17140.3 | Pleckstrin homology (PH) domain-containing protein | 2 | 0.90 | 0.447 | 1.000 | |
| AT4G17140.3 | Pleckstrin homology (PH) domain-containing protein | 3 | 0.95 | 0.338 | 1.000 | |
| AT4G17140.3 | Pleckstrin homology (PH) domain-containing protein | 4 | 0.99 | 0.357 | 1.000 | |
| AT4G17140.3 | Pleckstrin homology (PH) domain-containing protein | 6 | 1.10 | 0.398 | 1.000 | |
| AT4G17140.3 | Pleckstrin homology (PH) domain-containing protein | 7 | 1.10 | 0.401 | 1.000 | |
| AT4G17140.3 | Pleckstrin homology (PH) domain-containing protein | 8 | 0.98 | 0.414 | 1.000 | |
| AT4G17140.3 | Pleckstrin homology (PH) domain-containing protein | 12 | 0.52 | 0.332 | 0.999 | |
| AT4G17140.3 | Pleckstrin homology (PH) domain-containing protein | 13 | 0.77 | 0.309 | 1.000 | |
| AT4G17140.3 | Pleckstrin homology (PH) domain-containing protein | 14 | 1.18 | 0.284 | 1.000 | |
| AT4G17140.3 | Pleckstrin homology (PH) domain-containing protein | 15 | 1.47 | 0.260 | 1.000 | |

| | | | | | |
|---|---|---|---|---|---|
| AT4G17140.3 | Pleckstrin homology (PH) domain-containing protein | 16 | 1.51 | 0.246 | 1.000 |
| AT4G17140.3 | Pleckstrin homology (PH) domain-containing protein | 17 | 1.48 | 0.242 | 1.000 |
| AT4G17140.3 | Pleckstrin homology (PH) domain-containing protein | 18 | 1.55 | 0.245 | 1.000 |
| AT4G17140.3 | Pleckstrin homology (PH) domain-containing protein | 19 | 1.70 | 0.250 | 0.998 |
| AT4G17140.3 | Pleckstrin homology (PH) domain-containing protein | 20 | 1.81 | 0.255 | 0.995 |
| AT4G17140.3 | Pleckstrin homology (PH) domain-containing protein | 21 | 1.89 | 0.258 | 0.990 |
| AT4G17140.3 | Pleckstrin homology (PH) domain-containing protein | 22 | 1.92 | 0.258 | 0.987 |
| AT4G17140.3 | Pleckstrin homology (PH) domain-containing protein | 23 | 1.75 | 0.256 | 0.997 |
| AT4G17140.3 | Pleckstrin homology (PH) domain-containing protein | 24 | 1.50 | 0.252 | 1.000 |
| AT4G17140.3 | Pleckstrin homology (PH) domain-containing protein | 25 | 1.12 | 0.251 | 1.000 |
| AT4G17140.3 | Pleckstrin homology (PH) domain-containing protein | 26 | 0.66 | 0.253 | 1.000 |
| AT4G17140.3 | Pleckstrin homology (PH) domain-containing protein | 27 | 0.34 | 0.265 | 0.687 |
| AT4G17140.3 | Pleckstrin homology (PH) domain-containing protein | 28 | 0.24 | 0.288 | 0.405 |
| AT4G17140.3 | Pleckstrin homology (PH) domain-containing protein | 29 | 0.31 | 0.308 | 0.775 |
| AT4G17140.3 | Pleckstrin homology (PH) domain-containing protein | 30 | 0.52 | 0.334 | 0.999 |
| AT4G17140.3 | Pleckstrin homology (PH) domain-containing protein | 31 | 0.82 | 0.382 | 1.000 |
| AT4G17140.3 | Pleckstrin homology (PH) domain-containing protein | 32 | 1.20 | 0.516 | 1.000 |
| AT4G17140.3 | Pleckstrin homology (PH) domain-containing protein | 46 | 233.12 | 5.169 | 1.000 |
| AT4G17140.3 | Pleckstrin homology (PH) domain-containing protein | 47 | 341.09 | 5.691 | 1.000 |
| AT4G17170.1 | RAB GTPase homolog B1B (RAB-B1B) | 2 | 1.01 | 0.331 | 1.000 |
| AT4G17170.1 | RAB GTPase homolog B1B (RAB-B1B) | 3 | 1.05 | 0.252 | 1.000 |
| AT4G17170.1 | RAB GTPase homolog B1B (RAB-B1B) | 4 | 1.10 | 0.206 | 1.000 |
| AT4G17170.1 | RAB GTPase homolog B1B (RAB-B1B) | 5 | 1.15 | 0.195 | 1.000 |
| AT4G17170.1 | RAB GTPase homolog B1B (RAB-B1B) | 6 | 1.20 | 0.208 | 1.000 |
| AT4G17170.1 | RAB GTPase homolog B1B (RAB-B1B) | 7 | 1.24 | 0.234 | 1.000 |
| AT4G17170.1 | RAB GTPase homolog B1B (RAB-B1B) | 8 | 1.26 | 0.265 | 1.000 |
| AT4G17170.1 | RAB GTPase homolog B1B (RAB-B1B) | 17 | 1.28 | 0.294 | 1.000 |
| AT4G17170.1 | RAB GTPase homolog B1B (RAB-B1B) | 18 | 1.29 | 0.266 | 1.000 |
| AT4G17170.1 | RAB GTPase homolog B1B (RAB-B1B) | 19 | 1.30 | 0.242 | 1.000 |
| AT4G17170.1 | RAB GTPase homolog B1B (RAB-B1B) | 20 | 1.31 | 0.223 | 1.000 |
| AT4G17170.1 | RAB GTPase homolog B1B (RAB-B1B) | 21 | 1.31 | 0.210 | 1.000 |

| | | | | | |
|---|---|---|---|---|---|
| AT4G17170.1 | RAB GTPase homolog B1B (RAB-B1B) | 22 | 1.28 | 0.201 | 1.000 |
| AT4G17170.1 | RAB GTPase homolog B1B (RAB-B1B) | 23 | 1.23 | 0.196 | 1.000 |
| AT4G17170.1 | RAB GTPase homolog B1B (RAB-B1B) | 24 | 1.17 | 0.191 | 1.000 |
| AT4G17170.1 | RAB GTPase homolog B1B (RAB-B1B) | 25 | 1.09 | 0.186 | 1.000 |
| AT4G17170.1 | RAB GTPase homolog B1B (RAB-B1B) | 26 | 1.03 | 0.181 | 1.000 |
| AT4G17170.1 | RAB GTPase homolog B1B (RAB-B1B) | 27 | 0.97 | 0.176 | 1.000 |
| AT4G17170.1 | RAB GTPase homolog B1B (RAB-B1B) | 28 | 0.92 | 0.171 | 1.000 |
| AT4G17170.1 | RAB GTPase homolog B1B (RAB-B1B) | 29 | 0.85 | 0.167 | 1.000 |
| AT4G17170.1 | RAB GTPase homolog B1B (RAB-B1B) | 30 | 0.77 | 0.166 | 1.000 |
| AT4G17170.1 | RAB GTPase homolog B1B (RAB-B1B) | 31 | 0.69 | 0.167 | 0.989 |
| AT4G17170.1 | RAB GTPase homolog B1B (RAB-B1B) | 32 | 0.63 | 0.170 | 0.959 |
| AT4G17170.1 | RAB GTPase homolog B1B (RAB-B1B) | 33 | 0.61 | 0.178 | 0.951 |
| AT4G17170.1 | RAB GTPase homolog B1B (RAB-B1B) | 34 | 0.61 | 0.190 | 0.968 |
| AT4G17170.1 | RAB GTPase homolog B1B (RAB-B1B) | 35 | 0.63 | 0.206 | 0.988 |
| AT4G17170.1 | RAB GTPase homolog B1B (RAB-B1B) | 36 | 0.66 | 0.223 | 0.998 |
| AT4G17170.1 | RAB GTPase homolog B1B (RAB-B1B) | 39 | 0.89 | 0.249 | 1.000 |
| AT4G17170.1 | RAB GTPase homolog B1B (RAB-B1B) | 40 | 0.99 | 0.251 | 1.000 |
| AT4G17170.1 | RAB GTPase homolog B1B (RAB-B1B) | 41 | 1.11 | 0.249 | 1.000 |
| AT4G17170.1 | RAB GTPase homolog B1B (RAB-B1B) | 42 | 1.23 | 0.249 | 1.000 |
| AT4G17170.1 | RAB GTPase homolog B1B (RAB-B1B) | 43 | 1.35 | 0.254 | 1.000 |
| AT4G17170.1 | RAB GTPase homolog B1B (RAB-B1B) | 44 | 1.48 | 0.274 | 1.000 |
| AT4G17170.1 | RAB GTPase homolog B1B (RAB-B1B) | 45 | 1.62 | 0.313 | 1.000 |
| AT4G17170.1 | RAB GTPase homolog B1B (RAB-B1B) | 46 | 1.76 | 0.379 | 1.000 |
| AT4G17170.1 | RAB GTPase homolog B1B (RAB-B1B) | 47 | 1.91 | 0.472 | 1.000 |
| AT4G17300.1 | Asparaginyl-tRNA synthetase (ATNS1) / OVA8 | 2 | 0.71 | 0.644 | 1.000 |
| AT4G17300.1 | Asparaginyl-tRNA synthetase (ATNS1) / OVA8 | 3 | 0.62 | 0.619 | 1.000 |
| AT4G17300.1 | Asparaginyl-tRNA synthetase (ATNS1) / OVA8 | 33 | 0.56 | 0.439 | 0.998 |
| AT4G17300.1 | Asparaginyl-tRNA synthetase (ATNS1) / OVA8 | 34 | 0.72 | 0.343 | 1.000 |
| AT4G17300.1 | Asparaginyl-tRNA synthetase (ATNS1) / OVA8 | 35 | 1.01 | 0.299 | 1.000 |
| AT4G17300.1 | Asparaginyl-tRNA synthetase (ATNS1) / OVA8 | 36 | 1.47 | 0.284 | 0.998 |
| AT4G17300.1 | Asparaginyl-tRNA synthetase (ATNS1) / OVA8 | 37 | 2.01 | 0.279 | 0.910 |

| | | | | | |
|---|---|---|---|---|---|
| AT4G17300.1 | Asparaginyl-tRNA synthetase (ATNS1) / OVA8 | 38 | 2.40 | 0.278 | 0.757 |
| AT4G17300.1 | Asparaginyl-tRNA synthetase (ATNS1) / OVA8 | 39 | 2.49 | 0.277 | 0.711 |
| AT4G17300.1 | Asparaginyl-tRNA synthetase (ATNS1) / OVA8 | 40 | 2.29 | 0.277 | 0.801 |
| AT4G17300.1 | Asparaginyl-tRNA synthetase (ATNS1) / OVA8 | 41 | 1.85 | 0.278 | 0.953 |
| AT4G17300.1 | Asparaginyl-tRNA synthetase (ATNS1) / OVA8 | 42 | 1.45 | 0.281 | 0.999 |
| AT4G17300.1 | Asparaginyl-tRNA synthetase (ATNS1) / OVA8 | 43 | 1.11 | 0.282 | 1.000 |
| AT4G17300.1 | Asparaginyl-tRNA synthetase (ATNS1) / OVA8 | 44 | 0.82 | 0.281 | 1.000 |
| AT4G17300.1 | Asparaginyl-tRNA synthetase (ATNS1) / OVA8 | 45 | 0.55 | 0.290 | 0.969 |
| AT4G17300.1 | Asparaginyl-tRNA synthetase (ATNS1) / OVA8 | 46 | 0.32 | 0.343 | 0.711 |
| AT4G17300.1 | Asparaginyl-tRNA synthetase (ATNS1) / OVA8 | 47 | 0.17 | 0.492 | 0.612 |
| AT4G17360.1 | Putative 10-formyl tetrahydrofolate deformylase (PURU2) | 3 | 0.81 | 0.231 | 1.000 |
| AT4G17360.1 | Putative 10-formyl tetrahydrofolate deformylase (PURU2) | 4 | 0.90 | 0.199 | 1.000 |
| AT4G17360.1 | Putative 10-formyl tetrahydrofolate deformylase (PURU2) | 5 | 0.98 | 0.192 | 1.000 |
| AT4G17360.1 | Putative 10-formyl tetrahydrofolate deformylase (PURU2) | 6 | 1.05 | 0.192 | 1.000 |
| AT4G17360.1 | Putative 10-formyl tetrahydrofolate deformylase (PURU2) | 7 | 1.11 | 0.192 | 1.000 |
| AT4G17360.1 | Putative 10-formyl tetrahydrofolate deformylase (PURU2) | 8 | 1.14 | 0.190 | 1.000 |
| AT4G17360.1 | Putative 10-formyl tetrahydrofolate deformylase (PURU2) | 9 | 1.16 | 0.189 | 1.000 |
| AT4G17360.1 | Putative 10-formyl tetrahydrofolate deformylase (PURU2) | 10 | 1.15 | 0.188 | 1.000 |
| AT4G17360.1 | Putative 10-formyl tetrahydrofolate deformylase (PURU2) | 11 | 1.11 | 0.187 | 1.000 |
| AT4G17360.1 | Putative 10-formyl tetrahydrofolate deformylase (PURU2) | 12 | 1.05 | 0.187 | 1.000 |
| AT4G17360.1 | Putative 10-formyl tetrahydrofolate deformylase (PURU2) | 13 | 0.97 | 0.186 | 1.000 |
| AT4G17360.1 | Putative 10-formyl tetrahydrofolate deformylase (PURU2) | 14 | 0.90 | 0.185 | 1.000 |
| AT4G17360.1 | Putative 10-formyl tetrahydrofolate deformylase (PURU2) | 15 | 0.83 | 0.184 | 1.000 |
| AT4G17360.1 | Putative 10-formyl tetrahydrofolate deformylase (PURU2) | 16 | 0.79 | 0.183 | 1.000 |

| | | | | | |
|---|---|---|---|---|---|
| AT4G17360.1 | Putative 10-formyl tetrahydrofolate deformylase (PURU2) | 17 | 0.77 | 0.184 | 1.000 |
| AT4G17360.1 | Putative 10-formyl tetrahydrofolate deformylase (PURU2) | 18 | 0.80 | 0.185 | 1.000 |
| AT4G17360.1 | Putative 10-formyl tetrahydrofolate deformylase (PURU2) | 19 | 0.88 | 0.188 | 1.000 |
| AT4G17360.1 | Putative 10-formyl tetrahydrofolate deformylase (PURU2) | 20 | 1.04 | 0.190 | 1.000 |
| AT4G17360.1 | Putative 10-formyl tetrahydrofolate deformylase (PURU2) | 21 | 1.31 | 0.191 | 1.000 |
| AT4G17360.1 | Putative 10-formyl tetrahydrofolate deformylase (PURU2) | 22 | 1.63 | 0.191 | 0.987 |
| AT4G17360.1 | Putative 10-formyl tetrahydrofolate deformylase (PURU2) | 23 | 1.86 | 0.190 | 0.906 |
| AT4G17360.1 | Putative 10-formyl tetrahydrofolate deformylase (PURU2) | 24 | 1.92 | 0.189 | 0.864 |
| AT4G17360.1 | Putative 10-formyl tetrahydrofolate deformylase (PURU2) | 25 | 1.84 | 0.187 | 0.905 |
| AT4G17360.1 | Putative 10-formyl tetrahydrofolate deformylase (PURU2) | 26 | 1.72 | 0.186 | 0.960 |
| AT4G17360.1 | Putative 10-formyl tetrahydrofolate deformylase (PURU2) | 27 | 1.60 | 0.186 | 0.990 |
| AT4G17360.1 | Putative 10-formyl tetrahydrofolate deformylase (PURU2) | 28 | 1.44 | 0.186 | 0.999 |
| AT4G17360.1 | Putative 10-formyl tetrahydrofolate deformylase (PURU2) | 29 | 1.27 | 0.186 | 1.000 |
| AT4G17360.1 | Putative 10-formyl tetrahydrofolate deformylase (PURU2) | 30 | 1.10 | 0.186 | 1.000 |
| AT4G17360.1 | Putative 10-formyl tetrahydrofolate deformylase (PURU2) | 31 | 0.96 | 0.187 | 1.000 |
| AT4G17360.1 | Putative 10-formyl tetrahydrofolate deformylase (PURU2) | 32 | 0.83 | 0.187 | 1.000 |
| AT4G17360.1 | Putative 10-formyl tetrahydrofolate deformylase (PURU2) | 33 | 0.74 | 0.187 | 1.000 |
| AT4G17360.1 | Putative 10-formyl tetrahydrofolate deformylase (PURU2) | 34 | 0.69 | 0.187 | 0.999 |
| AT4G17360.1 | Putative 10-formyl tetrahydrofolate deformylase (PURU2) | 35 | 0.68 | 0.187 | 0.999 |
| AT4G17360.1 | Putative 10-formyl tetrahydrofolate deformylase (PURU2) | 36 | 0.69 | 0.187 | 0.999 |

| | | | | | | |
|---|---|---|---|---|---|---|
| AT4G17360.1 | Putative 10-formyl tetrahydrofolate deformylase (PURU2) | 37 | 0.73 | 0.187 | 1.000 | |
| AT4G17360.1 | Putative 10-formyl tetrahydrofolate deformylase (PURU2) | 38 | 0.82 | 0.187 | 1.000 | |
| AT4G17360.1 | Putative 10-formyl tetrahydrofolate deformylase (PURU2) | 39 | 0.97 | 0.187 | 1.000 | |
| AT4G17360.1 | Putative 10-formyl tetrahydrofolate deformylase (PURU2) | 40 | 1.18 | 0.188 | 1.000 | |
| AT4G17360.1 | Putative 10-formyl tetrahydrofolate deformylase (PURU2) | 41 | 1.43 | 0.190 | 1.000 | |
| AT4G17360.1 | Putative 10-formyl tetrahydrofolate deformylase (PURU2) | 42 | 1.63 | 0.191 | 0.988 | |
| AT4G17360.1 | Putative 10-formyl tetrahydrofolate deformylase (PURU2) | 43 | 1.70 | 0.191 | 0.974 | |
| AT4G17360.1 | Putative 10-formyl tetrahydrofolate deformylase (PURU2) | 44 | 1.60 | 0.192 | 0.992 | |
| AT4G17360.1 | Putative 10-formyl tetrahydrofolate deformylase (PURU2) | 45 | 1.35 | 0.202 | 1.000 | |
| AT4G17360.1 | Putative 10-formyl tetrahydrofolate deformylase (PURU2) | 46 | 1.09 | 0.243 | 1.000 | |
| AT4G17360.1 | Putative 10-formyl tetrahydrofolate deformylase (PURU2) | 47 | 0.88 | 0.334 | 1.000 | |
| AT4G17360.1 | Putative 10-formyl tetrahydrofolate deformylase (PURU2) | 48 | 0.71 | 0.459 | 1.000 | |
| AT4G18100.1 | 60S ribosomal protein L32-1 (RPL32A) | 2 | 0.41 | 0.157 | 0.087 | . |
| AT4G18100.1 | 60S ribosomal protein L32-1 (RPL32A) | 3 | 0.46 | 0.138 | 0.105 | |
| AT4G18100.1 | 60S ribosomal protein L32-1 (RPL32A) | 4 | 0.53 | 0.124 | 0.153 | |
| AT4G18100.1 | 60S ribosomal protein L32-1 (RPL32A) | 5 | 0.60 | 0.113 | 0.265 | |
| AT4G18100.1 | 60S ribosomal protein L32-1 (RPL32A) | 6 | 0.68 | 0.105 | 0.493 | |
| AT4G18100.1 | 60S ribosomal protein L32-1 (RPL32A) | 7 | 0.77 | 0.100 | 0.804 | |
| AT4G18100.1 | 60S ribosomal protein L32-1 (RPL32A) | 8 | 0.87 | 0.096 | 0.986 | |
| AT4G18100.1 | 60S ribosomal protein L32-1 (RPL32A) | 9 | 0.98 | 0.094 | 1.000 | |
| AT4G18100.1 | 60S ribosomal protein L32-1 (RPL32A) | 10 | 1.09 | 0.092 | 0.999 | |
| AT4G18100.1 | 60S ribosomal protein L32-1 (RPL32A) | 11 | 1.20 | 0.091 | 0.922 | |
| AT4G18100.1 | 60S ribosomal protein L32-1 (RPL32A) | 12 | 1.31 | 0.091 | 0.693 | |
| AT4G18100.1 | 60S ribosomal protein L32-1 (RPL32A) | 13 | 1.42 | 0.092 | 0.447 | |
| AT4G18100.1 | 60S ribosomal protein L32-1 (RPL32A) | 14 | 1.52 | 0.093 | 0.272 | |

| | | | | | | |
|---|---|---|---|---|---|---|
| AT4G18100.1 | 60S ribosomal protein L32-1 (RPL32A) | 15 | 1.61 | 0.095 | 0.171 | |
| AT4G18100.1 | 60S ribosomal protein L32-1 (RPL32A) | 16 | 1.71 | 0.098 | 0.116 | |
| AT4G18100.1 | 60S ribosomal protein L32-1 (RPL32A) | 17 | 1.79 | 0.103 | 0.092 | . |
| AT4G18100.1 | 60S ribosomal protein L32-1 (RPL32A) | 18 | 1.87 | 0.108 | 0.085 | . |
| AT4G18100.1 | 60S ribosomal protein L32-1 (RPL32A) | 19 | 1.93 | 0.115 | 0.090 | . |
| AT4G18100.1 | 60S ribosomal protein L32-1 (RPL32A) | 22 | 2.04 | 0.141 | 0.167 | |
| AT4G18100.1 | 60S ribosomal protein L32-1 (RPL32A) | 23 | 2.06 | 0.151 | 0.215 | |
| AT4G18100.1 | 60S ribosomal protein L32-1 (RPL32A) | 24 | 2.06 | 0.161 | 0.276 | |
| AT4G18100.1 | 60S ribosomal protein L32-1 (RPL32A) | 41 | 1.23 | 0.289 | 1.000 | |
| AT4G18100.1 | 60S ribosomal protein L32-1 (RPL32A) | 42 | 1.17 | 0.303 | 1.000 | |
| AT4G18100.1 | 60S ribosomal protein L32-1 (RPL32A) | 43 | 1.12 | 0.319 | 1.000 | |
| AT4G18100.1 | 60S ribosomal protein L32-1 (RPL32A) | 46 | 0.97 | 0.389 | 1.000 | |
| AT4G18100.1 | 60S ribosomal protein L32-1 (RPL32A) | 47 | 0.93 | 0.418 | 1.000 | |
| AT4G20150.1 | NADH:ubiquinone oxidoreductase subunit (B14.5b) | 2 | 0.74 | 0.610 | 1.000 | |
| AT4G20150.1 | NADH:ubiquinone oxidoreductase subunit (B14.5b) | 3 | 0.76 | 0.546 | 1.000 | |
| AT4G20150.1 | NADH:ubiquinone oxidoreductase subunit (B14.5b) | 14 | 0.94 | 0.354 | 1.000 | |
| AT4G20150.1 | NADH:ubiquinone oxidoreductase subunit (B14.5b) | 15 | 0.86 | 0.302 | 1.000 | |
| AT4G20150.1 | NADH:ubiquinone oxidoreductase subunit (B14.5b) | 16 | 0.72 | 0.274 | 1.000 | |
| AT4G20150.1 | NADH:ubiquinone oxidoreductase subunit (B14.5b) | 17 | 0.58 | 0.251 | 0.998 | |
| AT4G20150.1 | NADH:ubiquinone oxidoreductase subunit (B14.5b) | 18 | 0.56 | 0.244 | 0.995 | |
| AT4G20150.1 | NADH:ubiquinone oxidoreductase subunit (B14.5b) | 19 | 0.66 | 0.248 | 1.000 | |
| AT4G20150.1 | NADH:ubiquinone oxidoreductase subunit (B14.5b) | 20 | 0.81 | 0.253 | 1.000 | |
| AT4G20150.1 | NADH:ubiquinone oxidoreductase subunit (B14.5b) | 21 | 0.99 | 0.255 | 1.000 | |
| AT4G20150.1 | NADH:ubiquinone oxidoreductase subunit (B14.5b) | 22 | 1.24 | 0.255 | 1.000 | |
| AT4G20150.1 | NADH:ubiquinone oxidoreductase subunit (B14.5b) | 23 | 1.47 | 0.253 | 1.000 | |

| | | | | | |
|---|---|---|---|---|---|
| AT4G20150.1 | NADH:ubiquinone oxidoreductase subunit (B14.5b) | 24 | 1.50 | 0.249 | 1.000 |
| AT4G20150.1 | NADH:ubiquinone oxidoreductase subunit (B14.5b) | 25 | 1.30 | 0.245 | 1.000 |
| AT4G20150.1 | NADH:ubiquinone oxidoreductase subunit (B14.5b) | 26 | 1.14 | 0.244 | 1.000 |
| AT4G20150.1 | NADH:ubiquinone oxidoreductase subunit (B14.5b) | 27 | 1.07 | 0.244 | 1.000 |
| AT4G20150.1 | NADH:ubiquinone oxidoreductase subunit (B14.5b) | 28 | 0.93 | 0.245 | 1.000 |
| AT4G20150.1 | NADH:ubiquinone oxidoreductase subunit (B14.5b) | 29 | 0.73 | 0.245 | 1.000 |
| AT4G20150.1 | NADH:ubiquinone oxidoreductase subunit (B14.5b) | 30 | 0.64 | 0.247 | 1.000 |
| AT4G20150.1 | NADH:ubiquinone oxidoreductase subunit (B14.5b) | 31 | 0.68 | 0.254 | 1.000 |
| AT4G20150.1 | NADH:ubiquinone oxidoreductase subunit (B14.5b) | 32 | 0.72 | 0.275 | 1.000 |
| AT4G20150.1 | NADH:ubiquinone oxidoreductase subunit (B14.5b) | 33 | 0.59 | 0.320 | 1.000 |
| AT4G20150.1 | NADH:ubiquinone oxidoreductase subunit (B14.5b) | 34 | 0.49 | 0.371 | 1.000 |
| AT4G20150.1 | NADH:ubiquinone oxidoreductase subunit (B14.5b) | 38 | 0.47 | 0.380 | 0.999 |
| AT4G20150.1 | NADH:ubiquinone oxidoreductase subunit (B14.5b) | 39 | 0.46 | 0.335 | 0.996 |
| AT4G20150.1 | NADH:ubiquinone oxidoreductase subunit (B14.5b) | 40 | 0.42 | 0.359 | 0.994 |
| AT4G20150.1 | NADH:ubiquinone oxidoreductase subunit (B14.5b) | 41 | 0.37 | 0.487 | 0.999 |
| AT4G20150.1 | NADH:ubiquinone oxidoreductase subunit (B14.5b) | 46 | 0.37 | 1.885 | 1.000 |
| AT4G20150.1 | NADH:ubiquinone oxidoreductase subunit (B14.5b) | 47 | 0.39 | 2.302 | 1.000 |
| AT4G20760.1 | Putative NAD- or NADP-dependent oxidoreductase | 2 | 0.68 | 0.492 | 1.000 |
| AT4G20760.1 | Putative NAD- or NADP-dependent oxidoreductase | 3 | 0.69 | 0.413 | 1.000 |
| AT4G20760.1 | Putative NAD- or NADP-dependent oxidoreductase | 4 | 0.72 | 0.408 | 1.000 |
| AT4G20760.1 | Putative NAD- or NADP-dependent oxidoreductase | 7 | 0.72 | 0.381 | 1.000 |
| AT4G20760.1 | Putative NAD- or NADP-dependent oxidoreductase | 8 | 0.66 | 0.347 | 1.000 |

| | | | | | | |
|---|---|---|---|---|---|---|
| AT4G20760.1 | Putative NAD- or NADP-dependent oxidoreductase | 9 | 0.62 | 0.330 | 1.000 | |
| AT4G20760.1 | Putative NAD- or NADP-dependent oxidoreductase | 10 | 0.71 | 0.328 | 1.000 | |
| AT4G20760.1 | Putative NAD- or NADP-dependent oxidoreductase | 11 | 1.06 | 0.336 | 1.000 | |
| AT4G20760.1 | Putative NAD- or NADP-dependent oxidoreductase | 12 | 1.78 | 0.351 | 0.999 | |
| AT4G20760.1 | Putative NAD- or NADP-dependent oxidoreductase | 13 | 2.80 | 0.375 | 0.914 | |
| AT4G20760.1 | Putative NAD- or NADP-dependent oxidoreductase | 14 | 3.96 | 0.430 | 0.807 | |
| AT4G20760.1 | Putative NAD- or NADP-dependent oxidoreductase | 31 | 1.94 | 0.415 | 0.999 | |
| AT4G20760.1 | Putative NAD- or NADP-dependent oxidoreductase | 32 | 1.98 | 0.349 | 0.993 | |
| AT4G20760.1 | Putative NAD- or NADP-dependent oxidoreductase | 33 | 1.96 | 0.323 | 0.987 | |
| AT4G20760.1 | Putative NAD- or NADP-dependent oxidoreductase | 34 | 1.72 | 0.312 | 0.998 | |
| AT4G20760.1 | Putative NAD- or NADP-dependent oxidoreductase | 35 | 1.35 | 0.308 | 1.000 | |
| AT4G20760.1 | Putative NAD- or NADP-dependent oxidoreductase | 36 | 1.04 | 0.306 | 1.000 | |
| AT4G20760.1 | Putative NAD- or NADP-dependent oxidoreductase | 37 | 0.92 | 0.305 | 1.000 | |
| AT4G20760.1 | Putative NAD- or NADP-dependent oxidoreductase | 38 | 1.07 | 0.305 | 1.000 | |
| AT4G20760.1 | Putative NAD- or NADP-dependent oxidoreductase | 39 | 1.56 | 0.305 | 1.000 | |
| AT4G20760.1 | Putative NAD- or NADP-dependent oxidoreductase | 40 | 2.37 | 0.305 | 0.900 | |
| AT4G20760.1 | Putative NAD- or NADP-dependent oxidoreductase | 41 | 3.09 | 0.306 | 0.661 | |
| AT4G20760.1 | Putative NAD- or NADP-dependent oxidoreductase | 42 | 3.02 | 0.307 | 0.688 | |
| AT4G20760.1 | Putative NAD- or NADP-dependent oxidoreductase | 43 | 2.17 | 0.308 | 0.951 | |
| AT4G20760.1 | Putative NAD- or NADP-dependent oxidoreductase | 44 | 1.15 | 0.311 | 1.000 | |
| AT4G20760.1 | Putative NAD- or NADP-dependent oxidoreductase | 45 | 0.46 | 0.319 | 0.961 | |
| AT4G20760.1 | Putative NAD- or NADP-dependent oxidoreductase | 46 | 0.15 | 0.357 | 0.192 | |
| AT4G20760.1 | Putative NAD- or NADP-dependent oxidoreductase | 47 | 0.04 | 0.469 | 0.042 | * |
| AT4G21105.1 | Cytochrome c:O2 oxidoreductase subunit (putative) | 2 | 0.76 | 0.369 | 1.000 | |
| AT4G21105.1 | Cytochrome c:O2 oxidoreductase subunit (putative) | 3 | 0.79 | 0.215 | 1.000 | |
| AT4G21105.1 | Cytochrome c:O2 oxidoreductase subunit (putative) | 4 | 1.04 | 0.204 | 1.000 | |
| AT4G21105.1 | Cytochrome c:O2 oxidoreductase subunit (putative) | 5 | 1.72 | 0.203 | 0.999 | |
| AT4G21105.1 | Cytochrome c:O2 oxidoreductase subunit (putative) | 6 | 2.72 | 0.202 | 0.612 | |

| | | | | | |
|---|---|---|---|---|---|
| AT4G21105.1 | Cytochrome c:O2 oxidoreductase subunit (putative) | 7 | 3.12 | 0.203 | 0.376 |
| AT4G21105.1 | Cytochrome c:O2 oxidoreductase subunit (putative) | 8 | 2.98 | 0.199 | 0.417 |
| AT4G21105.1 | Cytochrome c:O2 oxidoreductase subunit (putative) | 9 | 2.69 | 0.197 | 0.580 |
| AT4G21105.1 | Cytochrome c:O2 oxidoreductase subunit (putative) | 10 | 2.41 | 0.198 | 0.791 |
| AT4G21105.1 | Cytochrome c:O2 oxidoreductase subunit (putative) | 11 | 2.42 | 0.200 | 0.801 |
| AT4G21105.1 | Cytochrome c:O2 oxidoreductase subunit (putative) | 12 | 2.52 | 0.199 | 0.724 |
| AT4G21105.1 | Cytochrome c:O2 oxidoreductase subunit (putative) | 13 | 2.27 | 0.197 | 0.865 |
| AT4G21105.1 | Cytochrome c:O2 oxidoreductase subunit (putative) | 14 | 1.97 | 0.193 | 0.974 |
| AT4G21105.1 | Cytochrome c:O2 oxidoreductase subunit (putative) | 15 | 1.79 | 0.192 | 0.995 |
| AT4G21105.1 | Cytochrome c:O2 oxidoreductase subunit (putative) | 16 | 1.84 | 0.190 | 0.991 |
| AT4G21105.1 | Cytochrome c:O2 oxidoreductase subunit (putative) | 17 | 2.21 | 0.190 | 0.867 |
| AT4G21105.1 | Cytochrome c:O2 oxidoreductase subunit (putative) | 18 | 2.49 | 0.192 | 0.690 |
| AT4G21105.1 | Cytochrome c:O2 oxidoreductase subunit (putative) | 19 | 2.11 | 0.197 | 0.943 |
| AT4G21105.1 | Cytochrome c:O2 oxidoreductase subunit (putative) | 20 | 1.94 | 0.203 | 0.988 |
| AT4G21105.1 | Cytochrome c:O2 oxidoreductase subunit (putative) | 21 | 2.33 | 0.205 | 0.881 |
| AT4G21105.1 | Cytochrome c:O2 oxidoreductase subunit (putative) | 22 | 2.90 | 0.206 | 0.530 |
| AT4G21105.1 | Cytochrome c:O2 oxidoreductase subunit (putative) | 23 | 2.70 | 0.204 | 0.645 |
| AT4G21105.1 | Cytochrome c:O2 oxidoreductase subunit (putative) | 24 | 1.70 | 0.198 | 0.999 |
| AT4G21105.1 | Cytochrome c:O2 oxidoreductase subunit (putative) | 25 | 1.04 | 0.196 | 1.000 |
| AT4G21105.1 | Cytochrome c:O2 oxidoreductase subunit (putative) | 26 | 1.07 | 0.196 | 1.000 |

| | | | | | | |
|---|---|---|---|---|---|---|
| AT4G21105.1 | Cytochrome c:O2 oxidoreductase subunit (putative) | 27 | 1.48 | 0.197 | 1.000 | |
| AT4G21105.1 | Cytochrome c:O2 oxidoreductase subunit (putative) | 28 | 1.72 | 0.197 | 0.999 | |
| AT4G21105.1 | Cytochrome c:O2 oxidoreductase subunit (putative) | 29 | 1.86 | 0.196 | 0.993 | |
| AT4G21105.1 | Cytochrome c:O2 oxidoreductase subunit (putative) | 30 | 1.83 | 0.196 | 0.994 | |
| AT4G21105.1 | Cytochrome c:O2 oxidoreductase subunit (putative) | 31 | 1.54 | 0.197 | 1.000 | |
| AT4G21105.1 | Cytochrome c:O2 oxidoreductase subunit (putative) | 32 | 1.02 | 0.198 | 1.000 | |
| AT4G21105.1 | Cytochrome c:O2 oxidoreductase subunit (putative) | 33 | 0.52 | 0.202 | 0.988 | |
| AT4G21105.1 | Cytochrome c:O2 oxidoreductase subunit (putative) | 34 | 0.29 | 0.216 | 0.328 | |
| AT4G21105.1 | Cytochrome c:O2 oxidoreductase subunit (putative) | 35 | 0.42 | 0.254 | 0.981 | |
| AT4G21105.1 | Cytochrome c:O2 oxidoreductase subunit (putative) | 36 | 0.93 | 0.364 | 1.000 | |
| AT4G21105.1 | Cytochrome c:O2 oxidoreductase subunit (putative) | 41 | 3.34 | 0.361 | 0.985 | |
| AT4G21105.1 | Cytochrome c:O2 oxidoreductase subunit (putative) | 42 | 2.63 | 0.338 | 0.998 | |
| AT4G21105.1 | Cytochrome c:O2 oxidoreductase subunit (putative) | 43 | 1.81 | 0.440 | 1.000 | |
| AT4G21105.1 | Cytochrome c:O2 oxidoreductase subunit (putative) | 46 | 0.34 | 1.587 | 1.000 | |
| AT4G21105.1 | Cytochrome c:O2 oxidoreductase subunit (putative) | 47 | 0.18 | 2.081 | 1.000 | |
| AT4G22220.1 | ISU1 | 2 | 0.98 | 0.451 | 1.000 | |
| AT4G22220.1 | ISU1 | 3 | 0.85 | 0.433 | 1.000 | |
| AT4G22220.1 | ISU1 | 28 | 0.84 | 0.320 | 1.000 | |
| AT4G22220.1 | ISU1 | 29 | 1.09 | 0.230 | 1.000 | |
| AT4G22220.1 | ISU1 | 30 | 1.35 | 0.204 | 1.000 | |
| AT4G22220.1 | ISU1 | 31 | 1.64 | 0.195 | 0.985 | |
| AT4G22220.1 | ISU1 | 32 | 1.91 | 0.191 | 0.874 | |
| AT4G22220.1 | ISU1 | 33 | 1.76 | 0.190 | 0.946 | |
| AT4G22220.1 | ISU1 | 34 | 1.23 | 0.190 | 1.000 | |
| AT4G22220.1 | ISU1 | 35 | 0.80 | 0.190 | 1.000 | |
| AT4G22220.1 | ISU1 | 36 | 0.62 | 0.189 | 0.987 | |
| AT4G22220.1 | ISU1 | 37 | 0.63 | 0.189 | 0.989 | |
| AT4G22220.1 | ISU1 | 38 | 0.73 | 0.188 | 1.000 | |
| AT4G22220.1 | ISU1 | 39 | 1.07 | 0.188 | 1.000 | |
| AT4G22220.1 | ISU1 | 40 | 1.98 | 0.188 | 0.812 | |
| AT4G22220.1 | ISU1 | 41 | 3.66 | 0.189 | 0.055 | . |
| AT4G22220.1 | ISU1 | 42 | 5.53 | 0.192 | 0.003 | ** |

| | | | | | | |
|---|---|---|---|---|---|---|
| AT4G22220.1 | ISU1 | 43 | 5.70 | 0.194 | 0.003 | ** |
| AT4G22220.1 | ISU1 | 44 | 3.86 | 0.194 | 0.049 | * |
| AT4G22220.1 | ISU1 | 45 | 1.97 | 0.194 | 0.852 | |
| AT4G22220.1 | ISU1 | 46 | 0.86 | 0.234 | 1.000 | |
| AT4G22220.1 | ISU1 | 47 | 0.35 | 0.419 | 0.987 | |
| AT4G22310.1 | ? | 2 | 0.79 | 0.291 | 1.000 | |
| AT4G22310.1 | ? | 3 | 0.69 | 0.182 | 1.000 | |
| AT4G22310.1 | ? | 4 | 0.68 | 0.166 | 1.000 | |
| AT4G22310.1 | ? | 5 | 0.79 | 0.167 | 1.000 | |
| AT4G22310.1 | ? | 6 | 0.89 | 0.167 | 1.000 | |
| AT4G22310.1 | ? | 7 | 0.91 | 0.166 | 1.000 | |
| AT4G22310.1 | ? | 8 | 0.95 | 0.163 | 1.000 | |
| AT4G22310.1 | ? | 9 | 1.02 | 0.161 | 1.000 | |
| AT4G22310.1 | ? | 10 | 1.02 | 0.161 | 1.000 | |
| AT4G22310.1 | ? | 11 | 1.00 | 0.161 | 1.000 | |
| AT4G22310.1 | ? | 12 | 0.99 | 0.161 | 1.000 | |
| AT4G22310.1 | ? | 13 | 0.99 | 0.160 | 1.000 | |
| AT4G22310.1 | ? | 14 | 0.99 | 0.158 | 1.000 | |
| AT4G22310.1 | ? | 15 | 0.94 | 0.157 | 1.000 | |
| AT4G22310.1 | ? | 16 | 0.88 | 0.156 | 1.000 | |
| AT4G22310.1 | ? | 17 | 0.94 | 0.156 | 1.000 | |
| AT4G22310.1 | ? | 18 | 1.18 | 0.158 | 1.000 | |
| AT4G22310.1 | ? | 19 | 1.64 | 0.162 | 0.994 | |
| AT4G22310.1 | ? | 20 | 2.34 | 0.166 | 0.538 | |
| AT4G22310.1 | ? | 21 | 3.04 | 0.168 | 0.133 | |
| AT4G22310.1 | ? | 22 | 3.17 | 0.169 | 0.100 | |
| AT4G22310.1 | ? | 23 | 2.79 | 0.170 | 0.249 | |
| AT4G22310.1 | ? | 24 | 2.65 | 0.175 | 0.385 | |
| AT4G22310.1 | ? | 25 | 2.58 | 0.187 | 0.557 | |
| AT4G22310.1 | ? | 26 | 2.53 | 0.192 | 0.647 | |
| AT4G22310.1 | ? | 27 | 2.61 | 0.194 | 0.604 | |
| AT4G22310.1 | ? | 28 | 2.95 | 0.180 | 0.254 | |
| AT4G22310.1 | ? | 29 | 3.50 | 0.171 | 0.053 | . |
| AT4G22310.1 | ? | 30 | 4.15 | 0.163 | 0.007 | ** |
| AT4G22310.1 | ? | 31 | 5.17 | 0.161 | 0.000 | *** |
| AT4G22310.1 | ? | 32 | 6.08 | 0.161 | 0.000 | *** |
| AT4G22310.1 | ? | 33 | 5.53 | 0.161 | 0.000 | *** |
| AT4G22310.1 | ? | 34 | 4.21 | 0.161 | 0.005 | ** |
| AT4G22310.1 | ? | 35 | 3.12 | 0.161 | 0.075 | . |
| AT4G22310.1 | ? | 36 | 2.42 | 0.160 | 0.407 | |
| AT4G22310.1 | ? | 37 | 2.09 | 0.160 | 0.727 | |
| AT4G22310.1 | ? | 38 | 2.14 | 0.161 | 0.684 | |
| AT4G22310.1 | ? | 39 | 2.30 | 0.161 | 0.520 | |
| AT4G22310.1 | ? | 40 | 2.16 | 0.162 | 0.671 | |
| AT4G22310.1 | ? | 41 | 1.65 | 0.162 | 0.994 | |
| AT4G22310.1 | ? | 42 | 1.16 | 0.164 | 1.000 | |
| AT4G22310.1 | ? | 43 | 0.82 | 0.166 | 1.000 | |
| AT4G22310.1 | ? | 44 | 0.52 | 0.166 | 0.915 | |
| AT4G22310.1 | ? | 45 | 0.26 | 0.166 | 0.020 | * |
| AT4G22310.1 | ? | 46 | 0.10 | 0.196 | 0.000 | *** |
| AT4G22310.1 | ? | 47 | 0.03 | 0.335 | 0.000 | *** |
| AT4G23660.2 | Polyprenyltransferase 1 (PPT1) | 2 | 0.95 | 0.566 | 1.000 | |
| AT4G23660.2 | Polyprenyltransferase 1 (PPT1) | 3 | 1.20 | 0.518 | 1.000 | |
| AT4G23660.2 | Polyprenyltransferase 1 (PPT1) | 38 | 16.90 | 0.383 | 0.018 | * |
| AT4G23660.2 | Polyprenyltransferase 1 (PPT1) | 39 | 6.09 | 0.261 | 0.032 | * |
| AT4G23660.2 | Polyprenyltransferase 1 (PPT1) | 40 | 1.95 | 0.217 | 0.745 | |
| AT4G23660.2 | Polyprenyltransferase 1 (PPT1) | 41 | 0.72 | 0.206 | 0.988 | |
| AT4G23660.2 | Polyprenyltransferase 1 (PPT1) | 42 | 0.56 | 0.214 | 0.830 | |
| AT4G23660.2 | Polyprenyltransferase 1 (PPT1) | 43 | 0.77 | 0.253 | 0.999 | |
| AT4G23660.2 | Polyprenyltransferase 1 (PPT1) | 44 | 1.25 | 0.420 | 1.000 | |
| AT4G23660.2 | Polyprenyltransferase 1 (PPT1) | 46 | 3.10 | 1.168 | 1.000 | |
| AT4G23660.2 | Polyprenyltransferase 1 (PPT1) | 47 | 4.82 | 1.677 | 1.000 | |

| | | | | | |
|---|---|---|---|---|---|
| AT4G23900.1 | Nucleoside diphosphate kinase 4 (NDPK4) | 2 | 1.35 | 0.286 | 1.000 |
| AT4G23900.1 | Nucleoside diphosphate kinase 4 (NDPK4) | 3 | 1.35 | 0.214 | 1.000 |
| AT4G23900.1 | Nucleoside diphosphate kinase 4 (NDPK4) | 4 | 1.32 | 0.184 | 1.000 |
| AT4G23900.1 | Nucleoside diphosphate kinase 4 (NDPK4) | 5 | 1.26 | 0.177 | 1.000 |
| AT4G23900.1 | Nucleoside diphosphate kinase 4 (NDPK4) | 6 | 1.20 | 0.177 | 1.000 |
| AT4G23900.1 | Nucleoside diphosphate kinase 4 (NDPK4) | 7 | 1.17 | 0.177 | 1.000 |
| AT4G23900.1 | Nucleoside diphosphate kinase 4 (NDPK4) | 8 | 1.22 | 0.176 | 1.000 |
| AT4G23900.1 | Nucleoside diphosphate kinase 4 (NDPK4) | 9 | 1.35 | 0.174 | 0.999 |
| AT4G23900.1 | Nucleoside diphosphate kinase 4 (NDPK4) | 10 | 1.53 | 0.173 | 0.982 |
| AT4G23900.1 | Nucleoside diphosphate kinase 4 (NDPK4) | 11 | 1.68 | 0.173 | 0.920 |
| AT4G23900.1 | Nucleoside diphosphate kinase 4 (NDPK4) | 12 | 1.68 | 0.172 | 0.918 |
| AT4G23900.1 | Nucleoside diphosphate kinase 4 (NDPK4) | 13 | 1.48 | 0.172 | 0.990 |
| AT4G23900.1 | Nucleoside diphosphate kinase 4 (NDPK4) | 14 | 1.20 | 0.171 | 1.000 |
| AT4G23900.1 | Nucleoside diphosphate kinase 4 (NDPK4) | 15 | 0.99 | 0.170 | 1.000 |
| AT4G23900.1 | Nucleoside diphosphate kinase 4 (NDPK4) | 16 | 0.89 | 0.170 | 1.000 |
| AT4G23900.1 | Nucleoside diphosphate kinase 4 (NDPK4) | 17 | 0.86 | 0.170 | 1.000 |
| AT4G23900.1 | Nucleoside diphosphate kinase 4 (NDPK4) | 18 | 0.84 | 0.171 | 1.000 |
| AT4G23900.1 | Nucleoside diphosphate kinase 4 (NDPK4) | 19 | 0.84 | 0.173 | 1.000 |
| AT4G23900.1 | Nucleoside diphosphate kinase 4 (NDPK4) | 20 | 0.88 | 0.175 | 1.000 |
| AT4G23900.1 | Nucleoside diphosphate kinase 4 (NDPK4) | 21 | 0.95 | 0.177 | 1.000 |
| AT4G23900.1 | Nucleoside diphosphate kinase 4 (NDPK4) | 22 | 1.02 | 0.177 | 1.000 |
| AT4G23900.1 | Nucleoside diphosphate kinase 4 (NDPK4) | 23 | 1.04 | 0.176 | 1.000 |
| AT4G23900.1 | Nucleoside diphosphate kinase 4 (NDPK4) | 24 | 1.00 | 0.176 | 1.000 |
| AT4G23900.1 | Nucleoside diphosphate kinase 4 (NDPK4) | 25 | 0.92 | 0.177 | 1.000 |
| AT4G23900.1 | Nucleoside diphosphate kinase 4 (NDPK4) | 26 | 0.83 | 0.181 | 1.000 |
| AT4G23900.1 | Nucleoside diphosphate kinase 4 (NDPK4) | 27 | 0.75 | 0.187 | 1.000 |
| AT4G23900.1 | Nucleoside diphosphate kinase 4 (NDPK4) | 28 | 0.67 | 0.193 | 0.997 |
| AT4G23900.1 | Nucleoside diphosphate kinase 4 (NDPK4) | 29 | 0.64 | 0.198 | 0.991 |
| AT4G23900.1 | Nucleoside diphosphate kinase 4 (NDPK4) | 30 | 0.63 | 0.202 | 0.991 |
| AT4G23900.1 | Nucleoside diphosphate kinase 4 (NDPK4) | 31 | 0.64 | 0.202 | 0.993 |
| AT4G23900.1 | Nucleoside diphosphate kinase 4 (NDPK4) | 32 | 0.66 | 0.206 | 0.997 |

| | | | | | | |
|---|---|---|---|---|---|---|
| AT4G23900.1 | Nucleoside diphosphate kinase 4 (NDPK4) | 33 | 0.71 | 0.217 | 1.000 | |
| AT4G23900.1 | Nucleoside diphosphate kinase 4 (NDPK4) | 34 | 0.77 | 0.245 | 1.000 | |
| AT4G23900.1 | Nucleoside diphosphate kinase 4 (NDPK4) | 35 | 0.86 | 0.300 | 1.000 | |
| AT4G23900.1 | Nucleoside diphosphate kinase 4 (NDPK4) | 46 | 3.69 | 2.051 | 1.000 | |
| AT4G23900.1 | Nucleoside diphosphate kinase 4 (NDPK4) | 47 | 4.25 | 2.289 | 1.000 | |
| AT4G24520.2 | P450 reductase 1 | 2 | 1.27 | 0.439 | 1.000 | |
| AT4G24520.2 | P450 reductase 1 | 3 | 1.19 | 0.416 | 1.000 | |
| AT4G24520.2 | P450 reductase 1 | 4 | 1.12 | 0.414 | 1.000 | |
| AT4G24520.2 | P450 reductase 1 | 22 | 0.75 | 0.396 | 0.999 | |
| AT4G24520.2 | P450 reductase 1 | 23 | 0.79 | 0.390 | 1.000 | |
| AT4G24520.2 | P450 reductase 1 | 24 | 0.85 | 0.385 | 1.000 | |
| AT4G24520.2 | P450 reductase 1 | 39 | 1.71 | 0.244 | 0.872 | |
| AT4G24520.2 | P450 reductase 1 | 40 | 1.53 | 0.224 | 0.926 | |
| AT4G24520.2 | P450 reductase 1 | 41 | 1.31 | 0.212 | 0.988 | |
| AT4G24520.2 | P450 reductase 1 | 42 | 1.06 | 0.209 | 1.000 | |
| AT4G24520.2 | P450 reductase 1 | 43 | 0.82 | 0.215 | 0.998 | |
| AT4G24520.2 | P450 reductase 1 | 44 | 0.62 | 0.232 | 0.896 | |
| AT4G24520.2 | P450 reductase 1 | 45 | 0.46 | 0.258 | 0.655 | |
| AT4G24520.2 | P450 reductase 1 | 46 | 0.34 | 0.295 | 0.461 | |
| AT4G24520.2 | P450 reductase 1 | 47 | 0.25 | 0.344 | 0.360 | |
| AT4G25130.1 | Methionine sulfoxide reductase A4 (MSRA4) | 2 | 1.00 | 0.320 | 1.000 | |
| AT4G25130.1 | Methionine sulfoxide reductase A4 (MSRA4) | 3 | 1.01 | 0.271 | 1.000 | |
| AT4G25130.1 | Methionine sulfoxide reductase A4 (MSRA4) | 9 | 1.09 | 0.167 | 1.000 | |
| AT4G25130.1 | Methionine sulfoxide reductase A4 (MSRA4) | 10 | 1.11 | 0.143 | 1.000 | |
| AT4G25130.1 | Methionine sulfoxide reductase A4 (MSRA4) | 11 | 1.13 | 0.125 | 1.000 | |
| AT4G25130.1 | Methionine sulfoxide reductase A4 (MSRA4) | 12 | 1.17 | 0.115 | 1.000 | |
| AT4G25130.1 | Methionine sulfoxide reductase A4 (MSRA4) | 13 | 1.23 | 0.109 | 1.000 | |
| AT4G25130.1 | Methionine sulfoxide reductase A4 (MSRA4) | 14 | 1.36 | 0.106 | 0.984 | |
| AT4G25130.1 | Methionine sulfoxide reductase A4 (MSRA4) | 15 | 1.58 | 0.105 | 0.695 | |
| AT4G25130.1 | Methionine sulfoxide reductase A4 (MSRA4) | 16 | 1.88 | 0.106 | 0.195 | |
| AT4G25130.1 | Methionine sulfoxide reductase A4 (MSRA4) | 17 | 2.24 | 0.108 | 0.032 | * |
| AT4G25130.1 | Methionine sulfoxide reductase A4 (MSRA4) | 18 | 2.58 | 0.113 | 0.008 | ** |
| AT4G25130.1 | Methionine sulfoxide reductase A4 (MSRA4) | 19 | 2.84 | 0.119 | 0.004 | ** |
| AT4G25130.1 | Methionine sulfoxide reductase A4 (MSRA4) | 20 | 2.95 | 0.126 | 0.006 | ** |
| AT4G25130.1 | Methionine sulfoxide reductase A4 (MSRA4) | 21 | 2.82 | 0.128 | 0.013 | * |
| AT4G25130.1 | Methionine sulfoxide reductase A4 (MSRA4) | 22 | 2.45 | 0.128 | 0.058 | . |
| AT4G25130.1 | Methionine sulfoxide reductase A4 (MSRA4) | 23 | 1.96 | 0.122 | 0.300 | |
| AT4G25130.1 | Methionine sulfoxide reductase A4 (MSRA4) | 24 | 1.52 | 0.115 | 0.890 | |

| | | | | | |
|---|---|---|---|---|---|
| AT4G25130.1 | Methionine sulfoxide reductase A4 (MSRA4) | 25 | 1.18 | 0.110 | 1.000 |
| AT4G25130.1 | Methionine sulfoxide reductase A4 (MSRA4) | 26 | 0.95 | 0.107 | 1.000 |
| AT4G25130.1 | Methionine sulfoxide reductase A4 (MSRA4) | 27 | 0.82 | 0.106 | 1.000 |
| AT4G25130.1 | Methionine sulfoxide reductase A4 (MSRA4) | 28 | 0.74 | 0.106 | 0.988 |
| AT4G25130.1 | Methionine sulfoxide reductase A4 (MSRA4) | 29 | 0.69 | 0.106 | 0.921 |
| AT4G25130.1 | Methionine sulfoxide reductase A4 (MSRA4) | 30 | 0.65 | 0.106 | 0.761 |
| AT4G25130.1 | Methionine sulfoxide reductase A4 (MSRA4) | 31 | 0.63 | 0.106 | 0.689 |
| AT4G25130.1 | Methionine sulfoxide reductase A4 (MSRA4) | 32 | 0.66 | 0.106 | 0.822 |
| AT4G25130.1 | Methionine sulfoxide reductase A4 (MSRA4) | 33 | 0.72 | 0.107 | 0.971 |
| AT4G25130.1 | Methionine sulfoxide reductase A4 (MSRA4) | 34 | 0.80 | 0.107 | 1.000 |
| AT4G25130.1 | Methionine sulfoxide reductase A4 (MSRA4) | 35 | 0.89 | 0.107 | 1.000 |
| AT4G25130.1 | Methionine sulfoxide reductase A4 (MSRA4) | 36 | 0.96 | 0.107 | 1.000 |
| AT4G25130.1 | Methionine sulfoxide reductase A4 (MSRA4) | 37 | 1.01 | 0.106 | 1.000 |
| AT4G25130.1 | Methionine sulfoxide reductase A4 (MSRA4) | 38 | 1.08 | 0.106 | 1.000 |
| AT4G25130.1 | Methionine sulfoxide reductase A4 (MSRA4) | 39 | 1.18 | 0.107 | 1.000 |
| AT4G25130.1 | Methionine sulfoxide reductase A4 (MSRA4) | 40 | 1.28 | 0.107 | 0.999 |
| AT4G25130.1 | Methionine sulfoxide reductase A4 (MSRA4) | 41 | 1.34 | 0.108 | 0.991 |
| AT4G25130.1 | Methionine sulfoxide reductase A4 (MSRA4) | 42 | 1.32 | 0.109 | 0.995 |
| AT4G25130.1 | Methionine sulfoxide reductase A4 (MSRA4) | 43 | 1.20 | 0.110 | 1.000 |
| AT4G25130.1 | Methionine sulfoxide reductase A4 (MSRA4) | 44 | 0.96 | 0.111 | 1.000 |
| AT4G25130.1 | Methionine sulfoxide reductase A4 (MSRA4) | 45 | 0.67 | 0.120 | 0.942 |
| AT4G25130.1 | Methionine sulfoxide reductase A4 (MSRA4) | 46 | 0.42 | 0.152 | 0.246 |
| AT4G25130.1 | Methionine sulfoxide reductase A4 (MSRA4) | 47 | 0.24 | 0.221 | 0.123 |
| AT4G25315.2 | ? | 2 | 0.87 | 0.315 | 1.000 |
| AT4G25315.2 | ? | 3 | 0.85 | 0.295 | 1.000 |
| AT4G25315.2 | ? | 4 | 0.82 | 0.341 | 1.000 |
| AT4G25315.2 | ? | 23 | 0.89 | 0.223 | 1.000 |
| AT4G25315.2 | ? | 24 | 0.95 | 0.191 | 1.000 |
| AT4G25315.2 | ? | 25 | 0.99 | 0.182 | 1.000 |
| AT4G25315.2 | ? | 26 | 0.99 | 0.186 | 1.000 |
| AT4G25315.2 | ? | 29 | 0.79 | 0.186 | 1.000 |
| AT4G25315.2 | ? | 30 | 0.73 | 0.178 | 0.996 |
| AT4G25315.2 | ? | 31 | 0.71 | 0.171 | 0.989 |
| AT4G25315.2 | ? | 32 | 0.72 | 0.163 | 0.988 |
| AT4G25315.2 | ? | 33 | 0.77 | 0.160 | 0.998 |
| AT4G25315.2 | ? | 34 | 0.86 | 0.161 | 1.000 |
| AT4G25315.2 | ? | 35 | 0.98 | 0.164 | 1.000 |
| AT4G25315.2 | ? | 37 | 1.38 | 0.155 | 0.986 |
| AT4G25315.2 | ? | 38 | 1.67 | 0.146 | 0.731 |

| | | | | | | |
|---|---|---|---|---|---|---|
| AT4G25315.2 | ? | 39 | 1.98 | 0.139 | 0.312 | |
| AT4G25315.2 | ? | 40 | 2.17 | 0.136 | 0.155 | |
| AT4G25315.2 | ? | 41 | 2.04 | 0.137 | 0.242 | |
| AT4G25315.2 | ? | 42 | 1.61 | 0.140 | 0.766 | |
| AT4G25315.2 | ? | 43 | 1.13 | 0.144 | 1.000 | |
| AT4G25315.2 | ? | 44 | 0.75 | 0.152 | 0.994 | |
| AT4G25315.2 | ? | 45 | 0.50 | 0.167 | 0.544 | |
| AT4G25315.2 | ? | 46 | 0.34 | 0.198 | 0.197 | |
| AT4G25315.2 | ? | 47 | 0.23 | 0.253 | 0.146 | |
| AT4G26180.1 | Mitochondrial substrate carrier family protein | 2 | 0.92 | 0.482 | 1.000 | |
| AT4G26180.1 | Mitochondrial substrate carrier family protein | 3 | 1.46 | 0.492 | 0.998 | |
| AT4G26180.1 | Mitochondrial substrate carrier family protein | 40 | 3.40 | 0.346 | 0.472 | |
| AT4G26180.1 | Mitochondrial substrate carrier family protein | 41 | 1.31 | 0.250 | 0.990 | |
| AT4G26180.1 | Mitochondrial substrate carrier family protein | 42 | 0.46 | 0.205 | 0.417 | |
| AT4G26180.1 | Mitochondrial substrate carrier family protein | 43 | 0.16 | 0.236 | 0.012 | * |
| AT4G26180.1 | Mitochondrial substrate carrier family protein | 44 | 0.06 | 0.323 | 0.003 | ** |
| AT4G26180.1 | Mitochondrial substrate carrier family protein | 46 | 0.01 | 0.578 | 0.006 | ** |
| AT4G26180.1 | Mitochondrial substrate carrier family protein | 47 | 0.00 | 0.738 | 0.009 | ** |
| AT4G26210.2 | ATP synthase subunit (ATPG) | 2 | 0.90 | 0.551 | 1.000 | |
| AT4G26210.2 | ATP synthase subunit (ATPG) | 3 | 0.86 | 0.517 | 1.000 | |
| AT4G26210.2 | ATP synthase subunit (ATPG) | 22 | 0.52 | 0.382 | 0.999 | |
| AT4G26210.2 | ATP synthase subunit (ATPG) | 23 | 0.64 | 0.297 | 1.000 | |
| AT4G26210.2 | ATP synthase subunit (ATPG) | 24 | 0.91 | 0.248 | 1.000 | |
| AT4G26210.2 | ATP synthase subunit (ATPG) | 25 | 1.45 | 0.245 | 1.000 | |
| AT4G26210.2 | ATP synthase subunit (ATPG) | 26 | 2.27 | 0.290 | 0.950 | |
| AT4G26210.2 | ATP synthase subunit (ATPG) | 27 | 3.18 | 0.372 | 0.907 | |
| AT4G26210.2 | ATP synthase subunit (ATPG) | 32 | 1.76 | 0.418 | 1.000 | |
| AT4G26210.2 | ATP synthase subunit (ATPG) | 33 | 1.38 | 0.395 | 1.000 | |
| AT4G26210.2 | ATP synthase subunit (ATPG) | 34 | 1.16 | 0.402 | 1.000 | |
| AT4G26210.2 | ATP synthase subunit (ATPG) | 38 | 2.33 | 0.335 | 0.978 | |
| AT4G26210.2 | ATP synthase subunit (ATPG) | 39 | 3.27 | 0.275 | 0.570 | |
| AT4G26210.2 | ATP synthase subunit (ATPG) | 40 | 3.89 | 0.234 | 0.170 | |
| AT4G26210.2 | ATP synthase subunit (ATPG) | 41 | 3.64 | 0.222 | 0.166 | |
| AT4G26210.2 | ATP synthase subunit (ATPG) | 42 | 2.81 | 0.223 | 0.459 | |
| AT4G26210.2 | ATP synthase subunit (ATPG) | 43 | 1.78 | 0.225 | 0.974 | |
| AT4G26210.2 | ATP synthase subunit (ATPG) | 44 | 0.91 | 0.224 | 1.000 | |
| AT4G26210.2 | ATP synthase subunit (ATPG) | 45 | 0.42 | 0.225 | 0.721 | |
| AT4G26210.2 | ATP synthase subunit (ATPG) | 46 | 0.20 | 0.276 | 0.157 | |
| AT4G26210.2 | ATP synthase subunit (ATPG) | 47 | 0.09 | 0.474 | 0.343 | |
| AT4G26300.1 | Arginyl-tRNA synthetase, class Ic | 2 | 0.97 | 0.374 | 1.000 | |
| AT4G26300.1 | Arginyl-tRNA synthetase, class Ic | 3 | 0.97 | 0.363 | 1.000 | |
| AT4G26300.1 | Arginyl-tRNA synthetase, class Ic | 32 | 1.49 | 0.270 | 1.000 | |
| AT4G26300.1 | Arginyl-tRNA synthetase, class Ic | 33 | 1.51 | 0.207 | 0.997 | |
| AT4G26300.1 | Arginyl-tRNA synthetase, class Ic | 34 | 1.47 | 0.183 | 0.994 | |
| AT4G26300.1 | Arginyl-tRNA synthetase, class Ic | 35 | 1.71 | 0.172 | 0.900 | |
| AT4G26300.1 | Arginyl-tRNA synthetase, class Ic | 36 | 2.25 | 0.168 | 0.392 | |

| | | | | | | |
|---|---|---|---|---|---|---|
| AT4G26300.1 | Arginyl-tRNA synthetase, class Ic | 37 | 2.78 | 0.168 | 0.123 | |
| AT4G26300.1 | Arginyl-tRNA synthetase, class Ic | 38 | 3.03 | 0.168 | 0.068 | . |
| AT4G26300.1 | Arginyl-tRNA synthetase, class Ic | 39 | 2.98 | 0.168 | 0.074 | . |
| AT4G26300.1 | Arginyl-tRNA synthetase, class Ic | 40 | 2.28 | 0.167 | 0.356 | |
| AT4G26300.1 | Arginyl-tRNA synthetase, class Ic | 41 | 1.39 | 0.168 | 0.997 | |
| AT4G26300.1 | Arginyl-tRNA synthetase, class Ic | 42 | 0.93 | 0.171 | 1.000 | |
| AT4G26300.1 | Arginyl-tRNA synthetase, class Ic | 43 | 0.71 | 0.172 | 0.997 | |
| AT4G26300.1 | Arginyl-tRNA synthetase, class Ic | 44 | 0.51 | 0.172 | 0.687 | |
| AT4G26300.1 | Arginyl-tRNA synthetase, class Ic | 45 | 0.36 | 0.171 | 0.133 | |
| AT4G26300.1 | Arginyl-tRNA synthetase, class Ic | 46 | 0.27 | 0.199 | 0.070 | . |
| AT4G26300.1 | Arginyl-tRNA synthetase, class Ic | 47 | 0.22 | 0.379 | 0.666 | |
| AT4G26410.1 | RGS1-HXK1 interacting protein 1 (RHIP1) | 2 | 1.10 | 0.304 | 1.000 | |
| AT4G26410.1 | RGS1-HXK1 interacting protein 1 (RHIP1) | 3 | 0.86 | 0.194 | 1.000 | |
| AT4G26410.1 | RGS1-HXK1 interacting protein 1 (RHIP1) | 4 | 0.71 | 0.167 | 1.000 | |
| AT4G26410.1 | RGS1-HXK1 interacting protein 1 (RHIP1) | 5 | 0.66 | 0.160 | 1.000 | |
| AT4G26410.1 | RGS1-HXK1 interacting protein 1 (RHIP1) | 6 | 0.67 | 0.157 | 1.000 | |
| AT4G26410.1 | RGS1-HXK1 interacting protein 1 (RHIP1) | 7 | 0.70 | 0.155 | 1.000 | |
| AT4G26410.1 | RGS1-HXK1 interacting protein 1 (RHIP1) | 8 | 0.83 | 0.152 | 1.000 | |
| AT4G26410.1 | RGS1-HXK1 interacting protein 1 (RHIP1) | 9 | 1.05 | 0.150 | 1.000 | |
| AT4G26410.1 | RGS1-HXK1 interacting protein 1 (RHIP1) | 10 | 1.24 | 0.150 | 1.000 | |
| AT4G26410.1 | RGS1-HXK1 interacting protein 1 (RHIP1) | 11 | 1.47 | 0.150 | 1.000 | |
| AT4G26410.1 | RGS1-HXK1 interacting protein 1 (RHIP1) | 12 | 1.81 | 0.151 | 0.924 | |
| AT4G26410.1 | RGS1-HXK1 interacting protein 1 (RHIP1) | 13 | 2.23 | 0.149 | 0.468 | |
| AT4G26410.1 | RGS1-HXK1 interacting protein 1 (RHIP1) | 14 | 2.60 | 0.147 | 0.161 | |
| AT4G26410.1 | RGS1-HXK1 interacting protein 1 (RHIP1) | 15 | 2.64 | 0.146 | 0.134 | |
| AT4G26410.1 | RGS1-HXK1 interacting protein 1 (RHIP1) | 16 | 2.35 | 0.145 | 0.299 | |
| AT4G26410.1 | RGS1-HXK1 interacting protein 1 (RHIP1) | 17 | 2.09 | 0.144 | 0.573 | |
| AT4G26410.1 | RGS1-HXK1 interacting protein 1 (RHIP1) | 18 | 1.84 | 0.147 | 0.880 | |
| AT4G26410.1 | RGS1-HXK1 interacting protein 1 (RHIP1) | 19 | 1.68 | 0.152 | 0.984 | |
| AT4G26410.1 | RGS1-HXK1 interacting protein 1 (RHIP1) | 20 | 1.74 | 0.155 | 0.975 | |
| AT4G26410.1 | RGS1-HXK1 interacting protein 1 (RHIP1) | 21 | 1.90 | 0.156 | 0.889 | |

| | | | | | |
|---|---|---|---|---|---|
| AT4G26410.1 | RGS1-HXK1 interacting protein 1 (RHIP1) | 22 | 1.74 | 0.156 | 0.976 |
| AT4G26410.1 | RGS1-HXK1 interacting protein 1 (RHIP1) | 23 | 1.44 | 0.154 | 1.000 |
| AT4G26410.1 | RGS1-HXK1 interacting protein 1 (RHIP1) | 24 | 1.29 | 0.151 | 1.000 |
| AT4G26410.1 | RGS1-HXK1 interacting protein 1 (RHIP1) | 25 | 1.29 | 0.149 | 1.000 |
| AT4G26410.1 | RGS1-HXK1 interacting protein 1 (RHIP1) | 26 | 1.55 | 0.148 | 0.998 |
| AT4G26410.1 | RGS1-HXK1 interacting protein 1 (RHIP1) | 27 | 1.93 | 0.148 | 0.810 |
| AT4G26410.1 | RGS1-HXK1 interacting protein 1 (RHIP1) | 28 | 2.04 | 0.149 | 0.697 |
| AT4G26410.1 | RGS1-HXK1 interacting protein 1 (RHIP1) | 29 | 1.98 | 0.150 | 0.769 |
| AT4G26410.1 | RGS1-HXK1 interacting protein 1 (RHIP1) | 30 | 1.72 | 0.150 | 0.971 |
| AT4G26410.1 | RGS1-HXK1 interacting protein 1 (RHIP1) | 31 | 1.28 | 0.151 | 1.000 |
| AT4G26410.1 | RGS1-HXK1 interacting protein 1 (RHIP1) | 32 | 0.84 | 0.151 | 1.000 |
| AT4G26410.1 | RGS1-HXK1 interacting protein 1 (RHIP1) | 33 | 0.58 | 0.151 | 0.965 |
| AT4G26410.1 | RGS1-HXK1 interacting protein 1 (RHIP1) | 34 | 0.54 | 0.151 | 0.890 |
| AT4G26410.1 | RGS1-HXK1 interacting protein 1 (RHIP1) | 35 | 0.73 | 0.151 | 1.000 |
| AT4G26410.1 | RGS1-HXK1 interacting protein 1 (RHIP1) | 36 | 1.03 | 0.151 | 1.000 |
| AT4G26410.1 | RGS1-HXK1 interacting protein 1 (RHIP1) | 37 | 1.25 | 0.151 | 1.000 |
| AT4G26410.1 | RGS1-HXK1 interacting protein 1 (RHIP1) | 38 | 1.39 | 0.151 | 1.000 |
| AT4G26410.1 | RGS1-HXK1 interacting protein 1 (RHIP1) | 39 | 1.50 | 0.150 | 1.000 |
| AT4G26410.1 | RGS1-HXK1 interacting protein 1 (RHIP1) | 40 | 1.56 | 0.150 | 0.998 |
| AT4G26410.1 | RGS1-HXK1 interacting protein 1 (RHIP1) | 41 | 1.42 | 0.150 | 1.000 |
| AT4G26410.1 | RGS1-HXK1 interacting protein 1 (RHIP1) | 42 | 1.35 | 0.153 | 1.000 |
| AT4G26410.1 | RGS1-HXK1 interacting protein 1 (RHIP1) | 43 | 1.32 | 0.155 | 1.000 |
| AT4G26410.1 | RGS1-HXK1 interacting protein 1 (RHIP1) | 44 | 1.03 | 0.154 | 1.000 |
| AT4G26410.1 | RGS1-HXK1 interacting protein 1 (RHIP1) | 45 | 0.63 | 0.154 | 0.997 |
| AT4G26410.1 | RGS1-HXK1 interacting protein 1 (RHIP1) | 46 | 0.30 | 0.181 | 0.131 |
| AT4G26410.1 | RGS1-HXK1 interacting protein 1 (RHIP1) | 47 | 0.12 | 0.322 | 0.155 |
| AT4G26620.1 | Sucrase/ferredoxin-like family protein | 2 | 0.89 | 0.305 | 1.000 |
| AT4G26620.1 | Sucrase/ferredoxin-like family protein | 3 | 0.76 | 0.292 | 1.000 |
| AT4G26620.1 | Sucrase/ferredoxin-like family protein | 25 | 0.47 | 0.248 | 0.969 |
| AT4G26620.1 | Sucrase/ferredoxin-like family protein | 26 | 0.60 | 0.172 | 0.980 |
| AT4G26620.1 | Sucrase/ferredoxin-like family protein | 27 | 0.83 | 0.153 | 1.000 |

| | | | | | | |
|---|---|---|---|---|---|---|
| AT4G26620.1 | Sucrase/ferredoxin-like family protein | 28 | 1.09 | 0.144 | 1.000 | |
| AT4G26620.1 | Sucrase/ferredoxin-like family protein | 29 | 1.23 | 0.144 | 1.000 | |
| AT4G26620.1 | Sucrase/ferredoxin-like family protein | 30 | 1.10 | 0.146 | 1.000 | |
| AT4G26620.1 | Sucrase/ferredoxin-like family protein | 31 | 0.95 | 0.146 | 1.000 | |
| AT4G26620.1 | Sucrase/ferredoxin-like family protein | 32 | 1.05 | 0.147 | 1.000 | |
| AT4G26620.1 | Sucrase/ferredoxin-like family protein | 33 | 1.21 | 0.146 | 1.000 | |
| AT4G26620.1 | Sucrase/ferredoxin-like family protein | 34 | 1.28 | 0.147 | 1.000 | |
| AT4G26620.1 | Sucrase/ferredoxin-like family protein | 35 | 1.25 | 0.147 | 1.000 | |
| AT4G26620.1 | Sucrase/ferredoxin-like family protein | 36 | 1.07 | 0.145 | 1.000 | |
| AT4G26620.1 | Sucrase/ferredoxin-like family protein | 37 | 0.89 | 0.144 | 1.000 | |
| AT4G26620.1 | Sucrase/ferredoxin-like family protein | 38 | 0.79 | 0.144 | 1.000 | |
| AT4G26620.1 | Sucrase/ferredoxin-like family protein | 39 | 0.94 | 0.144 | 1.000 | |
| AT4G26620.1 | Sucrase/ferredoxin-like family protein | 40 | 1.47 | 0.144 | 0.992 | |
| AT4G26620.1 | Sucrase/ferredoxin-like family protein | 41 | 2.26 | 0.146 | 0.274 | |
| AT4G26620.1 | Sucrase/ferredoxin-like family protein | 42 | 3.07 | 0.149 | 0.027 | * |
| AT4G26620.1 | Sucrase/ferredoxin-like family protein | 43 | 3.11 | 0.149 | 0.023 | * |
| AT4G26620.1 | Sucrase/ferredoxin-like family protein | 44 | 2.53 | 0.148 | 0.137 | |
| AT4G26620.1 | Sucrase/ferredoxin-like family protein | 45 | 1.58 | 0.149 | 0.970 | |
| AT4G26620.1 | Sucrase/ferredoxin-like family protein | 46 | 0.66 | 0.166 | 0.996 | |
| AT4G26620.1 | Sucrase/ferredoxin-like family protein | 47 | 0.21 | 0.331 | 0.563 | |
| AT4G26780.1 | Mitochondrial GRPE2 (MGE2) | 2 | 1.45 | 0.504 | 1.000 | |
| AT4G26780.1 | Mitochondrial GRPE2 (MGE2) | 3 | 0.83 | 0.369 | 1.000 | |
| AT4G26780.1 | Mitochondrial GRPE2 (MGE2) | 4 | 0.49 | 0.737 | 1.000 | |
| AT4G26780.1 | Mitochondrial GRPE2 (MGE2) | 41 | 3.21 | 0.579 | 0.975 | |
| AT4G26780.1 | Mitochondrial GRPE2 (MGE2) | 42 | 3.30 | 0.322 | 0.606 | |
| AT4G26780.1 | Mitochondrial GRPE2 (MGE2) | 43 | 1.70 | 0.267 | 0.977 | |
| AT4G26780.1 | Mitochondrial GRPE2 (MGE2) | 44 | 0.87 | 0.255 | 1.000 | |
| AT4G26780.1 | Mitochondrial GRPE2 (MGE2) | 45 | 1.20 | 0.255 | 1.000 | |
| AT4G26780.1 | Mitochondrial GRPE2 (MGE2) | 46 | 1.43 | 0.274 | 0.999 | |
| AT4G26780.1 | Mitochondrial GRPE2 (MGE2) | 47 | 1.08 | 0.644 | 1.000 | |
| AT4G26860.1 | Pyridoxal phosphate homeostasis protein | 2 | 1.18 | 0.521 | 1.000 | |
| AT4G26860.1 | Pyridoxal phosphate homeostasis protein | 3 | 1.09 | 0.499 | 1.000 | |
| AT4G26860.1 | Pyridoxal phosphate homeostasis protein | 25 | 0.57 | 0.319 | 0.929 | |
| AT4G26860.1 | Pyridoxal phosphate homeostasis protein | 26 | 0.62 | 0.298 | 0.952 | |
| AT4G26860.1 | Pyridoxal phosphate homeostasis protein | 27 | 0.68 | 0.283 | 0.978 | |
| AT4G26860.1 | Pyridoxal phosphate homeostasis protein | 28 | 0.75 | 0.273 | 0.995 | |

| | | | | | |
|---|---|---|---|---|---|
| AT4G26860.1 | Pyridoxal phosphate homeostasis protein | 29 | 0.84 | 0.268 | 1.000 |
| AT4G26860.1 | Pyridoxal phosphate homeostasis protein | 30 | 0.94 | 0.265 | 1.000 |
| AT4G26860.1 | Pyridoxal phosphate homeostasis protein | 31 | 1.05 | 0.264 | 1.000 |
| AT4G26860.1 | Pyridoxal phosphate homeostasis protein | 32 | 1.17 | 0.264 | 1.000 |
| AT4G26860.1 | Pyridoxal phosphate homeostasis protein | 33 | 1.30 | 0.264 | 0.996 |
| AT4G26860.1 | Pyridoxal phosphate homeostasis protein | 34 | 1.43 | 0.264 | 0.978 |
| AT4G26860.1 | Pyridoxal phosphate homeostasis protein | 35 | 1.56 | 0.264 | 0.939 |
| AT4G26860.1 | Pyridoxal phosphate homeostasis protein | 36 | 1.67 | 0.264 | 0.890 |
| AT4G26860.1 | Pyridoxal phosphate homeostasis protein | 37 | 1.76 | 0.264 | 0.843 |
| AT4G26860.1 | Pyridoxal phosphate homeostasis protein | 38 | 1.82 | 0.264 | 0.812 |
| AT4G26860.1 | Pyridoxal phosphate homeostasis protein | 39 | 1.83 | 0.265 | 0.803 |
| AT4G26860.1 | Pyridoxal phosphate homeostasis protein | 40 | 1.81 | 0.265 | 0.818 |
| AT4G26860.1 | Pyridoxal phosphate homeostasis protein | 41 | 1.74 | 0.265 | 0.856 |
| AT4G26860.1 | Pyridoxal phosphate homeostasis protein | 42 | 1.63 | 0.265 | 0.910 |
| AT4G26860.1 | Pyridoxal phosphate homeostasis protein | 43 | 1.50 | 0.267 | 0.962 |
| AT4G26860.1 | Pyridoxal phosphate homeostasis protein | 44 | 1.35 | 0.272 | 0.993 |
| AT4G26860.1 | Pyridoxal phosphate homeostasis protein | 45 | 1.20 | 0.283 | 1.000 |
| AT4G26860.1 | Pyridoxal phosphate homeostasis protein | 46 | 1.06 | 0.304 | 1.000 |
| AT4G26860.1 | Pyridoxal phosphate homeostasis protein | 47 | 0.93 | 0.337 | 1.000 |
| AT4G26910.3 | Dihydrolipoamide succinyltransferase (OGDCE2) | 2 | 0.73 | 0.747 | 1.000 |
| AT4G26910.3 | Dihydrolipoamide succinyltransferase (OGDCE2) | 3 | 0.95 | 0.610 | 1.000 |
| AT4G26910.3 | Dihydrolipoamide succinyltransferase (OGDCE2) | 8 | 2.25 | 0.446 | 0.995 |
| AT4G26910.3 | Dihydrolipoamide succinyltransferase (OGDCE2) | 9 | 2.29 | 0.371 | 0.976 |
| AT4G26910.3 | Dihydrolipoamide succinyltransferase (OGDCE2) | 10 | 2.22 | 0.323 | 0.955 |
| AT4G26910.3 | Dihydrolipoamide succinyltransferase (OGDCE2) | 11 | 2.04 | 0.296 | 0.962 |
| AT4G26910.3 | Dihydrolipoamide succinyltransferase (OGDCE2) | 12 | 1.50 | 0.285 | 0.999 |
| AT4G26910.3 | Dihydrolipoamide succinyltransferase (OGDCE2) | 13 | 0.87 | 0.280 | 1.000 |
| AT4G26910.3 | Dihydrolipoamide succinyltransferase (OGDCE2) | 14 | 0.53 | 0.279 | 0.972 |
| AT4G26910.3 | Dihydrolipoamide succinyltransferase (OGDCE2) | 15 | 0.44 | 0.280 | 0.882 |
| AT4G26910.3 | Dihydrolipoamide succinyltransferase (OGDCE2) | 16 | 0.64 | 0.282 | 0.998 |
| AT4G26910.3 | Dihydrolipoamide succinyltransferase (OGDCE2) | 17 | 1.54 | 0.292 | 0.999 |

| | | | | | |
|---|---|---|---|---|---|
| AT4G26910.3 | Dihydrolipoamide succinyltransferase (OGDCE2) | 18 | 4.09 | 0.322 | 0.459 |
| AT4G26910.3 | Dihydrolipoamide succinyltransferase (OGDCE2) | 19 | 7.88 | 0.400 | 0.249 |
| AT4G26910.3 | Dihydrolipoamide succinyltransferase (OGDCE2) | 20 | 11.87 | 0.573 | 0.477 |
| AT4G26910.3 | Dihydrolipoamide succinyltransferase (OGDCE2) | 46 | ########### | 24.065 | 1.000 |
| AT4G26910.3 | Dihydrolipoamide succinyltransferase (OGDCE2) | 47 | ########### | 25.414 | 1.000 |
| AT4G26970.1 | Aconitase 2 (ACO2) | 2 | 0.61 | 0.327 | 1.000 |
| AT4G26970.1 | Aconitase 2 (ACO2) | 3 | 0.53 | 0.262 | 0.994 |
| AT4G26970.1 | Aconitase 2 (ACO2) | 4 | 0.47 | 0.232 | 0.922 |
| AT4G26970.1 | Aconitase 2 (ACO2) | 6 | 0.43 | 0.211 | 0.745 |
| AT4G26970.1 | Aconitase 2 (ACO2) | 7 | 0.47 | 0.199 | 0.786 |
| AT4G26970.1 | Aconitase 2 (ACO2) | 8 | 0.55 | 0.185 | 0.933 |
| AT4G26970.1 | Aconitase 2 (ACO2) | 9 | 0.71 | 0.176 | 1.000 |
| AT4G26970.1 | Aconitase 2 (ACO2) | 10 | 0.95 | 0.170 | 1.000 |
| AT4G26970.1 | Aconitase 2 (ACO2) | 11 | 1.27 | 0.168 | 1.000 |
| AT4G26970.1 | Aconitase 2 (ACO2) | 12 | 1.67 | 0.166 | 0.948 |
| AT4G26970.1 | Aconitase 2 (ACO2) | 13 | 2.08 | 0.165 | 0.580 |
| AT4G26970.1 | Aconitase 2 (ACO2) | 14 | 2.42 | 0.164 | 0.288 |
| AT4G26970.1 | Aconitase 2 (ACO2) | 15 | 2.56 | 0.163 | 0.201 |
| AT4G26970.1 | Aconitase 2 (ACO2) | 16 | 2.52 | 0.162 | 0.219 |
| AT4G26970.1 | Aconitase 2 (ACO2) | 17 | 2.36 | 0.163 | 0.322 |
| AT4G26970.1 | Aconitase 2 (ACO2) | 18 | 2.05 | 0.164 | 0.611 |
| AT4G26970.1 | Aconitase 2 (ACO2) | 19 | 1.70 | 0.167 | 0.935 |
| AT4G26970.1 | Aconitase 2 (ACO2) | 20 | 1.42 | 0.169 | 0.999 |
| AT4G26970.1 | Aconitase 2 (ACO2) | 21 | 1.23 | 0.171 | 1.000 |
| AT4G26970.1 | Aconitase 2 (ACO2) | 22 | 1.12 | 0.171 | 1.000 |
| AT4G26970.1 | Aconitase 2 (ACO2) | 23 | 1.04 | 0.170 | 1.000 |
| AT4G26970.1 | Aconitase 2 (ACO2) | 24 | 0.98 | 0.168 | 1.000 |
| AT4G26970.1 | Aconitase 2 (ACO2) | 25 | 0.93 | 0.166 | 1.000 |
| AT4G26970.1 | Aconitase 2 (ACO2) | 26 | 0.92 | 0.165 | 1.000 |
| AT4G26970.1 | Aconitase 2 (ACO2) | 27 | 0.98 | 0.165 | 1.000 |
| AT4G26970.1 | Aconitase 2 (ACO2) | 28 | 1.06 | 0.165 | 1.000 |
| AT4G26970.1 | Aconitase 2 (ACO2) | 29 | 1.12 | 0.165 | 1.000 |
| AT4G26970.1 | Aconitase 2 (ACO2) | 30 | 1.14 | 0.166 | 1.000 |
| AT4G26970.1 | Aconitase 2 (ACO2) | 31 | 1.13 | 0.166 | 1.000 |
| AT4G26970.1 | Aconitase 2 (ACO2) | 32 | 1.11 | 0.166 | 1.000 |
| AT4G26970.1 | Aconitase 2 (ACO2) | 33 | 1.09 | 0.166 | 1.000 |
| AT4G26970.1 | Aconitase 2 (ACO2) | 34 | 1.08 | 0.166 | 1.000 |
| AT4G26970.1 | Aconitase 2 (ACO2) | 35 | 1.09 | 0.166 | 1.000 |
| AT4G26970.1 | Aconitase 2 (ACO2) | 36 | 1.11 | 0.166 | 1.000 |
| AT4G26970.1 | Aconitase 2 (ACO2) | 37 | 1.14 | 0.166 | 1.000 |
| AT4G26970.1 | Aconitase 2 (ACO2) | 38 | 1.18 | 0.166 | 1.000 |
| AT4G26970.1 | Aconitase 2 (ACO2) | 39 | 1.22 | 0.167 | 1.000 |
| AT4G26970.1 | Aconitase 2 (ACO2) | 40 | 1.24 | 0.167 | 1.000 |
| AT4G26970.1 | Aconitase 2 (ACO2) | 41 | 1.23 | 0.169 | 1.000 |
| AT4G26970.1 | Aconitase 2 (ACO2) | 42 | 1.16 | 0.170 | 1.000 |
| AT4G26970.1 | Aconitase 2 (ACO2) | 43 | 1.05 | 0.170 | 1.000 |
| AT4G26970.1 | Aconitase 2 (ACO2) | 44 | 0.90 | 0.172 | 1.000 |
| AT4G26970.1 | Aconitase 2 (ACO2) | 45 | 0.73 | 0.184 | 1.000 |
| AT4G26970.1 | Aconitase 2 (ACO2) | 46 | 0.57 | 0.223 | 0.990 |
| AT4G26970.1 | Aconitase 2 (ACO2) | 47 | 0.43 | 0.303 | 0.977 |
| AT4G27520.1 | Early nodulin-like protein 2 (ENODL2) | 2 | 0.92 | 0.318 | 1.000 |
| AT4G27520.1 | Early nodulin-like protein 2 (ENODL2) | 3 | 0.69 | 0.402 | 1.000 |
| AT4G27520.1 | Early nodulin-like protein 2 (ENODL2) | 28 | 1.28 | 0.213 | 1.000 |
| AT4G27520.1 | Early nodulin-like protein 2 (ENODL2) | 29 | 1.15 | 0.152 | 1.000 |

| | | | | | |
|---|---|---|---|---|---|
| AT4G27520.1 | Early nodulin-like protein 2 (ENODL2) | 30 | 1.18 | 0.147 | 1.000 |
| AT4G27520.1 | Early nodulin-like protein 2 (ENODL2) | 31 | 1.42 | 0.138 | 0.927 |
| AT4G27520.1 | Early nodulin-like protein 2 (ENODL2) | 32 | 1.31 | 0.139 | 0.986 |
| AT4G27520.1 | Early nodulin-like protein 2 (ENODL2) | 33 | 1.00 | 0.149 | 1.000 |
| AT4G27520.1 | Early nodulin-like protein 2 (ENODL2) | 34 | 0.87 | 0.183 | 1.000 |
| AT4G27520.1 | Early nodulin-like protein 2 (ENODL2) | 35 | 1.25 | 0.288 | 1.000 |
| AT4G27520.1 | Early nodulin-like protein 2 (ENODL2) | 46 | 904375.24 | 5.185 | 0.913 |
| AT4G27520.1 | Early nodulin-like protein 2 (ENODL2) | 47 | 4543971.17 | 5.872 | 0.919 |
| AT4G27585.1 | Stomatin-like protein 1 (SLP1) | 2 | 0.75 | 0.428 | 1.000 |
| AT4G27585.1 | Stomatin-like protein 1 (SLP1) | 3 | 0.70 | 0.281 | 1.000 |
| AT4G27585.1 | Stomatin-like protein 1 (SLP1) | 4 | 0.62 | 0.253 | 1.000 |
| AT4G27585.1 | Stomatin-like protein 1 (SLP1) | 5 | 0.56 | 0.253 | 0.999 |
| AT4G27585.1 | Stomatin-like protein 1 (SLP1) | 6 | 0.56 | 0.253 | 0.999 |
| AT4G27585.1 | Stomatin-like protein 1 (SLP1) | 7 | 0.61 | 0.251 | 1.000 |
| AT4G27585.1 | Stomatin-like protein 1 (SLP1) | 8 | 0.69 | 0.248 | 1.000 |
| AT4G27585.1 | Stomatin-like protein 1 (SLP1) | 9 | 0.79 | 0.246 | 1.000 |
| AT4G27585.1 | Stomatin-like protein 1 (SLP1) | 10 | 0.92 | 0.245 | 1.000 |
| AT4G27585.1 | Stomatin-like protein 1 (SLP1) | 11 | 1.04 | 0.245 | 1.000 |
| AT4G27585.1 | Stomatin-like protein 1 (SLP1) | 12 | 1.01 | 0.245 | 1.000 |
| AT4G27585.1 | Stomatin-like protein 1 (SLP1) | 13 | 0.81 | 0.244 | 1.000 |
| AT4G27585.1 | Stomatin-like protein 1 (SLP1) | 14 | 0.63 | 0.242 | 1.000 |
| AT4G27585.1 | Stomatin-like protein 1 (SLP1) | 15 | 0.53 | 0.241 | 0.992 |
| AT4G27585.1 | Stomatin-like protein 1 (SLP1) | 16 | 0.48 | 0.239 | 0.964 |
| AT4G27585.1 | Stomatin-like protein 1 (SLP1) | 17 | 0.50 | 0.239 | 0.980 |
| AT4G27585.1 | Stomatin-like protein 1 (SLP1) | 18 | 0.67 | 0.241 | 1.000 |
| AT4G27585.1 | Stomatin-like protein 1 (SLP1) | 19 | 1.21 | 0.246 | 1.000 |
| AT4G27585.1 | Stomatin-like protein 1 (SLP1) | 20 | 2.34 | 0.251 | 0.927 |
| AT4G27585.1 | Stomatin-like protein 1 (SLP1) | 21 | 3.39 | 0.253 | 0.502 |
| AT4G27585.1 | Stomatin-like protein 1 (SLP1) | 22 | 3.11 | 0.253 | 0.620 |
| AT4G27585.1 | Stomatin-like protein 1 (SLP1) | 23 | 2.12 | 0.251 | 0.974 |
| AT4G27585.1 | Stomatin-like protein 1 (SLP1) | 24 | 1.41 | 0.248 | 1.000 |
| AT4G27585.1 | Stomatin-like protein 1 (SLP1) | 25 | 1.01 | 0.245 | 1.000 |
| AT4G27585.1 | Stomatin-like protein 1 (SLP1) | 26 | 0.70 | 0.244 | 1.000 |
| AT4G27585.1 | Stomatin-like protein 1 (SLP1) | 27 | 0.45 | 0.244 | 0.941 |
| AT4G27585.1 | Stomatin-like protein 1 (SLP1) | 28 | 0.35 | 0.245 | 0.697 |
| AT4G27585.1 | Stomatin-like protein 1 (SLP1) | 29 | 0.41 | 0.246 | 0.882 |
| AT4G27585.1 | Stomatin-like protein 1 (SLP1) | 30 | 0.65 | 0.247 | 1.000 |
| AT4G27585.1 | Stomatin-like protein 1 (SLP1) | 31 | 1.23 | 0.255 | 1.000 |
| AT4G27585.1 | Stomatin-like protein 1 (SLP1) | 32 | 2.11 | 0.277 | 0.991 |
| AT4G27585.1 | Stomatin-like protein 1 (SLP1) | 33 | 2.68 | 0.325 | 0.971 |
| AT4G27585.1 | Stomatin-like protein 1 (SLP1) | 34 | 3.05 | 0.445 | 0.996 |
| AT4G27585.1 | Stomatin-like protein 1 (SLP1) | 46 | 27.48 | 6.092 | 1.000 |
| AT4G27585.1 | Stomatin-like protein 1 (SLP1) | 47 | 33.97 | 6.801 | 1.000 |
| AT4G27740.1 | Yippee family putative zinc-binding protein | 2 | 0.88 | 0.435 | 1.000 |
| AT4G27740.1 | Yippee family putative zinc-binding protein | 3 | 1.00 | 0.366 | 1.000 |
| AT4G27740.1 | Yippee family putative zinc-binding protein | 4 | 1.13 | 0.355 | 1.000 |
| AT4G27740.1 | Yippee family putative zinc-binding protein | 6 | 1.37 | 0.350 | 1.000 |
| AT4G27740.1 | Yippee family putative zinc-binding protein | 7 | 1.40 | 0.326 | 1.000 |

| | | | | | |
|---|---|---|---|---|---|
| AT4G27740.1 | Yippee family putative zinc-binding protein | 8 | 1.36 | 0.308 | 1.000 |
| AT4G27740.1 | Yippee family putative zinc-binding protein | 9 | 1.30 | 0.292 | 1.000 |
| AT4G27740.1 | Yippee family putative zinc-binding protein | 10 | 1.21 | 0.285 | 1.000 |
| AT4G27740.1 | Yippee family putative zinc-binding protein | 11 | 1.12 | 0.277 | 1.000 |
| AT4G27740.1 | Yippee family putative zinc-binding protein | 12 | 1.05 | 0.268 | 1.000 |
| AT4G27740.1 | Yippee family putative zinc-binding protein | 13 | 1.03 | 0.260 | 1.000 |
| AT4G27740.1 | Yippee family putative zinc-binding protein | 14 | 1.07 | 0.254 | 1.000 |
| AT4G27740.1 | Yippee family putative zinc-binding protein | 15 | 1.08 | 0.250 | 1.000 |
| AT4G27740.1 | Yippee family putative zinc-binding protein | 16 | 1.04 | 0.249 | 1.000 |
| AT4G27740.1 | Yippee family putative zinc-binding protein | 17 | 1.04 | 0.252 | 1.000 |
| AT4G27740.1 | Yippee family putative zinc-binding protein | 18 | 1.07 | 0.256 | 1.000 |
| AT4G27740.1 | Yippee family putative zinc-binding protein | 19 | 1.09 | 0.258 | 1.000 |
| AT4G27740.1 | Yippee family putative zinc-binding protein | 20 | 1.05 | 0.260 | 1.000 |
| AT4G27740.1 | Yippee family putative zinc-binding protein | 21 | 0.93 | 0.266 | 1.000 |
| AT4G27740.1 | Yippee family putative zinc-binding protein | 22 | 0.76 | 0.270 | 1.000 |
| AT4G27740.1 | Yippee family putative zinc-binding protein | 23 | 0.61 | 0.267 | 1.000 |
| AT4G27740.1 | Yippee family putative zinc-binding protein | 24 | 0.48 | 0.261 | 0.969 |
| AT4G27740.1 | Yippee family putative zinc-binding protein | 25 | 0.41 | 0.259 | 0.849 |
| AT4G27740.1 | Yippee family putative zinc-binding protein | 26 | 0.41 | 0.256 | 0.858 |
| AT4G27740.1 | Yippee family putative zinc-binding protein | 27 | 0.52 | 0.251 | 0.983 |
| AT4G27740.1 | Yippee family putative zinc-binding protein | 28 | 0.77 | 0.247 | 1.000 |
| AT4G27740.1 | Yippee family putative zinc-binding protein | 29 | 1.16 | 0.244 | 1.000 |
| AT4G27740.1 | Yippee family putative zinc-binding protein | 30 | 1.56 | 0.244 | 1.000 |
| AT4G27740.1 | Yippee family putative zinc-binding protein | 31 | 1.89 | 0.244 | 0.983 |
| AT4G27740.1 | Yippee family putative zinc-binding protein | 32 | 1.97 | 0.244 | 0.969 |
| AT4G27740.1 | Yippee family putative zinc-binding protein | 33 | 1.65 | 0.244 | 0.999 |
| AT4G27740.1 | Yippee family putative zinc-binding protein | 34 | 1.10 | 0.244 | 1.000 |
| AT4G27740.1 | Yippee family putative zinc-binding protein | 35 | 0.66 | 0.243 | 1.000 |
| AT4G27740.1 | Yippee family putative zinc-binding protein | 36 | 0.49 | 0.243 | 0.953 |
| AT4G27740.1 | Yippee family putative zinc-binding protein | 37 | 0.53 | 0.243 | 0.982 |
| AT4G27740.1 | Yippee family putative zinc-binding protein | 38 | 0.75 | 0.243 | 1.000 |

| | | | | | | |
|---|---|---|---|---|---|---|
| AT4G27740.1 | Yippee family putative zinc-binding protein | 39 | 1.24 | 0.244 | 1.000 | |
| AT4G27740.1 | Yippee family putative zinc-binding protein | 40 | 2.14 | 0.245 | 0.929 | |
| AT4G27740.1 | Yippee family putative zinc-binding protein | 41 | 3.34 | 0.247 | 0.406 | |
| AT4G27740.1 | Yippee family putative zinc-binding protein | 42 | 4.04 | 0.251 | 0.229 | |
| AT4G27740.1 | Yippee family putative zinc-binding protein | 43 | 3.41 | 0.256 | 0.428 | |
| AT4G27740.1 | Yippee family putative zinc-binding protein | 44 | 1.91 | 0.260 | 0.989 | |
| AT4G27740.1 | Yippee family putative zinc-binding protein | 45 | 0.74 | 0.268 | 1.000 | |
| AT4G27740.1 | Yippee family putative zinc-binding protein | 46 | 0.22 | 0.298 | 0.346 | |
| AT4G27740.1 | Yippee family putative zinc-binding protein | 47 | 0.06 | 0.391 | 0.039 | * |
| AT4G28070.1 | AFG1-like ATPase family protein | 2 | 1.08 | 0.552 | 1.000 | |
| AT4G28070.1 | AFG1-like ATPase family protein | 3 | 1.16 | 0.535 | 1.000 | |
| AT4G28070.1 | AFG1-like ATPase family protein | 34 | 2.94 | 0.242 | 0.200 | |
| AT4G28070.1 | AFG1-like ATPase family protein | 35 | 2.77 | 0.229 | 0.202 | |
| AT4G28070.1 | AFG1-like ATPase family protein | 36 | 2.58 | 0.218 | 0.219 | |
| AT4G28070.1 | AFG1-like ATPase family protein | 37 | 2.38 | 0.209 | 0.258 | |
| AT4G28070.1 | AFG1-like ATPase family protein | 42 | 1.38 | 0.185 | 0.889 | |
| AT4G28070.1 | AFG1-like ATPase family protein | 43 | 1.21 | 0.186 | 0.981 | |
| AT4G28070.1 | AFG1-like ATPase family protein | 44 | 1.07 | 0.191 | 1.000 | |
| AT4G28070.1 | AFG1-like ATPase family protein | 45 | 0.93 | 0.201 | 1.000 | |
| AT4G28070.1 | AFG1-like ATPase family protein | 46 | 0.82 | 0.217 | 0.987 | |
| AT4G28070.1 | AFG1-like ATPase family protein | 47 | 0.71 | 0.239 | 0.940 | |
| AT4G28220.1 | Alternative NAD(P)H dehydrogenase B1 (NDB1) | 2 | 1.05 | 0.324 | 1.000 | |
| AT4G28220.1 | Alternative NAD(P)H dehydrogenase B1 (NDB1) | 3 | 1.13 | 0.282 | 1.000 | |
| AT4G28220.1 | Alternative NAD(P)H dehydrogenase B1 (NDB1) | 11 | 1.78 | 0.270 | 1.000 | |
| AT4G28220.1 | Alternative NAD(P)H dehydrogenase B1 (NDB1) | 12 | 1.84 | 0.235 | 0.996 | |
| AT4G28220.1 | Alternative NAD(P)H dehydrogenase B1 (NDB1) | 13 | 1.88 | 0.225 | 0.991 | |
| AT4G28220.1 | Alternative NAD(P)H dehydrogenase B1 (NDB1) | 16 | 1.96 | 0.190 | 0.919 | |
| AT4G28220.1 | Alternative NAD(P)H dehydrogenase B1 (NDB1) | 17 | 2.09 | 0.176 | 0.758 | |
| AT4G28220.1 | Alternative NAD(P)H dehydrogenase B1 (NDB1) | 18 | 2.17 | 0.163 | 0.555 | |
| AT4G28220.1 | Alternative NAD(P)H dehydrogenase B1 (NDB1) | 19 | 2.28 | 0.159 | 0.422 | |
| AT4G28220.1 | Alternative NAD(P)H dehydrogenase B1 (NDB1) | 20 | 2.36 | 0.176 | 0.517 | |

| | | | | | |
|---|---|---|---|---|---|
| AT4G28220.1 | Alternative NAD(P)H dehydrogenase B1 (NDB1) | 21 | 2.26 | 0.204 | 0.816 |
| AT4G28220.1 | Alternative NAD(P)H dehydrogenase B1 (NDB1) | 22 | 1.95 | 0.213 | 0.971 |
| AT4G28220.1 | Alternative NAD(P)H dehydrogenase B1 (NDB1) | 23 | 1.49 | 0.225 | 1.000 |
| AT4G28220.1 | Alternative NAD(P)H dehydrogenase B1 (NDB1) | 24 | 0.99 | 0.219 | 1.000 |
| AT4G28220.1 | Alternative NAD(P)H dehydrogenase B1 (NDB1) | 26 | 0.46 | 0.195 | 0.803 |
| AT4G28220.1 | Alternative NAD(P)H dehydrogenase B1 (NDB1) | 27 | 0.43 | 0.169 | 0.501 |
| AT4G28220.1 | Alternative NAD(P)H dehydrogenase B1 (NDB1) | 28 | 0.61 | 0.155 | 0.962 |
| AT4G28220.1 | Alternative NAD(P)H dehydrogenase B1 (NDB1) | 29 | 1.00 | 0.148 | 1.000 |
| AT4G28220.1 | Alternative NAD(P)H dehydrogenase B1 (NDB1) | 30 | 1.38 | 0.146 | 1.000 |
| AT4G28220.1 | Alternative NAD(P)H dehydrogenase B1 (NDB1) | 31 | 1.59 | 0.146 | 0.967 |
| AT4G28220.1 | Alternative NAD(P)H dehydrogenase B1 (NDB1) | 32 | 1.61 | 0.147 | 0.961 |
| AT4G28220.1 | Alternative NAD(P)H dehydrogenase B1 (NDB1) | 33 | 1.44 | 0.147 | 0.998 |
| AT4G28220.1 | Alternative NAD(P)H dehydrogenase B1 (NDB1) | 34 | 1.25 | 0.147 | 1.000 |
| AT4G28220.1 | Alternative NAD(P)H dehydrogenase B1 (NDB1) | 35 | 1.17 | 0.147 | 1.000 |
| AT4G28220.1 | Alternative NAD(P)H dehydrogenase B1 (NDB1) | 36 | 1.17 | 0.147 | 1.000 |
| AT4G28220.1 | Alternative NAD(P)H dehydrogenase B1 (NDB1) | 37 | 1.20 | 0.146 | 1.000 |
| AT4G28220.1 | Alternative NAD(P)H dehydrogenase B1 (NDB1) | 38 | 1.14 | 0.146 | 1.000 |
| AT4G28220.1 | Alternative NAD(P)H dehydrogenase B1 (NDB1) | 39 | 0.95 | 0.147 | 1.000 |
| AT4G28220.1 | Alternative NAD(P)H dehydrogenase B1 (NDB1) | 40 | 0.73 | 0.146 | 1.000 |
| AT4G28220.1 | Alternative NAD(P)H dehydrogenase B1 (NDB1) | 41 | 0.57 | 0.147 | 0.862 |
| AT4G28220.1 | Alternative NAD(P)H dehydrogenase B1 (NDB1) | 42 | 0.53 | 0.149 | 0.743 |
| AT4G28220.1 | Alternative NAD(P)H dehydrogenase B1 (NDB1) | 43 | 0.65 | 0.150 | 0.987 |
| AT4G28220.1 | Alternative NAD(P)H dehydrogenase B1 (NDB1) | 44 | 0.78 | 0.150 | 1.000 |
| AT4G28220.1 | Alternative NAD(P)H dehydrogenase B1 (NDB1) | 45 | 0.74 | 0.149 | 1.000 |
| AT4G28220.1 | Alternative NAD(P)H dehydrogenase B1 (NDB1) | 46 | 0.62 | 0.173 | 0.992 |
| AT4G28220.1 | Alternative NAD(P)H dehydrogenase B1 (NDB1) | 47 | 0.52 | 0.292 | 0.999 |
| AT4G28360.1 | Ribosomal L22 family protein, putative | 2 | 1.11 | 0.612 | 1.000 |
| AT4G28360.1 | Ribosomal L22 family protein, putative | 3 | 1.32 | 0.542 | 1.000 |
| AT4G28360.1 | Ribosomal L22 family protein, putative | 12 | 3.23 | 0.329 | 0.742 |
| AT4G28360.1 | Ribosomal L22 family protein, putative | 13 | 3.02 | 0.288 | 0.656 |
| AT4G28360.1 | Ribosomal L22 family protein, putative | 14 | 2.64 | 0.259 | 0.684 |

| | | | | | |
|---|---|---|---|---|---|
| AT4G28360.1 | Ribosomal L22 family protein, putative | 15 | 2.14 | 0.239 | 0.846 |
| AT4G28360.1 | Ribosomal L22 family protein, putative | 16 | 1.72 | 0.228 | 0.973 |
| AT4G28360.1 | Ribosomal L22 family protein, putative | 17 | 1.50 | 0.224 | 0.997 |
| AT4G28360.1 | Ribosomal L22 family protein, putative | 18 | 1.38 | 0.225 | 1.000 |
| AT4G28360.1 | Ribosomal L22 family protein, putative | 19 | 1.29 | 0.229 | 1.000 |
| AT4G28360.1 | Ribosomal L22 family protein, putative | 20 | 1.20 | 0.236 | 1.000 |
| AT4G28360.1 | Ribosomal L22 family protein, putative | 21 | 1.08 | 0.247 | 1.000 |
| AT4G28360.1 | Ribosomal L22 family protein, putative | 22 | 0.94 | 0.264 | 1.000 |
| AT4G28360.1 | Ribosomal L22 family protein, putative | 23 | 0.81 | 0.288 | 1.000 |
| AT4G28360.1 | Ribosomal L22 family protein, putative | 24 | 0.69 | 0.314 | 1.000 |
| AT4G28360.1 | Ribosomal L22 family protein, putative | 29 | 0.34 | 0.303 | 0.730 |
| AT4G28360.1 | Ribosomal L22 family protein, putative | 30 | 0.34 | 0.275 | 0.619 |
| AT4G28360.1 | Ribosomal L22 family protein, putative | 31 | 0.37 | 0.256 | 0.644 |
| AT4G28360.1 | Ribosomal L22 family protein, putative | 32 | 0.44 | 0.251 | 0.828 |
| AT4G28360.1 | Ribosomal L22 family protein, putative | 33 | 0.56 | 0.263 | 0.983 |
| AT4G28360.1 | Ribosomal L22 family protein, putative | 34 | 0.73 | 0.293 | 1.000 |
| AT4G28360.1 | Ribosomal L22 family protein, putative | 35 | 0.96 | 0.332 | 1.000 |
| AT4G28360.1 | Ribosomal L22 family protein, putative | 41 | 3.64 | 0.423 | 0.875 |
| AT4G28360.1 | Ribosomal L22 family protein, putative | 42 | 3.83 | 0.445 | 0.884 |
| AT4G28360.1 | Ribosomal L22 family protein, putative | 43 | 3.69 | 0.493 | 0.945 |
| AT4G28360.1 | Ribosomal L22 family protein, putative | 46 | 2.30 | 0.878 | 1.000 |
| AT4G28360.1 | Ribosomal L22 family protein, putative | 47 | 1.86 | 1.070 | 1.000 |
| AT4G28390.1 | ADP/ATP carrier 3 (AAC3) | 2 | 0.93 | 0.529 | 1.000 |
| AT4G28390.1 | ADP/ATP carrier 3 (AAC3) | 3 | 0.99 | 0.392 | 1.000 |
| AT4G28390.1 | ADP/ATP carrier 3 (AAC3) | 5 | 1.10 | 0.328 | 1.000 |
| AT4G28390.1 | ADP/ATP carrier 3 (AAC3) | 6 | 1.14 | 0.297 | 1.000 |
| AT4G28390.1 | ADP/ATP carrier 3 (AAC3) | 7 | 1.13 | 0.272 | 1.000 |
| AT4G28390.1 | ADP/ATP carrier 3 (AAC3) | 8 | 1.11 | 0.257 | 1.000 |
| AT4G28390.1 | ADP/ATP carrier 3 (AAC3) | 9 | 1.09 | 0.248 | 1.000 |
| AT4G28390.1 | ADP/ATP carrier 3 (AAC3) | 10 | 1.11 | 0.241 | 1.000 |
| AT4G28390.1 | ADP/ATP carrier 3 (AAC3) | 11 | 1.23 | 0.234 | 1.000 |
| AT4G28390.1 | ADP/ATP carrier 3 (AAC3) | 12 | 1.43 | 0.230 | 1.000 |
| AT4G28390.1 | ADP/ATP carrier 3 (AAC3) | 13 | 1.64 | 0.229 | 0.999 |
| AT4G28390.1 | ADP/ATP carrier 3 (AAC3) | 14 | 1.84 | 0.233 | 0.990 |
| AT4G28390.1 | ADP/ATP carrier 3 (AAC3) | 15 | 2.24 | 0.238 | 0.910 |
| AT4G28390.1 | ADP/ATP carrier 3 (AAC3) | 16 | 2.81 | 0.248 | 0.704 |
| AT4G28390.1 | ADP/ATP carrier 3 (AAC3) | 17 | 3.90 | 0.281 | 0.475 |
| AT4G28390.1 | ADP/ATP carrier 3 (AAC3) | 18 | 5.38 | 0.371 | 0.580 |
| AT4G28390.1 | ADP/ATP carrier 3 (AAC3) | 33 | 4.31 | 0.364 | 0.753 |
| AT4G28390.1 | ADP/ATP carrier 3 (AAC3) | 34 | 5.49 | 0.284 | 0.171 |

| | | | | | | |
|---|---|---|---|---|---|---|
| AT4G28390.1 | ADP/ATP carrier 3 (AAC3) | 35 | 6.28 | 0.248 | 0.031 | * |
| AT4G28390.1 | ADP/ATP carrier 3 (AAC3) | 36 | 5.97 | 0.233 | 0.022 | * |
| AT4G28390.1 | ADP/ATP carrier 3 (AAC3) | 37 | 5.07 | 0.229 | 0.049 | * |
| AT4G28390.1 | ADP/ATP carrier 3 (AAC3) | 38 | 3.93 | 0.228 | 0.173 | |
| AT4G28390.1 | ADP/ATP carrier 3 (AAC3) | 39 | 2.71 | 0.228 | 0.633 | |
| AT4G28390.1 | ADP/ATP carrier 3 (AAC3) | 40 | 1.67 | 0.229 | 0.998 | |
| AT4G28390.1 | ADP/ATP carrier 3 (AAC3) | 41 | 1.14 | 0.230 | 1.000 | |
| AT4G28390.1 | ADP/ATP carrier 3 (AAC3) | 42 | 1.15 | 0.233 | 1.000 | |
| AT4G28390.1 | ADP/ATP carrier 3 (AAC3) | 43 | 1.39 | 0.236 | 1.000 | |
| AT4G28390.1 | ADP/ATP carrier 3 (AAC3) | 44 | 1.48 | 0.241 | 1.000 | |
| AT4G28390.1 | ADP/ATP carrier 3 (AAC3) | 45 | 1.56 | 0.281 | 1.000 | |
| AT4G28390.1 | ADP/ATP carrier 3 (AAC3) | 46 | 1.84 | 0.425 | 1.000 | |
| AT4G28390.1 | ADP/ATP carrier 3 (AAC3) | 47 | 2.31 | 0.690 | 1.000 | |
| AT4G28510.1 | Prohibitin 1 (PHB1) | 2 | 1.05 | 0.389 | 1.000 | |
| AT4G28510.1 | Prohibitin 1 (PHB1) | 3 | 0.79 | 0.250 | 1.000 | |
| AT4G28510.1 | Prohibitin 1 (PHB1) | 4 | 0.60 | 0.219 | 0.999 | |
| AT4G28510.1 | Prohibitin 1 (PHB1) | 5 | 0.52 | 0.218 | 0.982 | |
| AT4G28510.1 | Prohibitin 1 (PHB1) | 6 | 0.55 | 0.219 | 0.993 | |
| AT4G28510.1 | Prohibitin 1 (PHB1) | 7 | 0.66 | 0.217 | 1.000 | |
| AT4G28510.1 | Prohibitin 1 (PHB1) | 8 | 0.85 | 0.214 | 1.000 | |
| AT4G28510.1 | Prohibitin 1 (PHB1) | 9 | 1.12 | 0.211 | 1.000 | |
| AT4G28510.1 | Prohibitin 1 (PHB1) | 10 | 1.46 | 0.210 | 1.000 | |
| AT4G28510.1 | Prohibitin 1 (PHB1) | 11 | 1.93 | 0.211 | 0.974 | |
| AT4G28510.1 | Prohibitin 1 (PHB1) | 12 | 2.41 | 0.211 | 0.768 | |
| AT4G28510.1 | Prohibitin 1 (PHB1) | 13 | 2.46 | 0.209 | 0.721 | |
| AT4G28510.1 | Prohibitin 1 (PHB1) | 14 | 1.89 | 0.207 | 0.976 | |
| AT4G28510.1 | Prohibitin 1 (PHB1) | 15 | 1.17 | 0.206 | 1.000 | |
| AT4G28510.1 | Prohibitin 1 (PHB1) | 16 | 0.74 | 0.204 | 1.000 | |
| AT4G28510.1 | Prohibitin 1 (PHB1) | 17 | 0.63 | 0.204 | 0.999 | |
| AT4G28510.1 | Prohibitin 1 (PHB1) | 18 | 0.69 | 0.206 | 1.000 | |
| AT4G28510.1 | Prohibitin 1 (PHB1) | 19 | 0.82 | 0.211 | 1.000 | |
| AT4G28510.1 | Prohibitin 1 (PHB1) | 20 | 0.95 | 0.216 | 1.000 | |
| AT4G28510.1 | Prohibitin 1 (PHB1) | 21 | 1.10 | 0.218 | 1.000 | |
| AT4G28510.1 | Prohibitin 1 (PHB1) | 22 | 1.35 | 0.219 | 1.000 | |
| AT4G28510.1 | Prohibitin 1 (PHB1) | 23 | 1.81 | 0.217 | 0.993 | |
| AT4G28510.1 | Prohibitin 1 (PHB1) | 24 | 2.33 | 0.213 | 0.828 | |
| AT4G28510.1 | Prohibitin 1 (PHB1) | 25 | 2.14 | 0.210 | 0.909 | |
| AT4G28510.1 | Prohibitin 1 (PHB1) | 26 | 1.29 | 0.209 | 1.000 | |
| AT4G28510.1 | Prohibitin 1 (PHB1) | 27 | 0.70 | 0.209 | 1.000 | |
| AT4G28510.1 | Prohibitin 1 (PHB1) | 28 | 0.46 | 0.209 | 0.898 | |
| AT4G28510.1 | Prohibitin 1 (PHB1) | 29 | 0.49 | 0.210 | 0.948 | |
| AT4G28510.1 | Prohibitin 1 (PHB1) | 30 | 0.85 | 0.211 | 1.000 | |
| AT4G28510.1 | Prohibitin 1 (PHB1) | 31 | 1.74 | 0.214 | 0.996 | |
| AT4G28510.1 | Prohibitin 1 (PHB1) | 32 | 2.97 | 0.220 | 0.502 | |
| AT4G28510.1 | Prohibitin 1 (PHB1) | 33 | 3.49 | 0.240 | 0.411 | |
| AT4G28510.1 | Prohibitin 1 (PHB1) | 34 | 3.19 | 0.293 | 0.832 | |
| AT4G28510.1 | Prohibitin 1 (PHB1) | 35 | 2.80 | 0.415 | 0.998 | |
| AT4G28510.1 | Prohibitin 1 (PHB1) | 46 | 16.23 | 4.295 | 1.000 | |
| AT4G28510.1 | Prohibitin 1 (PHB1) | 47 | 21.94 | 4.826 | 1.000 | |
| AT4G28630.1 | ABC transporter of the mitochondrion 1 (ATM1) | 2 | 1.42 | 0.790 | 0.999 | |
| AT4G28630.1 | ABC transporter of the mitochondrion 1 (ATM1) | 3 | 1.54 | 0.763 | 0.998 | |
| AT4G28630.1 | ABC transporter of the mitochondrion 1 (ATM1) | 32 | 1.86 | 0.318 | 0.834 | |
| AT4G28630.1 | ABC transporter of the mitochondrion 1 (ATM1) | 33 | 1.60 | 0.302 | 0.917 | |
| AT4G28630.1 | ABC transporter of the mitochondrion 1 (ATM1) | 34 | 1.35 | 0.296 | 0.981 | |
| AT4G28630.1 | ABC transporter of the mitochondrion 1 (ATM1) | 35 | 1.12 | 0.298 | 1.000 | |
| AT4G28630.1 | ABC transporter of the mitochondrion 1 (ATM1) | 36 | 0.90 | 0.307 | 1.000 | |

| | | | | | | |
|---|---|---|---|---|---|---|
| AT4G28630.1 | ABC transporter of the mitochondrion 1 (ATM1) | 37 | 0.72 | 0.322 | 0.981 | |
| AT4G28630.1 | ABC transporter of the mitochondrion 1 (ATM1) | 46 | 0.06 | 0.501 | 0.071 | . |
| AT4G28630.1 | ABC transporter of the mitochondrion 1 (ATM1) | 47 | 0.05 | 0.537 | 0.060 | . |
| AT4G28750.1 | Photosystem I subunit (PSAE1) | 2 | 0.89 | 0.260 | 1.000 | |
| AT4G28750.1 | Photosystem I subunit (PSAE1) | 3 | 0.74 | 0.158 | 1.000 | |
| AT4G28750.1 | Photosystem I subunit (PSAE1) | 4 | 0.77 | 0.144 | 1.000 | |
| AT4G28750.1 | Photosystem I subunit (PSAE1) | 5 | 0.89 | 0.144 | 1.000 | |
| AT4G28750.1 | Photosystem I subunit (PSAE1) | 6 | 0.93 | 0.144 | 1.000 | |
| AT4G28750.1 | Photosystem I subunit (PSAE1) | 7 | 0.86 | 0.145 | 1.000 | |
| AT4G28750.1 | Photosystem I subunit (PSAE1) | 8 | 0.85 | 0.141 | 1.000 | |
| AT4G28750.1 | Photosystem I subunit (PSAE1) | 9 | 0.95 | 0.138 | 1.000 | |
| AT4G28750.1 | Photosystem I subunit (PSAE1) | 10 | 1.02 | 0.138 | 1.000 | |
| AT4G28750.1 | Photosystem I subunit (PSAE1) | 11 | 1.08 | 0.140 | 1.000 | |
| AT4G28750.1 | Photosystem I subunit (PSAE1) | 12 | 1.11 | 0.140 | 1.000 | |
| AT4G28750.1 | Photosystem I subunit (PSAE1) | 13 | 1.10 | 0.137 | 1.000 | |
| AT4G28750.1 | Photosystem I subunit (PSAE1) | 14 | 1.12 | 0.135 | 1.000 | |
| AT4G28750.1 | Photosystem I subunit (PSAE1) | 15 | 1.06 | 0.135 | 1.000 | |
| AT4G28750.1 | Photosystem I subunit (PSAE1) | 16 | 1.00 | 0.134 | 1.000 | |
| AT4G28750.1 | Photosystem I subunit (PSAE1) | 17 | 1.04 | 0.133 | 1.000 | |
| AT4G28750.1 | Photosystem I subunit (PSAE1) | 18 | 1.16 | 0.134 | 1.000 | |
| AT4G28750.1 | Photosystem I subunit (PSAE1) | 19 | 1.30 | 0.138 | 1.000 | |
| AT4G28750.1 | Photosystem I subunit (PSAE1) | 20 | 1.27 | 0.143 | 1.000 | |
| AT4G28750.1 | Photosystem I subunit (PSAE1) | 21 | 1.04 | 0.145 | 1.000 | |
| AT4G28750.1 | Photosystem I subunit (PSAE1) | 22 | 0.79 | 0.145 | 1.000 | |
| AT4G28750.1 | Photosystem I subunit (PSAE1) | 23 | 0.62 | 0.145 | 0.971 | |
| AT4G28750.1 | Photosystem I subunit (PSAE1) | 24 | 0.53 | 0.140 | 0.719 | |
| AT4G28750.1 | Photosystem I subunit (PSAE1) | 25 | 0.60 | 0.137 | 0.919 | |
| AT4G28750.1 | Photosystem I subunit (PSAE1) | 26 | 0.93 | 0.137 | 1.000 | |
| AT4G28750.1 | Photosystem I subunit (PSAE1) | 27 | 1.28 | 0.137 | 1.000 | |
| AT4G28750.1 | Photosystem I subunit (PSAE1) | 28 | 1.48 | 0.138 | 0.996 | |
| AT4G28750.1 | Photosystem I subunit (PSAE1) | 29 | 1.69 | 0.139 | 0.909 | |
| AT4G28750.1 | Photosystem I subunit (PSAE1) | 30 | 1.21 | 0.141 | 1.000 | |
| AT4G28750.1 | Photosystem I subunit (PSAE1) | 31 | 0.43 | 0.144 | 0.243 | |
| AT4G28750.1 | Photosystem I subunit (PSAE1) | 32 | 0.16 | 0.154 | 0.000 | *** |
| AT4G28750.1 | Photosystem I subunit (PSAE1) | 33 | 0.12 | 0.195 | 0.000 | *** |
| AT4G28750.1 | Photosystem I subunit (PSAE1) | 34 | 0.14 | 0.315 | 0.168 | |
| AT4G28750.1 | Photosystem I subunit (PSAE1) | 46 | 1081764.61 | 2.777 | 0.529 | |
| AT4G28750.1 | Photosystem I subunit (PSAE1) | 47 | 7112385.64 | 3.094 | 0.494 | |
| AT4G29120.1 | Probable 3-hydroxyisobutyrate dehydrogenase-like 1 | 2 | 0.88 | 0.805 | 1.000 | |
| AT4G29120.1 | Probable 3-hydroxyisobutyrate dehydrogenase-like 1 | 3 | 0.99 | 1.428 | 1.000 | |
| AT4G29120.1 | Probable 3-hydroxyisobutyrate dehydrogenase-like 1 | 33 | 0.15 | 0.614 | 0.830 | |
| AT4G29120.1 | Probable 3-hydroxyisobutyrate dehydrogenase-like 1 | 34 | 0.23 | 0.440 | 0.784 | |
| AT4G29120.1 | Probable 3-hydroxyisobutyrate dehydrogenase-like 1 | 35 | 0.46 | 0.413 | 0.993 | |
| AT4G29120.1 | Probable 3-hydroxyisobutyrate dehydrogenase-like 1 | 36 | 1.69 | 0.368 | 0.999 | |
| AT4G29120.1 | Probable 3-hydroxyisobutyrate dehydrogenase-like 1 | 37 | 5.45 | 0.341 | 0.289 | |
| AT4G29120.1 | Probable 3-hydroxyisobutyrate dehydrogenase-like 1 | 38 | 6.93 | 0.333 | 0.134 | |
| AT4G29120.1 | Probable 3-hydroxyisobutyrate dehydrogenase-like 1 | 39 | 2.59 | 0.331 | 0.888 | |
| AT4G29120.1 | Probable 3-hydroxyisobutyrate dehydrogenase-like 1 | 40 | 0.43 | 0.351 | 0.960 | |

| | | | | | |
|---|---|---|---|---|---|
| AT4G29120.1 | Probable 3-hydroxyisobutyrate dehydrogenase-like 1 | 41 | 0.11 | 0.484 | 0.389 |
| AT4G29120.1 | Probable 3-hydroxyisobutyrate dehydrogenase-like 1 | 42 | 0.07 | 0.778 | 0.764 |
| AT4G29120.1 | Probable 3-hydroxyisobutyrate dehydrogenase-like 1 | 46 | 7.12 | 1.766 | 1.000 |
| AT4G29120.1 | Probable 3-hydroxyisobutyrate dehydrogenase-like 1 | 47 | 46.74 | 2.204 | 0.996 |
| AT4G29130.1 | Hexokinase 1 (HXK1) | 3 | 0.55 | 0.211 | 0.988 |
| AT4G29130.1 | Hexokinase 1 (HXK1) | 4 | 0.96 | 0.181 | 1.000 |
| AT4G29130.1 | Hexokinase 1 (HXK1) | 5 | 1.53 | 0.178 | 0.998 |
| AT4G29130.1 | Hexokinase 1 (HXK1) | 6 | 2.11 | 0.178 | 0.731 |
| AT4G29130.1 | Hexokinase 1 (HXK1) | 7 | 2.54 | 0.178 | 0.373 |
| AT4G29130.1 | Hexokinase 1 (HXK1) | 8 | 2.79 | 0.175 | 0.219 |
| AT4G29130.1 | Hexokinase 1 (HXK1) | 9 | 2.80 | 0.173 | 0.201 |
| AT4G29130.1 | Hexokinase 1 (HXK1) | 10 | 2.67 | 0.173 | 0.258 |
| AT4G29130.1 | Hexokinase 1 (HXK1) | 11 | 2.52 | 0.173 | 0.341 |
| AT4G29130.1 | Hexokinase 1 (HXK1) | 12 | 2.38 | 0.172 | 0.447 |
| AT4G29130.1 | Hexokinase 1 (HXK1) | 13 | 2.21 | 0.171 | 0.574 |
| AT4G29130.1 | Hexokinase 1 (HXK1) | 14 | 2.08 | 0.170 | 0.689 |
| AT4G29130.1 | Hexokinase 1 (HXK1) | 15 | 2.02 | 0.169 | 0.738 |
| AT4G29130.1 | Hexokinase 1 (HXK1) | 16 | 2.11 | 0.168 | 0.650 |
| AT4G29130.1 | Hexokinase 1 (HXK1) | 17 | 2.41 | 0.168 | 0.384 |
| AT4G29130.1 | Hexokinase 1 (HXK1) | 18 | 2.77 | 0.170 | 0.192 |
| AT4G29130.1 | Hexokinase 1 (HXK1) | 19 | 2.84 | 0.173 | 0.183 |
| AT4G29130.1 | Hexokinase 1 (HXK1) | 20 | 2.44 | 0.176 | 0.428 |
| AT4G29130.1 | Hexokinase 1 (HXK1) | 21 | 1.83 | 0.178 | 0.928 |
| AT4G29130.1 | Hexokinase 1 (HXK1) | 22 | 1.32 | 0.178 | 1.000 |
| AT4G29130.1 | Hexokinase 1 (HXK1) | 23 | 1.00 | 0.177 | 1.000 |
| AT4G29130.1 | Hexokinase 1 (HXK1) | 24 | 0.81 | 0.175 | 1.000 |
| AT4G29130.1 | Hexokinase 1 (HXK1) | 25 | 0.73 | 0.173 | 1.000 |
| AT4G29130.1 | Hexokinase 1 (HXK1) | 26 | 0.73 | 0.172 | 1.000 |
| AT4G29130.1 | Hexokinase 1 (HXK1) | 27 | 0.79 | 0.171 | 1.000 |
| AT4G29130.1 | Hexokinase 1 (HXK1) | 28 | 0.86 | 0.171 | 1.000 |
| AT4G29130.1 | Hexokinase 1 (HXK1) | 29 | 0.91 | 0.172 | 1.000 |
| AT4G29130.1 | Hexokinase 1 (HXK1) | 30 | 0.94 | 0.172 | 1.000 |
| AT4G29130.1 | Hexokinase 1 (HXK1) | 31 | 0.95 | 0.172 | 1.000 |
| AT4G29130.1 | Hexokinase 1 (HXK1) | 32 | 0.96 | 0.172 | 1.000 |
| AT4G29130.1 | Hexokinase 1 (HXK1) | 33 | 0.94 | 0.173 | 1.000 |
| AT4G29130.1 | Hexokinase 1 (HXK1) | 34 | 0.93 | 0.173 | 1.000 |
| AT4G29130.1 | Hexokinase 1 (HXK1) | 35 | 0.93 | 0.173 | 1.000 |
| AT4G29130.1 | Hexokinase 1 (HXK1) | 36 | 0.96 | 0.172 | 1.000 |
| AT4G29130.1 | Hexokinase 1 (HXK1) | 37 | 1.00 | 0.172 | 1.000 |
| AT4G29130.1 | Hexokinase 1 (HXK1) | 38 | 1.06 | 0.172 | 1.000 |
| AT4G29130.1 | Hexokinase 1 (HXK1) | 39 | 1.14 | 0.173 | 1.000 |
| AT4G29130.1 | Hexokinase 1 (HXK1) | 40 | 1.24 | 0.173 | 1.000 |
| AT4G29130.1 | Hexokinase 1 (HXK1) | 41 | 1.31 | 0.174 | 1.000 |
| AT4G29130.1 | Hexokinase 1 (HXK1) | 42 | 1.31 | 0.176 | 1.000 |
| AT4G29130.1 | Hexokinase 1 (HXK1) | 43 | 1.21 | 0.177 | 1.000 |
| AT4G29130.1 | Hexokinase 1 (HXK1) | 44 | 1.00 | 0.177 | 1.000 |
| AT4G29130.1 | Hexokinase 1 (HXK1) | 45 | 0.73 | 0.184 | 1.000 |
| AT4G29130.1 | Hexokinase 1 (HXK1) | 46 | 0.48 | 0.223 | 0.948 |
| AT4G29130.1 | Hexokinase 1 (HXK1) | 47 | 0.30 | 0.325 | 0.872 |
| AT4G29130.1 | Hexokinase 1 (HXK1) | 48 | 0.19 | 0.470 | 0.899 |
| AT4G29480.1 | ATP synthase subunit (ATPG) | 2 | 0.87 | 0.486 | 1.000 |
| AT4G29480.1 | ATP synthase subunit (ATPG) | 3 | 0.88 | 0.333 | 1.000 |
| AT4G29480.1 | ATP synthase subunit (ATPG) | 4 | 0.87 | 0.322 | 1.000 |
| AT4G29480.1 | ATP synthase subunit (ATPG) | 5 | 0.81 | 0.290 | 1.000 |
| AT4G29480.1 | ATP synthase subunit (ATPG) | 6 | 0.73 | 0.276 | 1.000 |
| AT4G29480.1 | ATP synthase subunit (ATPG) | 7 | 0.69 | 0.248 | 1.000 |
| AT4G29480.1 | ATP synthase subunit (ATPG) | 8 | 0.90 | 0.233 | 1.000 |
| AT4G29480.1 | ATP synthase subunit (ATPG) | 9 | 1.54 | 0.224 | 1.000 |
| AT4G29480.1 | ATP synthase subunit (ATPG) | 10 | 2.35 | 0.221 | 0.937 |

| | | | | | | |
|---|---|---|---|---|---|---|
| AT4G29480.1 | ATP synthase subunit (ATPG) | 11 | 2.92 | 0.226 | 0.705 | |
| AT4G29480.1 | ATP synthase subunit (ATPG) | 12 | 3.08 | 0.234 | 0.682 | |
| AT4G29480.1 | ATP synthase subunit (ATPG) | 13 | 3.14 | 0.235 | 0.664 | |
| AT4G29480.1 | ATP synthase subunit (ATPG) | 14 | 3.38 | 0.235 | 0.542 | |
| AT4G29480.1 | ATP synthase subunit (ATPG) | 15 | 3.08 | 0.237 | 0.707 | |
| AT4G29480.1 | ATP synthase subunit (ATPG) | 16 | 2.22 | 0.243 | 0.990 | |
| AT4G29480.1 | ATP synthase subunit (ATPG) | 17 | 1.59 | 0.245 | 1.000 | |
| AT4G29480.1 | ATP synthase subunit (ATPG) | 18 | 1.48 | 0.234 | 1.000 | |
| AT4G29480.1 | ATP synthase subunit (ATPG) | 19 | 2.26 | 0.228 | 0.973 | |
| AT4G29480.1 | ATP synthase subunit (ATPG) | 20 | 4.80 | 0.228 | 0.101 | |
| AT4G29480.1 | ATP synthase subunit (ATPG) | 21 | 7.33 | 0.230 | 0.008 | ** |
| AT4G29480.1 | ATP synthase subunit (ATPG) | 22 | 6.09 | 0.230 | 0.027 | * |
| AT4G29480.1 | ATP synthase subunit (ATPG) | 23 | 3.44 | 0.229 | 0.461 | |
| AT4G29480.1 | ATP synthase subunit (ATPG) | 24 | 1.77 | 0.223 | 1.000 | |
| AT4G29480.1 | ATP synthase subunit (ATPG) | 25 | 1.14 | 0.219 | 1.000 | |
| AT4G29480.1 | ATP synthase subunit (ATPG) | 26 | 1.00 | 0.218 | 1.000 | |
| AT4G29480.1 | ATP synthase subunit (ATPG) | 27 | 0.93 | 0.219 | 1.000 | |
| AT4G29480.1 | ATP synthase subunit (ATPG) | 28 | 0.92 | 0.220 | 1.000 | |
| AT4G29480.1 | ATP synthase subunit (ATPG) | 29 | 1.13 | 0.220 | 1.000 | |
| AT4G29480.1 | ATP synthase subunit (ATPG) | 30 | 1.55 | 0.220 | 1.000 | |
| AT4G29480.1 | ATP synthase subunit (ATPG) | 31 | 1.84 | 0.220 | 0.999 | |
| AT4G29480.1 | ATP synthase subunit (ATPG) | 32 | 1.67 | 0.220 | 1.000 | |
| AT4G29480.1 | ATP synthase subunit (ATPG) | 33 | 1.46 | 0.220 | 1.000 | |
| AT4G29480.1 | ATP synthase subunit (ATPG) | 34 | 1.42 | 0.221 | 1.000 | |
| AT4G29480.1 | ATP synthase subunit (ATPG) | 35 | 1.57 | 0.222 | 1.000 | |
| AT4G29480.1 | ATP synthase subunit (ATPG) | 36 | 1.83 | 0.221 | 0.999 | |
| AT4G29480.1 | ATP synthase subunit (ATPG) | 37 | 2.31 | 0.221 | 0.950 | |
| AT4G29480.1 | ATP synthase subunit (ATPG) | 38 | 3.71 | 0.222 | 0.296 | |
| AT4G29480.1 | ATP synthase subunit (ATPG) | 39 | 7.55 | 0.221 | 0.003 | ** |
| AT4G29480.1 | ATP synthase subunit (ATPG) | 40 | 11.63 | 0.220 | 0.000 | *** |
| AT4G29480.1 | ATP synthase subunit (ATPG) | 41 | 7.48 | 0.221 | 0.004 | ** |
| AT4G29480.1 | ATP synthase subunit (ATPG) | 42 | 3.14 | 0.225 | 0.576 | |
| AT4G29480.1 | ATP synthase subunit (ATPG) | 43 | 1.35 | 0.226 | 1.000 | |
| AT4G29480.1 | ATP synthase subunit (ATPG) | 44 | 0.61 | 0.225 | 1.000 | |
| AT4G29480.1 | ATP synthase subunit (ATPG) | 45 | 0.33 | 0.225 | 0.650 | |
| AT4G29480.1 | ATP synthase subunit (ATPG) | 46 | 0.24 | 0.258 | 0.412 | |
| AT4G29480.1 | ATP synthase subunit (ATPG) | 47 | 0.18 | 0.492 | 0.983 | |
| AT4G29660.1 | ? | 2 | 0.97 | 0.446 | 1.000 | |
| AT4G29660.1 | ? | 3 | 1.08 | 0.423 | 1.000 | |
| AT4G29660.1 | ? | 22 | 1.22 | 0.344 | 1.000 | |
| AT4G29660.1 | ? | 23 | 1.10 | 0.266 | 1.000 | |
| AT4G29660.1 | ? | 24 | 1.02 | 0.226 | 1.000 | |
| AT4G29660.1 | ? | 25 | 1.16 | 0.208 | 1.000 | |
| AT4G29660.1 | ? | 26 | 1.84 | 0.208 | 0.887 | |
| AT4G29660.1 | ? | 27 | 2.55 | 0.207 | 0.421 | |
| AT4G29660.1 | ? | 28 | 1.49 | 0.213 | 0.994 | |
| AT4G29660.1 | ? | 29 | 0.70 | 0.227 | 0.999 | |
| AT4G29660.1 | ? | 30 | 0.51 | 0.285 | 0.968 | |
| AT4G29660.1 | ? | 31 | 0.48 | 0.472 | 0.999 | |
| AT4G29660.1 | ? | 46 | 581.02 | 10.467 | 1.000 | |
| AT4G29660.1 | ? | 47 | 1195.38 | 11.514 | 1.000 | |
| AT4G30000.1 | Bifunctional hydroxymethyl-dihydropterin-pyrophosphokinase & dihydropteroate synthase | 2 | 1.05 | 0.317 | 1.000 | |
| AT4G30000.1 | Bifunctional hydroxymethyl-dihydropterin-pyrophosphokinase & dihydropteroate synthase | 3 | 0.96 | 0.282 | 1.000 | |
| AT4G30000.1 | Bifunctional hydroxymethyl-dihydropterin-pyrophosphokinase & dihydropteroate synthase | 13 | 0.77 | 0.214 | 1.000 | |

| | | | | | |
|---|---|---|---|---|---|
| AT4G30000.1 | Bifunctional hydroxymethyl-dihydropterin-pyrophosphokinase & dihydropteroate synthase | 14 | 0.91 | 0.204 | 1.000 |
| AT4G30000.1 | Bifunctional hydroxymethyl-dihydropterin-pyrophosphokinase & dihydropteroate synthase | 15 | 1.17 | 0.206 | 1.000 |
| AT4G30000.1 | Bifunctional hydroxymethyl-dihydropterin-pyrophosphokinase & dihydropteroate synthase | 16 | 1.62 | 0.213 | 0.999 |
| AT4G30000.1 | Bifunctional hydroxymethyl-dihydropterin-pyrophosphokinase & dihydropteroate synthase | 25 | 0.33 | 0.222 | 0.456 |
| AT4G30000.1 | Bifunctional hydroxymethyl-dihydropterin-pyrophosphokinase & dihydropteroate synthase | 26 | 0.48 | 0.166 | 0.660 |
| AT4G30000.1 | Bifunctional hydroxymethyl-dihydropterin-pyrophosphokinase & dihydropteroate synthase | 27 | 0.89 | 0.147 | 1.000 |
| AT4G30000.1 | Bifunctional hydroxymethyl-dihydropterin-pyrophosphokinase & dihydropteroate synthase | 28 | 1.46 | 0.139 | 0.991 |
| AT4G30000.1 | Bifunctional hydroxymethyl-dihydropterin-pyrophosphokinase & dihydropteroate synthase | 29 | 1.83 | 0.137 | 0.648 |
| AT4G30000.1 | Bifunctional hydroxymethyl-dihydropterin-pyrophosphokinase & dihydropteroate synthase | 30 | 1.72 | 0.136 | 0.791 |
| AT4G30000.1 | Bifunctional hydroxymethyl-dihydropterin-pyrophosphokinase & dihydropteroate synthase | 31 | 1.46 | 0.137 | 0.988 |
| AT4G30000.1 | Bifunctional hydroxymethyl-dihydropterin-pyrophosphokinase & dihydropteroate synthase | 32 | 1.24 | 0.138 | 1.000 |
| AT4G30000.1 | Bifunctional hydroxymethyl-dihydropterin-pyrophosphokinase & dihydropteroate synthase | 33 | 1.02 | 0.138 | 1.000 |
| AT4G30000.1 | Bifunctional hydroxymethyl-dihydropterin-pyrophosphokinase & dihydropteroate synthase | 34 | 0.85 | 0.138 | 1.000 |
| AT4G30000.1 | Bifunctional hydroxymethyl-dihydropterin-pyrophosphokinase & dihydropteroate synthase | 35 | 0.75 | 0.138 | 1.000 |
| AT4G30000.1 | Bifunctional hydroxymethyl-dihydropterin-pyrophosphokinase & dihydropteroate synthase | 36 | 0.74 | 0.137 | 0.999 |

| | | | | | | |
|---|---|---|---|---|---|---|
| AT4G30000.1 | Bifunctional hydroxymethyl-dihydropterin-pyrophosphokinase & dihydropteroate synthase | 37 | 0.85 | 0.137 | 1.000 | |
| AT4G30000.1 | Bifunctional hydroxymethyl-dihydropterin-pyrophosphokinase & dihydropteroate synthase | 38 | 1.07 | 0.137 | 1.000 | |
| AT4G30000.1 | Bifunctional hydroxymethyl-dihydropterin-pyrophosphokinase & dihydropteroate synthase | 39 | 1.33 | 0.138 | 1.000 | |
| AT4G30000.1 | Bifunctional hydroxymethyl-dihydropterin-pyrophosphokinase & dihydropteroate synthase | 40 | 1.45 | 0.137 | 0.990 | |
| AT4G30000.1 | Bifunctional hydroxymethyl-dihydropterin-pyrophosphokinase & dihydropteroate synthase | 41 | 1.35 | 0.138 | 0.999 | |
| AT4G30000.1 | Bifunctional hydroxymethyl-dihydropterin-pyrophosphokinase & dihydropteroate synthase | 42 | 1.14 | 0.139 | 1.000 | |
| AT4G30000.1 | Bifunctional hydroxymethyl-dihydropterin-pyrophosphokinase & dihydropteroate synthase | 43 | 0.94 | 0.140 | 1.000 | |
| AT4G30000.1 | Bifunctional hydroxymethyl-dihydropterin-pyrophosphokinase & dihydropteroate synthase | 44 | 0.70 | 0.140 | 0.996 | |
| AT4G30000.1 | Bifunctional hydroxymethyl-dihydropterin-pyrophosphokinase & dihydropteroate synthase | 45 | 0.48 | 0.141 | 0.371 | |
| AT4G30000.1 | Bifunctional hydroxymethyl-dihydropterin-pyrophosphokinase & dihydropteroate synthase | 46 | 0.28 | 0.165 | 0.021 | * |
| AT4G30000.1 | Bifunctional hydroxymethyl-dihydropterin-pyrophosphokinase & dihydropteroate synthase | 47 | 0.14 | 0.282 | 0.053 | . |
| AT4G30010.1 | ATP synthase subunit (ATP17) | 2 | 0.87 | 0.461 | 1.000 | |
| AT4G30010.1 | ATP synthase subunit (ATP17) | 3 | 0.86 | 0.311 | 1.000 | |
| AT4G30010.1 | ATP synthase subunit (ATP17) | 4 | 0.93 | 0.289 | 1.000 | |
| AT4G30010.1 | ATP synthase subunit (ATP17) | 5 | 1.12 | 0.289 | 1.000 | |
| AT4G30010.1 | ATP synthase subunit (ATP17) | 6 | 1.34 | 0.289 | 1.000 | |
| AT4G30010.1 | ATP synthase subunit (ATP17) | 7 | 1.39 | 0.289 | 1.000 | |
| AT4G30010.1 | ATP synthase subunit (ATP17) | 8 | 1.41 | 0.285 | 1.000 | |
| AT4G30010.1 | ATP synthase subunit (ATP17) | 9 | 1.55 | 0.282 | 1.000 | |
| AT4G30010.1 | ATP synthase subunit (ATP17) | 10 | 1.79 | 0.282 | 1.000 | |
| AT4G30010.1 | ATP synthase subunit (ATP17) | 11 | 2.08 | 0.287 | 0.998 | |
| AT4G30010.1 | ATP synthase subunit (ATP17) | 12 | 2.58 | 0.293 | 0.965 | |
| AT4G30010.1 | ATP synthase subunit (ATP17) | 13 | 3.34 | 0.297 | 0.793 | |
| AT4G30010.1 | ATP synthase subunit (ATP17) | 14 | 3.99 | 0.298 | 0.602 | |
| AT4G30010.1 | ATP synthase subunit (ATP17) | 15 | 3.99 | 0.302 | 0.621 | |
| AT4G30010.1 | ATP synthase subunit (ATP17) | 16 | 3.40 | 0.304 | 0.802 | |
| AT4G30010.1 | ATP synthase subunit (ATP17) | 17 | 2.50 | 0.301 | 0.981 | |
| AT4G30010.1 | ATP synthase subunit (ATP17) | 18 | 1.72 | 0.292 | 1.000 | |
| AT4G30010.1 | ATP synthase subunit (ATP17) | 19 | 1.38 | 0.287 | 1.000 | |

| | | | | | |
|---|---|---|---|---|---|
| AT4G30010.1 | ATP synthase subunit (ATP17) | 20 | 1.20 | 0.289 | 1.000 |
| AT4G30010.1 | ATP synthase subunit (ATP17) | 21 | 0.97 | 0.290 | 1.000 |
| AT4G30010.1 | ATP synthase subunit (ATP17) | 22 | 0.81 | 0.290 | 1.000 |
| AT4G30010.1 | ATP synthase subunit (ATP17) | 23 | 0.81 | 0.289 | 1.000 |
| AT4G30010.1 | ATP synthase subunit (ATP17) | 24 | 1.09 | 0.284 | 1.000 |
| AT4G30010.1 | ATP synthase subunit (ATP17) | 25 | 1.49 | 0.281 | 1.000 |
| AT4G30010.1 | ATP synthase subunit (ATP17) | 26 | 1.32 | 0.280 | 1.000 |
| AT4G30010.1 | ATP synthase subunit (ATP17) | 27 | 0.70 | 0.280 | 1.000 |
| AT4G30010.1 | ATP synthase subunit (ATP17) | 28 | 0.44 | 0.282 | 0.990 |
| AT4G30010.1 | ATP synthase subunit (ATP17) | 29 | 0.52 | 0.282 | 1.000 |
| AT4G30010.1 | ATP synthase subunit (ATP17) | 30 | 0.70 | 0.282 | 1.000 |
| AT4G30010.1 | ATP synthase subunit (ATP17) | 31 | 0.72 | 0.282 | 1.000 |
| AT4G30010.1 | ATP synthase subunit (ATP17) | 32 | 0.56 | 0.283 | 1.000 |
| AT4G30010.1 | ATP synthase subunit (ATP17) | 33 | 0.41 | 0.283 | 0.977 |
| AT4G30010.1 | ATP synthase subunit (ATP17) | 34 | 0.36 | 0.284 | 0.907 |
| AT4G30010.1 | ATP synthase subunit (ATP17) | 35 | 0.34 | 0.285 | 0.866 |
| AT4G30010.1 | ATP synthase subunit (ATP17) | 36 | 0.42 | 0.291 | 0.985 |
| AT4G30010.1 | ATP synthase subunit (ATP17) | 37 | 0.72 | 0.300 | 1.000 |
| AT4G30010.1 | ATP synthase subunit (ATP17) | 38 | 1.29 | 0.307 | 1.000 |
| AT4G30010.1 | ATP synthase subunit (ATP17) | 39 | 2.03 | 0.298 | 0.999 |
| AT4G30010.1 | ATP synthase subunit (ATP17) | 40 | 2.59 | 0.289 | 0.959 |
| AT4G30010.1 | ATP synthase subunit (ATP17) | 41 | 2.46 | 0.285 | 0.972 |
| AT4G30010.1 | ATP synthase subunit (ATP17) | 42 | 1.88 | 0.286 | 1.000 |
| AT4G30010.1 | ATP synthase subunit (ATP17) | 43 | 1.63 | 0.288 | 1.000 |
| AT4G30010.1 | ATP synthase subunit (ATP17) | 44 | 1.88 | 0.288 | 1.000 |
| AT4G30010.1 | ATP synthase subunit (ATP17) | 45 | 2.37 | 0.289 | 0.985 |
| AT4G30010.1 | ATP synthase subunit (ATP17) | 46 | 2.62 | 0.329 | 0.988 |
| AT4G30010.1 | ATP synthase subunit (ATP17) | 47 | 2.58 | 0.520 | 1.000 |
| AT4G30490.1 | AFG1-like ATPase family protein | 2 | 1.25 | 0.437 | 1.000 |
| AT4G30490.1 | AFG1-like ATPase family protein | 3 | 0.97 | 0.420 | 1.000 |
| AT4G30490.1 | AFG1-like ATPase family protein | 30 | 0.51 | 0.345 | 0.997 |
| AT4G30490.1 | AFG1-like ATPase family protein | 31 | 0.76 | 0.265 | 1.000 |
| AT4G30490.1 | AFG1-like ATPase family protein | 32 | 1.02 | 0.235 | 1.000 |
| AT4G30490.1 | AFG1-like ATPase family protein | 33 | 1.16 | 0.216 | 1.000 |
| AT4G30490.1 | AFG1-like ATPase family protein | 34 | 1.11 | 0.204 | 1.000 |
| AT4G30490.1 | AFG1-like ATPase family protein | 35 | 0.98 | 0.196 | 1.000 |
| AT4G30490.1 | AFG1-like ATPase family protein | 36 | 1.05 | 0.193 | 1.000 |
| AT4G30490.1 | AFG1-like ATPase family protein | 37 | 1.59 | 0.191 | 0.983 |
| AT4G30490.1 | AFG1-like ATPase family protein | 38 | 2.31 | 0.192 | 0.543 |
| AT4G30490.1 | AFG1-like ATPase family protein | 39 | 2.33 | 0.192 | 0.530 |
| AT4G30490.1 | AFG1-like ATPase family protein | 40 | 1.82 | 0.192 | 0.900 |
| AT4G30490.1 | AFG1-like ATPase family protein | 41 | 1.29 | 0.192 | 1.000 |
| AT4G30490.1 | AFG1-like ATPase family protein | 42 | 0.99 | 0.195 | 1.000 |
| AT4G30490.1 | AFG1-like ATPase family protein | 43 | 0.91 | 0.197 | 1.000 |
| AT4G30490.1 | AFG1-like ATPase family protein | 44 | 0.92 | 0.197 | 1.000 |

| | | | | | |
|---|---|---|---|---|---|
| AT4G30490.1 | AFG1-like ATPase family protein | 45 | 0.91 | 0.196 | 1.000 |
| AT4G30490.1 | AFG1-like ATPase family protein | 46 | 0.76 | 0.231 | 1.000 |
| AT4G30490.1 | AFG1-like ATPase family protein | 47 | 0.55 | 0.417 | 1.000 |
| AT4G30920.1 | Leucyl aminopeptidase 2 (LAP2) | 2 | 1.12 | 0.825 | 1.000 |
| AT4G30920.1 | Leucyl aminopeptidase 2 (LAP2) | 3 | 0.88 | 0.611 | 1.000 |
| AT4G30920.1 | Leucyl aminopeptidase 2 (LAP2) | 4 | 0.72 | 1.170 | 1.000 |
| AT4G30920.1 | Leucyl aminopeptidase 2 (LAP2) | 27 | 0.24 | 0.638 | 0.998 |
| AT4G30920.1 | Leucyl aminopeptidase 2 (LAP2) | 28 | 0.22 | 0.457 | 0.931 |
| AT4G30920.1 | Leucyl aminopeptidase 2 (LAP2) | 29 | 0.28 | 0.408 | 0.950 |
| AT4G30920.1 | Leucyl aminopeptidase 2 (LAP2) | 30 | 0.68 | 0.385 | 1.000 |
| AT4G30920.1 | Leucyl aminopeptidase 2 (LAP2) | 31 | 4.06 | 0.380 | 0.848 |
| AT4G30920.1 | Leucyl aminopeptidase 2 (LAP2) | 32 | 9.04 | 0.383 | 0.218 |
| AT4G30920.1 | Leucyl aminopeptidase 2 (LAP2) | 33 | 4.55 | 0.381 | 0.768 |
| AT4G30920.1 | Leucyl aminopeptidase 2 (LAP2) | 34 | 2.06 | 0.381 | 1.000 |
| AT4G30920.1 | Leucyl aminopeptidase 2 (LAP2) | 35 | 1.18 | 0.381 | 1.000 |
| AT4G30920.1 | Leucyl aminopeptidase 2 (LAP2) | 36 | 0.47 | 0.385 | 1.000 |
| AT4G30920.1 | Leucyl aminopeptidase 2 (LAP2) | 37 | 0.25 | 0.399 | 0.893 |
| AT4G30920.1 | Leucyl aminopeptidase 2 (LAP2) | 38 | 0.27 | 0.419 | 0.955 |
| AT4G30920.1 | Leucyl aminopeptidase 2 (LAP2) | 39 | 0.60 | 0.437 | 1.000 |
| AT4G30920.1 | Leucyl aminopeptidase 2 (LAP2) | 40 | 1.68 | 0.418 | 1.000 |
| AT4G30920.1 | Leucyl aminopeptidase 2 (LAP2) | 41 | 2.78 | 0.393 | 0.990 |
| AT4G30920.1 | Leucyl aminopeptidase 2 (LAP2) | 42 | 1.65 | 0.390 | 1.000 |
| AT4G30920.1 | Leucyl aminopeptidase 2 (LAP2) | 43 | 0.69 | 0.402 | 1.000 |
| AT4G30920.1 | Leucyl aminopeptidase 2 (LAP2) | 44 | 0.47 | 0.528 | 1.000 |
| AT4G30920.1 | Leucyl aminopeptidase 2 (LAP2) | 45 | 0.67 | 0.771 | 1.000 |
| AT4G30920.1 | Leucyl aminopeptidase 2 (LAP2) | 46 | 1.61 | 1.156 | 1.000 |
| AT4G30920.1 | Leucyl aminopeptidase 2 (LAP2) | 47 | 4.85 | 1.770 | 1.000 |
| AT4G30930.1 | 50S ribosomal protein L21 (RPL21M) | 2 | 0.81 | 0.626 | 1.000 |
| AT4G30930.1 | 50S ribosomal protein L21 (RPL21M) | 3 | 0.93 | 0.549 | 1.000 |
| AT4G30930.1 | 50S ribosomal protein L21 (RPL21M) | 12 | 2.87 | 0.345 | 0.939 |
| AT4G30930.1 | 50S ribosomal protein L21 (RPL21M) | 13 | 3.05 | 0.291 | 0.763 |

| | | | | | | |
|---|---|---|---|---|---|---|
| AT4G30930.1 | 50S ribosomal protein L21 (RPL21M) | 14 | 2.98 | 0.258 | 0.640 | |
| AT4G30930.1 | 50S ribosomal protein L21 (RPL21M) | 15 | 2.55 | 0.236 | 0.728 | |
| AT4G30930.1 | 50S ribosomal protein L21 (RPL21M) | 16 | 2.03 | 0.224 | 0.922 | |
| AT4G30930.1 | 50S ribosomal protein L21 (RPL21M) | 17 | 1.74 | 0.219 | 0.987 | |
| AT4G30930.1 | 50S ribosomal protein L21 (RPL21M) | 18 | 1.61 | 0.221 | 0.997 | |
| AT4G30930.1 | 50S ribosomal protein L21 (RPL21M) | 19 | 1.57 | 0.226 | 0.999 | |
| AT4G30930.1 | 50S ribosomal protein L21 (RPL21M) | 20 | 1.59 | 0.230 | 0.998 | |
| AT4G30930.1 | 50S ribosomal protein L21 (RPL21M) | 21 | 1.55 | 0.234 | 0.999 | |
| AT4G30930.1 | 50S ribosomal protein L21 (RPL21M) | 22 | 1.33 | 0.238 | 1.000 | |
| AT4G30930.1 | 50S ribosomal protein L21 (RPL21M) | 23 | 1.01 | 0.242 | 1.000 | |
| AT4G30930.1 | 50S ribosomal protein L21 (RPL21M) | 24 | 0.72 | 0.245 | 1.000 | |
| AT4G30930.1 | 50S ribosomal protein L21 (RPL21M) | 25 | 0.50 | 0.247 | 0.968 | |
| AT4G30930.1 | 50S ribosomal protein L21 (RPL21M) | 26 | 0.36 | 0.248 | 0.692 | |
| AT4G30930.1 | 50S ribosomal protein L21 (RPL21M) | 27 | 0.29 | 0.248 | 0.378 | |
| AT4G30930.1 | 50S ribosomal protein L21 (RPL21M) | 28 | 0.24 | 0.250 | 0.222 | |
| AT4G30930.1 | 50S ribosomal protein L21 (RPL21M) | 29 | 0.22 | 0.253 | 0.151 | |
| AT4G30930.1 | 50S ribosomal protein L21 (RPL21M) | 30 | 0.21 | 0.262 | 0.153 | |
| AT4G30930.1 | 50S ribosomal protein L21 (RPL21M) | 31 | 0.22 | 0.276 | 0.251 | |
| AT4G30930.1 | 50S ribosomal protein L21 (RPL21M) | 32 | 0.25 | 0.306 | 0.550 | |
| AT4G30930.1 | 50S ribosomal protein L21 (RPL21M) | 33 | 0.31 | 0.364 | 0.912 | |
| AT4G30930.1 | 50S ribosomal protein L21 (RPL21M) | 34 | 0.38 | 0.444 | 0.996 | |
| AT4G30930.1 | 50S ribosomal protein L21 (RPL21M) | 44 | 0.88 | 0.475 | 1.000 | |
| AT4G30930.1 | 50S ribosomal protein L21 (RPL21M) | 45 | 0.78 | 0.337 | 1.000 | |
| AT4G30930.1 | 50S ribosomal protein L21 (RPL21M) | 46 | 0.65 | 0.319 | 1.000 | |
| AT4G30930.1 | 50S ribosomal protein L21 (RPL21M) | 47 | 0.53 | 0.502 | 1.000 | |
| AT4G31210.1 | DNA topoisomerase, type IA, core | 2 | 1.25 | 0.319 | 1.000 | |
| AT4G31210.1 | DNA topoisomerase, type IA, core | 3 | 0.97 | 0.293 | 1.000 | |
| AT4G31210.1 | DNA topoisomerase, type IA, core | 4 | 0.74 | 0.408 | 1.000 | |
| AT4G31210.1 | DNA topoisomerase, type IA, core | 28 | 0.19 | 0.222 | 0.014 | * |
| AT4G31210.1 | DNA topoisomerase, type IA, core | 29 | 0.29 | 0.173 | 0.022 | * |
| AT4G31210.1 | DNA topoisomerase, type IA, core | 30 | 0.47 | 0.156 | 0.261 | |

| | | | | | | |
|---|---|---|---|---|---|---|
| AT4G31210.1 | DNA topoisomerase, type IA, core | 31 | 0.79 | 0.155 | 0.992 | |
| AT4G31210.1 | DNA topoisomerase, type IA, core | 32 | 1.20 | 0.160 | 0.999 | |
| AT4G31210.1 | DNA topoisomerase, type IA, core | 33 | 1.50 | 0.177 | 0.920 | |
| AT4G31210.1 | DNA topoisomerase, type IA, core | 34 | 1.57 | 0.216 | 0.950 | |
| AT4G31210.1 | DNA topoisomerase, type IA, core | 35 | 1.49 | 0.280 | 0.994 | |
| AT4G31210.1 | DNA topoisomerase, type IA, core | 46 | 0.05 | 0.286 | 0.000 | *** |
| AT4G31210.1 | DNA topoisomerase, type IA, core | 47 | 0.03 | 0.307 | 0.000 | *** |
| AT4G31990.4 | Aspartate aminotransferase 5 (ASP5) | 2 | 0.98 | 0.421 | 1.000 | |
| AT4G31990.4 | Aspartate aminotransferase 5 (ASP5) | 3 | 1.05 | 0.345 | 1.000 | |
| AT4G31990.4 | Aspartate aminotransferase 5 (ASP5) | 4 | 1.13 | 0.315 | 1.000 | |
| AT4G31990.4 | Aspartate aminotransferase 5 (ASP5) | 6 | 1.30 | 0.320 | 1.000 | |
| AT4G31990.4 | Aspartate aminotransferase 5 (ASP5) | 7 | 1.38 | 0.334 | 1.000 | |
| AT4G31990.4 | Aspartate aminotransferase 5 (ASP5) | 8 | 1.46 | 0.341 | 1.000 | |
| AT4G31990.4 | Aspartate aminotransferase 5 (ASP5) | 13 | 1.31 | 0.269 | 1.000 | |
| AT4G31990.4 | Aspartate aminotransferase 5 (ASP5) | 14 | 1.17 | 0.249 | 1.000 | |
| AT4G31990.4 | Aspartate aminotransferase 5 (ASP5) | 15 | 1.03 | 0.239 | 1.000 | |
| AT4G31990.4 | Aspartate aminotransferase 5 (ASP5) | 16 | 0.93 | 0.247 | 1.000 | |
| AT4G31990.4 | Aspartate aminotransferase 5 (ASP5) | 17 | 0.84 | 0.279 | 1.000 | |
| AT4G31990.4 | Aspartate aminotransferase 5 (ASP5) | 18 | 0.77 | 0.336 | 1.000 | |
| AT4G31990.4 | Aspartate aminotransferase 5 (ASP5) | 40 | 1.32 | 0.355 | 1.000 | |
| AT4G31990.4 | Aspartate aminotransferase 5 (ASP5) | 41 | 1.32 | 0.283 | 1.000 | |
| AT4G31990.4 | Aspartate aminotransferase 5 (ASP5) | 42 | 1.26 | 0.232 | 1.000 | |
| AT4G31990.4 | Aspartate aminotransferase 5 (ASP5) | 43 | 1.12 | 0.210 | 1.000 | |
| AT4G31990.4 | Aspartate aminotransferase 5 (ASP5) | 44 | 0.98 | 0.225 | 1.000 | |
| AT4G31990.4 | Aspartate aminotransferase 5 (ASP5) | 45 | 0.88 | 0.280 | 1.000 | |
| AT4G31990.4 | Aspartate aminotransferase 5 (ASP5) | 46 | 0.80 | 0.373 | 1.000 | |
| AT4G31990.4 | Aspartate aminotransferase 5 (ASP5) | 47 | 0.73 | 0.499 | 1.000 | |
| AT4G32260.1 | ATP synthase subunit (ATPG) | 2 | 0.59 | 0.421 | 1.000 | |
| AT4G32260.1 | ATP synthase subunit (ATPG) | 3 | 0.48 | 0.284 | 0.993 | |
| AT4G32260.1 | ATP synthase subunit (ATPG) | 4 | 0.40 | 0.267 | 0.912 | |
| AT4G32260.1 | ATP synthase subunit (ATPG) | 5 | 0.39 | 0.267 | 0.892 | |
| AT4G32260.1 | ATP synthase subunit (ATPG) | 6 | 0.42 | 0.268 | 0.939 | |
| AT4G32260.1 | ATP synthase subunit (ATPG) | 7 | 0.43 | 0.268 | 0.956 | |
| AT4G32260.1 | ATP synthase subunit (ATPG) | 8 | 0.50 | 0.263 | 0.991 | |
| AT4G32260.1 | ATP synthase subunit (ATPG) | 9 | 0.62 | 0.261 | 1.000 | |

| | | | | | | |
|---|---|---|---|---|---|---|
| AT4G32260.1 | ATP synthase subunit (ATPG) | 10 | 0.74 | 0.261 | 1.000 | |
| AT4G32260.1 | ATP synthase subunit (ATPG) | 11 | 0.84 | 0.262 | 1.000 | |
| AT4G32260.1 | ATP synthase subunit (ATPG) | 12 | 1.00 | 0.262 | 1.000 | |
| AT4G32260.1 | ATP synthase subunit (ATPG) | 13 | 1.19 | 0.259 | 1.000 | |
| AT4G32260.1 | ATP synthase subunit (ATPG) | 14 | 1.23 | 0.257 | 1.000 | |
| AT4G32260.1 | ATP synthase subunit (ATPG) | 15 | 0.99 | 0.257 | 1.000 | |
| AT4G32260.1 | ATP synthase subunit (ATPG) | 16 | 0.72 | 0.256 | 1.000 | |
| AT4G32260.1 | ATP synthase subunit (ATPG) | 17 | 0.62 | 0.255 | 1.000 | |
| AT4G32260.1 | ATP synthase subunit (ATPG) | 18 | 0.67 | 0.256 | 1.000 | |
| AT4G32260.1 | ATP synthase subunit (ATPG) | 19 | 0.76 | 0.261 | 1.000 | |
| AT4G32260.1 | ATP synthase subunit (ATPG) | 20 | 0.82 | 0.267 | 1.000 | |
| AT4G32260.1 | ATP synthase subunit (ATPG) | 21 | 0.84 | 0.269 | 1.000 | |
| AT4G32260.1 | ATP synthase subunit (ATPG) | 22 | 0.86 | 0.269 | 1.000 | |
| AT4G32260.1 | ATP synthase subunit (ATPG) | 23 | 0.94 | 0.268 | 1.000 | |
| AT4G32260.1 | ATP synthase subunit (ATPG) | 24 | 0.85 | 0.263 | 1.000 | |
| AT4G32260.1 | ATP synthase subunit (ATPG) | 25 | 0.70 | 0.259 | 1.000 | |
| AT4G32260.1 | ATP synthase subunit (ATPG) | 26 | 1.01 | 0.259 | 1.000 | |
| AT4G32260.1 | ATP synthase subunit (ATPG) | 27 | 1.71 | 0.259 | 1.000 | |
| AT4G32260.1 | ATP synthase subunit (ATPG) | 28 | 1.88 | 0.260 | 0.997 | |
| AT4G32260.1 | ATP synthase subunit (ATPG) | 29 | 2.02 | 0.261 | 0.989 | |
| AT4G32260.1 | ATP synthase subunit (ATPG) | 30 | 2.30 | 0.262 | 0.946 | |
| AT4G32260.1 | ATP synthase subunit (ATPG) | 31 | 1.69 | 0.264 | 1.000 | |
| AT4G32260.1 | ATP synthase subunit (ATPG) | 32 | 0.58 | 0.268 | 1.000 | |
| AT4G32260.1 | ATP synthase subunit (ATPG) | 33 | 0.15 | 0.280 | 0.067 | . |
| AT4G32260.1 | ATP synthase subunit (ATPG) | 34 | 0.07 | 0.332 | 0.019 | * |
| AT4G32260.1 | ATP synthase subunit (ATPG) | 35 | 0.06 | 0.489 | 0.253 | |
| AT4G32260.1 | ATP synthase subunit (ATPG) | 46 | 316393.13 | 4.030 | 0.952 | |
| AT4G32260.1 | ATP synthase subunit (ATPG) | 47 | 2697372.54 | 4.527 | 0.934 | |
| AT4G32470.1 | Ubiquinol:cytochrome C oxidoreductase subunit (14kDa) | 2 | 0.58 | 0.412 | 1.000 | |
| AT4G32470.1 | Ubiquinol:cytochrome C oxidoreductase subunit (14kDa) | 3 | 0.70 | 0.242 | 1.000 | |
| AT4G32470.1 | Ubiquinol:cytochrome C oxidoreductase subunit (14kDa) | 4 | 0.74 | 0.234 | 1.000 | |
| AT4G32470.1 | Ubiquinol:cytochrome C oxidoreductase subunit (14kDa) | 5 | 0.75 | 0.232 | 1.000 | |
| AT4G32470.1 | Ubiquinol:cytochrome C oxidoreductase subunit (14kDa) | 6 | 0.71 | 0.231 | 1.000 | |
| AT4G32470.1 | Ubiquinol:cytochrome C oxidoreductase subunit (14kDa) | 7 | 0.77 | 0.235 | 1.000 | |
| AT4G32470.1 | Ubiquinol:cytochrome C oxidoreductase subunit (14kDa) | 8 | 1.16 | 0.228 | 1.000 | |
| AT4G32470.1 | Ubiquinol:cytochrome C oxidoreductase subunit (14kDa) | 9 | 1.52 | 0.224 | 1.000 | |
| AT4G32470.1 | Ubiquinol:cytochrome C oxidoreductase subunit (14kDa) | 10 | 1.48 | 0.227 | 1.000 | |
| AT4G32470.1 | Ubiquinol:cytochrome C oxidoreductase subunit (14kDa) | 11 | 1.42 | 0.233 | 1.000 | |
| AT4G32470.1 | Ubiquinol:cytochrome C oxidoreductase subunit (14kDa) | 12 | 1.45 | 0.230 | 1.000 | |
| AT4G32470.1 | Ubiquinol:cytochrome C oxidoreductase subunit (14kDa) | 13 | 1.46 | 0.226 | 1.000 | |
| AT4G32470.1 | Ubiquinol:cytochrome C oxidoreductase subunit (14kDa) | 14 | 1.73 | 0.221 | 1.000 | |
| AT4G32470.1 | Ubiquinol:cytochrome C oxidoreductase subunit (14kDa) | 15 | 1.73 | 0.220 | 1.000 | |
| AT4G32470.1 | Ubiquinol:cytochrome C oxidoreductase subunit (14kDa) | 16 | 1.22 | 0.218 | 1.000 | |
| AT4G32470.1 | Ubiquinol:cytochrome C oxidoreductase subunit (14kDa) | 17 | 0.76 | 0.219 | 1.000 | |
| AT4G32470.1 | Ubiquinol:cytochrome C oxidoreductase subunit (14kDa) | 18 | 0.62 | 0.220 | 1.000 | |

| | | | | | |
|---|---|---|---|---|---|
| AT4G32470.1 | Ubiquinol:cytochrome C oxidoreductase subunit (14kDa) | 19 | 0.78 | 0.225 | 1.000 |
| AT4G32470.1 | Ubiquinol:cytochrome C oxidoreductase subunit (14kDa) | 20 | 1.15 | 0.233 | 1.000 |
| AT4G32470.1 | Ubiquinol:cytochrome C oxidoreductase subunit (14kDa) | 21 | 1.75 | 0.236 | 1.000 |
| AT4G32470.1 | Ubiquinol:cytochrome C oxidoreductase subunit (14kDa) | 22 | 1.82 | 0.235 | 1.000 |
| AT4G32470.1 | Ubiquinol:cytochrome C oxidoreductase subunit (14kDa) | 23 | 0.85 | 0.235 | 1.000 |
| AT4G32470.1 | Ubiquinol:cytochrome C oxidoreductase subunit (14kDa) | 24 | 0.41 | 0.227 | 0.936 |
| AT4G32470.1 | Ubiquinol:cytochrome C oxidoreductase subunit (14kDa) | 25 | 0.51 | 0.223 | 0.997 |
| AT4G32470.1 | Ubiquinol:cytochrome C oxidoreductase subunit (14kDa) | 26 | 1.35 | 0.225 | 1.000 |
| AT4G32470.1 | Ubiquinol:cytochrome C oxidoreductase subunit (14kDa) | 27 | 3.57 | 0.227 | 0.395 |
| AT4G32470.1 | Ubiquinol:cytochrome C oxidoreductase subunit (14kDa) | 28 | 2.12 | 0.227 | 0.991 |
| AT4G32470.1 | Ubiquinol:cytochrome C oxidoreductase subunit (14kDa) | 29 | 0.42 | 0.225 | 0.939 |
| AT4G32470.1 | Ubiquinol:cytochrome C oxidoreductase subunit (14kDa) | 30 | 0.28 | 0.224 | 0.369 |
| AT4G32470.1 | Ubiquinol:cytochrome C oxidoreductase subunit (14kDa) | 31 | 0.71 | 0.225 | 1.000 |
| AT4G32470.1 | Ubiquinol:cytochrome C oxidoreductase subunit (14kDa) | 32 | 1.76 | 0.226 | 1.000 |
| AT4G32470.1 | Ubiquinol:cytochrome C oxidoreductase subunit (14kDa) | 33 | 1.73 | 0.227 | 1.000 |
| AT4G32470.1 | Ubiquinol:cytochrome C oxidoreductase subunit (14kDa) | 34 | 0.69 | 0.232 | 1.000 |
| AT4G32470.1 | Ubiquinol:cytochrome C oxidoreductase subunit (14kDa) | 35 | 0.23 | 0.243 | 0.274 |
| AT4G32470.1 | Ubiquinol:cytochrome C oxidoreductase subunit (14kDa) | 36 | 0.26 | 0.283 | 0.688 |
| AT4G32470.1 | Ubiquinol:cytochrome C oxidoreductase subunit (14kDa) | 37 | 0.53 | 0.427 | 1.000 |
| AT4G32470.1 | Ubiquinol:cytochrome C oxidoreductase subunit (14kDa) | 41 | 3.51 | 0.364 | 0.985 |
| AT4G32470.1 | Ubiquinol:cytochrome C oxidoreductase subunit (14kDa) | 42 | 3.93 | 0.369 | 0.964 |
| AT4G32470.1 | Ubiquinol:cytochrome C oxidoreductase subunit (14kDa) | 43 | 3.77 | 0.471 | 0.999 |
| AT4G32470.1 | Ubiquinol:cytochrome C oxidoreductase subunit (14kDa) | 46 | 1.80 | 1.538 | 1.000 |
| AT4G32470.1 | Ubiquinol:cytochrome C oxidoreductase subunit (14kDa) | 47 | 1.28 | 2.017 | 1.000 |
| AT4G32915.1 | Glutamyl-tRNA (Gln) amidotransferase subunit C | 2 | 1.10 | 0.535 | 1.000 |
| AT4G32915.1 | Glutamyl-tRNA (Gln) amidotransferase subunit C | 3 | 0.95 | 0.588 | 1.000 |
| AT4G32915.1 | Glutamyl-tRNA (Gln) amidotransferase subunit C | 33 | 0.49 | 0.362 | 0.952 |
| AT4G32915.1 | Glutamyl-tRNA (Gln) amidotransferase subunit C | 34 | 0.61 | 0.311 | 0.987 |
| AT4G32915.1 | Glutamyl-tRNA (Gln) amidotransferase subunit C | 35 | 0.79 | 0.294 | 1.000 |
| AT4G32915.1 | Glutamyl-tRNA (Gln) amidotransferase subunit C | 38 | 1.74 | 0.235 | 0.895 |
| AT4G32915.1 | Glutamyl-tRNA (Gln) amidotransferase subunit C | 39 | 2.23 | 0.210 | 0.495 |

| | | | | | | |
|---|---|---|---|---|---|---|
| AT4G32915.1 | Glutamyl-tRNA (Gln) amidotransferase subunit C | 40 | 2.49 | 0.201 | 0.308 | |
| AT4G32915.1 | Glutamyl-tRNA (Gln) amidotransferase subunit C | 41 | 2.17 | 0.216 | 0.571 | |
| AT4G32915.1 | Glutamyl-tRNA (Gln) amidotransferase subunit C | 42 | 1.53 | 0.251 | 0.979 | |
| AT4G32915.1 | Glutamyl-tRNA (Gln) amidotransferase subunit C | 43 | 0.97 | 0.291 | 1.000 | |
| AT4G32915.1 | Glutamyl-tRNA (Gln) amidotransferase subunit C | 46 | 0.16 | 0.361 | 0.200 | |
| AT4G32915.1 | Glutamyl-tRNA (Gln) amidotransferase subunit C | 47 | 0.08 | 0.424 | 0.094 | . |
| AT4G33010.1 | Glycine decarboxylase subunit (GDCP1) | 3 | 0.67 | 0.188 | 1.000 | |
| AT4G33010.1 | Glycine decarboxylase subunit (GDCP1) | 4 | 0.80 | 0.175 | 1.000 | |
| AT4G33010.1 | Glycine decarboxylase subunit (GDCP1) | 5 | 0.97 | 0.176 | 1.000 | |
| AT4G33010.1 | Glycine decarboxylase subunit (GDCP1) | 6 | 1.09 | 0.176 | 1.000 | |
| AT4G33010.1 | Glycine decarboxylase subunit (GDCP1) | 7 | 1.13 | 0.175 | 1.000 | |
| AT4G33010.1 | Glycine decarboxylase subunit (GDCP1) | 8 | 1.18 | 0.173 | 1.000 | |
| AT4G33010.1 | Glycine decarboxylase subunit (GDCP1) | 9 | 1.25 | 0.171 | 1.000 | |
| AT4G33010.1 | Glycine decarboxylase subunit (GDCP1) | 10 | 1.29 | 0.171 | 1.000 | |
| AT4G33010.1 | Glycine decarboxylase subunit (GDCP1) | 11 | 1.34 | 0.171 | 1.000 | |
| AT4G33010.1 | Glycine decarboxylase subunit (GDCP1) | 12 | 1.40 | 0.171 | 1.000 | |
| AT4G33010.1 | Glycine decarboxylase subunit (GDCP1) | 13 | 1.42 | 0.170 | 1.000 | |
| AT4G33010.1 | Glycine decarboxylase subunit (GDCP1) | 14 | 1.42 | 0.168 | 1.000 | |
| AT4G33010.1 | Glycine decarboxylase subunit (GDCP1) | 15 | 1.41 | 0.168 | 1.000 | |
| AT4G33010.1 | Glycine decarboxylase subunit (GDCP1) | 16 | 1.47 | 0.167 | 1.000 | |
| AT4G33010.1 | Glycine decarboxylase subunit (GDCP1) | 17 | 1.69 | 0.166 | 0.982 | |
| AT4G33010.1 | Glycine decarboxylase subunit (GDCP1) | 18 | 1.93 | 0.169 | 0.879 | |
| AT4G33010.1 | Glycine decarboxylase subunit (GDCP1) | 19 | 1.94 | 0.172 | 0.887 | |
| AT4G33010.1 | Glycine decarboxylase subunit (GDCP1) | 20 | 1.77 | 0.175 | 0.975 | |
| AT4G33010.1 | Glycine decarboxylase subunit (GDCP1) | 21 | 1.50 | 0.177 | 1.000 | |
| AT4G33010.1 | Glycine decarboxylase subunit (GDCP1) | 22 | 1.24 | 0.177 | 1.000 | |
| AT4G33010.1 | Glycine decarboxylase subunit (GDCP1) | 23 | 1.08 | 0.176 | 1.000 | |
| AT4G33010.1 | Glycine decarboxylase subunit (GDCP1) | 24 | 1.06 | 0.173 | 1.000 | |
| AT4G33010.1 | Glycine decarboxylase subunit (GDCP1) | 25 | 1.09 | 0.171 | 1.000 | |
| AT4G33010.1 | Glycine decarboxylase subunit (GDCP1) | 26 | 1.17 | 0.170 | 1.000 | |
| AT4G33010.1 | Glycine decarboxylase subunit (GDCP1) | 27 | 1.26 | 0.170 | 1.000 | |

| | | | | | | |
|---|---|---|---|---|---|---|
| AT4G33010.1 | Glycine decarboxylase subunit (GDCP1) | 28 | 1.30 | 0.170 | 1.000 | |
| AT4G33010.1 | Glycine decarboxylase subunit (GDCP1) | 29 | 1.36 | 0.170 | 1.000 | |
| AT4G33010.1 | Glycine decarboxylase subunit (GDCP1) | 30 | 1.48 | 0.170 | 1.000 | |
| AT4G33010.1 | Glycine decarboxylase subunit (GDCP1) | 31 | 1.55 | 0.171 | 0.999 | |
| AT4G33010.1 | Glycine decarboxylase subunit (GDCP1) | 32 | 1.46 | 0.171 | 1.000 | |
| AT4G33010.1 | Glycine decarboxylase subunit (GDCP1) | 33 | 1.36 | 0.171 | 1.000 | |
| AT4G33010.1 | Glycine decarboxylase subunit (GDCP1) | 34 | 1.39 | 0.171 | 1.000 | |
| AT4G33010.1 | Glycine decarboxylase subunit (GDCP1) | 35 | 1.50 | 0.171 | 1.000 | |
| AT4G33010.1 | Glycine decarboxylase subunit (GDCP1) | 36 | 1.52 | 0.170 | 0.999 | |
| AT4G33010.1 | Glycine decarboxylase subunit (GDCP1) | 37 | 1.43 | 0.170 | 1.000 | |
| AT4G33010.1 | Glycine decarboxylase subunit (GDCP1) | 38 | 1.21 | 0.171 | 1.000 | |
| AT4G33010.1 | Glycine decarboxylase subunit (GDCP1) | 39 | 1.02 | 0.171 | 1.000 | |
| AT4G33010.1 | Glycine decarboxylase subunit (GDCP1) | 40 | 1.17 | 0.171 | 1.000 | |
| AT4G33010.1 | Glycine decarboxylase subunit (GDCP1) | 41 | 1.87 | 0.172 | 0.931 | |
| AT4G33010.1 | Glycine decarboxylase subunit (GDCP1) | 42 | 3.11 | 0.174 | 0.126 | |
| AT4G33010.1 | Glycine decarboxylase subunit (GDCP1) | 43 | 4.47 | 0.175 | 0.008 | ** |
| AT4G33010.1 | Glycine decarboxylase subunit (GDCP1) | 44 | 4.50 | 0.175 | 0.008 | ** |
| AT4G33010.1 | Glycine decarboxylase subunit (GDCP1) | 45 | 3.12 | 0.175 | 0.128 | |
| AT4G33010.1 | Glycine decarboxylase subunit (GDCP1) | 46 | 1.71 | 0.200 | 0.998 | |
| AT4G33010.1 | Glycine decarboxylase subunit (GDCP1) | 47 | 0.84 | 0.340 | 1.000 | |
| AT4G33010.1 | Glycine decarboxylase subunit (GDCP1) | 48 | 0.41 | 0.558 | 1.000 | |
| AT4G33520.3 | Heavy metal ATPase 6 (HMAC6) | 2 | 0.93 | 0.265 | 1.000 | |
| AT4G33520.3 | Heavy metal ATPase 6 (HMAC6) | 3 | 0.93 | 0.278 | 1.000 | |
| AT4G33520.3 | Heavy metal ATPase 6 (HMAC6) | 28 | 1.53 | 0.205 | 0.996 | |
| AT4G33520.3 | Heavy metal ATPase 6 (HMAC6) | 29 | 1.44 | 0.155 | 0.989 | |
| AT4G33520.3 | Heavy metal ATPase 6 (HMAC6) | 30 | 1.25 | 0.142 | 1.000 | |
| AT4G33520.3 | Heavy metal ATPase 6 (HMAC6) | 31 | 1.52 | 0.134 | 0.911 | |
| AT4G33520.3 | Heavy metal ATPase 6 (HMAC6) | 32 | 2.04 | 0.134 | 0.265 | |
| AT4G33520.3 | Heavy metal ATPase 6 (HMAC6) | 33 | 1.54 | 0.134 | 0.896 | |
| AT4G33520.3 | Heavy metal ATPase 6 (HMAC6) | 34 | 1.09 | 0.135 | 1.000 | |
| AT4G33520.3 | Heavy metal ATPase 6 (HMAC6) | 35 | 1.00 | 0.134 | 1.000 | |
| AT4G33520.3 | Heavy metal ATPase 6 (HMAC6) | 36 | 1.16 | 0.134 | 1.000 | |
| AT4G33520.3 | Heavy metal ATPase 6 (HMAC6) | 37 | 0.76 | 0.135 | 0.997 | |
| AT4G33520.3 | Heavy metal ATPase 6 (HMAC6) | 38 | 0.28 | 0.140 | 0.002 | ** |
| AT4G33520.3 | Heavy metal ATPase 6 (HMAC6) | 39 | 0.17 | 0.171 | 0.000 | *** |
| AT4G33520.3 | Heavy metal ATPase 6 (HMAC6) | 40 | 0.23 | 0.321 | 0.462 | |
| AT4G33520.3 | Heavy metal ATPase 6 (HMAC6) | 46 | 464.16 | 1.888 | 0.884 | |
| AT4G33520.3 | Heavy metal ATPase 6 (HMAC6) | 47 | 2669.29 | 2.298 | 0.842 | |
| AT4G33760.1 | Putative aspartyl-tRNA synthetase (AspRS/SYDM) | 2 | 0.96 | 0.337 | 1.000 | |

| | | | | | | |
|---|---|---|---|---|---|---|
| AT4G33760.1 | Putative aspartyl-tRNA synthetase (AspRS/SYDM) | 3 | 1.20 | 0.329 | 1.000 | |
| AT4G33760.1 | Putative aspartyl-tRNA synthetase (AspRS/SYDM) | 28 | 1.03 | 0.259 | 1.000 | |
| AT4G33760.1 | Putative aspartyl-tRNA synthetase (AspRS/SYDM) | 29 | 0.67 | 0.191 | 0.998 | |
| AT4G33760.1 | Putative aspartyl-tRNA synthetase (AspRS/SYDM) | 30 | 0.49 | 0.176 | 0.687 | |
| AT4G33760.1 | Putative aspartyl-tRNA synthetase (AspRS/SYDM) | 31 | 0.57 | 0.171 | 0.892 | |
| AT4G33760.1 | Putative aspartyl-tRNA synthetase (AspRS/SYDM) | 32 | 0.76 | 0.170 | 1.000 | |
| AT4G33760.1 | Putative aspartyl-tRNA synthetase (AspRS/SYDM) | 33 | 0.71 | 0.170 | 0.998 | |
| AT4G33760.1 | Putative aspartyl-tRNA synthetase (AspRS/SYDM) | 34 | 0.71 | 0.170 | 0.998 | |
| AT4G33760.1 | Putative aspartyl-tRNA synthetase (AspRS/SYDM) | 35 | 0.95 | 0.170 | 1.000 | |
| AT4G33760.1 | Putative aspartyl-tRNA synthetase (AspRS/SYDM) | 36 | 1.46 | 0.169 | 0.995 | |
| AT4G33760.1 | Putative aspartyl-tRNA synthetase (AspRS/SYDM) | 37 | 1.93 | 0.169 | 0.732 | |
| AT4G33760.1 | Putative aspartyl-tRNA synthetase (AspRS/SYDM) | 38 | 1.43 | 0.168 | 0.997 | |
| AT4G33760.1 | Putative aspartyl-tRNA synthetase (AspRS/SYDM) | 39 | 0.73 | 0.167 | 0.999 | |
| AT4G33760.1 | Putative aspartyl-tRNA synthetase (AspRS/SYDM) | 40 | 0.52 | 0.167 | 0.723 | |
| AT4G33760.1 | Putative aspartyl-tRNA synthetase (AspRS/SYDM) | 41 | 0.60 | 0.169 | 0.942 | |
| AT4G33760.1 | Putative aspartyl-tRNA synthetase (AspRS/SYDM) | 42 | 1.03 | 0.171 | 1.000 | |
| AT4G33760.1 | Putative aspartyl-tRNA synthetase (AspRS/SYDM) | 43 | 1.79 | 0.171 | 0.872 | |
| AT4G33760.1 | Putative aspartyl-tRNA synthetase (AspRS/SYDM) | 44 | 1.77 | 0.171 | 0.882 | |
| AT4G33760.1 | Putative aspartyl-tRNA synthetase (AspRS/SYDM) | 45 | 0.93 | 0.173 | 1.000 | |
| AT4G33760.1 | Putative aspartyl-tRNA synthetase (AspRS/SYDM) | 46 | 0.31 | 0.192 | 0.132 | |
| AT4G33760.1 | Putative aspartyl-tRNA synthetase (AspRS/SYDM) | 47 | 0.08 | 0.323 | 0.014 | * |
| AT4G34030.1 | 3-methylcrotonyl-coenzyme A carboxylase complex subunit (MCCB) | 2 | 1.43 | 0.366 | 1.000 | |
| AT4G34030.1 | 3-methylcrotonyl-coenzyme A carboxylase complex subunit (MCCB) | 3 | 0.81 | 0.264 | 1.000 | |
| AT4G34030.1 | 3-methylcrotonyl-coenzyme A carboxylase complex subunit (MCCB) | 4 | 0.52 | 0.454 | 1.000 | |
| AT4G34030.1 | 3-methylcrotonyl-coenzyme A carboxylase complex subunit (MCCB) | 19 | 2.65 | 0.239 | 0.848 | |
| AT4G34030.1 | 3-methylcrotonyl-coenzyme A carboxylase complex subunit (MCCB) | 20 | 2.88 | 0.222 | 0.627 | |
| AT4G34030.1 | 3-methylcrotonyl-coenzyme A carboxylase complex subunit (MCCB) | 21 | 2.51 | 0.192 | 0.615 | |

| | | | | | |
|---|---|---|---|---|---|
| AT4G34030.1 | 3-methylcrotonyl-coenzyme A carboxylase complex subunit (MCCB) | 22 | 1.50 | 0.188 | 1.000 |
| AT4G34030.1 | 3-methylcrotonyl-coenzyme A carboxylase complex subunit (MCCB) | 23 | 0.82 | 0.186 | 1.000 |
| AT4G34030.1 | 3-methylcrotonyl-coenzyme A carboxylase complex subunit (MCCB) | 24 | 0.69 | 0.180 | 1.000 |
| AT4G34030.1 | 3-methylcrotonyl-coenzyme A carboxylase complex subunit (MCCB) | 25 | 0.75 | 0.177 | 1.000 |
| AT4G34030.1 | 3-methylcrotonyl-coenzyme A carboxylase complex subunit (MCCB) | 26 | 0.86 | 0.177 | 1.000 |
| AT4G34030.1 | 3-methylcrotonyl-coenzyme A carboxylase complex subunit (MCCB) | 27 | 0.92 | 0.177 | 1.000 |
| AT4G34030.1 | 3-methylcrotonyl-coenzyme A carboxylase complex subunit (MCCB) | 28 | 0.87 | 0.179 | 1.000 |
| AT4G34030.1 | 3-methylcrotonyl-coenzyme A carboxylase complex subunit (MCCB) | 29 | 0.89 | 0.180 | 1.000 |
| AT4G34030.1 | 3-methylcrotonyl-coenzyme A carboxylase complex subunit (MCCB) | 30 | 0.98 | 0.179 | 1.000 |
| AT4G34030.1 | 3-methylcrotonyl-coenzyme A carboxylase complex subunit (MCCB) | 31 | 1.07 | 0.180 | 1.000 |
| AT4G34030.1 | 3-methylcrotonyl-coenzyme A carboxylase complex subunit (MCCB) | 32 | 1.10 | 0.180 | 1.000 |
| AT4G34030.1 | 3-methylcrotonyl-coenzyme A carboxylase complex subunit (MCCB) | 33 | 1.05 | 0.180 | 1.000 |
| AT4G34030.1 | 3-methylcrotonyl-coenzyme A carboxylase complex subunit (MCCB) | 34 | 0.94 | 0.181 | 1.000 |
| AT4G34030.1 | 3-methylcrotonyl-coenzyme A carboxylase complex subunit (MCCB) | 35 | 0.85 | 0.180 | 1.000 |
| AT4G34030.1 | 3-methylcrotonyl-coenzyme A carboxylase complex subunit (MCCB) | 36 | 0.90 | 0.178 | 1.000 |
| AT4G34030.1 | 3-methylcrotonyl-coenzyme A carboxylase complex subunit (MCCB) | 37 | 1.29 | 0.177 | 1.000 |
| AT4G34030.1 | 3-methylcrotonyl-coenzyme A carboxylase complex subunit (MCCB) | 38 | 1.91 | 0.178 | 0.938 |
| AT4G34030.1 | 3-methylcrotonyl-coenzyme A carboxylase complex subunit (MCCB) | 39 | 2.48 | 0.178 | 0.504 |
| AT4G34030.1 | 3-methylcrotonyl-coenzyme A carboxylase complex subunit (MCCB) | 40 | 2.34 | 0.179 | 0.641 |
| AT4G34030.1 | 3-methylcrotonyl-coenzyme A carboxylase complex subunit (MCCB) | 41 | 1.25 | 0.180 | 1.000 |

| | | | | | |
|---|---|---|---|---|---|
| AT4G34030.1 | 3-methylcrotonyl-coenzyme A carboxylase complex subunit (MCCB) | 42 | 0.54 | 0.184 | 0.973 |
| AT4G34030.1 | 3-methylcrotonyl-coenzyme A carboxylase complex subunit (MCCB) | 43 | 0.44 | 0.185 | 0.756 |
| AT4G34030.1 | 3-methylcrotonyl-coenzyme A carboxylase complex subunit (MCCB) | 44 | 0.69 | 0.184 | 1.000 |
| AT4G34030.1 | 3-methylcrotonyl-coenzyme A carboxylase complex subunit (MCCB) | 45 | 1.07 | 0.185 | 1.000 |
| AT4G34030.1 | 3-methylcrotonyl-coenzyme A carboxylase complex subunit (MCCB) | 46 | 1.16 | 0.205 | 1.000 |
| AT4G34030.1 | 3-methylcrotonyl-coenzyme A carboxylase complex subunit (MCCB) | 47 | 1.08 | 0.433 | 1.000 |
| AT4G34310.1 | Alpha/beta-Hydrolases superfamily protein | 2 | 0.89 | 0.331 | 1.000 |
| AT4G34310.1 | Alpha/beta-Hydrolases superfamily protein | 3 | 0.89 | 0.321 | 1.000 |
| AT4G34310.1 | Alpha/beta-Hydrolases superfamily protein | 26 | 0.62 | 0.239 | 0.984 |
| AT4G34310.1 | Alpha/beta-Hydrolases superfamily protein | 27 | 0.60 | 0.169 | 0.848 |
| AT4G34310.1 | Alpha/beta-Hydrolases superfamily protein | 28 | 0.65 | 0.144 | 0.840 |
| AT4G34310.1 | Alpha/beta-Hydrolases superfamily protein | 29 | 0.80 | 0.135 | 0.996 |
| AT4G34310.1 | Alpha/beta-Hydrolases superfamily protein | 30 | 1.01 | 0.133 | 1.000 |
| AT4G34310.1 | Alpha/beta-Hydrolases superfamily protein | 31 | 1.32 | 0.133 | 0.979 |
| AT4G34310.1 | Alpha/beta-Hydrolases superfamily protein | 32 | 1.61 | 0.134 | 0.699 |
| AT4G34310.1 | Alpha/beta-Hydrolases superfamily protein | 33 | 1.60 | 0.137 | 0.738 |
| AT4G34310.1 | Alpha/beta-Hydrolases superfamily protein | 34 | 1.40 | 0.148 | 0.966 |
| AT4G34310.1 | Alpha/beta-Hydrolases superfamily protein | 35 | 1.24 | 0.189 | 1.000 |
| AT4G34310.1 | Alpha/beta-Hydrolases superfamily protein | 36 | 1.19 | 0.287 | 1.000 |
| AT4G34310.1 | Alpha/beta-Hydrolases superfamily protein | 46 | 15.75 | 2.156 | 0.999 |
| AT4G34310.1 | Alpha/beta-Hydrolases superfamily protein | 47 | 23.69 | 2.442 | 0.999 |
| AT4G34570.1 | Thymidylate synthase 2 (THY2) | 2 | 1.23 | 0.477 | 1.000 |
| AT4G34570.1 | Thymidylate synthase 2 (THY2) | 3 | 1.20 | 0.419 | 1.000 |
| AT4G34570.1 | Thymidylate synthase 2 (THY2) | 6 | 1.11 | 0.369 | 1.000 |
| AT4G34570.1 | Thymidylate synthase 2 (THY2) | 7 | 1.09 | 0.379 | 1.000 |
| AT4G34570.1 | Thymidylate synthase 2 (THY2) | 8 | 1.08 | 0.416 | 1.000 |
| AT4G34570.1 | Thymidylate synthase 2 (THY2) | 22 | 0.60 | 0.405 | 1.000 |
| AT4G34570.1 | Thymidylate synthase 2 (THY2) | 23 | 0.52 | 0.373 | 0.998 |
| AT4G34570.1 | Thymidylate synthase 2 (THY2) | 24 | 0.45 | 0.353 | 0.985 |
| AT4G34570.1 | Thymidylate synthase 2 (THY2) | 30 | 0.29 | 0.275 | 0.484 |
| AT4G34570.1 | Thymidylate synthase 2 (THY2) | 31 | 0.35 | 0.224 | 0.413 |
| AT4G34570.1 | Thymidylate synthase 2 (THY2) | 32 | 0.47 | 0.194 | 0.680 |
| AT4G34570.1 | Thymidylate synthase 2 (THY2) | 33 | 0.74 | 0.182 | 0.999 |
| AT4G34570.1 | Thymidylate synthase 2 (THY2) | 34 | 1.31 | 0.178 | 1.000 |
| AT4G34570.1 | Thymidylate synthase 2 (THY2) | 35 | 2.28 | 0.175 | 0.410 |

| | | | | | | |
|---|---|---|---|---|---|---|
| AT4G34570.1 | Thymidylate synthase 2 (THY2) | 36 | 3.30 | 0.174 | 0.047 | * |
| AT4G34570.1 | Thymidylate synthase 2 (THY2) | 37 | 3.44 | 0.173 | 0.034 | * |
| AT4G34570.1 | Thymidylate synthase 2 (THY2) | 38 | 2.48 | 0.173 | 0.269 | |
| AT4G34570.1 | Thymidylate synthase 2 (THY2) | 39 | 1.38 | 0.176 | 0.997 | |
| AT4G34570.1 | Thymidylate synthase 2 (THY2) | 40 | 0.66 | 0.184 | 0.986 | |
| AT4G34570.1 | Thymidylate synthase 2 (THY2) | 41 | 0.32 | 0.206 | 0.212 | |
| AT4G34570.1 | Thymidylate synthase 2 (THY2) | 42 | 0.18 | 0.247 | 0.050 | . |
| AT4G34570.1 | Thymidylate synthase 2 (THY2) | 43 | 0.13 | 0.291 | 0.035 | * |
| AT4G34570.1 | Thymidylate synthase 2 (THY2) | 46 | 0.08 | 0.333 | 0.024 | * |
| AT4G34570.1 | Thymidylate synthase 2 (THY2) | 47 | 0.08 | 0.396 | 0.094 | . |
| AT4G34700.1 | NADH:ubiquinone oxidoreductase subunit (B22) | 2 | 0.95 | 0.439 | 1.000 | |
| AT4G34700.1 | NADH:ubiquinone oxidoreductase subunit (B22) | 3 | 0.85 | 0.305 | 1.000 | |
| AT4G34700.1 | NADH:ubiquinone oxidoreductase subunit (B22) | 4 | 0.79 | 0.274 | 1.000 | |
| AT4G34700.1 | NADH:ubiquinone oxidoreductase subunit (B22) | 5 | 0.81 | 0.273 | 1.000 | |
| AT4G34700.1 | NADH:ubiquinone oxidoreductase subunit (B22) | 6 | 0.92 | 0.275 | 1.000 | |
| AT4G34700.1 | NADH:ubiquinone oxidoreductase subunit (B22) | 7 | 1.17 | 0.277 | 1.000 | |
| AT4G34700.1 | NADH:ubiquinone oxidoreductase subunit (B22) | 8 | 1.46 | 0.279 | 1.000 | |
| AT4G34700.1 | NADH:ubiquinone oxidoreductase subunit (B22) | 9 | 1.57 | 0.279 | 1.000 | |
| AT4G34700.1 | NADH:ubiquinone oxidoreductase subunit (B22) | 10 | 1.37 | 0.277 | 1.000 | |
| AT4G34700.1 | NADH:ubiquinone oxidoreductase subunit (B22) | 11 | 1.11 | 0.278 | 1.000 | |
| AT4G34700.1 | NADH:ubiquinone oxidoreductase subunit (B22) | 12 | 1.02 | 0.278 | 1.000 | |
| AT4G34700.1 | NADH:ubiquinone oxidoreductase subunit (B22) | 13 | 1.03 | 0.272 | 1.000 | |
| AT4G34700.1 | NADH:ubiquinone oxidoreductase subunit (B22) | 14 | 0.96 | 0.265 | 1.000 | |
| AT4G34700.1 | NADH:ubiquinone oxidoreductase subunit (B22) | 15 | 0.75 | 0.261 | 1.000 | |
| AT4G34700.1 | NADH:ubiquinone oxidoreductase subunit (B22) | 16 | 0.54 | 0.260 | 0.999 | |
| AT4G34700.1 | NADH:ubiquinone oxidoreductase subunit (B22) | 17 | 0.45 | 0.260 | 0.972 | |
| AT4G34700.1 | NADH:ubiquinone oxidoreductase subunit (B22) | 18 | 0.48 | 0.262 | 0.989 | |
| AT4G34700.1 | NADH:ubiquinone oxidoreductase subunit (B22) | 19 | 0.61 | 0.267 | 1.000 | |
| AT4G34700.1 | NADH:ubiquinone oxidoreductase subunit (B22) | 20 | 0.80 | 0.272 | 1.000 | |
| AT4G34700.1 | NADH:ubiquinone oxidoreductase subunit (B22) | 21 | 1.05 | 0.273 | 1.000 | |
| AT4G34700.1 | NADH:ubiquinone oxidoreductase subunit (B22) | 22 | 1.32 | 0.274 | 1.000 | |
| AT4G34700.1 | NADH:ubiquinone oxidoreductase subunit (B22) | 23 | 1.44 | 0.272 | 1.000 | |
| AT4G34700.1 | NADH:ubiquinone oxidoreductase subunit (B22) | 24 | 1.21 | 0.268 | 1.000 | |
| AT4G34700.1 | NADH:ubiquinone oxidoreductase subunit (B22) | 25 | 0.84 | 0.265 | 1.000 | |
| AT4G34700.1 | NADH:ubiquinone oxidoreductase subunit (B22) | 26 | 0.58 | 0.264 | 1.000 | |
| AT4G34700.1 | NADH:ubiquinone oxidoreductase subunit (B22) | 27 | 0.50 | 0.264 | 0.994 | |

| | | | | | | |
|---|---|---|---|---|---|---|
| AT4G34700.1 | NADH:ubiquinone oxidoreductase subunit (B22) | 28 | 0.57 | 0.264 | 1.000 | |
| AT4G34700.1 | NADH:ubiquinone oxidoreductase subunit (B22) | 29 | 0.82 | 0.265 | 1.000 | |
| AT4G34700.1 | NADH:ubiquinone oxidoreductase subunit (B22) | 30 | 1.17 | 0.266 | 1.000 | |
| AT4G34700.1 | NADH:ubiquinone oxidoreductase subunit (B22) | 31 | 1.47 | 0.266 | 1.000 | |
| AT4G34700.1 | NADH:ubiquinone oxidoreductase subunit (B22) | 32 | 1.74 | 0.267 | 1.000 | |
| AT4G34700.1 | NADH:ubiquinone oxidoreductase subunit (B22) | 33 | 2.05 | 0.267 | 0.993 | |
| AT4G34700.1 | NADH:ubiquinone oxidoreductase subunit (B22) | 34 | 2.28 | 0.267 | 0.971 | |
| AT4G34700.1 | NADH:ubiquinone oxidoreductase subunit (B22) | 35 | 2.44 | 0.267 | 0.941 | |
| AT4G34700.1 | NADH:ubiquinone oxidoreductase subunit (B22) | 36 | 2.71 | 0.266 | 0.858 | |
| AT4G34700.1 | NADH:ubiquinone oxidoreductase subunit (B22) | 37 | 3.14 | 0.266 | 0.691 | |
| AT4G34700.1 | NADH:ubiquinone oxidoreductase subunit (B22) | 38 | 3.83 | 0.266 | 0.437 | |
| AT4G34700.1 | NADH:ubiquinone oxidoreductase subunit (B22) | 39 | 4.67 | 0.266 | 0.232 | |
| AT4G34700.1 | NADH:ubiquinone oxidoreductase subunit (B22) | 40 | 5.35 | 0.266 | 0.138 | |
| AT4G34700.1 | NADH:ubiquinone oxidoreductase subunit (B22) | 41 | 6.16 | 0.267 | 0.079 | . |
| AT4G34700.1 | NADH:ubiquinone oxidoreductase subunit (B22) | 42 | 7.62 | 0.270 | 0.031 | * |
| AT4G34700.1 | NADH:ubiquinone oxidoreductase subunit (B22) | 43 | 8.62 | 0.272 | 0.019 | * |
| AT4G34700.1 | NADH:ubiquinone oxidoreductase subunit (B22) | 44 | 7.91 | 0.272 | 0.027 | * |
| AT4G34700.1 | NADH:ubiquinone oxidoreductase subunit (B22) | 45 | 6.23 | 0.275 | 0.094 | . |
| AT4G34700.1 | NADH:ubiquinone oxidoreductase subunit (B22) | 46 | 4.84 | 0.322 | 0.484 | |
| AT4G34700.1 | NADH:ubiquinone oxidoreductase subunit (B22) | 47 | 3.87 | 0.486 | 0.990 | |
| AT4G34870.1 | Rotamase cyclophilin 5 (ROC5) | 2 | 1.07 | 0.360 | 1.000 | |
| AT4G34870.1 | Rotamase cyclophilin 5 (ROC5) | 3 | 1.25 | 0.285 | 1.000 | |
| AT4G34870.1 | Rotamase cyclophilin 5 (ROC5) | 4 | 1.48 | 0.241 | 1.000 | |
| AT4G34870.1 | Rotamase cyclophilin 5 (ROC5) | 5 | 1.76 | 0.221 | 0.989 | |
| AT4G34870.1 | Rotamase cyclophilin 5 (ROC5) | 6 | 2.14 | 0.211 | 0.856 | |
| AT4G34870.1 | Rotamase cyclophilin 5 (ROC5) | 7 | 2.65 | 0.206 | 0.505 | |
| AT4G34870.1 | Rotamase cyclophilin 5 (ROC5) | 8 | 3.36 | 0.201 | 0.163 | |
| AT4G34870.1 | Rotamase cyclophilin 5 (ROC5) | 9 | 4.31 | 0.194 | 0.026 | * |
| AT4G34870.1 | Rotamase cyclophilin 5 (ROC5) | 10 | 5.42 | 0.186 | 0.003 | ** |
| AT4G34870.1 | Rotamase cyclophilin 5 (ROC5) | 11 | 6.42 | 0.178 | 0.000 | *** |
| AT4G34870.1 | Rotamase cyclophilin 5 (ROC5) | 12 | 7.02 | 0.170 | 0.000 | *** |
| AT4G34870.1 | Rotamase cyclophilin 5 (ROC5) | 13 | 7.06 | 0.163 | 0.000 | *** |
| AT4G34870.1 | Rotamase cyclophilin 5 (ROC5) | 14 | 6.62 | 0.159 | 0.000 | *** |
| AT4G34870.1 | Rotamase cyclophilin 5 (ROC5) | 15 | 5.95 | 0.157 | 0.000 | *** |
| AT4G34870.1 | Rotamase cyclophilin 5 (ROC5) | 16 | 5.29 | 0.157 | 0.000 | *** |
| AT4G34870.1 | Rotamase cyclophilin 5 (ROC5) | 17 | 4.73 | 0.159 | 0.001 | *** |
| AT4G34870.1 | Rotamase cyclophilin 5 (ROC5) | 18 | 4.29 | 0.163 | 0.003 | ** |
| AT4G34870.1 | Rotamase cyclophilin 5 (ROC5) | 19 | 3.99 | 0.167 | 0.009 | ** |
| AT4G34870.1 | Rotamase cyclophilin 5 (ROC5) | 20 | 3.78 | 0.172 | 0.019 | * |
| AT4G34870.1 | Rotamase cyclophilin 5 (ROC5) | 21 | 3.60 | 0.174 | 0.034 | * |
| AT4G34870.1 | Rotamase cyclophilin 5 (ROC5) | 22 | 3.38 | 0.173 | 0.050 | . |
| AT4G34870.1 | Rotamase cyclophilin 5 (ROC5) | 23 | 3.10 | 0.170 | 0.082 | . |

| | | | | | |
|---|---|---|---|---|---|
| AT4G34870.1 | Rotamase cyclophilin 5 (ROC5) | 24 | 2.74 | 0.166 | 0.156 |
| AT4G34870.1 | Rotamase cyclophilin 5 (ROC5) | 25 | 2.34 | 0.162 | 0.343 |
| AT4G34870.1 | Rotamase cyclophilin 5 (ROC5) | 26 | 1.97 | 0.160 | 0.668 |
| AT4G34870.1 | Rotamase cyclophilin 5 (ROC5) | 27 | 1.66 | 0.158 | 0.932 |
| AT4G34870.1 | Rotamase cyclophilin 5 (ROC5) | 28 | 1.44 | 0.158 | 0.996 |
| AT4G34870.1 | Rotamase cyclophilin 5 (ROC5) | 29 | 1.29 | 0.158 | 1.000 |
| AT4G34870.1 | Rotamase cyclophilin 5 (ROC5) | 30 | 1.18 | 0.158 | 1.000 |
| AT4G34870.1 | Rotamase cyclophilin 5 (ROC5) | 31 | 1.07 | 0.159 | 1.000 |
| AT4G34870.1 | Rotamase cyclophilin 5 (ROC5) | 32 | 0.96 | 0.159 | 1.000 |
| AT4G34870.1 | Rotamase cyclophilin 5 (ROC5) | 33 | 0.86 | 0.160 | 1.000 |
| AT4G34870.1 | Rotamase cyclophilin 5 (ROC5) | 34 | 0.79 | 0.162 | 1.000 |
| AT4G34870.1 | Rotamase cyclophilin 5 (ROC5) | 35 | 0.75 | 0.165 | 1.000 |
| AT4G34870.1 | Rotamase cyclophilin 5 (ROC5) | 36 | 0.75 | 0.168 | 1.000 |
| AT4G34870.1 | Rotamase cyclophilin 5 (ROC5) | 37 | 0.74 | 0.171 | 1.000 |
| AT4G34870.1 | Rotamase cyclophilin 5 (ROC5) | 38 | 0.73 | 0.173 | 1.000 |
| AT4G34870.1 | Rotamase cyclophilin 5 (ROC5) | 39 | 0.68 | 0.173 | 0.998 |
| AT4G34870.1 | Rotamase cyclophilin 5 (ROC5) | 40 | 0.60 | 0.171 | 0.962 |
| AT4G34870.1 | Rotamase cyclophilin 5 (ROC5) | 41 | 0.50 | 0.171 | 0.718 |
| AT4G34870.1 | Rotamase cyclophilin 5 (ROC5) | 42 | 0.40 | 0.176 | 0.365 |
| AT4G34870.1 | Rotamase cyclophilin 5 (ROC5) | 43 | 0.33 | 0.195 | 0.224 |
| AT4G34870.1 | Rotamase cyclophilin 5 (ROC5) | 44 | 0.27 | 0.238 | 0.273 |
| AT4G34870.1 | Rotamase cyclophilin 5 (ROC5) | 45 | 0.23 | 0.306 | 0.460 |
| AT4G34870.1 | Rotamase cyclophilin 5 (ROC5) | 46 | 0.19 | 0.401 | 0.699 |
| AT4G34870.1 | Rotamase cyclophilin 5 (ROC5) | 47 | 0.16 | 0.519 | 0.870 |
| AT4G35000.1 | Ascorbate peroxidase 3 (APX3) | 2 | 0.96 | 0.447 | 1.000 |
| AT4G35000.1 | Ascorbate peroxidase 3 (APX3) | 3 | 1.01 | 0.354 | 1.000 |
| AT4G35000.1 | Ascorbate peroxidase 3 (APX3) | 4 | 1.05 | 0.307 | 1.000 |
| AT4G35000.1 | Ascorbate peroxidase 3 (APX3) | 5 | 1.06 | 0.293 | 1.000 |
| AT4G35000.1 | Ascorbate peroxidase 3 (APX3) | 6 | 1.03 | 0.299 | 1.000 |
| AT4G35000.1 | Ascorbate peroxidase 3 (APX3) | 7 | 0.96 | 0.314 | 1.000 |
| AT4G35000.1 | Ascorbate peroxidase 3 (APX3) | 8 | 0.86 | 0.331 | 1.000 |
| AT4G35000.1 | Ascorbate peroxidase 3 (APX3) | 12 | 0.54 | 0.311 | 0.997 |
| AT4G35000.1 | Ascorbate peroxidase 3 (APX3) | 13 | 0.53 | 0.293 | 0.992 |
| AT4G35000.1 | Ascorbate peroxidase 3 (APX3) | 14 | 0.54 | 0.277 | 0.992 |
| AT4G35000.1 | Ascorbate peroxidase 3 (APX3) | 15 | 0.60 | 0.265 | 0.998 |
| AT4G35000.1 | Ascorbate peroxidase 3 (APX3) | 16 | 0.71 | 0.256 | 1.000 |
| AT4G35000.1 | Ascorbate peroxidase 3 (APX3) | 17 | 0.88 | 0.252 | 1.000 |
| AT4G35000.1 | Ascorbate peroxidase 3 (APX3) | 18 | 1.13 | 0.252 | 1.000 |
| AT4G35000.1 | Ascorbate peroxidase 3 (APX3) | 19 | 1.45 | 0.254 | 1.000 |
| AT4G35000.1 | Ascorbate peroxidase 3 (APX3) | 20 | 1.79 | 0.257 | 0.989 |
| AT4G35000.1 | Ascorbate peroxidase 3 (APX3) | 21 | 2.04 | 0.259 | 0.951 |
| AT4G35000.1 | Ascorbate peroxidase 3 (APX3) | 22 | 2.07 | 0.259 | 0.943 |
| AT4G35000.1 | Ascorbate peroxidase 3 (APX3) | 23 | 1.89 | 0.258 | 0.978 |
| AT4G35000.1 | Ascorbate peroxidase 3 (APX3) | 24 | 1.56 | 0.256 | 0.999 |
| AT4G35000.1 | Ascorbate peroxidase 3 (APX3) | 25 | 1.19 | 0.254 | 1.000 |
| AT4G35000.1 | Ascorbate peroxidase 3 (APX3) | 26 | 0.91 | 0.253 | 1.000 |
| AT4G35000.1 | Ascorbate peroxidase 3 (APX3) | 27 | 0.71 | 0.252 | 1.000 |
| AT4G35000.1 | Ascorbate peroxidase 3 (APX3) | 28 | 0.58 | 0.252 | 0.992 |
| AT4G35000.1 | Ascorbate peroxidase 3 (APX3) | 29 | 0.51 | 0.252 | 0.964 |
| AT4G35000.1 | Ascorbate peroxidase 3 (APX3) | 30 | 0.51 | 0.253 | 0.964 |
| AT4G35000.1 | Ascorbate peroxidase 3 (APX3) | 31 | 0.58 | 0.253 | 0.993 |
| AT4G35000.1 | Ascorbate peroxidase 3 (APX3) | 32 | 0.75 | 0.254 | 1.000 |
| AT4G35000.1 | Ascorbate peroxidase 3 (APX3) | 33 | 1.08 | 0.255 | 1.000 |
| AT4G35000.1 | Ascorbate peroxidase 3 (APX3) | 34 | 1.61 | 0.259 | 0.999 |
| AT4G35000.1 | Ascorbate peroxidase 3 (APX3) | 35 | 2.24 | 0.268 | 0.910 |
| AT4G35000.1 | Ascorbate peroxidase 3 (APX3) | 36 | 2.79 | 0.285 | 0.766 |
| AT4G35000.1 | Ascorbate peroxidase 3 (APX3) | 37 | 3.14 | 0.310 | 0.737 |
| AT4G35000.1 | Ascorbate peroxidase 3 (APX3) | 38 | 3.38 | 0.334 | 0.753 |
| AT4G35000.1 | Ascorbate peroxidase 3 (APX3) | 42 | 3.78 | 0.358 | 0.731 |
| AT4G35000.1 | Ascorbate peroxidase 3 (APX3) | 43 | 3.82 | 0.348 | 0.689 |
| AT4G35000.1 | Ascorbate peroxidase 3 (APX3) | 44 | 3.79 | 0.348 | 0.696 |
| AT4G35000.1 | Ascorbate peroxidase 3 (APX3) | 45 | 3.63 | 0.373 | 0.808 |
| AT4G35000.1 | Ascorbate peroxidase 3 (APX3) | 46 | 3.28 | 0.436 | 0.954 |

| | | | | | | |
|---|---|---|---|---|---|---|
| AT4G35000.1 | Ascorbate peroxidase 3 (APX3) | 47 | 2.87 | 0.543 | 0.998 | |
| AT4G35090.1 | Catalase 2 (Cat2) | 2 | 0.59 | 0.573 | 1.000 | |
| AT4G35090.1 | Catalase 2 (Cat2) | 3 | 0.47 | 0.461 | 0.999 | |
| AT4G35090.1 | Catalase 2 (Cat2) | 4 | 0.37 | 0.431 | 0.967 | |
| AT4G35090.1 | Catalase 2 (Cat2) | 5 | 0.29 | 0.428 | 0.856 | |
| AT4G35090.1 | Catalase 2 (Cat2) | 6 | 0.22 | 0.429 | 0.672 | |
| AT4G35090.1 | Catalase 2 (Cat2) | 7 | 0.17 | 0.428 | 0.454 | |
| AT4G35090.1 | Catalase 2 (Cat2) | 8 | 0.12 | 0.425 | 0.251 | |
| AT4G35090.1 | Catalase 2 (Cat2) | 9 | 0.08 | 0.423 | 0.110 | |
| AT4G35090.1 | Catalase 2 (Cat2) | 10 | 0.06 | 0.423 | 0.043 | * |
| AT4G35090.1 | Catalase 2 (Cat2) | 11 | 0.04 | 0.423 | 0.018 | * |
| AT4G35090.1 | Catalase 2 (Cat2) | 12 | 0.03 | 0.423 | 0.008 | ** |
| AT4G35090.1 | Catalase 2 (Cat2) | 13 | 0.03 | 0.422 | 0.005 | ** |
| AT4G35090.1 | Catalase 2 (Cat2) | 14 | 0.03 | 0.421 | 0.004 | ** |
| AT4G35090.1 | Catalase 2 (Cat2) | 15 | 0.03 | 0.421 | 0.005 | ** |
| AT4G35090.1 | Catalase 2 (Cat2) | 16 | 0.04 | 0.421 | 0.009 | ** |
| AT4G35090.1 | Catalase 2 (Cat2) | 17 | 0.04 | 0.421 | 0.020 | * |
| AT4G35090.1 | Catalase 2 (Cat2) | 18 | 0.05 | 0.423 | 0.038 | * |
| AT4G35090.1 | Catalase 2 (Cat2) | 19 | 0.06 | 0.427 | 0.059 | . |
| AT4G35090.1 | Catalase 2 (Cat2) | 20 | 0.07 | 0.430 | 0.071 | . |
| AT4G35090.1 | Catalase 2 (Cat2) | 21 | 0.06 | 0.432 | 0.063 | . |
| AT4G35090.1 | Catalase 2 (Cat2) | 22 | 0.06 | 0.433 | 0.049 | * |
| AT4G35090.1 | Catalase 2 (Cat2) | 23 | 0.05 | 0.431 | 0.045 | * |
| AT4G35090.1 | Catalase 2 (Cat2) | 24 | 0.07 | 0.428 | 0.067 | . |
| AT4G35090.1 | Catalase 2 (Cat2) | 25 | 0.10 | 0.425 | 0.180 | |
| AT4G35090.1 | Catalase 2 (Cat2) | 26 | 0.19 | 0.424 | 0.550 | |
| AT4G35090.1 | Catalase 2 (Cat2) | 27 | 0.40 | 0.423 | 0.980 | |
| AT4G35090.1 | Catalase 2 (Cat2) | 28 | 0.71 | 0.423 | 1.000 | |
| AT4G35090.1 | Catalase 2 (Cat2) | 29 | 1.02 | 0.423 | 1.000 | |
| AT4G35090.1 | Catalase 2 (Cat2) | 30 | 1.34 | 0.423 | 1.000 | |
| AT4G35090.1 | Catalase 2 (Cat2) | 31 | 1.73 | 0.423 | 1.000 | |
| AT4G35090.1 | Catalase 2 (Cat2) | 32 | 2.04 | 0.424 | 0.998 | |
| AT4G35090.1 | Catalase 2 (Cat2) | 33 | 1.98 | 0.424 | 0.999 | |
| AT4G35090.1 | Catalase 2 (Cat2) | 34 | 1.73 | 0.425 | 1.000 | |
| AT4G35090.1 | Catalase 2 (Cat2) | 35 | 1.44 | 0.430 | 1.000 | |
| AT4G35090.1 | Catalase 2 (Cat2) | 36 | 1.11 | 0.444 | 1.000 | |
| AT4G35090.1 | Catalase 2 (Cat2) | 37 | 0.87 | 0.469 | 1.000 | |
| AT4G35090.1 | Catalase 2 (Cat2) | 38 | 0.76 | 0.502 | 1.000 | |
| AT4G35090.1 | Catalase 2 (Cat2) | 41 | 1.03 | 0.494 | 1.000 | |
| AT4G35090.1 | Catalase 2 (Cat2) | 42 | 1.38 | 0.483 | 1.000 | |
| AT4G35090.1 | Catalase 2 (Cat2) | 43 | 1.93 | 0.505 | 1.000 | |
| AT4G35090.1 | Catalase 2 (Cat2) | 44 | 2.71 | 0.568 | 0.998 | |
| AT4G35090.1 | Catalase 2 (Cat2) | 46 | 5.17 | 0.864 | 0.994 | |
| AT4G35090.1 | Catalase 2 (Cat2) | 47 | 7.11 | 1.103 | 0.997 | |
| AT4G35100.2 | Plasma membrane intrinsic protein 2-7 (PIP2-7) | 2 | 0.33 | 0.246 | 0.508 | |
| AT4G35100.2 | Plasma membrane intrinsic protein 2-7 (PIP2-7) | 3 | 0.46 | 0.186 | 0.630 | |
| AT4G35100.2 | Plasma membrane intrinsic protein 2-7 (PIP2-7) | 4 | 0.65 | 0.153 | 0.956 | |
| AT4G35100.2 | Plasma membrane intrinsic protein 2-7 (PIP2-7) | 5 | 0.90 | 0.141 | 1.000 | |
| AT4G35100.2 | Plasma membrane intrinsic protein 2-7 (PIP2-7) | 6 | 1.19 | 0.139 | 1.000 | |
| AT4G35100.2 | Plasma membrane intrinsic protein 2-7 (PIP2-7) | 7 | 1.44 | 0.139 | 0.974 | |
| AT4G35100.2 | Plasma membrane intrinsic protein 2-7 (PIP2-7) | 8 | 1.62 | 0.139 | 0.840 | |
| AT4G35100.2 | Plasma membrane intrinsic protein 2-7 (PIP2-7) | 9 | 1.71 | 0.140 | 0.734 | |
| AT4G35100.2 | Plasma membrane intrinsic protein 2-7 (PIP2-7) | 10 | 1.72 | 0.143 | 0.743 | |

| | | | | | |
|---|---|---|---|---|---|
| AT4G35100.2 | Plasma membrane intrinsic protein 2-7 (PIP2-7) | 11 | 1.67 | 0.145 | 0.821 |
| AT4G35100.2 | Plasma membrane intrinsic protein 2-7 (PIP2-7) | 12 | 1.59 | 0.148 | 0.908 |
| AT4G35100.2 | Plasma membrane intrinsic protein 2-7 (PIP2-7) | 13 | 1.49 | 0.150 | 0.973 |
| AT4G35100.2 | Plasma membrane intrinsic protein 2-7 (PIP2-7) | 14 | 1.36 | 0.152 | 0.998 |
| AT4G35100.2 | Plasma membrane intrinsic protein 2-7 (PIP2-7) | 15 | 1.22 | 0.155 | 1.000 |
| AT4G35100.2 | Plasma membrane intrinsic protein 2-7 (PIP2-7) | 16 | 1.08 | 0.158 | 1.000 |
| AT4G35100.2 | Plasma membrane intrinsic protein 2-7 (PIP2-7) | 17 | 0.96 | 0.161 | 1.000 |
| AT4G35100.2 | Plasma membrane intrinsic protein 2-7 (PIP2-7) | 18 | 0.87 | 0.165 | 1.000 |
| AT4G35100.2 | Plasma membrane intrinsic protein 2-7 (PIP2-7) | 19 | 0.80 | 0.171 | 1.000 |
| AT4G35100.2 | Plasma membrane intrinsic protein 2-7 (PIP2-7) | 20 | 0.78 | 0.179 | 1.000 |
| AT4G35100.2 | Plasma membrane intrinsic protein 2-7 (PIP2-7) | 22 | 0.79 | 0.187 | 1.000 |
| AT4G35100.2 | Plasma membrane intrinsic protein 2-7 (PIP2-7) | 23 | 0.81 | 0.185 | 1.000 |
| AT4G35100.2 | Plasma membrane intrinsic protein 2-7 (PIP2-7) | 24 | 0.85 | 0.176 | 1.000 |
| AT4G35100.2 | Plasma membrane intrinsic protein 2-7 (PIP2-7) | 25 | 0.88 | 0.165 | 1.000 |
| AT4G35100.2 | Plasma membrane intrinsic protein 2-7 (PIP2-7) | 26 | 0.91 | 0.154 | 1.000 |
| AT4G35100.2 | Plasma membrane intrinsic protein 2-7 (PIP2-7) | 27 | 0.93 | 0.145 | 1.000 |
| AT4G35100.2 | Plasma membrane intrinsic protein 2-7 (PIP2-7) | 28 | 0.96 | 0.138 | 1.000 |
| AT4G35100.2 | Plasma membrane intrinsic protein 2-7 (PIP2-7) | 29 | 1.00 | 0.135 | 1.000 |
| AT4G35100.2 | Plasma membrane intrinsic protein 2-7 (PIP2-7) | 30 | 1.03 | 0.135 | 1.000 |
| AT4G35100.2 | Plasma membrane intrinsic protein 2-7 (PIP2-7) | 31 | 1.06 | 0.136 | 1.000 |
| AT4G35100.2 | Plasma membrane intrinsic protein 2-7 (PIP2-7) | 32 | 1.08 | 0.140 | 1.000 |
| AT4G35100.2 | Plasma membrane intrinsic protein 2-7 (PIP2-7) | 33 | 1.07 | 0.148 | 1.000 |
| AT4G35100.2 | Plasma membrane intrinsic protein 2-7 (PIP2-7) | 34 | 1.03 | 0.161 | 1.000 |
| AT4G35100.2 | Plasma membrane intrinsic protein 2-7 (PIP2-7) | 35 | 0.99 | 0.180 | 1.000 |
| AT4G35100.2 | Plasma membrane intrinsic protein 2-7 (PIP2-7) | 36 | 0.96 | 0.202 | 1.000 |
| AT4G35100.2 | Plasma membrane intrinsic protein 2-7 (PIP2-7) | 41 | 1.03 | 0.264 | 1.000 |
| AT4G35100.2 | Plasma membrane intrinsic protein 2-7 (PIP2-7) | 42 | 1.12 | 0.286 | 1.000 |
| AT4G35100.2 | Plasma membrane intrinsic protein 2-7 (PIP2-7) | 43 | 1.25 | 0.318 | 1.000 |
| AT4G35100.2 | Plasma membrane intrinsic protein 2-7 (PIP2-7) | 46 | 1.97 | 0.522 | 1.000 |
| AT4G35100.2 | Plasma membrane intrinsic protein 2-7 (PIP2-7) | 47 | 2.33 | 0.622 | 1.000 |
| AT4G35260.1 | NAD isocitrate dehydrogenase subunit (IDH1, regulatory) | 2 | 0.74 | 0.287 | 1.000 |

| | | | | | |
|---|---|---|---|---|---|
| AT4G35260.1 | NAD isocitrate dehydrogenase subunit (IDH1, regulatory) | 3 | 0.61 | 0.217 | 0.999 |
| AT4G35260.1 | NAD isocitrate dehydrogenase subunit (IDH1, regulatory) | 4 | 0.52 | 0.182 | 0.906 |
| AT4G35260.1 | NAD isocitrate dehydrogenase subunit (IDH1, regulatory) | 5 | 0.46 | 0.168 | 0.616 |
| AT4G35260.1 | NAD isocitrate dehydrogenase subunit (IDH1, regulatory) | 6 | 0.44 | 0.159 | 0.441 |
| AT4G35260.1 | NAD isocitrate dehydrogenase subunit (IDH1, regulatory) | 7 | 0.46 | 0.150 | 0.414 |
| AT4G35260.1 | NAD isocitrate dehydrogenase subunit (IDH1, regulatory) | 8 | 0.49 | 0.143 | 0.483 |
| AT4G35260.1 | NAD isocitrate dehydrogenase subunit (IDH1, regulatory) | 9 | 0.55 | 0.137 | 0.687 |
| AT4G35260.1 | NAD isocitrate dehydrogenase subunit (IDH1, regulatory) | 10 | 0.62 | 0.135 | 0.921 |
| AT4G35260.1 | NAD isocitrate dehydrogenase subunit (IDH1, regulatory) | 11 | 0.69 | 0.134 | 0.993 |
| AT4G35260.1 | NAD isocitrate dehydrogenase subunit (IDH1, regulatory) | 12 | 0.76 | 0.133 | 1.000 |
| AT4G35260.1 | NAD isocitrate dehydrogenase subunit (IDH1, regulatory) | 13 | 0.81 | 0.132 | 1.000 |
| AT4G35260.1 | NAD isocitrate dehydrogenase subunit (IDH1, regulatory) | 14 | 0.87 | 0.130 | 1.000 |
| AT4G35260.1 | NAD isocitrate dehydrogenase subunit (IDH1, regulatory) | 15 | 0.95 | 0.129 | 1.000 |
| AT4G35260.1 | NAD isocitrate dehydrogenase subunit (IDH1, regulatory) | 16 | 1.04 | 0.129 | 1.000 |
| AT4G35260.1 | NAD isocitrate dehydrogenase subunit (IDH1, regulatory) | 17 | 1.11 | 0.129 | 1.000 |
| AT4G35260.1 | NAD isocitrate dehydrogenase subunit (IDH1, regulatory) | 18 | 1.08 | 0.131 | 1.000 |
| AT4G35260.1 | NAD isocitrate dehydrogenase subunit (IDH1, regulatory) | 19 | 0.97 | 0.134 | 1.000 |
| AT4G35260.1 | NAD isocitrate dehydrogenase subunit (IDH1, regulatory) | 20 | 0.82 | 0.136 | 1.000 |
| AT4G35260.1 | NAD isocitrate dehydrogenase subunit (IDH1, regulatory) | 21 | 0.70 | 0.138 | 0.996 |
| AT4G35260.1 | NAD isocitrate dehydrogenase subunit (IDH1, regulatory) | 22 | 0.59 | 0.138 | 0.868 |
| AT4G35260.1 | NAD isocitrate dehydrogenase subunit (IDH1, regulatory) | 23 | 0.51 | 0.137 | 0.498 |
| AT4G35260.1 | NAD isocitrate dehydrogenase subunit (IDH1, regulatory) | 24 | 0.45 | 0.135 | 0.229 |
| AT4G35260.1 | NAD isocitrate dehydrogenase subunit (IDH1, regulatory) | 25 | 0.44 | 0.134 | 0.167 |
| AT4G35260.1 | NAD isocitrate dehydrogenase subunit (IDH1, regulatory) | 26 | 0.48 | 0.133 | 0.299 |
| AT4G35260.1 | NAD isocitrate dehydrogenase subunit (IDH1, regulatory) | 27 | 0.56 | 0.132 | 0.701 |
| AT4G35260.1 | NAD isocitrate dehydrogenase subunit (IDH1, regulatory) | 28 | 0.65 | 0.132 | 0.961 |
| AT4G35260.1 | NAD isocitrate dehydrogenase subunit (IDH1, regulatory) | 29 | 0.73 | 0.132 | 0.998 |
| AT4G35260.1 | NAD isocitrate dehydrogenase subunit (IDH1, regulatory) | 30 | 0.78 | 0.133 | 1.000 |
| AT4G35260.1 | NAD isocitrate dehydrogenase subunit (IDH1, regulatory) | 31 | 0.81 | 0.133 | 1.000 |
| AT4G35260.1 | NAD isocitrate dehydrogenase subunit (IDH1, regulatory) | 32 | 0.81 | 0.133 | 1.000 |
| AT4G35260.1 | NAD isocitrate dehydrogenase subunit (IDH1, regulatory) | 33 | 0.80 | 0.133 | 1.000 |

| | | | | | |
|---|---|---|---|---|---|
| AT4G35260.1 | NAD isocitrate dehydrogenase subunit (IDH1, regulatory) | 34 | 0.81 | 0.133 | 1.000 |
| AT4G35260.1 | NAD isocitrate dehydrogenase subunit (IDH1, regulatory) | 35 | 0.87 | 0.133 | 1.000 |
| AT4G35260.1 | NAD isocitrate dehydrogenase subunit (IDH1, regulatory) | 36 | 0.96 | 0.133 | 1.000 |
| AT4G35260.1 | NAD isocitrate dehydrogenase subunit (IDH1, regulatory) | 37 | 1.09 | 0.133 | 1.000 |
| AT4G35260.1 | NAD isocitrate dehydrogenase subunit (IDH1, regulatory) | 38 | 1.21 | 0.133 | 1.000 |
| AT4G35260.1 | NAD isocitrate dehydrogenase subunit (IDH1, regulatory) | 39 | 1.31 | 0.134 | 1.000 |
| AT4G35260.1 | NAD isocitrate dehydrogenase subunit (IDH1, regulatory) | 40 | 1.37 | 0.134 | 0.999 |
| AT4G35260.1 | NAD isocitrate dehydrogenase subunit (IDH1, regulatory) | 41 | 1.39 | 0.136 | 0.998 |
| AT4G35260.1 | NAD isocitrate dehydrogenase subunit (IDH1, regulatory) | 42 | 1.37 | 0.137 | 0.999 |
| AT4G35260.1 | NAD isocitrate dehydrogenase subunit (IDH1, regulatory) | 43 | 1.30 | 0.137 | 1.000 |
| AT4G35260.1 | NAD isocitrate dehydrogenase subunit (IDH1, regulatory) | 44 | 1.17 | 0.138 | 1.000 |
| AT4G35260.1 | NAD isocitrate dehydrogenase subunit (IDH1, regulatory) | 45 | 0.99 | 0.150 | 1.000 |
| AT4G35260.1 | NAD isocitrate dehydrogenase subunit (IDH1, regulatory) | 46 | 0.79 | 0.190 | 1.000 |
| AT4G35260.1 | NAD isocitrate dehydrogenase subunit (IDH1, regulatory) | 47 | 0.62 | 0.273 | 1.000 |
| AT4G35460.1 | NADPH-dependent thioredoxin reductase B (NTRB) | 2 | 0.75 | 0.387 | 1.000 |
| AT4G35460.1 | NADPH-dependent thioredoxin reductase B (NTRB) | 3 | 0.55 | 0.389 | 0.988 |
| AT4G35460.1 | NADPH-dependent thioredoxin reductase B (NTRB) | 42 | 1.31 | 0.465 | 1.000 |
| AT4G35460.1 | NADPH-dependent thioredoxin reductase B (NTRB) | 43 | 2.43 | 0.247 | 0.572 |
| AT4G35460.1 | NADPH-dependent thioredoxin reductase B (NTRB) | 44 | 2.93 | 0.206 | 0.177 |
| AT4G35460.1 | NADPH-dependent thioredoxin reductase B (NTRB) | 45 | 1.53 | 0.204 | 0.943 |
| AT4G35460.1 | NADPH-dependent thioredoxin reductase B (NTRB) | 46 | 0.66 | 0.216 | 0.960 |
| AT4G35460.1 | NADPH-dependent thioredoxin reductase B (NTRB) | 47 | 0.24 | 0.490 | 0.778 |
| AT4G35850.1 | Pentatricopeptide repeat (PPR) superfamily protein | 2 | 1.11 | 0.383 | 1.000 |
| AT4G35850.1 | Pentatricopeptide repeat (PPR) superfamily protein | 3 | 0.84 | 0.287 | 1.000 |
| AT4G35850.1 | Pentatricopeptide repeat (PPR) superfamily protein | 4 | 0.69 | 0.439 | 1.000 |
| AT4G35850.1 | Pentatricopeptide repeat (PPR) superfamily protein | 15 | 1.59 | 0.260 | 1.000 |
| AT4G35850.1 | Pentatricopeptide repeat (PPR) superfamily protein | 16 | 1.16 | 0.223 | 1.000 |
| AT4G35850.1 | Pentatricopeptide repeat (PPR) superfamily protein | 17 | 0.85 | 0.205 | 1.000 |
| AT4G35850.1 | Pentatricopeptide repeat (PPR) superfamily protein | 18 | 0.88 | 0.193 | 1.000 |
| AT4G35850.1 | Pentatricopeptide repeat (PPR) superfamily protein | 19 | 1.33 | 0.201 | 1.000 |
| AT4G35850.1 | Pentatricopeptide repeat (PPR) superfamily protein | 20 | 2.05 | 0.246 | 0.983 |

| | | | | | | |
|---|---|---|---|---|---|---|
| AT4G35850.1 | Pentatricopeptide repeat (PPR) superfamily protein | 21 | 2.30 | 0.325 | 0.996 | |
| AT4G35850.1 | Pentatricopeptide repeat (PPR) superfamily protein | 25 | 0.46 | 0.279 | 0.990 | |
| AT4G35850.1 | Pentatricopeptide repeat (PPR) superfamily protein | 26 | 0.48 | 0.221 | 0.951 | |
| AT4G35850.1 | Pentatricopeptide repeat (PPR) superfamily protein | 27 | 0.74 | 0.194 | 1.000 | |
| AT4G35850.1 | Pentatricopeptide repeat (PPR) superfamily protein | 28 | 1.19 | 0.182 | 1.000 | |
| AT4G35850.1 | Pentatricopeptide repeat (PPR) superfamily protein | 29 | 1.53 | 0.181 | 0.998 | |
| AT4G35850.1 | Pentatricopeptide repeat (PPR) superfamily protein | 30 | 1.54 | 0.180 | 0.998 | |
| AT4G35850.1 | Pentatricopeptide repeat (PPR) superfamily protein | 31 | 1.45 | 0.181 | 1.000 | |
| AT4G35850.1 | Pentatricopeptide repeat (PPR) superfamily protein | 32 | 1.38 | 0.182 | 1.000 | |
| AT4G35850.1 | Pentatricopeptide repeat (PPR) superfamily protein | 33 | 1.31 | 0.182 | 1.000 | |
| AT4G35850.1 | Pentatricopeptide repeat (PPR) superfamily protein | 34 | 1.29 | 0.182 | 1.000 | |
| AT4G35850.1 | Pentatricopeptide repeat (PPR) superfamily protein | 35 | 1.25 | 0.181 | 1.000 | |
| AT4G35850.1 | Pentatricopeptide repeat (PPR) superfamily protein | 36 | 1.00 | 0.182 | 1.000 | |
| AT4G35850.1 | Pentatricopeptide repeat (PPR) superfamily protein | 37 | 0.59 | 0.183 | 0.986 | |
| AT4G35850.1 | Pentatricopeptide repeat (PPR) superfamily protein | 38 | 0.29 | 0.184 | 0.086 | . |
| AT4G35850.1 | Pentatricopeptide repeat (PPR) superfamily protein | 39 | 0.20 | 0.196 | 0.011 | * |
| AT4G35850.1 | Pentatricopeptide repeat (PPR) superfamily protein | 40 | 0.32 | 0.255 | 0.641 | |
| AT4G35850.1 | Pentatricopeptide repeat (PPR) superfamily protein | 41 | 0.89 | 0.391 | 1.000 | |
| AT4G35850.1 | Pentatricopeptide repeat (PPR) superfamily protein | 46 | 2596.44 | 1.605 | 0.496 | |
| AT4G35850.1 | Pentatricopeptide repeat (PPR) superfamily protein | 47 | 16701.71 | 1.979 | 0.491 | |
| AT4G36250.1 | Aldehyde dehydrogenase 3F1 (ALDH3F1) | 2 | 1.29 | 0.422 | 1.000 | |
| AT4G36250.1 | Aldehyde dehydrogenase 3F1 (ALDH3F1) | 3 | 1.18 | 0.404 | 1.000 | |
| AT4G36250.1 | Aldehyde dehydrogenase 3F1 (ALDH3F1) | 28 | 0.80 | 0.177 | 0.998 | |
| AT4G36250.1 | Aldehyde dehydrogenase 3F1 (ALDH3F1) | 29 | 0.93 | 0.145 | 1.000 | |
| AT4G36250.1 | Aldehyde dehydrogenase 3F1 (ALDH3F1) | 30 | 1.10 | 0.125 | 1.000 | |
| AT4G36250.1 | Aldehyde dehydrogenase 3F1 (ALDH3F1) | 31 | 1.33 | 0.115 | 0.900 | |
| AT4G36250.1 | Aldehyde dehydrogenase 3F1 (ALDH3F1) | 32 | 1.55 | 0.112 | 0.494 | |
| AT4G36250.1 | Aldehyde dehydrogenase 3F1 (ALDH3F1) | 33 | 1.71 | 0.112 | 0.252 | |
| AT4G36250.1 | Aldehyde dehydrogenase 3F1 (ALDH3F1) | 34 | 1.77 | 0.112 | 0.199 | |
| AT4G36250.1 | Aldehyde dehydrogenase 3F1 (ALDH3F1) | 35 | 1.71 | 0.114 | 0.279 | |
| AT4G36250.1 | Aldehyde dehydrogenase 3F1 (ALDH3F1) | 36 | 1.55 | 0.117 | 0.535 | |

| | | | | | | |
|---|---|---|---|---|---|---|
| AT4G36250.1 | Aldehyde dehydrogenase 3F1 (ALDH3F1) | 37 | 1.34 | 0.119 | 0.901 | |
| AT4G36250.1 | Aldehyde dehydrogenase 3F1 (ALDH3F1) | 38 | 1.11 | 0.122 | 1.000 | |
| AT4G36250.1 | Aldehyde dehydrogenase 3F1 (ALDH3F1) | 39 | 0.90 | 0.125 | 1.000 | |
| AT4G36250.1 | Aldehyde dehydrogenase 3F1 (ALDH3F1) | 40 | 0.73 | 0.131 | 0.914 | |
| AT4G36250.1 | Aldehyde dehydrogenase 3F1 (ALDH3F1) | 41 | 0.59 | 0.140 | 0.549 | |
| AT4G36250.1 | Aldehyde dehydrogenase 3F1 (ALDH3F1) | 42 | 0.47 | 0.153 | 0.237 | |
| AT4G36250.1 | Aldehyde dehydrogenase 3F1 (ALDH3F1) | 43 | 0.37 | 0.165 | 0.082 | . |
| AT4G36250.1 | Aldehyde dehydrogenase 3F1 (ALDH3F1) | 46 | 0.17 | 0.218 | 0.005 | ** |
| AT4G36250.1 | Aldehyde dehydrogenase 3F1 (ALDH3F1) | 47 | 0.13 | 0.255 | 0.006 | ** |
| AT4G36400.2 | D-2-hydroxyglutarate dehydrogenase (D2HGDH) | 2 | 0.75 | 0.854 | 1.000 | |
| AT4G36400.2 | D-2-hydroxyglutarate dehydrogenase (D2HGDH) | 3 | 0.85 | 0.824 | 1.000 | |
| AT4G36400.2 | D-2-hydroxyglutarate dehydrogenase (D2HGDH) | 41 | 10.74 | 0.635 | 0.411 | |
| AT4G36400.2 | D-2-hydroxyglutarate dehydrogenase (D2HGDH) | 42 | 10.06 | 0.470 | 0.168 | |
| AT4G36400.2 | D-2-hydroxyglutarate dehydrogenase (D2HGDH) | 43 | 8.62 | 0.367 | 0.067 | . |
| AT4G36400.2 | D-2-hydroxyglutarate dehydrogenase (D2HGDH) | 44 | 6.13 | 0.322 | 0.088 | . |
| AT4G36400.2 | D-2-hydroxyglutarate dehydrogenase (D2HGDH) | 45 | 3.50 | 0.337 | 0.415 | |
| AT4G36400.2 | D-2-hydroxyglutarate dehydrogenase (D2HGDH) | 46 | 1.73 | 0.430 | 0.979 | |
| AT4G36400.2 | D-2-hydroxyglutarate dehydrogenase (D2HGDH) | 47 | 0.82 | 0.620 | 1.000 | |
| AT4G36680.1 | Pentatricopeptide repeat (PPR) superfamily protein | 2 | 1.19 | 0.604 | 1.000 | |
| AT4G36680.1 | Pentatricopeptide repeat (PPR) superfamily protein | 3 | 1.08 | 0.542 | 1.000 | |
| AT4G36680.1 | Pentatricopeptide repeat (PPR) superfamily protein | 14 | 2.42 | 0.359 | 0.976 | |
| AT4G36680.1 | Pentatricopeptide repeat (PPR) superfamily protein | 15 | 2.55 | 0.309 | 0.904 | |
| AT4G36680.1 | Pentatricopeptide repeat (PPR) superfamily protein | 16 | 2.08 | 0.281 | 0.963 | |
| AT4G36680.1 | Pentatricopeptide repeat (PPR) superfamily protein | 17 | 1.39 | 0.261 | 1.000 | |
| AT4G36680.1 | Pentatricopeptide repeat (PPR) superfamily protein | 18 | 0.97 | 0.259 | 1.000 | |
| AT4G36680.1 | Pentatricopeptide repeat (PPR) superfamily protein | 19 | 0.80 | 0.269 | 1.000 | |
| AT4G36680.1 | Pentatricopeptide repeat (PPR) superfamily protein | 20 | 0.81 | 0.290 | 1.000 | |
| AT4G36680.1 | Pentatricopeptide repeat (PPR) superfamily protein | 21 | 0.88 | 0.357 | 1.000 | |
| AT4G36680.1 | Pentatricopeptide repeat (PPR) superfamily protein | 22 | 0.95 | 0.501 | 1.000 | |
| AT4G36680.1 | Pentatricopeptide repeat (PPR) superfamily protein | 38 | 0.94 | 0.437 | 1.000 | |
| AT4G36680.1 | Pentatricopeptide repeat (PPR) superfamily protein | 39 | 1.34 | 0.320 | 1.000 | |

| | | | | | | |
|---|---|---|---|---|---|---|
| AT4G36680.1 | Pentatricopeptide repeat (PPR) superfamily protein | 40 | 1.99 | 0.271 | 0.970 | |
| AT4G36680.1 | Pentatricopeptide repeat (PPR) superfamily protein | 41 | 2.71 | 0.259 | 0.690 | |
| AT4G36680.1 | Pentatricopeptide repeat (PPR) superfamily protein | 42 | 2.96 | 0.259 | 0.579 | |
| AT4G36680.1 | Pentatricopeptide repeat (PPR) superfamily protein | 43 | 2.65 | 0.261 | 0.724 | |
| AT4G36680.1 | Pentatricopeptide repeat (PPR) superfamily protein | 44 | 1.87 | 0.260 | 0.979 | |
| AT4G36680.1 | Pentatricopeptide repeat (PPR) superfamily protein | 45 | 1.07 | 0.262 | 1.000 | |
| AT4G36680.1 | Pentatricopeptide repeat (PPR) superfamily protein | 46 | 0.53 | 0.313 | 0.995 | |
| AT4G36680.1 | Pentatricopeptide repeat (PPR) superfamily protein | 47 | 0.24 | 0.497 | 0.934 | |
| AT4G37660.1 | Ribosomal L12 family protein, putative | 2 | 1.07 | 0.489 | 1.000 | |
| AT4G37660.1 | Ribosomal L12 family protein, putative | 3 | 0.98 | 0.482 | 1.000 | |
| AT4G37660.1 | Ribosomal L12 family protein, putative | 32 | 0.59 | 0.270 | 0.959 | |
| AT4G37660.1 | Ribosomal L12 family protein, putative | 33 | 0.67 | 0.220 | 0.973 | |
| AT4G37660.1 | Ribosomal L12 family protein, putative | 34 | 0.77 | 0.192 | 0.996 | |
| AT4G37660.1 | Ribosomal L12 family protein, putative | 35 | 0.91 | 0.176 | 1.000 | |
| AT4G37660.1 | Ribosomal L12 family protein, putative | 36 | 1.10 | 0.165 | 1.000 | |
| AT4G37660.1 | Ribosomal L12 family protein, putative | 37 | 1.36 | 0.157 | 0.961 | |
| AT4G37660.1 | Ribosomal L12 family protein, putative | 38 | 1.62 | 0.153 | 0.706 | |
| AT4G37660.1 | Ribosomal L12 family protein, putative | 39 | 1.73 | 0.155 | 0.582 | |
| AT4G37660.1 | Ribosomal L12 family protein, putative | 40 | 1.56 | 0.166 | 0.829 | |
| AT4G37660.1 | Ribosomal L12 family protein, putative | 41 | 1.18 | 0.188 | 1.000 | |
| AT4G37660.1 | Ribosomal L12 family protein, putative | 42 | 0.80 | 0.220 | 0.999 | |
| AT4G37660.1 | Ribosomal L12 family protein, putative | 43 | 0.52 | 0.254 | 0.852 | |
| AT4G37660.1 | Ribosomal L12 family protein, putative | 46 | 0.12 | 0.354 | 0.079 | . |
| AT4G37660.1 | Ribosomal L12 family protein, putative | 47 | 0.07 | 0.416 | 0.055 | . |
| AT4G37830.1 | Cytochrome c:O2 oxidoreductase subunit (COXX6A) | 2 | 1.56 | 0.404 | 1.000 | |
| AT4G37830.1 | Cytochrome c:O2 oxidoreductase subunit (COXX6A) | 3 | 1.11 | 0.263 | 1.000 | |
| AT4G37830.1 | Cytochrome c:O2 oxidoreductase subunit (COXX6A) | 4 | 0.93 | 0.253 | 1.000 | |
| AT4G37830.1 | Cytochrome c:O2 oxidoreductase subunit (COXX6A) | 5 | 0.85 | 0.244 | 1.000 | |

| | | | | | |
|---|---|---|---|---|---|
| AT4G37830.1 | Cytochrome c:O2 oxidoreductase subunit (COXX6A) | 6 | 0.69 | 0.235 | 1.000 |
| AT4G37830.1 | Cytochrome c:O2 oxidoreductase subunit (COXX6A) | 7 | 0.51 | 0.233 | 0.994 |
| AT4G37830.1 | Cytochrome c:O2 oxidoreductase subunit (COXX6A) | 8 | 0.52 | 0.225 | 0.995 |
| AT4G37830.1 | Cytochrome c:O2 oxidoreductase subunit (COXX6A) | 9 | 0.66 | 0.222 | 1.000 |
| AT4G37830.1 | Cytochrome c:O2 oxidoreductase subunit (COXX6A) | 10 | 0.87 | 0.221 | 1.000 |
| AT4G37830.1 | Cytochrome c:O2 oxidoreductase subunit (COXX6A) | 11 | 1.00 | 0.224 | 1.000 |
| AT4G37830.1 | Cytochrome c:O2 oxidoreductase subunit (COXX6A) | 12 | 1.02 | 0.224 | 1.000 |
| AT4G37830.1 | Cytochrome c:O2 oxidoreductase subunit (COXX6A) | 13 | 1.06 | 0.222 | 1.000 |
| AT4G37830.1 | Cytochrome c:O2 oxidoreductase subunit (COXX6A) | 14 | 1.02 | 0.219 | 1.000 |
| AT4G37830.1 | Cytochrome c:O2 oxidoreductase subunit (COXX6A) | 15 | 0.89 | 0.220 | 1.000 |
| AT4G37830.1 | Cytochrome c:O2 oxidoreductase subunit (COXX6A) | 16 | 0.67 | 0.217 | 1.000 |
| AT4G37830.1 | Cytochrome c:O2 oxidoreductase subunit (COXX6A) | 17 | 0.64 | 0.215 | 1.000 |
| AT4G37830.1 | Cytochrome c:O2 oxidoreductase subunit (COXX6A) | 18 | 0.86 | 0.218 | 1.000 |
| AT4G37830.1 | Cytochrome c:O2 oxidoreductase subunit (COXX6A) | 19 | 1.30 | 0.227 | 1.000 |
| AT4G37830.1 | Cytochrome c:O2 oxidoreductase subunit (COXX6A) | 20 | 2.51 | 0.232 | 0.865 |
| AT4G37830.1 | Cytochrome c:O2 oxidoreductase subunit (COXX6A) | 21 | 4.23 | 0.234 | 0.188 |
| AT4G37830.1 | Cytochrome c:O2 oxidoreductase subunit (COXX6A) | 22 | 4.35 | 0.233 | 0.156 |
| AT4G37830.1 | Cytochrome c:O2 oxidoreductase subunit (COXX6A) | 23 | 2.43 | 0.231 | 0.894 |
| AT4G37830.1 | Cytochrome c:O2 oxidoreductase subunit (COXX6A) | 24 | 1.11 | 0.227 | 1.000 |
| AT4G37830.1 | Cytochrome c:O2 oxidoreductase subunit (COXX6A) | 25 | 0.76 | 0.223 | 1.000 |

| | | | | | | |
|---|---|---|---|---|---|---|
| AT4G37830.1 | Cytochrome c:O2 oxidoreductase subunit (COXX6A) | 26 | 1.04 | 0.222 | 1.000 | |
| AT4G37830.1 | Cytochrome c:O2 oxidoreductase subunit (COXX6A) | 27 | 1.78 | 0.221 | 0.999 | |
| AT4G37830.1 | Cytochrome c:O2 oxidoreductase subunit (COXX6A) | 28 | 1.93 | 0.221 | 0.992 | |
| AT4G37830.1 | Cytochrome c:O2 oxidoreductase subunit (COXX6A) | 29 | 1.86 | 0.221 | 0.996 | |
| AT4G37830.1 | Cytochrome c:O2 oxidoreductase subunit (COXX6A) | 30 | 1.78 | 0.221 | 0.999 | |
| AT4G37830.1 | Cytochrome c:O2 oxidoreductase subunit (COXX6A) | 31 | 1.63 | 0.223 | 1.000 | |
| AT4G37830.1 | Cytochrome c:O2 oxidoreductase subunit (COXX6A) | 32 | 1.24 | 0.225 | 1.000 | |
| AT4G37830.1 | Cytochrome c:O2 oxidoreductase subunit (COXX6A) | 33 | 0.58 | 0.229 | 1.000 | |
| AT4G37830.1 | Cytochrome c:O2 oxidoreductase subunit (COXX6A) | 34 | 0.19 | 0.241 | 0.092 | . |
| AT4G37830.1 | Cytochrome c:O2 oxidoreductase subunit (COXX6A) | 35 | 0.14 | 0.280 | 0.080 | . |
| AT4G37830.1 | Cytochrome c:O2 oxidoreductase subunit (COXX6A) | 36 | 0.22 | 0.450 | 0.970 | |
| AT4G37830.1 | Cytochrome c:O2 oxidoreductase subunit (COXX6A) | 46 | ########### | 4.938 | 0.740 | |
| AT4G37830.1 | Cytochrome c:O2 oxidoreductase subunit (COXX6A) | 47 | ########### | 5.624 | 0.727 | |
| AT4G37910.1 | Heat shock protein 70-1 (HSP70-1) | 2 | 0.87 | 0.272 | 1.000 | |
| AT4G37910.1 | Heat shock protein 70-1 (HSP70-1) | 3 | 0.69 | 0.198 | 1.000 | |
| AT4G37910.1 | Heat shock protein 70-1 (HSP70-1) | 4 | 0.56 | 0.182 | 0.979 | |
| AT4G37910.1 | Heat shock protein 70-1 (HSP70-1) | 5 | 0.50 | 0.178 | 0.858 | |
| AT4G37910.1 | Heat shock protein 70-1 (HSP70-1) | 6 | 0.50 | 0.167 | 0.793 | |
| AT4G37910.1 | Heat shock protein 70-1 (HSP70-1) | 7 | 0.58 | 0.156 | 0.954 | |
| AT4G37910.1 | Heat shock protein 70-1 (HSP70-1) | 8 | 0.79 | 0.148 | 1.000 | |
| AT4G37910.1 | Heat shock protein 70-1 (HSP70-1) | 9 | 1.09 | 0.144 | 1.000 | |
| AT4G37910.1 | Heat shock protein 70-1 (HSP70-1) | 10 | 1.33 | 0.143 | 1.000 | |
| AT4G37910.1 | Heat shock protein 70-1 (HSP70-1) | 11 | 1.47 | 0.143 | 0.997 | |
| AT4G37910.1 | Heat shock protein 70-1 (HSP70-1) | 12 | 1.50 | 0.143 | 0.994 | |

| | | | | | |
|---|---|---|---|---|---|
| AT4G37910.1 | Heat shock protein 70-1 (HSP70-1) | 13 | 1.42 | 0.142 | 0.999 |
| AT4G37910.1 | Heat shock protein 70-1 (HSP70-1) | 14 | 1.36 | 0.140 | 1.000 |
| AT4G37910.1 | Heat shock protein 70-1 (HSP70-1) | 15 | 1.34 | 0.139 | 1.000 |
| AT4G37910.1 | Heat shock protein 70-1 (HSP70-1) | 16 | 1.35 | 0.139 | 1.000 |
| AT4G37910.1 | Heat shock protein 70-1 (HSP70-1) | 17 | 1.34 | 0.139 | 1.000 |
| AT4G37910.1 | Heat shock protein 70-1 (HSP70-1) | 18 | 1.30 | 0.140 | 1.000 |
| AT4G37910.1 | Heat shock protein 70-1 (HSP70-1) | 19 | 1.29 | 0.143 | 1.000 |
| AT4G37910.1 | Heat shock protein 70-1 (HSP70-1) | 20 | 1.36 | 0.147 | 1.000 |
| AT4G37910.1 | Heat shock protein 70-1 (HSP70-1) | 21 | 1.31 | 0.148 | 1.000 |
| AT4G37910.1 | Heat shock protein 70-1 (HSP70-1) | 22 | 1.17 | 0.148 | 1.000 |
| AT4G37910.1 | Heat shock protein 70-1 (HSP70-1) | 23 | 0.99 | 0.147 | 1.000 |
| AT4G37910.1 | Heat shock protein 70-1 (HSP70-1) | 24 | 0.82 | 0.145 | 1.000 |
| AT4G37910.1 | Heat shock protein 70-1 (HSP70-1) | 25 | 0.71 | 0.143 | 0.999 |
| AT4G37910.1 | Heat shock protein 70-1 (HSP70-1) | 26 | 0.70 | 0.142 | 0.999 |
| AT4G37910.1 | Heat shock protein 70-1 (HSP70-1) | 27 | 0.77 | 0.142 | 1.000 |
| AT4G37910.1 | Heat shock protein 70-1 (HSP70-1) | 28 | 0.86 | 0.142 | 1.000 |
| AT4G37910.1 | Heat shock protein 70-1 (HSP70-1) | 29 | 0.91 | 0.142 | 1.000 |
| AT4G37910.1 | Heat shock protein 70-1 (HSP70-1) | 30 | 0.94 | 0.142 | 1.000 |
| AT4G37910.1 | Heat shock protein 70-1 (HSP70-1) | 31 | 0.94 | 0.142 | 1.000 |
| AT4G37910.1 | Heat shock protein 70-1 (HSP70-1) | 32 | 0.93 | 0.143 | 1.000 |
| AT4G37910.1 | Heat shock protein 70-1 (HSP70-1) | 33 | 0.88 | 0.143 | 1.000 |
| AT4G37910.1 | Heat shock protein 70-1 (HSP70-1) | 34 | 0.83 | 0.143 | 1.000 |
| AT4G37910.1 | Heat shock protein 70-1 (HSP70-1) | 35 | 0.80 | 0.143 | 1.000 |
| AT4G37910.1 | Heat shock protein 70-1 (HSP70-1) | 36 | 0.80 | 0.142 | 1.000 |
| AT4G37910.1 | Heat shock protein 70-1 (HSP70-1) | 37 | 0.88 | 0.142 | 1.000 |
| AT4G37910.1 | Heat shock protein 70-1 (HSP70-1) | 38 | 1.04 | 0.142 | 1.000 |
| AT4G37910.1 | Heat shock protein 70-1 (HSP70-1) | 39 | 1.26 | 0.143 | 1.000 |
| AT4G37910.1 | Heat shock protein 70-1 (HSP70-1) | 40 | 1.47 | 0.143 | 0.997 |
| AT4G37910.1 | Heat shock protein 70-1 (HSP70-1) | 41 | 1.55 | 0.144 | 0.987 |
| AT4G37910.1 | Heat shock protein 70-1 (HSP70-1) | 42 | 1.46 | 0.145 | 0.998 |
| AT4G37910.1 | Heat shock protein 70-1 (HSP70-1) | 43 | 1.20 | 0.146 | 1.000 |

| | | | | | | |
|---|---|---|---|---|---|---|
| AT4G37910.1 | Heat shock protein 70-1 (HSP70-1) | 44 | 0.80 | 0.146 | 1.000 | |
| AT4G37910.1 | Heat shock protein 70-1 (HSP70-1) | 45 | 0.43 | 0.149 | 0.285 | |
| AT4G37910.1 | Heat shock protein 70-1 (HSP70-1) | 46 | 0.18 | 0.178 | 0.001 | *** |
| AT4G37910.1 | Heat shock protein 70-1 (HSP70-1) | 47 | 0.06 | 0.275 | 0.001 | *** |
| AT4G37930.1 | Serine hydroxymethyltransferase 1 (SHMT1) | 3 | 0.85 | 0.127 | 1.000 | |
| AT4G37930.1 | Serine hydroxymethyltransferase 1 (SHMT1) | 4 | 0.85 | 0.115 | 1.000 | |
| AT4G37930.1 | Serine hydroxymethyltransferase 1 (SHMT1) | 5 | 0.87 | 0.114 | 1.000 | |
| AT4G37930.1 | Serine hydroxymethyltransferase 1 (SHMT1) | 6 | 0.90 | 0.114 | 1.000 | |
| AT4G37930.1 | Serine hydroxymethyltransferase 1 (SHMT1) | 7 | 0.93 | 0.114 | 1.000 | |
| AT4G37930.1 | Serine hydroxymethyltransferase 1 (SHMT1) | 8 | 0.96 | 0.113 | 1.000 | |
| AT4G37930.1 | Serine hydroxymethyltransferase 1 (SHMT1) | 9 | 0.99 | 0.112 | 1.000 | |
| AT4G37930.1 | Serine hydroxymethyltransferase 1 (SHMT1) | 10 | 1.02 | 0.112 | 1.000 | |
| AT4G37930.1 | Serine hydroxymethyltransferase 1 (SHMT1) | 11 | 1.03 | 0.112 | 1.000 | |
| AT4G37930.1 | Serine hydroxymethyltransferase 1 (SHMT1) | 12 | 1.04 | 0.112 | 1.000 | |
| AT4G37930.1 | Serine hydroxymethyltransferase 1 (SHMT1) | 13 | 1.04 | 0.111 | 1.000 | |
| AT4G37930.1 | Serine hydroxymethyltransferase 1 (SHMT1) | 14 | 1.03 | 0.111 | 1.000 | |
| AT4G37930.1 | Serine hydroxymethyltransferase 1 (SHMT1) | 15 | 1.02 | 0.110 | 1.000 | |
| AT4G37930.1 | Serine hydroxymethyltransferase 1 (SHMT1) | 16 | 1.01 | 0.110 | 1.000 | |
| AT4G37930.1 | Serine hydroxymethyltransferase 1 (SHMT1) | 17 | 1.00 | 0.110 | 1.000 | |
| AT4G37930.1 | Serine hydroxymethyltransferase 1 (SHMT1) | 18 | 0.99 | 0.111 | 1.000 | |
| AT4G37930.1 | Serine hydroxymethyltransferase 1 (SHMT1) | 19 | 0.98 | 0.112 | 1.000 | |
| AT4G37930.1 | Serine hydroxymethyltransferase 1 (SHMT1) | 20 | 0.97 | 0.113 | 1.000 | |

| | | | | | |
|---|---|---|---|---|---|
| AT4G37930.1 | Serine hydroxymethyltransferase 1 (SHMT1) | 21 | 0.93 | 0.114 | 1.000 |
| AT4G37930.1 | Serine hydroxymethyltransferase 1 (SHMT1) | 22 | 0.89 | 0.114 | 1.000 |
| AT4G37930.1 | Serine hydroxymethyltransferase 1 (SHMT1) | 23 | 0.85 | 0.114 | 1.000 |
| AT4G37930.1 | Serine hydroxymethyltransferase 1 (SHMT1) | 24 | 0.83 | 0.113 | 1.000 |
| AT4G37930.1 | Serine hydroxymethyltransferase 1 (SHMT1) | 25 | 0.85 | 0.112 | 1.000 |
| AT4G37930.1 | Serine hydroxymethyltransferase 1 (SHMT1) | 26 | 0.90 | 0.112 | 1.000 |
| AT4G37930.1 | Serine hydroxymethyltransferase 1 (SHMT1) | 27 | 0.98 | 0.111 | 1.000 |
| AT4G37930.1 | Serine hydroxymethyltransferase 1 (SHMT1) | 28 | 1.05 | 0.111 | 1.000 |
| AT4G37930.1 | Serine hydroxymethyltransferase 1 (SHMT1) | 29 | 1.09 | 0.112 | 1.000 |
| AT4G37930.1 | Serine hydroxymethyltransferase 1 (SHMT1) | 30 | 1.11 | 0.112 | 1.000 |
| AT4G37930.1 | Serine hydroxymethyltransferase 1 (SHMT1) | 31 | 1.10 | 0.112 | 1.000 |
| AT4G37930.1 | Serine hydroxymethyltransferase 1 (SHMT1) | 32 | 1.10 | 0.112 | 1.000 |
| AT4G37930.1 | Serine hydroxymethyltransferase 1 (SHMT1) | 33 | 1.10 | 0.112 | 1.000 |
| AT4G37930.1 | Serine hydroxymethyltransferase 1 (SHMT1) | 34 | 1.11 | 0.112 | 1.000 |
| AT4G37930.1 | Serine hydroxymethyltransferase 1 (SHMT1) | 35 | 1.12 | 0.112 | 1.000 |
| AT4G37930.1 | Serine hydroxymethyltransferase 1 (SHMT1) | 36 | 1.13 | 0.112 | 1.000 |
| AT4G37930.1 | Serine hydroxymethyltransferase 1 (SHMT1) | 37 | 1.11 | 0.112 | 1.000 |
| AT4G37930.1 | Serine hydroxymethyltransferase 1 (SHMT1) | 38 | 1.11 | 0.112 | 1.000 |
| AT4G37930.1 | Serine hydroxymethyltransferase 1 (SHMT1) | 39 | 1.15 | 0.112 | 1.000 |
| AT4G37930.1 | Serine hydroxymethyltransferase 1 (SHMT1) | 40 | 1.19 | 0.112 | 1.000 |

| | | | | | | |
|---|---|---|---|---|---|---|
| AT4G37930.1 | Serine hydroxymethyltransferase 1 (SHMT1) | 41 | 1.19 | 0.113 | 1.000 | |
| AT4G37930.1 | Serine hydroxymethyltransferase 1 (SHMT1) | 42 | 1.12 | 0.113 | 1.000 | |
| AT4G37930.1 | Serine hydroxymethyltransferase 1 (SHMT1) | 43 | 1.01 | 0.114 | 1.000 | |
| AT4G37930.1 | Serine hydroxymethyltransferase 1 (SHMT1) | 44 | 0.87 | 0.114 | 1.000 | |
| AT4G37930.1 | Serine hydroxymethyltransferase 1 (SHMT1) | 45 | 0.74 | 0.116 | 0.961 | |
| AT4G37930.1 | Serine hydroxymethyltransferase 1 (SHMT1) | 46 | 0.61 | 0.133 | 0.670 | |
| AT4G37930.1 | Serine hydroxymethyltransferase 1 (SHMT1) | 47 | 0.49 | 0.177 | 0.589 | |
| AT4G37930.1 | Serine hydroxymethyltransferase 1 (SHMT1) | 48 | 0.40 | 0.244 | 0.664 | |
| AT4G38740.1 | Rotamase cyclophilin 1 (ROC1) | 2 | 1.41 | 0.501 | 1.000 | |
| AT4G38740.1 | Rotamase cyclophilin 1 (ROC1) | 3 | 1.45 | 0.471 | 0.999 | |
| AT4G38740.1 | Rotamase cyclophilin 1 (ROC1) | 18 | 1.88 | 0.176 | 0.472 | |
| AT4G38740.1 | Rotamase cyclophilin 1 (ROC1) | 19 | 1.88 | 0.163 | 0.393 | |
| AT4G38740.1 | Rotamase cyclophilin 1 (ROC1) | 20 | 1.87 | 0.151 | 0.327 | |
| AT4G38740.1 | Rotamase cyclophilin 1 (ROC1) | 21 | 1.84 | 0.141 | 0.281 | |
| AT4G38740.1 | Rotamase cyclophilin 1 (ROC1) | 22 | 1.81 | 0.133 | 0.259 | |
| AT4G38740.1 | Rotamase cyclophilin 1 (ROC1) | 23 | 1.76 | 0.127 | 0.262 | |
| AT4G38740.1 | Rotamase cyclophilin 1 (ROC1) | 24 | 1.69 | 0.123 | 0.295 | |
| AT4G38740.1 | Rotamase cyclophilin 1 (ROC1) | 25 | 1.61 | 0.121 | 0.373 | |
| AT4G38740.1 | Rotamase cyclophilin 1 (ROC1) | 26 | 1.52 | 0.120 | 0.503 | |
| AT4G38740.1 | Rotamase cyclophilin 1 (ROC1) | 27 | 1.42 | 0.121 | 0.686 | |
| AT4G38740.1 | Rotamase cyclophilin 1 (ROC1) | 28 | 1.31 | 0.124 | 0.877 | |
| AT4G38740.1 | Rotamase cyclophilin 1 (ROC1) | 29 | 1.19 | 0.129 | 0.985 | |
| AT4G38740.1 | Rotamase cyclophilin 1 (ROC1) | 30 | 1.06 | 0.136 | 1.000 | |
| AT4G38740.1 | Rotamase cyclophilin 1 (ROC1) | 31 | 0.94 | 0.144 | 1.000 | |
| AT4G38740.1 | Rotamase cyclophilin 1 (ROC1) | 32 | 0.83 | 0.153 | 0.989 | |
| AT4G38740.1 | Rotamase cyclophilin 1 (ROC1) | 33 | 0.72 | 0.164 | 0.906 | |
| AT4G38740.1 | Rotamase cyclophilin 1 (ROC1) | 34 | 0.62 | 0.176 | 0.743 | |
| AT4G38740.1 | Rotamase cyclophilin 1 (ROC1) | 35 | 0.54 | 0.189 | 0.564 | |
| AT4G38740.1 | Rotamase cyclophilin 1 (ROC1) | 36 | 0.46 | 0.202 | 0.410 | |
| AT4G38740.1 | Rotamase cyclophilin 1 (ROC1) | 46 | 0.10 | 0.356 | 0.029 | * |
| AT4G38740.1 | Rotamase cyclophilin 1 (ROC1) | 47 | 0.08 | 0.380 | 0.028 | * |
| AT4G38970.1 | Fructose-bisphosphate aldolase 2 (FBA2) | 2 | 0.89 | 0.625 | 1.000 | |
| AT4G38970.1 | Fructose-bisphosphate aldolase 2 (FBA2) | 3 | 1.16 | 0.641 | 1.000 | |
| AT4G38970.1 | Fructose-bisphosphate aldolase 2 (FBA2) | 33 | 1.13 | 0.470 | 1.000 | |
| AT4G38970.1 | Fructose-bisphosphate aldolase 2 (FBA2) | 34 | 0.74 | 0.359 | 1.000 | |
| AT4G38970.1 | Fructose-bisphosphate aldolase 2 (FBA2) | 35 | 0.56 | 0.320 | 0.994 | |
| AT4G38970.1 | Fructose-bisphosphate aldolase 2 (FBA2) | 36 | 0.55 | 0.299 | 0.988 | |
| AT4G38970.1 | Fructose-bisphosphate aldolase 2 (FBA2) | 37 | 0.86 | 0.297 | 1.000 | |

| | | | | | | |
|---|---|---|---|---|---|---|
| AT4G38970.1 | Fructose-bisphosphate aldolase 2 (FBA2) | 38 | 1.60 | 0.296 | 0.998 | |
| AT4G38970.1 | Fructose-bisphosphate aldolase 2 (FBA2) | 39 | 1.47 | 0.301 | 1.000 | |
| AT4G38970.1 | Fructose-bisphosphate aldolase 2 (FBA2) | 40 | 0.76 | 0.310 | 1.000 | |
| AT4G38970.1 | Fructose-bisphosphate aldolase 2 (FBA2) | 41 | 0.52 | 0.411 | 0.998 | |
| AT4G38970.1 | Fructose-bisphosphate aldolase 2 (FBA2) | 42 | 0.73 | 0.693 | 1.000 | |
| AT4G38970.1 | Fructose-bisphosphate aldolase 2 (FBA2) | 46 | 540.39 | 2.607 | 0.961 | |
| AT4G38970.1 | Fructose-bisphosphate aldolase 2 (FBA2) | 47 | 5238.77 | 3.360 | 0.946 | |
| AT4G39080.1 | Vacuolar ATP synthase subunit A3 (VHA-A3) | 2 | 1.70 | 0.284 | 0.922 | |
| AT4G39080.1 | Vacuolar ATP synthase subunit A3 (VHA-A3) | 3 | 1.81 | 0.257 | 0.832 | |
| AT4G39080.1 | Vacuolar ATP synthase subunit A3 (VHA-A3) | 4 | 1.92 | 0.236 | 0.710 | |
| AT4G39080.1 | Vacuolar ATP synthase subunit A3 (VHA-A3) | 5 | 2.04 | 0.223 | 0.582 | |
| AT4G39080.1 | Vacuolar ATP synthase subunit A3 (VHA-A3) | 6 | 2.15 | 0.216 | 0.475 | |
| AT4G39080.1 | Vacuolar ATP synthase subunit A3 (VHA-A3) | 7 | 2.26 | 0.214 | 0.401 | |
| AT4G39080.1 | Vacuolar ATP synthase subunit A3 (VHA-A3) | 8 | 2.38 | 0.216 | 0.353 | |
| AT4G39080.1 | Vacuolar ATP synthase subunit A3 (VHA-A3) | 9 | 2.49 | 0.221 | 0.322 | |
| AT4G39080.1 | Vacuolar ATP synthase subunit A3 (VHA-A3) | 21 | 2.74 | 0.211 | 0.192 | |
| AT4G39080.1 | Vacuolar ATP synthase subunit A3 (VHA-A3) | 22 | 2.57 | 0.207 | 0.229 | |
| AT4G39080.1 | Vacuolar ATP synthase subunit A3 (VHA-A3) | 23 | 2.37 | 0.205 | 0.300 | |
| AT4G39080.1 | Vacuolar ATP synthase subunit A3 (VHA-A3) | 24 | 2.15 | 0.205 | 0.421 | |
| AT4G39080.1 | Vacuolar ATP synthase subunit A3 (VHA-A3) | 25 | 1.92 | 0.207 | 0.596 | |
| AT4G39080.1 | Vacuolar ATP synthase subunit A3 (VHA-A3) | 26 | 1.69 | 0.213 | 0.794 | |
| AT4G39080.1 | Vacuolar ATP synthase subunit A3 (VHA-A3) | 27 | 1.47 | 0.221 | 0.942 | |
| AT4G39080.1 | Vacuolar ATP synthase subunit A3 (VHA-A3) | 28 | 1.27 | 0.232 | 0.996 | |
| AT4G39080.1 | Vacuolar ATP synthase subunit A3 (VHA-A3) | 29 | 1.08 | 0.244 | 1.000 | |
| AT4G39080.1 | Vacuolar ATP synthase subunit A3 (VHA-A3) | 30 | 0.91 | 0.259 | 1.000 | |
| AT4G39080.1 | Vacuolar ATP synthase subunit A3 (VHA-A3) | 31 | 0.76 | 0.275 | 0.996 | |
| AT4G39080.1 | Vacuolar ATP synthase subunit A3 (VHA-A3) | 46 | 0.03 | 0.481 | 0.017 | * |
| AT4G39080.1 | Vacuolar ATP synthase subunit A3 (VHA-A3) | 47 | 0.03 | 0.504 | 0.014 | * |
| AT4G39460.2 | S-adenosylmethionine carrier 1 (SAMC1) | 2 | 0.63 | 0.423 | 1.000 | |
| AT4G39460.2 | S-adenosylmethionine carrier 1 (SAMC1) | 3 | 0.73 | 0.309 | 1.000 | |
| AT4G39460.2 | S-adenosylmethionine carrier 1 (SAMC1) | 5 | 1.06 | 0.290 | 1.000 | |

| | | | | | | |
|---|---|---|---|---|---|---|
| AT4G39460.2 | S-adenosylmethionine carrier 1 (SAMC1) | 6 | 1.31 | 0.242 | 1.000 | |
| AT4G39460.2 | S-adenosylmethionine carrier 1 (SAMC1) | 7 | 1.38 | 0.219 | 1.000 | |
| AT4G39460.2 | S-adenosylmethionine carrier 1 (SAMC1) | 8 | 1.21 | 0.209 | 1.000 | |
| AT4G39460.2 | S-adenosylmethionine carrier 1 (SAMC1) | 9 | 1.05 | 0.205 | 1.000 | |
| AT4G39460.2 | S-adenosylmethionine carrier 1 (SAMC1) | 10 | 1.00 | 0.205 | 1.000 | |
| AT4G39460.2 | S-adenosylmethionine carrier 1 (SAMC1) | 11 | 1.03 | 0.206 | 1.000 | |
| AT4G39460.2 | S-adenosylmethionine carrier 1 (SAMC1) | 12 | 1.05 | 0.205 | 1.000 | |
| AT4G39460.2 | S-adenosylmethionine carrier 1 (SAMC1) | 13 | 0.82 | 0.203 | 1.000 | |
| AT4G39460.2 | S-adenosylmethionine carrier 1 (SAMC1) | 14 | 0.61 | 0.201 | 0.996 | |
| AT4G39460.2 | S-adenosylmethionine carrier 1 (SAMC1) | 15 | 0.50 | 0.202 | 0.899 | |
| AT4G39460.2 | S-adenosylmethionine carrier 1 (SAMC1) | 16 | 0.45 | 0.201 | 0.769 | |
| AT4G39460.2 | S-adenosylmethionine carrier 1 (SAMC1) | 17 | 0.54 | 0.203 | 0.960 | |
| AT4G39460.2 | S-adenosylmethionine carrier 1 (SAMC1) | 18 | 0.76 | 0.209 | 1.000 | |
| AT4G39460.2 | S-adenosylmethionine carrier 1 (SAMC1) | 19 | 1.10 | 0.221 | 1.000 | |
| AT4G39460.2 | S-adenosylmethionine carrier 1 (SAMC1) | 20 | 1.34 | 0.266 | 1.000 | |
| AT4G39460.2 | S-adenosylmethionine carrier 1 (SAMC1) | 21 | 1.47 | 0.409 | 1.000 | |
| AT4G39460.2 | S-adenosylmethionine carrier 1 (SAMC1) | 39 | 24.40 | 0.349 | 0.002 | ** |
| AT4G39460.2 | S-adenosylmethionine carrier 1 (SAMC1) | 40 | 21.51 | 0.237 | 0.000 | *** |
| AT4G39460.2 | S-adenosylmethionine carrier 1 (SAMC1) | 41 | 13.17 | 0.212 | 0.000 | *** |
| AT4G39460.2 | S-adenosylmethionine carrier 1 (SAMC1) | 42 | 4.00 | 0.208 | 0.083 | . |
| AT4G39460.2 | S-adenosylmethionine carrier 1 (SAMC1) | 43 | 1.16 | 0.208 | 1.000 | |
| AT4G39460.2 | S-adenosylmethionine carrier 1 (SAMC1) | 44 | 0.35 | 0.208 | 0.410 | |
| AT4G39460.2 | S-adenosylmethionine carrier 1 (SAMC1) | 45 | 0.15 | 0.207 | 0.003 | ** |
| AT4G39460.2 | S-adenosylmethionine carrier 1 (SAMC1) | 46 | 0.11 | 0.230 | 0.001 | *** |
| AT4G39460.2 | S-adenosylmethionine carrier 1 (SAMC1) | 47 | 0.09 | 0.419 | 0.214 | |
| AT4G39660.1 | Alanine:glyoxylate aminotransferase 2 (AGT2) | 2 | 1.16 | 0.535 | 1.000 | |
| AT4G39660.1 | Alanine:glyoxylate aminotransferase 2 (AGT2) | 3 | 1.07 | 0.465 | 1.000 | |
| AT4G39660.1 | Alanine:glyoxylate aminotransferase 2 (AGT2) | 10 | 1.06 | 0.343 | 1.000 | |
| AT4G39660.1 | Alanine:glyoxylate aminotransferase 2 (AGT2) | 11 | 1.21 | 0.312 | 1.000 | |
| AT4G39660.1 | Alanine:glyoxylate aminotransferase 2 (AGT2) | 12 | 1.33 | 0.292 | 1.000 | |
| AT4G39660.1 | Alanine:glyoxylate aminotransferase 2 (AGT2) | 13 | 1.44 | 0.294 | 1.000 | |

| | | | | | |
|---|---|---|---|---|---|
| AT4G39660.1 | Alanine:glyoxylate aminotransferase 2 (AGT2) | 14 | 1.53 | 0.293 | 1.000 |
| AT4G39660.1 | Alanine:glyoxylate aminotransferase 2 (AGT2) | 15 | 1.35 | 0.289 | 1.000 |
| AT4G39660.1 | Alanine:glyoxylate aminotransferase 2 (AGT2) | 18 | 0.88 | 0.326 | 1.000 |
| AT4G39660.1 | Alanine:glyoxylate aminotransferase 2 (AGT2) | 19 | 1.15 | 0.285 | 1.000 |
| AT4G39660.1 | Alanine:glyoxylate aminotransferase 2 (AGT2) | 20 | 1.55 | 0.267 | 1.000 |
| AT4G39660.1 | Alanine:glyoxylate aminotransferase 2 (AGT2) | 21 | 1.73 | 0.261 | 1.000 |
| AT4G39660.1 | Alanine:glyoxylate aminotransferase 2 (AGT2) | 22 | 1.73 | 0.258 | 1.000 |
| AT4G39660.1 | Alanine:glyoxylate aminotransferase 2 (AGT2) | 23 | 1.69 | 0.258 | 1.000 |
| AT4G39660.1 | Alanine:glyoxylate aminotransferase 2 (AGT2) | 24 | 1.04 | 0.249 | 1.000 |
| AT4G39660.1 | Alanine:glyoxylate aminotransferase 2 (AGT2) | 25 | 0.40 | 0.244 | 0.923 |
| AT4G39660.1 | Alanine:glyoxylate aminotransferase 2 (AGT2) | 26 | 0.24 | 0.244 | 0.283 |
| AT4G39660.1 | Alanine:glyoxylate aminotransferase 2 (AGT2) | 27 | 0.38 | 0.246 | 0.901 |
| AT4G39660.1 | Alanine:glyoxylate aminotransferase 2 (AGT2) | 28 | 0.85 | 0.247 | 1.000 |
| AT4G39660.1 | Alanine:glyoxylate aminotransferase 2 (AGT2) | 29 | 1.04 | 0.247 | 1.000 |
| AT4G39660.1 | Alanine:glyoxylate aminotransferase 2 (AGT2) | 30 | 0.57 | 0.246 | 1.000 |
| AT4G39660.1 | Alanine:glyoxylate aminotransferase 2 (AGT2) | 31 | 0.34 | 0.246 | 0.773 |
| AT4G39660.1 | Alanine:glyoxylate aminotransferase 2 (AGT2) | 32 | 0.45 | 0.247 | 0.985 |
| AT4G39660.1 | Alanine:glyoxylate aminotransferase 2 (AGT2) | 33 | 1.24 | 0.247 | 1.000 |
| AT4G39660.1 | Alanine:glyoxylate aminotransferase 2 (AGT2) | 34 | 3.65 | 0.248 | 0.476 |
| AT4G39660.1 | Alanine:glyoxylate aminotransferase 2 (AGT2) | 35 | 4.04 | 0.248 | 0.343 |
| AT4G39660.1 | Alanine:glyoxylate aminotransferase 2 (AGT2) | 36 | 1.49 | 0.248 | 1.000 |
| AT4G39660.1 | Alanine:glyoxylate aminotransferase 2 (AGT2) | 37 | 0.59 | 0.248 | 1.000 |
| AT4G39660.1 | Alanine:glyoxylate aminotransferase 2 (AGT2) | 38 | 0.56 | 0.248 | 1.000 |
| AT4G39660.1 | Alanine:glyoxylate aminotransferase 2 (AGT2) | 39 | 0.98 | 0.246 | 1.000 |
| AT4G39660.1 | Alanine:glyoxylate aminotransferase 2 (AGT2) | 40 | 1.61 | 0.245 | 1.000 |
| AT4G39660.1 | Alanine:glyoxylate aminotransferase 2 (AGT2) | 41 | 1.90 | 0.247 | 0.999 |
| AT4G39660.1 | Alanine:glyoxylate aminotransferase 2 (AGT2) | 42 | 1.95 | 0.250 | 0.999 |
| AT4G39660.1 | Alanine:glyoxylate aminotransferase 2 (AGT2) | 43 | 2.14 | 0.251 | 0.993 |
| AT4G39660.1 | Alanine:glyoxylate aminotransferase 2 (AGT2) | 44 | 2.17 | 0.251 | 0.991 |
| AT4G39660.1 | Alanine:glyoxylate aminotransferase 2 (AGT2) | 45 | 1.49 | 0.254 | 1.000 |
| AT4G39660.1 | Alanine:glyoxylate aminotransferase 2 (AGT2) | 46 | 0.51 | 0.287 | 1.000 |

| | | | | | |
|---|---|---|---|---|---|
| AT4G39660.1 | Alanine:glyoxylate aminotransferase 2 (AGT2) | 47 | 0.09 | 0.503 | 0.664 |
| AT4G39690.1 | MICOS complex subunit (Mic60-like) | 3 | 0.53 | 0.192 | 0.989 |
| AT4G39690.1 | MICOS complex subunit (Mic60-like) | 4 | 0.38 | 0.171 | 0.357 |
| AT4G39690.1 | MICOS complex subunit (Mic60-like) | 5 | 0.43 | 0.171 | 0.643 |
| AT4G39690.1 | MICOS complex subunit (Mic60-like) | 6 | 0.63 | 0.171 | 0.999 |
| AT4G39690.1 | MICOS complex subunit (Mic60-like) | 7 | 0.86 | 0.171 | 1.000 |
| AT4G39690.1 | MICOS complex subunit (Mic60-like) | 8 | 1.09 | 0.167 | 1.000 |
| AT4G39690.1 | MICOS complex subunit (Mic60-like) | 9 | 1.25 | 0.164 | 1.000 |
| AT4G39690.1 | MICOS complex subunit (Mic60-like) | 10 | 1.30 | 0.164 | 1.000 |
| AT4G39690.1 | MICOS complex subunit (Mic60-like) | 11 | 1.36 | 0.165 | 1.000 |
| AT4G39690.1 | MICOS complex subunit (Mic60-like) | 12 | 1.49 | 0.165 | 1.000 |
| AT4G39690.1 | MICOS complex subunit (Mic60-like) | 13 | 1.71 | 0.163 | 0.990 |
| AT4G39690.1 | MICOS complex subunit (Mic60-like) | 14 | 1.90 | 0.160 | 0.914 |
| AT4G39690.1 | MICOS complex subunit (Mic60-like) | 15 | 1.87 | 0.159 | 0.927 |
| AT4G39690.1 | MICOS complex subunit (Mic60-like) | 16 | 1.70 | 0.158 | 0.986 |
| AT4G39690.1 | MICOS complex subunit (Mic60-like) | 17 | 1.62 | 0.158 | 0.996 |
| AT4G39690.1 | MICOS complex subunit (Mic60-like) | 18 | 1.58 | 0.161 | 0.998 |
| AT4G39690.1 | MICOS complex subunit (Mic60-like) | 19 | 1.58 | 0.165 | 0.999 |
| AT4G39690.1 | MICOS complex subunit (Mic60-like) | 20 | 1.57 | 0.170 | 1.000 |
| AT4G39690.1 | MICOS complex subunit (Mic60-like) | 21 | 1.53 | 0.173 | 1.000 |
| AT4G39690.1 | MICOS complex subunit (Mic60-like) | 22 | 1.38 | 0.173 | 1.000 |
| AT4G39690.1 | MICOS complex subunit (Mic60-like) | 23 | 1.09 | 0.172 | 1.000 |
| AT4G39690.1 | MICOS complex subunit (Mic60-like) | 24 | 0.76 | 0.168 | 1.000 |
| AT4G39690.1 | MICOS complex subunit (Mic60-like) | 25 | 0.50 | 0.164 | 0.877 |
| AT4G39690.1 | MICOS complex subunit (Mic60-like) | 26 | 0.42 | 0.163 | 0.481 |
| AT4G39690.1 | MICOS complex subunit (Mic60-like) | 27 | 0.48 | 0.163 | 0.791 |
| AT4G39690.1 | MICOS complex subunit (Mic60-like) | 28 | 0.59 | 0.163 | 0.993 |
| AT4G39690.1 | MICOS complex subunit (Mic60-like) | 29 | 0.70 | 0.163 | 1.000 |
| AT4G39690.1 | MICOS complex subunit (Mic60-like) | 30 | 0.80 | 0.163 | 1.000 |
| AT4G39690.1 | MICOS complex subunit (Mic60-like) | 31 | 0.97 | 0.164 | 1.000 |
| AT4G39690.1 | MICOS complex subunit (Mic60-like) | 32 | 1.30 | 0.164 | 1.000 |

| | | | | | |
|---|---|---|---|---|---|
| AT4G39690.1 | MICOS complex subunit (Mic60-like) | 33 | 1.72 | 0.165 | 0.989 |
| AT4G39690.1 | MICOS complex subunit (Mic60-like) | 34 | 1.96 | 0.165 | 0.895 |
| AT4G39690.1 | MICOS complex subunit (Mic60-like) | 35 | 1.79 | 0.164 | 0.975 |
| AT4G39690.1 | MICOS complex subunit (Mic60-like) | 36 | 1.24 | 0.164 | 1.000 |
| AT4G39690.1 | MICOS complex subunit (Mic60-like) | 37 | 0.68 | 0.164 | 1.000 |
| AT4G39690.1 | MICOS complex subunit (Mic60-like) | 38 | 0.39 | 0.164 | 0.333 |
| AT4G39690.1 | MICOS complex subunit (Mic60-like) | 39 | 0.33 | 0.164 | 0.113 |
| AT4G39690.1 | MICOS complex subunit (Mic60-like) | 40 | 0.48 | 0.165 | 0.810 |
| AT4G39690.1 | MICOS complex subunit (Mic60-like) | 41 | 0.89 | 0.166 | 1.000 |
| AT4G39690.1 | MICOS complex subunit (Mic60-like) | 42 | 1.56 | 0.168 | 1.000 |
| AT4G39690.1 | MICOS complex subunit (Mic60-like) | 43 | 2.46 | 0.169 | 0.493 |
| AT4G39690.1 | MICOS complex subunit (Mic60-like) | 44 | 2.85 | 0.170 | 0.229 |
| AT4G39690.1 | MICOS complex subunit (Mic60-like) | 45 | 2.10 | 0.172 | 0.843 |
| AT4G39690.1 | MICOS complex subunit (Mic60-like) | 46 | 1.16 | 0.207 | 1.000 |
| AT4G39690.1 | MICOS complex subunit (Mic60-like) | 47 | 0.62 | 0.348 | 1.000 |
| AT4G39690.1 | MICOS complex subunit (Mic60-like) | 48 | 0.34 | 0.568 | 1.000 |
| AT4G39740.1 | Cytochrome c:O2 oxidoreductase assembly protein (HCC2) | 2 | 0.61 | 0.517 | 1.000 |
| AT4G39740.1 | Cytochrome c:O2 oxidoreductase assembly protein (HCC2) | 3 | 0.65 | 0.417 | 1.000 |
| AT4G39740.1 | Cytochrome c:O2 oxidoreductase assembly protein (HCC2) | 6 | 0.78 | 0.272 | 1.000 |
| AT4G39740.1 | Cytochrome c:O2 oxidoreductase assembly protein (HCC2) | 7 | 0.86 | 0.240 | 1.000 |
| AT4G39740.1 | Cytochrome c:O2 oxidoreductase assembly protein (HCC2) | 8 | 0.97 | 0.214 | 1.000 |
| AT4G39740.1 | Cytochrome c:O2 oxidoreductase assembly protein (HCC2) | 9 | 1.14 | 0.196 | 1.000 |
| AT4G39740.1 | Cytochrome c:O2 oxidoreductase assembly protein (HCC2) | 10 | 1.35 | 0.186 | 1.000 |
| AT4G39740.1 | Cytochrome c:O2 oxidoreductase assembly protein (HCC2) | 11 | 1.55 | 0.180 | 0.993 |
| AT4G39740.1 | Cytochrome c:O2 oxidoreductase assembly protein (HCC2) | 12 | 1.69 | 0.178 | 0.961 |
| AT4G39740.1 | Cytochrome c:O2 oxidoreductase assembly protein (HCC2) | 13 | 1.74 | 0.177 | 0.939 |

| | | | | | | |
|---|---|---|---|---|---|---|
| AT4G39740.1 | Cytochrome c:O2 oxidoreductase assembly protein (HCC2) | 14 | 1.71 | 0.176 | 0.947 | |
| AT4G39740.1 | Cytochrome c:O2 oxidoreductase assembly protein (HCC2) | 15 | 1.67 | 0.175 | 0.960 | |
| AT4G39740.1 | Cytochrome c:O2 oxidoreductase assembly protein (HCC2) | 16 | 1.69 | 0.174 | 0.953 | |
| AT4G39740.1 | Cytochrome c:O2 oxidoreductase assembly protein (HCC2) | 17 | 1.90 | 0.174 | 0.830 | |
| AT4G39740.1 | Cytochrome c:O2 oxidoreductase assembly protein (HCC2) | 18 | 2.41 | 0.177 | 0.413 | |
| AT4G39740.1 | Cytochrome c:O2 oxidoreductase assembly protein (HCC2) | 19 | 3.32 | 0.182 | 0.083 | . |
| AT4G39740.1 | Cytochrome c:O2 oxidoreductase assembly protein (HCC2) | 20 | 4.64 | 0.190 | 0.011 | * |
| AT4G39740.1 | Cytochrome c:O2 oxidoreductase assembly protein (HCC2) | 21 | 6.15 | 0.202 | 0.003 | ** |
| AT4G39740.1 | Cytochrome c:O2 oxidoreductase assembly protein (HCC2) | 22 | 7.46 | 0.221 | 0.002 | ** |
| AT4G39740.1 | Cytochrome c:O2 oxidoreductase assembly protein (HCC2) | 23 | 8.17 | 0.247 | 0.006 | ** |
| AT4G39740.1 | Cytochrome c:O2 oxidoreductase assembly protein (HCC2) | 24 | 8.00 | 0.278 | 0.027 | * |
| AT4G39740.1 | Cytochrome c:O2 oxidoreductase assembly protein (HCC2) | 30 | 1.55 | 0.268 | 1.000 | |
| AT4G39740.1 | Cytochrome c:O2 oxidoreductase assembly protein (HCC2) | 31 | 1.12 | 0.237 | 1.000 | |
| AT4G39740.1 | Cytochrome c:O2 oxidoreductase assembly protein (HCC2) | 32 | 0.88 | 0.215 | 1.000 | |
| AT4G39740.1 | Cytochrome c:O2 oxidoreductase assembly protein (HCC2) | 33 | 0.75 | 0.207 | 1.000 | |
| AT4G39740.1 | Cytochrome c:O2 oxidoreductase assembly protein (HCC2) | 34 | 0.67 | 0.213 | 1.000 | |
| AT4G39740.1 | Cytochrome c:O2 oxidoreductase assembly protein (HCC2) | 35 | 0.65 | 0.232 | 1.000 | |
| AT4G39740.1 | Cytochrome c:O2 oxidoreductase assembly protein (HCC2) | 36 | 0.65 | 0.252 | 1.000 | |
| AT4G39740.1 | Cytochrome c:O2 oxidoreductase assembly protein (HCC2) | 39 | 0.73 | 0.286 | 1.000 | |
| AT4G39740.1 | Cytochrome c:O2 oxidoreductase assembly protein (HCC2) | 40 | 0.79 | 0.281 | 1.000 | |

| | | | | | |
|---|---|---|---|---|---|
| AT4G39740.1 | Cytochrome c:O2 oxidoreductase assembly protein (HCC2) | 41 | 0.86 | 0.267 | 1.000 |
| AT4G39740.1 | Cytochrome c:O2 oxidoreductase assembly protein (HCC2) | 42 | 0.95 | 0.246 | 1.000 |
| AT4G39740.1 | Cytochrome c:O2 oxidoreductase assembly protein (HCC2) | 43 | 1.02 | 0.224 | 1.000 |
| AT4G39740.1 | Cytochrome c:O2 oxidoreductase assembly protein (HCC2) | 44 | 1.04 | 0.206 | 1.000 |
| AT4G39740.1 | Cytochrome c:O2 oxidoreductase assembly protein (HCC2) | 45 | 0.97 | 0.213 | 1.000 |
| AT4G39740.1 | Cytochrome c:O2 oxidoreductase assembly protein (HCC2) | 46 | 0.84 | 0.270 | 1.000 |
| AT4G39740.1 | Cytochrome c:O2 oxidoreductase assembly protein (HCC2) | 47 | 0.71 | 0.383 | 1.000 |
| AT5G01530.1 | Photosystem II light harvesting complex subunit (LHCB4.1) | 2 | 1.21 | 0.317 | 1.000 |
| AT5G01530.1 | Photosystem II light harvesting complex subunit (LHCB4.1) | 3 | 0.84 | 0.190 | 1.000 |
| AT5G01530.1 | Photosystem II light harvesting complex subunit (LHCB4.1) | 4 | 0.80 | 0.177 | 1.000 |
| AT5G01530.1 | Photosystem II light harvesting complex subunit (LHCB4.1) | 5 | 1.01 | 0.177 | 1.000 |
| AT5G01530.1 | Photosystem II light harvesting complex subunit (LHCB4.1) | 6 | 1.19 | 0.177 | 1.000 |
| AT5G01530.1 | Photosystem II light harvesting complex subunit (LHCB4.1) | 7 | 1.18 | 0.178 | 1.000 |
| AT5G01530.1 | Photosystem II light harvesting complex subunit (LHCB4.1) | 8 | 1.12 | 0.174 | 1.000 |
| AT5G01530.1 | Photosystem II light harvesting complex subunit (LHCB4.1) | 9 | 1.05 | 0.172 | 1.000 |
| AT5G01530.1 | Photosystem II light harvesting complex subunit (LHCB4.1) | 10 | 0.97 | 0.172 | 1.000 |
| AT5G01530.1 | Photosystem II light harvesting complex subunit (LHCB4.1) | 11 | 1.00 | 0.172 | 1.000 |
| AT5G01530.1 | Photosystem II light harvesting complex subunit (LHCB4.1) | 12 | 1.15 | 0.173 | 1.000 |
| AT5G01530.1 | Photosystem II light harvesting complex subunit (LHCB4.1) | 13 | 1.28 | 0.170 | 1.000 |
| AT5G01530.1 | Photosystem II light harvesting complex subunit (LHCB4.1) | 14 | 1.32 | 0.168 | 1.000 |
| AT5G01530.1 | Photosystem II light harvesting complex subunit (LHCB4.1) | 15 | 1.23 | 0.167 | 1.000 |
| AT5G01530.1 | Photosystem II light harvesting complex subunit (LHCB4.1) | 16 | 1.09 | 0.165 | 1.000 |
| AT5G01530.1 | Photosystem II light harvesting complex subunit (LHCB4.1) | 17 | 1.05 | 0.164 | 1.000 |
| AT5G01530.1 | Photosystem II light harvesting complex subunit (LHCB4.1) | 18 | 1.14 | 0.167 | 1.000 |
| AT5G01530.1 | Photosystem II light harvesting complex subunit (LHCB4.1) | 19 | 1.26 | 0.174 | 1.000 |
| AT5G01530.1 | Photosystem II light harvesting complex subunit (LHCB4.1) | 20 | 1.35 | 0.178 | 1.000 |
| AT5G01530.1 | Photosystem II light harvesting complex subunit (LHCB4.1) | 21 | 1.40 | 0.178 | 1.000 |

| | | | | | | |
|---|---|---|---|---|---|---|
| AT5G01530.1 | Photosystem II light harvesting complex subunit (LHCB4.1) | 22 | 1.41 | 0.178 | 1.000 | |
| AT5G01530.1 | Photosystem II light harvesting complex subunit (LHCB4.1) | 23 | 1.29 | 0.177 | 1.000 | |
| AT5G01530.1 | Photosystem II light harvesting complex subunit (LHCB4.1) | 24 | 1.13 | 0.172 | 1.000 | |
| AT5G01530.1 | Photosystem II light harvesting complex subunit (LHCB4.1) | 25 | 1.04 | 0.169 | 1.000 | |
| AT5G01530.1 | Photosystem II light harvesting complex subunit (LHCB4.1) | 26 | 1.11 | 0.169 | 1.000 | |
| AT5G01530.1 | Photosystem II light harvesting complex subunit (LHCB4.1) | 27 | 1.19 | 0.169 | 1.000 | |
| AT5G01530.1 | Photosystem II light harvesting complex subunit (LHCB4.1) | 28 | 0.98 | 0.170 | 1.000 | |
| AT5G01530.1 | Photosystem II light harvesting complex subunit (LHCB4.1) | 29 | 0.75 | 0.171 | 1.000 | |
| AT5G01530.1 | Photosystem II light harvesting complex subunit (LHCB4.1) | 30 | 0.77 | 0.171 | 1.000 | |
| AT5G01530.1 | Photosystem II light harvesting complex subunit (LHCB4.1) | 31 | 1.11 | 0.172 | 1.000 | |
| AT5G01530.1 | Photosystem II light harvesting complex subunit (LHCB4.1) | 32 | 1.01 | 0.172 | 1.000 | |
| AT5G01530.1 | Photosystem II light harvesting complex subunit (LHCB4.1) | 33 | 0.41 | 0.172 | 0.561 | |
| AT5G01530.1 | Photosystem II light harvesting complex subunit (LHCB4.1) | 34 | 0.17 | 0.173 | 1.000 | |
| AT5G01530.1 | Photosystem II light harvesting complex subunit (LHCB4.1) | 35 | 0.12 | 0.172 | 0.000 | *** |
| AT5G01530.1 | Photosystem II light harvesting complex subunit (LHCB4.1) | 36 | 0.16 | 0.172 | 1.000 | |
| AT5G01530.1 | Photosystem II light harvesting complex subunit (LHCB4.1) | 37 | 0.27 | 0.172 | 0.037 | * |
| AT5G01530.1 | Photosystem II light harvesting complex subunit (LHCB4.1) | 38 | 0.45 | 0.172 | 0.758 | |
| AT5G01530.1 | Photosystem II light harvesting complex subunit (LHCB4.1) | 39 | 0.68 | 0.172 | 1.000 | |
| AT5G01530.1 | Photosystem II light harvesting complex subunit (LHCB4.1) | 40 | 0.81 | 0.172 | 1.000 | |
| AT5G01530.1 | Photosystem II light harvesting complex subunit (LHCB4.1) | 41 | 0.55 | 0.172 | 0.986 | |
| AT5G01530.1 | Photosystem II light harvesting complex subunit (LHCB4.1) | 42 | 0.24 | 0.176 | 0.021 | * |
| AT5G01530.1 | Photosystem II light harvesting complex subunit (LHCB4.1) | 43 | 0.13 | 0.178 | 0.000 | *** |
| AT5G01530.1 | Photosystem II light harvesting complex subunit (LHCB4.1) | 44 | 0.11 | 0.180 | 0.000 | *** |
| AT5G01530.1 | Photosystem II light harvesting complex subunit (LHCB4.1) | 45 | 0.16 | 0.215 | 1.000 | |
| AT5G01530.1 | Photosystem II light harvesting complex subunit (LHCB4.1) | 46 | 0.32 | 0.335 | 0.987 | |
| AT5G01530.1 | Photosystem II light harvesting complex subunit (LHCB4.1) | 47 | 0.69 | 0.586 | 1.000 | |
| AT5G02500.2 | Heat shock cognate protein 70-1 (HSC70-1) | 2 | 0.48 | 0.406 | 1.000 | |
| AT5G02500.2 | Heat shock cognate protein 70-1 (HSC70-1) | 3 | 0.37 | 0.273 | 0.921 | |
| AT5G02500.2 | Heat shock cognate protein 70-1 (HSC70-1) | 4 | 0.29 | 0.243 | 0.494 | |
| AT5G02500.2 | Heat shock cognate protein 70-1 (HSC70-1) | 5 | 0.26 | 0.243 | 1.000 | |
| AT5G02500.2 | Heat shock cognate protein 70-1 (HSC70-1) | 6 | 0.26 | 0.245 | 0.347 | |

| | | | | | |
|---|---|---|---|---|---|
| AT5G02500.2 | Heat shock cognate protein 70-1 (HSC70-1) | 7 | 0.28 | 0.247 | 0.438 |
| AT5G02500.2 | Heat shock cognate protein 70-1 (HSC70-1) | 8 | 0.30 | 0.251 | 0.555 |
| AT5G02500.2 | Heat shock cognate protein 70-1 (HSC70-1) | 9 | 0.33 | 0.259 | 0.756 |
| AT5G02500.2 | Heat shock cognate protein 70-1 (HSC70-1) | 10 | 0.39 | 0.265 | 0.927 |
| AT5G02500.2 | Heat shock cognate protein 70-1 (HSC70-1) | 11 | 0.43 | 0.272 | 0.983 |
| AT5G02500.2 | Heat shock cognate protein 70-1 (HSC70-1) | 12 | 0.46 | 0.278 | 0.994 |
| AT5G02500.2 | Heat shock cognate protein 70-1 (HSC70-1) | 13 | 0.47 | 0.268 | 0.994 |
| AT5G02500.2 | Heat shock cognate protein 70-1 (HSC70-1) | 14 | 0.48 | 0.259 | 0.993 |
| AT5G02500.2 | Heat shock cognate protein 70-1 (HSC70-1) | 15 | 0.51 | 0.252 | 0.996 |
| AT5G02500.2 | Heat shock cognate protein 70-1 (HSC70-1) | 16 | 0.58 | 0.244 | 1.000 |
| AT5G02500.2 | Heat shock cognate protein 70-1 (HSC70-1) | 17 | 0.73 | 0.236 | 1.000 |
| AT5G02500.2 | Heat shock cognate protein 70-1 (HSC70-1) | 18 | 1.00 | 0.234 | 1.000 |
| AT5G02500.2 | Heat shock cognate protein 70-1 (HSC70-1) | 19 | 1.48 | 0.236 | 1.000 |
| AT5G02500.2 | Heat shock cognate protein 70-1 (HSC70-1) | 20 | 2.14 | 0.241 | 0.978 |
| AT5G02500.2 | Heat shock cognate protein 70-1 (HSC70-1) | 21 | 2.73 | 0.243 | 0.799 |
| AT5G02500.2 | Heat shock cognate protein 70-1 (HSC70-1) | 22 | 2.83 | 0.244 | 0.756 |
| AT5G02500.2 | Heat shock cognate protein 70-1 (HSC70-1) | 23 | 2.41 | 0.242 | 0.919 |
| AT5G02500.2 | Heat shock cognate protein 70-1 (HSC70-1) | 24 | 1.85 | 0.238 | 0.998 |
| AT5G02500.2 | Heat shock cognate protein 70-1 (HSC70-1) | 25 | 1.24 | 0.235 | 1.000 |
| AT5G02500.2 | Heat shock cognate protein 70-1 (HSC70-1) | 26 | 0.78 | 0.234 | 1.000 |
| AT5G02500.2 | Heat shock cognate protein 70-1 (HSC70-1) | 27 | 0.63 | 0.234 | 1.000 |
| AT5G02500.2 | Heat shock cognate protein 70-1 (HSC70-1) | 28 | 0.76 | 0.235 | 1.000 |
| AT5G02500.2 | Heat shock cognate protein 70-1 (HSC70-1) | 29 | 1.06 | 0.235 | 1.000 |
| AT5G02500.2 | Heat shock cognate protein 70-1 (HSC70-1) | 30 | 1.29 | 0.235 | 1.000 |
| AT5G02500.2 | Heat shock cognate protein 70-1 (HSC70-1) | 31 | 1.46 | 0.236 | 1.000 |
| AT5G02500.2 | Heat shock cognate protein 70-1 (HSC70-1) | 32 | 1.54 | 0.236 | 1.000 |
| AT5G02500.2 | Heat shock cognate protein 70-1 (HSC70-1) | 33 | 1.37 | 0.236 | 1.000 |
| AT5G02500.2 | Heat shock cognate protein 70-1 (HSC70-1) | 34 | 1.06 | 0.237 | 1.000 |
| AT5G02500.2 | Heat shock cognate protein 70-1 (HSC70-1) | 35 | 0.85 | 0.236 | 1.000 |
| AT5G02500.2 | Heat shock cognate protein 70-1 (HSC70-1) | 36 | 0.73 | 0.236 | 1.000 |
| AT5G02500.2 | Heat shock cognate protein 70-1 (HSC70-1) | 37 | 0.60 | 0.235 | 1.000 |

| | | | | | |
|---|---|---|---|---|---|
| AT5G02500.2 | Heat shock cognate protein 70-1 (HSC70-1) | 38 | 0.48 | 0.235 | 0.979 |
| AT5G02500.2 | Heat shock cognate protein 70-1 (HSC70-1) | 39 | 0.49 | 0.235 | 0.985 |
| AT5G02500.2 | Heat shock cognate protein 70-1 (HSC70-1) | 40 | 0.70 | 0.236 | 1.000 |
| AT5G02500.2 | Heat shock cognate protein 70-1 (HSC70-1) | 41 | 1.09 | 0.237 | 1.000 |
| AT5G02500.2 | Heat shock cognate protein 70-1 (HSC70-1) | 42 | 1.26 | 0.240 | 1.000 |
| AT5G02500.2 | Heat shock cognate protein 70-1 (HSC70-1) | 43 | 1.05 | 0.242 | 1.000 |
| AT5G02500.2 | Heat shock cognate protein 70-1 (HSC70-1) | 44 | 0.77 | 0.241 | 1.000 |
| AT5G02500.2 | Heat shock cognate protein 70-1 (HSC70-1) | 45 | 0.63 | 0.245 | 1.000 |
| AT5G02500.2 | Heat shock cognate protein 70-1 (HSC70-1) | 46 | 0.65 | 0.290 | 1.000 |
| AT5G02500.2 | Heat shock cognate protein 70-1 (HSC70-1) | 47 | 0.82 | 0.452 | 1.000 |
| AT5G02740.1 | Ribosomal S24e family protein | 2 | 1.07 | 0.464 | 1.000 |
| AT5G02740.1 | Ribosomal S24e family protein | 3 | 1.17 | 0.405 | 1.000 |
| AT5G02740.1 | Ribosomal S24e family protein | 11 | 1.32 | 0.309 | 1.000 |
| AT5G02740.1 | Ribosomal S24e family protein | 12 | 1.17 | 0.264 | 1.000 |
| AT5G02740.1 | Ribosomal S24e family protein | 13 | 1.01 | 0.228 | 1.000 |
| AT5G02740.1 | Ribosomal S24e family protein | 14 | 0.86 | 0.209 | 1.000 |
| AT5G02740.1 | Ribosomal S24e family protein | 15 | 0.73 | 0.197 | 1.000 |
| AT5G02740.1 | Ribosomal S24e family protein | 16 | 0.64 | 0.189 | 0.996 |
| AT5G02740.1 | Ribosomal S24e family protein | 17 | 0.64 | 0.187 | 0.995 |
| AT5G02740.1 | Ribosomal S24e family protein | 18 | 0.78 | 0.190 | 1.000 |
| AT5G02740.1 | Ribosomal S24e family protein | 19 | 1.00 | 0.201 | 1.000 |
| AT5G02740.1 | Ribosomal S24e family protein | 20 | 1.17 | 0.220 | 1.000 |
| AT5G02740.1 | Ribosomal S24e family protein | 21 | 1.20 | 0.239 | 1.000 |
| AT5G02740.1 | Ribosomal S24e family protein | 22 | 1.14 | 0.266 | 1.000 |
| AT5G02740.1 | Ribosomal S24e family protein | 23 | 1.08 | 0.282 | 1.000 |
| AT5G02740.1 | Ribosomal S24e family protein | 26 | 1.11 | 0.258 | 1.000 |
| AT5G02740.1 | Ribosomal S24e family protein | 27 | 1.18 | 0.225 | 1.000 |
| AT5G02740.1 | Ribosomal S24e family protein | 28 | 1.31 | 0.202 | 1.000 |
| AT5G02740.1 | Ribosomal S24e family protein | 29 | 1.52 | 0.190 | 0.998 |
| AT5G02740.1 | Ribosomal S24e family protein | 30 | 1.57 | 0.187 | 0.995 |
| AT5G02740.1 | Ribosomal S24e family protein | 31 | 1.51 | 0.186 | 0.998 |
| AT5G02740.1 | Ribosomal S24e family protein | 32 | 1.44 | 0.187 | 1.000 |
| AT5G02740.1 | Ribosomal S24e family protein | 33 | 1.34 | 0.187 | 1.000 |
| AT5G02740.1 | Ribosomal S24e family protein | 34 | 1.23 | 0.188 | 1.000 |
| AT5G02740.1 | Ribosomal S24e family protein | 35 | 1.10 | 0.188 | 1.000 |
| AT5G02740.1 | Ribosomal S24e family protein | 36 | 0.89 | 0.189 | 1.000 |
| AT5G02740.1 | Ribosomal S24e family protein | 37 | 0.64 | 0.192 | 0.996 |
| AT5G02740.1 | Ribosomal S24e family protein | 38 | 0.46 | 0.207 | 0.814 |
| AT5G02740.1 | Ribosomal S24e family protein | 39 | 0.37 | 0.258 | 0.808 |
| AT5G02740.1 | Ribosomal S24e family protein | 40 | 0.37 | 0.361 | 0.984 |
| AT5G02740.1 | Ribosomal S24e family protein | 46 | 4.25 | 1.565 | 1.000 |
| AT5G02740.1 | Ribosomal S24e family protein | 47 | 7.76 | 1.869 | 1.000 |
| AT5G03290.1 | NAD isocitrate dehydrogenase subunit (IDH5, catalytic) | 2 | 0.64 | 0.264 | 1.000 |
| AT5G03290.1 | NAD isocitrate dehydrogenase subunit (IDH5, catalytic) | 3 | 0.60 | 0.199 | 0.992 |
| AT5G03290.1 | NAD isocitrate dehydrogenase subunit (IDH5, catalytic) | 4 | 0.56 | 0.164 | 0.895 |
| AT5G03290.1 | NAD isocitrate dehydrogenase subunit (IDH5, catalytic) | 5 | 0.55 | 0.150 | 0.761 |
| AT5G03290.1 | NAD isocitrate dehydrogenase subunit (IDH5, catalytic) | 6 | 0.56 | 0.143 | 0.767 |

| | | | | | | |
|---|---|---|---|---|---|---|
| AT5G03290.1 | NAD isocitrate dehydrogenase subunit (IDH5, catalytic) | 7 | 0.61 | 0.140 | 0.890 | |
| AT5G03290.1 | NAD isocitrate dehydrogenase subunit (IDH5, catalytic) | 8 | 0.69 | 0.137 | 0.987 | |
| AT5G03290.1 | NAD isocitrate dehydrogenase subunit (IDH5, catalytic) | 9 | 0.79 | 0.135 | 1.000 | |
| AT5G03290.1 | NAD isocitrate dehydrogenase subunit (IDH5, catalytic) | 10 | 0.88 | 0.134 | 1.000 | |
| AT5G03290.1 | NAD isocitrate dehydrogenase subunit (IDH5, catalytic) | 11 | 0.98 | 0.133 | 1.000 | |
| AT5G03290.1 | NAD isocitrate dehydrogenase subunit (IDH5, catalytic) | 12 | 1.08 | 0.132 | 1.000 | |
| AT5G03290.1 | NAD isocitrate dehydrogenase subunit (IDH5, catalytic) | 13 | 1.17 | 0.131 | 1.000 | |
| AT5G03290.1 | NAD isocitrate dehydrogenase subunit (IDH5, catalytic) | 14 | 1.20 | 0.130 | 1.000 | |
| AT5G03290.1 | NAD isocitrate dehydrogenase subunit (IDH5, catalytic) | 15 | 1.12 | 0.129 | 1.000 | |
| AT5G03290.1 | NAD isocitrate dehydrogenase subunit (IDH5, catalytic) | 16 | 0.98 | 0.129 | 1.000 | |
| AT5G03290.1 | NAD isocitrate dehydrogenase subunit (IDH5, catalytic) | 17 | 0.84 | 0.129 | 1.000 | |
| AT5G03290.1 | NAD isocitrate dehydrogenase subunit (IDH5, catalytic) | 18 | 0.73 | 0.131 | 0.996 | |
| AT5G03290.1 | NAD isocitrate dehydrogenase subunit (IDH5, catalytic) | 19 | 0.65 | 0.133 | 0.936 | |
| AT5G03290.1 | NAD isocitrate dehydrogenase subunit (IDH5, catalytic) | 20 | 0.59 | 0.135 | 0.788 | |
| AT5G03290.1 | NAD isocitrate dehydrogenase subunit (IDH5, catalytic) | 21 | 0.52 | 0.136 | 0.515 | |
| AT5G03290.1 | NAD isocitrate dehydrogenase subunit (IDH5, catalytic) | 22 | 0.44 | 0.137 | 0.195 | |
| AT5G03290.1 | NAD isocitrate dehydrogenase subunit (IDH5, catalytic) | 23 | 0.38 | 0.136 | 0.047 | * |
| AT5G03290.1 | NAD isocitrate dehydrogenase subunit (IDH5, catalytic) | 24 | 0.35 | 0.134 | 0.019 | * |
| AT5G03290.1 | NAD isocitrate dehydrogenase subunit (IDH5, catalytic) | 25 | 0.36 | 0.133 | 0.023 | * |
| AT5G03290.1 | NAD isocitrate dehydrogenase subunit (IDH5, catalytic) | 26 | 0.41 | 0.132 | 0.068 | . |
| AT5G03290.1 | NAD isocitrate dehydrogenase subunit (IDH5, catalytic) | 27 | 0.48 | 0.132 | 0.269 | |
| AT5G03290.1 | NAD isocitrate dehydrogenase subunit (IDH5, catalytic) | 28 | 0.57 | 0.132 | 0.686 | |
| AT5G03290.1 | NAD isocitrate dehydrogenase subunit (IDH5, catalytic) | 29 | 0.67 | 0.132 | 0.961 | |
| AT5G03290.1 | NAD isocitrate dehydrogenase subunit (IDH5, catalytic) | 30 | 0.74 | 0.132 | 0.998 | |
| AT5G03290.1 | NAD isocitrate dehydrogenase subunit (IDH5, catalytic) | 31 | 0.78 | 0.132 | 1.000 | |
| AT5G03290.1 | NAD isocitrate dehydrogenase subunit (IDH5, catalytic) | 32 | 0.77 | 0.133 | 1.000 | |
| AT5G03290.1 | NAD isocitrate dehydrogenase subunit (IDH5, catalytic) | 33 | 0.75 | 0.133 | 0.999 | |
| AT5G03290.1 | NAD isocitrate dehydrogenase subunit (IDH5, catalytic) | 34 | 0.76 | 0.133 | 0.999 | |
| AT5G03290.1 | NAD isocitrate dehydrogenase subunit (IDH5, catalytic) | 35 | 0.81 | 0.133 | 1.000 | |
| AT5G03290.1 | NAD isocitrate dehydrogenase subunit (IDH5, catalytic) | 36 | 0.90 | 0.133 | 1.000 | |
| AT5G03290.1 | NAD isocitrate dehydrogenase subunit (IDH5, catalytic) | 37 | 1.01 | 0.133 | 1.000 | |

| | | | | | |
|---|---|---|---|---|---|
| AT5G03290.1 | NAD isocitrate dehydrogenase subunit (IDH5, catalytic) | 38 | 1.13 | 0.133 | 1.000 |
| AT5G03290.1 | NAD isocitrate dehydrogenase subunit (IDH5, catalytic) | 39 | 1.22 | 0.133 | 1.000 |
| AT5G03290.1 | NAD isocitrate dehydrogenase subunit (IDH5, catalytic) | 40 | 1.28 | 0.134 | 1.000 |
| AT5G03290.1 | NAD isocitrate dehydrogenase subunit (IDH5, catalytic) | 41 | 1.31 | 0.135 | 1.000 |
| AT5G03290.1 | NAD isocitrate dehydrogenase subunit (IDH5, catalytic) | 42 | 1.34 | 0.136 | 0.999 |
| AT5G03290.1 | NAD isocitrate dehydrogenase subunit (IDH5, catalytic) | 43 | 1.33 | 0.136 | 0.999 |
| AT5G03290.1 | NAD isocitrate dehydrogenase subunit (IDH5, catalytic) | 44 | 1.27 | 0.139 | 1.000 |
| AT5G03290.1 | NAD isocitrate dehydrogenase subunit (IDH5, catalytic) | 45 | 1.15 | 0.152 | 1.000 |
| AT5G03290.1 | NAD isocitrate dehydrogenase subunit (IDH5, catalytic) | 46 | 0.99 | 0.188 | 1.000 |
| AT5G03290.1 | NAD isocitrate dehydrogenase subunit (IDH5, catalytic) | 47 | 0.84 | 0.257 | 1.000 |
| AT5G03905.1 | Iron-sulfur assembly protein IscA-like 2 | 2 | 1.14 | 0.306 | 0.983 |
| AT5G03905.1 | Iron-sulfur assembly protein IscA-like 2 | 3 | 1.13 | 0.297 | 0.984 |
| AT5G03905.1 | Iron-sulfur assembly protein IscA-like 2 | 26 | 1.00 | 0.124 | 1.000 |
| AT5G03905.1 | Iron-sulfur assembly protein IscA-like 2 | 27 | 0.99 | 0.119 | 1.000 |
| AT5G03905.1 | Iron-sulfur assembly protein IscA-like 2 | 28 | 0.99 | 0.114 | 0.999 |
| AT5G03905.1 | Iron-sulfur assembly protein IscA-like 2 | 29 | 0.98 | 0.110 | 0.998 |
| AT5G03905.1 | Iron-sulfur assembly protein IscA-like 2 | 30 | 0.98 | 0.106 | 0.996 |
| AT5G03905.1 | Iron-sulfur assembly protein IscA-like 2 | 31 | 0.97 | 0.102 | 0.993 |
| AT5G03905.1 | Iron-sulfur assembly protein IscA-like 2 | 32 | 0.97 | 0.100 | 0.990 |
| AT5G03905.1 | Iron-sulfur assembly protein IscA-like 2 | 33 | 0.96 | 0.098 | 0.986 |
| AT5G03905.1 | Iron-sulfur assembly protein IscA-like 2 | 34 | 0.96 | 0.097 | 0.981 |
| AT5G03905.1 | Iron-sulfur assembly protein IscA-like 2 | 35 | 0.95 | 0.096 | 0.976 |
| AT5G03905.1 | Iron-sulfur assembly protein IscA-like 2 | 36 | 0.95 | 0.097 | 0.971 |
| AT5G03905.1 | Iron-sulfur assembly protein IscA-like 2 | 37 | 0.94 | 0.098 | 0.965 |
| AT5G03905.1 | Iron-sulfur assembly protein IscA-like 2 | 40 | 0.93 | 0.107 | 0.953 |
| AT5G03905.1 | Iron-sulfur assembly protein IscA-like 2 | 41 | 0.92 | 0.111 | 0.950 |
| AT5G03905.1 | Iron-sulfur assembly protein IscA-like 2 | 42 | 0.92 | 0.115 | 0.948 |
| AT5G03905.1 | Iron-sulfur assembly protein IscA-like 2 | 43 | 0.91 | 0.120 | 0.946 |
| AT5G03905.1 | Iron-sulfur assembly protein IscA-like 2 | 44 | 0.91 | 0.126 | 0.944 |
| AT5G03905.1 | Iron-sulfur assembly protein IscA-like 2 | 45 | 0.90 | 0.131 | 0.943 |
| AT5G03905.1 | Iron-sulfur assembly protein IscA-like 2 | 46 | 0.90 | 0.138 | 0.943 |

| | | | | | | |
|---|---|---|---|---|---|---|
| AT5G03905.1 | Iron-sulfur assembly protein IscA-like 2 | 47 | 0.89 | 0.144 | 0.943 | |
| AT5G04140.1 | Glutamate synthase 1 (GLU1) | 2 | 0.98 | 0.749 | 1.000 | |
| AT5G04140.1 | Glutamate synthase 1 (GLU1) | 3 | 1.08 | 0.718 | 1.000 | |
| AT5G04140.1 | Glutamate synthase 1 (GLU1) | 25 | 5.84 | 0.338 | 0.118 | |
| AT5G04140.1 | Glutamate synthase 1 (GLU1) | 26 | 5.99 | 0.324 | 0.089 | . |
| AT5G04140.1 | Glutamate synthase 1 (GLU1) | 27 | 6.07 | 0.315 | 0.071 | . |
| AT5G04140.1 | Glutamate synthase 1 (GLU1) | 28 | 6.06 | 0.309 | 0.065 | . |
| AT5G04140.1 | Glutamate synthase 1 (GLU1) | 38 | 2.84 | 0.273 | 0.372 | |
| AT5G04140.1 | Glutamate synthase 1 (GLU1) | 39 | 2.44 | 0.270 | 0.516 | |
| AT5G04140.1 | Glutamate synthase 1 (GLU1) | 40 | 2.05 | 0.270 | 0.708 | |
| AT5G04140.1 | Glutamate synthase 1 (GLU1) | 41 | 1.69 | 0.276 | 0.895 | |
| AT5G04140.1 | Glutamate synthase 1 (GLU1) | 42 | 1.36 | 0.287 | 0.990 | |
| AT5G04140.1 | Glutamate synthase 1 (GLU1) | 43 | 1.07 | 0.304 | 1.000 | |
| AT5G04140.1 | Glutamate synthase 1 (GLU1) | 46 | 0.50 | 0.390 | 0.914 | |
| AT5G04140.1 | Glutamate synthase 1 (GLU1) | 47 | 0.38 | 0.431 | 0.820 | |
| AT5G04710.1 | M18 aspartyl aminopeptidase 2 (DAP2) | 2 | 1.11 | 0.783 | 1.000 | |
| AT5G04710.1 | M18 aspartyl aminopeptidase 2 (DAP2) | 3 | 0.54 | 0.906 | 1.000 | |
| AT5G04710.1 | M18 aspartyl aminopeptidase 2 (DAP2) | 25 | 0.09 | 0.443 | 0.142 | |
| AT5G04710.1 | M18 aspartyl aminopeptidase 2 (DAP2) | 26 | 0.31 | 0.334 | 0.564 | |
| AT5G04710.1 | M18 aspartyl aminopeptidase 2 (DAP2) | 27 | 1.15 | 0.307 | 1.000 | |
| AT5G04710.1 | M18 aspartyl aminopeptidase 2 (DAP2) | 28 | 3.50 | 0.333 | 0.481 | |
| AT5G04710.1 | M18 aspartyl aminopeptidase 2 (DAP2) | 29 | 6.68 | 0.423 | 0.294 | |
| AT5G04710.1 | M18 aspartyl aminopeptidase 2 (DAP2) | 30 | 8.95 | 0.585 | 0.486 | |
| AT5G04710.1 | M18 aspartyl aminopeptidase 2 (DAP2) | 46 | 0.01 | 0.618 | 0.018 | * |
| AT5G04710.1 | M18 aspartyl aminopeptidase 2 (DAP2) | 47 | 0.01 | 0.645 | 0.007 | ** |
| AT5G05370.1 | Ubiquinol:cytochrome c oxidoreductase subunit (QCR8) | 2 | 1.29 | 0.621 | 1.000 | |
| AT5G05370.1 | Ubiquinol:cytochrome c oxidoreductase subunit (QCR8) | 3 | 1.37 | 0.554 | 1.000 | |
| AT5G05370.1 | Ubiquinol:cytochrome c oxidoreductase subunit (QCR8) | 4 | 1.48 | 0.520 | 1.000 | |
| AT5G05370.1 | Ubiquinol:cytochrome c oxidoreductase subunit (QCR8) | 5 | 1.61 | 0.506 | 1.000 | |
| AT5G05370.1 | Ubiquinol:cytochrome c oxidoreductase subunit (QCR8) | 6 | 1.76 | 0.501 | 0.999 | |
| AT5G05370.1 | Ubiquinol:cytochrome c oxidoreductase subunit (QCR8) | 7 | 1.92 | 0.500 | 0.997 | |
| AT5G05370.1 | Ubiquinol:cytochrome c oxidoreductase subunit (QCR8) | 8 | 2.09 | 0.499 | 0.992 | |
| AT5G05370.1 | Ubiquinol:cytochrome c oxidoreductase subunit (QCR8) | 9 | 2.24 | 0.499 | 0.984 | |
| AT5G05370.1 | Ubiquinol:cytochrome c oxidoreductase subunit (QCR8) | 10 | 2.38 | 0.498 | 0.975 | |
| AT5G05370.1 | Ubiquinol:cytochrome c oxidoreductase subunit (QCR8) | 11 | 2.51 | 0.498 | 0.962 | |
| AT5G05370.1 | Ubiquinol:cytochrome c oxidoreductase subunit (QCR8) | 12 | 2.69 | 0.497 | 0.943 | |
| AT5G05370.1 | Ubiquinol:cytochrome c oxidoreductase subunit (QCR8) | 13 | 2.92 | 0.496 | 0.910 | |
| AT5G05370.1 | Ubiquinol:cytochrome c oxidoreductase subunit (QCR8) | 14 | 3.26 | 0.495 | 0.859 | |

| | | | | | |
|---|---|---|---|---|---|
| AT5G05370.1 | Ubiquinol:cytochrome c oxidoreductase subunit (QCR8) | 15 | 3.71 | 0.495 | 0.787 |
| AT5G05370.1 | Ubiquinol:cytochrome c oxidoreductase subunit (QCR8) | 16 | 4.34 | 0.495 | 0.689 |
| AT5G05370.1 | Ubiquinol:cytochrome c oxidoreductase subunit (QCR8) | 17 | 5.24 | 0.495 | 0.569 |
| AT5G05370.1 | Ubiquinol:cytochrome c oxidoreductase subunit (QCR8) | 18 | 6.42 | 0.496 | 0.447 |
| AT5G05370.1 | Ubiquinol:cytochrome c oxidoreductase subunit (QCR8) | 19 | 7.75 | 0.497 | 0.348 |
| AT5G05370.1 | Ubiquinol:cytochrome c oxidoreductase subunit (QCR8) | 20 | 8.76 | 0.498 | 0.292 |
| AT5G05370.1 | Ubiquinol:cytochrome c oxidoreductase subunit (QCR8) | 21 | 8.88 | 0.499 | 0.291 |
| AT5G05370.1 | Ubiquinol:cytochrome c oxidoreductase subunit (QCR8) | 22 | 8.00 | 0.499 | 0.339 |
| AT5G05370.1 | Ubiquinol:cytochrome c oxidoreductase subunit (QCR8) | 23 | 6.74 | 0.498 | 0.427 |
| AT5G05370.1 | Ubiquinol:cytochrome c oxidoreductase subunit (QCR8) | 24 | 5.94 | 0.497 | 0.497 |
| AT5G05370.1 | Ubiquinol:cytochrome c oxidoreductase subunit (QCR8) | 25 | 5.98 | 0.496 | 0.491 |
| AT5G05370.1 | Ubiquinol:cytochrome c oxidoreductase subunit (QCR8) | 26 | 6.75 | 0.495 | 0.418 |
| AT5G05370.1 | Ubiquinol:cytochrome c oxidoreductase subunit (QCR8) | 27 | 7.71 | 0.495 | 0.348 |
| AT5G05370.1 | Ubiquinol:cytochrome c oxidoreductase subunit (QCR8) | 28 | 7.89 | 0.495 | 0.338 |
| AT5G05370.1 | Ubiquinol:cytochrome c oxidoreductase subunit (QCR8) | 29 | 6.96 | 0.495 | 0.403 |
| AT5G05370.1 | Ubiquinol:cytochrome c oxidoreductase subunit (QCR8) | 30 | 5.81 | 0.496 | 0.507 |
| AT5G05370.1 | Ubiquinol:cytochrome c oxidoreductase subunit (QCR8) | 31 | 4.86 | 0.497 | 0.620 |
| AT5G05370.1 | Ubiquinol:cytochrome c oxidoreductase subunit (QCR8) | 32 | 4.11 | 0.498 | 0.729 |
| AT5G05370.1 | Ubiquinol:cytochrome c oxidoreductase subunit (QCR8) | 33 | 3.51 | 0.500 | 0.826 |
| AT5G05370.1 | Ubiquinol:cytochrome c oxidoreductase subunit (QCR8) | 34 | 3.03 | 0.503 | 0.902 |
| AT5G05370.1 | Ubiquinol:cytochrome c oxidoreductase subunit (QCR8) | 35 | 2.70 | 0.510 | 0.949 |
| AT5G05370.1 | Ubiquinol:cytochrome c oxidoreductase subunit (QCR8) | 36 | 2.56 | 0.522 | 0.968 |
| AT5G05370.1 | Ubiquinol:cytochrome c oxidoreductase subunit (QCR8) | 37 | 2.58 | 0.540 | 0.973 |
| AT5G05370.1 | Ubiquinol:cytochrome c oxidoreductase subunit (QCR8) | 38 | 2.71 | 0.560 | 0.970 |
| AT5G05370.1 | Ubiquinol:cytochrome c oxidoreductase subunit (QCR8) | 41 | 3.49 | 0.646 | 0.951 |
| AT5G05370.1 | Ubiquinol:cytochrome c oxidoreductase subunit (QCR8) | 42 | 3.74 | 0.690 | 0.954 |
| AT5G05370.1 | Ubiquinol:cytochrome c oxidoreductase subunit (QCR8) | 43 | 3.91 | 0.739 | 0.963 |
| AT5G05370.1 | Ubiquinol:cytochrome c oxidoreductase subunit (QCR8) | 46 | 3.99 | 0.987 | 0.995 |
| AT5G05370.1 | Ubiquinol:cytochrome c oxidoreductase subunit (QCR8) | 47 | 3.95 | 1.111 | 0.998 |
| AT5G05520.1 | Outer membrane OMP85 family protein | 2 | 0.84 | 0.261 | 1.000 |
| AT5G05520.1 | Outer membrane OMP85 family protein | 3 | 0.89 | 0.163 | 1.000 |

| | | | | | |
|---|---|---|---|---|---|
| AT5G05520.1 | Outer membrane OMP85 family protein | 4 | 1.00 | 0.158 | 1.000 |
| AT5G05520.1 | Outer membrane OMP85 family protein | 5 | 1.11 | 0.166 | 1.000 |
| AT5G05520.1 | Outer membrane OMP85 family protein | 6 | 1.14 | 0.171 | 1.000 |
| AT5G05520.1 | Outer membrane OMP85 family protein | 7 | 1.21 | 0.163 | 1.000 |
| AT5G05520.1 | Outer membrane OMP85 family protein | 8 | 1.27 | 0.154 | 1.000 |
| AT5G05520.1 | Outer membrane OMP85 family protein | 9 | 1.17 | 0.149 | 1.000 |
| AT5G05520.1 | Outer membrane OMP85 family protein | 10 | 0.90 | 0.147 | 1.000 |
| AT5G05520.1 | Outer membrane OMP85 family protein | 11 | 0.70 | 0.148 | 1.000 |
| AT5G05520.1 | Outer membrane OMP85 family protein | 12 | 0.69 | 0.148 | 0.999 |
| AT5G05520.1 | Outer membrane OMP85 family protein | 13 | 0.90 | 0.146 | 1.000 |
| AT5G05520.1 | Outer membrane OMP85 family protein | 14 | 1.41 | 0.144 | 1.000 |
| AT5G05520.1 | Outer membrane OMP85 family protein | 15 | 1.77 | 0.144 | 0.899 |
| AT5G05520.1 | Outer membrane OMP85 family protein | 16 | 1.63 | 0.143 | 0.973 |
| AT5G05520.1 | Outer membrane OMP85 family protein | 17 | 1.38 | 0.142 | 1.000 |
| AT5G05520.1 | Outer membrane OMP85 family protein | 18 | 1.23 | 0.143 | 1.000 |
| AT5G05520.1 | Outer membrane OMP85 family protein | 19 | 1.21 | 0.147 | 1.000 |
| AT5G05520.1 | Outer membrane OMP85 family protein | 20 | 1.25 | 0.151 | 1.000 |
| AT5G05520.1 | Outer membrane OMP85 family protein | 21 | 1.25 | 0.152 | 1.000 |
| AT5G05520.1 | Outer membrane OMP85 family protein | 22 | 1.22 | 0.153 | 1.000 |
| AT5G05520.1 | Outer membrane OMP85 family protein | 23 | 1.13 | 0.151 | 1.000 |
| AT5G05520.1 | Outer membrane OMP85 family protein | 24 | 0.94 | 0.148 | 1.000 |
| AT5G05520.1 | Outer membrane OMP85 family protein | 25 | 0.80 | 0.146 | 1.000 |
| AT5G05520.1 | Outer membrane OMP85 family protein | 26 | 0.88 | 0.146 | 1.000 |
| AT5G05520.1 | Outer membrane OMP85 family protein | 27 | 1.13 | 0.147 | 1.000 |
| AT5G05520.1 | Outer membrane OMP85 family protein | 28 | 1.27 | 0.147 | 1.000 |
| AT5G05520.1 | Outer membrane OMP85 family protein | 29 | 1.15 | 0.147 | 1.000 |
| AT5G05520.1 | Outer membrane OMP85 family protein | 30 | 0.90 | 0.146 | 1.000 |
| AT5G05520.1 | Outer membrane OMP85 family protein | 31 | 0.77 | 0.147 | 1.000 |
| AT5G05520.1 | Outer membrane OMP85 family protein | 32 | 0.86 | 0.147 | 1.000 |
| AT5G05520.1 | Outer membrane OMP85 family protein | 33 | 1.12 | 0.147 | 1.000 |
| AT5G05520.1 | Outer membrane OMP85 family protein | 34 | 1.43 | 0.147 | 1.000 |

| | | | | | | |
|---|---|---|---|---|---|---|
| AT5G05520.1 | Outer membrane OMP85 family protein | 35 | 1.58 | 0.147 | 0.992 | |
| AT5G05520.1 | Outer membrane OMP85 family protein | 36 | 1.40 | 0.146 | 1.000 | |
| AT5G05520.1 | Outer membrane OMP85 family protein | 37 | 1.14 | 0.147 | 1.000 | |
| AT5G05520.1 | Outer membrane OMP85 family protein | 38 | 1.05 | 0.148 | 1.000 | |
| AT5G05520.1 | Outer membrane OMP85 family protein | 39 | 0.82 | 0.149 | 1.000 | |
| AT5G05520.1 | Outer membrane OMP85 family protein | 40 | 0.45 | 0.157 | 0.532 | |
| AT5G05520.1 | Outer membrane OMP85 family protein | 41 | 0.26 | 0.204 | 0.129 | |
| AT5G05520.1 | Outer membrane OMP85 family protein | 42 | 0.21 | 0.297 | 0.457 | |
| AT5G05520.1 | Outer membrane OMP85 family protein | 46 | 0.85 | 1.001 | 1.000 | |
| AT5G05520.1 | Outer membrane OMP85 family protein | 47 | 1.59 | 1.271 | 1.000 | |
| AT5G06240.1 | ? | 2 | 1.17 | 0.421 | 0.999 | |
| AT5G06240.1 | ? | 3 | 1.18 | 0.402 | 0.999 | |
| AT5G06240.1 | ? | 4 | 1.18 | 0.387 | 0.999 | |
| AT5G06240.1 | ? | 25 | 0.74 | 0.271 | 0.959 | |
| AT5G06240.1 | ? | 26 | 0.68 | 0.273 | 0.919 | |
| AT5G06240.1 | ? | 27 | 0.63 | 0.278 | 0.864 | |
| AT5G06240.1 | ? | 28 | 0.57 | 0.286 | 0.796 | |
| AT5G06240.1 | ? | 29 | 0.52 | 0.296 | 0.721 | |
| AT5G06240.1 | ? | 46 | 0.05 | 0.532 | 0.051 | . |
| AT5G06240.1 | ? | 47 | 0.04 | 0.553 | 0.048 | * |
| AT5G06580.1 | D-lactate dehydrogenase (D-LDH) | 2 | 1.00 | 0.418 | 1.000 | |
| AT5G06580.1 | D-lactate dehydrogenase (D-LDH) | 3 | 0.76 | 0.412 | 1.000 | |
| AT5G06580.1 | D-lactate dehydrogenase (D-LDH) | 31 | 0.34 | 0.322 | 0.767 | |
| AT5G06580.1 | D-lactate dehydrogenase (D-LDH) | 32 | 0.55 | 0.254 | 0.970 | |
| AT5G06580.1 | D-lactate dehydrogenase (D-LDH) | 33 | 0.96 | 0.236 | 1.000 | |
| AT5G06580.1 | D-lactate dehydrogenase (D-LDH) | 34 | 1.02 | 0.230 | 1.000 | |
| AT5G06580.1 | D-lactate dehydrogenase (D-LDH) | 35 | 0.63 | 0.228 | 0.991 | |
| AT5G06580.1 | D-lactate dehydrogenase (D-LDH) | 36 | 0.51 | 0.227 | 0.871 | |
| AT5G06580.1 | D-lactate dehydrogenase (D-LDH) | 37 | 0.67 | 0.227 | 0.998 | |
| AT5G06580.1 | D-lactate dehydrogenase (D-LDH) | 38 | 1.07 | 0.227 | 1.000 | |
| AT5G06580.1 | D-lactate dehydrogenase (D-LDH) | 39 | 1.57 | 0.226 | 0.992 | |
| AT5G06580.1 | D-lactate dehydrogenase (D-LDH) | 40 | 1.93 | 0.225 | 0.883 | |
| AT5G06580.1 | D-lactate dehydrogenase (D-LDH) | 41 | 1.79 | 0.227 | 0.946 | |
| AT5G06580.1 | D-lactate dehydrogenase (D-LDH) | 42 | 1.83 | 0.228 | 0.935 | |
| AT5G06580.1 | D-lactate dehydrogenase (D-LDH) | 43 | 2.52 | 0.229 | 0.555 | |
| AT5G06580.1 | D-lactate dehydrogenase (D-LDH) | 44 | 3.16 | 0.229 | 0.272 | |

| | | | | | |
|---|---|---|---|---|---|
| AT5G06580.1 | D-lactate dehydrogenase (D-LDH) | 45 | 2.10 | 0.231 | 0.813 |
| AT5G06580.1 | D-lactate dehydrogenase (D-LDH) | 46 | 0.60 | 0.253 | 0.991 |
| AT5G06580.1 | D-lactate dehydrogenase (D-LDH) | 47 | 0.10 | 0.385 | 0.106 |
| AT5G07090.2 | 40S ribosomal protein S4-2 (RPS4B) | 2 | 0.75 | 0.190 | 0.801 |
| AT5G07090.2 | 40S ribosomal protein S4-2 (RPS4B) | 3 | 0.78 | 0.181 | 0.827 |
| AT5G07090.2 | 40S ribosomal protein S4-2 (RPS4B) | 4 | 0.80 | 0.172 | 0.855 |
| AT5G07090.2 | 40S ribosomal protein S4-2 (RPS4B) | 5 | 0.83 | 0.163 | 0.884 |
| AT5G07090.2 | 40S ribosomal protein S4-2 (RPS4B) | 6 | 0.86 | 0.155 | 0.915 |
| AT5G07090.2 | 40S ribosomal protein S4-2 (RPS4B) | 7 | 0.89 | 0.149 | 0.944 |
| AT5G07090.2 | 40S ribosomal protein S4-2 (RPS4B) | 8 | 0.92 | 0.143 | 0.970 |
| AT5G07090.2 | 40S ribosomal protein S4-2 (RPS4B) | 9 | 0.96 | 0.138 | 0.990 |
| AT5G07090.2 | 40S ribosomal protein S4-2 (RPS4B) | 10 | 0.99 | 0.134 | 0.999 |
| AT5G07090.2 | 40S ribosomal protein S4-2 (RPS4B) | 11 | 1.02 | 0.132 | 0.997 |
| AT5G07090.2 | 40S ribosomal protein S4-2 (RPS4B) | 12 | 1.06 | 0.130 | 0.980 |
| AT5G07090.2 | 40S ribosomal protein S4-2 (RPS4B) | 13 | 1.10 | 0.131 | 0.952 |
| AT5G07090.2 | 40S ribosomal protein S4-2 (RPS4B) | 14 | 1.14 | 0.133 | 0.913 |
| AT5G07090.2 | 40S ribosomal protein S4-2 (RPS4B) | 15 | 1.18 | 0.136 | 0.870 |
| AT5G07090.2 | 40S ribosomal protein S4-2 (RPS4B) | 16 | 1.22 | 0.140 | 0.824 |
| AT5G07090.2 | 40S ribosomal protein S4-2 (RPS4B) | 17 | 1.26 | 0.145 | 0.781 |
| AT5G07090.2 | 40S ribosomal protein S4-2 (RPS4B) | 18 | 1.30 | 0.152 | 0.741 |
| AT5G07090.2 | 40S ribosomal protein S4-2 (RPS4B) | 19 | 1.35 | 0.159 | 0.707 |
| AT5G07090.2 | 40S ribosomal protein S4-2 (RPS4B) | 20 | 1.40 | 0.167 | 0.677 |
| AT5G07090.2 | 40S ribosomal protein S4-2 (RPS4B) | 21 | 1.45 | 0.176 | 0.651 |
| AT5G07090.2 | 40S ribosomal protein S4-2 (RPS4B) | 22 | 1.50 | 0.185 | 0.630 |
| AT5G07090.2 | 40S ribosomal protein S4-2 (RPS4B) | 46 | 3.43 | 0.475 | 0.518 |
| AT5G07090.2 | 40S ribosomal protein S4-2 (RPS4B) | 47 | 3.55 | 0.488 | 0.519 |
| AT5G07320.1 | ATP-Mg/Pi transporter | 2 | 1.03 | 0.348 | 1.000 |
| AT5G07320.1 | ATP-Mg/Pi transporter | 3 | 1.12 | 0.256 | 1.000 |
| AT5G07320.1 | ATP-Mg/Pi transporter | 5 | 1.37 | 0.208 | 1.000 |
| AT5G07320.1 | ATP-Mg/Pi transporter | 6 | 1.56 | 0.181 | 0.999 |
| AT5G07320.1 | ATP-Mg/Pi transporter | 7 | 1.79 | 0.163 | 0.940 |
| AT5G07320.1 | ATP-Mg/Pi transporter | 8 | 2.03 | 0.152 | 0.650 |
| AT5G07320.1 | ATP-Mg/Pi transporter | 9 | 2.29 | 0.147 | 0.321 |
| AT5G07320.1 | ATP-Mg/Pi transporter | 10 | 2.38 | 0.146 | 0.240 |
| AT5G07320.1 | ATP-Mg/Pi transporter | 11 | 2.08 | 0.146 | 0.519 |
| AT5G07320.1 | ATP-Mg/Pi transporter | 12 | 1.74 | 0.146 | 0.904 |

| | | | | | |
|---|---|---|---|---|---|
| AT5G07320.1 | ATP-Mg/Pi transporter | 13 | 1.52 | 0.145 | 0.994 |
| AT5G07320.1 | ATP-Mg/Pi transporter | 14 | 1.34 | 0.144 | 1.000 |
| AT5G07320.1 | ATP-Mg/Pi transporter | 15 | 1.19 | 0.143 | 1.000 |
| AT5G07320.1 | ATP-Mg/Pi transporter | 16 | 1.12 | 0.142 | 1.000 |
| AT5G07320.1 | ATP-Mg/Pi transporter | 17 | 1.24 | 0.142 | 1.000 |
| AT5G07320.1 | ATP-Mg/Pi transporter | 18 | 1.60 | 0.144 | 0.975 |
| AT5G07320.1 | ATP-Mg/Pi transporter | 19 | 2.05 | 0.148 | 0.573 |
| AT5G07320.1 | ATP-Mg/Pi transporter | 20 | 2.13 | 0.151 | 0.523 |
| AT5G07320.1 | ATP-Mg/Pi transporter | 21 | 1.75 | 0.153 | 0.927 |
| AT5G07320.1 | ATP-Mg/Pi transporter | 22 | 1.40 | 0.156 | 1.000 |
| AT5G07320.1 | ATP-Mg/Pi transporter | 23 | 1.21 | 0.165 | 1.000 |
| AT5G07320.1 | ATP-Mg/Pi transporter | 24 | 1.13 | 0.181 | 1.000 |
| AT5G07320.1 | ATP-Mg/Pi transporter | 25 | 1.11 | 0.202 | 1.000 |
| AT5G07320.1 | ATP-Mg/Pi transporter | 28 | 1.22 | 0.201 | 1.000 |
| AT5G07320.1 | ATP-Mg/Pi transporter | 29 | 1.24 | 0.175 | 1.000 |
| AT5G07320.1 | ATP-Mg/Pi transporter | 30 | 1.25 | 0.159 | 1.000 |
| AT5G07320.1 | ATP-Mg/Pi transporter | 31 | 1.23 | 0.150 | 1.000 |
| AT5G07320.1 | ATP-Mg/Pi transporter | 32 | 1.17 | 0.147 | 1.000 |
| AT5G07320.1 | ATP-Mg/Pi transporter | 33 | 1.09 | 0.147 | 1.000 |
| AT5G07320.1 | ATP-Mg/Pi transporter | 34 | 1.05 | 0.147 | 1.000 |
| AT5G07320.1 | ATP-Mg/Pi transporter | 35 | 1.08 | 0.147 | 1.000 |
| AT5G07320.1 | ATP-Mg/Pi transporter | 36 | 1.23 | 0.147 | 1.000 |
| AT5G07320.1 | ATP-Mg/Pi transporter | 37 | 1.46 | 0.146 | 0.998 |
| AT5G07320.1 | ATP-Mg/Pi transporter | 38 | 1.50 | 0.146 | 0.996 |
| AT5G07320.1 | ATP-Mg/Pi transporter | 39 | 1.20 | 0.146 | 1.000 |
| AT5G07320.1 | ATP-Mg/Pi transporter | 40 | 0.83 | 0.146 | 1.000 |
| AT5G07320.1 | ATP-Mg/Pi transporter | 41 | 0.58 | 0.147 | 0.915 |
| AT5G07320.1 | ATP-Mg/Pi transporter | 42 | 0.46 | 0.149 | 0.447 |
| AT5G07320.1 | ATP-Mg/Pi transporter | 43 | 0.43 | 0.151 | 0.328 |
| AT5G07320.1 | ATP-Mg/Pi transporter | 44 | 0.44 | 0.150 | 0.369 |
| AT5G07320.1 | ATP-Mg/Pi transporter | 45 | 0.49 | 0.151 | 0.636 |
| AT5G07320.1 | ATP-Mg/Pi transporter | 46 | 0.62 | 0.184 | 0.998 |
| AT5G07320.1 | ATP-Mg/Pi transporter | 47 | 0.80 | 0.303 | 1.000 |
| AT5G07440.2 | Glutamate dehydrogenase 2 (GDH2) | 2 | 0.73 | 0.570 | 1.000 |
| AT5G07440.2 | Glutamate dehydrogenase 2 (GDH2) | 3 | 0.57 | 0.434 | 1.000 |
| AT5G07440.2 | Glutamate dehydrogenase 2 (GDH2) | 4 | 0.46 | 0.425 | 1.000 |
| AT5G07440.2 | Glutamate dehydrogenase 2 (GDH2) | 6 | 0.39 | 0.423 | 0.999 |
| AT5G07440.2 | Glutamate dehydrogenase 2 (GDH2) | 7 | 0.44 | 0.375 | 0.999 |
| AT5G07440.2 | Glutamate dehydrogenase 2 (GDH2) | 8 | 0.54 | 0.341 | 1.000 |
| AT5G07440.2 | Glutamate dehydrogenase 2 (GDH2) | 9 | 0.66 | 0.323 | 1.000 |
| AT5G07440.2 | Glutamate dehydrogenase 2 (GDH2) | 10 | 0.77 | 0.316 | 1.000 |
| AT5G07440.2 | Glutamate dehydrogenase 2 (GDH2) | 11 | 0.87 | 0.316 | 1.000 |
| AT5G07440.2 | Glutamate dehydrogenase 2 (GDH2) | 12 | 1.02 | 0.315 | 1.000 |
| AT5G07440.2 | Glutamate dehydrogenase 2 (GDH2) | 13 | 1.14 | 0.312 | 1.000 |
| AT5G07440.2 | Glutamate dehydrogenase 2 (GDH2) | 14 | 1.28 | 0.310 | 1.000 |
| AT5G07440.2 | Glutamate dehydrogenase 2 (GDH2) | 15 | 1.44 | 0.308 | 1.000 |
| AT5G07440.2 | Glutamate dehydrogenase 2 (GDH2) | 16 | 1.45 | 0.307 | 1.000 |

| | | | | | |
|---|---|---|---|---|---|
| AT5G07440.2 | Glutamate dehydrogenase 2 (GDH2) | 17 | 1.35 | 0.306 | 1.000 |
| AT5G07440.2 | Glutamate dehydrogenase 2 (GDH2) | 18 | 1.18 | 0.310 | 1.000 |
| AT5G07440.2 | Glutamate dehydrogenase 2 (GDH2) | 19 | 1.18 | 0.316 | 1.000 |
| AT5G07440.2 | Glutamate dehydrogenase 2 (GDH2) | 20 | 1.36 | 0.321 | 1.000 |
| AT5G07440.2 | Glutamate dehydrogenase 2 (GDH2) | 21 | 1.73 | 0.324 | 1.000 |
| AT5G07440.2 | Glutamate dehydrogenase 2 (GDH2) | 22 | 2.29 | 0.324 | 0.994 |
| AT5G07440.2 | Glutamate dehydrogenase 2 (GDH2) | 23 | 2.66 | 0.323 | 0.967 |
| AT5G07440.2 | Glutamate dehydrogenase 2 (GDH2) | 24 | 2.62 | 0.317 | 0.967 |
| AT5G07440.2 | Glutamate dehydrogenase 2 (GDH2) | 25 | 2.10 | 0.313 | 0.998 |
| AT5G07440.2 | Glutamate dehydrogenase 2 (GDH2) | 26 | 1.74 | 0.312 | 1.000 |
| AT5G07440.2 | Glutamate dehydrogenase 2 (GDH2) | 27 | 1.54 | 0.312 | 1.000 |
| AT5G07440.2 | Glutamate dehydrogenase 2 (GDH2) | 28 | 1.24 | 0.313 | 1.000 |
| AT5G07440.2 | Glutamate dehydrogenase 2 (GDH2) | 29 | 0.99 | 0.313 | 1.000 |
| AT5G07440.2 | Glutamate dehydrogenase 2 (GDH2) | 30 | 0.96 | 0.313 | 1.000 |
| AT5G07440.2 | Glutamate dehydrogenase 2 (GDH2) | 31 | 1.34 | 0.314 | 1.000 |
| AT5G07440.2 | Glutamate dehydrogenase 2 (GDH2) | 32 | 2.29 | 0.314 | 0.991 |
| AT5G07440.2 | Glutamate dehydrogenase 2 (GDH2) | 33 | 2.71 | 0.315 | 0.952 |
| AT5G07440.2 | Glutamate dehydrogenase 2 (GDH2) | 34 | 1.85 | 0.316 | 1.000 |
| AT5G07440.2 | Glutamate dehydrogenase 2 (GDH2) | 35 | 1.09 | 0.318 | 1.000 |
| AT5G07440.2 | Glutamate dehydrogenase 2 (GDH2) | 36 | 0.89 | 0.321 | 1.000 |
| AT5G07440.2 | Glutamate dehydrogenase 2 (GDH2) | 37 | 0.99 | 0.342 | 1.000 |
| AT5G07440.2 | Glutamate dehydrogenase 2 (GDH2) | 38 | 1.14 | 0.414 | 1.000 |
| AT5G07440.2 | Glutamate dehydrogenase 2 (GDH2) | 39 | 1.58 | 0.567 | 1.000 |
| AT5G07440.2 | Glutamate dehydrogenase 2 (GDH2) | 46 | 4472.72 | 2.717 | 0.961 |
| AT5G07440.2 | Glutamate dehydrogenase 2 (GDH2) | 47 | 20608.38 | 3.185 | 0.958 |
| AT5G08040.1 | Translocase of the outer mitochondrial membrane 5 (TOM5, putative) | 2 | 1.12 | 0.524 | 1.000 |
| AT5G08040.1 | Translocase of the outer mitochondrial membrane 5 (TOM5, putative) | 3 | 1.35 | 0.390 | 1.000 |
| AT5G08040.1 | Translocase of the outer mitochondrial membrane 5 (TOM5, putative) | 4 | 1.43 | 0.417 | 1.000 |
| AT5G08040.1 | Translocase of the outer mitochondrial membrane 5 (TOM5, putative) | 6 | 1.62 | 0.472 | 1.000 |

| | | | | | |
|---|---|---|---|---|---|
| AT5G08040.1 | Translocase of the outer mitochondrial membrane 5 (TOM5, putative) | 7 | 1.92 | 0.470 | 1.000 |
| AT5G08040.1 | Translocase of the outer mitochondrial membrane 5 (TOM5, putative) | 8 | 2.27 | 0.531 | 1.000 |
| AT5G08040.1 | Translocase of the outer mitochondrial membrane 5 (TOM5, putative) | 15 | 1.38 | 0.369 | 1.000 |
| AT5G08040.1 | Translocase of the outer mitochondrial membrane 5 (TOM5, putative) | 16 | 1.49 | 0.330 | 1.000 |
| AT5G08040.1 | Translocase of the outer mitochondrial membrane 5 (TOM5, putative) | 17 | 1.41 | 0.296 | 1.000 |
| AT5G08040.1 | Translocase of the outer mitochondrial membrane 5 (TOM5, putative) | 18 | 1.62 | 0.279 | 1.000 |
| AT5G08040.1 | Translocase of the outer mitochondrial membrane 5 (TOM5, putative) | 19 | 2.36 | 0.279 | 0.974 |
| AT5G08040.1 | Translocase of the outer mitochondrial membrane 5 (TOM5, putative) | 20 | 3.24 | 0.284 | 0.770 |
| AT5G08040.1 | Translocase of the outer mitochondrial membrane 5 (TOM5, putative) | 21 | 3.67 | 0.288 | 0.642 |
| AT5G08040.1 | Translocase of the outer mitochondrial membrane 5 (TOM5, putative) | 22 | 3.47 | 0.287 | 0.704 |
| AT5G08040.1 | Translocase of the outer mitochondrial membrane 5 (TOM5, putative) | 23 | 2.86 | 0.286 | 0.889 |
| AT5G08040.1 | Translocase of the outer mitochondrial membrane 5 (TOM5, putative) | 24 | 1.90 | 0.278 | 0.999 |
| AT5G08040.1 | Translocase of the outer mitochondrial membrane 5 (TOM5, putative) | 25 | 1.28 | 0.274 | 1.000 |
| AT5G08040.1 | Translocase of the outer mitochondrial membrane 5 (TOM5, putative) | 26 | 1.08 | 0.273 | 1.000 |
| AT5G08040.1 | Translocase of the outer mitochondrial membrane 5 (TOM5, putative) | 27 | 0.98 | 0.273 | 1.000 |
| AT5G08040.1 | Translocase of the outer mitochondrial membrane 5 (TOM5, putative) | 28 | 1.09 | 0.274 | 1.000 |
| AT5G08040.1 | Translocase of the outer mitochondrial membrane 5 (TOM5, putative) | 29 | 1.72 | 0.274 | 1.000 |
| AT5G08040.1 | Translocase of the outer mitochondrial membrane 5 (TOM5, putative) | 30 | 2.67 | 0.274 | 0.907 |
| AT5G08040.1 | Translocase of the outer mitochondrial membrane 5 (TOM5, putative) | 31 | 2.36 | 0.276 | 0.972 |
| AT5G08040.1 | Translocase of the outer mitochondrial membrane 5 (TOM5, putative) | 32 | 0.98 | 0.277 | 1.000 |

| | | | | | | |
|---|---|---|---|---|---|---|
| AT5G08040.1 | Translocase of the outer mitochondrial membrane 5 (TOM5, putative) | 33 | 0.25 | 0.281 | 0.492 | |
| AT5G08040.1 | Translocase of the outer mitochondrial membrane 5 (TOM5, putative) | 34 | 0.09 | 0.299 | 0.012 | * |
| AT5G08040.1 | Translocase of the outer mitochondrial membrane 5 (TOM5, putative) | 35 | 0.09 | 0.338 | 0.042 | * |
| AT5G08040.1 | Translocase of the outer mitochondrial membrane 5 (TOM5, putative) | 36 | 0.14 | 0.377 | 0.379 | |
| AT5G08040.1 | Translocase of the outer mitochondrial membrane 5 (TOM5, putative) | 38 | 0.54 | 0.382 | 1.000 | |
| AT5G08040.1 | Translocase of the outer mitochondrial membrane 5 (TOM5, putative) | 39 | 1.13 | 0.343 | 1.000 | |
| AT5G08040.1 | Translocase of the outer mitochondrial membrane 5 (TOM5, putative) | 40 | 1.90 | 0.394 | 1.000 | |
| AT5G08040.1 | Translocase of the outer mitochondrial membrane 5 (TOM5, putative) | 41 | 2.25 | 0.569 | 1.000 | |
| AT5G08040.1 | Translocase of the outer mitochondrial membrane 5 (TOM5, putative) | 46 | 0.12 | 1.826 | 1.000 | |
| AT5G08040.1 | Translocase of the outer mitochondrial membrane 5 (TOM5, putative) | 47 | 0.05 | 2.227 | 1.000 | |
| AT5G08060.1 | NADH:ubiquinone oxidoreductase subunit (B14.5a) | 2 | 0.91 | 0.479 | 1.000 | |
| AT5G08060.1 | NADH:ubiquinone oxidoreductase subunit (B14.5a) | 3 | 0.88 | 0.338 | 1.000 | |
| AT5G08060.1 | NADH:ubiquinone oxidoreductase subunit (B14.5a) | 4 | 0.86 | 0.302 | 1.000 | |
| AT5G08060.1 | NADH:ubiquinone oxidoreductase subunit (B14.5a) | 5 | 0.87 | 0.296 | 1.000 | |
| AT5G08060.1 | NADH:ubiquinone oxidoreductase subunit (B14.5a) | 6 | 0.91 | 0.288 | 1.000 | |
| AT5G08060.1 | NADH:ubiquinone oxidoreductase subunit (B14.5a) | 7 | 0.94 | 0.286 | 1.000 | |
| AT5G08060.1 | NADH:ubiquinone oxidoreductase subunit (B14.5a) | 8 | 1.00 | 0.286 | 1.000 | |
| AT5G08060.1 | NADH:ubiquinone oxidoreductase subunit (B14.5a) | 9 | 1.11 | 0.286 | 1.000 | |
| AT5G08060.1 | NADH:ubiquinone oxidoreductase subunit (B14.5a) | 10 | 1.21 | 0.284 | 1.000 | |
| AT5G08060.1 | NADH:ubiquinone oxidoreductase subunit (B14.5a) | 11 | 1.23 | 0.285 | 1.000 | |

| | | | | | |
|---|---|---|---|---|---|
| AT5G08060.1 | NADH:ubiquinone oxidoreductase subunit (B14.5a) | 12 | 1.24 | 0.284 | 1.000 |
| AT5G08060.1 | NADH:ubiquinone oxidoreductase subunit (B14.5a) | 13 | 1.24 | 0.277 | 1.000 |
| AT5G08060.1 | NADH:ubiquinone oxidoreductase subunit (B14.5a) | 14 | 1.13 | 0.269 | 1.000 |
| AT5G08060.1 | NADH:ubiquinone oxidoreductase subunit (B14.5a) | 15 | 0.87 | 0.266 | 1.000 |
| AT5G08060.1 | NADH:ubiquinone oxidoreductase subunit (B14.5a) | 16 | 0.56 | 0.265 | 1.000 |
| AT5G08060.1 | NADH:ubiquinone oxidoreductase subunit (B14.5a) | 17 | 0.39 | 0.265 | 0.915 |
| AT5G08060.1 | NADH:ubiquinone oxidoreductase subunit (B14.5a) | 18 | 0.39 | 0.267 | 0.917 |
| AT5G08060.1 | NADH:ubiquinone oxidoreductase subunit (B14.5a) | 19 | 0.52 | 0.271 | 0.999 |
| AT5G08060.1 | NADH:ubiquinone oxidoreductase subunit (B14.5a) | 20 | 0.74 | 0.277 | 1.000 |
| AT5G08060.1 | NADH:ubiquinone oxidoreductase subunit (B14.5a) | 21 | 1.09 | 0.279 | 1.000 |
| AT5G08060.1 | NADH:ubiquinone oxidoreductase subunit (B14.5a) | 22 | 1.50 | 0.279 | 1.000 |
| AT5G08060.1 | NADH:ubiquinone oxidoreductase subunit (B14.5a) | 23 | 1.63 | 0.278 | 1.000 |
| AT5G08060.1 | NADH:ubiquinone oxidoreductase subunit (B14.5a) | 24 | 1.19 | 0.273 | 1.000 |
| AT5G08060.1 | NADH:ubiquinone oxidoreductase subunit (B14.5a) | 25 | 0.62 | 0.270 | 1.000 |
| AT5G08060.1 | NADH:ubiquinone oxidoreductase subunit (B14.5a) | 26 | 0.37 | 0.269 | 0.878 |
| AT5G08060.1 | NADH:ubiquinone oxidoreductase subunit (B14.5a) | 27 | 0.32 | 0.269 | 0.745 |
| AT5G08060.1 | NADH:ubiquinone oxidoreductase subunit (B14.5a) | 28 | 0.39 | 0.270 | 0.921 |
| AT5G08060.1 | NADH:ubiquinone oxidoreductase subunit (B14.5a) | 29 | 0.56 | 0.270 | 1.000 |
| AT5G08060.1 | NADH:ubiquinone oxidoreductase subunit (B14.5a) | 30 | 0.84 | 0.271 | 1.000 |
| AT5G08060.1 | NADH:ubiquinone oxidoreductase subunit (B14.5a) | 31 | 1.20 | 0.271 | 1.000 |

| | | | | | |
|---|---|---|---|---|---|
| AT5G08060.1 | NADH:ubiquinone oxidoreductase subunit (B14.5a) | 32 | 1.50 | 0.271 | 1.000 |
| AT5G08060.1 | NADH:ubiquinone oxidoreductase subunit (B14.5a) | 33 | 1.63 | 0.272 | 1.000 |
| AT5G08060.1 | NADH:ubiquinone oxidoreductase subunit (B14.5a) | 34 | 1.78 | 0.273 | 1.000 |
| AT5G08060.1 | NADH:ubiquinone oxidoreductase subunit (B14.5a) | 35 | 2.06 | 0.275 | 0.996 |
| AT5G08060.1 | NADH:ubiquinone oxidoreductase subunit (B14.5a) | 36 | 2.48 | 0.280 | 0.961 |
| AT5G08060.1 | NADH:ubiquinone oxidoreductase subunit (B14.5a) | 37 | 3.15 | 0.288 | 0.811 |
| AT5G08060.1 | NADH:ubiquinone oxidoreductase subunit (B14.5a) | 38 | 4.07 | 0.294 | 0.544 |
| AT5G08060.1 | NADH:ubiquinone oxidoreductase subunit (B14.5a) | 39 | 4.71 | 0.286 | 0.340 |
| AT5G08060.1 | NADH:ubiquinone oxidoreductase subunit (B14.5a) | 40 | 4.89 | 0.278 | 0.263 |
| AT5G08060.1 | NADH:ubiquinone oxidoreductase subunit (B14.5a) | 41 | 4.45 | 0.275 | 0.332 |
| AT5G08060.1 | NADH:ubiquinone oxidoreductase subunit (B14.5a) | 42 | 3.75 | 0.276 | 0.536 |
| AT5G08060.1 | NADH:ubiquinone oxidoreductase subunit (B14.5a) | 43 | 3.31 | 0.277 | 0.702 |
| AT5G08060.1 | NADH:ubiquinone oxidoreductase subunit (B14.5a) | 44 | 2.79 | 0.277 | 0.884 |
| AT5G08060.1 | NADH:ubiquinone oxidoreductase subunit (B14.5a) | 45 | 2.12 | 0.280 | 0.995 |
| AT5G08060.1 | NADH:ubiquinone oxidoreductase subunit (B14.5a) | 46 | 1.46 | 0.326 | 1.000 |
| AT5G08060.1 | NADH:ubiquinone oxidoreductase subunit (B14.5a) | 47 | 0.91 | 0.499 | 1.000 |
| AT5G08300.1 | Succinyl-CoA ligase subunit (SCL alpha) | 2 | 1.01 | 0.483 | 1.000 |
| AT5G08300.1 | Succinyl-CoA ligase subunit (SCL alpha) | 3 | 0.64 | 0.361 | 1.000 |
| AT5G08300.1 | Succinyl-CoA ligase subunit (SCL alpha) | 4 | 0.44 | 0.356 | 0.999 |
| AT5G08300.1 | Succinyl-CoA ligase subunit (SCL alpha) | 6 | 0.31 | 0.358 | 0.963 |
| AT5G08300.1 | Succinyl-CoA ligase subunit (SCL alpha) | 7 | 0.35 | 0.317 | 0.958 |
| AT5G08300.1 | Succinyl-CoA ligase subunit (SCL alpha) | 8 | 0.46 | 0.287 | 0.996 |
| AT5G08300.1 | Succinyl-CoA ligase subunit (SCL alpha) | 9 | 0.69 | 0.271 | 1.000 |

| | | | | | |
|---|---|---|---|---|---|
| AT5G08300.1 | Succinyl-CoA ligase subunit (SCL alpha) | 10 | 1.01 | 0.266 | 1.000 |
| AT5G08300.1 | Succinyl-CoA ligase subunit (SCL alpha) | 11 | 1.23 | 0.266 | 1.000 |
| AT5G08300.1 | Succinyl-CoA ligase subunit (SCL alpha) | 12 | 1.25 | 0.265 | 1.000 |
| AT5G08300.1 | Succinyl-CoA ligase subunit (SCL alpha) | 13 | 1.16 | 0.265 | 1.000 |
| AT5G08300.1 | Succinyl-CoA ligase subunit (SCL alpha) | 14 | 1.20 | 0.267 | 1.000 |
| AT5G08300.1 | Succinyl-CoA ligase subunit (SCL alpha) | 15 | 1.49 | 0.271 | 1.000 |
| AT5G08300.1 | Succinyl-CoA ligase subunit (SCL alpha) | 16 | 2.01 | 0.275 | 0.998 |
| AT5G08300.1 | Succinyl-CoA ligase subunit (SCL alpha) | 17 | 2.70 | 0.291 | 0.942 |
| AT5G08300.1 | Succinyl-CoA ligase subunit (SCL alpha) | 18 | 3.18 | 0.313 | 0.892 |
| AT5G08300.1 | Succinyl-CoA ligase subunit (SCL alpha) | 19 | 3.06 | 0.338 | 0.957 |
| AT5G08300.1 | Succinyl-CoA ligase subunit (SCL alpha) | 20 | 2.41 | 0.352 | 0.998 |
| AT5G08300.1 | Succinyl-CoA ligase subunit (SCL alpha) | 21 | 1.68 | 0.368 | 1.000 |
| AT5G08300.1 | Succinyl-CoA ligase subunit (SCL alpha) | 22 | 1.13 | 0.354 | 1.000 |
| AT5G08300.1 | Succinyl-CoA ligase subunit (SCL alpha) | 23 | 0.77 | 0.343 | 1.000 |
| AT5G08300.1 | Succinyl-CoA ligase subunit (SCL alpha) | 24 | 0.57 | 0.312 | 1.000 |
| AT5G08300.1 | Succinyl-CoA ligase subunit (SCL alpha) | 25 | 0.51 | 0.290 | 0.999 |
| AT5G08300.1 | Succinyl-CoA ligase subunit (SCL alpha) | 26 | 0.57 | 0.274 | 1.000 |
| AT5G08300.1 | Succinyl-CoA ligase subunit (SCL alpha) | 27 | 0.72 | 0.266 | 1.000 |
| AT5G08300.1 | Succinyl-CoA ligase subunit (SCL alpha) | 28 | 0.82 | 0.263 | 1.000 |
| AT5G08300.1 | Succinyl-CoA ligase subunit (SCL alpha) | 29 | 0.89 | 0.263 | 1.000 |
| AT5G08300.1 | Succinyl-CoA ligase subunit (SCL alpha) | 30 | 1.01 | 0.264 | 1.000 |
| AT5G08300.1 | Succinyl-CoA ligase subunit (SCL alpha) | 31 | 1.39 | 0.264 | 1.000 |
| AT5G08300.1 | Succinyl-CoA ligase subunit (SCL alpha) | 32 | 1.89 | 0.264 | 0.999 |
| AT5G08300.1 | Succinyl-CoA ligase subunit (SCL alpha) | 33 | 1.98 | 0.264 | 0.997 |
| AT5G08300.1 | Succinyl-CoA ligase subunit (SCL alpha) | 34 | 1.63 | 0.265 | 1.000 |
| AT5G08300.1 | Succinyl-CoA ligase subunit (SCL alpha) | 35 | 1.28 | 0.265 | 1.000 |
| AT5G08300.1 | Succinyl-CoA ligase subunit (SCL alpha) | 36 | 1.19 | 0.265 | 1.000 |
| AT5G08300.1 | Succinyl-CoA ligase subunit (SCL alpha) | 37 | 1.37 | 0.264 | 1.000 |
| AT5G08300.1 | Succinyl-CoA ligase subunit (SCL alpha) | 38 | 1.78 | 0.264 | 1.000 |
| AT5G08300.1 | Succinyl-CoA ligase subunit (SCL alpha) | 39 | 2.20 | 0.264 | 0.984 |
| AT5G08300.1 | Succinyl-CoA ligase subunit (SCL alpha) | 40 | 2.41 | 0.265 | 0.955 |

| | | | | | |
|---|---|---|---|---|---|
| AT5G08300.1 | Succinyl-CoA ligase subunit (SCL alpha) | 41 | 2.22 | 0.265 | 0.983 |
| AT5G08300.1 | Succinyl-CoA ligase subunit (SCL alpha) | 42 | 1.75 | 0.268 | 1.000 |
| AT5G08300.1 | Succinyl-CoA ligase subunit (SCL alpha) | 43 | 1.18 | 0.270 | 1.000 |
| AT5G08300.1 | Succinyl-CoA ligase subunit (SCL alpha) | 44 | 0.75 | 0.270 | 1.000 |
| AT5G08300.1 | Succinyl-CoA ligase subunit (SCL alpha) | 45 | 0.53 | 0.271 | 0.999 |
| AT5G08300.1 | Succinyl-CoA ligase subunit (SCL alpha) | 46 | 0.41 | 0.318 | 0.993 |
| AT5G08300.1 | Succinyl-CoA ligase subunit (SCL alpha) | 47 | 0.32 | 0.502 | 1.000 |
| AT5G08530.1 | NADH:ubiquinone oxidoreductase subunit (51kDa) | 3 | 0.83 | 0.248 | 1.000 |
| AT5G08530.1 | NADH:ubiquinone oxidoreductase subunit (51kDa) | 4 | 0.83 | 0.229 | 1.000 |
| AT5G08530.1 | NADH:ubiquinone oxidoreductase subunit (51kDa) | 5 | 0.99 | 0.229 | 1.000 |
| AT5G08530.1 | NADH:ubiquinone oxidoreductase subunit (51kDa) | 6 | 1.09 | 0.230 | 1.000 |
| AT5G08530.1 | NADH:ubiquinone oxidoreductase subunit (51kDa) | 7 | 1.04 | 0.230 | 1.000 |
| AT5G08530.1 | NADH:ubiquinone oxidoreductase subunit (51kDa) | 8 | 1.06 | 0.225 | 1.000 |
| AT5G08530.1 | NADH:ubiquinone oxidoreductase subunit (51kDa) | 9 | 1.20 | 0.222 | 1.000 |
| AT5G08530.1 | NADH:ubiquinone oxidoreductase subunit (51kDa) | 10 | 1.38 | 0.222 | 1.000 |
| AT5G08530.1 | NADH:ubiquinone oxidoreductase subunit (51kDa) | 11 | 1.63 | 0.223 | 1.000 |
| AT5G08530.1 | NADH:ubiquinone oxidoreductase subunit (51kDa) | 12 | 1.85 | 0.223 | 0.999 |
| AT5G08530.1 | NADH:ubiquinone oxidoreductase subunit (51kDa) | 13 | 1.80 | 0.220 | 0.999 |
| AT5G08530.1 | NADH:ubiquinone oxidoreductase subunit (51kDa) | 14 | 1.40 | 0.217 | 1.000 |
| AT5G08530.1 | NADH:ubiquinone oxidoreductase subunit (51kDa) | 15 | 0.88 | 0.216 | 1.000 |
| AT5G08530.1 | NADH:ubiquinone oxidoreductase subunit (51kDa) | 16 | 0.51 | 0.215 | 0.994 |
| AT5G08530.1 | NADH:ubiquinone oxidoreductase subunit (51kDa) | 17 | 0.38 | 0.215 | 0.752 |
| AT5G08530.1 | NADH:ubiquinone oxidoreductase subunit (51kDa) | 18 | 0.43 | 0.218 | 0.927 |
| AT5G08530.1 | NADH:ubiquinone oxidoreductase subunit (51kDa) | 19 | 0.63 | 0.224 | 1.000 |
| AT5G08530.1 | NADH:ubiquinone oxidoreductase subunit (51kDa) | 20 | 0.86 | 0.229 | 1.000 |
| AT5G08530.1 | NADH:ubiquinone oxidoreductase subunit (51kDa) | 21 | 1.16 | 0.232 | 1.000 |
| AT5G08530.1 | NADH:ubiquinone oxidoreductase subunit (51kDa) | 22 | 1.59 | 0.232 | 1.000 |
| AT5G08530.1 | NADH:ubiquinone oxidoreductase subunit (51kDa) | 23 | 1.83 | 0.231 | 0.999 |
| AT5G08530.1 | NADH:ubiquinone oxidoreductase subunit (51kDa) | 24 | 1.38 | 0.226 | 1.000 |
| AT5G08530.1 | NADH:ubiquinone oxidoreductase subunit (51kDa) | 25 | 0.70 | 0.222 | 1.000 |
| AT5G08530.1 | NADH:ubiquinone oxidoreductase subunit (51kDa) | 26 | 0.41 | 0.221 | 0.892 |

| | | | | | |
|---|---|---|---|---|---|
| AT5G08530.1 | NADH:ubiquinone oxidoreductase subunit (51kDa) | 27 | 0.39 | 0.221 | 0.850 |
| AT5G08530.1 | NADH:ubiquinone oxidoreductase subunit (51kDa) | 28 | 0.54 | 0.221 | 0.998 |
| AT5G08530.1 | NADH:ubiquinone oxidoreductase subunit (51kDa) | 29 | 0.82 | 0.221 | 1.000 |
| AT5G08530.1 | NADH:ubiquinone oxidoreductase subunit (51kDa) | 30 | 1.06 | 0.221 | 1.000 |
| AT5G08530.1 | NADH:ubiquinone oxidoreductase subunit (51kDa) | 31 | 1.11 | 0.221 | 1.000 |
| AT5G08530.1 | NADH:ubiquinone oxidoreductase subunit (51kDa) | 32 | 1.13 | 0.222 | 1.000 |
| AT5G08530.1 | NADH:ubiquinone oxidoreductase subunit (51kDa) | 33 | 1.21 | 0.222 | 1.000 |
| AT5G08530.1 | NADH:ubiquinone oxidoreductase subunit (51kDa) | 34 | 1.30 | 0.223 | 1.000 |
| AT5G08530.1 | NADH:ubiquinone oxidoreductase subunit (51kDa) | 35 | 1.30 | 0.222 | 1.000 |
| AT5G08530.1 | NADH:ubiquinone oxidoreductase subunit (51kDa) | 36 | 1.18 | 0.221 | 1.000 |
| AT5G08530.1 | NADH:ubiquinone oxidoreductase subunit (51kDa) | 37 | 0.98 | 0.221 | 1.000 |
| AT5G08530.1 | NADH:ubiquinone oxidoreductase subunit (51kDa) | 38 | 0.82 | 0.222 | 1.000 |
| AT5G08530.1 | NADH:ubiquinone oxidoreductase subunit (51kDa) | 39 | 0.82 | 0.222 | 1.000 |
| AT5G08530.1 | NADH:ubiquinone oxidoreductase subunit (51kDa) | 40 | 1.03 | 0.223 | 1.000 |
| AT5G08530.1 | NADH:ubiquinone oxidoreductase subunit (51kDa) | 41 | 1.39 | 0.224 | 1.000 |
| AT5G08530.1 | NADH:ubiquinone oxidoreductase subunit (51kDa) | 42 | 1.70 | 0.226 | 1.000 |
| AT5G08530.1 | NADH:ubiquinone oxidoreductase subunit (51kDa) | 43 | 1.72 | 0.228 | 1.000 |
| AT5G08530.1 | NADH:ubiquinone oxidoreductase subunit (51kDa) | 44 | 1.32 | 0.228 | 1.000 |
| AT5G08530.1 | NADH:ubiquinone oxidoreductase subunit (51kDa) | 45 | 0.69 | 0.229 | 1.000 |
| AT5G08530.1 | NADH:ubiquinone oxidoreductase subunit (51kDa) | 46 | 0.23 | 0.266 | 0.392 |
| AT5G08530.1 | NADH:ubiquinone oxidoreductase subunit (51kDa) | 47 | 0.06 | 0.456 | 0.206 |
| AT5G08530.1 | NADH:ubiquinone oxidoreductase subunit (51kDa) | 48 | 0.01 | 0.757 | 0.376 |
| AT5G08690.1 | ATP synthase subunit (ATPbeta) | 3 | 0.93 | 0.161 | 1.000 |
| AT5G08690.1 | ATP synthase subunit (ATPbeta) | 4 | 0.93 | 0.153 | 1.000 |
| AT5G08690.1 | ATP synthase subunit (ATPbeta) | 5 | 1.09 | 0.153 | 1.000 |
| AT5G08690.1 | ATP synthase subunit (ATPbeta) | 6 | 1.17 | 0.154 | 1.000 |
| AT5G08690.1 | ATP synthase subunit (ATPbeta) | 7 | 1.14 | 0.154 | 1.000 |
| AT5G08690.1 | ATP synthase subunit (ATPbeta) | 8 | 1.18 | 0.151 | 1.000 |
| AT5G08690.1 | ATP synthase subunit (ATPbeta) | 9 | 1.29 | 0.149 | 1.000 |
| AT5G08690.1 | ATP synthase subunit (ATPbeta) | 10 | 1.34 | 0.149 | 1.000 |
| AT5G08690.1 | ATP synthase subunit (ATPbeta) | 11 | 1.42 | 0.150 | 1.000 |

| | | | | | | |
|---|---|---|---|---|---|---|
| AT5G08690.1 | ATP synthase subunit (ATPbeta) | 12 | 1.54 | 0.150 | 0.997 | |
| AT5G08690.1 | ATP synthase subunit (ATPbeta) | 13 | 1.68 | 0.148 | 0.965 | |
| AT5G08690.1 | ATP synthase subunit (ATPbeta) | 14 | 1.79 | 0.146 | 0.885 | |
| AT5G08690.1 | ATP synthase subunit (ATPbeta) | 15 | 1.68 | 0.146 | 0.960 | |
| AT5G08690.1 | ATP synthase subunit (ATPbeta) | 16 | 1.49 | 0.145 | 0.999 | |
| AT5G08690.1 | ATP synthase subunit (ATPbeta) | 17 | 1.37 | 0.145 | 1.000 | |
| AT5G08690.1 | ATP synthase subunit (ATPbeta) | 18 | 1.30 | 0.147 | 1.000 | |
| AT5G08690.1 | ATP synthase subunit (ATPbeta) | 19 | 1.39 | 0.150 | 1.000 | |
| AT5G08690.1 | ATP synthase subunit (ATPbeta) | 20 | 1.48 | 0.154 | 1.000 | |
| AT5G08690.1 | ATP synthase subunit (ATPbeta) | 21 | 1.29 | 0.156 | 1.000 | |
| AT5G08690.1 | ATP synthase subunit (ATPbeta) | 22 | 1.03 | 0.155 | 1.000 | |
| AT5G08690.1 | ATP synthase subunit (ATPbeta) | 23 | 0.93 | 0.155 | 1.000 | |
| AT5G08690.1 | ATP synthase subunit (ATPbeta) | 24 | 1.15 | 0.152 | 1.000 | |
| AT5G08690.1 | ATP synthase subunit (ATPbeta) | 25 | 1.59 | 0.149 | 0.991 | |
| AT5G08690.1 | ATP synthase subunit (ATPbeta) | 26 | 1.54 | 0.149 | 0.997 | |
| AT5G08690.1 | ATP synthase subunit (ATPbeta) | 27 | 0.92 | 0.149 | 1.000 | |
| AT5G08690.1 | ATP synthase subunit (ATPbeta) | 28 | 0.71 | 0.149 | 1.000 | |
| AT5G08690.1 | ATP synthase subunit (ATPbeta) | 29 | 1.10 | 0.148 | 1.000 | |
| AT5G08690.1 | ATP synthase subunit (ATPbeta) | 30 | 2.12 | 0.148 | 0.529 | |
| AT5G08690.1 | ATP synthase subunit (ATPbeta) | 31 | 2.72 | 0.149 | 0.108 | |
| AT5G08690.1 | ATP synthase subunit (ATPbeta) | 32 | 2.04 | 0.149 | 0.630 | |
| AT5G08690.1 | ATP synthase subunit (ATPbeta) | 33 | 1.41 | 0.149 | 1.000 | |
| AT5G08690.1 | ATP synthase subunit (ATPbeta) | 34 | 1.32 | 0.150 | 1.000 | |
| AT5G08690.1 | ATP synthase subunit (ATPbeta) | 35 | 1.51 | 0.149 | 0.998 | |
| AT5G08690.1 | ATP synthase subunit (ATPbeta) | 36 | 1.87 | 0.149 | 0.827 | |
| AT5G08690.1 | ATP synthase subunit (ATPbeta) | 37 | 2.28 | 0.148 | 0.367 | |
| AT5G08690.1 | ATP synthase subunit (ATPbeta) | 38 | 2.68 | 0.149 | 0.125 | |
| AT5G08690.1 | ATP synthase subunit (ATPbeta) | 39 | 3.19 | 0.149 | 0.029 | * |
| AT5G08690.1 | ATP synthase subunit (ATPbeta) | 40 | 3.72 | 0.150 | 0.006 | ** |
| AT5G08690.1 | ATP synthase subunit (ATPbeta) | 41 | 3.85 | 0.150 | 0.004 | ** |
| AT5G08690.1 | ATP synthase subunit (ATPbeta) | 42 | 3.48 | 0.152 | 0.014 | * |

| | | | | | | |
|---|---|---|---|---|---|---|
| AT5G08690.1 | ATP synthase subunit (ATPbeta) | 43 | 3.29 | 0.153 | 0.027 | * |
| AT5G08690.1 | ATP synthase subunit (ATPbeta) | 44 | 2.73 | 0.153 | 0.131 | |
| AT5G08690.1 | ATP synthase subunit (ATPbeta) | 45 | 1.84 | 0.153 | 0.888 | |
| AT5G08690.1 | ATP synthase subunit (ATPbeta) | 46 | 1.06 | 0.170 | 1.000 | |
| AT5G08690.1 | ATP synthase subunit (ATPbeta) | 47 | 0.58 | 0.294 | 1.000 | |
| AT5G08690.1 | ATP synthase subunit (ATPbeta) | 48 | 0.31 | 0.504 | 1.000 | |
| AT5G09270.2 | ? | 2 | 1.00 | 0.594 | 1.000 | |
| AT5G09270.2 | ? | 3 | 1.19 | 0.539 | 1.000 | |
| AT5G09270.2 | ? | 10 | 1.86 | 0.431 | 1.000 | |
| AT5G09270.2 | ? | 11 | 1.91 | 0.404 | 1.000 | |
| AT5G09270.2 | ? | 12 | 2.21 | 0.379 | 1.000 | |
| AT5G09270.2 | ? | 13 | 2.82 | 0.377 | 0.995 | |
| AT5G09270.2 | ? | 14 | 3.62 | 0.365 | 0.936 | |
| AT5G09270.2 | ? | 17 | 4.30 | 0.368 | 0.847 | |
| AT5G09270.2 | ? | 18 | 3.72 | 0.375 | 0.940 | |
| AT5G09270.2 | ? | 19 | 3.03 | 0.346 | 0.973 | |
| AT5G09270.2 | ? | 20 | 2.00 | 0.400 | 1.000 | |
| AT5G09270.2 | ? | 21 | 0.92 | 0.464 | 1.000 | |
| AT5G09270.2 | ? | 25 | 0.17 | 0.390 | 0.650 | |
| AT5G09270.2 | ? | 26 | 0.54 | 0.299 | 1.000 | |
| AT5G09270.2 | ? | 27 | 2.29 | 0.265 | 0.979 | |
| AT5G09270.2 | ? | 28 | 2.80 | 0.251 | 0.811 | |
| AT5G09270.2 | ? | 29 | 0.94 | 0.248 | 1.000 | |
| AT5G09270.2 | ? | 30 | 0.38 | 0.247 | 0.848 | |
| AT5G09270.2 | ? | 31 | 0.41 | 0.249 | 0.934 | |
| AT5G09270.2 | ? | 32 | 0.69 | 0.249 | 1.000 | |
| AT5G09270.2 | ? | 33 | 1.12 | 0.250 | 1.000 | |
| AT5G09270.2 | ? | 34 | 1.40 | 0.250 | 1.000 | |
| AT5G09270.2 | ? | 35 | 0.99 | 0.249 | 1.000 | |
| AT5G09270.2 | ? | 36 | 0.86 | 0.249 | 1.000 | |
| AT5G09270.2 | ? | 37 | 1.58 | 0.248 | 1.000 | |
| AT5G09270.2 | ? | 38 | 2.68 | 0.250 | 0.854 | |
| AT5G09270.2 | ? | 39 | 1.84 | 0.249 | 0.999 | |
| AT5G09270.2 | ? | 40 | 0.75 | 0.251 | 1.000 | |
| AT5G09270.2 | ? | 41 | 0.27 | 0.269 | 0.529 | |
| AT5G09270.2 | ? | 42 | 0.09 | 0.386 | 0.158 | |
| AT5G09270.2 | ? | 43 | 0.05 | 0.576 | 0.377 | |
| AT5G09270.2 | ? | 46 | 0.04 | 1.090 | 0.989 | |
| AT5G09270.2 | ? | 47 | 0.05 | 1.395 | 1.000 | |
| AT5G09300.2 | Branched chain keto acid dehydrogenase subunit (BCKDHE1alpha) | 2 | 0.99 | 0.550 | 1.000 | |
| AT5G09300.2 | Branched chain keto acid dehydrogenase subunit (BCKDHE1alpha) | 3 | 0.94 | 0.588 | 1.000 | |
| AT5G09300.2 | Branched chain keto acid dehydrogenase subunit (BCKDHE1alpha) | 33 | 0.74 | 0.344 | 1.000 | |
| AT5G09300.2 | Branched chain keto acid dehydrogenase subunit (BCKDHE1alpha) | 34 | 0.79 | 0.284 | 1.000 | |
| AT5G09300.2 | Branched chain keto acid dehydrogenase subunit (BCKDHE1alpha) | 35 | 0.85 | 0.259 | 1.000 | |

| | | | | | | |
|---|---|---|---|---|---|---|
| AT5G09300.2 | Branched chain keto acid dehydrogenase subunit (BCKDHE1alpha) | 36 | 0.91 | 0.251 | 1.000 | |
| AT5G09300.2 | Branched chain keto acid dehydrogenase subunit (BCKDHE1alpha) | 38 | 1.05 | 0.226 | 1.000 | |
| AT5G09300.2 | Branched chain keto acid dehydrogenase subunit (BCKDHE1alpha) | 39 | 1.06 | 0.209 | 1.000 | |
| AT5G09300.2 | Branched chain keto acid dehydrogenase subunit (BCKDHE1alpha) | 40 | 0.96 | 0.204 | 1.000 | |
| AT5G09300.2 | Branched chain keto acid dehydrogenase subunit (BCKDHE1alpha) | 41 | 0.71 | 0.224 | 0.990 | |
| AT5G09300.2 | Branched chain keto acid dehydrogenase subunit (BCKDHE1alpha) | 42 | 0.43 | 0.268 | 0.716 | |
| AT5G09300.2 | Branched chain keto acid dehydrogenase subunit (BCKDHE1alpha) | 43 | 0.23 | 0.325 | 0.309 | |
| AT5G09300.2 | Branched chain keto acid dehydrogenase subunit (BCKDHE1alpha) | 46 | 0.02 | 0.508 | 0.013 | * |
| AT5G09300.2 | Branched chain keto acid dehydrogenase subunit (BCKDHE1alpha) | 47 | 0.01 | 0.610 | 0.011 | * |
| AT5G09420.1 | Outer membrane protein 64 (OM64) | 2 | 0.87 | 0.448 | 1.000 | |
| AT5G09420.1 | Outer membrane protein 64 (OM64) | 3 | 0.77 | 0.408 | 1.000 | |
| AT5G09420.1 | Outer membrane protein 64 (OM64) | 15 | 0.88 | 0.190 | 1.000 | |
| AT5G09420.1 | Outer membrane protein 64 (OM64) | 16 | 1.11 | 0.167 | 1.000 | |
| AT5G09420.1 | Outer membrane protein 64 (OM64) | 17 | 1.38 | 0.152 | 0.996 | |
| AT5G09420.1 | Outer membrane protein 64 (OM64) | 18 | 1.71 | 0.144 | 0.779 | |
| AT5G09420.1 | Outer membrane protein 64 (OM64) | 19 | 2.04 | 0.142 | 0.359 | |
| AT5G09420.1 | Outer membrane protein 64 (OM64) | 20 | 2.26 | 0.142 | 0.185 | |
| AT5G09420.1 | Outer membrane protein 64 (OM64) | 21 | 2.27 | 0.142 | 0.179 | |
| AT5G09420.1 | Outer membrane protein 64 (OM64) | 22 | 2.11 | 0.141 | 0.286 | |
| AT5G09420.1 | Outer membrane protein 64 (OM64) | 23 | 1.89 | 0.139 | 0.497 | |
| AT5G09420.1 | Outer membrane protein 64 (OM64) | 24 | 1.70 | 0.136 | 0.727 | |
| AT5G09420.1 | Outer membrane protein 64 (OM64) | 25 | 1.63 | 0.134 | 0.796 | |
| AT5G09420.1 | Outer membrane protein 64 (OM64) | 26 | 1.79 | 0.133 | 0.567 | |
| AT5G09420.1 | Outer membrane protein 64 (OM64) | 27 | 2.11 | 0.132 | 0.203 | |
| AT5G09420.1 | Outer membrane protein 64 (OM64) | 28 | 2.41 | 0.132 | 0.070 | . |
| AT5G09420.1 | Outer membrane protein 64 (OM64) | 29 | 2.43 | 0.132 | 0.065 | . |

| | | | | | |
|---|---|---|---|---|---|
| AT5G09420.1 | Outer membrane protein 64 (OM64) | 30 | 2.15 | 0.133 | 0.181 |
| AT5G09420.1 | Outer membrane protein 64 (OM64) | 31 | 1.74 | 0.133 | 0.634 |
| AT5G09420.1 | Outer membrane protein 64 (OM64) | 32 | 1.34 | 0.133 | 0.995 |
| AT5G09420.1 | Outer membrane protein 64 (OM64) | 33 | 1.05 | 0.133 | 1.000 |
| AT5G09420.1 | Outer membrane protein 64 (OM64) | 34 | 0.87 | 0.133 | 1.000 |
| AT5G09420.1 | Outer membrane protein 64 (OM64) | 35 | 0.79 | 0.133 | 0.999 |
| AT5G09420.1 | Outer membrane protein 64 (OM64) | 36 | 0.79 | 0.133 | 0.999 |
| AT5G09420.1 | Outer membrane protein 64 (OM64) | 37 | 0.85 | 0.133 | 1.000 |
| AT5G09420.1 | Outer membrane protein 64 (OM64) | 38 | 0.98 | 0.133 | 1.000 |
| AT5G09420.1 | Outer membrane protein 64 (OM64) | 39 | 1.16 | 0.134 | 1.000 |
| AT5G09420.1 | Outer membrane protein 64 (OM64) | 40 | 1.39 | 0.135 | 0.988 |
| AT5G09420.1 | Outer membrane protein 64 (OM64) | 41 | 1.64 | 0.138 | 0.819 |
| AT5G09420.1 | Outer membrane protein 64 (OM64) | 42 | 1.87 | 0.141 | 0.539 |
| AT5G09420.1 | Outer membrane protein 64 (OM64) | 43 | 2.04 | 0.145 | 0.394 |
| AT5G09420.1 | Outer membrane protein 64 (OM64) | 44 | 1.99 | 0.150 | 0.500 |
| AT5G09420.1 | Outer membrane protein 64 (OM64) | 45 | 1.69 | 0.161 | 0.890 |
| AT5G09420.1 | Outer membrane protein 64 (OM64) | 46 | 1.29 | 0.190 | 1.000 |
| AT5G09420.1 | Outer membrane protein 64 (OM64) | 47 | 0.93 | 0.251 | 1.000 |
| AT5G09450.1 | Pentatricopeptide repeat (PPR) superfamily protein | 2 | 1.00 | 0.345 | 1.000 |
| AT5G09450.1 | Pentatricopeptide repeat (PPR) superfamily protein | 3 | 0.98 | 0.333 | 1.000 |
| AT5G09450.1 | Pentatricopeptide repeat (PPR) superfamily protein | 29 | 1.29 | 0.257 | 1.000 |
| AT5G09450.1 | Pentatricopeptide repeat (PPR) superfamily protein | 30 | 1.37 | 0.200 | 1.000 |
| AT5G09450.1 | Pentatricopeptide repeat (PPR) superfamily protein | 31 | 1.41 | 0.183 | 0.999 |
| AT5G09450.1 | Pentatricopeptide repeat (PPR) superfamily protein | 32 | 1.47 | 0.175 | 0.994 |
| AT5G09450.1 | Pentatricopeptide repeat (PPR) superfamily protein | 33 | 1.29 | 0.173 | 1.000 |
| AT5G09450.1 | Pentatricopeptide repeat (PPR) superfamily protein | 34 | 0.84 | 0.173 | 1.000 |
| AT5G09450.1 | Pentatricopeptide repeat (PPR) superfamily protein | 35 | 0.51 | 0.173 | 0.701 |
| AT5G09450.1 | Pentatricopeptide repeat (PPR) superfamily protein | 36 | 0.39 | 0.173 | 0.238 |
| AT5G09450.1 | Pentatricopeptide repeat (PPR) superfamily protein | 37 | 0.50 | 0.172 | 0.661 |
| AT5G09450.1 | Pentatricopeptide repeat (PPR) superfamily protein | 38 | 0.82 | 0.172 | 1.000 |
| AT5G09450.1 | Pentatricopeptide repeat (PPR) superfamily protein | 39 | 1.31 | 0.171 | 1.000 |

| | | | | | |
|---|---|---|---|---|---|
| AT5G09450.1 | Pentatricopeptide repeat (PPR) superfamily protein | 40 | 2.09 | 0.171 | 0.566 |
| AT5G09450.1 | Pentatricopeptide repeat (PPR) superfamily protein | 41 | 2.73 | 0.172 | 0.163 |
| AT5G09450.1 | Pentatricopeptide repeat (PPR) superfamily protein | 42 | 2.44 | 0.174 | 0.314 |
| AT5G09450.1 | Pentatricopeptide repeat (PPR) superfamily protein | 43 | 1.76 | 0.175 | 0.882 |
| AT5G09450.1 | Pentatricopeptide repeat (PPR) superfamily protein | 44 | 1.10 | 0.175 | 1.000 |
| AT5G09450.1 | Pentatricopeptide repeat (PPR) superfamily protein | 45 | 0.61 | 0.175 | 0.953 |
| AT5G09450.1 | Pentatricopeptide repeat (PPR) superfamily protein | 46 | 0.32 | 0.195 | 0.153 |
| AT5G09450.1 | Pentatricopeptide repeat (PPR) superfamily protein | 47 | 0.16 | 0.350 | 0.272 |
| AT5G09590.1 | Heat shock cognate 70 (HSC70-5) | 2 | 0.73 | 0.314 | 1.000 |
| AT5G09590.1 | Heat shock cognate 70 (HSC70-5) | 3 | 0.54 | 0.223 | 0.995 |
| AT5G09590.1 | Heat shock cognate 70 (HSC70-5) | 4 | 0.41 | 0.208 | 0.738 |
| AT5G09590.1 | Heat shock cognate 70 (HSC70-5) | 5 | 0.34 | 0.213 | 0.476 |
| AT5G09590.1 | Heat shock cognate 70 (HSC70-5) | 6 | 0.34 | 0.206 | 0.381 |
| AT5G09590.1 | Heat shock cognate 70 (HSC70-5) | 7 | 0.37 | 0.199 | 0.491 |
| AT5G09590.1 | Heat shock cognate 70 (HSC70-5) | 8 | 0.46 | 0.187 | 0.765 |
| AT5G09590.1 | Heat shock cognate 70 (HSC70-5) | 9 | 0.60 | 0.181 | 0.992 |
| AT5G09590.1 | Heat shock cognate 70 (HSC70-5) | 10 | 0.73 | 0.180 | 1.000 |
| AT5G09590.1 | Heat shock cognate 70 (HSC70-5) | 11 | 0.87 | 0.179 | 1.000 |
| AT5G09590.1 | Heat shock cognate 70 (HSC70-5) | 12 | 1.02 | 0.179 | 1.000 |
| AT5G09590.1 | Heat shock cognate 70 (HSC70-5) | 13 | 1.18 | 0.177 | 1.000 |
| AT5G09590.1 | Heat shock cognate 70 (HSC70-5) | 14 | 1.35 | 0.175 | 1.000 |
| AT5G09590.1 | Heat shock cognate 70 (HSC70-5) | 15 | 1.54 | 0.174 | 0.998 |
| AT5G09590.1 | Heat shock cognate 70 (HSC70-5) | 16 | 1.74 | 0.174 | 0.971 |
| AT5G09590.1 | Heat shock cognate 70 (HSC70-5) | 17 | 1.91 | 0.174 | 0.884 |
| AT5G09590.1 | Heat shock cognate 70 (HSC70-5) | 18 | 2.01 | 0.175 | 0.818 |
| AT5G09590.1 | Heat shock cognate 70 (HSC70-5) | 19 | 2.13 | 0.179 | 0.744 |
| AT5G09590.1 | Heat shock cognate 70 (HSC70-5) | 20 | 2.19 | 0.183 | 0.723 |
| AT5G09590.1 | Heat shock cognate 70 (HSC70-5) | 21 | 2.13 | 0.184 | 0.786 |
| AT5G09590.1 | Heat shock cognate 70 (HSC70-5) | 22 | 1.86 | 0.184 | 0.950 |
| AT5G09590.1 | Heat shock cognate 70 (HSC70-5) | 23 | 1.37 | 0.183 | 1.000 |
| AT5G09590.1 | Heat shock cognate 70 (HSC70-5) | 24 | 0.90 | 0.180 | 1.000 |

| | | | | | | |
|---|---|---|---|---|---|---|
| AT5G09590.1 | Heat shock cognate 70 (HSC70-5) | 25 | 0.67 | 0.178 | 1.000 | |
| AT5G09590.1 | Heat shock cognate 70 (HSC70-5) | 26 | 0.70 | 0.177 | 1.000 | |
| AT5G09590.1 | Heat shock cognate 70 (HSC70-5) | 27 | 0.81 | 0.177 | 1.000 | |
| AT5G09590.1 | Heat shock cognate 70 (HSC70-5) | 28 | 0.84 | 0.177 | 1.000 | |
| AT5G09590.1 | Heat shock cognate 70 (HSC70-5) | 29 | 0.80 | 0.177 | 1.000 | |
| AT5G09590.1 | Heat shock cognate 70 (HSC70-5) | 30 | 0.72 | 0.177 | 1.000 | |
| AT5G09590.1 | Heat shock cognate 70 (HSC70-5) | 31 | 0.67 | 0.178 | 1.000 | |
| AT5G09590.1 | Heat shock cognate 70 (HSC70-5) | 32 | 0.68 | 0.178 | 1.000 | |
| AT5G09590.1 | Heat shock cognate 70 (HSC70-5) | 33 | 0.78 | 0.178 | 1.000 | |
| AT5G09590.1 | Heat shock cognate 70 (HSC70-5) | 34 | 0.93 | 0.178 | 1.000 | |
| AT5G09590.1 | Heat shock cognate 70 (HSC70-5) | 35 | 0.99 | 0.178 | 1.000 | |
| AT5G09590.1 | Heat shock cognate 70 (HSC70-5) | 36 | 0.86 | 0.178 | 1.000 | |
| AT5G09590.1 | Heat shock cognate 70 (HSC70-5) | 37 | 0.75 | 0.178 | 1.000 | |
| AT5G09590.1 | Heat shock cognate 70 (HSC70-5) | 38 | 0.78 | 0.178 | 1.000 | |
| AT5G09590.1 | Heat shock cognate 70 (HSC70-5) | 39 | 0.92 | 0.178 | 1.000 | |
| AT5G09590.1 | Heat shock cognate 70 (HSC70-5) | 40 | 1.15 | 0.178 | 1.000 | |
| AT5G09590.1 | Heat shock cognate 70 (HSC70-5) | 41 | 1.44 | 0.179 | 1.000 | |
| AT5G09590.1 | Heat shock cognate 70 (HSC70-5) | 42 | 1.77 | 0.181 | 0.974 | |
| AT5G09590.1 | Heat shock cognate 70 (HSC70-5) | 43 | 1.88 | 0.182 | 0.934 | |
| AT5G09590.1 | Heat shock cognate 70 (HSC70-5) | 44 | 1.40 | 0.182 | 1.000 | |
| AT5G09590.1 | Heat shock cognate 70 (HSC70-5) | 45 | 0.72 | 0.184 | 1.000 | |
| AT5G09590.1 | Heat shock cognate 70 (HSC70-5) | 46 | 0.29 | 0.211 | 0.225 | |
| AT5G09590.1 | Heat shock cognate 70 (HSC70-5) | 47 | 0.10 | 0.326 | 0.053 | . |
| AT5G09600.2 | Succinate dehydrogenase subunit (SDH3-1) | 2 | 0.95 | 0.506 | 1.000 | |
| AT5G09600.2 | Succinate dehydrogenase subunit (SDH3-1) | 3 | 1.10 | 0.392 | 1.000 | |
| AT5G09600.2 | Succinate dehydrogenase subunit (SDH3-1) | 6 | 1.29 | 0.408 | 1.000 | |
| AT5G09600.2 | Succinate dehydrogenase subunit (SDH3-1) | 7 | 1.16 | 0.327 | 1.000 | |
| AT5G09600.2 | Succinate dehydrogenase subunit (SDH3-1) | 8 | 1.07 | 0.293 | 1.000 | |
| AT5G09600.2 | Succinate dehydrogenase subunit (SDH3-1) | 9 | 1.10 | 0.270 | 1.000 | |
| AT5G09600.2 | Succinate dehydrogenase subunit (SDH3-1) | 10 | 1.24 | 0.259 | 1.000 | |
| AT5G09600.2 | Succinate dehydrogenase subunit (SDH3-1) | 11 | 1.59 | 0.252 | 1.000 | |

| | | | | | | |
|---|---|---|---|---|---|---|
| AT5G09600.2 | Succinate dehydrogenase subunit (SDH3-1) | 12 | 2.00 | 0.249 | 0.996 | |
| AT5G09600.2 | Succinate dehydrogenase subunit (SDH3-1) | 13 | 1.84 | 0.246 | 0.999 | |
| AT5G09600.2 | Succinate dehydrogenase subunit (SDH3-1) | 14 | 1.39 | 0.245 | 1.000 | |
| AT5G09600.2 | Succinate dehydrogenase subunit (SDH3-1) | 15 | 1.64 | 0.244 | 1.000 | |
| AT5G09600.2 | Succinate dehydrogenase subunit (SDH3-1) | 16 | 2.34 | 0.241 | 0.946 | |
| AT5G09600.2 | Succinate dehydrogenase subunit (SDH3-1) | 17 | 3.19 | 0.242 | 0.591 | |
| AT5G09600.2 | Succinate dehydrogenase subunit (SDH3-1) | 18 | 4.24 | 0.249 | 0.266 | |
| AT5G09600.2 | Succinate dehydrogenase subunit (SDH3-1) | 19 | 4.57 | 0.258 | 0.245 | |
| AT5G09600.2 | Succinate dehydrogenase subunit (SDH3-1) | 20 | 3.72 | 0.271 | 0.571 | |
| AT5G09600.2 | Succinate dehydrogenase subunit (SDH3-1) | 21 | 2.51 | 0.306 | 0.990 | |
| AT5G09600.2 | Succinate dehydrogenase subunit (SDH3-1) | 22 | 1.88 | 0.374 | 1.000 | |
| AT5G09600.2 | Succinate dehydrogenase subunit (SDH3-1) | 27 | 2.33 | 0.355 | 1.000 | |
| AT5G09600.2 | Succinate dehydrogenase subunit (SDH3-1) | 28 | 2.90 | 0.289 | 0.920 | |
| AT5G09600.2 | Succinate dehydrogenase subunit (SDH3-1) | 29 | 3.41 | 0.259 | 0.612 | |
| AT5G09600.2 | Succinate dehydrogenase subunit (SDH3-1) | 30 | 4.19 | 0.251 | 0.290 | |
| AT5G09600.2 | Succinate dehydrogenase subunit (SDH3-1) | 31 | 5.93 | 0.249 | 0.060 | . |
| AT5G09600.2 | Succinate dehydrogenase subunit (SDH3-1) | 32 | 7.23 | 0.248 | 0.019 | * |
| AT5G09600.2 | Succinate dehydrogenase subunit (SDH3-1) | 33 | 5.31 | 0.248 | 0.100 | . |
| AT5G09600.2 | Succinate dehydrogenase subunit (SDH3-1) | 34 | 2.76 | 0.249 | 0.829 | |
| AT5G09600.2 | Succinate dehydrogenase subunit (SDH3-1) | 35 | 1.78 | 0.248 | 1.000 | |
| AT5G09600.2 | Succinate dehydrogenase subunit (SDH3-1) | 36 | 2.00 | 0.246 | 0.996 | |
| AT5G09600.2 | Succinate dehydrogenase subunit (SDH3-1) | 37 | 3.44 | 0.247 | 0.517 | |
| AT5G09600.2 | Succinate dehydrogenase subunit (SDH3-1) | 38 | 6.87 | 0.249 | 0.027 | * |
| AT5G09600.2 | Succinate dehydrogenase subunit (SDH3-1) | 39 | 11.48 | 0.248 | 0.001 | *** |
| AT5G09600.2 | Succinate dehydrogenase subunit (SDH3-1) | 40 | 10.52 | 0.248 | 0.002 | ** |
| AT5G09600.2 | Succinate dehydrogenase subunit (SDH3-1) | 41 | 4.92 | 0.247 | 0.139 | |
| AT5G09600.2 | Succinate dehydrogenase subunit (SDH3-1) | 42 | 1.94 | 0.251 | 0.998 | |
| AT5G09600.2 | Succinate dehydrogenase subunit (SDH3-1) | 43 | 1.12 | 0.252 | 1.000 | |
| AT5G09600.2 | Succinate dehydrogenase subunit (SDH3-1) | 44 | 0.78 | 0.251 | 1.000 | |
| AT5G09600.2 | Succinate dehydrogenase subunit (SDH3-1) | 45 | 0.57 | 0.251 | 1.000 | |
| AT5G09600.2 | Succinate dehydrogenase subunit (SDH3-1) | 46 | 0.44 | 0.276 | 0.992 | |

| | | | | | |
|---|---|---|---|---|---|
| AT5G09600.2 | Succinate dehydrogenase subunit (SDH3-1) | 47 | 0.35 | 0.522 | 1.000 |
| AT5G09660.2 | Peroxisomal NAD-malate dehydrogenase 2 ( PMDH2) | 2 | 0.82 | 0.325 | 1.000 |
| AT5G09660.2 | Peroxisomal NAD-malate dehydrogenase 2 ( PMDH2) | 3 | 1.02 | 0.309 | 1.000 |
| AT5G09660.2 | Peroxisomal NAD-malate dehydrogenase 2 ( PMDH2) | 24 | 1.15 | 0.253 | 1.000 |
| AT5G09660.2 | Peroxisomal NAD-malate dehydrogenase 2 ( PMDH2) | 25 | 0.78 | 0.176 | 1.000 |
| AT5G09660.2 | Peroxisomal NAD-malate dehydrogenase 2 ( PMDH2) | 26 | 0.61 | 0.152 | 0.927 |
| AT5G09660.2 | Peroxisomal NAD-malate dehydrogenase 2 ( PMDH2) | 27 | 0.65 | 0.144 | 0.961 |
| AT5G09660.2 | Peroxisomal NAD-malate dehydrogenase 2 ( PMDH2) | 28 | 0.87 | 0.142 | 1.000 |
| AT5G09660.2 | Peroxisomal NAD-malate dehydrogenase 2 ( PMDH2) | 29 | 1.13 | 0.143 | 1.000 |
| AT5G09660.2 | Peroxisomal NAD-malate dehydrogenase 2 ( PMDH2) | 30 | 1.22 | 0.143 | 1.000 |
| AT5G09660.2 | Peroxisomal NAD-malate dehydrogenase 2 ( PMDH2) | 31 | 1.24 | 0.143 | 1.000 |
| AT5G09660.2 | Peroxisomal NAD-malate dehydrogenase 2 ( PMDH2) | 32 | 1.26 | 0.144 | 1.000 |
| AT5G09660.2 | Peroxisomal NAD-malate dehydrogenase 2 ( PMDH2) | 33 | 1.25 | 0.144 | 1.000 |
| AT5G09660.2 | Peroxisomal NAD-malate dehydrogenase 2 ( PMDH2) | 34 | 1.14 | 0.144 | 1.000 |
| AT5G09660.2 | Peroxisomal NAD-malate dehydrogenase 2 ( PMDH2) | 35 | 0.85 | 0.144 | 1.000 |
| AT5G09660.2 | Peroxisomal NAD-malate dehydrogenase 2 ( PMDH2) | 36 | 0.63 | 0.143 | 0.932 |
| AT5G09660.2 | Peroxisomal NAD-malate dehydrogenase 2 ( PMDH2) | 37 | 0.63 | 0.142 | 0.929 |
| AT5G09660.2 | Peroxisomal NAD-malate dehydrogenase 2 ( PMDH2) | 38 | 0.81 | 0.142 | 1.000 |
| AT5G09660.2 | Peroxisomal NAD-malate dehydrogenase 2 ( PMDH2) | 39 | 1.10 | 0.142 | 1.000 |
| AT5G09660.2 | Peroxisomal NAD-malate dehydrogenase 2 ( PMDH2) | 40 | 1.41 | 0.143 | 0.996 |
| AT5G09660.2 | Peroxisomal NAD-malate dehydrogenase 2 ( PMDH2) | 41 | 1.62 | 0.144 | 0.911 |
| AT5G09660.2 | Peroxisomal NAD-malate dehydrogenase 2 ( PMDH2) | 42 | 1.75 | 0.146 | 0.805 |
| AT5G09660.2 | Peroxisomal NAD-malate dehydrogenase 2 ( PMDH2) | 43 | 1.76 | 0.147 | 0.800 |
| AT5G09660.2 | Peroxisomal NAD-malate dehydrogenase 2 ( PMDH2) | 44 | 1.48 | 0.147 | 0.988 |
| AT5G09660.2 | Peroxisomal NAD-malate dehydrogenase 2 ( PMDH2) | 45 | 0.99 | 0.148 | 1.000 |
| AT5G09660.2 | Peroxisomal NAD-malate dehydrogenase 2 ( PMDH2) | 46 | 0.55 | 0.172 | 0.896 |
| AT5G09660.2 | Peroxisomal NAD-malate dehydrogenase 2 ( PMDH2) | 47 | 0.27 | 0.296 | 0.621 |
| AT5G09840.1 | Mitochondrial nuclease 2 (MNU2) | 2 | 0.70 | 0.358 | 1.000 |
| AT5G09840.1 | Mitochondrial nuclease 2 (MNU2) | 3 | 0.52 | 0.231 | 0.950 |
| AT5G09840.1 | Mitochondrial nuclease 2 (MNU2) | 4 | 0.44 | 0.207 | 0.668 |
| AT5G09840.1 | Mitochondrial nuclease 2 (MNU2) | 5 | 0.45 | 0.207 | 0.721 |

| | | | | | | |
|---|---|---|---|---|---|---|
| AT5G09840.1 | Mitochondrial nuclease 2 (MNU2) | 6 | 0.53 | 0.208 | 0.915 | |
| AT5G09840.1 | Mitochondrial nuclease 2 (MNU2) | 7 | 0.66 | 0.206 | 0.996 | |
| AT5G09840.1 | Mitochondrial nuclease 2 (MNU2) | 8 | 0.85 | 0.203 | 1.000 | |
| AT5G09840.1 | Mitochondrial nuclease 2 (MNU2) | 9 | 1.06 | 0.201 | 1.000 | |
| AT5G09840.1 | Mitochondrial nuclease 2 (MNU2) | 10 | 1.20 | 0.201 | 1.000 | |
| AT5G09840.1 | Mitochondrial nuclease 2 (MNU2) | 11 | 1.31 | 0.203 | 1.000 | |
| AT5G09840.1 | Mitochondrial nuclease 2 (MNU2) | 12 | 1.47 | 0.208 | 0.999 | |
| AT5G09840.1 | Mitochondrial nuclease 2 (MNU2) | 13 | 1.83 | 0.215 | 0.952 | |
| AT5G09840.1 | Mitochondrial nuclease 2 (MNU2) | 14 | 2.49 | 0.222 | 0.634 | |
| AT5G09840.1 | Mitochondrial nuclease 2 (MNU2) | 15 | 3.28 | 0.238 | 0.347 | |
| AT5G09840.1 | Mitochondrial nuclease 2 (MNU2) | 16 | 3.69 | 0.288 | 0.487 | |
| AT5G09840.1 | Mitochondrial nuclease 2 (MNU2) | 32 | 1.37 | 0.404 | 1.000 | |
| AT5G09840.1 | Mitochondrial nuclease 2 (MNU2) | 33 | 1.21 | 0.321 | 1.000 | |
| AT5G09840.1 | Mitochondrial nuclease 2 (MNU2) | 34 | 1.00 | 0.298 | 1.000 | |
| AT5G09840.1 | Mitochondrial nuclease 2 (MNU2) | 35 | 0.76 | 0.321 | 1.000 | |
| AT5G09840.1 | Mitochondrial nuclease 2 (MNU2) | 36 | 0.55 | 0.430 | 1.000 | |
| AT5G09840.1 | Mitochondrial nuclease 2 (MNU2) | 46 | 0.01 | 3.620 | 1.000 | |
| AT5G09840.1 | Mitochondrial nuclease 2 (MNU2) | 47 | 0.01 | 4.101 | 1.000 | |
| AT5G10360.1 | 40S ribosomal protein S6-2 (RPS6B) | 2 | 0.19 | 0.363 | 0.312 | |
| AT5G10360.1 | 40S ribosomal protein S6-2 (RPS6B) | 3 | 0.41 | 0.286 | 0.725 | |
| AT5G10360.1 | 40S ribosomal protein S6-2 (RPS6B) | 4 | 0.85 | 0.254 | 1.000 | |
| AT5G10360.1 | 40S ribosomal protein S6-2 (RPS6B) | 5 | 1.66 | 0.243 | 0.956 | |
| AT5G10360.1 | 40S ribosomal protein S6-2 (RPS6B) | 6 | 2.74 | 0.238 | 0.393 | |
| AT5G10360.1 | 40S ribosomal protein S6-2 (RPS6B) | 7 | 3.66 | 0.232 | 0.125 | |
| AT5G10360.1 | 40S ribosomal protein S6-2 (RPS6B) | 8 | 3.95 | 0.226 | 0.075 | . |
| AT5G10360.1 | 40S ribosomal protein S6-2 (RPS6B) | 9 | 3.49 | 0.221 | 0.119 | |
| AT5G10360.1 | 40S ribosomal protein S6-2 (RPS6B) | 10 | 2.70 | 0.220 | 0.321 | |
| AT5G10360.1 | 40S ribosomal protein S6-2 (RPS6B) | 11 | 2.05 | 0.220 | 0.690 | |
| AT5G10360.1 | 40S ribosomal protein S6-2 (RPS6B) | 12 | 1.69 | 0.221 | 0.916 | |
| AT5G10360.1 | 40S ribosomal protein S6-2 (RPS6B) | 13 | 1.59 | 0.223 | 0.954 | |
| AT5G10360.1 | 40S ribosomal protein S6-2 (RPS6B) | 14 | 1.70 | 0.227 | 0.919 | |

| | | | | | | |
|---|---|---|---|---|---|---|
| AT5G10360.1 | 40S ribosomal protein S6-2 (RPS6B) | 15 | 1.85 | 0.234 | 0.862 | |
| AT5G10360.1 | 40S ribosomal protein S6-2 (RPS6B) | 16 | 1.90 | 0.246 | 0.864 | |
| AT5G10360.1 | 40S ribosomal protein S6-2 (RPS6B) | 17 | 1.80 | 0.269 | 0.944 | |
| AT5G10360.1 | 40S ribosomal protein S6-2 (RPS6B) | 18 | 1.58 | 0.319 | 0.996 | |
| AT5G10360.1 | 40S ribosomal protein S6-2 (RPS6B) | 19 | 1.35 | 0.405 | 1.000 | |
| AT5G10360.1 | 40S ribosomal protein S6-2 (RPS6B) | 46 | 19.32 | 0.732 | 0.448 | |
| AT5G10360.1 | 40S ribosomal protein S6-2 (RPS6B) | 47 | 24.18 | 0.754 | 0.396 | |
| AT5G10730.1 | NAD(P)-binding Rossmann-fold superfamily protein | 2 | 1.07 | 0.393 | 1.000 | |
| AT5G10730.1 | NAD(P)-binding Rossmann-fold superfamily protein | 3 | 0.66 | 0.324 | 1.000 | |
| AT5G10730.1 | NAD(P)-binding Rossmann-fold superfamily protein | 4 | 0.43 | 0.387 | 0.988 | |
| AT5G10730.1 | NAD(P)-binding Rossmann-fold superfamily protein | 27 | 2.76 | 0.304 | 0.822 | |
| AT5G10730.1 | NAD(P)-binding Rossmann-fold superfamily protein | 28 | 3.22 | 0.235 | 0.327 | |
| AT5G10730.1 | NAD(P)-binding Rossmann-fold superfamily protein | 29 | 3.59 | 0.195 | 0.067 | . |
| AT5G10730.1 | NAD(P)-binding Rossmann-fold superfamily protein | 30 | 3.92 | 0.176 | 0.014 | * |
| AT5G10730.1 | NAD(P)-binding Rossmann-fold superfamily protein | 31 | 4.50 | 0.170 | 0.002 | ** |
| AT5G10730.1 | NAD(P)-binding Rossmann-fold superfamily protein | 32 | 5.41 | 0.168 | 0.000 | *** |
| AT5G10730.1 | NAD(P)-binding Rossmann-fold superfamily protein | 33 | 6.27 | 0.168 | 0.000 | *** |
| AT5G10730.1 | NAD(P)-binding Rossmann-fold superfamily protein | 34 | 6.60 | 0.167 | 0.000 | *** |
| AT5G10730.1 | NAD(P)-binding Rossmann-fold superfamily protein | 35 | 6.09 | 0.167 | 0.000 | *** |
| AT5G10730.1 | NAD(P)-binding Rossmann-fold superfamily protein | 36 | 5.10 | 0.166 | 0.000 | *** |
| AT5G10730.1 | NAD(P)-binding Rossmann-fold superfamily protein | 37 | 4.14 | 0.166 | 0.005 | ** |
| AT5G10730.1 | NAD(P)-binding Rossmann-fold superfamily protein | 38 | 3.19 | 0.166 | 0.038 | * |
| AT5G10730.1 | NAD(P)-binding Rossmann-fold superfamily protein | 39 | 2.32 | 0.167 | 0.306 | |
| AT5G10730.1 | NAD(P)-binding Rossmann-fold superfamily protein | 40 | 1.71 | 0.168 | 0.855 | |
| AT5G10730.1 | NAD(P)-binding Rossmann-fold superfamily protein | 41 | 1.35 | 0.170 | 0.998 | |
| AT5G10730.1 | NAD(P)-binding Rossmann-fold superfamily protein | 42 | 1.15 | 0.171 | 1.000 | |
| AT5G10730.1 | NAD(P)-binding Rossmann-fold superfamily protein | 43 | 1.03 | 0.171 | 1.000 | |
| AT5G10730.1 | NAD(P)-binding Rossmann-fold superfamily protein | 44 | 0.90 | 0.173 | 1.000 | |
| AT5G10730.1 | NAD(P)-binding Rossmann-fold superfamily protein | 45 | 0.74 | 0.189 | 0.999 | |
| AT5G10730.1 | NAD(P)-binding Rossmann-fold superfamily protein | 46 | 0.57 | 0.244 | 0.983 | |
| AT5G10730.1 | NAD(P)-binding Rossmann-fold superfamily protein | 47 | 0.43 | 0.350 | 0.976 | |

| | | | | | | |
|---|---|---|---|---|---|---|
| AT5G10860.1 | Cystathionine beta-synthase domain-containing protein (CBSX3) | 2 | 1.04 | 0.293 | 1.000 | |
| AT5G10860.1 | Cystathionine beta-synthase domain-containing protein (CBSX3) | 3 | 0.94 | 0.276 | 1.000 | |
| AT5G10860.1 | Cystathionine beta-synthase domain-containing protein (CBSX3) | 4 | 0.85 | 0.268 | 1.000 | |
| AT5G10860.1 | Cystathionine beta-synthase domain-containing protein (CBSX3) | 21 | 0.36 | 0.164 | 0.057 | . |
| AT5G10860.1 | Cystathionine beta-synthase domain-containing protein (CBSX3) | 22 | 0.37 | 0.149 | 0.040 | * |
| AT5G10860.1 | Cystathionine beta-synthase domain-containing protein (CBSX3) | 23 | 0.40 | 0.137 | 0.034 | * |
| AT5G10860.1 | Cystathionine beta-synthase domain-containing protein (CBSX3) | 24 | 0.43 | 0.127 | 0.037 | * |
| AT5G10860.1 | Cystathionine beta-synthase domain-containing protein (CBSX3) | 25 | 0.47 | 0.120 | 0.057 | . |
| AT5G10860.1 | Cystathionine beta-synthase domain-containing protein (CBSX3) | 26 | 0.51 | 0.116 | 0.104 | |
| AT5G10860.1 | Cystathionine beta-synthase domain-containing protein (CBSX3) | 27 | 0.56 | 0.113 | 0.206 | |
| AT5G10860.1 | Cystathionine beta-synthase domain-containing protein (CBSX3) | 28 | 0.62 | 0.111 | 0.384 | |
| AT5G10860.1 | Cystathionine beta-synthase domain-containing protein (CBSX3) | 29 | 0.68 | 0.110 | 0.623 | |
| AT5G10860.1 | Cystathionine beta-synthase domain-containing protein (CBSX3) | 30 | 0.74 | 0.109 | 0.847 | |
| AT5G10860.1 | Cystathionine beta-synthase domain-containing protein (CBSX3) | 31 | 0.81 | 0.109 | 0.974 | |
| AT5G10860.1 | Cystathionine beta-synthase domain-containing protein (CBSX3) | 32 | 0.89 | 0.108 | 0.999 | |
| AT5G10860.1 | Cystathionine beta-synthase domain-containing protein (CBSX3) | 33 | 0.97 | 0.108 | 1.000 | |
| AT5G10860.1 | Cystathionine beta-synthase domain-containing protein (CBSX3) | 34 | 1.06 | 0.108 | 1.000 | |
| AT5G10860.1 | Cystathionine beta-synthase domain-containing protein (CBSX3) | 35 | 1.14 | 0.109 | 0.999 | |
| AT5G10860.1 | Cystathionine beta-synthase domain-containing protein (CBSX3) | 36 | 1.23 | 0.109 | 0.974 | |
| AT5G10860.1 | Cystathionine beta-synthase domain-containing protein (CBSX3) | 37 | 1.32 | 0.110 | 0.887 | |

| | | | | | | |
|---|---|---|---|---|---|---|
| AT5G10860.1 | Cystathionine beta-synthase domain-containing protein (CBSX3) | 38 | 1.41 | 0.111 | 0.754 | |
| AT5G10860.1 | Cystathionine beta-synthase domain-containing protein (CBSX3) | 39 | 1.48 | 0.112 | 0.628 | |
| AT5G10860.1 | Cystathionine beta-synthase domain-containing protein (CBSX3) | 40 | 1.52 | 0.114 | 0.563 | |
| AT5G10860.1 | Cystathionine beta-synthase domain-containing protein (CBSX3) | 41 | 1.53 | 0.116 | 0.578 | |
| AT5G10860.1 | Cystathionine beta-synthase domain-containing protein (CBSX3) | 42 | 1.49 | 0.119 | 0.676 | |
| AT5G10860.1 | Cystathionine beta-synthase domain-containing protein (CBSX3) | 43 | 1.41 | 0.124 | 0.835 | |
| AT5G10860.1 | Cystathionine beta-synthase domain-containing protein (CBSX3) | 44 | 1.30 | 0.133 | 0.967 | |
| AT5G10860.1 | Cystathionine beta-synthase domain-containing protein (CBSX3) | 45 | 1.18 | 0.146 | 0.999 | |
| AT5G10860.1 | Cystathionine beta-synthase domain-containing protein (CBSX3) | 46 | 1.04 | 0.167 | 1.000 | |
| AT5G10860.1 | Cystathionine beta-synthase domain-containing protein (CBSX3) | 47 | 0.92 | 0.195 | 1.000 | |
| AT5G10980.1 | Histone H3-like 8 (HTR8) | 2 | 0.40 | 0.229 | 0.275 | |
| AT5G10980.1 | Histone H3-like 8 (HTR8) | 3 | 0.40 | 0.218 | 0.225 | |
| AT5G10980.1 | Histone H3-like 8 (HTR8) | 4 | 0.39 | 0.209 | 0.184 | |
| AT5G10980.1 | Histone H3-like 8 (HTR8) | 5 | 0.39 | 0.201 | 0.152 | |
| AT5G10980.1 | Histone H3-like 8 (HTR8) | 6 | 0.38 | 0.195 | 0.125 | |
| AT5G10980.1 | Histone H3-like 8 (HTR8) | 7 | 0.38 | 0.190 | 0.106 | |
| AT5G10980.1 | Histone H3-like 8 (HTR8) | 8 | 0.38 | 0.185 | 0.092 | . |
| AT5G10980.1 | Histone H3-like 8 (HTR8) | 9 | 0.37 | 0.182 | 0.080 | . |
| AT5G10980.1 | Histone H3-like 8 (HTR8) | 10 | 0.37 | 0.180 | 0.070 | . |
| AT5G10980.1 | Histone H3-like 8 (HTR8) | 11 | 0.37 | 0.178 | 0.063 | . |
| AT5G10980.1 | Histone H3-like 8 (HTR8) | 12 | 0.36 | 0.177 | 0.059 | . |
| AT5G10980.1 | Histone H3-like 8 (HTR8) | 13 | 0.36 | 0.176 | 0.055 | . |
| AT5G10980.1 | Histone H3-like 8 (HTR8) | 14 | 0.36 | 0.175 | 0.053 | . |
| AT5G10980.1 | Histone H3-like 8 (HTR8) | 15 | 0.36 | 0.175 | 0.053 | . |
| AT5G10980.1 | Histone H3-like 8 (HTR8) | 16 | 0.36 | 0.175 | 0.054 | . |
| AT5G10980.1 | Histone H3-like 8 (HTR8) | 17 | 0.37 | 0.175 | 0.058 | . |
| AT5G10980.1 | Histone H3-like 8 (HTR8) | 18 | 0.37 | 0.175 | 0.064 | . |
| AT5G10980.1 | Histone H3-like 8 (HTR8) | 19 | 0.38 | 0.176 | 0.074 | . |
| AT5G10980.1 | Histone H3-like 8 (HTR8) | 20 | 0.39 | 0.176 | 0.087 | . |
| AT5G10980.1 | Histone H3-like 8 (HTR8) | 21 | 0.40 | 0.177 | 0.104 | |
| AT5G10980.1 | Histone H3-like 8 (HTR8) | 22 | 0.42 | 0.178 | 0.128 | |
| AT5G10980.1 | Histone H3-like 8 (HTR8) | 23 | 0.43 | 0.179 | 0.157 | |
| AT5G10980.1 | Histone H3-like 8 (HTR8) | 24 | 0.45 | 0.180 | 0.196 | |
| AT5G10980.1 | Histone H3-like 8 (HTR8) | 25 | 0.47 | 0.181 | 0.243 | |
| AT5G10980.1 | Histone H3-like 8 (HTR8) | 26 | 0.49 | 0.183 | 0.299 | |
| AT5G10980.1 | Histone H3-like 8 (HTR8) | 27 | 0.51 | 0.184 | 0.367 | |
| AT5G10980.1 | Histone H3-like 8 (HTR8) | 28 | 0.54 | 0.186 | 0.441 | |
| AT5G10980.1 | Histone H3-like 8 (HTR8) | 29 | 0.56 | 0.188 | 0.523 | |
| AT5G10980.1 | Histone H3-like 8 (HTR8) | 30 | 0.59 | 0.190 | 0.608 | |
| AT5G10980.1 | Histone H3-like 8 (HTR8) | 31 | 0.62 | 0.192 | 0.690 | |
| AT5G10980.1 | Histone H3-like 8 (HTR8) | 32 | 0.65 | 0.194 | 0.771 | |
| AT5G10980.1 | Histone H3-like 8 (HTR8) | 33 | 0.68 | 0.196 | 0.843 | |

| | | | | | |
|---|---|---|---|---|---|
| AT5G10980.1 | Histone H3-like 8 (HTR8) | 38 | 0.88 | 0.208 | 1.000 |
| AT5G10980.1 | Histone H3-like 8 (HTR8) | 39 | 0.93 | 0.211 | 1.000 |
| AT5G10980.1 | Histone H3-like 8 (HTR8) | 40 | 0.98 | 0.214 | 1.000 |
| AT5G10980.1 | Histone H3-like 8 (HTR8) | 41 | 1.04 | 0.218 | 1.000 |
| AT5G10980.1 | Histone H3-like 8 (HTR8) | 42 | 1.11 | 0.223 | 1.000 |
| AT5G10980.1 | Histone H3-like 8 (HTR8) | 43 | 1.17 | 0.229 | 0.998 |
| AT5G10980.1 | Histone H3-like 8 (HTR8) | 44 | 1.25 | 0.236 | 0.991 |
| AT5G10980.1 | Histone H3-like 8 (HTR8) | 45 | 1.33 | 0.244 | 0.976 |
| AT5G10980.1 | Histone H3-like 8 (HTR8) | 46 | 1.42 | 0.253 | 0.952 |
| AT5G10980.1 | Histone H3-like 8 (HTR8) | 47 | 1.52 | 0.264 | 0.923 |
| AT5G11350.1 | DNAse I-like superfamily protein | 2 | 0.93 | 0.801 | 1.000 |
| AT5G11350.1 | DNAse I-like superfamily protein | 3 | 0.95 | 0.759 | 1.000 |
| AT5G11350.1 | DNAse I-like superfamily protein | 27 | 1.04 | 0.461 | 1.000 |
| AT5G11350.1 | DNAse I-like superfamily protein | 28 | 1.02 | 0.362 | 1.000 |
| AT5G11350.1 | DNAse I-like superfamily protein | 29 | 1.02 | 0.299 | 1.000 |
| AT5G11350.1 | DNAse I-like superfamily protein | 30 | 1.03 | 0.268 | 1.000 |
| AT5G11350.1 | DNAse I-like superfamily protein | 31 | 1.02 | 0.257 | 1.000 |
| AT5G11350.1 | DNAse I-like superfamily protein | 32 | 0.98 | 0.254 | 1.000 |
| AT5G11350.1 | DNAse I-like superfamily protein | 33 | 0.90 | 0.254 | 1.000 |
| AT5G11350.1 | DNAse I-like superfamily protein | 34 | 0.89 | 0.254 | 1.000 |
| AT5G11350.1 | DNAse I-like superfamily protein | 35 | 0.98 | 0.256 | 1.000 |
| AT5G11350.1 | DNAse I-like superfamily protein | 36 | 1.19 | 0.260 | 1.000 |
| AT5G11350.1 | DNAse I-like superfamily protein | 37 | 1.61 | 0.272 | 0.989 |
| AT5G11350.1 | DNAse I-like superfamily protein | 38 | 2.29 | 0.303 | 0.871 |
| AT5G11350.1 | DNAse I-like superfamily protein | 39 | 3.44 | 0.368 | 0.708 |
| AT5G11350.1 | DNAse I-like superfamily protein | 40 | 5.34 | 0.477 | 0.662 |
| AT5G11350.1 | DNAse I-like superfamily protein | 46 | 77.71 | 1.873 | 0.942 |
| AT5G11350.1 | DNAse I-like superfamily protein | 47 | 121.67 | 2.205 | 0.959 |
| AT5G11520.1 | Aspartate aminotransferase 3 (ASP3) | 2 | 0.60 | 0.399 | 1.000 |
| AT5G11520.1 | Aspartate aminotransferase 3 (ASP3) | 3 | 0.68 | 0.322 | 1.000 |
| AT5G11520.1 | Aspartate aminotransferase 3 (ASP3) | 4 | 0.79 | 0.286 | 1.000 |
| AT5G11520.1 | Aspartate aminotransferase 3 (ASP3) | 5 | 0.94 | 0.273 | 1.000 |
| AT5G11520.1 | Aspartate aminotransferase 3 (ASP3) | 6 | 1.10 | 0.271 | 1.000 |
| AT5G11520.1 | Aspartate aminotransferase 3 (ASP3) | 7 | 1.27 | 0.270 | 1.000 |
| AT5G11520.1 | Aspartate aminotransferase 3 (ASP3) | 8 | 1.43 | 0.269 | 1.000 |
| AT5G11520.1 | Aspartate aminotransferase 3 (ASP3) | 9 | 1.57 | 0.268 | 0.997 |

| | | | | | | |
|---|---|---|---|---|---|---|
| AT5G11520.1 | Aspartate aminotransferase 3 (ASP3) | 10 | 1.69 | 0.268 | 0.990 | |
| AT5G11520.1 | Aspartate aminotransferase 3 (ASP3) | 11 | 1.79 | 0.267 | 0.977 | |
| AT5G11520.1 | Aspartate aminotransferase 3 (ASP3) | 12 | 1.83 | 0.267 | 0.970 | |
| AT5G11520.1 | Aspartate aminotransferase 3 (ASP3) | 13 | 1.77 | 0.266 | 0.979 | |
| AT5G11520.1 | Aspartate aminotransferase 3 (ASP3) | 14 | 1.66 | 0.265 | 0.992 | |
| AT5G11520.1 | Aspartate aminotransferase 3 (ASP3) | 15 | 1.51 | 0.264 | 0.999 | |
| AT5G11520.1 | Aspartate aminotransferase 3 (ASP3) | 16 | 1.31 | 0.264 | 1.000 | |
| AT5G11520.1 | Aspartate aminotransferase 3 (ASP3) | 17 | 1.02 | 0.264 | 1.000 | |
| AT5G11520.1 | Aspartate aminotransferase 3 (ASP3) | 18 | 0.71 | 0.266 | 1.000 | |
| AT5G11520.1 | Aspartate aminotransferase 3 (ASP3) | 19 | 0.48 | 0.267 | 0.908 | |
| AT5G11520.1 | Aspartate aminotransferase 3 (ASP3) | 20 | 0.34 | 0.270 | 0.540 | |
| AT5G11520.1 | Aspartate aminotransferase 3 (ASP3) | 21 | 0.26 | 0.272 | 0.263 | |
| AT5G11520.1 | Aspartate aminotransferase 3 (ASP3) | 22 | 0.21 | 0.275 | 0.142 | |
| AT5G11520.1 | Aspartate aminotransferase 3 (ASP3) | 23 | 0.19 | 0.277 | 0.097 | . |
| AT5G11520.1 | Aspartate aminotransferase 3 (ASP3) | 24 | 0.18 | 0.279 | 0.083 | . |
| AT5G11520.1 | Aspartate aminotransferase 3 (ASP3) | 25 | 0.18 | 0.283 | 0.091 | . |
| AT5G11520.1 | Aspartate aminotransferase 3 (ASP3) | 26 | 0.19 | 0.291 | 0.136 | |
| AT5G11520.1 | Aspartate aminotransferase 3 (ASP3) | 27 | 0.22 | 0.310 | 0.280 | |
| AT5G11520.1 | Aspartate aminotransferase 3 (ASP3) | 28 | 0.27 | 0.345 | 0.592 | |
| AT5G11520.1 | Aspartate aminotransferase 3 (ASP3) | 29 | 0.34 | 0.391 | 0.894 | |
| AT5G11520.1 | Aspartate aminotransferase 3 (ASP3) | 41 | 1.65 | 0.423 | 1.000 | |
| AT5G11520.1 | Aspartate aminotransferase 3 (ASP3) | 42 | 1.42 | 0.389 | 1.000 | |
| AT5G11520.1 | Aspartate aminotransferase 3 (ASP3) | 43 | 1.15 | 0.373 | 1.000 | |
| AT5G11520.1 | Aspartate aminotransferase 3 (ASP3) | 44 | 0.89 | 0.391 | 1.000 | |
| AT5G11520.1 | Aspartate aminotransferase 3 (ASP3) | 45 | 0.68 | 0.440 | 1.000 | |
| AT5G11520.1 | Aspartate aminotransferase 3 (ASP3) | 46 | 0.52 | 0.525 | 1.000 | |
| AT5G11520.1 | Aspartate aminotransferase 3 (ASP3) | 47 | 0.40 | 0.645 | 0.999 | |
| AT5G11670.1 | NADP malic enzyme 2 (NADP-ME2) | 2 | 0.99 | 0.342 | 1.000 | |
| AT5G11670.1 | NADP malic enzyme 2 (NADP-ME2) | 3 | 1.01 | 0.307 | 1.000 | |
| AT5G11670.1 | NADP malic enzyme 2 (NADP-ME2) | 4 | 1.03 | 0.283 | 1.000 | |
| AT5G11670.1 | NADP malic enzyme 2 (NADP-ME2) | 6 | 1.07 | 0.268 | 1.000 | |

| | | | | | |
|---|---|---|---|---|---|
| AT5G11670.1 | NADP malic enzyme 2 (NADP-ME2) | 7 | 1.09 | 0.272 | 1.000 |
| AT5G11670.1 | NADP malic enzyme 2 (NADP-ME2) | 8 | 1.11 | 0.282 | 1.000 |
| AT5G11670.1 | NADP malic enzyme 2 (NADP-ME2) | 22 | 1.06 | 0.284 | 1.000 |
| AT5G11670.1 | NADP malic enzyme 2 (NADP-ME2) | 23 | 1.03 | 0.270 | 1.000 |
| AT5G11670.1 | NADP malic enzyme 2 (NADP-ME2) | 24 | 0.99 | 0.256 | 1.000 |
| AT5G11670.1 | NADP malic enzyme 2 (NADP-ME2) | 29 | 0.84 | 0.187 | 1.000 |
| AT5G11670.1 | NADP malic enzyme 2 (NADP-ME2) | 30 | 0.84 | 0.177 | 0.999 |
| AT5G11670.1 | NADP malic enzyme 2 (NADP-ME2) | 31 | 0.86 | 0.169 | 1.000 |
| AT5G11670.1 | NADP malic enzyme 2 (NADP-ME2) | 32 | 0.89 | 0.164 | 1.000 |
| AT5G11670.1 | NADP malic enzyme 2 (NADP-ME2) | 33 | 0.94 | 0.162 | 1.000 |
| AT5G11670.1 | NADP malic enzyme 2 (NADP-ME2) | 34 | 1.00 | 0.162 | 1.000 |
| AT5G11670.1 | NADP malic enzyme 2 (NADP-ME2) | 35 | 1.07 | 0.164 | 1.000 |
| AT5G11670.1 | NADP malic enzyme 2 (NADP-ME2) | 37 | 1.24 | 0.174 | 0.997 |
| AT5G11670.1 | NADP malic enzyme 2 (NADP-ME2) | 38 | 1.33 | 0.183 | 0.988 |
| AT5G11670.1 | NADP malic enzyme 2 (NADP-ME2) | 39 | 1.42 | 0.197 | 0.974 |
| AT5G11670.1 | NADP malic enzyme 2 (NADP-ME2) | 40 | 1.51 | 0.216 | 0.962 |
| AT5G11670.1 | NADP malic enzyme 2 (NADP-ME2) | 41 | 1.61 | 0.241 | 0.957 |
| AT5G11670.1 | NADP malic enzyme 2 (NADP-ME2) | 42 | 1.70 | 0.274 | 0.960 |
| AT5G11670.1 | NADP malic enzyme 2 (NADP-ME2) | 43 | 1.80 | 0.314 | 0.967 |
| AT5G11670.1 | NADP malic enzyme 2 (NADP-ME2) | 46 | 2.11 | 0.471 | 0.986 |
| AT5G11670.1 | NADP malic enzyme 2 (NADP-ME2) | 47 | 2.23 | 0.535 | 0.990 |
| AT5G11690.1 | Translocase of the inner mitochondrial membrane 17 (TIM17-3) | 2 | 0.96 | 0.511 | 1.000 |
| AT5G11690.1 | Translocase of the inner mitochondrial membrane 17 (TIM17-3) | 3 | 0.98 | 0.462 | 1.000 |
| AT5G11690.1 | Translocase of the inner mitochondrial membrane 17 (TIM17-3) | 17 | 1.95 | 0.364 | 0.999 |
| AT5G11690.1 | Translocase of the inner mitochondrial membrane 17 (TIM17-3) | 18 | 2.18 | 0.319 | 0.986 |
| AT5G11690.1 | Translocase of the inner mitochondrial membrane 17 (TIM17-3) | 19 | 2.45 | 0.295 | 0.928 |
| AT5G11690.1 | Translocase of the inner mitochondrial membrane 17 (TIM17-3) | 20 | 2.65 | 0.315 | 0.920 |

| | | | | | | |
|---|---|---|---|---|---|---|
| AT5G11690.1 | Translocase of the inner mitochondrial membrane 17 (TIM17-3) | 21 | 2.60 | 0.343 | 0.961 | |
| AT5G11690.1 | Translocase of the inner mitochondrial membrane 17 (TIM17-3) | 22 | 2.31 | 0.398 | 0.996 | |
| AT5G11690.1 | Translocase of the inner mitochondrial membrane 17 (TIM17-3) | 32 | 0.96 | 0.308 | 1.000 | |
| AT5G11690.1 | Translocase of the inner mitochondrial membrane 17 (TIM17-3) | 33 | 1.36 | 0.244 | 1.000 | |
| AT5G11690.1 | Translocase of the inner mitochondrial membrane 17 (TIM17-3) | 34 | 1.88 | 0.216 | 0.945 | |
| AT5G11690.1 | Translocase of the inner mitochondrial membrane 17 (TIM17-3) | 35 | 2.34 | 0.200 | 0.601 | |
| AT5G11690.1 | Translocase of the inner mitochondrial membrane 17 (TIM17-3) | 36 | 3.20 | 0.194 | 0.142 | |
| AT5G11690.1 | Translocase of the inner mitochondrial membrane 17 (TIM17-3) | 37 | 4.88 | 0.193 | 0.009 | ** |
| AT5G11690.1 | Translocase of the inner mitochondrial membrane 17 (TIM17-3) | 38 | 5.51 | 0.193 | 0.004 | ** |
| AT5G11690.1 | Translocase of the inner mitochondrial membrane 17 (TIM17-3) | 39 | 3.55 | 0.193 | 0.076 | . |
| AT5G11690.1 | Translocase of the inner mitochondrial membrane 17 (TIM17-3) | 40 | 1.61 | 0.194 | 0.986 | |
| AT5G11690.1 | Translocase of the inner mitochondrial membrane 17 (TIM17-3) | 41 | 0.76 | 0.196 | 1.000 | |
| AT5G11690.1 | Translocase of the inner mitochondrial membrane 17 (TIM17-3) | 42 | 0.63 | 0.203 | 0.992 | |
| AT5G11690.1 | Translocase of the inner mitochondrial membrane 17 (TIM17-3) | 43 | 1.04 | 0.211 | 1.000 | |
| AT5G11690.1 | Translocase of the inner mitochondrial membrane 17 (TIM17-3) | 44 | 2.27 | 0.249 | 0.884 | |
| AT5G11690.1 | Translocase of the inner mitochondrial membrane 17 (TIM17-3) | 45 | 5.07 | 0.398 | 0.663 | |
| AT5G11690.1 | Translocase of the inner mitochondrial membrane 17 (TIM17-3) | 46 | 11.09 | 0.671 | 0.812 | |
| AT5G11690.1 | Translocase of the inner mitochondrial membrane 17 (TIM17-3) | 47 | 23.98 | 1.033 | 0.923 | |
| AT5G11770.1 | NADH:ubiquinone oxidoreductase subunit (PSST) | 2 | 0.72 | 0.614 | 1.000 | |
| AT5G11770.1 | NADH:ubiquinone oxidoreductase subunit (PSST) | 3 | 0.78 | 0.530 | 1.000 | |
| AT5G11770.1 | NADH:ubiquinone oxidoreductase subunit (PSST) | 10 | 1.24 | 0.395 | 1.000 | |
| AT5G11770.1 | NADH:ubiquinone oxidoreductase subunit (PSST) | 11 | 1.26 | 0.344 | 1.000 | |

| | | | | | |
|---|---|---|---|---|---|
| AT5G11770.1 | NADH:ubiquinone oxidoreductase subunit (PSST) | 12 | 1.19 | 0.315 | 1.000 |
| AT5G11770.1 | NADH:ubiquinone oxidoreductase subunit (PSST) | 13 | 1.11 | 0.298 | 1.000 |
| AT5G11770.1 | NADH:ubiquinone oxidoreductase subunit (PSST) | 14 | 1.06 | 0.286 | 1.000 |
| AT5G11770.1 | NADH:ubiquinone oxidoreductase subunit (PSST) | 15 | 0.86 | 0.282 | 1.000 |
| AT5G11770.1 | NADH:ubiquinone oxidoreductase subunit (PSST) | 16 | 0.59 | 0.280 | 1.000 |
| AT5G11770.1 | NADH:ubiquinone oxidoreductase subunit (PSST) | 17 | 0.45 | 0.280 | 0.984 |
| AT5G11770.1 | NADH:ubiquinone oxidoreductase subunit (PSST) | 18 | 0.48 | 0.283 | 0.995 |
| AT5G11770.1 | NADH:ubiquinone oxidoreductase subunit (PSST) | 19 | 0.66 | 0.289 | 1.000 |
| AT5G11770.1 | NADH:ubiquinone oxidoreductase subunit (PSST) | 20 | 0.89 | 0.295 | 1.000 |
| AT5G11770.1 | NADH:ubiquinone oxidoreductase subunit (PSST) | 21 | 1.19 | 0.297 | 1.000 |
| AT5G11770.1 | NADH:ubiquinone oxidoreductase subunit (PSST) | 22 | 1.59 | 0.297 | 1.000 |
| AT5G11770.1 | NADH:ubiquinone oxidoreductase subunit (PSST) | 23 | 1.77 | 0.295 | 1.000 |
| AT5G11770.1 | NADH:ubiquinone oxidoreductase subunit (PSST) | 24 | 1.32 | 0.291 | 1.000 |
| AT5G11770.1 | NADH:ubiquinone oxidoreductase subunit (PSST) | 25 | 0.70 | 0.287 | 1.000 |
| AT5G11770.1 | NADH:ubiquinone oxidoreductase subunit (PSST) | 26 | 0.42 | 0.286 | 0.974 |
| AT5G11770.1 | NADH:ubiquinone oxidoreductase subunit (PSST) | 27 | 0.37 | 0.286 | 0.906 |
| AT5G11770.1 | NADH:ubiquinone oxidoreductase subunit (PSST) | 28 | 0.46 | 0.286 | 0.993 |
| AT5G11770.1 | NADH:ubiquinone oxidoreductase subunit (PSST) | 29 | 0.73 | 0.286 | 1.000 |
| AT5G11770.1 | NADH:ubiquinone oxidoreductase subunit (PSST) | 30 | 1.02 | 0.286 | 1.000 |
| AT5G11770.1 | NADH:ubiquinone oxidoreductase subunit (PSST) | 31 | 1.21 | 0.287 | 1.000 |
| AT5G11770.1 | NADH:ubiquinone oxidoreductase subunit (PSST) | 32 | 1.36 | 0.287 | 1.000 |
| AT5G11770.1 | NADH:ubiquinone oxidoreductase subunit (PSST) | 33 | 1.34 | 0.287 | 1.000 |
| AT5G11770.1 | NADH:ubiquinone oxidoreductase subunit (PSST) | 34 | 1.25 | 0.288 | 1.000 |
| AT5G11770.1 | NADH:ubiquinone oxidoreductase subunit (PSST) | 35 | 1.06 | 0.287 | 1.000 |
| AT5G11770.1 | NADH:ubiquinone oxidoreductase subunit (PSST) | 36 | 0.95 | 0.286 | 1.000 |
| AT5G11770.1 | NADH:ubiquinone oxidoreductase subunit (PSST) | 37 | 0.94 | 0.286 | 1.000 |
| AT5G11770.1 | NADH:ubiquinone oxidoreductase subunit (PSST) | 38 | 0.95 | 0.287 | 1.000 |
| AT5G11770.1 | NADH:ubiquinone oxidoreductase subunit (PSST) | 39 | 1.05 | 0.287 | 1.000 |
| AT5G11770.1 | NADH:ubiquinone oxidoreductase subunit (PSST) | 40 | 1.30 | 0.288 | 1.000 |
| AT5G11770.1 | NADH:ubiquinone oxidoreductase subunit (PSST) | 41 | 1.55 | 0.289 | 1.000 |
| AT5G11770.1 | NADH:ubiquinone oxidoreductase subunit (PSST) | 42 | 1.68 | 0.291 | 1.000 |

| | | | | | | |
|---|---|---|---|---|---|---|
| AT5G11770.1 | NADH:ubiquinone oxidoreductase subunit (PSST) | 43 | 1.51 | 0.293 | 1.000 | |
| AT5G11770.1 | NADH:ubiquinone oxidoreductase subunit (PSST) | 44 | 1.02 | 0.293 | 1.000 | |
| AT5G11770.1 | NADH:ubiquinone oxidoreductase subunit (PSST) | 45 | 0.51 | 0.295 | 0.999 | |
| AT5G11770.1 | NADH:ubiquinone oxidoreductase subunit (PSST) | 46 | 0.20 | 0.338 | 0.509 | |
| AT5G11770.1 | NADH:ubiquinone oxidoreductase subunit (PSST) | 47 | 0.07 | 0.529 | 0.468 | |
| AT5G12040.1 | Putative omega-amidase (NLP3) | 2 | 0.90 | 0.342 | 1.000 | |
| AT5G12040.1 | Putative omega-amidase (NLP3) | 3 | 0.78 | 0.338 | 1.000 | |
| AT5G12040.1 | Putative omega-amidase (NLP3) | 40 | 0.55 | 0.330 | 0.987 | |
| AT5G12040.1 | Putative omega-amidase (NLP3) | 41 | 0.79 | 0.209 | 0.999 | |
| AT5G12040.1 | Putative omega-amidase (NLP3) | 42 | 1.59 | 0.180 | 0.901 | |
| AT5G12040.1 | Putative omega-amidase (NLP3) | 43 | 2.87 | 0.178 | 0.100 | . |
| AT5G12040.1 | Putative omega-amidase (NLP3) | 44 | 2.20 | 0.174 | 0.354 | |
| AT5G12040.1 | Putative omega-amidase (NLP3) | 45 | 1.15 | 0.174 | 1.000 | |
| AT5G12040.1 | Putative omega-amidase (NLP3) | 46 | 0.61 | 0.189 | 0.899 | |
| AT5G12040.1 | Putative omega-amidase (NLP3) | 47 | 0.26 | 0.405 | 0.725 | |
| AT5G12290.1 | DGD1 suppressor 1 (DGS1) | 2 | 0.92 | 0.560 | 1.000 | |
| AT5G12290.1 | DGD1 suppressor 1 (DGS1) | 3 | 0.89 | 0.512 | 1.000 | |
| AT5G12290.1 | DGD1 suppressor 1 (DGS1) | 17 | 1.35 | 0.369 | 1.000 | |
| AT5G12290.1 | DGD1 suppressor 1 (DGS1) | 18 | 1.59 | 0.314 | 1.000 | |
| AT5G12290.1 | DGD1 suppressor 1 (DGS1) | 19 | 1.86 | 0.292 | 0.996 | |
| AT5G12290.1 | DGD1 suppressor 1 (DGS1) | 20 | 1.94 | 0.285 | 0.991 | |
| AT5G12290.1 | DGD1 suppressor 1 (DGS1) | 21 | 1.78 | 0.283 | 0.998 | |
| AT5G12290.1 | DGD1 suppressor 1 (DGS1) | 22 | 1.51 | 0.282 | 1.000 | |
| AT5G12290.1 | DGD1 suppressor 1 (DGS1) | 23 | 1.15 | 0.281 | 1.000 | |
| AT5G12290.1 | DGD1 suppressor 1 (DGS1) | 24 | 0.86 | 0.275 | 1.000 | |
| AT5G12290.1 | DGD1 suppressor 1 (DGS1) | 25 | 0.75 | 0.273 | 1.000 | |
| AT5G12290.1 | DGD1 suppressor 1 (DGS1) | 26 | 0.90 | 0.272 | 1.000 | |
| AT5G12290.1 | DGD1 suppressor 1 (DGS1) | 27 | 1.32 | 0.271 | 1.000 | |
| AT5G12290.1 | DGD1 suppressor 1 (DGS1) | 28 | 1.83 | 0.272 | 0.994 | |
| AT5G12290.1 | DGD1 suppressor 1 (DGS1) | 29 | 1.76 | 0.273 | 0.997 | |
| AT5G12290.1 | DGD1 suppressor 1 (DGS1) | 30 | 0.96 | 0.273 | 1.000 | |
| AT5G12290.1 | DGD1 suppressor 1 (DGS1) | 31 | 0.44 | 0.274 | 0.939 | |
| AT5G12290.1 | DGD1 suppressor 1 (DGS1) | 32 | 0.32 | 0.280 | 0.667 | |
| AT5G12290.1 | DGD1 suppressor 1 (DGS1) | 33 | 0.44 | 0.294 | 0.957 | |
| AT5G12290.1 | DGD1 suppressor 1 (DGS1) | 34 | 0.72 | 0.352 | 1.000 | |
| AT5G12290.1 | DGD1 suppressor 1 (DGS1) | 35 | 1.20 | 0.538 | 1.000 | |
| AT5G12290.1 | DGD1 suppressor 1 (DGS1) | 46 | 45091.29 | 5.942 | 0.999 | |
| AT5G12290.1 | DGD1 suppressor 1 (DGS1) | 47 | 145770.44 | 6.698 | 0.999 | |
| AT5G12470.1 | Reticula-related 4 (RER4) | 2 | 1.00 | 0.335 | 1.000 | |
| AT5G12470.1 | Reticula-related 4 (RER4) | 3 | 1.10 | 0.322 | 1.000 | |
| AT5G12470.1 | Reticula-related 4 (RER4) | 4 | 1.20 | 0.352 | 0.999 | |
| AT5G12470.1 | Reticula-related 4 (RER4) | 42 | 0.74 | 0.252 | 0.975 | |
| AT5G12470.1 | Reticula-related 4 (RER4) | 43 | 0.58 | 0.214 | 0.712 | |
| AT5G12470.1 | Reticula-related 4 (RER4) | 44 | 0.44 | 0.193 | 0.268 | |
| AT5G12470.1 | Reticula-related 4 (RER4) | 45 | 0.32 | 0.194 | 0.065 | . |
| AT5G12470.1 | Reticula-related 4 (RER4) | 46 | 0.23 | 0.219 | 0.024 | * |
| AT5G12470.1 | Reticula-related 4 (RER4) | 47 | 0.16 | 0.266 | 0.021 | * |

| | | | | | | |
|---|---|---|---|---|---|---|
| AT5G12860.2 | Dicarboxylate transporter 1 (DIT1) | 2 | 1.11 | 0.580 | 1.000 | |
| AT5G12860.2 | Dicarboxylate transporter 1 (DIT1) | 3 | 1.16 | 0.564 | 1.000 | |
| AT5G12860.2 | Dicarboxylate transporter 1 (DIT1) | 33 | 1.53 | 0.245 | 0.902 | |
| AT5G12860.2 | Dicarboxylate transporter 1 (DIT1) | 34 | 1.44 | 0.225 | 0.922 | |
| AT5G12860.2 | Dicarboxylate transporter 1 (DIT1) | 35 | 1.34 | 0.217 | 0.961 | |
| AT5G12860.2 | Dicarboxylate transporter 1 (DIT1) | 36 | 1.20 | 0.220 | 0.994 | |
| AT5G12860.2 | Dicarboxylate transporter 1 (DIT1) | 37 | 1.05 | 0.231 | 1.000 | |
| AT5G12860.2 | Dicarboxylate transporter 1 (DIT1) | 38 | 0.88 | 0.247 | 0.999 | |
| AT5G12860.2 | Dicarboxylate transporter 1 (DIT1) | 46 | 0.11 | 0.361 | 0.048 | * |
| AT5G12860.2 | Dicarboxylate transporter 1 (DIT1) | 47 | 0.08 | 0.390 | 0.035 | * |
| AT5G13030.1 | Selenoprotein O | 2 | 0.92 | 0.317 | 1.000 | |
| AT5G13030.1 | Selenoprotein O | 3 | 0.62 | 0.305 | 1.000 | |
| AT5G13030.1 | Selenoprotein O | 27 | 0.18 | 0.239 | 0.041 | * |
| AT5G13030.1 | Selenoprotein O | 28 | 0.33 | 0.171 | 0.097 | . |
| AT5G13030.1 | Selenoprotein O | 29 | 0.58 | 0.153 | 0.848 | |
| AT5G13030.1 | Selenoprotein O | 30 | 0.86 | 0.146 | 1.000 | |
| AT5G13030.1 | Selenoprotein O | 31 | 1.03 | 0.144 | 1.000 | |
| AT5G13030.1 | Selenoprotein O | 32 | 0.90 | 0.145 | 1.000 | |
| AT5G13030.1 | Selenoprotein O | 33 | 0.61 | 0.145 | 0.881 | |
| AT5G13030.1 | Selenoprotein O | 34 | 0.48 | 0.145 | 0.352 | |
| AT5G13030.1 | Selenoprotein O | 35 | 0.50 | 0.145 | 0.456 | |
| AT5G13030.1 | Selenoprotein O | 36 | 0.69 | 0.144 | 0.983 | |
| AT5G13030.1 | Selenoprotein O | 37 | 1.08 | 0.144 | 1.000 | |
| AT5G13030.1 | Selenoprotein O | 38 | 1.44 | 0.143 | 0.985 | |
| AT5G13030.1 | Selenoprotein O | 39 | 1.64 | 0.143 | 0.861 | |
| AT5G13030.1 | Selenoprotein O | 40 | 1.70 | 0.143 | 0.800 | |
| AT5G13030.1 | Selenoprotein O | 41 | 1.55 | 0.144 | 0.940 | |
| AT5G13030.1 | Selenoprotein O | 42 | 1.41 | 0.146 | 0.993 | |
| AT5G13030.1 | Selenoprotein O | 43 | 1.34 | 0.147 | 0.999 | |
| AT5G13030.1 | Selenoprotein O | 44 | 1.14 | 0.147 | 1.000 | |
| AT5G13030.1 | Selenoprotein O | 45 | 0.80 | 0.148 | 1.000 | |
| AT5G13030.1 | Selenoprotein O | 46 | 0.43 | 0.171 | 0.385 | |
| AT5G13030.1 | Selenoprotein O | 47 | 0.18 | 0.295 | 0.186 | |
| AT5G13430.1 | Ubiquinol:cytochrome c oxidoreductase subunit (iron-sulfur subunit) | 2 | 0.86 | 0.440 | 1.000 | |
| AT5G13430.1 | Ubiquinol:cytochrome c oxidoreductase subunit (iron-sulfur subunit) | 3 | 0.65 | 0.269 | 1.000 | |
| AT5G13430.1 | Ubiquinol:cytochrome c oxidoreductase subunit (iron-sulfur subunit) | 4 | 0.62 | 0.253 | 1.000 | |
| AT5G13430.1 | Ubiquinol:cytochrome c oxidoreductase subunit (iron-sulfur subunit) | 5 | 0.67 | 0.253 | 1.000 | |
| AT5G13430.1 | Ubiquinol:cytochrome c oxidoreductase subunit (iron-sulfur subunit) | 6 | 0.70 | 0.254 | 1.000 | |
| AT5G13430.1 | Ubiquinol:cytochrome c oxidoreductase subunit (iron-sulfur subunit) | 7 | 0.70 | 0.255 | 1.000 | |

| | | | | | |
|---|---|---|---|---|---|
| AT5G13430.1 | Ubiquinol:cytochrome c oxidoreductase subunit (iron-sulfur subunit) | 8 | 0.77 | 0.249 | 1.000 |
| AT5G13430.1 | Ubiquinol:cytochrome c oxidoreductase subunit (iron-sulfur subunit) | 9 | 0.85 | 0.245 | 1.000 |
| AT5G13430.1 | Ubiquinol:cytochrome c oxidoreductase subunit (iron-sulfur subunit) | 10 | 0.86 | 0.245 | 1.000 |
| AT5G13430.1 | Ubiquinol:cytochrome c oxidoreductase subunit (iron-sulfur subunit) | 11 | 0.90 | 0.246 | 1.000 |
| AT5G13430.1 | Ubiquinol:cytochrome c oxidoreductase subunit (iron-sulfur subunit) | 12 | 0.96 | 0.246 | 1.000 |
| AT5G13430.1 | Ubiquinol:cytochrome c oxidoreductase subunit (iron-sulfur subunit) | 13 | 1.06 | 0.243 | 1.000 |
| AT5G13430.1 | Ubiquinol:cytochrome c oxidoreductase subunit (iron-sulfur subunit) | 14 | 1.18 | 0.239 | 1.000 |
| AT5G13430.1 | Ubiquinol:cytochrome c oxidoreductase subunit (iron-sulfur subunit) | 15 | 1.07 | 0.239 | 1.000 |
| AT5G13430.1 | Ubiquinol:cytochrome c oxidoreductase subunit (iron-sulfur subunit) | 16 | 0.78 | 0.237 | 1.000 |
| AT5G13430.1 | Ubiquinol:cytochrome c oxidoreductase subunit (iron-sulfur subunit) | 17 | 0.55 | 0.237 | 1.000 |
| AT5G13430.1 | Ubiquinol:cytochrome c oxidoreductase subunit (iron-sulfur subunit) | 18 | 0.54 | 0.240 | 1.000 |
| AT5G13430.1 | Ubiquinol:cytochrome c oxidoreductase subunit (iron-sulfur subunit) | 19 | 0.73 | 0.247 | 1.000 |
| AT5G13430.1 | Ubiquinol:cytochrome c oxidoreductase subunit (iron-sulfur subunit) | 20 | 1.11 | 0.253 | 1.000 |
| AT5G13430.1 | Ubiquinol:cytochrome c oxidoreductase subunit (iron-sulfur subunit) | 21 | 1.60 | 0.257 | 1.000 |
| AT5G13430.1 | Ubiquinol:cytochrome c oxidoreductase subunit (iron-sulfur subunit) | 22 | 1.51 | 0.256 | 1.000 |
| AT5G13430.1 | Ubiquinol:cytochrome c oxidoreductase subunit (iron-sulfur subunit) | 23 | 0.76 | 0.257 | 1.000 |
| AT5G13430.1 | Ubiquinol:cytochrome c oxidoreductase subunit (iron-sulfur subunit) | 24 | 0.41 | 0.250 | 0.977 |
| AT5G13430.1 | Ubiquinol:cytochrome c oxidoreductase subunit (iron-sulfur subunit) | 25 | 0.47 | 0.244 | 0.997 |
| AT5G13430.1 | Ubiquinol:cytochrome c oxidoreductase subunit (iron-sulfur subunit) | 26 | 0.93 | 0.243 | 1.000 |
| AT5G13430.1 | Ubiquinol:cytochrome c oxidoreductase subunit (iron-sulfur subunit) | 27 | 1.76 | 0.243 | 1.000 |

| | | | | | | |
|---|---|---|---|---|---|---|
| AT5G13430.1 | Ubiquinol:cytochrome c oxidoreductase subunit (iron-sulfur subunit) | 28 | 1.43 | 0.243 | 1.000 | |
| AT5G13430.1 | Ubiquinol:cytochrome c oxidoreductase subunit (iron-sulfur subunit) | 29 | 0.63 | 0.243 | 1.000 | |
| AT5G13430.1 | Ubiquinol:cytochrome c oxidoreductase subunit (iron-sulfur subunit) | 30 | 0.49 | 0.243 | 0.998 | |
| AT5G13430.1 | Ubiquinol:cytochrome c oxidoreductase subunit (iron-sulfur subunit) | 31 | 0.79 | 0.244 | 1.000 | |
| AT5G13430.1 | Ubiquinol:cytochrome c oxidoreductase subunit (iron-sulfur subunit) | 32 | 1.30 | 0.245 | 1.000 | |
| AT5G13430.1 | Ubiquinol:cytochrome c oxidoreductase subunit (iron-sulfur subunit) | 33 | 1.45 | 0.245 | 1.000 | |
| AT5G13430.1 | Ubiquinol:cytochrome c oxidoreductase subunit (iron-sulfur subunit) | 34 | 1.19 | 0.247 | 1.000 | |
| AT5G13430.1 | Ubiquinol:cytochrome c oxidoreductase subunit (iron-sulfur subunit) | 35 | 0.83 | 0.250 | 1.000 | |
| AT5G13430.1 | Ubiquinol:cytochrome c oxidoreductase subunit (iron-sulfur subunit) | 36 | 0.75 | 0.261 | 1.000 | |
| AT5G13430.1 | Ubiquinol:cytochrome c oxidoreductase subunit (iron-sulfur subunit) | 37 | 1.15 | 0.298 | 1.000 | |
| AT5G13430.1 | Ubiquinol:cytochrome c oxidoreductase subunit (iron-sulfur subunit) | 38 | 2.37 | 0.364 | 1.000 | |
| AT5G13430.1 | Ubiquinol:cytochrome c oxidoreductase subunit (iron-sulfur subunit) | 40 | 7.48 | 0.331 | 0.252 | |
| AT5G13430.1 | Ubiquinol:cytochrome c oxidoreductase subunit (iron-sulfur subunit) | 41 | 7.44 | 0.288 | 0.088 | . |
| AT5G13430.1 | Ubiquinol:cytochrome c oxidoreductase subunit (iron-sulfur subunit) | 42 | 4.92 | 0.260 | 0.240 | |
| AT5G13430.1 | Ubiquinol:cytochrome c oxidoreductase subunit (iron-sulfur subunit) | 43 | 3.47 | 0.253 | 0.646 | |
| AT5G13430.1 | Ubiquinol:cytochrome c oxidoreductase subunit (iron-sulfur subunit) | 44 | 1.95 | 0.252 | 1.000 | |
| AT5G13430.1 | Ubiquinol:cytochrome c oxidoreductase subunit (iron-sulfur subunit) | 45 | 0.79 | 0.253 | 1.000 | |
| AT5G13430.1 | Ubiquinol:cytochrome c oxidoreductase subunit (iron-sulfur subunit) | 46 | 0.27 | 0.288 | 0.760 | |
| AT5G13430.1 | Ubiquinol:cytochrome c oxidoreductase subunit (iron-sulfur subunit) | 47 | 0.09 | 0.514 | 0.703 | |
| AT5G13450.1 | ATP synthase subunit (ATPO) | 2 | 1.97 | 0.439 | 1.000 | |
| AT5G13450.1 | ATP synthase subunit (ATPO) | 3 | 1.52 | 0.307 | 1.000 | |
| AT5G13450.1 | ATP synthase subunit (ATPO) | 4 | 1.19 | 0.278 | 1.000 | |
| AT5G13450.1 | ATP synthase subunit (ATPO) | 5 | 0.98 | 0.278 | 1.000 | |
| AT5G13450.1 | ATP synthase subunit (ATPO) | 6 | 0.86 | 0.278 | 1.000 | |

| | | | | | | |
|---|---|---|---|---|---|---|
| AT5G13450.1 | ATP synthase subunit (ATPO) | 7 | 0.82 | 0.277 | 1.000 | |
| AT5G13450.1 | ATP synthase subunit (ATPO) | 8 | 0.83 | 0.274 | 1.000 | |
| AT5G13450.1 | ATP synthase subunit (ATPO) | 9 | 0.87 | 0.272 | 1.000 | |
| AT5G13450.1 | ATP synthase subunit (ATPO) | 10 | 0.92 | 0.271 | 1.000 | |
| AT5G13450.1 | ATP synthase subunit (ATPO) | 11 | 1.03 | 0.271 | 1.000 | |
| AT5G13450.1 | ATP synthase subunit (ATPO) | 12 | 1.17 | 0.271 | 1.000 | |
| AT5G13450.1 | ATP synthase subunit (ATPO) | 13 | 1.41 | 0.269 | 1.000 | |
| AT5G13450.1 | ATP synthase subunit (ATPO) | 14 | 1.76 | 0.267 | 1.000 | |
| AT5G13450.1 | ATP synthase subunit (ATPO) | 15 | 2.02 | 0.266 | 0.993 | |
| AT5G13450.1 | ATP synthase subunit (ATPO) | 16 | 1.91 | 0.265 | 0.997 | |
| AT5G13450.1 | ATP synthase subunit (ATPO) | 17 | 1.61 | 0.265 | 1.000 | |
| AT5G13450.1 | ATP synthase subunit (ATPO) | 18 | 1.29 | 0.268 | 1.000 | |
| AT5G13450.1 | ATP synthase subunit (ATPO) | 19 | 1.10 | 0.272 | 1.000 | |
| AT5G13450.1 | ATP synthase subunit (ATPO) | 20 | 1.05 | 0.276 | 1.000 | |
| AT5G13450.1 | ATP synthase subunit (ATPO) | 21 | 1.02 | 0.278 | 1.000 | |
| AT5G13450.1 | ATP synthase subunit (ATPO) | 22 | 0.98 | 0.279 | 1.000 | |
| AT5G13450.1 | ATP synthase subunit (ATPO) | 23 | 0.96 | 0.277 | 1.000 | |
| AT5G13450.1 | ATP synthase subunit (ATPO) | 24 | 1.00 | 0.274 | 1.000 | |
| AT5G13450.1 | ATP synthase subunit (ATPO) | 25 | 1.07 | 0.271 | 1.000 | |
| AT5G13450.1 | ATP synthase subunit (ATPO) | 26 | 1.08 | 0.270 | 1.000 | |
| AT5G13450.1 | ATP synthase subunit (ATPO) | 27 | 1.00 | 0.269 | 1.000 | |
| AT5G13450.1 | ATP synthase subunit (ATPO) | 28 | 1.07 | 0.269 | 1.000 | |
| AT5G13450.1 | ATP synthase subunit (ATPO) | 29 | 1.55 | 0.270 | 1.000 | |
| AT5G13450.1 | ATP synthase subunit (ATPO) | 30 | 2.44 | 0.270 | 0.932 | |
| AT5G13450.1 | ATP synthase subunit (ATPO) | 31 | 3.04 | 0.271 | 0.728 | |
| AT5G13450.1 | ATP synthase subunit (ATPO) | 32 | 2.89 | 0.271 | 0.787 | |
| AT5G13450.1 | ATP synthase subunit (ATPO) | 33 | 2.46 | 0.271 | 0.929 | |
| AT5G13450.1 | ATP synthase subunit (ATPO) | 34 | 1.95 | 0.272 | 0.997 | |
| AT5G13450.1 | ATP synthase subunit (ATPO) | 35 | 1.62 | 0.272 | 1.000 | |
| AT5G13450.1 | ATP synthase subunit (ATPO) | 36 | 1.66 | 0.271 | 1.000 | |
| AT5G13450.1 | ATP synthase subunit (ATPO) | 37 | 2.09 | 0.271 | 0.990 | |
| AT5G13450.1 | ATP synthase subunit (ATPO) | 38 | 2.88 | 0.272 | 0.792 | |
| AT5G13450.1 | ATP synthase subunit (ATPO) | 39 | 4.09 | 0.272 | 0.372 | |
| AT5G13450.1 | ATP synthase subunit (ATPO) | 40 | 5.45 | 0.273 | 0.144 | |
| AT5G13450.1 | ATP synthase subunit (ATPO) | 41 | 6.27 | 0.276 | 0.088 | . |
| AT5G13450.1 | ATP synthase subunit (ATPO) | 42 | 5.88 | 0.293 | 0.168 | |
| AT5G13450.1 | ATP synthase subunit (ATPO) | 43 | 4.72 | 0.348 | 0.607 | |
| AT5G13450.1 | ATP synthase subunit (ATPO) | 44 | 3.48 | 0.454 | 0.988 | |
| AT5G13450.1 | ATP synthase subunit (ATPO) | 46 | 1.61 | 0.887 | 1.000 | |
| AT5G13450.1 | ATP synthase subunit (ATPO) | 47 | 1.06 | 1.200 | 1.000 | |
| AT5G13490.2 | ADP/ATP carrier 2 (AAC2) | 3 | 0.56 | 0.290 | 1.000 | |
| AT5G13490.2 | ADP/ATP carrier 2 (AAC2) | 4 | 0.52 | 0.250 | 0.993 | |
| AT5G13490.2 | ADP/ATP carrier 2 (AAC2) | 5 | 0.58 | 0.245 | 0.999 | |
| AT5G13490.2 | ADP/ATP carrier 2 (AAC2) | 6 | 0.77 | 0.245 | 1.000 | |
| AT5G13490.2 | ADP/ATP carrier 2 (AAC2) | 7 | 1.17 | 0.244 | 1.000 | |
| AT5G13490.2 | ADP/ATP carrier 2 (AAC2) | 8 | 1.73 | 0.242 | 0.999 | |
| AT5G13490.2 | ADP/ATP carrier 2 (AAC2) | 9 | 2.05 | 0.240 | 0.969 | |
| AT5G13490.2 | ADP/ATP carrier 2 (AAC2) | 10 | 1.91 | 0.238 | 0.988 | |
| AT5G13490.2 | ADP/ATP carrier 2 (AAC2) | 11 | 1.56 | 0.238 | 1.000 | |
| AT5G13490.2 | ADP/ATP carrier 2 (AAC2) | 12 | 1.21 | 0.238 | 1.000 | |
| AT5G13490.2 | ADP/ATP carrier 2 (AAC2) | 13 | 0.93 | 0.236 | 1.000 | |
| AT5G13490.2 | ADP/ATP carrier 2 (AAC2) | 14 | 0.76 | 0.235 | 1.000 | |
| AT5G13490.2 | ADP/ATP carrier 2 (AAC2) | 15 | 0.70 | 0.234 | 1.000 | |
| AT5G13490.2 | ADP/ATP carrier 2 (AAC2) | 16 | 0.75 | 0.233 | 1.000 | |
| AT5G13490.2 | ADP/ATP carrier 2 (AAC2) | 17 | 0.95 | 0.233 | 1.000 | |
| AT5G13490.2 | ADP/ATP carrier 2 (AAC2) | 18 | 1.24 | 0.235 | 1.000 | |
| AT5G13490.2 | ADP/ATP carrier 2 (AAC2) | 19 | 1.53 | 0.239 | 1.000 | |
| AT5G13490.2 | ADP/ATP carrier 2 (AAC2) | 20 | 1.79 | 0.242 | 0.997 | |
| AT5G13490.2 | ADP/ATP carrier 2 (AAC2) | 21 | 2.07 | 0.245 | 0.971 | |
| AT5G13490.2 | ADP/ATP carrier 2 (AAC2) | 22 | 2.28 | 0.245 | 0.919 | |
| AT5G13490.2 | ADP/ATP carrier 2 (AAC2) | 23 | 2.39 | 0.243 | 0.876 | |
| AT5G13490.2 | ADP/ATP carrier 2 (AAC2) | 24 | 2.34 | 0.241 | 0.887 | |
| AT5G13490.2 | ADP/ATP carrier 2 (AAC2) | 25 | 2.15 | 0.238 | 0.944 | |

| | | | | | | |
|---|---|---|---|---|---|---|
| AT5G13490.2 | ADP/ATP carrier 2 (AAC2) | 26 | 1.93 | 0.237 | 0.985 | |
| AT5G13490.2 | ADP/ATP carrier 2 (AAC2) | 27 | 1.74 | 0.237 | 0.998 | |
| AT5G13490.2 | ADP/ATP carrier 2 (AAC2) | 28 | 1.61 | 0.237 | 1.000 | |
| AT5G13490.2 | ADP/ATP carrier 2 (AAC2) | 29 | 1.57 | 0.237 | 1.000 | |
| AT5G13490.2 | ADP/ATP carrier 2 (AAC2) | 30 | 1.61 | 0.237 | 1.000 | |
| AT5G13490.2 | ADP/ATP carrier 2 (AAC2) | 31 | 1.72 | 0.238 | 0.998 | |
| AT5G13490.2 | ADP/ATP carrier 2 (AAC2) | 32 | 1.81 | 0.238 | 0.995 | |
| AT5G13490.2 | ADP/ATP carrier 2 (AAC2) | 33 | 1.81 | 0.238 | 0.995 | |
| AT5G13490.2 | ADP/ATP carrier 2 (AAC2) | 34 | 1.80 | 0.238 | 0.996 | |
| AT5G13490.2 | ADP/ATP carrier 2 (AAC2) | 35 | 1.76 | 0.238 | 0.997 | |
| AT5G13490.2 | ADP/ATP carrier 2 (AAC2) | 36 | 1.65 | 0.238 | 0.999 | |
| AT5G13490.2 | ADP/ATP carrier 2 (AAC2) | 37 | 1.50 | 0.238 | 1.000 | |
| AT5G13490.2 | ADP/ATP carrier 2 (AAC2) | 38 | 1.36 | 0.238 | 1.000 | |
| AT5G13490.2 | ADP/ATP carrier 2 (AAC2) | 39 | 1.20 | 0.238 | 1.000 | |
| AT5G13490.2 | ADP/ATP carrier 2 (AAC2) | 40 | 0.99 | 0.239 | 1.000 | |
| AT5G13490.2 | ADP/ATP carrier 2 (AAC2) | 41 | 0.78 | 0.241 | 1.000 | |
| AT5G13490.2 | ADP/ATP carrier 2 (AAC2) | 42 | 0.62 | 0.243 | 1.000 | |
| AT5G13490.2 | ADP/ATP carrier 2 (AAC2) | 43 | 0.52 | 0.244 | 0.990 | |
| AT5G13490.2 | ADP/ATP carrier 2 (AAC2) | 44 | 0.45 | 0.244 | 0.935 | |
| AT5G13490.2 | ADP/ATP carrier 2 (AAC2) | 45 | 0.40 | 0.253 | 0.855 | |
| AT5G13490.2 | ADP/ATP carrier 2 (AAC2) | 46 | 0.34 | 0.307 | 0.899 | |
| AT5G13490.2 | ADP/ATP carrier 2 (AAC2) | 47 | 0.29 | 0.443 | 0.983 | |
| AT5G13490.2 | ADP/ATP carrier 2 (AAC2) | 48 | 0.24 | 0.635 | 0.999 | |
| AT5G13610.1 | Involved in ABI4-mediated mitochondrial retrograde signalling | 2 | 1.01 | 0.485 | 1.000 | |
| AT5G13610.1 | Involved in ABI4-mediated mitochondrial retrograde signalling | 3 | 0.79 | 0.481 | 1.000 | |
| AT5G13610.1 | Involved in ABI4-mediated mitochondrial retrograde signalling | 27 | 0.58 | 0.290 | 0.966 | |
| AT5G13610.1 | Involved in ABI4-mediated mitochondrial retrograde signalling | 28 | 0.88 | 0.219 | 1.000 | |
| AT5G13610.1 | Involved in ABI4-mediated mitochondrial retrograde signalling | 29 | 1.35 | 0.186 | 0.986 | |
| AT5G13610.1 | Involved in ABI4-mediated mitochondrial retrograde signalling | 30 | 1.97 | 0.172 | 0.473 | |
| AT5G13610.1 | Involved in ABI4-mediated mitochondrial retrograde signalling | 31 | 2.69 | 0.167 | 0.087 | . |
| AT5G13610.1 | Involved in ABI4-mediated mitochondrial retrograde signalling | 32 | 3.37 | 0.172 | 0.024 | * |
| AT5G13610.1 | Involved in ABI4-mediated mitochondrial retrograde signalling | 33 | 3.51 | 0.196 | 0.052 | . |
| AT5G13610.1 | Involved in ABI4-mediated mitochondrial retrograde signalling | 34 | 2.82 | 0.254 | 0.430 | |
| AT5G13610.1 | Involved in ABI4-mediated mitochondrial retrograde signalling | 35 | 1.92 | 0.347 | 0.969 | |
| AT5G13610.1 | Involved in ABI4-mediated mitochondrial retrograde signalling | 46 | 0.00 | 1.163 | 0.040 | * |
| AT5G13610.1 | Involved in ABI4-mediated mitochondrial retrograde signalling | 47 | 0.00 | 1.282 | 0.035 | * |

| | | | | | | |
|---|---|---|---|---|---|---|
| AT5G14040.1 | Mitochondrial phosphate transporter 3 (MPT3) | 3 | 0.88 | 0.110 | 1.000 | |
| AT5G14040.1 | Mitochondrial phosphate transporter 3 (MPT3) | 4 | 0.90 | 0.104 | 1.000 | |
| AT5G14040.1 | Mitochondrial phosphate transporter 3 (MPT3) | 5 | 0.95 | 0.104 | 1.000 | |
| AT5G14040.1 | Mitochondrial phosphate transporter 3 (MPT3) | 6 | 0.97 | 0.104 | 1.000 | |
| AT5G14040.1 | Mitochondrial phosphate transporter 3 (MPT3) | 7 | 0.95 | 0.104 | 1.000 | |
| AT5G14040.1 | Mitochondrial phosphate transporter 3 (MPT3) | 8 | 0.98 | 0.102 | 1.000 | |
| AT5G14040.1 | Mitochondrial phosphate transporter 3 (MPT3) | 9 | 1.04 | 0.101 | 1.000 | |
| AT5G14040.1 | Mitochondrial phosphate transporter 3 (MPT3) | 10 | 1.09 | 0.101 | 1.000 | |
| AT5G14040.1 | Mitochondrial phosphate transporter 3 (MPT3) | 11 | 1.16 | 0.101 | 1.000 | |
| AT5G14040.1 | Mitochondrial phosphate transporter 3 (MPT3) | 12 | 1.18 | 0.101 | 1.000 | |
| AT5G14040.1 | Mitochondrial phosphate transporter 3 (MPT3) | 13 | 1.14 | 0.100 | 1.000 | |
| AT5G14040.1 | Mitochondrial phosphate transporter 3 (MPT3) | 14 | 1.09 | 0.099 | 1.000 | |
| AT5G14040.1 | Mitochondrial phosphate transporter 3 (MPT3) | 15 | 1.02 | 0.098 | 1.000 | |
| AT5G14040.1 | Mitochondrial phosphate transporter 3 (MPT3) | 16 | 0.98 | 0.098 | 1.000 | |
| AT5G14040.1 | Mitochondrial phosphate transporter 3 (MPT3) | 17 | 1.11 | 0.097 | 1.000 | |
| AT5G14040.1 | Mitochondrial phosphate transporter 3 (MPT3) | 18 | 1.45 | 0.099 | 0.948 | |
| AT5G14040.1 | Mitochondrial phosphate transporter 3 (MPT3) | 19 | 1.97 | 0.102 | 0.128 | |
| AT5G14040.1 | Mitochondrial phosphate transporter 3 (MPT3) | 20 | 2.41 | 0.104 | 0.011 | * |
| AT5G14040.1 | Mitochondrial phosphate transporter 3 (MPT3) | 21 | 2.33 | 0.105 | 0.020 | * |
| AT5G14040.1 | Mitochondrial phosphate transporter 3 (MPT3) | 22 | 1.80 | 0.105 | 0.381 | |
| AT5G14040.1 | Mitochondrial phosphate transporter 3 (MPT3) | 23 | 1.33 | 0.104 | 0.999 | |
| AT5G14040.1 | Mitochondrial phosphate transporter 3 (MPT3) | 24 | 1.12 | 0.102 | 1.000 | |
| AT5G14040.1 | Mitochondrial phosphate transporter 3 (MPT3) | 25 | 1.18 | 0.101 | 1.000 | |
| AT5G14040.1 | Mitochondrial phosphate transporter 3 (MPT3) | 26 | 1.33 | 0.100 | 0.999 | |
| AT5G14040.1 | Mitochondrial phosphate transporter 3 (MPT3) | 27 | 1.27 | 0.100 | 1.000 | |
| AT5G14040.1 | Mitochondrial phosphate transporter 3 (MPT3) | 28 | 1.11 | 0.100 | 1.000 | |
| AT5G14040.1 | Mitochondrial phosphate transporter 3 (MPT3) | 29 | 1.03 | 0.100 | 1.000 | |
| AT5G14040.1 | Mitochondrial phosphate transporter 3 (MPT3) | 30 | 1.02 | 0.100 | 1.000 | |
| AT5G14040.1 | Mitochondrial phosphate transporter 3 (MPT3) | 31 | 1.15 | 0.100 | 1.000 | |
| AT5G14040.1 | Mitochondrial phosphate transporter 3 (MPT3) | 32 | 1.32 | 0.101 | 0.999 | |
| AT5G14040.1 | Mitochondrial phosphate transporter 3 (MPT3) | 33 | 1.34 | 0.101 | 0.998 | |

| | | | | | | |
|---|---|---|---|---|---|---|
| AT5G14040.1 | Mitochondrial phosphate transporter 3 (MPT3) | 34 | 1.30 | 0.101 | 1.000 | |
| AT5G14040.1 | Mitochondrial phosphate transporter 3 (MPT3) | 35 | 1.33 | 0.101 | 0.999 | |
| AT5G14040.1 | Mitochondrial phosphate transporter 3 (MPT3) | 36 | 1.42 | 0.100 | 0.974 | |
| AT5G14040.1 | Mitochondrial phosphate transporter 3 (MPT3) | 37 | 1.59 | 0.100 | 0.735 | |
| AT5G14040.1 | Mitochondrial phosphate transporter 3 (MPT3) | 38 | 1.90 | 0.101 | 0.178 | |
| AT5G14040.1 | Mitochondrial phosphate transporter 3 (MPT3) | 39 | 2.45 | 0.101 | 0.005 | ** |
| AT5G14040.1 | Mitochondrial phosphate transporter 3 (MPT3) | 40 | 3.10 | 0.101 | 0.000 | *** |
| AT5G14040.1 | Mitochondrial phosphate transporter 3 (MPT3) | 41 | 2.95 | 0.101 | 0.000 | *** |
| AT5G14040.1 | Mitochondrial phosphate transporter 3 (MPT3) | 42 | 2.07 | 0.103 | 0.075 | . |
| AT5G14040.1 | Mitochondrial phosphate transporter 3 (MPT3) | 43 | 1.32 | 0.104 | 0.999 | |
| AT5G14040.1 | Mitochondrial phosphate transporter 3 (MPT3) | 44 | 0.82 | 0.104 | 1.000 | |
| AT5G14040.1 | Mitochondrial phosphate transporter 3 (MPT3) | 45 | 0.51 | 0.103 | 0.151 | |
| AT5G14040.1 | Mitochondrial phosphate transporter 3 (MPT3) | 46 | 0.31 | 0.118 | 0.001 | *** |
| AT5G14040.1 | Mitochondrial phosphate transporter 3 (MPT3) | 47 | 0.18 | 0.215 | 0.019 | * |
| AT5G14040.1 | Mitochondrial phosphate transporter 3 (MPT3) | 48 | 0.10 | 0.367 | 0.202 | |
| AT5G14590.1 | Putative NADP-dependent isocitrate dehydrogenase | 2 | 1.04 | 0.419 | 1.000 | |
| AT5G14590.1 | Putative NADP-dependent isocitrate dehydrogenase | 3 | 1.08 | 0.407 | 1.000 | |
| AT5G14590.1 | Putative NADP-dependent isocitrate dehydrogenase | 32 | 0.68 | 0.315 | 1.000 | |
| AT5G14590.1 | Putative NADP-dependent isocitrate dehydrogenase | 33 | 0.62 | 0.255 | 0.997 | |
| AT5G14590.1 | Putative NADP-dependent isocitrate dehydrogenase | 34 | 0.61 | 0.230 | 0.991 | |
| AT5G14590.1 | Putative NADP-dependent isocitrate dehydrogenase | 35 | 0.75 | 0.226 | 1.000 | |
| AT5G14590.1 | Putative NADP-dependent isocitrate dehydrogenase | 36 | 1.37 | 0.229 | 1.000 | |
| AT5G14590.1 | Putative NADP-dependent isocitrate dehydrogenase | 37 | 2.99 | 0.232 | 0.390 | |
| AT5G14590.1 | Putative NADP-dependent isocitrate dehydrogenase | 38 | 5.38 | 0.228 | 0.024 | * |
| AT5G14590.1 | Putative NADP-dependent isocitrate dehydrogenase | 39 | 6.68 | 0.217 | 0.003 | ** |
| AT5G14590.1 | Putative NADP-dependent isocitrate dehydrogenase | 40 | 4.89 | 0.213 | 0.022 | * |
| AT5G14590.1 | Putative NADP-dependent isocitrate dehydrogenase | 41 | 2.63 | 0.213 | 0.443 | |
| AT5G14590.1 | Putative NADP-dependent isocitrate dehydrogenase | 42 | 1.57 | 0.216 | 0.992 | |
| AT5G14590.1 | Putative NADP-dependent isocitrate dehydrogenase | 43 | 1.24 | 0.216 | 1.000 | |
| AT5G14590.1 | Putative NADP-dependent isocitrate dehydrogenase | 44 | 1.10 | 0.216 | 1.000 | |
| AT5G14590.1 | Putative NADP-dependent isocitrate dehydrogenase | 45 | 0.94 | 0.215 | 1.000 | |

| | | | | | | |
|---|---|---|---|---|---|---|
| AT5G14590.1 | Putative NADP-dependent isocitrate dehydrogenase | 46 | 0.67 | 0.238 | 0.999 | |
| AT5G14590.1 | Putative NADP-dependent isocitrate dehydrogenase | 47 | 0.38 | 0.439 | 0.988 | |
| AT5G14780.1 | Formate dehydrogenase (FDH1) | 3 | 0.97 | 0.304 | 1.000 | |
| AT5G14780.1 | Formate dehydrogenase (FDH1) | 4 | 1.01 | 0.283 | 1.000 | |
| AT5G14780.1 | Formate dehydrogenase (FDH1) | 5 | 0.98 | 0.284 | 1.000 | |
| AT5G14780.1 | Formate dehydrogenase (FDH1) | 6 | 1.00 | 0.284 | 1.000 | |
| AT5G14780.1 | Formate dehydrogenase (FDH1) | 7 | 1.14 | 0.284 | 1.000 | |
| AT5G14780.1 | Formate dehydrogenase (FDH1) | 8 | 1.47 | 0.280 | 1.000 | |
| AT5G14780.1 | Formate dehydrogenase (FDH1) | 9 | 1.68 | 0.277 | 1.000 | |
| AT5G14780.1 | Formate dehydrogenase (FDH1) | 10 | 1.68 | 0.277 | 1.000 | |
| AT5G14780.1 | Formate dehydrogenase (FDH1) | 11 | 1.87 | 0.277 | 1.000 | |
| AT5G14780.1 | Formate dehydrogenase (FDH1) | 12 | 2.26 | 0.277 | 0.988 | |
| AT5G14780.1 | Formate dehydrogenase (FDH1) | 13 | 2.80 | 0.275 | 0.883 | |
| AT5G14780.1 | Formate dehydrogenase (FDH1) | 14 | 3.69 | 0.272 | 0.547 | |
| AT5G14780.1 | Formate dehydrogenase (FDH1) | 15 | 4.62 | 0.272 | 0.287 | |
| AT5G14780.1 | Formate dehydrogenase (FDH1) | 16 | 5.17 | 0.270 | 0.187 | |
| AT5G14780.1 | Formate dehydrogenase (FDH1) | 17 | 5.56 | 0.270 | 0.139 | |
| AT5G14780.1 | Formate dehydrogenase (FDH1) | 18 | 5.62 | 0.273 | 0.145 | |
| AT5G14780.1 | Formate dehydrogenase (FDH1) | 19 | 5.37 | 0.278 | 0.195 | |
| AT5G14780.1 | Formate dehydrogenase (FDH1) | 20 | 4.52 | 0.283 | 0.374 | |
| AT5G14780.1 | Formate dehydrogenase (FDH1) | 21 | 3.38 | 0.286 | 0.735 | |
| AT5G14780.1 | Formate dehydrogenase (FDH1) | 22 | 2.30 | 0.286 | 0.989 | |
| AT5G14780.1 | Formate dehydrogenase (FDH1) | 23 | 1.77 | 0.285 | 1.000 | |
| AT5G14780.1 | Formate dehydrogenase (FDH1) | 24 | 1.85 | 0.280 | 1.000 | |
| AT5G14780.1 | Formate dehydrogenase (FDH1) | 25 | 2.35 | 0.277 | 0.980 | |
| AT5G14780.1 | Formate dehydrogenase (FDH1) | 26 | 3.02 | 0.276 | 0.813 | |
| AT5G14780.1 | Formate dehydrogenase (FDH1) | 27 | 3.88 | 0.276 | 0.503 | |
| AT5G14780.1 | Formate dehydrogenase (FDH1) | 28 | 4.67 | 0.276 | 0.296 | |
| AT5G14780.1 | Formate dehydrogenase (FDH1) | 29 | 5.84 | 0.276 | 0.133 | |
| AT5G14780.1 | Formate dehydrogenase (FDH1) | 30 | 6.88 | 0.275 | 0.067 | . |
| AT5G14780.1 | Formate dehydrogenase (FDH1) | 31 | 6.88 | 0.276 | 0.068 | . |

| | | | | | |
|---|---|---|---|---|---|
| AT5G14780.1 | Formate dehydrogenase (FDH1) | 32 | 4.60 | 0.277 | 0.318 |
| AT5G14780.1 | Formate dehydrogenase (FDH1) | 33 | 1.89 | 0.277 | 1.000 |
| AT5G14780.1 | Formate dehydrogenase (FDH1) | 34 | 0.79 | 0.277 | 1.000 |
| AT5G14780.1 | Formate dehydrogenase (FDH1) | 35 | 0.58 | 0.277 | 1.000 |
| AT5G14780.1 | Formate dehydrogenase (FDH1) | 36 | 0.69 | 0.276 | 1.000 |
| AT5G14780.1 | Formate dehydrogenase (FDH1) | 37 | 1.03 | 0.276 | 1.000 |
| AT5G14780.1 | Formate dehydrogenase (FDH1) | 38 | 1.59 | 0.277 | 1.000 |
| AT5G14780.1 | Formate dehydrogenase (FDH1) | 39 | 2.02 | 0.277 | 0.998 |
| AT5G14780.1 | Formate dehydrogenase (FDH1) | 40 | 1.78 | 0.277 | 1.000 |
| AT5G14780.1 | Formate dehydrogenase (FDH1) | 41 | 1.20 | 0.278 | 1.000 |
| AT5G14780.1 | Formate dehydrogenase (FDH1) | 42 | 0.84 | 0.281 | 1.000 |
| AT5G14780.1 | Formate dehydrogenase (FDH1) | 43 | 0.74 | 0.283 | 1.000 |
| AT5G14780.1 | Formate dehydrogenase (FDH1) | 44 | 0.74 | 0.283 | 1.000 |
| AT5G14780.1 | Formate dehydrogenase (FDH1) | 45 | 0.72 | 0.283 | 1.000 |
| AT5G14780.1 | Formate dehydrogenase (FDH1) | 46 | 0.59 | 0.321 | 1.000 |
| AT5G14780.1 | Formate dehydrogenase (FDH1) | 47 | 0.40 | 0.519 | 1.000 |
| AT5G14780.1 | Formate dehydrogenase (FDH1) | 48 | 0.27 | 0.836 | 1.000 |
| AT5G15090.2 | Voltage-dependent anion channel 3 (VDAC3) | 3 | 0.89 | 0.134 | 1.000 |
| AT5G15090.2 | Voltage-dependent anion channel 3 (VDAC3) | 4 | 0.94 | 0.123 | 1.000 |
| AT5G15090.2 | Voltage-dependent anion channel 3 (VDAC3) | 5 | 1.14 | 0.123 | 1.000 |
| AT5G15090.2 | Voltage-dependent anion channel 3 (VDAC3) | 6 | 1.34 | 0.123 | 0.999 |
| AT5G15090.2 | Voltage-dependent anion channel 3 (VDAC3) | 7 | 1.42 | 0.123 | 0.991 |
| AT5G15090.2 | Voltage-dependent anion channel 3 (VDAC3) | 8 | 1.40 | 0.121 | 0.993 |
| AT5G15090.2 | Voltage-dependent anion channel 3 (VDAC3) | 9 | 1.34 | 0.120 | 0.999 |
| AT5G15090.2 | Voltage-dependent anion channel 3 (VDAC3) | 10 | 1.28 | 0.120 | 1.000 |
| AT5G15090.2 | Voltage-dependent anion channel 3 (VDAC3) | 11 | 1.25 | 0.120 | 1.000 |
| AT5G15090.2 | Voltage-dependent anion channel 3 (VDAC3) | 12 | 1.27 | 0.120 | 1.000 |
| AT5G15090.2 | Voltage-dependent anion channel 3 (VDAC3) | 13 | 1.28 | 0.119 | 1.000 |
| AT5G15090.2 | Voltage-dependent anion channel 3 (VDAC3) | 14 | 1.25 | 0.118 | 1.000 |
| AT5G15090.2 | Voltage-dependent anion channel 3 (VDAC3) | 15 | 1.13 | 0.118 | 1.000 |
| AT5G15090.2 | Voltage-dependent anion channel 3 (VDAC3) | 16 | 0.94 | 0.117 | 1.000 |

| | | | | | | |
|---|---|---|---|---|---|---|
| AT5G15090.2 | Voltage-dependent anion channel 3 (VDAC3) | 17 | 0.85 | 0.117 | 1.000 | |
| AT5G15090.2 | Voltage-dependent anion channel 3 (VDAC3) | 18 | 0.88 | 0.119 | 1.000 | |
| AT5G15090.2 | Voltage-dependent anion channel 3 (VDAC3) | 19 | 0.95 | 0.121 | 1.000 | |
| AT5G15090.2 | Voltage-dependent anion channel 3 (VDAC3) | 20 | 1.02 | 0.123 | 1.000 | |
| AT5G15090.2 | Voltage-dependent anion channel 3 (VDAC3) | 21 | 1.06 | 0.124 | 1.000 | |
| AT5G15090.2 | Voltage-dependent anion channel 3 (VDAC3) | 22 | 1.02 | 0.124 | 1.000 | |
| AT5G15090.2 | Voltage-dependent anion channel 3 (VDAC3) | 23 | 0.93 | 0.123 | 1.000 | |
| AT5G15090.2 | Voltage-dependent anion channel 3 (VDAC3) | 24 | 0.86 | 0.121 | 1.000 | |
| AT5G15090.2 | Voltage-dependent anion channel 3 (VDAC3) | 25 | 0.84 | 0.120 | 1.000 | |
| AT5G15090.2 | Voltage-dependent anion channel 3 (VDAC3) | 26 | 0.86 | 0.120 | 1.000 | |
| AT5G15090.2 | Voltage-dependent anion channel 3 (VDAC3) | 27 | 0.87 | 0.120 | 1.000 | |
| AT5G15090.2 | Voltage-dependent anion channel 3 (VDAC3) | 28 | 0.90 | 0.119 | 1.000 | |
| AT5G15090.2 | Voltage-dependent anion channel 3 (VDAC3) | 29 | 0.96 | 0.119 | 1.000 | |
| AT5G15090.2 | Voltage-dependent anion channel 3 (VDAC3) | 30 | 1.07 | 0.119 | 1.000 | |
| AT5G15090.2 | Voltage-dependent anion channel 3 (VDAC3) | 31 | 1.11 | 0.120 | 1.000 | |
| AT5G15090.2 | Voltage-dependent anion channel 3 (VDAC3) | 32 | 1.07 | 0.120 | 1.000 | |
| AT5G15090.2 | Voltage-dependent anion channel 3 (VDAC3) | 33 | 1.02 | 0.120 | 1.000 | |
| AT5G15090.2 | Voltage-dependent anion channel 3 (VDAC3) | 34 | 1.02 | 0.120 | 1.000 | |
| AT5G15090.2 | Voltage-dependent anion channel 3 (VDAC3) | 35 | 1.06 | 0.120 | 1.000 | |
| AT5G15090.2 | Voltage-dependent anion channel 3 (VDAC3) | 36 | 1.08 | 0.120 | 1.000 | |
| AT5G15090.2 | Voltage-dependent anion channel 3 (VDAC3) | 37 | 1.20 | 0.120 | 1.000 | |
| AT5G15090.2 | Voltage-dependent anion channel 3 (VDAC3) | 38 | 1.49 | 0.120 | 0.955 | |
| AT5G15090.2 | Voltage-dependent anion channel 3 (VDAC3) | 39 | 1.98 | 0.120 | 0.275 | |
| AT5G15090.2 | Voltage-dependent anion channel 3 (VDAC3) | 40 | 2.50 | 0.120 | 0.030 | * |
| AT5G15090.2 | Voltage-dependent anion channel 3 (VDAC3) | 41 | 2.41 | 0.121 | 0.045 | * |
| AT5G15090.2 | Voltage-dependent anion channel 3 (VDAC3) | 42 | 1.64 | 0.122 | 0.794 | |
| AT5G15090.2 | Voltage-dependent anion channel 3 (VDAC3) | 43 | 0.93 | 0.123 | 1.000 | |
| AT5G15090.2 | Voltage-dependent anion channel 3 (VDAC3) | 44 | 0.54 | 0.123 | 0.470 | |
| AT5G15090.2 | Voltage-dependent anion channel 3 (VDAC3) | 45 | 0.38 | 0.123 | 0.021 | * |
| AT5G15090.2 | Voltage-dependent anion channel 3 (VDAC3) | 46 | 0.31 | 0.142 | 0.012 | * |
| AT5G15090.2 | Voltage-dependent anion channel 3 (VDAC3) | 47 | 0.26 | 0.232 | 0.240 | |

| | | | | | |
|---|---|---|---|---|---|
| AT5G15090.2 | Voltage-dependent anion channel 3 (VDAC3) | 48 | 0.22 | 0.370 | 0.773 |
| AT5G15320.2 | ATP synthase subunit (E chain) | 2 | 1.18 | 0.576 | 1.000 |
| AT5G15320.2 | ATP synthase subunit (E chain) | 3 | 0.94 | 0.480 | 1.000 |
| AT5G15320.2 | ATP synthase subunit (E chain) | 8 | 0.59 | 0.392 | 1.000 |
| AT5G15320.2 | ATP synthase subunit (E chain) | 9 | 0.69 | 0.313 | 1.000 |
| AT5G15320.2 | ATP synthase subunit (E chain) | 10 | 0.88 | 0.281 | 1.000 |
| AT5G15320.2 | ATP synthase subunit (E chain) | 11 | 1.30 | 0.274 | 1.000 |
| AT5G15320.2 | ATP synthase subunit (E chain) | 12 | 2.23 | 0.278 | 0.996 |
| AT5G15320.2 | ATP synthase subunit (E chain) | 13 | 3.75 | 0.292 | 0.726 |
| AT5G15320.2 | ATP synthase subunit (E chain) | 14 | 4.98 | 0.295 | 0.400 |
| AT5G15320.2 | ATP synthase subunit (E chain) | 15 | 5.57 | 0.326 | 0.458 |
| AT5G15320.2 | ATP synthase subunit (E chain) | 16 | 5.09 | 0.393 | 0.847 |
| AT5G15320.2 | ATP synthase subunit (E chain) | 22 | 0.13 | 0.420 | 0.626 |
| AT5G15320.2 | ATP synthase subunit (E chain) | 23 | 0.21 | 0.318 | 0.570 |
| AT5G15320.2 | ATP synthase subunit (E chain) | 24 | 0.62 | 0.271 | 1.000 |
| AT5G15320.2 | ATP synthase subunit (E chain) | 25 | 1.71 | 0.251 | 1.000 |
| AT5G15320.2 | ATP synthase subunit (E chain) | 26 | 1.40 | 0.249 | 1.000 |
| AT5G15320.2 | ATP synthase subunit (E chain) | 27 | 0.66 | 0.250 | 1.000 |
| AT5G15320.2 | ATP synthase subunit (E chain) | 28 | 0.60 | 0.249 | 1.000 |
| AT5G15320.2 | ATP synthase subunit (E chain) | 29 | 0.76 | 0.249 | 1.000 |
| AT5G15320.2 | ATP synthase subunit (E chain) | 30 | 0.80 | 0.251 | 1.000 |
| AT5G15320.2 | ATP synthase subunit (E chain) | 31 | 0.84 | 0.251 | 1.000 |
| AT5G15320.2 | ATP synthase subunit (E chain) | 32 | 0.88 | 0.251 | 1.000 |
| AT5G15320.2 | ATP synthase subunit (E chain) | 33 | 1.06 | 0.252 | 1.000 |
| AT5G15320.2 | ATP synthase subunit (E chain) | 34 | 1.19 | 0.257 | 1.000 |
| AT5G15320.2 | ATP synthase subunit (E chain) | 35 | 1.20 | 0.273 | 1.000 |
| AT5G15320.2 | ATP synthase subunit (E chain) | 36 | 1.37 | 0.314 | 1.000 |
| AT5G15320.2 | ATP synthase subunit (E chain) | 37 | 1.87 | 0.357 | 1.000 |
| AT5G15320.2 | ATP synthase subunit (E chain) | 39 | 4.18 | 0.365 | 0.900 |
| AT5G15320.2 | ATP synthase subunit (E chain) | 40 | 5.35 | 0.296 | 0.330 |
| AT5G15320.2 | ATP synthase subunit (E chain) | 41 | 4.73 | 0.263 | 0.261 |
| AT5G15320.2 | ATP synthase subunit (E chain) | 42 | 3.19 | 0.257 | 0.735 |
| AT5G15320.2 | ATP synthase subunit (E chain) | 43 | 1.96 | 0.258 | 0.999 |
| AT5G15320.2 | ATP synthase subunit (E chain) | 44 | 1.05 | 0.257 | 1.000 |
| AT5G15320.2 | ATP synthase subunit (E chain) | 45 | 0.48 | 0.257 | 0.997 |
| AT5G15320.2 | ATP synthase subunit (E chain) | 46 | 0.19 | 0.295 | 0.313 |
| AT5G15320.2 | ATP synthase subunit (E chain) | 47 | 0.05 | 0.548 | 0.447 |
| AT5G15450.1 | Caseinolytic protease B3 (CLPB3) | 2 | 0.77 | 0.374 | 1.000 |
| AT5G15450.1 | Caseinolytic protease B3 (CLPB3) | 3 | 0.74 | 0.287 | 1.000 |
| AT5G15450.1 | Caseinolytic protease B3 (CLPB3) | 4 | 0.71 | 0.256 | 0.999 |
| AT5G15450.1 | Caseinolytic protease B3 (CLPB3) | 6 | 0.66 | 0.293 | 0.999 |
| AT5G15450.1 | Caseinolytic protease B3 (CLPB3) | 7 | 0.63 | 0.328 | 0.999 |
| AT5G15450.1 | Caseinolytic protease B3 (CLPB3) | 8 | 0.61 | 0.378 | 0.999 |
| AT5G15450.1 | Caseinolytic protease B3 (CLPB3) | 30 | 1.33 | 0.250 | 1.000 |
| AT5G15450.1 | Caseinolytic protease B3 (CLPB3) | 31 | 1.19 | 0.204 | 1.000 |
| AT5G15450.1 | Caseinolytic protease B3 (CLPB3) | 32 | 1.01 | 0.177 | 1.000 |
| AT5G15450.1 | Caseinolytic protease B3 (CLPB3) | 33 | 0.82 | 0.163 | 1.000 |
| AT5G15450.1 | Caseinolytic protease B3 (CLPB3) | 34 | 0.65 | 0.158 | 0.898 |
| AT5G15450.1 | Caseinolytic protease B3 (CLPB3) | 35 | 0.52 | 0.156 | 0.512 |

| | | | | | |
|---|---|---|---|---|---|
| AT5G15450.1 | Caseinolytic protease B3 (CLPB3) | 36 | 0.48 | 0.157 | 0.351 |
| AT5G15450.1 | Caseinolytic protease B3 (CLPB3) | 37 | 0.52 | 0.159 | 0.523 |
| AT5G15450.1 | Caseinolytic protease B3 (CLPB3) | 38 | 0.64 | 0.165 | 0.911 |
| AT5G15450.1 | Caseinolytic protease B3 (CLPB3) | 39 | 0.85 | 0.181 | 1.000 |
| AT5G15450.1 | Caseinolytic protease B3 (CLPB3) | 40 | 1.17 | 0.220 | 1.000 |
| AT5G15450.1 | Caseinolytic protease B3 (CLPB3) | 41 | 1.61 | 0.294 | 0.997 |
| AT5G15450.1 | Caseinolytic protease B3 (CLPB3) | 46 | 5.11 | 1.096 | 0.998 |
| AT5G15450.1 | Caseinolytic protease B3 (CLPB3) | 47 | 6.16 | 1.321 | 0.999 |
| AT5G15640.1 | Mitochondrial substrate carrier family protein | 2 | 1.06 | 0.361 | 1.000 |
| AT5G15640.1 | Mitochondrial substrate carrier family protein | 3 | 1.27 | 0.223 | 1.000 |
| AT5G15640.1 | Mitochondrial substrate carrier family protein | 4 | 1.48 | 0.212 | 1.000 |
| AT5G15640.1 | Mitochondrial substrate carrier family protein | 5 | 1.64 | 0.212 | 1.000 |
| AT5G15640.1 | Mitochondrial substrate carrier family protein | 6 | 1.52 | 0.212 | 1.000 |
| AT5G15640.1 | Mitochondrial substrate carrier family protein | 7 | 1.20 | 0.213 | 1.000 |
| AT5G15640.1 | Mitochondrial substrate carrier family protein | 8 | 1.05 | 0.209 | 1.000 |
| AT5G15640.1 | Mitochondrial substrate carrier family protein | 9 | 1.06 | 0.206 | 1.000 |
| AT5G15640.1 | Mitochondrial substrate carrier family protein | 10 | 1.06 | 0.206 | 1.000 |
| AT5G15640.1 | Mitochondrial substrate carrier family protein | 11 | 1.11 | 0.206 | 1.000 |
| AT5G15640.1 | Mitochondrial substrate carrier family protein | 12 | 1.20 | 0.206 | 1.000 |
| AT5G15640.1 | Mitochondrial substrate carrier family protein | 13 | 1.27 | 0.204 | 1.000 |
| AT5G15640.1 | Mitochondrial substrate carrier family protein | 14 | 1.25 | 0.201 | 1.000 |
| AT5G15640.1 | Mitochondrial substrate carrier family protein | 15 | 1.16 | 0.202 | 1.000 |
| AT5G15640.1 | Mitochondrial substrate carrier family protein | 16 | 1.18 | 0.200 | 1.000 |
| AT5G15640.1 | Mitochondrial substrate carrier family protein | 17 | 1.52 | 0.200 | 1.000 |
| AT5G15640.1 | Mitochondrial substrate carrier family protein | 18 | 2.36 | 0.203 | 0.832 |
| AT5G15640.1 | Mitochondrial substrate carrier family protein | 19 | 3.23 | 0.211 | 0.377 |
| AT5G15640.1 | Mitochondrial substrate carrier family protein | 20 | 2.96 | 0.220 | 0.594 |
| AT5G15640.1 | Mitochondrial substrate carrier family protein | 21 | 1.49 | 0.229 | 1.000 |
| AT5G15640.1 | Mitochondrial substrate carrier family protein | 22 | 0.54 | 0.260 | 1.000 |
| AT5G15640.1 | Mitochondrial substrate carrier family protein | 23 | 0.29 | 0.335 | 0.939 |
| AT5G15640.1 | Mitochondrial substrate carrier family protein | 28 | 1.51 | 0.303 | 1.000 |

| | | | | | | |
|---|---|---|---|---|---|---|
| AT5G15640.1 | Mitochondrial substrate carrier family protein | 29 | 2.13 | 0.237 | 0.989 | |
| AT5G15640.1 | Mitochondrial substrate carrier family protein | 30 | 2.35 | 0.218 | 0.909 | |
| AT5G15640.1 | Mitochondrial substrate carrier family protein | 31 | 2.22 | 0.209 | 0.930 | |
| AT5G15640.1 | Mitochondrial substrate carrier family protein | 32 | 2.01 | 0.206 | 0.979 | |
| AT5G15640.1 | Mitochondrial substrate carrier family protein | 33 | 1.97 | 0.206 | 0.984 | |
| AT5G15640.1 | Mitochondrial substrate carrier family protein | 34 | 3.10 | 0.206 | 0.395 | |
| AT5G15640.1 | Mitochondrial substrate carrier family protein | 35 | 4.56 | 0.206 | 0.048 | * |
| AT5G15640.1 | Mitochondrial substrate carrier family protein | 36 | 3.22 | 0.205 | 0.327 | |
| AT5G15640.1 | Mitochondrial substrate carrier family protein | 37 | 2.20 | 0.205 | 0.921 | |
| AT5G15640.1 | Mitochondrial substrate carrier family protein | 38 | 3.23 | 0.206 | 0.328 | |
| AT5G15640.1 | Mitochondrial substrate carrier family protein | 39 | 5.45 | 0.206 | 0.014 | * |
| AT5G15640.1 | Mitochondrial substrate carrier family protein | 40 | 4.61 | 0.207 | 0.048 | * |
| AT5G15640.1 | Mitochondrial substrate carrier family protein | 41 | 2.18 | 0.207 | 0.936 | |
| AT5G15640.1 | Mitochondrial substrate carrier family protein | 42 | 0.85 | 0.210 | 1.000 | |
| AT5G15640.1 | Mitochondrial substrate carrier family protein | 43 | 0.37 | 0.212 | 0.658 | |
| AT5G15640.1 | Mitochondrial substrate carrier family protein | 44 | 0.18 | 0.212 | 0.014 | * |
| AT5G15640.1 | Mitochondrial substrate carrier family protein | 45 | 0.10 | 0.211 | 0.000 | *** |
| AT5G15640.1 | Mitochondrial substrate carrier family protein | 46 | 0.08 | 0.237 | 0.000 | *** |
| AT5G15640.1 | Mitochondrial substrate carrier family protein | 47 | 0.08 | 0.434 | 0.314 | |
| AT5G15910.1 | NAD(P)-binding Rossmann-fold superfamily protein | 2 | 1.15 | 0.379 | 1.000 | |
| AT5G15910.1 | NAD(P)-binding Rossmann-fold superfamily protein | 3 | 1.16 | 0.310 | 1.000 | |
| AT5G15910.1 | NAD(P)-binding Rossmann-fold superfamily protein | 6 | 1.13 | 0.246 | 1.000 | |
| AT5G15910.1 | NAD(P)-binding Rossmann-fold superfamily protein | 7 | 1.11 | 0.227 | 1.000 | |
| AT5G15910.1 | NAD(P)-binding Rossmann-fold superfamily protein | 8 | 1.08 | 0.205 | 1.000 | |
| AT5G15910.1 | NAD(P)-binding Rossmann-fold superfamily protein | 9 | 1.08 | 0.191 | 1.000 | |
| AT5G15910.1 | NAD(P)-binding Rossmann-fold superfamily protein | 10 | 1.13 | 0.183 | 1.000 | |
| AT5G15910.1 | NAD(P)-binding Rossmann-fold superfamily protein | 11 | 1.25 | 0.180 | 1.000 | |
| AT5G15910.1 | NAD(P)-binding Rossmann-fold superfamily protein | 12 | 1.45 | 0.185 | 0.994 | |
| AT5G15910.1 | NAD(P)-binding Rossmann-fold superfamily protein | 13 | 1.67 | 0.196 | 0.957 | |
| AT5G15910.1 | NAD(P)-binding Rossmann-fold superfamily protein | 14 | 1.81 | 0.210 | 0.928 | |
| AT5G15910.1 | NAD(P)-binding Rossmann-fold superfamily protein | 15 | 1.84 | 0.220 | 0.937 | |

| | | | | | | |
|---|---|---|---|---|---|---|
| AT5G15910.1 | NAD(P)-binding Rossmann-fold superfamily protein | 16 | 1.82 | 0.246 | 0.974 | |
| AT5G15910.1 | NAD(P)-binding Rossmann-fold superfamily protein | 34 | 2.19 | 0.320 | 0.972 | |
| AT5G15910.1 | NAD(P)-binding Rossmann-fold superfamily protein | 35 | 2.07 | 0.289 | 0.967 | |
| AT5G15910.1 | NAD(P)-binding Rossmann-fold superfamily protein | 36 | 2.17 | 0.280 | 0.938 | |
| AT5G15910.1 | NAD(P)-binding Rossmann-fold superfamily protein | 40 | 6.81 | 0.234 | 0.008 | ** |
| AT5G15910.1 | NAD(P)-binding Rossmann-fold superfamily protein | 41 | 9.35 | 0.201 | 0.000 | *** |
| AT5G15910.1 | NAD(P)-binding Rossmann-fold superfamily protein | 42 | 11.42 | 0.176 | 0.000 | *** |
| AT5G15910.1 | NAD(P)-binding Rossmann-fold superfamily protein | 43 | 11.35 | 0.165 | 0.000 | *** |
| AT5G15910.1 | NAD(P)-binding Rossmann-fold superfamily protein | 44 | 8.45 | 0.163 | 0.000 | *** |
| AT5G15910.1 | NAD(P)-binding Rossmann-fold superfamily protein | 45 | 4.54 | 0.172 | 0.003 | ** |
| AT5G15910.1 | NAD(P)-binding Rossmann-fold superfamily protein | 46 | 1.84 | 0.215 | 0.926 | |
| AT5G15910.1 | NAD(P)-binding Rossmann-fold superfamily protein | 47 | 0.69 | 0.320 | 1.000 | |
| AT5G15980.1 | Pentatricopeptide repeat (PPR) superfamily protein | 2 | 1.15 | 0.415 | 1.000 | |
| AT5G15980.1 | Pentatricopeptide repeat (PPR) superfamily protein | 3 | 1.22 | 0.400 | 1.000 | |
| AT5G15980.1 | Pentatricopeptide repeat (PPR) superfamily protein | 30 | 0.94 | 0.303 | 1.000 | |
| AT5G15980.1 | Pentatricopeptide repeat (PPR) superfamily protein | 31 | 0.80 | 0.237 | 1.000 | |
| AT5G15980.1 | Pentatricopeptide repeat (PPR) superfamily protein | 32 | 0.62 | 0.215 | 0.989 | |
| AT5G15980.1 | Pentatricopeptide repeat (PPR) superfamily protein | 33 | 0.51 | 0.206 | 0.844 | |
| AT5G15980.1 | Pentatricopeptide repeat (PPR) superfamily protein | 34 | 0.48 | 0.203 | 0.749 | |
| AT5G15980.1 | Pentatricopeptide repeat (PPR) superfamily protein | 35 | 0.55 | 0.202 | 0.918 | |
| AT5G15980.1 | Pentatricopeptide repeat (PPR) superfamily protein | 36 | 0.83 | 0.202 | 1.000 | |
| AT5G15980.1 | Pentatricopeptide repeat (PPR) superfamily protein | 37 | 1.25 | 0.201 | 1.000 | |
| AT5G15980.1 | Pentatricopeptide repeat (PPR) superfamily protein | 38 | 1.41 | 0.201 | 0.999 | |
| AT5G15980.1 | Pentatricopeptide repeat (PPR) superfamily protein | 39 | 1.33 | 0.201 | 1.000 | |
| AT5G15980.1 | Pentatricopeptide repeat (PPR) superfamily protein | 40 | 1.31 | 0.200 | 1.000 | |
| AT5G15980.1 | Pentatricopeptide repeat (PPR) superfamily protein | 41 | 1.25 | 0.201 | 1.000 | |
| AT5G15980.1 | Pentatricopeptide repeat (PPR) superfamily protein | 42 | 1.10 | 0.204 | 1.000 | |
| AT5G15980.1 | Pentatricopeptide repeat (PPR) superfamily protein | 43 | 0.88 | 0.205 | 1.000 | |
| AT5G15980.1 | Pentatricopeptide repeat (PPR) superfamily protein | 44 | 0.60 | 0.205 | 0.971 | |
| AT5G15980.1 | Pentatricopeptide repeat (PPR) superfamily protein | 45 | 0.37 | 0.205 | 0.342 | |
| AT5G15980.1 | Pentatricopeptide repeat (PPR) superfamily protein | 46 | 0.24 | 0.233 | 0.103 | |

| | | | | | | |
|---|---|---|---|---|---|---|
| AT5G15980.1 | Pentatricopeptide repeat (PPR) superfamily protein | 47 | 0.16 | 0.400 | 0.455 | |
| AT5G16060.1 | Cytochrome c:O2 oxidoreductase assembly protein (CMC1-like) | 2 | 0.92 | 0.565 | 1.000 | |
| AT5G16060.1 | Cytochrome c:O2 oxidoreductase assembly protein (CMC1-like) | 3 | 0.79 | 0.662 | 1.000 | |
| AT5G16060.1 | Cytochrome c:O2 oxidoreductase assembly protein (CMC1-like) | 32 | 5.64 | 0.440 | 0.573 | |
| AT5G16060.1 | Cytochrome c:O2 oxidoreductase assembly protein (CMC1-like) | 33 | 7.26 | 0.312 | 0.076 | . |
| AT5G16060.1 | Cytochrome c:O2 oxidoreductase assembly protein (CMC1-like) | 34 | 8.23 | 0.248 | 0.005 | ** |
| AT5G16060.1 | Cytochrome c:O2 oxidoreductase assembly protein (CMC1-like) | 35 | 7.13 | 0.226 | 0.004 | ** |
| AT5G16060.1 | Cytochrome c:O2 oxidoreductase assembly protein (CMC1-like) | 36 | 5.03 | 0.233 | 0.039 | * |
| AT5G16060.1 | Cytochrome c:O2 oxidoreductase assembly protein (CMC1-like) | 37 | 3.12 | 0.269 | 0.482 | |
| AT5G16060.1 | Cytochrome c:O2 oxidoreductase assembly protein (CMC1-like) | 38 | 1.88 | 0.323 | 0.986 | |
| AT5G16060.1 | Cytochrome c:O2 oxidoreductase assembly protein (CMC1-like) | 41 | 0.73 | 0.375 | 1.000 | |
| AT5G16060.1 | Cytochrome c:O2 oxidoreductase assembly protein (CMC1-like) | 42 | 0.79 | 0.360 | 1.000 | |
| AT5G16060.1 | Cytochrome c:O2 oxidoreductase assembly protein (CMC1-like) | 43 | 0.99 | 0.346 | 1.000 | |
| AT5G16060.1 | Cytochrome c:O2 oxidoreductase assembly protein (CMC1-like) | 44 | 1.28 | 0.329 | 1.000 | |
| AT5G16060.1 | Cytochrome c:O2 oxidoreductase assembly protein (CMC1-like) | 45 | 1.63 | 0.341 | 0.999 | |
| AT5G16060.1 | Cytochrome c:O2 oxidoreductase assembly protein (CMC1-like) | 46 | 1.99 | 0.433 | 0.997 | |
| AT5G16060.1 | Cytochrome c:O2 oxidoreductase assembly protein (CMC1-like) | 47 | 2.38 | 0.632 | 0.999 | |
| AT5G16715.1 | Putative valyl-tRNA synthetase (ValRS) / EMB2247 | 2 | 0.99 | 0.442 | 1.000 | |
| AT5G16715.1 | Putative valyl-tRNA synthetase (ValRS) / EMB2247 | 3 | 1.07 | 0.429 | 1.000 | |
| AT5G16715.1 | Putative valyl-tRNA synthetase (ValRS) / EMB2247 | 33 | 1.55 | 0.262 | 0.987 | |
| AT5G16715.1 | Putative valyl-tRNA synthetase (ValRS) / EMB2247 | 34 | 1.46 | 0.199 | 0.973 | |
| AT5G16715.1 | Putative valyl-tRNA synthetase (ValRS) / EMB2247 | 35 | 1.46 | 0.168 | 0.932 | |
| AT5G16715.1 | Putative valyl-tRNA synthetase (ValRS) / EMB2247 | 36 | 1.55 | 0.154 | 0.812 | |

| | | | | | | |
|---|---|---|---|---|---|---|
| AT5G16715.1 | Putative valyl-tRNA synthetase (ValRS) / EMB2247 | 37 | 1.68 | 0.148 | 0.624 | |
| AT5G16715.1 | Putative valyl-tRNA synthetase (ValRS) / EMB2247 | 38 | 1.82 | 0.145 | 0.430 | |
| AT5G16715.1 | Putative valyl-tRNA synthetase (ValRS) / EMB2247 | 39 | 1.97 | 0.146 | 0.294 | |
| AT5G16715.1 | Putative valyl-tRNA synthetase (ValRS) / EMB2247 | 40 | 1.95 | 0.155 | 0.386 | |
| AT5G16715.1 | Putative valyl-tRNA synthetase (ValRS) / EMB2247 | 41 | 1.56 | 0.180 | 0.898 | |
| AT5G16715.1 | Putative valyl-tRNA synthetase (ValRS) / EMB2247 | 42 | 1.00 | 0.223 | 1.000 | |
| AT5G16715.1 | Putative valyl-tRNA synthetase (ValRS) / EMB2247 | 43 | 0.55 | 0.269 | 0.934 | |
| AT5G16715.1 | Putative valyl-tRNA synthetase (ValRS) / EMB2247 | 46 | 0.05 | 0.399 | 0.012 | * |
| AT5G16715.1 | Putative valyl-tRNA synthetase (ValRS) / EMB2247 | 47 | 0.02 | 0.479 | 0.005 | ** |
| AT5G16930.1 | AAA-type ATPase family protein | 2 | 1.25 | 0.561 | 1.000 | |
| AT5G16930.1 | AAA-type ATPase family protein | 3 | 1.19 | 0.480 | 1.000 | |
| AT5G16930.1 | AAA-type ATPase family protein | 10 | 1.00 | 0.340 | 1.000 | |
| AT5G16930.1 | AAA-type ATPase family protein | 11 | 1.00 | 0.290 | 1.000 | |
| AT5G16930.1 | AAA-type ATPase family protein | 12 | 0.96 | 0.260 | 1.000 | |
| AT5G16930.1 | AAA-type ATPase family protein | 13 | 0.90 | 0.243 | 1.000 | |
| AT5G16930.1 | AAA-type ATPase family protein | 14 | 0.86 | 0.230 | 1.000 | |
| AT5G16930.1 | AAA-type ATPase family protein | 15 | 0.82 | 0.226 | 1.000 | |
| AT5G16930.1 | AAA-type ATPase family protein | 16 | 0.85 | 0.225 | 1.000 | |
| AT5G16930.1 | AAA-type ATPase family protein | 17 | 1.04 | 0.225 | 1.000 | |
| AT5G16930.1 | AAA-type ATPase family protein | 18 | 1.36 | 0.227 | 1.000 | |
| AT5G16930.1 | AAA-type ATPase family protein | 19 | 1.65 | 0.232 | 1.000 | |
| AT5G16930.1 | AAA-type ATPase family protein | 20 | 1.72 | 0.238 | 1.000 | |
| AT5G16930.1 | AAA-type ATPase family protein | 21 | 1.65 | 0.240 | 1.000 | |
| AT5G16930.1 | AAA-type ATPase family protein | 22 | 1.53 | 0.240 | 1.000 | |
| AT5G16930.1 | AAA-type ATPase family protein | 23 | 1.35 | 0.239 | 1.000 | |
| AT5G16930.1 | AAA-type ATPase family protein | 24 | 1.14 | 0.234 | 1.000 | |
| AT5G16930.1 | AAA-type ATPase family protein | 25 | 0.90 | 0.230 | 1.000 | |
| AT5G16930.1 | AAA-type ATPase family protein | 26 | 0.84 | 0.229 | 1.000 | |
| AT5G16930.1 | AAA-type ATPase family protein | 27 | 0.89 | 0.229 | 1.000 | |
| AT5G16930.1 | AAA-type ATPase family protein | 28 | 0.99 | 0.230 | 1.000 | |
| AT5G16930.1 | AAA-type ATPase family protein | 29 | 1.18 | 0.231 | 1.000 | |
| AT5G16930.1 | AAA-type ATPase family protein | 30 | 1.36 | 0.231 | 1.000 | |
| AT5G16930.1 | AAA-type ATPase family protein | 31 | 1.40 | 0.231 | 1.000 | |
| AT5G16930.1 | AAA-type ATPase family protein | 32 | 1.10 | 0.232 | 1.000 | |
| AT5G16930.1 | AAA-type ATPase family protein | 33 | 0.60 | 0.232 | 1.000 | |
| AT5G16930.1 | AAA-type ATPase family protein | 34 | 0.29 | 0.232 | 0.380 | |
| AT5G16930.1 | AAA-type ATPase family protein | 35 | 0.19 | 0.232 | 0.052 | . |
| AT5G16930.1 | AAA-type ATPase family protein | 36 | 0.22 | 0.231 | 0.117 | |
| AT5G16930.1 | AAA-type ATPase family protein | 37 | 0.42 | 0.231 | 0.897 | |
| AT5G16930.1 | AAA-type ATPase family protein | 38 | 0.99 | 0.231 | 1.000 | |
| AT5G16930.1 | AAA-type ATPase family protein | 39 | 1.94 | 0.231 | 0.991 | |
| AT5G16930.1 | AAA-type ATPase family protein | 40 | 2.48 | 0.232 | 0.853 | |
| AT5G16930.1 | AAA-type ATPase family protein | 41 | 2.09 | 0.234 | 0.975 | |
| AT5G16930.1 | AAA-type ATPase family protein | 42 | 1.40 | 0.236 | 1.000 | |
| AT5G16930.1 | AAA-type ATPase family protein | 43 | 0.88 | 0.239 | 1.000 | |
| AT5G16930.1 | AAA-type ATPase family protein | 44 | 0.58 | 0.248 | 1.000 | |
| AT5G16930.1 | AAA-type ATPase family protein | 45 | 0.42 | 0.293 | 0.986 | |
| AT5G16930.1 | AAA-type ATPase family protein | 46 | 0.29 | 0.404 | 0.981 | |
| AT5G16930.1 | AAA-type ATPase family protein | 47 | 0.19 | 0.620 | 0.997 | |
| AT5G16950.1 | Krueppel-like factor | 2 | 0.95 | 0.620 | 1.000 | |
| AT5G16950.1 | Krueppel-like factor | 3 | 0.45 | 0.641 | 1.000 | |
| AT5G16950.1 | Krueppel-like factor | 25 | 0.09 | 0.478 | 0.380 | |
| AT5G16950.1 | Krueppel-like factor | 26 | 0.45 | 0.329 | 0.993 | |
| AT5G16950.1 | Krueppel-like factor | 27 | 2.26 | 0.292 | 0.976 | |

| | | | | | | |
|---|---|---|---|---|---|---|
| AT5G16950.1 | Krueppel-like factor | 28 | 3.76 | 0.293 | 0.567 | |
| AT5G16950.1 | Krueppel-like factor | 29 | 1.71 | 0.328 | 1.000 | |
| AT5G16950.1 | Krueppel-like factor | 30 | 0.49 | 0.355 | 0.999 | |
| AT5G16950.1 | Krueppel-like factor | 31 | 0.24 | 0.343 | 0.692 | |
| AT5G16950.1 | Krueppel-like factor | 32 | 0.27 | 0.323 | 0.735 | |
| AT5G16950.1 | Krueppel-like factor | 33 | 0.53 | 0.315 | 0.999 | |
| AT5G16950.1 | Krueppel-like factor | 34 | 1.13 | 0.313 | 1.000 | |
| AT5G16950.1 | Krueppel-like factor | 35 | 2.26 | 0.293 | 0.977 | |
| AT5G16950.1 | Krueppel-like factor | 36 | 3.84 | 0.277 | 0.452 | |
| AT5G16950.1 | Krueppel-like factor | 37 | 4.76 | 0.270 | 0.205 | |
| AT5G16950.1 | Krueppel-like factor | 38 | 3.52 | 0.268 | 0.503 | |
| AT5G16950.1 | Krueppel-like factor | 39 | 1.92 | 0.268 | 0.994 | |
| AT5G16950.1 | Krueppel-like factor | 40 | 1.01 | 0.269 | 1.000 | |
| AT5G16950.1 | Krueppel-like factor | 41 | 0.50 | 0.271 | 0.990 | |
| AT5G16950.1 | Krueppel-like factor | 42 | 0.42 | 0.288 | 0.956 | |
| AT5G16950.1 | Krueppel-like factor | 43 | 0.60 | 0.402 | 1.000 | |
| AT5G16950.1 | Krueppel-like factor | 44 | 1.19 | 0.582 | 1.000 | |
| AT5G16950.1 | Krueppel-like factor | 46 | 7.62 | 0.977 | 0.999 | |
| AT5G16950.1 | Krueppel-like factor | 47 | 21.71 | 1.332 | 0.996 | |
| AT5G17400.1 | Endoplasmic reticulum-adenine nucleotide transporter 1 (ER-ANT1) | 2 | 0.99 | 0.432 | 1.000 | |
| AT5G17400.1 | Endoplasmic reticulum-adenine nucleotide transporter 1 (ER-ANT1) | 3 | 0.84 | 0.371 | 1.000 | |
| AT5G17400.1 | Endoplasmic reticulum-adenine nucleotide transporter 1 (ER-ANT1) | 11 | 1.46 | 0.355 | 1.000 | |
| AT5G17400.1 | Endoplasmic reticulum-adenine nucleotide transporter 1 (ER-ANT1) | 12 | 1.88 | 0.309 | 0.991 | |
| AT5G17400.1 | Endoplasmic reticulum-adenine nucleotide transporter 1 (ER-ANT1) | 13 | 2.39 | 0.336 | 0.952 | |
| AT5G17400.1 | Endoplasmic reticulum-adenine nucleotide transporter 1 (ER-ANT1) | 38 | 4.78 | 0.361 | 0.503 | |
| AT5G17400.1 | Endoplasmic reticulum-adenine nucleotide transporter 1 (ER-ANT1) | 39 | 5.61 | 0.247 | 0.039 | * |
| AT5G17400.1 | Endoplasmic reticulum-adenine nucleotide transporter 1 (ER-ANT1) | 40 | 4.86 | 0.215 | 0.025 | * |
| AT5G17400.1 | Endoplasmic reticulum-adenine nucleotide transporter 1 (ER-ANT1) | 41 | 1.85 | 0.208 | 0.893 | |
| AT5G17400.1 | Endoplasmic reticulum-adenine nucleotide transporter 1 (ER-ANT1) | 42 | 0.45 | 0.211 | 0.674 | |
| AT5G17400.1 | Endoplasmic reticulum-adenine nucleotide transporter 1 (ER-ANT1) | 43 | 0.14 | 0.213 | 0.002 | ** |
| AT5G17400.1 | Endoplasmic reticulum-adenine nucleotide transporter 1 (ER-ANT1) | 44 | 0.09 | 0.213 | 0.000 | *** |
| AT5G17400.1 | Endoplasmic reticulum-adenine nucleotide transporter 1 (ER-ANT1) | 45 | 0.17 | 0.253 | 0.038 | * |
| AT5G17400.1 | Endoplasmic reticulum-adenine nucleotide transporter 1 (ER-ANT1) | 46 | 0.56 | 0.421 | 1.000 | |

| | | | | | | |
|---|---|---|---|---|---|---|
| AT5G17400.1 | Endoplasmic reticulum-adenine nucleotide transporter 1 (ER-ANT1) | 47 | 2.40 | 0.809 | 1.000 | |
| AT5G17770.1 | NADH:cytochrome (Cyt) b5 reductase (CBR1) | 2 | 0.97 | 0.487 | 1.000 | |
| AT5G17770.1 | NADH:cytochrome (Cyt) b5 reductase (CBR1) | 3 | 1.03 | 0.451 | 1.000 | |
| AT5G17770.1 | NADH:cytochrome (Cyt) b5 reductase (CBR1) | 18 | 1.48 | 0.269 | 1.000 | |
| AT5G17770.1 | NADH:cytochrome (Cyt) b5 reductase (CBR1) | 19 | 1.43 | 0.240 | 1.000 | |
| AT5G17770.1 | NADH:cytochrome (Cyt) b5 reductase (CBR1) | 20 | 1.38 | 0.220 | 1.000 | |
| AT5G17770.1 | NADH:cytochrome (Cyt) b5 reductase (CBR1) | 21 | 1.30 | 0.209 | 1.000 | |
| AT5G17770.1 | NADH:cytochrome (Cyt) b5 reductase (CBR1) | 22 | 1.16 | 0.203 | 1.000 | |
| AT5G17770.1 | NADH:cytochrome (Cyt) b5 reductase (CBR1) | 23 | 0.97 | 0.199 | 1.000 | |
| AT5G17770.1 | NADH:cytochrome (Cyt) b5 reductase (CBR1) | 24 | 0.78 | 0.194 | 1.000 | |
| AT5G17770.1 | NADH:cytochrome (Cyt) b5 reductase (CBR1) | 25 | 0.60 | 0.191 | 0.935 | |
| AT5G17770.1 | NADH:cytochrome (Cyt) b5 reductase (CBR1) | 26 | 0.47 | 0.188 | 0.552 | |
| AT5G17770.1 | NADH:cytochrome (Cyt) b5 reductase (CBR1) | 27 | 0.38 | 0.186 | 0.225 | |
| AT5G17770.1 | NADH:cytochrome (Cyt) b5 reductase (CBR1) | 28 | 0.33 | 0.186 | 0.110 | |
| AT5G17770.1 | NADH:cytochrome (Cyt) b5 reductase (CBR1) | 29 | 0.33 | 0.185 | 0.118 | |
| AT5G17770.1 | NADH:cytochrome (Cyt) b5 reductase (CBR1) | 30 | 0.39 | 0.185 | 0.266 | |
| AT5G17770.1 | NADH:cytochrome (Cyt) b5 reductase (CBR1) | 31 | 0.51 | 0.185 | 0.697 | |
| AT5G17770.1 | NADH:cytochrome (Cyt) b5 reductase (CBR1) | 32 | 0.74 | 0.186 | 0.999 | |
| AT5G17770.1 | NADH:cytochrome (Cyt) b5 reductase (CBR1) | 33 | 1.17 | 0.186 | 1.000 | |
| AT5G17770.1 | NADH:cytochrome (Cyt) b5 reductase (CBR1) | 34 | 1.91 | 0.186 | 0.736 | |
| AT5G17770.1 | NADH:cytochrome (Cyt) b5 reductase (CBR1) | 35 | 2.97 | 0.186 | 0.128 | |
| AT5G17770.1 | NADH:cytochrome (Cyt) b5 reductase (CBR1) | 36 | 4.05 | 0.186 | 0.017 | * |
| AT5G17770.1 | NADH:cytochrome (Cyt) b5 reductase (CBR1) | 37 | 4.65 | 0.186 | 0.006 | ** |
| AT5G17770.1 | NADH:cytochrome (Cyt) b5 reductase (CBR1) | 38 | 4.48 | 0.186 | 0.008 | ** |
| AT5G17770.1 | NADH:cytochrome (Cyt) b5 reductase (CBR1) | 39 | 3.86 | 0.187 | 0.025 | * |
| AT5G17770.1 | NADH:cytochrome (Cyt) b5 reductase (CBR1) | 40 | 3.21 | 0.187 | 0.086 | . |
| AT5G17770.1 | NADH:cytochrome (Cyt) b5 reductase (CBR1) | 41 | 2.60 | 0.189 | 0.268 | |
| AT5G17770.1 | NADH:cytochrome (Cyt) b5 reductase (CBR1) | 42 | 2.03 | 0.191 | 0.667 | |
| AT5G17770.1 | NADH:cytochrome (Cyt) b5 reductase (CBR1) | 43 | 1.51 | 0.193 | 0.986 | |
| AT5G17770.1 | NADH:cytochrome (Cyt) b5 reductase (CBR1) | 44 | 1.06 | 0.197 | 1.000 | |

| | | | | | |
|---|---|---|---|---|---|
| AT5G17770.1 | NADH:cytochrome (Cyt) b5 reductase (CBR1) | 45 | 0.72 | 0.211 | 0.999 |
| AT5G17770.1 | NADH:cytochrome (Cyt) b5 reductase (CBR1) | 46 | 0.49 | 0.251 | 0.903 |
| AT5G17770.1 | NADH:cytochrome (Cyt) b5 reductase (CBR1) | 47 | 0.33 | 0.331 | 0.786 |
| AT5G17920.2 | Methionine synthase 1 (MS1) | 2 | 0.92 | 0.407 | 1.000 |
| AT5G17920.2 | Methionine synthase 1 (MS1) | 3 | 1.06 | 0.399 | 1.000 |
| AT5G17920.2 | Methionine synthase 1 (MS1) | 39 | 1.95 | 0.283 | 0.901 |
| AT5G17920.2 | Methionine synthase 1 (MS1) | 40 | 1.64 | 0.206 | 0.891 |
| AT5G17920.2 | Methionine synthase 1 (MS1) | 41 | 1.61 | 0.174 | 0.822 |
| AT5G17920.2 | Methionine synthase 1 (MS1) | 42 | 1.86 | 0.162 | 0.496 |
| AT5G17920.2 | Methionine synthase 1 (MS1) | 43 | 2.25 | 0.157 | 0.181 |
| AT5G17920.2 | Methionine synthase 1 (MS1) | 44 | 2.13 | 0.159 | 0.258 |
| AT5G17920.2 | Methionine synthase 1 (MS1) | 45 | 1.29 | 0.168 | 0.990 |
| AT5G17920.2 | Methionine synthase 1 (MS1) | 46 | 0.54 | 0.198 | 0.726 |
| AT5G17920.2 | Methionine synthase 1 (MS1) | 47 | 0.20 | 0.306 | 0.155 |
| AT5G18170.1 | NADH glutamate dehydrogenase subunit (GDH1) | 2 | 0.84 | 0.571 | 1.000 |
| AT5G18170.1 | NADH glutamate dehydrogenase subunit (GDH1) | 3 | 0.67 | 0.414 | 1.000 |
| AT5G18170.1 | NADH glutamate dehydrogenase subunit (GDH1) | 4 | 0.55 | 0.443 | 1.000 |
| AT5G18170.1 | NADH glutamate dehydrogenase subunit (GDH1) | 6 | 0.52 | 0.470 | 1.000 |
| AT5G18170.1 | NADH glutamate dehydrogenase subunit (GDH1) | 7 | 0.59 | 0.404 | 1.000 |
| AT5G18170.1 | NADH glutamate dehydrogenase subunit (GDH1) | 8 | 0.67 | 0.358 | 1.000 |
| AT5G18170.1 | NADH glutamate dehydrogenase subunit (GDH1) | 9 | 0.71 | 0.328 | 1.000 |
| AT5G18170.1 | NADH glutamate dehydrogenase subunit (GDH1) | 10 | 0.73 | 0.311 | 1.000 |
| AT5G18170.1 | NADH glutamate dehydrogenase subunit (GDH1) | 11 | 0.87 | 0.300 | 1.000 |
| AT5G18170.1 | NADH glutamate dehydrogenase subunit (GDH1) | 12 | 1.29 | 0.294 | 1.000 |
| AT5G18170.1 | NADH glutamate dehydrogenase subunit (GDH1) | 13 | 1.84 | 0.290 | 1.000 |
| AT5G18170.1 | NADH glutamate dehydrogenase subunit (GDH1) | 14 | 2.30 | 0.291 | 0.996 |
| AT5G18170.1 | NADH glutamate dehydrogenase subunit (GDH1) | 15 | 2.63 | 0.291 | 0.976 |
| AT5G18170.1 | NADH glutamate dehydrogenase subunit (GDH1) | 16 | 2.80 | 0.290 | 0.953 |
| AT5G18170.1 | NADH glutamate dehydrogenase subunit (GDH1) | 17 | 2.95 | 0.297 | 0.943 |
| AT5G18170.1 | NADH glutamate dehydrogenase subunit (GDH1) | 18 | 3.04 | 0.306 | 0.945 |
| AT5G18170.1 | NADH glutamate dehydrogenase subunit (GDH1) | 19 | 3.20 | 0.310 | 0.926 |
| AT5G18170.1 | NADH glutamate dehydrogenase subunit (GDH1) | 20 | 3.17 | 0.309 | 0.929 |
| AT5G18170.1 | NADH glutamate dehydrogenase subunit (GDH1) | 21 | 2.99 | 0.306 | 0.951 |
| AT5G18170.1 | NADH glutamate dehydrogenase subunit (GDH1) | 22 | 3.15 | 0.304 | 0.923 |
| AT5G18170.1 | NADH glutamate dehydrogenase subunit (GDH1) | 23 | 3.40 | 0.301 | 0.860 |
| AT5G18170.1 | NADH glutamate dehydrogenase subunit (GDH1) | 24 | 2.60 | 0.295 | 0.982 |

| | | | | | |
|---|---|---|---|---|---|
| AT5G18170.1 | NADH glutamate dehydrogenase subunit (GDH1) | 25 | 1.48 | 0.288 | 1.000 |
| AT5G18170.1 | NADH glutamate dehydrogenase subunit (GDH1) | 26 | 1.26 | 0.288 | 1.000 |
| AT5G18170.1 | NADH glutamate dehydrogenase subunit (GDH1) | 27 | 1.37 | 0.288 | 1.000 |
| AT5G18170.1 | NADH glutamate dehydrogenase subunit (GDH1) | 28 | 1.13 | 0.288 | 1.000 |
| AT5G18170.1 | NADH glutamate dehydrogenase subunit (GDH1) | 29 | 0.88 | 0.288 | 1.000 |
| AT5G18170.1 | NADH glutamate dehydrogenase subunit (GDH1) | 30 | 1.09 | 0.290 | 1.000 |
| AT5G18170.1 | NADH glutamate dehydrogenase subunit (GDH1) | 31 | 2.46 | 0.290 | 0.989 |
| AT5G18170.1 | NADH glutamate dehydrogenase subunit (GDH1) | 32 | 4.96 | 0.290 | 0.374 |
| AT5G18170.1 | NADH glutamate dehydrogenase subunit (GDH1) | 33 | 4.16 | 0.290 | 0.589 |
| AT5G18170.1 | NADH glutamate dehydrogenase subunit (GDH1) | 34 | 1.84 | 0.292 | 1.000 |
| AT5G18170.1 | NADH glutamate dehydrogenase subunit (GDH1) | 35 | 1.05 | 0.292 | 1.000 |
| AT5G18170.1 | NADH glutamate dehydrogenase subunit (GDH1) | 36 | 1.07 | 0.295 | 1.000 |
| AT5G18170.1 | NADH glutamate dehydrogenase subunit (GDH1) | 37 | 1.24 | 0.313 | 1.000 |
| AT5G18170.1 | NADH glutamate dehydrogenase subunit (GDH1) | 38 | 1.62 | 0.361 | 1.000 |
| AT5G18170.1 | NADH glutamate dehydrogenase subunit (GDH1) | 39 | 2.54 | 0.432 | 1.000 |
| AT5G18170.1 | NADH glutamate dehydrogenase subunit (GDH1) | 41 | 5.68 | 0.443 | 0.893 |
| AT5G18170.1 | NADH glutamate dehydrogenase subunit (GDH1) | 42 | 7.02 | 0.515 | 0.921 |
| AT5G18170.1 | NADH glutamate dehydrogenase subunit (GDH1) | 43 | 7.17 | 0.641 | 0.990 |
| AT5G18170.1 | NADH glutamate dehydrogenase subunit (GDH1) | 46 | 3.42 | 1.627 | 1.000 |
| AT5G18170.1 | NADH glutamate dehydrogenase subunit (GDH1) | 47 | 2.36 | 2.105 | 1.000 |
| AT5G18800.2 | NADH:ubiquinone oxidoreductase subunit (NDUFA8) | 2 | 1.04 | 0.466 | 1.000 |
| AT5G18800.2 | NADH:ubiquinone oxidoreductase subunit (NDUFA8) | 3 | 0.95 | 0.430 | 1.000 |
| AT5G18800.2 | NADH:ubiquinone oxidoreductase subunit (NDUFA8) | 15 | 0.54 | 0.305 | 0.991 |
| AT5G18800.2 | NADH:ubiquinone oxidoreductase subunit (NDUFA8) | 16 | 0.41 | 0.260 | 0.777 |
| AT5G18800.2 | NADH:ubiquinone oxidoreductase subunit (NDUFA8) | 17 | 0.31 | 0.234 | 0.284 |
| AT5G18800.2 | NADH:ubiquinone oxidoreductase subunit (NDUFA8) | 18 | 0.34 | 0.218 | 0.321 |
| AT5G18800.2 | NADH:ubiquinone oxidoreductase subunit (NDUFA8) | 19 | 0.60 | 0.221 | 0.978 |

| | | | | | |
|---|---|---|---|---|---|
| AT5G18800.2 | NADH:ubiquinone oxidoreductase subunit (NDUFA8) | 20 | 0.93 | 0.225 | 1.000 |
| AT5G18800.2 | NADH:ubiquinone oxidoreductase subunit (NDUFA8) | 21 | 1.20 | 0.227 | 1.000 |
| AT5G18800.2 | NADH:ubiquinone oxidoreductase subunit (NDUFA8) | 22 | 1.64 | 0.229 | 0.985 |
| AT5G18800.2 | NADH:ubiquinone oxidoreductase subunit (NDUFA8) | 23 | 2.21 | 0.233 | 0.785 |
| AT5G18800.2 | NADH:ubiquinone oxidoreductase subunit (NDUFA8) | 24 | 1.77 | 0.235 | 0.968 |
| AT5G18800.2 | NADH:ubiquinone oxidoreductase subunit (NDUFA8) | 25 | 0.87 | 0.276 | 1.000 |
| AT5G18800.2 | NADH:ubiquinone oxidoreductase subunit (NDUFA8) | 26 | 0.55 | 0.445 | 1.000 |
| AT5G18800.2 | NADH:ubiquinone oxidoreductase subunit (NDUFA8) | 46 | ########### | 11.375 | 0.934 |
| AT5G18800.2 | NADH:ubiquinone oxidoreductase subunit (NDUFA8) | 47 | ########### | 12.197 | 0.930 |
| AT5G19150.2 | ATP-dependent (S)-NAD(P)H-hydrate dehydratase | 2 | 1.09 | 0.632 | 1.000 |
| AT5G19150.2 | ATP-dependent (S)-NAD(P)H-hydrate dehydratase | 3 | 1.30 | 0.631 | 1.000 |
| AT5G19150.2 | ATP-dependent (S)-NAD(P)H-hydrate dehydratase | 40 | 2.41 | 0.316 | 0.619 |
| AT5G19150.2 | ATP-dependent (S)-NAD(P)H-hydrate dehydratase | 41 | 1.74 | 0.283 | 0.843 |
| AT5G19150.2 | ATP-dependent (S)-NAD(P)H-hydrate dehydratase | 42 | 1.20 | 0.270 | 0.997 |
| AT5G19150.2 | ATP-dependent (S)-NAD(P)H-hydrate dehydratase | 43 | 0.79 | 0.275 | 0.992 |
| AT5G19150.2 | ATP-dependent (S)-NAD(P)H-hydrate dehydratase | 44 | 0.51 | 0.292 | 0.747 |
| AT5G19150.2 | ATP-dependent (S)-NAD(P)H-hydrate dehydratase | 46 | 0.19 | 0.355 | 0.185 |
| AT5G19150.2 | ATP-dependent (S)-NAD(P)H-hydrate dehydratase | 47 | 0.12 | 0.408 | 0.111 |
| AT5G19370.1 | Rhodanese-like/PpiC domain-containing protein 12 (PIN3) | 2 | 1.00 | 0.414 | 1.000 |
| AT5G19370.1 | Rhodanese-like/PpiC domain-containing protein 12 (PIN3) | 3 | 1.05 | 0.359 | 1.000 |
| AT5G19370.1 | Rhodanese-like/PpiC domain-containing protein 12 (PIN3) | 9 | 1.41 | 0.232 | 1.000 |
| AT5G19370.1 | Rhodanese-like/PpiC domain-containing protein 12 (PIN3) | 10 | 1.48 | 0.226 | 0.998 |
| AT5G19370.1 | Rhodanese-like/PpiC domain-containing protein 12 (PIN3) | 11 | 1.55 | 0.226 | 0.996 |
| AT5G19370.1 | Rhodanese-like/PpiC domain-containing protein 12 (PIN3) | 12 | 1.60 | 0.234 | 0.994 |
| AT5G19370.1 | Rhodanese-like/PpiC domain-containing protein 12 (PIN3) | 22 | 0.82 | 0.256 | 1.000 |
| AT5G19370.1 | Rhodanese-like/PpiC domain-containing protein 12 (PIN3) | 23 | 0.73 | 0.235 | 1.000 |

| | | | | | | |
|---|---|---|---|---|---|---|
| AT5G19370.1 | Rhodanese-like/PpiC domain-containing protein 12 (PIN3) | 24 | 0.64 | 0.213 | 0.992 | |
| AT5G19370.1 | Rhodanese-like/PpiC domain-containing protein 12 (PIN3) | 26 | 0.54 | 0.174 | 0.772 | |
| AT5G19370.1 | Rhodanese-like/PpiC domain-containing protein 12 (PIN3) | 27 | 0.52 | 0.157 | 0.580 | |
| AT5G19370.1 | Rhodanese-like/PpiC domain-containing protein 12 (PIN3) | 28 | 0.52 | 0.143 | 0.452 | |
| AT5G19370.1 | Rhodanese-like/PpiC domain-containing protein 12 (PIN3) | 29 | 0.54 | 0.134 | 0.418 | |
| AT5G19370.1 | Rhodanese-like/PpiC domain-containing protein 12 (PIN3) | 30 | 0.57 | 0.129 | 0.526 | |
| AT5G19370.1 | Rhodanese-like/PpiC domain-containing protein 12 (PIN3) | 31 | 0.65 | 0.127 | 0.812 | |
| AT5G19370.1 | Rhodanese-like/PpiC domain-containing protein 12 (PIN3) | 32 | 0.78 | 0.127 | 0.994 | |
| AT5G19370.1 | Rhodanese-like/PpiC domain-containing protein 12 (PIN3) | 33 | 0.95 | 0.128 | 1.000 | |
| AT5G19370.1 | Rhodanese-like/PpiC domain-containing protein 12 (PIN3) | 34 | 1.18 | 0.130 | 1.000 | |
| AT5G19370.1 | Rhodanese-like/PpiC domain-containing protein 12 (PIN3) | 35 | 1.48 | 0.133 | 0.911 | |
| AT5G19370.1 | Rhodanese-like/PpiC domain-containing protein 12 (PIN3) | 36 | 1.85 | 0.138 | 0.482 | |
| AT5G19370.1 | Rhodanese-like/PpiC domain-containing protein 12 (PIN3) | 37 | 2.21 | 0.143 | 0.192 | |
| AT5G19370.1 | Rhodanese-like/PpiC domain-containing protein 12 (PIN3) | 38 | 2.49 | 0.146 | 0.092 | . |
| AT5G19370.1 | Rhodanese-like/PpiC domain-containing protein 12 (PIN3) | 39 | 2.62 | 0.146 | 0.062 | . |
| AT5G19370.1 | Rhodanese-like/PpiC domain-containing protein 12 (PIN3) | 40 | 2.55 | 0.143 | 0.067 | . |
| AT5G19370.1 | Rhodanese-like/PpiC domain-containing protein 12 (PIN3) | 41 | 2.28 | 0.139 | 0.132 | |
| AT5G19370.1 | Rhodanese-like/PpiC domain-containing protein 12 (PIN3) | 42 | 1.86 | 0.135 | 0.430 | |
| AT5G19370.1 | Rhodanese-like/PpiC domain-containing protein 12 (PIN3) | 43 | 1.41 | 0.133 | 0.963 | |
| AT5G19370.1 | Rhodanese-like/PpiC domain-containing protein 12 (PIN3) | 44 | 1.00 | 0.138 | 1.000 | |
| AT5G19370.1 | Rhodanese-like/PpiC domain-containing protein 12 (PIN3) | 45 | 0.71 | 0.155 | 0.985 | |
| AT5G19370.1 | Rhodanese-like/PpiC domain-containing protein 12 (PIN3) | 46 | 0.50 | 0.193 | 0.748 | |
| AT5G19370.1 | Rhodanese-like/PpiC domain-containing protein 12 (PIN3) | 47 | 0.35 | 0.254 | 0.591 | |
| AT5G19760.1 | Dicarboxylate and tricarboxylate carrier (DTC) | 3 | 0.86 | 0.131 | 1.000 | |
| AT5G19760.1 | Dicarboxylate and tricarboxylate carrier (DTC) | 4 | 0.91 | 0.128 | 1.000 | |
| AT5G19760.1 | Dicarboxylate and tricarboxylate carrier (DTC) | 5 | 1.06 | 0.127 | 1.000 | |
| AT5G19760.1 | Dicarboxylate and tricarboxylate carrier (DTC) | 6 | 1.04 | 0.127 | 1.000 | |
| AT5G19760.1 | Dicarboxylate and tricarboxylate carrier (DTC) | 7 | 0.93 | 0.129 | 1.000 | |
| AT5G19760.1 | Dicarboxylate and tricarboxylate carrier (DTC) | 8 | 1.00 | 0.126 | 1.000 | |
| AT5G19760.1 | Dicarboxylate and tricarboxylate carrier (DTC) | 9 | 1.12 | 0.124 | 1.000 | |
| AT5G19760.1 | Dicarboxylate and tricarboxylate carrier (DTC) | 10 | 1.15 | 0.123 | 1.000 | |

| | | | | | | |
|---|---|---|---|---|---|---|
| AT5G19760.1 | Dicarboxylate and tricarboxylate carrier (DTC) | 11 | 1.15 | 0.124 | 1.000 | |
| AT5G19760.1 | Dicarboxylate and tricarboxylate carrier (DTC) | 12 | 1.13 | 0.124 | 1.000 | |
| AT5G19760.1 | Dicarboxylate and tricarboxylate carrier (DTC) | 13 | 1.16 | 0.123 | 1.000 | |
| AT5G19760.1 | Dicarboxylate and tricarboxylate carrier (DTC) | 14 | 1.23 | 0.121 | 1.000 | |
| AT5G19760.1 | Dicarboxylate and tricarboxylate carrier (DTC) | 15 | 1.17 | 0.121 | 1.000 | |
| AT5G19760.1 | Dicarboxylate and tricarboxylate carrier (DTC) | 16 | 1.02 | 0.120 | 1.000 | |
| AT5G19760.1 | Dicarboxylate and tricarboxylate carrier (DTC) | 17 | 1.09 | 0.120 | 1.000 | |
| AT5G19760.1 | Dicarboxylate and tricarboxylate carrier (DTC) | 18 | 1.34 | 0.122 | 1.000 | |
| AT5G19760.1 | Dicarboxylate and tricarboxylate carrier (DTC) | 19 | 1.62 | 0.125 | 0.942 | |
| AT5G19760.1 | Dicarboxylate and tricarboxylate carrier (DTC) | 20 | 1.76 | 0.128 | 0.820 | |
| AT5G19760.1 | Dicarboxylate and tricarboxylate carrier (DTC) | 21 | 1.75 | 0.130 | 0.844 | |
| AT5G19760.1 | Dicarboxylate and tricarboxylate carrier (DTC) | 22 | 1.67 | 0.129 | 0.925 | |
| AT5G19760.1 | Dicarboxylate and tricarboxylate carrier (DTC) | 23 | 1.46 | 0.129 | 0.999 | |
| AT5G19760.1 | Dicarboxylate and tricarboxylate carrier (DTC) | 24 | 1.09 | 0.126 | 1.000 | |
| AT5G19760.1 | Dicarboxylate and tricarboxylate carrier (DTC) | 25 | 0.80 | 0.124 | 1.000 | |
| AT5G19760.1 | Dicarboxylate and tricarboxylate carrier (DTC) | 26 | 0.84 | 0.123 | 1.000 | |
| AT5G19760.1 | Dicarboxylate and tricarboxylate carrier (DTC) | 27 | 1.26 | 0.123 | 1.000 | |
| AT5G19760.1 | Dicarboxylate and tricarboxylate carrier (DTC) | 28 | 1.69 | 0.123 | 0.855 | |
| AT5G19760.1 | Dicarboxylate and tricarboxylate carrier (DTC) | 29 | 1.95 | 0.123 | 0.450 | |
| AT5G19760.1 | Dicarboxylate and tricarboxylate carrier (DTC) | 30 | 1.87 | 0.122 | 0.574 | |
| AT5G19760.1 | Dicarboxylate and tricarboxylate carrier (DTC) | 31 | 1.71 | 0.123 | 0.834 | |
| AT5G19760.1 | Dicarboxylate and tricarboxylate carrier (DTC) | 32 | 1.52 | 0.123 | 0.988 | |
| AT5G19760.1 | Dicarboxylate and tricarboxylate carrier (DTC) | 33 | 1.45 | 0.123 | 0.998 | |
| AT5G19760.1 | Dicarboxylate and tricarboxylate carrier (DTC) | 34 | 1.61 | 0.124 | 0.942 | |
| AT5G19760.1 | Dicarboxylate and tricarboxylate carrier (DTC) | 35 | 1.96 | 0.124 | 0.454 | |
| AT5G19760.1 | Dicarboxylate and tricarboxylate carrier (DTC) | 36 | 2.21 | 0.123 | 0.172 | |
| AT5G19760.1 | Dicarboxylate and tricarboxylate carrier (DTC) | 37 | 2.30 | 0.123 | 0.118 | |
| AT5G19760.1 | Dicarboxylate and tricarboxylate carrier (DTC) | 38 | 2.62 | 0.123 | 0.029 | * |
| AT5G19760.1 | Dicarboxylate and tricarboxylate carrier (DTC) | 39 | 3.98 | 0.124 | 0.000 | *** |
| AT5G19760.1 | Dicarboxylate and tricarboxylate carrier (DTC) | 40 | 6.45 | 0.125 | 0.000 | *** |
| AT5G19760.1 | Dicarboxylate and tricarboxylate carrier (DTC) | 41 | 6.18 | 0.125 | 0.000 | *** |

| | | | | | | |
|---|---|---|---|---|---|---|
| AT5G19760.1 | Dicarboxylate and tricarboxylate carrier (DTC) | 42 | 3.22 | 0.126 | 0.003 | ** |
| AT5G19760.1 | Dicarboxylate and tricarboxylate carrier (DTC) | 43 | 1.35 | 0.127 | 1.000 | |
| AT5G19760.1 | Dicarboxylate and tricarboxylate carrier (DTC) | 44 | 0.56 | 0.127 | 0.776 | |
| AT5G19760.1 | Dicarboxylate and tricarboxylate carrier (DTC) | 45 | 0.27 | 0.127 | 0.000 | *** |
| AT5G19760.1 | Dicarboxylate and tricarboxylate carrier (DTC) | 46 | 0.14 | 0.138 | 0.000 | *** |
| AT5G19760.1 | Dicarboxylate and tricarboxylate carrier (DTC) | 47 | 0.06 | 0.266 | 0.000 | *** |
| AT5G19760.1 | Dicarboxylate and tricarboxylate carrier (DTC) | 48 | 0.03 | 0.482 | 0.038 | * |
| AT5G20080.1 | NADH:cytochrome (Cyt) b5 reductase (CBR2) | 3 | 0.80 | 0.454 | 1.000 | |
| AT5G20080.1 | NADH:cytochrome (Cyt) b5 reductase (CBR2) | 4 | 0.82 | 0.439 | 1.000 | |
| AT5G20080.1 | NADH:cytochrome (Cyt) b5 reductase (CBR2) | 5 | 0.85 | 0.435 | 1.000 | |
| AT5G20080.1 | NADH:cytochrome (Cyt) b5 reductase (CBR2) | 6 | 0.88 | 0.435 | 1.000 | |
| AT5G20080.1 | NADH:cytochrome (Cyt) b5 reductase (CBR2) | 7 | 0.91 | 0.435 | 1.000 | |
| AT5G20080.1 | NADH:cytochrome (Cyt) b5 reductase (CBR2) | 8 | 0.93 | 0.435 | 1.000 | |
| AT5G20080.1 | NADH:cytochrome (Cyt) b5 reductase (CBR2) | 9 | 0.92 | 0.434 | 1.000 | |
| AT5G20080.1 | NADH:cytochrome (Cyt) b5 reductase (CBR2) | 10 | 0.89 | 0.433 | 1.000 | |
| AT5G20080.1 | NADH:cytochrome (Cyt) b5 reductase (CBR2) | 11 | 0.86 | 0.433 | 1.000 | |
| AT5G20080.1 | NADH:cytochrome (Cyt) b5 reductase (CBR2) | 12 | 0.83 | 0.433 | 1.000 | |
| AT5G20080.1 | NADH:cytochrome (Cyt) b5 reductase (CBR2) | 13 | 0.81 | 0.433 | 1.000 | |
| AT5G20080.1 | NADH:cytochrome (Cyt) b5 reductase (CBR2) | 14 | 0.80 | 0.432 | 1.000 | |
| AT5G20080.1 | NADH:cytochrome (Cyt) b5 reductase (CBR2) | 15 | 0.79 | 0.432 | 1.000 | |
| AT5G20080.1 | NADH:cytochrome (Cyt) b5 reductase (CBR2) | 16 | 0.79 | 0.431 | 1.000 | |
| AT5G20080.1 | NADH:cytochrome (Cyt) b5 reductase (CBR2) | 17 | 0.79 | 0.432 | 1.000 | |
| AT5G20080.1 | NADH:cytochrome (Cyt) b5 reductase (CBR2) | 18 | 0.80 | 0.432 | 1.000 | |
| AT5G20080.1 | NADH:cytochrome (Cyt) b5 reductase (CBR2) | 19 | 0.81 | 0.433 | 1.000 | |
| AT5G20080.1 | NADH:cytochrome (Cyt) b5 reductase (CBR2) | 20 | 0.82 | 0.434 | 1.000 | |
| AT5G20080.1 | NADH:cytochrome (Cyt) b5 reductase (CBR2) | 21 | 0.80 | 0.435 | 1.000 | |
| AT5G20080.1 | NADH:cytochrome (Cyt) b5 reductase (CBR2) | 22 | 0.74 | 0.435 | 1.000 | |
| AT5G20080.1 | NADH:cytochrome (Cyt) b5 reductase (CBR2) | 23 | 0.68 | 0.435 | 1.000 | |
| AT5G20080.1 | NADH:cytochrome (Cyt) b5 reductase (CBR2) | 24 | 0.62 | 0.434 | 0.998 | |
| AT5G20080.1 | NADH:cytochrome (Cyt) b5 reductase (CBR2) | 25 | 0.57 | 0.433 | 0.990 | |
| AT5G20080.1 | NADH:cytochrome (Cyt) b5 reductase (CBR2) | 26 | 0.55 | 0.433 | 0.984 | |

| | | | | | |
|---|---|---|---|---|---|
| AT5G20080.1 | NADH:cytochrome (Cyt) b5 reductase (CBR2) | 27 | 0.56 | 0.433 | 0.989 |
| AT5G20080.1 | NADH:cytochrome (Cyt) b5 reductase (CBR2) | 28 | 0.59 | 0.433 | 0.995 |
| AT5G20080.1 | NADH:cytochrome (Cyt) b5 reductase (CBR2) | 29 | 0.63 | 0.433 | 0.999 |
| AT5G20080.1 | NADH:cytochrome (Cyt) b5 reductase (CBR2) | 30 | 0.68 | 0.433 | 1.000 |
| AT5G20080.1 | NADH:cytochrome (Cyt) b5 reductase (CBR2) | 31 | 0.74 | 0.433 | 1.000 |
| AT5G20080.1 | NADH:cytochrome (Cyt) b5 reductase (CBR2) | 32 | 0.77 | 0.433 | 1.000 |
| AT5G20080.1 | NADH:cytochrome (Cyt) b5 reductase (CBR2) | 33 | 0.74 | 0.433 | 1.000 |
| AT5G20080.1 | NADH:cytochrome (Cyt) b5 reductase (CBR2) | 34 | 0.71 | 0.433 | 1.000 |
| AT5G20080.1 | NADH:cytochrome (Cyt) b5 reductase (CBR2) | 35 | 0.72 | 0.433 | 1.000 |
| AT5G20080.1 | NADH:cytochrome (Cyt) b5 reductase (CBR2) | 36 | 0.79 | 0.433 | 1.000 |
| AT5G20080.1 | NADH:cytochrome (Cyt) b5 reductase (CBR2) | 37 | 0.96 | 0.433 | 1.000 |
| AT5G20080.1 | NADH:cytochrome (Cyt) b5 reductase (CBR2) | 38 | 1.24 | 0.433 | 1.000 |
| AT5G20080.1 | NADH:cytochrome (Cyt) b5 reductase (CBR2) | 39 | 1.64 | 0.433 | 0.997 |
| AT5G20080.1 | NADH:cytochrome (Cyt) b5 reductase (CBR2) | 40 | 2.02 | 0.434 | 0.953 |
| AT5G20080.1 | NADH:cytochrome (Cyt) b5 reductase (CBR2) | 41 | 2.18 | 0.434 | 0.917 |
| AT5G20080.1 | NADH:cytochrome (Cyt) b5 reductase (CBR2) | 42 | 2.10 | 0.435 | 0.938 |
| AT5G20080.1 | NADH:cytochrome (Cyt) b5 reductase (CBR2) | 43 | 1.80 | 0.435 | 0.987 |
| AT5G20080.1 | NADH:cytochrome (Cyt) b5 reductase (CBR2) | 44 | 1.33 | 0.435 | 1.000 |
| AT5G20080.1 | NADH:cytochrome (Cyt) b5 reductase (CBR2) | 45 | 0.85 | 0.440 | 1.000 |
| AT5G20080.1 | NADH:cytochrome (Cyt) b5 reductase (CBR2) | 46 | 0.49 | 0.460 | 0.968 |
| AT5G20080.1 | NADH:cytochrome (Cyt) b5 reductase (CBR2) | 47 | 0.27 | 0.513 | 0.704 |
| AT5G20080.1 | NADH:cytochrome (Cyt) b5 reductase (CBR2) | 48 | 0.15 | 0.601 | 0.513 |
| AT5G20090.3 | Mitochondrial pyruvate carrier 1 (MPC1) | 3 | 1.46 | 0.198 | 1.000 |
| AT5G20090.3 | Mitochondrial pyruvate carrier 1 (MPC1) | 4 | 1.24 | 0.175 | 1.000 |
| AT5G20090.3 | Mitochondrial pyruvate carrier 1 (MPC1) | 5 | 1.14 | 0.174 | 1.000 |
| AT5G20090.3 | Mitochondrial pyruvate carrier 1 (MPC1) | 6 | 1.04 | 0.175 | 1.000 |
| AT5G20090.3 | Mitochondrial pyruvate carrier 1 (MPC1) | 7 | 0.94 | 0.174 | 1.000 |
| AT5G20090.3 | Mitochondrial pyruvate carrier 1 (MPC1) | 8 | 0.88 | 0.172 | 1.000 |
| AT5G20090.3 | Mitochondrial pyruvate carrier 1 (MPC1) | 9 | 0.82 | 0.170 | 1.000 |
| AT5G20090.3 | Mitochondrial pyruvate carrier 1 (MPC1) | 10 | 0.78 | 0.169 | 1.000 |
| AT5G20090.3 | Mitochondrial pyruvate carrier 1 (MPC1) | 11 | 0.80 | 0.169 | 1.000 |

| | | | | | |
|---|---|---|---|---|---|
| AT5G20090.3 | Mitochondrial pyruvate carrier 1 (MPC1) | 12 | 0.85 | 0.169 | 1.000 |
| AT5G20090.3 | Mitochondrial pyruvate carrier 1 (MPC1) | 13 | 0.91 | 0.168 | 1.000 |
| AT5G20090.3 | Mitochondrial pyruvate carrier 1 (MPC1) | 14 | 0.92 | 0.166 | 1.000 |
| AT5G20090.3 | Mitochondrial pyruvate carrier 1 (MPC1) | 15 | 0.89 | 0.164 | 1.000 |
| AT5G20090.3 | Mitochondrial pyruvate carrier 1 (MPC1) | 16 | 0.87 | 0.164 | 1.000 |
| AT5G20090.3 | Mitochondrial pyruvate carrier 1 (MPC1) | 17 | 0.90 | 0.164 | 1.000 |
| AT5G20090.3 | Mitochondrial pyruvate carrier 1 (MPC1) | 18 | 0.98 | 0.166 | 1.000 |
| AT5G20090.3 | Mitochondrial pyruvate carrier 1 (MPC1) | 19 | 1.07 | 0.169 | 1.000 |
| AT5G20090.3 | Mitochondrial pyruvate carrier 1 (MPC1) | 20 | 1.09 | 0.173 | 1.000 |
| AT5G20090.3 | Mitochondrial pyruvate carrier 1 (MPC1) | 21 | 1.02 | 0.175 | 1.000 |
| AT5G20090.3 | Mitochondrial pyruvate carrier 1 (MPC1) | 22 | 0.88 | 0.175 | 1.000 |
| AT5G20090.3 | Mitochondrial pyruvate carrier 1 (MPC1) | 23 | 0.70 | 0.173 | 1.000 |
| AT5G20090.3 | Mitochondrial pyruvate carrier 1 (MPC1) | 24 | 0.52 | 0.171 | 0.869 |
| AT5G20090.3 | Mitochondrial pyruvate carrier 1 (MPC1) | 25 | 0.39 | 0.169 | 0.316 |
| AT5G20090.3 | Mitochondrial pyruvate carrier 1 (MPC1) | 26 | 0.36 | 0.168 | 0.172 |
| AT5G20090.3 | Mitochondrial pyruvate carrier 1 (MPC1) | 27 | 0.42 | 0.169 | 0.426 |
| AT5G20090.3 | Mitochondrial pyruvate carrier 1 (MPC1) | 28 | 0.59 | 0.169 | 0.972 |
| AT5G20090.3 | Mitochondrial pyruvate carrier 1 (MPC1) | 29 | 0.89 | 0.169 | 1.000 |
| AT5G20090.3 | Mitochondrial pyruvate carrier 1 (MPC1) | 30 | 1.23 | 0.168 | 1.000 |
| AT5G20090.3 | Mitochondrial pyruvate carrier 1 (MPC1) | 31 | 1.50 | 0.168 | 0.999 |
| AT5G20090.3 | Mitochondrial pyruvate carrier 1 (MPC1) | 32 | 1.66 | 0.168 | 0.982 |
| AT5G20090.3 | Mitochondrial pyruvate carrier 1 (MPC1) | 33 | 1.65 | 0.168 | 0.984 |
| AT5G20090.3 | Mitochondrial pyruvate carrier 1 (MPC1) | 34 | 1.72 | 0.168 | 0.967 |
| AT5G20090.3 | Mitochondrial pyruvate carrier 1 (MPC1) | 35 | 2.04 | 0.168 | 0.737 |
| AT5G20090.3 | Mitochondrial pyruvate carrier 1 (MPC1) | 36 | 2.44 | 0.168 | 0.372 |
| AT5G20090.3 | Mitochondrial pyruvate carrier 1 (MPC1) | 37 | 2.61 | 0.168 | 0.265 |
| AT5G20090.3 | Mitochondrial pyruvate carrier 1 (MPC1) | 38 | 2.60 | 0.168 | 0.272 |
| AT5G20090.3 | Mitochondrial pyruvate carrier 1 (MPC1) | 39 | 2.47 | 0.168 | 0.355 |
| AT5G20090.3 | Mitochondrial pyruvate carrier 1 (MPC1) | 40 | 2.20 | 0.169 | 0.593 |
| AT5G20090.3 | Mitochondrial pyruvate carrier 1 (MPC1) | 41 | 1.81 | 0.170 | 0.930 |
| AT5G20090.3 | Mitochondrial pyruvate carrier 1 (MPC1) | 42 | 1.37 | 0.172 | 1.000 |

| | | | | | | |
|---|---|---|---|---|---|---|
| AT5G20090.3 | Mitochondrial pyruvate carrier 1 (MPC1) | 43 | 0.97 | 0.173 | 1.000 | |
| AT5G20090.3 | Mitochondrial pyruvate carrier 1 (MPC1) | 44 | 0.60 | 0.173 | 0.988 | |
| AT5G20090.3 | Mitochondrial pyruvate carrier 1 (MPC1) | 45 | 0.33 | 0.178 | 0.161 | |
| AT5G20090.3 | Mitochondrial pyruvate carrier 1 (MPC1) | 46 | 0.17 | 0.213 | 0.011 | * |
| AT5G20090.3 | Mitochondrial pyruvate carrier 1 (MPC1) | 47 | 0.08 | 0.320 | 0.026 | * |
| AT5G20090.3 | Mitochondrial pyruvate carrier 1 (MPC1) | 48 | 0.04 | 0.479 | 0.104 | |
| AT5G20290.1 | 40S ribosomal protein S8-1 (RPS8A) | 2 | 0.71 | 0.166 | 0.667 | |
| AT5G20290.1 | 40S ribosomal protein S8-1 (RPS8A) | 3 | 0.76 | 0.155 | 0.729 | |
| AT5G20290.1 | 40S ribosomal protein S8-1 (RPS8A) | 4 | 0.80 | 0.144 | 0.800 | |
| AT5G20290.1 | 40S ribosomal protein S8-1 (RPS8A) | 5 | 0.85 | 0.134 | 0.874 | |
| AT5G20290.1 | 40S ribosomal protein S8-1 (RPS8A) | 6 | 0.91 | 0.125 | 0.943 | |
| AT5G20290.1 | 40S ribosomal protein S8-1 (RPS8A) | 7 | 0.96 | 0.117 | 0.991 | |
| AT5G20290.1 | 40S ribosomal protein S8-1 (RPS8A) | 8 | 1.03 | 0.110 | 0.995 | |
| AT5G20290.1 | 40S ribosomal protein S8-1 (RPS8A) | 9 | 1.09 | 0.105 | 0.937 | |
| AT5G20290.1 | 40S ribosomal protein S8-1 (RPS8A) | 10 | 1.16 | 0.101 | 0.817 | |
| AT5G20290.1 | 40S ribosomal protein S8-1 (RPS8A) | 11 | 1.23 | 0.100 | 0.662 | |
| AT5G20290.1 | 40S ribosomal protein S8-1 (RPS8A) | 13 | 1.39 | 0.104 | 0.384 | |
| AT5G20290.1 | 40S ribosomal protein S8-1 (RPS8A) | 14 | 1.48 | 0.109 | 0.294 | |
| AT5G20290.1 | 40S ribosomal protein S8-1 (RPS8A) | 15 | 1.57 | 0.116 | 0.234 | |
| AT5G20290.1 | 40S ribosomal protein S8-1 (RPS8A) | 16 | 1.67 | 0.124 | 0.194 | |
| AT5G20290.1 | 40S ribosomal protein S8-1 (RPS8A) | 17 | 1.78 | 0.133 | 0.168 | |
| AT5G20290.1 | 40S ribosomal protein S8-1 (RPS8A) | 18 | 1.89 | 0.143 | 0.152 | |
| AT5G20290.1 | 40S ribosomal protein S8-1 (RPS8A) | 19 | 2.01 | 0.153 | 0.142 | |
| AT5G20290.1 | 40S ribosomal protein S8-1 (RPS8A) | 20 | 2.13 | 0.165 | 0.135 | |
| AT5G20290.1 | 40S ribosomal protein S8-1 (RPS8A) | 46 | 10.46 | 0.521 | 0.147 | |
| AT5G20290.1 | 40S ribosomal protein S8-1 (RPS8A) | 47 | 11.11 | 0.536 | 0.148 | |
| AT5G20720.3 | Chaperonin 20 (CPN20) | 2 | 1.03 | 0.325 | 1.000 | |
| AT5G20720.3 | Chaperonin 20 (CPN20) | 3 | 1.32 | 0.327 | 0.992 | |
| AT5G20720.3 | Chaperonin 20 (CPN20) | 43 | 2.49 | 0.199 | 0.199 | |
| AT5G20720.3 | Chaperonin 20 (CPN20) | 44 | 1.52 | 0.152 | 0.633 | |
| AT5G20720.3 | Chaperonin 20 (CPN20) | 45 | 0.88 | 0.135 | 0.990 | |
| AT5G20720.3 | Chaperonin 20 (CPN20) | 46 | 0.48 | 0.150 | 0.150 | |
| AT5G20720.3 | Chaperonin 20 (CPN20) | 47 | 0.25 | 0.202 | 0.022 | * |
| AT5G20980.2 | Methionine synthase 3 (MS3) | 2 | 1.29 | 0.423 | 1.000 | |
| AT5G20980.2 | Methionine synthase 3 (MS3) | 3 | 0.81 | 0.327 | 1.000 | |
| AT5G20980.2 | Methionine synthase 3 (MS3) | 4 | 0.55 | 0.523 | 1.000 | |

| | | | | | | |
|---|---|---|---|---|---|---|
| AT5G20980.2 | Methionine synthase 3 (MS3) | 31 | 1.14 | 0.326 | 1.000 | |
| AT5G20980.2 | Methionine synthase 3 (MS3) | 32 | 0.91 | 0.252 | 1.000 | |
| AT5G20980.2 | Methionine synthase 3 (MS3) | 33 | 0.79 | 0.224 | 1.000 | |
| AT5G20980.2 | Methionine synthase 3 (MS3) | 34 | 0.80 | 0.214 | 1.000 | |
| AT5G20980.2 | Methionine synthase 3 (MS3) | 35 | 0.99 | 0.209 | 1.000 | |
| AT5G20980.2 | Methionine synthase 3 (MS3) | 36 | 1.47 | 0.208 | 0.997 | |
| AT5G20980.2 | Methionine synthase 3 (MS3) | 37 | 2.38 | 0.207 | 0.565 | |
| AT5G20980.2 | Methionine synthase 3 (MS3) | 38 | 3.48 | 0.207 | 0.120 | |
| AT5G20980.2 | Methionine synthase 3 (MS3) | 39 | 3.91 | 0.207 | 0.062 | . |
| AT5G20980.2 | Methionine synthase 3 (MS3) | 40 | 3.15 | 0.207 | 0.193 | |
| AT5G20980.2 | Methionine synthase 3 (MS3) | 41 | 1.99 | 0.208 | 0.833 | |
| AT5G20980.2 | Methionine synthase 3 (MS3) | 42 | 1.23 | 0.210 | 1.000 | |
| AT5G20980.2 | Methionine synthase 3 (MS3) | 43 | 0.80 | 0.212 | 1.000 | |
| AT5G20980.2 | Methionine synthase 3 (MS3) | 44 | 0.54 | 0.212 | 0.920 | |
| AT5G20980.2 | Methionine synthase 3 (MS3) | 45 | 0.41 | 0.213 | 0.573 | |
| AT5G20980.2 | Methionine synthase 3 (MS3) | 46 | 0.36 | 0.253 | 0.627 | |
| AT5G20980.2 | Methionine synthase 3 (MS3) | 47 | 0.33 | 0.406 | 0.946 | |
| AT5G22350.1 | Elongated mitochondria 1 (ELM1) | 2 | 0.84 | 0.277 | 1.000 | |
| AT5G22350.1 | Elongated mitochondria 1 (ELM1) | 3 | 1.02 | 0.283 | 1.000 | |
| AT5G22350.1 | Elongated mitochondria 1 (ELM1) | 40 | 1.51 | 0.263 | 0.997 | |
| AT5G22350.1 | Elongated mitochondria 1 (ELM1) | 41 | 1.15 | 0.177 | 1.000 | |
| AT5G22350.1 | Elongated mitochondria 1 (ELM1) | 42 | 1.53 | 0.155 | 0.887 | |
| AT5G22350.1 | Elongated mitochondria 1 (ELM1) | 43 | 6.13 | 0.154 | 0.000 | *** |
| AT5G22350.1 | Elongated mitochondria 1 (ELM1) | 44 | 5.38 | 0.150 | 0.000 | *** |
| AT5G22350.1 | Elongated mitochondria 1 (ELM1) | 45 | 1.30 | 0.148 | 0.991 | |
| AT5G22350.1 | Elongated mitochondria 1 (ELM1) | 46 | 0.24 | 0.168 | 0.005 | ** |
| AT5G22350.1 | Elongated mitochondria 1 (ELM1) | 47 | 0.02 | 0.331 | 0.000 | *** |
| AT5G22800.1 | Alanyl-tRNA synthetase 2 (ALATS2) | 2 | 0.83 | 0.364 | 1.000 | |
| AT5G22800.1 | Alanyl-tRNA synthetase 2 (ALATS2) | 3 | 0.86 | 0.349 | 1.000 | |
| AT5G22800.1 | Alanyl-tRNA synthetase 2 (ALATS2) | 4 | 0.89 | 0.337 | 1.000 | |
| AT5G22800.1 | Alanyl-tRNA synthetase 2 (ALATS2) | 31 | 1.52 | 0.196 | 0.848 | |
| AT5G22800.1 | Alanyl-tRNA synthetase 2 (ALATS2) | 32 | 1.49 | 0.182 | 0.835 | |
| AT5G22800.1 | Alanyl-tRNA synthetase 2 (ALATS2) | 33 | 1.45 | 0.169 | 0.827 | |
| AT5G22800.1 | Alanyl-tRNA synthetase 2 (ALATS2) | 34 | 1.42 | 0.156 | 0.827 | |
| AT5G22800.1 | Alanyl-tRNA synthetase 2 (ALATS2) | 35 | 1.37 | 0.146 | 0.837 | |
| AT5G22800.1 | Alanyl-tRNA synthetase 2 (ALATS2) | 36 | 1.33 | 0.136 | 0.858 | |
| AT5G22800.1 | Alanyl-tRNA synthetase 2 (ALATS2) | 37 | 1.28 | 0.129 | 0.890 | |
| AT5G22800.1 | Alanyl-tRNA synthetase 2 (ALATS2) | 38 | 1.23 | 0.123 | 0.929 | |
| AT5G22800.1 | Alanyl-tRNA synthetase 2 (ALATS2) | 39 | 1.18 | 0.119 | 0.967 | |

| | | | | | |
|---|---|---|---|---|---|
| AT5G22800.1 | Alanyl-tRNA synthetase 2 (ALATS2) | 40 | 1.13 | 0.118 | 0.991 |
| AT5G22800.1 | Alanyl-tRNA synthetase 2 (ALATS2) | 41 | 1.08 | 0.120 | 0.999 |
| AT5G22800.1 | Alanyl-tRNA synthetase 2 (ALATS2) | 42 | 1.03 | 0.125 | 1.000 |
| AT5G22800.1 | Alanyl-tRNA synthetase 2 (ALATS2) | 43 | 0.98 | 0.134 | 1.000 |
| AT5G22800.1 | Alanyl-tRNA synthetase 2 (ALATS2) | 44 | 0.93 | 0.148 | 1.000 |
| AT5G22800.1 | Alanyl-tRNA synthetase 2 (ALATS2) | 45 | 0.89 | 0.166 | 0.999 |
| AT5G22800.1 | Alanyl-tRNA synthetase 2 (ALATS2) | 46 | 0.84 | 0.187 | 0.995 |
| AT5G22800.1 | Alanyl-tRNA synthetase 2 (ALATS2) | 47 | 0.80 | 0.213 | 0.991 |
| AT5G23060.1 | Calcium sensing receptor (CAS) | 2 | 1.12 | 0.452 | 1.000 |
| AT5G23060.1 | Calcium sensing receptor (CAS) | 3 | 0.89 | 0.369 | 1.000 |
| AT5G23060.1 | Calcium sensing receptor (CAS) | 4 | 0.73 | 0.322 | 1.000 |
| AT5G23060.1 | Calcium sensing receptor (CAS) | 5 | 0.62 | 0.300 | 0.999 |
| AT5G23060.1 | Calcium sensing receptor (CAS) | 6 | 0.57 | 0.290 | 1.000 |
| AT5G23060.1 | Calcium sensing receptor (CAS) | 7 | 0.54 | 0.286 | 1.000 |
| AT5G23060.1 | Calcium sensing receptor (CAS) | 8 | 0.55 | 0.284 | 1.000 |
| AT5G23060.1 | Calcium sensing receptor (CAS) | 9 | 0.61 | 0.283 | 0.997 |
| AT5G23060.1 | Calcium sensing receptor (CAS) | 10 | 0.72 | 0.281 | 1.000 |
| AT5G23060.1 | Calcium sensing receptor (CAS) | 11 | 0.88 | 0.280 | 1.000 |
| AT5G23060.1 | Calcium sensing receptor (CAS) | 12 | 1.07 | 0.279 | 1.000 |
| AT5G23060.1 | Calcium sensing receptor (CAS) | 13 | 1.28 | 0.279 | 1.000 |
| AT5G23060.1 | Calcium sensing receptor (CAS) | 14 | 1.48 | 0.278 | 1.000 |
| AT5G23060.1 | Calcium sensing receptor (CAS) | 15 | 1.63 | 0.277 | 0.997 |
| AT5G23060.1 | Calcium sensing receptor (CAS) | 16 | 1.72 | 0.277 | 1.000 |
| AT5G23060.1 | Calcium sensing receptor (CAS) | 17 | 1.79 | 0.278 | 1.000 |
| AT5G23060.1 | Calcium sensing receptor (CAS) | 18 | 1.84 | 0.279 | 1.000 |
| AT5G23060.1 | Calcium sensing receptor (CAS) | 19 | 1.81 | 0.281 | 1.000 |
| AT5G23060.1 | Calcium sensing receptor (CAS) | 20 | 1.68 | 0.284 | 0.996 |
| AT5G23060.1 | Calcium sensing receptor (CAS) | 21 | 1.44 | 0.287 | 1.000 |
| AT5G23060.1 | Calcium sensing receptor (CAS) | 22 | 1.15 | 0.289 | 1.000 |
| AT5G23060.1 | Calcium sensing receptor (CAS) | 23 | 0.89 | 0.291 | 1.000 |
| AT5G23060.1 | Calcium sensing receptor (CAS) | 24 | 0.68 | 0.291 | 1.000 |
| AT5G23060.1 | Calcium sensing receptor (CAS) | 25 | 0.52 | 0.289 | 1.000 |
| AT5G23060.1 | Calcium sensing receptor (CAS) | 26 | 0.42 | 0.286 | 0.839 |
| AT5G23060.1 | Calcium sensing receptor (CAS) | 27 | 0.36 | 0.284 | 0.678 |
| AT5G23060.1 | Calcium sensing receptor (CAS) | 28 | 0.34 | 0.281 | 0.595 |
| AT5G23060.1 | Calcium sensing receptor (CAS) | 29 | 0.34 | 0.281 | 0.596 |
| AT5G23060.1 | Calcium sensing receptor (CAS) | 30 | 0.36 | 0.282 | 0.671 |
| AT5G23060.1 | Calcium sensing receptor (CAS) | 31 | 0.41 | 0.286 | 0.817 |
| AT5G23060.1 | Calcium sensing receptor (CAS) | 32 | 0.48 | 0.295 | 1.000 |
| AT5G23060.1 | Calcium sensing receptor (CAS) | 33 | 0.58 | 0.311 | 0.997 |
| AT5G23060.1 | Calcium sensing receptor (CAS) | 34 | 0.71 | 0.336 | 1.000 |
| AT5G23060.1 | Calcium sensing receptor (CAS) | 35 | 0.84 | 0.371 | 1.000 |
| AT5G23060.1 | Calcium sensing receptor (CAS) | 43 | 0.89 | 0.387 | 1.000 |
| AT5G23060.1 | Calcium sensing receptor (CAS) | 44 | 0.85 | 0.348 | 1.000 |
| AT5G23060.1 | Calcium sensing receptor (CAS) | 45 | 0.82 | 0.328 | 1.000 |
| AT5G23060.1 | Calcium sensing receptor (CAS) | 46 | 0.81 | 0.360 | 1.000 |
| AT5G23060.1 | Calcium sensing receptor (CAS) | 47 | 0.80 | 0.454 | 1.000 |
| AT5G23140.1 | Caseinolytic protease P7 (CLPP7) | 2 | 0.94 | 0.512 | 1.000 |
| AT5G23140.1 | Caseinolytic protease P7 (CLPP7) | 3 | 0.90 | 0.463 | 1.000 |
| AT5G23140.1 | Caseinolytic protease P7 (CLPP7) | 15 | 1.62 | 0.306 | 1.000 |
| AT5G23140.1 | Caseinolytic protease P7 (CLPP7) | 16 | 1.69 | 0.279 | 1.000 |

| | | | | | |
|---|---|---|---|---|---|
| AT5G23140.1 | Caseinolytic protease P7 (CLPP7) | 17 | 1.42 | 0.249 | 1.000 |
| AT5G23140.1 | Caseinolytic protease P7 (CLPP7) | 18 | 1.07 | 0.234 | 1.000 |
| AT5G23140.1 | Caseinolytic protease P7 (CLPP7) | 19 | 0.94 | 0.236 | 1.000 |
| AT5G23140.1 | Caseinolytic protease P7 (CLPP7) | 20 | 0.90 | 0.241 | 1.000 |
| AT5G23140.1 | Caseinolytic protease P7 (CLPP7) | 21 | 0.85 | 0.243 | 1.000 |
| AT5G23140.1 | Caseinolytic protease P7 (CLPP7) | 22 | 0.94 | 0.242 | 1.000 |
| AT5G23140.1 | Caseinolytic protease P7 (CLPP7) | 23 | 1.11 | 0.241 | 1.000 |
| AT5G23140.1 | Caseinolytic protease P7 (CLPP7) | 24 | 1.33 | 0.236 | 1.000 |
| AT5G23140.1 | Caseinolytic protease P7 (CLPP7) | 25 | 1.43 | 0.231 | 1.000 |
| AT5G23140.1 | Caseinolytic protease P7 (CLPP7) | 26 | 1.06 | 0.230 | 1.000 |
| AT5G23140.1 | Caseinolytic protease P7 (CLPP7) | 27 | 0.67 | 0.230 | 1.000 |
| AT5G23140.1 | Caseinolytic protease P7 (CLPP7) | 28 | 0.69 | 0.230 | 1.000 |
| AT5G23140.1 | Caseinolytic protease P7 (CLPP7) | 29 | 1.20 | 0.230 | 1.000 |
| AT5G23140.1 | Caseinolytic protease P7 (CLPP7) | 30 | 2.75 | 0.232 | 0.693 |
| AT5G23140.1 | Caseinolytic protease P7 (CLPP7) | 31 | 4.09 | 0.233 | 0.186 |
| AT5G23140.1 | Caseinolytic protease P7 (CLPP7) | 32 | 2.34 | 0.233 | 0.902 |
| AT5G23140.1 | Caseinolytic protease P7 (CLPP7) | 33 | 0.91 | 0.233 | 1.000 |
| AT5G23140.1 | Caseinolytic protease P7 (CLPP7) | 34 | 0.69 | 0.234 | 1.000 |
| AT5G23140.1 | Caseinolytic protease P7 (CLPP7) | 35 | 1.00 | 0.233 | 1.000 |
| AT5G23140.1 | Caseinolytic protease P7 (CLPP7) | 36 | 1.56 | 0.232 | 1.000 |
| AT5G23140.1 | Caseinolytic protease P7 (CLPP7) | 37 | 1.88 | 0.231 | 0.994 |
| AT5G23140.1 | Caseinolytic protease P7 (CLPP7) | 38 | 1.66 | 0.231 | 1.000 |
| AT5G23140.1 | Caseinolytic protease P7 (CLPP7) | 39 | 1.42 | 0.231 | 1.000 |
| AT5G23140.1 | Caseinolytic protease P7 (CLPP7) | 40 | 1.41 | 0.233 | 1.000 |
| AT5G23140.1 | Caseinolytic protease P7 (CLPP7) | 41 | 1.45 | 0.243 | 1.000 |
| AT5G23140.1 | Caseinolytic protease P7 (CLPP7) | 42 | 1.42 | 0.265 | 1.000 |
| AT5G23140.1 | Caseinolytic protease P7 (CLPP7) | 43 | 1.68 | 0.266 | 1.000 |
| AT5G23140.1 | Caseinolytic protease P7 (CLPP7) | 44 | 2.24 | 0.255 | 0.969 |
| AT5G23140.1 | Caseinolytic protease P7 (CLPP7) | 45 | 2.63 | 0.244 | 0.822 |
| AT5G23140.1 | Caseinolytic protease P7 (CLPP7) | 46 | 2.12 | 0.275 | 0.993 |
| AT5G23140.1 | Caseinolytic protease P7 (CLPP7) | 47 | 1.39 | 0.453 | 1.000 |

| | | | | | |
|---|---|---|---|---|---|
| AT5G23250.1 | Succinyl-CoA ligase subunit (SCLalpha) | 2 | 0.70 | 0.403 | 1.000 |
| AT5G23250.1 | Succinyl-CoA ligase subunit (SCLalpha) | 3 | 0.78 | 0.391 | 1.000 |
| AT5G23250.1 | Succinyl-CoA ligase subunit (SCLalpha) | 27 | 1.18 | 0.361 | 1.000 |
| AT5G23250.1 | Succinyl-CoA ligase subunit (SCLalpha) | 28 | 0.97 | 0.288 | 1.000 |
| AT5G23250.1 | Succinyl-CoA ligase subunit (SCLalpha) | 29 | 0.72 | 0.272 | 1.000 |
| AT5G23250.1 | Succinyl-CoA ligase subunit (SCLalpha) | 30 | 0.54 | 0.255 | 0.984 |
| AT5G23250.1 | Succinyl-CoA ligase subunit (SCLalpha) | 31 | 0.63 | 0.250 | 0.999 |
| AT5G23250.1 | Succinyl-CoA ligase subunit (SCLalpha) | 32 | 0.86 | 0.245 | 1.000 |
| AT5G23250.1 | Succinyl-CoA ligase subunit (SCLalpha) | 33 | 0.85 | 0.242 | 1.000 |
| AT5G23250.1 | Succinyl-CoA ligase subunit (SCLalpha) | 34 | 0.79 | 0.243 | 1.000 |
| AT5G23250.1 | Succinyl-CoA ligase subunit (SCLalpha) | 35 | 0.84 | 0.242 | 1.000 |
| AT5G23250.1 | Succinyl-CoA ligase subunit (SCLalpha) | 36 | 1.15 | 0.240 | 1.000 |
| AT5G23250.1 | Succinyl-CoA ligase subunit (SCLalpha) | 37 | 1.26 | 0.240 | 1.000 |
| AT5G23250.1 | Succinyl-CoA ligase subunit (SCLalpha) | 38 | 1.06 | 0.242 | 1.000 |
| AT5G23250.1 | Succinyl-CoA ligase subunit (SCLalpha) | 39 | 1.61 | 0.243 | 0.998 |
| AT5G23250.1 | Succinyl-CoA ligase subunit (SCLalpha) | 40 | 3.35 | 0.242 | 0.330 |
| AT5G23250.1 | Succinyl-CoA ligase subunit (SCLalpha) | 41 | 4.31 | 0.241 | 0.123 |
| AT5G23250.1 | Succinyl-CoA ligase subunit (SCLalpha) | 42 | 1.98 | 0.244 | 0.951 |
| AT5G23250.1 | Succinyl-CoA ligase subunit (SCLalpha) | 43 | 0.76 | 0.244 | 1.000 |
| AT5G23250.1 | Succinyl-CoA ligase subunit (SCLalpha) | 44 | 0.58 | 0.243 | 0.993 |
| AT5G23250.1 | Succinyl-CoA ligase subunit (SCLalpha) | 45 | 0.60 | 0.244 | 0.996 |
| AT5G23250.1 | Succinyl-CoA ligase subunit (SCLalpha) | 46 | 0.54 | 0.258 | 0.987 |
| AT5G23250.1 | Succinyl-CoA ligase subunit (SCLalpha) | 47 | 0.39 | 0.454 | 0.996 |
| AT5G23300.1 | Pyrimidine D (PYRD) | 2 | 1.08 | 0.290 | 1.000 |
| AT5G23300.1 | Pyrimidine D (PYRD) | 3 | 0.98 | 0.196 | 1.000 |
| AT5G23300.1 | Pyrimidine D (PYRD) | 4 | 0.89 | 0.170 | 1.000 |
| AT5G23300.1 | Pyrimidine D (PYRD) | 5 | 0.81 | 0.147 | 1.000 |
| AT5G23300.1 | Pyrimidine D (PYRD) | 6 | 0.74 | 0.136 | 1.000 |
| AT5G23300.1 | Pyrimidine D (PYRD) | 7 | 0.69 | 0.132 | 0.999 |
| AT5G23300.1 | Pyrimidine D (PYRD) | 8 | 0.72 | 0.127 | 1.000 |
| AT5G23300.1 | Pyrimidine D (PYRD) | 9 | 0.81 | 0.125 | 1.000 |
| AT5G23300.1 | Pyrimidine D (PYRD) | 10 | 0.84 | 0.125 | 1.000 |
| AT5G23300.1 | Pyrimidine D (PYRD) | 11 | 0.84 | 0.125 | 1.000 |
| AT5G23300.1 | Pyrimidine D (PYRD) | 12 | 0.81 | 0.125 | 1.000 |
| AT5G23300.1 | Pyrimidine D (PYRD) | 13 | 0.78 | 0.123 | 1.000 |
| AT5G23300.1 | Pyrimidine D (PYRD) | 14 | 0.79 | 0.122 | 1.000 |
| AT5G23300.1 | Pyrimidine D (PYRD) | 15 | 0.81 | 0.122 | 1.000 |
| AT5G23300.1 | Pyrimidine D (PYRD) | 16 | 0.81 | 0.121 | 1.000 |
| AT5G23300.1 | Pyrimidine D (PYRD) | 17 | 0.89 | 0.121 | 1.000 |

| | | | | | |
|---|---|---|---|---|---|
| AT5G23300.1 | Pyrimidine D (PYRD) | 18 | 1.07 | 0.122 | 1.000 |
| AT5G23300.1 | Pyrimidine D (PYRD) | 19 | 1.29 | 0.126 | 1.000 |
| AT5G23300.1 | Pyrimidine D (PYRD) | 20 | 1.38 | 0.129 | 1.000 |
| AT5G23300.1 | Pyrimidine D (PYRD) | 21 | 1.30 | 0.131 | 1.000 |
| AT5G23300.1 | Pyrimidine D (PYRD) | 22 | 1.14 | 0.131 | 1.000 |
| AT5G23300.1 | Pyrimidine D (PYRD) | 23 | 0.96 | 0.130 | 1.000 |
| AT5G23300.1 | Pyrimidine D (PYRD) | 24 | 0.79 | 0.127 | 1.000 |
| AT5G23300.1 | Pyrimidine D (PYRD) | 25 | 0.62 | 0.125 | 0.937 |
| AT5G23300.1 | Pyrimidine D (PYRD) | 26 | 0.53 | 0.124 | 0.555 |
| AT5G23300.1 | Pyrimidine D (PYRD) | 27 | 0.57 | 0.124 | 0.772 |
| AT5G23300.1 | Pyrimidine D (PYRD) | 28 | 0.66 | 0.124 | 0.988 |
| AT5G23300.1 | Pyrimidine D (PYRD) | 29 | 0.68 | 0.124 | 0.995 |
| AT5G23300.1 | Pyrimidine D (PYRD) | 30 | 0.65 | 0.124 | 0.979 |
| AT5G23300.1 | Pyrimidine D (PYRD) | 31 | 0.68 | 0.125 | 0.996 |
| AT5G23300.1 | Pyrimidine D (PYRD) | 32 | 0.73 | 0.125 | 1.000 |
| AT5G23300.1 | Pyrimidine D (PYRD) | 33 | 0.77 | 0.125 | 1.000 |
| AT5G23300.1 | Pyrimidine D (PYRD) | 34 | 0.75 | 0.125 | 1.000 |
| AT5G23300.1 | Pyrimidine D (PYRD) | 35 | 0.74 | 0.125 | 1.000 |
| AT5G23300.1 | Pyrimidine D (PYRD) | 36 | 0.76 | 0.125 | 1.000 |
| AT5G23300.1 | Pyrimidine D (PYRD) | 37 | 0.85 | 0.124 | 1.000 |
| AT5G23300.1 | Pyrimidine D (PYRD) | 38 | 0.97 | 0.125 | 1.000 |
| AT5G23300.1 | Pyrimidine D (PYRD) | 39 | 1.09 | 0.125 | 1.000 |
| AT5G23300.1 | Pyrimidine D (PYRD) | 40 | 1.16 | 0.125 | 1.000 |
| AT5G23300.1 | Pyrimidine D (PYRD) | 41 | 1.25 | 0.126 | 1.000 |
| AT5G23300.1 | Pyrimidine D (PYRD) | 42 | 1.44 | 0.128 | 0.998 |
| AT5G23300.1 | Pyrimidine D (PYRD) | 43 | 1.74 | 0.129 | 0.851 |
| AT5G23300.1 | Pyrimidine D (PYRD) | 44 | 1.74 | 0.129 | 0.852 |
| AT5G23300.1 | Pyrimidine D (PYRD) | 45 | 1.21 | 0.129 | 1.000 |
| AT5G23300.1 | Pyrimidine D (PYRD) | 46 | 0.61 | 0.153 | 0.993 |
| AT5G23300.1 | Pyrimidine D (PYRD) | 47 | 0.26 | 0.267 | 0.573 |
| AT5G23535.1 | 50S ribosomal protein L24 | 2 | 0.95 | 0.377 | 1.000 |
| AT5G23535.1 | 50S ribosomal protein L24 | 3 | 0.99 | 0.295 | 1.000 |
| AT5G23535.1 | 50S ribosomal protein L24 | 4 | 1.03 | 0.280 | 1.000 |
| AT5G23535.1 | 50S ribosomal protein L24 | 6 | 1.15 | 0.306 | 1.000 |
| AT5G23535.1 | 50S ribosomal protein L24 | 7 | 1.21 | 0.306 | 1.000 |
| AT5G23535.1 | 50S ribosomal protein L24 | 8 | 1.27 | 0.294 | 1.000 |
| AT5G23535.1 | 50S ribosomal protein L24 | 10 | 1.28 | 0.260 | 1.000 |
| AT5G23535.1 | 50S ribosomal protein L24 | 11 | 1.18 | 0.242 | 1.000 |
| AT5G23535.1 | 50S ribosomal protein L24 | 12 | 1.00 | 0.223 | 1.000 |
| AT5G23535.1 | 50S ribosomal protein L24 | 13 | 0.79 | 0.205 | 1.000 |
| AT5G23535.1 | 50S ribosomal protein L24 | 14 | 0.59 | 0.191 | 0.976 |
| AT5G23535.1 | 50S ribosomal protein L24 | 15 | 0.46 | 0.183 | 0.668 |
| AT5G23535.1 | 50S ribosomal protein L24 | 16 | 0.42 | 0.179 | 0.442 |
| AT5G23535.1 | 50S ribosomal protein L24 | 17 | 0.45 | 0.178 | 0.563 |
| AT5G23535.1 | 50S ribosomal protein L24 | 18 | 0.53 | 0.180 | 0.856 |
| AT5G23535.1 | 50S ribosomal protein L24 | 19 | 0.63 | 0.183 | 0.986 |
| AT5G23535.1 | 50S ribosomal protein L24 | 20 | 0.72 | 0.187 | 1.000 |
| AT5G23535.1 | 50S ribosomal protein L24 | 21 | 0.80 | 0.191 | 1.000 |
| AT5G23535.1 | 50S ribosomal protein L24 | 22 | 0.86 | 0.196 | 1.000 |
| AT5G23535.1 | 50S ribosomal protein L24 | 23 | 0.86 | 0.204 | 1.000 |
| AT5G23535.1 | 50S ribosomal protein L24 | 24 | 0.85 | 0.217 | 1.000 |
| AT5G23535.1 | 50S ribosomal protein L24 | 25 | 0.84 | 0.235 | 1.000 |
| AT5G23535.1 | 50S ribosomal protein L24 | 26 | 0.86 | 0.254 | 1.000 |
| AT5G23535.1 | 50S ribosomal protein L24 | 27 | 0.87 | 0.264 | 1.000 |
| AT5G23535.1 | 50S ribosomal protein L24 | 28 | 0.88 | 0.267 | 1.000 |
| AT5G23535.1 | 50S ribosomal protein L24 | 30 | 0.86 | 0.253 | 1.000 |
| AT5G23535.1 | 50S ribosomal protein L24 | 31 | 0.85 | 0.244 | 1.000 |
| AT5G23535.1 | 50S ribosomal protein L24 | 32 | 0.87 | 0.242 | 1.000 |
| AT5G23535.1 | 50S ribosomal protein L24 | 33 | 0.92 | 0.258 | 1.000 |
| AT5G23535.1 | 50S ribosomal protein L24 | 34 | 0.98 | 0.298 | 1.000 |
| AT5G23535.1 | 50S ribosomal protein L24 | 41 | 1.05 | 0.401 | 1.000 |
| AT5G23535.1 | 50S ribosomal protein L24 | 42 | 1.03 | 0.406 | 1.000 |
| AT5G23535.1 | 50S ribosomal protein L24 | 43 | 1.02 | 0.454 | 1.000 |

| | | | | | |
|---|---|---|---|---|---|
| AT5G23535.1 | 50S ribosomal protein L24 | 46 | 1.01 | 0.893 | 1.000 |
| AT5G23535.1 | 50S ribosomal protein L24 | 47 | 1.01 | 1.110 | 1.000 |
| AT5G24165.1 | ? | 2 | 0.79 | 0.409 | 1.000 |
| AT5G24165.1 | ? | 3 | 0.83 | 0.271 | 1.000 |
| AT5G24165.1 | ? | 4 | 0.86 | 0.241 | 1.000 |
| AT5G24165.1 | ? | 5 | 0.88 | 0.242 | 1.000 |
| AT5G24165.1 | ? | 6 | 0.90 | 0.245 | 1.000 |
| AT5G24165.1 | ? | 7 | 0.96 | 0.249 | 1.000 |
| AT5G24165.1 | ? | 8 | 1.06 | 0.248 | 1.000 |
| AT5G24165.1 | ? | 9 | 1.16 | 0.248 | 1.000 |
| AT5G24165.1 | ? | 10 | 1.33 | 0.248 | 1.000 |
| AT5G24165.1 | ? | 11 | 1.51 | 0.246 | 1.000 |
| AT5G24165.1 | ? | 12 | 1.42 | 0.240 | 1.000 |
| AT5G24165.1 | ? | 13 | 1.06 | 0.234 | 1.000 |
| AT5G24165.1 | ? | 14 | 0.69 | 0.230 | 1.000 |
| AT5G24165.1 | ? | 15 | 0.50 | 0.229 | 0.988 |
| AT5G24165.1 | ? | 16 | 0.57 | 0.227 | 0.999 |
| AT5G24165.1 | ? | 17 | 1.10 | 0.226 | 1.000 |
| AT5G24165.1 | ? | 18 | 2.31 | 0.229 | 0.928 |
| AT5G24165.1 | ? | 19 | 3.53 | 0.235 | 0.397 |
| AT5G24165.1 | ? | 20 | 3.53 | 0.239 | 0.420 |
| AT5G24165.1 | ? | 21 | 2.53 | 0.240 | 0.881 |
| AT5G24165.1 | ? | 22 | 1.62 | 0.240 | 1.000 |
| AT5G24165.1 | ? | 23 | 1.28 | 0.238 | 1.000 |
| AT5G24165.1 | ? | 24 | 1.31 | 0.235 | 1.000 |
| AT5G24165.1 | ? | 25 | 1.52 | 0.233 | 1.000 |
| AT5G24165.1 | ? | 26 | 1.78 | 0.232 | 0.999 |
| AT5G24165.1 | ? | 27 | 1.94 | 0.231 | 0.994 |
| AT5G24165.1 | ? | 28 | 1.81 | 0.231 | 0.999 |
| AT5G24165.1 | ? | 29 | 1.56 | 0.232 | 1.000 |
| AT5G24165.1 | ? | 30 | 1.43 | 0.233 | 1.000 |
| AT5G24165.1 | ? | 31 | 1.36 | 0.233 | 1.000 |
| AT5G24165.1 | ? | 32 | 1.25 | 0.234 | 1.000 |
| AT5G24165.1 | ? | 33 | 1.09 | 0.234 | 1.000 |
| AT5G24165.1 | ? | 34 | 0.98 | 0.234 | 1.000 |
| AT5G24165.1 | ? | 35 | 0.94 | 0.234 | 1.000 |
| AT5G24165.1 | ? | 36 | 0.92 | 0.234 | 1.000 |
| AT5G24165.1 | ? | 37 | 0.95 | 0.233 | 1.000 |
| AT5G24165.1 | ? | 38 | 1.02 | 0.234 | 1.000 |
| AT5G24165.1 | ? | 39 | 1.12 | 0.234 | 1.000 |
| AT5G24165.1 | ? | 40 | 1.19 | 0.234 | 1.000 |
| AT5G24165.1 | ? | 41 | 1.14 | 0.234 | 1.000 |
| AT5G24165.1 | ? | 42 | 1.00 | 0.238 | 1.000 |
| AT5G24165.1 | ? | 43 | 0.87 | 0.241 | 1.000 |
| AT5G24165.1 | ? | 44 | 0.74 | 0.239 | 1.000 |
| AT5G24165.1 | ? | 45 | 0.61 | 0.241 | 1.000 |
| AT5G24165.1 | ? | 46 | 0.50 | 0.291 | 1.000 |
| AT5G24165.1 | ? | 47 | 0.41 | 0.473 | 1.000 |
| AT5G24970.1 | Protein kinase superfamily protein | 2 | 1.03 | 0.378 | 1.000 |
| AT5G24970.1 | Protein kinase superfamily protein | 3 | 0.95 | 0.293 | 1.000 |
| AT5G24970.1 | Protein kinase superfamily protein | 4 | 0.90 | 0.538 | 1.000 |
| AT5G24970.1 | Protein kinase superfamily protein | 28 | 1.63 | 0.266 | 0.998 |
| AT5G24970.1 | Protein kinase superfamily protein | 29 | 1.55 | 0.197 | 0.990 |
| AT5G24970.1 | Protein kinase superfamily protein | 30 | 1.55 | 0.177 | 0.978 |
| AT5G24970.1 | Protein kinase superfamily protein | 31 | 1.58 | 0.168 | 0.952 |

| | | | | | |
|---|---|---|---|---|---|
| AT5G24970.1 | Protein kinase superfamily protein | 32 | 1.49 | 0.165 | 0.980 |
| AT5G24970.1 | Protein kinase superfamily protein | 33 | 1.28 | 0.165 | 1.000 |
| AT5G24970.1 | Protein kinase superfamily protein | 34 | 1.07 | 0.165 | 1.000 |
| AT5G24970.1 | Protein kinase superfamily protein | 35 | 0.94 | 0.165 | 1.000 |
| AT5G24970.1 | Protein kinase superfamily protein | 36 | 0.91 | 0.165 | 1.000 |
| AT5G24970.1 | Protein kinase superfamily protein | 37 | 0.97 | 0.168 | 1.000 |
| AT5G24970.1 | Protein kinase superfamily protein | 38 | 0.97 | 0.180 | 1.000 |
| AT5G24970.1 | Protein kinase superfamily protein | 39 | 0.92 | 0.229 | 1.000 |
| AT5G24970.1 | Protein kinase superfamily protein | 40 | 1.07 | 0.351 | 1.000 |
| AT5G24970.1 | Protein kinase superfamily protein | 46 | 328.15 | 2.001 | 0.931 |
| AT5G24970.1 | Protein kinase superfamily protein | 47 | 1248.35 | 2.423 | 0.924 |
| AT5G25100.1 | Putative TM9-type membrane protein (EMP2/TMN9) | 2 | 0.91 | 0.515 | 1.000 |
| AT5G25100.1 | Putative TM9-type membrane protein (EMP2/TMN9) | 3 | 0.98 | 0.380 | 1.000 |
| AT5G25100.1 | Putative TM9-type membrane protein (EMP2/TMN9) | 4 | 1.06 | 0.343 | 1.000 |
| AT5G25100.1 | Putative TM9-type membrane protein (EMP2/TMN9) | 5 | 1.15 | 0.369 | 1.000 |
| AT5G25100.1 | Putative TM9-type membrane protein (EMP2/TMN9) | 6 | 1.26 | 0.433 | 1.000 |
| AT5G25100.1 | Putative TM9-type membrane protein (EMP2/TMN9) | 7 | 1.36 | 0.552 | 1.000 |
| AT5G25100.1 | Putative TM9-type membrane protein (EMP2/TMN9) | 30 | 0.67 | 0.379 | 1.000 |
| AT5G25100.1 | Putative TM9-type membrane protein (EMP2/TMN9) | 31 | 0.68 | 0.287 | 1.000 |
| AT5G25100.1 | Putative TM9-type membrane protein (EMP2/TMN9) | 32 | 0.73 | 0.248 | 1.000 |
| AT5G25100.1 | Putative TM9-type membrane protein (EMP2/TMN9) | 33 | 0.87 | 0.235 | 1.000 |
| AT5G25100.1 | Putative TM9-type membrane protein (EMP2/TMN9) | 34 | 1.08 | 0.231 | 1.000 |
| AT5G25100.1 | Putative TM9-type membrane protein (EMP2/TMN9) | 35 | 1.19 | 0.229 | 1.000 |
| AT5G25100.1 | Putative TM9-type membrane protein (EMP2/TMN9) | 36 | 0.99 | 0.228 | 1.000 |
| AT5G25100.1 | Putative TM9-type membrane protein (EMP2/TMN9) | 37 | 0.63 | 0.228 | 0.988 |
| AT5G25100.1 | Putative TM9-type membrane protein (EMP2/TMN9) | 38 | 0.37 | 0.235 | 0.518 |
| AT5G25100.1 | Putative TM9-type membrane protein (EMP2/TMN9) | 39 | 0.23 | 0.265 | 0.191 |
| AT5G25100.1 | Putative TM9-type membrane protein (EMP2/TMN9) | 40 | 0.17 | 0.340 | 0.253 |
| AT5G25100.1 | Putative TM9-type membrane protein (EMP2/TMN9) | 41 | 0.15 | 0.458 | 0.525 |
| AT5G25100.1 | Putative TM9-type membrane protein (EMP2/TMN9) | 46 | 0.22 | 1.226 | 1.000 |
| AT5G25100.1 | Putative TM9-type membrane protein (EMP2/TMN9) | 47 | 0.28 | 1.464 | 1.000 |

| | | | | | |
|---|---|---|---|---|---|
| AT5G25940.1 | Early nodulin-related | 2 | 0.88 | 0.409 | 1.000 |
| AT5G25940.1 | Early nodulin-related | 3 | 0.92 | 0.274 | 1.000 |
| AT5G25940.1 | Early nodulin-related | 4 | 1.03 | 0.252 | 1.000 |
| AT5G25940.1 | Early nodulin-related | 5 | 1.31 | 0.218 | 1.000 |
| AT5G25940.1 | Early nodulin-related | 6 | 1.61 | 0.201 | 1.000 |
| AT5G25940.1 | Early nodulin-related | 7 | 1.78 | 0.195 | 0.998 |
| AT5G25940.1 | Early nodulin-related | 8 | 1.89 | 0.189 | 0.987 |
| AT5G25940.1 | Early nodulin-related | 9 | 1.83 | 0.186 | 0.992 |
| AT5G25940.1 | Early nodulin-related | 10 | 1.59 | 0.186 | 1.000 |
| AT5G25940.1 | Early nodulin-related | 11 | 1.43 | 0.187 | 1.000 |
| AT5G25940.1 | Early nodulin-related | 12 | 1.38 | 0.188 | 1.000 |
| AT5G25940.1 | Early nodulin-related | 13 | 1.37 | 0.185 | 1.000 |
| AT5G25940.1 | Early nodulin-related | 14 | 1.39 | 0.183 | 1.000 |
| AT5G25940.1 | Early nodulin-related | 15 | 1.31 | 0.183 | 1.000 |
| AT5G25940.1 | Early nodulin-related | 16 | 1.18 | 0.182 | 1.000 |
| AT5G25940.1 | Early nodulin-related | 17 | 1.25 | 0.180 | 1.000 |
| AT5G25940.1 | Early nodulin-related | 18 | 1.67 | 0.181 | 0.999 |
| AT5G25940.1 | Early nodulin-related | 19 | 2.53 | 0.187 | 0.629 |
| AT5G25940.1 | Early nodulin-related | 20 | 3.52 | 0.192 | 0.152 |
| AT5G25940.1 | Early nodulin-related | 21 | 3.59 | 0.194 | 0.143 |
| AT5G25940.1 | Early nodulin-related | 22 | 2.89 | 0.194 | 0.440 |
| AT5G25940.1 | Early nodulin-related | 23 | 2.31 | 0.192 | 0.831 |
| AT5G25940.1 | Early nodulin-related | 24 | 1.94 | 0.187 | 0.976 |
| AT5G25940.1 | Early nodulin-related | 25 | 1.58 | 0.185 | 1.000 |
| AT5G25940.1 | Early nodulin-related | 26 | 1.43 | 0.185 | 1.000 |
| AT5G25940.1 | Early nodulin-related | 27 | 1.55 | 0.186 | 1.000 |
| AT5G25940.1 | Early nodulin-related | 28 | 1.66 | 0.187 | 0.999 |
| AT5G25940.1 | Early nodulin-related | 29 | 1.66 | 0.187 | 0.999 |
| AT5G25940.1 | Early nodulin-related | 30 | 1.32 | 0.187 | 1.000 |
| AT5G25940.1 | Early nodulin-related | 31 | 0.97 | 0.187 | 1.000 |
| AT5G25940.1 | Early nodulin-related | 32 | 0.86 | 0.187 | 1.000 |
| AT5G25940.1 | Early nodulin-related | 33 | 0.71 | 0.187 | 1.000 |
| AT5G25940.1 | Early nodulin-related | 34 | 0.56 | 0.187 | 0.997 |
| AT5G25940.1 | Early nodulin-related | 35 | 0.58 | 0.189 | 0.998 |
| AT5G25940.1 | Early nodulin-related | 36 | 0.78 | 0.194 | 1.000 |
| AT5G25940.1 | Early nodulin-related | 37 | 1.32 | 0.202 | 1.000 |
| AT5G25940.1 | Early nodulin-related | 38 | 2.13 | 0.210 | 0.972 |
| AT5G25940.1 | Early nodulin-related | 39 | 2.07 | 0.203 | 0.972 |
| AT5G25940.1 | Early nodulin-related | 40 | 1.47 | 0.193 | 1.000 |
| AT5G25940.1 | Early nodulin-related | 41 | 1.09 | 0.189 | 1.000 |
| AT5G25940.1 | Early nodulin-related | 42 | 1.11 | 0.191 | 1.000 |
| AT5G25940.1 | Early nodulin-related | 43 | 1.61 | 0.192 | 1.000 |
| AT5G25940.1 | Early nodulin-related | 44 | 2.23 | 0.192 | 0.879 |
| AT5G25940.1 | Early nodulin-related | 45 | 2.31 | 0.192 | 0.832 |
| AT5G25940.1 | Early nodulin-related | 46 | 1.87 | 0.218 | 0.999 |
| AT5G25940.1 | Early nodulin-related | 47 | 1.56 | 0.399 | 1.000 |
| AT5G26000.2 | Thioglucoside glucohydrolase 1 (TGG1) | 2 | 0.97 | 0.598 | 1.000 |
| AT5G26000.2 | Thioglucoside glucohydrolase 1 (TGG1) | 3 | 0.98 | 0.589 | 1.000 |
| AT5G26000.2 | Thioglucoside glucohydrolase 1 (TGG1) | 28 | 0.78 | 0.435 | 1.000 |
| AT5G26000.2 | Thioglucoside glucohydrolase 1 (TGG1) | 29 | 0.70 | 0.313 | 1.000 |
| AT5G26000.2 | Thioglucoside glucohydrolase 1 (TGG1) | 30 | 0.62 | 0.281 | 1.000 |
| AT5G26000.2 | Thioglucoside glucohydrolase 1 (TGG1) | 31 | 0.65 | 0.268 | 1.000 |
| AT5G26000.2 | Thioglucoside glucohydrolase 1 (TGG1) | 32 | 0.81 | 0.268 | 1.000 |
| AT5G26000.2 | Thioglucoside glucohydrolase 1 (TGG1) | 33 | 1.01 | 0.285 | 1.000 |

| | | | | | |
|---|---|---|---|---|---|
| AT5G26000.2 | Thioglucoside glucohydrolase 1 (TGG1) | 34 | 0.95 | 0.329 | 1.000 |
| AT5G26000.2 | Thioglucoside glucohydrolase 1 (TGG1) | 35 | 0.76 | 0.368 | 1.000 |
| AT5G26000.2 | Thioglucoside glucohydrolase 1 (TGG1) | 37 | 0.71 | 0.362 | 1.000 |
| AT5G26000.2 | Thioglucoside glucohydrolase 1 (TGG1) | 38 | 1.06 | 0.312 | 1.000 |
| AT5G26000.2 | Thioglucoside glucohydrolase 1 (TGG1) | 39 | 1.89 | 0.275 | 0.992 |
| AT5G26000.2 | Thioglucoside glucohydrolase 1 (TGG1) | 40 | 2.22 | 0.261 | 0.933 |
| AT5G26000.2 | Thioglucoside glucohydrolase 1 (TGG1) | 41 | 1.34 | 0.262 | 1.000 |
| AT5G26000.2 | Thioglucoside glucohydrolase 1 (TGG1) | 42 | 0.78 | 0.266 | 1.000 |
| AT5G26000.2 | Thioglucoside glucohydrolase 1 (TGG1) | 43 | 0.79 | 0.268 | 1.000 |
| AT5G26000.2 | Thioglucoside glucohydrolase 1 (TGG1) | 44 | 1.06 | 0.273 | 1.000 |
| AT5G26000.2 | Thioglucoside glucohydrolase 1 (TGG1) | 45 | 1.35 | 0.333 | 1.000 |
| AT5G26000.2 | Thioglucoside glucohydrolase 1 (TGG1) | 46 | 1.86 | 0.495 | 1.000 |
| AT5G26000.2 | Thioglucoside glucohydrolase 1 (TGG1) | 47 | 2.83 | 0.832 | 1.000 |
| AT5G26780.1 | Serine hydroxymethyltransferase 2 (SHMT2) | 3 | 0.73 | 0.176 | 1.000 |
| AT5G26780.1 | Serine hydroxymethyltransferase 2 (SHMT2) | 4 | 0.75 | 0.153 | 0.999 |
| AT5G26780.1 | Serine hydroxymethyltransferase 2 (SHMT2) | 5 | 0.76 | 0.148 | 1.000 |
| AT5G26780.1 | Serine hydroxymethyltransferase 2 (SHMT2) | 6 | 0.78 | 0.148 | 1.000 |
| AT5G26780.1 | Serine hydroxymethyltransferase 2 (SHMT2) | 7 | 0.80 | 0.148 | 1.000 |
| AT5G26780.1 | Serine hydroxymethyltransferase 2 (SHMT2) | 8 | 0.85 | 0.146 | 1.000 |
| AT5G26780.1 | Serine hydroxymethyltransferase 2 (SHMT2) | 9 | 0.94 | 0.145 | 1.000 |
| AT5G26780.1 | Serine hydroxymethyltransferase 2 (SHMT2) | 10 | 1.04 | 0.145 | 1.000 |
| AT5G26780.1 | Serine hydroxymethyltransferase 2 (SHMT2) | 11 | 1.13 | 0.144 | 1.000 |
| AT5G26780.1 | Serine hydroxymethyltransferase 2 (SHMT2) | 12 | 1.17 | 0.144 | 1.000 |
| AT5G26780.1 | Serine hydroxymethyltransferase 2 (SHMT2) | 13 | 1.14 | 0.143 | 1.000 |
| AT5G26780.1 | Serine hydroxymethyltransferase 2 (SHMT2) | 14 | 1.07 | 0.142 | 1.000 |

| | | | | | |
|---|---|---|---|---|---|
| AT5G26780.1 | Serine hydroxymethyltransferase 2 (SHMT2) | 15 | 0.99 | 0.142 | 1.000 |
| AT5G26780.1 | Serine hydroxymethyltransferase 2 (SHMT2) | 16 | 0.91 | 0.141 | 1.000 |
| AT5G26780.1 | Serine hydroxymethyltransferase 2 (SHMT2) | 17 | 0.85 | 0.142 | 1.000 |
| AT5G26780.1 | Serine hydroxymethyltransferase 2 (SHMT2) | 18 | 0.83 | 0.143 | 1.000 |
| AT5G26780.1 | Serine hydroxymethyltransferase 2 (SHMT2) | 19 | 0.84 | 0.144 | 1.000 |
| AT5G26780.1 | Serine hydroxymethyltransferase 2 (SHMT2) | 20 | 0.85 | 0.146 | 1.000 |
| AT5G26780.1 | Serine hydroxymethyltransferase 2 (SHMT2) | 21 | 0.84 | 0.147 | 1.000 |
| AT5G26780.1 | Serine hydroxymethyltransferase 2 (SHMT2) | 22 | 0.83 | 0.147 | 1.000 |
| AT5G26780.1 | Serine hydroxymethyltransferase 2 (SHMT2) | 23 | 0.82 | 0.147 | 1.000 |
| AT5G26780.1 | Serine hydroxymethyltransferase 2 (SHMT2) | 24 | 0.83 | 0.146 | 1.000 |
| AT5G26780.1 | Serine hydroxymethyltransferase 2 (SHMT2) | 25 | 0.85 | 0.144 | 1.000 |
| AT5G26780.1 | Serine hydroxymethyltransferase 2 (SHMT2) | 26 | 0.89 | 0.144 | 1.000 |
| AT5G26780.1 | Serine hydroxymethyltransferase 2 (SHMT2) | 27 | 0.94 | 0.143 | 1.000 |
| AT5G26780.1 | Serine hydroxymethyltransferase 2 (SHMT2) | 28 | 0.97 | 0.143 | 1.000 |
| AT5G26780.1 | Serine hydroxymethyltransferase 2 (SHMT2) | 29 | 0.98 | 0.144 | 1.000 |
| AT5G26780.1 | Serine hydroxymethyltransferase 2 (SHMT2) | 30 | 0.96 | 0.144 | 1.000 |
| AT5G26780.1 | Serine hydroxymethyltransferase 2 (SHMT2) | 31 | 0.96 | 0.144 | 1.000 |
| AT5G26780.1 | Serine hydroxymethyltransferase 2 (SHMT2) | 32 | 0.99 | 0.144 | 1.000 |
| AT5G26780.1 | Serine hydroxymethyltransferase 2 (SHMT2) | 33 | 1.05 | 0.144 | 1.000 |
| AT5G26780.1 | Serine hydroxymethyltransferase 2 (SHMT2) | 34 | 1.14 | 0.144 | 1.000 |

| | | | | | | |
|---|---|---|---|---|---|---|
| AT5G26780.1 | Serine hydroxymethyltransferase 2 (SHMT2) | 35 | 1.22 | 0.144 | 1.000 | |
| AT5G26780.1 | Serine hydroxymethyltransferase 2 (SHMT2) | 36 | 1.23 | 0.144 | 1.000 | |
| AT5G26780.1 | Serine hydroxymethyltransferase 2 (SHMT2) | 37 | 1.17 | 0.144 | 1.000 | |
| AT5G26780.1 | Serine hydroxymethyltransferase 2 (SHMT2) | 38 | 1.10 | 0.144 | 1.000 | |
| AT5G26780.1 | Serine hydroxymethyltransferase 2 (SHMT2) | 39 | 1.07 | 0.144 | 1.000 | |
| AT5G26780.1 | Serine hydroxymethyltransferase 2 (SHMT2) | 40 | 1.07 | 0.145 | 1.000 | |
| AT5G26780.1 | Serine hydroxymethyltransferase 2 (SHMT2) | 41 | 1.03 | 0.146 | 1.000 | |
| AT5G26780.1 | Serine hydroxymethyltransferase 2 (SHMT2) | 42 | 0.92 | 0.146 | 1.000 | |
| AT5G26780.1 | Serine hydroxymethyltransferase 2 (SHMT2) | 43 | 0.74 | 0.147 | 0.998 | |
| AT5G26780.1 | Serine hydroxymethyltransferase 2 (SHMT2) | 44 | 0.55 | 0.148 | 0.665 | |
| AT5G26780.1 | Serine hydroxymethyltransferase 2 (SHMT2) | 45 | 0.38 | 0.156 | 0.112 | |
| AT5G26780.1 | Serine hydroxymethyltransferase 2 (SHMT2) | 46 | 0.24 | 0.183 | 0.018 | * |
| AT5G26780.1 | Serine hydroxymethyltransferase 2 (SHMT2) | 47 | 0.15 | 0.244 | 0.018 | * |
| AT5G26780.1 | Serine hydroxymethyltransferase 2 (SHMT2) | 48 | 0.10 | 0.331 | 0.039 | * |
| AT5G26800.1 | XAA-Pro aminopeptidase P | 2 | 1.72 | 0.592 | 1.000 | |
| AT5G26800.1 | XAA-Pro aminopeptidase P | 3 | 1.86 | 0.520 | 0.997 | |
| AT5G26800.1 | XAA-Pro aminopeptidase P | 11 | 1.29 | 0.306 | 1.000 | |
| AT5G26800.1 | XAA-Pro aminopeptidase P | 12 | 1.05 | 0.269 | 1.000 | |
| AT5G26800.1 | XAA-Pro aminopeptidase P | 13 | 0.88 | 0.240 | 1.000 | |
| AT5G26800.1 | XAA-Pro aminopeptidase P | 14 | 0.81 | 0.219 | 0.999 | |
| AT5G26800.1 | XAA-Pro aminopeptidase P | 15 | 0.87 | 0.207 | 1.000 | |
| AT5G26800.1 | XAA-Pro aminopeptidase P | 16 | 1.11 | 0.204 | 1.000 | |
| AT5G26800.1 | XAA-Pro aminopeptidase P | 17 | 1.63 | 0.209 | 0.900 | |
| AT5G26800.1 | XAA-Pro aminopeptidase P | 18 | 2.52 | 0.223 | 0.402 | |
| AT5G26800.1 | XAA-Pro aminopeptidase P | 19 | 3.87 | 0.250 | 0.137 | |
| AT5G26800.1 | XAA-Pro aminopeptidase P | 20 | 5.91 | 0.297 | 0.077 | . |
| AT5G26800.1 | XAA-Pro aminopeptidase P | 21 | 9.04 | 0.378 | 0.090 | . |
| AT5G26800.1 | XAA-Pro aminopeptidase P | 46 | 236847.99 | 8.311 | 0.990 | |
| AT5G26800.1 | XAA-Pro aminopeptidase P | 47 | 352788.01 | 8.773 | 0.991 | |
| AT5G26830.1 | Dual-targeted threonyl-tRNA synthetase of chloroplasts and mitochondria | 2 | 0.88 | 0.457 | 1.000 | |

| | | | | | |
|---|---|---|---|---|---|
| AT5G26830.1 | Dual-targeted threonyl-tRNA synthetase of chloroplasts and mitochondria | 3 | 0.91 | 0.449 | 1.000 |
| AT5G26830.1 | Dual-targeted threonyl-tRNA synthetase of chloroplasts and mitochondria | 29 | 1.06 | 0.322 | 1.000 |
| AT5G26830.1 | Dual-targeted threonyl-tRNA synthetase of chloroplasts and mitochondria | 30 | 1.07 | 0.235 | 1.000 |
| AT5G26830.1 | Dual-targeted threonyl-tRNA synthetase of chloroplasts and mitochondria | 31 | 1.25 | 0.211 | 1.000 |
| AT5G26830.1 | Dual-targeted threonyl-tRNA synthetase of chloroplasts and mitochondria | 32 | 1.68 | 0.201 | 0.925 |
| AT5G26830.1 | Dual-targeted threonyl-tRNA synthetase of chloroplasts and mitochondria | 33 | 2.09 | 0.196 | 0.629 |
| AT5G26830.1 | Dual-targeted threonyl-tRNA synthetase of chloroplasts and mitochondria | 34 | 2.17 | 0.196 | 0.568 |
| AT5G26830.1 | Dual-targeted threonyl-tRNA synthetase of chloroplasts and mitochondria | 35 | 1.83 | 0.200 | 0.840 |
| AT5G26830.1 | Dual-targeted threonyl-tRNA synthetase of chloroplasts and mitochondria | 36 | 1.39 | 0.212 | 0.997 |
| AT5G26830.1 | Dual-targeted threonyl-tRNA synthetase of chloroplasts and mitochondria | 37 | 1.26 | 0.266 | 1.000 |
| AT5G26830.1 | Dual-targeted threonyl-tRNA synthetase of chloroplasts and mitochondria | 38 | 1.55 | 0.419 | 1.000 |
| AT5G26830.1 | Dual-targeted threonyl-tRNA synthetase of chloroplasts and mitochondria | 46 | 517.91 | 3.154 | 0.985 |
| AT5G26830.1 | Dual-targeted threonyl-tRNA synthetase of chloroplasts and mitochondria | 47 | 1370.95 | 3.675 | 0.986 |
| AT5G26860.1 | Lon protease 1 (LON1) | 2 | 0.91 | 0.486 | 1.000 |
| AT5G26860.1 | Lon protease 1 (LON1) | 3 | 0.84 | 0.432 | 1.000 |
| AT5G26860.1 | Lon protease 1 (LON1) | 13 | 1.55 | 0.289 | 1.000 |
| AT5G26860.1 | Lon protease 1 (LON1) | 14 | 1.79 | 0.256 | 0.999 |
| AT5G26860.1 | Lon protease 1 (LON1) | 15 | 1.64 | 0.233 | 1.000 |
| AT5G26860.1 | Lon protease 1 (LON1) | 16 | 1.21 | 0.215 | 1.000 |
| AT5G26860.1 | Lon protease 1 (LON1) | 17 | 0.92 | 0.206 | 1.000 |
| AT5G26860.1 | Lon protease 1 (LON1) | 18 | 0.89 | 0.206 | 1.000 |
| AT5G26860.1 | Lon protease 1 (LON1) | 19 | 0.97 | 0.210 | 1.000 |
| AT5G26860.1 | Lon protease 1 (LON1) | 20 | 1.03 | 0.215 | 1.000 |
| AT5G26860.1 | Lon protease 1 (LON1) | 21 | 1.01 | 0.217 | 1.000 |
| AT5G26860.1 | Lon protease 1 (LON1) | 22 | 0.96 | 0.217 | 1.000 |
| AT5G26860.1 | Lon protease 1 (LON1) | 23 | 0.90 | 0.215 | 1.000 |
| AT5G26860.1 | Lon protease 1 (LON1) | 24 | 0.85 | 0.211 | 1.000 |
| AT5G26860.1 | Lon protease 1 (LON1) | 25 | 0.82 | 0.209 | 1.000 |
| AT5G26860.1 | Lon protease 1 (LON1) | 26 | 0.88 | 0.208 | 1.000 |
| AT5G26860.1 | Lon protease 1 (LON1) | 27 | 1.00 | 0.209 | 1.000 |
| AT5G26860.1 | Lon protease 1 (LON1) | 28 | 1.04 | 0.209 | 1.000 |
| AT5G26860.1 | Lon protease 1 (LON1) | 29 | 0.99 | 0.209 | 1.000 |
| AT5G26860.1 | Lon protease 1 (LON1) | 30 | 0.94 | 0.209 | 1.000 |
| AT5G26860.1 | Lon protease 1 (LON1) | 31 | 0.96 | 0.209 | 1.000 |
| AT5G26860.1 | Lon protease 1 (LON1) | 32 | 1.04 | 0.210 | 1.000 |
| AT5G26860.1 | Lon protease 1 (LON1) | 33 | 1.09 | 0.210 | 1.000 |

| | | | | | | |
|---|---|---|---|---|---|---|
| AT5G26860.1 | Lon protease 1 (LON1) | 34 | 1.10 | 0.211 | 1.000 | |
| AT5G26860.1 | Lon protease 1 (LON1) | 35 | 1.08 | 0.210 | 1.000 | |
| AT5G26860.1 | Lon protease 1 (LON1) | 36 | 1.03 | 0.210 | 1.000 | |
| AT5G26860.1 | Lon protease 1 (LON1) | 37 | 0.97 | 0.210 | 1.000 | |
| AT5G26860.1 | Lon protease 1 (LON1) | 38 | 0.90 | 0.209 | 1.000 | |
| AT5G26860.1 | Lon protease 1 (LON1) | 39 | 0.76 | 0.209 | 1.000 | |
| AT5G26860.1 | Lon protease 1 (LON1) | 40 | 0.55 | 0.210 | 0.986 | |
| AT5G26860.1 | Lon protease 1 (LON1) | 41 | 0.38 | 0.212 | 0.624 | |
| AT5G26860.1 | Lon protease 1 (LON1) | 42 | 0.35 | 0.215 | 0.490 | |
| AT5G26860.1 | Lon protease 1 (LON1) | 43 | 0.42 | 0.227 | 0.845 | |
| AT5G26860.1 | Lon protease 1 (LON1) | 44 | 0.59 | 0.273 | 1.000 | |
| AT5G26860.1 | Lon protease 1 (LON1) | 45 | 1.00 | 0.365 | 1.000 | |
| AT5G26860.1 | Lon protease 1 (LON1) | 46 | 1.94 | 0.532 | 1.000 | |
| AT5G26860.1 | Lon protease 1 (LON1) | 47 | 4.00 | 0.773 | 1.000 | |
| AT5G27540.2 | MIRO-related GTPase (MIRO1) | 3 | 0.78 | 0.166 | 1.000 | |
| AT5G27540.2 | MIRO-related GTPase (MIRO1) | 4 | 0.85 | 0.146 | 1.000 | |
| AT5G27540.2 | MIRO-related GTPase (MIRO1) | 5 | 0.94 | 0.143 | 1.000 | |
| AT5G27540.2 | MIRO-related GTPase (MIRO1) | 6 | 1.04 | 0.143 | 1.000 | |
| AT5G27540.2 | MIRO-related GTPase (MIRO1) | 7 | 1.18 | 0.143 | 1.000 | |
| AT5G27540.2 | MIRO-related GTPase (MIRO1) | 8 | 1.42 | 0.141 | 0.997 | |
| AT5G27540.2 | MIRO-related GTPase (MIRO1) | 9 | 1.69 | 0.140 | 0.847 | |
| AT5G27540.2 | MIRO-related GTPase (MIRO1) | 10 | 1.94 | 0.139 | 0.515 | |
| AT5G27540.2 | MIRO-related GTPase (MIRO1) | 11 | 2.24 | 0.140 | 0.232 | |
| AT5G27540.2 | MIRO-related GTPase (MIRO1) | 12 | 2.48 | 0.139 | 0.105 | |
| AT5G27540.2 | MIRO-related GTPase (MIRO1) | 13 | 2.53 | 0.138 | 0.083 | . |
| AT5G27540.2 | MIRO-related GTPase (MIRO1) | 14 | 2.39 | 0.137 | 0.122 | |
| AT5G27540.2 | MIRO-related GTPase (MIRO1) | 15 | 2.08 | 0.136 | 0.317 | |
| AT5G27540.2 | MIRO-related GTPase (MIRO1) | 16 | 1.84 | 0.136 | 0.613 | |
| AT5G27540.2 | MIRO-related GTPase (MIRO1) | 17 | 1.76 | 0.136 | 0.725 | |
| AT5G27540.2 | MIRO-related GTPase (MIRO1) | 18 | 1.75 | 0.137 | 0.749 | |
| AT5G27540.2 | MIRO-related GTPase (MIRO1) | 19 | 1.68 | 0.140 | 0.857 | |
| AT5G27540.2 | MIRO-related GTPase (MIRO1) | 20 | 1.49 | 0.142 | 0.988 | |
| AT5G27540.2 | MIRO-related GTPase (MIRO1) | 21 | 1.24 | 0.144 | 1.000 | |
| AT5G27540.2 | MIRO-related GTPase (MIRO1) | 22 | 1.02 | 0.144 | 1.000 | |
| AT5G27540.2 | MIRO-related GTPase (MIRO1) | 23 | 0.87 | 0.143 | 1.000 | |
| AT5G27540.2 | MIRO-related GTPase (MIRO1) | 24 | 0.75 | 0.141 | 1.000 | |
| AT5G27540.2 | MIRO-related GTPase (MIRO1) | 25 | 0.68 | 0.139 | 0.989 | |
| AT5G27540.2 | MIRO-related GTPase (MIRO1) | 26 | 0.70 | 0.138 | 0.996 | |
| AT5G27540.2 | MIRO-related GTPase (MIRO1) | 27 | 0.80 | 0.138 | 1.000 | |
| AT5G27540.2 | MIRO-related GTPase (MIRO1) | 28 | 0.90 | 0.138 | 1.000 | |
| AT5G27540.2 | MIRO-related GTPase (MIRO1) | 29 | 0.96 | 0.138 | 1.000 | |
| AT5G27540.2 | MIRO-related GTPase (MIRO1) | 30 | 1.00 | 0.139 | 1.000 | |
| AT5G27540.2 | MIRO-related GTPase (MIRO1) | 31 | 1.03 | 0.139 | 1.000 | |
| AT5G27540.2 | MIRO-related GTPase (MIRO1) | 32 | 1.06 | 0.139 | 1.000 | |
| AT5G27540.2 | MIRO-related GTPase (MIRO1) | 33 | 1.08 | 0.139 | 1.000 | |
| AT5G27540.2 | MIRO-related GTPase (MIRO1) | 34 | 1.08 | 0.139 | 1.000 | |
| AT5G27540.2 | MIRO-related GTPase (MIRO1) | 35 | 1.06 | 0.139 | 1.000 | |
| AT5G27540.2 | MIRO-related GTPase (MIRO1) | 36 | 1.02 | 0.139 | 1.000 | |
| AT5G27540.2 | MIRO-related GTPase (MIRO1) | 37 | 0.99 | 0.139 | 1.000 | |
| AT5G27540.2 | MIRO-related GTPase (MIRO1) | 38 | 1.01 | 0.139 | 1.000 | |
| AT5G27540.2 | MIRO-related GTPase (MIRO1) | 39 | 1.10 | 0.139 | 1.000 | |
| AT5G27540.2 | MIRO-related GTPase (MIRO1) | 40 | 1.28 | 0.140 | 1.000 | |
| AT5G27540.2 | MIRO-related GTPase (MIRO1) | 41 | 1.47 | 0.140 | 0.990 | |
| AT5G27540.2 | MIRO-related GTPase (MIRO1) | 42 | 1.54 | 0.141 | 0.970 | |
| AT5G27540.2 | MIRO-related GTPase (MIRO1) | 43 | 1.37 | 0.142 | 0.999 | |
| AT5G27540.2 | MIRO-related GTPase (MIRO1) | 44 | 0.96 | 0.143 | 1.000 | |
| AT5G27540.2 | MIRO-related GTPase (MIRO1) | 45 | 0.50 | 0.148 | 0.554 | |
| AT5G27540.2 | MIRO-related GTPase (MIRO1) | 46 | 0.20 | 0.174 | 0.002 | ** |
| AT5G27540.2 | MIRO-related GTPase (MIRO1) | 47 | 0.07 | 0.249 | 0.000 | *** |
| AT5G27540.2 | MIRO-related GTPase (MIRO1) | 48 | 0.02 | 0.364 | 0.000 | *** |
| AT5G27760.1 | Cytochrome c:O2 oxidoreductase subunit (COXX1) | 2 | 0.89 | 0.499 | 1.000 | |

| | | | | | | |
|---|---|---|---|---|---|---|
| AT5G27760.1 | Cytochrome c:O2 oxidoreductase subunit (COXX1) | 3 | 0.90 | 0.385 | 1.000 | |
| AT5G27760.1 | Cytochrome c:O2 oxidoreductase subunit (COXX1) | 6 | 1.05 | 0.352 | 1.000 | |
| AT5G27760.1 | Cytochrome c:O2 oxidoreductase subunit (COXX1) | 7 | 1.23 | 0.273 | 1.000 | |
| AT5G27760.1 | Cytochrome c:O2 oxidoreductase subunit (COXX1) | 8 | 1.57 | 0.235 | 1.000 | |
| AT5G27760.1 | Cytochrome c:O2 oxidoreductase subunit (COXX1) | 9 | 1.96 | 0.222 | 0.986 | |
| AT5G27760.1 | Cytochrome c:O2 oxidoreductase subunit (COXX1) | 10 | 2.01 | 0.221 | 0.977 | |
| AT5G27760.1 | Cytochrome c:O2 oxidoreductase subunit (COXX1) | 11 | 1.93 | 0.220 | 0.987 | |
| AT5G27760.1 | Cytochrome c:O2 oxidoreductase subunit (COXX1) | 12 | 2.21 | 0.220 | 0.925 | |
| AT5G27760.1 | Cytochrome c:O2 oxidoreductase subunit (COXX1) | 13 | 3.10 | 0.216 | 0.422 | |
| AT5G27760.1 | Cytochrome c:O2 oxidoreductase subunit (COXX1) | 14 | 3.86 | 0.213 | 0.147 | |
| AT5G27760.1 | Cytochrome c:O2 oxidoreductase subunit (COXX1) | 15 | 3.31 | 0.213 | 0.304 | |
| AT5G27760.1 | Cytochrome c:O2 oxidoreductase subunit (COXX1) | 16 | 2.53 | 0.210 | 0.717 | |
| AT5G27760.1 | Cytochrome c:O2 oxidoreductase subunit (COXX1) | 17 | 2.20 | 0.212 | 0.906 | |
| AT5G27760.1 | Cytochrome c:O2 oxidoreductase subunit (COXX1) | 18 | 2.15 | 0.218 | 0.940 | |
| AT5G27760.1 | Cytochrome c:O2 oxidoreductase subunit (COXX1) | 19 | 2.04 | 0.228 | 0.980 | |
| AT5G27760.1 | Cytochrome c:O2 oxidoreductase subunit (COXX1) | 20 | 1.95 | 0.241 | 0.995 | |
| AT5G27760.1 | Cytochrome c:O2 oxidoreductase subunit (COXX1) | 21 | 1.59 | 0.284 | 1.000 | |
| AT5G27760.1 | Cytochrome c:O2 oxidoreductase subunit (COXX1) | 22 | 1.06 | 0.381 | 1.000 | |
| AT5G27760.1 | Cytochrome c:O2 oxidoreductase subunit (COXX1) | 25 | 0.43 | 0.320 | 0.997 | |
| AT5G27760.1 | Cytochrome c:O2 oxidoreductase subunit (COXX1) | 26 | 0.50 | 0.258 | 0.996 | |
| AT5G27760.1 | Cytochrome c:O2 oxidoreductase subunit (COXX1) | 27 | 0.81 | 0.231 | 1.000 | |
| AT5G27760.1 | Cytochrome c:O2 oxidoreductase subunit (COXX1) | 28 | 1.60 | 0.219 | 1.000 | |
| AT5G27760.1 | Cytochrome c:O2 oxidoreductase subunit (COXX1) | 29 | 2.45 | 0.218 | 0.811 | |
| AT5G27760.1 | Cytochrome c:O2 oxidoreductase subunit (COXX1) | 30 | 2.45 | 0.218 | 0.811 | |
| AT5G27760.1 | Cytochrome c:O2 oxidoreductase subunit (COXX1) | 31 | 1.74 | 0.219 | 0.998 | |
| AT5G27760.1 | Cytochrome c:O2 oxidoreductase subunit (COXX1) | 32 | 0.91 | 0.224 | 1.000 | |
| AT5G27760.1 | Cytochrome c:O2 oxidoreductase subunit (COXX1) | 33 | 0.26 | 0.237 | 0.307 | |
| AT5G27760.1 | Cytochrome c:O2 oxidoreductase subunit (COXX1) | 34 | 0.08 | 0.293 | 0.008 | ** |
| AT5G27760.1 | Cytochrome c:O2 oxidoreductase subunit (COXX1) | 35 | 0.04 | 0.488 | 0.127 | |
| AT5G27760.1 | Cytochrome c:O2 oxidoreductase subunit (COXX1) | 46 | 18400.72 | 5.030 | 1.000 | |
| AT5G27760.1 | Cytochrome c:O2 oxidoreductase subunit (COXX1) | 47 | 146735.90 | 5.669 | 1.000 | |

| | | | | | |
|---|---|---|---|---|---|
| AT5G27850.1 | 60S ribosomal protein L18-3 (RPL18C) | 2 | 0.53 | 0.200 | 0.384 |
| AT5G27850.1 | 60S ribosomal protein L18-3 (RPL18C) | 3 | 0.56 | 0.189 | 0.405 |
| AT5G27850.1 | 60S ribosomal protein L18-3 (RPL18C) | 4 | 0.59 | 0.179 | 0.433 |
| AT5G27850.1 | 60S ribosomal protein L18-3 (RPL18C) | 5 | 0.62 | 0.169 | 0.469 |
| AT5G27850.1 | 60S ribosomal protein L18-3 (RPL18C) | 6 | 0.66 | 0.160 | 0.513 |
| AT5G27850.1 | 60S ribosomal protein L18-3 (RPL18C) | 7 | 0.69 | 0.152 | 0.568 |
| AT5G27850.1 | 60S ribosomal protein L18-3 (RPL18C) | 8 | 0.73 | 0.145 | 0.635 |
| AT5G27850.1 | 60S ribosomal protein L18-3 (RPL18C) | 9 | 0.77 | 0.139 | 0.713 |
| AT5G27850.1 | 60S ribosomal protein L18-3 (RPL18C) | 10 | 0.81 | 0.135 | 0.796 |
| AT5G27850.1 | 60S ribosomal protein L18-3 (RPL18C) | 11 | 0.86 | 0.132 | 0.878 |
| AT5G27850.1 | 60S ribosomal protein L18-3 (RPL18C) | 12 | 0.90 | 0.132 | 0.945 |
| AT5G27850.1 | 60S ribosomal protein L18-3 (RPL18C) | 13 | 0.95 | 0.132 | 0.988 |
| AT5G27850.1 | 60S ribosomal protein L18-3 (RPL18C) | 14 | 1.01 | 0.135 | 1.000 |
| AT5G27850.1 | 60S ribosomal protein L18-3 (RPL18C) | 15 | 1.06 | 0.139 | 0.983 |
| AT5G27850.1 | 60S ribosomal protein L18-3 (RPL18C) | 16 | 1.12 | 0.145 | 0.943 |
| AT5G27850.1 | 60S ribosomal protein L18-3 (RPL18C) | 17 | 1.18 | 0.152 | 0.891 |
| AT5G27850.1 | 60S ribosomal protein L18-3 (RPL18C) | 18 | 1.24 | 0.160 | 0.835 |
| AT5G27850.1 | 60S ribosomal protein L18-3 (RPL18C) | 19 | 1.31 | 0.169 | 0.779 |
| AT5G27850.1 | 60S ribosomal protein L18-3 (RPL18C) | 20 | 1.38 | 0.179 | 0.727 |
| AT5G27850.1 | 60S ribosomal protein L18-3 (RPL18C) | 21 | 1.46 | 0.190 | 0.681 |
| AT5G27850.1 | 60S ribosomal protein L18-3 (RPL18C) | 22 | 1.54 | 0.201 | 0.642 |
| AT5G27850.1 | 60S ribosomal protein L18-3 (RPL18C) | 46 | 5.52 | 0.533 | 0.372 |
| AT5G27850.1 | 60S ribosomal protein L18-3 (RPL18C) | 47 | 5.82 | 0.548 | 0.369 |
| AT5G33320.1 | Phosphoenolpyruvate/phosphate translocator (PPT) | 2 | 0.97 | 0.565 | 1.000 |
| AT5G33320.1 | Phosphoenolpyruvate/phosphate translocator (PPT) | 3 | 1.38 | 0.517 | 1.000 |
| AT5G33320.1 | Phosphoenolpyruvate/phosphate translocator (PPT) | 40 | 12.27 | 0.454 | 0.137 |
| AT5G33320.1 | Phosphoenolpyruvate/phosphate translocator (PPT) | 41 | 2.31 | 0.277 | 0.745 |
| AT5G33320.1 | Phosphoenolpyruvate/phosphate translocator (PPT) | 42 | 0.61 | 0.233 | 0.935 |

| | | | | | |
|---|---|---|---|---|---|
| AT5G33320.1 | Phosphoenolpyruvate/phosphate translocator (PPT) | 43 | 0.44 | 0.227 | 0.561 |
| AT5G33320.1 | Phosphoenolpyruvate/phosphate translocator (PPT) | 44 | 0.89 | 0.248 | 1.000 |
| AT5G33320.1 | Phosphoenolpyruvate/phosphate translocator (PPT) | 45 | 2.71 | 0.486 | 0.948 |
| AT5G33320.1 | Phosphoenolpyruvate/phosphate translocator (PPT) | 46 | 8.27 | 0.976 | 0.933 |
| AT5G33320.1 | Phosphoenolpyruvate/phosphate translocator (PPT) | 47 | 25.10 | 1.621 | 0.956 |
| AT5G35630.3 | Glutamine synthetase 2 (GS2) | 2 | 0.89 | 0.541 | 1.000 |
| AT5G35630.3 | Glutamine synthetase 2 (GS2) | 3 | 0.74 | 0.564 | 1.000 |
| AT5G35630.3 | Glutamine synthetase 2 (GS2) | 28 | 1.63 | 0.352 | 1.000 |
| AT5G35630.3 | Glutamine synthetase 2 (GS2) | 29 | 1.78 | 0.259 | 0.984 |
| AT5G35630.3 | Glutamine synthetase 2 (GS2) | 30 | 1.88 | 0.230 | 0.934 |
| AT5G35630.3 | Glutamine synthetase 2 (GS2) | 31 | 1.91 | 0.217 | 0.894 |
| AT5G35630.3 | Glutamine synthetase 2 (GS2) | 32 | 1.95 | 0.213 | 0.863 |
| AT5G35630.3 | Glutamine synthetase 2 (GS2) | 33 | 1.86 | 0.211 | 0.905 |
| AT5G35630.3 | Glutamine synthetase 2 (GS2) | 34 | 1.45 | 0.212 | 0.998 |
| AT5G35630.3 | Glutamine synthetase 2 (GS2) | 35 | 1.05 | 0.216 | 1.000 |
| AT5G35630.3 | Glutamine synthetase 2 (GS2) | 36 | 1.00 | 0.227 | 1.000 |
| AT5G35630.3 | Glutamine synthetase 2 (GS2) | 37 | 1.20 | 0.247 | 1.000 |
| AT5G35630.3 | Glutamine synthetase 2 (GS2) | 38 | 1.47 | 0.270 | 1.000 |
| AT5G35630.3 | Glutamine synthetase 2 (GS2) | 39 | 1.66 | 0.280 | 0.997 |
| AT5G35630.3 | Glutamine synthetase 2 (GS2) | 40 | 1.74 | 0.284 | 0.995 |
| AT5G35630.3 | Glutamine synthetase 2 (GS2) | 41 | 1.71 | 0.291 | 0.996 |
| AT5G35630.3 | Glutamine synthetase 2 (GS2) | 42 | 1.64 | 0.345 | 1.000 |
| AT5G35630.3 | Glutamine synthetase 2 (GS2) | 43 | 1.61 | 0.467 | 1.000 |
| AT5G35630.3 | Glutamine synthetase 2 (GS2) | 46 | 1.84 | 1.204 | 1.000 |
| AT5G35630.3 | Glutamine synthetase 2 (GS2) | 47 | 1.98 | 1.558 | 1.000 |
| AT5G36950.1 | DEGP protease 10 (DEGP10) | 2 | 0.94 | 0.499 | 1.000 |
| AT5G36950.1 | DEGP protease 10 (DEGP10) | 3 | 0.84 | 0.825 | 1.000 |
| AT5G36950.1 | DEGP protease 10 (DEGP10) | 24 | 0.24 | 0.361 | 0.624 |
| AT5G36950.1 | DEGP protease 10 (DEGP10) | 25 | 0.33 | 0.246 | 0.432 |
| AT5G36950.1 | DEGP protease 10 (DEGP10) | 26 | 0.65 | 0.214 | 0.990 |
| AT5G36950.1 | DEGP protease 10 (DEGP10) | 27 | 1.66 | 0.204 | 0.958 |
| AT5G36950.1 | DEGP protease 10 (DEGP10) | 28 | 2.48 | 0.205 | 0.460 |
| AT5G36950.1 | DEGP protease 10 (DEGP10) | 29 | 1.64 | 0.220 | 0.980 |
| AT5G36950.1 | DEGP protease 10 (DEGP10) | 30 | 0.66 | 0.264 | 0.999 |
| AT5G36950.1 | DEGP protease 10 (DEGP10) | 31 | 0.34 | 0.325 | 0.795 |
| AT5G36950.1 | DEGP protease 10 (DEGP10) | 33 | 0.41 | 0.326 | 0.925 |
| AT5G36950.1 | DEGP protease 10 (DEGP10) | 34 | 0.69 | 0.347 | 1.000 |
| AT5G36950.1 | DEGP protease 10 (DEGP10) | 35 | 1.28 | 0.521 | 1.000 |
| AT5G36950.1 | DEGP protease 10 (DEGP10) | 46 | 216.77 | 3.026 | 0.997 |
| AT5G36950.1 | DEGP protease 10 (DEGP10) | 47 | 315.79 | 3.390 | 0.998 |
| AT5G37510.1 | NADH:ubiquinone oxidoreductase intermediate-associated protein | 3 | 0.89 | 0.224 | 1.000 |
| AT5G37510.1 | NADH:ubiquinone oxidoreductase intermediate-associated protein | 4 | 1.05 | 0.207 | 1.000 |
| AT5G37510.1 | NADH:ubiquinone oxidoreductase intermediate-associated protein | 5 | 1.36 | 0.207 | 1.000 |
| AT5G37510.1 | NADH:ubiquinone oxidoreductase intermediate-associated protein | 6 | 1.63 | 0.208 | 1.000 |

| | | | | | |
|---|---|---|---|---|---|
| AT5G37510.1 | NADH:ubiquinone oxidoreductase intermediate-associated protein | 7 | 1.73 | 0.208 | 0.999 |
| AT5G37510.1 | NADH:ubiquinone oxidoreductase intermediate-associated protein | 8 | 1.79 | 0.204 | 0.997 |
| AT5G37510.1 | NADH:ubiquinone oxidoreductase intermediate-associated protein | 9 | 1.80 | 0.201 | 0.997 |
| AT5G37510.1 | NADH:ubiquinone oxidoreductase intermediate-associated protein | 10 | 1.72 | 0.201 | 0.999 |
| AT5G37510.1 | NADH:ubiquinone oxidoreductase intermediate-associated protein | 11 | 1.74 | 0.202 | 0.999 |
| AT5G37510.1 | NADH:ubiquinone oxidoreductase intermediate-associated protein | 12 | 1.73 | 0.202 | 0.999 |
| AT5G37510.1 | NADH:ubiquinone oxidoreductase intermediate-associated protein | 13 | 1.52 | 0.200 | 1.000 |
| AT5G37510.1 | NADH:ubiquinone oxidoreductase intermediate-associated protein | 14 | 1.20 | 0.197 | 1.000 |
| AT5G37510.1 | NADH:ubiquinone oxidoreductase intermediate-associated protein | 15 | 0.85 | 0.196 | 1.000 |
| AT5G37510.1 | NADH:ubiquinone oxidoreductase intermediate-associated protein | 16 | 0.59 | 0.195 | 0.999 |
| AT5G37510.1 | NADH:ubiquinone oxidoreductase intermediate-associated protein | 17 | 0.47 | 0.195 | 0.918 |
| AT5G37510.1 | NADH:ubiquinone oxidoreductase intermediate-associated protein | 18 | 0.52 | 0.198 | 0.982 |
| AT5G37510.1 | NADH:ubiquinone oxidoreductase intermediate-associated protein | 19 | 0.69 | 0.203 | 1.000 |
| AT5G37510.1 | NADH:ubiquinone oxidoreductase intermediate-associated protein | 20 | 0.90 | 0.207 | 1.000 |
| AT5G37510.1 | NADH:ubiquinone oxidoreductase intermediate-associated protein | 21 | 1.20 | 0.210 | 1.000 |
| AT5G37510.1 | NADH:ubiquinone oxidoreductase intermediate-associated protein | 22 | 1.64 | 0.210 | 1.000 |
| AT5G37510.1 | NADH:ubiquinone oxidoreductase intermediate-associated protein | 23 | 1.84 | 0.209 | 0.996 |
| AT5G37510.1 | NADH:ubiquinone oxidoreductase intermediate-associated protein | 24 | 1.32 | 0.205 | 1.000 |
| AT5G37510.1 | NADH:ubiquinone oxidoreductase intermediate-associated protein | 25 | 0.63 | 0.201 | 1.000 |
| AT5G37510.1 | NADH:ubiquinone oxidoreductase intermediate-associated protein | 26 | 0.36 | 0.200 | 0.542 |

| | | | | | | |
|---|---|---|---|---|---|---|
| AT5G37510.1 | NADH:ubiquinone oxidoreductase intermediate-associated protein | 27 | 0.35 | 0.200 | 0.487 | |
| AT5G37510.1 | NADH:ubiquinone oxidoreductase intermediate-associated protein | 28 | 0.51 | 0.200 | 0.981 | |
| AT5G37510.1 | NADH:ubiquinone oxidoreductase intermediate-associated protein | 29 | 0.89 | 0.200 | 1.000 | |
| AT5G37510.1 | NADH:ubiquinone oxidoreductase intermediate-associated protein | 30 | 1.24 | 0.200 | 1.000 | |
| AT5G37510.1 | NADH:ubiquinone oxidoreductase intermediate-associated protein | 31 | 1.36 | 0.200 | 1.000 | |
| AT5G37510.1 | NADH:ubiquinone oxidoreductase intermediate-associated protein | 32 | 1.43 | 0.201 | 1.000 | |
| AT5G37510.1 | NADH:ubiquinone oxidoreductase intermediate-associated protein | 33 | 1.45 | 0.201 | 1.000 | |
| AT5G37510.1 | NADH:ubiquinone oxidoreductase intermediate-associated protein | 34 | 1.27 | 0.202 | 1.000 | |
| AT5G37510.1 | NADH:ubiquinone oxidoreductase intermediate-associated protein | 35 | 0.90 | 0.201 | 1.000 | |
| AT5G37510.1 | NADH:ubiquinone oxidoreductase intermediate-associated protein | 36 | 0.67 | 0.200 | 1.000 | |
| AT5G37510.1 | NADH:ubiquinone oxidoreductase intermediate-associated protein | 37 | 0.63 | 0.200 | 1.000 | |
| AT5G37510.1 | NADH:ubiquinone oxidoreductase intermediate-associated protein | 38 | 0.73 | 0.201 | 1.000 | |
| AT5G37510.1 | NADH:ubiquinone oxidoreductase intermediate-associated protein | 39 | 0.94 | 0.201 | 1.000 | |
| AT5G37510.1 | NADH:ubiquinone oxidoreductase intermediate-associated protein | 40 | 1.18 | 0.202 | 1.000 | |
| AT5G37510.1 | NADH:ubiquinone oxidoreductase intermediate-associated protein | 41 | 1.20 | 0.203 | 1.000 | |
| AT5G37510.1 | NADH:ubiquinone oxidoreductase intermediate-associated protein | 42 | 1.07 | 0.205 | 1.000 | |
| AT5G37510.1 | NADH:ubiquinone oxidoreductase intermediate-associated protein | 43 | 0.87 | 0.206 | 1.000 | |
| AT5G37510.1 | NADH:ubiquinone oxidoreductase intermediate-associated protein | 44 | 0.52 | 0.207 | 0.989 | |
| AT5G37510.1 | NADH:ubiquinone oxidoreductase intermediate-associated protein | 45 | 0.25 | 0.207 | 0.128 | |
| AT5G37510.1 | NADH:ubiquinone oxidoreductase intermediate-associated protein | 46 | 0.13 | 0.239 | 0.010 | ** |

| | | | | | | |
|---|---|---|---|---|---|---|
| AT5G37510.1 | NADH:ubiquinone oxidoreductase intermediate-associated protein | 47 | 0.07 | 0.404 | 0.156 | |
| AT5G37510.1 | NADH:ubiquinone oxidoreductase intermediate-associated protein | 48 | 0.04 | 0.668 | 0.669 | |
| AT5G37530.2 | NAD(P)-binding Rossmann-fold superfamily protein; | 2 | 1.02 | 0.492 | 1.000 | |
| AT5G37530.2 | NAD(P)-binding Rossmann-fold superfamily protein; | 3 | 1.12 | 0.459 | 1.000 | |
| AT5G37530.2 | NAD(P)-binding Rossmann-fold superfamily protein; | 4 | 1.23 | 0.440 | 1.000 | |
| AT5G37530.2 | NAD(P)-binding Rossmann-fold superfamily protein; | 22 | 3.51 | 0.304 | 0.401 | |
| AT5G37530.2 | NAD(P)-binding Rossmann-fold superfamily protein; | 23 | 3.43 | 0.279 | 0.328 | |
| AT5G37530.2 | NAD(P)-binding Rossmann-fold superfamily protein; | 24 | 3.30 | 0.257 | 0.275 | |
| AT5G37530.2 | NAD(P)-binding Rossmann-fold superfamily protein; | 25 | 3.11 | 0.238 | 0.247 | |
| AT5G37530.2 | NAD(P)-binding Rossmann-fold superfamily protein; | 26 | 2.89 | 0.223 | 0.251 | |
| AT5G37530.2 | NAD(P)-binding Rossmann-fold superfamily protein; | 27 | 2.64 | 0.212 | 0.291 | |
| AT5G37530.2 | NAD(P)-binding Rossmann-fold superfamily protein; | 28 | 2.37 | 0.205 | 0.380 | |
| AT5G37530.2 | NAD(P)-binding Rossmann-fold superfamily protein; | 29 | 2.09 | 0.200 | 0.536 | |
| AT5G37530.2 | NAD(P)-binding Rossmann-fold superfamily protein; | 30 | 1.80 | 0.198 | 0.750 | |
| AT5G37530.2 | NAD(P)-binding Rossmann-fold superfamily protein; | 31 | 1.53 | 0.197 | 0.933 | |
| AT5G37530.2 | NAD(P)-binding Rossmann-fold superfamily protein; | 32 | 1.29 | 0.197 | 0.997 | |
| AT5G37530.2 | NAD(P)-binding Rossmann-fold superfamily protein; | 33 | 1.08 | 0.198 | 1.000 | |
| AT5G37530.2 | NAD(P)-binding Rossmann-fold superfamily protein; | 34 | 0.92 | 0.200 | 1.000 | |
| AT5G37530.2 | NAD(P)-binding Rossmann-fold superfamily protein; | 35 | 0.78 | 0.201 | 0.998 | |
| AT5G37530.2 | NAD(P)-binding Rossmann-fold superfamily protein; | 36 | 0.67 | 0.203 | 0.956 | |
| AT5G37530.2 | NAD(P)-binding Rossmann-fold superfamily protein; | 37 | 0.57 | 0.204 | 0.814 | |
| AT5G37530.2 | NAD(P)-binding Rossmann-fold superfamily protein; | 38 | 0.49 | 0.204 | 0.584 | |
| AT5G37530.2 | NAD(P)-binding Rossmann-fold superfamily protein; | 39 | 0.41 | 0.205 | 0.351 | |
| AT5G37530.2 | NAD(P)-binding Rossmann-fold superfamily protein; | 40 | 0.35 | 0.206 | 0.185 | |
| AT5G37530.2 | NAD(P)-binding Rossmann-fold superfamily protein; | 41 | 0.30 | 0.209 | 0.096 | . |
| AT5G37530.2 | NAD(P)-binding Rossmann-fold superfamily protein; | 42 | 0.26 | 0.216 | 0.057 | . |
| AT5G37530.2 | NAD(P)-binding Rossmann-fold superfamily protein; | 43 | 0.23 | 0.229 | 0.045 | * |
| AT5G37530.2 | NAD(P)-binding Rossmann-fold superfamily protein; | 44 | 0.20 | 0.250 | 0.051 | . |
| AT5G37530.2 | NAD(P)-binding Rossmann-fold superfamily protein; | 45 | 0.18 | 0.280 | 0.072 | . |
| AT5G37530.2 | NAD(P)-binding Rossmann-fold superfamily protein; | 46 | 0.17 | 0.321 | 0.117 | |

| | | | | | | |
|---|---|---|---|---|---|---|
| AT5G37530.2 | NAD(P)-binding Rossmann-fold superfamily protein; | 47 | 0.15 | 0.372 | 0.191 | |
| AT5G37590.1 | Tetratricopeptide repeat (TPR)-like superfamily protein | 2 | 0.89 | 0.288 | 1.000 | |
| AT5G37590.1 | Tetratricopeptide repeat (TPR)-like superfamily protein | 3 | 1.32 | 0.301 | 1.000 | |
| AT5G37590.1 | Tetratricopeptide repeat (TPR)-like superfamily protein | 41 | 0.90 | 0.256 | 1.000 | |
| AT5G37590.1 | Tetratricopeptide repeat (TPR)-like superfamily protein | 42 | 0.58 | 0.177 | 0.742 | |
| AT5G37590.1 | Tetratricopeptide repeat (TPR)-like superfamily protein | 43 | 1.30 | 0.153 | 0.984 | |
| AT5G37590.1 | Tetratricopeptide repeat (TPR)-like superfamily protein | 44 | 2.27 | 0.142 | 0.110 | |
| AT5G37590.1 | Tetratricopeptide repeat (TPR)-like superfamily protein | 45 | 1.44 | 0.144 | 0.884 | |
| AT5G37590.1 | Tetratricopeptide repeat (TPR)-like superfamily protein | 46 | 0.45 | 0.164 | 0.242 | |
| AT5G37590.1 | Tetratricopeptide repeat (TPR)-like superfamily protein | 47 | 0.08 | 0.284 | 0.003 | ** |
| AT5G38420.1 | RUBISCO small subunit 2B (RBCS2B) | 2 | 0.77 | 0.863 | 1.000 | |
| AT5G38420.1 | RUBISCO small subunit 2B (RBCS2B) | 3 | 1.24 | 0.843 | 1.000 | |
| AT5G38420.1 | RUBISCO small subunit 2B (RBCS2B) | 24 | 6.39 | 0.710 | 0.947 | |
| AT5G38420.1 | RUBISCO small subunit 2B (RBCS2B) | 25 | 3.58 | 0.529 | 0.968 | |
| AT5G38420.1 | RUBISCO small subunit 2B (RBCS2B) | 26 | 2.11 | 0.428 | 0.997 | |
| AT5G38420.1 | RUBISCO small subunit 2B (RBCS2B) | 27 | 1.95 | 0.389 | 0.997 | |
| AT5G38420.1 | RUBISCO small subunit 2B (RBCS2B) | 28 | 3.90 | 0.369 | 0.701 | |
| AT5G38420.1 | RUBISCO small subunit 2B (RBCS2B) | 29 | 9.26 | 0.356 | 0.088 | . |
| AT5G38420.1 | RUBISCO small subunit 2B (RBCS2B) | 30 | 8.41 | 0.352 | 0.110 | |
| AT5G38420.1 | RUBISCO small subunit 2B (RBCS2B) | 31 | 3.02 | 0.353 | 0.854 | |
| AT5G38420.1 | RUBISCO small subunit 2B (RBCS2B) | 32 | 1.32 | 0.356 | 1.000 | |
| AT5G38420.1 | RUBISCO small subunit 2B (RBCS2B) | 33 | 1.01 | 0.365 | 1.000 | |
| AT5G38420.1 | RUBISCO small subunit 2B (RBCS2B) | 34 | 0.92 | 0.391 | 1.000 | |
| AT5G38420.1 | RUBISCO small subunit 2B (RBCS2B) | 35 | 0.87 | 0.499 | 1.000 | |
| AT5G38420.1 | RUBISCO small subunit 2B (RBCS2B) | 36 | 0.91 | 0.786 | 1.000 | |
| AT5G38420.1 | RUBISCO small subunit 2B (RBCS2B) | 46 | 554.56 | 5.714 | 1.000 | |
| AT5G38420.1 | RUBISCO small subunit 2B (RBCS2B) | 47 | 1412.37 | 6.475 | 1.000 | |
| AT5G38430.1 | RUBISCO small subunit 1B (RBCS1B) | 2 | 0.71 | 0.804 | 1.000 | |
| AT5G38430.1 | RUBISCO small subunit 1B (RBCS1B) | 3 | 0.77 | 0.769 | 1.000 | |
| AT5G38430.1 | RUBISCO small subunit 1B (RBCS1B) | 4 | 0.83 | 0.747 | 1.000 | |
| AT5G38430.1 | RUBISCO small subunit 1B (RBCS1B) | 27 | 0.76 | 0.434 | 0.998 | |

| | | | | | | |
|---|---|---|---|---|---|---|
| AT5G38430.1 | RUBISCO small subunit 1B (RBCS1B) | 28 | 0.65 | 0.419 | 0.983 | |
| AT5G38430.1 | RUBISCO small subunit 1B (RBCS1B) | 29 | 0.54 | 0.412 | 0.933 | |
| AT5G38430.1 | RUBISCO small subunit 1B (RBCS1B) | 30 | 0.43 | 0.416 | 0.830 | |
| AT5G38430.1 | RUBISCO small subunit 1B (RBCS1B) | 31 | 0.34 | 0.428 | 0.689 | |
| AT5G38430.1 | RUBISCO small subunit 1B (RBCS1B) | 32 | 0.26 | 0.447 | 0.547 | |
| AT5G38430.1 | RUBISCO small subunit 1B (RBCS1B) | 33 | 0.20 | 0.472 | 0.425 | |
| AT5G38430.1 | RUBISCO small subunit 1B (RBCS1B) | 46 | 0.00 | 0.821 | 0.011 | * |
| AT5G38430.1 | RUBISCO small subunit 1B (RBCS1B) | 47 | 0.00 | 0.867 | 0.010 | * |
| AT5G39410.1 | Saccharopine dehydrogenase | 2 | 1.00 | 0.330 | 1.000 | |
| AT5G39410.1 | Saccharopine dehydrogenase | 3 | 1.05 | 0.331 | 1.000 | |
| AT5G39410.1 | Saccharopine dehydrogenase | 28 | 2.12 | 0.282 | 0.917 | |
| AT5G39410.1 | Saccharopine dehydrogenase | 29 | 2.15 | 0.257 | 0.857 | |
| AT5G39410.1 | Saccharopine dehydrogenase | 30 | 2.20 | 0.319 | 0.945 | |
| AT5G39410.1 | Saccharopine dehydrogenase | 39 | 3.67 | 0.237 | 0.179 | |
| AT5G39410.1 | Saccharopine dehydrogenase | 40 | 3.84 | 0.171 | 0.011 | * |
| AT5G39410.1 | Saccharopine dehydrogenase | 41 | 3.81 | 0.146 | 0.002 | ** |
| AT5G39410.1 | Saccharopine dehydrogenase | 42 | 3.52 | 0.140 | 0.002 | ** |
| AT5G39410.1 | Saccharopine dehydrogenase | 43 | 2.93 | 0.140 | 0.015 | * |
| AT5G39410.1 | Saccharopine dehydrogenase | 44 | 1.82 | 0.140 | 0.467 | |
| AT5G39410.1 | Saccharopine dehydrogenase | 45 | 0.82 | 0.141 | 0.999 | |
| AT5G39410.1 | Saccharopine dehydrogenase | 46 | 0.29 | 0.168 | 0.022 | * |
| AT5G39410.1 | Saccharopine dehydrogenase | 47 | 0.09 | 0.289 | 0.005 | ** |
| AT5G40610.1 | Glycerol-3-phosphate dehydrogenase (GPDHP) | 2 | 1.55 | 0.584 | 1.000 | |
| AT5G40610.1 | Glycerol-3-phosphate dehydrogenase (GPDHP) | 3 | 1.53 | 0.556 | 1.000 | |
| AT5G40610.1 | Glycerol-3-phosphate dehydrogenase (GPDHP) | 22 | 0.84 | 0.420 | 1.000 | |
| AT5G40610.1 | Glycerol-3-phosphate dehydrogenase (GPDHP) | 23 | 0.78 | 0.377 | 1.000 | |
| AT5G40610.1 | Glycerol-3-phosphate dehydrogenase (GPDHP) | 24 | 0.71 | 0.353 | 1.000 | |
| AT5G40610.1 | Glycerol-3-phosphate dehydrogenase (GPDHP) | 25 | 0.64 | 0.341 | 0.997 | |
| AT5G40610.1 | Glycerol-3-phosphate dehydrogenase (GPDHP) | 26 | 0.60 | 0.337 | 0.989 | |
| AT5G40610.1 | Glycerol-3-phosphate dehydrogenase (GPDHP) | 27 | 0.59 | 0.336 | 0.987 | |
| AT5G40610.1 | Glycerol-3-phosphate dehydrogenase (GPDHP) | 28 | 0.62 | 0.335 | 0.994 | |
| AT5G40610.1 | Glycerol-3-phosphate dehydrogenase (GPDHP) | 29 | 0.70 | 0.335 | 1.000 | |
| AT5G40610.1 | Glycerol-3-phosphate dehydrogenase (GPDHP) | 30 | 0.81 | 0.336 | 1.000 | |
| AT5G40610.1 | Glycerol-3-phosphate dehydrogenase (GPDHP) | 31 | 0.90 | 0.336 | 1.000 | |
| AT5G40610.1 | Glycerol-3-phosphate dehydrogenase (GPDHP) | 32 | 0.93 | 0.336 | 1.000 | |
| AT5G40610.1 | Glycerol-3-phosphate dehydrogenase (GPDHP) | 33 | 0.88 | 0.336 | 1.000 | |
| AT5G40610.1 | Glycerol-3-phosphate dehydrogenase (GPDHP) | 34 | 0.83 | 0.336 | 1.000 | |
| AT5G40610.1 | Glycerol-3-phosphate dehydrogenase (GPDHP) | 35 | 0.85 | 0.336 | 1.000 | |

| | | | | | |
|---|---|---|---|---|---|
| AT5G40610.1 | Glycerol-3-phosphate dehydrogenase (GPDHP) | 36 | 0.94 | 0.336 | 1.000 |
| AT5G40610.1 | Glycerol-3-phosphate dehydrogenase (GPDHP) | 37 | 1.09 | 0.336 | 1.000 |
| AT5G40610.1 | Glycerol-3-phosphate dehydrogenase (GPDHP) | 38 | 1.25 | 0.336 | 1.000 |
| AT5G40610.1 | Glycerol-3-phosphate dehydrogenase (GPDHP) | 39 | 1.44 | 0.336 | 0.999 |
| AT5G40610.1 | Glycerol-3-phosphate dehydrogenase (GPDHP) | 40 | 1.67 | 0.336 | 0.990 |
| AT5G40610.1 | Glycerol-3-phosphate dehydrogenase (GPDHP) | 41 | 1.91 | 0.337 | 0.954 |
| AT5G40610.1 | Glycerol-3-phosphate dehydrogenase (GPDHP) | 42 | 2.04 | 0.338 | 0.923 |
| AT5G40610.1 | Glycerol-3-phosphate dehydrogenase (GPDHP) | 43 | 1.96 | 0.338 | 0.943 |
| AT5G40610.1 | Glycerol-3-phosphate dehydrogenase (GPDHP) | 44 | 1.59 | 0.339 | 0.996 |
| AT5G40610.1 | Glycerol-3-phosphate dehydrogenase (GPDHP) | 45 | 1.04 | 0.347 | 1.000 |
| AT5G40610.1 | Glycerol-3-phosphate dehydrogenase (GPDHP) | 46 | 0.57 | 0.375 | 0.990 |
| AT5G40610.1 | Glycerol-3-phosphate dehydrogenase (GPDHP) | 47 | 0.29 | 0.446 | 0.743 |
| AT5G40650.1 | Succinate dehydrogenase subunit (SDH2-2, Fe-S subunit) | 2 | 0.64 | 0.616 | 1.000 |
| AT5G40650.1 | Succinate dehydrogenase subunit (SDH2-2, Fe-S subunit) | 3 | 0.89 | 0.440 | 1.000 |
| AT5G40650.1 | Succinate dehydrogenase subunit (SDH2-2, Fe-S subunit) | 4 | 1.11 | 0.414 | 1.000 |
| AT5G40650.1 | Succinate dehydrogenase subunit (SDH2-2, Fe-S subunit) | 5 | 1.14 | 0.372 | 1.000 |
| AT5G40650.1 | Succinate dehydrogenase subunit (SDH2-2, Fe-S subunit) | 6 | 1.09 | 0.350 | 1.000 |
| AT5G40650.1 | Succinate dehydrogenase subunit (SDH2-2, Fe-S subunit) | 7 | 1.11 | 0.343 | 1.000 |
| AT5G40650.1 | Succinate dehydrogenase subunit (SDH2-2, Fe-S subunit) | 8 | 1.23 | 0.337 | 1.000 |
| AT5G40650.1 | Succinate dehydrogenase subunit (SDH2-2, Fe-S subunit) | 9 | 1.32 | 0.334 | 1.000 |
| AT5G40650.1 | Succinate dehydrogenase subunit (SDH2-2, Fe-S subunit) | 10 | 1.30 | 0.335 | 1.000 |
| AT5G40650.1 | Succinate dehydrogenase subunit (SDH2-2, Fe-S subunit) | 11 | 1.13 | 0.337 | 1.000 |
| AT5G40650.1 | Succinate dehydrogenase subunit (SDH2-2, Fe-S subunit) | 12 | 1.06 | 0.336 | 1.000 |
| AT5G40650.1 | Succinate dehydrogenase subunit (SDH2-2, Fe-S subunit) | 13 | 1.33 | 0.334 | 1.000 |
| AT5G40650.1 | Succinate dehydrogenase subunit (SDH2-2, Fe-S subunit) | 14 | 1.77 | 0.331 | 1.000 |
| AT5G40650.1 | Succinate dehydrogenase subunit (SDH2-2, Fe-S subunit) | 15 | 2.13 | 0.330 | 0.999 |
| AT5G40650.1 | Succinate dehydrogenase subunit (SDH2-2, Fe-S subunit) | 16 | 2.38 | 0.328 | 0.996 |
| AT5G40650.1 | Succinate dehydrogenase subunit (SDH2-2, Fe-S subunit) | 17 | 3.09 | 0.329 | 0.927 |
| AT5G40650.1 | Succinate dehydrogenase subunit (SDH2-2, Fe-S subunit) | 18 | 4.35 | 0.334 | 0.650 |
| AT5G40650.1 | Succinate dehydrogenase subunit (SDH2-2, Fe-S subunit) | 19 | 4.84 | 0.339 | 0.562 |
| AT5G40650.1 | Succinate dehydrogenase subunit (SDH2-2, Fe-S subunit) | 20 | 3.95 | 0.347 | 0.799 |

| | | | | | |
|---|---|---|---|---|---|
| AT5G40650.1 | Succinate dehydrogenase subunit (SDH2-2, Fe-S subunit) | 21 | 2.68 | 0.362 | 0.994 |
| AT5G40650.1 | Succinate dehydrogenase subunit (SDH2-2, Fe-S subunit) | 22 | 1.97 | 0.386 | 1.000 |
| AT5G40650.1 | Succinate dehydrogenase subunit (SDH2-2, Fe-S subunit) | 23 | 1.78 | 0.380 | 1.000 |
| AT5G40650.1 | Succinate dehydrogenase subunit (SDH2-2, Fe-S subunit) | 24 | 1.79 | 0.394 | 1.000 |
| AT5G40650.1 | Succinate dehydrogenase subunit (SDH2-2, Fe-S subunit) | 25 | 1.86 | 0.426 | 1.000 |
| AT5G40650.1 | Succinate dehydrogenase subunit (SDH2-2, Fe-S subunit) | 27 | 2.26 | 0.403 | 1.000 |
| AT5G40650.1 | Succinate dehydrogenase subunit (SDH2-2, Fe-S subunit) | 28 | 2.89 | 0.373 | 0.989 |
| AT5G40650.1 | Succinate dehydrogenase subunit (SDH2-2, Fe-S subunit) | 29 | 3.77 | 0.350 | 0.844 |
| AT5G40650.1 | Succinate dehydrogenase subunit (SDH2-2, Fe-S subunit) | 30 | 3.93 | 0.338 | 0.768 |
| AT5G40650.1 | Succinate dehydrogenase subunit (SDH2-2, Fe-S subunit) | 31 | 4.91 | 0.335 | 0.527 |
| AT5G40650.1 | Succinate dehydrogenase subunit (SDH2-2, Fe-S subunit) | 32 | 6.44 | 0.335 | 0.285 |
| AT5G40650.1 | Succinate dehydrogenase subunit (SDH2-2, Fe-S subunit) | 33 | 6.12 | 0.334 | 0.323 |
| AT5G40650.1 | Succinate dehydrogenase subunit (SDH2-2, Fe-S subunit) | 34 | 4.48 | 0.335 | 0.627 |
| AT5G40650.1 | Succinate dehydrogenase subunit (SDH2-2, Fe-S subunit) | 35 | 2.62 | 0.336 | 0.988 |
| AT5G40650.1 | Succinate dehydrogenase subunit (SDH2-2, Fe-S subunit) | 36 | 1.07 | 0.335 | 1.000 |
| AT5G40650.1 | Succinate dehydrogenase subunit (SDH2-2, Fe-S subunit) | 37 | 0.48 | 0.335 | 1.000 |
| AT5G40650.1 | Succinate dehydrogenase subunit (SDH2-2, Fe-S subunit) | 38 | 0.52 | 0.336 | 1.000 |
| AT5G40650.1 | Succinate dehydrogenase subunit (SDH2-2, Fe-S subunit) | 39 | 1.07 | 0.335 | 1.000 |
| AT5G40650.1 | Succinate dehydrogenase subunit (SDH2-2, Fe-S subunit) | 40 | 1.97 | 0.334 | 1.000 |
| AT5G40650.1 | Succinate dehydrogenase subunit (SDH2-2, Fe-S subunit) | 41 | 2.46 | 0.335 | 0.995 |
| AT5G40650.1 | Succinate dehydrogenase subunit (SDH2-2, Fe-S subunit) | 42 | 2.73 | 0.338 | 0.982 |
| AT5G40650.1 | Succinate dehydrogenase subunit (SDH2-2, Fe-S subunit) | 43 | 3.69 | 0.339 | 0.829 |
| AT5G40650.1 | Succinate dehydrogenase subunit (SDH2-2, Fe-S subunit) | 44 | 4.99 | 0.339 | 0.531 |
| AT5G40650.1 | Succinate dehydrogenase subunit (SDH2-2, Fe-S subunit) | 45 | 3.89 | 0.339 | 0.784 |
| AT5G40650.1 | Succinate dehydrogenase subunit (SDH2-2, Fe-S subunit) | 46 | 1.41 | 0.370 | 1.000 |
| AT5G40650.1 | Succinate dehydrogenase subunit (SDH2-2, Fe-S subunit) | 47 | 0.31 | 0.609 | 1.000 |
| AT5G40660.1 | ATP synthase subunit (ATP12) | 2 | 0.89 | 0.455 | 1.000 |
| AT5G40660.1 | ATP synthase subunit (ATP12) | 3 | 0.88 | 0.443 | 1.000 |
| AT5G40660.1 | ATP synthase subunit (ATP12) | 39 | 1.27 | 0.330 | 0.998 |
| AT5G40660.1 | ATP synthase subunit (ATP12) | 40 | 1.32 | 0.273 | 0.992 |
| AT5G40660.1 | ATP synthase subunit (ATP12) | 41 | 1.38 | 0.223 | 0.963 |
| AT5G40660.1 | ATP synthase subunit (ATP12) | 42 | 1.44 | 0.185 | 0.885 |
| AT5G40660.1 | ATP synthase subunit (ATP12) | 43 | 1.48 | 0.160 | 0.766 |
| AT5G40660.1 | ATP synthase subunit (ATP12) | 44 | 1.48 | 0.153 | 0.741 |
| AT5G40660.1 | ATP synthase subunit (ATP12) | 45 | 1.42 | 0.172 | 0.874 |
| AT5G40660.1 | ATP synthase subunit (ATP12) | 46 | 1.30 | 0.218 | 0.982 |

| | | | | | |
|---|---|---|---|---|---|
| AT5G40660.1 | ATP synthase subunit (ATP12) | 47 | 1.18 | 0.289 | 1.000 |
| AT5G40770.1 | Prohibitin 3 (PHB3) | 2 | 0.43 | 0.349 | 0.998 |
| AT5G40770.1 | Prohibitin 3 (PHB3) | 3 | 0.41 | 0.233 | 0.839 |
| AT5G40770.1 | Prohibitin 3 (PHB3) | 4 | 0.42 | 0.204 | 0.726 |
| AT5G40770.1 | Prohibitin 3 (PHB3) | 5 | 0.47 | 0.203 | 0.878 |
| AT5G40770.1 | Prohibitin 3 (PHB3) | 6 | 0.55 | 0.204 | 0.981 |
| AT5G40770.1 | Prohibitin 3 (PHB3) | 7 | 0.64 | 0.203 | 0.999 |
| AT5G40770.1 | Prohibitin 3 (PHB3) | 8 | 0.76 | 0.199 | 1.000 |
| AT5G40770.1 | Prohibitin 3 (PHB3) | 9 | 0.93 | 0.197 | 1.000 |
| AT5G40770.1 | Prohibitin 3 (PHB3) | 10 | 1.12 | 0.197 | 1.000 |
| AT5G40770.1 | Prohibitin 3 (PHB3) | 11 | 1.30 | 0.197 | 1.000 |
| AT5G40770.1 | Prohibitin 3 (PHB3) | 12 | 1.48 | 0.197 | 1.000 |
| AT5G40770.1 | Prohibitin 3 (PHB3) | 13 | 1.57 | 0.195 | 0.999 |
| AT5G40770.1 | Prohibitin 3 (PHB3) | 14 | 1.40 | 0.193 | 1.000 |
| AT5G40770.1 | Prohibitin 3 (PHB3) | 15 | 1.04 | 0.192 | 1.000 |
| AT5G40770.1 | Prohibitin 3 (PHB3) | 16 | 0.78 | 0.191 | 1.000 |
| AT5G40770.1 | Prohibitin 3 (PHB3) | 17 | 0.71 | 0.191 | 1.000 |
| AT5G40770.1 | Prohibitin 3 (PHB3) | 18 | 0.76 | 0.193 | 1.000 |
| AT5G40770.1 | Prohibitin 3 (PHB3) | 19 | 0.88 | 0.197 | 1.000 |
| AT5G40770.1 | Prohibitin 3 (PHB3) | 20 | 1.04 | 0.202 | 1.000 |
| AT5G40770.1 | Prohibitin 3 (PHB3) | 21 | 1.24 | 0.204 | 1.000 |
| AT5G40770.1 | Prohibitin 3 (PHB3) | 22 | 1.48 | 0.204 | 1.000 |
| AT5G40770.1 | Prohibitin 3 (PHB3) | 23 | 1.60 | 0.203 | 0.999 |
| AT5G40770.1 | Prohibitin 3 (PHB3) | 24 | 1.39 | 0.199 | 1.000 |
| AT5G40770.1 | Prohibitin 3 (PHB3) | 25 | 0.92 | 0.196 | 1.000 |
| AT5G40770.1 | Prohibitin 3 (PHB3) | 26 | 0.56 | 0.195 | 0.980 |
| AT5G40770.1 | Prohibitin 3 (PHB3) | 27 | 0.41 | 0.195 | 0.597 |
| AT5G40770.1 | Prohibitin 3 (PHB3) | 28 | 0.39 | 0.195 | 0.510 |
| AT5G40770.1 | Prohibitin 3 (PHB3) | 29 | 0.49 | 0.196 | 0.893 |
| AT5G40770.1 | Prohibitin 3 (PHB3) | 30 | 0.78 | 0.197 | 1.000 |
| AT5G40770.1 | Prohibitin 3 (PHB3) | 31 | 1.36 | 0.198 | 1.000 |
| AT5G40770.1 | Prohibitin 3 (PHB3) | 32 | 2.00 | 0.200 | 0.922 |
| AT5G40770.1 | Prohibitin 3 (PHB3) | 33 | 2.06 | 0.204 | 0.908 |
| AT5G40770.1 | Prohibitin 3 (PHB3) | 34 | 1.62 | 0.220 | 0.999 |
| AT5G40770.1 | Prohibitin 3 (PHB3) | 35 | 1.21 | 0.273 | 1.000 |
| AT5G40770.1 | Prohibitin 3 (PHB3) | 36 | 1.04 | 0.385 | 1.000 |
| AT5G40770.1 | Prohibitin 3 (PHB3) | 46 | 805.31 | 2.605 | 0.995 |
| AT5G40770.1 | Prohibitin 3 (PHB3) | 47 | 2330.11 | 2.942 | 0.994 |
| AT5G40810.2 | Ubiquinol:cytochrome c oxidoreductase subunit (cyt c1-1) | 3 | 0.88 | 0.279 | 1.000 |
| AT5G40810.2 | Ubiquinol:cytochrome c oxidoreductase subunit (cyt c1-1) | 4 | 0.86 | 0.262 | 1.000 |
| AT5G40810.2 | Ubiquinol:cytochrome c oxidoreductase subunit (cyt c1-1) | 5 | 0.94 | 0.261 | 1.000 |
| AT5G40810.2 | Ubiquinol:cytochrome c oxidoreductase subunit (cyt c1-1) | 6 | 0.83 | 0.263 | 1.000 |
| AT5G40810.2 | Ubiquinol:cytochrome c oxidoreductase subunit (cyt c1-1) | 7 | 0.72 | 0.266 | 1.000 |
| AT5G40810.2 | Ubiquinol:cytochrome c oxidoreductase subunit (cyt c1-1) | 8 | 0.93 | 0.257 | 1.000 |
| AT5G40810.2 | Ubiquinol:cytochrome c oxidoreductase subunit (cyt c1-1) | 9 | 1.16 | 0.252 | 1.000 |
| AT5G40810.2 | Ubiquinol:cytochrome c oxidoreductase subunit (cyt c1-1) | 10 | 1.06 | 0.253 | 1.000 |

| | | | | | | |
|---|---|---|---|---|---|---|
| AT5G40810.2 | Ubiquinol:cytochrome c oxidoreductase subunit (cyt c1-1) | 11 | 0.83 | 0.254 | 1.000 | |
| AT5G40810.2 | Ubiquinol:cytochrome c oxidoreductase subunit (cyt c1-1) | 12 | 0.81 | 0.255 | 1.000 | |
| AT5G40810.2 | Ubiquinol:cytochrome c oxidoreductase subunit (cyt c1-1) | 13 | 1.11 | 0.251 | 1.000 | |
| AT5G40810.2 | Ubiquinol:cytochrome c oxidoreductase subunit (cyt c1-1) | 14 | 1.45 | 0.246 | 1.000 | |
| AT5G40810.2 | Ubiquinol:cytochrome c oxidoreductase subunit (cyt c1-1) | 15 | 1.32 | 0.246 | 1.000 | |
| AT5G40810.2 | Ubiquinol:cytochrome c oxidoreductase subunit (cyt c1-1) | 16 | 0.92 | 0.244 | 1.000 | |
| AT5G40810.2 | Ubiquinol:cytochrome c oxidoreductase subunit (cyt c1-1) | 17 | 0.66 | 0.245 | 1.000 | |
| AT5G40810.2 | Ubiquinol:cytochrome c oxidoreductase subunit (cyt c1-1) | 18 | 0.69 | 0.248 | 1.000 | |
| AT5G40810.2 | Ubiquinol:cytochrome c oxidoreductase subunit (cyt c1-1) | 19 | 0.99 | 0.255 | 1.000 | |
| AT5G40810.2 | Ubiquinol:cytochrome c oxidoreductase subunit (cyt c1-1) | 20 | 1.41 | 0.263 | 1.000 | |
| AT5G40810.2 | Ubiquinol:cytochrome c oxidoreductase subunit (cyt c1-1) | 21 | 1.42 | 0.267 | 1.000 | |
| AT5G40810.2 | Ubiquinol:cytochrome c oxidoreductase subunit (cyt c1-1) | 22 | 0.72 | 0.266 | 1.000 | |
| AT5G40810.2 | Ubiquinol:cytochrome c oxidoreductase subunit (cyt c1-1) | 23 | 0.21 | 0.266 | 0.309 | |
| AT5G40810.2 | Ubiquinol:cytochrome c oxidoreductase subunit (cyt c1-1) | 24 | 0.11 | 0.259 | 0.008 | ** |
| AT5G40810.2 | Ubiquinol:cytochrome c oxidoreductase subunit (cyt c1-1) | 25 | 0.18 | 0.252 | 0.111 | |
| AT5G40810.2 | Ubiquinol:cytochrome c oxidoreductase subunit (cyt c1-1) | 26 | 0.75 | 0.252 | 1.000 | |
| AT5G40810.2 | Ubiquinol:cytochrome c oxidoreductase subunit (cyt c1-1) | 27 | 2.69 | 0.252 | 0.938 | |
| AT5G40810.2 | Ubiquinol:cytochrome c oxidoreductase subunit (cyt c1-1) | 28 | 2.17 | 0.251 | 0.997 | |
| AT5G40810.2 | Ubiquinol:cytochrome c oxidoreductase subunit (cyt c1-1) | 29 | 0.69 | 0.251 | 1.000 | |
| AT5G40810.2 | Ubiquinol:cytochrome c oxidoreductase subunit (cyt c1-1) | 30 | 0.46 | 0.251 | 0.997 | |

| | | | | | |
|---|---|---|---|---|---|
| AT5G40810.2 | Ubiquinol:cytochrome c oxidoreductase subunit (cyt c1-1) | 31 | 0.75 | 0.252 | 1.000 |
| AT5G40810.2 | Ubiquinol:cytochrome c oxidoreductase subunit (cyt c1-1) | 32 | 1.42 | 0.253 | 1.000 |
| AT5G40810.2 | Ubiquinol:cytochrome c oxidoreductase subunit (cyt c1-1) | 33 | 1.68 | 0.253 | 1.000 |
| AT5G40810.2 | Ubiquinol:cytochrome c oxidoreductase subunit (cyt c1-1) | 34 | 1.03 | 0.253 | 1.000 |
| AT5G40810.2 | Ubiquinol:cytochrome c oxidoreductase subunit (cyt c1-1) | 35 | 0.51 | 0.253 | 1.000 |
| AT5G40810.2 | Ubiquinol:cytochrome c oxidoreductase subunit (cyt c1-1) | 36 | 0.69 | 0.252 | 1.000 |
| AT5G40810.2 | Ubiquinol:cytochrome c oxidoreductase subunit (cyt c1-1) | 37 | 1.74 | 0.251 | 1.000 |
| AT5G40810.2 | Ubiquinol:cytochrome c oxidoreductase subunit (cyt c1-1) | 38 | 2.61 | 0.252 | 0.954 |
| AT5G40810.2 | Ubiquinol:cytochrome c oxidoreductase subunit (cyt c1-1) | 39 | 3.18 | 0.253 | 0.779 |
| AT5G40810.2 | Ubiquinol:cytochrome c oxidoreductase subunit (cyt c1-1) | 40 | 4.28 | 0.254 | 0.367 |
| AT5G40810.2 | Ubiquinol:cytochrome c oxidoreductase subunit (cyt c1-1) | 41 | 4.56 | 0.255 | 0.296 |
| AT5G40810.2 | Ubiquinol:cytochrome c oxidoreductase subunit (cyt c1-1) | 42 | 3.49 | 0.258 | 0.685 |
| AT5G40810.2 | Ubiquinol:cytochrome c oxidoreductase subunit (cyt c1-1) | 43 | 3.60 | 0.260 | 0.654 |
| AT5G40810.2 | Ubiquinol:cytochrome c oxidoreductase subunit (cyt c1-1) | 44 | 3.00 | 0.260 | 0.879 |
| AT5G40810.2 | Ubiquinol:cytochrome c oxidoreductase subunit (cyt c1-1) | 45 | 1.33 | 0.263 | 1.000 |
| AT5G40810.2 | Ubiquinol:cytochrome c oxidoreductase subunit (cyt c1-1) | 46 | 0.40 | 0.297 | 0.996 |
| AT5G40810.2 | Ubiquinol:cytochrome c oxidoreductase subunit (cyt c1-1) | 47 | 0.13 | 0.514 | 0.941 |
| AT5G40810.2 | Ubiquinol:cytochrome c oxidoreductase subunit (cyt c1-1) | 48 | 0.05 | 0.898 | 0.988 |
| AT5G40930.1 | Translocase of the outer mitochondrial membrane 20 (TOM20-4) | 2 | 0.95 | 0.554 | 1.000 |
| AT5G40930.1 | Translocase of the outer mitochondrial membrane 20 (TOM20-4) | 3 | 1.07 | 0.510 | 1.000 |

| | | | | | |
|---|---|---|---|---|---|
| AT5G40930.1 | Translocase of the outer mitochondrial membrane 20 (TOM20-4) | 16 | 0.62 | 0.372 | 1.000 |
| AT5G40930.1 | Translocase of the outer mitochondrial membrane 20 (TOM20-4) | 17 | 0.68 | 0.325 | 1.000 |
| AT5G40930.1 | Translocase of the outer mitochondrial membrane 20 (TOM20-4) | 18 | 0.99 | 0.301 | 1.000 |
| AT5G40930.1 | Translocase of the outer mitochondrial membrane 20 (TOM20-4) | 19 | 1.70 | 0.293 | 0.999 |
| AT5G40930.1 | Translocase of the outer mitochondrial membrane 20 (TOM20-4) | 20 | 2.57 | 0.295 | 0.895 |
| AT5G40930.1 | Translocase of the outer mitochondrial membrane 20 (TOM20-4) | 21 | 3.07 | 0.296 | 0.737 |
| AT5G40930.1 | Translocase of the outer mitochondrial membrane 20 (TOM20-4) | 22 | 2.45 | 0.296 | 0.926 |
| AT5G40930.1 | Translocase of the outer mitochondrial membrane 20 (TOM20-4) | 23 | 1.45 | 0.295 | 1.000 |
| AT5G40930.1 | Translocase of the outer mitochondrial membrane 20 (TOM20-4) | 24 | 0.74 | 0.290 | 1.000 |
| AT5G40930.1 | Translocase of the outer mitochondrial membrane 20 (TOM20-4) | 25 | 0.45 | 0.286 | 0.961 |
| AT5G40930.1 | Translocase of the outer mitochondrial membrane 20 (TOM20-4) | 26 | 0.51 | 0.284 | 0.988 |
| AT5G40930.1 | Translocase of the outer mitochondrial membrane 20 (TOM20-4) | 27 | 0.64 | 0.284 | 1.000 |
| AT5G40930.1 | Translocase of the outer mitochondrial membrane 20 (TOM20-4) | 28 | 0.85 | 0.286 | 1.000 |
| AT5G40930.1 | Translocase of the outer mitochondrial membrane 20 (TOM20-4) | 29 | 1.68 | 0.287 | 0.999 |
| AT5G40930.1 | Translocase of the outer mitochondrial membrane 20 (TOM20-4) | 30 | 3.36 | 0.286 | 0.590 |
| AT5G40930.1 | Translocase of the outer mitochondrial membrane 20 (TOM20-4) | 31 | 2.84 | 0.287 | 0.786 |
| AT5G40930.1 | Translocase of the outer mitochondrial membrane 20 (TOM20-4) | 32 | 0.87 | 0.292 | 1.000 |
| AT5G40930.1 | Translocase of the outer mitochondrial membrane 20 (TOM20-4) | 33 | 0.22 | 0.305 | 0.366 |
| AT5G40930.1 | Translocase of the outer mitochondrial membrane 20 (TOM20-4) | 34 | 0.12 | 0.360 | 0.160 |
| AT5G40930.1 | Translocase of the outer mitochondrial membrane 20 (TOM20-4) | 35 | 0.14 | 0.564 | 0.819 |

| | | | | | | |
|---|---|---|---|---|---|---|
| AT5G40930.1 | Translocase of the outer mitochondrial membrane 20 (TOM20-4) | 46 | ########### | 5.115 | 0.767 | |
| AT5G40930.1 | Translocase of the outer mitochondrial membrane 20 (TOM20-4) | 47 | ########### | 5.759 | 0.747 | |
| AT5G41685.1 | Translocase of the outer mitochondrial membrane 7 (TOM7-1) | 2 | 1.04 | 0.486 | 1.000 | |
| AT5G41685.1 | Translocase of the outer mitochondrial membrane 7 (TOM7-1) | 3 | 1.00 | 0.448 | 1.000 | |
| AT5G41685.1 | Translocase of the outer mitochondrial membrane 7 (TOM7-1) | 20 | 2.80 | 0.341 | 0.926 | |
| AT5G41685.1 | Translocase of the outer mitochondrial membrane 7 (TOM7-1) | 21 | 2.69 | 0.283 | 0.821 | |
| AT5G41685.1 | Translocase of the outer mitochondrial membrane 7 (TOM7-1) | 22 | 2.10 | 0.249 | 0.929 | |
| AT5G41685.1 | Translocase of the outer mitochondrial membrane 7 (TOM7-1) | 23 | 1.51 | 0.239 | 0.999 | |
| AT5G41685.1 | Translocase of the outer mitochondrial membrane 7 (TOM7-1) | 24 | 1.03 | 0.230 | 1.000 | |
| AT5G41685.1 | Translocase of the outer mitochondrial membrane 7 (TOM7-1) | 25 | 0.89 | 0.224 | 1.000 | |
| AT5G41685.1 | Translocase of the outer mitochondrial membrane 7 (TOM7-1) | 26 | 0.95 | 0.224 | 1.000 | |
| AT5G41685.1 | Translocase of the outer mitochondrial membrane 7 (TOM7-1) | 27 | 0.99 | 0.225 | 1.000 | |
| AT5G41685.1 | Translocase of the outer mitochondrial membrane 7 (TOM7-1) | 28 | 1.06 | 0.224 | 1.000 | |
| AT5G41685.1 | Translocase of the outer mitochondrial membrane 7 (TOM7-1) | 29 | 1.62 | 0.225 | 0.994 | |
| AT5G41685.1 | Translocase of the outer mitochondrial membrane 7 (TOM7-1) | 30 | 2.51 | 0.226 | 0.648 | |
| AT5G41685.1 | Translocase of the outer mitochondrial membrane 7 (TOM7-1) | 31 | 1.94 | 0.227 | 0.940 | |
| AT5G41685.1 | Translocase of the outer mitochondrial membrane 7 (TOM7-1) | 32 | 0.77 | 0.232 | 1.000 | |
| AT5G41685.1 | Translocase of the outer mitochondrial membrane 7 (TOM7-1) | 33 | 0.26 | 0.245 | 0.239 | |
| AT5G41685.1 | Translocase of the outer mitochondrial membrane 7 (TOM7-1) | 34 | 0.15 | 0.300 | 0.089 | . |
| AT5G41685.1 | Translocase of the outer mitochondrial membrane 7 (TOM7-1) | 35 | 0.14 | 0.489 | 0.664 | |

| | | | | | | |
|---|---|---|---|---|---|---|
| AT5G41685.1 | Translocase of the outer mitochondrial membrane 7 (TOM7-1) | 46 | 2289303.51 | 4.960 | 0.934 | |
| AT5G41685.1 | Translocase of the outer mitochondrial membrane 7 (TOM7-1) | 47 | 22030839.97 | 5.589 | 0.923 | |
| AT5G42270.1 | FTSH protease 5 (FTSH5) | 2 | 0.77 | 0.510 | 1.000 | |
| AT5G42270.1 | FTSH protease 5 (FTSH5) | 3 | 0.57 | 0.391 | 1.000 | |
| AT5G42270.1 | FTSH protease 5 (FTSH5) | 4 | 0.44 | 0.352 | 0.989 | |
| AT5G42270.1 | FTSH protease 5 (FTSH5) | 5 | 0.35 | 0.347 | 0.893 | |
| AT5G42270.1 | FTSH protease 5 (FTSH5) | 6 | 0.26 | 0.348 | 0.660 | |
| AT5G42270.1 | FTSH protease 5 (FTSH5) | 7 | 0.21 | 0.347 | 0.430 | |
| AT5G42270.1 | FTSH protease 5 (FTSH5) | 8 | 0.18 | 0.344 | 0.312 | |
| AT5G42270.1 | FTSH protease 5 (FTSH5) | 9 | 0.17 | 0.342 | 0.273 | |
| AT5G42270.1 | FTSH protease 5 (FTSH5) | 10 | 0.18 | 0.342 | 0.320 | |
| AT5G42270.1 | FTSH protease 5 (FTSH5) | 11 | 0.23 | 0.342 | 0.496 | |
| AT5G42270.1 | FTSH protease 5 (FTSH5) | 12 | 0.32 | 0.341 | 0.817 | |
| AT5G42270.1 | FTSH protease 5 (FTSH5) | 13 | 0.44 | 0.340 | 0.985 | |
| AT5G42270.1 | FTSH protease 5 (FTSH5) | 14 | 0.51 | 0.338 | 0.998 | |
| AT5G42270.1 | FTSH protease 5 (FTSH5) | 15 | 0.46 | 0.338 | 0.988 | |
| AT5G42270.1 | FTSH protease 5 (FTSH5) | 16 | 0.35 | 0.338 | 0.888 | |
| AT5G42270.1 | FTSH protease 5 (FTSH5) | 17 | 0.30 | 0.340 | 0.755 | |
| AT5G42270.1 | FTSH protease 5 (FTSH5) | 18 | 0.31 | 0.344 | 0.798 | |
| AT5G42270.1 | FTSH protease 5 (FTSH5) | 19 | 0.36 | 0.352 | 0.923 | |
| AT5G42270.1 | FTSH protease 5 (FTSH5) | 20 | 0.40 | 0.364 | 0.977 | |
| AT5G42270.1 | FTSH protease 5 (FTSH5) | 21 | 0.39 | 0.376 | 0.976 | |
| AT5G42270.1 | FTSH protease 5 (FTSH5) | 22 | 0.33 | 0.381 | 0.918 | |
| AT5G42270.1 | FTSH protease 5 (FTSH5) | 23 | 0.27 | 0.385 | 0.812 | |
| AT5G42270.1 | FTSH protease 5 (FTSH5) | 24 | 0.25 | 0.382 | 0.724 | |
| AT5G42270.1 | FTSH protease 5 (FTSH5) | 25 | 0.24 | 0.374 | 0.656 | |
| AT5G42270.1 | FTSH protease 5 (FTSH5) | 26 | 0.23 | 0.362 | 0.576 | |
| AT5G42270.1 | FTSH protease 5 (FTSH5) | 27 | 0.21 | 0.350 | 0.467 | |
| AT5G42270.1 | FTSH protease 5 (FTSH5) | 28 | 0.19 | 0.344 | 0.347 | |
| AT5G42270.1 | FTSH protease 5 (FTSH5) | 29 | 0.16 | 0.341 | 0.230 | |
| AT5G42270.1 | FTSH protease 5 (FTSH5) | 30 | 0.14 | 0.341 | 0.153 | |
| AT5G42270.1 | FTSH protease 5 (FTSH5) | 31 | 0.13 | 0.344 | 0.125 | |
| AT5G42270.1 | FTSH protease 5 (FTSH5) | 32 | 0.13 | 0.349 | 0.142 | |
| AT5G42270.1 | FTSH protease 5 (FTSH5) | 33 | 0.14 | 0.359 | 0.201 | |
| AT5G42270.1 | FTSH protease 5 (FTSH5) | 34 | 0.15 | 0.369 | 0.263 | |
| AT5G42270.1 | FTSH protease 5 (FTSH5) | 35 | 0.15 | 0.378 | 0.294 | |
| AT5G42270.1 | FTSH protease 5 (FTSH5) | 36 | 0.14 | 0.372 | 0.260 | |
| AT5G42270.1 | FTSH protease 5 (FTSH5) | 37 | 0.14 | 0.365 | 0.222 | |
| AT5G42270.1 | FTSH protease 5 (FTSH5) | 38 | 0.15 | 0.353 | 0.220 | |
| AT5G42270.1 | FTSH protease 5 (FTSH5) | 39 | 0.18 | 0.346 | 0.320 | |
| AT5G42270.1 | FTSH protease 5 (FTSH5) | 40 | 0.24 | 0.344 | 0.579 | |
| AT5G42270.1 | FTSH protease 5 (FTSH5) | 41 | 0.35 | 0.345 | 0.887 | |
| AT5G42270.1 | FTSH protease 5 (FTSH5) | 42 | 0.42 | 0.346 | 0.975 | |
| AT5G42270.1 | FTSH protease 5 (FTSH5) | 43 | 0.39 | 0.347 | 0.955 | |
| AT5G42270.1 | FTSH protease 5 (FTSH5) | 44 | 0.30 | 0.346 | 0.793 | |
| AT5G42270.1 | FTSH protease 5 (FTSH5) | 45 | 0.20 | 0.355 | 0.442 | |
| AT5G42270.1 | FTSH protease 5 (FTSH5) | 46 | 0.12 | 0.403 | 0.252 | |
| AT5G42270.1 | FTSH protease 5 (FTSH5) | 47 | 0.07 | 0.540 | 0.321 | |
| AT5G42650.1 | Allene oxide synthase (AOS) | 2 | 0.92 | 0.327 | 1.000 | |
| AT5G42650.1 | Allene oxide synthase (AOS) | 3 | 0.35 | 0.326 | 0.708 | |
| AT5G42650.1 | Allene oxide synthase (AOS) | 42 | 2.69 | 0.423 | 0.921 | |
| AT5G42650.1 | Allene oxide synthase (AOS) | 43 | 12.38 | 0.220 | 0.000 | *** |
| AT5G42650.1 | Allene oxide synthase (AOS) | 44 | 3.73 | 0.189 | 0.031 | * |
| AT5G42650.1 | Allene oxide synthase (AOS) | 45 | 0.43 | 0.186 | 0.336 | |
| AT5G42650.1 | Allene oxide synthase (AOS) | 46 | 0.19 | 0.191 | 0.004 | ** |
| AT5G42650.1 | Allene oxide synthase (AOS) | 47 | 0.18 | 0.578 | 0.787 | |
| AT5G43430.1 | Electron transfer flavoprotein complex subunit (ETF beta) | 2 | 0.99 | 0.330 | 1.000 | |

| | | | | | | |
|---|---|---|---|---|---|---|
| AT5G43430.1 | Electron transfer flavoprotein complex subunit (ETF beta) | 3 | 0.92 | 0.268 | 1.000 | |
| AT5G43430.1 | Electron transfer flavoprotein complex subunit (ETF beta) | 4 | 0.85 | 0.274 | 1.000 | |
| AT5G43430.1 | Electron transfer flavoprotein complex subunit (ETF beta) | 9 | 0.67 | 0.281 | 1.000 | |
| AT5G43430.1 | Electron transfer flavoprotein complex subunit (ETF beta) | 10 | 0.69 | 0.252 | 1.000 | |
| AT5G43430.1 | Electron transfer flavoprotein complex subunit (ETF beta) | 11 | 0.73 | 0.222 | 1.000 | |
| AT5G43430.1 | Electron transfer flavoprotein complex subunit (ETF beta) | 12 | 0.82 | 0.200 | 1.000 | |
| AT5G43430.1 | Electron transfer flavoprotein complex subunit (ETF beta) | 13 | 0.94 | 0.188 | 1.000 | |
| AT5G43430.1 | Electron transfer flavoprotein complex subunit (ETF beta) | 14 | 1.08 | 0.182 | 1.000 | |
| AT5G43430.1 | Electron transfer flavoprotein complex subunit (ETF beta) | 15 | 1.23 | 0.182 | 1.000 | |
| AT5G43430.1 | Electron transfer flavoprotein complex subunit (ETF beta) | 16 | 1.35 | 0.191 | 1.000 | |
| AT5G43430.1 | Electron transfer flavoprotein complex subunit (ETF beta) | 17 | 1.44 | 0.209 | 1.000 | |
| AT5G43430.1 | Electron transfer flavoprotein complex subunit (ETF beta) | 18 | 1.47 | 0.237 | 1.000 | |
| AT5G43430.1 | Electron transfer flavoprotein complex subunit (ETF beta) | 22 | 1.22 | 0.294 | 1.000 | |
| AT5G43430.1 | Electron transfer flavoprotein complex subunit (ETF beta) | 23 | 1.15 | 0.284 | 1.000 | |
| AT5G43430.1 | Electron transfer flavoprotein complex subunit (ETF beta) | 24 | 1.10 | 0.261 | 1.000 | |
| AT5G43430.1 | Electron transfer flavoprotein complex subunit (ETF beta) | 26 | 1.07 | 0.224 | 1.000 | |
| AT5G43430.1 | Electron transfer flavoprotein complex subunit (ETF beta) | 27 | 1.07 | 0.201 | 1.000 | |
| AT5G43430.1 | Electron transfer flavoprotein complex subunit (ETF beta) | 28 | 1.05 | 0.183 | 1.000 | |
| AT5G43430.1 | Electron transfer flavoprotein complex subunit (ETF beta) | 29 | 0.97 | 0.172 | 1.000 | |
| AT5G43430.1 | Electron transfer flavoprotein complex subunit (ETF beta) | 30 | 0.85 | 0.166 | 1.000 | |
| AT5G43430.1 | Electron transfer flavoprotein complex subunit (ETF beta) | 31 | 0.72 | 0.165 | 1.000 | |
| AT5G43430.1 | Electron transfer flavoprotein complex subunit (ETF beta) | 32 | 0.62 | 0.165 | 0.974 | |
| AT5G43430.1 | Electron transfer flavoprotein complex subunit (ETF beta) | 33 | 0.58 | 0.165 | 0.917 | |
| AT5G43430.1 | Electron transfer flavoprotein complex subunit (ETF beta) | 34 | 0.63 | 0.165 | 0.982 | |
| AT5G43430.1 | Electron transfer flavoprotein complex subunit (ETF beta) | 35 | 0.81 | 0.165 | 1.000 | |
| AT5G43430.1 | Electron transfer flavoprotein complex subunit (ETF beta) | 36 | 1.10 | 0.165 | 1.000 | |
| AT5G43430.1 | Electron transfer flavoprotein complex subunit (ETF beta) | 37 | 1.43 | 0.164 | 0.998 | |
| AT5G43430.1 | Electron transfer flavoprotein complex subunit (ETF beta) | 38 | 1.80 | 0.164 | 0.868 | |
| AT5G43430.1 | Electron transfer flavoprotein complex subunit (ETF beta) | 39 | 2.23 | 0.164 | 0.458 | |
| AT5G43430.1 | Electron transfer flavoprotein complex subunit (ETF beta) | 40 | 2.75 | 0.165 | 0.152 | |
| AT5G43430.1 | Electron transfer flavoprotein complex subunit (ETF beta) | 41 | 3.19 | 0.166 | 0.058 | . |

| | | | | | | |
|---|---|---|---|---|---|---|
| AT5G43430.1 | Electron transfer flavoprotein complex subunit (ETF beta) | 42 | 3.22 | 0.168 | 0.057 | . |
| AT5G43430.1 | Electron transfer flavoprotein complex subunit (ETF beta) | 43 | 2.72 | 0.169 | 0.187 | |
| AT5G43430.1 | Electron transfer flavoprotein complex subunit (ETF beta) | 44 | 1.84 | 0.169 | 0.859 | |
| AT5G43430.1 | Electron transfer flavoprotein complex subunit (ETF beta) | 45 | 1.04 | 0.176 | 1.000 | |
| AT5G43430.1 | Electron transfer flavoprotein complex subunit (ETF beta) | 46 | 0.52 | 0.216 | 0.956 | |
| AT5G43430.1 | Electron transfer flavoprotein complex subunit (ETF beta) | 47 | 0.24 | 0.314 | 0.579 | |
| AT5G43970.1 | Translocase of the outer mitochondrial membrane 22 (TOM22-2) | 2 | 0.64 | 0.318 | 1.000 | |
| AT5G43970.1 | Translocase of the outer mitochondrial membrane 22 (TOM22-2) | 3 | 0.73 | 0.204 | 1.000 | |
| AT5G43970.1 | Translocase of the outer mitochondrial membrane 22 (TOM22-2) | 4 | 0.93 | 0.189 | 1.000 | |
| AT5G43970.1 | Translocase of the outer mitochondrial membrane 22 (TOM22-2) | 5 | 1.29 | 0.189 | 1.000 | |
| AT5G43970.1 | Translocase of the outer mitochondrial membrane 22 (TOM22-2) | 6 | 1.55 | 0.189 | 1.000 | |
| AT5G43970.1 | Translocase of the outer mitochondrial membrane 22 (TOM22-2) | 7 | 1.62 | 0.189 | 0.999 | |
| AT5G43970.1 | Translocase of the outer mitochondrial membrane 22 (TOM22-2) | 8 | 1.58 | 0.186 | 0.999 | |
| AT5G43970.1 | Translocase of the outer mitochondrial membrane 22 (TOM22-2) | 9 | 1.57 | 0.184 | 0.999 | |
| AT5G43970.1 | Translocase of the outer mitochondrial membrane 22 (TOM22-2) | 10 | 1.54 | 0.183 | 1.000 | |
| AT5G43970.1 | Translocase of the outer mitochondrial membrane 22 (TOM22-2) | 11 | 1.70 | 0.185 | 0.994 | |
| AT5G43970.1 | Translocase of the outer mitochondrial membrane 22 (TOM22-2) | 12 | 2.19 | 0.185 | 0.780 | |
| AT5G43970.1 | Translocase of the outer mitochondrial membrane 22 (TOM22-2) | 13 | 2.93 | 0.183 | 0.250 | |
| AT5G43970.1 | Translocase of the outer mitochondrial membrane 22 (TOM22-2) | 14 | 3.69 | 0.181 | 0.054 | . |
| AT5G43970.1 | Translocase of the outer mitochondrial membrane 22 (TOM22-2) | 15 | 3.49 | 0.181 | 0.078 | . |
| AT5G43970.1 | Translocase of the outer mitochondrial membrane 22 (TOM22-2) | 16 | 2.51 | 0.179 | 0.470 | |
| AT5G43970.1 | Translocase of the outer mitochondrial membrane 22 (TOM22-2) | 17 | 1.90 | 0.178 | 0.936 | |

| | | | | | |
|---|---|---|---|---|---|
| AT5G43970.1 | Translocase of the outer mitochondrial membrane 22 (TOM22-2) | 18 | 1.88 | 0.180 | 0.948 |
| AT5G43970.1 | Translocase of the outer mitochondrial membrane 22 (TOM22-2) | 19 | 2.12 | 0.186 | 0.840 |
| AT5G43970.1 | Translocase of the outer mitochondrial membrane 22 (TOM22-2) | 20 | 2.23 | 0.190 | 0.783 |
| AT5G43970.1 | Translocase of the outer mitochondrial membrane 22 (TOM22-2) | 21 | 2.07 | 0.190 | 0.890 |
| AT5G43970.1 | Translocase of the outer mitochondrial membrane 22 (TOM22-2) | 22 | 1.78 | 0.191 | 0.989 |
| AT5G43970.1 | Translocase of the outer mitochondrial membrane 22 (TOM22-2) | 23 | 1.54 | 0.189 | 1.000 |
| AT5G43970.1 | Translocase of the outer mitochondrial membrane 22 (TOM22-2) | 24 | 1.31 | 0.186 | 1.000 |
| AT5G43970.1 | Translocase of the outer mitochondrial membrane 22 (TOM22-2) | 25 | 1.10 | 0.184 | 1.000 |
| AT5G43970.1 | Translocase of the outer mitochondrial membrane 22 (TOM22-2) | 26 | 1.00 | 0.183 | 1.000 |
| AT5G43970.1 | Translocase of the outer mitochondrial membrane 22 (TOM22-2) | 27 | 0.97 | 0.183 | 1.000 |
| AT5G43970.1 | Translocase of the outer mitochondrial membrane 22 (TOM22-2) | 28 | 1.09 | 0.183 | 1.000 |
| AT5G43970.1 | Translocase of the outer mitochondrial membrane 22 (TOM22-2) | 29 | 1.49 | 0.183 | 1.000 |
| AT5G43970.1 | Translocase of the outer mitochondrial membrane 22 (TOM22-2) | 30 | 1.93 | 0.183 | 0.936 |
| AT5G43970.1 | Translocase of the outer mitochondrial membrane 22 (TOM22-2) | 31 | 1.67 | 0.184 | 0.996 |
| AT5G43970.1 | Translocase of the outer mitochondrial membrane 22 (TOM22-2) | 32 | 0.96 | 0.184 | 1.000 |
| AT5G43970.1 | Translocase of the outer mitochondrial membrane 22 (TOM22-2) | 33 | 0.50 | 0.184 | 0.907 |
| AT5G43970.1 | Translocase of the outer mitochondrial membrane 22 (TOM22-2) | 34 | 0.32 | 0.185 | 0.205 |
| AT5G43970.1 | Translocase of the outer mitochondrial membrane 22 (TOM22-2) | 35 | 0.36 | 0.187 | 0.378 |
| AT5G43970.1 | Translocase of the outer mitochondrial membrane 22 (TOM22-2) | 36 | 0.61 | 0.192 | 0.999 |
| AT5G43970.1 | Translocase of the outer mitochondrial membrane 22 (TOM22-2) | 37 | 0.98 | 0.199 | 1.000 |

| | | | | | | |
|---|---|---|---|---|---|---|
| AT5G43970.1 | Translocase of the outer mitochondrial membrane 22 (TOM22-2) | 38 | 1.26 | 0.206 | 1.000 | |
| AT5G43970.1 | Translocase of the outer mitochondrial membrane 22 (TOM22-2) | 39 | 1.27 | 0.213 | 1.000 | |
| AT5G43970.1 | Translocase of the outer mitochondrial membrane 22 (TOM22-2) | 40 | 1.12 | 0.252 | 1.000 | |
| AT5G43970.1 | Translocase of the outer mitochondrial membrane 22 (TOM22-2) | 41 | 1.08 | 0.353 | 1.000 | |
| AT5G43970.1 | Translocase of the outer mitochondrial membrane 22 (TOM22-2) | 46 | 4.15 | 1.453 | 1.000 | |
| AT5G43970.1 | Translocase of the outer mitochondrial membrane 22 (TOM22-2) | 47 | 6.30 | 1.788 | 1.000 | |
| AT5G44070.1 | Phytochelatin synthase 1 (PCS1) | 2 | 1.06 | 0.418 | 1.000 | |
| AT5G44070.1 | Phytochelatin synthase 1 (PCS1) | 3 | 0.87 | 0.384 | 1.000 | |
| AT5G44070.1 | Phytochelatin synthase 1 (PCS1) | 4 | 0.70 | 0.565 | 1.000 | |
| AT5G44070.1 | Phytochelatin synthase 1 (PCS1) | 37 | 0.45 | 0.275 | 0.789 | |
| AT5G44070.1 | Phytochelatin synthase 1 (PCS1) | 38 | 0.80 | 0.213 | 0.999 | |
| AT5G44070.1 | Phytochelatin synthase 1 (PCS1) | 39 | 1.45 | 0.193 | 0.965 | |
| AT5G44070.1 | Phytochelatin synthase 1 (PCS1) | 40 | 2.46 | 0.191 | 0.273 | |
| AT5G44070.1 | Phytochelatin synthase 1 (PCS1) | 41 | 3.29 | 0.197 | 0.080 | . |
| AT5G44070.1 | Phytochelatin synthase 1 (PCS1) | 42 | 3.12 | 0.213 | 0.157 | |
| AT5G44070.1 | Phytochelatin synthase 1 (PCS1) | 43 | 2.29 | 0.241 | 0.626 | |
| AT5G44070.1 | Phytochelatin synthase 1 (PCS1) | 44 | 1.47 | 0.268 | 0.993 | |
| AT5G44070.1 | Phytochelatin synthase 1 (PCS1) | 46 | 0.49 | 0.282 | 0.881 | |
| AT5G44070.1 | Phytochelatin synthase 1 (PCS1) | 47 | 0.27 | 0.341 | 0.516 | |
| AT5G44140.1 | Prohibitin 7 (PHB7) | 2 | 1.07 | 0.501 | 1.000 | |
| AT5G44140.1 | Prohibitin 7 (PHB7) | 3 | 0.93 | 0.462 | 1.000 | |
| AT5G44140.1 | Prohibitin 7 (PHB7) | 16 | 0.44 | 0.193 | 0.326 | |
| AT5G44140.1 | Prohibitin 7 (PHB7) | 17 | 0.50 | 0.170 | 0.388 | |
| AT5G44140.1 | Prohibitin 7 (PHB7) | 18 | 0.61 | 0.157 | 0.656 | |
| AT5G44140.1 | Prohibitin 7 (PHB7) | 19 | 0.78 | 0.152 | 0.973 | |
| AT5G44140.1 | Prohibitin 7 (PHB7) | 20 | 1.03 | 0.153 | 1.000 | |
| AT5G44140.1 | Prohibitin 7 (PHB7) | 21 | 1.38 | 0.159 | 0.922 | |
| AT5G44140.1 | Prohibitin 7 (PHB7) | 22 | 1.86 | 0.171 | 0.497 | |
| AT5G44140.1 | Prohibitin 7 (PHB7) | 23 | 2.37 | 0.193 | 0.283 | |
| AT5G44140.1 | Prohibitin 7 (PHB7) | 24 | 2.81 | 0.230 | 0.275 | |
| AT5G44140.1 | Prohibitin 7 (PHB7) | 25 | 3.16 | 0.281 | 0.373 | |
| AT5G44140.1 | Prohibitin 7 (PHB7) | 46 | 0.02 | 0.493 | 0.006 | ** |
| AT5G44140.1 | Prohibitin 7 (PHB7) | 47 | 0.01 | 0.513 | 0.003 | ** |
| AT5G44710.1 | 37S ribosomal protein S27 (RPS27) | 2 | 1.49 | 0.631 | 1.000 | |
| AT5G44710.1 | 37S ribosomal protein S27 (RPS27) | 3 | 1.74 | 0.573 | 1.000 | |
| AT5G44710.1 | 37S ribosomal protein S27 (RPS27) | 11 | 4.15 | 0.301 | 0.268 | |
| AT5G44710.1 | 37S ribosomal protein S27 (RPS27) | 12 | 4.19 | 0.274 | 0.172 | |
| AT5G44710.1 | 37S ribosomal protein S27 (RPS27) | 13 | 4.10 | 0.251 | 0.116 | |
| AT5G44710.1 | 37S ribosomal protein S27 (RPS27) | 14 | 3.84 | 0.232 | 0.097 | . |
| AT5G44710.1 | 37S ribosomal protein S27 (RPS27) | 15 | 3.47 | 0.218 | 0.107 | |
| AT5G44710.1 | 37S ribosomal protein S27 (RPS27) | 16 | 3.02 | 0.210 | 0.165 | |

| | | | | | | |
|---|---|---|---|---|---|---|
| AT5G44710.1 | 37S ribosomal protein S27 (RPS27) | 17 | 2.52 | 0.206 | 0.322 | |
| AT5G44710.1 | 37S ribosomal protein S27 (RPS27) | 18 | 2.02 | 0.206 | 0.636 | |
| AT5G44710.1 | 37S ribosomal protein S27 (RPS27) | 19 | 1.58 | 0.210 | 0.937 | |
| AT5G44710.1 | 37S ribosomal protein S27 (RPS27) | 20 | 1.22 | 0.215 | 1.000 | |
| AT5G44710.1 | 37S ribosomal protein S27 (RPS27) | 21 | 0.95 | 0.222 | 1.000 | |
| AT5G44710.1 | 37S ribosomal protein S27 (RPS27) | 22 | 0.76 | 0.229 | 0.998 | |
| AT5G44710.1 | 37S ribosomal protein S27 (RPS27) | 23 | 0.61 | 0.234 | 0.948 | |
| AT5G44710.1 | 37S ribosomal protein S27 (RPS27) | 24 | 0.51 | 0.238 | 0.804 | |
| AT5G44710.1 | 37S ribosomal protein S27 (RPS27) | 25 | 0.44 | 0.242 | 0.630 | |
| AT5G44710.1 | 37S ribosomal protein S27 (RPS27) | 26 | 0.39 | 0.247 | 0.497 | |
| AT5G44710.1 | 37S ribosomal protein S27 (RPS27) | 27 | 0.35 | 0.255 | 0.429 | |
| AT5G44710.1 | 37S ribosomal protein S27 (RPS27) | 28 | 0.33 | 0.271 | 0.424 | |
| AT5G44710.1 | 37S ribosomal protein S27 (RPS27) | 29 | 0.31 | 0.294 | 0.471 | |
| AT5G44710.1 | 37S ribosomal protein S27 (RPS27) | 30 | 0.30 | 0.325 | 0.557 | |
| AT5G44710.1 | 37S ribosomal protein S27 (RPS27) | 31 | 0.30 | 0.364 | 0.662 | |
| AT5G44710.1 | 37S ribosomal protein S27 (RPS27) | 46 | 0.52 | 0.962 | 1.000 | |
| AT5G44710.1 | 37S ribosomal protein S27 (RPS27) | 47 | 0.55 | 1.026 | 1.000 | |
| AT5G44730.2 | Haloacid dehalogenase-like hydrolase (HAD) superfamily protein | 2 | 1.20 | 0.276 | 1.000 | |
| AT5G44730.2 | Haloacid dehalogenase-like hydrolase (HAD) superfamily protein | 3 | 0.64 | 0.221 | 0.992 | |
| AT5G44730.2 | Haloacid dehalogenase-like hydrolase (HAD) superfamily protein | 4 | 0.38 | 0.240 | 0.560 | |
| AT5G44730.2 | Haloacid dehalogenase-like hydrolase (HAD) superfamily protein | 28 | 2.18 | 0.193 | 0.580 | |
| AT5G44730.2 | Haloacid dehalogenase-like hydrolase (HAD) superfamily protein | 29 | 2.30 | 0.159 | 0.242 | |
| AT5G44730.2 | Haloacid dehalogenase-like hydrolase (HAD) superfamily protein | 30 | 2.37 | 0.140 | 0.093 | . |
| AT5G44730.2 | Haloacid dehalogenase-like hydrolase (HAD) superfamily protein | 31 | 2.42 | 0.129 | 0.042 | * |
| AT5G44730.2 | Haloacid dehalogenase-like hydrolase (HAD) superfamily protein | 32 | 2.54 | 0.124 | 0.017 | * |
| AT5G44730.2 | Haloacid dehalogenase-like hydrolase (HAD) superfamily protein | 33 | 2.88 | 0.121 | 0.003 | ** |

| | | | | | | |
|---|---|---|---|---|---|---|
| AT5G44730.2 | Haloacid dehalogenase-like hydrolase (HAD) superfamily protein | 34 | 3.57 | 0.121 | 0.000 | *** |
| AT5G44730.2 | Haloacid dehalogenase-like hydrolase (HAD) superfamily protein | 35 | 4.69 | 0.123 | 0.000 | *** |
| AT5G44730.2 | Haloacid dehalogenase-like hydrolase (HAD) superfamily protein | 36 | 5.99 | 0.126 | 0.000 | *** |
| AT5G44730.2 | Haloacid dehalogenase-like hydrolase (HAD) superfamily protein | 37 | 6.95 | 0.131 | 0.000 | *** |
| AT5G44730.2 | Haloacid dehalogenase-like hydrolase (HAD) superfamily protein | 38 | 7.08 | 0.136 | 0.000 | *** |
| AT5G44730.2 | Haloacid dehalogenase-like hydrolase (HAD) superfamily protein | 39 | 6.30 | 0.137 | 0.000 | *** |
| AT5G44730.2 | Haloacid dehalogenase-like hydrolase (HAD) superfamily protein | 40 | 4.99 | 0.134 | 0.000 | *** |
| AT5G44730.2 | Haloacid dehalogenase-like hydrolase (HAD) superfamily protein | 41 | 3.64 | 0.129 | 0.000 | *** |
| AT5G44730.2 | Haloacid dehalogenase-like hydrolase (HAD) superfamily protein | 42 | 2.49 | 0.126 | 0.026 | * |
| AT5G44730.2 | Haloacid dehalogenase-like hydrolase (HAD) superfamily protein | 43 | 1.60 | 0.126 | 0.672 | |
| AT5G44730.2 | Haloacid dehalogenase-like hydrolase (HAD) superfamily protein | 44 | 1.01 | 0.127 | 1.000 | |
| AT5G44730.2 | Haloacid dehalogenase-like hydrolase (HAD) superfamily protein | 45 | 0.69 | 0.135 | 0.918 | |
| AT5G44730.2 | Haloacid dehalogenase-like hydrolase (HAD) superfamily protein | 46 | 0.50 | 0.167 | 0.540 | |
| AT5G44730.2 | Haloacid dehalogenase-like hydrolase (HAD) superfamily protein | 47 | 0.37 | 0.240 | 0.543 | |
| AT5G46110.2 | Acclimation of photosynthesis to environment 2 (APE2) | 2 | 1.02 | 0.537 | 1.000 | |
| AT5G46110.2 | Acclimation of photosynthesis to environment 2 (APE2) | 3 | 0.69 | 0.350 | 1.000 | |
| AT5G46110.2 | Acclimation of photosynthesis to environment 2 (APE2) | 4 | 0.57 | 0.340 | 1.000 | |
| AT5G46110.2 | Acclimation of photosynthesis to environment 2 (APE2) | 5 | 0.70 | 0.334 | 1.000 | |
| AT5G46110.2 | Acclimation of photosynthesis to environment 2 (APE2) | 6 | 0.99 | 0.322 | 1.000 | |
| AT5G46110.2 | Acclimation of photosynthesis to environment 2 (APE2) | 7 | 1.13 | 0.315 | 1.000 | |
| AT5G46110.2 | Acclimation of photosynthesis to environment 2 (APE2) | 8 | 1.42 | 0.309 | 1.000 | |
| AT5G46110.2 | Acclimation of photosynthesis to environment 2 (APE2) | 9 | 1.57 | 0.305 | 1.000 | |
| AT5G46110.2 | Acclimation of photosynthesis to environment 2 (APE2) | 10 | 1.22 | 0.304 | 1.000 | |
| AT5G46110.2 | Acclimation of photosynthesis to environment 2 (APE2) | 11 | 0.84 | 0.306 | 1.000 | |

| | | | | | | |
|---|---|---|---|---|---|---|
| AT5G46110.2 | Acclimation of photosynthesis to environment 2 (APE2) | 12 | 0.72 | 0.307 | 1.000 | |
| AT5G46110.2 | Acclimation of photosynthesis to environment 2 (APE2) | 13 | 0.83 | 0.305 | 1.000 | |
| AT5G46110.2 | Acclimation of photosynthesis to environment 2 (APE2) | 14 | 1.10 | 0.304 | 1.000 | |
| AT5G46110.2 | Acclimation of photosynthesis to environment 2 (APE2) | 15 | 1.31 | 0.305 | 1.000 | |
| AT5G46110.2 | Acclimation of photosynthesis to environment 2 (APE2) | 16 | 1.32 | 0.306 | 1.000 | |
| AT5G46110.2 | Acclimation of photosynthesis to environment 2 (APE2) | 17 | 1.40 | 0.311 | 1.000 | |
| AT5G46110.2 | Acclimation of photosynthesis to environment 2 (APE2) | 18 | 1.33 | 0.317 | 1.000 | |
| AT5G46110.2 | Acclimation of photosynthesis to environment 2 (APE2) | 19 | 0.91 | 0.377 | 1.000 | |
| AT5G46110.2 | Acclimation of photosynthesis to environment 2 (APE2) | 20 | 0.49 | 0.530 | 1.000 | |
| AT5G46110.2 | Acclimation of photosynthesis to environment 2 (APE2) | 31 | 0.52 | 0.423 | 1.000 | |
| AT5G46110.2 | Acclimation of photosynthesis to environment 2 (APE2) | 32 | 1.23 | 0.331 | 1.000 | |
| AT5G46110.2 | Acclimation of photosynthesis to environment 2 (APE2) | 33 | 3.12 | 0.329 | 0.942 | |
| AT5G46110.2 | Acclimation of photosynthesis to environment 2 (APE2) | 34 | 5.65 | 0.313 | 0.312 | |
| AT5G46110.2 | Acclimation of photosynthesis to environment 2 (APE2) | 35 | 4.06 | 0.310 | 0.648 | |
| AT5G46110.2 | Acclimation of photosynthesis to environment 2 (APE2) | 36 | 2.14 | 0.308 | 0.999 | |
| AT5G46110.2 | Acclimation of photosynthesis to environment 2 (APE2) | 37 | 1.73 | 0.307 | 1.000 | |
| AT5G46110.2 | Acclimation of photosynthesis to environment 2 (APE2) | 38 | 3.31 | 0.305 | 0.845 | |
| AT5G46110.2 | Acclimation of photosynthesis to environment 2 (APE2) | 39 | 9.93 | 0.305 | 0.032 | * |
| AT5G46110.2 | Acclimation of photosynthesis to environment 2 (APE2) | 40 | 12.88 | 0.305 | 0.009 | ** |
| AT5G46110.2 | Acclimation of photosynthesis to environment 2 (APE2) | 41 | 3.22 | 0.306 | 0.872 | |
| AT5G46110.2 | Acclimation of photosynthesis to environment 2 (APE2) | 42 | 0.53 | 0.312 | 1.000 | |
| AT5G46110.2 | Acclimation of photosynthesis to environment 2 (APE2) | 43 | 0.16 | 0.312 | 0.254 | |
| AT5G46110.2 | Acclimation of photosynthesis to environment 2 (APE2) | 44 | 0.18 | 0.311 | 0.343 | |
| AT5G46110.2 | Acclimation of photosynthesis to environment 2 (APE2) | 45 | 0.47 | 0.312 | 0.999 | |
| AT5G46110.2 | Acclimation of photosynthesis to environment 2 (APE2) | 46 | 0.98 | 0.336 | 1.000 | |
| AT5G46110.2 | Acclimation of photosynthesis to environment 2 (APE2) | 47 | 1.88 | 0.652 | 1.000 | |
| AT5G46160.2 | 50S ribosomal protein (HLP) | 2 | 1.26 | 0.754 | 1.000 | |
| AT5G46160.2 | 50S ribosomal protein (HLP) | 3 | 1.46 | 0.671 | 1.000 | |
| AT5G46160.2 | 50S ribosomal protein (HLP) | 12 | 2.63 | 0.427 | 0.977 | |
| AT5G46160.2 | 50S ribosomal protein (HLP) | 13 | 2.36 | 0.370 | 0.973 | |
| AT5G46160.2 | 50S ribosomal protein (HLP) | 14 | 2.01 | 0.331 | 0.986 | |
| AT5G46160.2 | 50S ribosomal protein (HLP) | 15 | 1.65 | 0.307 | 0.998 | |
| AT5G46160.2 | 50S ribosomal protein (HLP) | 16 | 1.41 | 0.293 | 1.000 | |
| AT5G46160.2 | 50S ribosomal protein (HLP) | 17 | 1.31 | 0.289 | 1.000 | |
| AT5G46160.2 | 50S ribosomal protein (HLP) | 18 | 1.29 | 0.291 | 1.000 | |
| AT5G46160.2 | 50S ribosomal protein (HLP) | 19 | 1.33 | 0.297 | 1.000 | |

| | | | | | |
|---|---|---|---|---|---|
| AT5G46160.2 | 50S ribosomal protein (HLP) | 20 | 1.39 | 0.304 | 1.000 |
| AT5G46160.2 | 50S ribosomal protein (HLP) | 21 | 1.39 | 0.314 | 1.000 |
| AT5G46160.2 | 50S ribosomal protein (HLP) | 22 | 1.20 | 0.327 | 1.000 |
| AT5G46160.2 | 50S ribosomal protein (HLP) | 23 | 0.90 | 0.347 | 1.000 |
| AT5G46160.2 | 50S ribosomal protein (HLP) | 24 | 0.60 | 0.374 | 1.000 |
| AT5G46160.2 | 50S ribosomal protein (HLP) | 25 | 0.38 | 0.404 | 0.968 |
| AT5G46160.2 | 50S ribosomal protein (HLP) | 26 | 0.26 | 0.444 | 0.860 |
| AT5G46160.2 | 50S ribosomal protein (HLP) | 27 | 0.19 | 0.474 | 0.738 |
| AT5G46160.2 | 50S ribosomal protein (HLP) | 32 | 0.18 | 0.527 | 0.798 |
| AT5G46160.2 | 50S ribosomal protein (HLP) | 33 | 0.23 | 0.548 | 0.931 |
| AT5G46160.2 | 50S ribosomal protein (HLP) | 34 | 0.31 | 0.631 | 0.995 |
| AT5G46160.2 | 50S ribosomal protein (HLP) | 46 | 3.62 | 4.302 | 1.000 |
| AT5G46160.2 | 50S ribosomal protein (HLP) | 47 | 4.21 | 4.767 | 1.000 |
| AT5G46180.1 | Ornithine delta-aminotransferase (OAT1) | 2 | 1.15 | 0.356 | 1.000 |
| AT5G46180.1 | Ornithine delta-aminotransferase (OAT1) | 3 | 0.82 | 0.271 | 1.000 |
| AT5G46180.1 | Ornithine delta-aminotransferase (OAT1) | 4 | 0.61 | 0.490 | 1.000 |
| AT5G46180.1 | Ornithine delta-aminotransferase (OAT1) | 28 | 1.28 | 0.277 | 1.000 |
| AT5G46180.1 | Ornithine delta-aminotransferase (OAT1) | 29 | 1.35 | 0.203 | 1.000 |
| AT5G46180.1 | Ornithine delta-aminotransferase (OAT1) | 30 | 1.48 | 0.186 | 0.998 |
| AT5G46180.1 | Ornithine delta-aminotransferase (OAT1) | 31 | 1.77 | 0.179 | 0.918 |
| AT5G46180.1 | Ornithine delta-aminotransferase (OAT1) | 32 | 2.13 | 0.176 | 0.617 |
| AT5G46180.1 | Ornithine delta-aminotransferase (OAT1) | 33 | 2.38 | 0.176 | 0.406 |
| AT5G46180.1 | Ornithine delta-aminotransferase (OAT1) | 34 | 2.59 | 0.176 | 0.273 |
| AT5G46180.1 | Ornithine delta-aminotransferase (OAT1) | 35 | 2.64 | 0.176 | 0.245 |
| AT5G46180.1 | Ornithine delta-aminotransferase (OAT1) | 36 | 2.16 | 0.176 | 0.588 |
| AT5G46180.1 | Ornithine delta-aminotransferase (OAT1) | 37 | 1.79 | 0.175 | 0.896 |
| AT5G46180.1 | Ornithine delta-aminotransferase (OAT1) | 38 | 1.79 | 0.175 | 0.892 |
| AT5G46180.1 | Ornithine delta-aminotransferase (OAT1) | 39 | 1.96 | 0.174 | 0.753 |
| AT5G46180.1 | Ornithine delta-aminotransferase (OAT1) | 40 | 2.24 | 0.174 | 0.500 |
| AT5G46180.1 | Ornithine delta-aminotransferase (OAT1) | 41 | 2.27 | 0.175 | 0.489 |
| AT5G46180.1 | Ornithine delta-aminotransferase (OAT1) | 42 | 2.25 | 0.178 | 0.523 |
| AT5G46180.1 | Ornithine delta-aminotransferase (OAT1) | 43 | 2.30 | 0.179 | 0.490 |
| AT5G46180.1 | Ornithine delta-aminotransferase (OAT1) | 44 | 1.83 | 0.179 | 0.882 |
| AT5G46180.1 | Ornithine delta-aminotransferase (OAT1) | 45 | 1.04 | 0.179 | 1.000 |
| AT5G46180.1 | Ornithine delta-aminotransferase (OAT1) | 46 | 0.46 | 0.204 | 0.788 |
| AT5G46180.1 | Ornithine delta-aminotransferase (OAT1) | 47 | 0.17 | 0.352 | 0.380 |
| AT5G46800.1 | A bout de souffle (BOU) | 3 | 0.98 | 0.123 | 1.000 |
| AT5G46800.1 | A bout de souffle (BOU) | 4 | 0.98 | 0.118 | 1.000 |
| AT5G46800.1 | A bout de souffle (BOU) | 5 | 1.14 | 0.118 | 1.000 |

| | | | | | | |
|---|---|---|---|---|---|---|
| AT5G46800.1 | A bout de souffle (BOU) | 6 | 1.21 | 0.118 | 1.000 | |
| AT5G46800.1 | A bout de souffle (BOU) | 7 | 1.14 | 0.119 | 1.000 | |
| AT5G46800.1 | A bout de souffle (BOU) | 8 | 1.11 | 0.117 | 1.000 | |
| AT5G46800.1 | A bout de souffle (BOU) | 9 | 1.10 | 0.115 | 1.000 | |
| AT5G46800.1 | A bout de souffle (BOU) | 10 | 1.03 | 0.115 | 1.000 | |
| AT5G46800.1 | A bout de souffle (BOU) | 11 | 1.02 | 0.115 | 1.000 | |
| AT5G46800.1 | A bout de souffle (BOU) | 12 | 1.06 | 0.115 | 1.000 | |
| AT5G46800.1 | A bout de souffle (BOU) | 13 | 1.11 | 0.114 | 1.000 | |
| AT5G46800.1 | A bout de souffle (BOU) | 14 | 1.16 | 0.113 | 1.000 | |
| AT5G46800.1 | A bout de souffle (BOU) | 15 | 1.13 | 0.113 | 1.000 | |
| AT5G46800.1 | A bout de souffle (BOU) | 16 | 1.08 | 0.112 | 1.000 | |
| AT5G46800.1 | A bout de souffle (BOU) | 17 | 1.22 | 0.111 | 1.000 | |
| AT5G46800.1 | A bout de souffle (BOU) | 18 | 1.60 | 0.113 | 0.867 | |
| AT5G46800.1 | A bout de souffle (BOU) | 19 | 1.98 | 0.116 | 0.279 | |
| AT5G46800.1 | A bout de souffle (BOU) | 20 | 2.05 | 0.118 | 0.240 | |
| AT5G46800.1 | A bout de souffle (BOU) | 21 | 1.76 | 0.120 | 0.686 | |
| AT5G46800.1 | A bout de souffle (BOU) | 22 | 1.40 | 0.120 | 0.999 | |
| AT5G46800.1 | A bout de souffle (BOU) | 23 | 1.13 | 0.119 | 1.000 | |
| AT5G46800.1 | A bout de souffle (BOU) | 24 | 1.05 | 0.117 | 1.000 | |
| AT5G46800.1 | A bout de souffle (BOU) | 25 | 0.99 | 0.115 | 1.000 | |
| AT5G46800.1 | A bout de souffle (BOU) | 26 | 0.90 | 0.114 | 1.000 | |
| AT5G46800.1 | A bout de souffle (BOU) | 27 | 1.00 | 0.115 | 1.000 | |
| AT5G46800.1 | A bout de souffle (BOU) | 28 | 1.16 | 0.114 | 1.000 | |
| AT5G46800.1 | A bout de souffle (BOU) | 29 | 1.25 | 0.114 | 1.000 | |
| AT5G46800.1 | A bout de souffle (BOU) | 30 | 1.22 | 0.114 | 1.000 | |
| AT5G46800.1 | A bout de souffle (BOU) | 31 | 1.19 | 0.114 | 1.000 | |
| AT5G46800.1 | A bout de souffle (BOU) | 32 | 1.20 | 0.115 | 1.000 | |
| AT5G46800.1 | A bout de souffle (BOU) | 33 | 1.27 | 0.115 | 1.000 | |
| AT5G46800.1 | A bout de souffle (BOU) | 34 | 1.48 | 0.115 | 0.980 | |
| AT5G46800.1 | A bout de souffle (BOU) | 35 | 1.70 | 0.115 | 0.725 | |
| AT5G46800.1 | A bout de souffle (BOU) | 36 | 1.61 | 0.114 | 0.851 | |
| AT5G46800.1 | A bout de souffle (BOU) | 37 | 1.45 | 0.114 | 0.989 | |
| AT5G46800.1 | A bout de souffle (BOU) | 38 | 1.56 | 0.115 | 0.919 | |
| AT5G46800.1 | A bout de souffle (BOU) | 39 | 2.21 | 0.115 | 0.098 | . |
| AT5G46800.1 | A bout de souffle (BOU) | 40 | 3.08 | 0.116 | 0.001 | ** |
| AT5G46800.1 | A bout de souffle (BOU) | 41 | 2.75 | 0.116 | 0.006 | ** |
| AT5G46800.1 | A bout de souffle (BOU) | 42 | 1.64 | 0.117 | 0.838 | |
| AT5G46800.1 | A bout de souffle (BOU) | 43 | 0.90 | 0.118 | 1.000 | |
| AT5G46800.1 | A bout de souffle (BOU) | 44 | 0.53 | 0.118 | 0.461 | |
| AT5G46800.1 | A bout de souffle (BOU) | 45 | 0.36 | 0.118 | 0.008 | ** |
| AT5G46800.1 | A bout de souffle (BOU) | 46 | 0.27 | 0.130 | 0.001 | *** |
| AT5G46800.1 | A bout de souffle (BOU) | 47 | 0.21 | 0.242 | 0.144 | |
| AT5G46800.1 | A bout de souffle (BOU) | 48 | 0.16 | 0.425 | 0.804 | |
| AT5G47030.1 | ATP synthase subunit (ATPdelta) | 2 | 0.79 | 0.418 | 1.000 | |
| AT5G47030.1 | ATP synthase subunit (ATPdelta) | 3 | 0.70 | 0.273 | 1.000 | |
| AT5G47030.1 | ATP synthase subunit (ATPdelta) | 4 | 0.67 | 0.247 | 1.000 | |
| AT5G47030.1 | ATP synthase subunit (ATPdelta) | 5 | 0.75 | 0.247 | 1.000 | |
| AT5G47030.1 | ATP synthase subunit (ATPdelta) | 6 | 1.05 | 0.247 | 1.000 | |
| AT5G47030.1 | ATP synthase subunit (ATPdelta) | 7 | 1.50 | 0.246 | 1.000 | |
| AT5G47030.1 | ATP synthase subunit (ATPdelta) | 8 | 1.93 | 0.243 | 0.997 | |
| AT5G47030.1 | ATP synthase subunit (ATPdelta) | 9 | 2.21 | 0.241 | 0.974 | |
| AT5G47030.1 | ATP synthase subunit (ATPdelta) | 10 | 2.22 | 0.241 | 0.972 | |
| AT5G47030.1 | ATP synthase subunit (ATPdelta) | 11 | 2.20 | 0.241 | 0.974 | |

| | | | | | |
|---|---|---|---|---|---|
| AT5G47030.1 | ATP synthase subunit (ATPdelta) | 12 | 2.47 | 0.241 | 0.911 |
| AT5G47030.1 | ATP synthase subunit (ATPdelta) | 13 | 3.08 | 0.238 | 0.626 |
| AT5G47030.1 | ATP synthase subunit (ATPdelta) | 14 | 3.65 | 0.236 | 0.367 |
| AT5G47030.1 | ATP synthase subunit (ATPdelta) | 15 | 3.74 | 0.235 | 0.330 |
| AT5G47030.1 | ATP synthase subunit (ATPdelta) | 16 | 3.32 | 0.234 | 0.483 |
| AT5G47030.1 | ATP synthase subunit (ATPdelta) | 17 | 2.45 | 0.233 | 0.890 |
| AT5G47030.1 | ATP synthase subunit (ATPdelta) | 18 | 1.64 | 0.236 | 1.000 |
| AT5G47030.1 | ATP synthase subunit (ATPdelta) | 19 | 1.22 | 0.242 | 1.000 |
| AT5G47030.1 | ATP synthase subunit (ATPdelta) | 20 | 1.02 | 0.247 | 1.000 |
| AT5G47030.1 | ATP synthase subunit (ATPdelta) | 21 | 0.86 | 0.249 | 1.000 |
| AT5G47030.1 | ATP synthase subunit (ATPdelta) | 22 | 0.78 | 0.249 | 1.000 |
| AT5G47030.1 | ATP synthase subunit (ATPdelta) | 23 | 0.81 | 0.247 | 1.000 |
| AT5G47030.1 | ATP synthase subunit (ATPdelta) | 24 | 0.98 | 0.243 | 1.000 |
| AT5G47030.1 | ATP synthase subunit (ATPdelta) | 25 | 1.23 | 0.239 | 1.000 |
| AT5G47030.1 | ATP synthase subunit (ATPdelta) | 26 | 1.28 | 0.239 | 1.000 |
| AT5G47030.1 | ATP synthase subunit (ATPdelta) | 27 | 1.06 | 0.239 | 1.000 |
| AT5G47030.1 | ATP synthase subunit (ATPdelta) | 28 | 0.97 | 0.239 | 1.000 |
| AT5G47030.1 | ATP synthase subunit (ATPdelta) | 29 | 1.25 | 0.239 | 1.000 |
| AT5G47030.1 | ATP synthase subunit (ATPdelta) | 30 | 1.96 | 0.239 | 0.996 |
| AT5G47030.1 | ATP synthase subunit (ATPdelta) | 31 | 2.61 | 0.241 | 0.861 |
| AT5G47030.1 | ATP synthase subunit (ATPdelta) | 32 | 2.34 | 0.247 | 0.958 |
| AT5G47030.1 | ATP synthase subunit (ATPdelta) | 33 | 1.69 | 0.259 | 1.000 |
| AT5G47030.1 | ATP synthase subunit (ATPdelta) | 34 | 1.42 | 0.276 | 1.000 |
| AT5G47030.1 | ATP synthase subunit (ATPdelta) | 35 | 1.42 | 0.290 | 1.000 |
| AT5G47030.1 | ATP synthase subunit (ATPdelta) | 36 | 1.57 | 0.297 | 1.000 |
| AT5G47030.1 | ATP synthase subunit (ATPdelta) | 37 | 1.78 | 0.283 | 1.000 |
| AT5G47030.1 | ATP synthase subunit (ATPdelta) | 38 | 2.30 | 0.275 | 0.989 |
| AT5G47030.1 | ATP synthase subunit (ATPdelta) | 39 | 3.48 | 0.270 | 0.663 |
| AT5G47030.1 | ATP synthase subunit (ATPdelta) | 40 | 4.93 | 0.261 | 0.202 |
| AT5G47030.1 | ATP synthase subunit (ATPdelta) | 41 | 5.78 | 0.265 | 0.116 |
| AT5G47030.1 | ATP synthase subunit (ATPdelta) | 42 | 6.07 | 0.298 | 0.214 |

| | | | | | |
|---|---|---|---|---|---|
| AT5G47030.1 | ATP synthase subunit (ATPdelta) | 43 | 6.46 | 0.323 | 0.282 |
| AT5G47030.1 | ATP synthase subunit (ATPdelta) | 44 | 6.32 | 0.325 | 0.311 |
| AT5G47030.1 | ATP synthase subunit (ATPdelta) | 45 | 4.76 | 0.346 | 0.701 |
| AT5G47030.1 | ATP synthase subunit (ATPdelta) | 46 | 2.46 | 0.439 | 1.000 |
| AT5G47030.1 | ATP synthase subunit (ATPdelta) | 47 | 1.04 | 0.674 | 1.000 |
| AT5G47320.1 | 40S ribosomal protein S19 (RPS19) | 2 | 1.59 | 0.456 | 1.000 |
| AT5G47320.1 | 40S ribosomal protein S19 (RPS19) | 3 | 1.62 | 0.414 | 0.999 |
| AT5G47320.1 | 40S ribosomal protein S19 (RPS19) | 10 | 1.76 | 0.241 | 0.910 |
| AT5G47320.1 | 40S ribosomal protein S19 (RPS19) | 11 | 1.76 | 0.226 | 0.879 |
| AT5G47320.1 | 40S ribosomal protein S19 (RPS19) | 12 | 1.75 | 0.214 | 0.853 |
| AT5G47320.1 | 40S ribosomal protein S19 (RPS19) | 13 | 1.73 | 0.204 | 0.835 |
| AT5G47320.1 | 40S ribosomal protein S19 (RPS19) | 14 | 1.69 | 0.196 | 0.837 |
| AT5G47320.1 | 40S ribosomal protein S19 (RPS19) | 15 | 1.64 | 0.191 | 0.858 |
| AT5G47320.1 | 40S ribosomal protein S19 (RPS19) | 16 | 1.57 | 0.187 | 0.896 |
| AT5G47320.1 | 40S ribosomal protein S19 (RPS19) | 17 | 1.48 | 0.185 | 0.946 |
| AT5G47320.1 | 40S ribosomal protein S19 (RPS19) | 18 | 1.37 | 0.185 | 0.986 |
| AT5G47320.1 | 40S ribosomal protein S19 (RPS19) | 19 | 1.24 | 0.185 | 0.999 |
| AT5G47320.1 | 40S ribosomal protein S19 (RPS19) | 20 | 1.12 | 0.186 | 1.000 |
| AT5G47320.1 | 40S ribosomal protein S19 (RPS19) | 21 | 1.01 | 0.187 | 1.000 |
| AT5G47320.1 | 40S ribosomal protein S19 (RPS19) | 22 | 0.92 | 0.188 | 1.000 |
| AT5G47320.1 | 40S ribosomal protein S19 (RPS19) | 23 | 0.85 | 0.188 | 1.000 |
| AT5G47320.1 | 40S ribosomal protein S19 (RPS19) | 24 | 0.81 | 0.189 | 0.999 |
| AT5G47320.1 | 40S ribosomal protein S19 (RPS19) | 25 | 0.77 | 0.190 | 0.997 |
| AT5G47320.1 | 40S ribosomal protein S19 (RPS19) | 26 | 0.75 | 0.190 | 0.994 |
| AT5G47320.1 | 40S ribosomal protein S19 (RPS19) | 27 | 0.75 | 0.191 | 0.993 |
| AT5G47320.1 | 40S ribosomal protein S19 (RPS19) | 28 | 0.75 | 0.191 | 0.994 |
| AT5G47320.1 | 40S ribosomal protein S19 (RPS19) | 29 | 0.76 | 0.192 | 0.996 |
| AT5G47320.1 | 40S ribosomal protein S19 (RPS19) | 30 | 0.79 | 0.193 | 0.998 |
| AT5G47320.1 | 40S ribosomal protein S19 (RPS19) | 31 | 0.82 | 0.194 | 1.000 |
| AT5G47320.1 | 40S ribosomal protein S19 (RPS19) | 32 | 0.86 | 0.195 | 1.000 |
| AT5G47320.1 | 40S ribosomal protein S19 (RPS19) | 33 | 0.91 | 0.197 | 1.000 |

| | | | | | |
|---|---|---|---|---|---|
| AT5G47320.1 | 40S ribosomal protein S19 (RPS19) | 34 | 0.96 | 0.199 | 1.000 |
| AT5G47320.1 | 40S ribosomal protein S19 (RPS19) | 35 | 1.02 | 0.202 | 1.000 |
| AT5G47320.1 | 40S ribosomal protein S19 (RPS19) | 36 | 1.07 | 0.204 | 1.000 |
| AT5G47320.1 | 40S ribosomal protein S19 (RPS19) | 37 | 1.13 | 0.205 | 1.000 |
| AT5G47320.1 | 40S ribosomal protein S19 (RPS19) | 38 | 1.19 | 0.205 | 1.000 |
| AT5G47320.1 | 40S ribosomal protein S19 (RPS19) | 39 | 1.24 | 0.205 | 1.000 |
| AT5G47320.1 | 40S ribosomal protein S19 (RPS19) | 40 | 1.29 | 0.205 | 0.998 |
| AT5G47320.1 | 40S ribosomal protein S19 (RPS19) | 41 | 1.34 | 0.205 | 0.995 |
| AT5G47320.1 | 40S ribosomal protein S19 (RPS19) | 42 | 1.39 | 0.206 | 0.990 |
| AT5G47320.1 | 40S ribosomal protein S19 (RPS19) | 43 | 1.44 | 0.209 | 0.982 |
| AT5G47320.1 | 40S ribosomal protein S19 (RPS19) | 44 | 1.49 | 0.217 | 0.977 |
| AT5G47320.1 | 40S ribosomal protein S19 (RPS19) | 45 | 1.52 | 0.232 | 0.977 |
| AT5G47320.1 | 40S ribosomal protein S19 (RPS19) | 46 | 1.55 | 0.255 | 0.984 |
| AT5G47320.1 | 40S ribosomal protein S19 (RPS19) | 47 | 1.58 | 0.287 | 0.990 |
| AT5G47435.2 | Formyltetrahydrofolate deformylase, putative (PURU1) | 3 | 1.07 | 0.210 | 1.000 |
| AT5G47435.2 | Formyltetrahydrofolate deformylase, putative (PURU1) | 4 | 1.14 | 0.178 | 1.000 |
| AT5G47435.2 | Formyltetrahydrofolate deformylase, putative (PURU1) | 5 | 1.20 | 0.171 | 1.000 |
| AT5G47435.2 | Formyltetrahydrofolate deformylase, putative (PURU1) | 6 | 1.23 | 0.171 | 1.000 |
| AT5G47435.2 | Formyltetrahydrofolate deformylase, putative (PURU1) | 7 | 1.22 | 0.171 | 1.000 |
| AT5G47435.2 | Formyltetrahydrofolate deformylase, putative (PURU1) | 8 | 1.19 | 0.169 | 1.000 |
| AT5G47435.2 | Formyltetrahydrofolate deformylase, putative (PURU1) | 9 | 1.14 | 0.167 | 1.000 |
| AT5G47435.2 | Formyltetrahydrofolate deformylase, putative (PURU1) | 10 | 1.08 | 0.166 | 1.000 |
| AT5G47435.2 | Formyltetrahydrofolate deformylase, putative (PURU1) | 11 | 1.01 | 0.166 | 1.000 |
| AT5G47435.2 | Formyltetrahydrofolate deformylase, putative (PURU1) | 12 | 0.94 | 0.165 | 1.000 |
| AT5G47435.2 | Formyltetrahydrofolate deformylase, putative (PURU1) | 13 | 0.87 | 0.164 | 1.000 |
| AT5G47435.2 | Formyltetrahydrofolate deformylase, putative (PURU1) | 14 | 0.81 | 0.163 | 1.000 |
| AT5G47435.2 | Formyltetrahydrofolate deformylase, putative (PURU1) | 15 | 0.76 | 0.162 | 1.000 |
| AT5G47435.2 | Formyltetrahydrofolate deformylase, putative (PURU1) | 16 | 0.70 | 0.161 | 0.999 |
| AT5G47435.2 | Formyltetrahydrofolate deformylase, putative (PURU1) | 17 | 0.61 | 0.162 | 0.965 |
| AT5G47435.2 | Formyltetrahydrofolate deformylase, putative (PURU1) | 18 | 0.51 | 0.163 | 0.733 |
| AT5G47435.2 | Formyltetrahydrofolate deformylase, putative (PURU1) | 19 | 0.42 | 0.166 | 0.357 |

| | | | | | |
|---|---|---|---|---|---|
| AT5G47435.2 | Formyltetrahydrofolate deformylase, putative (PURU1) | 20 | 0.35 | 0.169 | 0.156 |
| AT5G47435.2 | Formyltetrahydrofolate deformylase, putative (PURU1) | 21 | 0.34 | 0.171 | 0.117 |
| AT5G47435.2 | Formyltetrahydrofolate deformylase, putative (PURU1) | 22 | 0.37 | 0.171 | 0.231 |
| AT5G47435.2 | Formyltetrahydrofolate deformylase, putative (PURU1) | 23 | 0.49 | 0.170 | 0.693 |
| AT5G47435.2 | Formyltetrahydrofolate deformylase, putative (PURU1) | 24 | 0.66 | 0.168 | 0.996 |
| AT5G47435.2 | Formyltetrahydrofolate deformylase, putative (PURU1) | 25 | 0.87 | 0.166 | 1.000 |
| AT5G47435.2 | Formyltetrahydrofolate deformylase, putative (PURU1) | 26 | 1.08 | 0.165 | 1.000 |
| AT5G47435.2 | Formyltetrahydrofolate deformylase, putative (PURU1) | 27 | 1.23 | 0.164 | 1.000 |
| AT5G47435.2 | Formyltetrahydrofolate deformylase, putative (PURU1) | 28 | 1.28 | 0.164 | 1.000 |
| AT5G47435.2 | Formyltetrahydrofolate deformylase, putative (PURU1) | 29 | 1.23 | 0.165 | 1.000 |
| AT5G47435.2 | Formyltetrahydrofolate deformylase, putative (PURU1) | 30 | 1.10 | 0.165 | 1.000 |
| AT5G47435.2 | Formyltetrahydrofolate deformylase, putative (PURU1) | 31 | 0.96 | 0.165 | 1.000 |
| AT5G47435.2 | Formyltetrahydrofolate deformylase, putative (PURU1) | 32 | 0.85 | 0.165 | 1.000 |
| AT5G47435.2 | Formyltetrahydrofolate deformylase, putative (PURU1) | 33 | 0.78 | 0.166 | 1.000 |
| AT5G47435.2 | Formyltetrahydrofolate deformylase, putative (PURU1) | 34 | 0.79 | 0.166 | 1.000 |
| AT5G47435.2 | Formyltetrahydrofolate deformylase, putative (PURU1) | 35 | 0.89 | 0.166 | 1.000 |
| AT5G47435.2 | Formyltetrahydrofolate deformylase, putative (PURU1) | 36 | 1.12 | 0.165 | 1.000 |
| AT5G47435.2 | Formyltetrahydrofolate deformylase, putative (PURU1) | 37 | 1.45 | 0.165 | 0.999 |
| AT5G47435.2 | Formyltetrahydrofolate deformylase, putative (PURU1) | 38 | 1.77 | 0.165 | 0.908 |
| AT5G47435.2 | Formyltetrahydrofolate deformylase, putative (PURU1) | 39 | 1.97 | 0.166 | 0.738 |
| AT5G47435.2 | Formyltetrahydrofolate deformylase, putative (PURU1) | 40 | 2.04 | 0.167 | 0.677 |
| AT5G47435.2 | Formyltetrahydrofolate deformylase, putative (PURU1) | 41 | 1.95 | 0.168 | 0.774 |
| AT5G47435.2 | Formyltetrahydrofolate deformylase, putative (PURU1) | 42 | 1.70 | 0.169 | 0.958 |
| AT5G47435.2 | Formyltetrahydrofolate deformylase, putative (PURU1) | 43 | 1.36 | 0.170 | 1.000 |
| AT5G47435.2 | Formyltetrahydrofolate deformylase, putative (PURU1) | 44 | 1.00 | 0.171 | 1.000 |
| AT5G47435.2 | Formyltetrahydrofolate deformylase, putative (PURU1) | 45 | 0.68 | 0.181 | 0.999 |
| AT5G47435.2 | Formyltetrahydrofolate deformylase, putative (PURU1) | 46 | 0.45 | 0.222 | 0.878 |
| AT5G47435.2 | Formyltetrahydrofolate deformylase, putative (PURU1) | 47 | 0.29 | 0.310 | 0.789 |
| AT5G47435.2 | Formyltetrahydrofolate deformylase, putative (PURU1) | 48 | 0.19 | 0.433 | 0.829 |
| AT5G47570.1 | NADH:ubiquinone oxidoreductase subunit (ASHI) | 2 | 0.90 | 0.596 | 1.000 |
| AT5G47570.1 | NADH:ubiquinone oxidoreductase subunit (ASHI) | 3 | 0.84 | 0.542 | 1.000 |

| | | | | | |
|---|---|---|---|---|---|
| AT5G47570.1 | NADH:ubiquinone oxidoreductase subunit (ASHI) | 14 | 0.42 | 0.361 | 0.988 |
| AT5G47570.1 | NADH:ubiquinone oxidoreductase subunit (ASHI) | 15 | 0.47 | 0.317 | 0.988 |
| AT5G47570.1 | NADH:ubiquinone oxidoreductase subunit (ASHI) | 16 | 0.47 | 0.283 | 0.968 |
| AT5G47570.1 | NADH:ubiquinone oxidoreductase subunit (ASHI) | 17 | 0.40 | 0.256 | 0.798 |
| AT5G47570.1 | NADH:ubiquinone oxidoreductase subunit (ASHI) | 18 | 0.43 | 0.250 | 0.849 |
| AT5G47570.1 | NADH:ubiquinone oxidoreductase subunit (ASHI) | 19 | 0.58 | 0.257 | 0.996 |
| AT5G47570.1 | NADH:ubiquinone oxidoreductase subunit (ASHI) | 20 | 0.73 | 0.263 | 1.000 |
| AT5G47570.1 | NADH:ubiquinone oxidoreductase subunit (ASHI) | 21 | 0.97 | 0.266 | 1.000 |
| AT5G47570.1 | NADH:ubiquinone oxidoreductase subunit (ASHI) | 22 | 1.56 | 0.272 | 1.000 |
| AT5G47570.1 | NADH:ubiquinone oxidoreductase subunit (ASHI) | 23 | 2.71 | 0.286 | 0.831 |
| AT5G47570.1 | NADH:ubiquinone oxidoreductase subunit (ASHI) | 24 | 3.79 | 0.316 | 0.608 |
| AT5G47570.1 | NADH:ubiquinone oxidoreductase subunit (ASHI) | 25 | 3.24 | 0.359 | 0.883 |
| AT5G47570.1 | NADH:ubiquinone oxidoreductase subunit (ASHI) | 27 | 0.79 | 0.344 | 1.000 |
| AT5G47570.1 | NADH:ubiquinone oxidoreductase subunit (ASHI) | 28 | 0.49 | 0.313 | 0.991 |
| AT5G47570.1 | NADH:ubiquinone oxidoreductase subunit (ASHI) | 29 | 0.41 | 0.306 | 0.942 |
| AT5G47570.1 | NADH:ubiquinone oxidoreductase subunit (ASHI) | 30 | 0.29 | 0.365 | 0.850 |
| AT5G47570.1 | NADH:ubiquinone oxidoreductase subunit (ASHI) | 31 | 0.19 | 0.550 | 0.923 |
| AT5G47570.1 | NADH:ubiquinone oxidoreductase subunit (ASHI) | 46 | 0.01 | 8.315 | 1.000 |
| AT5G47570.1 | NADH:ubiquinone oxidoreductase subunit (ASHI) | 47 | 0.01 | 9.121 | 1.000 |
| AT5G47890.1 | NADH:ubiquinone oxidoreductase subunit (B8) | 3 | 0.94 | 0.244 | 1.000 |
| AT5G47890.1 | NADH:ubiquinone oxidoreductase subunit (B8) | 4 | 0.85 | 0.230 | 1.000 |
| AT5G47890.1 | NADH:ubiquinone oxidoreductase subunit (B8) | 5 | 0.90 | 0.229 | 1.000 |
| AT5G47890.1 | NADH:ubiquinone oxidoreductase subunit (B8) | 6 | 0.95 | 0.231 | 1.000 |
| AT5G47890.1 | NADH:ubiquinone oxidoreductase subunit (B8) | 7 | 0.88 | 0.233 | 1.000 |
| AT5G47890.1 | NADH:ubiquinone oxidoreductase subunit (B8) | 8 | 0.85 | 0.226 | 1.000 |
| AT5G47890.1 | NADH:ubiquinone oxidoreductase subunit (B8) | 9 | 0.87 | 0.221 | 1.000 |
| AT5G47890.1 | NADH:ubiquinone oxidoreductase subunit (B8) | 10 | 1.03 | 0.222 | 1.000 |
| AT5G47890.1 | NADH:ubiquinone oxidoreductase subunit (B8) | 11 | 1.36 | 0.223 | 1.000 |
| AT5G47890.1 | NADH:ubiquinone oxidoreductase subunit (B8) | 12 | 1.50 | 0.223 | 1.000 |
| AT5G47890.1 | NADH:ubiquinone oxidoreductase subunit (B8) | 13 | 1.27 | 0.220 | 1.000 |
| AT5G47890.1 | NADH:ubiquinone oxidoreductase subunit (B8) | 14 | 1.25 | 0.215 | 1.000 |

| | | | | | | |
|---|---|---|---|---|---|---|
| AT5G47890.1 | NADH:ubiquinone oxidoreductase subunit (B8) | 15 | 1.12 | 0.215 | 1.000 | |
| AT5G47890.1 | NADH:ubiquinone oxidoreductase subunit (B8) | 16 | 0.83 | 0.213 | 1.000 | |
| AT5G47890.1 | NADH:ubiquinone oxidoreductase subunit (B8) | 17 | 0.59 | 0.213 | 1.000 | |
| AT5G47890.1 | NADH:ubiquinone oxidoreductase subunit (B8) | 18 | 0.56 | 0.217 | 1.000 | |
| AT5G47890.1 | NADH:ubiquinone oxidoreductase subunit (B8) | 19 | 0.69 | 0.224 | 1.000 | |
| AT5G47890.1 | NADH:ubiquinone oxidoreductase subunit (B8) | 20 | 0.78 | 0.231 | 1.000 | |
| AT5G47890.1 | NADH:ubiquinone oxidoreductase subunit (B8) | 21 | 0.97 | 0.235 | 1.000 | |
| AT5G47890.1 | NADH:ubiquinone oxidoreductase subunit (B8) | 22 | 1.63 | 0.234 | 1.000 | |
| AT5G47890.1 | NADH:ubiquinone oxidoreductase subunit (B8) | 23 | 2.31 | 0.233 | 0.982 | |
| AT5G47890.1 | NADH:ubiquinone oxidoreductase subunit (B8) | 24 | 1.29 | 0.227 | 1.000 | |
| AT5G47890.1 | NADH:ubiquinone oxidoreductase subunit (B8) | 25 | 0.27 | 0.221 | 0.294 | |
| AT5G47890.1 | NADH:ubiquinone oxidoreductase subunit (B8) | 26 | 0.13 | 0.221 | 0.003 | ** |
| AT5G47890.1 | NADH:ubiquinone oxidoreductase subunit (B8) | 27 | 0.32 | 0.221 | 0.602 | |
| AT5G47890.1 | NADH:ubiquinone oxidoreductase subunit (B8) | 28 | 0.82 | 0.220 | 1.000 | |
| AT5G47890.1 | NADH:ubiquinone oxidoreductase subunit (B8) | 29 | 1.48 | 0.220 | 1.000 | |
| AT5G47890.1 | NADH:ubiquinone oxidoreductase subunit (B8) | 30 | 1.37 | 0.220 | 1.000 | |
| AT5G47890.1 | NADH:ubiquinone oxidoreductase subunit (B8) | 31 | 0.89 | 0.220 | 1.000 | |
| AT5G47890.1 | NADH:ubiquinone oxidoreductase subunit (B8) | 32 | 0.84 | 0.221 | 1.000 | |
| AT5G47890.1 | NADH:ubiquinone oxidoreductase subunit (B8) | 33 | 1.23 | 0.222 | 1.000 | |
| AT5G47890.1 | NADH:ubiquinone oxidoreductase subunit (B8) | 34 | 1.53 | 0.222 | 1.000 | |
| AT5G47890.1 | NADH:ubiquinone oxidoreductase subunit (B8) | 35 | 1.23 | 0.221 | 1.000 | |
| AT5G47890.1 | NADH:ubiquinone oxidoreductase subunit (B8) | 36 | 1.02 | 0.220 | 1.000 | |
| AT5G47890.1 | NADH:ubiquinone oxidoreductase subunit (B8) | 37 | 1.12 | 0.220 | 1.000 | |
| AT5G47890.1 | NADH:ubiquinone oxidoreductase subunit (B8) | 38 | 1.80 | 0.221 | 1.000 | |
| AT5G47890.1 | NADH:ubiquinone oxidoreductase subunit (B8) | 39 | 4.04 | 0.222 | 0.219 | |
| AT5G47890.1 | NADH:ubiquinone oxidoreductase subunit (B8) | 40 | 5.71 | 0.223 | 0.031 | * |
| AT5G47890.1 | NADH:ubiquinone oxidoreductase subunit (B8) | 41 | 3.81 | 0.224 | 0.303 | |
| AT5G47890.1 | NADH:ubiquinone oxidoreductase subunit (B8) | 42 | 3.07 | 0.226 | 0.671 | |
| AT5G47890.1 | NADH:ubiquinone oxidoreductase subunit (B8) | 43 | 3.31 | 0.229 | 0.560 | |
| AT5G47890.1 | NADH:ubiquinone oxidoreductase subunit (B8) | 44 | 1.75 | 0.228 | 1.000 | |
| AT5G47890.1 | NADH:ubiquinone oxidoreductase subunit (B8) | 45 | 0.93 | 0.230 | 1.000 | |

| | | | | | |
|---|---|---|---|---|---|
| AT5G47890.1 | NADH:ubiquinone oxidoreductase subunit (B8) | 46 | 0.79 | 0.261 | 1.000 |
| AT5G47890.1 | NADH:ubiquinone oxidoreductase subunit (B8) | 47 | 0.78 | 0.475 | 1.000 |
| AT5G47890.1 | NADH:ubiquinone oxidoreductase subunit (B8) | 48 | 0.78 | 0.841 | 1.000 |
| AT5G48300.1 | ADP glucose pyrophosphorylase small subunit 1 (APS1) | 2 | 1.08 | 0.409 | 1.000 |
| AT5G48300.1 | ADP glucose pyrophosphorylase small subunit 1 (APS1) | 3 | 1.12 | 0.394 | 1.000 |
| AT5G48300.1 | ADP glucose pyrophosphorylase small subunit 1 (APS1) | 4 | 1.17 | 0.392 | 1.000 |
| AT5G48300.1 | ADP glucose pyrophosphorylase small subunit 1 (APS1) | 34 | 1.83 | 0.232 | 0.690 |
| AT5G48300.1 | ADP glucose pyrophosphorylase small subunit 1 (APS1) | 35 | 1.76 | 0.216 | 0.685 |
| AT5G48300.1 | ADP glucose pyrophosphorylase small subunit 1 (APS1) | 36 | 1.68 | 0.208 | 0.720 |
| AT5G48300.1 | ADP glucose pyrophosphorylase small subunit 1 (APS1) | 37 | 1.57 | 0.206 | 0.804 |
| AT5G48300.1 | ADP glucose pyrophosphorylase small subunit 1 (APS1) | 38 | 1.43 | 0.211 | 0.912 |
| AT5G48300.1 | ADP glucose pyrophosphorylase small subunit 1 (APS1) | 39 | 1.27 | 0.220 | 0.985 |
| AT5G48300.1 | ADP glucose pyrophosphorylase small subunit 1 (APS1) | 40 | 1.10 | 0.233 | 1.000 |
| AT5G48300.1 | ADP glucose pyrophosphorylase small subunit 1 (APS1) | 46 | 0.35 | 0.340 | 0.553 |
| AT5G48300.1 | ADP glucose pyrophosphorylase small subunit 1 (APS1) | 47 | 0.29 | 0.370 | 0.462 |
| AT5G48970.1 | Mitochondrial substrate carrier family protein | 2 | 0.97 | 0.401 | 1.000 |
| AT5G48970.1 | Mitochondrial substrate carrier family protein | 3 | 0.97 | 0.333 | 1.000 |
| AT5G48970.1 | Mitochondrial substrate carrier family protein | 8 | 1.03 | 0.261 | 1.000 |
| AT5G48970.1 | Mitochondrial substrate carrier family protein | 9 | 1.07 | 0.211 | 1.000 |
| AT5G48970.1 | Mitochondrial substrate carrier family protein | 10 | 1.09 | 0.187 | 1.000 |
| AT5G48970.1 | Mitochondrial substrate carrier family protein | 11 | 1.13 | 0.175 | 1.000 |
| AT5G48970.1 | Mitochondrial substrate carrier family protein | 12 | 1.32 | 0.174 | 1.000 |
| AT5G48970.1 | Mitochondrial substrate carrier family protein | 13 | 1.69 | 0.179 | 0.982 |
| AT5G48970.1 | Mitochondrial substrate carrier family protein | 14 | 1.98 | 0.183 | 0.877 |
| AT5G48970.1 | Mitochondrial substrate carrier family protein | 15 | 2.06 | 0.190 | 0.857 |

| | | | | | | |
|---|---|---|---|---|---|---|
| AT5G48970.1 | Mitochondrial substrate carrier family protein | 16 | 1.89 | 0.207 | 0.974 | |
| AT5G48970.1 | Mitochondrial substrate carrier family protein | 18 | 1.74 | 0.244 | 0.999 | |
| AT5G48970.1 | Mitochondrial substrate carrier family protein | 19 | 1.79 | 0.291 | 1.000 | |
| AT5G48970.1 | Mitochondrial substrate carrier family protein | 20 | 1.84 | 0.360 | 1.000 | |
| AT5G48970.1 | Mitochondrial substrate carrier family protein | 31 | 1.02 | 0.324 | 1.000 | |
| AT5G48970.1 | Mitochondrial substrate carrier family protein | 32 | 0.84 | 0.267 | 1.000 | |
| AT5G48970.1 | Mitochondrial substrate carrier family protein | 33 | 0.77 | 0.264 | 1.000 | |
| AT5G48970.1 | Mitochondrial substrate carrier family protein | 35 | 1.42 | 0.234 | 1.000 | |
| AT5G48970.1 | Mitochondrial substrate carrier family protein | 36 | 2.60 | 0.219 | 0.693 | |
| AT5G48970.1 | Mitochondrial substrate carrier family protein | 37 | 4.47 | 0.198 | 0.030 | * |
| AT5G48970.1 | Mitochondrial substrate carrier family protein | 38 | 6.43 | 0.191 | 1.000 | |
| AT5G48970.1 | Mitochondrial substrate carrier family protein | 39 | 6.23 | 0.186 | 1.000 | |
| AT5G48970.1 | Mitochondrial substrate carrier family protein | 40 | 4.09 | 0.174 | 0.013 | * |
| AT5G48970.1 | Mitochondrial substrate carrier family protein | 41 | 1.88 | 0.169 | 0.873 | |
| AT5G48970.1 | Mitochondrial substrate carrier family protein | 42 | 0.81 | 0.171 | 1.000 | |
| AT5G48970.1 | Mitochondrial substrate carrier family protein | 43 | 0.46 | 0.172 | 0.647 | |
| AT5G48970.1 | Mitochondrial substrate carrier family protein | 44 | 0.35 | 0.172 | 0.167 | |
| AT5G48970.1 | Mitochondrial substrate carrier family protein | 45 | 0.31 | 0.171 | 0.072 | . |
| AT5G48970.1 | Mitochondrial substrate carrier family protein | 46 | 0.33 | 0.201 | 0.304 | |
| AT5G48970.1 | Mitochondrial substrate carrier family protein | 47 | 0.39 | 0.376 | 0.996 | |
| AT5G49030.1 | tRNA synthase class I family protein (OVA2) | 2 | 0.82 | 0.433 | 1.000 | |
| AT5G49030.1 | tRNA synthase class I family protein (OVA2) | 3 | 0.62 | 0.427 | 1.000 | |
| AT5G49030.1 | tRNA synthase class I family protein (OVA2) | 27 | 0.44 | 0.327 | 0.986 | |
| AT5G49030.1 | tRNA synthase class I family protein (OVA2) | 28 | 0.75 | 0.233 | 1.000 | |
| AT5G49030.1 | tRNA synthase class I family protein (OVA2) | 29 | 1.28 | 0.207 | 1.000 | |
| AT5G49030.1 | tRNA synthase class I family protein (OVA2) | 30 | 1.76 | 0.196 | 0.958 | |
| AT5G49030.1 | tRNA synthase class I family protein (OVA2) | 31 | 2.16 | 0.192 | 0.704 | |
| AT5G49030.1 | tRNA synthase class I family protein (OVA2) | 32 | 2.16 | 0.192 | 0.699 | |
| AT5G49030.1 | tRNA synthase class I family protein (OVA2) | 33 | 1.65 | 0.193 | 0.981 | |
| AT5G49030.1 | tRNA synthase class I family protein (OVA2) | 34 | 1.11 | 0.193 | 1.000 | |
| AT5G49030.1 | tRNA synthase class I family protein (OVA2) | 35 | 0.76 | 0.192 | 1.000 | |

| | | | | | |
|---|---|---|---|---|---|
| AT5G49030.1 | tRNA synthase class I family protein (OVA2) | 36 | 0.60 | 0.191 | 0.973 |
| AT5G49030.1 | tRNA synthase class I family protein (OVA2) | 37 | 0.60 | 0.192 | 0.978 |
| AT5G49030.1 | tRNA synthase class I family protein (OVA2) | 38 | 0.78 | 0.193 | 1.000 |
| AT5G49030.1 | tRNA synthase class I family protein (OVA2) | 39 | 1.04 | 0.193 | 1.000 |
| AT5G49030.1 | tRNA synthase class I family protein (OVA2) | 40 | 0.81 | 0.193 | 1.000 |
| AT5G49030.1 | tRNA synthase class I family protein (OVA2) | 41 | 0.38 | 0.205 | 0.477 |
| AT5G49030.1 | tRNA synthase class I family protein (OVA2) | 42 | 0.22 | 0.291 | 0.311 |
| AT5G49030.1 | tRNA synthase class I family protein (OVA2) | 43 | 0.24 | 0.436 | 0.904 |
| AT5G49030.1 | tRNA synthase class I family protein (OVA2) | 46 | 6.69 | 1.084 | 1.000 |
| AT5G49030.1 | tRNA synthase class I family protein (OVA2) | 47 | 32.24 | 1.417 | 0.989 |
| AT5G49555.1 | FAD/NAD(P)-binding oxidoreductase family protein | 2 | 1.11 | 0.295 | 1.000 |
| AT5G49555.1 | FAD/NAD(P)-binding oxidoreductase family protein | 3 | 0.77 | 0.221 | 1.000 |
| AT5G49555.1 | FAD/NAD(P)-binding oxidoreductase family protein | 4 | 0.58 | 0.281 | 1.000 |
| AT5G49555.1 | FAD/NAD(P)-binding oxidoreductase family protein | 11 | 0.94 | 0.214 | 1.000 |
| AT5G49555.1 | FAD/NAD(P)-binding oxidoreductase family protein | 12 | 1.32 | 0.190 | 1.000 |
| AT5G49555.1 | FAD/NAD(P)-binding oxidoreductase family protein | 13 | 1.82 | 0.174 | 0.949 |
| AT5G49555.1 | FAD/NAD(P)-binding oxidoreductase family protein | 14 | 2.33 | 0.164 | 0.458 |
| AT5G49555.1 | FAD/NAD(P)-binding oxidoreductase family protein | 15 | 2.62 | 0.160 | 0.205 |
| AT5G49555.1 | FAD/NAD(P)-binding oxidoreductase family protein | 16 | 2.58 | 0.164 | 0.267 |
| AT5G49555.1 | FAD/NAD(P)-binding oxidoreductase family protein | 17 | 2.30 | 0.174 | 0.591 |
| AT5G49555.1 | FAD/NAD(P)-binding oxidoreductase family protein | 18 | 1.99 | 0.192 | 0.929 |
| AT5G49555.1 | FAD/NAD(P)-binding oxidoreductase family protein | 19 | 1.81 | 0.216 | 0.996 |
| AT5G49555.1 | FAD/NAD(P)-binding oxidoreductase family protein | 20 | 1.72 | 0.243 | 1.000 |
| AT5G49555.1 | FAD/NAD(P)-binding oxidoreductase family protein | 24 | 1.36 | 0.220 | 1.000 |
| AT5G49555.1 | FAD/NAD(P)-binding oxidoreductase family protein | 25 | 1.20 | 0.184 | 1.000 |
| AT5G49555.1 | FAD/NAD(P)-binding oxidoreductase family protein | 26 | 1.03 | 0.157 | 1.000 |
| AT5G49555.1 | FAD/NAD(P)-binding oxidoreductase family protein | 27 | 0.86 | 0.148 | 1.000 |
| AT5G49555.1 | FAD/NAD(P)-binding oxidoreductase family protein | 28 | 0.85 | 0.146 | 1.000 |
| AT5G49555.1 | FAD/NAD(P)-binding oxidoreductase family protein | 29 | 1.08 | 0.146 | 1.000 |
| AT5G49555.1 | FAD/NAD(P)-binding oxidoreductase family protein | 30 | 1.28 | 0.146 | 1.000 |
| AT5G49555.1 | FAD/NAD(P)-binding oxidoreductase family protein | 31 | 1.28 | 0.147 | 1.000 |

| | | | | | |
|---|---|---|---|---|---|
| AT5G49555.1 | FAD/NAD(P)-binding oxidoreductase family protein | 32 | 1.17 | 0.147 | 1.000 |
| AT5G49555.1 | FAD/NAD(P)-binding oxidoreductase family protein | 33 | 1.06 | 0.147 | 1.000 |
| AT5G49555.1 | FAD/NAD(P)-binding oxidoreductase family protein | 34 | 1.02 | 0.147 | 1.000 |
| AT5G49555.1 | FAD/NAD(P)-binding oxidoreductase family protein | 35 | 1.10 | 0.147 | 1.000 |
| AT5G49555.1 | FAD/NAD(P)-binding oxidoreductase family protein | 36 | 1.30 | 0.147 | 1.000 |
| AT5G49555.1 | FAD/NAD(P)-binding oxidoreductase family protein | 37 | 1.67 | 0.147 | 0.945 |
| AT5G49555.1 | FAD/NAD(P)-binding oxidoreductase family protein | 38 | 2.05 | 0.146 | 0.547 |
| AT5G49555.1 | FAD/NAD(P)-binding oxidoreductase family protein | 39 | 2.15 | 0.146 | 0.426 |
| AT5G49555.1 | FAD/NAD(P)-binding oxidoreductase family protein | 40 | 1.91 | 0.147 | 0.720 |
| AT5G49555.1 | FAD/NAD(P)-binding oxidoreductase family protein | 41 | 1.45 | 0.147 | 0.998 |
| AT5G49555.1 | FAD/NAD(P)-binding oxidoreductase family protein | 42 | 1.03 | 0.149 | 1.000 |
| AT5G49555.1 | FAD/NAD(P)-binding oxidoreductase family protein | 43 | 0.75 | 0.151 | 1.000 |
| AT5G49555.1 | FAD/NAD(P)-binding oxidoreductase family protein | 44 | 0.62 | 0.150 | 0.978 |
| AT5G49555.1 | FAD/NAD(P)-binding oxidoreductase family protein | 45 | 0.55 | 0.150 | 0.854 |
| AT5G49555.1 | FAD/NAD(P)-binding oxidoreductase family protein | 46 | 0.44 | 0.180 | 0.667 |
| AT5G49555.1 | FAD/NAD(P)-binding oxidoreductase family protein | 47 | 0.31 | 0.310 | 0.890 |
| AT5G49840.1 | Caseinolytic protease X2 (CLPX2) | 2 | 1.16 | 0.444 | 1.000 |
| AT5G49840.1 | Caseinolytic protease X2 (CLPX2) | 3 | 1.24 | 0.409 | 1.000 |
| AT5G49840.1 | Caseinolytic protease X2 (CLPX2) | 19 | 0.81 | 0.278 | 1.000 |
| AT5G49840.1 | Caseinolytic protease X2 (CLPX2) | 20 | 0.69 | 0.243 | 1.000 |
| AT5G49840.1 | Caseinolytic protease X2 (CLPX2) | 21 | 0.63 | 0.217 | 0.999 |
| AT5G49840.1 | Caseinolytic protease X2 (CLPX2) | 22 | 0.71 | 0.210 | 1.000 |
| AT5G49840.1 | Caseinolytic protease X2 (CLPX2) | 23 | 1.15 | 0.207 | 1.000 |
| AT5G49840.1 | Caseinolytic protease X2 (CLPX2) | 24 | 2.03 | 0.208 | 0.912 |
| AT5G49840.1 | Caseinolytic protease X2 (CLPX2) | 25 | 2.40 | 0.213 | 0.730 |
| AT5G49840.1 | Caseinolytic protease X2 (CLPX2) | 26 | 1.63 | 0.212 | 0.998 |
| AT5G49840.1 | Caseinolytic protease X2 (CLPX2) | 27 | 1.02 | 0.207 | 1.000 |
| AT5G49840.1 | Caseinolytic protease X2 (CLPX2) | 28 | 0.89 | 0.203 | 1.000 |
| AT5G49840.1 | Caseinolytic protease X2 (CLPX2) | 29 | 1.04 | 0.198 | 1.000 |
| AT5G49840.1 | Caseinolytic protease X2 (CLPX2) | 30 | 1.23 | 0.198 | 1.000 |
| AT5G49840.1 | Caseinolytic protease X2 (CLPX2) | 31 | 1.33 | 0.198 | 1.000 |

| | | | | | |
|---|---|---|---|---|---|
| AT5G49840.1 | Caseinolytic protease X2 (CLPX2) | 32 | 1.23 | 0.198 | 1.000 |
| AT5G49840.1 | Caseinolytic protease X2 (CLPX2) | 33 | 0.95 | 0.198 | 1.000 |
| AT5G49840.1 | Caseinolytic protease X2 (CLPX2) | 34 | 0.67 | 0.198 | 1.000 |
| AT5G49840.1 | Caseinolytic protease X2 (CLPX2) | 35 | 0.51 | 0.200 | 0.921 |
| AT5G49840.1 | Caseinolytic protease X2 (CLPX2) | 36 | 0.47 | 0.205 | 0.847 |
| AT5G49840.1 | Caseinolytic protease X2 (CLPX2) | 37 | 0.48 | 0.212 | 0.903 |
| AT5G49840.1 | Caseinolytic protease X2 (CLPX2) | 38 | 0.48 | 0.216 | 0.917 |
| AT5G49840.1 | Caseinolytic protease X2 (CLPX2) | 39 | 0.48 | 0.212 | 0.898 |
| AT5G49840.1 | Caseinolytic protease X2 (CLPX2) | 40 | 0.54 | 0.203 | 0.966 |
| AT5G49840.1 | Caseinolytic protease X2 (CLPX2) | 41 | 0.75 | 0.199 | 1.000 |
| AT5G49840.1 | Caseinolytic protease X2 (CLPX2) | 42 | 1.11 | 0.200 | 1.000 |
| AT5G49840.1 | Caseinolytic protease X2 (CLPX2) | 43 | 1.33 | 0.201 | 1.000 |
| AT5G49840.1 | Caseinolytic protease X2 (CLPX2) | 44 | 1.19 | 0.201 | 1.000 |
| AT5G49840.1 | Caseinolytic protease X2 (CLPX2) | 45 | 0.86 | 0.203 | 1.000 |
| AT5G49840.1 | Caseinolytic protease X2 (CLPX2) | 46 | 0.54 | 0.237 | 0.991 |
| AT5G49840.1 | Caseinolytic protease X2 (CLPX2) | 47 | 0.31 | 0.384 | 0.963 |
| AT5G49970.1 | Pyridoxin (pyrodoxamine) 5'-phosphate oxidase (PPOX) | 2 | 0.92 | 0.671 | 1.000 |
| AT5G49970.1 | Pyridoxin (pyrodoxamine) 5'-phosphate oxidase (PPOX) | 3 | 1.00 | 0.668 | 1.000 |
| AT5G49970.1 | Pyridoxin (pyrodoxamine) 5'-phosphate oxidase (PPOX) | 32 | 1.16 | 0.496 | 1.000 |
| AT5G49970.1 | Pyridoxin (pyrodoxamine) 5'-phosphate oxidase (PPOX) | 33 | 1.37 | 0.392 | 1.000 |
| AT5G49970.1 | Pyridoxin (pyrodoxamine) 5'-phosphate oxidase (PPOX) | 34 | 1.88 | 0.375 | 0.999 |
| AT5G49970.1 | Pyridoxin (pyrodoxamine) 5'-phosphate oxidase (PPOX) | 35 | 1.64 | 0.405 | 1.000 |
| AT5G49970.1 | Pyridoxin (pyrodoxamine) 5'-phosphate oxidase (PPOX) | 36 | 1.67 | 0.488 | 1.000 |
| AT5G49970.1 | Pyridoxin (pyrodoxamine) 5'-phosphate oxidase (PPOX) | 39 | 3.02 | 0.485 | 0.986 |
| AT5G49970.1 | Pyridoxin (pyrodoxamine) 5'-phosphate oxidase (PPOX) | 40 | 3.58 | 0.385 | 0.842 |
| AT5G49970.1 | Pyridoxin (pyrodoxamine) 5'-phosphate oxidase (PPOX) | 41 | 3.54 | 0.325 | 0.674 |
| AT5G49970.1 | Pyridoxin (pyrodoxamine) 5'-phosphate oxidase (PPOX) | 42 | 1.66 | 0.320 | 0.999 |
| AT5G49970.1 | Pyridoxin (pyrodoxamine) 5'-phosphate oxidase (PPOX) | 43 | 0.66 | 0.322 | 1.000 |
| AT5G49970.1 | Pyridoxin (pyrodoxamine) 5'-phosphate oxidase (PPOX) | 44 | 0.38 | 0.321 | 0.899 |
| AT5G49970.1 | Pyridoxin (pyrodoxamine) 5'-phosphate oxidase (PPOX) | 45 | 0.36 | 0.319 | 0.863 |
| AT5G49970.1 | Pyridoxin (pyrodoxamine) 5'-phosphate oxidase (PPOX) | 46 | 0.45 | 0.358 | 0.989 |

| | | | | | | |
|---|---|---|---|---|---|---|
| AT5G49970.1 | Pyridoxin (pyrodoxamine) 5'-phosphate oxidase (PPOX) | 47 | 0.67 | 0.737 | 1.000 | |
| AT5G50370.1 | Adenylate kinase family protein (AMK3 / ADK2) | 2 | 1.31 | 0.437 | 1.000 | |
| AT5G50370.1 | Adenylate kinase family protein (AMK3 / ADK2) | 3 | 1.15 | 0.395 | 1.000 | |
| AT5G50370.1 | Adenylate kinase family protein (AMK3 / ADK2) | 11 | 1.35 | 0.329 | 1.000 | |
| AT5G50370.1 | Adenylate kinase family protein (AMK3 / ADK2) | 12 | 1.61 | 0.287 | 1.000 | |
| AT5G50370.1 | Adenylate kinase family protein (AMK3 / ADK2) | 13 | 1.97 | 0.260 | 0.988 | |
| AT5G50370.1 | Adenylate kinase family protein (AMK3 / ADK2) | 14 | 2.32 | 0.254 | 0.914 | |
| AT5G50370.1 | Adenylate kinase family protein (AMK3 / ADK2) | 16 | 2.52 | 0.252 | 0.836 | |
| AT5G50370.1 | Adenylate kinase family protein (AMK3 / ADK2) | 17 | 2.41 | 0.255 | 0.886 | |
| AT5G50370.1 | Adenylate kinase family protein (AMK3 / ADK2) | 18 | 2.37 | 0.260 | 0.910 | |
| AT5G50370.1 | Adenylate kinase family protein (AMK3 / ADK2) | 19 | 2.40 | 0.281 | 0.944 | |
| AT5G50370.1 | Adenylate kinase family protein (AMK3 / ADK2) | 20 | 2.30 | 0.318 | 0.987 | |
| AT5G50370.1 | Adenylate kinase family protein (AMK3 / ADK2) | 26 | 1.23 | 0.342 | 1.000 | |
| AT5G50370.1 | Adenylate kinase family protein (AMK3 / ADK2) | 27 | 1.24 | 0.293 | 1.000 | |
| AT5G50370.1 | Adenylate kinase family protein (AMK3 / ADK2) | 28 | 1.33 | 0.262 | 1.000 | |
| AT5G50370.1 | Adenylate kinase family protein (AMK3 / ADK2) | 29 | 1.54 | 0.236 | 1.000 | |
| AT5G50370.1 | Adenylate kinase family protein (AMK3 / ADK2) | 30 | 1.82 | 0.221 | 0.983 | |
| AT5G50370.1 | Adenylate kinase family protein (AMK3 / ADK2) | 31 | 2.22 | 0.210 | 0.797 | |
| AT5G50370.1 | Adenylate kinase family protein (AMK3 / ADK2) | 32 | 2.76 | 0.202 | 0.393 | |
| AT5G50370.1 | Adenylate kinase family protein (AMK3 / ADK2) | 33 | 3.09 | 0.198 | 0.222 | |
| AT5G50370.1 | Adenylate kinase family protein (AMK3 / ADK2) | 34 | 3.15 | 0.198 | 0.202 | |
| AT5G50370.1 | Adenylate kinase family protein (AMK3 / ADK2) | 35 | 3.37 | 0.199 | 0.150 | |
| AT5G50370.1 | Adenylate kinase family protein (AMK3 / ADK2) | 36 | 4.05 | 0.202 | 0.059 | . |
| AT5G50370.1 | Adenylate kinase family protein (AMK3 / ADK2) | 37 | 5.05 | 0.208 | 0.019 | * |
| AT5G50370.1 | Adenylate kinase family protein (AMK3 / ADK2) | 38 | 5.62 | 0.212 | 0.011 | * |
| AT5G50370.1 | Adenylate kinase family protein (AMK3 / ADK2) | 39 | 4.95 | 0.209 | 0.023 | * |
| AT5G50370.1 | Adenylate kinase family protein (AMK3 / ADK2) | 40 | 3.44 | 0.202 | 0.144 | |
| AT5G50370.1 | Adenylate kinase family protein (AMK3 / ADK2) | 41 | 1.81 | 0.197 | 0.958 | |
| AT5G50370.1 | Adenylate kinase family protein (AMK3 / ADK2) | 42 | 0.91 | 0.198 | 1.000 | |
| AT5G50370.1 | Adenylate kinase family protein (AMK3 / ADK2) | 43 | 0.55 | 0.200 | 0.958 | |
| AT5G50370.1 | Adenylate kinase family protein (AMK3 / ADK2) | 44 | 0.41 | 0.200 | 0.588 | |

| | | | | | |
|---|---|---|---|---|---|
| AT5G50370.1 | Adenylate kinase family protein (AMK3 / ADK2) | 45 | 0.38 | 0.201 | 0.475 |
| AT5G50370.1 | Adenylate kinase family protein (AMK3 / ADK2) | 46 | 0.41 | 0.238 | 0.811 |
| AT5G50370.1 | Adenylate kinase family protein (AMK3 / ADK2) | 47 | 0.44 | 0.375 | 0.998 |
| AT5G50850.1 | Pyruvate dehydrogenase subunit (PDCE1 beta) | 3 | 0.77 | 0.204 | 1.000 |
| AT5G50850.1 | Pyruvate dehydrogenase subunit (PDCE1 beta) | 4 | 0.64 | 0.186 | 1.000 |
| AT5G50850.1 | Pyruvate dehydrogenase subunit (PDCE1 beta) | 5 | 0.56 | 0.186 | 0.994 |
| AT5G50850.1 | Pyruvate dehydrogenase subunit (PDCE1 beta) | 6 | 0.48 | 0.186 | 0.913 |
| AT5G50850.1 | Pyruvate dehydrogenase subunit (PDCE1 beta) | 7 | 0.42 | 0.186 | 0.696 |
| AT5G50850.1 | Pyruvate dehydrogenase subunit (PDCE1 beta) | 8 | 0.39 | 0.183 | 0.513 |
| AT5G50850.1 | Pyruvate dehydrogenase subunit (PDCE1 beta) | 9 | 0.39 | 0.180 | 0.473 |
| AT5G50850.1 | Pyruvate dehydrogenase subunit (PDCE1 beta) | 10 | 0.41 | 0.180 | 0.594 |
| AT5G50850.1 | Pyruvate dehydrogenase subunit (PDCE1 beta) | 11 | 0.53 | 0.180 | 0.967 |
| AT5G50850.1 | Pyruvate dehydrogenase subunit (PDCE1 beta) | 12 | 0.78 | 0.180 | 1.000 |
| AT5G50850.1 | Pyruvate dehydrogenase subunit (PDCE1 beta) | 13 | 1.05 | 0.179 | 1.000 |
| AT5G50850.1 | Pyruvate dehydrogenase subunit (PDCE1 beta) | 14 | 1.18 | 0.176 | 1.000 |
| AT5G50850.1 | Pyruvate dehydrogenase subunit (PDCE1 beta) | 15 | 1.16 | 0.176 | 1.000 |
| AT5G50850.1 | Pyruvate dehydrogenase subunit (PDCE1 beta) | 16 | 1.06 | 0.174 | 1.000 |
| AT5G50850.1 | Pyruvate dehydrogenase subunit (PDCE1 beta) | 17 | 0.97 | 0.174 | 1.000 |
| AT5G50850.1 | Pyruvate dehydrogenase subunit (PDCE1 beta) | 18 | 0.91 | 0.177 | 1.000 |
| AT5G50850.1 | Pyruvate dehydrogenase subunit (PDCE1 beta) | 19 | 0.83 | 0.181 | 1.000 |
| AT5G50850.1 | Pyruvate dehydrogenase subunit (PDCE1 beta) | 20 | 0.70 | 0.186 | 1.000 |
| AT5G50850.1 | Pyruvate dehydrogenase subunit (PDCE1 beta) | 21 | 0.56 | 0.188 | 0.993 |
| AT5G50850.1 | Pyruvate dehydrogenase subunit (PDCE1 beta) | 22 | 0.50 | 0.188 | 0.944 |
| AT5G50850.1 | Pyruvate dehydrogenase subunit (PDCE1 beta) | 23 | 0.52 | 0.186 | 0.971 |
| AT5G50850.1 | Pyruvate dehydrogenase subunit (PDCE1 beta) | 24 | 0.62 | 0.183 | 0.999 |
| AT5G50850.1 | Pyruvate dehydrogenase subunit (PDCE1 beta) | 25 | 0.72 | 0.180 | 1.000 |
| AT5G50850.1 | Pyruvate dehydrogenase subunit (PDCE1 beta) | 26 | 0.76 | 0.179 | 1.000 |
| AT5G50850.1 | Pyruvate dehydrogenase subunit (PDCE1 beta) | 27 | 0.76 | 0.179 | 1.000 |
| AT5G50850.1 | Pyruvate dehydrogenase subunit (PDCE1 beta) | 28 | 0.76 | 0.179 | 1.000 |
| AT5G50850.1 | Pyruvate dehydrogenase subunit (PDCE1 beta) | 29 | 0.76 | 0.179 | 1.000 |
| AT5G50850.1 | Pyruvate dehydrogenase subunit (PDCE1 beta) | 30 | 0.75 | 0.179 | 1.000 |

| | | | | | |
|---|---|---|---|---|---|
| AT5G50850.1 | Pyruvate dehydrogenase subunit (PDCE1 beta) | 31 | 0.75 | 0.180 | 1.000 |
| AT5G50850.1 | Pyruvate dehydrogenase subunit (PDCE1 beta) | 32 | 0.80 | 0.180 | 1.000 |
| AT5G50850.1 | Pyruvate dehydrogenase subunit (PDCE1 beta) | 33 | 0.86 | 0.180 | 1.000 |
| AT5G50850.1 | Pyruvate dehydrogenase subunit (PDCE1 beta) | 34 | 0.94 | 0.181 | 1.000 |
| AT5G50850.1 | Pyruvate dehydrogenase subunit (PDCE1 beta) | 35 | 1.06 | 0.180 | 1.000 |
| AT5G50850.1 | Pyruvate dehydrogenase subunit (PDCE1 beta) | 36 | 1.20 | 0.179 | 1.000 |
| AT5G50850.1 | Pyruvate dehydrogenase subunit (PDCE1 beta) | 37 | 1.33 | 0.179 | 1.000 |
| AT5G50850.1 | Pyruvate dehydrogenase subunit (PDCE1 beta) | 38 | 1.47 | 0.180 | 1.000 |
| AT5G50850.1 | Pyruvate dehydrogenase subunit (PDCE1 beta) | 39 | 1.35 | 0.180 | 1.000 |
| AT5G50850.1 | Pyruvate dehydrogenase subunit (PDCE1 beta) | 40 | 0.97 | 0.181 | 1.000 |
| AT5G50850.1 | Pyruvate dehydrogenase subunit (PDCE1 beta) | 41 | 0.71 | 0.181 | 1.000 |
| AT5G50850.1 | Pyruvate dehydrogenase subunit (PDCE1 beta) | 42 | 0.65 | 0.184 | 1.000 |
| AT5G50850.1 | Pyruvate dehydrogenase subunit (PDCE1 beta) | 43 | 0.69 | 0.186 | 1.000 |
| AT5G50850.1 | Pyruvate dehydrogenase subunit (PDCE1 beta) | 44 | 0.73 | 0.185 | 1.000 |
| AT5G50850.1 | Pyruvate dehydrogenase subunit (PDCE1 beta) | 45 | 0.72 | 0.185 | 1.000 |
| AT5G50850.1 | Pyruvate dehydrogenase subunit (PDCE1 beta) | 46 | 0.69 | 0.219 | 1.000 |
| AT5G50850.1 | Pyruvate dehydrogenase subunit (PDCE1 beta) | 47 | 0.67 | 0.376 | 1.000 |
| AT5G50850.1 | Pyruvate dehydrogenase subunit (PDCE1 beta) | 48 | 0.66 | 0.616 | 1.000 |
| AT5G50920.1 | Caseinolytic protease C1 (CLPC1) | 2 | 0.81 | 0.516 | 1.000 |
| AT5G50920.1 | Caseinolytic protease C1 (CLPC1) | 3 | 0.98 | 0.497 | 1.000 |
| AT5G50920.1 | Caseinolytic protease C1 (CLPC1) | 28 | 1.35 | 0.409 | 1.000 |
| AT5G50920.1 | Caseinolytic protease C1 (CLPC1) | 29 | 1.07 | 0.314 | 1.000 |
| AT5G50920.1 | Caseinolytic protease C1 (CLPC1) | 30 | 1.01 | 0.271 | 1.000 |
| AT5G50920.1 | Caseinolytic protease C1 (CLPC1) | 31 | 1.18 | 0.250 | 1.000 |
| AT5G50920.1 | Caseinolytic protease C1 (CLPC1) | 32 | 1.45 | 0.238 | 0.999 |
| AT5G50920.1 | Caseinolytic protease C1 (CLPC1) | 33 | 1.64 | 0.232 | 0.986 |
| AT5G50920.1 | Caseinolytic protease C1 (CLPC1) | 34 | 1.68 | 0.229 | 0.978 |
| AT5G50920.1 | Caseinolytic protease C1 (CLPC1) | 35 | 1.73 | 0.228 | 0.966 |
| AT5G50920.1 | Caseinolytic protease C1 (CLPC1) | 36 | 1.80 | 0.227 | 0.949 |
| AT5G50920.1 | Caseinolytic protease C1 (CLPC1) | 37 | 1.78 | 0.227 | 0.954 |
| AT5G50920.1 | Caseinolytic protease C1 (CLPC1) | 38 | 1.65 | 0.228 | 0.982 |

| | | | | | | |
|---|---|---|---|---|---|---|
| AT5G50920.1 | Caseinolytic protease C1 (CLPC1) | 39 | 1.39 | 0.231 | 1.000 | |
| AT5G50920.1 | Caseinolytic protease C1 (CLPC1) | 40 | 1.15 | 0.246 | 1.000 | |
| AT5G50920.1 | Caseinolytic protease C1 (CLPC1) | 41 | 1.06 | 0.309 | 1.000 | |
| AT5G50920.1 | Caseinolytic protease C1 (CLPC1) | 42 | 1.26 | 0.436 | 1.000 | |
| AT5G50920.1 | Caseinolytic protease C1 (CLPC1) | 46 | 17.42 | 1.391 | 0.989 | |
| AT5G50920.1 | Caseinolytic protease C1 (CLPC1) | 47 | 42.39 | 1.756 | 0.986 | |
| AT5G51050.1 | ATP/Pi carrier 2 (APC2) | 2 | 1.00 | 0.683 | 1.000 | |
| AT5G51050.1 | ATP/Pi carrier 2 (APC2) | 3 | 0.96 | 0.732 | 1.000 | |
| AT5G51050.1 | ATP/Pi carrier 2 (APC2) | 33 | 1.42 | 0.421 | 1.000 | |
| AT5G51050.1 | ATP/Pi carrier 2 (APC2) | 34 | 1.53 | 0.327 | 0.998 | |
| AT5G51050.1 | ATP/Pi carrier 2 (APC2) | 35 | 1.70 | 0.287 | 0.980 | |
| AT5G51050.1 | ATP/Pi carrier 2 (APC2) | 36 | 2.07 | 0.267 | 0.853 | |
| AT5G51050.1 | ATP/Pi carrier 2 (APC2) | 37 | 2.79 | 0.249 | 0.450 | |
| AT5G51050.1 | ATP/Pi carrier 2 (APC2) | 38 | 3.59 | 0.235 | 0.154 | |
| AT5G51050.1 | ATP/Pi carrier 2 (APC2) | 39 | 3.30 | 0.230 | 0.196 | |
| AT5G51050.1 | ATP/Pi carrier 2 (APC2) | 40 | 1.88 | 0.241 | 0.877 | |
| AT5G51050.1 | ATP/Pi carrier 2 (APC2) | 41 | 0.76 | 0.279 | 1.000 | |
| AT5G51050.1 | ATP/Pi carrier 2 (APC2) | 42 | 0.28 | 0.347 | 0.594 | |
| AT5G51050.1 | ATP/Pi carrier 2 (APC2) | 43 | 0.11 | 0.422 | 0.207 | |
| AT5G51050.1 | ATP/Pi carrier 2 (APC2) | 46 | 0.01 | 0.606 | 0.020 | * |
| AT5G51050.1 | ATP/Pi carrier 2 (APC2) | 47 | 0.01 | 0.726 | 0.029 | * |
| AT5G51220.1 | Ubiquinol:cytochrome c oxidoreductase assembly protein | 2 | 1.51 | 0.465 | 1.000 | |
| AT5G51220.1 | Ubiquinol:cytochrome c oxidoreductase assembly protein | 3 | 1.18 | 0.357 | 1.000 | |
| AT5G51220.1 | Ubiquinol:cytochrome c oxidoreductase assembly protein | 6 | 0.65 | 0.322 | 1.000 | |
| AT5G51220.1 | Ubiquinol:cytochrome c oxidoreductase assembly protein | 7 | 0.59 | 0.262 | 1.000 | |
| AT5G51220.1 | Ubiquinol:cytochrome c oxidoreductase assembly protein | 8 | 0.61 | 0.230 | 1.000 | |
| AT5G51220.1 | Ubiquinol:cytochrome c oxidoreductase assembly protein | 9 | 0.70 | 0.217 | 1.000 | |
| AT5G51220.1 | Ubiquinol:cytochrome c oxidoreductase assembly protein | 10 | 0.86 | 0.212 | 1.000 | |
| AT5G51220.1 | Ubiquinol:cytochrome c oxidoreductase assembly protein | 11 | 0.98 | 0.213 | 1.000 | |
| AT5G51220.1 | Ubiquinol:cytochrome c oxidoreductase assembly protein | 12 | 1.08 | 0.213 | 1.000 | |
| AT5G51220.1 | Ubiquinol:cytochrome c oxidoreductase assembly protein | 13 | 1.13 | 0.211 | 1.000 | |
| AT5G51220.1 | Ubiquinol:cytochrome c oxidoreductase assembly protein | 14 | 1.17 | 0.210 | 1.000 | |

| | | | | | |
|---|---|---|---|---|---|
| AT5G51220.1 | Ubiquinol:cytochrome c oxidoreductase assembly protein | 15 | 1.21 | 0.209 | 1.000 |
| AT5G51220.1 | Ubiquinol:cytochrome c oxidoreductase assembly protein | 16 | 1.24 | 0.211 | 1.000 |
| AT5G51220.1 | Ubiquinol:cytochrome c oxidoreductase assembly protein | 17 | 1.64 | 0.217 | 1.000 |
| AT5G51220.1 | Ubiquinol:cytochrome c oxidoreductase assembly protein | 18 | 2.87 | 0.229 | 0.659 |
| AT5G51220.1 | Ubiquinol:cytochrome c oxidoreductase assembly protein | 19 | 4.82 | 0.267 | 0.242 |
| AT5G51220.1 | Ubiquinol:cytochrome c oxidoreductase assembly protein | 20 | 5.92 | 0.343 | 0.452 |
| AT5G51220.1 | Ubiquinol:cytochrome c oxidoreductase assembly protein | 26 | 1.16 | 0.360 | 1.000 |
| AT5G51220.1 | Ubiquinol:cytochrome c oxidoreductase assembly protein | 27 | 1.30 | 0.284 | 1.000 |
| AT5G51220.1 | Ubiquinol:cytochrome c oxidoreductase assembly protein | 28 | 1.67 | 0.251 | 1.000 |
| AT5G51220.1 | Ubiquinol:cytochrome c oxidoreductase assembly protein | 29 | 1.91 | 0.230 | 0.995 |
| AT5G51220.1 | Ubiquinol:cytochrome c oxidoreductase assembly protein | 30 | 1.59 | 0.222 | 1.000 |
| AT5G51220.1 | Ubiquinol:cytochrome c oxidoreductase assembly protein | 31 | 1.48 | 0.216 | 1.000 |
| AT5G51220.1 | Ubiquinol:cytochrome c oxidoreductase assembly protein | 32 | 1.74 | 0.213 | 0.998 |
| AT5G51220.1 | Ubiquinol:cytochrome c oxidoreductase assembly protein | 33 | 1.92 | 0.213 | 0.987 |
| AT5G51220.1 | Ubiquinol:cytochrome c oxidoreductase assembly protein | 34 | 2.11 | 0.213 | 0.950 |
| AT5G51220.1 | Ubiquinol:cytochrome c oxidoreductase assembly protein | 35 | 2.21 | 0.213 | 0.912 |
| AT5G51220.1 | Ubiquinol:cytochrome c oxidoreductase assembly protein | 36 | 2.22 | 0.211 | 0.900 |
| AT5G51220.1 | Ubiquinol:cytochrome c oxidoreductase assembly protein | 37 | 2.06 | 0.212 | 0.959 |
| AT5G51220.1 | Ubiquinol:cytochrome c oxidoreductase assembly protein | 38 | 1.27 | 0.213 | 1.000 |
| AT5G51220.1 | Ubiquinol:cytochrome c oxidoreductase assembly protein | 39 | 0.51 | 0.213 | 0.982 |

| | | | | | |
|---|---|---|---|---|---|
| AT5G51220.1 | Ubiquinol:cytochrome c oxidoreductase assembly protein | 40 | 0.25 | 0.212 | 0.112 |
| AT5G51220.1 | Ubiquinol:cytochrome c oxidoreductase assembly protein | 41 | 0.26 | 0.212 | 0.165 |
| AT5G51220.1 | Ubiquinol:cytochrome c oxidoreductase assembly protein | 42 | 0.66 | 0.215 | 1.000 |
| AT5G51220.1 | Ubiquinol:cytochrome c oxidoreductase assembly protein | 43 | 1.61 | 0.217 | 1.000 |
| AT5G51220.1 | Ubiquinol:cytochrome c oxidoreductase assembly protein | 44 | 2.11 | 0.216 | 0.955 |
| AT5G51220.1 | Ubiquinol:cytochrome c oxidoreductase assembly protein | 45 | 1.56 | 0.216 | 1.000 |
| AT5G51220.1 | Ubiquinol:cytochrome c oxidoreductase assembly protein | 46 | 0.87 | 0.247 | 1.000 |
| AT5G51220.1 | Ubiquinol:cytochrome c oxidoreductase assembly protein | 47 | 0.44 | 0.445 | 1.000 |
| AT5G51540.1 | Mitochondrial ATP-independend protease (OCT1) | 2 | 0.94 | 0.503 | 1.000 |
| AT5G51540.1 | Mitochondrial ATP-independend protease (OCT1) | 3 | 1.00 | 0.479 | 1.000 |
| AT5G51540.1 | Mitochondrial ATP-independend protease (OCT1) | 29 | 1.49 | 0.327 | 1.000 |
| AT5G51540.1 | Mitochondrial ATP-independend protease (OCT1) | 30 | 1.39 | 0.254 | 1.000 |
| AT5G51540.1 | Mitochondrial ATP-independend protease (OCT1) | 31 | 1.23 | 0.219 | 1.000 |
| AT5G51540.1 | Mitochondrial ATP-independend protease (OCT1) | 32 | 1.05 | 0.206 | 1.000 |
| AT5G51540.1 | Mitochondrial ATP-independend protease (OCT1) | 33 | 0.91 | 0.201 | 1.000 |
| AT5G51540.1 | Mitochondrial ATP-independend protease (OCT1) | 34 | 0.90 | 0.199 | 1.000 |
| AT5G51540.1 | Mitochondrial ATP-independend protease (OCT1) | 35 | 1.08 | 0.198 | 1.000 |
| AT5G51540.1 | Mitochondrial ATP-independend protease (OCT1) | 36 | 1.52 | 0.197 | 0.989 |
| AT5G51540.1 | Mitochondrial ATP-independend protease (OCT1) | 37 | 2.28 | 0.197 | 0.544 |
| AT5G51540.1 | Mitochondrial ATP-independend protease (OCT1) | 38 | 2.91 | 0.198 | 0.211 |
| AT5G51540.1 | Mitochondrial ATP-independend protease (OCT1) | 39 | 2.63 | 0.198 | 0.333 |
| AT5G51540.1 | Mitochondrial ATP-independend protease (OCT1) | 40 | 1.79 | 0.198 | 0.898 |
| AT5G51540.1 | Mitochondrial ATP-independend protease (OCT1) | 41 | 1.05 | 0.200 | 1.000 |
| AT5G51540.1 | Mitochondrial ATP-independend protease (OCT1) | 42 | 0.61 | 0.203 | 0.965 |
| AT5G51540.1 | Mitochondrial ATP-independend protease (OCT1) | 43 | 0.43 | 0.205 | 0.567 |
| AT5G51540.1 | Mitochondrial ATP-independend protease (OCT1) | 44 | 0.39 | 0.212 | 0.441 |
| AT5G51540.1 | Mitochondrial ATP-independend protease (OCT1) | 45 | 0.40 | 0.253 | 0.714 |

| | | | | | |
|---|---|---|---|---|---|
| AT5G51540.1 | Mitochondrial ATP-independend protease (OCT1) | 46 | 0.42 | 0.373 | 0.978 |
| AT5G51540.1 | Mitochondrial ATP-independend protease (OCT1) | 47 | 0.46 | 0.576 | 1.000 |
| AT5G51740.1 | Mitochondrial ATP-independend protease (OMA1) | 2 | 0.97 | 0.451 | 1.000 |
| AT5G51740.1 | Mitochondrial ATP-independend protease (OMA1) | 3 | 0.97 | 0.385 | 1.000 |
| AT5G51740.1 | Mitochondrial ATP-independend protease (OMA1) | 10 | 1.45 | 0.279 | 1.000 |
| AT5G51740.1 | Mitochondrial ATP-independend protease (OMA1) | 11 | 1.71 | 0.240 | 1.000 |
| AT5G51740.1 | Mitochondrial ATP-independend protease (OMA1) | 12 | 2.08 | 0.217 | 0.983 |
| AT5G51740.1 | Mitochondrial ATP-independend protease (OMA1) | 13 | 2.46 | 0.203 | 0.793 |
| AT5G51740.1 | Mitochondrial ATP-independend protease (OMA1) | 14 | 2.55 | 0.190 | 0.621 |
| AT5G51740.1 | Mitochondrial ATP-independend protease (OMA1) | 15 | 1.94 | 0.187 | 0.972 |
| AT5G51740.1 | Mitochondrial ATP-independend protease (OMA1) | 16 | 1.13 | 0.186 | 1.000 |
| AT5G51740.1 | Mitochondrial ATP-independend protease (OMA1) | 17 | 0.77 | 0.185 | 1.000 |
| AT5G51740.1 | Mitochondrial ATP-independend protease (OMA1) | 18 | 0.71 | 0.189 | 1.000 |
| AT5G51740.1 | Mitochondrial ATP-independend protease (OMA1) | 19 | 0.78 | 0.195 | 1.000 |
| AT5G51740.1 | Mitochondrial ATP-independend protease (OMA1) | 20 | 0.94 | 0.201 | 1.000 |
| AT5G51740.1 | Mitochondrial ATP-independend protease (OMA1) | 21 | 1.17 | 0.204 | 1.000 |
| AT5G51740.1 | Mitochondrial ATP-independend protease (OMA1) | 22 | 1.51 | 0.203 | 1.000 |
| AT5G51740.1 | Mitochondrial ATP-independend protease (OMA1) | 23 | 1.63 | 0.202 | 1.000 |
| AT5G51740.1 | Mitochondrial ATP-independend protease (OMA1) | 24 | 1.27 | 0.197 | 1.000 |
| AT5G51740.1 | Mitochondrial ATP-independend protease (OMA1) | 25 | 0.85 | 0.193 | 1.000 |
| AT5G51740.1 | Mitochondrial ATP-independend protease (OMA1) | 26 | 0.91 | 0.192 | 1.000 |
| AT5G51740.1 | Mitochondrial ATP-independend protease (OMA1) | 27 | 1.41 | 0.193 | 1.000 |
| AT5G51740.1 | Mitochondrial ATP-independend protease (OMA1) | 28 | 1.85 | 0.192 | 0.991 |
| AT5G51740.1 | Mitochondrial ATP-independend protease (OMA1) | 29 | 2.28 | 0.192 | 0.837 |
| AT5G51740.1 | Mitochondrial ATP-independend protease (OMA1) | 30 | 2.29 | 0.192 | 0.825 |
| AT5G51740.1 | Mitochondrial ATP-independend protease (OMA1) | 31 | 2.00 | 0.192 | 0.964 |
| AT5G51740.1 | Mitochondrial ATP-independend protease (OMA1) | 32 | 1.77 | 0.193 | 0.997 |
| AT5G51740.1 | Mitochondrial ATP-independend protease (OMA1) | 33 | 1.52 | 0.193 | 1.000 |
| AT5G51740.1 | Mitochondrial ATP-independend protease (OMA1) | 34 | 1.41 | 0.194 | 1.000 |
| AT5G51740.1 | Mitochondrial ATP-independend protease (OMA1) | 35 | 1.50 | 0.193 | 1.000 |
| AT5G51740.1 | Mitochondrial ATP-independend protease (OMA1) | 36 | 1.77 | 0.192 | 0.997 |

| | | | | | |
|---|---|---|---|---|---|
| AT5G51740.1 | Mitochondrial ATP-independend protease (OMA1) | 37 | 2.21 | 0.192 | 0.878 |
| AT5G51740.1 | Mitochondrial ATP-independend protease (OMA1) | 38 | 2.49 | 0.193 | 0.689 |
| AT5G51740.1 | Mitochondrial ATP-independend protease (OMA1) | 39 | 2.46 | 0.193 | 0.719 |
| AT5G51740.1 | Mitochondrial ATP-independend protease (OMA1) | 40 | 2.03 | 0.194 | 0.960 |
| AT5G51740.1 | Mitochondrial ATP-independend protease (OMA1) | 41 | 1.40 | 0.195 | 1.000 |
| AT5G51740.1 | Mitochondrial ATP-independend protease (OMA1) | 42 | 1.09 | 0.197 | 1.000 |
| AT5G51740.1 | Mitochondrial ATP-independend protease (OMA1) | 43 | 1.08 | 0.199 | 1.000 |
| AT5G51740.1 | Mitochondrial ATP-independend protease (OMA1) | 44 | 1.06 | 0.200 | 1.000 |
| AT5G51740.1 | Mitochondrial ATP-independend protease (OMA1) | 45 | 0.80 | 0.199 | 1.000 |
| AT5G51740.1 | Mitochondrial ATP-independend protease (OMA1) | 46 | 0.45 | 0.231 | 0.980 |
| AT5G51740.1 | Mitochondrial ATP-independend protease (OMA1) | 47 | 0.21 | 0.430 | 0.965 |
| AT5G52310.1 | Cold regulated 78 (COR78) | 2 | 1.27 | 0.771 | 1.000 |
| AT5G52310.1 | Cold regulated 78 (COR78) | 3 | 1.37 | 0.745 | 0.999 |
| AT5G52310.1 | Cold regulated 78 (COR78) | 31 | 1.99 | 0.343 | 0.806 |
| AT5G52310.1 | Cold regulated 78 (COR78) | 32 | 1.78 | 0.334 | 0.869 |
| AT5G52310.1 | Cold regulated 78 (COR78) | 33 | 1.57 | 0.333 | 0.938 |
| AT5G52310.1 | Cold regulated 78 (COR78) | 34 | 1.36 | 0.340 | 0.985 |
| AT5G52310.1 | Cold regulated 78 (COR78) | 35 | 1.15 | 0.352 | 1.000 |
| AT5G52310.1 | Cold regulated 78 (COR78) | 46 | 0.08 | 0.540 | 0.154 |
| AT5G52310.1 | Cold regulated 78 (COR78) | 47 | 0.06 | 0.572 | 0.126 |
| AT5G52520.1 | Prolyl-tRNA synthetase Organellar (PRORS-ORG / PRORS1) | 2 | 1.02 | 0.307 | 1.000 |
| AT5G52520.1 | Prolyl-tRNA synthetase Organellar (PRORS-ORG / PRORS1) | 3 | 1.02 | 0.304 | 1.000 |
| AT5G52520.1 | Prolyl-tRNA synthetase Organellar (PRORS-ORG / PRORS1) | 33 | 1.55 | 0.235 | 0.997 |
| AT5G52520.1 | Prolyl-tRNA synthetase Organellar (PRORS-ORG / PRORS1) | 34 | 1.50 | 0.192 | 0.990 |
| AT5G52520.1 | Prolyl-tRNA synthetase Organellar (PRORS-ORG / PRORS1) | 35 | 1.29 | 0.181 | 1.000 |
| AT5G52520.1 | Prolyl-tRNA synthetase Organellar (PRORS-ORG / PRORS1) | 36 | 1.11 | 0.187 | 1.000 |
| AT5G52520.1 | Prolyl-tRNA synthetase Organellar (PRORS-ORG / PRORS1) | 37 | 1.06 | 0.184 | 1.000 |
| AT5G52520.1 | Prolyl-tRNA synthetase Organellar (PRORS-ORG / PRORS1) | 38 | 1.25 | 0.164 | 1.000 |
| AT5G52520.1 | Prolyl-tRNA synthetase Organellar (PRORS-ORG / PRORS1) | 39 | 1.69 | 0.150 | 0.779 |
| AT5G52520.1 | Prolyl-tRNA synthetase Organellar (PRORS-ORG / PRORS1) | 40 | 2.25 | 0.145 | 0.192 |

| | | | | | |
|---|---|---|---|---|---|
| AT5G52520.1 | Prolyl-tRNA synthetase Organellar (PRORS-ORG / PRORS1) | 41 | 2.38 | 0.145 | 0.124 |
| AT5G52520.1 | Prolyl-tRNA synthetase Organellar (PRORS-ORG / PRORS1) | 42 | 1.80 | 0.147 | 0.625 |
| AT5G52520.1 | Prolyl-tRNA synthetase Organellar (PRORS-ORG / PRORS1) | 43 | 1.24 | 0.148 | 1.000 |
| AT5G52520.1 | Prolyl-tRNA synthetase Organellar (PRORS-ORG / PRORS1) | 44 | 0.89 | 0.147 | 1.000 |
| AT5G52520.1 | Prolyl-tRNA synthetase Organellar (PRORS-ORG / PRORS1) | 45 | 0.62 | 0.147 | 0.851 |
| AT5G52520.1 | Prolyl-tRNA synthetase Organellar (PRORS-ORG / PRORS1) | 46 | 0.36 | 0.167 | 0.117 |
| AT5G52520.1 | Prolyl-tRNA synthetase Organellar (PRORS-ORG / PRORS1) | 47 | 0.17 | 0.304 | 0.140 |
| AT5G52840.1 | NADH:ubiquinone oxidoreductase subunit (B13) | 3 | 0.72 | 0.212 | 1.000 |
| AT5G52840.1 | NADH:ubiquinone oxidoreductase subunit (B13) | 4 | 0.73 | 0.197 | 1.000 |
| AT5G52840.1 | NADH:ubiquinone oxidoreductase subunit (B13) | 5 | 0.82 | 0.197 | 1.000 |
| AT5G52840.1 | NADH:ubiquinone oxidoreductase subunit (B13) | 6 | 0.93 | 0.198 | 1.000 |
| AT5G52840.1 | NADH:ubiquinone oxidoreductase subunit (B13) | 7 | 0.97 | 0.198 | 1.000 |
| AT5G52840.1 | NADH:ubiquinone oxidoreductase subunit (B13) | 8 | 0.98 | 0.194 | 1.000 |
| AT5G52840.1 | NADH:ubiquinone oxidoreductase subunit (B13) | 9 | 0.96 | 0.191 | 1.000 |
| AT5G52840.1 | NADH:ubiquinone oxidoreductase subunit (B13) | 10 | 0.97 | 0.191 | 1.000 |
| AT5G52840.1 | NADH:ubiquinone oxidoreductase subunit (B13) | 11 | 1.04 | 0.192 | 1.000 |
| AT5G52840.1 | NADH:ubiquinone oxidoreductase subunit (B13) | 12 | 1.16 | 0.192 | 1.000 |
| AT5G52840.1 | NADH:ubiquinone oxidoreductase subunit (B13) | 13 | 1.24 | 0.190 | 1.000 |
| AT5G52840.1 | NADH:ubiquinone oxidoreductase subunit (B13) | 14 | 1.15 | 0.187 | 1.000 |
| AT5G52840.1 | NADH:ubiquinone oxidoreductase subunit (B13) | 15 | 0.95 | 0.186 | 1.000 |
| AT5G52840.1 | NADH:ubiquinone oxidoreductase subunit (B13) | 16 | 0.70 | 0.185 | 1.000 |
| AT5G52840.1 | NADH:ubiquinone oxidoreductase subunit (B13) | 17 | 0.54 | 0.185 | 0.988 |
| AT5G52840.1 | NADH:ubiquinone oxidoreductase subunit (B13) | 18 | 0.56 | 0.188 | 0.995 |
| AT5G52840.1 | NADH:ubiquinone oxidoreductase subunit (B13) | 19 | 0.72 | 0.192 | 1.000 |
| AT5G52840.1 | NADH:ubiquinone oxidoreductase subunit (B13) | 20 | 0.88 | 0.197 | 1.000 |
| AT5G52840.1 | NADH:ubiquinone oxidoreductase subunit (B13) | 21 | 1.09 | 0.200 | 1.000 |
| AT5G52840.1 | NADH:ubiquinone oxidoreductase subunit (B13) | 22 | 1.42 | 0.200 | 1.000 |

| | | | | | |
|---|---|---|---|---|---|
| AT5G52840.1 | NADH:ubiquinone oxidoreductase subunit (B13) | 23 | 1.65 | 0.199 | 1.000 |
| AT5G52840.1 | NADH:ubiquinone oxidoreductase subunit (B13) | 24 | 1.32 | 0.195 | 1.000 |
| AT5G52840.1 | NADH:ubiquinone oxidoreductase subunit (B13) | 25 | 0.74 | 0.191 | 1.000 |
| AT5G52840.1 | NADH:ubiquinone oxidoreductase subunit (B13) | 26 | 0.52 | 0.190 | 0.976 |
| AT5G52840.1 | NADH:ubiquinone oxidoreductase subunit (B13) | 27 | 0.58 | 0.190 | 0.999 |
| AT5G52840.1 | NADH:ubiquinone oxidoreductase subunit (B13) | 28 | 0.83 | 0.190 | 1.000 |
| AT5G52840.1 | NADH:ubiquinone oxidoreductase subunit (B13) | 29 | 1.24 | 0.190 | 1.000 |
| AT5G52840.1 | NADH:ubiquinone oxidoreductase subunit (B13) | 30 | 1.61 | 0.190 | 1.000 |
| AT5G52840.1 | NADH:ubiquinone oxidoreductase subunit (B13) | 31 | 1.77 | 0.190 | 0.997 |
| AT5G52840.1 | NADH:ubiquinone oxidoreductase subunit (B13) | 32 | 1.75 | 0.191 | 0.997 |
| AT5G52840.1 | NADH:ubiquinone oxidoreductase subunit (B13) | 33 | 1.62 | 0.191 | 1.000 |
| AT5G52840.1 | NADH:ubiquinone oxidoreductase subunit (B13) | 34 | 1.52 | 0.191 | 1.000 |
| AT5G52840.1 | NADH:ubiquinone oxidoreductase subunit (B13) | 35 | 1.53 | 0.191 | 1.000 |
| AT5G52840.1 | NADH:ubiquinone oxidoreductase subunit (B13) | 36 | 1.60 | 0.190 | 1.000 |
| AT5G52840.1 | NADH:ubiquinone oxidoreductase subunit (B13) | 37 | 1.60 | 0.190 | 1.000 |
| AT5G52840.1 | NADH:ubiquinone oxidoreductase subunit (B13) | 38 | 1.69 | 0.191 | 0.999 |
| AT5G52840.1 | NADH:ubiquinone oxidoreductase subunit (B13) | 39 | 2.13 | 0.191 | 0.915 |
| AT5G52840.1 | NADH:ubiquinone oxidoreductase subunit (B13) | 40 | 2.54 | 0.192 | 0.646 |
| AT5G52840.1 | NADH:ubiquinone oxidoreductase subunit (B13) | 41 | 2.49 | 0.192 | 0.685 |
| AT5G52840.1 | NADH:ubiquinone oxidoreductase subunit (B13) | 42 | 2.15 | 0.195 | 0.917 |
| AT5G52840.1 | NADH:ubiquinone oxidoreductase subunit (B13) | 43 | 1.78 | 0.196 | 0.997 |
| AT5G52840.1 | NADH:ubiquinone oxidoreductase subunit (B13) | 44 | 1.31 | 0.197 | 1.000 |
| AT5G52840.1 | NADH:ubiquinone oxidoreductase subunit (B13) | 45 | 0.79 | 0.197 | 1.000 |
| AT5G52840.1 | NADH:ubiquinone oxidoreductase subunit (B13) | 46 | 0.35 | 0.227 | 0.744 |
| AT5G52840.1 | NADH:ubiquinone oxidoreductase subunit (B13) | 47 | 0.12 | 0.395 | 0.482 |
| AT5G52840.1 | NADH:ubiquinone oxidoreductase subunit (B13) | 48 | 0.04 | 0.663 | 0.669 |
| AT5G53350.1 | Caseinolytic protease X1 (CLPX1) | 2 | 0.97 | 0.413 | 1.000 |
| AT5G53350.1 | Caseinolytic protease X1 (CLPX1) | 3 | 0.98 | 0.374 | 1.000 |
| AT5G53350.1 | Caseinolytic protease X1 (CLPX1) | 16 | 1.16 | 0.251 | 1.000 |
| AT5G53350.1 | Caseinolytic protease X1 (CLPX1) | 17 | 1.04 | 0.230 | 1.000 |
| AT5G53350.1 | Caseinolytic protease X1 (CLPX1) | 18 | 0.82 | 0.205 | 1.000 |

| | | | | | |
|---|---|---|---|---|---|
| AT5G53350.1 | Caseinolytic protease X1 (CLPX1) | 19 | 0.76 | 0.199 | 1.000 |
| AT5G53350.1 | Caseinolytic protease X1 (CLPX1) | 20 | 0.76 | 0.201 | 1.000 |
| AT5G53350.1 | Caseinolytic protease X1 (CLPX1) | 21 | 0.77 | 0.200 | 1.000 |
| AT5G53350.1 | Caseinolytic protease X1 (CLPX1) | 22 | 0.87 | 0.201 | 1.000 |
| AT5G53350.1 | Caseinolytic protease X1 (CLPX1) | 23 | 1.04 | 0.199 | 1.000 |
| AT5G53350.1 | Caseinolytic protease X1 (CLPX1) | 24 | 1.23 | 0.194 | 1.000 |
| AT5G53350.1 | Caseinolytic protease X1 (CLPX1) | 25 | 1.32 | 0.192 | 1.000 |
| AT5G53350.1 | Caseinolytic protease X1 (CLPX1) | 26 | 0.90 | 0.192 | 1.000 |
| AT5G53350.1 | Caseinolytic protease X1 (CLPX1) | 27 | 0.47 | 0.193 | 0.865 |
| AT5G53350.1 | Caseinolytic protease X1 (CLPX1) | 28 | 0.44 | 0.193 | 0.769 |
| AT5G53350.1 | Caseinolytic protease X1 (CLPX1) | 29 | 0.68 | 0.193 | 1.000 |
| AT5G53350.1 | Caseinolytic protease X1 (CLPX1) | 30 | 1.16 | 0.193 | 1.000 |
| AT5G53350.1 | Caseinolytic protease X1 (CLPX1) | 31 | 1.73 | 0.193 | 0.992 |
| AT5G53350.1 | Caseinolytic protease X1 (CLPX1) | 32 | 1.90 | 0.193 | 0.960 |
| AT5G53350.1 | Caseinolytic protease X1 (CLPX1) | 33 | 1.89 | 0.193 | 0.964 |
| AT5G53350.1 | Caseinolytic protease X1 (CLPX1) | 34 | 1.91 | 0.193 | 0.959 |
| AT5G53350.1 | Caseinolytic protease X1 (CLPX1) | 35 | 2.10 | 0.193 | 0.874 |
| AT5G53350.1 | Caseinolytic protease X1 (CLPX1) | 36 | 2.41 | 0.192 | 0.648 |
| AT5G53350.1 | Caseinolytic protease X1 (CLPX1) | 37 | 2.42 | 0.192 | 0.638 |
| AT5G53350.1 | Caseinolytic protease X1 (CLPX1) | 38 | 1.86 | 0.193 | 0.971 |
| AT5G53350.1 | Caseinolytic protease X1 (CLPX1) | 39 | 1.44 | 0.193 | 1.000 |
| AT5G53350.1 | Caseinolytic protease X1 (CLPX1) | 40 | 1.70 | 0.194 | 0.995 |
| AT5G53350.1 | Caseinolytic protease X1 (CLPX1) | 41 | 2.20 | 0.194 | 0.820 |
| AT5G53350.1 | Caseinolytic protease X1 (CLPX1) | 42 | 2.51 | 0.197 | 0.606 |
| AT5G53350.1 | Caseinolytic protease X1 (CLPX1) | 43 | 2.67 | 0.198 | 0.511 |
| AT5G53350.1 | Caseinolytic protease X1 (CLPX1) | 44 | 2.33 | 0.198 | 0.749 |
| AT5G53350.1 | Caseinolytic protease X1 (CLPX1) | 45 | 1.55 | 0.198 | 1.000 |
| AT5G53350.1 | Caseinolytic protease X1 (CLPX1) | 46 | 0.79 | 0.224 | 1.000 |
| AT5G53350.1 | Caseinolytic protease X1 (CLPX1) | 47 | 0.33 | 0.396 | 0.995 |
| AT5G53540.1 | 26S proteasome regulatory subunit (RPT6-like) | 2 | 1.29 | 0.512 | 1.000 |
| AT5G53540.1 | 26S proteasome regulatory subunit (RPT6-like) | 3 | 1.28 | 0.471 | 1.000 |

| | | | | | |
|---|---|---|---|---|---|
| AT5G53540.1 | 26S proteasome regulatory subunit (RPT6-like) | 14 | 1.18 | 0.207 | 1.000 |
| AT5G53540.1 | 26S proteasome regulatory subunit (RPT6-like) | 15 | 1.18 | 0.193 | 1.000 |
| AT5G53540.1 | 26S proteasome regulatory subunit (RPT6-like) | 16 | 1.17 | 0.180 | 1.000 |
| AT5G53540.1 | 26S proteasome regulatory subunit (RPT6-like) | 17 | 1.16 | 0.170 | 1.000 |
| AT5G53540.1 | 26S proteasome regulatory subunit (RPT6-like) | 18 | 1.15 | 0.162 | 1.000 |
| AT5G53540.1 | 26S proteasome regulatory subunit (RPT6-like) | 19 | 1.13 | 0.157 | 1.000 |
| AT5G53540.1 | 26S proteasome regulatory subunit (RPT6-like) | 20 | 1.11 | 0.153 | 1.000 |
| AT5G53540.1 | 26S proteasome regulatory subunit (RPT6-like) | 21 | 1.09 | 0.151 | 1.000 |
| AT5G53540.1 | 26S proteasome regulatory subunit (RPT6-like) | 22 | 1.07 | 0.149 | 1.000 |
| AT5G53540.1 | 26S proteasome regulatory subunit (RPT6-like) | 23 | 1.05 | 0.148 | 1.000 |
| AT5G53540.1 | 26S proteasome regulatory subunit (RPT6-like) | 24 | 1.04 | 0.148 | 1.000 |
| AT5G53540.1 | 26S proteasome regulatory subunit (RPT6-like) | 25 | 1.02 | 0.147 | 1.000 |
| AT5G53540.1 | 26S proteasome regulatory subunit (RPT6-like) | 26 | 1.01 | 0.147 | 1.000 |
| AT5G53540.1 | 26S proteasome regulatory subunit (RPT6-like) | 27 | 0.98 | 0.148 | 1.000 |
| AT5G53540.1 | 26S proteasome regulatory subunit (RPT6-like) | 28 | 0.96 | 0.149 | 1.000 |
| AT5G53540.1 | 26S proteasome regulatory subunit (RPT6-like) | 29 | 0.93 | 0.151 | 1.000 |
| AT5G53540.1 | 26S proteasome regulatory subunit (RPT6-like) | 30 | 0.90 | 0.154 | 1.000 |
| AT5G53540.1 | 26S proteasome regulatory subunit (RPT6-like) | 31 | 0.87 | 0.158 | 1.000 |
| AT5G53540.1 | 26S proteasome regulatory subunit (RPT6-like) | 32 | 0.84 | 0.165 | 0.999 |
| AT5G53540.1 | 26S proteasome regulatory subunit (RPT6-like) | 33 | 0.81 | 0.175 | 0.998 |
| AT5G53540.1 | 26S proteasome regulatory subunit (RPT6-like) | 34 | 0.79 | 0.187 | 0.997 |
| AT5G53540.1 | 26S proteasome regulatory subunit (RPT6-like) | 35 | 0.77 | 0.204 | 0.996 |
| AT5G53540.1 | 26S proteasome regulatory subunit (RPT6-like) | 36 | 0.75 | 0.225 | 0.996 |
| AT5G53540.1 | 26S proteasome regulatory subunit (RPT6-like) | 37 | 0.73 | 0.250 | 0.997 |
| AT5G53540.1 | 26S proteasome regulatory subunit (RPT6-like) | 46 | 0.56 | 0.620 | 1.000 |
| AT5G53540.1 | 26S proteasome regulatory subunit (RPT6-like) | 47 | 0.55 | 0.677 | 1.000 |
| AT5G53650.1 | ABC transporter A family protein | 2 | 1.05 | 0.284 | 1.000 |
| AT5G53650.1 | ABC transporter A family protein | 3 | 0.73 | 0.179 | 1.000 |
| AT5G53650.1 | ABC transporter A family protein | 4 | 0.55 | 0.152 | 0.935 |
| AT5G53650.1 | ABC transporter A family protein | 5 | 0.49 | 0.145 | 0.656 |
| AT5G53650.1 | ABC transporter A family protein | 6 | 0.53 | 0.142 | 0.820 |

| | | | | | | |
|---|---|---|---|---|---|---|
| AT5G53650.1 | ABC transporter A family protein | 7 | 0.65 | 0.140 | 0.997 | |
| AT5G53650.1 | ABC transporter A family protein | 8 | 0.81 | 0.137 | 1.000 | |
| AT5G53650.1 | ABC transporter A family protein | 9 | 0.91 | 0.135 | 1.000 | |
| AT5G53650.1 | ABC transporter A family protein | 10 | 0.93 | 0.135 | 1.000 | |
| AT5G53650.1 | ABC transporter A family protein | 11 | 1.04 | 0.135 | 1.000 | |
| AT5G53650.1 | ABC transporter A family protein | 12 | 1.31 | 0.136 | 1.000 | |
| AT5G53650.1 | ABC transporter A family protein | 13 | 1.83 | 0.134 | 0.792 | |
| AT5G53650.1 | ABC transporter A family protein | 14 | 2.47 | 0.132 | 0.104 | |
| AT5G53650.1 | ABC transporter A family protein | 15 | 2.64 | 0.131 | 0.049 | * |
| AT5G53650.1 | ABC transporter A family protein | 16 | 2.31 | 0.130 | 0.167 | |
| AT5G53650.1 | ABC transporter A family protein | 17 | 2.11 | 0.129 | 0.342 | |
| AT5G53650.1 | ABC transporter A family protein | 18 | 2.02 | 0.132 | 0.492 | |
| AT5G53650.1 | ABC transporter A family protein | 19 | 1.91 | 0.137 | 0.724 | |
| AT5G53650.1 | ABC transporter A family protein | 20 | 1.89 | 0.140 | 0.785 | |
| AT5G53650.1 | ABC transporter A family protein | 21 | 2.04 | 0.141 | 0.600 | |
| AT5G53650.1 | ABC transporter A family protein | 22 | 2.12 | 0.141 | 0.506 | |
| AT5G53650.1 | ABC transporter A family protein | 23 | 1.82 | 0.139 | 0.851 | |
| AT5G53650.1 | ABC transporter A family protein | 24 | 1.28 | 0.136 | 1.000 | |
| AT5G53650.1 | ABC transporter A family protein | 25 | 0.86 | 0.134 | 1.000 | |
| AT5G53650.1 | ABC transporter A family protein | 26 | 0.73 | 0.133 | 1.000 | |
| AT5G53650.1 | ABC transporter A family protein | 27 | 0.87 | 0.133 | 1.000 | |
| AT5G53650.1 | ABC transporter A family protein | 28 | 1.13 | 0.134 | 1.000 | |
| AT5G53650.1 | ABC transporter A family protein | 29 | 1.31 | 0.135 | 1.000 | |
| AT5G53650.1 | ABC transporter A family protein | 30 | 1.34 | 0.135 | 1.000 | |
| AT5G53650.1 | ABC transporter A family protein | 31 | 1.24 | 0.135 | 1.000 | |
| AT5G53650.1 | ABC transporter A family protein | 32 | 1.05 | 0.136 | 1.000 | |
| AT5G53650.1 | ABC transporter A family protein | 33 | 0.90 | 0.136 | 1.000 | |
| AT5G53650.1 | ABC transporter A family protein | 34 | 0.88 | 0.136 | 1.000 | |
| AT5G53650.1 | ABC transporter A family protein | 35 | 0.93 | 0.136 | 1.000 | |
| AT5G53650.1 | ABC transporter A family protein | 36 | 0.97 | 0.135 | 1.000 | |
| AT5G53650.1 | ABC transporter A family protein | 37 | 0.94 | 0.135 | 1.000 | |

| | | | | | |
|---|---|---|---|---|---|
| AT5G53650.1 | ABC transporter A family protein | 38 | 0.91 | 0.135 | 1.000 |
| AT5G53650.1 | ABC transporter A family protein | 39 | 0.95 | 0.135 | 1.000 |
| AT5G53650.1 | ABC transporter A family protein | 40 | 1.01 | 0.135 | 1.000 |
| AT5G53650.1 | ABC transporter A family protein | 41 | 0.94 | 0.135 | 1.000 |
| AT5G53650.1 | ABC transporter A family protein | 42 | 0.79 | 0.138 | 1.000 |
| AT5G53650.1 | ABC transporter A family protein | 43 | 0.69 | 0.139 | 1.000 |
| AT5G53650.1 | ABC transporter A family protein | 44 | 0.58 | 0.139 | 0.929 |
| AT5G53650.1 | ABC transporter A family protein | 45 | 0.47 | 0.139 | 0.458 |
| AT5G53650.1 | ABC transporter A family protein | 46 | 0.39 | 0.166 | 0.362 |
| AT5G53650.1 | ABC transporter A family protein | 47 | 0.32 | 0.294 | 0.942 |
| AT5G54100.1 | Stomatin-like protein 2 (SLP2) | 2 | 1.11 | 0.481 | 1.000 |
| AT5G54100.1 | Stomatin-like protein 2 (SLP2) | 3 | 0.95 | 0.326 | 1.000 |
| AT5G54100.1 | Stomatin-like protein 2 (SLP2) | 4 | 0.78 | 0.322 | 1.000 |
| AT5G54100.1 | Stomatin-like protein 2 (SLP2) | 5 | 0.64 | 0.326 | 1.000 |
| AT5G54100.1 | Stomatin-like protein 2 (SLP2) | 6 | 0.56 | 0.288 | 0.999 |
| AT5G54100.1 | Stomatin-like protein 2 (SLP2) | 7 | 0.54 | 0.261 | 0.994 |
| AT5G54100.1 | Stomatin-like protein 2 (SLP2) | 8 | 0.53 | 0.248 | 0.988 |
| AT5G54100.1 | Stomatin-like protein 2 (SLP2) | 9 | 0.50 | 0.241 | 0.970 |
| AT5G54100.1 | Stomatin-like protein 2 (SLP2) | 10 | 0.51 | 0.241 | 0.975 |
| AT5G54100.1 | Stomatin-like protein 2 (SLP2) | 11 | 0.64 | 0.241 | 1.000 |
| AT5G54100.1 | Stomatin-like protein 2 (SLP2) | 12 | 0.78 | 0.241 | 1.000 |
| AT5G54100.1 | Stomatin-like protein 2 (SLP2) | 13 | 0.74 | 0.238 | 1.000 |
| AT5G54100.1 | Stomatin-like protein 2 (SLP2) | 14 | 0.59 | 0.235 | 0.997 |
| AT5G54100.1 | Stomatin-like protein 2 (SLP2) | 15 | 0.49 | 0.234 | 0.948 |
| AT5G54100.1 | Stomatin-like protein 2 (SLP2) | 16 | 0.46 | 0.231 | 0.905 |
| AT5G54100.1 | Stomatin-like protein 2 (SLP2) | 17 | 0.54 | 0.231 | 0.983 |
| AT5G54100.1 | Stomatin-like protein 2 (SLP2) | 18 | 0.91 | 0.235 | 1.000 |
| AT5G54100.1 | Stomatin-like protein 2 (SLP2) | 19 | 2.06 | 0.248 | 0.966 |
| AT5G54100.1 | Stomatin-like protein 2 (SLP2) | 20 | 2.84 | 0.260 | 0.730 |
| AT5G54100.1 | Stomatin-like protein 2 (SLP2) | 21 | 1.53 | 0.271 | 1.000 |
| AT5G54100.1 | Stomatin-like protein 2 (SLP2) | 22 | 0.43 | 0.332 | 0.989 |
| AT5G54100.1 | Stomatin-like protein 2 (SLP2) | 23 | 0.11 | 0.543 | 0.722 |
| AT5G54100.1 | Stomatin-like protein 2 (SLP2) | 46 | 154613.65 | 16.468 | 1.000 |
| AT5G54100.1 | Stomatin-like protein 2 (SLP2) | 47 | 756325.36 | 17.506 | 1.000 |
| AT5G54270.1 | Photosystem II light-harvesting complex subunit (LHCB3) | 2 | 0.96 | 0.265 | 1.000 |
| AT5G54270.1 | Photosystem II light-harvesting complex subunit (LHCB3) | 3 | 1.04 | 0.189 | 1.000 |
| AT5G54270.1 | Photosystem II light-harvesting complex subunit (LHCB3) | 4 | 1.13 | 0.154 | 1.000 |
| AT5G54270.1 | Photosystem II light-harvesting complex subunit (LHCB3) | 5 | 1.18 | 0.145 | 1.000 |
| AT5G54270.1 | Photosystem II light-harvesting complex subunit (LHCB3) | 6 | 1.18 | 0.144 | 1.000 |
| AT5G54270.1 | Photosystem II light-harvesting complex subunit (LHCB3) | 7 | 1.16 | 0.144 | 1.000 |
| AT5G54270.1 | Photosystem II light-harvesting complex subunit (LHCB3) | 8 | 1.13 | 0.142 | 1.000 |
| AT5G54270.1 | Photosystem II light-harvesting complex subunit (LHCB3) | 9 | 1.09 | 0.140 | 1.000 |
| AT5G54270.1 | Photosystem II light-harvesting complex subunit (LHCB3) | 10 | 1.04 | 0.139 | 1.000 |

| | | | | | |
|---|---|---|---|---|---|
| AT5G54270.1 | Photosystem II light-harvesting complex subunit (LHCB3) | 11 | 1.00 | 0.139 | 1.000 |
| AT5G54270.1 | Photosystem II light-harvesting complex subunit (LHCB3) | 12 | 0.97 | 0.138 | 1.000 |
| AT5G54270.1 | Photosystem II light-harvesting complex subunit (LHCB3) | 13 | 0.96 | 0.137 | 1.000 |
| AT5G54270.1 | Photosystem II light-harvesting complex subunit (LHCB3) | 14 | 0.95 | 0.135 | 1.000 |
| AT5G54270.1 | Photosystem II light-harvesting complex subunit (LHCB3) | 15 | 0.93 | 0.134 | 1.000 |
| AT5G54270.1 | Photosystem II light-harvesting complex subunit (LHCB3) | 16 | 0.92 | 0.134 | 1.000 |
| AT5G54270.1 | Photosystem II light-harvesting complex subunit (LHCB3) | 17 | 0.96 | 0.134 | 1.000 |
| AT5G54270.1 | Photosystem II light-harvesting complex subunit (LHCB3) | 18 | 1.06 | 0.136 | 1.000 |
| AT5G54270.1 | Photosystem II light-harvesting complex subunit (LHCB3) | 19 | 1.19 | 0.139 | 1.000 |
| AT5G54270.1 | Photosystem II light-harvesting complex subunit (LHCB3) | 20 | 1.32 | 0.142 | 1.000 |
| AT5G54270.1 | Photosystem II light-harvesting complex subunit (LHCB3) | 21 | 1.40 | 0.143 | 0.998 |
| AT5G54270.1 | Photosystem II light-harvesting complex subunit (LHCB3) | 22 | 1.38 | 0.144 | 0.999 |
| AT5G54270.1 | Photosystem II light-harvesting complex subunit (LHCB3) | 23 | 1.27 | 0.143 | 1.000 |
| AT5G54270.1 | Photosystem II light-harvesting complex subunit (LHCB3) | 24 | 1.11 | 0.140 | 1.000 |
| AT5G54270.1 | Photosystem II light-harvesting complex subunit (LHCB3) | 25 | 0.97 | 0.139 | 1.000 |
| AT5G54270.1 | Photosystem II light-harvesting complex subunit (LHCB3) | 26 | 0.91 | 0.138 | 1.000 |
| AT5G54270.1 | Photosystem II light-harvesting complex subunit (LHCB3) | 27 | 0.91 | 0.137 | 1.000 |
| AT5G54270.1 | Photosystem II light-harvesting complex subunit (LHCB3) | 28 | 0.94 | 0.137 | 1.000 |
| AT5G54270.1 | Photosystem II light-harvesting complex subunit (LHCB3) | 29 | 1.01 | 0.138 | 1.000 |
| AT5G54270.1 | Photosystem II light-harvesting complex subunit (LHCB3) | 30 | 1.13 | 0.138 | 1.000 |
| AT5G54270.1 | Photosystem II light-harvesting complex subunit (LHCB3) | 31 | 1.29 | 0.138 | 1.000 |
| AT5G54270.1 | Photosystem II light-harvesting complex subunit (LHCB3) | 32 | 1.41 | 0.139 | 0.996 |
| AT5G54270.1 | Photosystem II light-harvesting complex subunit (LHCB3) | 33 | 1.45 | 0.139 | 0.993 |
| AT5G54270.1 | Photosystem II light-harvesting complex subunit (LHCB3) | 34 | 1.48 | 0.142 | 0.988 |
| AT5G54270.1 | Photosystem II light-harvesting complex subunit (LHCB3) | 35 | 1.53 | 0.148 | 0.983 |
| AT5G54270.1 | Photosystem II light-harvesting complex subunit (LHCB3) | 36 | 1.53 | 0.160 | 0.992 |
| AT5G54270.1 | Photosystem II light-harvesting complex subunit (LHCB3) | 37 | 1.55 | 0.179 | 0.997 |
| AT5G54270.1 | Photosystem II light-harvesting complex subunit (LHCB3) | 38 | 1.64 | 0.198 | 0.996 |
| AT5G54270.1 | Photosystem II light-harvesting complex subunit (LHCB3) | 41 | 2.06 | 0.194 | 0.868 |
| AT5G54270.1 | Photosystem II light-harvesting complex subunit (LHCB3) | 42 | 2.08 | 0.183 | 0.794 |
| AT5G54270.1 | Photosystem II light-harvesting complex subunit (LHCB3) | 43 | 1.97 | 0.181 | 0.863 |

| | | | | | |
|---|---|---|---|---|---|
| AT5G54270.1 | Photosystem II light-harvesting complex subunit (LHCB3) | 44 | 1.76 | 0.204 | 0.989 |
| AT5G54270.1 | Photosystem II light-harvesting complex subunit (LHCB3) | 45 | 1.51 | 0.253 | 1.000 |
| AT5G54270.1 | Photosystem II light-harvesting complex subunit (LHCB3) | 46 | 1.27 | 0.332 | 1.000 |
| AT5G54270.1 | Photosystem II light-harvesting complex subunit (LHCB3) | 47 | 1.06 | 0.440 | 1.000 |
| AT5G55070.1 | Dihydrolipoamide succinyltransferase (OGDCE2) | 2 | 0.54 | 0.622 | 1.000 |
| AT5G55070.1 | Dihydrolipoamide succinyltransferase (OGDCE2) | 3 | 0.61 | 0.418 | 1.000 |
| AT5G55070.1 | Dihydrolipoamide succinyltransferase (OGDCE2) | 4 | 0.67 | 0.374 | 1.000 |
| AT5G55070.1 | Dihydrolipoamide succinyltransferase (OGDCE2) | 5 | 0.77 | 0.364 | 1.000 |
| AT5G55070.1 | Dihydrolipoamide succinyltransferase (OGDCE2) | 6 | 0.97 | 0.351 | 1.000 |
| AT5G55070.1 | Dihydrolipoamide succinyltransferase (OGDCE2) | 7 | 1.19 | 0.340 | 1.000 |
| AT5G55070.1 | Dihydrolipoamide succinyltransferase (OGDCE2) | 8 | 1.43 | 0.332 | 1.000 |
| AT5G55070.1 | Dihydrolipoamide succinyltransferase (OGDCE2) | 9 | 1.71 | 0.327 | 1.000 |
| AT5G55070.1 | Dihydrolipoamide succinyltransferase (OGDCE2) | 10 | 1.78 | 0.326 | 1.000 |
| AT5G55070.1 | Dihydrolipoamide succinyltransferase (OGDCE2) | 11 | 1.57 | 0.327 | 1.000 |
| AT5G55070.1 | Dihydrolipoamide succinyltransferase (OGDCE2) | 12 | 1.15 | 0.326 | 1.000 |
| AT5G55070.1 | Dihydrolipoamide succinyltransferase (OGDCE2) | 13 | 0.74 | 0.324 | 1.000 |
| AT5G55070.1 | Dihydrolipoamide succinyltransferase (OGDCE2) | 14 | 0.47 | 0.321 | 0.998 |
| AT5G55070.1 | Dihydrolipoamide succinyltransferase (OGDCE2) | 15 | 0.35 | 0.321 | 0.955 |
| AT5G55070.1 | Dihydrolipoamide succinyltransferase (OGDCE2) | 16 | 0.43 | 0.320 | 0.994 |
| AT5G55070.1 | Dihydrolipoamide succinyltransferase (OGDCE2) | 17 | 0.96 | 0.323 | 1.000 |
| AT5G55070.1 | Dihydrolipoamide succinyltransferase (OGDCE2) | 18 | 2.35 | 0.335 | 0.996 |
| AT5G55070.1 | Dihydrolipoamide succinyltransferase (OGDCE2) | 19 | 3.92 | 0.351 | 0.829 |
| AT5G55070.1 | Dihydrolipoamide succinyltransferase (OGDCE2) | 20 | 4.46 | 0.377 | 0.806 |
| AT5G55070.1 | Dihydrolipoamide succinyltransferase (OGDCE2) | 21 | 3.98 | 0.407 | 0.933 |
| AT5G55070.1 | Dihydrolipoamide succinyltransferase (OGDCE2) | 22 | 2.93 | 0.405 | 0.994 |
| AT5G55070.1 | Dihydrolipoamide succinyltransferase (OGDCE2) | 23 | 1.76 | 0.397 | 1.000 |
| AT5G55070.1 | Dihydrolipoamide succinyltransferase (OGDCE2) | 24 | 0.87 | 0.371 | 1.000 |
| AT5G55070.1 | Dihydrolipoamide succinyltransferase (OGDCE2) | 25 | 0.42 | 0.364 | 0.998 |
| AT5G55070.1 | Dihydrolipoamide succinyltransferase (OGDCE2) | 26 | 0.25 | 0.370 | 0.875 |
| AT5G55070.1 | Dihydrolipoamide succinyltransferase (OGDCE2) | 27 | 0.21 | 0.376 | 0.767 |
| AT5G55070.1 | Dihydrolipoamide succinyltransferase (OGDCE2) | 28 | 0.24 | 0.371 | 0.842 |

| | | | | | |
|---|---|---|---|---|---|
| AT5G55070.1 | Dihydrolipoamide succinyltransferase (OGDCE2) | 29 | 0.33 | 0.355 | 0.963 |
| AT5G55070.1 | Dihydrolipoamide succinyltransferase (OGDCE2) | 30 | 0.51 | 0.344 | 1.000 |
| AT5G55070.1 | Dihydrolipoamide succinyltransferase (OGDCE2) | 31 | 1.06 | 0.335 | 1.000 |
| AT5G55070.1 | Dihydrolipoamide succinyltransferase (OGDCE2) | 32 | 2.31 | 0.339 | 0.997 |
| AT5G55070.1 | Dihydrolipoamide succinyltransferase (OGDCE2) | 33 | 3.53 | 0.366 | 0.925 |
| AT5G55070.1 | Dihydrolipoamide succinyltransferase (OGDCE2) | 34 | 3.60 | 0.438 | 0.982 |
| AT5G55070.1 | Dihydrolipoamide succinyltransferase (OGDCE2) | 35 | 3.21 | 0.606 | 1.000 |
| AT5G55070.1 | Dihydrolipoamide succinyltransferase (OGDCE2) | 46 | 15.94 | 7.332 | 1.000 |
| AT5G55070.1 | Dihydrolipoamide succinyltransferase (OGDCE2) | 47 | 21.08 | 8.256 | 1.000 |
| AT5G55610.1 | Isopentenyl-diphosphate delta-isomerase | 2 | 0.68 | 0.347 | 1.000 |
| AT5G55610.1 | Isopentenyl-diphosphate delta-isomerase | 3 | 0.94 | 0.259 | 1.000 |
| AT5G55610.1 | Isopentenyl-diphosphate delta-isomerase | 4 | 1.31 | 0.220 | 1.000 |
| AT5G55610.1 | Isopentenyl-diphosphate delta-isomerase | 5 | 1.86 | 0.207 | 0.972 |
| AT5G55610.1 | Isopentenyl-diphosphate delta-isomerase | 6 | 2.55 | 0.200 | 0.542 |
| AT5G55610.1 | Isopentenyl-diphosphate delta-isomerase | 7 | 3.17 | 0.196 | 0.200 |
| AT5G55610.1 | Isopentenyl-diphosphate delta-isomerase | 8 | 3.40 | 0.192 | 0.122 |
| AT5G55610.1 | Isopentenyl-diphosphate delta-isomerase | 9 | 3.12 | 0.190 | 0.180 |
| AT5G55610.1 | Isopentenyl-diphosphate delta-isomerase | 10 | 2.61 | 0.189 | 0.412 |
| AT5G55610.1 | Isopentenyl-diphosphate delta-isomerase | 11 | 2.12 | 0.189 | 0.772 |
| AT5G55610.1 | Isopentenyl-diphosphate delta-isomerase | 12 | 1.80 | 0.188 | 0.959 |
| AT5G55610.1 | Isopentenyl-diphosphate delta-isomerase | 13 | 1.67 | 0.186 | 0.987 |
| AT5G55610.1 | Isopentenyl-diphosphate delta-isomerase | 14 | 1.77 | 0.184 | 0.960 |
| AT5G55610.1 | Isopentenyl-diphosphate delta-isomerase | 15 | 1.91 | 0.183 | 0.891 |
| AT5G55610.1 | Isopentenyl-diphosphate delta-isomerase | 16 | 2.09 | 0.183 | 0.760 |
| AT5G55610.1 | Isopentenyl-diphosphate delta-isomerase | 17 | 2.37 | 0.183 | 0.524 |
| AT5G55610.1 | Isopentenyl-diphosphate delta-isomerase | 18 | 2.68 | 0.185 | 0.333 |
| AT5G55610.1 | Isopentenyl-diphosphate delta-isomerase | 19 | 2.70 | 0.188 | 0.349 |
| AT5G55610.1 | Isopentenyl-diphosphate delta-isomerase | 20 | 2.22 | 0.191 | 0.718 |
| AT5G55610.1 | Isopentenyl-diphosphate delta-isomerase | 21 | 1.58 | 0.193 | 0.998 |
| AT5G55610.1 | Isopentenyl-diphosphate delta-isomerase | 22 | 1.10 | 0.193 | 1.000 |
| AT5G55610.1 | Isopentenyl-diphosphate delta-isomerase | 23 | 0.81 | 0.192 | 1.000 |

| | | | | | |
|---|---|---|---|---|---|
| AT5G55610.1 | Isopentenyl-diphosphate delta-isomerase | 24 | 0.66 | 0.189 | 0.999 |
| AT5G55610.1 | Isopentenyl-diphosphate delta-isomerase | 25 | 0.57 | 0.187 | 0.975 |
| AT5G55610.1 | Isopentenyl-diphosphate delta-isomerase | 26 | 0.59 | 0.186 | 0.986 |
| AT5G55610.1 | Isopentenyl-diphosphate delta-isomerase | 27 | 0.76 | 0.186 | 1.000 |
| AT5G55610.1 | Isopentenyl-diphosphate delta-isomerase | 28 | 0.96 | 0.186 | 1.000 |
| AT5G55610.1 | Isopentenyl-diphosphate delta-isomerase | 29 | 1.05 | 0.186 | 1.000 |
| AT5G55610.1 | Isopentenyl-diphosphate delta-isomerase | 30 | 1.00 | 0.187 | 1.000 |
| AT5G55610.1 | Isopentenyl-diphosphate delta-isomerase | 31 | 0.90 | 0.187 | 1.000 |
| AT5G55610.1 | Isopentenyl-diphosphate delta-isomerase | 32 | 0.83 | 0.188 | 1.000 |
| AT5G55610.1 | Isopentenyl-diphosphate delta-isomerase | 33 | 0.79 | 0.188 | 1.000 |
| AT5G55610.1 | Isopentenyl-diphosphate delta-isomerase | 34 | 0.75 | 0.188 | 1.000 |
| AT5G55610.1 | Isopentenyl-diphosphate delta-isomerase | 35 | 0.69 | 0.188 | 1.000 |
| AT5G55610.1 | Isopentenyl-diphosphate delta-isomerase | 36 | 0.65 | 0.187 | 0.998 |
| AT5G55610.1 | Isopentenyl-diphosphate delta-isomerase | 37 | 0.69 | 0.187 | 1.000 |
| AT5G55610.1 | Isopentenyl-diphosphate delta-isomerase | 38 | 0.84 | 0.188 | 1.000 |
| AT5G55610.1 | Isopentenyl-diphosphate delta-isomerase | 39 | 1.09 | 0.188 | 1.000 |
| AT5G55610.1 | Isopentenyl-diphosphate delta-isomerase | 40 | 1.39 | 0.188 | 1.000 |
| AT5G55610.1 | Isopentenyl-diphosphate delta-isomerase | 41 | 1.63 | 0.189 | 0.993 |
| AT5G55610.1 | Isopentenyl-diphosphate delta-isomerase | 42 | 1.76 | 0.191 | 0.974 |
| AT5G55610.1 | Isopentenyl-diphosphate delta-isomerase | 43 | 1.70 | 0.192 | 0.987 |
| AT5G55610.1 | Isopentenyl-diphosphate delta-isomerase | 44 | 1.37 | 0.194 | 1.000 |
| AT5G55610.1 | Isopentenyl-diphosphate delta-isomerase | 45 | 0.90 | 0.205 | 1.000 |
| AT5G55610.1 | Isopentenyl-diphosphate delta-isomerase | 46 | 0.50 | 0.244 | 0.985 |
| AT5G55610.1 | Isopentenyl-diphosphate delta-isomerase | 47 | 0.26 | 0.340 | 0.793 |
| AT5G56090.1 | Cytochrome c:O2 oxidoreductase assembly protein (COX15) | 2 | 0.82 | 0.400 | 1.000 |
| AT5G56090.1 | Cytochrome c:O2 oxidoreductase assembly protein (COX15) | 3 | 0.97 | 0.314 | 1.000 |
| AT5G56090.1 | Cytochrome c:O2 oxidoreductase assembly protein (COX15) | 6 | 1.34 | 0.337 | 1.000 |
| AT5G56090.1 | Cytochrome c:O2 oxidoreductase assembly protein (COX15) | 7 | 1.33 | 0.270 | 1.000 |

| | | | | | | |
|---|---|---|---|---|---|---|
| AT5G56090.1 | Cytochrome c:O2 oxidoreductase assembly protein (COX15) | 8 | 1.30 | 0.242 | 1.000 | |
| AT5G56090.1 | Cytochrome c:O2 oxidoreductase assembly protein (COX15) | 9 | 1.30 | 0.232 | 1.000 | |
| AT5G56090.1 | Cytochrome c:O2 oxidoreductase assembly protein (COX15) | 10 | 1.42 | 0.221 | 1.000 | |
| AT5G56090.1 | Cytochrome c:O2 oxidoreductase assembly protein (COX15) | 11 | 1.75 | 0.216 | 0.991 | |
| AT5G56090.1 | Cytochrome c:O2 oxidoreductase assembly protein (COX15) | 12 | 2.22 | 0.211 | 0.815 | |
| AT5G56090.1 | Cytochrome c:O2 oxidoreductase assembly protein (COX15) | 13 | 3.45 | 0.209 | 0.179 | |
| AT5G56090.1 | Cytochrome c:O2 oxidoreductase assembly protein (COX15) | 14 | 4.98 | 0.207 | 0.020 | * |
| AT5G56090.1 | Cytochrome c:O2 oxidoreductase assembly protein (COX15) | 15 | 4.60 | 0.207 | 0.034 | * |
| AT5G56090.1 | Cytochrome c:O2 oxidoreductase assembly protein (COX15) | 16 | 3.46 | 0.206 | 0.163 | |
| AT5G56090.1 | Cytochrome c:O2 oxidoreductase assembly protein (COX15) | 17 | 3.54 | 0.208 | 0.154 | |
| AT5G56090.1 | Cytochrome c:O2 oxidoreductase assembly protein (COX15) | 18 | 3.61 | 0.216 | 0.176 | |
| AT5G56090.1 | Cytochrome c:O2 oxidoreductase assembly protein (COX15) | 19 | 3.35 | 0.225 | 0.300 | |
| AT5G56090.1 | Cytochrome c:O2 oxidoreductase assembly protein (COX15) | 20 | 3.37 | 0.258 | 0.508 | |
| AT5G56090.1 | Cytochrome c:O2 oxidoreductase assembly protein (COX15) | 21 | 3.56 | 0.378 | 0.914 | |
| AT5G56090.1 | Cytochrome c:O2 oxidoreductase assembly protein (COX15) | 39 | 16.02 | 0.348 | 0.014 | * |
| AT5G56090.1 | Cytochrome c:O2 oxidoreductase assembly protein (COX15) | 40 | 9.89 | 0.244 | 0.002 | ** |
| AT5G56090.1 | Cytochrome c:O2 oxidoreductase assembly protein (COX15) | 41 | 3.31 | 0.215 | 0.257 | |
| AT5G56090.1 | Cytochrome c:O2 oxidoreductase assembly protein (COX15) | 42 | 1.01 | 0.214 | 1.000 | |
| AT5G56090.1 | Cytochrome c:O2 oxidoreductase assembly protein (COX15) | 43 | 0.35 | 0.214 | 0.455 | |
| AT5G56090.1 | Cytochrome c:O2 oxidoreductase assembly protein (COX15) | 44 | 0.13 | 0.212 | 0.001 | *** |

| | | | | | | |
|---|---|---|---|---|---|---|
| AT5G56090.1 | Cytochrome c:O2 oxidoreductase assembly protein (COX15) | 45 | 0.09 | 0.212 | 0.000 | *** |
| AT5G56090.1 | Cytochrome c:O2 oxidoreductase assembly protein (COX15) | 46 | 0.26 | 0.226 | 0.170 | |
| AT5G56090.1 | Cytochrome c:O2 oxidoreductase assembly protein (COX15) | 47 | 1.65 | 0.432 | 1.000 | |
| AT5G56450.1 | Plasma membrane located adenine nucleotide transporter (PM-ANT) | 2 | 1.14 | 0.635 | 1.000 | |
| AT5G56450.1 | Plasma membrane located adenine nucleotide transporter (PM-ANT) | 3 | 0.24 | 1.812 | 1.000 | |
| AT5G56450.1 | Plasma membrane located adenine nucleotide transporter (PM-ANT) | 38 | 20.68 | 0.483 | 0.078 | . |
| AT5G56450.1 | Plasma membrane located adenine nucleotide transporter (PM-ANT) | 39 | 21.04 | 0.319 | 0.001 | ** |
| AT5G56450.1 | Plasma membrane located adenine nucleotide transporter (PM-ANT) | 40 | 8.19 | 0.297 | 0.032 | * |
| AT5G56450.1 | Plasma membrane located adenine nucleotide transporter (PM-ANT) | 41 | 0.84 | 0.287 | 1.000 | |
| AT5G56450.1 | Plasma membrane located adenine nucleotide transporter (PM-ANT) | 42 | 0.10 | 0.301 | 0.013 | * |
| AT5G56450.1 | Plasma membrane located adenine nucleotide transporter (PM-ANT) | 43 | 0.05 | 0.398 | 0.020 | * |
| AT5G56450.1 | Plasma membrane located adenine nucleotide transporter (PM-ANT) | 44 | 0.23 | 0.750 | 0.984 | |
| AT5G56450.1 | Plasma membrane located adenine nucleotide transporter (PM-ANT) | 46 | 82.72 | 2.142 | 0.978 | |
| AT5G56450.1 | Plasma membrane located adenine nucleotide transporter (PM-ANT) | 47 | 3012.54 | 3.234 | 0.936 | |
| AT5G56730.1 | Insulinase (Peptidase family M16) family protein | 2 | 0.87 | 0.315 | 1.000 | |
| AT5G56730.1 | Insulinase (Peptidase family M16) family protein | 3 | 0.83 | 0.305 | 1.000 | |
| AT5G56730.1 | Insulinase (Peptidase family M16) family protein | 29 | 0.57 | 0.219 | 0.965 | |
| AT5G56730.1 | Insulinase (Peptidase family M16) family protein | 30 | 0.63 | 0.161 | 0.927 | |
| AT5G56730.1 | Insulinase (Peptidase family M16) family protein | 31 | 0.80 | 0.142 | 0.999 | |
| AT5G56730.1 | Insulinase (Peptidase family M16) family protein | 32 | 1.06 | 0.135 | 1.000 | |
| AT5G56730.1 | Insulinase (Peptidase family M16) family protein | 33 | 1.20 | 0.131 | 1.000 | |
| AT5G56730.1 | Insulinase (Peptidase family M16) family protein | 34 | 1.19 | 0.131 | 1.000 | |
| AT5G56730.1 | Insulinase (Peptidase family M16) family protein | 35 | 1.15 | 0.130 | 1.000 | |
| AT5G56730.1 | Insulinase (Peptidase family M16) family protein | 36 | 1.17 | 0.130 | 1.000 | |

| | | | | | |
|---|---|---|---|---|---|
| AT5G56730.1 | Insulinase (Peptidase family M16) family protein | 37 | 1.27 | 0.129 | 0.997 |
| AT5G56730.1 | Insulinase (Peptidase family M16) family protein | 38 | 1.30 | 0.130 | 0.994 |
| AT5G56730.1 | Insulinase (Peptidase family M16) family protein | 39 | 1.12 | 0.131 | 1.000 |
| AT5G56730.1 | Insulinase (Peptidase family M16) family protein | 40 | 0.89 | 0.131 | 1.000 |
| AT5G56730.1 | Insulinase (Peptidase family M16) family protein | 41 | 0.69 | 0.143 | 0.966 |
| AT5G56730.1 | Insulinase (Peptidase family M16) family protein | 42 | 0.65 | 0.194 | 0.989 |
| AT5G56730.1 | Insulinase (Peptidase family M16) family protein | 43 | 0.75 | 0.283 | 1.000 |
| AT5G56730.1 | Insulinase (Peptidase family M16) family protein | 46 | 2.39 | 0.769 | 1.000 |
| AT5G56730.1 | Insulinase (Peptidase family M16) family protein | 47 | 3.94 | 1.005 | 1.000 |
| AT5G56940.1 | 30S ribosomal protein S16-2 (RPS16B) | 2 | 1.30 | 0.705 | 1.000 |
| AT5G56940.1 | 30S ribosomal protein S16-2 (RPS16B) | 3 | 1.35 | 0.622 | 1.000 |
| AT5G56940.1 | 30S ribosomal protein S16-2 (RPS16B) | 11 | 1.72 | 0.333 | 0.984 |
| AT5G56940.1 | 30S ribosomal protein S16-2 (RPS16B) | 12 | 1.70 | 0.297 | 0.974 |
| AT5G56940.1 | 30S ribosomal protein S16-2 (RPS16B) | 13 | 1.61 | 0.268 | 0.975 |
| AT5G56940.1 | 30S ribosomal protein S16-2 (RPS16B) | 14 | 1.43 | 0.247 | 0.992 |
| AT5G56940.1 | 30S ribosomal protein S16-2 (RPS16B) | 15 | 1.23 | 0.234 | 1.000 |
| AT5G56940.1 | 30S ribosomal protein S16-2 (RPS16B) | 16 | 1.06 | 0.227 | 1.000 |
| AT5G56940.1 | 30S ribosomal protein S16-2 (RPS16B) | 17 | 0.91 | 0.225 | 1.000 |
| AT5G56940.1 | 30S ribosomal protein S16-2 (RPS16B) | 18 | 0.75 | 0.228 | 0.996 |
| AT5G56940.1 | 30S ribosomal protein S16-2 (RPS16B) | 19 | 0.59 | 0.238 | 0.922 |
| AT5G56940.1 | 30S ribosomal protein S16-2 (RPS16B) | 20 | 0.45 | 0.258 | 0.724 |
| AT5G56940.1 | 30S ribosomal protein S16-2 (RPS16B) | 21 | 0.35 | 0.297 | 0.573 |
| AT5G56940.1 | 30S ribosomal protein S16-2 (RPS16B) | 22 | 0.27 | 0.359 | 0.559 |
| AT5G56940.1 | 30S ribosomal protein S16-2 (RPS16B) | 23 | 0.22 | 0.445 | 0.639 |
| AT5G56940.1 | 30S ribosomal protein S16-2 (RPS16B) | 46 | 3.19 | 3.793 | 1.000 |
| AT5G56940.1 | 30S ribosomal protein S16-2 (RPS16B) | 47 | 4.18 | 4.007 | 1.000 |
| AT5G57230.1 | Thioredoxin superfamily protein | 2 | 0.89 | 0.525 | 1.000 |
| AT5G57230.1 | Thioredoxin superfamily protein | 3 | 1.51 | 0.580 | 1.000 |
| AT5G57230.1 | Thioredoxin superfamily protein | 22 | 0.92 | 0.405 | 1.000 |
| AT5G57230.1 | Thioredoxin superfamily protein | 23 | 0.64 | 0.297 | 0.999 |
| AT5G57230.1 | Thioredoxin superfamily protein | 24 | 0.57 | 0.258 | 0.979 |

| | | | | | |
|---|---|---|---|---|---|
| AT5G57230.1 | Thioredoxin superfamily protein | 25 | 1.14 | 0.241 | 1.000 |
| AT5G57230.1 | Thioredoxin superfamily protein | 26 | 2.71 | 0.239 | 0.513 |
| AT5G57230.1 | Thioredoxin superfamily protein | 27 | 2.65 | 0.244 | 0.568 |
| AT5G57230.1 | Thioredoxin superfamily protein | 28 | 0.86 | 0.262 | 1.000 |
| AT5G57230.1 | Thioredoxin superfamily protein | 29 | 0.28 | 0.326 | 0.594 |
| AT5G57230.1 | Thioredoxin superfamily protein | 30 | 0.18 | 0.557 | 0.837 |
| AT5G57230.1 | Thioredoxin superfamily protein | 46 | 886055.34 | 10.135 | 0.999 |
| AT5G57230.1 | Thioredoxin superfamily protein | 47 | 4321726.21 | 11.075 | 0.999 |
| AT5G57490.1 | Voltage-dependent anion channel 4 (VDAC4) | 3 | 0.71 | 0.181 | 1.000 |
| AT5G57490.1 | Voltage-dependent anion channel 4 (VDAC4) | 4 | 0.87 | 0.172 | 1.000 |
| AT5G57490.1 | Voltage-dependent anion channel 4 (VDAC4) | 5 | 1.20 | 0.172 | 1.000 |
| AT5G57490.1 | Voltage-dependent anion channel 4 (VDAC4) | 6 | 1.47 | 0.172 | 1.000 |
| AT5G57490.1 | Voltage-dependent anion channel 4 (VDAC4) | 7 | 1.61 | 0.173 | 0.998 |
| AT5G57490.1 | Voltage-dependent anion channel 4 (VDAC4) | 8 | 1.70 | 0.170 | 0.991 |
| AT5G57490.1 | Voltage-dependent anion channel 4 (VDAC4) | 9 | 1.69 | 0.168 | 0.991 |
| AT5G57490.1 | Voltage-dependent anion channel 4 (VDAC4) | 10 | 1.59 | 0.167 | 0.998 |
| AT5G57490.1 | Voltage-dependent anion channel 4 (VDAC4) | 11 | 1.64 | 0.168 | 0.996 |
| AT5G57490.1 | Voltage-dependent anion channel 4 (VDAC4) | 12 | 1.81 | 0.168 | 0.963 |
| AT5G57490.1 | Voltage-dependent anion channel 4 (VDAC4) | 13 | 1.87 | 0.167 | 0.928 |
| AT5G57490.1 | Voltage-dependent anion channel 4 (VDAC4) | 14 | 1.68 | 0.165 | 0.989 |
| AT5G57490.1 | Voltage-dependent anion channel 4 (VDAC4) | 15 | 1.30 | 0.164 | 1.000 |
| AT5G57490.1 | Voltage-dependent anion channel 4 (VDAC4) | 16 | 0.95 | 0.163 | 1.000 |
| AT5G57490.1 | Voltage-dependent anion channel 4 (VDAC4) | 17 | 0.89 | 0.163 | 1.000 |
| AT5G57490.1 | Voltage-dependent anion channel 4 (VDAC4) | 18 | 0.98 | 0.165 | 1.000 |
| AT5G57490.1 | Voltage-dependent anion channel 4 (VDAC4) | 19 | 1.05 | 0.169 | 1.000 |
| AT5G57490.1 | Voltage-dependent anion channel 4 (VDAC4) | 20 | 1.12 | 0.172 | 1.000 |
| AT5G57490.1 | Voltage-dependent anion channel 4 (VDAC4) | 21 | 1.11 | 0.175 | 1.000 |
| AT5G57490.1 | Voltage-dependent anion channel 4 (VDAC4) | 22 | 1.07 | 0.174 | 1.000 |
| AT5G57490.1 | Voltage-dependent anion channel 4 (VDAC4) | 23 | 1.04 | 0.173 | 1.000 |
| AT5G57490.1 | Voltage-dependent anion channel 4 (VDAC4) | 24 | 1.03 | 0.170 | 1.000 |
| AT5G57490.1 | Voltage-dependent anion channel 4 (VDAC4) | 25 | 0.97 | 0.168 | 1.000 |

| | | | | | | |
|---|---|---|---|---|---|---|
| AT5G57490.1 | Voltage-dependent anion channel 4 (VDAC4) | 26 | 0.94 | 0.167 | 1.000 | |
| AT5G57490.1 | Voltage-dependent anion channel 4 (VDAC4) | 27 | 0.92 | 0.167 | 1.000 | |
| AT5G57490.1 | Voltage-dependent anion channel 4 (VDAC4) | 28 | 0.95 | 0.167 | 1.000 | |
| AT5G57490.1 | Voltage-dependent anion channel 4 (VDAC4) | 29 | 1.19 | 0.167 | 1.000 | |
| AT5G57490.1 | Voltage-dependent anion channel 4 (VDAC4) | 30 | 1.49 | 0.167 | 1.000 | |
| AT5G57490.1 | Voltage-dependent anion channel 4 (VDAC4) | 31 | 1.35 | 0.167 | 1.000 | |
| AT5G57490.1 | Voltage-dependent anion channel 4 (VDAC4) | 32 | 1.10 | 0.168 | 1.000 | |
| AT5G57490.1 | Voltage-dependent anion channel 4 (VDAC4) | 33 | 1.06 | 0.168 | 1.000 | |
| AT5G57490.1 | Voltage-dependent anion channel 4 (VDAC4) | 34 | 1.12 | 0.168 | 1.000 | |
| AT5G57490.1 | Voltage-dependent anion channel 4 (VDAC4) | 35 | 1.10 | 0.168 | 1.000 | |
| AT5G57490.1 | Voltage-dependent anion channel 4 (VDAC4) | 36 | 1.03 | 0.167 | 1.000 | |
| AT5G57490.1 | Voltage-dependent anion channel 4 (VDAC4) | 37 | 1.17 | 0.167 | 1.000 | |
| AT5G57490.1 | Voltage-dependent anion channel 4 (VDAC4) | 38 | 1.66 | 0.168 | 0.994 | |
| AT5G57490.1 | Voltage-dependent anion channel 4 (VDAC4) | 39 | 2.56 | 0.168 | 0.354 | |
| AT5G57490.1 | Voltage-dependent anion channel 4 (VDAC4) | 40 | 3.61 | 0.168 | 0.034 | * |
| AT5G57490.1 | Voltage-dependent anion channel 4 (VDAC4) | 41 | 3.36 | 0.169 | 0.062 | . |
| AT5G57490.1 | Voltage-dependent anion channel 4 (VDAC4) | 42 | 1.92 | 0.171 | 0.918 | |
| AT5G57490.1 | Voltage-dependent anion channel 4 (VDAC4) | 43 | 0.88 | 0.172 | 1.000 | |
| AT5G57490.1 | Voltage-dependent anion channel 4 (VDAC4) | 44 | 0.41 | 0.172 | 0.481 | |
| AT5G57490.1 | Voltage-dependent anion channel 4 (VDAC4) | 45 | 0.24 | 0.171 | 0.011 | * |
| AT5G57490.1 | Voltage-dependent anion channel 4 (VDAC4) | 46 | 0.19 | 0.192 | 0.009 | ** |
| AT5G57490.1 | Voltage-dependent anion channel 4 (VDAC4) | 47 | 0.19 | 0.342 | 0.635 | |
| AT5G57490.1 | Voltage-dependent anion channel 4 (VDAC4) | 48 | 0.20 | 0.584 | 0.999 | |
| AT5G57590.1 | Biotin synthase 1 (BIO1) | 2 | 1.05 | 0.892 | 1.000 | |
| AT5G57590.1 | Biotin synthase 1 (BIO1) | 3 | 1.16 | 0.985 | 1.000 | |
| AT5G57590.1 | Biotin synthase 1 (BIO1) | 29 | 1.26 | 0.669 | 1.000 | |
| AT5G57590.1 | Biotin synthase 1 (BIO1) | 30 | 1.03 | 0.609 | 1.000 | |
| AT5G57590.1 | Biotin synthase 1 (BIO1) | 31 | 0.86 | 0.614 | 1.000 | |
| AT5G57590.1 | Biotin synthase 1 (BIO1) | 36 | 1.13 | 0.510 | 1.000 | |
| AT5G57590.1 | Biotin synthase 1 (BIO1) | 37 | 1.66 | 0.454 | 0.999 | |
| AT5G57590.1 | Biotin synthase 1 (BIO1) | 38 | 2.60 | 0.411 | 0.930 | |
| AT5G57590.1 | Biotin synthase 1 (BIO1) | 39 | 3.65 | 0.378 | 0.665 | |
| AT5G57590.1 | Biotin synthase 1 (BIO1) | 40 | 3.53 | 0.363 | 0.647 | |
| AT5G57590.1 | Biotin synthase 1 (BIO1) | 41 | 2.07 | 0.385 | 0.977 | |
| AT5G57590.1 | Biotin synthase 1 (BIO1) | 42 | 0.84 | 0.454 | 1.000 | |
| AT5G57590.1 | Biotin synthase 1 (BIO1) | 43 | 0.29 | 0.534 | 0.931 | |
| AT5G57590.1 | Biotin synthase 1 (BIO1) | 46 | 0.01 | 0.746 | 0.059 | . |
| AT5G57590.1 | Biotin synthase 1 (BIO1) | 47 | 0.00 | 0.880 | 0.037 | * |

| | | | | | |
|---|---|---|---|---|---|
| AT5G58270.1 | ABC transporter of the mitochondrion 3 (ATM3, STA1) | 2 | 1.02 | 0.573 | 1.000 |
| AT5G58270.1 | ABC transporter of the mitochondrion 3 (ATM3, STA1) | 3 | 0.94 | 0.403 | 1.000 |
| AT5G58270.1 | ABC transporter of the mitochondrion 3 (ATM3, STA1) | 4 | 0.89 | 0.498 | 1.000 |
| AT5G58270.1 | ABC transporter of the mitochondrion 3 (ATM3, STA1) | 6 | 1.04 | 0.498 | 1.000 |
| AT5G58270.1 | ABC transporter of the mitochondrion 3 (ATM3, STA1) | 7 | 1.13 | 0.478 | 1.000 |
| AT5G58270.1 | ABC transporter of the mitochondrion 3 (ATM3, STA1) | 8 | 1.12 | 0.419 | 1.000 |
| AT5G58270.1 | ABC transporter of the mitochondrion 3 (ATM3, STA1) | 9 | 1.02 | 0.402 | 1.000 |
| AT5G58270.1 | ABC transporter of the mitochondrion 3 (ATM3, STA1) | 10 | 0.92 | 0.395 | 1.000 |
| AT5G58270.1 | ABC transporter of the mitochondrion 3 (ATM3, STA1) | 11 | 0.94 | 0.374 | 1.000 |
| AT5G58270.1 | ABC transporter of the mitochondrion 3 (ATM3, STA1) | 12 | 1.08 | 0.384 | 1.000 |
| AT5G58270.1 | ABC transporter of the mitochondrion 3 (ATM3, STA1) | 13 | 1.46 | 0.415 | 1.000 |
| AT5G58270.1 | ABC transporter of the mitochondrion 3 (ATM3, STA1) | 14 | 2.32 | 0.409 | 0.999 |
| AT5G58270.1 | ABC transporter of the mitochondrion 3 (ATM3, STA1) | 28 | 0.48 | 0.465 | 1.000 |
| AT5G58270.1 | ABC transporter of the mitochondrion 3 (ATM3, STA1) | 29 | 0.49 | 0.342 | 0.999 |
| AT5G58270.1 | ABC transporter of the mitochondrion 3 (ATM3, STA1) | 30 | 0.62 | 0.307 | 1.000 |
| AT5G58270.1 | ABC transporter of the mitochondrion 3 (ATM3, STA1) | 31 | 1.04 | 0.294 | 1.000 |
| AT5G58270.1 | ABC transporter of the mitochondrion 3 (ATM3, STA1) | 32 | 2.03 | 0.288 | 0.995 |
| AT5G58270.1 | ABC transporter of the mitochondrion 3 (ATM3, STA1) | 33 | 3.82 | 0.288 | 0.542 |
| AT5G58270.1 | ABC transporter of the mitochondrion 3 (ATM3, STA1) | 34 | 5.79 | 0.290 | 0.164 |
| AT5G58270.1 | ABC transporter of the mitochondrion 3 (ATM3, STA1) | 35 | 3.70 | 0.295 | 0.618 |
| AT5G58270.1 | ABC transporter of the mitochondrion 3 (ATM3, STA1) | 36 | 1.10 | 0.311 | 1.000 |
| AT5G58270.1 | ABC transporter of the mitochondrion 3 (ATM3, STA1) | 37 | 0.38 | 0.387 | 0.994 |
| AT5G58270.1 | ABC transporter of the mitochondrion 3 (ATM3, STA1) | 38 | 0.34 | 0.659 | 1.000 |
| AT5G58270.1 | ABC transporter of the mitochondrion 3 (ATM3, STA1) | 46 | 7663117.42 | 4.792 | 0.927 |
| AT5G58270.1 | ABC transporter of the mitochondrion 3 (ATM3, STA1) | 47 | ########### | 5.593 | 0.911 |
| AT5G58920.1 | Homeobox prospero protein | 2 | 0.86 | 0.446 | 1.000 |
| AT5G58920.1 | Homeobox prospero protein | 3 | 1.88 | 1.295 | 1.000 |
| AT5G58920.1 | Homeobox prospero protein | 23 | 0.40 | 0.317 | 0.858 |
| AT5G58920.1 | Homeobox prospero protein | 24 | 0.27 | 0.227 | 0.137 |
| AT5G58920.1 | Homeobox prospero protein | 25 | 0.35 | 0.209 | 0.260 |
| AT5G58920.1 | Homeobox prospero protein | 26 | 1.28 | 0.207 | 1.000 |
| AT5G58920.1 | Homeobox prospero protein | 27 | 1.20 | 0.219 | 1.000 |
| AT5G58920.1 | Homeobox prospero protein | 28 | 0.20 | 0.272 | 0.106 |
| AT5G58920.1 | Homeobox prospero protein | 29 | 0.13 | 0.463 | 0.436 |
| AT5G58920.1 | Homeobox prospero protein | 46 | 6875057.82 | 8.591 | 0.991 |
| AT5G58920.1 | Homeobox prospero protein | 47 | 26613413.10 | 9.341 | 0.991 |
| AT5G59370.2 | Actin 4 (ACT4) | 2 | 0.61 | 0.446 | 0.997 |

| | | | | | |
|---|---|---|---|---|---|
| AT5G59370.2 | Actin 4 (ACT4) | 3 | 0.59 | 0.412 | 0.991 |
| AT5G59370.2 | Actin 4 (ACT4) | 4 | 0.57 | 0.390 | 0.982 |
| AT5G59370.2 | Actin 4 (ACT4) | 5 | 0.56 | 0.377 | 0.971 |
| AT5G59370.2 | Actin 4 (ACT4) | 6 | 0.55 | 0.370 | 0.961 |
| AT5G59370.2 | Actin 4 (ACT4) | 7 | 0.54 | 0.366 | 0.954 |
| AT5G59370.2 | Actin 4 (ACT4) | 8 | 0.54 | 0.365 | 0.949 |
| AT5G59370.2 | Actin 4 (ACT4) | 9 | 0.53 | 0.364 | 0.945 |
| AT5G59370.2 | Actin 4 (ACT4) | 10 | 0.53 | 0.363 | 0.942 |
| AT5G59370.2 | Actin 4 (ACT4) | 11 | 0.52 | 0.363 | 0.937 |
| AT5G59370.2 | Actin 4 (ACT4) | 12 | 0.52 | 0.362 | 0.932 |
| AT5G59370.2 | Actin 4 (ACT4) | 13 | 0.52 | 0.362 | 0.927 |
| AT5G59370.2 | Actin 4 (ACT4) | 14 | 0.51 | 0.362 | 0.925 |
| AT5G59370.2 | Actin 4 (ACT4) | 15 | 0.51 | 0.362 | 0.921 |
| AT5G59370.2 | Actin 4 (ACT4) | 16 | 0.50 | 0.362 | 0.915 |
| AT5G59370.2 | Actin 4 (ACT4) | 17 | 0.49 | 0.363 | 0.899 |
| AT5G59370.2 | Actin 4 (ACT4) | 18 | 0.46 | 0.365 | 0.867 |
| AT5G59370.2 | Actin 4 (ACT4) | 19 | 0.43 | 0.367 | 0.816 |
| AT5G59370.2 | Actin 4 (ACT4) | 20 | 0.39 | 0.370 | 0.749 |
| AT5G59370.2 | Actin 4 (ACT4) | 21 | 0.36 | 0.372 | 0.672 |
| AT5G59370.2 | Actin 4 (ACT4) | 22 | 0.33 | 0.374 | 0.600 |
| AT5G59370.2 | Actin 4 (ACT4) | 23 | 0.30 | 0.375 | 0.542 |
| AT5G59370.2 | Actin 4 (ACT4) | 24 | 0.29 | 0.375 | 0.500 |
| AT5G59370.2 | Actin 4 (ACT4) | 25 | 0.28 | 0.374 | 0.480 |
| AT5G59370.2 | Actin 4 (ACT4) | 26 | 0.28 | 0.373 | 0.477 |
| AT5G59370.2 | Actin 4 (ACT4) | 27 | 0.29 | 0.372 | 0.489 |
| AT5G59370.2 | Actin 4 (ACT4) | 28 | 0.29 | 0.373 | 0.505 |
| AT5G59370.2 | Actin 4 (ACT4) | 29 | 0.30 | 0.374 | 0.521 |
| AT5G59370.2 | Actin 4 (ACT4) | 30 | 0.30 | 0.378 | 0.529 |
| AT5G59370.2 | Actin 4 (ACT4) | 31 | 0.29 | 0.383 | 0.524 |
| AT5G59370.2 | Actin 4 (ACT4) | 32 | 0.28 | 0.388 | 0.513 |
| AT5G59370.2 | Actin 4 (ACT4) | 33 | 0.27 | 0.395 | 0.497 |
| AT5G59370.2 | Actin 4 (ACT4) | 34 | 0.26 | 0.400 | 0.476 |
| AT5G59370.2 | Actin 4 (ACT4) | 35 | 0.25 | 0.405 | 0.458 |
| AT5G59370.2 | Actin 4 (ACT4) | 39 | 0.24 | 0.403 | 0.432 |
| AT5G59370.2 | Actin 4 (ACT4) | 40 | 0.25 | 0.400 | 0.451 |
| AT5G59370.2 | Actin 4 (ACT4) | 41 | 0.27 | 0.395 | 0.489 |
| AT5G59370.2 | Actin 4 (ACT4) | 42 | 0.30 | 0.391 | 0.559 |
| AT5G59370.2 | Actin 4 (ACT4) | 43 | 0.34 | 0.389 | 0.669 |
| AT5G59370.2 | Actin 4 (ACT4) | 44 | 0.41 | 0.393 | 0.813 |
| AT5G59370.2 | Actin 4 (ACT4) | 45 | 0.50 | 0.403 | 0.943 |
| AT5G59370.2 | Actin 4 (ACT4) | 46 | 0.62 | 0.425 | 0.996 |
| AT5G59370.2 | Actin 4 (ACT4) | 47 | 0.78 | 0.460 | 1.000 |
| AT5G59613.2 | ATP synthase subunit (ATP6kDa) | 2 | 0.62 | 0.392 | 1.000 |
| AT5G59613.2 | ATP synthase subunit (ATP6kDa) | 3 | 0.85 | 0.249 | 1.000 |
| AT5G59613.2 | ATP synthase subunit (ATP6kDa) | 4 | 1.24 | 0.223 | 1.000 |
| AT5G59613.2 | ATP synthase subunit (ATP6kDa) | 5 | 1.87 | 0.223 | 0.996 |
| AT5G59613.2 | ATP synthase subunit (ATP6kDa) | 6 | 2.52 | 0.223 | 0.834 |
| AT5G59613.2 | ATP synthase subunit (ATP6kDa) | 7 | 2.70 | 0.222 | 0.736 |
| AT5G59613.2 | ATP synthase subunit (ATP6kDa) | 8 | 2.37 | 0.219 | 0.882 |
| AT5G59613.2 | ATP synthase subunit (ATP6kDa) | 9 | 2.04 | 0.216 | 0.976 |
| AT5G59613.2 | ATP synthase subunit (ATP6kDa) | 10 | 2.04 | 0.216 | 0.977 |
| AT5G59613.2 | ATP synthase subunit (ATP6kDa) | 11 | 2.33 | 0.218 | 0.894 |

| | | | | | | |
|---|---|---|---|---|---|---|
| AT5G59613.2 | ATP synthase subunit (ATP6kDa) | 12 | 2.76 | 0.217 | 0.663 | |
| AT5G59613.2 | ATP synthase subunit (ATP6kDa) | 13 | 3.35 | 0.215 | 0.334 | |
| AT5G59613.2 | ATP synthase subunit (ATP6kDa) | 14 | 4.36 | 0.212 | 0.079 | . |
| AT5G59613.2 | ATP synthase subunit (ATP6kDa) | 15 | 5.16 | 0.210 | 0.025 | * |
| AT5G59613.2 | ATP synthase subunit (ATP6kDa) | 16 | 5.07 | 0.209 | 0.027 | * |
| AT5G59613.2 | ATP synthase subunit (ATP6kDa) | 17 | 4.28 | 0.209 | 0.078 | . |
| AT5G59613.2 | ATP synthase subunit (ATP6kDa) | 18 | 3.13 | 0.212 | 0.412 | |
| AT5G59613.2 | ATP synthase subunit (ATP6kDa) | 19 | 2.21 | 0.217 | 0.940 | |
| AT5G59613.2 | ATP synthase subunit (ATP6kDa) | 20 | 1.64 | 0.222 | 1.000 | |
| AT5G59613.2 | ATP synthase subunit (ATP6kDa) | 21 | 1.25 | 0.224 | 1.000 | |
| AT5G59613.2 | ATP synthase subunit (ATP6kDa) | 22 | 1.02 | 0.225 | 1.000 | |
| AT5G59613.2 | ATP synthase subunit (ATP6kDa) | 23 | 0.94 | 0.223 | 1.000 | |
| AT5G59613.2 | ATP synthase subunit (ATP6kDa) | 24 | 1.06 | 0.219 | 1.000 | |
| AT5G59613.2 | ATP synthase subunit (ATP6kDa) | 25 | 1.31 | 0.215 | 1.000 | |
| AT5G59613.2 | ATP synthase subunit (ATP6kDa) | 26 | 1.35 | 0.214 | 1.000 | |
| AT5G59613.2 | ATP synthase subunit (ATP6kDa) | 27 | 1.07 | 0.214 | 1.000 | |
| AT5G59613.2 | ATP synthase subunit (ATP6kDa) | 28 | 0.92 | 0.215 | 1.000 | |
| AT5G59613.2 | ATP synthase subunit (ATP6kDa) | 29 | 1.03 | 0.215 | 1.000 | |
| AT5G59613.2 | ATP synthase subunit (ATP6kDa) | 30 | 1.23 | 0.215 | 1.000 | |
| AT5G59613.2 | ATP synthase subunit (ATP6kDa) | 31 | 1.19 | 0.216 | 1.000 | |
| AT5G59613.2 | ATP synthase subunit (ATP6kDa) | 32 | 0.89 | 0.216 | 1.000 | |
| AT5G59613.2 | ATP synthase subunit (ATP6kDa) | 33 | 0.64 | 0.216 | 1.000 | |
| AT5G59613.2 | ATP synthase subunit (ATP6kDa) | 34 | 0.56 | 0.218 | 0.999 | |
| AT5G59613.2 | ATP synthase subunit (ATP6kDa) | 35 | 0.59 | 0.221 | 1.000 | |
| AT5G59613.2 | ATP synthase subunit (ATP6kDa) | 36 | 0.80 | 0.235 | 1.000 | |
| AT5G59613.2 | ATP synthase subunit (ATP6kDa) | 37 | 1.39 | 0.266 | 1.000 | |
| AT5G59613.2 | ATP synthase subunit (ATP6kDa) | 38 | 2.68 | 0.306 | 0.982 | |
| AT5G59613.2 | ATP synthase subunit (ATP6kDa) | 40 | 7.24 | 0.296 | 0.109 | |
| AT5G59613.2 | ATP synthase subunit (ATP6kDa) | 41 | 7.81 | 0.279 | 0.046 | * |
| AT5G59613.2 | ATP synthase subunit (ATP6kDa) | 42 | 6.03 | 0.326 | 0.370 | |
| AT5G59613.2 | ATP synthase subunit (ATP6kDa) | 43 | 4.07 | 0.445 | 0.985 | |

| | | | | | | |
|---|---|---|---|---|---|---|
| AT5G59613.2 | ATP synthase subunit (ATP6kDa) | 46 | 0.86 | 1.155 | 1.000 | |
| AT5G59613.2 | ATP synthase subunit (ATP6kDa) | 47 | 0.48 | 1.498 | 1.000 | |
| AT5G59970.1 | Histone H4, putative | 2 | 0.68 | 0.316 | 1.000 | |
| AT5G59970.1 | Histone H4, putative | 3 | 0.62 | 0.272 | 0.990 | |
| AT5G59970.1 | Histone H4, putative | 4 | 0.56 | 0.244 | 0.932 | |
| AT5G59970.1 | Histone H4, putative | 5 | 0.50 | 0.228 | 0.786 | |
| AT5G59970.1 | Histone H4, putative | 6 | 0.44 | 0.219 | 0.566 | |
| AT5G59970.1 | Histone H4, putative | 7 | 0.38 | 0.214 | 0.331 | |
| AT5G59970.1 | Histone H4, putative | 8 | 0.32 | 0.212 | 0.158 | |
| AT5G59970.1 | Histone H4, putative | 9 | 0.26 | 0.210 | 0.063 | . |
| AT5G59970.1 | Histone H4, putative | 10 | 0.22 | 0.209 | 0.021 | * |
| AT5G59970.1 | Histone H4, putative | 11 | 0.19 | 0.208 | 0.007 | ** |
| AT5G59970.1 | Histone H4, putative | 12 | 0.16 | 0.207 | 0.002 | ** |
| AT5G59970.1 | Histone H4, putative | 13 | 0.15 | 0.207 | 0.001 | *** |
| AT5G59970.1 | Histone H4, putative | 14 | 0.14 | 0.207 | 0.001 | *** |
| AT5G59970.1 | Histone H4, putative | 15 | 0.14 | 0.206 | 0.001 | *** |
| AT5G59970.1 | Histone H4, putative | 16 | 0.15 | 0.206 | 0.001 | ** |
| AT5G59970.1 | Histone H4, putative | 17 | 0.17 | 0.207 | 0.003 | ** |
| AT5G59970.1 | Histone H4, putative | 18 | 0.21 | 0.207 | 0.014 | * |
| AT5G59970.1 | Histone H4, putative | 19 | 0.26 | 0.209 | 0.054 | . |
| AT5G59970.1 | Histone H4, putative | 20 | 0.32 | 0.210 | 0.165 | |
| AT5G59970.1 | Histone H4, putative | 21 | 0.39 | 0.211 | 0.353 | |
| AT5G59970.1 | Histone H4, putative | 22 | 0.45 | 0.212 | 0.546 | |
| AT5G59970.1 | Histone H4, putative | 23 | 0.48 | 0.213 | 0.665 | |
| AT5G59970.1 | Histone H4, putative | 24 | 0.50 | 0.213 | 0.709 | |
| AT5G59970.1 | Histone H4, putative | 25 | 0.49 | 0.213 | 0.694 | |
| AT5G59970.1 | Histone H4, putative | 26 | 0.47 | 0.213 | 0.632 | |
| AT5G59970.1 | Histone H4, putative | 27 | 0.44 | 0.212 | 0.538 | |
| AT5G59970.1 | Histone H4, putative | 28 | 0.41 | 0.212 | 0.436 | |
| AT5G59970.1 | Histone H4, putative | 29 | 0.38 | 0.214 | 0.342 | |
| AT5G59970.1 | Histone H4, putative | 30 | 0.35 | 0.216 | 0.263 | |
| AT5G59970.1 | Histone H4, putative | 31 | 0.33 | 0.219 | 0.211 | |
| AT5G59970.1 | Histone H4, putative | 32 | 0.31 | 0.224 | 0.183 | |
| AT5G59970.1 | Histone H4, putative | 33 | 0.30 | 0.231 | 0.184 | |
| AT5G59970.1 | Histone H4, putative | 34 | 0.30 | 0.239 | 0.216 | |
| AT5G59970.1 | Histone H4, putative | 35 | 0.31 | 0.248 | 0.283 | |
| AT5G59970.1 | Histone H4, putative | 36 | 0.33 | 0.259 | 0.391 | |
| AT5G59970.1 | Histone H4, putative | 39 | 0.43 | 0.307 | 0.861 | |
| AT5G59970.1 | Histone H4, putative | 40 | 0.48 | 0.339 | 0.963 | |
| AT5G59970.1 | Histone H4, putative | 41 | 0.55 | 0.380 | 0.996 | |
| AT5G59970.1 | Histone H4, putative | 46 | 1.08 | 0.779 | 1.000 | |
| AT5G59970.1 | Histone H4, putative | 47 | 1.25 | 0.889 | 1.000 | |
| AT5G60960.1 | Pentatricopeptide repeat (PPR) superfamily protein (PNM1) | 2 | 1.39 | 0.453 | 1.000 | |
| AT5G60960.1 | Pentatricopeptide repeat (PPR) superfamily protein (PNM1) | 3 | 1.41 | 0.407 | 1.000 | |
| AT5G60960.1 | Pentatricopeptide repeat (PPR) superfamily protein (PNM1) | 13 | 3.07 | 0.279 | 1.000 | |
| AT5G60960.1 | Pentatricopeptide repeat (PPR) superfamily protein (PNM1) | 14 | 3.15 | 0.242 | 0.474 | |
| AT5G60960.1 | Pentatricopeptide repeat (PPR) superfamily protein (PNM1) | 15 | 2.97 | 0.224 | 0.432 | |
| AT5G60960.1 | Pentatricopeptide repeat (PPR) superfamily protein (PNM1) | 16 | 2.76 | 0.212 | 0.456 | |
| AT5G60960.1 | Pentatricopeptide repeat (PPR) superfamily protein (PNM1) | 17 | 2.56 | 0.210 | 0.555 | |
| AT5G60960.1 | Pentatricopeptide repeat (PPR) superfamily protein (PNM1) | 18 | 2.10 | 0.222 | 0.894 | |
| AT5G60960.1 | Pentatricopeptide repeat (PPR) superfamily protein (PNM1) | 19 | 1.51 | 0.262 | 1.000 | |

| | | | | | |
|---|---|---|---|---|---|
| AT5G60960.1 | Pentatricopeptide repeat (PPR) superfamily protein (PNM1) | 20 | 1.03 | 0.333 | 1.000 |
| AT5G60960.1 | Pentatricopeptide repeat (PPR) superfamily protein (PNM1) | 27 | 0.40 | 0.330 | 0.974 |
| AT5G60960.1 | Pentatricopeptide repeat (PPR) superfamily protein (PNM1) | 28 | 0.50 | 0.296 | 0.995 |
| AT5G60960.1 | Pentatricopeptide repeat (PPR) superfamily protein (PNM1) | 29 | 0.69 | 0.259 | 1.000 |
| AT5G60960.1 | Pentatricopeptide repeat (PPR) superfamily protein (PNM1) | 30 | 1.01 | 0.243 | 1.000 |
| AT5G60960.1 | Pentatricopeptide repeat (PPR) superfamily protein (PNM1) | 31 | 1.34 | 0.224 | 1.000 |
| AT5G60960.1 | Pentatricopeptide repeat (PPR) superfamily protein (PNM1) | 32 | 1.68 | 0.214 | 0.992 |
| AT5G60960.1 | Pentatricopeptide repeat (PPR) superfamily protein (PNM1) | 33 | 2.00 | 0.210 | 0.905 |
| AT5G60960.1 | Pentatricopeptide repeat (PPR) superfamily protein (PNM1) | 34 | 2.07 | 0.213 | 0.880 |
| AT5G60960.1 | Pentatricopeptide repeat (PPR) superfamily protein (PNM1) | 35 | 1.90 | 0.221 | 0.961 |
| AT5G60960.1 | Pentatricopeptide repeat (PPR) superfamily protein (PNM1) | 36 | 1.77 | 0.230 | 0.990 |
| AT5G60960.1 | Pentatricopeptide repeat (PPR) superfamily protein (PNM1) | 37 | 1.72 | 0.233 | 0.995 |
| AT5G60960.1 | Pentatricopeptide repeat (PPR) superfamily protein (PNM1) | 38 | 1.67 | 0.228 | 0.996 |
| AT5G60960.1 | Pentatricopeptide repeat (PPR) superfamily protein (PNM1) | 39 | 1.50 | 0.217 | 1.000 |
| AT5G60960.1 | Pentatricopeptide repeat (PPR) superfamily protein (PNM1) | 40 | 1.33 | 0.208 | 1.000 |
| AT5G60960.1 | Pentatricopeptide repeat (PPR) superfamily protein (PNM1) | 41 | 1.20 | 0.204 | 1.000 |
| AT5G60960.1 | Pentatricopeptide repeat (PPR) superfamily protein (PNM1) | 42 | 1.18 | 0.205 | 1.000 |
| AT5G60960.1 | Pentatricopeptide repeat (PPR) superfamily protein (PNM1) | 43 | 1.22 | 0.206 | 1.000 |
| AT5G60960.1 | Pentatricopeptide repeat (PPR) superfamily protein (PNM1) | 44 | 1.11 | 0.206 | 1.000 |
| AT5G60960.1 | Pentatricopeptide repeat (PPR) superfamily protein (PNM1) | 45 | 0.75 | 0.208 | 1.000 |
| AT5G60960.1 | Pentatricopeptide repeat (PPR) superfamily protein (PNM1) | 46 | 0.35 | 0.243 | 0.596 |
| AT5G60960.1 | Pentatricopeptide repeat (PPR) superfamily protein (PNM1) | 47 | 0.13 | 0.375 | 0.263 |
| AT5G61220.1 | LYR family of Fe/S cluster biogenesis protein | 2 | 1.12 | 0.648 | 1.000 |
| AT5G61220.1 | LYR family of Fe/S cluster biogenesis protein | 3 | 1.26 | 0.606 | 1.000 |
| AT5G61220.1 | LYR family of Fe/S cluster biogenesis protein | 22 | 1.51 | 0.531 | 1.000 |
| AT5G61220.1 | LYR family of Fe/S cluster biogenesis protein | 23 | 1.26 | 0.441 | 1.000 |
| AT5G61220.1 | LYR family of Fe/S cluster biogenesis protein | 24 | 1.09 | 0.395 | 1.000 |
| AT5G61220.1 | LYR family of Fe/S cluster biogenesis protein | 26 | 1.14 | 0.324 | 1.000 |
| AT5G61220.1 | LYR family of Fe/S cluster biogenesis protein | 27 | 1.43 | 0.289 | 1.000 |
| AT5G61220.1 | LYR family of Fe/S cluster biogenesis protein | 28 | 1.88 | 0.261 | 0.988 |
| AT5G61220.1 | LYR family of Fe/S cluster biogenesis protein | 29 | 2.39 | 0.247 | 0.833 |

| | | | | | |
|---|---|---|---|---|---|
| AT5G61220.1 | LYR family of Fe/S cluster biogenesis protein | 30 | 2.47 | 0.241 | 0.773 |
| AT5G61220.1 | LYR family of Fe/S cluster biogenesis protein | 31 | 2.18 | 0.240 | 0.897 |
| AT5G61220.1 | LYR family of Fe/S cluster biogenesis protein | 32 | 1.71 | 0.240 | 0.994 |
| AT5G61220.1 | LYR family of Fe/S cluster biogenesis protein | 33 | 1.37 | 0.240 | 1.000 |
| AT5G61220.1 | LYR family of Fe/S cluster biogenesis protein | 34 | 1.20 | 0.241 | 1.000 |
| AT5G61220.1 | LYR family of Fe/S cluster biogenesis protein | 35 | 1.14 | 0.241 | 1.000 |
| AT5G61220.1 | LYR family of Fe/S cluster biogenesis protein | 36 | 1.09 | 0.240 | 1.000 |
| AT5G61220.1 | LYR family of Fe/S cluster biogenesis protein | 37 | 1.01 | 0.239 | 1.000 |
| AT5G61220.1 | LYR family of Fe/S cluster biogenesis protein | 38 | 0.87 | 0.239 | 1.000 |
| AT5G61220.1 | LYR family of Fe/S cluster biogenesis protein | 39 | 0.68 | 0.240 | 1.000 |
| AT5G61220.1 | LYR family of Fe/S cluster biogenesis protein | 40 | 0.49 | 0.241 | 0.947 |
| AT5G61220.1 | LYR family of Fe/S cluster biogenesis protein | 41 | 0.39 | 0.242 | 0.745 |
| AT5G61220.1 | LYR family of Fe/S cluster biogenesis protein | 42 | 0.44 | 0.245 | 0.867 |
| AT5G61220.1 | LYR family of Fe/S cluster biogenesis protein | 43 | 0.64 | 0.247 | 0.999 |
| AT5G61220.1 | LYR family of Fe/S cluster biogenesis protein | 44 | 0.98 | 0.247 | 1.000 |
| AT5G61220.1 | LYR family of Fe/S cluster biogenesis protein | 45 | 1.42 | 0.257 | 1.000 |
| AT5G61220.1 | LYR family of Fe/S cluster biogenesis protein | 46 | 1.93 | 0.320 | 0.997 |
| AT5G61220.1 | LYR family of Fe/S cluster biogenesis protein | 47 | 2.48 | 0.490 | 0.999 |
| AT5G61310.4 | Cytochrome c:O2 oxidoreductase subunit (COX5C) | 2 | 1.03 | 0.416 | 1.000 |
| AT5G61310.4 | Cytochrome c:O2 oxidoreductase subunit (COX5C) | 3 | 1.08 | 0.327 | 1.000 |
| AT5G61310.4 | Cytochrome c:O2 oxidoreductase subunit (COX5C) | 6 | 1.23 | 0.340 | 1.000 |
| AT5G61310.4 | Cytochrome c:O2 oxidoreductase subunit (COX5C) | 7 | 1.26 | 0.287 | 1.000 |
| AT5G61310.4 | Cytochrome c:O2 oxidoreductase subunit (COX5C) | 8 | 1.18 | 0.262 | 1.000 |
| AT5G61310.4 | Cytochrome c:O2 oxidoreductase subunit (COX5C) | 9 | 1.01 | 0.229 | 1.000 |
| AT5G61310.4 | Cytochrome c:O2 oxidoreductase subunit (COX5C) | 10 | 0.90 | 0.220 | 1.000 |
| AT5G61310.4 | Cytochrome c:O2 oxidoreductase subunit (COX5C) | 11 | 0.93 | 0.211 | 1.000 |
| AT5G61310.4 | Cytochrome c:O2 oxidoreductase subunit (COX5C) | 12 | 1.01 | 0.198 | 1.000 |
| AT5G61310.4 | Cytochrome c:O2 oxidoreductase subunit (COX5C) | 13 | 1.06 | 0.190 | 1.000 |
| AT5G61310.4 | Cytochrome c:O2 oxidoreductase subunit (COX5C) | 14 | 1.16 | 0.187 | 1.000 |
| AT5G61310.4 | Cytochrome c:O2 oxidoreductase subunit (COX5C) | 15 | 1.02 | 0.186 | 1.000 |
| AT5G61310.4 | Cytochrome c:O2 oxidoreductase subunit (COX5C) | 16 | 0.65 | 0.185 | 0.999 |

| | | | | | | |
|---|---|---|---|---|---|---|
| AT5G61310.4 | Cytochrome c:O2 oxidoreductase subunit (COX5C) | 17 | 0.50 | 0.185 | 0.883 | |
| AT5G61310.4 | Cytochrome c:O2 oxidoreductase subunit (COX5C) | 18 | 0.56 | 0.187 | 0.980 | |
| AT5G61310.4 | Cytochrome c:O2 oxidoreductase subunit (COX5C) | 19 | 0.69 | 0.192 | 1.000 | |
| AT5G61310.4 | Cytochrome c:O2 oxidoreductase subunit (COX5C) | 20 | 0.76 | 0.197 | 1.000 | |
| AT5G61310.4 | Cytochrome c:O2 oxidoreductase subunit (COX5C) | 21 | 0.69 | 0.204 | 1.000 | |
| AT5G61310.4 | Cytochrome c:O2 oxidoreductase subunit (COX5C) | 22 | 0.51 | 0.213 | 0.976 | |
| AT5G61310.4 | Cytochrome c:O2 oxidoreductase subunit (COX5C) | 23 | 0.41 | 0.237 | 0.892 | |
| AT5G61310.4 | Cytochrome c:O2 oxidoreductase subunit (COX5C) | 24 | 0.34 | 0.275 | 0.842 | |
| AT5G61310.4 | Cytochrome c:O2 oxidoreductase subunit (COX5C) | 26 | 0.27 | 0.255 | 0.453 | |
| AT5G61310.4 | Cytochrome c:O2 oxidoreductase subunit (COX5C) | 27 | 0.42 | 0.218 | 0.825 | |
| AT5G61310.4 | Cytochrome c:O2 oxidoreductase subunit (COX5C) | 28 | 0.89 | 0.201 | 1.000 | |
| AT5G61310.4 | Cytochrome c:O2 oxidoreductase subunit (COX5C) | 29 | 1.55 | 0.193 | 0.999 | |
| AT5G61310.4 | Cytochrome c:O2 oxidoreductase subunit (COX5C) | 30 | 1.76 | 0.190 | 0.986 | |
| AT5G61310.4 | Cytochrome c:O2 oxidoreductase subunit (COX5C) | 31 | 1.47 | 0.191 | 1.000 | |
| AT5G61310.4 | Cytochrome c:O2 oxidoreductase subunit (COX5C) | 32 | 0.98 | 0.193 | 1.000 | |
| AT5G61310.4 | Cytochrome c:O2 oxidoreductase subunit (COX5C) | 33 | 0.54 | 0.196 | 0.976 | |
| AT5G61310.4 | Cytochrome c:O2 oxidoreductase subunit (COX5C) | 34 | 0.27 | 0.205 | 0.146 | |
| AT5G61310.4 | Cytochrome c:O2 oxidoreductase subunit (COX5C) | 35 | 0.18 | 0.249 | 0.081 | . |
| AT5G61310.4 | Cytochrome c:O2 oxidoreductase subunit (COX5C) | 36 | 0.17 | 0.405 | 0.704 | |
| AT5G61310.4 | Cytochrome c:O2 oxidoreductase subunit (COX5C) | 46 | 792.44 | 3.610 | 1.000 | |
| AT5G61310.4 | Cytochrome c:O2 oxidoreductase subunit (COX5C) | 47 | 2891.14 | 4.107 | 1.000 | |
| AT5G61790.1 | Calnexin 1 (CNX1) | 2 | 1.64 | 0.534 | 1.000 | |
| AT5G61790.1 | Calnexin 1 (CNX1) | 3 | 1.06 | 0.390 | 1.000 | |
| AT5G61790.1 | Calnexin 1 (CNX1) | 4 | 0.73 | 0.674 | 1.000 | |
| AT5G61790.1 | Calnexin 1 (CNX1) | 18 | 0.54 | 0.376 | 1.000 | |
| AT5G61790.1 | Calnexin 1 (CNX1) | 19 | 0.56 | 0.373 | 1.000 | |
| AT5G61790.1 | Calnexin 1 (CNX1) | 20 | 0.61 | 0.375 | 1.000 | |
| AT5G61790.1 | Calnexin 1 (CNX1) | 22 | 0.84 | 0.402 | 1.000 | |
| AT5G61790.1 | Calnexin 1 (CNX1) | 23 | 0.95 | 0.411 | 1.000 | |
| AT5G61790.1 | Calnexin 1 (CNX1) | 24 | 0.98 | 0.341 | 1.000 | |
| AT5G61790.1 | Calnexin 1 (CNX1) | 25 | 0.91 | 0.316 | 1.000 | |
| AT5G61790.1 | Calnexin 1 (CNX1) | 26 | 0.77 | 0.285 | 1.000 | |
| AT5G61790.1 | Calnexin 1 (CNX1) | 27 | 0.64 | 0.263 | 1.000 | |
| AT5G61790.1 | Calnexin 1 (CNX1) | 28 | 0.58 | 0.253 | 1.000 | |
| AT5G61790.1 | Calnexin 1 (CNX1) | 29 | 0.52 | 0.253 | 0.997 | |
| AT5G61790.1 | Calnexin 1 (CNX1) | 30 | 0.35 | 0.253 | 0.787 | |
| AT5G61790.1 | Calnexin 1 (CNX1) | 31 | 0.28 | 0.257 | 0.526 | |
| AT5G61790.1 | Calnexin 1 (CNX1) | 32 | 0.27 | 0.257 | 0.467 | |
| AT5G61790.1 | Calnexin 1 (CNX1) | 33 | 0.27 | 0.258 | 0.463 | |
| AT5G61790.1 | Calnexin 1 (CNX1) | 34 | 0.23 | 0.262 | 0.299 | |
| AT5G61790.1 | Calnexin 1 (CNX1) | 35 | 0.24 | 0.271 | 0.402 | |

| | | | | | | |
|---|---|---|---|---|---|---|
| AT5G61790.1 | Calnexin 1 (CNX1) | 36 | 0.38 | 0.277 | 0.934 | |
| AT5G61790.1 | Calnexin 1 (CNX1) | 37 | 1.03 | 0.283 | 1.000 | |
| AT5G61790.1 | Calnexin 1 (CNX1) | 38 | 4.31 | 0.276 | 0.392 | |
| AT5G61790.1 | Calnexin 1 (CNX1) | 39 | 8.95 | 0.264 | 0.011 | * |
| AT5G61790.1 | Calnexin 1 (CNX1) | 40 | 2.50 | 0.257 | 0.921 | |
| AT5G61790.1 | Calnexin 1 (CNX1) | 41 | 0.23 | 0.258 | 0.259 | |
| AT5G61790.1 | Calnexin 1 (CNX1) | 42 | 0.04 | 0.264 | 0.000 | *** |
| AT5G61790.1 | Calnexin 1 (CNX1) | 43 | 0.05 | 0.268 | 0.000 | *** |
| AT5G61790.1 | Calnexin 1 (CNX1) | 44 | 0.26 | 0.333 | 0.815 | |
| AT5G61790.1 | Calnexin 1 (CNX1) | 45 | 2.15 | 0.521 | 1.000 | |
| AT5G61790.1 | Calnexin 1 (CNX1) | 46 | 23.13 | 0.858 | 0.903 | |
| AT5G61790.1 | Calnexin 1 (CNX1) | 47 | 280.24 | 1.388 | 0.806 | |
| AT5G61810.1 | ATP/Pi carrier 1 (APC1) | 2 | 0.93 | 0.286 | 1.000 | |
| AT5G61810.1 | ATP/Pi carrier 1 (APC1) | 3 | 1.07 | 0.188 | 1.000 | |
| AT5G61810.1 | ATP/Pi carrier 1 (APC1) | 4 | 1.19 | 0.170 | 1.000 | |
| AT5G61810.1 | ATP/Pi carrier 1 (APC1) | 5 | 1.22 | 0.171 | 1.000 | |
| AT5G61810.1 | ATP/Pi carrier 1 (APC1) | 6 | 1.15 | 0.171 | 1.000 | |
| AT5G61810.1 | ATP/Pi carrier 1 (APC1) | 7 | 1.08 | 0.170 | 1.000 | |
| AT5G61810.1 | ATP/Pi carrier 1 (APC1) | 8 | 1.07 | 0.168 | 1.000 | |
| AT5G61810.1 | ATP/Pi carrier 1 (APC1) | 9 | 1.09 | 0.166 | 1.000 | |
| AT5G61810.1 | ATP/Pi carrier 1 (APC1) | 10 | 1.09 | 0.165 | 1.000 | |
| AT5G61810.1 | ATP/Pi carrier 1 (APC1) | 11 | 1.09 | 0.166 | 1.000 | |
| AT5G61810.1 | ATP/Pi carrier 1 (APC1) | 12 | 1.11 | 0.166 | 1.000 | |
| AT5G61810.1 | ATP/Pi carrier 1 (APC1) | 13 | 1.10 | 0.164 | 1.000 | |
| AT5G61810.1 | ATP/Pi carrier 1 (APC1) | 14 | 1.04 | 0.163 | 1.000 | |
| AT5G61810.1 | ATP/Pi carrier 1 (APC1) | 15 | 0.96 | 0.162 | 1.000 | |
| AT5G61810.1 | ATP/Pi carrier 1 (APC1) | 16 | 0.92 | 0.161 | 1.000 | |
| AT5G61810.1 | ATP/Pi carrier 1 (APC1) | 17 | 1.01 | 0.161 | 1.000 | |
| AT5G61810.1 | ATP/Pi carrier 1 (APC1) | 18 | 1.30 | 0.163 | 1.000 | |
| AT5G61810.1 | ATP/Pi carrier 1 (APC1) | 19 | 1.81 | 0.167 | 0.944 | |
| AT5G61810.1 | ATP/Pi carrier 1 (APC1) | 20 | 2.27 | 0.171 | 0.597 | |
| AT5G61810.1 | ATP/Pi carrier 1 (APC1) | 21 | 2.29 | 0.175 | 0.622 | |
| AT5G61810.1 | ATP/Pi carrier 1 (APC1) | 22 | 1.92 | 0.180 | 0.934 | |
| AT5G61810.1 | ATP/Pi carrier 1 (APC1) | 23 | 1.49 | 0.186 | 1.000 | |
| AT5G61810.1 | ATP/Pi carrier 1 (APC1) | 24 | 1.20 | 0.191 | 1.000 | |
| AT5G61810.1 | ATP/Pi carrier 1 (APC1) | 25 | 1.06 | 0.194 | 1.000 | |
| AT5G61810.1 | ATP/Pi carrier 1 (APC1) | 26 | 1.01 | 0.185 | 1.000 | |
| AT5G61810.1 | ATP/Pi carrier 1 (APC1) | 27 | 1.01 | 0.178 | 1.000 | |
| AT5G61810.1 | ATP/Pi carrier 1 (APC1) | 28 | 1.09 | 0.172 | 1.000 | |
| AT5G61810.1 | ATP/Pi carrier 1 (APC1) | 29 | 1.23 | 0.167 | 1.000 | |
| AT5G61810.1 | ATP/Pi carrier 1 (APC1) | 30 | 1.17 | 0.165 | 1.000 | |
| AT5G61810.1 | ATP/Pi carrier 1 (APC1) | 31 | 1.11 | 0.165 | 1.000 | |
| AT5G61810.1 | ATP/Pi carrier 1 (APC1) | 32 | 1.18 | 0.165 | 1.000 | |
| AT5G61810.1 | ATP/Pi carrier 1 (APC1) | 33 | 1.29 | 0.166 | 1.000 | |
| AT5G61810.1 | ATP/Pi carrier 1 (APC1) | 34 | 1.36 | 0.166 | 1.000 | |
| AT5G61810.1 | ATP/Pi carrier 1 (APC1) | 35 | 1.41 | 0.166 | 1.000 | |
| AT5G61810.1 | ATP/Pi carrier 1 (APC1) | 36 | 1.49 | 0.165 | 1.000 | |
| AT5G61810.1 | ATP/Pi carrier 1 (APC1) | 37 | 1.61 | 0.165 | 0.994 | |
| AT5G61810.1 | ATP/Pi carrier 1 (APC1) | 38 | 1.63 | 0.165 | 0.992 | |
| AT5G61810.1 | ATP/Pi carrier 1 (APC1) | 39 | 1.39 | 0.166 | 1.000 | |
| AT5G61810.1 | ATP/Pi carrier 1 (APC1) | 40 | 1.03 | 0.166 | 1.000 | |
| AT5G61810.1 | ATP/Pi carrier 1 (APC1) | 41 | 0.72 | 0.166 | 1.000 | |
| AT5G61810.1 | ATP/Pi carrier 1 (APC1) | 42 | 0.54 | 0.169 | 0.921 | |
| AT5G61810.1 | ATP/Pi carrier 1 (APC1) | 43 | 0.46 | 0.170 | 0.670 | |
| AT5G61810.1 | ATP/Pi carrier 1 (APC1) | 44 | 0.43 | 0.170 | 0.513 | |
| AT5G61810.1 | ATP/Pi carrier 1 (APC1) | 45 | 0.43 | 0.170 | 0.539 | |
| AT5G61810.1 | ATP/Pi carrier 1 (APC1) | 46 | 0.48 | 0.201 | 0.934 | |
| AT5G61810.1 | ATP/Pi carrier 1 (APC1) | 47 | 0.56 | 0.328 | 1.000 | |
| AT5G61990.1 | Pentatricopeptide repeat (PPR) superfamily protein | 2 | 1.13 | 0.431 | 1.000 | |
| AT5G61990.1 | Pentatricopeptide repeat (PPR) superfamily protein | 3 | 1.01 | 0.392 | 1.000 | |

| | | | | | | |
|---|---|---|---|---|---|---|
| AT5G61990.1 | Pentatricopeptide repeat (PPR) superfamily protein | 16 | 0.69 | 0.293 | 1.000 | |
| AT5G61990.1 | Pentatricopeptide repeat (PPR) superfamily protein | 17 | 0.75 | 0.258 | 1.000 | |
| AT5G61990.1 | Pentatricopeptide repeat (PPR) superfamily protein | 18 | 0.81 | 0.241 | 1.000 | |
| AT5G61990.1 | Pentatricopeptide repeat (PPR) superfamily protein | 19 | 0.89 | 0.241 | 1.000 | |
| AT5G61990.1 | Pentatricopeptide repeat (PPR) superfamily protein | 20 | 0.95 | 0.244 | 1.000 | |
| AT5G61990.1 | Pentatricopeptide repeat (PPR) superfamily protein | 22 | 0.86 | 0.237 | 1.000 | |
| AT5G61990.1 | Pentatricopeptide repeat (PPR) superfamily protein | 23 | 0.74 | 0.224 | 1.000 | |
| AT5G61990.1 | Pentatricopeptide repeat (PPR) superfamily protein | 24 | 0.66 | 0.206 | 0.996 | |
| AT5G61990.1 | Pentatricopeptide repeat (PPR) superfamily protein | 25 | 0.69 | 0.193 | 0.997 | |
| AT5G61990.1 | Pentatricopeptide repeat (PPR) superfamily protein | 26 | 0.92 | 0.187 | 1.000 | |
| AT5G61990.1 | Pentatricopeptide repeat (PPR) superfamily protein | 27 | 1.41 | 0.181 | 0.998 | |
| AT5G61990.1 | Pentatricopeptide repeat (PPR) superfamily protein | 28 | 2.03 | 0.175 | 0.646 | |
| AT5G61990.1 | Pentatricopeptide repeat (PPR) superfamily protein | 29 | 2.50 | 0.173 | 0.269 | |
| AT5G61990.1 | Pentatricopeptide repeat (PPR) superfamily protein | 30 | 2.36 | 0.173 | 0.350 | |
| AT5G61990.1 | Pentatricopeptide repeat (PPR) superfamily protein | 31 | 1.83 | 0.174 | 0.810 | |
| AT5G61990.1 | Pentatricopeptide repeat (PPR) superfamily protein | 32 | 1.36 | 0.174 | 0.999 | |
| AT5G61990.1 | Pentatricopeptide repeat (PPR) superfamily protein | 33 | 1.07 | 0.175 | 1.000 | |
| AT5G61990.1 | Pentatricopeptide repeat (PPR) superfamily protein | 34 | 0.97 | 0.176 | 1.000 | |
| AT5G61990.1 | Pentatricopeptide repeat (PPR) superfamily protein | 35 | 1.01 | 0.181 | 1.000 | |
| AT5G61990.1 | Pentatricopeptide repeat (PPR) superfamily protein | 36 | 1.33 | 0.195 | 1.000 | |
| AT5G61990.1 | Pentatricopeptide repeat (PPR) superfamily protein | 37 | 2.20 | 0.233 | 0.839 | |
| AT5G61990.1 | Pentatricopeptide repeat (PPR) superfamily protein | 38 | 4.20 | 0.314 | 0.466 | |
| AT5G61990.1 | Pentatricopeptide repeat (PPR) superfamily protein | 46 | 2713.75 | 2.113 | 0.737 | |
| AT5G61990.1 | Pentatricopeptide repeat (PPR) superfamily protein | 47 | 6551.97 | 2.448 | 0.782 | |
| AT5G62050.1 | Oxidase assembly 1 (OXA1) | 2 | 1.20 | 0.283 | 1.000 | |
| AT5G62050.1 | Oxidase assembly 1 (OXA1) | 3 | 1.24 | 0.265 | 1.000 | |
| AT5G62050.1 | Oxidase assembly 1 (OXA1) | 4 | 1.28 | 0.250 | 0.999 | |
| AT5G62050.1 | Oxidase assembly 1 (OXA1) | 12 | 1.67 | 0.188 | 0.761 | |
| AT5G62050.1 | Oxidase assembly 1 (OXA1) | 13 | 1.72 | 0.184 | 0.692 | |
| AT5G62050.1 | Oxidase assembly 1 (OXA1) | 14 | 1.78 | 0.179 | 0.617 | |
| AT5G62050.1 | Oxidase assembly 1 (OXA1) | 15 | 1.83 | 0.175 | 0.537 | |
| AT5G62050.1 | Oxidase assembly 1 (OXA1) | 16 | 1.89 | 0.170 | 0.460 | |
| AT5G62050.1 | Oxidase assembly 1 (OXA1) | 21 | 2.09 | 0.147 | 0.170 | |
| AT5G62050.1 | Oxidase assembly 1 (OXA1) | 22 | 2.11 | 0.143 | 0.140 | |
| AT5G62050.1 | Oxidase assembly 1 (OXA1) | 23 | 2.12 | 0.139 | 0.118 | |
| AT5G62050.1 | Oxidase assembly 1 (OXA1) | 24 | 2.11 | 0.135 | 0.105 | |
| AT5G62050.1 | Oxidase assembly 1 (OXA1) | 25 | 2.09 | 0.132 | 0.098 | . |
| AT5G62050.1 | Oxidase assembly 1 (OXA1) | 26 | 2.06 | 0.130 | 0.100 | . |

| | | | | | |
|---|---|---|---|---|---|
| AT5G62050.1 | Oxidase assembly 1 (OXA1) | 27 | 2.02 | 0.128 | 0.110 |
| AT5G62050.1 | Oxidase assembly 1 (OXA1) | 28 | 1.96 | 0.127 | 0.129 |
| AT5G62050.1 | Oxidase assembly 1 (OXA1) | 30 | 1.82 | 0.125 | 0.211 |
| AT5G62050.1 | Oxidase assembly 1 (OXA1) | 31 | 1.73 | 0.124 | 0.284 |
| AT5G62050.1 | Oxidase assembly 1 (OXA1) | 32 | 1.64 | 0.124 | 0.387 |
| AT5G62050.1 | Oxidase assembly 1 (OXA1) | 33 | 1.55 | 0.124 | 0.518 |
| AT5G62050.1 | Oxidase assembly 1 (OXA1) | 34 | 1.46 | 0.124 | 0.671 |
| AT5G62050.1 | Oxidase assembly 1 (OXA1) | 35 | 1.37 | 0.125 | 0.821 |
| AT5G62050.1 | Oxidase assembly 1 (OXA1) | 36 | 1.27 | 0.125 | 0.936 |
| AT5G62050.1 | Oxidase assembly 1 (OXA1) | 37 | 1.18 | 0.126 | 0.990 |
| AT5G62050.1 | Oxidase assembly 1 (OXA1) | 38 | 1.09 | 0.128 | 1.000 |
| AT5G62050.1 | Oxidase assembly 1 (OXA1) | 39 | 1.00 | 0.129 | 1.000 |
| AT5G62050.1 | Oxidase assembly 1 (OXA1) | 40 | 0.91 | 0.132 | 1.000 |
| AT5G62050.1 | Oxidase assembly 1 (OXA1) | 41 | 0.81 | 0.135 | 0.979 |
| AT5G62050.1 | Oxidase assembly 1 (OXA1) | 42 | 0.72 | 0.139 | 0.862 |
| AT5G62050.1 | Oxidase assembly 1 (OXA1) | 43 | 0.64 | 0.146 | 0.652 |
| AT5G62050.1 | Oxidase assembly 1 (OXA1) | 44 | 0.56 | 0.154 | 0.443 |
| AT5G62050.1 | Oxidase assembly 1 (OXA1) | 45 | 0.49 | 0.166 | 0.298 |
| AT5G62050.1 | Oxidase assembly 1 (OXA1) | 46 | 0.42 | 0.180 | 0.211 |
| AT5G62050.1 | Oxidase assembly 1 (OXA1) | 47 | 0.37 | 0.198 | 0.166 |
| AT5G62270.2 | ? | 2 | 0.82 | 0.680 | 1.000 |
| AT5G62270.2 | ? | 3 | 0.75 | 0.592 | 1.000 |
| AT5G62270.2 | ? | 10 | 0.89 | 0.352 | 1.000 |
| AT5G62270.2 | ? | 11 | 1.11 | 0.312 | 1.000 |
| AT5G62270.2 | ? | 12 | 1.39 | 0.283 | 1.000 |
| AT5G62270.2 | ? | 13 | 1.60 | 0.263 | 0.997 |
| AT5G62270.2 | ? | 14 | 1.72 | 0.250 | 0.987 |
| AT5G62270.2 | ? | 15 | 1.82 | 0.243 | 0.968 |
| AT5G62270.2 | ? | 16 | 1.94 | 0.241 | 0.933 |
| AT5G62270.2 | ? | 17 | 1.95 | 0.240 | 0.927 |
| AT5G62270.2 | ? | 18 | 1.76 | 0.243 | 0.977 |
| AT5G62270.2 | ? | 19 | 1.43 | 0.247 | 1.000 |
| AT5G62270.2 | ? | 20 | 1.06 | 0.251 | 1.000 |
| AT5G62270.2 | ? | 21 | 0.74 | 0.256 | 1.000 |
| AT5G62270.2 | ? | 22 | 0.52 | 0.262 | 0.963 |
| AT5G62270.2 | ? | 23 | 0.40 | 0.269 | 0.780 |
| AT5G62270.2 | ? | 24 | 0.35 | 0.276 | 0.664 |
| AT5G62270.2 | ? | 25 | 0.36 | 0.280 | 0.709 |
| AT5G62270.2 | ? | 26 | 0.42 | 0.277 | 0.863 |
| AT5G62270.2 | ? | 27 | 0.56 | 0.269 | 0.988 |
| AT5G62270.2 | ? | 28 | 0.81 | 0.261 | 1.000 |
| AT5G62270.2 | ? | 29 | 1.13 | 0.255 | 1.000 |
| AT5G62270.2 | ? | 30 | 1.44 | 0.254 | 1.000 |
| AT5G62270.2 | ? | 31 | 1.66 | 0.264 | 0.995 |
| AT5G62270.2 | ? | 32 | 1.76 | 0.295 | 0.995 |
| AT5G62270.2 | ? | 33 | 1.79 | 0.358 | 0.999 |
| AT5G62270.2 | ? | 34 | 1.83 | 0.451 | 1.000 |
| AT5G62270.2 | ? | 46 | 4.24 | 1.926 | 1.000 |
| AT5G62270.2 | ? | 47 | 4.65 | 2.115 | 1.000 |
| AT5G62530.1 | Delta-pyrroline-5-carboxylate dehydrogenase (5CDH) | 2 | 1.04 | 0.575 | 1.000 |
| AT5G62530.1 | Delta-pyrroline-5-carboxylate dehydrogenase (5CDH) | 3 | 0.80 | 0.426 | 1.000 |
| AT5G62530.1 | Delta-pyrroline-5-carboxylate dehydrogenase (5CDH) | 4 | 0.66 | 0.424 | 1.000 |
| AT5G62530.1 | Delta-pyrroline-5-carboxylate dehydrogenase (5CDH) | 6 | 0.65 | 0.430 | 1.000 |
| AT5G62530.1 | Delta-pyrroline-5-carboxylate dehydrogenase (5CDH) | 7 | 0.79 | 0.379 | 1.000 |
| AT5G62530.1 | Delta-pyrroline-5-carboxylate dehydrogenase (5CDH) | 8 | 1.01 | 0.343 | 1.000 |
| AT5G62530.1 | Delta-pyrroline-5-carboxylate dehydrogenase (5CDH) | 9 | 1.22 | 0.324 | 1.000 |

| | | | | | |
|---|---|---|---|---|---|
| AT5G62530.1 | Delta-pyrroline-5-carboxylate dehydrogenase (5CDH) | 10 | 1.42 | 0.317 | 1.000 |
| AT5G62530.1 | Delta-pyrroline-5-carboxylate dehydrogenase (5CDH) | 11 | 1.72 | 0.318 | 1.000 |
| AT5G62530.1 | Delta-pyrroline-5-carboxylate dehydrogenase (5CDH) | 12 | 2.23 | 0.319 | 0.998 |
| AT5G62530.1 | Delta-pyrroline-5-carboxylate dehydrogenase (5CDH) | 13 | 2.74 | 0.322 | 0.976 |
| AT5G62530.1 | Delta-pyrroline-5-carboxylate dehydrogenase (5CDH) | 14 | 3.16 | 0.326 | 0.925 |
| AT5G62530.1 | Delta-pyrroline-5-carboxylate dehydrogenase (5CDH) | 15 | 3.52 | 0.327 | 0.859 |
| AT5G62530.1 | Delta-pyrroline-5-carboxylate dehydrogenase (5CDH) | 16 | 3.54 | 0.332 | 0.869 |
| AT5G62530.1 | Delta-pyrroline-5-carboxylate dehydrogenase (5CDH) | 17 | 3.05 | 0.344 | 0.966 |
| AT5G62530.1 | Delta-pyrroline-5-carboxylate dehydrogenase (5CDH) | 18 | 2.34 | 0.352 | 0.999 |
| AT5G62530.1 | Delta-pyrroline-5-carboxylate dehydrogenase (5CDH) | 19 | 1.81 | 0.347 | 1.000 |
| AT5G62530.1 | Delta-pyrroline-5-carboxylate dehydrogenase (5CDH) | 20 | 1.53 | 0.338 | 1.000 |
| AT5G62530.1 | Delta-pyrroline-5-carboxylate dehydrogenase (5CDH) | 21 | 1.44 | 0.330 | 1.000 |
| AT5G62530.1 | Delta-pyrroline-5-carboxylate dehydrogenase (5CDH) | 22 | 1.44 | 0.326 | 1.000 |
| AT5G62530.1 | Delta-pyrroline-5-carboxylate dehydrogenase (5CDH) | 23 | 1.53 | 0.323 | 1.000 |
| AT5G62530.1 | Delta-pyrroline-5-carboxylate dehydrogenase (5CDH) | 24 | 1.74 | 0.318 | 1.000 |
| AT5G62530.1 | Delta-pyrroline-5-carboxylate dehydrogenase (5CDH) | 25 | 1.86 | 0.316 | 1.000 |
| AT5G62530.1 | Delta-pyrroline-5-carboxylate dehydrogenase (5CDH) | 26 | 1.89 | 0.315 | 1.000 |
| AT5G62530.1 | Delta-pyrroline-5-carboxylate dehydrogenase (5CDH) | 27 | 2.01 | 0.315 | 1.000 |
| AT5G62530.1 | Delta-pyrroline-5-carboxylate dehydrogenase (5CDH) | 28 | 2.15 | 0.314 | 0.999 |
| AT5G62530.1 | Delta-pyrroline-5-carboxylate dehydrogenase (5CDH) | 29 | 2.09 | 0.314 | 0.999 |
| AT5G62530.1 | Delta-pyrroline-5-carboxylate dehydrogenase (5CDH) | 30 | 1.82 | 0.316 | 1.000 |
| AT5G62530.1 | Delta-pyrroline-5-carboxylate dehydrogenase (5CDH) | 31 | 1.65 | 0.316 | 1.000 |
| AT5G62530.1 | Delta-pyrroline-5-carboxylate dehydrogenase (5CDH) | 32 | 1.68 | 0.316 | 1.000 |
| AT5G62530.1 | Delta-pyrroline-5-carboxylate dehydrogenase (5CDH) | 33 | 1.68 | 0.317 | 1.000 |
| AT5G62530.1 | Delta-pyrroline-5-carboxylate dehydrogenase (5CDH) | 34 | 1.64 | 0.317 | 1.000 |
| AT5G62530.1 | Delta-pyrroline-5-carboxylate dehydrogenase (5CDH) | 35 | 1.74 | 0.316 | 1.000 |
| AT5G62530.1 | Delta-pyrroline-5-carboxylate dehydrogenase (5CDH) | 36 | 2.01 | 0.316 | 1.000 |
| AT5G62530.1 | Delta-pyrroline-5-carboxylate dehydrogenase (5CDH) | 37 | 2.21 | 0.316 | 0.998 |
| AT5G62530.1 | Delta-pyrroline-5-carboxylate dehydrogenase (5CDH) | 38 | 2.18 | 0.316 | 0.999 |
| AT5G62530.1 | Delta-pyrroline-5-carboxylate dehydrogenase (5CDH) | 39 | 2.40 | 0.316 | 0.994 |
| AT5G62530.1 | Delta-pyrroline-5-carboxylate dehydrogenase (5CDH) | 40 | 3.42 | 0.316 | 0.849 |

| | | | | | |
|---|---|---|---|---|---|
| AT5G62530.1 | Delta-pyrroline-5-carboxylate dehydrogenase (5CDH) | 41 | 4.89 | 0.319 | 0.488 |
| AT5G62530.1 | Delta-pyrroline-5-carboxylate dehydrogenase (5CDH) | 42 | 5.87 | 0.325 | 0.340 |
| AT5G62530.1 | Delta-pyrroline-5-carboxylate dehydrogenase (5CDH) | 43 | 5.43 | 0.338 | 0.478 |
| AT5G62530.1 | Delta-pyrroline-5-carboxylate dehydrogenase (5CDH) | 44 | 4.05 | 0.395 | 0.925 |
| AT5G62530.1 | Delta-pyrroline-5-carboxylate dehydrogenase (5CDH) | 45 | 3.15 | 0.545 | 1.000 |
| AT5G62530.1 | Delta-pyrroline-5-carboxylate dehydrogenase (5CDH) | 46 | 2.64 | 0.830 | 1.000 |
| AT5G62530.1 | Delta-pyrroline-5-carboxylate dehydrogenase (5CDH) | 47 | 2.29 | 1.231 | 1.000 |
| AT5G62575.1 | Succinate dehydrogenase subunit (SDH7-2) | 2 | 0.93 | 0.396 | 1.000 |
| AT5G62575.1 | Succinate dehydrogenase subunit (SDH7-2) | 3 | 0.99 | 0.304 | 1.000 |
| AT5G62575.1 | Succinate dehydrogenase subunit (SDH7-2) | 6 | 1.40 | 0.309 | 1.000 |
| AT5G62575.1 | Succinate dehydrogenase subunit (SDH7-2) | 7 | 1.70 | 0.246 | 1.000 |
| AT5G62575.1 | Succinate dehydrogenase subunit (SDH7-2) | 8 | 1.98 | 0.218 | 0.993 |
| AT5G62575.1 | Succinate dehydrogenase subunit (SDH7-2) | 9 | 2.07 | 0.205 | 0.968 |
| AT5G62575.1 | Succinate dehydrogenase subunit (SDH7-2) | 10 | 2.10 | 0.196 | 0.941 |
| AT5G62575.1 | Succinate dehydrogenase subunit (SDH7-2) | 11 | 2.31 | 0.186 | 0.763 |
| AT5G62575.1 | Succinate dehydrogenase subunit (SDH7-2) | 12 | 2.81 | 0.181 | 0.344 |
| AT5G62575.1 | Succinate dehydrogenase subunit (SDH7-2) | 13 | 3.00 | 0.182 | 0.244 |
| AT5G62575.1 | Succinate dehydrogenase subunit (SDH7-2) | 14 | 2.78 | 0.175 | 0.295 |
| AT5G62575.1 | Succinate dehydrogenase subunit (SDH7-2) | 15 | 2.54 | 0.169 | 0.404 |
| AT5G62575.1 | Succinate dehydrogenase subunit (SDH7-2) | 16 | 2.52 | 0.169 | 0.416 |
| AT5G62575.1 | Succinate dehydrogenase subunit (SDH7-2) | 17 | 2.80 | 0.169 | 0.238 |
| AT5G62575.1 | Succinate dehydrogenase subunit (SDH7-2) | 18 | 3.17 | 0.176 | 0.142 |
| AT5G62575.1 | Succinate dehydrogenase subunit (SDH7-2) | 19 | 3.39 | 0.195 | 0.194 |
| AT5G62575.1 | Succinate dehydrogenase subunit (SDH7-2) | 20 | 3.08 | 0.207 | 0.429 |
| AT5G62575.1 | Succinate dehydrogenase subunit (SDH7-2) | 21 | 2.56 | 0.224 | 0.859 |
| AT5G62575.1 | Succinate dehydrogenase subunit (SDH7-2) | 22 | 2.13 | 0.231 | 0.987 |
| AT5G62575.1 | Succinate dehydrogenase subunit (SDH7-2) | 23 | 1.89 | 0.269 | 1.000 |
| AT5G62575.1 | Succinate dehydrogenase subunit (SDH7-2) | 24 | 1.84 | 0.293 | 1.000 |
| AT5G62575.1 | Succinate dehydrogenase subunit (SDH7-2) | 28 | 2.28 | 0.251 | 0.986 |
| AT5G62575.1 | Succinate dehydrogenase subunit (SDH7-2) | 29 | 2.33 | 0.204 | 0.870 |
| AT5G62575.1 | Succinate dehydrogenase subunit (SDH7-2) | 30 | 2.37 | 0.184 | 0.707 |

| | | | | | | |
|---|---|---|---|---|---|---|
| AT5G62575.1 | Succinate dehydrogenase subunit (SDH7-2) | 31 | 2.28 | 0.174 | 0.684 | |
| AT5G62575.1 | Succinate dehydrogenase subunit (SDH7-2) | 32 | 2.02 | 0.171 | 0.882 | |
| AT5G62575.1 | Succinate dehydrogenase subunit (SDH7-2) | 33 | 2.04 | 0.171 | 0.865 | |
| AT5G62575.1 | Succinate dehydrogenase subunit (SDH7-2) | 34 | 2.09 | 0.172 | 0.833 | |
| AT5G62575.1 | Succinate dehydrogenase subunit (SDH7-2) | 35 | 1.81 | 0.172 | 0.976 | |
| AT5G62575.1 | Succinate dehydrogenase subunit (SDH7-2) | 36 | 1.10 | 0.170 | 1.000 | |
| AT5G62575.1 | Succinate dehydrogenase subunit (SDH7-2) | 37 | 0.78 | 0.170 | 1.000 | |
| AT5G62575.1 | Succinate dehydrogenase subunit (SDH7-2) | 38 | 1.15 | 0.172 | 1.000 | |
| AT5G62575.1 | Succinate dehydrogenase subunit (SDH7-2) | 39 | 2.14 | 0.171 | 0.785 | |
| AT5G62575.1 | Succinate dehydrogenase subunit (SDH7-2) | 40 | 3.17 | 0.170 | 0.106 | |
| AT5G62575.1 | Succinate dehydrogenase subunit (SDH7-2) | 41 | 3.18 | 0.171 | 0.113 | |
| AT5G62575.1 | Succinate dehydrogenase subunit (SDH7-2) | 42 | 2.16 | 0.174 | 0.795 | |
| AT5G62575.1 | Succinate dehydrogenase subunit (SDH7-2) | 43 | 1.16 | 0.175 | 1.000 | |
| AT5G62575.1 | Succinate dehydrogenase subunit (SDH7-2) | 44 | 0.55 | 0.175 | 0.981 | |
| AT5G62575.1 | Succinate dehydrogenase subunit (SDH7-2) | 45 | 0.30 | 0.174 | 0.094 | . |
| AT5G62575.1 | Succinate dehydrogenase subunit (SDH7-2) | 46 | 0.23 | 0.200 | 0.051 | . |
| AT5G62575.1 | Succinate dehydrogenase subunit (SDH7-2) | 47 | 0.22 | 0.386 | 0.914 | |
| AT5G63000.1 | Translocase of the inner mitochondrial membrane Tim17/Tim22/Tim23 family protein | 2 | 0.58 | 0.337 | 0.999 | |
| AT5G63000.1 | Translocase of the inner mitochondrial membrane Tim17/Tim22/Tim23 family protein | 3 | 0.60 | 0.295 | 0.998 | |
| AT5G63000.1 | Translocase of the inner mitochondrial membrane Tim17/Tim22/Tim23 family protein | 9 | 0.89 | 0.155 | 1.000 | |
| AT5G63000.1 | Translocase of the inner mitochondrial membrane Tim17/Tim22/Tim23 family protein | 10 | 1.01 | 0.138 | 1.000 | |
| AT5G63000.1 | Translocase of the inner mitochondrial membrane Tim17/Tim22/Tim23 family protein | 11 | 1.17 | 0.125 | 1.000 | |
| AT5G63000.1 | Translocase of the inner mitochondrial membrane Tim17/Tim22/Tim23 family protein | 12 | 1.37 | 0.116 | 0.930 | |
| AT5G63000.1 | Translocase of the inner mitochondrial membrane Tim17/Tim22/Tim23 family protein | 13 | 1.60 | 0.110 | 0.505 | |

| | | | | | | |
|---|---|---|---|---|---|---|
| AT5G63000.1 | Translocase of the inner mitochondrial membrane Tim17/Tim22/Tim23 family protein | 14 | 1.82 | 0.105 | 0.162 | |
| AT5G63000.1 | Translocase of the inner mitochondrial membrane Tim17/Tim22/Tim23 family protein | 15 | 1.99 | 0.103 | 0.053 | . |
| AT5G63000.1 | Translocase of the inner mitochondrial membrane Tim17/Tim22/Tim23 family protein | 16 | 2.11 | 0.103 | 0.026 | * |
| AT5G63000.1 | Translocase of the inner mitochondrial membrane Tim17/Tim22/Tim23 family protein | 17 | 2.17 | 0.105 | 0.021 | * |
| AT5G63000.1 | Translocase of the inner mitochondrial membrane Tim17/Tim22/Tim23 family protein | 18 | 2.20 | 0.110 | 0.029 | * |
| AT5G63000.1 | Translocase of the inner mitochondrial membrane Tim17/Tim22/Tim23 family protein | 19 | 2.18 | 0.117 | 0.053 | . |
| AT5G63000.1 | Translocase of the inner mitochondrial membrane Tim17/Tim22/Tim23 family protein | 20 | 2.14 | 0.126 | 0.109 | |
| AT5G63000.1 | Translocase of the inner mitochondrial membrane Tim17/Tim22/Tim23 family protein | 21 | 2.09 | 0.134 | 0.193 | |
| AT5G63000.1 | Translocase of the inner mitochondrial membrane Tim17/Tim22/Tim23 family protein | 25 | 1.89 | 0.133 | 0.360 | |
| AT5G63000.1 | Translocase of the inner mitochondrial membrane Tim17/Tim22/Tim23 family protein | 26 | 1.86 | 0.129 | 0.346 | |
| AT5G63000.1 | Translocase of the inner mitochondrial membrane Tim17/Tim22/Tim23 family protein | 27 | 1.82 | 0.129 | 0.396 | |
| AT5G63000.1 | Translocase of the inner mitochondrial membrane Tim17/Tim22/Tim23 family protein | 28 | 1.71 | 0.132 | 0.573 | |
| AT5G63000.1 | Translocase of the inner mitochondrial membrane Tim17/Tim22/Tim23 family protein | 29 | 1.57 | 0.136 | 0.802 | |
| AT5G63000.1 | Translocase of the inner mitochondrial membrane Tim17/Tim22/Tim23 family protein | 32 | 1.29 | 0.144 | 0.997 | |
| AT5G63000.1 | Translocase of the inner mitochondrial membrane Tim17/Tim22/Tim23 family protein | 33 | 1.29 | 0.145 | 0.997 | |

| | | | | | | |
|---|---|---|---|---|---|---|
| AT5G63000.1 | Translocase of the inner mitochondrial membrane Tim17/Tim22/Tim23 family protein | 34 | 1.34 | 0.147 | 0.993 | |
| AT5G63000.1 | Translocase of the inner mitochondrial membrane Tim17/Tim22/Tim23 family protein | 35 | 1.42 | 0.149 | 0.974 | |
| AT5G63000.1 | Translocase of the inner mitochondrial membrane Tim17/Tim22/Tim23 family protein | 38 | 1.65 | 0.150 | 0.801 | |
| AT5G63000.1 | Translocase of the inner mitochondrial membrane Tim17/Tim22/Tim23 family protein | 39 | 1.61 | 0.150 | 0.841 | |
| AT5G63000.1 | Translocase of the inner mitochondrial membrane Tim17/Tim22/Tim23 family protein | 40 | 1.48 | 0.155 | 0.957 | |
| AT5G63000.1 | Translocase of the inner mitochondrial membrane Tim17/Tim22/Tim23 family protein | 41 | 1.27 | 0.163 | 1.000 | |
| AT5G63000.1 | Translocase of the inner mitochondrial membrane Tim17/Tim22/Tim23 family protein | 42 | 1.04 | 0.174 | 1.000 | |
| AT5G63000.1 | Translocase of the inner mitochondrial membrane Tim17/Tim22/Tim23 family protein | 46 | 0.35 | 0.226 | 0.385 | |
| AT5G63000.1 | Translocase of the inner mitochondrial membrane Tim17/Tim22/Tim23 family protein | 47 | 0.26 | 0.257 | 0.228 | |
| AT5G63400.1 | Adenylate kinase 1 (ADK1) | 2 | 0.76 | 0.391 | 1.000 | |
| AT5G63400.1 | Adenylate kinase 1 (ADK1) | 3 | 0.67 | 0.330 | 1.000 | |
| AT5G63400.1 | Adenylate kinase 1 (ADK1) | 4 | 0.60 | 0.349 | 1.000 | |
| AT5G63400.1 | Adenylate kinase 1 (ADK1) | 9 | 0.67 | 0.278 | 1.000 | |
| AT5G63400.1 | Adenylate kinase 1 (ADK1) | 10 | 0.86 | 0.243 | 1.000 | |
| AT5G63400.1 | Adenylate kinase 1 (ADK1) | 11 | 1.16 | 0.224 | 1.000 | |
| AT5G63400.1 | Adenylate kinase 1 (ADK1) | 12 | 1.55 | 0.213 | 1.000 | |
| AT5G63400.1 | Adenylate kinase 1 (ADK1) | 13 | 1.94 | 0.204 | 0.944 | |
| AT5G63400.1 | Adenylate kinase 1 (ADK1) | 14 | 2.31 | 0.199 | 0.697 | |
| AT5G63400.1 | Adenylate kinase 1 (ADK1) | 15 | 2.73 | 0.197 | 0.394 | |
| AT5G63400.1 | Adenylate kinase 1 (ADK1) | 16 | 3.12 | 0.196 | 0.213 | |
| AT5G63400.1 | Adenylate kinase 1 (ADK1) | 17 | 3.42 | 0.196 | 0.133 | |
| AT5G63400.1 | Adenylate kinase 1 (ADK1) | 18 | 3.77 | 0.197 | 0.080 | . |
| AT5G63400.1 | Adenylate kinase 1 (ADK1) | 19 | 4.03 | 0.201 | 0.063 | . |
| AT5G63400.1 | Adenylate kinase 1 (ADK1) | 20 | 3.65 | 0.206 | 0.133 | |
| AT5G63400.1 | Adenylate kinase 1 (ADK1) | 21 | 2.66 | 0.210 | 0.541 | |
| AT5G63400.1 | Adenylate kinase 1 (ADK1) | 22 | 1.72 | 0.214 | 0.994 | |
| AT5G63400.1 | Adenylate kinase 1 (ADK1) | 23 | 1.02 | 0.220 | 1.000 | |
| AT5G63400.1 | Adenylate kinase 1 (ADK1) | 24 | 0.56 | 0.229 | 0.994 | |
| AT5G63400.1 | Adenylate kinase 1 (ADK1) | 25 | 0.32 | 0.236 | 0.492 | |
| AT5G63400.1 | Adenylate kinase 1 (ADK1) | 26 | 0.22 | 0.228 | 0.097 | . |
| AT5G63400.1 | Adenylate kinase 1 (ADK1) | 27 | 0.22 | 0.217 | 0.054 | . |
| AT5G63400.1 | Adenylate kinase 1 (ADK1) | 28 | 0.30 | 0.208 | 0.219 | |
| AT5G63400.1 | Adenylate kinase 1 (ADK1) | 29 | 0.53 | 0.203 | 0.953 | |
| AT5G63400.1 | Adenylate kinase 1 (ADK1) | 30 | 0.98 | 0.202 | 1.000 | |
| AT5G63400.1 | Adenylate kinase 1 (ADK1) | 31 | 1.66 | 0.201 | 0.995 | |

| | | | | | |
|---|---|---|---|---|---|
| AT5G63400.1 | Adenylate kinase 1 (ADK1) | 32 | 2.34 | 0.201 | 0.691 |
| AT5G63400.1 | Adenylate kinase 1 (ADK1) | 33 | 2.51 | 0.201 | 0.572 |
| AT5G63400.1 | Adenylate kinase 1 (ADK1) | 34 | 2.07 | 0.201 | 0.872 |
| AT5G63400.1 | Adenylate kinase 1 (ADK1) | 35 | 1.42 | 0.201 | 1.000 |
| AT5G63400.1 | Adenylate kinase 1 (ADK1) | 36 | 0.93 | 0.201 | 1.000 |
| AT5G63400.1 | Adenylate kinase 1 (ADK1) | 37 | 0.72 | 0.201 | 1.000 |
| AT5G63400.1 | Adenylate kinase 1 (ADK1) | 38 | 0.77 | 0.201 | 1.000 |
| AT5G63400.1 | Adenylate kinase 1 (ADK1) | 39 | 1.10 | 0.201 | 1.000 |
| AT5G63400.1 | Adenylate kinase 1 (ADK1) | 40 | 1.65 | 0.201 | 0.995 |
| AT5G63400.1 | Adenylate kinase 1 (ADK1) | 41 | 2.18 | 0.202 | 0.810 |
| AT5G63400.1 | Adenylate kinase 1 (ADK1) | 42 | 2.33 | 0.203 | 0.714 |
| AT5G63400.1 | Adenylate kinase 1 (ADK1) | 43 | 2.03 | 0.206 | 0.909 |
| AT5G63400.1 | Adenylate kinase 1 (ADK1) | 44 | 1.43 | 0.211 | 1.000 |
| AT5G63400.1 | Adenylate kinase 1 (ADK1) | 45 | 0.81 | 0.222 | 1.000 |
| AT5G63400.1 | Adenylate kinase 1 (ADK1) | 46 | 0.39 | 0.257 | 0.854 |
| AT5G63400.1 | Adenylate kinase 1 (ADK1) | 47 | 0.17 | 0.357 | 0.446 |
| AT5G63510.1 | NADH:ubiquinone oxidoreductase subunit (gamma CAL1) | 2 | 0.69 | 0.355 | 1.000 |
| AT5G63510.1 | NADH:ubiquinone oxidoreductase subunit (gamma CAL1) | 3 | 0.69 | 0.209 | 1.000 |
| AT5G63510.1 | NADH:ubiquinone oxidoreductase subunit (gamma CAL1) | 4 | 0.77 | 0.201 | 1.000 |
| AT5G63510.1 | NADH:ubiquinone oxidoreductase subunit (gamma CAL1) | 5 | 1.01 | 0.200 | 1.000 |
| AT5G63510.1 | NADH:ubiquinone oxidoreductase subunit (gamma CAL1) | 6 | 1.23 | 0.201 | 1.000 |
| AT5G63510.1 | NADH:ubiquinone oxidoreductase subunit (gamma CAL1) | 7 | 1.29 | 0.202 | 1.000 |
| AT5G63510.1 | NADH:ubiquinone oxidoreductase subunit (gamma CAL1) | 8 | 1.39 | 0.196 | 1.000 |
| AT5G63510.1 | NADH:ubiquinone oxidoreductase subunit (gamma CAL1) | 9 | 1.32 | 0.193 | 1.000 |
| AT5G63510.1 | NADH:ubiquinone oxidoreductase subunit (gamma CAL1) | 10 | 1.11 | 0.193 | 1.000 |
| AT5G63510.1 | NADH:ubiquinone oxidoreductase subunit (gamma CAL1) | 11 | 1.10 | 0.195 | 1.000 |
| AT5G63510.1 | NADH:ubiquinone oxidoreductase subunit (gamma CAL1) | 12 | 1.16 | 0.196 | 1.000 |
| AT5G63510.1 | NADH:ubiquinone oxidoreductase subunit (gamma CAL1) | 13 | 1.23 | 0.193 | 1.000 |
| AT5G63510.1 | NADH:ubiquinone oxidoreductase subunit (gamma CAL1) | 14 | 1.37 | 0.190 | 1.000 |
| AT5G63510.1 | NADH:ubiquinone oxidoreductase subunit (gamma CAL1) | 15 | 1.12 | 0.191 | 1.000 |
| AT5G63510.1 | NADH:ubiquinone oxidoreductase subunit (gamma CAL1) | 16 | 0.63 | 0.189 | 1.000 |

| | | | | | |
|---|---|---|---|---|---|
| AT5G63510.1 | NADH:ubiquinone oxidoreductase subunit (gamma CAL1) | 17 | 0.41 | 0.187 | 0.732 |
| AT5G63510.1 | NADH:ubiquinone oxidoreductase subunit (gamma CAL1) | 18 | 0.48 | 0.188 | 0.949 |
| AT5G63510.1 | NADH:ubiquinone oxidoreductase subunit (gamma CAL1) | 19 | 0.72 | 0.195 | 1.000 |
| AT5G63510.1 | NADH:ubiquinone oxidoreductase subunit (gamma CAL1) | 20 | 0.85 | 0.201 | 1.000 |
| AT5G63510.1 | NADH:ubiquinone oxidoreductase subunit (gamma CAL1) | 21 | 1.00 | 0.202 | 1.000 |
| AT5G63510.1 | NADH:ubiquinone oxidoreductase subunit (gamma CAL1) | 22 | 1.36 | 0.202 | 1.000 |
| AT5G63510.1 | NADH:ubiquinone oxidoreductase subunit (gamma CAL1) | 23 | 1.76 | 0.201 | 0.999 |
| AT5G63510.1 | NADH:ubiquinone oxidoreductase subunit (gamma CAL1) | 24 | 1.49 | 0.195 | 1.000 |
| AT5G63510.1 | NADH:ubiquinone oxidoreductase subunit (gamma CAL1) | 25 | 0.85 | 0.193 | 1.000 |
| AT5G63510.1 | NADH:ubiquinone oxidoreductase subunit (gamma CAL1) | 26 | 0.64 | 0.193 | 1.000 |
| AT5G63510.1 | NADH:ubiquinone oxidoreductase subunit (gamma CAL1) | 27 | 0.82 | 0.195 | 1.000 |
| AT5G63510.1 | NADH:ubiquinone oxidoreductase subunit (gamma CAL1) | 28 | 0.98 | 0.195 | 1.000 |
| AT5G63510.1 | NADH:ubiquinone oxidoreductase subunit (gamma CAL1) | 29 | 0.93 | 0.194 | 1.000 |
| AT5G63510.1 | NADH:ubiquinone oxidoreductase subunit (gamma CAL1) | 30 | 0.94 | 0.194 | 1.000 |
| AT5G63510.1 | NADH:ubiquinone oxidoreductase subunit (gamma CAL1) | 31 | 1.10 | 0.194 | 1.000 |
| AT5G63510.1 | NADH:ubiquinone oxidoreductase subunit (gamma CAL1) | 32 | 1.14 | 0.195 | 1.000 |
| AT5G63510.1 | NADH:ubiquinone oxidoreductase subunit (gamma CAL1) | 33 | 0.77 | 0.194 | 1.000 |
| AT5G63510.1 | NADH:ubiquinone oxidoreductase subunit (gamma CAL1) | 34 | 0.49 | 0.194 | 0.974 |
| AT5G63510.1 | NADH:ubiquinone oxidoreductase subunit (gamma CAL1) | 35 | 0.53 | 0.194 | 0.994 |
| AT5G63510.1 | NADH:ubiquinone oxidoreductase subunit (gamma CAL1) | 36 | 0.86 | 0.193 | 1.000 |

| | | | | | | |
|---|---|---|---|---|---|---|
| AT5G63510.1 | NADH:ubiquinone oxidoreductase subunit (gamma CAL1) | 37 | 1.46 | 0.193 | 1.000 | |
| AT5G63510.1 | NADH:ubiquinone oxidoreductase subunit (gamma CAL1) | 38 | 1.88 | 0.194 | 0.995 | |
| AT5G63510.1 | NADH:ubiquinone oxidoreductase subunit (gamma CAL1) | 39 | 1.26 | 0.196 | 1.000 | |
| AT5G63510.1 | NADH:ubiquinone oxidoreductase subunit (gamma CAL1) | 40 | 0.53 | 0.196 | 0.995 | |
| AT5G63510.1 | NADH:ubiquinone oxidoreductase subunit (gamma CAL1) | 41 | 0.23 | 0.203 | 0.071 | . |
| AT5G63510.1 | NADH:ubiquinone oxidoreductase subunit (gamma CAL1) | 42 | 0.23 | 0.234 | 0.233 | |
| AT5G63510.1 | NADH:ubiquinone oxidoreductase subunit (gamma CAL1) | 43 | 0.60 | 0.237 | 1.000 | |
| AT5G63510.1 | NADH:ubiquinone oxidoreductase subunit (gamma CAL1) | 44 | 2.03 | 0.281 | 1.000 | |
| AT5G63510.1 | NADH:ubiquinone oxidoreductase subunit (gamma CAL1) | 45 | 5.34 | 0.387 | 0.870 | |
| AT5G63510.1 | NADH:ubiquinone oxidoreductase subunit (gamma CAL1) | 46 | 11.28 | 0.651 | 0.971 | |
| AT5G63510.1 | NADH:ubiquinone oxidoreductase subunit (gamma CAL1) | 47 | 21.65 | 1.026 | 0.999 | |
| AT5G64040.1 | Photosystem I subunit (PSAN) | 2 | 0.73 | 0.275 | 1.000 | |
| AT5G64040.1 | Photosystem I subunit (PSAN) | 3 | 0.77 | 0.211 | 1.000 | |
| AT5G64040.1 | Photosystem I subunit (PSAN) | 4 | 0.82 | 0.177 | 1.000 | |
| AT5G64040.1 | Photosystem I subunit (PSAN) | 5 | 0.88 | 0.164 | 1.000 | |
| AT5G64040.1 | Photosystem I subunit (PSAN) | 6 | 0.94 | 0.161 | 1.000 | |
| AT5G64040.1 | Photosystem I subunit (PSAN) | 7 | 0.99 | 0.160 | 1.000 | |
| AT5G64040.1 | Photosystem I subunit (PSAN) | 8 | 1.04 | 0.159 | 1.000 | |
| AT5G64040.1 | Photosystem I subunit (PSAN) | 9 | 1.09 | 0.158 | 1.000 | |
| AT5G64040.1 | Photosystem I subunit (PSAN) | 10 | 1.15 | 0.157 | 1.000 | |
| AT5G64040.1 | Photosystem I subunit (PSAN) | 11 | 1.20 | 0.156 | 1.000 | |
| AT5G64040.1 | Photosystem I subunit (PSAN) | 12 | 1.22 | 0.155 | 1.000 | |
| AT5G64040.1 | Photosystem I subunit (PSAN) | 13 | 1.20 | 0.154 | 1.000 | |
| AT5G64040.1 | Photosystem I subunit (PSAN) | 14 | 1.15 | 0.154 | 1.000 | |
| AT5G64040.1 | Photosystem I subunit (PSAN) | 15 | 1.12 | 0.153 | 1.000 | |
| AT5G64040.1 | Photosystem I subunit (PSAN) | 16 | 1.12 | 0.153 | 1.000 | |
| AT5G64040.1 | Photosystem I subunit (PSAN) | 17 | 1.17 | 0.153 | 1.000 | |
| AT5G64040.1 | Photosystem I subunit (PSAN) | 18 | 1.25 | 0.154 | 1.000 | |
| AT5G64040.1 | Photosystem I subunit (PSAN) | 19 | 1.32 | 0.156 | 0.998 | |
| AT5G64040.1 | Photosystem I subunit (PSAN) | 20 | 1.33 | 0.158 | 0.998 | |
| AT5G64040.1 | Photosystem I subunit (PSAN) | 21 | 1.24 | 0.159 | 1.000 | |
| AT5G64040.1 | Photosystem I subunit (PSAN) | 22 | 1.12 | 0.159 | 1.000 | |
| AT5G64040.1 | Photosystem I subunit (PSAN) | 23 | 1.04 | 0.158 | 1.000 | |
| AT5G64040.1 | Photosystem I subunit (PSAN) | 24 | 1.03 | 0.158 | 1.000 | |
| AT5G64040.1 | Photosystem I subunit (PSAN) | 25 | 1.11 | 0.157 | 1.000 | |
| AT5G64040.1 | Photosystem I subunit (PSAN) | 26 | 1.33 | 0.158 | 0.998 | |
| AT5G64040.1 | Photosystem I subunit (PSAN) | 27 | 1.74 | 0.162 | 0.807 | |
| AT5G64040.1 | Photosystem I subunit (PSAN) | 28 | 2.30 | 0.171 | 0.359 | |
| AT5G64040.1 | Photosystem I subunit (PSAN) | 29 | 2.99 | 0.188 | 0.148 | |
| AT5G64040.1 | Photosystem I subunit (PSAN) | 30 | 3.85 | 0.213 | 0.089 | . |

| | | | | | | |
|---|---|---|---|---|---|---|
| AT5G64040.1 | Photosystem I subunit (PSAN) | 31 | 4.80 | 0.251 | 0.095 | . |
| AT5G64040.1 | Photosystem I subunit (PSAN) | 45 | 1.49 | 0.318 | 1.000 | |
| AT5G64040.1 | Photosystem I subunit (PSAN) | 46 | 1.13 | 0.304 | 1.000 | |
| AT5G64040.1 | Photosystem I subunit (PSAN) | 47 | 0.85 | 0.348 | 1.000 | |
| AT5G64050.1 | Glutamate tRNA synthetase (OVA3) | 2 | 1.16 | 0.604 | 1.000 | |
| AT5G64050.1 | Glutamate tRNA synthetase (OVA3) | 3 | 1.21 | 0.579 | 1.000 | |
| AT5G64050.1 | Glutamate tRNA synthetase (OVA3) | 32 | 3.06 | 0.407 | 0.895 | |
| AT5G64050.1 | Glutamate tRNA synthetase (OVA3) | 33 | 2.97 | 0.332 | 0.769 | |
| AT5G64050.1 | Glutamate tRNA synthetase (OVA3) | 34 | 2.73 | 0.303 | 0.758 | |
| AT5G64050.1 | Glutamate tRNA synthetase (OVA3) | 35 | 2.44 | 0.297 | 0.842 | |
| AT5G64050.1 | Glutamate tRNA synthetase (OVA3) | 36 | 2.13 | 0.298 | 0.931 | |
| AT5G64050.1 | Glutamate tRNA synthetase (OVA3) | 37 | 1.80 | 0.287 | 0.981 | |
| AT5G64050.1 | Glutamate tRNA synthetase (OVA3) | 38 | 1.43 | 0.270 | 0.999 | |
| AT5G64050.1 | Glutamate tRNA synthetase (OVA3) | 39 | 1.12 | 0.252 | 1.000 | |
| AT5G64050.1 | Glutamate tRNA synthetase (OVA3) | 40 | 0.90 | 0.237 | 1.000 | |
| AT5G64050.1 | Glutamate tRNA synthetase (OVA3) | 41 | 0.72 | 0.230 | 0.999 | |
| AT5G64050.1 | Glutamate tRNA synthetase (OVA3) | 42 | 0.56 | 0.231 | 0.936 | |
| AT5G64050.1 | Glutamate tRNA synthetase (OVA3) | 43 | 0.47 | 0.231 | 0.769 | |
| AT5G64050.1 | Glutamate tRNA synthetase (OVA3) | 44 | 0.44 | 0.230 | 0.676 | |
| AT5G64050.1 | Glutamate tRNA synthetase (OVA3) | 45 | 0.45 | 0.238 | 0.758 | |
| AT5G64050.1 | Glutamate tRNA synthetase (OVA3) | 46 | 0.46 | 0.301 | 0.928 | |
| AT5G64050.1 | Glutamate tRNA synthetase (OVA3) | 47 | 0.46 | 0.466 | 0.995 | |
| AT5G64290.1 | Dicarboxylate transport 2.1 (DCT / DIT2.1) | 2 | 1.09 | 0.590 | 1.000 | |
| AT5G64290.1 | Dicarboxylate transport 2.1 (DCT / DIT2.1) | 3 | 1.19 | 0.573 | 1.000 | |
| AT5G64290.1 | Dicarboxylate transport 2.1 (DCT / DIT2.1) | 33 | 2.60 | 0.245 | 0.323 | |
| AT5G64290.1 | Dicarboxylate transport 2.1 (DCT / DIT2.1) | 34 | 2.33 | 0.228 | 0.366 | |
| AT5G64290.1 | Dicarboxylate transport 2.1 (DCT / DIT2.1) | 35 | 2.05 | 0.221 | 0.491 | |
| AT5G64290.1 | Dicarboxylate transport 2.1 (DCT / DIT2.1) | 36 | 1.75 | 0.223 | 0.705 | |
| AT5G64290.1 | Dicarboxylate transport 2.1 (DCT / DIT2.1) | 37 | 1.46 | 0.233 | 0.918 | |
| AT5G64290.1 | Dicarboxylate transport 2.1 (DCT / DIT2.1) | 38 | 1.19 | 0.246 | 0.997 | |
| AT5G64290.1 | Dicarboxylate transport 2.1 (DCT / DIT2.1) | 46 | 0.16 | 0.366 | 0.125 | |
| AT5G64290.1 | Dicarboxylate transport 2.1 (DCT / DIT2.1) | 47 | 0.12 | 0.396 | 0.092 | . |
| AT5G64670.1 | Ribosomal L15 family protein | 2 | 1.00 | 0.647 | 1.000 | |
| AT5G64670.1 | Ribosomal L15 family protein | 3 | 0.87 | 0.572 | 1.000 | |

| | | | | | | |
|---|---|---|---|---|---|---|
| AT5G64670.1 | Ribosomal L15 family protein | 11 | 0.67 | 0.315 | 1.000 | |
| AT5G64670.1 | Ribosomal L15 family protein | 12 | 0.80 | 0.276 | 1.000 | |
| AT5G64670.1 | Ribosomal L15 family protein | 13 | 1.00 | 0.247 | 1.000 | |
| AT5G64670.1 | Ribosomal L15 family protein | 14 | 1.32 | 0.226 | 1.000 | |
| AT5G64670.1 | Ribosomal L15 family protein | 15 | 1.80 | 0.212 | 0.965 | |
| AT5G64670.1 | Ribosomal L15 family protein | 16 | 2.44 | 0.203 | 0.569 | |
| AT5G64670.1 | Ribosomal L15 family protein | 17 | 3.00 | 0.201 | 0.246 | |
| AT5G64670.1 | Ribosomal L15 family protein | 18 | 2.98 | 0.203 | 0.270 | |
| AT5G64670.1 | Ribosomal L15 family protein | 19 | 2.29 | 0.210 | 0.716 | |
| AT5G64670.1 | Ribosomal L15 family protein | 20 | 1.46 | 0.223 | 1.000 | |
| AT5G64670.1 | Ribosomal L15 family protein | 21 | 0.85 | 0.242 | 1.000 | |
| AT5G64670.1 | Ribosomal L15 family protein | 22 | 0.50 | 0.261 | 0.977 | |
| AT5G64670.1 | Ribosomal L15 family protein | 23 | 0.32 | 0.274 | 0.644 | |
| AT5G64670.1 | Ribosomal L15 family protein | 24 | 0.23 | 0.280 | 0.284 | |
| AT5G64670.1 | Ribosomal L15 family protein | 26 | 0.15 | 0.255 | 0.030 | * |
| AT5G64670.1 | Ribosomal L15 family protein | 27 | 0.15 | 0.237 | 0.011 | * |
| AT5G64670.1 | Ribosomal L15 family protein | 28 | 0.15 | 0.221 | 0.006 | ** |
| AT5G64670.1 | Ribosomal L15 family protein | 29 | 0.17 | 0.212 | 0.007 | ** |
| AT5G64670.1 | Ribosomal L15 family protein | 30 | 0.20 | 0.208 | 0.018 | * |
| AT5G64670.1 | Ribosomal L15 family protein | 31 | 0.25 | 0.208 | 0.064 | . |
| AT5G64670.1 | Ribosomal L15 family protein | 32 | 0.31 | 0.212 | 0.237 | |
| AT5G64670.1 | Ribosomal L15 family protein | 33 | 0.40 | 0.222 | 0.639 | |
| AT5G64670.1 | Ribosomal L15 family protein | 34 | 0.50 | 0.240 | 0.951 | |
| AT5G64670.1 | Ribosomal L15 family protein | 35 | 0.60 | 0.263 | 0.999 | |
| AT5G64670.1 | Ribosomal L15 family protein | 36 | 0.71 | 0.290 | 1.000 | |
| AT5G64670.1 | Ribosomal L15 family protein | 37 | 0.83 | 0.308 | 1.000 | |
| AT5G64670.1 | Ribosomal L15 family protein | 41 | 1.24 | 0.285 | 1.000 | |
| AT5G64670.1 | Ribosomal L15 family protein | 42 | 1.21 | 0.258 | 1.000 | |
| AT5G64670.1 | Ribosomal L15 family protein | 43 | 1.05 | 0.239 | 1.000 | |
| AT5G64670.1 | Ribosomal L15 family protein | 44 | 0.79 | 0.235 | 1.000 | |
| AT5G64670.1 | Ribosomal L15 family protein | 45 | 0.53 | 0.251 | 0.983 | |
| AT5G64670.1 | Ribosomal L15 family protein | 46 | 0.35 | 0.301 | 0.842 | |
| AT5G64670.1 | Ribosomal L15 family protein | 47 | 0.23 | 0.401 | 0.794 | |
| AT5G65720.1 | Nitrogen fixation S (NIFS)-like 1 (NFS1) | 2 | 0.83 | 0.322 | 1.000 | |
| AT5G65720.1 | Nitrogen fixation S (NIFS)-like 1 (NFS1) | 3 | 0.69 | 0.256 | 1.000 | |
| AT5G65720.1 | Nitrogen fixation S (NIFS)-like 1 (NFS1) | 4 | 0.61 | 0.247 | 1.000 | |
| AT5G65720.1 | Nitrogen fixation S (NIFS)-like 1 (NFS1) | 9 | 0.78 | 0.221 | 1.000 | |
| AT5G65720.1 | Nitrogen fixation S (NIFS)-like 1 (NFS1) | 10 | 0.94 | 0.191 | 1.000 | |
| AT5G65720.1 | Nitrogen fixation S (NIFS)-like 1 (NFS1) | 11 | 1.09 | 0.170 | 1.000 | |
| AT5G65720.1 | Nitrogen fixation S (NIFS)-like 1 (NFS1) | 12 | 1.19 | 0.155 | 1.000 | |
| AT5G65720.1 | Nitrogen fixation S (NIFS)-like 1 (NFS1) | 13 | 1.21 | 0.148 | 1.000 | |
| AT5G65720.1 | Nitrogen fixation S (NIFS)-like 1 (NFS1) | 14 | 1.17 | 0.145 | 1.000 | |
| AT5G65720.1 | Nitrogen fixation S (NIFS)-like 1 (NFS1) | 15 | 1.15 | 0.145 | 1.000 | |
| AT5G65720.1 | Nitrogen fixation S (NIFS)-like 1 (NFS1) | 16 | 1.25 | 0.145 | 1.000 | |
| AT5G65720.1 | Nitrogen fixation S (NIFS)-like 1 (NFS1) | 17 | 1.49 | 0.148 | 0.993 | |
| AT5G65720.1 | Nitrogen fixation S (NIFS)-like 1 (NFS1) | 18 | 1.85 | 0.151 | 0.793 | |
| AT5G65720.1 | Nitrogen fixation S (NIFS)-like 1 (NFS1) | 19 | 2.21 | 0.153 | 0.422 | |

| | | | | | |
|---|---|---|---|---|---|
| AT5G65720.1 | Nitrogen fixation S (NIFS)-like 1 (NFS1) | 20 | 2.38 | 0.153 | 0.280 |
| AT5G65720.1 | Nitrogen fixation S (NIFS)-like 1 (NFS1) | 21 | 2.27 | 0.152 | 0.357 |
| AT5G65720.1 | Nitrogen fixation S (NIFS)-like 1 (NFS1) | 22 | 1.96 | 0.150 | 0.667 |
| AT5G65720.1 | Nitrogen fixation S (NIFS)-like 1 (NFS1) | 23 | 1.67 | 0.148 | 0.930 |
| AT5G65720.1 | Nitrogen fixation S (NIFS)-like 1 (NFS1) | 24 | 1.56 | 0.146 | 0.977 |
| AT5G65720.1 | Nitrogen fixation S (NIFS)-like 1 (NFS1) | 25 | 1.53 | 0.144 | 0.982 |
| AT5G65720.1 | Nitrogen fixation S (NIFS)-like 1 (NFS1) | 26 | 1.49 | 0.143 | 0.991 |
| AT5G65720.1 | Nitrogen fixation S (NIFS)-like 1 (NFS1) | 27 | 1.38 | 0.142 | 0.999 |
| AT5G65720.1 | Nitrogen fixation S (NIFS)-like 1 (NFS1) | 28 | 1.26 | 0.142 | 1.000 |
| AT5G65720.1 | Nitrogen fixation S (NIFS)-like 1 (NFS1) | 29 | 1.17 | 0.143 | 1.000 |
| AT5G65720.1 | Nitrogen fixation S (NIFS)-like 1 (NFS1) | 30 | 1.12 | 0.143 | 1.000 |
| AT5G65720.1 | Nitrogen fixation S (NIFS)-like 1 (NFS1) | 31 | 1.10 | 0.144 | 1.000 |
| AT5G65720.1 | Nitrogen fixation S (NIFS)-like 1 (NFS1) | 32 | 1.10 | 0.144 | 1.000 |
| AT5G65720.1 | Nitrogen fixation S (NIFS)-like 1 (NFS1) | 33 | 1.10 | 0.144 | 1.000 |
| AT5G65720.1 | Nitrogen fixation S (NIFS)-like 1 (NFS1) | 34 | 1.11 | 0.144 | 1.000 |
| AT5G65720.1 | Nitrogen fixation S (NIFS)-like 1 (NFS1) | 35 | 1.12 | 0.144 | 1.000 |
| AT5G65720.1 | Nitrogen fixation S (NIFS)-like 1 (NFS1) | 36 | 1.12 | 0.144 | 1.000 |
| AT5G65720.1 | Nitrogen fixation S (NIFS)-like 1 (NFS1) | 37 | 1.12 | 0.143 | 1.000 |
| AT5G65720.1 | Nitrogen fixation S (NIFS)-like 1 (NFS1) | 38 | 1.11 | 0.143 | 1.000 |
| AT5G65720.1 | Nitrogen fixation S (NIFS)-like 1 (NFS1) | 39 | 1.08 | 0.144 | 1.000 |
| AT5G65720.1 | Nitrogen fixation S (NIFS)-like 1 (NFS1) | 40 | 1.05 | 0.145 | 1.000 |
| AT5G65720.1 | Nitrogen fixation S (NIFS)-like 1 (NFS1) | 41 | 1.01 | 0.146 | 1.000 |
| AT5G65720.1 | Nitrogen fixation S (NIFS)-like 1 (NFS1) | 42 | 0.95 | 0.148 | 1.000 |
| AT5G65720.1 | Nitrogen fixation S (NIFS)-like 1 (NFS1) | 43 | 0.86 | 0.149 | 1.000 |
| AT5G65720.1 | Nitrogen fixation S (NIFS)-like 1 (NFS1) | 44 | 0.71 | 0.149 | 0.999 |
| AT5G65720.1 | Nitrogen fixation S (NIFS)-like 1 (NFS1) | 45 | 0.53 | 0.160 | 0.829 |
| AT5G65720.1 | Nitrogen fixation S (NIFS)-like 1 (NFS1) | 46 | 0.37 | 0.206 | 0.528 |
| AT5G65720.1 | Nitrogen fixation S (NIFS)-like 1 (NFS1) | 47 | 0.24 | 0.306 | 0.602 |
| AT5G65750.1 | 2-oxoglutarate dehydrogenase subunit (OGDCE1) | 2 | 1.04 | 0.648 | 1.000 |
| AT5G65750.1 | 2-oxoglutarate dehydrogenase subunit (OGDCE1) | 3 | 0.80 | 0.460 | 1.000 |
| AT5G65750.1 | 2-oxoglutarate dehydrogenase subunit (OGDCE1) | 4 | 0.63 | 0.481 | 1.000 |

| | | | | | |
|---|---|---|---|---|---|
| AT5G65750.1 | 2-oxoglutarate dehydrogenase subunit (OGDCE1) | 6 | 0.58 | 0.518 | 1.000 |
| AT5G65750.1 | 2-oxoglutarate dehydrogenase subunit (OGDCE1) | 7 | 0.71 | 0.447 | 1.000 |
| AT5G65750.1 | 2-oxoglutarate dehydrogenase subunit (OGDCE1) | 8 | 0.90 | 0.395 | 1.000 |
| AT5G65750.1 | 2-oxoglutarate dehydrogenase subunit (OGDCE1) | 9 | 1.02 | 0.359 | 1.000 |
| AT5G65750.1 | 2-oxoglutarate dehydrogenase subunit (OGDCE1) | 10 | 1.06 | 0.337 | 1.000 |
| AT5G65750.1 | 2-oxoglutarate dehydrogenase subunit (OGDCE1) | 11 | 1.03 | 0.325 | 1.000 |
| AT5G65750.1 | 2-oxoglutarate dehydrogenase subunit (OGDCE1) | 12 | 0.91 | 0.317 | 1.000 |
| AT5G65750.1 | 2-oxoglutarate dehydrogenase subunit (OGDCE1) | 13 | 0.72 | 0.311 | 1.000 |
| AT5G65750.1 | 2-oxoglutarate dehydrogenase subunit (OGDCE1) | 14 | 0.52 | 0.308 | 1.000 |
| AT5G65750.1 | 2-oxoglutarate dehydrogenase subunit (OGDCE1) | 15 | 0.46 | 0.309 | 0.999 |
| AT5G65750.1 | 2-oxoglutarate dehydrogenase subunit (OGDCE1) | 16 | 0.77 | 0.312 | 1.000 |
| AT5G65750.1 | 2-oxoglutarate dehydrogenase subunit (OGDCE1) | 17 | 1.97 | 0.319 | 1.000 |
| AT5G65750.1 | 2-oxoglutarate dehydrogenase subunit (OGDCE1) | 18 | 3.97 | 0.339 | 0.834 |
| AT5G65750.1 | 2-oxoglutarate dehydrogenase subunit (OGDCE1) | 19 | 4.13 | 0.385 | 0.921 |
| AT5G65750.1 | 2-oxoglutarate dehydrogenase subunit (OGDCE1) | 20 | 2.95 | 0.434 | 0.999 |
| AT5G65750.1 | 2-oxoglutarate dehydrogenase subunit (OGDCE1) | 22 | 1.40 | 0.456 | 1.000 |
| AT5G65750.1 | 2-oxoglutarate dehydrogenase subunit (OGDCE1) | 23 | 1.19 | 0.400 | 1.000 |
| AT5G65750.1 | 2-oxoglutarate dehydrogenase subunit (OGDCE1) | 24 | 1.07 | 0.344 | 1.000 |
| AT5G65750.1 | 2-oxoglutarate dehydrogenase subunit (OGDCE1) | 25 | 0.87 | 0.319 | 1.000 |
| AT5G65750.1 | 2-oxoglutarate dehydrogenase subunit (OGDCE1) | 26 | 0.74 | 0.312 | 1.000 |
| AT5G65750.1 | 2-oxoglutarate dehydrogenase subunit (OGDCE1) | 27 | 0.81 | 0.311 | 1.000 |
| AT5G65750.1 | 2-oxoglutarate dehydrogenase subunit (OGDCE1) | 28 | 1.06 | 0.311 | 1.000 |
| AT5G65750.1 | 2-oxoglutarate dehydrogenase subunit (OGDCE1) | 29 | 1.48 | 0.312 | 1.000 |
| AT5G65750.1 | 2-oxoglutarate dehydrogenase subunit (OGDCE1) | 30 | 1.80 | 0.312 | 1.000 |
| AT5G65750.1 | 2-oxoglutarate dehydrogenase subunit (OGDCE1) | 31 | 1.93 | 0.313 | 1.000 |
| AT5G65750.1 | 2-oxoglutarate dehydrogenase subunit (OGDCE1) | 32 | 2.13 | 0.315 | 0.999 |
| AT5G65750.1 | 2-oxoglutarate dehydrogenase subunit (OGDCE1) | 33 | 2.37 | 0.316 | 0.996 |
| AT5G65750.1 | 2-oxoglutarate dehydrogenase subunit (OGDCE1) | 34 | 2.26 | 0.319 | 0.998 |
| AT5G65750.1 | 2-oxoglutarate dehydrogenase subunit (OGDCE1) | 35 | 1.62 | 0.324 | 1.000 |
| AT5G65750.1 | 2-oxoglutarate dehydrogenase subunit (OGDCE1) | 36 | 0.98 | 0.346 | 1.000 |
| AT5G65750.1 | 2-oxoglutarate dehydrogenase subunit (OGDCE1) | 37 | 0.79 | 0.434 | 1.000 |

| | | | | | | |
|---|---|---|---|---|---|---|
| AT5G65750.1 | 2-oxoglutarate dehydrogenase subunit (OGDCE1) | 38 | 1.07 | 0.657 | 1.000 | |
| AT5G65750.1 | 2-oxoglutarate dehydrogenase subunit (OGDCE1) | 46 | 3508351.28 | 4.994 | 0.988 | |
| AT5G65750.1 | 2-oxoglutarate dehydrogenase subunit (OGDCE1) | 47 | 48802220.23 | 5.811 | 0.987 | |
| AT5G65880.1 | ? | 2 | 0.63 | 0.592 | 1.000 | |
| AT5G65880.1 | ? | 3 | 0.79 | 0.470 | 1.000 | |
| AT5G65880.1 | ? | 6 | 1.44 | 0.327 | 1.000 | |
| AT5G65880.1 | ? | 7 | 1.62 | 0.284 | 1.000 | |
| AT5G65880.1 | ? | 8 | 1.73 | 0.251 | 0.999 | |
| AT5G65880.1 | ? | 9 | 1.77 | 0.229 | 0.995 | |
| AT5G65880.1 | ? | 10 | 1.78 | 0.217 | 0.990 | |
| AT5G65880.1 | ? | 11 | 1.76 | 0.213 | 0.990 | |
| AT5G65880.1 | ? | 12 | 1.69 | 0.211 | 0.995 | |
| AT5G65880.1 | ? | 13 | 1.60 | 0.210 | 0.999 | |
| AT5G65880.1 | ? | 14 | 1.54 | 0.209 | 0.999 | |
| AT5G65880.1 | ? | 15 | 1.46 | 0.208 | 1.000 | |
| AT5G65880.1 | ? | 16 | 1.30 | 0.207 | 1.000 | |
| AT5G65880.1 | ? | 17 | 1.26 | 0.209 | 1.000 | |
| AT5G65880.1 | ? | 18 | 1.51 | 0.217 | 1.000 | |
| AT5G65880.1 | ? | 19 | 2.33 | 0.232 | 0.867 | |
| AT5G65880.1 | ? | 20 | 4.27 | 0.258 | 0.257 | |
| AT5G65880.1 | ? | 21 | 8.03 | 0.301 | 0.062 | . |
| AT5G65880.1 | ? | 22 | 13.42 | 0.360 | 0.042 | * |
| AT5G65880.1 | ? | 31 | 15.59 | 0.340 | 0.013 | * |
| AT5G65880.1 | ? | 32 | 12.28 | 0.284 | 0.004 | ** |
| AT5G65880.1 | ? | 33 | 9.54 | 0.247 | 0.002 | ** |
| AT5G65880.1 | ? | 34 | 6.73 | 0.226 | 0.007 | ** |
| AT5G65880.1 | ? | 35 | 4.31 | 0.217 | 0.076 | . |
| AT5G65880.1 | ? | 36 | 2.67 | 0.213 | 0.570 | |
| AT5G65880.1 | ? | 37 | 1.70 | 0.213 | 0.995 | |
| AT5G65880.1 | ? | 38 | 1.23 | 0.213 | 1.000 | |
| AT5G65880.1 | ? | 39 | 1.19 | 0.213 | 1.000 | |
| AT5G65880.1 | ? | 40 | 1.49 | 0.214 | 1.000 | |
| AT5G65880.1 | ? | 41 | 1.98 | 0.216 | 0.953 | |
| AT5G65880.1 | ? | 42 | 2.52 | 0.219 | 0.707 | |
| AT5G65880.1 | ? | 43 | 2.99 | 0.220 | 0.443 | |
| AT5G65880.1 | ? | 44 | 3.54 | 0.220 | 0.229 | |
| AT5G65880.1 | ? | 45 | 4.40 | 0.236 | 0.133 | |
| AT5G65880.1 | ? | 46 | 5.65 | 0.307 | 0.254 | |
| AT5G65880.1 | ? | 47 | 7.28 | 0.461 | 0.676 | |
| AT5G66380.1 | Chloroplast folate transporter | 2 | 0.88 | 0.440 | 1.000 | |
| AT5G66380.1 | Chloroplast folate transporter | 3 | 0.99 | 0.381 | 1.000 | |
| AT5G66380.1 | Chloroplast folate transporter | 4 | 1.11 | 0.498 | 1.000 | |
| AT5G66380.1 | Chloroplast folate transporter | 39 | 3.15 | 0.359 | 0.639 | |
| AT5G66380.1 | Chloroplast folate transporter | 40 | 2.44 | 0.276 | 0.626 | |
| AT5G66380.1 | Chloroplast folate transporter | 41 | 1.72 | 0.223 | 0.846 | |
| AT5G66380.1 | Chloroplast folate transporter | 42 | 1.00 | 0.204 | 1.000 | |
| AT5G66380.1 | Chloroplast folate transporter | 43 | 0.51 | 0.221 | 0.689 | |
| AT5G66380.1 | Chloroplast folate transporter | 44 | 0.25 | 0.270 | 0.157 | |
| AT5G66380.1 | Chloroplast folate transporter | 45 | 0.11 | 0.343 | 0.054 | . |
| AT5G66380.1 | Chloroplast folate transporter | 46 | 0.05 | 0.439 | 0.034 | * |
| AT5G66380.1 | Chloroplast folate transporter | 47 | 0.02 | 0.563 | 0.039 | * |
| AT5G66510.1 | NADH:ubiquinone oxidoreductase subunit (gamma CA3) | 2 | 0.91 | 0.428 | 1.000 | |
| AT5G66510.1 | NADH:ubiquinone oxidoreductase subunit (gamma CA3) | 3 | 0.79 | 0.255 | 1.000 | |
| AT5G66510.1 | NADH:ubiquinone oxidoreductase subunit (gamma CA3) | 4 | 0.75 | 0.244 | 1.000 | |

| | | | | | |
|---|---|---|---|---|---|
| AT5G66510.1 | NADH:ubiquinone oxidoreductase subunit (gamma CA3) | 5 | 0.82 | 0.243 | 1.000 |
| AT5G66510.1 | NADH:ubiquinone oxidoreductase subunit (gamma CA3) | 6 | 0.79 | 0.244 | 1.000 |
| AT5G66510.1 | NADH:ubiquinone oxidoreductase subunit (gamma CA3) | 7 | 0.76 | 0.246 | 1.000 |
| AT5G66510.1 | NADH:ubiquinone oxidoreductase subunit (gamma CA3) | 8 | 0.89 | 0.239 | 1.000 |
| AT5G66510.1 | NADH:ubiquinone oxidoreductase subunit (gamma CA3) | 9 | 0.90 | 0.235 | 1.000 |
| AT5G66510.1 | NADH:ubiquinone oxidoreductase subunit (gamma CA3) | 10 | 0.78 | 0.235 | 1.000 |
| AT5G66510.1 | NADH:ubiquinone oxidoreductase subunit (gamma CA3) | 11 | 0.81 | 0.236 | 1.000 |
| AT5G66510.1 | NADH:ubiquinone oxidoreductase subunit (gamma CA3) | 12 | 1.01 | 0.236 | 1.000 |
| AT5G66510.1 | NADH:ubiquinone oxidoreductase subunit (gamma CA3) | 13 | 1.07 | 0.233 | 1.000 |
| AT5G66510.1 | NADH:ubiquinone oxidoreductase subunit (gamma CA3) | 14 | 0.93 | 0.229 | 1.000 |
| AT5G66510.1 | NADH:ubiquinone oxidoreductase subunit (gamma CA3) | 15 | 0.74 | 0.230 | 1.000 |
| AT5G66510.1 | NADH:ubiquinone oxidoreductase subunit (gamma CA3) | 16 | 0.51 | 0.228 | 0.998 |
| AT5G66510.1 | NADH:ubiquinone oxidoreductase subunit (gamma CA3) | 17 | 0.38 | 0.228 | 0.883 |
| AT5G66510.1 | NADH:ubiquinone oxidoreductase subunit (gamma CA3) | 18 | 0.45 | 0.231 | 0.982 |
| AT5G66510.1 | NADH:ubiquinone oxidoreductase subunit (gamma CA3) | 19 | 0.65 | 0.238 | 1.000 |
| AT5G66510.1 | NADH:ubiquinone oxidoreductase subunit (gamma CA3) | 20 | 0.81 | 0.244 | 1.000 |
| AT5G66510.1 | NADH:ubiquinone oxidoreductase subunit (gamma CA3) | 21 | 1.02 | 0.247 | 1.000 |
| AT5G66510.1 | NADH:ubiquinone oxidoreductase subunit (gamma CA3) | 22 | 1.43 | 0.247 | 1.000 |
| AT5G66510.1 | NADH:ubiquinone oxidoreductase subunit (gamma CA3) | 23 | 1.75 | 0.248 | 1.000 |
| AT5G66510.1 | NADH:ubiquinone oxidoreductase subunit (gamma CA3) | 24 | 1.44 | 0.241 | 1.000 |

| | | | | | | |
|---|---|---|---|---|---|---|
| AT5G66510.1 | NADH:ubiquinone oxidoreductase subunit (gamma CA3) | 25 | 0.82 | 0.234 | 1.000 | |
| AT5G66510.1 | NADH:ubiquinone oxidoreductase subunit (gamma CA3) | 26 | 0.64 | 0.233 | 1.000 | |
| AT5G66510.1 | NADH:ubiquinone oxidoreductase subunit (gamma CA3) | 27 | 0.72 | 0.233 | 1.000 | |
| AT5G66510.1 | NADH:ubiquinone oxidoreductase subunit (gamma CA3) | 28 | 0.79 | 0.233 | 1.000 | |
| AT5G66510.1 | NADH:ubiquinone oxidoreductase subunit (gamma CA3) | 29 | 0.78 | 0.233 | 1.000 | |
| AT5G66510.1 | NADH:ubiquinone oxidoreductase subunit (gamma CA3) | 30 | 0.85 | 0.233 | 1.000 | |
| AT5G66510.1 | NADH:ubiquinone oxidoreductase subunit (gamma CA3) | 31 | 1.07 | 0.235 | 1.000 | |
| AT5G66510.1 | NADH:ubiquinone oxidoreductase subunit (gamma CA3) | 32 | 1.13 | 0.235 | 1.000 | |
| AT5G66510.1 | NADH:ubiquinone oxidoreductase subunit (gamma CA3) | 33 | 0.73 | 0.236 | 1.000 | |
| AT5G66510.1 | NADH:ubiquinone oxidoreductase subunit (gamma CA3) | 34 | 0.37 | 0.237 | 0.891 | |
| AT5G66510.1 | NADH:ubiquinone oxidoreductase subunit (gamma CA3) | 35 | 0.33 | 0.237 | 0.742 | |
| AT5G66510.1 | NADH:ubiquinone oxidoreductase subunit (gamma CA3) | 36 | 0.69 | 0.237 | 1.000 | |
| AT5G66510.1 | NADH:ubiquinone oxidoreductase subunit (gamma CA3) | 37 | 1.87 | 0.237 | 1.000 | |
| AT5G66510.1 | NADH:ubiquinone oxidoreductase subunit (gamma CA3) | 38 | 3.49 | 0.238 | 0.526 | |
| AT5G66510.1 | NADH:ubiquinone oxidoreductase subunit (gamma CA3) | 39 | 2.51 | 0.238 | 0.946 | |
| AT5G66510.1 | NADH:ubiquinone oxidoreductase subunit (gamma CA3) | 40 | 0.92 | 0.239 | 1.000 | |
| AT5G66510.1 | NADH:ubiquinone oxidoreductase subunit (gamma CA3) | 41 | 0.24 | 0.248 | 0.368 | |
| AT5G66510.1 | NADH:ubiquinone oxidoreductase subunit (gamma CA3) | 42 | 0.08 | 0.342 | 0.045 | * |
| AT5G66510.1 | NADH:ubiquinone oxidoreductase subunit (gamma CA3) | 43 | 0.05 | 0.520 | 0.374 | |
| AT5G66510.1 | NADH:ubiquinone oxidoreductase subunit (gamma CA3) | 46 | 0.50 | 1.613 | 1.000 | |

| | | | | | |
|---|---|---|---|---|---|
| AT5G66510.1 | NADH:ubiquinone oxidoreductase subunit (gamma CA3) | 47 | 1.83 | 2.144 | 1.000 |
| AT5G66570.1 | Photosystem II subunit (PSBO1) | 3 | 0.50 | 0.206 | 0.900 |
| AT5G66570.1 | Photosystem II subunit (PSBO1) | 4 | 0.58 | 0.174 | 0.936 |
| AT5G66570.1 | Photosystem II subunit (PSBO1) | 5 | 0.66 | 0.165 | 0.991 |
| AT5G66570.1 | Photosystem II subunit (PSBO1) | 6 | 0.73 | 0.165 | 1.000 |
| AT5G66570.1 | Photosystem II subunit (PSBO1) | 7 | 0.79 | 0.165 | 1.000 |
| AT5G66570.1 | Photosystem II subunit (PSBO1) | 8 | 0.85 | 0.163 | 1.000 |
| AT5G66570.1 | Photosystem II subunit (PSBO1) | 9 | 0.90 | 0.162 | 1.000 |
| AT5G66570.1 | Photosystem II subunit (PSBO1) | 10 | 0.96 | 0.161 | 1.000 |
| AT5G66570.1 | Photosystem II subunit (PSBO1) | 11 | 1.00 | 0.160 | 1.000 |
| AT5G66570.1 | Photosystem II subunit (PSBO1) | 12 | 1.04 | 0.159 | 1.000 |
| AT5G66570.1 | Photosystem II subunit (PSBO1) | 13 | 1.07 | 0.159 | 1.000 |
| AT5G66570.1 | Photosystem II subunit (PSBO1) | 14 | 1.08 | 0.157 | 1.000 |
| AT5G66570.1 | Photosystem II subunit (PSBO1) | 15 | 1.08 | 0.157 | 1.000 |
| AT5G66570.1 | Photosystem II subunit (PSBO1) | 16 | 1.07 | 0.156 | 1.000 |
| AT5G66570.1 | Photosystem II subunit (PSBO1) | 17 | 1.07 | 0.157 | 1.000 |
| AT5G66570.1 | Photosystem II subunit (PSBO1) | 18 | 1.09 | 0.158 | 1.000 |
| AT5G66570.1 | Photosystem II subunit (PSBO1) | 19 | 1.10 | 0.160 | 1.000 |
| AT5G66570.1 | Photosystem II subunit (PSBO1) | 20 | 1.09 | 0.163 | 1.000 |
| AT5G66570.1 | Photosystem II subunit (PSBO1) | 21 | 1.03 | 0.164 | 1.000 |
| AT5G66570.1 | Photosystem II subunit (PSBO1) | 22 | 0.93 | 0.164 | 1.000 |
| AT5G66570.1 | Photosystem II subunit (PSBO1) | 23 | 0.82 | 0.163 | 1.000 |
| AT5G66570.1 | Photosystem II subunit (PSBO1) | 24 | 0.72 | 0.162 | 0.999 |
| AT5G66570.1 | Photosystem II subunit (PSBO1) | 25 | 0.68 | 0.160 | 0.995 |
| AT5G66570.1 | Photosystem II subunit (PSBO1) | 26 | 0.72 | 0.159 | 0.999 |
| AT5G66570.1 | Photosystem II subunit (PSBO1) | 27 | 0.81 | 0.159 | 1.000 |
| AT5G66570.1 | Photosystem II subunit (PSBO1) | 28 | 0.91 | 0.159 | 1.000 |
| AT5G66570.1 | Photosystem II subunit (PSBO1) | 29 | 1.02 | 0.159 | 1.000 |
| AT5G66570.1 | Photosystem II subunit (PSBO1) | 30 | 1.07 | 0.159 | 1.000 |
| AT5G66570.1 | Photosystem II subunit (PSBO1) | 31 | 1.00 | 0.160 | 1.000 |
| AT5G66570.1 | Photosystem II subunit (PSBO1) | 32 | 0.90 | 0.160 | 1.000 |
| AT5G66570.1 | Photosystem II subunit (PSBO1) | 33 | 0.83 | 0.160 | 1.000 |
| AT5G66570.1 | Photosystem II subunit (PSBO1) | 34 | 0.81 | 0.160 | 1.000 |
| AT5G66570.1 | Photosystem II subunit (PSBO1) | 35 | 0.82 | 0.160 | 1.000 |
| AT5G66570.1 | Photosystem II subunit (PSBO1) | 36 | 0.84 | 0.160 | 1.000 |
| AT5G66570.1 | Photosystem II subunit (PSBO1) | 37 | 0.82 | 0.160 | 1.000 |
| AT5G66570.1 | Photosystem II subunit (PSBO1) | 38 | 0.78 | 0.160 | 1.000 |
| AT5G66570.1 | Photosystem II subunit (PSBO1) | 39 | 0.72 | 0.160 | 0.999 |
| AT5G66570.1 | Photosystem II subunit (PSBO1) | 40 | 0.67 | 0.161 | 0.993 |
| AT5G66570.1 | Photosystem II subunit (PSBO1) | 41 | 0.63 | 0.162 | 0.971 |
| AT5G66570.1 | Photosystem II subunit (PSBO1) | 42 | 0.60 | 0.163 | 0.940 |
| AT5G66570.1 | Photosystem II subunit (PSBO1) | 43 | 0.60 | 0.164 | 0.943 |
| AT5G66570.1 | Photosystem II subunit (PSBO1) | 44 | 0.63 | 0.165 | 0.972 |
| AT5G66570.1 | Photosystem II subunit (PSBO1) | 45 | 0.66 | 0.178 | 0.995 |
| AT5G66570.1 | Photosystem II subunit (PSBO1) | 46 | 0.69 | 0.216 | 1.000 |
| AT5G66570.1 | Photosystem II subunit (PSBO1) | 47 | 0.71 | 0.293 | 1.000 |
| AT5G66570.1 | Photosystem II subunit (PSBO1) | 48 | 0.73 | 0.398 | 1.000 |
| AT5G66760.1 | Succinate dehydrogenase subunit (SDH1-1, FAD subunit) | 3 | 0.36 | 0.251 | 0.901 |
| AT5G66760.1 | Succinate dehydrogenase subunit (SDH1-1, FAD subunit) | 4 | 0.37 | 0.234 | 0.853 |
| AT5G66760.1 | Succinate dehydrogenase subunit (SDH1-1, FAD subunit) | 5 | 0.47 | 0.234 | 0.993 |
| AT5G66760.1 | Succinate dehydrogenase subunit (SDH1-1, FAD subunit) | 6 | 0.61 | 0.234 | 1.000 |
| AT5G66760.1 | Succinate dehydrogenase subunit (SDH1-1, FAD subunit) | 7 | 0.73 | 0.235 | 1.000 |
| AT5G66760.1 | Succinate dehydrogenase subunit (SDH1-1, FAD subunit) | 8 | 0.90 | 0.230 | 1.000 |
| AT5G66760.1 | Succinate dehydrogenase subunit (SDH1-1, FAD subunit) | 9 | 1.00 | 0.226 | 1.000 |

| | | | | | | |
|---|---|---|---|---|---|---|
| AT5G66760.1 | Succinate dehydrogenase subunit (SDH1-1, FAD subunit) | 10 | 0.93 | 0.226 | 1.000 | |
| AT5G66760.1 | Succinate dehydrogenase subunit (SDH1-1, FAD subunit) | 11 | 0.86 | 0.227 | 1.000 | |
| AT5G66760.1 | Succinate dehydrogenase subunit (SDH1-1, FAD subunit) | 12 | 0.84 | 0.227 | 1.000 | |
| AT5G66760.1 | Succinate dehydrogenase subunit (SDH1-1, FAD subunit) | 13 | 0.85 | 0.224 | 1.000 | |
| AT5G66760.1 | Succinate dehydrogenase subunit (SDH1-1, FAD subunit) | 14 | 0.84 | 0.221 | 1.000 | |
| AT5G66760.1 | Succinate dehydrogenase subunit (SDH1-1, FAD subunit) | 15 | 0.77 | 0.220 | 1.000 | |
| AT5G66760.1 | Succinate dehydrogenase subunit (SDH1-1, FAD subunit) | 16 | 0.77 | 0.218 | 1.000 | |
| AT5G66760.1 | Succinate dehydrogenase subunit (SDH1-1, FAD subunit) | 17 | 1.04 | 0.218 | 1.000 | |
| AT5G66760.1 | Succinate dehydrogenase subunit (SDH1-1, FAD subunit) | 18 | 1.85 | 0.222 | 0.999 | |
| AT5G66760.1 | Succinate dehydrogenase subunit (SDH1-1, FAD subunit) | 19 | 3.61 | 0.228 | 0.381 | |
| AT5G66760.1 | Succinate dehydrogenase subunit (SDH1-1, FAD subunit) | 20 | 5.26 | 0.234 | 0.076 | . |
| AT5G66760.1 | Succinate dehydrogenase subunit (SDH1-1, FAD subunit) | 21 | 4.55 | 0.237 | 0.181 | |
| AT5G66760.1 | Succinate dehydrogenase subunit (SDH1-1, FAD subunit) | 22 | 2.57 | 0.237 | 0.919 | |
| AT5G66760.1 | Succinate dehydrogenase subunit (SDH1-1, FAD subunit) | 23 | 1.25 | 0.236 | 1.000 | |
| AT5G66760.1 | Succinate dehydrogenase subunit (SDH1-1, FAD subunit) | 24 | 0.81 | 0.230 | 1.000 | |
| AT5G66760.1 | Succinate dehydrogenase subunit (SDH1-1, FAD subunit) | 25 | 0.75 | 0.226 | 1.000 | |
| AT5G66760.1 | Succinate dehydrogenase subunit (SDH1-1, FAD subunit) | 26 | 0.88 | 0.225 | 1.000 | |
| AT5G66760.1 | Succinate dehydrogenase subunit (SDH1-1, FAD subunit) | 27 | 1.38 | 0.225 | 1.000 | |
| AT5G66760.1 | Succinate dehydrogenase subunit (SDH1-1, FAD subunit) | 28 | 1.72 | 0.225 | 1.000 | |
| AT5G66760.1 | Succinate dehydrogenase subunit (SDH1-1, FAD subunit) | 29 | 1.65 | 0.225 | 1.000 | |
| AT5G66760.1 | Succinate dehydrogenase subunit (SDH1-1, FAD subunit) | 30 | 1.40 | 0.224 | 1.000 | |
| AT5G66760.1 | Succinate dehydrogenase subunit (SDH1-1, FAD subunit) | 31 | 1.36 | 0.225 | 1.000 | |
| AT5G66760.1 | Succinate dehydrogenase subunit (SDH1-1, FAD subunit) | 32 | 1.65 | 0.226 | 1.000 | |
| AT5G66760.1 | Succinate dehydrogenase subunit (SDH1-1, FAD subunit) | 33 | 2.10 | 0.226 | 0.991 | |
| AT5G66760.1 | Succinate dehydrogenase subunit (SDH1-1, FAD subunit) | 34 | 2.55 | 0.227 | 0.890 | |
| AT5G66760.1 | Succinate dehydrogenase subunit (SDH1-1, FAD subunit) | 35 | 2.40 | 0.226 | 0.939 | |
| AT5G66760.1 | Succinate dehydrogenase subunit (SDH1-1, FAD subunit) | 36 | 1.37 | 0.225 | 1.000 | |
| AT5G66760.1 | Succinate dehydrogenase subunit (SDH1-1, FAD subunit) | 37 | 0.66 | 0.225 | 1.000 | |
| AT5G66760.1 | Succinate dehydrogenase subunit (SDH1-1, FAD subunit) | 38 | 0.50 | 0.226 | 0.997 | |
| AT5G66760.1 | Succinate dehydrogenase subunit (SDH1-1, FAD subunit) | 39 | 0.70 | 0.226 | 1.000 | |
| AT5G66760.1 | Succinate dehydrogenase subunit (SDH1-1, FAD subunit) | 40 | 1.13 | 0.227 | 1.000 | |

| | | | | | |
|---|---|---|---|---|---|
| AT5G66760.1 | Succinate dehydrogenase subunit (SDH1-1, FAD subunit) | 41 | 1.52 | 0.228 | 1.000 |
| AT5G66760.1 | Succinate dehydrogenase subunit (SDH1-1, FAD subunit) | 42 | 1.79 | 0.231 | 1.000 |
| AT5G66760.1 | Succinate dehydrogenase subunit (SDH1-1, FAD subunit) | 43 | 2.09 | 0.233 | 0.994 |
| AT5G66760.1 | Succinate dehydrogenase subunit (SDH1-1, FAD subunit) | 44 | 2.07 | 0.233 | 0.995 |
| AT5G66760.1 | Succinate dehydrogenase subunit (SDH1-1, FAD subunit) | 45 | 1.29 | 0.233 | 1.000 |
| AT5G66760.1 | Succinate dehydrogenase subunit (SDH1-1, FAD subunit) | 46 | 0.45 | 0.270 | 0.997 |
| AT5G66760.1 | Succinate dehydrogenase subunit (SDH1-1, FAD subunit) | 47 | 0.10 | 0.479 | 0.653 |
| AT5G66760.1 | Succinate dehydrogenase subunit (SDH1-1, FAD subunit) | 48 | 0.02 | 0.811 | 0.664 |
| AT5G66860.1 | Ribosomal L25 family protein /GLN-tRNA synthetase | 2 | 0.78 | 0.446 | 1.000 |
| AT5G66860.1 | Ribosomal L25 family protein /GLN-tRNA synthetase | 3 | 0.69 | 0.391 | 1.000 |
| AT5G66860.1 | Ribosomal L25 family protein /GLN-tRNA synthetase | 4 | 0.62 | 0.370 | 1.000 |
| AT5G66860.1 | Ribosomal L25 family protein /GLN-tRNA synthetase | 10 | 0.55 | 0.291 | 0.994 |
| AT5G66860.1 | Ribosomal L25 family protein /GLN-tRNA synthetase | 11 | 0.63 | 0.268 | 0.999 |
| AT5G66860.1 | Ribosomal L25 family protein /GLN-tRNA synthetase | 12 | 0.75 | 0.248 | 1.000 |
| AT5G66860.1 | Ribosomal L25 family protein /GLN-tRNA synthetase | 13 | 0.95 | 0.233 | 1.000 |
| AT5G66860.1 | Ribosomal L25 family protein /GLN-tRNA synthetase | 14 | 1.26 | 0.222 | 1.000 |
| AT5G66860.1 | Ribosomal L25 family protein /GLN-tRNA synthetase | 15 | 1.77 | 0.215 | 0.959 |
| AT5G66860.1 | Ribosomal L25 family protein /GLN-tRNA synthetase | 16 | 2.50 | 0.212 | 0.524 |
| AT5G66860.1 | Ribosomal L25 family protein /GLN-tRNA synthetase | 17 | 3.19 | 0.210 | 0.205 |
| AT5G66860.1 | Ribosomal L25 family protein /GLN-tRNA synthetase | 18 | 3.45 | 0.211 | 0.147 |
| AT5G66860.1 | Ribosomal L25 family protein /GLN-tRNA synthetase | 19 | 3.18 | 0.213 | 0.224 |
| AT5G66860.1 | Ribosomal L25 family protein /GLN-tRNA synthetase | 20 | 2.57 | 0.216 | 0.512 |
| AT5G66860.1 | Ribosomal L25 family protein /GLN-tRNA synthetase | 21 | 1.87 | 0.218 | 0.930 |
| AT5G66860.1 | Ribosomal L25 family protein /GLN-tRNA synthetase | 22 | 1.29 | 0.221 | 1.000 |
| AT5G66860.1 | Ribosomal L25 family protein /GLN-tRNA synthetase | 23 | 0.88 | 0.222 | 1.000 |
| AT5G66860.1 | Ribosomal L25 family protein /GLN-tRNA synthetase | 24 | 0.63 | 0.223 | 0.995 |
| AT5G66860.1 | Ribosomal L25 family protein /GLN-tRNA synthetase | 25 | 0.50 | 0.224 | 0.897 |
| AT5G66860.1 | Ribosomal L25 family protein /GLN-tRNA synthetase | 26 | 0.44 | 0.222 | 0.748 |
| AT5G66860.1 | Ribosomal L25 family protein /GLN-tRNA synthetase | 27 | 0.43 | 0.220 | 0.675 |
| AT5G66860.1 | Ribosomal L25 family protein /GLN-tRNA synthetase | 28 | 0.43 | 0.218 | 0.676 |
| AT5G66860.1 | Ribosomal L25 family protein /GLN-tRNA synthetase | 29 | 0.44 | 0.216 | 0.704 |

| | | | | | |
|---|---|---|---|---|---|
| AT5G66860.1 | Ribosomal L25 family protein /GLN-tRNA synthetase | 30 | 0.45 | 0.216 | 0.729 |
| AT5G66860.1 | Ribosomal L25 family protein /GLN-tRNA synthetase | 31 | 0.45 | 0.217 | 0.745 |
| AT5G66860.1 | Ribosomal L25 family protein /GLN-tRNA synthetase | 32 | 0.44 | 0.220 | 0.723 |
| AT5G66860.1 | Ribosomal L25 family protein /GLN-tRNA synthetase | 33 | 0.42 | 0.225 | 0.677 |
| AT5G66860.1 | Ribosomal L25 family protein /GLN-tRNA synthetase | 34 | 0.40 | 0.231 | 0.637 |
| AT5G66860.1 | Ribosomal L25 family protein /GLN-tRNA synthetase | 35 | 0.38 | 0.237 | 0.616 |
| AT5G66860.1 | Ribosomal L25 family protein /GLN-tRNA synthetase | 36 | 0.38 | 0.240 | 0.609 |
| AT5G66860.1 | Ribosomal L25 family protein /GLN-tRNA synthetase | 37 | 0.38 | 0.243 | 0.625 |
| AT5G66860.1 | Ribosomal L25 family protein /GLN-tRNA synthetase | 38 | 0.38 | 0.245 | 0.656 |
| AT5G66860.1 | Ribosomal L25 family protein /GLN-tRNA synthetase | 39 | 0.39 | 0.247 | 0.707 |
| AT5G66860.1 | Ribosomal L25 family protein /GLN-tRNA synthetase | 40 | 0.42 | 0.246 | 0.780 |
| AT5G66860.1 | Ribosomal L25 family protein /GLN-tRNA synthetase | 41 | 0.47 | 0.242 | 0.877 |
| AT5G66860.1 | Ribosomal L25 family protein /GLN-tRNA synthetase | 42 | 0.55 | 0.238 | 0.971 |
| AT5G66860.1 | Ribosomal L25 family protein /GLN-tRNA synthetase | 43 | 0.68 | 0.234 | 1.000 |
| AT5G66860.1 | Ribosomal L25 family protein /GLN-tRNA synthetase | 44 | 0.88 | 0.237 | 1.000 |
| AT5G66860.1 | Ribosomal L25 family protein /GLN-tRNA synthetase | 45 | 1.18 | 0.252 | 1.000 |
| AT5G66860.1 | Ribosomal L25 family protein /GLN-tRNA synthetase | 46 | 1.61 | 0.291 | 1.000 |
| AT5G66860.1 | Ribosomal L25 family protein /GLN-tRNA synthetase | 47 | 2.21 | 0.366 | 0.992 |
| AT5G67500.1 | Voltage dependent anion channel 2 (VDAC2) | 3 | 0.97 | 0.155 | 1.000 |
| AT5G67500.1 | Voltage dependent anion channel 2 (VDAC2) | 4 | 1.23 | 0.140 | 1.000 |
| AT5G67500.1 | Voltage dependent anion channel 2 (VDAC2) | 5 | 1.55 | 0.140 | 0.961 |
| AT5G67500.1 | Voltage dependent anion channel 2 (VDAC2) | 6 | 1.73 | 0.141 | 0.799 |
| AT5G67500.1 | Voltage dependent anion channel 2 (VDAC2) | 7 | 1.73 | 0.140 | 0.796 |
| AT5G67500.1 | Voltage dependent anion channel 2 (VDAC2) | 8 | 1.66 | 0.138 | 0.864 |
| AT5G67500.1 | Voltage dependent anion channel 2 (VDAC2) | 9 | 1.59 | 0.137 | 0.924 |
| AT5G67500.1 | Voltage dependent anion channel 2 (VDAC2) | 10 | 1.54 | 0.137 | 0.960 |
| AT5G67500.1 | Voltage dependent anion channel 2 (VDAC2) | 11 | 1.55 | 0.137 | 0.952 |
| AT5G67500.1 | Voltage dependent anion channel 2 (VDAC2) | 12 | 1.62 | 0.137 | 0.898 |
| AT5G67500.1 | Voltage dependent anion channel 2 (VDAC2) | 13 | 1.68 | 0.136 | 0.822 |
| AT5G67500.1 | Voltage dependent anion channel 2 (VDAC2) | 14 | 1.67 | 0.135 | 0.830 |
| AT5G67500.1 | Voltage dependent anion channel 2 (VDAC2) | 15 | 1.53 | 0.134 | 0.957 |

| | | | | | | |
|---|---|---|---|---|---|---|
| AT5G67500.1 | Voltage dependent anion channel 2 (VDAC2) | 16 | 1.37 | 0.134 | 0.998 | |
| AT5G67500.1 | Voltage dependent anion channel 2 (VDAC2) | 17 | 1.32 | 0.134 | 1.000 | |
| AT5G67500.1 | Voltage dependent anion channel 2 (VDAC2) | 18 | 1.36 | 0.135 | 0.999 | |
| AT5G67500.1 | Voltage dependent anion channel 2 (VDAC2) | 19 | 1.45 | 0.137 | 0.990 | |
| AT5G67500.1 | Voltage dependent anion channel 2 (VDAC2) | 20 | 1.48 | 0.140 | 0.987 | |
| AT5G67500.1 | Voltage dependent anion channel 2 (VDAC2) | 21 | 1.38 | 0.141 | 0.999 | |
| AT5G67500.1 | Voltage dependent anion channel 2 (VDAC2) | 22 | 1.23 | 0.141 | 1.000 | |
| AT5G67500.1 | Voltage dependent anion channel 2 (VDAC2) | 23 | 1.09 | 0.140 | 1.000 | |
| AT5G67500.1 | Voltage dependent anion channel 2 (VDAC2) | 24 | 0.98 | 0.138 | 1.000 | |
| AT5G67500.1 | Voltage dependent anion channel 2 (VDAC2) | 25 | 0.90 | 0.137 | 1.000 | |
| AT5G67500.1 | Voltage dependent anion channel 2 (VDAC2) | 26 | 0.89 | 0.136 | 1.000 | |
| AT5G67500.1 | Voltage dependent anion channel 2 (VDAC2) | 27 | 0.93 | 0.136 | 1.000 | |
| AT5G67500.1 | Voltage dependent anion channel 2 (VDAC2) | 28 | 1.02 | 0.136 | 1.000 | |
| AT5G67500.1 | Voltage dependent anion channel 2 (VDAC2) | 29 | 1.18 | 0.136 | 1.000 | |
| AT5G67500.1 | Voltage dependent anion channel 2 (VDAC2) | 30 | 1.28 | 0.136 | 1.000 | |
| AT5G67500.1 | Voltage dependent anion channel 2 (VDAC2) | 31 | 1.20 | 0.137 | 1.000 | |
| AT5G67500.1 | Voltage dependent anion channel 2 (VDAC2) | 32 | 1.08 | 0.137 | 1.000 | |
| AT5G67500.1 | Voltage dependent anion channel 2 (VDAC2) | 33 | 1.06 | 0.137 | 1.000 | |
| AT5G67500.1 | Voltage dependent anion channel 2 (VDAC2) | 34 | 1.12 | 0.137 | 1.000 | |
| AT5G67500.1 | Voltage dependent anion channel 2 (VDAC2) | 35 | 1.15 | 0.137 | 1.000 | |
| AT5G67500.1 | Voltage dependent anion channel 2 (VDAC2) | 36 | 1.14 | 0.137 | 1.000 | |
| AT5G67500.1 | Voltage dependent anion channel 2 (VDAC2) | 37 | 1.21 | 0.137 | 1.000 | |
| AT5G67500.1 | Voltage dependent anion channel 2 (VDAC2) | 38 | 1.37 | 0.137 | 0.999 | |
| AT5G67500.1 | Voltage dependent anion channel 2 (VDAC2) | 39 | 1.65 | 0.137 | 0.860 | |
| AT5G67500.1 | Voltage dependent anion channel 2 (VDAC2) | 40 | 2.07 | 0.137 | 0.336 | |
| AT5G67500.1 | Voltage dependent anion channel 2 (VDAC2) | 41 | 2.31 | 0.138 | 0.163 | |
| AT5G67500.1 | Voltage dependent anion channel 2 (VDAC2) | 42 | 2.01 | 0.139 | 0.417 | |
| AT5G67500.1 | Voltage dependent anion channel 2 (VDAC2) | 43 | 1.37 | 0.140 | 0.999 | |
| AT5G67500.1 | Voltage dependent anion channel 2 (VDAC2) | 44 | 0.75 | 0.140 | 1.000 | |
| AT5G67500.1 | Voltage dependent anion channel 2 (VDAC2) | 45 | 0.35 | 0.141 | 0.035 | * |
| AT5G67500.1 | Voltage dependent anion channel 2 (VDAC2) | 46 | 0.15 | 0.163 | 0.000 | *** |

| | | | | | | |
|---|---|---|---|---|---|---|
| AT5G67500.1 | Voltage dependent anion channel 2 (VDAC2) | 47 | 0.06 | 0.244 | 0.000 | *** |
| AT5G67500.1 | Voltage dependent anion channel 2 (VDAC2) | 48 | 0.02 | 0.371 | 0.000 | *** |
| AT5G67590.1 | NADH:ubiquinone oxidoreductase subunit (FRO1) | 2 | 1.05 | 0.577 | 1.000 | |
| AT5G67590.1 | NADH:ubiquinone oxidoreductase subunit (FRO1) | 3 | 0.89 | 0.392 | 1.000 | |
| AT5G67590.1 | NADH:ubiquinone oxidoreductase subunit (FRO1) | 4 | 0.81 | 0.356 | 1.000 | |
| AT5G67590.1 | NADH:ubiquinone oxidoreductase subunit (FRO1) | 5 | 0.88 | 0.353 | 1.000 | |
| AT5G67590.1 | NADH:ubiquinone oxidoreductase subunit (FRO1) | 6 | 1.16 | 0.354 | 1.000 | |
| AT5G67590.1 | NADH:ubiquinone oxidoreductase subunit (FRO1) | 7 | 1.45 | 0.378 | 1.000 | |
| AT5G67590.1 | NADH:ubiquinone oxidoreductase subunit (FRO1) | 8 | 1.44 | 0.446 | 1.000 | |
| AT5G67590.1 | NADH:ubiquinone oxidoreductase subunit (FRO1) | 11 | 0.65 | 0.374 | 1.000 | |
| AT5G67590.1 | NADH:ubiquinone oxidoreductase subunit (FRO1) | 12 | 0.78 | 0.343 | 1.000 | |
| AT5G67590.1 | NADH:ubiquinone oxidoreductase subunit (FRO1) | 13 | 0.98 | 0.317 | 1.000 | |
| AT5G67590.1 | NADH:ubiquinone oxidoreductase subunit (FRO1) | 14 | 0.92 | 0.293 | 1.000 | |
| AT5G67590.1 | NADH:ubiquinone oxidoreductase subunit (FRO1) | 15 | 0.61 | 0.279 | 1.000 | |
| AT5G67590.1 | NADH:ubiquinone oxidoreductase subunit (FRO1) | 16 | 0.43 | 0.276 | 0.981 | |
| AT5G67590.1 | NADH:ubiquinone oxidoreductase subunit (FRO1) | 17 | 0.36 | 0.273 | 0.891 | |
| AT5G67590.1 | NADH:ubiquinone oxidoreductase subunit (FRO1) | 18 | 0.42 | 0.277 | 0.978 | |
| AT5G67590.1 | NADH:ubiquinone oxidoreductase subunit (FRO1) | 19 | 0.57 | 0.286 | 1.000 | |
| AT5G67590.1 | NADH:ubiquinone oxidoreductase subunit (FRO1) | 20 | 0.70 | 0.294 | 1.000 | |
| AT5G67590.1 | NADH:ubiquinone oxidoreductase subunit (FRO1) | 21 | 1.07 | 0.296 | 1.000 | |
| AT5G67590.1 | NADH:ubiquinone oxidoreductase subunit (FRO1) | 22 | 2.13 | 0.296 | 0.998 | |
| AT5G67590.1 | NADH:ubiquinone oxidoreductase subunit (FRO1) | 23 | 3.18 | 0.295 | 0.857 | |
| AT5G67590.1 | NADH:ubiquinone oxidoreductase subunit (FRO1) | 24 | 1.60 | 0.288 | 1.000 | |
| AT5G67590.1 | NADH:ubiquinone oxidoreductase subunit (FRO1) | 25 | 0.27 | 0.283 | 0.634 | |
| AT5G67590.1 | NADH:ubiquinone oxidoreductase subunit (FRO1) | 26 | 0.07 | 0.280 | 0.002 | ** |
| AT5G67590.1 | NADH:ubiquinone oxidoreductase subunit (FRO1) | 27 | 0.08 | 0.281 | 0.003 | ** |
| AT5G67590.1 | NADH:ubiquinone oxidoreductase subunit (FRO1) | 28 | 0.23 | 0.283 | 0.429 | |
| AT5G67590.1 | NADH:ubiquinone oxidoreductase subunit (FRO1) | 29 | 0.81 | 0.284 | 1.000 | |
| AT5G67590.1 | NADH:ubiquinone oxidoreductase subunit (FRO1) | 30 | 1.16 | 0.285 | 1.000 | |
| AT5G67590.1 | NADH:ubiquinone oxidoreductase subunit (FRO1) | 31 | 1.13 | 0.291 | 1.000 | |
| AT5G67590.1 | NADH:ubiquinone oxidoreductase subunit (FRO1) | 32 | 1.21 | 0.309 | 1.000 | |

| | | | | | | |
|---|---|---|---|---|---|---|
| AT5G67590.1 | NADH:ubiquinone oxidoreductase subunit (FRO1) | 33 | 1.26 | 0.390 | 1.000 | |
| AT5G67590.1 | NADH:ubiquinone oxidoreductase subunit (FRO1) | 34 | 1.25 | 0.647 | 1.000 | |
| AT5G67590.1 | NADH:ubiquinone oxidoreductase subunit (FRO1) | 46 | 84.86 | 5.313 | 1.000 | |
| AT5G67590.1 | NADH:ubiquinone oxidoreductase subunit (FRO1) | 47 | 144.26 | 5.921 | 1.000 | |
| ATCG00020.1 | Photosystem II subunit (PSBA) | 2 | 0.50 | 0.256 | 0.998 | |
| ATCG00020.1 | Photosystem II subunit (PSBA) | 3 | 0.71 | 0.161 | 1.000 | |
| ATCG00020.1 | Photosystem II subunit (PSBA) | 4 | 0.97 | 0.142 | 1.000 | |
| ATCG00020.1 | Photosystem II subunit (PSBA) | 5 | 1.19 | 0.141 | 1.000 | |
| ATCG00020.1 | Photosystem II subunit (PSBA) | 6 | 1.17 | 0.142 | 1.000 | |
| ATCG00020.1 | Photosystem II subunit (PSBA) | 7 | 1.02 | 0.142 | 1.000 | |
| ATCG00020.1 | Photosystem II subunit (PSBA) | 8 | 0.94 | 0.138 | 1.000 | |
| ATCG00020.1 | Photosystem II subunit (PSBA) | 9 | 0.89 | 0.135 | 1.000 | |
| ATCG00020.1 | Photosystem II subunit (PSBA) | 10 | 0.85 | 0.136 | 1.000 | |
| ATCG00020.1 | Photosystem II subunit (PSBA) | 11 | 0.87 | 0.137 | 1.000 | |
| ATCG00020.1 | Photosystem II subunit (PSBA) | 12 | 0.93 | 0.136 | 1.000 | |
| ATCG00020.1 | Photosystem II subunit (PSBA) | 13 | 0.98 | 0.134 | 1.000 | |
| ATCG00020.1 | Photosystem II subunit (PSBA) | 14 | 1.01 | 0.131 | 1.000 | |
| ATCG00020.1 | Photosystem II subunit (PSBA) | 15 | 0.96 | 0.131 | 1.000 | |
| ATCG00020.1 | Photosystem II subunit (PSBA) | 16 | 0.90 | 0.131 | 1.000 | |
| ATCG00020.1 | Photosystem II subunit (PSBA) | 17 | 0.88 | 0.131 | 1.000 | |
| ATCG00020.1 | Photosystem II subunit (PSBA) | 18 | 0.93 | 0.132 | 1.000 | |
| ATCG00020.1 | Photosystem II subunit (PSBA) | 19 | 1.08 | 0.135 | 1.000 | |
| ATCG00020.1 | Photosystem II subunit (PSBA) | 20 | 1.29 | 0.140 | 1.000 | |
| ATCG00020.1 | Photosystem II subunit (PSBA) | 21 | 1.49 | 0.142 | 0.996 | |
| ATCG00020.1 | Photosystem II subunit (PSBA) | 22 | 1.51 | 0.143 | 0.995 | |
| ATCG00020.1 | Photosystem II subunit (PSBA) | 23 | 1.20 | 0.142 | 1.000 | |
| ATCG00020.1 | Photosystem II subunit (PSBA) | 24 | 0.82 | 0.138 | 1.000 | |
| ATCG00020.1 | Photosystem II subunit (PSBA) | 25 | 0.57 | 0.134 | 0.825 | |
| ATCG00020.1 | Photosystem II subunit (PSBA) | 26 | 0.55 | 0.134 | 0.715 | |
| ATCG00020.1 | Photosystem II subunit (PSBA) | 27 | 0.63 | 0.134 | 0.964 | |
| ATCG00020.1 | Photosystem II subunit (PSBA) | 28 | 0.70 | 0.134 | 0.997 | |
| ATCG00020.1 | Photosystem II subunit (PSBA) | 29 | 1.01 | 0.134 | 1.000 | |
| ATCG00020.1 | Photosystem II subunit (PSBA) | 30 | 1.63 | 0.136 | 0.943 | |
| ATCG00020.1 | Photosystem II subunit (PSBA) | 31 | 1.74 | 0.137 | 0.855 | |
| ATCG00020.1 | Photosystem II subunit (PSBA) | 32 | 1.16 | 0.137 | 1.000 | |
| ATCG00020.1 | Photosystem II subunit (PSBA) | 33 | 0.64 | 0.139 | 0.981 | |
| ATCG00020.1 | Photosystem II subunit (PSBA) | 34 | 0.35 | 0.141 | 0.043 | * |
| ATCG00020.1 | Photosystem II subunit (PSBA) | 35 | 0.26 | 0.152 | 0.004 | ** |
| ATCG00020.1 | Photosystem II subunit (PSBA) | 36 | 0.29 | 0.196 | 0.156 | |
| ATCG00020.1 | Photosystem II subunit (PSBA) | 37 | 0.42 | 0.300 | 0.994 | |
| ATCG00020.1 | Photosystem II subunit (PSBA) | 46 | 6227.77 | 1.511 | 0.282 | |
| ATCG00020.1 | Photosystem II subunit (PSBA) | 47 | 25252.94 | 1.721 | 0.254 | |
| ATCG00120.1 | ATP synthase subunit (ATPalpha) | 3 | 0.24 | 0.262 | 0.379 | |
| ATCG00120.1 | ATP synthase subunit (ATPalpha) | 4 | 0.20 | 0.245 | 0.117 | |
| ATCG00120.1 | ATP synthase subunit (ATPalpha) | 5 | 0.25 | 0.245 | 0.312 | |
| ATCG00120.1 | ATP synthase subunit (ATPalpha) | 6 | 0.33 | 0.246 | 0.721 | |
| ATCG00120.1 | ATP synthase subunit (ATPalpha) | 7 | 0.40 | 0.246 | 0.925 | |
| ATCG00120.1 | ATP synthase subunit (ATPalpha) | 8 | 0.52 | 0.242 | 0.998 | |
| ATCG00120.1 | ATP synthase subunit (ATPalpha) | 9 | 0.68 | 0.239 | 1.000 | |
| ATCG00120.1 | ATP synthase subunit (ATPalpha) | 10 | 0.74 | 0.239 | 1.000 | |

| | | | | | |
|---|---|---|---|---|---|
| ATCG00120.1 | ATP synthase subunit (ATPalpha) | 11 | 0.73 | 0.240 | 1.000 |
| ATCG00120.1 | ATP synthase subunit (ATPalpha) | 12 | 0.73 | 0.240 | 1.000 |
| ATCG00120.1 | ATP synthase subunit (ATPalpha) | 13 | 0.76 | 0.237 | 1.000 |
| ATCG00120.1 | ATP synthase subunit (ATPalpha) | 14 | 0.80 | 0.234 | 1.000 |
| ATCG00120.1 | ATP synthase subunit (ATPalpha) | 15 | 0.73 | 0.234 | 1.000 |
| ATCG00120.1 | ATP synthase subunit (ATPalpha) | 16 | 0.64 | 0.232 | 1.000 |
| ATCG00120.1 | ATP synthase subunit (ATPalpha) | 17 | 0.67 | 0.232 | 1.000 |
| ATCG00120.1 | ATP synthase subunit (ATPalpha) | 18 | 0.87 | 0.235 | 1.000 |
| ATCG00120.1 | ATP synthase subunit (ATPalpha) | 19 | 1.19 | 0.240 | 1.000 |
| ATCG00120.1 | ATP synthase subunit (ATPalpha) | 20 | 1.15 | 0.246 | 1.000 |
| ATCG00120.1 | ATP synthase subunit (ATPalpha) | 21 | 0.70 | 0.249 | 1.000 |
| ATCG00120.1 | ATP synthase subunit (ATPalpha) | 22 | 0.39 | 0.248 | 0.902 |
| ATCG00120.1 | ATP synthase subunit (ATPalpha) | 23 | 0.27 | 0.247 | 0.406 |
| ATCG00120.1 | ATP synthase subunit (ATPalpha) | 24 | 0.25 | 0.243 | 0.297 |
| ATCG00120.1 | ATP synthase subunit (ATPalpha) | 25 | 0.33 | 0.239 | 0.677 |
| ATCG00120.1 | ATP synthase subunit (ATPalpha) | 26 | 0.69 | 0.238 | 1.000 |
| ATCG00120.1 | ATP synthase subunit (ATPalpha) | 27 | 1.34 | 0.238 | 1.000 |
| ATCG00120.1 | ATP synthase subunit (ATPalpha) | 28 | 1.58 | 0.238 | 1.000 |
| ATCG00120.1 | ATP synthase subunit (ATPalpha) | 29 | 1.87 | 0.238 | 0.999 |
| ATCG00120.1 | ATP synthase subunit (ATPalpha) | 30 | 2.28 | 0.238 | 0.959 |
| ATCG00120.1 | ATP synthase subunit (ATPalpha) | 31 | 2.07 | 0.238 | 0.991 |
| ATCG00120.1 | ATP synthase subunit (ATPalpha) | 32 | 1.28 | 0.239 | 1.000 |
| ATCG00120.1 | ATP synthase subunit (ATPalpha) | 33 | 0.78 | 0.239 | 1.000 |
| ATCG00120.1 | ATP synthase subunit (ATPalpha) | 34 | 0.69 | 0.239 | 1.000 |
| ATCG00120.1 | ATP synthase subunit (ATPalpha) | 35 | 0.82 | 0.239 | 1.000 |
| ATCG00120.1 | ATP synthase subunit (ATPalpha) | 36 | 1.12 | 0.238 | 1.000 |
| ATCG00120.1 | ATP synthase subunit (ATPalpha) | 37 | 1.52 | 0.238 | 1.000 |
| ATCG00120.1 | ATP synthase subunit (ATPalpha) | 38 | 1.74 | 0.239 | 1.000 |
| ATCG00120.1 | ATP synthase subunit (ATPalpha) | 39 | 1.75 | 0.239 | 1.000 |
| ATCG00120.1 | ATP synthase subunit (ATPalpha) | 40 | 1.82 | 0.240 | 0.999 |
| ATCG00120.1 | ATP synthase subunit (ATPalpha) | 41 | 1.92 | 0.240 | 0.998 |

| | | | | | |
|---|---|---|---|---|---|
| ATCG00120.1 | ATP synthase subunit (ATPalpha) | 42 | 1.93 | 0.243 | 0.998 |
| ATCG00120.1 | ATP synthase subunit (ATPalpha) | 43 | 1.93 | 0.245 | 0.998 |
| ATCG00120.1 | ATP synthase subunit (ATPalpha) | 44 | 1.74 | 0.245 | 1.000 |
| ATCG00120.1 | ATP synthase subunit (ATPalpha) | 45 | 1.40 | 0.245 | 1.000 |
| ATCG00120.1 | ATP synthase subunit (ATPalpha) | 46 | 1.02 | 0.278 | 1.000 |
| ATCG00120.1 | ATP synthase subunit (ATPalpha) | 47 | 0.71 | 0.464 | 1.000 |
| ATCG00120.1 | ATP synthase subunit (ATPalpha) | 48 | 0.49 | 0.766 | 1.000 |
| ATCG00130.E | ATP synthase subunit (ATPF) | 2 | 0.34 | 0.373 | 0.987 |
| ATCG00130.E | ATP synthase subunit (ATPF) | 3 | 0.40 | 0.241 | 0.878 |
| ATCG00130.E | ATP synthase subunit (ATPF) | 4 | 0.53 | 0.222 | 0.988 |
| ATCG00130.E | ATP synthase subunit (ATPF) | 5 | 0.74 | 0.221 | 1.000 |
| ATCG00130.E | ATP synthase subunit (ATPF) | 6 | 0.83 | 0.222 | 1.000 |
| ATCG00130.E | ATP synthase subunit (ATPF) | 7 | 0.81 | 0.222 | 1.000 |
| ATCG00130.E | ATP synthase subunit (ATPF) | 8 | 0.86 | 0.218 | 1.000 |
| ATCG00130.E | ATP synthase subunit (ATPF) | 9 | 0.94 | 0.215 | 1.000 |
| ATCG00130.E | ATP synthase subunit (ATPF) | 10 | 0.95 | 0.215 | 1.000 |
| ATCG00130.E | ATP synthase subunit (ATPF) | 11 | 0.96 | 0.216 | 1.000 |
| ATCG00130.E | ATP synthase subunit (ATPF) | 12 | 1.02 | 0.216 | 1.000 |
| ATCG00130.E | ATP synthase subunit (ATPF) | 13 | 1.12 | 0.213 | 1.000 |
| ATCG00130.E | ATP synthase subunit (ATPF) | 14 | 1.13 | 0.211 | 1.000 |
| ATCG00130.E | ATP synthase subunit (ATPF) | 15 | 0.94 | 0.211 | 1.000 |
| ATCG00130.E | ATP synthase subunit (ATPF) | 16 | 0.76 | 0.210 | 1.000 |
| ATCG00130.E | ATP synthase subunit (ATPF) | 17 | 0.76 | 0.209 | 1.000 |
| ATCG00130.E | ATP synthase subunit (ATPF) | 18 | 0.96 | 0.211 | 1.000 |
| ATCG00130.E | ATP synthase subunit (ATPF) | 19 | 1.26 | 0.215 | 1.000 |
| ATCG00130.E | ATP synthase subunit (ATPF) | 20 | 1.30 | 0.221 | 1.000 |
| ATCG00130.E | ATP synthase subunit (ATPF) | 21 | 1.05 | 0.223 | 1.000 |
| ATCG00130.E | ATP synthase subunit (ATPF) | 22 | 0.83 | 0.223 | 1.000 |
| ATCG00130.E | ATP synthase subunit (ATPF) | 23 | 0.71 | 0.222 | 1.000 |
| ATCG00130.E | ATP synthase subunit (ATPF) | 24 | 0.60 | 0.217 | 0.999 |
| ATCG00130.E | ATP synthase subunit (ATPF) | 25 | 0.55 | 0.214 | 0.993 |
| ATCG00130.E | ATP synthase subunit (ATPF) | 26 | 0.77 | 0.213 | 1.000 |
| ATCG00130.E | ATP synthase subunit (ATPF) | 27 | 1.23 | 0.213 | 1.000 |
| ATCG00130.E | ATP synthase subunit (ATPF) | 28 | 1.52 | 0.214 | 1.000 |
| ATCG00130.E | ATP synthase subunit (ATPF) | 29 | 1.89 | 0.215 | 0.984 |
| ATCG00130.E | ATP synthase subunit (ATPF) | 30 | 2.37 | 0.216 | 0.814 |
| ATCG00130.E | ATP synthase subunit (ATPF) | 31 | 1.96 | 0.218 | 0.977 |
| ATCG00130.E | ATP synthase subunit (ATPF) | 32 | 0.84 | 0.222 | 1.000 |
| ATCG00130.E | ATP synthase subunit (ATPF) | 33 | 0.28 | 0.235 | 0.360 |
| ATCG00130.E | ATP synthase subunit (ATPF) | 34 | 0.17 | 0.289 | 0.168 |
| ATCG00130.E | ATP synthase subunit (ATPF) | 35 | 0.17 | 0.436 | 0.786 |
| ATCG00130.E | ATP synthase subunit (ATPF) | 46 | 3291278.16 | 3.543 | 0.743 |
| ATCG00130.E | ATP synthase subunit (ATPF) | 47 | 31970140.85 | 3.978 | 0.706 |
| ATCG00140.1 | ATP synthase subunit (ATPH) | 2 | 1.07 | 0.304 | 1.000 |
| ATCG00140.1 | ATP synthase subunit (ATPH) | 3 | 0.95 | 0.238 | 1.000 |
| ATCG00140.1 | ATP synthase subunit (ATPH) | 4 | 0.84 | 0.202 | 1.000 |
| ATCG00140.1 | ATP synthase subunit (ATPH) | 5 | 0.74 | 0.190 | 0.999 |
| ATCG00140.1 | ATP synthase subunit (ATPH) | 6 | 0.66 | 0.191 | 0.991 |
| ATCG00140.1 | ATP synthase subunit (ATPH) | 7 | 0.61 | 0.197 | 0.969 |
| ATCG00140.1 | ATP synthase subunit (ATPH) | 8 | 0.58 | 0.203 | 0.949 |
| ATCG00140.1 | ATP synthase subunit (ATPH) | 9 | 0.56 | 0.208 | 0.941 |
| ATCG00140.1 | ATP synthase subunit (ATPH) | 10 | 0.56 | 0.212 | 0.945 |
| ATCG00140.1 | ATP synthase subunit (ATPH) | 11 | 0.57 | 0.216 | 0.959 |
| ATCG00140.1 | ATP synthase subunit (ATPH) | 12 | 0.59 | 0.219 | 0.979 |
| ATCG00140.1 | ATP synthase subunit (ATPH) | 13 | 0.63 | 0.220 | 0.993 |
| ATCG00140.1 | ATP synthase subunit (ATPH) | 14 | 0.68 | 0.221 | 0.999 |

| | | | | | | |
|---|---|---|---|---|---|---|
| ATCG00140.1 | ATP synthase subunit (ATPH) | 15 | 0.74 | 0.222 | 1.000 | |
| ATCG00140.1 | ATP synthase subunit (ATPH) | 16 | 0.82 | 0.222 | 1.000 | |
| ATCG00140.1 | ATP synthase subunit (ATPH) | 17 | 0.90 | 0.223 | 1.000 | |
| ATCG00140.1 | ATP synthase subunit (ATPH) | 18 | 0.97 | 0.223 | 1.000 | |
| ATCG00140.1 | ATP synthase subunit (ATPH) | 19 | 1.03 | 0.223 | 1.000 | |
| ATCG00140.1 | ATP synthase subunit (ATPH) | 21 | 1.07 | 0.215 | 1.000 | |
| ATCG00140.1 | ATP synthase subunit (ATPH) | 22 | 1.06 | 0.206 | 1.000 | |
| ATCG00140.1 | ATP synthase subunit (ATPH) | 23 | 1.02 | 0.197 | 1.000 | |
| ATCG00140.1 | ATP synthase subunit (ATPH) | 24 | 0.97 | 0.187 | 1.000 | |
| ATCG00140.1 | ATP synthase subunit (ATPH) | 25 | 0.91 | 0.179 | 1.000 | |
| ATCG00140.1 | ATP synthase subunit (ATPH) | 26 | 0.86 | 0.172 | 1.000 | |
| ATCG00140.1 | ATP synthase subunit (ATPH) | 27 | 0.82 | 0.167 | 1.000 | |
| ATCG00140.1 | ATP synthase subunit (ATPH) | 28 | 0.80 | 0.165 | 1.000 | |
| ATCG00140.1 | ATP synthase subunit (ATPH) | 29 | 0.79 | 0.166 | 1.000 | |
| ATCG00140.1 | ATP synthase subunit (ATPH) | 30 | 0.80 | 0.171 | 1.000 | |
| ATCG00140.1 | ATP synthase subunit (ATPH) | 31 | 0.84 | 0.181 | 1.000 | |
| ATCG00140.1 | ATP synthase subunit (ATPH) | 32 | 0.89 | 0.199 | 1.000 | |
| ATCG00140.1 | ATP synthase subunit (ATPH) | 33 | 0.94 | 0.223 | 1.000 | |
| ATCG00140.1 | ATP synthase subunit (ATPH) | 34 | 0.98 | 0.251 | 1.000 | |
| ATCG00140.1 | ATP synthase subunit (ATPH) | 41 | 1.26 | 0.352 | 1.000 | |
| ATCG00140.1 | ATP synthase subunit (ATPH) | 42 | 1.33 | 0.372 | 1.000 | |
| ATCG00140.1 | ATP synthase subunit (ATPH) | 43 | 1.42 | 0.412 | 1.000 | |
| ATCG00140.1 | ATP synthase subunit (ATPH) | 46 | 1.84 | 0.667 | 1.000 | |
| ATCG00140.1 | ATP synthase subunit (ATPH) | 47 | 2.02 | 0.790 | 1.000 | |
| ATCG00150.1 | ATP synthase subunit (ATPI) | 2 | 1.98 | 0.360 | 0.966 | |
| ATCG00150.1 | ATP synthase subunit (ATPI) | 3 | 1.78 | 0.331 | 0.979 | |
| ATCG00150.1 | ATP synthase subunit (ATPI) | 4 | 1.61 | 0.311 | 0.991 | |
| ATCG00150.1 | ATP synthase subunit (ATPI) | 12 | 0.79 | 0.217 | 0.999 | |
| ATCG00150.1 | ATP synthase subunit (ATPI) | 13 | 0.75 | 0.202 | 0.993 | |
| ATCG00150.1 | ATP synthase subunit (ATPI) | 14 | 0.71 | 0.190 | 0.974 | |
| ATCG00150.1 | ATP synthase subunit (ATPI) | 15 | 0.68 | 0.181 | 0.939 | |
| ATCG00150.1 | ATP synthase subunit (ATPI) | 16 | 0.66 | 0.176 | 0.898 | |
| ATCG00150.1 | ATP synthase subunit (ATPI) | 17 | 0.64 | 0.175 | 0.866 | |
| ATCG00150.1 | ATP synthase subunit (ATPI) | 18 | 0.63 | 0.178 | 0.854 | |
| ATCG00150.1 | ATP synthase subunit (ATPI) | 19 | 0.63 | 0.183 | 0.860 | |
| ATCG00150.1 | ATP synthase subunit (ATPI) | 25 | 0.73 | 0.173 | 0.975 | |
| ATCG00150.1 | ATP synthase subunit (ATPI) | 26 | 0.78 | 0.166 | 0.993 | |
| ATCG00150.1 | ATP synthase subunit (ATPI) | 27 | 0.84 | 0.161 | 0.999 | |
| ATCG00150.1 | ATP synthase subunit (ATPI) | 28 | 0.90 | 0.159 | 1.000 | |
| ATCG00150.1 | ATP synthase subunit (ATPI) | 29 | 0.94 | 0.162 | 1.000 | |
| ATCG00150.1 | ATP synthase subunit (ATPI) | 30 | 0.94 | 0.169 | 1.000 | |
| ATCG00150.1 | ATP synthase subunit (ATPI) | 31 | 0.89 | 0.181 | 1.000 | |
| ATCG00150.1 | ATP synthase subunit (ATPI) | 32 | 0.80 | 0.197 | 0.999 | |
| ATCG00150.1 | ATP synthase subunit (ATPI) | 33 | 0.69 | 0.218 | 0.983 | |
| ATCG00150.1 | ATP synthase subunit (ATPI) | 34 | 0.58 | 0.241 | 0.923 | |
| ATCG00150.1 | ATP synthase subunit (ATPI) | 46 | 0.03 | 0.381 | 0.001 | *** |
| ATCG00150.1 | ATP synthase subunit (ATPI) | 47 | 0.02 | 0.406 | 0.000 | *** |
| ATCG00270.1 | Photosystem II subunit (PSBA) | 2 | 1.29 | 0.238 | 1.000 | |
| ATCG00270.1 | Photosystem II subunit (PSBA) | 3 | 0.96 | 0.148 | 1.000 | |
| ATCG00270.1 | Photosystem II subunit (PSBA) | 4 | 0.86 | 0.131 | 1.000 | |
| ATCG00270.1 | Photosystem II subunit (PSBA) | 5 | 0.93 | 0.131 | 1.000 | |
| ATCG00270.1 | Photosystem II subunit (PSBA) | 6 | 1.02 | 0.131 | 1.000 | |
| ATCG00270.1 | Photosystem II subunit (PSBA) | 7 | 1.07 | 0.130 | 1.000 | |
| ATCG00270.1 | Photosystem II subunit (PSBA) | 8 | 1.08 | 0.128 | 1.000 | |
| ATCG00270.1 | Photosystem II subunit (PSBA) | 9 | 1.07 | 0.126 | 1.000 | |
| ATCG00270.1 | Photosystem II subunit (PSBA) | 10 | 1.02 | 0.127 | 1.000 | |
| ATCG00270.1 | Photosystem II subunit (PSBA) | 11 | 0.98 | 0.127 | 1.000 | |
| ATCG00270.1 | Photosystem II subunit (PSBA) | 12 | 0.98 | 0.127 | 1.000 | |
| ATCG00270.1 | Photosystem II subunit (PSBA) | 13 | 1.00 | 0.126 | 1.000 | |
| ATCG00270.1 | Photosystem II subunit (PSBA) | 14 | 1.04 | 0.124 | 1.000 | |
| ATCG00270.1 | Photosystem II subunit (PSBA) | 15 | 1.03 | 0.123 | 1.000 | |
| ATCG00270.1 | Photosystem II subunit (PSBA) | 16 | 0.98 | 0.122 | 1.000 | |
| ATCG00270.1 | Photosystem II subunit (PSBA) | 17 | 0.98 | 0.122 | 1.000 | |

| | | | | | |
|---|---|---|---|---|---|
| ATCG00270.1 | Photosystem II subunit (PSBA) | 18 | 1.05 | 0.124 | 1.000 |
| ATCG00270.1 | Photosystem II subunit (PSBA) | 19 | 1.13 | 0.128 | 1.000 |
| ATCG00270.1 | Photosystem II subunit (PSBA) | 20 | 1.19 | 0.131 | 1.000 |
| ATCG00270.1 | Photosystem II subunit (PSBA) | 21 | 1.21 | 0.132 | 1.000 |
| ATCG00270.1 | Photosystem II subunit (PSBA) | 22 | 1.18 | 0.132 | 1.000 |
| ATCG00270.1 | Photosystem II subunit (PSBA) | 23 | 1.06 | 0.131 | 1.000 |
| ATCG00270.1 | Photosystem II subunit (PSBA) | 24 | 0.93 | 0.128 | 1.000 |
| ATCG00270.1 | Photosystem II subunit (PSBA) | 25 | 0.87 | 0.126 | 1.000 |
| ATCG00270.1 | Photosystem II subunit (PSBA) | 26 | 0.89 | 0.125 | 1.000 |
| ATCG00270.1 | Photosystem II subunit (PSBA) | 27 | 0.89 | 0.126 | 1.000 |
| ATCG00270.1 | Photosystem II subunit (PSBA) | 28 | 0.84 | 0.126 | 1.000 |
| ATCG00270.1 | Photosystem II subunit (PSBA) | 29 | 0.92 | 0.126 | 1.000 |
| ATCG00270.1 | Photosystem II subunit (PSBA) | 30 | 1.18 | 0.126 | 1.000 |
| ATCG00270.1 | Photosystem II subunit (PSBA) | 31 | 1.34 | 0.126 | 1.000 |
| ATCG00270.1 | Photosystem II subunit (PSBA) | 32 | 1.28 | 0.127 | 1.000 |
| ATCG00270.1 | Photosystem II subunit (PSBA) | 33 | 1.18 | 0.127 | 1.000 |
| ATCG00270.1 | Photosystem II subunit (PSBA) | 34 | 1.15 | 0.128 | 1.000 |
| ATCG00270.1 | Photosystem II subunit (PSBA) | 35 | 1.26 | 0.130 | 1.000 |
| ATCG00270.1 | Photosystem II subunit (PSBA) | 36 | 1.55 | 0.139 | 0.988 |
| ATCG00270.1 | Photosystem II subunit (PSBA) | 37 | 1.94 | 0.158 | 0.836 |
| ATCG00270.1 | Photosystem II subunit (PSBA) | 38 | 2.28 | 0.183 | 0.743 |
| ATCG00270.1 | Photosystem II subunit (PSBA) | 40 | 2.22 | 0.179 | 0.755 |
| ATCG00270.1 | Photosystem II subunit (PSBA) | 41 | 1.73 | 0.168 | 0.983 |
| ATCG00270.1 | Photosystem II subunit (PSBA) | 42 | 1.20 | 0.196 | 1.000 |
| ATCG00270.1 | Photosystem II subunit (PSBA) | 43 | 0.88 | 0.271 | 1.000 |
| ATCG00270.1 | Photosystem II subunit (PSBA) | 46 | 0.51 | 0.739 | 1.000 |
| ATCG00270.1 | Photosystem II subunit (PSBA) | 47 | 0.45 | 0.962 | 1.000 |
| ATCG00280.1 | photosystem II subunit (PSBC) | 2 | 0.71 | 0.353 | 1.000 |
| ATCG00280.1 | photosystem II subunit (PSBC) | 3 | 0.64 | 0.208 | 1.000 |
| ATCG00280.1 | photosystem II subunit (PSBC) | 4 | 0.73 | 0.200 | 1.000 |
| ATCG00280.1 | photosystem II subunit (PSBC) | 5 | 0.96 | 0.199 | 1.000 |
| ATCG00280.1 | photosystem II subunit (PSBC) | 6 | 1.08 | 0.198 | 1.000 |
| ATCG00280.1 | photosystem II subunit (PSBC) | 7 | 1.02 | 0.199 | 1.000 |
| ATCG00280.1 | photosystem II subunit (PSBC) | 8 | 1.01 | 0.195 | 1.000 |
| ATCG00280.1 | photosystem II subunit (PSBC) | 9 | 1.02 | 0.192 | 1.000 |
| ATCG00280.1 | photosystem II subunit (PSBC) | 10 | 0.95 | 0.192 | 1.000 |
| ATCG00280.1 | photosystem II subunit (PSBC) | 11 | 0.91 | 0.194 | 1.000 |
| ATCG00280.1 | photosystem II subunit (PSBC) | 12 | 0.93 | 0.194 | 1.000 |
| ATCG00280.1 | photosystem II subunit (PSBC) | 13 | 0.97 | 0.193 | 1.000 |
| ATCG00280.1 | photosystem II subunit (PSBC) | 14 | 1.04 | 0.190 | 1.000 |
| ATCG00280.1 | photosystem II subunit (PSBC) | 15 | 1.02 | 0.189 | 1.000 |
| ATCG00280.1 | photosystem II subunit (PSBC) | 16 | 0.89 | 0.187 | 1.000 |
| ATCG00280.1 | photosystem II subunit (PSBC) | 17 | 0.90 | 0.185 | 1.000 |
| ATCG00280.1 | photosystem II subunit (PSBC) | 18 | 1.05 | 0.190 | 1.000 |
| ATCG00280.1 | photosystem II subunit (PSBC) | 19 | 1.26 | 0.196 | 1.000 |
| ATCG00280.1 | photosystem II subunit (PSBC) | 20 | 1.34 | 0.200 | 1.000 |
| ATCG00280.1 | photosystem II subunit (PSBC) | 21 | 1.29 | 0.203 | 1.000 |
| ATCG00280.1 | photosystem II subunit (PSBC) | 22 | 1.23 | 0.202 | 1.000 |
| ATCG00280.1 | photosystem II subunit (PSBC) | 23 | 1.02 | 0.203 | 1.000 |
| ATCG00280.1 | photosystem II subunit (PSBC) | 24 | 0.76 | 0.196 | 1.000 |
| ATCG00280.1 | photosystem II subunit (PSBC) | 25 | 0.56 | 0.191 | 0.998 |
| ATCG00280.1 | photosystem II subunit (PSBC) | 26 | 0.54 | 0.191 | 0.994 |
| ATCG00280.1 | photosystem II subunit (PSBC) | 27 | 0.64 | 0.191 | 1.000 |
| ATCG00280.1 | photosystem II subunit (PSBC) | 28 | 0.60 | 0.192 | 1.000 |
| ATCG00280.1 | photosystem II subunit (PSBC) | 29 | 0.76 | 0.194 | 1.000 |
| ATCG00280.1 | photosystem II subunit (PSBC) | 30 | 1.42 | 0.194 | 1.000 |
| ATCG00280.1 | photosystem II subunit (PSBC) | 31 | 1.25 | 0.194 | 1.000 |
| ATCG00280.1 | photosystem II subunit (PSBC) | 32 | 0.47 | 0.194 | 0.951 |
| ATCG00280.1 | photosystem II subunit (PSBC) | 33 | 0.27 | 0.194 | 0.130 |
| ATCG00280.1 | photosystem II subunit (PSBC) | 34 | 0.37 | 0.197 | 0.616 |
| ATCG00280.1 | photosystem II subunit (PSBC) | 35 | 0.49 | 0.206 | 0.988 |
| ATCG00280.1 | photosystem II subunit (PSBC) | 36 | 0.65 | 0.227 | 1.000 |
| ATCG00280.1 | photosystem II subunit (PSBC) | 37 | 1.12 | 0.250 | 1.000 |

| | | | | | | |
|---|---|---|---|---|---|---|
| ATCG00280.1 | photosystem II subunit (PSBC) | 38 | 1.76 | 0.246 | 1.000 | |
| ATCG00280.1 | photosystem II subunit (PSBC) | 39 | 1.82 | 0.225 | 1.000 | |
| ATCG00280.1 | photosystem II subunit (PSBC) | 40 | 1.01 | 0.201 | 1.000 | |
| ATCG00280.1 | photosystem II subunit (PSBC) | 41 | 0.39 | 0.196 | 0.700 | |
| ATCG00280.1 | photosystem II subunit (PSBC) | 42 | 0.17 | 0.200 | 0.006 | ** |
| ATCG00280.1 | photosystem II subunit (PSBC) | 43 | 0.16 | 0.207 | 0.007 | ** |
| ATCG00280.1 | photosystem II subunit (PSBC) | 44 | 0.40 | 0.257 | 0.979 | |
| ATCG00280.1 | photosystem II subunit (PSBC) | 45 | 1.83 | 0.389 | 1.000 | |
| ATCG00280.1 | photosystem II subunit (PSBC) | 46 | 12.71 | 0.663 | 0.956 | |
| ATCG00280.1 | photosystem II subunit (PSBC) | 47 | 105.26 | 1.050 | 0.833 | |
| ATCG00340.1 | Photosystem I subunit (PSAB) | 2 | 0.56 | 0.227 | 0.999 | |
| ATCG00340.1 | Photosystem I subunit (PSAB) | 3 | 0.59 | 0.139 | 0.917 | |
| ATCG00340.1 | Photosystem I subunit (PSAB) | 4 | 0.70 | 0.127 | 0.996 | |
| ATCG00340.1 | Photosystem I subunit (PSAB) | 5 | 0.88 | 0.127 | 1.000 | |
| ATCG00340.1 | Photosystem I subunit (PSAB) | 6 | 0.96 | 0.127 | 1.000 | |
| ATCG00340.1 | Photosystem I subunit (PSAB) | 7 | 0.95 | 0.127 | 1.000 | |
| ATCG00340.1 | Photosystem I subunit (PSAB) | 8 | 0.93 | 0.124 | 1.000 | |
| ATCG00340.1 | Photosystem I subunit (PSAB) | 9 | 0.91 | 0.122 | 1.000 | |
| ATCG00340.1 | Photosystem I subunit (PSAB) | 10 | 0.86 | 0.122 | 1.000 | |
| ATCG00340.1 | Photosystem I subunit (PSAB) | 11 | 0.82 | 0.123 | 1.000 | |
| ATCG00340.1 | Photosystem I subunit (PSAB) | 12 | 0.84 | 0.123 | 1.000 | |
| ATCG00340.1 | Photosystem I subunit (PSAB) | 13 | 0.88 | 0.121 | 1.000 | |
| ATCG00340.1 | Photosystem I subunit (PSAB) | 14 | 0.89 | 0.120 | 1.000 | |
| ATCG00340.1 | Photosystem I subunit (PSAB) | 15 | 0.86 | 0.119 | 1.000 | |
| ATCG00340.1 | Photosystem I subunit (PSAB) | 16 | 0.84 | 0.119 | 1.000 | |
| ATCG00340.1 | Photosystem I subunit (PSAB) | 17 | 0.89 | 0.118 | 1.000 | |
| ATCG00340.1 | Photosystem I subunit (PSAB) | 18 | 1.00 | 0.119 | 1.000 | |
| ATCG00340.1 | Photosystem I subunit (PSAB) | 19 | 1.13 | 0.123 | 1.000 | |
| ATCG00340.1 | Photosystem I subunit (PSAB) | 20 | 1.18 | 0.127 | 1.000 | |
| ATCG00340.1 | Photosystem I subunit (PSAB) | 21 | 1.10 | 0.128 | 1.000 | |
| ATCG00340.1 | Photosystem I subunit (PSAB) | 22 | 0.93 | 0.128 | 1.000 | |
| ATCG00340.1 | Photosystem I subunit (PSAB) | 23 | 0.76 | 0.127 | 1.000 | |
| ATCG00340.1 | Photosystem I subunit (PSAB) | 24 | 0.64 | 0.124 | 0.944 | |
| ATCG00340.1 | Photosystem I subunit (PSAB) | 25 | 0.66 | 0.122 | 0.965 | |
| ATCG00340.1 | Photosystem I subunit (PSAB) | 26 | 0.91 | 0.121 | 1.000 | |
| ATCG00340.1 | Photosystem I subunit (PSAB) | 27 | 1.21 | 0.121 | 1.000 | |
| ATCG00340.1 | Photosystem I subunit (PSAB) | 28 | 1.38 | 0.122 | 0.997 | |
| ATCG00340.1 | Photosystem I subunit (PSAB) | 29 | 1.65 | 0.123 | 0.836 | |
| ATCG00340.1 | Photosystem I subunit (PSAB) | 30 | 1.66 | 0.123 | 0.825 | |
| ATCG00340.1 | Photosystem I subunit (PSAB) | 31 | 1.18 | 0.124 | 1.000 | |
| ATCG00340.1 | Photosystem I subunit (PSAB) | 32 | 0.75 | 0.128 | 1.000 | |
| ATCG00340.1 | Photosystem I subunit (PSAB) | 33 | 0.49 | 0.136 | 0.422 | |
| ATCG00340.1 | Photosystem I subunit (PSAB) | 34 | 0.37 | 0.170 | 0.274 | |
| ATCG00340.1 | Photosystem I subunit (PSAB) | 35 | 0.36 | 0.264 | 0.892 | |
| ATCG00340.1 | Photosystem I subunit (PSAB) | 46 | 22794.24 | 2.421 | 0.819 | |
| ATCG00340.1 | Photosystem I subunit (PSAB) | 47 | 103279.77 | 2.721 | 0.789 | |
| ATCG00350.1 | Photosystem I subunit (PSAA) | 2 | 0.45 | 0.216 | 0.935 | |
| ATCG00350.1 | Photosystem I subunit (PSAA) | 3 | 0.42 | 0.130 | 0.113 | |
| ATCG00350.1 | Photosystem I subunit (PSAA) | 4 | 0.47 | 0.120 | 0.164 | |
| ATCG00350.1 | Photosystem I subunit (PSAA) | 5 | 0.57 | 0.120 | 0.652 | |
| ATCG00350.1 | Photosystem I subunit (PSAA) | 6 | 0.61 | 0.120 | 0.840 | |
| ATCG00350.1 | Photosystem I subunit (PSAA) | 7 | 0.60 | 0.120 | 0.813 | |
| ATCG00350.1 | Photosystem I subunit (PSAA) | 8 | 0.63 | 0.117 | 0.880 | |
| ATCG00350.1 | Photosystem I subunit (PSAA) | 9 | 0.65 | 0.116 | 0.920 | |
| ATCG00350.1 | Photosystem I subunit (PSAA) | 10 | 0.65 | 0.116 | 0.919 | |
| ATCG00350.1 | Photosystem I subunit (PSAA) | 11 | 0.68 | 0.117 | 0.972 | |
| ATCG00350.1 | Photosystem I subunit (PSAA) | 12 | 0.75 | 0.117 | 0.999 | |
| ATCG00350.1 | Photosystem I subunit (PSAA) | 13 | 0.82 | 0.115 | 1.000 | |
| ATCG00350.1 | Photosystem I subunit (PSAA) | 14 | 0.88 | 0.113 | 1.000 | |
| ATCG00350.1 | Photosystem I subunit (PSAA) | 15 | 0.85 | 0.113 | 1.000 | |
| ATCG00350.1 | Photosystem I subunit (PSAA) | 16 | 0.81 | 0.112 | 1.000 | |
| ATCG00350.1 | Photosystem I subunit (PSAA) | 17 | 0.87 | 0.112 | 1.000 | |
| ATCG00350.1 | Photosystem I subunit (PSAA) | 18 | 1.04 | 0.113 | 1.000 | |

| | | | | | | |
|---|---|---|---|---|---|---|
| ATCG00350.1 | Photosystem I subunit (PSAA) | 19 | 1.28 | 0.116 | 1.000 | |
| ATCG00350.1 | Photosystem I subunit (PSAA) | 20 | 1.49 | 0.120 | 0.975 | |
| ATCG00350.1 | Photosystem I subunit (PSAA) | 21 | 1.49 | 0.121 | 0.977 | |
| ATCG00350.1 | Photosystem I subunit (PSAA) | 22 | 1.25 | 0.121 | 1.000 | |
| ATCG00350.1 | Photosystem I subunit (PSAA) | 23 | 0.94 | 0.121 | 1.000 | |
| ATCG00350.1 | Photosystem I subunit (PSAA) | 24 | 0.71 | 0.117 | 0.995 | |
| ATCG00350.1 | Photosystem I subunit (PSAA) | 25 | 0.68 | 0.115 | 0.966 | |
| ATCG00350.1 | Photosystem I subunit (PSAA) | 26 | 0.91 | 0.115 | 1.000 | |
| ATCG00350.1 | Photosystem I subunit (PSAA) | 27 | 1.24 | 0.115 | 1.000 | |
| ATCG00350.1 | Photosystem I subunit (PSAA) | 28 | 1.49 | 0.115 | 0.959 | |
| ATCG00350.1 | Photosystem I subunit (PSAA) | 29 | 1.89 | 0.116 | 0.363 | |
| ATCG00350.1 | Photosystem I subunit (PSAA) | 30 | 1.97 | 0.116 | 0.264 | |
| ATCG00350.1 | Photosystem I subunit (PSAA) | 31 | 1.41 | 0.118 | 0.994 | |
| ATCG00350.1 | Photosystem I subunit (PSAA) | 32 | 0.95 | 0.121 | 1.000 | |
| ATCG00350.1 | Photosystem I subunit (PSAA) | 33 | 0.74 | 0.129 | 1.000 | |
| ATCG00350.1 | Photosystem I subunit (PSAA) | 34 | 0.65 | 0.159 | 0.998 | |
| ATCG00350.1 | Photosystem I subunit (PSAA) | 35 | 0.67 | 0.250 | 1.000 | |
| ATCG00350.1 | Photosystem I subunit (PSAA) | 46 | 3490.12 | 2.638 | 0.988 | |
| ATCG00350.1 | Photosystem I subunit (PSAA) | 47 | 10991.05 | 2.969 | 0.986 | |
| ATCG00470.1 | ATP synthase subunit (ATPepsilon) | 2 | 1.07 | 0.615 | 1.000 | |
| ATCG00470.1 | ATP synthase subunit (ATPepsilon) | 3 | 0.52 | 0.608 | 1.000 | |
| ATCG00470.1 | ATP synthase subunit (ATPepsilon) | 25 | 0.06 | 0.447 | 0.088 | . |
| ATCG00470.1 | ATP synthase subunit (ATPepsilon) | 26 | 0.20 | 0.312 | 0.257 | |
| ATCG00470.1 | ATP synthase subunit (ATPepsilon) | 27 | 0.66 | 0.268 | 0.998 | |
| ATCG00470.1 | ATP synthase subunit (ATPepsilon) | 28 | 1.50 | 0.252 | 0.997 | |
| ATCG00470.1 | ATP synthase subunit (ATPepsilon) | 29 | 3.44 | 0.248 | 0.281 | |
| ATCG00470.1 | ATP synthase subunit (ATPepsilon) | 30 | 5.22 | 0.247 | 0.048 | * |
| ATCG00470.1 | ATP synthase subunit (ATPepsilon) | 31 | 3.33 | 0.250 | 0.325 | |
| ATCG00470.1 | ATP synthase subunit (ATPepsilon) | 32 | 0.96 | 0.255 | 1.000 | |
| ATCG00470.1 | ATP synthase subunit (ATPepsilon) | 33 | 0.29 | 0.274 | 0.411 | |
| ATCG00470.1 | ATP synthase subunit (ATPepsilon) | 34 | 0.17 | 0.362 | 0.282 | |
| ATCG00470.1 | ATP synthase subunit (ATPepsilon) | 35 | 0.14 | 0.590 | 0.760 | |
| ATCG00470.1 | ATP synthase subunit (ATPepsilon) | 46 | 93.08 | 2.686 | 0.996 | |
| ATCG00470.1 | ATP synthase subunit (ATPepsilon) | 47 | 245.68 | 2.995 | 0.993 | |
| ATCG00480.1 | Beta subunit of ATP synthase | 3 | 0.33 | 0.309 | 0.929 | |
| ATCG00480.1 | Beta subunit of ATP synthase | 4 | 0.35 | 0.285 | 0.917 | |
| ATCG00480.1 | Beta subunit of ATP synthase | 5 | 0.41 | 0.285 | 0.983 | |
| ATCG00480.1 | Beta subunit of ATP synthase | 6 | 0.43 | 0.286 | 0.990 | |
| ATCG00480.1 | Beta subunit of ATP synthase | 7 | 0.39 | 0.286 | 0.964 | |
| ATCG00480.1 | Beta subunit of ATP synthase | 8 | 0.35 | 0.281 | 0.905 | |
| ATCG00480.1 | Beta subunit of ATP synthase | 9 | 0.35 | 0.278 | 0.886 | |
| ATCG00480.1 | Beta subunit of ATP synthase | 10 | 0.33 | 0.278 | 0.831 | |
| ATCG00480.1 | Beta subunit of ATP synthase | 11 | 0.36 | 0.279 | 0.906 | |
| ATCG00480.1 | Beta subunit of ATP synthase | 12 | 0.40 | 0.279 | 0.968 | |
| ATCG00480.1 | Beta subunit of ATP synthase | 13 | 0.41 | 0.276 | 0.975 | |
| ATCG00480.1 | Beta subunit of ATP synthase | 14 | 0.43 | 0.273 | 0.981 | |
| ATCG00480.1 | Beta subunit of ATP synthase | 15 | 0.42 | 0.272 | 0.974 | |

| | | | | | | |
|---|---|---|---|---|---|---|
| ATCG00480.1 | Beta subunit of ATP synthase | 16 | 0.40 | 0.271 | 0.955 | |
| ATCG00480.1 | Beta subunit of ATP synthase | 17 | 0.43 | 0.271 | 0.978 | |
| ATCG00480.1 | Beta subunit of ATP synthase | 18 | 0.46 | 0.274 | 0.993 | |
| ATCG00480.1 | Beta subunit of ATP synthase | 19 | 0.43 | 0.279 | 0.985 | |
| ATCG00480.1 | Beta subunit of ATP synthase | 20 | 0.36 | 0.285 | 0.922 | |
| ATCG00480.1 | Beta subunit of ATP synthase | 21 | 0.26 | 0.288 | 0.621 | |
| ATCG00480.1 | Beta subunit of ATP synthase | 22 | 0.19 | 0.288 | 0.265 | |
| ATCG00480.1 | Beta subunit of ATP synthase | 23 | 0.16 | 0.286 | 0.131 | |
| ATCG00480.1 | Beta subunit of ATP synthase | 24 | 0.19 | 0.282 | 0.234 | |
| ATCG00480.1 | Beta subunit of ATP synthase | 25 | 0.43 | 0.278 | 0.986 | |
| ATCG00480.1 | Beta subunit of ATP synthase | 26 | 1.19 | 0.276 | 1.000 | |
| ATCG00480.1 | Beta subunit of ATP synthase | 27 | 1.91 | 0.277 | 1.000 | |
| ATCG00480.1 | Beta subunit of ATP synthase | 28 | 1.89 | 0.276 | 1.000 | |
| ATCG00480.1 | Beta subunit of ATP synthase | 29 | 1.94 | 0.277 | 0.999 | |
| ATCG00480.1 | Beta subunit of ATP synthase | 30 | 2.14 | 0.276 | 0.996 | |
| ATCG00480.1 | Beta subunit of ATP synthase | 31 | 1.82 | 0.277 | 1.000 | |
| ATCG00480.1 | Beta subunit of ATP synthase | 32 | 1.08 | 0.278 | 1.000 | |
| ATCG00480.1 | Beta subunit of ATP synthase | 33 | 0.62 | 0.278 | 1.000 | |
| ATCG00480.1 | Beta subunit of ATP synthase | 34 | 0.52 | 0.278 | 1.000 | |
| ATCG00480.1 | Beta subunit of ATP synthase | 35 | 0.62 | 0.278 | 1.000 | |
| ATCG00480.1 | Beta subunit of ATP synthase | 36 | 0.84 | 0.277 | 1.000 | |
| ATCG00480.1 | Beta subunit of ATP synthase | 37 | 1.18 | 0.277 | 1.000 | |
| ATCG00480.1 | Beta subunit of ATP synthase | 38 | 1.62 | 0.277 | 1.000 | |
| ATCG00480.1 | Beta subunit of ATP synthase | 39 | 2.32 | 0.278 | 0.986 | |
| ATCG00480.1 | Beta subunit of ATP synthase | 40 | 3.08 | 0.278 | 0.821 | |
| ATCG00480.1 | Beta subunit of ATP synthase | 41 | 2.94 | 0.279 | 0.868 | |
| ATCG00480.1 | Beta subunit of ATP synthase | 42 | 2.19 | 0.282 | 0.995 | |
| ATCG00480.1 | Beta subunit of ATP synthase | 43 | 1.63 | 0.284 | 1.000 | |
| ATCG00480.1 | Beta subunit of ATP synthase | 44 | 1.41 | 0.284 | 1.000 | |
| ATCG00480.1 | Beta subunit of ATP synthase | 45 | 1.13 | 0.286 | 1.000 | |
| ATCG00480.1 | Beta subunit of ATP synthase | 46 | 0.66 | 0.327 | 1.000 | |
| ATCG00480.1 | Beta subunit of ATP synthase | 47 | 0.29 | 0.521 | 1.000 | |
| ATCG00480.1 | Beta subunit of ATP synthase | 48 | 0.13 | 0.833 | 0.999 | |
| ATCG00490.1 | RUBISCO large subunit (RBCL) | 2 | 0.06 | 0.470 | 1.000 | |
| ATCG00490.1 | RUBISCO large subunit (RBCL) | 3 | 0.06 | 0.347 | 0.011 | * |
| ATCG00490.1 | RUBISCO large subunit (RBCL) | 4 | 0.07 | 0.325 | 0.009 | ** |
| ATCG00490.1 | RUBISCO large subunit (RBCL) | 5 | 0.08 | 0.326 | 0.019 | * |
| ATCG00490.1 | RUBISCO large subunit (RBCL) | 6 | 0.09 | 0.326 | 0.033 | * |
| ATCG00490.1 | RUBISCO large subunit (RBCL) | 7 | 0.10 | 0.325 | 0.041 | * |
| ATCG00490.1 | RUBISCO large subunit (RBCL) | 8 | 0.10 | 0.322 | 0.040 | * |
| ATCG00490.1 | RUBISCO large subunit (RBCL) | 9 | 0.09 | 0.320 | 0.027 | * |
| ATCG00490.1 | RUBISCO large subunit (RBCL) | 10 | 0.09 | 0.320 | 0.020 | * |
| ATCG00490.1 | RUBISCO large subunit (RBCL) | 11 | 0.08 | 0.321 | 0.016 | * |
| ATCG00490.1 | RUBISCO large subunit (RBCL) | 12 | 0.07 | 0.320 | 0.010 | * |
| ATCG00490.1 | RUBISCO large subunit (RBCL) | 13 | 0.06 | 0.319 | 0.004 | ** |
| ATCG00490.1 | RUBISCO large subunit (RBCL) | 14 | 0.06 | 0.318 | 0.004 | ** |
| ATCG00490.1 | RUBISCO large subunit (RBCL) | 15 | 0.07 | 0.319 | 0.007 | ** |
| ATCG00490.1 | RUBISCO large subunit (RBCL) | 16 | 0.09 | 0.319 | 0.019 | * |
| ATCG00490.1 | RUBISCO large subunit (RBCL) | 17 | 0.12 | 0.320 | 0.069 | . |
| ATCG00490.1 | RUBISCO large subunit (RBCL) | 18 | 0.17 | 0.328 | 1.000 | |
| ATCG00490.1 | RUBISCO large subunit (RBCL) | 19 | 0.24 | 0.335 | 1.000 | |
| ATCG00490.1 | RUBISCO large subunit (RBCL) | 20 | 0.31 | 0.337 | 0.840 | |
| ATCG00490.1 | RUBISCO large subunit (RBCL) | 21 | 0.40 | 0.333 | 0.969 | |
| ATCG00490.1 | RUBISCO large subunit (RBCL) | 22 | 0.52 | 0.329 | 0.999 | |
| ATCG00490.1 | RUBISCO large subunit (RBCL) | 23 | 0.63 | 0.326 | 1.000 | |
| ATCG00490.1 | RUBISCO large subunit (RBCL) | 24 | 0.74 | 0.323 | 1.000 | |
| ATCG00490.1 | RUBISCO large subunit (RBCL) | 25 | 0.77 | 0.320 | 1.000 | |
| ATCG00490.1 | RUBISCO large subunit (RBCL) | 26 | 0.69 | 0.318 | 1.000 | |
| ATCG00490.1 | RUBISCO large subunit (RBCL) | 27 | 0.78 | 0.318 | 1.000 | |
| ATCG00490.1 | RUBISCO large subunit (RBCL) | 28 | 1.27 | 0.318 | 1.000 | |
| ATCG00490.1 | RUBISCO large subunit (RBCL) | 29 | 2.29 | 0.319 | 0.984 | |
| ATCG00490.1 | RUBISCO large subunit (RBCL) | 30 | 2.49 | 0.320 | 0.959 | |
| ATCG00490.1 | RUBISCO large subunit (RBCL) | 31 | 1.53 | 0.320 | 1.000 | |

| | | | | | |
|---|---|---|---|---|---|
| ATCG00490.1 | RUBISCO large subunit (RBCL) | 32 | 0.96 | 0.320 | 1.000 |
| ATCG00490.1 | RUBISCO large subunit (RBCL) | 33 | 0.77 | 0.320 | 1.000 |
| ATCG00490.1 | RUBISCO large subunit (RBCL) | 34 | 0.70 | 0.321 | 1.000 |
| ATCG00490.1 | RUBISCO large subunit (RBCL) | 35 | 0.69 | 0.320 | 1.000 |
| ATCG00490.1 | RUBISCO large subunit (RBCL) | 36 | 0.79 | 0.320 | 1.000 |
| ATCG00490.1 | RUBISCO large subunit (RBCL) | 37 | 1.04 | 0.322 | 1.000 |
| ATCG00490.1 | RUBISCO large subunit (RBCL) | 38 | 1.38 | 0.327 | 1.000 |
| ATCG00490.1 | RUBISCO large subunit (RBCL) | 39 | 1.77 | 0.335 | 1.000 |
| ATCG00490.1 | RUBISCO large subunit (RBCL) | 40 | 2.15 | 0.341 | 0.997 |
| ATCG00490.1 | RUBISCO large subunit (RBCL) | 41 | 2.33 | 0.350 | 0.993 |
| ATCG00490.1 | RUBISCO large subunit (RBCL) | 42 | 2.17 | 0.372 | 0.999 |
| ATCG00490.1 | RUBISCO large subunit (RBCL) | 43 | 1.81 | 0.386 | 1.000 |
| ATCG00490.1 | RUBISCO large subunit (RBCL) | 44 | 1.32 | 0.374 | 1.000 |
| ATCG00490.1 | RUBISCO large subunit (RBCL) | 45 | 0.81 | 0.354 | 1.000 |
| ATCG00490.1 | RUBISCO large subunit (RBCL) | 46 | 0.40 | 0.366 | 0.990 |
| ATCG00490.1 | RUBISCO large subunit (RBCL) | 47 | 0.20 | 0.522 | 0.916 |
| ATCG00540.1 | Cytochrome b6f complex subunit (PETA) | 2 | 0.45 | 0.411 | 1.000 |
| ATCG00540.1 | Cytochrome b6f complex subunit (PETA) | 3 | 0.57 | 0.276 | 1.000 |
| ATCG00540.1 | Cytochrome b6f complex subunit (PETA) | 4 | 0.73 | 0.257 | 1.000 |
| ATCG00540.1 | Cytochrome b6f complex subunit (PETA) | 5 | 0.95 | 0.257 | 1.000 |
| ATCG00540.1 | Cytochrome b6f complex subunit (PETA) | 6 | 1.04 | 0.258 | 1.000 |
| ATCG00540.1 | Cytochrome b6f complex subunit (PETA) | 7 | 0.93 | 0.259 | 1.000 |
| ATCG00540.1 | Cytochrome b6f complex subunit (PETA) | 8 | 0.85 | 0.254 | 1.000 |
| ATCG00540.1 | Cytochrome b6f complex subunit (PETA) | 9 | 0.84 | 0.250 | 1.000 |
| ATCG00540.1 | Cytochrome b6f complex subunit (PETA) | 10 | 0.83 | 0.251 | 1.000 |
| ATCG00540.1 | Cytochrome b6f complex subunit (PETA) | 11 | 0.83 | 0.252 | 1.000 |
| ATCG00540.1 | Cytochrome b6f complex subunit (PETA) | 12 | 0.82 | 0.252 | 1.000 |
| ATCG00540.1 | Cytochrome b6f complex subunit (PETA) | 13 | 0.83 | 0.249 | 1.000 |
| ATCG00540.1 | Cytochrome b6f complex subunit (PETA) | 14 | 0.84 | 0.245 | 1.000 |
| ATCG00540.1 | Cytochrome b6f complex subunit (PETA) | 15 | 0.81 | 0.245 | 1.000 |
| ATCG00540.1 | Cytochrome b6f complex subunit (PETA) | 16 | 0.84 | 0.244 | 1.000 |
| ATCG00540.1 | Cytochrome b6f complex subunit (PETA) | 17 | 1.02 | 0.244 | 1.000 |
| ATCG00540.1 | Cytochrome b6f complex subunit (PETA) | 18 | 1.22 | 0.247 | 1.000 |
| ATCG00540.1 | Cytochrome b6f complex subunit (PETA) | 19 | 1.42 | 0.252 | 1.000 |
| ATCG00540.1 | Cytochrome b6f complex subunit (PETA) | 20 | 1.47 | 0.257 | 1.000 |
| ATCG00540.1 | Cytochrome b6f complex subunit (PETA) | 21 | 1.32 | 0.260 | 1.000 |
| ATCG00540.1 | Cytochrome b6f complex subunit (PETA) | 22 | 1.02 | 0.260 | 1.000 |
| ATCG00540.1 | Cytochrome b6f complex subunit (PETA) | 23 | 0.75 | 0.259 | 1.000 |
| ATCG00540.1 | Cytochrome b6f complex subunit (PETA) | 24 | 0.58 | 0.254 | 1.000 |

| | | | | | | |
|---|---|---|---|---|---|---|
| ATCG00540.1 | Cytochrome b6f complex subunit (PETA) | 25 | 0.49 | 0.250 | 0.995 | |
| ATCG00540.1 | Cytochrome b6f complex subunit (PETA) | 26 | 0.52 | 0.249 | 0.998 | |
| ATCG00540.1 | Cytochrome b6f complex subunit (PETA) | 27 | 0.80 | 0.249 | 1.000 | |
| ATCG00540.1 | Cytochrome b6f complex subunit (PETA) | 28 | 1.10 | 0.249 | 1.000 | |
| ATCG00540.1 | Cytochrome b6f complex subunit (PETA) | 29 | 1.64 | 0.249 | 1.000 | |
| ATCG00540.1 | Cytochrome b6f complex subunit (PETA) | 30 | 2.83 | 0.249 | 0.778 | |
| ATCG00540.1 | Cytochrome b6f complex subunit (PETA) | 31 | 3.46 | 0.250 | 0.505 | |
| ATCG00540.1 | Cytochrome b6f complex subunit (PETA) | 32 | 1.41 | 0.250 | 1.000 | |
| ATCG00540.1 | Cytochrome b6f complex subunit (PETA) | 33 | 0.35 | 0.251 | 0.776 | |
| ATCG00540.1 | Cytochrome b6f complex subunit (PETA) | 34 | 0.17 | 0.251 | 0.071 | . |
| ATCG00540.1 | Cytochrome b6f complex subunit (PETA) | 35 | 0.22 | 0.251 | 0.188 | |
| ATCG00540.1 | Cytochrome b6f complex subunit (PETA) | 36 | 0.51 | 0.251 | 0.997 | |
| ATCG00540.1 | Cytochrome b6f complex subunit (PETA) | 37 | 1.01 | 0.251 | 1.000 | |
| ATCG00540.1 | Cytochrome b6f complex subunit (PETA) | 38 | 0.98 | 0.251 | 1.000 | |
| ATCG00540.1 | Cytochrome b6f complex subunit (PETA) | 39 | 0.65 | 0.251 | 1.000 | |
| ATCG00540.1 | Cytochrome b6f complex subunit (PETA) | 40 | 0.53 | 0.252 | 0.999 | |
| ATCG00540.1 | Cytochrome b6f complex subunit (PETA) | 41 | 0.54 | 0.264 | 1.000 | |
| ATCG00540.1 | Cytochrome b6f complex subunit (PETA) | 42 | 0.63 | 0.306 | 1.000 | |
| ATCG00540.1 | Cytochrome b6f complex subunit (PETA) | 43 | 0.83 | 0.322 | 1.000 | |
| ATCG00540.1 | Cytochrome b6f complex subunit (PETA) | 44 | 1.06 | 0.310 | 1.000 | |
| ATCG00540.1 | Cytochrome b6f complex subunit (PETA) | 45 | 1.11 | 0.285 | 1.000 | |
| ATCG00540.1 | Cytochrome b6f complex subunit (PETA) | 46 | 0.89 | 0.295 | 1.000 | |
| ATCG00540.1 | Cytochrome b6f complex subunit (PETA) | 47 | 0.66 | 0.457 | 1.000 | |
| ATCG00560.1 | Photosystem II subunit (PSBL) | 2 | 1.08 | 0.252 | 1.000 | |
| ATCG00560.1 | Photosystem II subunit (PSBL) | 3 | 1.30 | 0.172 | 1.000 | |
| ATCG00560.1 | Photosystem II subunit (PSBL) | 4 | 1.55 | 0.142 | 0.964 | |
| ATCG00560.1 | Photosystem II subunit (PSBL) | 5 | 1.74 | 0.137 | 0.775 | |
| ATCG00560.1 | Photosystem II subunit (PSBL) | 6 | 1.75 | 0.137 | 0.756 | |
| ATCG00560.1 | Photosystem II subunit (PSBL) | 7 | 1.63 | 0.136 | 0.886 | |
| ATCG00560.1 | Photosystem II subunit (PSBL) | 8 | 1.50 | 0.134 | 0.971 | |
| ATCG00560.1 | Photosystem II subunit (PSBL) | 9 | 1.42 | 0.132 | 0.992 | |
| ATCG00560.1 | Photosystem II subunit (PSBL) | 10 | 1.38 | 0.131 | 0.997 | |
| ATCG00560.1 | Photosystem II subunit (PSBL) | 11 | 1.35 | 0.132 | 0.998 | |
| ATCG00560.1 | Photosystem II subunit (PSBL) | 12 | 1.33 | 0.131 | 0.999 | |
| ATCG00560.1 | Photosystem II subunit (PSBL) | 13 | 1.28 | 0.130 | 1.000 | |
| ATCG00560.1 | Photosystem II subunit (PSBL) | 14 | 1.18 | 0.128 | 1.000 | |
| ATCG00560.1 | Photosystem II subunit (PSBL) | 15 | 1.07 | 0.127 | 1.000 | |
| ATCG00560.1 | Photosystem II subunit (PSBL) | 16 | 0.98 | 0.126 | 1.000 | |
| ATCG00560.1 | Photosystem II subunit (PSBL) | 17 | 0.95 | 0.127 | 1.000 | |

| | | | | | | |
|---|---|---|---|---|---|---|
| ATCG00560.1 | Photosystem II subunit (PSBL) | 18 | 1.01 | 0.128 | 1.000 | |
| ATCG00560.1 | Photosystem II subunit (PSBL) | 19 | 1.14 | 0.132 | 1.000 | |
| ATCG00560.1 | Photosystem II subunit (PSBL) | 20 | 1.26 | 0.135 | 1.000 | |
| ATCG00560.1 | Photosystem II subunit (PSBL) | 21 | 1.30 | 0.137 | 1.000 | |
| ATCG00560.1 | Photosystem II subunit (PSBL) | 22 | 1.25 | 0.137 | 1.000 | |
| ATCG00560.1 | Photosystem II subunit (PSBL) | 23 | 1.09 | 0.136 | 1.000 | |
| ATCG00560.1 | Photosystem II subunit (PSBL) | 24 | 0.88 | 0.133 | 1.000 | |
| ATCG00560.1 | Photosystem II subunit (PSBL) | 25 | 0.68 | 0.131 | 0.978 | |
| ATCG00560.1 | Photosystem II subunit (PSBL) | 26 | 0.56 | 0.130 | 0.638 | |
| ATCG00560.1 | Photosystem II subunit (PSBL) | 27 | 0.48 | 0.130 | 0.272 | |
| ATCG00560.1 | Photosystem II subunit (PSBL) | 28 | 0.43 | 0.130 | 0.117 | |
| ATCG00560.1 | Photosystem II subunit (PSBL) | 29 | 0.45 | 0.131 | 0.156 | |
| ATCG00560.1 | Photosystem II subunit (PSBL) | 30 | 0.48 | 0.133 | 0.315 | |
| ATCG00560.1 | Photosystem II subunit (PSBL) | 31 | 0.47 | 0.137 | 0.274 | |
| ATCG00560.1 | Photosystem II subunit (PSBL) | 32 | 0.39 | 0.143 | 0.089 | . |
| ATCG00560.1 | Photosystem II subunit (PSBL) | 33 | 0.31 | 0.156 | 1.000 | |
| ATCG00560.1 | Photosystem II subunit (PSBL) | 34 | 0.27 | 0.181 | 1.000 | |
| ATCG00560.1 | Photosystem II subunit (PSBL) | 35 | 0.27 | 0.230 | 0.251 | |
| ATCG00560.1 | Photosystem II subunit (PSBL) | 36 | 0.30 | 0.297 | 0.768 | |
| ATCG00560.1 | Photosystem II subunit (PSBL) | 46 | 60.43 | 1.338 | 0.967 | |
| ATCG00560.1 | Photosystem II subunit (PSBL) | 47 | 126.57 | 1.496 | 0.948 | |
| ATCG00580.E | Photosystem II subunit (PSBE) | 2 | 0.29 | 0.300 | 0.816 | |
| ATCG00580.E | Photosystem II subunit (PSBE) | 3 | 0.47 | 0.190 | 0.859 | |
| ATCG00580.E | Photosystem II subunit (PSBE) | 4 | 0.79 | 0.169 | 1.000 | |
| ATCG00580.E | Photosystem II subunit (PSBE) | 5 | 1.11 | 0.165 | 1.000 | |
| ATCG00580.E | Photosystem II subunit (PSBE) | 6 | 1.17 | 0.166 | 1.000 | |
| ATCG00580.E | Photosystem II subunit (PSBE) | 7 | 1.02 | 0.167 | 1.000 | |
| ATCG00580.E | Photosystem II subunit (PSBE) | 8 | 0.90 | 0.162 | 1.000 | |
| ATCG00580.E | Photosystem II subunit (PSBE) | 9 | 0.86 | 0.159 | 1.000 | |
| ATCG00580.E | Photosystem II subunit (PSBE) | 10 | 0.89 | 0.160 | 1.000 | |
| ATCG00580.E | Photosystem II subunit (PSBE) | 11 | 0.95 | 0.161 | 1.000 | |
| ATCG00580.E | Photosystem II subunit (PSBE) | 12 | 1.03 | 0.161 | 1.000 | |
| ATCG00580.E | Photosystem II subunit (PSBE) | 13 | 1.08 | 0.157 | 1.000 | |
| ATCG00580.E | Photosystem II subunit (PSBE) | 14 | 1.07 | 0.154 | 1.000 | |
| ATCG00580.E | Photosystem II subunit (PSBE) | 15 | 1.00 | 0.154 | 1.000 | |
| ATCG00580.E | Photosystem II subunit (PSBE) | 16 | 0.98 | 0.154 | 1.000 | |
| ATCG00580.E | Photosystem II subunit (PSBE) | 17 | 1.06 | 0.153 | 1.000 | |
| ATCG00580.E | Photosystem II subunit (PSBE) | 18 | 1.24 | 0.155 | 1.000 | |
| ATCG00580.E | Photosystem II subunit (PSBE) | 19 | 1.47 | 0.159 | 0.999 | |
| ATCG00580.E | Photosystem II subunit (PSBE) | 20 | 1.63 | 0.165 | 0.991 | |
| ATCG00580.E | Photosystem II subunit (PSBE) | 21 | 1.78 | 0.167 | 0.960 | |
| ATCG00580.E | Photosystem II subunit (PSBE) | 22 | 1.88 | 0.167 | 0.910 | |
| ATCG00580.E | Photosystem II subunit (PSBE) | 23 | 1.62 | 0.167 | 0.994 | |
| ATCG00580.E | Photosystem II subunit (PSBE) | 24 | 1.07 | 0.162 | 1.000 | |
| ATCG00580.E | Photosystem II subunit (PSBE) | 25 | 0.66 | 0.158 | 0.998 | |
| ATCG00580.E | Photosystem II subunit (PSBE) | 26 | 0.53 | 0.157 | 0.846 | |
| ATCG00580.E | Photosystem II subunit (PSBE) | 27 | 0.49 | 0.157 | 0.704 | |
| ATCG00580.E | Photosystem II subunit (PSBE) | 28 | 0.57 | 0.157 | 0.940 | |
| ATCG00580.E | Photosystem II subunit (PSBE) | 29 | 1.14 | 0.159 | 1.000 | |
| ATCG00580.E | Photosystem II subunit (PSBE) | 30 | 2.34 | 0.160 | 0.417 | |
| ATCG00580.E | Photosystem II subunit (PSBE) | 31 | 2.58 | 0.161 | 0.251 | |
| ATCG00580.E | Photosystem II subunit (PSBE) | 32 | 1.37 | 0.163 | 1.000 | |
| ATCG00580.E | Photosystem II subunit (PSBE) | 33 | 0.48 | 0.166 | 0.737 | |
| ATCG00580.E | Photosystem II subunit (PSBE) | 34 | 0.18 | 0.180 | 0.001 | ** |
| ATCG00580.E | Photosystem II subunit (PSBE) | 35 | 0.11 | 0.234 | 0.002 | ** |
| ATCG00580.E | Photosystem II subunit (PSBE) | 36 | 0.10 | 0.362 | 0.134 | |
| ATCG00580.E | Photosystem II subunit (PSBE) | 46 | 292.38 | 1.589 | 0.943 | |
| ATCG00580.E | Photosystem II subunit (PSBE) | 47 | 1021.43 | 1.785 | 0.888 | |
| ATCG00680.1 | Photosystem II subunit (PSBB / CP47) | 2 | 0.43 | 0.259 | 0.990 | |
| ATCG00680.1 | Photosystem II subunit (PSBB / CP47) | 3 | 0.55 | 0.159 | 0.935 | |

| | | | | | |
|---|---|---|---|---|---|
| ATCG00680.1 | Photosystem II subunit (PSBB / CP47) | 4 | 0.72 | 0.146 | 1.000 |
| ATCG00680.1 | Photosystem II subunit (PSBB / CP47) | 5 | 0.90 | 0.146 | 1.000 |
| ATCG00680.1 | Photosystem II subunit (PSBB / CP47) | 6 | 0.95 | 0.146 | 1.000 |
| ATCG00680.1 | Photosystem II subunit (PSBB / CP47) | 7 | 0.92 | 0.146 | 1.000 |
| ATCG00680.1 | Photosystem II subunit (PSBB / CP47) | 8 | 0.93 | 0.143 | 1.000 |
| ATCG00680.1 | Photosystem II subunit (PSBB / CP47) | 9 | 0.93 | 0.141 | 1.000 |
| ATCG00680.1 | Photosystem II subunit (PSBB / CP47) | 10 | 0.89 | 0.141 | 1.000 |
| ATCG00680.1 | Photosystem II subunit (PSBB / CP47) | 11 | 0.87 | 0.143 | 1.000 |
| ATCG00680.1 | Photosystem II subunit (PSBB / CP47) | 12 | 0.88 | 0.143 | 1.000 |
| ATCG00680.1 | Photosystem II subunit (PSBB / CP47) | 13 | 0.91 | 0.141 | 1.000 |
| ATCG00680.1 | Photosystem II subunit (PSBB / CP47) | 14 | 0.96 | 0.139 | 1.000 |
| ATCG00680.1 | Photosystem II subunit (PSBB / CP47) | 15 | 0.97 | 0.138 | 1.000 |
| ATCG00680.1 | Photosystem II subunit (PSBB / CP47) | 16 | 0.96 | 0.137 | 1.000 |
| ATCG00680.1 | Photosystem II subunit (PSBB / CP47) | 17 | 1.01 | 0.136 | 1.000 |
| ATCG00680.1 | Photosystem II subunit (PSBB / CP47) | 18 | 1.14 | 0.139 | 1.000 |
| ATCG00680.1 | Photosystem II subunit (PSBB / CP47) | 19 | 1.28 | 0.143 | 1.000 |
| ATCG00680.1 | Photosystem II subunit (PSBB / CP47) | 20 | 1.37 | 0.146 | 1.000 |
| ATCG00680.1 | Photosystem II subunit (PSBB / CP47) | 21 | 1.39 | 0.148 | 1.000 |
| ATCG00680.1 | Photosystem II subunit (PSBB / CP47) | 22 | 1.28 | 0.148 | 1.000 |
| ATCG00680.1 | Photosystem II subunit (PSBB / CP47) | 23 | 0.99 | 0.147 | 1.000 |
| ATCG00680.1 | Photosystem II subunit (PSBB / CP47) | 24 | 0.73 | 0.143 | 1.000 |
| ATCG00680.1 | Photosystem II subunit (PSBB / CP47) | 25 | 0.61 | 0.140 | 0.975 |
| ATCG00680.1 | Photosystem II subunit (PSBB / CP47) | 26 | 0.65 | 0.140 | 0.993 |
| ATCG00680.1 | Photosystem II subunit (PSBB / CP47) | 27 | 0.71 | 0.140 | 1.000 |
| ATCG00680.1 | Photosystem II subunit (PSBB / CP47) | 28 | 0.71 | 0.141 | 1.000 |
| ATCG00680.1 | Photosystem II subunit (PSBB / CP47) | 29 | 0.94 | 0.141 | 1.000 |
| ATCG00680.1 | Photosystem II subunit (PSBB / CP47) | 30 | 1.56 | 0.141 | 0.992 |
| ATCG00680.1 | Photosystem II subunit (PSBB / CP47) | 31 | 1.73 | 0.142 | 0.926 |
| ATCG00680.1 | Photosystem II subunit (PSBB / CP47) | 32 | 1.14 | 0.144 | 1.000 |
| ATCG00680.1 | Photosystem II subunit (PSBB / CP47) | 33 | 0.66 | 0.149 | 0.999 |
| ATCG00680.1 | Photosystem II subunit (PSBB / CP47) | 34 | 0.49 | 0.156 | 0.751 |

| | | | | | | |
|---|---|---|---|---|---|---|
| ATCG00680.1 | Photosystem II subunit (PSBB / CP47) | 35 | 0.55 | 0.160 | 0.944 | |
| ATCG00680.1 | Photosystem II subunit (PSBB / CP47) | 36 | 0.75 | 0.154 | 1.000 | |
| ATCG00680.1 | Photosystem II subunit (PSBB / CP47) | 37 | 0.91 | 0.146 | 1.000 | |
| ATCG00680.1 | Photosystem II subunit (PSBB / CP47) | 38 | 0.93 | 0.144 | 1.000 | |
| ATCG00680.1 | Photosystem II subunit (PSBB / CP47) | 39 | 0.90 | 0.150 | 1.000 | |
| ATCG00680.1 | Photosystem II subunit (PSBB / CP47) | 40 | 0.93 | 0.161 | 1.000 | |
| ATCG00680.1 | Photosystem II subunit (PSBB / CP47) | 41 | 0.90 | 0.176 | 1.000 | |
| ATCG00680.1 | Photosystem II subunit (PSBB / CP47) | 42 | 0.84 | 0.235 | 1.000 | |
| ATCG00680.1 | Photosystem II subunit (PSBB / CP47) | 43 | 0.82 | 0.318 | 1.000 | |
| ATCG00680.1 | Photosystem II subunit (PSBB / CP47) | 46 | 0.98 | 0.824 | 1.000 | |
| ATCG00680.1 | Photosystem II subunit (PSBB / CP47) | 47 | 1.07 | 1.071 | 1.000 | |
| ATCG00710.1 | Photosystem II subunit (PSBH) | 2 | 0.65 | 0.245 | 1.000 | |
| ATCG00710.1 | Photosystem II subunit (PSBH) | 3 | 0.77 | 0.155 | 1.000 | |
| ATCG00710.1 | Photosystem II subunit (PSBH) | 4 | 0.96 | 0.137 | 1.000 | |
| ATCG00710.1 | Photosystem II subunit (PSBH) | 5 | 1.18 | 0.135 | 1.000 | |
| ATCG00710.1 | Photosystem II subunit (PSBH) | 6 | 1.24 | 0.135 | 1.000 | |
| ATCG00710.1 | Photosystem II subunit (PSBH) | 7 | 1.18 | 0.136 | 1.000 | |
| ATCG00710.1 | Photosystem II subunit (PSBH) | 8 | 1.22 | 0.132 | 1.000 | |
| ATCG00710.1 | Photosystem II subunit (PSBH) | 9 | 1.35 | 0.130 | 1.000 | |
| ATCG00710.1 | Photosystem II subunit (PSBH) | 10 | 1.48 | 0.130 | 0.988 | |
| ATCG00710.1 | Photosystem II subunit (PSBH) | 11 | 1.63 | 0.131 | 0.907 | |
| ATCG00710.1 | Photosystem II subunit (PSBH) | 12 | 1.76 | 0.131 | 0.754 | |
| ATCG00710.1 | Photosystem II subunit (PSBH) | 13 | 1.77 | 0.128 | 0.706 | |
| ATCG00710.1 | Photosystem II subunit (PSBH) | 14 | 1.66 | 0.126 | 0.836 | |
| ATCG00710.1 | Photosystem II subunit (PSBH) | 15 | 1.48 | 0.126 | 0.981 | |
| ATCG00710.1 | Photosystem II subunit (PSBH) | 16 | 1.35 | 0.125 | 0.999 | |
| ATCG00710.1 | Photosystem II subunit (PSBH) | 17 | 1.29 | 0.125 | 1.000 | |
| ATCG00710.1 | Photosystem II subunit (PSBH) | 18 | 1.32 | 0.126 | 1.000 | |
| ATCG00710.1 | Photosystem II subunit (PSBH) | 19 | 1.44 | 0.130 | 0.995 | |
| ATCG00710.1 | Photosystem II subunit (PSBH) | 20 | 1.55 | 0.134 | 0.974 | |
| ATCG00710.1 | Photosystem II subunit (PSBH) | 21 | 1.54 | 0.136 | 0.979 | |
| ATCG00710.1 | Photosystem II subunit (PSBH) | 22 | 1.29 | 0.136 | 1.000 | |
| ATCG00710.1 | Photosystem II subunit (PSBH) | 23 | 0.85 | 0.136 | 1.000 | |
| ATCG00710.1 | Photosystem II subunit (PSBH) | 24 | 0.52 | 0.132 | 0.528 | |
| ATCG00710.1 | Photosystem II subunit (PSBH) | 25 | 0.38 | 0.129 | 0.032 | * |
| ATCG00710.1 | Photosystem II subunit (PSBH) | 26 | 0.38 | 0.128 | 0.029 | * |
| ATCG00710.1 | Photosystem II subunit (PSBH) | 27 | 0.46 | 0.128 | 0.201 | |
| ATCG00710.1 | Photosystem II subunit (PSBH) | 28 | 0.60 | 0.129 | 0.862 | |
| ATCG00710.1 | Photosystem II subunit (PSBH) | 29 | 0.88 | 0.130 | 1.000 | |
| ATCG00710.1 | Photosystem II subunit (PSBH) | 30 | 1.09 | 0.131 | 1.000 | |
| ATCG00710.1 | Photosystem II subunit (PSBH) | 31 | 0.75 | 0.133 | 1.000 | |
| ATCG00710.1 | Photosystem II subunit (PSBH) | 32 | 0.32 | 0.136 | 0.010 | * |
| ATCG00710.1 | Photosystem II subunit (PSBH) | 33 | 0.16 | 0.147 | 0.000 | *** |
| ATCG00710.1 | Photosystem II subunit (PSBH) | 34 | 0.12 | 0.192 | 0.000 | *** |
| ATCG00710.1 | Photosystem II subunit (PSBH) | 35 | 0.12 | 0.296 | 0.055 | . |
| ATCG00710.1 | Photosystem II subunit (PSBH) | 46 | 457.89 | 1.575 | 0.875 | |
| ATCG00710.1 | Photosystem II subunit (PSBH) | 47 | 1403.75 | 1.758 | 0.815 | |
| ATCG00720.1 | Cytochrome b6f complex subunit (PETB) | 2 | 1.17 | 0.433 | 1.000 | |
| ATCG00720.1 | Cytochrome b6f complex subunit (PETB) | 3 | 1.10 | 0.296 | 1.000 | |

| | | | | | |
|---|---|---|---|---|---|
| ATCG00720.1 | Cytochrome b6f complex subunit (PETB) | 4 | 1.11 | 0.273 | 1.000 |
| ATCG00720.1 | Cytochrome b6f complex subunit (PETB) | 5 | 1.24 | 0.271 | 1.000 |
| ATCG00720.1 | Cytochrome b6f complex subunit (PETB) | 6 | 1.35 | 0.272 | 1.000 |
| ATCG00720.1 | Cytochrome b6f complex subunit (PETB) | 7 | 1.38 | 0.273 | 1.000 |
| ATCG00720.1 | Cytochrome b6f complex subunit (PETB) | 8 | 1.37 | 0.269 | 1.000 |
| ATCG00720.1 | Cytochrome b6f complex subunit (PETB) | 9 | 1.26 | 0.266 | 1.000 |
| ATCG00720.1 | Cytochrome b6f complex subunit (PETB) | 10 | 1.17 | 0.267 | 1.000 |
| ATCG00720.1 | Cytochrome b6f complex subunit (PETB) | 11 | 1.12 | 0.268 | 1.000 |
| ATCG00720.1 | Cytochrome b6f complex subunit (PETB) | 12 | 1.11 | 0.267 | 1.000 |
| ATCG00720.1 | Cytochrome b6f complex subunit (PETB) | 13 | 1.16 | 0.264 | 1.000 |
| ATCG00720.1 | Cytochrome b6f complex subunit (PETB) | 14 | 1.34 | 0.261 | 1.000 |
| ATCG00720.1 | Cytochrome b6f complex subunit (PETB) | 15 | 1.23 | 0.261 | 1.000 |
| ATCG00720.1 | Cytochrome b6f complex subunit (PETB) | 16 | 1.07 | 0.260 | 1.000 |
| ATCG00720.1 | Cytochrome b6f complex subunit (PETB) | 17 | 1.07 | 0.259 | 1.000 |
| ATCG00720.1 | Cytochrome b6f complex subunit (PETB) | 18 | 1.19 | 0.261 | 1.000 |
| ATCG00720.1 | Cytochrome b6f complex subunit (PETB) | 19 | 1.35 | 0.267 | 1.000 |
| ATCG00720.1 | Cytochrome b6f complex subunit (PETB) | 20 | 1.27 | 0.273 | 1.000 |
| ATCG00720.1 | Cytochrome b6f complex subunit (PETB) | 21 | 0.90 | 0.276 | 1.000 |
| ATCG00720.1 | Cytochrome b6f complex subunit (PETB) | 22 | 0.62 | 0.278 | 1.000 |
| ATCG00720.1 | Cytochrome b6f complex subunit (PETB) | 23 | 0.50 | 0.284 | 0.998 |
| ATCG00720.1 | Cytochrome b6f complex subunit (PETB) | 24 | 0.44 | 0.286 | 0.983 |
| ATCG00720.1 | Cytochrome b6f complex subunit (PETB) | 25 | 0.33 | 0.288 | 0.831 |
| ATCG00720.1 | Cytochrome b6f complex subunit (PETB) | 26 | 0.25 | 0.280 | 0.440 |
| ATCG00720.1 | Cytochrome b6f complex subunit (PETB) | 27 | 0.22 | 0.272 | 0.279 |
| ATCG00720.1 | Cytochrome b6f complex subunit (PETB) | 28 | 0.31 | 0.267 | 0.658 |
| ATCG00720.1 | Cytochrome b6f complex subunit (PETB) | 29 | 0.79 | 0.266 | 1.000 |
| ATCG00720.1 | Cytochrome b6f complex subunit (PETB) | 30 | 2.38 | 0.266 | 0.947 |
| ATCG00720.1 | Cytochrome b6f complex subunit (PETB) | 31 | 4.01 | 0.270 | 0.393 |
| ATCG00720.1 | Cytochrome b6f complex subunit (PETB) | 32 | 1.68 | 0.285 | 1.000 |
| ATCG00720.1 | Cytochrome b6f complex subunit (PETB) | 33 | 0.44 | 0.332 | 0.997 |
| ATCG00720.1 | Cytochrome b6f complex subunit (PETB) | 34 | 0.21 | 0.422 | 0.848 |

| | | | | | |
|---|---|---|---|---|---|
| ATCG00720.1 | Cytochrome b6f complex subunit (PETB) | 38 | 1.04 | 0.389 | 1.000 |
| ATCG00720.1 | Cytochrome b6f complex subunit (PETB) | 39 | 2.45 | 0.422 | 1.000 |
| ATCG00720.1 | Cytochrome b6f complex subunit (PETB) | 40 | 5.28 | 0.491 | 0.925 |
| ATCG00720.1 | Cytochrome b6f complex subunit (PETB) | 46 | 81.36 | 1.581 | 0.988 |
| ATCG00720.1 | Cytochrome b6f complex subunit (PETB) | 47 | 110.46 | 1.865 | 0.996 |
| ATCG00730.1 | Cytochrome b6f complex subunit (PETD) | 2 | 1.17 | 0.547 | 1.000 |
| ATCG00730.1 | Cytochrome b6f complex subunit (PETD) | 3 | 1.22 | 0.456 | 1.000 |
| ATCG00730.1 | Cytochrome b6f complex subunit (PETD) | 6 | 1.19 | 0.359 | 1.000 |
| ATCG00730.1 | Cytochrome b6f complex subunit (PETD) | 7 | 1.10 | 0.328 | 1.000 |
| ATCG00730.1 | Cytochrome b6f complex subunit (PETD) | 8 | 0.98 | 0.307 | 1.000 |
| ATCG00730.1 | Cytochrome b6f complex subunit (PETD) | 9 | 0.87 | 0.291 | 1.000 |
| ATCG00730.1 | Cytochrome b6f complex subunit (PETD) | 10 | 0.79 | 0.282 | 1.000 |
| ATCG00730.1 | Cytochrome b6f complex subunit (PETD) | 11 | 0.77 | 0.277 | 1.000 |
| ATCG00730.1 | Cytochrome b6f complex subunit (PETD) | 12 | 0.79 | 0.276 | 1.000 |
| ATCG00730.1 | Cytochrome b6f complex subunit (PETD) | 13 | 0.84 | 0.285 | 1.000 |
| ATCG00730.1 | Cytochrome b6f complex subunit (PETD) | 14 | 0.90 | 0.301 | 1.000 |
| ATCG00730.1 | Cytochrome b6f complex subunit (PETD) | 15 | 0.95 | 0.321 | 1.000 |
| ATCG00730.1 | Cytochrome b6f complex subunit (PETD) | 20 | 0.92 | 0.367 | 1.000 |
| ATCG00730.1 | Cytochrome b6f complex subunit (PETD) | 21 | 0.84 | 0.374 | 1.000 |
| ATCG00730.1 | Cytochrome b6f complex subunit (PETD) | 22 | 0.71 | 0.388 | 1.000 |
| ATCG00730.1 | Cytochrome b6f complex subunit (PETD) | 27 | 0.43 | 0.297 | 0.949 |
| ATCG00730.1 | Cytochrome b6f complex subunit (PETD) | 28 | 0.63 | 0.250 | 0.999 |
| ATCG00730.1 | Cytochrome b6f complex subunit (PETD) | 29 | 1.14 | 0.225 | 1.000 |
| ATCG00730.1 | Cytochrome b6f complex subunit (PETD) | 30 | 2.27 | 0.220 | 0.759 |
| ATCG00730.1 | Cytochrome b6f complex subunit (PETD) | 31 | 3.99 | 0.234 | 0.155 |
| ATCG00730.1 | Cytochrome b6f complex subunit (PETD) | 32 | 4.55 | 0.267 | 0.195 |
| ATCG00730.1 | Cytochrome b6f complex subunit (PETD) | 33 | 3.55 | 0.306 | 0.629 |
| ATCG00730.1 | Cytochrome b6f complex subunit (PETD) | 36 | 0.90 | 0.356 | 1.000 |
| ATCG00730.1 | Cytochrome b6f complex subunit (PETD) | 37 | 0.62 | 0.397 | 1.000 |
| ATCG00730.1 | Cytochrome b6f complex subunit (PETD) | 38 | 0.49 | 0.475 | 1.000 |
| ATCG00730.1 | Cytochrome b6f complex subunit (PETD) | 46 | 3.56 | 1.231 | 1.000 |

| | | | | | |
|---|---|---|---|---|---|
| ATCG00730.1 | Cytochrome b6f complex subunit (PETD) | 47 | 5.69 | 1.384 | 1.000 |
| ATCG00800.1 | 30S ribosomal protein S3 (RPS3) | 2 | 0.21 | 0.361 | 0.467 |
| ATCG00800.1 | 30S ribosomal protein S3 (RPS3) | 3 | 0.21 | 0.286 | 0.190 |
| ATCG00800.1 | 30S ribosomal protein S3 (RPS3) | 4 | 0.22 | 0.247 | 0.104 |
| ATCG00800.1 | 30S ribosomal protein S3 (RPS3) | 5 | 0.26 | 0.231 | 0.137 |
| ATCG00800.1 | 30S ribosomal protein S3 (RPS3) | 6 | 0.34 | 0.226 | 0.343 |
| ATCG00800.1 | 30S ribosomal protein S3 (RPS3) | 7 | 0.49 | 0.224 | 0.834 |
| ATCG00800.1 | 30S ribosomal protein S3 (RPS3) | 8 | 0.77 | 0.222 | 1.000 |
| ATCG00800.1 | 30S ribosomal protein S3 (RPS3) | 9 | 1.19 | 0.221 | 1.000 |
| ATCG00800.1 | 30S ribosomal protein S3 (RPS3) | 10 | 1.72 | 0.220 | 0.960 |
| ATCG00800.1 | 30S ribosomal protein S3 (RPS3) | 11 | 2.24 | 0.219 | 0.685 |
| ATCG00800.1 | 30S ribosomal protein S3 (RPS3) | 12 | 2.66 | 0.218 | 0.426 |
| ATCG00800.1 | 30S ribosomal protein S3 (RPS3) | 13 | 2.89 | 0.218 | 0.321 |
| ATCG00800.1 | 30S ribosomal protein S3 (RPS3) | 14 | 2.83 | 0.218 | 0.343 |
| ATCG00800.1 | 30S ribosomal protein S3 (RPS3) | 15 | 2.51 | 0.219 | 0.511 |
| ATCG00800.1 | 30S ribosomal protein S3 (RPS3) | 16 | 2.03 | 0.221 | 0.824 |
| ATCG00800.1 | 30S ribosomal protein S3 (RPS3) | 17 | 1.54 | 0.226 | 0.993 |
| ATCG00800.1 | 30S ribosomal protein S3 (RPS3) | 18 | 1.17 | 0.235 | 1.000 |
| ATCG00800.1 | 30S ribosomal protein S3 (RPS3) | 19 | 0.94 | 0.250 | 1.000 |
| ATCG00800.1 | 30S ribosomal protein S3 (RPS3) | 20 | 0.83 | 0.272 | 1.000 |
| ATCG00800.1 | 30S ribosomal protein S3 (RPS3) | 21 | 0.80 | 0.299 | 1.000 |
| ATCG00800.1 | 30S ribosomal protein S3 (RPS3) | 27 | 1.51 | 0.381 | 1.000 |
| ATCG00800.1 | 30S ribosomal protein S3 (RPS3) | 28 | 1.72 | 0.385 | 1.000 |
| ATCG00800.1 | 30S ribosomal protein S3 (RPS3) | 29 | 1.94 | 0.382 | 0.997 |
| ATCG00800.1 | 30S ribosomal protein S3 (RPS3) | 35 | 3.39 | 0.397 | 0.854 |
| ATCG00800.1 | 30S ribosomal protein S3 (RPS3) | 36 | 3.72 | 0.398 | 0.797 |
| ATCG00800.1 | 30S ribosomal protein S3 (RPS3) | 37 | 4.03 | 0.410 | 0.768 |
| ATCG00800.1 | 30S ribosomal protein S3 (RPS3) | 44 | 3.35 | 0.383 | 0.833 |
| ATCG00800.1 | 30S ribosomal protein S3 (RPS3) | 45 | 3.05 | 0.395 | 0.907 |
| ATCG00800.1 | 30S ribosomal protein S3 (RPS3) | 46 | 2.76 | 0.450 | 0.978 |
| ATCG00800.1 | 30S ribosomal protein S3 (RPS3) | 47 | 2.49 | 0.559 | 0.998 |

| | | | | | |
|---|---|---|---|---|---|
| ATCG01060.1 | Photosystem I subunit (PSAC) | 2 | 0.69 | 0.264 | 1.000 |
| ATCG01060.1 | Photosystem I subunit (PSAC) | 3 | 0.62 | 0.179 | 0.988 |
| ATCG01060.1 | Photosystem I subunit (PSAC) | 4 | 0.58 | 0.147 | 0.853 |
| ATCG01060.1 | Photosystem I subunit (PSAC) | 5 | 0.58 | 0.143 | 0.809 |
| ATCG01060.1 | Photosystem I subunit (PSAC) | 6 | 0.60 | 0.143 | 0.867 |
| ATCG01060.1 | Photosystem I subunit (PSAC) | 7 | 0.63 | 0.142 | 0.932 |
| ATCG01060.1 | Photosystem I subunit (PSAC) | 8 | 0.68 | 0.140 | 0.980 |
| ATCG01060.1 | Photosystem I subunit (PSAC) | 9 | 0.73 | 0.138 | 0.998 |
| ATCG01060.1 | Photosystem I subunit (PSAC) | 10 | 0.80 | 0.137 | 1.000 |
| ATCG01060.1 | Photosystem I subunit (PSAC) | 11 | 0.87 | 0.137 | 1.000 |
| ATCG01060.1 | Photosystem I subunit (PSAC) | 12 | 0.94 | 0.137 | 1.000 |
| ATCG01060.1 | Photosystem I subunit (PSAC) | 13 | 1.00 | 0.136 | 1.000 |
| ATCG01060.1 | Photosystem I subunit (PSAC) | 14 | 1.04 | 0.135 | 1.000 |
| ATCG01060.1 | Photosystem I subunit (PSAC) | 15 | 1.07 | 0.134 | 1.000 |
| ATCG01060.1 | Photosystem I subunit (PSAC) | 16 | 1.11 | 0.133 | 1.000 |
| ATCG01060.1 | Photosystem I subunit (PSAC) | 17 | 1.19 | 0.133 | 1.000 |
| ATCG01060.1 | Photosystem I subunit (PSAC) | 18 | 1.32 | 0.135 | 0.999 |
| ATCG01060.1 | Photosystem I subunit (PSAC) | 19 | 1.41 | 0.138 | 0.992 |
| ATCG01060.1 | Photosystem I subunit (PSAC) | 20 | 1.43 | 0.140 | 0.991 |
| ATCG01060.1 | Photosystem I subunit (PSAC) | 21 | 1.37 | 0.142 | 0.998 |
| ATCG01060.1 | Photosystem I subunit (PSAC) | 22 | 1.24 | 0.142 | 1.000 |
| ATCG01060.1 | Photosystem I subunit (PSAC) | 23 | 1.11 | 0.141 | 1.000 |
| ATCG01060.1 | Photosystem I subunit (PSAC) | 24 | 1.02 | 0.139 | 1.000 |
| ATCG01060.1 | Photosystem I subunit (PSAC) | 25 | 0.97 | 0.138 | 1.000 |
| ATCG01060.1 | Photosystem I subunit (PSAC) | 26 | 0.97 | 0.137 | 1.000 |
| ATCG01060.1 | Photosystem I subunit (PSAC) | 27 | 1.05 | 0.138 | 1.000 |
| ATCG01060.1 | Photosystem I subunit (PSAC) | 28 | 1.23 | 0.140 | 1.000 |
| ATCG01060.1 | Photosystem I subunit (PSAC) | 29 | 1.59 | 0.147 | 0.945 |
| ATCG01060.1 | Photosystem I subunit (PSAC) | 30 | 1.90 | 0.165 | 0.782 |
| ATCG01060.1 | Photosystem I subunit (PSAC) | 31 | 2.05 | 0.201 | 0.868 |
| ATCG01060.1 | Photosystem I subunit (PSAC) | 32 | 2.22 | 0.268 | 0.964 |
| ATCG01060.1 | Photosystem I subunit (PSAC) | 46 | 2443.57 | 3.031 | 0.990 |
| ATCG01060.1 | Photosystem I subunit (PSAC) | 47 | 4961.66 | 3.328 | 0.991 |
| ATCG01090.1 | NAD(P)H dehydrogenase complex subunit (NDHI) | 2 | 1.01 | 0.373 | 1.000 |
| ATCG01090.1 | NAD(P)H dehydrogenase complex subunit (NDHI) | 3 | 0.98 | 0.290 | 1.000 |
| ATCG01090.1 | NAD(P)H dehydrogenase complex subunit (NDHI) | 4 | 0.94 | 0.237 | 1.000 |
| ATCG01090.1 | NAD(P)H dehydrogenase complex subunit (NDHI) | 5 | 0.90 | 0.207 | 1.000 |
| ATCG01090.1 | NAD(P)H dehydrogenase complex subunit (NDHI) | 6 | 0.83 | 0.190 | 1.000 |
| ATCG01090.1 | NAD(P)H dehydrogenase complex subunit (NDHI) | 7 | 0.75 | 0.180 | 0.999 |
| ATCG01090.1 | NAD(P)H dehydrogenase complex subunit (NDHI) | 8 | 0.67 | 0.172 | 0.975 |
| ATCG01090.1 | NAD(P)H dehydrogenase complex subunit (NDHI) | 9 | 0.61 | 0.167 | 0.884 |
| ATCG01090.1 | NAD(P)H dehydrogenase complex subunit (NDHI) | 10 | 0.58 | 0.163 | 0.799 |
| ATCG01090.1 | NAD(P)H dehydrogenase complex subunit (NDHI) | 11 | 0.58 | 0.161 | 0.781 |
| ATCG01090.1 | NAD(P)H dehydrogenase complex subunit (NDHI) | 12 | 0.59 | 0.160 | 0.812 |
| ATCG01090.1 | NAD(P)H dehydrogenase complex subunit (NDHI) | 13 | 0.61 | 0.159 | 0.852 |
| ATCG01090.1 | NAD(P)H dehydrogenase complex subunit (NDHI) | 14 | 0.62 | 0.158 | 0.870 |
| ATCG01090.1 | NAD(P)H dehydrogenase complex subunit (NDHI) | 15 | 0.60 | 0.157 | 0.832 |

| | | | | | | |
|---|---|---|---|---|---|---|
| ATCG01090.1 | NAD(P)H dehydrogenase complex subunit (NDHI) | 16 | 0.57 | 0.157 | 0.721 | |
| ATCG01090.1 | NAD(P)H dehydrogenase complex subunit (NDHI) | 17 | 0.55 | 0.159 | 0.676 | |
| ATCG01090.1 | NAD(P)H dehydrogenase complex subunit (NDHI) | 18 | 0.59 | 0.163 | 0.824 | |
| ATCG01090.1 | NAD(P)H dehydrogenase complex subunit (NDHI) | 19 | 0.69 | 0.172 | 0.985 | |
| ATCG01090.1 | NAD(P)H dehydrogenase complex subunit (NDHI) | 20 | 0.87 | 0.186 | 1.000 | |
| ATCG01090.1 | NAD(P)H dehydrogenase complex subunit (NDHI) | 21 | 1.10 | 0.207 | 1.000 | |
| ATCG01090.1 | NAD(P)H dehydrogenase complex subunit (NDHI) | 22 | 1.35 | 0.238 | 1.000 | |
| ATCG01090.1 | NAD(P)H dehydrogenase complex subunit (NDHI) | 23 | 1.60 | 0.278 | 0.998 | |
| ATCG01090.1 | NAD(P)H dehydrogenase complex subunit (NDHI) | 24 | 1.82 | 0.330 | 0.996 | |
| ATCG01090.1 | NAD(P)H dehydrogenase complex subunit (NDHI) | 41 | 1.10 | 0.403 | 1.000 | |
| ATCG01090.1 | NAD(P)H dehydrogenase complex subunit (NDHI) | 42 | 0.99 | 0.390 | 1.000 | |
| ATCG01090.1 | NAD(P)H dehydrogenase complex subunit (NDHI) | 43 | 0.88 | 0.421 | 1.000 | |
| ATCG01090.1 | NAD(P)H dehydrogenase complex subunit (NDHI) | 46 | 0.63 | 0.794 | 1.000 | |
| ATCG01090.1 | NAD(P)H dehydrogenase complex subunit (NDHI) | 47 | 0.57 | 0.969 | 1.000 | |
| ATCG01110.1 | NAD(P)H dehydrogenase complex subunit (NDHH) | 2 | 1.10 | 0.398 | 1.000 | |
| ATCG01110.1 | NAD(P)H dehydrogenase complex subunit (NDHH) | 3 | 1.06 | 0.316 | 1.000 | |
| ATCG01110.1 | NAD(P)H dehydrogenase complex subunit (NDHH) | 4 | 1.02 | 0.272 | 1.000 | |
| ATCG01110.1 | NAD(P)H dehydrogenase complex subunit (NDHH) | 5 | 0.95 | 0.252 | 1.000 | |
| ATCG01110.1 | NAD(P)H dehydrogenase complex subunit (NDHH) | 6 | 0.85 | 0.242 | 1.000 | |
| ATCG01110.1 | NAD(P)H dehydrogenase complex subunit (NDHH) | 7 | 0.71 | 0.236 | 0.999 | |
| ATCG01110.1 | NAD(P)H dehydrogenase complex subunit (NDHH) | 8 | 0.56 | 0.231 | 0.938 | |
| ATCG01110.1 | NAD(P)H dehydrogenase complex subunit (NDHH) | 9 | 0.42 | 0.226 | 0.595 | |
| ATCG01110.1 | NAD(P)H dehydrogenase complex subunit (NDHH) | 10 | 0.32 | 0.223 | 0.235 | |
| ATCG01110.1 | NAD(P)H dehydrogenase complex subunit (NDHH) | 11 | 0.25 | 0.221 | 0.080 | . |
| ATCG01110.1 | NAD(P)H dehydrogenase complex subunit (NDHH) | 12 | 0.21 | 0.220 | 0.032 | * |
| ATCG01110.1 | NAD(P)H dehydrogenase complex subunit (NDHH) | 13 | 0.19 | 0.220 | 0.014 | * |
| ATCG01110.1 | NAD(P)H dehydrogenase complex subunit (NDHH) | 14 | 0.17 | 0.219 | 0.009 | ** |
| ATCG01110.1 | NAD(P)H dehydrogenase complex subunit (NDHH) | 15 | 0.18 | 0.219 | 0.011 | * |
| ATCG01110.1 | NAD(P)H dehydrogenase complex subunit (NDHH) | 16 | 0.21 | 0.220 | 0.033 | * |
| ATCG01110.1 | NAD(P)H dehydrogenase complex subunit (NDHH) | 17 | 0.30 | 0.223 | 0.200 | |
| ATCG01110.1 | NAD(P)H dehydrogenase complex subunit (NDHH) | 18 | 0.49 | 0.228 | 0.811 | |

| | | | | | |
|---|---|---|---|---|---|
| ATCG01110.1 | NAD(P)H dehydrogenase complex subunit (NDHH) | 19 | 0.84 | 0.237 | 1.000 |
| ATCG01110.1 | NAD(P)H dehydrogenase complex subunit (NDHH) | 20 | 1.48 | 0.250 | 0.998 |
| ATCG01110.1 | NAD(P)H dehydrogenase complex subunit (NDHH) | 21 | 2.50 | 0.267 | 0.736 |
| ATCG01110.1 | NAD(P)H dehydrogenase complex subunit (NDHH) | 22 | 3.82 | 0.293 | 0.380 |
| ATCG01110.1 | NAD(P)H dehydrogenase complex subunit (NDHH) | 23 | 5.32 | 0.328 | 0.250 |
| ATCG01110.1 | NAD(P)H dehydrogenase complex subunit (NDHH) | 24 | 6.86 | 0.380 | 0.255 |
| ATCG01110.1 | NAD(P)H dehydrogenase complex subunit (NDHH) | 41 | 1.50 | 0.484 | 1.000 |
| ATCG01110.1 | NAD(P)H dehydrogenase complex subunit (NDHH) | 42 | 1.16 | 0.476 | 1.000 |
| ATCG01110.1 | NAD(P)H dehydrogenase complex subunit (NDHH) | 43 | 0.90 | 0.531 | 1.000 |
| ATCG01110.1 | NAD(P)H dehydrogenase complex subunit (NDHH) | 46 | 0.42 | 1.051 | 1.000 |
| ATCG01110.1 | NAD(P)H dehydrogenase complex subunit (NDHH) | 47 | 0.32 | 1.284 | 1.000 |
| ATMG00070.E | NADH:ubiquinone oxidoreductase subunit (NAD9) | 3 | 0.97 | 0.217 | 1.000 |
| ATMG00070.E | NADH:ubiquinone oxidoreductase subunit (NAD9) | 4 | 1.04 | 0.202 | 1.000 |
| ATMG00070.E | NADH:ubiquinone oxidoreductase subunit (NAD9) | 5 | 1.35 | 0.203 | 1.000 |
| ATMG00070.E | NADH:ubiquinone oxidoreductase subunit (NAD9) | 6 | 1.81 | 0.203 | 0.998 |
| ATMG00070.E | NADH:ubiquinone oxidoreductase subunit (NAD9) | 7 | 2.01 | 0.203 | 0.981 |
| ATMG00070.E | NADH:ubiquinone oxidoreductase subunit (NAD9) | 8 | 2.07 | 0.199 | 0.963 |
| ATMG00070.E | NADH:ubiquinone oxidoreductase subunit (NAD9) | 9 | 2.25 | 0.196 | 0.884 |
| ATMG00070.E | NADH:ubiquinone oxidoreductase subunit (NAD9) | 10 | 2.43 | 0.196 | 0.774 |
| ATMG00070.E | NADH:ubiquinone oxidoreductase subunit (NAD9) | 11 | 2.33 | 0.197 | 0.842 |
| ATMG00070.E | NADH:ubiquinone oxidoreductase subunit (NAD9) | 12 | 1.99 | 0.197 | 0.978 |
| ATMG00070.E | NADH:ubiquinone oxidoreductase subunit (NAD9) | 13 | 1.64 | 0.195 | 1.000 |
| ATMG00070.E | NADH:ubiquinone oxidoreductase subunit (NAD9) | 14 | 1.28 | 0.192 | 1.000 |
| ATMG00070.E | NADH:ubiquinone oxidoreductase subunit (NAD9) | 15 | 0.90 | 0.191 | 1.000 |
| ATMG00070.E | NADH:ubiquinone oxidoreductase subunit (NAD9) | 16 | 0.57 | 0.190 | 0.998 |
| ATMG00070.E | NADH:ubiquinone oxidoreductase subunit (NAD9) | 17 | 0.44 | 0.190 | 0.826 |
| ATMG00070.E | NADH:ubiquinone oxidoreductase subunit (NAD9) | 18 | 0.51 | 0.193 | 0.975 |
| ATMG00070.E | NADH:ubiquinone oxidoreductase subunit (NAD9) | 19 | 0.70 | 0.198 | 1.000 |
| ATMG00070.E | NADH:ubiquinone oxidoreductase subunit (NAD9) | 20 | 0.88 | 0.203 | 1.000 |
| ATMG00070.E | NADH:ubiquinone oxidoreductase subunit (NAD9) | 21 | 1.11 | 0.206 | 1.000 |
| ATMG00070.E | NADH:ubiquinone oxidoreductase subunit (NAD9) | 22 | 1.49 | 0.205 | 1.000 |

| | | | | | |
|---|---|---|---|---|---|
| ATMG00070.E | NADH:ubiquinone oxidoreductase subunit (NAD9) | 23 | 1.84 | 0.204 | 0.997 |
| ATMG00070.E | NADH:ubiquinone oxidoreductase subunit (NAD9) | 24 | 1.68 | 0.200 | 1.000 |
| ATMG00070.E | NADH:ubiquinone oxidoreductase subunit (NAD9) | 25 | 1.03 | 0.196 | 1.000 |
| ATMG00070.E | NADH:ubiquinone oxidoreductase subunit (NAD9) | 26 | 0.61 | 0.195 | 1.000 |
| ATMG00070.E | NADH:ubiquinone oxidoreductase subunit (NAD9) | 27 | 0.55 | 0.195 | 0.995 |
| ATMG00070.E | NADH:ubiquinone oxidoreductase subunit (NAD9) | 28 | 0.68 | 0.195 | 1.000 |
| ATMG00070.E | NADH:ubiquinone oxidoreductase subunit (NAD9) | 29 | 1.03 | 0.195 | 1.000 |
| ATMG00070.E | NADH:ubiquinone oxidoreductase subunit (NAD9) | 30 | 1.46 | 0.195 | 1.000 |
| ATMG00070.E | NADH:ubiquinone oxidoreductase subunit (NAD9) | 31 | 1.70 | 0.195 | 0.999 |
| ATMG00070.E | NADH:ubiquinone oxidoreductase subunit (NAD9) | 32 | 1.78 | 0.196 | 0.998 |
| ATMG00070.E | NADH:ubiquinone oxidoreductase subunit (NAD9) | 33 | 1.63 | 0.196 | 1.000 |
| ATMG00070.E | NADH:ubiquinone oxidoreductase subunit (NAD9) | 34 | 1.43 | 0.197 | 1.000 |
| ATMG00070.E | NADH:ubiquinone oxidoreductase subunit (NAD9) | 35 | 1.32 | 0.196 | 1.000 |
| ATMG00070.E | NADH:ubiquinone oxidoreductase subunit (NAD9) | 36 | 1.39 | 0.195 | 1.000 |
| ATMG00070.E | NADH:ubiquinone oxidoreductase subunit (NAD9) | 37 | 1.62 | 0.195 | 1.000 |
| ATMG00070.E | NADH:ubiquinone oxidoreductase subunit (NAD9) | 38 | 1.92 | 0.196 | 0.987 |
| ATMG00070.E | NADH:ubiquinone oxidoreductase subunit (NAD9) | 39 | 2.27 | 0.196 | 0.875 |
| ATMG00070.E | NADH:ubiquinone oxidoreductase subunit (NAD9) | 40 | 2.40 | 0.197 | 0.797 |
| ATMG00070.E | NADH:ubiquinone oxidoreductase subunit (NAD9) | 41 | 2.15 | 0.198 | 0.934 |
| ATMG00070.E | NADH:ubiquinone oxidoreductase subunit (NAD9) | 42 | 1.79 | 0.200 | 0.998 |
| ATMG00070.E | NADH:ubiquinone oxidoreductase subunit (NAD9) | 43 | 1.47 | 0.202 | 1.000 |
| ATMG00070.E | NADH:ubiquinone oxidoreductase subunit (NAD9) | 44 | 1.14 | 0.202 | 1.000 |
| ATMG00070.E | NADH:ubiquinone oxidoreductase subunit (NAD9) | 45 | 0.80 | 0.202 | 1.000 |
| ATMG00070.E | NADH:ubiquinone oxidoreductase subunit (NAD9) | 46 | 0.48 | 0.233 | 0.995 |
| ATMG00070.E | NADH:ubiquinone oxidoreductase subunit (NAD9) | 47 | 0.24 | 0.415 | 0.985 |
| ATMG00070.E | NADH:ubiquinone oxidoreductase subunit (NAD9) | 48 | 0.12 | 0.704 | 0.997 |
| ATMG00090.E | Ribosomal S3 family protein | 2 | 1.00 | 0.635 | 1.000 |
| ATMG00090.E | Ribosomal S3 family protein | 3 | 0.97 | 0.544 | 1.000 |
| ATMG00090.E | Ribosomal S3 family protein | 10 | 1.41 | 0.343 | 1.000 |
| ATMG00090.E | Ribosomal S3 family protein | 11 | 1.68 | 0.290 | 1.000 |
| ATMG00090.E | Ribosomal S3 family protein | 12 | 1.91 | 0.255 | 0.996 |
| ATMG00090.E | Ribosomal S3 family protein | 13 | 1.94 | 0.234 | 0.987 |
| ATMG00090.E | Ribosomal S3 family protein | 14 | 1.88 | 0.221 | 0.985 |
| ATMG00090.E | Ribosomal S3 family protein | 15 | 2.02 | 0.215 | 0.949 |
| ATMG00090.E | Ribosomal S3 family protein | 16 | 2.38 | 0.212 | 0.772 |
| ATMG00090.E | Ribosomal S3 family protein | 17 | 2.65 | 0.212 | 0.602 |

| | | | | | |
|---|---|---|---|---|---|
| ATMG00090.E | Ribosomal S3 family protein | 18 | 2.45 | 0.214 | 0.745 |
| ATMG00090.E | Ribosomal S3 family protein | 19 | 1.78 | 0.220 | 0.994 |
| ATMG00090.E | Ribosomal S3 family protein | 20 | 1.03 | 0.226 | 1.000 |
| ATMG00090.E | Ribosomal S3 family protein | 21 | 0.53 | 0.228 | 0.990 |
| ATMG00090.E | Ribosomal S3 family protein | 22 | 0.32 | 0.229 | 0.460 |
| ATMG00090.E | Ribosomal S3 family protein | 23 | 0.26 | 0.226 | 0.192 |
| ATMG00090.E | Ribosomal S3 family protein | 24 | 0.29 | 0.223 | 0.280 |
| ATMG00090.E | Ribosomal S3 family protein | 25 | 0.40 | 0.219 | 0.754 |
| ATMG00090.E | Ribosomal S3 family protein | 26 | 0.64 | 0.217 | 1.000 |
| ATMG00090.E | Ribosomal S3 family protein | 27 | 0.97 | 0.217 | 1.000 |
| ATMG00090.E | Ribosomal S3 family protein | 28 | 1.23 | 0.217 | 1.000 |
| ATMG00090.E | Ribosomal S3 family protein | 29 | 1.31 | 0.217 | 1.000 |
| ATMG00090.E | Ribosomal S3 family protein | 30 | 1.21 | 0.218 | 1.000 |
| ATMG00090.E | Ribosomal S3 family protein | 31 | 1.03 | 0.219 | 1.000 |
| ATMG00090.E | Ribosomal S3 family protein | 32 | 0.82 | 0.220 | 1.000 |
| ATMG00090.E | Ribosomal S3 family protein | 33 | 0.60 | 0.220 | 0.999 |
| ATMG00090.E | Ribosomal S3 family protein | 34 | 0.44 | 0.220 | 0.856 |
| ATMG00090.E | Ribosomal S3 family protein | 35 | 0.35 | 0.220 | 0.527 |
| ATMG00090.E | Ribosomal S3 family protein | 36 | 0.33 | 0.220 | 0.450 |
| ATMG00090.E | Ribosomal S3 family protein | 37 | 0.37 | 0.220 | 0.615 |
| ATMG00090.E | Ribosomal S3 family protein | 38 | 0.44 | 0.221 | 0.874 |
| ATMG00090.E | Ribosomal S3 family protein | 39 | 0.54 | 0.222 | 0.987 |
| ATMG00090.E | Ribosomal S3 family protein | 40 | 0.58 | 0.229 | 0.998 |
| ATMG00090.E | Ribosomal S3 family protein | 41 | 0.55 | 0.261 | 0.999 |
| ATMG00090.E | Ribosomal S3 family protein | 42 | 0.50 | 0.336 | 1.000 |
| ATMG00090.E | Ribosomal S3 family protein | 43 | 0.48 | 0.435 | 1.000 |
| ATMG00090.E | Ribosomal S3 family protein | 46 | 0.55 | 0.896 | 1.000 |
| ATMG00090.E | Ribosomal S3 family protein | 47 | 0.60 | 1.122 | 1.000 |
| ATMG00160.E | Cytochrome c:O2 oxidoreductase subunit (COX2) | 3 | 0.98 | 0.222 | 1.000 |
| ATMG00160.E | Cytochrome c:O2 oxidoreductase subunit (COX2) | 4 | 0.85 | 0.206 | 1.000 |
| ATMG00160.E | Cytochrome c:O2 oxidoreductase subunit (COX2) | 5 | 0.88 | 0.206 | 1.000 |
| ATMG00160.E | Cytochrome c:O2 oxidoreductase subunit (COX2) | 6 | 0.92 | 0.206 | 1.000 |
| ATMG00160.E | Cytochrome c:O2 oxidoreductase subunit (COX2) | 7 | 0.89 | 0.206 | 1.000 |
| ATMG00160.E | Cytochrome c:O2 oxidoreductase subunit (COX2) | 8 | 0.89 | 0.203 | 1.000 |
| ATMG00160.E | Cytochrome c:O2 oxidoreductase subunit (COX2) | 9 | 0.89 | 0.200 | 1.000 |
| ATMG00160.E | Cytochrome c:O2 oxidoreductase subunit (COX2) | 10 | 0.85 | 0.199 | 1.000 |
| ATMG00160.E | Cytochrome c:O2 oxidoreductase subunit (COX2) | 11 | 0.82 | 0.200 | 1.000 |
| ATMG00160.E | Cytochrome c:O2 oxidoreductase subunit (COX2) | 12 | 0.83 | 0.200 | 1.000 |
| ATMG00160.E | Cytochrome c:O2 oxidoreductase subunit (COX2) | 13 | 0.88 | 0.198 | 1.000 |
| ATMG00160.E | Cytochrome c:O2 oxidoreductase subunit (COX2) | 14 | 0.94 | 0.196 | 1.000 |
| ATMG00160.E | Cytochrome c:O2 oxidoreductase subunit (COX2) | 15 | 0.97 | 0.195 | 1.000 |
| ATMG00160.E | Cytochrome c:O2 oxidoreductase subunit (COX2) | 16 | 0.91 | 0.193 | 1.000 |
| ATMG00160.E | Cytochrome c:O2 oxidoreductase subunit (COX2) | 17 | 0.88 | 0.193 | 1.000 |
| ATMG00160.E | Cytochrome c:O2 oxidoreductase subunit (COX2) | 18 | 0.94 | 0.196 | 1.000 |
| ATMG00160.E | Cytochrome c:O2 oxidoreductase subunit (COX2) | 19 | 1.06 | 0.201 | 1.000 |

| | | | | | | |
|---|---|---|---|---|---|---|
| ATMG00160.E | Cytochrome c:O2 oxidoreductase subunit (COX2) | 20 | 1.33 | 0.206 | 1.000 | |
| ATMG00160.E | Cytochrome c:O2 oxidoreductase subunit (COX2) | 21 | 1.82 | 0.209 | 0.998 | |
| ATMG00160.E | Cytochrome c:O2 oxidoreductase subunit (COX2) | 22 | 2.07 | 0.208 | 0.975 | |
| ATMG00160.E | Cytochrome c:O2 oxidoreductase subunit (COX2) | 23 | 1.59 | 0.207 | 1.000 | |
| ATMG00160.E | Cytochrome c:O2 oxidoreductase subunit (COX2) | 24 | 1.10 | 0.203 | 1.000 | |
| ATMG00160.E | Cytochrome c:O2 oxidoreductase subunit (COX2) | 25 | 0.92 | 0.200 | 1.000 | |
| ATMG00160.E | Cytochrome c:O2 oxidoreductase subunit (COX2) | 26 | 1.01 | 0.199 | 1.000 | |
| ATMG00160.E | Cytochrome c:O2 oxidoreductase subunit (COX2) | 27 | 1.34 | 0.199 | 1.000 | |
| ATMG00160.E | Cytochrome c:O2 oxidoreductase subunit (COX2) | 28 | 1.65 | 0.199 | 1.000 | |
| ATMG00160.E | Cytochrome c:O2 oxidoreductase subunit (COX2) | 29 | 1.85 | 0.199 | 0.994 | |
| ATMG00160.E | Cytochrome c:O2 oxidoreductase subunit (COX2) | 30 | 1.80 | 0.198 | 0.997 | |
| ATMG00160.E | Cytochrome c:O2 oxidoreductase subunit (COX2) | 31 | 1.50 | 0.199 | 1.000 | |
| ATMG00160.E | Cytochrome c:O2 oxidoreductase subunit (COX2) | 32 | 1.15 | 0.200 | 1.000 | |
| ATMG00160.E | Cytochrome c:O2 oxidoreductase subunit (COX2) | 33 | 0.79 | 0.200 | 1.000 | |
| ATMG00160.E | Cytochrome c:O2 oxidoreductase subunit (COX2) | 34 | 0.55 | 0.200 | 0.997 | |
| ATMG00160.E | Cytochrome c:O2 oxidoreductase subunit (COX2) | 35 | 0.50 | 0.200 | 0.975 | |
| ATMG00160.E | Cytochrome c:O2 oxidoreductase subunit (COX2) | 36 | 0.60 | 0.199 | 1.000 | |
| ATMG00160.E | Cytochrome c:O2 oxidoreductase subunit (COX2) | 37 | 0.92 | 0.199 | 1.000 | |
| ATMG00160.E | Cytochrome c:O2 oxidoreductase subunit (COX2) | 38 | 1.69 | 0.200 | 0.999 | |
| ATMG00160.E | Cytochrome c:O2 oxidoreductase subunit (COX2) | 39 | 3.48 | 0.200 | 0.203 | |
| ATMG00160.E | Cytochrome c:O2 oxidoreductase subunit (COX2) | 40 | 5.11 | 0.200 | 0.017 | * |
| ATMG00160.E | Cytochrome c:O2 oxidoreductase subunit (COX2) | 41 | 5.10 | 0.201 | 0.018 | * |
| ATMG00160.E | Cytochrome c:O2 oxidoreductase subunit (COX2) | 42 | 4.63 | 0.204 | 0.043 | * |
| ATMG00160.E | Cytochrome c:O2 oxidoreductase subunit (COX2) | 43 | 4.49 | 0.206 | 0.057 | . |
| ATMG00160.E | Cytochrome c:O2 oxidoreductase subunit (COX2) | 44 | 3.94 | 0.206 | 0.126 | |
| ATMG00160.E | Cytochrome c:O2 oxidoreductase subunit (COX2) | 45 | 2.49 | 0.205 | 0.786 | |
| ATMG00160.E | Cytochrome c:O2 oxidoreductase subunit (COX2) | 46 | 1.04 | 0.239 | 1.000 | |
| ATMG00160.E | Cytochrome c:O2 oxidoreductase subunit (COX2) | 47 | 0.35 | 0.423 | 1.000 | |
| ATMG00160.E | Cytochrome c:O2 oxidoreductase subunit (COX2) | 48 | 0.11 | 0.708 | 0.995 | |
| ATMG00180.E | Cytochrome C biogenesis 452 (CCB452) | 2 | 1.04 | 0.497 | 1.000 | |
| ATMG00180.E | Cytochrome C biogenesis 452 (CCB452) | 3 | 0.96 | 0.398 | 1.000 | |

| | | | | | |
|---|---|---|---|---|---|
| ATMG00180.E | Cytochrome C biogenesis 452 (CCB452) | 4 | 0.88 | 0.700 | 1.000 |
| ATMG00180.E | Cytochrome C biogenesis 452 (CCB452) | 18 | 1.61 | 0.428 | 1.000 |
| ATMG00180.E | Cytochrome C biogenesis 452 (CCB452) | 19 | 1.57 | 0.387 | 1.000 |
| ATMG00180.E | Cytochrome C biogenesis 452 (CCB452) | 20 | 1.42 | 0.416 | 1.000 |
| ATMG00180.E | Cytochrome C biogenesis 452 (CCB452) | 22 | 0.95 | 0.366 | 1.000 |
| ATMG00180.E | Cytochrome C biogenesis 452 (CCB452) | 23 | 0.96 | 0.309 | 1.000 |
| ATMG00180.E | Cytochrome C biogenesis 452 (CCB452) | 24 | 1.07 | 0.266 | 1.000 |
| ATMG00180.E | Cytochrome C biogenesis 452 (CCB452) | 25 | 0.97 | 0.252 | 1.000 |
| ATMG00180.E | Cytochrome C biogenesis 452 (CCB452) | 26 | 1.15 | 0.249 | 1.000 |
| ATMG00180.E | Cytochrome C biogenesis 452 (CCB452) | 27 | 1.40 | 0.252 | 1.000 |
| ATMG00180.E | Cytochrome C biogenesis 452 (CCB452) | 28 | 0.68 | 0.270 | 1.000 |
| ATMG00180.E | Cytochrome C biogenesis 452 (CCB452) | 29 | 0.23 | 0.332 | 0.502 |
| ATMG00180.E | Cytochrome C biogenesis 452 (CCB452) | 30 | 0.13 | 0.529 | 0.676 |
| ATMG00180.E | Cytochrome C biogenesis 452 (CCB452) | 46 | ########### | 9.535 | 0.993 |
| ATMG00180.E | Cytochrome C biogenesis 452 (CCB452) | 47 | ########### | 10.415 | 0.992 |
| ATMG00210.E | Ribosomal L5 family protein | 2 | 1.00 | 0.377 | 1.000 |
| ATMG00210.E | Ribosomal L5 family protein | 3 | 0.86 | 0.376 | 1.000 |
| ATMG00210.E | Ribosomal L5 family protein | 33 | 0.55 | 0.305 | 0.991 |
| ATMG00210.E | Ribosomal L5 family protein | 34 | 0.85 | 0.263 | 1.000 |
| ATMG00210.E | Ribosomal L5 family protein | 35 | 1.59 | 0.259 | 0.996 |
| ATMG00210.E | Ribosomal L5 family protein | 36 | 2.42 | 0.300 | 0.881 |
| ATMG00210.E | Ribosomal L5 family protein | 39 | 1.60 | 0.305 | 0.999 |
| ATMG00210.E | Ribosomal L5 family protein | 40 | 1.06 | 0.240 | 1.000 |
| ATMG00210.E | Ribosomal L5 family protein | 41 | 0.79 | 0.214 | 1.000 |
| ATMG00210.E | Ribosomal L5 family protein | 42 | 0.90 | 0.212 | 1.000 |
| ATMG00210.E | Ribosomal L5 family protein | 43 | 0.93 | 0.212 | 1.000 |
| ATMG00210.E | Ribosomal L5 family protein | 44 | 0.75 | 0.212 | 1.000 |
| ATMG00210.E | Ribosomal L5 family protein | 45 | 0.62 | 0.211 | 0.976 |
| ATMG00210.E | Ribosomal L5 family protein | 46 | 0.57 | 0.224 | 0.952 |
| ATMG00210.E | Ribosomal L5 family protein | 47 | 0.65 | 0.422 | 1.000 |
| ATMG00220.E | Ubiquinol:cytochrome C oxidoreductase subunit (COB) | 2 | 1.03 | 0.414 | 1.000 |
| ATMG00220.E | Ubiquinol:cytochrome C oxidoreductase subunit (COB) | 3 | 0.84 | 0.245 | 1.000 |
| ATMG00220.E | Ubiquinol:cytochrome C oxidoreductase subunit (COB) | 4 | 0.84 | 0.233 | 1.000 |
| ATMG00220.E | Ubiquinol:cytochrome C oxidoreductase subunit (COB) | 5 | 1.01 | 0.232 | 1.000 |
| ATMG00220.E | Ubiquinol:cytochrome C oxidoreductase subunit (COB) | 6 | 1.10 | 0.234 | 1.000 |
| ATMG00220.E | Ubiquinol:cytochrome C oxidoreductase subunit (COB) | 7 | 0.98 | 0.237 | 1.000 |
| ATMG00220.E | Ubiquinol:cytochrome C oxidoreductase subunit (COB) | 8 | 0.93 | 0.229 | 1.000 |
| ATMG00220.E | Ubiquinol:cytochrome C oxidoreductase subunit (COB) | 9 | 0.97 | 0.226 | 1.000 |

| | | | | | |
|---|---|---|---|---|---|
| ATMG00220.E | Ubiquinol:cytochrome C oxidoreductase subunit (COB) | 10 | 1.06 | 0.227 | 1.000 |
| ATMG00220.E | Ubiquinol:cytochrome C oxidoreductase subunit (COB) | 11 | 1.23 | 0.228 | 1.000 |
| ATMG00220.E | Ubiquinol:cytochrome C oxidoreductase subunit (COB) | 12 | 1.34 | 0.228 | 1.000 |
| ATMG00220.E | Ubiquinol:cytochrome C oxidoreductase subunit (COB) | 13 | 1.45 | 0.224 | 1.000 |
| ATMG00220.E | Ubiquinol:cytochrome C oxidoreductase subunit (COB) | 14 | 1.65 | 0.220 | 1.000 |
| ATMG00220.E | Ubiquinol:cytochrome C oxidoreductase subunit (COB) | 15 | 1.63 | 0.219 | 1.000 |
| ATMG00220.E | Ubiquinol:cytochrome C oxidoreductase subunit (COB) | 16 | 1.22 | 0.217 | 1.000 |
| ATMG00220.E | Ubiquinol:cytochrome C oxidoreductase subunit (COB) | 17 | 0.83 | 0.215 | 1.000 |
| ATMG00220.E | Ubiquinol:cytochrome C oxidoreductase subunit (COB) | 18 | 0.78 | 0.218 | 1.000 |
| ATMG00220.E | Ubiquinol:cytochrome C oxidoreductase subunit (COB) | 19 | 0.94 | 0.228 | 1.000 |
| ATMG00220.E | Ubiquinol:cytochrome C oxidoreductase subunit (COB) | 20 | 1.11 | 0.235 | 1.000 |
| ATMG00220.E | Ubiquinol:cytochrome C oxidoreductase subunit (COB) | 21 | 1.08 | 0.233 | 1.000 |
| ATMG00220.E | Ubiquinol:cytochrome C oxidoreductase subunit (COB) | 22 | 0.85 | 0.234 | 1.000 |
| ATMG00220.E | Ubiquinol:cytochrome C oxidoreductase subunit (COB) | 23 | 0.48 | 0.233 | 0.997 |
| ATMG00220.E | Ubiquinol:cytochrome C oxidoreductase subunit (COB) | 24 | 0.38 | 0.225 | 0.891 |
| ATMG00220.E | Ubiquinol:cytochrome C oxidoreductase subunit (COB) | 25 | 0.64 | 0.221 | 1.000 |
| ATMG00220.E | Ubiquinol:cytochrome C oxidoreductase subunit (COB) | 26 | 1.43 | 0.222 | 1.000 |
| ATMG00220.E | Ubiquinol:cytochrome C oxidoreductase subunit (COB) | 27 | 2.71 | 0.222 | 0.833 |
| ATMG00220.E | Ubiquinol:cytochrome C oxidoreductase subunit (COB) | 28 | 1.52 | 0.223 | 1.000 |
| ATMG00220.E | Ubiquinol:cytochrome C oxidoreductase subunit (COB) | 29 | 0.39 | 0.225 | 0.905 |
| ATMG00220.E | Ubiquinol:cytochrome C oxidoreductase subunit (COB) | 30 | 0.29 | 0.226 | 0.450 |
| ATMG00220.E | Ubiquinol:cytochrome C oxidoreductase subunit (COB) | 31 | 0.70 | 0.227 | 1.000 |
| ATMG00220.E | Ubiquinol:cytochrome C oxidoreductase subunit (COB) | 32 | 1.58 | 0.227 | 1.000 |
| ATMG00220.E | Ubiquinol:cytochrome C oxidoreductase subunit (COB) | 33 | 1.90 | 0.227 | 1.000 |
| ATMG00220.E | Ubiquinol:cytochrome C oxidoreductase subunit (COB) | 34 | 1.50 | 0.228 | 1.000 |
| ATMG00220.E | Ubiquinol:cytochrome C oxidoreductase subunit (COB) | 35 | 1.34 | 0.227 | 1.000 |
| ATMG00220.E | Ubiquinol:cytochrome C oxidoreductase subunit (COB) | 36 | 1.37 | 0.227 | 1.000 |
| ATMG00220.E | Ubiquinol:cytochrome C oxidoreductase subunit (COB) | 37 | 1.25 | 0.227 | 1.000 |
| ATMG00220.E | Ubiquinol:cytochrome C oxidoreductase subunit (COB) | 38 | 1.25 | 0.227 | 1.000 |
| ATMG00220.E | Ubiquinol:cytochrome C oxidoreductase subunit (COB) | 39 | 1.39 | 0.226 | 1.000 |
| ATMG00220.E | Ubiquinol:cytochrome C oxidoreductase subunit (COB) | 40 | 1.40 | 0.226 | 1.000 |

| | | | | | |
|---|---|---|---|---|---|
| ATMG00220.E | Ubiquinol:cytochrome C oxidoreductase subunit (COB) | 41 | 1.31 | 0.226 | 1.000 |
| ATMG00220.E | Ubiquinol:cytochrome C oxidoreductase subunit (COB) | 42 | 1.64 | 0.231 | 1.000 |
| ATMG00220.E | Ubiquinol:cytochrome C oxidoreductase subunit (COB) | 43 | 2.63 | 0.233 | 0.917 |
| ATMG00220.E | Ubiquinol:cytochrome C oxidoreductase subunit (COB) | 44 | 2.94 | 0.235 | 0.806 |
| ATMG00220.E | Ubiquinol:cytochrome C oxidoreductase subunit (COB) | 45 | 2.34 | 0.288 | 0.999 |
| ATMG00220.E | Ubiquinol:cytochrome C oxidoreductase subunit (COB) | 46 | 1.95 | 0.420 | 1.000 |
| ATMG00220.E | Ubiquinol:cytochrome C oxidoreductase subunit (COB) | 47 | 1.70 | 0.728 | 1.000 |
| ATMG00270.E | NADH:ubiquinone oxidoreductase subunit (NAD6) | 2 | 0.92 | 0.752 | 1.000 |
| ATMG00270.E | NADH:ubiquinone oxidoreductase subunit (NAD6) | 3 | 1.11 | 0.676 | 1.000 |
| ATMG00270.E | NADH:ubiquinone oxidoreductase subunit (NAD6) | 15 | 1.44 | 0.396 | 1.000 |
| ATMG00270.E | NADH:ubiquinone oxidoreductase subunit (NAD6) | 16 | 1.20 | 0.332 | 1.000 |
| ATMG00270.E | NADH:ubiquinone oxidoreductase subunit (NAD6) | 17 | 1.00 | 0.294 | 1.000 |
| ATMG00270.E | NADH:ubiquinone oxidoreductase subunit (NAD6) | 18 | 0.86 | 0.273 | 1.000 |
| ATMG00270.E | NADH:ubiquinone oxidoreductase subunit (NAD6) | 19 | 0.78 | 0.268 | 1.000 |
| ATMG00270.E | NADH:ubiquinone oxidoreductase subunit (NAD6) | 20 | 0.70 | 0.270 | 1.000 |
| ATMG00270.E | NADH:ubiquinone oxidoreductase subunit (NAD6) | 21 | 0.62 | 0.272 | 0.996 |
| ATMG00270.E | NADH:ubiquinone oxidoreductase subunit (NAD6) | 22 | 0.59 | 0.271 | 0.992 |
| ATMG00270.E | NADH:ubiquinone oxidoreductase subunit (NAD6) | 23 | 0.63 | 0.269 | 0.997 |
| ATMG00270.E | NADH:ubiquinone oxidoreductase subunit (NAD6) | 24 | 0.71 | 0.266 | 1.000 |
| ATMG00270.E | NADH:ubiquinone oxidoreductase subunit (NAD6) | 25 | 0.75 | 0.264 | 1.000 |
| ATMG00270.E | NADH:ubiquinone oxidoreductase subunit (NAD6) | 26 | 0.75 | 0.265 | 1.000 |
| ATMG00270.E | NADH:ubiquinone oxidoreductase subunit (NAD6) | 27 | 0.71 | 0.276 | 1.000 |
| ATMG00270.E | NADH:ubiquinone oxidoreductase subunit (NAD6) | 28 | 0.64 | 0.310 | 0.999 |
| ATMG00270.E | NADH:ubiquinone oxidoreductase subunit (NAD6) | 29 | 0.58 | 0.385 | 0.999 |
| ATMG00270.E | NADH:ubiquinone oxidoreductase subunit (NAD6) | 30 | 0.58 | 0.520 | 1.000 |
| ATMG00270.E | NADH:ubiquinone oxidoreductase subunit (NAD6) | 46 | 3701.43 | 6.722 | 1.000 |
| ATMG00270.E | NADH:ubiquinone oxidoreductase subunit (NAD6) | 47 | 8369.89 | 7.309 | 1.000 |
| ATMG00290.1 | Ribosomal S4 family protein | 2 | 1.07 | 0.469 | 1.000 |
| ATMG00290.1 | Ribosomal S4 family protein | 3 | 1.06 | 0.411 | 1.000 |
| ATMG00290.1 | Ribosomal S4 family protein | 11 | 1.24 | 0.223 | 1.000 |
| ATMG00290.1 | Ribosomal S4 family protein | 12 | 1.30 | 0.194 | 1.000 |
| ATMG00290.1 | Ribosomal S4 family protein | 13 | 1.34 | 0.172 | 0.998 |
| ATMG00290.1 | Ribosomal S4 family protein | 14 | 1.36 | 0.156 | 0.994 |
| ATMG00290.1 | Ribosomal S4 family protein | 15 | 1.42 | 0.146 | 0.974 |
| ATMG00290.1 | Ribosomal S4 family protein | 16 | 1.53 | 0.141 | 0.894 |

| | | | | | | |
|---|---|---|---|---|---|---|
| ATMG00290.1 | Ribosomal S4 family protein | 17 | 1.64 | 0.140 | 0.764 | |
| ATMG00290.1 | Ribosomal S4 family protein | 18 | 1.68 | 0.141 | 0.714 | |
| ATMG00290.1 | Ribosomal S4 family protein | 19 | 1.64 | 0.143 | 0.784 | |
| ATMG00290.1 | Ribosomal S4 family protein | 20 | 1.51 | 0.145 | 0.921 | |
| ATMG00290.1 | Ribosomal S4 family protein | 21 | 1.33 | 0.147 | 0.995 | |
| ATMG00290.1 | Ribosomal S4 family protein | 22 | 1.13 | 0.147 | 1.000 | |
| ATMG00290.1 | Ribosomal S4 family protein | 23 | 0.94 | 0.147 | 1.000 | |
| ATMG00290.1 | Ribosomal S4 family protein | 24 | 0.81 | 0.145 | 1.000 | |
| ATMG00290.1 | Ribosomal S4 family protein | 25 | 0.72 | 0.144 | 0.984 | |
| ATMG00290.1 | Ribosomal S4 family protein | 26 | 0.68 | 0.143 | 0.946 | |
| ATMG00290.1 | Ribosomal S4 family protein | 27 | 0.67 | 0.143 | 0.921 | |
| ATMG00290.1 | Ribosomal S4 family protein | 28 | 0.65 | 0.144 | 0.892 | |
| ATMG00290.1 | Ribosomal S4 family protein | 29 | 0.60 | 0.148 | 0.794 | |
| ATMG00290.1 | Ribosomal S4 family protein | 30 | 0.53 | 0.159 | 0.611 | |
| ATMG00290.1 | Ribosomal S4 family protein | 31 | 0.44 | 0.183 | 0.460 | |
| ATMG00290.1 | Ribosomal S4 family protein | 32 | 0.36 | 0.225 | 0.432 | |
| ATMG00290.1 | Ribosomal S4 family protein | 33 | 0.29 | 0.285 | 0.495 | |
| ATMG00290.1 | Ribosomal S4 family protein | 46 | 0.02 | 1.581 | 0.975 | |
| ATMG00290.1 | Ribosomal S4 family protein | 47 | 0.02 | 1.731 | 0.982 | |
| ATMG00290.E | Ribosomal S4 family protein | 2 | 1.00 | 0.563 | 1.000 | |
| ATMG00290.E | Ribosomal S4 family protein | 3 | 0.92 | 0.472 | 1.000 | |
| ATMG00290.E | Ribosomal S4 family protein | 4 | 0.85 | 0.485 | 1.000 | |
| ATMG00290.E | Ribosomal S4 family protein | 10 | 1.09 | 0.408 | 1.000 | |
| ATMG00290.E | Ribosomal S4 family protein | 11 | 1.38 | 0.361 | 1.000 | |
| ATMG00290.E | Ribosomal S4 family protein | 12 | 1.67 | 0.327 | 1.000 | |
| ATMG00290.E | Ribosomal S4 family protein | 13 | 1.79 | 0.304 | 0.999 | |
| ATMG00290.E | Ribosomal S4 family protein | 14 | 1.78 | 0.290 | 0.998 | |
| ATMG00290.E | Ribosomal S4 family protein | 15 | 1.87 | 0.283 | 0.994 | |
| ATMG00290.E | Ribosomal S4 family protein | 16 | 2.11 | 0.281 | 0.970 | |
| ATMG00290.E | Ribosomal S4 family protein | 17 | 2.29 | 0.281 | 0.936 | |
| ATMG00290.E | Ribosomal S4 family protein | 18 | 2.22 | 0.284 | 0.954 | |
| ATMG00290.E | Ribosomal S4 family protein | 19 | 1.97 | 0.289 | 0.989 | |
| ATMG00290.E | Ribosomal S4 family protein | 20 | 1.59 | 0.293 | 1.000 | |
| ATMG00290.E | Ribosomal S4 family protein | 21 | 1.12 | 0.296 | 1.000 | |
| ATMG00290.E | Ribosomal S4 family protein | 22 | 0.70 | 0.296 | 1.000 | |
| ATMG00290.E | Ribosomal S4 family protein | 23 | 0.41 | 0.295 | 0.930 | |
| ATMG00290.E | Ribosomal S4 family protein | 24 | 0.27 | 0.291 | 0.488 | |
| ATMG00290.E | Ribosomal S4 family protein | 25 | 0.22 | 0.287 | 0.263 | |
| ATMG00290.E | Ribosomal S4 family protein | 26 | 0.24 | 0.286 | 0.366 | |
| ATMG00290.E | Ribosomal S4 family protein | 27 | 0.36 | 0.286 | 0.789 | |
| ATMG00290.E | Ribosomal S4 family protein | 28 | 0.57 | 0.287 | 0.998 | |
| ATMG00290.E | Ribosomal S4 family protein | 29 | 0.84 | 0.288 | 1.000 | |
| ATMG00290.E | Ribosomal S4 family protein | 30 | 1.03 | 0.289 | 1.000 | |
| ATMG00290.E | Ribosomal S4 family protein | 31 | 1.05 | 0.292 | 1.000 | |
| ATMG00290.E | Ribosomal S4 family protein | 32 | 0.91 | 0.302 | 1.000 | |
| ATMG00290.E | Ribosomal S4 family protein | 33 | 0.72 | 0.334 | 1.000 | |
| ATMG00290.E | Ribosomal S4 family protein | 34 | 0.57 | 0.411 | 1.000 | |
| ATMG00290.E | Ribosomal S4 family protein | 35 | 0.47 | 0.539 | 1.000 | |
| ATMG00290.E | Ribosomal S4 family protein | 46 | 0.86 | 2.521 | 1.000 | |
| ATMG00290.E | Ribosomal S4 family protein | 47 | 1.02 | 2.800 | 1.000 | |
| ATMG00480.1 | ATP synthase subunit (ATP8) | 2 | 0.73 | 0.392 | 1.000 | |
| ATMG00480.1 | ATP synthase subunit (ATP8) | 3 | 0.76 | 0.255 | 1.000 | |
| ATMG00480.1 | ATP synthase subunit (ATP8) | 4 | 0.85 | 0.233 | 1.000 | |
| ATMG00480.1 | ATP synthase subunit (ATP8) | 5 | 1.07 | 0.233 | 1.000 | |
| ATMG00480.1 | ATP synthase subunit (ATP8) | 6 | 1.18 | 0.233 | 1.000 | |
| ATMG00480.1 | ATP synthase subunit (ATP8) | 7 | 1.03 | 0.232 | 1.000 | |
| ATMG00480.1 | ATP synthase subunit (ATP8) | 8 | 0.93 | 0.229 | 1.000 | |
| ATMG00480.1 | ATP synthase subunit (ATP8) | 9 | 1.04 | 0.227 | 1.000 | |
| ATMG00480.1 | ATP synthase subunit (ATP8) | 10 | 1.26 | 0.227 | 1.000 | |
| ATMG00480.1 | ATP synthase subunit (ATP8) | 11 | 1.67 | 0.228 | 1.000 | |
| ATMG00480.1 | ATP synthase subunit (ATP8) | 12 | 2.54 | 0.228 | 0.819 | |
| ATMG00480.1 | ATP synthase subunit (ATP8) | 13 | 3.82 | 0.225 | 0.230 | |
| ATMG00480.1 | ATP synthase subunit (ATP8) | 14 | 4.70 | 0.223 | 0.076 | . |

| | | | | | | |
|---|---|---|---|---|---|---|
| ATMG00480.1 | ATP synthase subunit (ATP8) | 15 | 4.70 | 0.222 | 0.071 | . |
| ATMG00480.1 | ATP synthase subunit (ATP8) | 16 | 4.09 | 0.220 | 0.144 | |
| ATMG00480.1 | ATP synthase subunit (ATP8) | 17 | 3.23 | 0.220 | 0.400 | |
| ATMG00480.1 | ATP synthase subunit (ATP8) | 18 | 2.21 | 0.223 | 0.937 | |
| ATMG00480.1 | ATP synthase subunit (ATP8) | 19 | 1.69 | 0.228 | 1.000 | |
| ATMG00480.1 | ATP synthase subunit (ATP8) | 20 | 1.62 | 0.232 | 1.000 | |
| ATMG00480.1 | ATP synthase subunit (ATP8) | 21 | 1.60 | 0.234 | 1.000 | |
| ATMG00480.1 | ATP synthase subunit (ATP8) | 22 | 1.46 | 0.235 | 1.000 | |
| ATMG00480.1 | ATP synthase subunit (ATP8) | 23 | 1.32 | 0.233 | 1.000 | |
| ATMG00480.1 | ATP synthase subunit (ATP8) | 24 | 1.34 | 0.229 | 1.000 | |
| ATMG00480.1 | ATP synthase subunit (ATP8) | 25 | 1.47 | 0.226 | 1.000 | |
| ATMG00480.1 | ATP synthase subunit (ATP8) | 26 | 1.42 | 0.225 | 1.000 | |
| ATMG00480.1 | ATP synthase subunit (ATP8) | 27 | 1.14 | 0.225 | 1.000 | |
| ATMG00480.1 | ATP synthase subunit (ATP8) | 28 | 1.04 | 0.226 | 1.000 | |
| ATMG00480.1 | ATP synthase subunit (ATP8) | 29 | 1.30 | 0.226 | 1.000 | |
| ATMG00480.1 | ATP synthase subunit (ATP8) | 30 | 1.77 | 0.226 | 0.999 | |
| ATMG00480.1 | ATP synthase subunit (ATP8) | 31 | 1.97 | 0.226 | 0.989 | |
| ATMG00480.1 | ATP synthase subunit (ATP8) | 32 | 1.62 | 0.227 | 1.000 | |
| ATMG00480.1 | ATP synthase subunit (ATP8) | 33 | 1.13 | 0.227 | 1.000 | |
| ATMG00480.1 | ATP synthase subunit (ATP8) | 34 | 0.87 | 0.227 | 1.000 | |
| ATMG00480.1 | ATP synthase subunit (ATP8) | 35 | 0.81 | 0.228 | 1.000 | |
| ATMG00480.1 | ATP synthase subunit (ATP8) | 36 | 0.96 | 0.231 | 1.000 | |
| ATMG00480.1 | ATP synthase subunit (ATP8) | 37 | 1.42 | 0.244 | 1.000 | |
| ATMG00480.1 | ATP synthase subunit (ATP8) | 38 | 2.59 | 0.275 | 0.949 | |
| ATMG00480.1 | ATP synthase subunit (ATP8) | 39 | 4.66 | 0.314 | 0.544 | |
| ATMG00480.1 | ATP synthase subunit (ATP8) | 41 | 8.70 | 0.335 | 0.135 | |
| ATMG00480.1 | ATP synthase subunit (ATP8) | 42 | 8.51 | 0.384 | 0.325 | |
| ATMG00480.1 | ATP synthase subunit (ATP8) | 43 | 6.56 | 0.473 | 0.849 | |
| ATMG00480.1 | ATP synthase subunit (ATP8) | 46 | 1.16 | 1.229 | 1.000 | |
| ATMG00480.1 | ATP synthase subunit (ATP8) | 47 | 0.56 | 1.589 | 1.000 | |
| ATMG00510.E | NADH:ubiquinone oxidoreductase subunit (NAD7) | 3 | 0.63 | 0.166 | 0.999 | |
| ATMG00510.E | NADH:ubiquinone oxidoreductase subunit (NAD7) | 4 | 0.69 | 0.156 | 1.000 | |
| ATMG00510.E | NADH:ubiquinone oxidoreductase subunit (NAD7) | 5 | 0.86 | 0.156 | 1.000 | |
| ATMG00510.E | NADH:ubiquinone oxidoreductase subunit (NAD7) | 6 | 1.07 | 0.156 | 1.000 | |
| ATMG00510.E | NADH:ubiquinone oxidoreductase subunit (NAD7) | 7 | 1.17 | 0.157 | 1.000 | |
| ATMG00510.E | NADH:ubiquinone oxidoreductase subunit (NAD7) | 8 | 1.17 | 0.154 | 1.000 | |
| ATMG00510.E | NADH:ubiquinone oxidoreductase subunit (NAD7) | 9 | 1.18 | 0.151 | 1.000 | |
| ATMG00510.E | NADH:ubiquinone oxidoreductase subunit (NAD7) | 10 | 1.18 | 0.151 | 1.000 | |
| ATMG00510.E | NADH:ubiquinone oxidoreductase subunit (NAD7) | 11 | 1.29 | 0.152 | 1.000 | |
| ATMG00510.E | NADH:ubiquinone oxidoreductase subunit (NAD7) | 12 | 1.41 | 0.152 | 1.000 | |
| ATMG00510.E | NADH:ubiquinone oxidoreductase subunit (NAD7) | 13 | 1.39 | 0.150 | 1.000 | |
| ATMG00510.E | NADH:ubiquinone oxidoreductase subunit (NAD7) | 14 | 1.24 | 0.148 | 1.000 | |
| ATMG00510.E | NADH:ubiquinone oxidoreductase subunit (NAD7) | 15 | 0.96 | 0.147 | 1.000 | |
| ATMG00510.E | NADH:ubiquinone oxidoreductase subunit (NAD7) | 16 | 0.65 | 0.146 | 0.998 | |
| ATMG00510.E | NADH:ubiquinone oxidoreductase subunit (NAD7) | 17 | 0.48 | 0.146 | 0.599 | |
| ATMG00510.E | NADH:ubiquinone oxidoreductase subunit (NAD7) | 18 | 0.52 | 0.149 | 0.808 | |

| | | | | | |
|---|---|---|---|---|---|
| ATMG00510.E | NADH:ubiquinone oxidoreductase subunit (NAD7) | 19 | 0.71 | 0.153 | 1.000 |
| ATMG00510.E | NADH:ubiquinone oxidoreductase subunit (NAD7) | 20 | 0.89 | 0.156 | 1.000 |
| ATMG00510.E | NADH:ubiquinone oxidoreductase subunit (NAD7) | 21 | 1.09 | 0.159 | 1.000 |
| ATMG00510.E | NADH:ubiquinone oxidoreductase subunit (NAD7) | 22 | 1.41 | 0.158 | 1.000 |
| ATMG00510.E | NADH:ubiquinone oxidoreductase subunit (NAD7) | 23 | 1.67 | 0.158 | 0.991 |
| ATMG00510.E | NADH:ubiquinone oxidoreductase subunit (NAD7) | 24 | 1.51 | 0.154 | 1.000 |
| ATMG00510.E | NADH:ubiquinone oxidoreductase subunit (NAD7) | 25 | 1.00 | 0.151 | 1.000 |
| ATMG00510.E | NADH:ubiquinone oxidoreductase subunit (NAD7) | 26 | 0.71 | 0.150 | 1.000 |
| ATMG00510.E | NADH:ubiquinone oxidoreductase subunit (NAD7) | 27 | 0.65 | 0.151 | 0.998 |
| ATMG00510.E | NADH:ubiquinone oxidoreductase subunit (NAD7) | 28 | 0.74 | 0.150 | 1.000 |
| ATMG00510.E | NADH:ubiquinone oxidoreductase subunit (NAD7) | 29 | 1.00 | 0.150 | 1.000 |
| ATMG00510.E | NADH:ubiquinone oxidoreductase subunit (NAD7) | 30 | 1.23 | 0.150 | 1.000 |
| ATMG00510.E | NADH:ubiquinone oxidoreductase subunit (NAD7) | 31 | 1.35 | 0.151 | 1.000 |
| ATMG00510.E | NADH:ubiquinone oxidoreductase subunit (NAD7) | 32 | 1.50 | 0.151 | 1.000 |
| ATMG00510.E | NADH:ubiquinone oxidoreductase subunit (NAD7) | 33 | 1.59 | 0.151 | 0.996 |
| ATMG00510.E | NADH:ubiquinone oxidoreductase subunit (NAD7) | 34 | 1.54 | 0.152 | 0.999 |
| ATMG00510.E | NADH:ubiquinone oxidoreductase subunit (NAD7) | 35 | 1.44 | 0.151 | 1.000 |
| ATMG00510.E | NADH:ubiquinone oxidoreductase subunit (NAD7) | 36 | 1.44 | 0.151 | 1.000 |
| ATMG00510.E | NADH:ubiquinone oxidoreductase subunit (NAD7) | 37 | 1.60 | 0.150 | 0.995 |
| ATMG00510.E | NADH:ubiquinone oxidoreductase subunit (NAD7) | 38 | 1.86 | 0.151 | 0.889 |
| ATMG00510.E | NADH:ubiquinone oxidoreductase subunit (NAD7) | 39 | 2.14 | 0.151 | 0.602 |
| ATMG00510.E | NADH:ubiquinone oxidoreductase subunit (NAD7) | 40 | 2.33 | 0.152 | 0.402 |
| ATMG00510.E | NADH:ubiquinone oxidoreductase subunit (NAD7) | 41 | 2.21 | 0.152 | 0.529 |
| ATMG00510.E | NADH:ubiquinone oxidoreductase subunit (NAD7) | 42 | 1.89 | 0.154 | 0.885 |
| ATMG00510.E | NADH:ubiquinone oxidoreductase subunit (NAD7) | 43 | 1.60 | 0.156 | 0.997 |
| ATMG00510.E | NADH:ubiquinone oxidoreductase subunit (NAD7) | 44 | 1.29 | 0.156 | 1.000 |
| ATMG00510.E | NADH:ubiquinone oxidoreductase subunit (NAD7) | 45 | 0.91 | 0.156 | 1.000 |
| ATMG00510.E | NADH:ubiquinone oxidoreductase subunit (NAD7) | 46 | 0.50 | 0.178 | 0.931 |
| ATMG00510.E | NADH:ubiquinone oxidoreductase subunit (NAD7) | 47 | 0.21 | 0.320 | 0.638 |
| ATMG00510.E | NADH:ubiquinone oxidoreductase subunit (NAD7) | 48 | 0.08 | 0.545 | 0.767 |

| | | | | | |
|---|---|---|---|---|---|
| ATMG00513.E | NADH:ubiquinone oxidoreductase subunit (NAD5A) | 2 | 1.40 | 0.379 | 1.000 |
| ATMG00513.E | NADH:ubiquinone oxidoreductase subunit (NAD5A) | 3 | 1.79 | 0.230 | 0.999 |
| ATMG00513.E | NADH:ubiquinone oxidoreductase subunit (NAD5A) | 4 | 2.32 | 0.210 | 0.880 |
| ATMG00513.E | NADH:ubiquinone oxidoreductase subunit (NAD5A) | 5 | 2.82 | 0.210 | 0.588 |
| ATMG00513.E | NADH:ubiquinone oxidoreductase subunit (NAD5A) | 6 | 2.79 | 0.210 | 0.602 |
| ATMG00513.E | NADH:ubiquinone oxidoreductase subunit (NAD5A) | 7 | 2.32 | 0.209 | 0.878 |
| ATMG00513.E | NADH:ubiquinone oxidoreductase subunit (NAD5A) | 8 | 2.04 | 0.206 | 0.968 |
| ATMG00513.E | NADH:ubiquinone oxidoreductase subunit (NAD5A) | 9 | 2.14 | 0.203 | 0.934 |
| ATMG00513.E | NADH:ubiquinone oxidoreductase subunit (NAD5A) | 10 | 2.32 | 0.204 | 0.853 |
| ATMG00513.E | NADH:ubiquinone oxidoreductase subunit (NAD5A) | 11 | 2.39 | 0.206 | 0.827 |
| ATMG00513.E | NADH:ubiquinone oxidoreductase subunit (NAD5A) | 12 | 2.36 | 0.205 | 0.839 |
| ATMG00513.E | NADH:ubiquinone oxidoreductase subunit (NAD5A) | 13 | 1.96 | 0.202 | 0.979 |
| ATMG00513.E | NADH:ubiquinone oxidoreductase subunit (NAD5A) | 14 | 1.51 | 0.199 | 1.000 |
| ATMG00513.E | NADH:ubiquinone oxidoreductase subunit (NAD5A) | 15 | 1.16 | 0.198 | 1.000 |
| ATMG00513.E | NADH:ubiquinone oxidoreductase subunit (NAD5A) | 16 | 0.81 | 0.196 | 1.000 |
| ATMG00513.E | NADH:ubiquinone oxidoreductase subunit (NAD5A) | 17 | 0.62 | 0.196 | 1.000 |
| ATMG00513.E | NADH:ubiquinone oxidoreductase subunit (NAD5A) | 18 | 0.66 | 0.199 | 1.000 |
| ATMG00513.E | NADH:ubiquinone oxidoreductase subunit (NAD5A) | 19 | 0.84 | 0.204 | 1.000 |
| ATMG00513.E | NADH:ubiquinone oxidoreductase subunit (NAD5A) | 20 | 0.99 | 0.210 | 1.000 |
| ATMG00513.E | NADH:ubiquinone oxidoreductase subunit (NAD5A) | 21 | 1.16 | 0.212 | 1.000 |

| | | | | | |
|---|---|---|---|---|---|
| ATMG00513.E | NADH:ubiquinone oxidoreductase subunit (NAD5A) | 22 | 1.44 | 0.212 | 1.000 |
| ATMG00513.E | NADH:ubiquinone oxidoreductase subunit (NAD5A) | 23 | 1.65 | 0.210 | 1.000 |
| ATMG00513.E | NADH:ubiquinone oxidoreductase subunit (NAD5A) | 24 | 1.33 | 0.205 | 1.000 |
| ATMG00513.E | NADH:ubiquinone oxidoreductase subunit (NAD5A) | 25 | 0.73 | 0.203 | 1.000 |
| ATMG00513.E | NADH:ubiquinone oxidoreductase subunit (NAD5A) | 26 | 0.42 | 0.202 | 0.795 |
| ATMG00513.E | NADH:ubiquinone oxidoreductase subunit (NAD5A) | 27 | 0.35 | 0.203 | 0.503 |
| ATMG00513.E | NADH:ubiquinone oxidoreductase subunit (NAD5A) | 28 | 0.48 | 0.203 | 0.948 |
| ATMG00513.E | NADH:ubiquinone oxidoreductase subunit (NAD5A) | 29 | 0.82 | 0.202 | 1.000 |
| ATMG00513.E | NADH:ubiquinone oxidoreductase subunit (NAD5A) | 30 | 1.21 | 0.202 | 1.000 |
| ATMG00513.E | NADH:ubiquinone oxidoreductase subunit (NAD5A) | 31 | 1.56 | 0.203 | 1.000 |
| ATMG00513.E | NADH:ubiquinone oxidoreductase subunit (NAD5A) | 32 | 2.20 | 0.205 | 0.916 |
| ATMG00513.E | NADH:ubiquinone oxidoreductase subunit (NAD5A) | 33 | 3.38 | 0.209 | 0.286 |
| ATMG00513.E | NADH:ubiquinone oxidoreductase subunit (NAD5A) | 34 | 4.53 | 0.225 | 0.107 |
| ATMG00513.E | NADH:ubiquinone oxidoreductase subunit (NAD5A) | 35 | 4.84 | 0.269 | 0.272 |
| ATMG00513.E | NADH:ubiquinone oxidoreductase subunit (NAD5A) | 36 | 4.90 | 0.367 | 0.794 |
| ATMG00513.E | NADH:ubiquinone oxidoreductase subunit (NAD5A) | 41 | 5.74 | 0.367 | 0.651 |
| ATMG00513.E | NADH:ubiquinone oxidoreductase subunit (NAD5A) | 42 | 4.67 | 0.367 | 0.834 |
| ATMG00513.E | NADH:ubiquinone oxidoreductase subunit (NAD5A) | 43 | 3.27 | 0.446 | 0.999 |
| ATMG00513.E | NADH:ubiquinone oxidoreductase subunit (NAD5A) | 46 | 0.62 | 1.323 | 1.000 |
| ATMG00513.E | NADH:ubiquinone oxidoreductase subunit (NAD5A) | 47 | 0.32 | 1.716 | 1.000 |
| ATMG00560.1 | Ribosomal L2 family protein | 2 | 1.77 | 0.469 | 1.000 |
| ATMG00560.1 | Ribosomal L2 family protein | 3 | 1.70 | 0.429 | 1.000 |

| | | | | | |
|---|---|---|---|---|---|
| ATMG00560.1 | Ribosomal L2 family protein | 13 | 2.19 | 0.286 | 0.910 |
| ATMG00560.1 | Ribosomal L2 family protein | 14 | 2.31 | 0.264 | 0.803 |
| ATMG00560.1 | Ribosomal L2 family protein | 15 | 2.27 | 0.252 | 0.778 |
| ATMG00560.1 | Ribosomal L2 family protein | 16 | 2.16 | 0.244 | 0.806 |
| ATMG00560.1 | Ribosomal L2 family protein | 17 | 2.00 | 0.241 | 0.878 |
| ATMG00560.1 | Ribosomal L2 family protein | 18 | 1.68 | 0.242 | 0.985 |
| ATMG00560.1 | Ribosomal L2 family protein | 19 | 1.28 | 0.248 | 1.000 |
| ATMG00560.1 | Ribosomal L2 family protein | 20 | 0.94 | 0.262 | 1.000 |
| ATMG00560.1 | Ribosomal L2 family protein | 21 | 0.71 | 0.280 | 1.000 |
| ATMG00560.1 | Ribosomal L2 family protein | 22 | 0.57 | 0.285 | 0.993 |
| ATMG00560.1 | Ribosomal L2 family protein | 23 | 0.47 | 0.286 | 0.934 |
| ATMG00560.1 | Ribosomal L2 family protein | 24 | 0.42 | 0.276 | 0.805 |
| ATMG00560.1 | Ribosomal L2 family protein | 25 | 0.39 | 0.263 | 0.684 |
| ATMG00560.1 | Ribosomal L2 family protein | 26 | 0.41 | 0.251 | 0.676 |
| ATMG00560.1 | Ribosomal L2 family protein | 27 | 0.47 | 0.243 | 0.821 |
| ATMG00560.1 | Ribosomal L2 family protein | 28 | 0.59 | 0.239 | 0.983 |
| ATMG00560.1 | Ribosomal L2 family protein | 29 | 0.78 | 0.238 | 1.000 |
| ATMG00560.1 | Ribosomal L2 family protein | 30 | 0.99 | 0.237 | 1.000 |
| ATMG00560.1 | Ribosomal L2 family protein | 31 | 1.17 | 0.237 | 1.000 |
| ATMG00560.1 | Ribosomal L2 family protein | 32 | 1.21 | 0.237 | 1.000 |
| ATMG00560.1 | Ribosomal L2 family protein | 33 | 1.06 | 0.237 | 1.000 |
| ATMG00560.1 | Ribosomal L2 family protein | 34 | 0.87 | 0.237 | 1.000 |
| ATMG00560.1 | Ribosomal L2 family protein | 35 | 0.75 | 0.237 | 1.000 |
| ATMG00560.1 | Ribosomal L2 family protein | 36 | 0.69 | 0.237 | 0.999 |
| ATMG00560.1 | Ribosomal L2 family protein | 37 | 0.65 | 0.237 | 0.997 |
| ATMG00560.1 | Ribosomal L2 family protein | 38 | 0.62 | 0.237 | 0.992 |
| ATMG00560.1 | Ribosomal L2 family protein | 39 | 0.66 | 0.237 | 0.998 |
| ATMG00560.1 | Ribosomal L2 family protein | 40 | 0.79 | 0.237 | 1.000 |
| ATMG00560.1 | Ribosomal L2 family protein | 41 | 1.11 | 0.238 | 1.000 |
| ATMG00560.1 | Ribosomal L2 family protein | 42 | 1.65 | 0.238 | 0.987 |
| ATMG00560.1 | Ribosomal L2 family protein | 43 | 2.16 | 0.240 | 0.790 |
| ATMG00560.1 | Ribosomal L2 family protein | 44 | 2.06 | 0.242 | 0.854 |
| ATMG00560.1 | Ribosomal L2 family protein | 45 | 1.26 | 0.249 | 1.000 |
| ATMG00560.1 | Ribosomal L2 family protein | 46 | 0.52 | 0.272 | 0.966 |
| ATMG00560.1 | Ribosomal L2 family protein | 47 | 0.18 | 0.344 | 0.271 |
| ATMG00580.E | NADH:ubiquinone oxidoreductase subunit (NAD4) | 2 | 0.90 | 0.355 | 1.000 |
| ATMG00580.E | NADH:ubiquinone oxidoreductase subunit (NAD4) | 3 | 0.89 | 0.231 | 1.000 |
| ATMG00580.E | NADH:ubiquinone oxidoreductase subunit (NAD4) | 4 | 0.91 | 0.208 | 1.000 |
| ATMG00580.E | NADH:ubiquinone oxidoreductase subunit (NAD4) | 5 | 1.00 | 0.211 | 1.000 |
| ATMG00580.E | NADH:ubiquinone oxidoreductase subunit (NAD4) | 6 | 1.11 | 0.217 | 1.000 |
| ATMG00580.E | NADH:ubiquinone oxidoreductase subunit (NAD4) | 7 | 1.11 | 0.222 | 1.000 |
| ATMG00580.E | NADH:ubiquinone oxidoreductase subunit (NAD4) | 8 | 1.05 | 0.227 | 1.000 |
| ATMG00580.E | NADH:ubiquinone oxidoreductase subunit (NAD4) | 9 | 1.02 | 0.229 | 1.000 |
| ATMG00580.E | NADH:ubiquinone oxidoreductase subunit (NAD4) | 10 | 1.06 | 0.226 | 1.000 |
| ATMG00580.E | NADH:ubiquinone oxidoreductase subunit (NAD4) | 11 | 1.17 | 0.218 | 1.000 |
| ATMG00580.E | NADH:ubiquinone oxidoreductase subunit (NAD4) | 12 | 1.36 | 0.214 | 1.000 |
| ATMG00580.E | NADH:ubiquinone oxidoreductase subunit (NAD4) | 13 | 1.61 | 0.212 | 1.000 |
| ATMG00580.E | NADH:ubiquinone oxidoreductase subunit (NAD4) | 14 | 1.76 | 0.206 | 0.995 |

| | | | | | |
|---|---|---|---|---|---|
| ATMG00580.E | NADH:ubiquinone oxidoreductase subunit (NAD4) | 15 | 1.54 | 0.200 | 1.000 |
| ATMG00580.E | NADH:ubiquinone oxidoreductase subunit (NAD4) | 16 | 1.12 | 0.197 | 1.000 |
| ATMG00580.E | NADH:ubiquinone oxidoreductase subunit (NAD4) | 17 | 0.82 | 0.196 | 1.000 |
| ATMG00580.E | NADH:ubiquinone oxidoreductase subunit (NAD4) | 18 | 0.75 | 0.198 | 1.000 |
| ATMG00580.E | NADH:ubiquinone oxidoreductase subunit (NAD4) | 19 | 0.82 | 0.203 | 1.000 |
| ATMG00580.E | NADH:ubiquinone oxidoreductase subunit (NAD4) | 20 | 0.93 | 0.207 | 1.000 |
| ATMG00580.E | NADH:ubiquinone oxidoreductase subunit (NAD4) | 21 | 1.09 | 0.209 | 1.000 |
| ATMG00580.E | NADH:ubiquinone oxidoreductase subunit (NAD4) | 22 | 1.32 | 0.209 | 1.000 |
| ATMG00580.E | NADH:ubiquinone oxidoreductase subunit (NAD4) | 23 | 1.45 | 0.208 | 1.000 |
| ATMG00580.E | NADH:ubiquinone oxidoreductase subunit (NAD4) | 24 | 1.31 | 0.204 | 1.000 |
| ATMG00580.E | NADH:ubiquinone oxidoreductase subunit (NAD4) | 25 | 1.03 | 0.201 | 1.000 |
| ATMG00580.E | NADH:ubiquinone oxidoreductase subunit (NAD4) | 26 | 0.91 | 0.201 | 1.000 |
| ATMG00580.E | NADH:ubiquinone oxidoreductase subunit (NAD4) | 27 | 0.97 | 0.201 | 1.000 |
| ATMG00580.E | NADH:ubiquinone oxidoreductase subunit (NAD4) | 28 | 1.18 | 0.201 | 1.000 |
| ATMG00580.E | NADH:ubiquinone oxidoreductase subunit (NAD4) | 29 | 1.50 | 0.201 | 1.000 |
| ATMG00580.E | NADH:ubiquinone oxidoreductase subunit (NAD4) | 30 | 1.68 | 0.201 | 0.998 |
| ATMG00580.E | NADH:ubiquinone oxidoreductase subunit (NAD4) | 31 | 1.65 | 0.202 | 0.999 |
| ATMG00580.E | NADH:ubiquinone oxidoreductase subunit (NAD4) | 32 | 1.79 | 0.202 | 0.992 |
| ATMG00580.E | NADH:ubiquinone oxidoreductase subunit (NAD4) | 33 | 2.24 | 0.203 | 0.845 |
| ATMG00580.E | NADH:ubiquinone oxidoreductase subunit (NAD4) | 34 | 2.67 | 0.203 | 0.564 |
| ATMG00580.E | NADH:ubiquinone oxidoreductase subunit (NAD4) | 35 | 2.57 | 0.206 | 0.659 |
| ATMG00580.E | NADH:ubiquinone oxidoreductase subunit (NAD4) | 36 | 2.27 | 0.219 | 0.902 |
| ATMG00580.E | NADH:ubiquinone oxidoreductase subunit (NAD4) | 37 | 2.25 | 0.253 | 0.974 |
| ATMG00580.E | NADH:ubiquinone oxidoreductase subunit (NAD4) | 38 | 2.51 | 0.305 | 0.987 |
| ATMG00580.E | NADH:ubiquinone oxidoreductase subunit (NAD4) | 41 | 3.54 | 0.323 | 0.863 |
| ATMG00580.E | NADH:ubiquinone oxidoreductase subunit (NAD4) | 42 | 3.60 | 0.356 | 0.928 |
| ATMG00580.E | NADH:ubiquinone oxidoreductase subunit (NAD4) | 43 | 3.34 | 0.426 | 0.993 |
| ATMG00580.E | NADH:ubiquinone oxidoreductase subunit (NAD4) | 46 | 1.86 | 1.067 | 1.000 |
| ATMG00580.E | NADH:ubiquinone oxidoreductase subunit (NAD4) | 47 | 1.44 | 1.372 | 1.000 |
| ATMG00640.E | ATP synthase subunit (ATP4) | 2 | 1.35 | 0.369 | 1.000 |
| ATMG00640.E | ATP synthase subunit (ATP4) | 3 | 1.24 | 0.246 | 1.000 |
| ATMG00640.E | ATP synthase subunit (ATP4) | 4 | 1.22 | 0.225 | 1.000 |
| ATMG00640.E | ATP synthase subunit (ATP4) | 5 | 1.32 | 0.226 | 1.000 |

| | | | | | | |
|---|---|---|---|---|---|---|
| ATMG00640.E | ATP synthase subunit (ATP4) | 6 | 1.41 | 0.226 | 1.000 | |
| ATMG00640.E | ATP synthase subunit (ATP4) | 7 | 1.43 | 0.225 | 1.000 | |
| ATMG00640.E | ATP synthase subunit (ATP4) | 8 | 1.49 | 0.222 | 1.000 | |
| ATMG00640.E | ATP synthase subunit (ATP4) | 9 | 1.58 | 0.220 | 1.000 | |
| ATMG00640.E | ATP synthase subunit (ATP4) | 10 | 1.67 | 0.220 | 1.000 | |
| ATMG00640.E | ATP synthase subunit (ATP4) | 11 | 1.86 | 0.220 | 0.993 | |
| ATMG00640.E | ATP synthase subunit (ATP4) | 12 | 2.23 | 0.220 | 0.910 | |
| ATMG00640.E | ATP synthase subunit (ATP4) | 13 | 2.79 | 0.218 | 0.594 | |
| ATMG00640.E | ATP synthase subunit (ATP4) | 14 | 3.55 | 0.216 | 0.238 | |
| ATMG00640.E | ATP synthase subunit (ATP4) | 15 | 3.96 | 0.216 | 0.139 | |
| ATMG00640.E | ATP synthase subunit (ATP4) | 16 | 3.14 | 0.214 | 0.376 | |
| ATMG00640.E | ATP synthase subunit (ATP4) | 17 | 2.05 | 0.214 | 0.958 | |
| ATMG00640.E | ATP synthase subunit (ATP4) | 18 | 1.37 | 0.217 | 1.000 | |
| ATMG00640.E | ATP synthase subunit (ATP4) | 19 | 1.08 | 0.221 | 1.000 | |
| ATMG00640.E | ATP synthase subunit (ATP4) | 20 | 1.03 | 0.225 | 1.000 | |
| ATMG00640.E | ATP synthase subunit (ATP4) | 21 | 1.01 | 0.227 | 1.000 | |
| ATMG00640.E | ATP synthase subunit (ATP4) | 22 | 0.96 | 0.227 | 1.000 | |
| ATMG00640.E | ATP synthase subunit (ATP4) | 23 | 0.93 | 0.226 | 1.000 | |
| ATMG00640.E | ATP synthase subunit (ATP4) | 24 | 1.00 | 0.223 | 1.000 | |
| ATMG00640.E | ATP synthase subunit (ATP4) | 25 | 1.16 | 0.220 | 1.000 | |
| ATMG00640.E | ATP synthase subunit (ATP4) | 26 | 1.25 | 0.218 | 1.000 | |
| ATMG00640.E | ATP synthase subunit (ATP4) | 27 | 1.10 | 0.218 | 1.000 | |
| ATMG00640.E | ATP synthase subunit (ATP4) | 28 | 1.03 | 0.218 | 1.000 | |
| ATMG00640.E | ATP synthase subunit (ATP4) | 29 | 1.23 | 0.219 | 1.000 | |
| ATMG00640.E | ATP synthase subunit (ATP4) | 30 | 1.54 | 0.219 | 1.000 | |
| ATMG00640.E | ATP synthase subunit (ATP4) | 31 | 1.60 | 0.219 | 1.000 | |
| ATMG00640.E | ATP synthase subunit (ATP4) | 32 | 1.37 | 0.220 | 1.000 | |
| ATMG00640.E | ATP synthase subunit (ATP4) | 33 | 1.13 | 0.220 | 1.000 | |
| ATMG00640.E | ATP synthase subunit (ATP4) | 34 | 1.11 | 0.220 | 1.000 | |
| ATMG00640.E | ATP synthase subunit (ATP4) | 35 | 1.32 | 0.220 | 1.000 | |
| ATMG00640.E | ATP synthase subunit (ATP4) | 36 | 1.90 | 0.220 | 0.990 | |
| ATMG00640.E | ATP synthase subunit (ATP4) | 37 | 3.39 | 0.220 | 0.315 | |
| ATMG00640.E | ATP synthase subunit (ATP4) | 38 | 6.03 | 0.220 | 0.014 | * |
| ATMG00640.E | ATP synthase subunit (ATP4) | 39 | 8.34 | 0.220 | 0.001 | ** |
| ATMG00640.E | ATP synthase subunit (ATP4) | 40 | 8.26 | 0.221 | 0.002 | ** |
| ATMG00640.E | ATP synthase subunit (ATP4) | 41 | 6.42 | 0.222 | 0.010 | ** |
| ATMG00640.E | ATP synthase subunit (ATP4) | 42 | 4.21 | 0.226 | 0.141 | |
| ATMG00640.E | ATP synthase subunit (ATP4) | 43 | 2.59 | 0.235 | 0.811 | |
| ATMG00640.E | ATP synthase subunit (ATP4) | 44 | 1.36 | 0.276 | 1.000 | |
| ATMG00640.E | ATP synthase subunit (ATP4) | 45 | 0.68 | 0.377 | 1.000 | |
| ATMG00640.E | ATP synthase subunit (ATP4) | 46 | 0.33 | 0.575 | 1.000 | |
| ATMG00640.E | ATP synthase subunit (ATP4) | 47 | 0.16 | 0.850 | 1.000 | |
| ATMG00650.E | NADH:ubiquinone oxidoreductase subunit (NAD4L) | 2 | 0.91 | 0.507 | 1.000 | |
| ATMG00650.E | NADH:ubiquinone oxidoreductase subunit (NAD4L) | 3 | 0.91 | 0.458 | 1.000 | |
| ATMG00650.E | NADH:ubiquinone oxidoreductase subunit (NAD4L) | 15 | 0.65 | 0.280 | 1.000 | |
| ATMG00650.E | NADH:ubiquinone oxidoreductase subunit (NAD4L) | 16 | 0.58 | 0.238 | 0.985 | |
| ATMG00650.E | NADH:ubiquinone oxidoreductase subunit (NAD4L) | 17 | 0.54 | 0.213 | 0.923 | |
| ATMG00650.E | NADH:ubiquinone oxidoreductase subunit (NAD4L) | 18 | 0.60 | 0.198 | 0.970 | |

| | | | | | |
|---|---|---|---|---|---|
| ATMG00650.E | NADH:ubiquinone oxidoreductase subunit (NAD4L) | 19 | 0.83 | 0.196 | 1.000 |
| ATMG00650.E | NADH:ubiquinone oxidoreductase subunit (NAD4L) | 20 | 1.23 | 0.200 | 1.000 |
| ATMG00650.E | NADH:ubiquinone oxidoreductase subunit (NAD4L) | 21 | 1.89 | 0.202 | 0.881 |
| ATMG00650.E | NADH:ubiquinone oxidoreductase subunit (NAD4L) | 22 | 2.68 | 0.202 | 0.361 |
| ATMG00650.E | NADH:ubiquinone oxidoreductase subunit (NAD4L) | 23 | 2.84 | 0.200 | 0.274 |
| ATMG00650.E | NADH:ubiquinone oxidoreductase subunit (NAD4L) | 24 | 2.25 | 0.196 | 0.599 |
| ATMG00650.E | NADH:ubiquinone oxidoreductase subunit (NAD4L) | 25 | 1.50 | 0.192 | 0.993 |
| ATMG00650.E | NADH:ubiquinone oxidoreductase subunit (NAD4L) | 26 | 1.09 | 0.192 | 1.000 |
| ATMG00650.E | NADH:ubiquinone oxidoreductase subunit (NAD4L) | 27 | 0.97 | 0.193 | 1.000 |
| ATMG00650.E | NADH:ubiquinone oxidoreductase subunit (NAD4L) | 28 | 0.93 | 0.195 | 1.000 |
| ATMG00650.E | NADH:ubiquinone oxidoreductase subunit (NAD4L) | 29 | 0.87 | 0.200 | 1.000 |
| ATMG00650.E | NADH:ubiquinone oxidoreductase subunit (NAD4L) | 30 | 0.81 | 0.221 | 1.000 |
| ATMG00650.E | NADH:ubiquinone oxidoreductase subunit (NAD4L) | 31 | 0.75 | 0.284 | 1.000 |
| ATMG00650.E | NADH:ubiquinone oxidoreductase subunit (NAD4L) | 32 | 0.75 | 0.411 | 1.000 |
| ATMG00650.E | NADH:ubiquinone oxidoreductase subunit (NAD4L) | 46 | 88.25 | 3.476 | 1.000 |
| ATMG00650.E | NADH:ubiquinone oxidoreductase subunit (NAD4L) | 47 | 147.20 | 3.815 | 1.000 |
| ATMG00730.E | Cytochrome c:O2 oxidoreductase subunit (COX3) | 2 | 1.00 | 0.474 | 1.000 |
| ATMG00730.E | Cytochrome c:O2 oxidoreductase subunit (COX3) | 3 | 0.98 | 0.545 | 1.000 |
| ATMG00730.E | Cytochrome c:O2 oxidoreductase subunit (COX3) | 11 | 0.85 | 0.321 | 1.000 |
| ATMG00730.E | Cytochrome c:O2 oxidoreductase subunit (COX3) | 12 | 1.02 | 0.296 | 1.000 |
| ATMG00730.E | Cytochrome c:O2 oxidoreductase subunit (COX3) | 13 | 1.23 | 0.324 | 1.000 |
| ATMG00730.E | Cytochrome c:O2 oxidoreductase subunit (COX3) | 14 | 1.39 | 0.305 | 1.000 |
| ATMG00730.E | Cytochrome c:O2 oxidoreductase subunit (COX3) | 20 | 0.89 | 0.391 | 1.000 |

| | | | | | | |
|---|---|---|---|---|---|---|
| ATMG00730.E | Cytochrome c:O2 oxidoreductase subunit (COX3) | 21 | 0.86 | 0.352 | 1.000 | |
| ATMG00730.E | Cytochrome c:O2 oxidoreductase subunit (COX3) | 22 | 0.80 | 0.434 | 1.000 | |
| ATMG00730.E | Cytochrome c:O2 oxidoreductase subunit (COX3) | 27 | 0.49 | 0.308 | 0.991 | |
| ATMG00730.E | Cytochrome c:O2 oxidoreductase subunit (COX3) | 28 | 0.96 | 0.237 | 1.000 | |
| ATMG00730.E | Cytochrome c:O2 oxidoreductase subunit (COX3) | 29 | 1.88 | 0.210 | 0.937 | |
| ATMG00730.E | Cytochrome c:O2 oxidoreductase subunit (COX3) | 30 | 2.39 | 0.201 | 0.583 | |
| ATMG00730.E | Cytochrome c:O2 oxidoreductase subunit (COX3) | 31 | 2.02 | 0.199 | 0.834 | |
| ATMG00730.E | Cytochrome c:O2 oxidoreductase subunit (COX3) | 32 | 1.24 | 0.204 | 1.000 | |
| ATMG00730.E | Cytochrome c:O2 oxidoreductase subunit (COX3) | 33 | 0.45 | 0.216 | 0.802 | |
| ATMG00730.E | Cytochrome c:O2 oxidoreductase subunit (COX3) | 34 | 0.14 | 0.275 | 0.035 | * |
| ATMG00730.E | Cytochrome c:O2 oxidoreductase subunit (COX3) | 35 | 0.06 | 0.466 | 0.166 | |
| ATMG00730.E | Cytochrome c:O2 oxidoreductase subunit (COX3) | 46 | 1607.47 | 3.911 | 0.999 | |
| ATMG00730.E | Cytochrome c:O2 oxidoreductase subunit (COX3) | 47 | 8908.48 | 4.402 | 0.997 | |
| ATMG00960.E | [MISSING] | 2 | 0.88 | 0.497 | 1.000 | |
| ATMG00960.E | [MISSING] | 3 | 0.84 | 0.484 | 1.000 | |
| ATMG00960.E | [MISSING] | 23 | 0.58 | 0.324 | 0.982 | |
| ATMG00960.E | [MISSING] | 24 | 0.52 | 0.236 | 0.827 | |
| ATMG00960.E | [MISSING] | 25 | 0.50 | 0.199 | 0.616 | |
| ATMG00960.E | [MISSING] | 26 | 0.60 | 0.189 | 0.825 | |
| ATMG00960.E | [MISSING] | 27 | 0.91 | 0.207 | 1.000 | |
| ATMG00960.E | [MISSING] | 28 | 1.58 | 0.255 | 0.975 | |
| ATMG00960.E | [MISSING] | 29 | 3.21 | 0.342 | 0.635 | |
| ATMG00960.E | [MISSING] | 46 | 445923.31 | 7.602 | 0.981 | |
| ATMG00960.E | [MISSING] | 47 | 858021.01 | 8.256 | 0.984 | |
| ATMG00990.E | [MISSING] | 2 | 0.79 | 0.626 | 1.000 | |
| ATMG00990.E | [MISSING] | 3 | 0.91 | 0.460 | 1.000 | |
| ATMG00990.E | [MISSING] | 5 | 1.17 | 0.456 | 1.000 | |
| ATMG00990.E | [MISSING] | 6 | 1.29 | 0.448 | 1.000 | |
| ATMG00990.E | [MISSING] | 7 | 1.34 | 0.488 | 1.000 | |
| ATMG00990.E | [MISSING] | 13 | 0.35 | 0.378 | 0.980 | |
| ATMG00990.E | [MISSING] | 14 | 0.30 | 0.334 | 0.868 | |
| ATMG00990.E | [MISSING] | 15 | 0.34 | 0.304 | 0.883 | |
| ATMG00990.E | [MISSING] | 16 | 0.45 | 0.278 | 0.972 | |
| ATMG00990.E | [MISSING] | 17 | 0.51 | 0.266 | 0.992 | |
| ATMG00990.E | [MISSING] | 18 | 0.62 | 0.267 | 1.000 | |
| ATMG00990.E | [MISSING] | 19 | 0.81 | 0.274 | 1.000 | |
| ATMG00990.E | [MISSING] | 20 | 0.93 | 0.279 | 1.000 | |
| ATMG00990.E | [MISSING] | 21 | 0.99 | 0.282 | 1.000 | |
| ATMG00990.E | [MISSING] | 22 | 1.11 | 0.281 | 1.000 | |
| ATMG00990.E | [MISSING] | 23 | 1.43 | 0.278 | 1.000 | |
| ATMG00990.E | [MISSING] | 24 | 2.15 | 0.273 | 0.979 | |
| ATMG00990.E | [MISSING] | 25 | 2.80 | 0.271 | 0.810 | |
| ATMG00990.E | [MISSING] | 26 | 1.63 | 0.270 | 1.000 | |
| ATMG00990.E | [MISSING] | 27 | 0.49 | 0.270 | 0.987 | |
| ATMG00990.E | [MISSING] | 28 | 0.19 | 0.271 | 0.153 | |
| ATMG00990.E | [MISSING] | 29 | 0.20 | 0.272 | 0.172 | |
| ATMG00990.E | [MISSING] | 30 | 0.43 | 0.279 | 0.961 | |
| ATMG00990.E | [MISSING] | 31 | 1.04 | 0.306 | 1.000 | |
| ATMG00990.E | [MISSING] | 32 | 1.73 | 0.367 | 1.000 | |

| | | | | | | |
|---|---|---|---|---|---|---|
| ATMG00990.E | [MISSING] | 33 | 2.23 | 0.504 | 1.000 | |
| ATMG00990.E | [MISSING] | 46 | 65.53 | 8.912 | 1.000 | |
| ATMG00990.E | [MISSING] | 47 | 86.12 | 9.902 | 1.000 | |
| ATMG01080.E | [MISSING] | 2 | 2.82 | 0.679 | 1.000 | |
| ATMG01080.E | [MISSING] | 3 | 3.26 | 0.545 | 0.999 | |
| ATMG01080.E | [MISSING] | 6 | 5.45 | 0.470 | 0.868 | |
| ATMG01080.E | [MISSING] | 7 | 6.69 | 0.410 | 0.545 | |
| ATMG01080.E | [MISSING] | 8 | 7.95 | 0.369 | 0.254 | |
| ATMG01080.E | [MISSING] | 9 | 8.74 | 0.341 | 0.121 | |
| ATMG01080.E | [MISSING] | 10 | 8.52 | 0.328 | 0.099 | . |
| ATMG01080.E | [MISSING] | 11 | 6.92 | 0.323 | 0.178 | |
| ATMG01080.E | [MISSING] | 12 | 4.95 | 0.321 | 0.431 | |
| ATMG01080.E | [MISSING] | 13 | 3.25 | 0.319 | 0.845 | |
| ATMG01080.E | [MISSING] | 14 | 2.21 | 0.315 | 0.995 | |
| ATMG01080.E | [MISSING] | 15 | 1.75 | 0.314 | 1.000 | |
| ATMG01080.E | [MISSING] | 16 | 1.52 | 0.313 | 1.000 | |
| ATMG01080.E | [MISSING] | 17 | 1.52 | 0.307 | 1.000 | |
| ATMG01080.E | [MISSING] | 18 | 1.68 | 0.305 | 1.000 | |
| ATMG01080.E | [MISSING] | 19 | 1.86 | 0.310 | 1.000 | |
| ATMG01080.E | [MISSING] | 20 | 1.90 | 0.315 | 1.000 | |
| ATMG01080.E | [MISSING] | 21 | 1.71 | 0.318 | 1.000 | |
| ATMG01080.E | [MISSING] | 22 | 1.38 | 0.317 | 1.000 | |
| ATMG01080.E | [MISSING] | 23 | 1.34 | 0.315 | 1.000 | |
| ATMG01080.E | [MISSING] | 24 | 1.56 | 0.311 | 1.000 | |
| ATMG01080.E | [MISSING] | 25 | 1.89 | 0.307 | 1.000 | |
| ATMG01080.E | [MISSING] | 26 | 2.06 | 0.306 | 0.998 | |
| ATMG01080.E | [MISSING] | 27 | 1.96 | 0.307 | 0.999 | |
| ATMG01080.E | [MISSING] | 28 | 1.83 | 0.306 | 1.000 | |
| ATMG01080.E | [MISSING] | 29 | 2.01 | 0.307 | 0.998 | |
| ATMG01080.E | [MISSING] | 30 | 2.53 | 0.308 | 0.967 | |
| ATMG01080.E | [MISSING] | 31 | 2.86 | 0.308 | 0.909 | |
| ATMG01080.E | [MISSING] | 32 | 2.68 | 0.308 | 0.946 | |
| ATMG01080.E | [MISSING] | 33 | 2.33 | 0.308 | 0.987 | |
| ATMG01080.E | [MISSING] | 34 | 2.31 | 0.309 | 0.988 | |
| ATMG01080.E | [MISSING] | 35 | 2.74 | 0.309 | 0.935 | |
| ATMG01080.E | [MISSING] | 36 | 4.15 | 0.309 | 0.554 | |
| ATMG01080.E | [MISSING] | 37 | 7.41 | 0.309 | 0.105 | |
| ATMG01080.E | [MISSING] | 38 | 11.25 | 0.312 | 0.022 | * |
| ATMG01080.E | [MISSING] | 39 | 10.78 | 0.322 | 0.033 | * |
| ATMG01080.E | [MISSING] | 40 | 7.84 | 0.340 | 0.164 | |
| ATMG01080.E | [MISSING] | 41 | 5.22 | 0.379 | 0.637 | |
| ATMG01080.E | [MISSING] | 42 | 3.55 | 0.467 | 0.988 | |
| ATMG01080.E | [MISSING] | 43 | 2.82 | 0.588 | 1.000 | |
| ATMG01080.E | [MISSING] | 46 | 2.82 | 1.247 | 1.000 | |
| ATMG01080.E | [MISSING] | 47 | 3.15 | 1.574 | 1.000 | |
| ATMG01170.1 | ATP synthase subunit (ATPA2) | 2 | 0.87 | 0.442 | 1.000 | |
| ATMG01170.1 | ATP synthase subunit (ATPA2) | 3 | 0.81 | 0.319 | 1.000 | |
| ATMG01170.1 | ATP synthase subunit (ATPA2) | 4 | 0.77 | 0.356 | 1.000 | |
| ATMG01170.1 | ATP synthase subunit (ATPA2) | 6 | 0.91 | 0.363 | 1.000 | |
| ATMG01170.1 | ATP synthase subunit (ATPA2) | 7 | 1.19 | 0.317 | 1.000 | |
| ATMG01170.1 | ATP synthase subunit (ATPA2) | 8 | 1.66 | 0.274 | 1.000 | |
| ATMG01170.1 | ATP synthase subunit (ATPA2) | 9 | 2.03 | 0.253 | 0.997 | |
| ATMG01170.1 | ATP synthase subunit (ATPA2) | 10 | 2.03 | 0.243 | 0.995 | |
| ATMG01170.1 | ATP synthase subunit (ATPA2) | 11 | 2.34 | 0.245 | 0.962 | |
| ATMG01170.1 | ATP synthase subunit (ATPA2) | 12 | 3.02 | 0.260 | 0.806 | |
| ATMG01170.1 | ATP synthase subunit (ATPA2) | 13 | 3.29 | 0.274 | 0.774 | |
| ATMG01170.1 | ATP synthase subunit (ATPA2) | 14 | 2.54 | 0.291 | 0.984 | |
| ATMG01170.1 | ATP synthase subunit (ATPA2) | 18 | 0.78 | 0.309 | 1.000 | |
| ATMG01170.1 | ATP synthase subunit (ATPA2) | 19 | 1.18 | 0.268 | 1.000 | |
| ATMG01170.1 | ATP synthase subunit (ATPA2) | 20 | 1.37 | 0.248 | 1.000 | |
| ATMG01170.1 | ATP synthase subunit (ATPA2) | 21 | 0.96 | 0.240 | 1.000 | |
| ATMG01170.1 | ATP synthase subunit (ATPA2) | 22 | 0.61 | 0.236 | 1.000 | |
| ATMG01170.1 | ATP synthase subunit (ATPA2) | 23 | 0.64 | 0.234 | 1.000 | |

| | | | | | | |
|---|---|---|---|---|---|---|
| ATMG01170.1 | ATP synthase subunit (ATPA2) | 24 | 1.03 | 0.227 | 1.000 | |
| ATMG01170.1 | ATP synthase subunit (ATPA2) | 25 | 1.47 | 0.224 | 1.000 | |
| ATMG01170.1 | ATP synthase subunit (ATPA2) | 26 | 1.11 | 0.222 | 1.000 | |
| ATMG01170.1 | ATP synthase subunit (ATPA2) | 27 | 0.57 | 0.223 | 0.999 | |
| ATMG01170.1 | ATP synthase subunit (ATPA2) | 28 | 0.53 | 0.223 | 0.996 | |
| ATMG01170.1 | ATP synthase subunit (ATPA2) | 29 | 1.02 | 0.223 | 1.000 | |
| ATMG01170.1 | ATP synthase subunit (ATPA2) | 30 | 2.22 | 0.224 | 0.951 | |
| ATMG01170.1 | ATP synthase subunit (ATPA2) | 31 | 3.28 | 0.225 | 0.442 | |
| ATMG01170.1 | ATP synthase subunit (ATPA2) | 32 | 2.94 | 0.229 | 0.656 | |
| ATMG01170.1 | ATP synthase subunit (ATPA2) | 33 | 2.05 | 0.238 | 0.992 | |
| ATMG01170.1 | ATP synthase subunit (ATPA2) | 34 | 1.88 | 0.256 | 0.999 | |
| ATMG01170.1 | ATP synthase subunit (ATPA2) | 35 | 2.14 | 0.285 | 0.998 | |
| ATMG01170.1 | ATP synthase subunit (ATPA2) | 36 | 2.64 | 0.310 | 0.987 | |
| ATMG01170.1 | ATP synthase subunit (ATPA2) | 37 | 3.31 | 0.351 | 0.968 | |
| ATMG01170.1 | ATP synthase subunit (ATPA2) | 39 | 5.36 | 0.318 | 0.444 | |
| ATMG01170.1 | ATP synthase subunit (ATPA2) | 40 | 7.36 | 0.268 | 0.041 | * |
| ATMG01170.1 | ATP synthase subunit (ATPA2) | 41 | 9.16 | 0.251 | 0.005 | ** |
| ATMG01170.1 | ATP synthase subunit (ATPA2) | 42 | 7.49 | 0.335 | 0.232 | |
| ATMG01170.1 | ATP synthase subunit (ATPA2) | 43 | 4.24 | 0.498 | 0.995 | |
| ATMG01170.1 | ATP synthase subunit (ATPA2) | 46 | 0.20 | 1.231 | 1.000 | |
| ATMG01170.1 | ATP synthase subunit (ATPA2) | 47 | 0.06 | 1.604 | 1.000 | |
| ATMG01190.E | ATP synthase subunit (ATP1) | 3 | 0.90 | 0.167 | 1.000 | |
| ATMG01190.E | ATP synthase subunit (ATP1) | 4 | 0.98 | 0.160 | 1.000 | |
| ATMG01190.E | ATP synthase subunit (ATP1) | 5 | 1.18 | 0.160 | 1.000 | |
| ATMG01190.E | ATP synthase subunit (ATP1) | 6 | 1.33 | 0.160 | 1.000 | |
| ATMG01190.E | ATP synthase subunit (ATP1) | 7 | 1.33 | 0.161 | 1.000 | |
| ATMG01190.E | ATP synthase subunit (ATP1) | 8 | 1.38 | 0.158 | 1.000 | |
| ATMG01190.E | ATP synthase subunit (ATP1) | 9 | 1.48 | 0.156 | 0.999 | |
| ATMG01190.E | ATP synthase subunit (ATP1) | 10 | 1.49 | 0.156 | 0.999 | |
| ATMG01190.E | ATP synthase subunit (ATP1) | 11 | 1.55 | 0.156 | 0.997 | |
| ATMG01190.E | ATP synthase subunit (ATP1) | 12 | 1.73 | 0.157 | 0.956 | |
| ATMG01190.E | ATP synthase subunit (ATP1) | 13 | 1.92 | 0.155 | 0.800 | |
| ATMG01190.E | ATP synthase subunit (ATP1) | 14 | 2.05 | 0.153 | 0.639 | |
| ATMG01190.E | ATP synthase subunit (ATP1) | 15 | 1.96 | 0.153 | 0.740 | |
| ATMG01190.E | ATP synthase subunit (ATP1) | 16 | 1.75 | 0.152 | 0.927 | |
| ATMG01190.E | ATP synthase subunit (ATP1) | 17 | 1.60 | 0.152 | 0.989 | |
| ATMG01190.E | ATP synthase subunit (ATP1) | 18 | 1.45 | 0.154 | 1.000 | |
| ATMG01190.E | ATP synthase subunit (ATP1) | 19 | 1.46 | 0.157 | 1.000 | |
| ATMG01190.E | ATP synthase subunit (ATP1) | 20 | 1.54 | 0.160 | 0.998 | |
| ATMG01190.E | ATP synthase subunit (ATP1) | 21 | 1.40 | 0.162 | 1.000 | |
| ATMG01190.E | ATP synthase subunit (ATP1) | 22 | 1.14 | 0.162 | 1.000 | |
| ATMG01190.E | ATP synthase subunit (ATP1) | 23 | 1.00 | 0.162 | 1.000 | |
| ATMG01190.E | ATP synthase subunit (ATP1) | 24 | 1.16 | 0.159 | 1.000 | |
| ATMG01190.E | ATP synthase subunit (ATP1) | 25 | 1.53 | 0.156 | 0.998 | |
| ATMG01190.E | ATP synthase subunit (ATP1) | 26 | 1.53 | 0.156 | 0.998 | |
| ATMG01190.E | ATP synthase subunit (ATP1) | 27 | 0.97 | 0.156 | 1.000 | |
| ATMG01190.E | ATP synthase subunit (ATP1) | 28 | 0.75 | 0.155 | 1.000 | |
| ATMG01190.E | ATP synthase subunit (ATP1) | 29 | 1.12 | 0.155 | 1.000 | |
| ATMG01190.E | ATP synthase subunit (ATP1) | 30 | 2.15 | 0.155 | 0.546 | |
| ATMG01190.E | ATP synthase subunit (ATP1) | 31 | 2.80 | 0.156 | 0.118 | |
| ATMG01190.E | ATP synthase subunit (ATP1) | 32 | 2.09 | 0.156 | 0.623 | |
| ATMG01190.E | ATP synthase subunit (ATP1) | 33 | 1.42 | 0.156 | 1.000 | |
| ATMG01190.E | ATP synthase subunit (ATP1) | 34 | 1.29 | 0.156 | 1.000 | |
| ATMG01190.E | ATP synthase subunit (ATP1) | 35 | 1.44 | 0.156 | 1.000 | |
| ATMG01190.E | ATP synthase subunit (ATP1) | 36 | 1.85 | 0.155 | 0.868 | |
| ATMG01190.E | ATP synthase subunit (ATP1) | 37 | 2.44 | 0.155 | 0.292 | |
| ATMG01190.E | ATP synthase subunit (ATP1) | 38 | 3.06 | 0.156 | 0.061 | . |
| ATMG01190.E | ATP synthase subunit (ATP1) | 39 | 3.77 | 0.156 | 0.010 | * |
| ATMG01190.E | ATP synthase subunit (ATP1) | 40 | 4.47 | 0.157 | 0.002 | ** |
| ATMG01190.E | ATP synthase subunit (ATP1) | 41 | 4.48 | 0.157 | 0.002 | ** |
| ATMG01190.E | ATP synthase subunit (ATP1) | 42 | 3.69 | 0.159 | 0.013 | * |
| ATMG01190.E | ATP synthase subunit (ATP1) | 43 | 3.02 | 0.160 | 0.083 | . |
| ATMG01190.E | ATP synthase subunit (ATP1) | 44 | 2.24 | 0.160 | 0.498 | |

| | | | | | |
|---|---|---|---|---|---|
| ATMG01190.E | ATP synthase subunit (ATP1) | 45 | 1.49 | 0.160 | 1.000 |
| ATMG01190.E | ATP synthase subunit (ATP1) | 46 | 0.96 | 0.176 | 1.000 |
| ATMG01190.E | ATP synthase subunit (ATP1) | 47 | 0.64 | 0.301 | 1.000 |
| ATMG01190.E | ATP synthase subunit (ATP1) | 48 | 0.43 | 0.513 | 1.000 |
| ATMG01275.E | NADH:ubiquinone oxidoreductase subunit (NAD1) | 2 | 0.85 | 0.453 | 1.000 |
| ATMG01275.E | NADH:ubiquinone oxidoreductase subunit (NAD1) | 3 | 1.02 | 0.317 | 1.000 |
| ATMG01275.E | NADH:ubiquinone oxidoreductase subunit (NAD1) | 4 | 1.26 | 0.277 | 1.000 |
| ATMG01275.E | NADH:ubiquinone oxidoreductase subunit (NAD1) | 5 | 1.60 | 0.273 | 1.000 |
| ATMG01275.E | NADH:ubiquinone oxidoreductase subunit (NAD1) | 6 | 1.95 | 0.272 | 0.999 |
| ATMG01275.E | NADH:ubiquinone oxidoreductase subunit (NAD1) | 7 | 2.02 | 0.272 | 0.998 |
| ATMG01275.E | NADH:ubiquinone oxidoreductase subunit (NAD1) | 8 | 1.77 | 0.265 | 1.000 |
| ATMG01275.E | NADH:ubiquinone oxidoreductase subunit (NAD1) | 9 | 1.51 | 0.260 | 1.000 |
| ATMG01275.E | NADH:ubiquinone oxidoreductase subunit (NAD1) | 10 | 1.37 | 0.255 | 1.000 |
| ATMG01275.E | NADH:ubiquinone oxidoreductase subunit (NAD1) | 11 | 1.32 | 0.250 | 1.000 |
| ATMG01275.E | NADH:ubiquinone oxidoreductase subunit (NAD1) | 12 | 1.34 | 0.246 | 1.000 |
| ATMG01275.E | NADH:ubiquinone oxidoreductase subunit (NAD1) | 13 | 1.37 | 0.242 | 1.000 |
| ATMG01275.E | NADH:ubiquinone oxidoreductase subunit (NAD1) | 14 | 1.26 | 0.239 | 1.000 |
| ATMG01275.E | NADH:ubiquinone oxidoreductase subunit (NAD1) | 15 | 0.97 | 0.238 | 1.000 |
| ATMG01275.E | NADH:ubiquinone oxidoreductase subunit (NAD1) | 16 | 0.69 | 0.237 | 1.000 |
| ATMG01275.E | NADH:ubiquinone oxidoreductase subunit (NAD1) | 17 | 0.54 | 0.237 | 0.998 |
| ATMG01275.E | NADH:ubiquinone oxidoreductase subunit (NAD1) | 18 | 0.56 | 0.239 | 0.999 |
| ATMG01275.E | NADH:ubiquinone oxidoreductase subunit (NAD1) | 19 | 0.70 | 0.244 | 1.000 |
| ATMG01275.E | NADH:ubiquinone oxidoreductase subunit (NAD1) | 20 | 0.91 | 0.249 | 1.000 |
| ATMG01275.E | NADH:ubiquinone oxidoreductase subunit (NAD1) | 21 | 1.16 | 0.251 | 1.000 |
| ATMG01275.E | NADH:ubiquinone oxidoreductase subunit (NAD1) | 22 | 1.36 | 0.252 | 1.000 |
| ATMG01275.E | NADH:ubiquinone oxidoreductase subunit (NAD1) | 23 | 1.32 | 0.250 | 1.000 |
| ATMG01275.E | NADH:ubiquinone oxidoreductase subunit (NAD1) | 24 | 1.00 | 0.246 | 1.000 |
| ATMG01275.E | NADH:ubiquinone oxidoreductase subunit (NAD1) | 25 | 0.65 | 0.243 | 1.000 |
| ATMG01275.E | NADH:ubiquinone oxidoreductase subunit (NAD1) | 26 | 0.47 | 0.242 | 0.979 |
| ATMG01275.E | NADH:ubiquinone oxidoreductase subunit (NAD1) | 27 | 0.41 | 0.241 | 0.903 |
| ATMG01275.E | NADH:ubiquinone oxidoreductase subunit (NAD1) | 28 | 0.36 | 0.242 | 0.774 |
| ATMG01275.E | NADH:ubiquinone oxidoreductase subunit (NAD1) | 29 | 0.37 | 0.243 | 0.798 |
| ATMG01275.E | NADH:ubiquinone oxidoreductase subunit (NAD1) | 30 | 0.56 | 0.243 | 0.999 |

| | | | | | |
|---|---|---|---|---|---|
| ATMG01275.E | NADH:ubiquinone oxidoreductase subunit (NAD1) | 31 | 1.08 | 0.243 | 1.000 |
| ATMG01275.E | NADH:ubiquinone oxidoreductase subunit (NAD1) | 32 | 1.84 | 0.244 | 0.998 |
| ATMG01275.E | NADH:ubiquinone oxidoreductase subunit (NAD1) | 33 | 2.13 | 0.244 | 0.981 |
| ATMG01275.E | NADH:ubiquinone oxidoreductase subunit (NAD1) | 34 | 1.78 | 0.244 | 0.999 |
| ATMG01275.E | NADH:ubiquinone oxidoreductase subunit (NAD1) | 35 | 1.33 | 0.244 | 1.000 |
| ATMG01275.E | NADH:ubiquinone oxidoreductase subunit (NAD1) | 36 | 1.15 | 0.243 | 1.000 |
| ATMG01275.E | NADH:ubiquinone oxidoreductase subunit (NAD1) | 37 | 1.23 | 0.243 | 1.000 |
| ATMG01275.E | NADH:ubiquinone oxidoreductase subunit (NAD1) | 38 | 1.62 | 0.243 | 1.000 |
| ATMG01275.E | NADH:ubiquinone oxidoreductase subunit (NAD1) | 39 | 2.43 | 0.243 | 0.913 |
| ATMG01275.E | NADH:ubiquinone oxidoreductase subunit (NAD1) | 40 | 3.31 | 0.244 | 0.531 |
| ATMG01275.E | NADH:ubiquinone oxidoreductase subunit (NAD1) | 41 | 3.76 | 0.246 | 0.371 |
| ATMG01275.E | NADH:ubiquinone oxidoreductase subunit (NAD1) | 42 | 4.03 | 0.249 | 0.306 |
| ATMG01275.E | NADH:ubiquinone oxidoreductase subunit (NAD1) | 43 | 4.43 | 0.251 | 0.225 |
| ATMG01275.E | NADH:ubiquinone oxidoreductase subunit (NAD1) | 44 | 4.29 | 0.261 | 0.315 |
| ATMG01275.E | NADH:ubiquinone oxidoreductase subunit (NAD1) | 45 | 3.32 | 0.306 | 0.850 |
| ATMG01275.E | NADH:ubiquinone oxidoreductase subunit (NAD1) | 46 | 2.24 | 0.417 | 1.000 |
| ATMG01275.E | NADH:ubiquinone oxidoreductase subunit (NAD1) | 47 | 1.43 | 0.626 | 1.000 |
| ATMG01320.E | NADH:ubiquinone oxidoreductase subunit (NAD2) | 2 | 0.80 | 0.343 | 1.000 |
| ATMG01320.E | NADH:ubiquinone oxidoreductase subunit (NAD2) | 3 | 0.70 | 0.207 | 1.000 |
| ATMG01320.E | NADH:ubiquinone oxidoreductase subunit (NAD2) | 4 | 0.65 | 0.189 | 1.000 |
| ATMG01320.E | NADH:ubiquinone oxidoreductase subunit (NAD2) | 5 | 0.68 | 0.188 | 1.000 |
| ATMG01320.E | NADH:ubiquinone oxidoreductase subunit (NAD2) | 6 | 0.74 | 0.189 | 1.000 |
| ATMG01320.E | NADH:ubiquinone oxidoreductase subunit (NAD2) | 7 | 0.77 | 0.188 | 1.000 |
| ATMG01320.E | NADH:ubiquinone oxidoreductase subunit (NAD2) | 8 | 0.72 | 0.185 | 1.000 |
| ATMG01320.E | NADH:ubiquinone oxidoreductase subunit (NAD2) | 9 | 0.65 | 0.183 | 1.000 |
| ATMG01320.E | NADH:ubiquinone oxidoreductase subunit (NAD2) | 10 | 0.64 | 0.184 | 1.000 |
| ATMG01320.E | NADH:ubiquinone oxidoreductase subunit (NAD2) | 11 | 0.70 | 0.184 | 1.000 |
| ATMG01320.E | NADH:ubiquinone oxidoreductase subunit (NAD2) | 12 | 0.74 | 0.184 | 1.000 |
| ATMG01320.E | NADH:ubiquinone oxidoreductase subunit (NAD2) | 13 | 0.79 | 0.181 | 1.000 |
| ATMG01320.E | NADH:ubiquinone oxidoreductase subunit (NAD2) | 14 | 0.88 | 0.178 | 1.000 |
| ATMG01320.E | NADH:ubiquinone oxidoreductase subunit (NAD2) | 15 | 0.84 | 0.178 | 1.000 |

| | | | | | | |
|---|---|---|---|---|---|---|
| ATMG01320.E | NADH:ubiquinone oxidoreductase subunit (NAD2) | 16 | 0.69 | 0.177 | 1.000 | |
| ATMG01320.E | NADH:ubiquinone oxidoreductase subunit (NAD2) | 17 | 0.57 | 0.175 | 0.988 | |
| ATMG01320.E | NADH:ubiquinone oxidoreductase subunit (NAD2) | 18 | 0.58 | 0.178 | 0.995 | |
| ATMG01320.E | NADH:ubiquinone oxidoreductase subunit (NAD2) | 19 | 0.70 | 0.185 | 1.000 | |
| ATMG01320.E | NADH:ubiquinone oxidoreductase subunit (NAD2) | 20 | 0.84 | 0.191 | 1.000 | |
| ATMG01320.E | NADH:ubiquinone oxidoreductase subunit (NAD2) | 21 | 1.11 | 0.193 | 1.000 | |
| ATMG01320.E | NADH:ubiquinone oxidoreductase subunit (NAD2) | 22 | 1.58 | 0.193 | 1.000 | |
| ATMG01320.E | NADH:ubiquinone oxidoreductase subunit (NAD2) | 23 | 1.84 | 0.192 | 0.991 | |
| ATMG01320.E | NADH:ubiquinone oxidoreductase subunit (NAD2) | 24 | 1.31 | 0.187 | 1.000 | |
| ATMG01320.E | NADH:ubiquinone oxidoreductase subunit (NAD2) | 25 | 0.68 | 0.181 | 1.000 | |
| ATMG01320.E | NADH:ubiquinone oxidoreductase subunit (NAD2) | 26 | 0.50 | 0.180 | 0.922 | |
| ATMG01320.E | NADH:ubiquinone oxidoreductase subunit (NAD2) | 27 | 0.56 | 0.181 | 0.990 | |
| ATMG01320.E | NADH:ubiquinone oxidoreductase subunit (NAD2) | 28 | 0.72 | 0.182 | 1.000 | |
| ATMG01320.E | NADH:ubiquinone oxidoreductase subunit (NAD2) | 29 | 0.93 | 0.183 | 1.000 | |
| ATMG01320.E | NADH:ubiquinone oxidoreductase subunit (NAD2) | 30 | 1.24 | 0.182 | 1.000 | |
| ATMG01320.E | NADH:ubiquinone oxidoreductase subunit (NAD2) | 31 | 1.70 | 0.183 | 0.997 | |
| ATMG01320.E | NADH:ubiquinone oxidoreductase subunit (NAD2) | 32 | 1.75 | 0.184 | 0.995 | |
| ATMG01320.E | NADH:ubiquinone oxidoreductase subunit (NAD2) | 33 | 0.94 | 0.186 | 1.000 | |
| ATMG01320.E | NADH:ubiquinone oxidoreductase subunit (NAD2) | 34 | 0.37 | 0.194 | 0.528 | |
| ATMG01320.E | NADH:ubiquinone oxidoreductase subunit (NAD2) | 35 | 0.23 | 0.207 | 0.069 | . |
| ATMG01320.E | NADH:ubiquinone oxidoreductase subunit (NAD2) | 36 | 0.29 | 0.233 | 0.475 | |
| ATMG01320.E | NADH:ubiquinone oxidoreductase subunit (NAD2) | 37 | 0.57 | 0.272 | 1.000 | |
| ATMG01320.E | NADH:ubiquinone oxidoreductase subunit (NAD2) | 38 | 1.13 | 0.323 | 1.000 | |
| ATMG01320.E | NADH:ubiquinone oxidoreductase subunit (NAD2) | 41 | 3.80 | 0.302 | 0.788 | |
| ATMG01320.E | NADH:ubiquinone oxidoreductase subunit (NAD2) | 42 | 4.30 | 0.335 | 0.808 | |
| ATMG01320.E | NADH:ubiquinone oxidoreductase subunit (NAD2) | 43 | 4.28 | 0.404 | 0.962 | |
| ATMG01320.E | NADH:ubiquinone oxidoreductase subunit (NAD2) | 46 | 2.51 | 1.016 | 1.000 | |
| ATMG01320.E | NADH:ubiquinone oxidoreductase subunit (NAD2) | 47 | 1.94 | 1.308 | 1.000 | |
| ATMG01360.1 | Cytochrome c:O2 oxidoreductase subunit (COX1) | 2 | 1.11 | 0.455 | 1.000 | |
| ATMG01360.1 | Cytochrome c:O2 oxidoreductase subunit (COX1) | 3 | 1.23 | 0.293 | 1.000 | |
| ATMG01360.1 | Cytochrome c:O2 oxidoreductase subunit (COX1) | 4 | 1.37 | 0.291 | 1.000 | |

| | | | | | |
|---|---|---|---|---|---|
| ATMG01360.1 | Cytochrome c:O2 oxidoreductase subunit (COX1) | 5 | 1.42 | 0.299 | 1.000 |
| ATMG01360.1 | Cytochrome c:O2 oxidoreductase subunit (COX1) | 6 | 1.42 | 0.282 | 1.000 |
| ATMG01360.1 | Cytochrome c:O2 oxidoreductase subunit (COX1) | 7 | 1.36 | 0.267 | 1.000 |
| ATMG01360.1 | Cytochrome c:O2 oxidoreductase subunit (COX1) | 8 | 1.29 | 0.259 | 1.000 |
| ATMG01360.1 | Cytochrome c:O2 oxidoreductase subunit (COX1) | 9 | 1.26 | 0.255 | 1.000 |
| ATMG01360.1 | Cytochrome c:O2 oxidoreductase subunit (COX1) | 10 | 1.23 | 0.257 | 1.000 |
| ATMG01360.1 | Cytochrome c:O2 oxidoreductase subunit (COX1) | 11 | 1.16 | 0.257 | 1.000 |
| ATMG01360.1 | Cytochrome c:O2 oxidoreductase subunit (COX1) | 12 | 1.15 | 0.258 | 1.000 |
| ATMG01360.1 | Cytochrome c:O2 oxidoreductase subunit (COX1) | 13 | 1.10 | 0.255 | 1.000 |
| ATMG01360.1 | Cytochrome c:O2 oxidoreductase subunit (COX1) | 14 | 0.89 | 0.252 | 1.000 |
| ATMG01360.1 | Cytochrome c:O2 oxidoreductase subunit (COX1) | 15 | 0.73 | 0.253 | 1.000 |
| ATMG01360.1 | Cytochrome c:O2 oxidoreductase subunit (COX1) | 16 | 0.65 | 0.254 | 1.000 |
| ATMG01360.1 | Cytochrome c:O2 oxidoreductase subunit (COX1) | 17 | 0.55 | 0.252 | 1.000 |
| ATMG01360.1 | Cytochrome c:O2 oxidoreductase subunit (COX1) | 18 | 0.51 | 0.253 | 0.998 |
| ATMG01360.1 | Cytochrome c:O2 oxidoreductase subunit (COX1) | 19 | 0.49 | 0.264 | 0.998 |
| ATMG01360.1 | Cytochrome c:O2 oxidoreductase subunit (COX1) | 20 | 0.61 | 0.272 | 1.000 |
| ATMG01360.1 | Cytochrome c:O2 oxidoreductase subunit (COX1) | 21 | 0.82 | 0.281 | 1.000 |
| ATMG01360.1 | Cytochrome c:O2 oxidoreductase subunit (COX1) | 22 | 0.93 | 0.313 | 1.000 |
| ATMG01360.1 | Cytochrome c:O2 oxidoreductase subunit (COX1) | 23 | 0.84 | 0.362 | 1.000 |
| ATMG01360.1 | Cytochrome c:O2 oxidoreductase subunit (COX1) | 25 | 0.57 | 0.337 | 1.000 |
| ATMG01360.1 | Cytochrome c:O2 oxidoreductase subunit (COX1) | 26 | 0.55 | 0.290 | 1.000 |
| ATMG01360.1 | Cytochrome c:O2 oxidoreductase subunit (COX1) | 27 | 0.63 | 0.264 | 1.000 |
| ATMG01360.1 | Cytochrome c:O2 oxidoreductase subunit (COX1) | 28 | 0.84 | 0.254 | 1.000 |
| ATMG01360.1 | Cytochrome c:O2 oxidoreductase subunit (COX1) | 29 | 1.33 | 0.254 | 1.000 |
| ATMG01360.1 | Cytochrome c:O2 oxidoreductase subunit (COX1) | 30 | 2.02 | 0.254 | 0.996 |
| ATMG01360.1 | Cytochrome c:O2 oxidoreductase subunit (COX1) | 31 | 2.37 | 0.255 | 0.965 |
| ATMG01360.1 | Cytochrome c:O2 oxidoreductase subunit (COX1) | 32 | 1.95 | 0.256 | 0.998 |
| ATMG01360.1 | Cytochrome c:O2 oxidoreductase subunit (COX1) | 33 | 0.85 | 0.257 | 1.000 |
| ATMG01360.1 | Cytochrome c:O2 oxidoreductase subunit (COX1) | 34 | 0.28 | 0.261 | 0.579 |
| ATMG01360.1 | Cytochrome c:O2 oxidoreductase subunit (COX1) | 35 | 0.17 | 0.266 | 0.103 |
| ATMG01360.1 | Cytochrome c:O2 oxidoreductase subunit (COX1) | 36 | 0.32 | 0.281 | 0.845 |

| | | | | | |
|---|---|---|---|---|---|
| ATMG01360.1 | Cytochrome c:O2 oxidoreductase subunit (COX1) | 37 | 1.12 | 0.354 | 1.000 |
| ATMG01360.1 | Cytochrome c:O2 oxidoreductase subunit (COX1) | 38 | 4.07 | 0.554 | 0.999 |
| ATMG01360.1 | Cytochrome c:O2 oxidoreductase subunit (COX1) | 46 | 8784.92 | 3.902 | 1.000 |
| ATMG01360.1 | Cytochrome c:O2 oxidoreductase subunit (COX1) | 47 | 19519.12 | 4.516 | 1.000 |
